# Somatic and germline mutational processes across the tree of life

**DOI:** 10.64898/2025.12.28.691837

**Authors:** Sangjin Lee, Yichen Wang, Haynes Heaton, Emily Mitchell, Mark Maddison, Liam M. Crowley, Patrick Adkins, Nova Mieszkowska, Mark Blaxter, Darwin Tree of Life consortium, Richard Durbin, Peter J. Campbell

**Affiliations:** Wellcome Sanger Institute, Cancer Ageing and Somatic Mutations, Hinxton, Cambridgeshire, CB10 1SA, United Kingdom; Auburn University, Computer Science and Software Engineering, 345 W Magnolia Ave, Auburn, AL 36849; Department of Biology, University of Oxford, Oxford, OX1 3SZ, UK; Thomson Environmental Consultants, North Yorkshire, HG1 1HQ, United Kingdom; Marine Biological Association of the United Kingdom, Plymouth PL1 2PB, UK; University of Liverpool, Liverpool L69 3BX, UK; Wellcome Sanger Institute, Darwin Tree of Life, Hinxton, Cambridgeshire, CB10 1SA, United Kingdom; Department of Genetics, University of Cambridge, Downing Street, Cambridge, Cambridgeshire, CB2 3EH, United Kingdom; Wellcome-MRC Cambridge Stem Cell Institute, Cambridge Biomedical Campus, Cambridge, UK

**Author notes:** These authors contributed equally: Sangjin Lee, Yichen Wang. These authors jointly supervised this work: Richard Durbin and Peter J. Campbell.

## Abstract

The characterisation of mutational processes active in somatic and germline cells *in vivo* has predominantly focused on human cancers^1,2^, normal tissues^3–5^ and model organisms^6–8^. Beyond mammals^9^, little is known about mutational processes across the tree of life. We developed himut (high-fidelity mutation), an algorithm to identify somatic mutations from circular consensus long-read sequencing data, deploying it on 708 samples from 661 species from Britain and Ireland^10^. The spectra of somatic mutations, categorised by mutation type and local sequence context, showed considerable between-species divergence but within-species similarity. From normalised spectra across the dataset, we extracted 95 distinct patterns, or ‘signatures’, of somatic mutations, together with 18 signatures of germline mutational processes. Only two of the somatic signatures resembled mutational signatures extracted from human cancers^1,2^. Of the somatic signatures, 22 had significant clustering within the taxonomic classification, with some distributed across an entire kingdom or phylum, while others were restricted to a single family or species. Three somatic signatures were found only in water-dwelling species, with one distributed across multiple clades that might suggest it derives from a water-borne mutagen. Of germline signatures, eight were found in multiple species, showed significant clustering by taxonomic classification and had counterpart somatic signatures with matching mutational spectrum and species distribution. Thus, there is considerably greater diversity of mutational processes across the tree of life than found in human samples, likely reflecting either cell-intrinsic biological processes or environmental exposures (or both) operative within distinct clades.

## INTRODUCTION

*De novo* mutations result from the interplay of DNA damage with failed or incorrect DNA repair and error-prone DNA replication. DNA lesions, arising from endogenous^11,12^ or exogenous^13,14^ sources, vary in half-life, target nucleotide, local sequence context, influence of chromatin packaging and preferred pathways for repair. This variability leaves characteristic mutational ‘signatures’ in the genome^15^, which can be described as vectors of 96 numbers, reflecting the six different classes of base substitution and 16 combinations of the bases immediately 5’ and 3’ to the mutation. In normal human germ cells and somatic cells, most *de novo* mutations result from the activity of endogenous mutational processes, acting at constant rates throughout life^3–5^, with DNA repair defects and exposure to exogenous mutagens generating additional mutations in some tissues^5,16–19^. Across 16 mammalian species, somatic mutations in normal colon epithelium were accounted for by just three mutational signatures^9^ – it remains unknown whether this striking homogeneity of signatures in mammals extends to all organisms. On the one hand, most mutations derive from fundamental chemical reactions generating spontaneous lesions in DNA^11^, the universality of which would argue for conservation of signatures across the tree of life; on the other hand, the huge diversity of life cycles, environments and niches among species might introduce considerable variability in mutagen exposures and mutational processes.

The Earth BioGenome Project aims to generate reference genomes for all species on Earth^20^. Contributing to this effort, the goal of the Darwin Tree of Life (DToL) project is to sequence and assemble the genomes of the 70,000 eukaryotic species found in Britain and Ireland^10^. DToL has produced high-quality reference genomes for over 2,000 species to date. Many of these reference genomes were assembled from long-read sequencing on the PacBio platform, with high accuracy base calls derived through circular consensus sequencing (CCS)^21^. Short-read sequencing of both strands of the same DNA molecule via duplex sequencing enables accurate identification of somatic mutations^22–24^ – we reasoned that this concept could be extended to CCS sequences in which both strands of a native, unamplified DNA molecule are repeatedly sequenced. This would enable us to study germline and somatic mutational processes across the tree of life.

## RESULTS

### Single-molecule somatic mutation detection

We developed an algorithm, himut, to call somatic mutations in CCS sequencing data of single molecules of dsDNA from bulk tissue samples, agnostic of clonality or species (**Figure 1a**; **Extended Data Fig. 1, Methods**). The key features of the algorithm are: first, identification of non-chimeric primary reads with unambiguous alignment to the reference genome; second, detection of single-base substitution candidates from high-quality mismatches to the reference genome seen in single reads; third, optionally phasing to nearby heterozygous germline polymorphisms; and, fourth, filtering to remove false positives resulting from alignment and sequencing errors. Where available, a panel of normal samples can aid removal of false positives arising from recurrent sequencing artefacts.

**Fig. 1.**
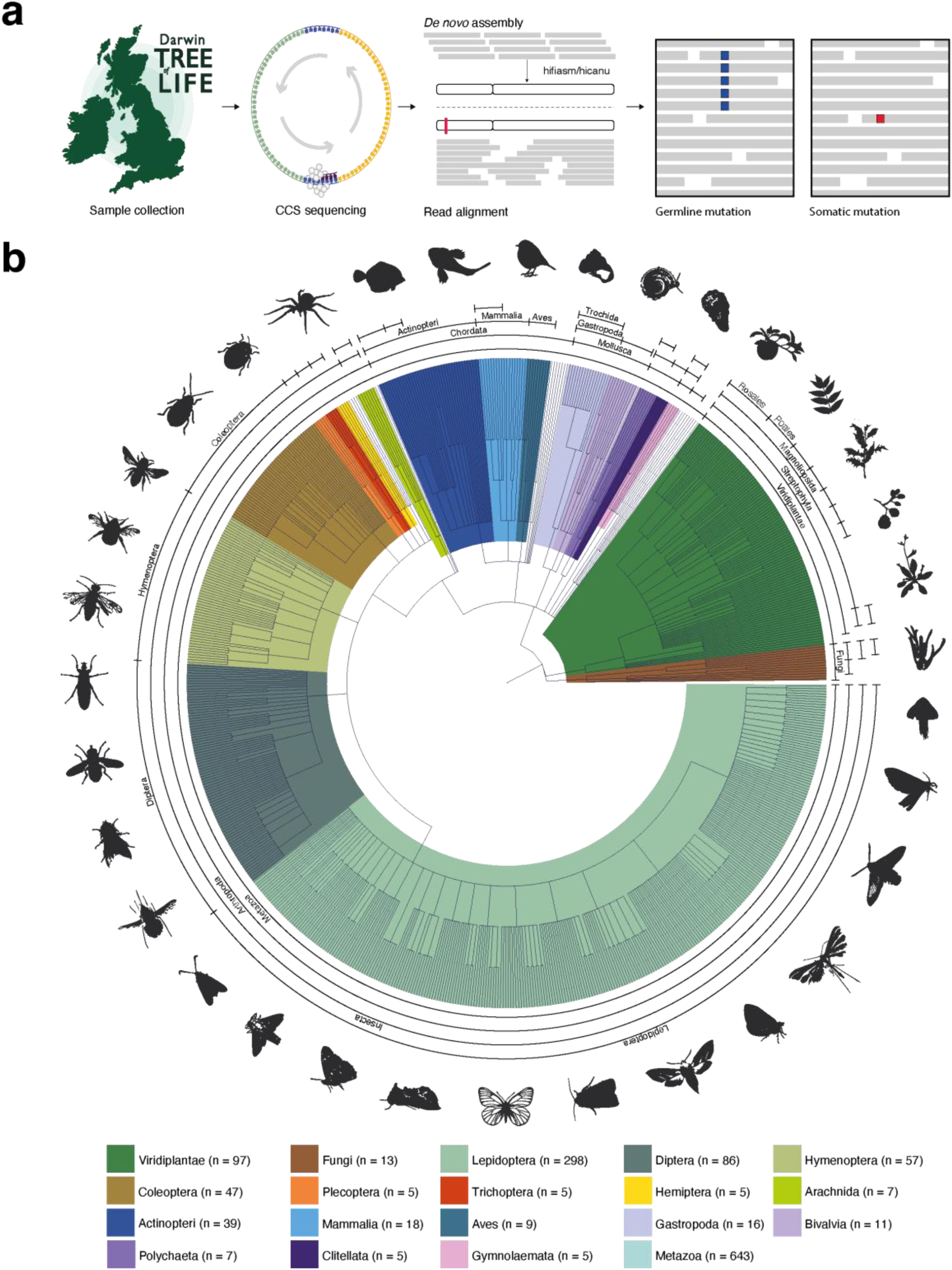
Experimental Design. **a**, Sample collection, preparation and processing. The DToL consortium acquired samples, performed library preparation and CCS sequencing, and de novo assembled high-quality reference genomes. CCS reads were aligned to publicly available reference genomes, and alignments were processed using deepvariant and himut to call germline and somatic mutations, respectively. Blue and red squares indicate a germline and a somatic mutation, respectively. **b**, Taxonomic tree constructed from the NCBI TaxonomyDB classification of samples with complete taxonomic information, with branch shading of major clades (also used across other figures). Labels follow the clockwise order in which the taxonomic classification appears in the tree. The Metazoa label is not used in this figure but serves as a reference in subsequent figures.

As positive controls to validate the algorithm, we performed CCS sequencing on two human cancer cell lines, BC-1 and HT-115, known to have characteristic mutational processes that remain active in culture^25^ and peripheral blood granulocytes from an 82-year-old woman^26^ (**Extended Data Fig. 2a-b**; **Supplementary Tables 1-3**). The BC-1 cell line exhibits on-going APOBEC-mediated mutagenesis *in vitro*^25^, and the somatic mutations we called from single molecules of DNA in this culture showed the expected pattern of C>T mutations in a TpC context (cosine similarity with previous BC-1 spectrum^25^, 0.971). The HT-115 cell line has a mutation in the proof-reading domain of POLE that generates a distinctive spectrum of C>A and T>G mutations^25^, which again was faithfully recaptured from the long-read sequencing (cosine similarity with HT-115 spectrum^25^, 0.988). Finally, for normal granulocytes, the expected spectrum of somatic mutations shows a flatter pattern with transitions dominating transversions^26,27^, and this is what was extracted from the long-read data (cosine similarity with spectrum from short-read sequencing of monoclonal blood colonies^26^, 0.946).

As a negative control, we sequenced umbilical cord blood granulocytes, which have only ∼50-60 somatic mutations/cell (6×10^9^ base-pairs)^26,27^. Here, the discordance between the expected mutational spectrum and that generated from long-read sequencing illustrates both the limits of resolution and residual error mode for single-molecule mutation calling. We estimated the true error rate of high-quality base calls to range between 1/10^6^ (phred quality 60) and 1/10^9^ calls (phred quality 90), depending on the substitution and trinucleotide sequence context (**Extended Data Fig. 3**; **Supplementary Table 3**).

Overall, these data demonstrate that error rates of circular consensus sequences from single molecules of dsDNA are sufficiently low and predictable to permit accurate identification of somatic mutation signatures when present at the levels seen in, for example, adult human blood.

### Single-molecule somatic mutation detection in non-mammalian samples

We selected 708 CCS datasets from individuals representing 661 species that had more than 20x mean coverage. Most sequenced individuals were presumed to be diploid, aside from a few exceptions (Methods). From these, we called germline heterozygous single nucleotide polymorphisms in the nuclear genome to study germline mutational spectra and, using himut, called somatic mutations (**Figure 1b**; **Supplementary Figs. 1-3**; **Supplementary Tables 4-7, Supplementary Information**). For 14 non-mammalian species, we had sequencing data from two or more individuals, enabling us to compare somatic mutational spectra within and between species **(Supplementary Table 8)**. While most of these biological replicates were obtained before the development of the himut algorithm, an early version of the algorithm had identified distinctive somatic mutation patterns in single samples from *Syritta pipiens* (thick-legged hoverfly), *Vespula vulgaris* (common wasp) and *Platycheirus albimanus* (white-footed hoverfly). To demonstrate the reproducibility of these patterns, we acquired and sequenced further individuals from these species (for totals of 9, 4 and 3 individuals respectively).

The raw spectrum of somatic mutations, comprising the number of called variants in each of the 96 categories, was generally concordant among the replicates within a species (**Figure 2a-c**; **Extended Data Fig. 2c-d**). In contrast, there was much greater variation between species, and none of these spectra (mostly from insects) were consistent with the signatures previously observed in mammals. These observations provide reassurance that the algorithm is identifying robust, reproducible patterns of somatic mutation, especially given that the repeated individuals within a species were sequenced in different batches at different stages of the project.

**Fig. 2.**
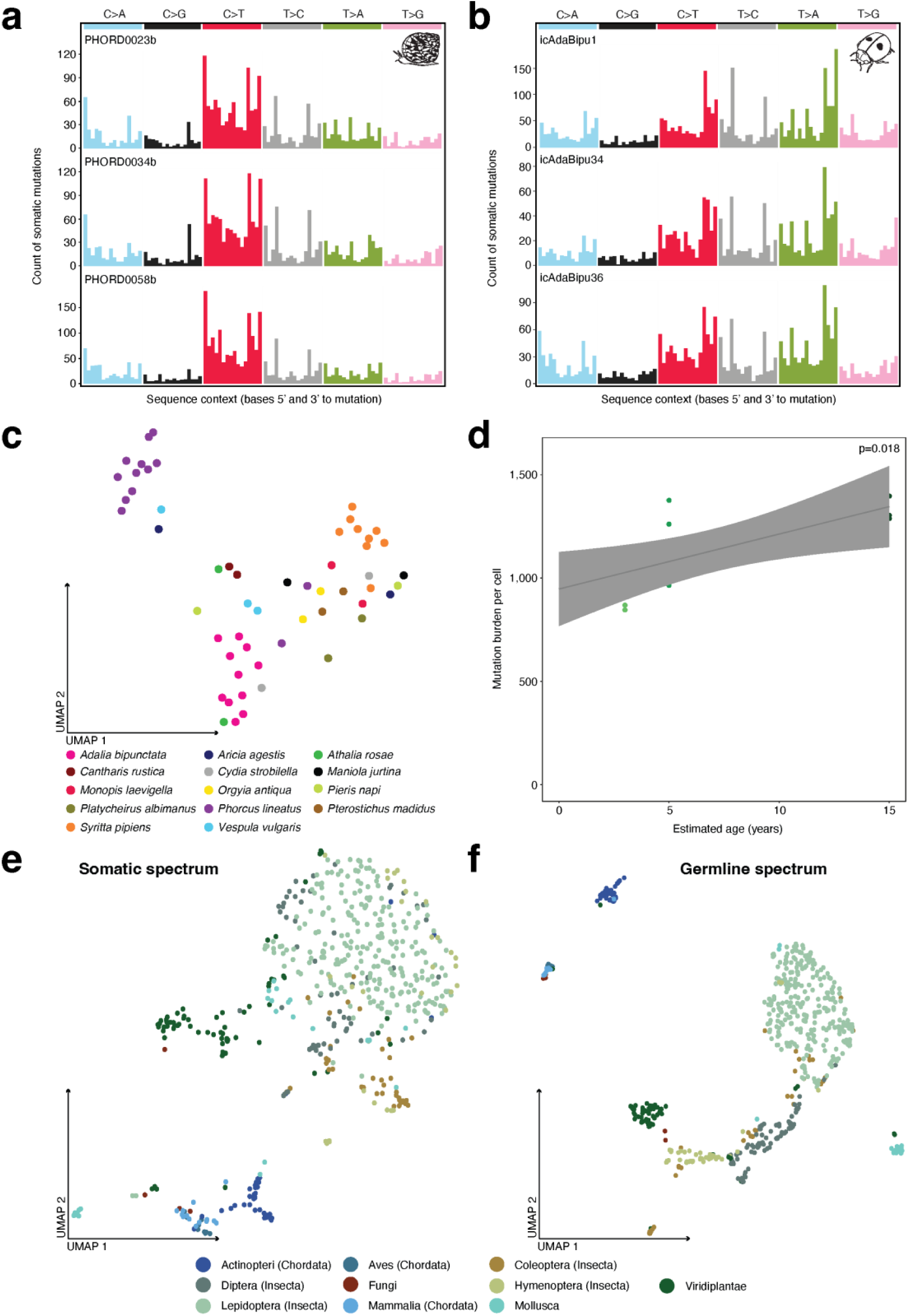
Robust and reproducible somatic mutational spectra across a phylogenetically diverse set of invertebrates species. **a-b**, Somatic mutational spectra for multiple individuals from two species: **a**, *Phorcus lineatus* (thick topshell); **b**, *Adalia bipunctata* (2-spot ladybird); **c**, Scatter plot showing the relationship between estimated age and mutation burden in *Phorcus lineatus*. The grey line represents a regression of age with mutation burden, with shading indicating the 95% confidence interval. **d**, UMAP projection of somatic mutational spectra from species with biological replicates. **e**, UMAP projection of somatic mutational spectra across 562 samples with >1,000 somatic mutations, excluding those with high artefactual signature contributions identified through manual inspection. **f,** UMAP projection of germline mutational spectra across 627 samples from lineages with adequate sample representation.

*Phorcus lineatus* (thick topshell; Mollusca) can be classified by age based on the maturity of its shell^28^, so we sequenced 10 individuals estimated to be 3, 5 or 15 years old to assess correlation of mutation burden with age (**Supplementary Tables 8-9**). We found that somatic mutation burden significantly increased with age at an estimated rate of 67 mutations/cell/year (9.58 x10^8^ base-pairs, CI_95%_=12-121; p=0.02; linear regression; **Figure 2d**). Notwithstanding inter-individual variability in mutation burden, this increase appeared linear with time, suggesting a constant mutation rate across life in these molluscs, as observed in normal cells from humans and other mammals^3,9,29–31^. Interestingly, there is a substantial extrapolated offset in the linear fit at age zero, which may reflect a combination of residual false positive errors and changes in exposure during life (topshell larvae go through an unsampled planktonic stage).

We then extended the analysis of mutational spectra in repeated samples from the same species to all 708 individuals in the dataset. Both somatic (**Figure 2e**) and germline (**Figure 2f**) spectra showed clustering by higher levels of the taxonomic spectrum. Both the clustering of mutational spectra by taxonomic classification and the correlation of inferred age with mutation burden in *Phorcus lineatus* imply that informative mutational signatures can be identified across the tree of life using the approach described here.

### Somatic mutation signatures across the Tree of Life

In the analysis of mutation patterns in human cancers, the overall spectrum of somatic mutations can be expressed as the summation of a small number of component ‘mutational signatures’, with each tumour combining these signatures in different proportions^15^. We adapted methods^32^ based on Bayesian hierarchical Dirichlet processes to extract mutational signatures from the DToL samples (**Methods**). Unlike human cancers, where the reference genome is equivalent in all patients, genomes across different species show markedly different trinucleotide frequencies (**Extended Data Fig. 4**) – this has the consequence that even if an identical mutational process was active in two species, the observed spectrum of mutations could vary in the relative proportions of the 96 mutation categories. We therefore normalised the counts in each of the 96 mutation classes by the number of callable bases in each reference genome prior to extracting signatures.

Applying this analysis to the 708 samples, we obtained 95 somatic mutational signatures (sToL1-95; **Supplementary Figures 4-98**; **Supplementary Tables 10, 11 and 13**), after exclusion of three artefact signatures (**Supplementary Figures 99-101, Supplementary Tables 12-13**). Only two sToL signatures showed any resemblance to the mutational signatures documented in human cancers (**Extended Data Fig. 5**). The signature sToL1 was distributed widely over species sampled in the plant and animal kingdoms, where it typically accounted for 4-79% of somatic mutations, but was much less prevalent in fungi (**Extended Data Fig. 6**). This signature was characterised by transitions (T>C and C>T) predominating over transversions (C>A, C>G, T>A and T>G), with minimal influence of local sequence context, a similar pattern to the SBS5 signature that is seen broadly across normal human tissues^3–5^, human cancers^1,2^ and other mammals^9^. The ubiquity of this signature across plants and animals likely reflects damage to DNA accrued during eukaryotic life from universal metabolic processes^11^. Likewise, sToL4, characterised by C>T mutations in a CpG context, closely resembles the SBS1 signature in human tissues, caused by spontaneous deamination of methylated cytosine to thymine^33^.

At the other end of the prevalence spectrum, 54 of the 95 somatic signatures were identified based on their presence in only one sample. While it is difficult to interpret these signatures in the absence of confirmatory data, many have distinctive spectra that diverge from the typical patterns of sequencing errors. As discussed in the previous section, we were able to reproduce one-off signatures from an earlier run of the algorithm in 3 species by sequencing more members of those species (**Figure 2a**; **Extended Data Fig. 2c-d**) – this suggests that many of the single-sample signatures reported here contain plausible biological signal.

### Taxonomic clustering of somatic mutational signatures

Many somatic mutational signatures were seen patchily across species, present in some but absent from others. One potential route to understanding their relevance is to identify those signatures that show clustering by taxonomy, since signatures co-occurring in closely related species may arise through shared cellular processes or environmental niches. We used Abouheif’s *C_mean_* statistic for this purpose^34,35^. This metric scores how correlated a quantitative measure is with the structure of a tree but not its branch lengths (since these are unknown here). It ranges from +1.0 (maximal positive correlation) through 0 (uncorrelated) to -1.0 (maximal inverse correlation).

After correction for multiple hypothesis testing, 22 somatic signatures showed significant clustering on the taxonomy (q<0.01; **Extended Data Fig. 7-8, Supplementary Table 14, Methods**). Intriguingly, this clustering occurred at all levels of the taxonomic classification, with some signatures prominent across an entire kingdom or phylum, while others clustered in specific families or genera (**Figure 3**; **Extended Data Fig. 9**). For example, three different signatures with a strong cytosine mutation component in different sequence contexts showed distinctive phylogenetic patterns: sToL5, characterised by C>A mutations especially in an R<u>C</u>R context, was widely distributed across the Viridiplantae (plant kingdom), but seen only sparsely in the animal kingdom; sToL10 with C>T mutations in CHG and CHH contexts was also largely restricted to plants (where H represents A, C or G); and sToL4, the C>T at NCG signature of methyl-cytosine deamination, was most pronounced in the Chordata phylum (vertebrates), while also present in some annelids, fungi and plants.

**Fig. 3.**
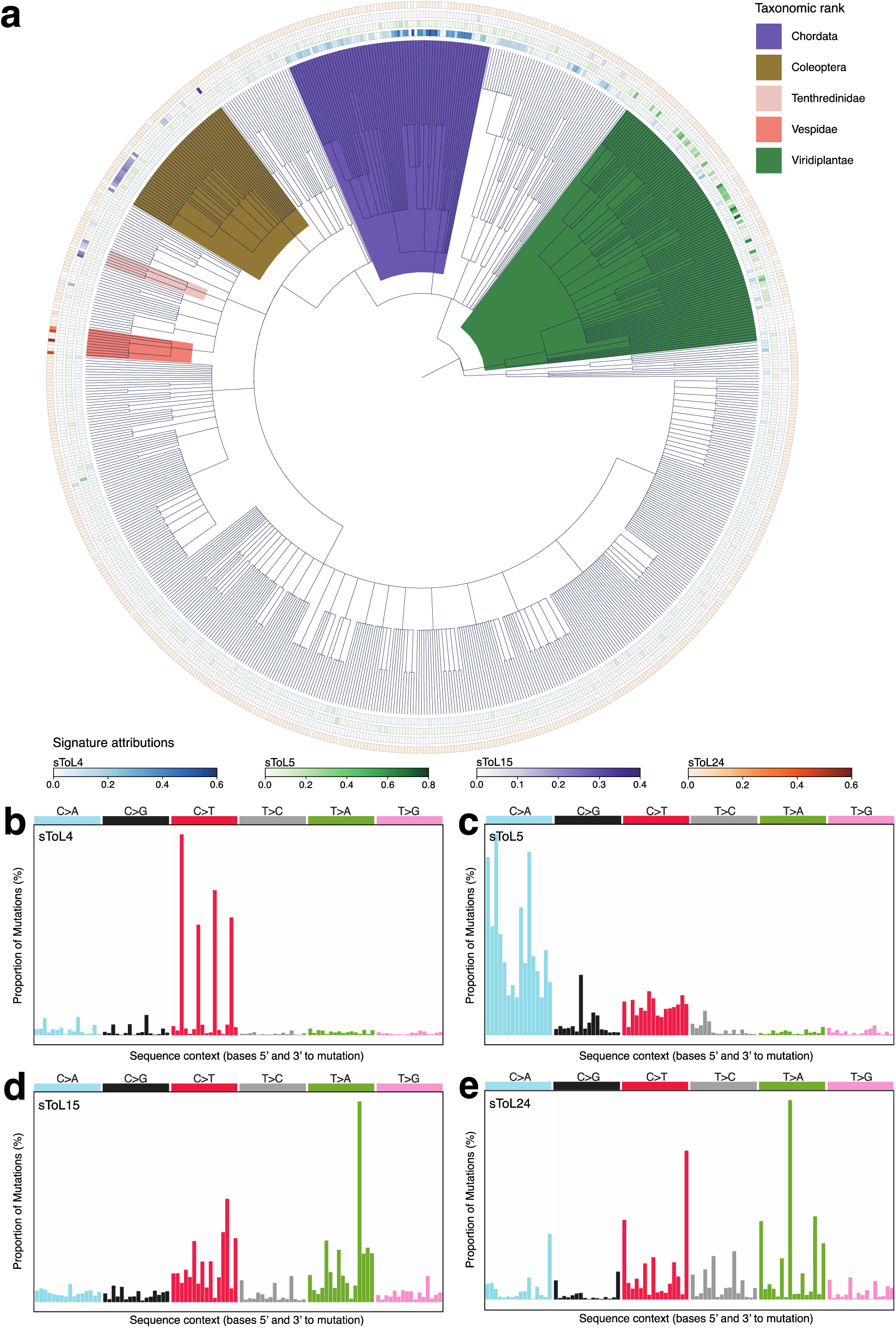
Somatic mutational processes detected across the tree of life. **a**, Taxonomic tree with branches shaded by relevant clade, and sample-level mutational signature attributions displayed at the tips of the tree. **b-e**, somatic mutational signatures: **b**, sToL4 signature, **c**, sToL5 signature, **d**, sToL15 signature, and **e**, sToL24 signature.

Some signatures had more restricted taxonomic distributions (**Figure 3**; **Extended Data Fig. 9**). To provide a few examples: sToL8 marked the molluscan class Gastropoda (snails and slugs); sToL15 was found predominantly in the arthropod order Coleoptera (beetles); sToL24 was restricted to the arthropod family Vespidae (wasps); and sToL41 was found only in the species *Platycheirus albimanus* (white-footed hoverfly). The spectra for these more restricted signatures show no clear resemblance to any of the known mutational signatures in human cancers, and it is unclear whether they arise from exogenous mutagens in shared environments or clade-specific biological processes.

### Shared somatic signatures in aquatic animals

Three somatic signatures were identified in multiple water-dwelling species (**Figure 4**). Of particular interest was sToL9, characterised by C>T and C>A mutations especially in a <u>C</u>pS context – this signature showed strongly significant clustering across the taxonomic tree (C_mean_=0.51; q<0.0001), but not in a single clade. Instead, we observed this signature in a variety of aquatic species, including molluscs, where it accounted for 6-40% of mutations identified; species from the phylum Echinodermata (sea urchins and starfish), comprising 6-25% of mutations; species from the order Actiniaria (sea anemones), ranging from 7-17% of mutations; and sporadically among the Actinopteri (fish), where 4-15% of mutations were attributed in positive samples. The distribution of sToL9 across different phyla suggests that it is most likely to be caused by a water-borne mutagen.

**Fig. 4.**
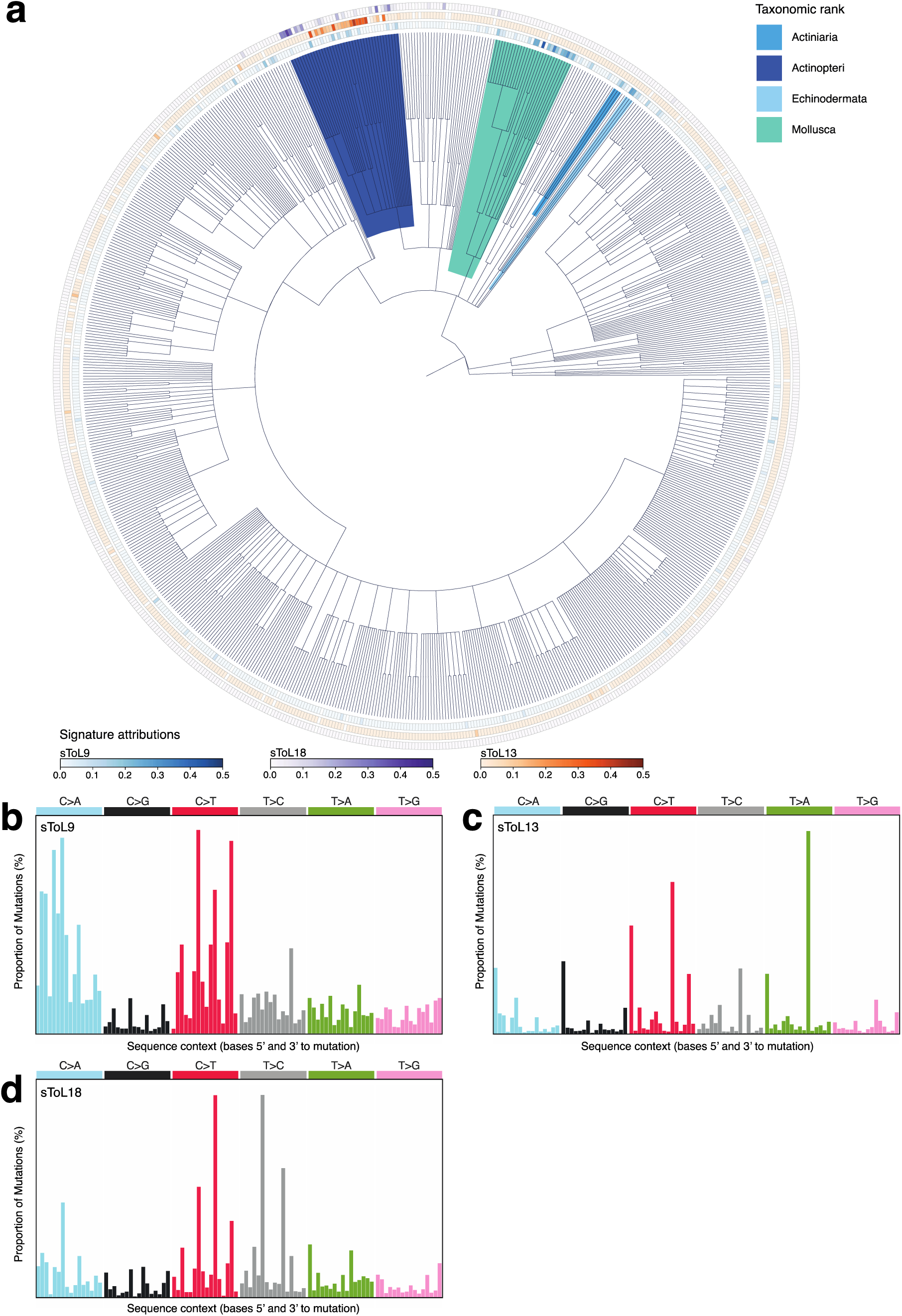
Somatic mutational processes shared across aquatic animals. **a**, Taxonomic tree shaded according to taxonomic classification, with sample-level mutational signature exposures annotated at the tips of the tree. **b-d**, somatic mutational signatures: **b**, sToL9 signature, **c**, sToL13 signature, and **d**, sToL 18 signature.

Fish species were notable for two other mutational signatures, sToL13 and sToL18, that were barely evident elsewhere on the taxonomy. sToL13 had a spiky spectrum, with cytosine mutations in an A<u>CA</u> context and T>G mutations in an G<u>T</u>G context – this was distributed widely across Actinopteri, where it accounted for 10-36% of somatic mutations in positive samples. Interestingly, sToL18 was anticorrelated with sToL13 among Actinopteri, and was especially prevalent in the Cypriniformes order (carp and minnows), species that are mostly freshwater in habitat, although it was also observed in some saltwater species too.

### Germline mutational signatures

We also identified germline variants as heterozygous variants in the diploid species within the sample set (**Methods**). Unlike somatic variants, where the direction of the mutation is known, we do not know which of the two alleles for a given heterozygous germline variant is ancestral and which is derived. This means that the 96 categories for somatic mutations collapse into 52 categories for germline variants (**Supplementary Table 15**).

Signature extraction identified 18 germline mutational signatures, only 8 of which were seen in more than 2 samples (gToL1-gToL18, **Figure 5**; **Extended Data Fig. 10**; **Supplementary Figs. 102-119**; **Supplementary Tables 16-18**). All 8 of these showed significant clustering by taxonomic classification (Abouheif’s C_mean_ range, 0.40-0.87; q<0.0001 for all; **Extended Figure 7b**; **Supplementary Table 19**). The most widespread signature, gToL1, followed a flat distribution where transitions (C<>T) predominated over transversions, but with little variation by sequence context – this signature accounted for a large fraction of polymorphisms across all clades studied, with the exception of fungi. The gToL3 signature, denoting the methylated CpG deamination signature of C/T polymorphisms in a CpG context, was largely restricted to Chordata. Viridiplantae showed a high rate of gToL4, which comprised C/T polymorphisms with some enrichment for the CpG context.

**Fig. 5.**
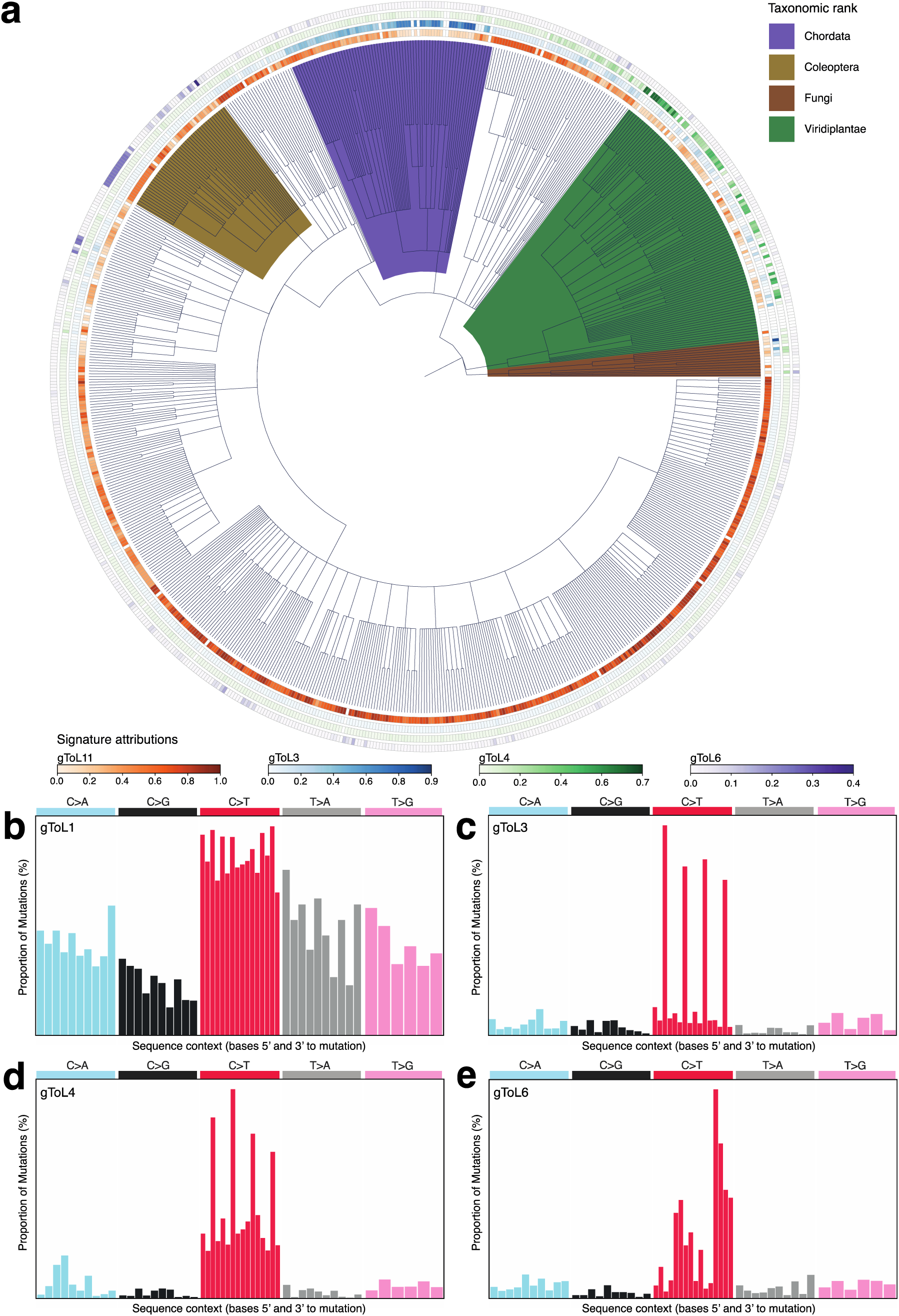
Germline mutational processes detected across the tree of life. **a**, Taxonomic tree coloured by taxonomic classification and annotated at the tips with sample-specific mutational signature attributions. **b-e**: germline mutational signatures, **b**, gToL1 signature, **c**, gToL3 signature, **d**, gToL4 signature, **e**, gToL6 signature.

We compared the 18 germline signatures to the 95 somatic signatures we extracted, both in terms of their spectrum (after collapsing the somatic signatures to the 52-channel spectrum) and their co-occurrence in the same species (**Extended Data Fig. 11**). With the exception of gToL5, the most widespread germline signatures generally each had a match in the somatic signatures. The most ubiquitous signatures among somatic and germline mutations, sToL1 and gToL1 respectively, overlapped in spectrum and species distribution, consistent with them capturing universal mutational processes. Of the more restricted signatures, sToL4 and gToL3 represented the methylated CpG deamination signature seen in vertebrates; plants showed high frequencies of sToL10 and gToL4, comprising mostly C>T mutations, consistent with the broader patterns of cytosine methylation in NCG, CHH and CHG sequence contexts in plants (where H can be A, C or G)^36^; and there were matching spectra for signatures seen in the soma and germline in Coleoptera (sToL15 and gToL6) and wasps (sToL24 and gToL8). The overall excess of somatic to germline signatures mirrors observations in humans where the repertoire of germline mutational processes represents only a subset of the processes observed in the soma, in keeping with strong evolutionary pressure to constrain new variants entering the inherited genome.

## Discussion

We identified a wide spectrum of mutational signatures in the soma and germlines of species across Metazoa, Fungi and Viridiplantae. It is striking how many of the somatic mutational signatures cluster at different levels of the Linnaean taxonomic classification. Signatures operative in entire kingdoms and phyla likely represent consequences of endogenous biological processes given the diversity of environments occupied at this level of taxonomy. The well-understood mutational process of deamination of methylated cytosines, sToL4 and gToL3 here (SBS1 in COSMIC), exemplifies this, seen across the whole phylum of Chordata. The two signatures present throughout the plant kingdom, sToL5 and sToL10, may perhaps be a consequence of life as a photosynthesising organism. Clustering of signatures was also abundant at more restricted levels of the taxonomy – this might result from exposure to a given mutagen in a family, genus or species that occupies a particular environmental niche or it may result from some quirk of cell-intrinsic metabolism and DNA repair in that clade.

Both marine and freshwater habitats suffer extensive pollution in the modern world. Our observation of somatic mutational signatures in aquatic species is therefore particularly interesting. While it is possible that these signatures may be the consequence of some endogenous adaptation to life in water, their distribution across taxa as diverse as fish, molluscs, sea anemones and starfish suggests that they are more likely to result from exposure to a water-borne mutagen. This exemplifies the exciting opportunity to use measurement of somatic mutations to monitor the health of ecosystems – for example, deploying these methods in environments affected by known chemical spills or unexplained die-offs of resident species could identify the presence of DNA-damaging chemicals in that ecosystem. Monitoring somatic mutation patterns in sentinel species that bioaccumulate chemicals in the environment would be a logical experimental design for this purpose.

Leveraging high-fidelity long-read sequencing, we have glimpsed the heterogeneity of mutational processes operative across a large cross-section of the tree of life, revealing a much richer landscape than that previously studied in humans. If this represents the diversity present in species resident on a few temperate rocky islands on the northwestern fringes of Europe, we can only wonder at what might await discovery planet-wide.

## Acknowledgements

The Wellcome Trust funded this research. We thank the staff of Wellcome Sanger Institute Cancer, Ageing Somatic Mutation (CASM) and Darwin Tree of Life (DTOL) programme for sample collection, management, and preparation and for the generation and public release of chromosome-level assemblies. We thank the staff of Wellcome Sanger Institute long-read sequencing team for CCS library preparation and sequencing. We sincerely thank M.M, H.H, Y.Z for their invaluable contributions in recovering germline and somatic mutation calls following the data incident at the Wellcome Sanger Institute. We thank M. Lawniczak for helpful introductions and discussions. We thank H. Lee for help with the illustrations. S.L, Y.W and H.H were supported by a Wellcome PhD Studentship. E. M is supported by a Wellcome Clinical PhD fellowship. The Darwin Tree of Life Project is funded by the Wellcome Trust through a Discretionary Award to the partnership (218328) and core funding to the Sanger Institute (206194), and by in-kind support from the partner institutions. Principal investigators at the Wellcome Sanger Institute were supported by a core grant from the Wellcome Trust or Wellcome Trust Award 220540/Z/20/A, Wellcome Sanger Institute Quinquennial Review 2021-2026. For the purpose of Open Access, the authors have applied a CC BY public copyright licence to any Author Accepted Manuscript version arising from this submission.

## Author contributions

P.J.C and R.D designed the experiments. S.L performed data collection, curation, and analysis. S.L developed the somatic mutation detection algorithm. S.L and H.H developed the haplotype phasing algorithm. S.L and Y.W performed mutational signature analysis across the Tree of Life. M.M cultured cancer cell lines. E.M, L.M, P.A, and N.M provided samples for evaluation of somatic mutation detection in human and non-human samples. S.L, R.D and P.J.C wrote the manuscript. All authors reviewed and edited the manuscript.

## Competing interests

S.L. is a shareholder and was previously an employee at Pacific Biosciences. P.J.C. is an employee and shareholder of Quotient Therapeutics Ltd.

## Data availability

The DToL consortium has made all genome assemblies and associated sequencing data publicly available through the accessions provided in **Supplementary Table 4**. PacBio HiFi datasets used for evaluation of somatic mutation detection in human and non-human samples are listed separately in **Supplementary Tables 3** and **8**.

## Code availability

Himut is available as an open-source software under the MIT license at https://github.com/sjin09/himut. Additionally, the code and documentation accompanying this manuscript for data preparation, mutational signature extraction and phylogentic analysis are available at https://github.com/sjin09/treeoflife.

## METHODS

### Human CCS library preparation and sequencing

BC-1 and HT-115 cell lines were cultured in RPMI and DMEM media with 5% foetal bovine serum and penicillin, respectively, at 37 °C with 5% CO_2_. Cambridge Blood and Stem Cell Biobank (CBSB) provided one peripheral blood sample (60mL; PD48473b, 82-year-old female) and one cord blood sample (PD47269d, newborn female), obtained with informed consent from patients at Addenbrooke’s Hospital (Cambridge East Ethics Committee reference 18/EE/0199). Whole blood and cord blood was diluted 1:1 with PBS, after which granulocytes were isolated using Lymphoprep^TM^ density gradient centrifugation (STEMCELL Technologies). Red cell lysis was performed on the granulocyte fraction using two 30-minute incubations at 20°C using RBC lysis buffer (BioLegend).

High molecular weight (HMW) DNA from these four samples was extracted using Qiagen MagAttract HMW DNA extraction kit (67563) and sheared into 16-20kb DNA fragments using Megaruptor 3 system (B06010003) at a speed setting 30. CCS libraries were then prepared following the standard CCS library preparation protocol v1.0 (100-222-300) and sequenced on a Sequel IIe instrument at the Wellcome Sanger Institute.

### DToL consortium sample selection

The DToL consortium coordinated the sample collection, CCS library preparation, sequencing, assembly construction, and the public release of *de novo* assemblies^10^. For most species, a single sample was sequenced and assembled. Biological replicates from the same species were sequenced in some cases to assess the consistency of mutational spectra. The tissue sources and NCBI taxonomic classifications for each sample, retrieved using the Genomes on A Tree^37^, are summarised in **Supplementary Table 4** and **Supplementary Table 7**, respectively.

The himut tool requires unbinned base quality (BQ) scores. The PacBio Sequel II/IIe instruments generate CCS reads with unbinned BQ scores (Q1-Q93), whereas Revio instruments generate CCS reads with binned BQ scores. CCS reads with unbinned BQ scores were collected from 770 samples sequenced on the Sequel II/IIe, representing 714 species.

Accurate detection of germline and somatic mutations depends on the ploidy of the sample and the models used to account for it. Both deepvariant and himut, which we used for germline and somatic mutation detection, assume that input samples have a diploid genome. As a result, samples with non-diploid genomes were identified and excluded from the analysis.

To determine ploidy, the DToL consortium relied on either known sex information or the k-mer coverage histogram. For instance, in insects of the Hymenoptera order, the sex determines the ploidy of the sample: maples are haploid, while females are diploid. If ploidy information was not available, samples were assumed to be diploid by default. Nineteen haploid and ten polyploid samples were removed from downstream sequence analysis.

### CCS read alignment and downstream sequence analysis

CCS reads with adapter sequences were identified using HiFiAdapterFilt^38^ and excluded from downstream analysis. CCS reads were aligned using minimap2^39^ (version 2.24-r1155-dirty, parameters: -ax map-hifi –cs=short) to the appropriate reference genome: human_g1k_v37_decoy.fasta for human samples and DToL reference genomes for non-human species. Primary alignments were selected, compressed, merged, and sorted with SAMtools (version 1.6)^40^. DToL samples with autosomal sequence coverage below 20, except for *Phorcus lineatus* sample PHORD060b, where additional sequencing could not be performed, were excluded from downstream sequence analysis. Germline and somatic mutations were detected using deepvariant (version 1.1.0)^41^ and himut, respectively. A total of 708 samples, representing 661 eukaryotic species, were analysed. Within this dataset, the analysis was limited to autosomes due to ploidy variation and the lower assembly quality of sex chromosomes.

### Haplotype phasing of germline mutations

Himut requires a phased VCF file of germline heterozygous variants. It generates this via a graph algorithm problem, where each heterozygous SNV (hetSNV) is represented as a node in a graph, with edges connecting pairs of hetSNVs that are spanned by at least one read.

A pair of hetSNVs is considered haplotype-consistent either if all their spanning reads support the cis configuration or if all reads support the trans configuration. Conversely, a pair is haplotype-inconsistent if there are reads that support both configurations. In a cis configuration, both sites on the read carry the reference allele or both sites carry the alternate allele. In contrast, a trans configuration occurs when one hetSNV carries the reference allele and the other carries the alternate allele.

Himut leverages the long read lengths of CCS reads (10 – 20 kb on average) to evaluate haplotype consistency across multiple hetSNV pairs and connect haplotype-consistent pairs to assemble haplotype blocks. It determines whether a given pair is haplotype-consistent based on the number of reads supporting the cis configuration and the total number of spanning reads (p < 0.0001, one-sided binomial test). HetSNVs that are consistent with at least 20% of their possible pairings are then connected using a breadth-first search algorithm to construct haplotype blocks.

### Overview of himut (high-fidelity mutation)

Accurate detection of single-molecule somatic mutation, where a single CCS read observation supports the alternate allele, requires the separation of germline mutations from somatic mutations, followed by the differentiation of somatic mutations from sequencing errors and non-canonical artefacts. To accomplish this, himut performs germline and somatic mutation identification in parallel, limiting somatic mutation detection to homozygous reference positions with a minimum genotype quality (GQ) score above 20. Consequently, somatic mutations called from heterozygous, heterozygous alternative and homozygous alternative positions are excluded from analysis. As a result, somatic reversions, and clonal somatic mutations, which could be misinterpreted as heterozygous alleles, are not called. Afterwards, himut leverages CCS base accuracy and applies a set base-level hard filters to remove sequencing errors and non-canonical artefacts, yielding a set of high-confidence single-molecule somatic mutations.

### Germline mutation detection using a Bayesian classifier

To call germline variants, himut employs a Bayesian classifier on the pileup of bases at each reference position, using an approach similar to GATK^42^, as follows:

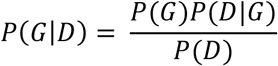

*P*(*D*|*G*) is the probability of observing the data given the genotype. *G* is the genotype and there are 10 possible genotypes (AA, CA, CC, CT, GA, GC, GG, GT, TA, and TT). *D* denotes the data, which includes the pileup of bases and corresponding BQ scores that represent sequencing error probabilities for each base. *P*(*G*) is the prior probability of observing the genotype and depends on whether the genotype is heterozygous, heterozygous alternative (tri-allelic), homozygous alternative or homozygous reference with respect to the reference base^43^, as follows:

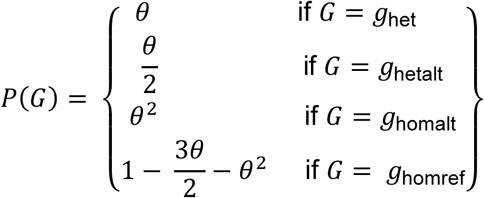

where *θ* is the expected germline SNP frequency, with a default value of 1 × 10^-3^, corresponding to the human germline SNP frequency. *P*(*D*) is a constant across all possible genotypes and is therefore ignored. For germline SNP detection in non-human samples, a deepvariant VCF file containing germline mutations must be provided to compute the *θ* for the given sample, unless *θ* is already known for the species in question.

The binomial likelihood is calculated for each genotype based on the pileup of bases and the associated base quality scores, assuming that sequencing errors and read sampling are independent and identically distributed, as follows:

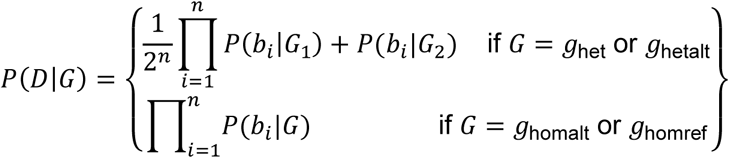

*P*(*b*_*i*_|*G*) represents the probability of observing a base given the genotype, defined as:

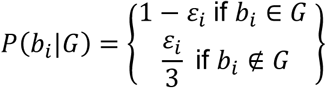

Here, *b* represents the base and *ε* denotes the sequencing error probability associated with the base. In practice, all calculations are performed on a logarithmic scale and the Phred-scaled likelihood (PL) is computed for all the possible genotypes.

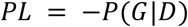

To normalise PL values, the lowest PL is subtracted from each PL. The genotype with the lowest PL is identified as the germline genotype and assigned a genotype quality (GQ) score. The GQ score is the second-lowest normalised PL and represents the relative likelihood that the true germline genotype is the one with the second-lowest PL. The GQ score is capped at a maximum value of 99.

### Single-molecule somatic mutation detection from bulk tissue

Somatic mutation candidates are initially identified as variants in high quality reads at sites that are homozygous reference according to the himut germline variant caller, where high quality reads are defined by meeting the following conditions:

- Minimum read quality (RQ) score (default = Q20)
- Minimum sequence identity (default = 0.99)
- Minimum mapping quality (MAPQ) score (default = Q60)
- Read length is within the pre-determined lower and upper bounds.

The RQ score is the average of the BQ scores, while the sequence identity is determined as the ratio of the matches to the total number of matches and mismatches. Lower and upper bounds for the read length are calculated as *u*_rl_ − 2*σ*_rl_ and *u*_rl_ + 2*σ*_rl_, respectively, where *u*_rl_ represents the mean read length and *σ*_rl_ denotes the standard deviation of read length.

Following initial selection of candidates, a series of filters are applied to differentiate somatic mutations from alignment and sequencing artefacts. The default base-level thresholds are as follows:

- Minimum trim proportion (default = 1%)
- Minimum window size (default = 40) and maximum number of mismatches within the window (default = 0)
- Minimum GQ score (default = 20)
- Minimum BQ score (default = 93)
- Maximum depth threshold (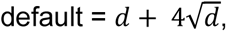 where *d* is the average read depth)
- Maximum number of overlapping insertion and deletions (default = 0)
- Minimum reference allele count (default = 3)
- Minimum alternate allele count (default = 1)

Previously, we observed that substitution error rates were highest at the ends of reads and progressively decreased toward their middle^44^. This result is consistent with earlier publications where end-repair and dA-tailing during sequencing library generation were pinpointed as major sources of technical artefacts for double-strand somatic variant detection^24,45^. As a result, somatic mutation candidates within a defined percentage of the 5’ and 3’ end of the read, defined as the minimum trim proportion, were excluded from further analysis. Furthermore, if the number of mismatches adjacent to a somatic mutation candidate exceeded the allowable maximum within a specified window, the candidate was discarded to reduce false positives arising from alignment errors. If the window extends beyond the start or end of the read, its size is adjusted to preserve the specified window size.

By default, himut processes a BAM file with read alignments as input and returns a VCF file with somatic mutations identified from homozygous reference positions. Here, we emphasize that himut exclusively uses Q93 CCS bases for somatic mutation detection, and therefore, requires CCS reads with unbinned base quality scores. To improve somatic mutation detection specificity, himut can optionally accept a VCF file with germline common SNPs (minor allele frequency >1%), a VCF file of himut calls on unrelated individuals (called as a panel of normals), and/or a VCF file with haplotype phased germline mutations. Filtering out sites found in these files can reduce false positives from genomic DNA contamination or non-canonical artefacts, while ensuring that somatic mutation detection is confined to haplotype-phased reads.

### Haplotype-phased somatic mutation detection

Although sequencing errors can impact both haplotypes, somatic mutations are transmitted exclusively through a single haplotype once established. Himut can leverage this biological characteristic to improve the specificity of somatic mutation detection.

To perform haplotype-phased somatic mutation detection, himut requires a VCF file containing haplotype-phased germline mutations. Himut fetches reads located within a haplotype block and uses those with an exact match to the consensus haplotype for somatic mutation detection. Consequently, reads that intersect a haplotype block, span two or more haplotype blocks, or have an inexact match to the consensus haplotype are not considered. Somatic mutation detection is performed as described above, but with the additional requirement that both haplotypes must be observed more than the minimum haplotype count (--min-hap-count is set to 3 by default).

### Panel of normals file preparation

To generate a panel of normals (PoN) VCF file, HiFi reads from eight normal unrelated individuals (**Supplementary Information**), publicly available through the Genome in a Bottle consortium^46^, were processed using the same workflow described above, but with more permissive filtering thresholds to maximise the detection of false positives arising from non-canonical artefacts, ensuring that the PoN captures recurrent technical artefacts. The changes in the hard filter thresholds are as follows:

- Minimum MAPQ score: 60 → 30
- Minimum sequence identity: 0.99 → 0.8
- Minimum proportion of bases to trim from end of reads: 1% → 0%
- Minimum GQ score: 20 → 10
- Minimum BQ score: 93 → 20

### SBS96 classification

The six types of substitution classes based on Watson-Crick base pairs are C:G>A:T, C:G>G:C, C:G>T:A, T:A>A:T, T:A>C:G, T:A>G:C. Both the SBS52 and the SBS96 classifications use the representation where the mutated base is the pyrimidine in the Watson-Crick base pair (C>A, C>G, C>T, T>A, T>C and T>G). The SBS96 classification uses the four bases upstream and downstream of the substitution to define the canonical 96 categories (four upstream bases × six substitution classes × four downstream bases), capturing both the substitution type and the local sequence context of somatic mutations^1^.

### SBS96 count correction

To enable single-molecule somatic mutation detection, himut identifies homozygous reference positions that are free of insertions and deletions, and leverages highly accurate bases from high-quality reads, referred to as callable reference positions and callable CCS bases, respectively. As a result of this selection process, the trinucleotide frequencies from callable reference positions and callable CCS bases deviate from the trinucleotide frequencies in the reference FASTA file. This deviation must be accounted for to obtain corrected SBS96 counts, which are essential for mutation burden calculation and mutational signature extraction.

First, trinucleotide frequencies are calculated from the reference FASTA file, callable reference positions and callable CCS bases.

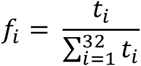

*t* denotes a trinucleotide, and there are 32 possible trinucleotides with a pyrimidine as the middle base (four upstream bases × two pyrimidine bases × four downstream bases).

Second, the ratio of trinucleotide frequencies derived from callable reference positions to those from reference FASTA file is calculated as follows:

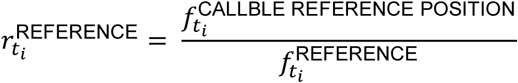

Additionally, the ratio of trinucleotide frequencies derived from callable CCS bases to those from callable reference positions is calculated as:

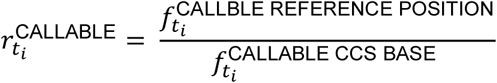

Third, the ratio of trinucleotide frequencies is leveraged to obtain the corrected SBS96 counts.

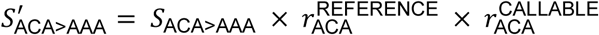

*S*_*ACA*>*AAA*_ represents the uncorrected count of a C>A substitution within the ACA trinucleotide context, and 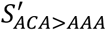 denotes the corrected SBS96 count.

### Cosine similarity calculation

The cosine similarity between the expected and the observed mutational spectrum is calculated as follows:

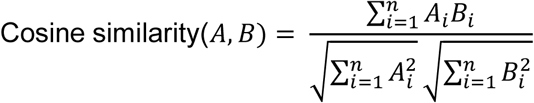

where *A* and *B* are vectors with SBS96 counts from two samples. Cosine similarity ranges from 0 to 1, where 0 indicates independence, 1 denotes identity and the values in between represent intermediate similarity.

### Mutation burden calculation

To estimate the average mutation burden per cell, the somatic mutation rate for each trinucleotide is determined as the ratio of the sum of corrected SBS96 counts for mutations away from that trinucleotide to the total number of callable trinucleotides in the aligned CCS reads, as follows:

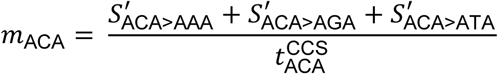

These rates are then multiplied by the number of corresponding trinucleotides in the reference FASTA file to obtain the mutation burden per cell:

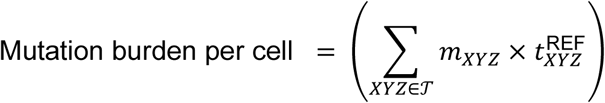

where *T* is the set of 32 trinucleotide contexts with a pyrimidine in the middle base, and 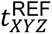 is calculated from both the forwards and reverse-complemented reference.

### Calculation of the CCS technical error rate using CCS reads from a negative control

Mismatches between reads and the reference genome arise from two processes: technical artefacts and mutational processes. Distinguishing somatic mutations from false positives introduced during library preparation and sequencing remains challenging. The former is dependent on the age and somatic mutation rate of the tissue, while the latter is effectively a constant. Here, blood granulocytes from cord blood with very few somatic mutations (50 – 60 somatic mutations per cell) were sequenced as a negative control sample. Additionally, because the mutational process and the mutation burden of cord blood have been characterised previously through single-cell-derived colonies^24^, the expected SBS96 counts for this sample can be derived from the number of callable trinucleotides and subtracted from the total observed SBS96 counts to calculate the SBS96 counts attributed to technical artefacts. This process was used to calculate the trinucleotide context dependent substitution error rates.

### *Phorcus lineatus* somatic mutation rate

Somatic mutations continue to accumulate throughout an individual’s lifetime, leading to higher expected numbers of somatic mutations in older individuals. To help confirm that the somatic mutations we were calling in non-human samples are real, *Phorcus lineatus* specimens of various ages (three-, five-, and fifteen-year-old) were analysed to calculate their mutation burdens and determine their correlation with age.

Staff scientists at the Marine Biological Association collected *Phorcus lineatus* samples from the rocky intertidal zone at Batten Bay, Plymouth (50.3570, -4.1278), and estimated the age of the specimens based on their shell rings. The samples were snap frozen, transferred to the Wellcome Sanger Institute, and stored at −80 °C. In a sterile environment, a bench-mounted vice was used to carefully crack open the shell, and the foot muscle tissue was isolated and HMW DNA was extracted and sequenced on a Sequel lIe instrument. To control for batch effects, all samples were processed using the same extraction and library preparation protocol.

To ensure uniform calling properties for these samples, the reads were truncated so that there were ten full-length subreads for each template proceeding to CCS base calling (dropping ZMWs with fewer than ten subreads). Somatic mutation detection and mutation burden calculation were performed as described above.

### Data preparation for somatic mutational signature extraction

In this study, a single sample from each species was sequenced and assembled. The CCS reads were then aligned back to the resulting primary reference genome to call germline and somatic mutations. To account for variation in trinucleotide frequencies across species during mutational signature extraction SBS96 counts were normalised, as shown below:

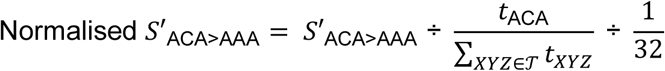

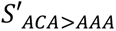 represents the corrected count of A[C>A]A substitutions. *t*_ACA_ indicates the number of ACA trinucleotides in the reference genome, and 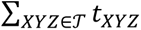 denotes the total number of trinucleotides in the reference genome.

### Data preparation for germline mutational signature extraction

We called germline mutations using deepvariant with standard settings for PacBio CCS, and further excluded samples with fewer than 1,000 germline substitutions. If both the reads and the reference genome originated from the same individual, as in the majority of cases, then homozygous alternate alleles are likely assembly errors so only heterozygous mutation calls were used in germline mutational signature extraction. If the reads and the reference genome originated from different individuals, both heterozygous and homozygous alternate alleles were used.

The SBS96 classification, developed for somatic mutations, depends on knowing the mutation’s direction. In contrast to somatic mutations, for which the ancestral allele is known, determining the ancestral allele and the direction of germline mutations requires comparison with an outgroup. Consequently, we developed the SBS52 classification germline mutation categorisation (**Supplementary Table 15**). This consolidates reciprocal pairs of mutations by selecting the SBS96 category with the higher number of pyrimidines. If both SBS96 categories under consideration have the same number of pyrimidines, the one listed first alphabetically is chosen as the representative category. For example, the germline substitution ACA>ATA is indistinguishable from ATA>ACA. In the SBS52 classification, ACA>ATA represents both ACA>ATA and ATA>ACA.

### Mutational signature extraction

Mutational signature extraction was conducted using hierarchical Dirichlet processes (HDP) v0.1.5^32^ without prior information about potential signatures. HDP extraction was performed with ten independent chains, each run for 120,000 iterations, with a burn-in of 20,000. SigProfiler^47^ was not used for mutational signature extraction because it does not support extraction with the SBS52 classification. Samples exhibiting a mutational signature attribution greater than 3.5% were classified as positive for that signature.

### Somatic mutational signature extraction

Before signature extraction, samples with fewer than 100 somatic substitutions were excluded, and the somatic mutation counts of the remaining samples were scaled to the median value of each dataset. To obtain the final somatic mutational signatures, two rounds of extraction were performed. The first round, based on corrected SBS96 counts, identified samples where signatures associated with technical artefacts greatly contributed to the observed mutations. Before the second round, samples with more than 80% cumulative attribution to artefactual mutational signatures were excluded. The identification and definition of artefact-associated mutational signatures are described below. The second round of extraction used trinucleotide normalised SBS96 counts to account for variation in trinucleotide frequencies across species.

### Germline mutational signature extraction

Before germline mutational signature extraction, samples with fewer than 10,000 germline substitutions were excluded. The extraction was performed using a dataset of trinucleotide normalised SBS52, scaled to 3,000 mutations per sample.

### Mutational signatures associated with library and sequencing artefacts

Single-molecule somatic mutations called in each sample are a collection of differences between the reference and the sample, which can result from library preparation and sequencing errors as well as from mutational processes. During library preparation, single-stranded and double-stranded DNA damage can be introduced to the template molecule. If this damage is improperly repaired and the template molecule is subsequently sequenced, it becomes indistinguishable from somatic mutations.

In this study, blood granulocytes from cord blood were sequenced as a negative control, as the mutation burden of this sample (50 – 60 mutations per cell) is among the lowest that has been well characterised in biological material^26^. Because the number of called mutations exceeded the expected number of somatic mutations in this sample, the derived mutational spectrum was dominated by errors and discordant with that expected for the sample. Consequently, we took this as reflecting the residual error profile, and mutational signatures with a cosine similarity greater than 0.85 to the PD47269d mutational spectrum were identified as artefactual (rToL1 and rToL2)

In addition, during our analysis, we identified multiple samples in which another suspected artefactual process appeared to be the primary source of observed somatic mutations (fTraTra1, gfFlaVelt1, icAdaBipu9, icElmAene2, icHalSede1, icLepMacu1, idCisGlob1, idSarSubv1, ihDrePlat2, iiAthCine2, ilBraVimi1, ilHypCost1, ilProPyga1, ilSynAndr1, ilTelLucu1, ilTheBrit1, ilTinPell1, ilXesSexs1, ilYpoCang5, ilYpoPade1 and xgPatVulg1). In these samples, the total number of observed somatic mutations was over ten times higher than found in both biological replicates (e.g., icAdaBipu9) and species with the same taxonomic classification. Additionally, the mutational spectra observed in these samples were distinct from those in biological replicates and related species but resembled each other, being dominated by a third mutational signature rToL3 that we also designated as artefactual. We suggest that this may have been caused by post-mortem DNA damage or by a library preparation process common to these samples.

### Phylogenetic matrix

We used a measure of phylogenetic proximity (Matrix *A*), derived from Abouheif’s statistic, to measure the relatedness between two species^34^. Unlike conventional distance matrices that depend on branch lengths of a phylogenetic tree, Matrix *A* is based on the topology of the tree (**Supplementary Table 7**). We note that seven samples with incomplete taxonomic classification were excluded from the phylogenetic analysis (paTetJugo1, pxEimPrae1, xgAniVort1, xgPatDepr1, xgPatPell1, xgPatUlys2 and xgPatVulg1). The proximity score *a*_*ij*_ for two species *i* and *j* is defined as the inverse of the product of the number of direct descendants at each interior node on the path connecting species *i* and *j*.

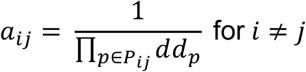

*P*_*ij*_ represents the set of interior nodes along the path connecting species *i* and *j*, while *dd*_*p*_ denotes the number of direct descents at each interior node. The diagonal elements *a*_*ii*_ are defined as 1 minus the sum of *a*_*ij*_ across all species.

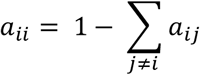

Hence, Matrix *A* is a square matrix composed of non-negative real numbers, where the sum of each row and column equals 1. The non-diagonal values represent the proximity between two species *i* and *j*, while the diagonal value capture the uniqueness of species *i* in relation to the root of the phylogeny.

### Phylogenetic signal analysis of germline and somatic mutational signatures

We calculated Abouheif’s C_mean_ to evaluate the phylogenetic signal of germline and somatic mutational signatures (i.e., the traits of interest). During mutational signature extraction, each species is assigned a mutational signature exposure, which quantifies the proportion of observed mutations that can be attributed to a specific mutational signature *x* = (*x*_1_, *x*_2_, …, *x*_*n*_). Before calculating C_mean_, the first signature extracted by HDP, the averaging component, was removed from the germline and somatic mutational signature attributions. Additionally, artefactual somatic signatures (rToL1, rToL2, and rToL3) were excluded from somatic mutational signature attributions. For each sample, mutational signature exposures were normalised by dividing by the total exposure excluding the aforementioned signatures.

Abouheif’s C_mean_ assesses how strongly phylogenetic relationships influence the trait observed in each species, with higher values indicating a stronger phylogenetic signal.

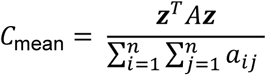

***z*** = (*z*_1_, *z*_2_ …, *z*_*n*_) represents the standardised mutational signature exposure across samples, obtained by subtracting the mean and dividing by the standard deviation.

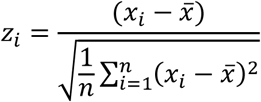

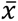 denotes the mean mutational signature exposure across samples.

To assess the statistical significance of the observed value of C_mean_, we performed a permutation test by randomly shuffling the mutational signature exposures across samples and recalculating C_mean_ over 1,000 iterations. The p-value was then estimated as the proportion of permuted C_mean_ values that exceeded the observed C_mean_. To account for multiple hypothesis testing, Benjamini-Hochberg correction was applied, and mutational signatures with a q-value below 0.01 were considered to exhibit a statistically significant phylogenetic signal.

---

**Extended Data Fig. 1.**
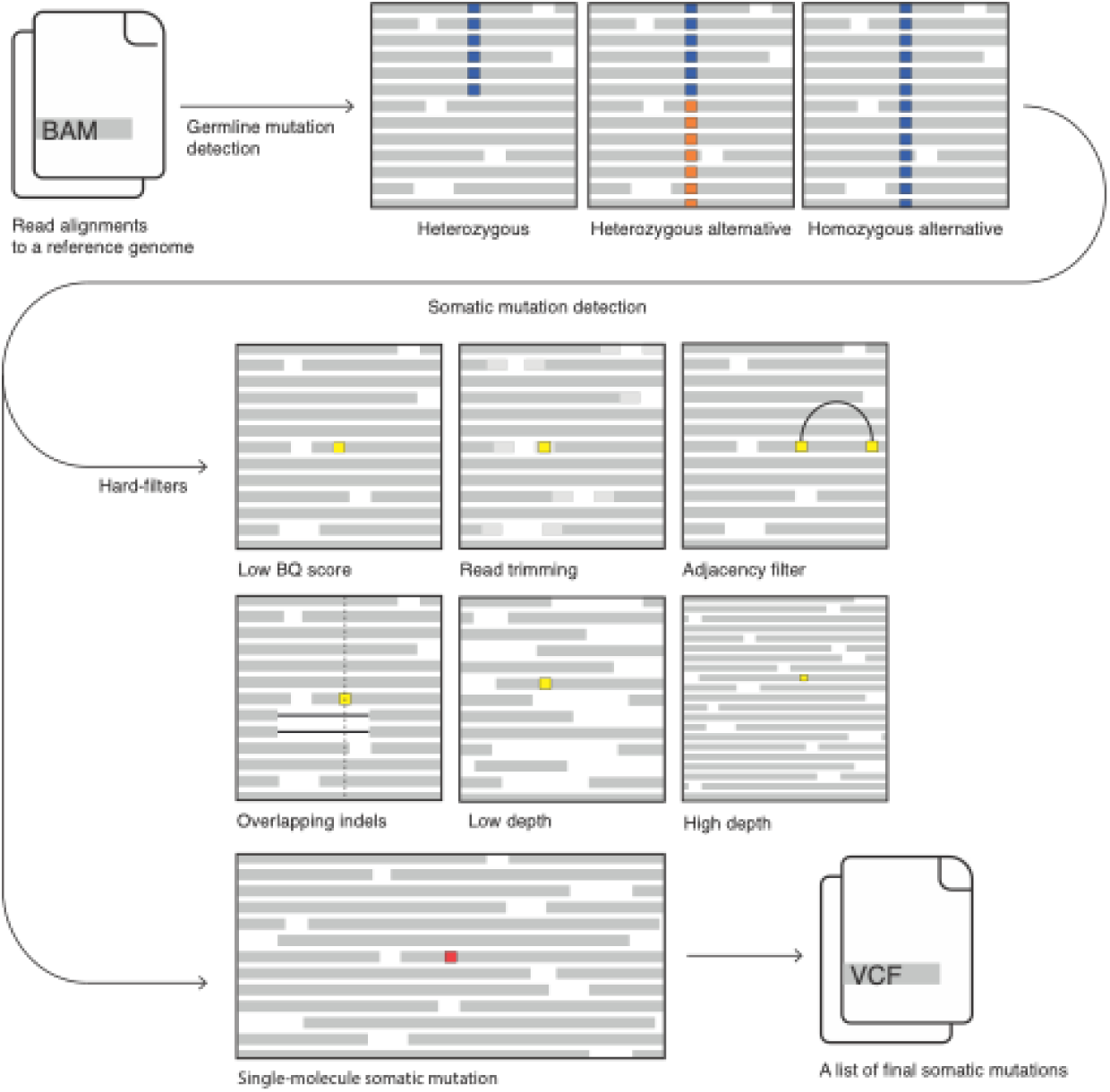
An overview of somatic mutation detection using himut. Himut takes as input a BAM file with primary read alignments and returns a VCF file containing a filtered list of proposed somatic mutations. To achieve this, it performs germline mutation detection first, then limits somatic mutation detection to homozygous reference positions (i.e., exclude heterozygous and homozygous alternative positions). Next, various base-level filters are applied as described in the Methods section. Blue and orange squares indicate germline variants, while yellow and red squares indicate candidates and a somatic mutation, respectively.

**Extended Data Fig. 2.**
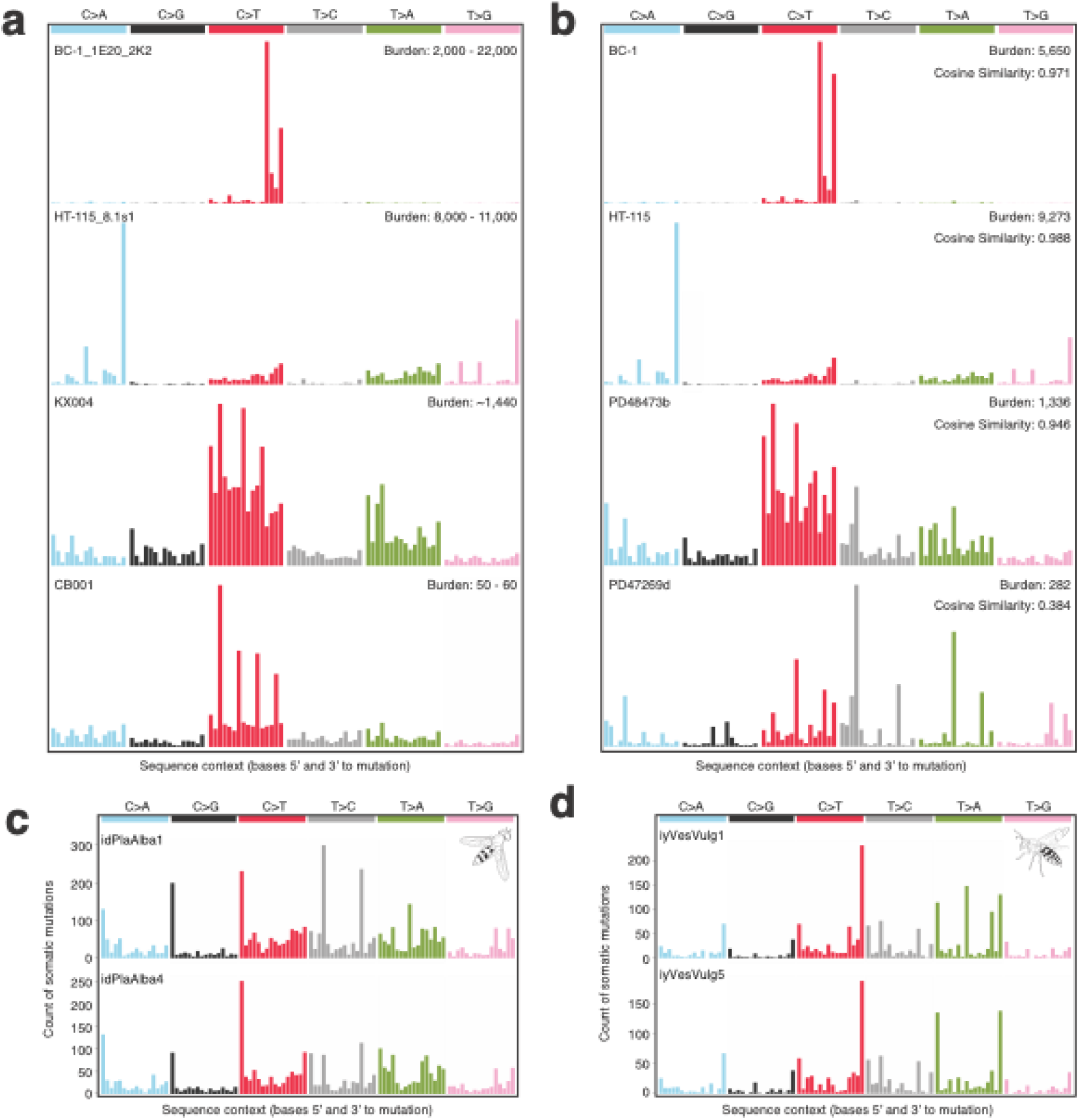
Evaluation of single-molecule somatic mutation calls in human and non-human samples. **a**, Expected mutational spectra derived from single-cell derived colonies^25,26^. CB001 and KX004 correspond to blood granulocyte samples from a neonate and a 77-year-old female, respectively. **b**, Observed Mutational spectra called by himut in experimentally matched cell lines (BC-1 and HT-115) and blood granulocytes of similar age (PD47269d: neonate and PD48473b: 82-year-old female). For these positive control samples, mutation burdens estimated from somatic mutation calls fall within the expected range relative to those from single-cell–derived colonies. For a conservative evaluation of somatic mutation detection, single-molecule somatic mutations supported by a single observation (alternate count of 1) were used in human samples. **c-d**, Somatic mutational spectra from himut for multiple individuals from two species: **c**, *Platycheirus albimanus* (white-footed hoverfly); **d**, *Vespula vulgaris* (common wasp).

**Extended Data Fig. 3.**
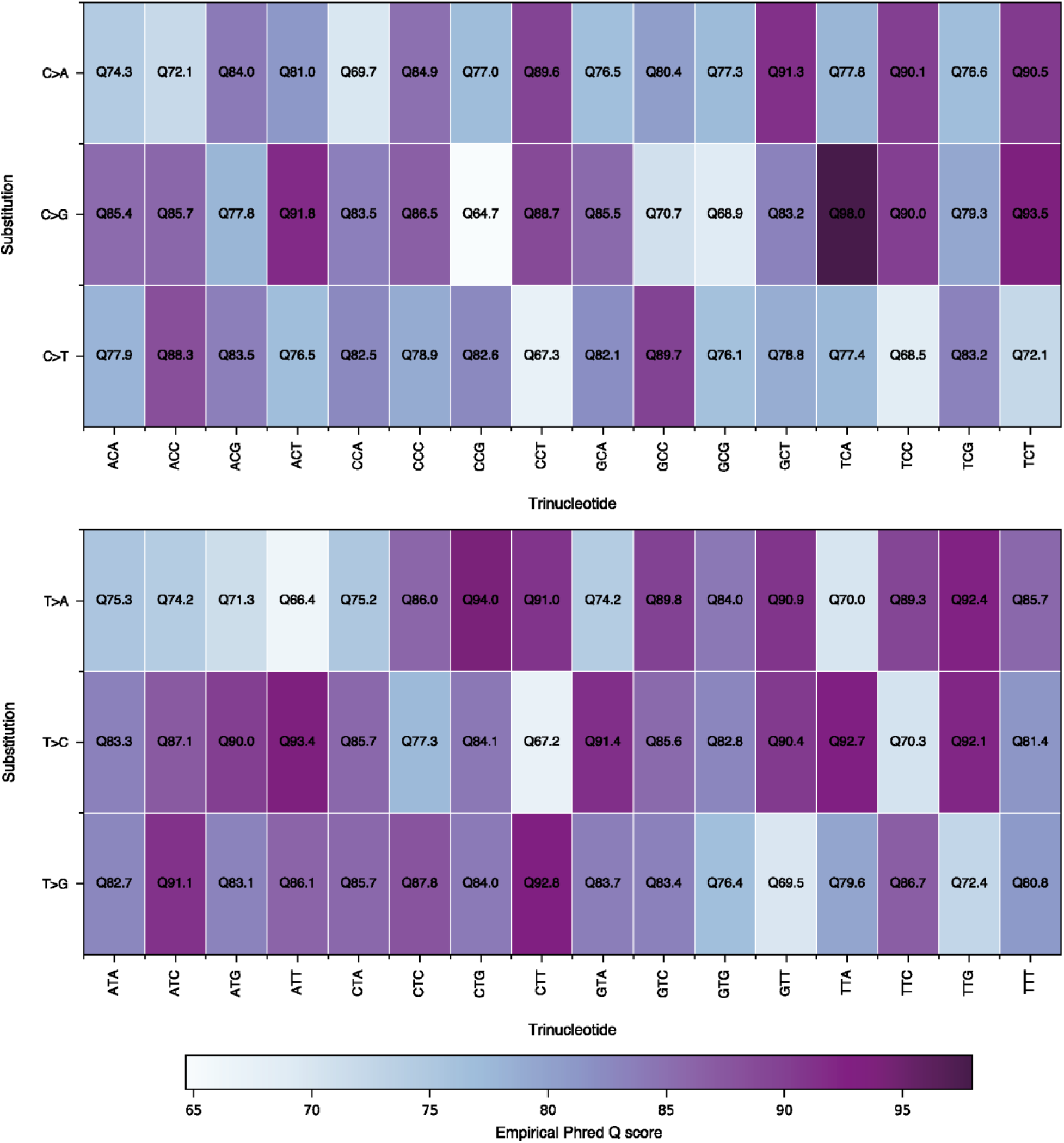
Heatmap showing substitution- and context-specific error rates in Q93 CCS bases, estimated from somatic mutation calls in the cord blood granulocyte sample (PD47269d). The T[C>G]A substitution has an empirical Phred quality score of Q98.0, which is not clearly visible in the figure.

**Extended Data Fig. 4.**
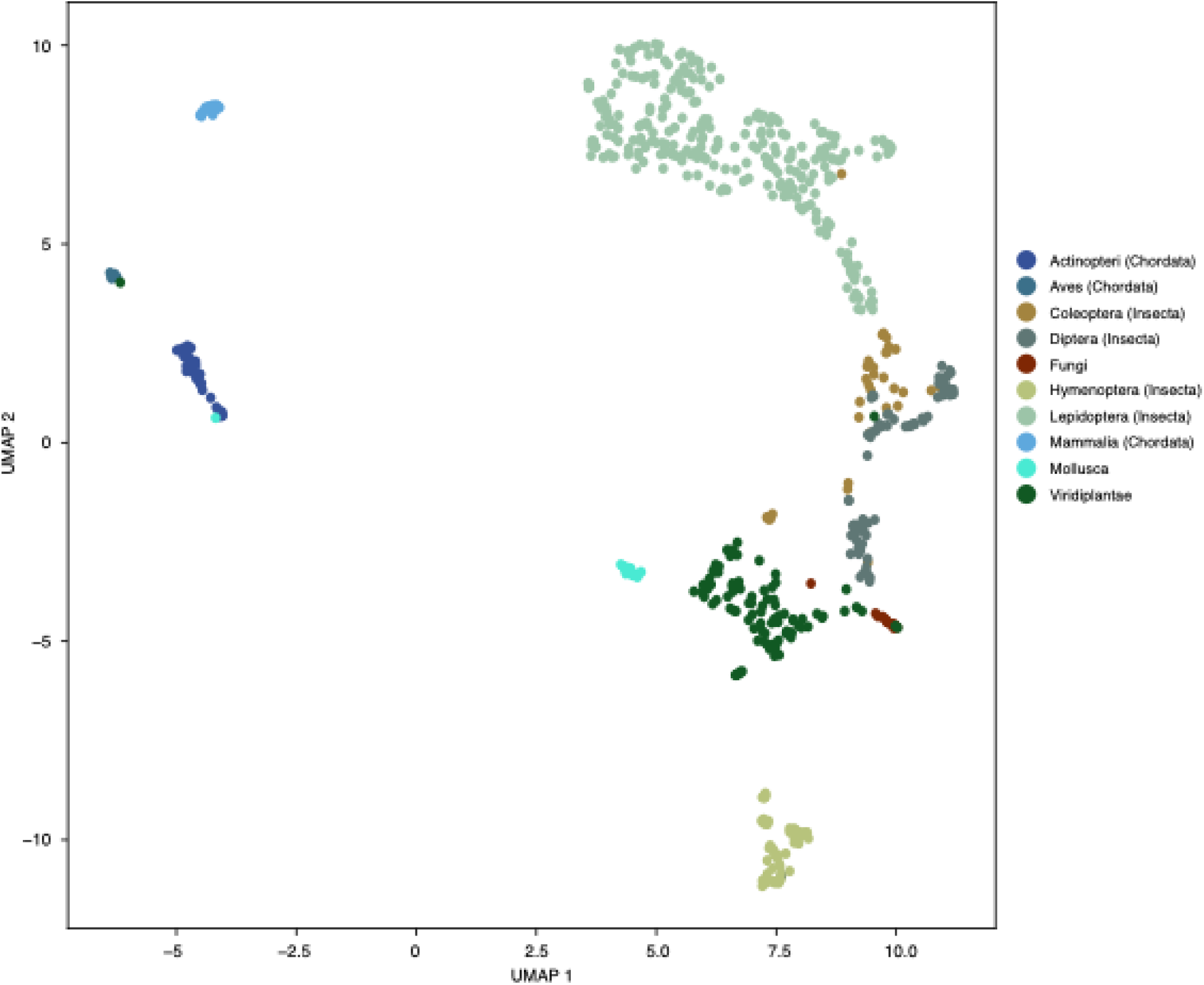
UMAP of genomic trinucleotide frequencies. Samples belonging to Fungi and Viridiplantae are coloured according to their kingdom-level taxonomic classification, whereas samples from the Metazoa kingdom are coloured by their lower-level taxonomic classification where sample sizes are sufficient.

**Extended Data Fig. 5.**
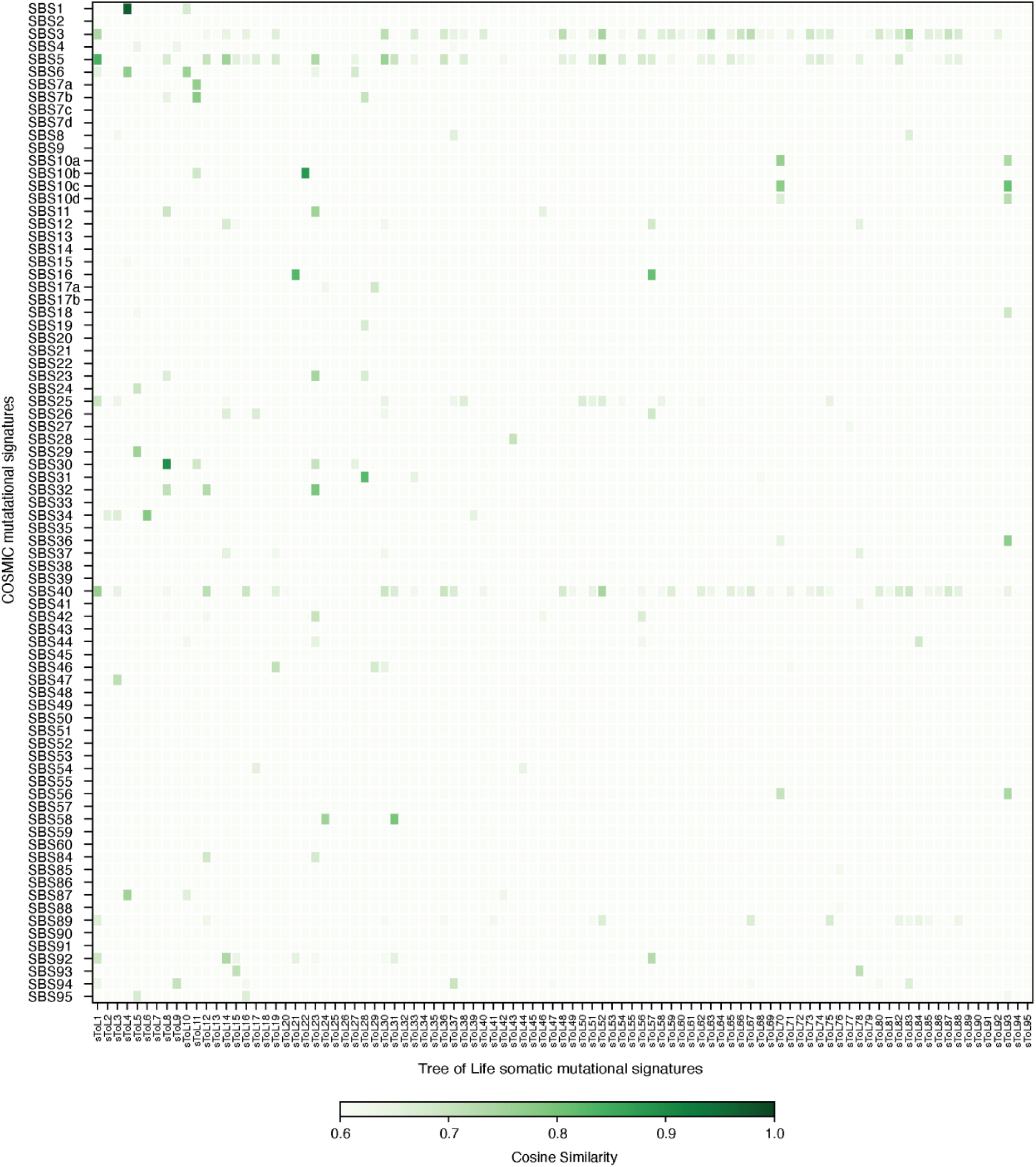
Heatmap of COSMIC and sToL somatic mutational signature similarities. Pairwise cosine similarities were calculated between somatic mutational signatures identified across the tree of life and known COSMIC reference signatures. High similarity values indicate mutational processes that are potentially shared between those operating across diverse species and those observed in human tissues.

**Extended Data Fig. 6.**
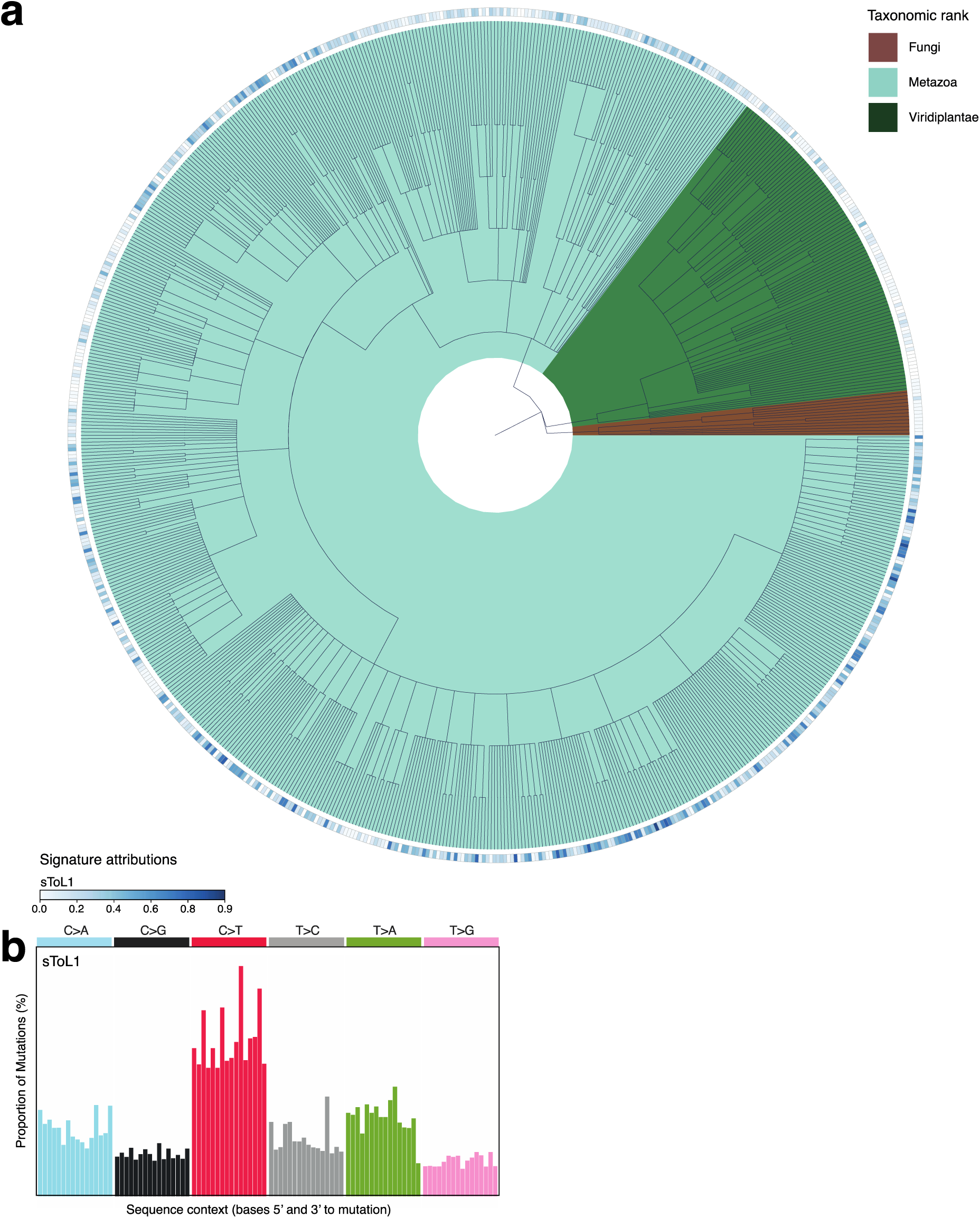
Universal somatic mutational process. **a**, Phylogenetic tree with branches shaded by Kingdom-level classification and tips annotated with sample-level sToL1 mutational signature attributions. **b**, sToL1 mutational signature.

**Extended Data Fig. 7.**
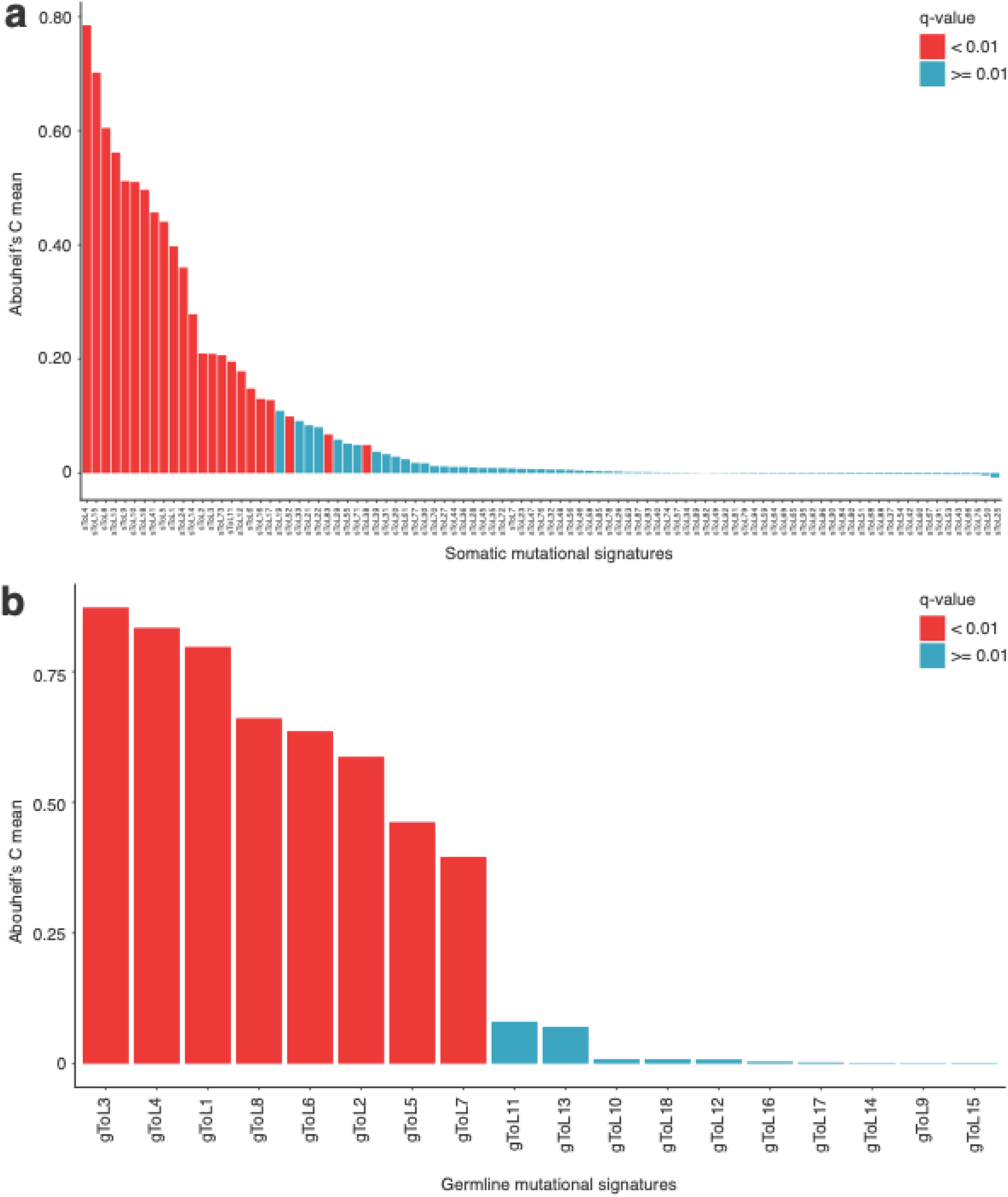
Mutational signatures with phylogenetic signals. **a**, Bar plot showing Abouheif’s C_mean_ values for somatic mutational signatures. **b**, Bar plot showing Abouheif’s C_mean_ values for germline mutational signatures. Bars are coloured to indicate the statistical significance of phylogenetic signal after correction for multiple testing.

**Extended Data Fig. 8.**
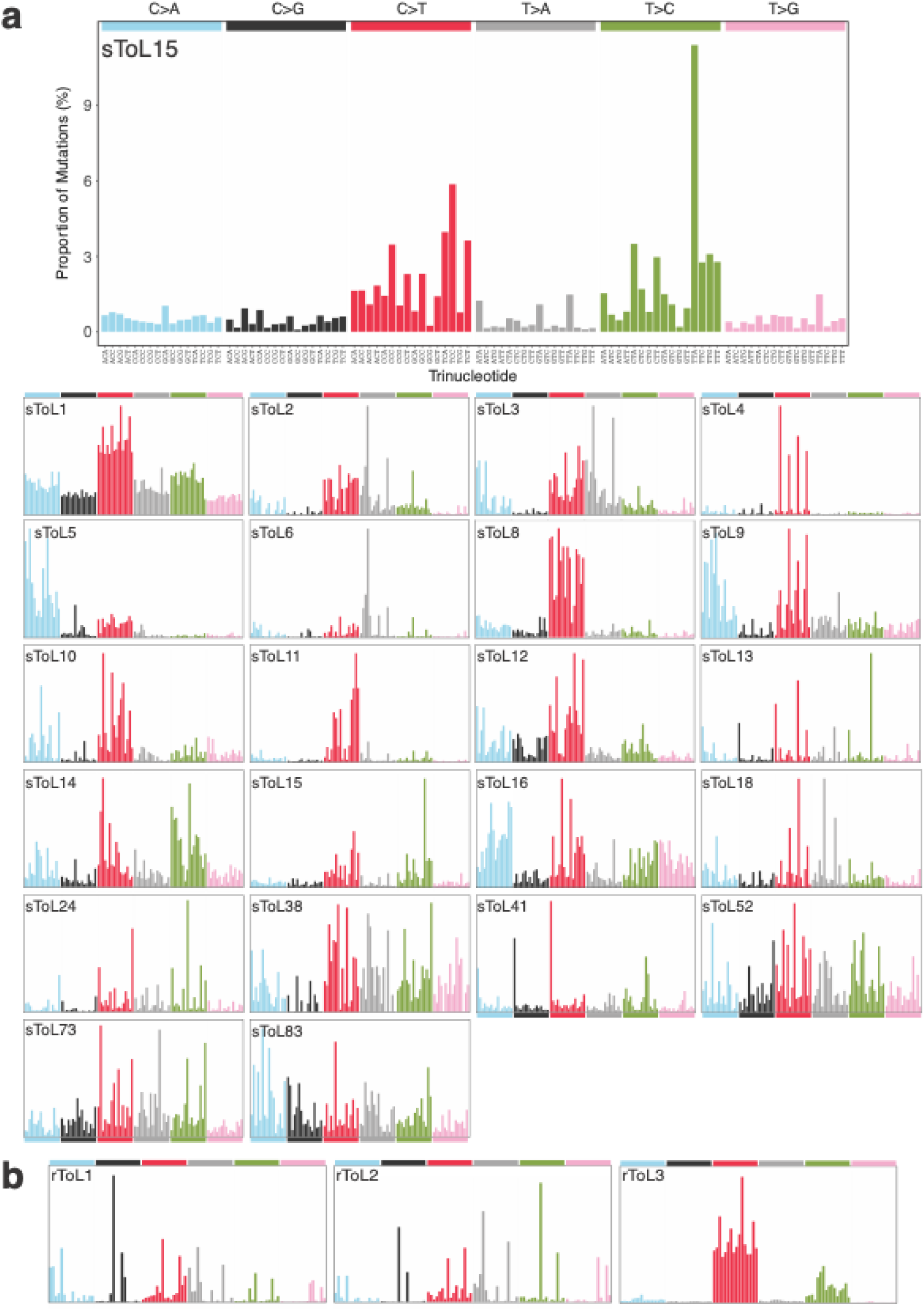
An overview of somatic mutational signatures detected across the tree of life. **a**, Somatic mutational signatures with statistically significant phylogenetic signals (q-value <0.01). **b**, Artefactual mutational signatures recurrently detected across the majority of samples.

**Extended Data Fig. 9.**
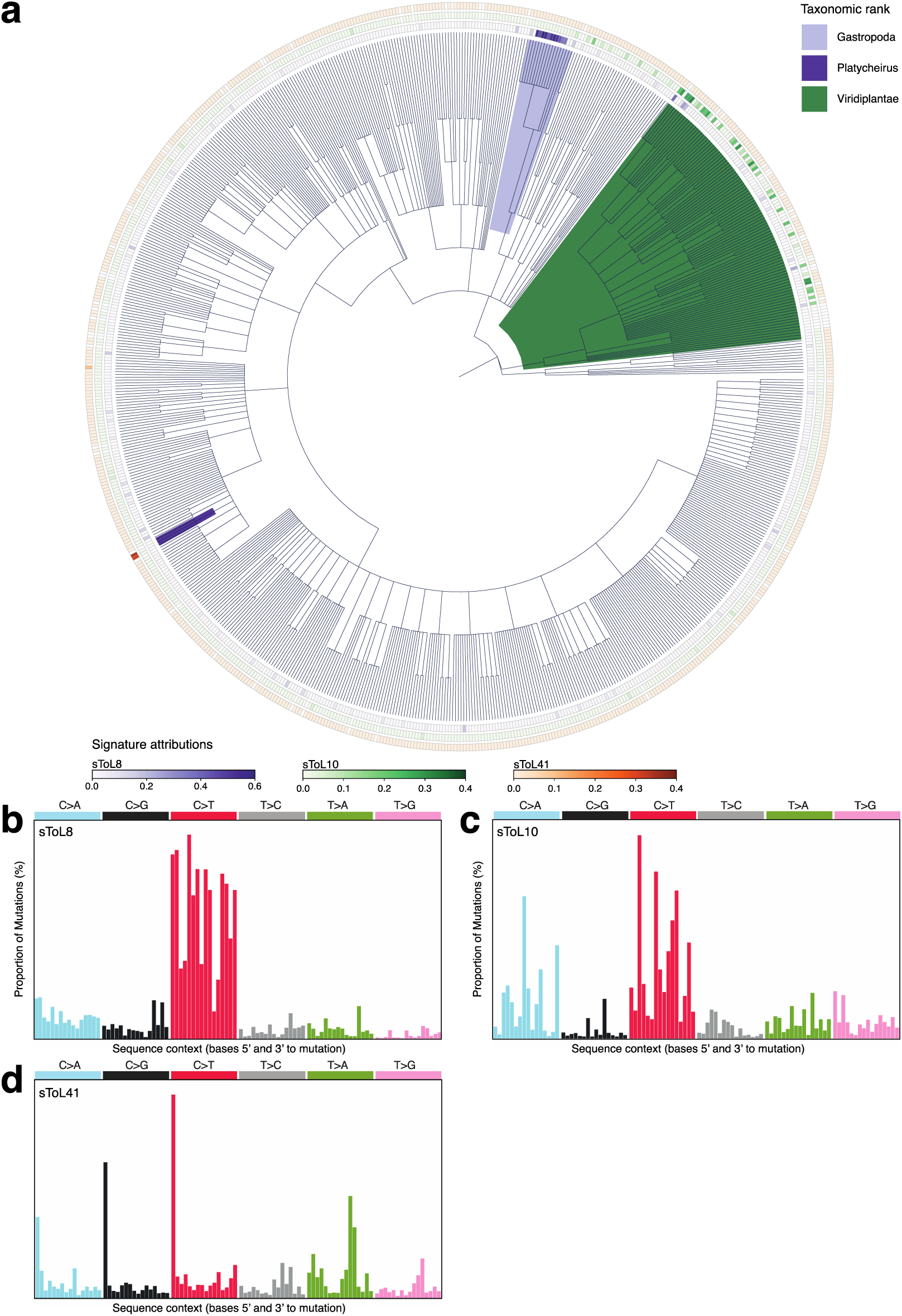
Clade-specific somatic mutational processes. **a**, Taxonomic tree with branches shaded to highlight clades in which the somatic mutational processes operate. **b-d**, Somatic mutational signatures: **b**, sToL8; **c**, sToL10; **d**, sToL41.

**Extended Data Fig. 10.**
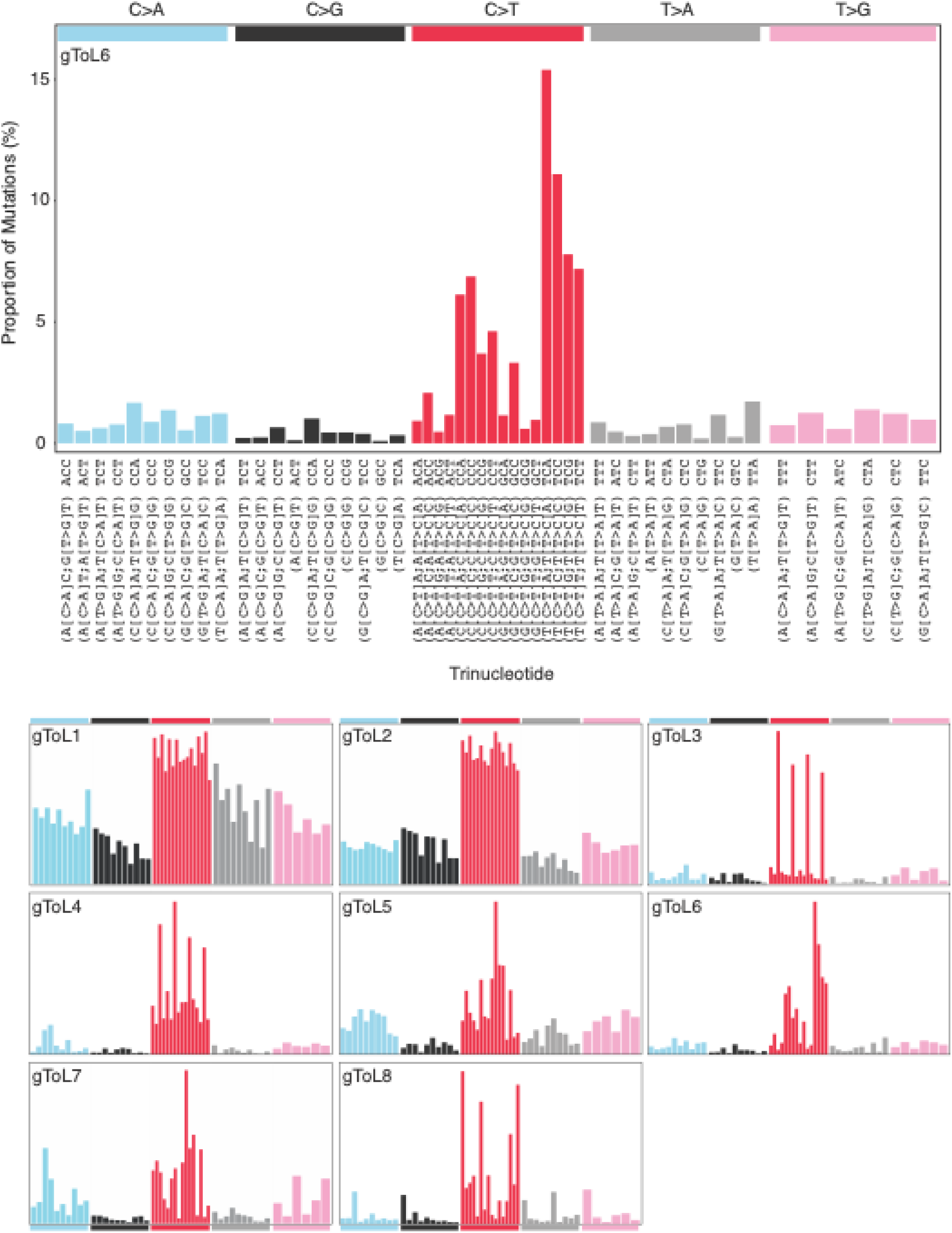
Germline mutational signatures with statistically significant phylogenetic signals (q-value < 0.01).

**Extended Data Fig. 11.**
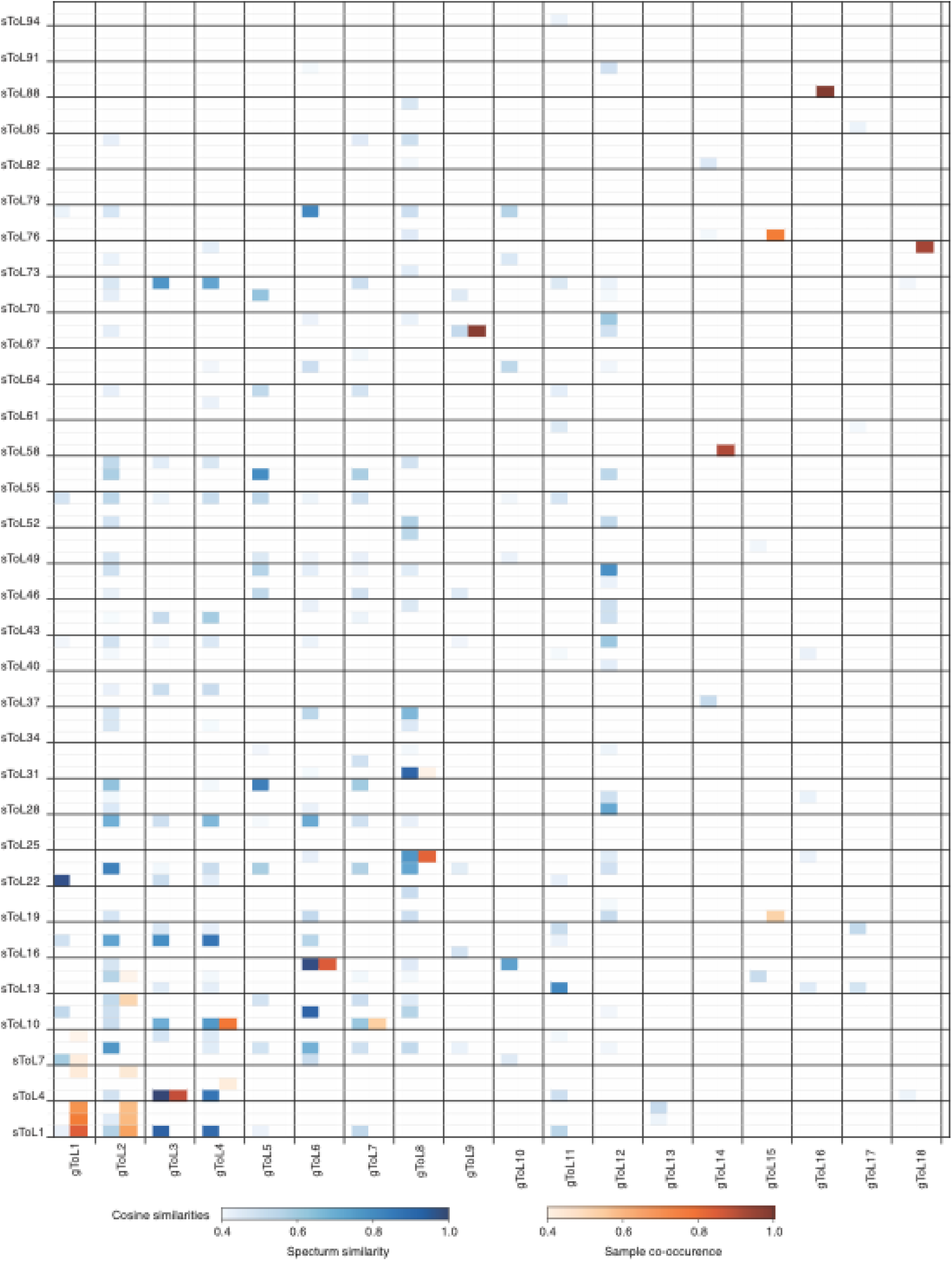
Heatmap showing similarities between gToL and sToL somatic mutational signatures and their attributions.

## References

1. Alexandrov, L. B. et al. The repertoire of mutational signatures in human cancer. Nature 578, 94–101 (2020).

2. Degasperi, A. et al. Substitution mutational signatures in whole-genome–sequenced cancers in the UK population. Science (1979) 376, (2022).

3. Moore, L. et al. The mutational landscape of human somatic and germline cells. Nature 597, 381–386 (2021).

4. Li, R. et al. A body map of somatic mutagenesis in morphologically normal human tissues. Nature 597, 398–403 (2021).

5. Kaplanis, J. et al. Genetic and chemotherapeutic influences on germline hypermutation. Nature 605, 503–508 (2022).

6. Serero, A., Jubin, C., Loeillet, S., Legoix-Né, P. & Nicolas, A. G. Mutational landscape of yeast mutator strains. Proc Natl Acad Sci U S A 111, 1897–1902 (2014).

7. Meier, B. et al. C. elegans whole genome sequencing reveals mutational signatures related to carcinogens and DNA repair deficiency. Genome Res 24, gr.175547.114 (2014).

8. Riva, L. et al. The mutational signature profile of known and suspected human carcinogens in mice. Nat Genet 52, 1189–1197 (2020).

9. Cagan, A. et al. Somatic mutation rates scale with lifespan across mammals. Nature 604, 517–524 (2022).

10. Darwin Tree of Life Consortium, Sequence locally, think globally: The Darwin Tree of Life Project. Proc. Natl Acad. Sci. USA 119(e), (2022).

11. Lindahl, T. Instability and decay of the primary structure of DNA. Nature 362, 709–715 (1993).

12. Garaycoechea, J. I. et al. Alcohol and endogenous aldehydes damage chromosomes and mutate stem cells. Nature 553, 171–177 (2018).

13. Pfeifer, G. P. et al. Tobacco smoke carcinogens, DNA damage and p53 mutations in smoking-associated cancers. Oncogene 21, 7435–7451 (2002).

14. Pfeifer, G. P., You, Y.-H. & Besaratinia, A. Mutations induced by ultraviolet light. Mutat Res 571, 19–31 (2005).

15. Alexandrov, L. B., Nik-Zainal, S., Wedge, D. C., Campbell, P. J. & Stratton, M. R. Deciphering Signatures of Mutational Processes Operative in Human Cancer. Cell Rep 3, 246–259 (2013).

16. Ng, S. W. K. et al. Convergent somatic mutations in metabolism genes in chronic liver disease. Nature 598, 473–478 (2021).

17. Yoshida, K. et al. Tobacco smoking and somatic mutations in human bronchial epithelium. Nature 578, 266–272 (2020).

18. Huang, Z. et al. Single-cell analysis of somatic mutations in human bronchial epithelial cells in relation to aging and smoking. Nat Genet 54, 492–498 (2022).

19. Lawson, A. R. J. et al. Extensive heterogeneity in somatic mutation and selection in the human bladder. Science (1979) 370, 75–82 (2020).

20. Lewin, H. A. et al. The Earth BioGenome Project 2020: Starting the clock. https://doi.org/10.1073/pnas.2115635118/-/DCSupplemental doi:10.1073/pnas.2115635118/-/DCSupplemental.

21. Wenger, A. M. et al. Accurate circular consensus long-read sequencing improves variant detection and assembly of a human genome. Nat Biotechnol 37, 1155–1162 (2019).

22. Schmitt, M. W. et al. Detection of ultra-rare mutations by next-generation sequencing. Proc Natl Acad Sci U S A 109, 14508–14513 (2012).

23. Hoang, M. L. et al. Genome-wide quantification of rare somatic mutations in normal human tissues using massively parallel sequencing. Proc Natl Acad Sci U S A 113, 9846–9851 (2016).

24. Abascal, F. et al. Somatic mutation landscapes at single-molecule resolution. Nature 593, 405–410 (2021).

25. Petljak, M. et al. Characterizing Mutational Signatures in Human Cancer Cell Lines Reveals Episodic APOBEC Mutagenesis. Cell 176, 1282–1294.e20 (2019).

26. Mitchell, E. et al. Clonal dynamics of haematopoiesis across the human lifespan. Nature 606, 343–350 (2022).

27. Osorio, F. G. et al. Somatic Mutations Reveal Lineage Relationships and Age-Related Mutagenesis in Human Hematopoiesis. Cell Rep 25, 2308–2316.e4 (2018).

28. Mieszkowska, N., Hawkins, S.J., Burrows, M.T. and Kendall, M.A. Long-term changes in the geographic distribution and population structures of *Osilinus lineatus* (Gastropoda: Trochidae) in Britain and Ireland. J. Mar. Biol. Assoc. U.K. 87, 537–545 (2007).

29. Blokzijl, F. et al. Tissue-specific mutation accumulation in human adult stem cells during life. Nature 538, 260–264 (2016).

30. Machado, H. E. et al. Diverse mutational landscapes in human lymphocytes. Nature 608, 724–732 (2022).

31. Alexandrov, L. B. et al. Clock-like mutational processes in human somatic cells. Nat Genet 47, 1402–1407 (2015).

32. Li, Y. et al. Patterns of somatic structural variation in human cancer genomes. Nature 578, 112–121 (2020).

33. Shen, J. cheng, Rideout, W. M. & Jones, P. A. The rate of hydrolytic deamination of 5-methylcytosine in double-stranded DNA. Nucleic Acids Res 22, 972–976 (1994).

34. Abouheif, E. A Method for Testing the Assumption of Phylogenetic Independence in Comparative Data. Evolutionary Ecology Research vol. 1 (1999).

35. Pavoine, S., Ollier, S., Pontier, D. & Chessel, D. Testing for phylogenetic signal in phenotypic traits: New matrices of phylogenetic proximities. Theor Popul Biol 73, 79–91 (2008).

36. Gallego-Bartolomé, J. DNA methylation in plants: mechanisms and tools for targeted manipulation. New Phytologist vol. 227 38–44 Preprint at 10.1111/nph.16529 (2020).

37. Challis, R., Kumar, S., Sotero-Caio, C., Brown, M. & Blaxter, M. Genomes on a Tree (GoaT): A versatile, scalable search engine for genomic and sequencing project metadata across the eukaryotic tree of life. Wellcome Open Res 8, (2023).

38. Sim, S. B., Corpuz, R. L., Simmonds, T. J. & Geib, S. M. HiFiAdapterFilt, a memory efficient read processing pipeline, prevents occurrence of adapter sequence in PacBio HiFi reads and their negative impacts on genome assembly. BMC Genomics 23, 157 (2022).

39. Li, H. Minimap2: Pairwise alignment for nucleotide sequences. Bioinformatics 34, 3094–3100 (2018).

40. Li, H. et al. The Sequence Alignment/Map format and SAMtools. Bioinformatics 25, 2078–2079 (2009).

41. Poplin, R. et al. A universal SNP and small-indel variant caller using deep neural networks. Nat Biotechnol 36, 983–987 (2018).

42. McKenna, A. et al. The Genome Analysis Toolkit: a MapReduce framework for analyzing next-generation DNA sequencing data. Genome Res 20, 1297–1303 (2010).

43. Kim, S. et al. Strelka2: fast and accurate calling of germline and somatic variants. Nat Methods 15, 591–594 (2018).

44. Schweiger, R. et al. Insights into non-crossover recombination from long-read sperm sequencing. BioRXiv https://doi.org/10.1101/2024.07.05.602249 (2024) doi:10.1101/2024.07.05.602249.

45. Xiong, K. et al. Duplex-Repair enables highly accurate sequencing, despite DNA damage. Nucleic Acids Res 50, (2022).

46. Wagner, J. et al. Benchmarking challenging small variants with linked and long reads. Cell Genomics 2, (2022).

47. Bergstrom, E. N. et al. SigProfilerMatrixGenerator: A tool for visualizing and exploring patterns of small mutational events. BMC Genomics 20, 1–12 (2019).

