## Supplementary Figures for "Somatic and germline mutational processes across the tree of life"

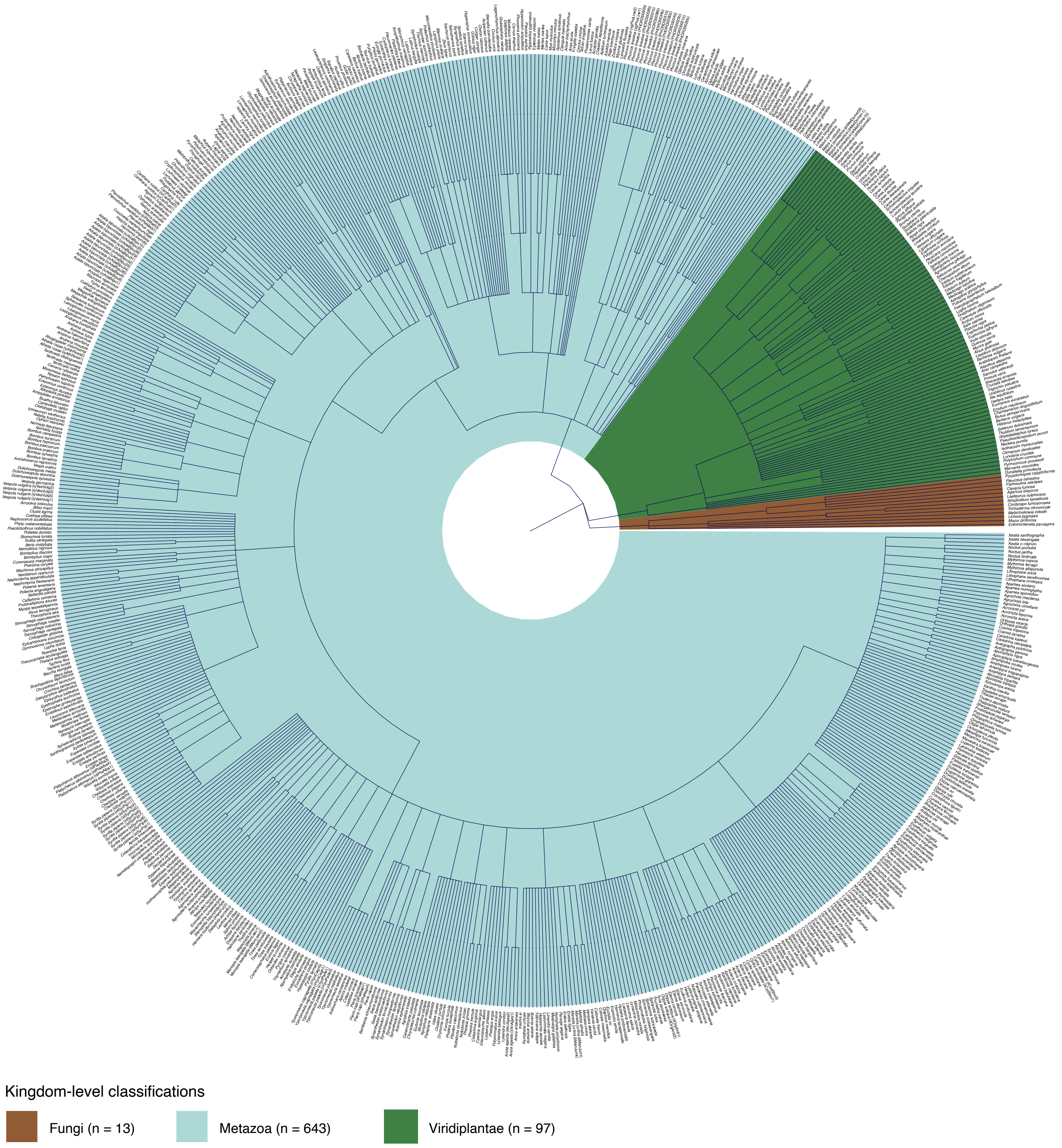


**Supplementary Fig. 1** | Phylogenetic tree constructed from samples with complete taxonomic classifications, shaded by their Kingdom-level classification. Seven samples with missing taxonomic information were excluded. Taxonomic classifications with fewer than five samples are not shaded.

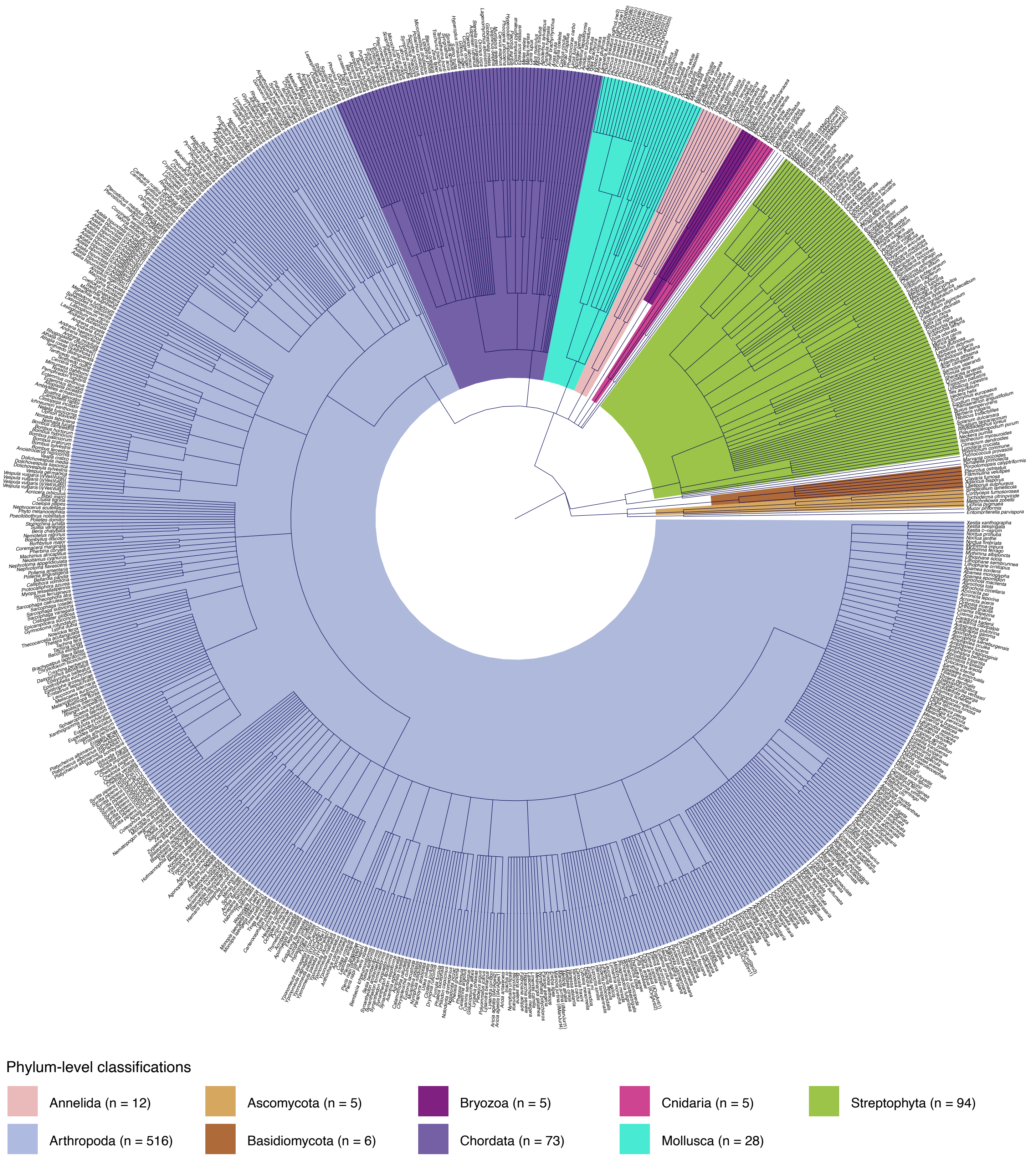


**Supplementary Fig. 2** | Phylogenetic tree constructed from samples with complete taxonomic classifications, shaded by their Phylum-level classification. Seven samples with missing taxonomic information were excluded. Taxonomic classifications with fewer than five samples are not shaded.

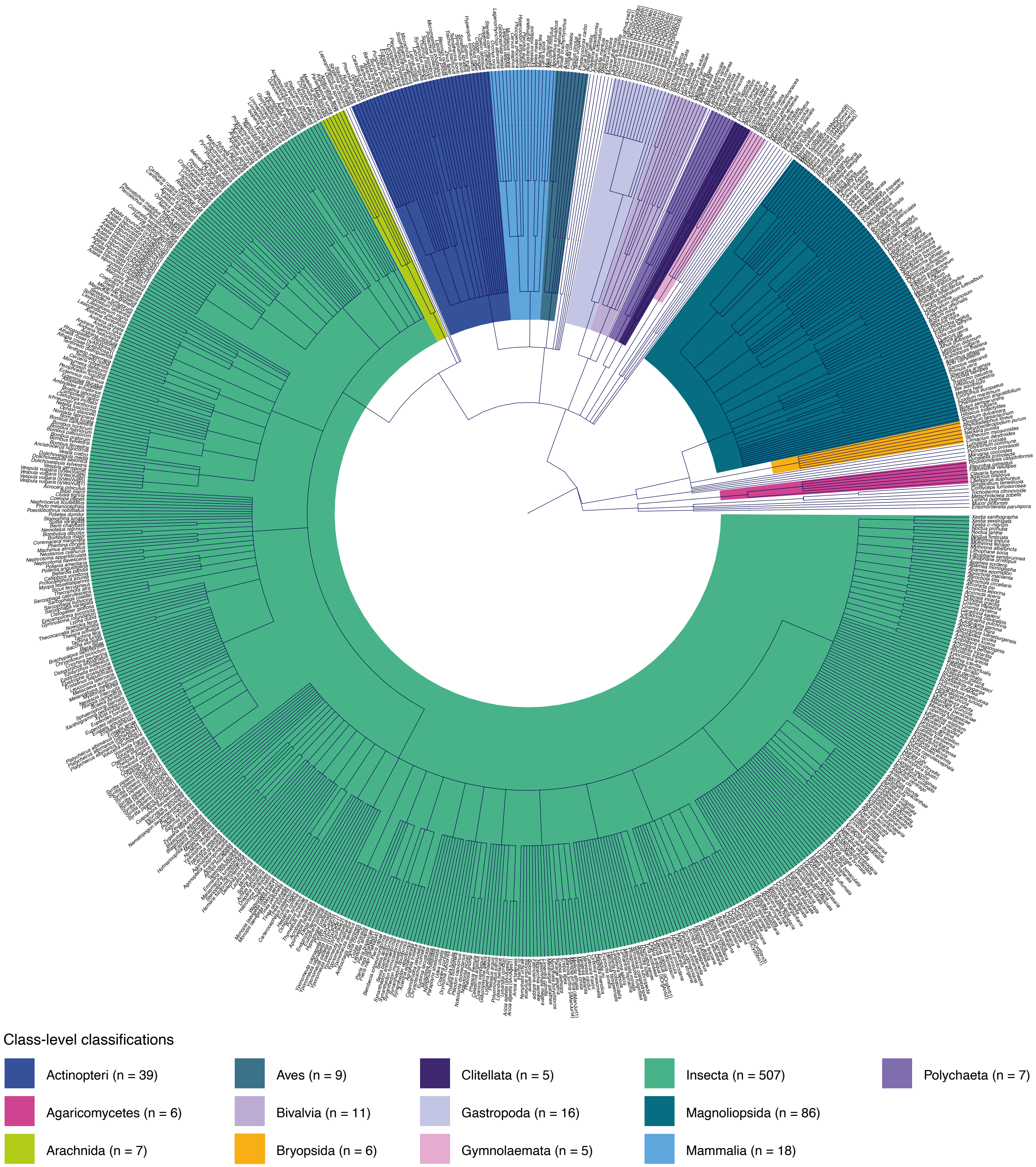


**Supplementary Fig. 3 |** Phylogenetic tree constructed from samples with complete taxonomic classifications, shaded by their Class-level classification. Seven samples with missing taxonomic information were excluded. Taxonomic classifications with fewer than five samples are not shaded.

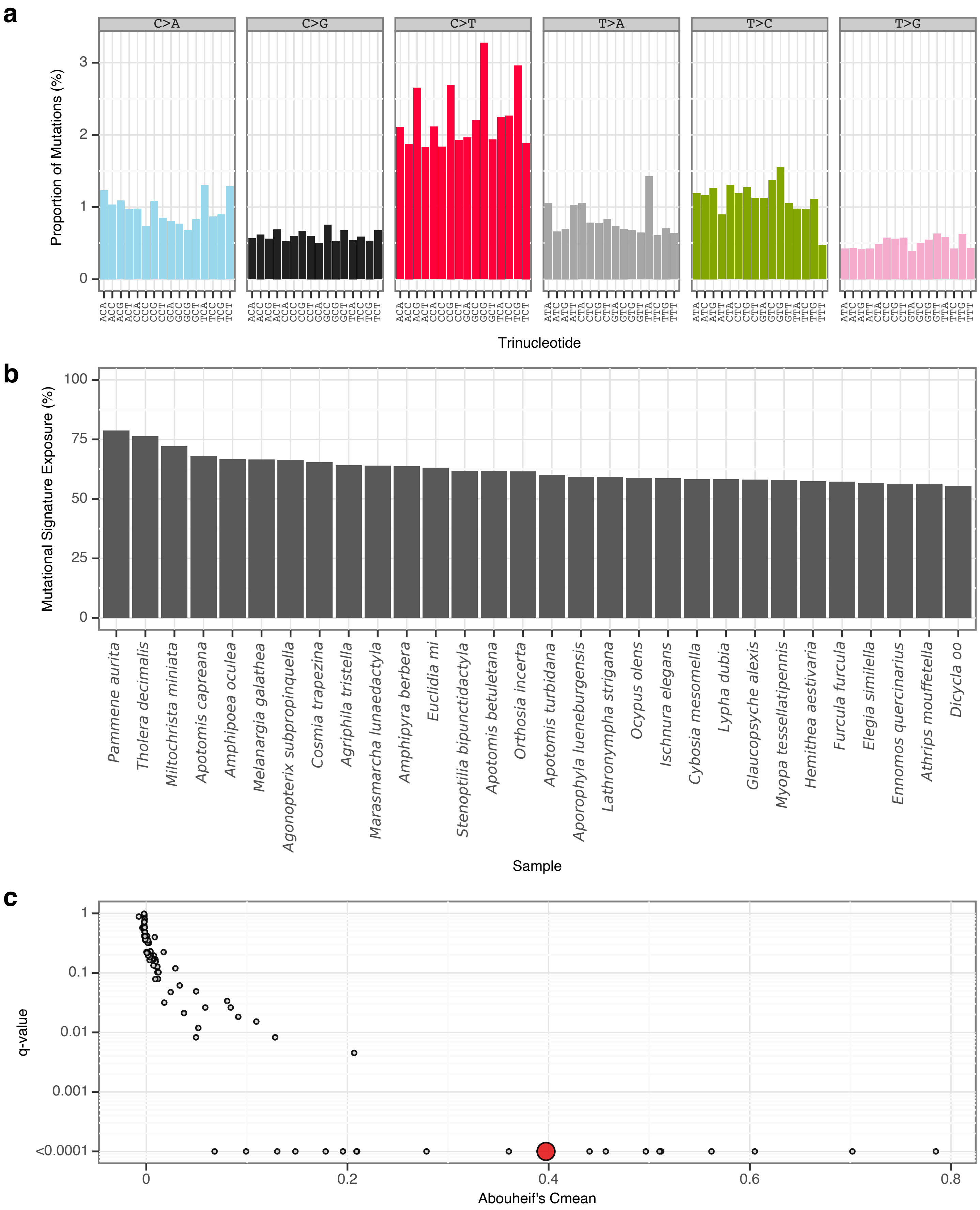


**Supplementary Fig. 4** | sToL1 mutational signature.
**a**, Mutational signature spectrum.
**b**, Bar plot displaying the 30 samples that exhibit the greatest exposure to the mutational signature.
**c**, Phylogenetic signal of the mutational signature.

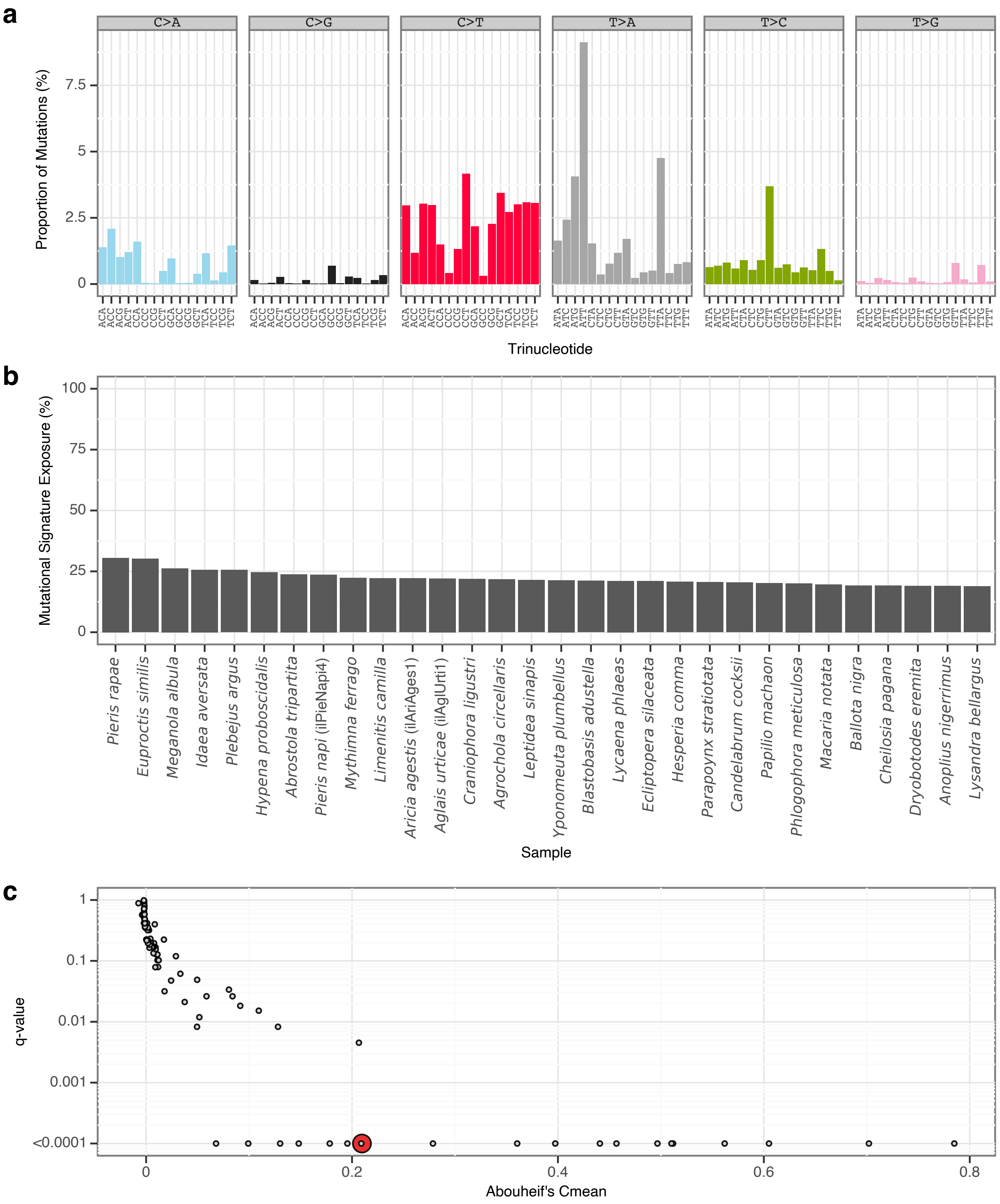


**Supplementary Fig. 5** | sToL2 mutational signature.

**a**, Mutational signature spectrum.

**b**, Bar plot displaying the 30 samples that exhibit the greatest exposure to the mutational signature.

**c**, Phylogenetic signal of the mutational signature.

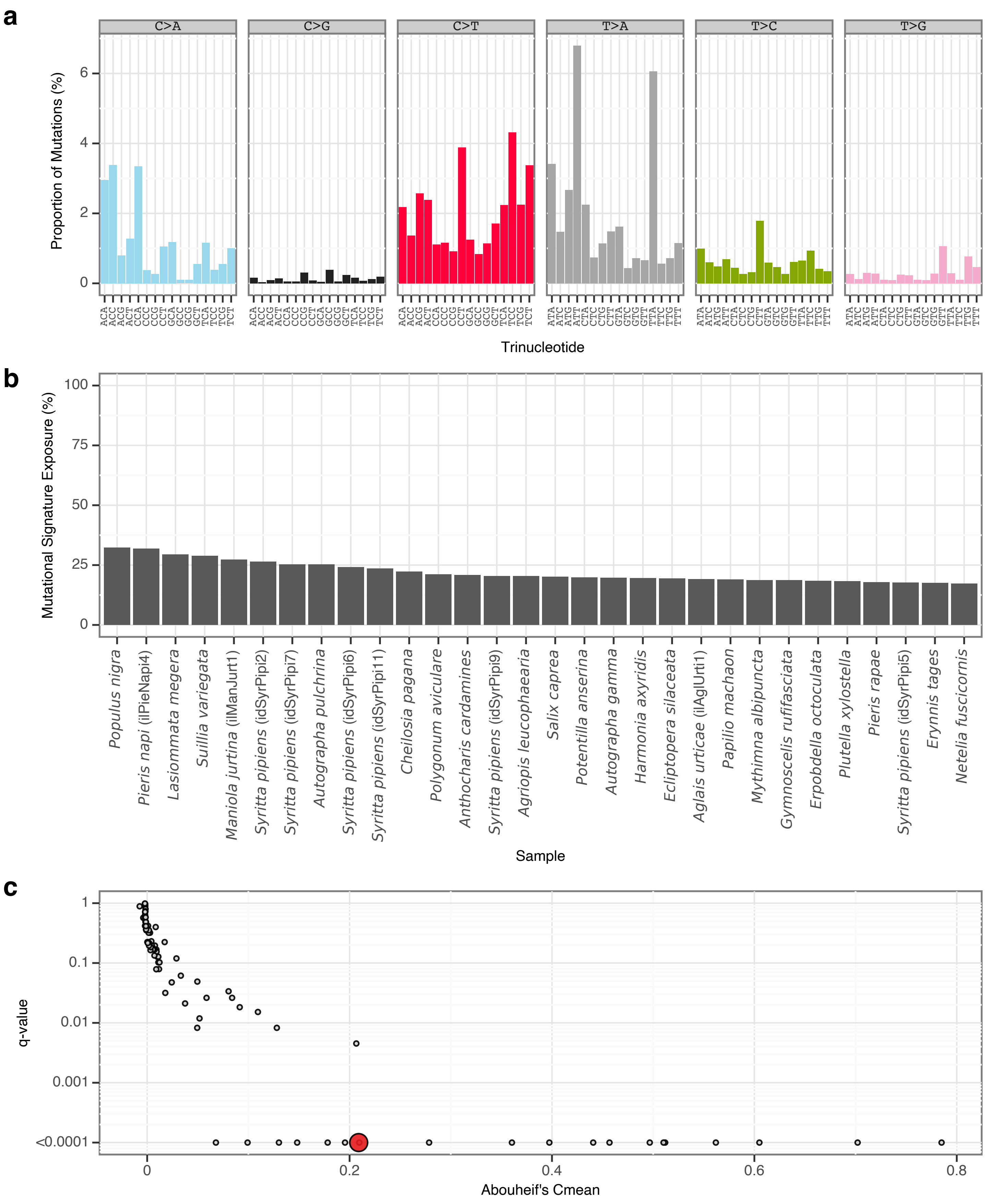


**Supplementary Fig. 6** | sToL3 mutational signature.

**a**, Mutational signature spectrum.

**b**, Bar plot displaying the 30 samples that exhibit the greatest exposure to the mutational signature.

**c**, Phylogenetic signal of the mutational signature.

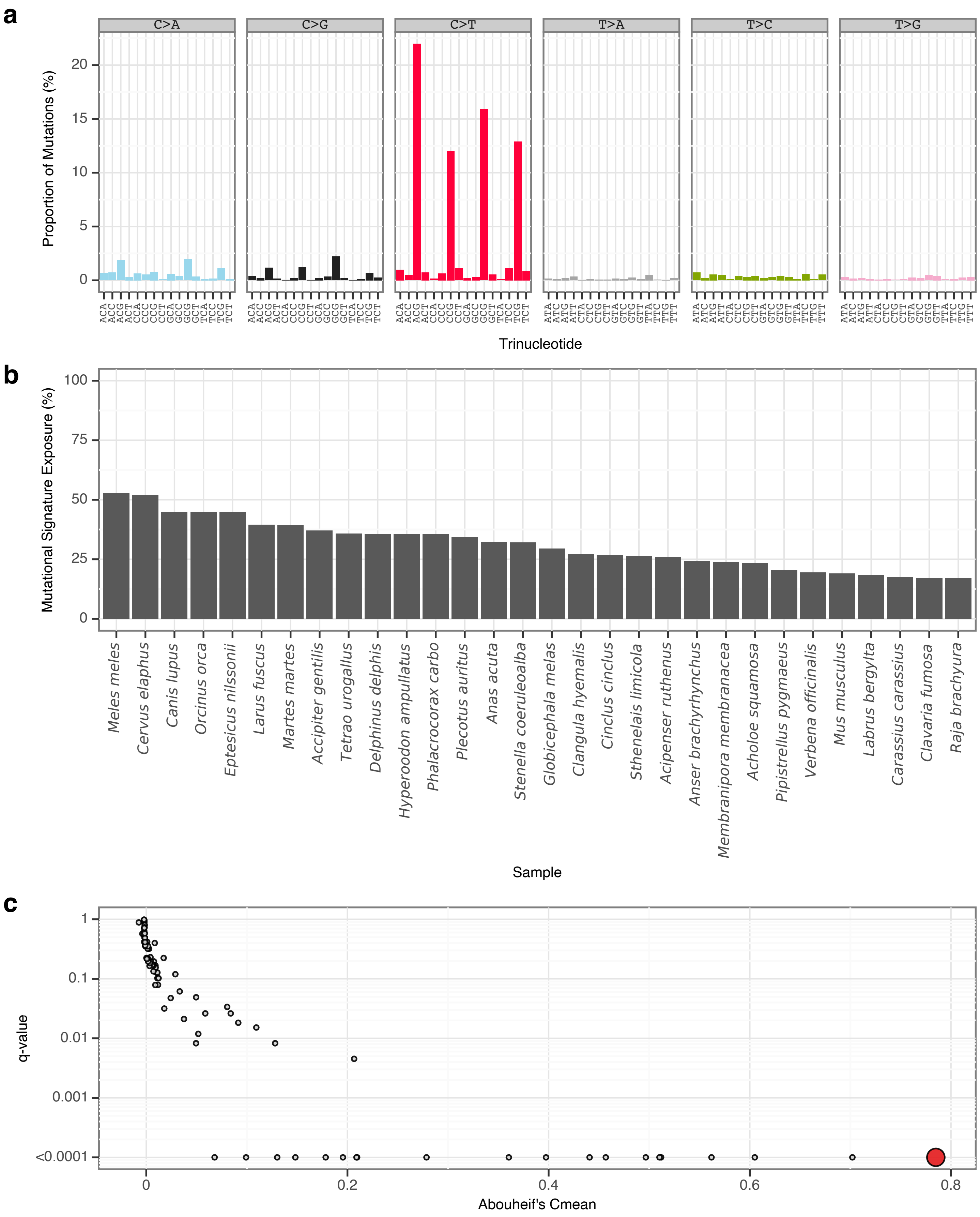


**Supplementary Fig. 7** | sToL4 mutational signature.

**a**, Mutational signature spectrum.

**b**, Bar plot displaying the 30 samples that exhibit the greatest exposure to the mutational signature.

**c**, Phylogenetic signal of the mutational signature.

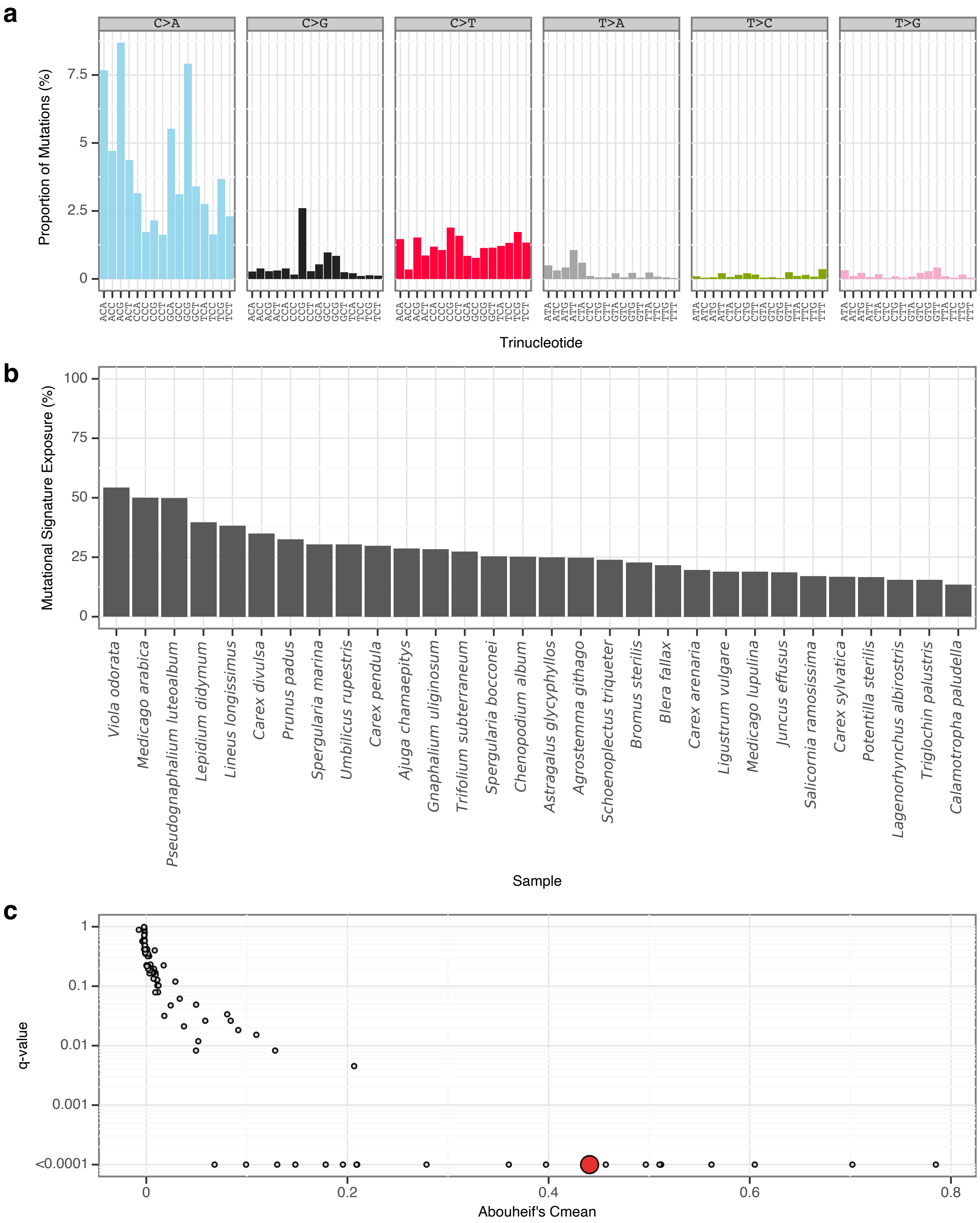


**Supplementary Fig. 8** | sToL5 mutational signature.
**a**, Mutational signature spectrum.
**b**, Bar plot displaying the 30 samples that exhibit the greatest exposure to the mutational signature.
**c**, Phylogenetic signal of the mutational signature.

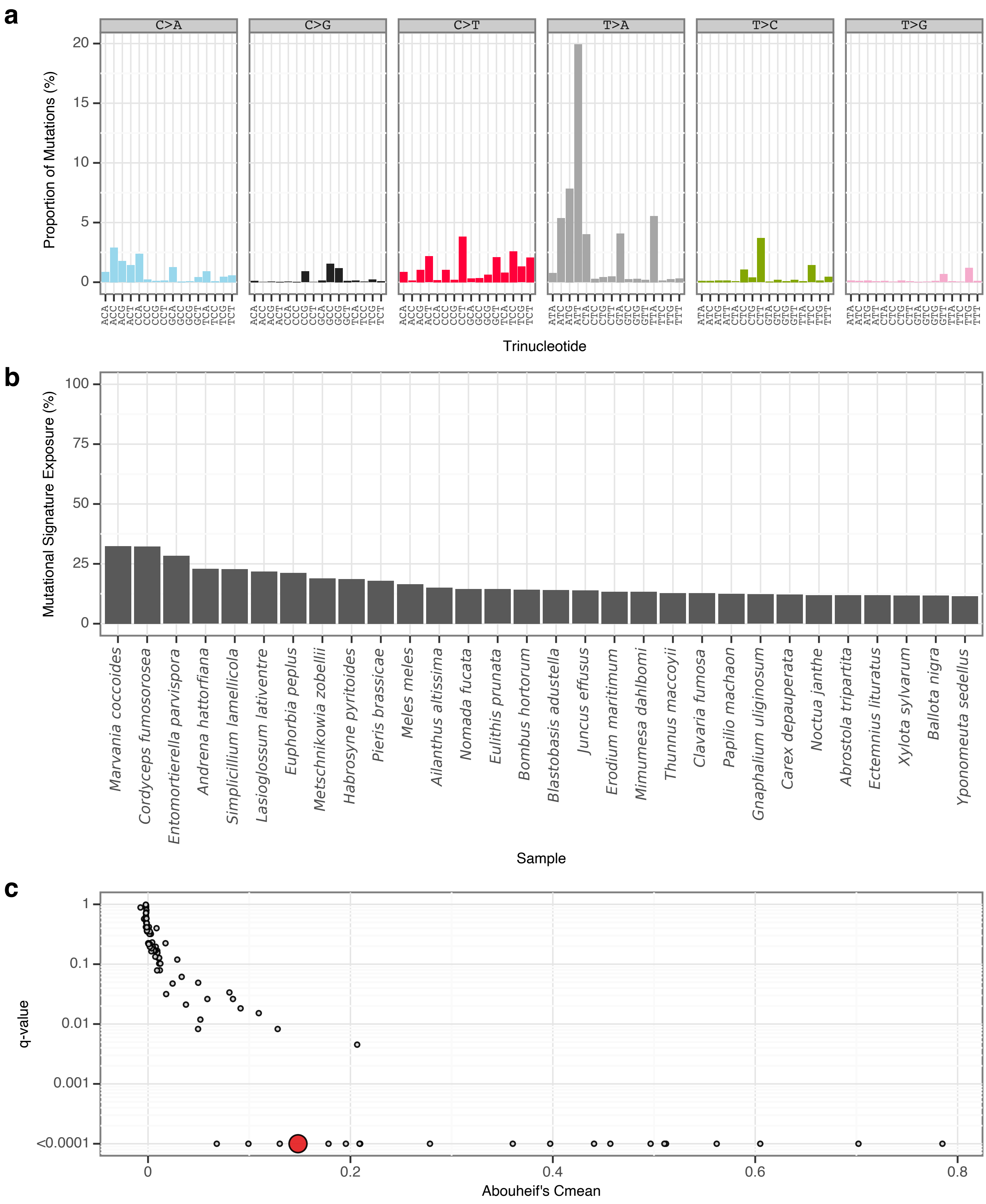


**Supplementary Fig. 9** | sToL6 mutational signature.
**a**, Mutational signature spectrum.
**b**, Bar plot displaying the 30 samples that exhibit the greatest exposure to the mutational signature.
**c**, Phylogenetic signal of the mutational signature.

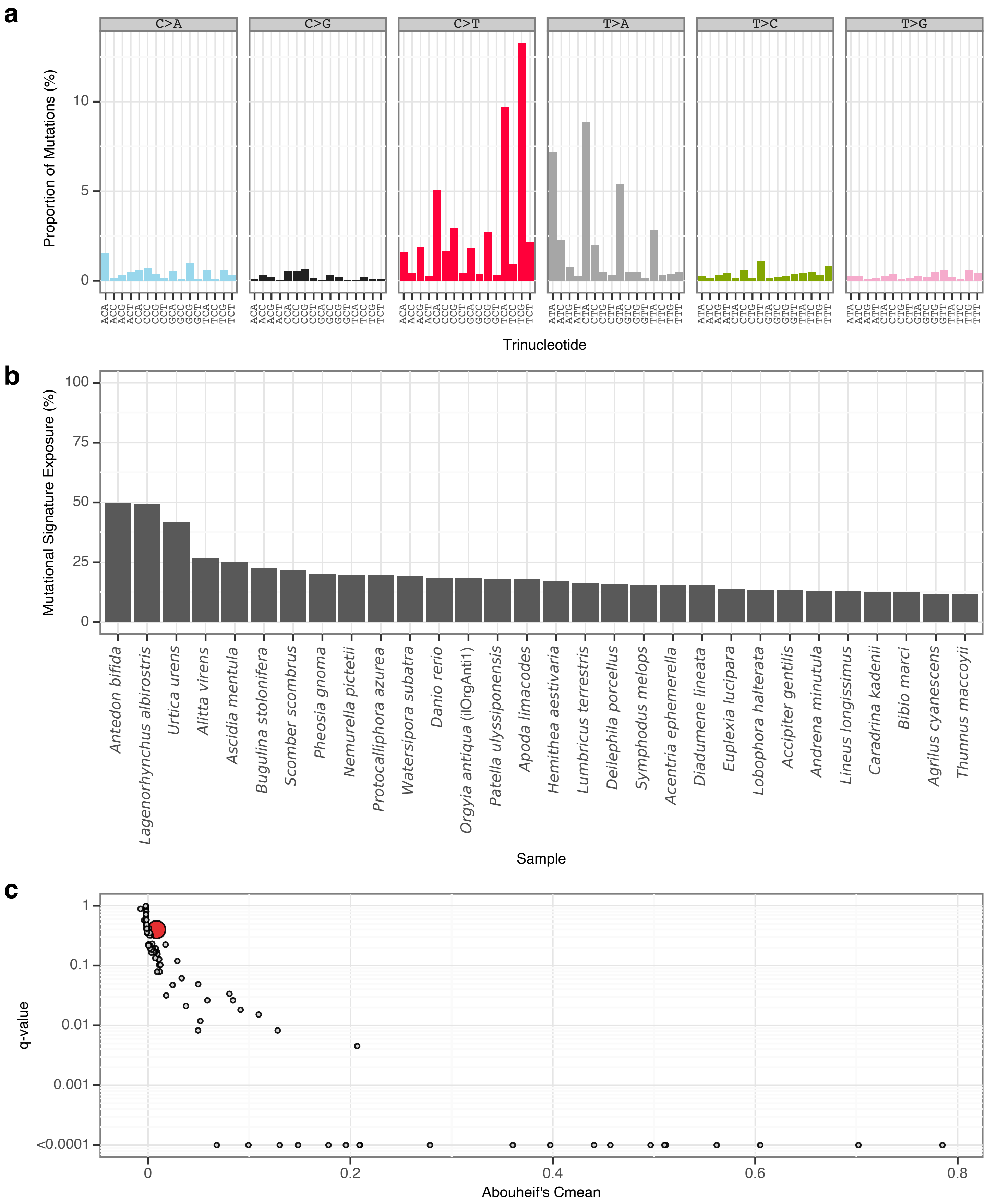


**Supplementary Fig. 10** | sToL7 mutational signature.

**a**, Mutational signature spectrum.

**b**, Bar plot displaying the 30 samples that exhibit the greatest exposure to the mutational signature.

**c**, Phylogenetic signal of the mutational signature.

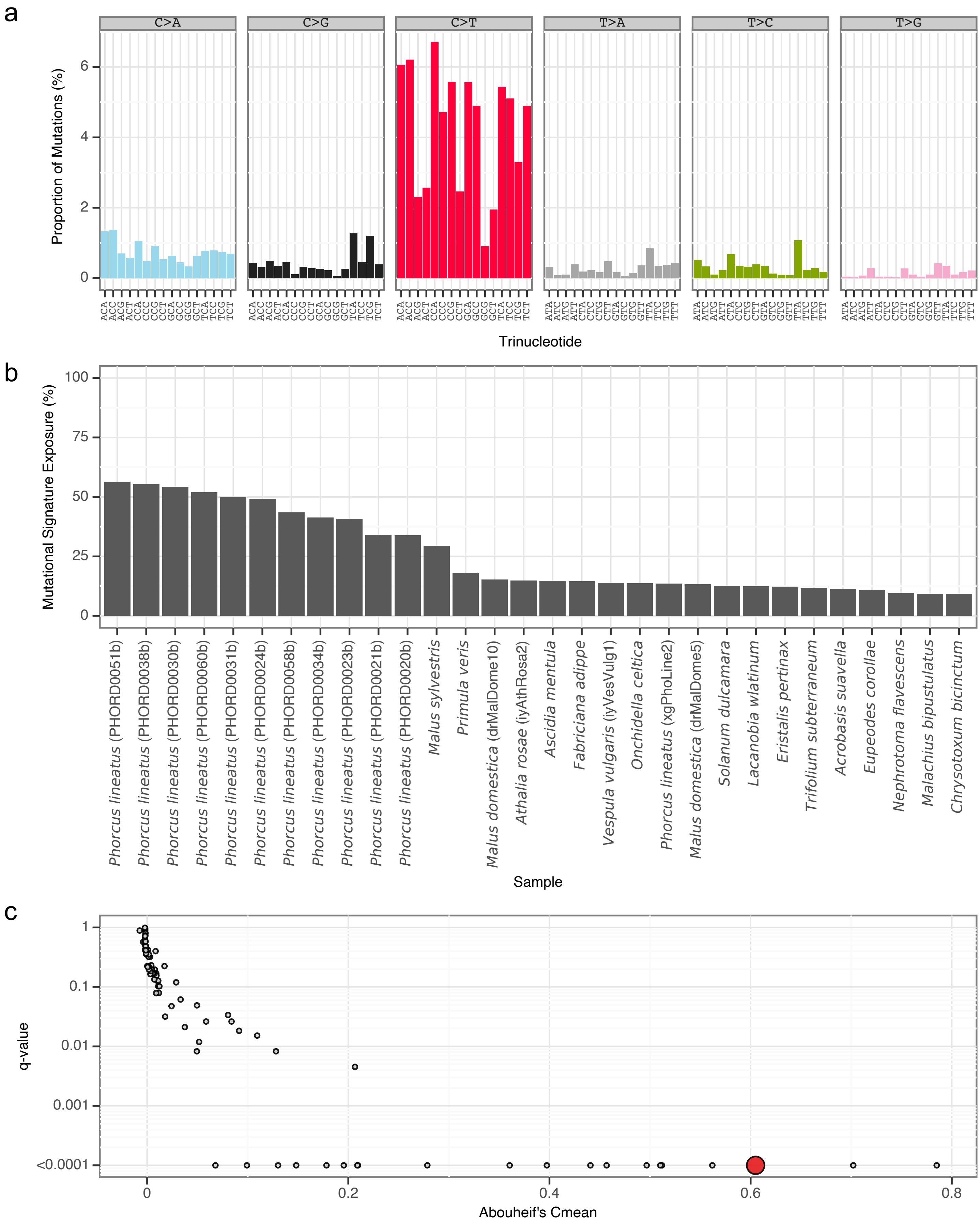


**Supplementary Fig. 11** I sToL8 mutational signature.
**a**, Mutational signature spectrum.
**b**, Bar plot displaying the 30 samples that exhibit the greatest exposure to the mutational signature.
**c**, Phylogenetic signal of the mutational signature.

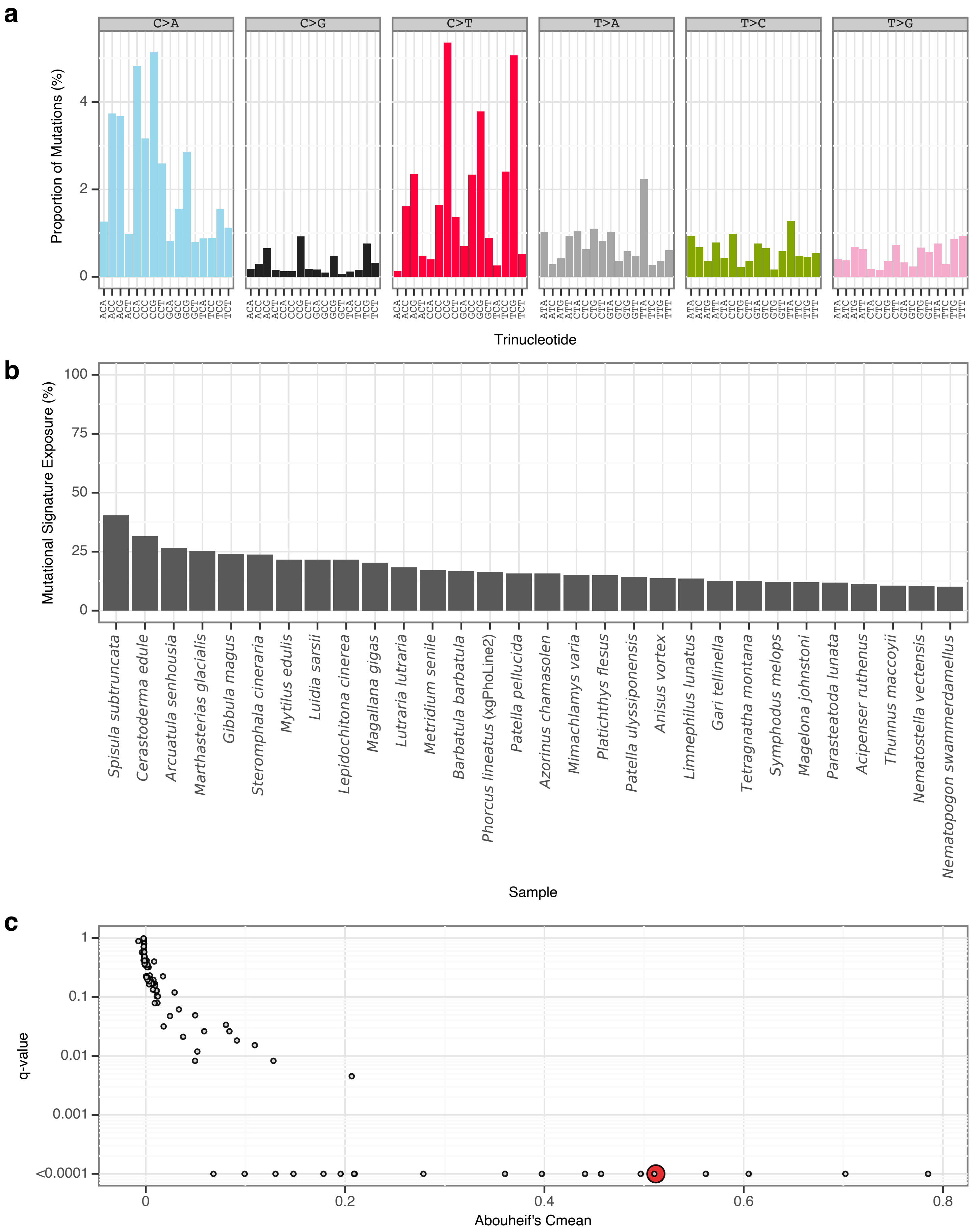


**Supplementary Fig. 12** | sToL9 mutational signature.

**a**, Mutational signature spectrum.

**b**, Bar plot displaying the 30 samples that exhibit the greatest exposure to the mutational signature.

**c**, Phylogenetic signal of the mutational signature.

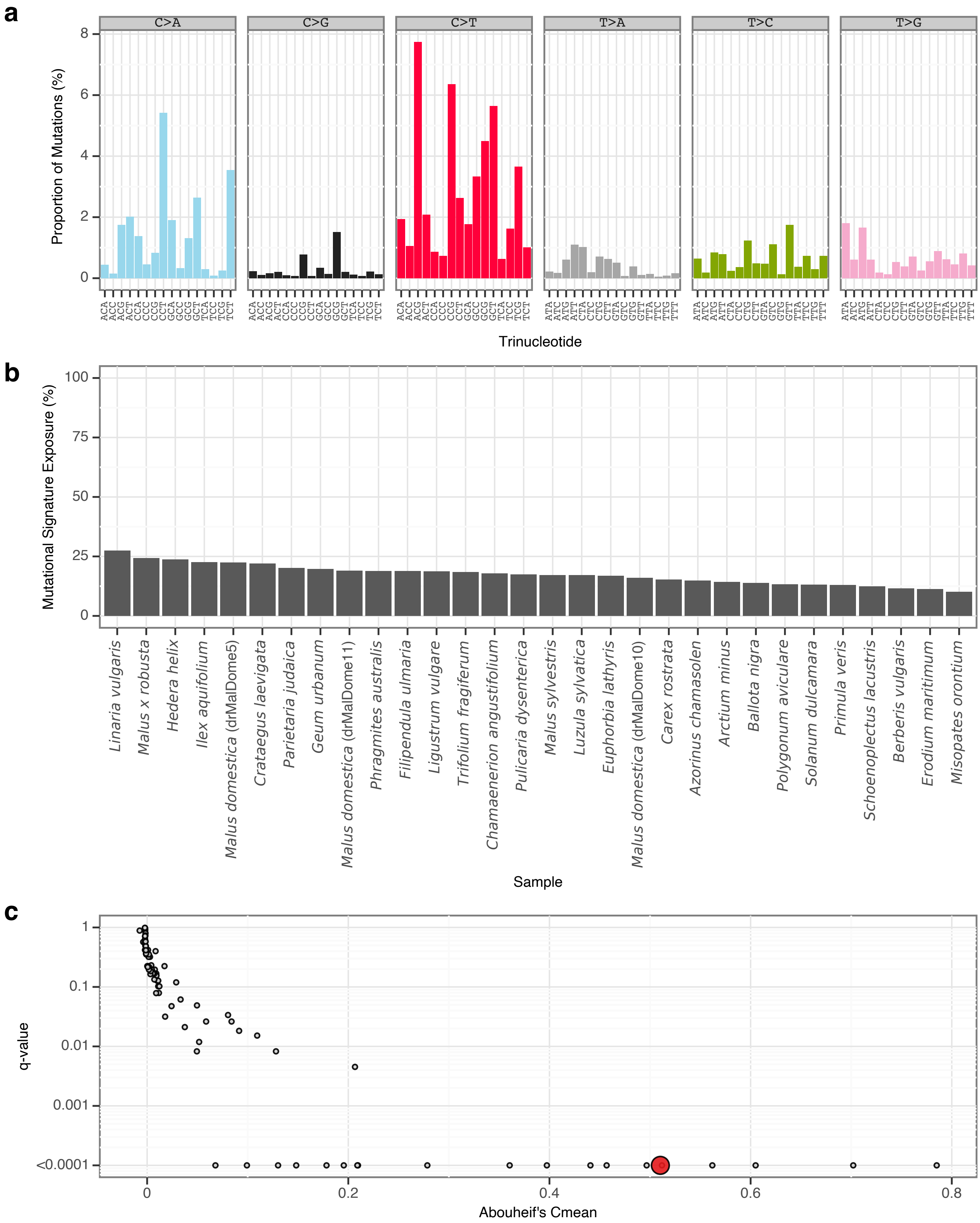


**Supplementary Fig. 13** l sToL10 mutational signature.

**a**, Mutational signature spectrum.

**b**, Bar plot displaying the 30 samples that exhibit the greatest exposure to the mutational signature.

**c**, Phylogenetic signal of the mutational signature.

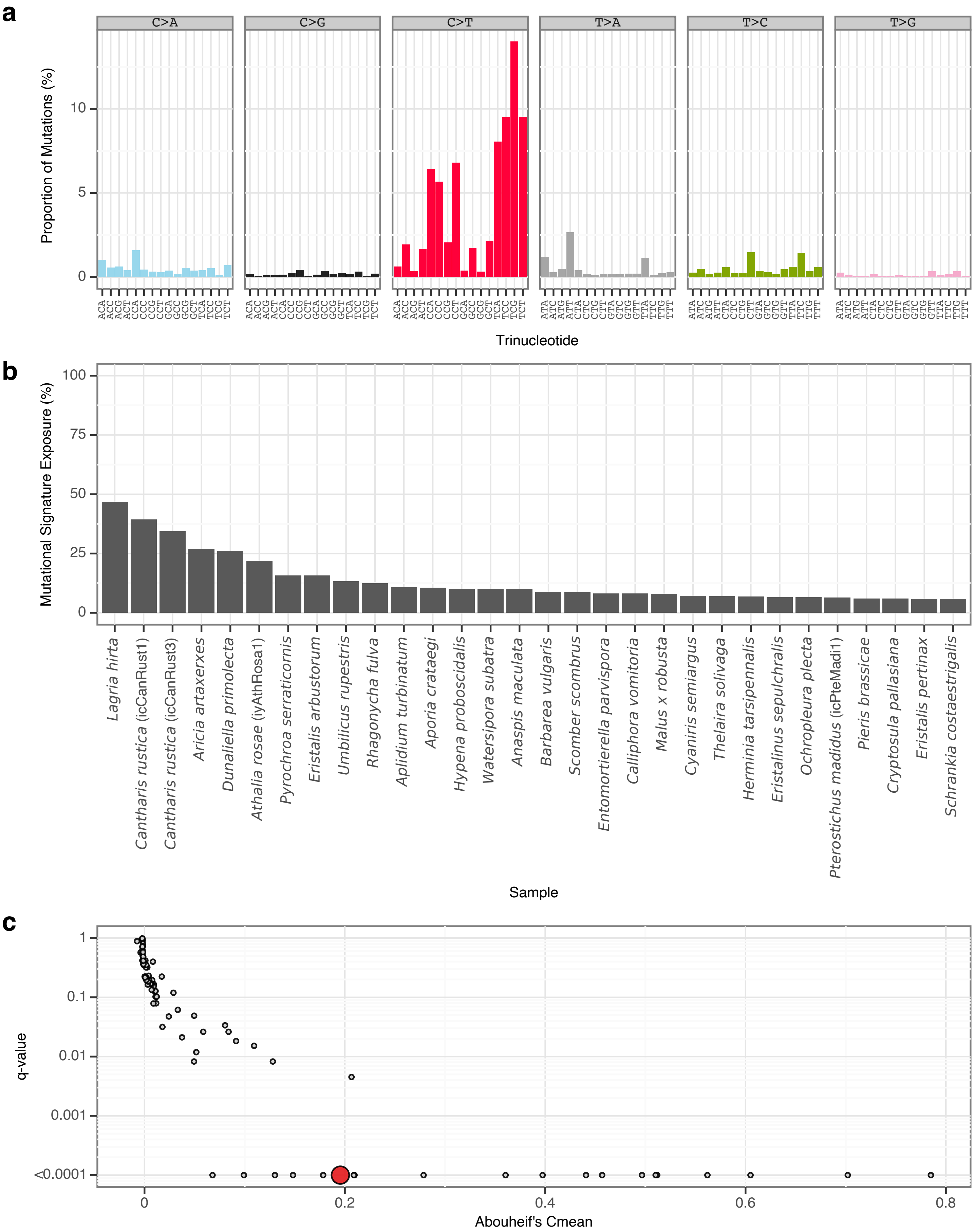


**Supplementary Fig. 14** | sToL11 mutational signature.
**a**, Mutational signature spectrum.
**b**, Bar plot displaying the 30 samples that exhibit the greatest exposure to the mutational signature.
**c**, Phylogenetic signal of the mutational signature.

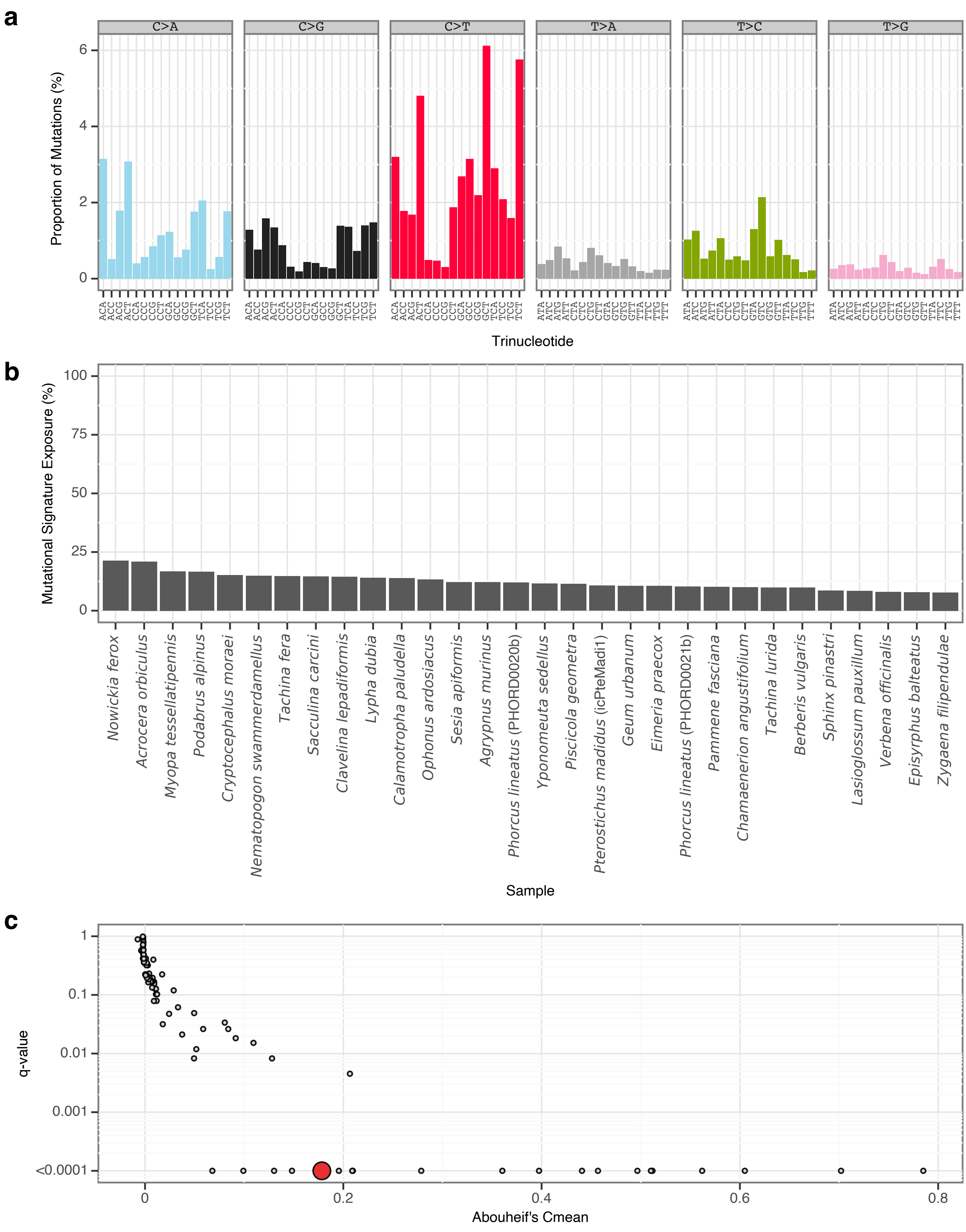


**Supplementary Fig. 15** | sToL12 mutational signature.

**a**, Mutational signature spectrum.

**b**, Bar plot displaying the 30 samples that exhibit the greatest exposure to the mutational signature.

**c**, Phylogenetic signal of the mutational signature.

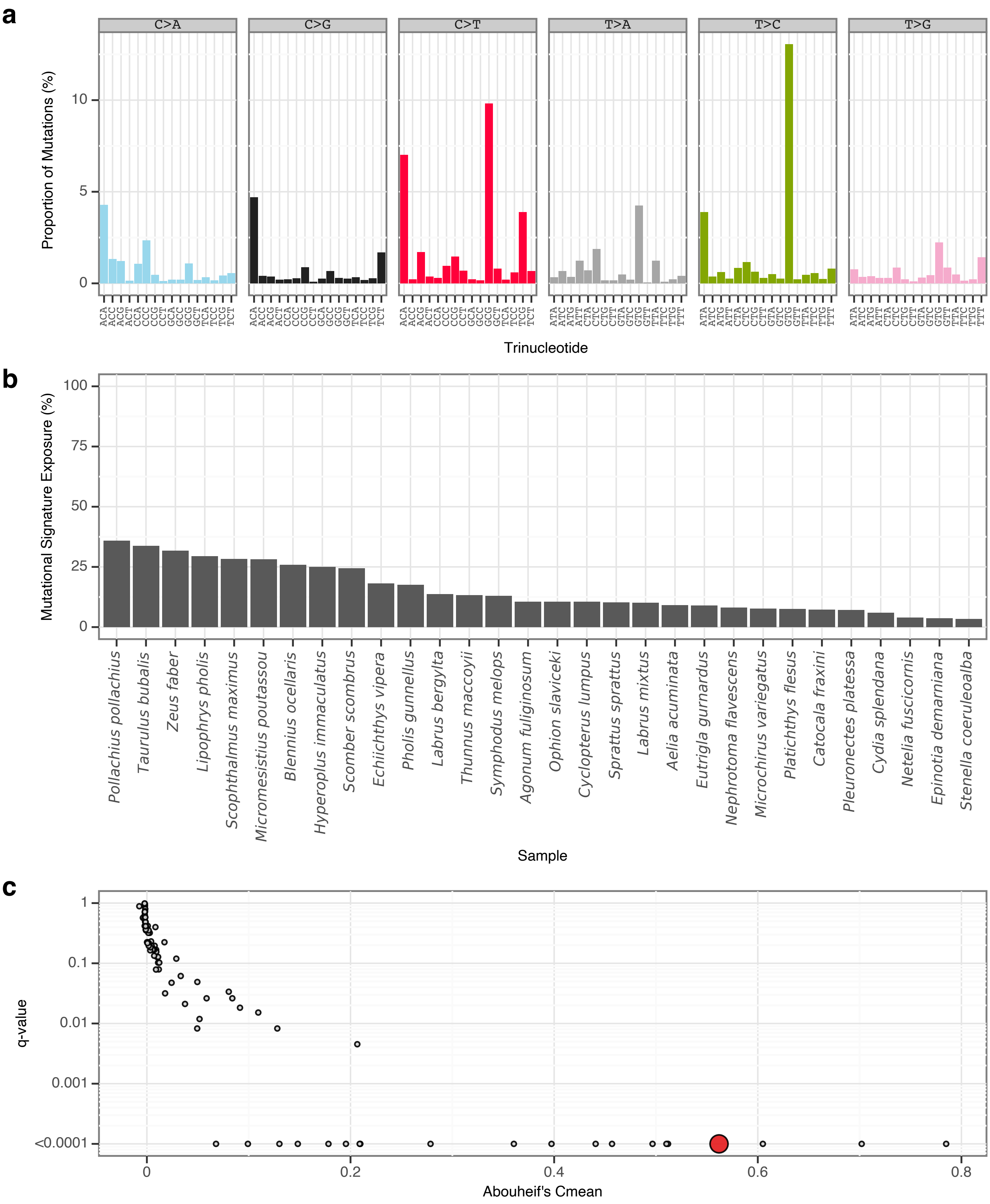


**Supplementary Fig. 16** | sToL13 mutational signature.
**a**, Mutational signature spectrum.
**b**, Bar plot displaying the 30 samples that exhibit the greatest exposure to the mutational signature.
**c**, Phylogenetic signal of the mutational signature.

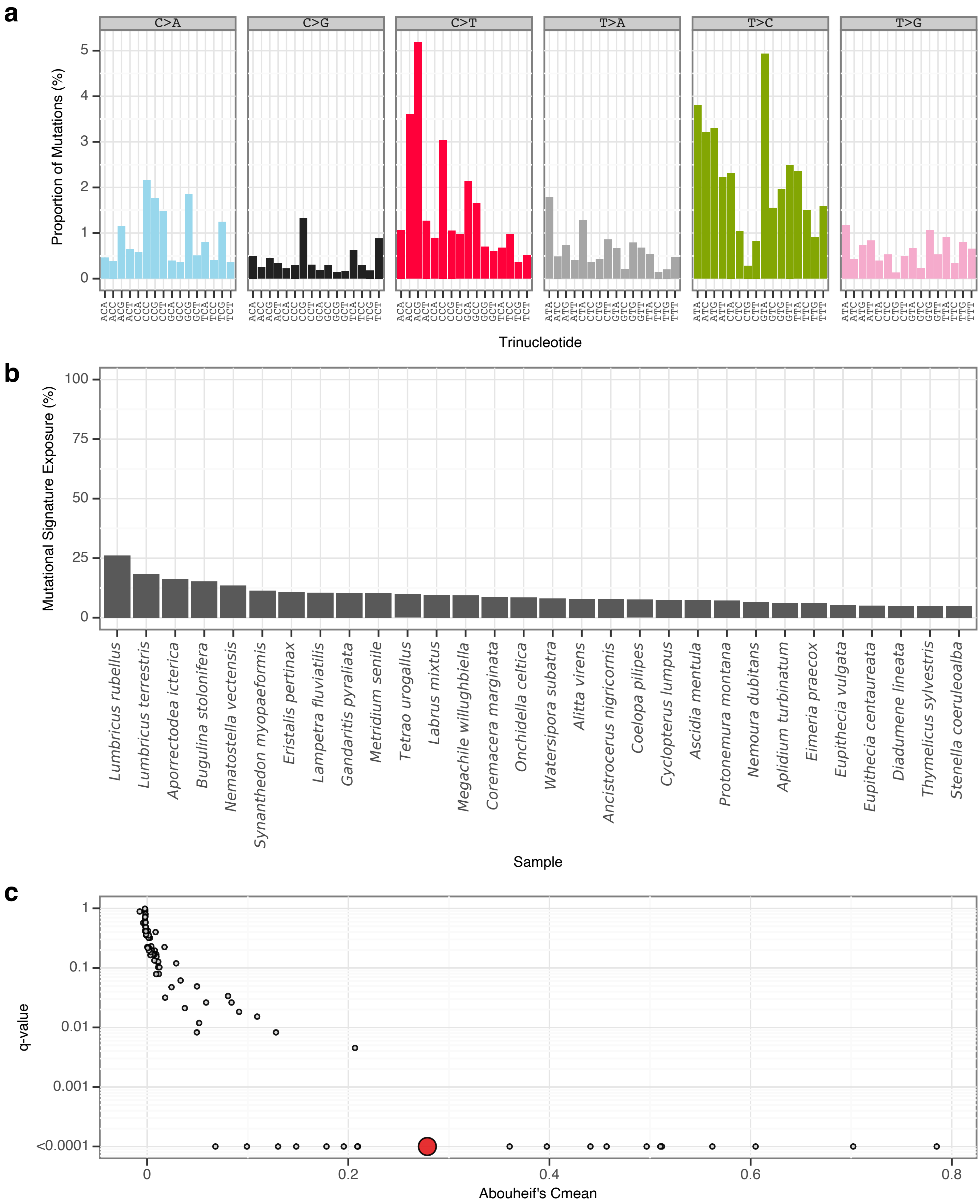


**Supplementary Fig. 17** I sToL14 mutational signature.
**a**, Mutational signature spectrum.
**b**, Bar plot displaying the 30 samples that exhibit the greatest exposure to the mutational signature.
**c**, Phylogenetic signal of the mutational signature.

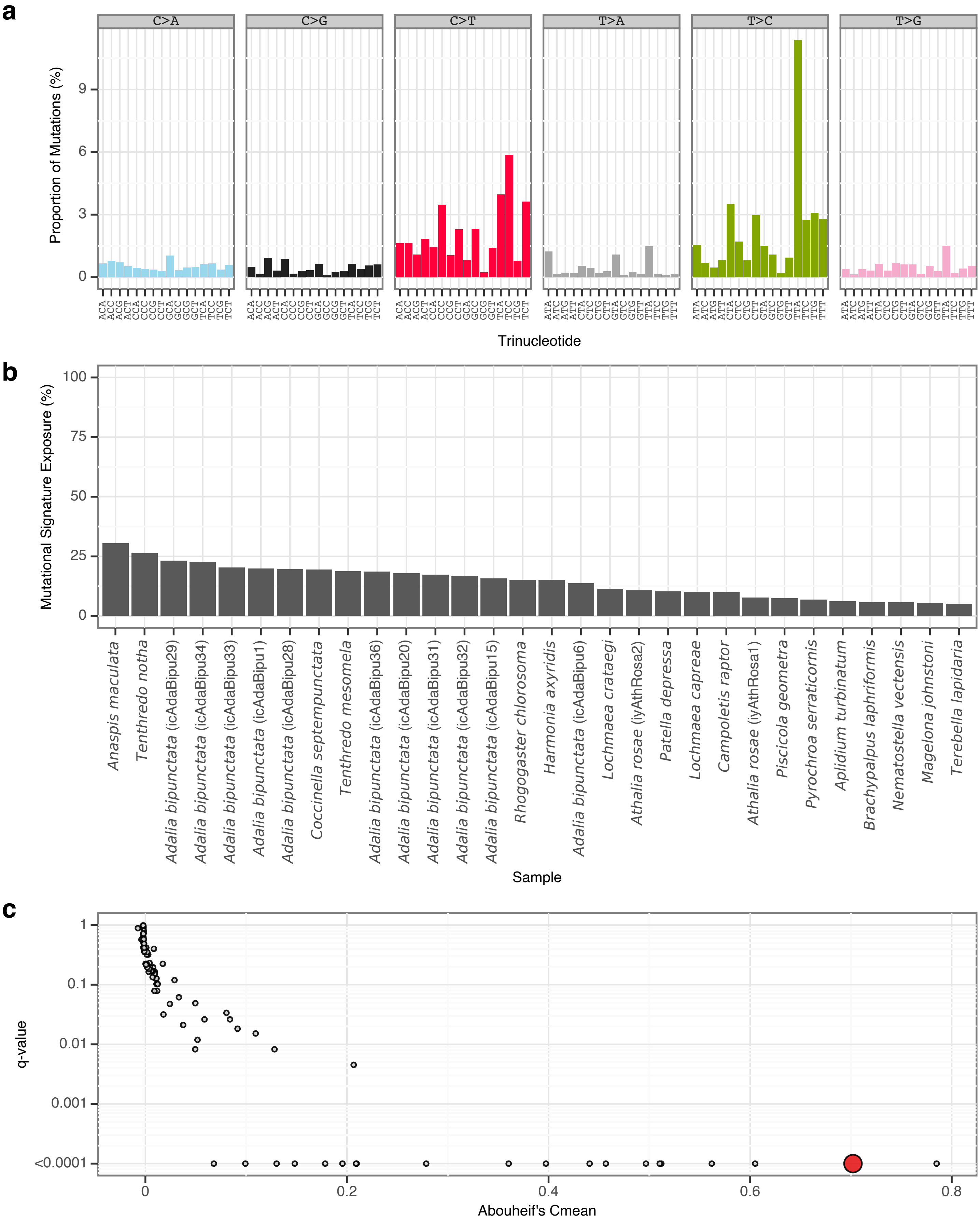


**Supplementary Fig. 18** | sToL15 mutational signature.

**a**, Mutational signature spectrum.

**b**, Bar plot displaying the 30 samples that exhibit the greatest exposure to the mutational signature.

**c**, Phylogenetic signal of the mutational signature.

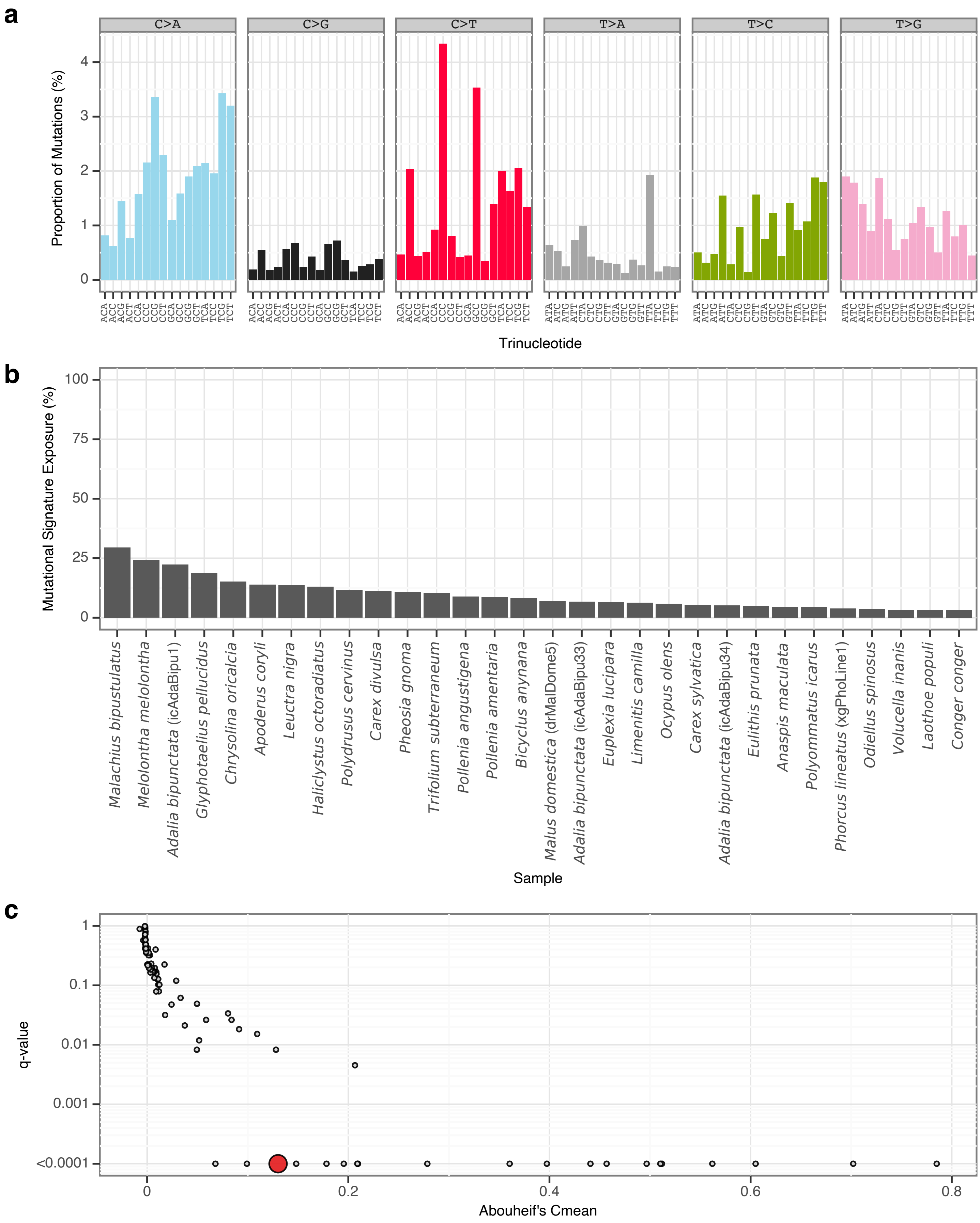


**Supplementary Fig. 19** I sToL16 mutational signature.

**a**, Mutational signature spectrum.

**b**, Bar plot displaying the 30 samples that exhibit the greatest exposure to the mutational signature.

**c**, Phylogenetic signal of the mutational signature.

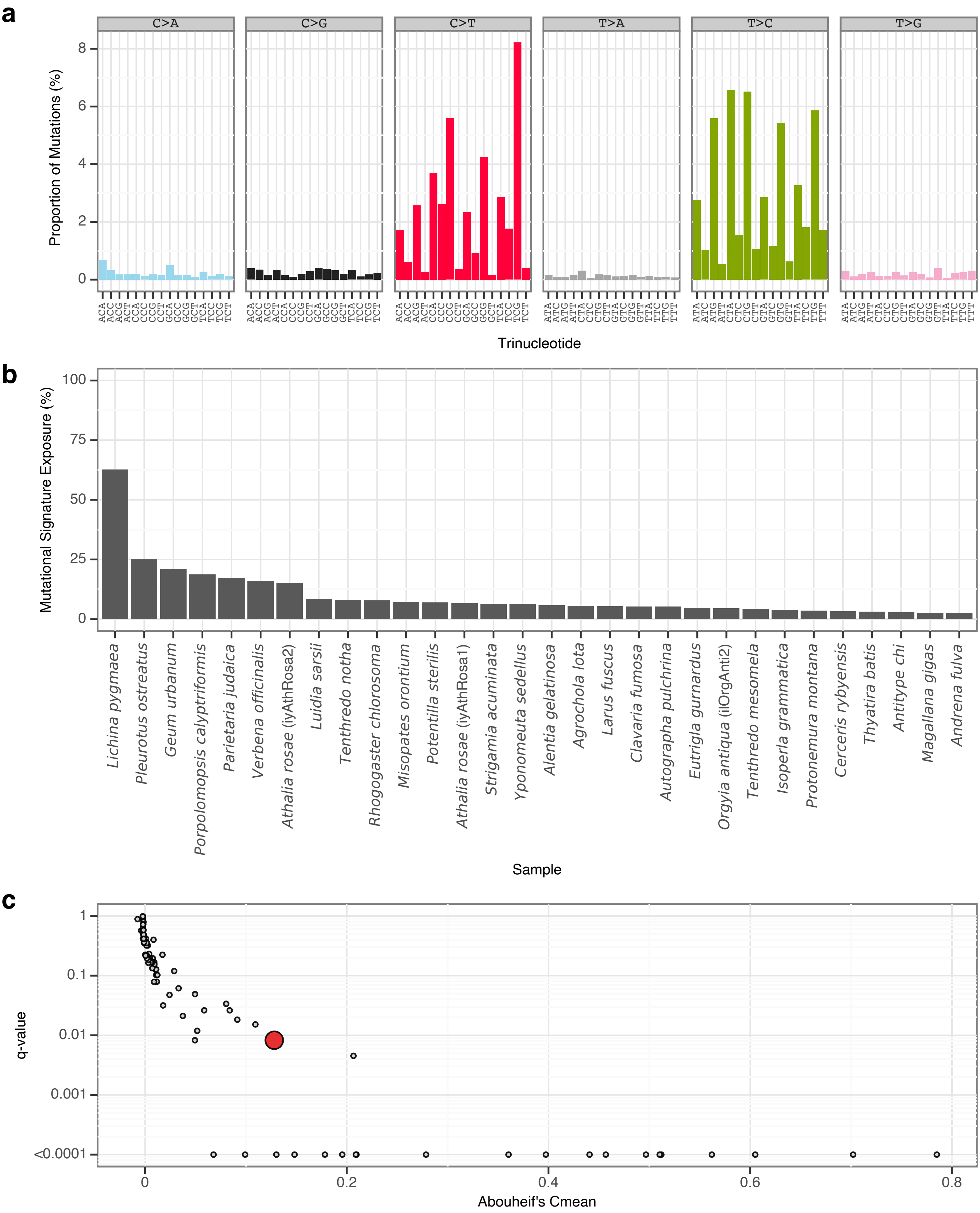


**Supplementary Fig. 20** | sToL17 mutational signature.
**a**, Mutational signature spectrum.
**b**, Bar plot displaying the 30 samples that exhibit the greatest exposure to the mutational signature.
**c**, Phylogenetic signal of the mutational signature.

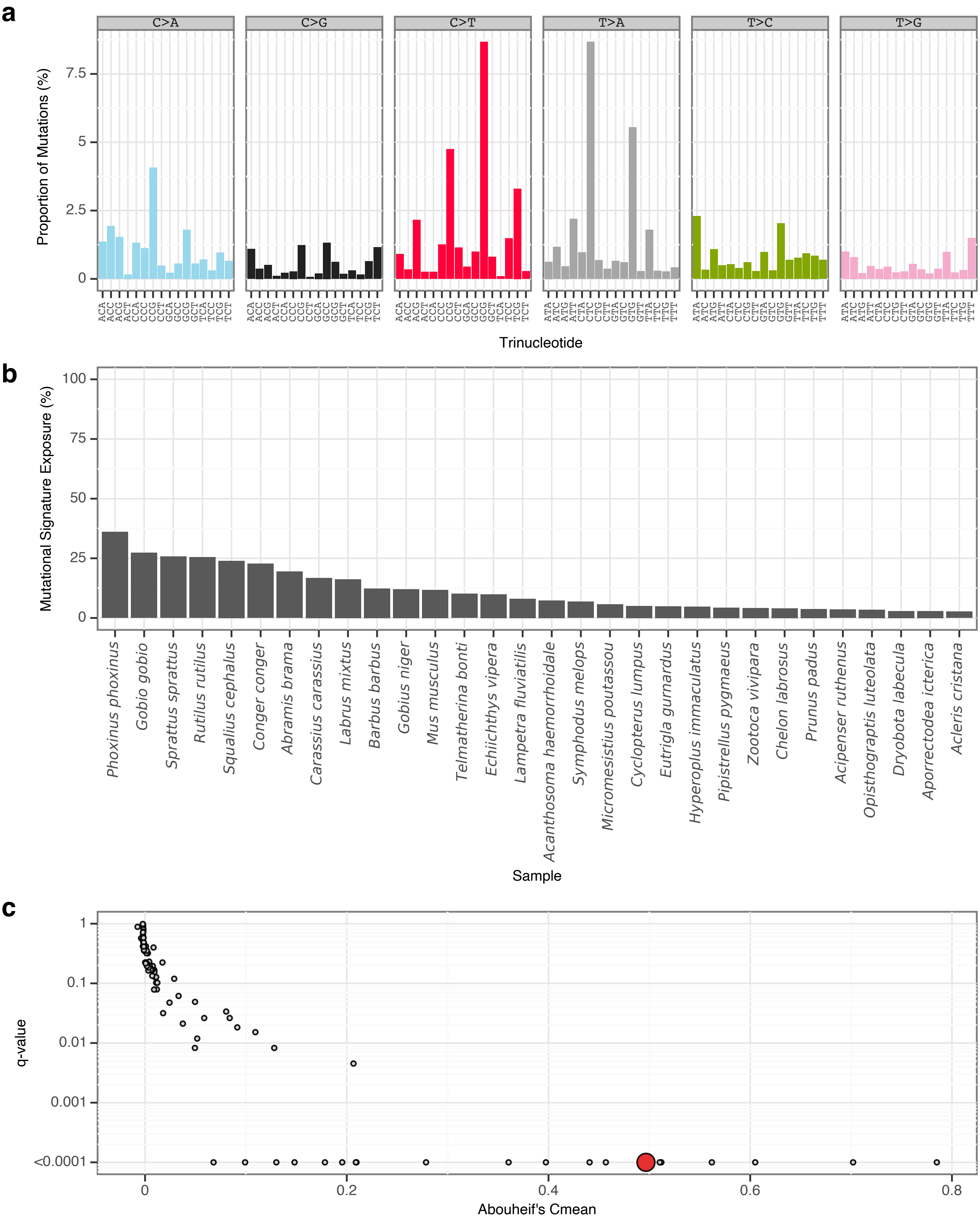


**Supplementary Fig. 21** | sToL18 mutational signature.

**a**, Mutational signature spectrum.

**b**, Bar plot displaying the 30 samples that exhibit the greatest exposure to the mutational signature.

**c**, Phylogenetic signal of the mutational signature.

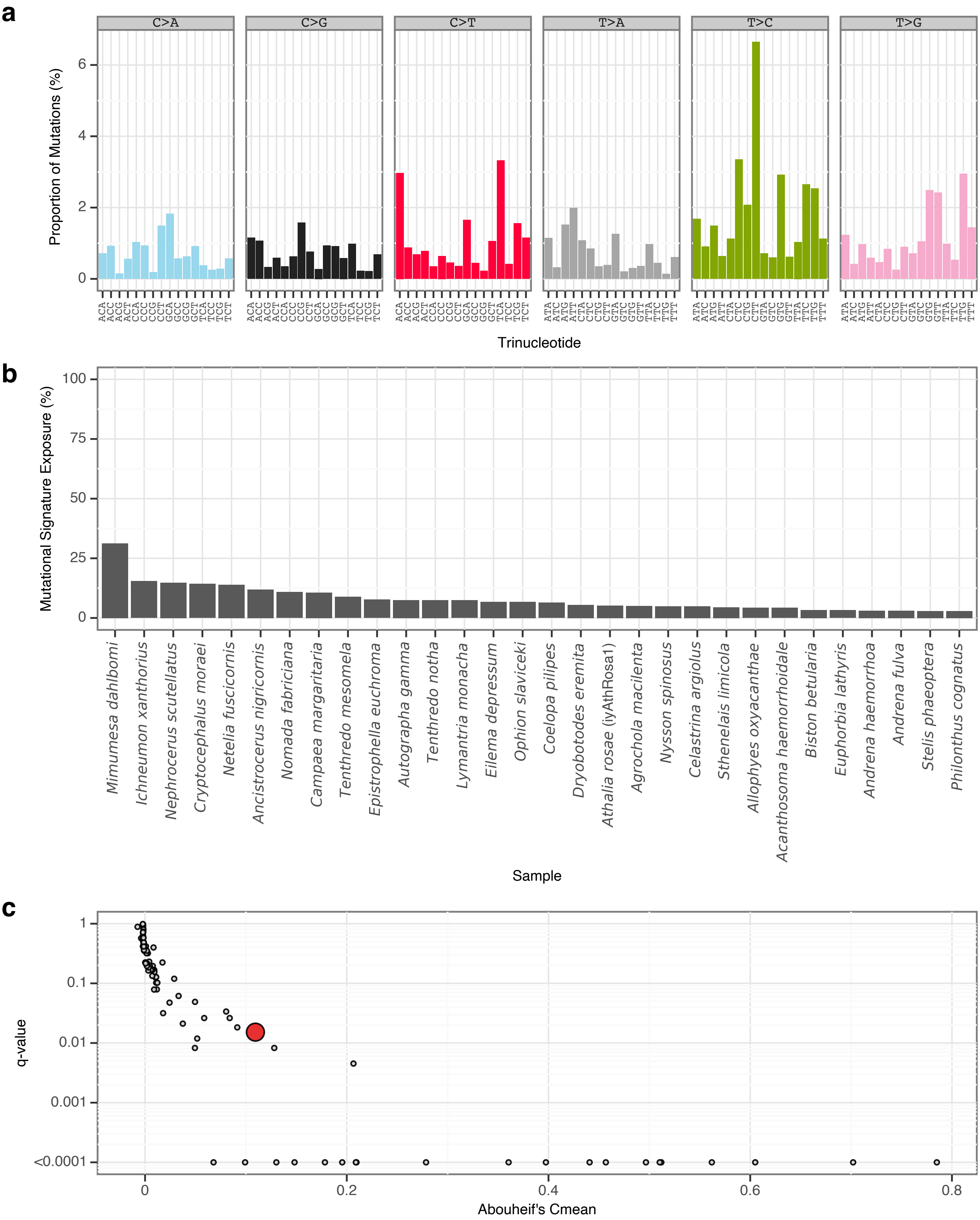


**Supplementary Fig. 22** | sToL19 mutational signature.

**a**, Mutational signature spectrum.

**b**, Bar plot displaying the 30 samples that exhibit the greatest exposure to the mutational signature.

**c**, Phylogenetic signal of the mutational signature.

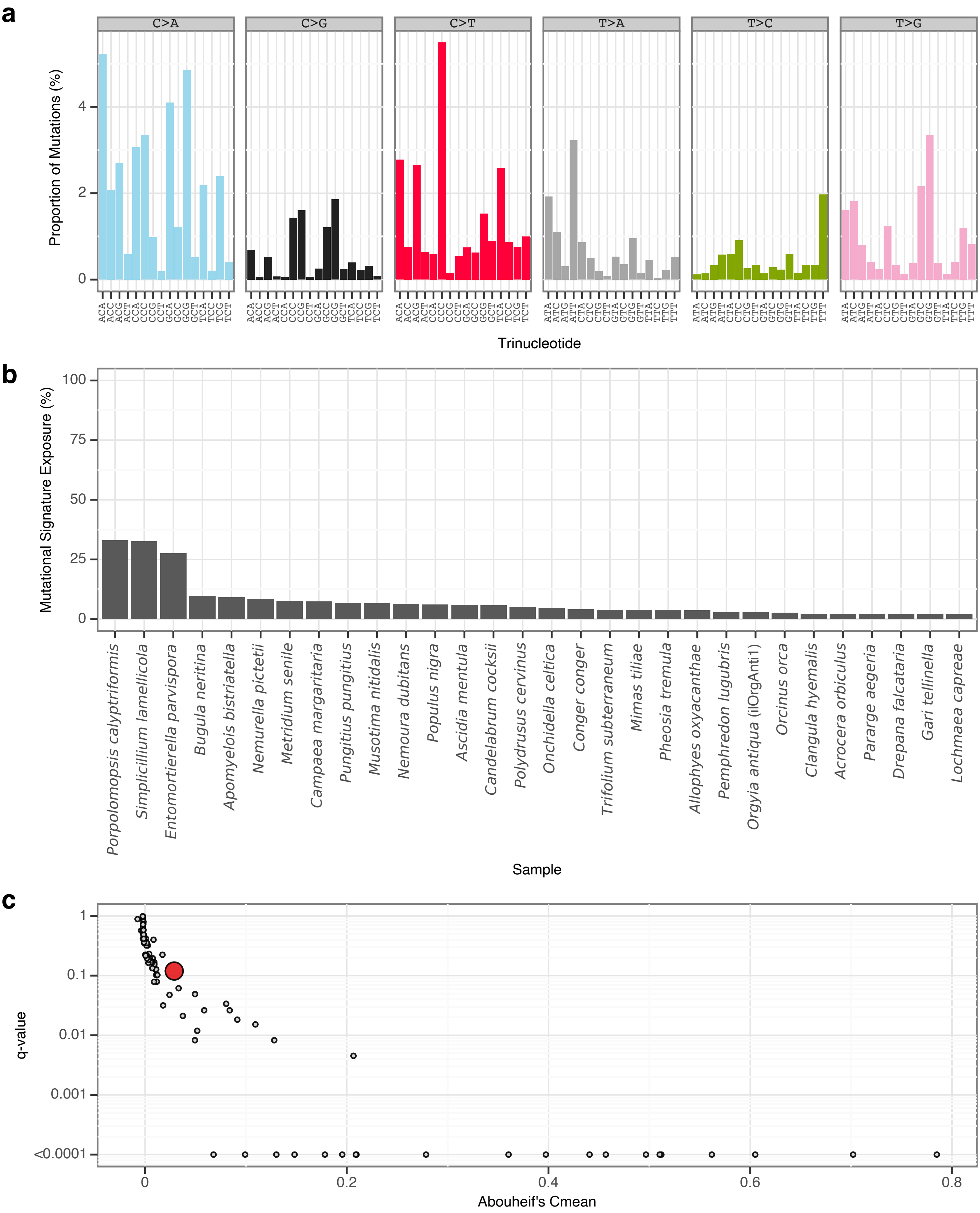


**Supplementary Fig. 23** | sToL20 mutational signature.

**a**, Mutational signature spectrum.

**b**, Bar plot displaying the 30 samples that exhibit the greatest exposure to the mutational signature.

**c**, Phylogenetic signal of the mutational signature.

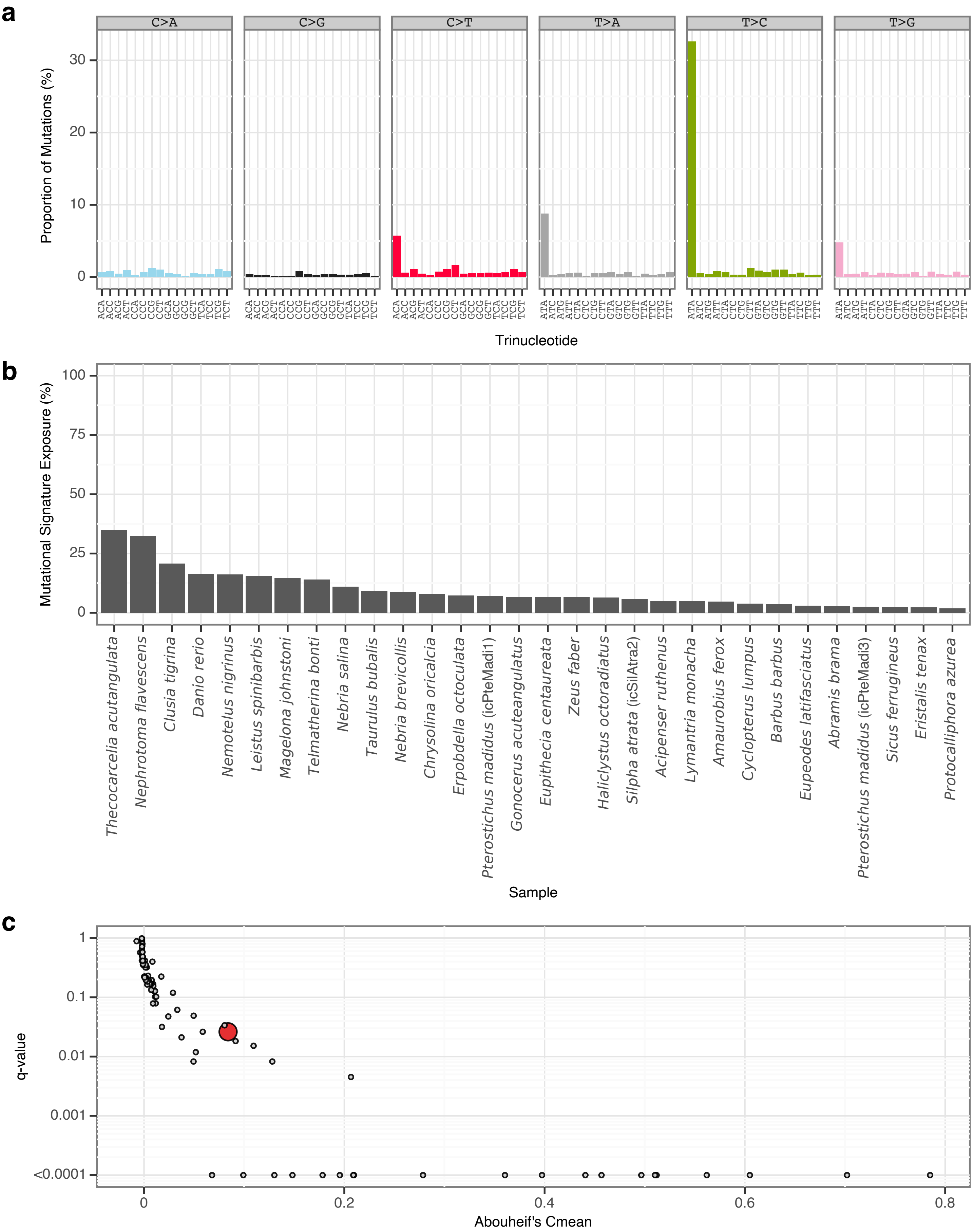


**Supplementary Fig. 24** | sToL21 mutational signature.
**a**, Mutational signature spectrum.
**b**, Bar plot displaying the 30 samples that exhibit the greatest exposure to the mutational signature.
**c**, Phylogenetic signal of the mutational signature.

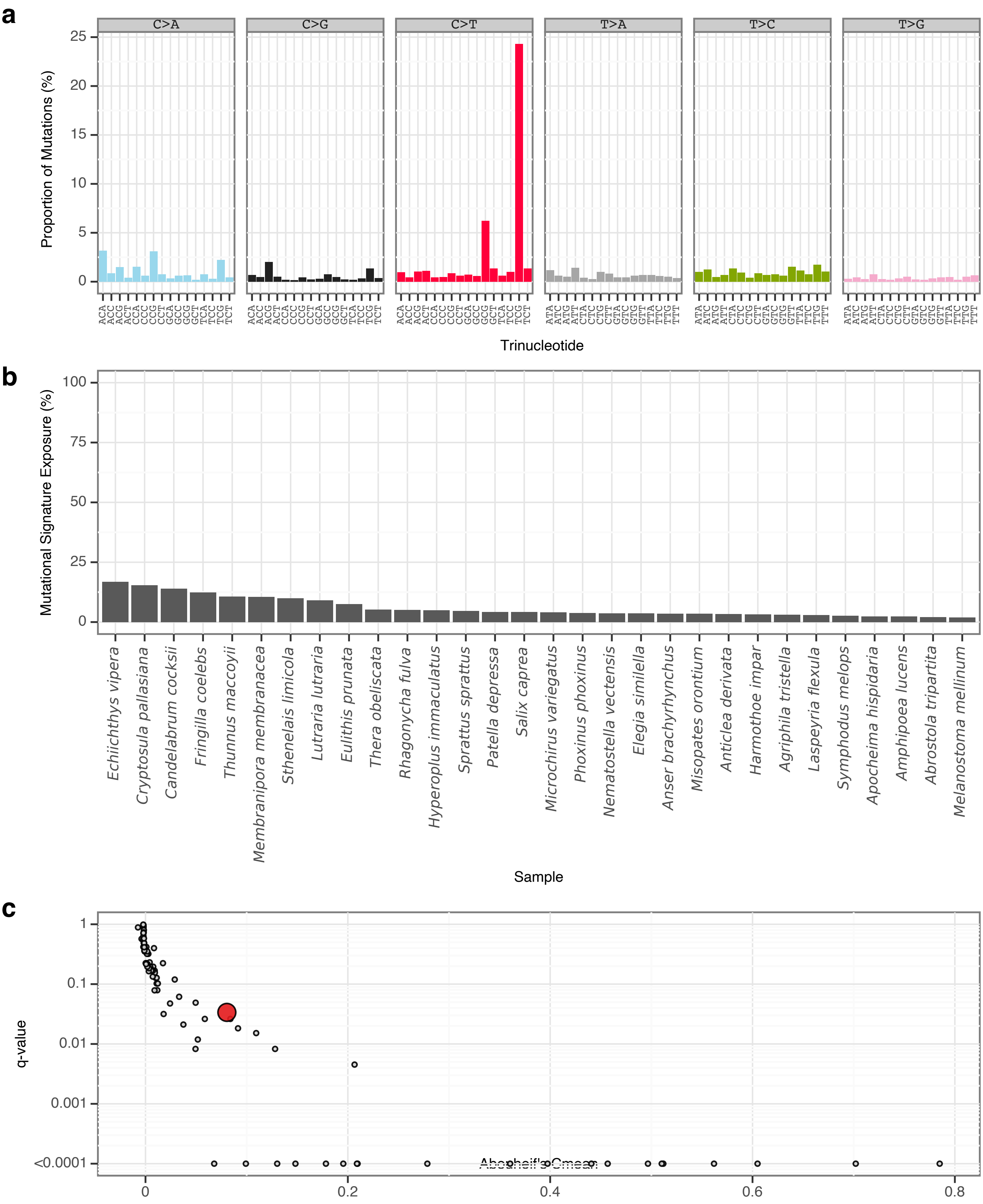


**Supplementary Fig. 25** | sToL22 mutational signature.
**a**, Mutational signature spectrum.
**b**, Bar plot displaying the 30 samples that exhibit the greatest exposure to the mutational signature.
**c**, Phylogenetic signal of the mutational signature.

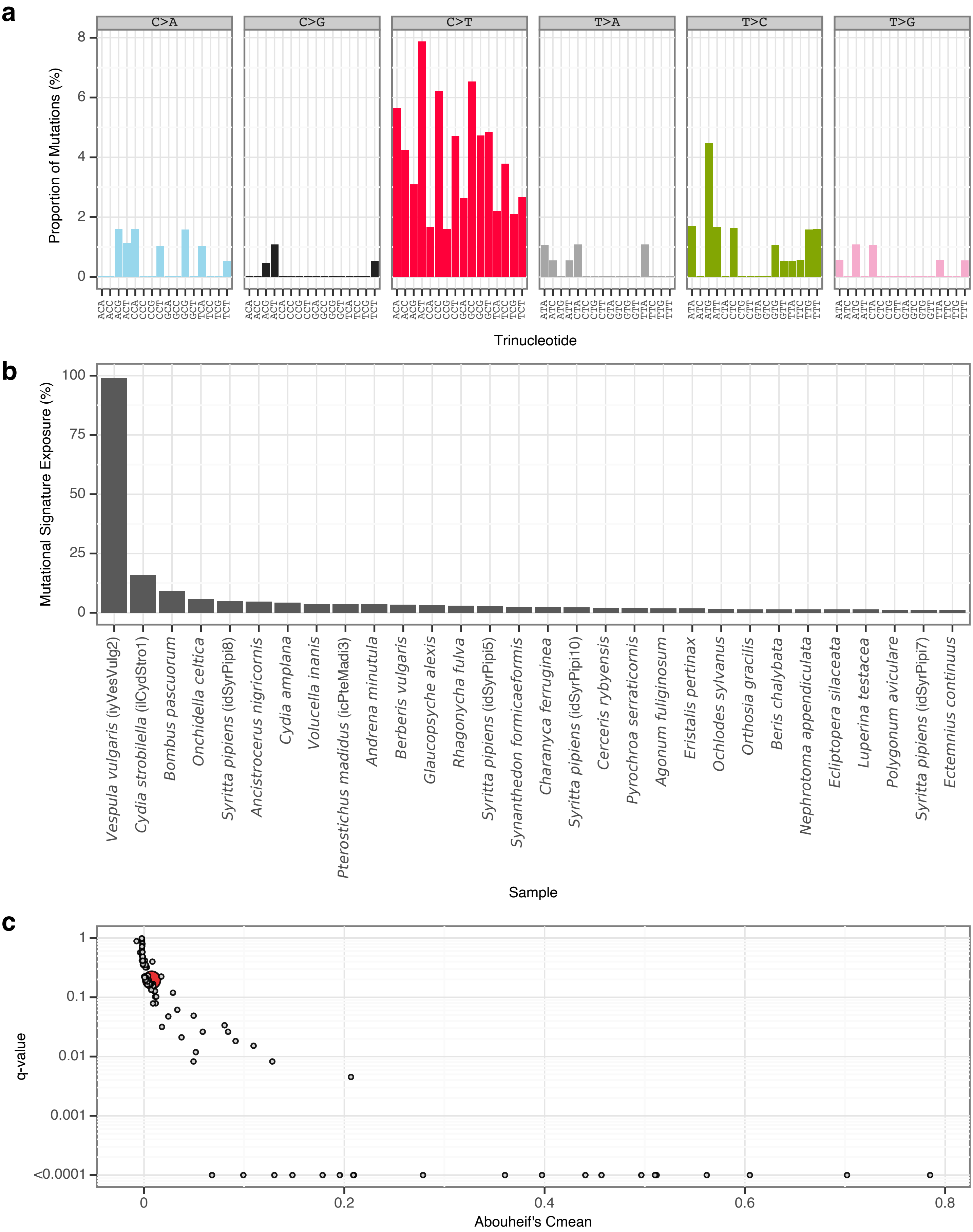


**Supplementary Fig. 26** | sToL23 mutational signature.

**a**, Mutational signature spectrum.

**b**, Bar plot displaying the 30 samples that exhibit the greatest exposure to the mutational signature.

**c**, Phylogenetic signal of the mutational signature.

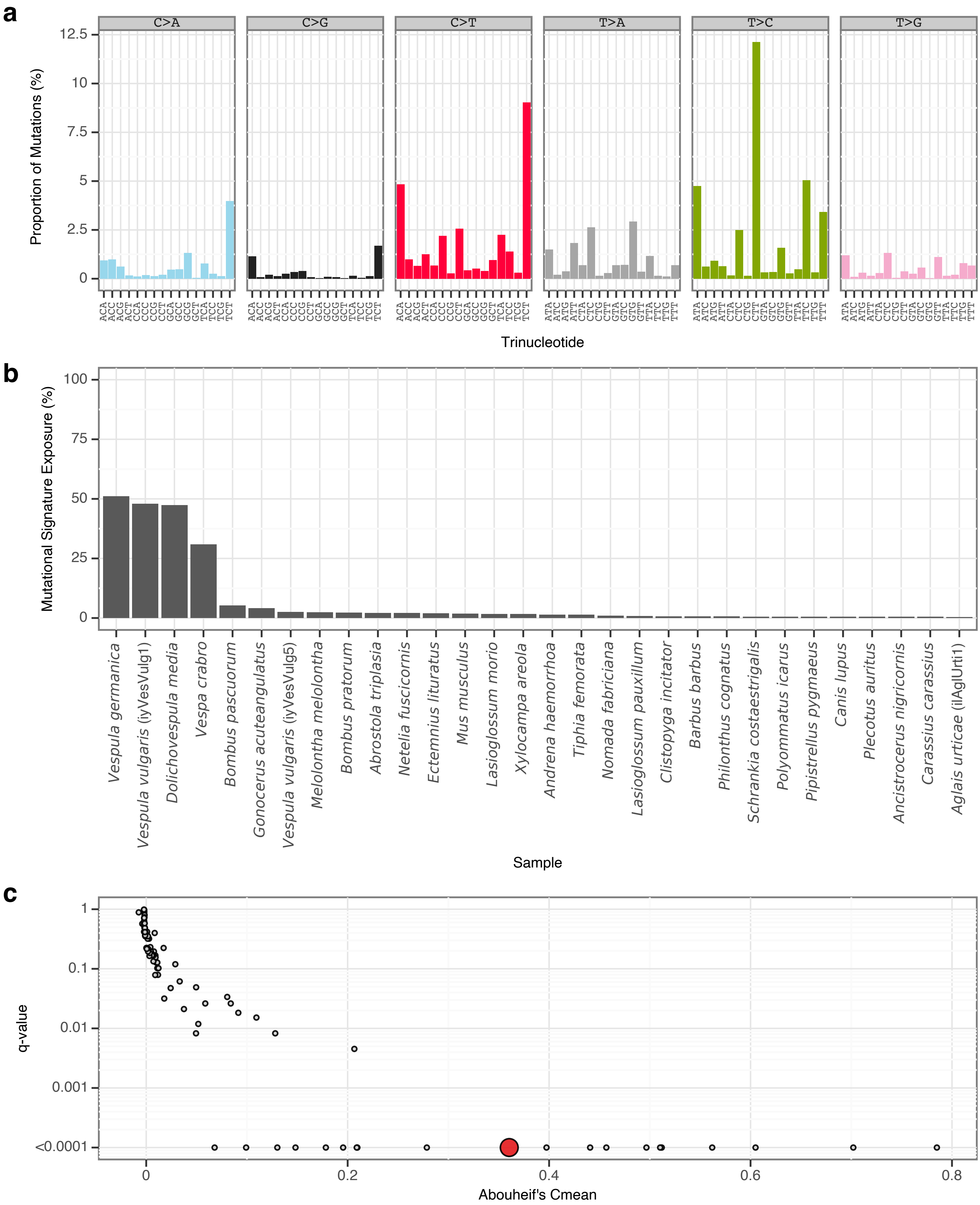


**Supplementary Fig. 27** | sToL24 mutational signature.

**a**, Mutational signature spectrum.

**b**, Bar plot displaying the 30 samples that exhibit the greatest exposure to the mutational signature.

**c**, Phylogenetic signal of the mutational signature.

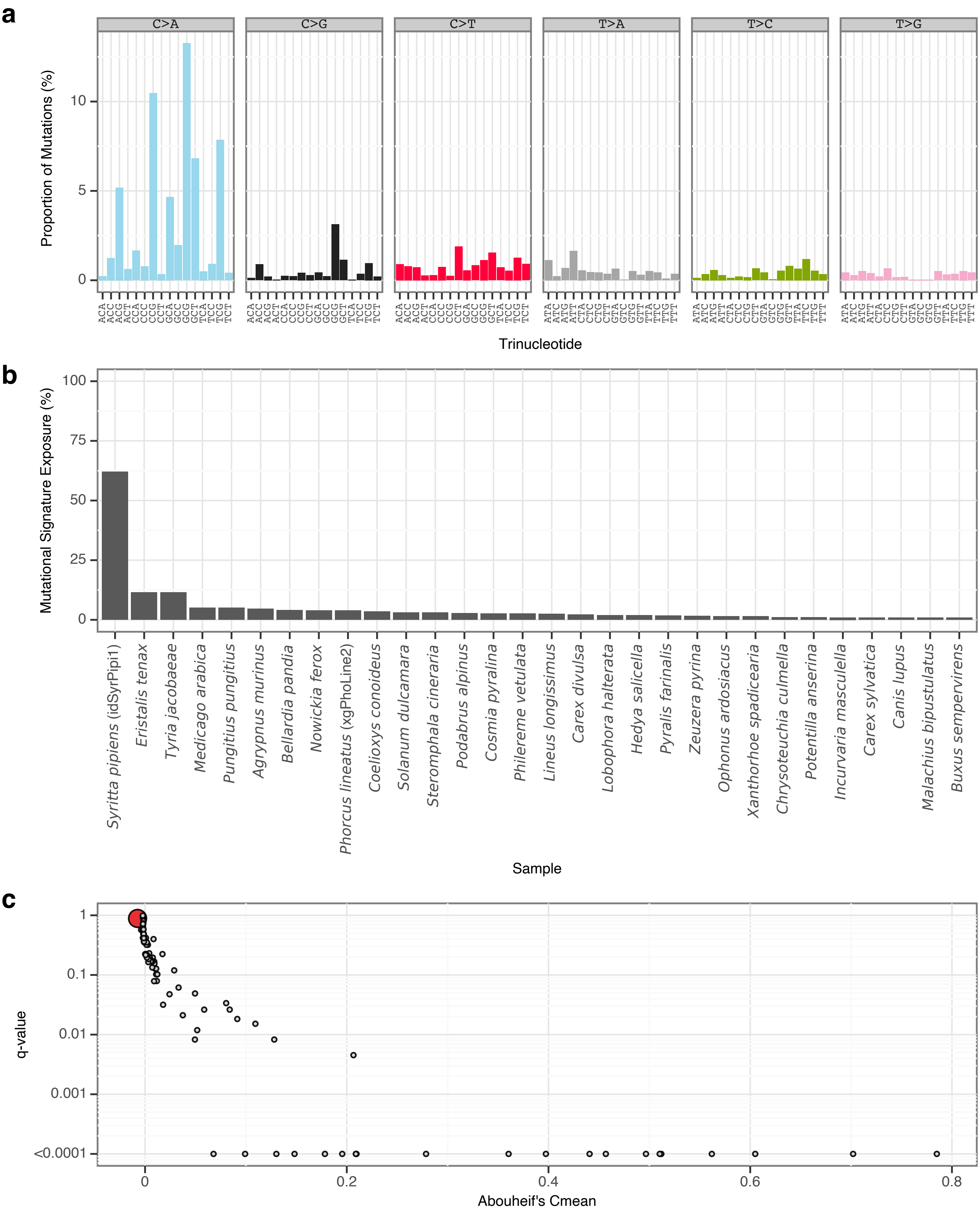


**Supplementary Fig. 28** | sToL25 mutational signature.

**a**, Mutational signature spectrum.

**b**, Bar plot displaying the 30 samples that exhibit the greatest exposure to the mutational signature.

**c**, Phylogenetic signal of the mutational signature.

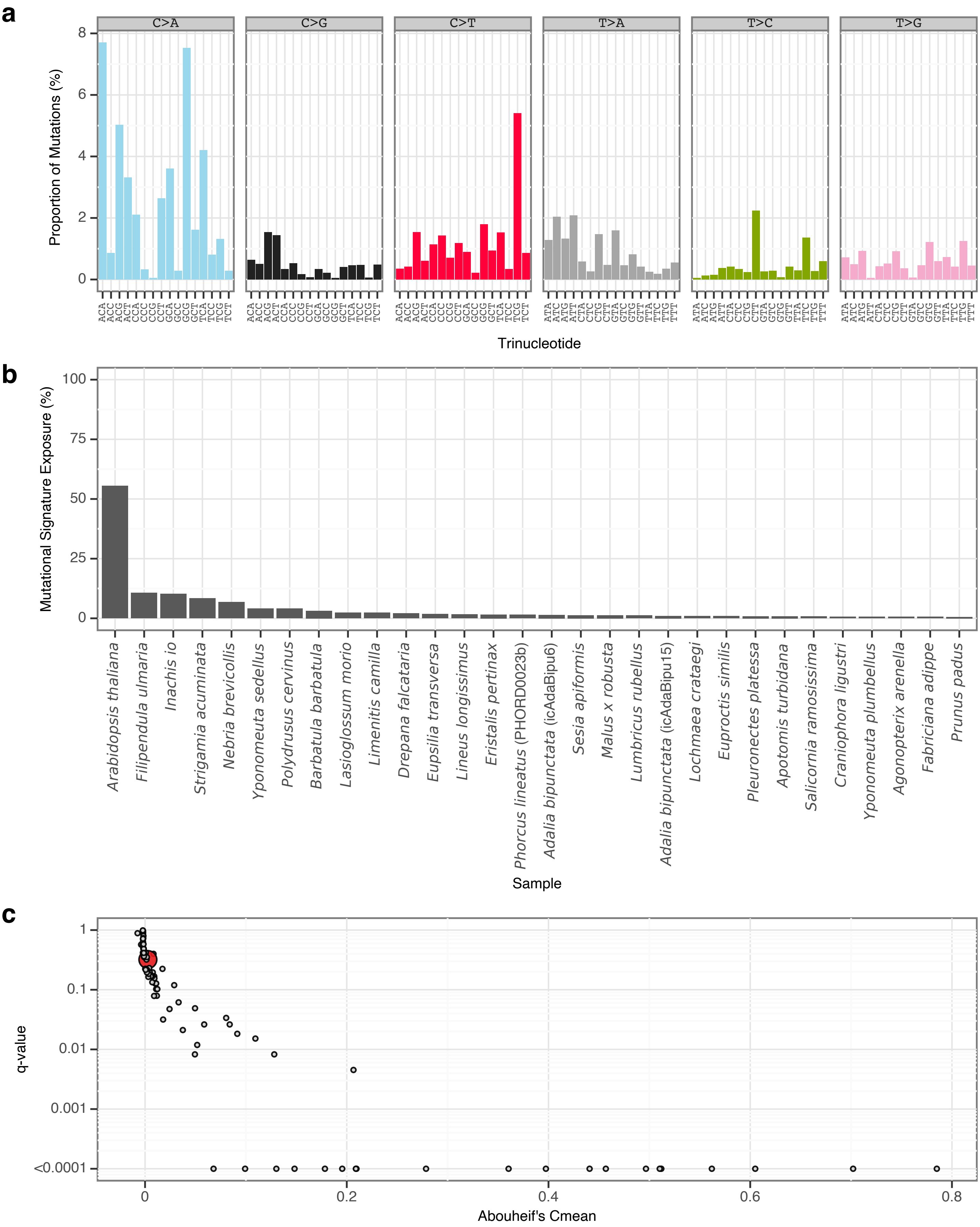


**Supplementary Fig. 29** | sToL26 mutational signature.

**a**, Mutational signature spectrum.

**b**, Bar plot displaying the 30 samples that exhibit the greatest exposure to the mutational signature.

**c**, Phylogenetic signal of the mutational signature.

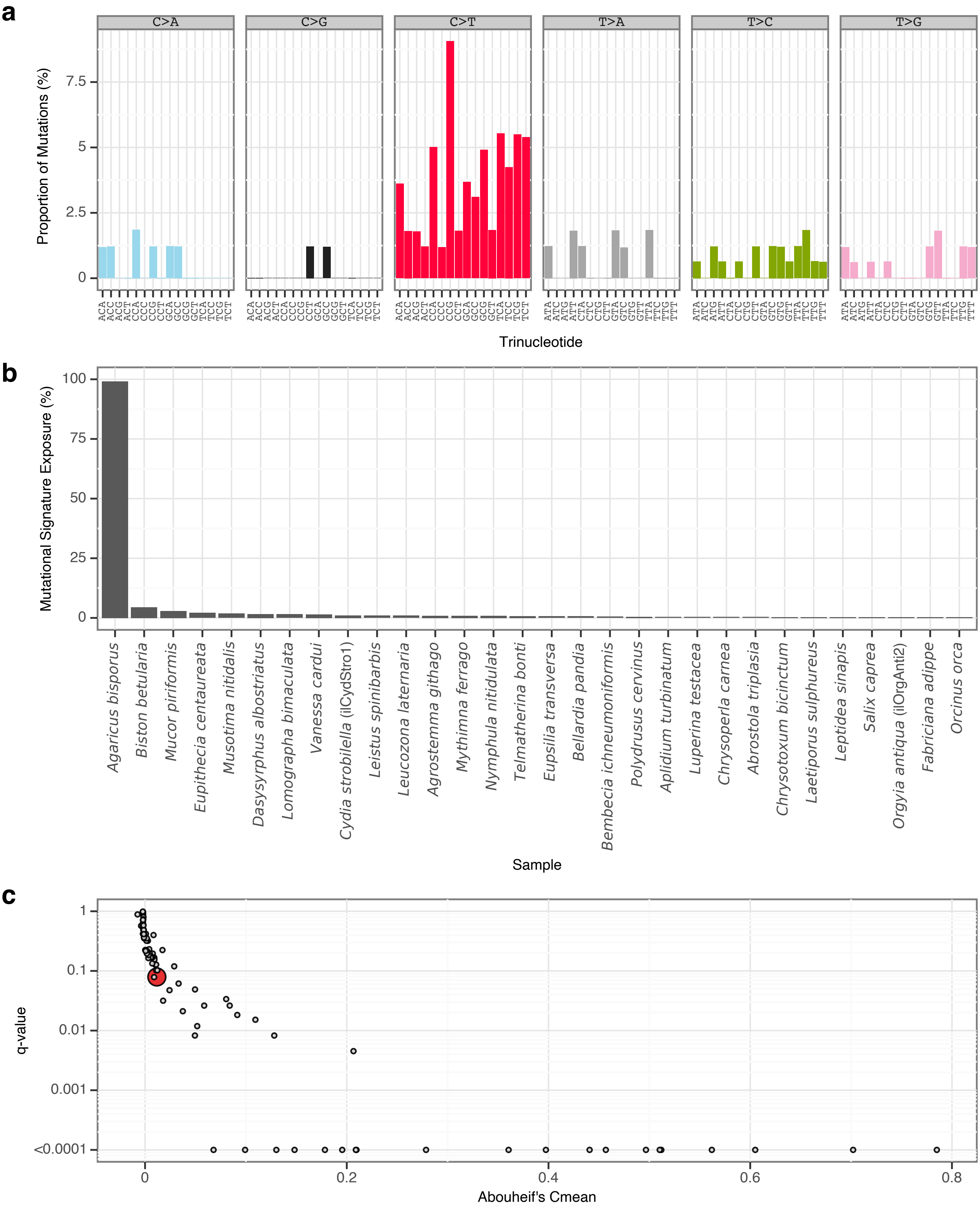


**Supplementary Fig. 30** | sToL27 mutational signature.

**a**, Mutational signature spectrum.

**b**, Bar plot displaying the 30 samples that exhibit the greatest exposure to the mutational signature.

**c**, Phylogenetic signal of the mutational signature.

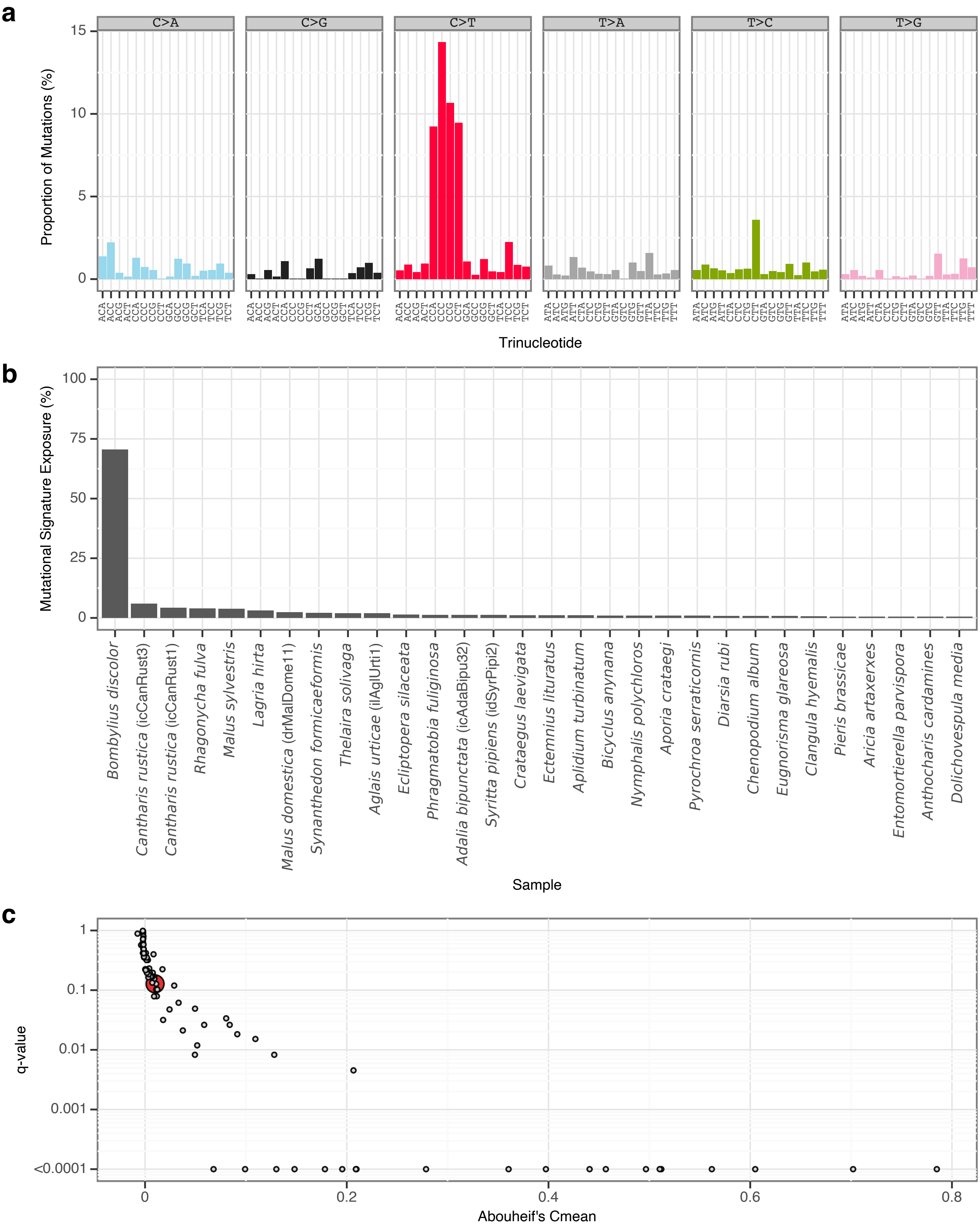


**Supplementary Fig. 31** | sToL28 mutational signature.

**a**, Mutational signature spectrum.

**b**, Bar plot displaying the 30 samples that exhibit the greatest exposure to the mutational signature.

**c**, Phylogenetic signal of the mutational signature.

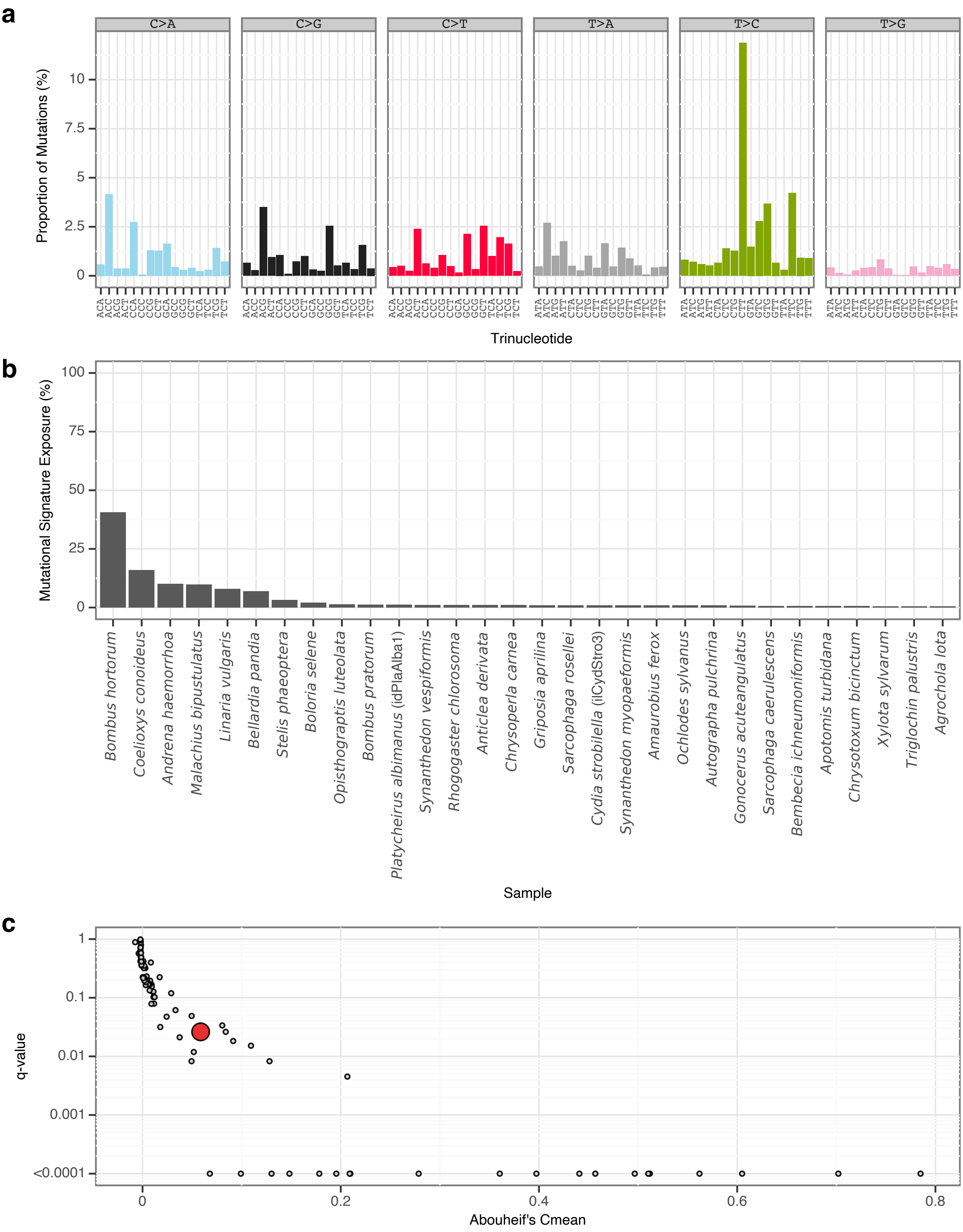


**Supplementary Fig. 32 |** sToL29 mutational signature.
**a**, Mutational signature spectrum.
**b**, Bar plot displaying the 30 samples that exhibit the greatest exposure to the mutational signature.
**c**, Phylogenetic signal of the mutational signature.

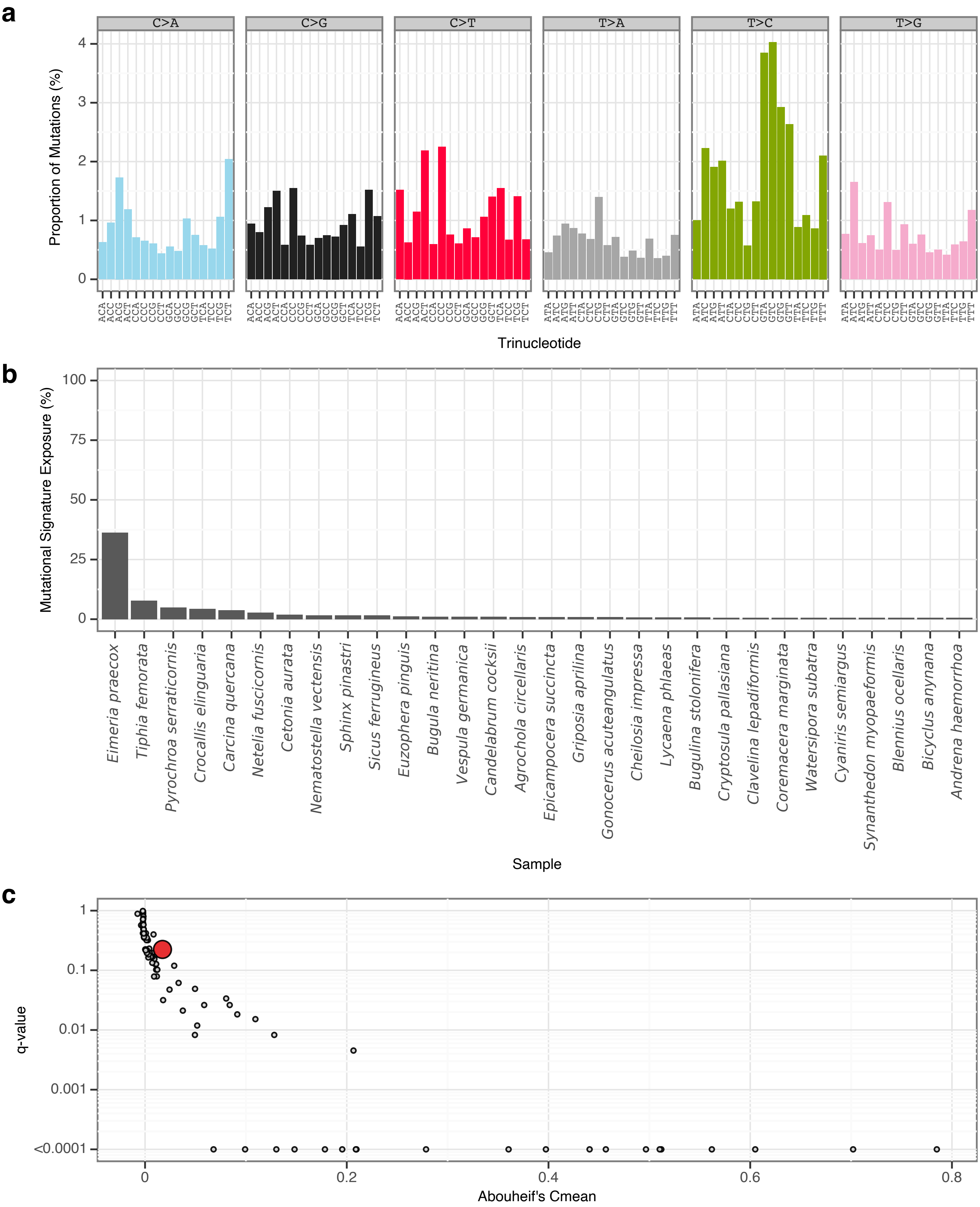


**Supplementary Fig. 33** | sToL30 mutational signature.

**a**, Mutational signature spectrum.

**b**, Bar plot displaying the 30 samples that exhibit the greatest exposure to the mutational signature.

**c**, Phylogenetic signal of the mutational signature.

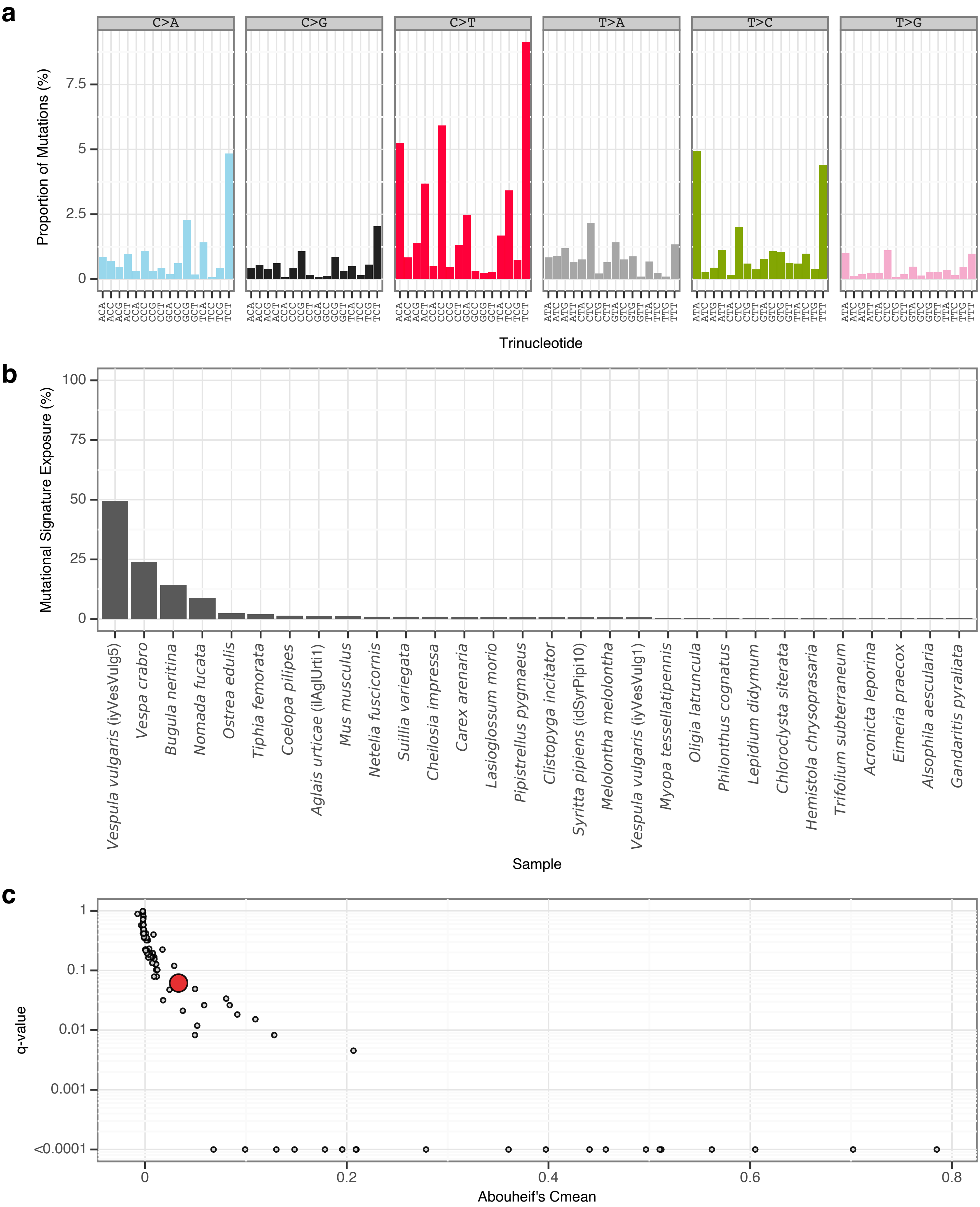


**Supplementary Fig. 34** | sToL31 mutational signature.
**a**, Mutational signature spectrum.
**b**, Bar plot displaying the 30 samples that exhibit the greatest exposure to the mutational signature.
**c**, Phylogenetic signal of the mutational signature.

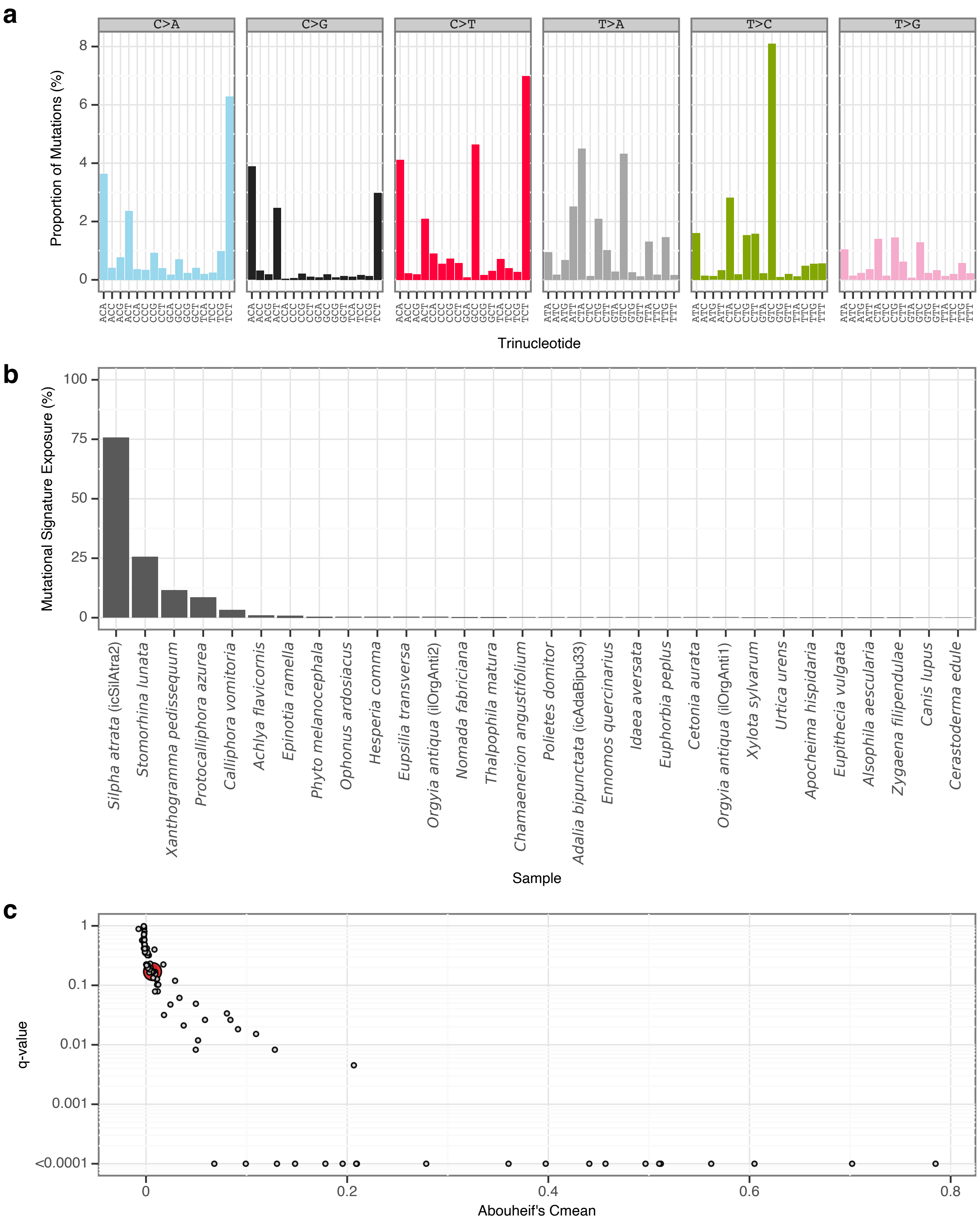


**Supplementary Fig. 35** | sToL32 mutational signature.

**a**, Mutational signature spectrum.

**b**, Bar plot displaying the 30 samples that exhibit the greatest exposure to the mutational signature.

**c**, Phylogenetic signal of the mutational signature.

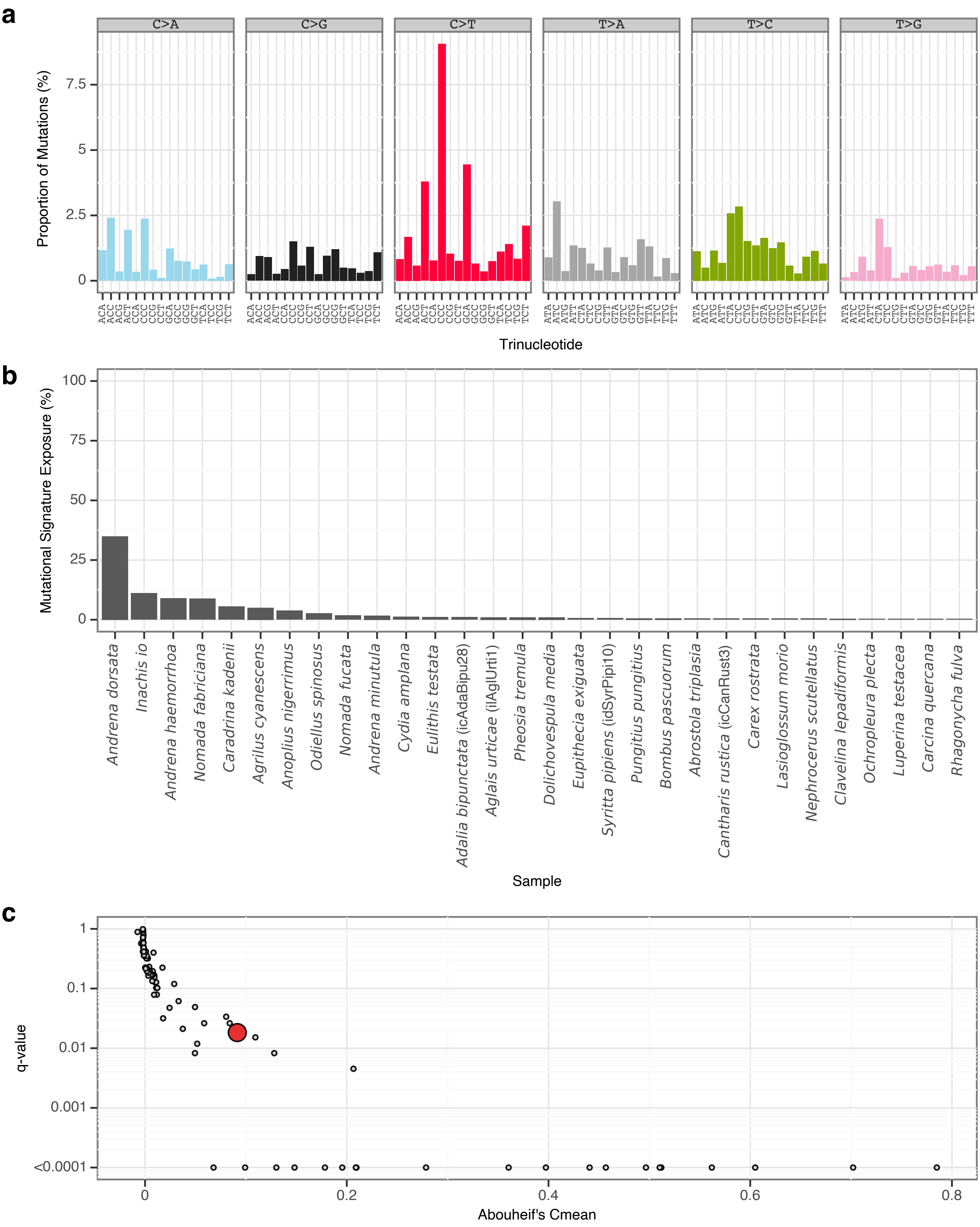


**Supplementary Fig. 36** | sToL33 mutational signature.
**a**, Mutational signature spectrum.
**b**, Bar plot displaying the 30 samples that exhibit the greatest exposure to the mutational signature.
**c**, Phylogenetic signal of the mutational signature.

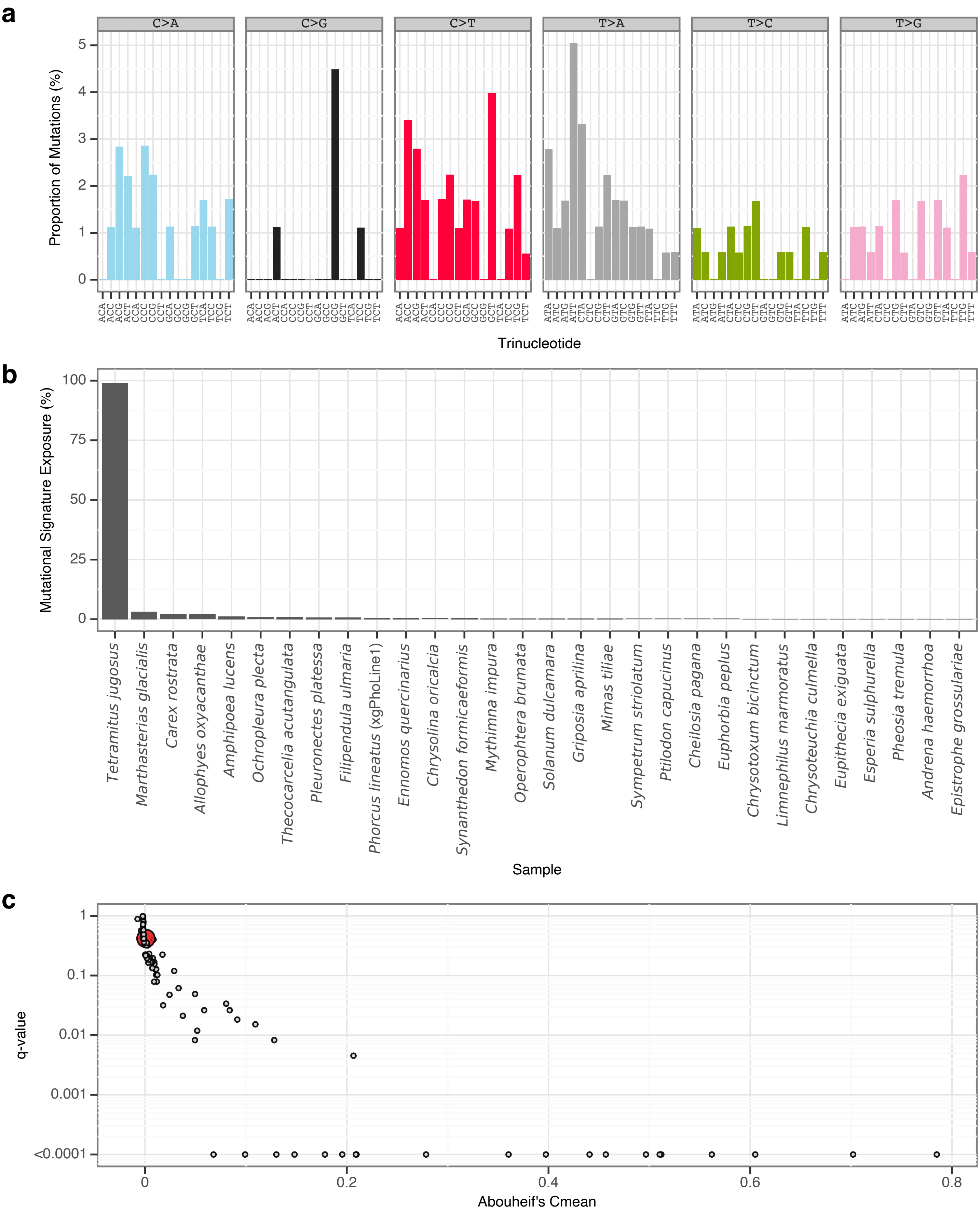


**Supplementary Fig. 37** l sToL34 mutational signature.

**a**, Mutational signature spectrum.

**b**, Bar plot displaying the 30 samples that exhibit the greatest exposure to the mutational signature.

**c**, Phylogenetic signal of the mutational signature.

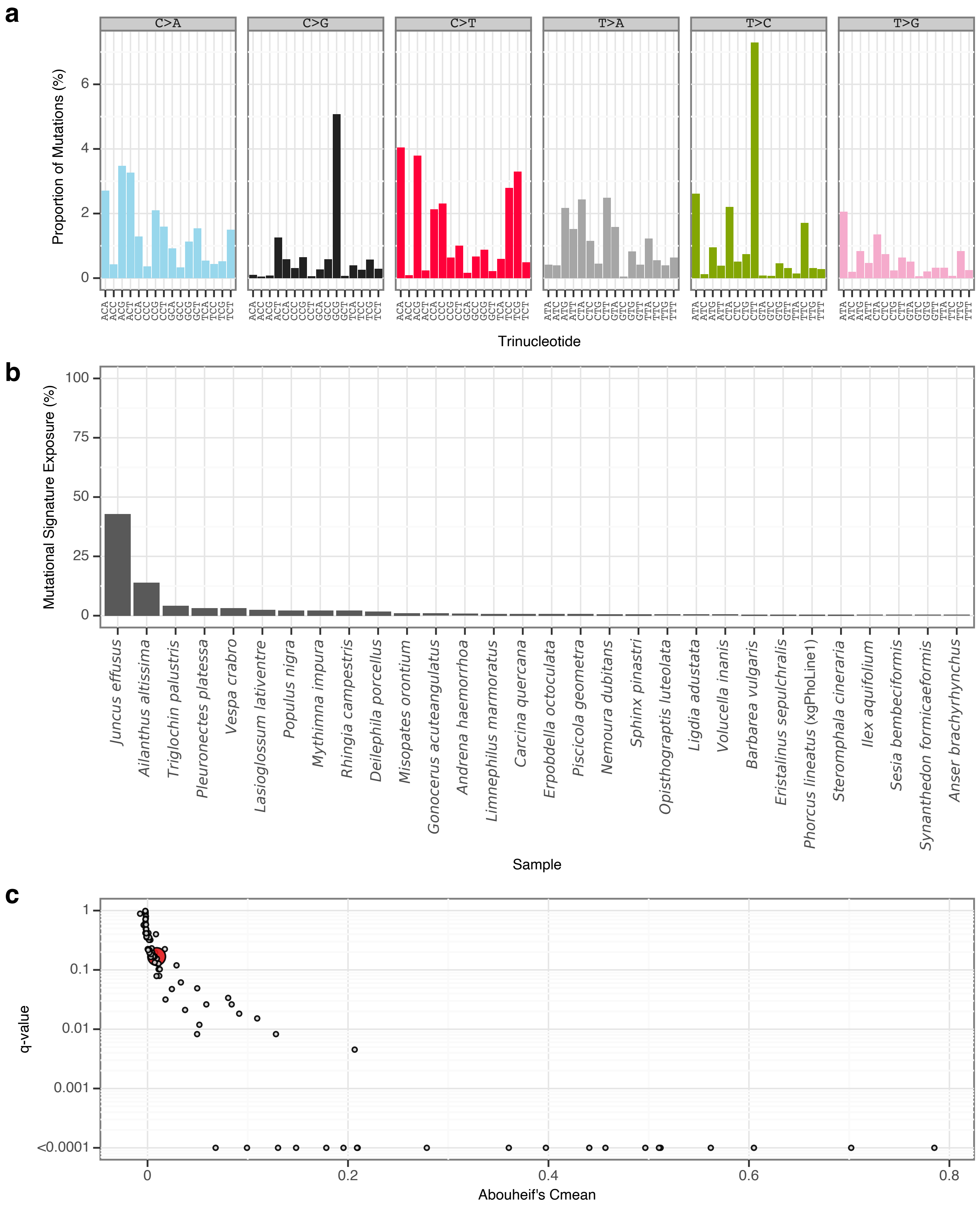


**Supplementary Fig. 38** | sToL35 mutational signature.

**a**, Mutational signature spectrum.

**b**, Bar plot displaying the 30 samples that exhibit the greatest exposure to the mutational signature.

**c**, Phylogenetic signal of the mutational signature.

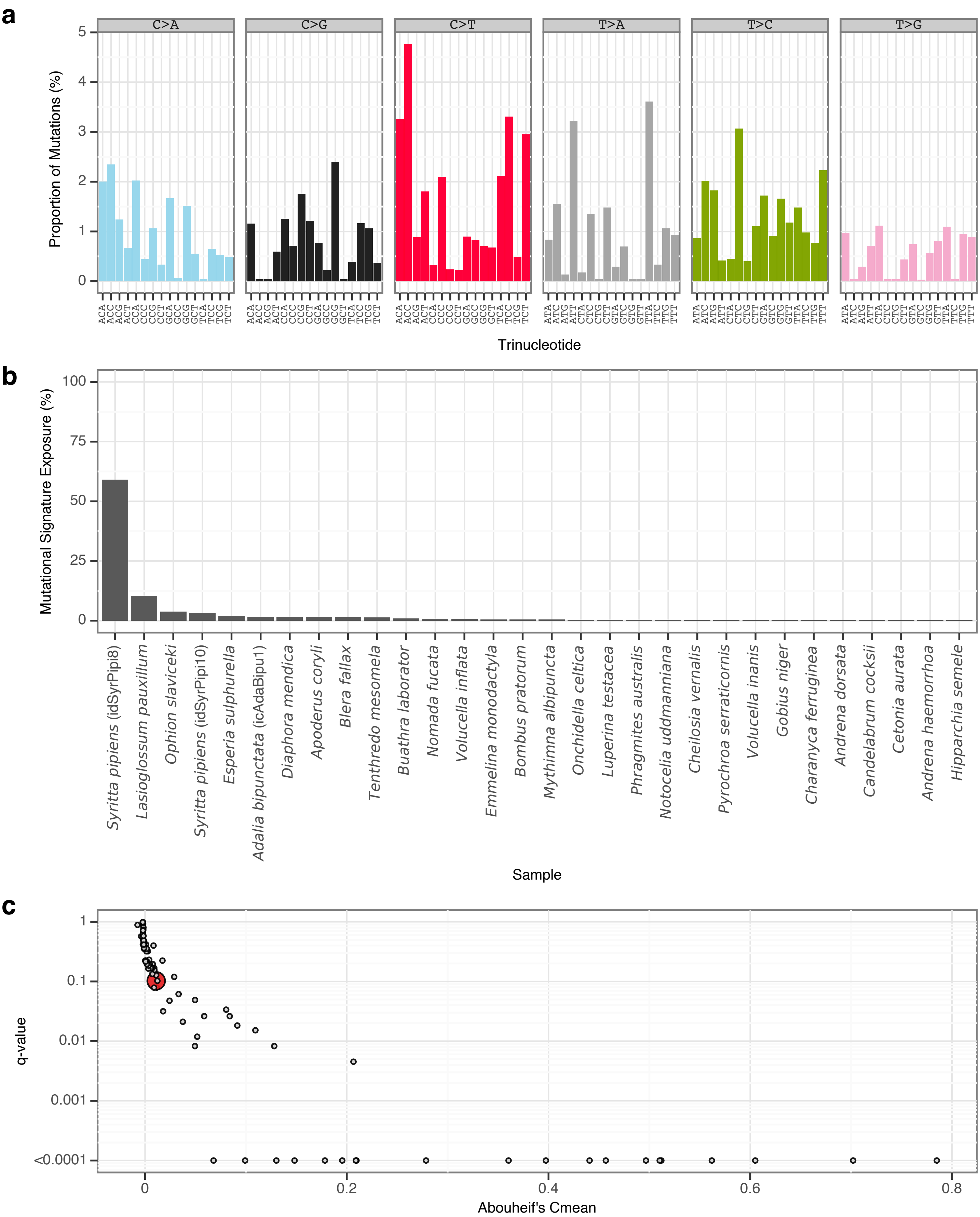


**Supplementary Fig. 39** | sToL36 mutational signature.

**a**, Mutational signature spectrum.

**b**, Bar plot displaying the 30 samples that exhibit the greatest exposure to the mutational signature.

**c**, Phylogenetic signal of the mutational signature.

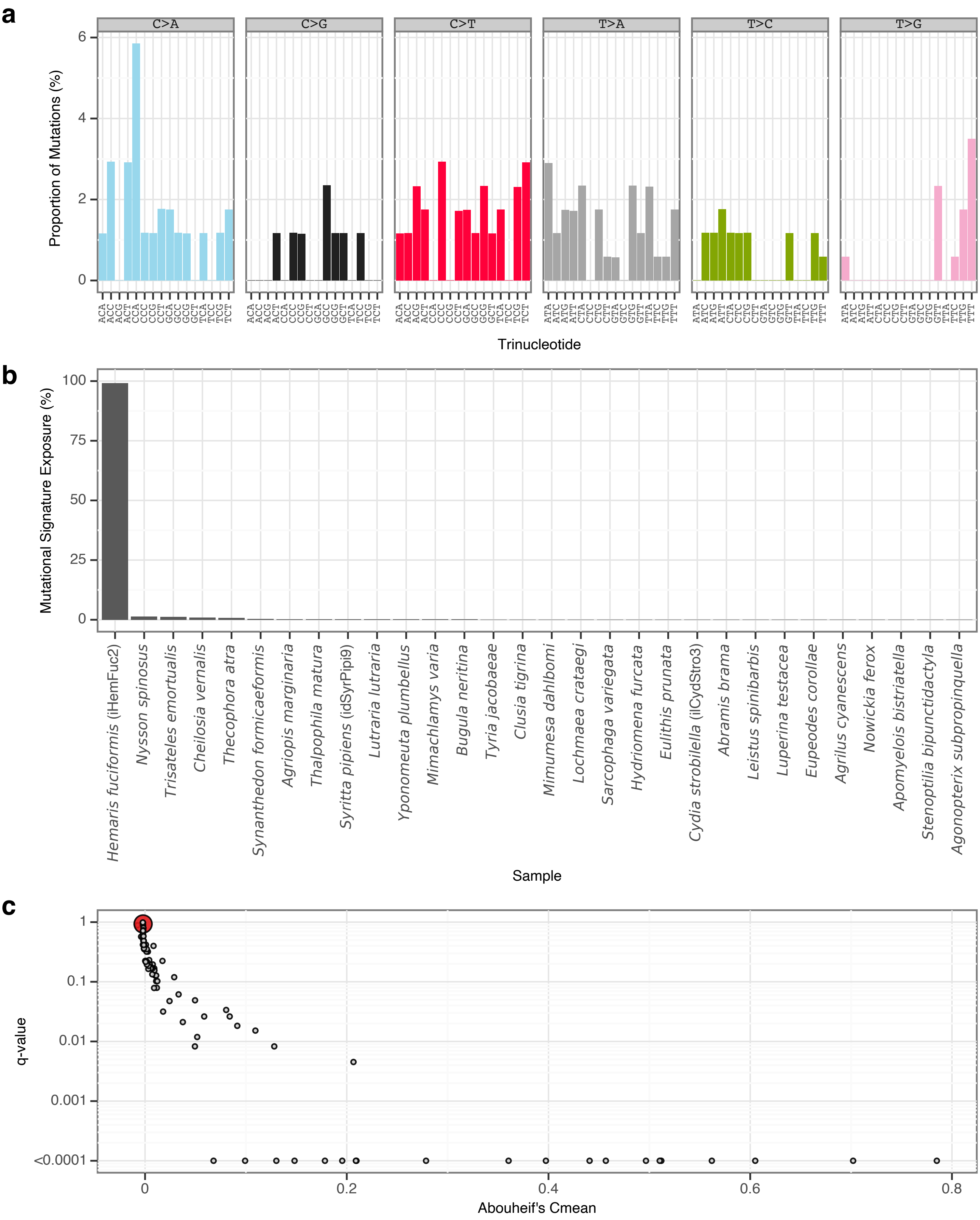


**Supplementary Fig. 40** | sToL37 mutational signature.
**a**, Mutational signature spectrum.
**b**, Bar plot displaying the 30 samples that exhibit the greatest exposure to the mutational signature.
**c**, Phylogenetic signal of the mutational signature.

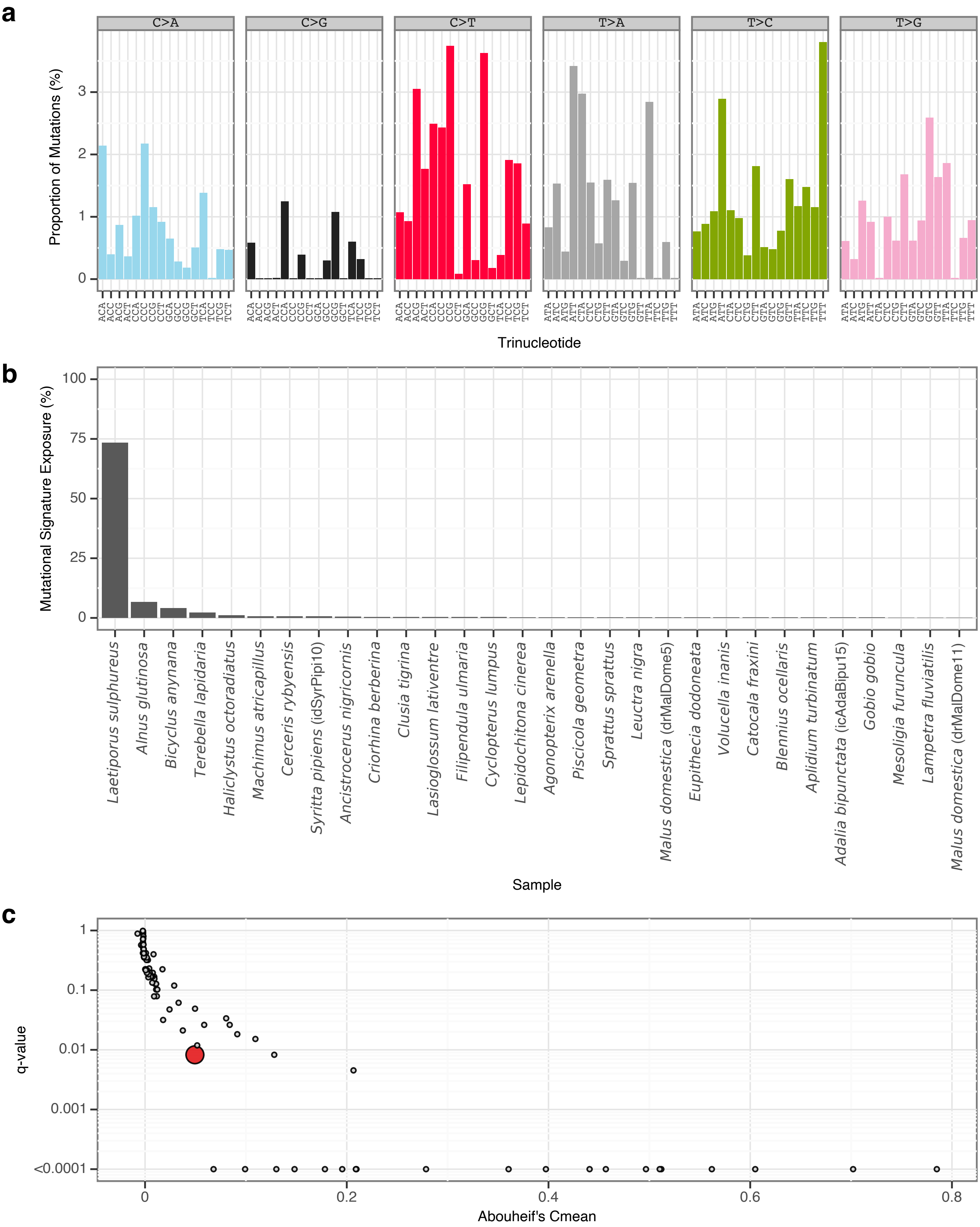


**Supplementary Fig. 41** | sToL38 mutational signature.

**a**, Mutational signature spectrum.

**b**, Bar plot displaying the 30 samples that exhibit the greatest exposure to the mutational signature.

**c**, Phylogenetic signal of the mutational signature.

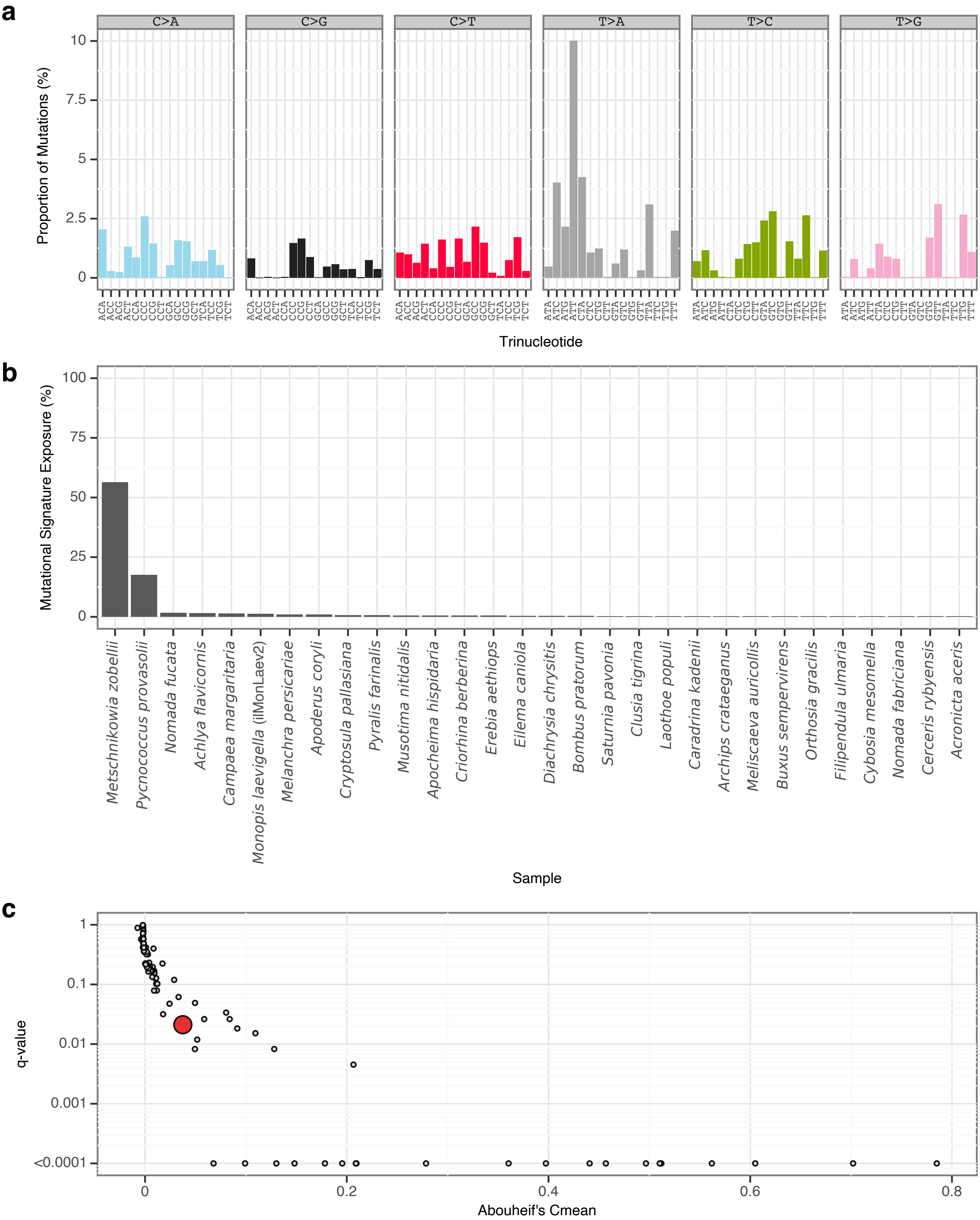


**Supplementary Fig. 42** | sToL39 mutational signature.

**a**, Mutational signature spectrum.

**b**, Bar plot displaying the 30 samples that exhibit the greatest exposure to the mutational signature.

**c**, Phylogenetic signal of the mutational signature.

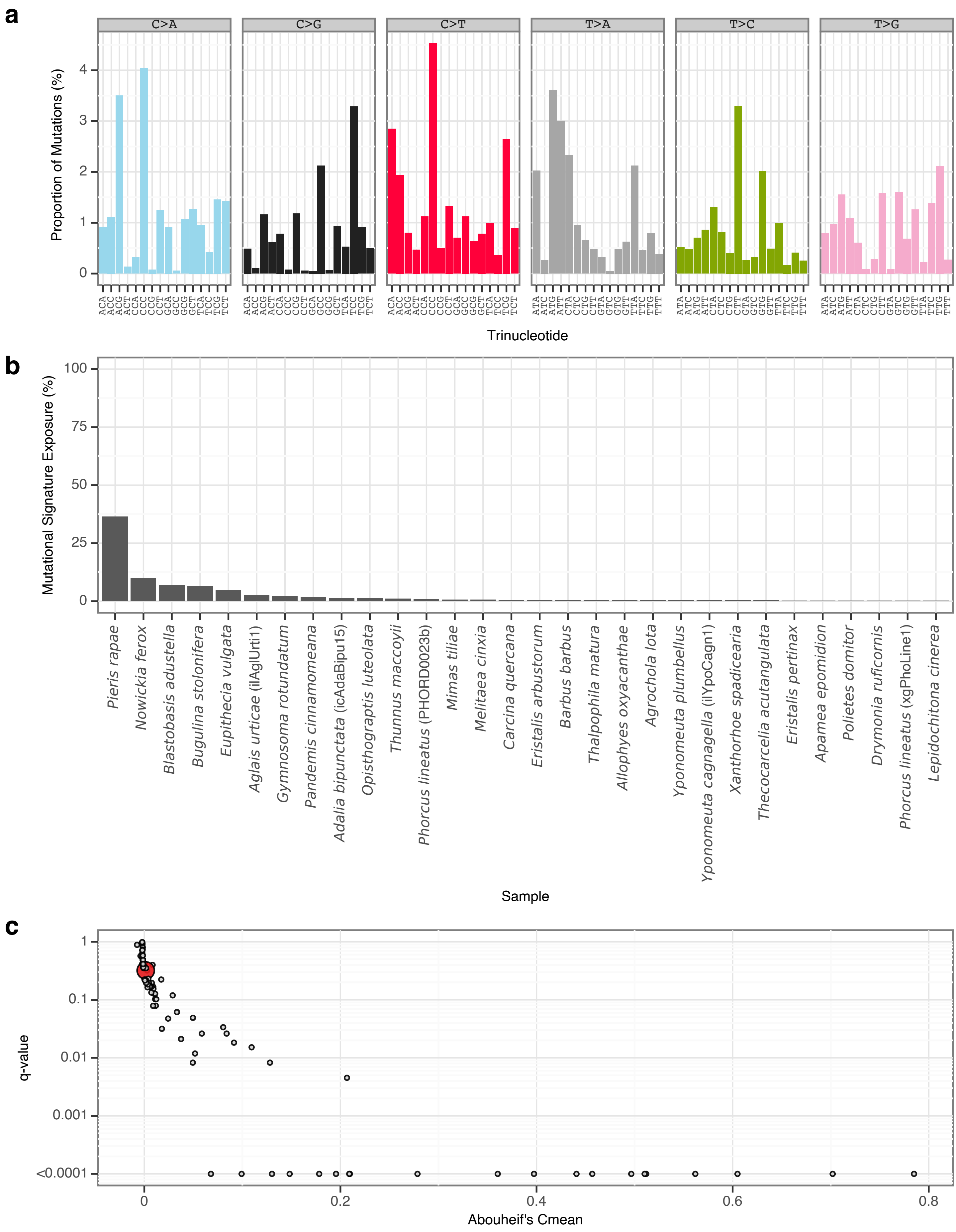


**Supplementary Fig. 43** | sToL40 mutational signature.
**a**, Mutational signature spectrum.
**b**, Bar plot displaying the 30 samples that exhibit the greatest exposure to the mutational signature.
**c**, Phylogenetic signal of the mutational signature.

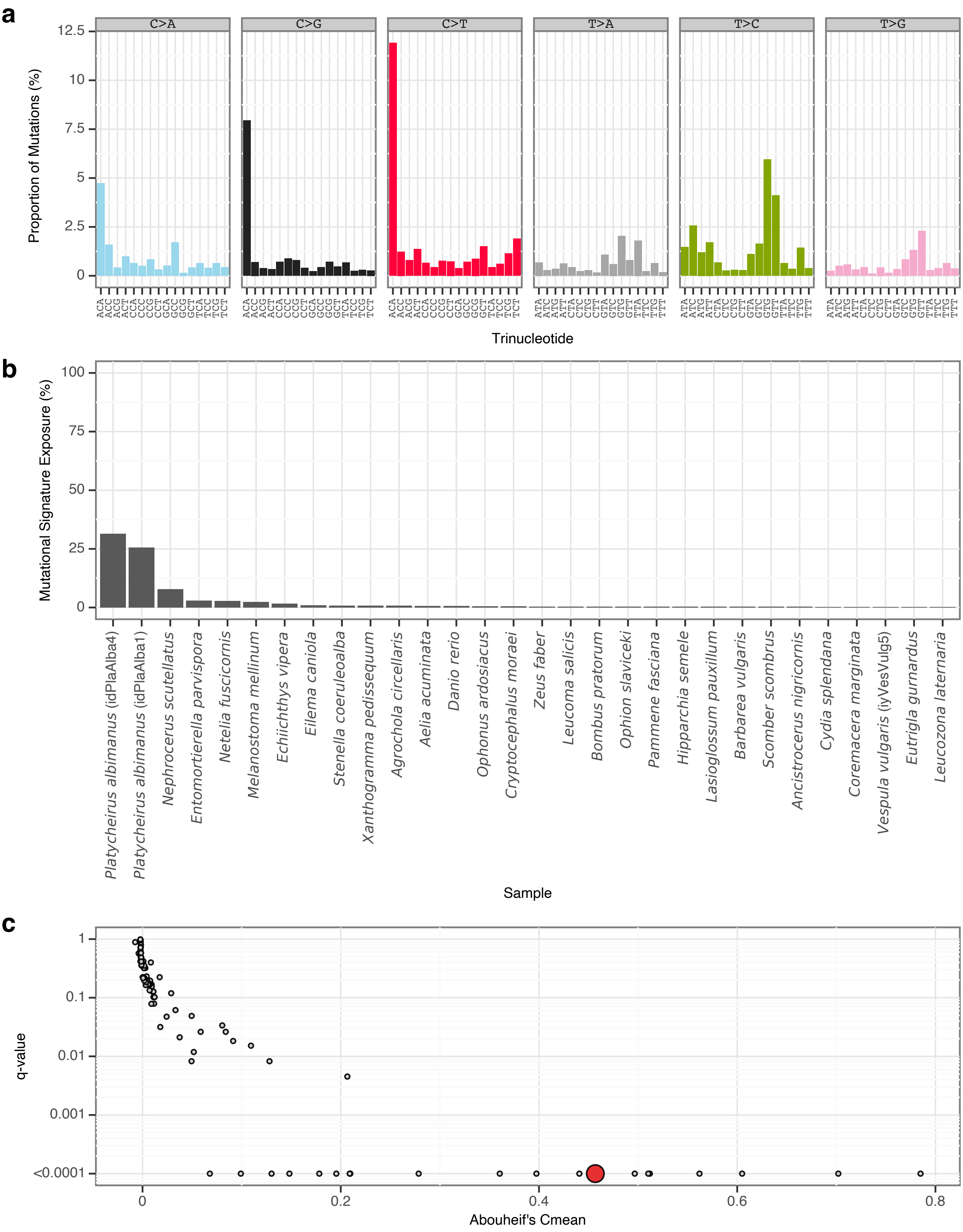


**Supplementary Fig. 44 | sToL41 mutational signature.**
**a**, Mutational signature spectrum.
**b**, Bar plot displaying the 30 samples that exhibit the greatest exposure to the mutational signature.
**c**, Phylogenetic signal of the mutational signature.

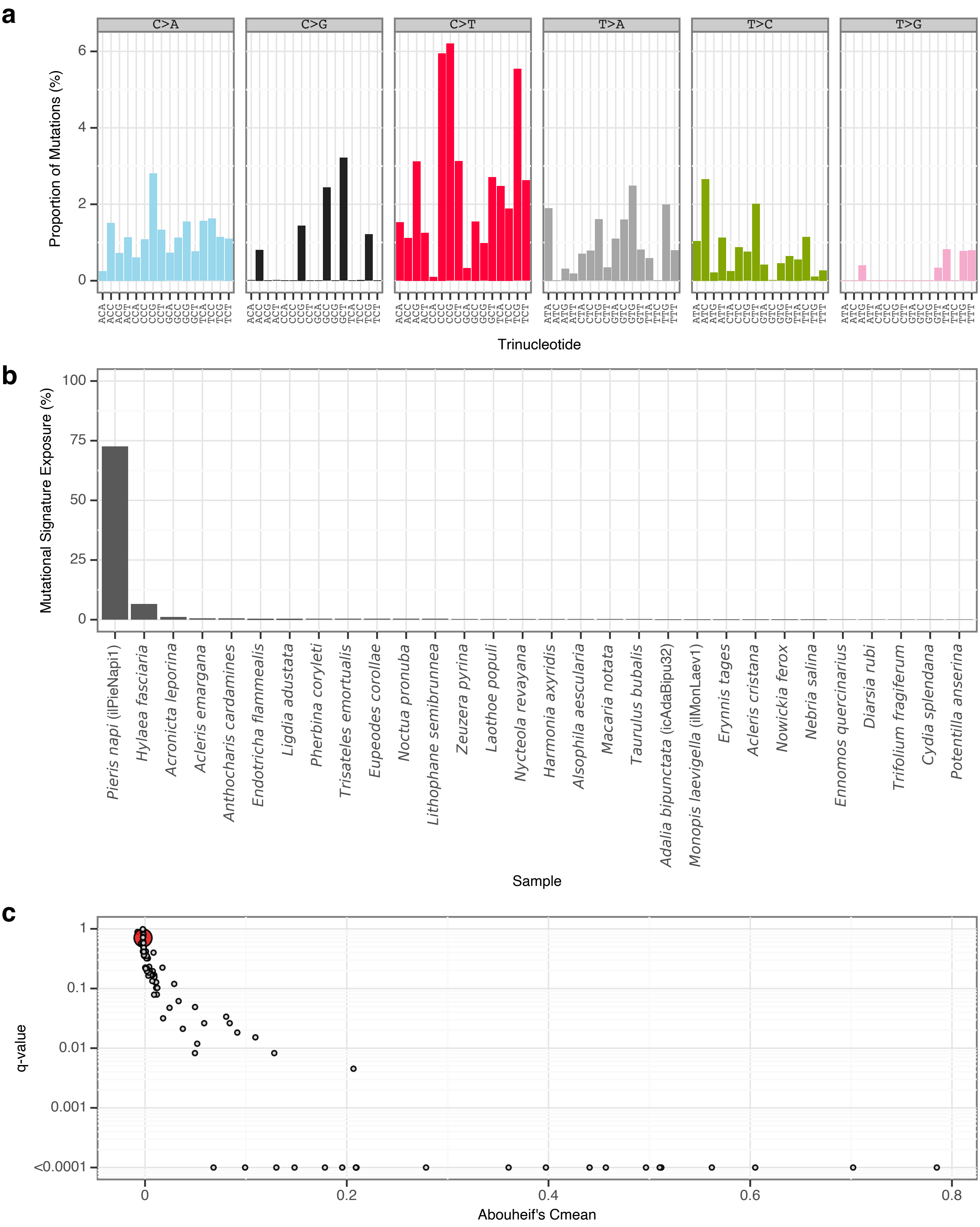


**Supplementary Fig. 45** | sToL42 mutational signature.

**a**, Mutational signature spectrum.

**b**, Bar plot displaying the 30 samples that exhibit the greatest exposure to the mutational signature.

**c**, Phylogenetic signal of the mutational signature.

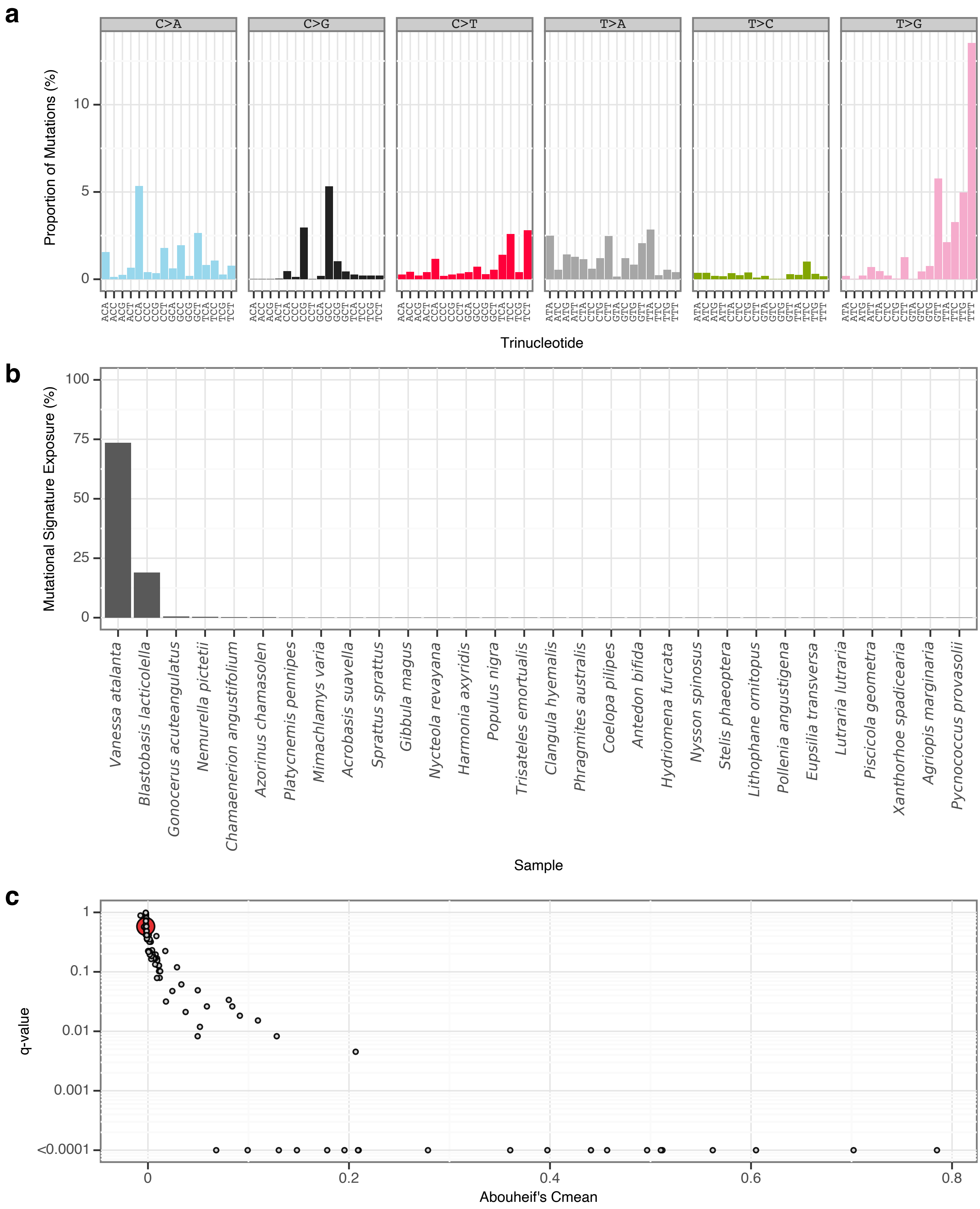


**Supplementary Fig. 46** | sToL43 mutational signature.

**a**, Mutational signature spectrum.

**b**, Bar plot displaying the 30 samples that exhibit the greatest exposure to the mutational signature.

**c**, Phylogenetic signal of the mutational signature.

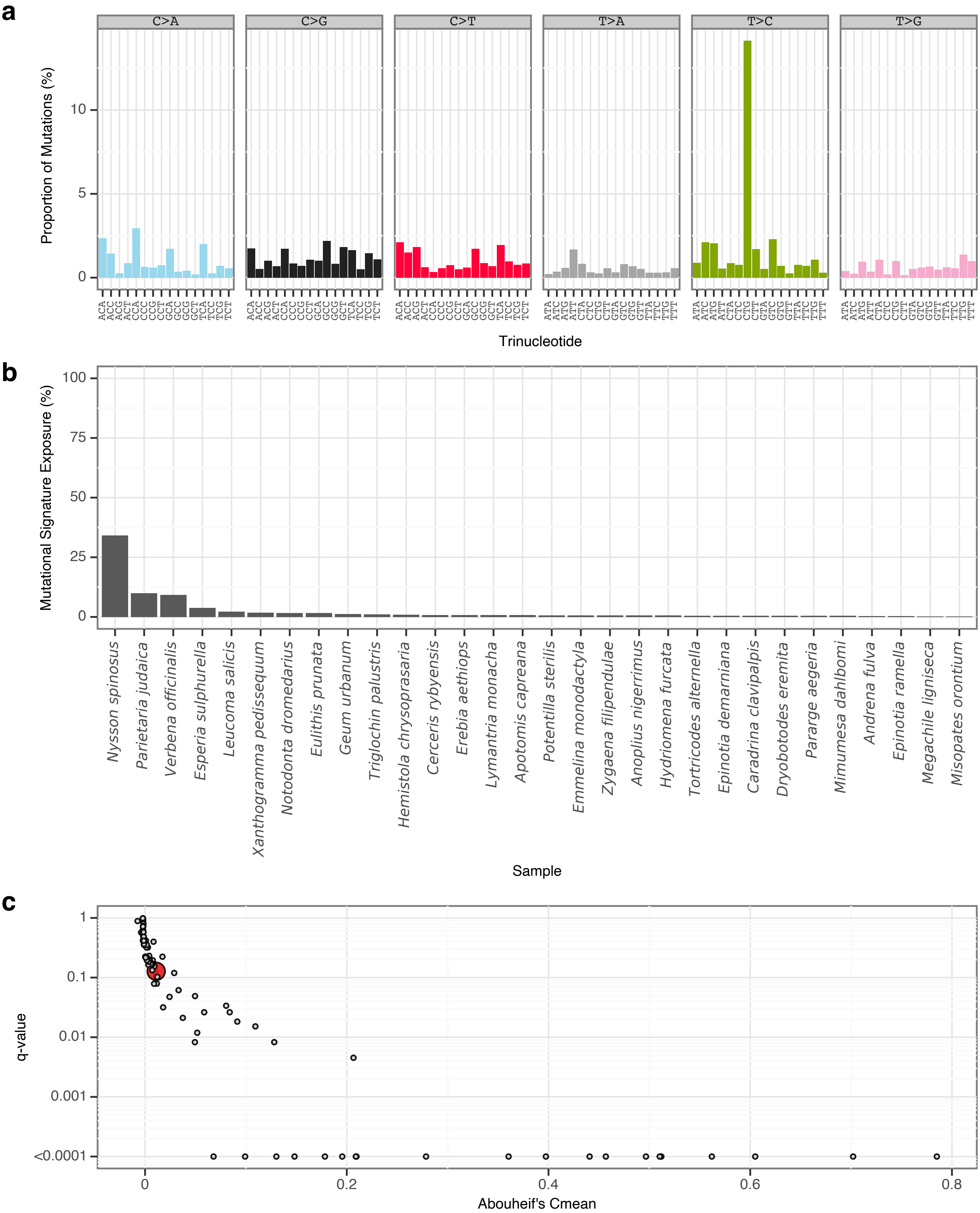


**Supplementary Fig. 47** | sToL44 mutational signature.

**a**, Mutational signature spectrum.

**b**, Bar plot displaying the 30 samples that exhibit the greatest exposure to the mutational signature.

**c**, Phylogenetic signal of the mutational signature.

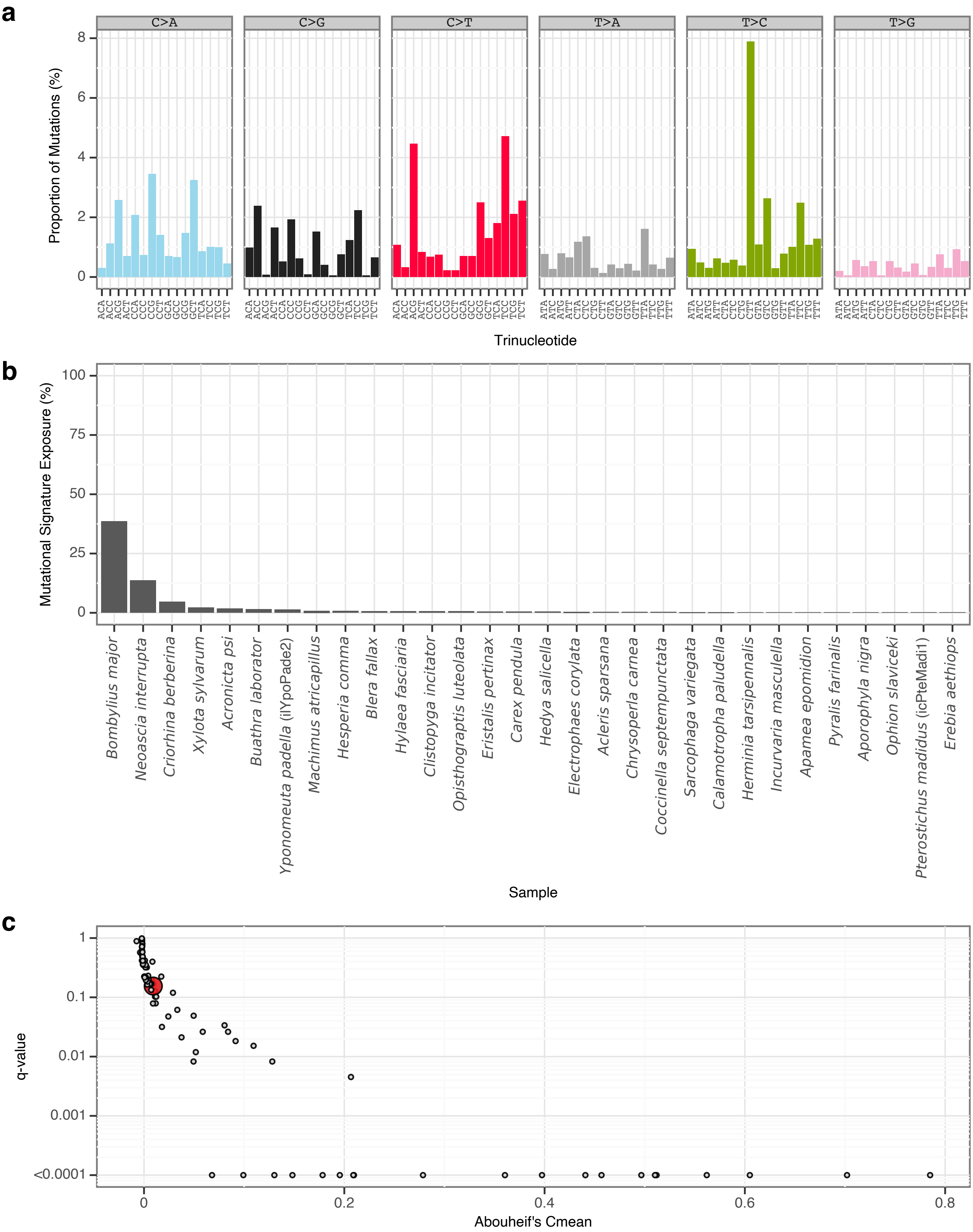


**Supplementary Fig. 48** | sToL45 mutational signature.

**a**, Mutational signature spectrum.

**b**, Bar plot displaying the 30 samples that exhibit the greatest exposure to the mutational signature.

**c**, Phylogenetic signal of the mutational signature.

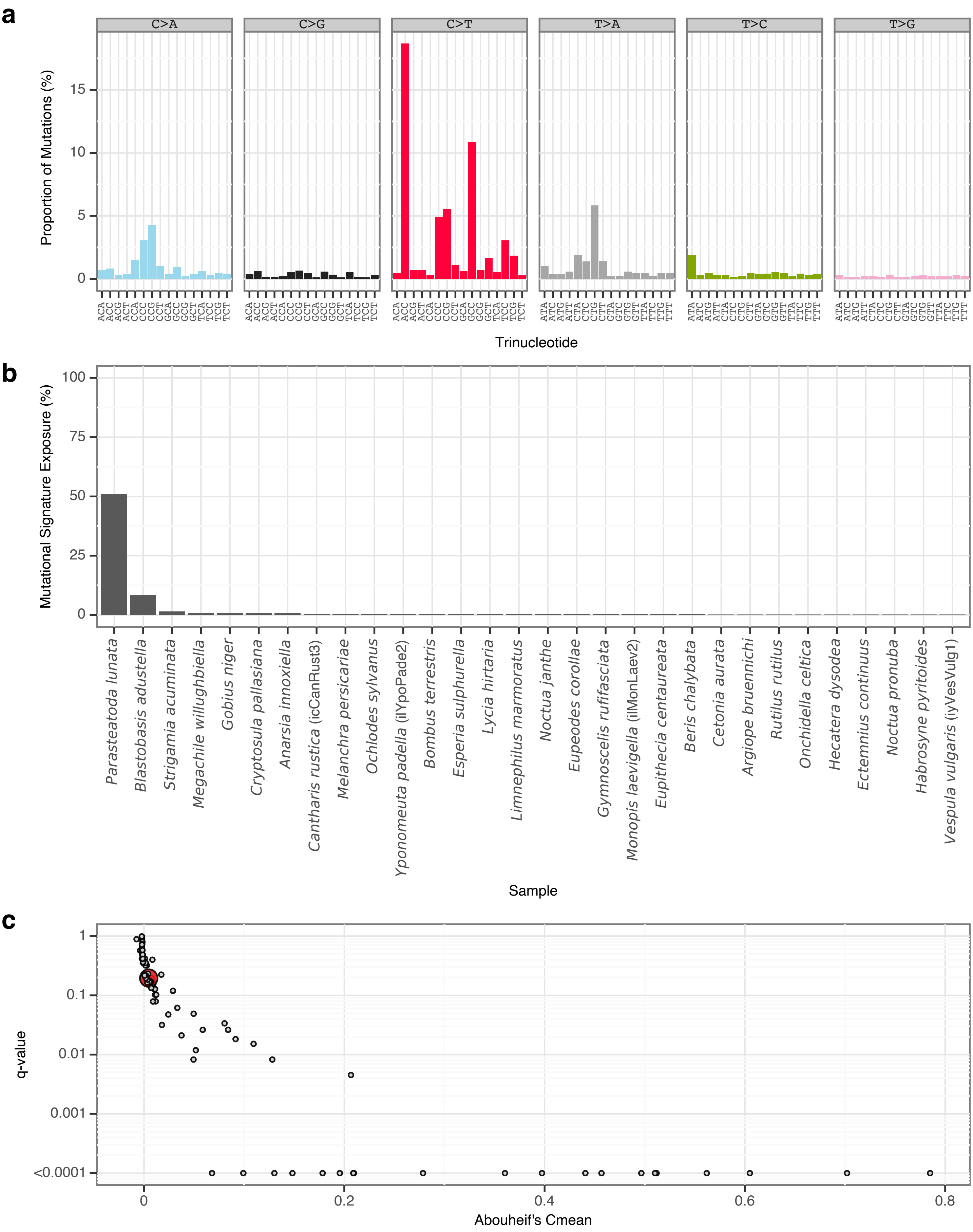


**Supplementary Fig. 49** | sToL46 mutational signature.
**a**, Mutational signature spectrum.
**b**, Bar plot displaying the 30 samples that exhibit the greatest exposure to the mutational signature.
**c**, Phylogenetic signal of the mutational signature.

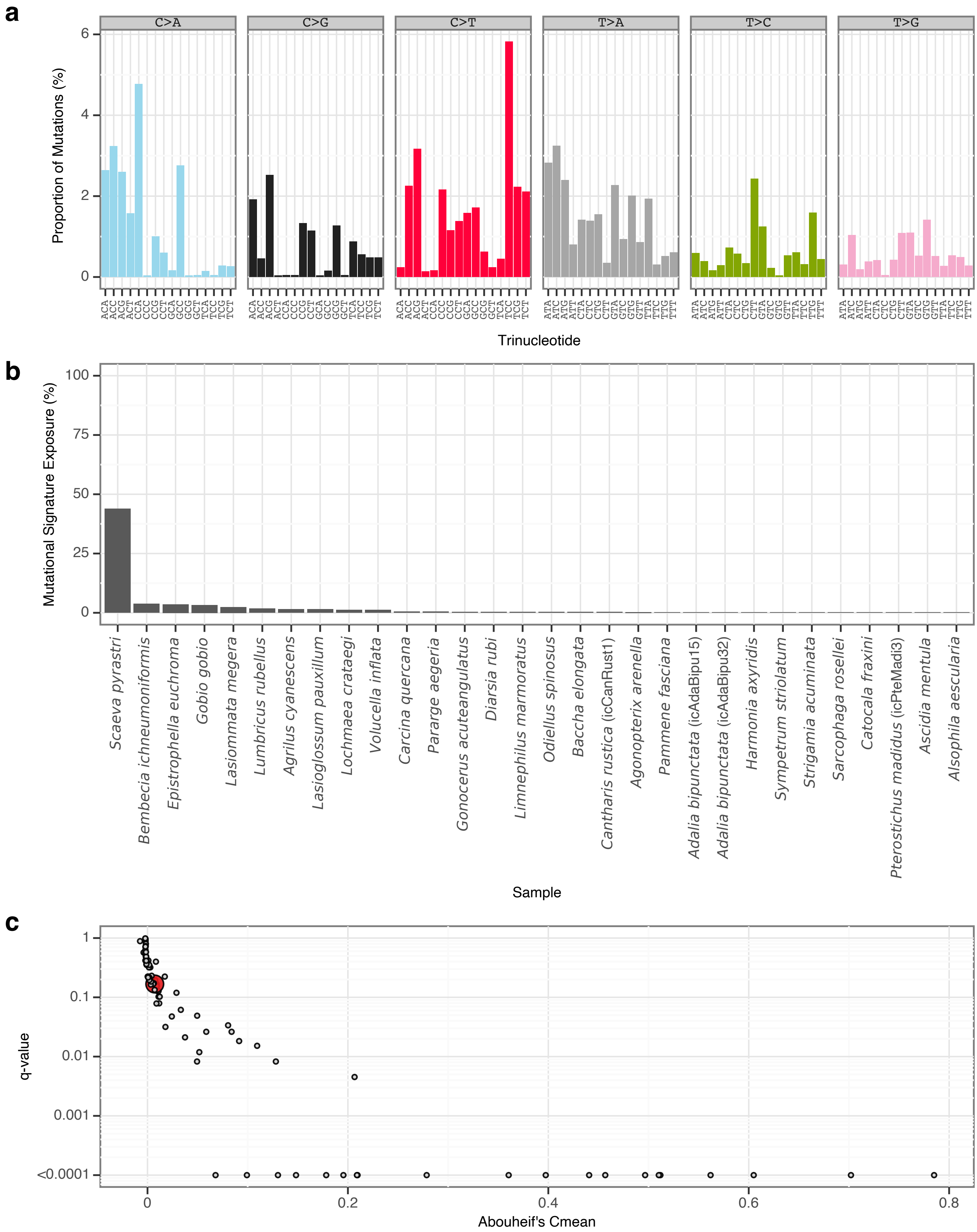


**Supplementary Fig. 50** | sToL47 mutational signature.

**a**, Mutational signature spectrum.

**b**, Bar plot displaying the 30 samples that exhibit the greatest exposure to the mutational signature.

**c**, Phylogenetic signal of the mutational signature.

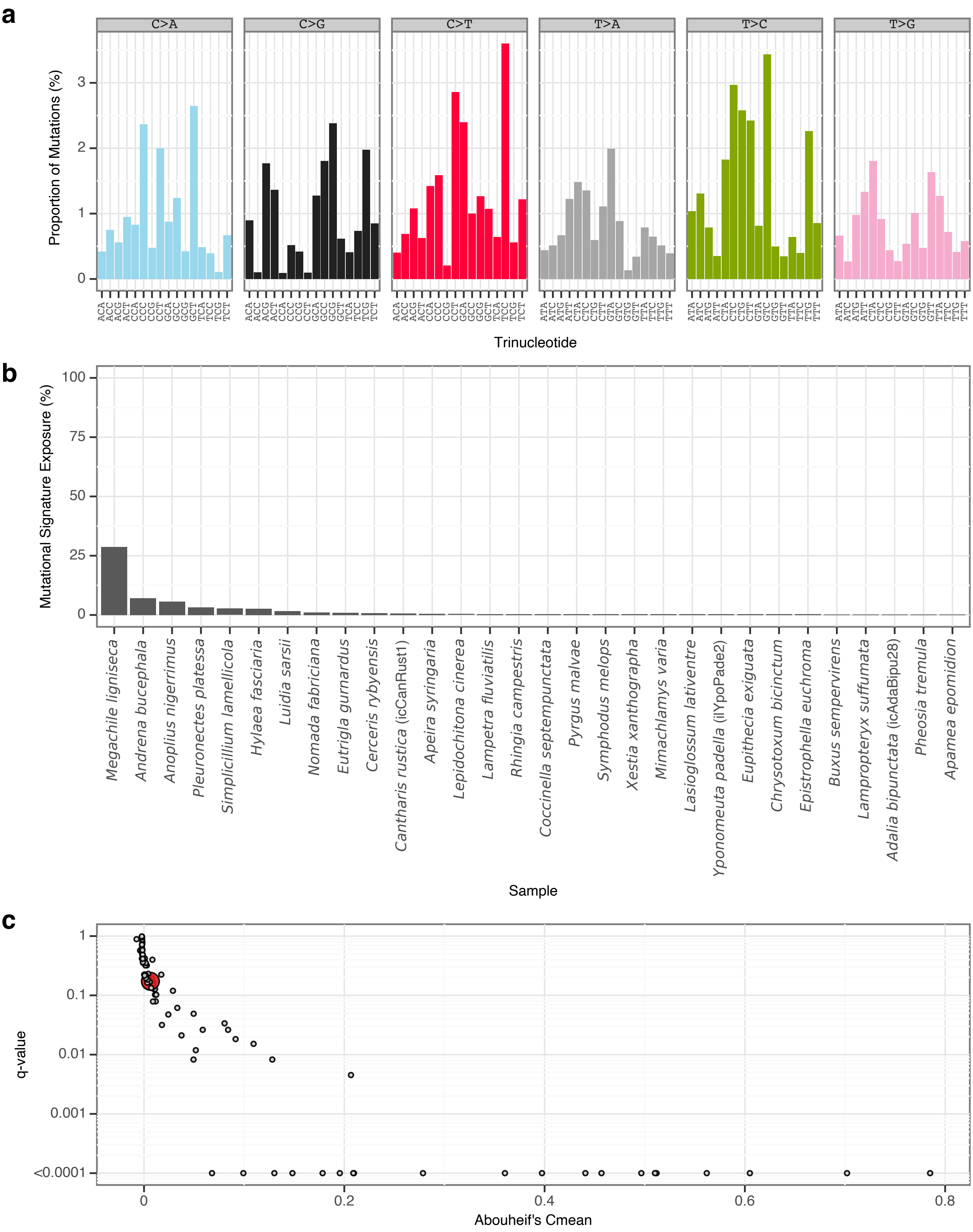


**Supplementary Fig. 51** | sToL48 mutational signature.

**a**, Mutational signature spectrum.

**b**, Bar plot displaying the 30 samples that exhibit the greatest exposure to the mutational signature.

**c**, Phylogenetic signal of the mutational signature.

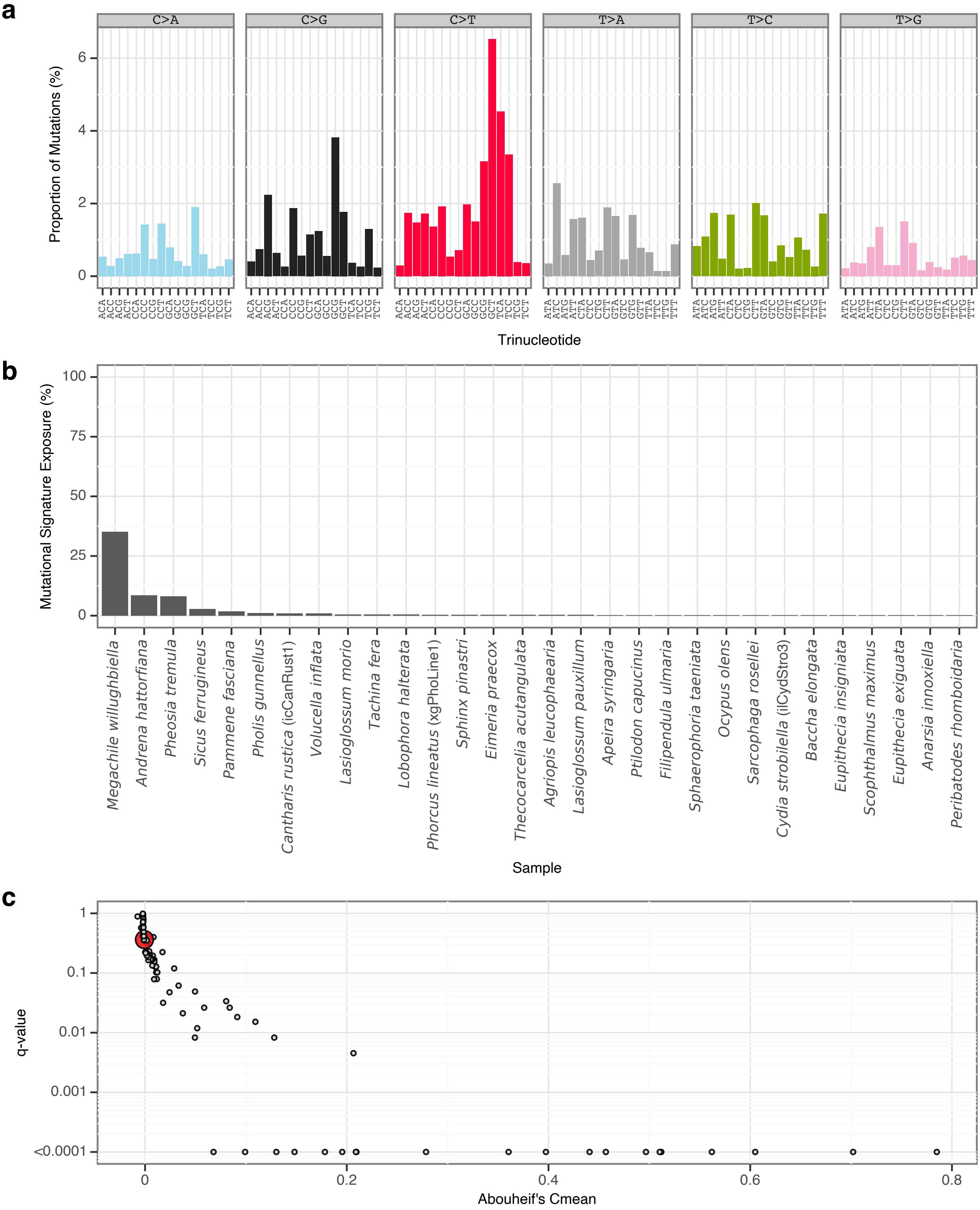


**Supplementary Fig. 52** | sToL49 mutational signature.

**a**, Mutational signature spectrum.

**b**, Bar plot displaying the 30 samples that exhibit the greatest exposure to the mutational signature.

**c**, Phylogenetic signal of the mutational signature.

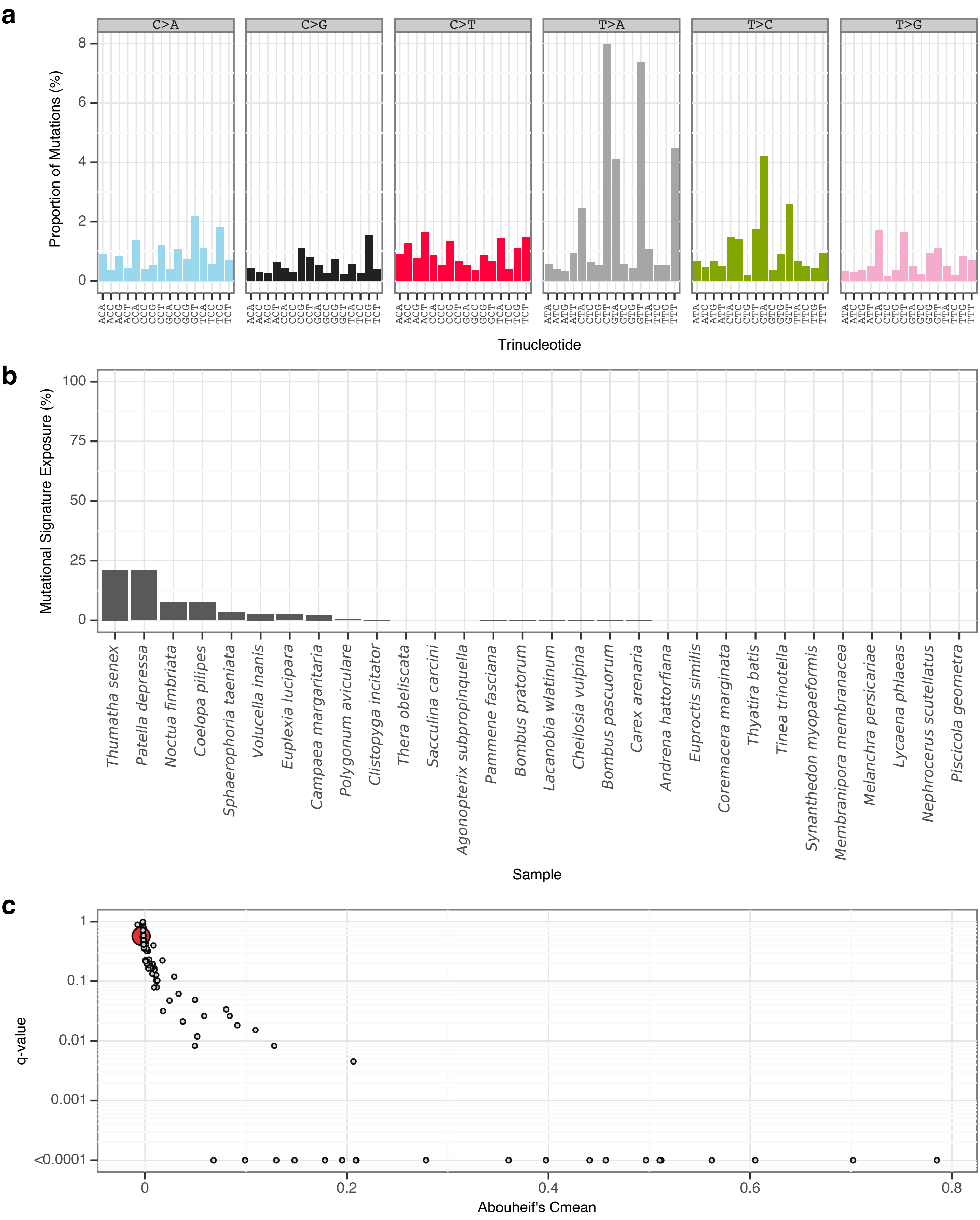


**Supplementary Fig. 53** | sToL50 mutational signature.

**a**, Mutational signature spectrum.

**b**, Bar plot displaying the 30 samples that exhibit the greatest exposure to the mutational signature.

**c**, Phylogenetic signal of the mutational signature.

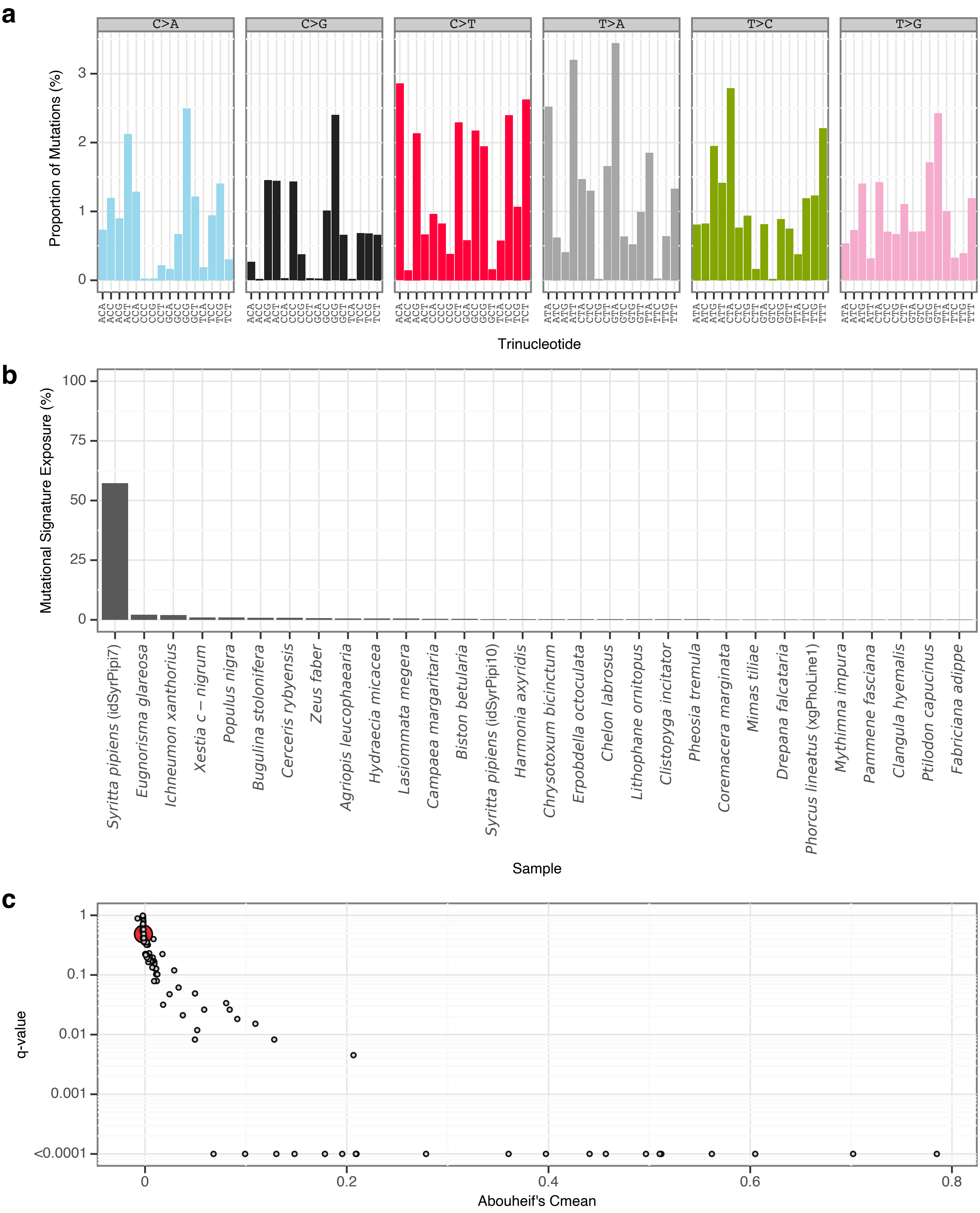


**Supplementary Fig. 54** | sToL51 mutational signature.

**a**, Mutational signature spectrum.

**b**, Bar plot displaying the 30 samples that exhibit the greatest exposure to the mutational signature.

**c**, Phylogenetic signal of the mutational signature.

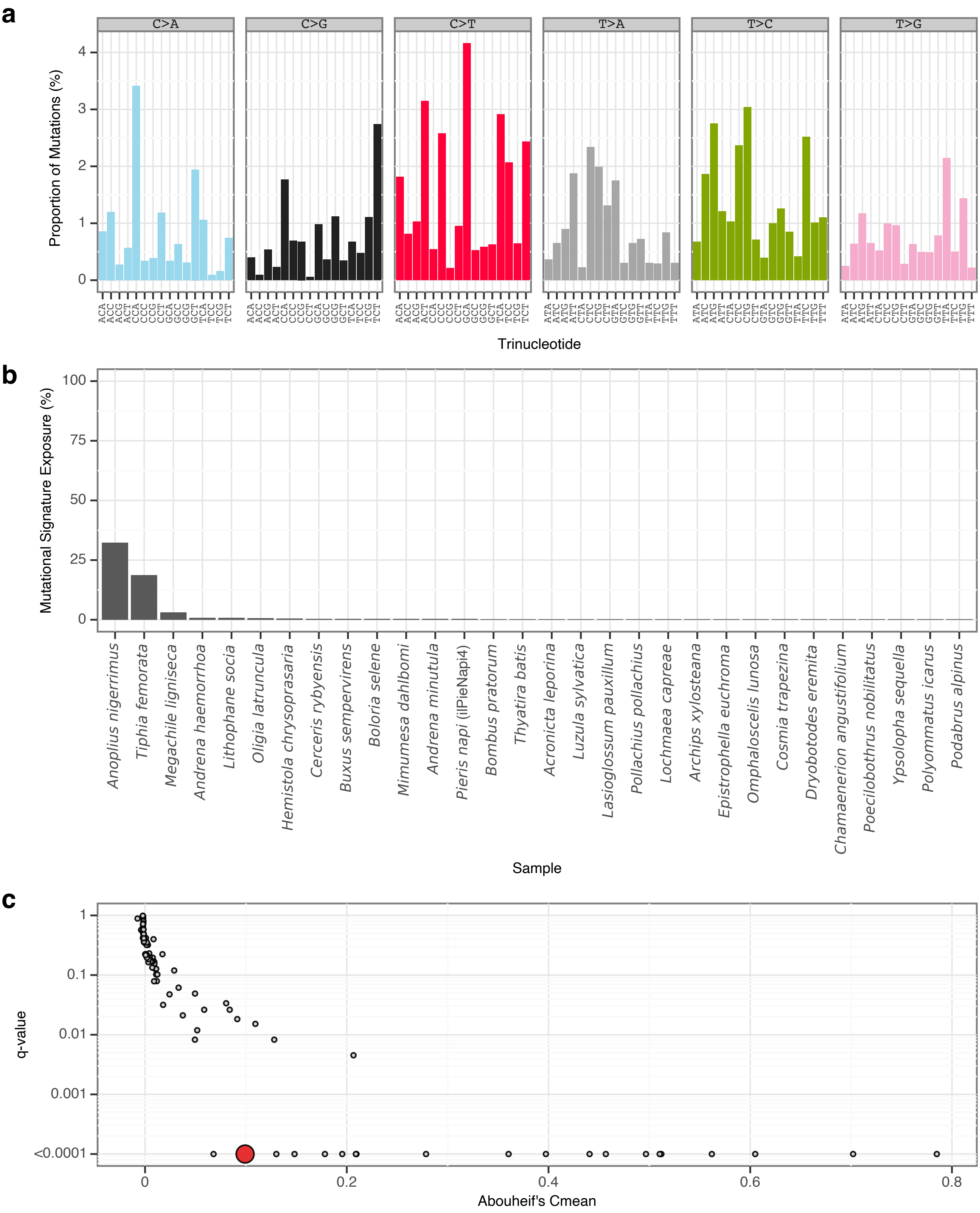


**Supplementary Fig. 55** | sToL52 mutational signature.

**a**, Mutational signature spectrum.

**b**, Bar plot displaying the 30 samples that exhibit the greatest exposure to the mutational signature.

**c**, Phylogenetic signal of the mutational signature.

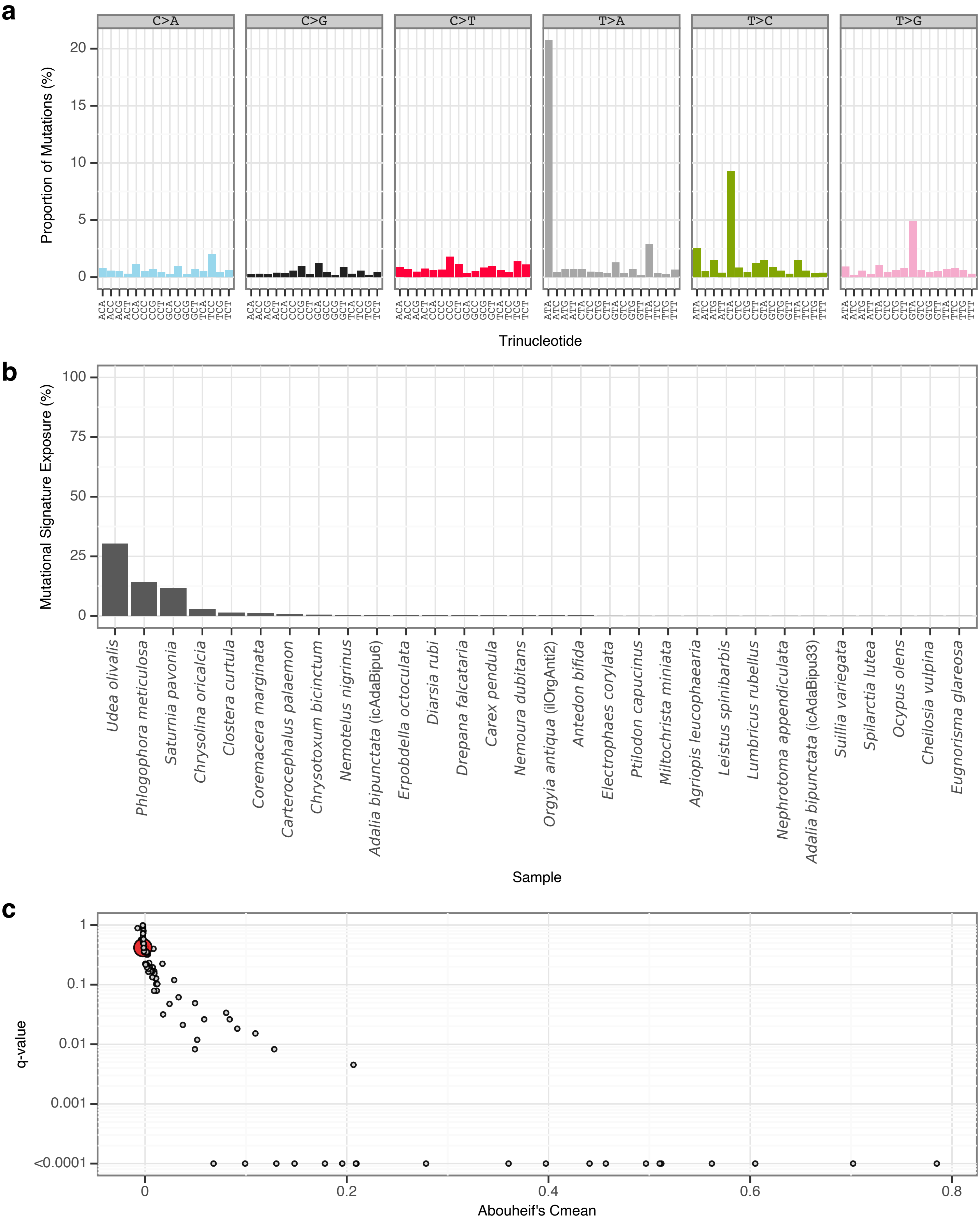


**Supplementary Fig. 56** | sToL53 mutational signature.

**a**, Mutational signature spectrum.

**b**, Bar plot displaying the 30 samples that exhibit the greatest exposure to the mutational signature.

**c**, Phylogenetic signal of the mutational signature.

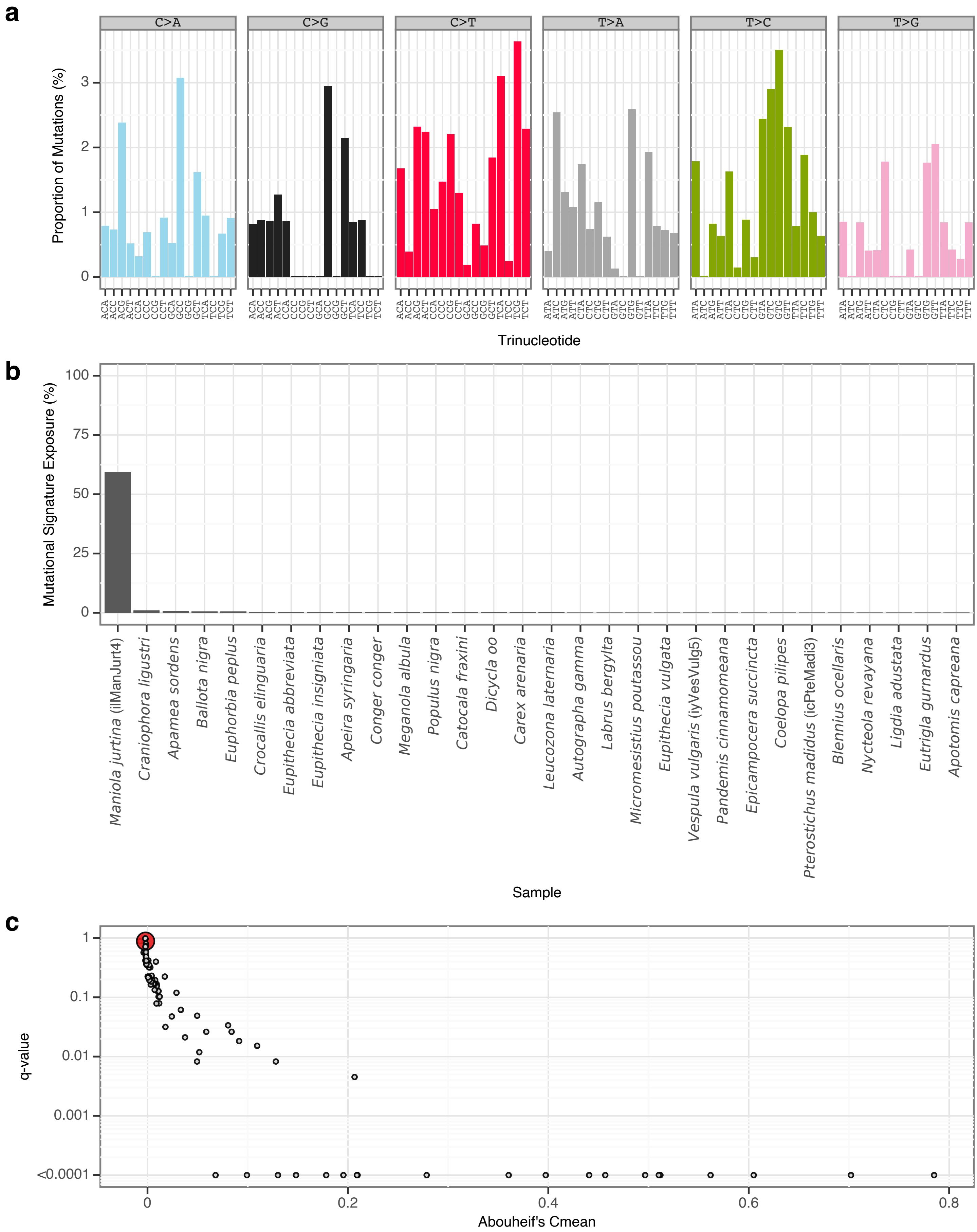


**Supplementary Fig. 57** | sToL54 mutational signature.
**a**, Mutational signature spectrum.
**b**, Bar plot displaying the 30 samples that exhibit the greatest exposure to the mutational signature.
**c**, Phylogenetic signal of the mutational signature.

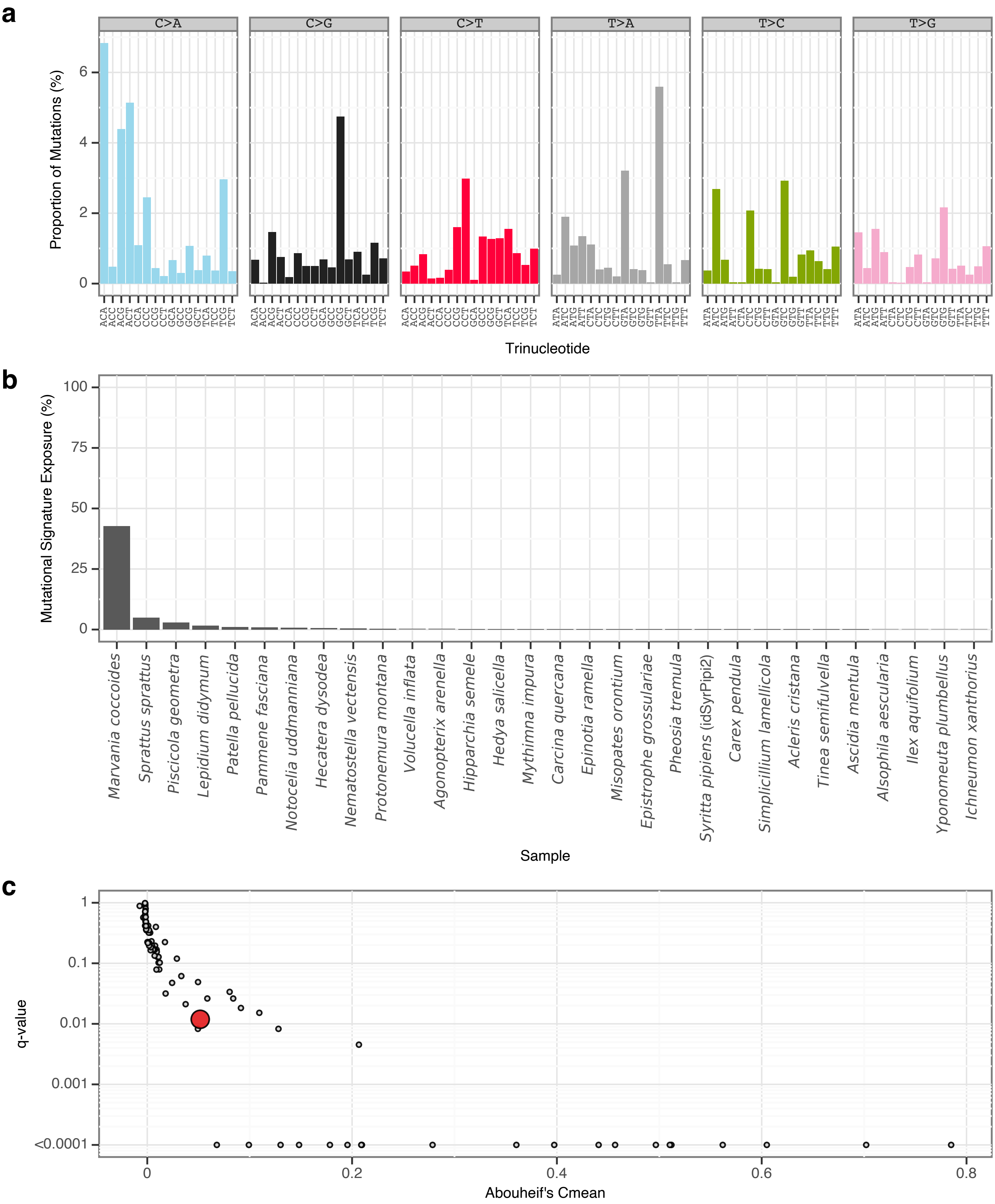


**Supplementary Fig. 58** | sToL55 mutational signature.

**a**, Mutational signature spectrum.

**b**, Bar plot displaying the 30 samples that exhibit the greatest exposure to the mutational signature.

**c**, Phylogenetic signal of the mutational signature.

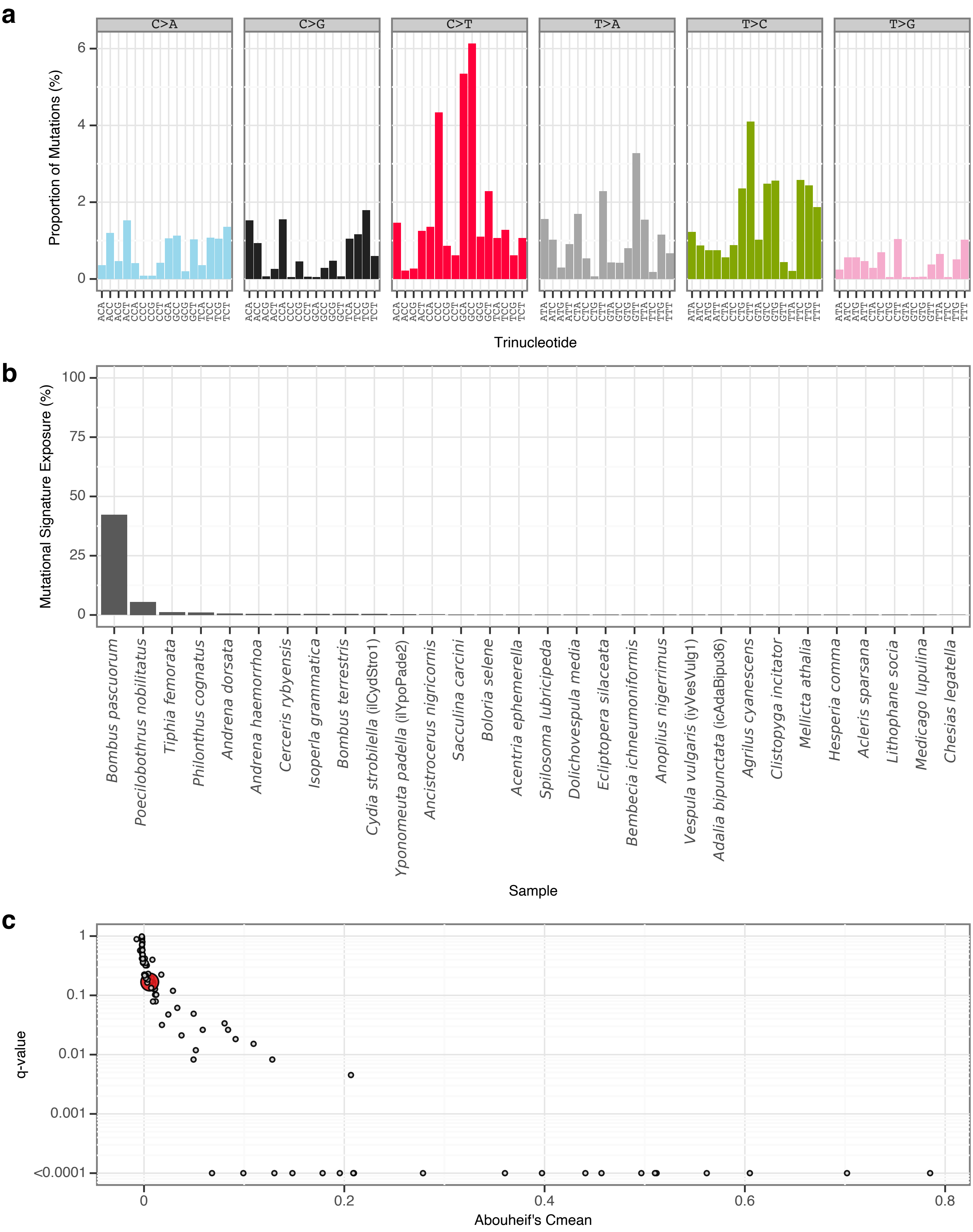


**Supplementary Fig. 59** | sToL56 mutational signature.

**a**, Mutational signature spectrum.

**b**, Bar plot displaying the 30 samples that exhibit the greatest exposure to the mutational signature.

**c**, Phylogenetic signal of the mutational signature.

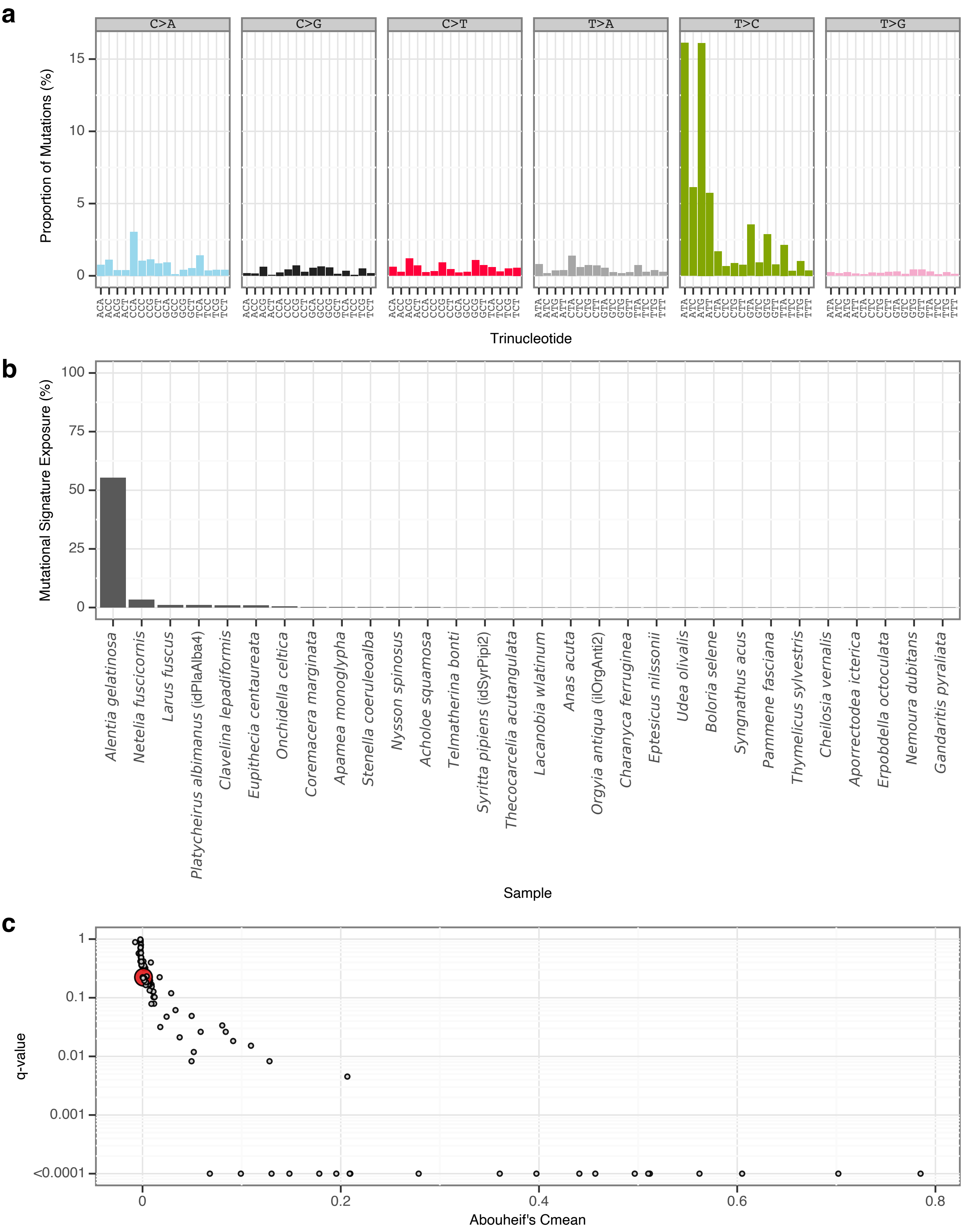


**Supplementary Fig. 60** | sToL57 mutational signature.

**a**, Mutational signature spectrum.

**b**, Bar plot displaying the 30 samples that exhibit the greatest exposure to the mutational signature.

**c**, Phylogenetic signal of the mutational signature.

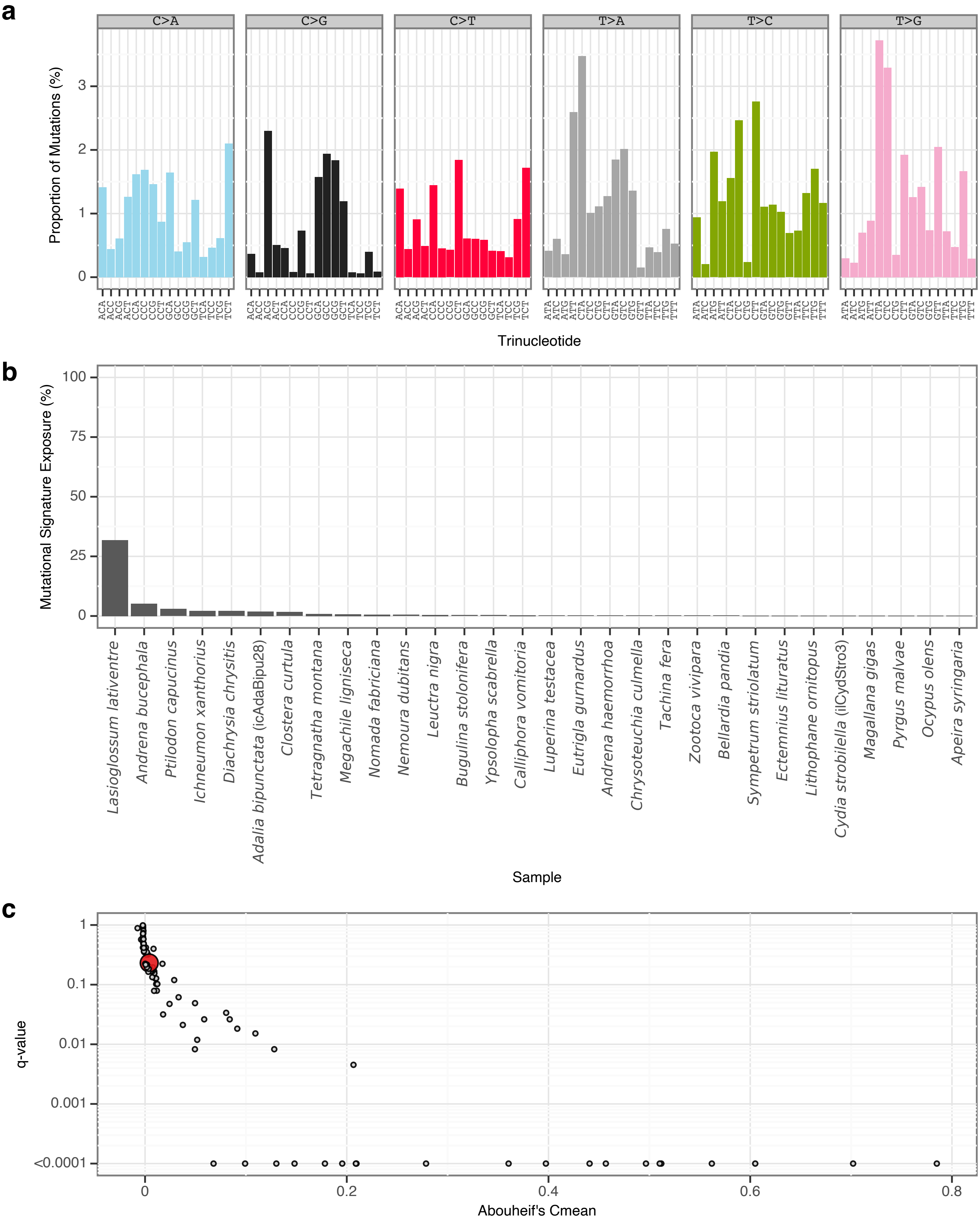


**Supplementary Fig. 61** | sToL58 mutational signature.

**a**, Mutational signature spectrum.

**b**, Bar plot displaying the 30 samples that exhibit the greatest exposure to the mutational signature.

**c**, Phylogenetic signal of the mutational signature.

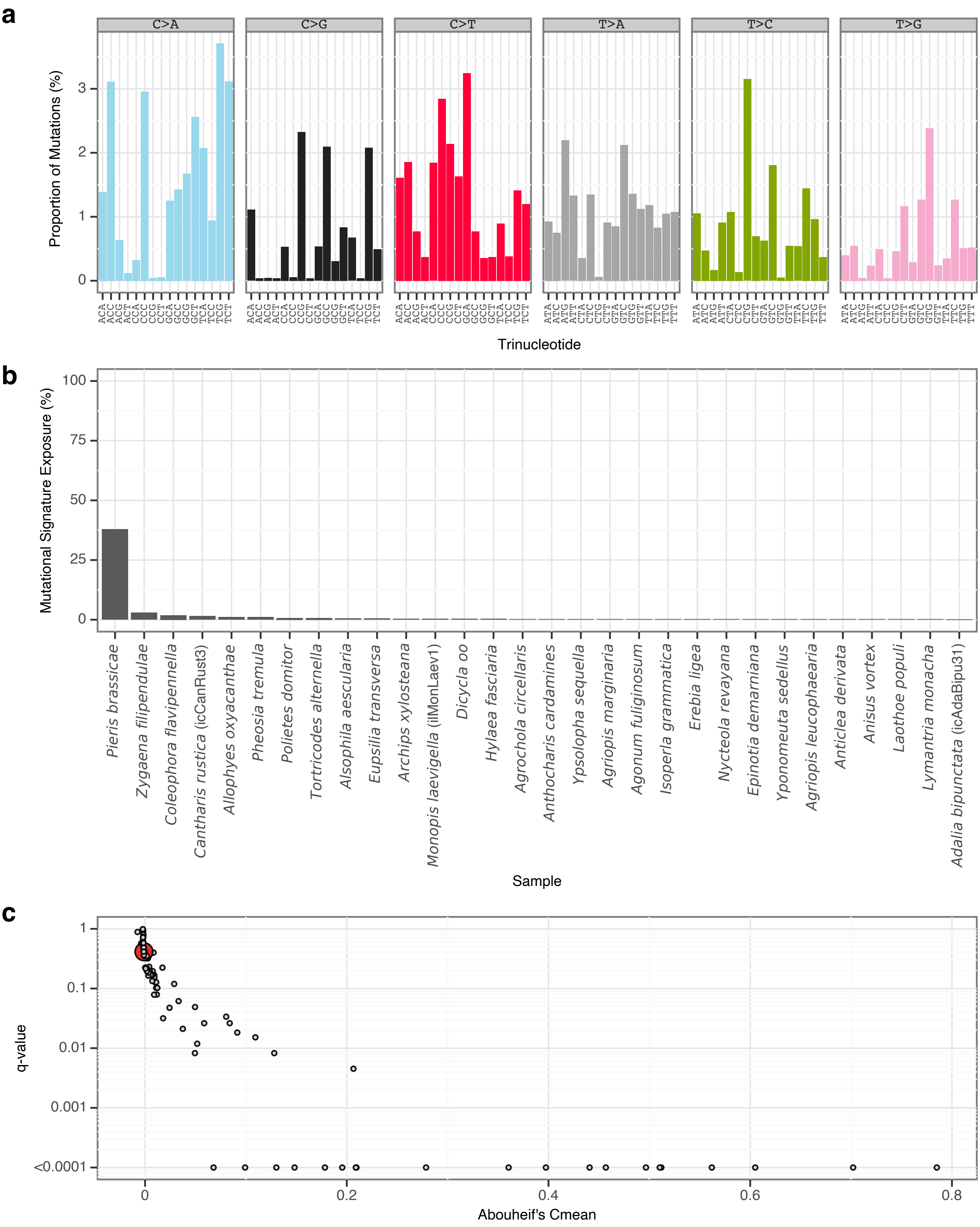


**Supplementary Fig. 62** | sToL59 mutational signature.
**a**, Mutational signature spectrum.
**b**, Bar plot displaying the 30 samples that exhibit the greatest exposure to the mutational signature.
**c**, Phylogenetic signal of the mutational signature.

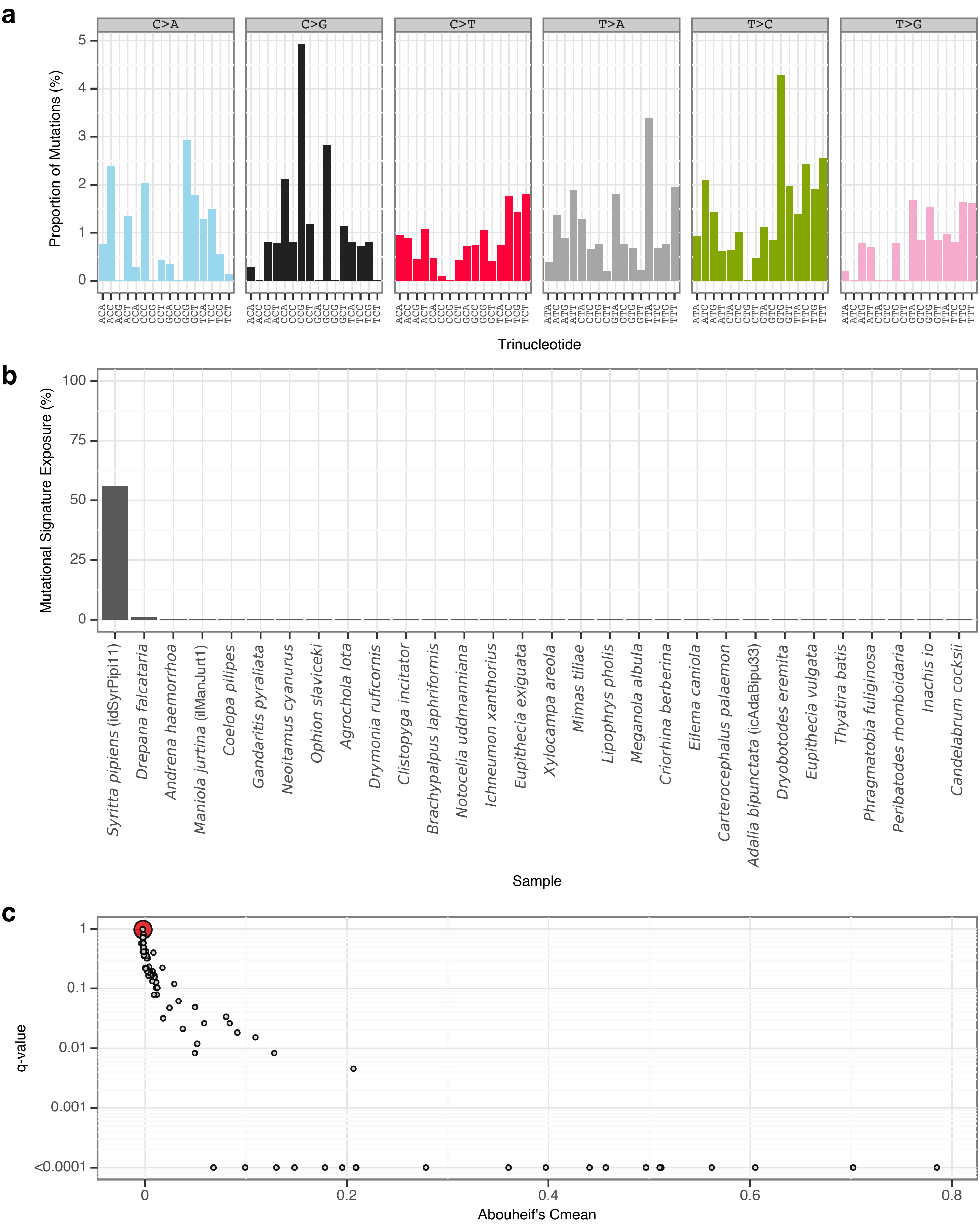


**Supplementary Fig. 63** | sToL60 mutational signature.

**a**, Mutational signature spectrum.

**b**, Bar plot displaying the 30 samples that exhibit the greatest exposure to the mutational signature.

**c**, Phylogenetic signal of the mutational signature.

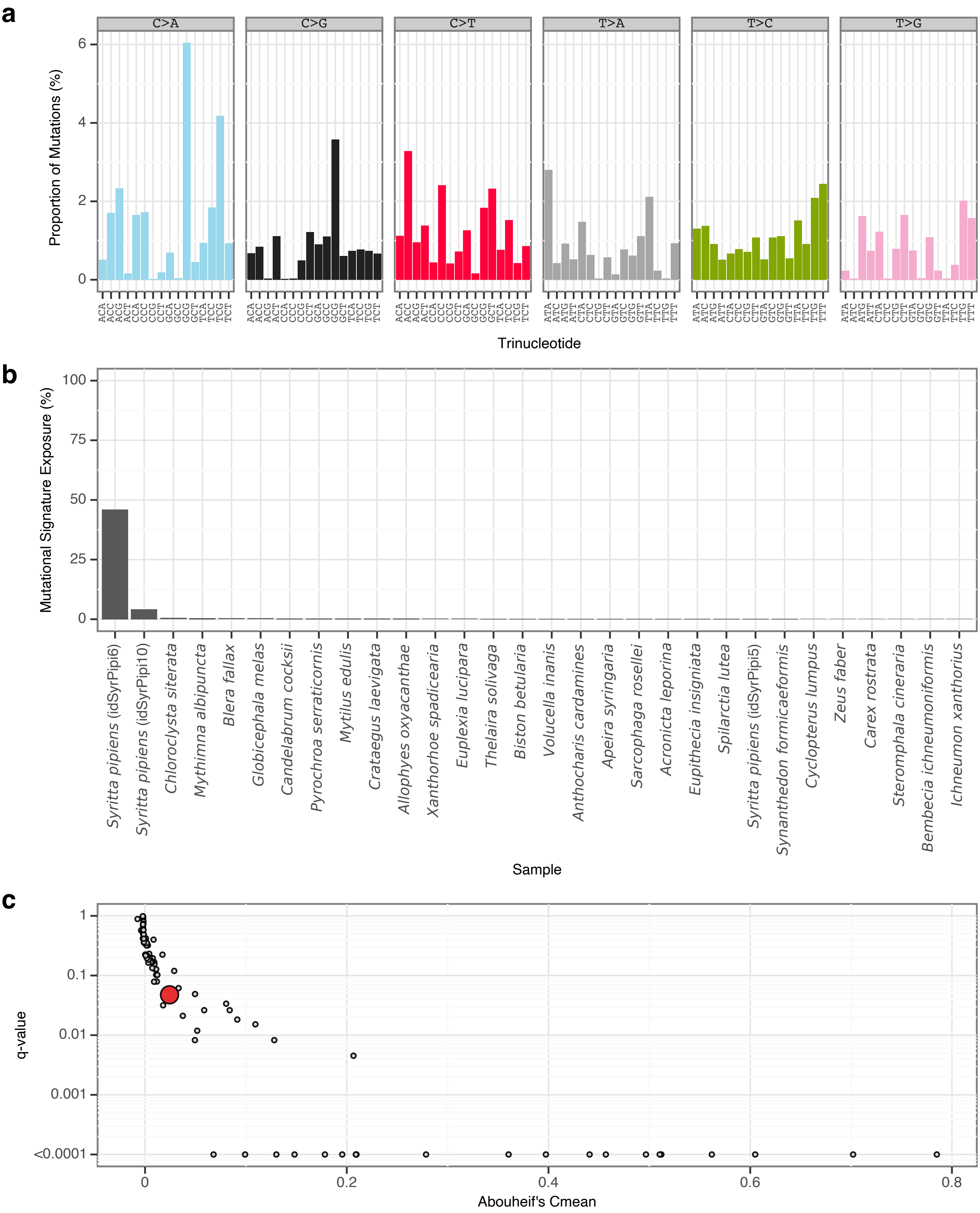


**Supplementary Fig. 64** | sToL61 mutational signature.
**a**, Mutational signature spectrum.
**b**, Bar plot displaying the 30 samples that exhibit the greatest exposure to the mutational signature.
**c**, Phylogenetic signal of the mutational signature.

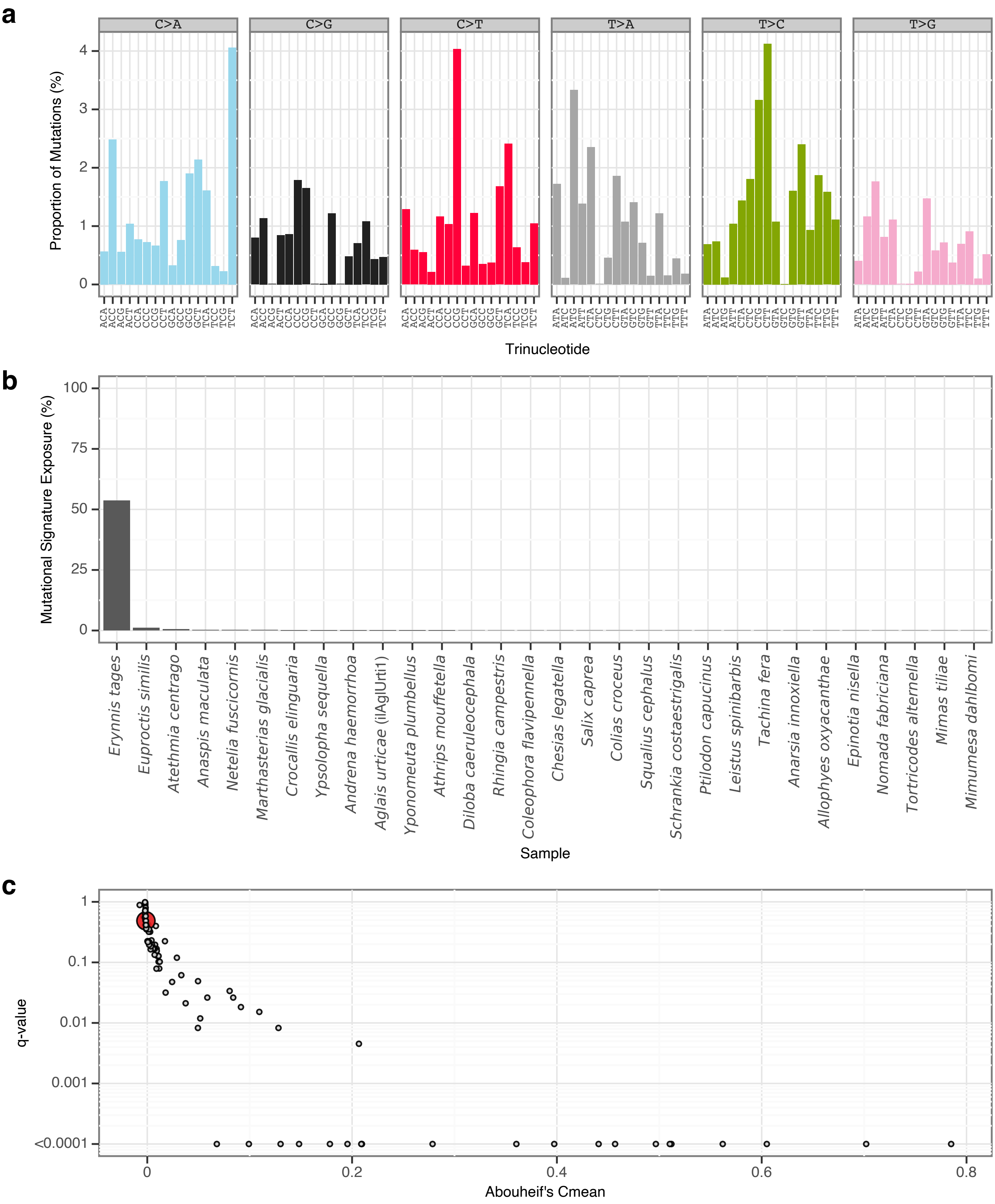


**Supplementary Fig. 65** | sToL62 mutational signature.
**a**, Mutational signature spectrum.
**b**, Bar plot displaying the 30 samples that exhibit the greatest exposure to the mutational signature.
**c**, Phylogenetic signal of the mutational signature.

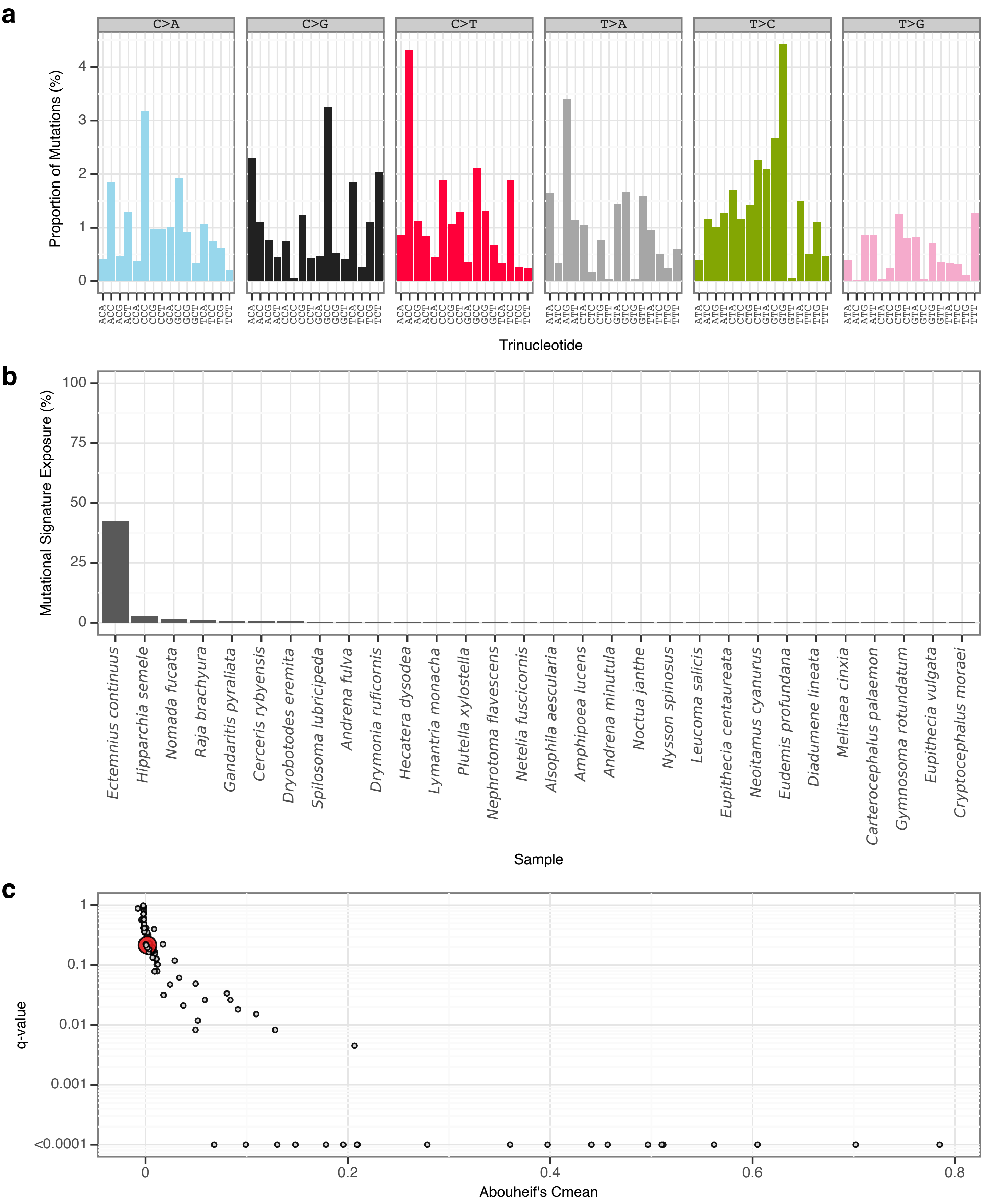


**Supplementary Fig. 66** | sToL63 mutational signature.

**a**, Mutational signature spectrum.

**b**, Bar plot displaying the 30 samples that exhibit the greatest exposure to the mutational signature.

**c**, Phylogenetic signal of the mutational signature.

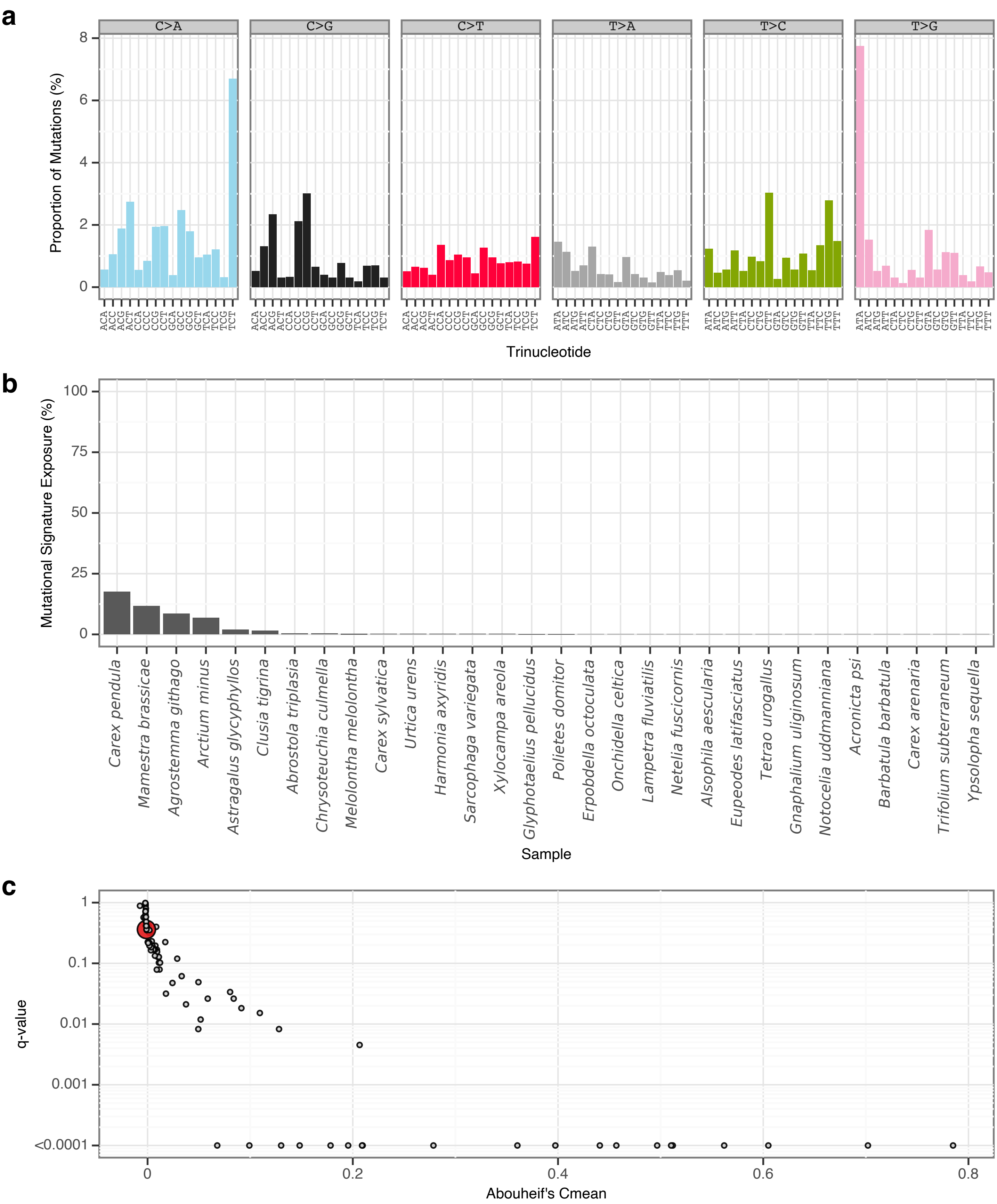


**Supplementary Fig. 67** I sToL64 mutational signature.
**a**, Mutational signature spectrum.
**b**, Bar plot displaying the 30 samples that exhibit the greatest exposure to the mutational signature.
**c**, Phylogenetic signal of the mutational signature.

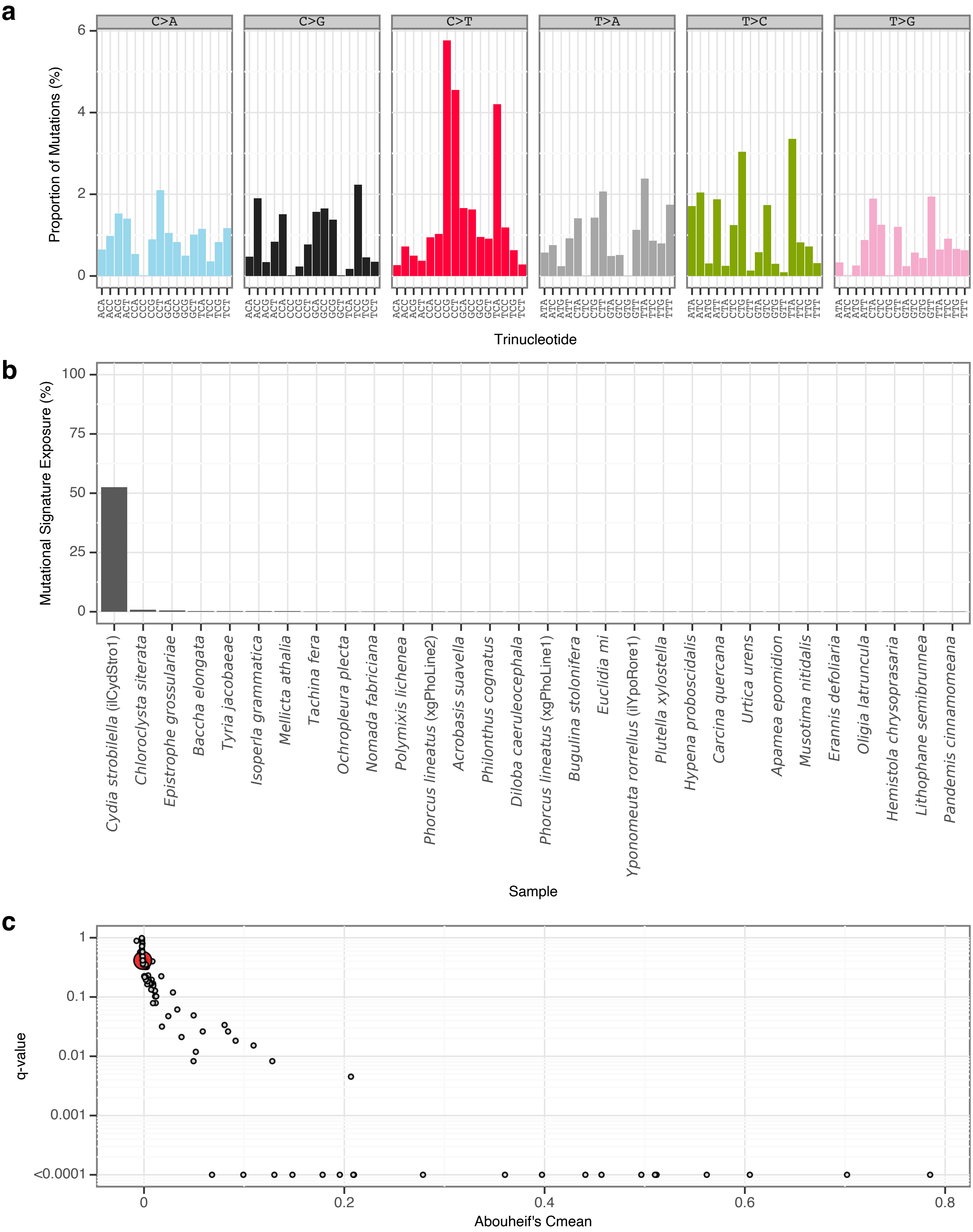


**Supplementary Fig. 68** | sToL65 mutational signature.

**a**, Mutational signature spectrum.

**b**, Bar plot displaying the 30 samples that exhibit the greatest exposure to the mutational signature.

**c**, Phylogenetic signal of the mutational signature.

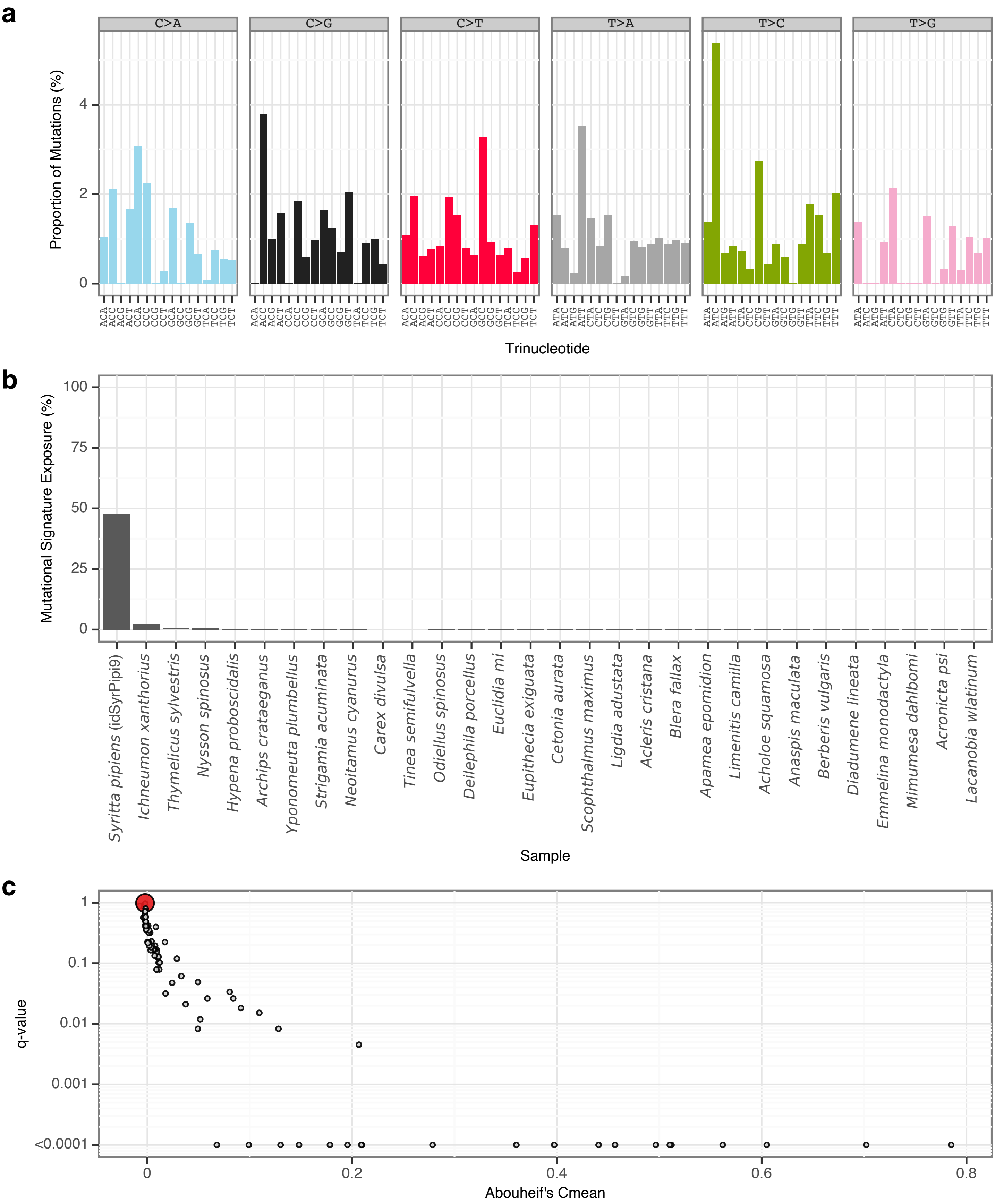


**Supplementary Fig. 69** | sToL66 mutational signature.

**a**, Mutational signature spectrum.

**b**, Bar plot displaying the 30 samples that exhibit the greatest exposure to the mutational signature.

**c**, Phylogenetic signal of the mutational signature.

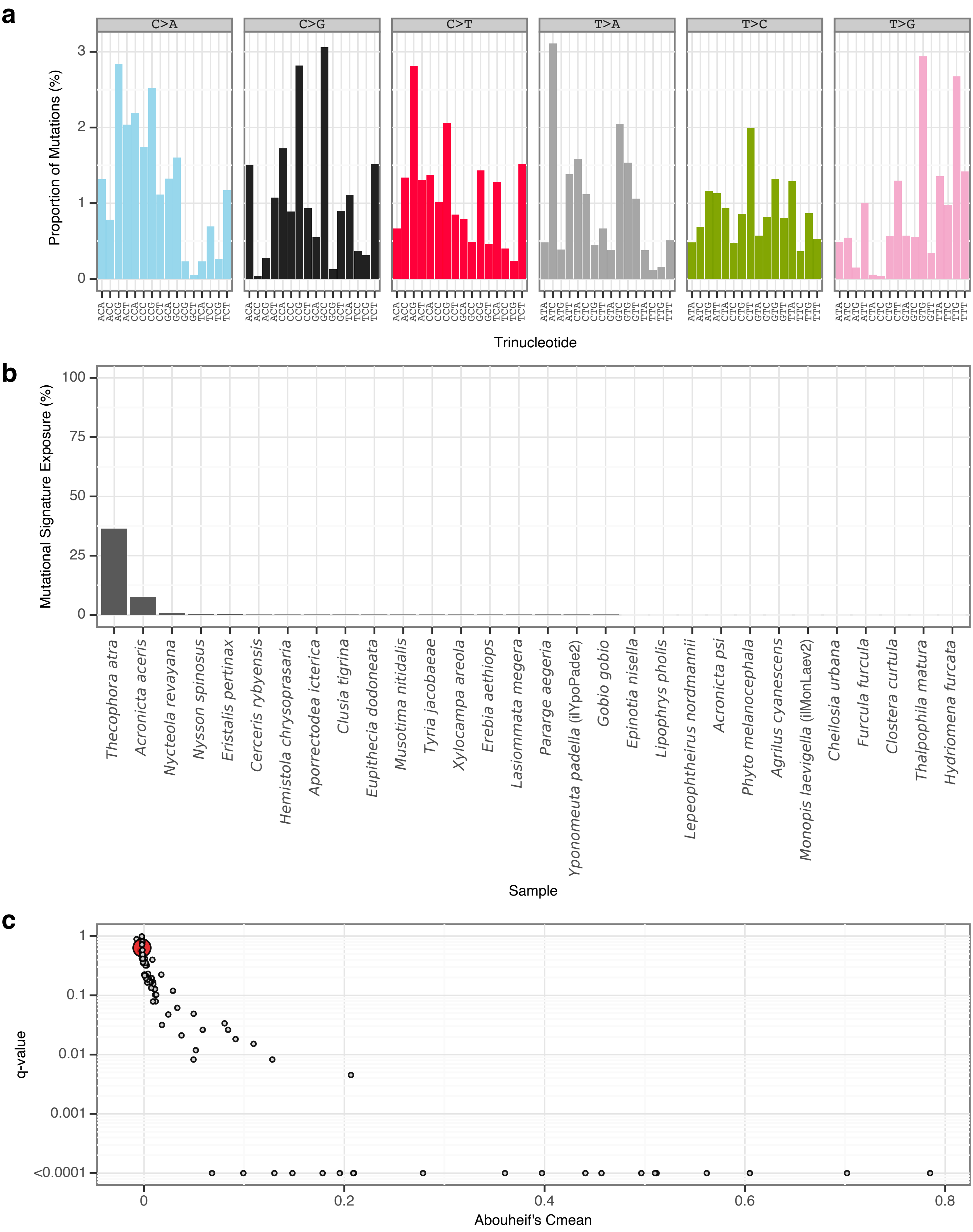


**Supplementary Fig. 70** | sToL67 mutational signature.

**a**, Mutational signature spectrum.

**b**, Bar plot displaying the 30 samples that exhibit the greatest exposure to the mutational signature.

**c**, Phylogenetic signal of the mutational signature.

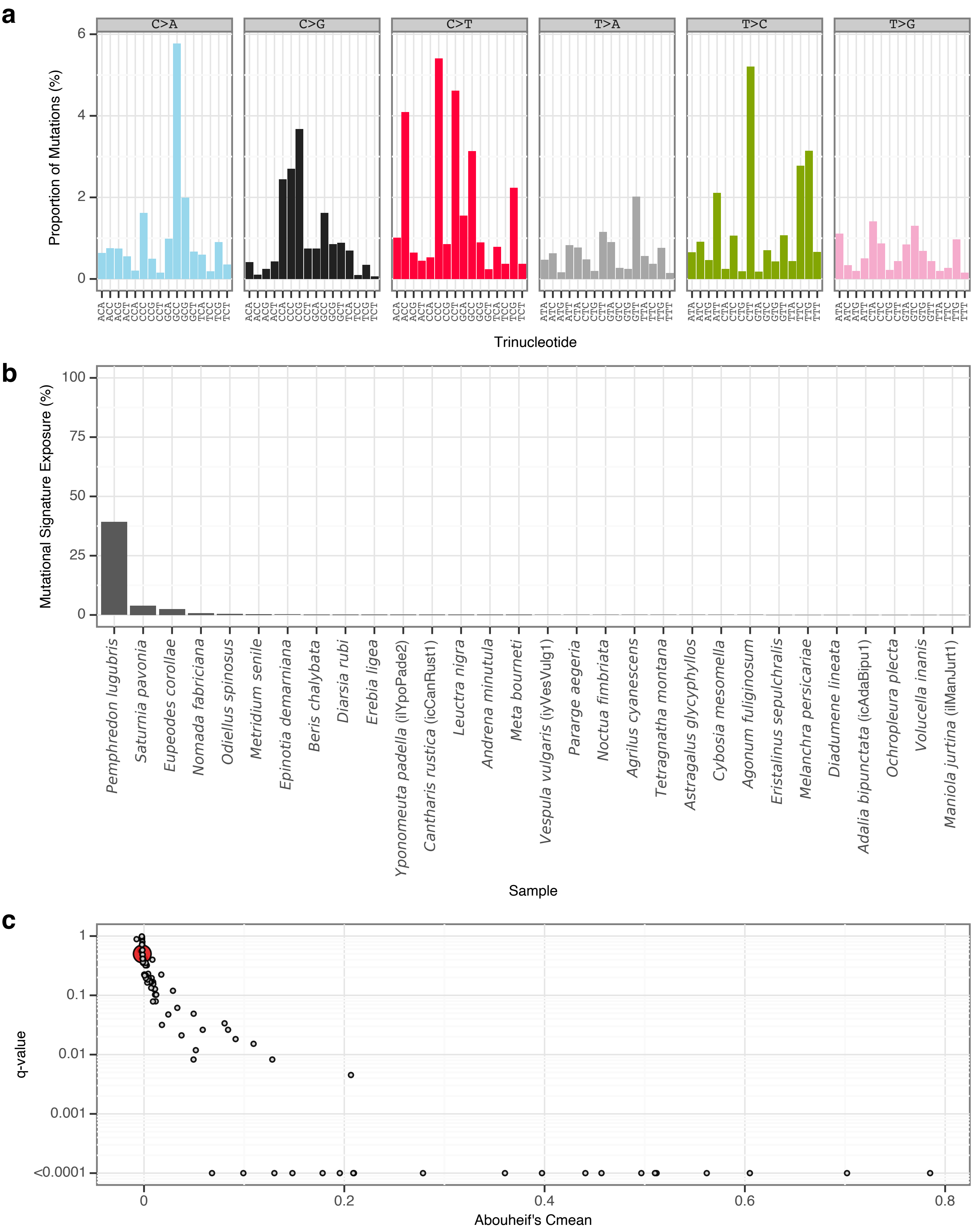


**Supplementary Fig. 71** | sToL68 mutational signature.

**a**, Mutational signature spectrum.

**b**, Bar plot displaying the 30 samples that exhibit the greatest exposure to the mutational signature.

**c**, Phylogenetic signal of the mutational signature.

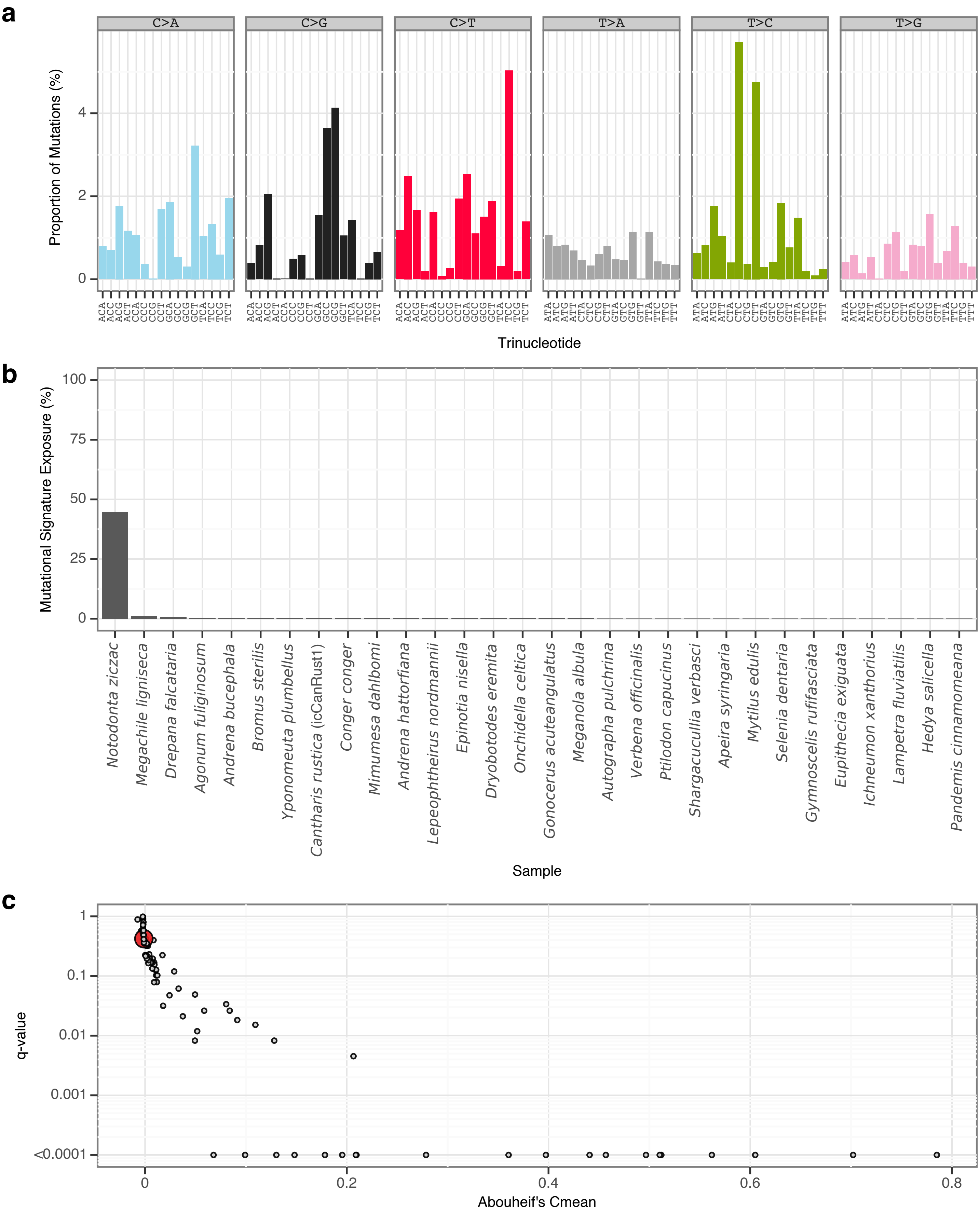


**Supplementary Fig. 72** | sToL69 mutational signature.

**a**, Mutational signature spectrum.

**b**, Bar plot displaying the 30 samples that exhibit the greatest exposure to the mutational signature.

**c**, Phylogenetic signal of the mutational signature.

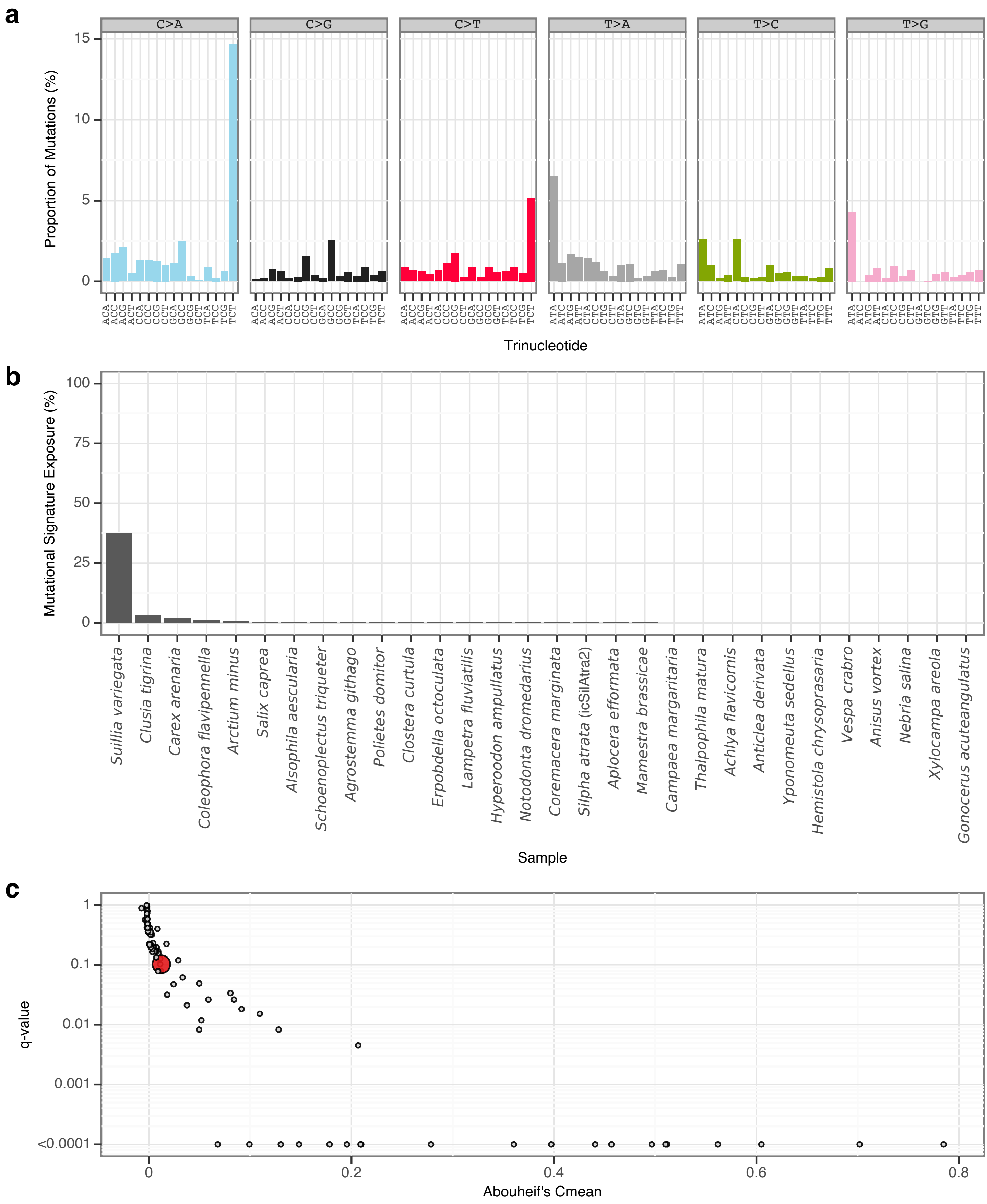


**Supplementary Fig. 73** | sToL70 mutational signature.

**a**, Mutational signature spectrum.

**b**, Bar plot displaying the 30 samples that exhibit the greatest exposure to the mutational signature.

**c**, Phylogenetic signal of the mutational signature.

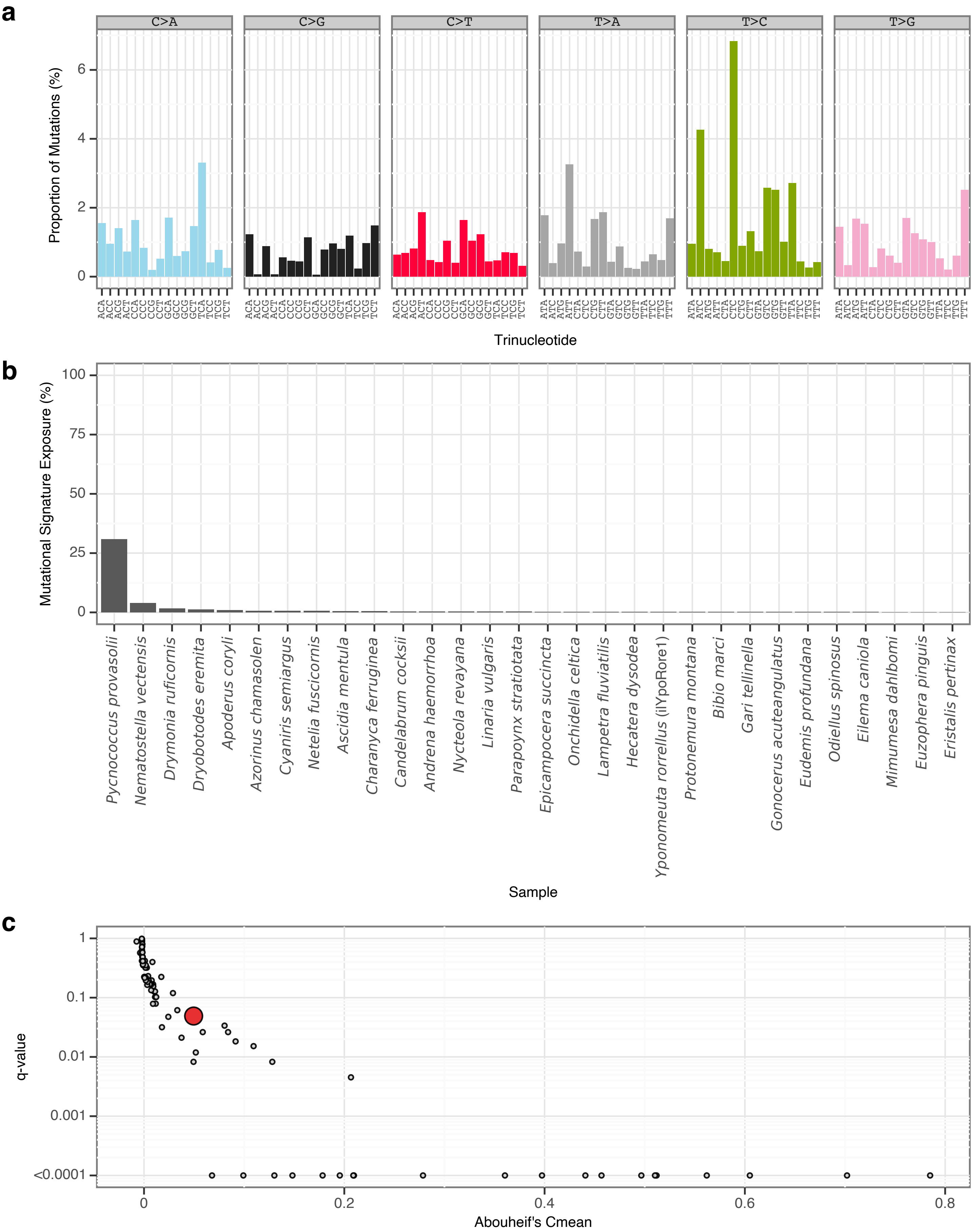


**Supplementary Fig. 74 | sToL71 mutational signature.**
**a**, Mutational signature spectrum.
**b**, Bar plot displaying the 30 samples that exhibit the greatest exposure to the mutational signature.
**c**, Phylogenetic signal of the mutational signature.

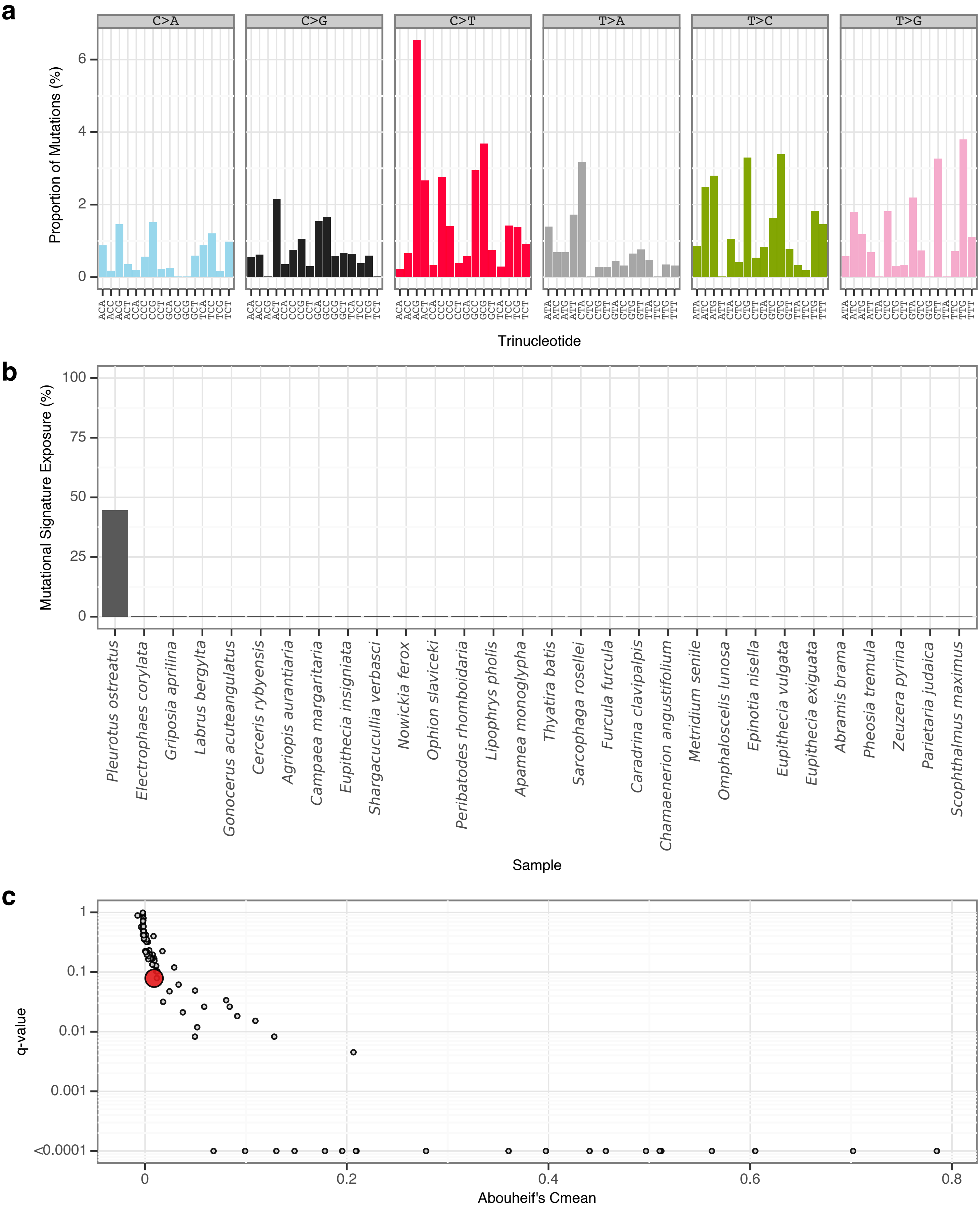


**Supplementary Fig. 75** | sToL72 mutational signature.
**a**, Mutational signature spectrum.
**b**, Bar plot displaying the 30 samples that exhibit the greatest exposure to the mutational signature.
**c**, Phylogenetic signal of the mutational signature.

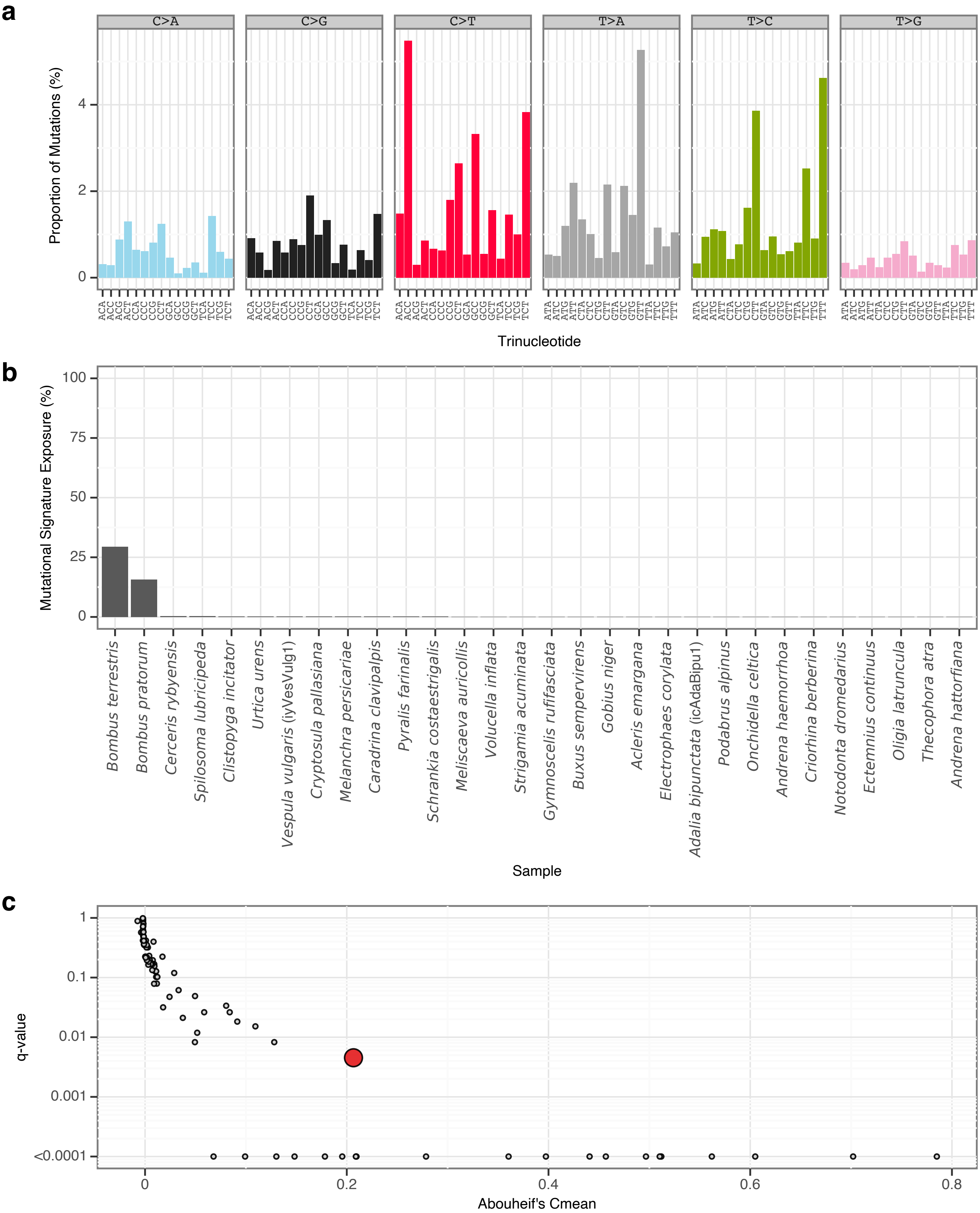


**Supplementary Fig. 76** | sToL73 mutational signature.
**a**, Mutational signature spectrum.
**b**, Bar plot displaying the 30 samples that exhibit the greatest exposure to the mutational signature.
**c**, Phylogenetic signal of the mutational signature.

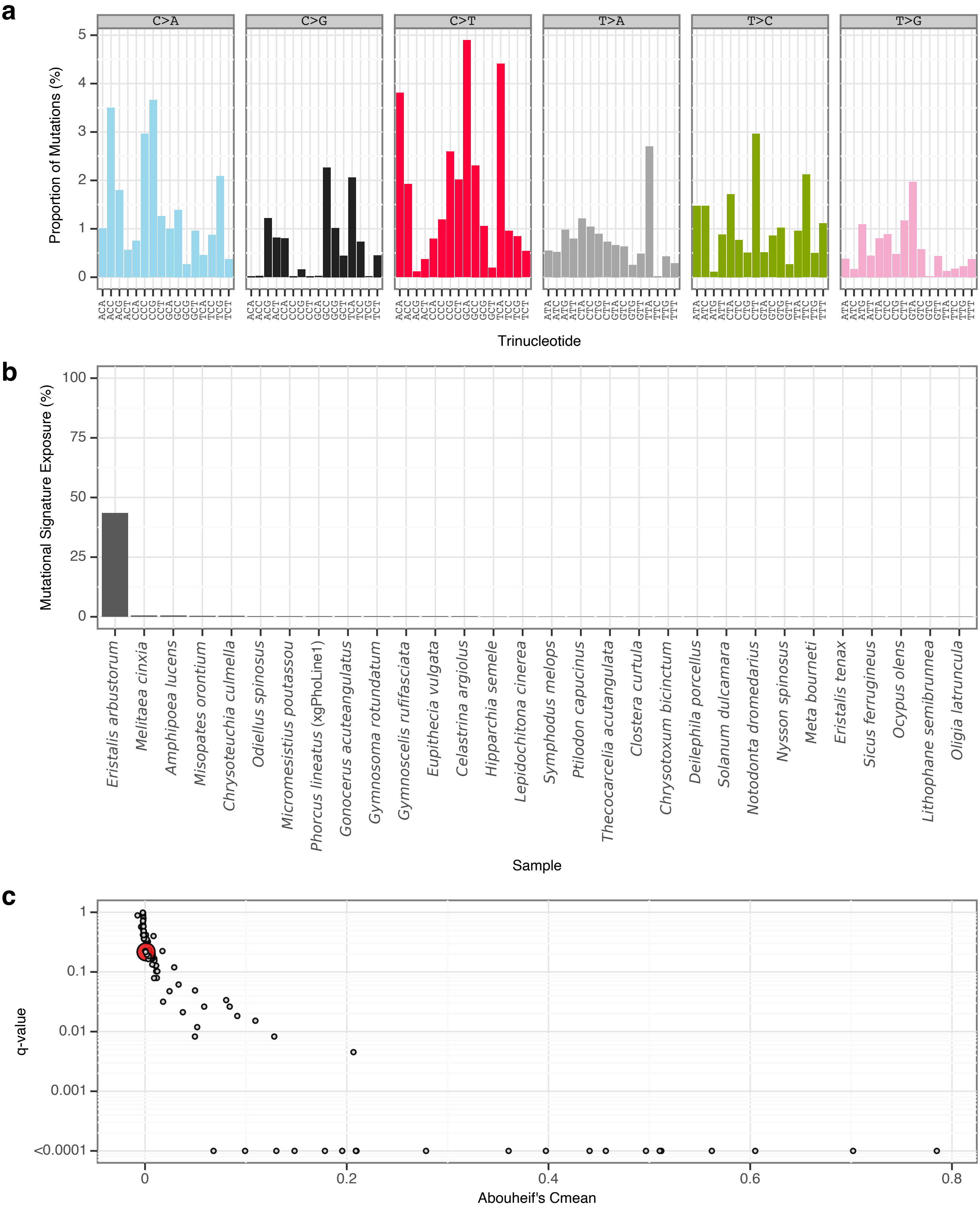


**Supplementary Fig. 77** I sToL74 mutational signature.

**a**, Mutational signature spectrum.

**b**, Bar plot displaying the 30 samples that exhibit the greatest exposure to the mutational signature.

**c**, Phylogenetic signal of the mutational signature.

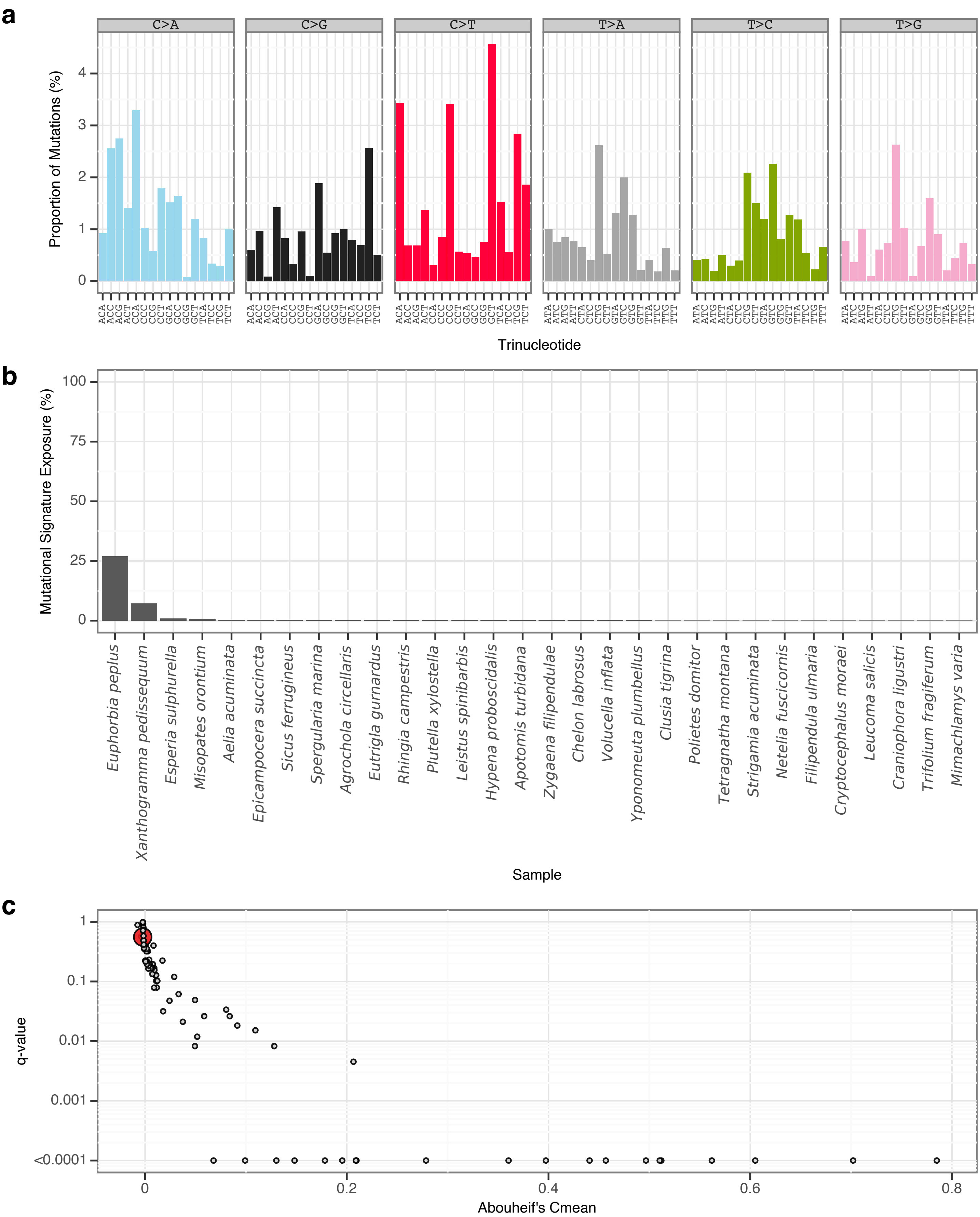


**Supplementary Fig. 78** | sToL75 mutational signature.

**a**, Mutational signature spectrum.

**b**, Bar plot displaying the 30 samples that exhibit the greatest exposure to the mutational signature.

**c**, Phylogenetic signal of the mutational signature.

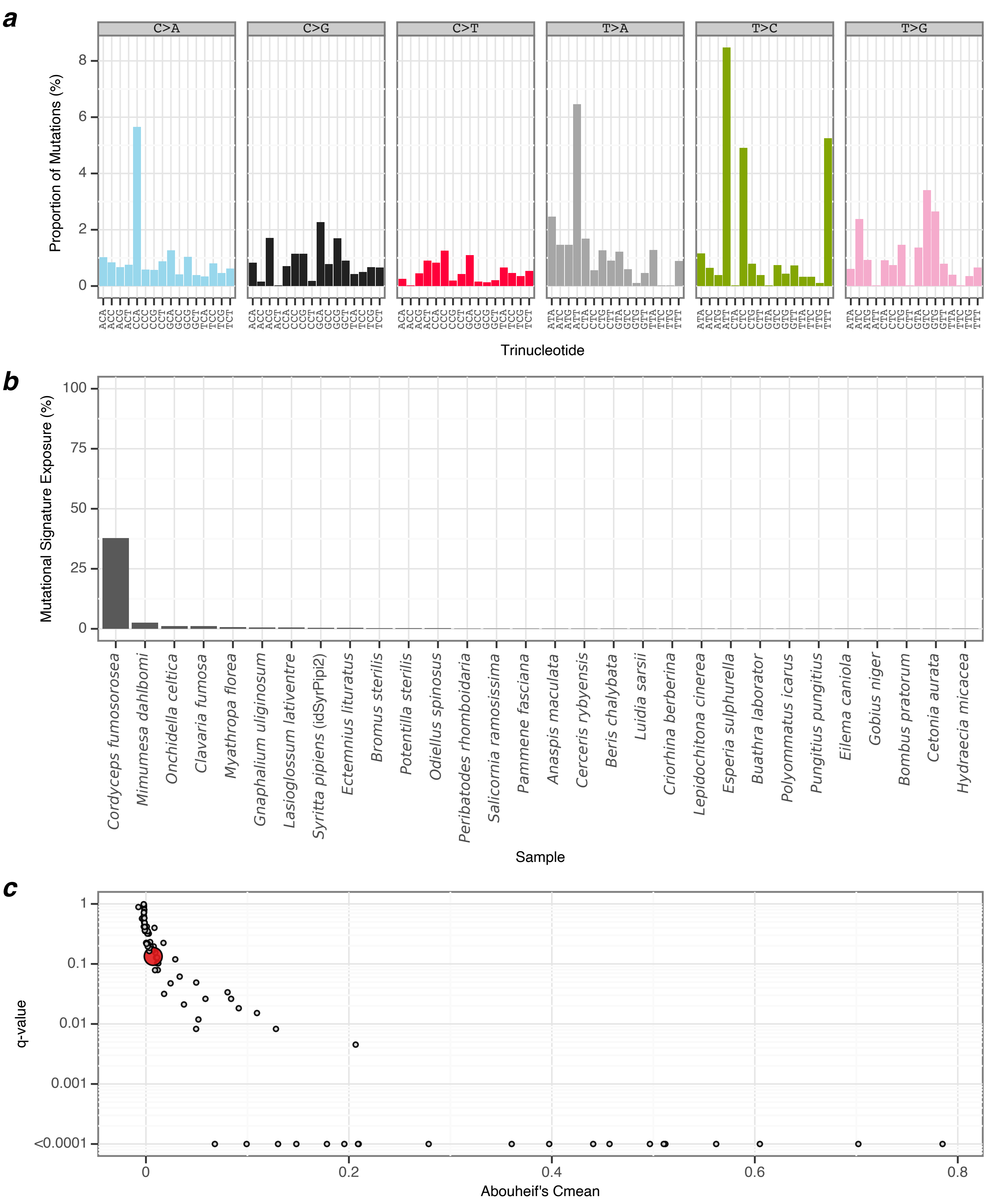


**Supplementary Fig. 79** | sToL76 mutational signature.
**a**, Mutational signature spectrum.
**b**, Bar plot displaying the 30 samples that exhibit the greatest exposure to the mutational signature.
**c**, Phylogenetic signal of the mutational signature.

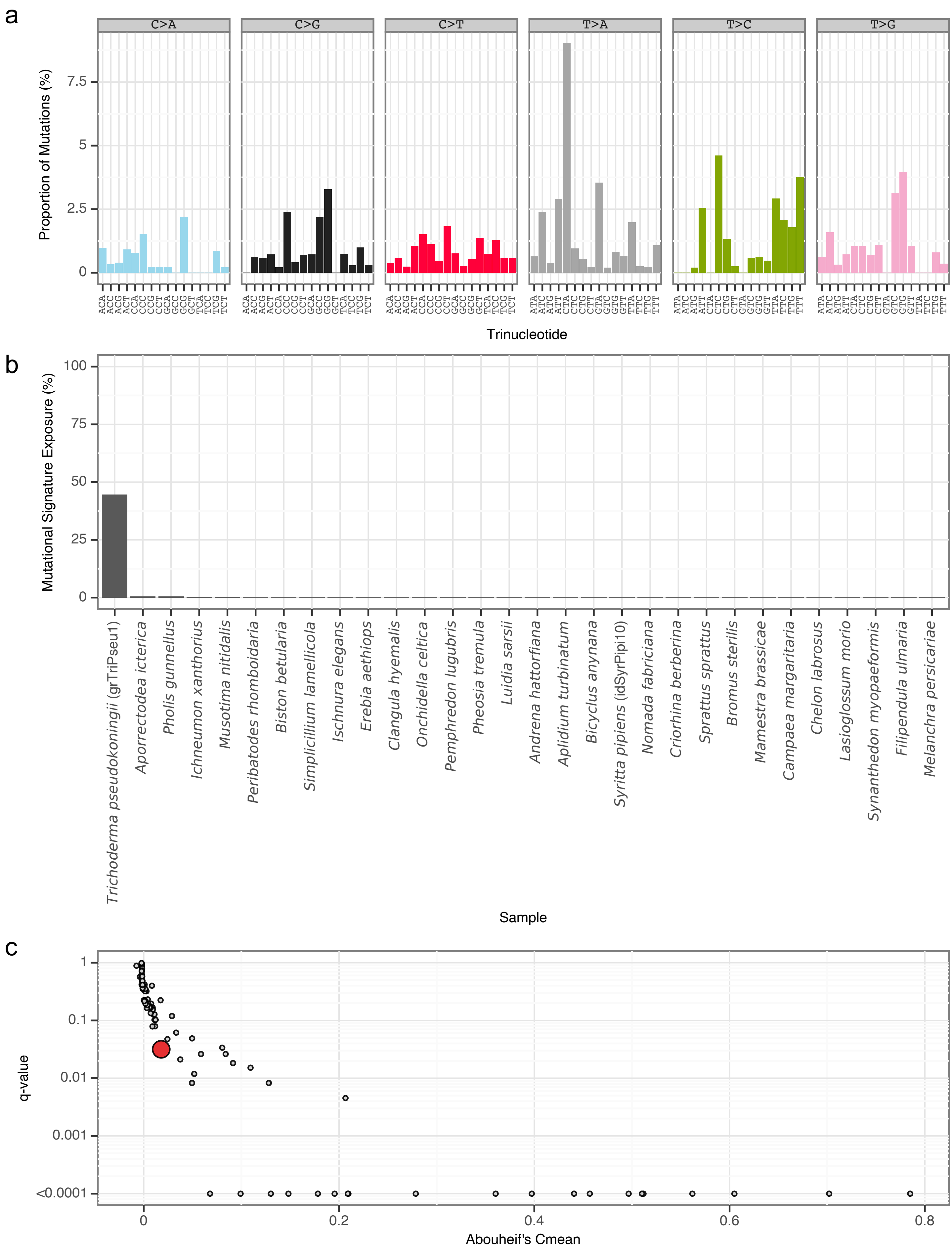


**Supplementary Fig. 80** | sToL77 mutational signature.
**a**, Mutational signature spectrum.
**b**, Bar plot displaying the 30 samples that exhibit the greatest exposure to the mutational signature.
**c**, Phylogenetic signal of the mutational signature.

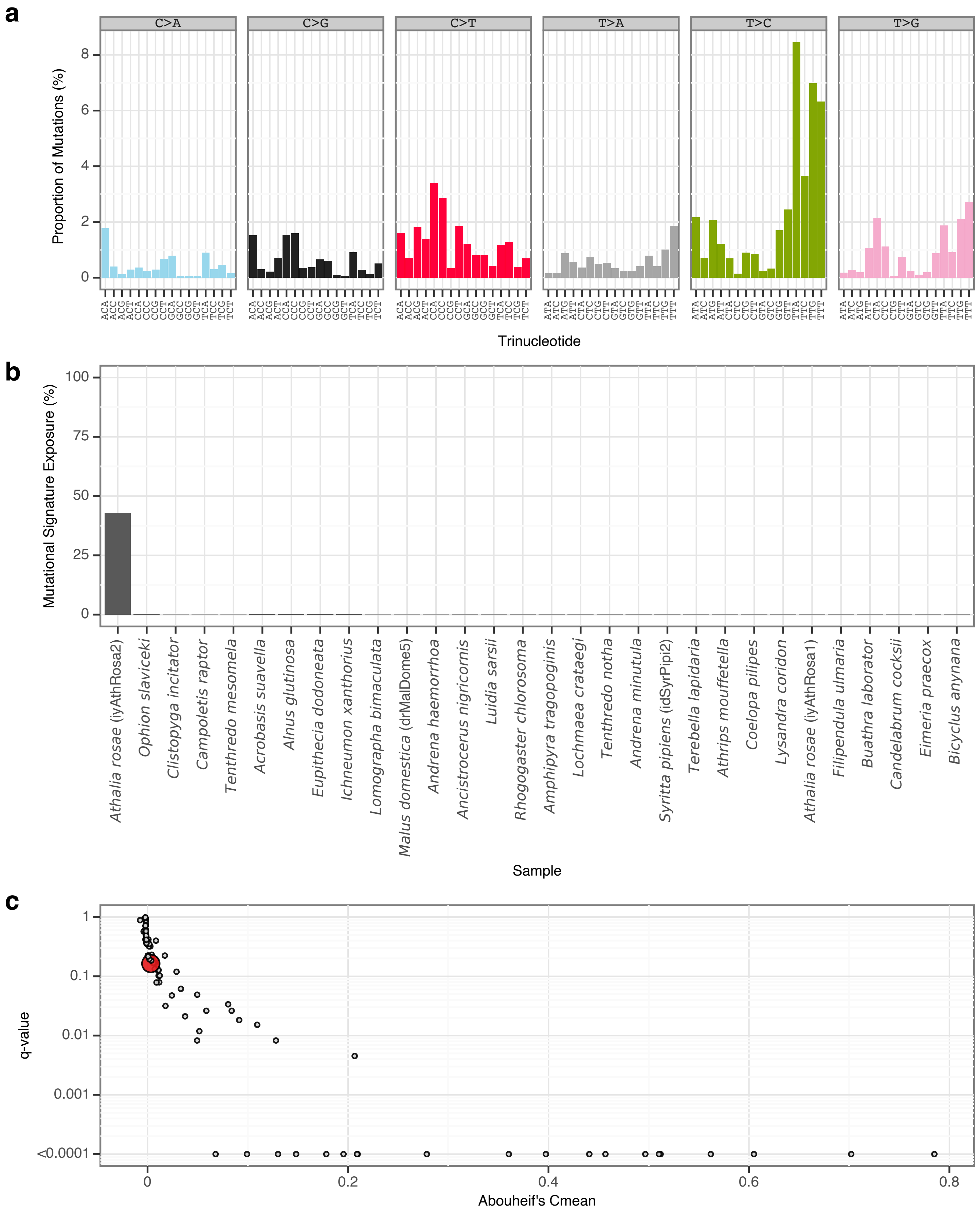


**Supplementary Fig. 81** | sToL78 mutational signature.

**a**, Mutational signature spectrum.

**b**, Bar plot displaying the 30 samples that exhibit the greatest exposure to the mutational signature.

**c**, Phylogenetic signal of the mutational signature.

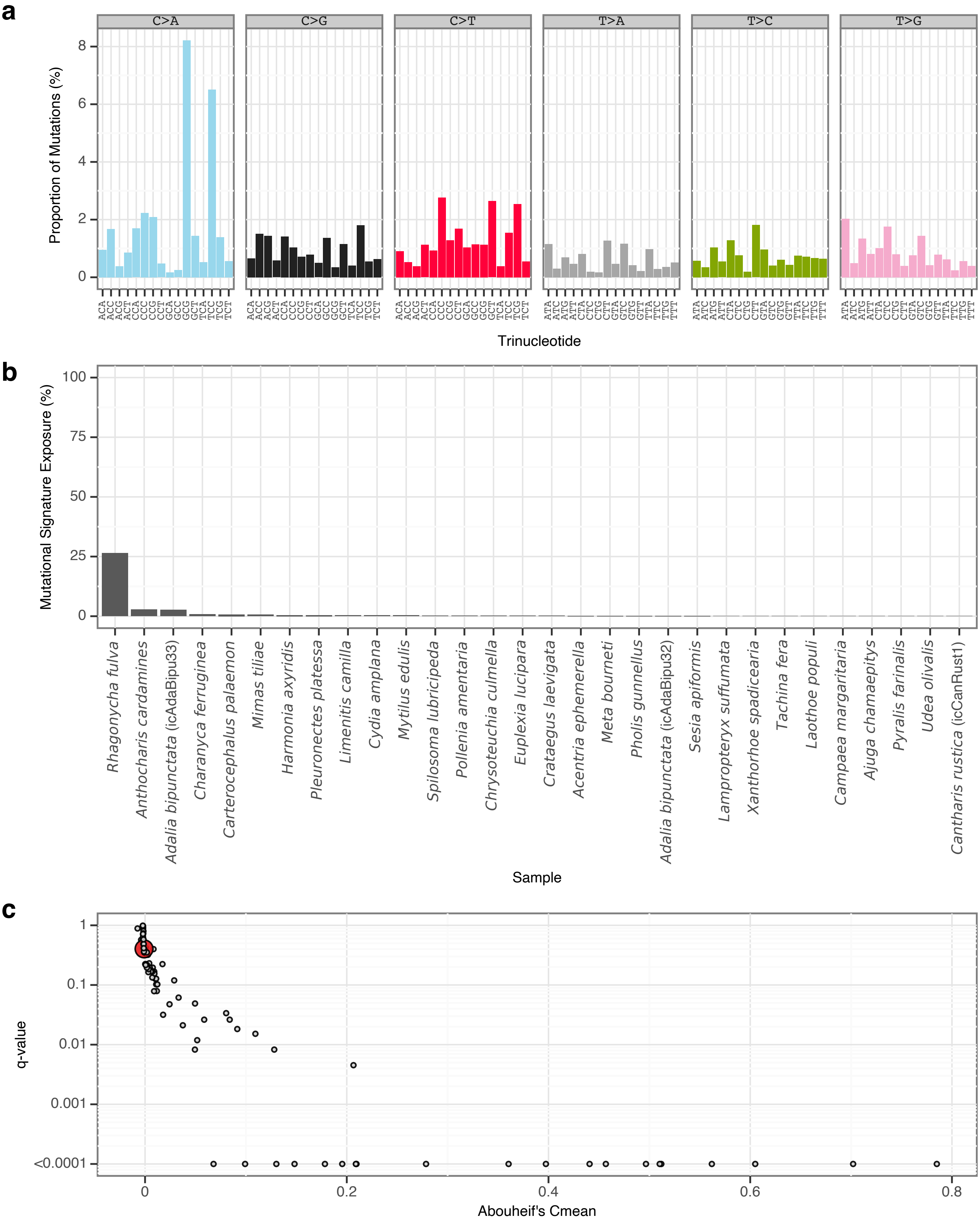


**Supplementary Fig. 82** | sToL79 mutational signature.

**a**, Mutational signature spectrum.

**b**, Bar plot displaying the 30 samples that exhibit the greatest exposure to the mutational signature.

**c**, Phylogenetic signal of the mutational signature.

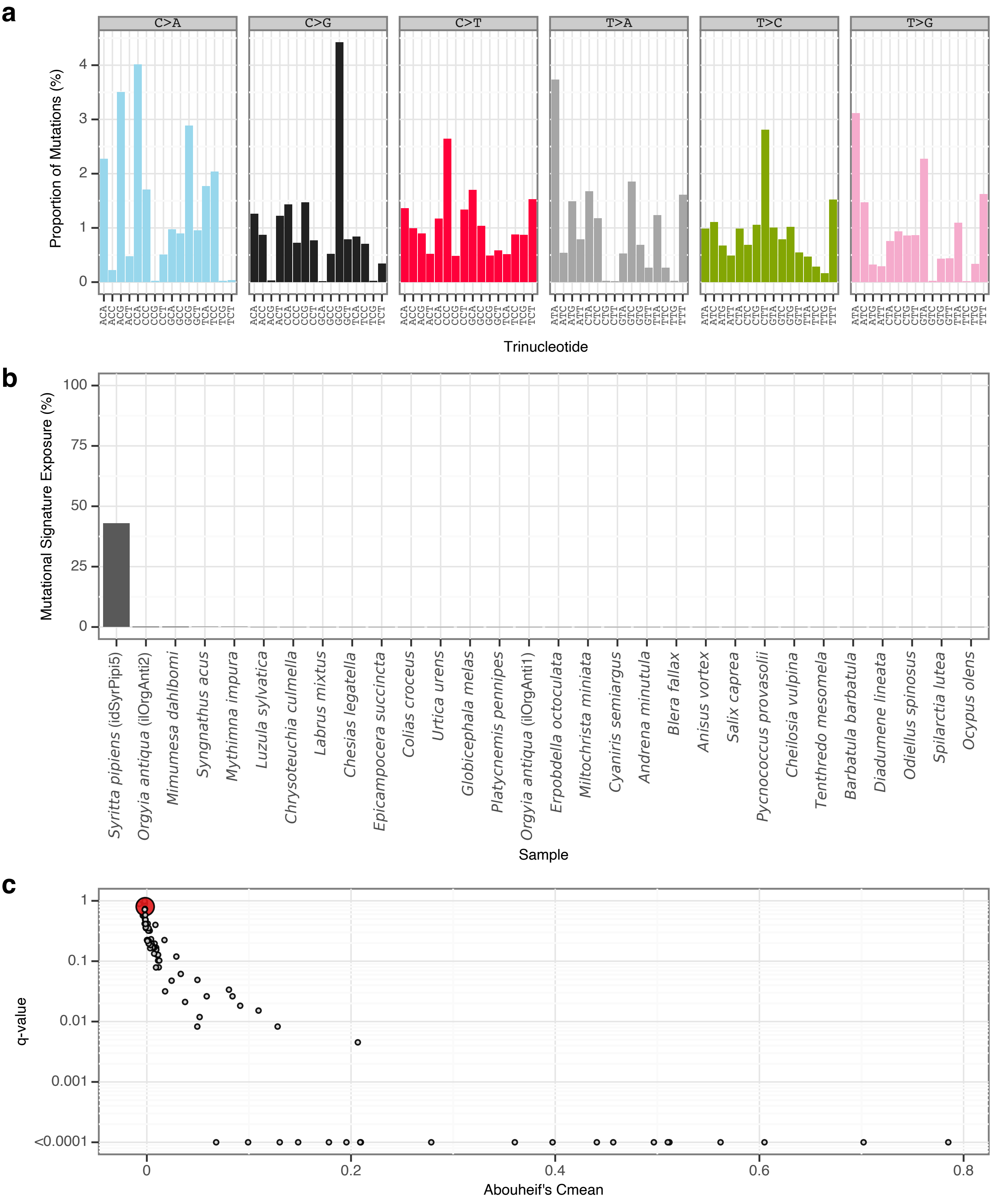


**Supplementary Fig. 83** | sToL80 mutational signature.

**a**, Mutational signature spectrum.

**b**, Bar plot displaying the 30 samples that exhibit the greatest exposure to the mutational signature.

**c**, Phylogenetic signal of the mutational signature.

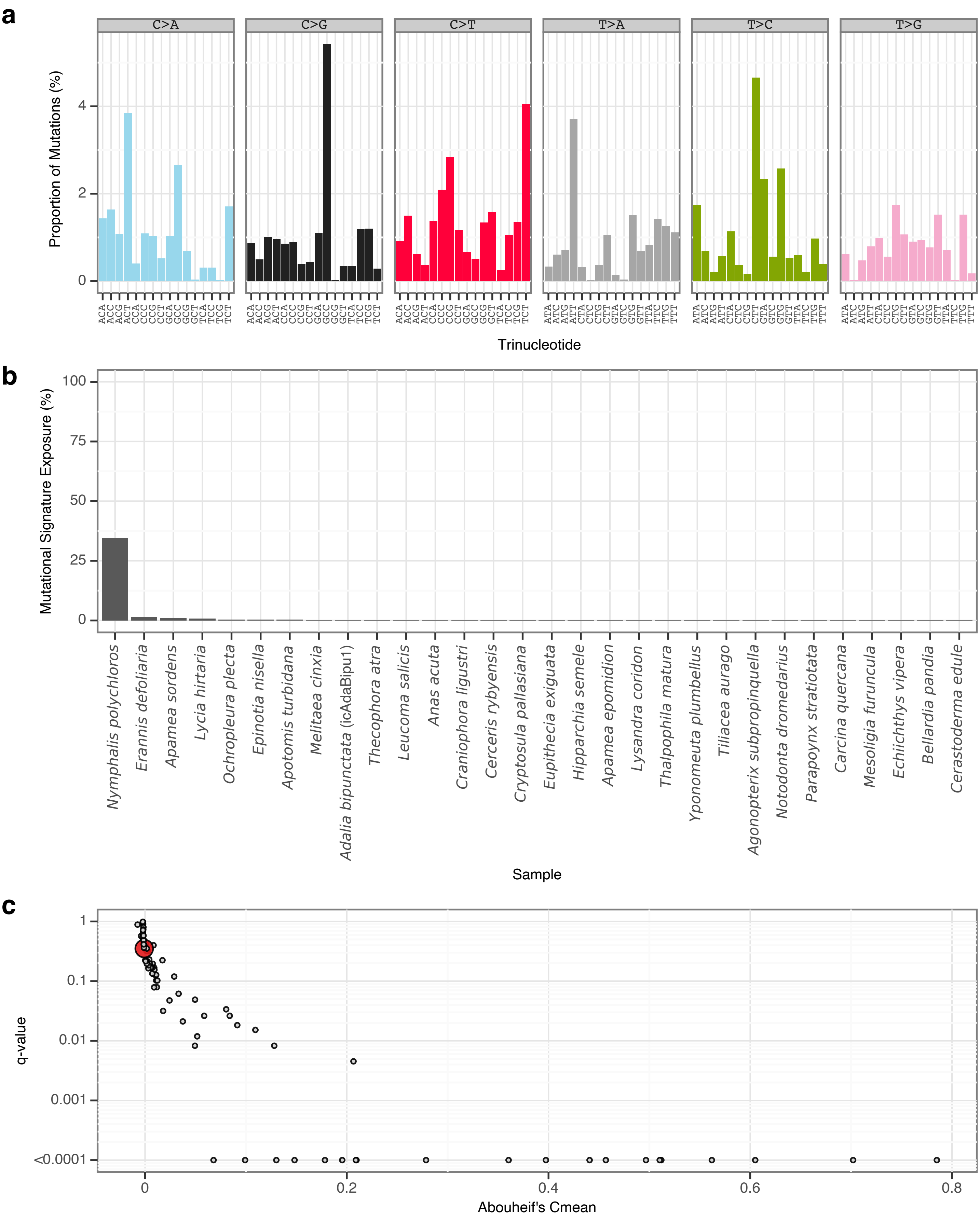


**Supplementary Fig. 84** | sToL81 mutational signature.

**a**, Mutational signature spectrum.

**b**, Bar plot displaying the 30 samples that exhibit the greatest exposure to the mutational signature.

**c**, Phylogenetic signal of the mutational signature.

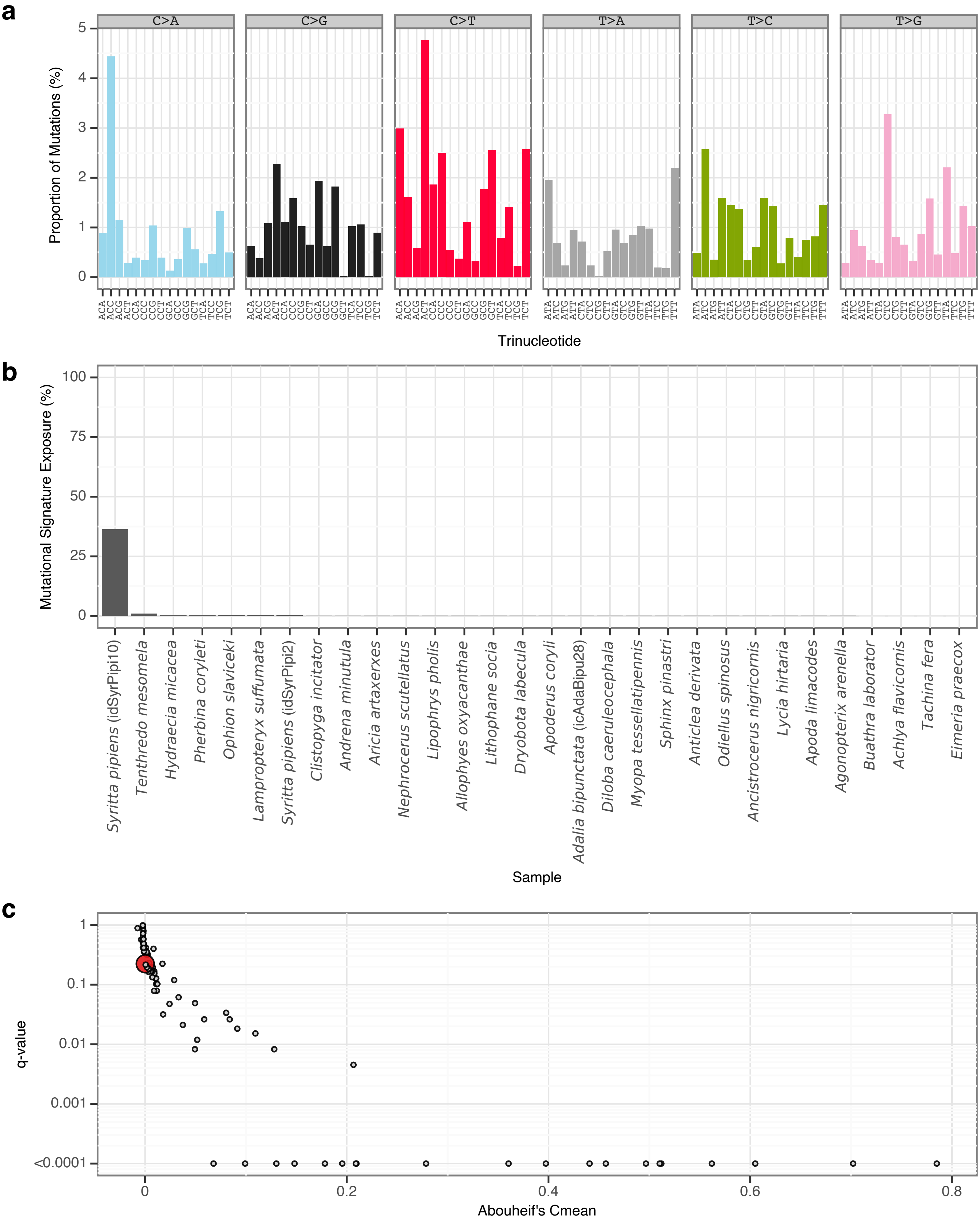


**Supplementary Fig. 85** | sToL82 mutational signature.

**a**, Mutational signature spectrum.

**b**, Bar plot displaying the 30 samples that exhibit the greatest exposure to the mutational signature.

**c**, Phylogenetic signal of the mutational signature.

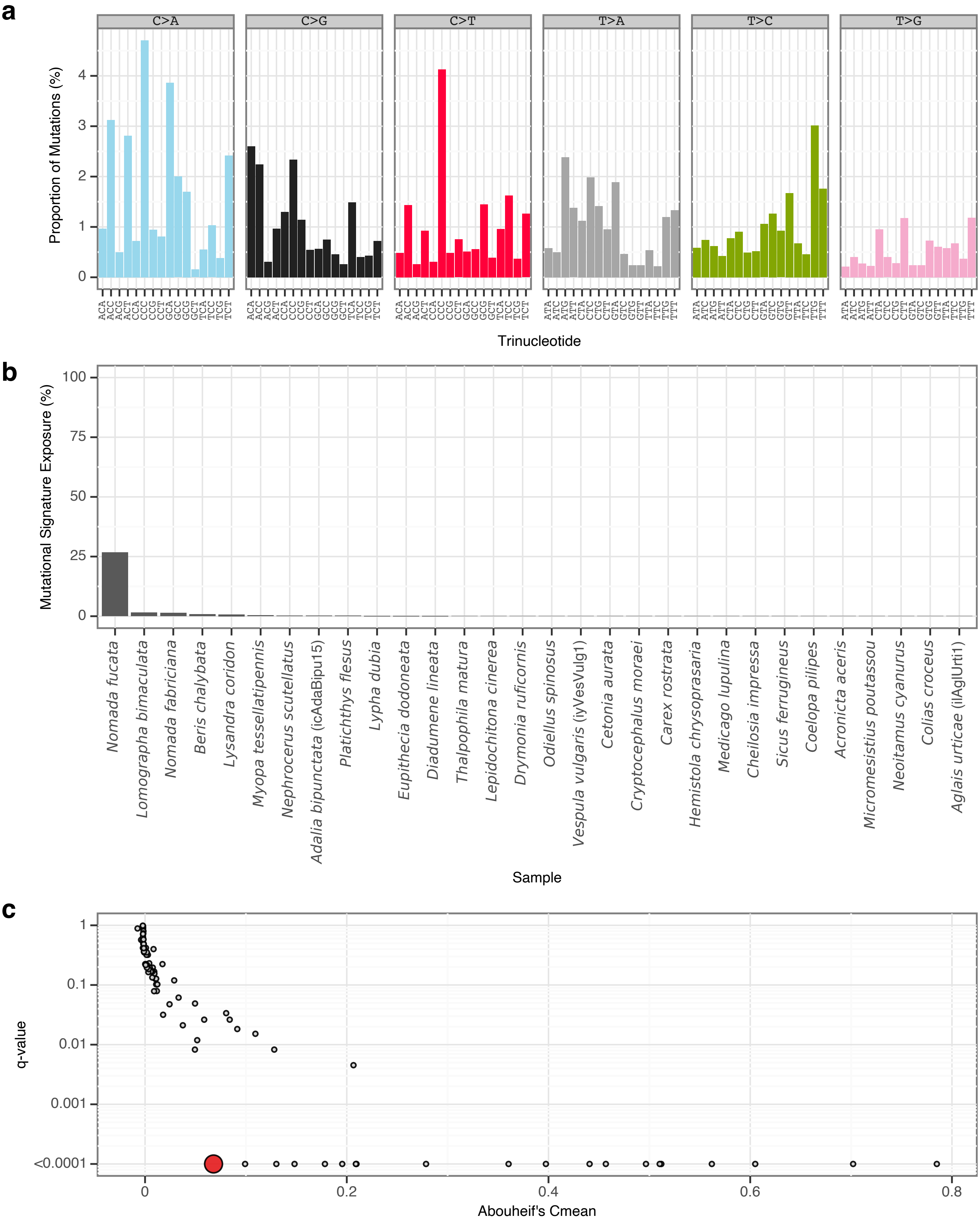


**Supplementary Fig. 86** | sToL83 mutational signature.

**a**, Mutational signature spectrum.

**b**, Bar plot displaying the 30 samples that exhibit the greatest exposure to the mutational signature.

**c**, Phylogenetic signal of the mutational signature.

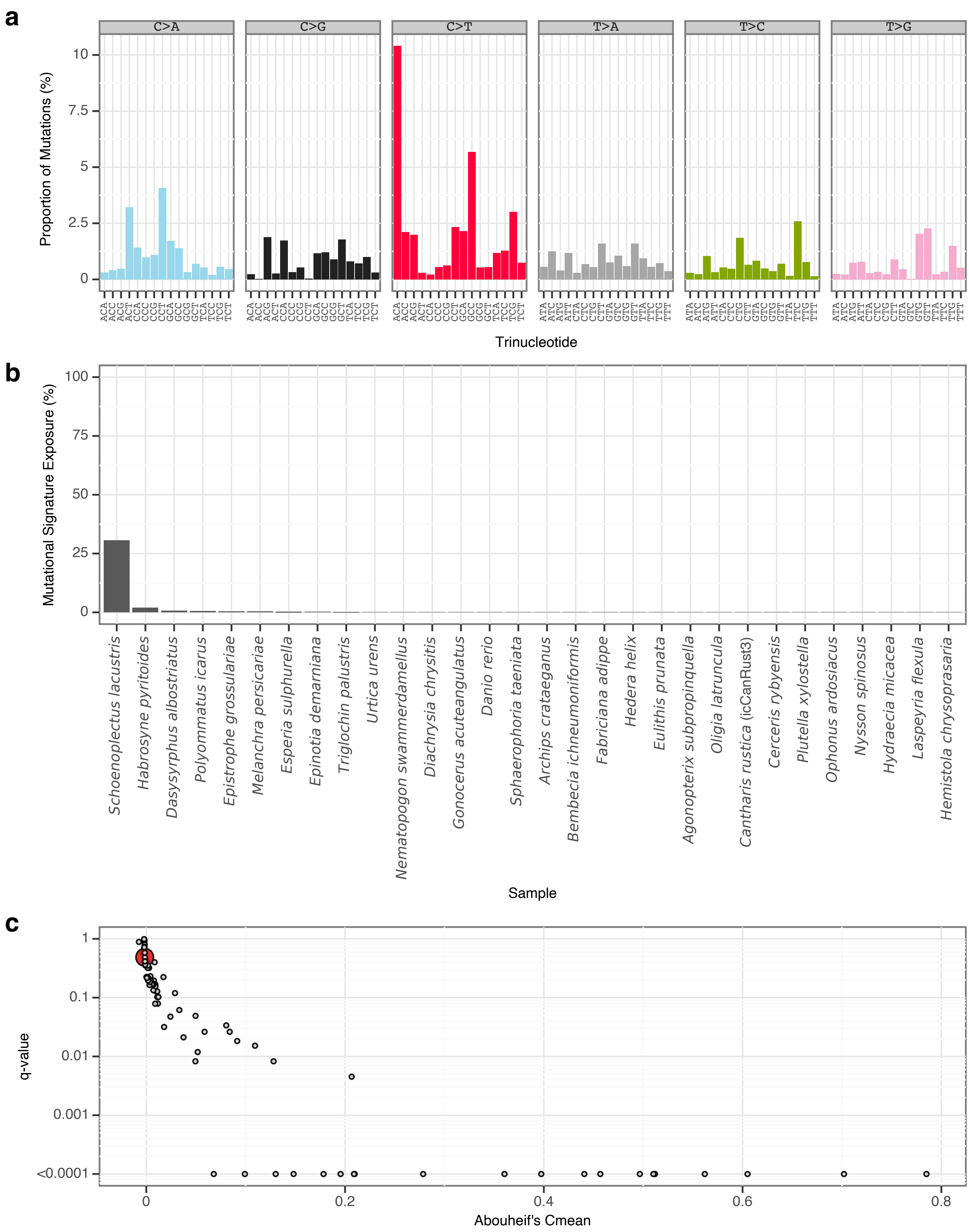


**Supplementary Fig. 87** l sToL84 mutational signature.

**a**, Mutational signature spectrum..

**b**, Bar plot displaying the 30 samples that exhibit the greatest exposure to the mutational signature.

**c**, Phylogenetic signal of the mutational signature.

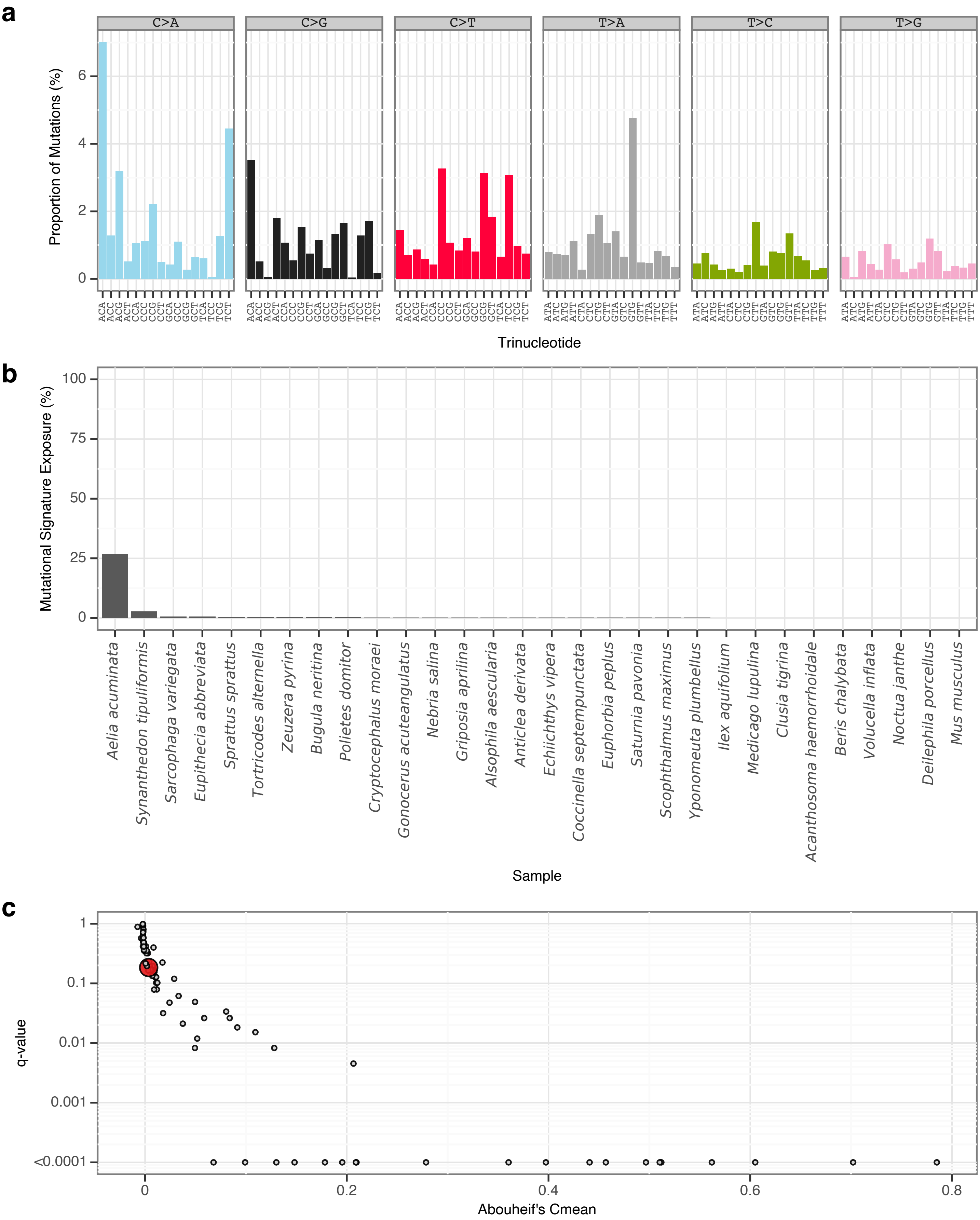


**Supplementary Fig. 88** | sToL85 mutational signature.
**a**, Mutational signature spectrum.
**b**, Bar plot displaying the 30 samples that exhibit the greatest exposure to the mutational signature.
**c**, Phylogenetic signal of the mutational signature.

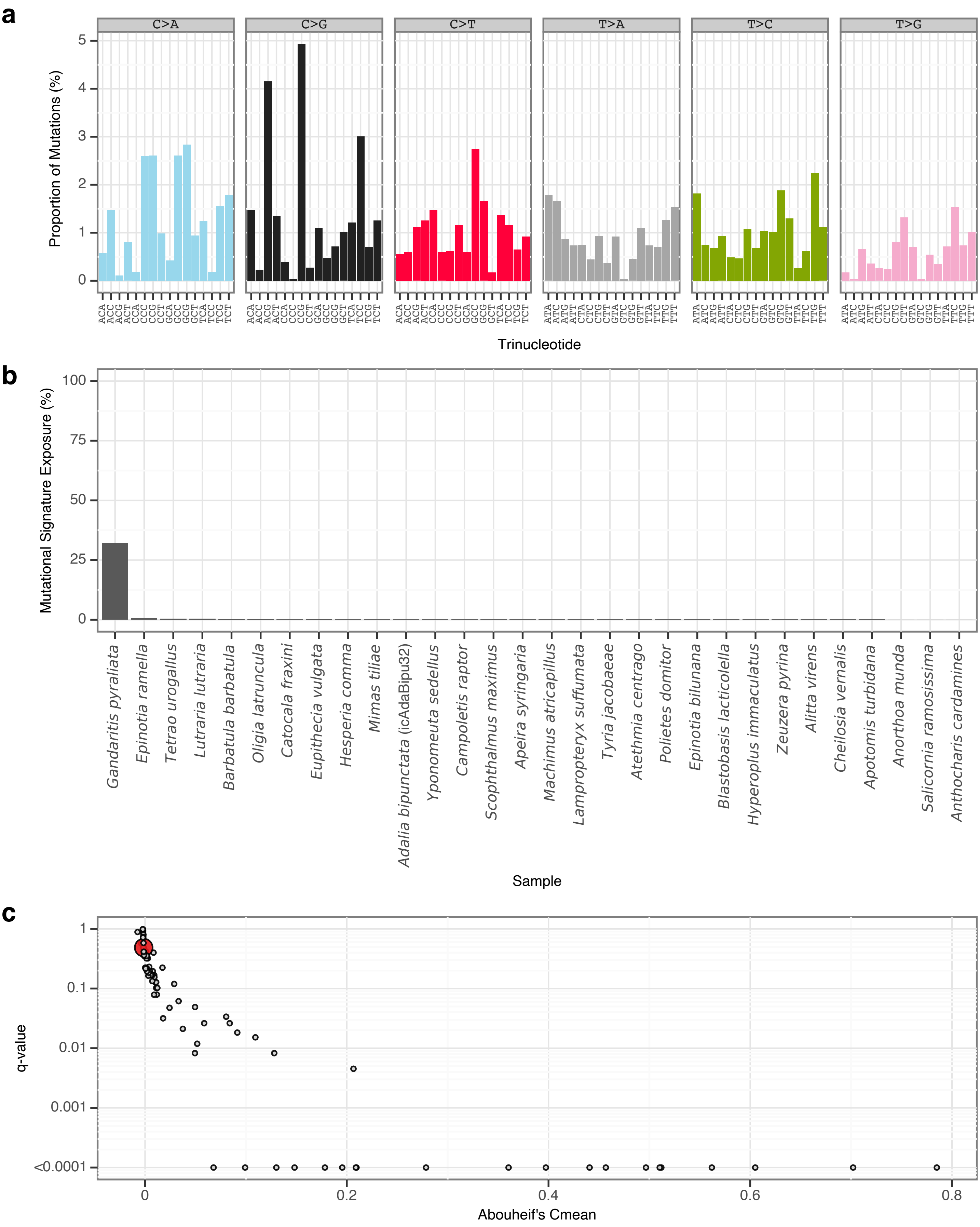


**Supplementary Fig. 89** | sToL86 mutational signature.

**a**, Mutational signature spectrum.

**b**, Bar plot displaying the 30 samples that exhibit the greatest exposure to the mutational signature.

**c**, Phylogenetic signal of the mutational signature.

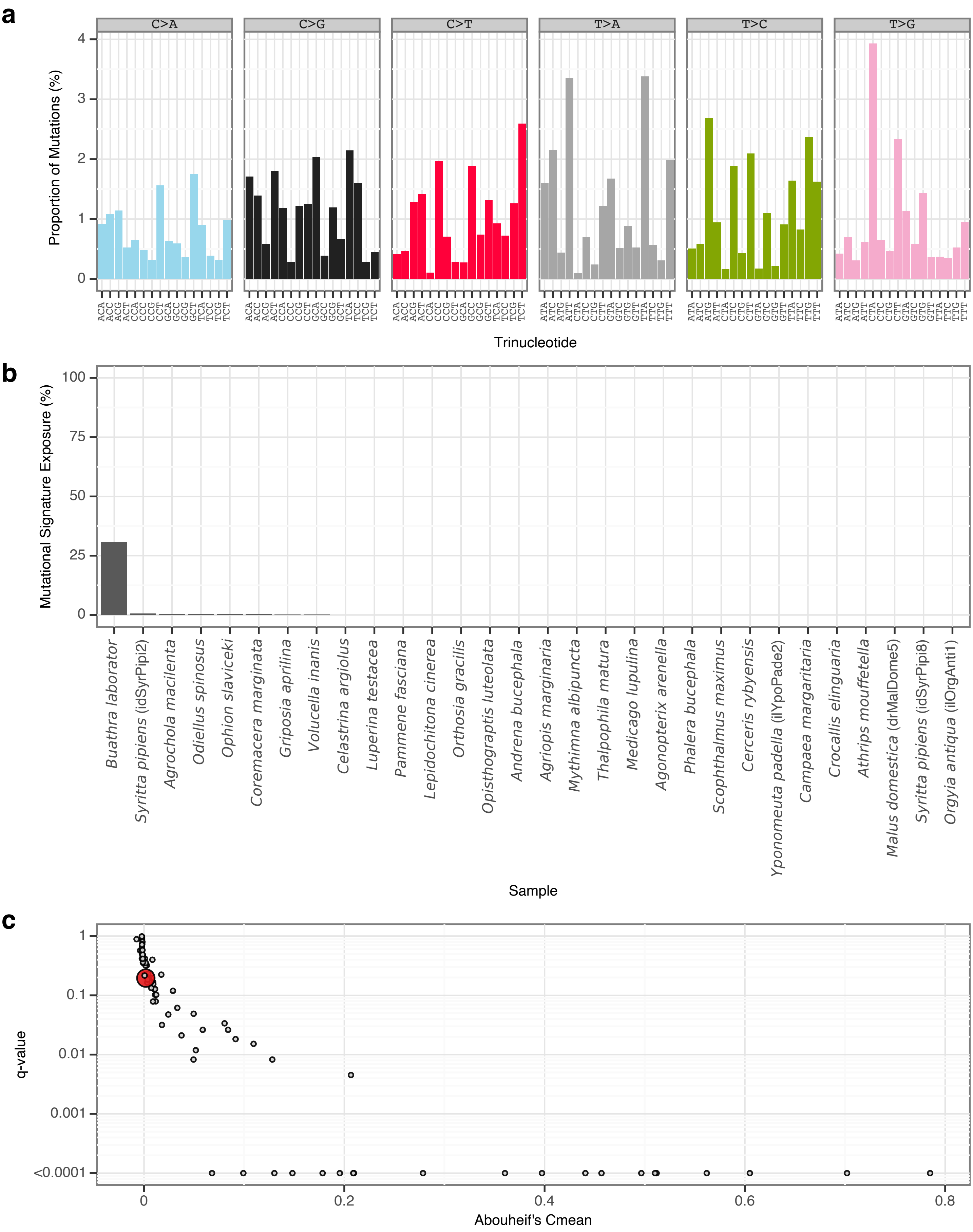


**Supplementary Fig. 90** | sToL87 mutational signature.
**a**, Mutational signature spectrum.
**b**, Bar plot displaying the 30 samples that exhibit the greatest exposure to the mutational signature.
**c**, Phylogenetic signal of the mutational signature.

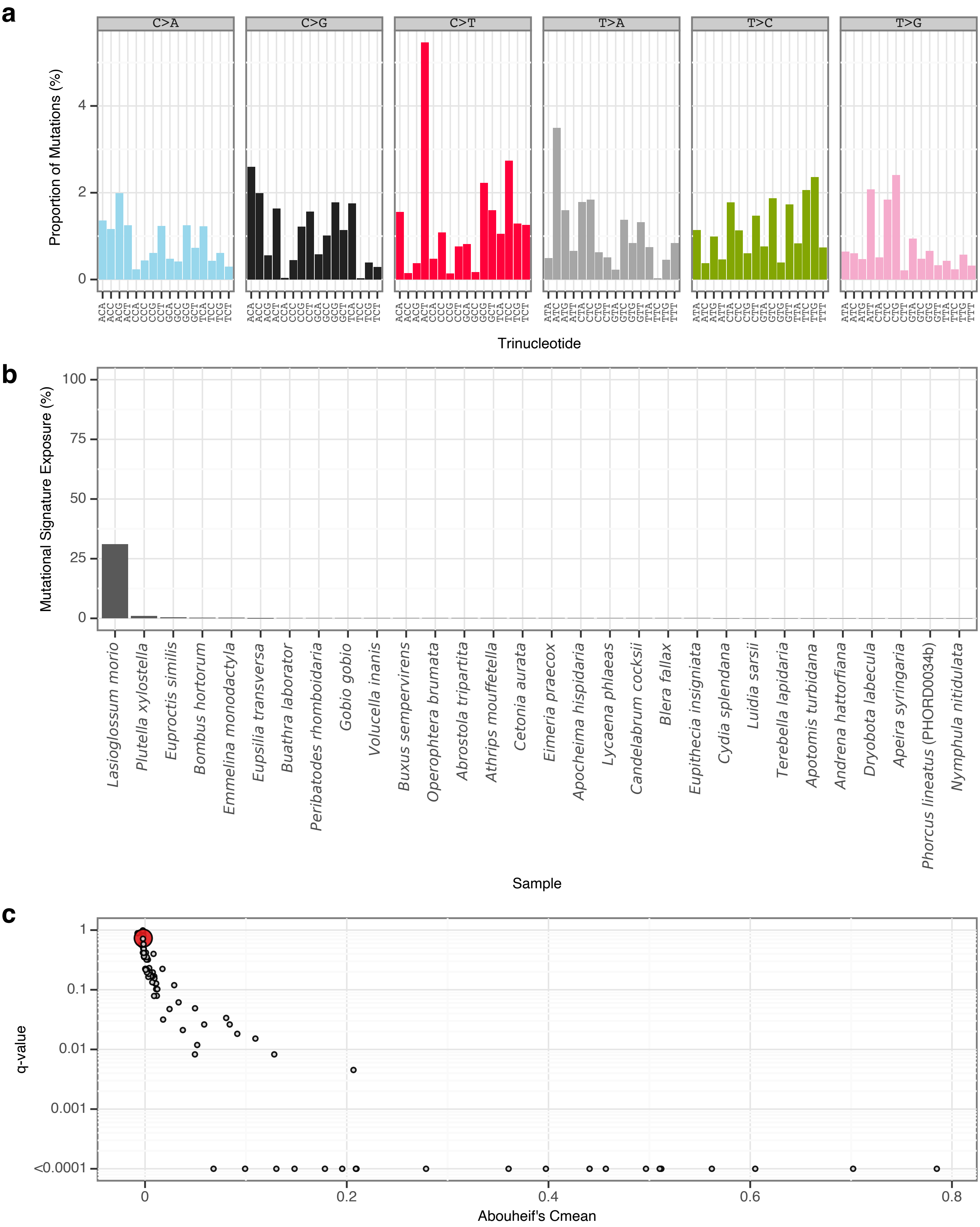


**Supplementary Fig. 91** | sToL88 mutational signature.

**a**, Mutational signature spectrum.

**b**, Bar plot displaying the 30 samples that exhibit the greatest exposure to the mutational signature.

**c**, Phylogenetic signal of the mutational signature.

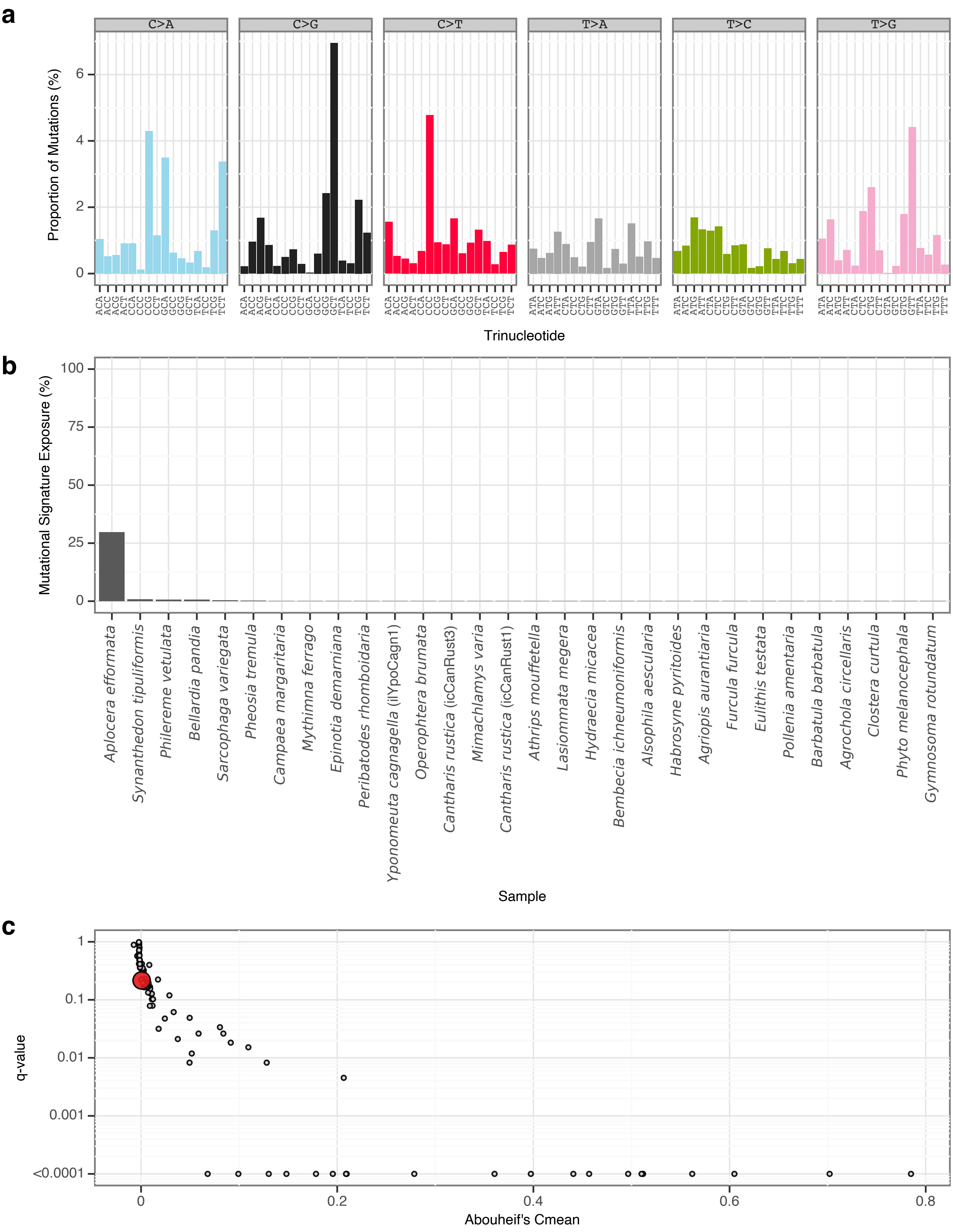


**Supplementary Fig. 92** | sToL89 mutational signature.
**a**, Mutational signature spectrum.
**b**, Bar plot displaying the 30 samples that exhibit the greatest exposure to the mutational signature.
**c**, Phylogenetic signal of the mutational signature.

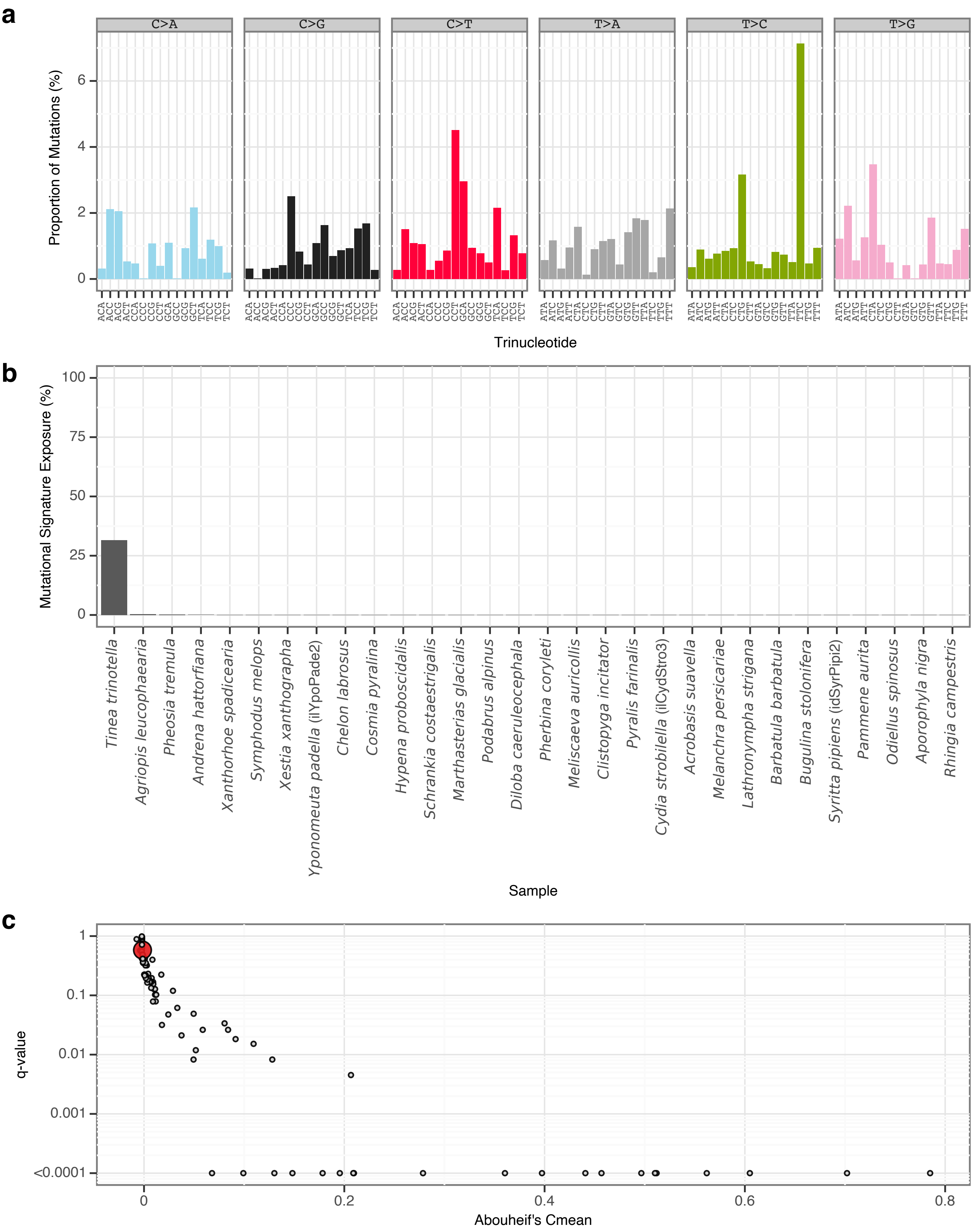


**Supplementary Fig. 93** | sToL90 mutational signature.

**a**, Mutational signature spectrum.

**b**, Bar plot displaying the 30 samples that exhibit the greatest exposure to the mutational signature.

**c**, Phylogenetic signal of the mutational signature.

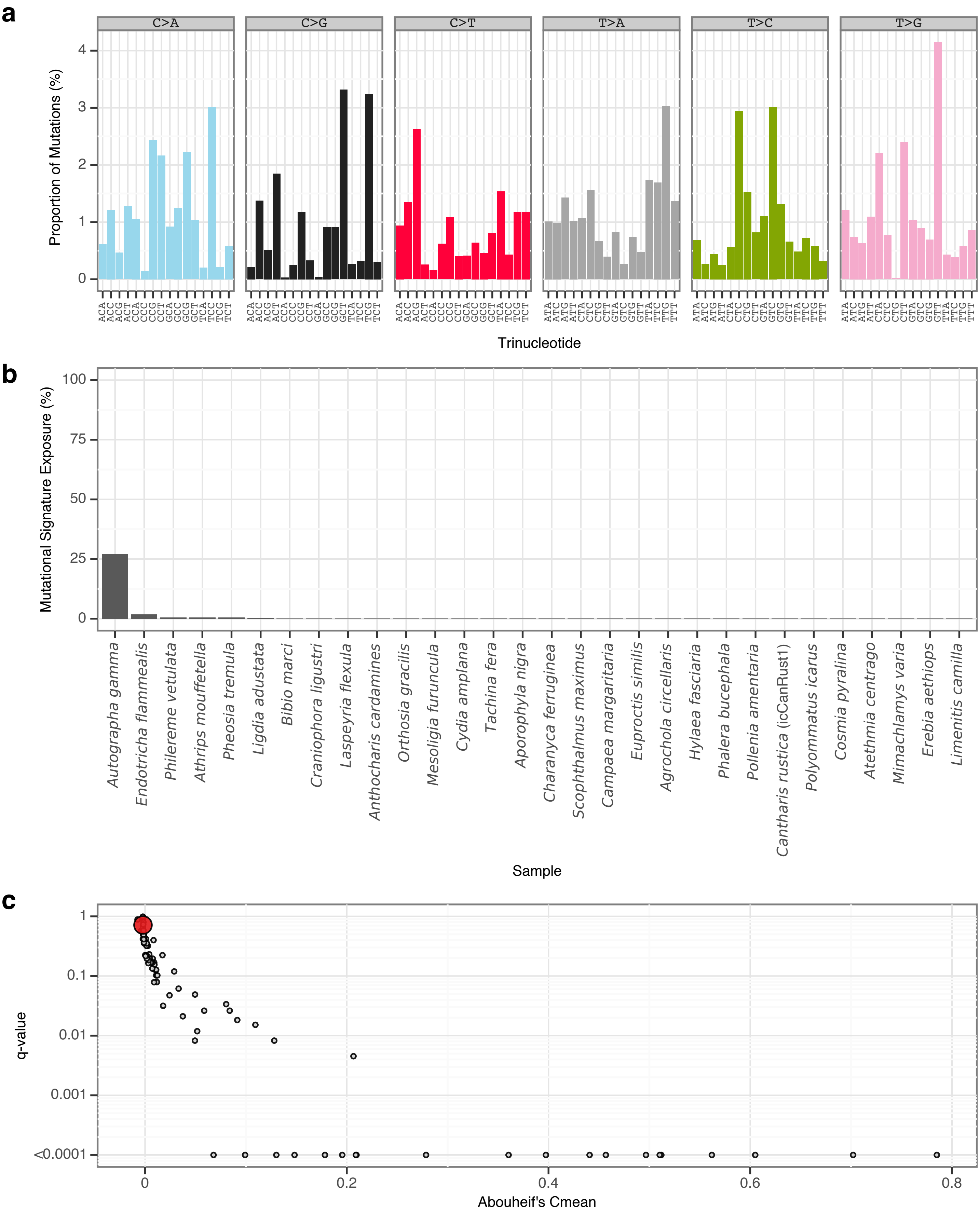


**Supplementary Fig. 94** | sToL91 mutational signature.

**a**, Mutational signature spectrum.

**b**, Bar plot displaying the 30 samples that exhibit the greatest exposure to the mutational signature.

**c**, Phylogenetic signal of the mutational signature.

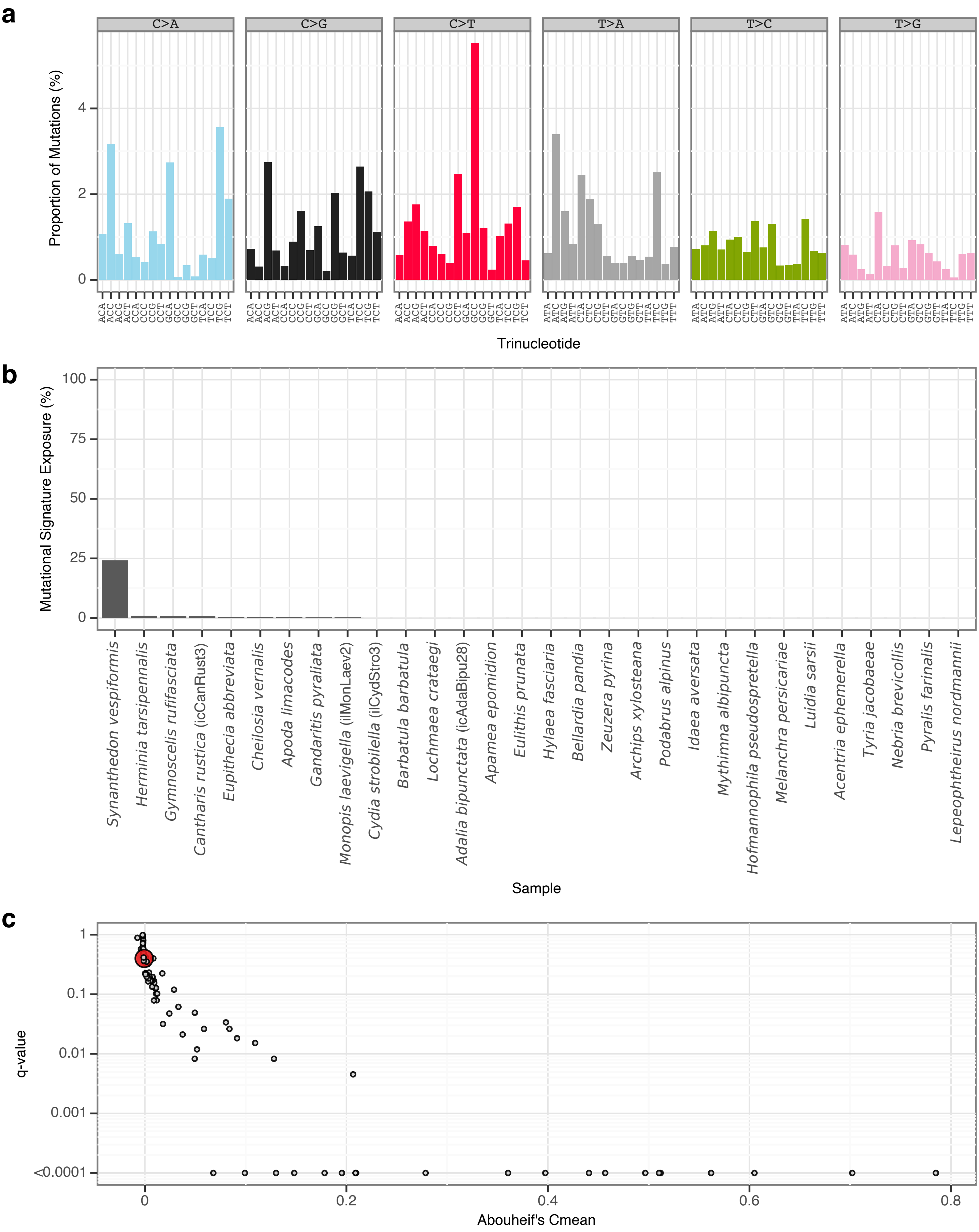


**Supplementary Fig. 95** | sToL92 mutational signature.

**a**, Mutational signature spectrum.

**b**, Bar plot displaying the 30 samples that exhibit the greatest exposure to the mutational signature.

**c**, Phylogenetic signal of the mutational signature.

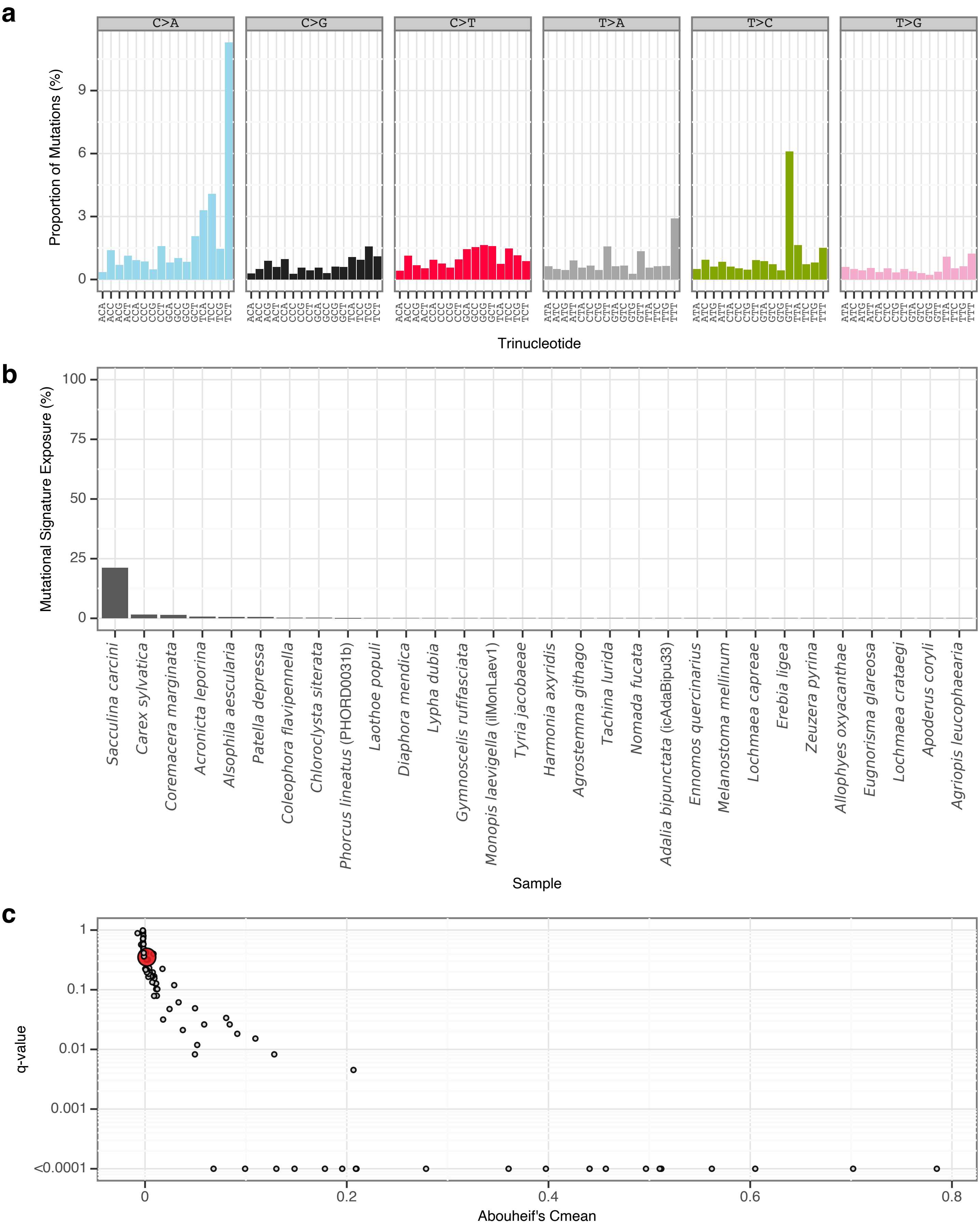


**Supplementary Fig. 96** | sToL93 mutational signature.
**a**, Mutational signature spectrum.
**b**, Bar plot displaying the 30 samples that exhibit the greatest exposure to the mutational signature.
**c**, Phylogenetic signal of the mutational signature.

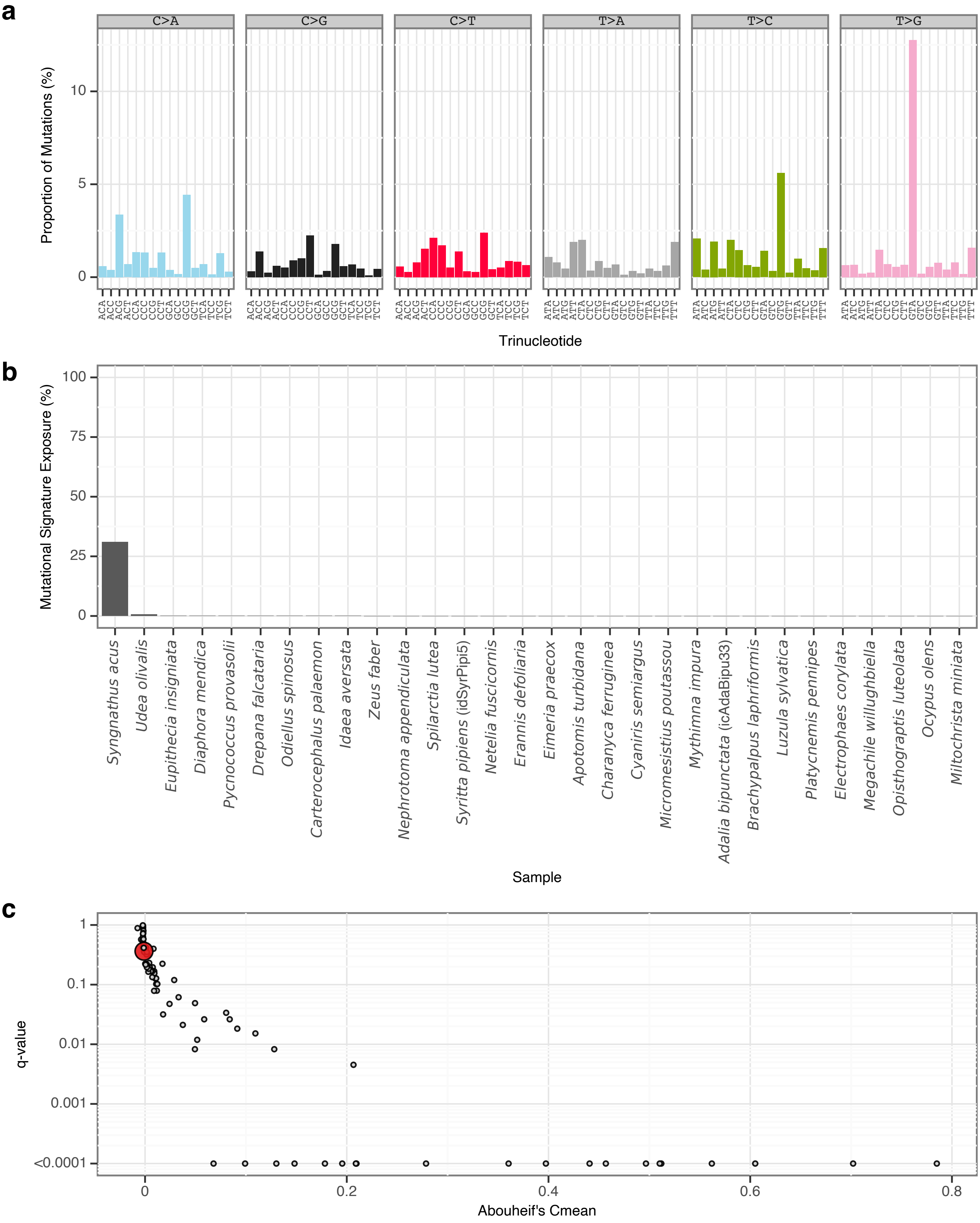


**Supplementary Fig. 97** | sToL94 mutational signature.

**a**, Mutational signature spectrum.

**b**, Bar plot displaying the 30 samples that exhibit the greatest exposure to the mutational signature.

**c**, Phylogenetic signal of the mutational signature.

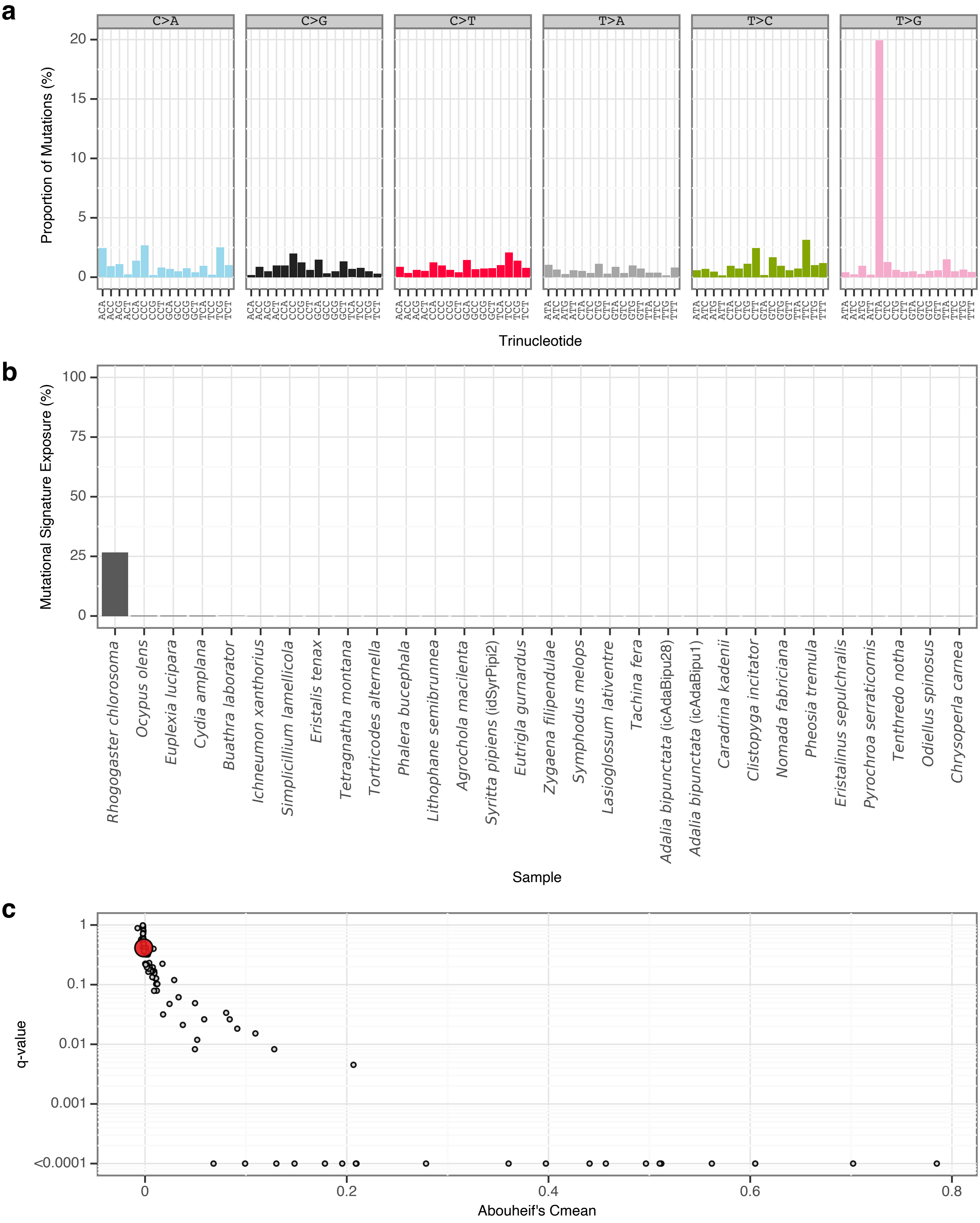


**Supplementary Fig. 98** | sToL95 mutational signature.

**a**, Mutational signature spectrum.

**b**, Bar plot displaying the 30 samples that exhibit the greatest exposure to the mutational signature.

**c**, Phylogenetic signal of the mutational signature.

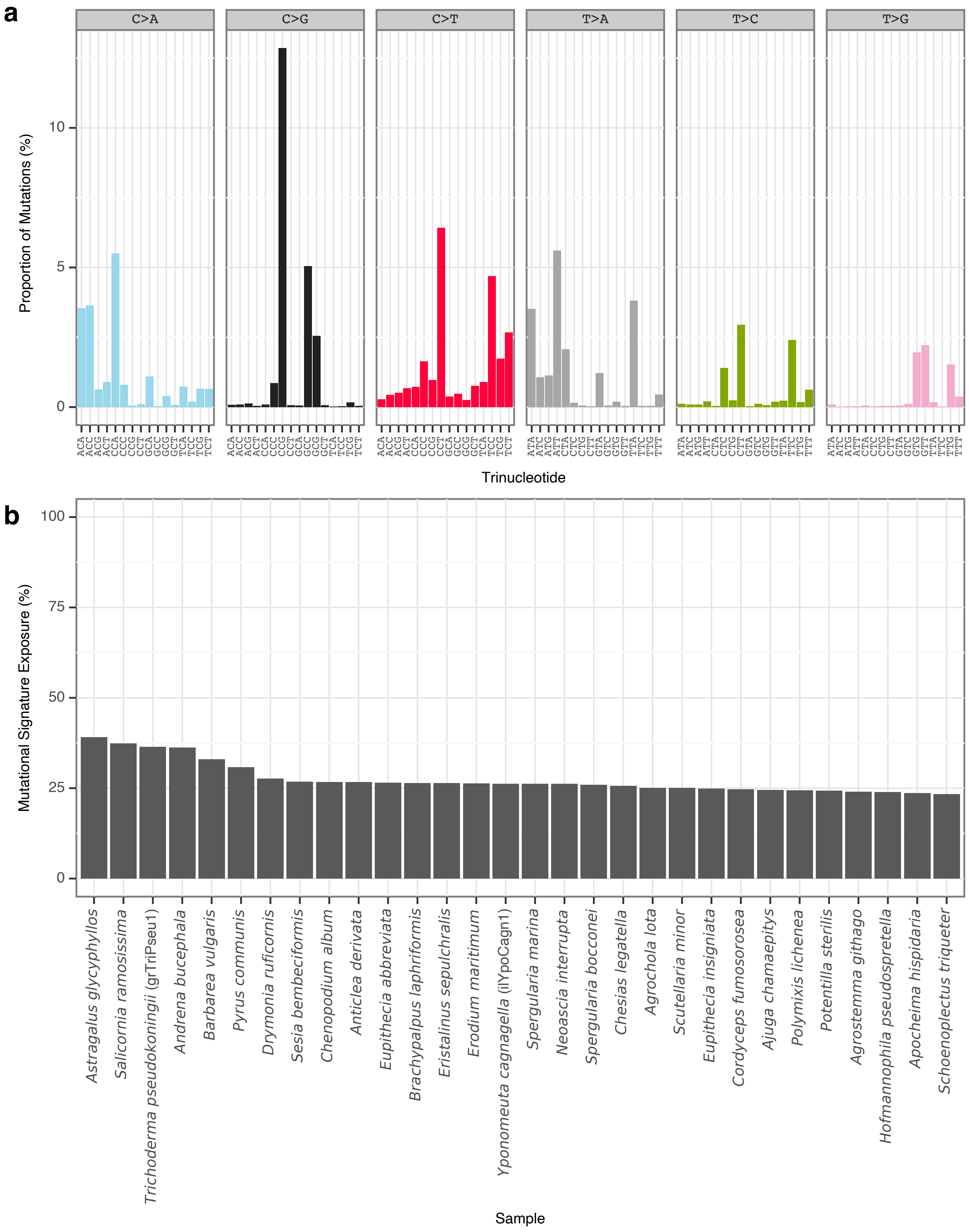


**Supplementary Fig. 99** | rToL1 mutational signature.
**a**, Mutational signature spectrum.
**b**, Bar plot displaying the 30 samples that exhibit the greatest exposure to the mutational signature.

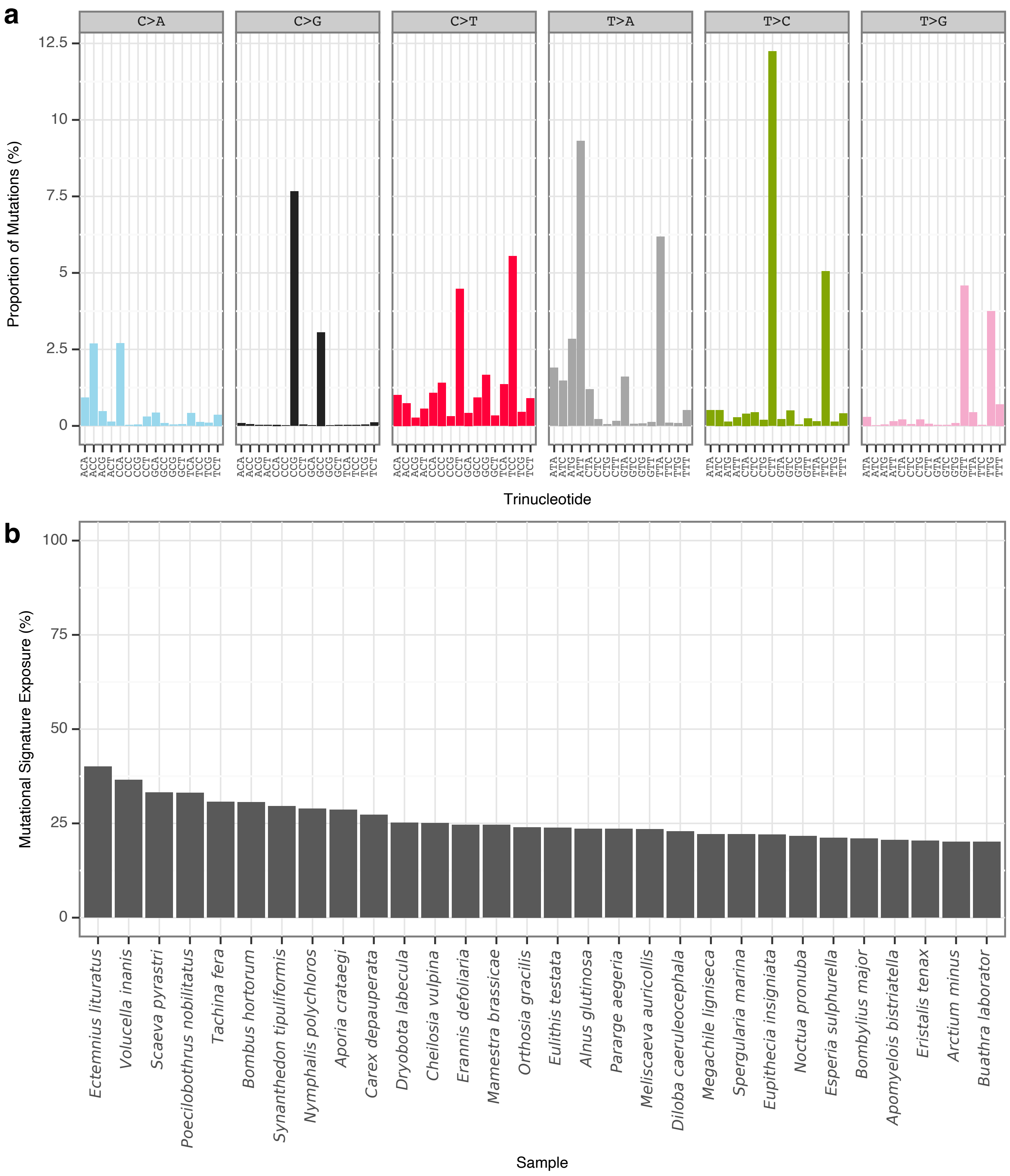


**Supplementary Fig. 100** | rToL2 mutational signature.
**a**, Mutational signature spectrum.
**b**, Bar plot displaying the 30 samples that exhibit the greatest exposure to the mutational signature.

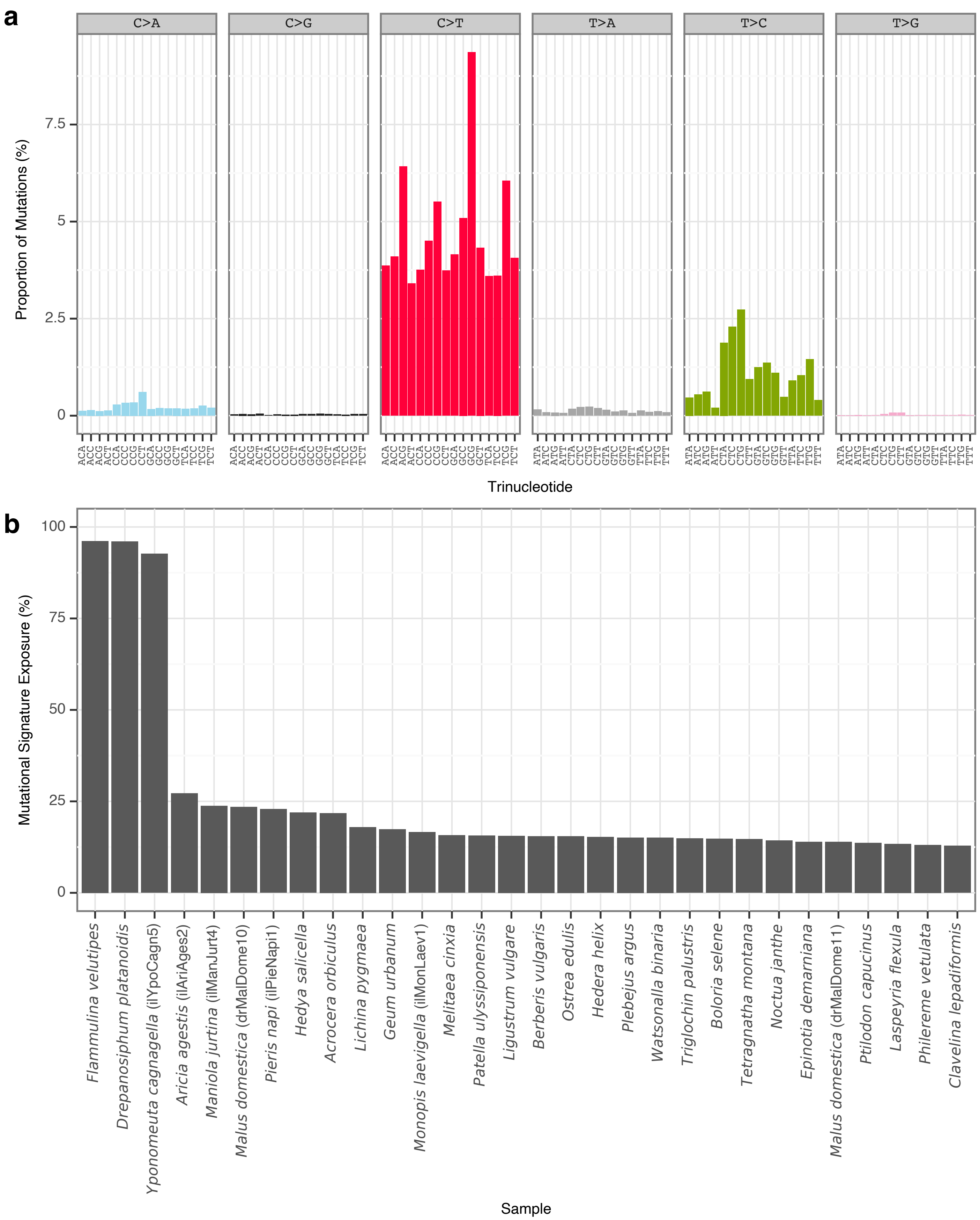


**Supplementary Fig. 101** | rToL3 mutational signature.
**a**, Mutational signature spectrum.
**b**, Bar plot displaying the 30 samples that exhibit the greatest exposure to the mutational signature.

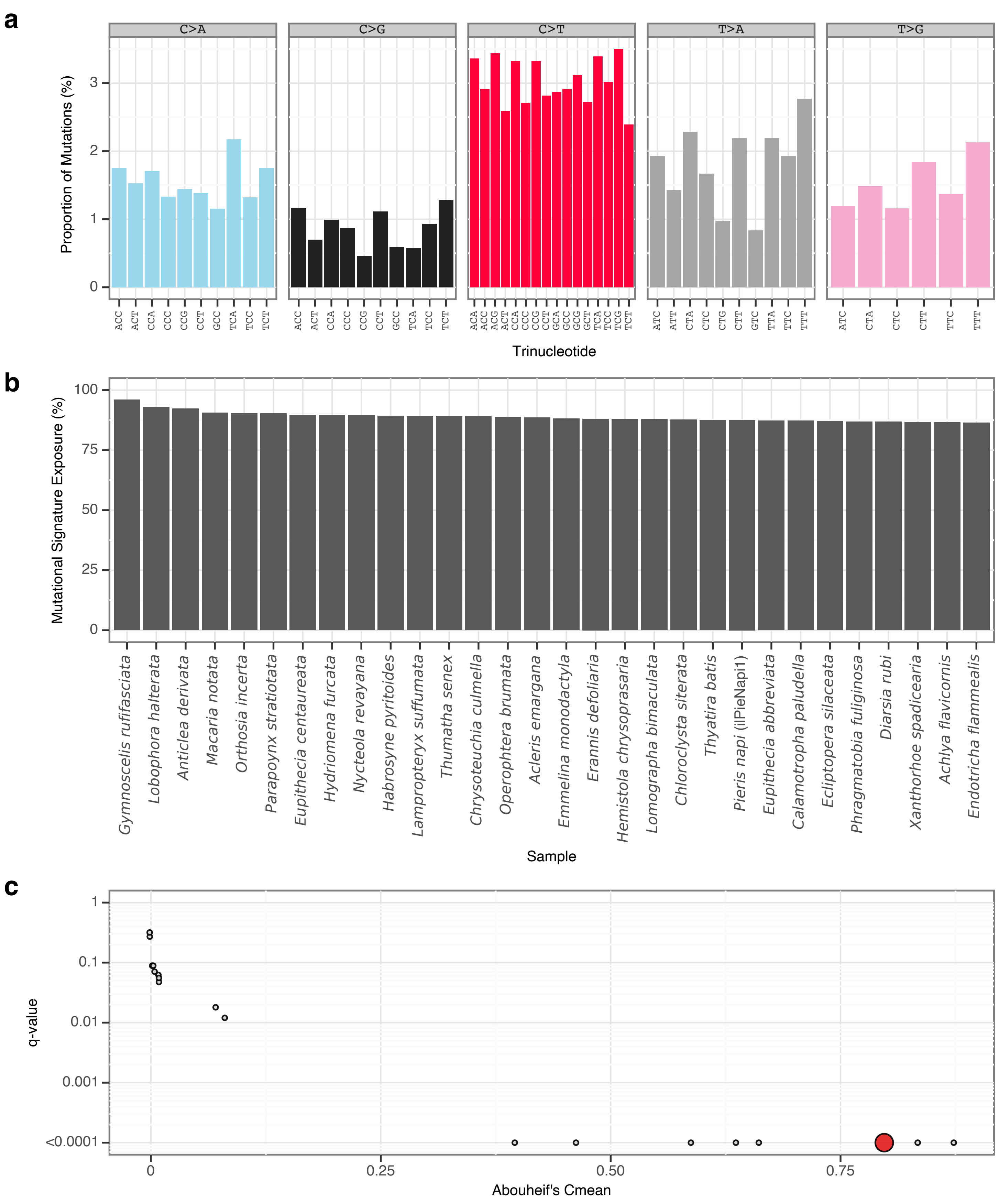


**Supplementary Fig. 102** | gToL1 mutational signature.

**a**, Mutational signature spectrum.

**b**, Bar plot displaying the 30 samples that exhibit the greatest exposure to the mutational signature.

**c**, Phylogenetic signal of the mutational signature.

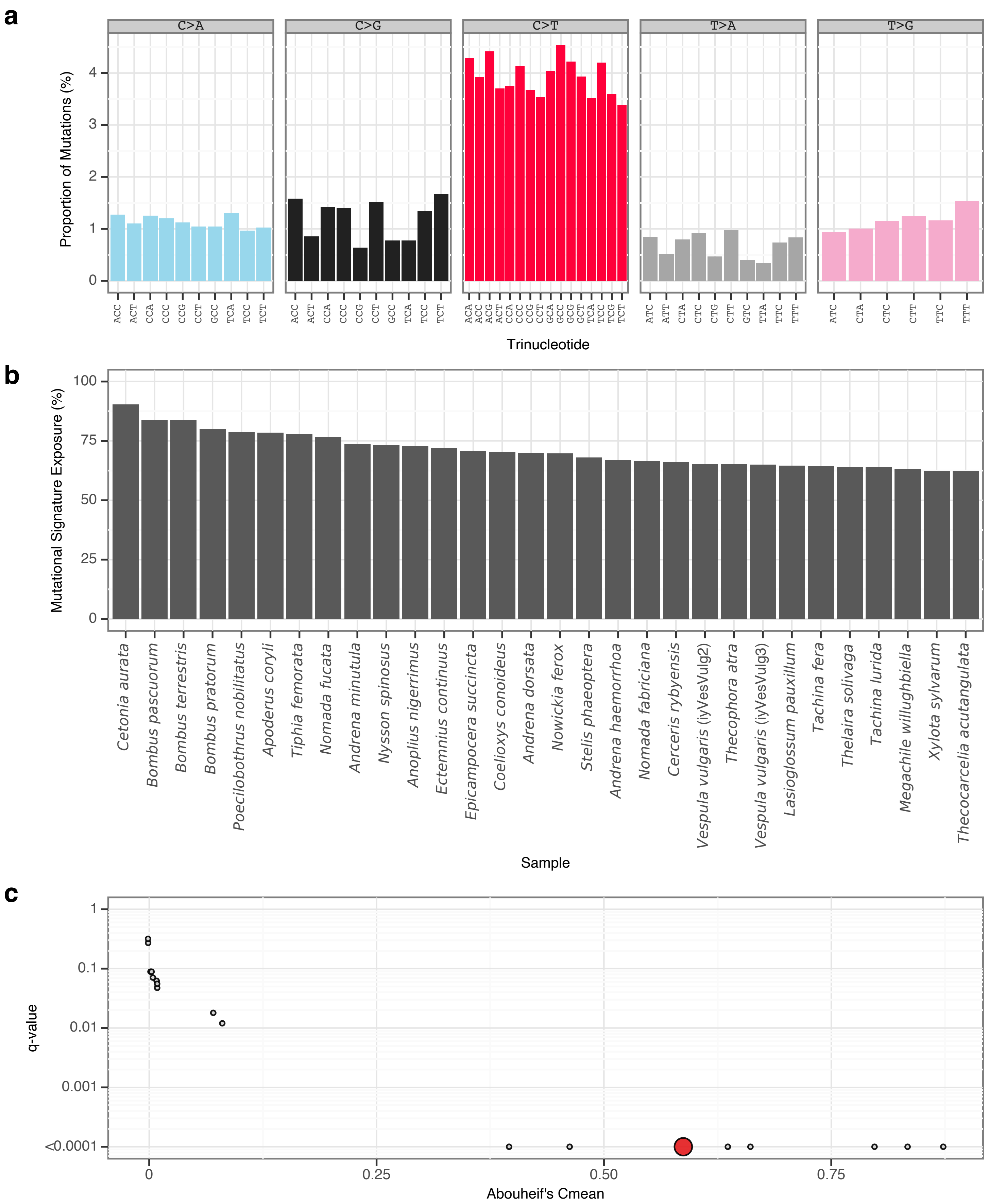


**Supplementary Fig. 103** | gToL2 mutational signature.

**a**, Mutational signature spectrum.

**b**, Bar plot displaying the 30 samples that exhibit the greatest exposure to the mutational signature.

**c**, Phylogenetic signal of the mutational signature.

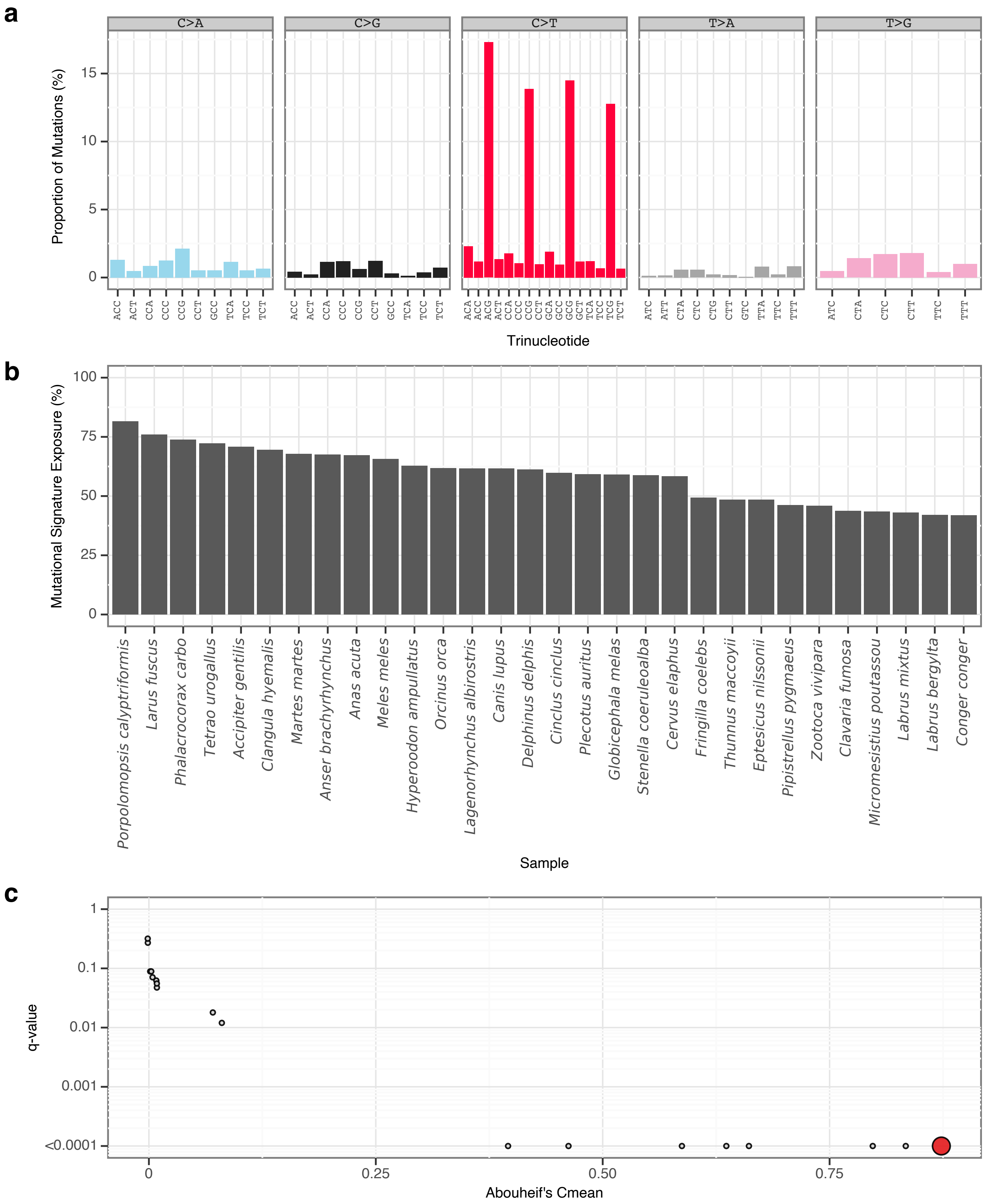


**Supplementary Fig. 104** | gToL3 mutational signature.
**a**, Mutational signature spectrum.
**b**, Bar plot displaying the 30 samples that exhibit the greatest exposure to the mutational signature.
**c**, Phylogenetic signal of the mutational signature.

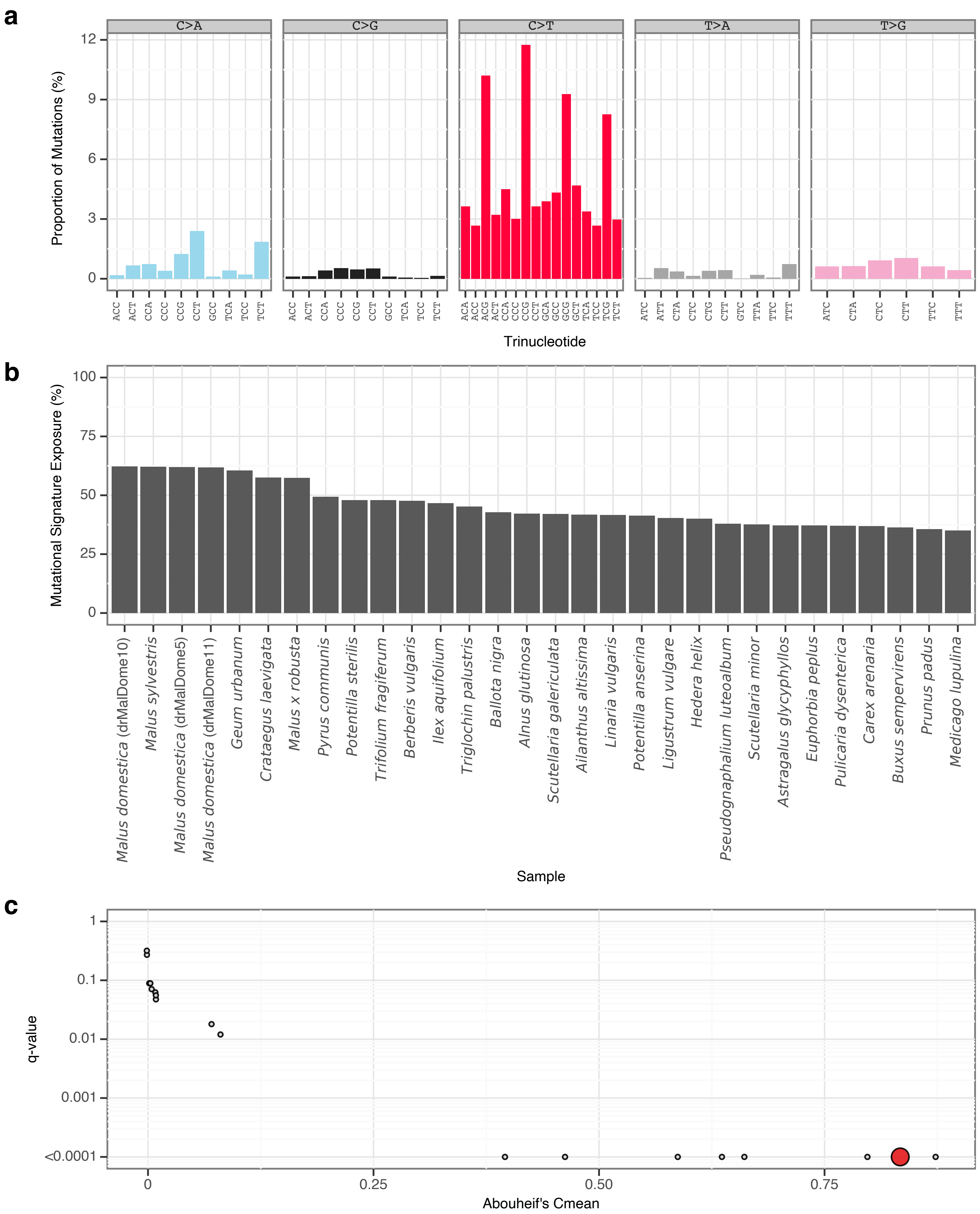


**Supplementary Fig. 105** | gToL4 mutational signature.

**a**, Mutational signature spectrum.

**b**, Bar plot displaying the 30 samples that exhibit the greatest exposure to the mutational signature.

**c**, Phylogenetic signal of the mutational signature.

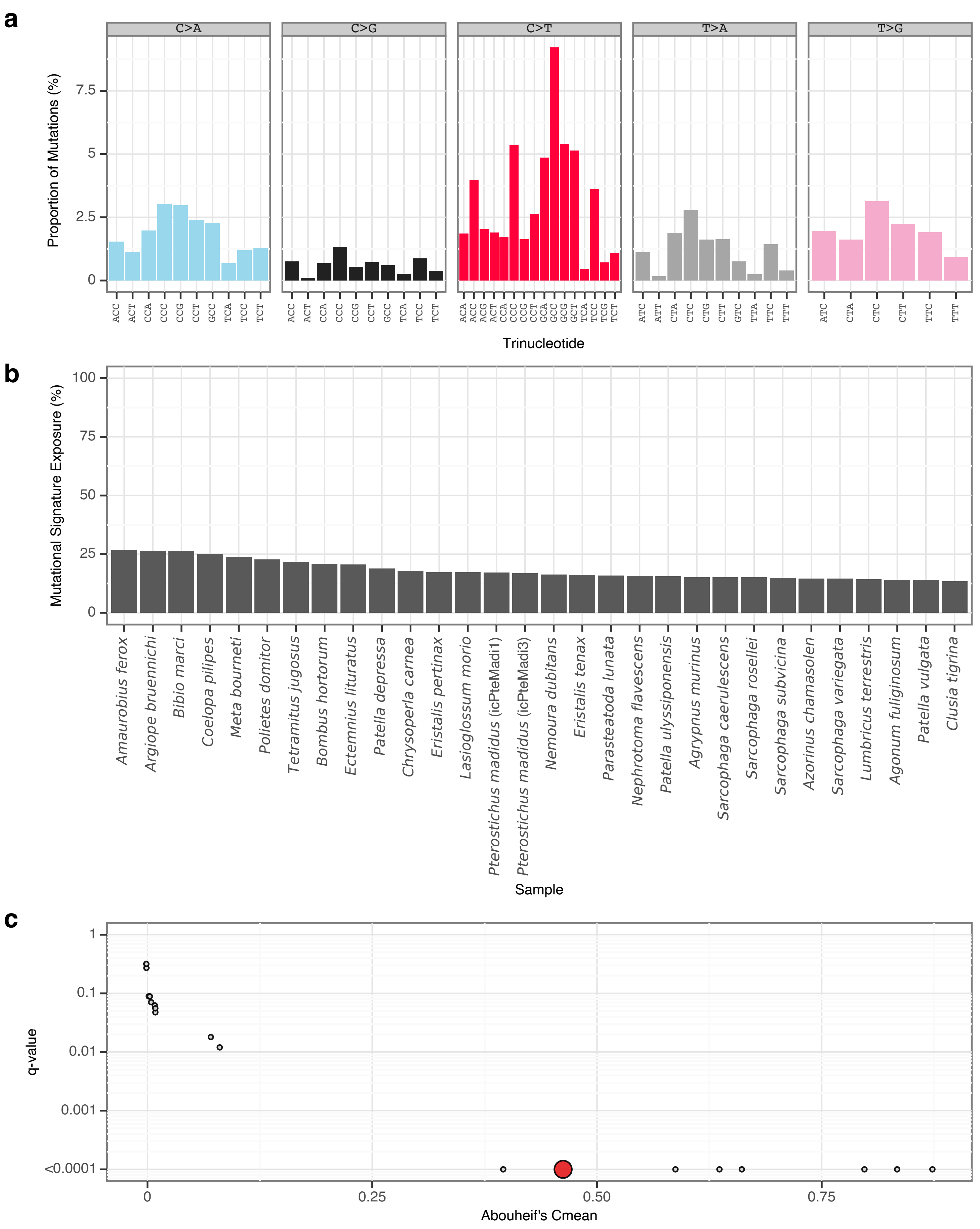


**Supplementary Fig. 106** | gToL5 mutational signature.

**a**, Mutational signature spectrum.

**b**, Bar plot displaying the 30 samples that exhibit the greatest exposure to the mutational signature.

**c**, Phylogenetic signal of the mutational signature.

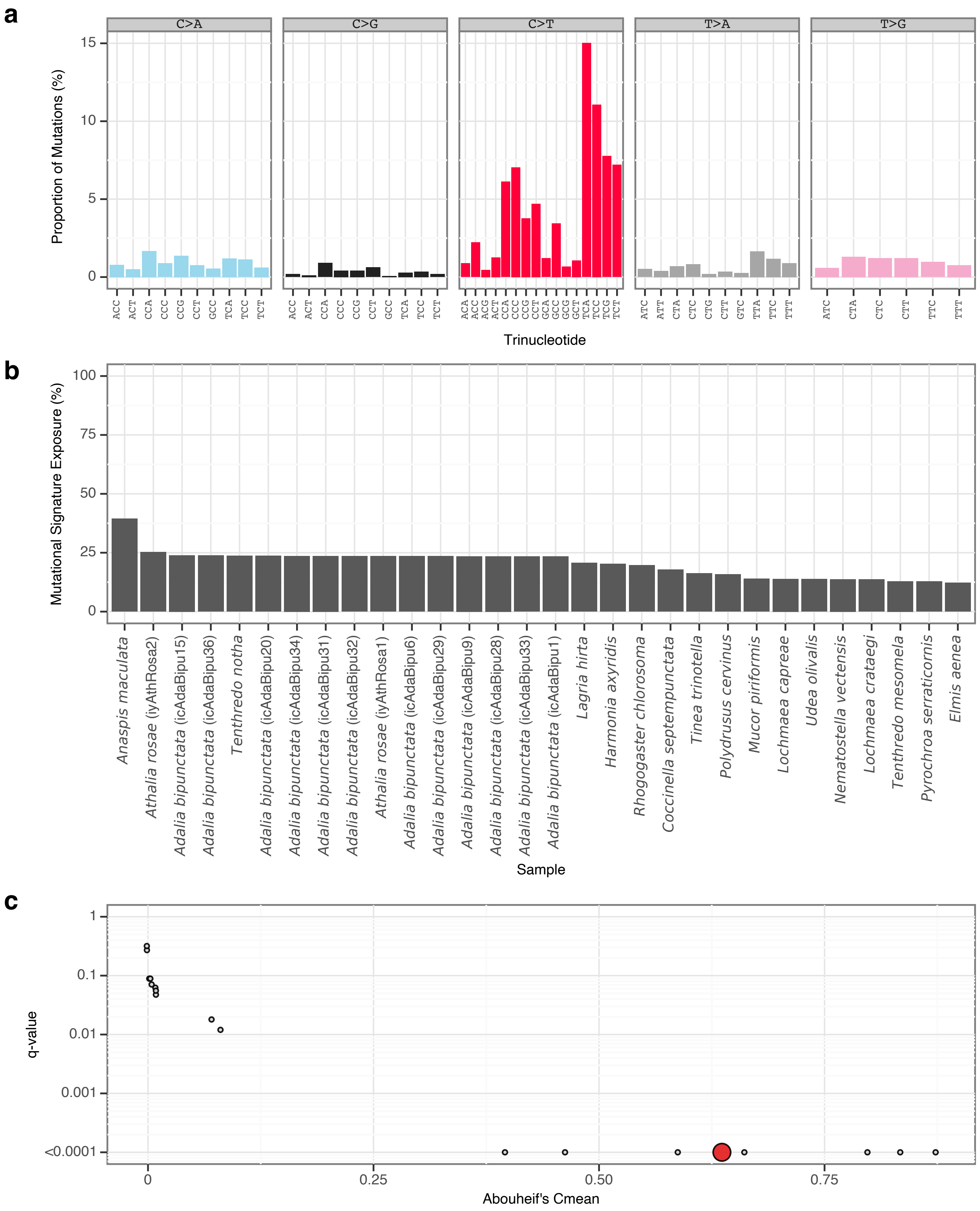


**Supplementary Fig. 107** | gToL6 mutational signature.
**a**, Mutational signature spectrum.
**b**, Bar plot displaying the 30 samples that exhibit the greatest exposure to the mutational signature.
**c**, Phylogenetic signal of the mutational signature.

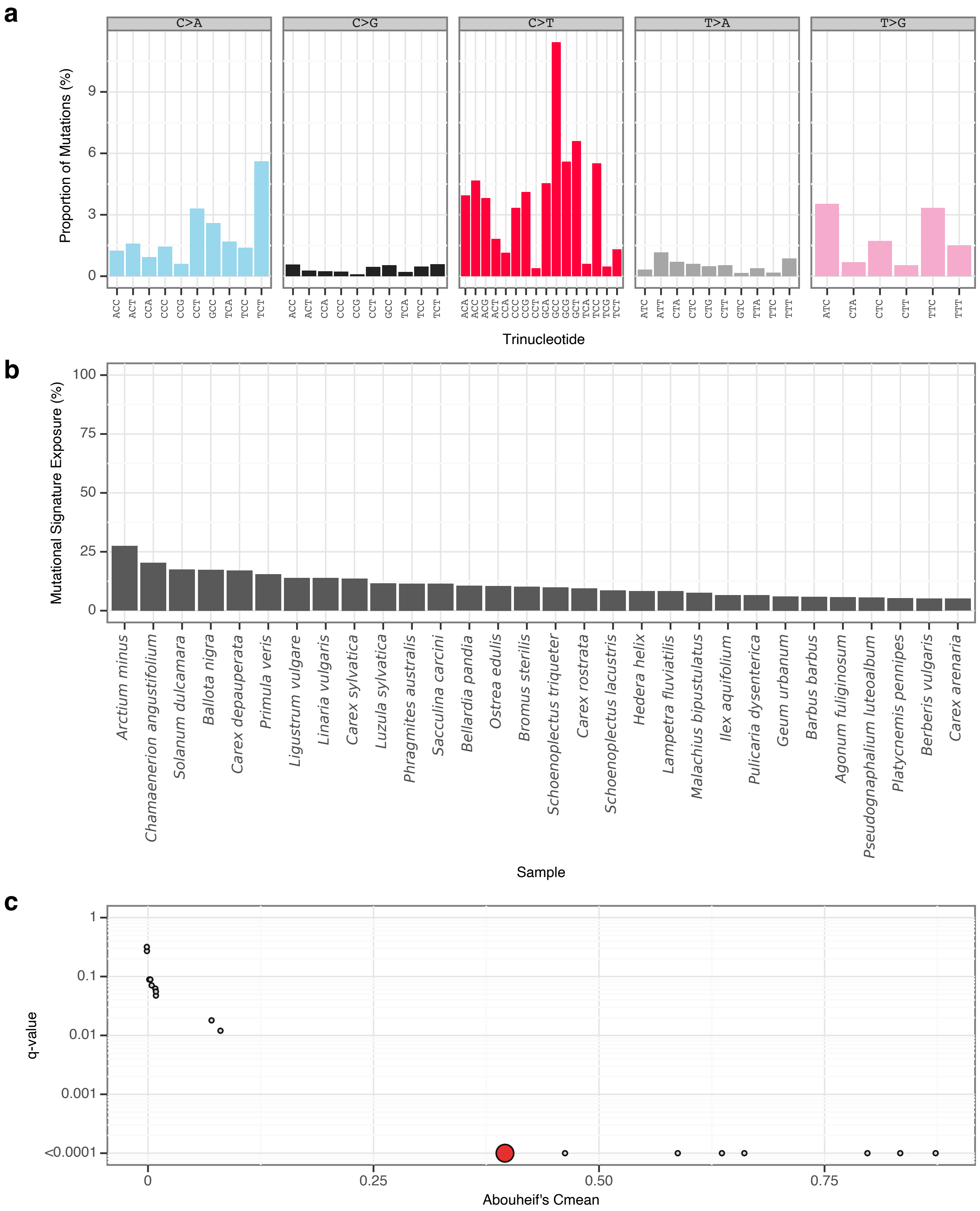


**Supplementary Fig. 108** | gToL7 mutational signature.
**a**, Mutational signature spectrum.
**b**, Bar plot displaying the 30 samples that exhibit the greatest exposure to the mutational signature.
**c**, Phylogenetic signal of the mutational signature.

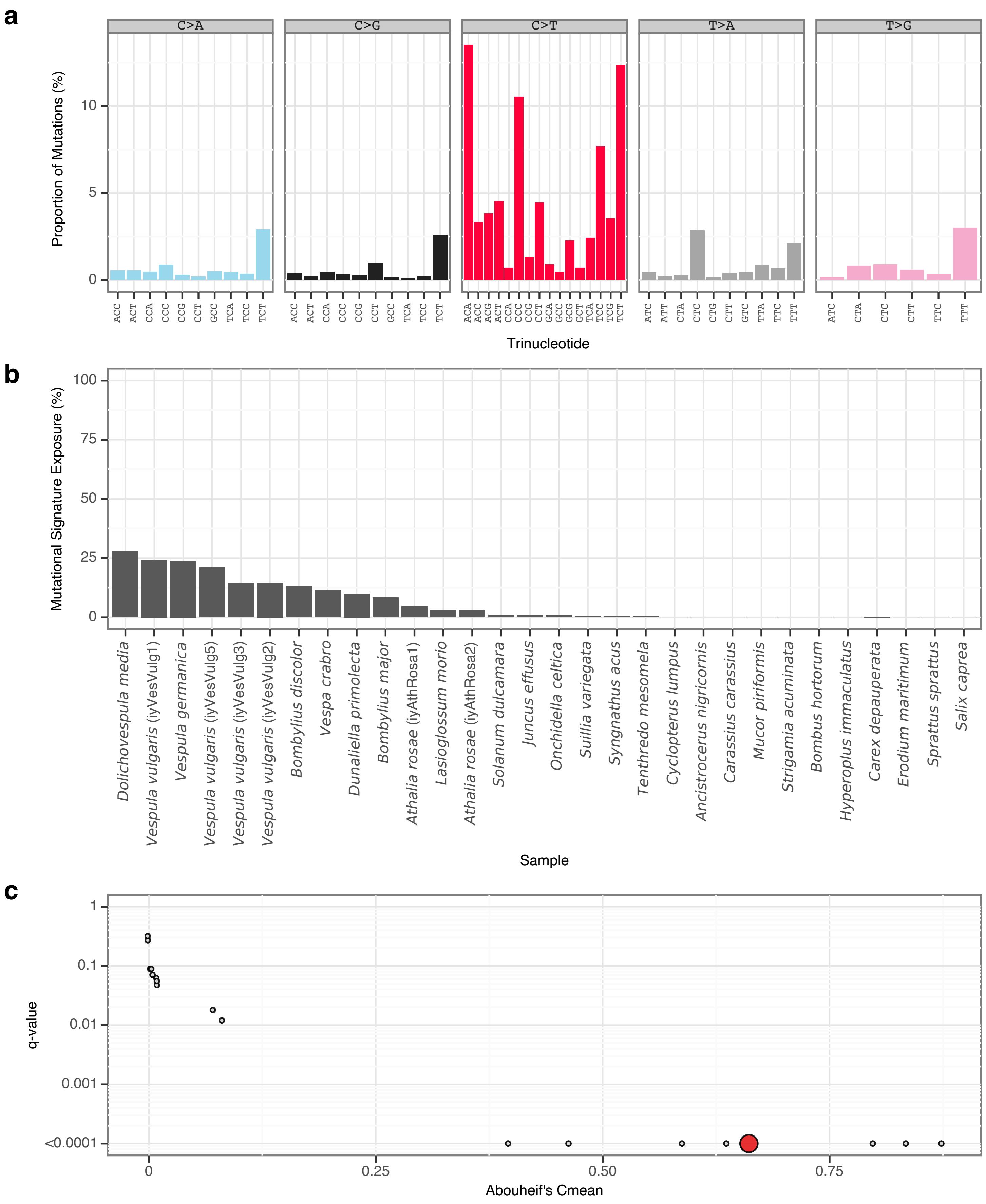


**Supplementary Fig. 109** | gToL8 mutational signature.

**a**, Mutational signature spectrum.

**b**, Bar plot displaying the 30 samples that exhibit the greatest exposure to the mutational signature.

**c**, Phylogenetic signal of the mutational signature.

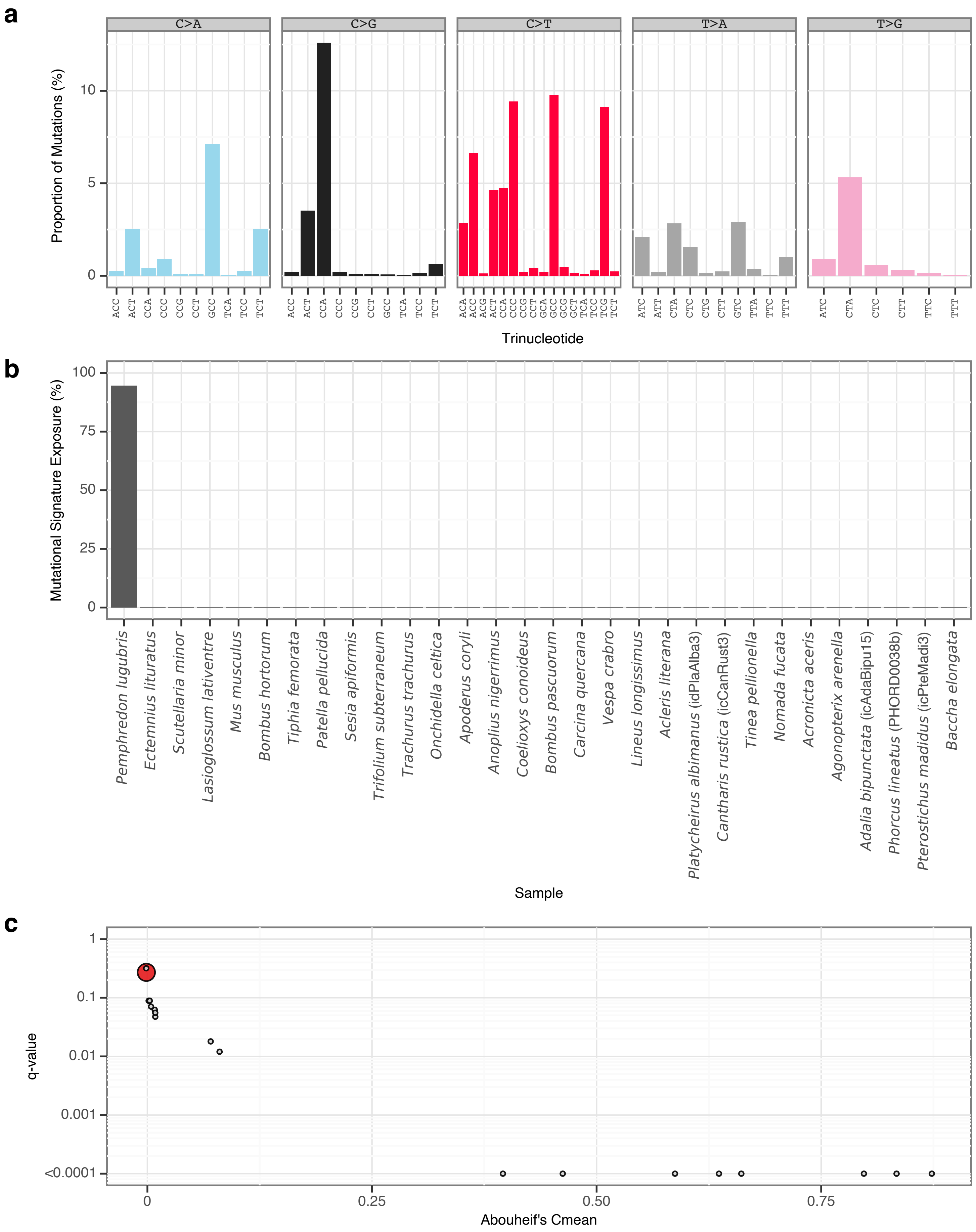


**Supplementary Fig. 110** | gToL9 mutational signature.

**a**, Mutational signature spectrum.

**b**, Bar plot displaying the 30 samples that exhibit the greatest exposure to the mutational signature.

**c**, Phylogenetic signal of the mutational signature.

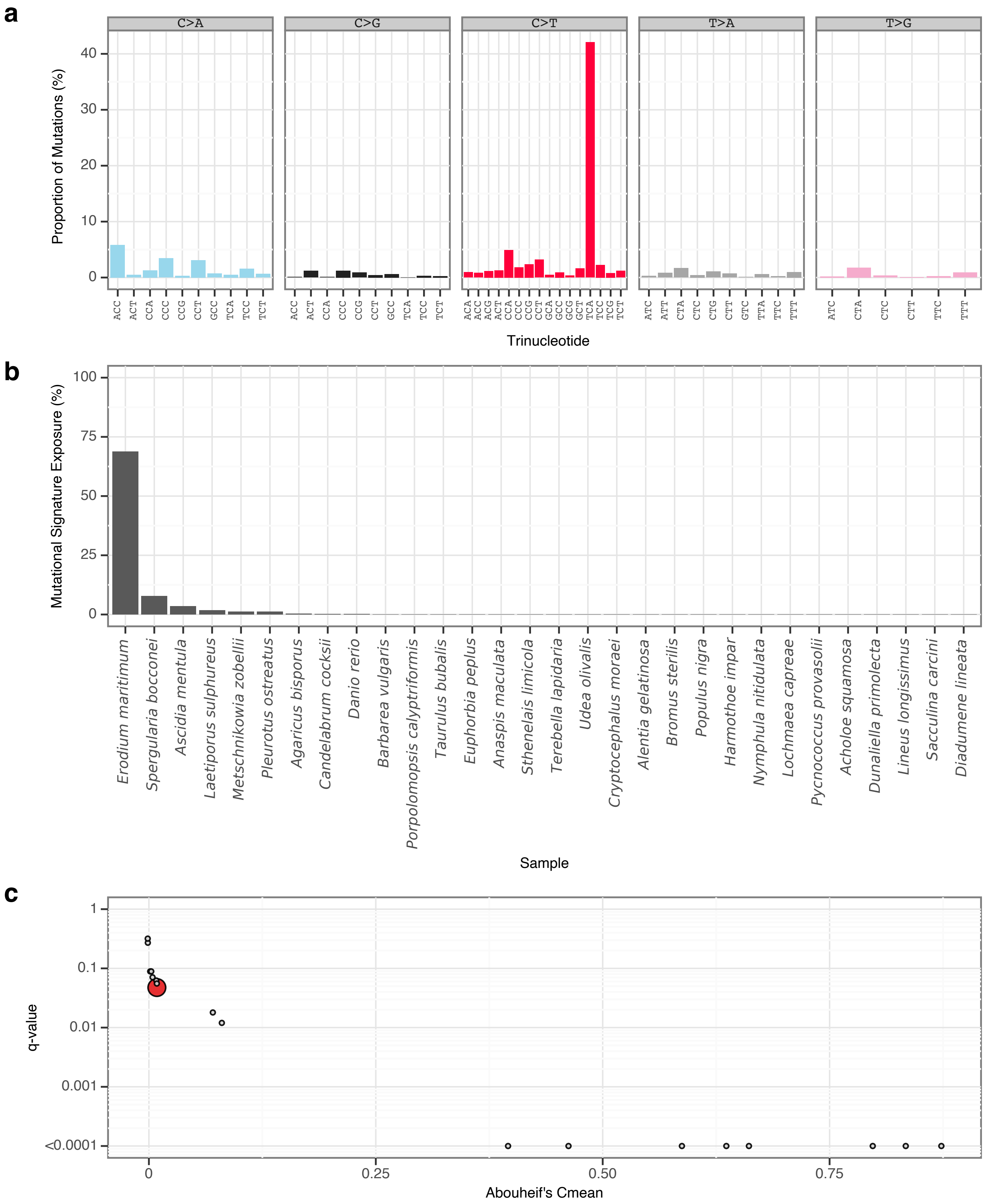


**Supplementary Fig. 111** | gToL10 mutational signature.
**a**, Mutational signature spectrum.
**b**, Bar plot displaying the 30 samples that exhibit the greatest exposure to the mutational signature.
**c**, Phylogenetic signal of the mutational signature.

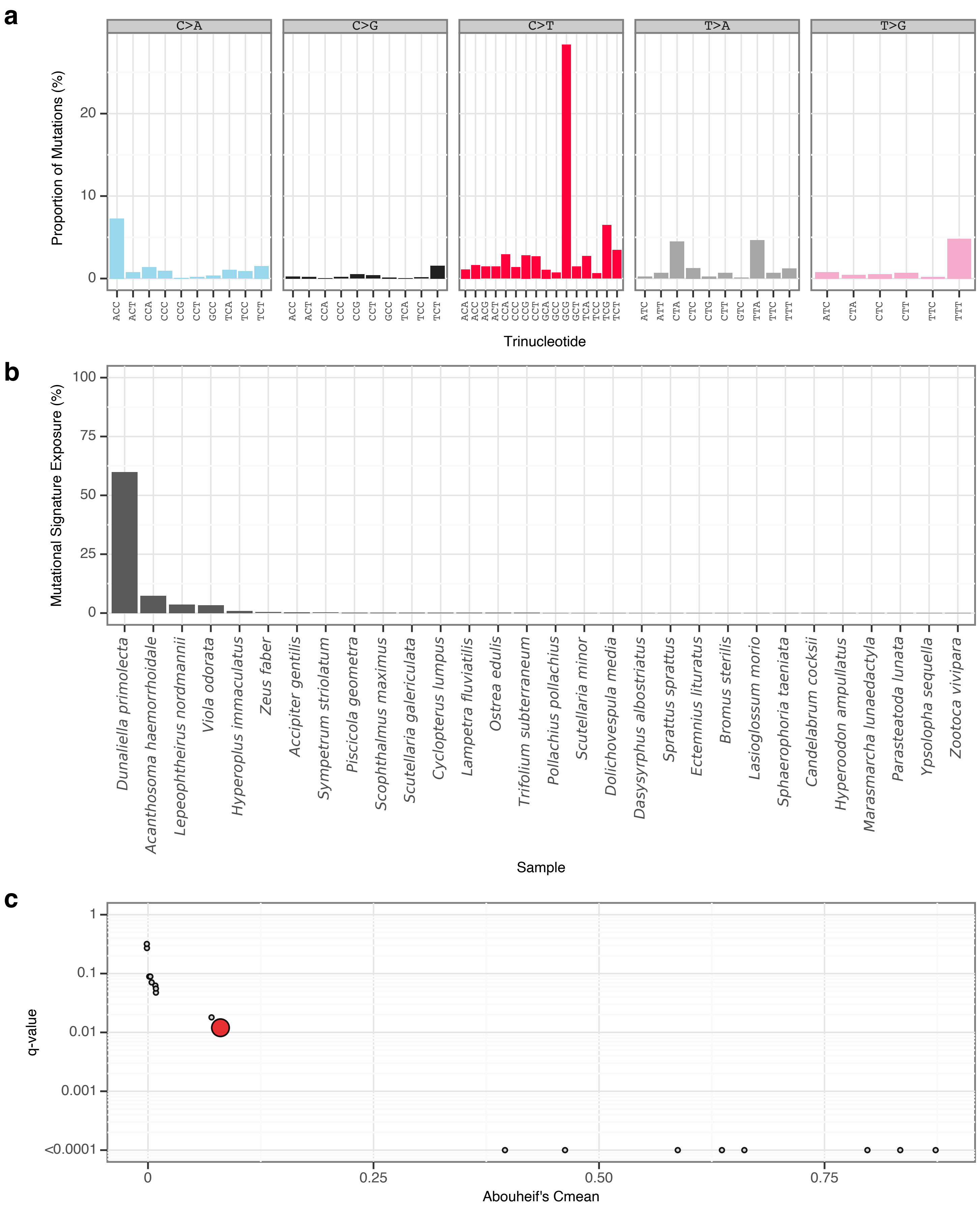


**Supplementary Fig. 112** | gToL11 mutational signature.

**a**, Mutational signature spectrum.

**b**, Bar plot displaying the 30 samples that exhibit the greatest exposure to the mutational signature.

**c**, Phylogenetic signal of the mutational signature.

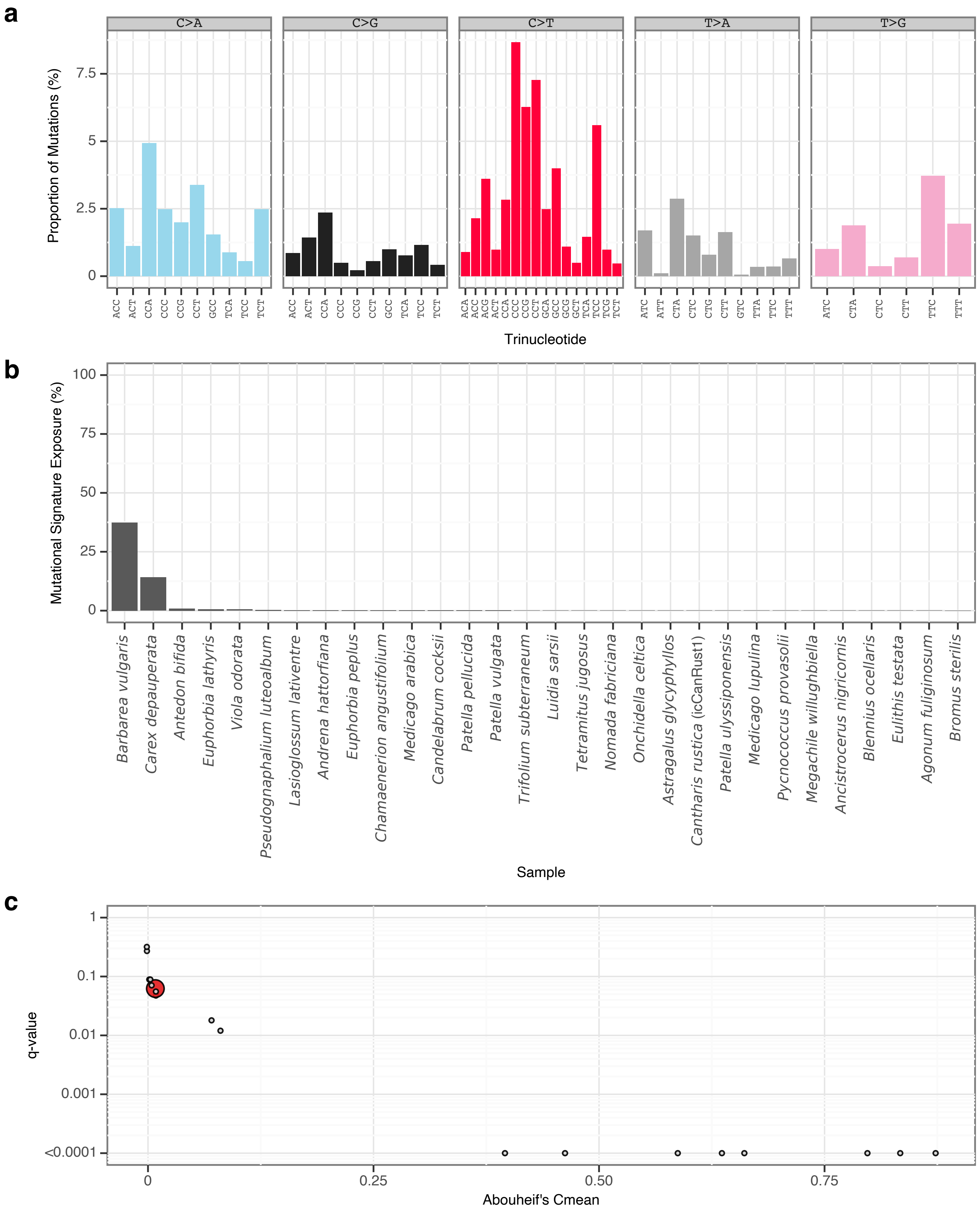


**Supplementary Fig. 113 |** gToL12 mutational signature.
**a**, Mutational signature spectrum.
**b**, Bar plot displaying the 30 samples that exhibit the greatest exposure to the mutational signature.
**c**, Phylogenetic signal of the mutational signature.

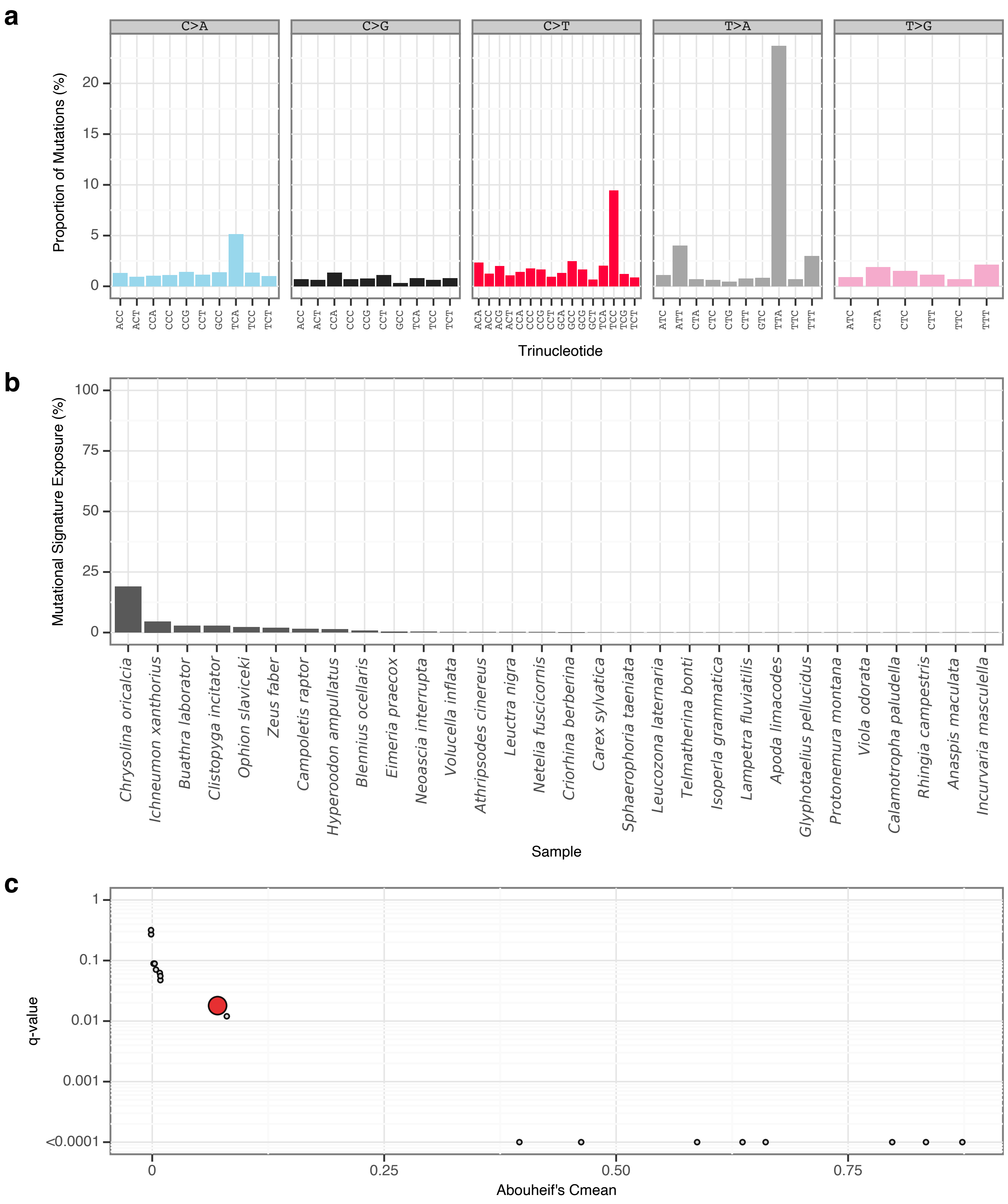


**Supplementary Fig. 114** | gToL13 mutational signature.

**a**, Mutational signature spectrum.

**b**, Bar plot displaying the 30 samples that exhibit the greatest exposure to the mutational signature.

**c**, Phylogenetic signal of the mutational signature.

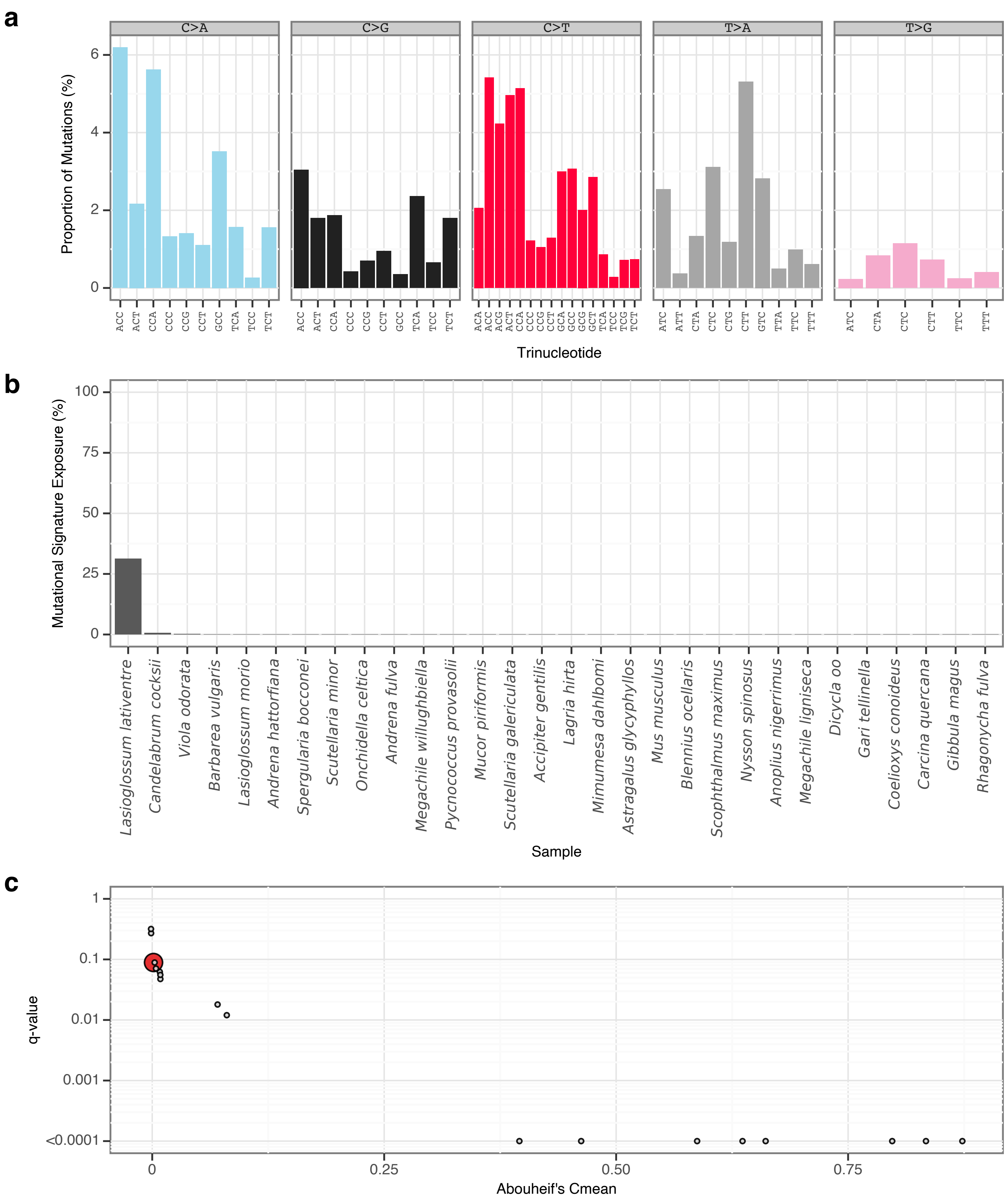


**Supplementary Fig. 115** | gToL14 mutational signature.

**a**, Mutational signature spectrum.

**b**, Bar plot displaying the 30 samples that exhibit the greatest exposure to the mutational signature.

**c**, Phylogenetic signal of the mutational signature.

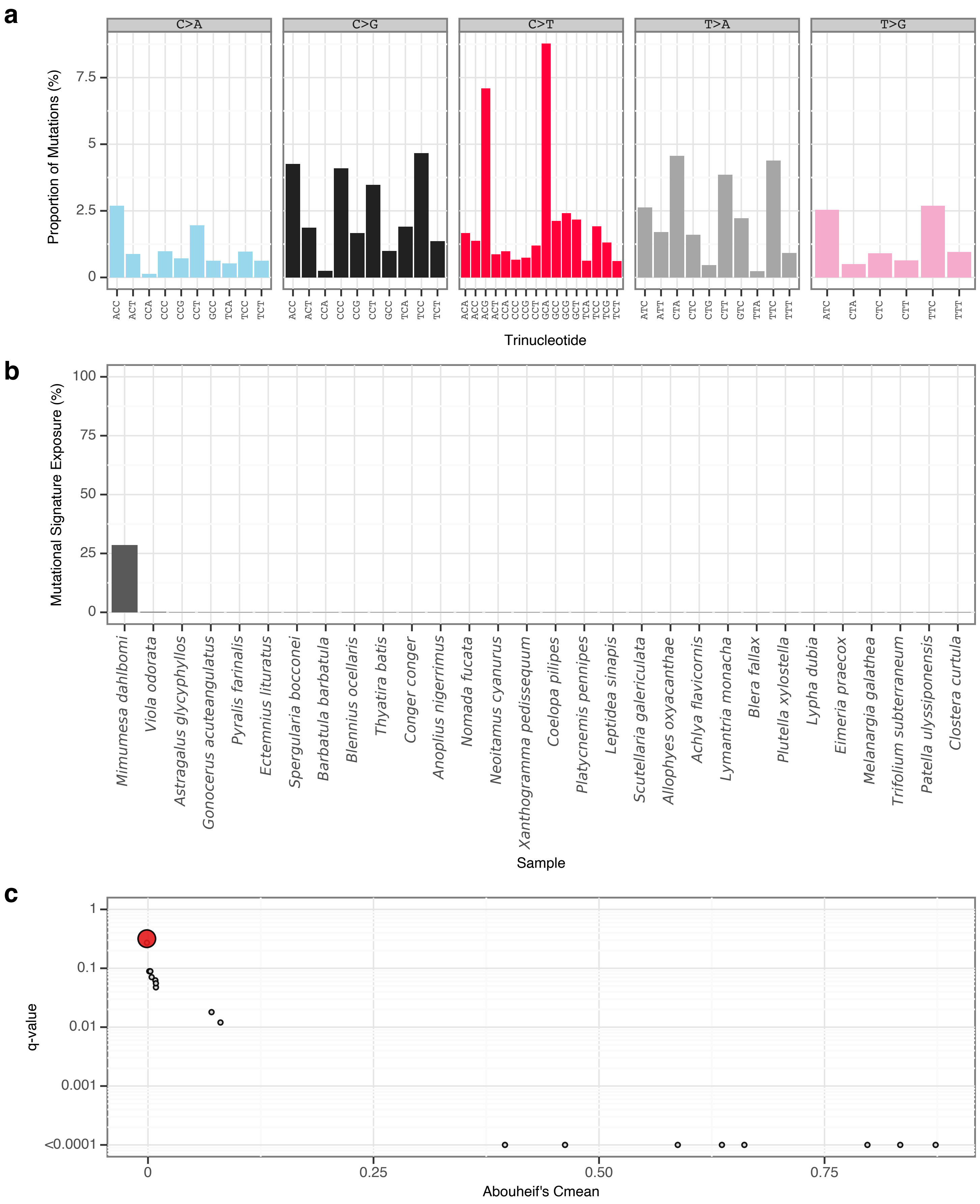


**Supplementary Fig. 116** | gToL15 mutational signature.

**a**, Mutational signature spectrum.

**b**, Bar plot displaying the 30 samples that exhibit the greatest exposure to the mutational signature.

**c**, Phylogenetic signal of the mutational signature.

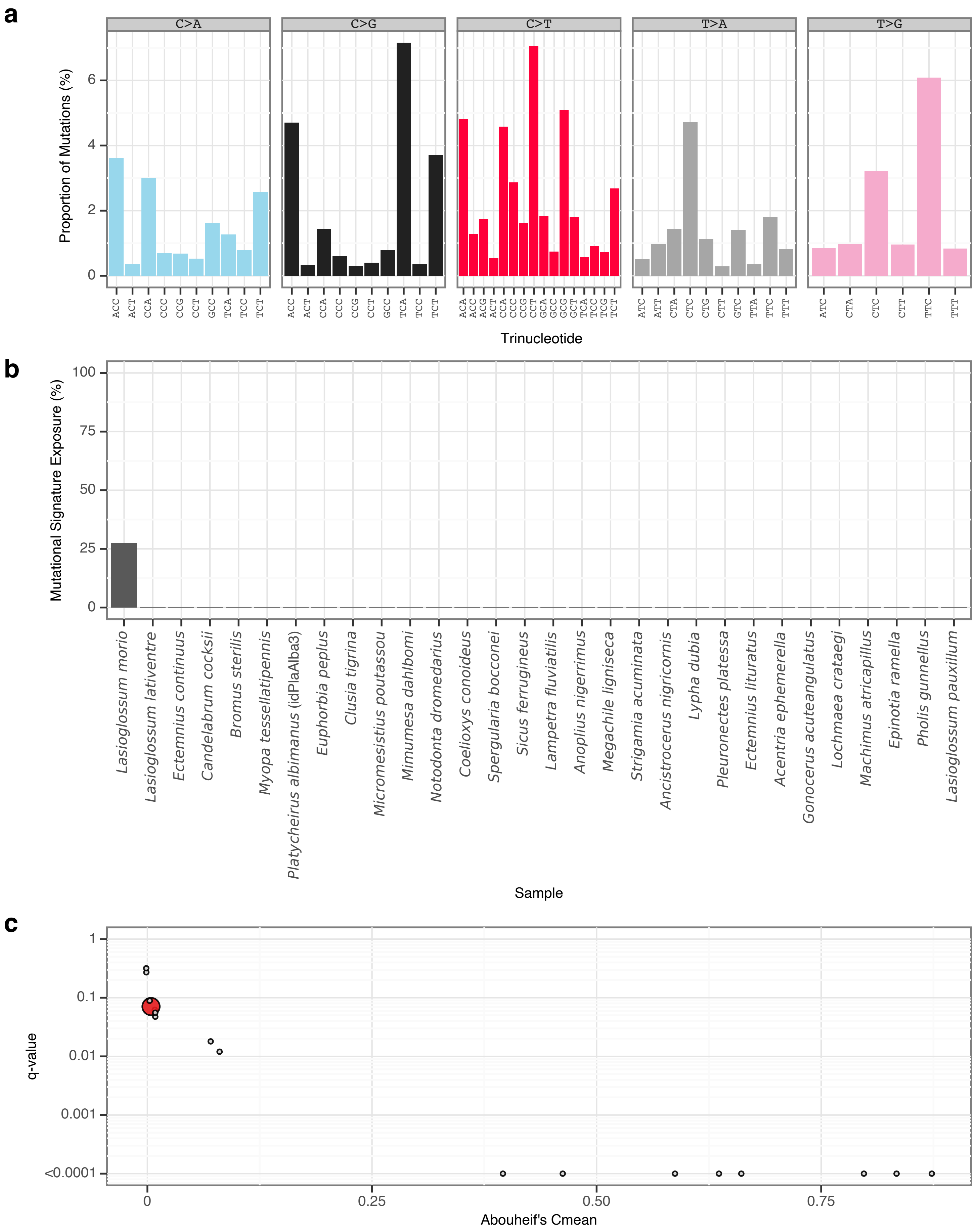


**Supplementary Fig. 117** | gToL16 mutational signature.

**a**, Mutational signature spectrum.

**b**, Bar plot displaying the 30 samples that exhibit the greatest exposure to the mutational signature.

**c**, Phylogenetic signal of the mutational signature.

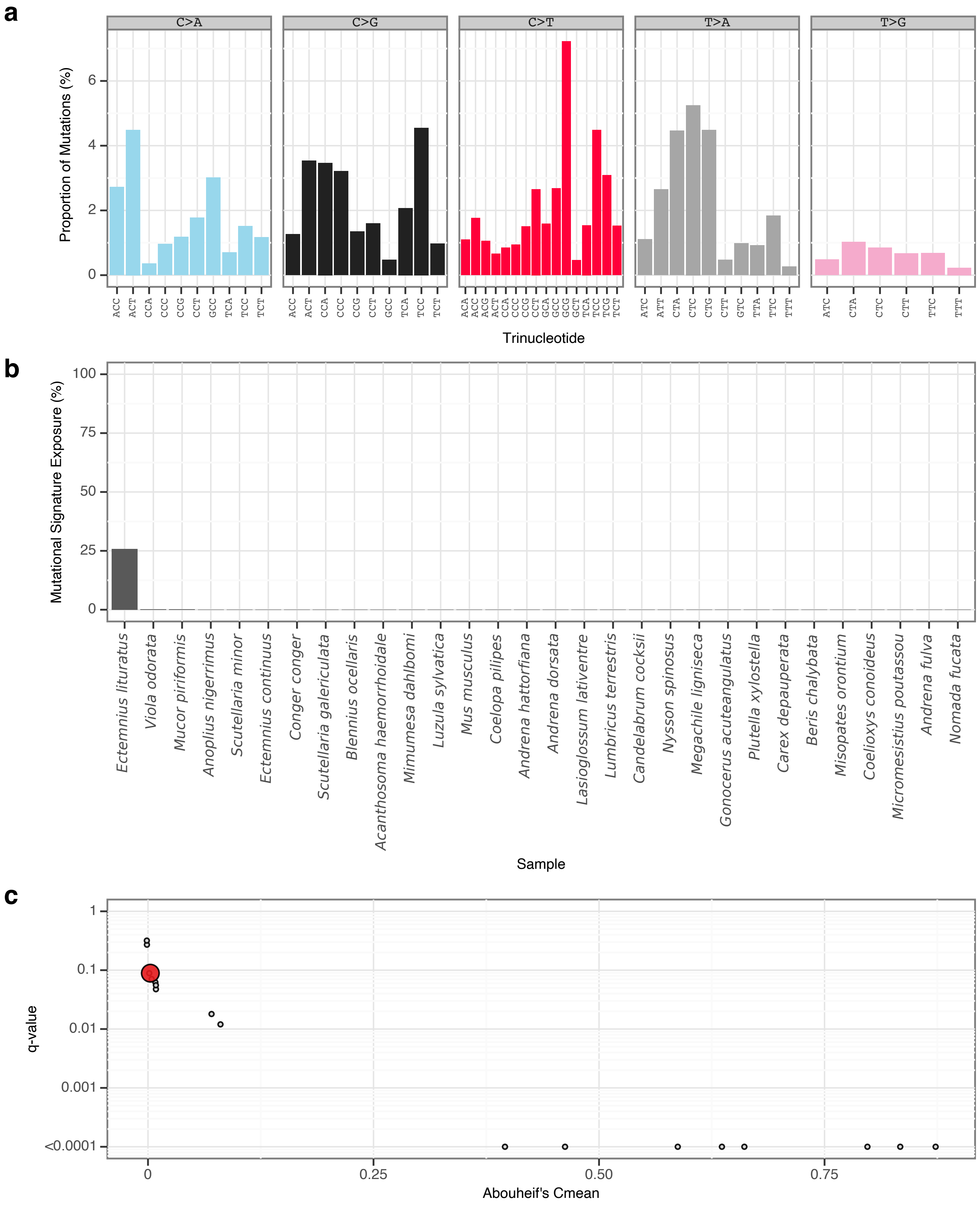


**Supplementary Fig. 118** | gToL17 mutational signature.
**a**, Mutational signature spectrum.
**b**, Bar plot displaying the 30 samples that exhibit the greatest exposure to the mutational signature.
**c**, Phylogenetic signal of the mutational signature.

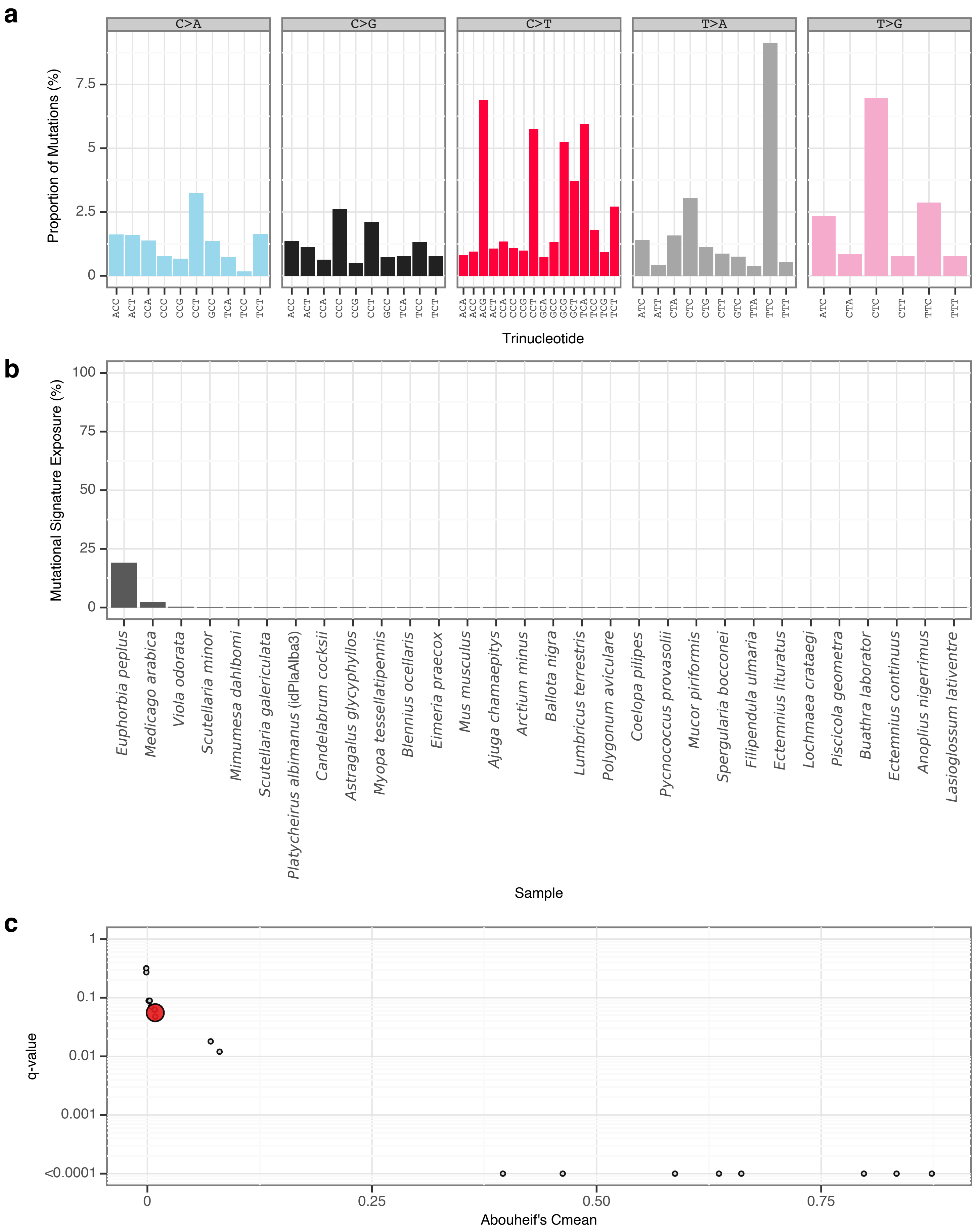


**Supplementary Fig. 119** | gToL18 mutational signature.

**a**, Mutational signature spectrum.

**b**, Bar plot displaying the 30 samples that exhibit the greatest exposure to the mutational signature.

**c**, Phylogenetic signal of the mutational signature.
