## Supplementary Information for "Somatic and germline mutational processes across the tree of life"

*Accipiter gentilis* (bAccGen1) germline mutational spectrum

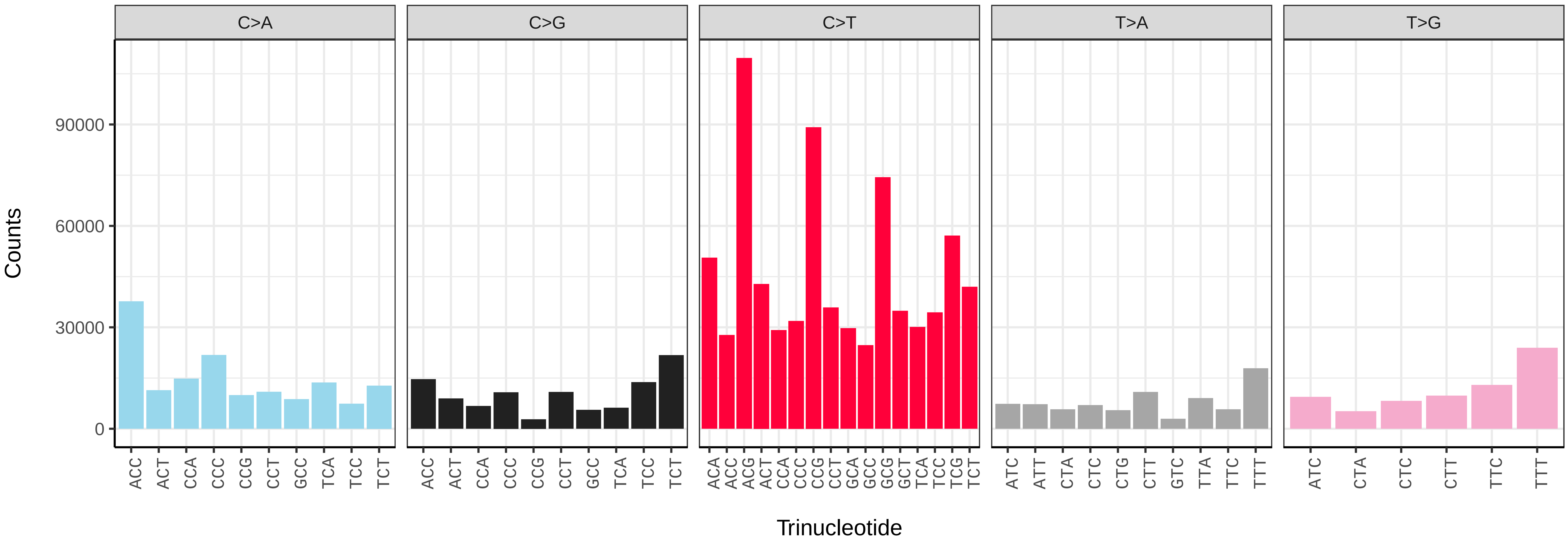


*Accipiter gentilis* (bAccGen1) somatic mutational spectrum (SBS96 classification)

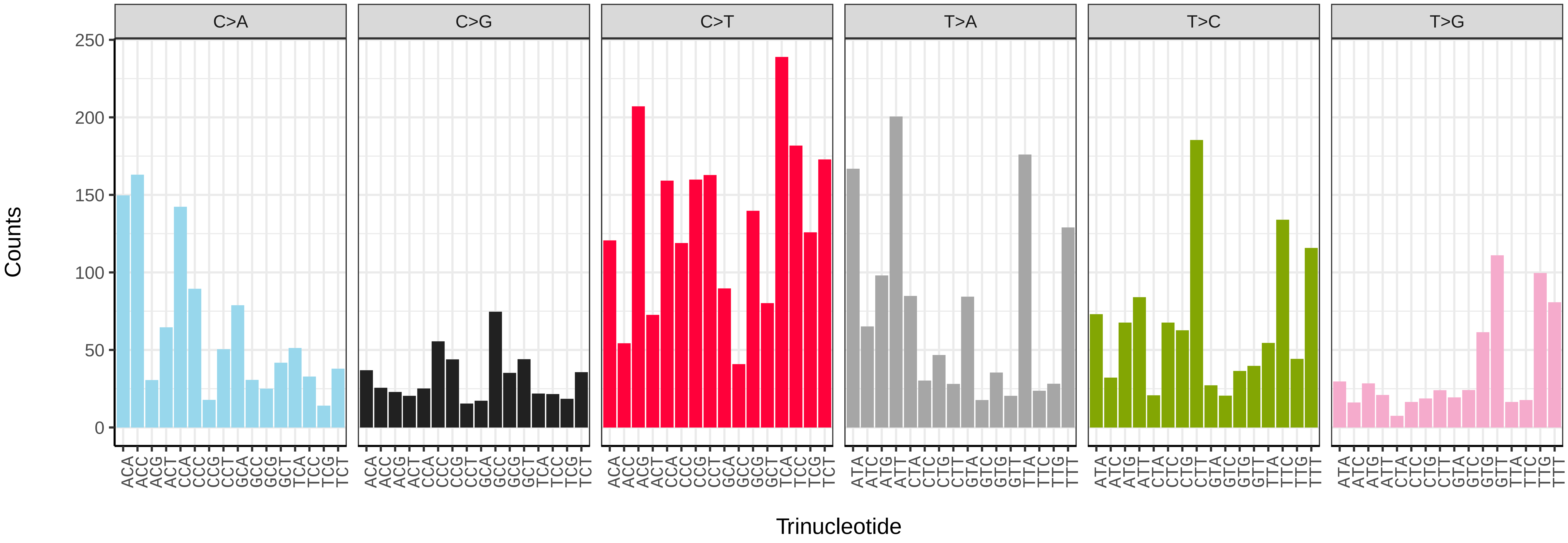


*Accipiter gentilis* (bAccGen1) somatic mutational spectrum (SBS52 classification)

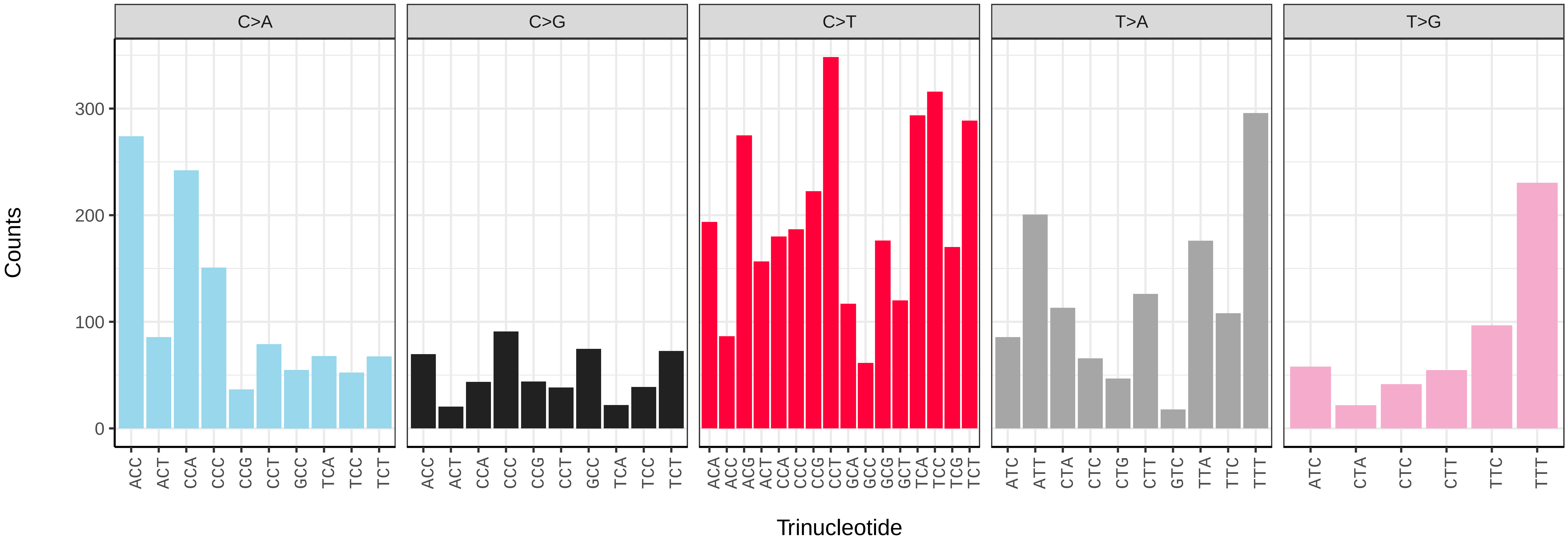

*Anas acuta* (bAnaAcu1) germline mutational spectrum

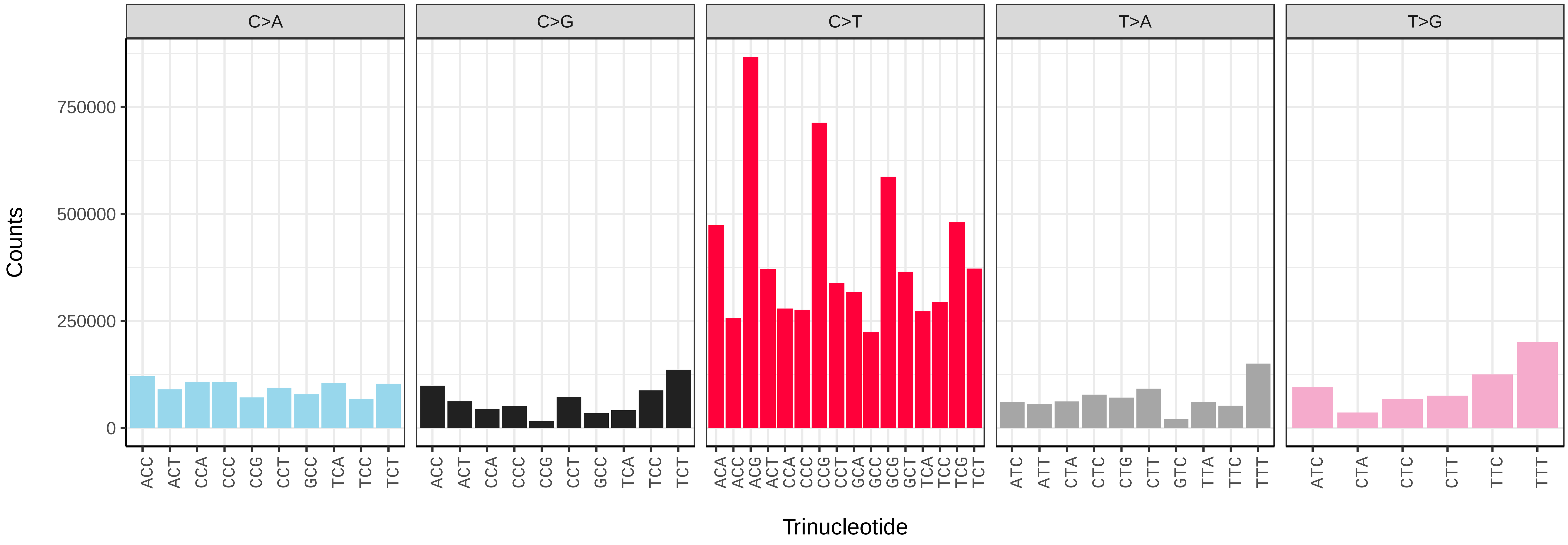


*Anas acuta* (bAnaAcu1) somatic mutational spectrum (SBS96 classification)

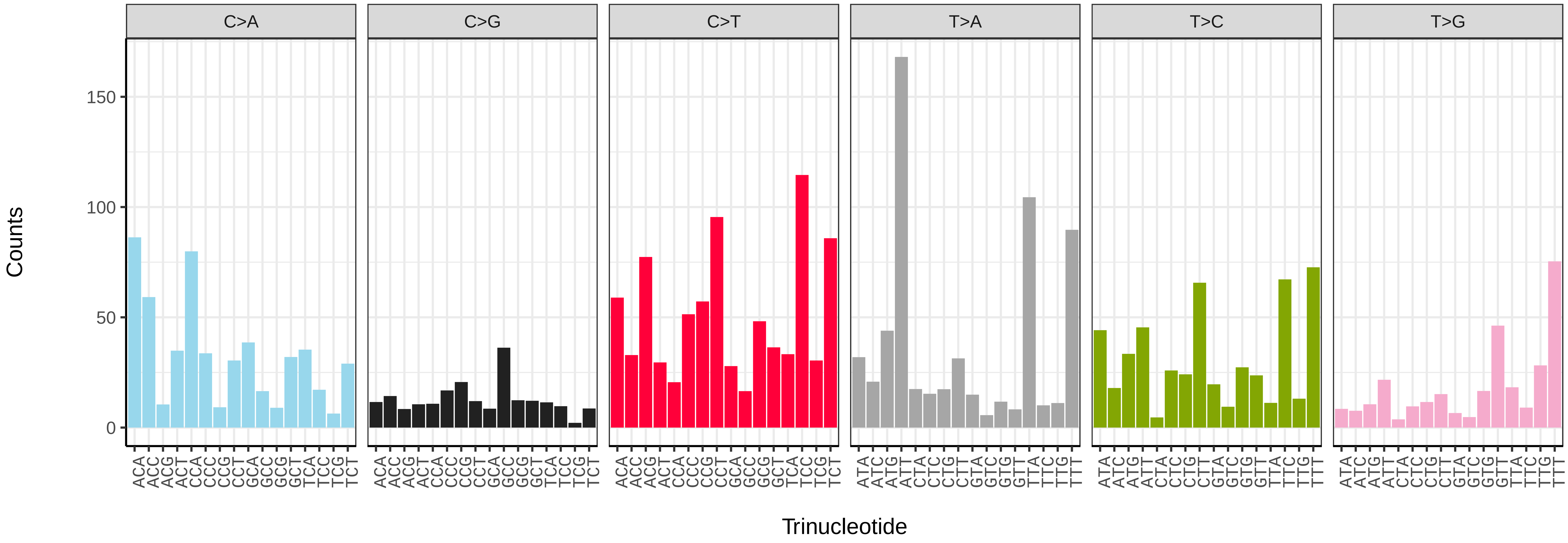


*Anas acuta* (bAnaAcu1) somatic mutational spectrum (SBS52 classification)

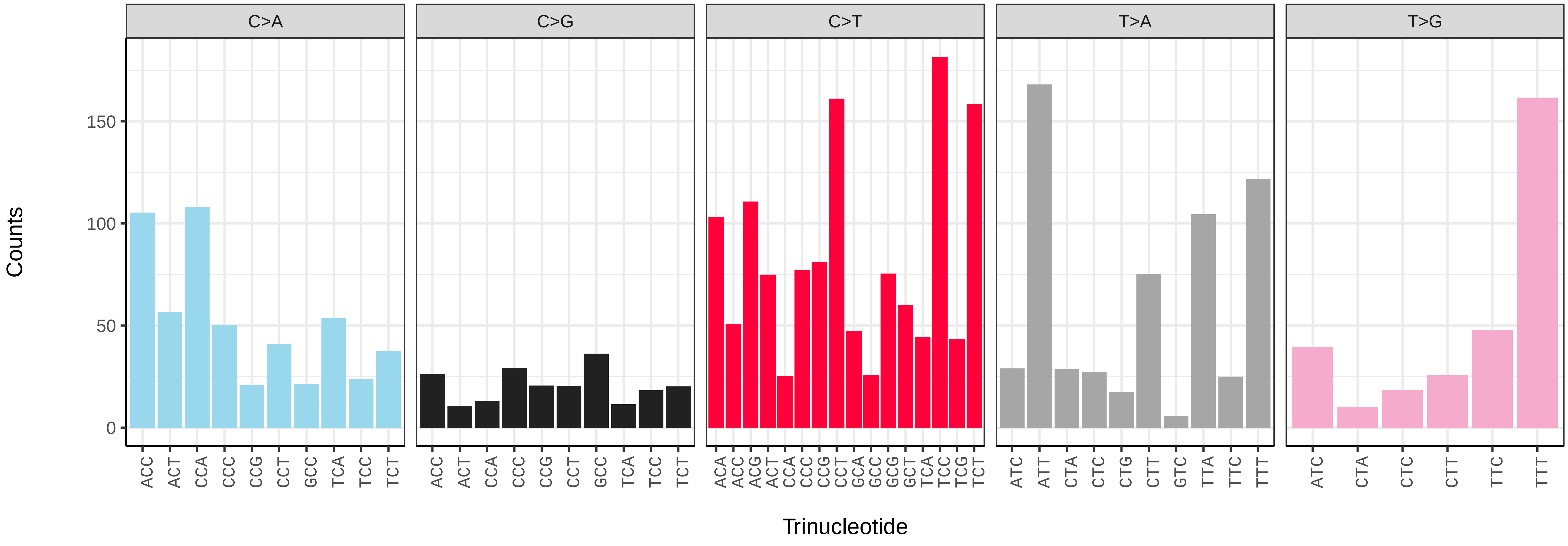

*Anser brachyrhynchus* (bAnsBra1) germline mutational spectrum

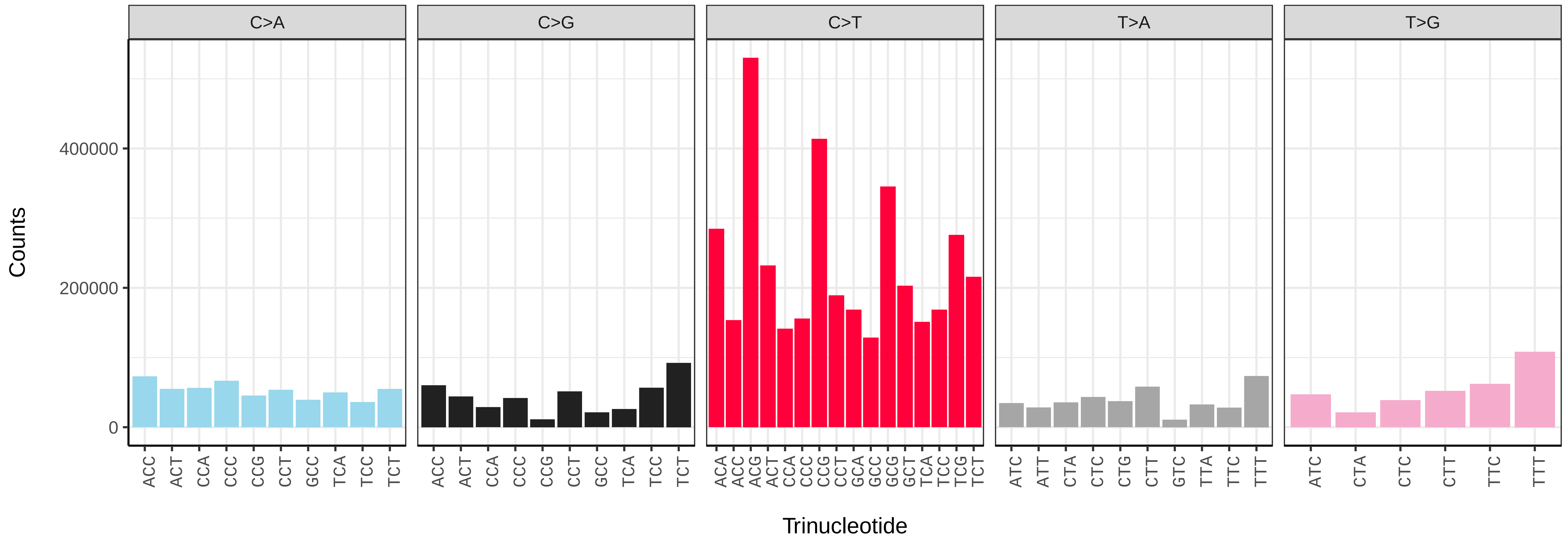


*Anser brachyrhynchus* (bAnsBra1) somatic mutational spectrum (SBS96 classification)

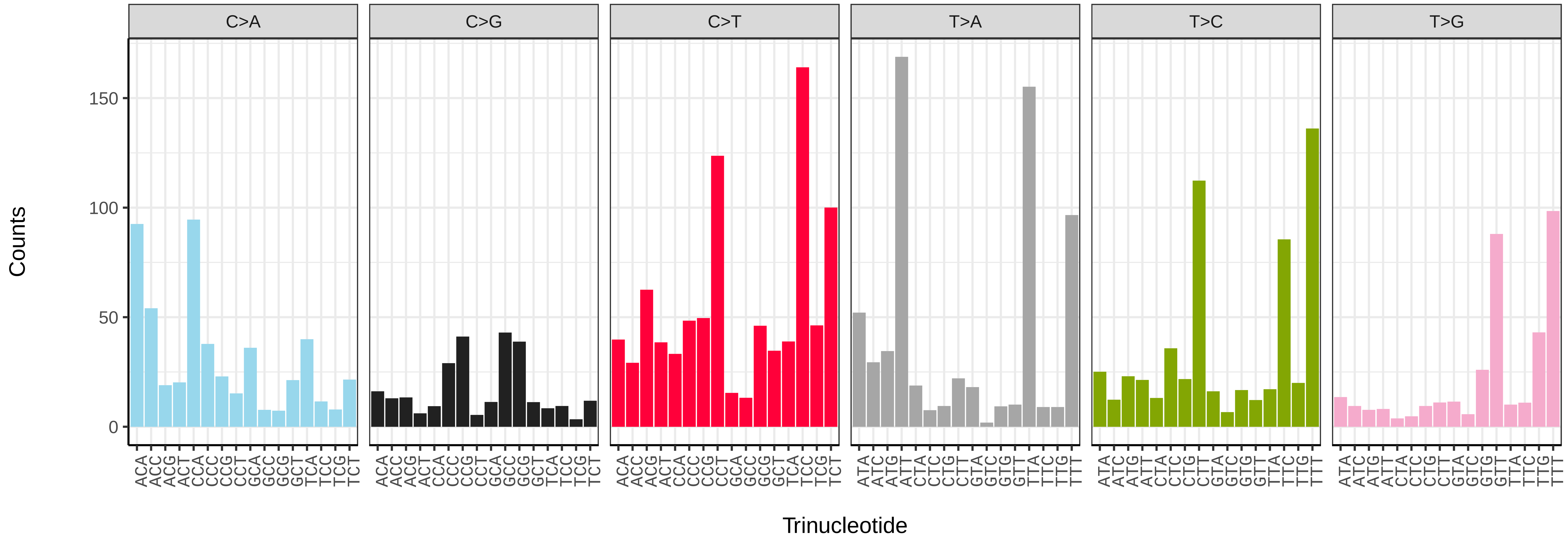


*Anser brachyrhynchus* (bAnsBra1) somatic mutational spectrum (SBS52 classification)

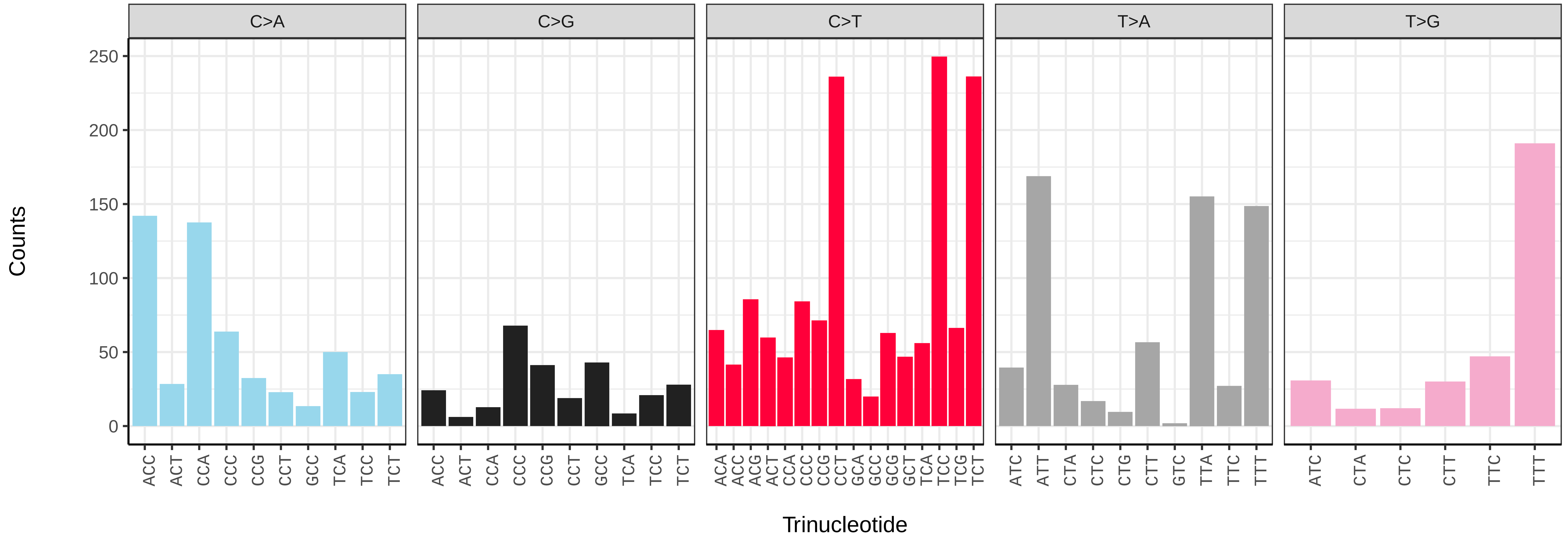

*Cinclus cinclus* (bCinCin1) germline mutational spectrum

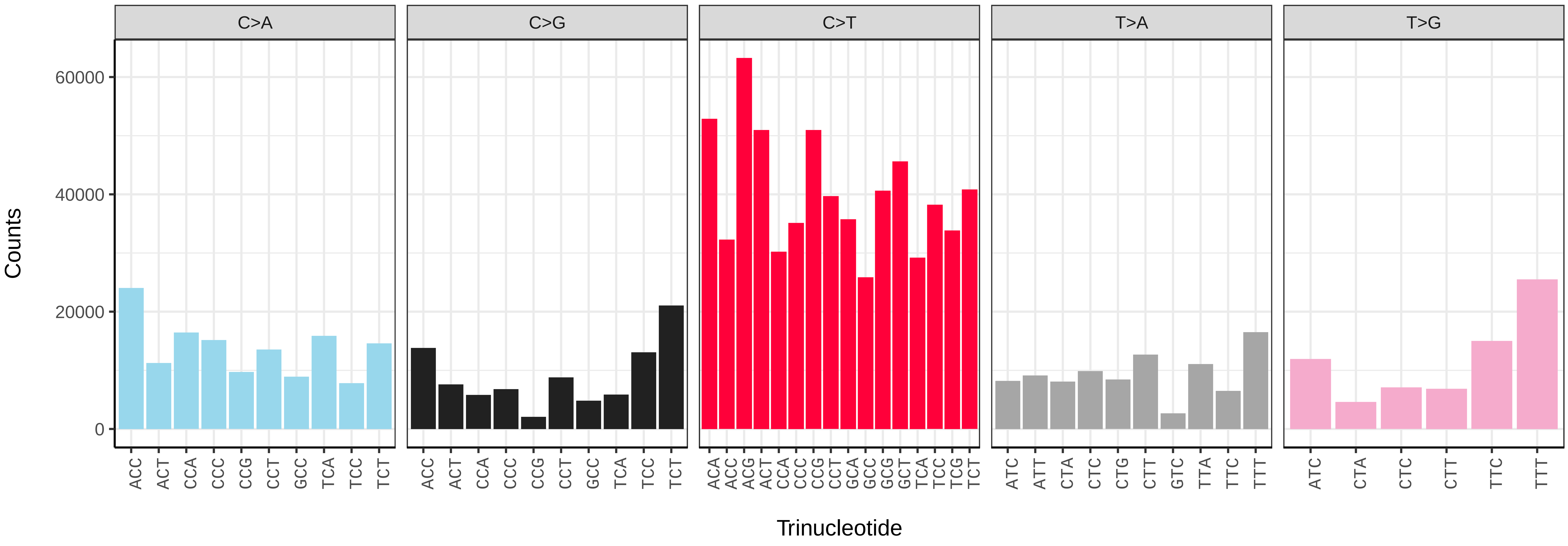


*Cinclus cinclus* (bCinCin1) somatic mutational spectrum (SBS96 classification)

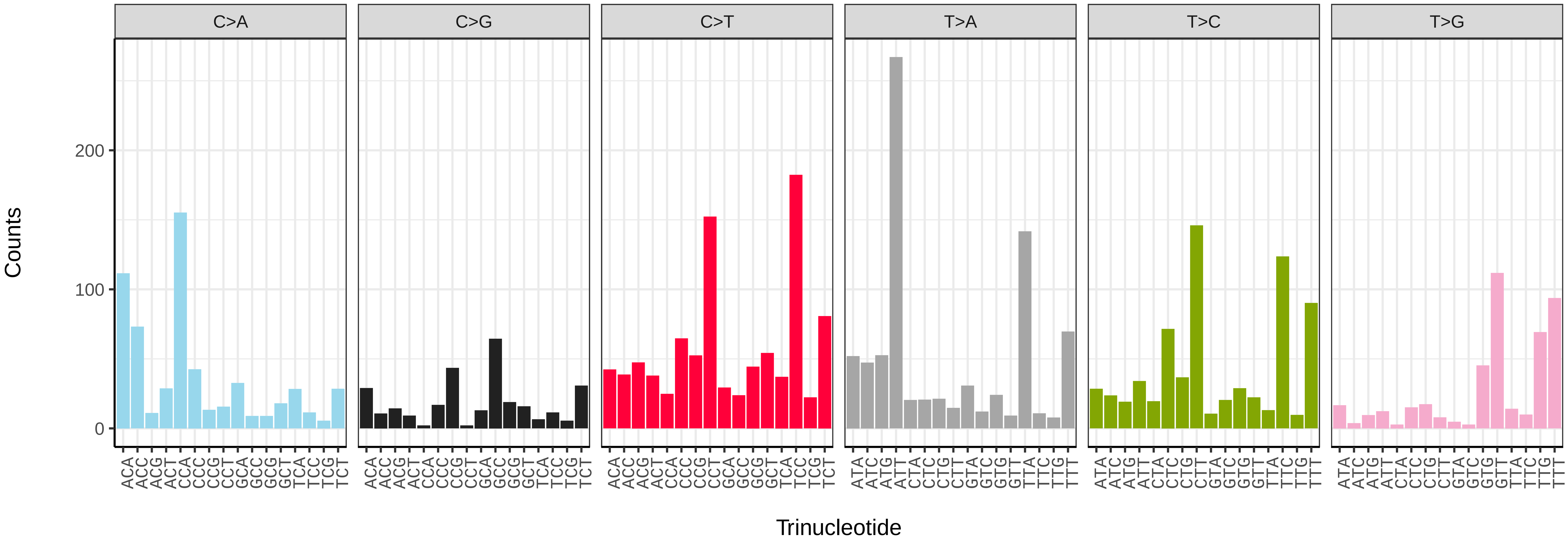


*Cinclus cinclus* (bCinCin1) somatic mutational spectrum (SBS52 classification)

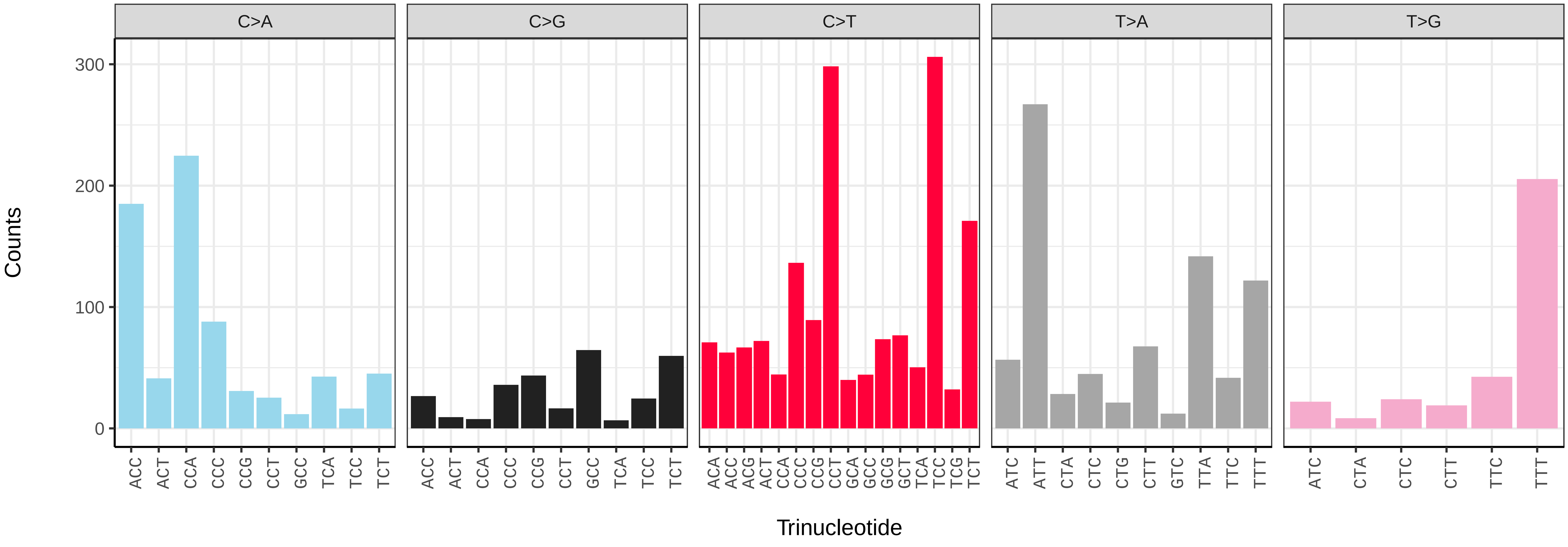

*Clangula hyemalis* (bClaHye2) germline mutational spectrum

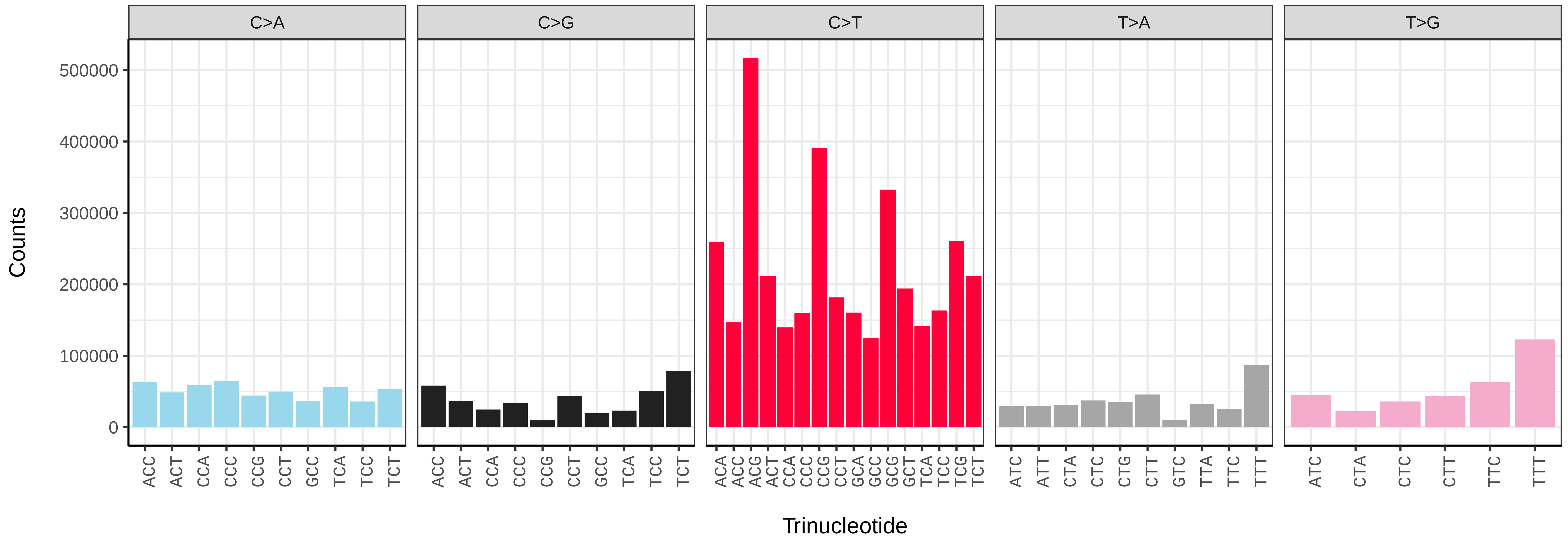


*Clangula hyemalis* (bClaHye2) somatic mutational spectrum (SBS96 classification)

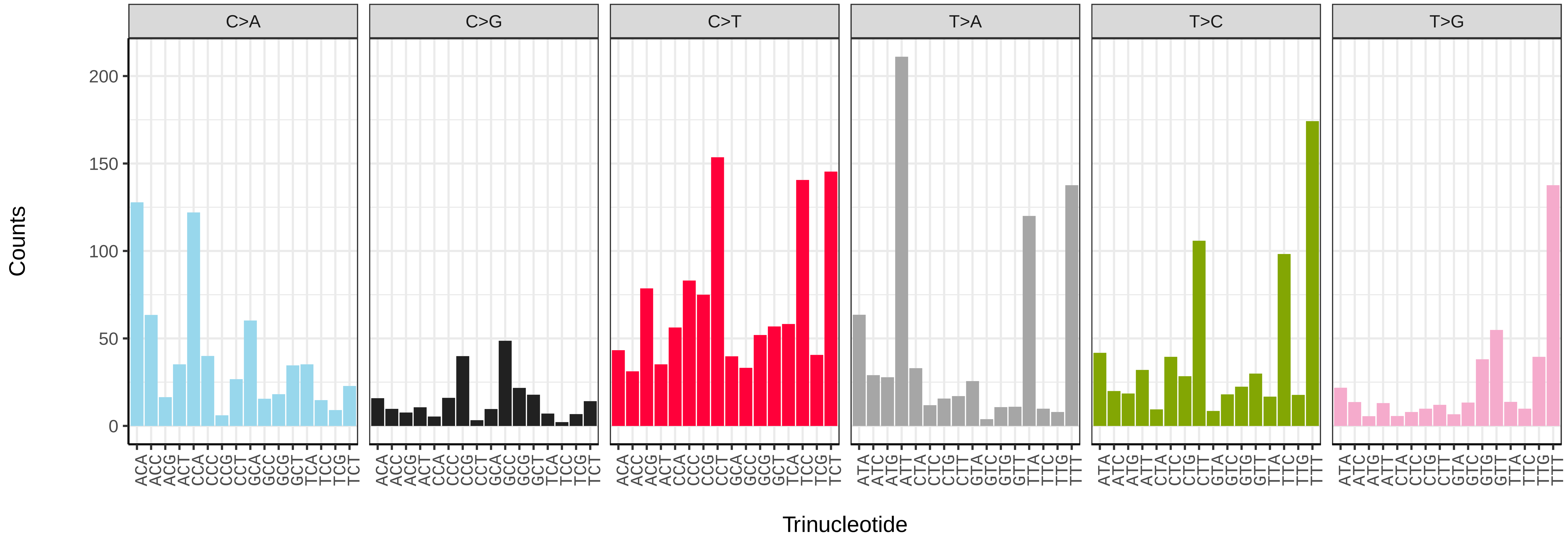


*Clangula hyemalis* (bClaHye2) somatic mutational spectrum (SBS52 classification)

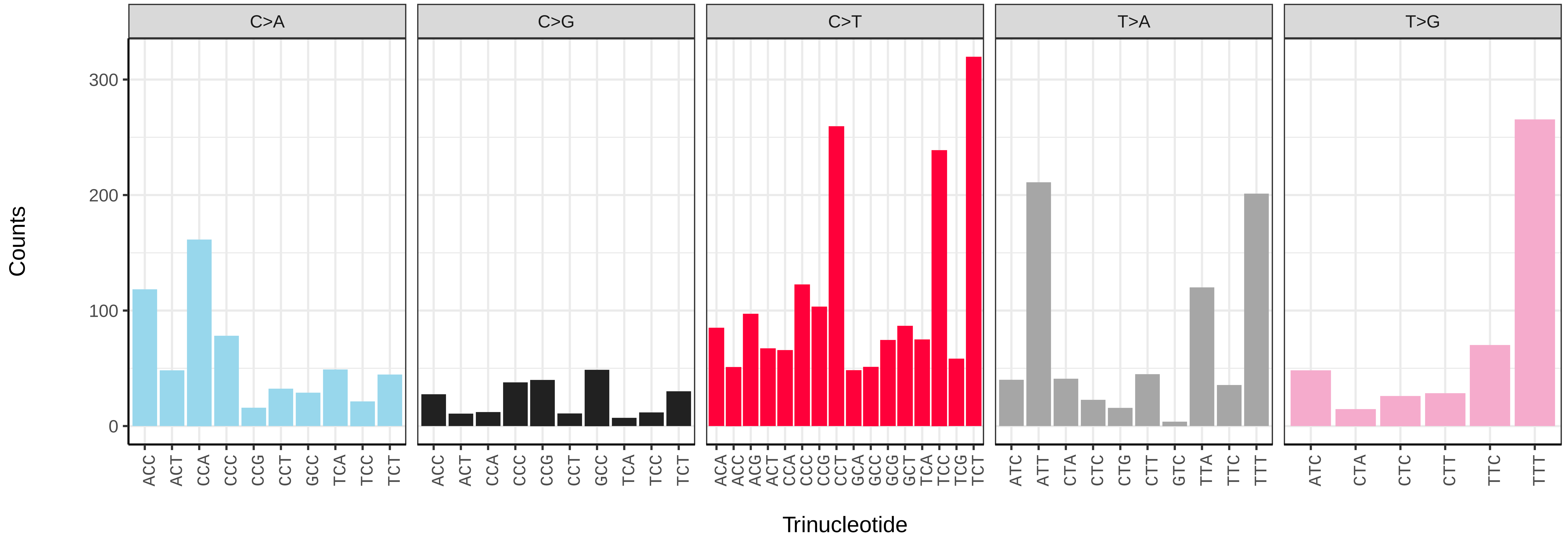

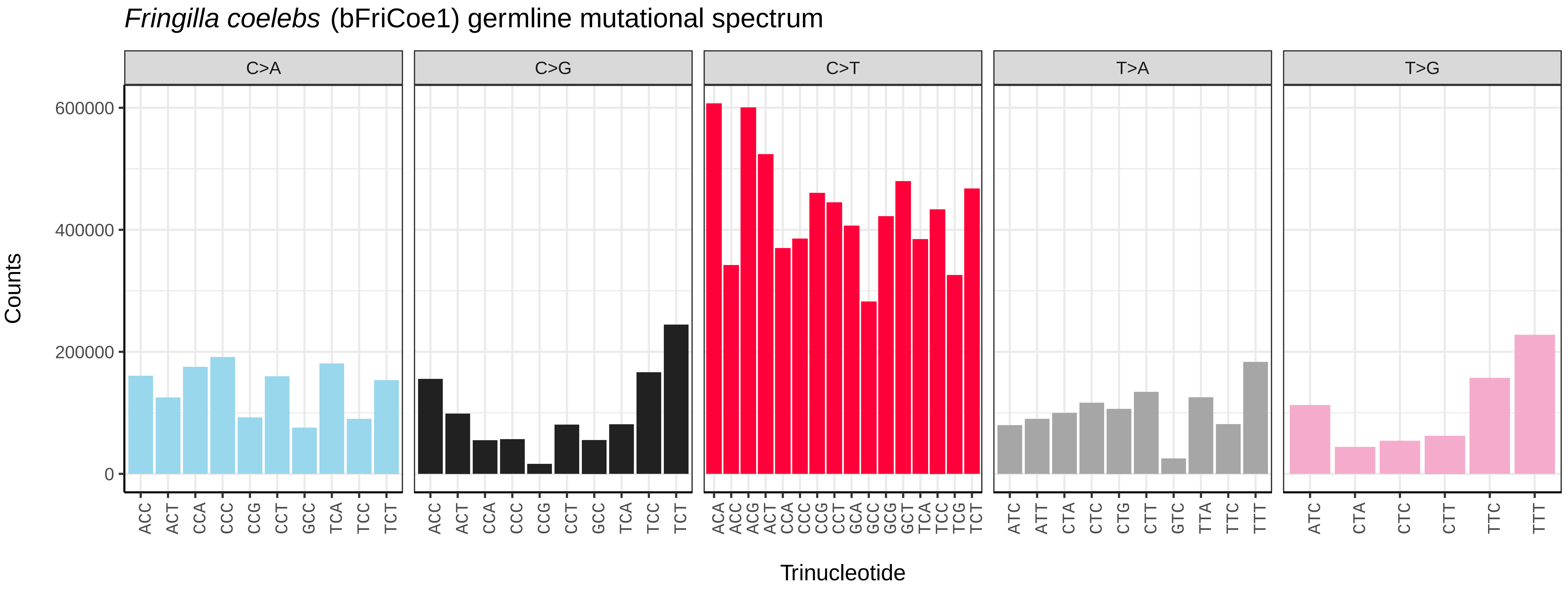
*Fringilla coelebs* (bFriCoe1) germline mutational spectrum
C>A
C>G
C>T
T>A
T>G
Counts
600000
400000
200000
0
ACC ACT CCA CCC CCG CCT GCC TCA TCC TCT
ACC ACT CCA CCC CCG CCT GCC TCA TCC TCT
ACA ACC ACG ACT CCA CCC CCG CCT GCA GCC GCG GCT TCA TCC TCG TCT
ATC ATT CTA CTC CTG CTT GTC TTA TTC TTT
ATC CTA CTC CTT TTC TTT
Trinucleotide

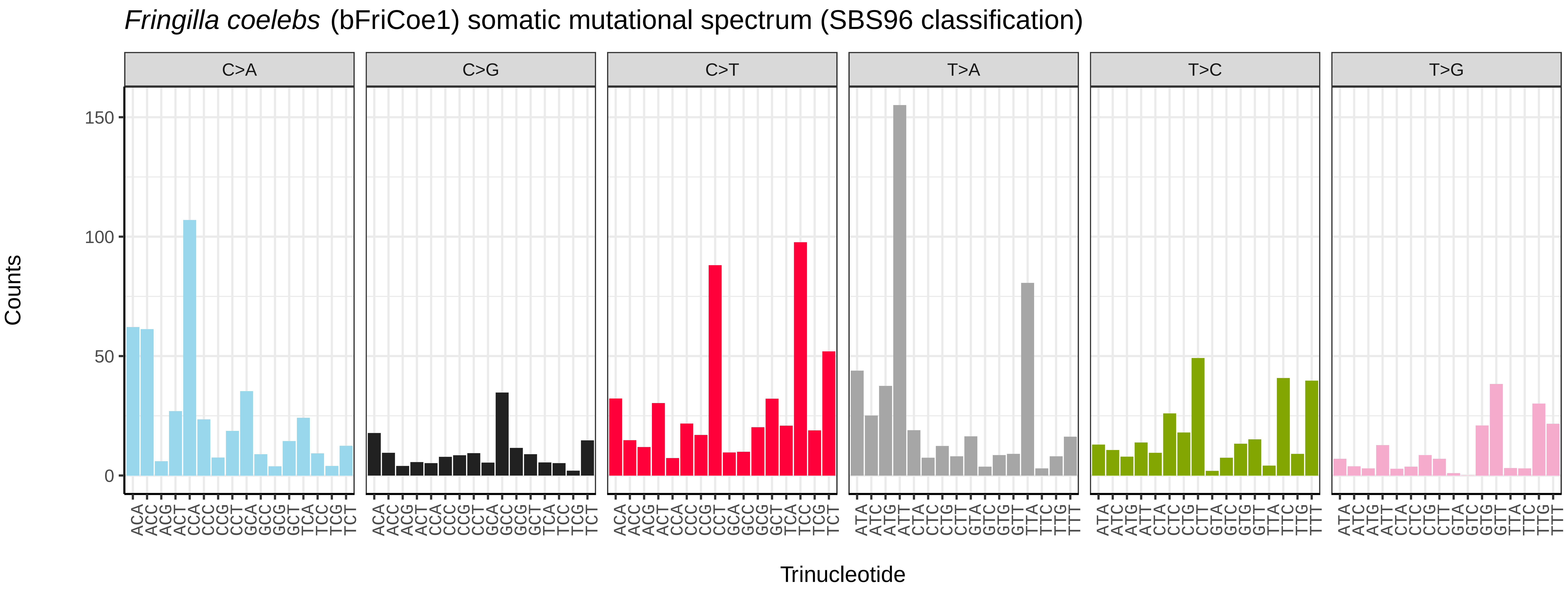
*Fringilla coelebs* (bFriCoe1) somatic mutational spectrum (SBS96 classification)
C>A
C>G
C>T
T>A
T>C
T>G
Counts
150
100
50
0
Trinucleotide

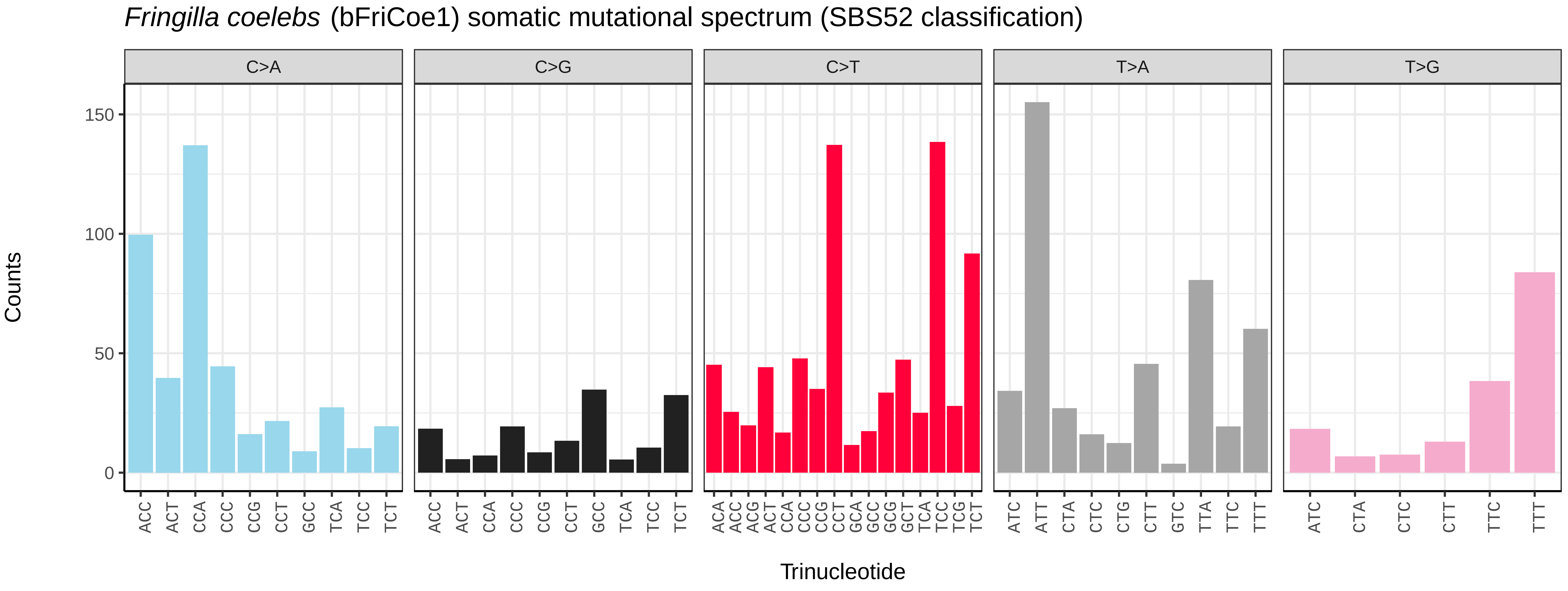
*Fringilla coelebs* (bFriCoe1) somatic mutational spectrum (SBS52 classification)
C>A
C>G
C>T
T>A
T>G
Counts
150
100
50
0
ACC ACT CCA CCC CCG CCT GCC TCA TCC TCT
ACC ACT CCA CCC CCG CCT GCC TCA TCC TCT
ACA ACC ACG ACT CCA CCC CCG CCT GCA GCC GCG GCT TCA TCC TCG TCT
ATC ATT CTA CTC CTG CTT GTC TTA TTC TTT
ATC CTA CTC CTT TTC TTT
Trinucleotide

*Larus fuscus* (bLarFus1) germline mutational spectrum

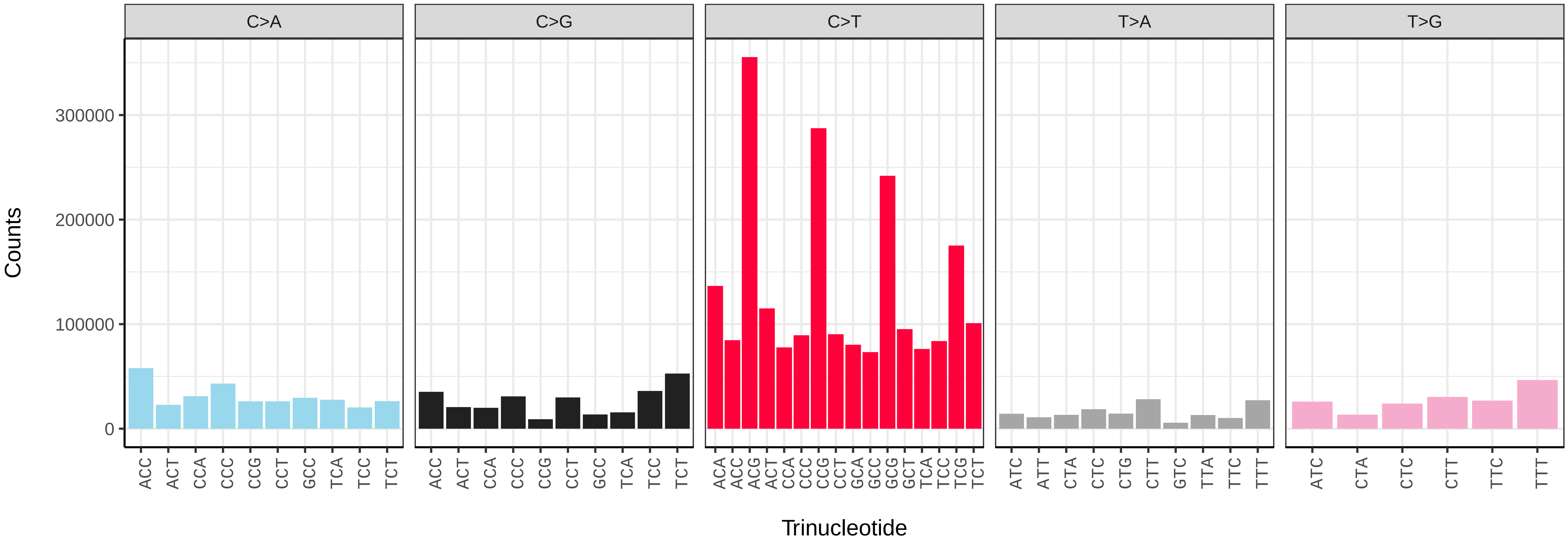


*Larus fuscus* (bLarFus1) somatic mutational spectrum (SBS96 classification)

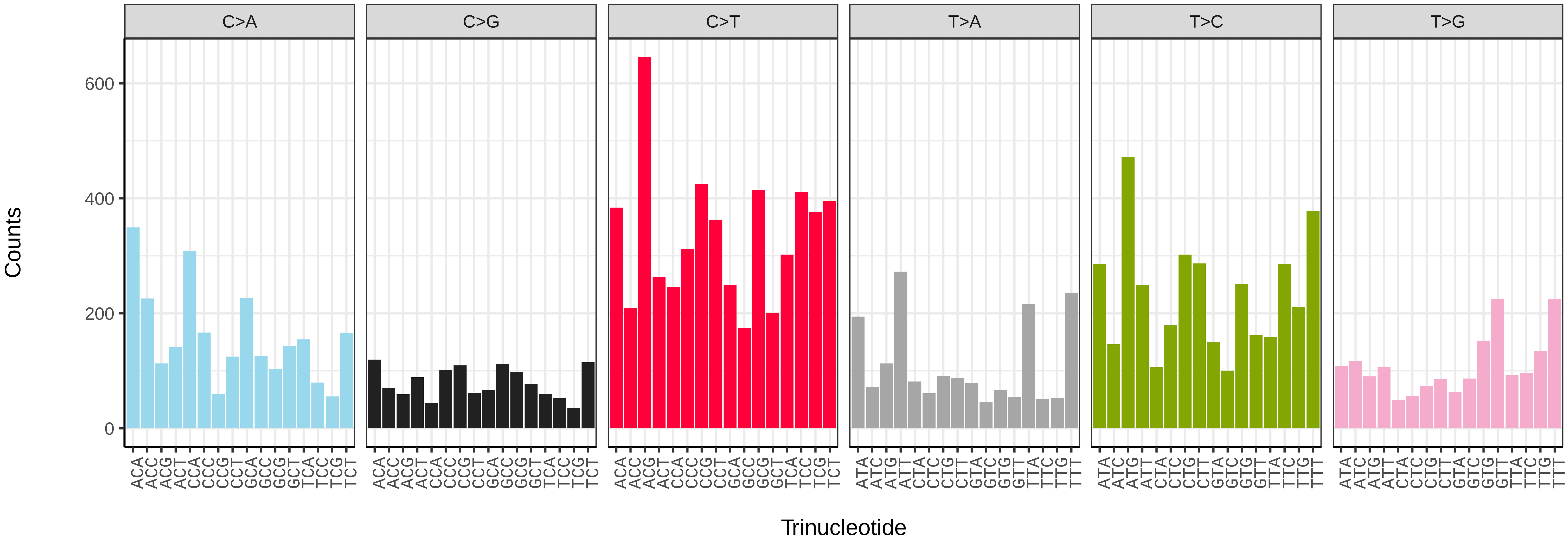


*Larus fuscus* (bLarFus1) somatic mutational spectrum (SBS52 classification)

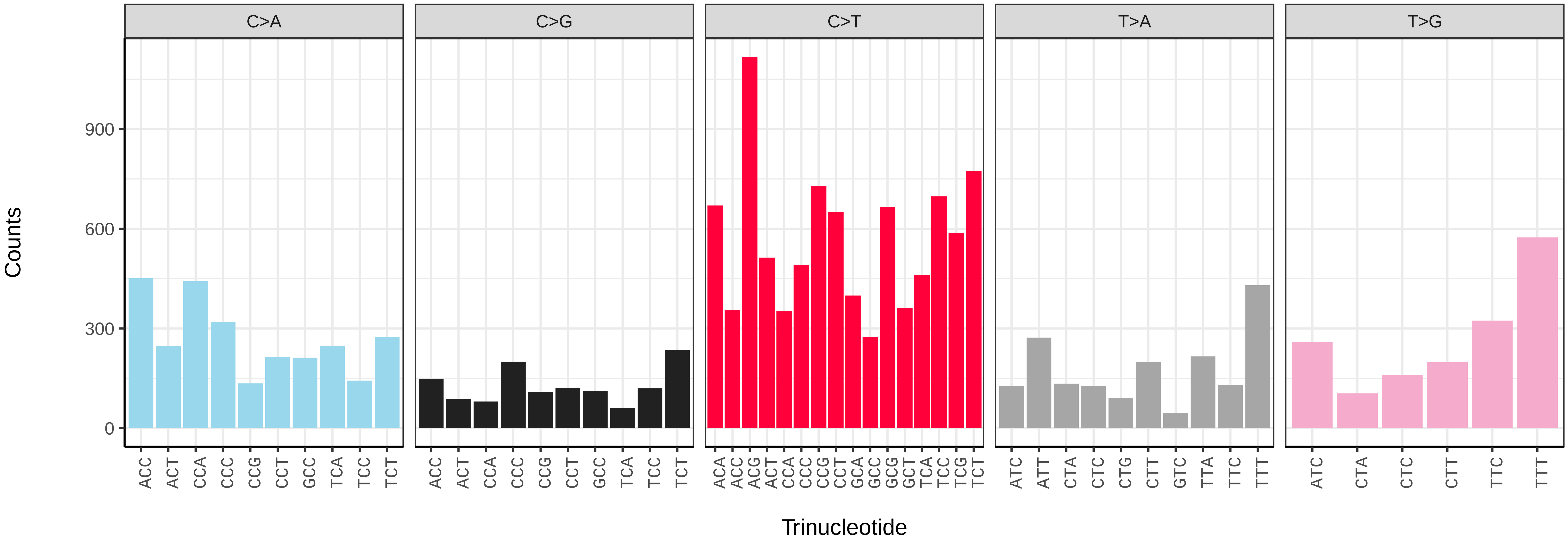

*Phalacrocorax carbo* (bPhaCar2) germline mutational spectrum

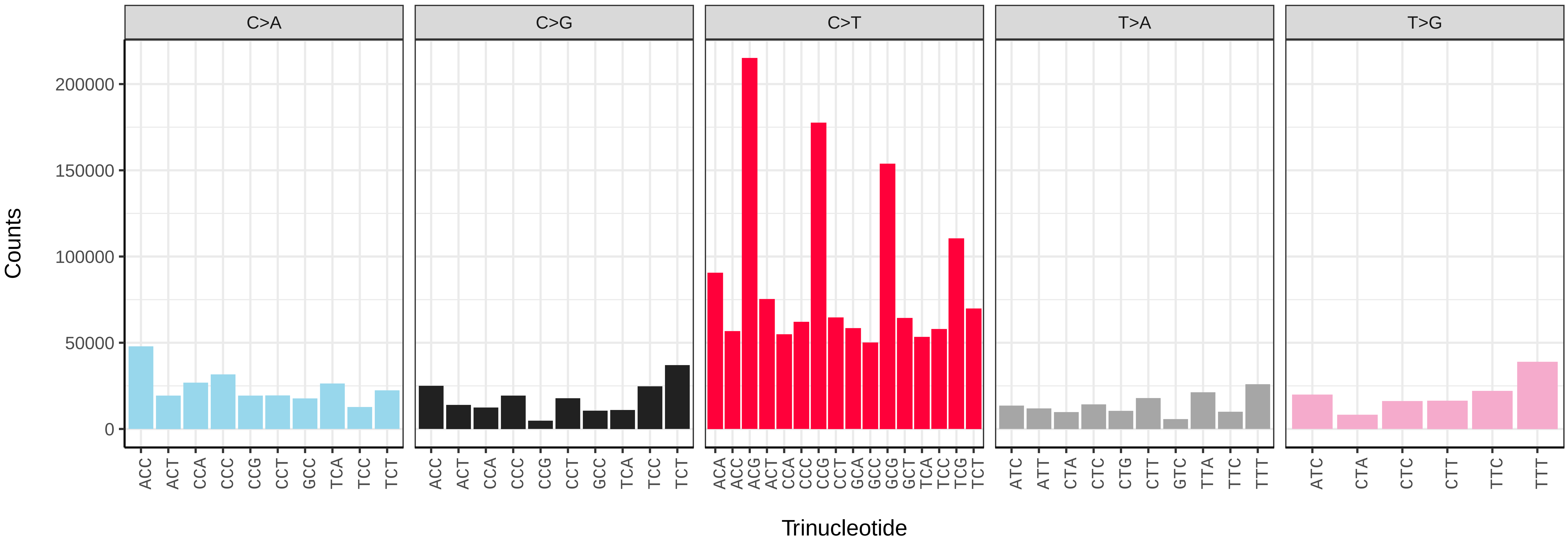


*Phalacrocorax carbo* (bPhaCar2) somatic mutational spectrum (SBS96 classification)

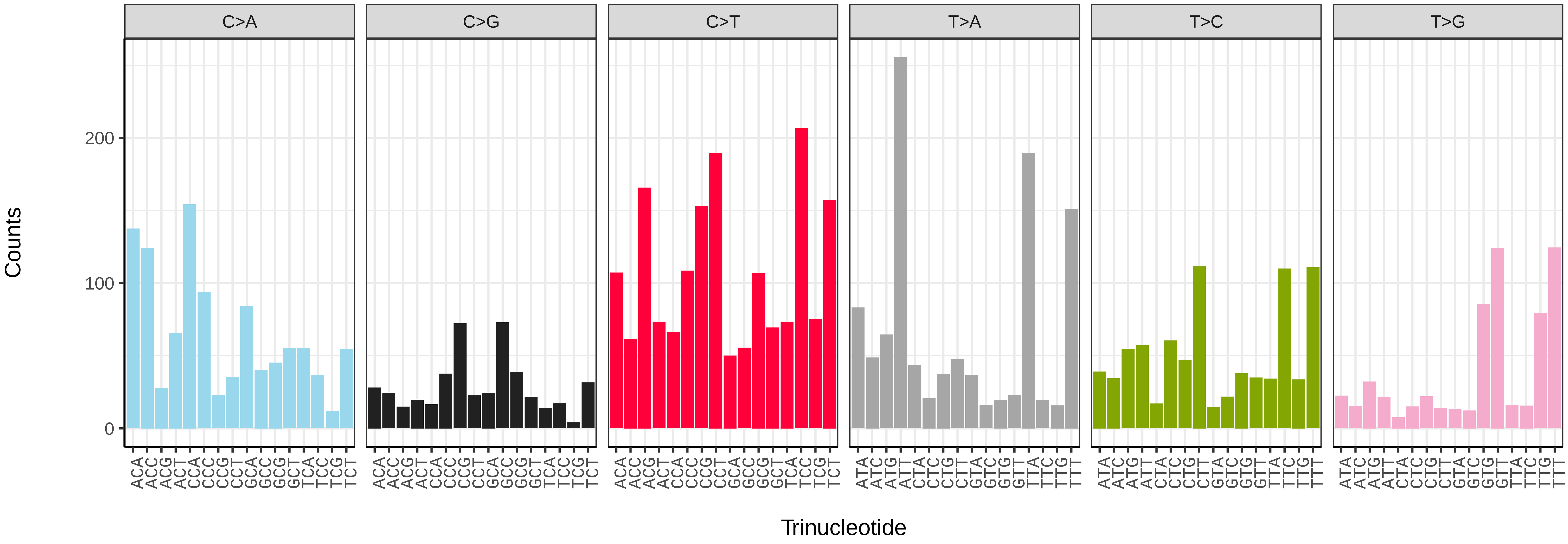


*Phalacrocorax carbo* (bPhaCar2) somatic mutational spectrum (SBS52 classification)

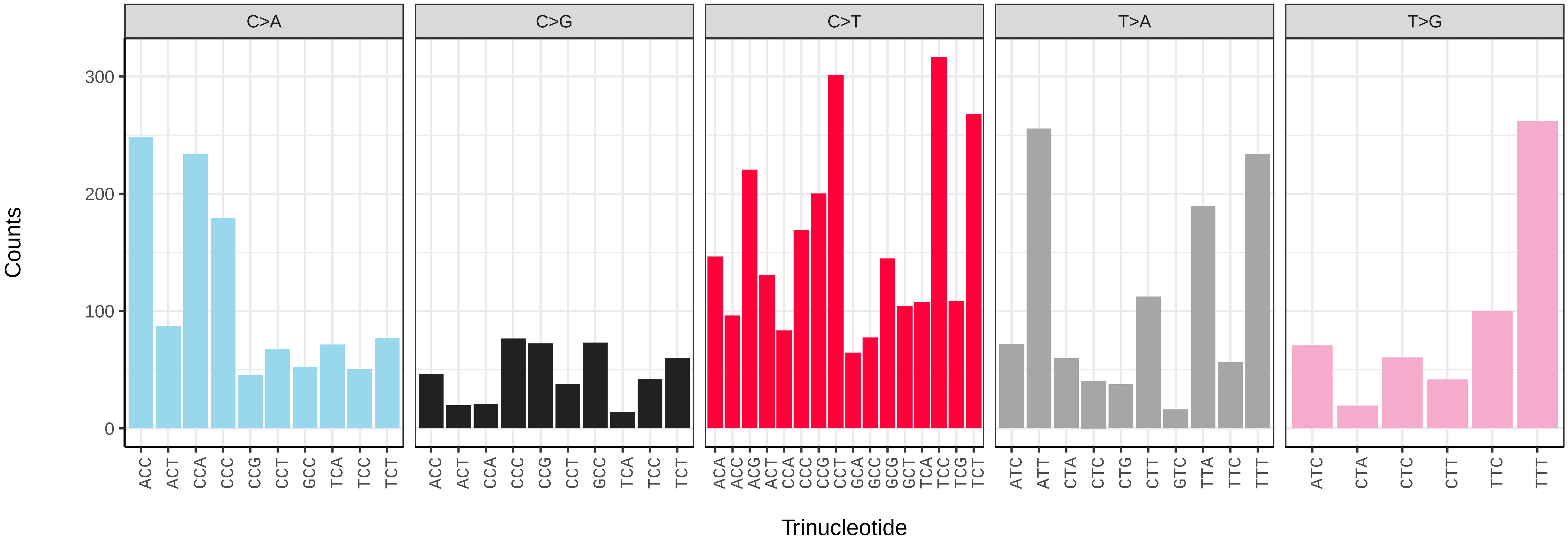

*Tetrao urogallus* (bTetUro1) germline mutational spectrum

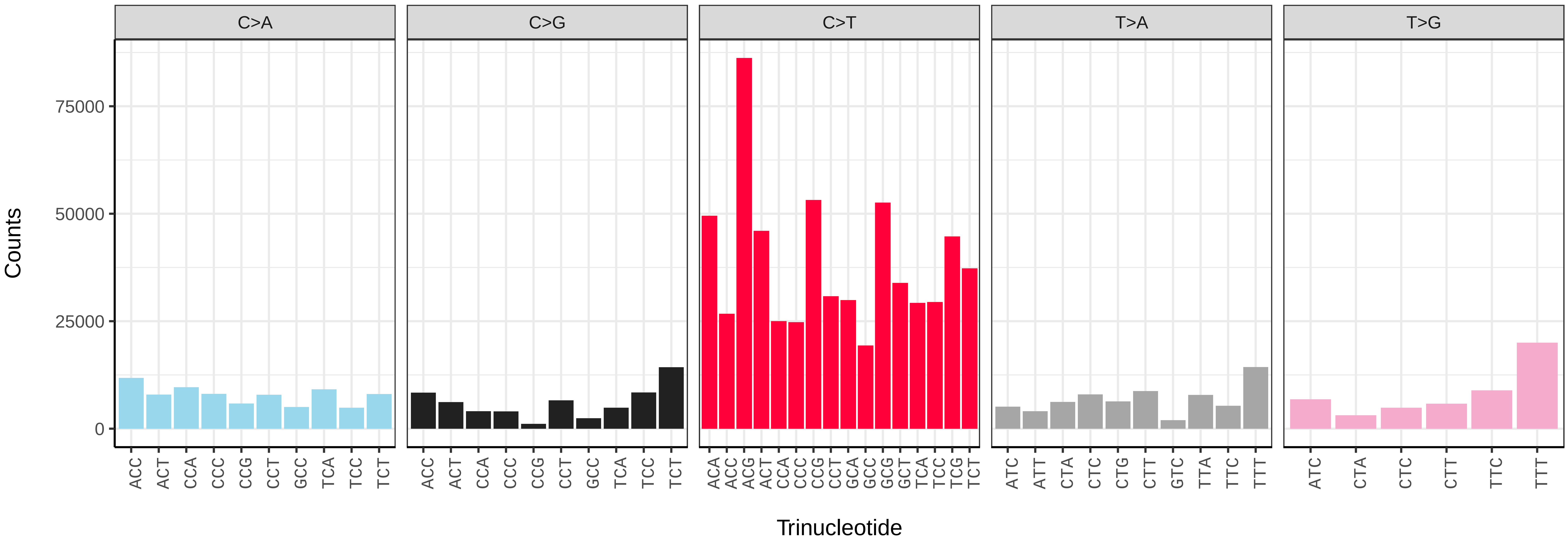


*Tetrao urogallus* (bTetUro1) somatic mutational spectrum (SBS96 classification)

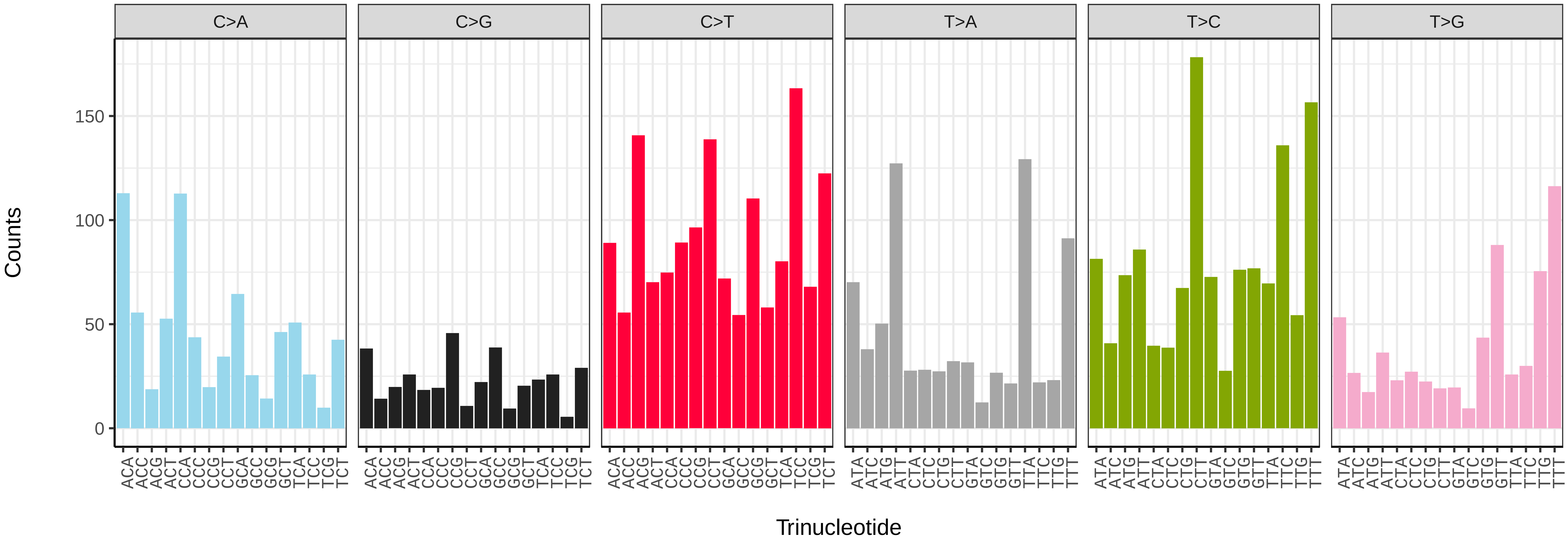


*Tetrao urogallus* (bTetUro1) somatic mutational spectrum (SBS52 classification)

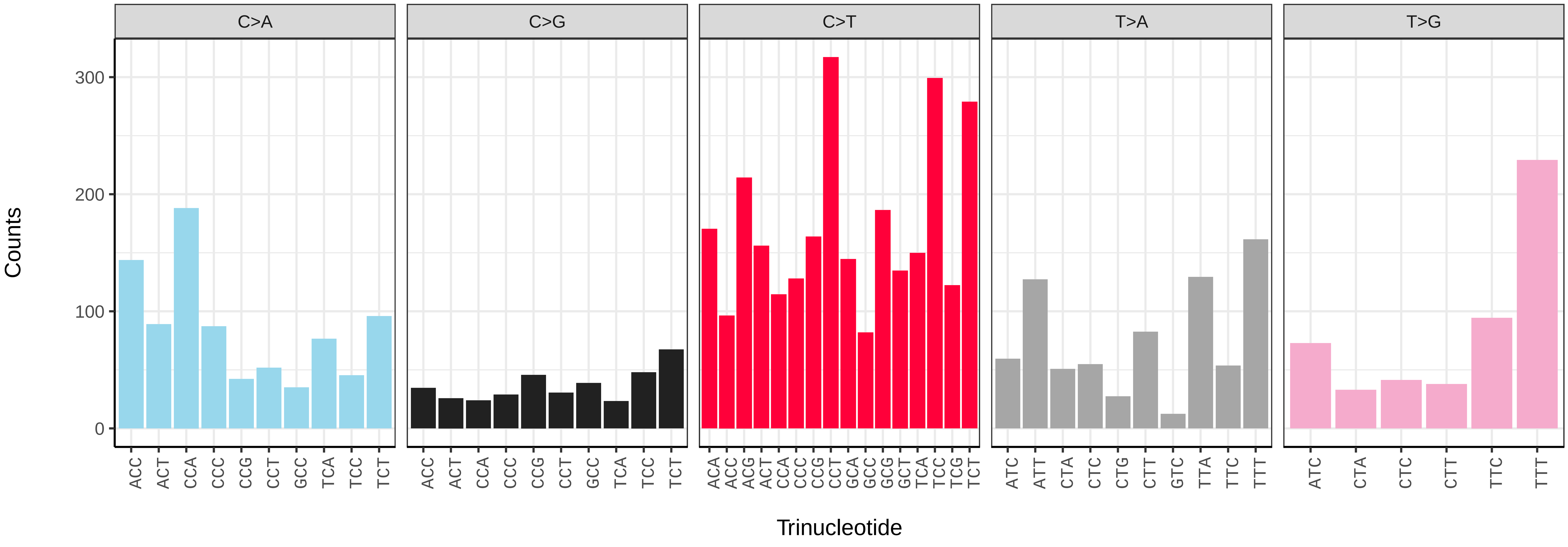

*Ajuga chamaepitys* (daAjuCham1) germline mutational spectrum

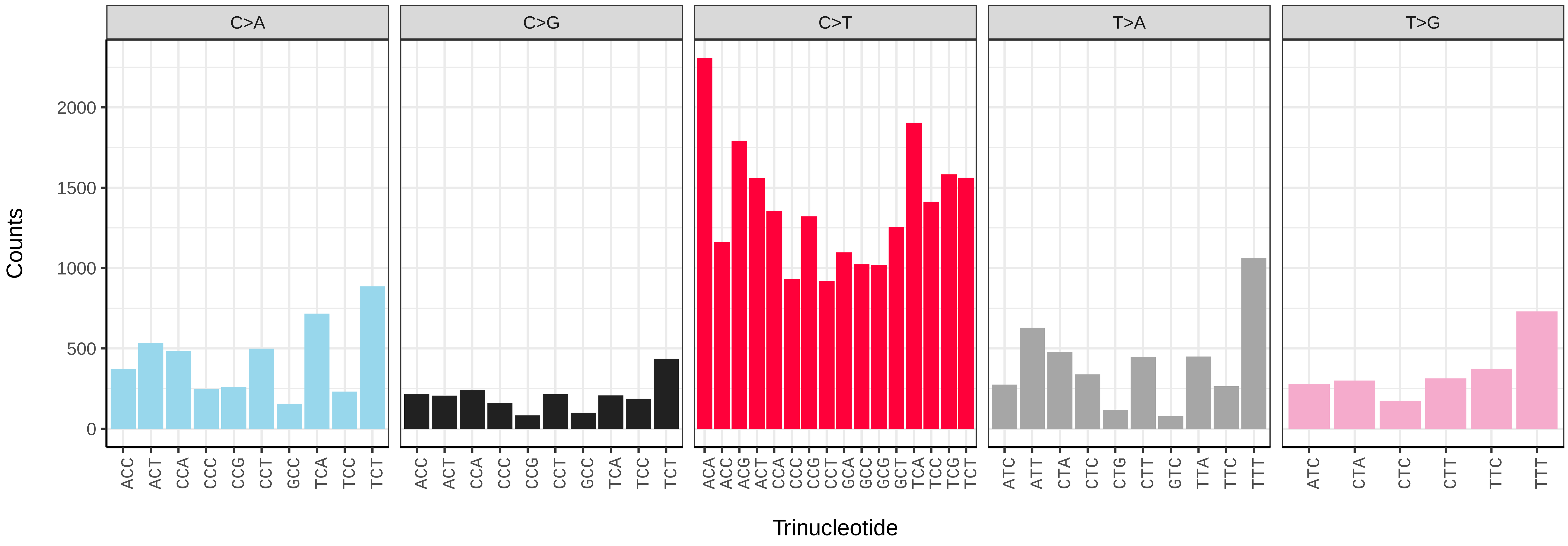


*Ajuga chamaepitys* (daAjuCham1) somatic mutational spectrum (SBS96 classification)

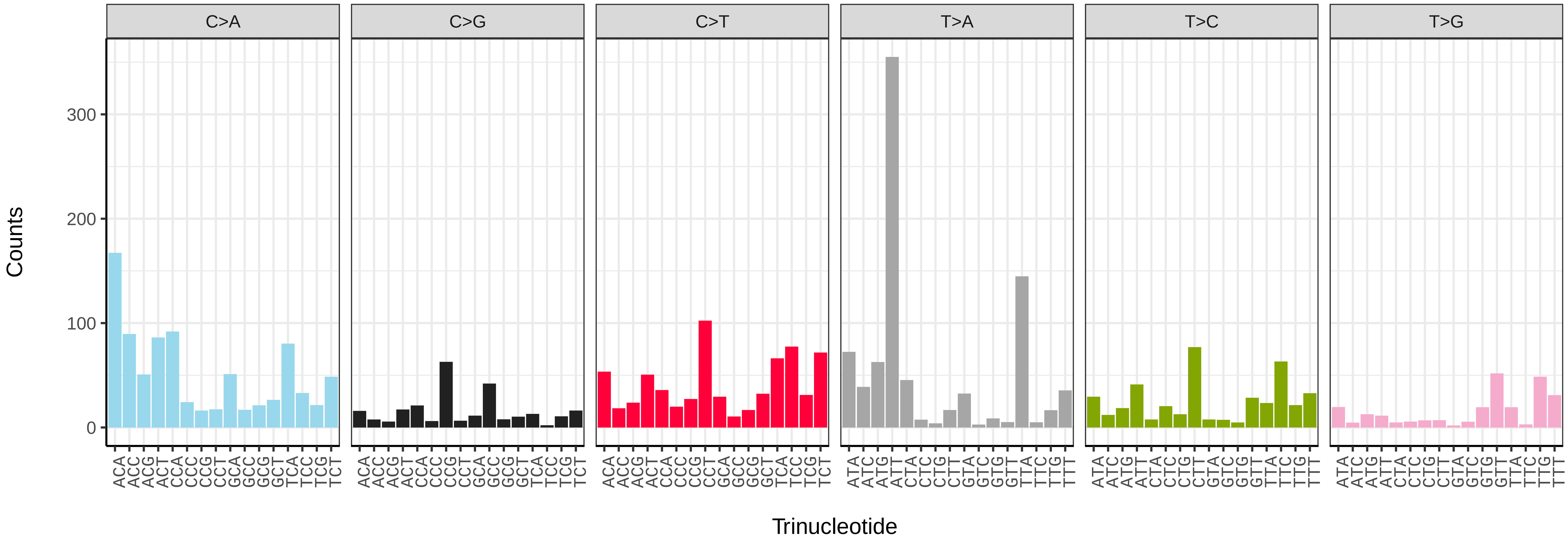


*Ajuga chamaepitys* (daAjuCham1) somatic mutational spectrum (SBS52 classification)

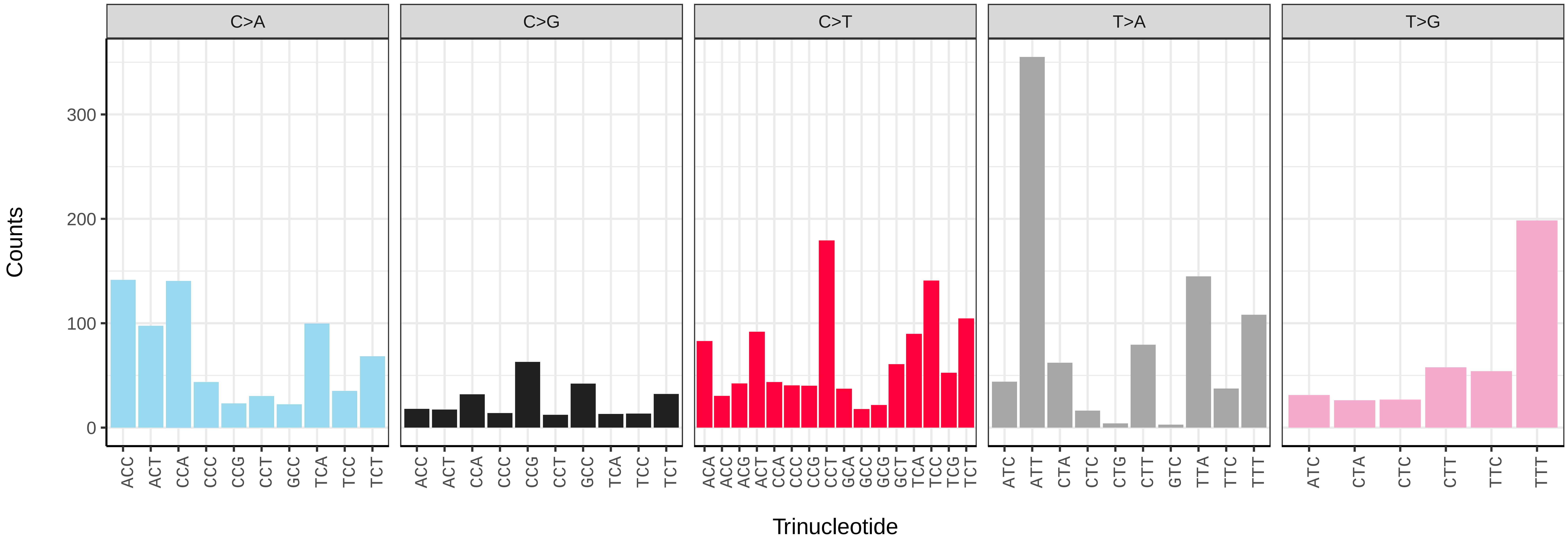

*Arctium minus* (daArcMinu1) germline mutational spectrum

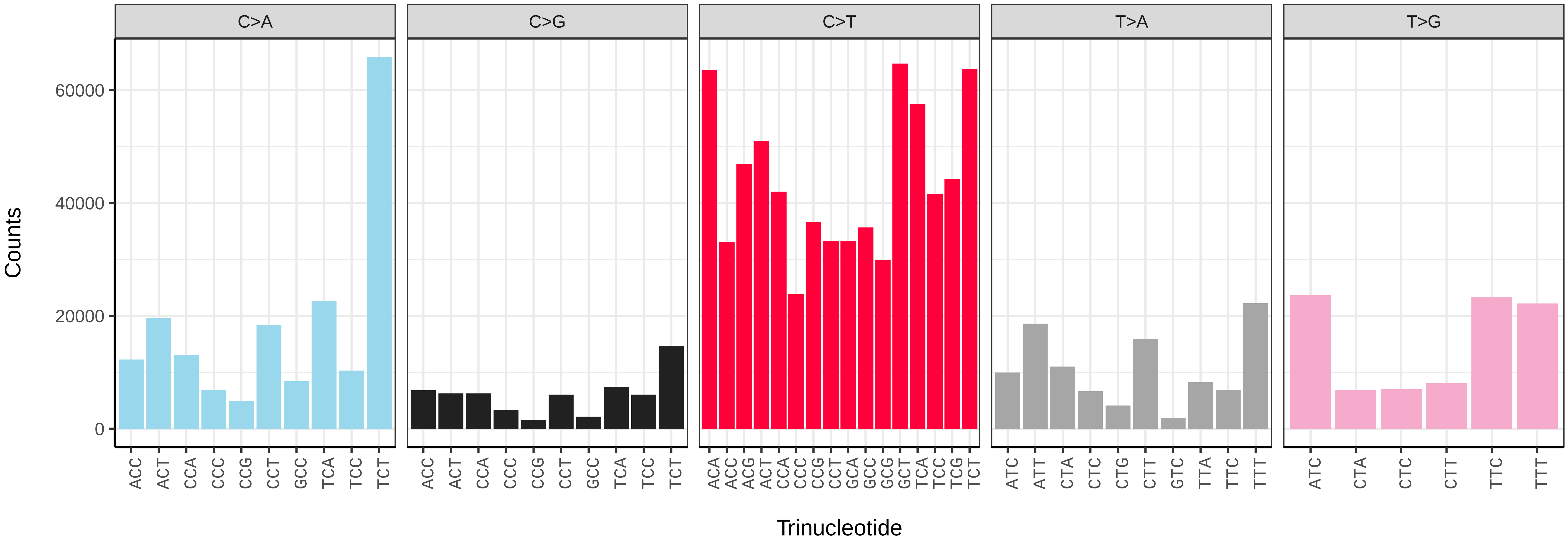


*Arctium minus* (daArcMinu1) somatic mutational spectrum (SBS96 classification)

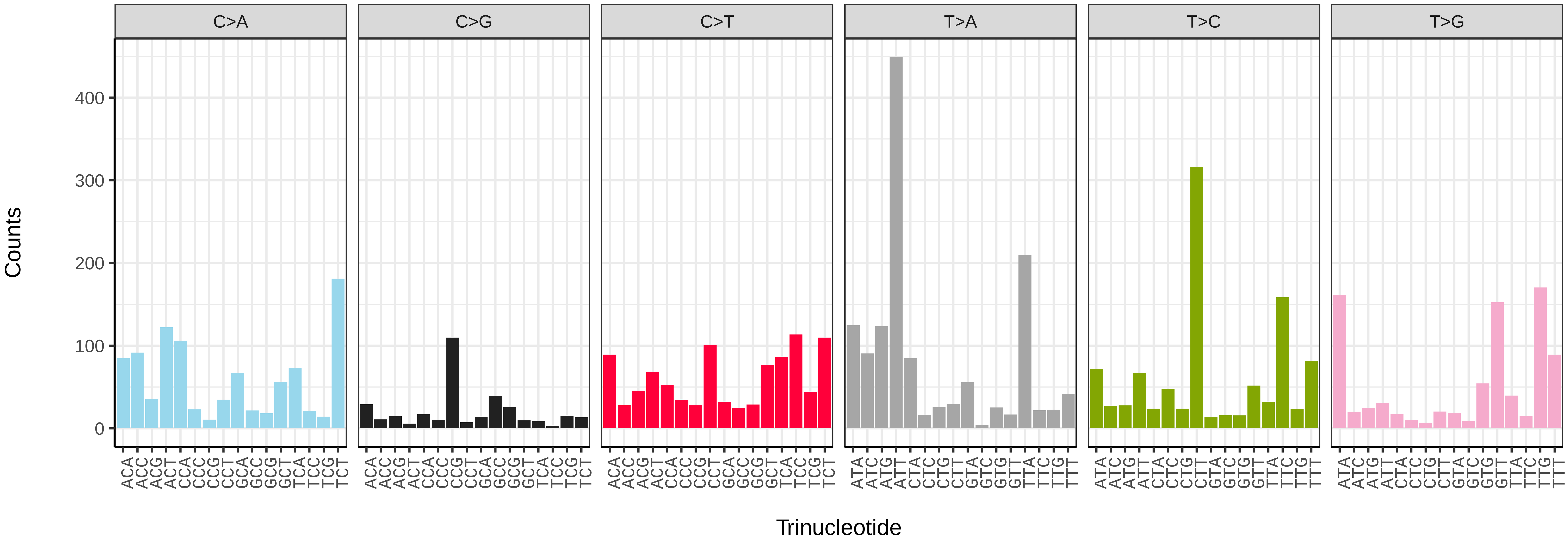


*Arctium minus* (daArcMinu1) somatic mutational spectrum (SBS52 classification)

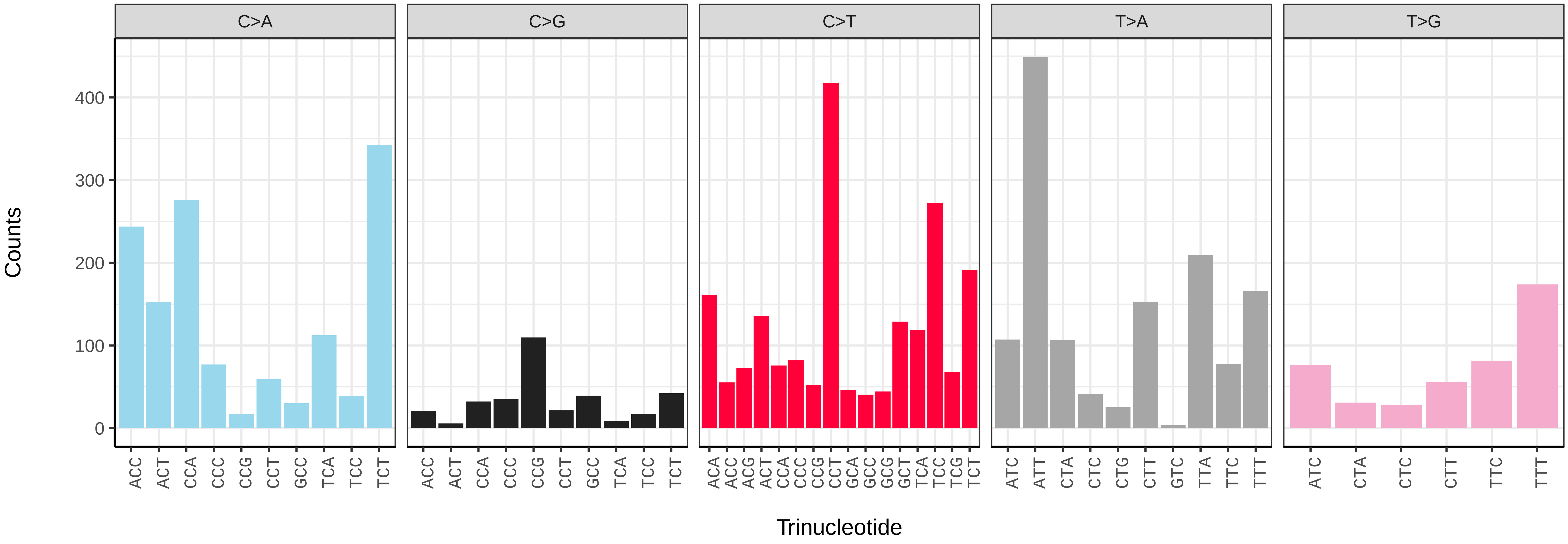

*Ballota nigra* (daBalNigr1) germline mutational spectrum

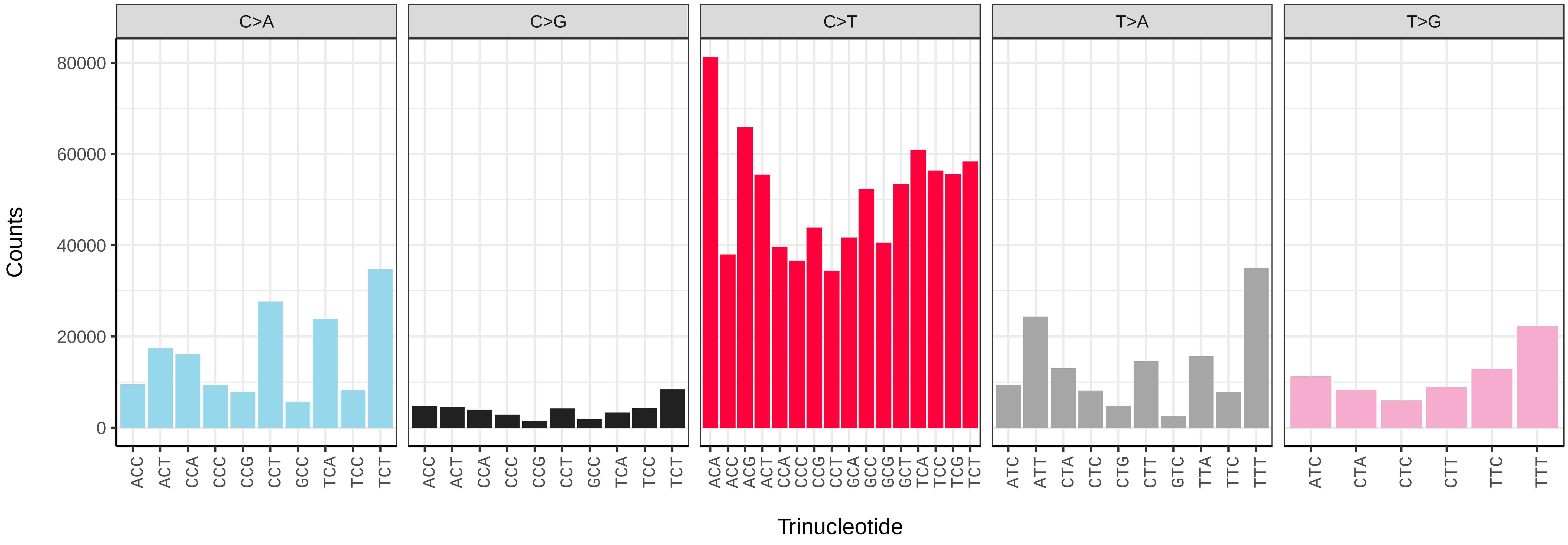


*Ballota nigra* (daBalNigr1) somatic mutational spectrum (SBS96 classification)

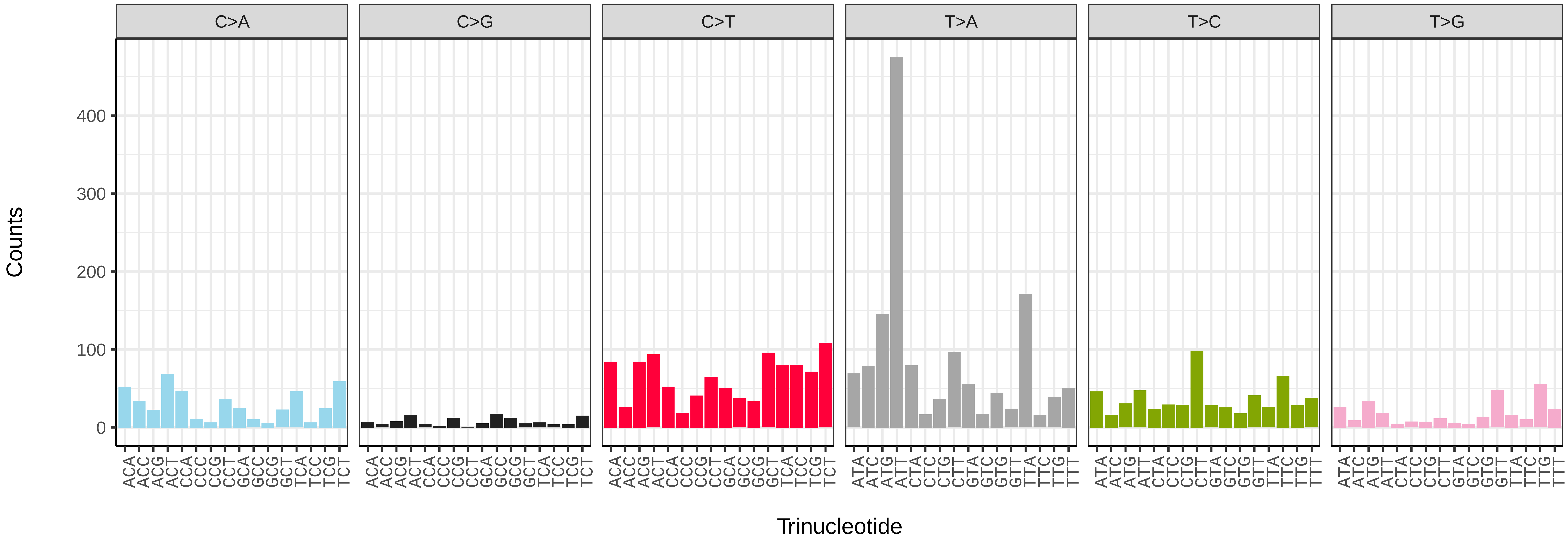


*Ballota nigra* (daBalNigr1) somatic mutational spectrum (SBS52 classification)

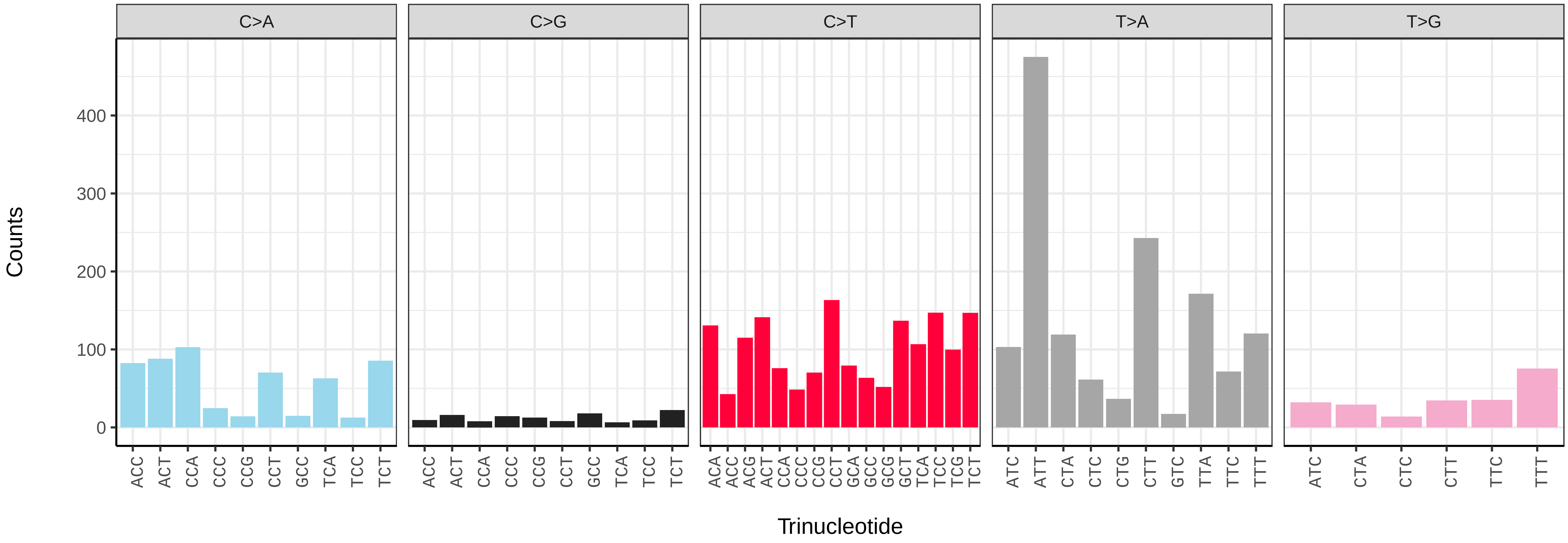

*Gnaphalium uliginosum* (daGnaUlig1) germline mutational spectrum

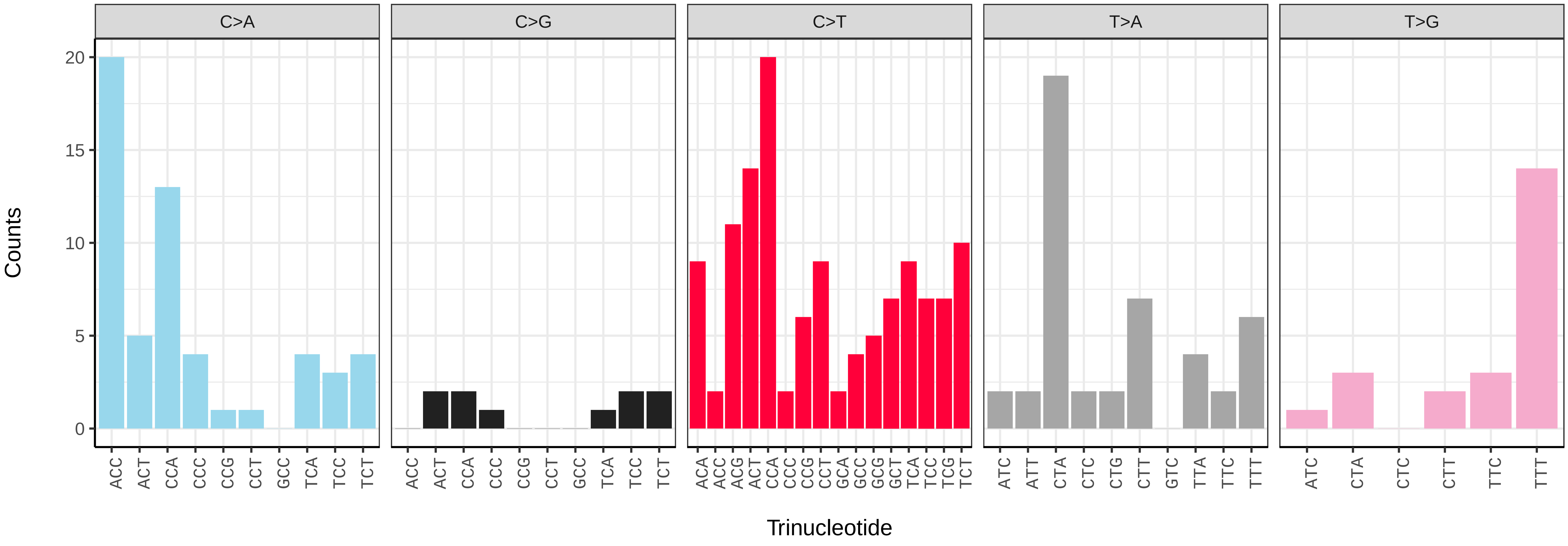


*Gnaphalium uliginosum* (daGnaUlig1) somatic mutational spectrum (SBS96 classification)

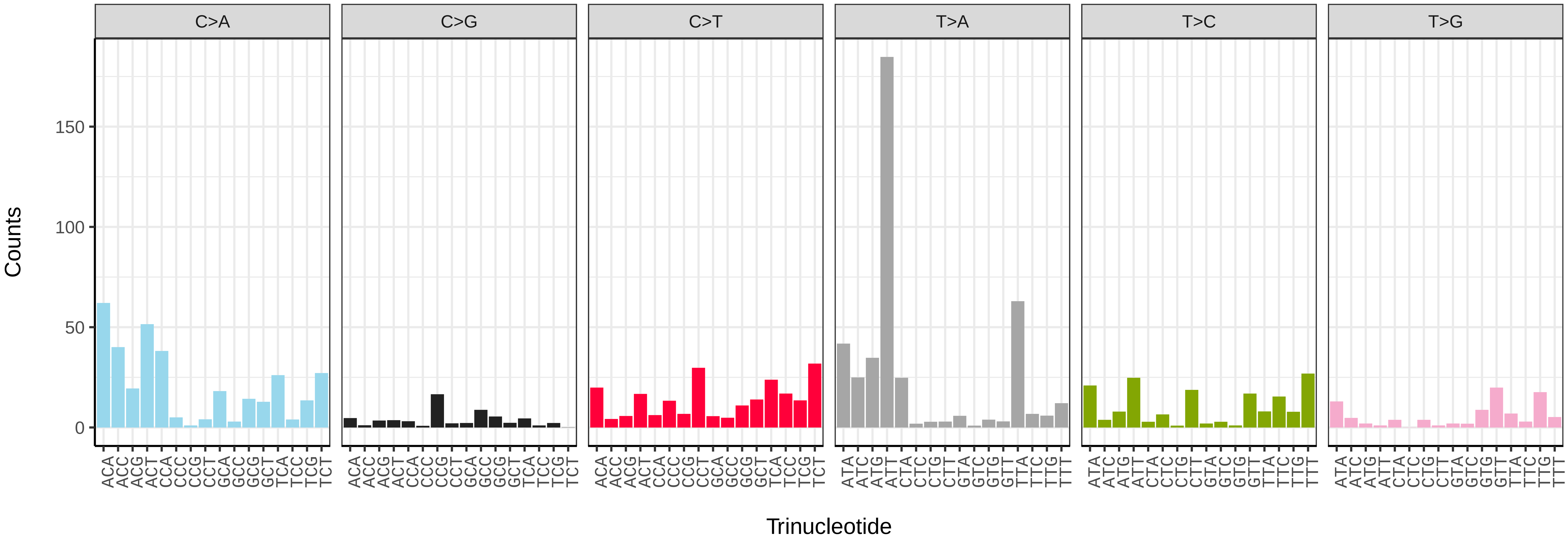


*Gnaphalium uliginosum* (daGnaUlig1) somatic mutational spectrum (SBS52 classification)

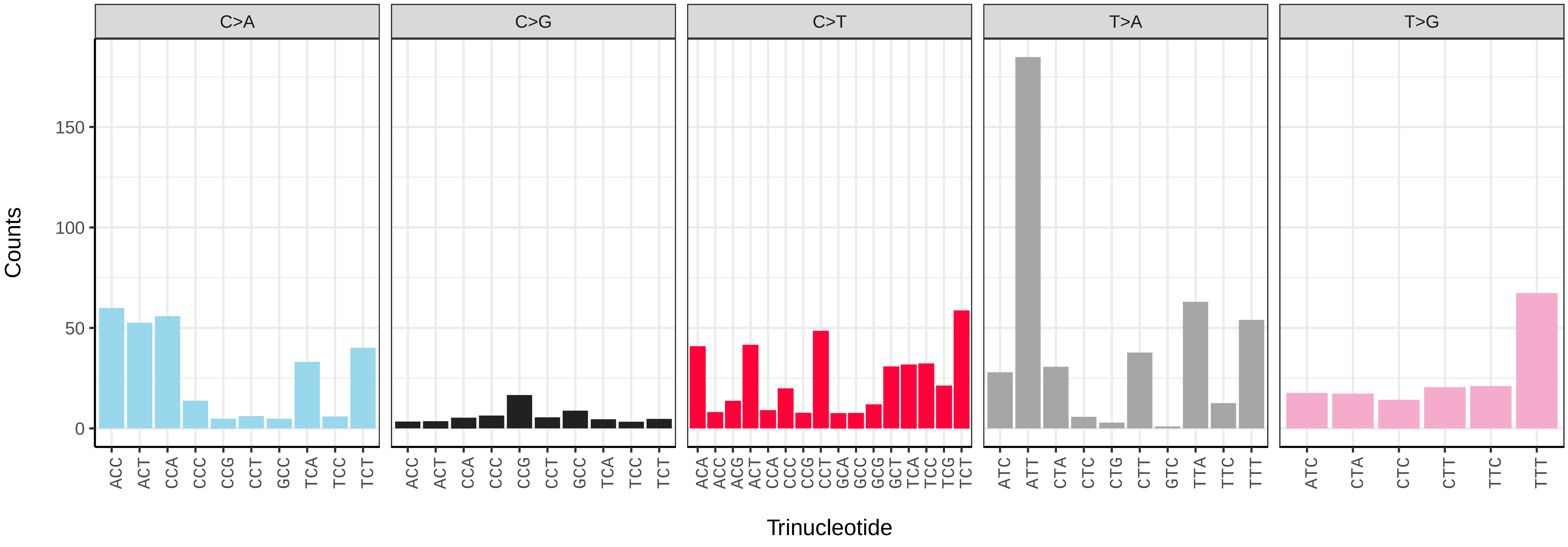

*Ligustrum vulgare* (daLigVulg1) germline mutational spectrum

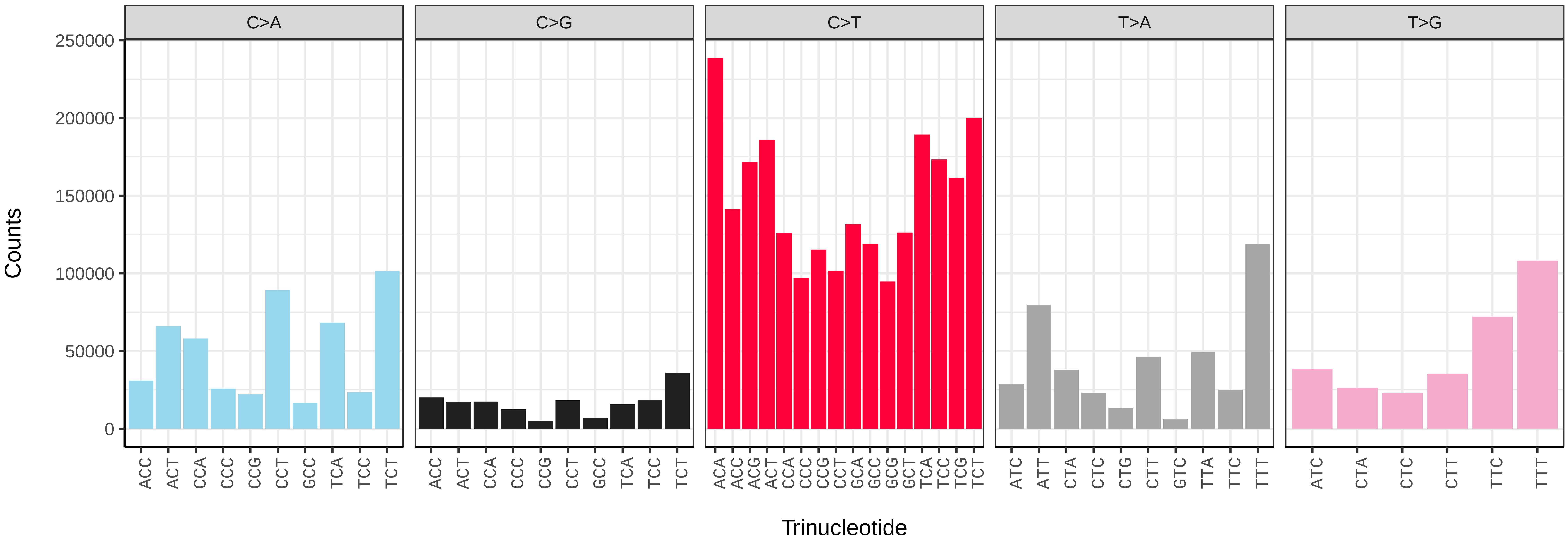


*Ligustrum vulgare* (daLigVulg1) somatic mutational spectrum (SBS96 classification)

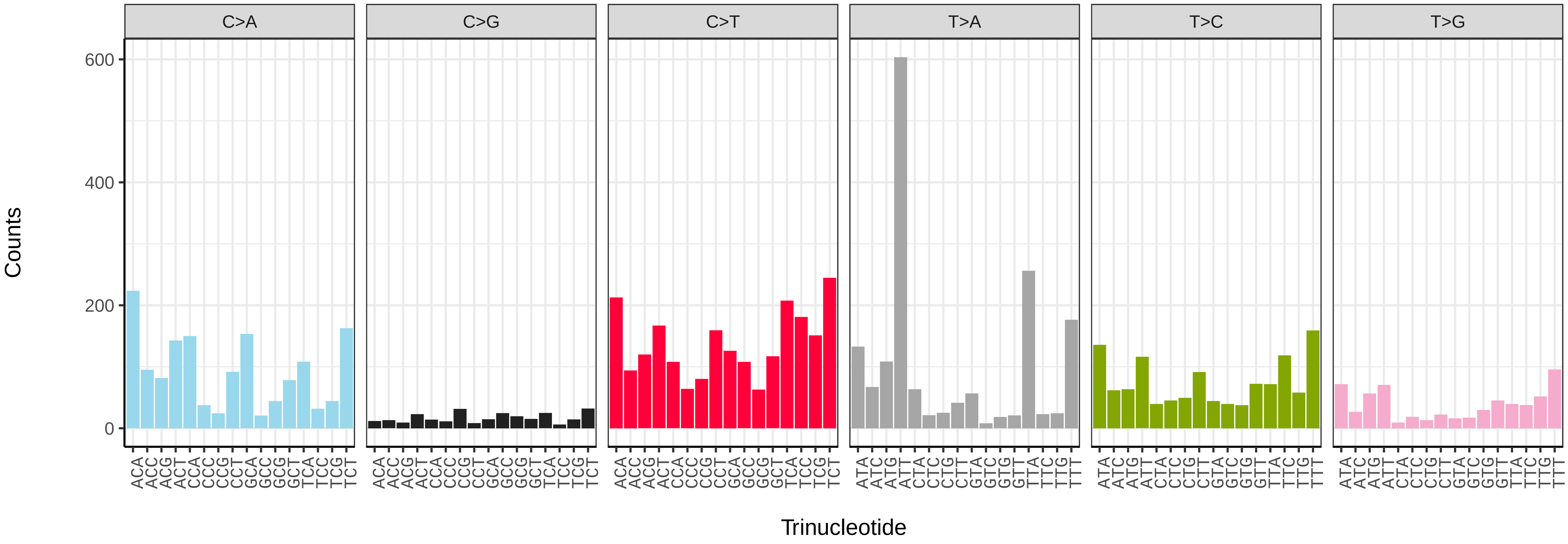


*Ligustrum vulgare* (daLigVulg1) somatic mutational spectrum (SBS52 classification)

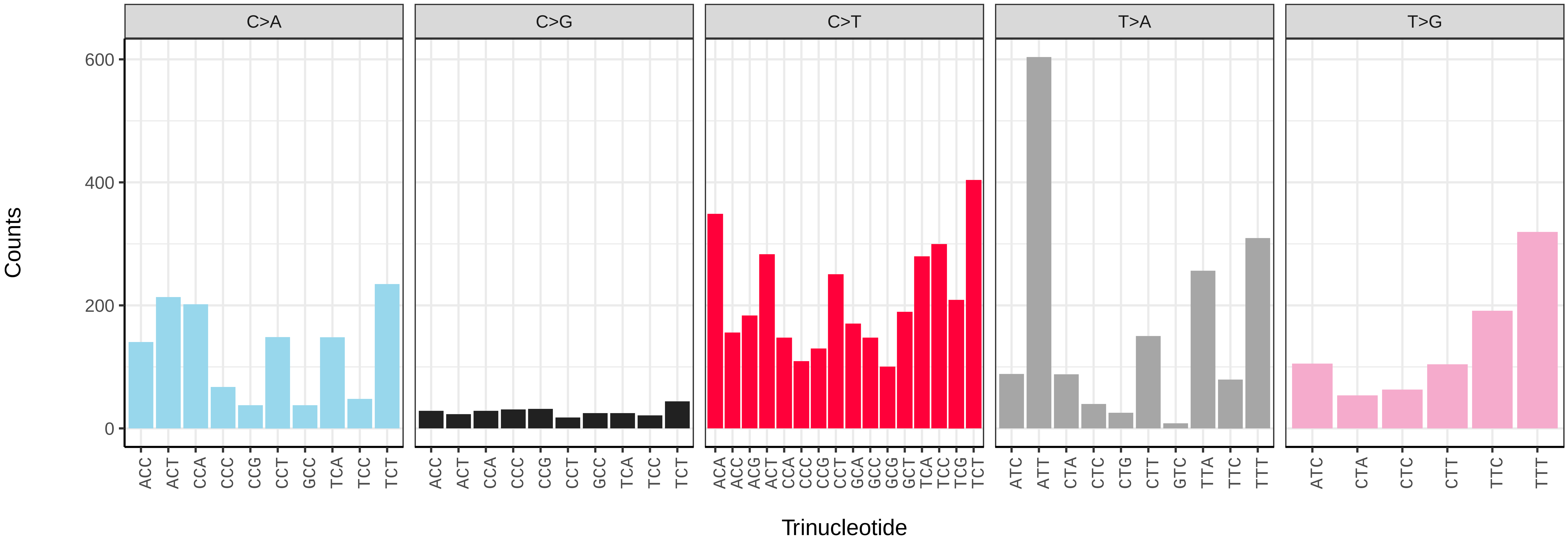

*Linaria vulgaris* (daLinVulg1) germline mutational spectrum

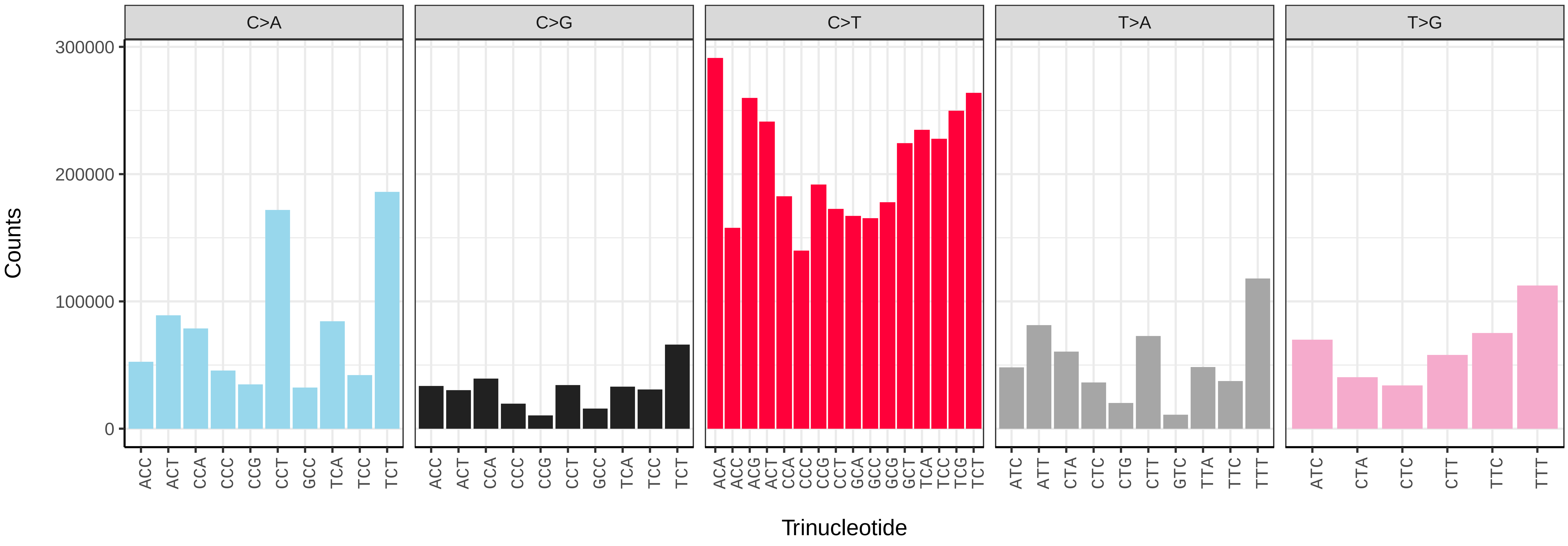


*Linaria vulgaris* (daLinVulg1) somatic mutational spectrum (SBS96 classification)

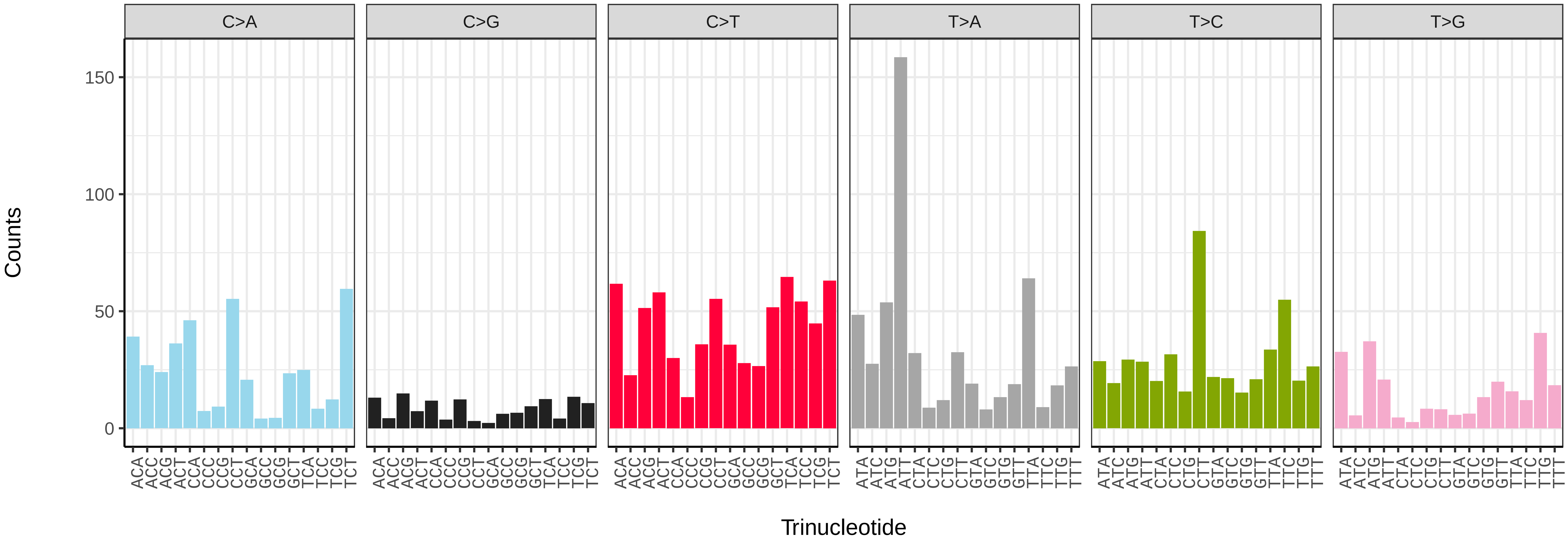


*Linaria vulgaris* (daLinVulg1) somatic mutational spectrum (SBS52 classification)

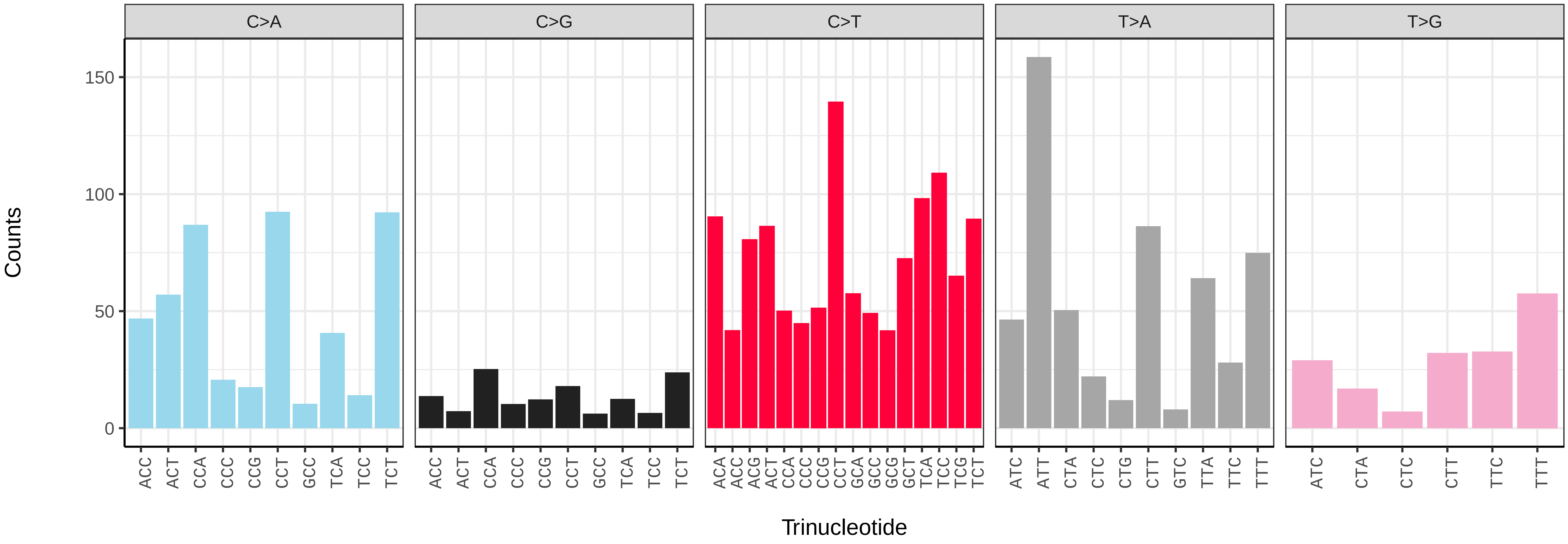

*Misopates orontium* (daMisOron1) germline mutational spectrum

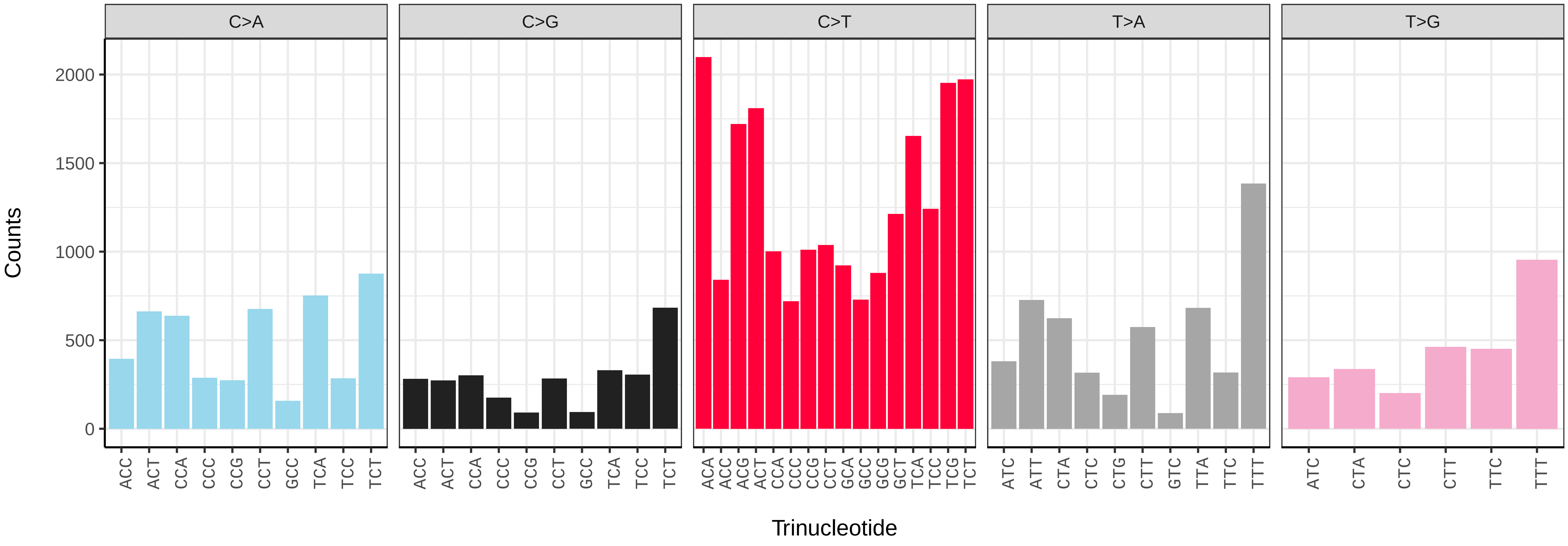


*Misopates orontium* (daMisOron1) somatic mutational spectrum (SBS96 classification)

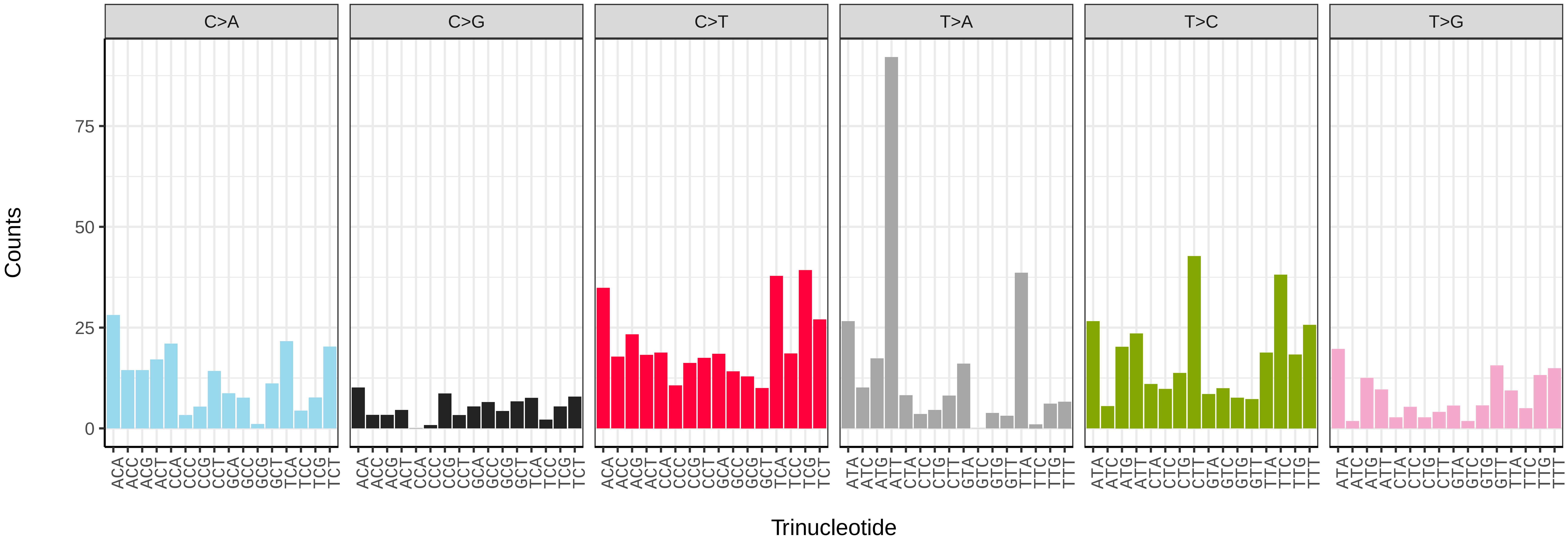


*Misopates orontium* (daMisOron1) somatic mutational spectrum (SBS52 classification)

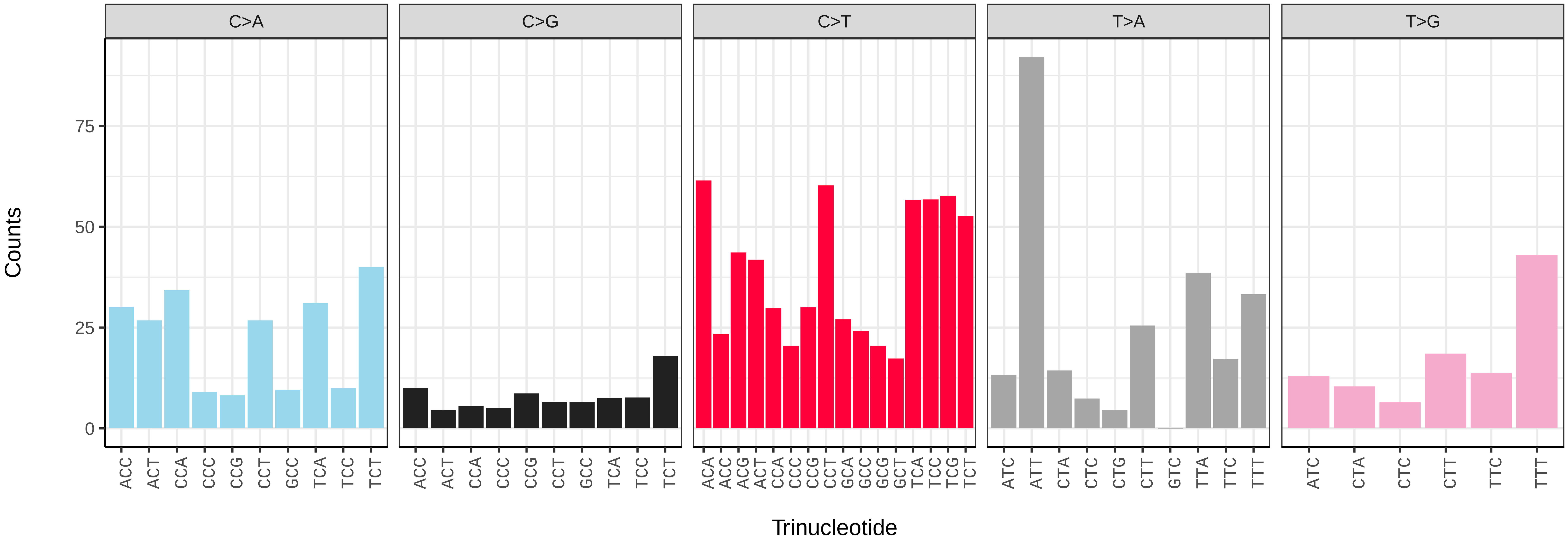

*Pseudognaphalium luteoalbum* (daPseLute1) germline mutational spectrum

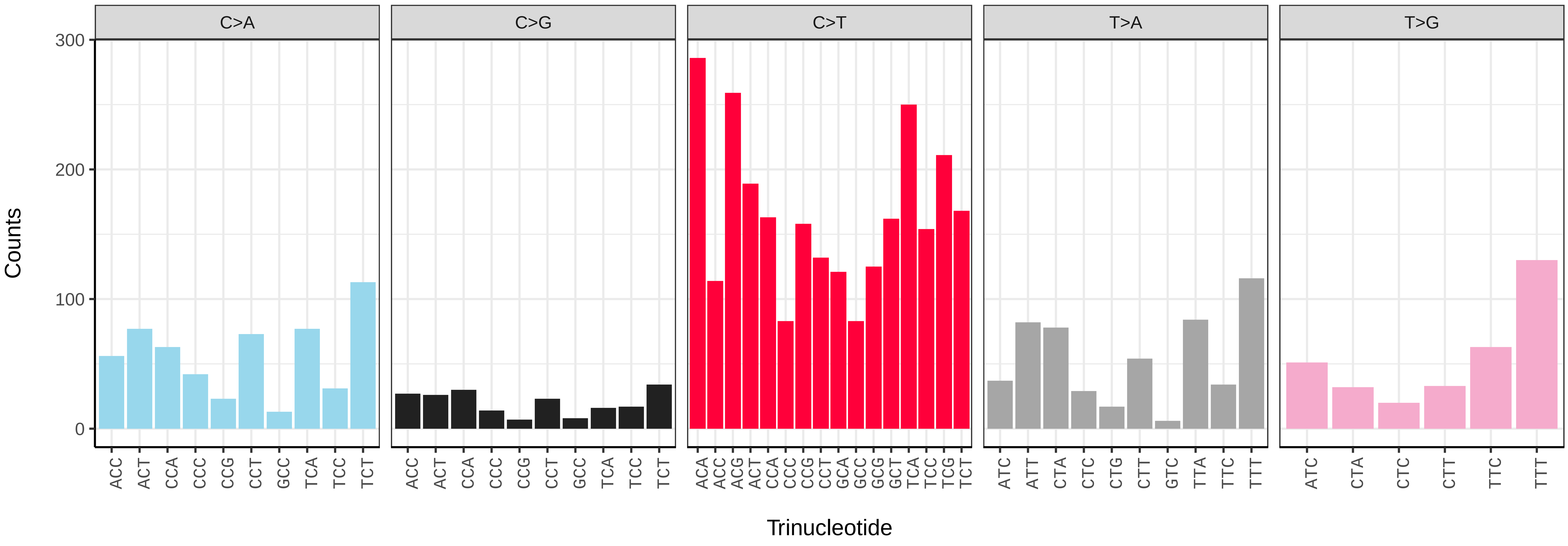


*Pseudognaphalium luteoalbum* (daPseLute1) somatic mutational spectrum (SBS96 classification)

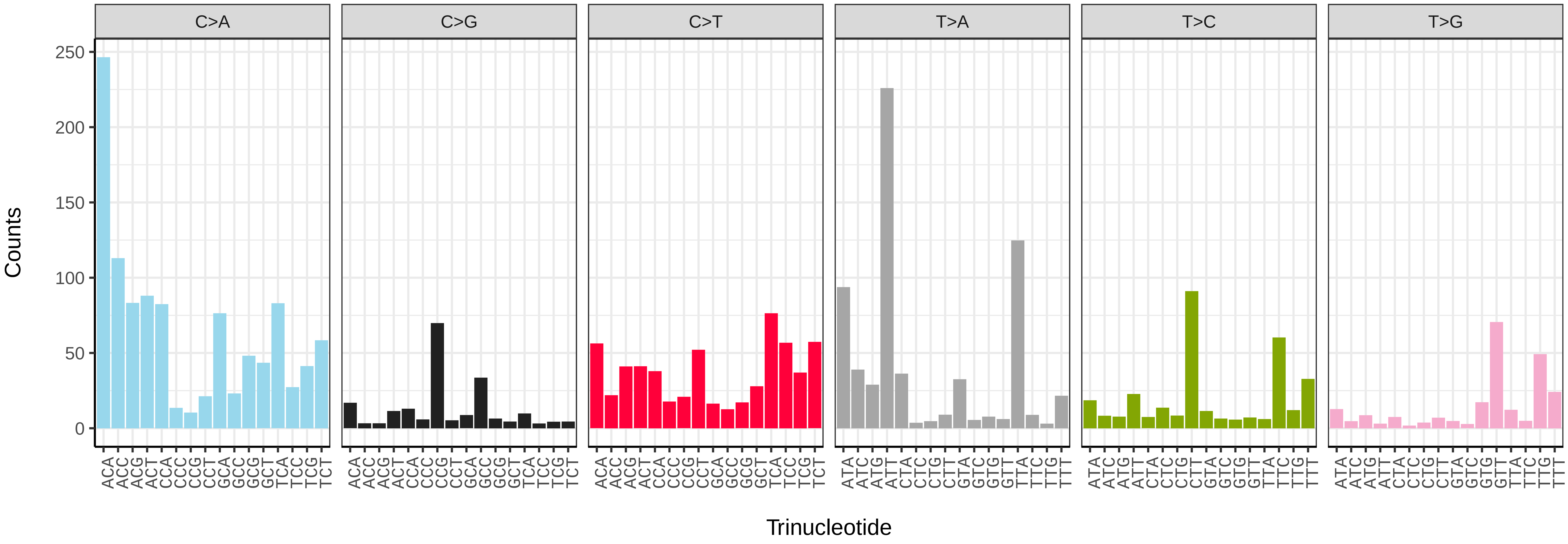


*Pseudognaphalium luteoalbum* (daPseLute1) somatic mutational spectrum (SBS52 classification)

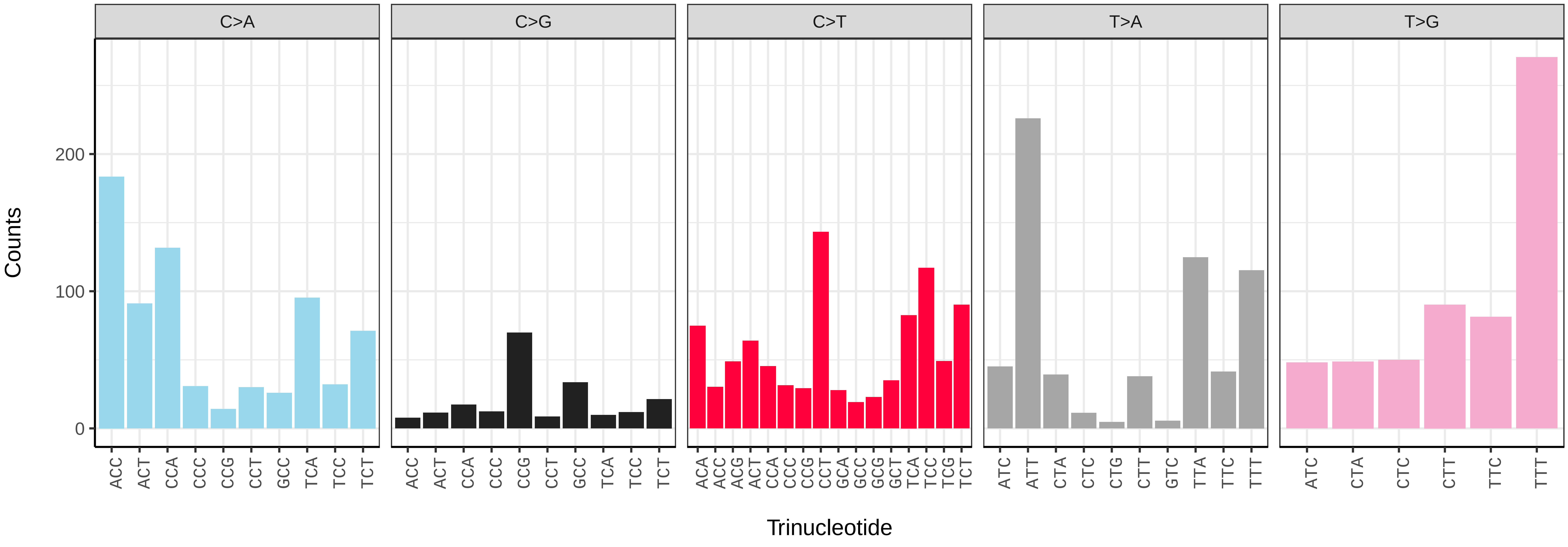

*Pulicaria dysenterica* (daPulDyse1) germline mutational spectrum

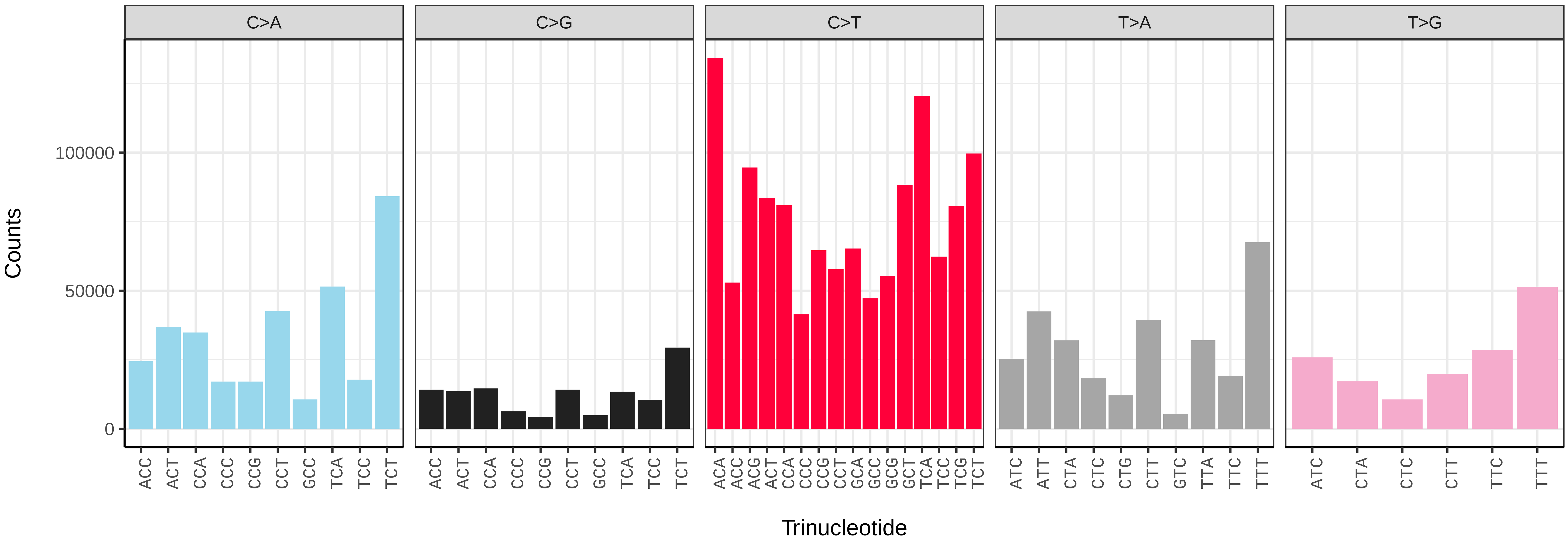


*Pulicaria dysenterica* (daPulDyse1) somatic mutational spectrum (SBS96 classification)

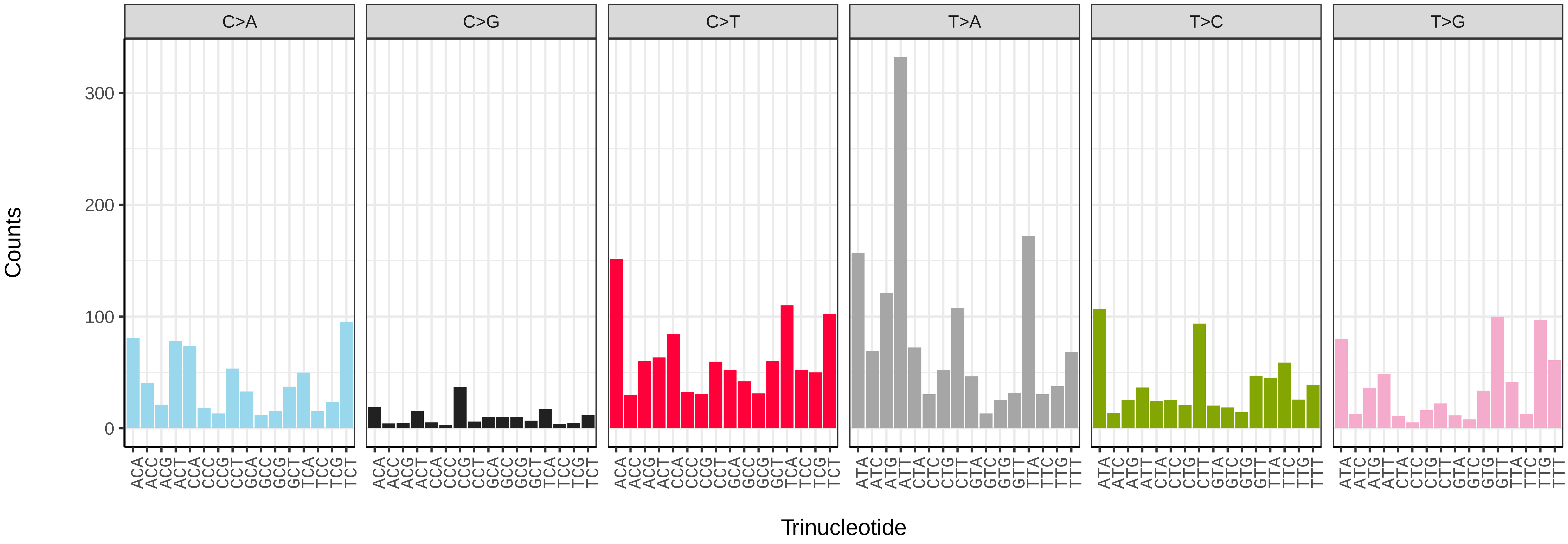


*Pulicaria dysenterica* (daPulDyse1) somatic mutational spectrum (SBS52 classification)

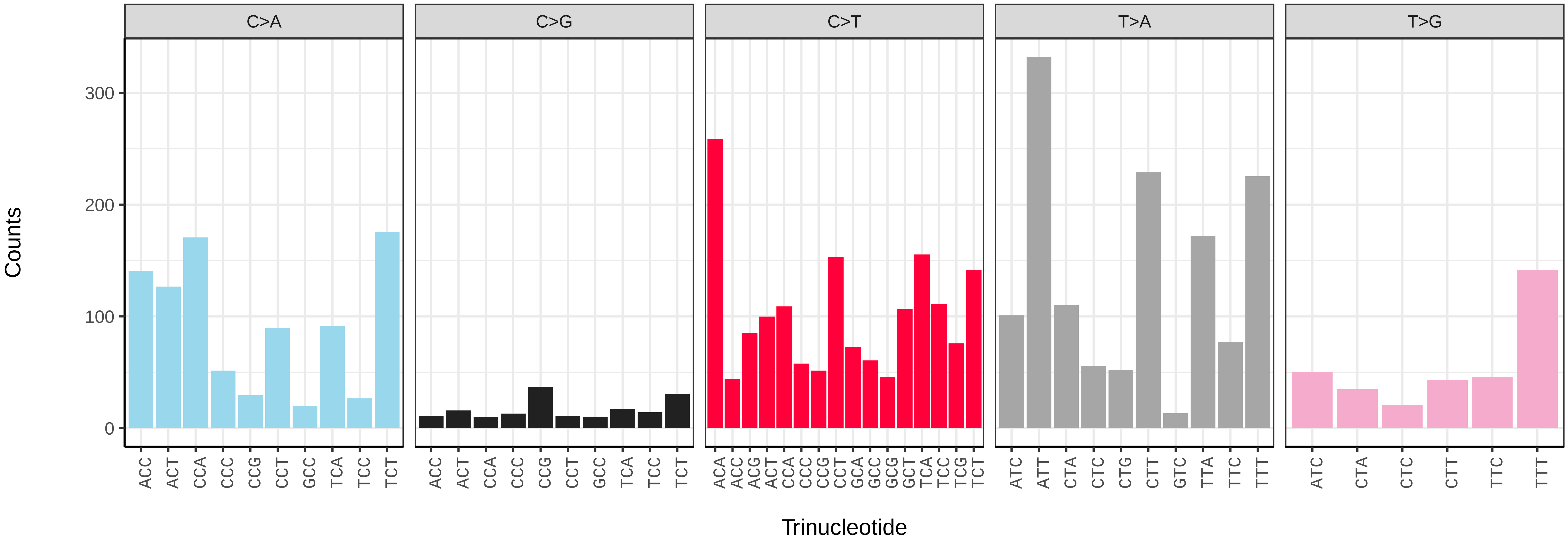

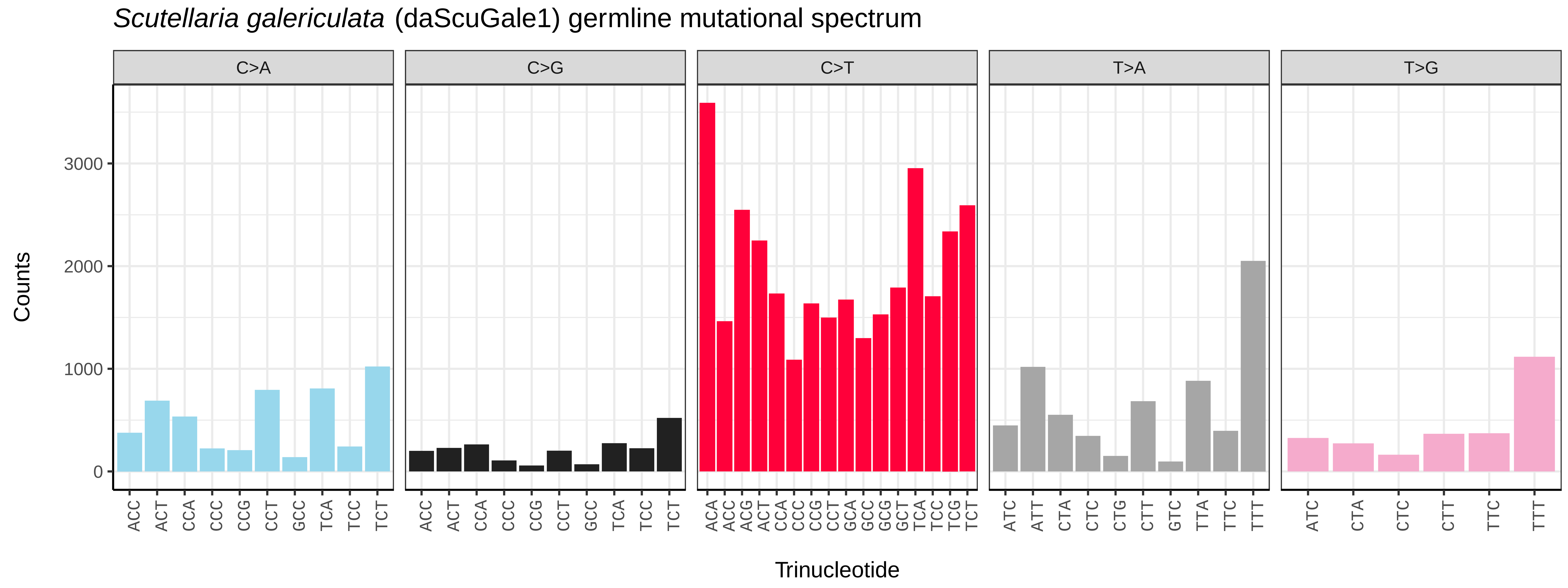

*Scutellaria galericulata* (daScuGale1) germline mutational spectrum
C>A
C>G
C>T
T>A
T>G
Counts
0
1000
2000
3000
ACC ACT CCA CCC CCG CCT GCC TCA TCC TCT
ACA ACC ACG ACT CCA CCC CCG CCT GCA GCC GCG GCT TCA TCC TCG TCT
ATC ATT CTA CTC CTG CTT GTC TTA TTC TTT
ATC CTA CTC CTT TTC TTT
Trinucleotide


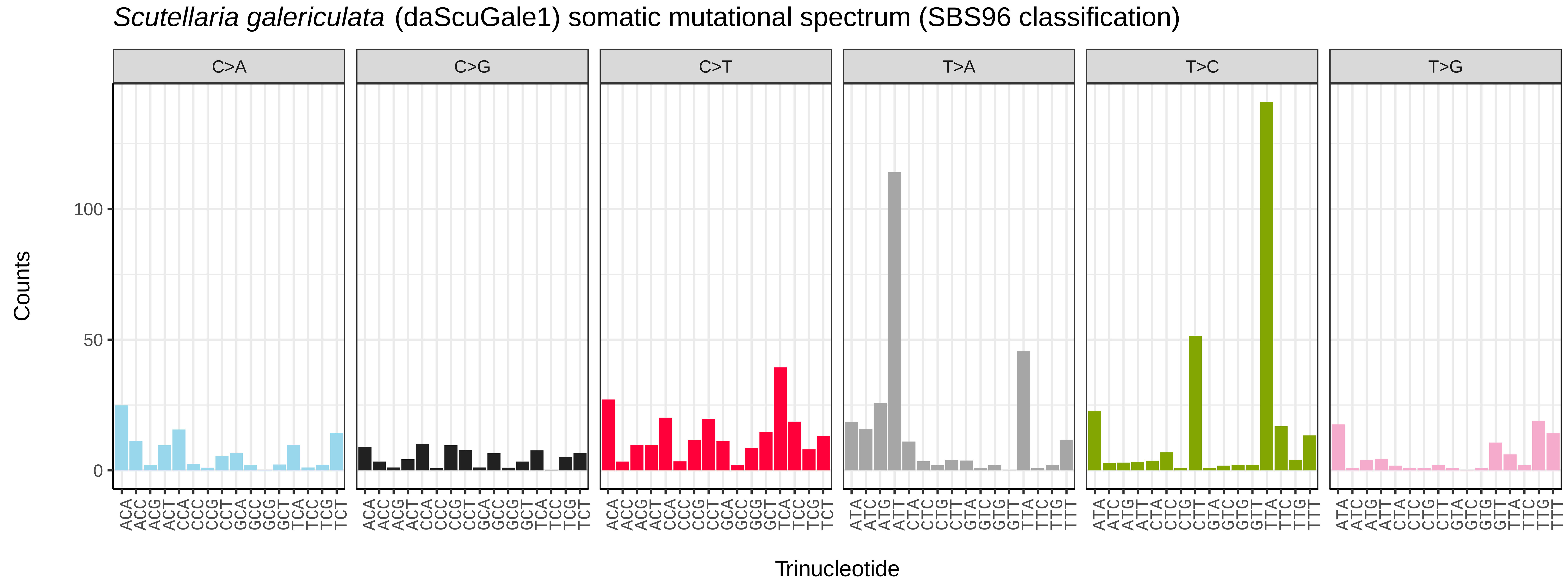

*Scutellaria galericulata* (daScuGale1) somatic mutational spectrum (SBS96 classification)
C>A
C>G
C>T
T>A
T>C
T>G
Counts
0
50
100
Trinucleotide


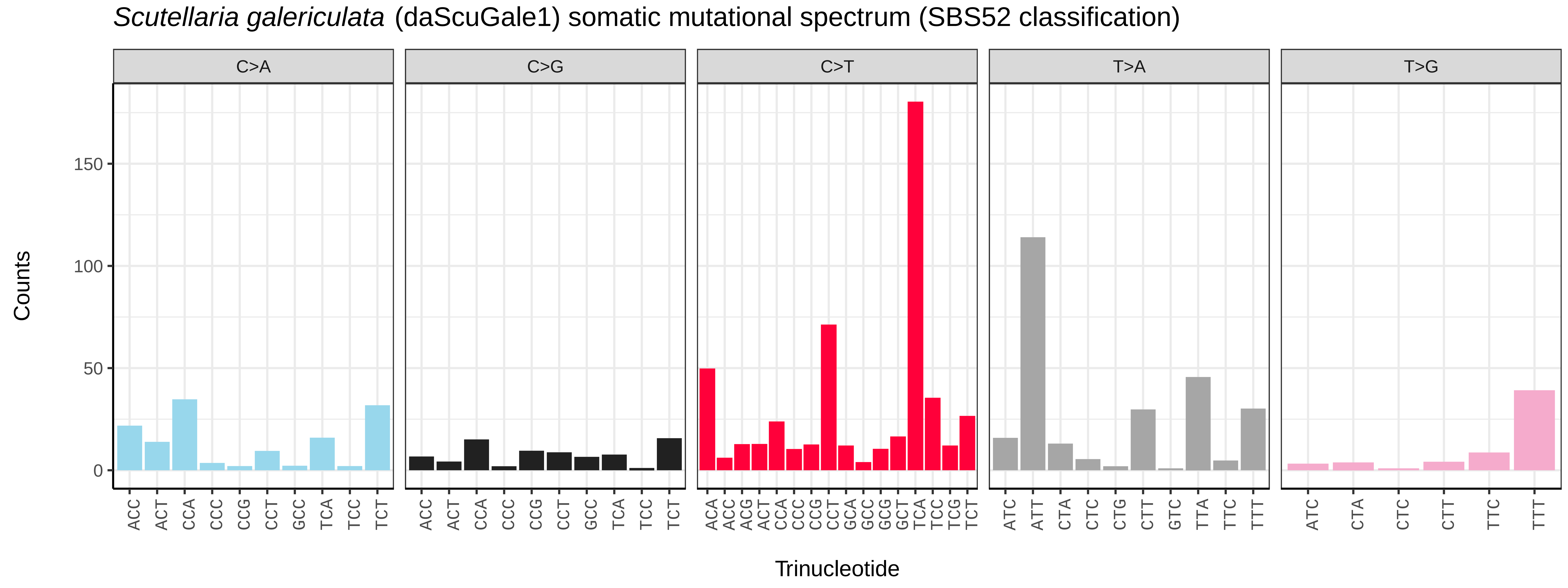

*Scutellaria galericulata* (daScuGale1) somatic mutational spectrum (SBS52 classification)
C>A
C>G
C>T
T>A
T>G
Counts
0
50
100
150
ACC ACT CCA CCC CCG CCT GCC TCA TCC TCT
ACA ACC ACG ACT CCA CCC CCG CCT GCA GCC GCG GCT TCA TCC TCG TCT
ATC ATT CTA CTC CTG CTT GTC TTA TTC TTT
ATC CTA CTC CTT TTC TTT
Trinucleotide

*Scutellaria minor* (daScuMino1) germline mutational spectrum

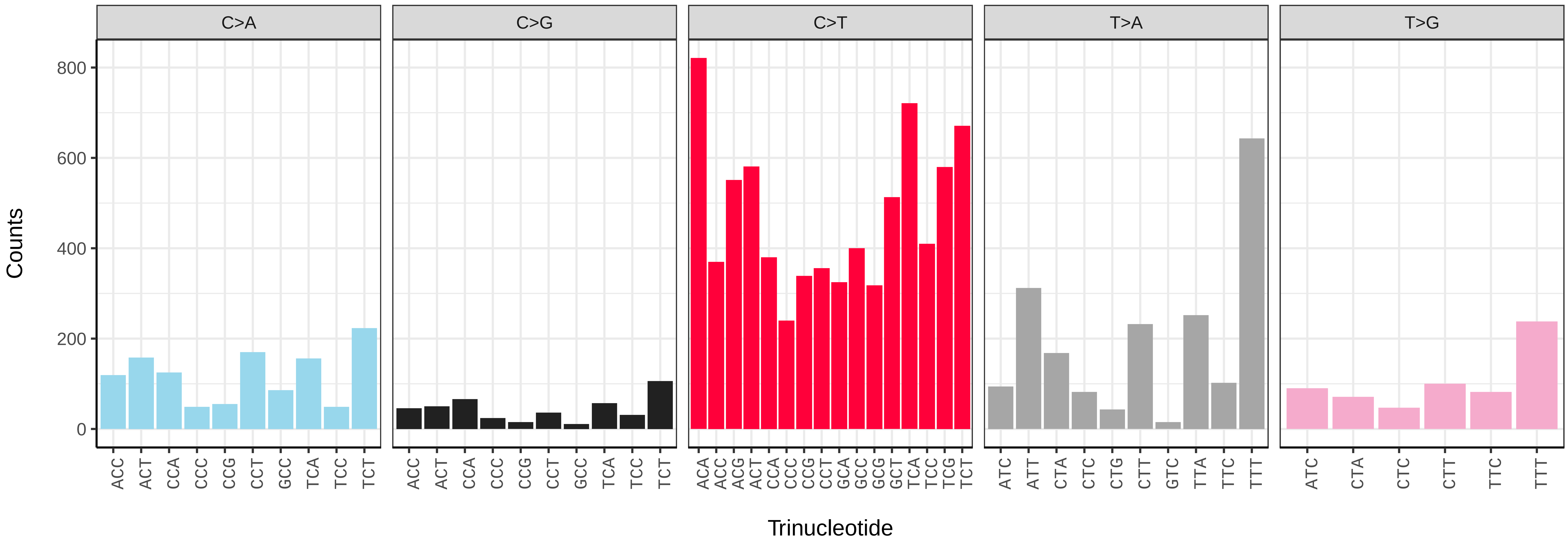


*Scutellaria minor* (daScuMino1) somatic mutational spectrum (SBS96 classification)

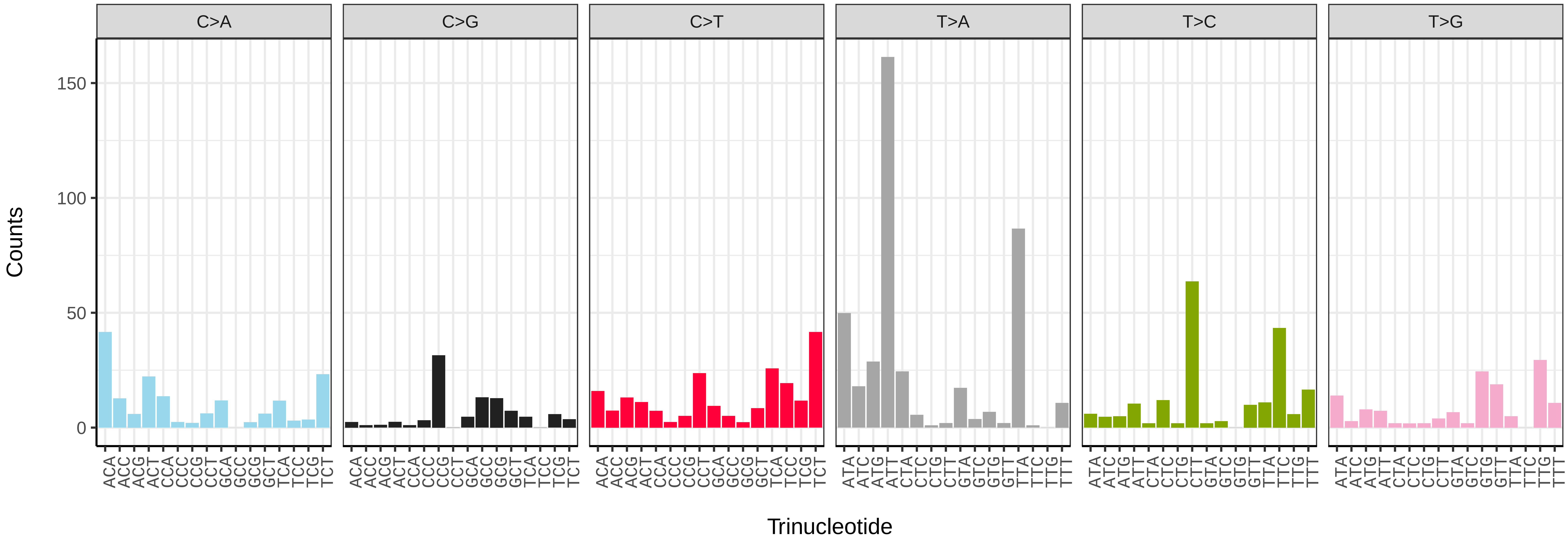


*Scutellaria minor* (daScuMino1) somatic mutational spectrum (SBS52 classification)

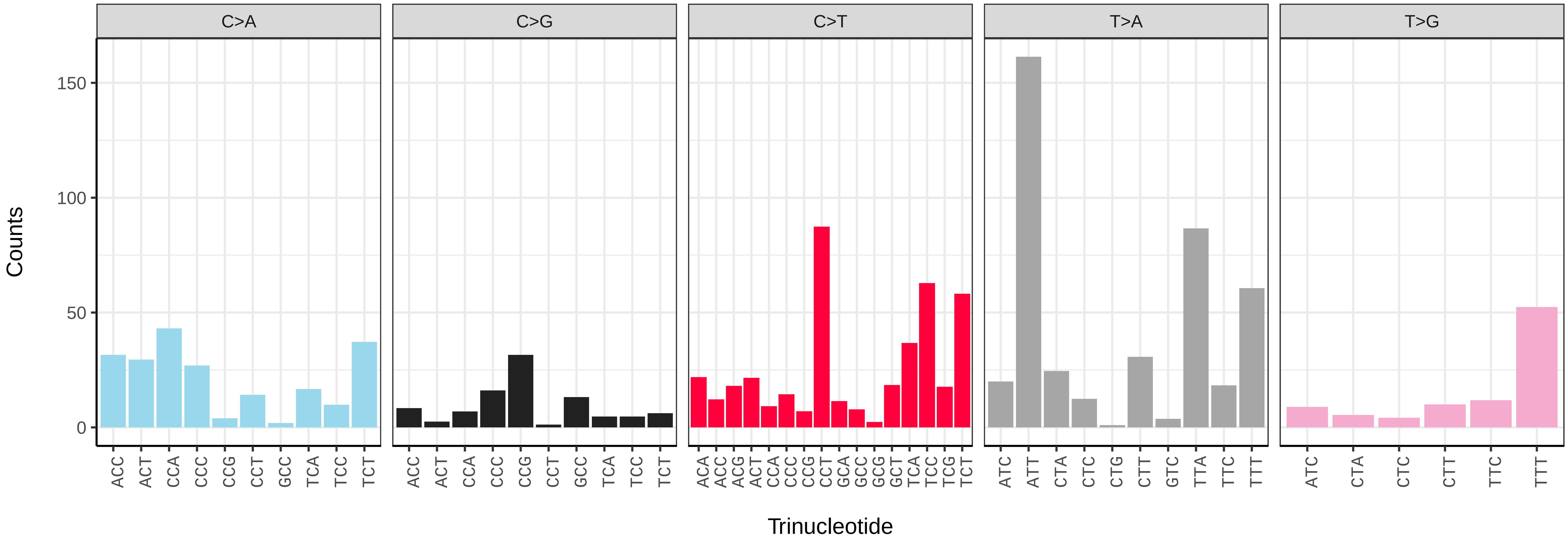

*Solanum dulcamara* (daSolDulc1) germline mutational spectrum

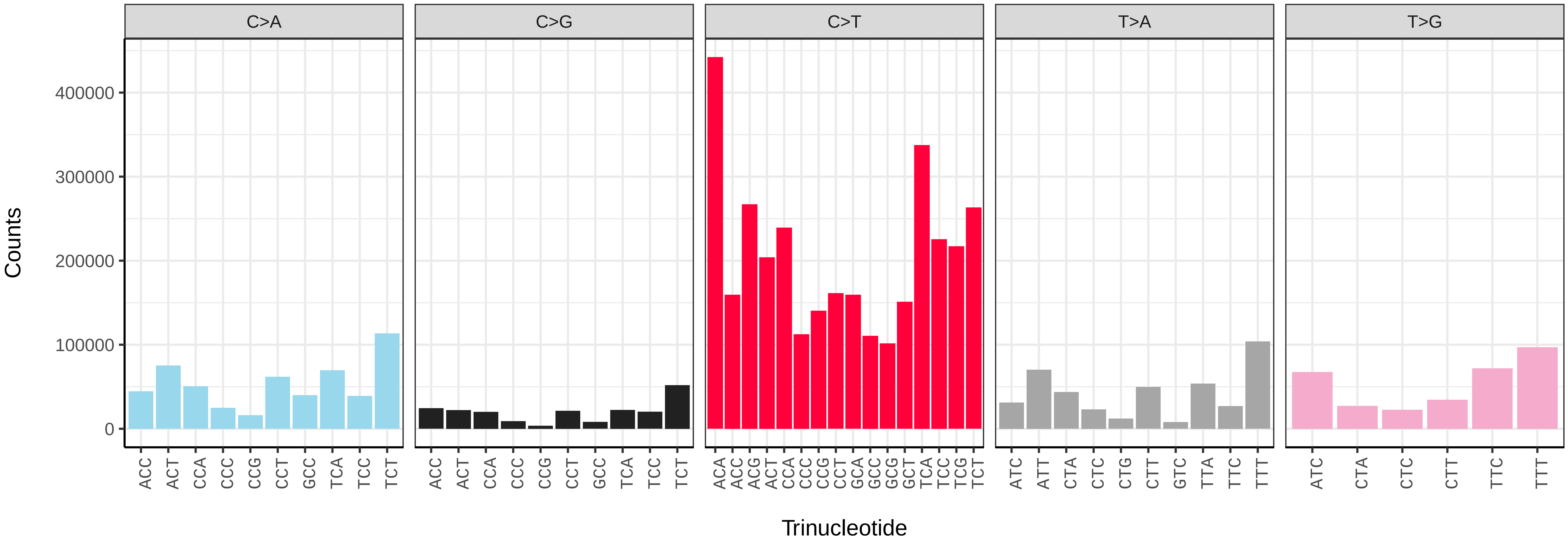


*Solanum dulcamara* (daSolDulc1) somatic mutational spectrum (SBS96 classification)

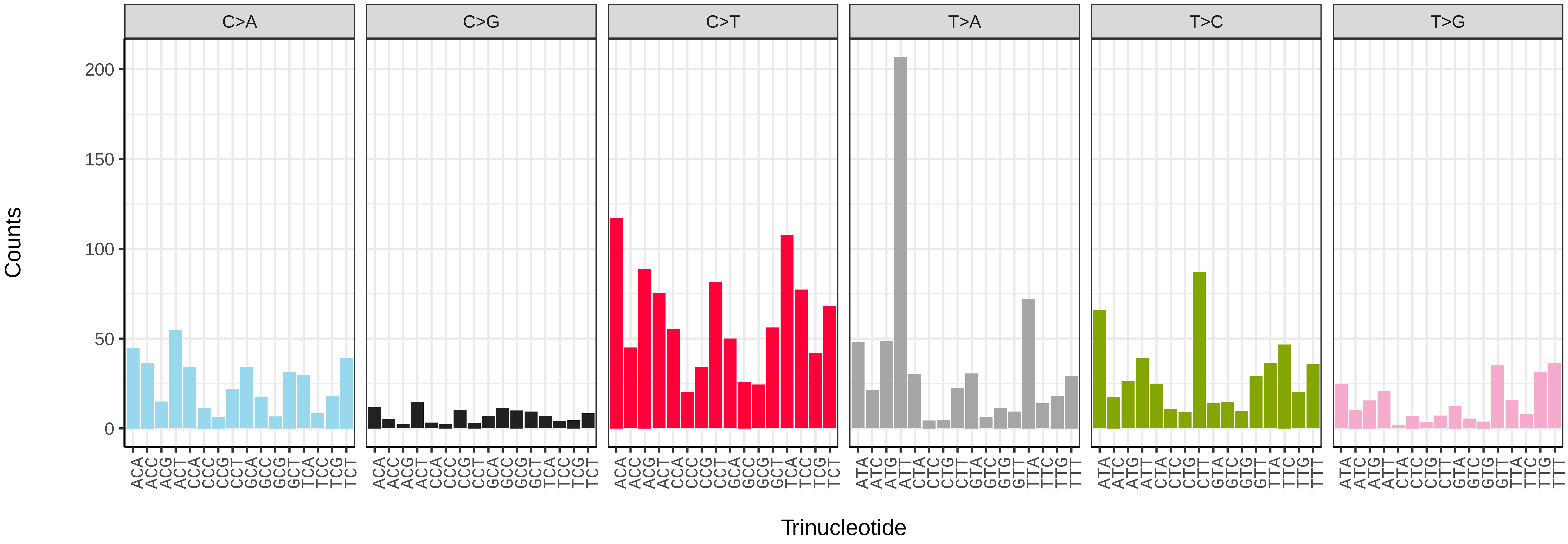


*Solanum dulcamara* (daSolDulc1) somatic mutational spectrum (SBS52 classification)

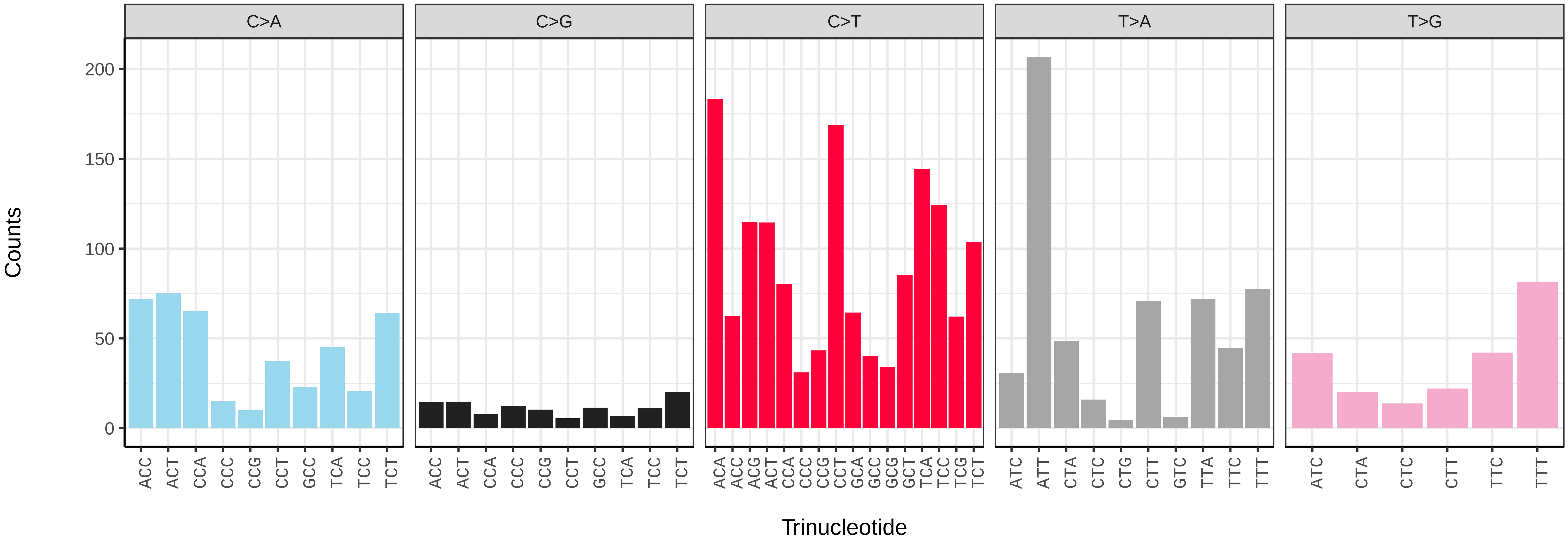

*Verbena officinalis* (daVerOffi1) germline mutational spectrum

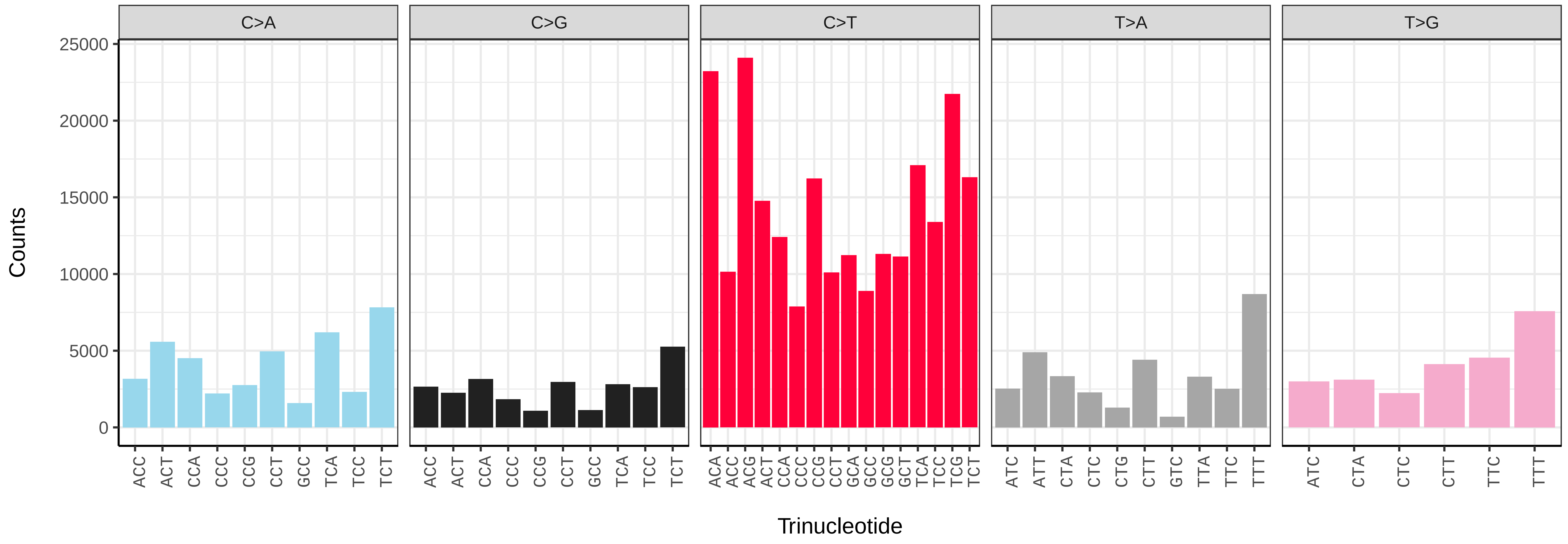


*Verbena officinalis* (daVerOffi1) somatic mutational spectrum (SBS96 classification)

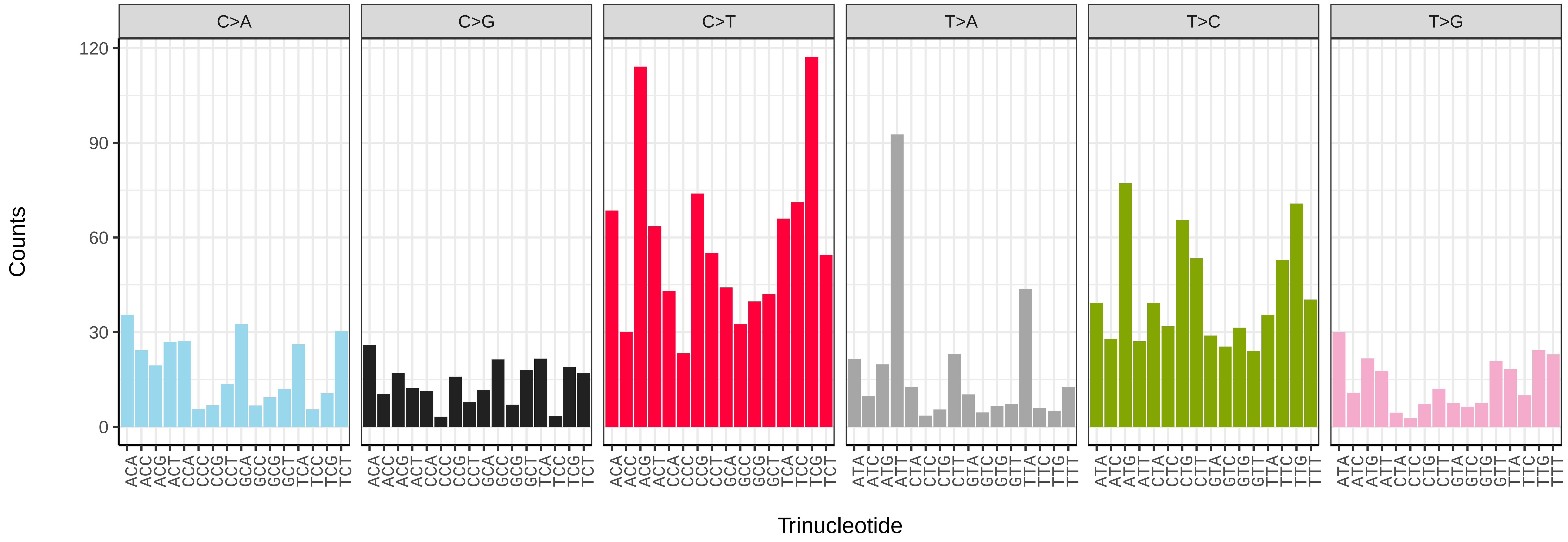


*Verbena officinalis* (daVerOffi1) somatic mutational spectrum (SBS52 classification)

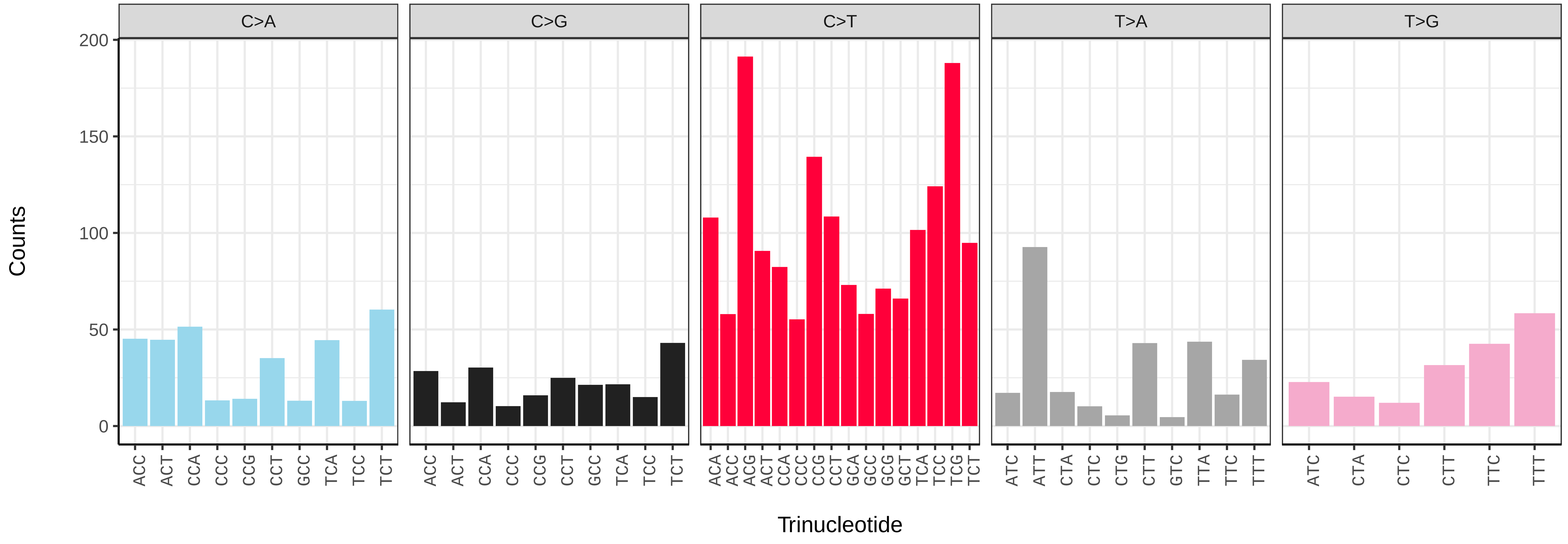

*Agrostemma githago* (dcAgrGith1) germline mutational spectrum

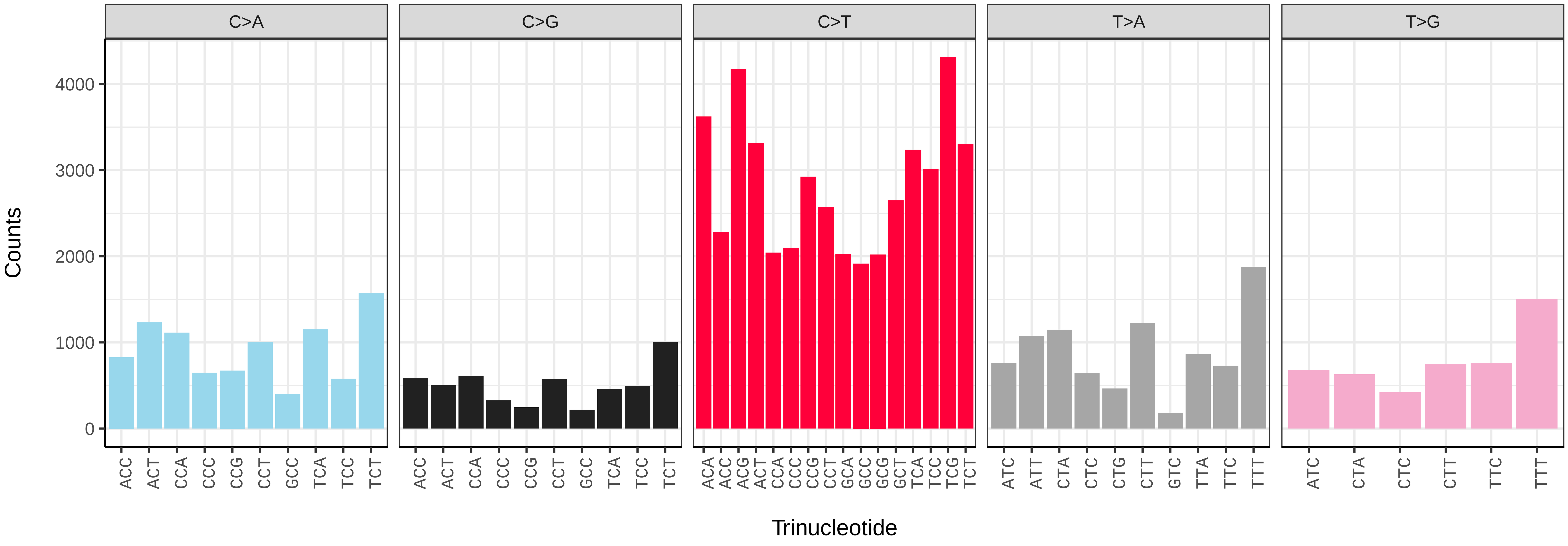


*Agrostemma githago* (dcAgrGith1) somatic mutational spectrum (SBS96 classification)

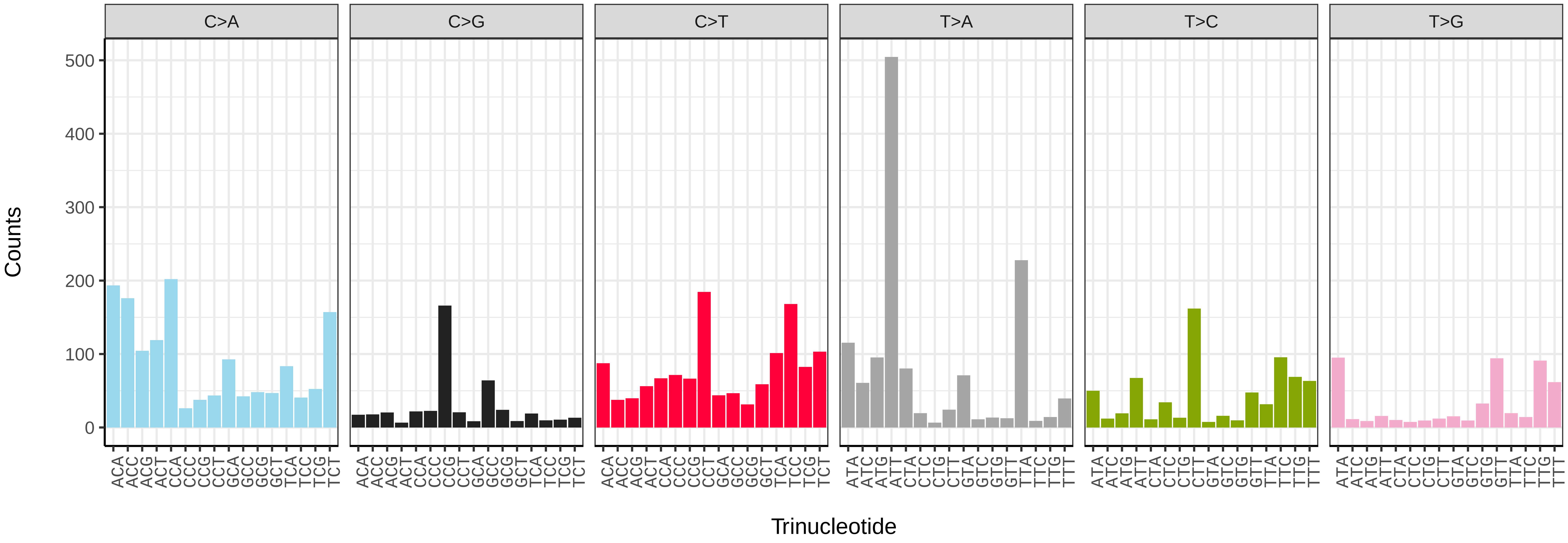


*Agrostemma githago* (dcAgrGith1) somatic mutational spectrum (SBS52 classification)

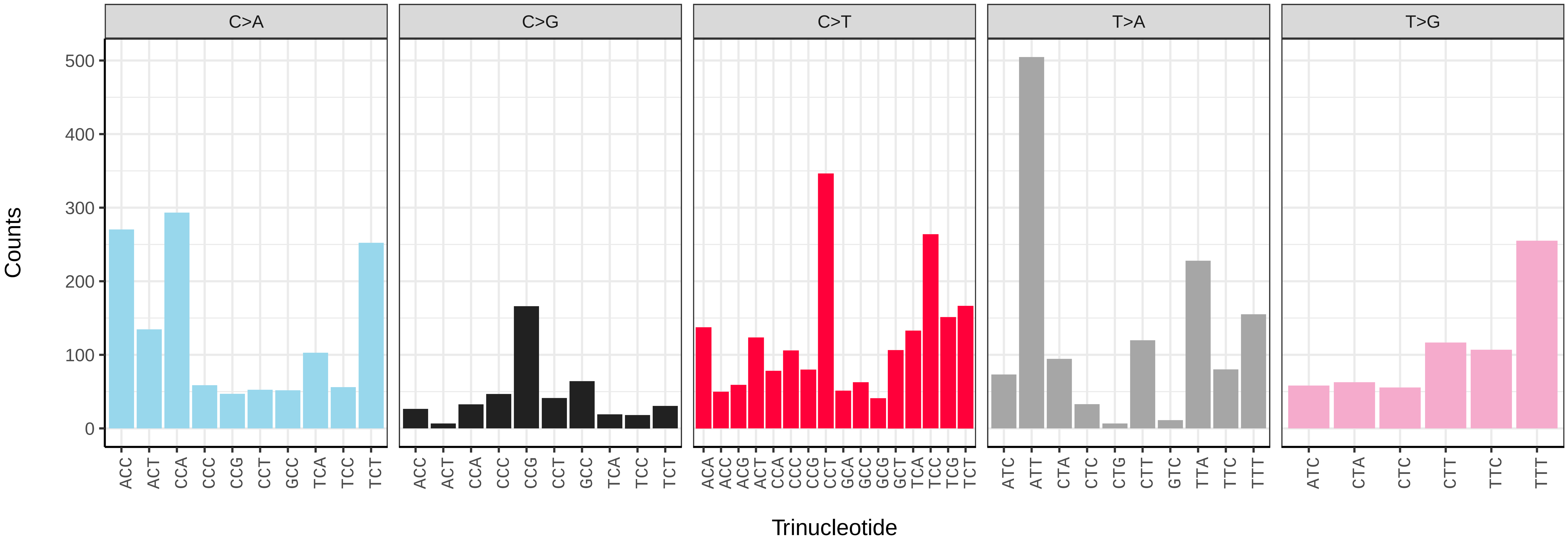

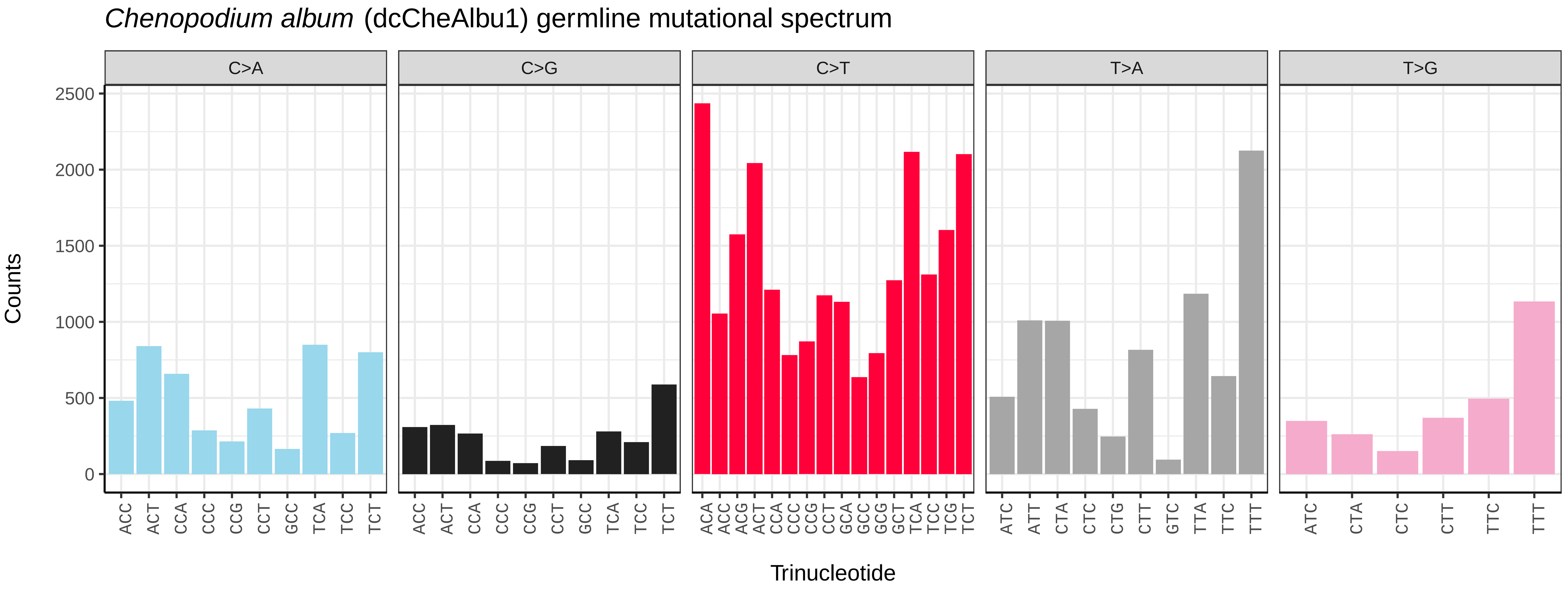
*Chenopodium album* (dcCheAlbu1) germline mutational spectrum
C>A
C>G
C>T
T>A
T>G
Counts
Trinucleotide

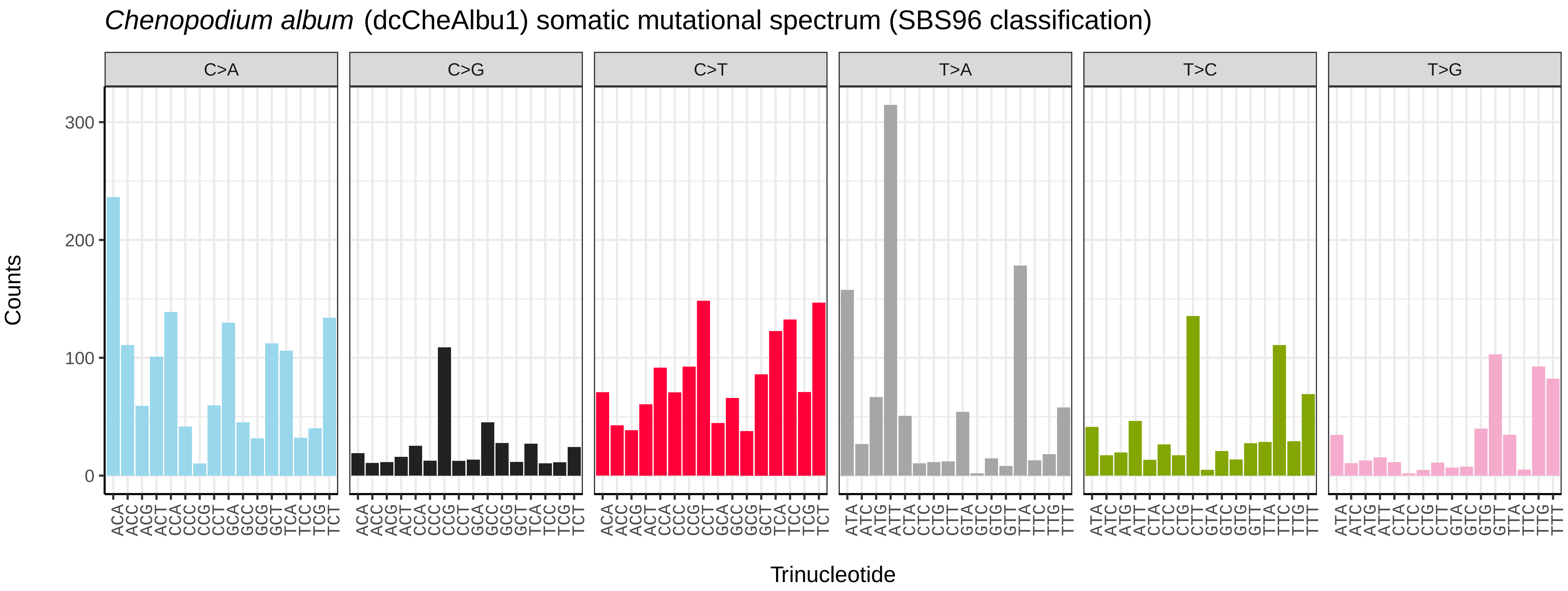
*Chenopodium album* (dcCheAlbu1) somatic mutational spectrum (SBS96 classification)
C>A
C>G
C>T
T>A
T>C
T>G
Counts
Trinucleotide

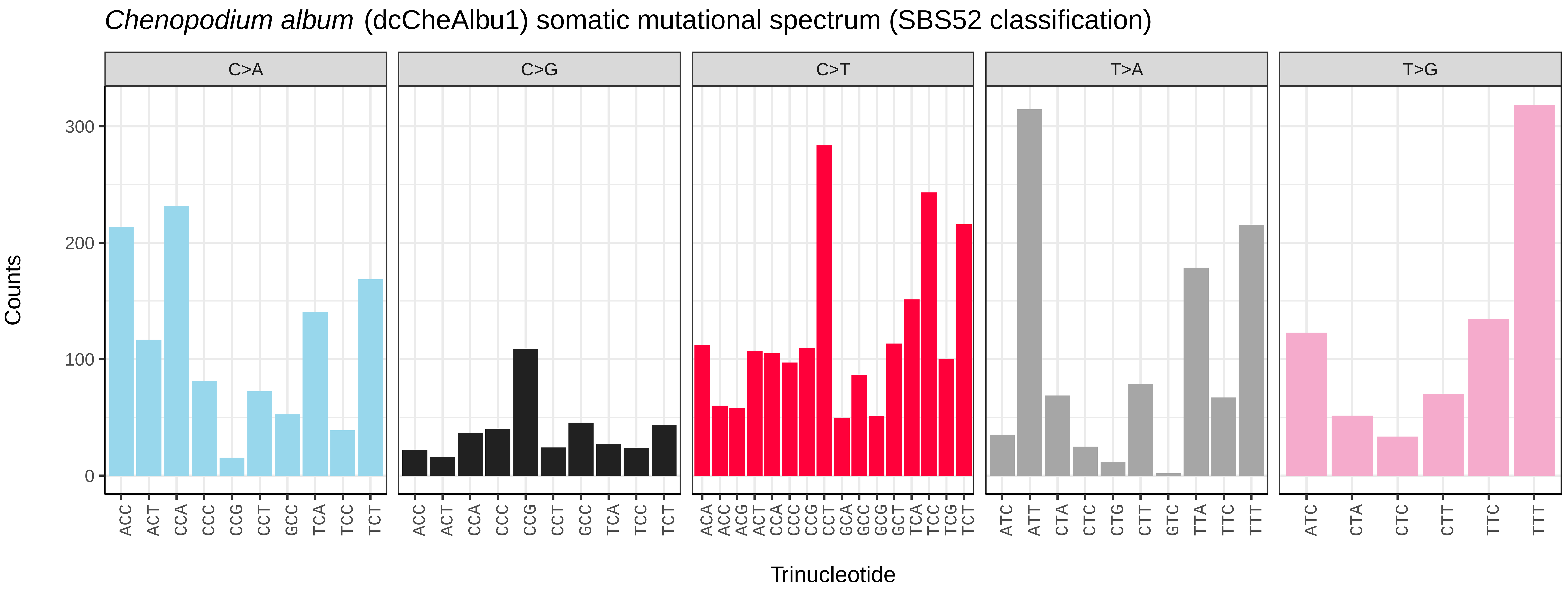
*Chenopodium album* (dcCheAlbu1) somatic mutational spectrum (SBS52 classification)
C>A
C>G
C>T
T>A
T>G
Counts
Trinucleotide

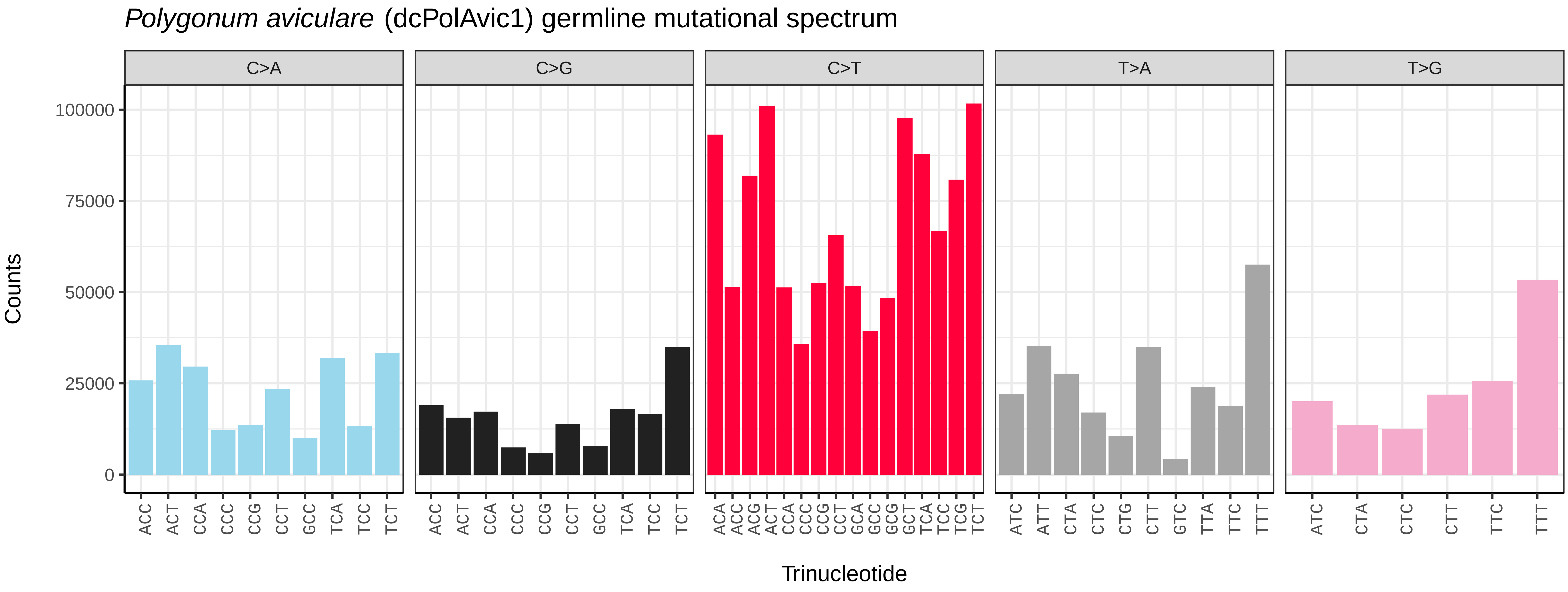
*Polygonum aviculare* (dcPolAvic1) germline mutational spectrum
C>A
C>G
C>T
T>A
T>G
Counts
0
25000
50000
75000
100000
ACC ACT CCA CCC CCG CCT GCC TCA TCC TCT
ACC ACT CCA CCC CCG CCT GCC TCA TCC TCT
ACA ACC ACG ACT CCA CCC CCG CCT GCA GCC GCG GCT TCA TCC TCG TCT
ATC ATT CTA CTC CTG CTT GTC TTA TTC TTT
ATC CTA CTC CTT TTC TTT
Trinucleotide

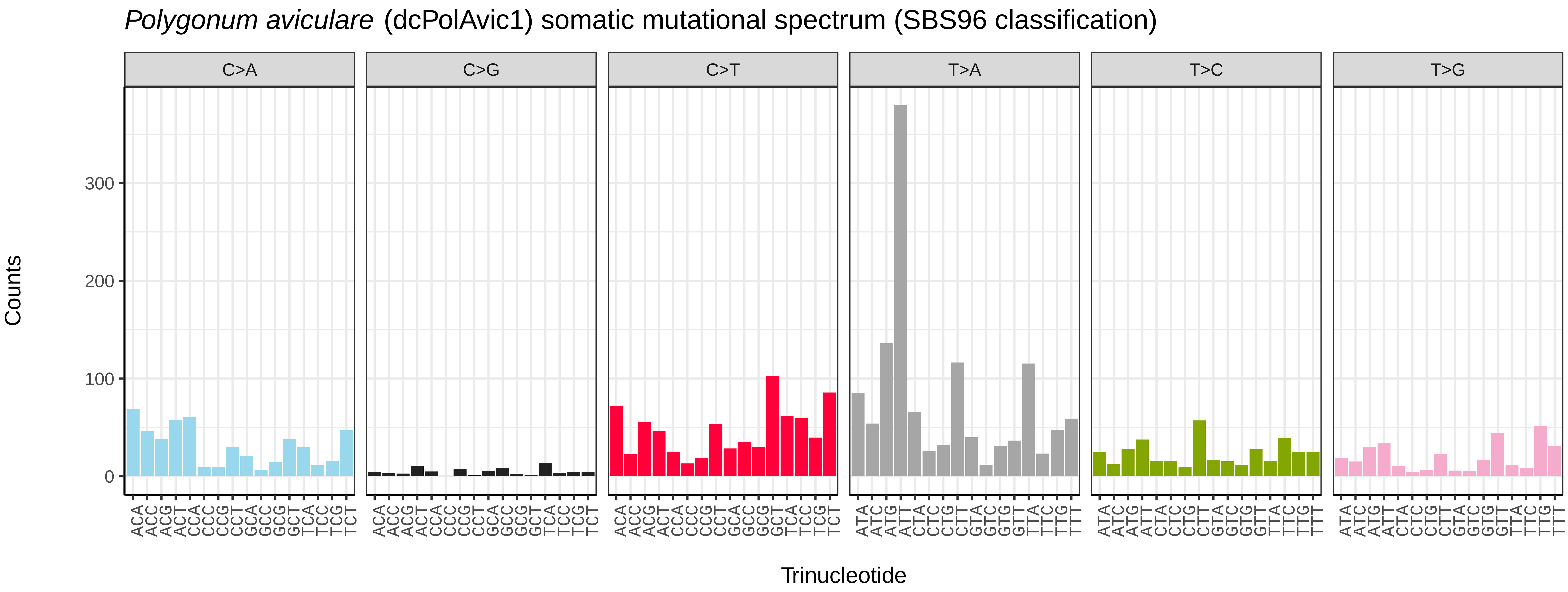
*Polygonum aviculare* (dcPolAvic1) somatic mutational spectrum (SBS96 classification)
C>A
C>G
C>T
T>A
T>C
T>G
Counts
0
100
200
300
ACA ACC ACG ACT CCA CCC CCG CCT GCA GCC GCG GCT TCA TCC TCG TCT
ACA ACC ACG ACT CCA CCC CCG CCT GCA GCC GCG GCT TCA TCC TCG TCT
ACA ACC ACG ACT CCA CCC CCG CCT GCA GCC GCG GCT TCA TCC TCG TCT
ATA ATC ATG ATT CTA CTC CTG CTT GTA GTC GTG GTT TTA TTC TTG TTT
ATA ATC ATG ATT CTA CTC CTG CTT GTA GTC GTG GTT TTA TTC TTG TTT
ATA ATC ATG ATT CTA CTC CTG CTT GTA GTC GTG GTT TTA TTC TTG TTT
Trinucleotide

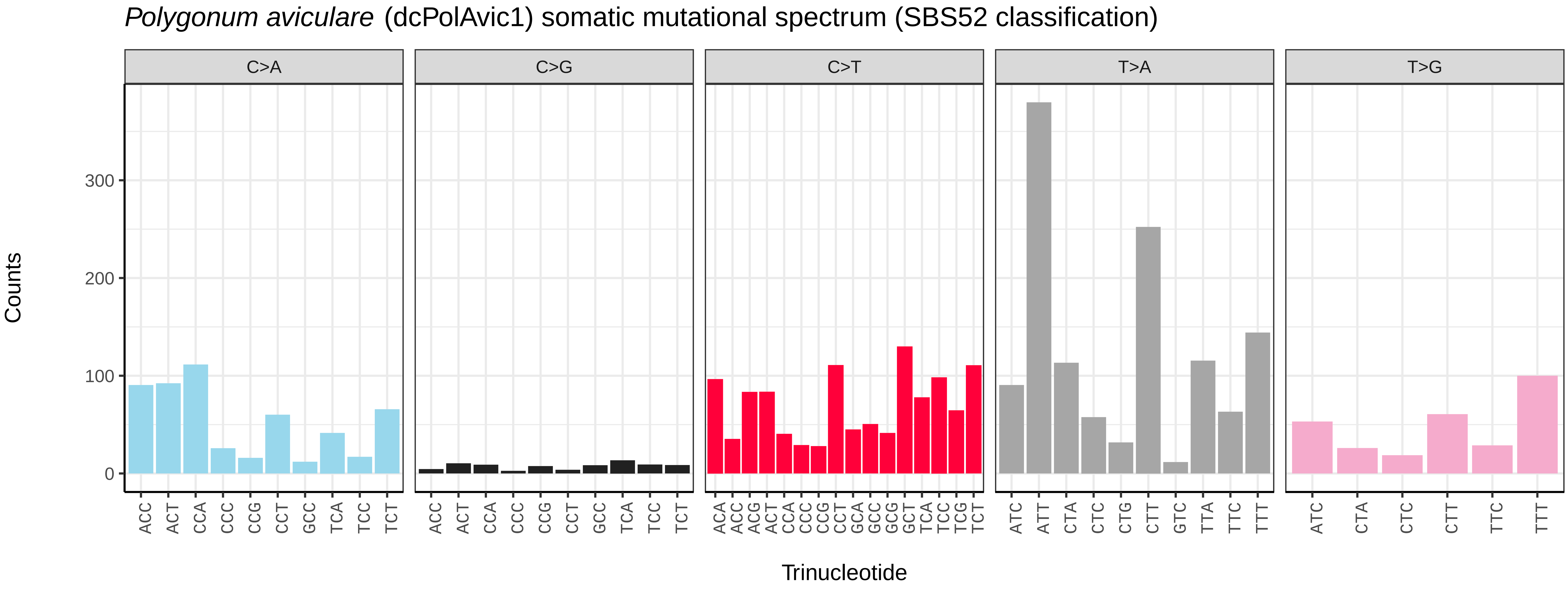
*Polygonum aviculare* (dcPolAvic1) somatic mutational spectrum (SBS52 classification)
C>A
C>G
C>T
T>A
T>G
Counts
0
100
200
300
ACC ACT CCA CCC CCG CCT GCC TCA TCC TCT
ACC ACT CCA CCC CCG CCT GCC TCA TCC TCT
ACA ACC ACG ACT CCA CCC CCG CCT GCA GCC GCG GCT TCA TCC TCG TCT
ATC ATT CTA CTC CTG CTT GTC TTA TTC TTT
ATC CTA CTC CTT TTC TTT
Trinucleotide

*Salicornia ramosissima* (dcSalRamo1) germline mutational spectrum

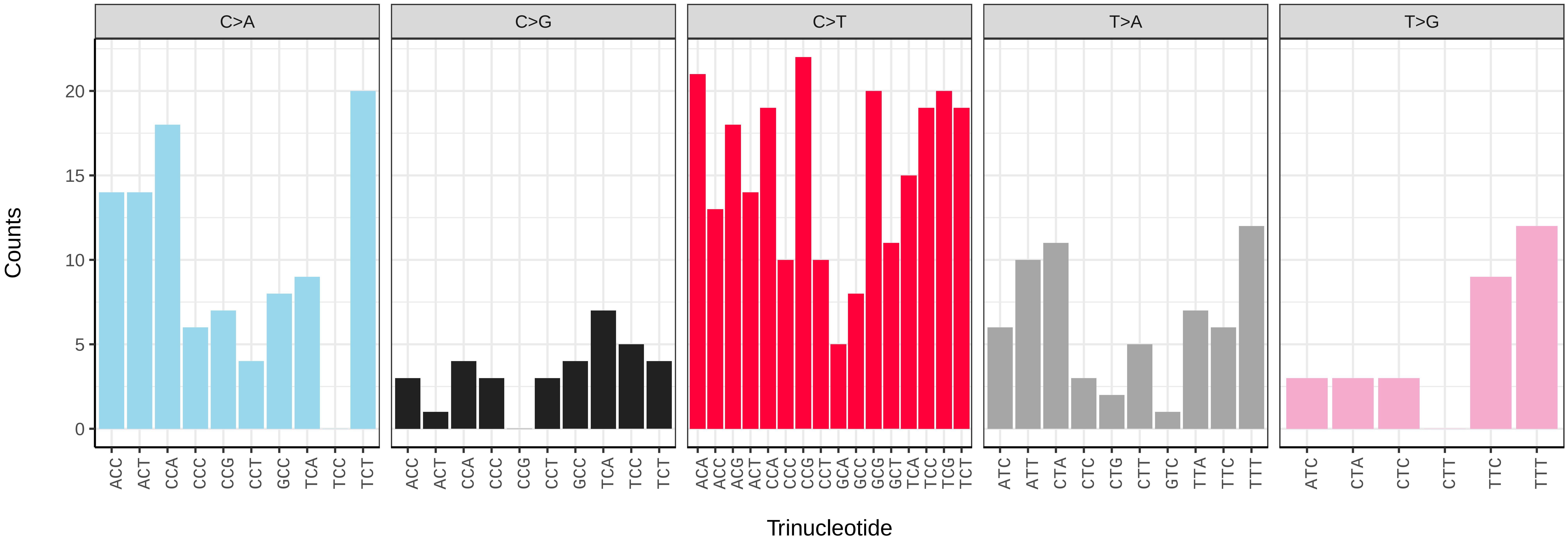


*Salicornia ramosissima* (dcSalRamo1) somatic mutational spectrum (SBS96 classification)

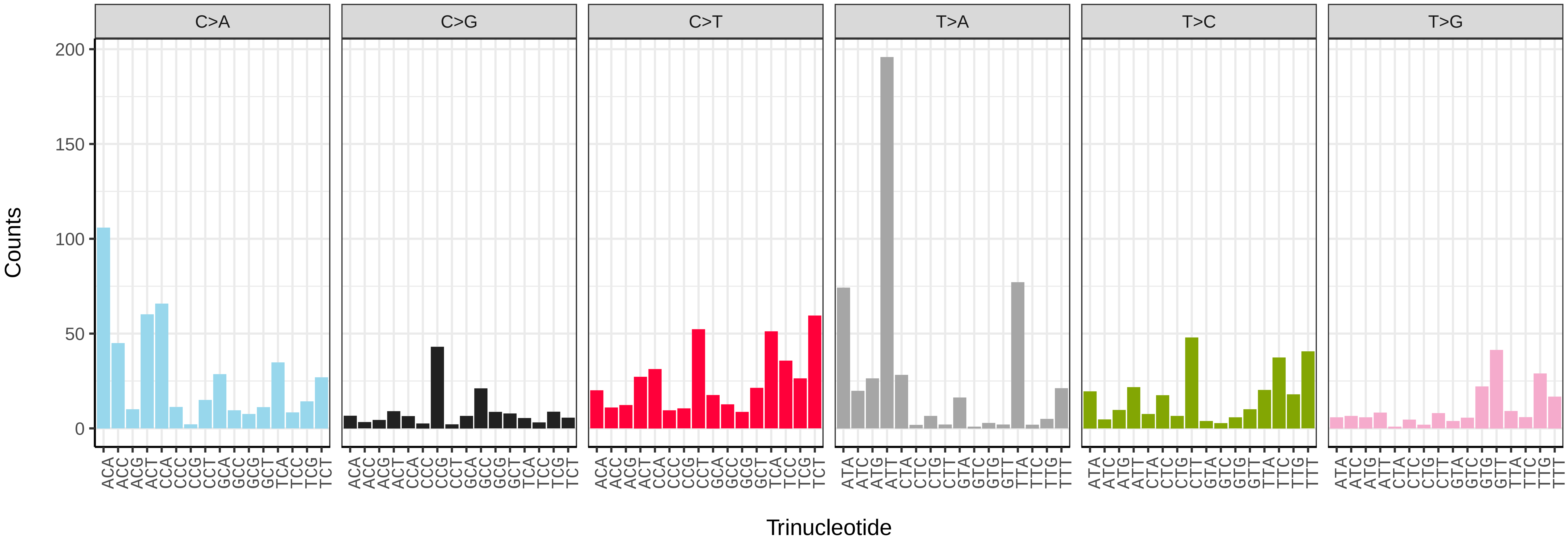


*Salicornia ramosissima* (dcSalRamo1) somatic mutational spectrum (SBS52 classification)

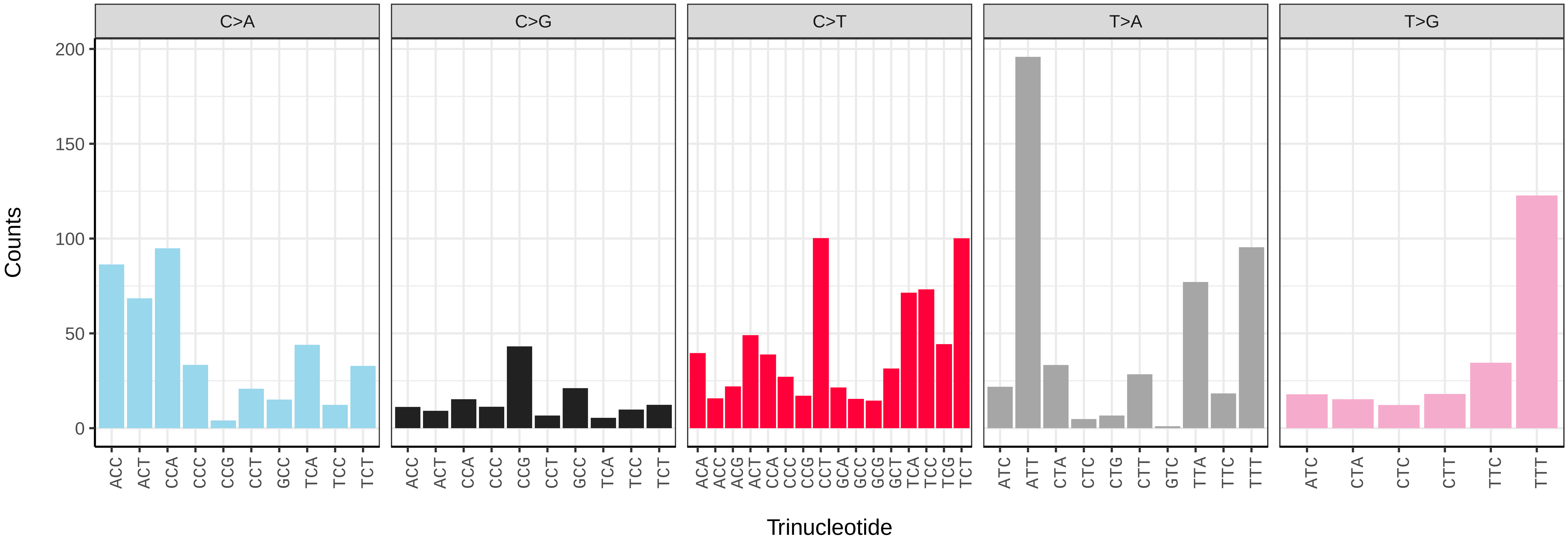

*Spergularia bocconei* (dcSpeBocc1) germline mutational spectrum

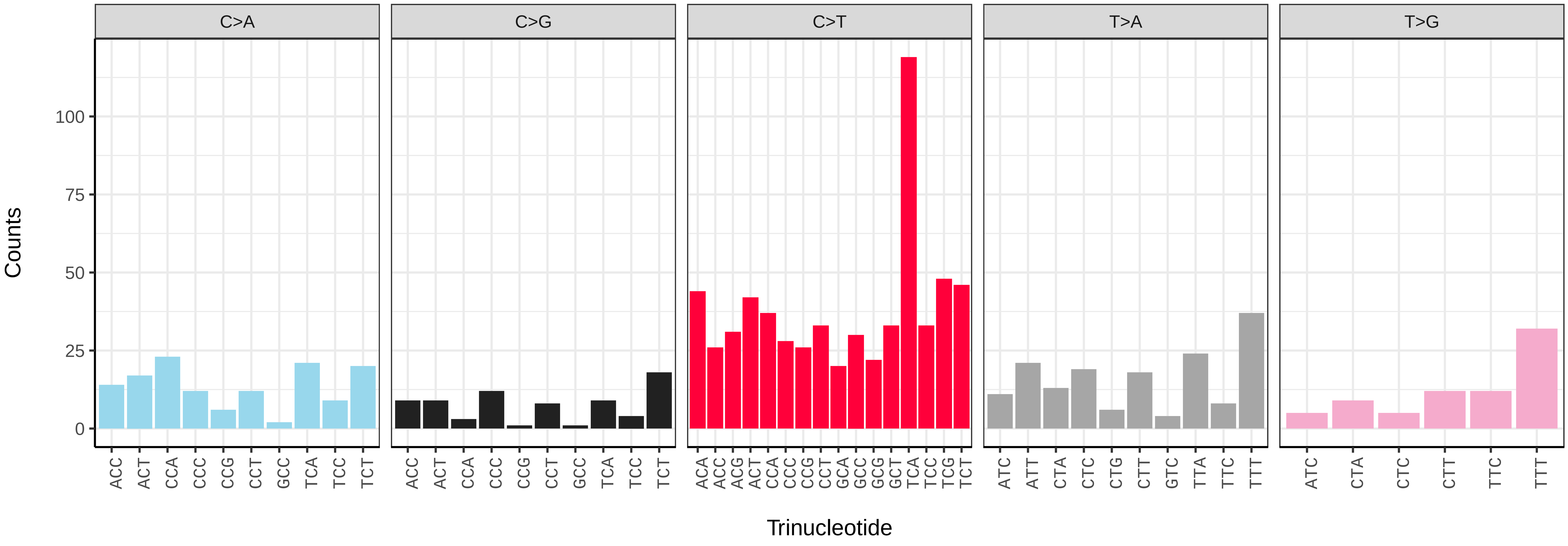


*Spergularia bocconei* (dcSpeBocc1) somatic mutational spectrum (SBS96 classification)

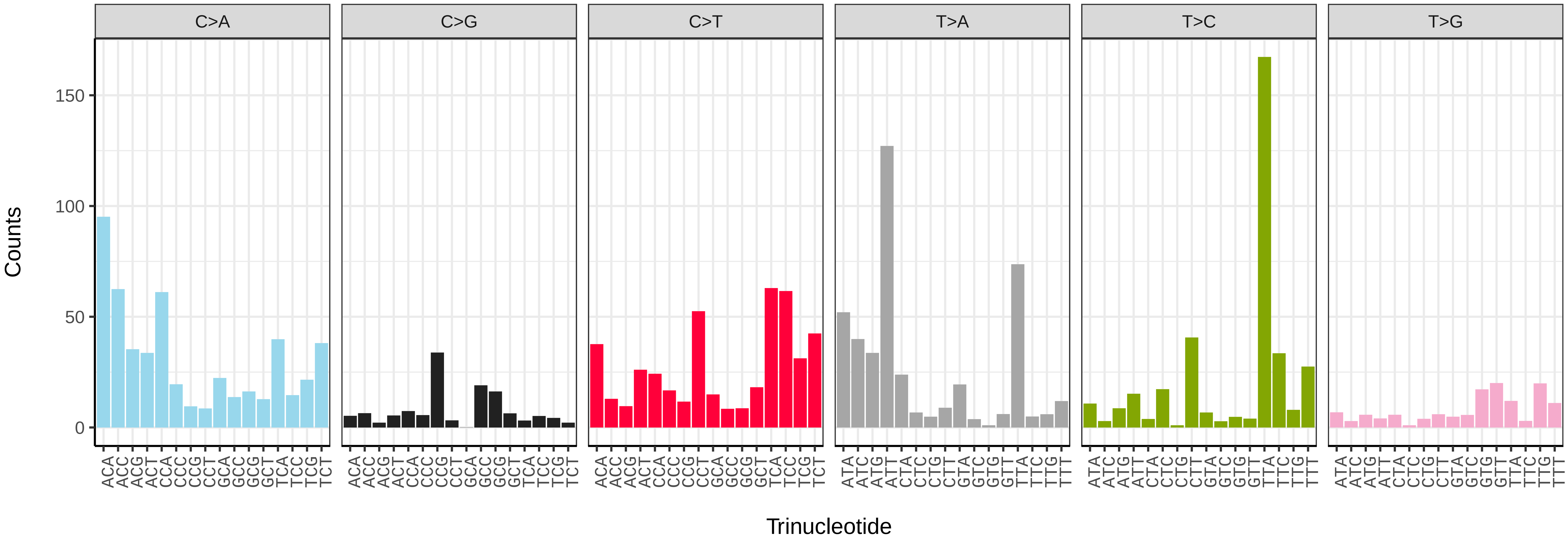


*Spergularia bocconei* (dcSpeBocc1) somatic mutational spectrum (SBS52 classification)

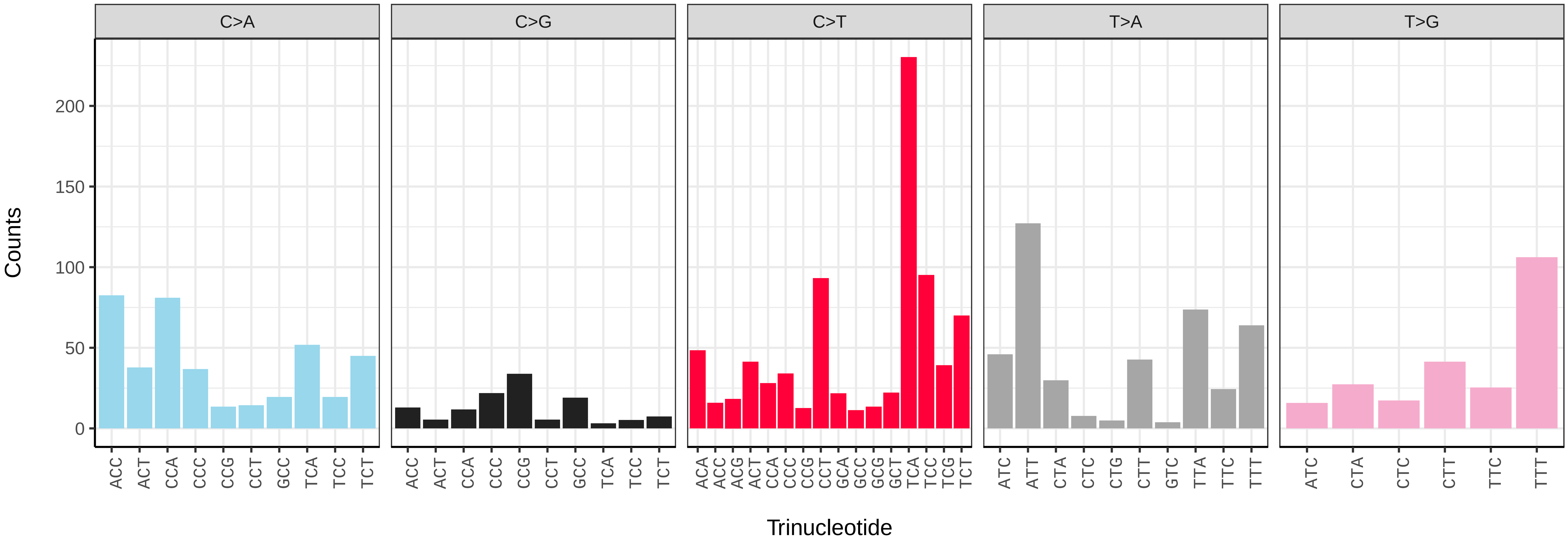

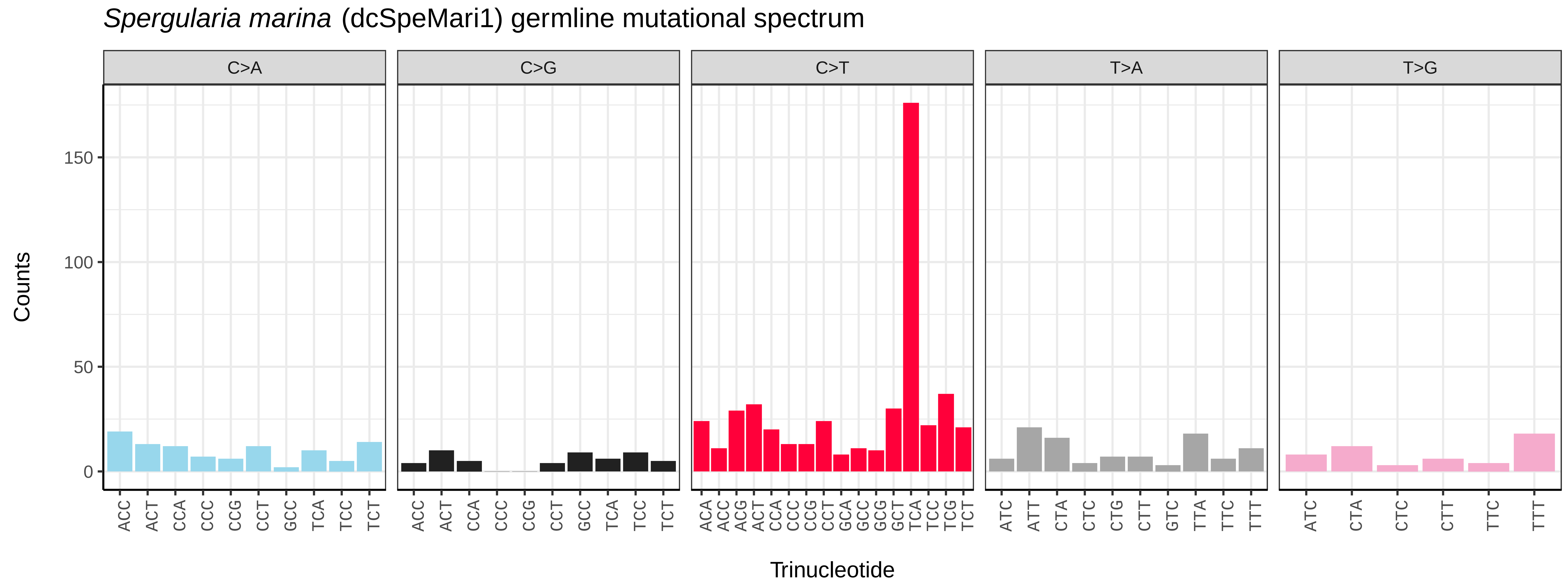
*Spergularia marina* (dcSpeMari1) germline mutational spectrum
C>A
C>G
C>T
T>A
T>G
Counts
0
50
100
150
ACC ACT CCA CCC CCG CCT GCC TCA TCC TCT
ACC ACT CCA CCC CCG CCT GCC TCA TCC TCT
ACA ACC ACG ACT CCA CCC CCG CCT GCA GCC GCG GCT TCA TCC TCG TCT
ATC ATT CTA CTC CTG CTT GTC TTA TTC TTT
ATC CTA CTC CTT TTC TTT
Trinucleotide

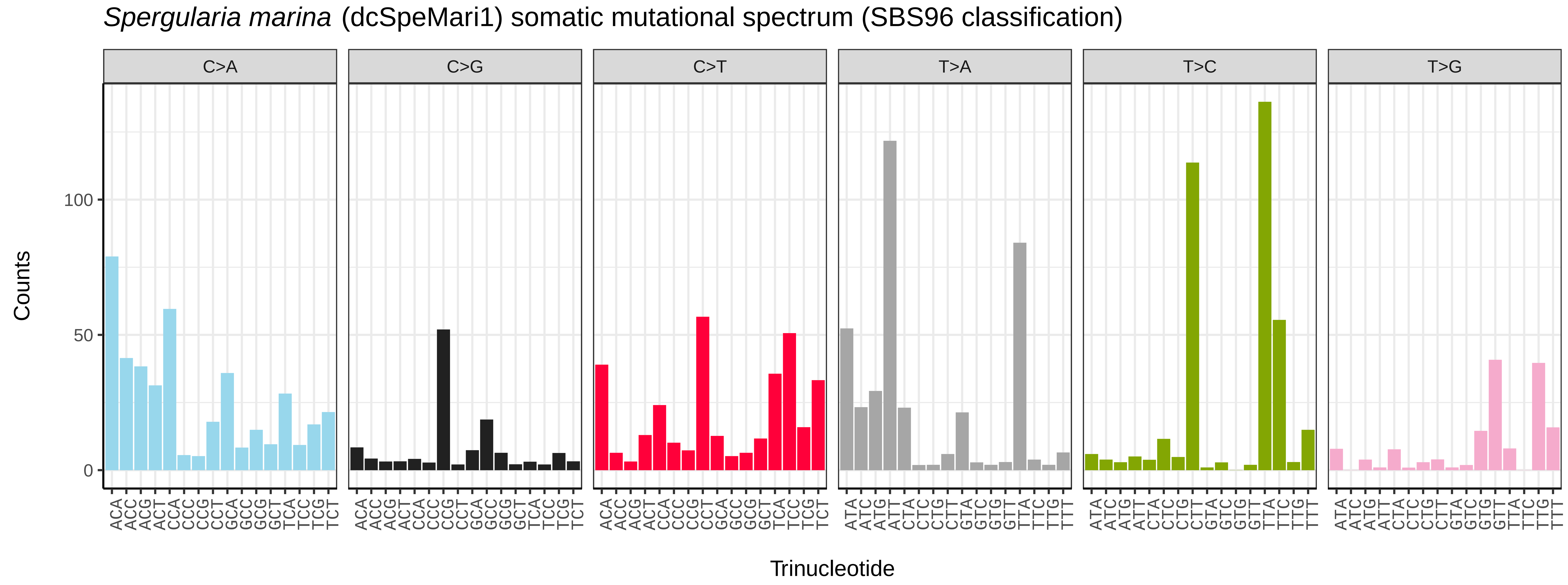
*Spergularia marina* (dcSpeMari1) somatic mutational spectrum (SBS96 classification)
C>A
C>G
C>T
T>A
T>C
T>G
Counts
0
50
100
ACA ACC ACG ACT CCA CCC CCG CCT GCA GCC GCG GCT TCA TCC TCG TCT
ACA ACC ACG ACT CCA CCC CCG CCT GCA GCC GCG GCT TCA TCC TCG TCT
ACA ACC ACG ACT CCA CCC CCG CCT GCA GCC GCG GCT TCA TCC TCG TCT
ATA ATC ATG ATT CTA CTC CTG CTT GTA GTC GTG GTT TTA TTC TTG TTT
ATA ATC ATG ATT CTA CTC CTG CTT GTA GTC GTG GTT TTA TTC TTG TTT
ATA ATC ATG ATT CTA CTC CTG CTT GTA GTC GTG GTT TTA TTC TTG TTT
Trinucleotide

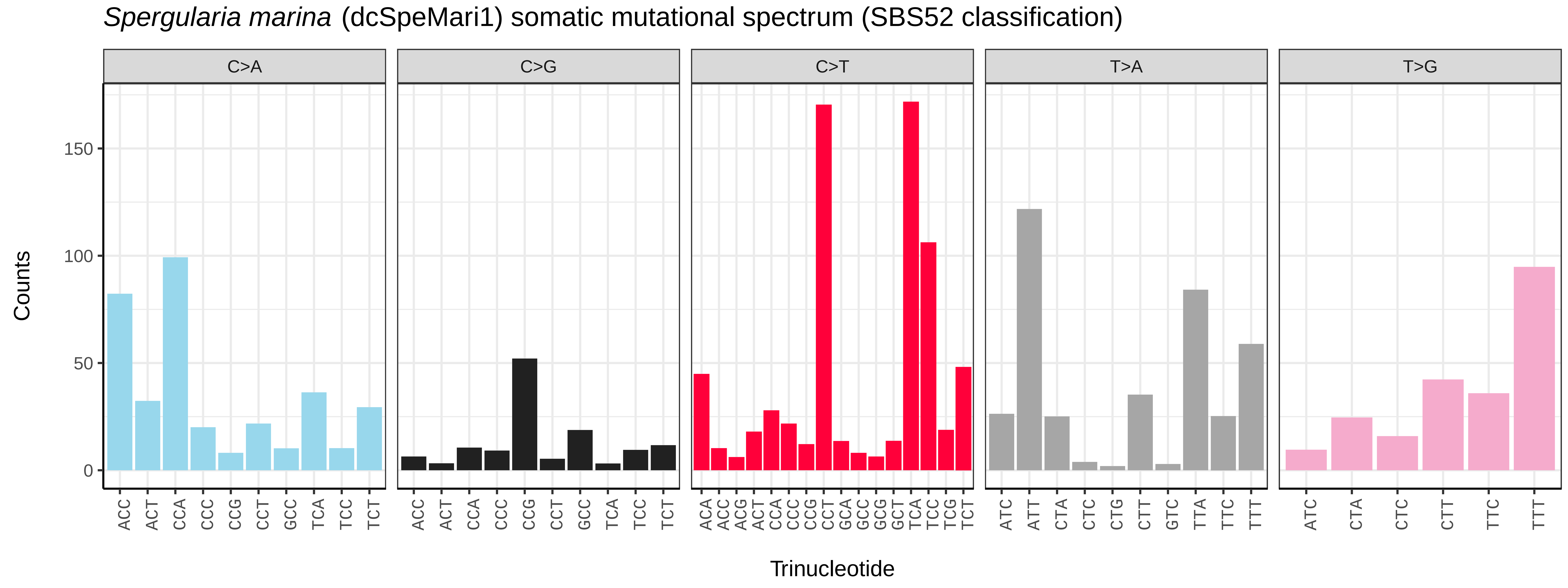
*Spergularia marina* (dcSpeMari1) somatic mutational spectrum (SBS52 classification)
C>A
C>G
C>T
T>A
T>G
Counts
0
50
100
150
ACC ACT CCA CCC CCG CCT GCC TCA TCC TCT
ACC ACT CCA CCC CCG CCT GCC TCA TCC TCT
ACA ACC ACG ACT CCA CCC CCG CCT GCA GCC GCG GCT TCA TCC TCG TCT
ATC ATT CTA CTC CTG CTT GTC TTA TTC TTT
ATC CTA CTC CTT TTC TTT
Trinucleotide

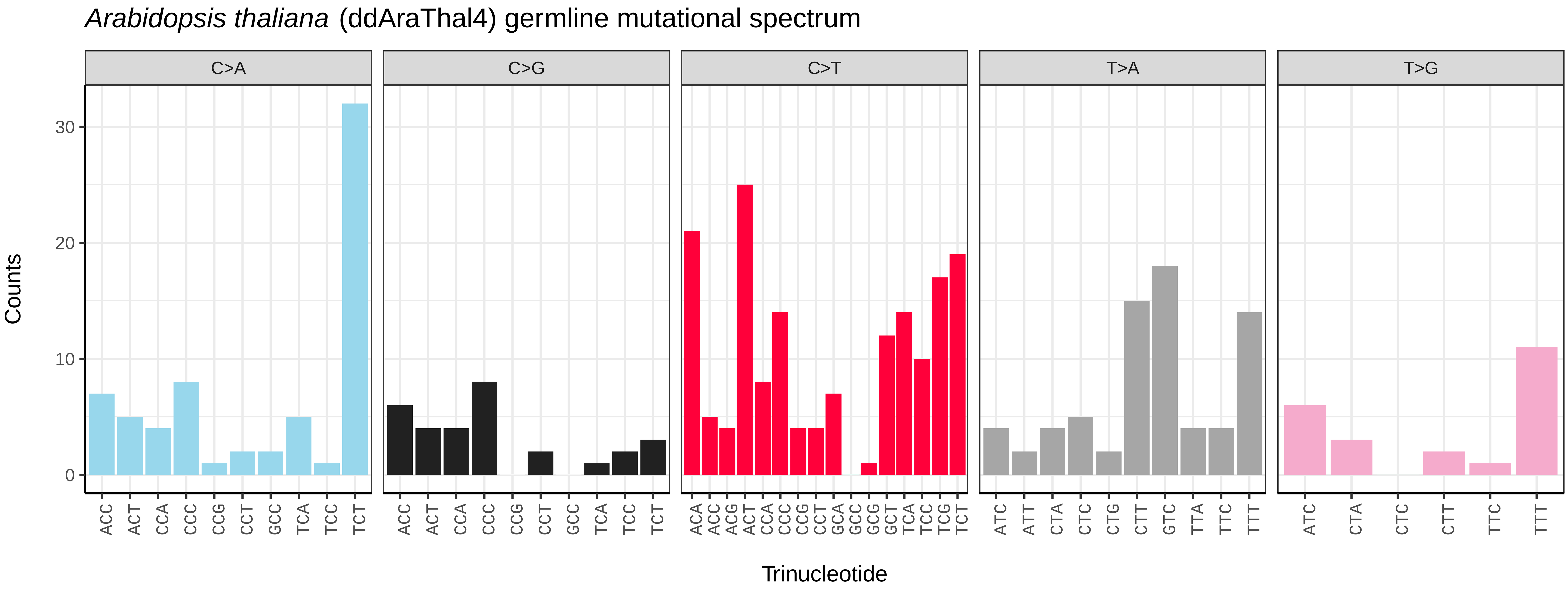
*Arabidopsis thaliana* (ddAraThal4) germline mutational spectrum
C>A
C>G
C>T
T>A
T>G
Counts
0
10
20
30
ACC ACT CCA CCC CCG CCT GCC TCA TCC TCT
ACA ACC ACG ACT CCA CCC CCG CCT GCA GCC GCG GCT TCA TCC TCG TCT
ATC ATT CTA CTC CTG CTT GTC TTA TTC TTT
ATC CTA CTC CTT TTC TTT
Trinucleotide

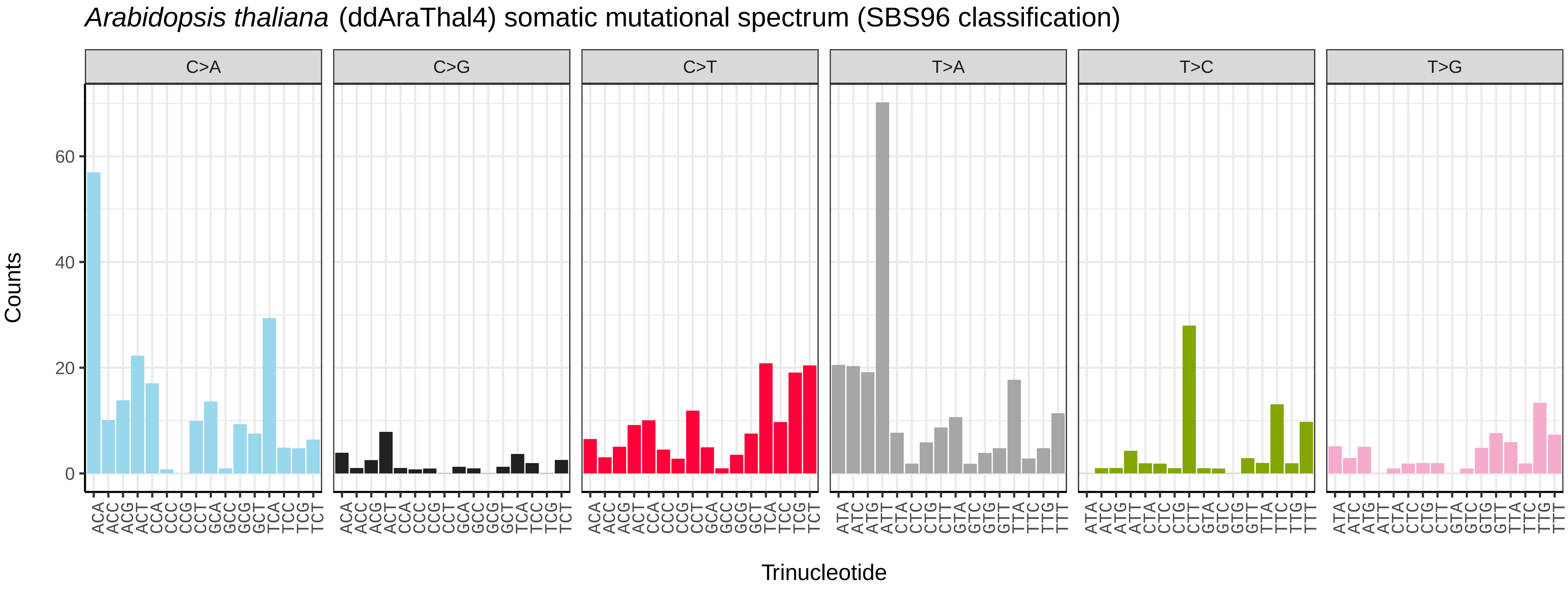
*Arabidopsis thaliana* (ddAraThal4) somatic mutational spectrum (SBS96 classification)
C>A
C>G
C>T
T>A
T>C
T>G
Counts
0
20
40
60
ACA ACC ACG ACT CCA CCC CCG CCT GCA GCC GCG GCT TCA TCC TCG TCT
ATA ATC ATG ATT CTA CTC CTG CTT GTA GTC GTG GTT TTA TTC TTG TTT
Trinucleotide

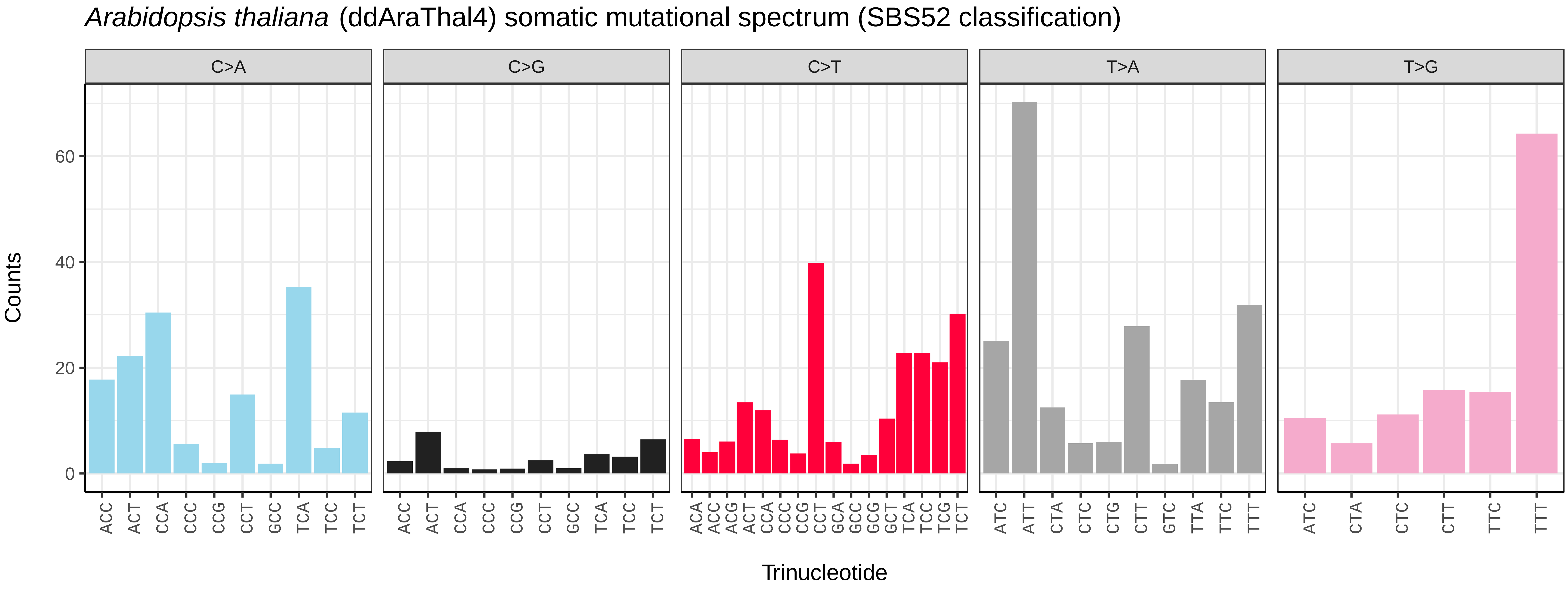
*Arabidopsis thaliana* (ddAraThal4) somatic mutational spectrum (SBS52 classification)
C>A
C>G
C>T
T>A
T>G
Counts
0
20
40
60
ACC ACT CCA CCC CCG CCT GCC TCA TCC TCT
ACA ACC ACG ACT CCA CCC CCG CCT GCA GCC GCG GCT TCA TCC TCG TCT
ATC ATT CTA CTC CTG CTT GTC TTA TTC TTT
ATC CTA CTC CTT TTC TTT
Trinucleotide

*Barbarea vulgaris* (ddBarVulg1) germline mutational spectrum

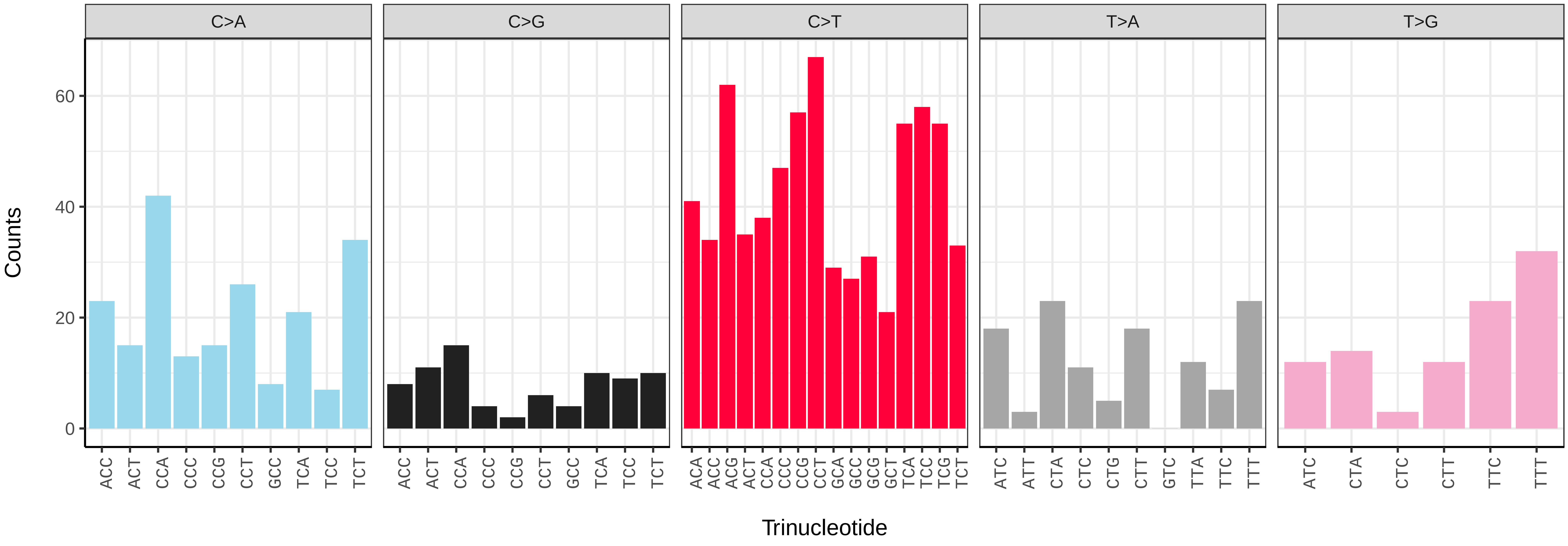


*Barbarea vulgaris* (ddBarVulg1) somatic mutational spectrum (SBS96 classification)

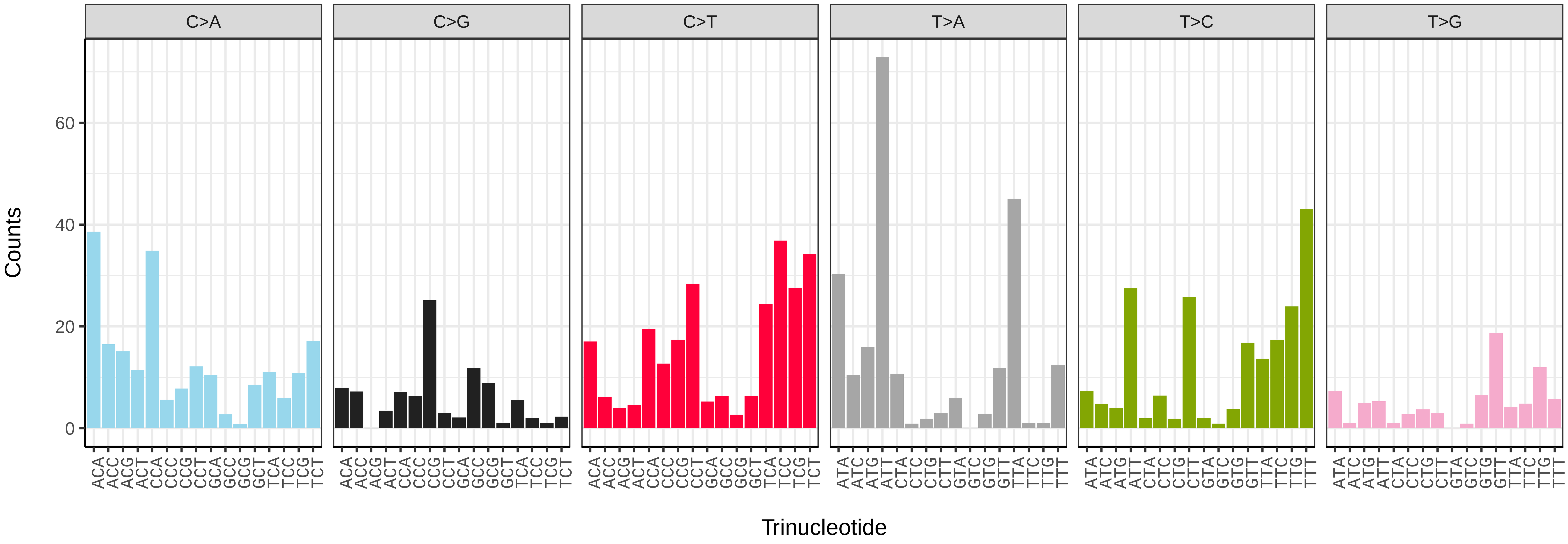


*Barbarea vulgaris* (ddBarVulg1) somatic mutational spectrum (SBS52 classification)

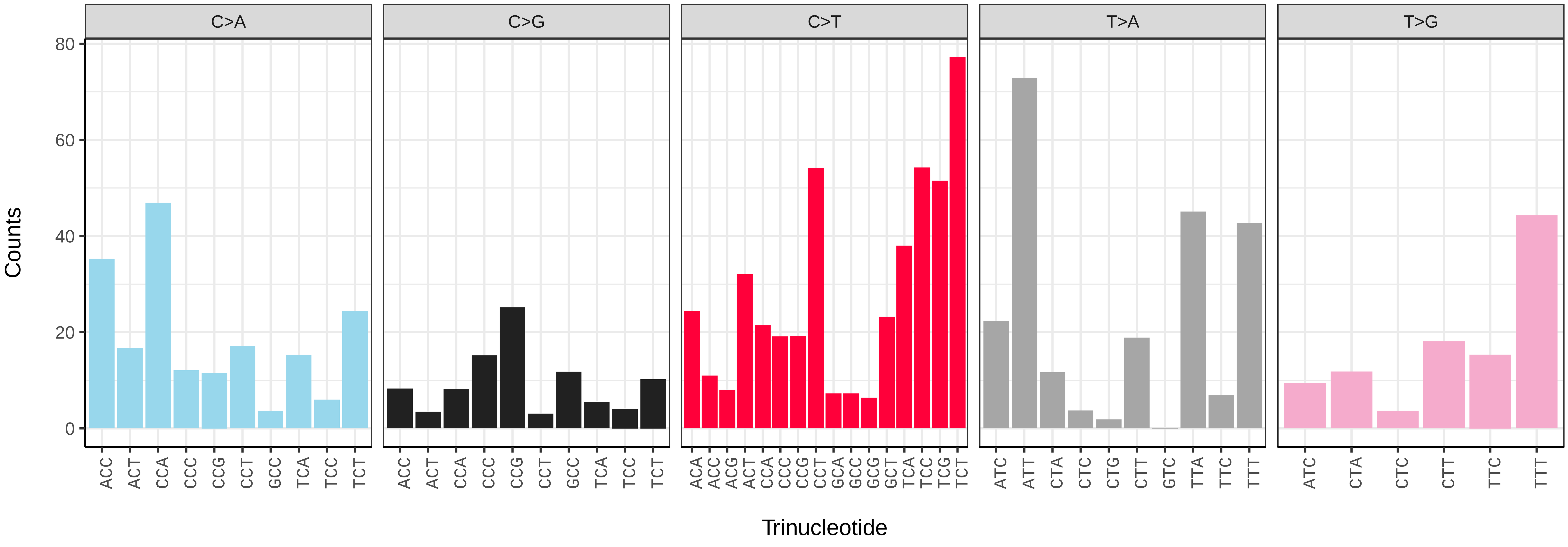

*Euphorbia lathyris* (ddEupLath1) germline mutational spectrum

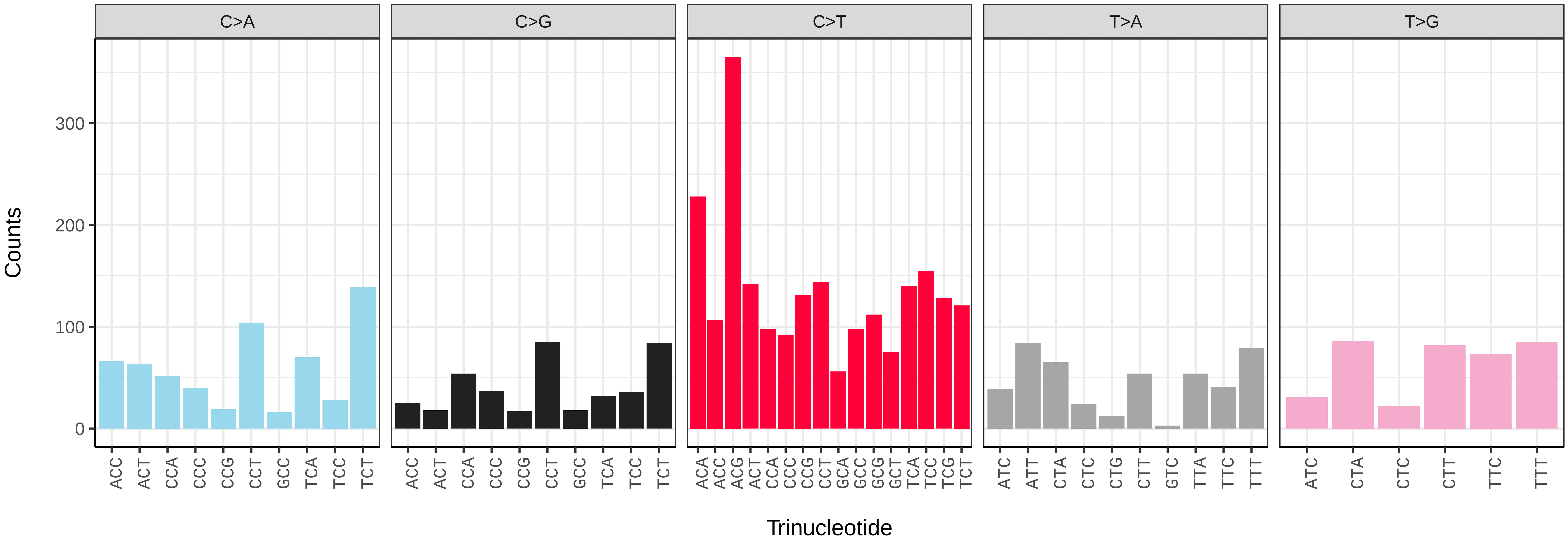


*Euphorbia lathyris* (ddEupLath1) somatic mutational spectrum (SBS96 classification)

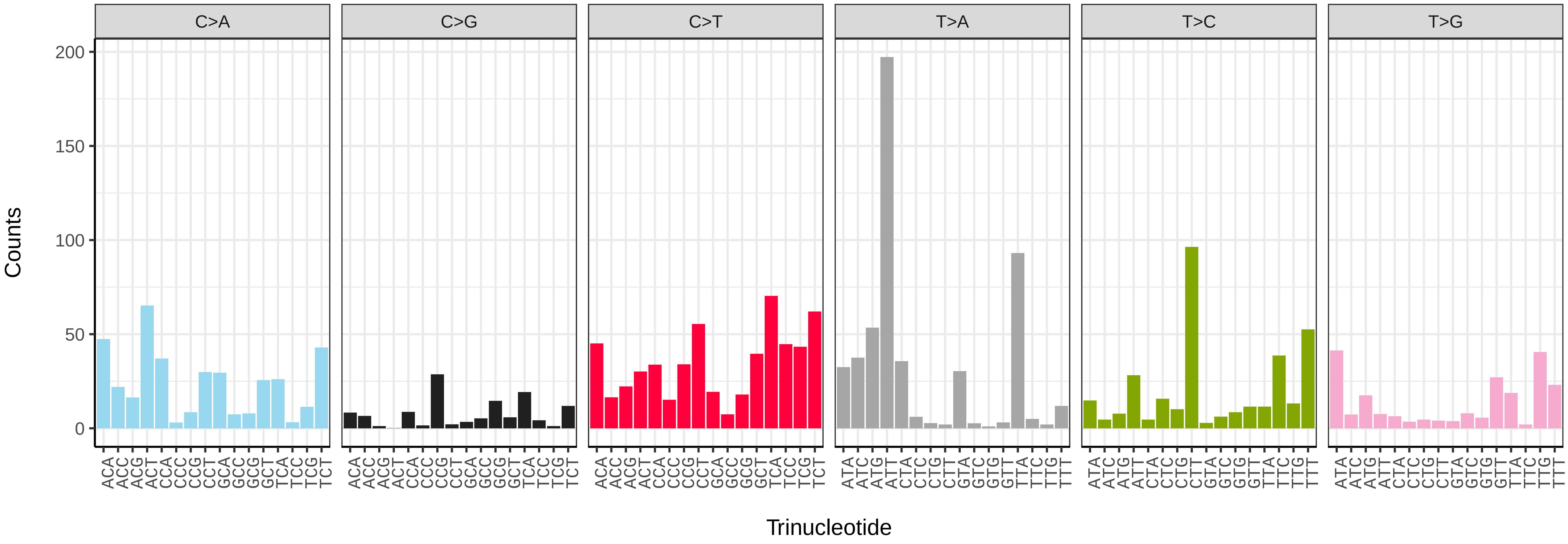


*Euphorbia lathyris* (ddEupLath1) somatic mutational spectrum (SBS52 classification)

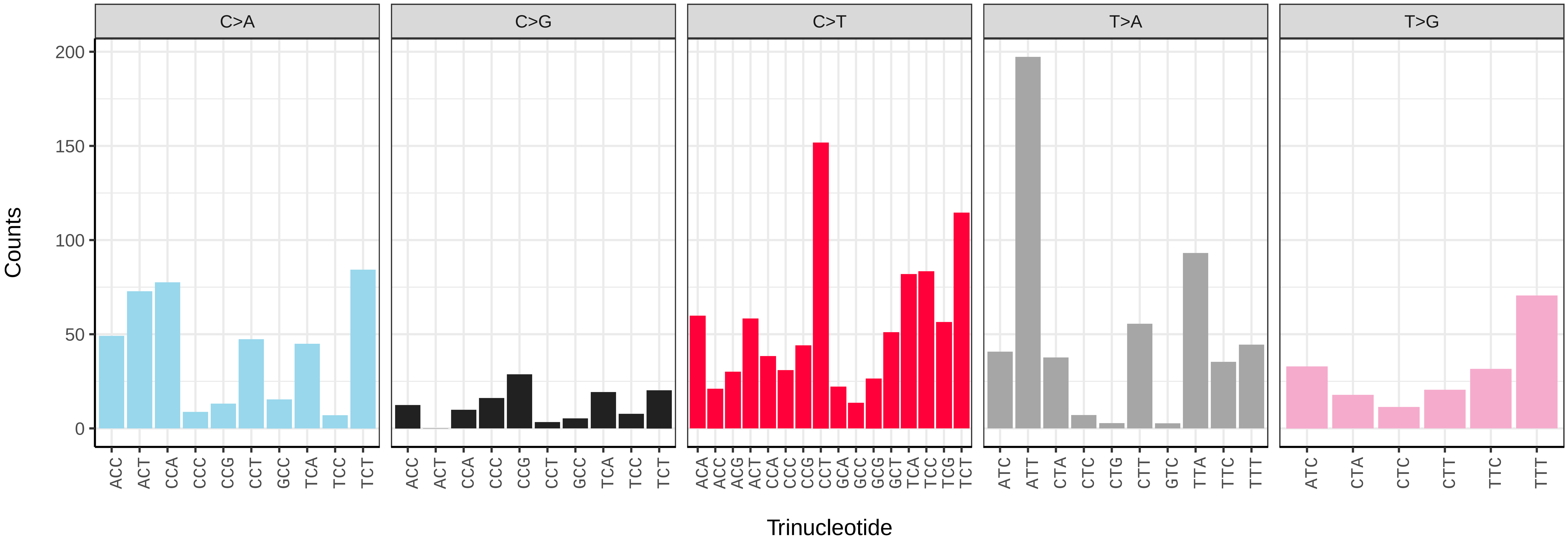

*Euphorbia peplus* (ddEupPepu3) germline mutational spectrum

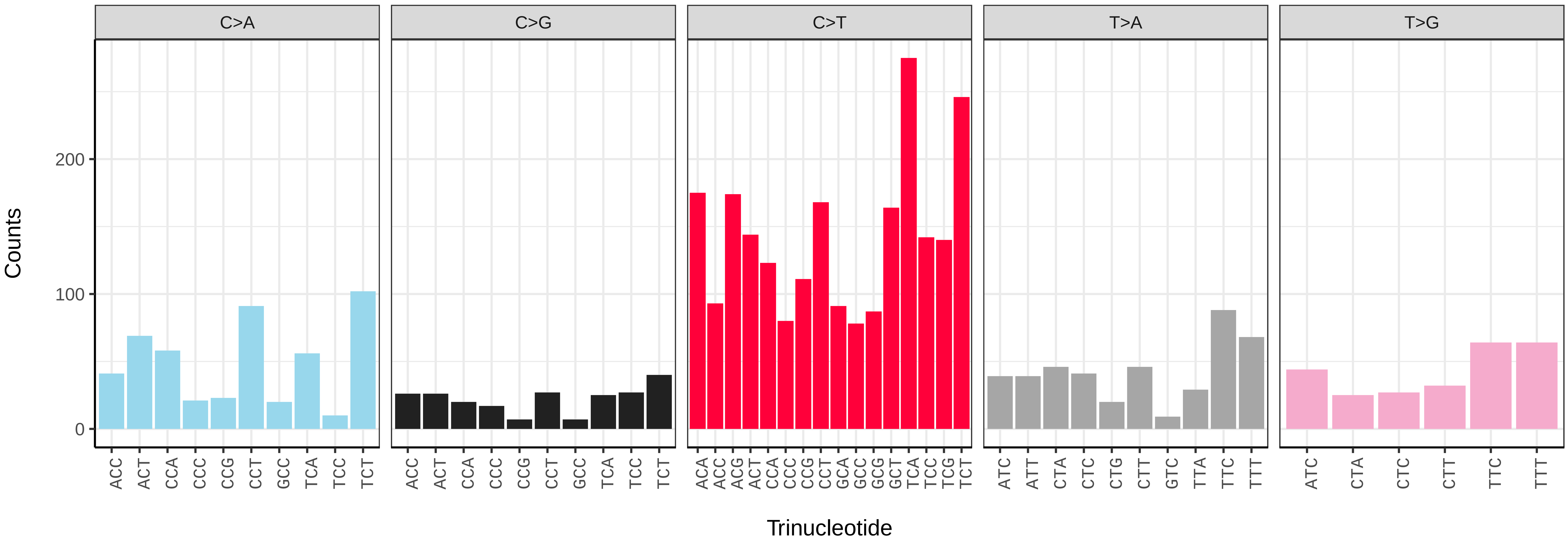


*Euphorbia peplus* (ddEupPepu3) somatic mutational spectrum (SBS96 classification)

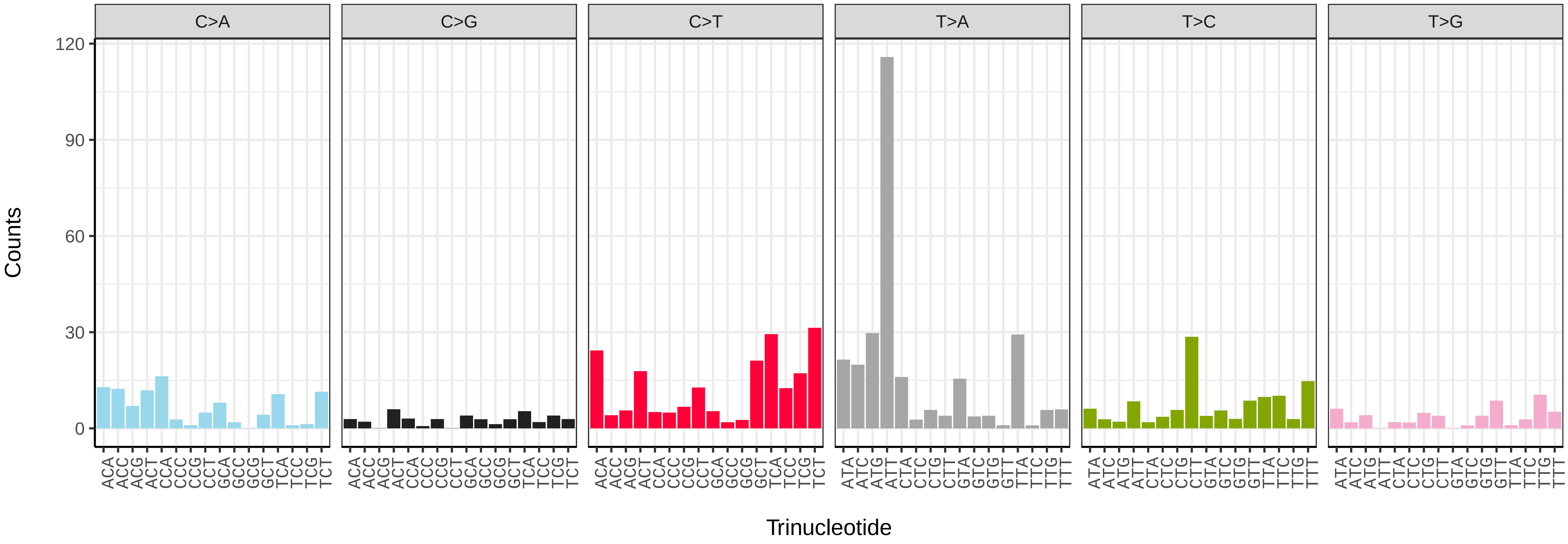


*Euphorbia peplus* (ddEupPepu3) somatic mutational spectrum (SBS52 classification)

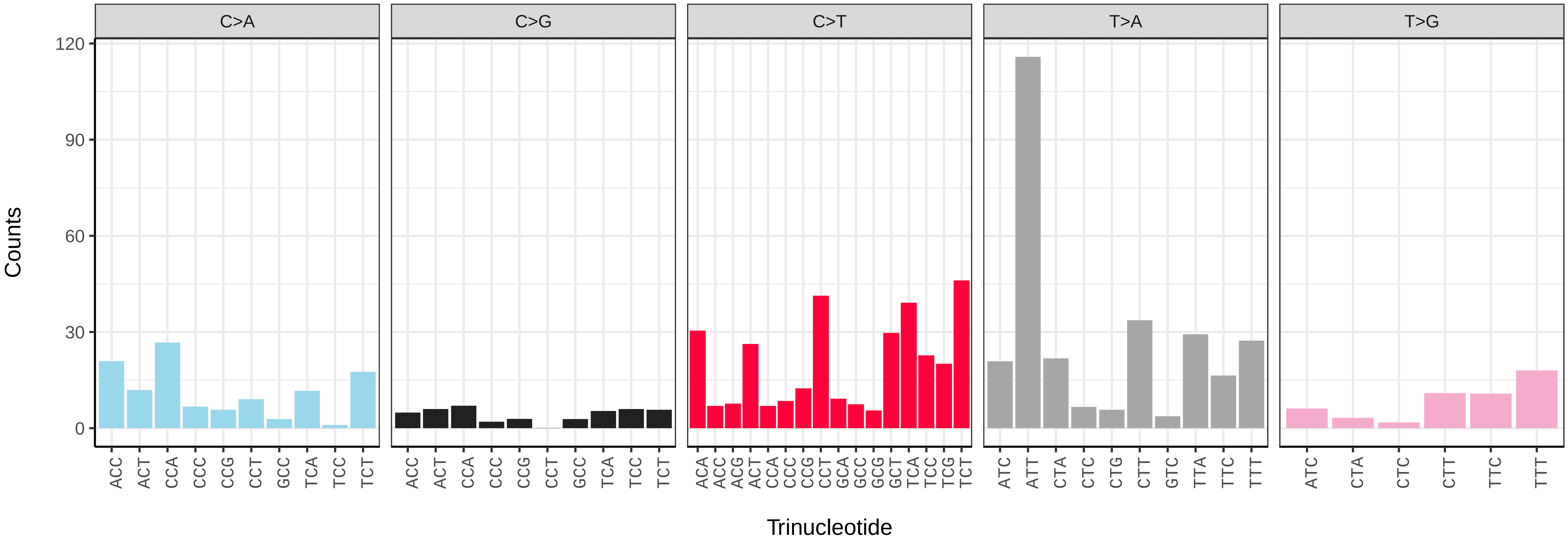

*Lepidium didymum* (ddLepDidy1) germline mutational spectrum

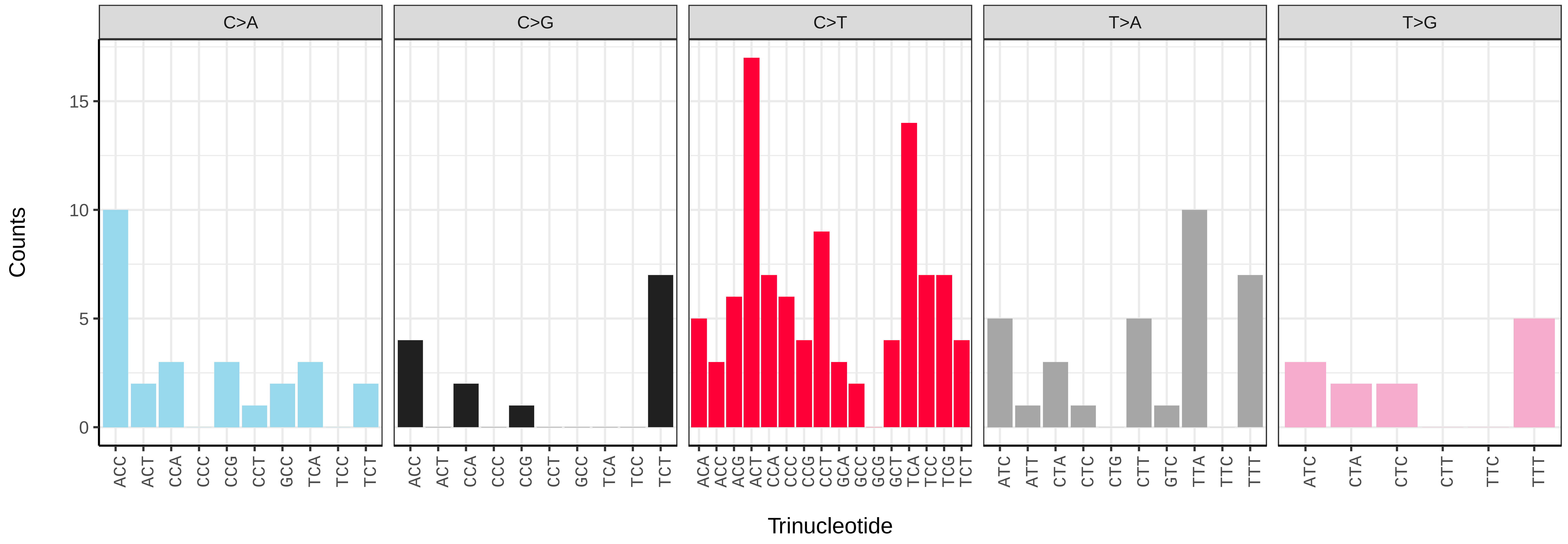


*Lepidium didymum* (ddLepDidy1) somatic mutational spectrum (SBS96 classification)

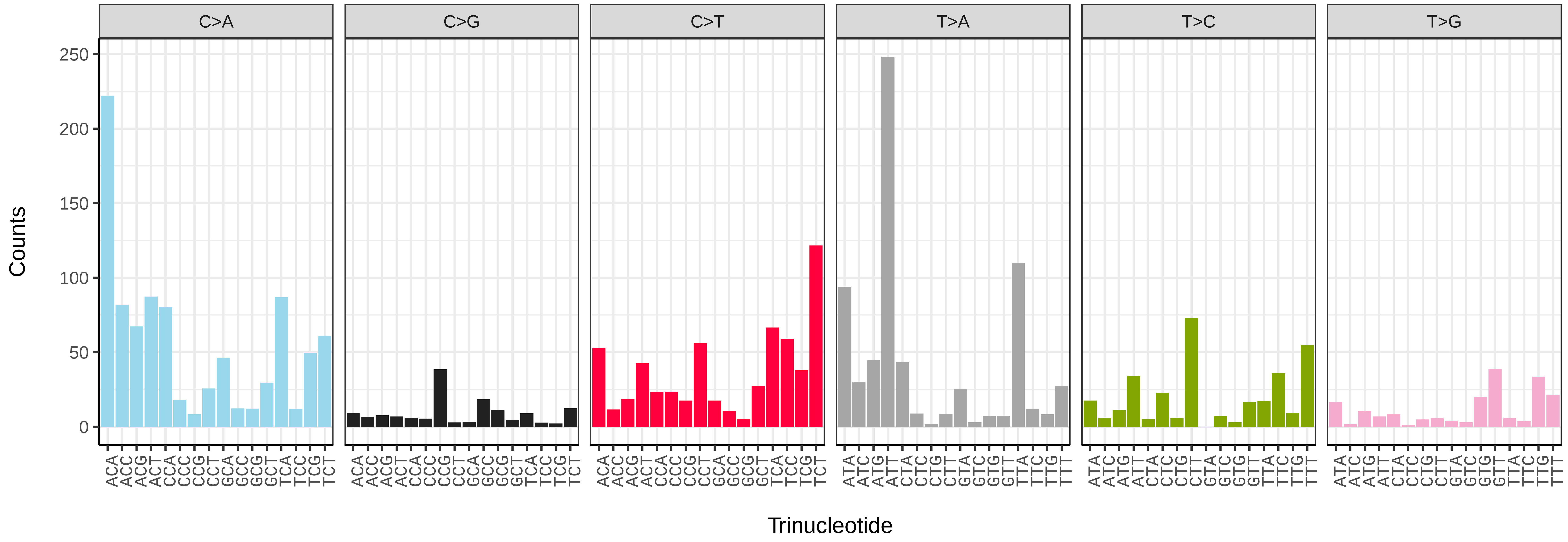


*Lepidium didymum* (ddLepDidy1) somatic mutational spectrum (SBS52 classification)

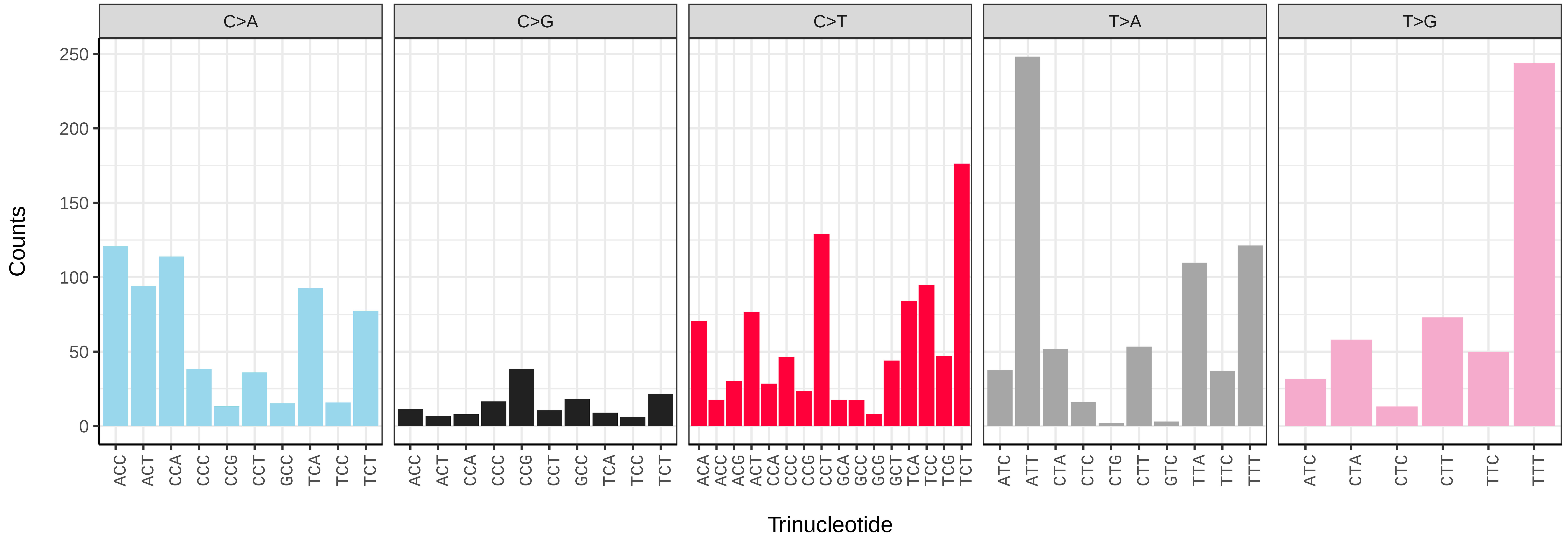

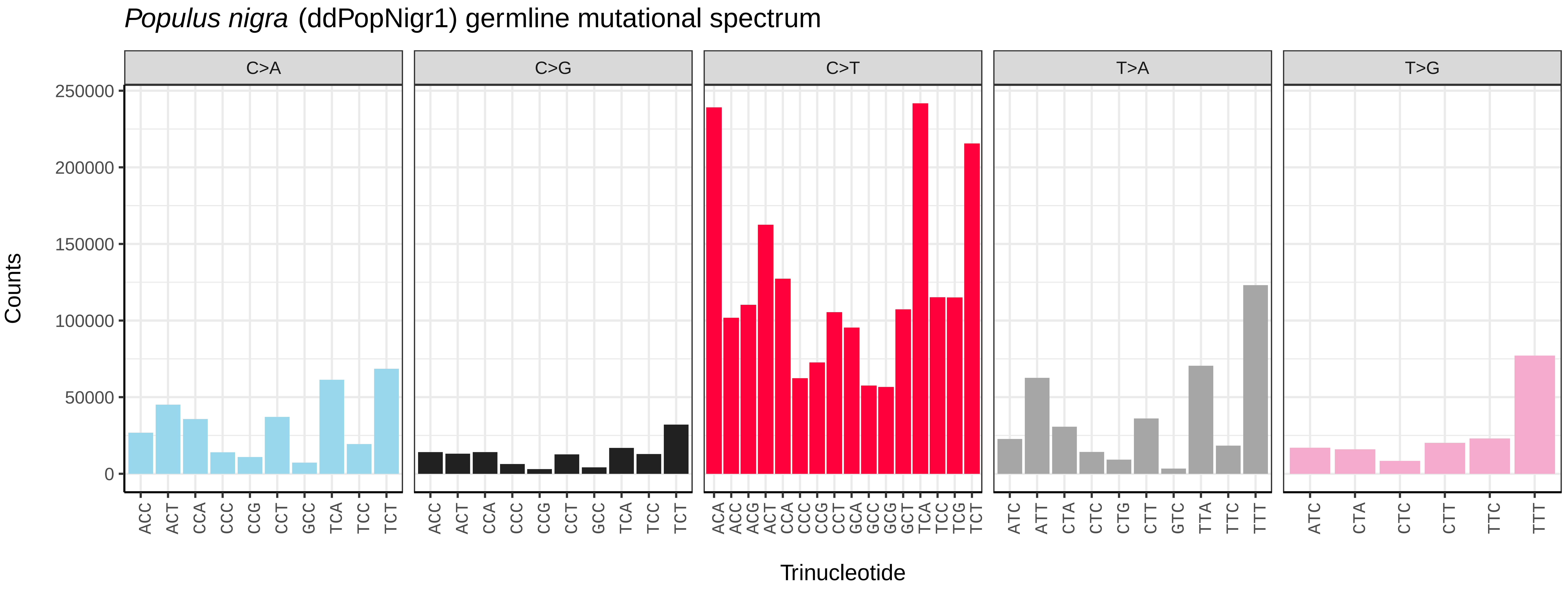
Populus nigra (ddPopNigr1) germline mutational spectrum
C>A
C>G
C>T
T>A
T>G
Counts
250000
200000
150000
100000
50000
0
ACC ACT CCA CCC CCG CCT GCC TCA TCC TCT
ACA ACC ACG ACT CCA CCC CCG CCT GCA GCC GCG GCT TCA TCC TCG TCT
ATC ATT CTA CTC CTG CTT GTC TTA TTC TTT
ATC CTA CTC CTT TTC TTT
Trinucleotide

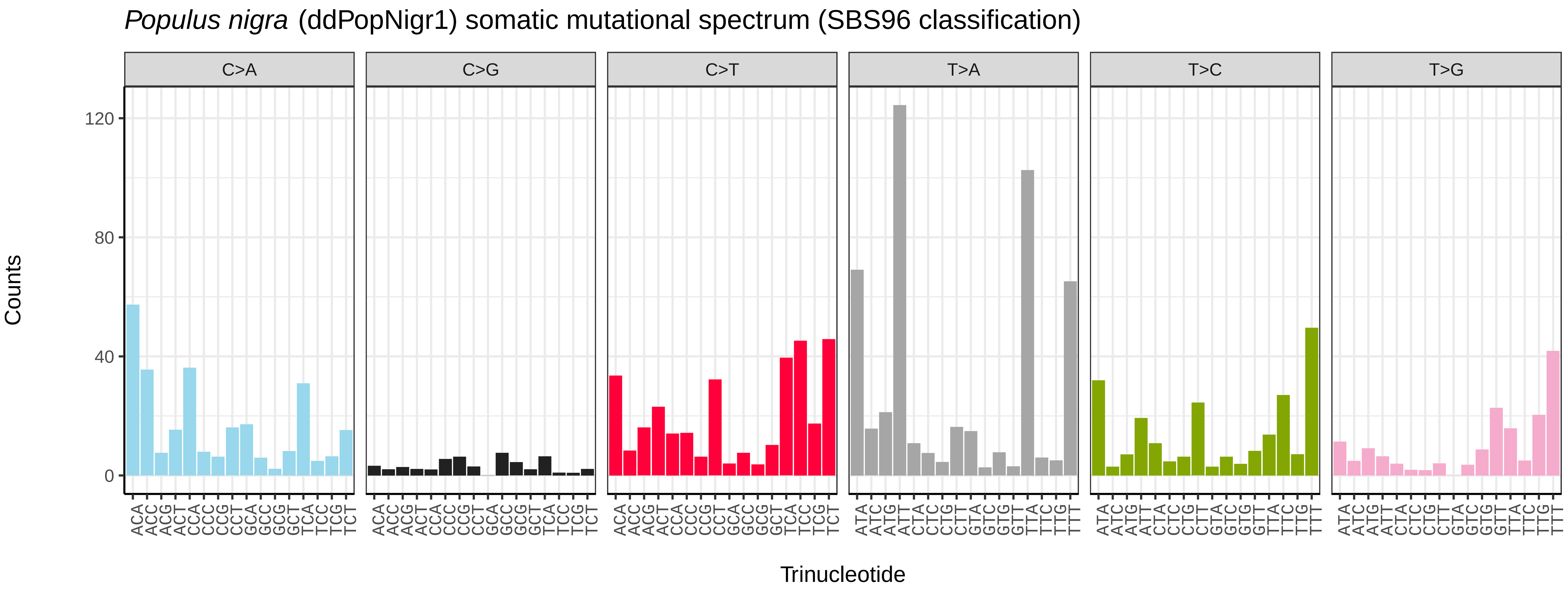
Populus nigra (ddPopNigr1) somatic mutational spectrum (SBS96 classification)
C>A
C>G
C>T
T>A
T>C
T>G
Counts
120
80
40
0
ACA ACC ACG ACT CCA CCC CCG CCT GCA GCC GCG GCT TCA TCC TCG TCT
ATA ATC ATG ATT CTA CTC CTG CTT GTA GTC GTG GTT TTA TTC TTG TTT
Trinucleotide

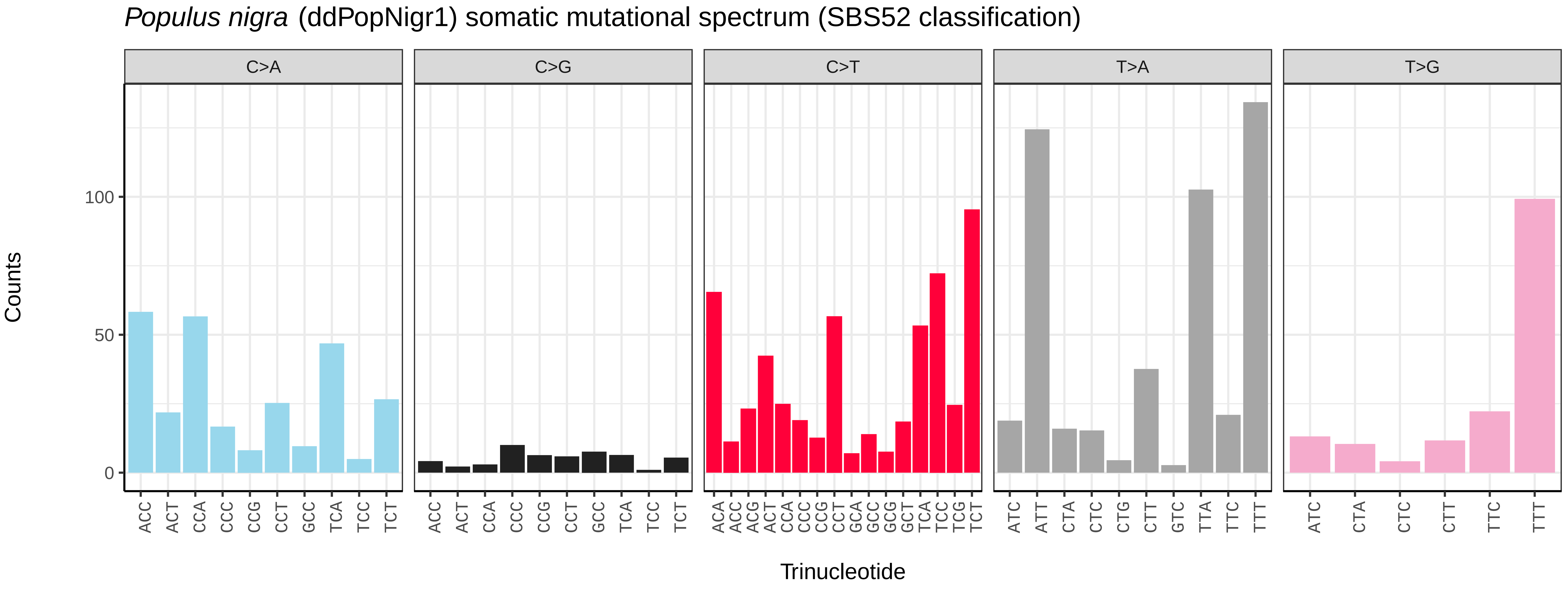
Populus nigra (ddPopNigr1) somatic mutational spectrum (SBS52 classification)
C>A
C>G
C>T
T>A
T>G
Counts
100
50
0
ACC ACT CCA CCC CCG CCT GCC TCA TCC TCT
ACA ACC ACG ACT CCA CCC CCG CCT GCA GCC GCG GCT TCA TCC TCG TCT
ATC ATT CTA CTC CTG CTT GTC TTA TTC TTT
ATC CTA CTC CTT TTC TTT
Trinucleotide

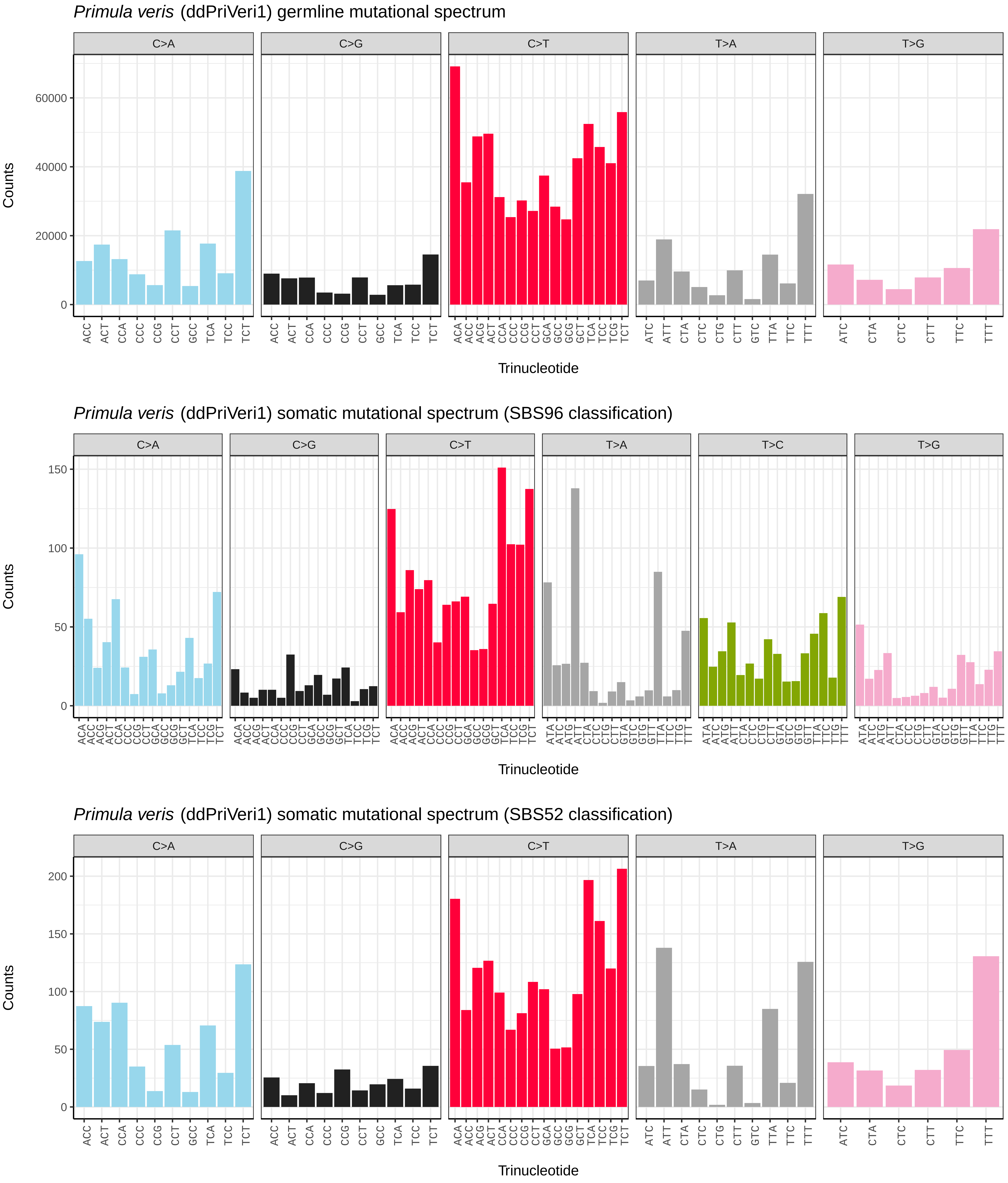
Primula veris (ddPriVeri1) germline mutational spectrum
C>A
C>G
C>T
T>A
T>G
Counts
Trinucleotide
Primula veris (ddPriVeri1) somatic mutational spectrum (SBS96 classification)
C>A
C>G
C>T
T>A
T>C
T>G
Counts
Trinucleotide
Primula veris (ddPriVeri1) somatic mutational spectrum (SBS52 classification)
C>A
C>G
C>T
T>A
T>G
Counts
Trinucleotide

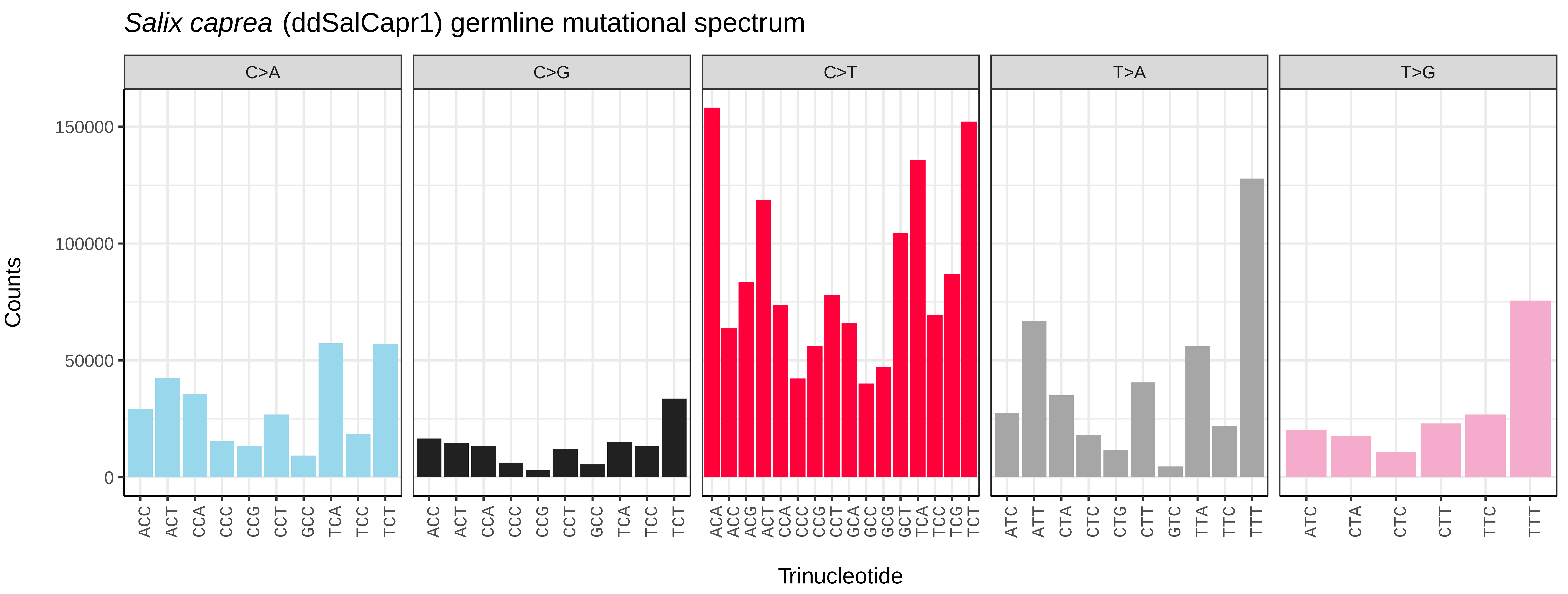
*Salix caprea* (ddSalCapr1) germline mutational spectrum
C>A
C>G
C>T
T>A
T>G
Counts
150000
100000
50000
0
ACC ACT CCA CCC CCG CCT GCC TCA TCC TCT
ACC ACT CCA CCC CCG CCT GCC TCA TCC TCT
ACA ACC ACG ACT CCA CCC CCG CCT GCA GCC GCG GCT TCA TCC TCG TCT
ATC ATT CTA CTC CTG CTT GTC TTA TTC TTT
ATC CTA CTC CTT TTC TTT
Trinucleotide

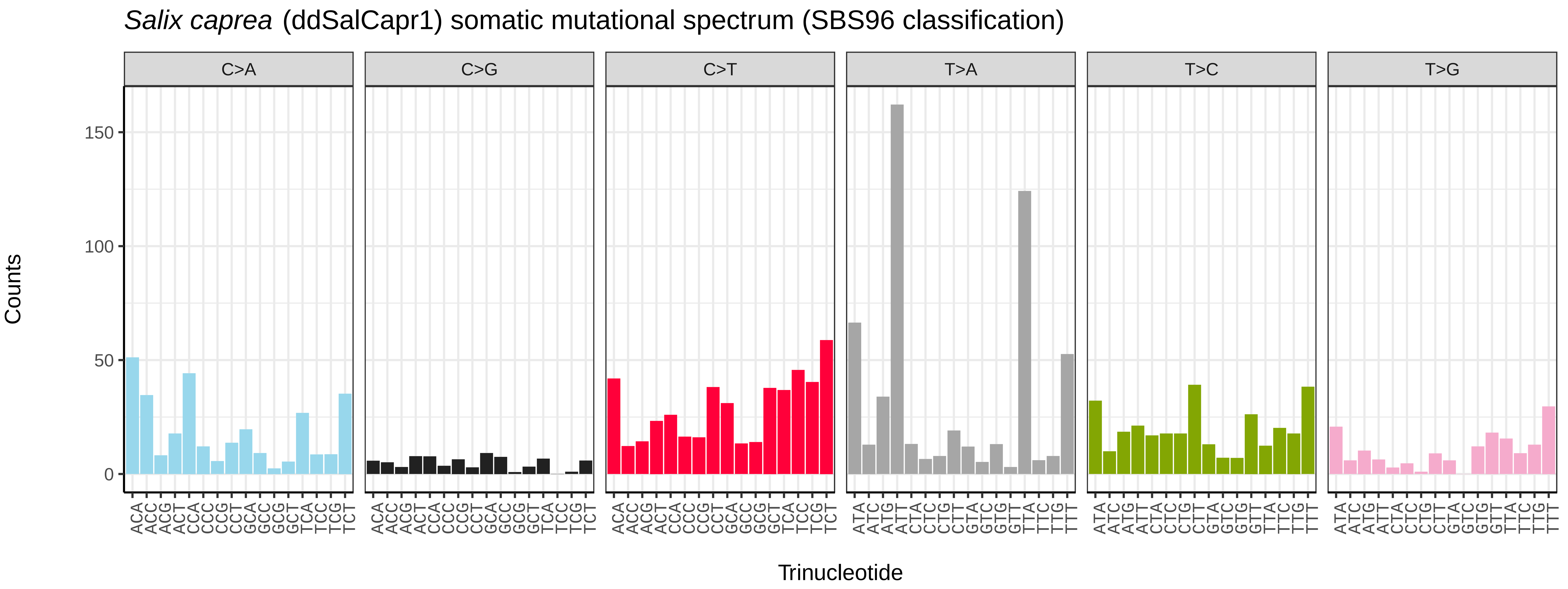
*Salix caprea* (ddSalCapr1) somatic mutational spectrum (SBS96 classification)
C>A
C>G
C>T
T>A
T>C
T>G
Counts
150
100
50
0
ACA ACC ACG ACT CCA CCC CCG CCT GCA GCC GCG GCT TCA TCC TCG TCT
ATA ATC ATG ATT CTA CTC CTG CTT GTA GTC GTG GTT TTA TTC TTG TTT
Trinucleotide

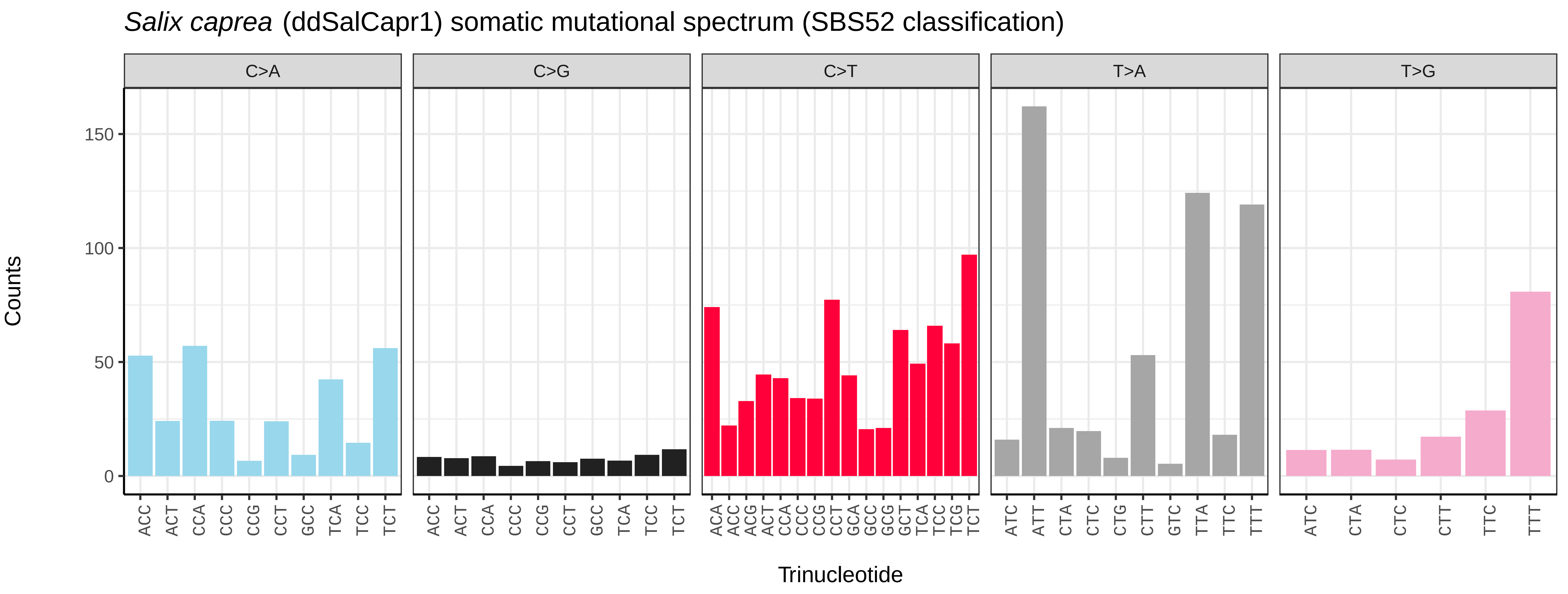
*Salix caprea* (ddSalCapr1) somatic mutational spectrum (SBS52 classification)
C>A
C>G
C>T
T>A
T>G
Counts
150
100
50
0
ACC ACT CCA CCC CCG CCT GCC TCA TCC TCT
ACC ACT CCA CCC CCG CCT GCC TCA TCC TCT
ACA ACC ACG ACT CCA CCC CCG CCT GCA GCC GCG GCT TCA TCC TCG TCT
ATC ATT CTA CTC CTG CTT GTC TTA TTC TTT
ATC CTA CTC CTT TTC TTT
Trinucleotide

*Viola odorata* (ddVioOdor1) germline mutational spectrum

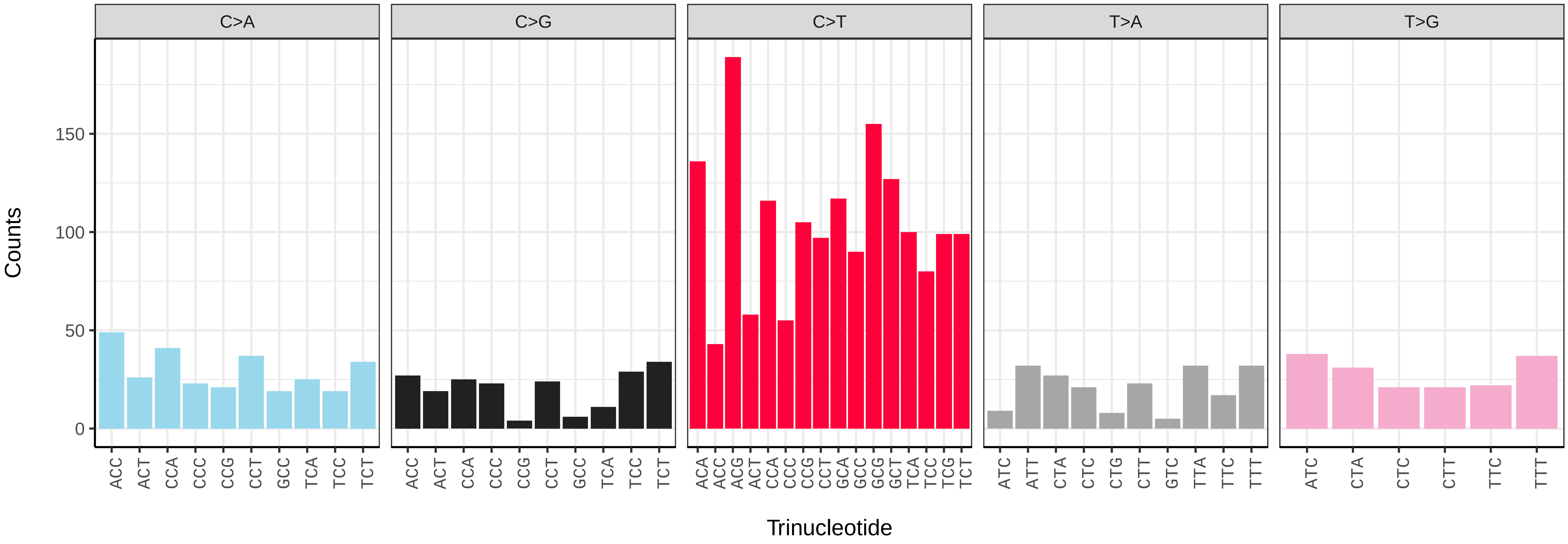


*Viola odorata* (ddVioOdor1) somatic mutational spectrum (SBS96 classification)

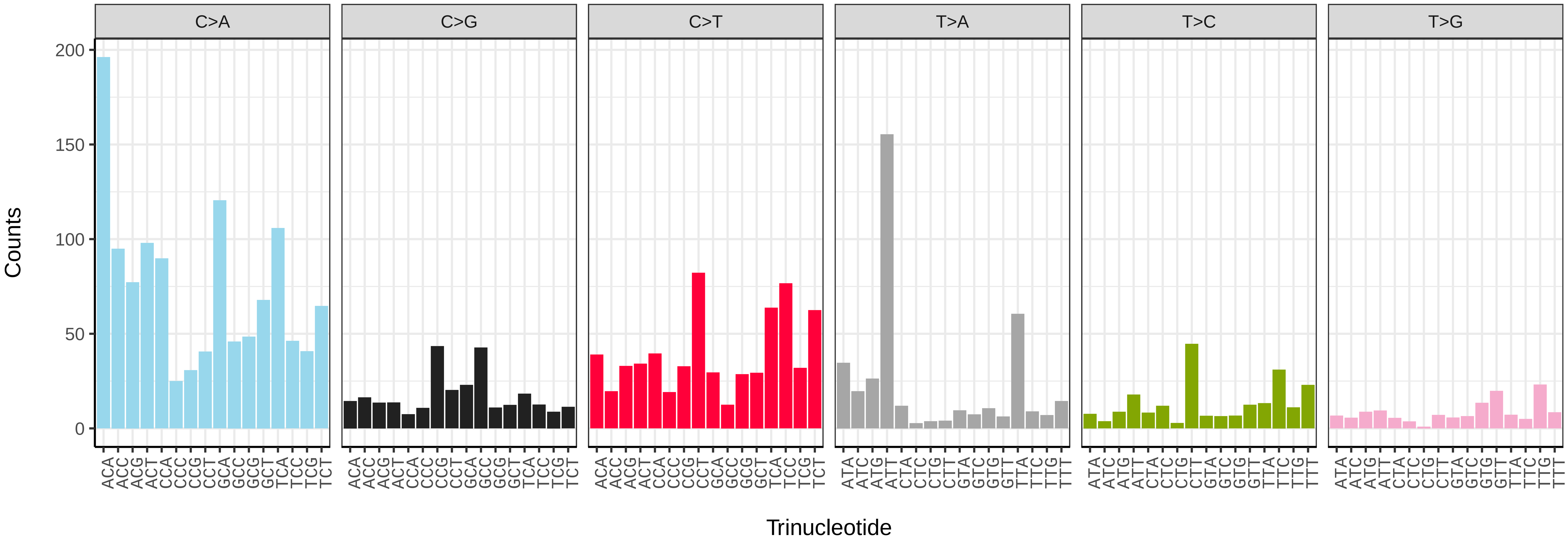


*Viola odorata* (ddVioOdor1) somatic mutational spectrum (SBS52 classification)

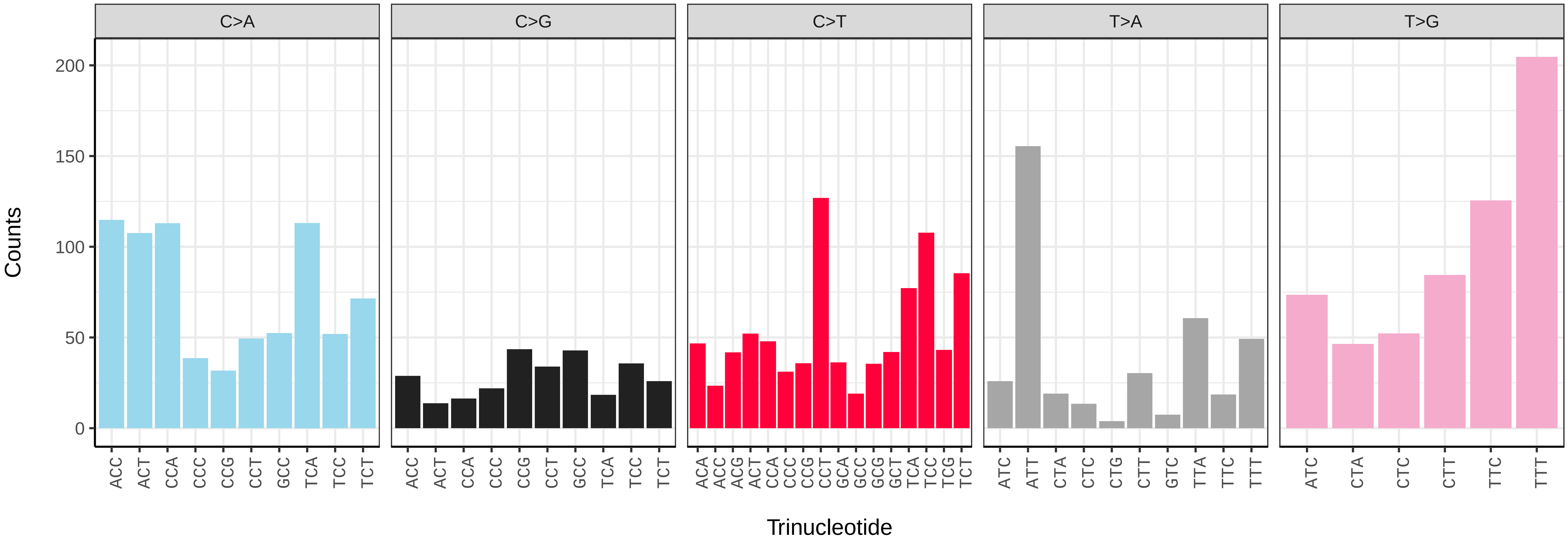

*Alnus glutinosa* (dhAlnGlut1) germline mutational spectrum

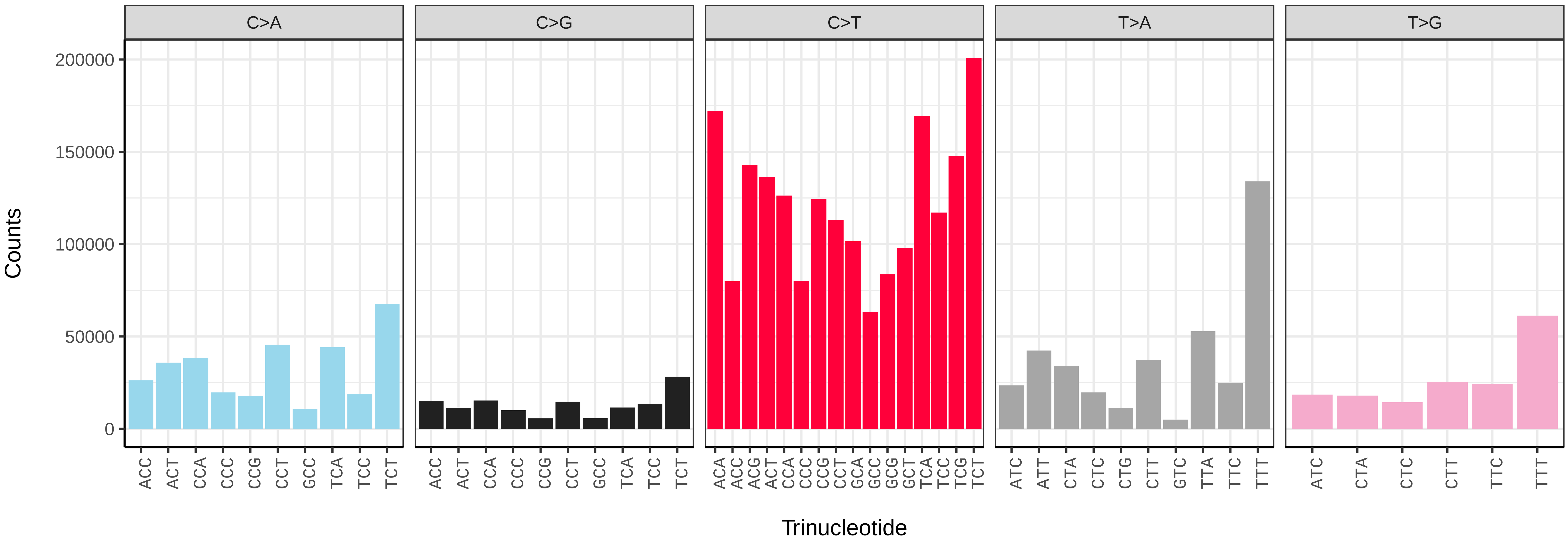


*Alnus glutinosa* (dhAlnGlut1) somatic mutational spectrum (SBS96 classification)

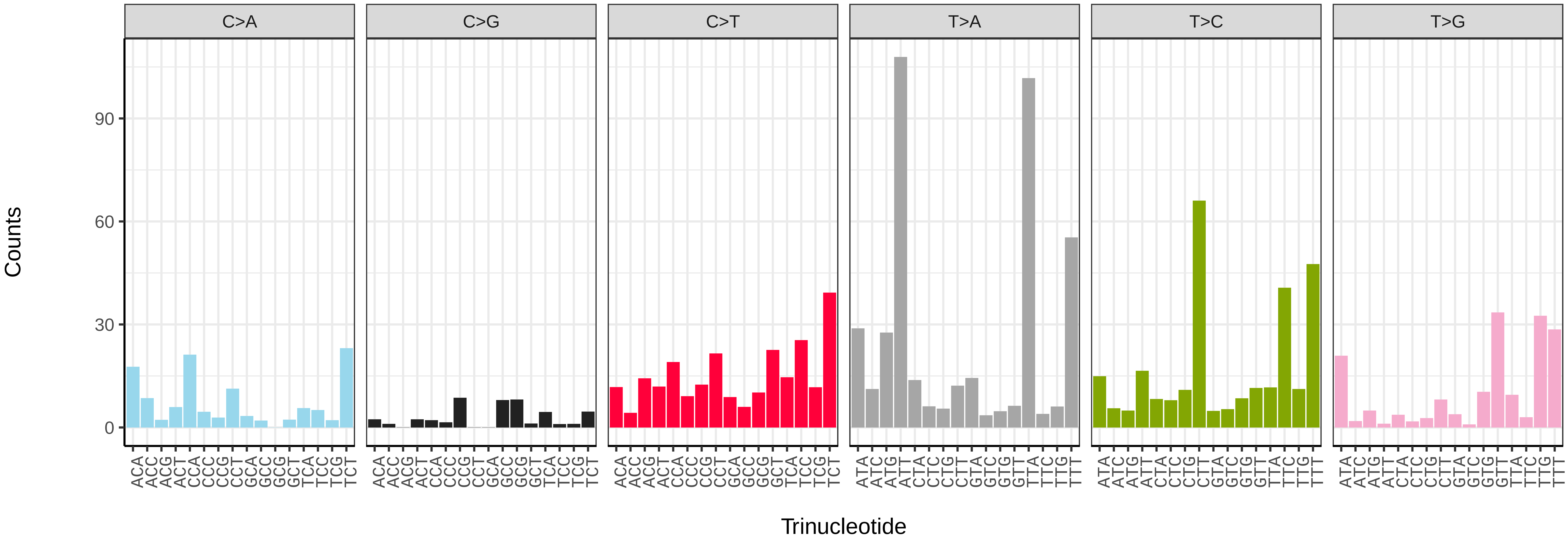


*Alnus glutinosa* (dhAlnGlut1) somatic mutational spectrum (SBS52 classification)

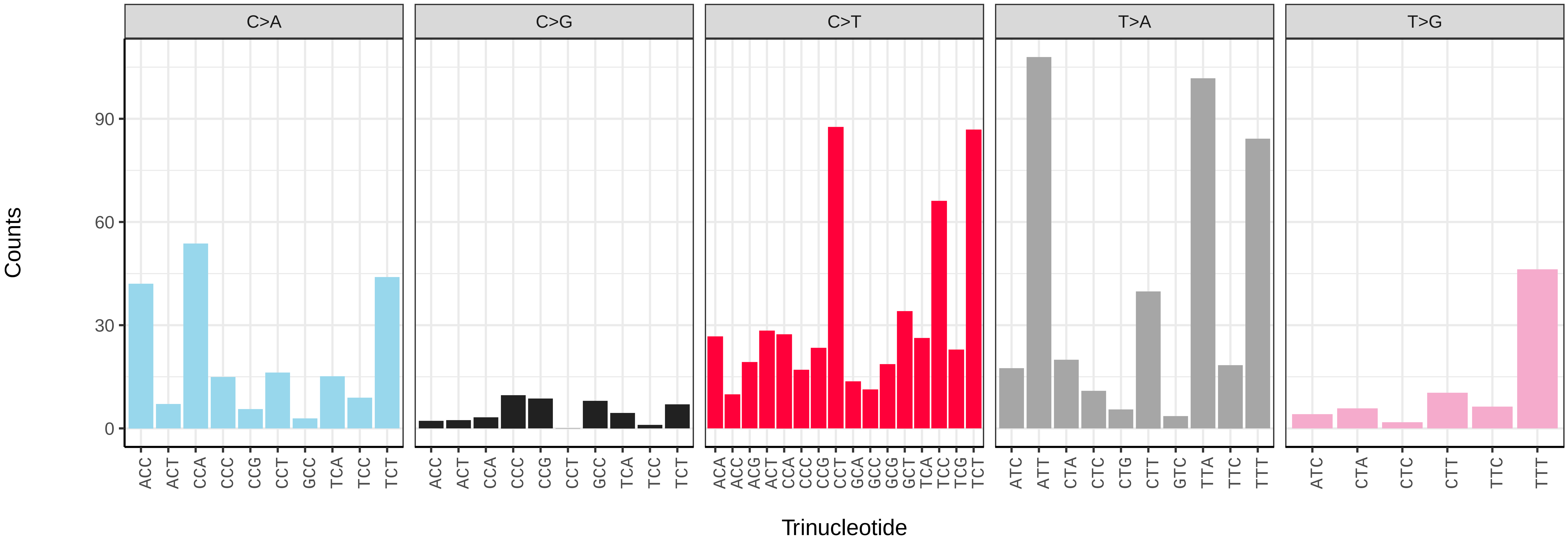

*Berberis vulgaris* (dmBerVulg1) germline mutational spectrum

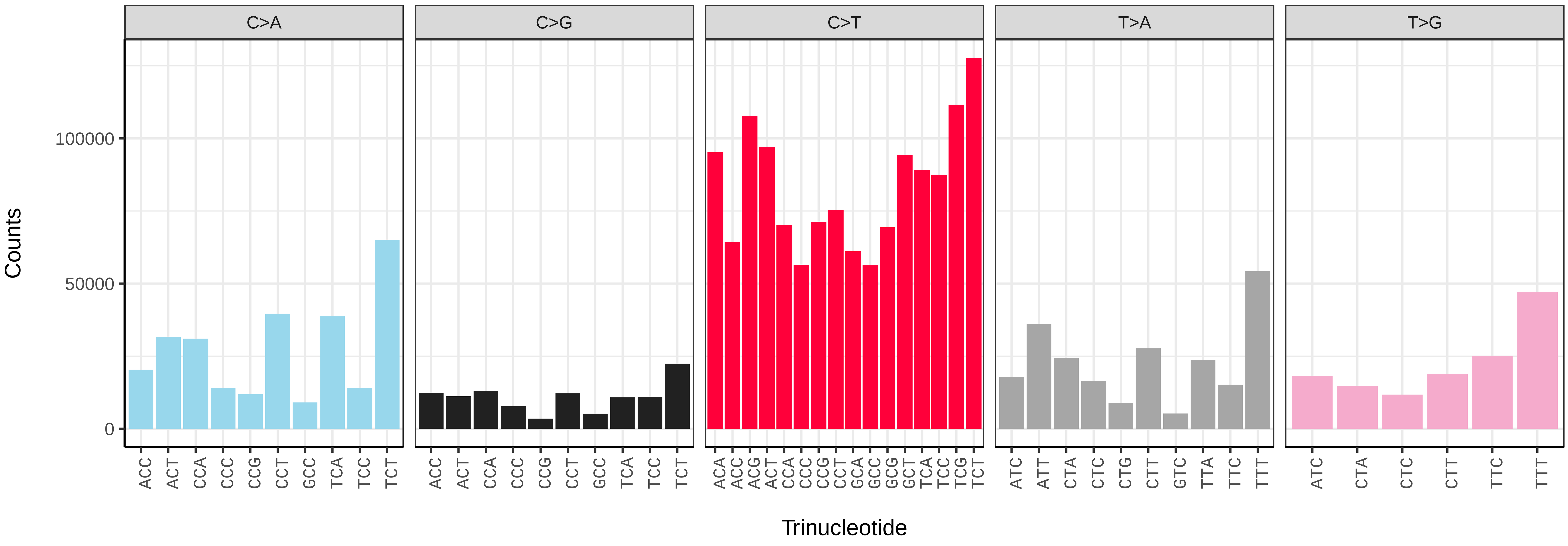


*Berberis vulgaris* (dmBerVulg1) somatic mutational spectrum (SBS96 classification)

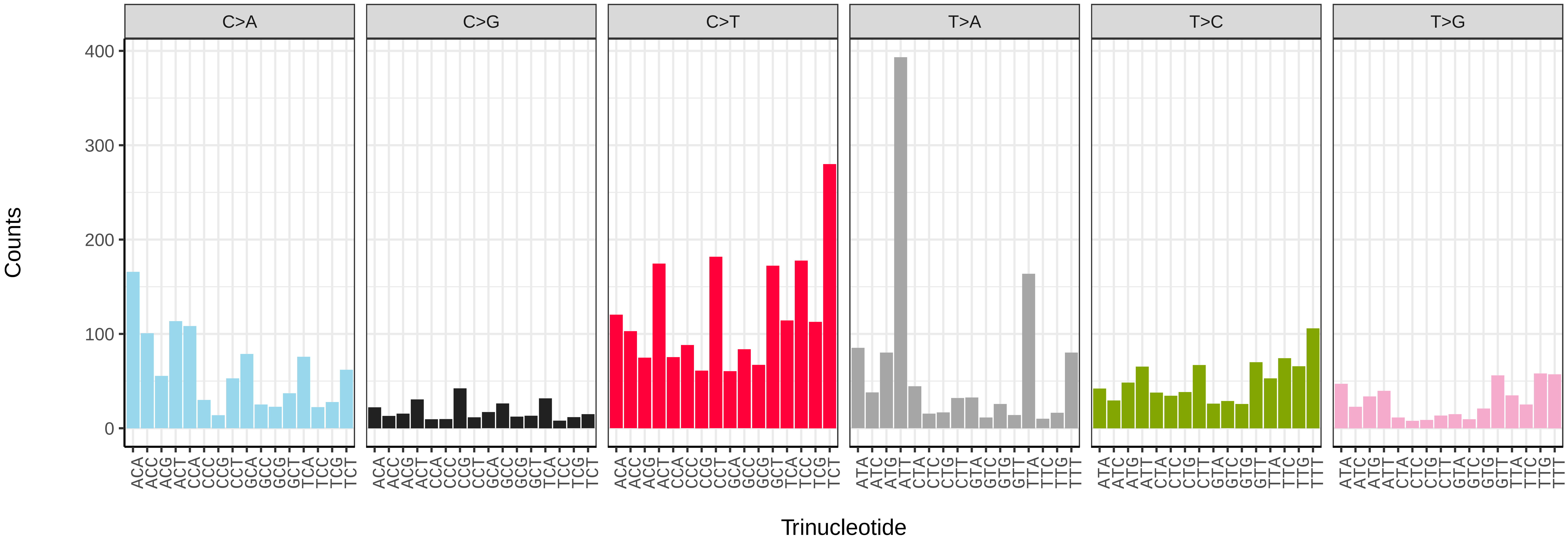


*Berberis vulgaris* (dmBerVulg1) somatic mutational spectrum (SBS52 classification)

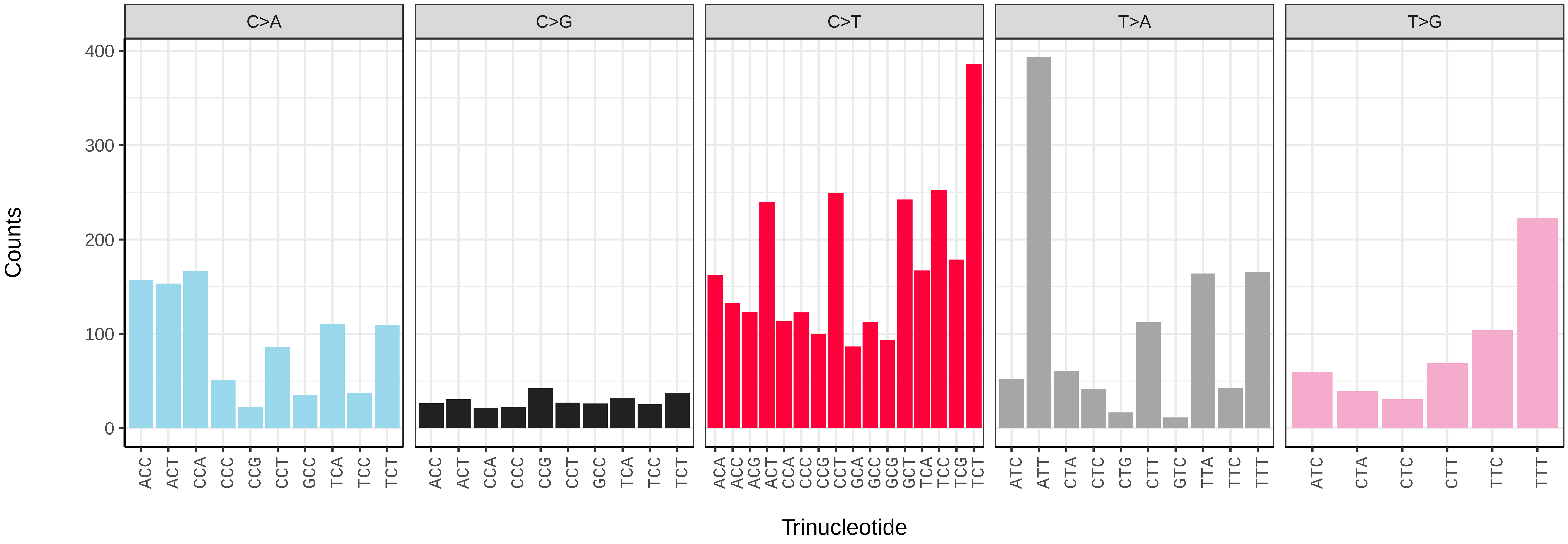

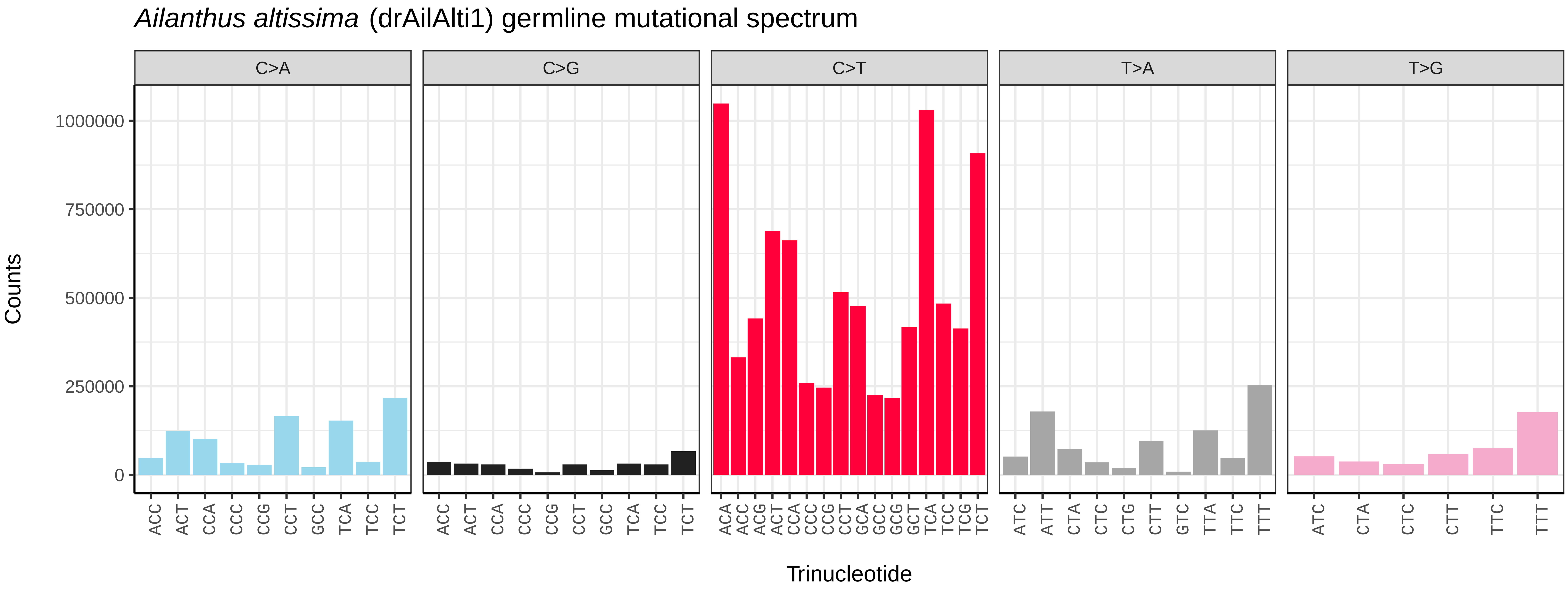
*Ailanthus altissima* (drAilAlti1) germline mutational spectrum
C>A
C>G
C>T
T>A
T>G
Counts
1000000
750000
500000
250000
0
Trinucleotide

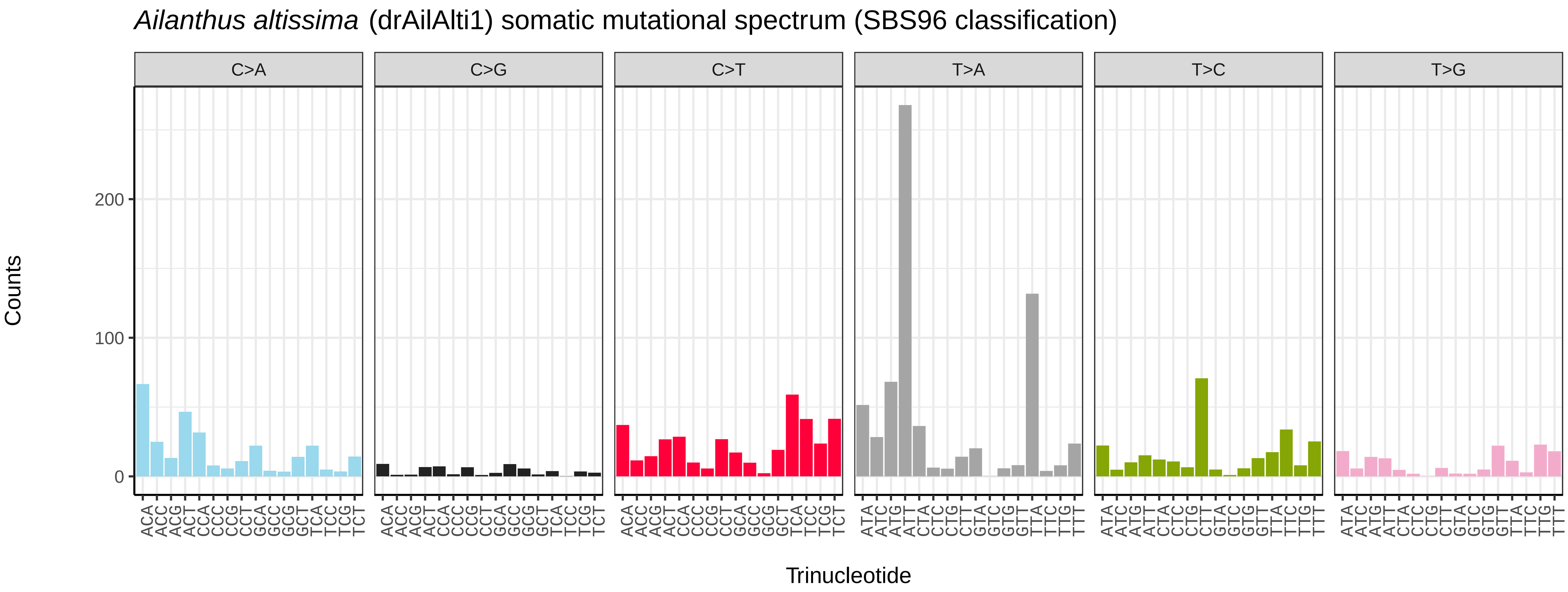
*Ailanthus altissima* (drAilAlti1) somatic mutational spectrum (SBS96 classification)
C>A
C>G
C>T
T>A
T>C
T>G
Counts
200
100
0
Trinucleotide

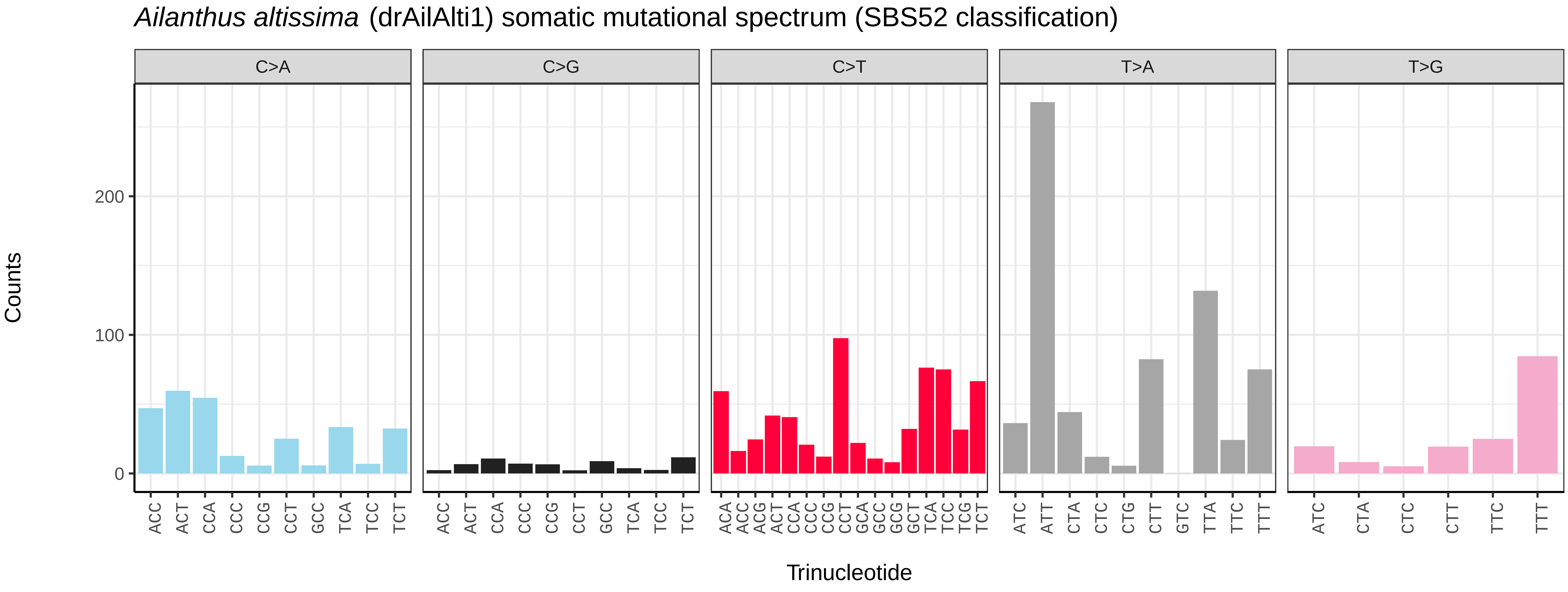
*Ailanthus altissima* (drAilAlti1) somatic mutational spectrum (SBS52 classification)
C>A
C>G
C>T
T>A
T>G
Counts
200
100
0
Trinucleotide

*Astragalus glycyphyllos* (drAstGlyc1) germline mutational spectrum

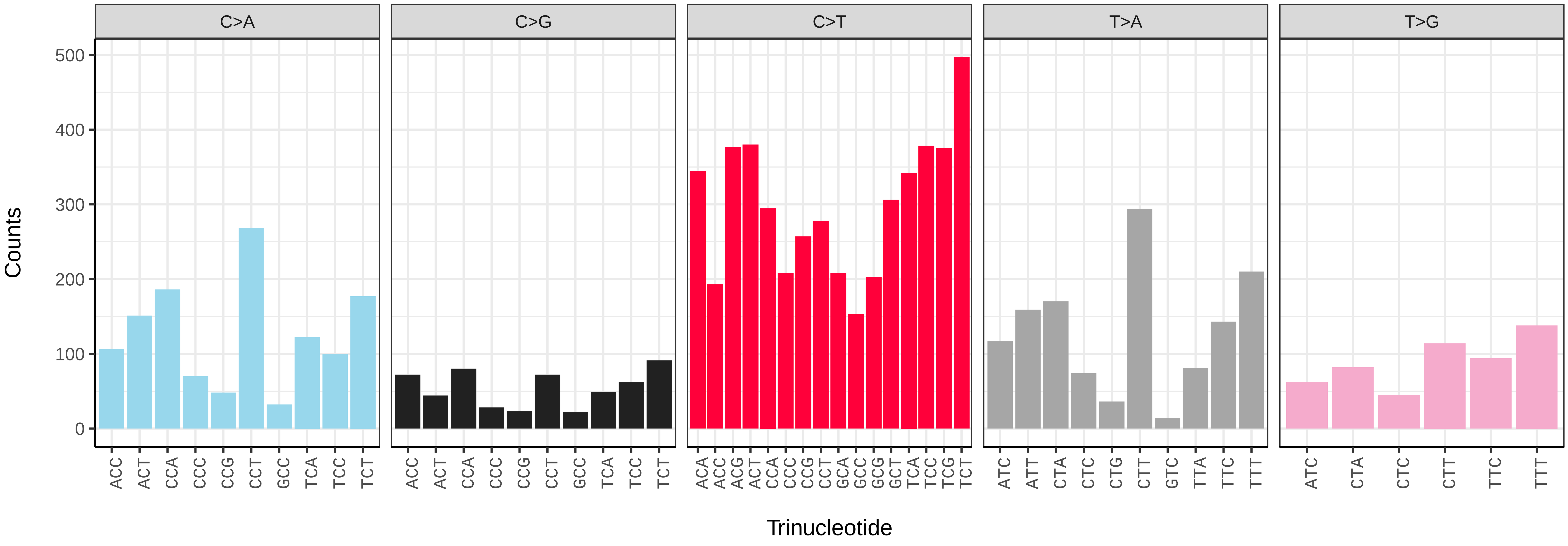


*Astragalus glycyphyllos* (drAstGlyc1) somatic mutational spectrum (SBS96 classification)

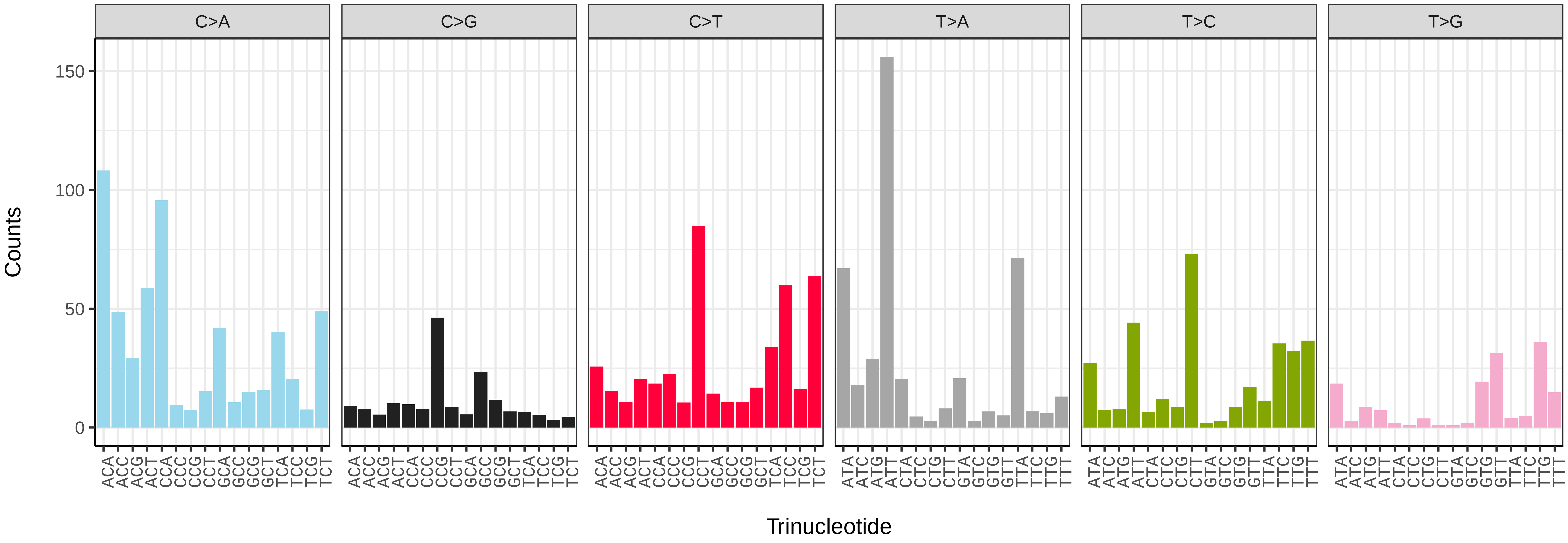


*Astragalus glycyphyllos* (drAstGlyc1) somatic mutational spectrum (SBS52 classification)

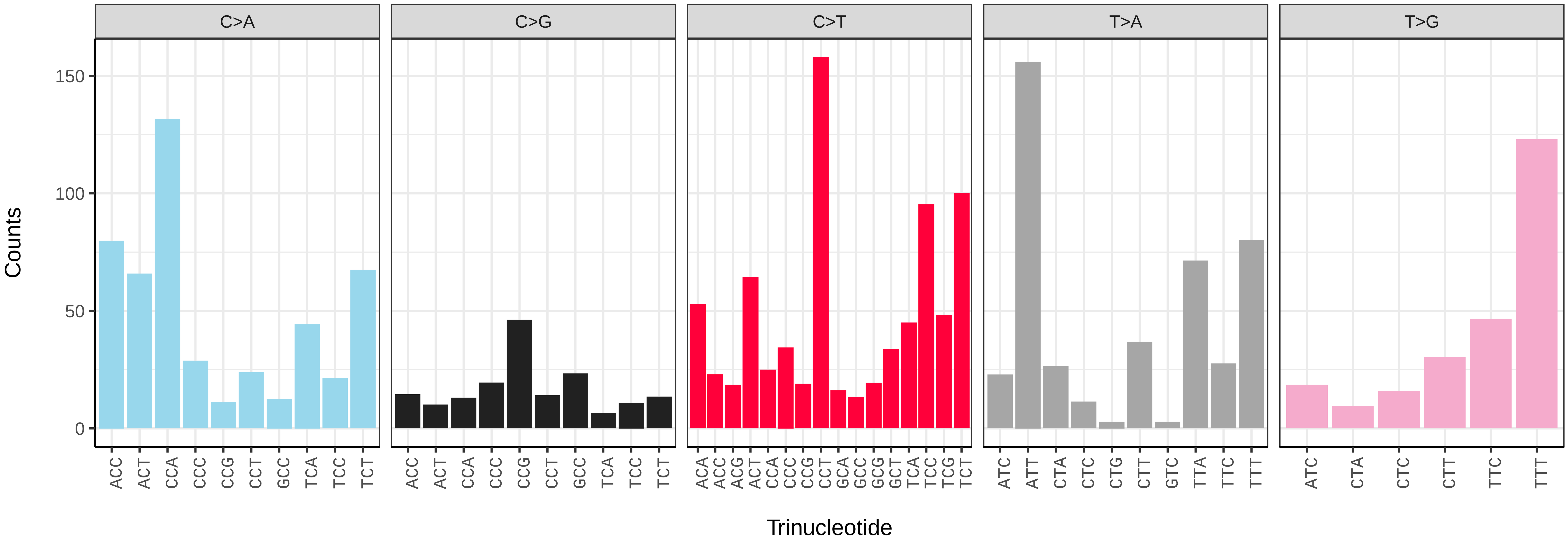

*Buxus sempervirens* (drBuxSemp1) germline mutational spectrum

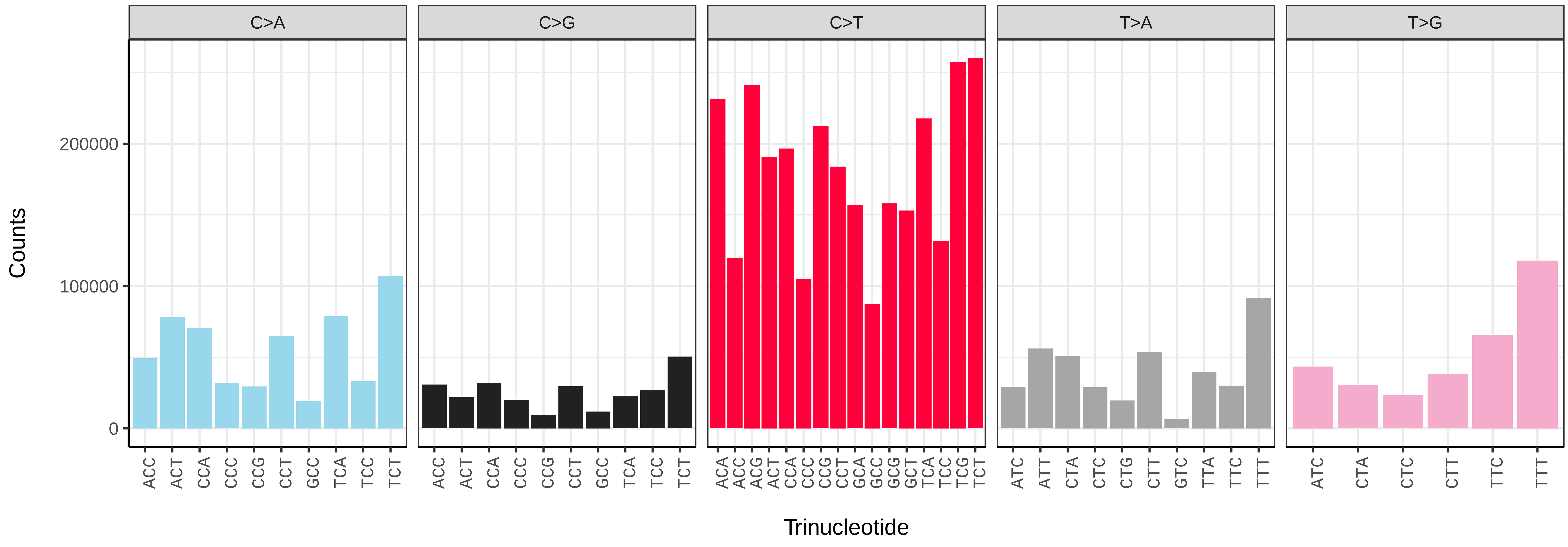


*Buxus sempervirens* (drBuxSemp1) somatic mutational spectrum (SBS96 classification)

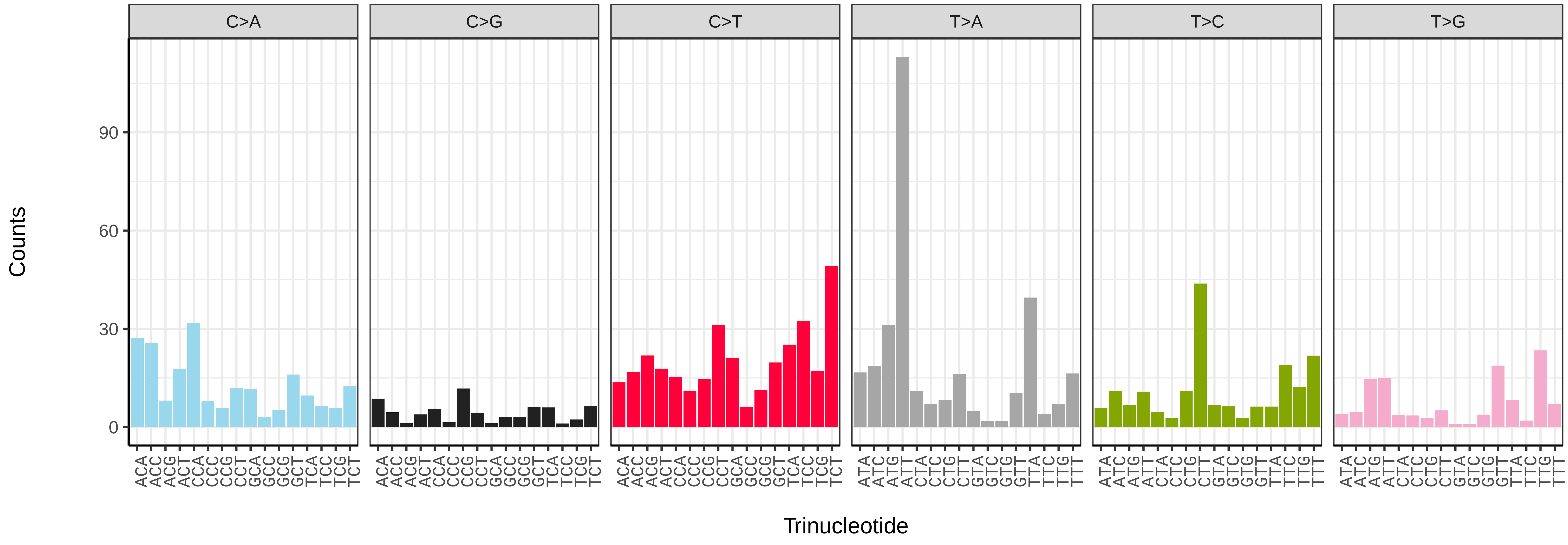


*Buxus sempervirens* (drBuxSemp1) somatic mutational spectrum (SBS52 classification)

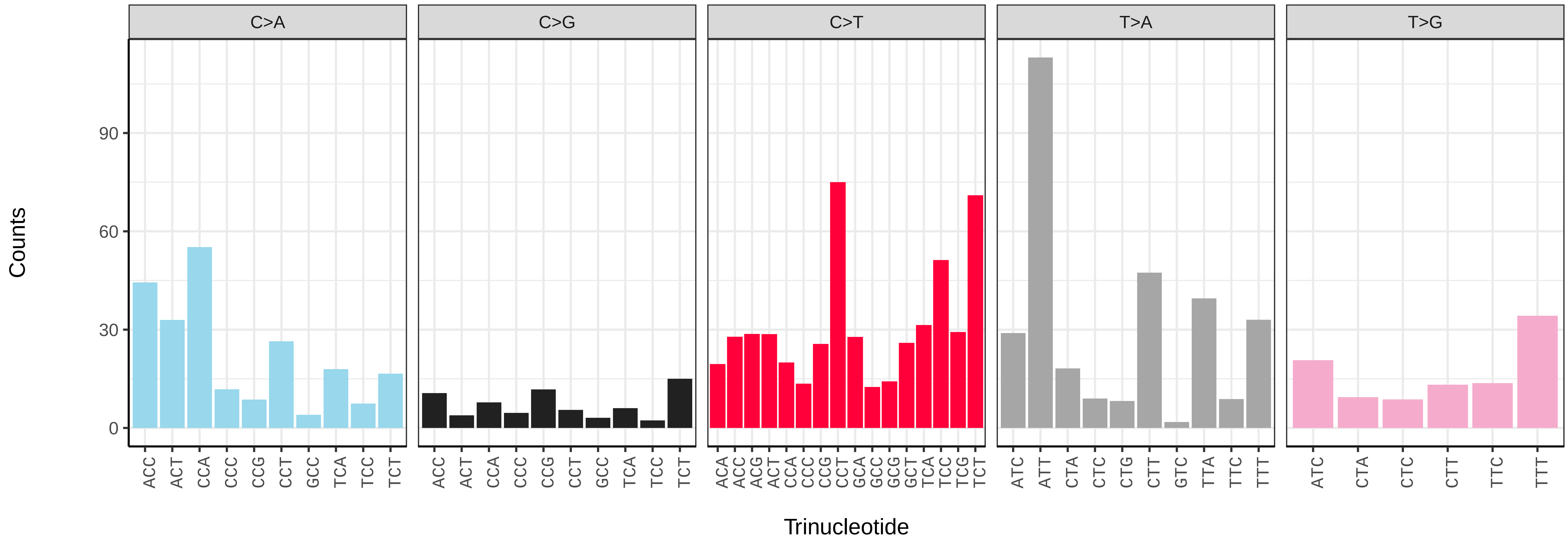

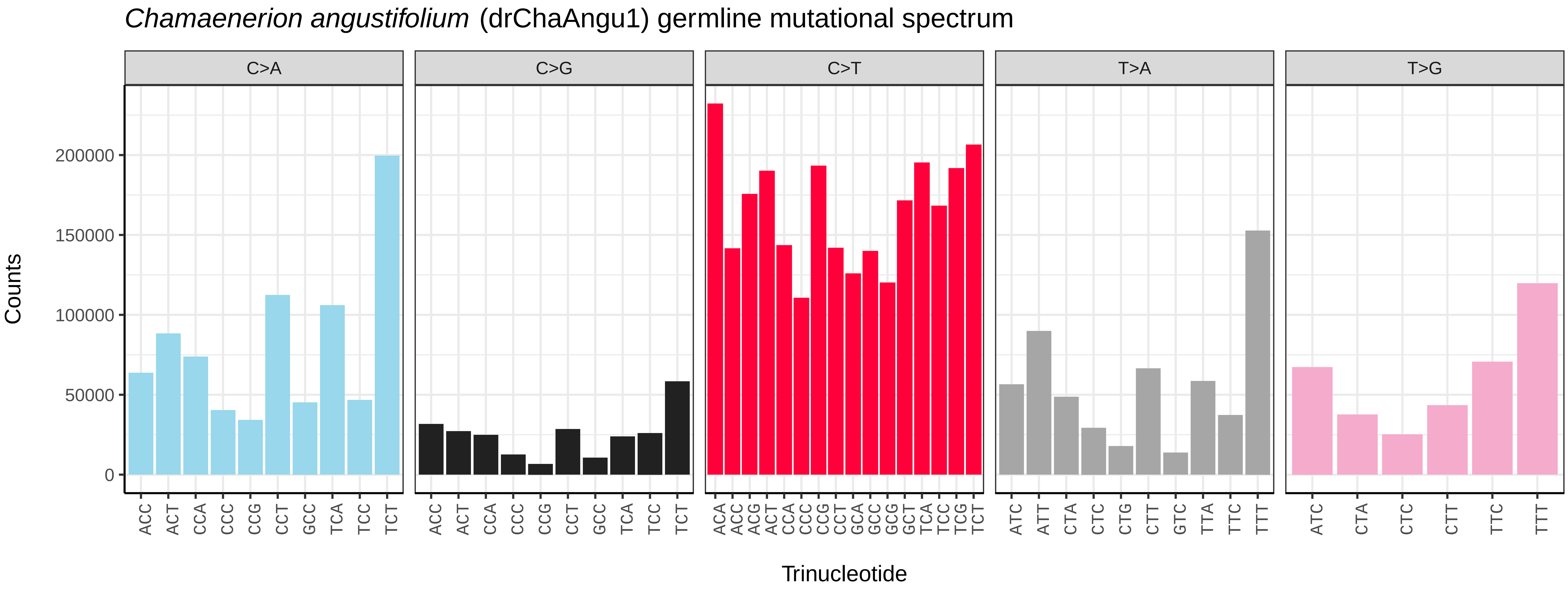
*Chamaenerion angustifolium* (drChaAngu1) germline mutational spectrum
C>A
C>G
C>T
T>A
T>G
Counts
0
50000
100000
150000
200000
Trinucleotide

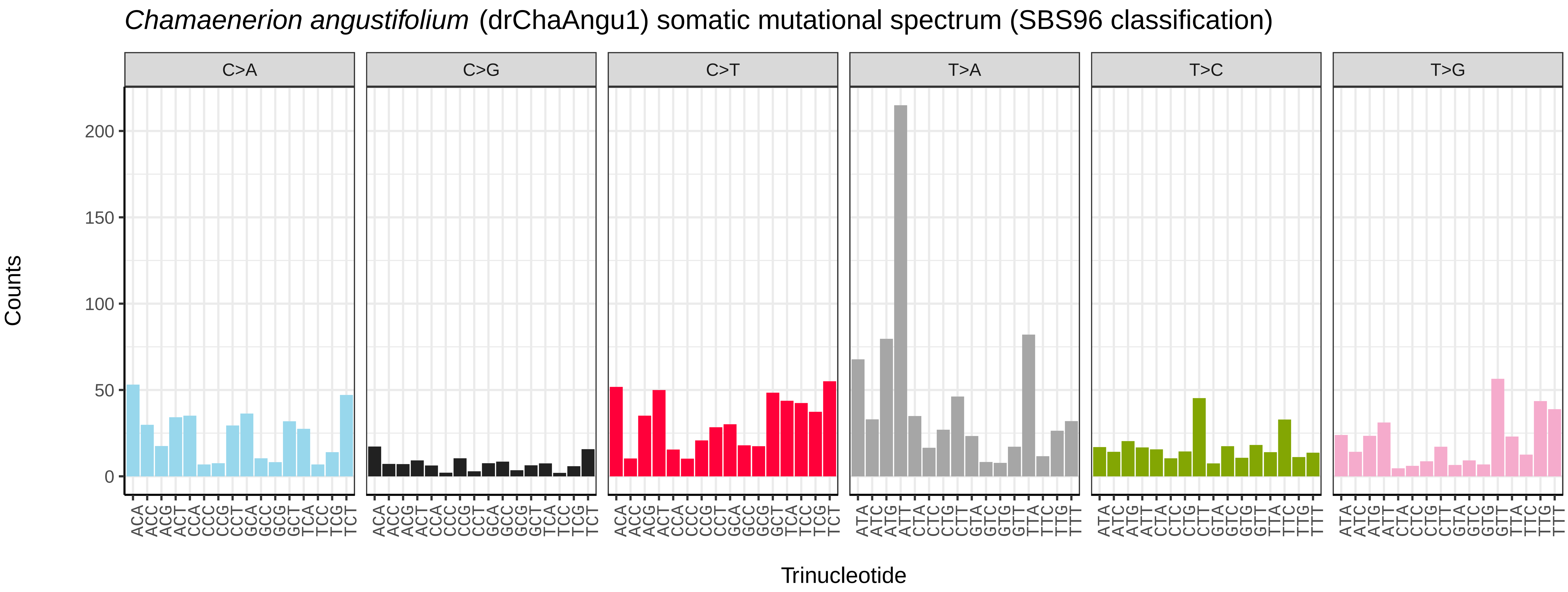
*Chamaenerion angustifolium* (drChaAngu1) somatic mutational spectrum (SBS96 classification)
C>A
C>G
C>T
T>A
T>C
T>G
Counts
0
50
100
150
200
Trinucleotide

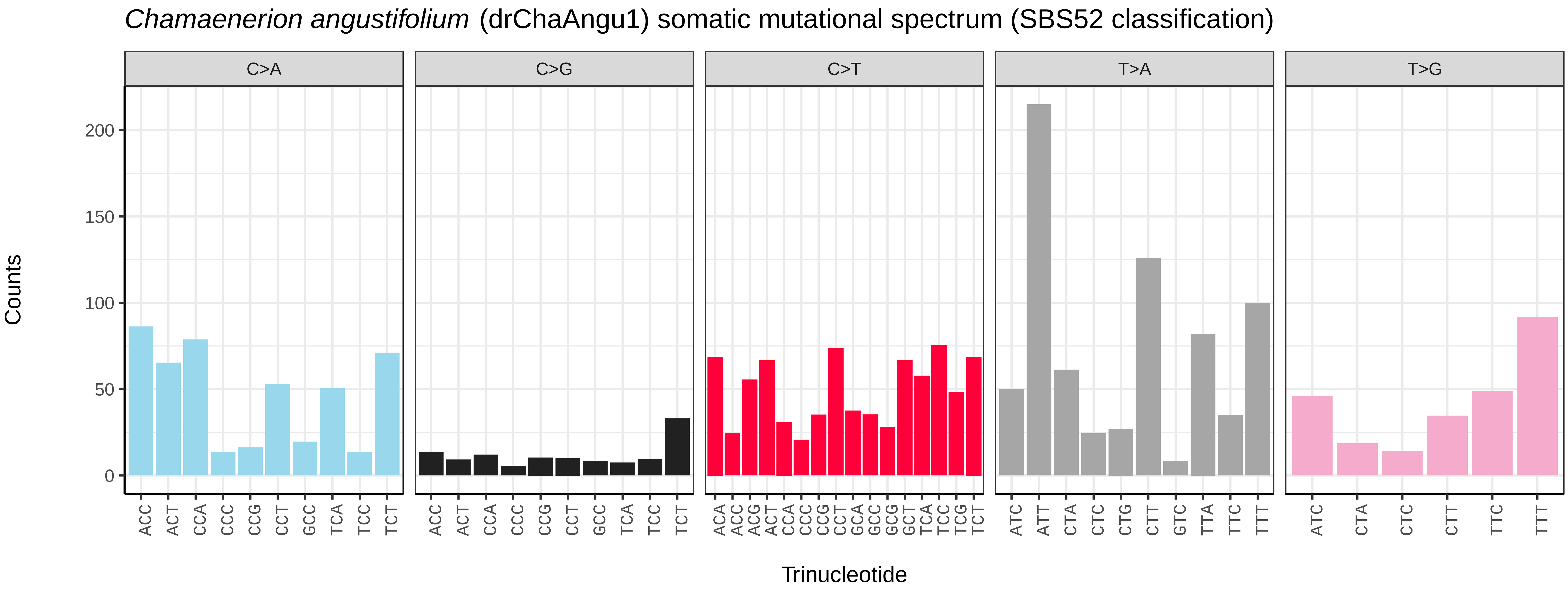
*Chamaenerion angustifolium* (drChaAngu1) somatic mutational spectrum (SBS52 classification)
C>A
C>G
C>T
T>A
T>G
Counts
0
50
100
150
200
Trinucleotide

*Crataegus laevigata* (drCraLaev2) germline mutational spectrum

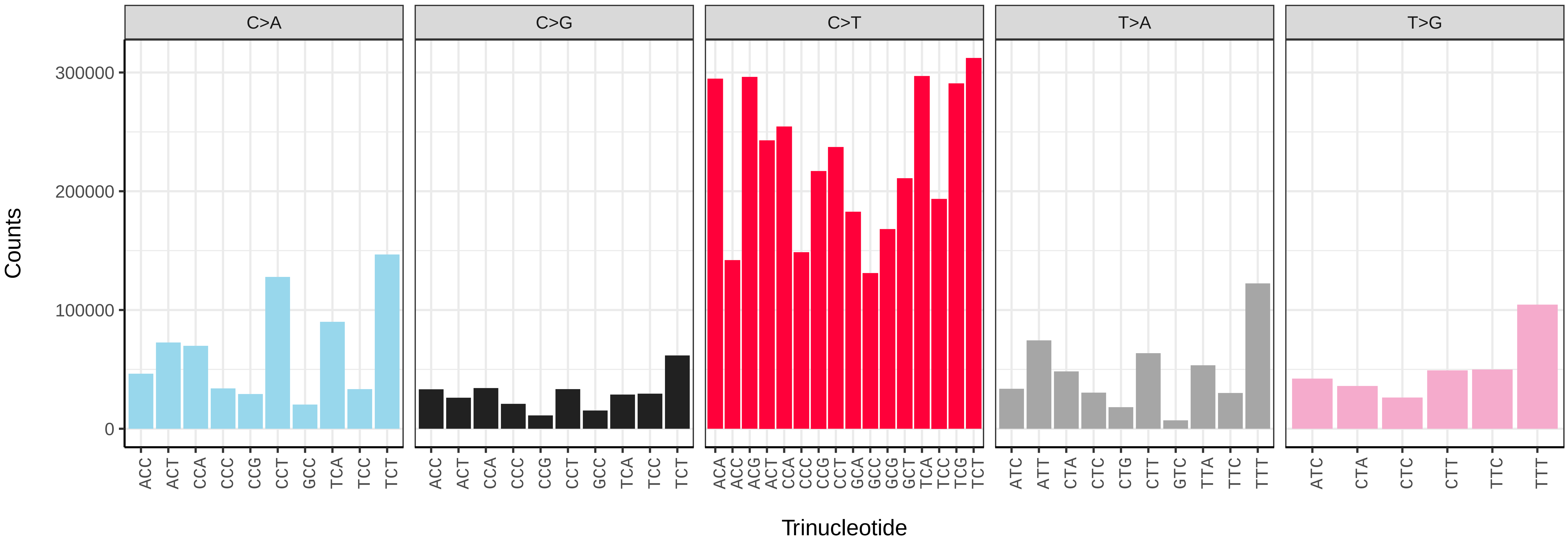


*Crataegus laevigata* (drCraLaev2) somatic mutational spectrum (SBS96 classification)

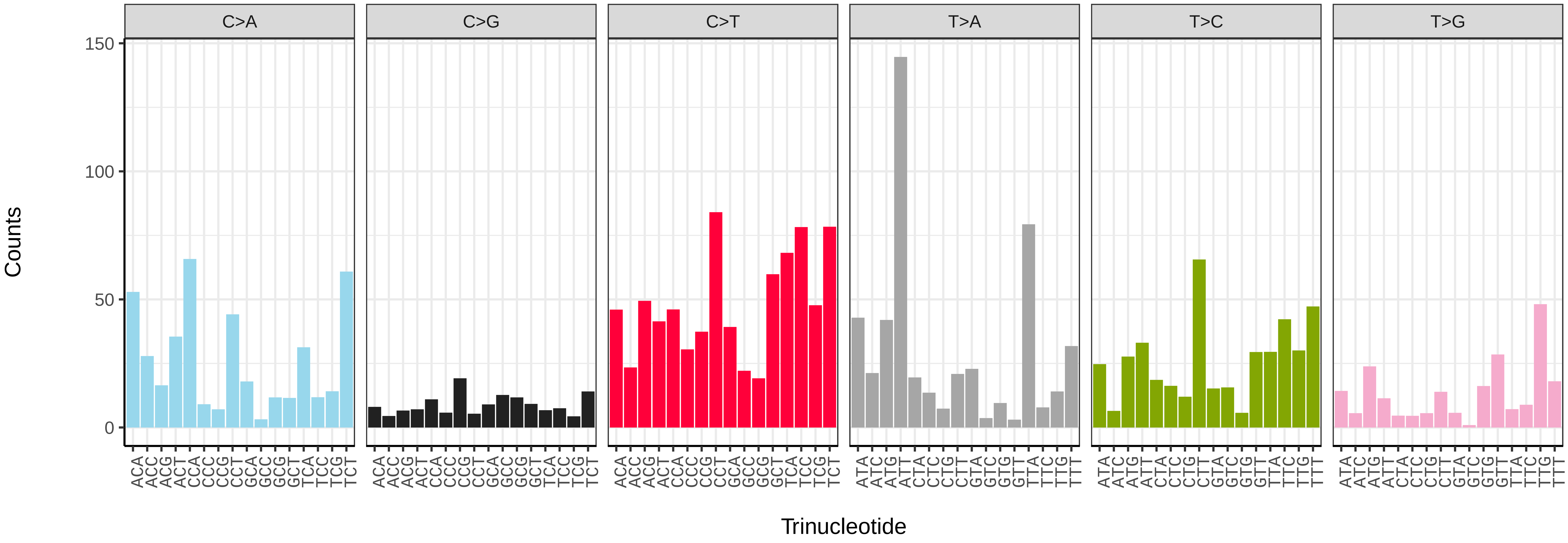


*Crataegus laevigata* (drCraLaev2) somatic mutational spectrum (SBS52 classification)

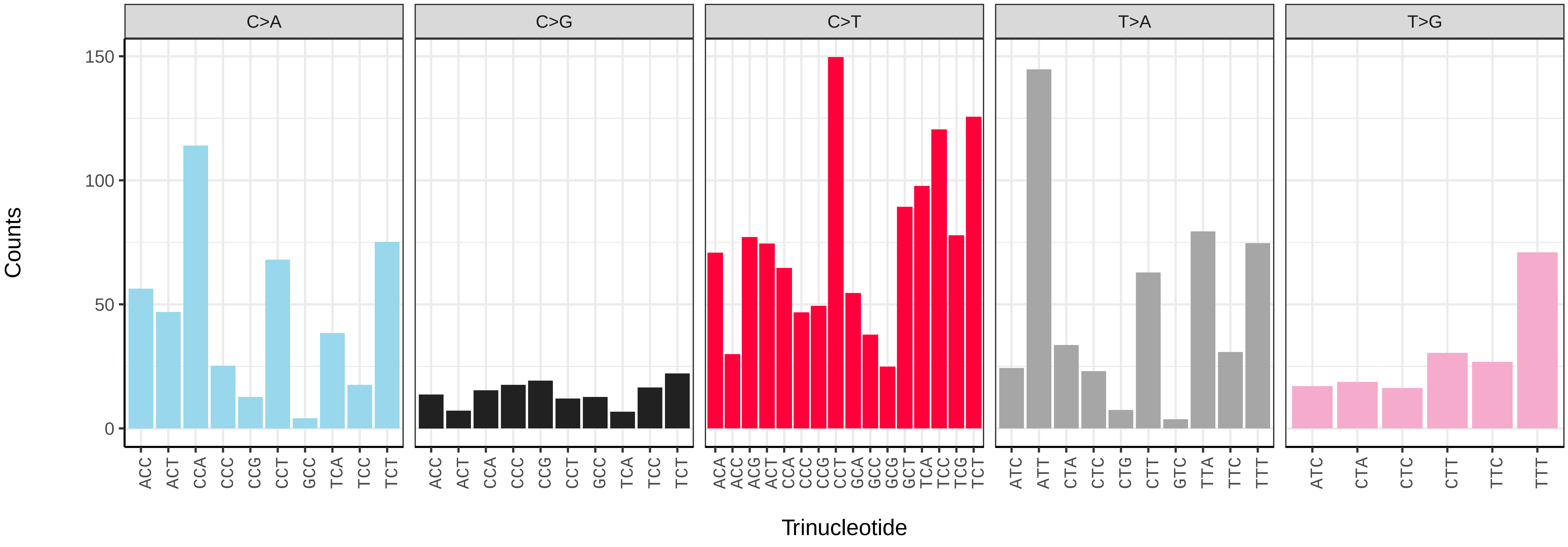

*Erodium maritimum* (drEroMari8) germline mutational spectrum

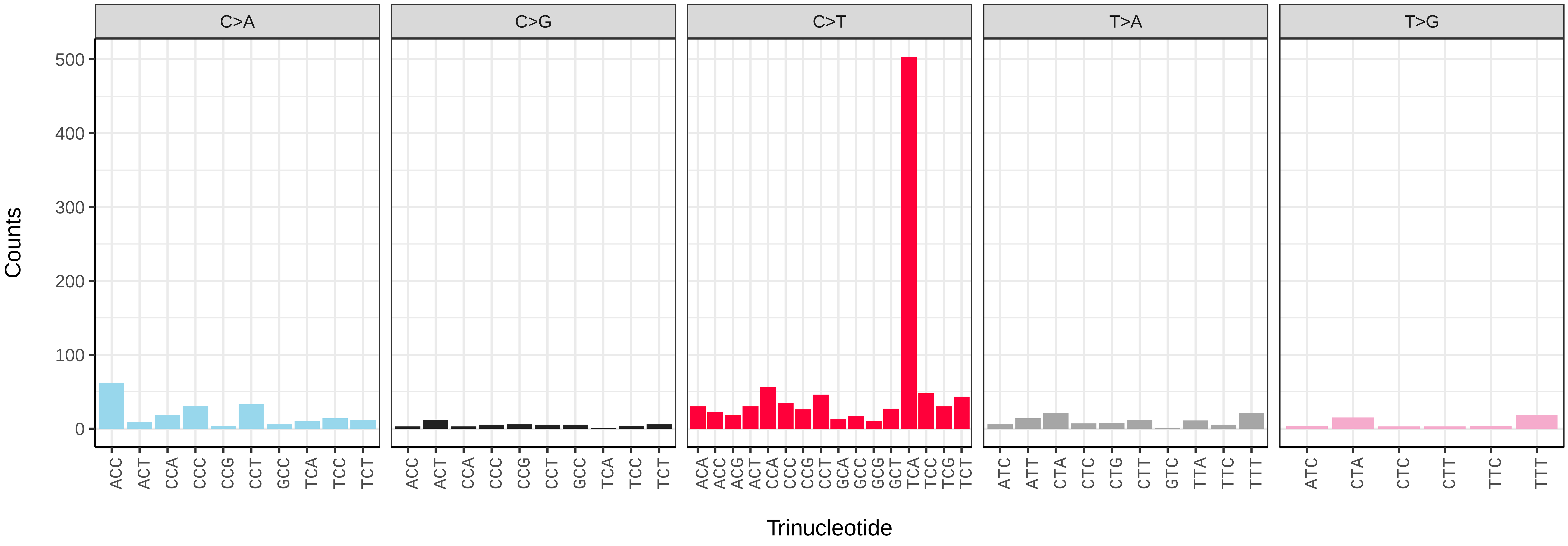


*Erodium maritimum* (drEroMari8) somatic mutational spectrum (SBS96 classification)

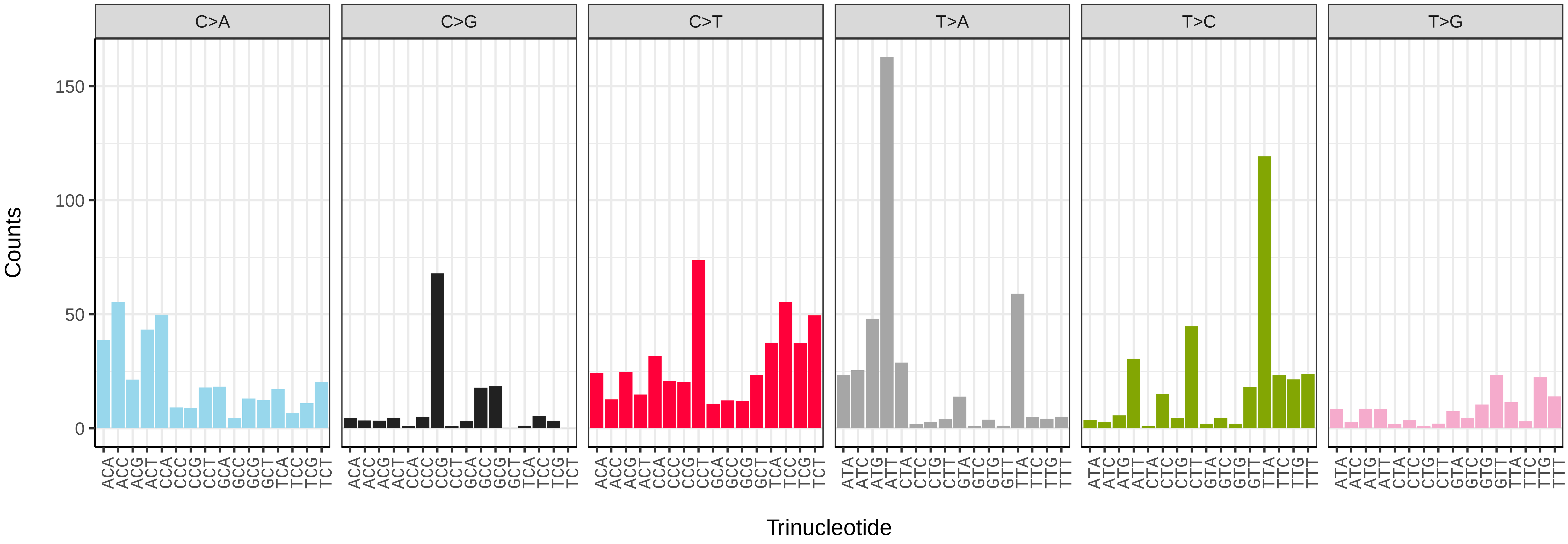


*Erodium maritimum* (drEroMari8) somatic mutational spectrum (SBS52 classification)

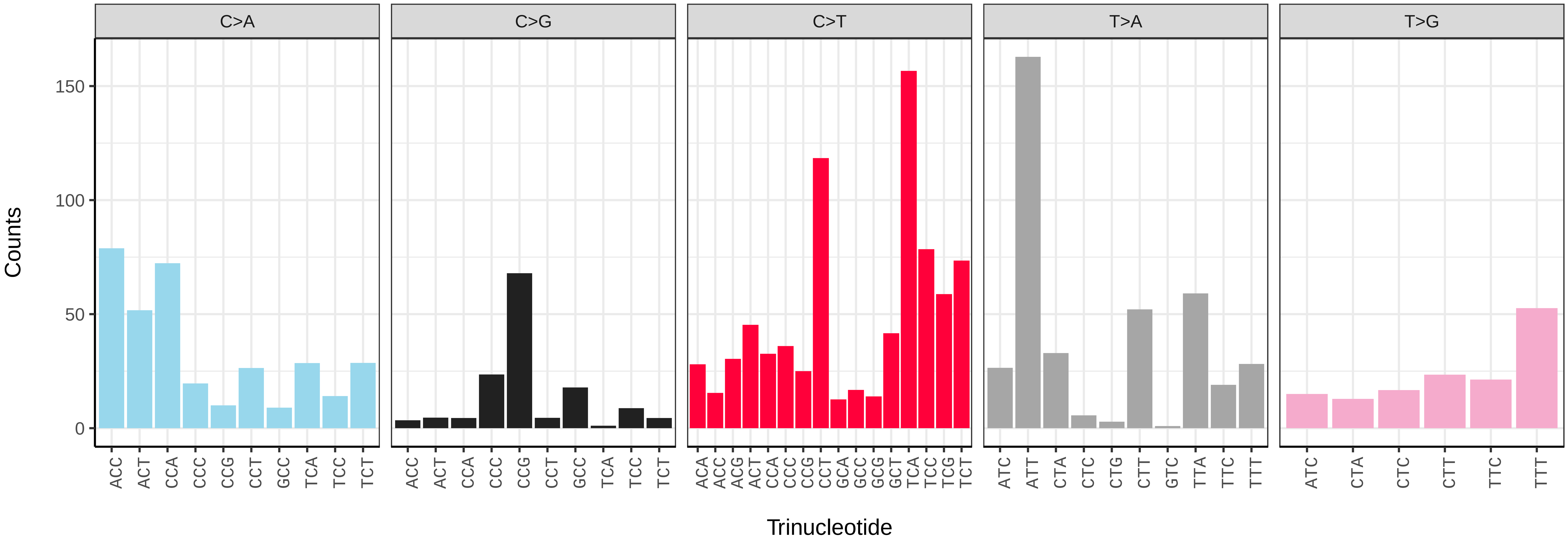

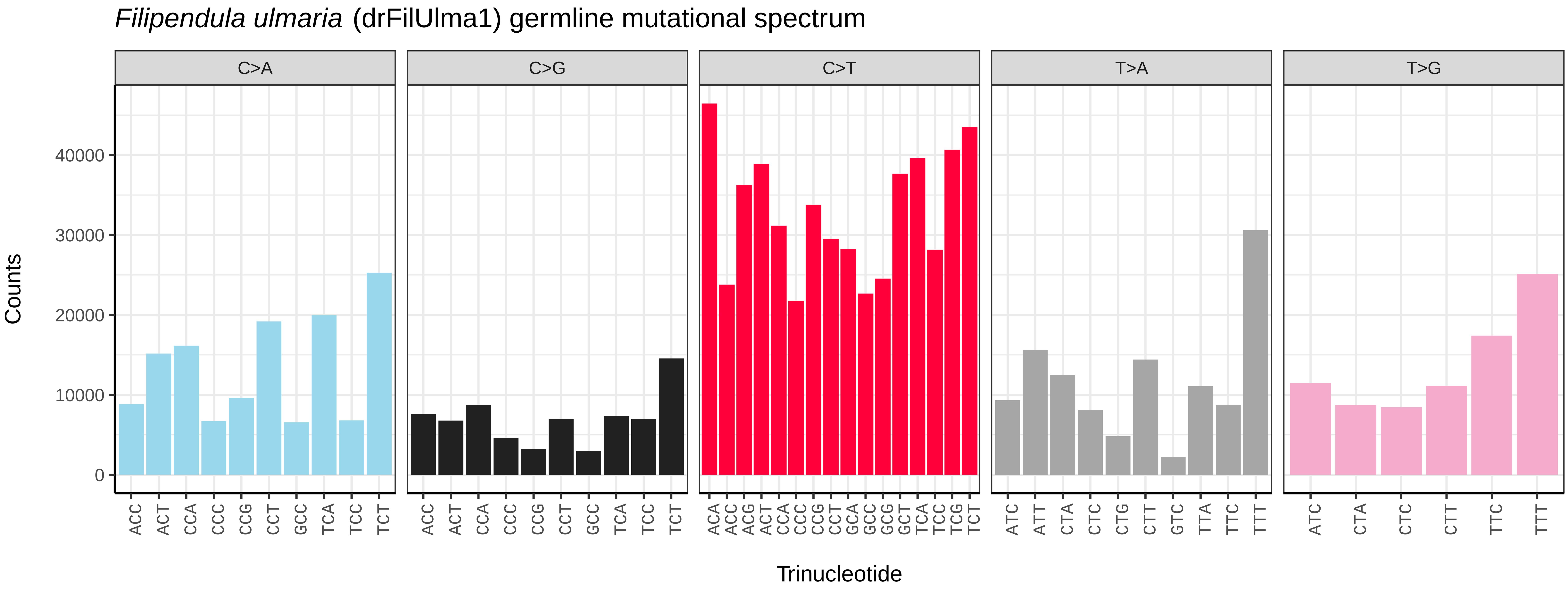
*Filipendula ulmaria* (drFilUlma1) germline mutational spectrum
C>A
C>G
C>T
T>A
T>G
Counts
40000
30000
20000
10000
0
Trinucleotide

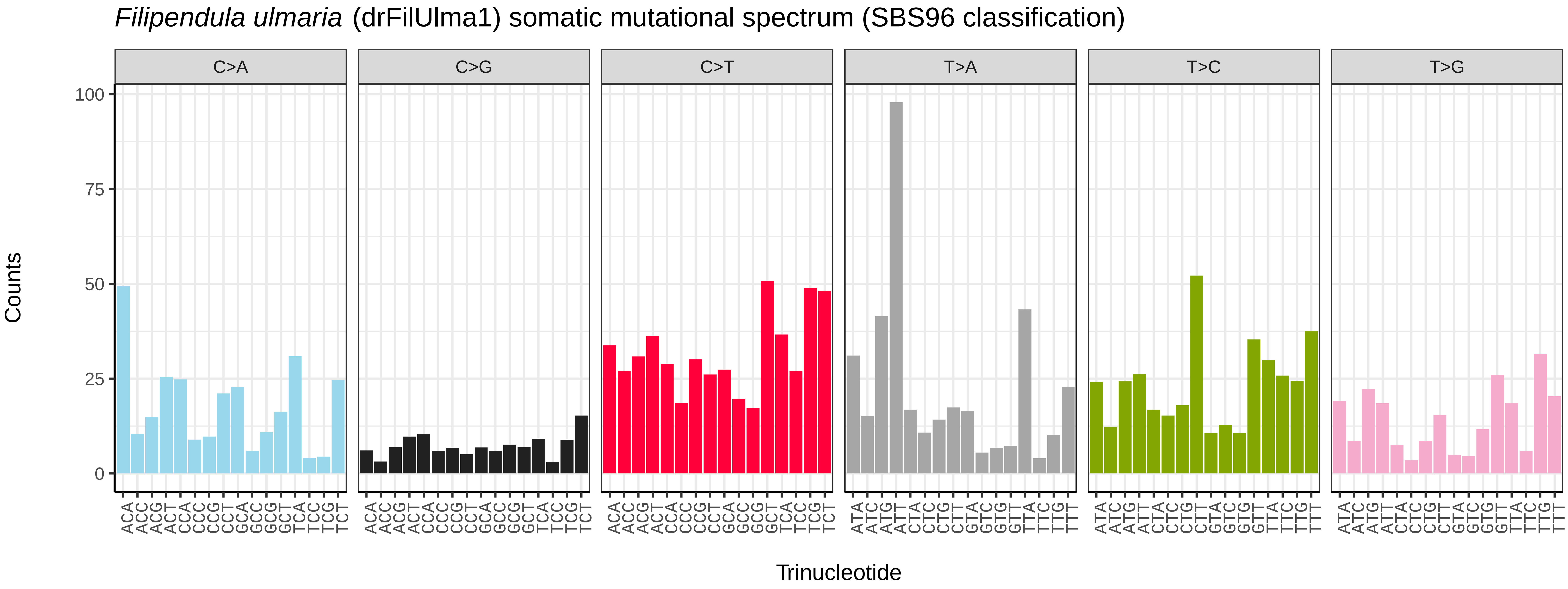
*Filipendula ulmaria* (drFilUlma1) somatic mutational spectrum (SBS96 classification)
C>A
C>G
C>T
T>A
T>C
T>G
Counts
100
75
50
25
0
Trinucleotide

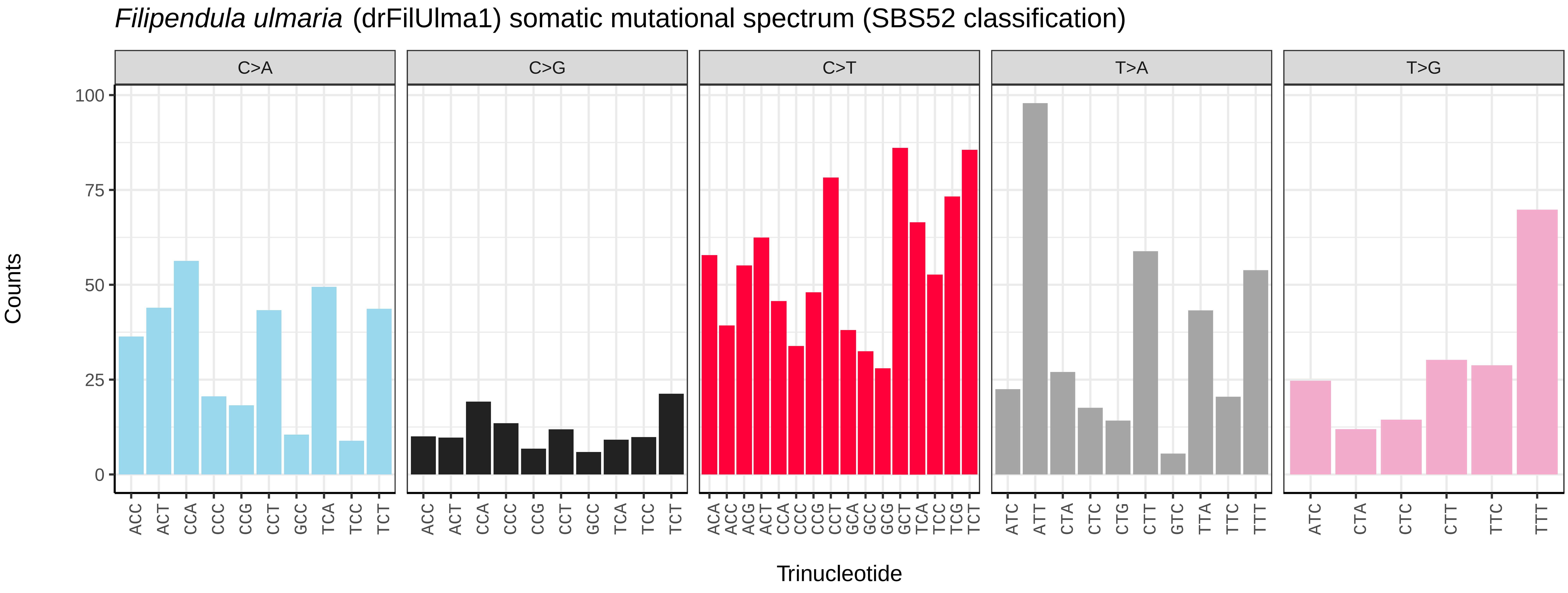
*Filipendula ulmaria* (drFilUlma1) somatic mutational spectrum (SBS52 classification)
C>A
C>G
C>T
T>A
T>G
Counts
100
75
50
25
0
Trinucleotide

*Geum urbanum* (drGeuUrba1) germline mutational spectrum

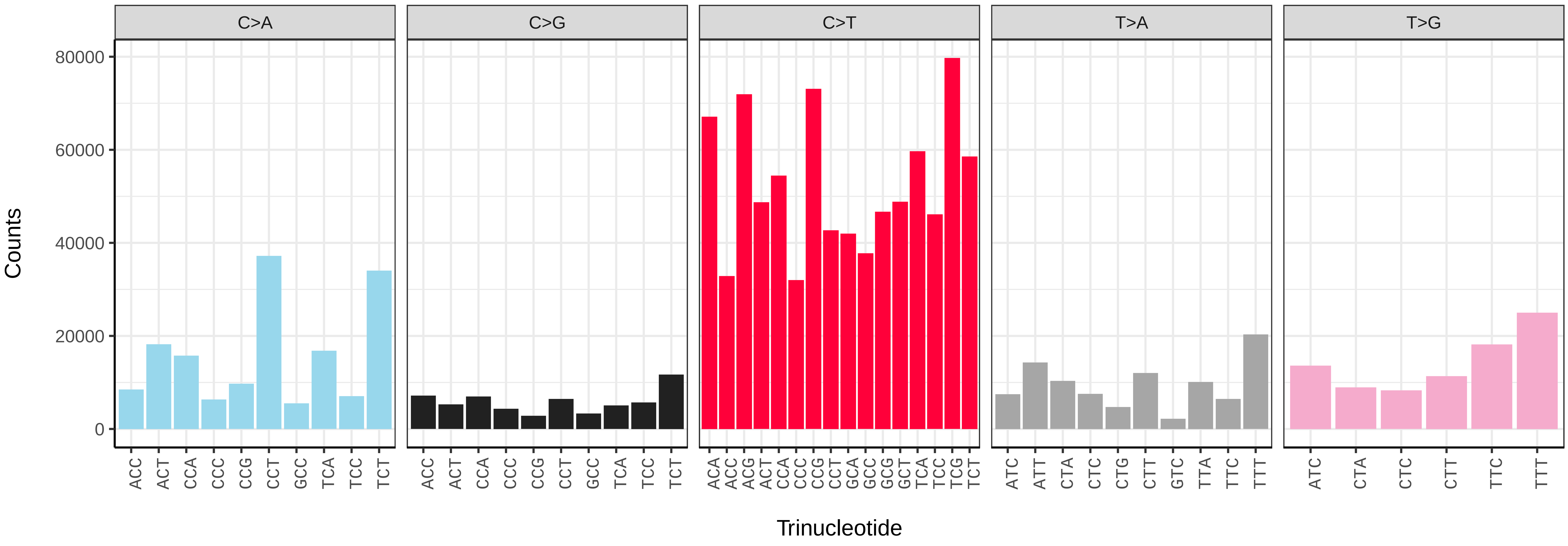


*Geum urbanum* (drGeuUrba1) somatic mutational spectrum (SBS96 classification)

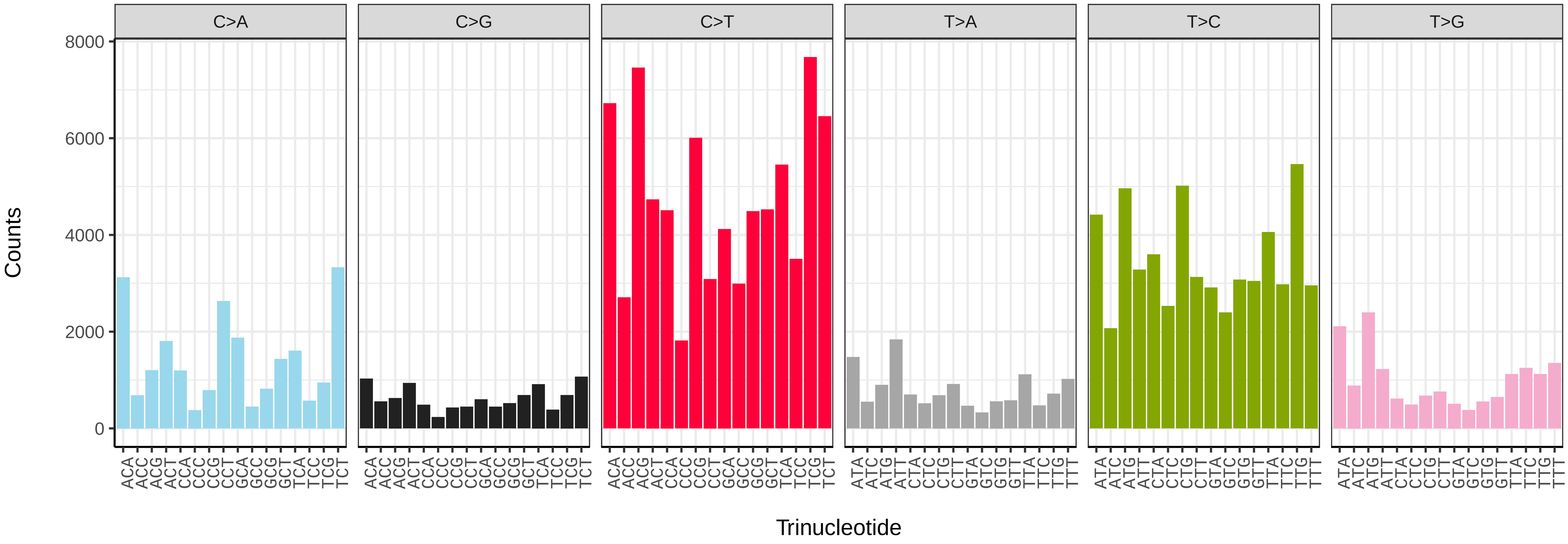


*Geum urbanum* (drGeuUrba1) somatic mutational spectrum (SBS52 classification)

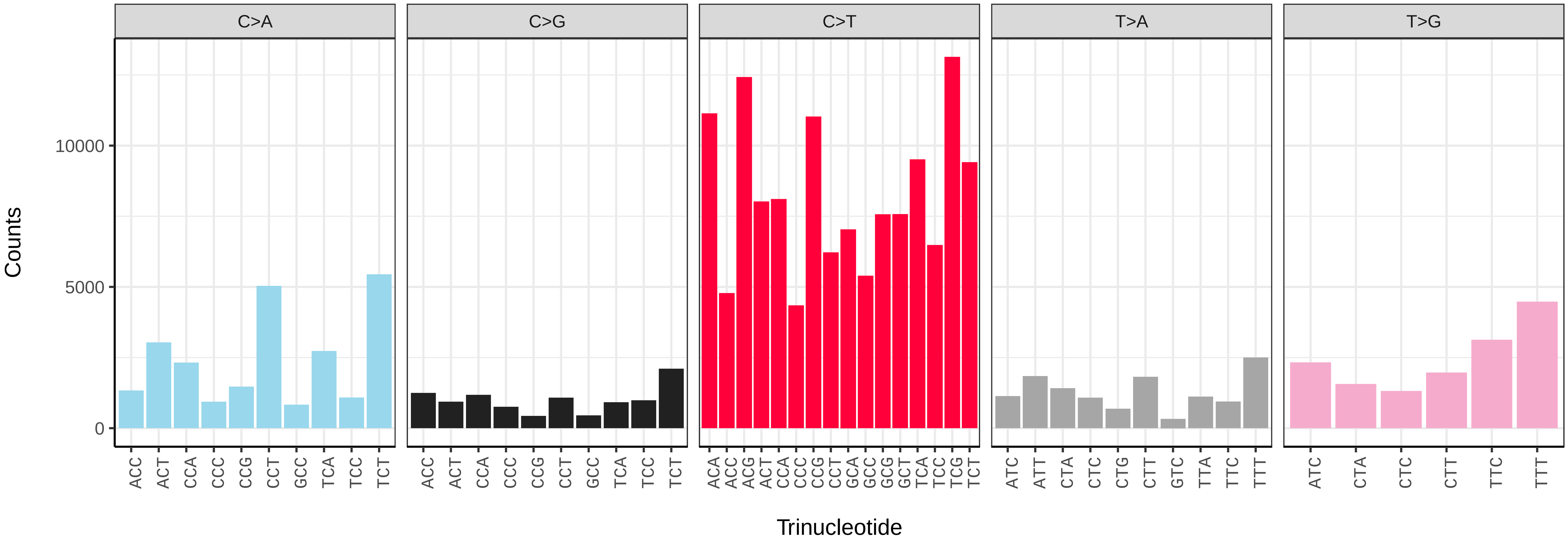

*Hedera helix* (drHedHeli1) germline mutational spectrum

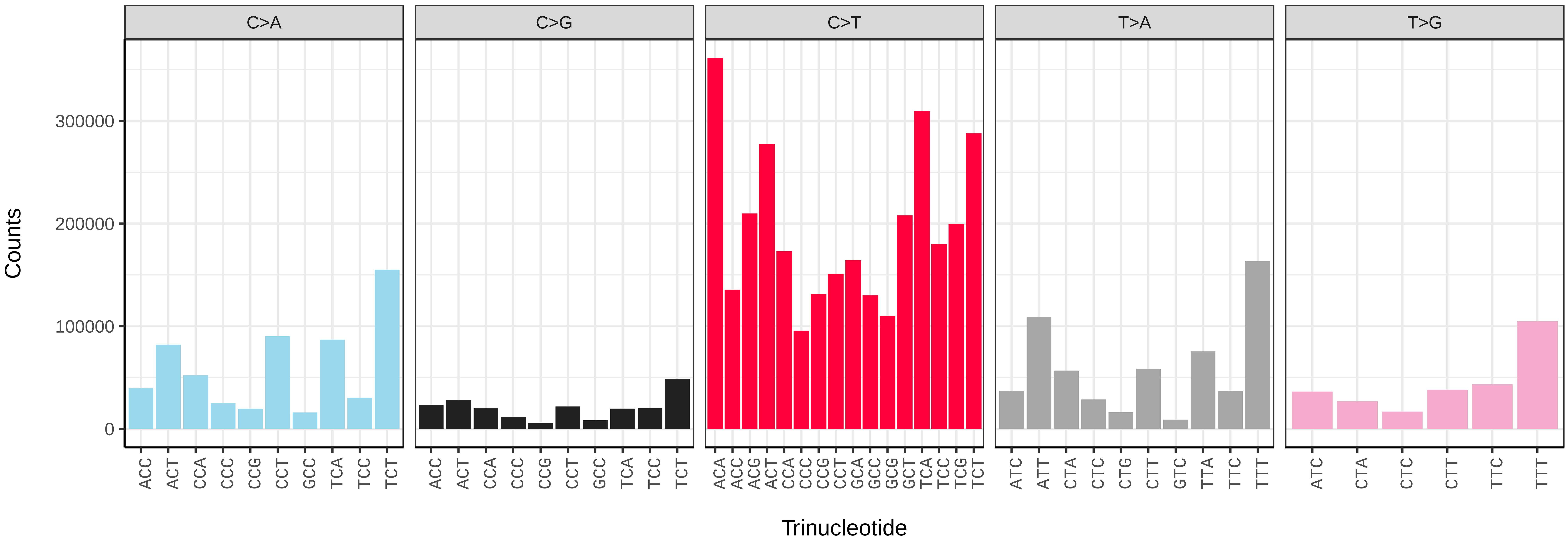


*Hedera helix* (drHedHeli1) somatic mutational spectrum (SBS96 classification)

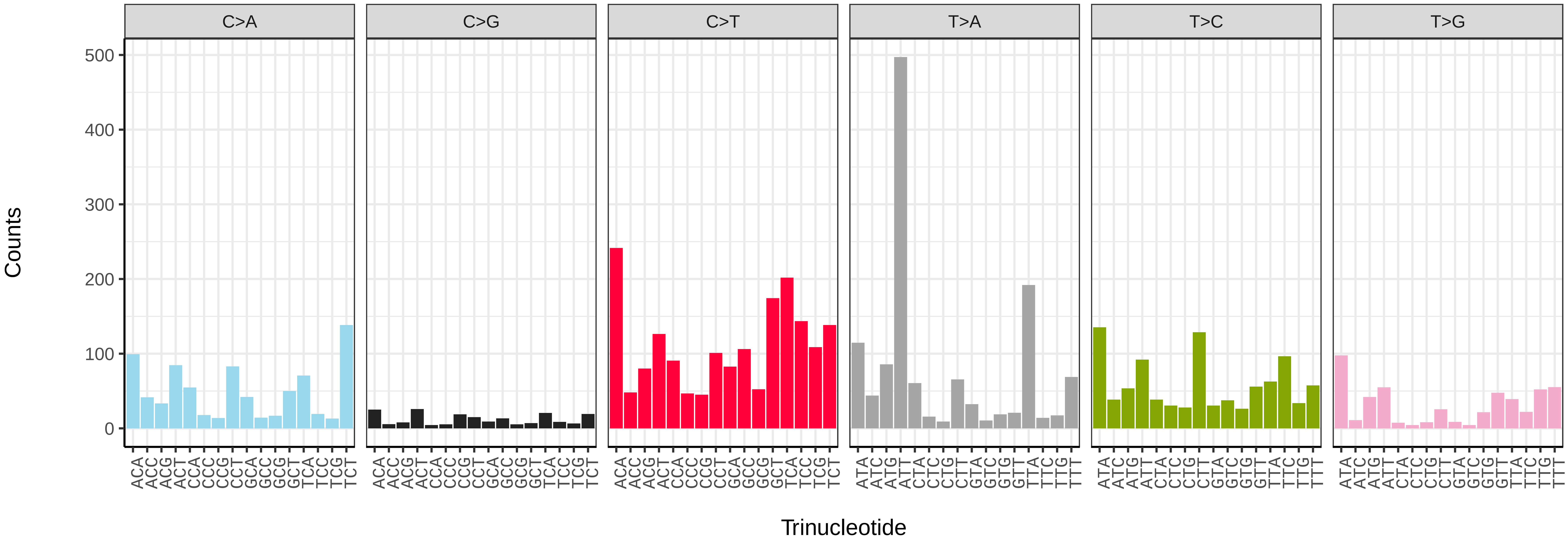


*Hedera helix* (drHedHeli1) somatic mutational spectrum (SBS52 classification)

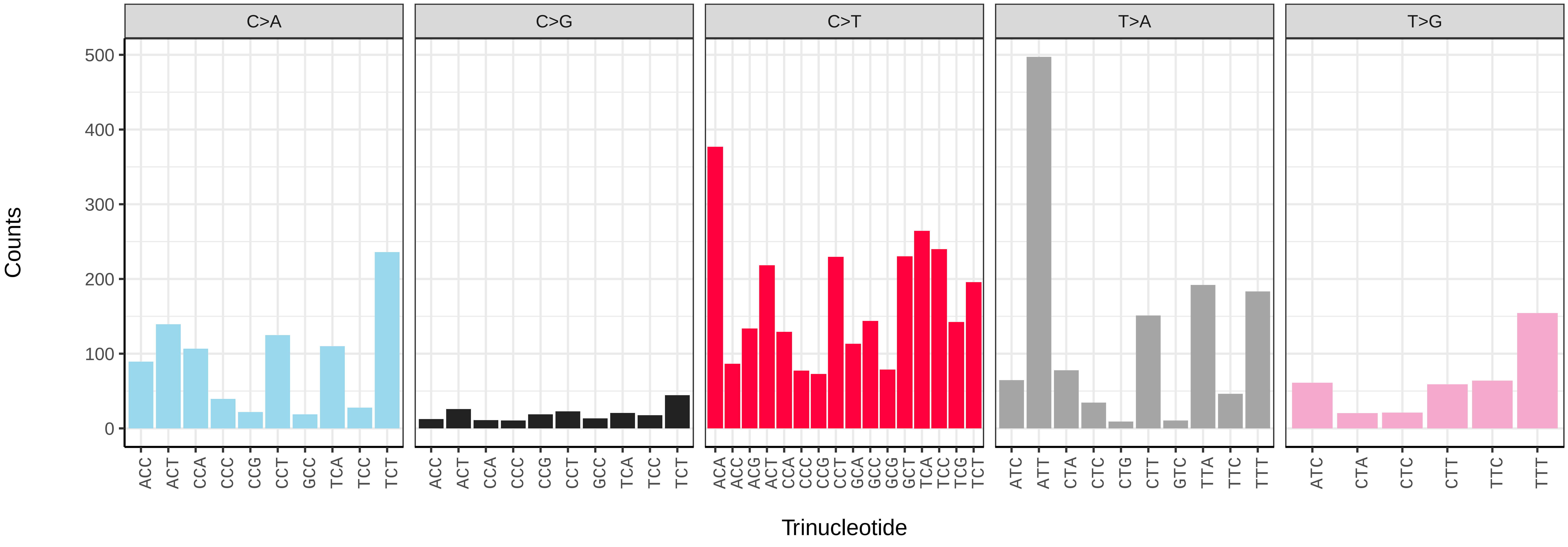

*Ilex aquifolium* (drIleAqui2) germline mutational spectrum

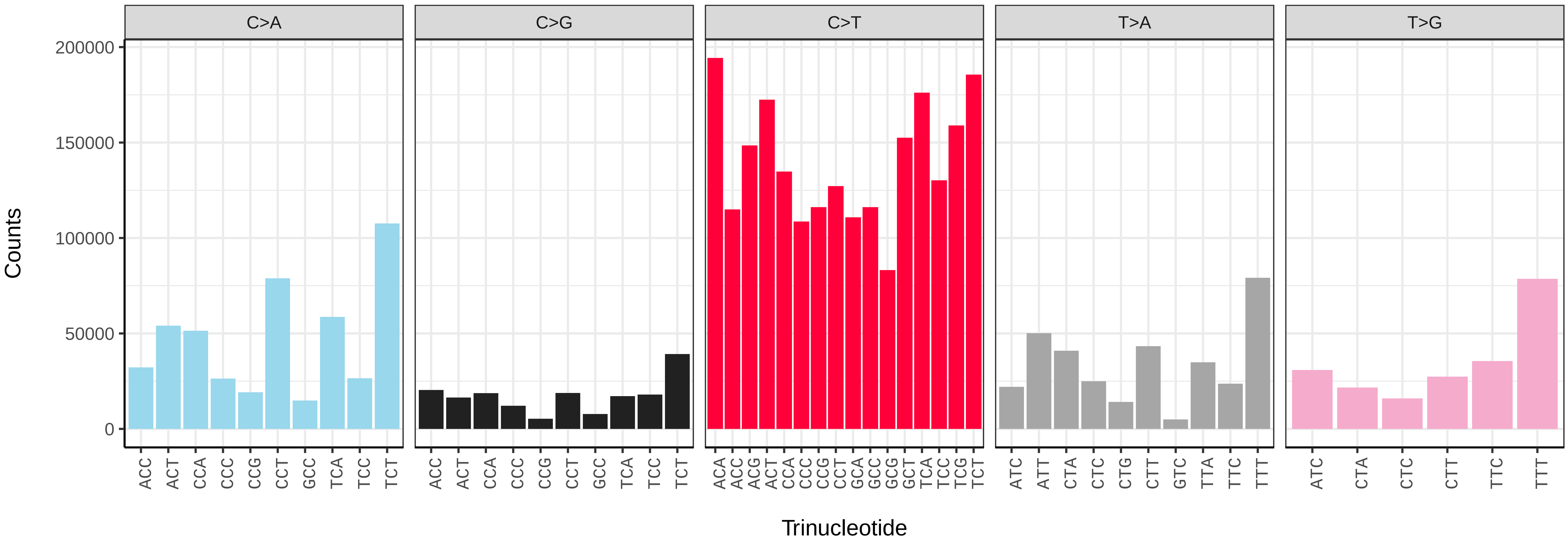


*Ilex aquifolium* (drIleAqui2) somatic mutational spectrum (SBS96 classification)

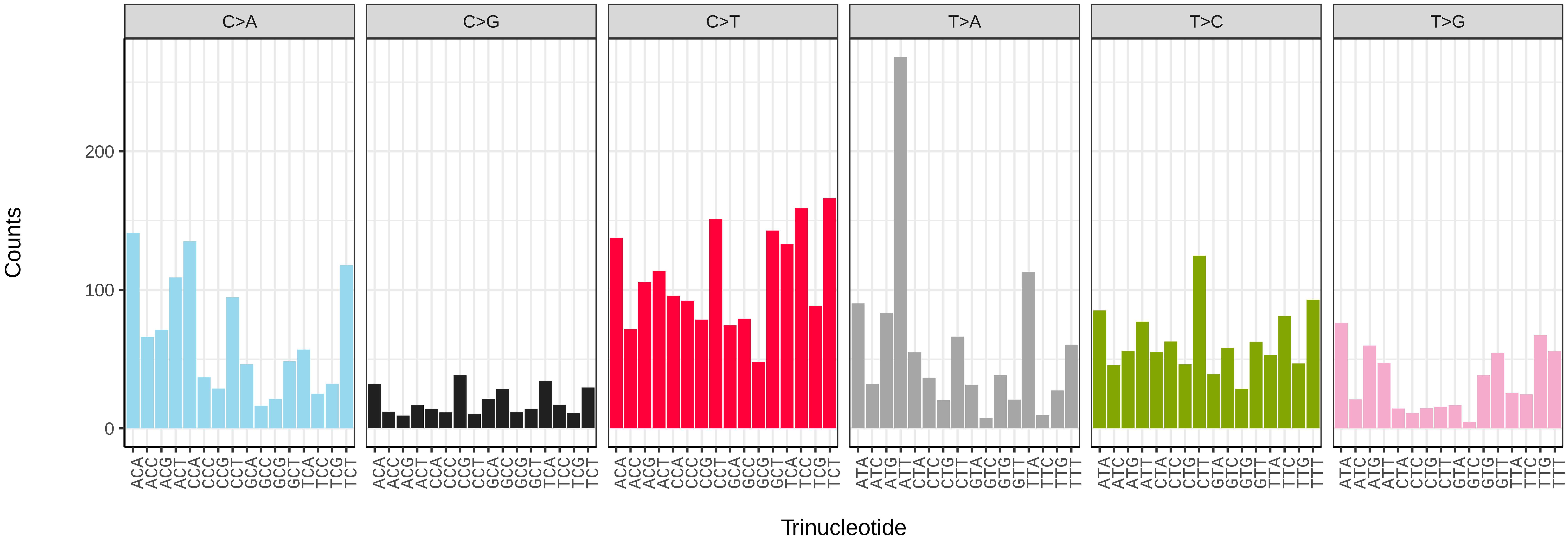


*Ilex aquifolium* (drIleAqui2) somatic mutational spectrum (SBS52 classification)

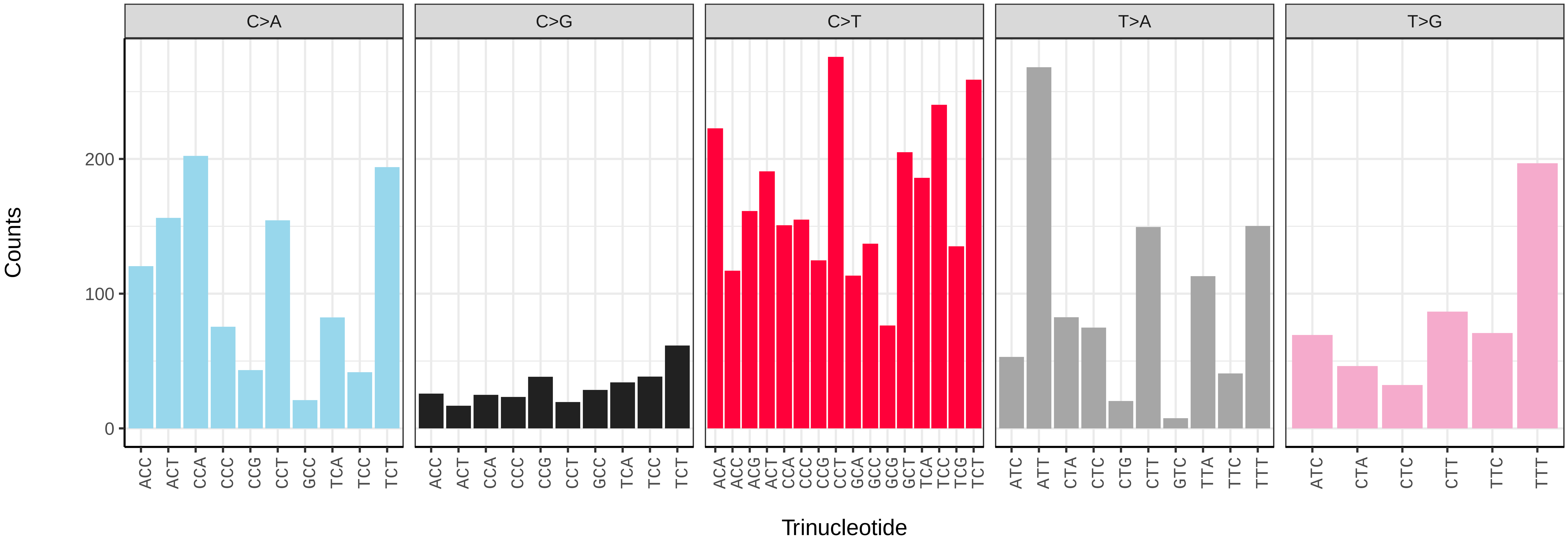

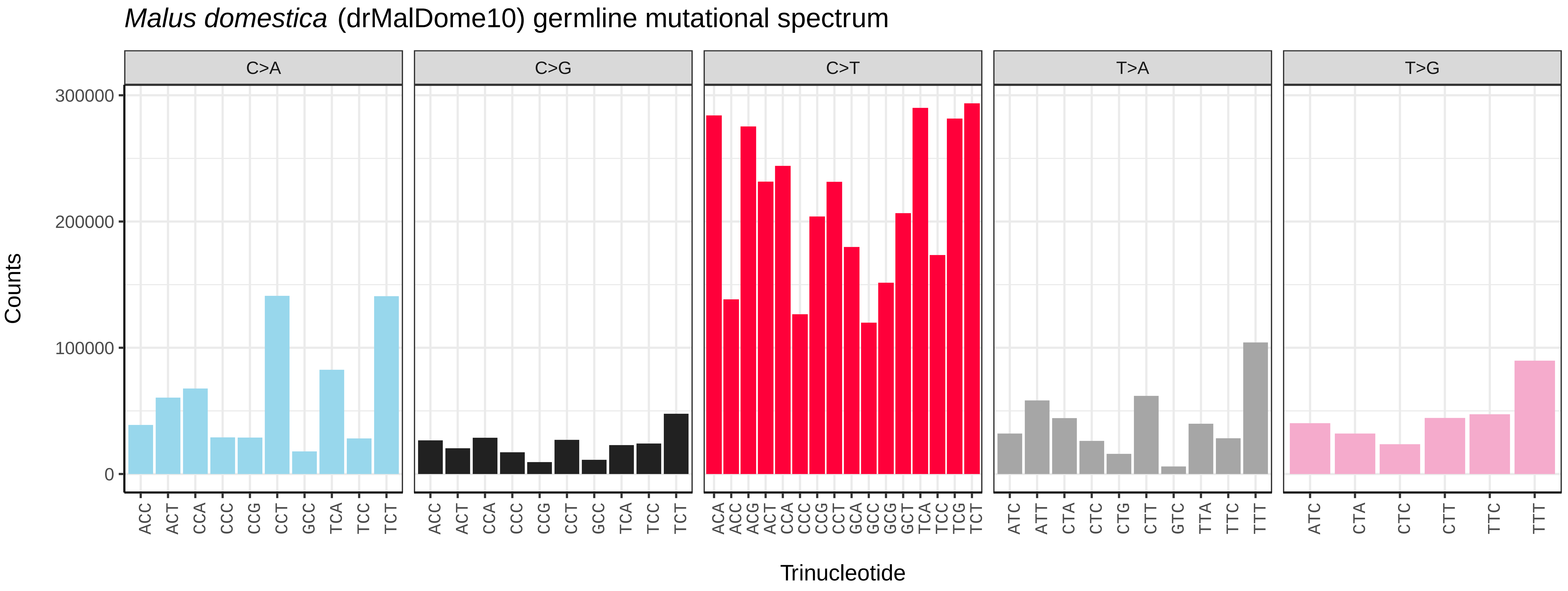

Malus domestica (drMalDome10) germline mutational spectrum
C>A
C>G
C>T
T>A
T>G
300000
200000
100000
0
Counts
ACC ACT CCA CCC CCG CCT GCC TCA TCC TCT
ACA ACC ACG ACT CCA CCC CCG CCT GCA GCC GCG GCT TCA TCC TCG TCT
ATC ATT CTA CTC CTG CTT GTC TTA TTC TTT
ATC CTA CTC CTT TTC TTT
Trinucleotide


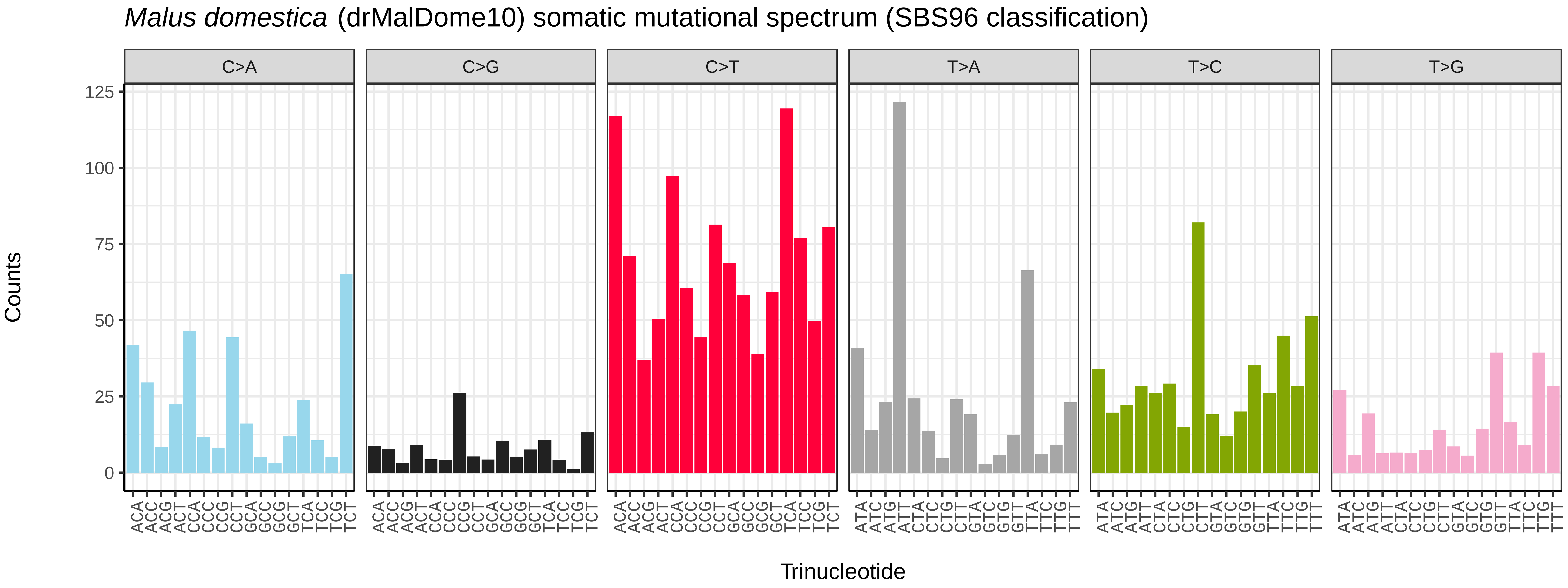

Malus domestica (drMalDome10) somatic mutational spectrum (SBS96 classification)
C>A
C>G
C>T
T>A
T>C
T>G
125
100
75
50
25
0
Counts
ACA ACC ACG ACT CCA CCC CCG CCT GCA GCC GCG GCT TCA TCC TCG TCT
ATA ATC ATG ATT CTA CTC CTG CTT GTA GTC GTG GTT TTA TTC TTG TTT
Trinucleotide


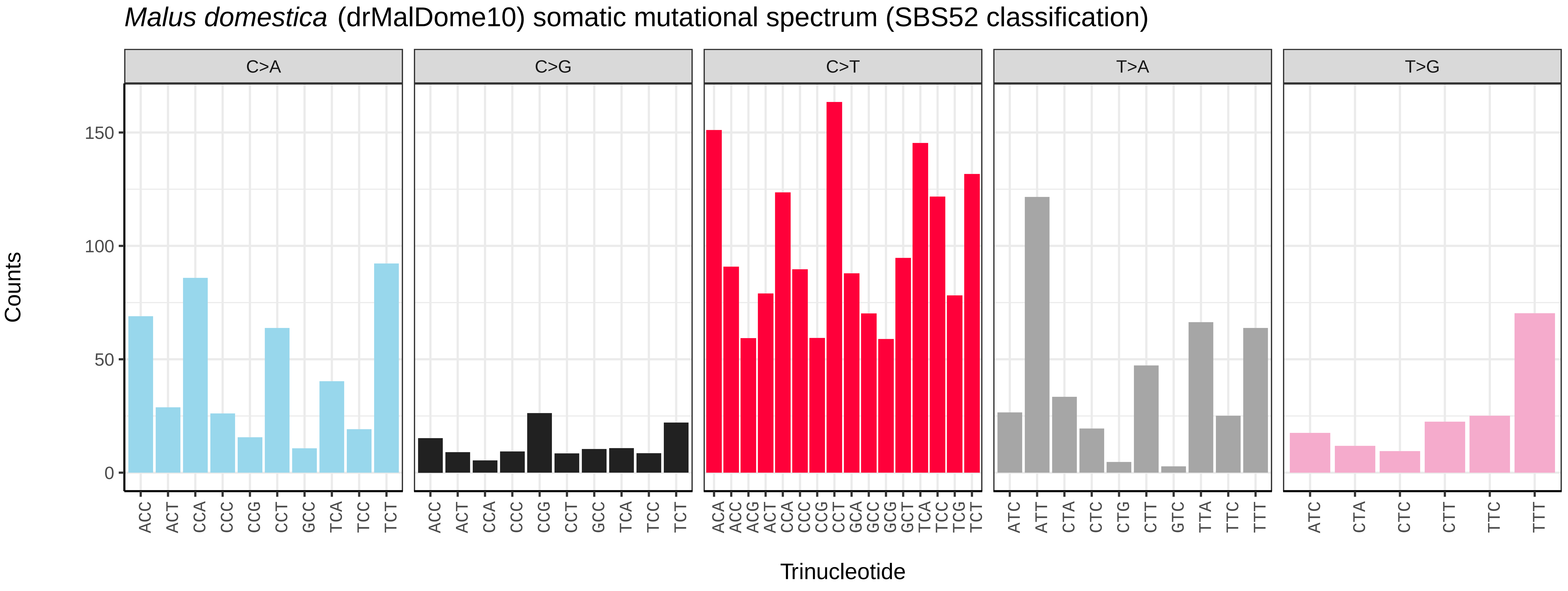

Malus domestica (drMalDome10) somatic mutational spectrum (SBS52 classification)
C>A
C>G
C>T
T>A
T>G
150
100
50
0
Counts
ACC ACT CCA CCC CCG CCT GCC TCA TCC TCT
ACA ACC ACG ACT CCA CCC CCG CCT GCA GCC GCG GCT TCA TCC TCG TCT
ATC ATT CTA CTC CTG CTT GTC TTA TTC TTT
ATC CTA CTC CTT TTC TTT
Trinucleotide

*Malus domestica* (drMalDome11) germline mutational spectrum

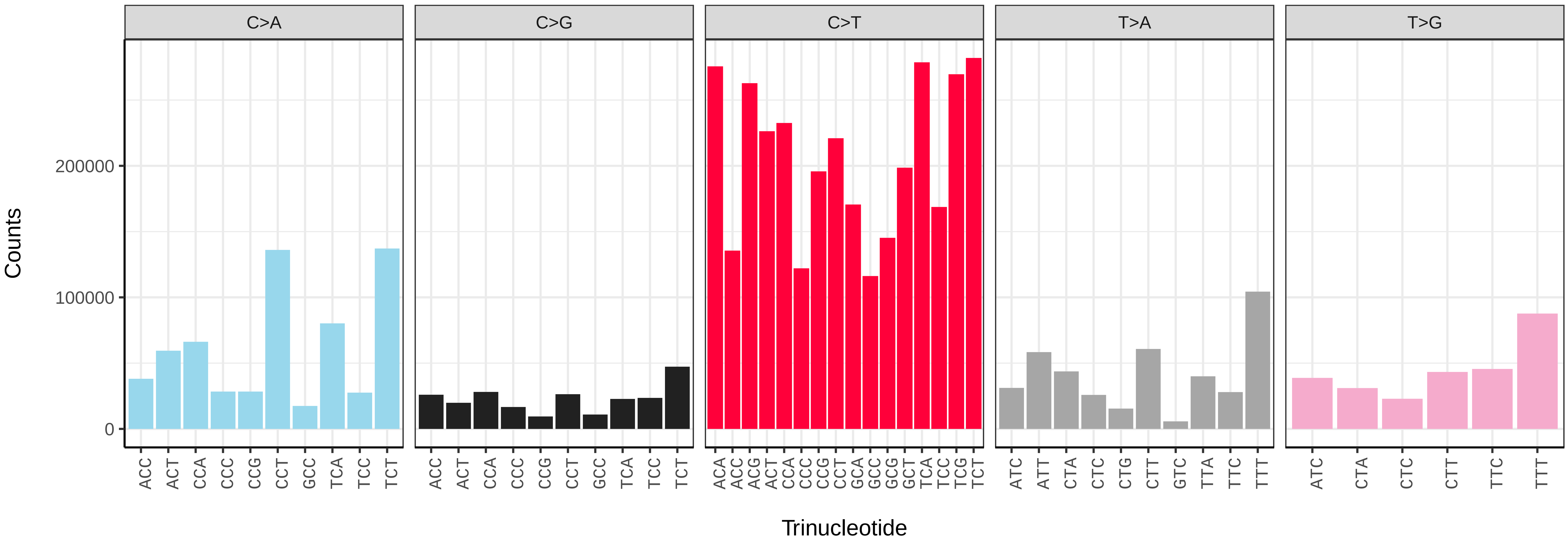


*Malus domestica* (drMalDome11) somatic mutational spectrum (SBS96 classification)

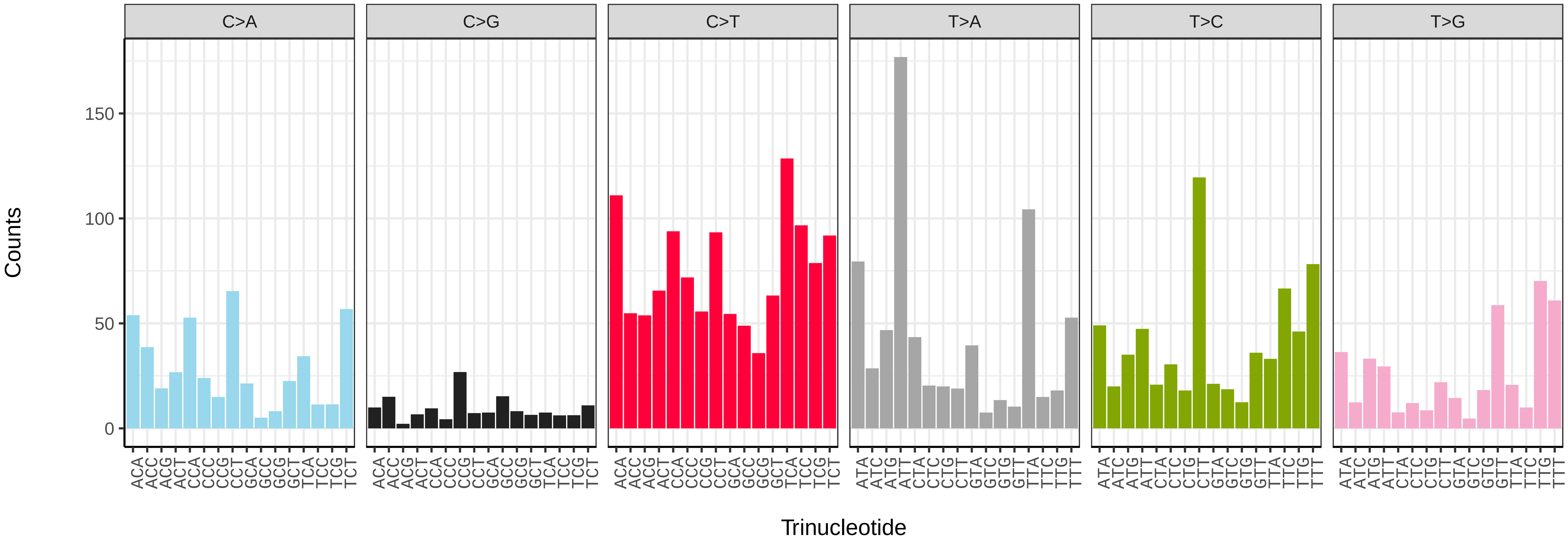


*Malus domestica* (drMalDome11) somatic mutational spectrum (SBS52 classification)

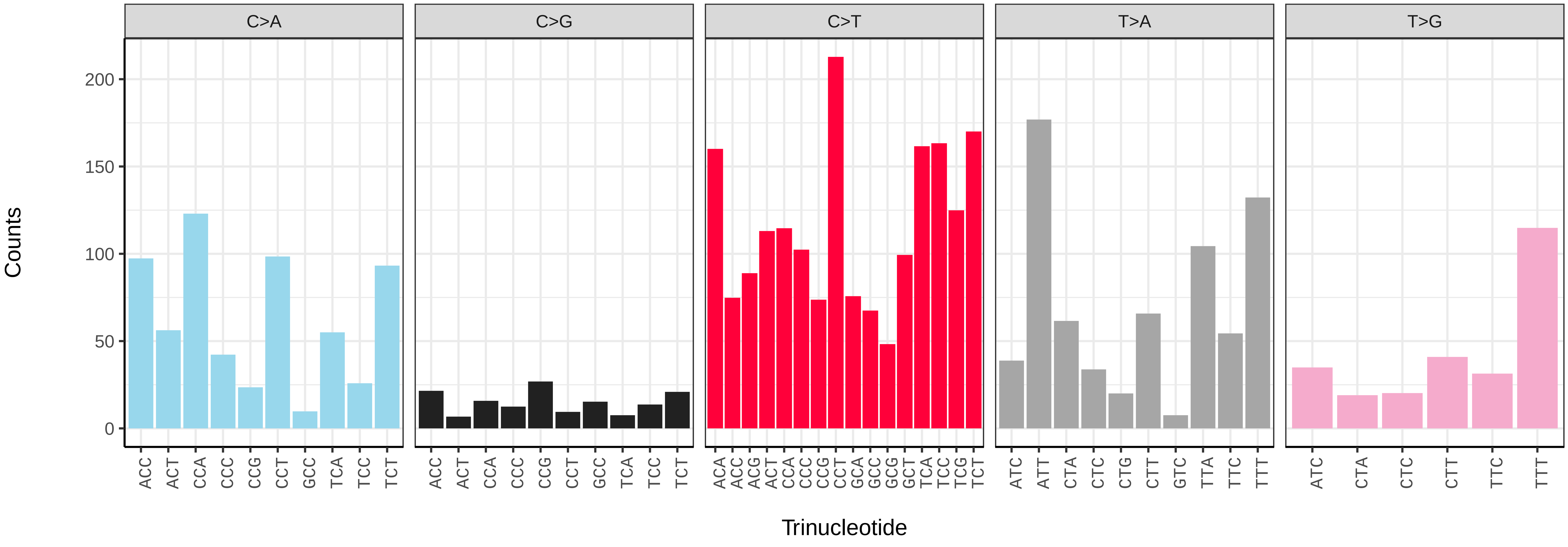

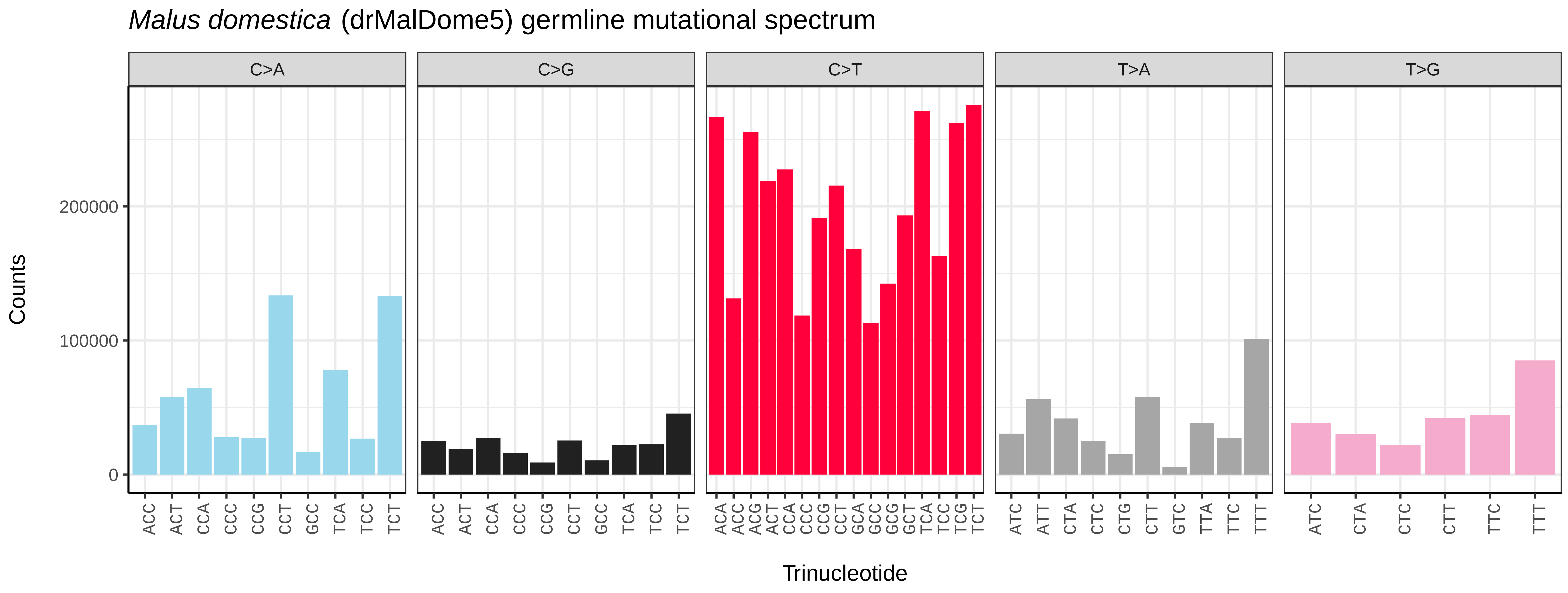
Malus domestica (drMalDome5) germline mutational spectrum
C>A
C>G
C>T
T>A
T>G
Counts
0
100000
200000
ACC ACT CCA CCC CCG CCT GCC TCA TCC TCT
ACA ACC ACG ACT CCA CCC CCG CCT GCA GCC GCG GCT TCA TCC TCG TCT
ATC ATT CTA CTC CTG CTT GTC TTA TTC TTT
ATC CTA CTC CTT TTC TTT
Trinucleotide

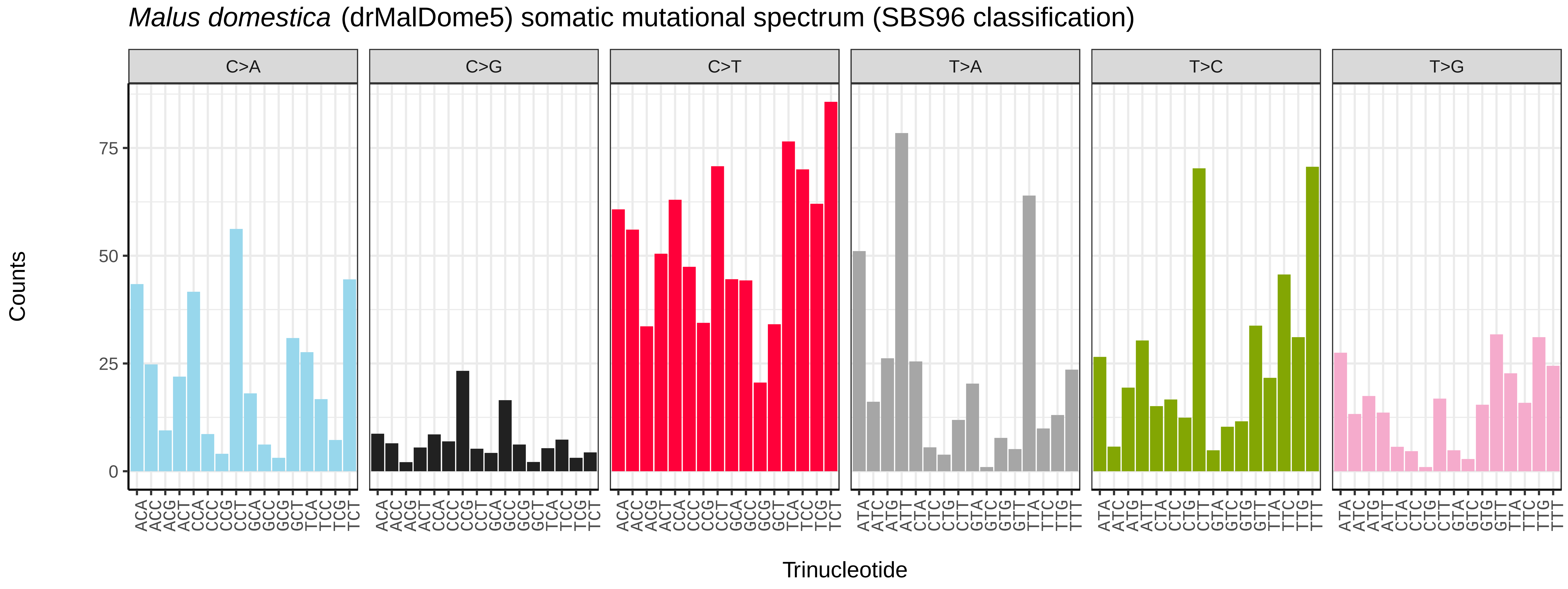
Malus domestica (drMalDome5) somatic mutational spectrum (SBS96 classification)
C>A
C>G
C>T
T>A
T>C
T>G
Counts
0
25
50
75
ACA ACC ACG ACT CCA CCC CCG CCT GCA GCC GCG GCT TCA TCC TCG TCT
ATA ATC ATG ATT CTA CTC CTG CTT GTA GTC GTG GTT TTA TTC TTG TTT
Trinucleotide

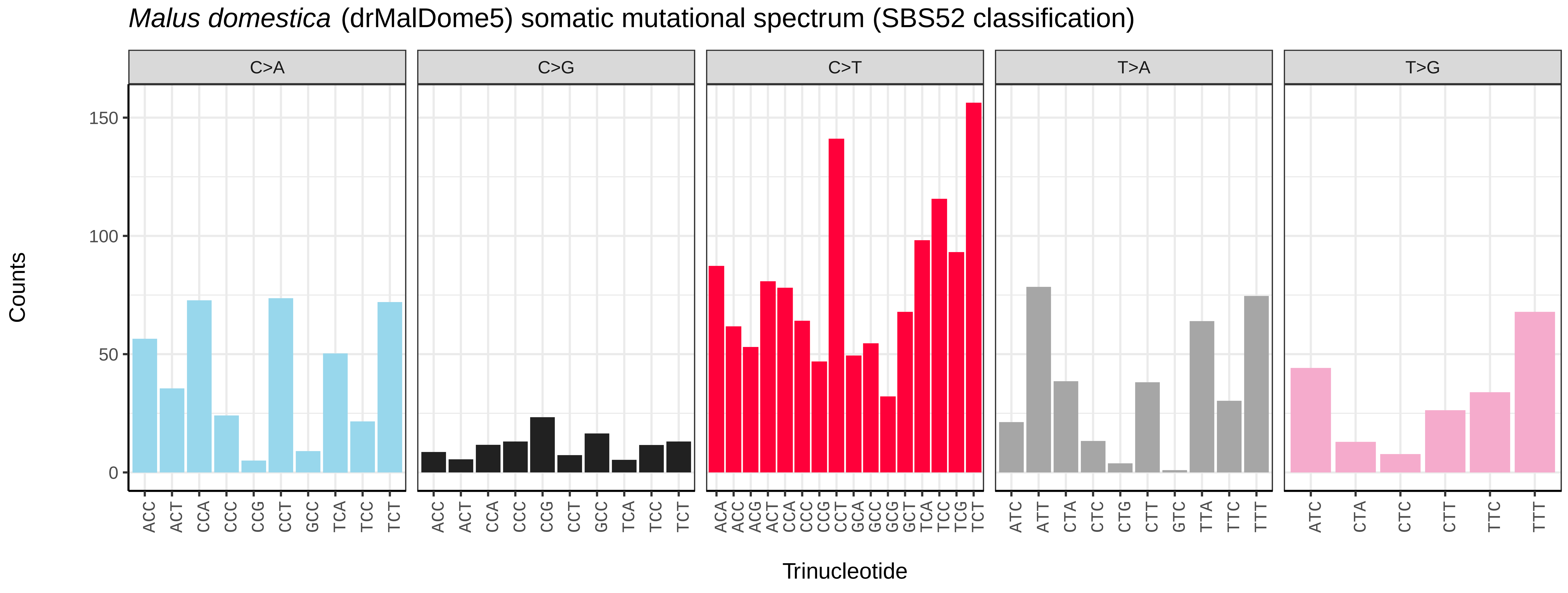
Malus domestica (drMalDome5) somatic mutational spectrum (SBS52 classification)
C>A
C>G
C>T
T>A
T>G
Counts
0
50
100
150
ACC ACT CCA CCC CCG CCT GCC TCA TCC TCT
ACA ACC ACG ACT CCA CCC CCG CCT GCA GCC GCG GCT TCA TCC TCG TCT
ATC ATT CTA CTC CTG CTT GTC TTA TTC TTT
ATC CTA CTC CTT TTC TTT
Trinucleotide

*Malus x robusta* (drMalRobu1) germline mutational spectrum

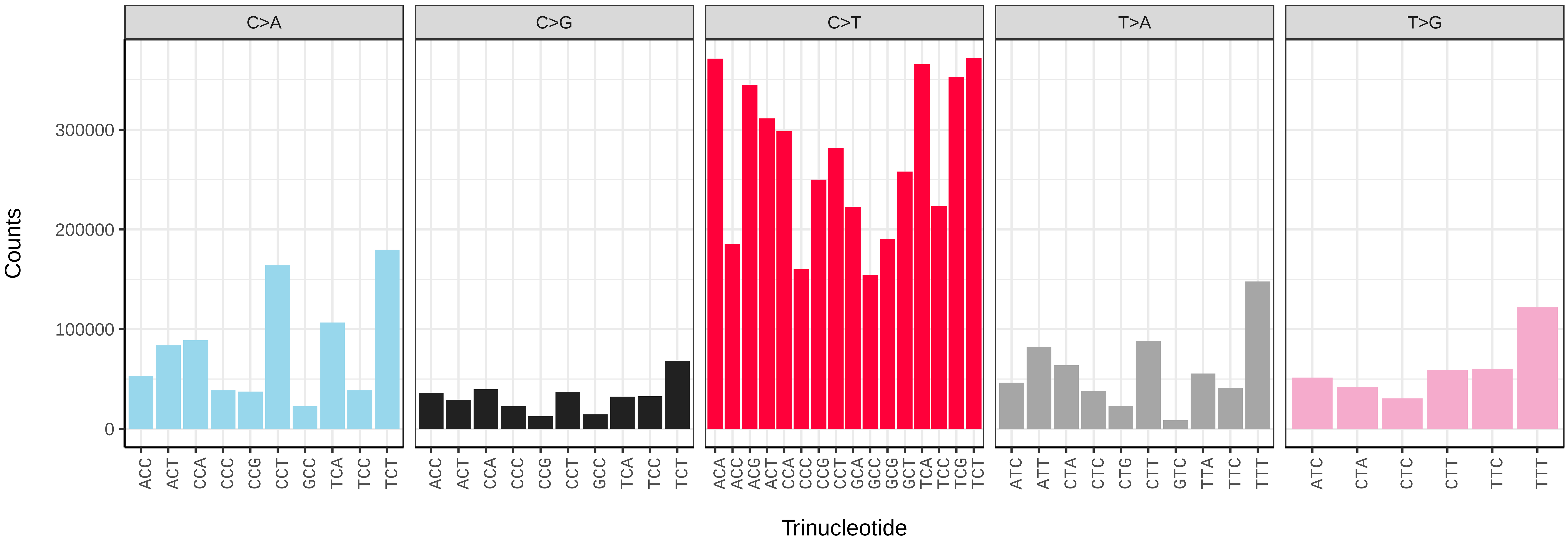


*Malus x robusta* (drMalRobu1) somatic mutational spectrum (SBS96 classification)

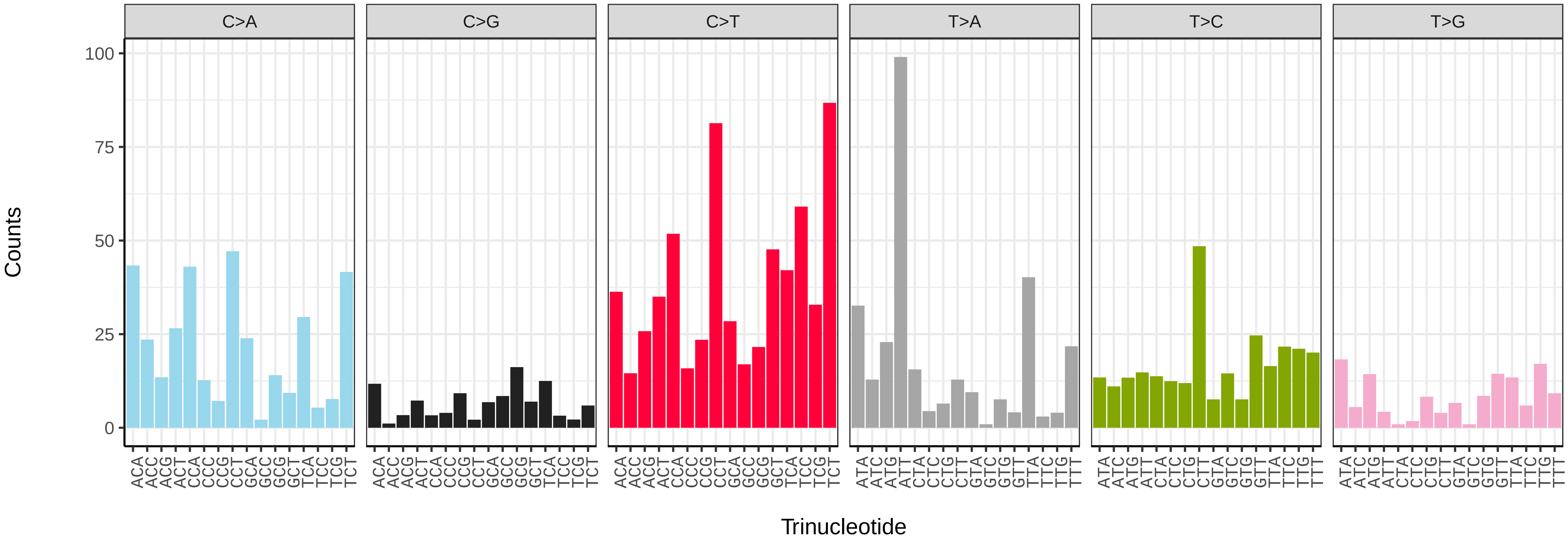


*Malus x robusta* (drMalRobu1) somatic mutational spectrum (SBS52 classification)

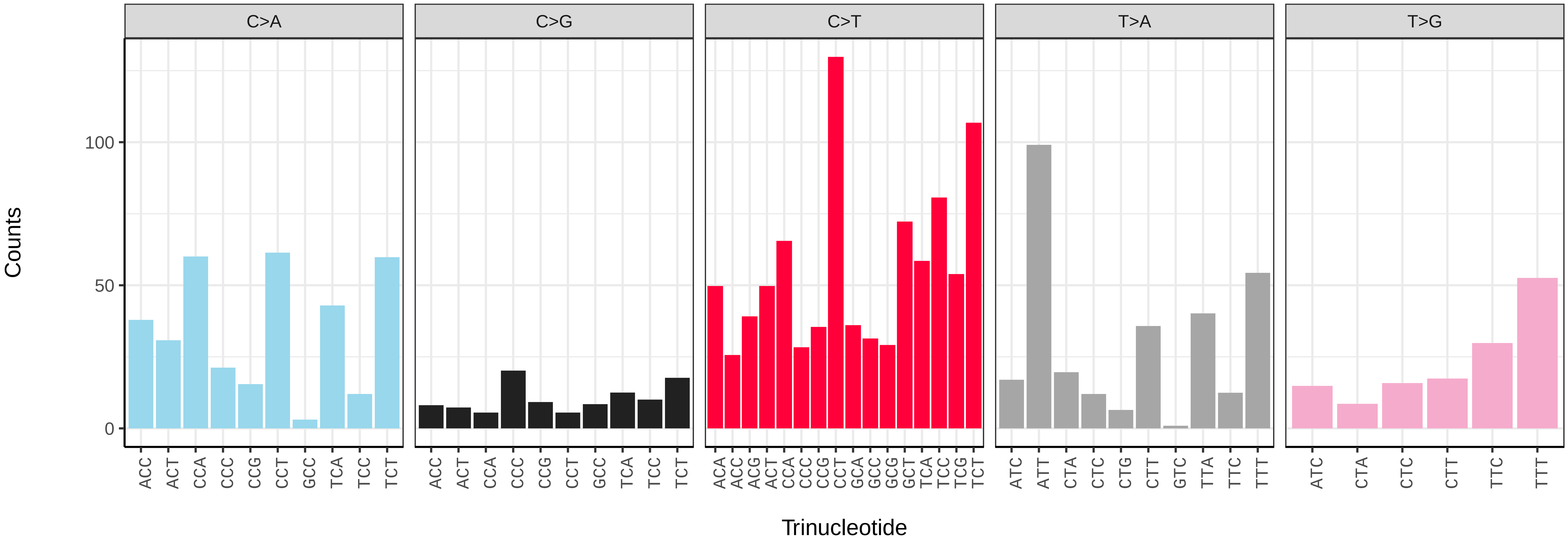

*Malus sylvestris* (drMalSylv7) germline mutational spectrum

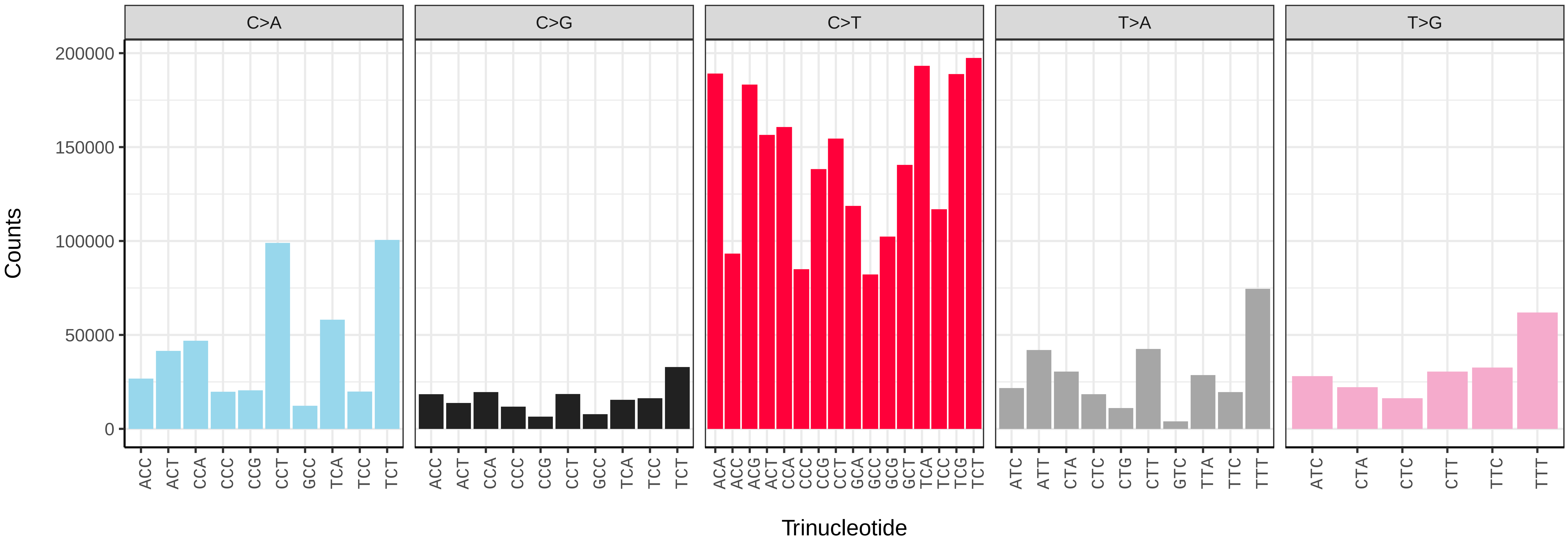


*Malus sylvestris* (drMalSylv7) somatic mutational spectrum (SBS96 classification)

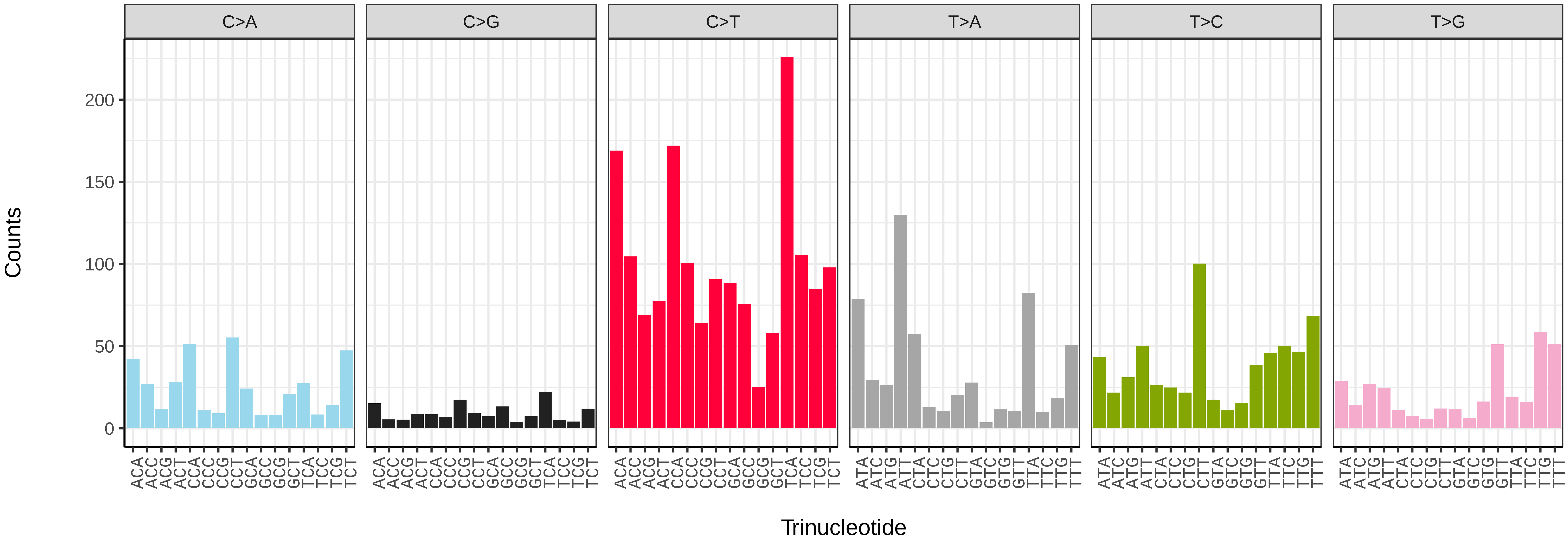


*Malus sylvestris* (drMalSylv7) somatic mutational spectrum (SBS52 classification)

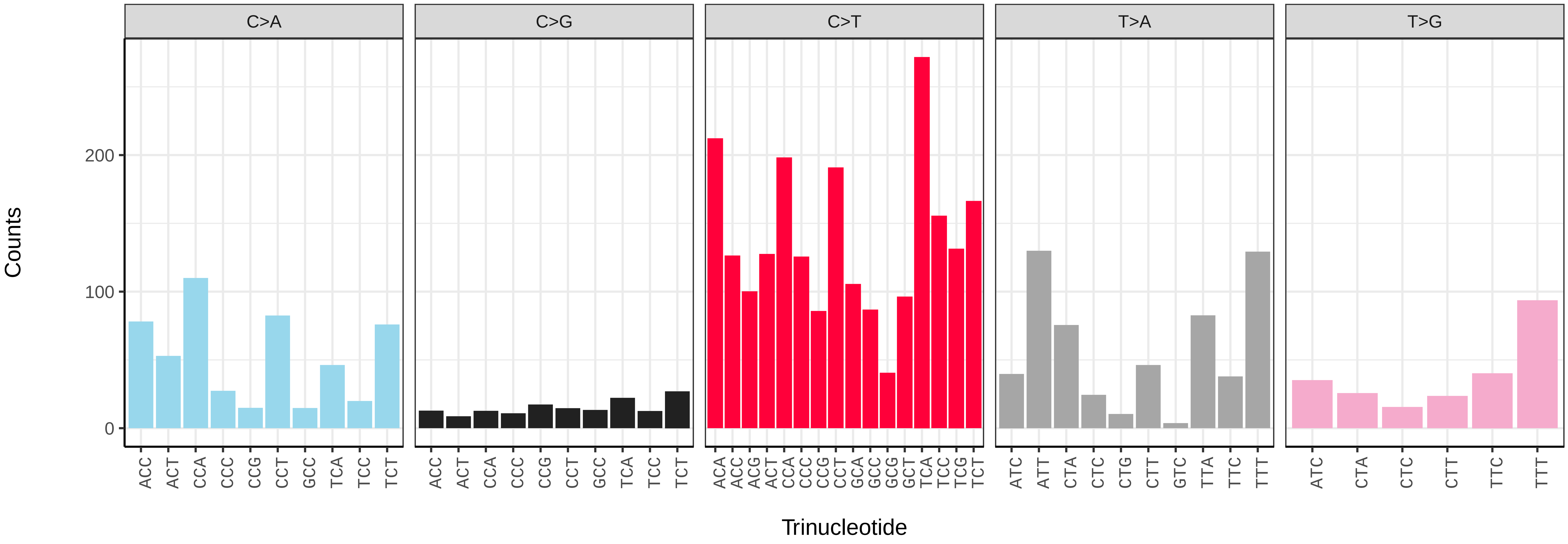

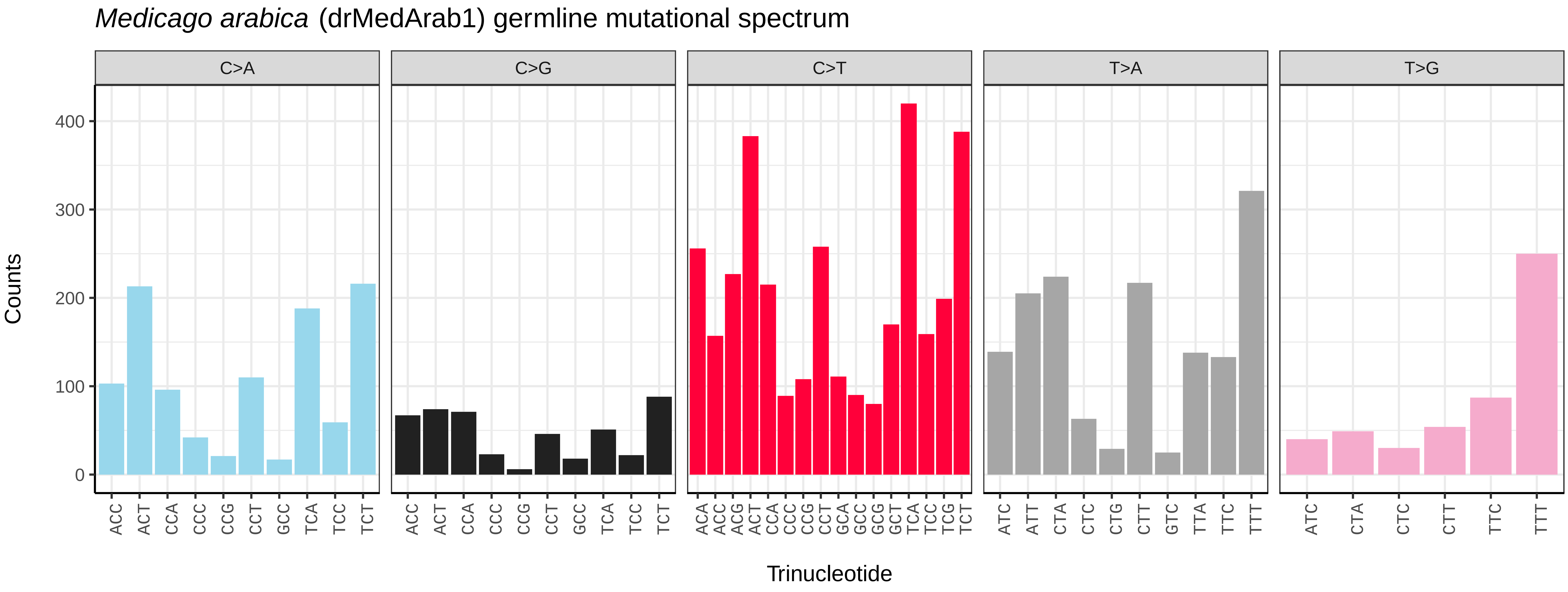

*Medicago arabica* (drMedArab1) germline mutational spectrum
C>A
C>G
C>T
T>A
T>G
Counts
0
100
200
300
400
ACC ACT CCA CCC CCG CCT GCC TCA TCC TCT
ACA ACC ACG ACT CCA CCC CCG CCT GCA GCC GCG GCT TCA TCC TCG TCT
ATC ATT CTA CTC CTG CTT GTC TTA TTC TTT
ATC CTA CTC CTT TTC TTT
Trinucleotide


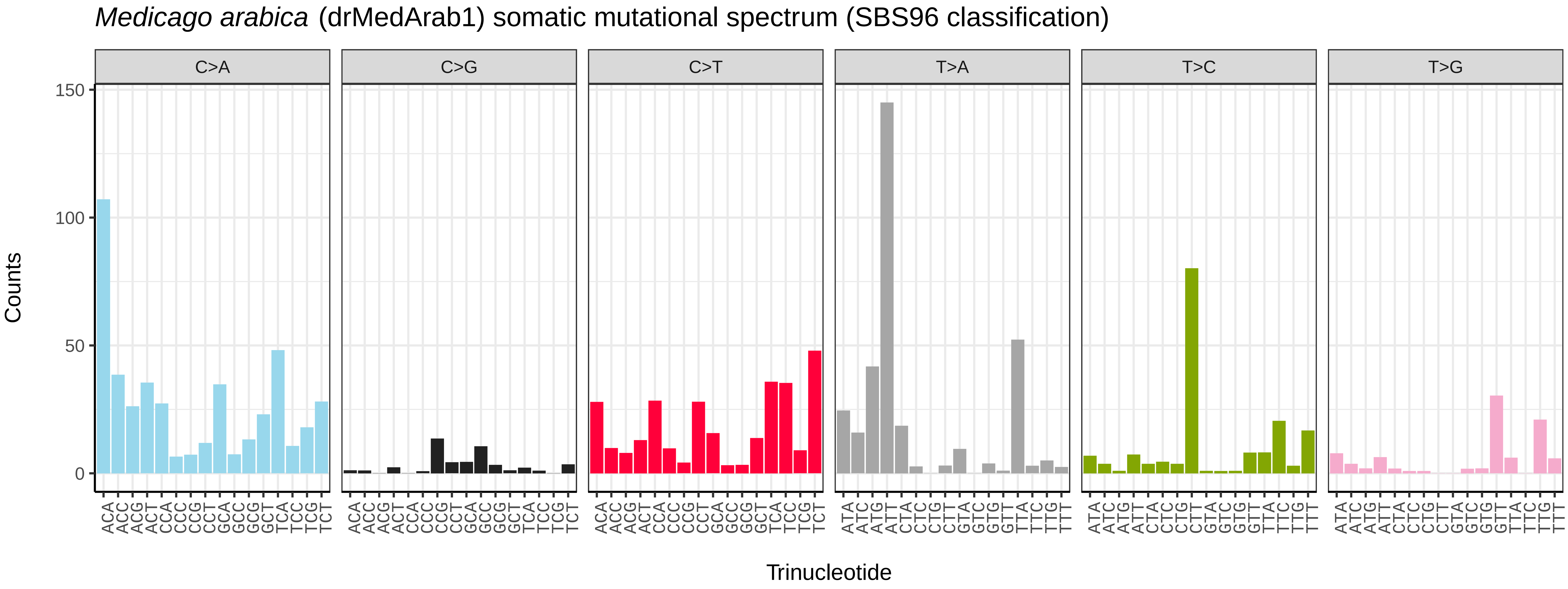

*Medicago arabica* (drMedArab1) somatic mutational spectrum (SBS96 classification)
C>A
C>G
C>T
T>A
T>C
T>G
Counts
0
50
100
150
ACA ACC ACG ACT CCA CCC CCG CCT GCA GCC GCG GCT TCA TCC TCG TCT
ATA ATC ATG ATT CTA CTC CTG CTT GTA GTC GTG GTT TTA TTC TTG TTT
Trinucleotide


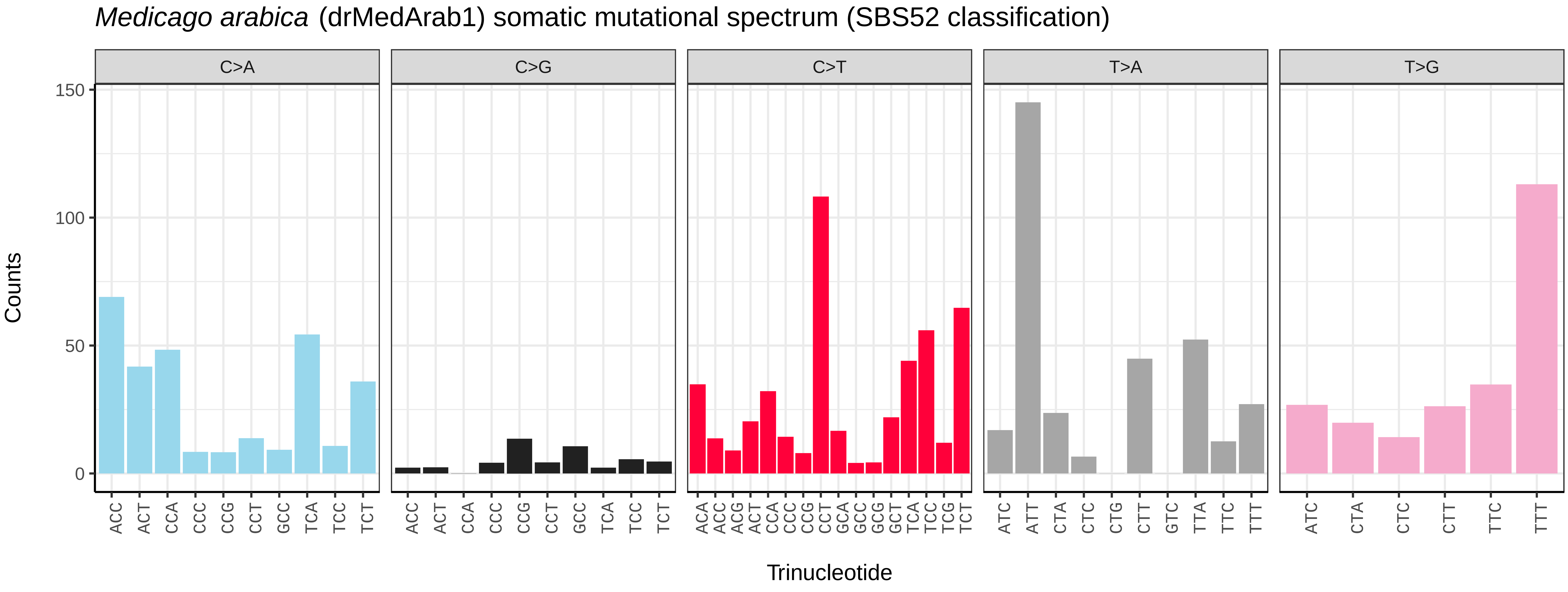

*Medicago arabica* (drMedArab1) somatic mutational spectrum (SBS52 classification)
C>A
C>G
C>T
T>A
T>G
Counts
0
50
100
150
ACC ACT CCA CCC CCG CCT GCC TCA TCC TCT
ACA ACC ACG ACT CCA CCC CCG CCT GCA GCC GCG GCT TCA TCC TCG TCT
ATC ATT CTA CTC CTG CTT GTC TTA TTC TTT
ATC CTA CTC CTT TTC TTT
Trinucleotide

*Medicago lupulina* (drMedLupu1) germline mutational spectrum

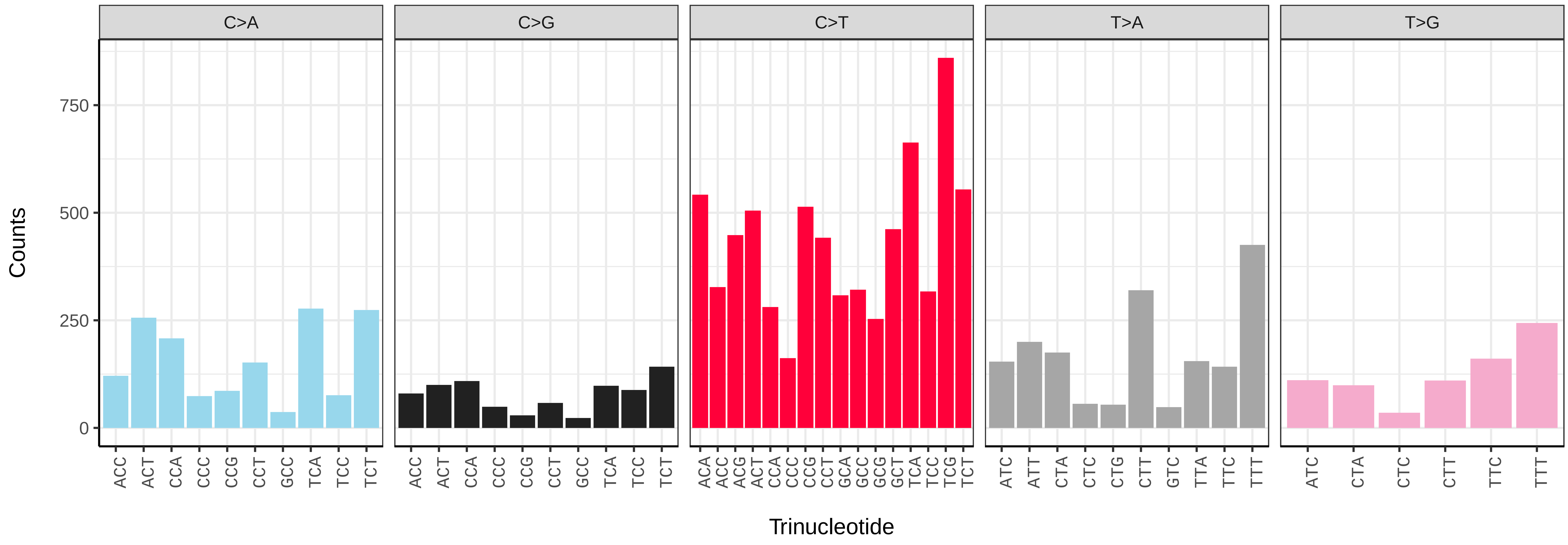


*Medicago lupulina* (drMedLupu1) somatic mutational spectrum (SBS96 classification)

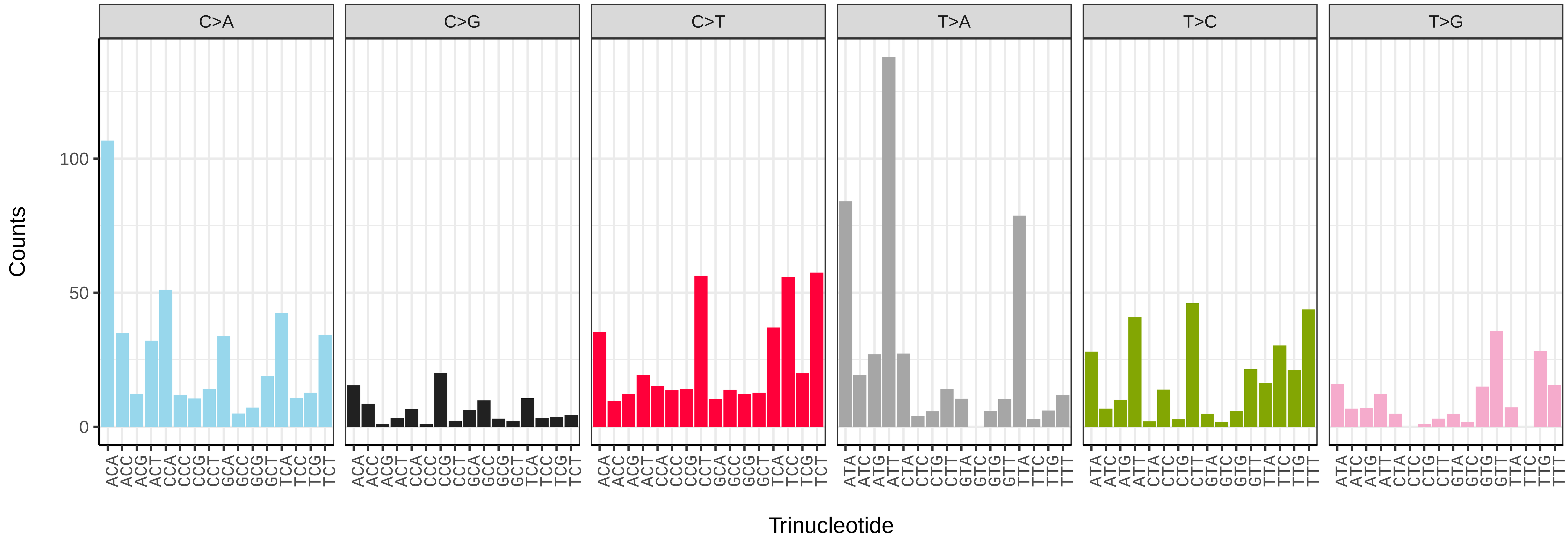


*Medicago lupulina* (drMedLupu1) somatic mutational spectrum (SBS52 classification)

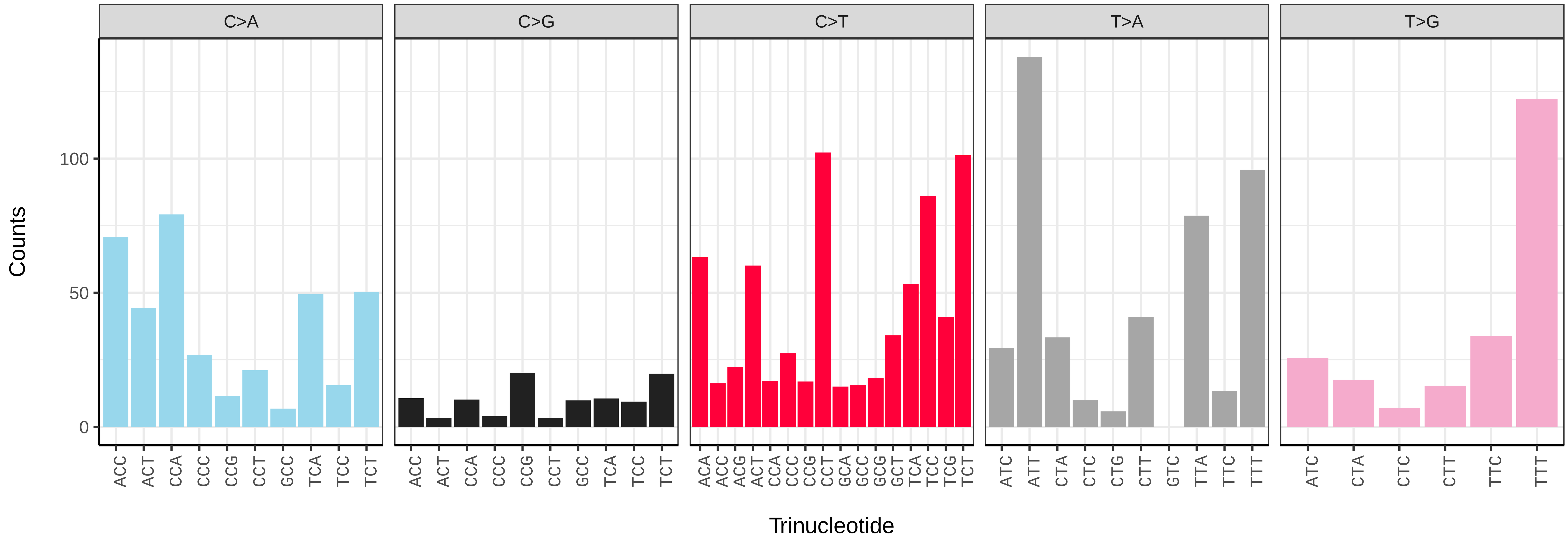

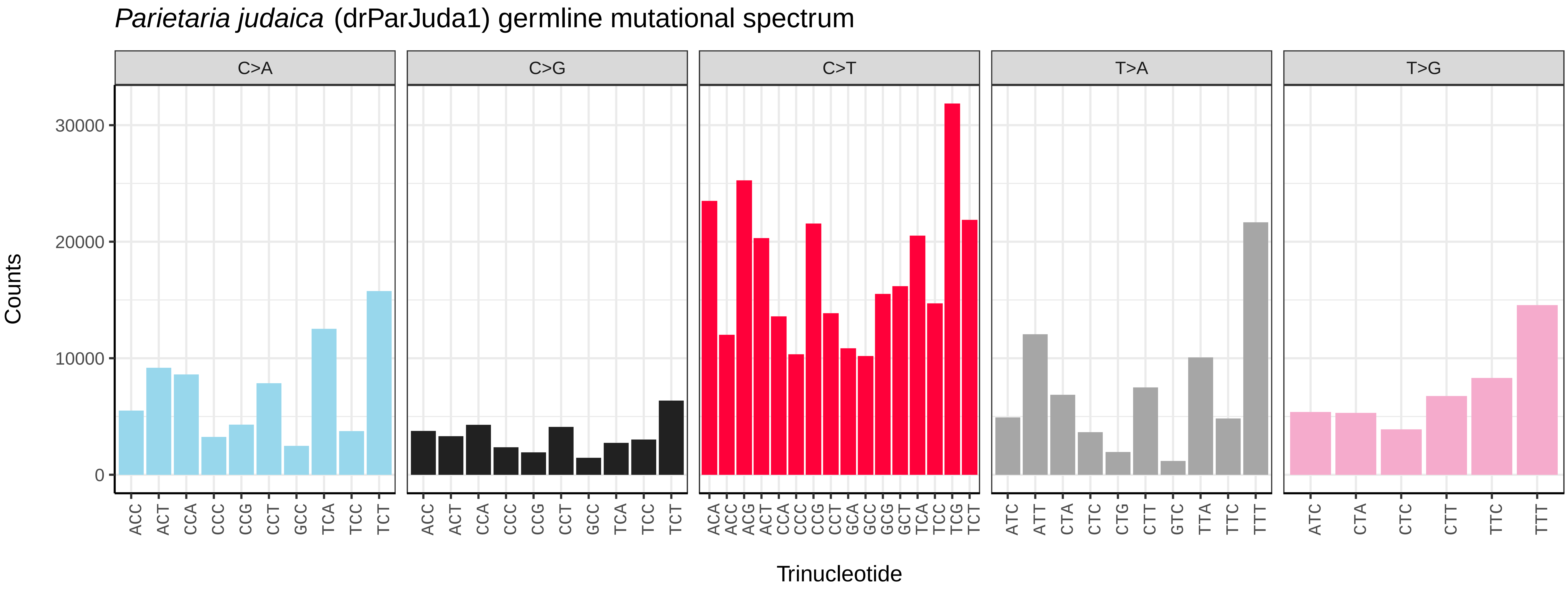
*Parietaria judaica* (drParJuda1) germline mutational spectrum
C>A
C>G
C>T
T>A
T>G
Counts
0
10000
20000
30000
ACC ACT CCA CCC CCG CCT GCC TCA TCC TCT
ACA ACC ACG ACT CCA CCC CCG CCT GCA GCC GCG GCT TCA TCC TCG TCT
ATC ATT CTA CTC CTG CTT GTC TTA TTC TTT
ATC CTA CTC CTT TTC TTT
Trinucleotide

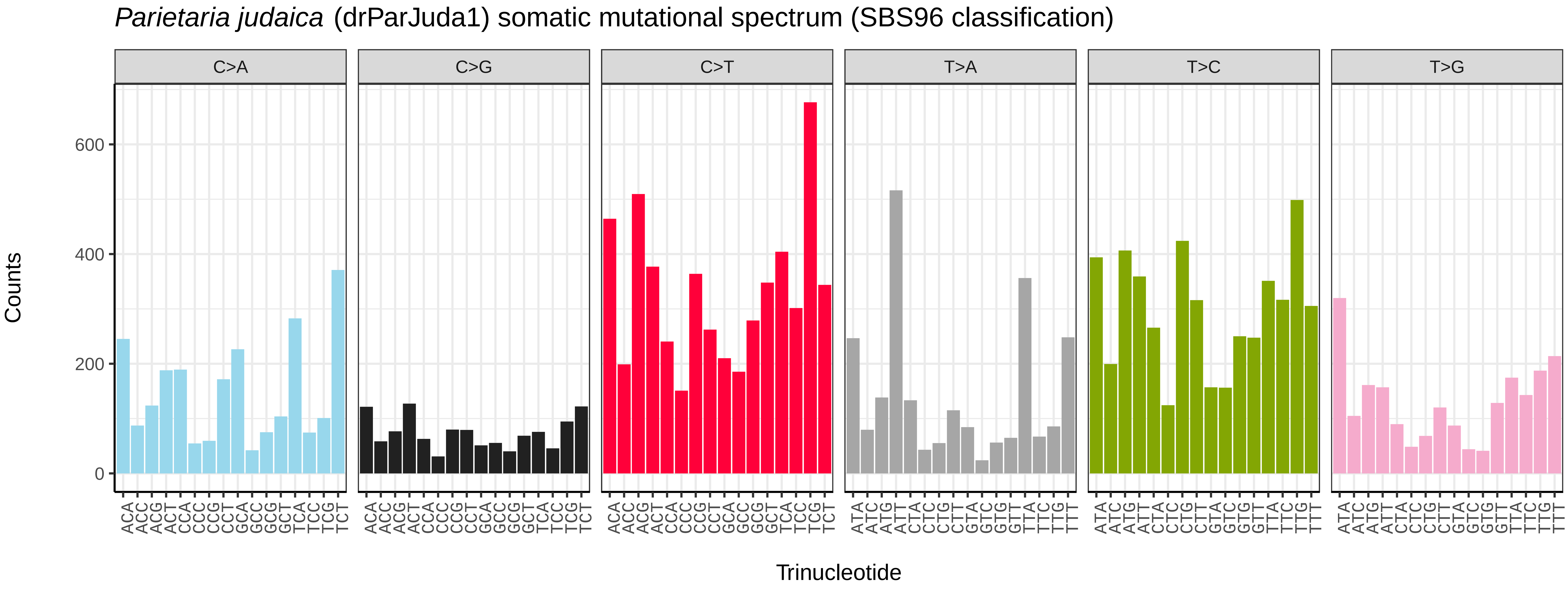
*Parietaria judaica* (drParJuda1) somatic mutational spectrum (SBS96 classification)
C>A
C>G
C>T
T>A
T>C
T>G
Counts
0
200
400
600
ACA ACC ACG ACT CCA CCC CCG CCT GCA GCC GCG GCT TCA TCC TCG TCT
ATA ATC ATG ATT CTA CTC CTG CTT GTA GTC GTG GTT TTA TTC TTG TTT
Trinucleotide

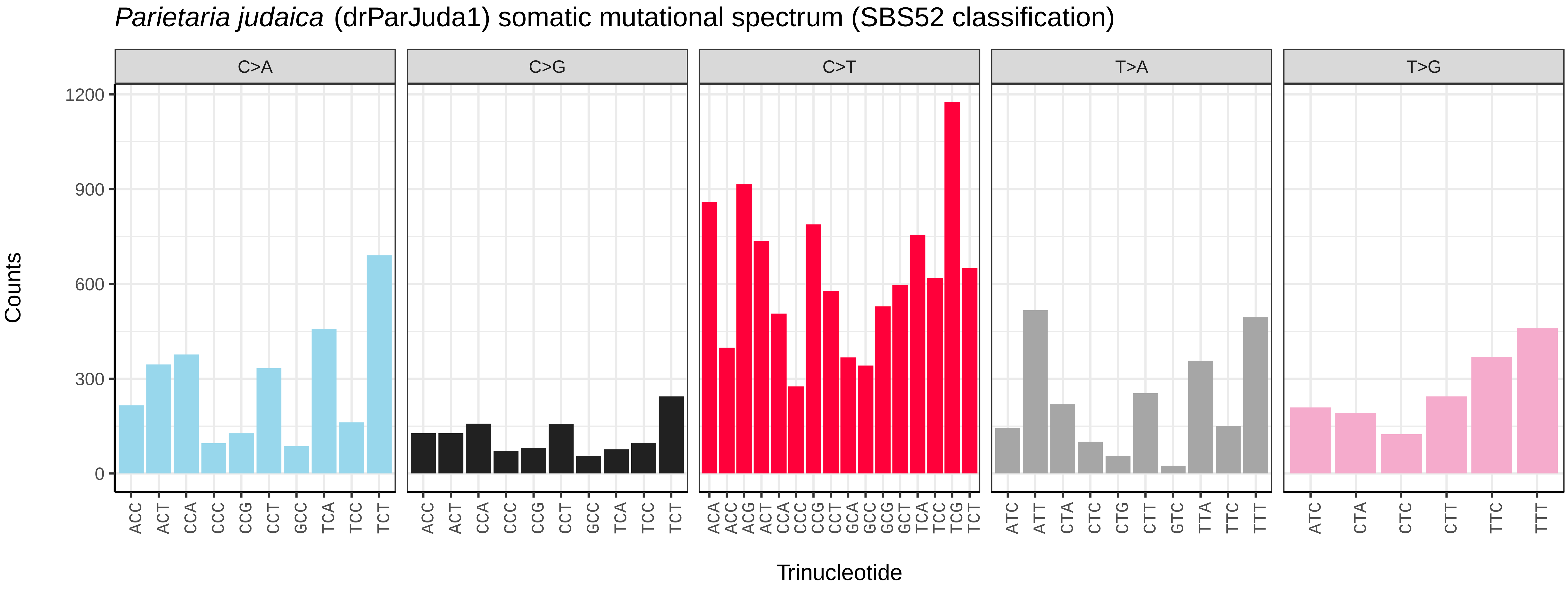
*Parietaria judaica* (drParJuda1) somatic mutational spectrum (SBS52 classification)
C>A
C>G
C>T
T>A
T>G
Counts
0
300
600
900
1200
ACC ACT CCA CCC CCG CCT GCC TCA TCC TCT
ACA ACC ACG ACT CCA CCC CCG CCT GCA GCC GCG GCT TCA TCC TCG TCT
ATC ATT CTA CTC CTG CTT GTC TTA TTC TTT
ATC CTA CTC CTT TTC TTT
Trinucleotide

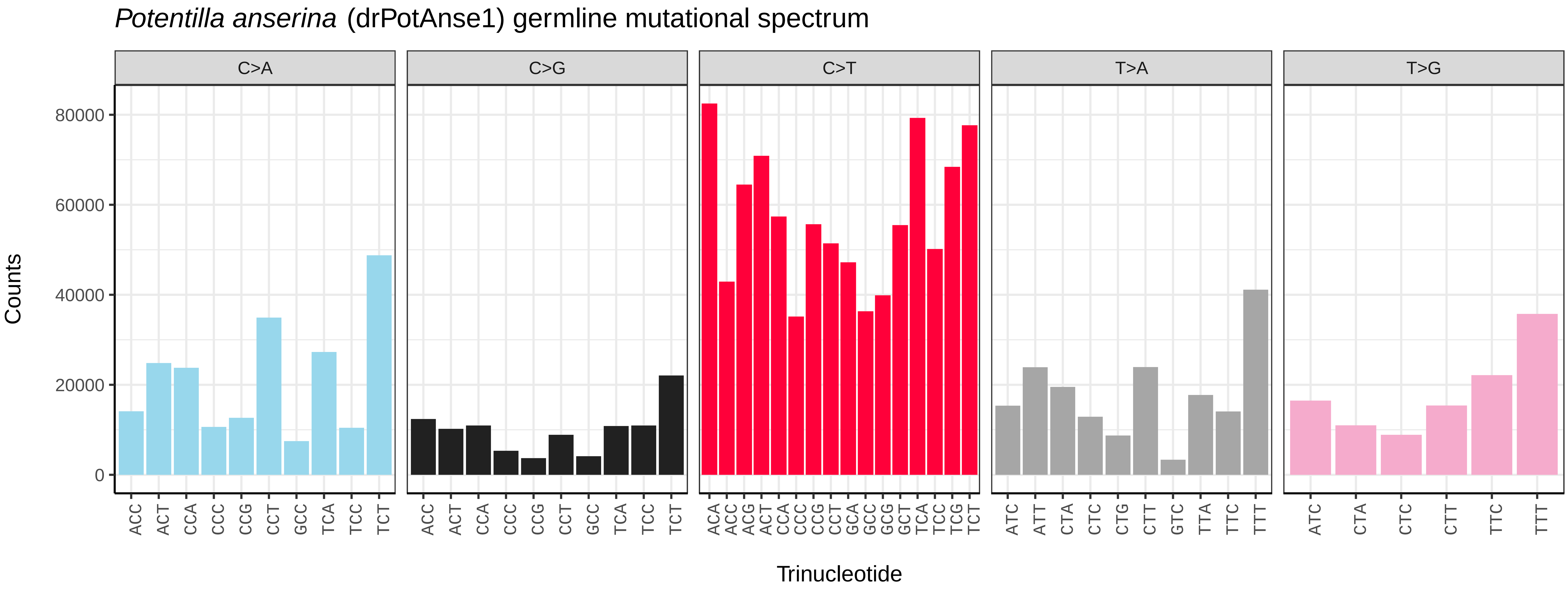
*Potentilla anserina* (drPotAnse1) germline mutational spectrum
C>A
C>G
C>T
T>A
T>G
Counts
0
20000
40000
60000
80000
Trinucleotide

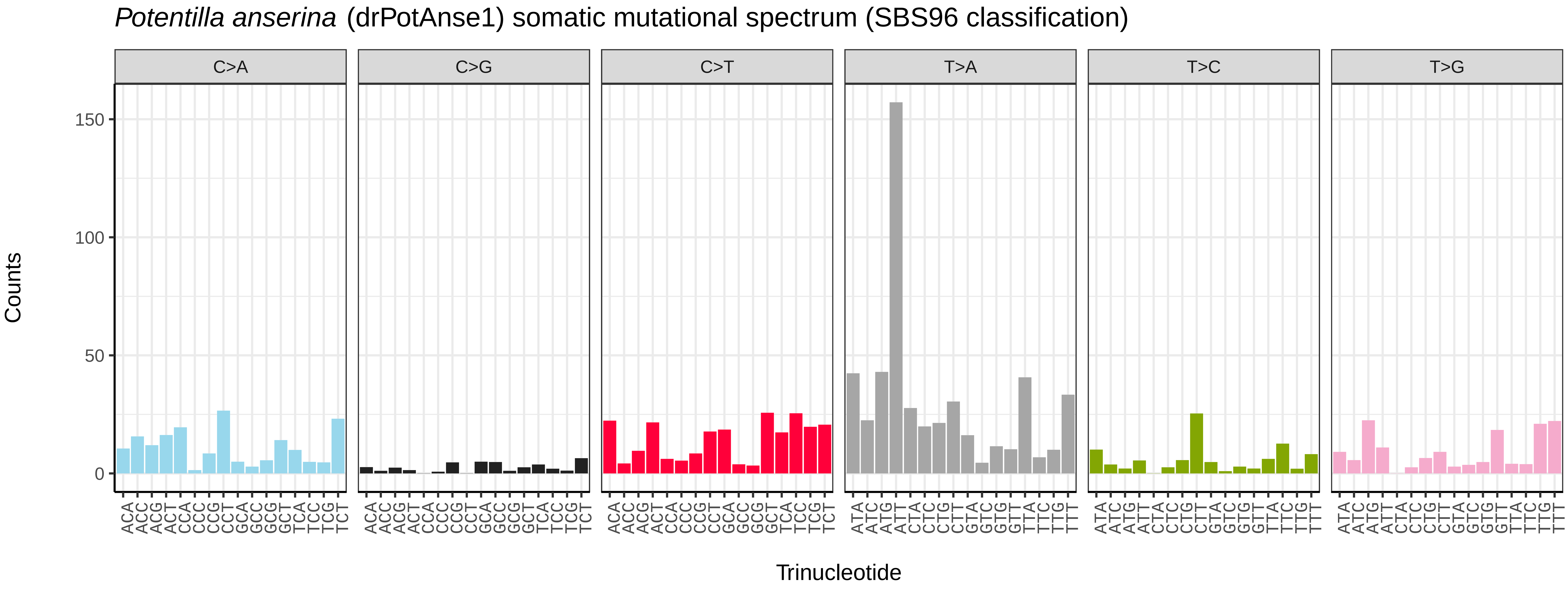
*Potentilla anserina* (drPotAnse1) somatic mutational spectrum (SBS96 classification)
C>A
C>G
C>T
T>A
T>C
T>G
Counts
0
50
100
150
Trinucleotide

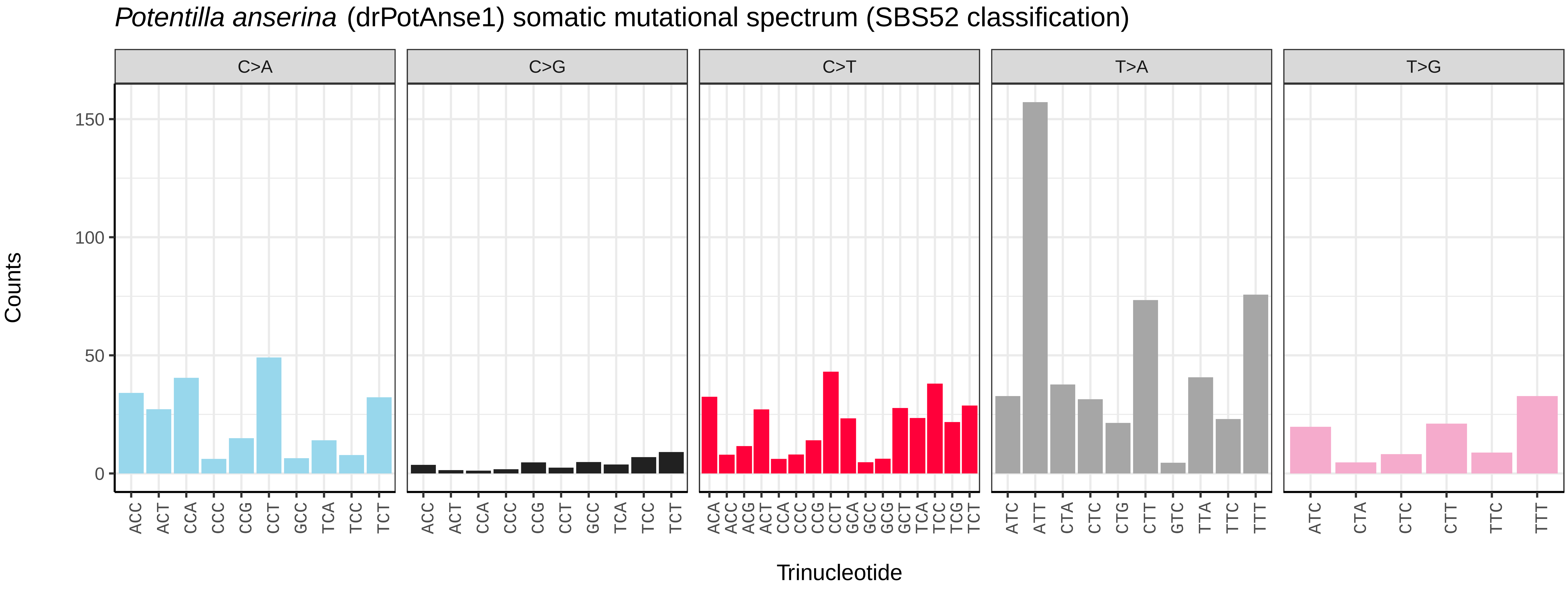
*Potentilla anserina* (drPotAnse1) somatic mutational spectrum (SBS52 classification)
C>A
C>G
C>T
T>A
T>G
Counts
0
50
100
150
Trinucleotide

*Potentilla sterilis* (drPotSter1) germline mutational spectrum

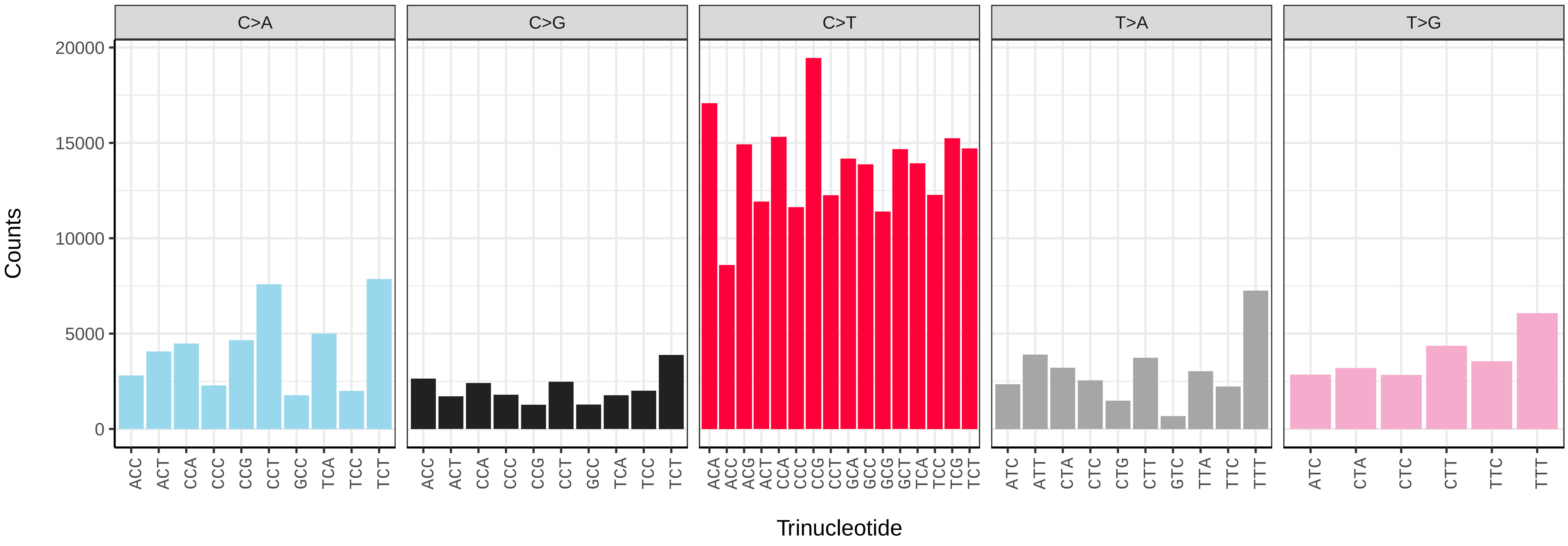


*Potentilla sterilis* (drPotSter1) somatic mutational spectrum (SBS96 classification)

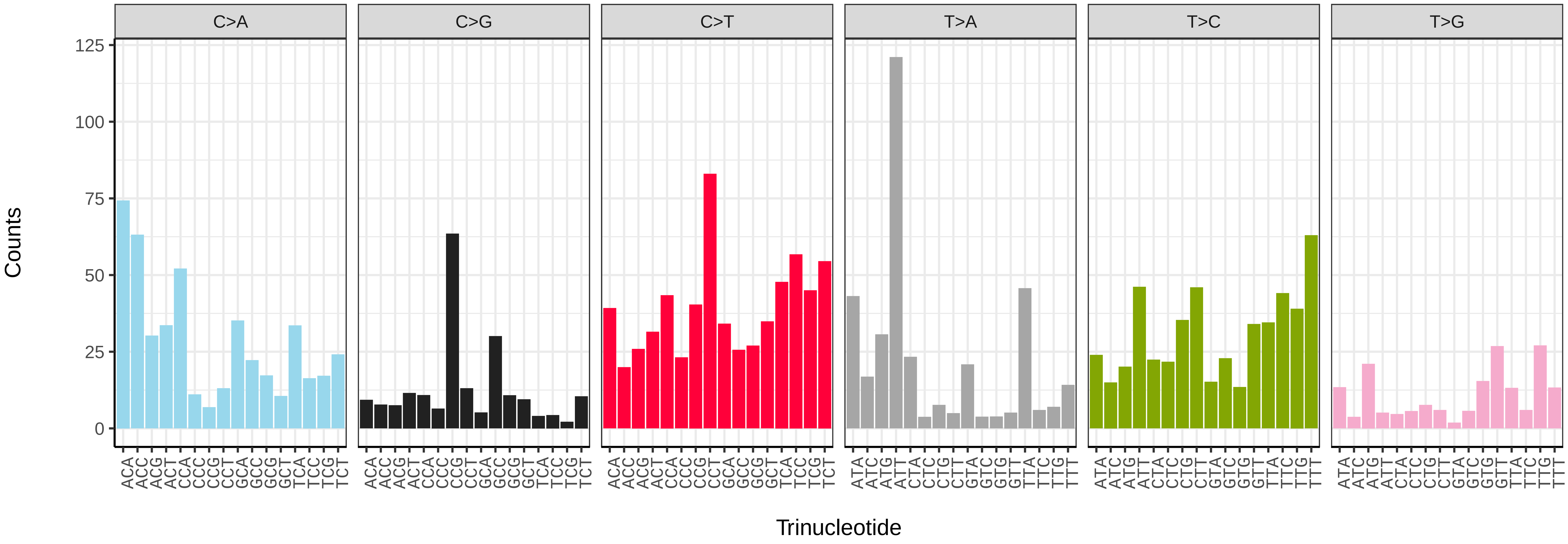


*Potentilla sterilis* (drPotSter1) somatic mutational spectrum (SBS52 classification)

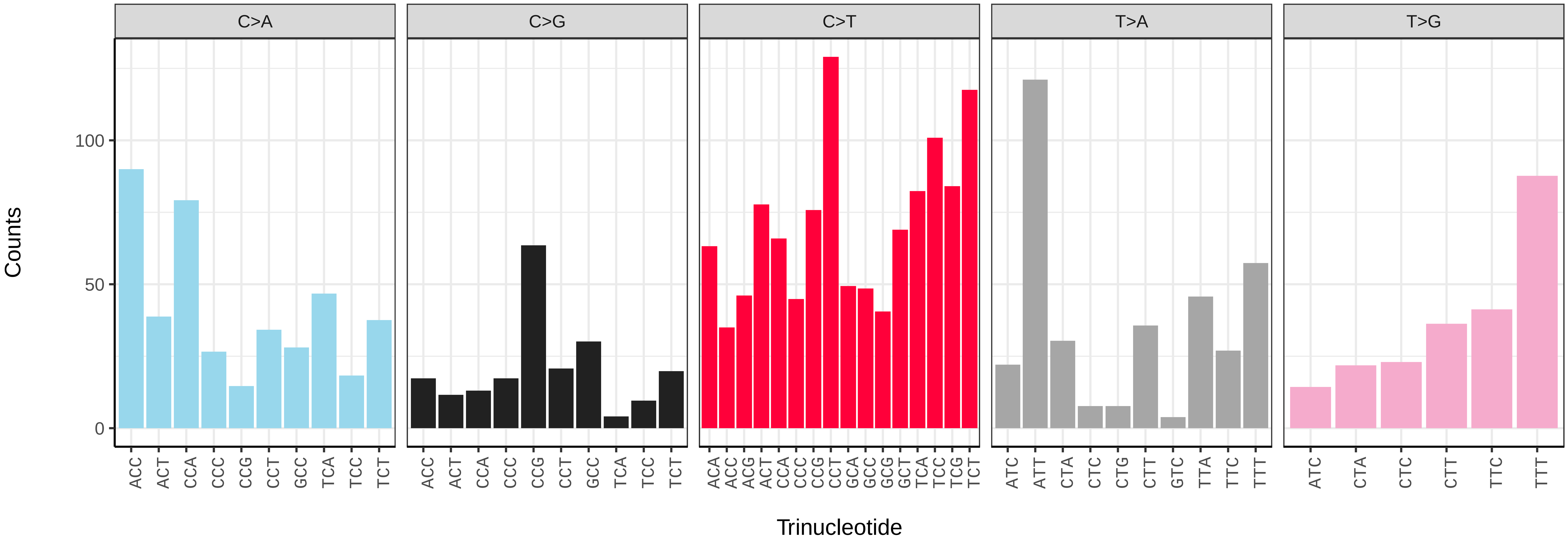

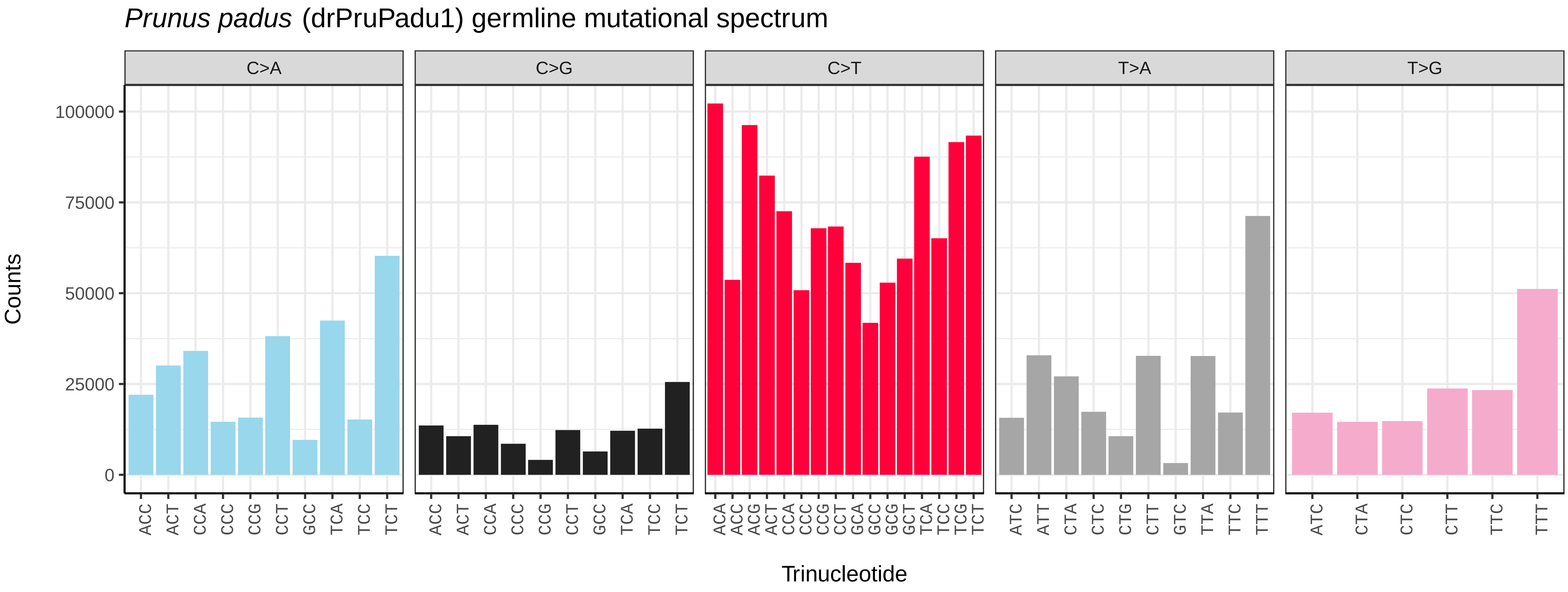
*Prunus padus* (drPruPadu1) germline mutational spectrum
C>A
C>G
C>T
T>A
T>G
Counts
Trinucleotide

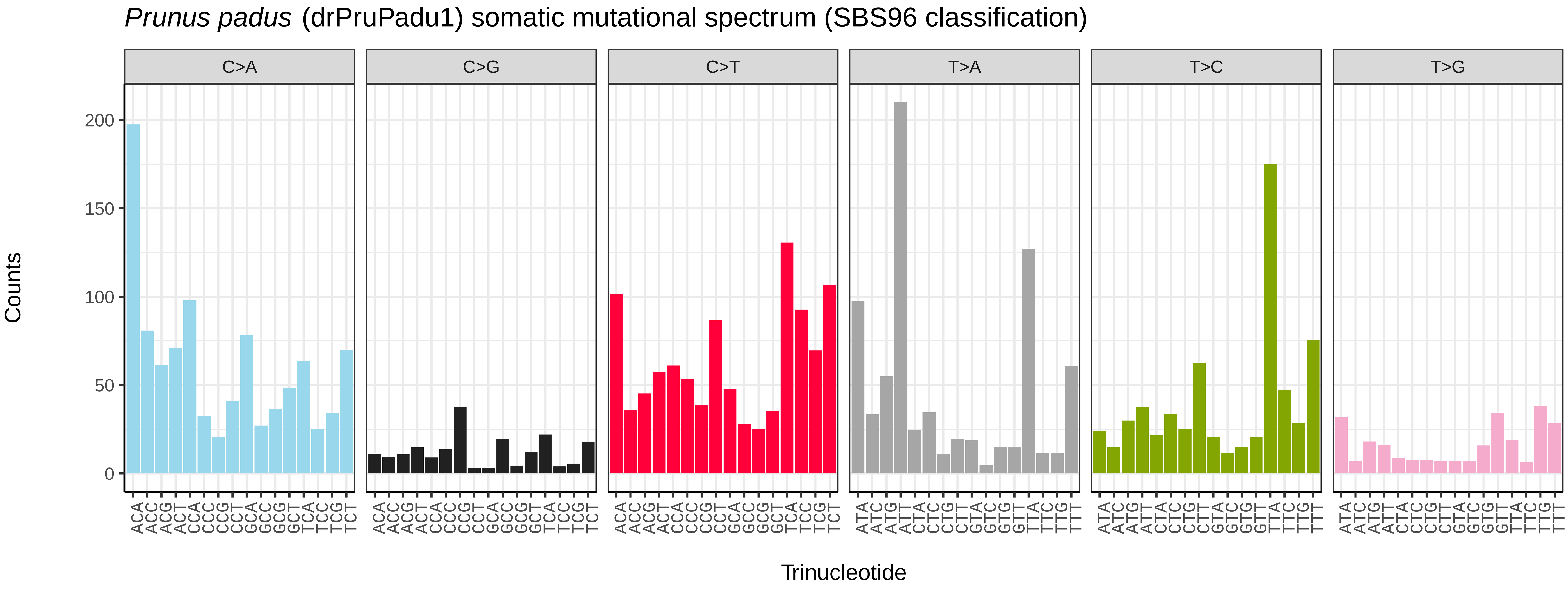
*Prunus padus* (drPruPadu1) somatic mutational spectrum (SBS96 classification)
C>A
C>G
C>T
T>A
T>C
T>G
Counts
Trinucleotide

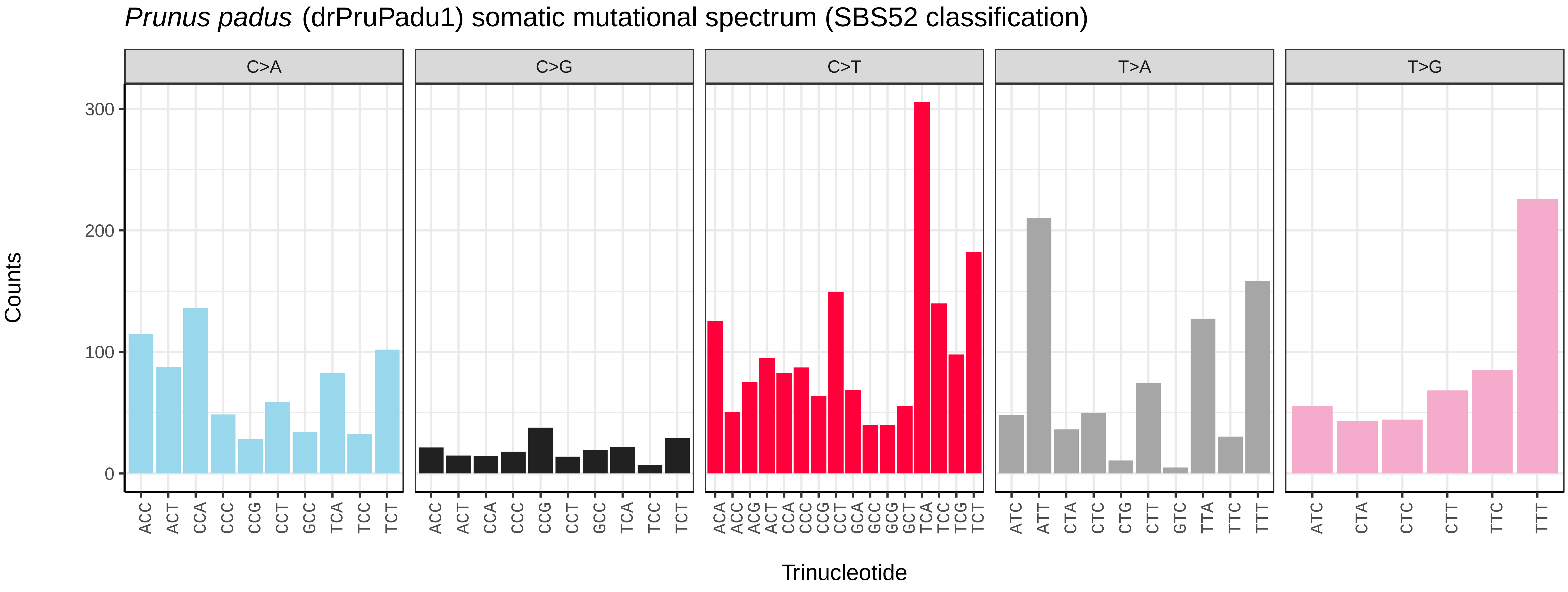
*Prunus padus* (drPruPadu1) somatic mutational spectrum (SBS52 classification)
C>A
C>G
C>T
T>A
T>G
Counts
Trinucleotide

*Pyrus communis* (drPyrComm1) germline mutational spectrum

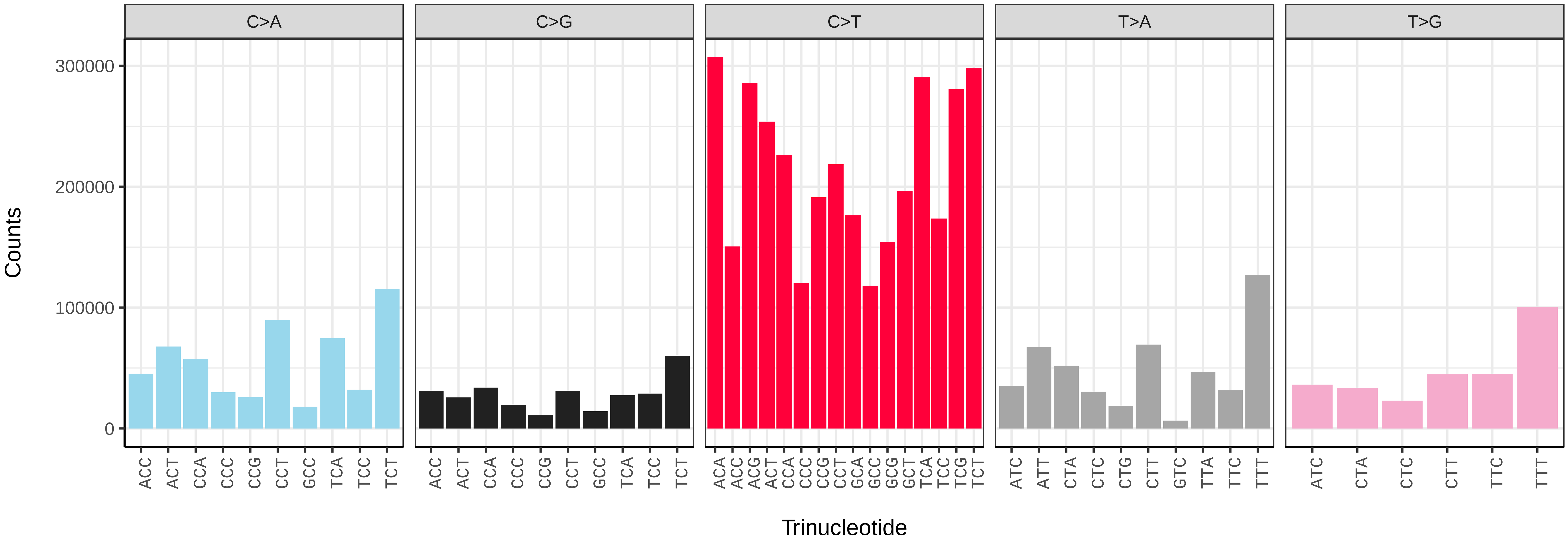


*Pyrus communis* (drPyrComm1) somatic mutational spectrum (SBS96 classification)

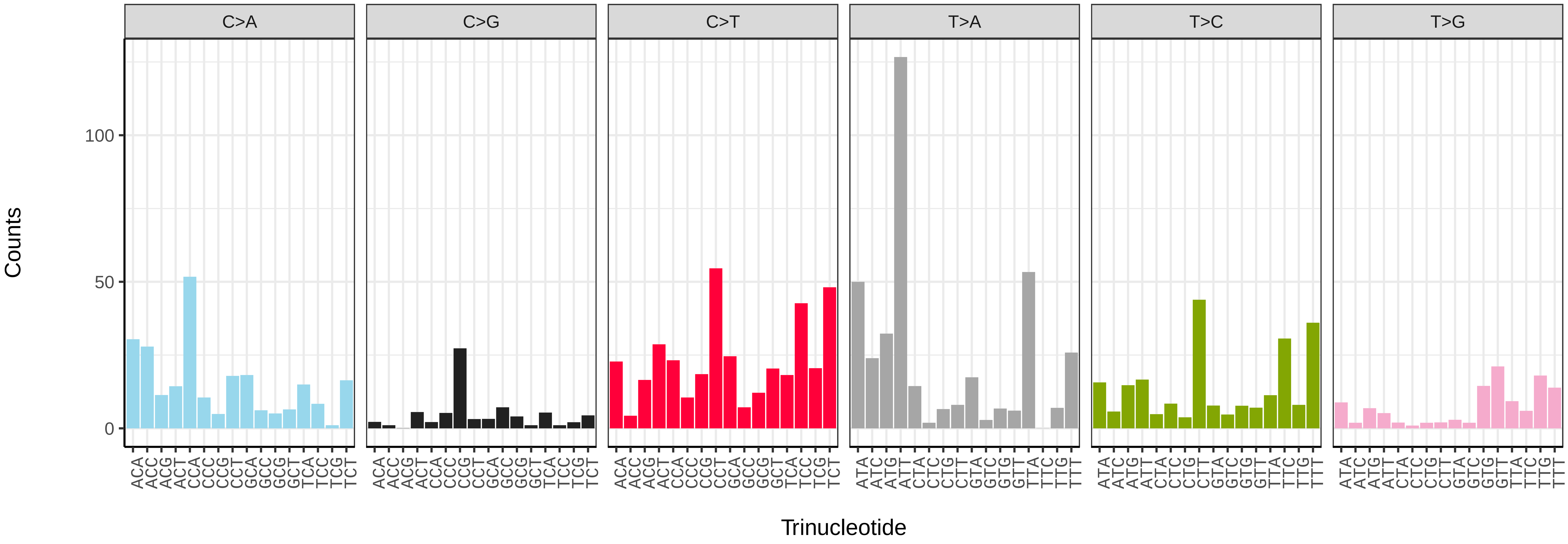


*Pyrus communis* (drPyrComm1) somatic mutational spectrum (SBS52 classification)

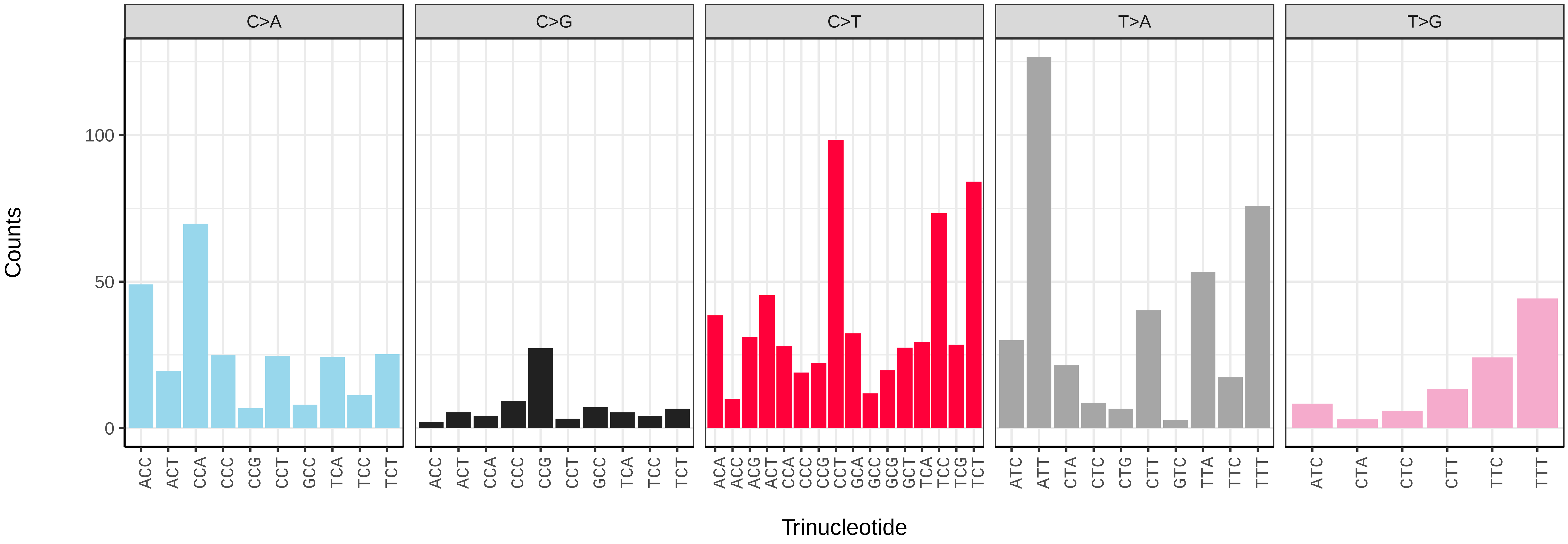

*Trifolium fragiferum* (drTriFrag1) germline mutational spectrum

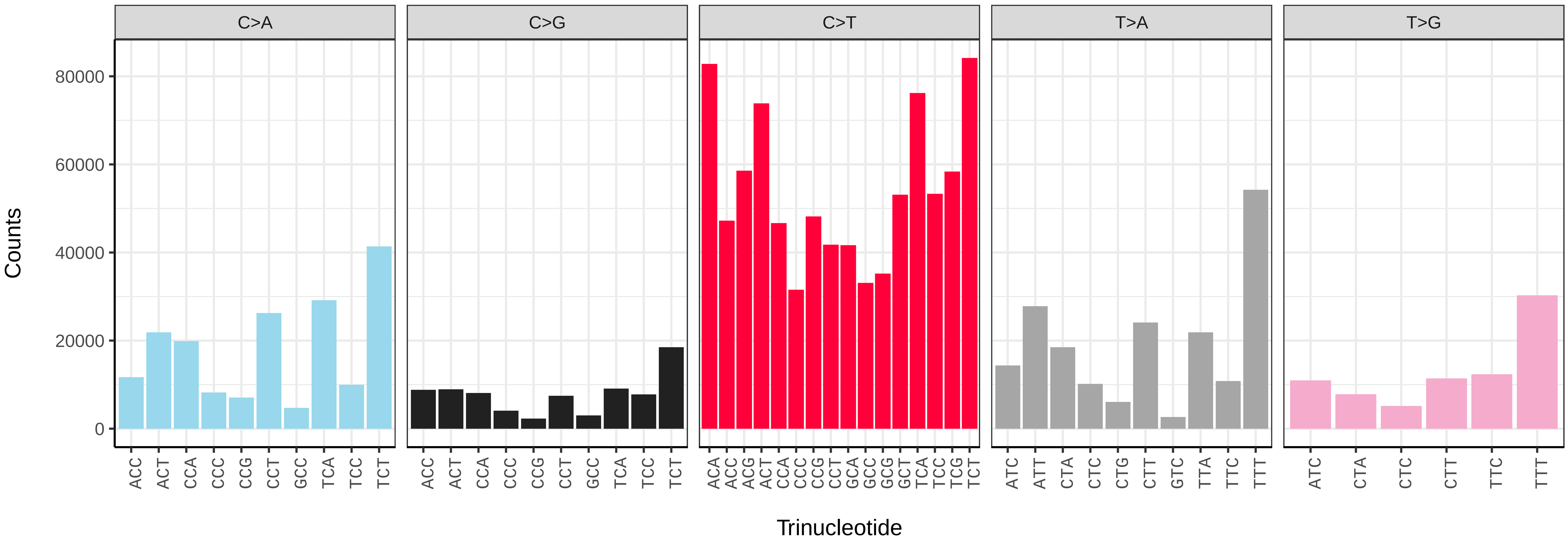


*Trifolium fragiferum* (drTriFrag1) somatic mutational spectrum (SBS96 classification)

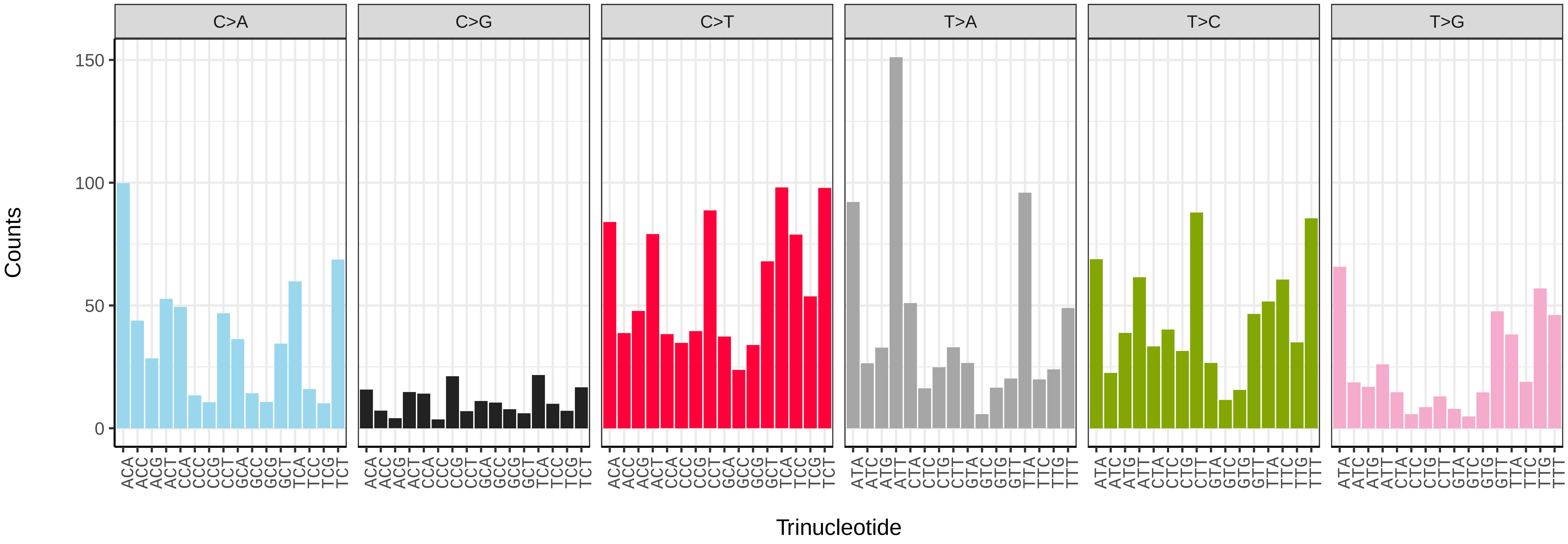


*Trifolium fragiferum* (drTriFrag1) somatic mutational spectrum (SBS52 classification)

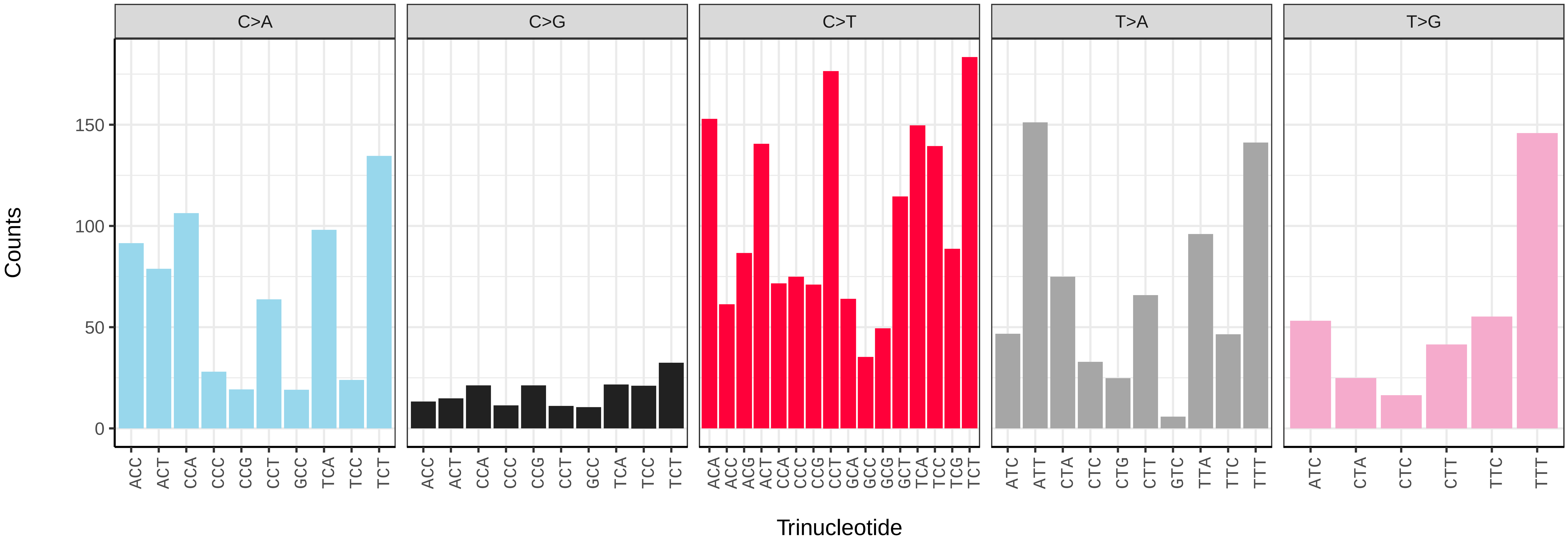

*Trifolium subterraneum* (drTriSubt1) germline mutational spectrum

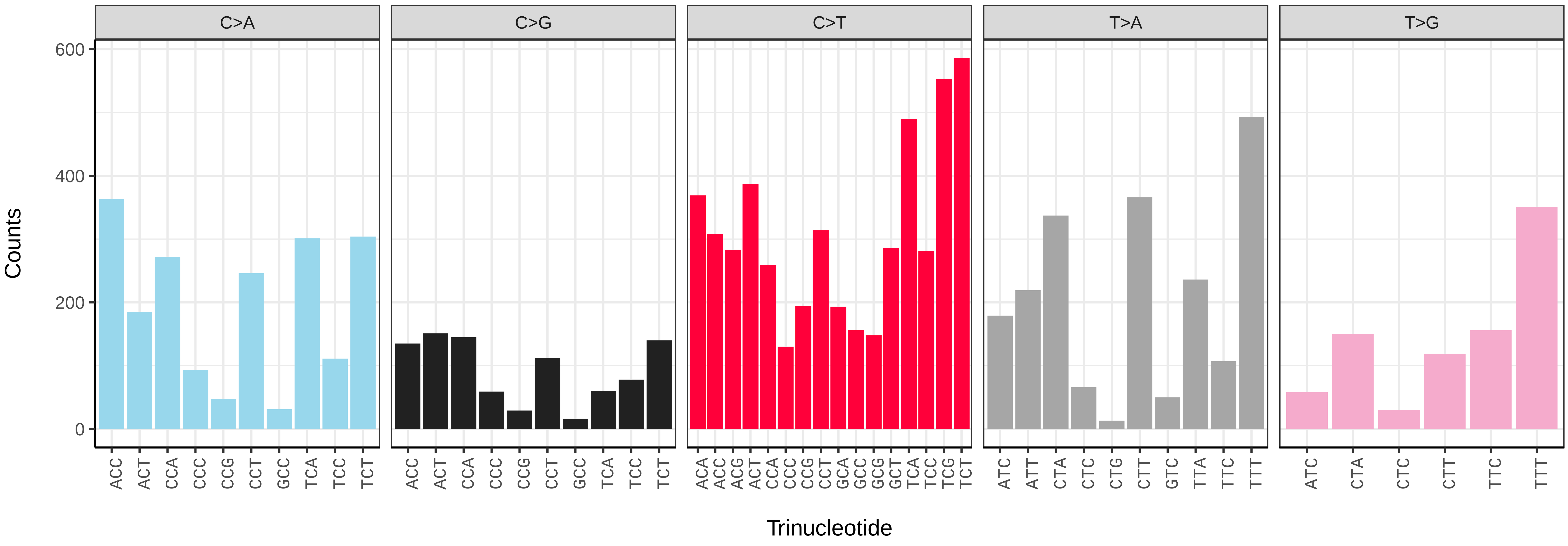


*Trifolium subterraneum* (drTriSubt1) somatic mutational spectrum (SBS96 classification)

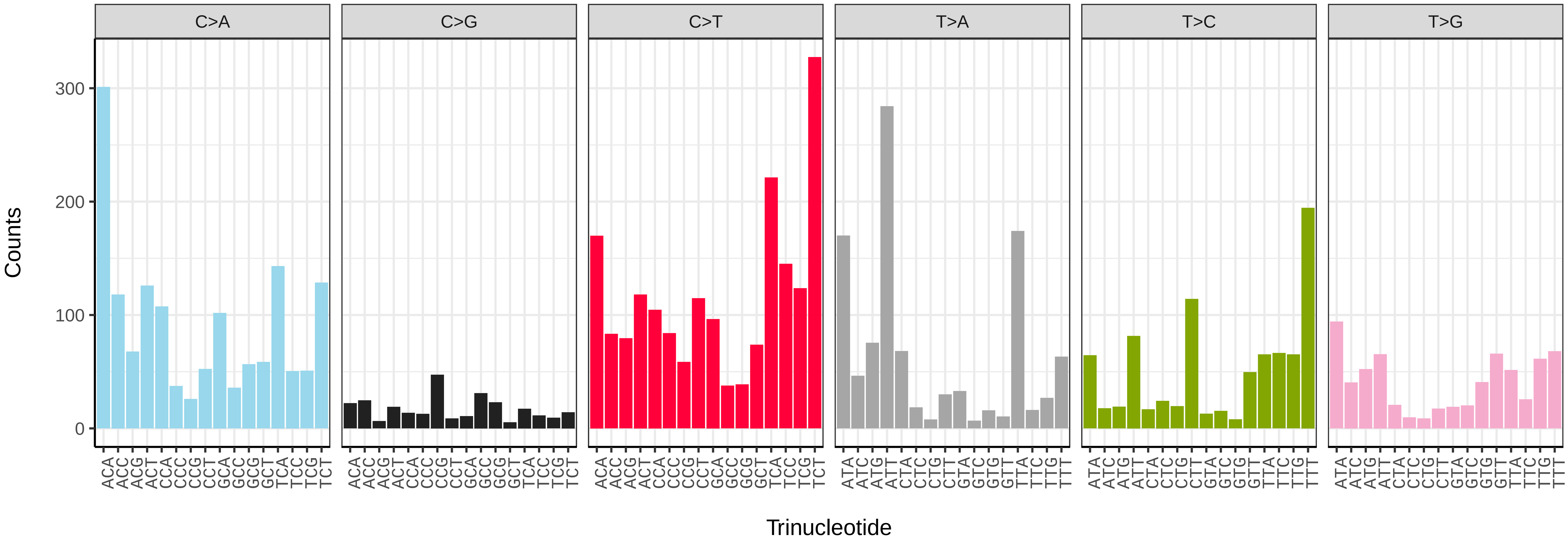


*Trifolium subterraneum* (drTriSubt1) somatic mutational spectrum (SBS52 classification)

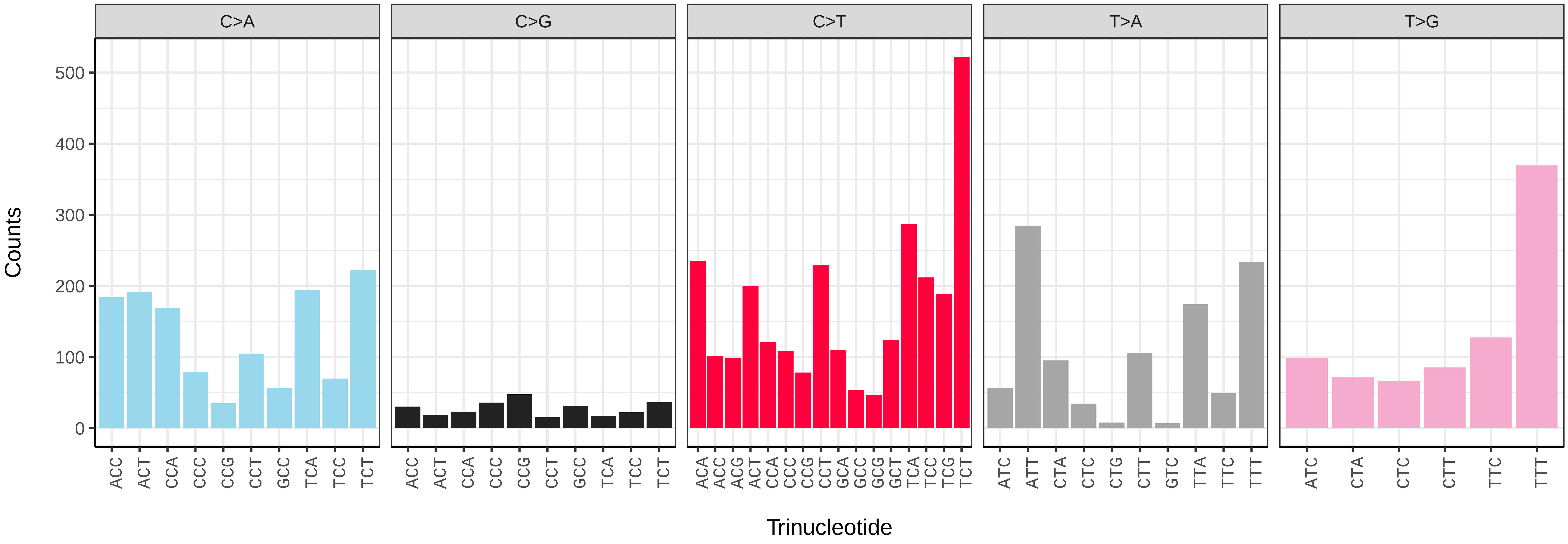

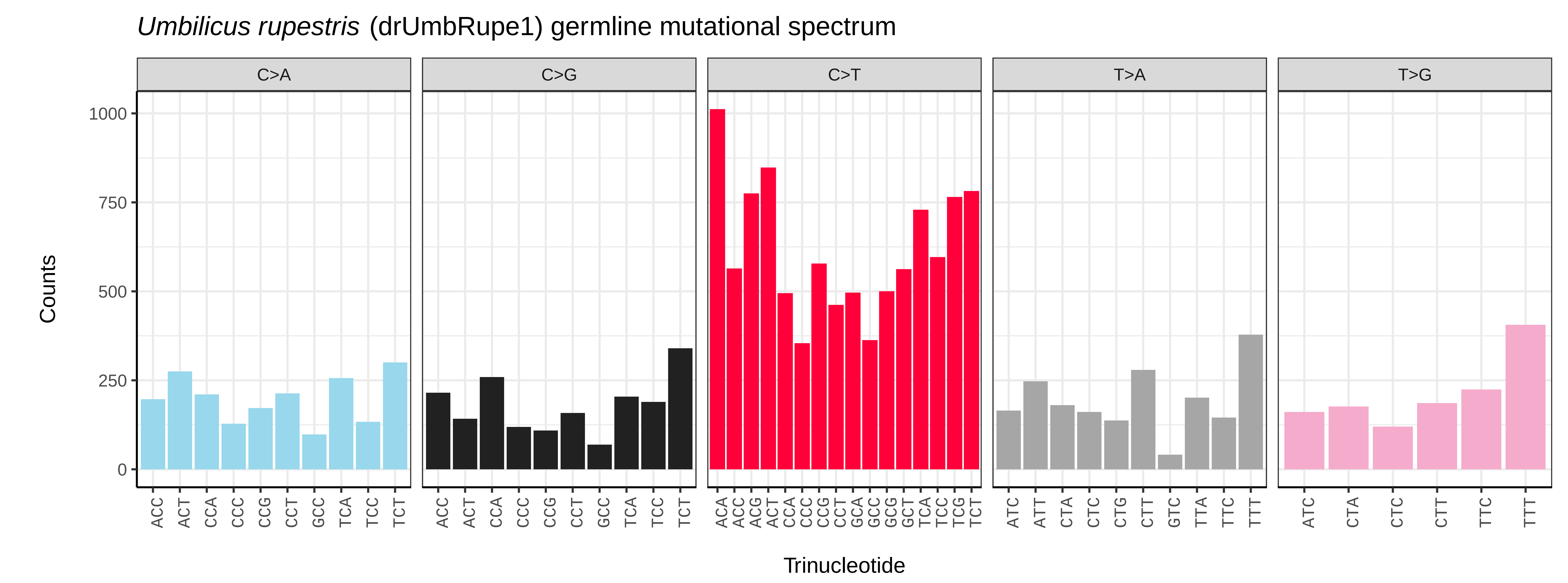
*Umbilicus rupestris* (drUmbRupe1) germline mutational spectrum
C>A
C>G
C>T
T>A
T>G
Counts
0
250
500
750
1000
Trinucleotide
ACC ACT CCA CCC CCG CCT GCC TCA TCC TCT
ACA ACC ACG ACT CCA CCC CCG CCT GCA GCC GCG GCT TCA TCC TCG TCT
ATC ATT CTA CTC CTG CTT GTC TTA TTC TTT
ATC CTA CTC CTT TTC TTT

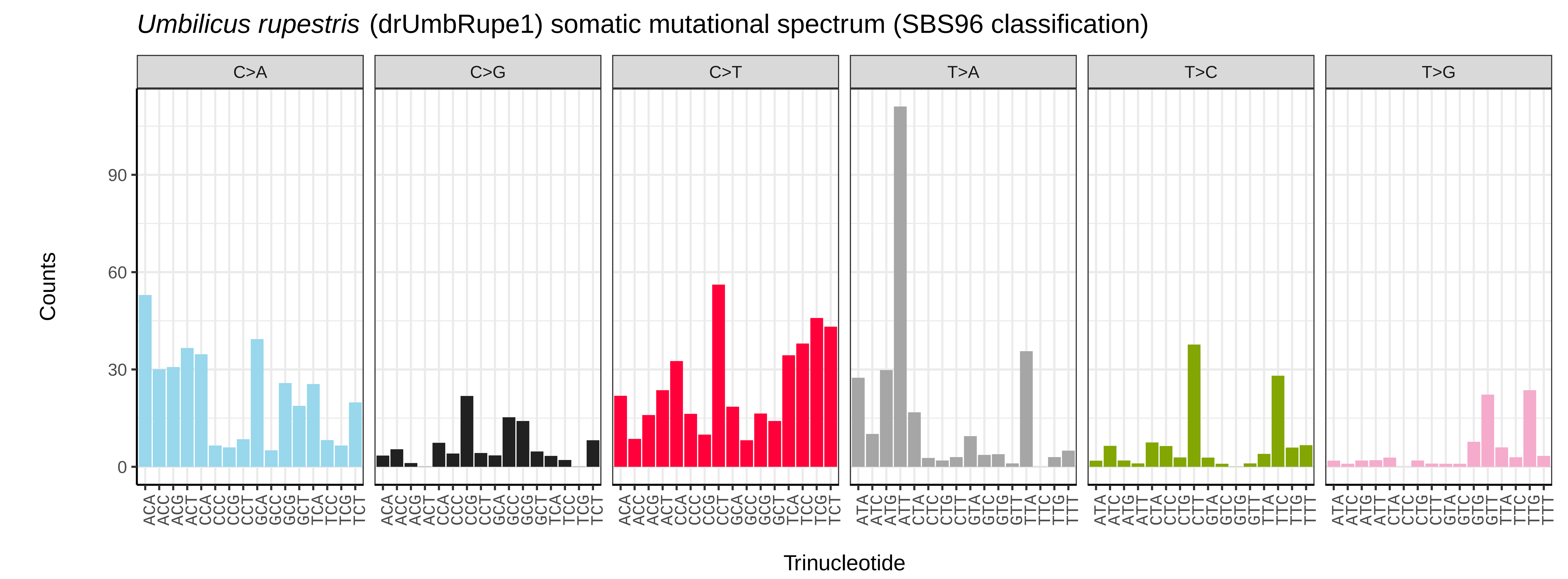
*Umbilicus rupestris* (drUmbRupe1) somatic mutational spectrum (SBS96 classification)
C>A
C>G
C>T
T>A
T>C
T>G
Counts
0
30
60
90
Trinucleotide
ACA ACC ACG ACT CCA CCC CCG CCT GCA GCC GCG GCT TCA TCC TCG TCT
ATA ATC ATG ATT CTA CTC CTG CTT GTA GTC GTG GTT TTA TTC TTG TTT

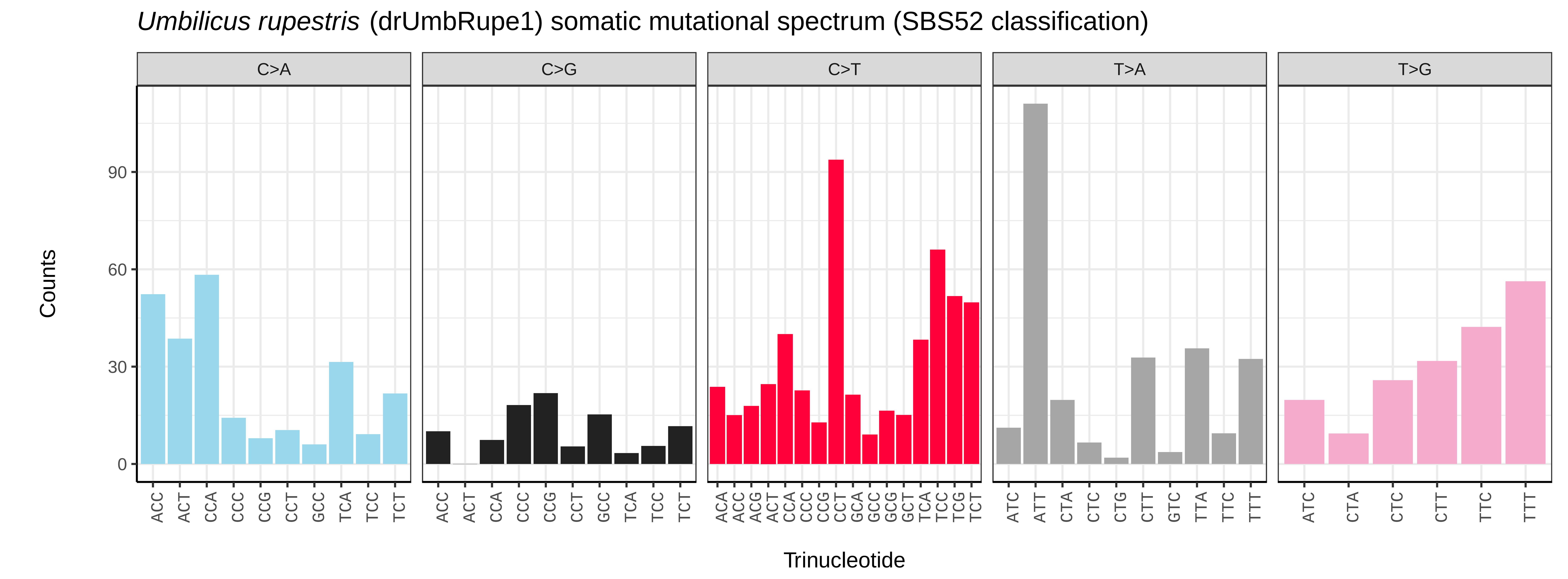
*Umbilicus rupestris* (drUmbRupe1) somatic mutational spectrum (SBS52 classification)
C>A
C>G
C>T
T>A
T>G
Counts
0
30
60
90
Trinucleotide
ACC ACT CCA CCC CCG CCT GCC TCA TCC TCT
ACA ACC ACG ACT CCA CCC CCG CCT GCA GCC GCG GCT TCA TCC TCG TCT
ATC ATT CTA CTC CTG CTT GTC TTA TTC TTT
ATC CTA CTC CTT TTC TTT

*Urtica urens* (drUrtUren1) germline mutational spectrum

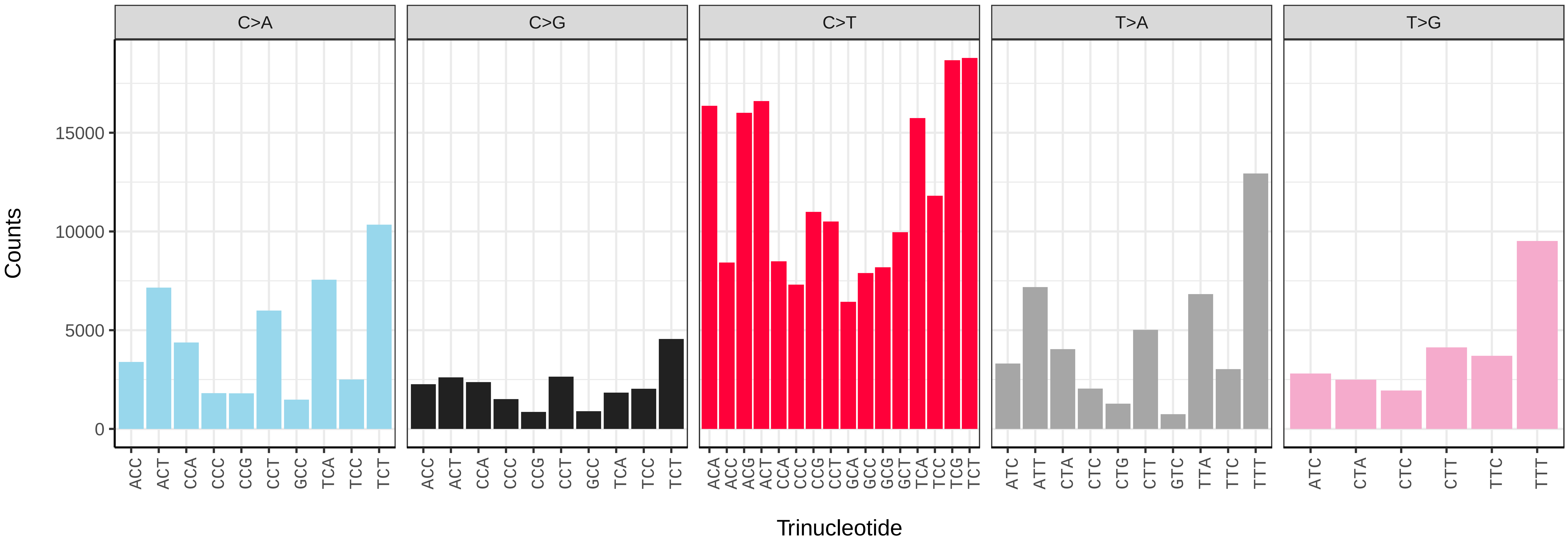


*Urtica urens* (drUrtUren1) somatic mutational spectrum (SBS96 classification)

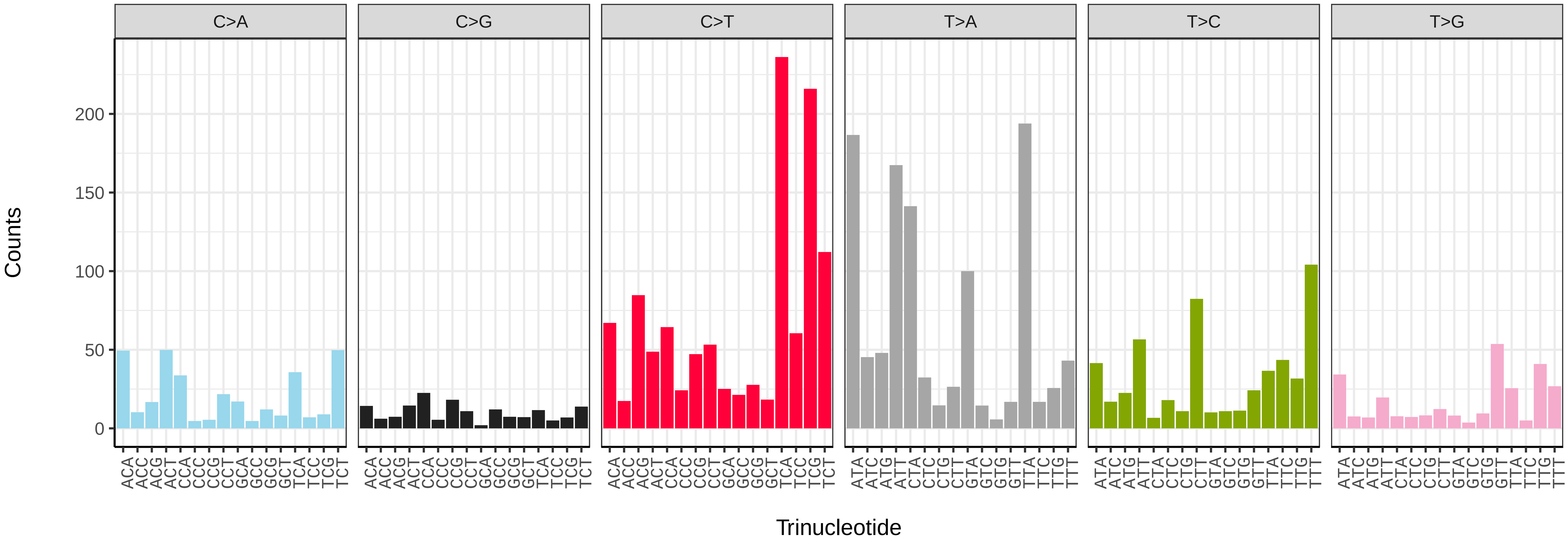


*Urtica urens* (drUrtUren1) somatic mutational spectrum (SBS52 classification)

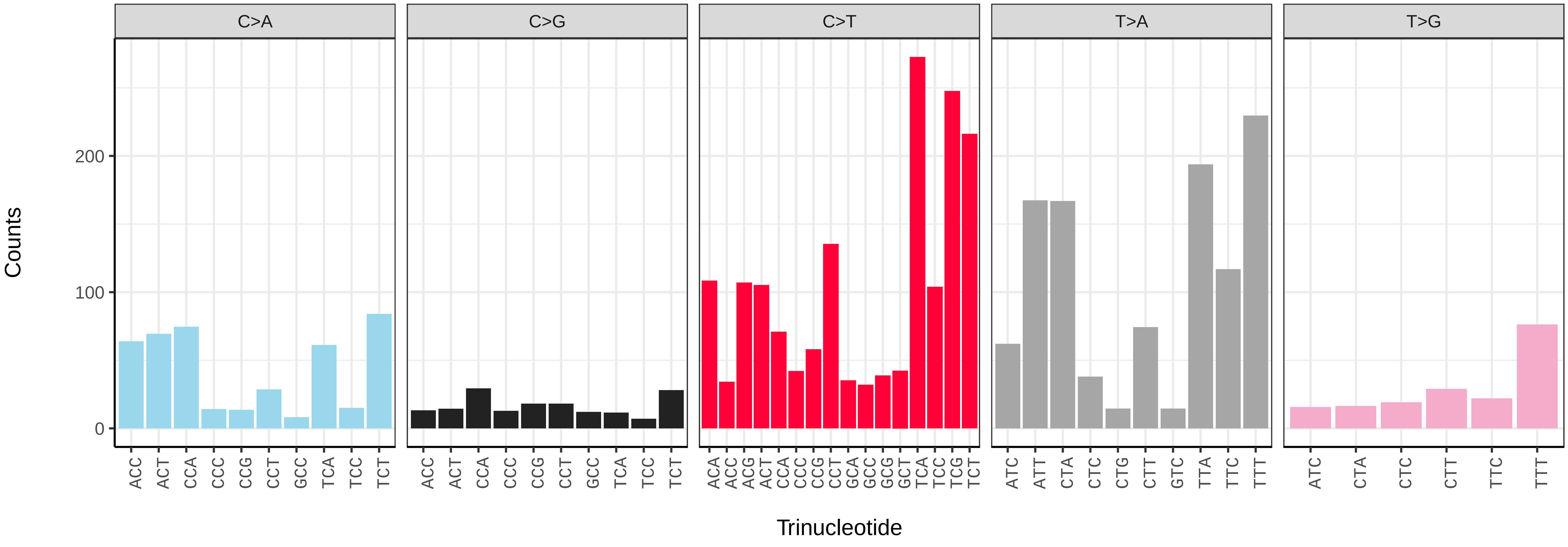

*Luidia sarsii* (eaLuiSars1) germline mutational spectrum

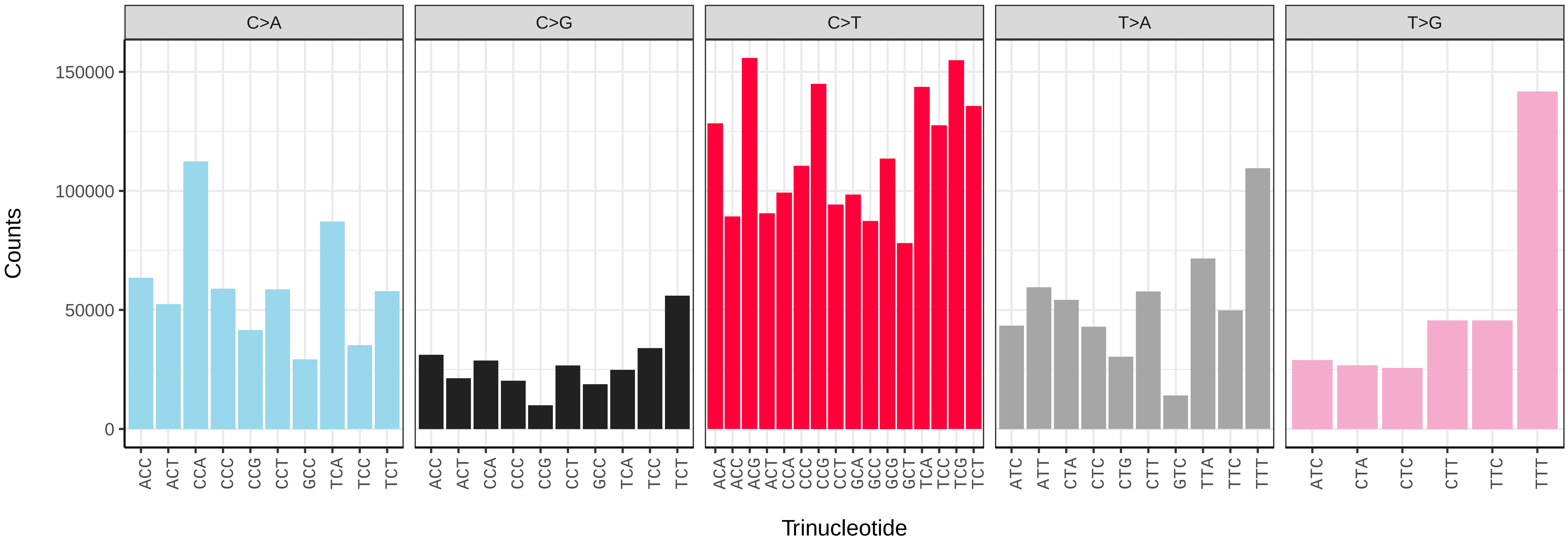


*Luidia sarsii* (eaLuiSars1) somatic mutational spectrum (SBS96 classification)

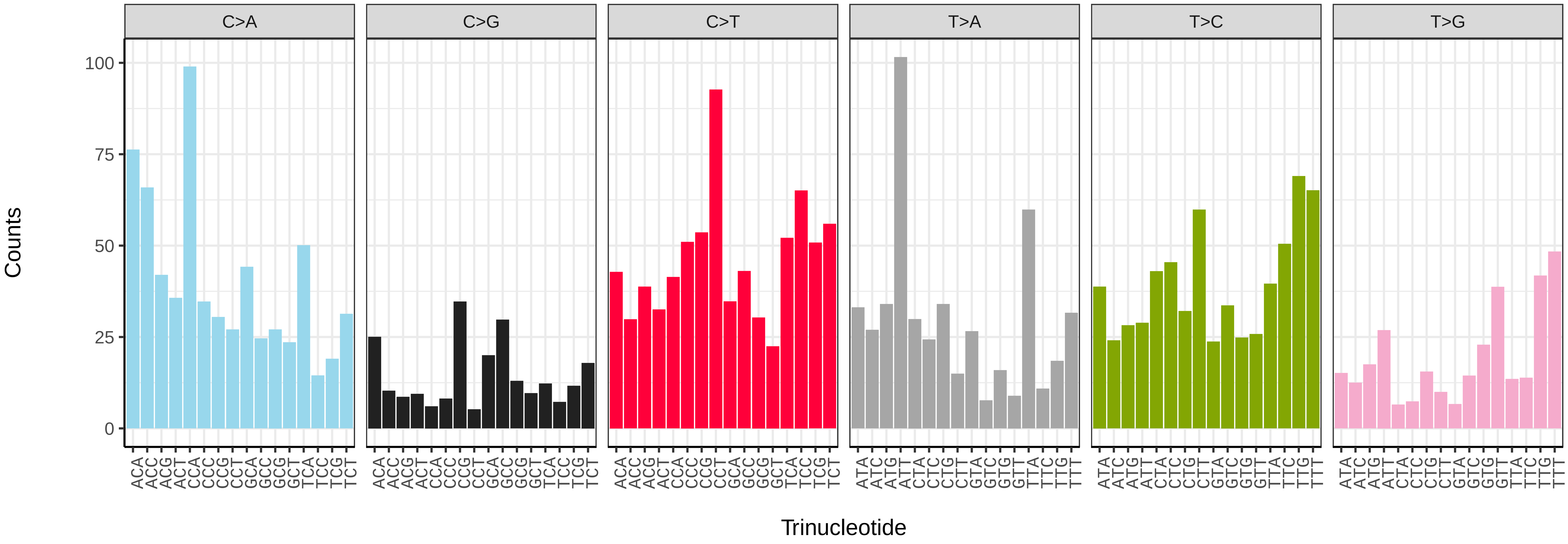


*Luidia sarsii* (eaLuiSars1) somatic mutational spectrum (SBS52 classification)

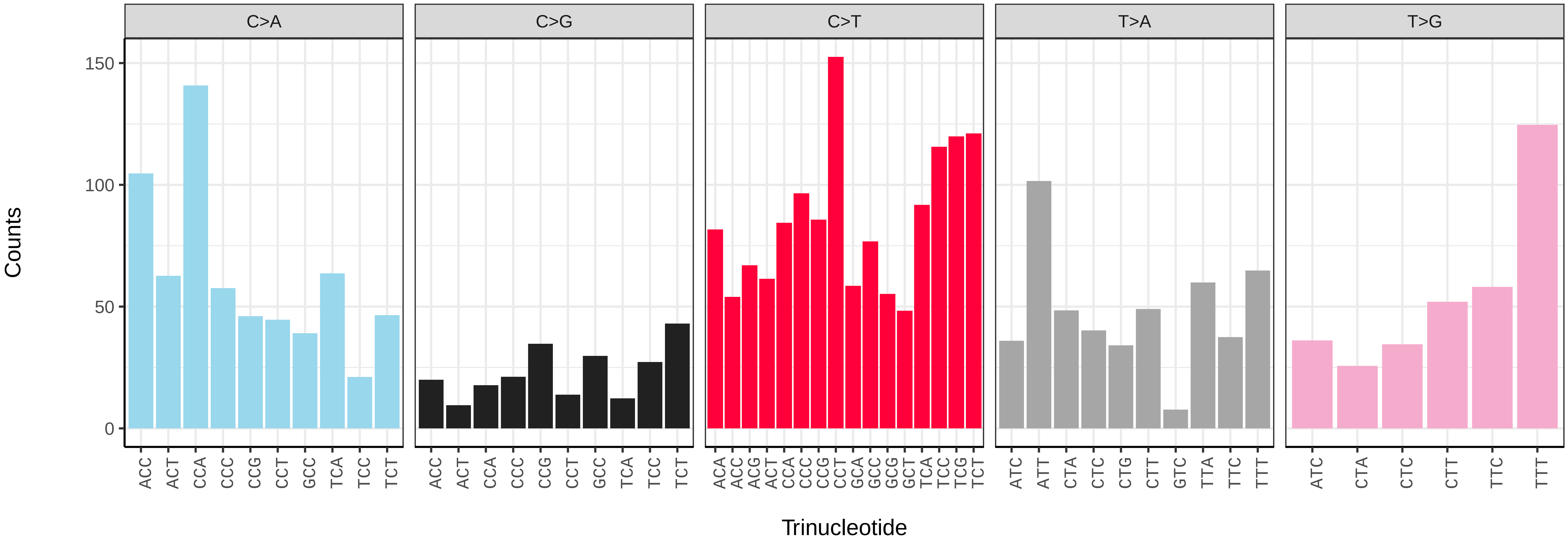

*Marthasterias glacialis* (eaMarGlac1) germline mutational spectrum

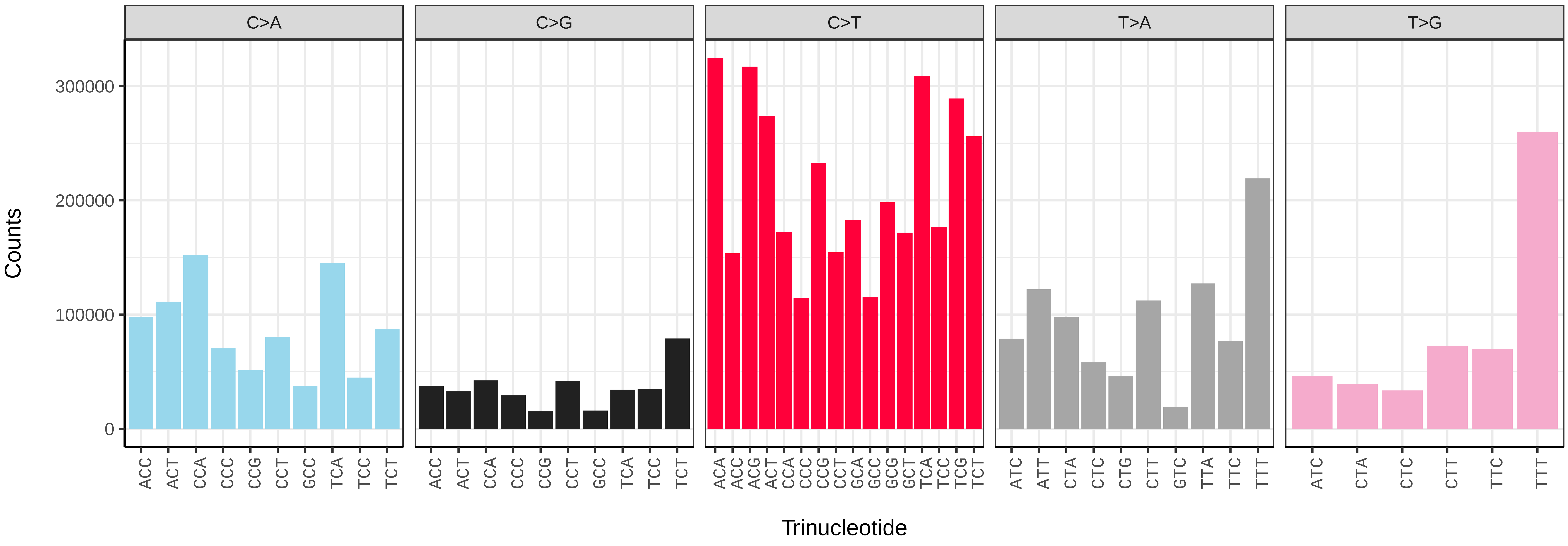


*Marthasterias glacialis* (eaMarGlac1) somatic mutational spectrum (SBS96 classification)

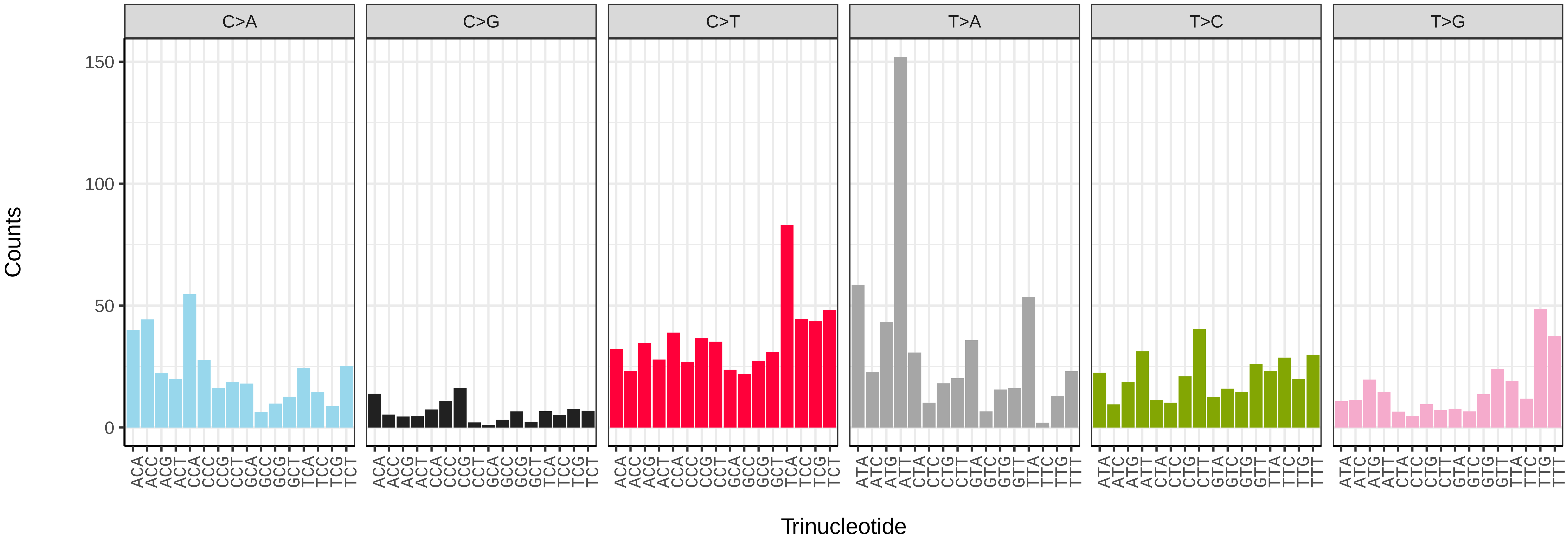


*Marthasterias glacialis* (eaMarGlac1) somatic mutational spectrum (SBS52 classification)

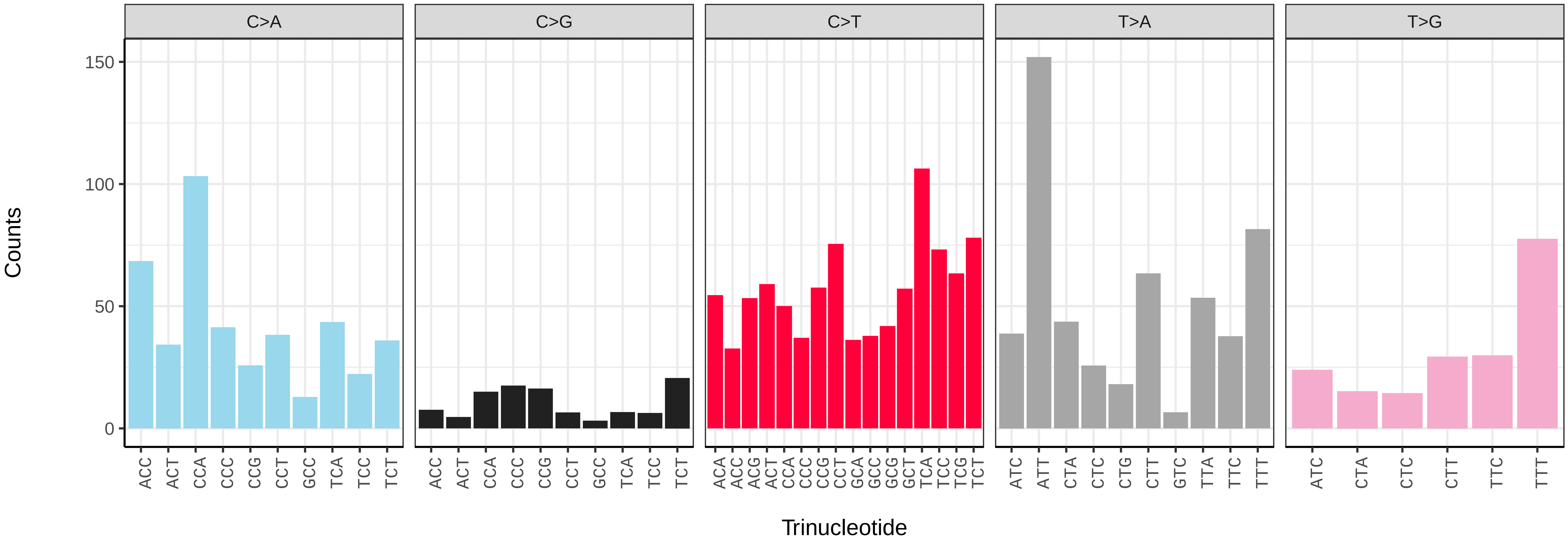

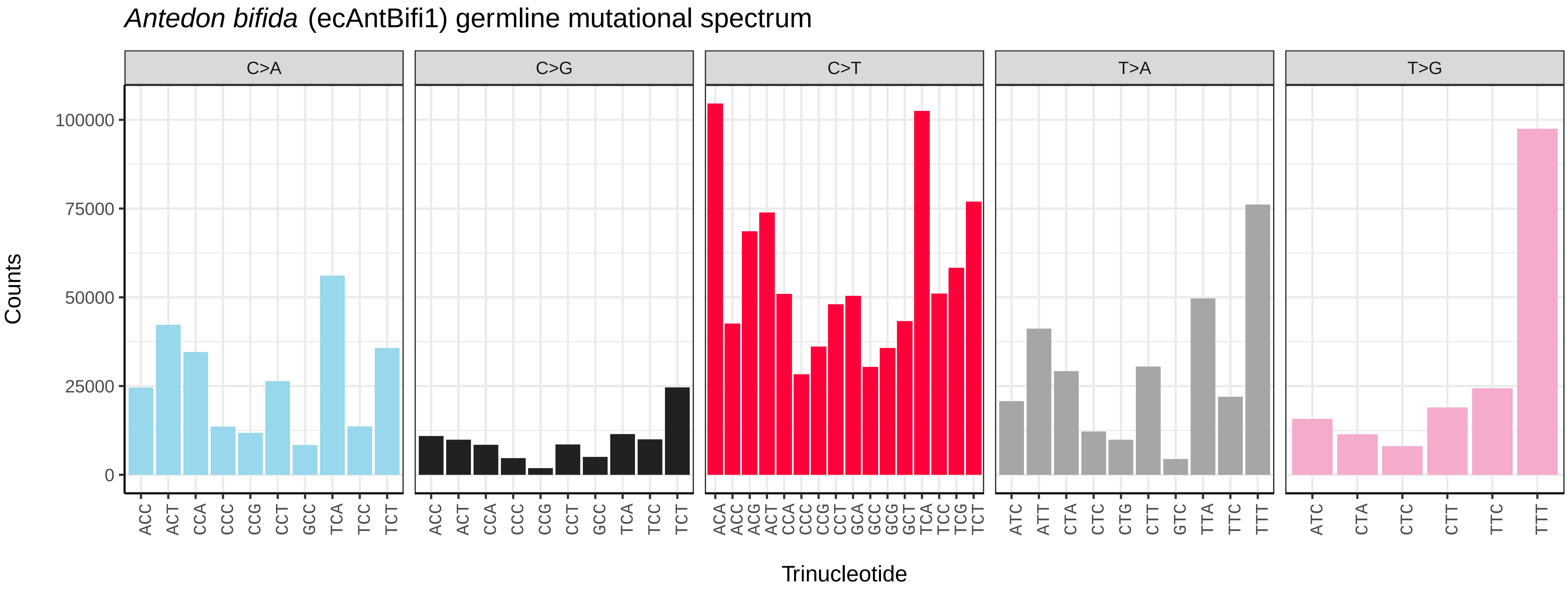
Antedon bifida (ecAntBifi1) germline mutational spectrum
C>A
C>G
C>T
T>A
T>G
Counts
0
25000
50000
75000
100000
Trinucleotide

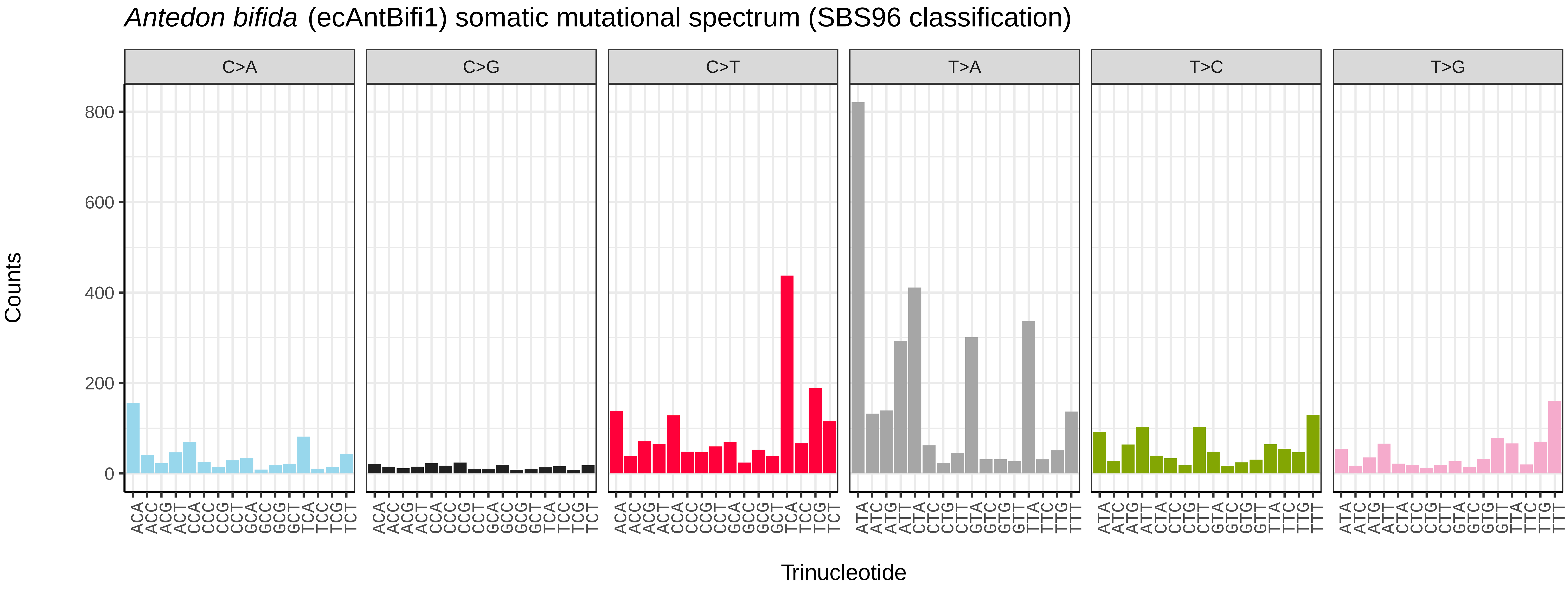
Antedon bifida (ecAntBifi1) somatic mutational spectrum (SBS96 classification)
C>A
C>G
C>T
T>A
T>C
T>G
Counts
0
200
400
600
800
Trinucleotide

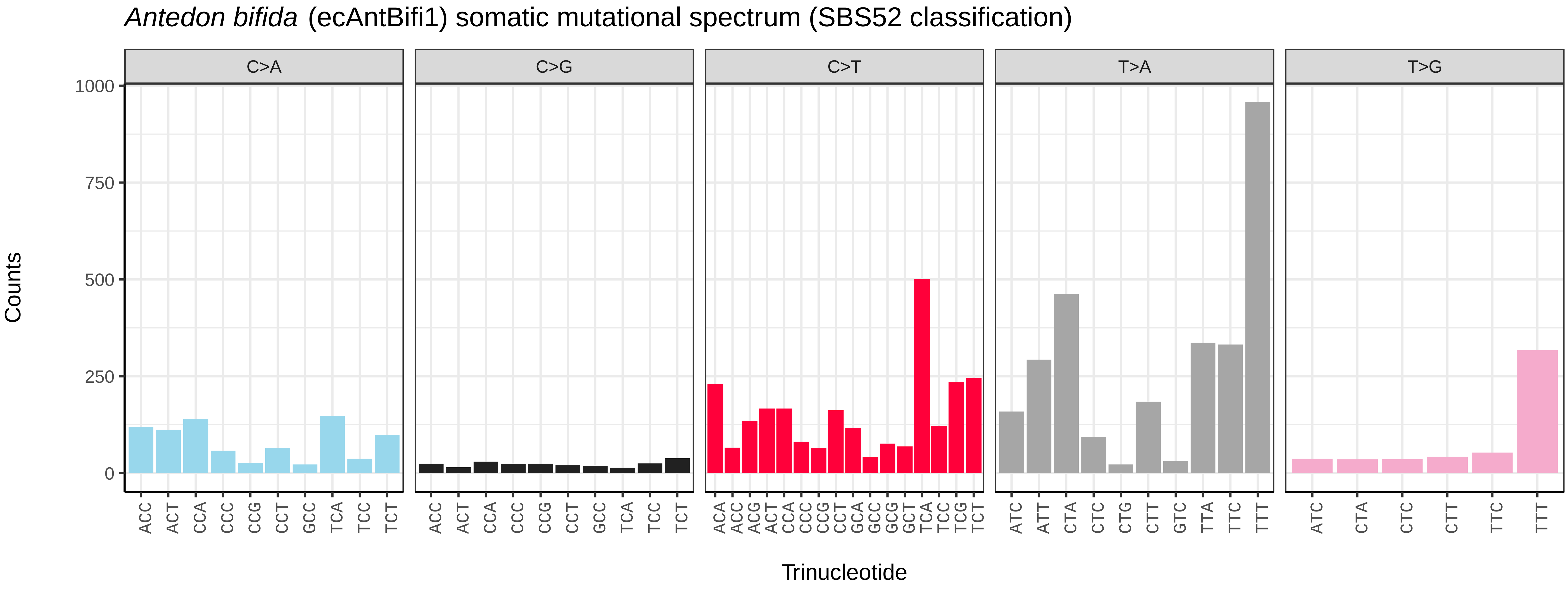
Antedon bifida (ecAntBifi1) somatic mutational spectrum (SBS52 classification)
C>A
C>G
C>T
T>A
T>G
Counts
0
250
500
750
1000
Trinucleotide

*Abramis brama* (fAbrBra2) germline mutational spectrum

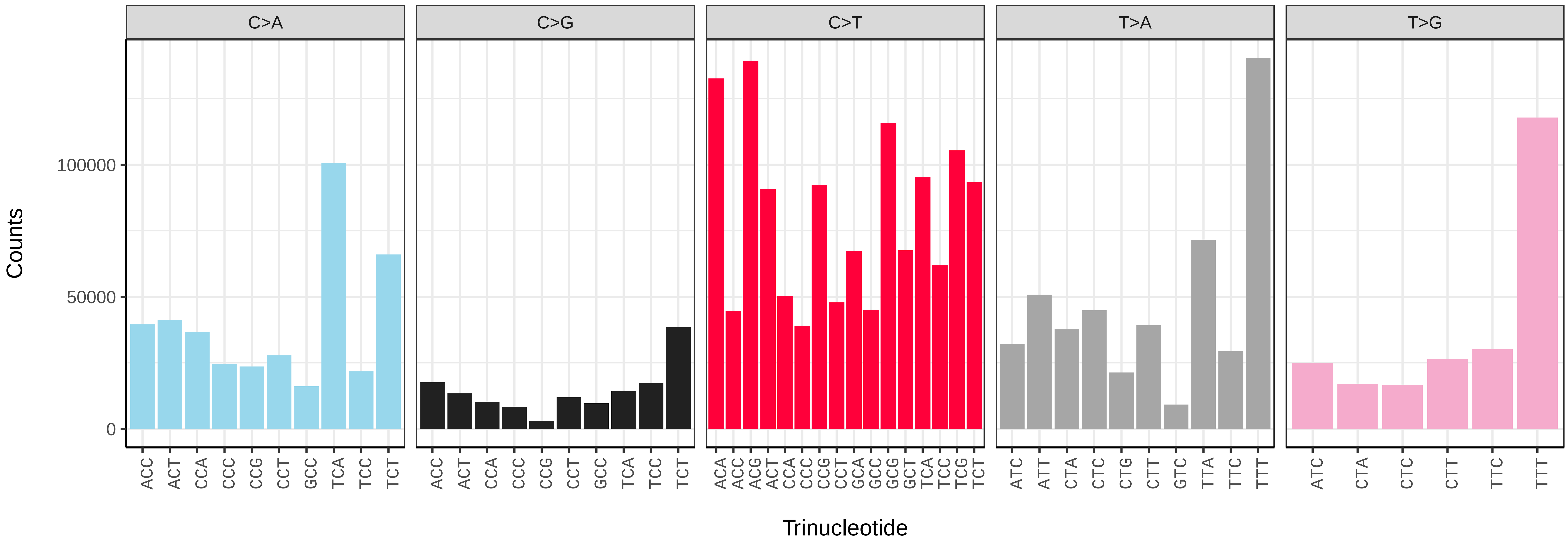


*Abramis brama* (fAbrBra2) somatic mutational spectrum (SBS96 classification)

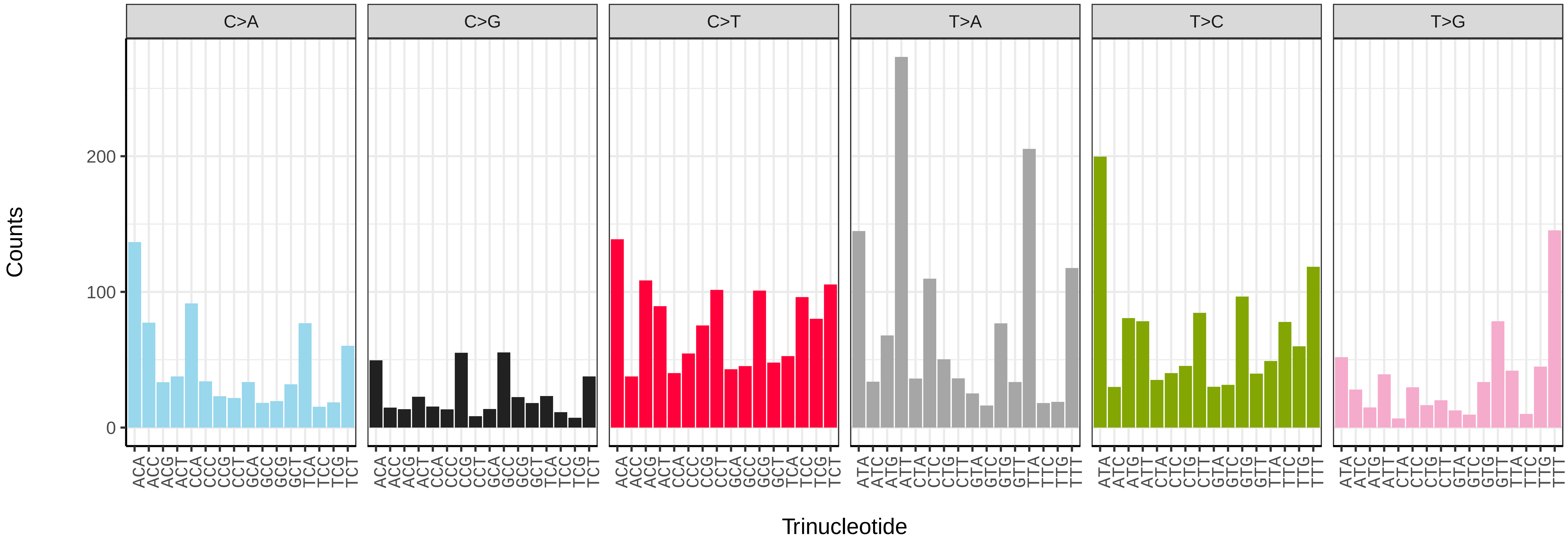


*Abramis brama* (fAbrBra2) somatic mutational spectrum (SBS52 classification)

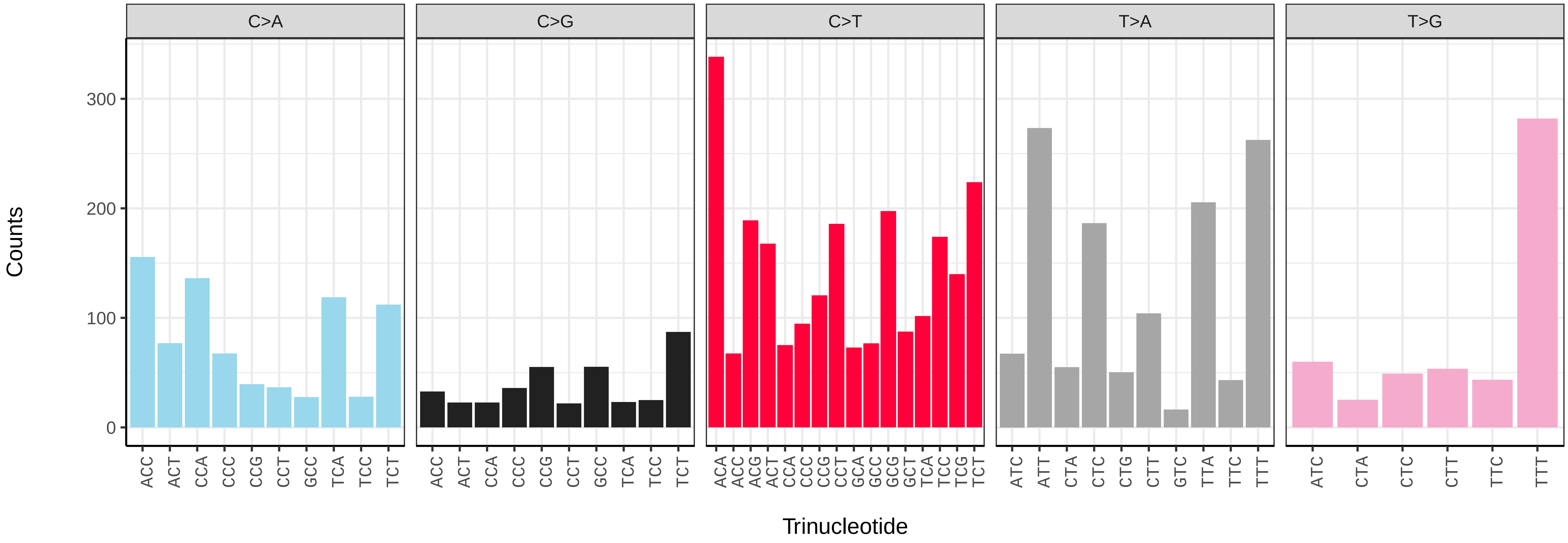

*Acipenser ruthenus* (fAciRut3) germline mutational spectrum

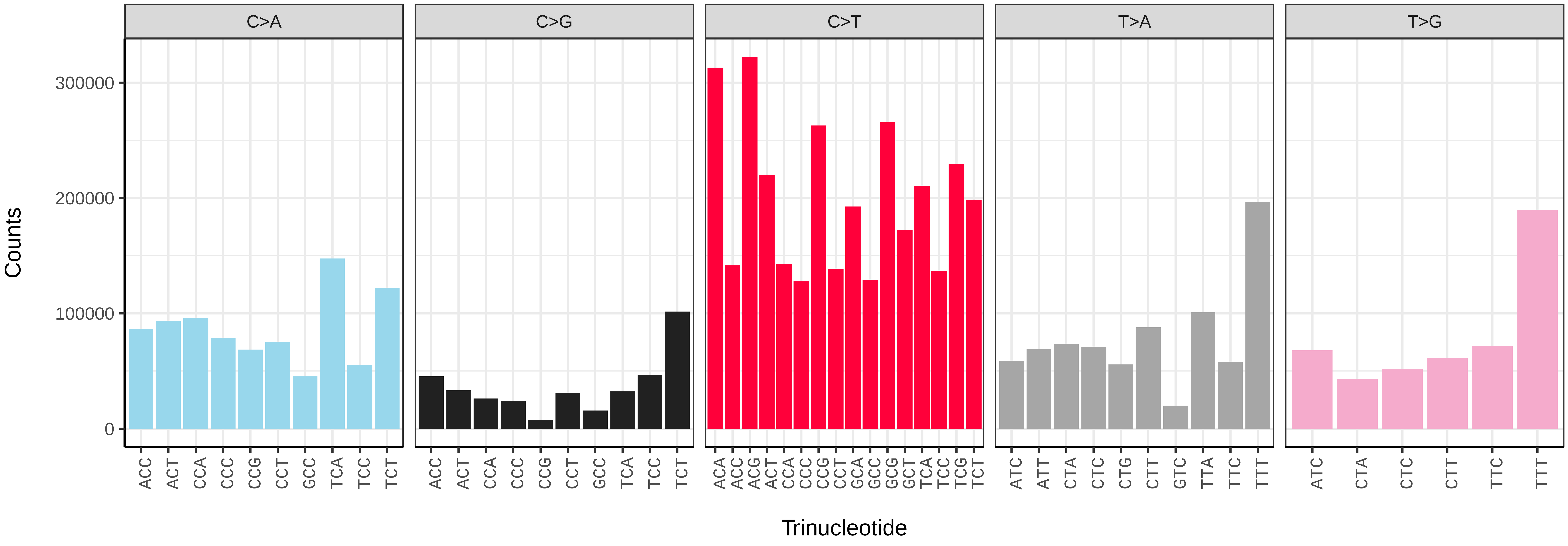


*Acipenser ruthenus* (fAciRut3) somatic mutational spectrum (SBS96 classification)

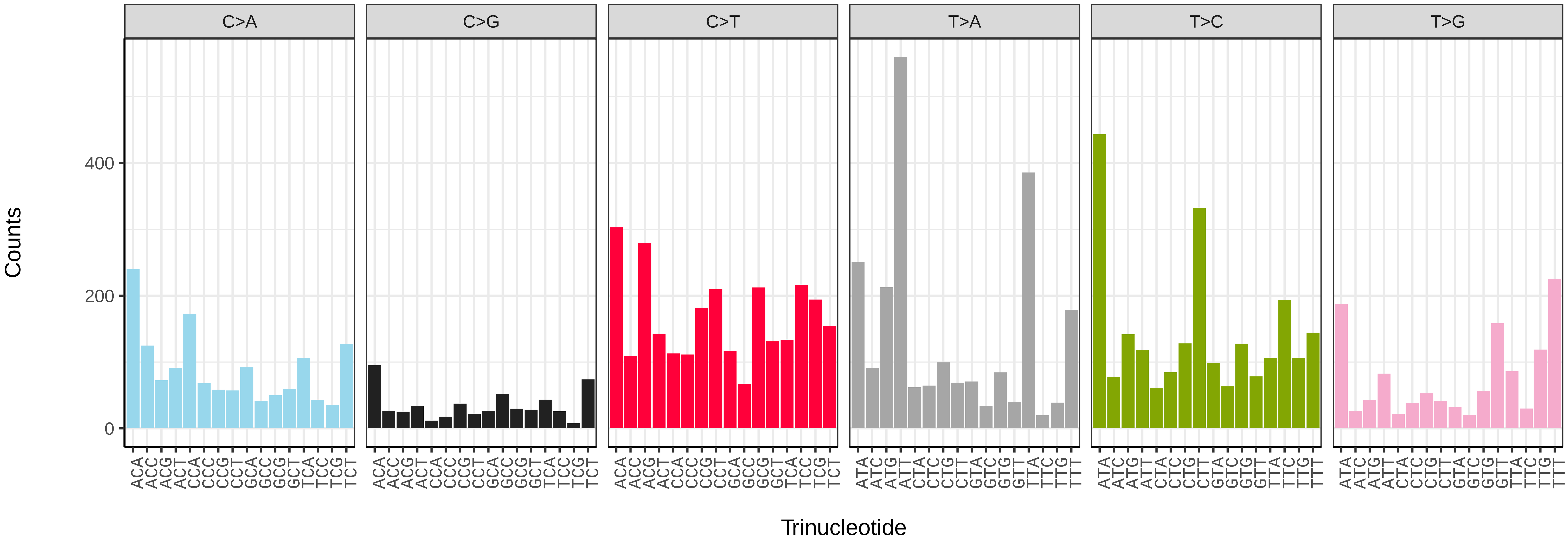


*Acipenser ruthenus* (fAciRut3) somatic mutational spectrum (SBS52 classification)

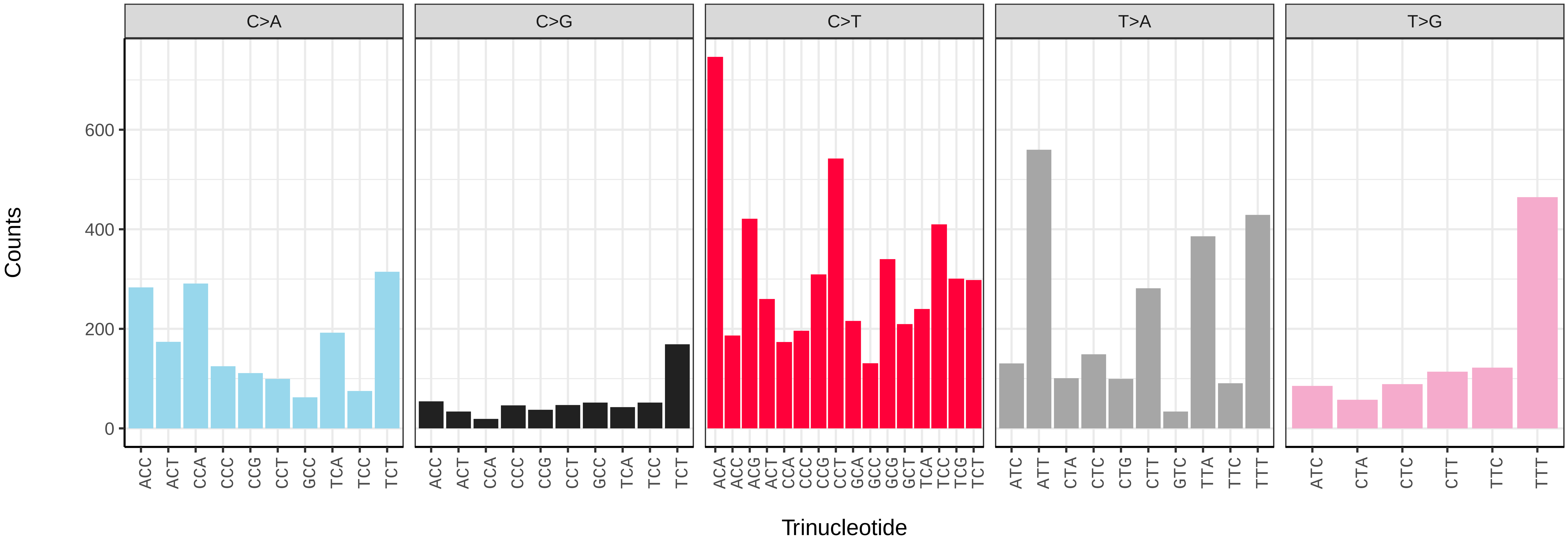

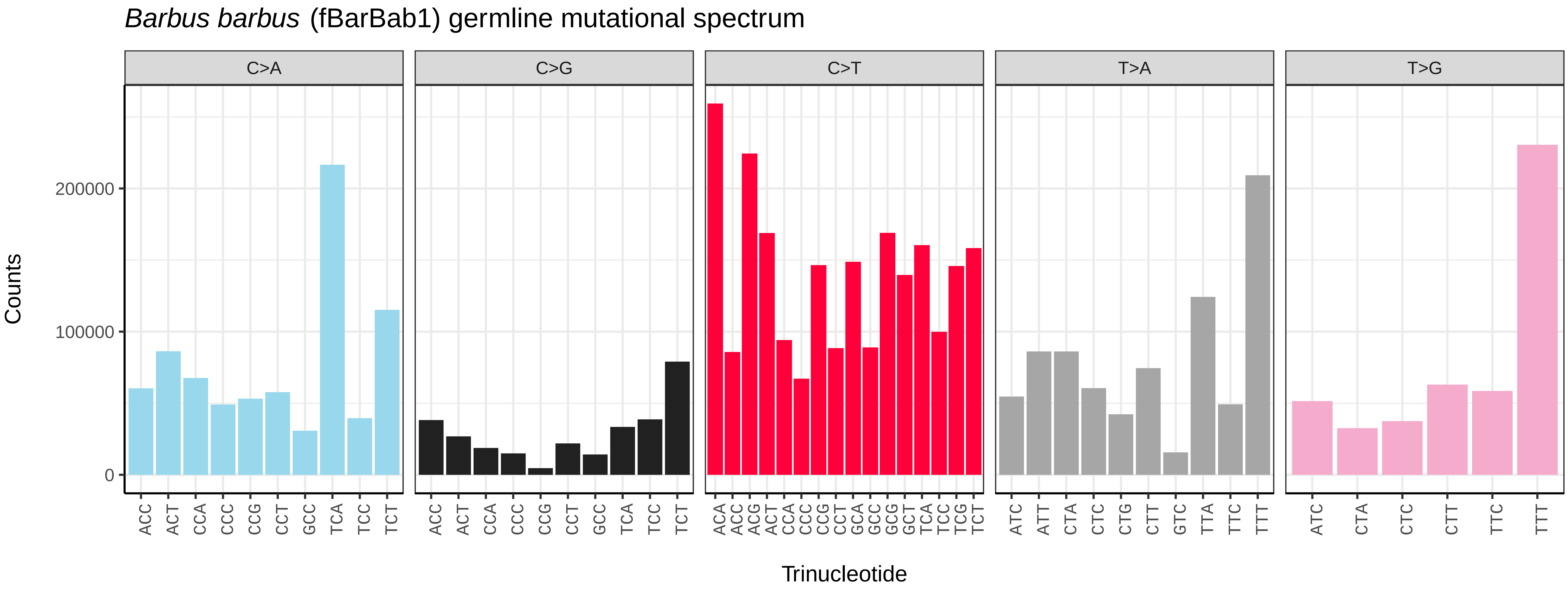
Barbus barbus (fBarBab1) germline mutational spectrum
C>A
C>G
C>T
T>A
T>G
Counts
200000
100000
0
Trinucleotide

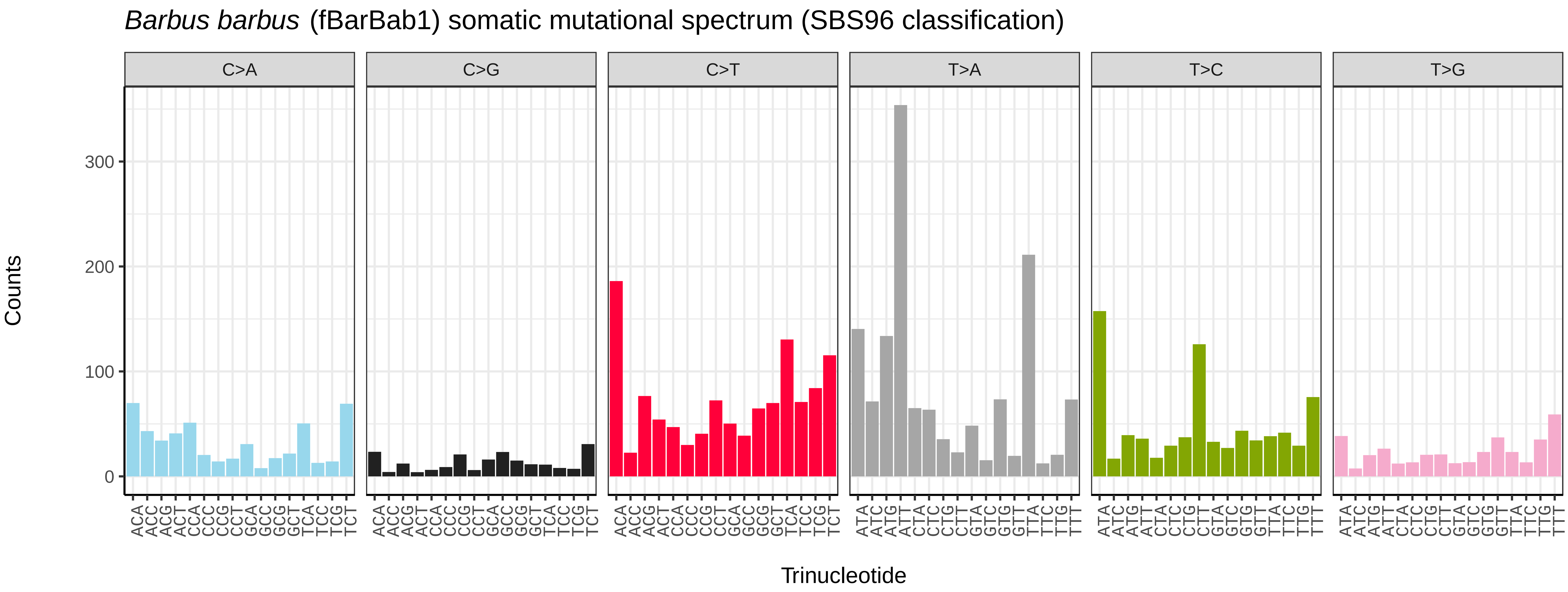
Barbus barbus (fBarBab1) somatic mutational spectrum (SBS96 classification)
C>A
C>G
C>T
T>A
T>C
T>G
Counts
300
200
100
0
Trinucleotide

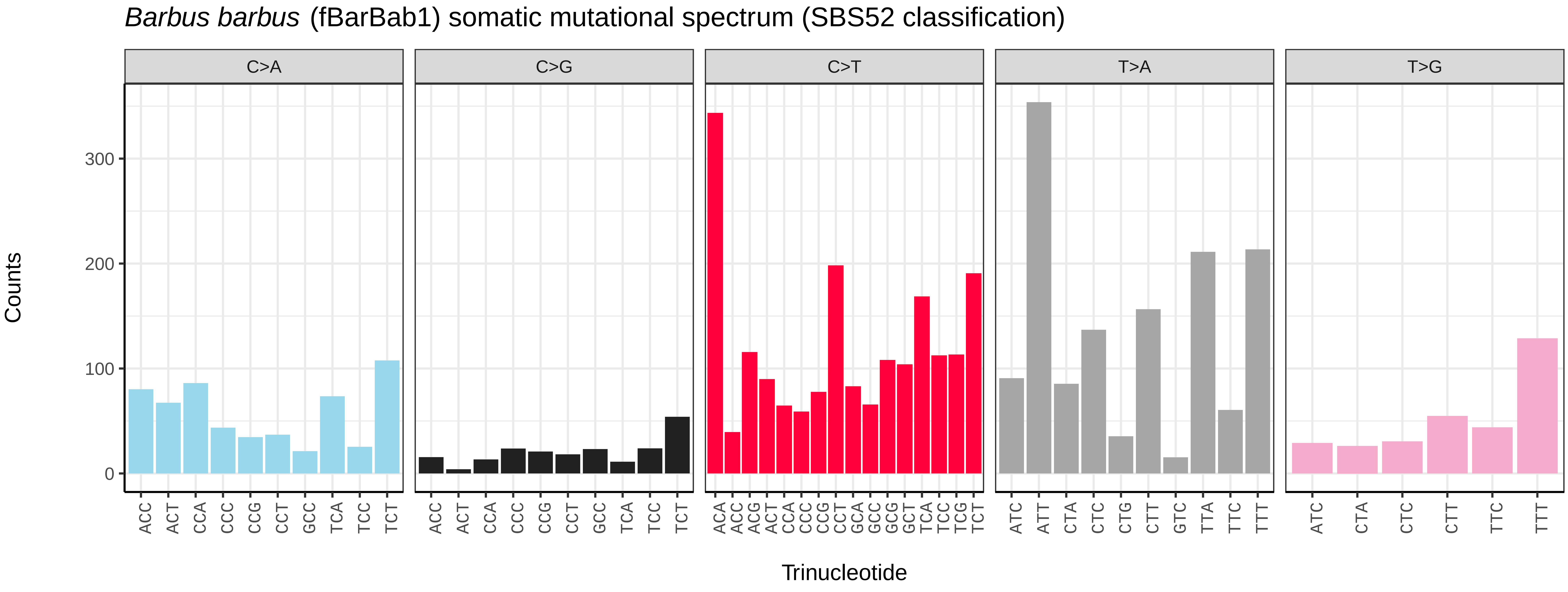
Barbus barbus (fBarBab1) somatic mutational spectrum (SBS52 classification)
C>A
C>G
C>T
T>A
T>G
Counts
300
200
100
0
Trinucleotide

*Barbatula barbatula* (fBarBar1) germline mutational spectrum

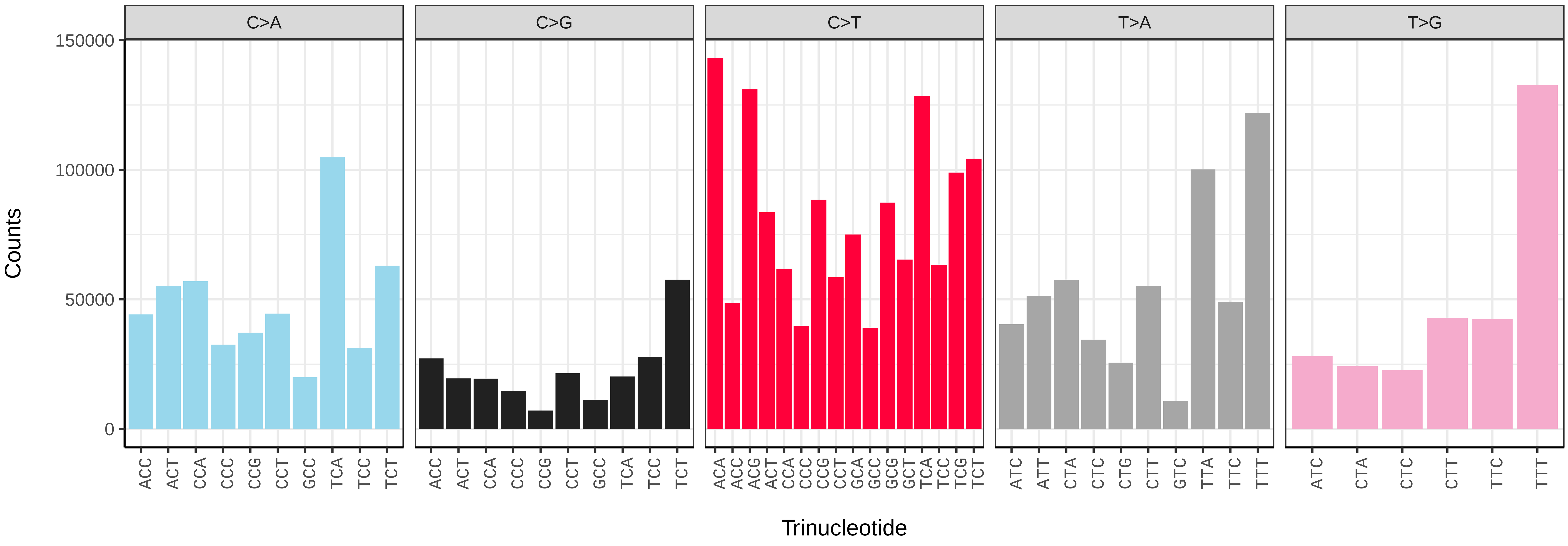


*Barbatula barbatula* (fBarBar1) somatic mutational spectrum (SBS96 classification)

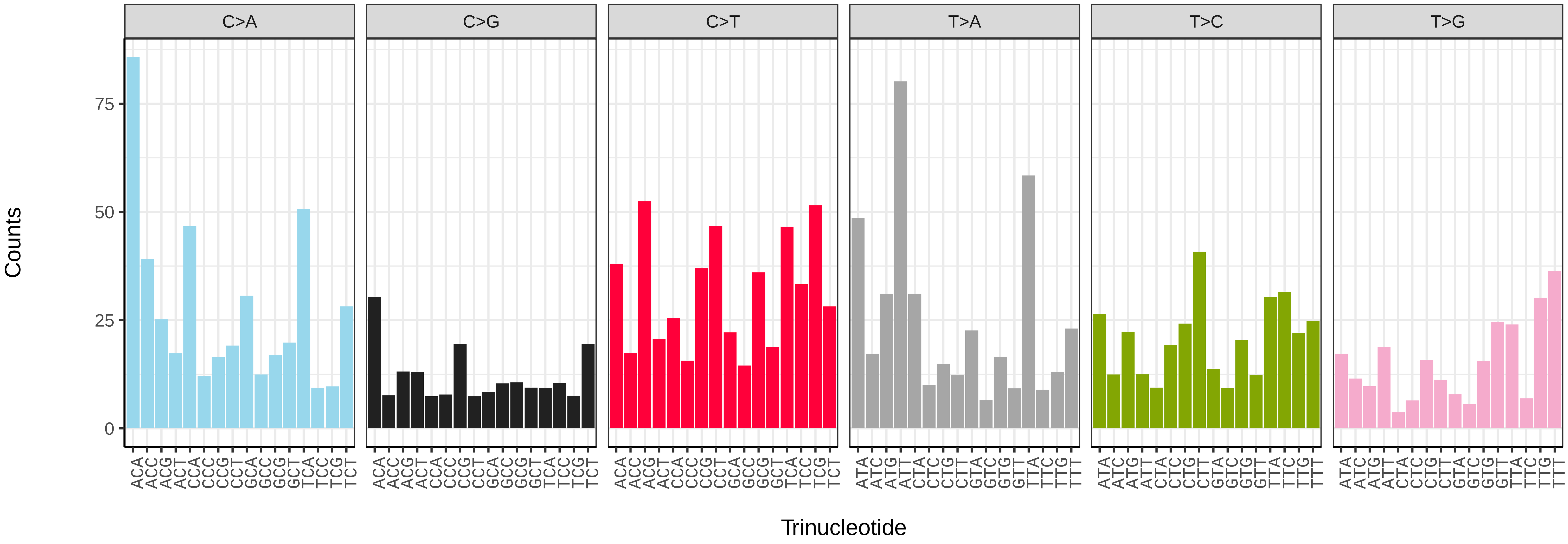


*Barbatula barbatula* (fBarBar1) somatic mutational spectrum (SBS52 classification)

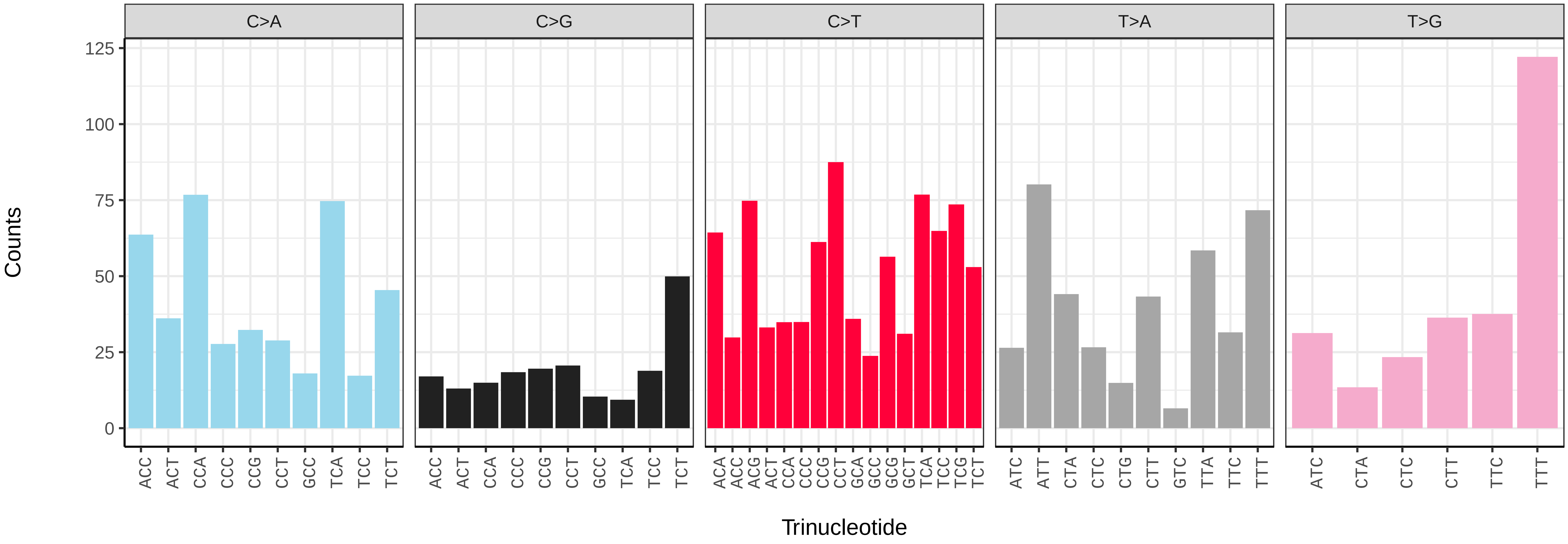

*Blennius ocellaris* (fBleOce1) germline mutational spectrum

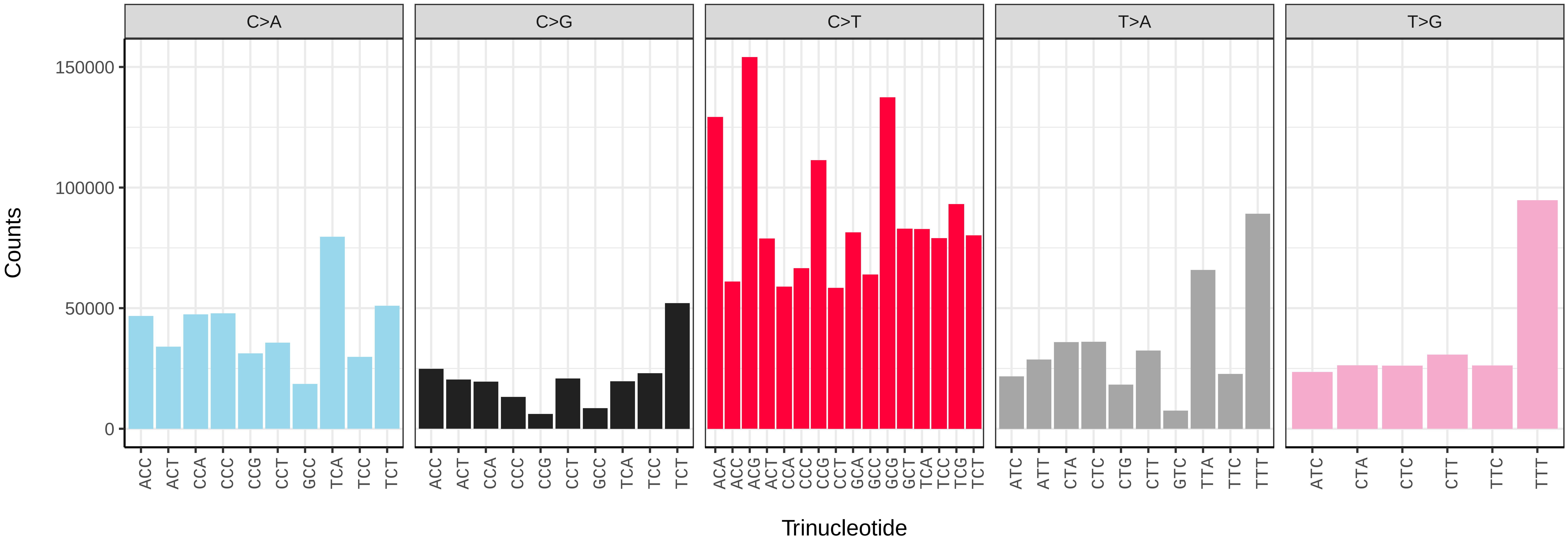


*Blennius ocellaris* (fBleOce1) somatic mutational spectrum (SBS96 classification)

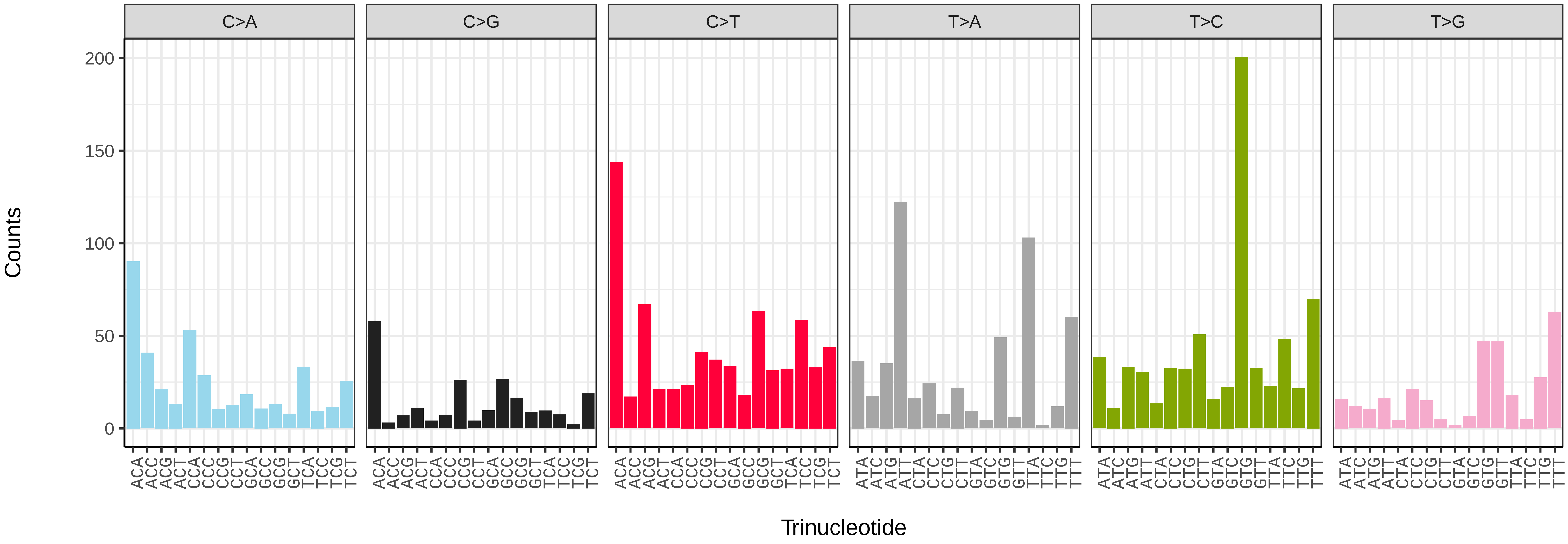


*Blennius ocellaris* (fBleOce1) somatic mutational spectrum (SBS52 classification)

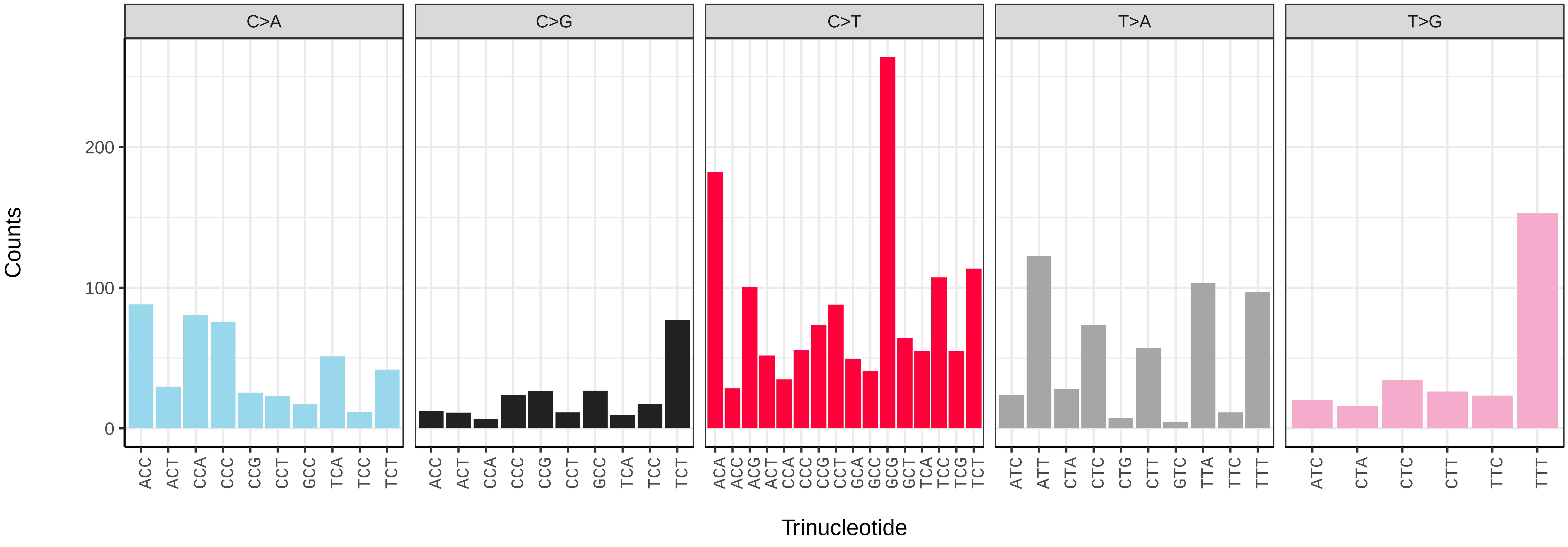

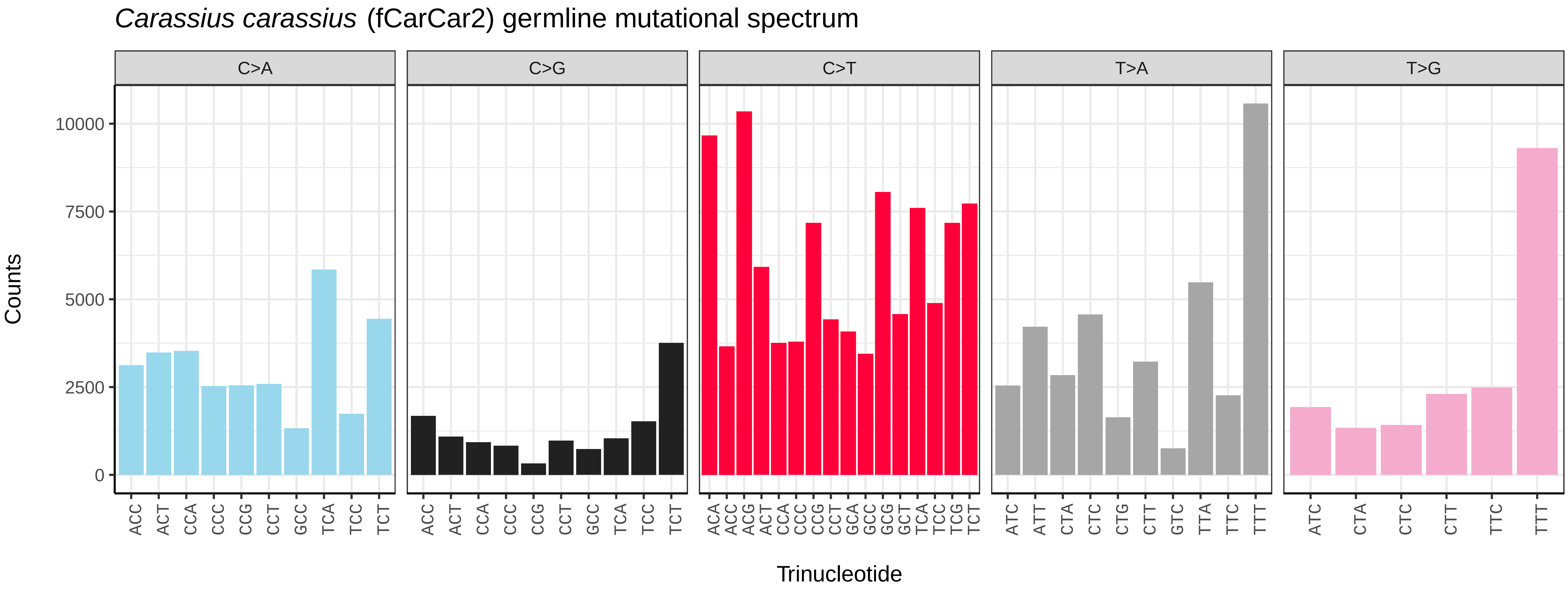
*Carassius carassius* (fCarCar2) germline mutational spectrum
C>A
C>G
C>T
T>A
T>G
Counts
0
2500
5000
7500
10000
ACC ACT CCA CCC CCG CCT GCC TCA TCC TCT
ACA ACC ACG ACT CCA CCC CCG CCT GCA GCC GCG GCT TCA TCC TCG TCT
ATC ATT CTA CTC CTG CTT GTC TTA TTC TTT
ATC CTA CTC CTT TTC TTT
Trinucleotide

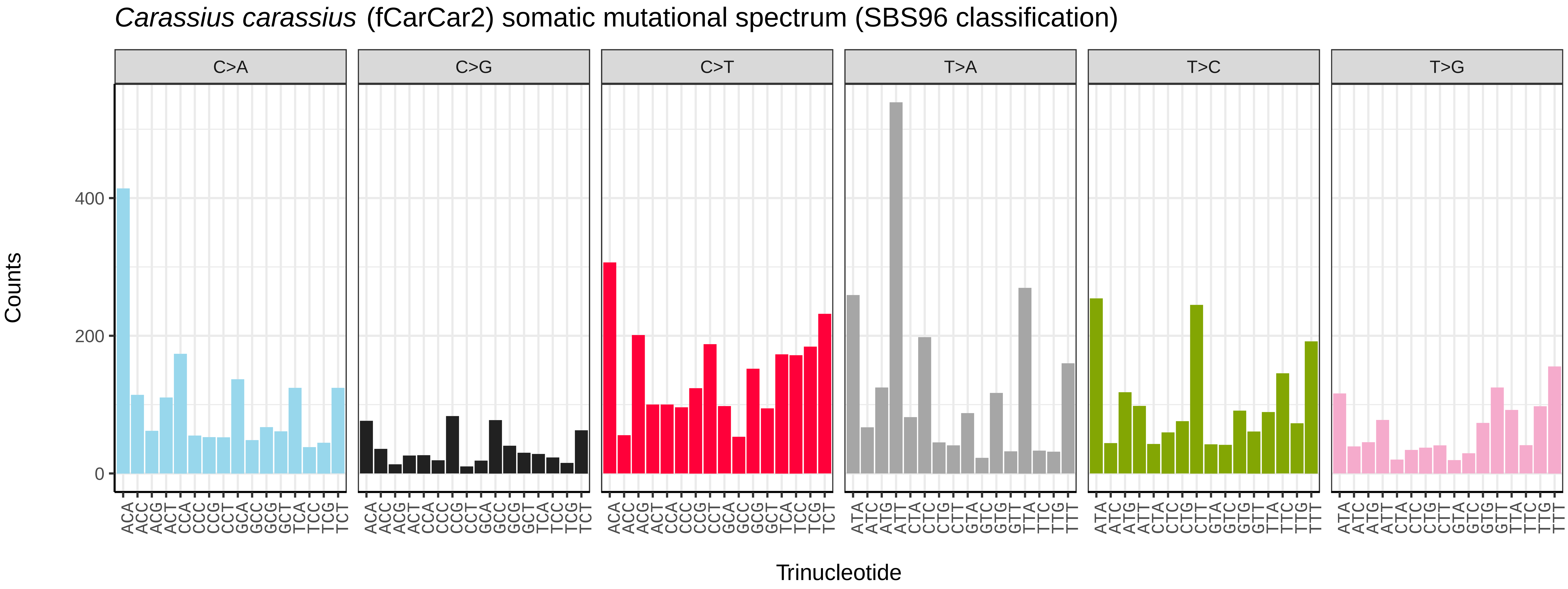
*Carassius carassius* (fCarCar2) somatic mutational spectrum (SBS96 classification)
C>A
C>G
C>T
T>A
T>C
T>G
Counts
0
200
400
ACA ACC ACG ACT CCA CCC CCG CCT GCA GCC GCG GCT TCA TCC TCG TCT
ATA ATC ATG ATT CTA CTC CTG CTT GTA GTC GTG GTT TTA TTC TTG TTT
Trinucleotide

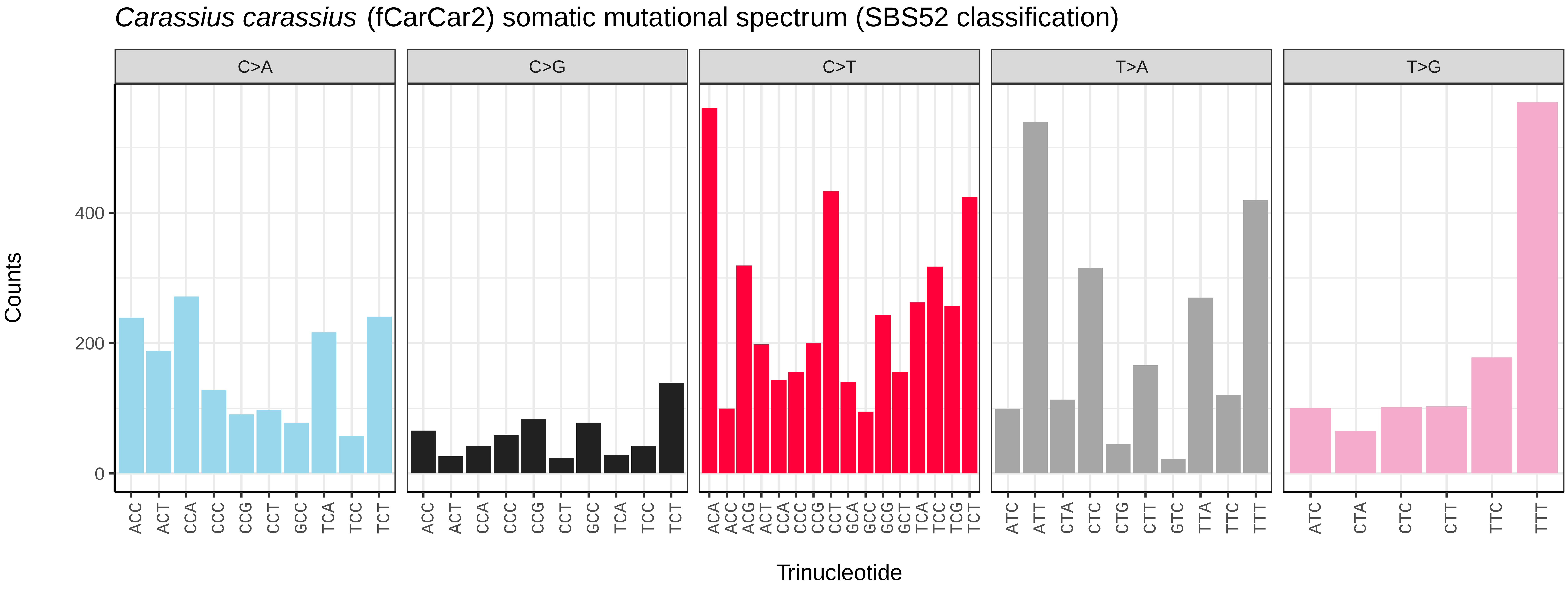
*Carassius carassius* (fCarCar2) somatic mutational spectrum (SBS52 classification)
C>A
C>G
C>T
T>A
T>G
Counts
0
200
400
ACC ACT CCA CCC CCG CCT GCC TCA TCC TCT
ACA ACC ACG ACT CCA CCC CCG CCT GCA GCC GCG GCT TCA TCC TCG TCT
ATC ATT CTA CTC CTG CTT GTC TTA TTC TTT
ATC CTA CTC CTT TTC TTT
Trinucleotide

*Chelon labrosus* (fCheLab1) germline mutational spectrum

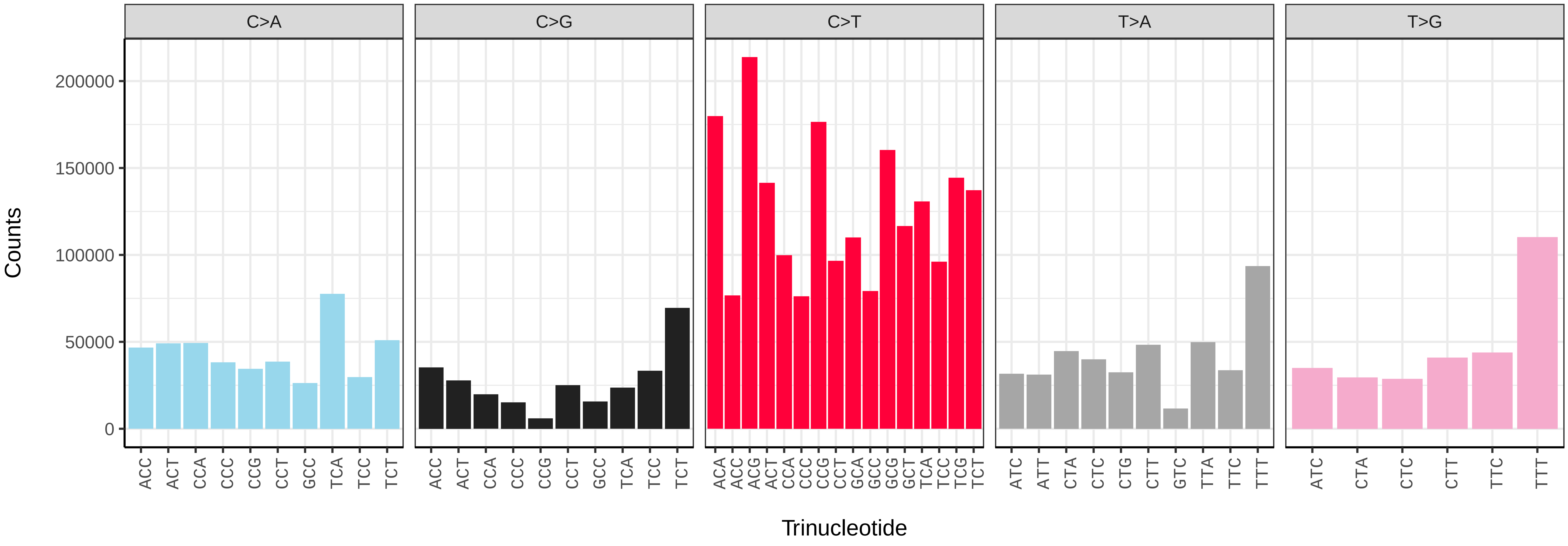


*Chelon labrosus* (fCheLab1) somatic mutational spectrum (SBS96 classification)

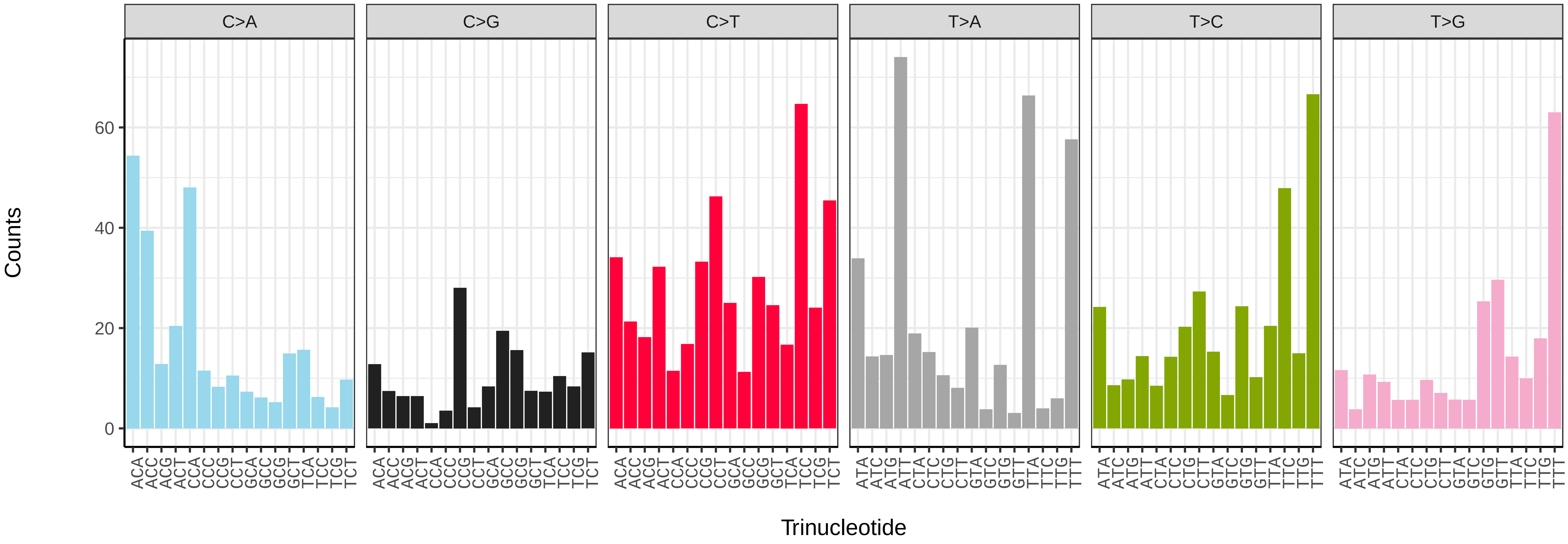


*Chelon labrosus* (fCheLab1) somatic mutational spectrum (SBS52 classification)

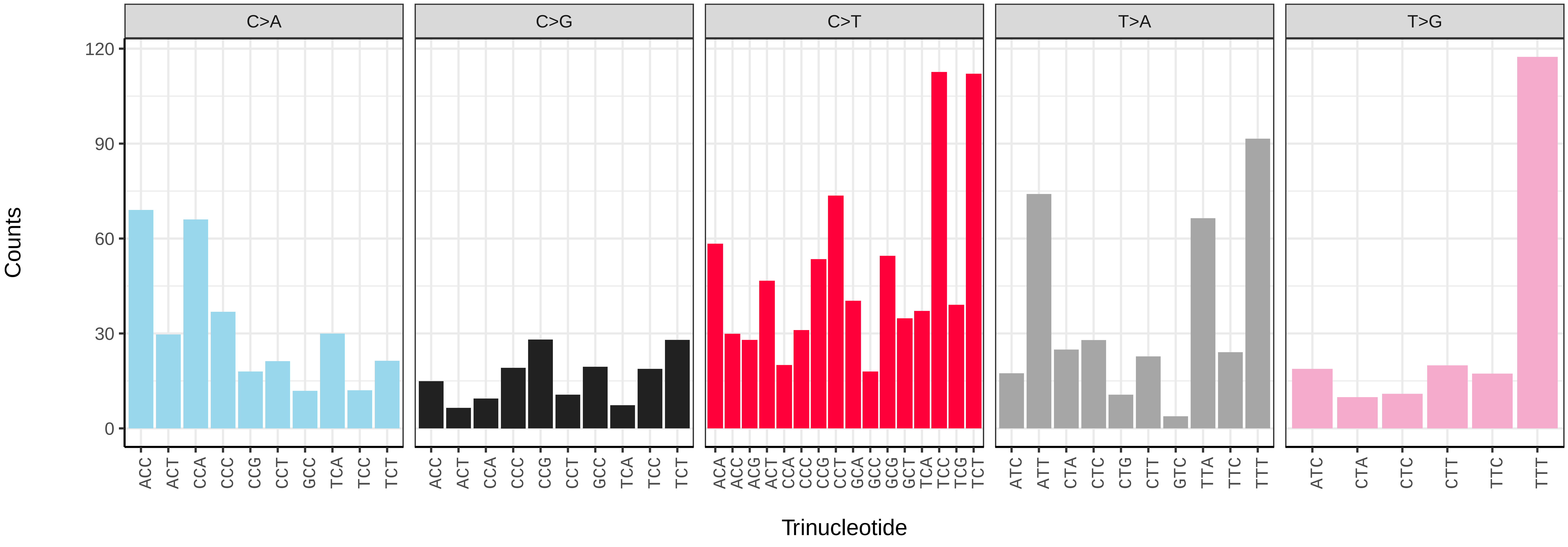

*Conger conger* (fConCon1) germline mutational spectrum

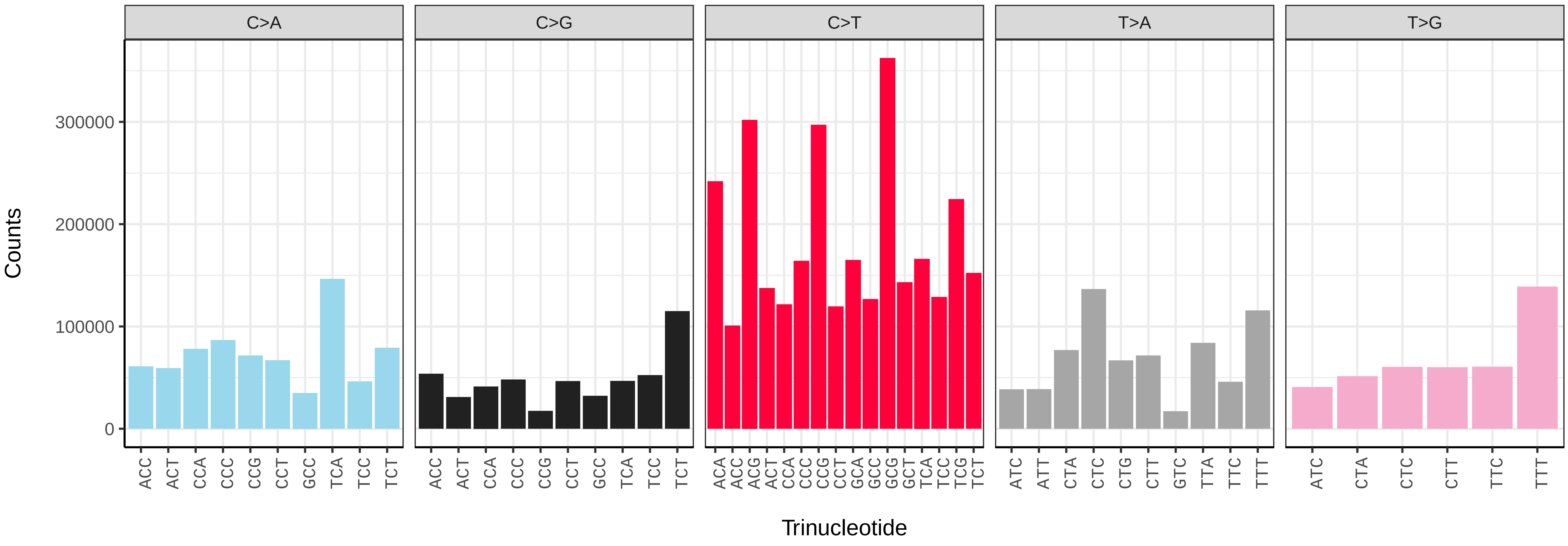


*Conger conger* (fConCon1) somatic mutational spectrum (SBS96 classification)

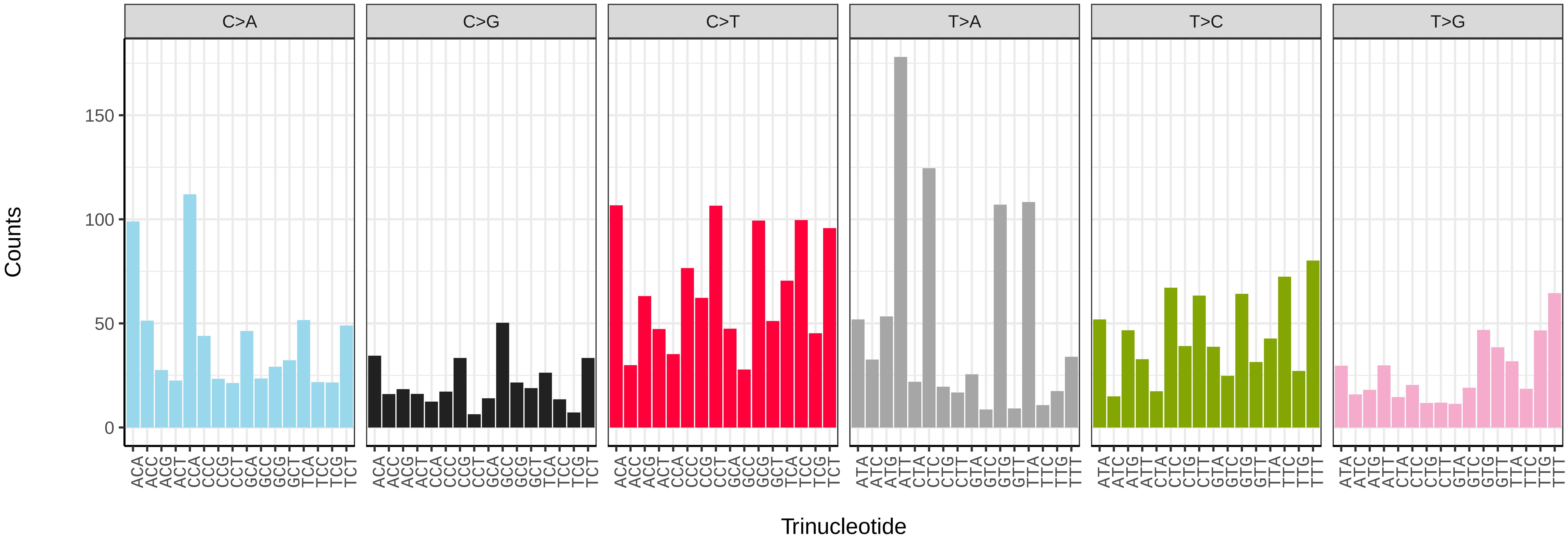


*Conger conger* (fConCon1) somatic mutational spectrum (SBS52 classification)

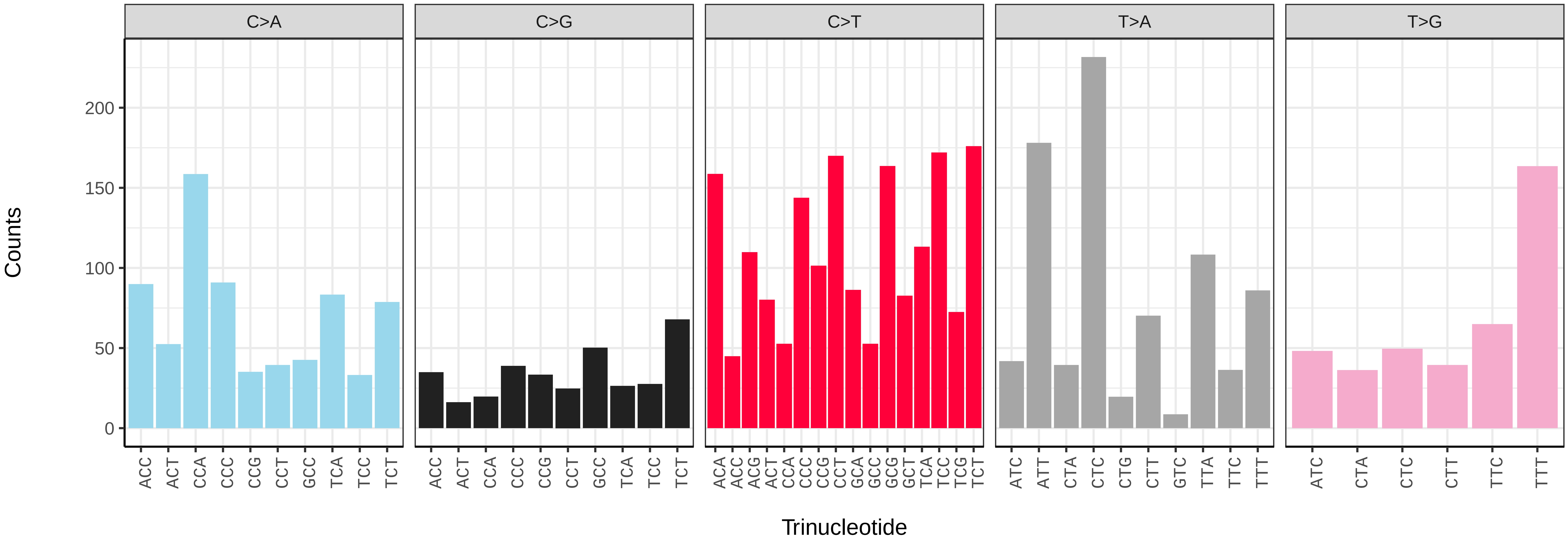

*Cyclopterus lumpus* (fCycLum2) germline mutational spectrum

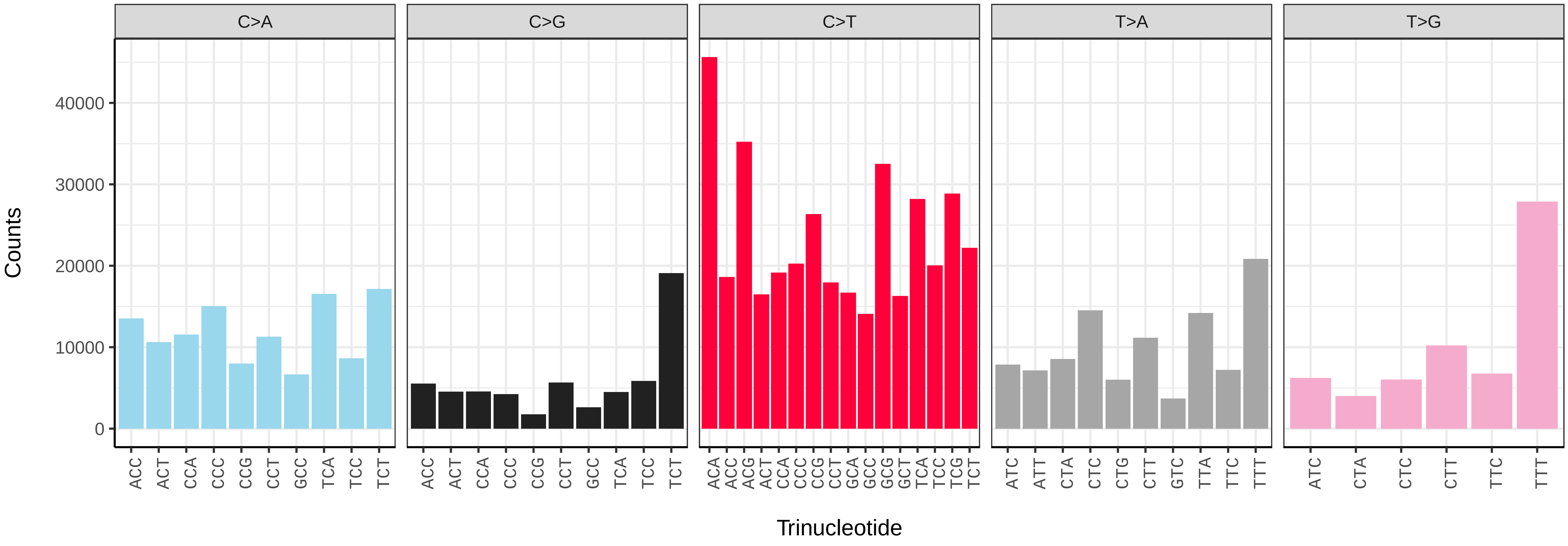


*Cyclopterus lumpus* (fCycLum2) somatic mutational spectrum (SBS96 classification)

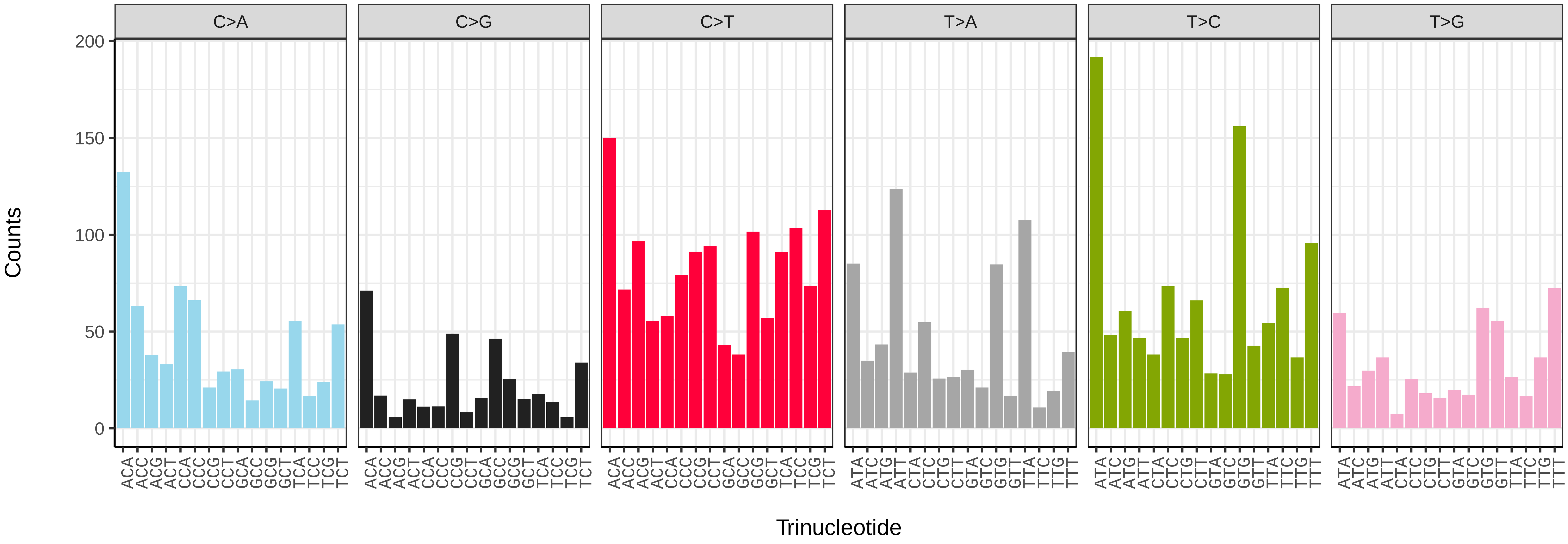


*Cyclopterus lumpus* (fCycLum2) somatic mutational spectrum (SBS52 classification)

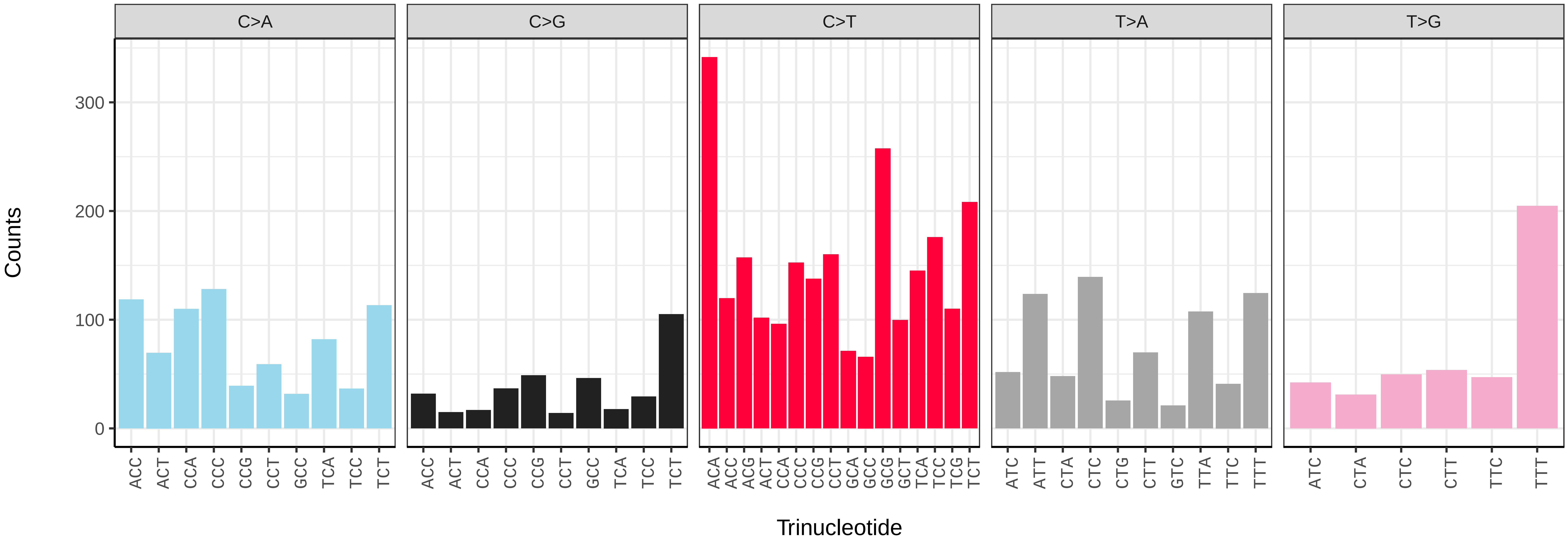

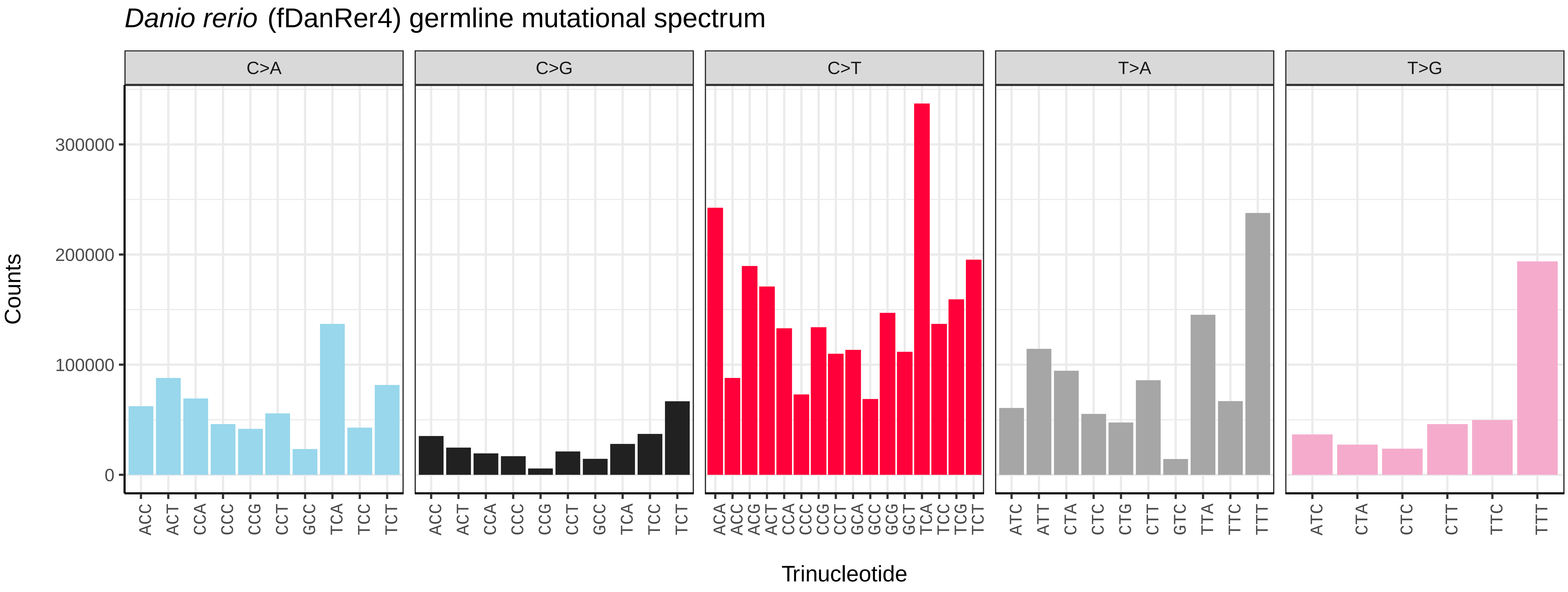
*Danio rerio* (fDanRer4) germline mutational spectrum
C>A
C>G
C>T
T>A
T>G
Counts
300000
200000
100000
0
ACC ACT CCA CCC CCG CCT GCC TCA TCC TCT
ACA ACC ACG ACT CCA CCC CCG CCT GCA GCC GCG GCT TCA TCC TCG TCT
ATC ATT CTA CTC CTG CTT GTC TTA TTC TTT
ATC CTA CTC CTT TTC TTT
Trinucleotide

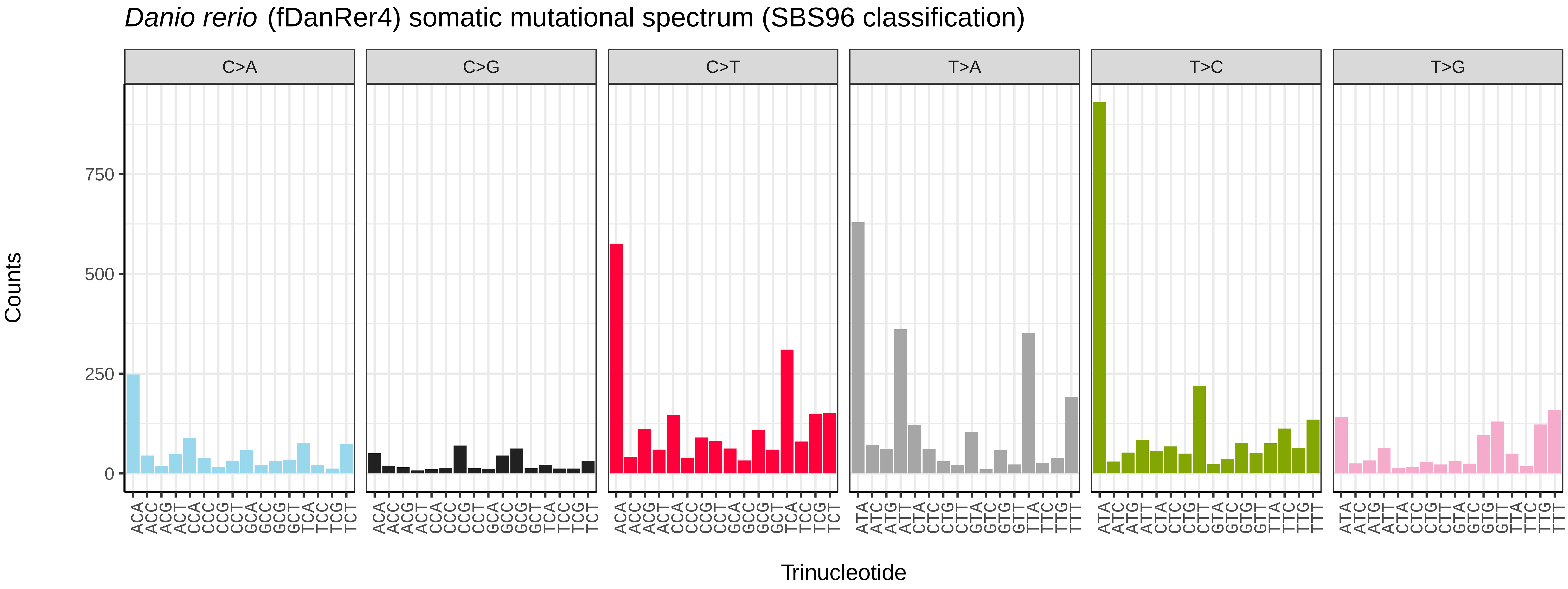
*Danio rerio* (fDanRer4) somatic mutational spectrum (SBS96 classification)
C>A
C>G
C>T
T>A
T>C
T>G
Counts
750
500
250
0
ACA ACC ACG ACT CCA CCC CCG CCT GCA GCC GCG GCT TCA TCC TCG TCT
ATA ATC ATG ATT CTA CTC CTG CTT GTA GTC GTG GTT TTA TTC TTG TTT
Trinucleotide

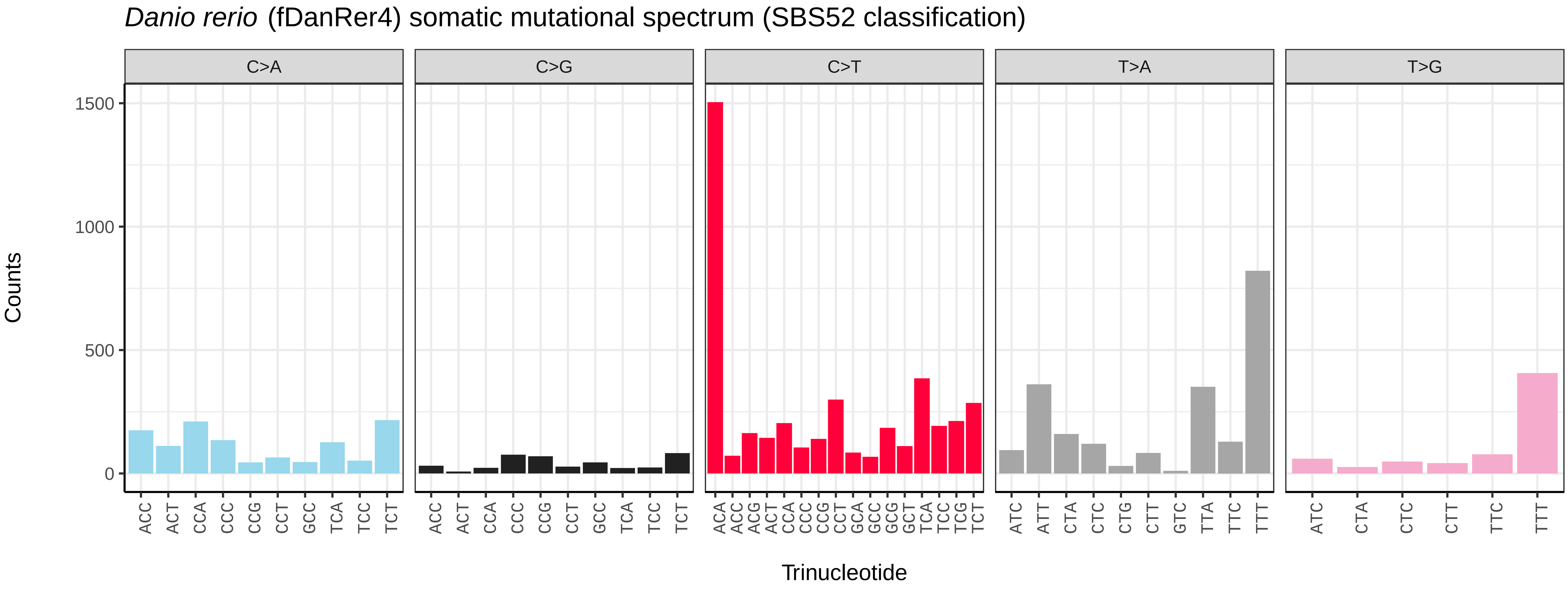
*Danio rerio* (fDanRer4) somatic mutational spectrum (SBS52 classification)
C>A
C>G
C>T
T>A
T>G
Counts
1500
1000
500
0
ACC ACT CCA CCC CCG CCT GCC TCA TCC TCT
ACA ACC ACG ACT CCA CCC CCG CCT GCA GCC GCG GCT TCA TCC TCG TCT
ATC ATT CTA CTC CTG CTT GTC TTA TTC TTT
ATC CTA CTC CTT TTC TTT
Trinucleotide

*Echiichthys vipera* (fEchVip8) germline mutational spectrum

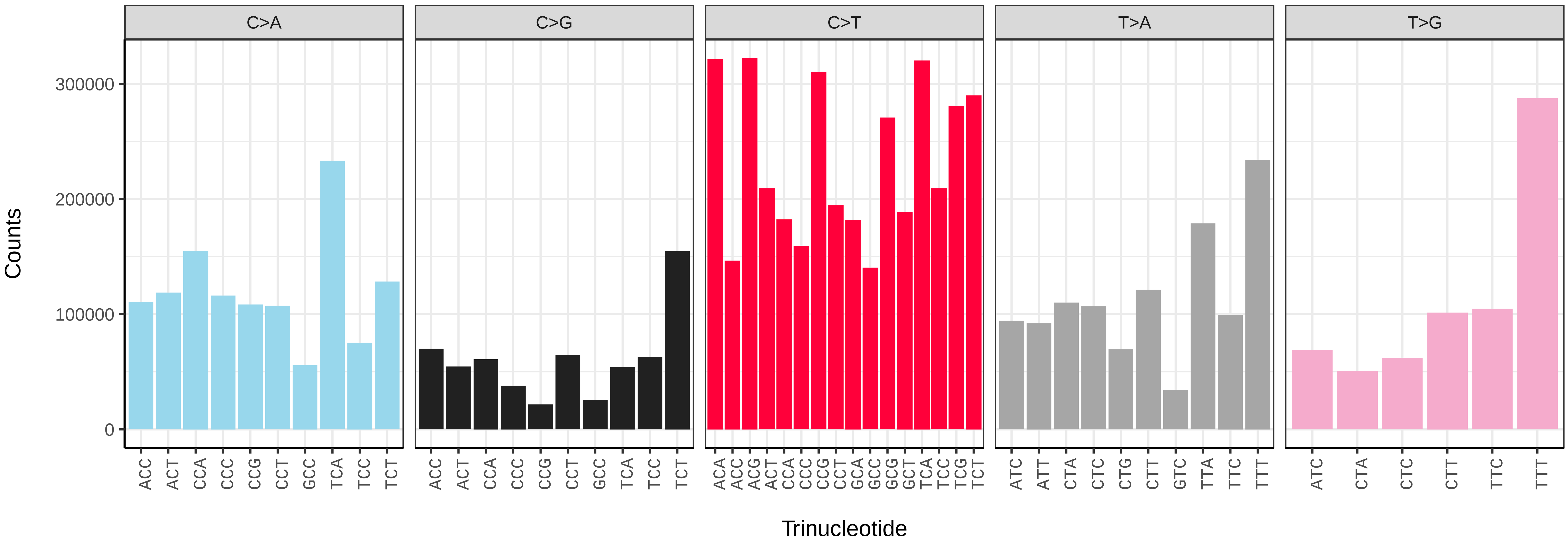


*Echiichthys vipera* (fEchVip8) somatic mutational spectrum (SBS96 classification)

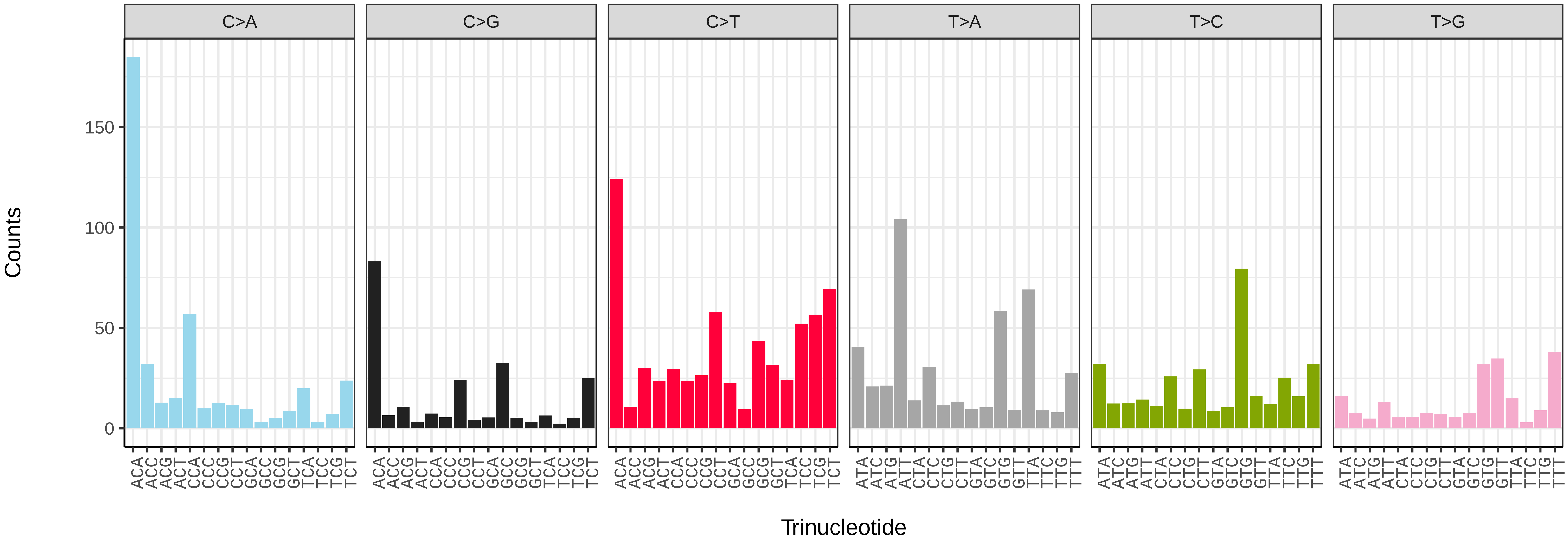


*Echiichthys vipera* (fEchVip8) somatic mutational spectrum (SBS52 classification)

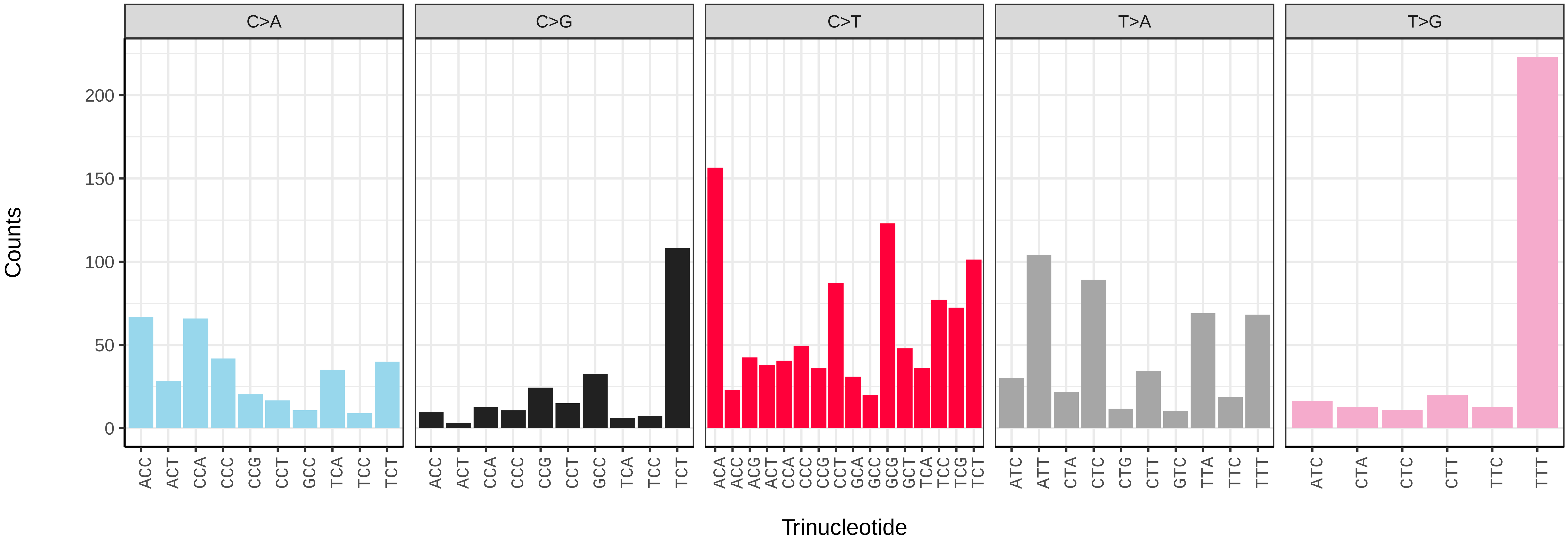

*Eutrigla gurnardus* (fEutGur1) germline mutational spectrum

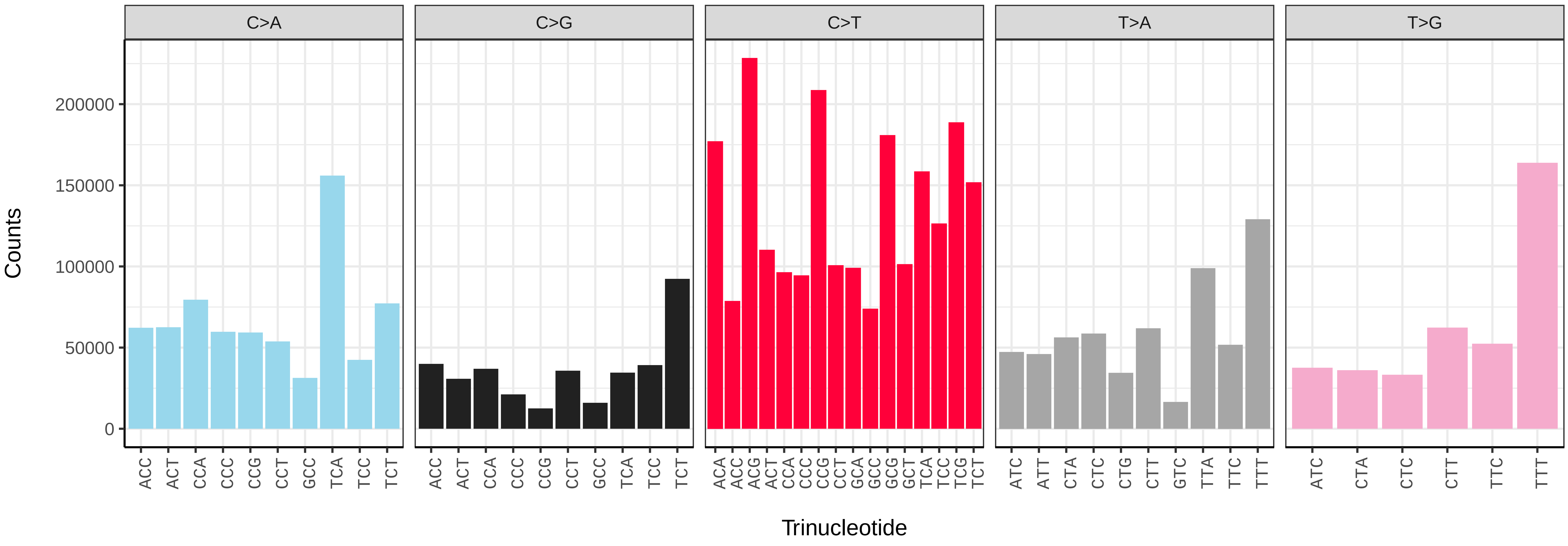


*Eutrigla gurnardus* (fEutGur1) somatic mutational spectrum (SBS96 classification)

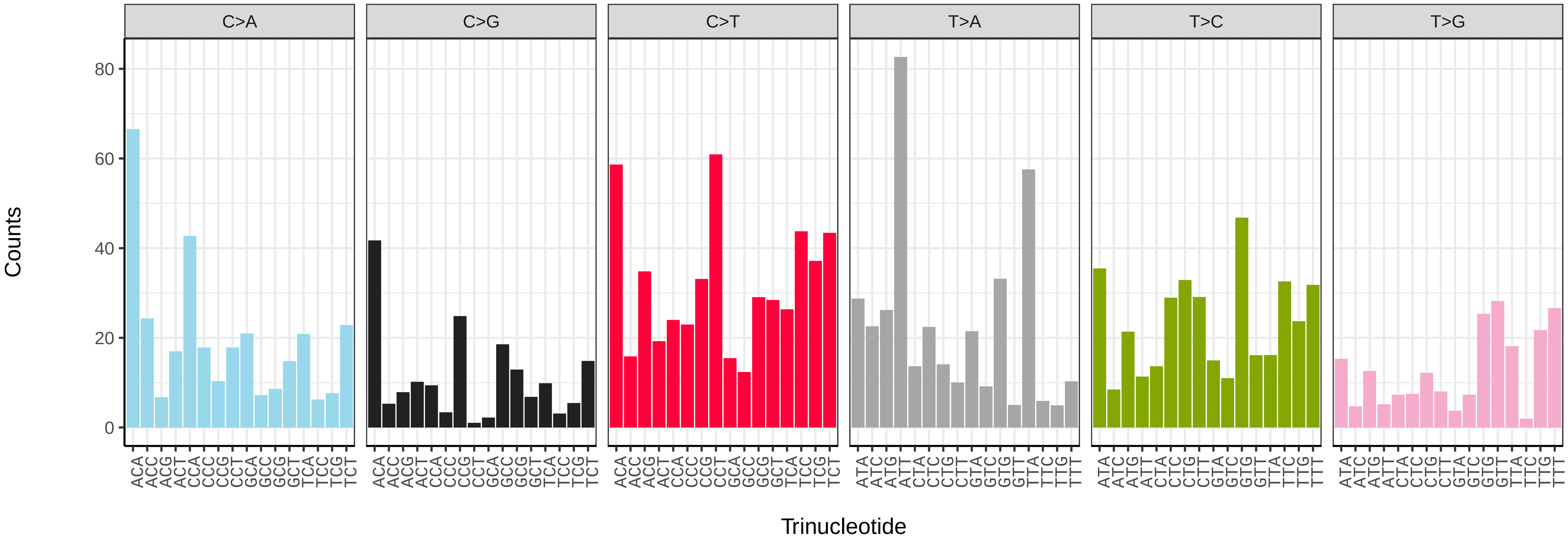


*Eutrigla gurnardus* (fEutGur1) somatic mutational spectrum (SBS52 classification)

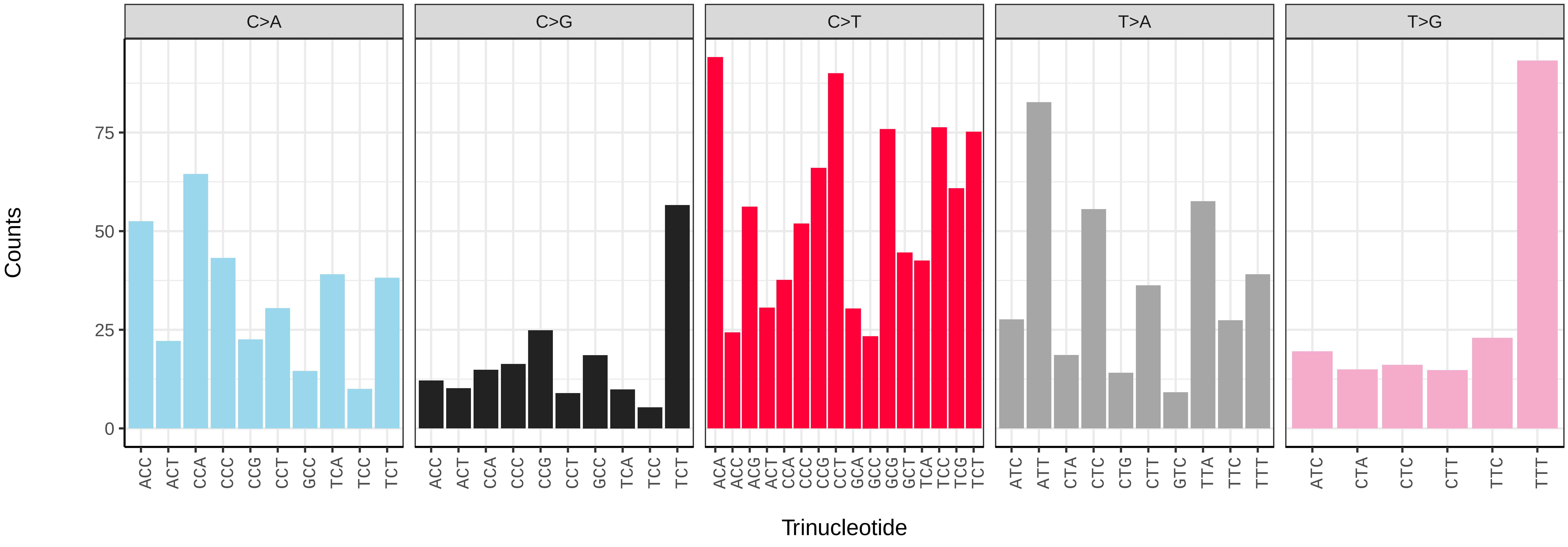

*Gobio gobio* (fGobGob1) germline mutational spectrum

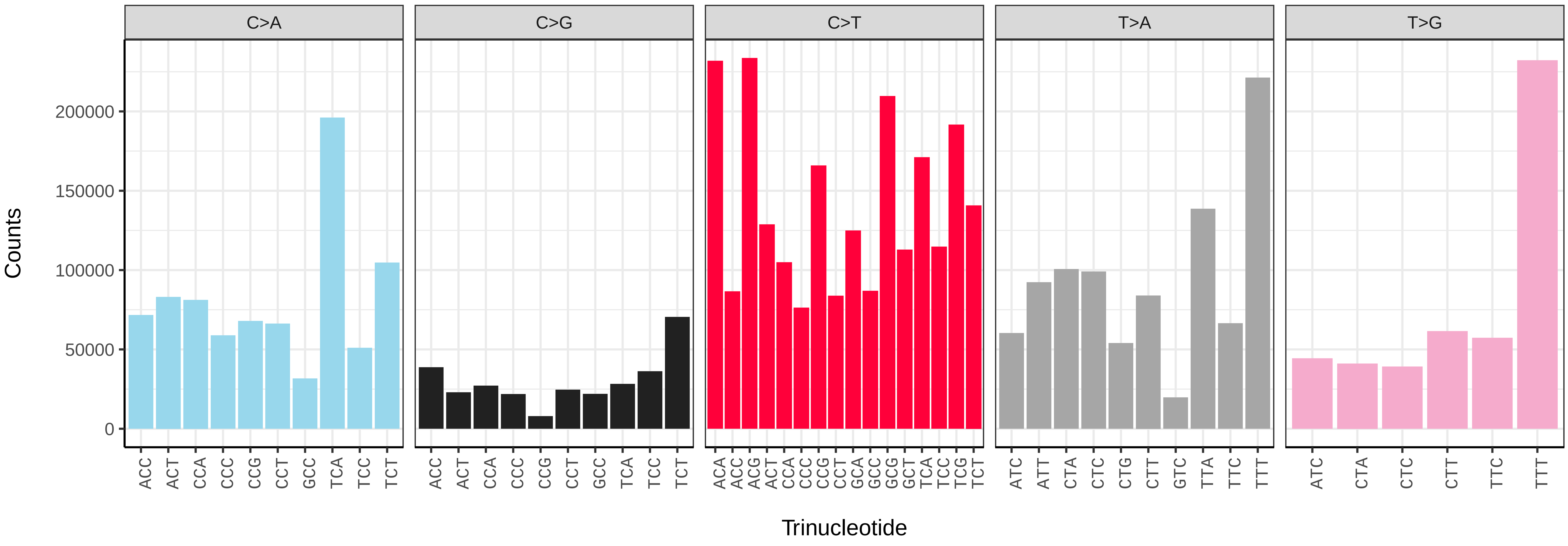


*Gobio gobio* (fGobGob1) somatic mutational spectrum (SBS96 classification)

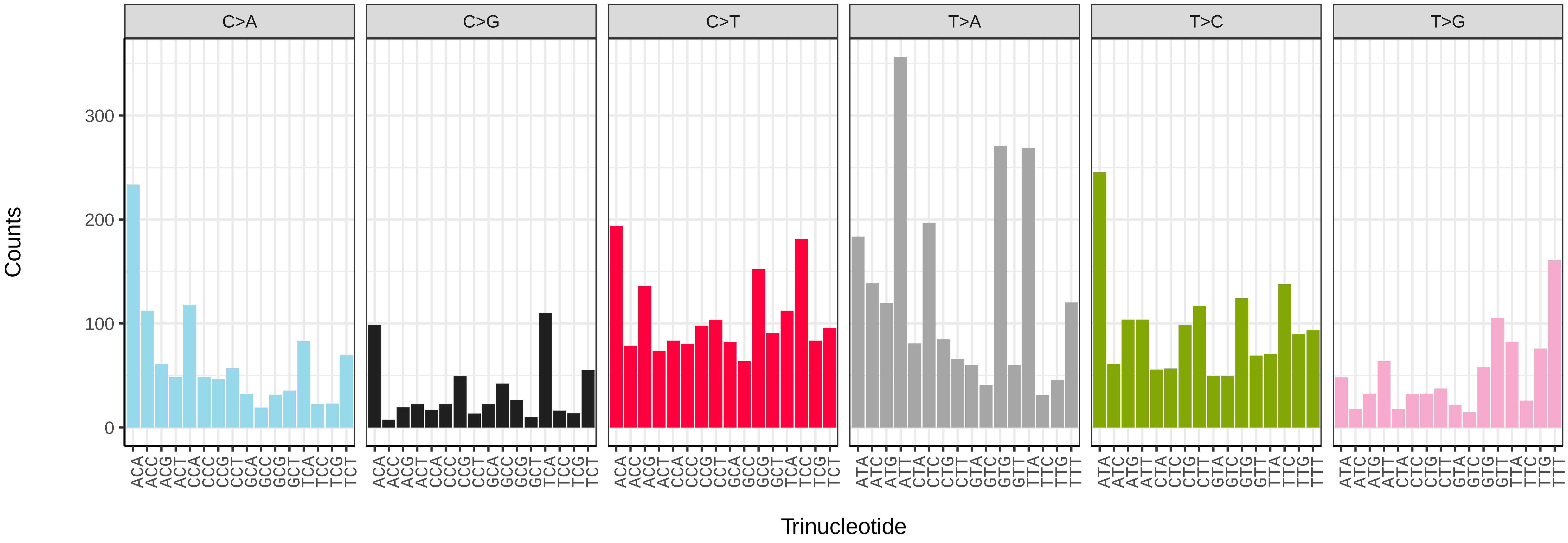


*Gobio gobio* (fGobGob1) somatic mutational spectrum (SBS52 classification)

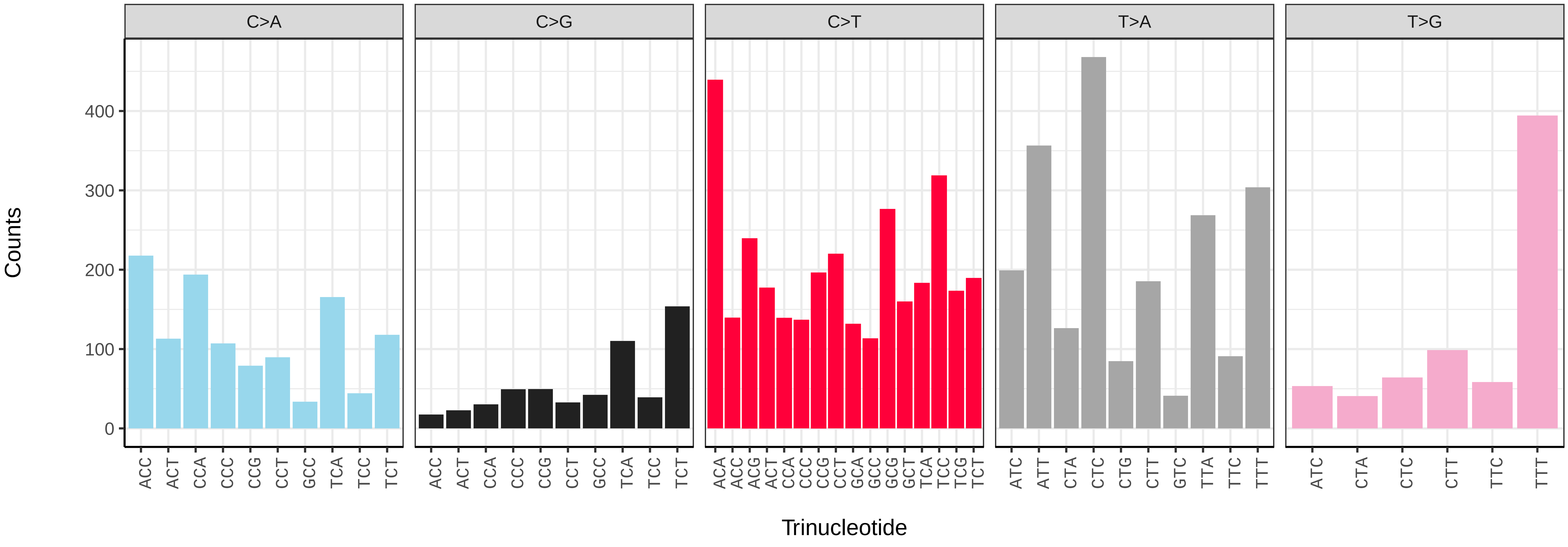

*Gobius niger* (fGobNig1) germline mutational spectrum

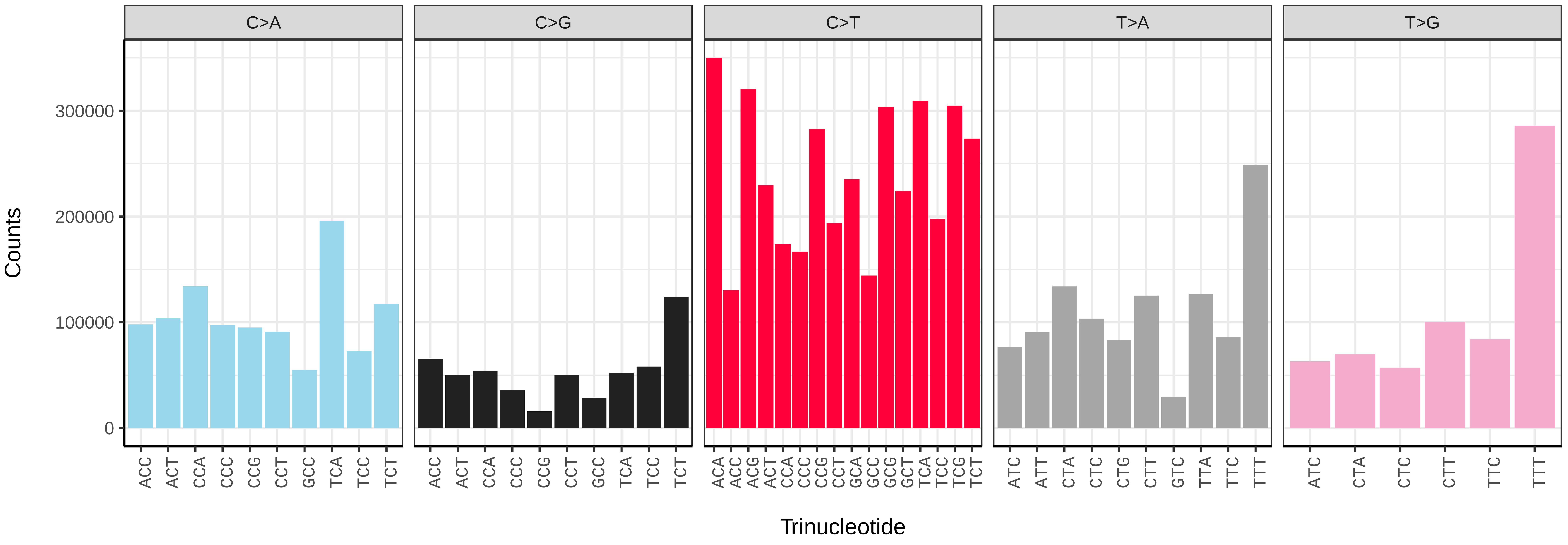


*Gobius niger* (fGobNig1) somatic mutational spectrum (SBS96 classification)

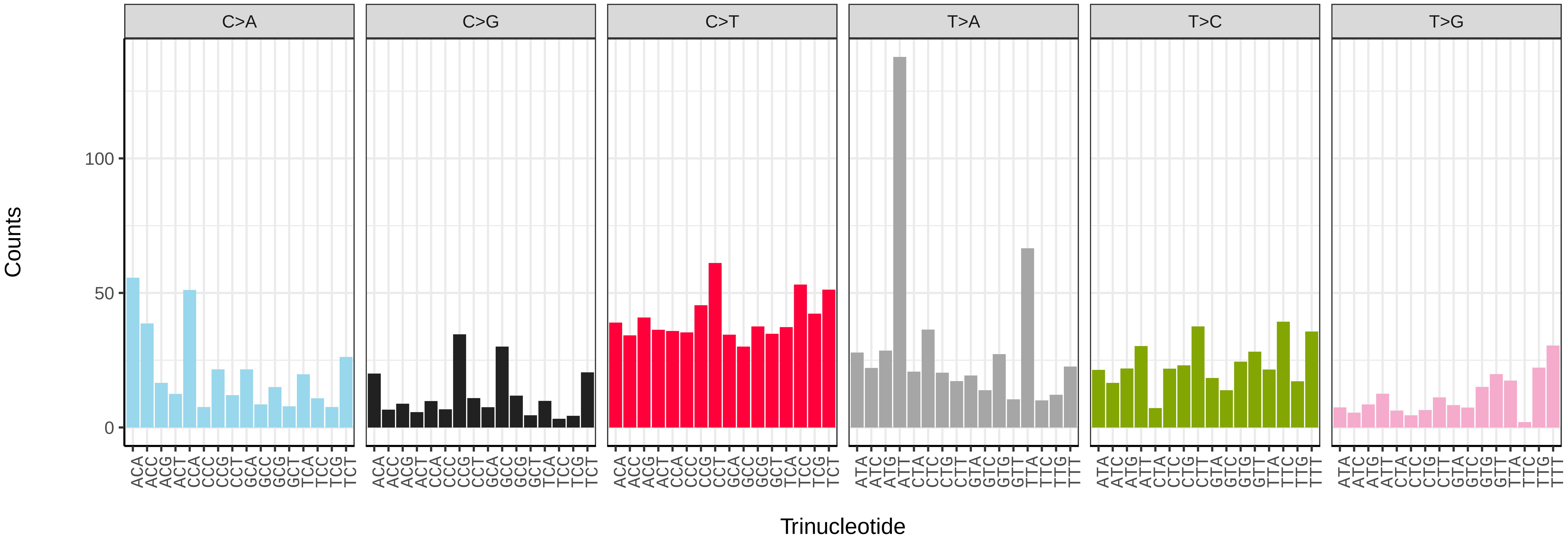


*Gobius niger* (fGobNig1) somatic mutational spectrum (SBS52 classification)

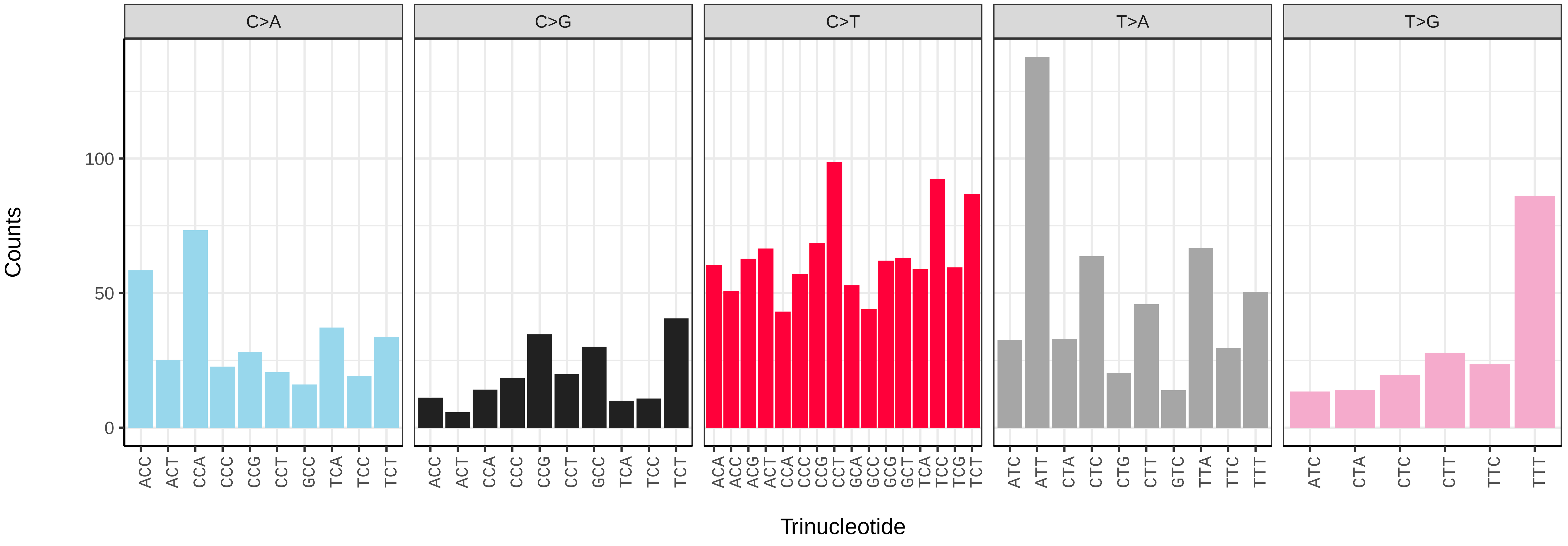

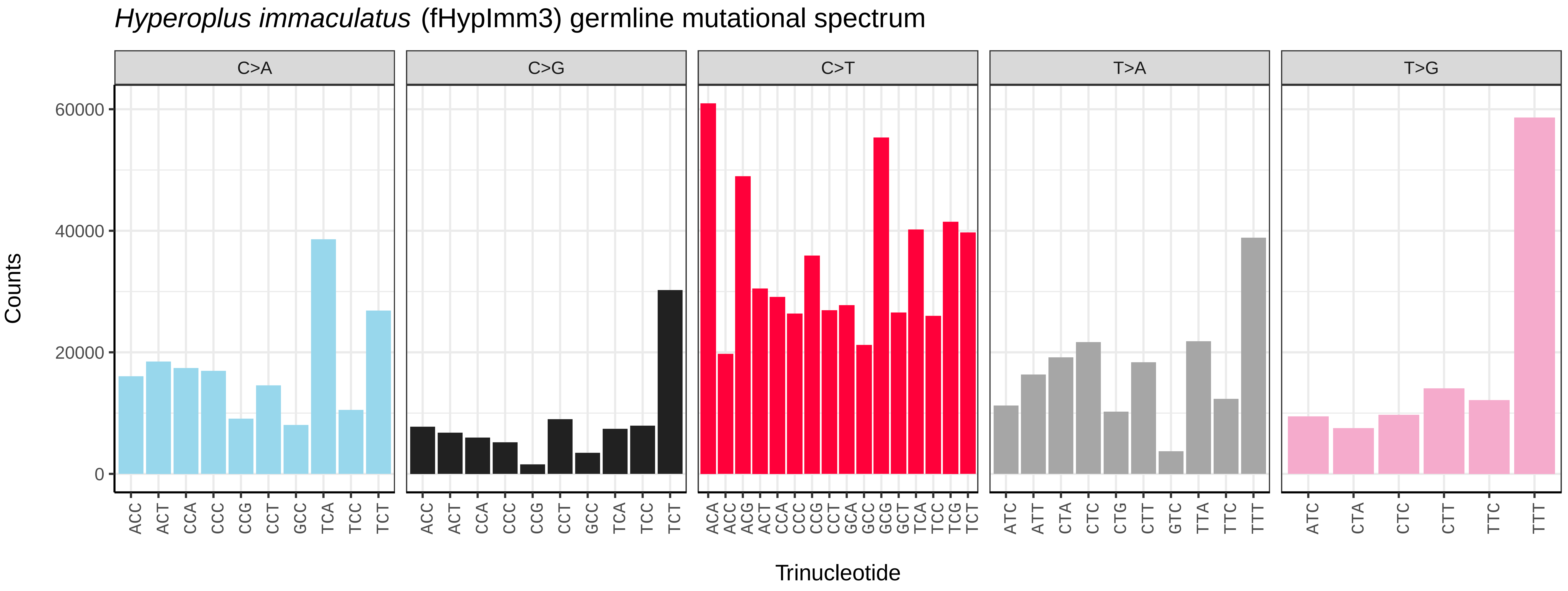
*Hyperoplus immaculatus* (fHypImm3) germline mutational spectrum
C>A
C>G
C>T
T>A
T>G
Counts
0
20000
40000
60000
Trinucleotide

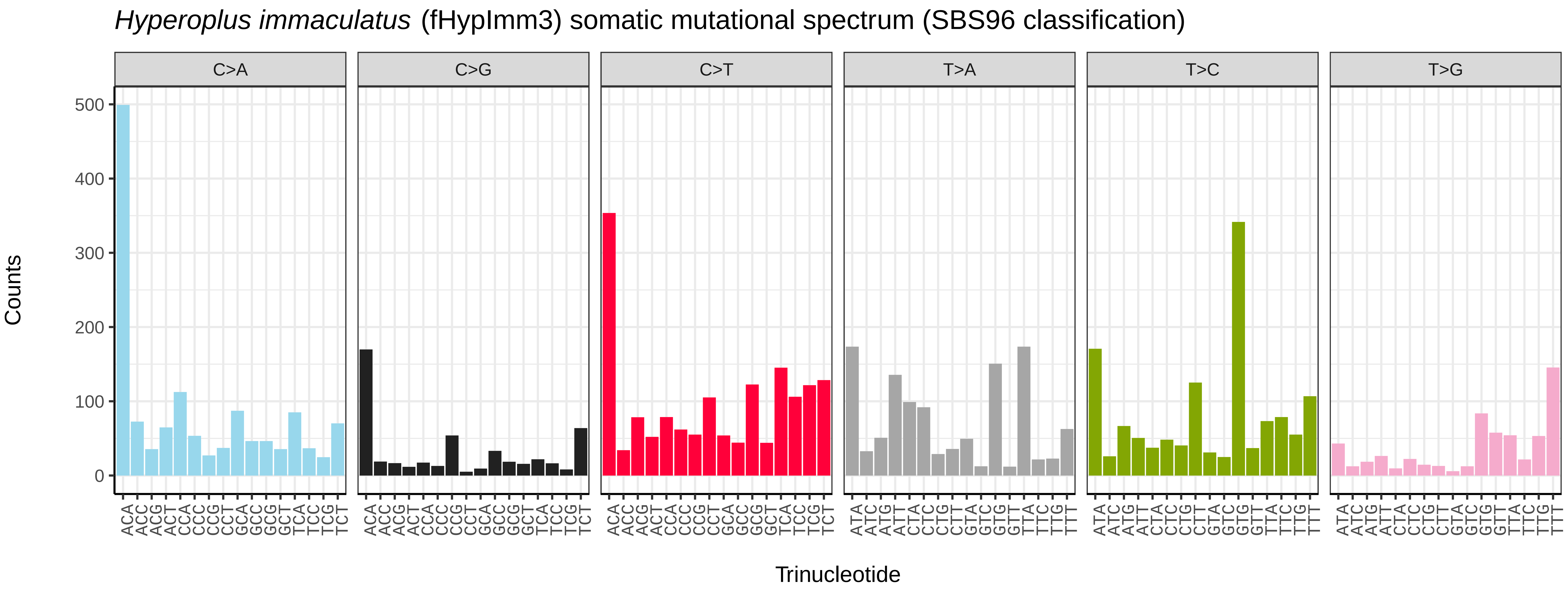
*Hyperoplus immaculatus* (fHypImm3) somatic mutational spectrum (SBS96 classification)
C>A
C>G
C>T
T>A
T>C
T>G
Counts
0
100
200
300
400
500
Trinucleotide

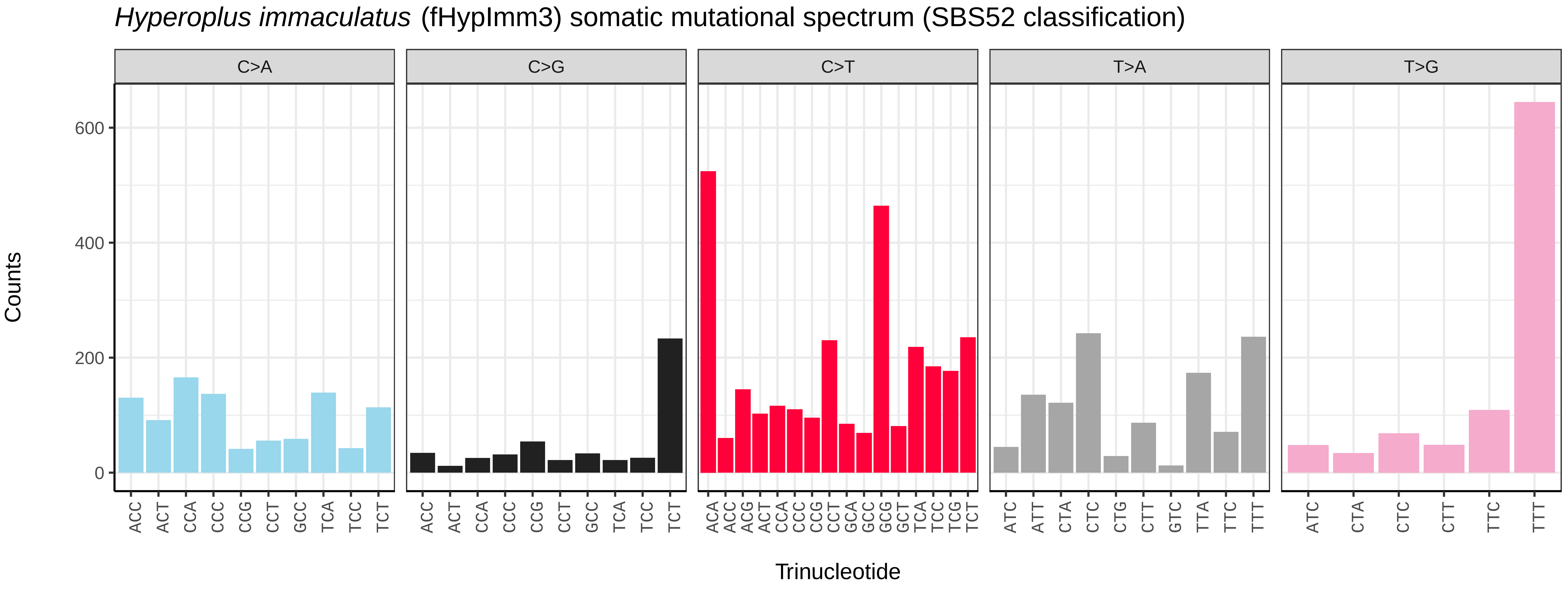
*Hyperoplus immaculatus* (fHypImm3) somatic mutational spectrum (SBS52 classification)
C>A
C>G
C>T
T>A
T>G
Counts
0
200
400
600
Trinucleotide

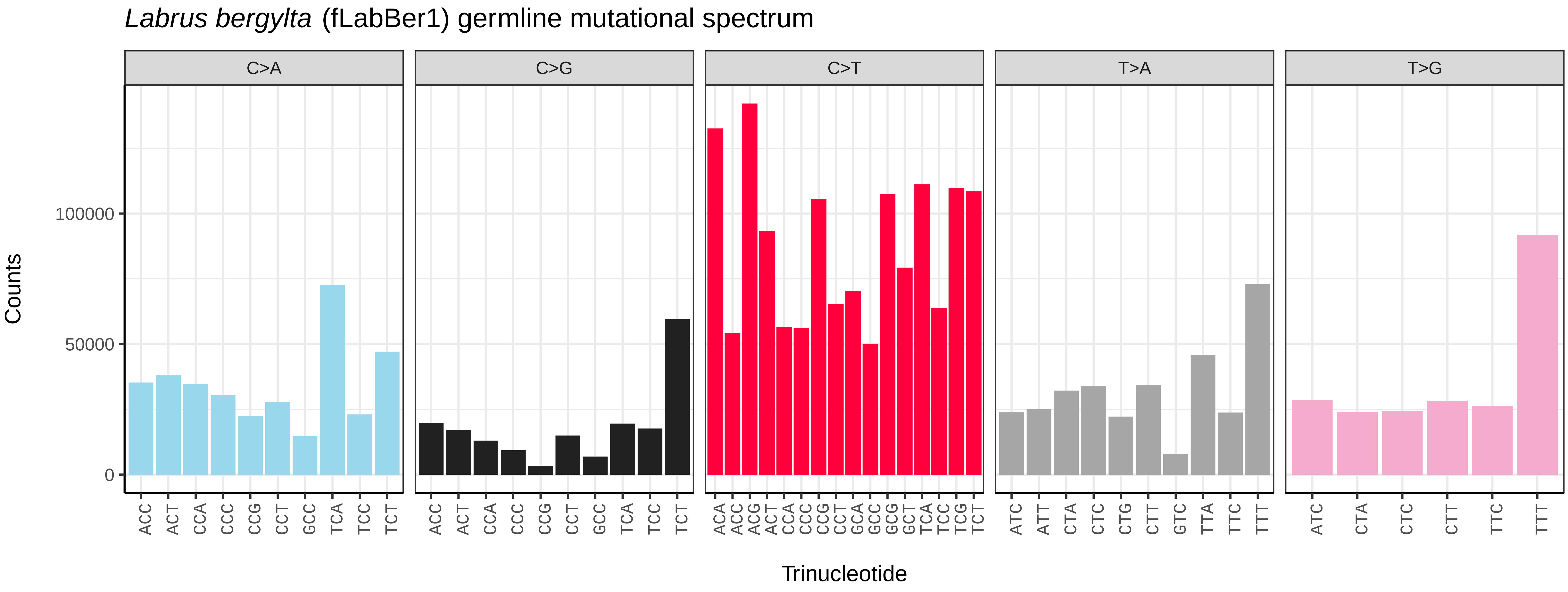

Labrus bergylta (fLabBer1) germline mutational spectrum
C>A
C>G
C>T
T>A
T>G
Counts
0
50000
100000
Trinucleotide
ACC ACT CCA CCC CCG CCT GCC TCA TCC TCT
ACA ACC ACG ACT CCA CCC CCG CCT GCA GCC GCG GCT TCA TCC TCG TCT
ATC ATT CTA CTC CTG CTT GTC TTA TTC TTT
ATC CTA CTC CTT TTC TTT


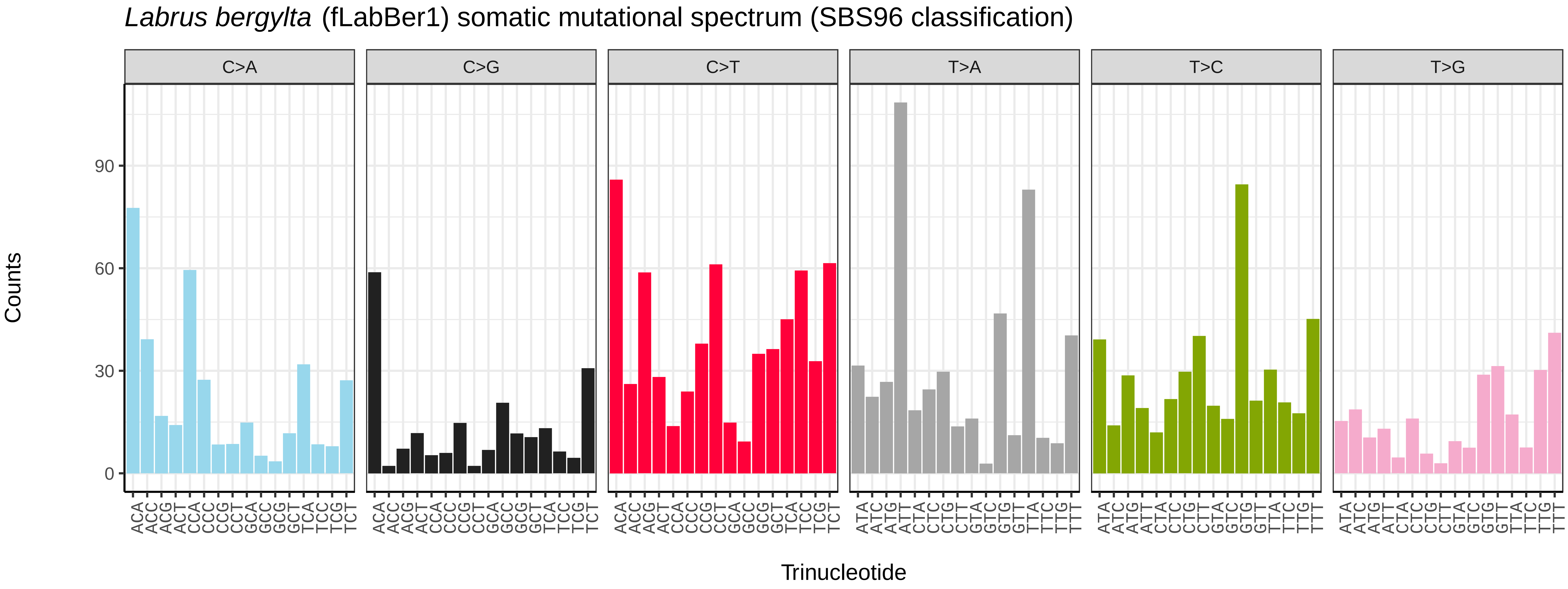

Labrus bergylta (fLabBer1) somatic mutational spectrum (SBS96 classification)
C>A
C>G
C>T
T>A
T>C
T>G
Counts
0
30
60
90
Trinucleotide
ACA ACC ACG ACT CCA CCC CCG CCT GCA GCC GCG GCT TCA TCC TCG TCT
ATA ATC ATG ATT CTA CTC CTG CTT GTA GTC GTG GTT TTA TTC TTG TTT


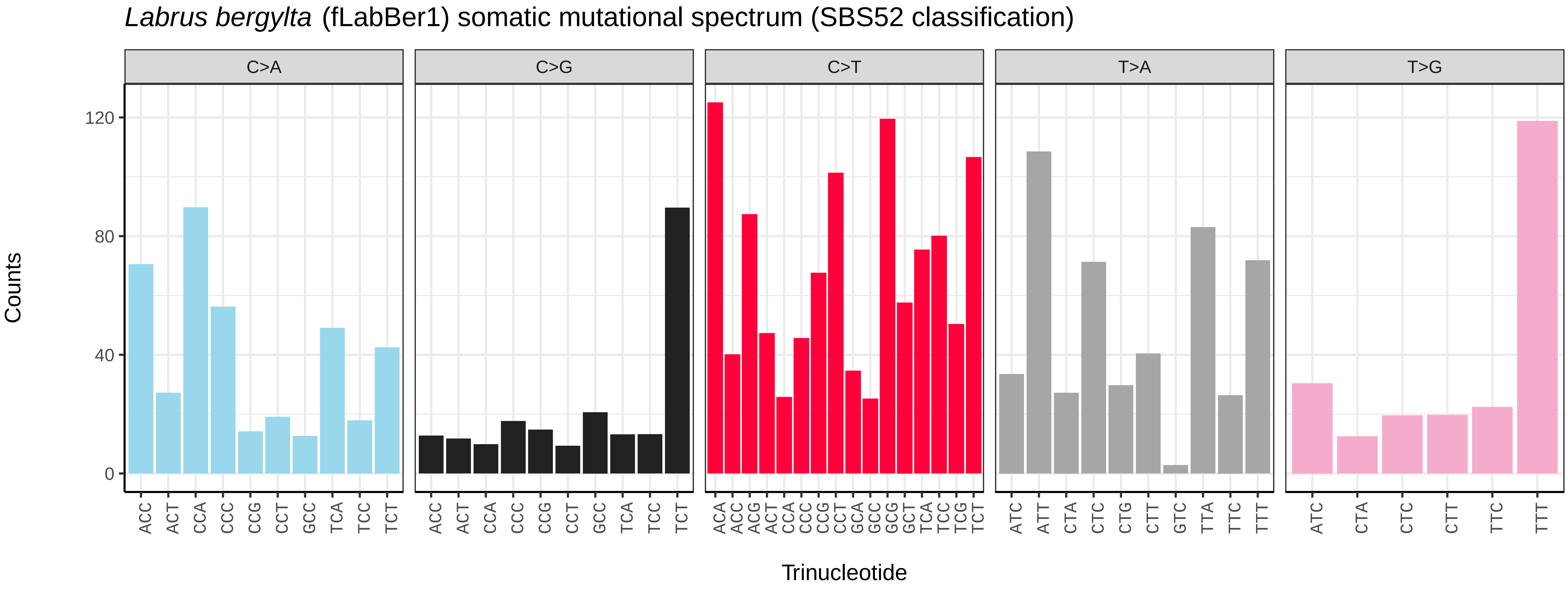

Labrus bergylta (fLabBer1) somatic mutational spectrum (SBS52 classification)
C>A
C>G
C>T
T>A
T>G
Counts
0
40
80
120
Trinucleotide
ACC ACT CCA CCC CCG CCT GCC TCA TCC TCT
ACA ACC ACG ACT CCA CCC CCG CCT GCA GCC GCG GCT TCA TCC TCG TCT
ATC ATT CTA CTC CTG CTT GTC TTA TTC TTT
ATC CTA CTC CTT TTC TTT

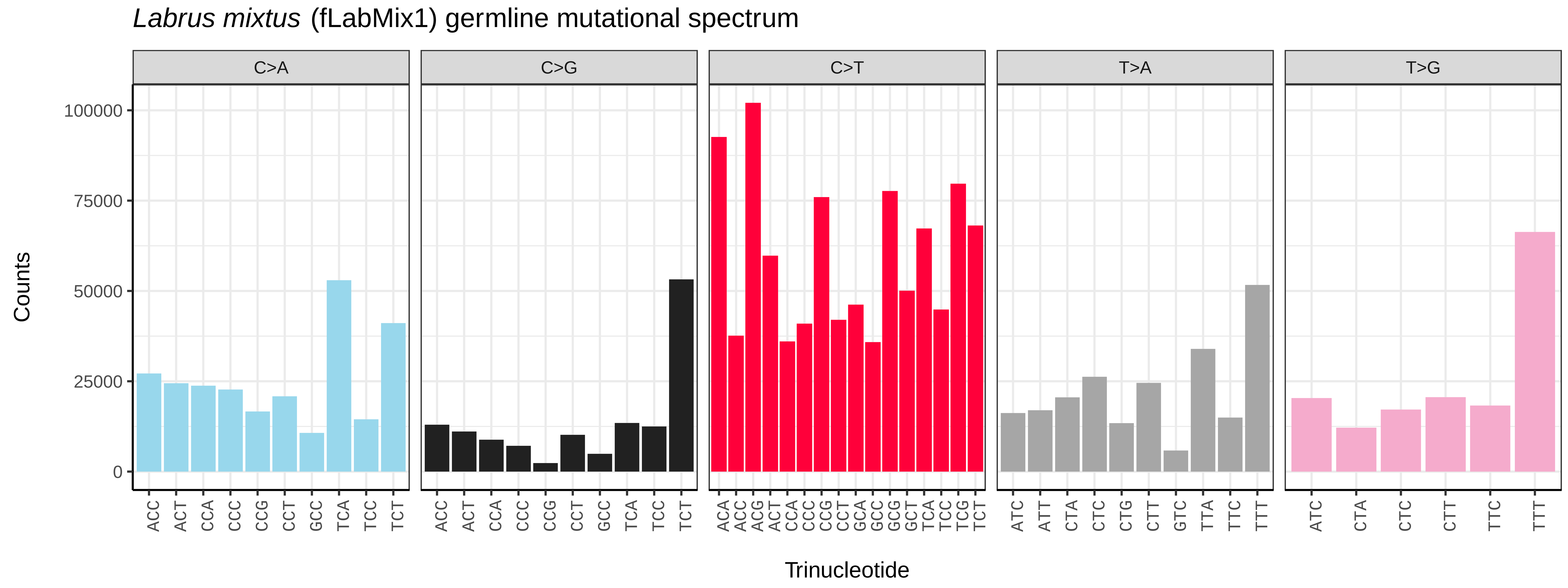
*Labrus mixtus* (fLabMix1) germline mutational spectrum
C>A
C>G
C>T
T>A
T>G
Counts
0
25000
50000
75000
100000
ACC ACT CCA CCC CCG CCT GCC TCA TCC TCT
ACC ACT CCA CCC CCG CCT GCC TCA TCC TCT
ACA ACC ACG ACT CCA CCC CCG CCT GCA GCC GCG GCT TCA TCC TCG TCT
ATC ATT CTA CTC CTG CTT GTC TTA TTC TTT
ATC CTA CTC CTT TTC TTT
Trinucleotide

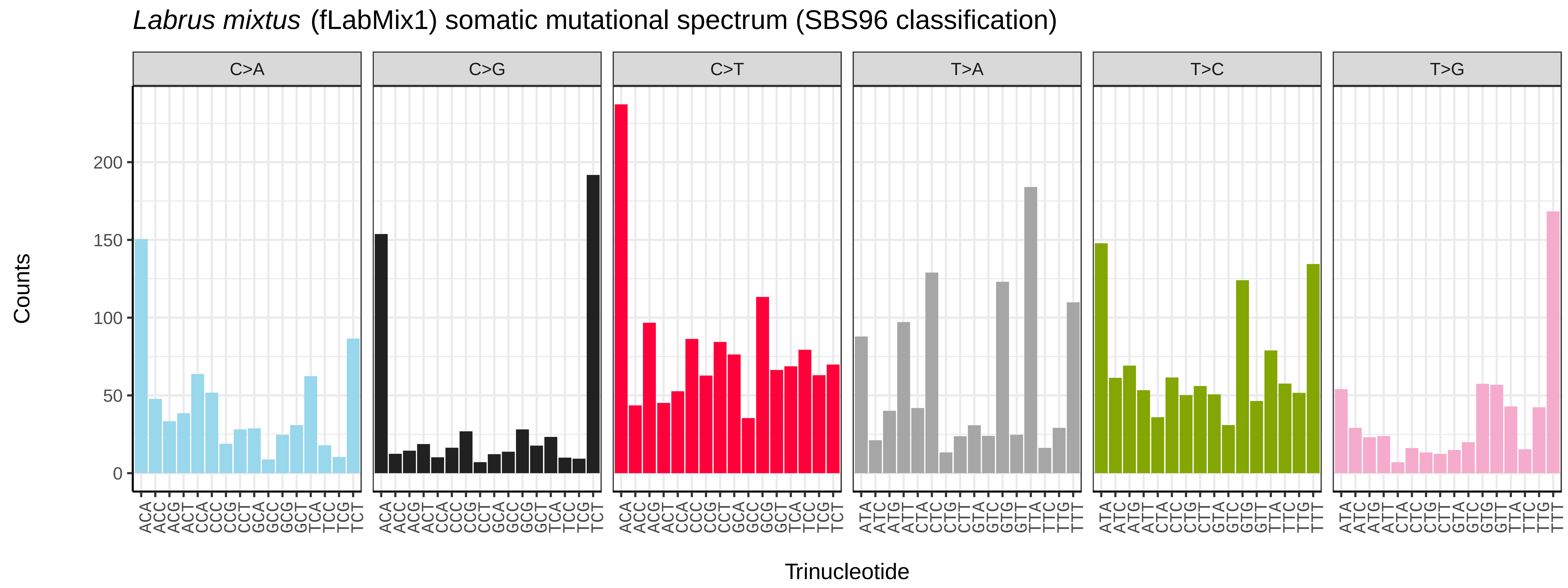
*Labrus mixtus* (fLabMix1) somatic mutational spectrum (SBS96 classification)
C>A
C>G
C>T
T>A
T>C
T>G
Counts
0
50
100
150
200
ACA ACC ACG ACT CCA CCC CCG CCT GCA GCC GCG GCT TCA TCC TCG TCT
ACA ACC ACG ACT CCA CCC CCG CCT GCA GCC GCG GCT TCA TCC TCG TCT
ACA ACC ACG ACT CCA CCC CCG CCT GCA GCC GCG GCT TCA TCC TCG TCT
ATA ATC ATG ATT CTA CTC CTG CTT GTA GTC GTG GTT TTA TTC TTG TTT
ATA ATC ATG ATT CTA CTC CTG CTT GTA GTC GTG GTT TTA TTC TTG TTT
ATA ATC ATG ATT CTA CTC CTG CTT GTA GTC GTG GTT TTA TTC TTG TTT
Trinucleotide

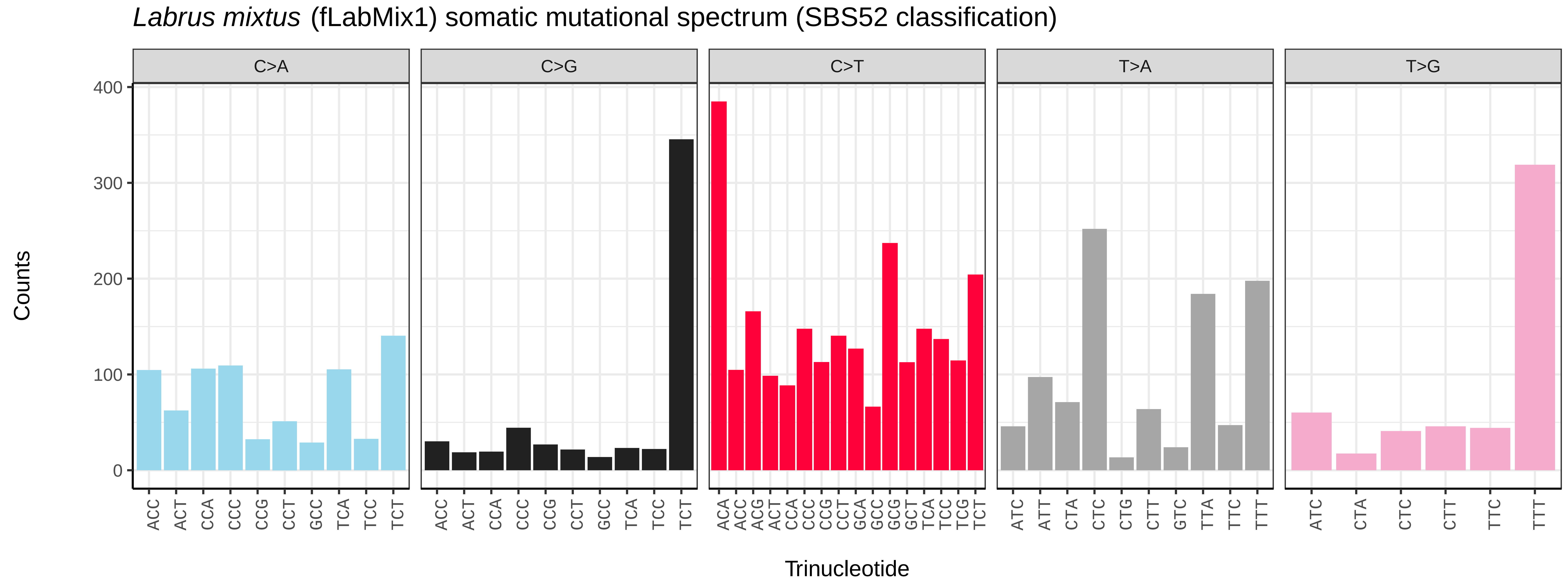
*Labrus mixtus* (fLabMix1) somatic mutational spectrum (SBS52 classification)
C>A
C>G
C>T
T>A
T>G
Counts
0
100
200
300
400
ACC ACT CCA CCC CCG CCT GCC TCA TCC TCT
ACC ACT CCA CCC CCG CCT GCC TCA TCC TCT
ACA ACC ACG ACT CCA CCC CCG CCT GCA GCC GCG GCT TCA TCC TCG TCT
ATC ATT CTA CTC CTG CTT GTC TTA TTC TTT
ATC CTA CTC CTT TTC TTT
Trinucleotide

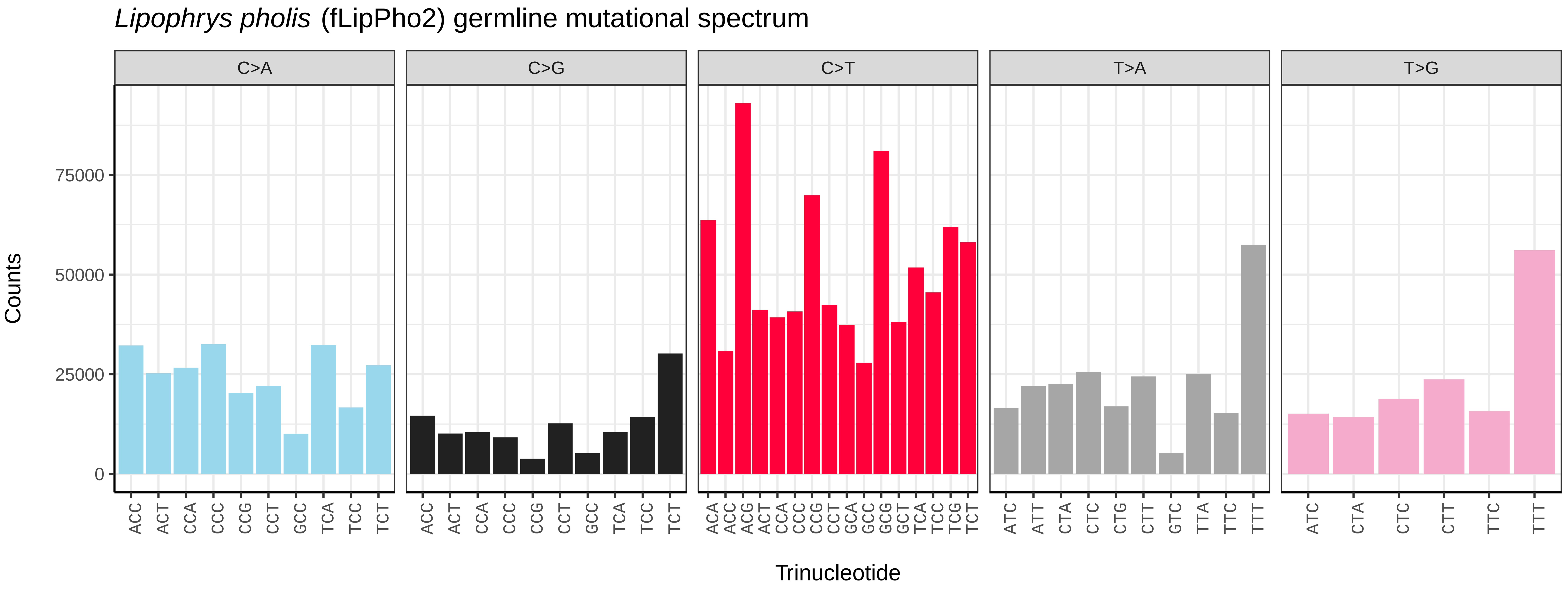
Lipophrys pholis (fLipPho2) germline mutational spectrum
C>A
C>G
C>T
T>A
T>G
Counts
Trinucleotide
0
25000
50000
75000

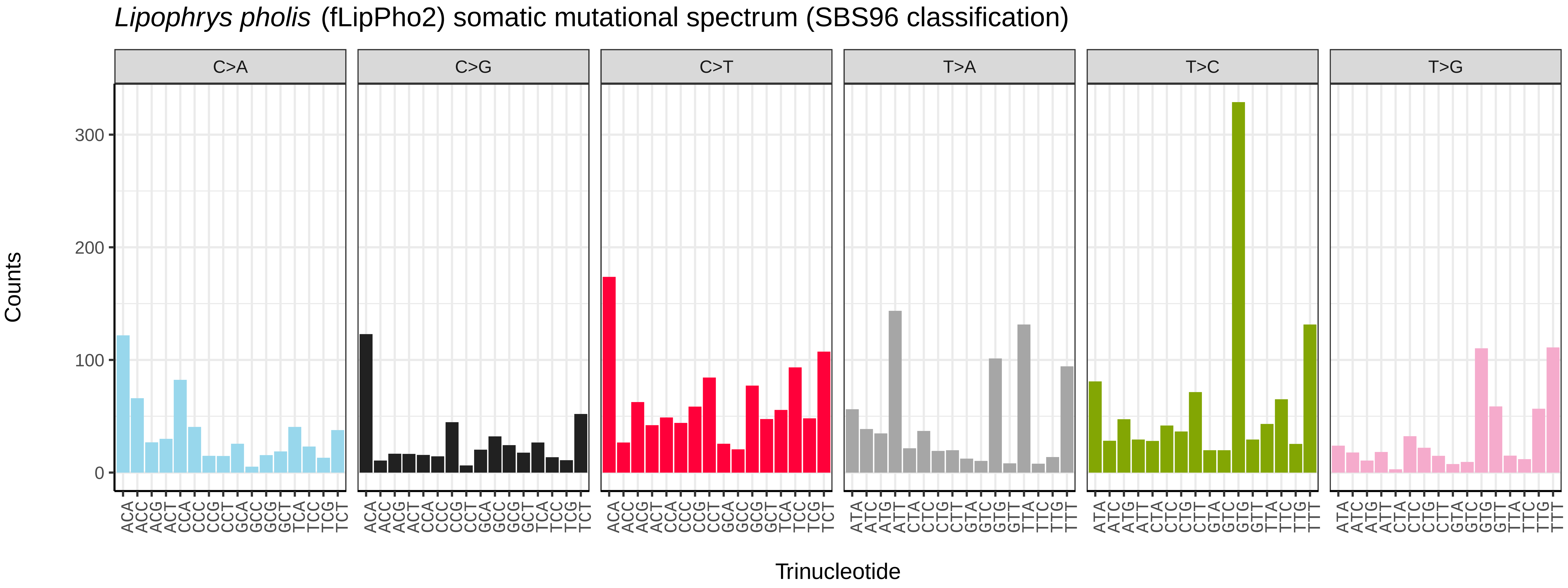
Lipophrys pholis (fLipPho2) somatic mutational spectrum (SBS96 classification)
C>A
C>G
C>T
T>A
T>C
T>G
Counts
Trinucleotide
0
100
200
300

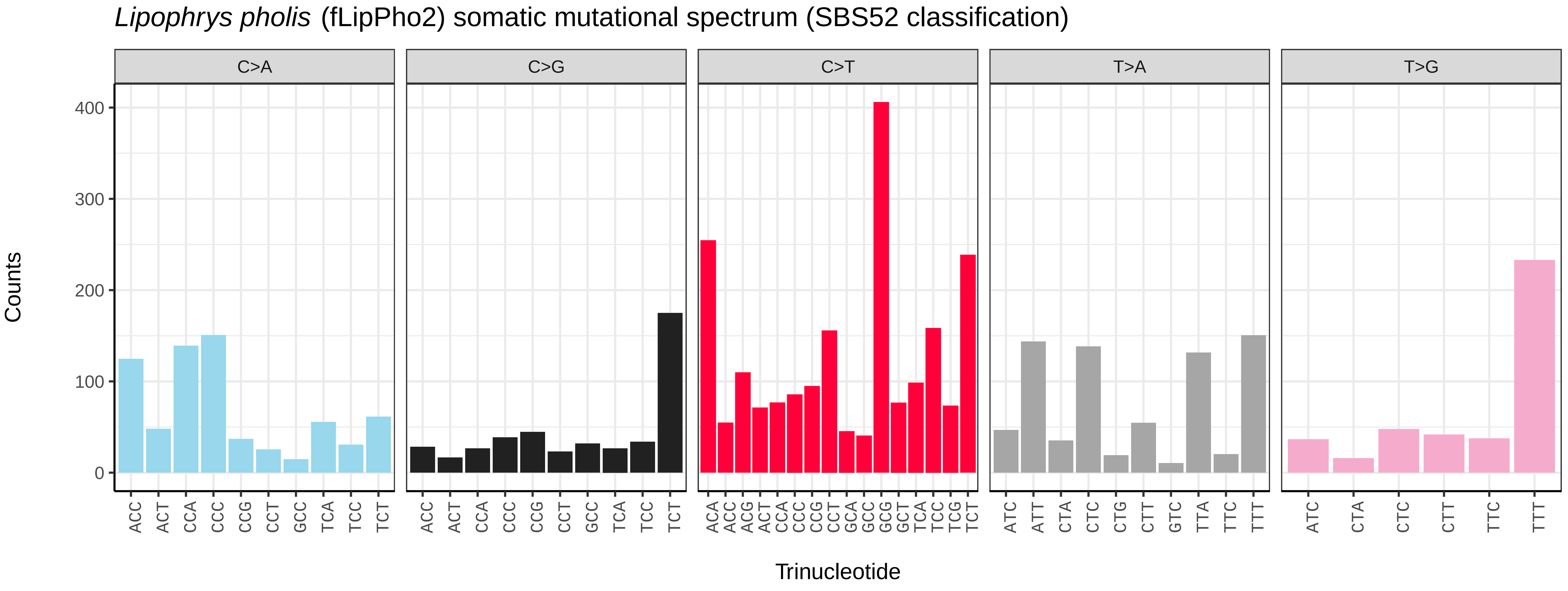
Lipophrys pholis (fLipPho2) somatic mutational spectrum (SBS52 classification)
C>A
C>G
C>T
T>A
T>G
Counts
Trinucleotide
0
100
200
300
400

*Micromesistius poutassou* (fMicPou1) germline mutational spectrum

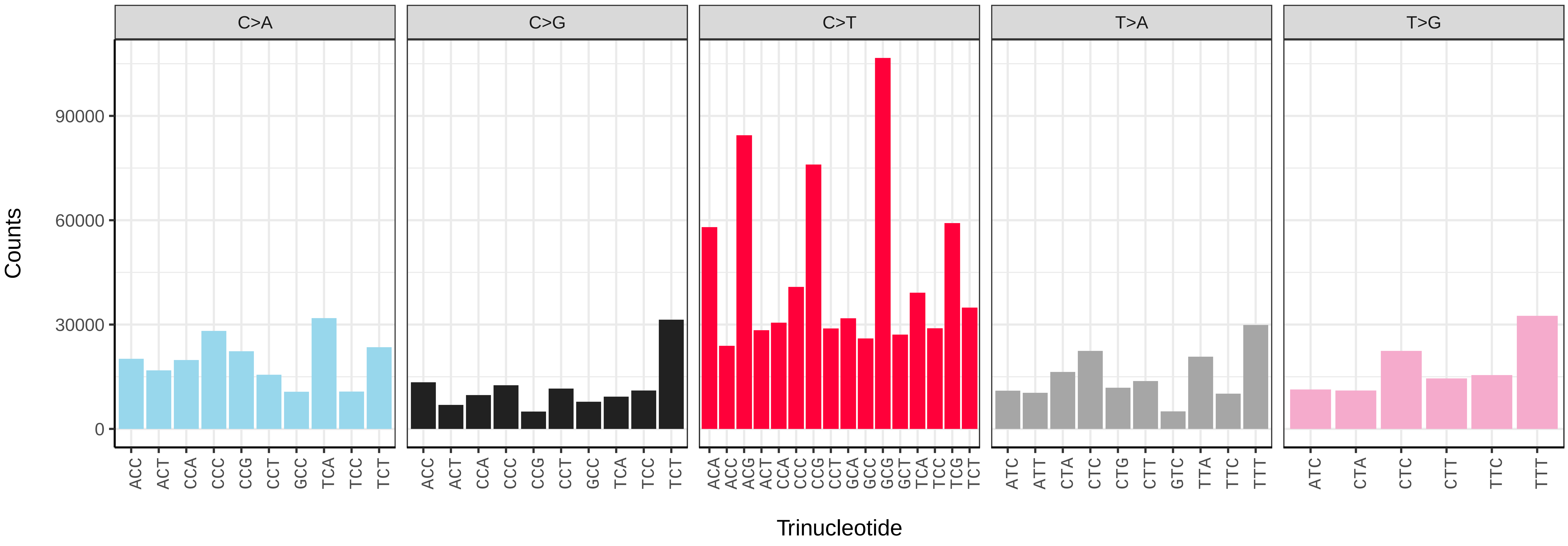


*Micromesistius poutassou* (fMicPou1) somatic mutational spectrum (SBS96 classification)

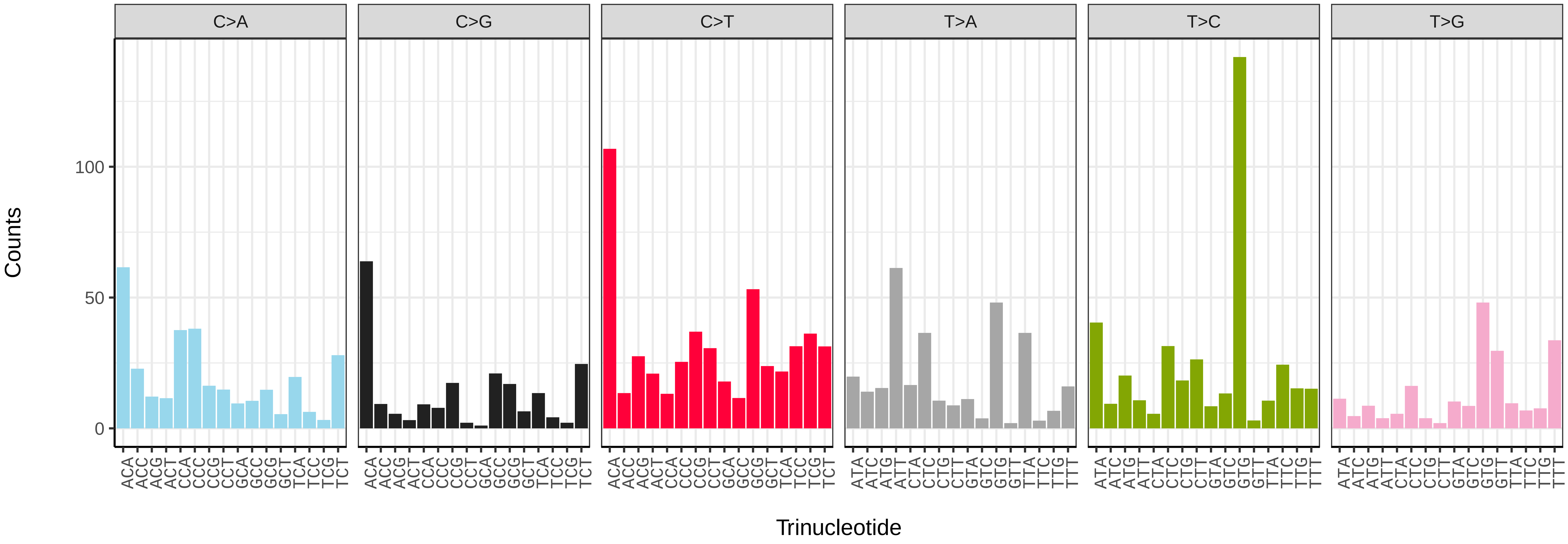


*Micromesistius poutassou* (fMicPou1) somatic mutational spectrum (SBS52 classification)

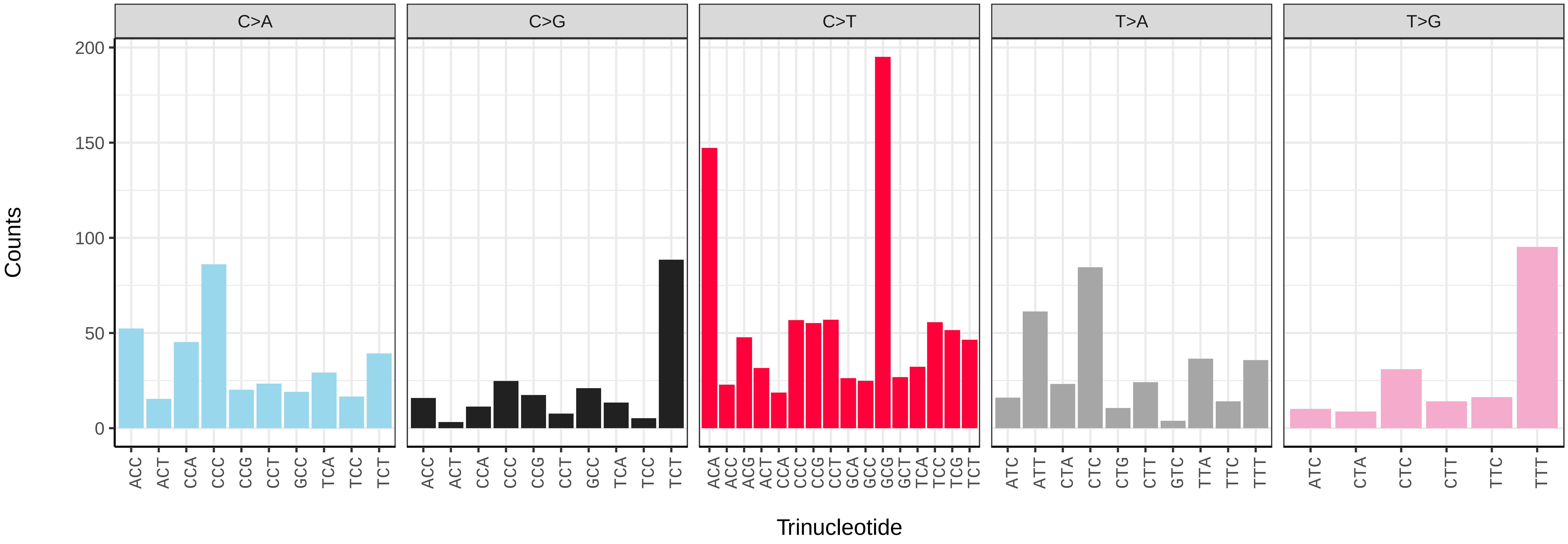

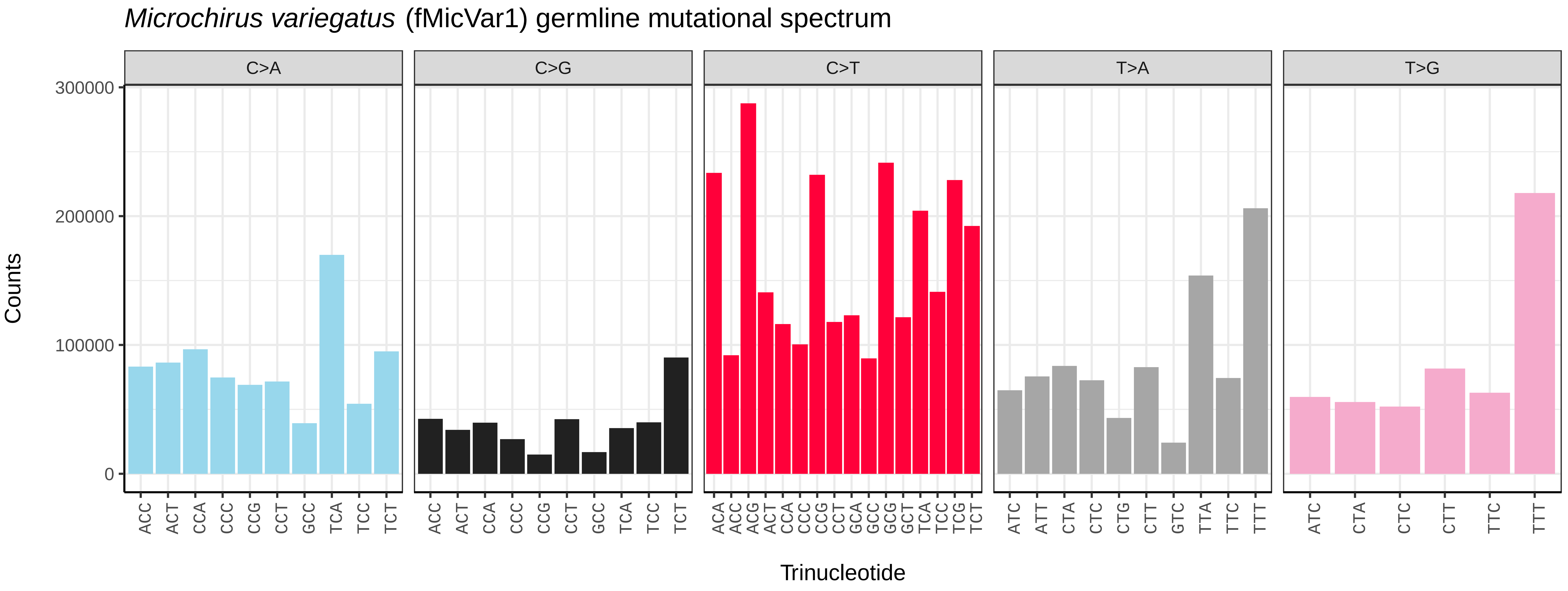
Microchirus variegatus (fMicVar1) germline mutational spectrum
C>A
C>G
C>T
T>A
T>G
Counts
300000
200000
100000
0
Trinucleotide

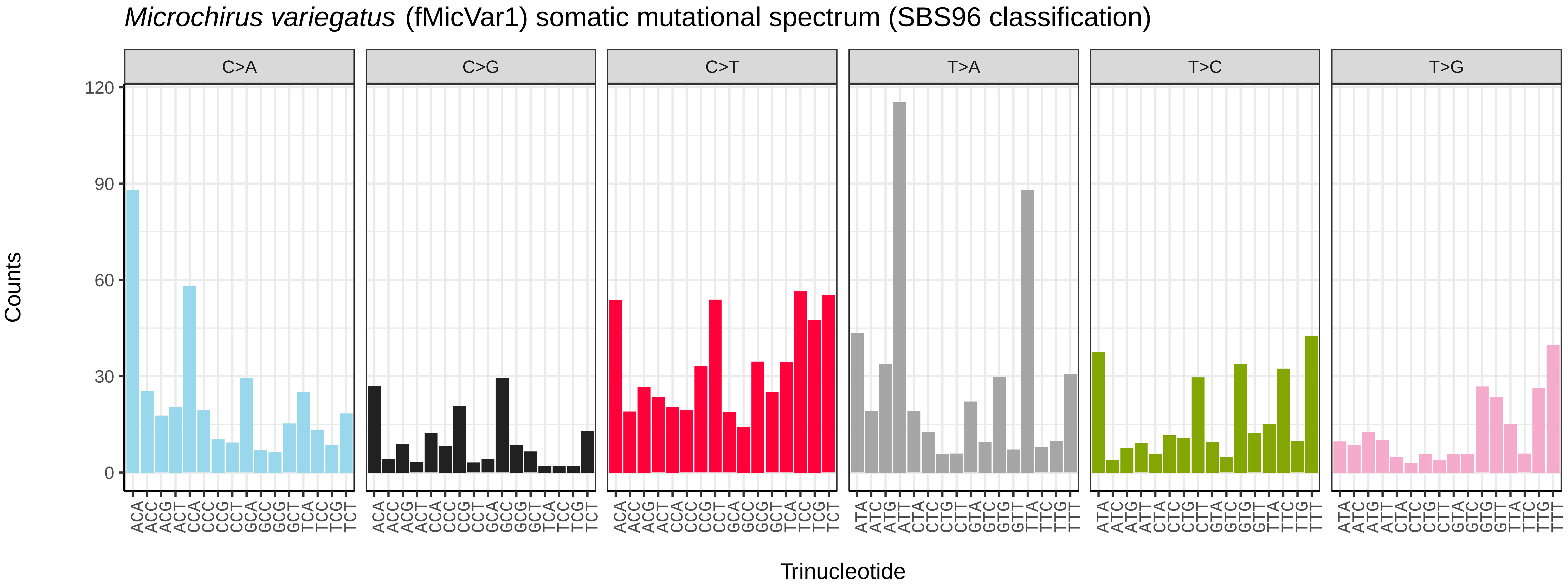
Microchirus variegatus (fMicVar1) somatic mutational spectrum (SBS96 classification)
C>A
C>G
C>T
T>A
T>C
T>G
Counts
120
90
60
30
0
Trinucleotide

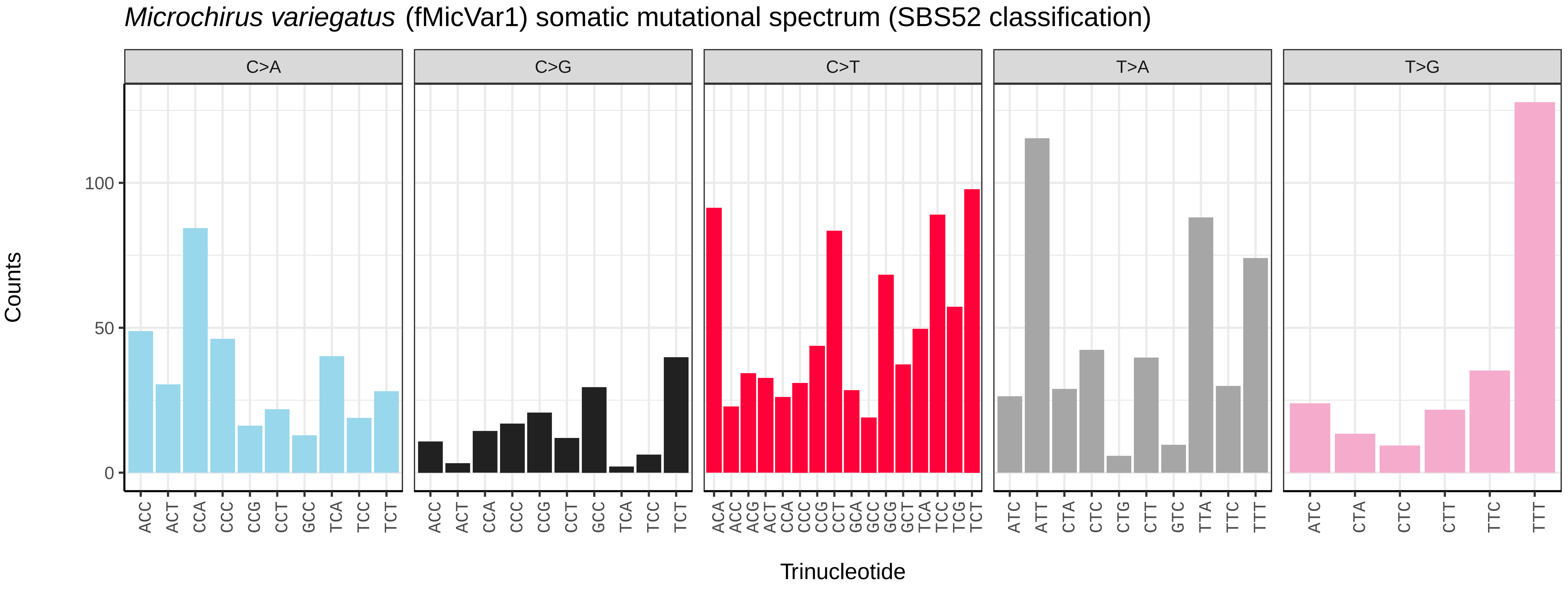
Microchirus variegatus (fMicVar1) somatic mutational spectrum (SBS52 classification)
C>A
C>G
C>T
T>A
T>G
Counts
100
50
0
Trinucleotide

*Pholis gunnellus* (fPhoGun1) germline mutational spectrum

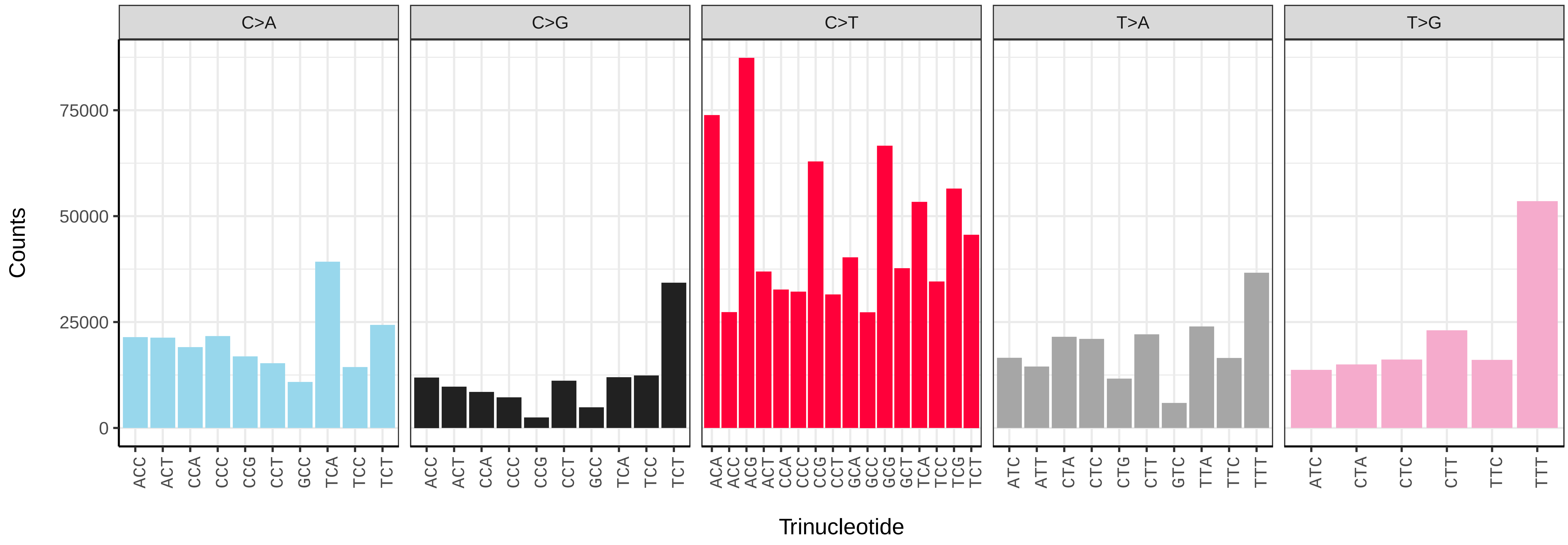


*Pholis gunnellus* (fPhoGun1) somatic mutational spectrum (SBS96 classification)

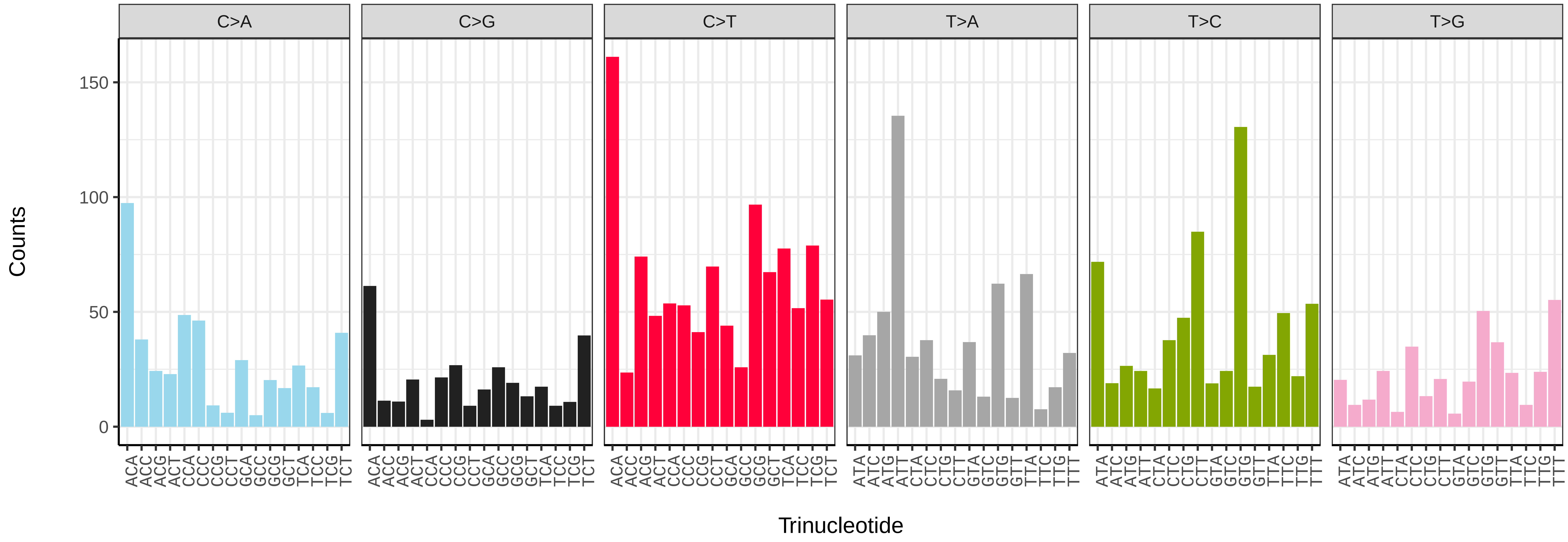


*Pholis gunnellus* (fPhoGun1) somatic mutational spectrum (SBS52 classification)

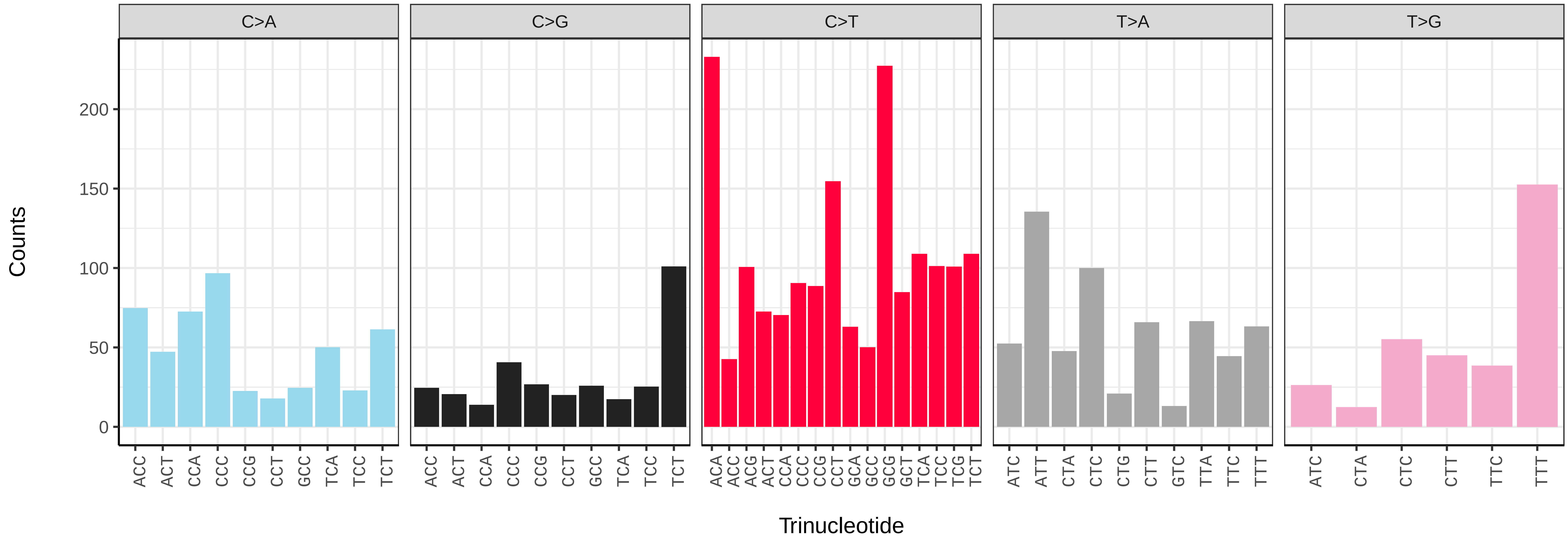

*Phoxinus phoxinus* (fPhoPho1) germline mutational spectrum

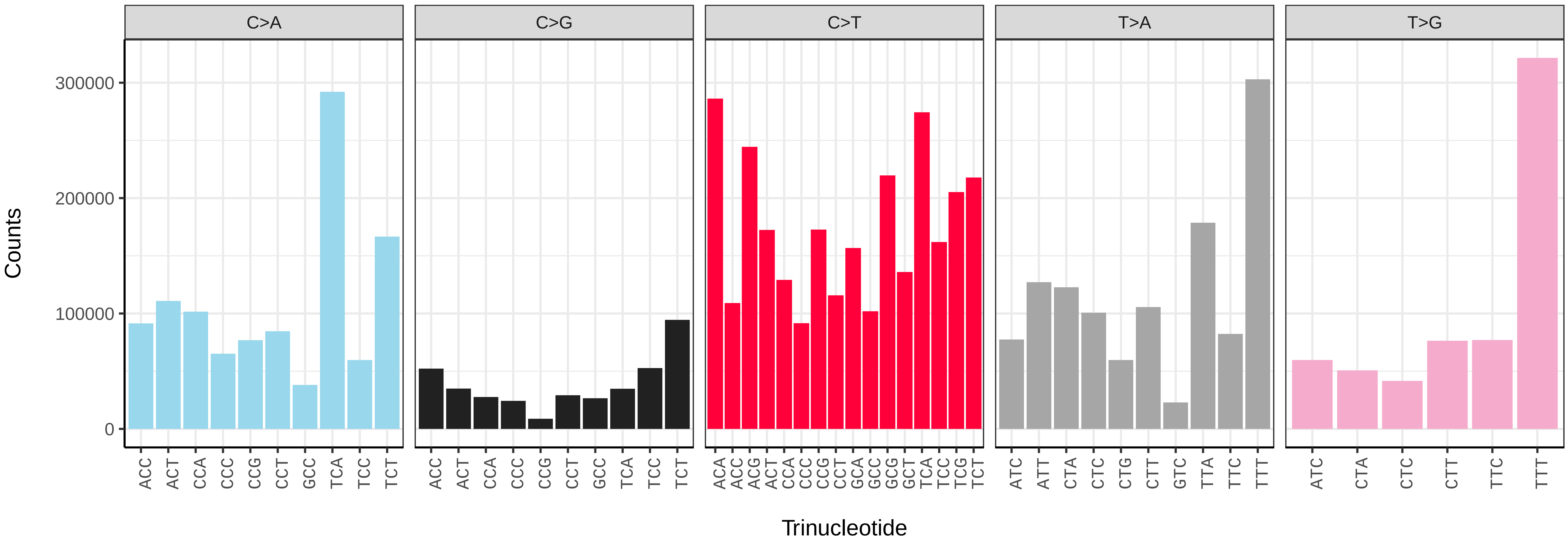


*Phoxinus phoxinus* (fPhoPho1) somatic mutational spectrum (SBS96 classification)

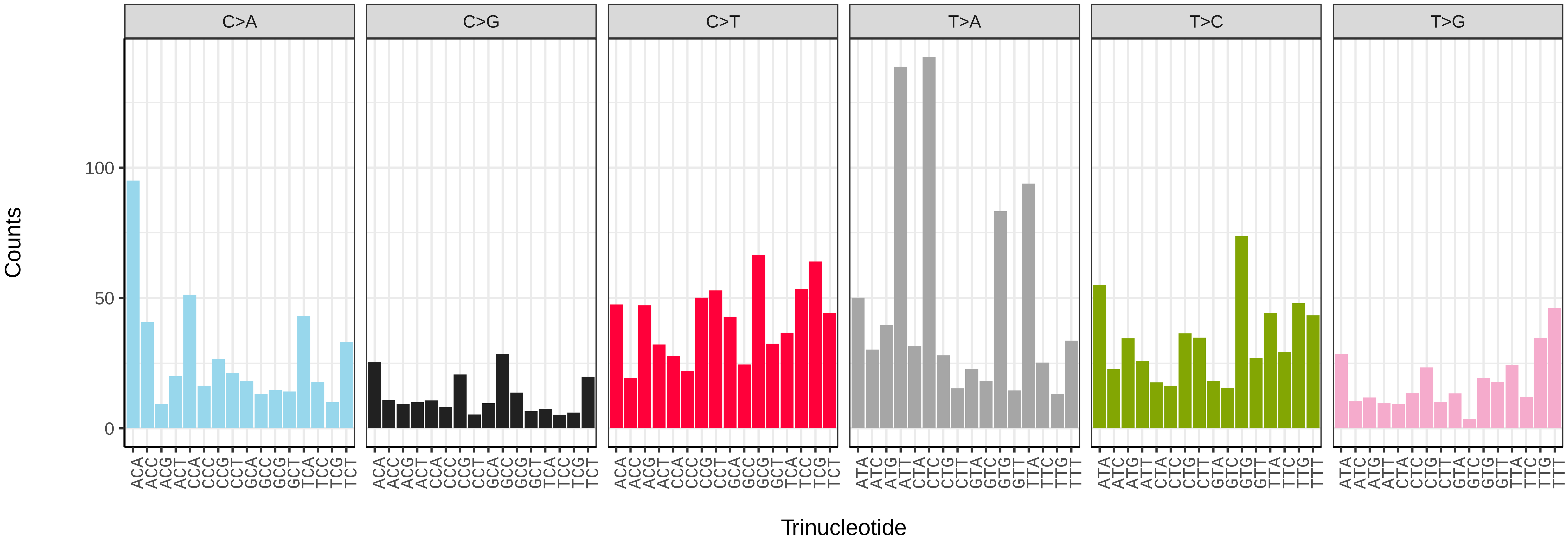


*Phoxinus phoxinus* (fPhoPho1) somatic mutational spectrum (SBS52 classification)

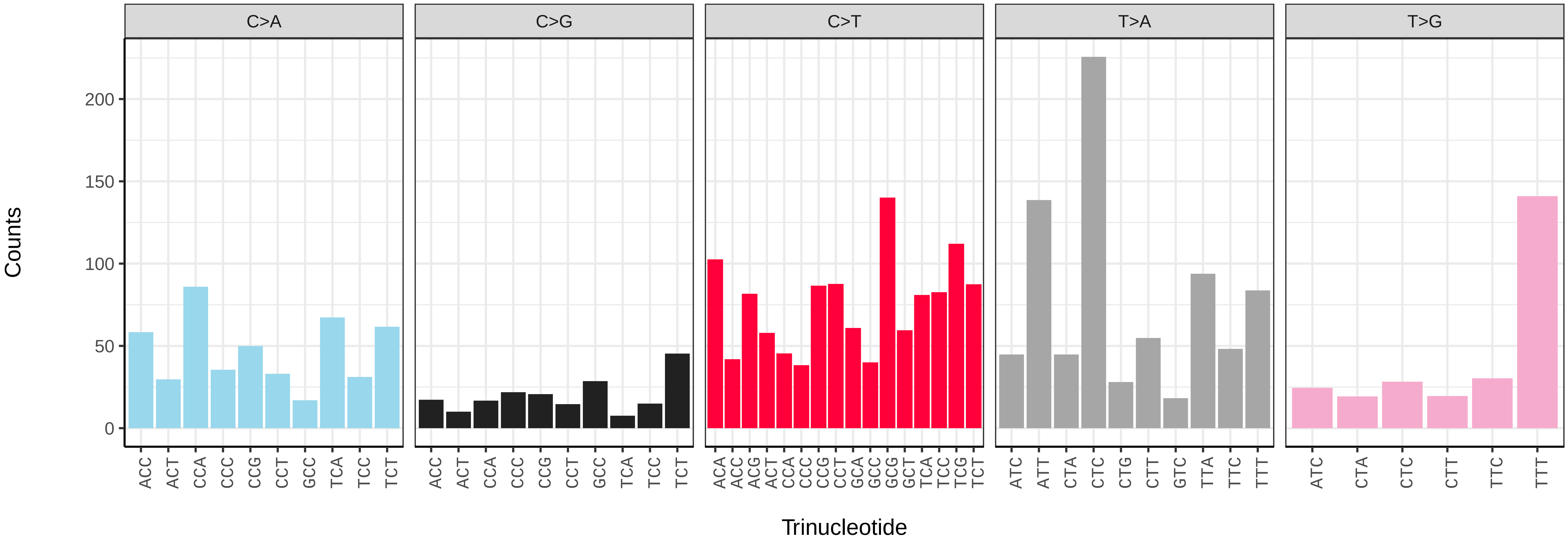

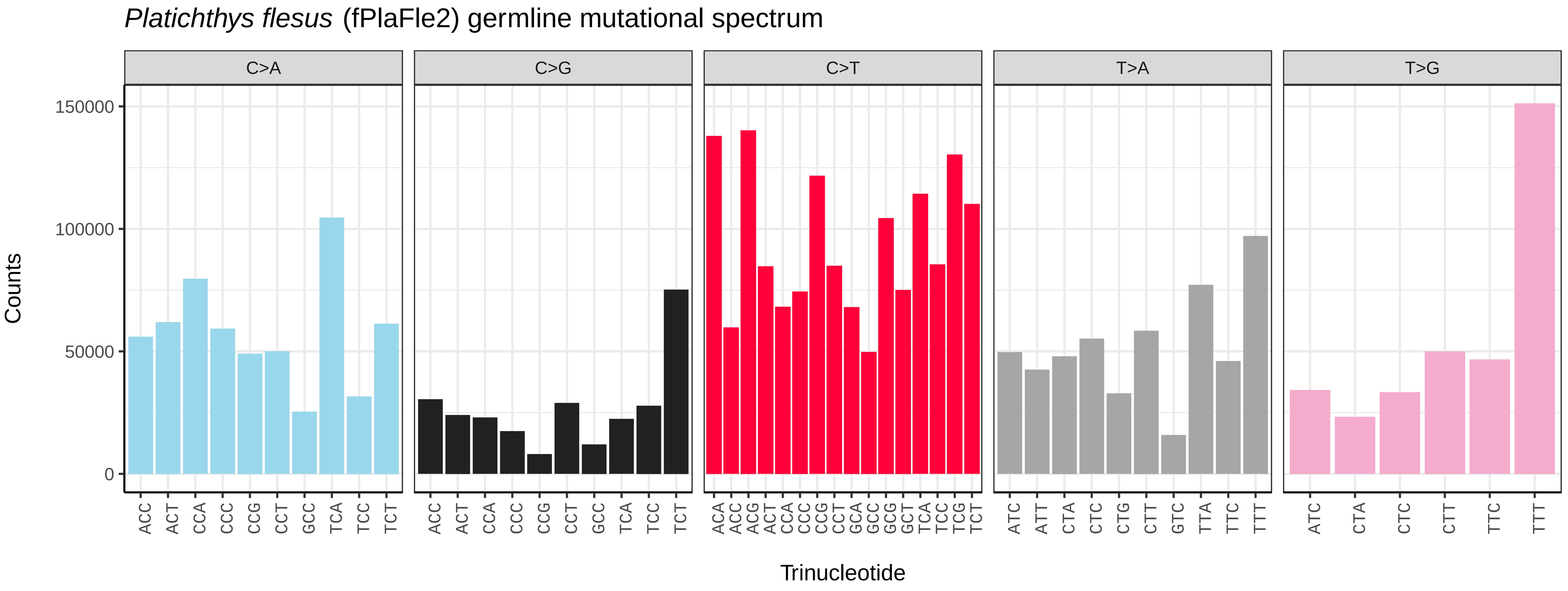
*Platichthys flesus* (fPlaFle2) germline mutational spectrum
C>A
C>G
C>T
T>A
T>G
Counts
150000
100000
50000
0
ACC ACT CCA CCC CCG CCT GCC TCA TCC TCT
ACA ACC ACG ACT CCA CCC CCG CCT GCA GCC GCG GCT TCA TCC TCG TCT
ATC ATT CTA CTC CTG CTT GTC TTA TTC TTT
ATC CTA CTC CTT TTC TTT
Trinucleotide

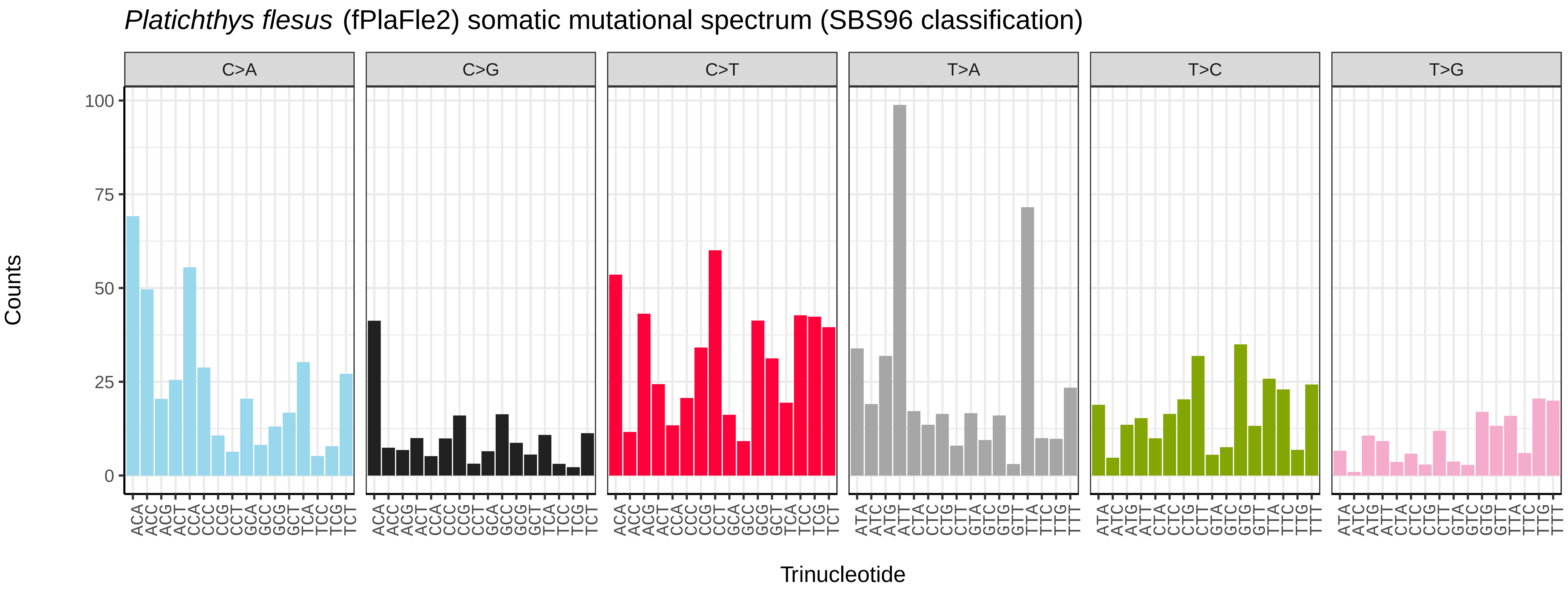
*Platichthys flesus* (fPlaFle2) somatic mutational spectrum (SBS96 classification)
C>A
C>G
C>T
T>A
T>C
T>G
Counts
100
75
50
25
0
ACA ACC ACG ACT CCA CCC CCG CCT GCA GCC GCG GCT TCA TCC TCG TCT
ATA ATC ATG ATT CTA CTC CTG CTT GTA GTC GTG GTT TTA TTC TTG TTT
Trinucleotide

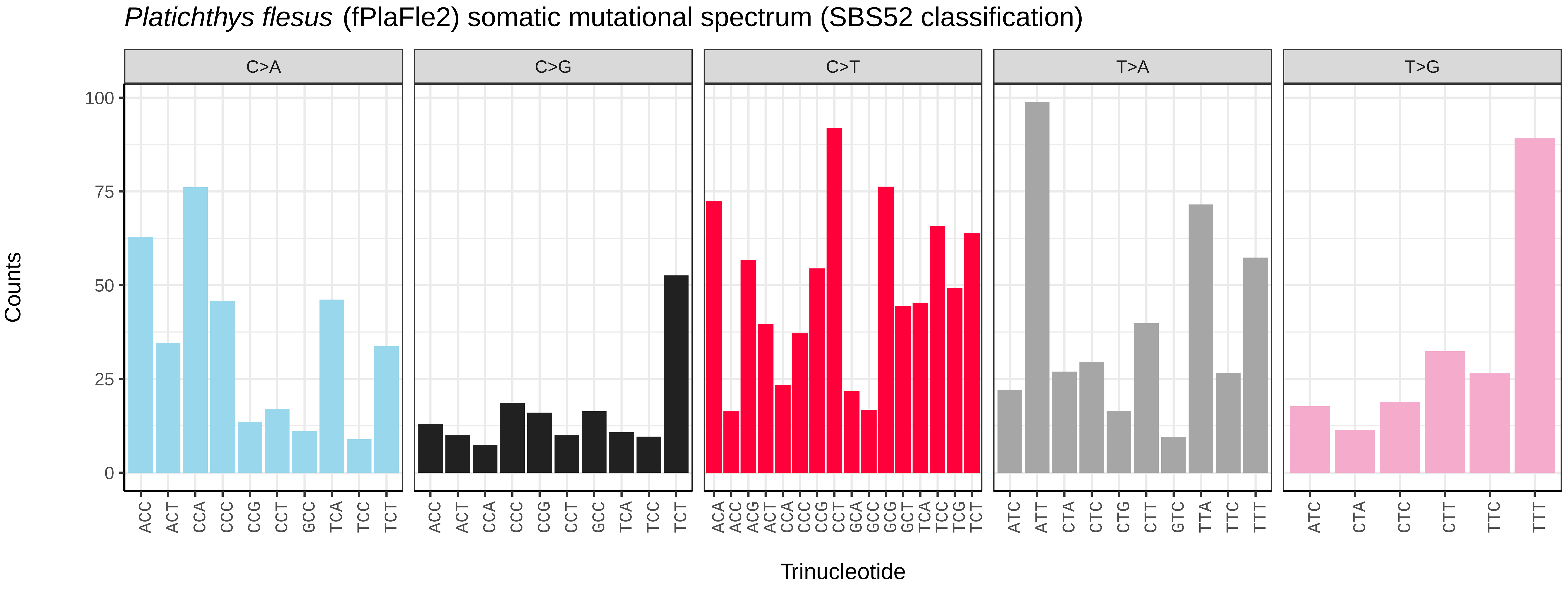
*Platichthys flesus* (fPlaFle2) somatic mutational spectrum (SBS52 classification)
C>A
C>G
C>T
T>A
T>G
Counts
100
75
50
25
0
ACC ACT CCA CCC CCG CCT GCC TCA TCC TCT
ACA ACC ACG ACT CCA CCC CCG CCT GCA GCC GCG GCT TCA TCC TCG TCT
ATC ATT CTA CTC CTG CTT GTC TTA TTC TTT
ATC CTA CTC CTT TTC TTT
Trinucleotide

*Pleuronectes platessa* (fPlePla1) germline mutational spectrum

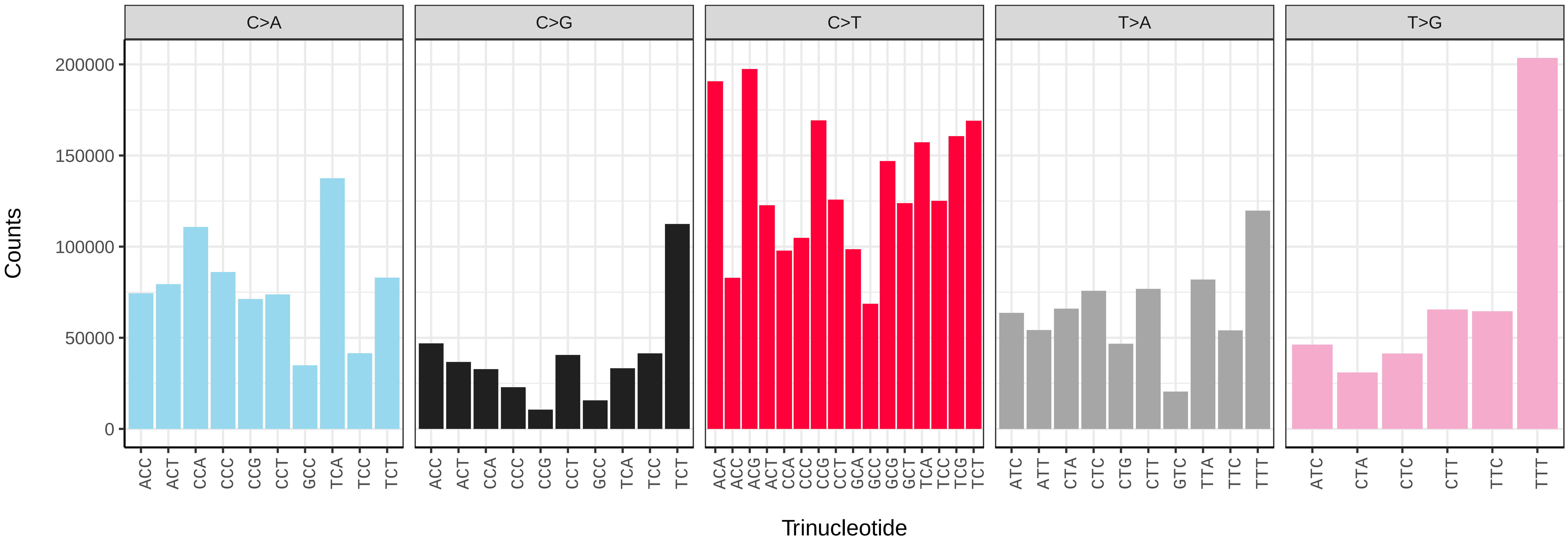


*Pleuronectes platessa* (fPlePla1) somatic mutational spectrum (SBS96 classification)

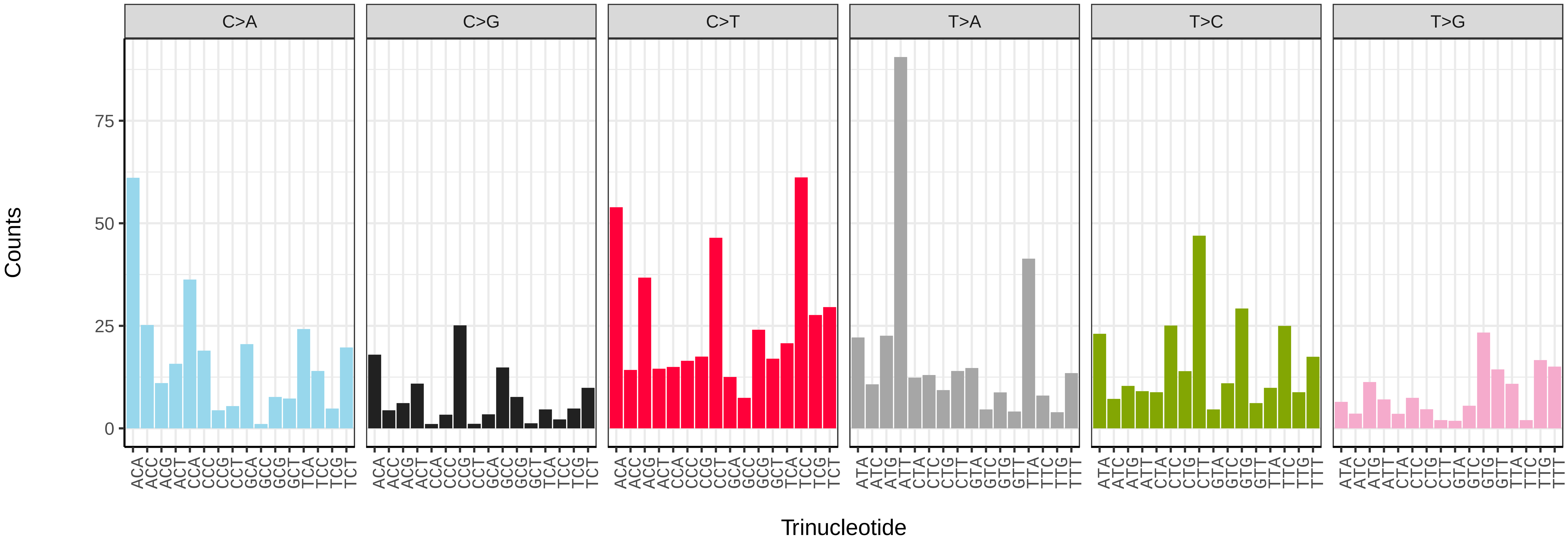


*Pleuronectes platessa* (fPlePla1) somatic mutational spectrum (SBS52 classification)

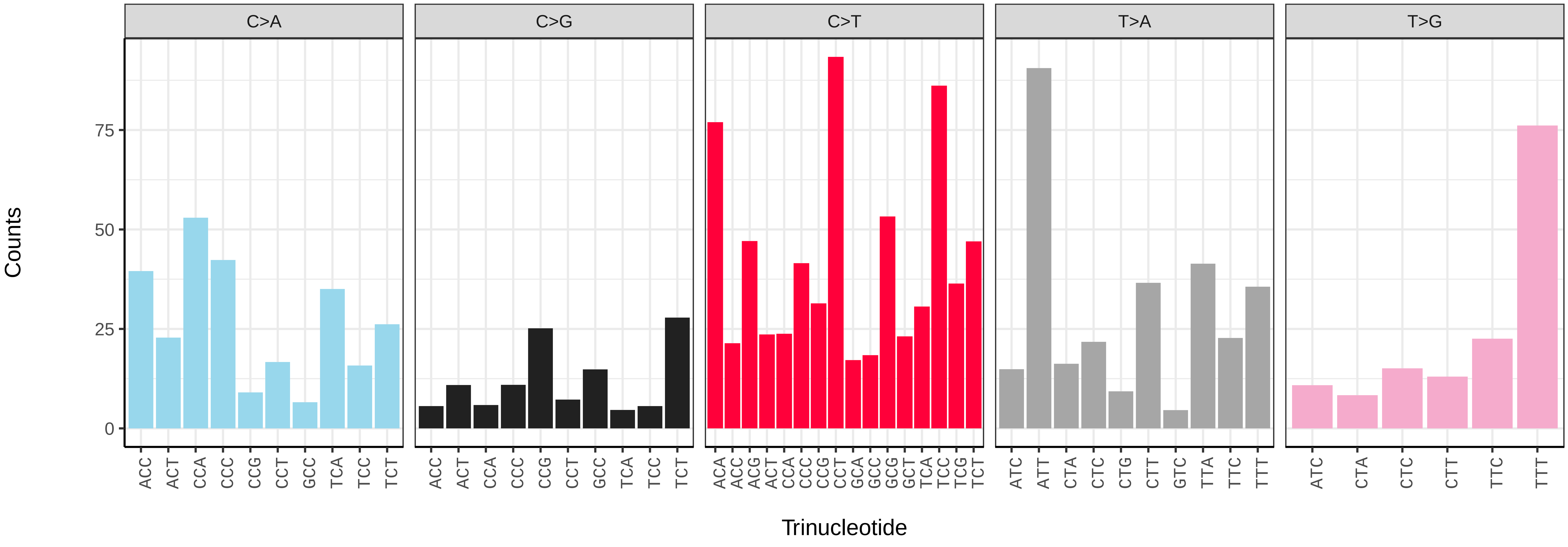

*Pollachius pollachius* (fPolPol2) germline mutational spectrum

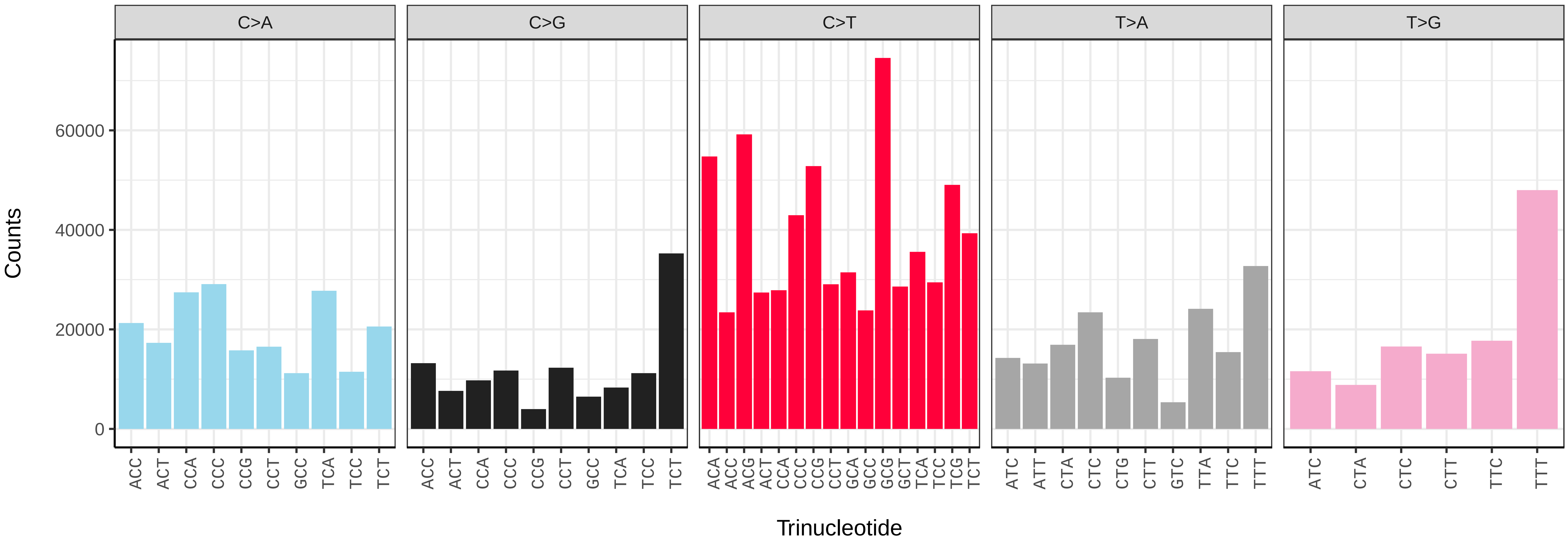


*Pollachius pollachius* (fPolPol2) somatic mutational spectrum (SBS96 classification)

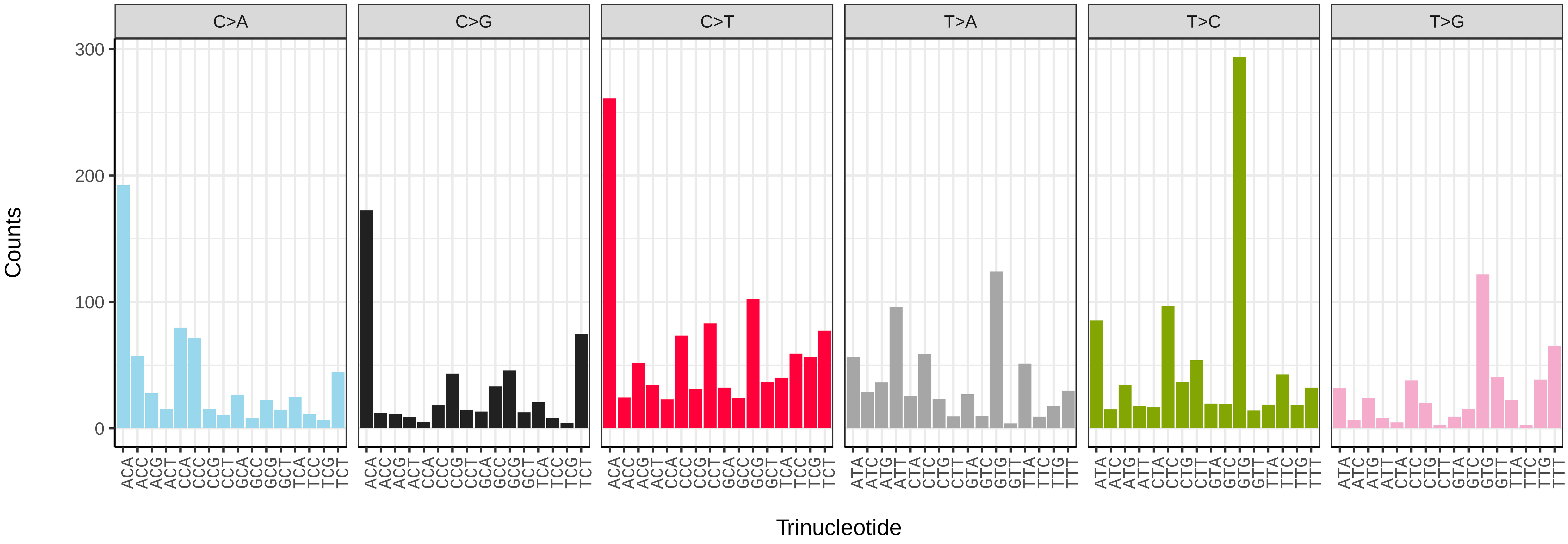


*Pollachius pollachius* (fPolPol2) somatic mutational spectrum (SBS52 classification)

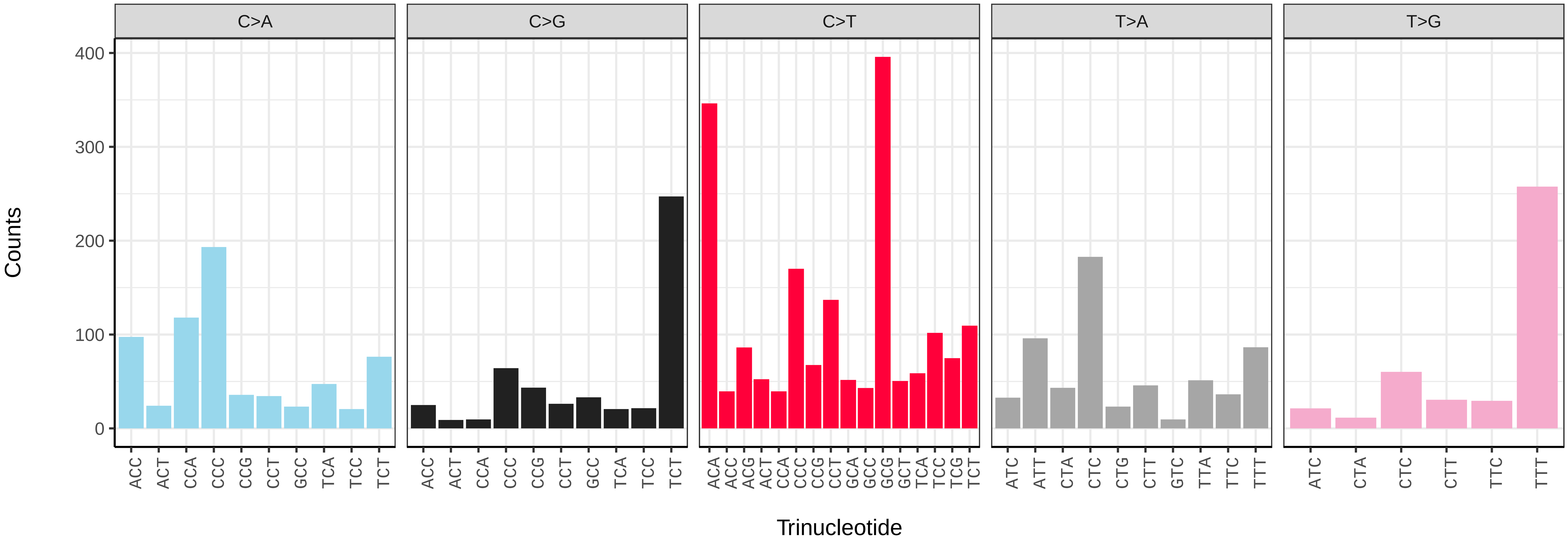

*Pungitius pungitius* (fPunPun2) germline mutational spectrum

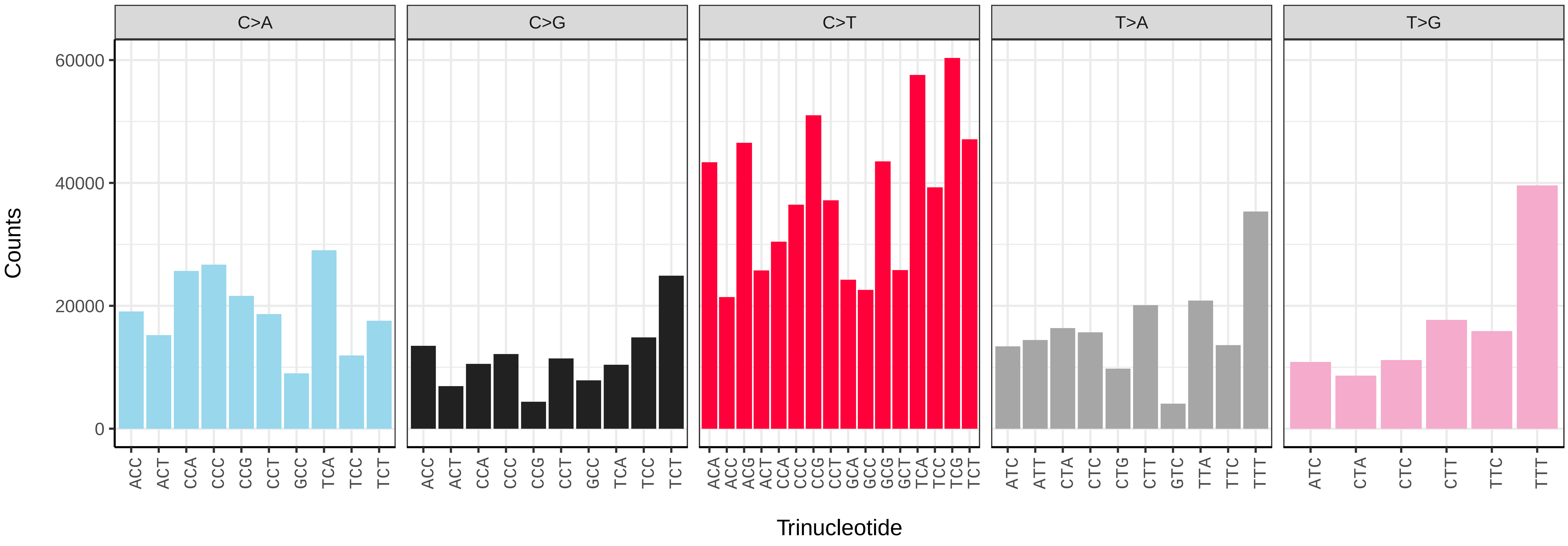


*Pungitius pungitius* (fPunPun2) somatic mutational spectrum (SBS96 classification)

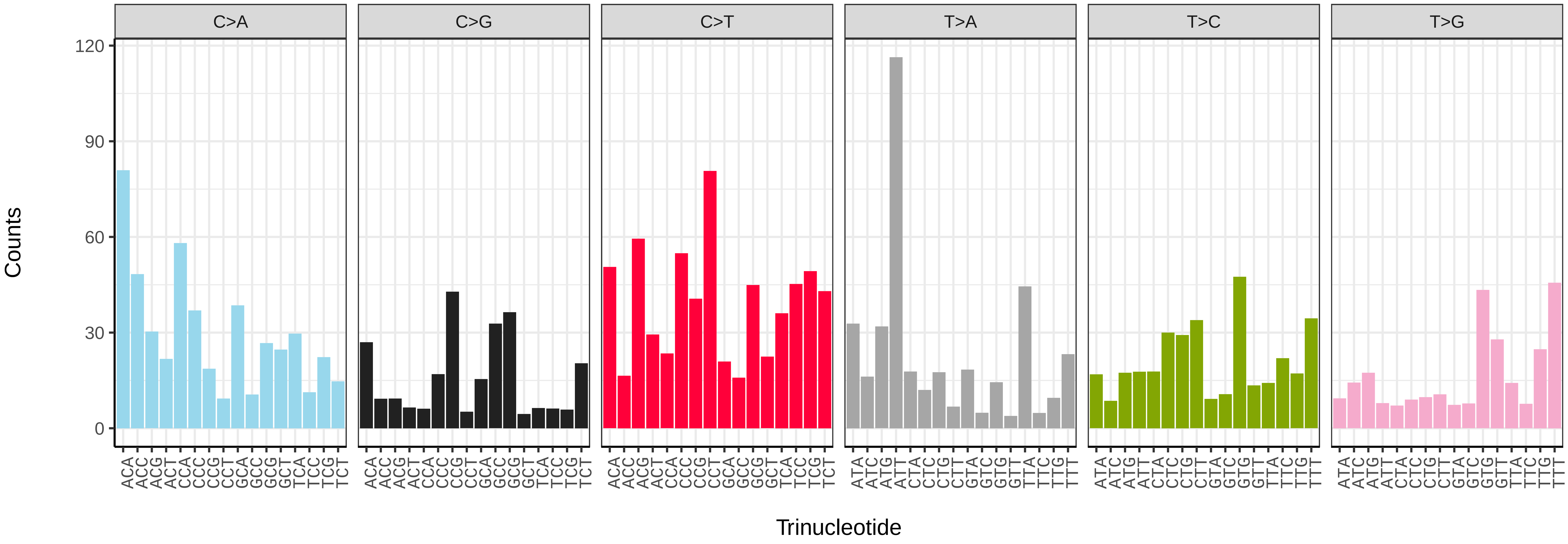


*Pungitius pungitius* (fPunPun2) somatic mutational spectrum (SBS52 classification)

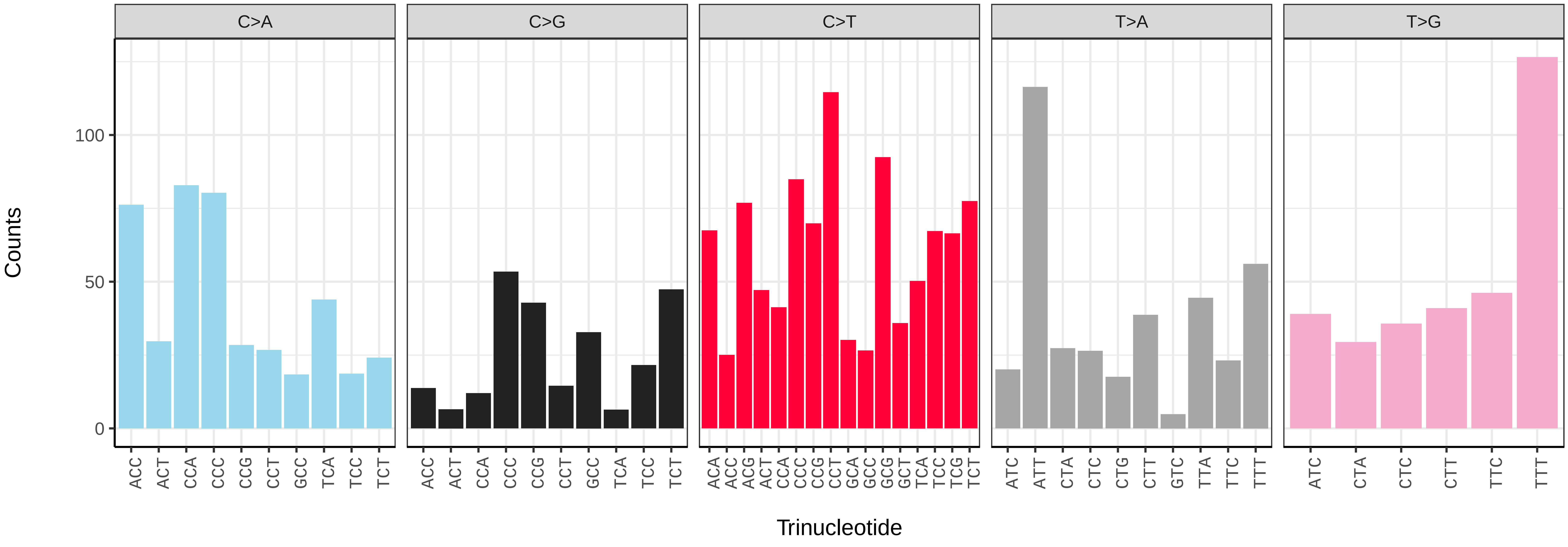

*Rutilus rutilus* (fRutRut2) germline mutational spectrum

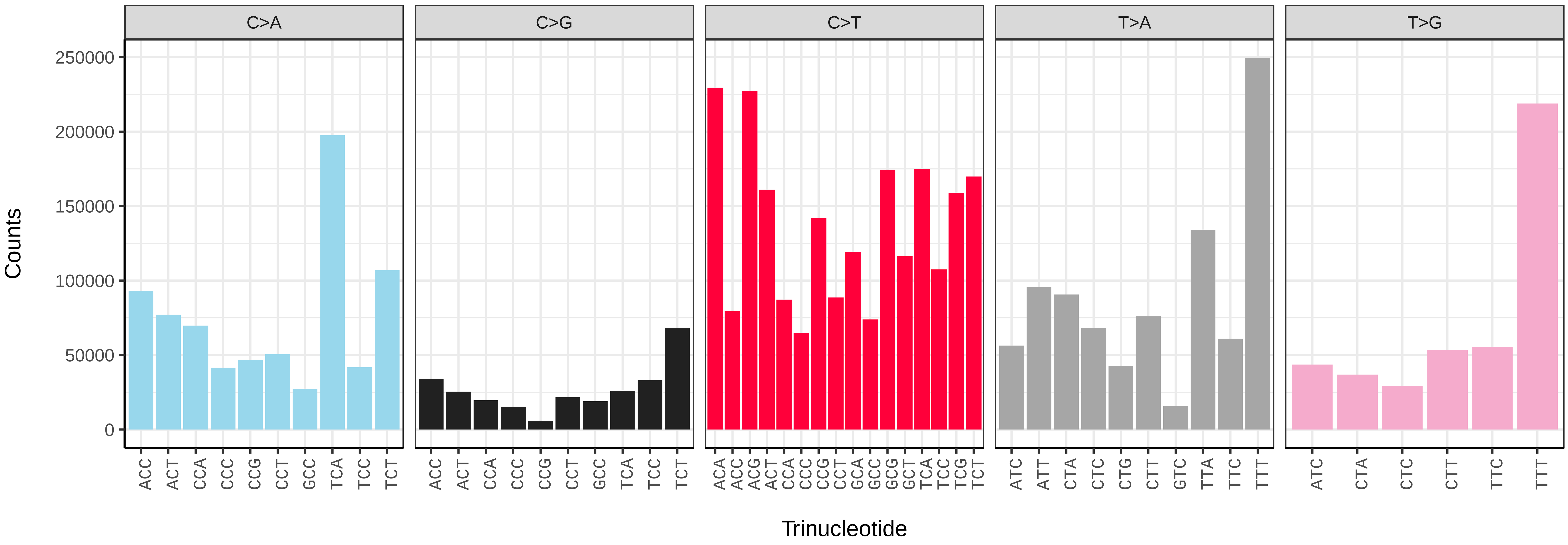


*Rutilus rutilus* (fRutRut2) somatic mutational spectrum (SBS96 classification)

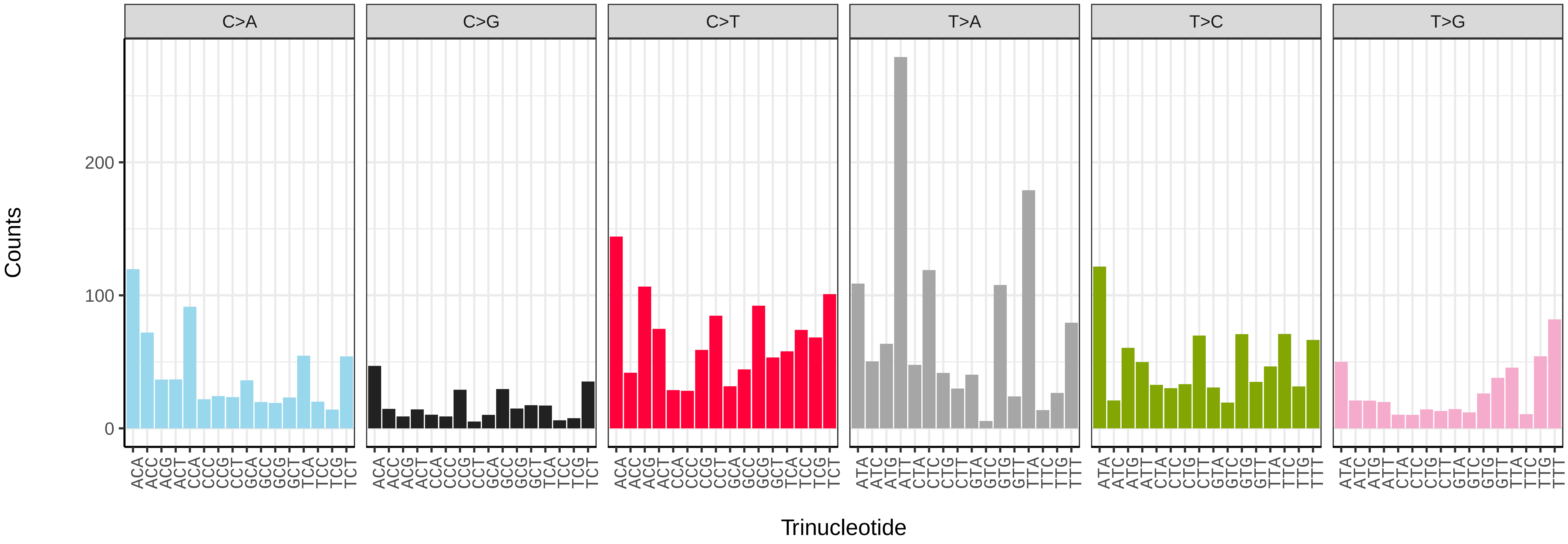


*Rutilus rutilus* (fRutRut2) somatic mutational spectrum (SBS52 classification)

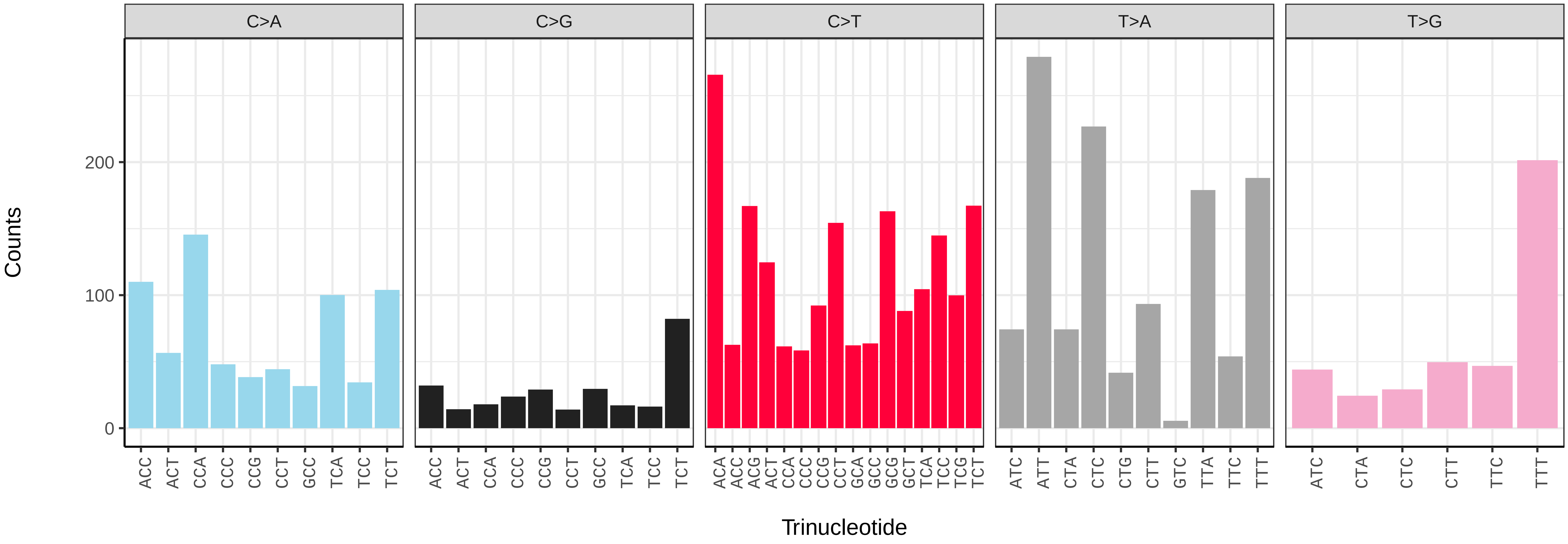

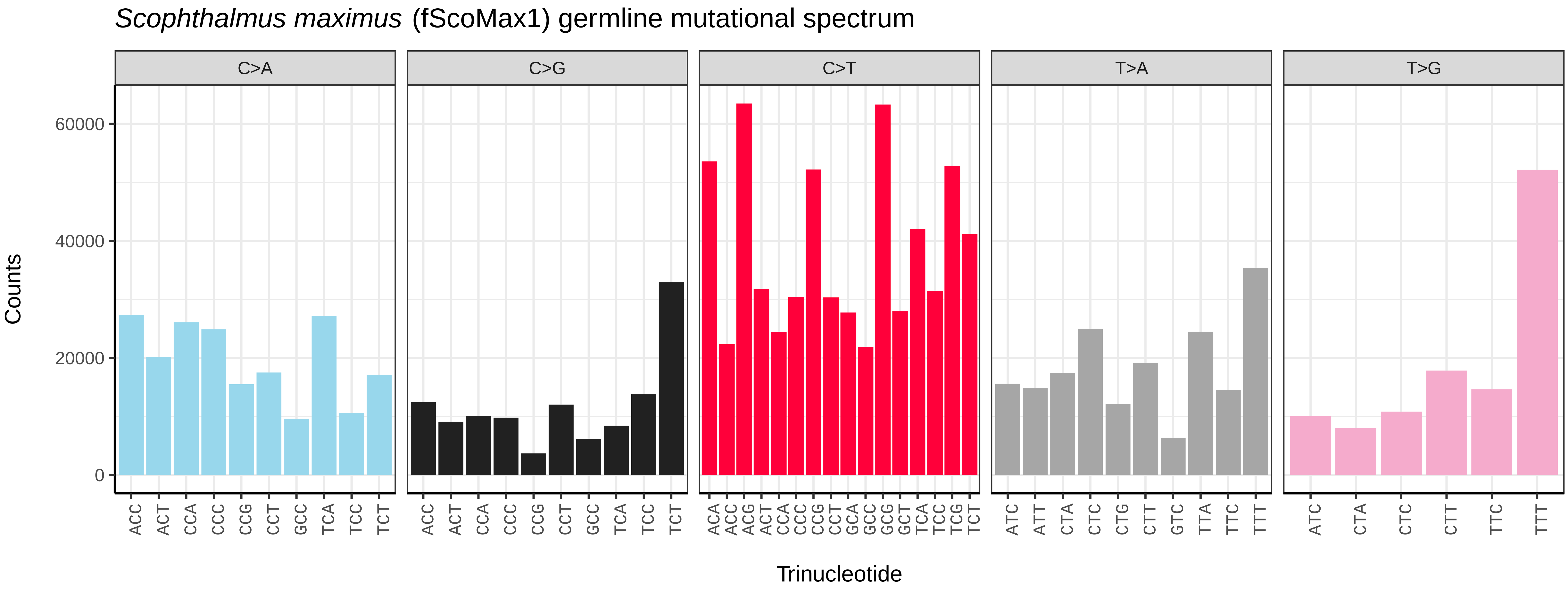
*Scophthalmus maximus* (fScoMax1) germline mutational spectrum
C>A
C>G
C>T
T>A
T>G
Counts
0
20000
40000
60000
Trinucleotide

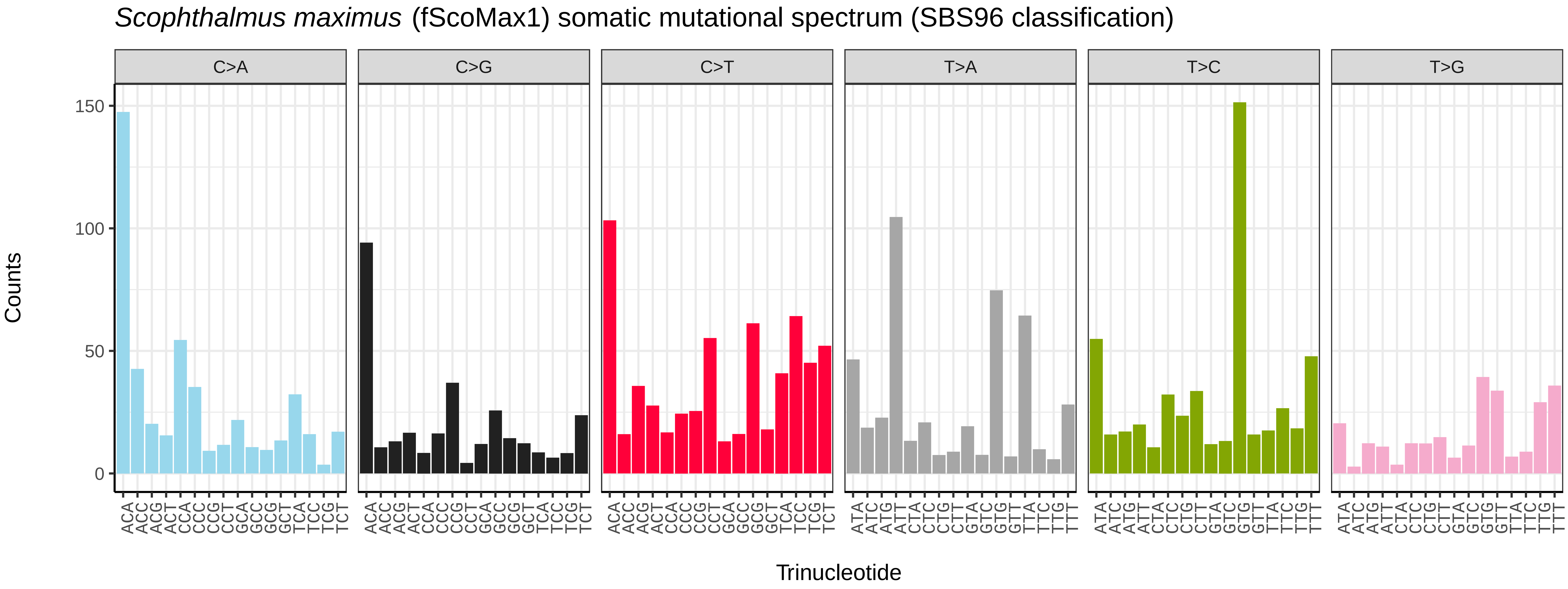
*Scophthalmus maximus* (fScoMax1) somatic mutational spectrum (SBS96 classification)
C>A
C>G
C>T
T>A
T>C
T>G
Counts
0
50
100
150
Trinucleotide

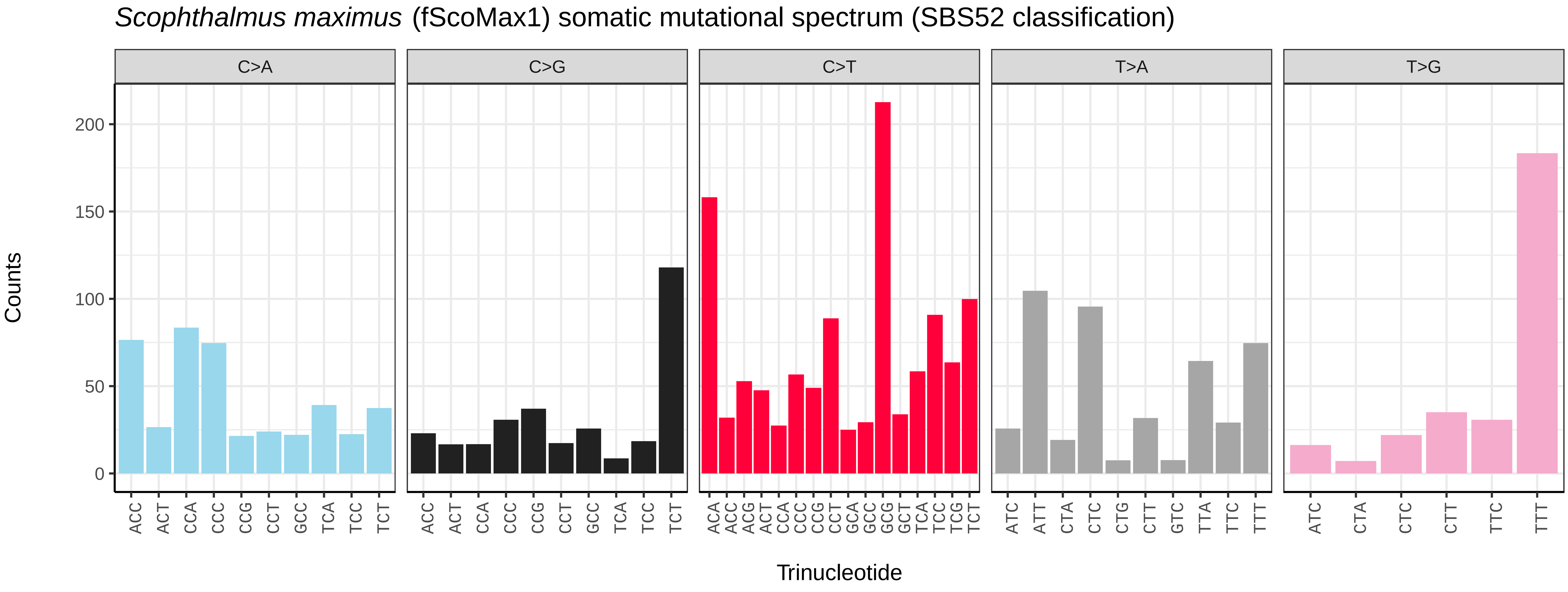
*Scophthalmus maximus* (fScoMax1) somatic mutational spectrum (SBS52 classification)
C>A
C>G
C>T
T>A
T>G
Counts
0
50
100
150
200
Trinucleotide

*Scomber scombrus* (fScoSco1) germline mutational spectrum

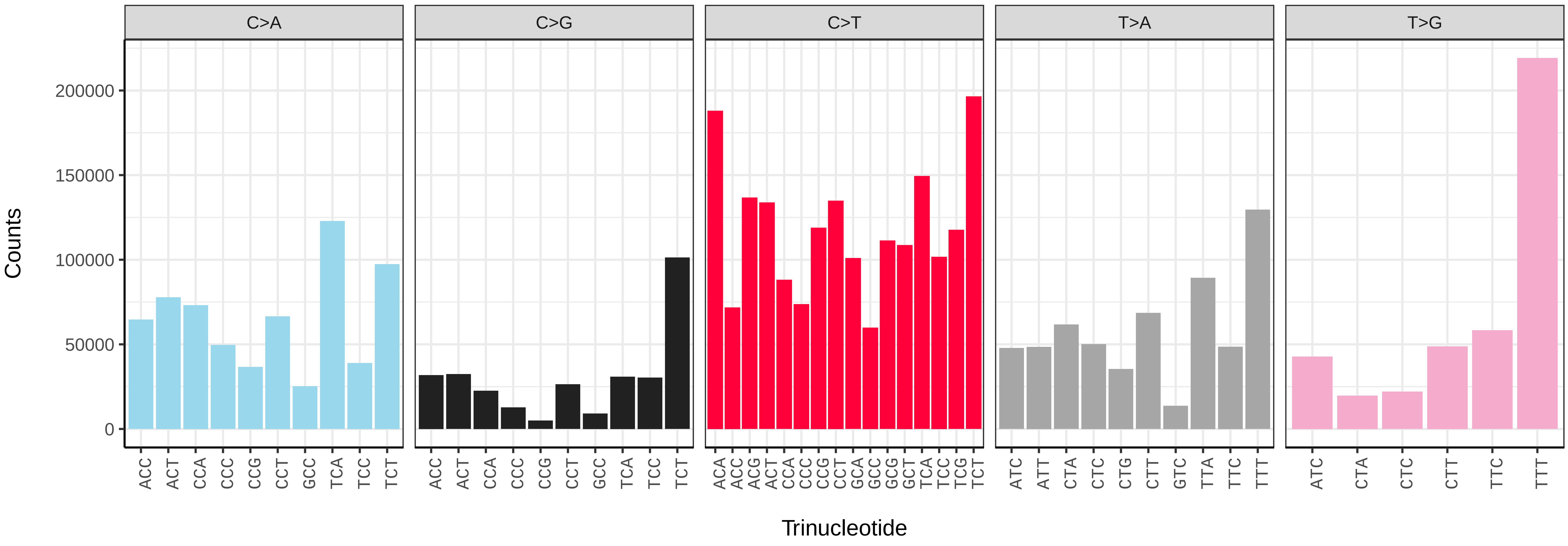


*Scomber scombrus* (fScoSco1) somatic mutational spectrum (SBS96 classification)

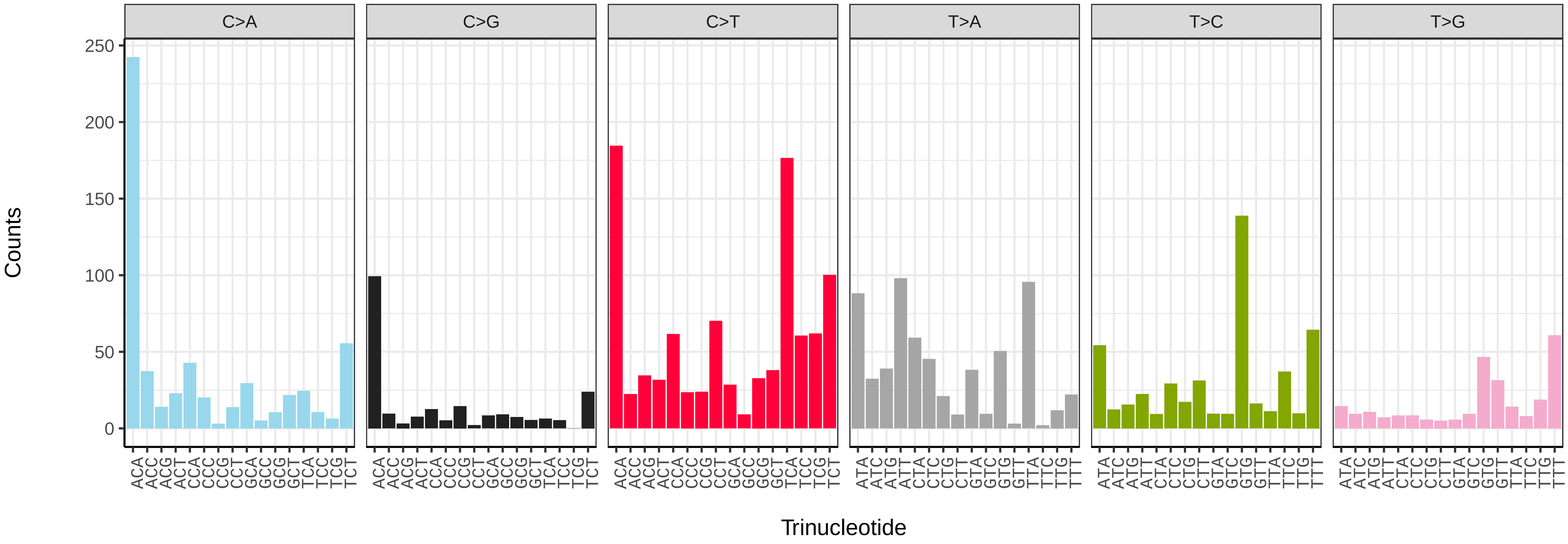


*Scomber scombrus* (fScoSco1) somatic mutational spectrum (SBS52 classification)

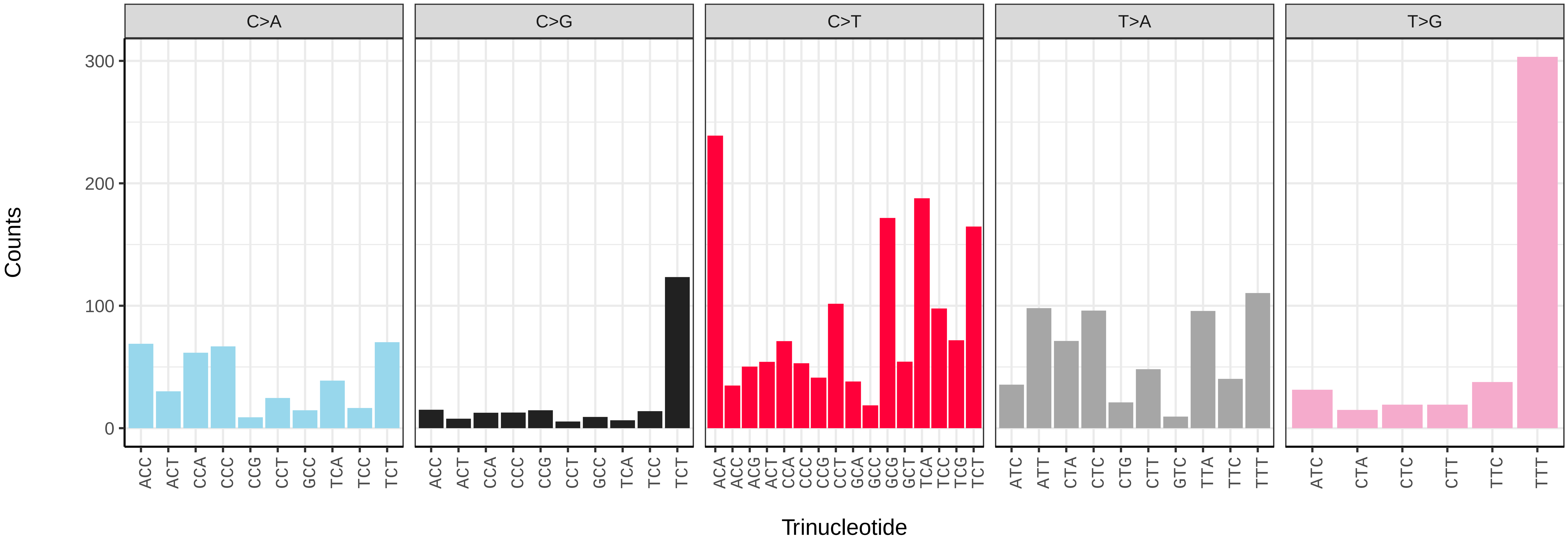

*Sprattus sprattus* (fSprSpr1) germline mutational spectrum

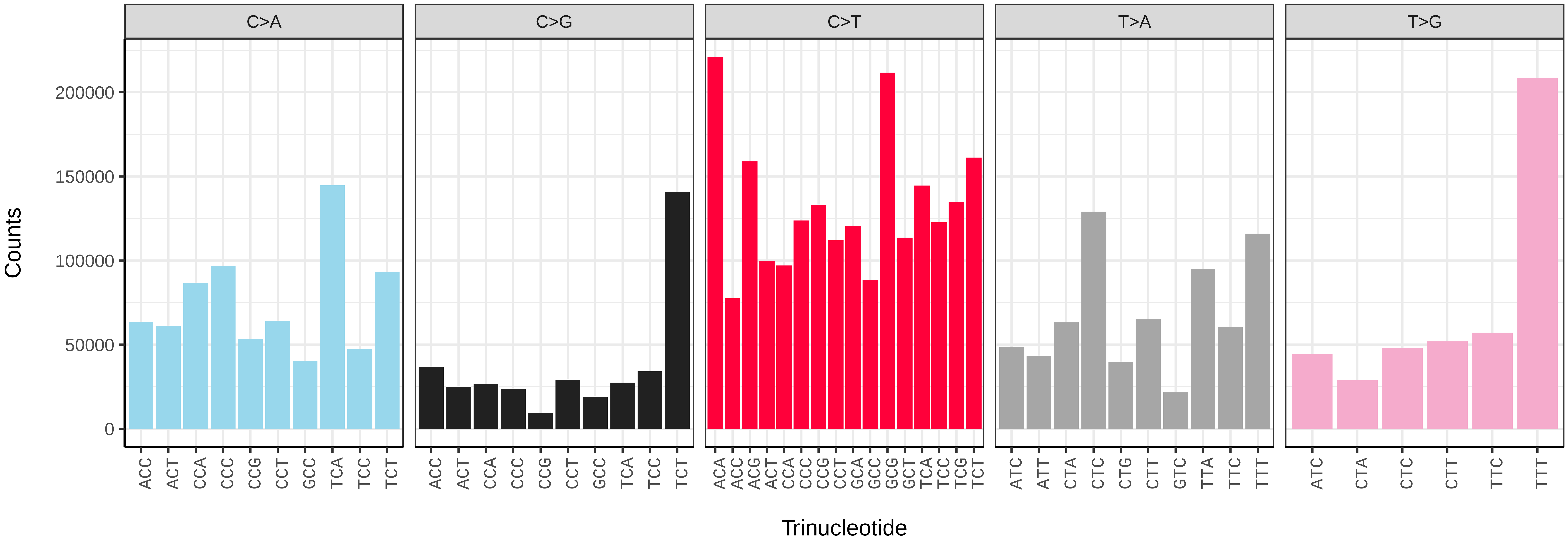


*Sprattus sprattus* (fSprSpr1) somatic mutational spectrum (SBS96 classification)

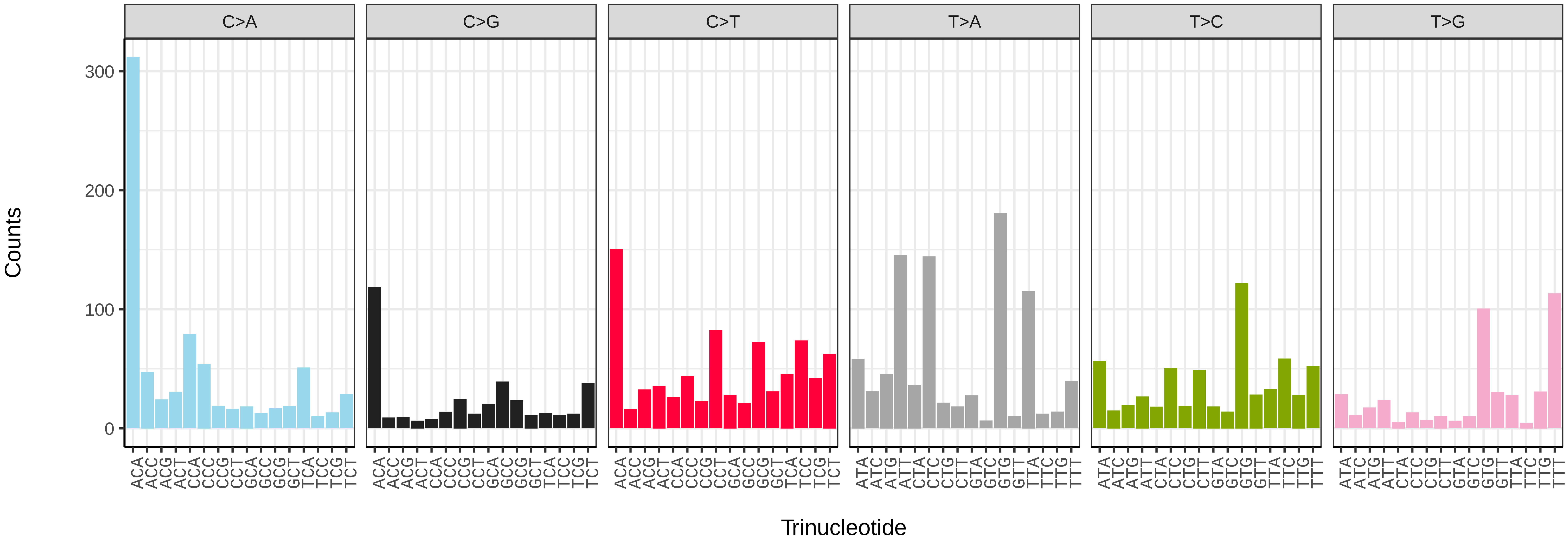


*Sprattus sprattus* (fSprSpr1) somatic mutational spectrum (SBS52 classification)

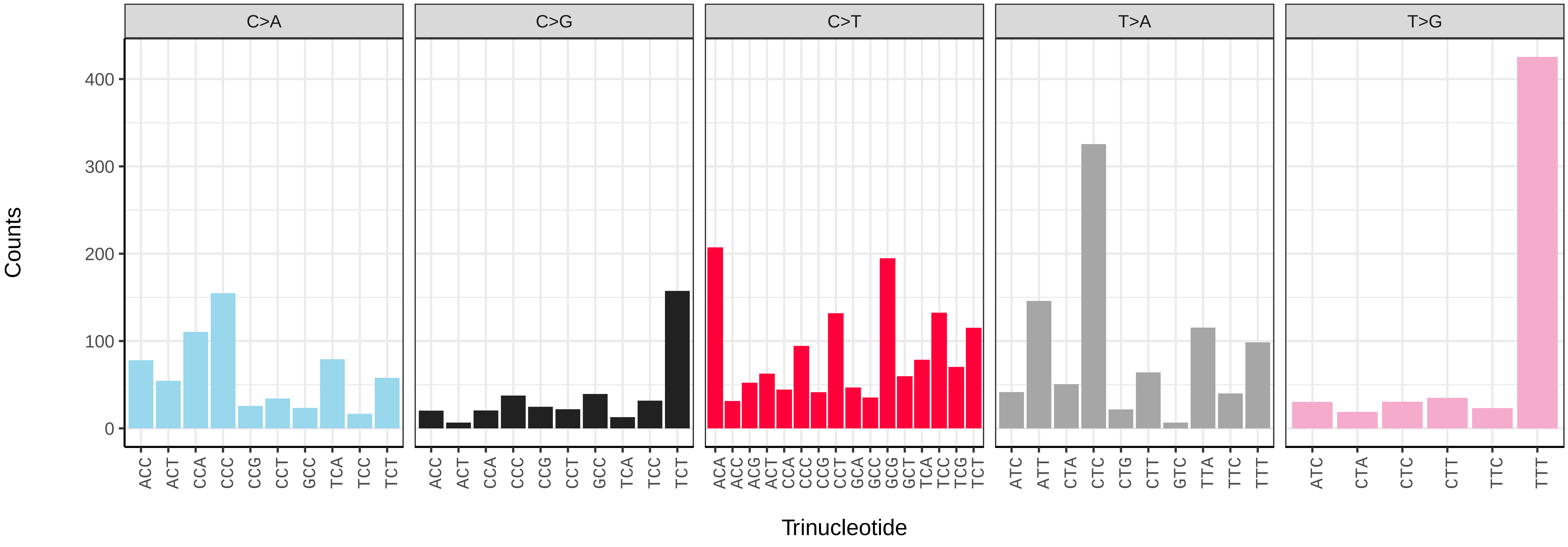

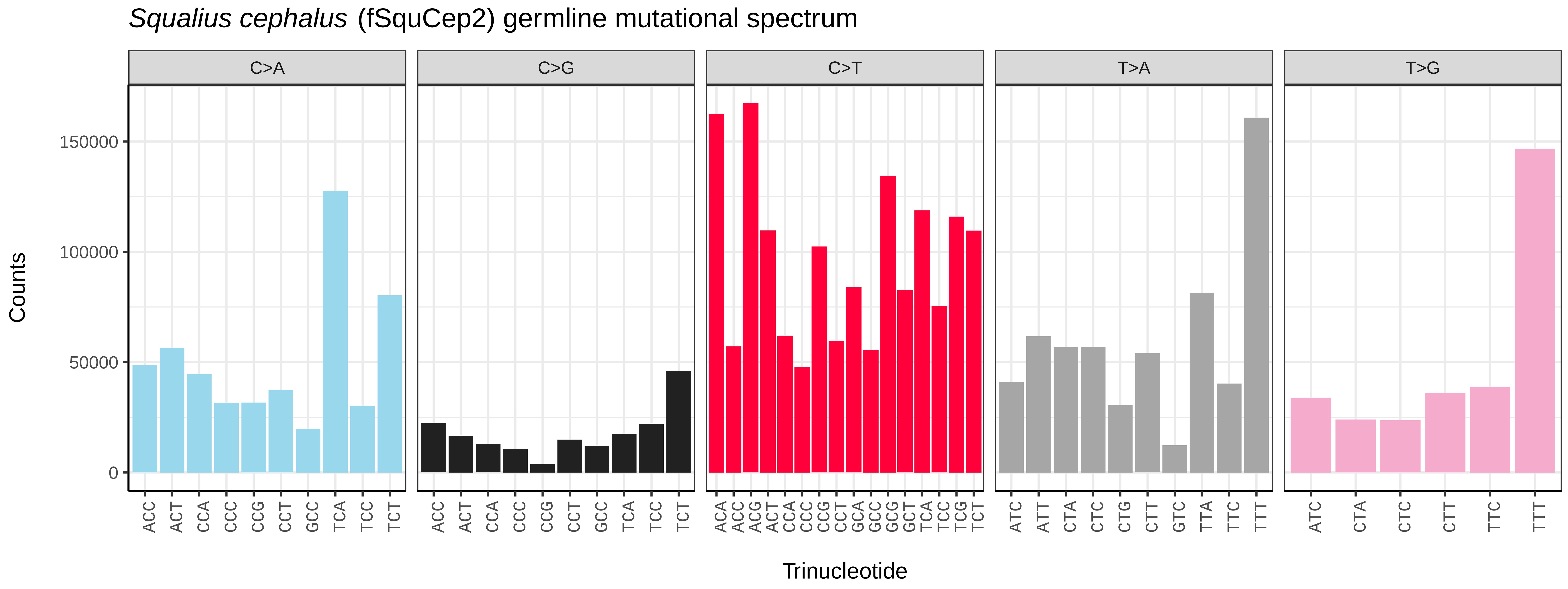
*Squalius cephalus* (fSquCep2) germline mutational spectrum
C>A
C>G
C>T
T>A
T>G
Counts
0
50000
100000
150000
Trinucleotide

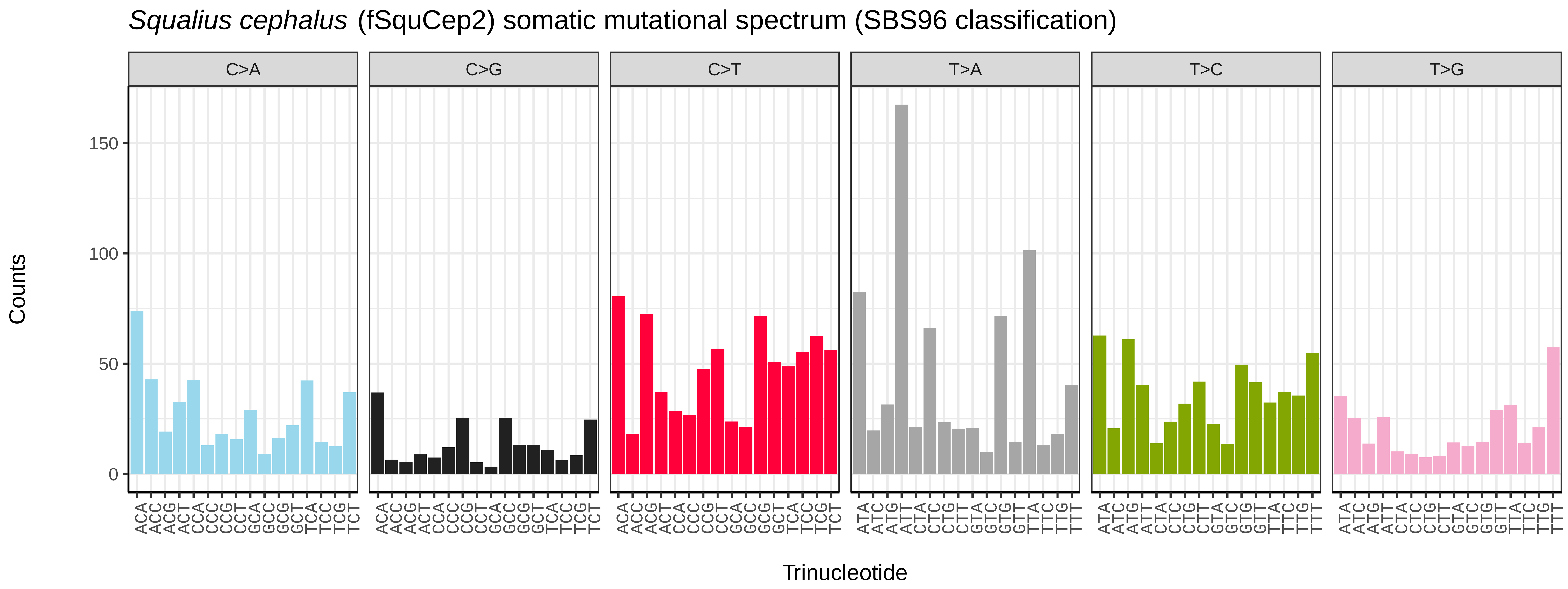
*Squalius cephalus* (fSquCep2) somatic mutational spectrum (SBS96 classification)
C>A
C>G
C>T
T>A
T>C
T>G
Counts
0
50
100
150
Trinucleotide

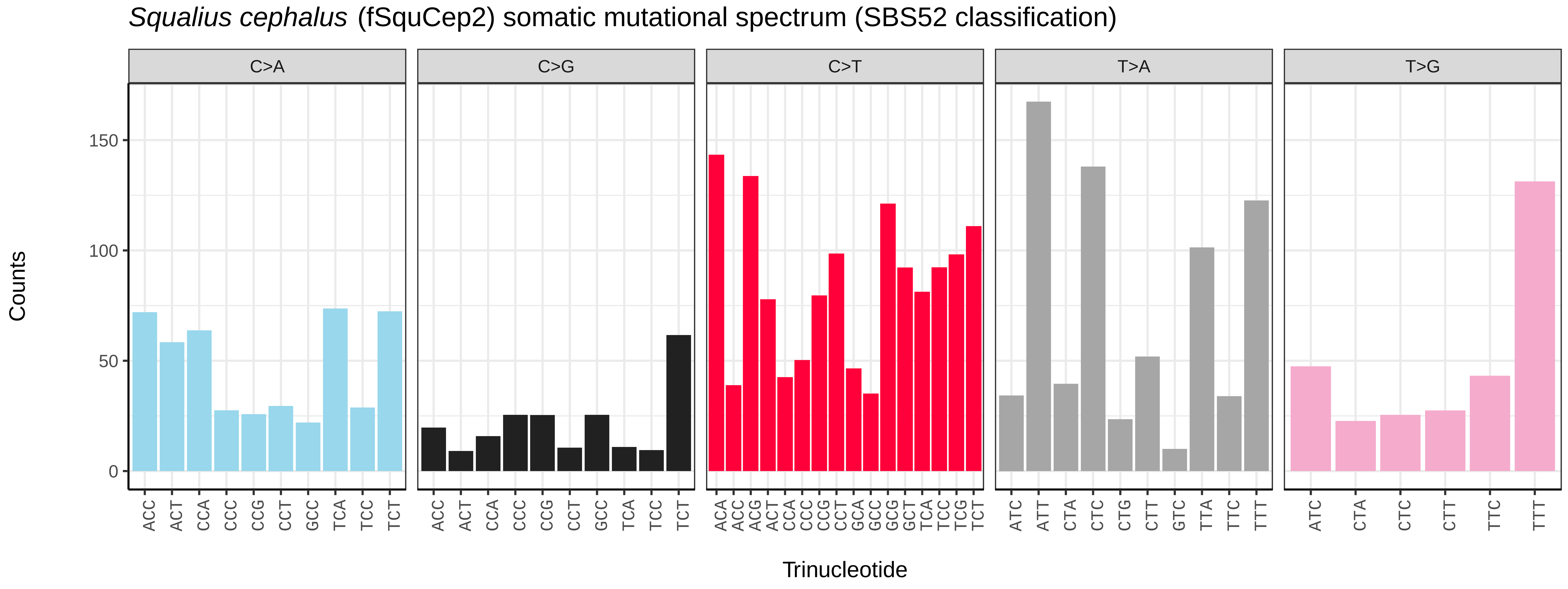
*Squalius cephalus* (fSquCep2) somatic mutational spectrum (SBS52 classification)
C>A
C>G
C>T
T>A
T>G
Counts
0
50
100
150
Trinucleotide

*Symphodus melops* (fSymMel2) germline mutational spectrum

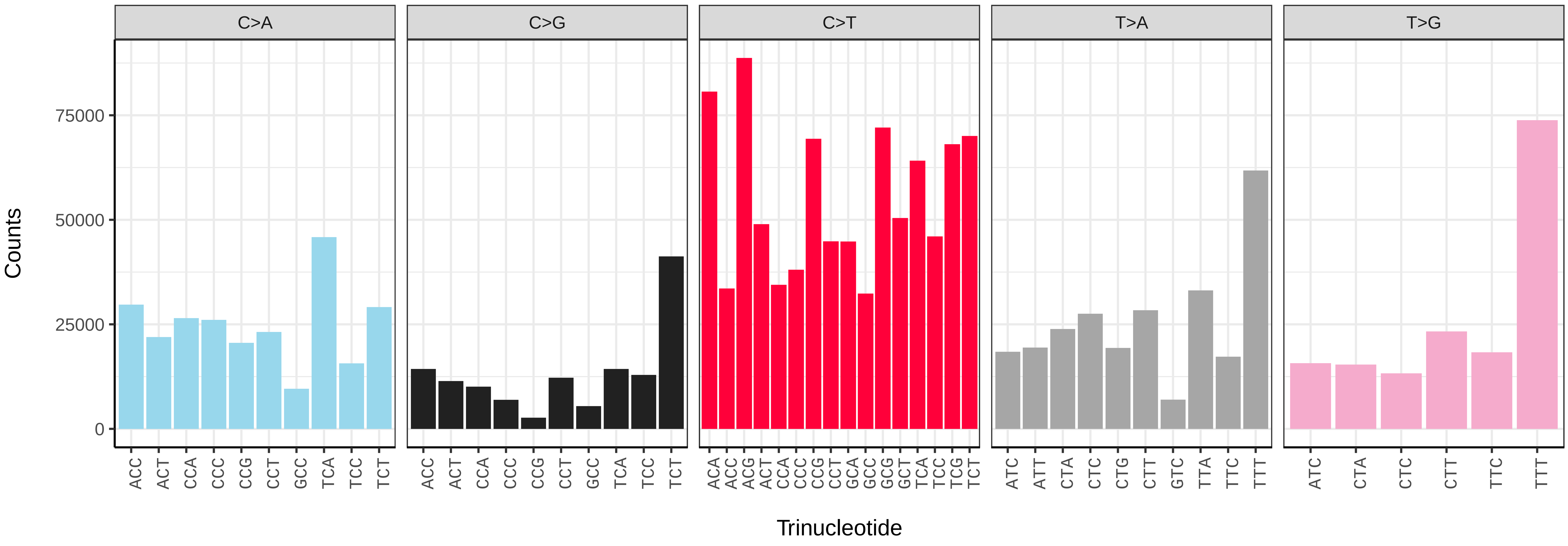


*Symphodus melops* (fSymMel2) somatic mutational spectrum (SBS96 classification)

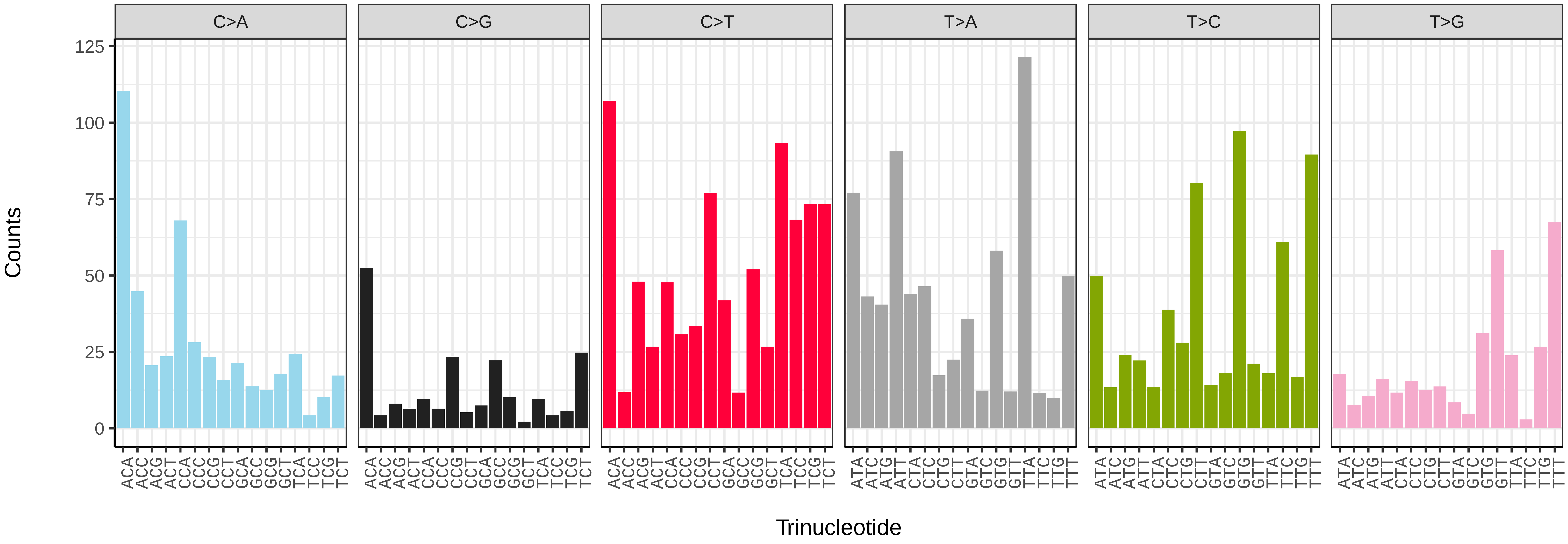


*Symphodus melops* (fSymMel2) somatic mutational spectrum (SBS52 classification)

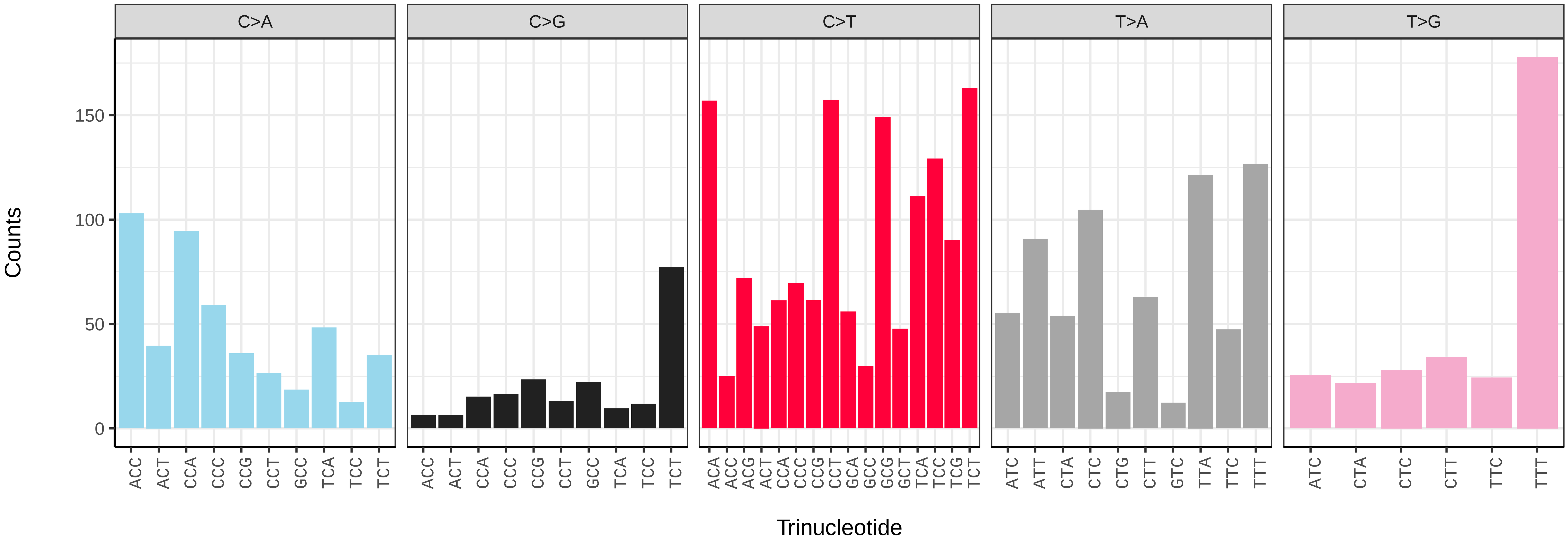

*Syngnathus acus* (fSynAcu2) germline mutational spectrum

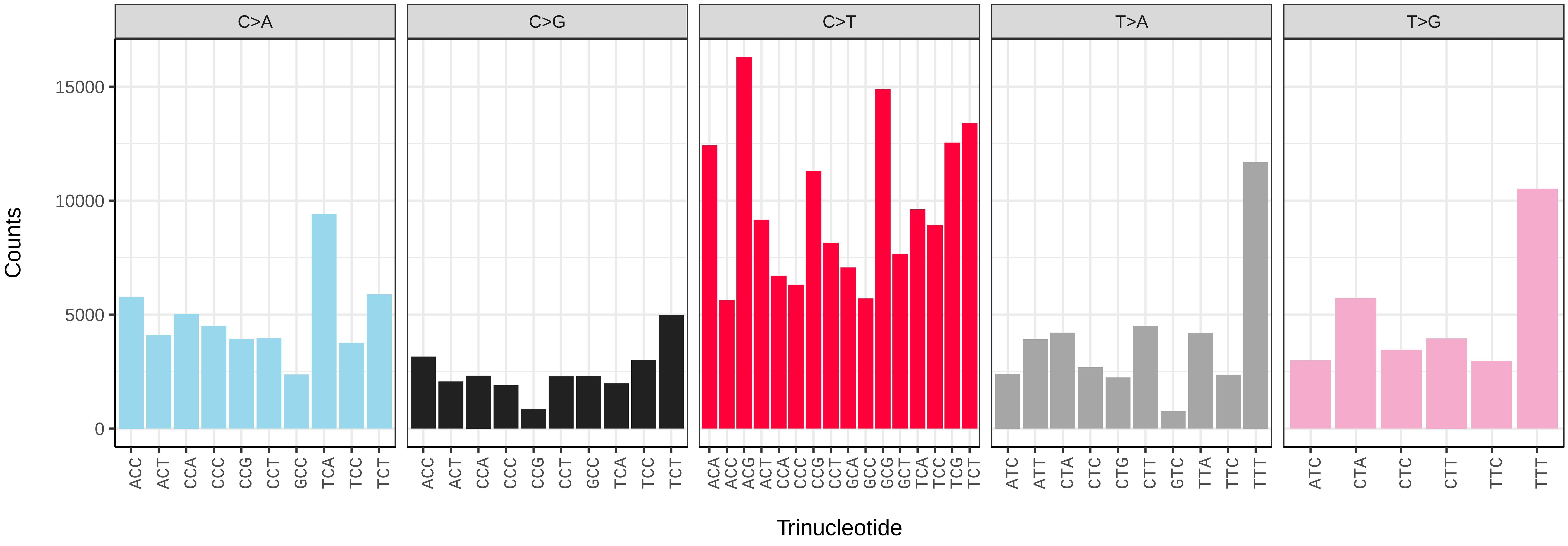


*Syngnathus acus* (fSynAcu2) somatic mutational spectrum (SBS96 classification)

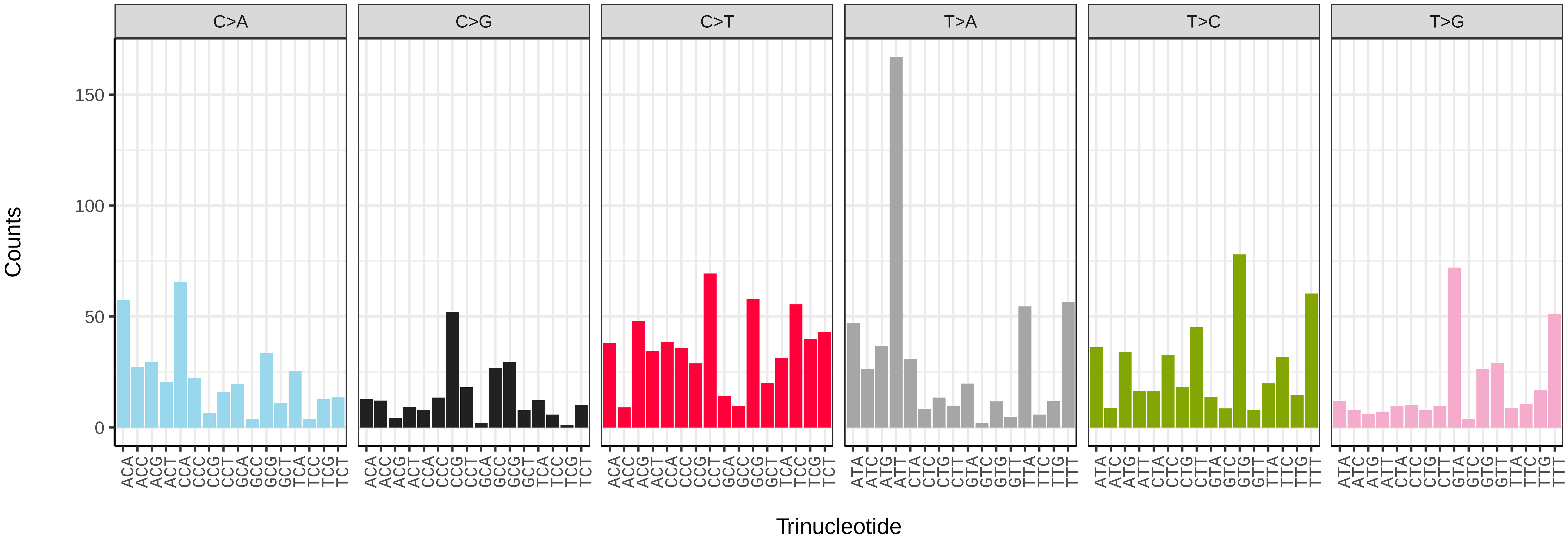


*Syngnathus acus* (fSynAcu2) somatic mutational spectrum (SBS52 classification)

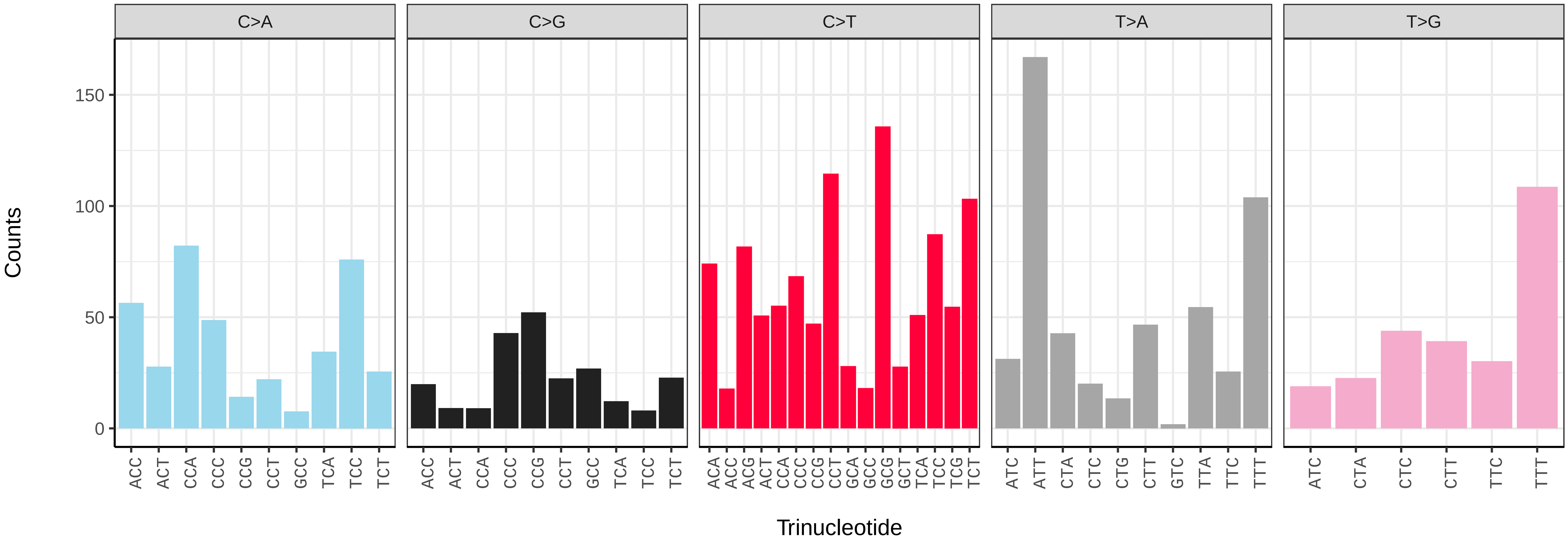

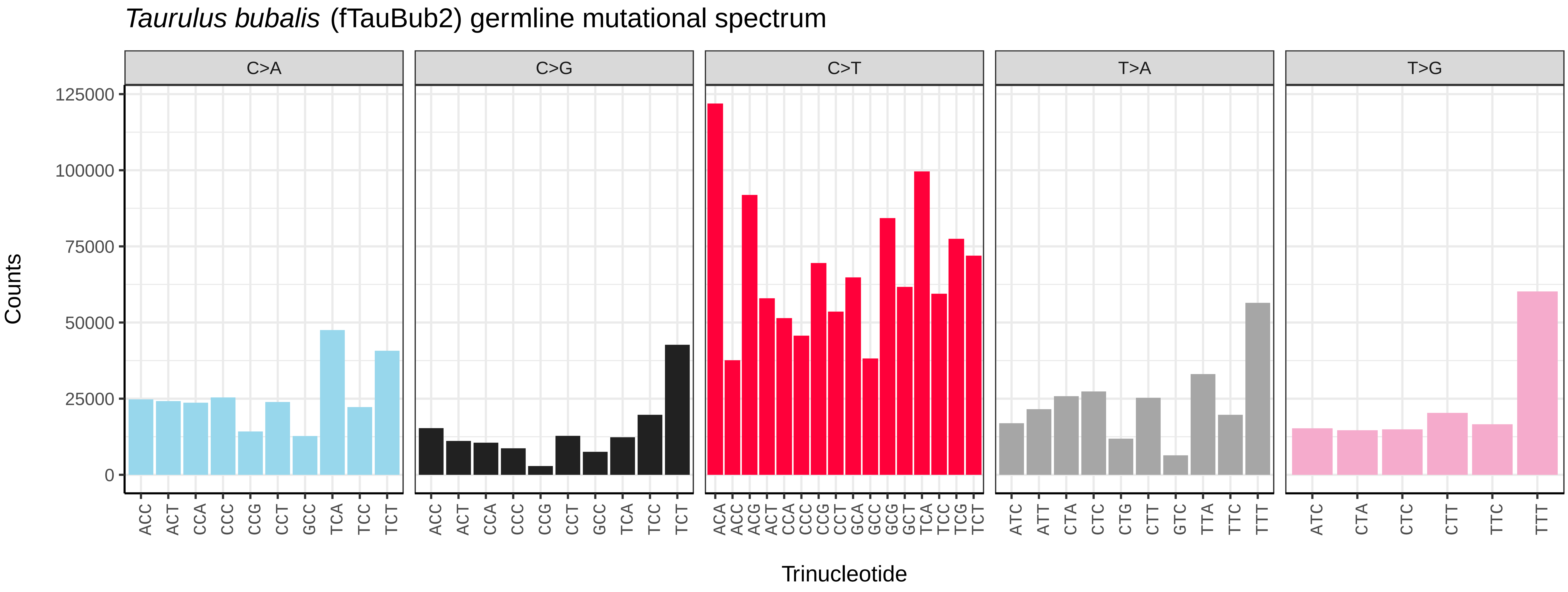
Taurulus bubalis (fTauBub2) germline mutational spectrum
C>A
C>G
C>T
T>A
T>G
Counts
0
25000
50000
75000
100000
125000
Trinucleotide

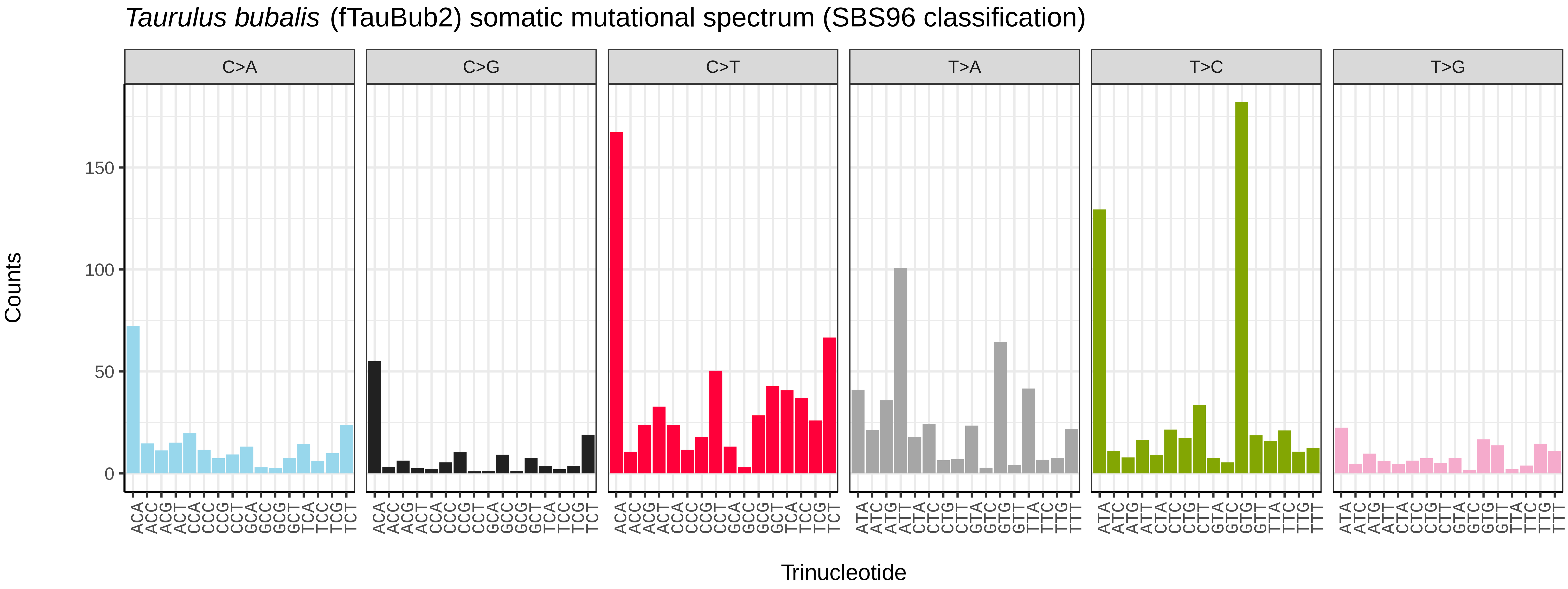
Taurulus bubalis (fTauBub2) somatic mutational spectrum (SBS96 classification)
C>A
C>G
C>T
T>A
T>C
T>G
Counts
0
50
100
150
Trinucleotide

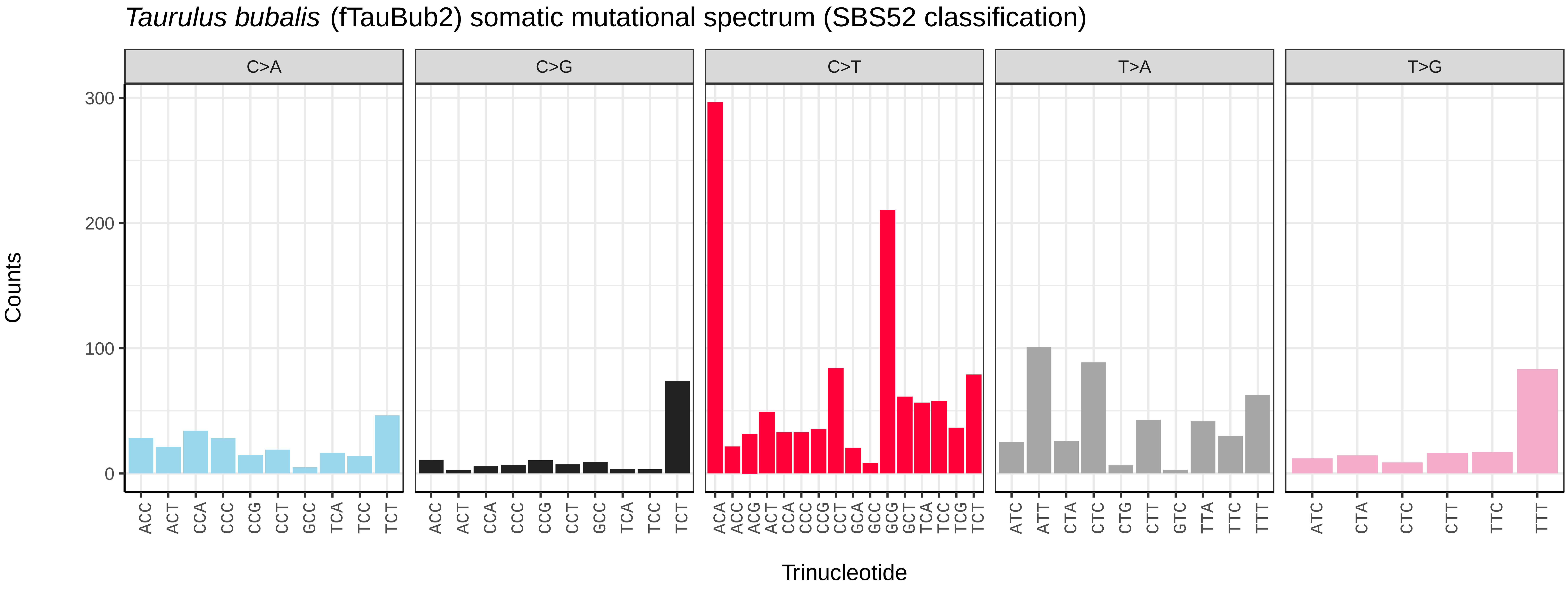
Taurulus bubalis (fTauBub2) somatic mutational spectrum (SBS52 classification)
C>A
C>G
C>T
T>A
T>G
Counts
0
100
200
300
Trinucleotide

*Telmatherina bonti* (fTelBon1) germline mutational spectrum

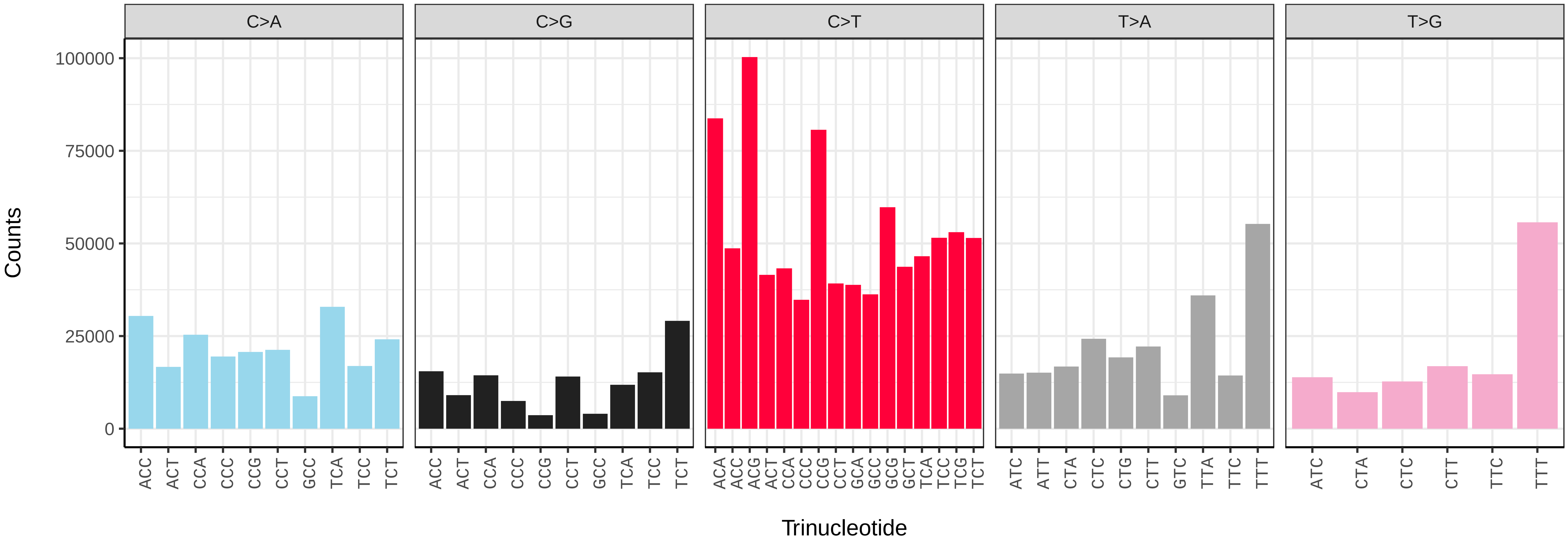


*Telmatherina bonti* (fTelBon1) somatic mutational spectrum (SBS96 classification)

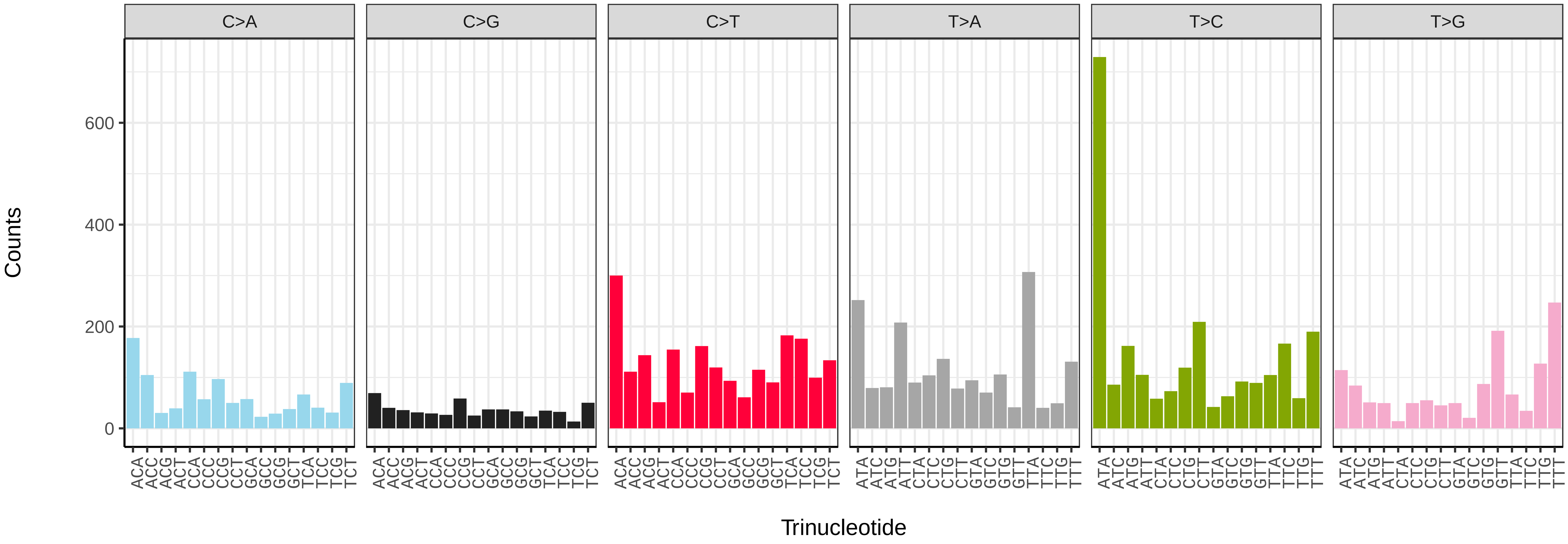


*Telmatherina bonti* (fTelBon1) somatic mutational spectrum (SBS52 classification)

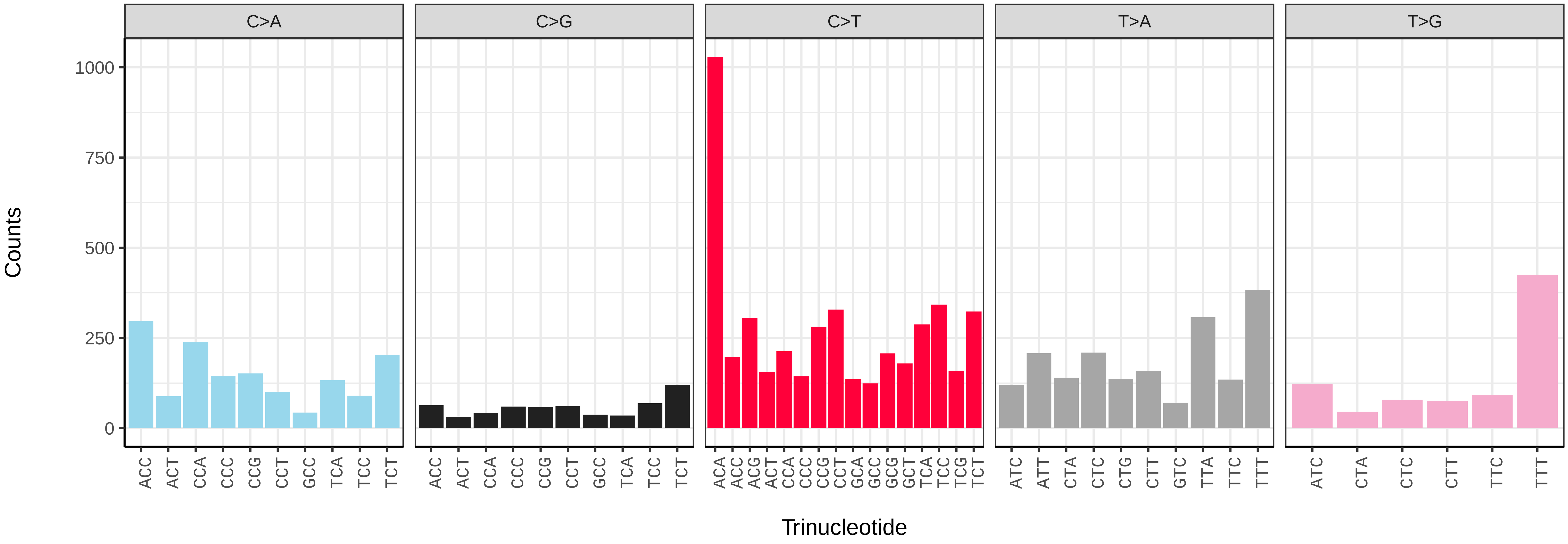

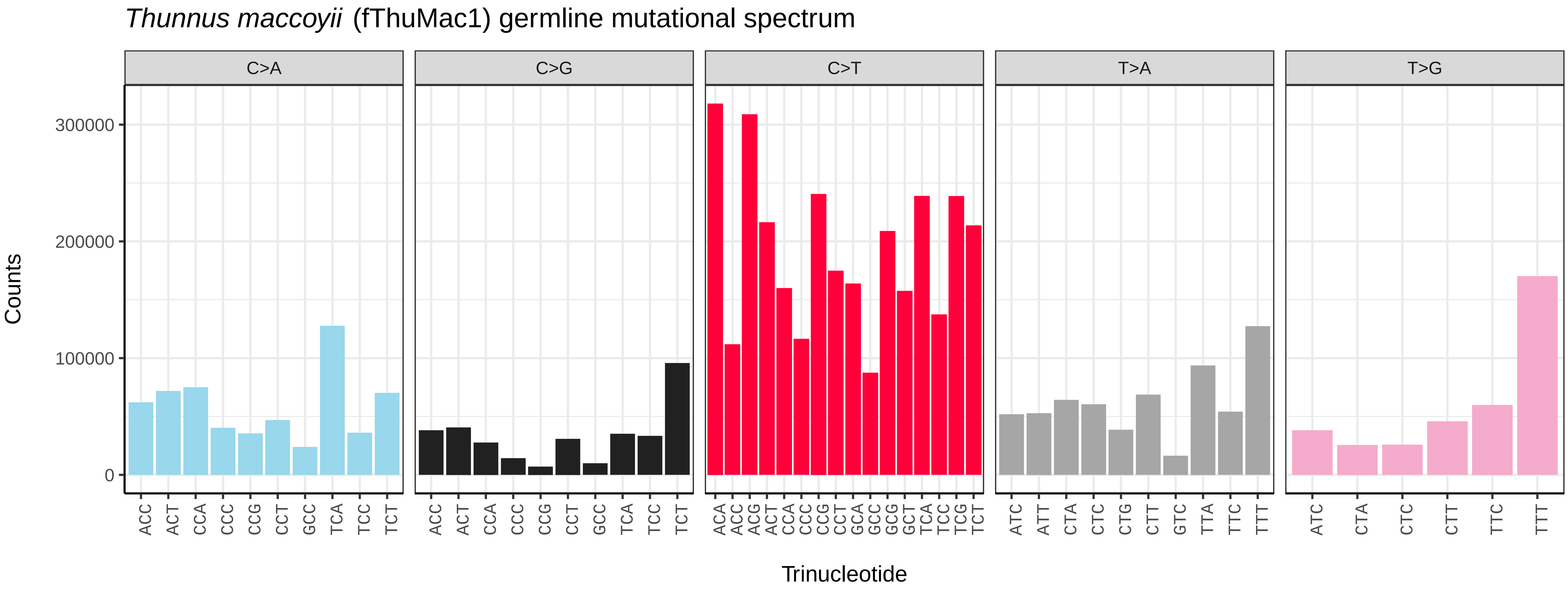
*Thunnus maccoyii* (fThuMac1) germline mutational spectrum
C>A
C>G
C>T
T>A
T>G
Counts
300000
200000
100000
0
ACC ACT CCA CCC CCG CCT GCC TCA TCC TCT
ACA ACC ACG ACT CCA CCC CCG CCT GCA GCC GCG GCT TCA TCC TCG TCT
ATC ATT CTA CTC CTG CTT GTC TTA TTC TTT
ATC CTA CTC CTT TTC TTT
Trinucleotide

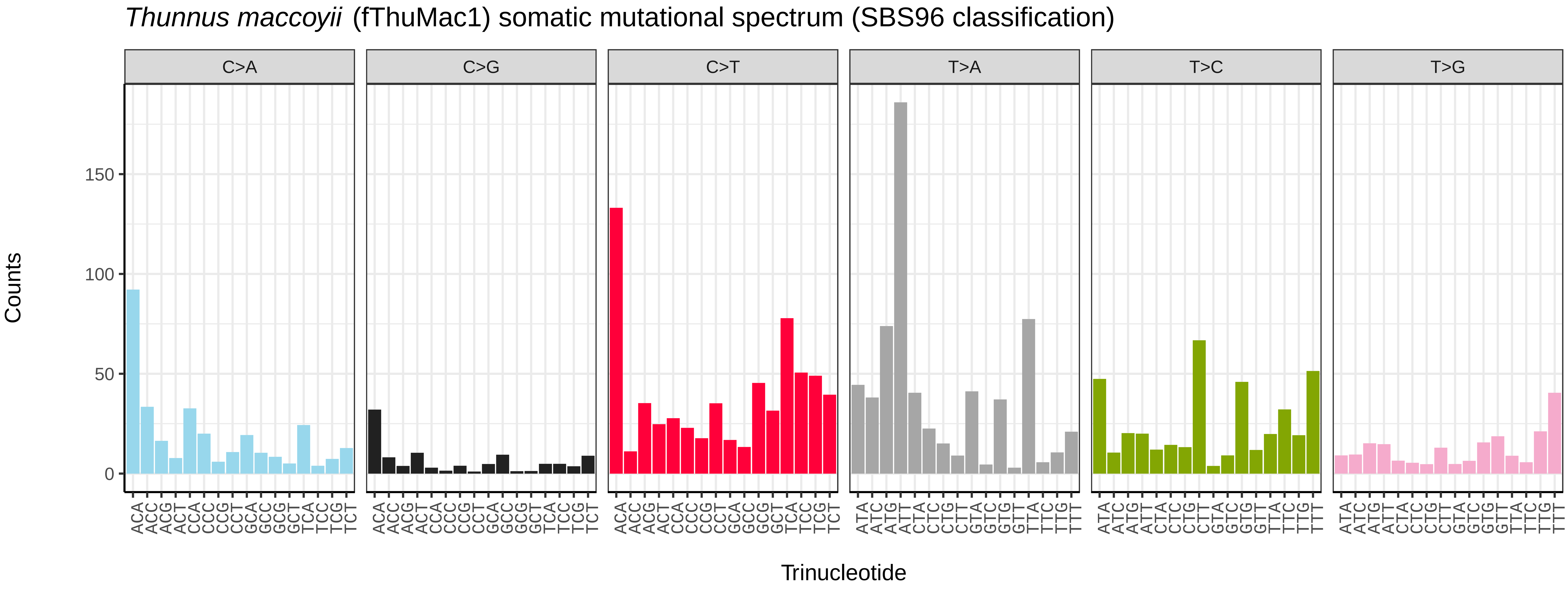
*Thunnus maccoyii* (fThuMac1) somatic mutational spectrum (SBS96 classification)
C>A
C>G
C>T
T>A
T>C
T>G
Counts
150
100
50
0
ACA ACC ACG ACT CCA CCC CCG CCT GCA GCC GCG GCT TCA TCC TCG TCT
ATA ATC ATG ATT CTA CTC CTG CTT GTA GTC GTG GTT TTA TTC TTG TTT
Trinucleotide

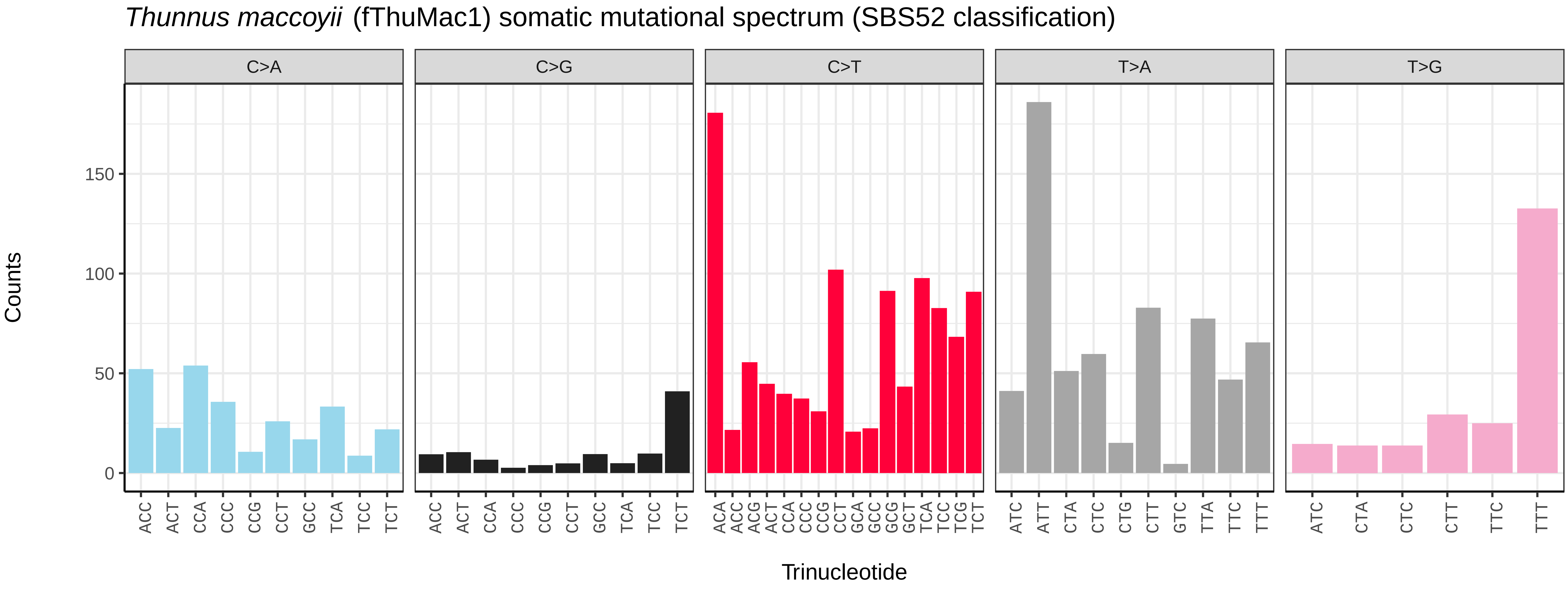
*Thunnus maccoyii* (fThuMac1) somatic mutational spectrum (SBS52 classification)
C>A
C>G
C>T
T>A
T>G
Counts
150
100
50
0
ACC ACT CCA CCC CCG CCT GCC TCA TCC TCT
ACA ACC ACG ACT CCA CCC CCG CCT GCA GCC GCG GCT TCA TCC TCG TCT
ATC ATT CTA CTC CTG CTT GTC TTA TTC TTT
ATC CTA CTC CTT TTC TTT
Trinucleotide

*Trachurus trachurus* (fTraTra1) germline mutational spectrum

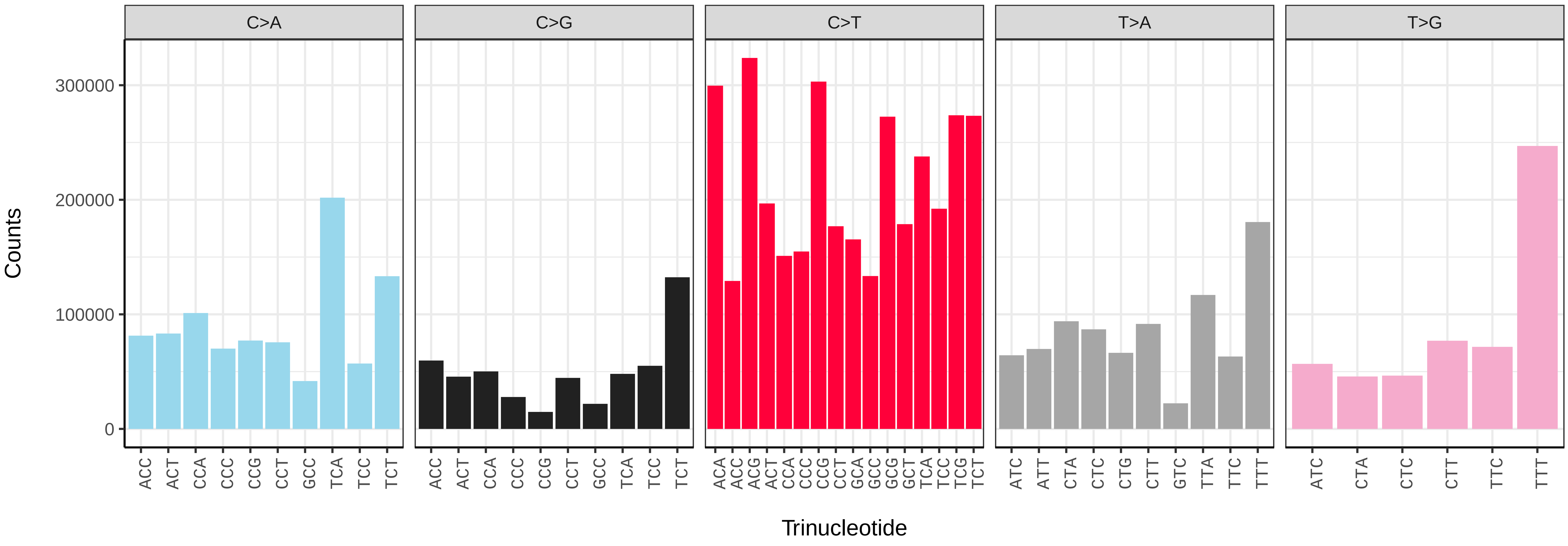


*Trachurus trachurus* (fTraTra1) somatic mutational spectrum (SBS96 classification)

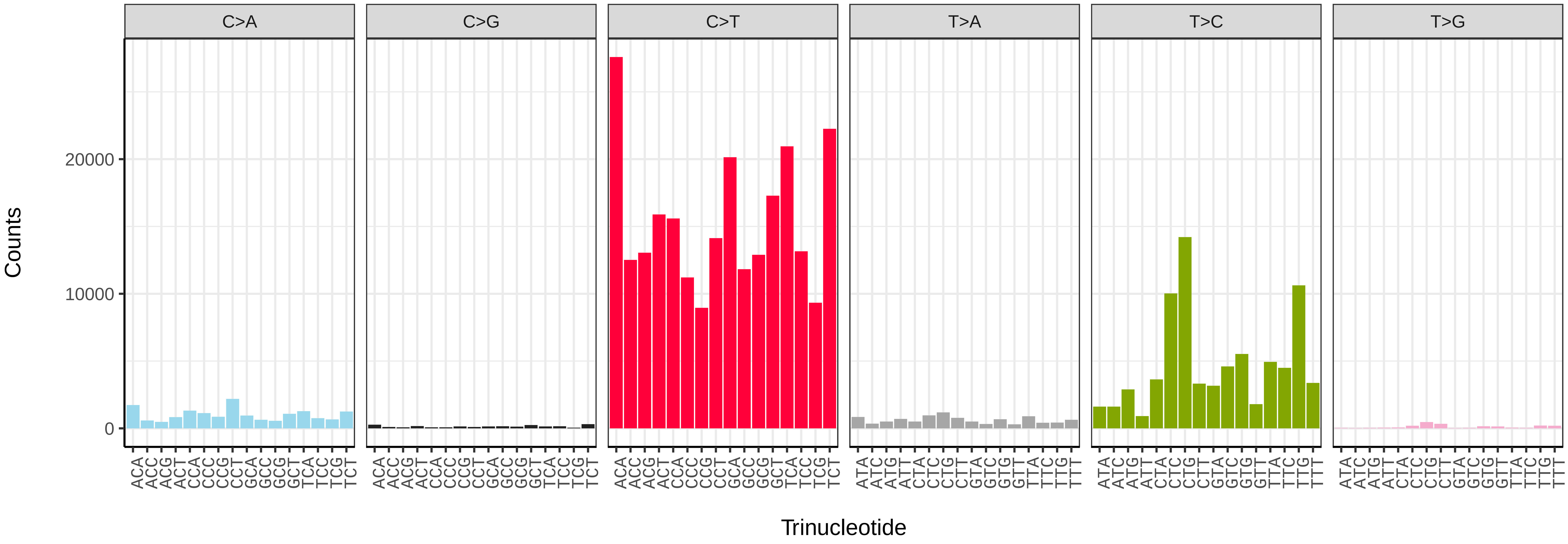


*Trachurus trachurus* (fTraTra1) somatic mutational spectrum (SBS52 classification)

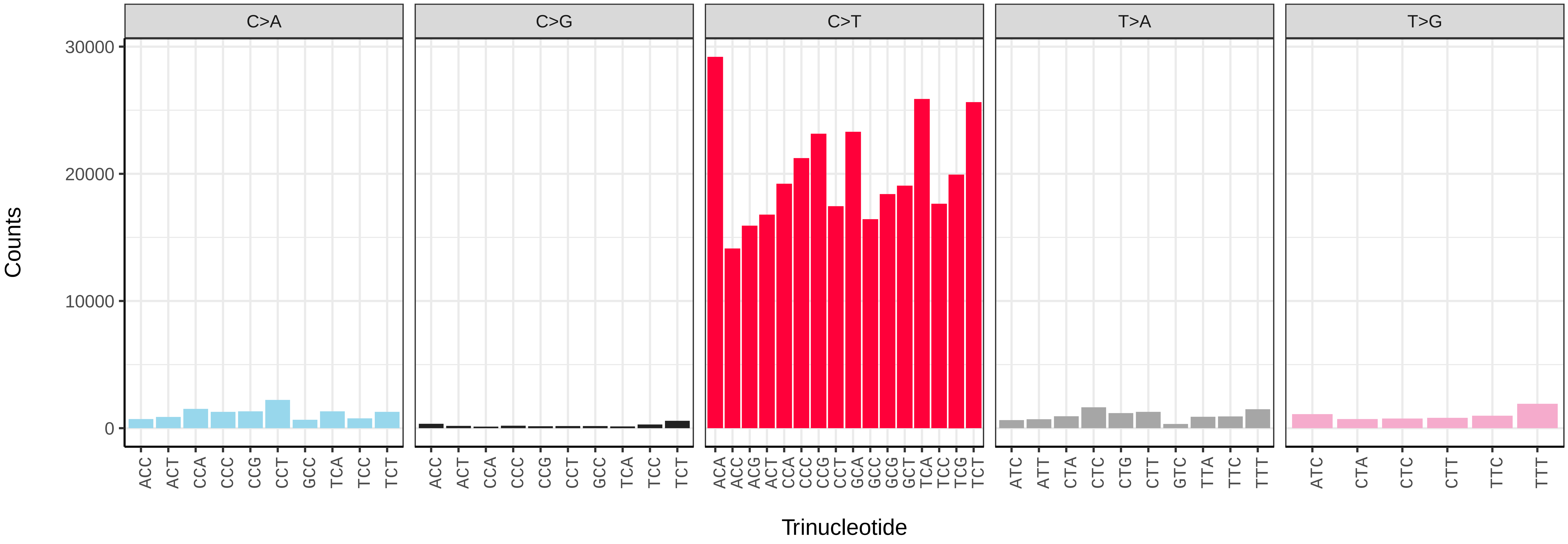

*Zeus faber* (fZeuFab8) germline mutational spectrum

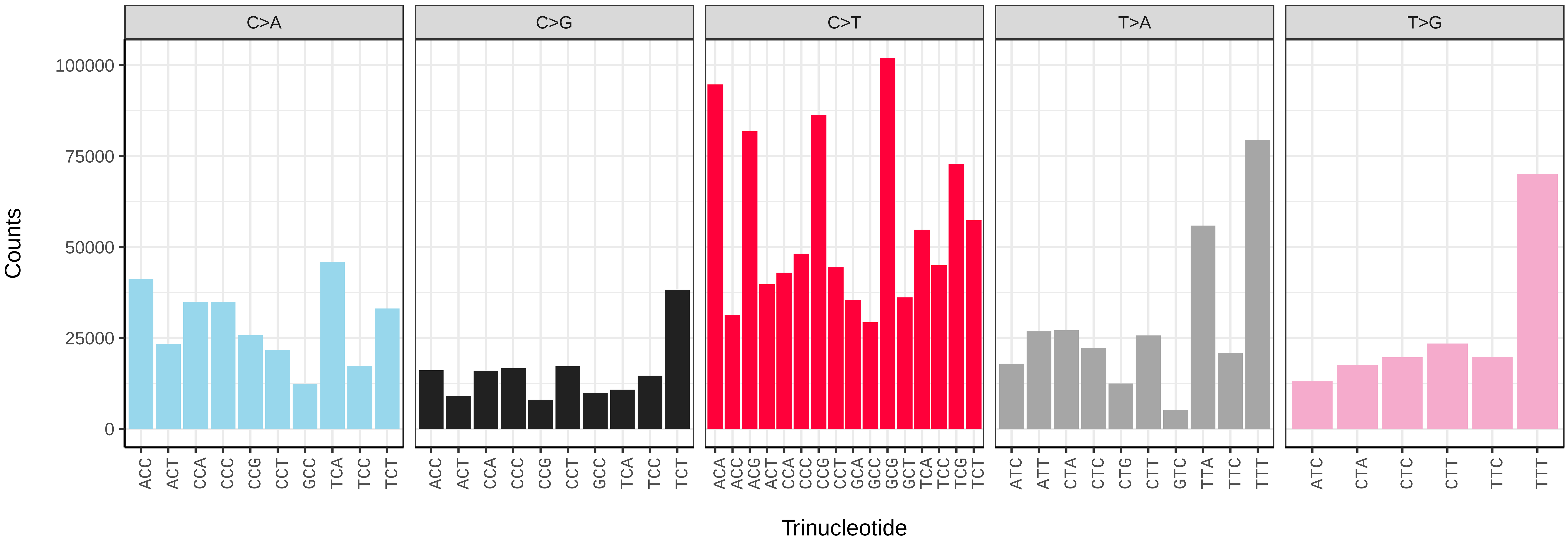


*Zeus faber* (fZeuFab8) somatic mutational spectrum (SBS96 classification)

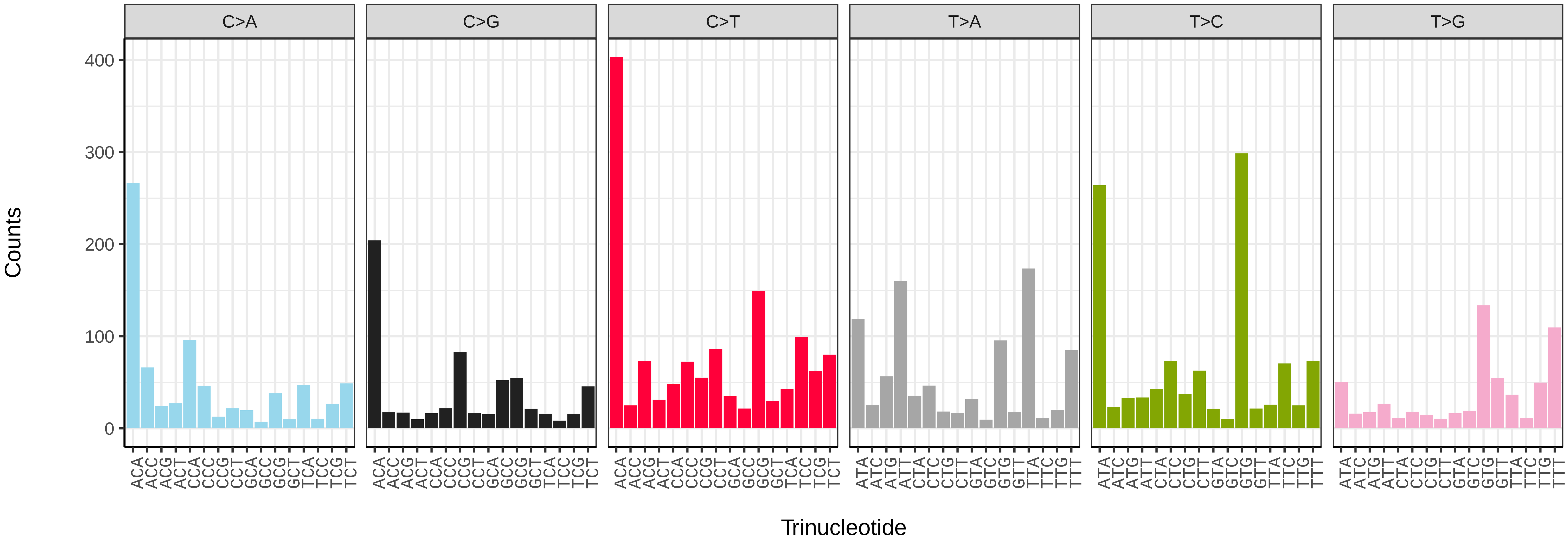


*Zeus faber* (fZeuFab8) somatic mutational spectrum (SBS52 classification)

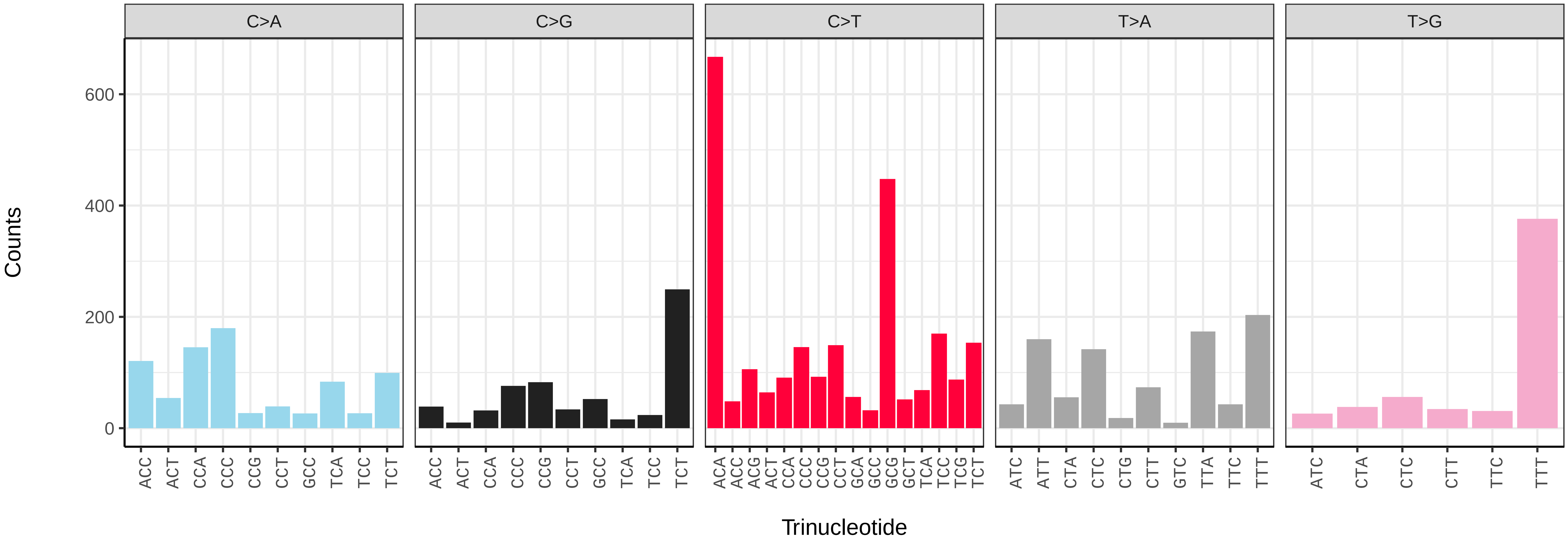

*Agaricus bisporus* (gfAgaBisp1) germline mutational spectrum

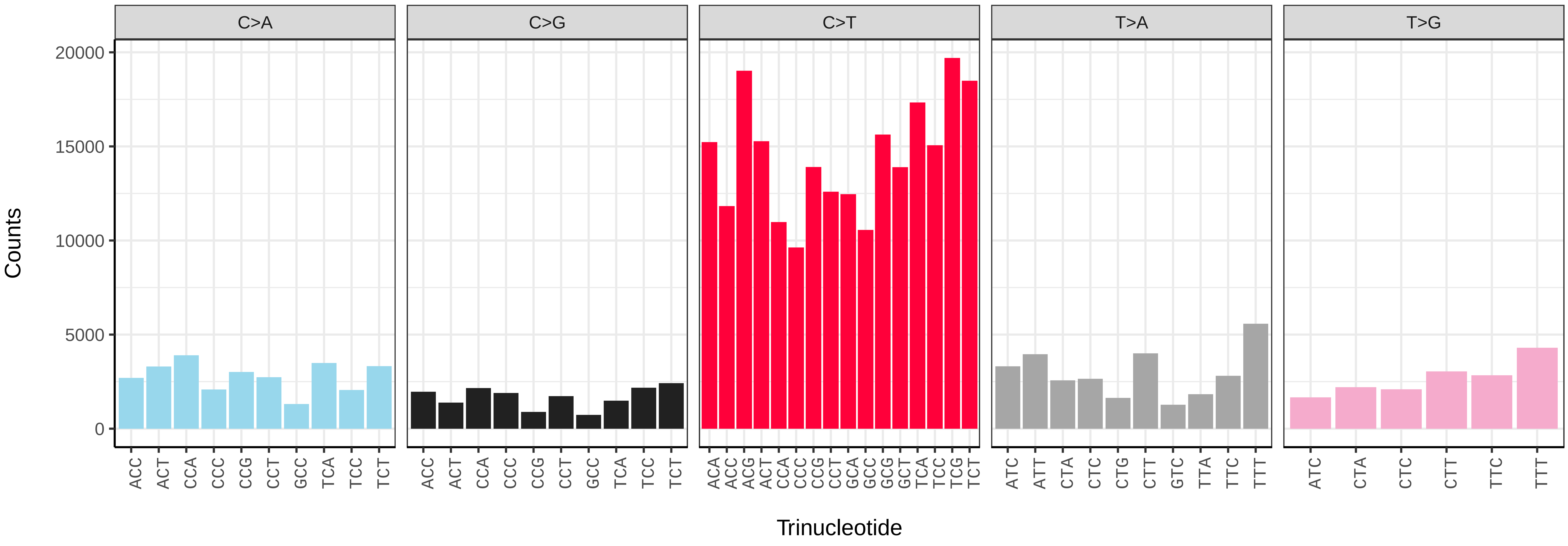


*Agaricus bisporus* (gfAgaBisp1) somatic mutational spectrum (SBS96 classification)

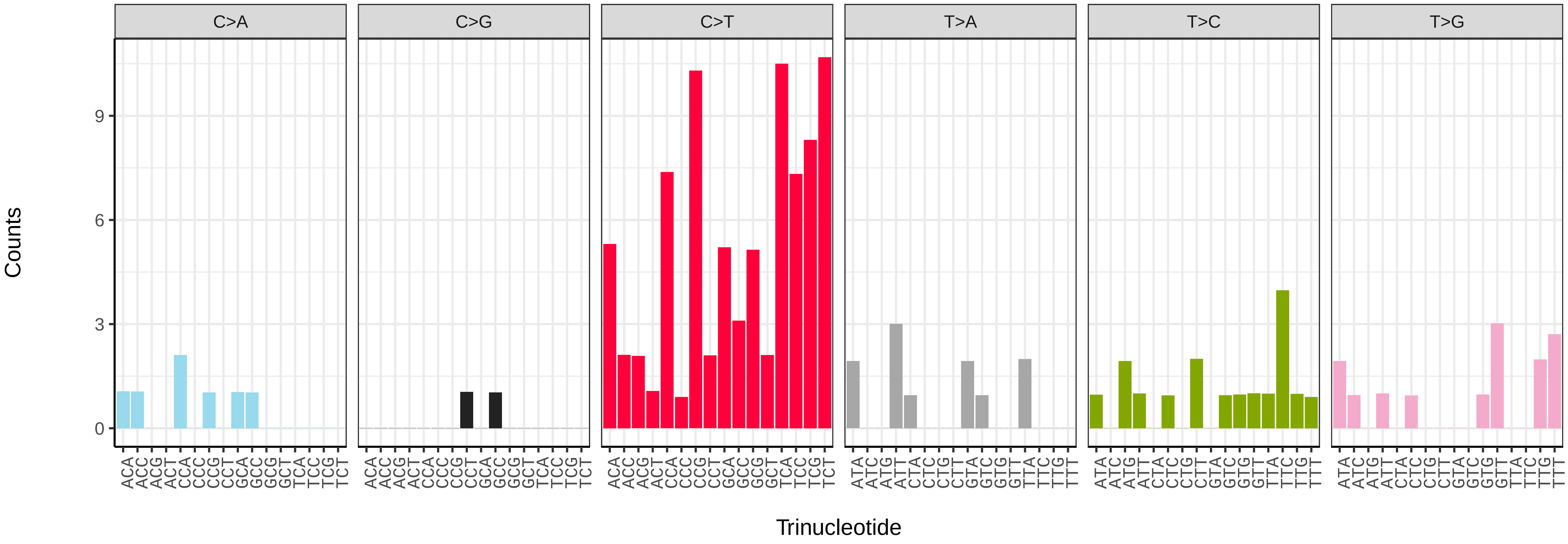


*Agaricus bisporus* (gfAgaBisp1) somatic mutational spectrum (SBS52 classification)

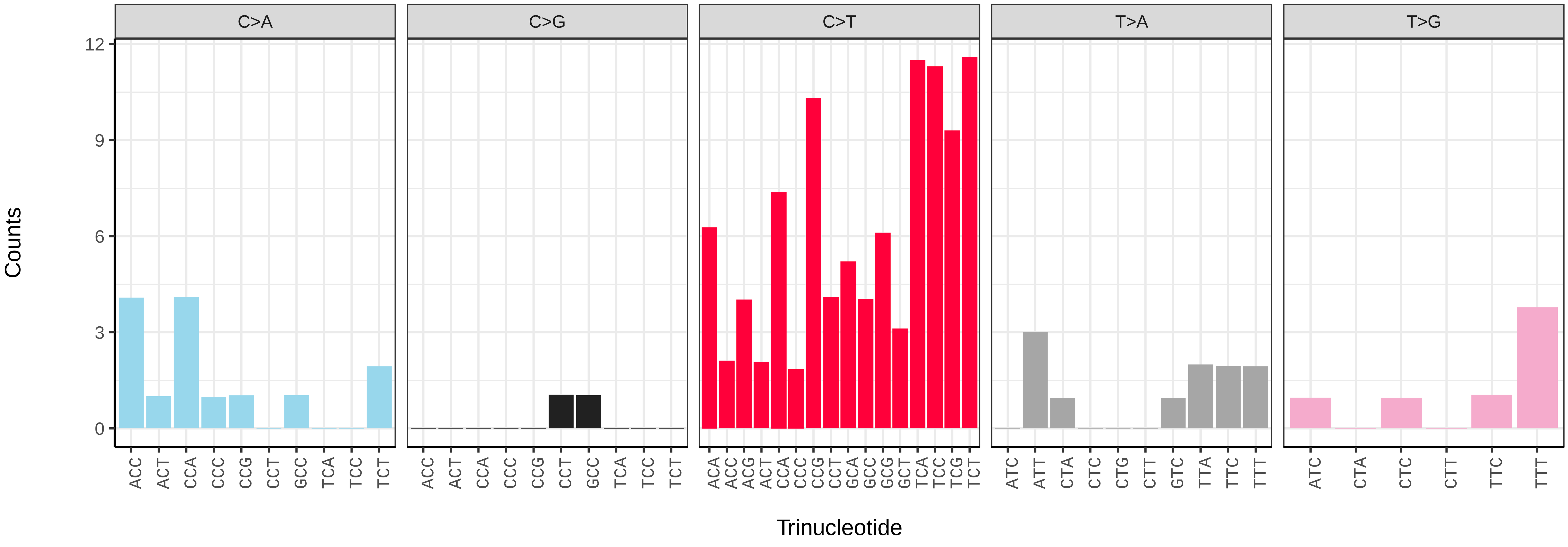

*Clavaria fumosa* (gfClaFumo1) germline mutational spectrum

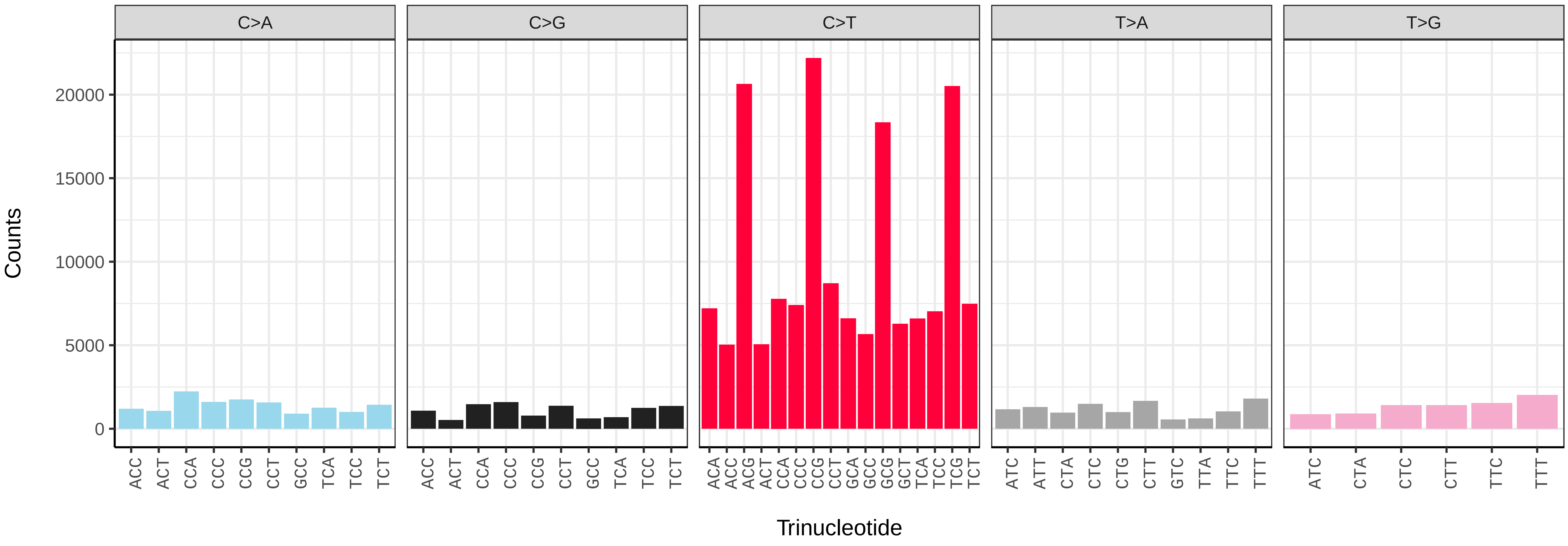


*Clavaria fumosa* (gfClaFumo1) somatic mutational spectrum (SBS96 classification)

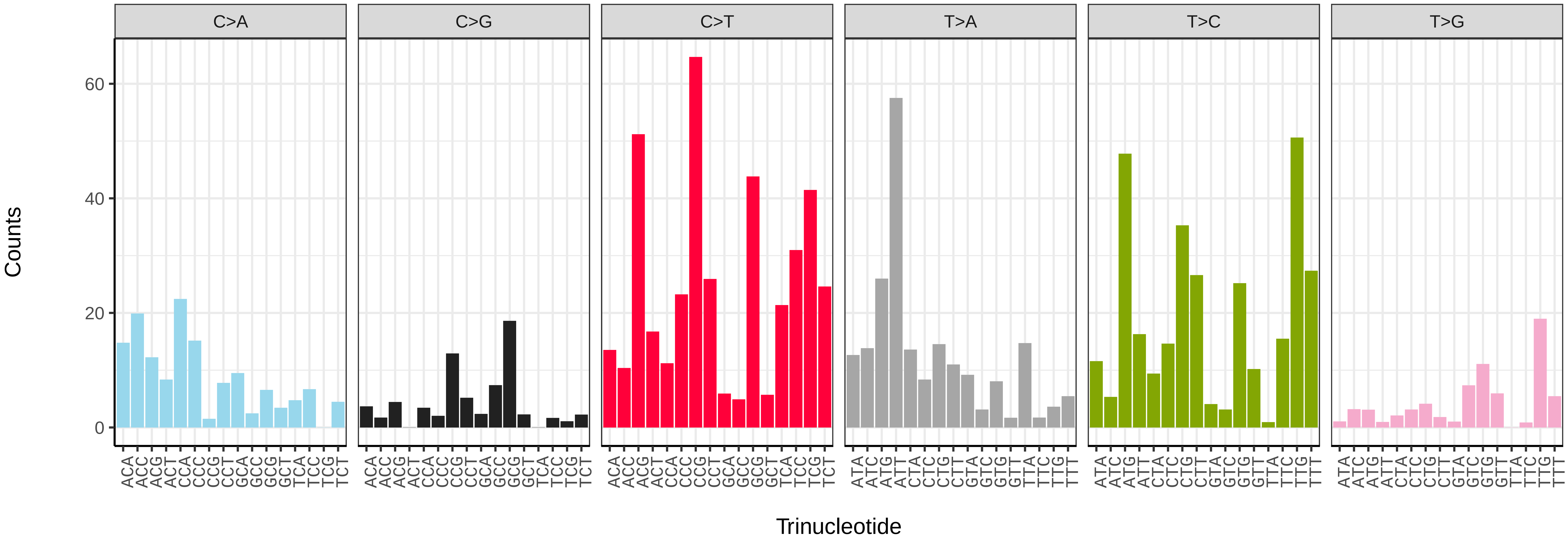


*Clavaria fumosa* (gfClaFumo1) somatic mutational spectrum (SBS52 classification)

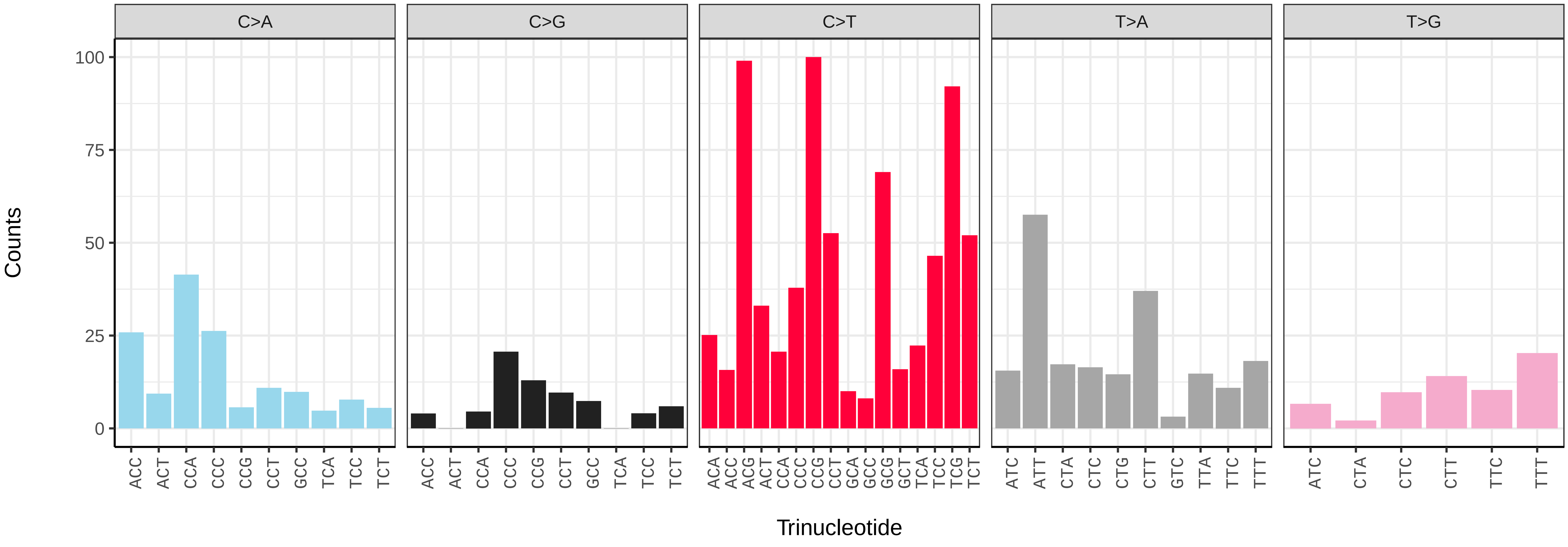

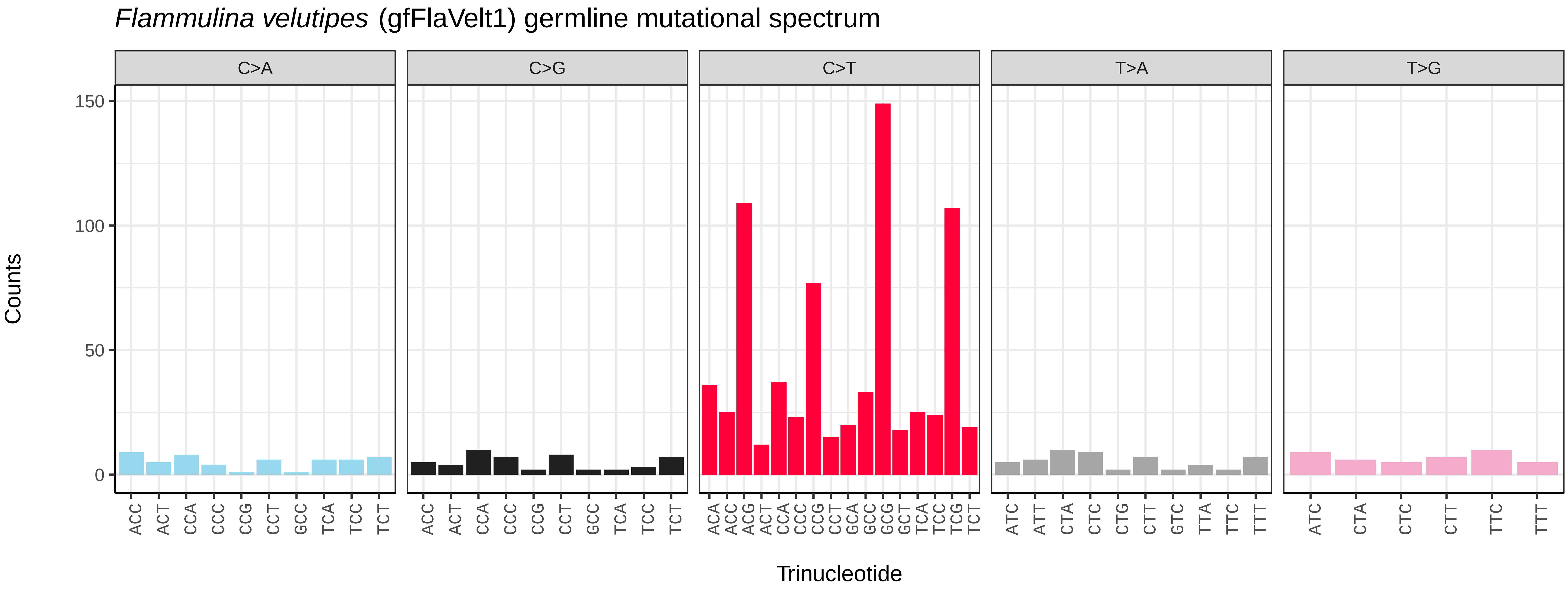

Flammulina velutipes (gfFlaVelt1) germline mutational spectrum
C>A
C>G
C>T
T>A
T>G
Counts
0
50
100
150
ACC ACT CCA CCC CCG CCT GCC TCA TCC TCT
ACC ACT CCA CCC CCG CCT GCC TCA TCC TCT
ACA ACC ACG ACT CCA CCC CCG CCT GCA GCC GCG GCT TCA TCC TCG TCT
ATC ATT CTA CTC CTG CTT GTC TTA TTC TTT
ATC CTA CTC CTT TTC TTT
Trinucleotide


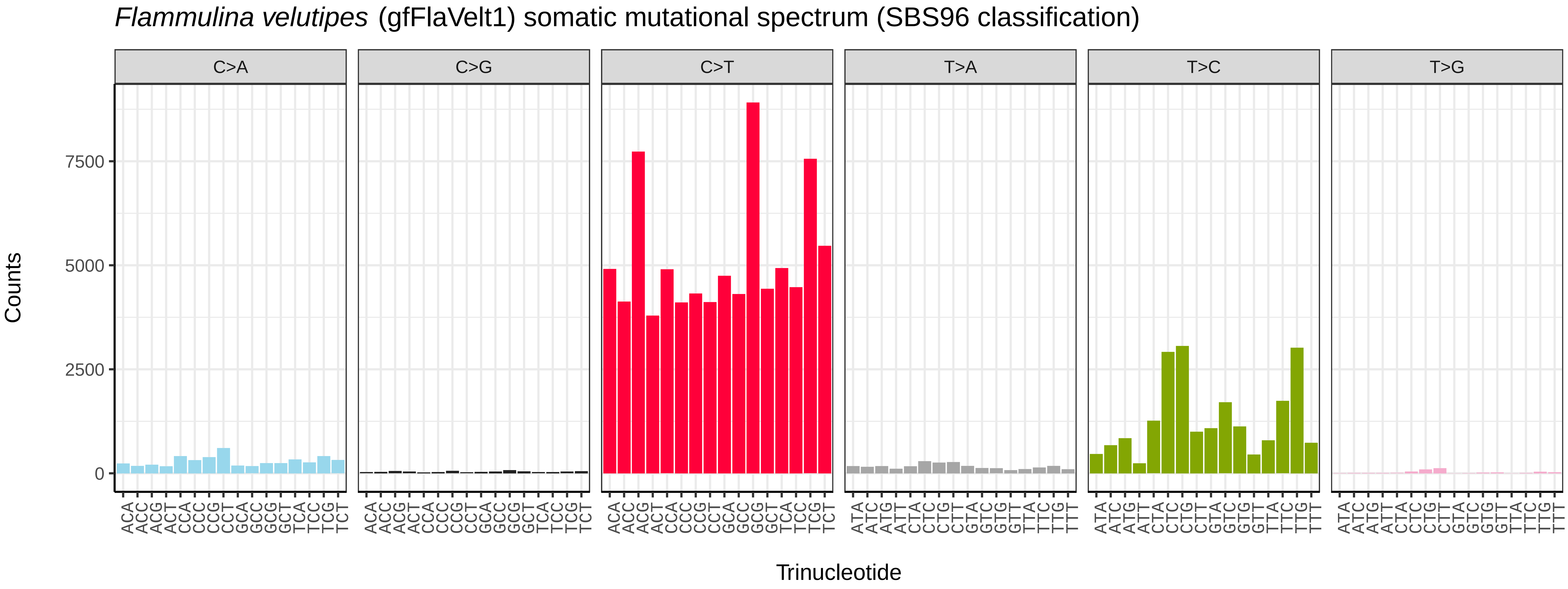

Flammulina velutipes (gfFlaVelt1) somatic mutational spectrum (SBS96 classification)
C>A
C>G
C>T
T>A
T>C
T>G
Counts
0
2500
5000
7500
Trinucleotide


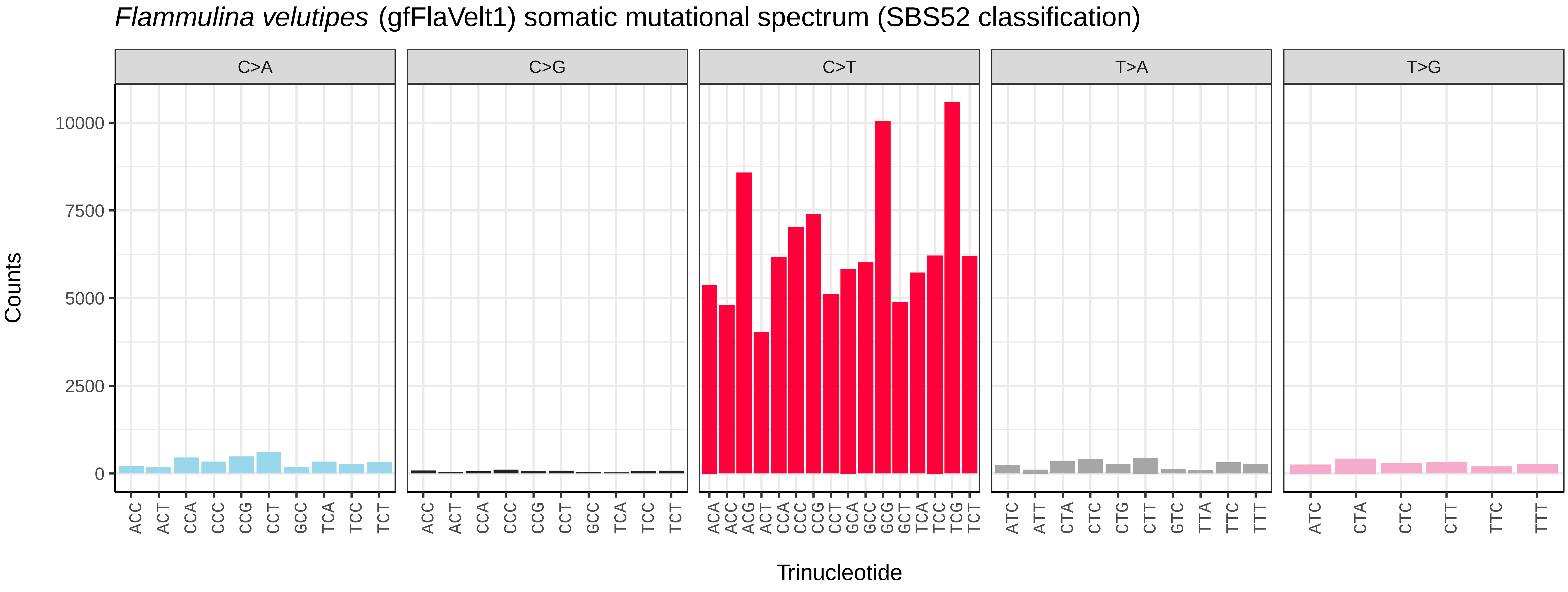

Flammulina velutipes (gfFlaVelt1) somatic mutational spectrum (SBS52 classification)
C>A
C>G
C>T
T>A
T>G
Counts
0
2500
5000
7500
10000
ACC ACT CCA CCC CCG CCT GCC TCA TCC TCT
ACC ACT CCA CCC CCG CCT GCC TCA TCC TCT
ACA ACC ACG ACT CCA CCC CCG CCT GCA GCC GCG GCT TCA TCC TCG TCT
ATC ATT CTA CTC CTG CTT GTC TTA TTC TTT
ATC CTA CTC CTT TTC TTT
Trinucleotide

*Laetiporus sulphureus* (gfLaeSulp1) germline mutational spectrum

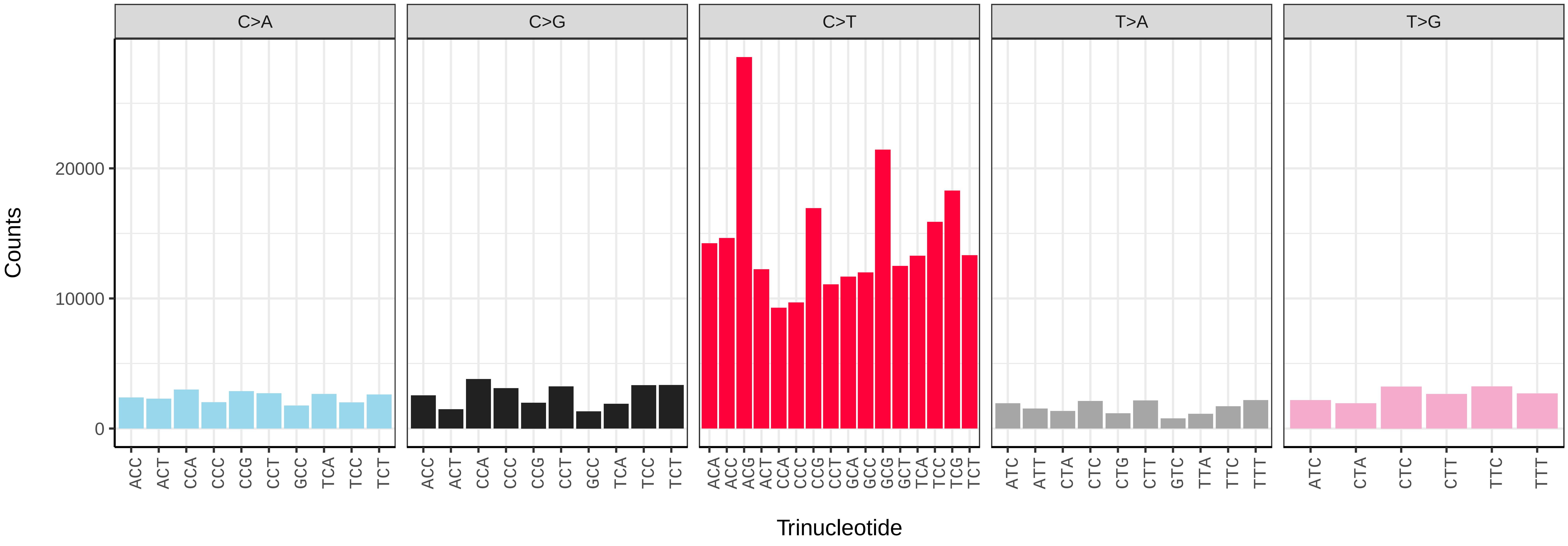


*Laetiporus sulphureus* (gfLaeSulp1) somatic mutational spectrum (SBS96 classification)

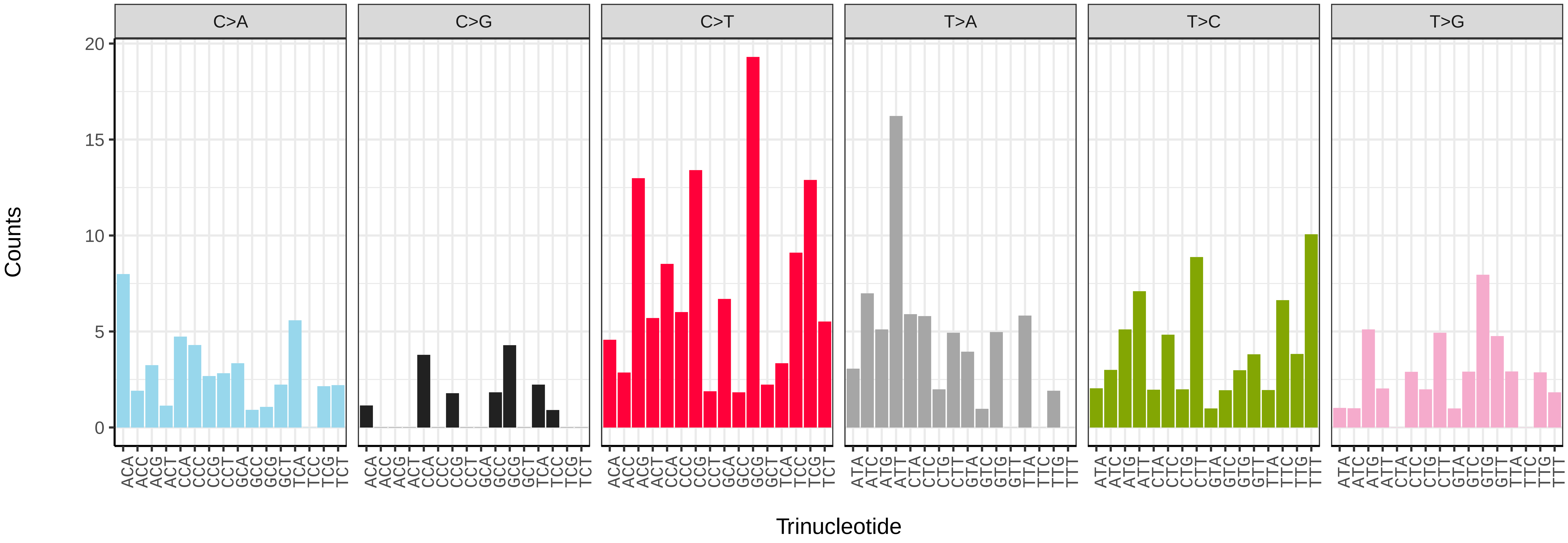


*Laetiporus sulphureus* (gfLaeSulp1) somatic mutational spectrum (SBS52 classification)

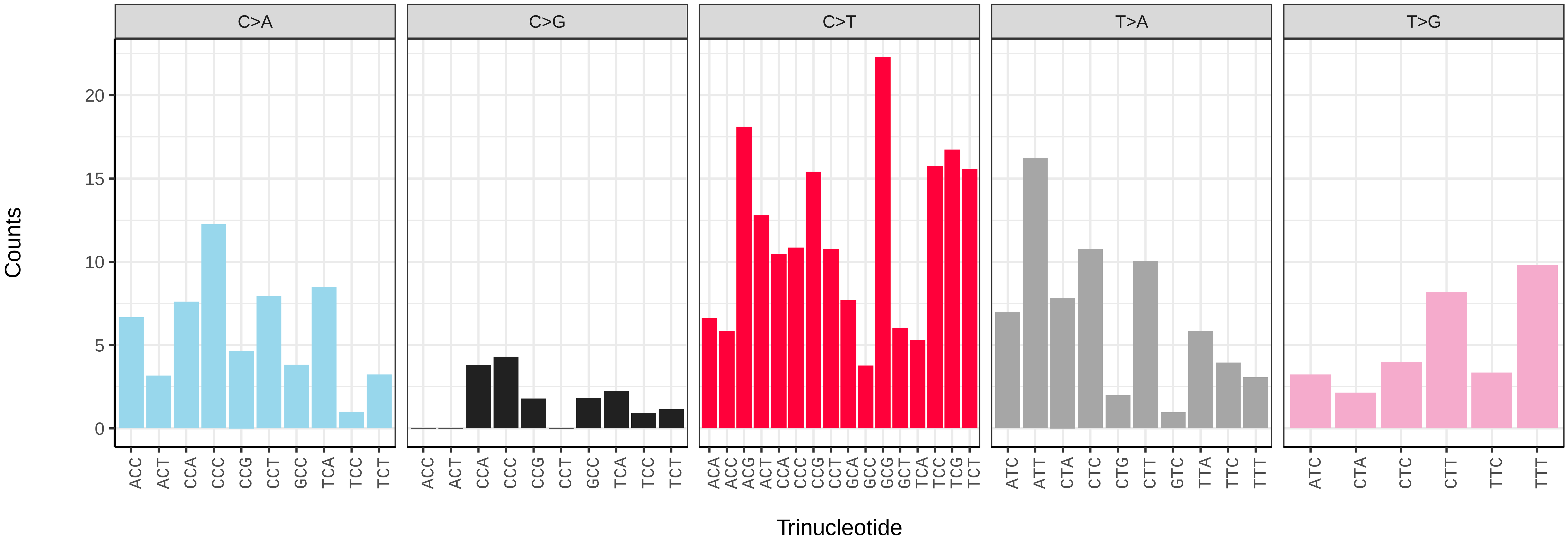

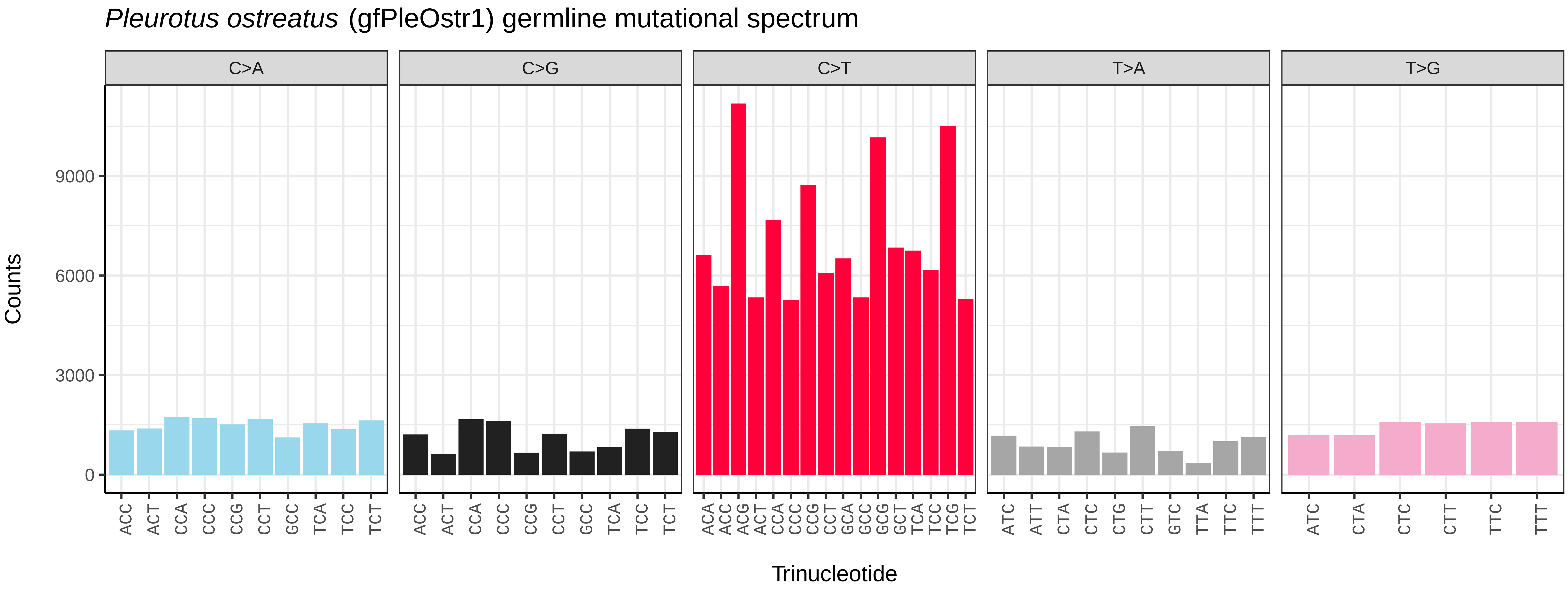
Pleurotus ostreatus (gfPleOstr1) germline mutational spectrum
C>A
C>G
C>T
T>A
T>G
Counts
0
3000
6000
9000
Trinucleotide

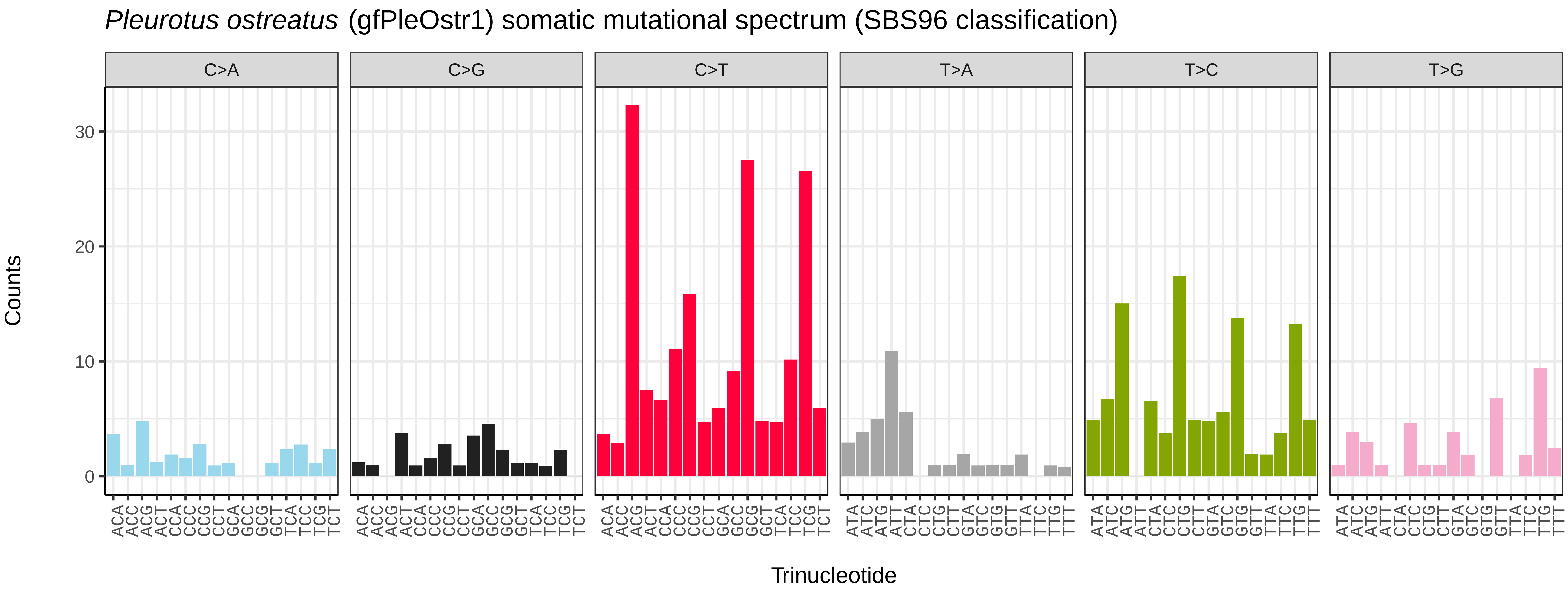
Pleurotus ostreatus (gfPleOstr1) somatic mutational spectrum (SBS96 classification)
C>A
C>G
C>T
T>A
T>C
T>G
Counts
0
10
20
30
Trinucleotide

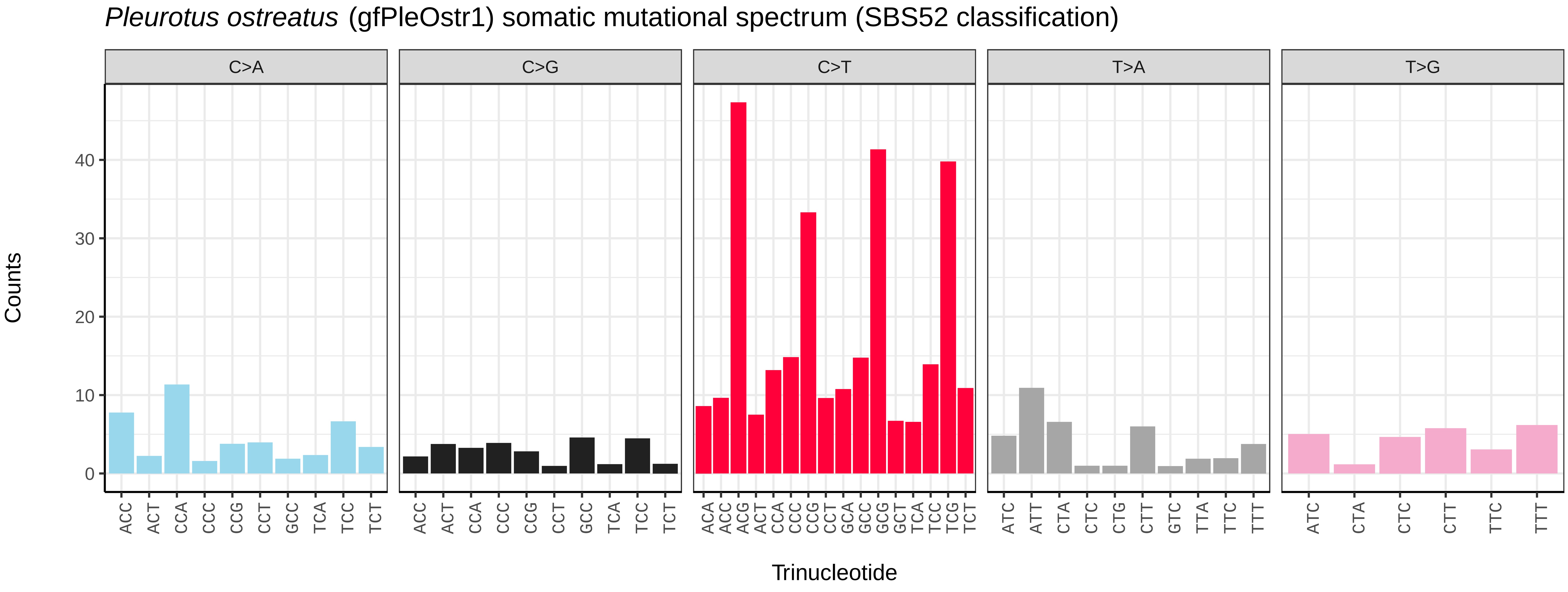
Pleurotus ostreatus (gfPleOstr1) somatic mutational spectrum (SBS52 classification)
C>A
C>G
C>T
T>A
T>G
Counts
0
10
20
30
40
Trinucleotide

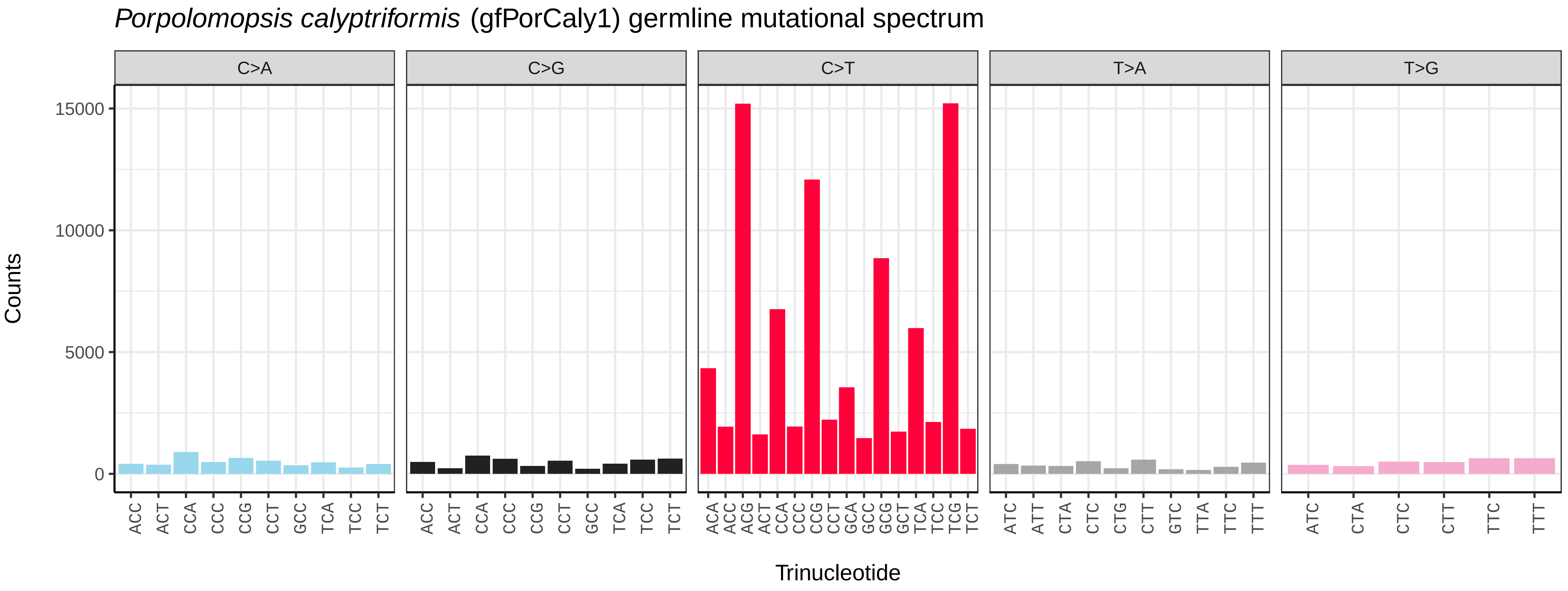
Porpolomopsis calyptriformis (gfPorCaly1) germline mutational spectrum
C>A
C>G
C>T
T>A
T>G
Counts
Trinucleotide

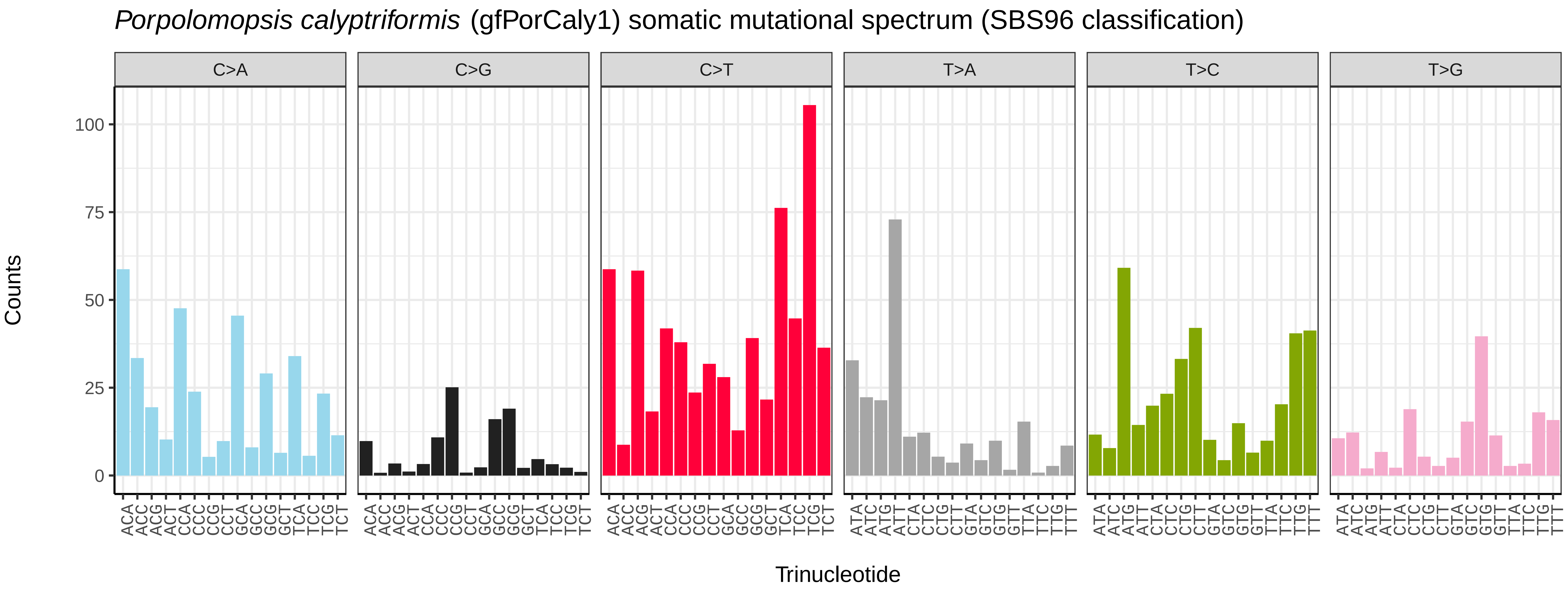
Porpolomopsis calyptriformis (gfPorCaly1) somatic mutational spectrum (SBS96 classification)
C>A
C>G
C>T
T>A
T>C
T>G
Counts
Trinucleotide

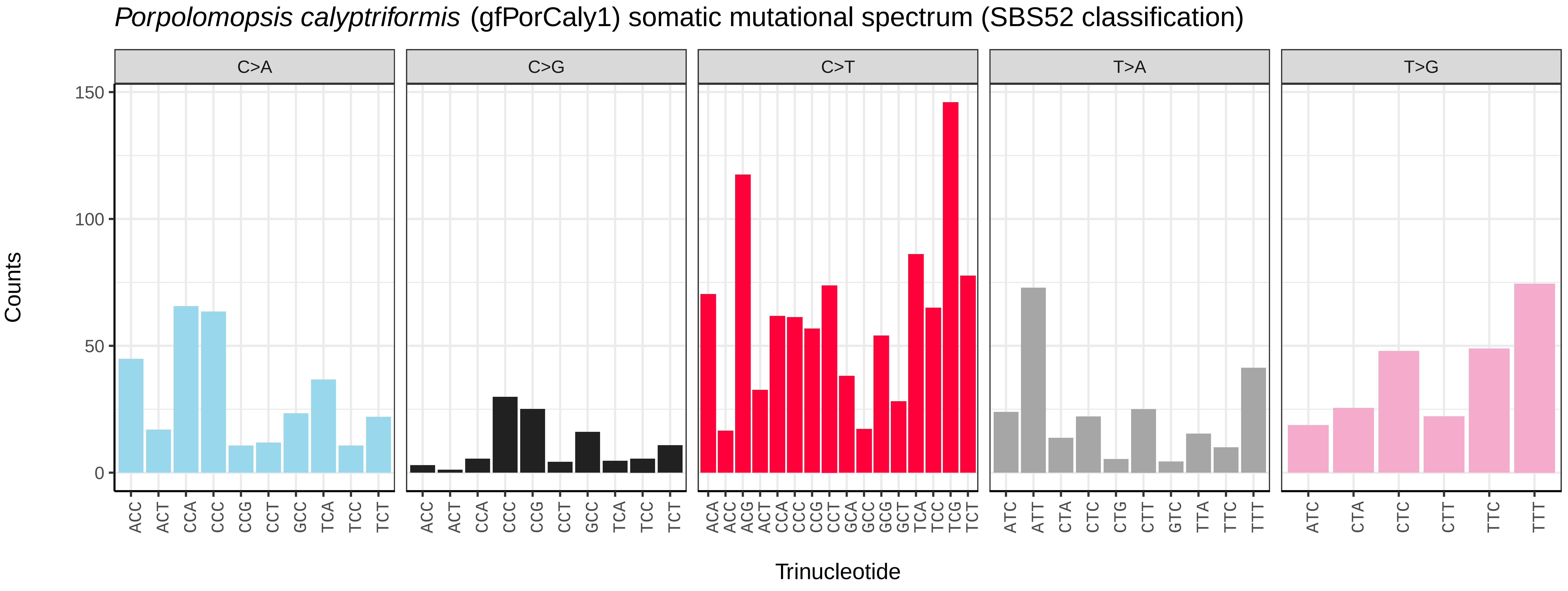
Porpolomopsis calyptriformis (gfPorCaly1) somatic mutational spectrum (SBS52 classification)
C>A
C>G
C>T
T>A
T>G
Counts
Trinucleotide

*Lichina pygmaea* (glLicPygm2) germline mutational spectrum

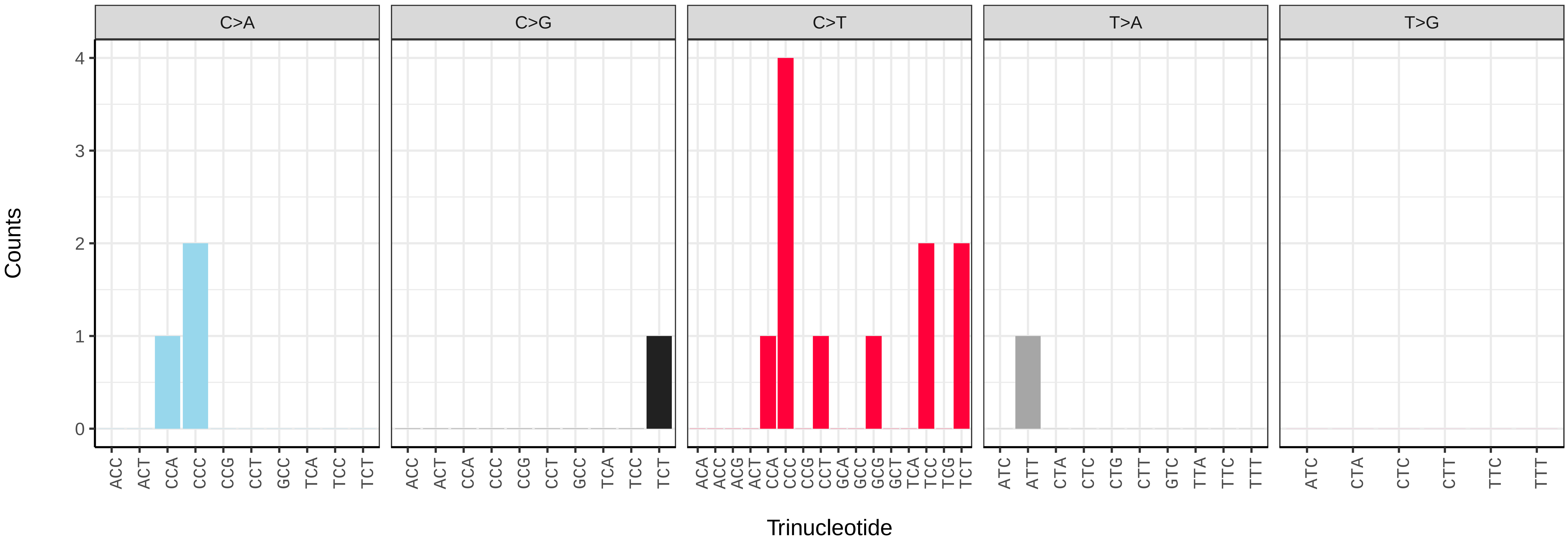


*Lichina pygmaea* (glLicPygm2) somatic mutational spectrum (SBS96 classification)

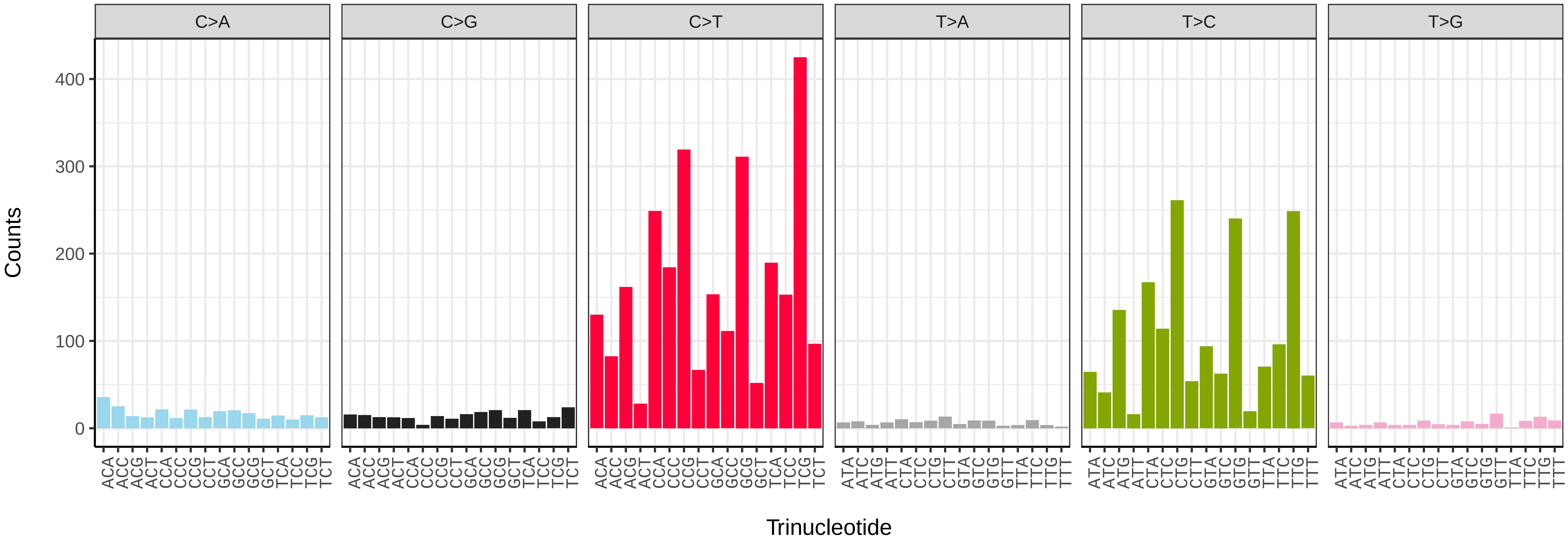


*Lichina pygmaea* (glLicPygm2) somatic mutational spectrum (SBS52 classification)

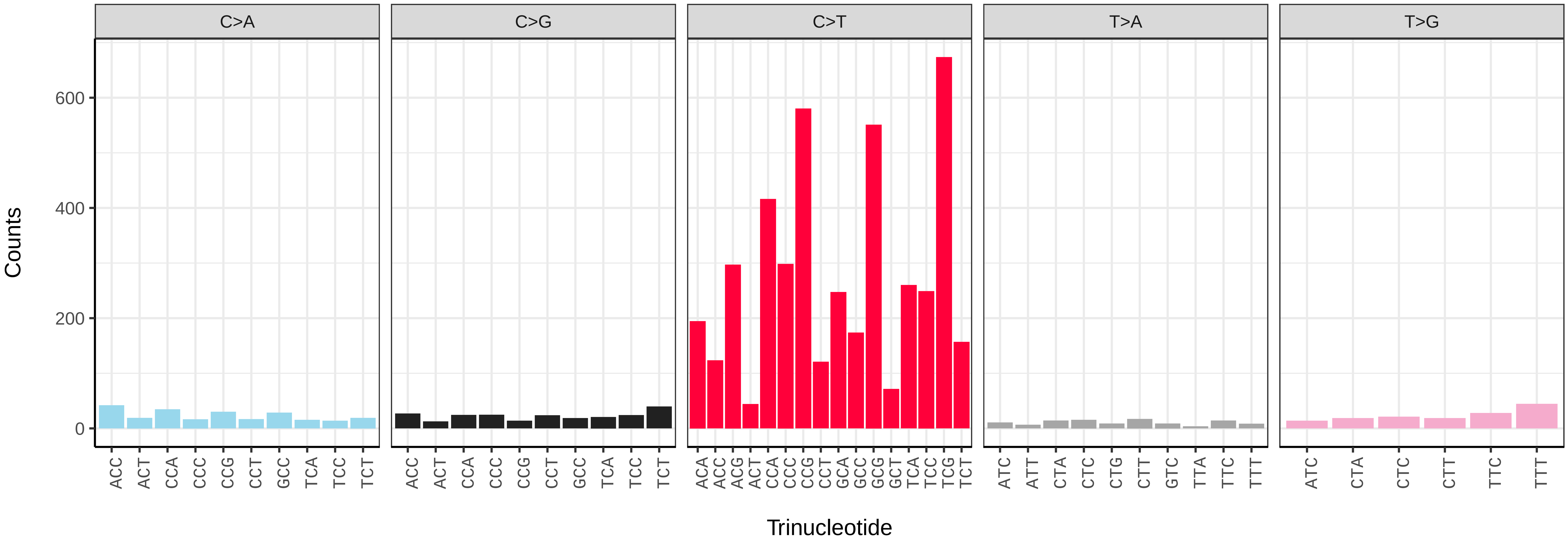

*Cordyceps fumosorosea* (grCorFumo1) germline mutational spectrum

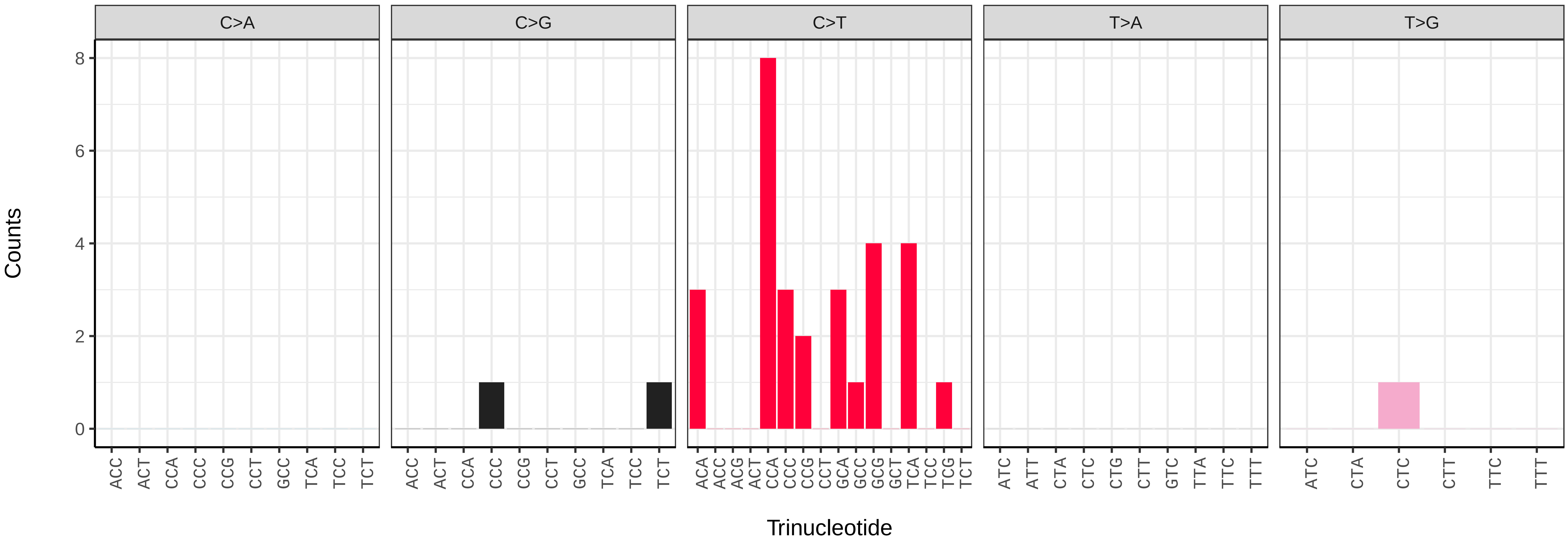


*Cordyceps fumosorosea* (grCorFumo1) somatic mutational spectrum (SBS96 classification)

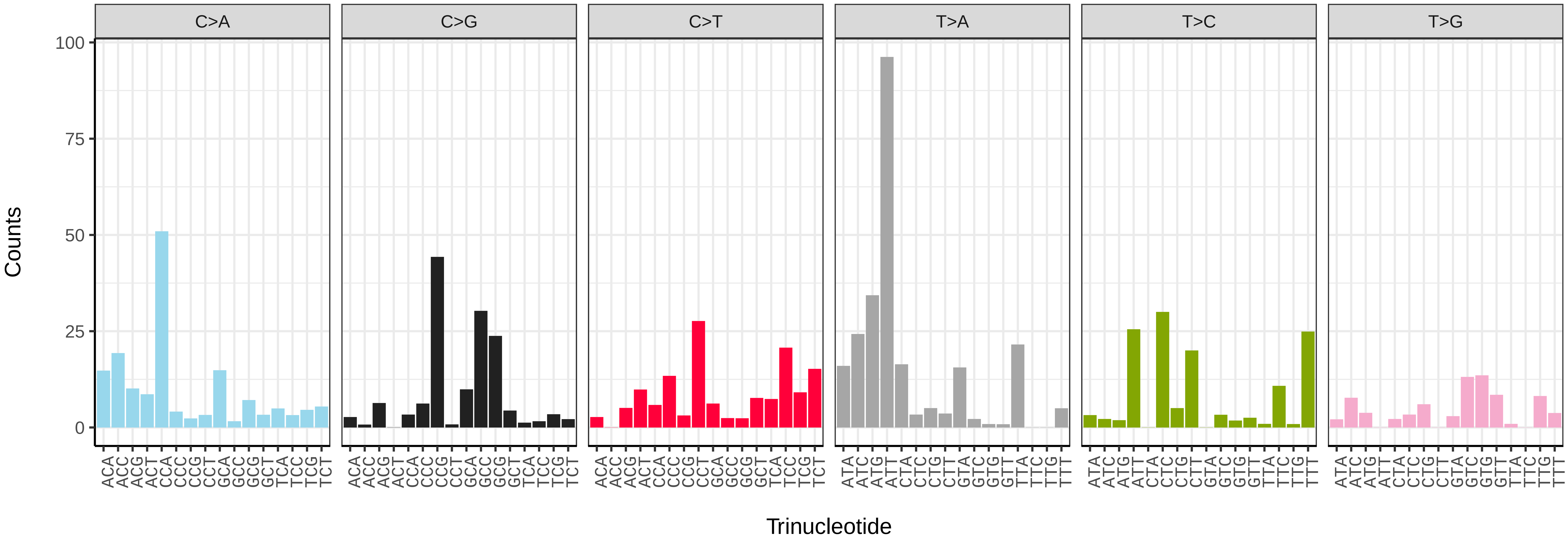


*Cordyceps fumosorosea* (grCorFumo1) somatic mutational spectrum (SBS52 classification)

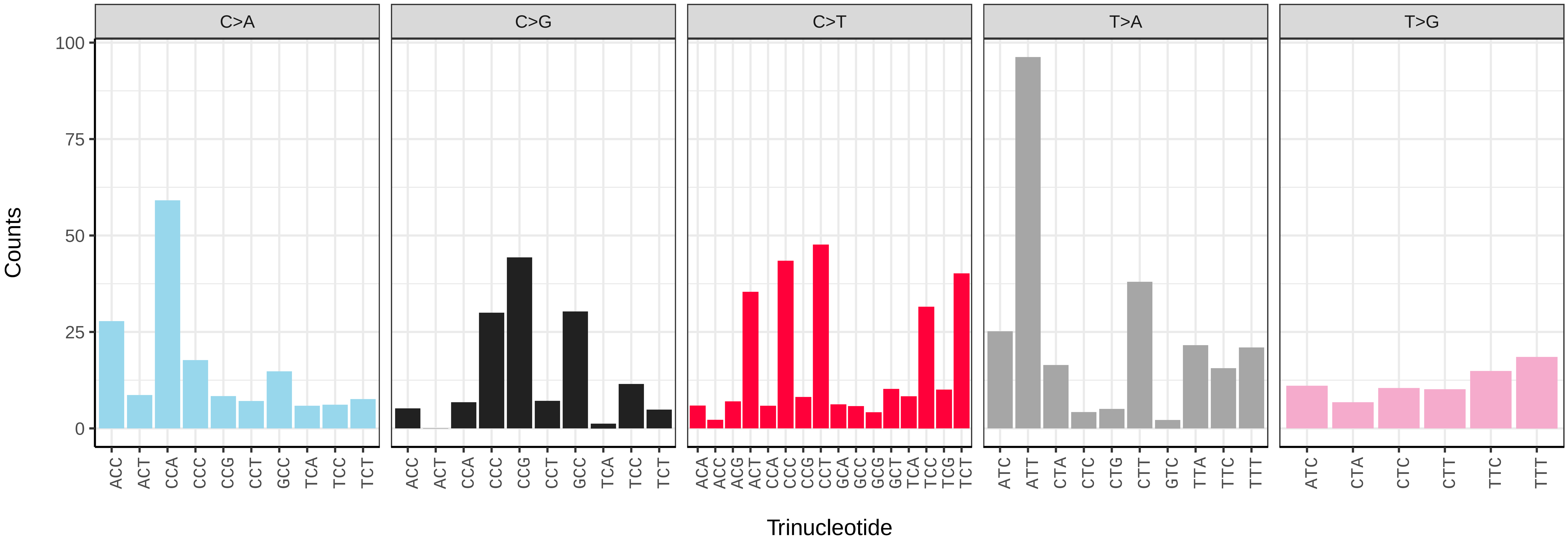

*Simplicillium lamellicola* (grSimLame1) germline mutational spectrum

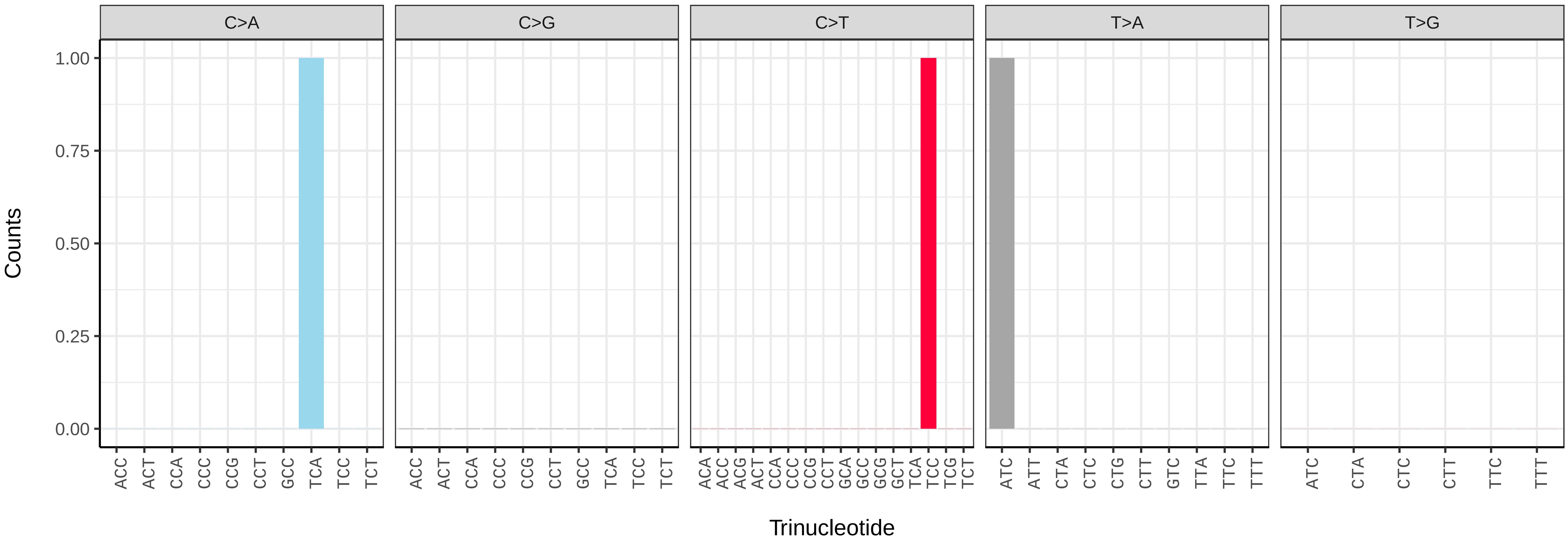


*Simplicillium lamellicola* (grSimLame1) somatic mutational spectrum (SBS96 classification)

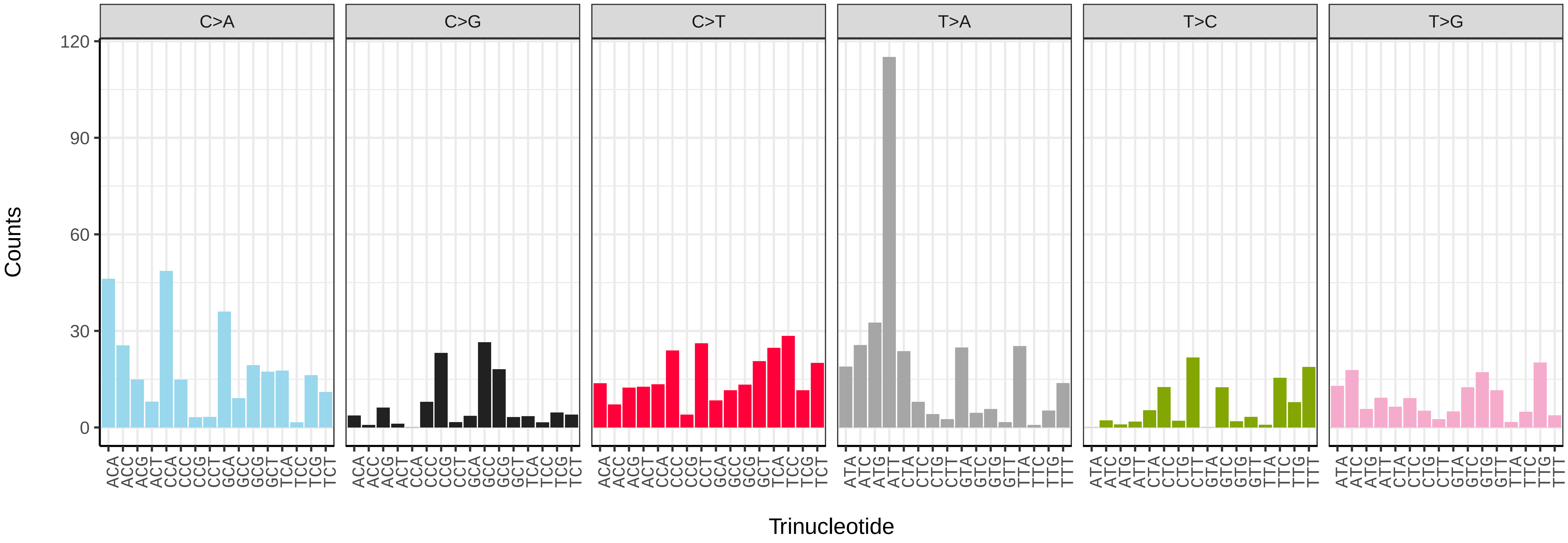


*Simplicillium lamellicola* (grSimLame1) somatic mutational spectrum (SBS52 classification)

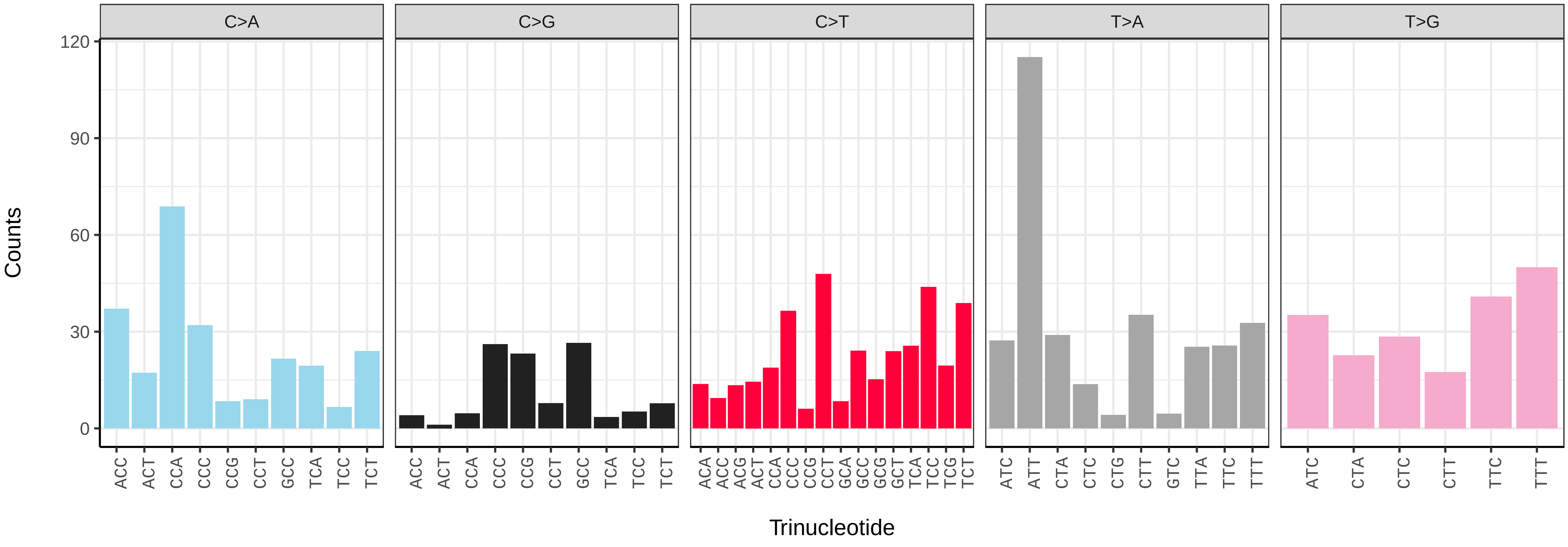

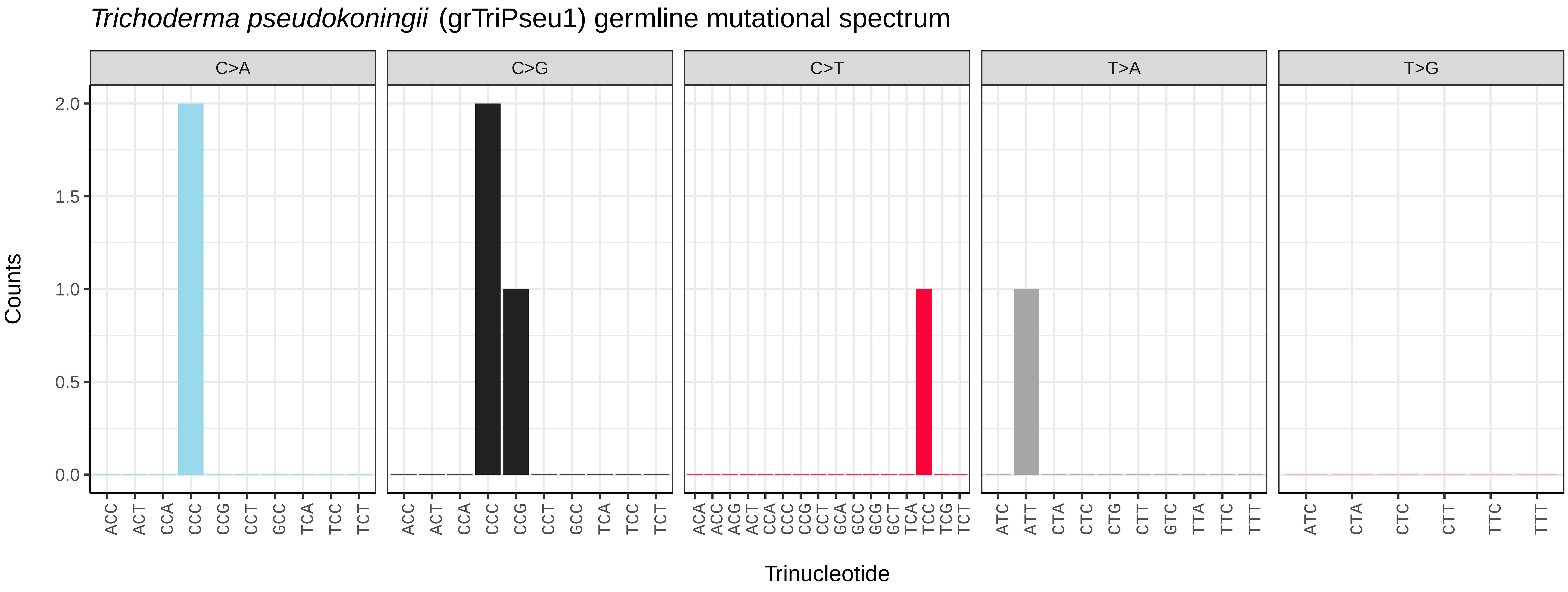
Trichoderma pseudokoningii (grTriPseu1) germline mutational spectrum
C>A
C>G
C>T
T>A
T>G
Counts
Trinucleotide
0.0
0.5
1.0
1.5
2.0

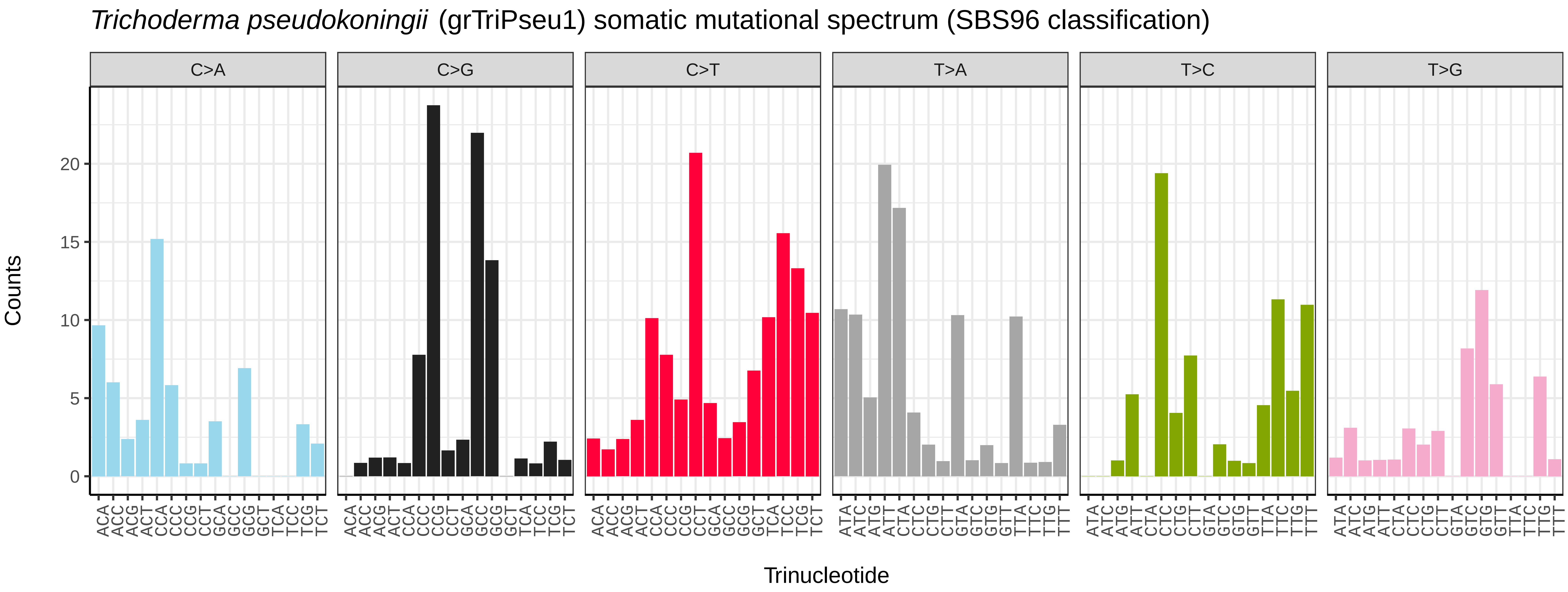
Trichoderma pseudokoningii (grTriPseu1) somatic mutational spectrum (SBS96 classification)
C>A
C>G
C>T
T>A
T>C
T>G
Counts
Trinucleotide
0
5
10
15
20

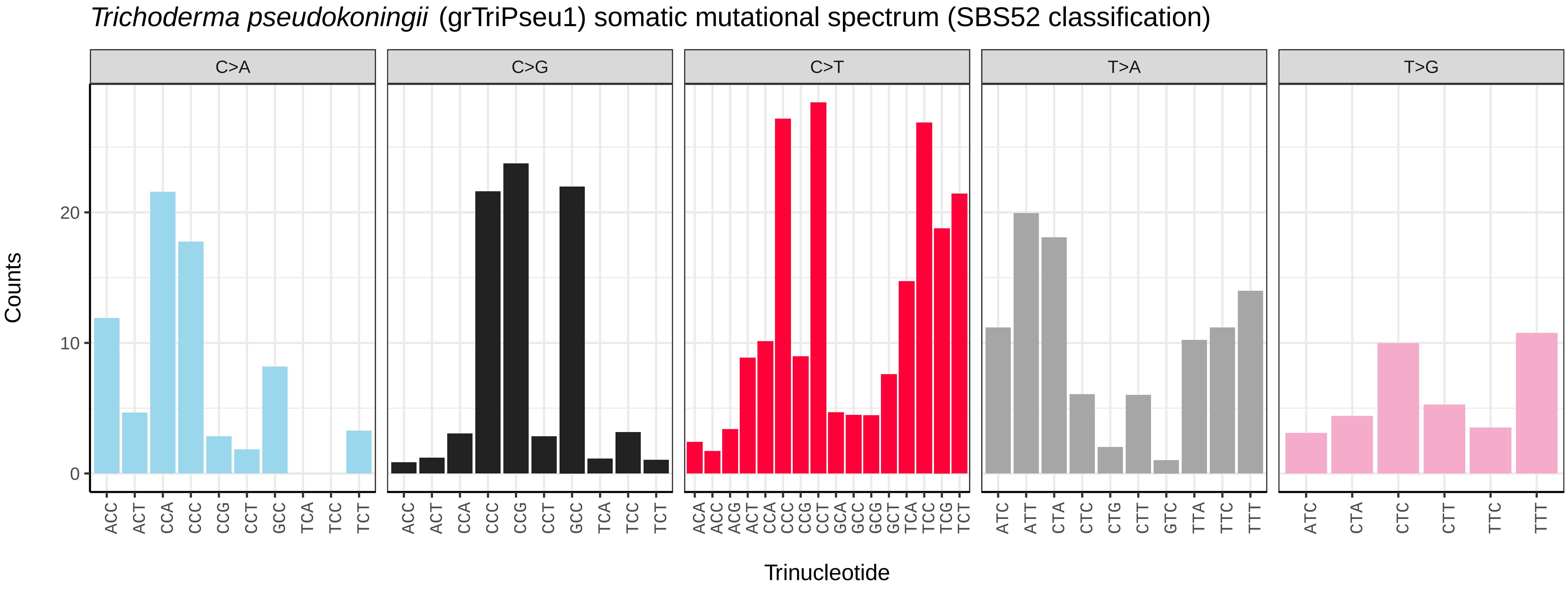
Trichoderma pseudokoningii (grTriPseu1) somatic mutational spectrum (SBS52 classification)
C>A
C>G
C>T
T>A
T>G
Counts
Trinucleotide
0
10
20

*Metschnikowia zobellii* (gsMetZobe1) germline mutational spectrum

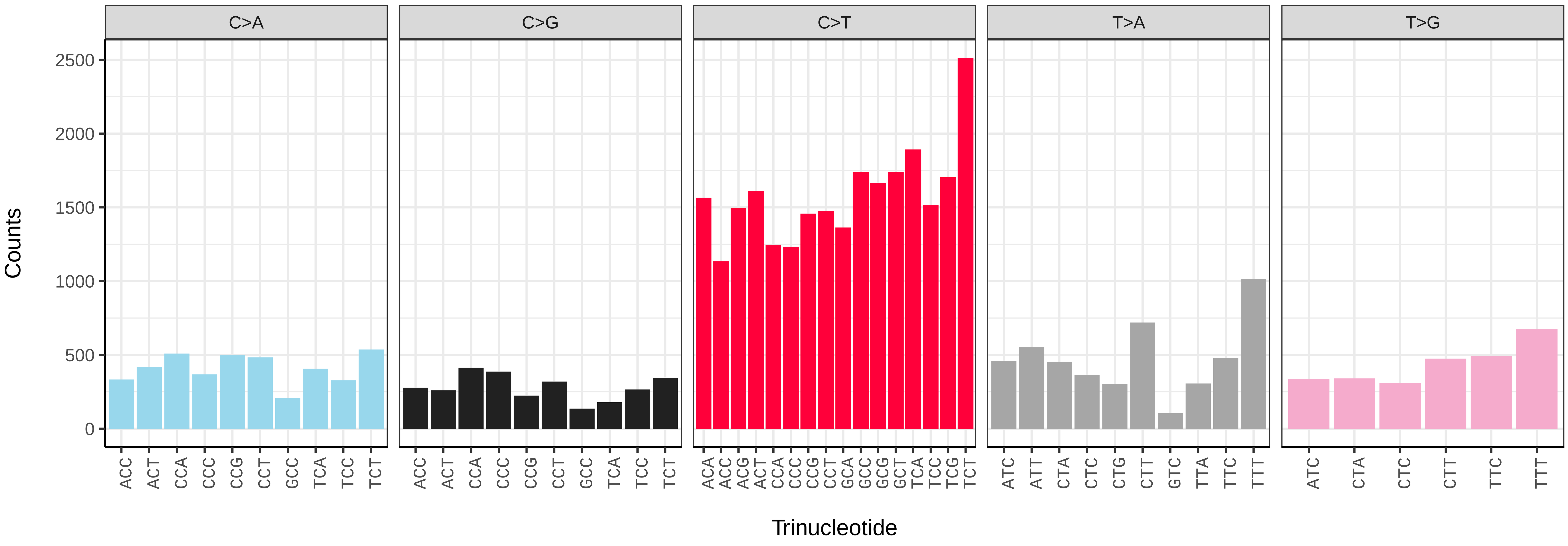


*Metschnikowia zobellii* (gsMetZobe1) somatic mutational spectrum (SBS96 classification)

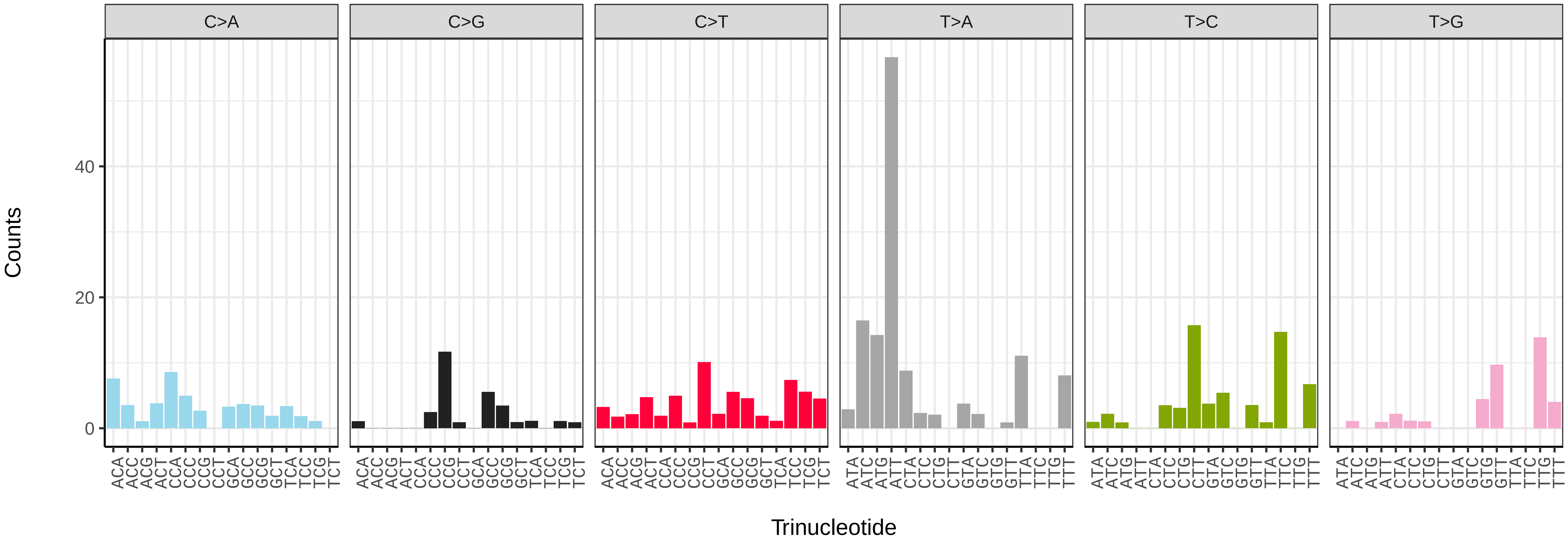


*Metschnikowia zobellii* (gsMetZobe1) somatic mutational spectrum (SBS52 classification)

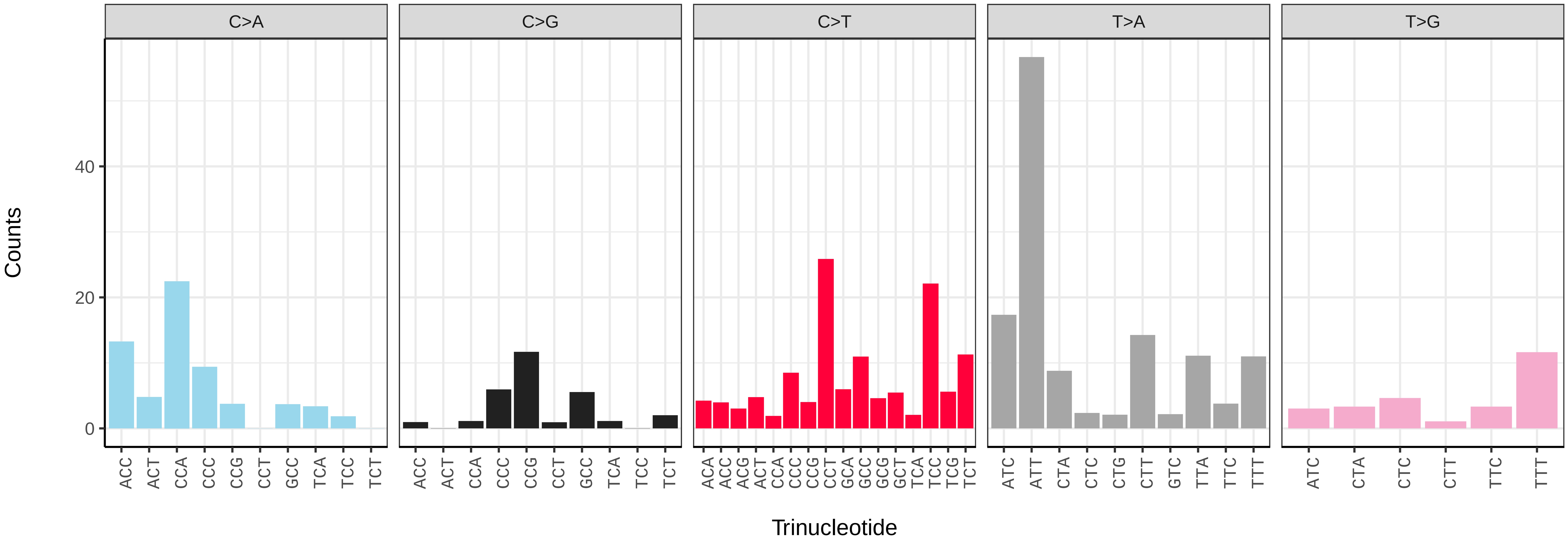

*Entomortierella parvispora* (gzEntParv1) germline mutational spectrum

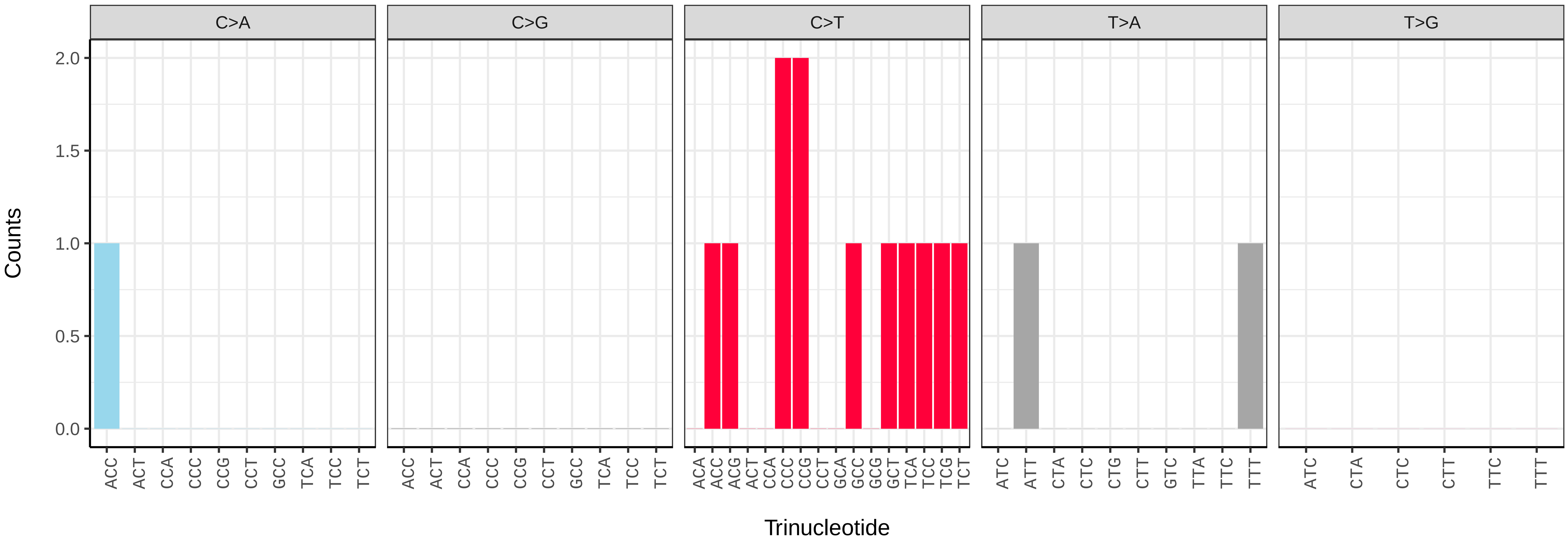


*Entomortierella parvispora* (gzEntParv1) somatic mutational spectrum (SBS96 classification)

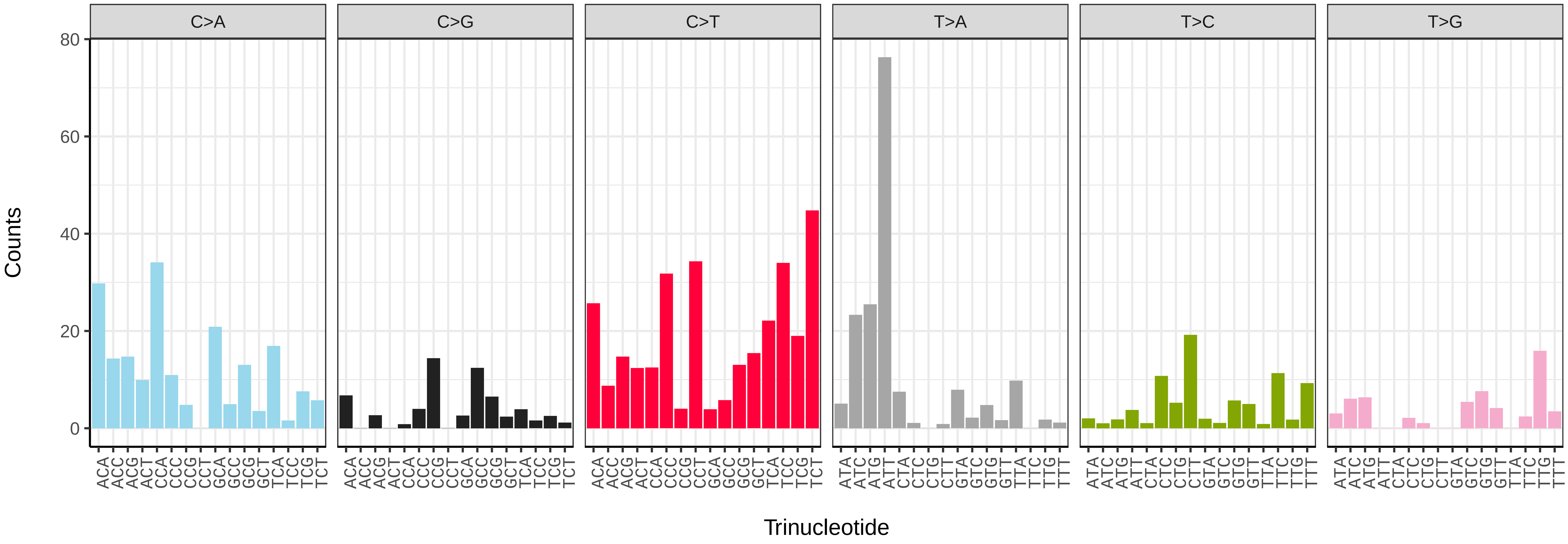


*Entomortierella parvispora* (gzEntParv1) somatic mutational spectrum (SBS52 classification)

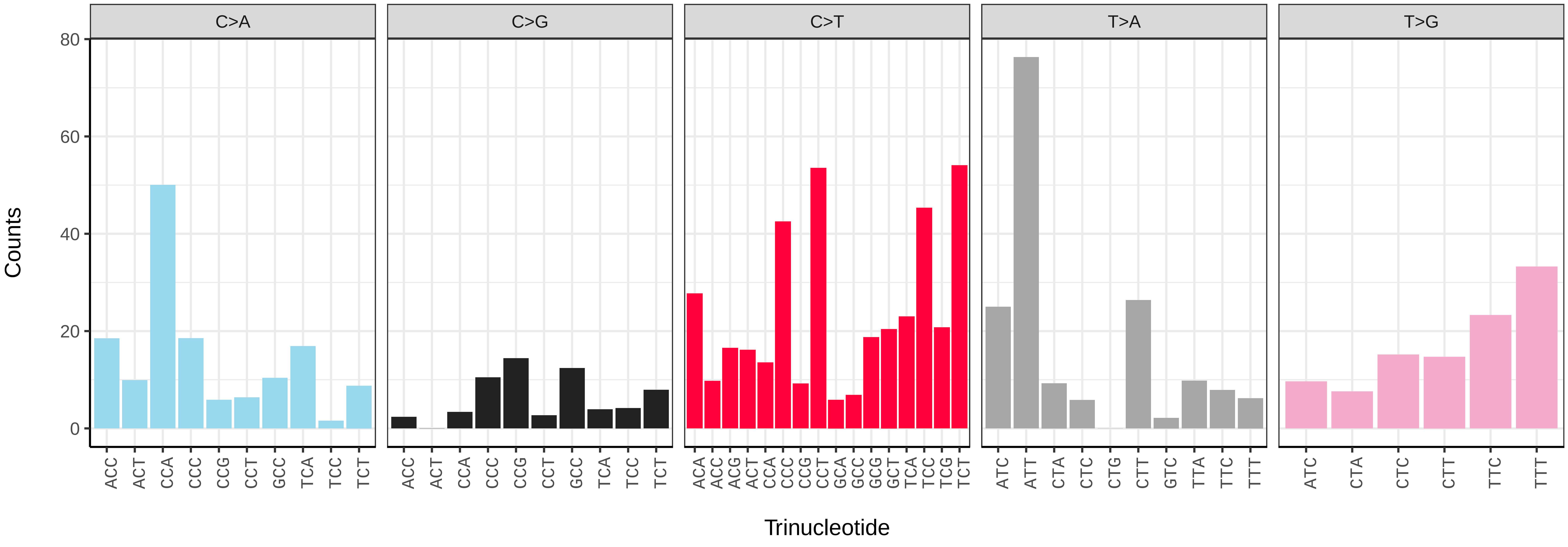

*Mucor piriformis* (gzMucPiri1) germline mutational spectrum

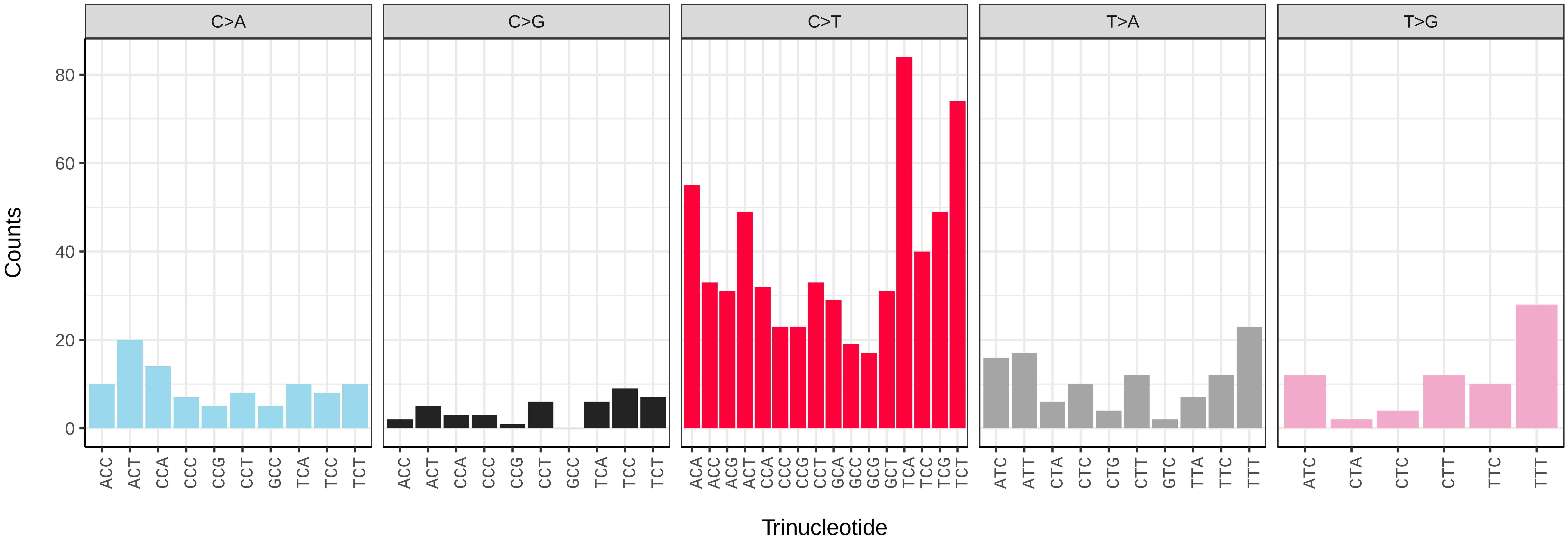


*Mucor piriformis* (gzMucPiri1) somatic mutational spectrum (SBS96 classification)

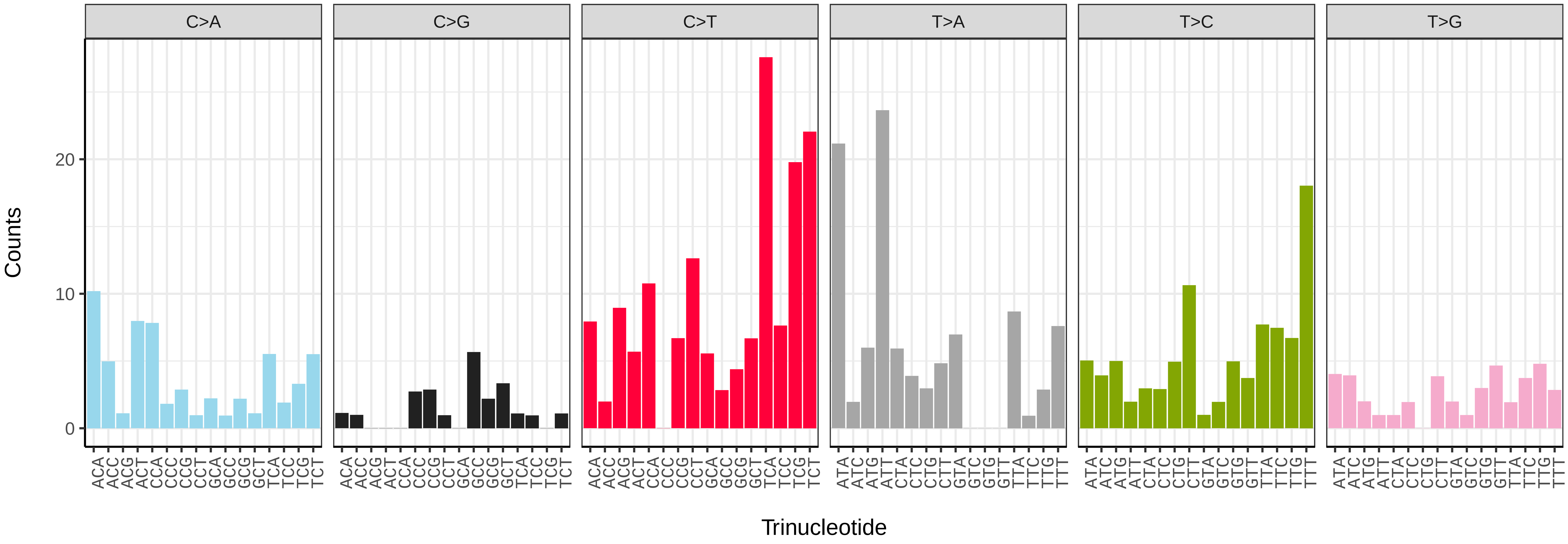


*Mucor piriformis* (gzMucPiri1) somatic mutational spectrum (SBS52 classification)

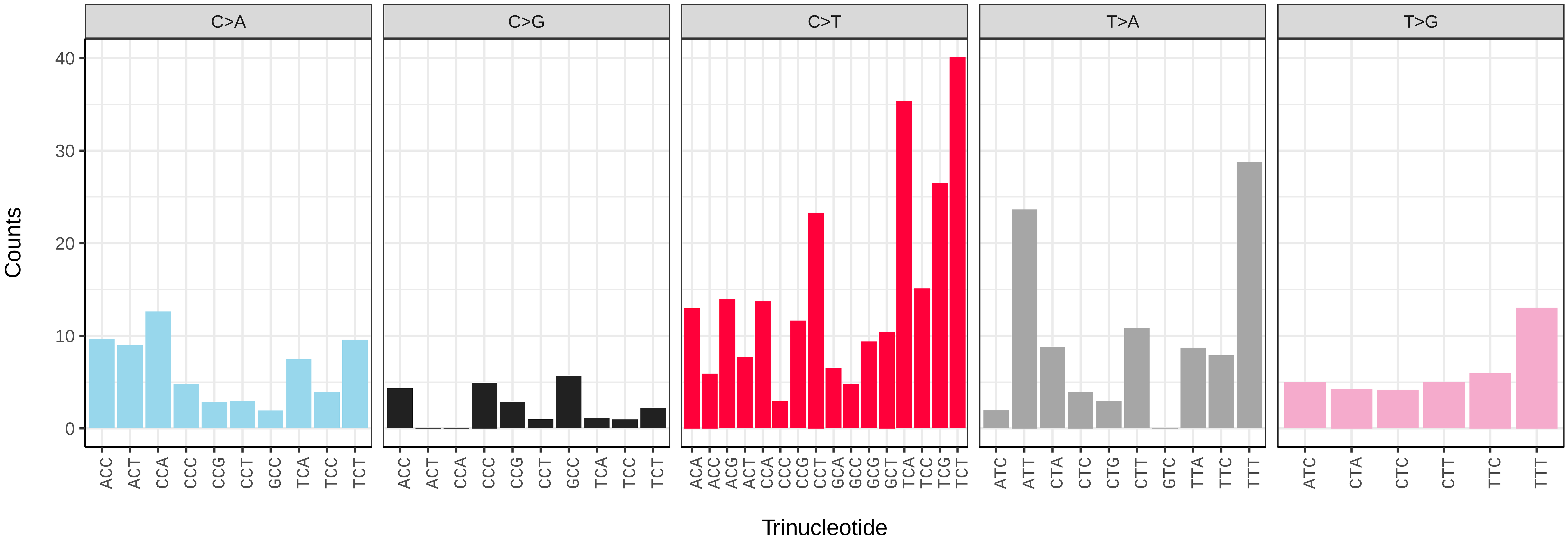

*Adalia bipunctata* (icAdaBipu1) germline mutational spectrum

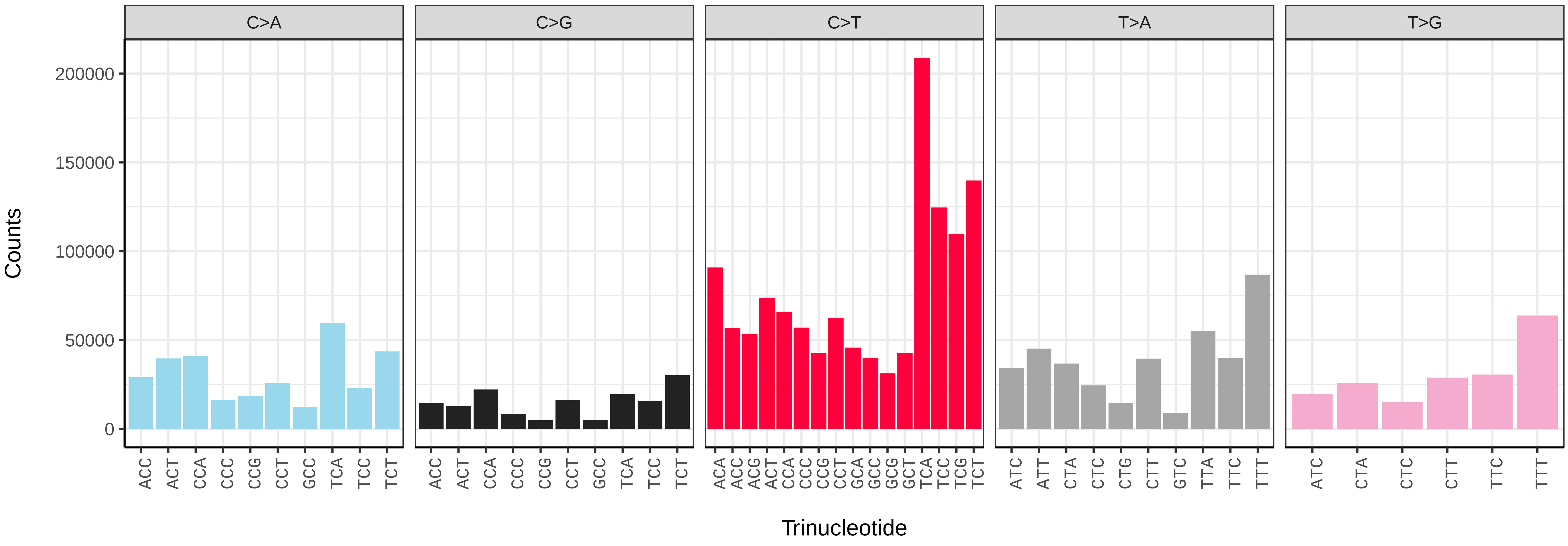


*Adalia bipunctata* (icAdaBipu1) somatic mutational spectrum (SBS96 classification)

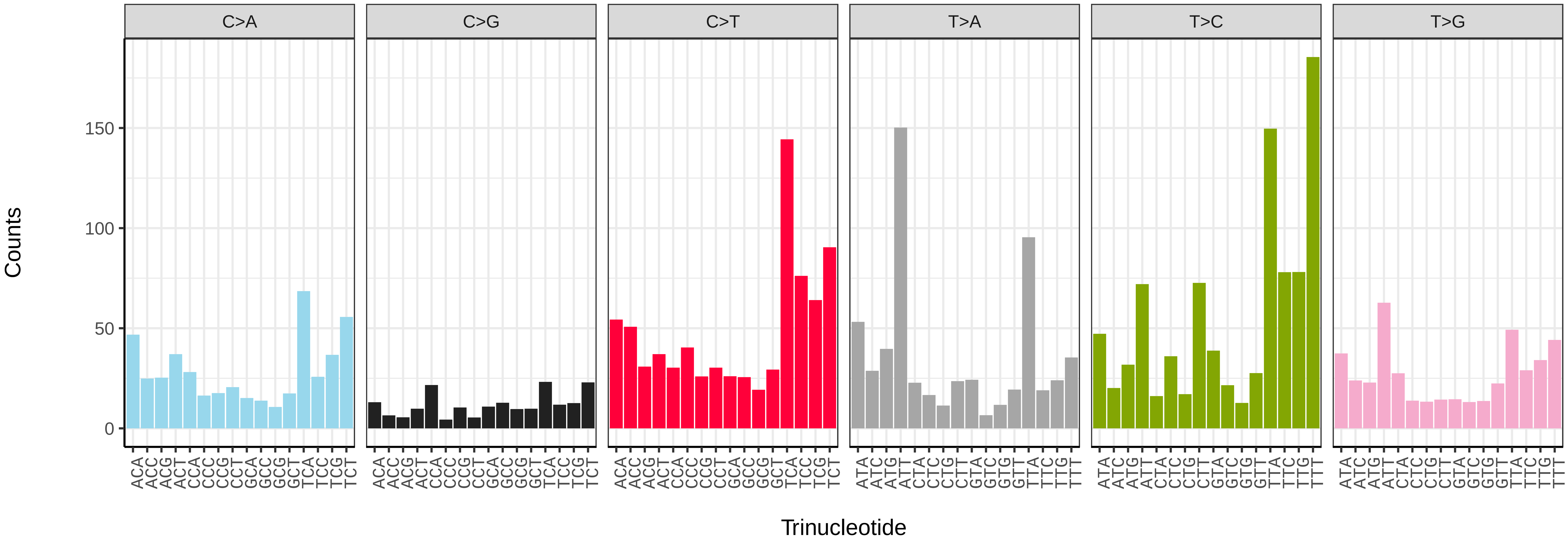


*Adalia bipunctata* (icAdaBipu1) somatic mutational spectrum (SBS52 classification)

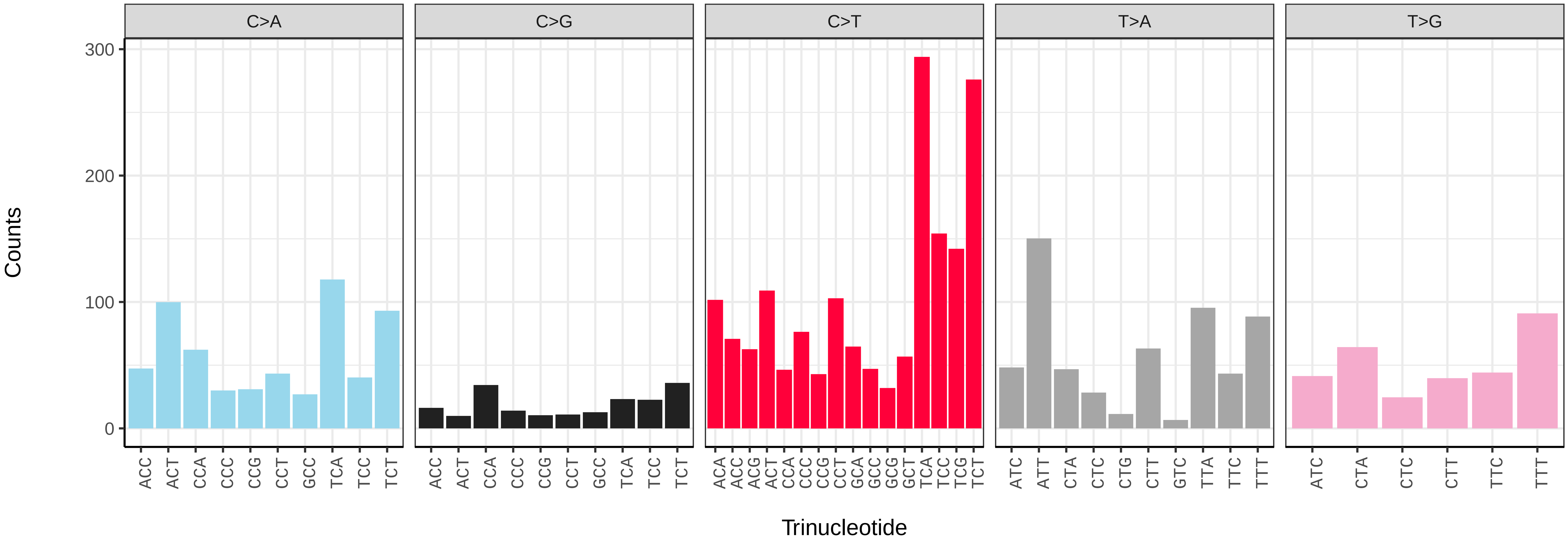

*Adalia bipunctata* (icAdaBipu1.icAdaBipu15) germline mutational spectrum

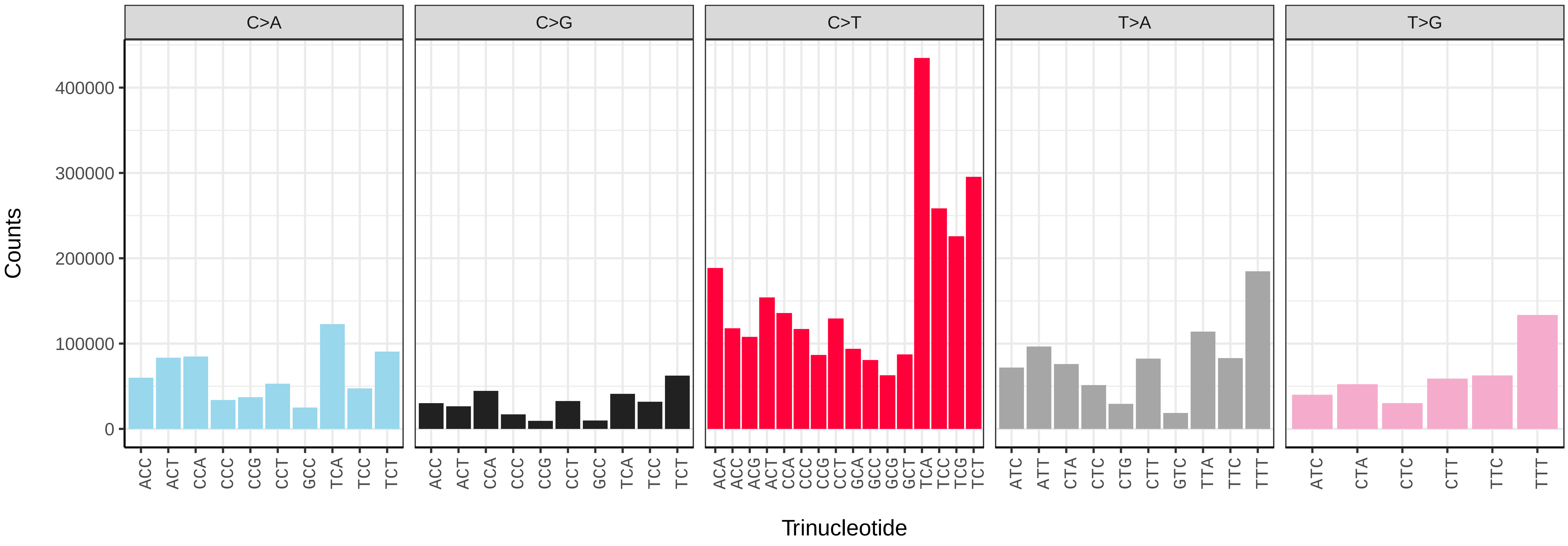


*Adalia bipunctata* (icAdaBipu1.icAdaBipu15) somatic mutational spectrum (SBS96 classification)

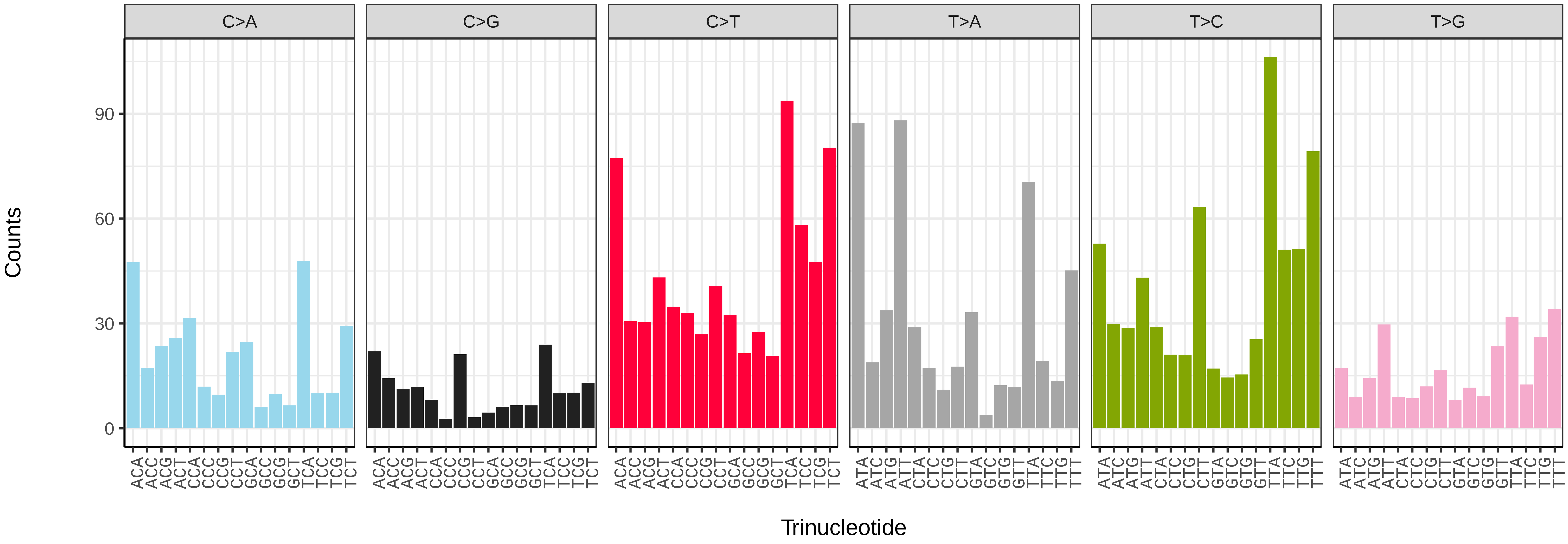


*Adalia bipunctata* (icAdaBipu1.icAdaBipu15) somatic mutational spectrum (SBS52 classification)

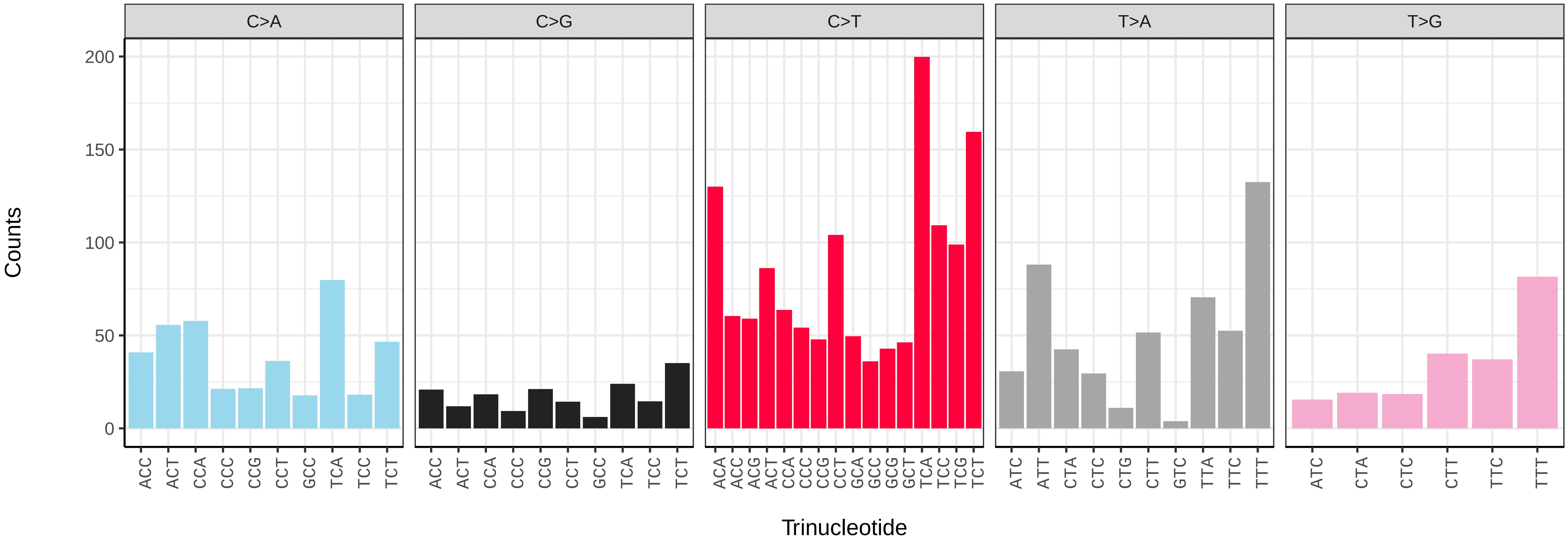

*Adalia bipunctata* (icAdaBipu1.icAdaBipu20) germline mutational spectrum

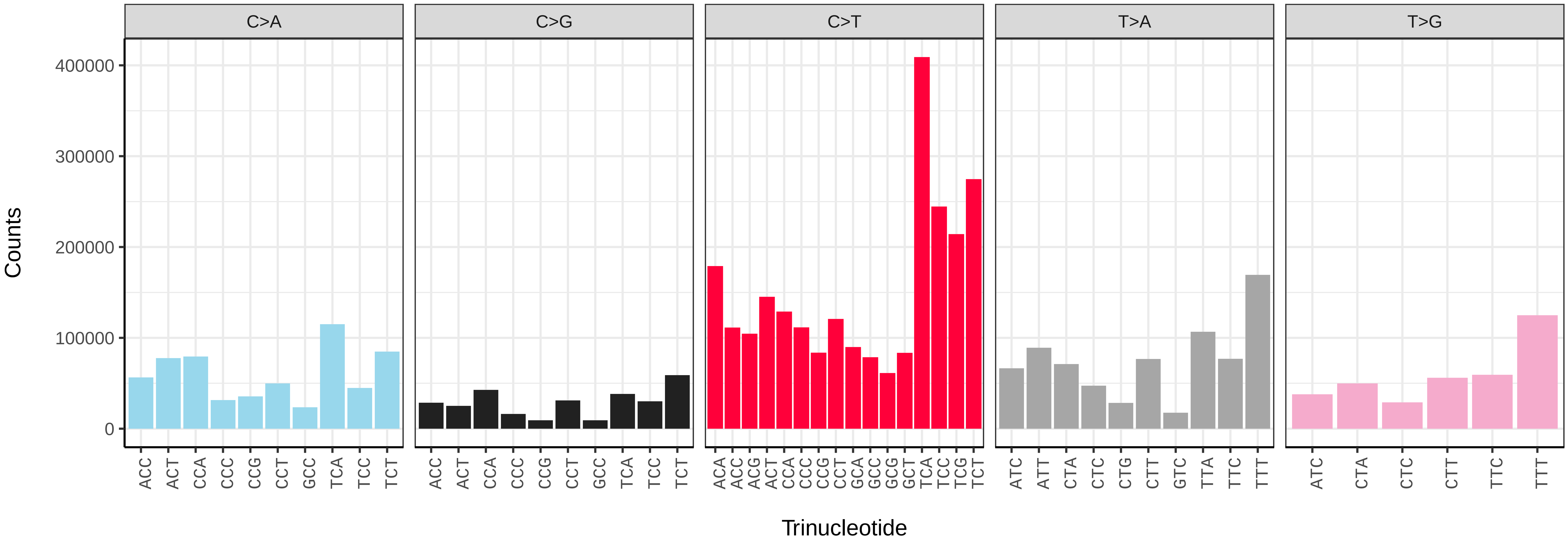


*Adalia bipunctata* (icAdaBipu1.icAdaBipu20) somatic mutational spectrum (SBS96 classification)

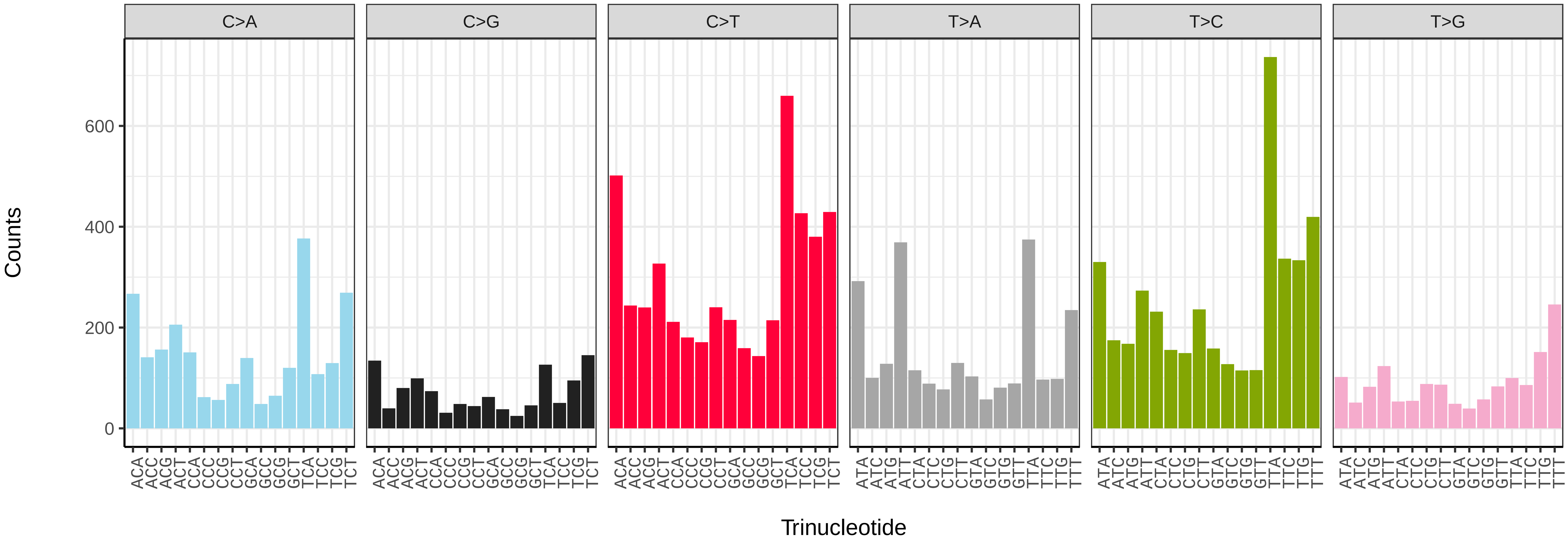


*Adalia bipunctata* (icAdaBipu1.icAdaBipu20) somatic mutational spectrum (SBS52 classification)

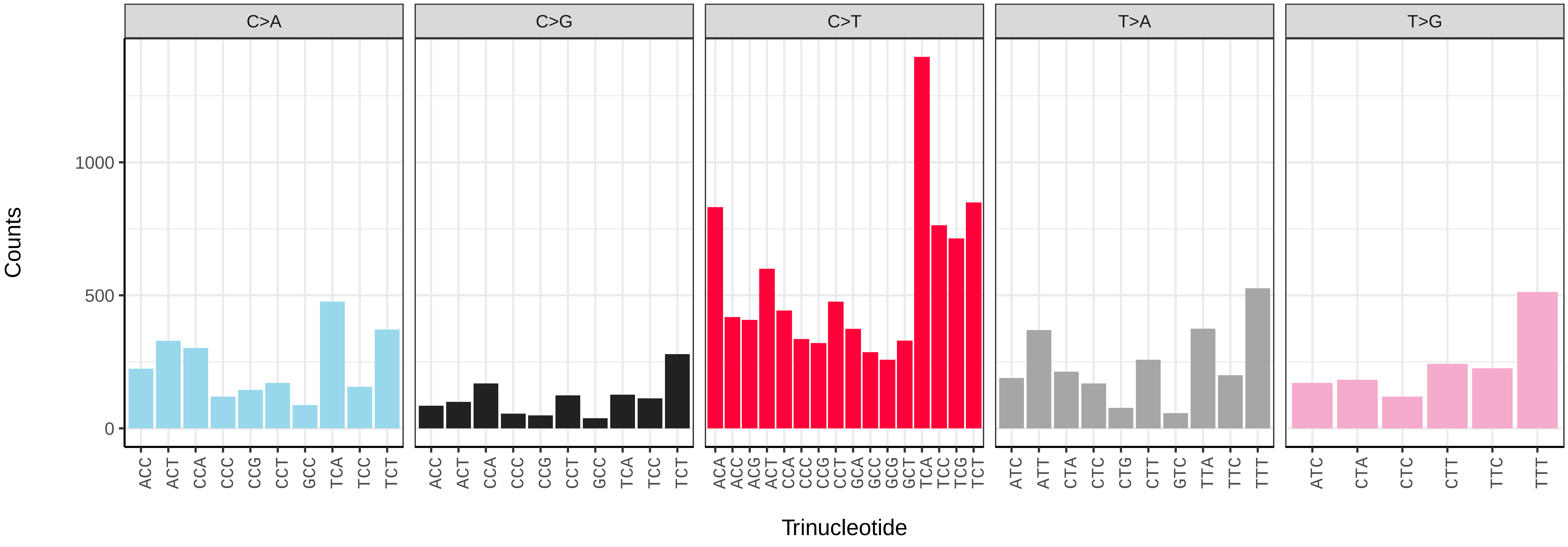

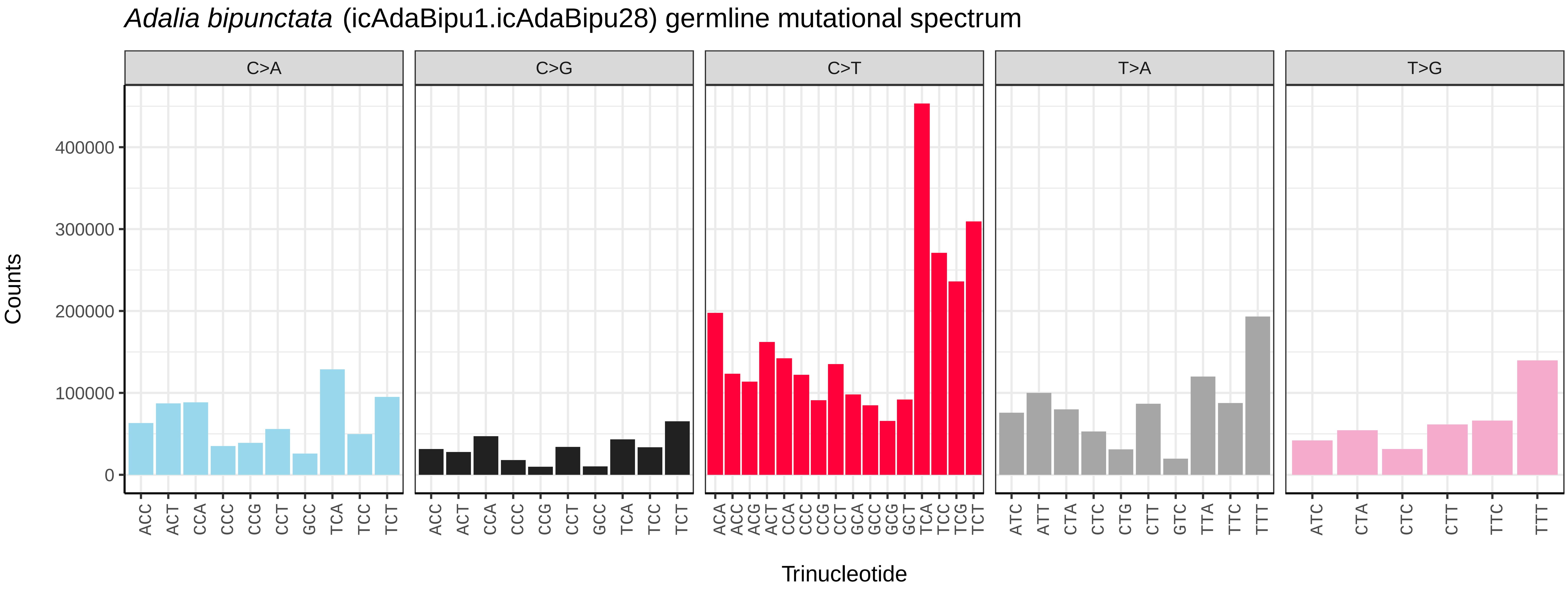
*Adalia bipunctata* (icAdaBipu1.icAdaBipu28) germline mutational spectrum
C>A
C>G
C>T
T>A
T>G
Counts
0
100000
200000
300000
400000
ACC ACT CCA CCC CCG CCT GCC TCA TCC TCT
ACC ACT CCA CCC CCG CCT GCC TCA TCC TCT
ACA ACC ACG ACT CCA CCC CCG CCT GCA GCC GCG GCT TCA TCC TCG TCT
ATC ATT CTA CTC CTG CTT GTC TTA TTC TTT
ATC CTA CTC CTT TTC TTT
Trinucleotide

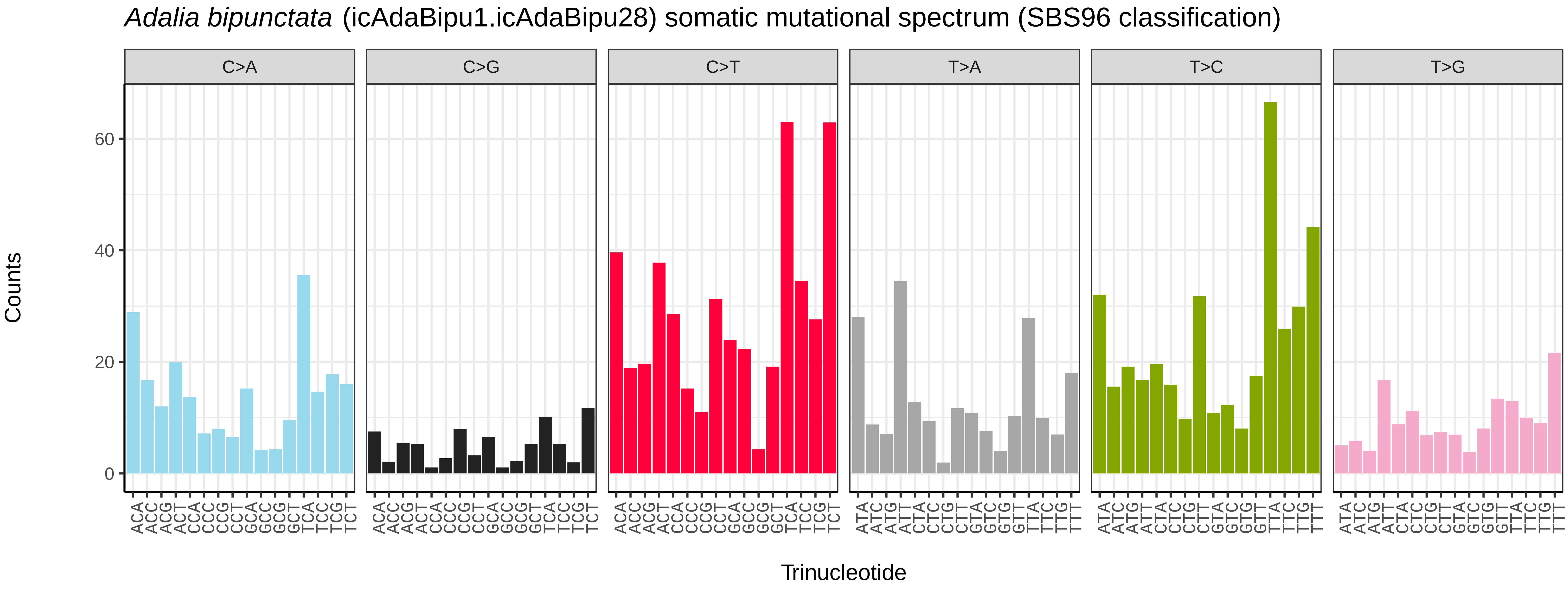
*Adalia bipunctata* (icAdaBipu1.icAdaBipu28) somatic mutational spectrum (SBS96 classification)
C>A
C>G
C>T
T>A
T>C
T>G
Counts
0
20
40
60
ACA ACC ACG ACT CCA CCC CCG CCT GCA GCC GCG GCT TCA TCC TCG TCT
ACA ACC ACG ACT CCA CCC CCG CCT GCA GCC GCG GCT TCA TCC TCG TCT
ACA ACC ACG ACT CCA CCC CCG CCT GCA GCC GCG GCT TCA TCC TCG TCT
ATA ATC ATG ATT CTA CTC CTG CTT GTA GTC GTG GTT TTA TTC TTG TTT
ATA ATC ATG ATT CTA CTC CTG CTT GTA GTC GTG GTT TTA TTC TTG TTT
ATA ATC ATG ATT CTA CTC CTG CTT GTA GTC GTG GTT TTA TTC TTG TTT
Trinucleotide

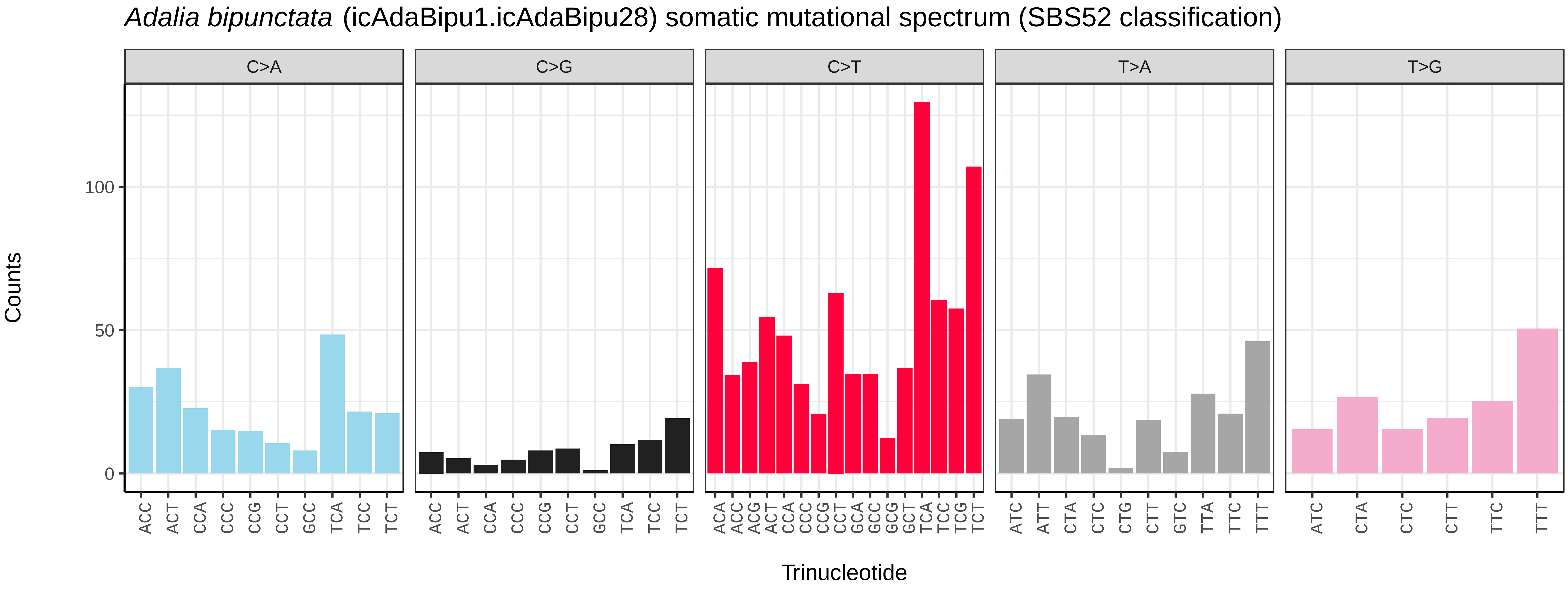
*Adalia bipunctata* (icAdaBipu1.icAdaBipu28) somatic mutational spectrum (SBS52 classification)
C>A
C>G
C>T
T>A
T>G
Counts
0
50
100
ACC ACT CCA CCC CCG CCT GCC TCA TCC TCT
ACC ACT CCA CCC CCG CCT GCC TCA TCC TCT
ACA ACC ACG ACT CCA CCC CCG CCT GCA GCC GCG GCT TCA TCC TCG TCT
ATC ATT CTA CTC CTG CTT GTC TTA TTC TTT
ATC CTA CTC CTT TTC TTT
Trinucleotide

*Adalia bipunctata* (icAdaBipu1.icAdaBipu29) germline mutational spectrum

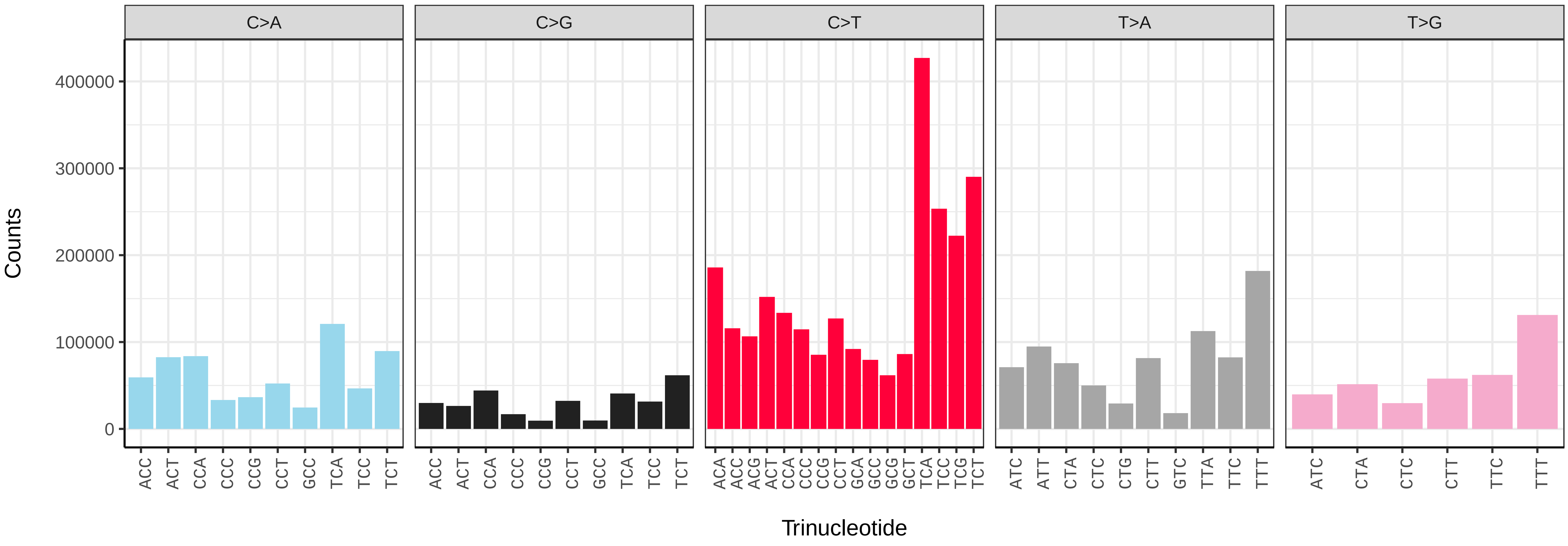


*Adalia bipunctata* (icAdaBipu1.icAdaBipu29) somatic mutational spectrum (SBS96 classification)

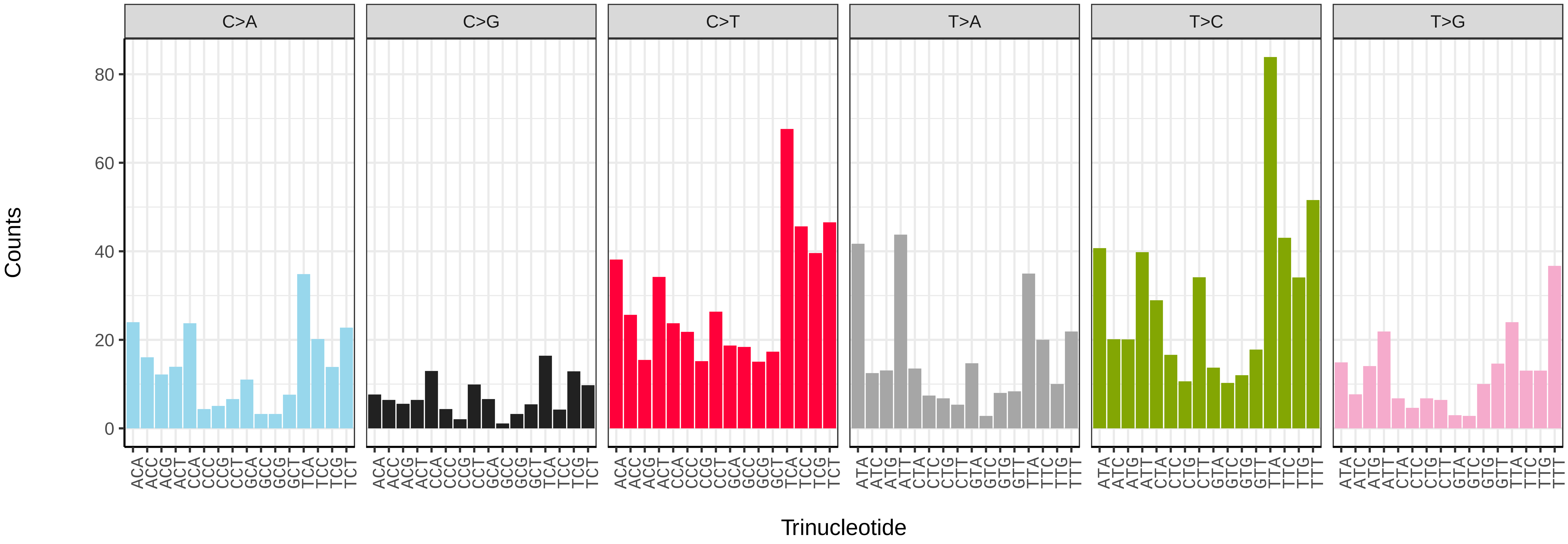


*Adalia bipunctata* (icAdaBipu1.icAdaBipu29) somatic mutational spectrum (SBS52 classification)

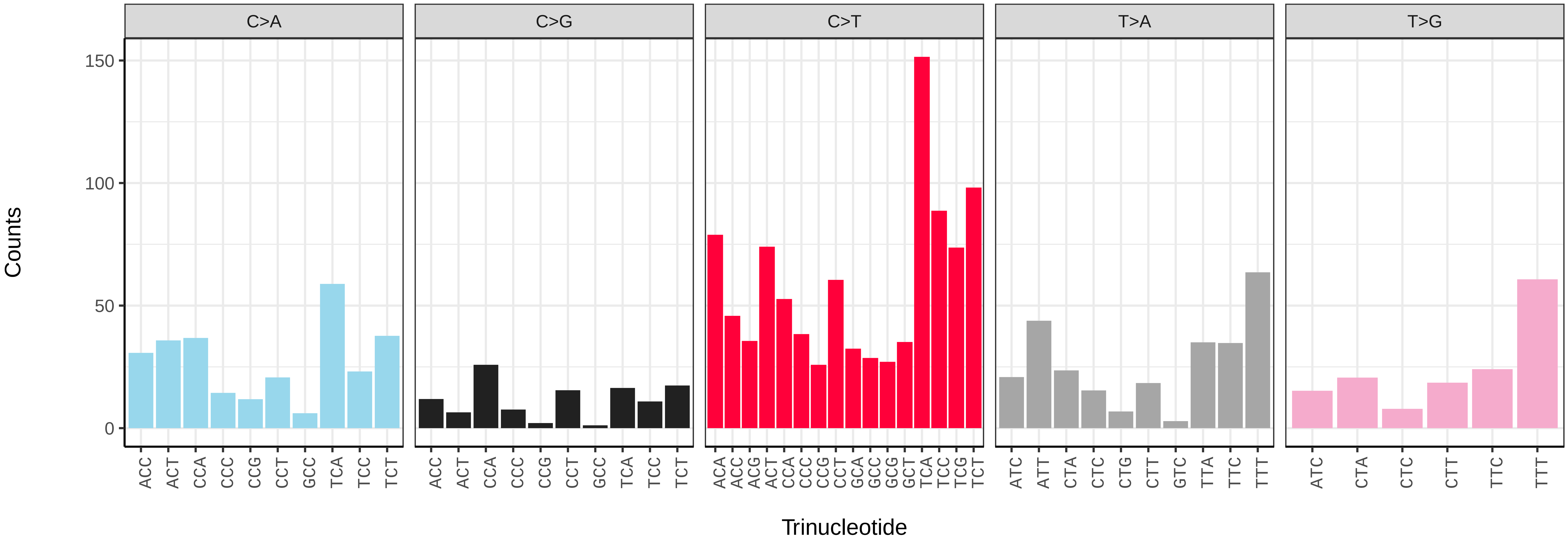

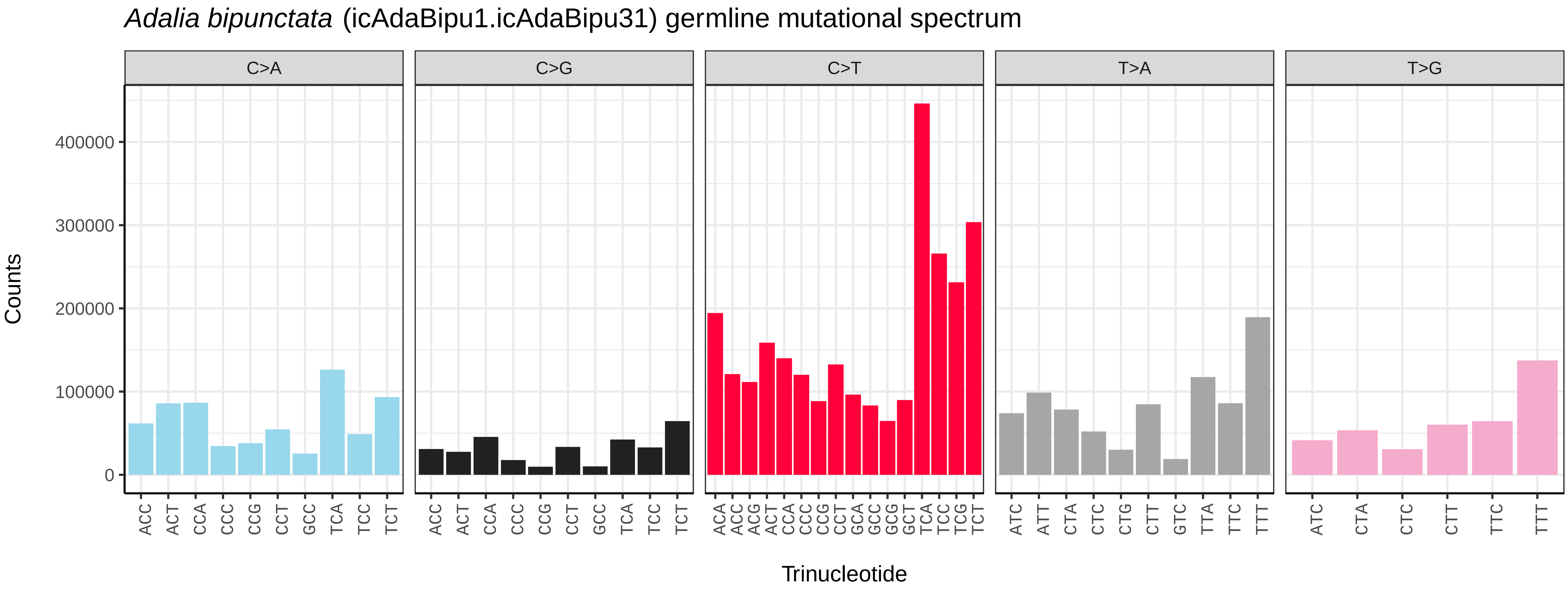
Adalia bipunctata (icAdaBipu1.icAdaBipu31) germline mutational spectrum
C>A
C>G
C>T
T>A
T>G
Counts
400000
300000
200000
100000
0
ACC ACT CCA CCC CCG CCT GCC TCA TCC TCT
ACC ACT CCA CCC CCG CCT GCC TCA TCC TCT
ACA ACC ACG ACT CCA CCC CCG CCT GCA GCC GCG GCT TCA TCC TCG TCT
ATC ATT CTA CTC CTG CTT GTC TTA TTC TTT
ATC CTA CTC CTT TTC TTT
Trinucleotide

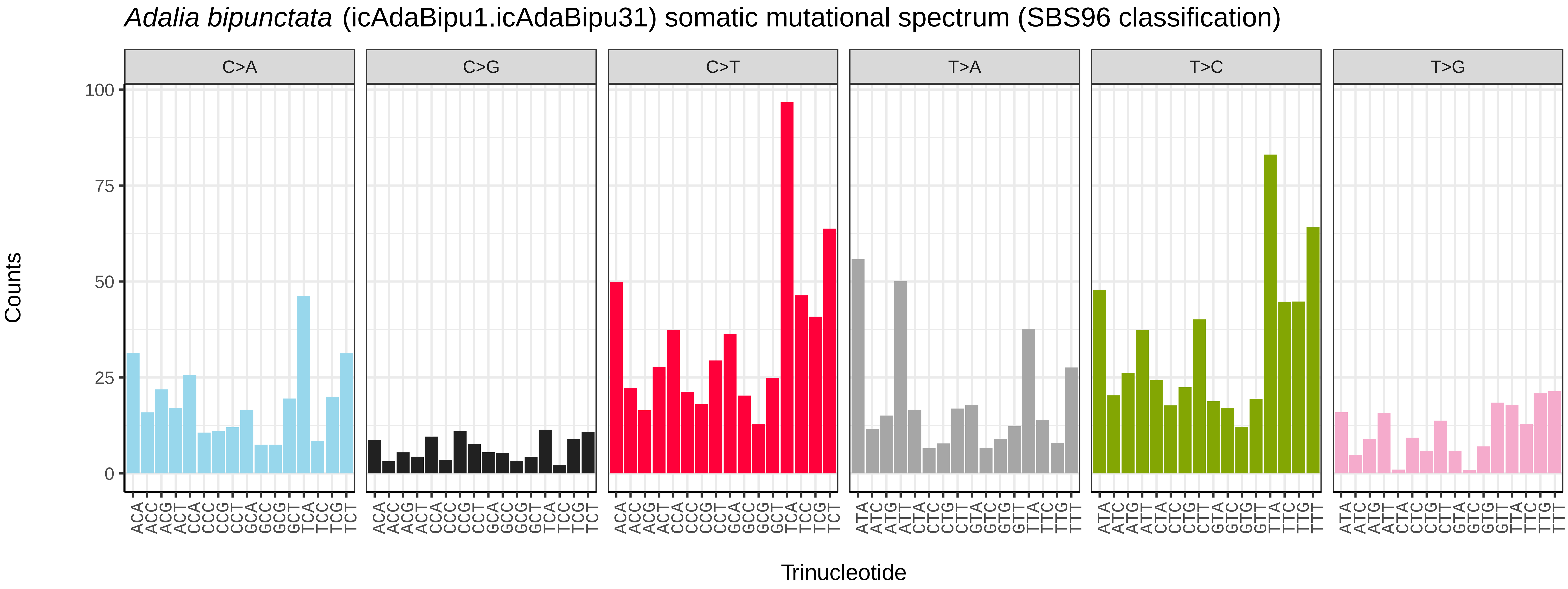
Adalia bipunctata (icAdaBipu1.icAdaBipu31) somatic mutational spectrum (SBS96 classification)
C>A
C>G
C>T
T>A
T>C
T>G
Counts
100
75
50
25
0
ACA ACC ACG ACT CCA CCC CCG CCT GCA GCC GCG GCT TCA TCC TCG TCT
ACA ACC ACG ACT CCA CCC CCG CCT GCA GCC GCG GCT TCA TCC TCG TCT
ACA ACC ACG ACT CCA CCC CCG CCT GCA GCC GCG GCT TCA TCC TCG TCT
ATA ATC ATG ATT CTA CTC CTG CTT GTA GTC GTG GTT TTA TTC TTG TTT
ATA ATC ATG ATT CTA CTC CTG CTT GTA GTC GTG GTT TTA TTC TTG TTT
ATA ATC ATG ATT CTA CTC CTG CTT GTA GTC GTG GTT TTA TTC TTG TTT
Trinucleotide

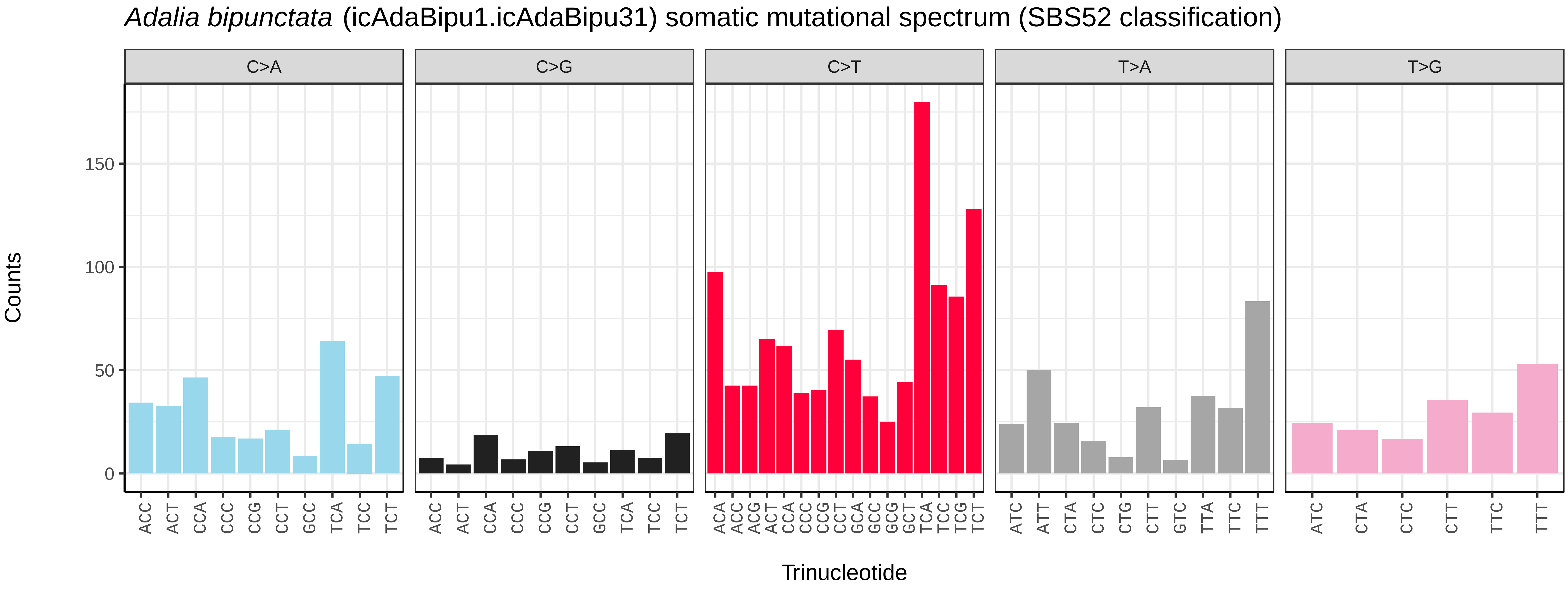
Adalia bipunctata (icAdaBipu1.icAdaBipu31) somatic mutational spectrum (SBS52 classification)
C>A
C>G
C>T
T>A
T>G
Counts
150
100
50
0
ACC ACT CCA CCC CCG CCT GCC TCA TCC TCT
ACC ACT CCA CCC CCG CCT GCC TCA TCC TCT
ACA ACC ACG ACT CCA CCC CCG CCT GCA GCC GCG GCT TCA TCC TCG TCT
ATC ATT CTA CTC CTG CTT GTC TTA TTC TTT
ATC CTA CTC CTT TTC TTT
Trinucleotide

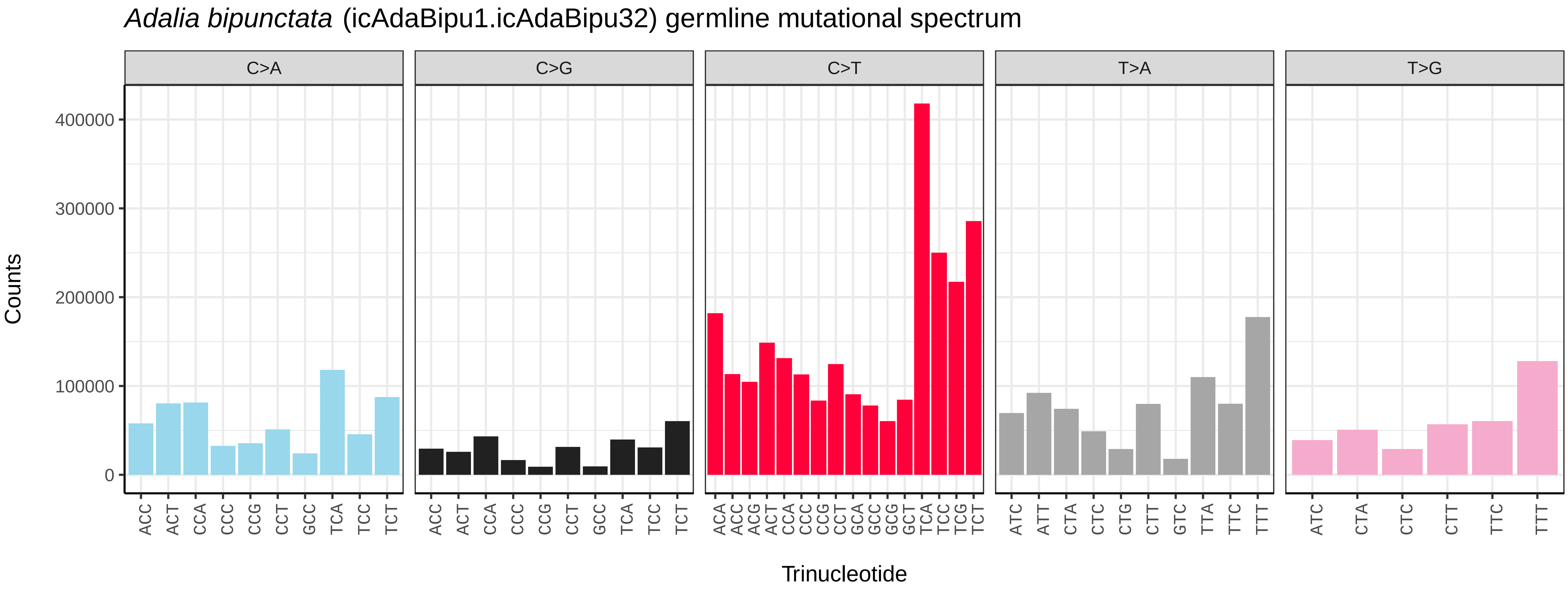
*Adalia bipunctata* (icAdaBipu1.icAdaBipu32) germline mutational spectrum
C>A
C>G
C>T
T>A
T>G
Counts
0
100000
200000
300000
400000
Trinucleotide
ACC ACT CCA CCC CCG CCT GCC TCA TCC TCT
ACA ACC ACG ACT CCA CCC CCG CCT GCA GCC GCG GCT TCA TCC TCG TCT
ATC ATT CTA CTC CTG CTT GTC TTA TTC TTT
ATC CTA CTC CTT TTC TTT

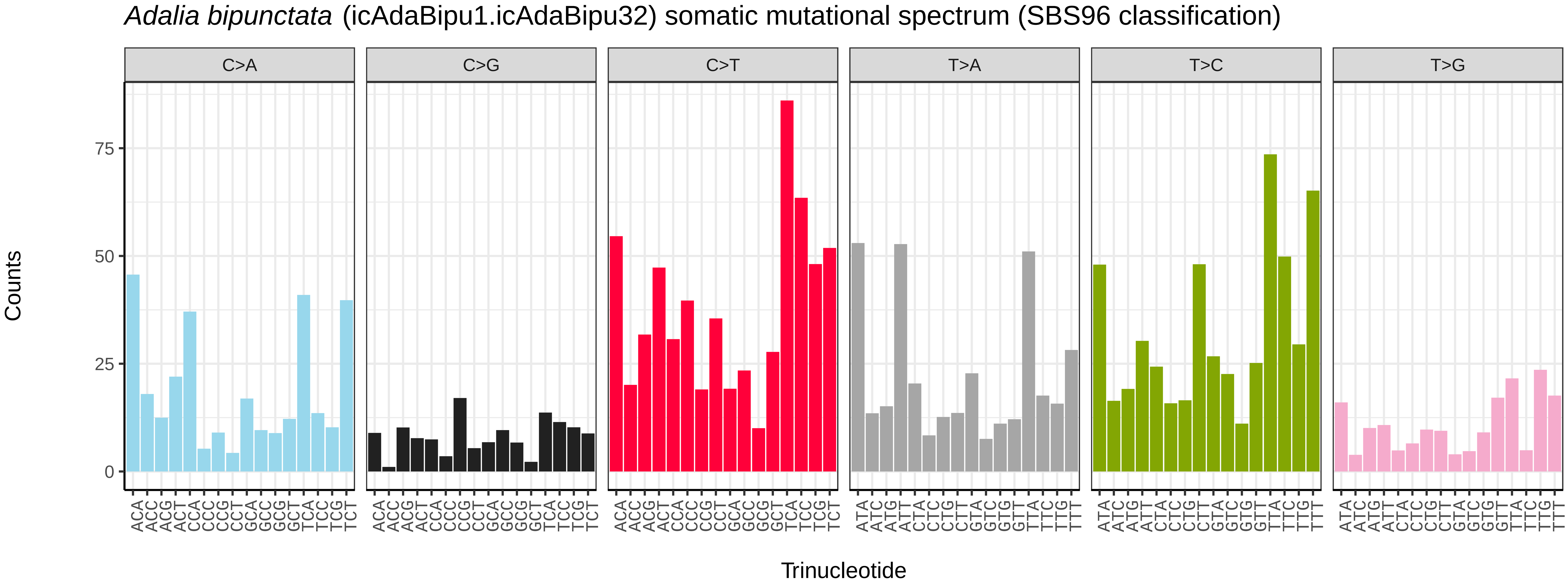
*Adalia bipunctata* (icAdaBipu1.icAdaBipu32) somatic mutational spectrum (SBS96 classification)
C>A
C>G
C>T
T>A
T>C
T>G
Counts
0
25
50
75
Trinucleotide
ACA ACC ACG ACT CCA CCC CCG CCT GCA GCC GCG GCT TCA TCC TCG TCT
ATA ATC ATG ATT CTA CTC CTG CTT GTA GTC GTG GTT TTA TTC TTG TTT

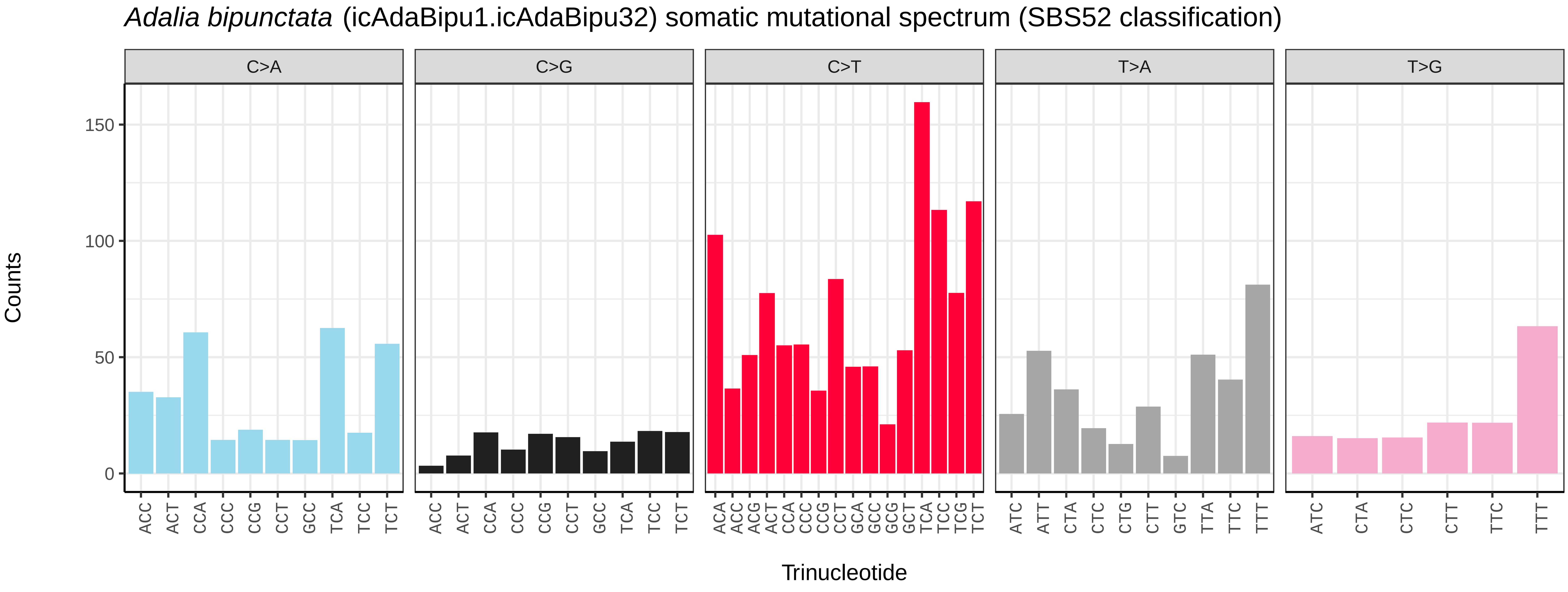
*Adalia bipunctata* (icAdaBipu1.icAdaBipu32) somatic mutational spectrum (SBS52 classification)
C>A
C>G
C>T
T>A
T>G
Counts
0
50
100
150
Trinucleotide
ACC ACT CCA CCC CCG CCT GCC TCA TCC TCT
ACA ACC ACG ACT CCA CCC CCG CCT GCA GCC GCG GCT TCA TCC TCG TCT
ATC ATT CTA CTC CTG CTT GTC TTA TTC TTT
ATC CTA CTC CTT TTC TTT

*Adalia bipunctata* (icAdaBipu1.icAdaBipu33) germline mutational spectrum

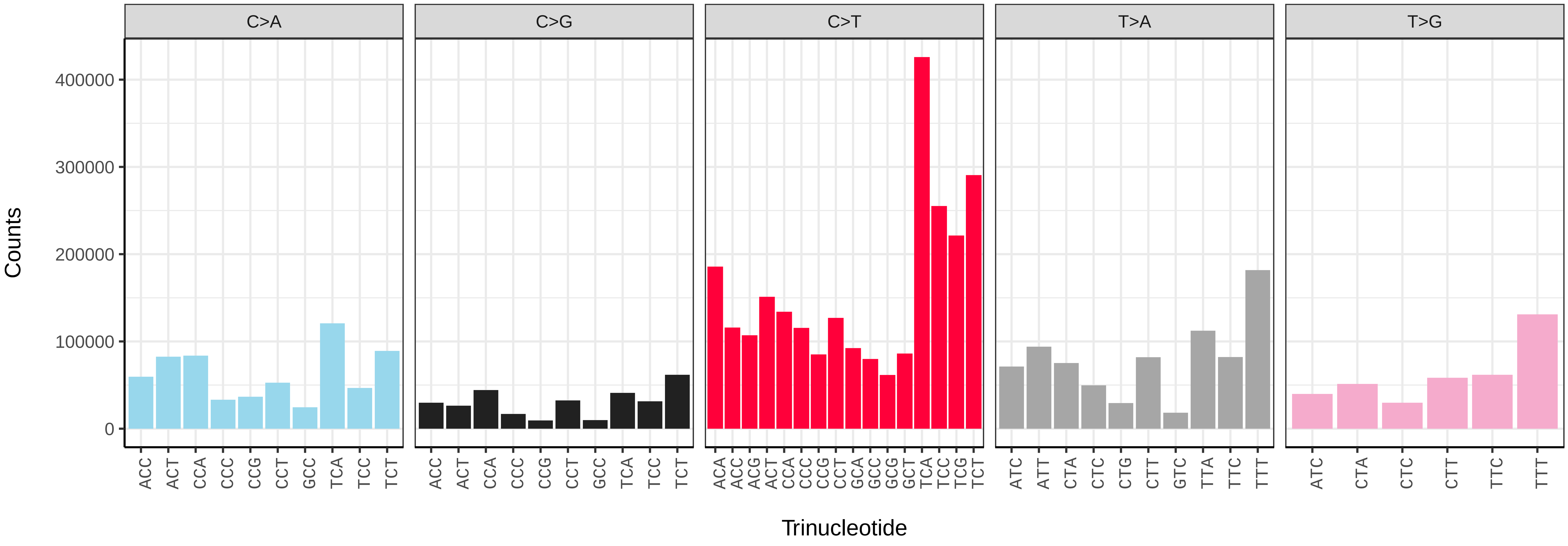


*Adalia bipunctata* (icAdaBipu1.icAdaBipu33) somatic mutational spectrum (SBS96 classification)

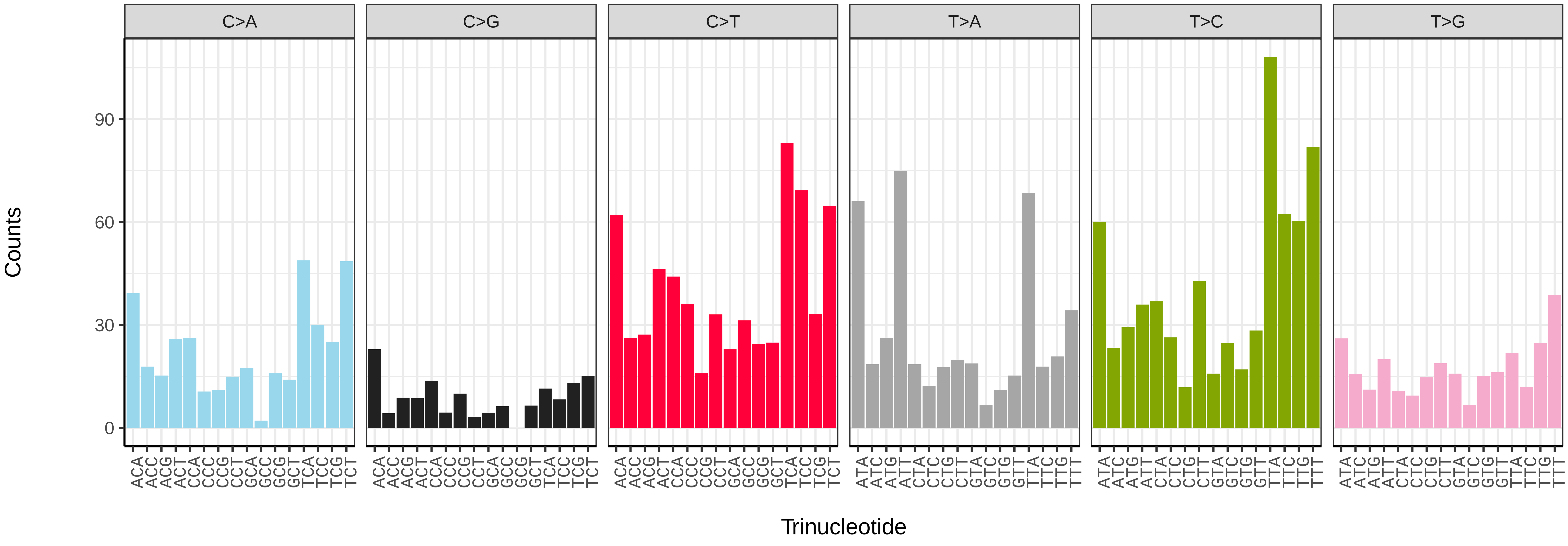


*Adalia bipunctata* (icAdaBipu1.icAdaBipu33) somatic mutational spectrum (SBS52 classification)

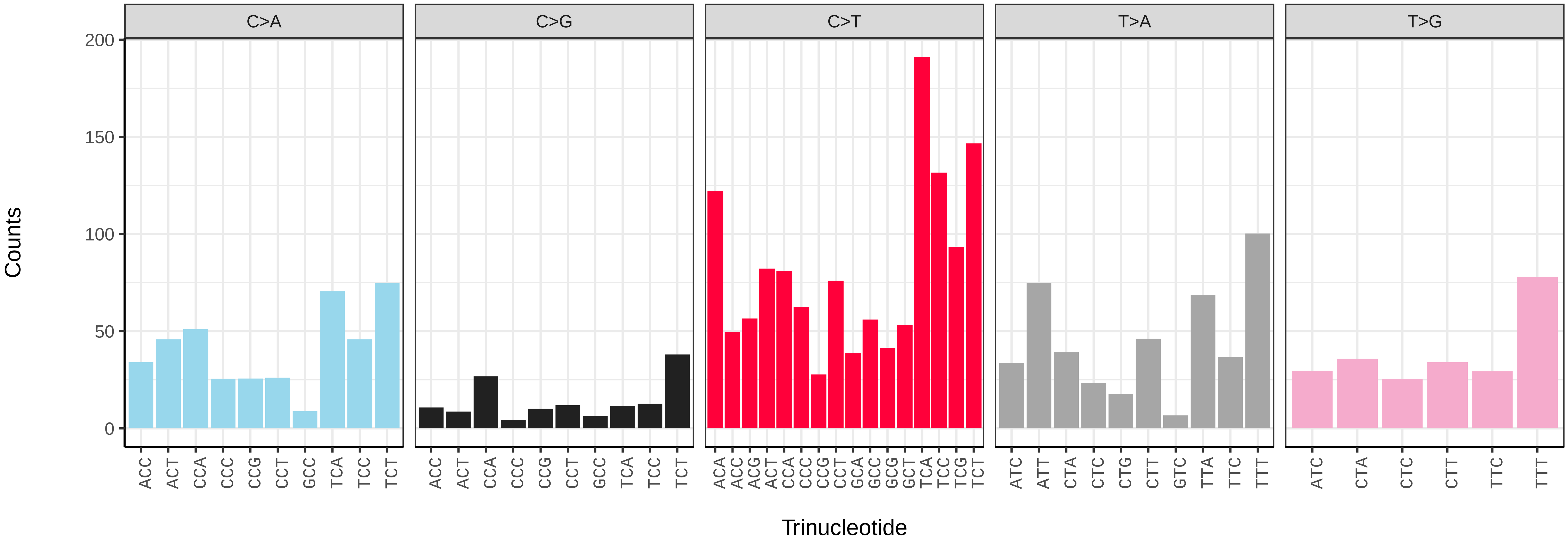

*Adalia bipunctata* (icAdaBipu1.icAdaBipu34) germline mutational spectrum

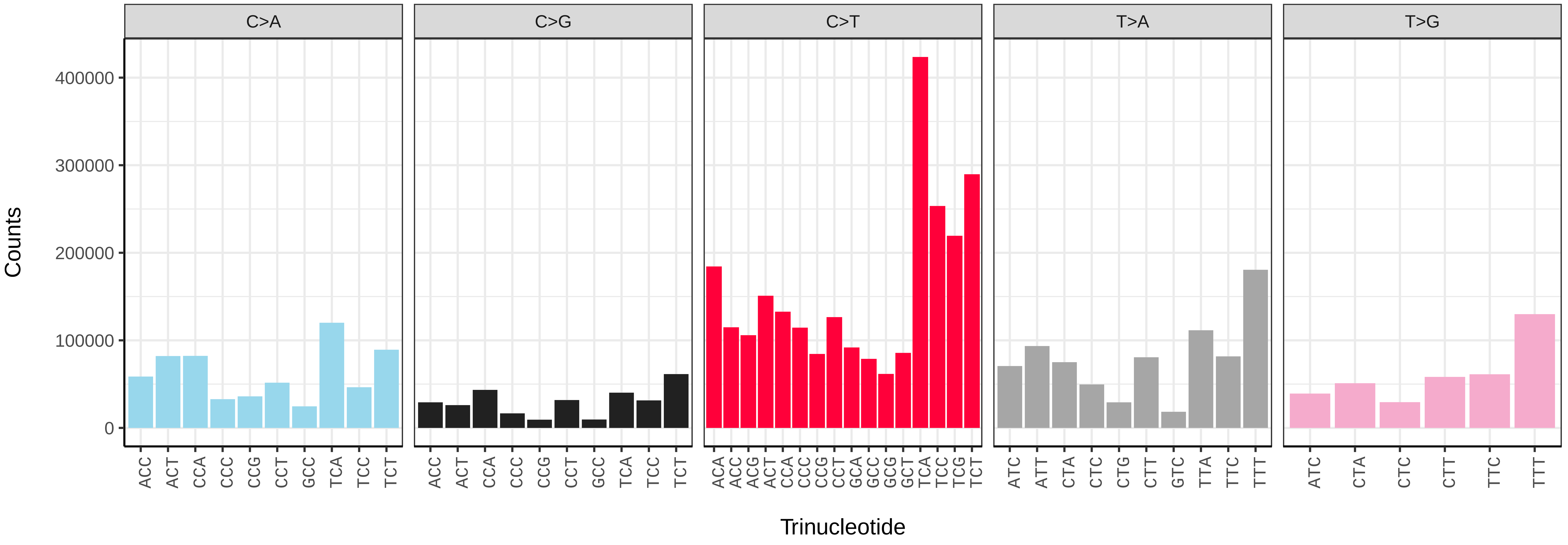


*Adalia bipunctata* (icAdaBipu1.icAdaBipu34) somatic mutational spectrum (SBS96 classification)

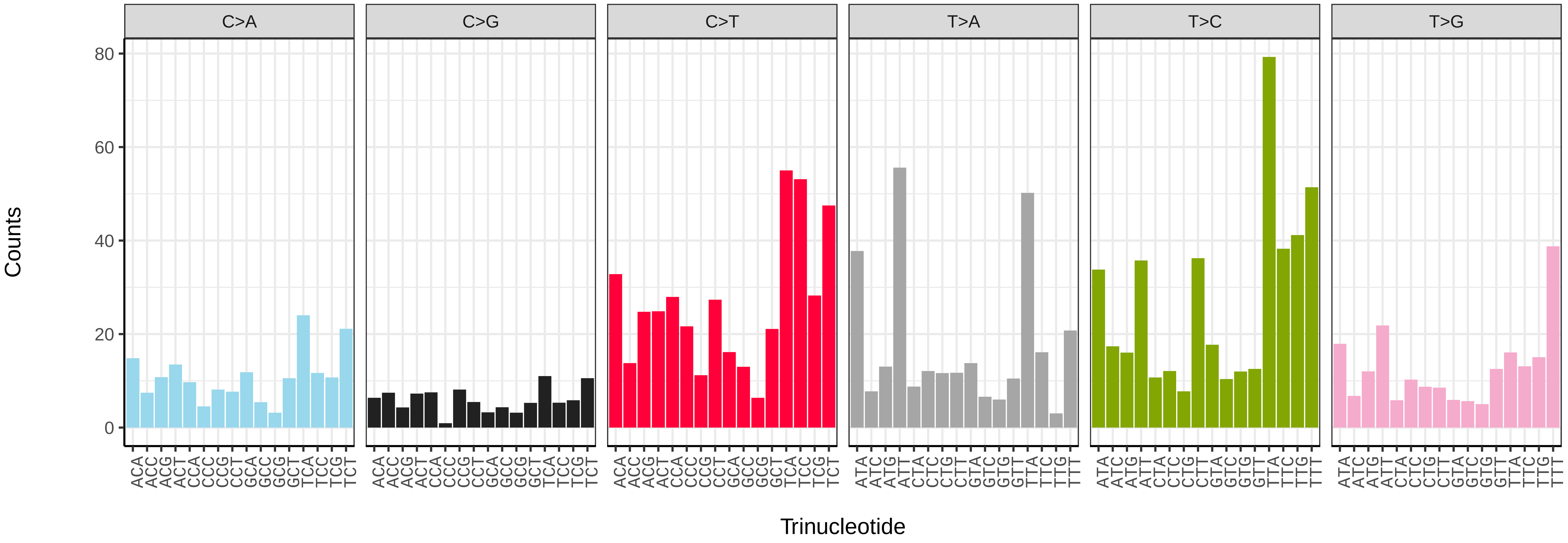


*Adalia bipunctata* (icAdaBipu1.icAdaBipu34) somatic mutational spectrum (SBS52 classification)

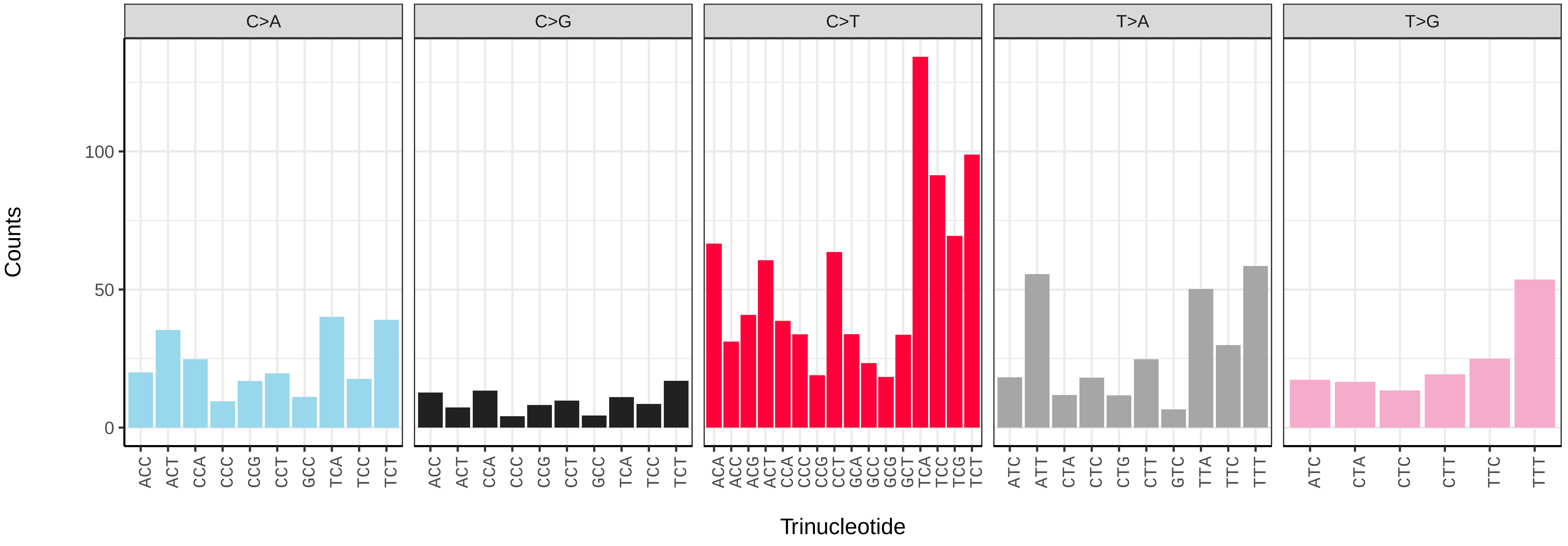

*Adalia bipunctata* (icAdaBipu1.icAdaBipu36) germline mutational spectrum

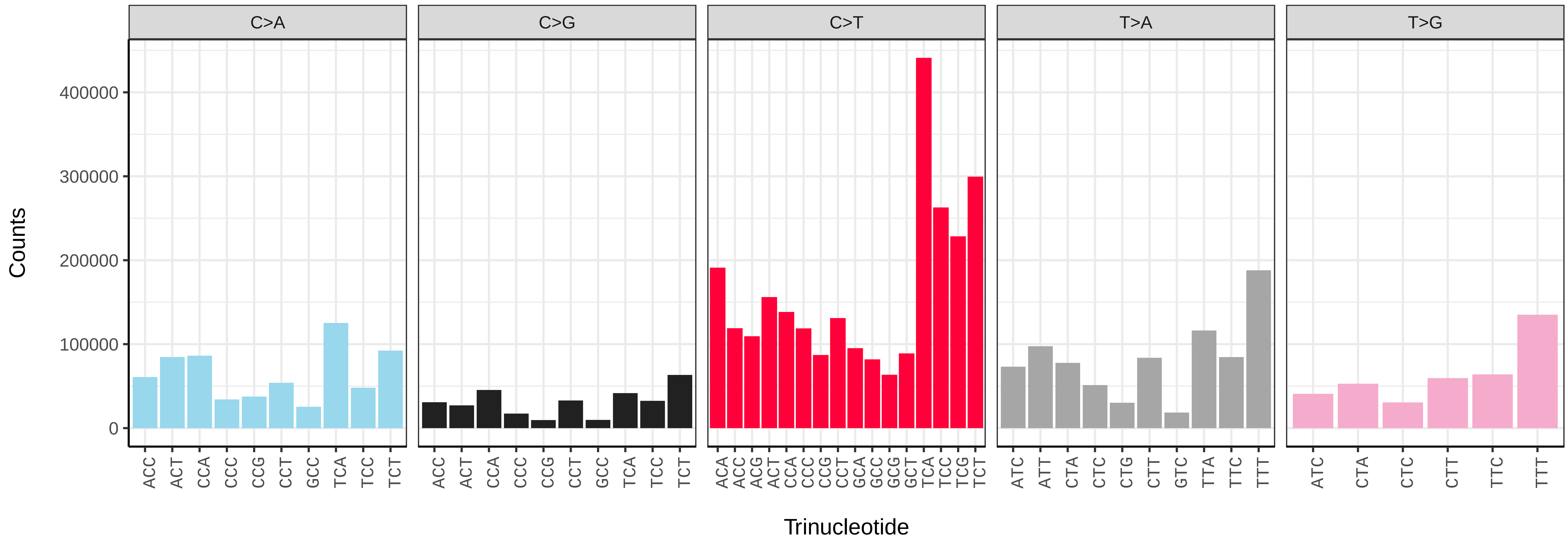


*Adalia bipunctata* (icAdaBipu1.icAdaBipu36) somatic mutational spectrum (SBS96 classification)

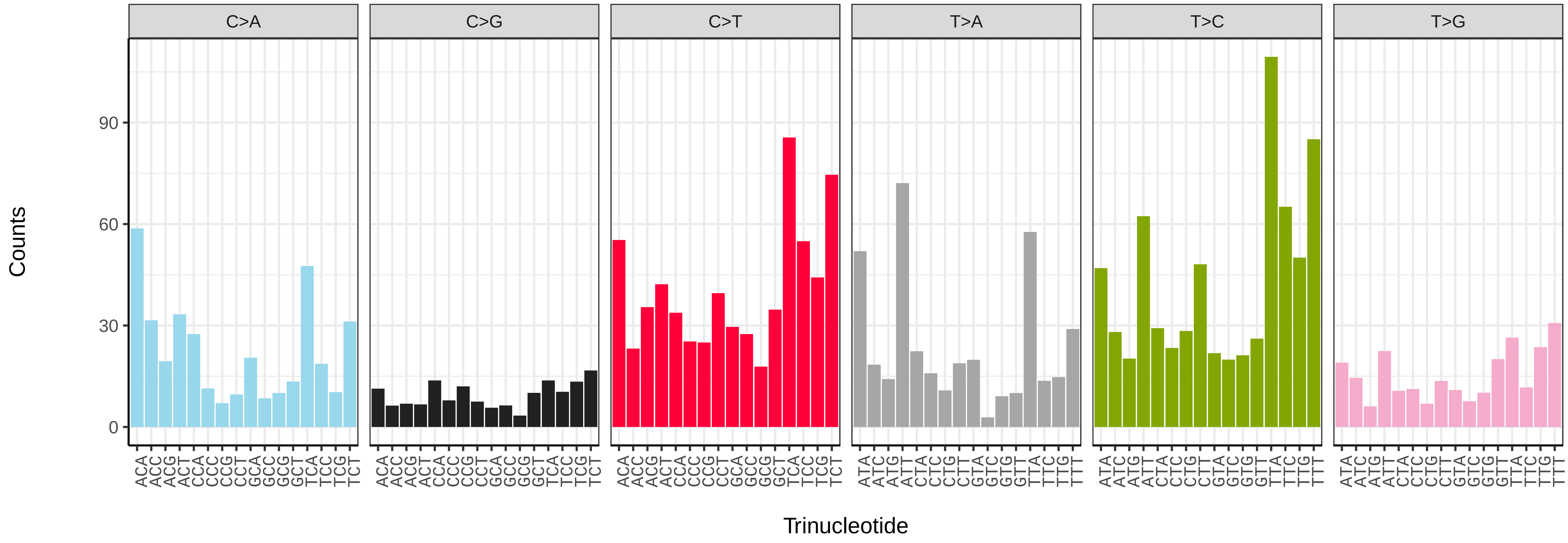


*Adalia bipunctata* (icAdaBipu1.icAdaBipu36) somatic mutational spectrum (SBS52 classification)

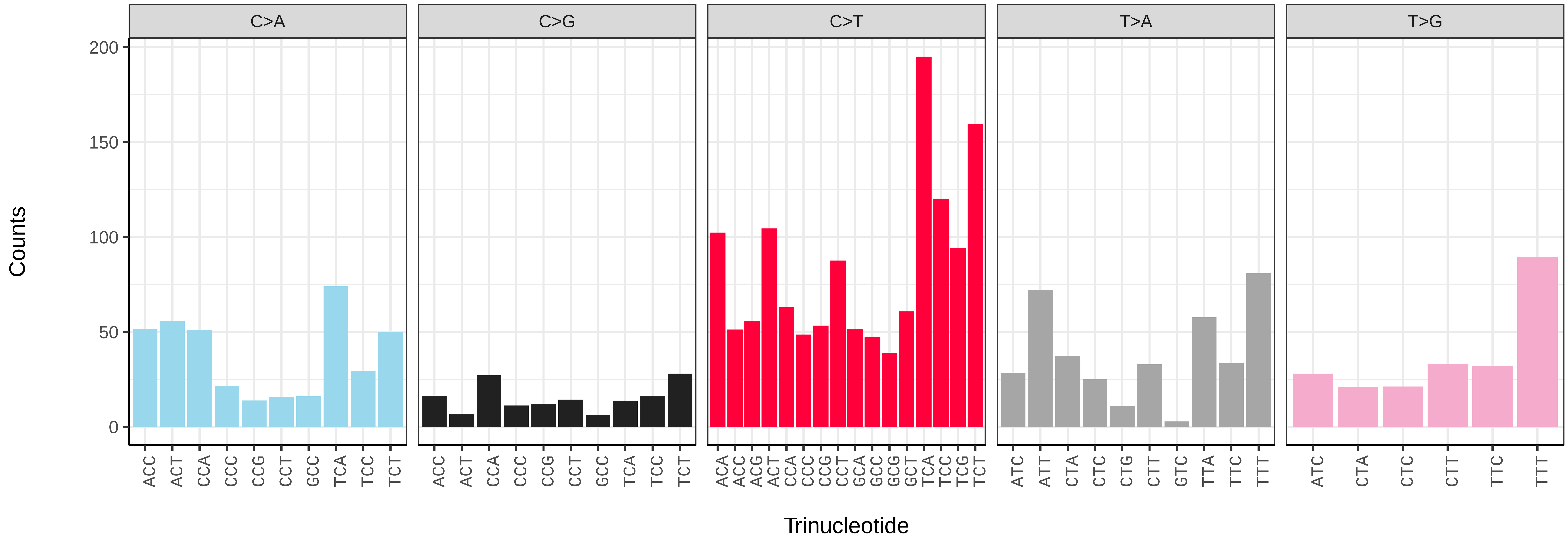

*Adalia bipunctata* (icAdaBipu1.icAdaBipu6) germline mutational spectrum

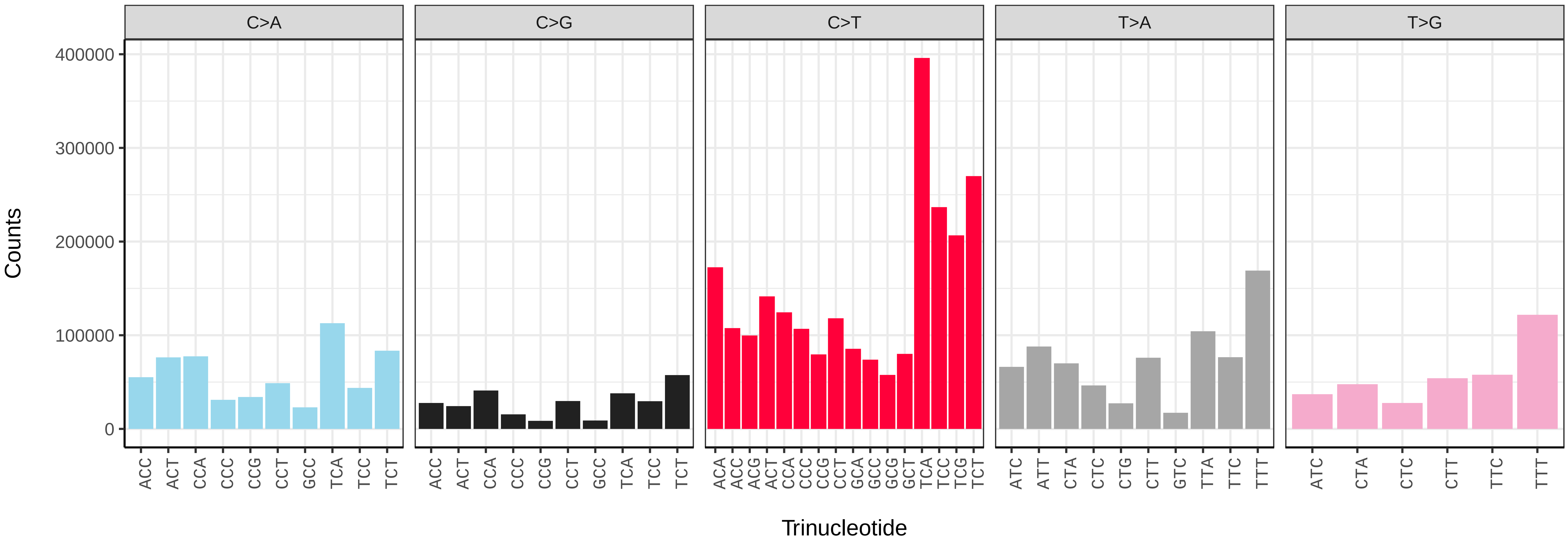


*Adalia bipunctata* (icAdaBipu1.icAdaBipu6) somatic mutational spectrum (SBS96 classification)

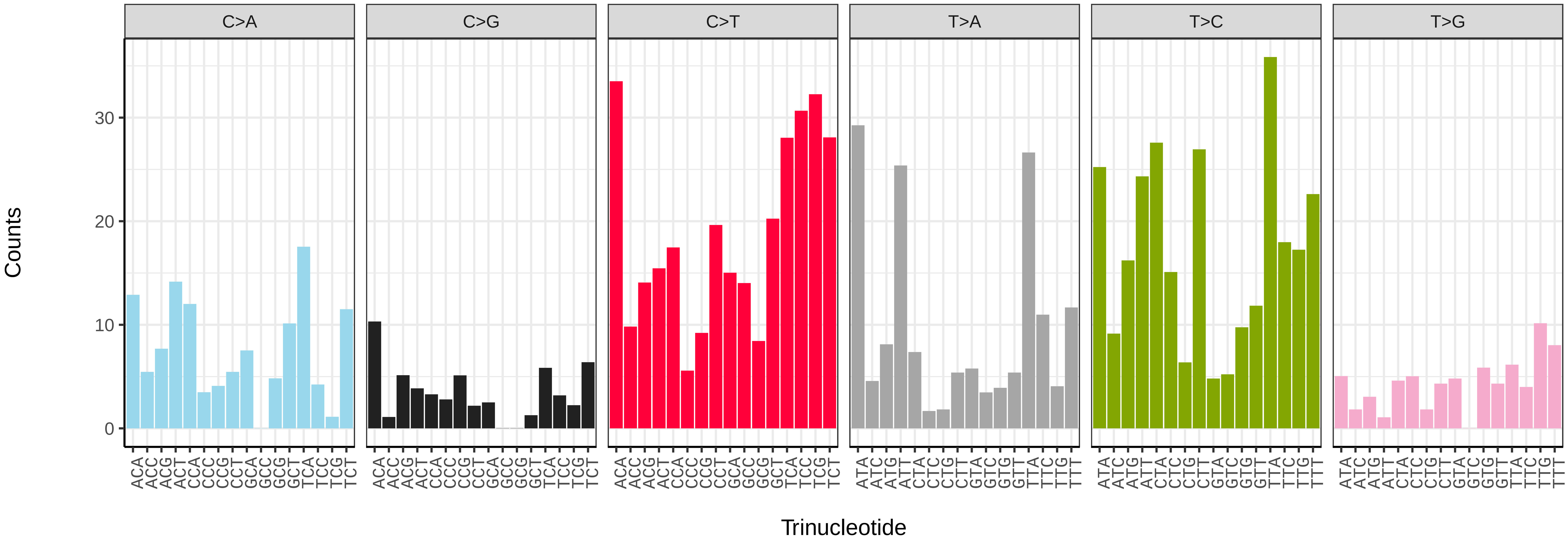


*Adalia bipunctata* (icAdaBipu1.icAdaBipu6) somatic mutational spectrum (SBS52 classification)

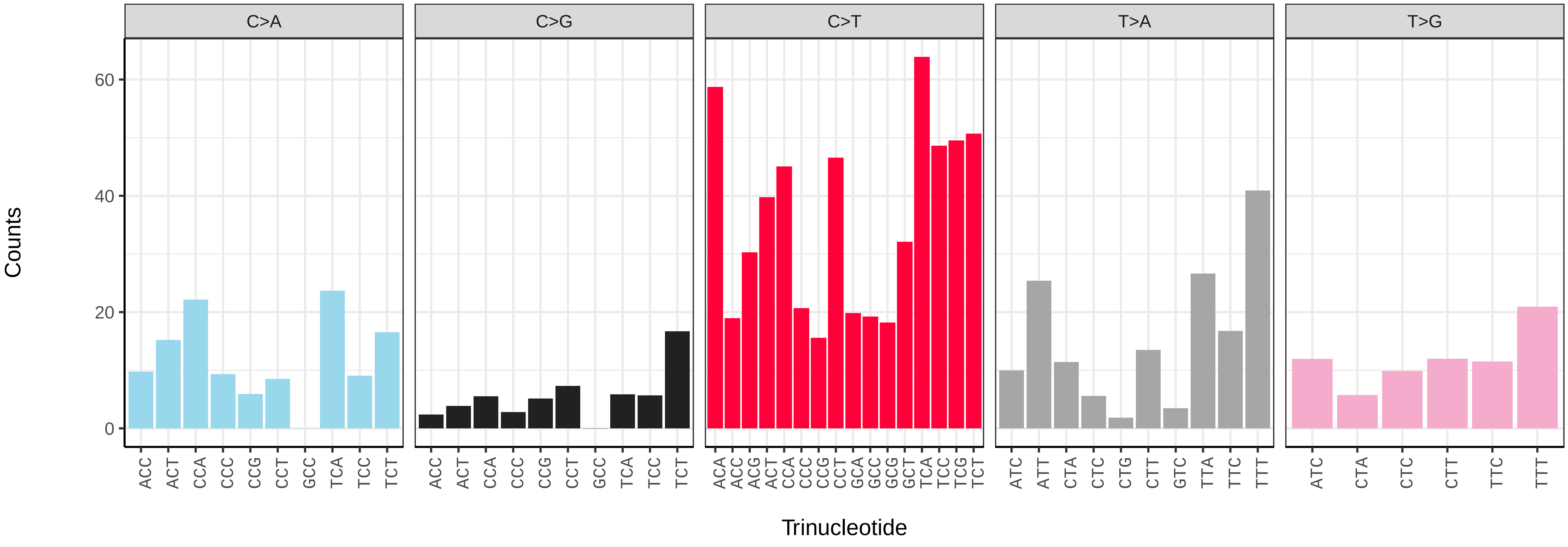

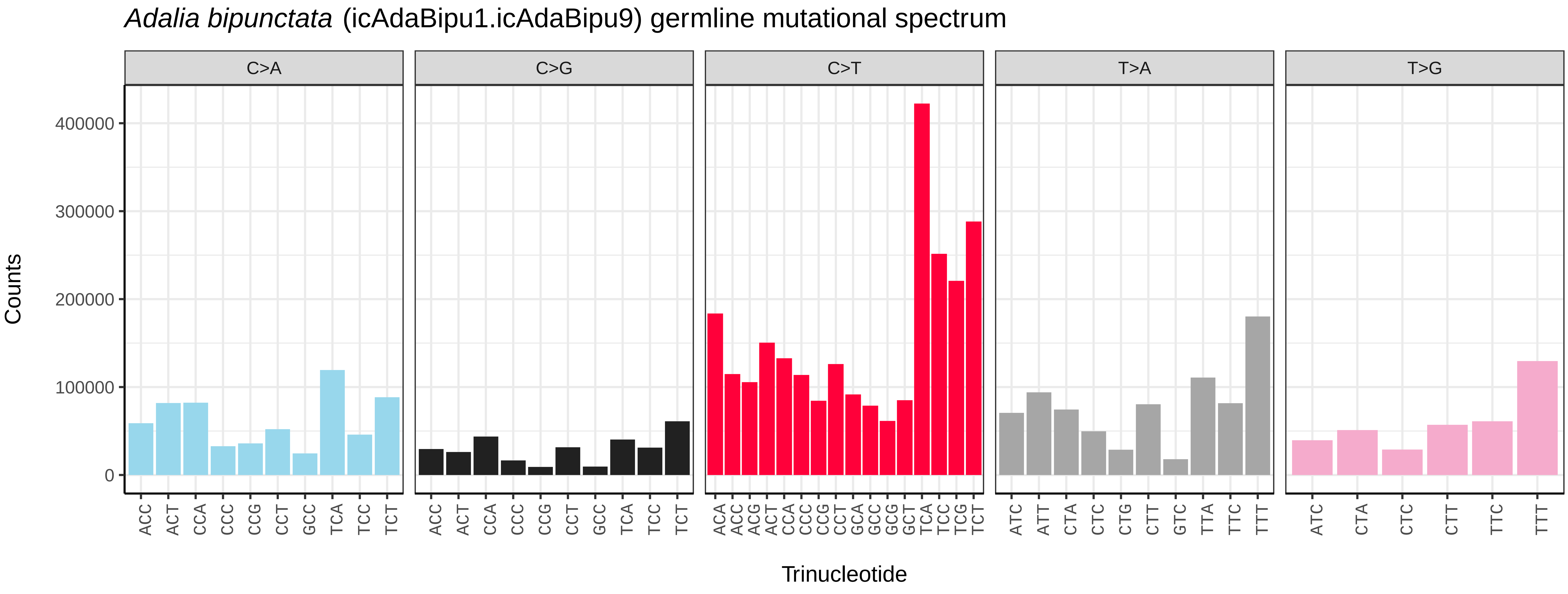
*Adalia bipunctata* (icAdaBipu1.icAdaBipu9) germline mutational spectrum
C>A
C>G
C>T
T>A
T>G
Counts
400000
300000
200000
100000
0
Trinucleotide

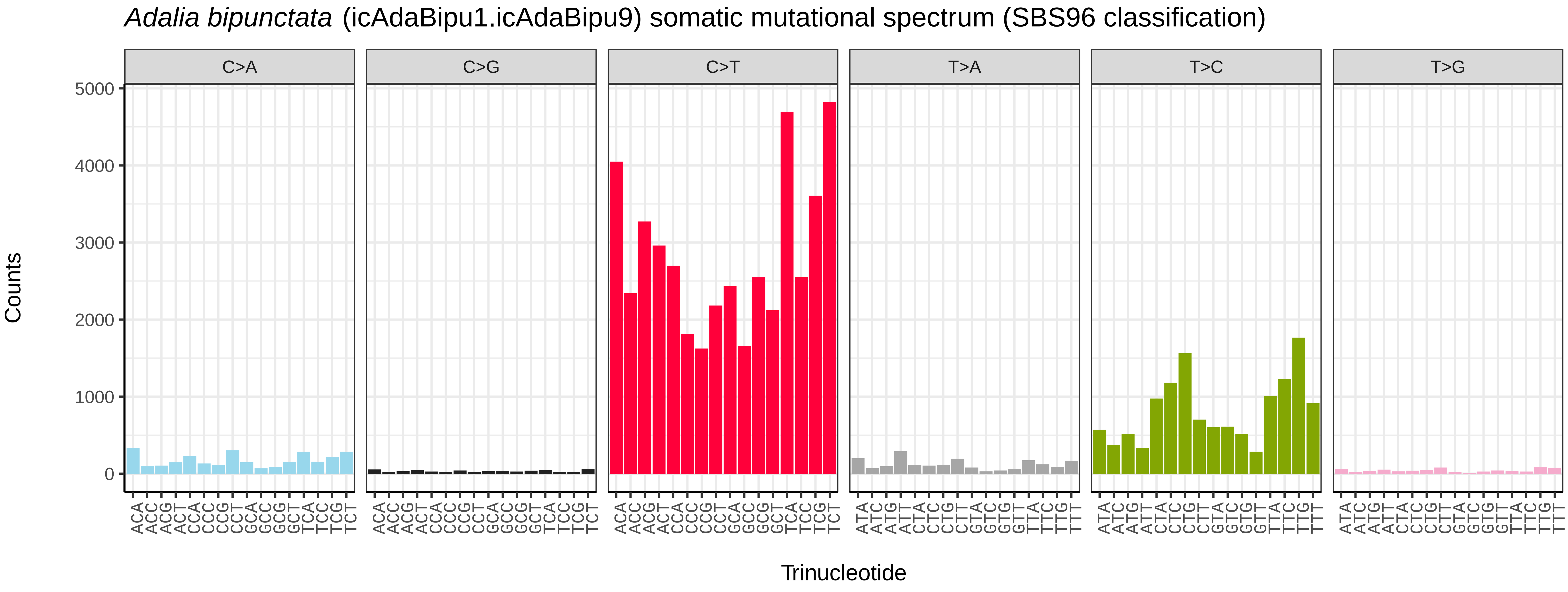
*Adalia bipunctata* (icAdaBipu1.icAdaBipu9) somatic mutational spectrum (SBS96 classification)
C>A
C>G
C>T
T>A
T>C
T>G
Counts
5000
4000
3000
2000
1000
0
Trinucleotide

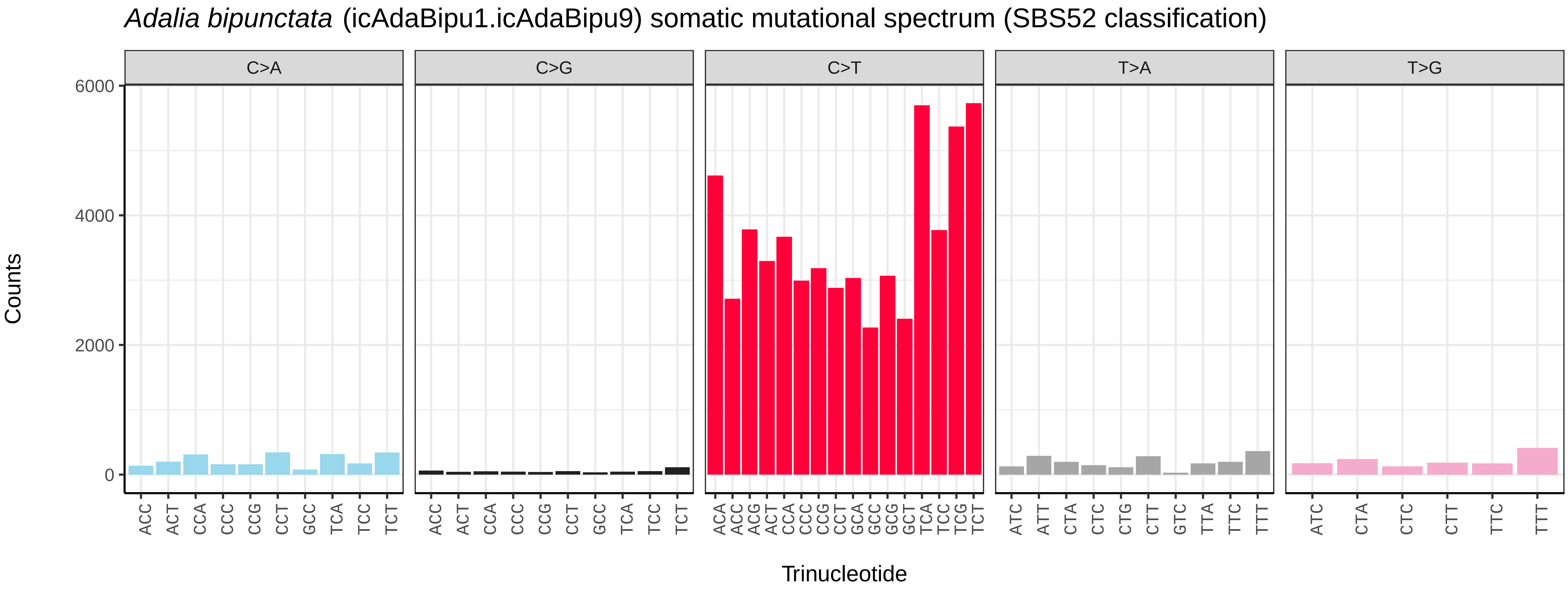
*Adalia bipunctata* (icAdaBipu1.icAdaBipu9) somatic mutational spectrum (SBS52 classification)
C>A
C>G
C>T
T>A
T>G
Counts
6000
4000
2000
0
Trinucleotide

*Agonum fuliginosum* (icAgoFuli1) germline mutational spectrum

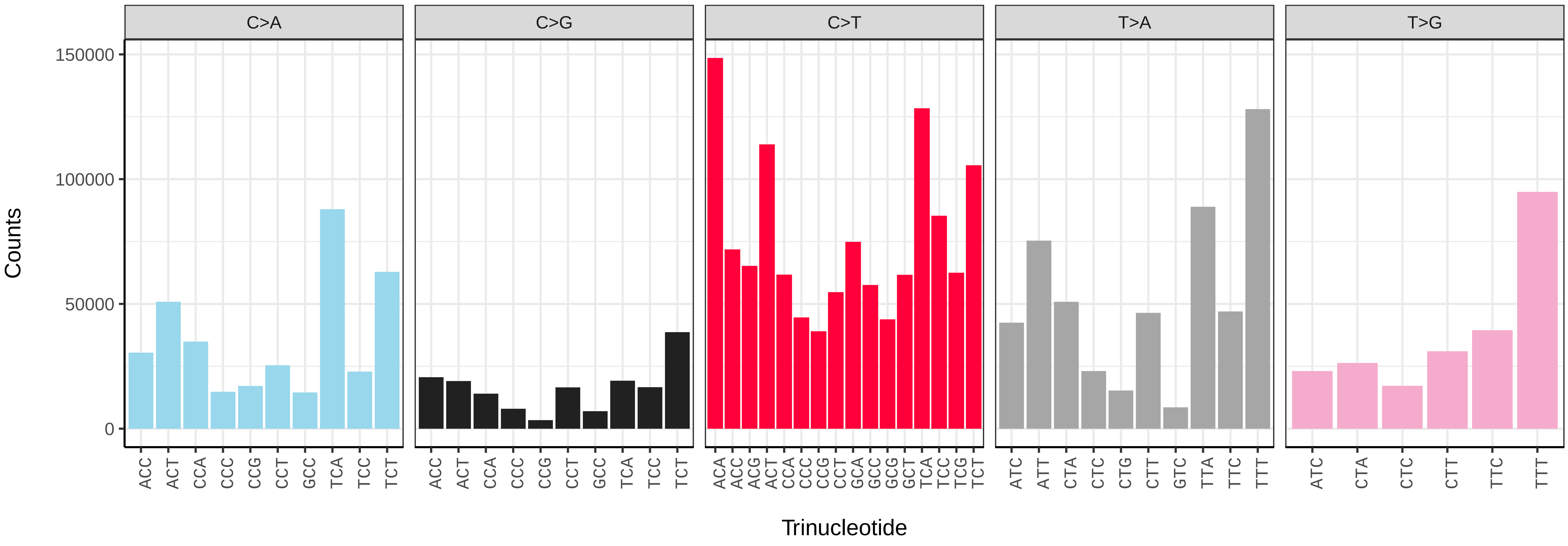


*Agonum fuliginosum* (icAgoFuli1) somatic mutational spectrum (SBS96 classification)

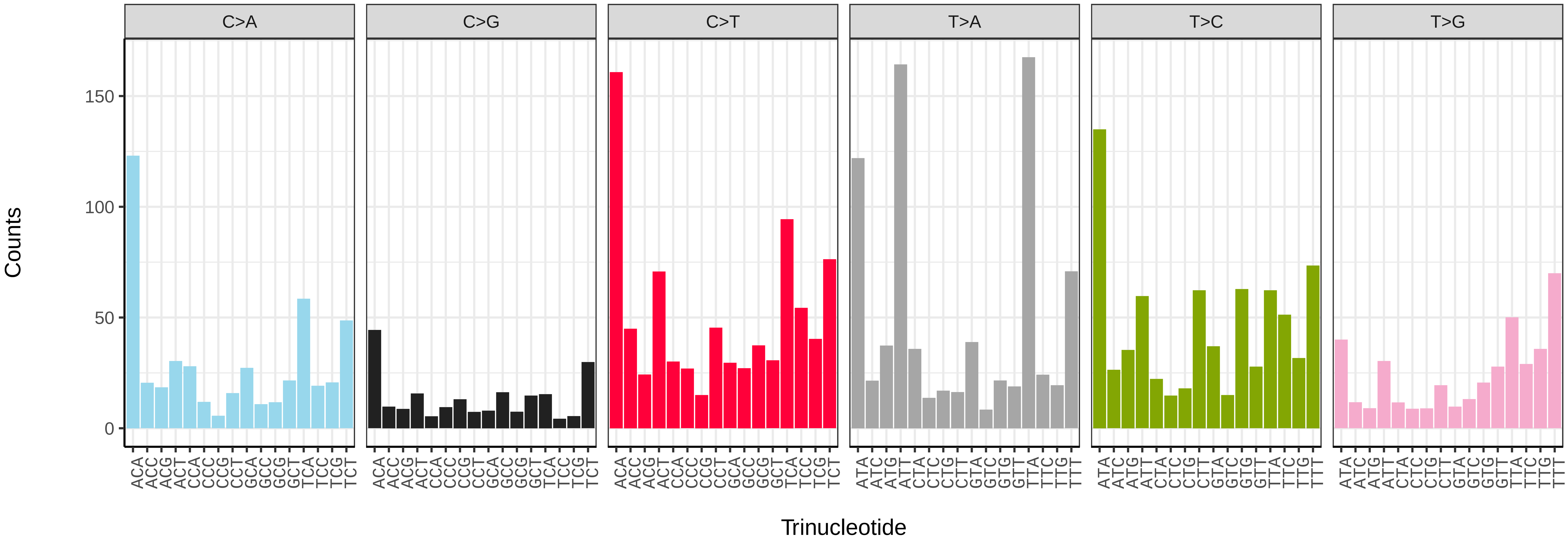


*Agonum fuliginosum* (icAgoFuli1) somatic mutational spectrum (SBS52 classification)

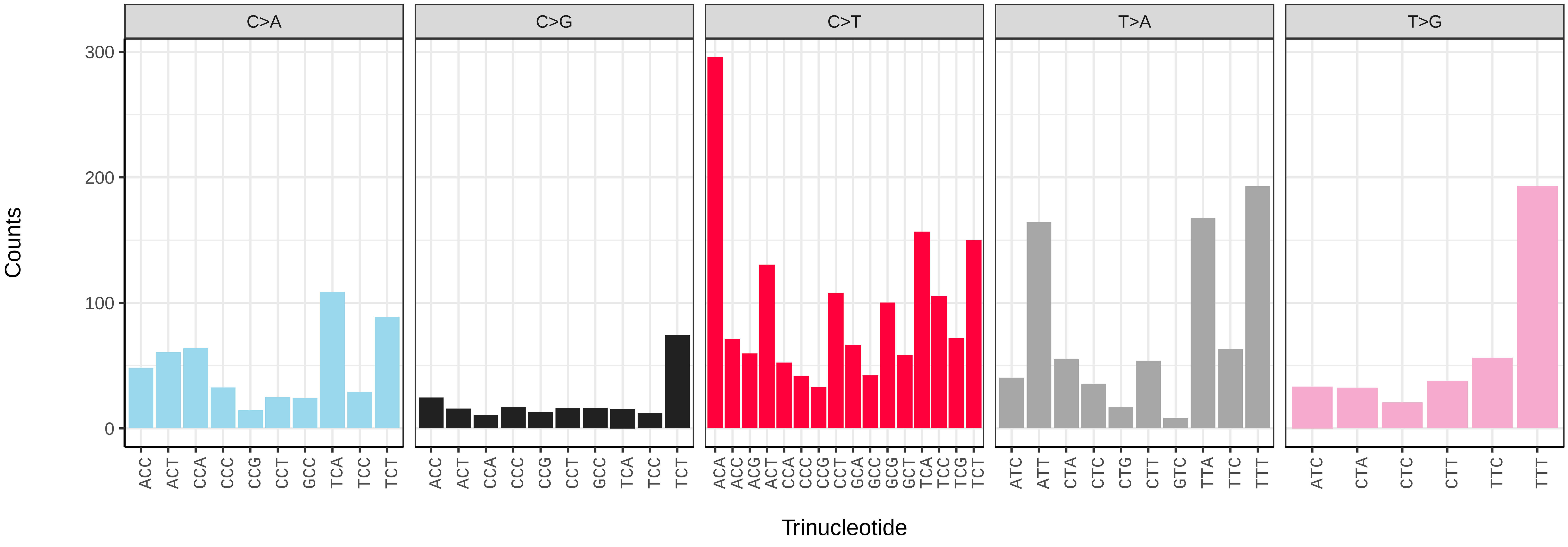

*Agrilus cyanescens* (icAgrCyan1) germline mutational spectrum

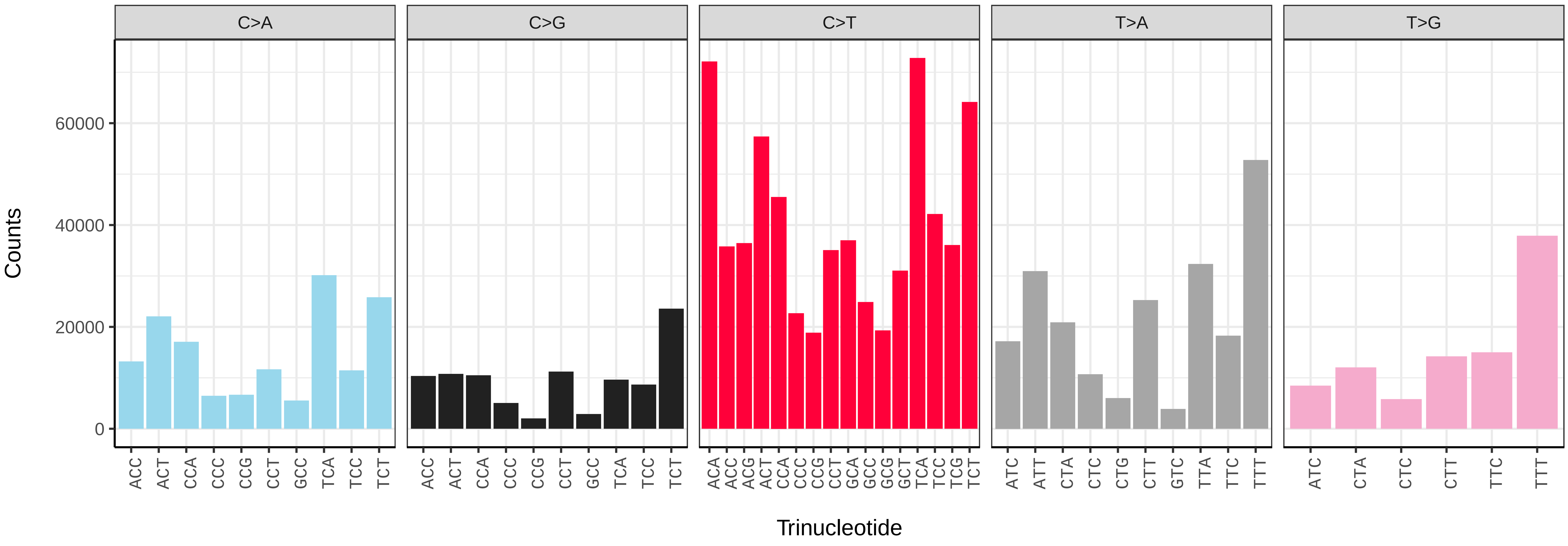


*Agrilus cyanescens* (icAgrCyan1) somatic mutational spectrum (SBS96 classification)

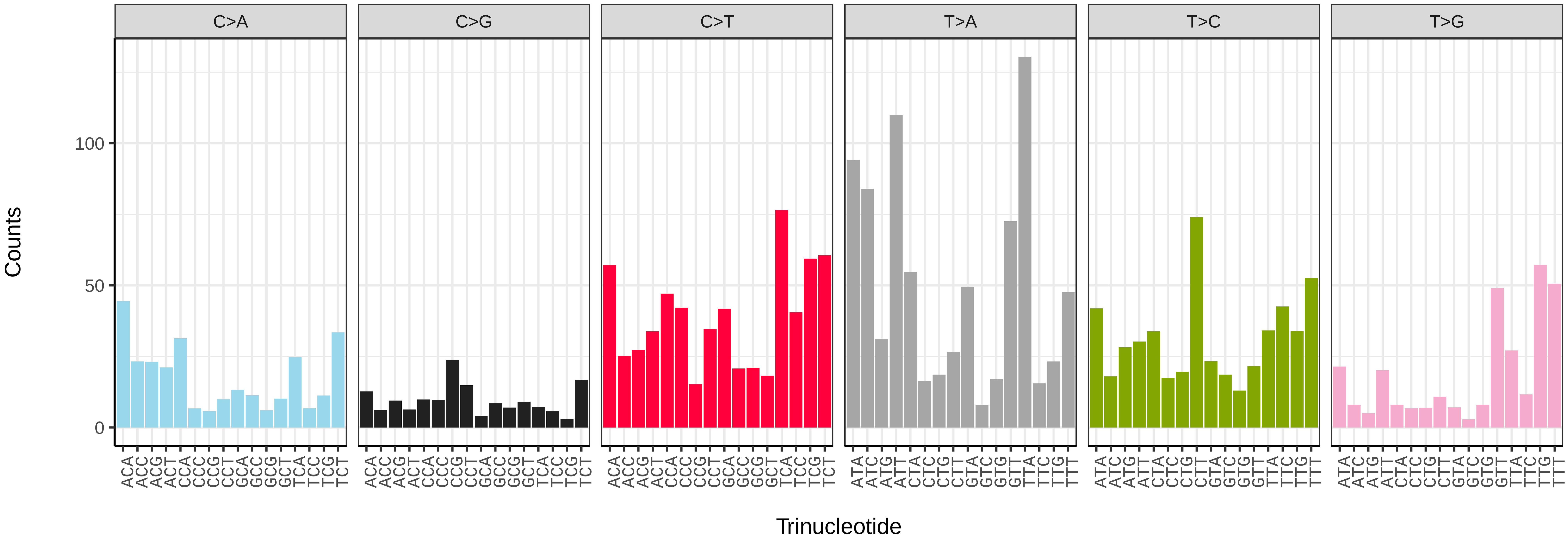


*Agrilus cyanescens* (icAgrCyan1) somatic mutational spectrum (SBS52 classification)

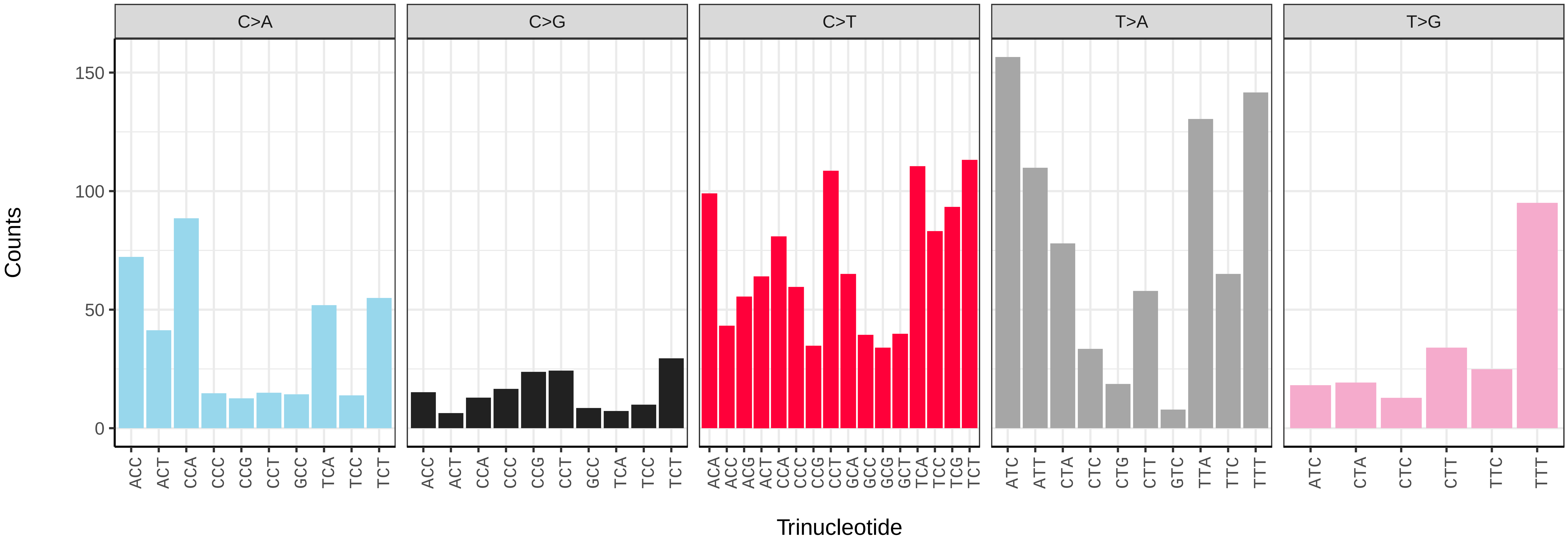

*Agrypnus murinus* (icAgrMuri1) germline mutational spectrum

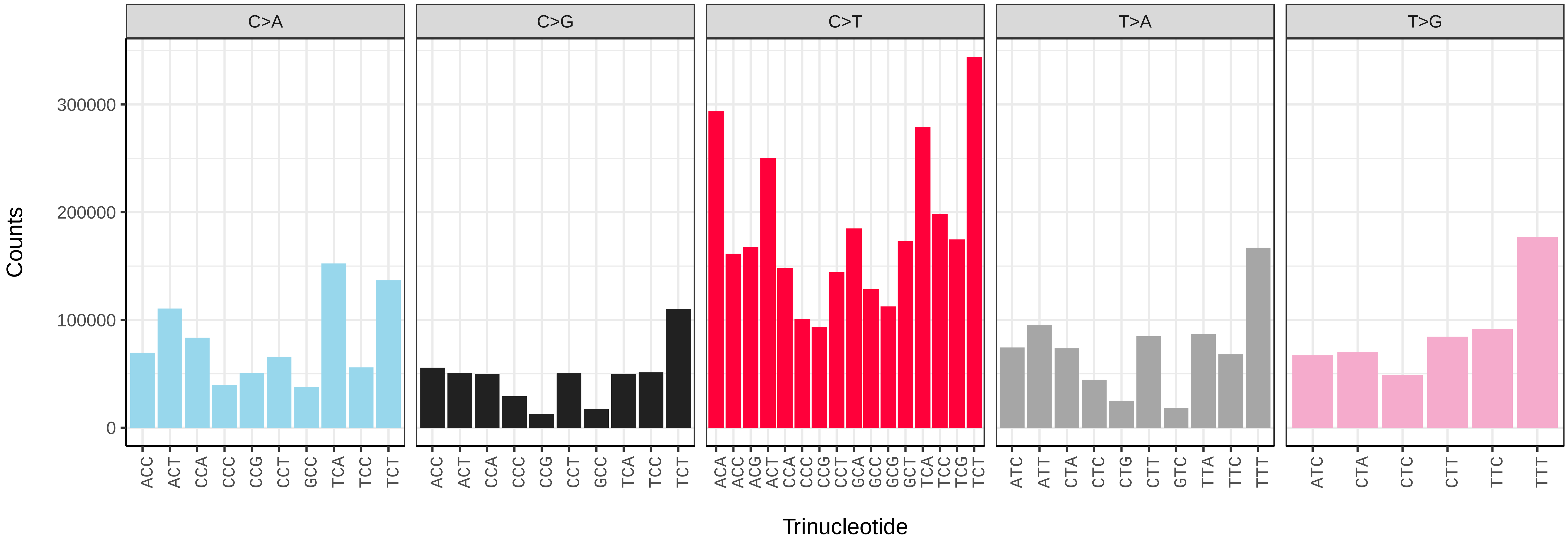


*Agrypnus murinus* (icAgrMuri1) somatic mutational spectrum (SBS96 classification)

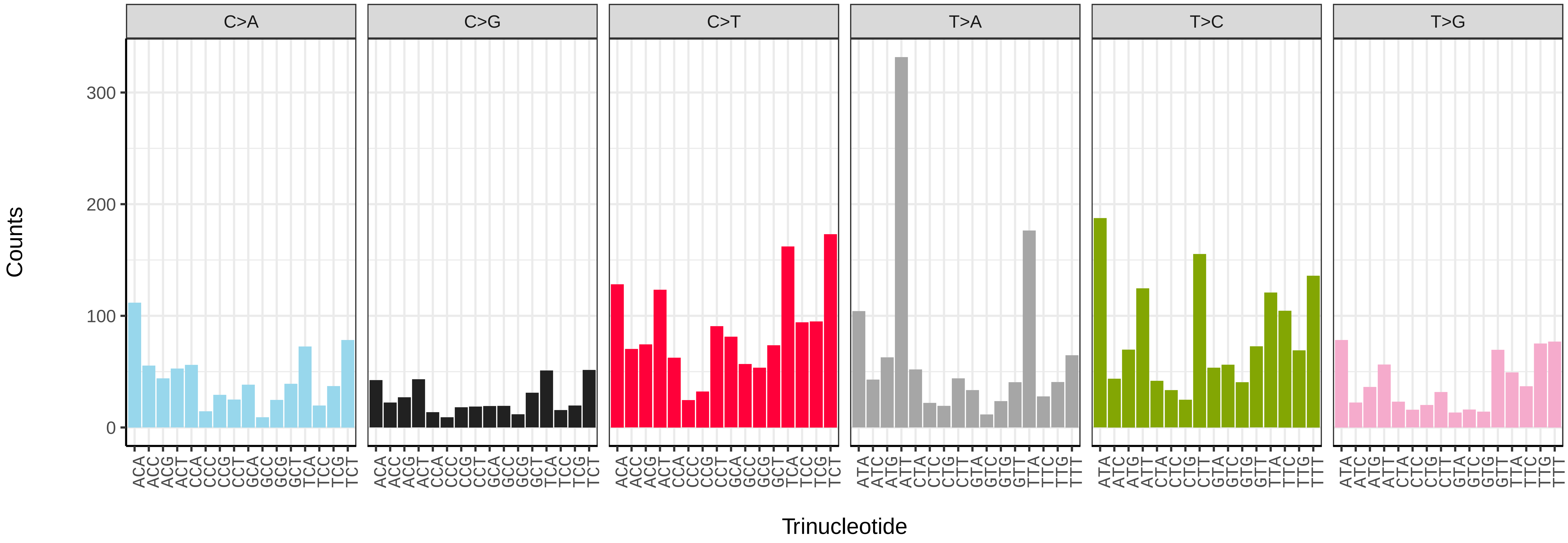


*Agrypnus murinus* (icAgrMuri1) somatic mutational spectrum (SBS52 classification)

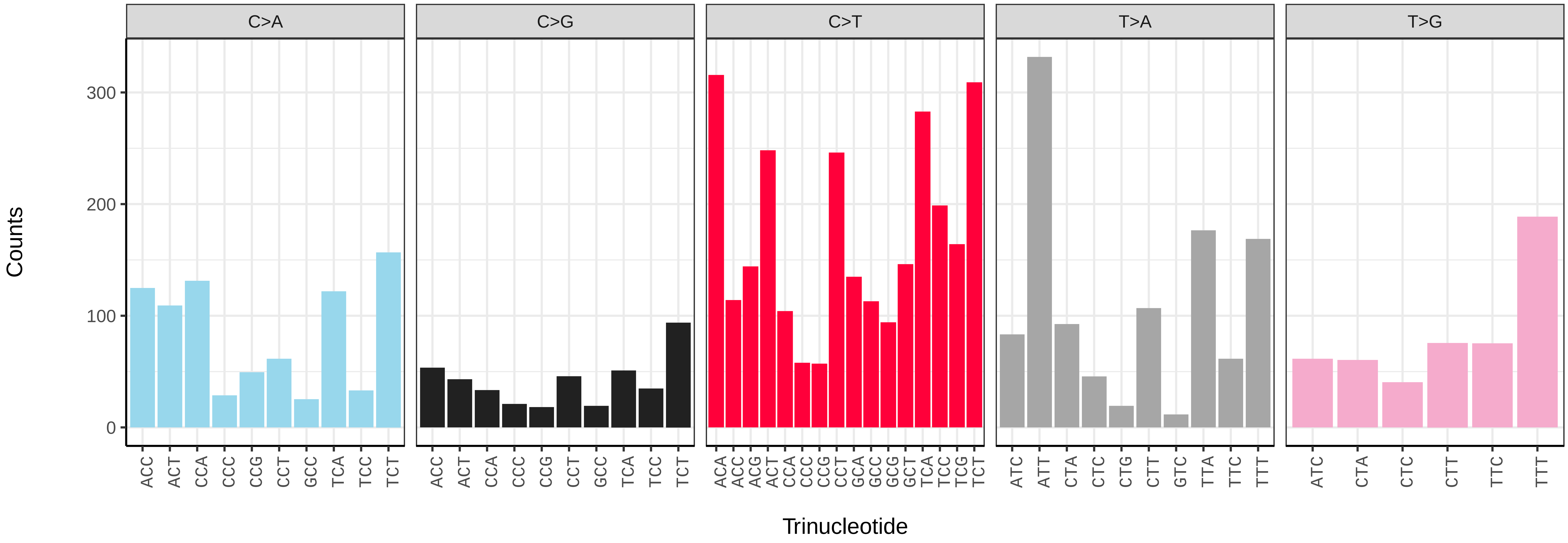

*Anaspis maculata* (icAnaMacu3) germline mutational spectrum

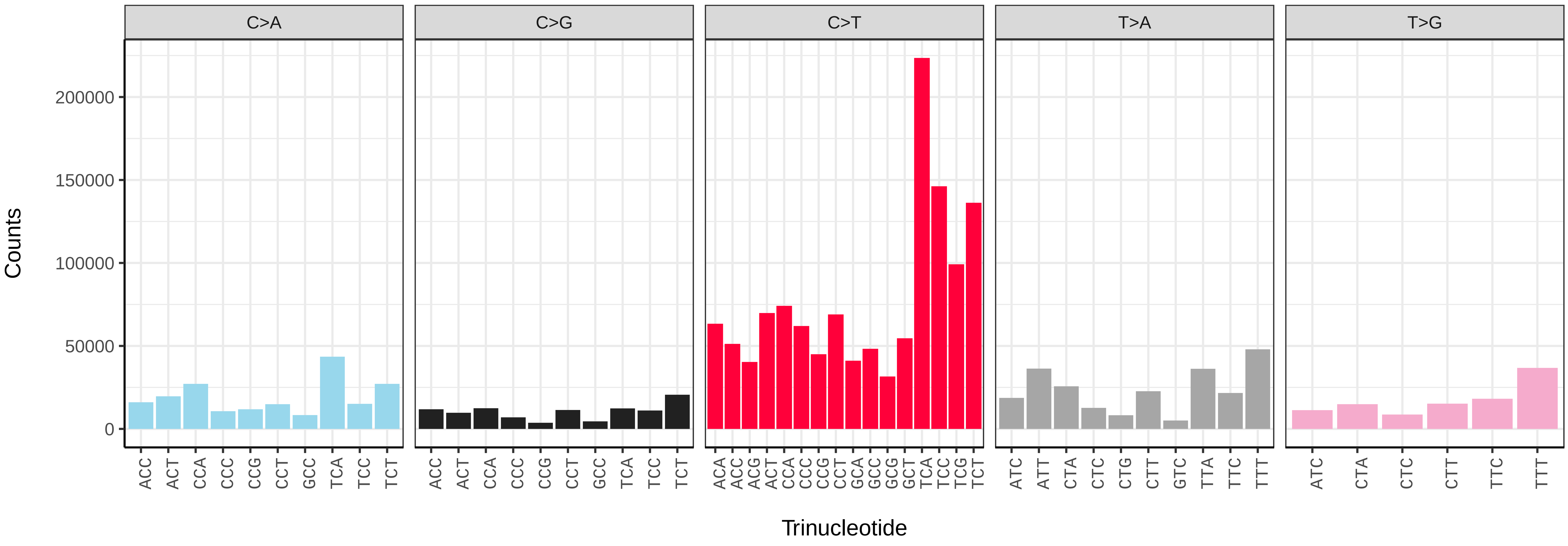


*Anaspis maculata* (icAnaMacu3) somatic mutational spectrum (SBS96 classification)

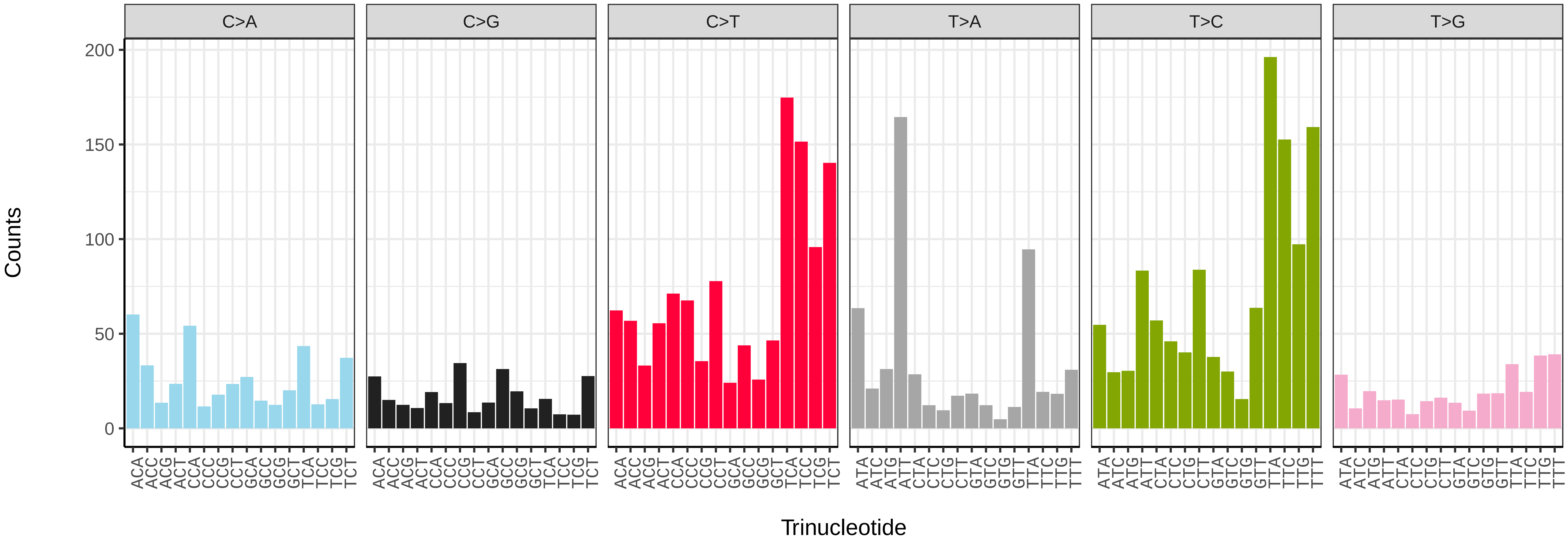


*Anaspis maculata* (icAnaMacu3) somatic mutational spectrum (SBS52 classification)

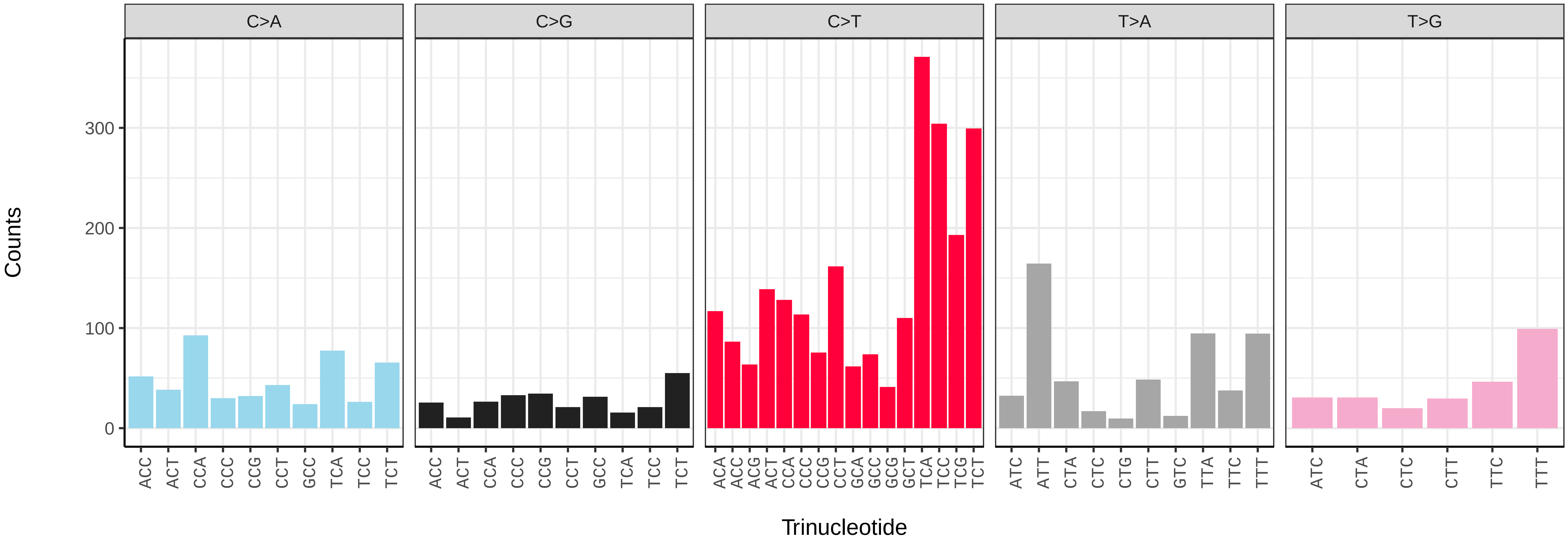

*Apoderus coryli* (icApoCory1) germline mutational spectrum

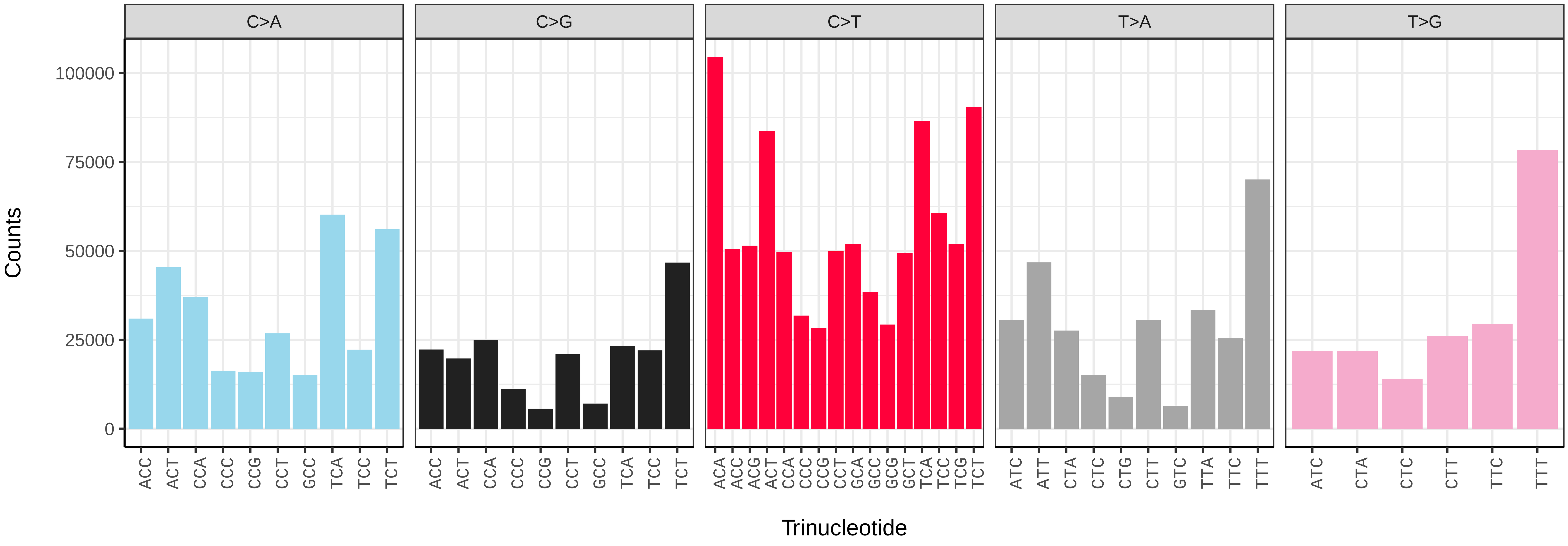


*Apoderus coryli* (icApoCory1) somatic mutational spectrum (SBS96 classification)

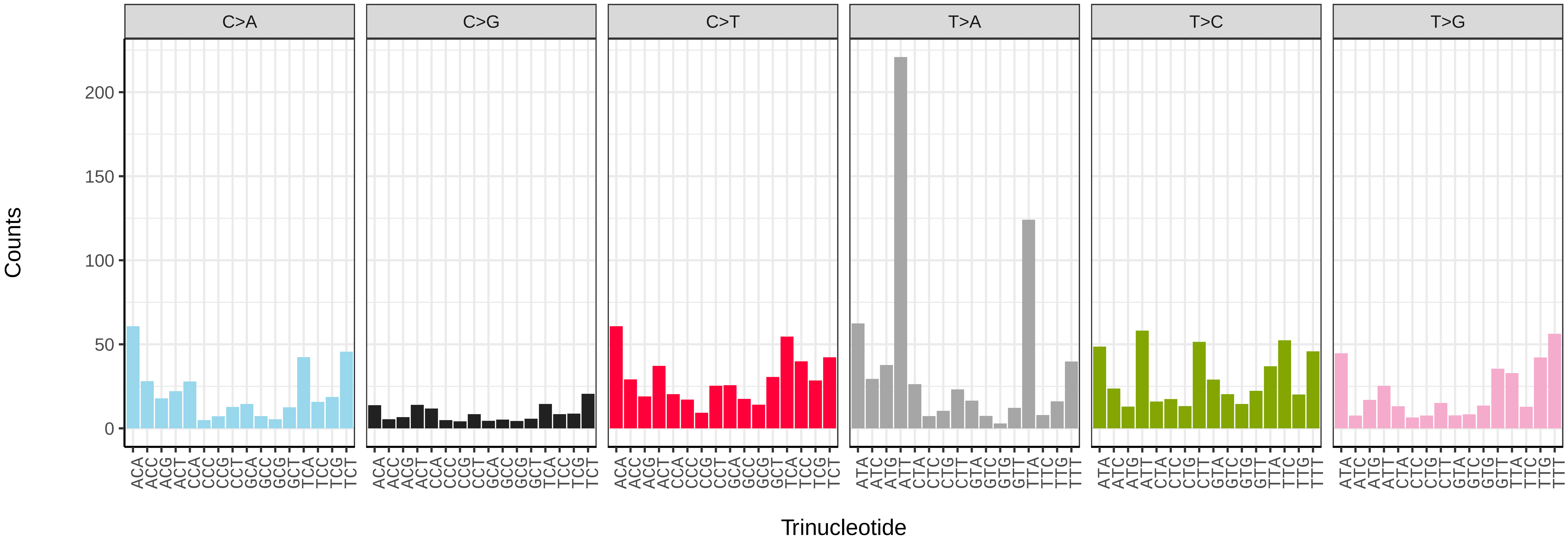


*Apoderus coryli* (icApoCory1) somatic mutational spectrum (SBS52 classification)

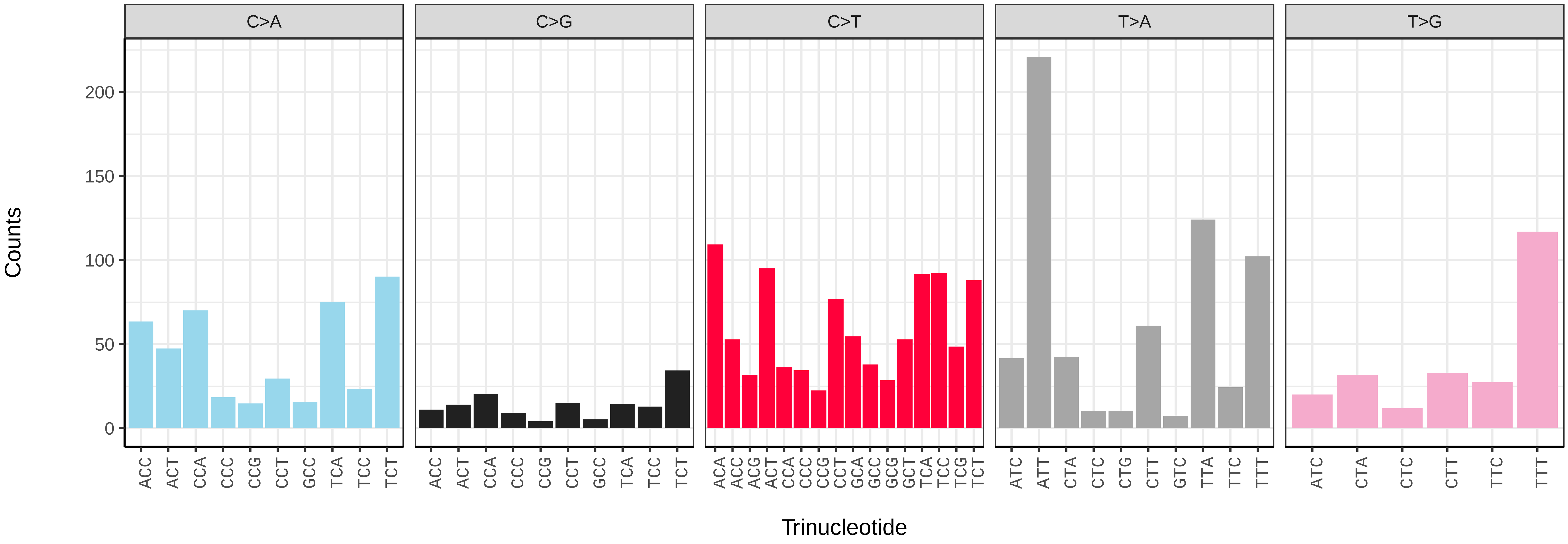

*Cantharis rustica* (icCanRust1) germline mutational spectrum

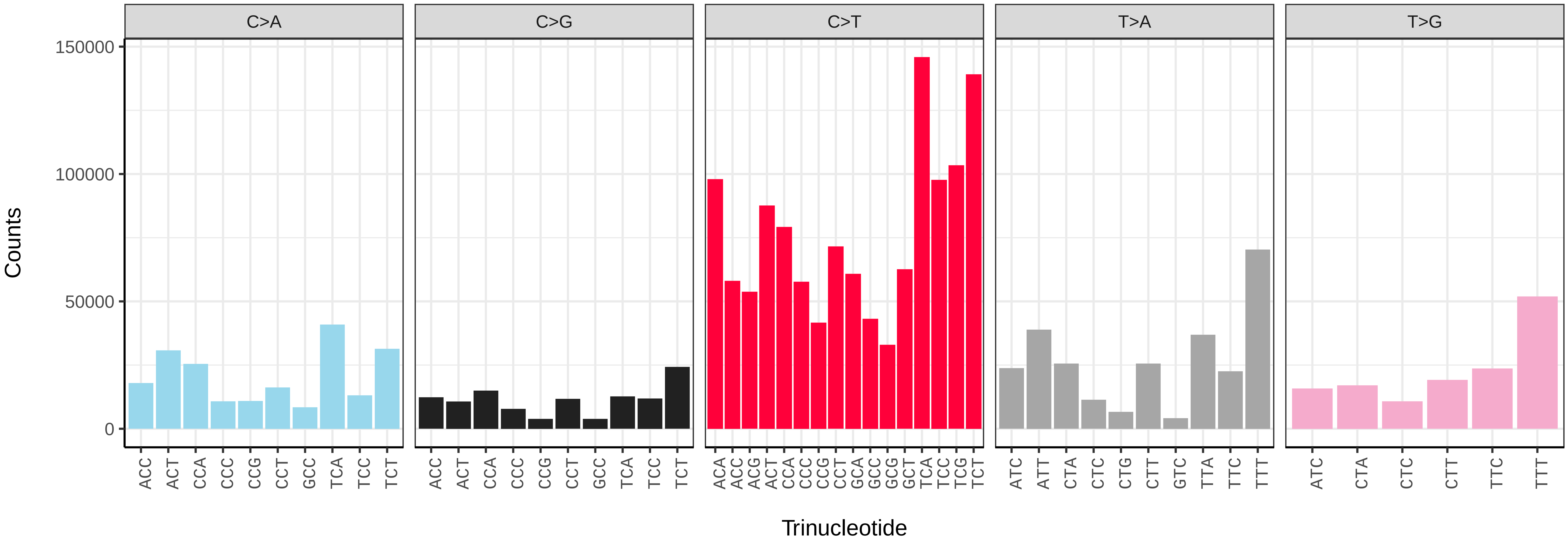


*Cantharis rustica* (icCanRust1) somatic mutational spectrum (SBS96 classification)

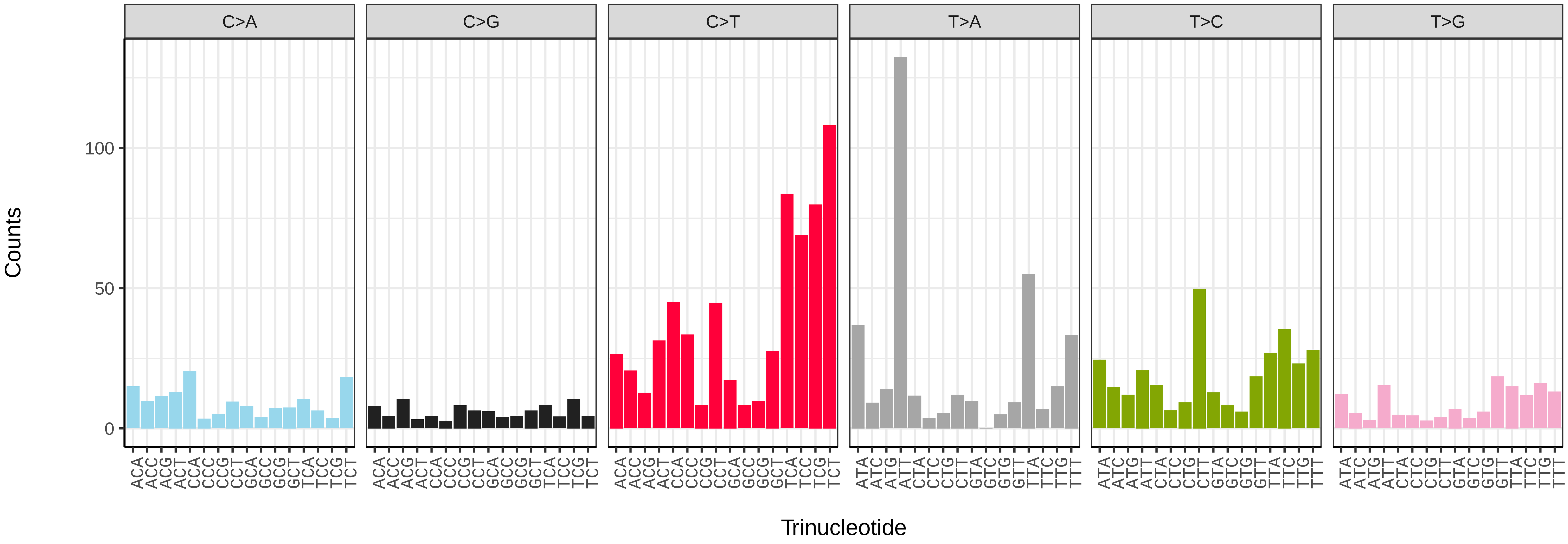


*Cantharis rustica* (icCanRust1) somatic mutational spectrum (SBS52 classification)

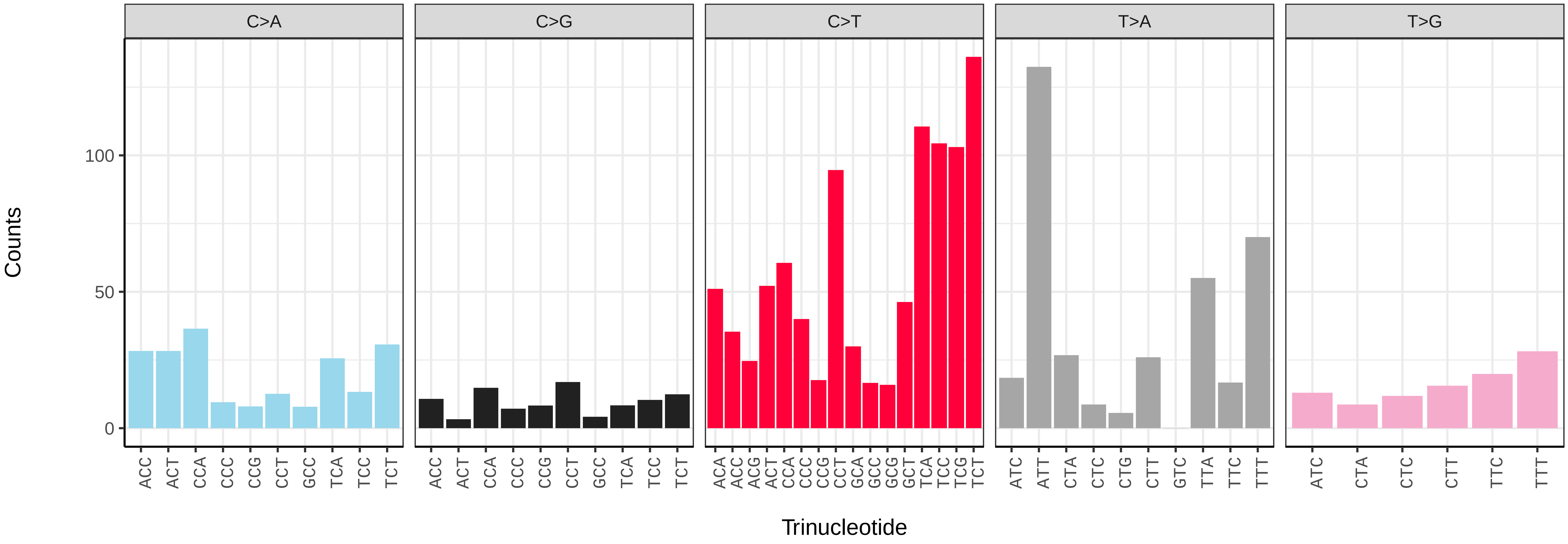

*Cantharis rustica* (icCanRust1.icCanRust3) germline mutational spectrum

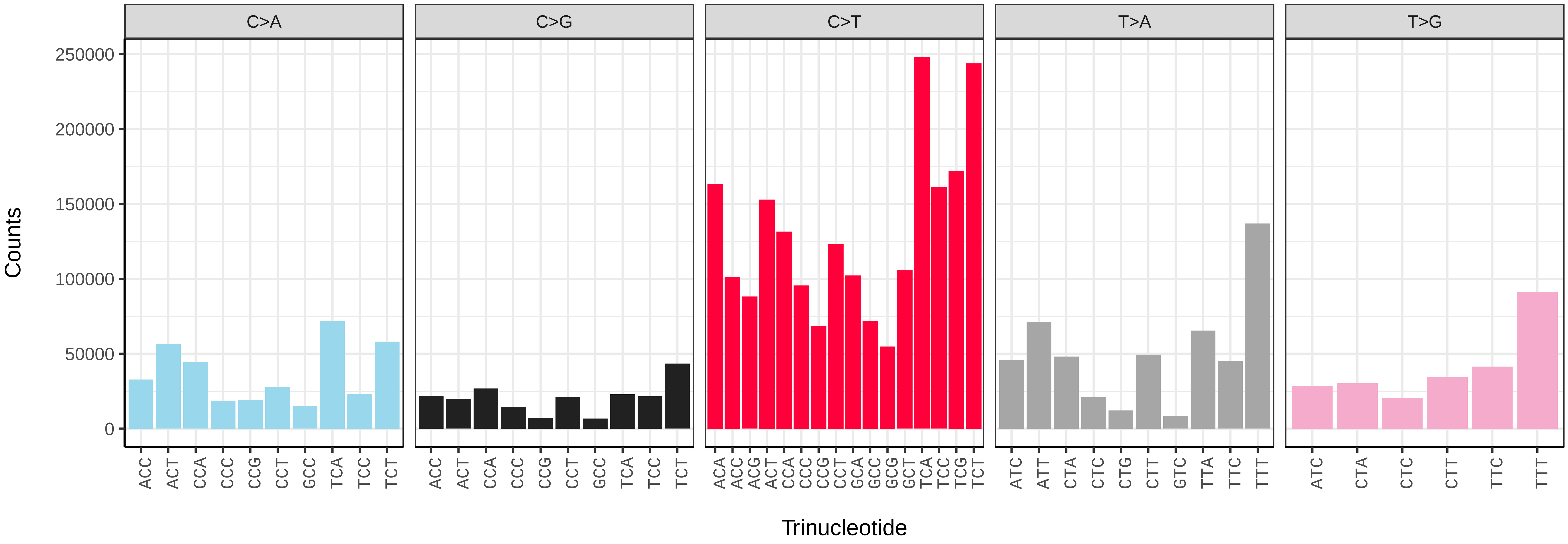


*Cantharis rustica* (icCanRust1.icCanRust3) somatic mutational spectrum (SBS96 classification)

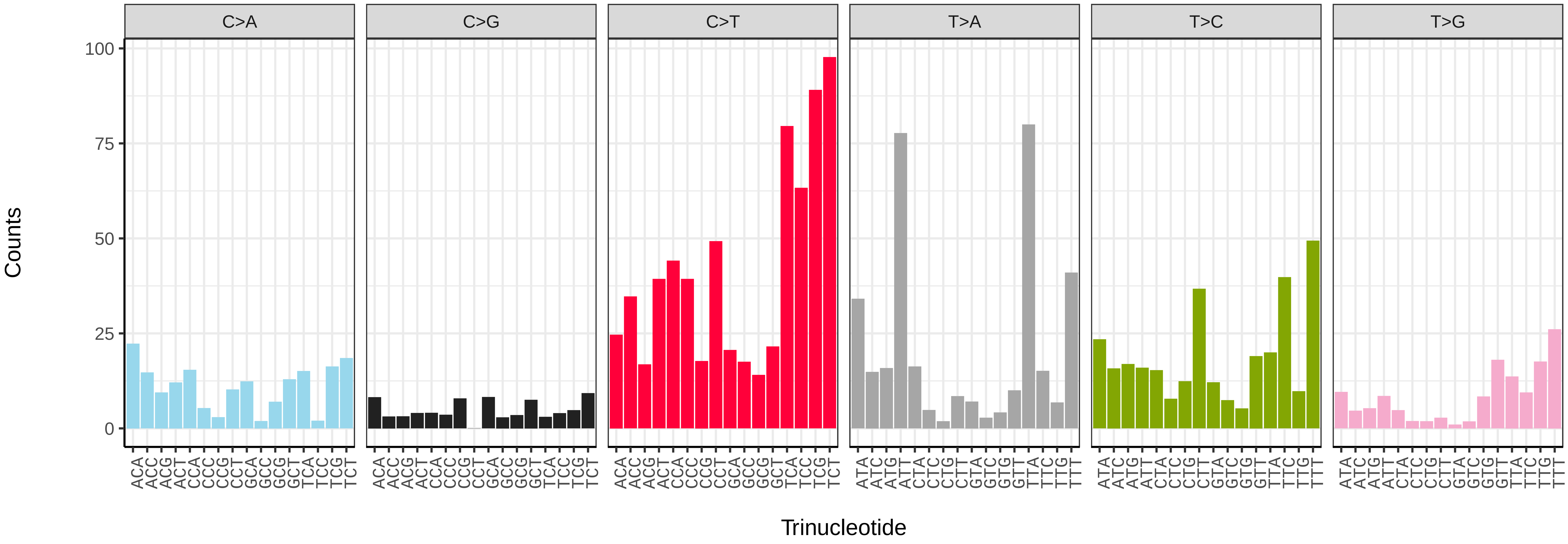


*Cantharis rustica* (icCanRust1.icCanRust3) somatic mutational spectrum (SBS52 classification)

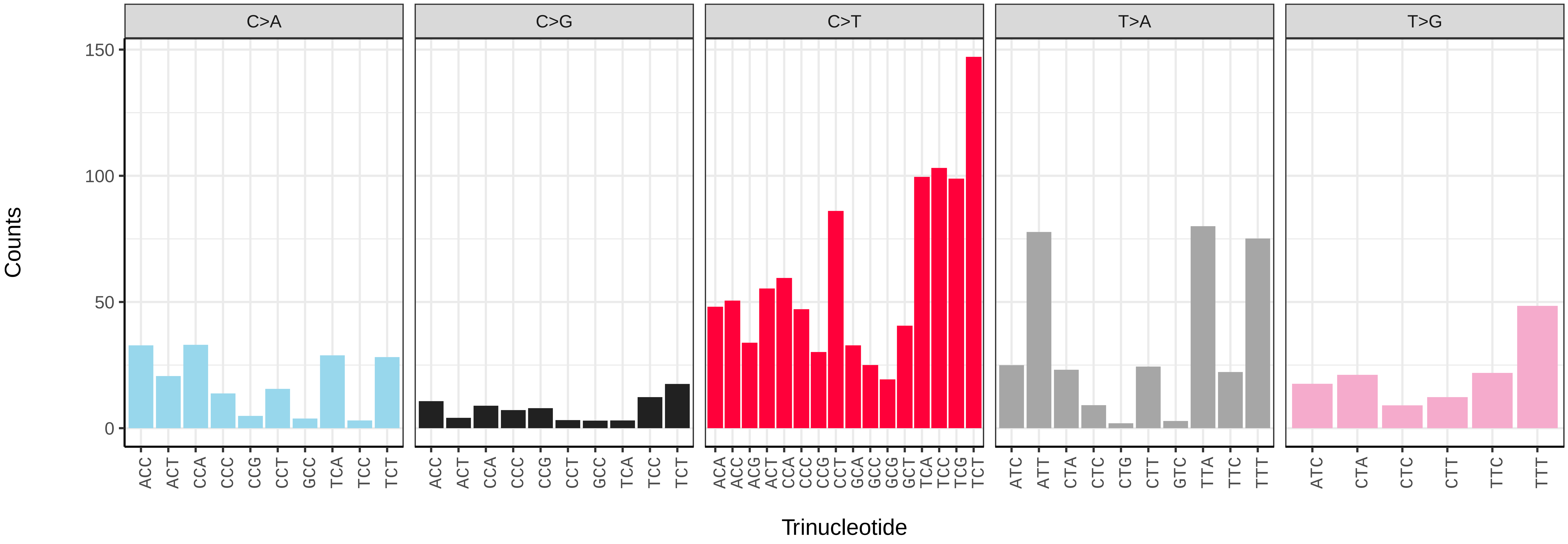

*Cetonia aurata* (icCetAura1) germline mutational spectrum

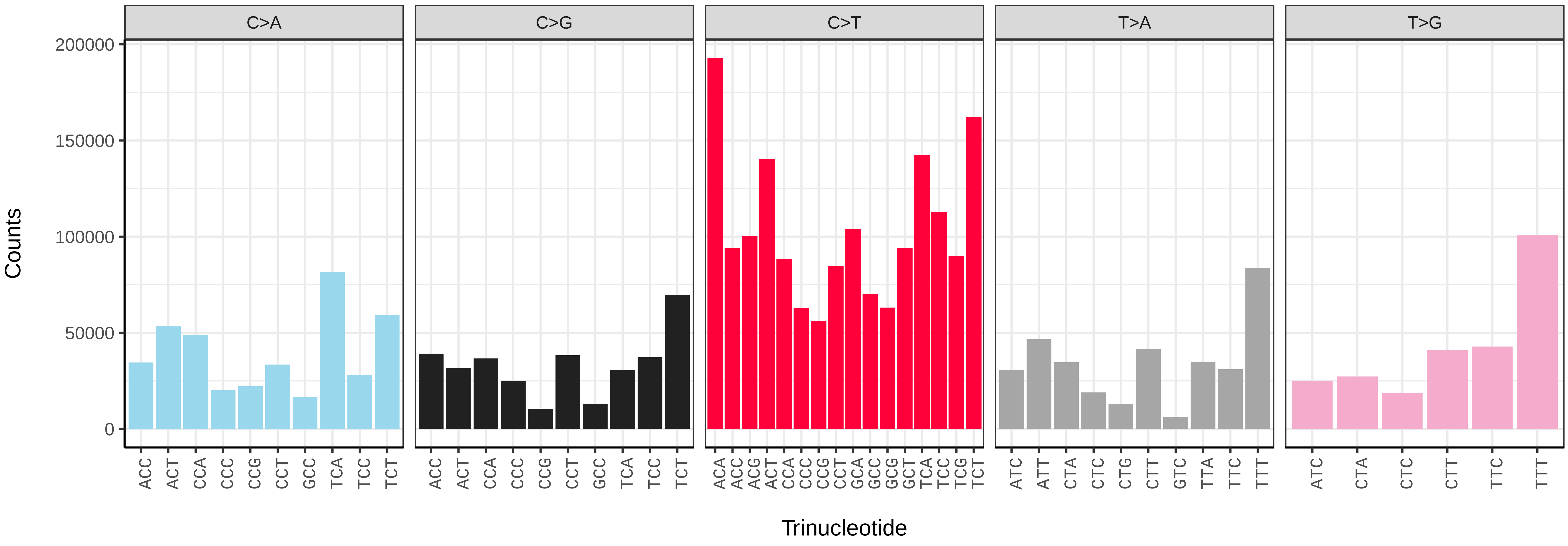


*Cetonia aurata* (icCetAura1) somatic mutational spectrum (SBS96 classification)

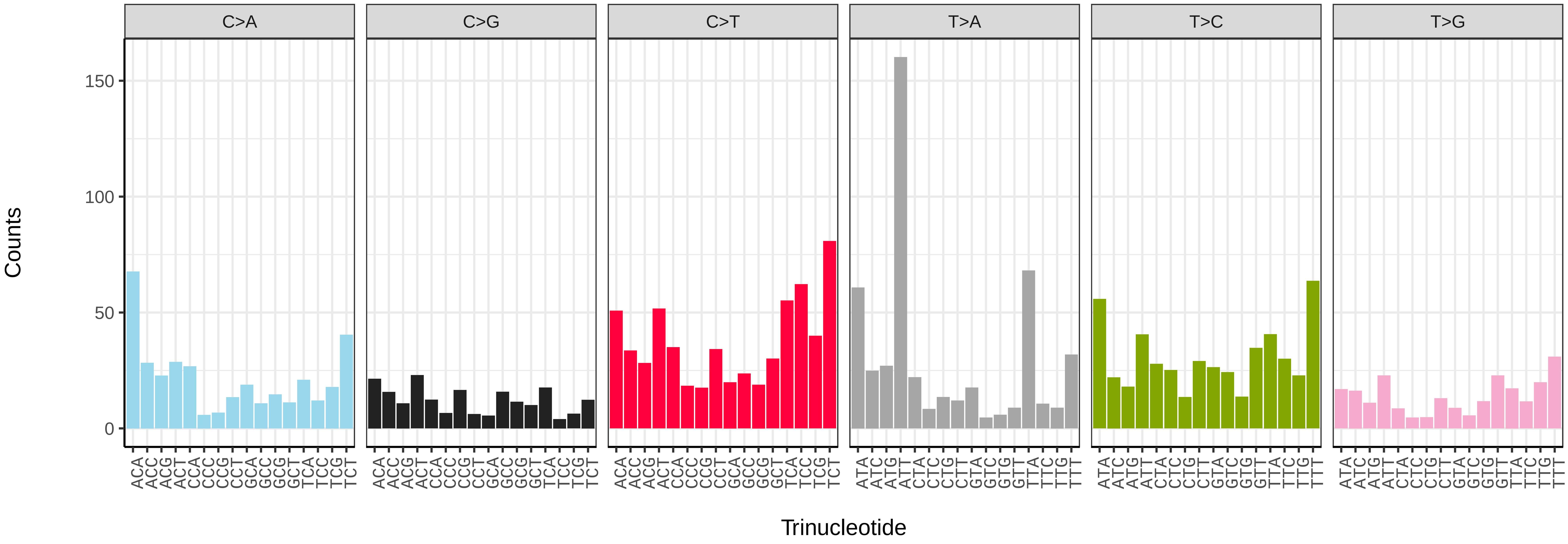


*Cetonia aurata* (icCetAura1) somatic mutational spectrum (SBS52 classification)

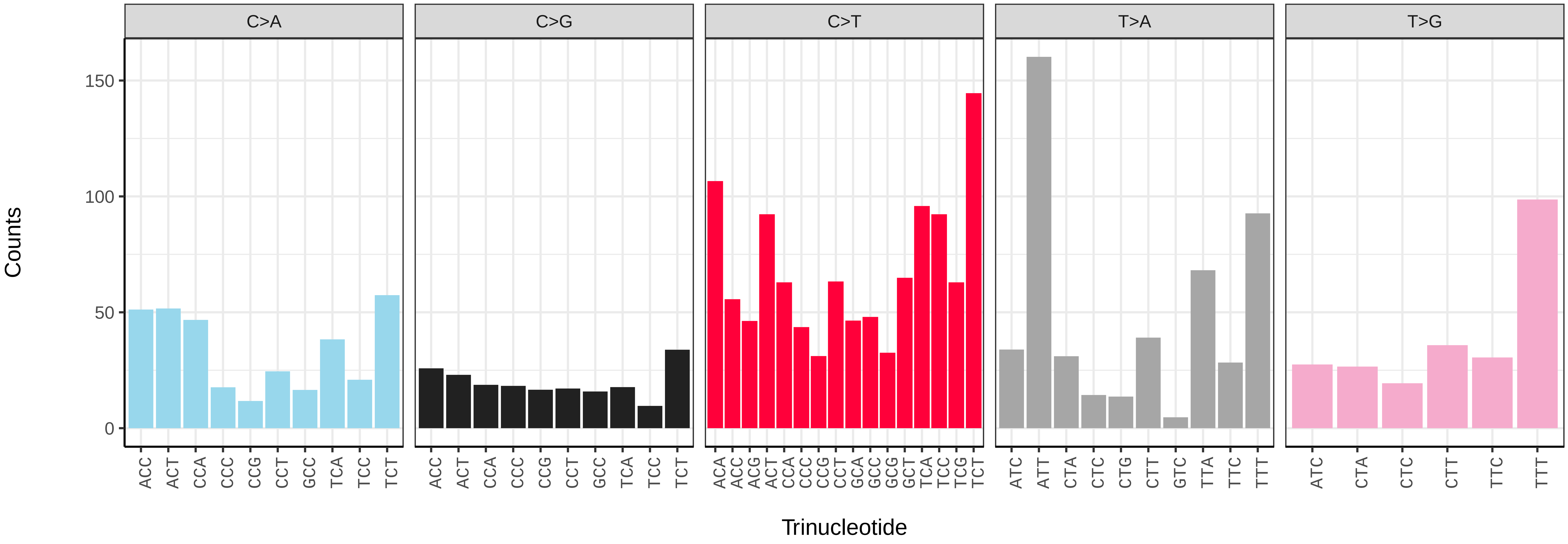

*Chrysolina oricalcia* (icChrOric1) germline mutational spectrum

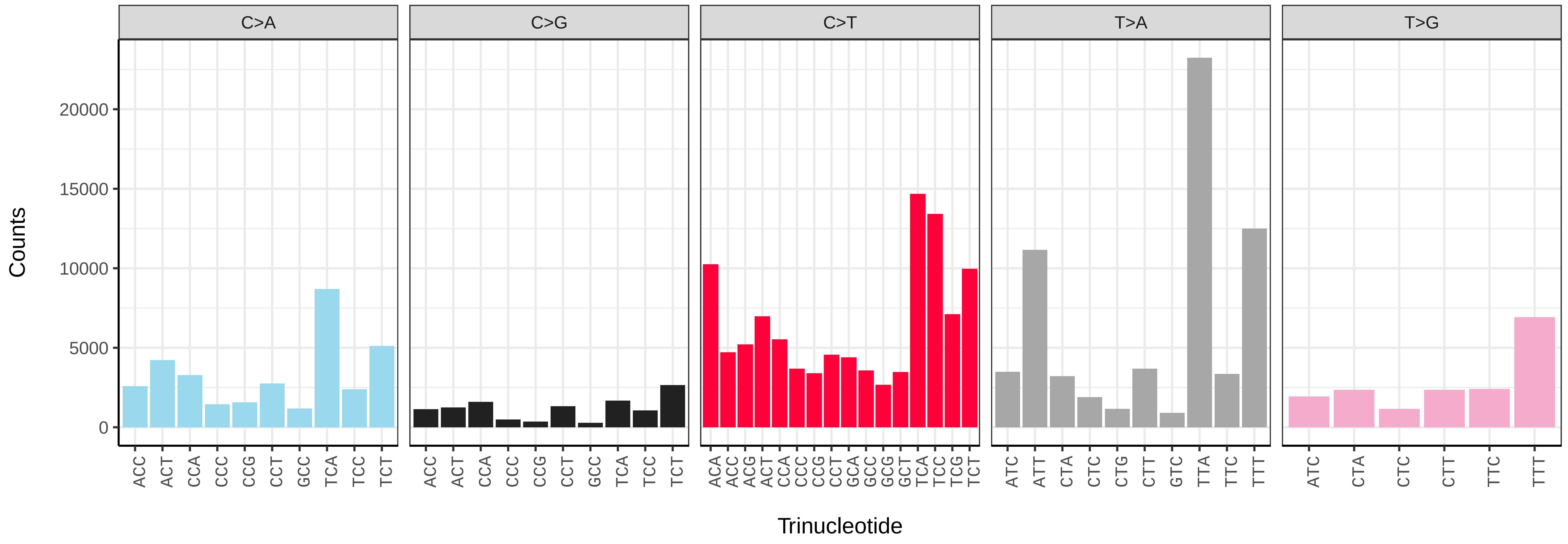


*Chrysolina oricalcia* (icChrOric1) somatic mutational spectrum (SBS96 classification)

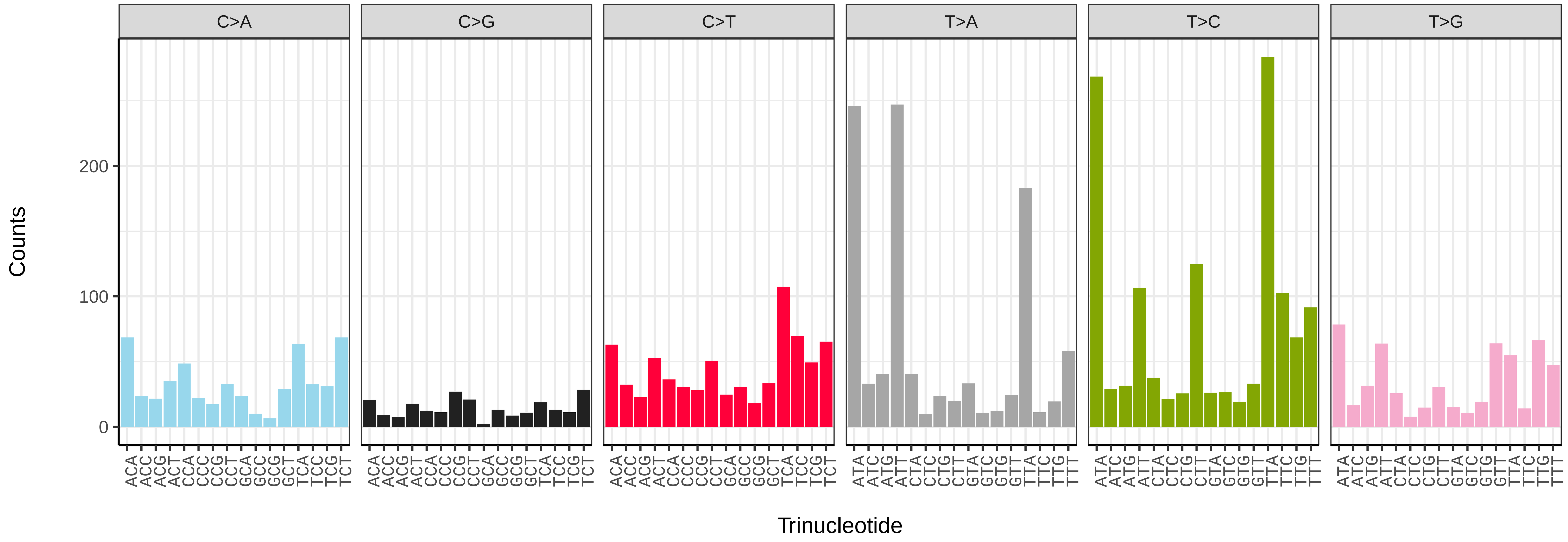


*Chrysolina oricalcia* (icChrOric1) somatic mutational spectrum (SBS52 classification)

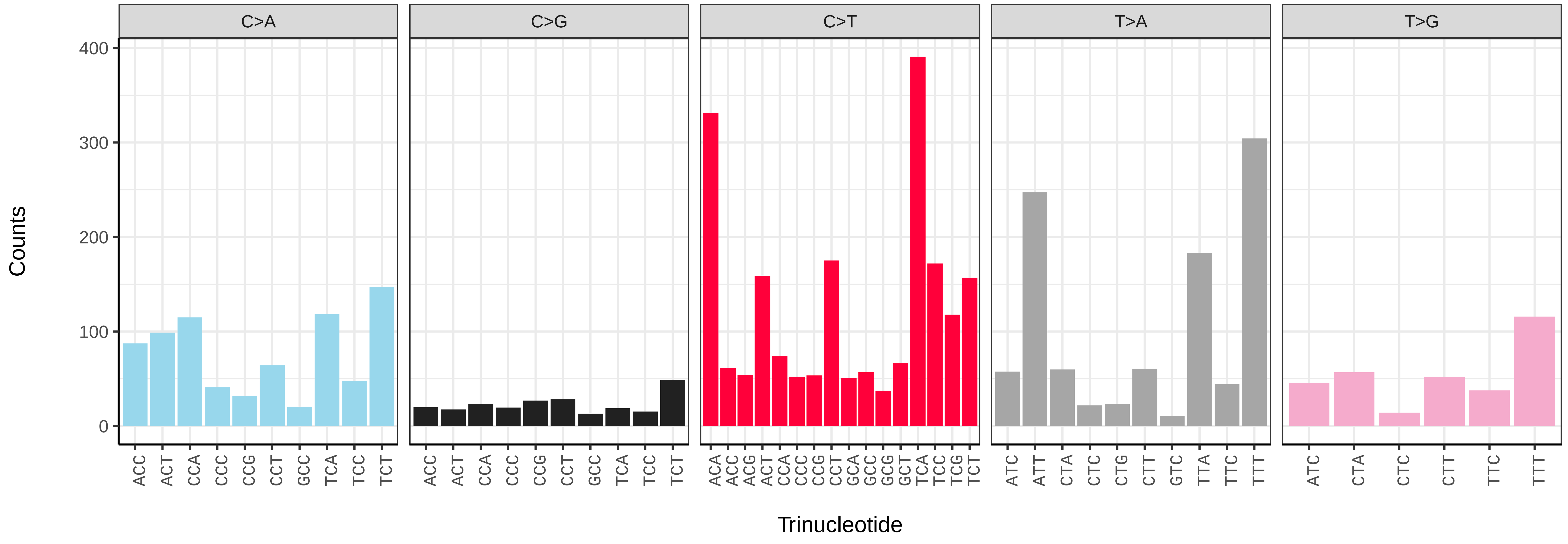

*Coccinella septempunctata* (icCocSept1) germline mutational spectrum

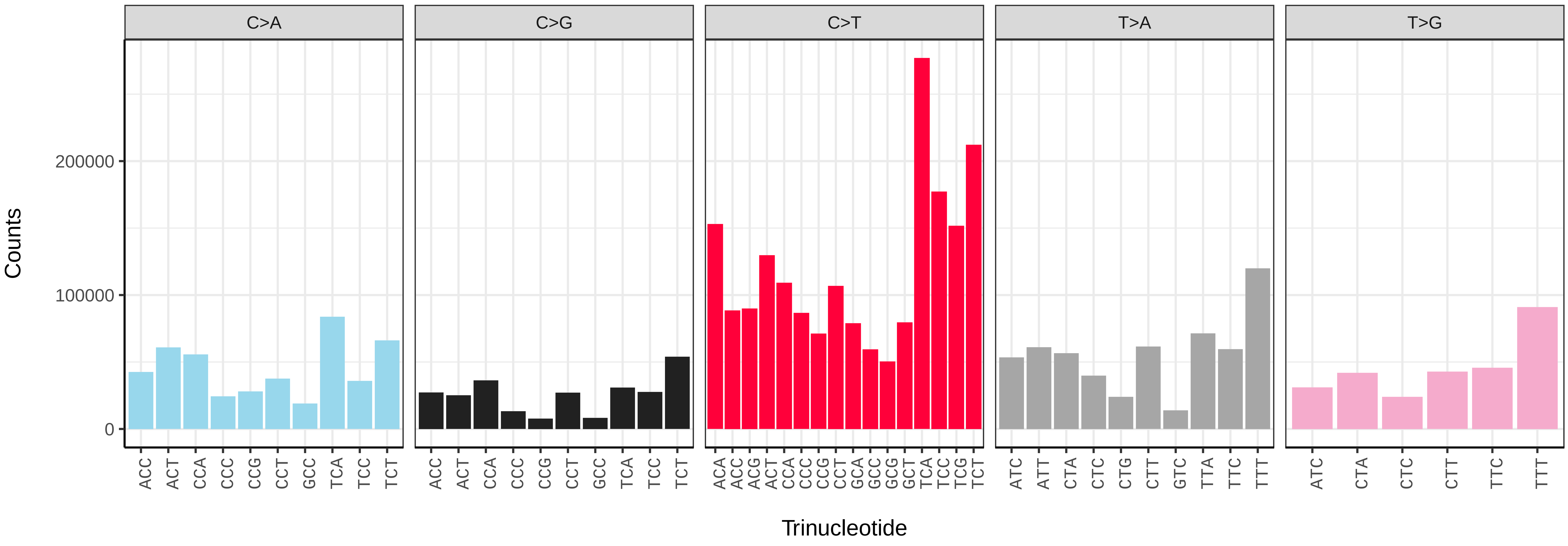


*Coccinella septempunctata* (icCocSept1) somatic mutational spectrum (SBS96 classification)

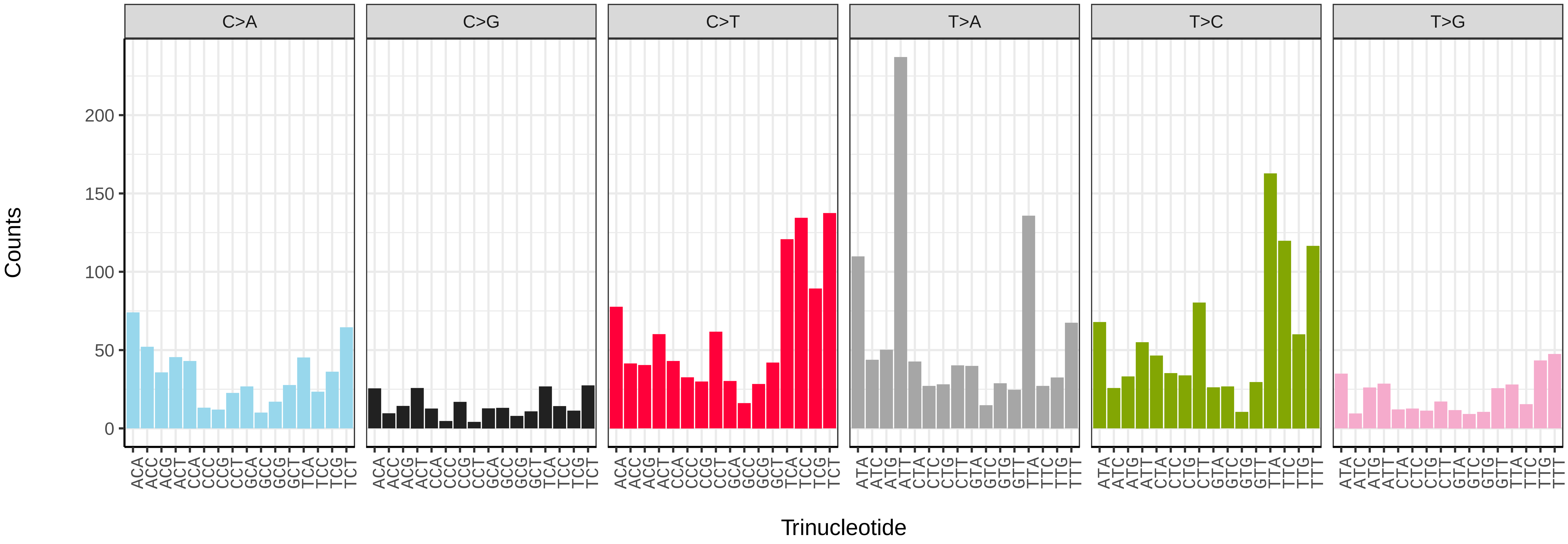


*Coccinella septempunctata* (icCocSept1) somatic mutational spectrum (SBS52 classification)

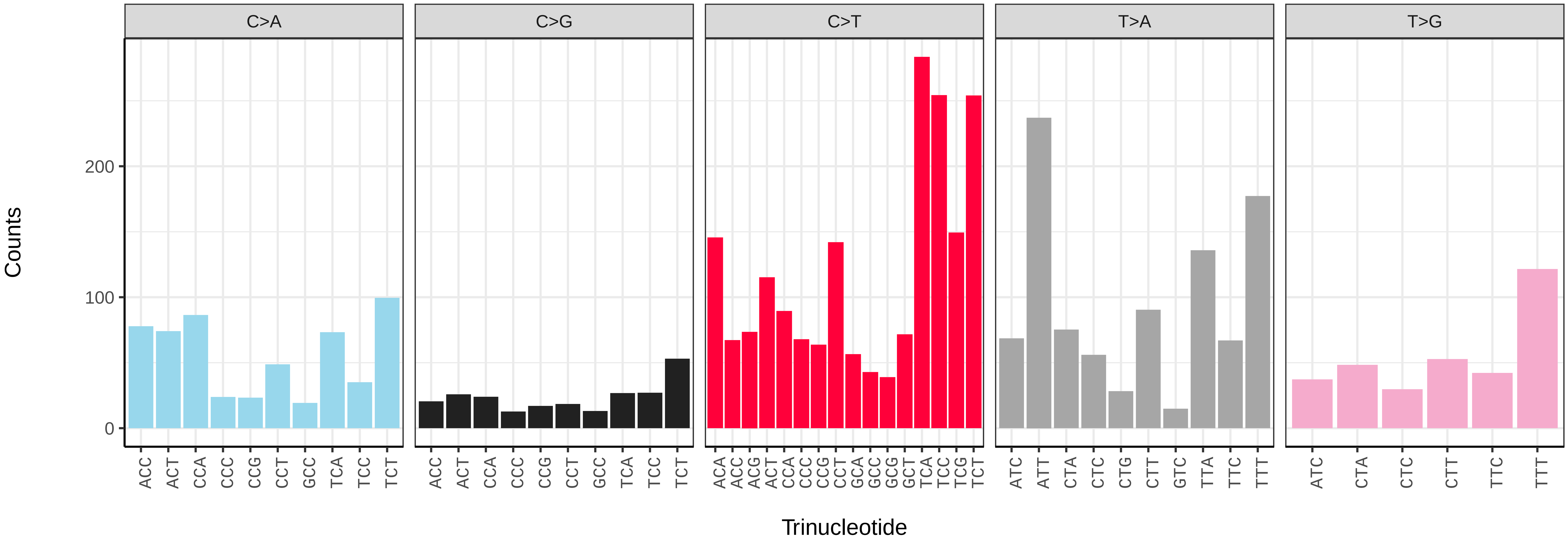

*Cryptocephalus moraei* (icCryMora2) germline mutational spectrum

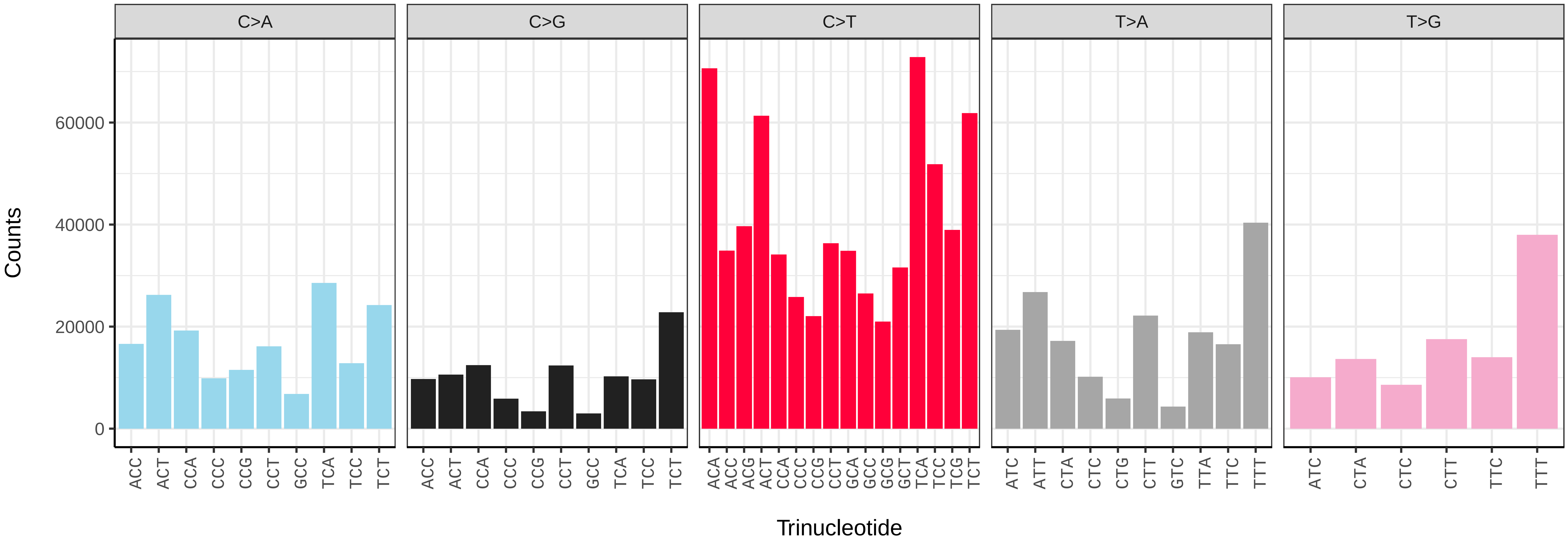


*Cryptocephalus moraei* (icCryMora2) somatic mutational spectrum (SBS96 classification)

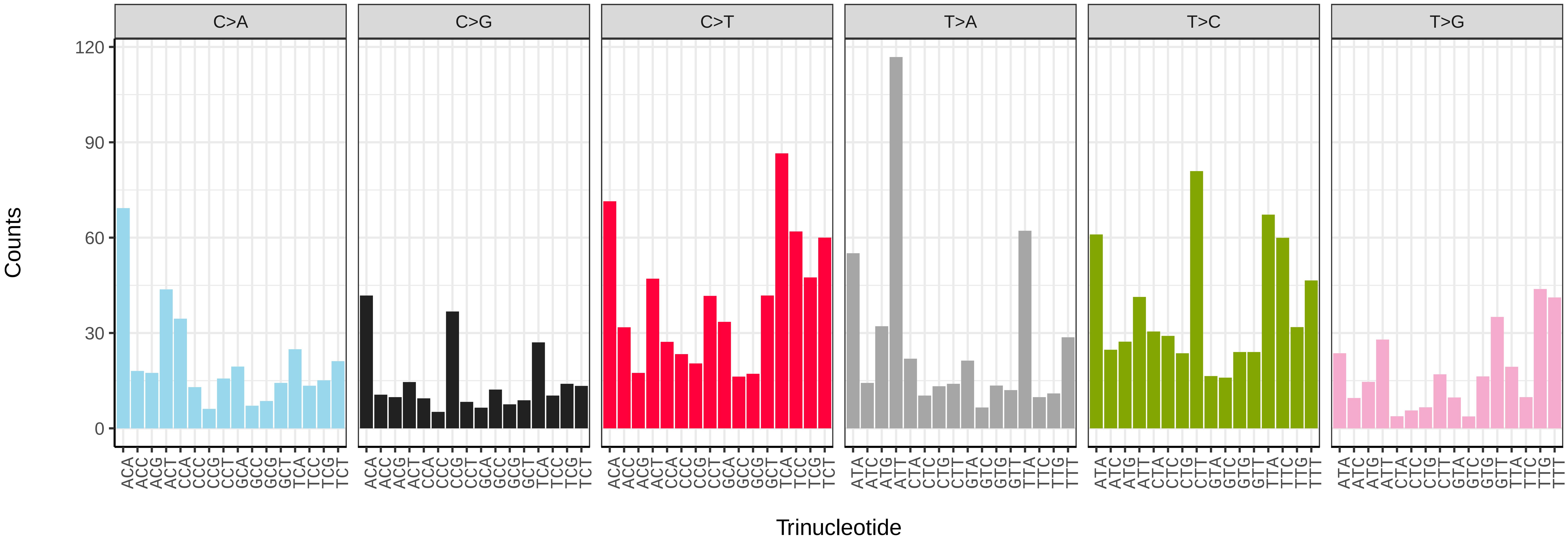


*Cryptocephalus moraei* (icCryMora2) somatic mutational spectrum (SBS52 classification)

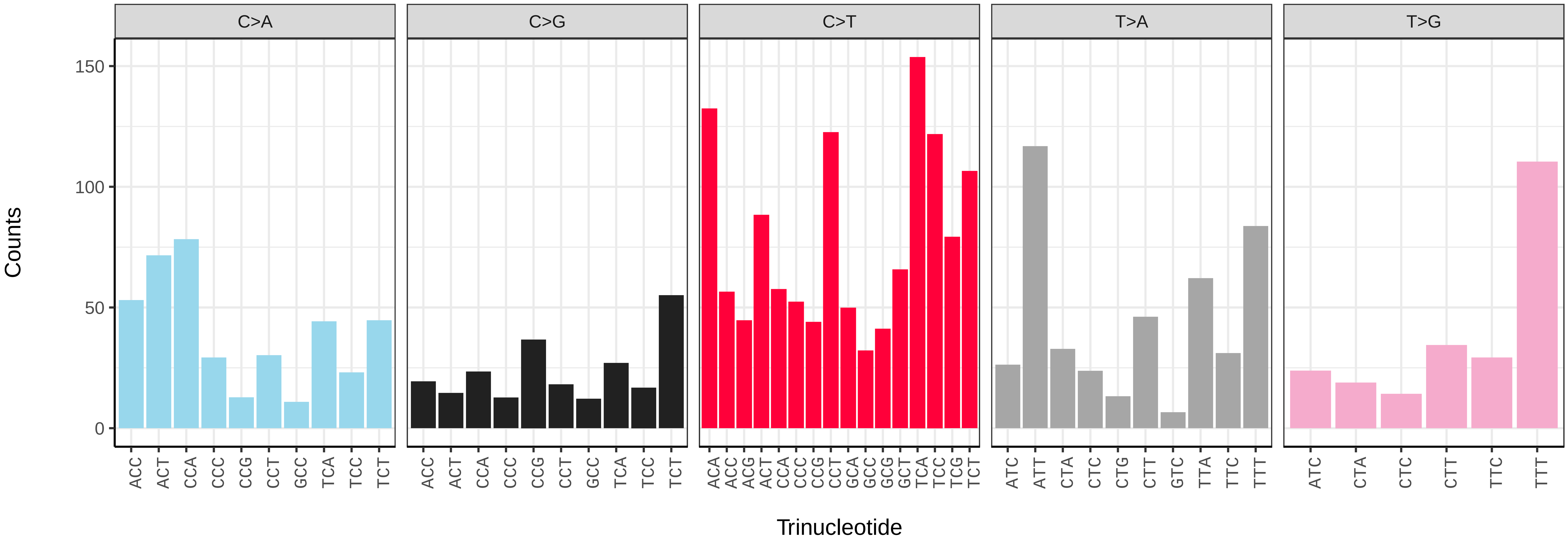

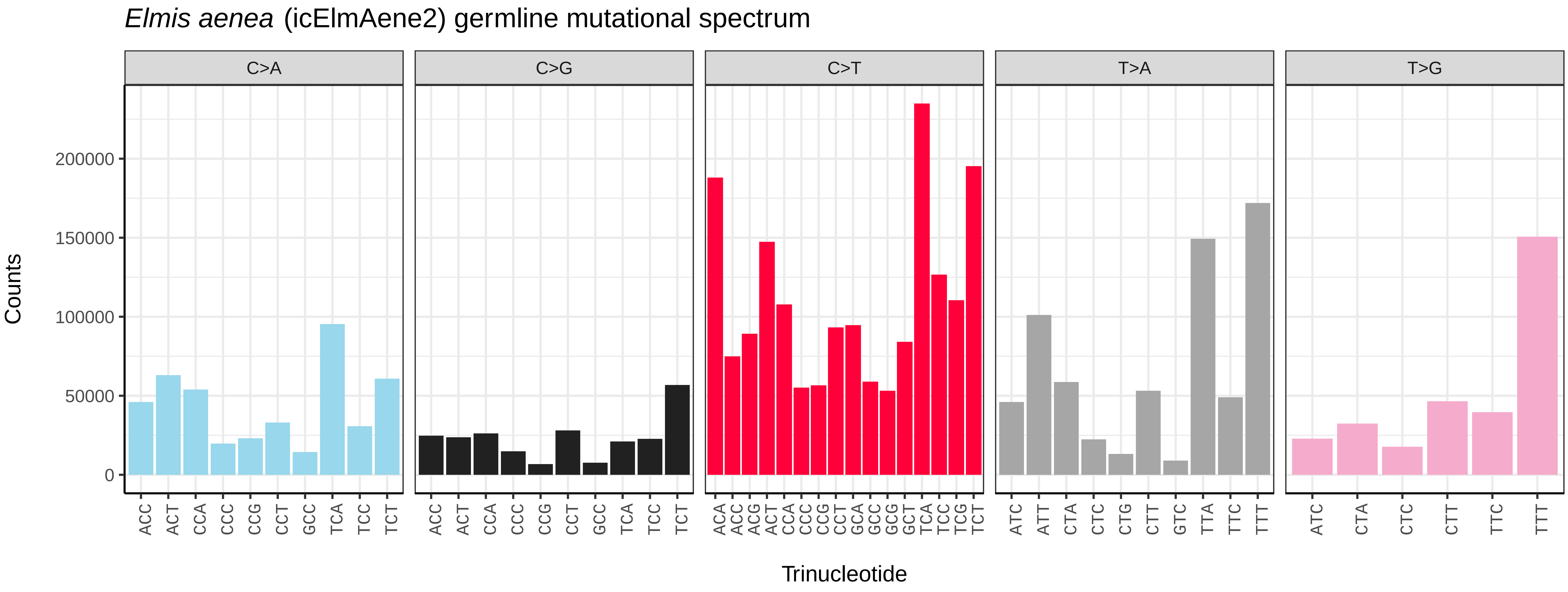
Elmis aenea (icElmAene2) germline mutational spectrum
C>A
C>G
C>T
T>A
T>G
Counts
0
50000
100000
150000
200000
Trinucleotide

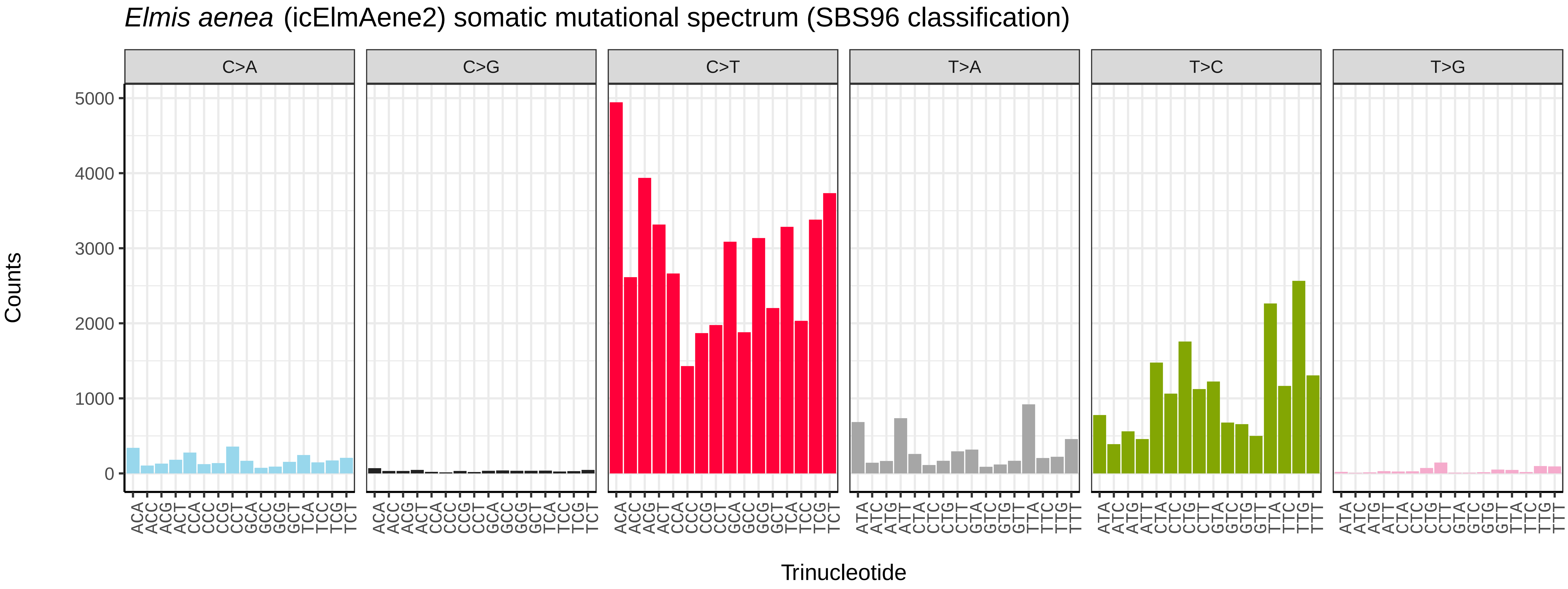
Elmis aenea (icElmAene2) somatic mutational spectrum (SBS96 classification)
C>A
C>G
C>T
T>A
T>C
T>G
Counts
0
1000
2000
3000
4000
5000
Trinucleotide

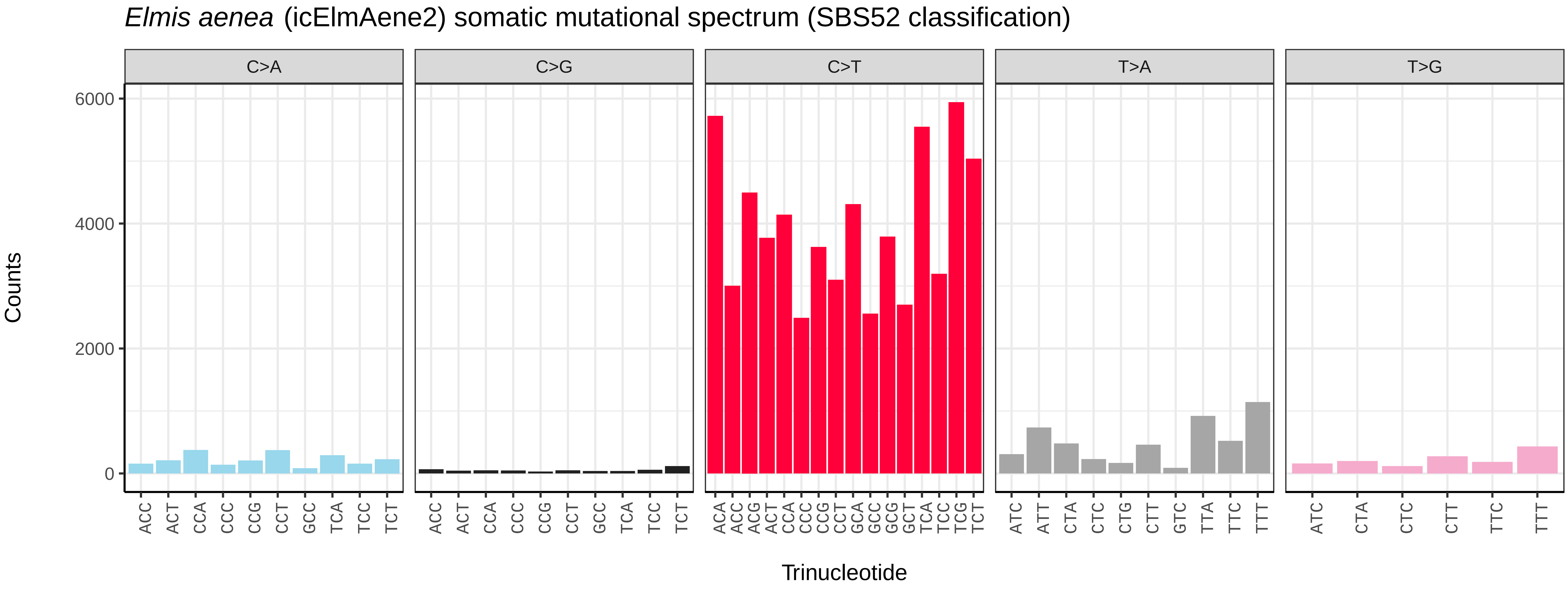
Elmis aenea (icElmAene2) somatic mutational spectrum (SBS52 classification)
C>A
C>G
C>T
T>A
T>G
Counts
0
2000
4000
6000
Trinucleotide

*Halyzia sedecimguttata* (icHalSede1) germline mutational spectrum

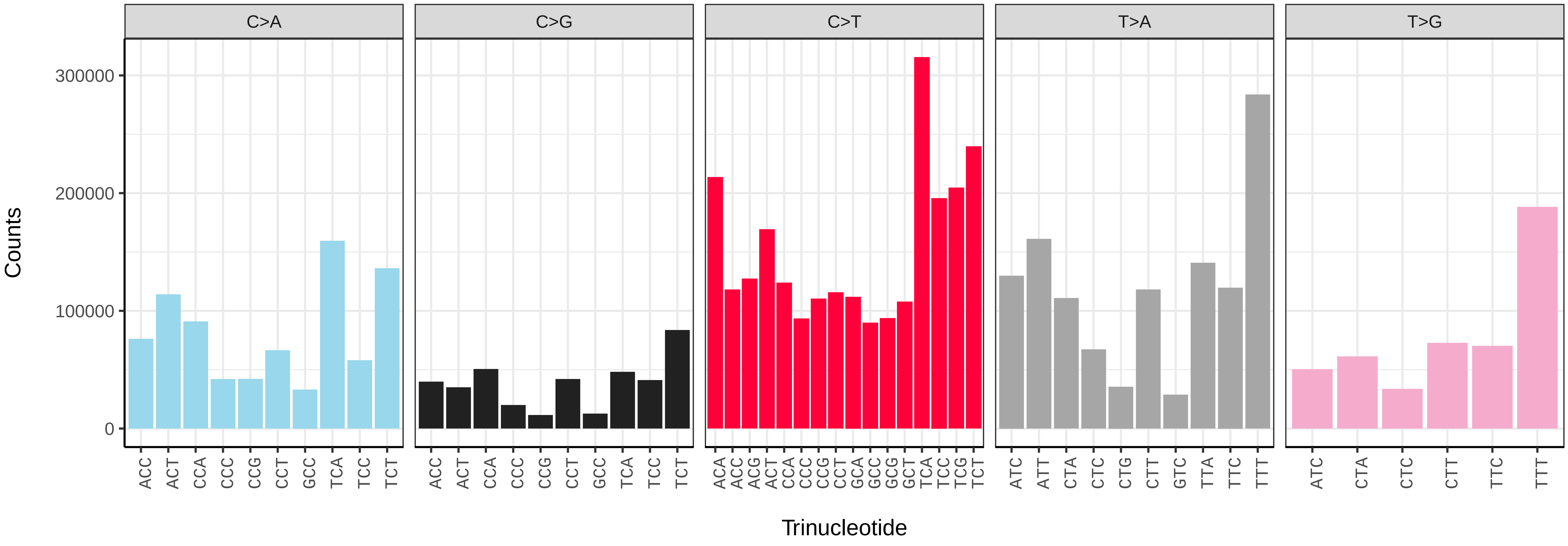


*Halyzia sedecimguttata* (icHalSede1) somatic mutational spectrum (SBS96 classification)

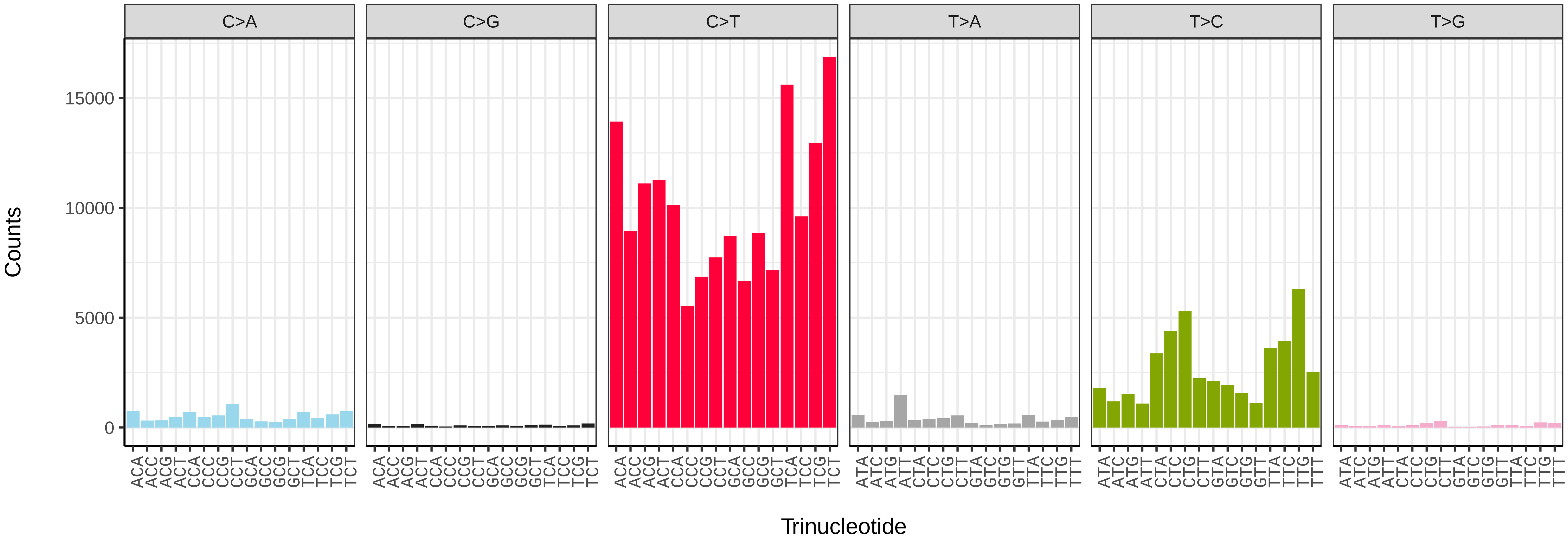


*Halyzia sedecimguttata* (icHalSede1) somatic mutational spectrum (SBS52 classification)

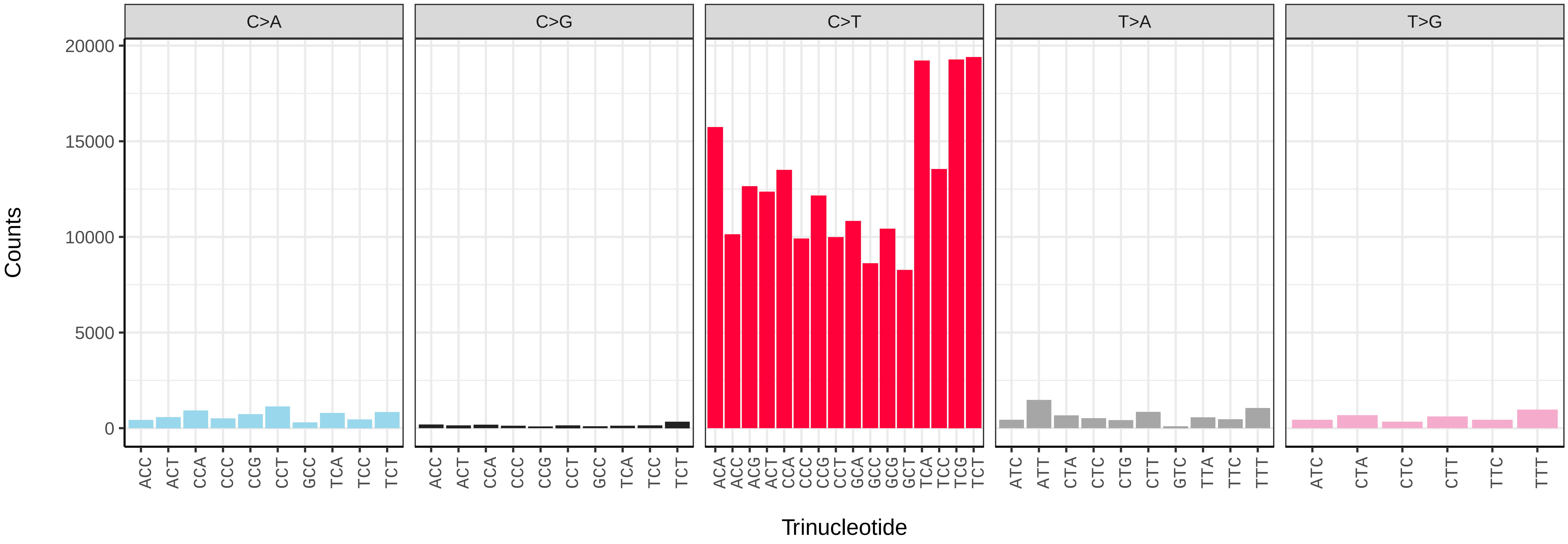

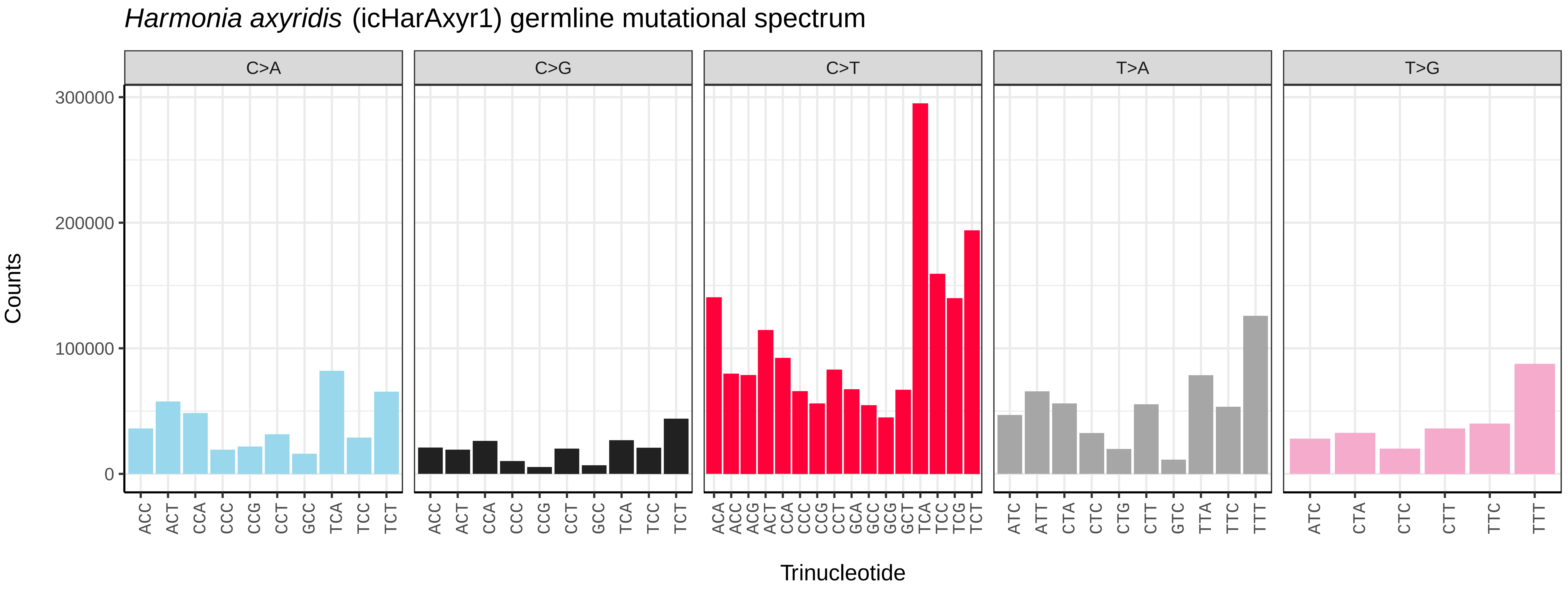
*Harmonia axyridis* (icHarAxyr1) germline mutational spectrum
C>A
C>G
C>T
T>A
T>G
Counts
300000
200000
100000
0
ACC ACT CCA CCC CCG CCT GCC TCA TCC TCT
ACA ACC ACG ACT CCA CCC CCG CCT GCA GCC GCG GCT TCA TCC TCG TCT
ATC ATT CTA CTC CTG CTT GTC TTA TTC TTT
ATC CTA CTC CTT TTC TTT
Trinucleotide

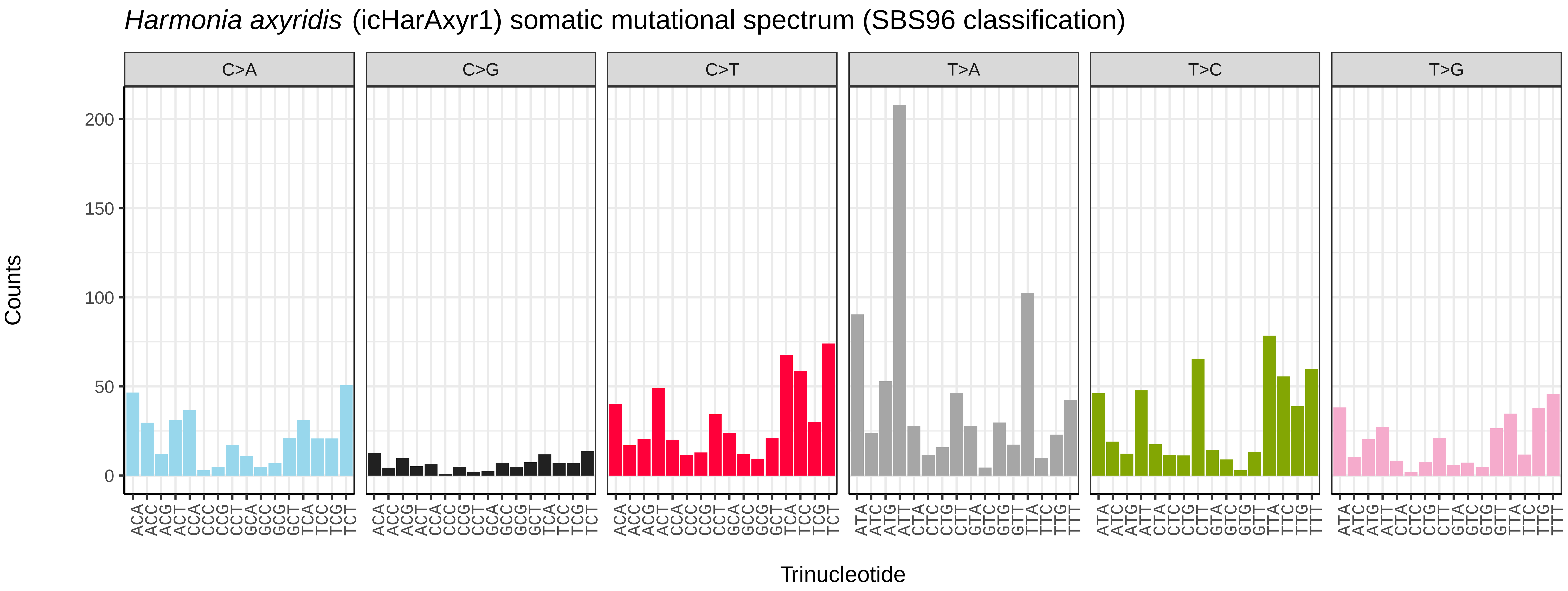
*Harmonia axyridis* (icHarAxyr1) somatic mutational spectrum (SBS96 classification)
C>A
C>G
C>T
T>A
T>C
T>G
Counts
200
150
100
50
0
ACA ACC ACG ACT CCA CCC CCG CCT GCA GCC GCG GCT TCA TCC TCG TCT
ATA ATC ATG ATT CTA CTC CTG CTT GTA GTC GTG GTT TTA TTC TTG TTT
Trinucleotide

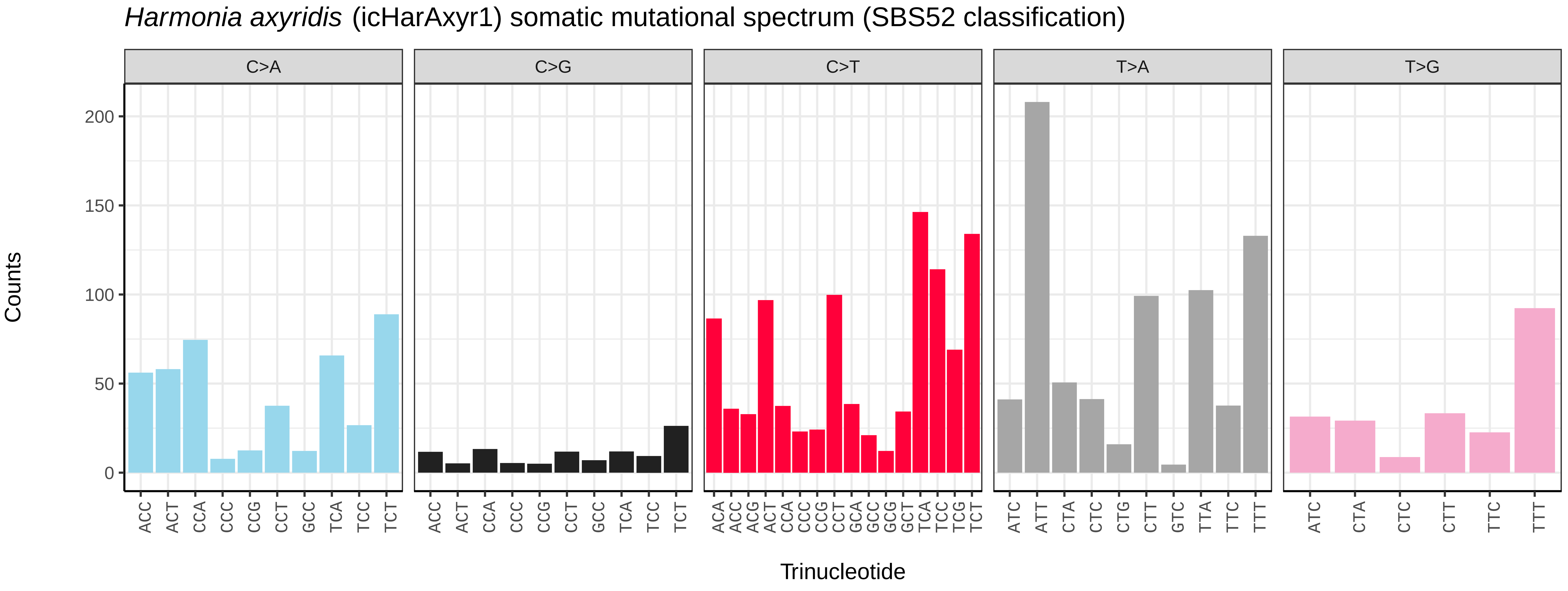
*Harmonia axyridis* (icHarAxyr1) somatic mutational spectrum (SBS52 classification)
C>A
C>G
C>T
T>A
T>G
Counts
200
150
100
50
0
ACC ACT CCA CCC CCG CCT GCC TCA TCC TCT
ACA ACC ACG ACT CCA CCC CCG CCT GCA GCC GCG GCT TCA TCC TCG TCT
ATC ATT CTA CTC CTG CTT GTC TTA TTC TTT
ATC CTA CTC CTT TTC TTT
Trinucleotide

*Lagria hirta* (icLagHirt1) germline mutational spectrum

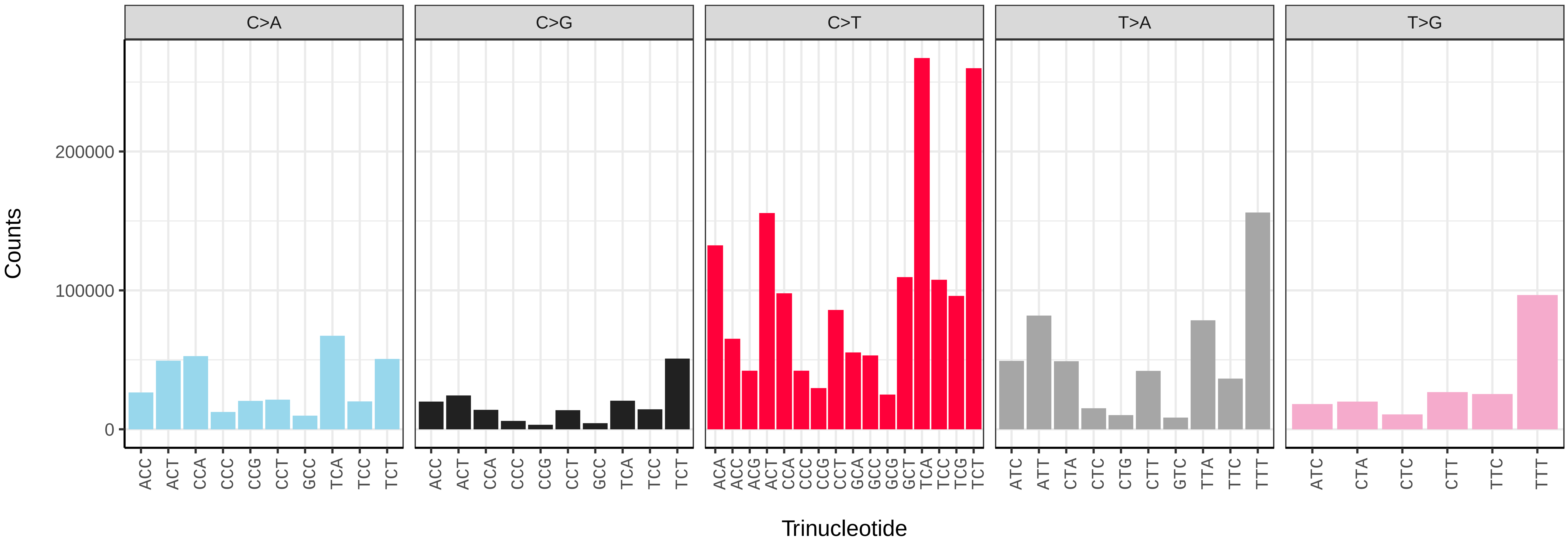


*Lagria hirta* (icLagHirt1) somatic mutational spectrum (SBS96 classification)

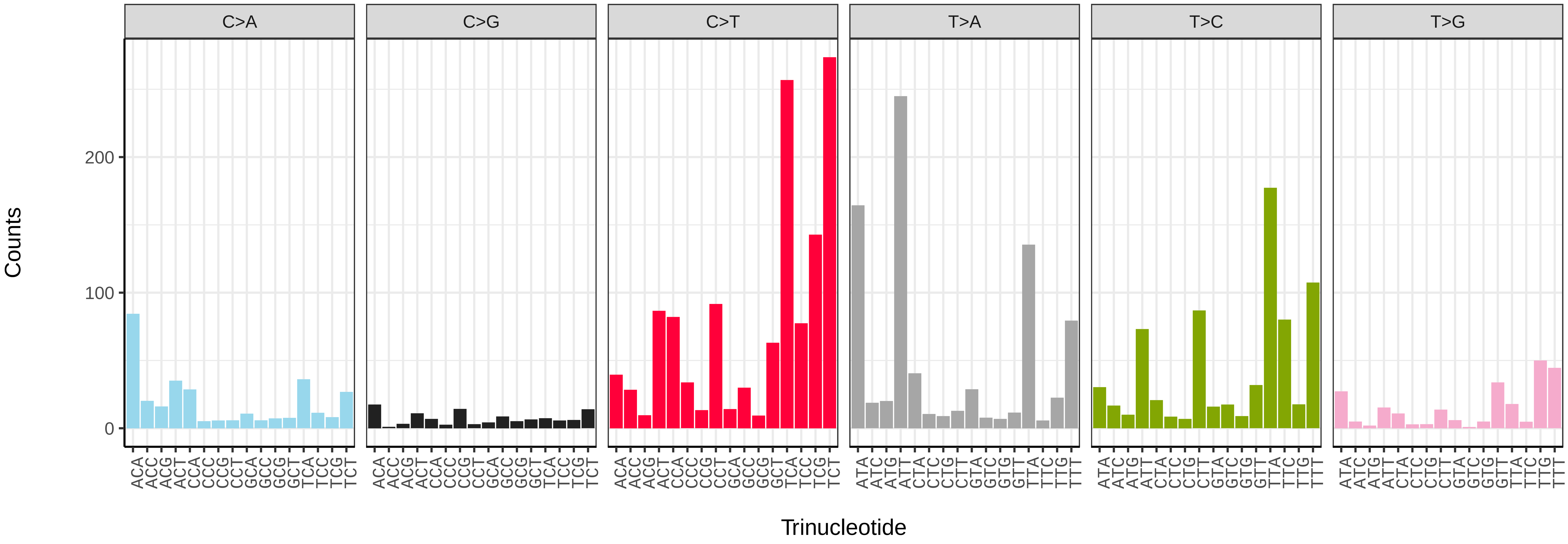


*Lagria hirta* (icLagHirt1) somatic mutational spectrum (SBS52 classification)

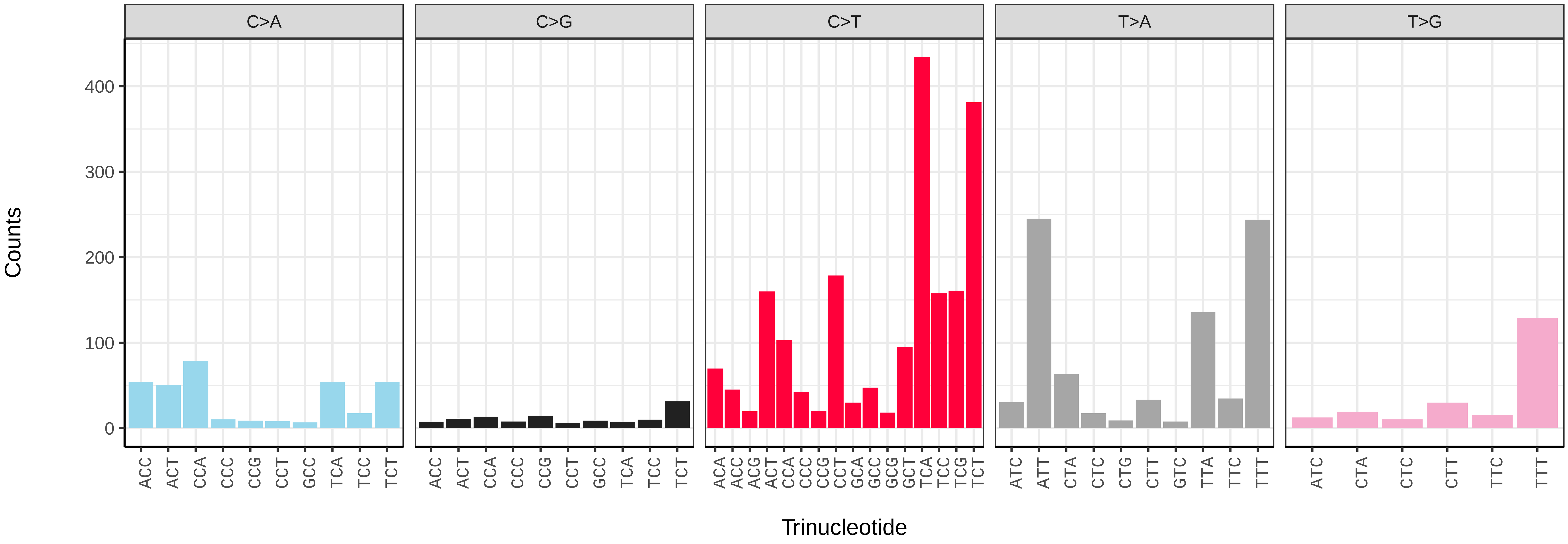

*Leistus spinibarbis* (icLeiSpin1) germline mutational spectrum

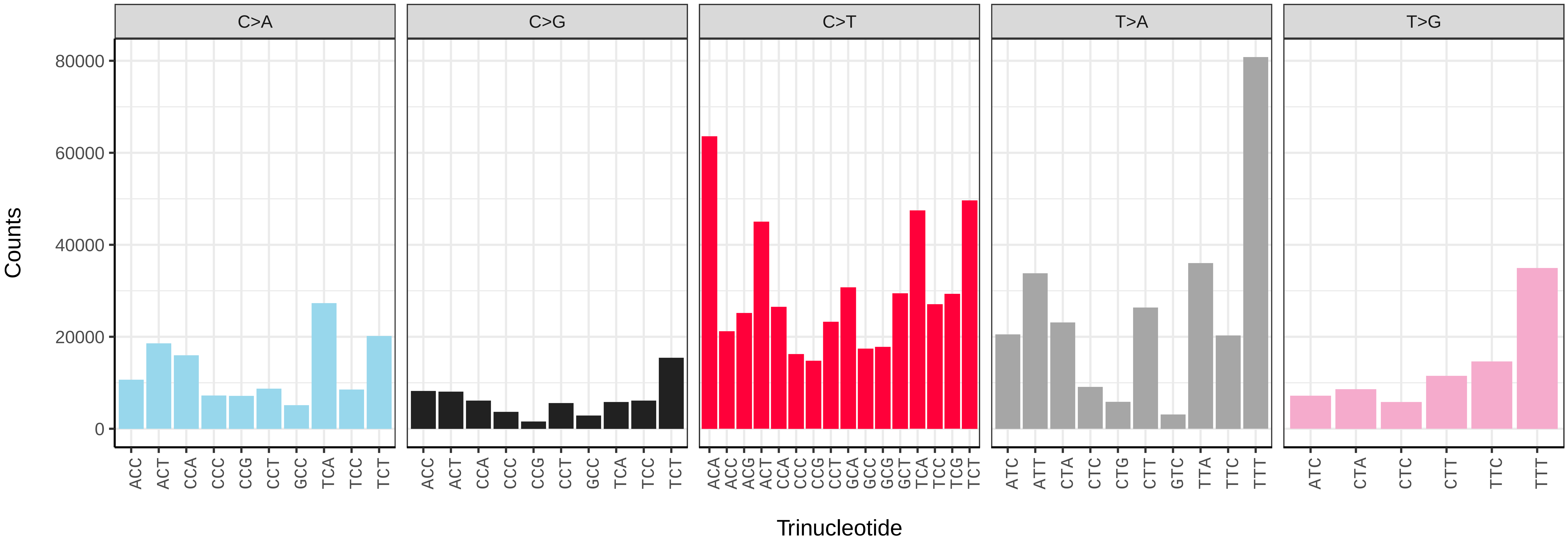


*Leistus spinibarbis* (icLeiSpin1) somatic mutational spectrum (SBS96 classification)

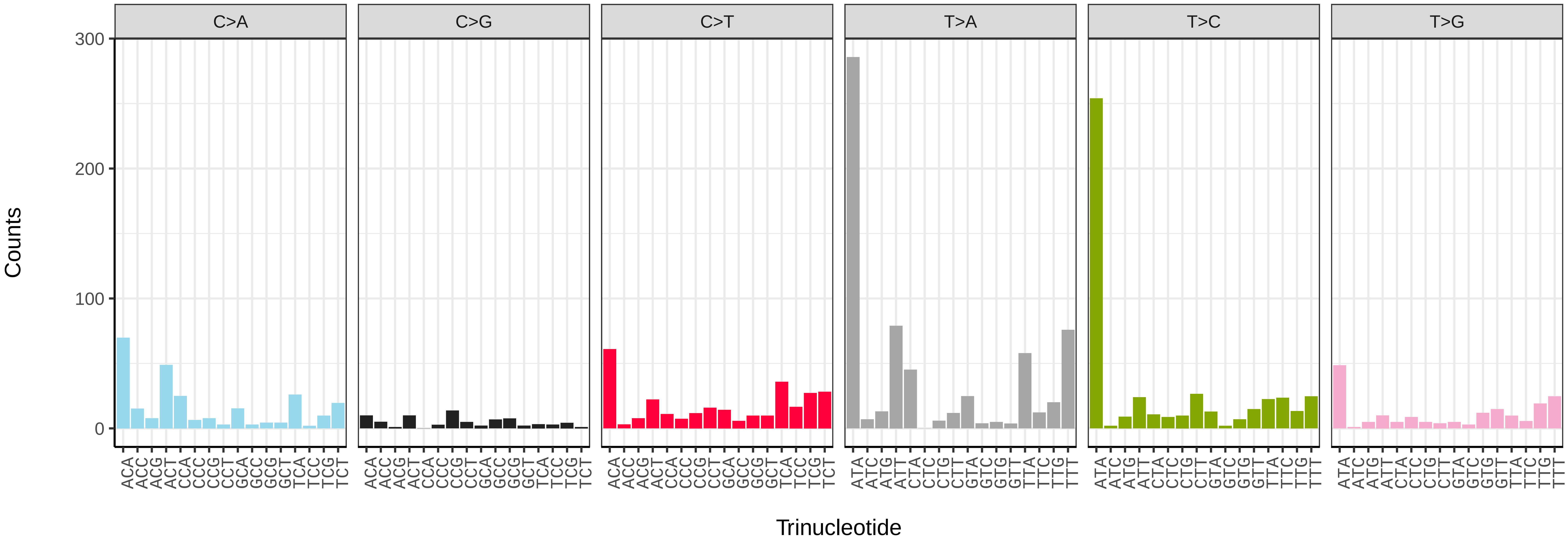


*Leistus spinibarbis* (icLeiSpin1) somatic mutational spectrum (SBS52 classification)

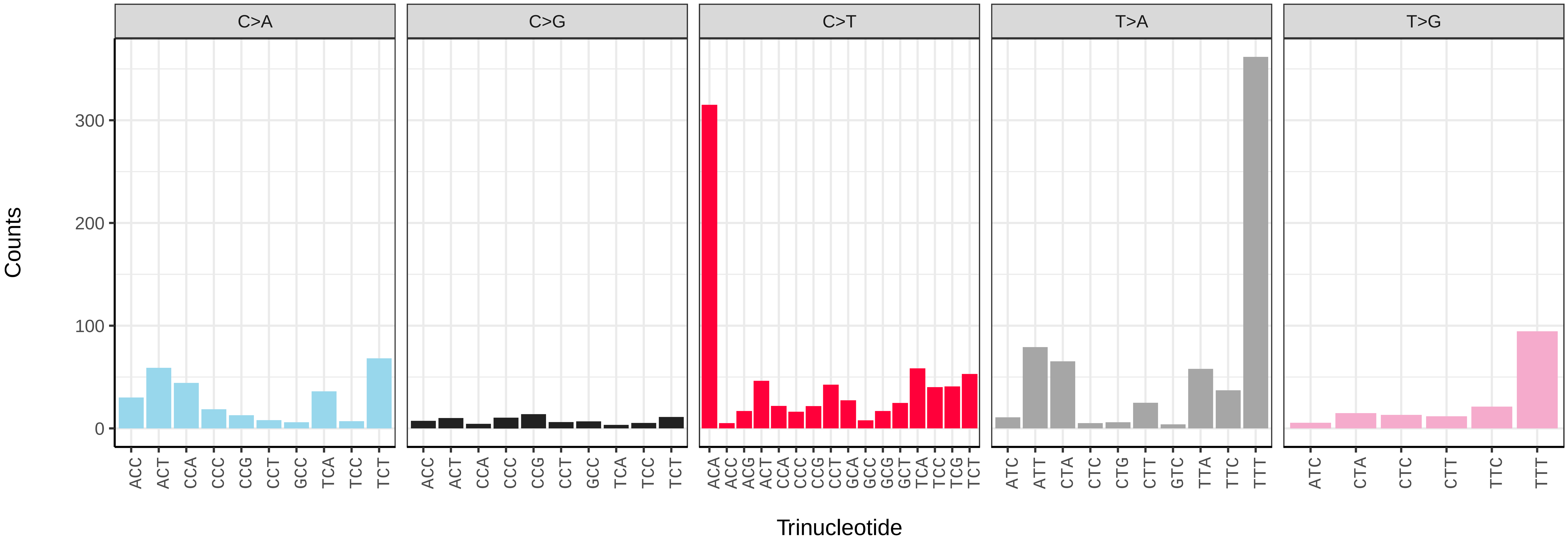

*Rutpela maculata* (icLepMacu1) germline mutational spectrum

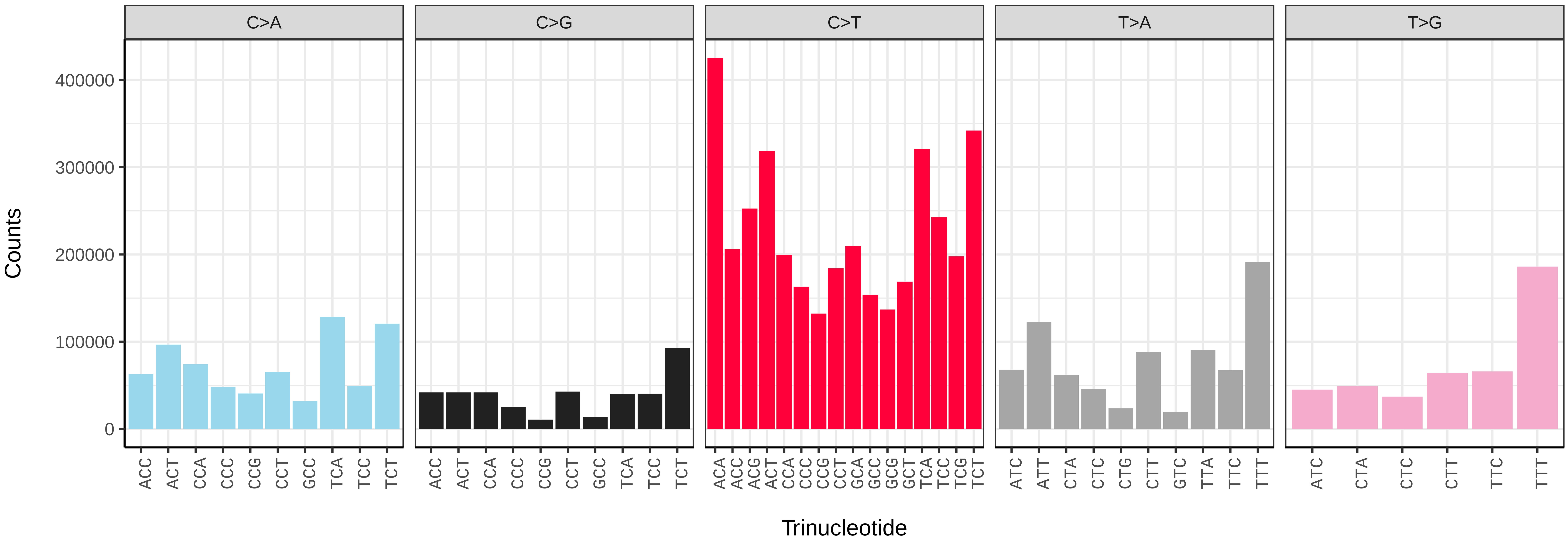


*Rutpela maculata* (icLepMacu1) somatic mutational spectrum (SBS96 classification)

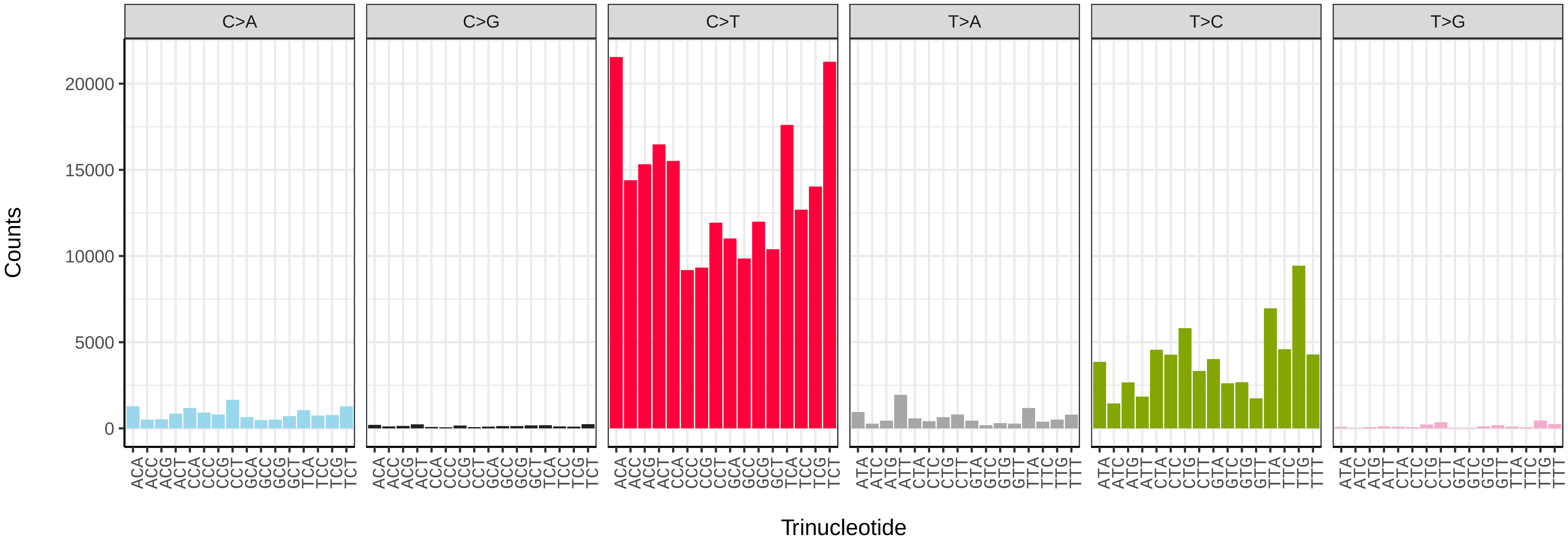


*Rutpela maculata* (icLepMacu1) somatic mutational spectrum (SBS52 classification)

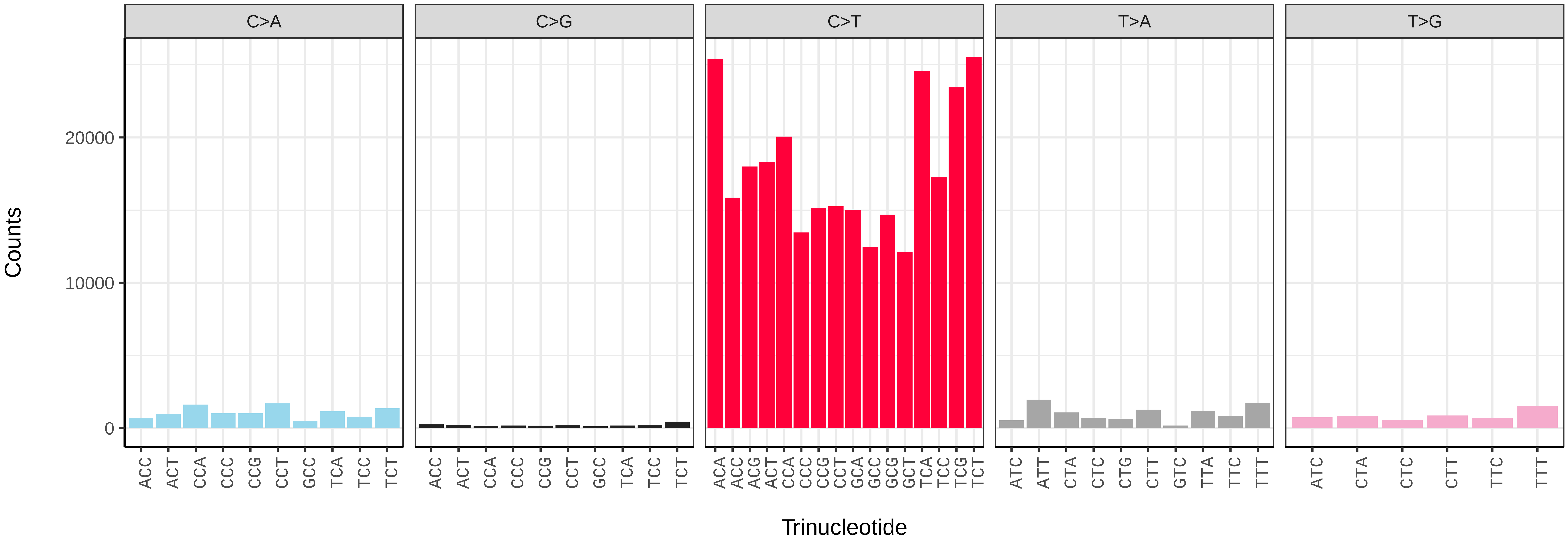

*Lochmaea capreae* (icLocCapr1) germline mutational spectrum

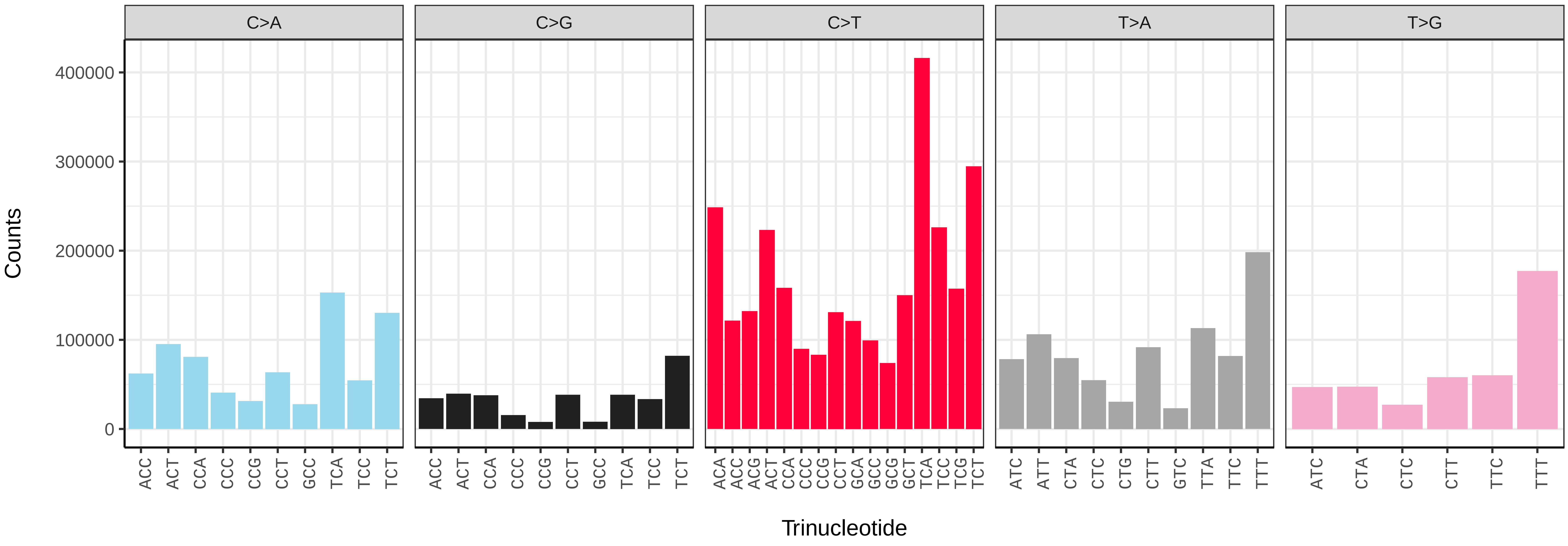


*Lochmaea capreae* (icLocCapr1) somatic mutational spectrum (SBS96 classification)

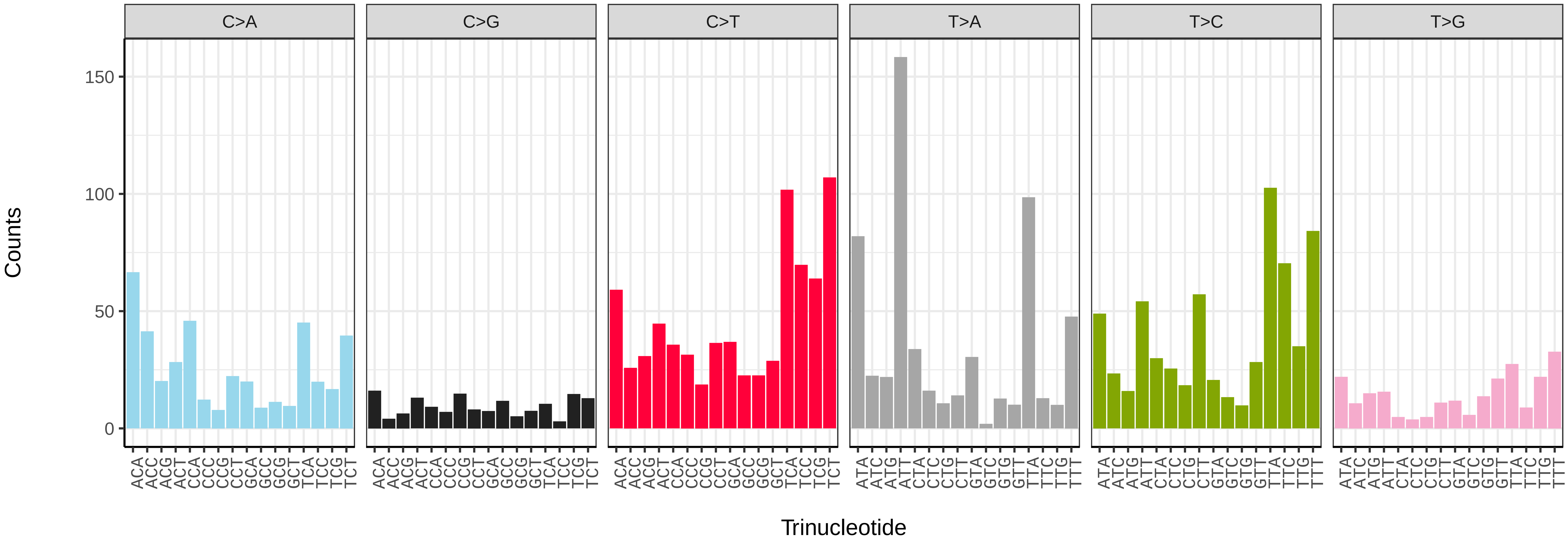


*Lochmaea capreae* (icLocCapr1) somatic mutational spectrum (SBS52 classification)

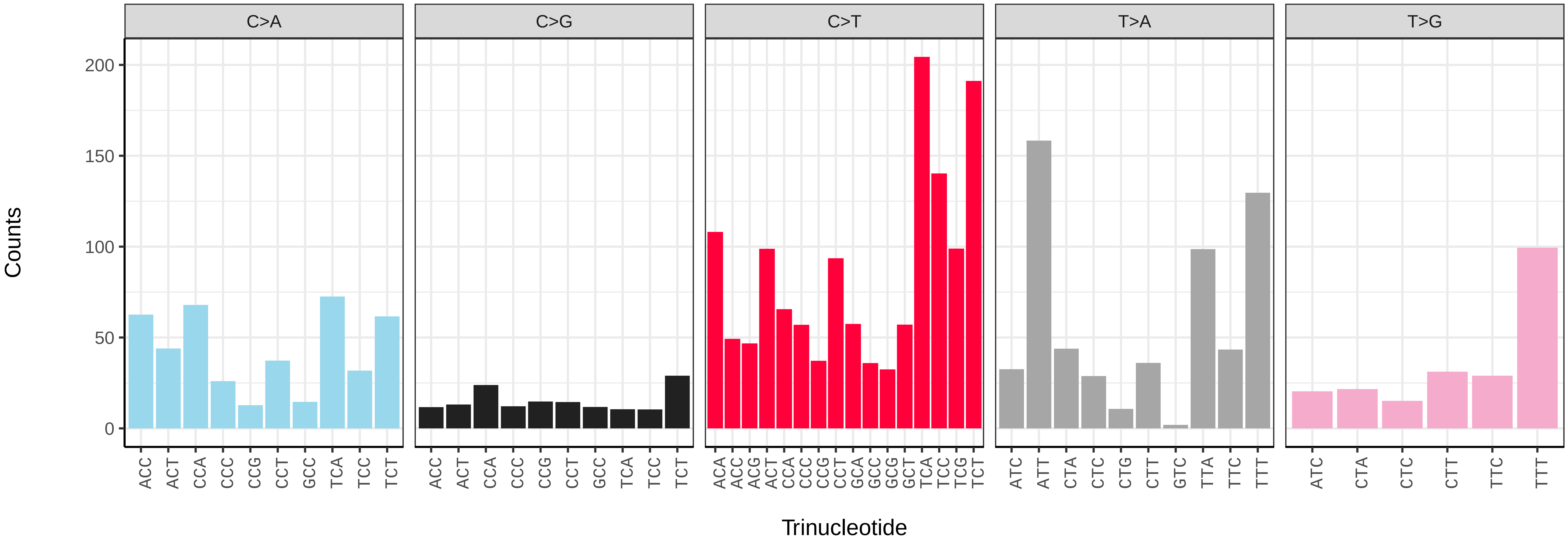

*Lochmaea crataegi* (icLocCrat2) germline mutational spectrum

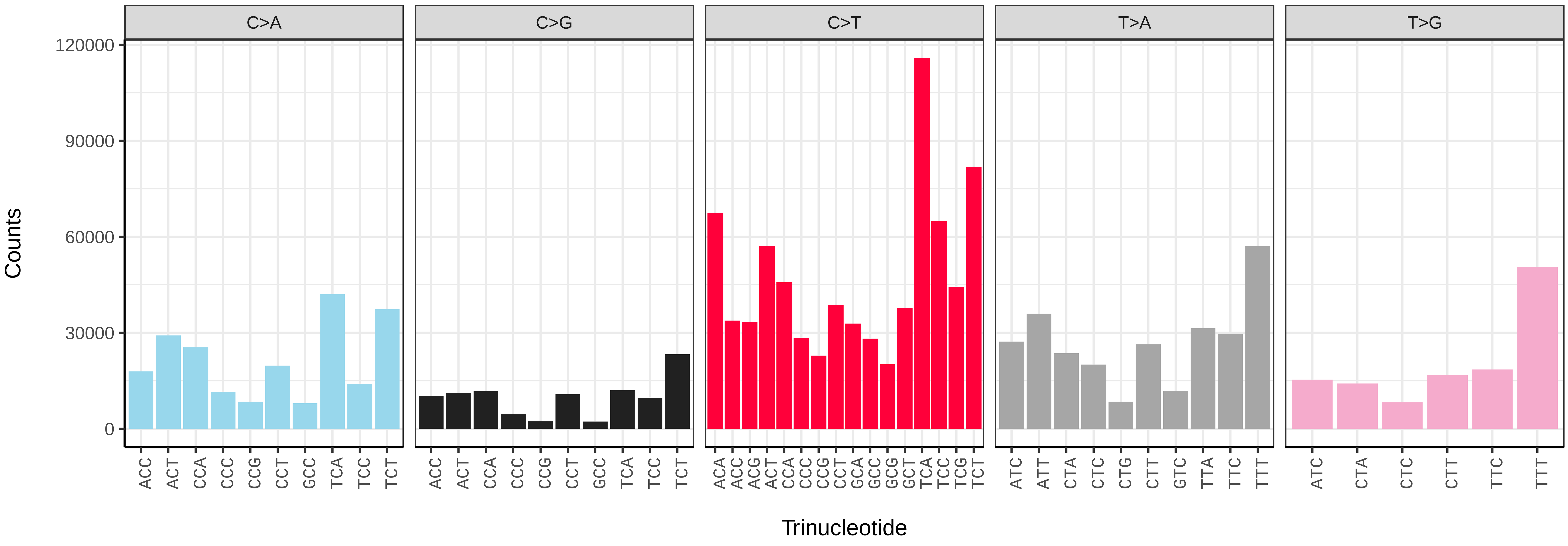


*Lochmaea crataegi* (icLocCrat2) somatic mutational spectrum (SBS96 classification)

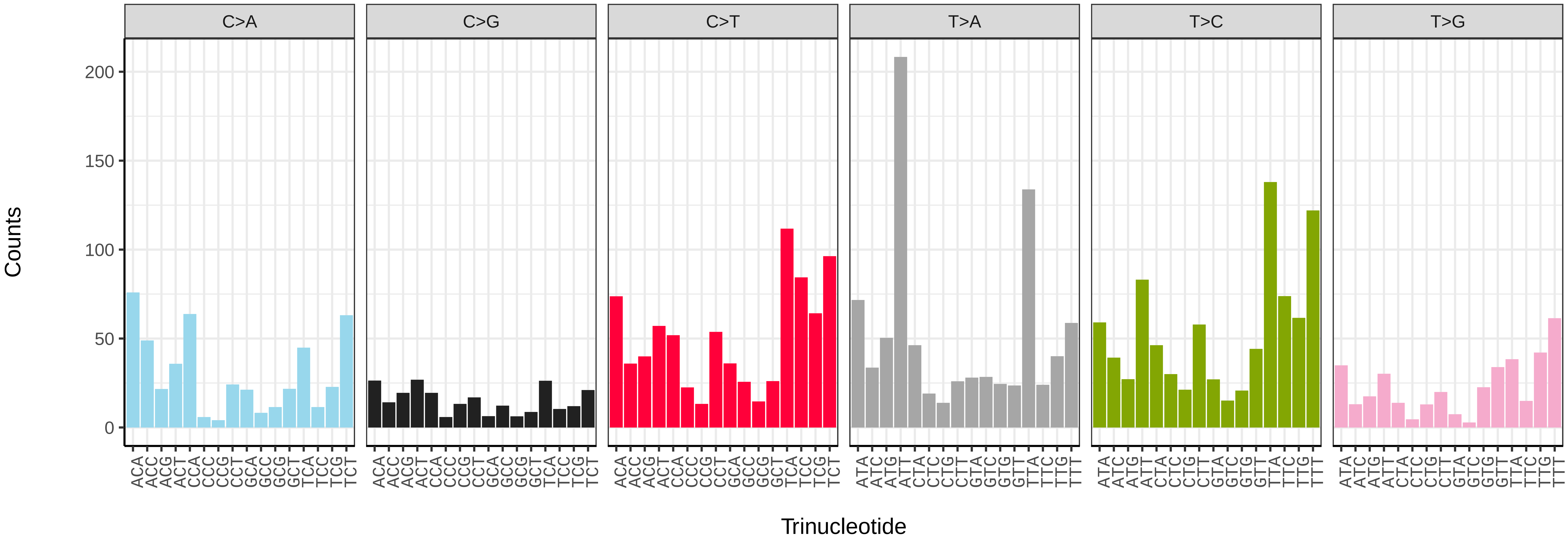


*Lochmaea crataegi* (icLocCrat2) somatic mutational spectrum (SBS52 classification)

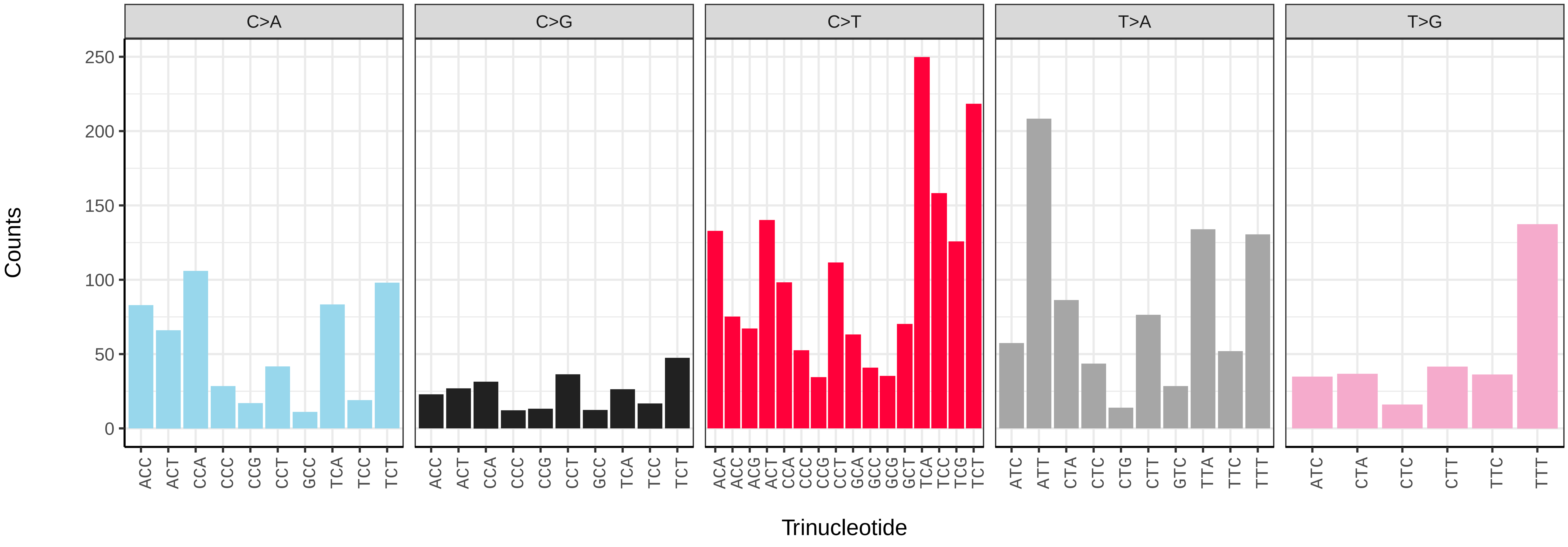

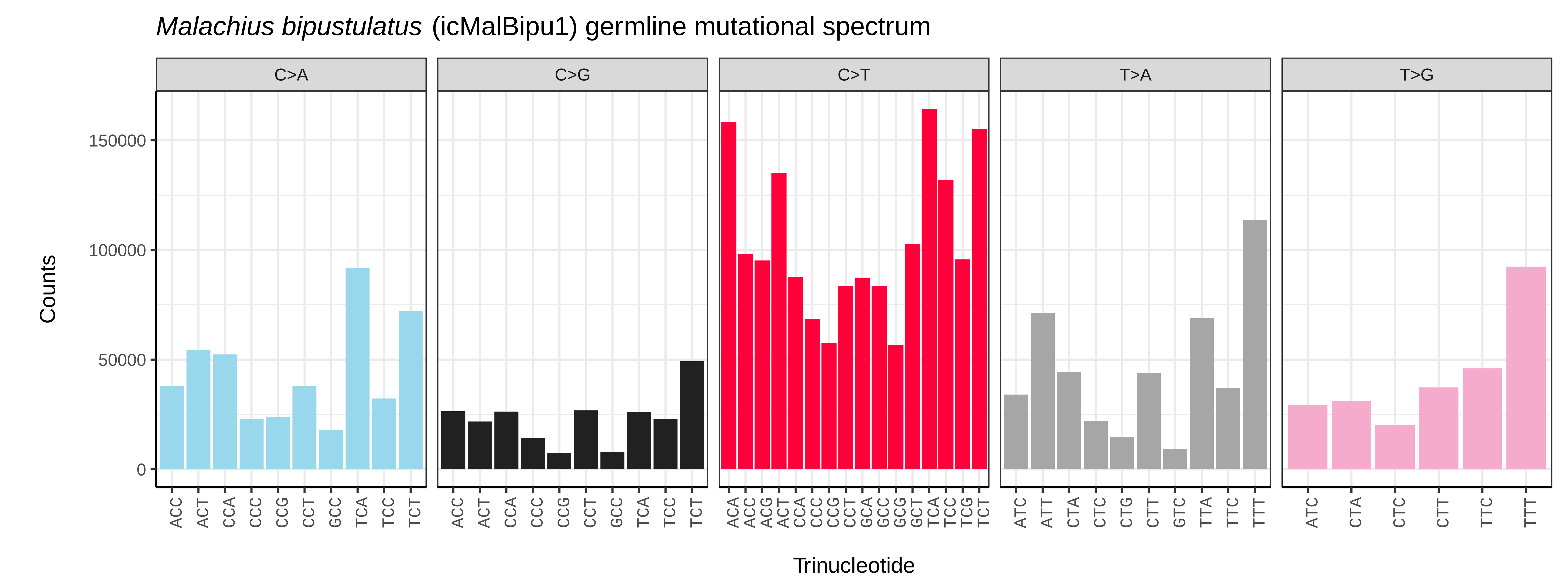
*Malachius bipustulatus* (icMalBipu1) germline mutational spectrum
C>A
C>G
C>T
T>A
T>G
Counts
150000
100000
50000
0
Trinucleotide

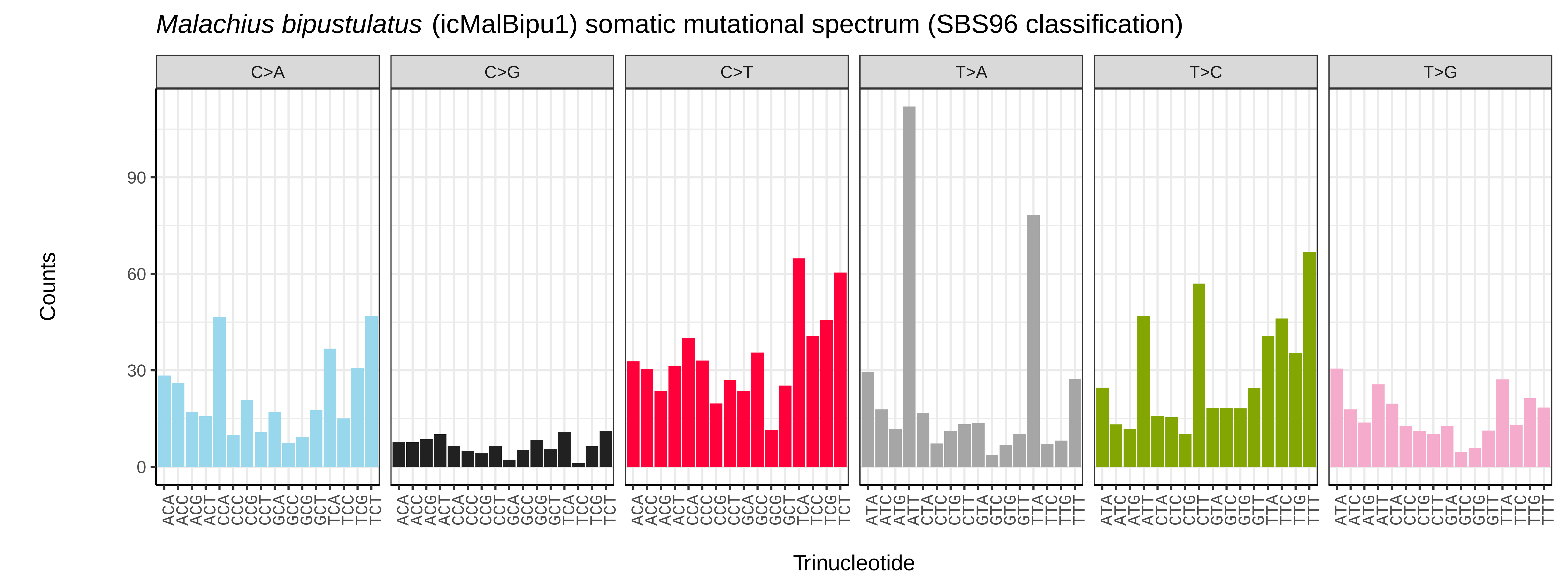
*Malachius bipustulatus* (icMalBipu1) somatic mutational spectrum (SBS96 classification)
C>A
C>G
C>T
T>A
T>C
T>G
Counts
90
60
30
0
Trinucleotide

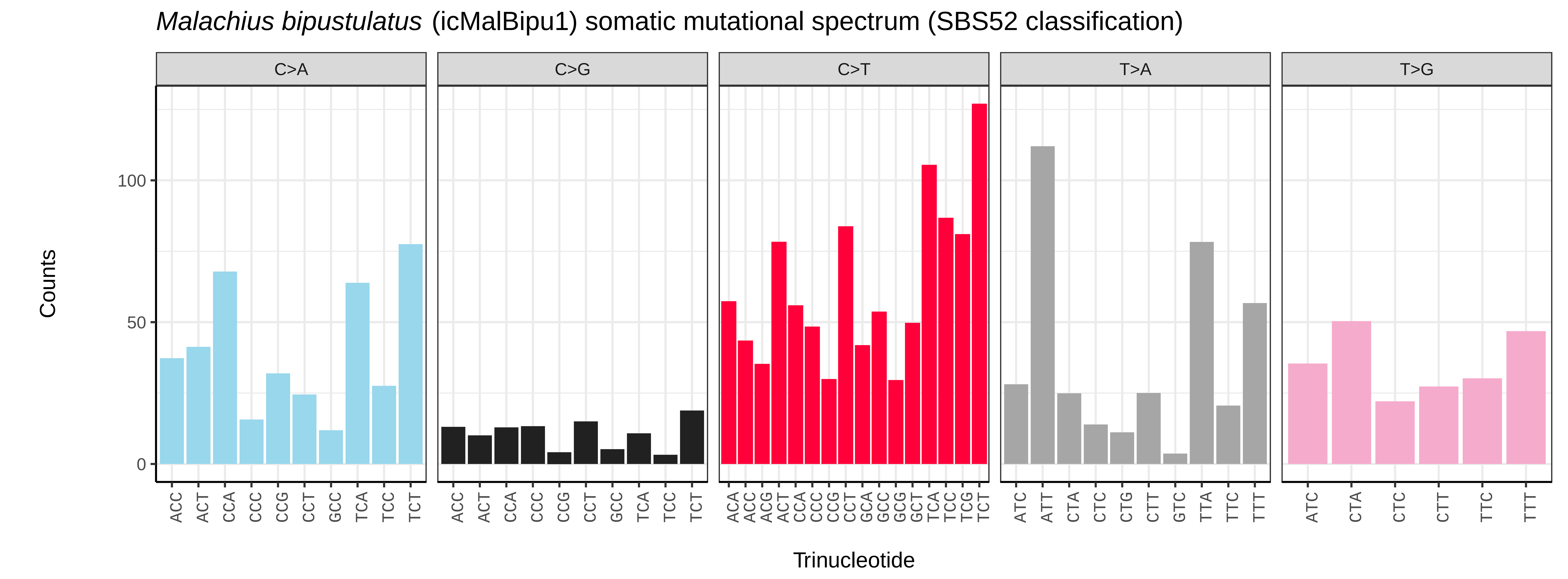
*Malachius bipustulatus* (icMalBipu1) somatic mutational spectrum (SBS52 classification)
C>A
C>G
C>T
T>A
T>G
Counts
100
50
0
Trinucleotide

*Melolontha melolontha* (icMelMelo1) germline mutational spectrum

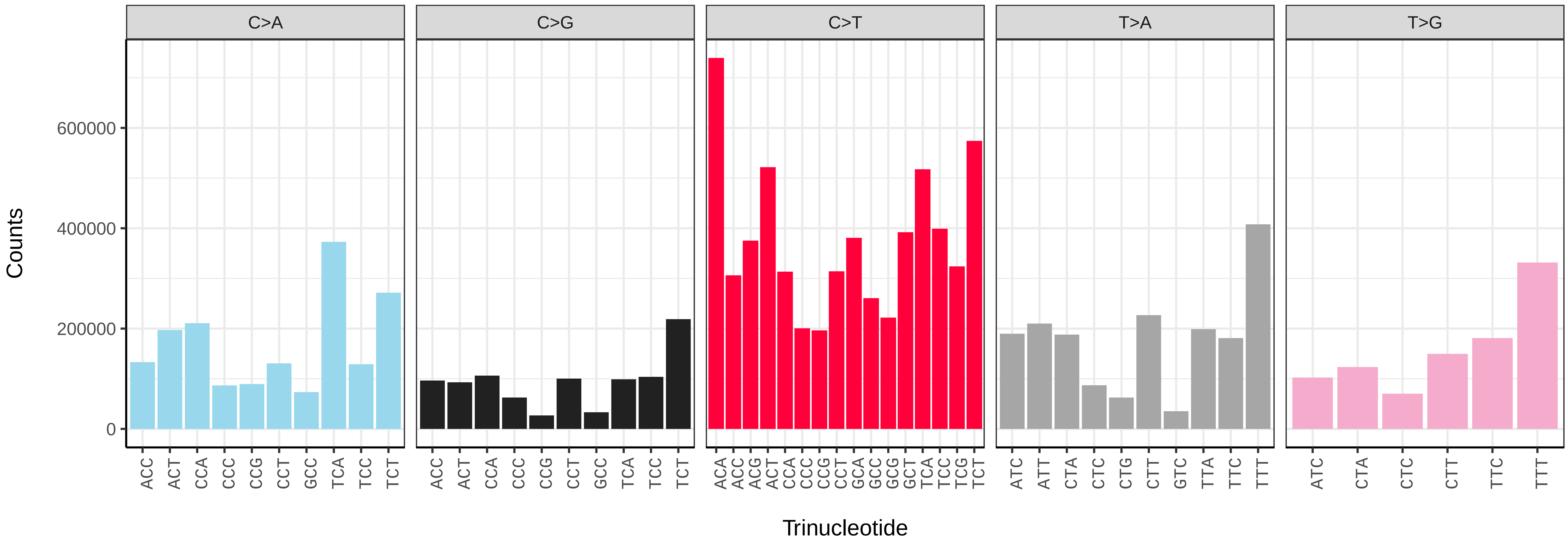


*Melolontha melolontha* (icMelMelo1) somatic mutational spectrum (SBS96 classification)

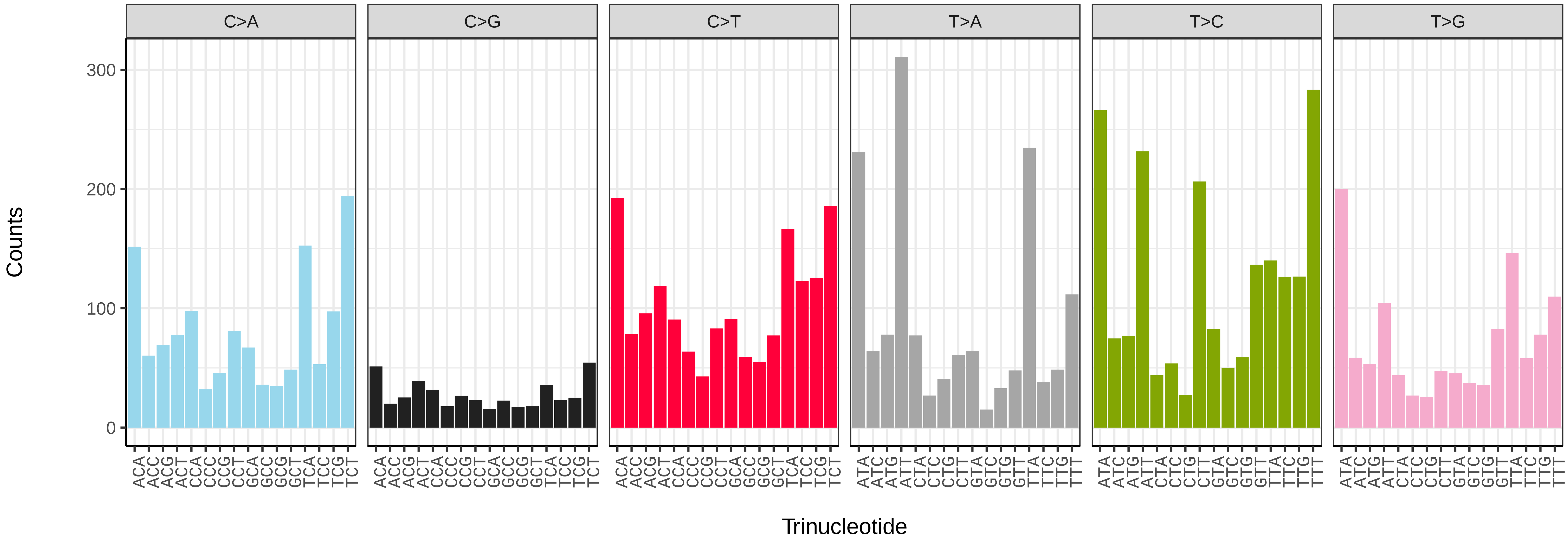


*Melolontha melolontha* (icMelMelo1) somatic mutational spectrum (SBS52 classification)

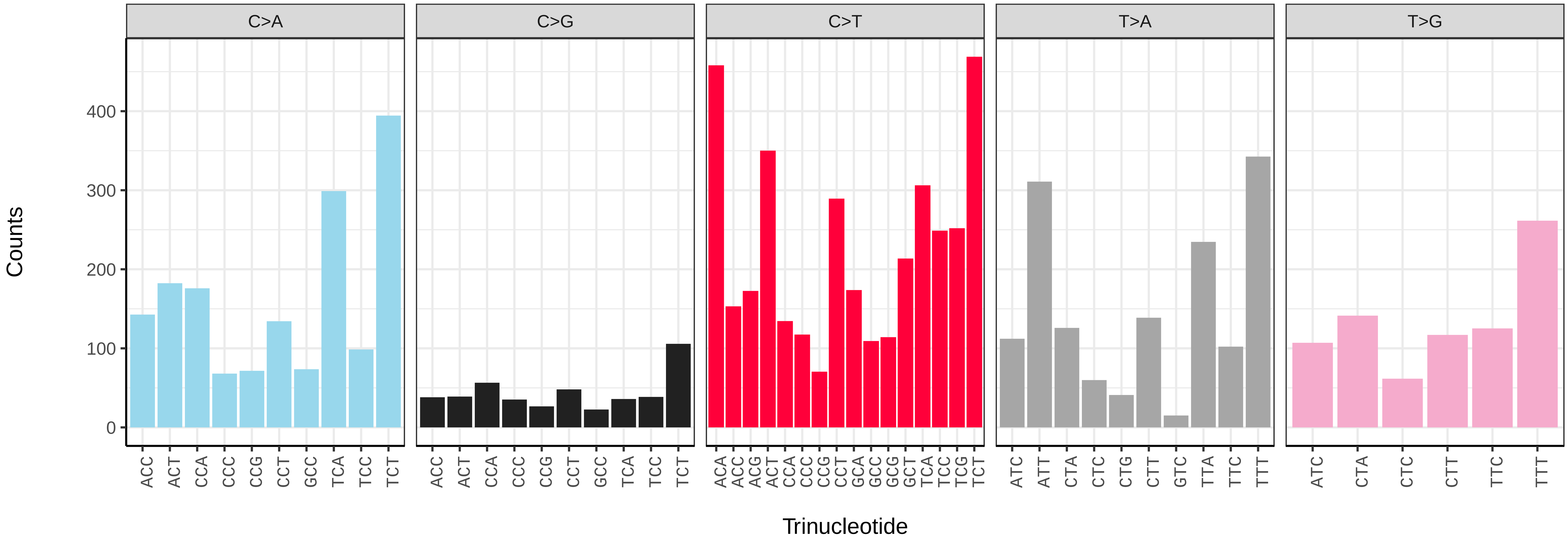

*Nebria brevicollis* (icNebBrev1) germline mutational spectrum

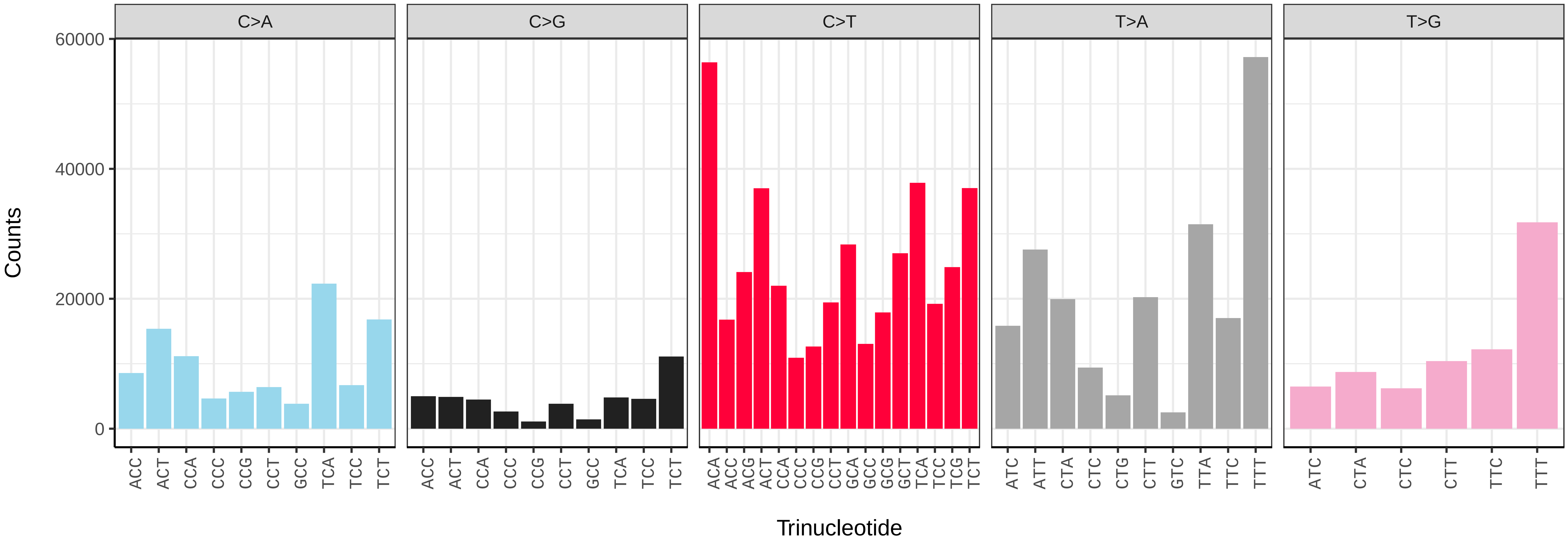


*Nebria brevicollis* (icNebBrev1) somatic mutational spectrum (SBS96 classification)

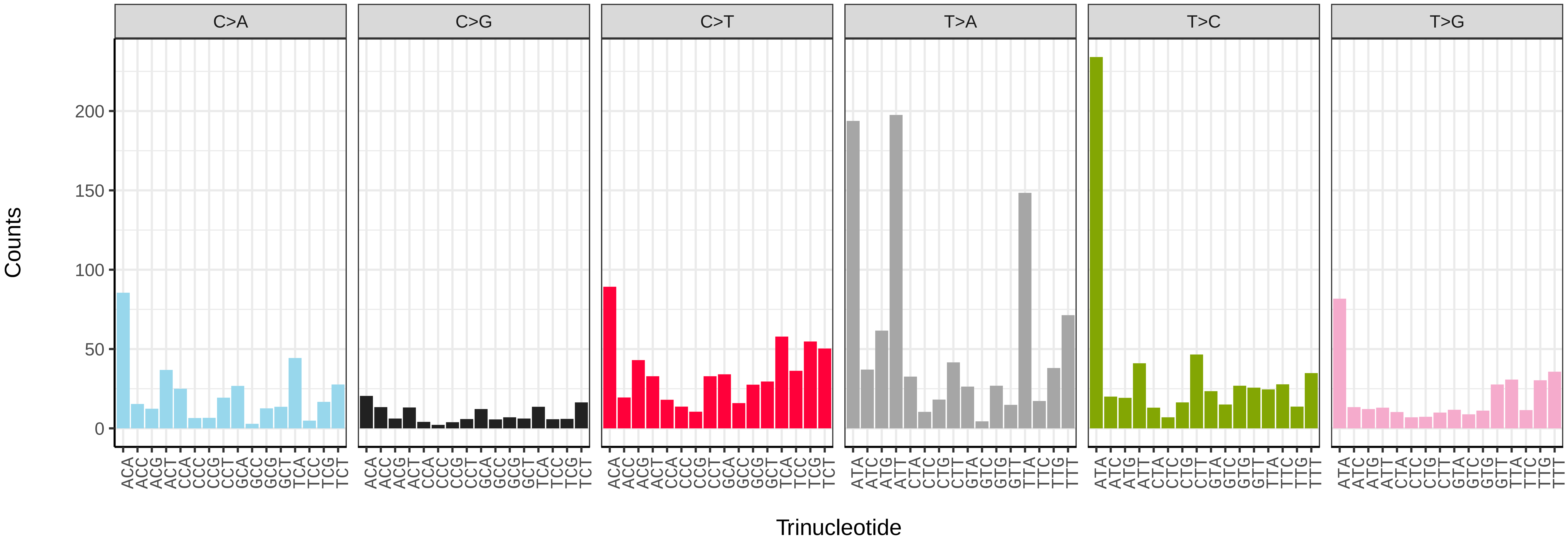


*Nebria brevicollis* (icNebBrev1) somatic mutational spectrum (SBS52 classification)

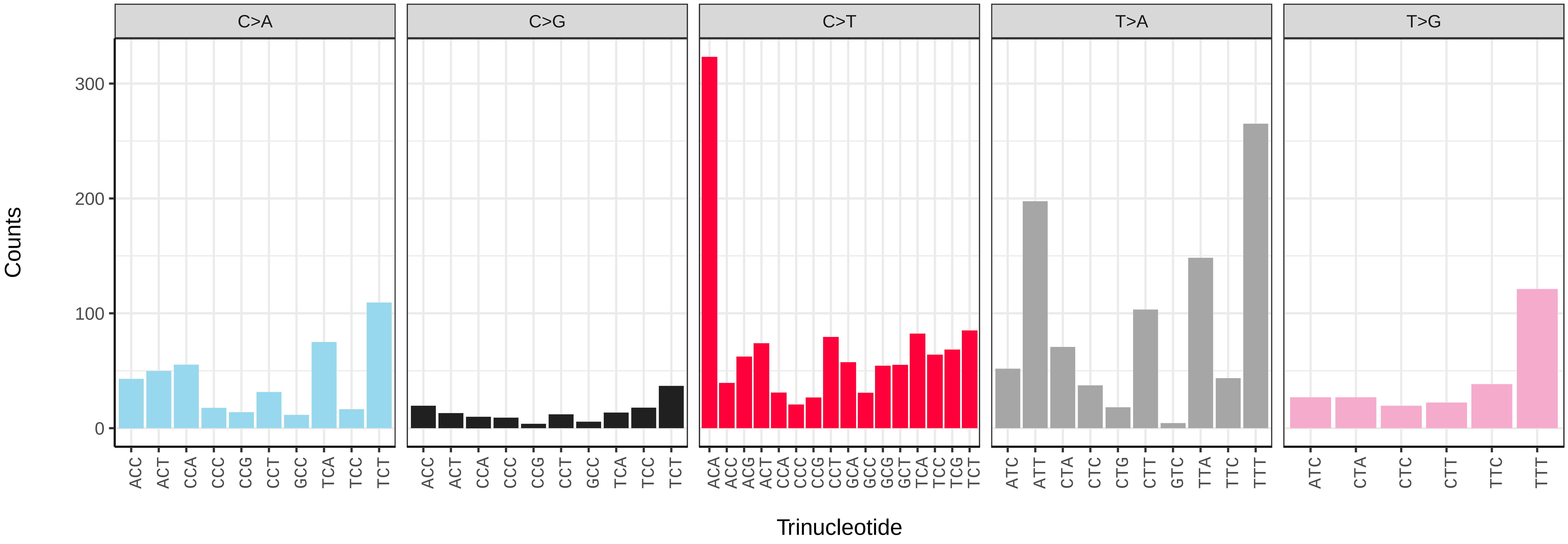

*Nebria salina* (icNebSali1) germline mutational spectrum

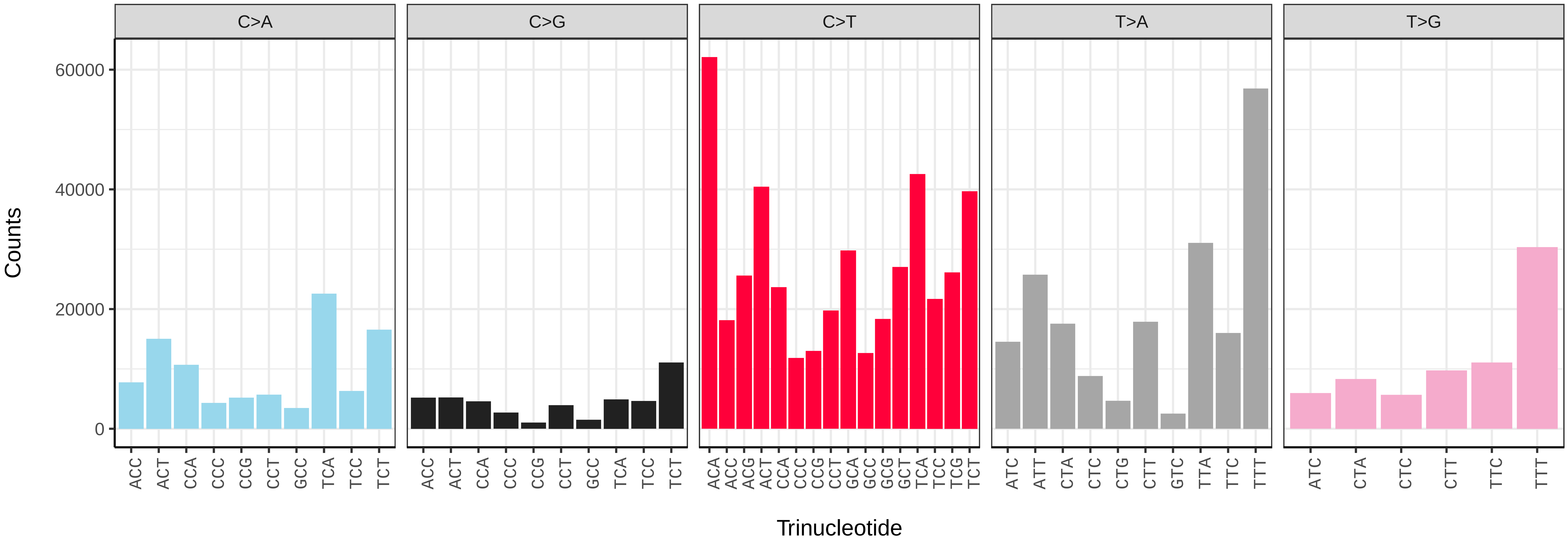


*Nebria salina* (icNebSali1) somatic mutational spectrum (SBS96 classification)

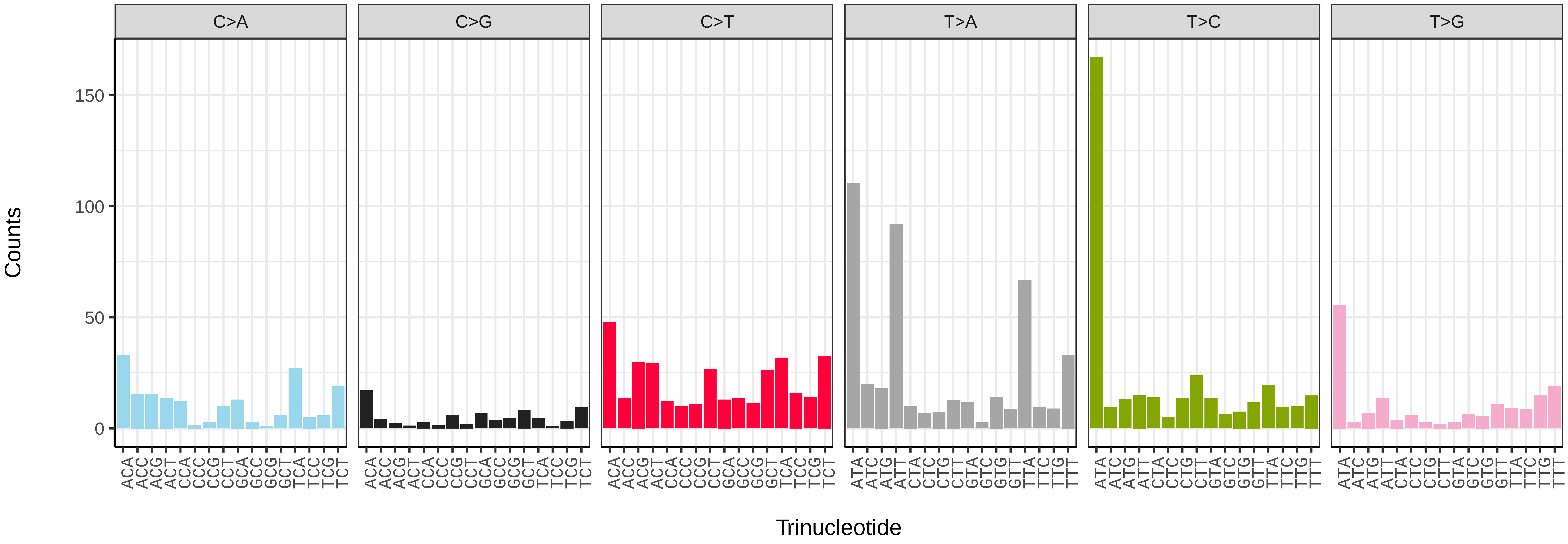


*Nebria salina* (icNebSali1) somatic mutational spectrum (SBS52 classification)

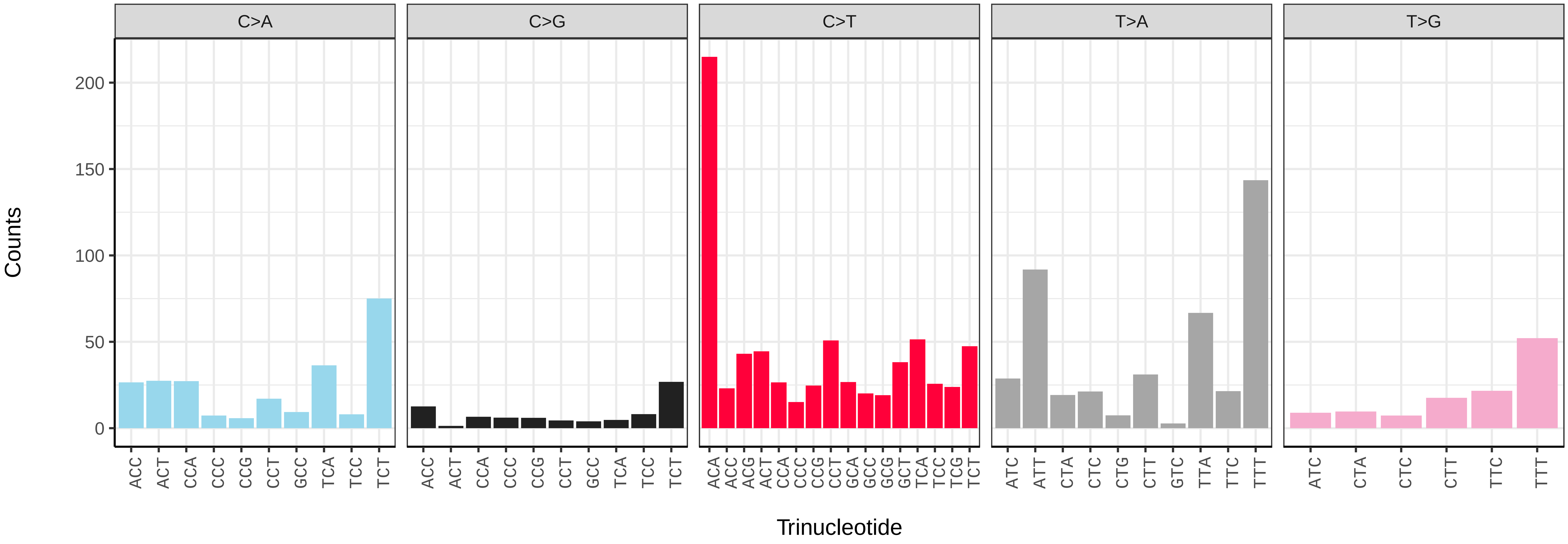

*Ocypus olens* (icOcyOlen1) germline mutational spectrum

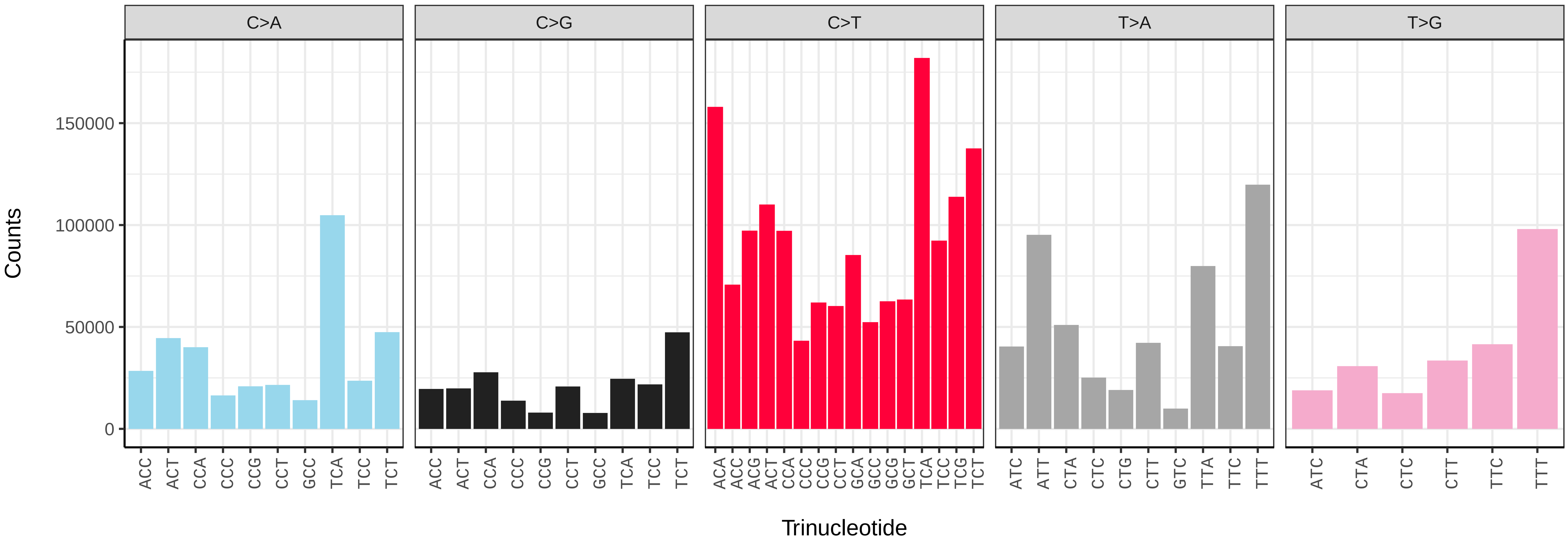


*Ocypus olens* (icOcyOlen1) somatic mutational spectrum (SBS96 classification)

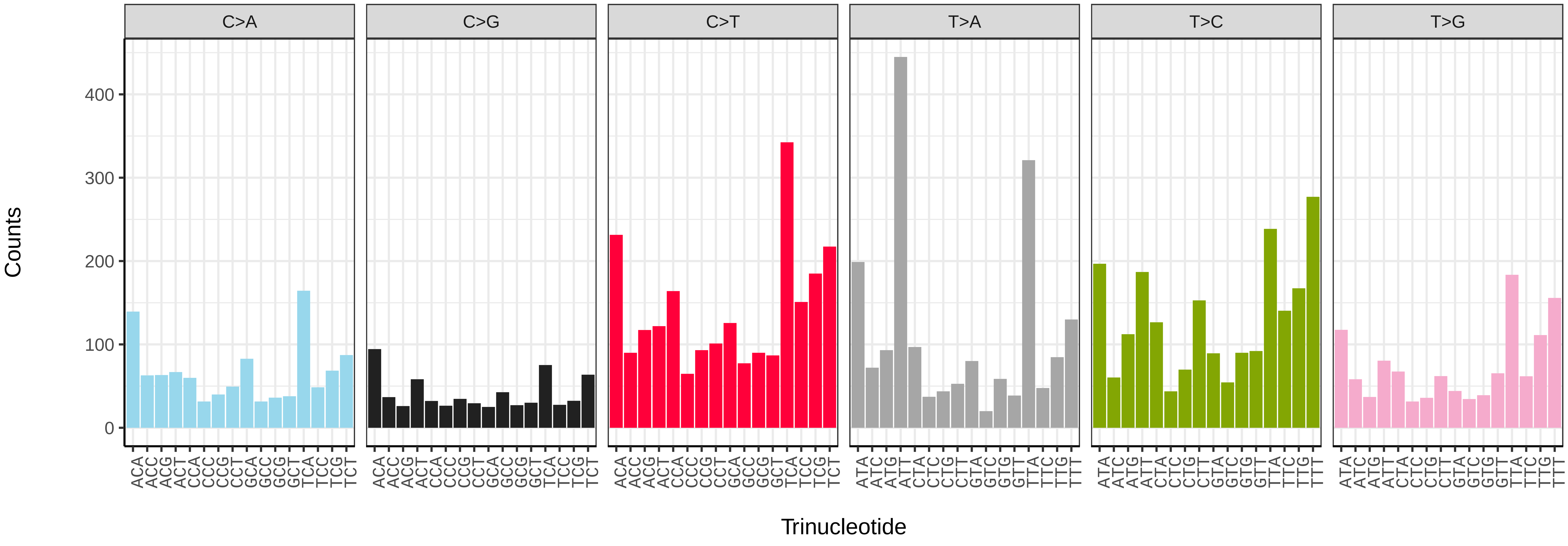


*Ocypus olens* (icOcyOlen1) somatic mutational spectrum (SBS52 classification)

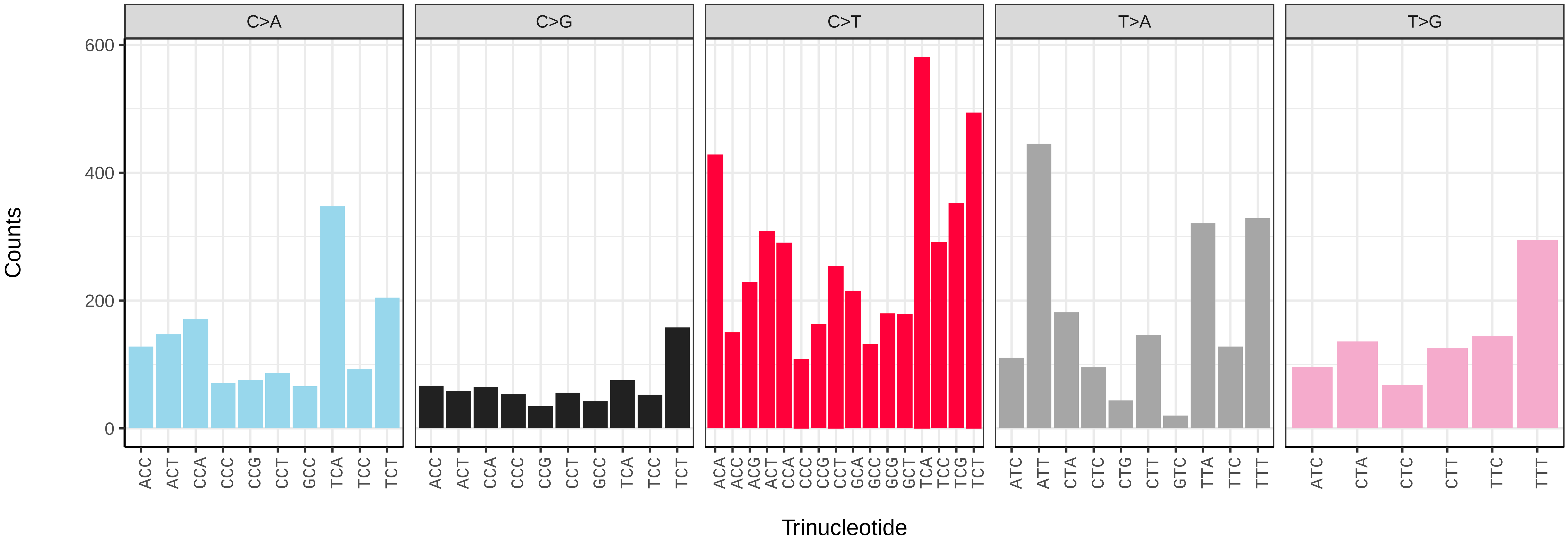

*Ophonus ardosiacus* (icOphArdo1) germline mutational spectrum

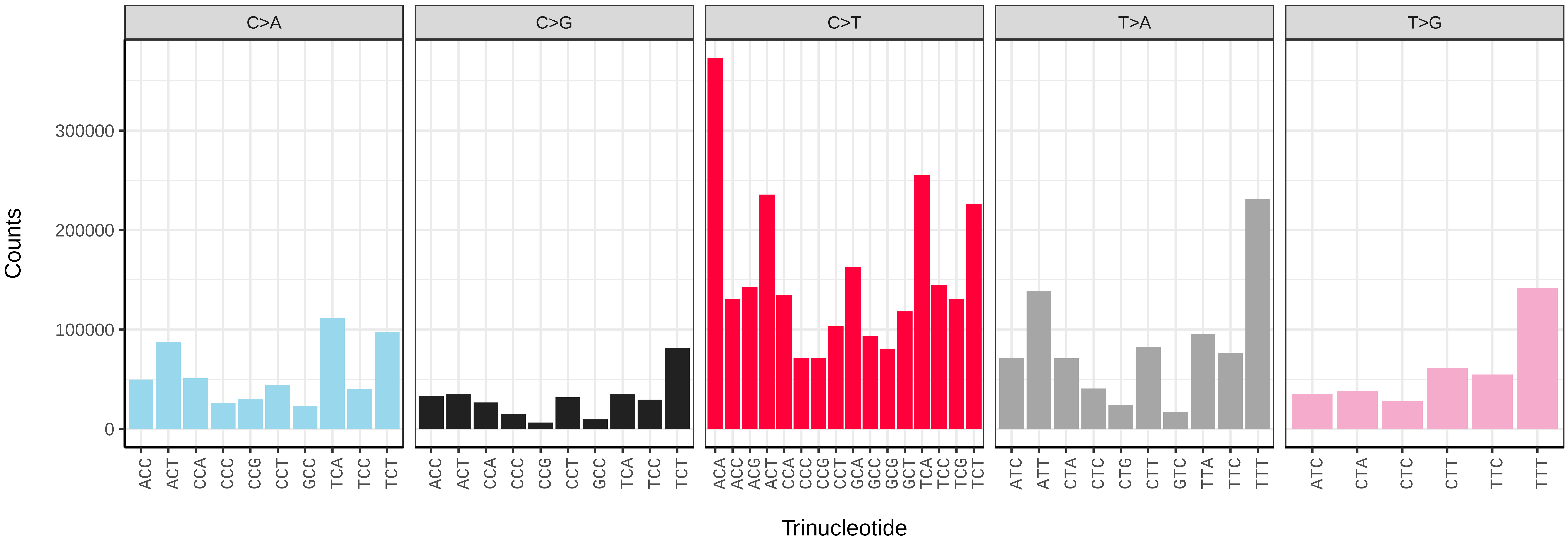


*Ophonus ardosiacus* (icOphArdo1) somatic mutational spectrum (SBS96 classification)

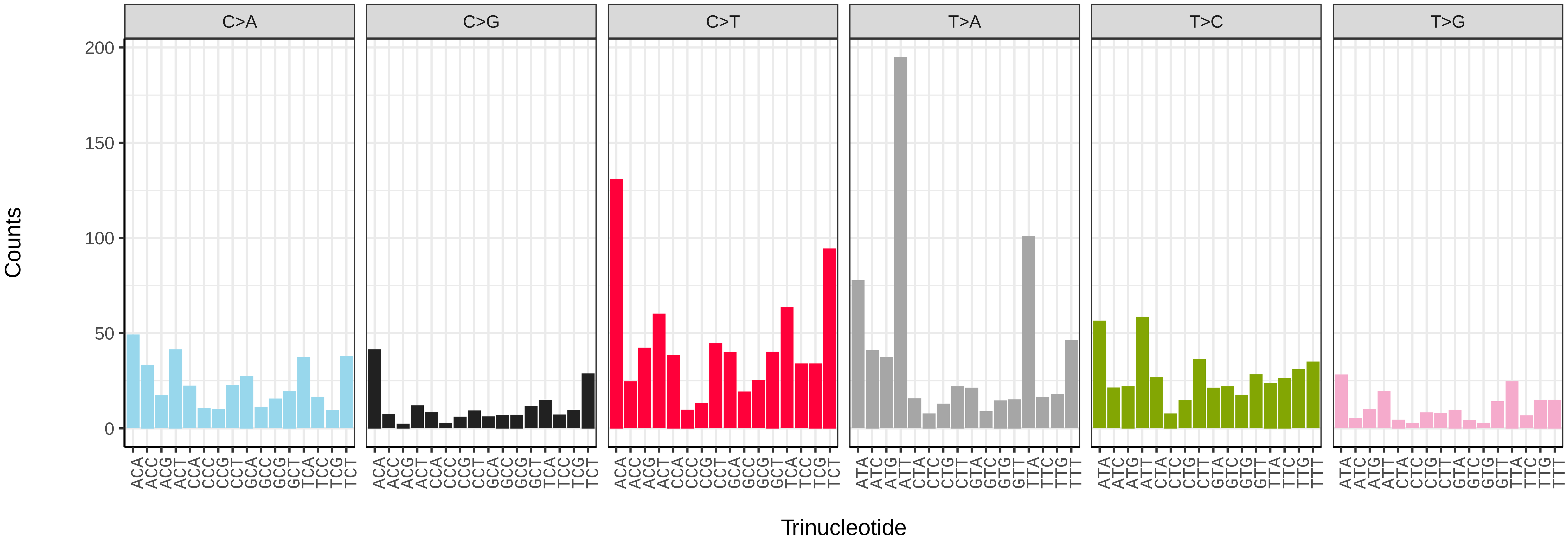


*Ophonus ardosiacus* (icOphArdo1) somatic mutational spectrum (SBS52 classification)

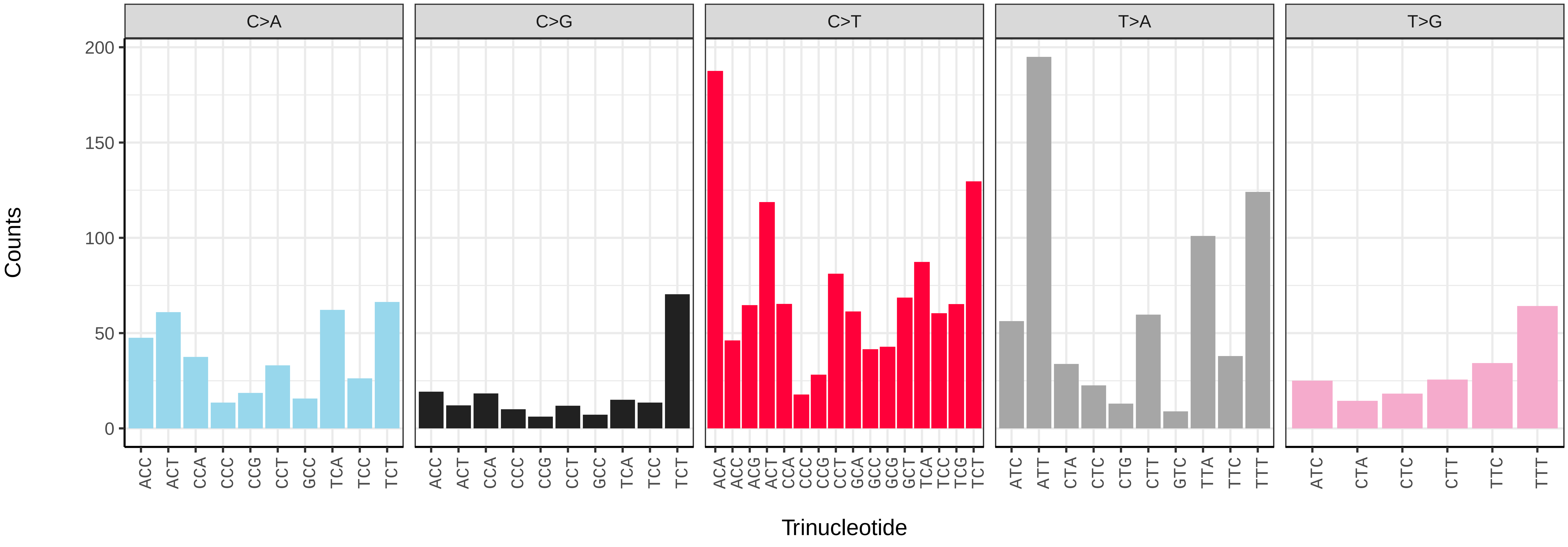

*Philonthus cognatus* (icPhiCogn1) germline mutational spectrum

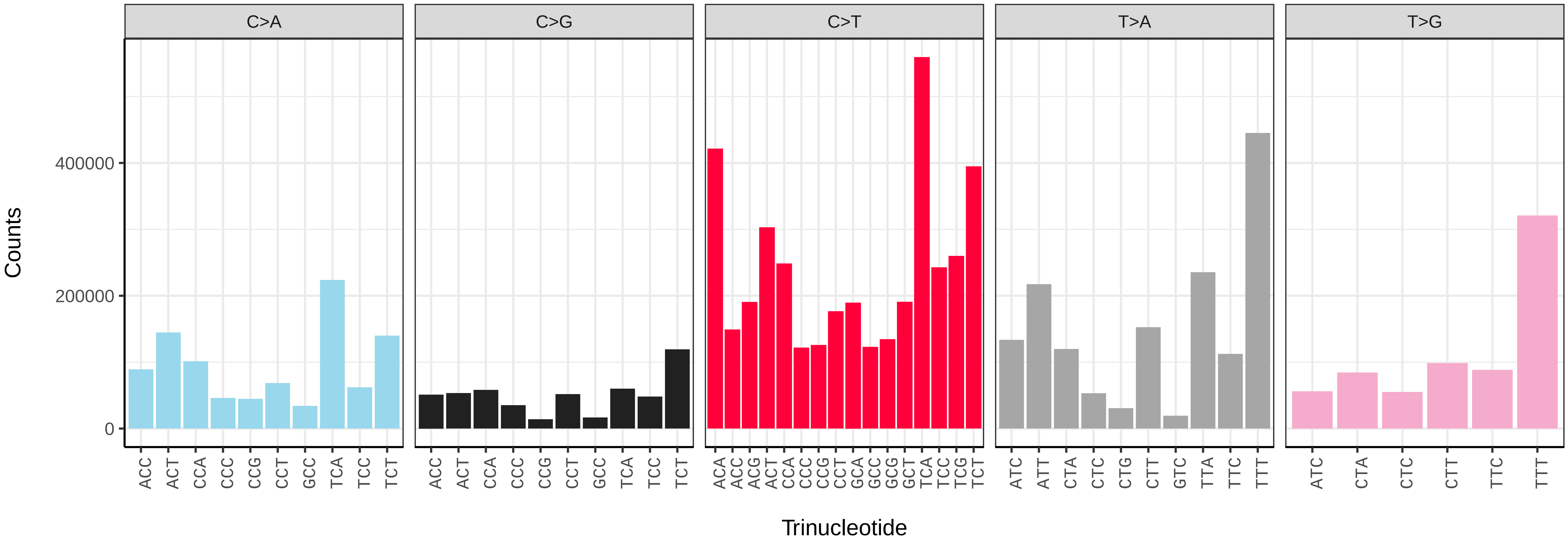


*Philonthus cognatus* (icPhiCogn1) somatic mutational spectrum (SBS96 classification)

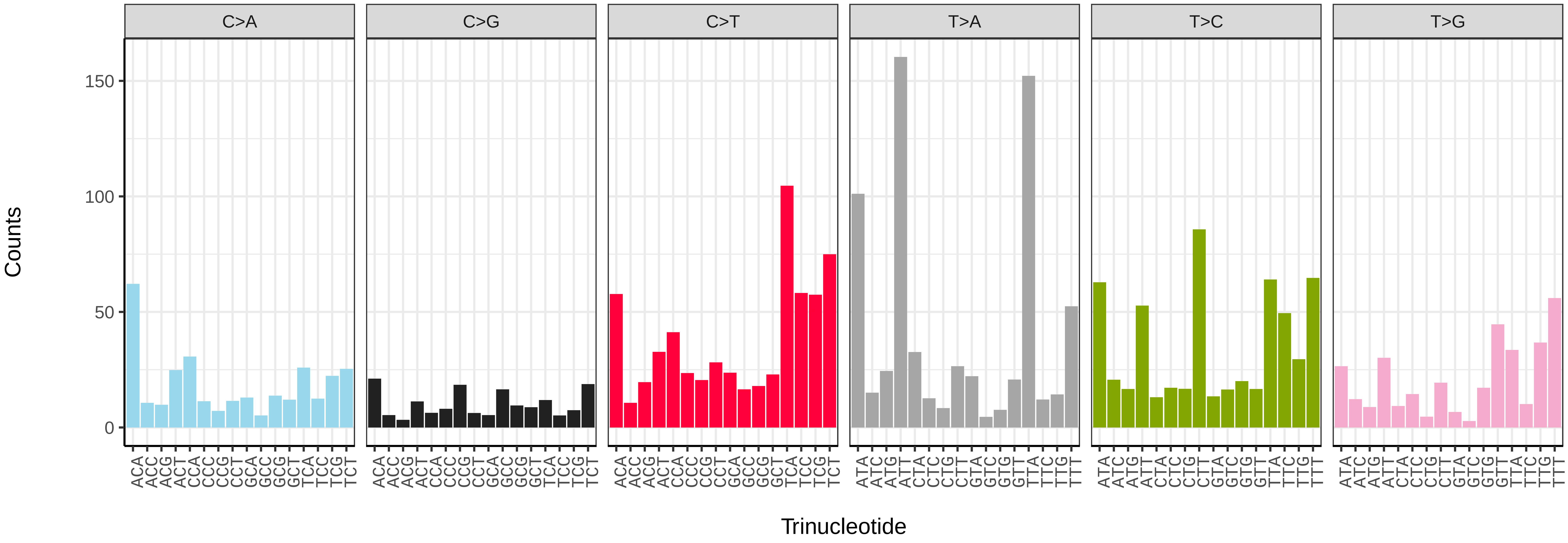


*Philonthus cognatus* (icPhiCogn1) somatic mutational spectrum (SBS52 classification)

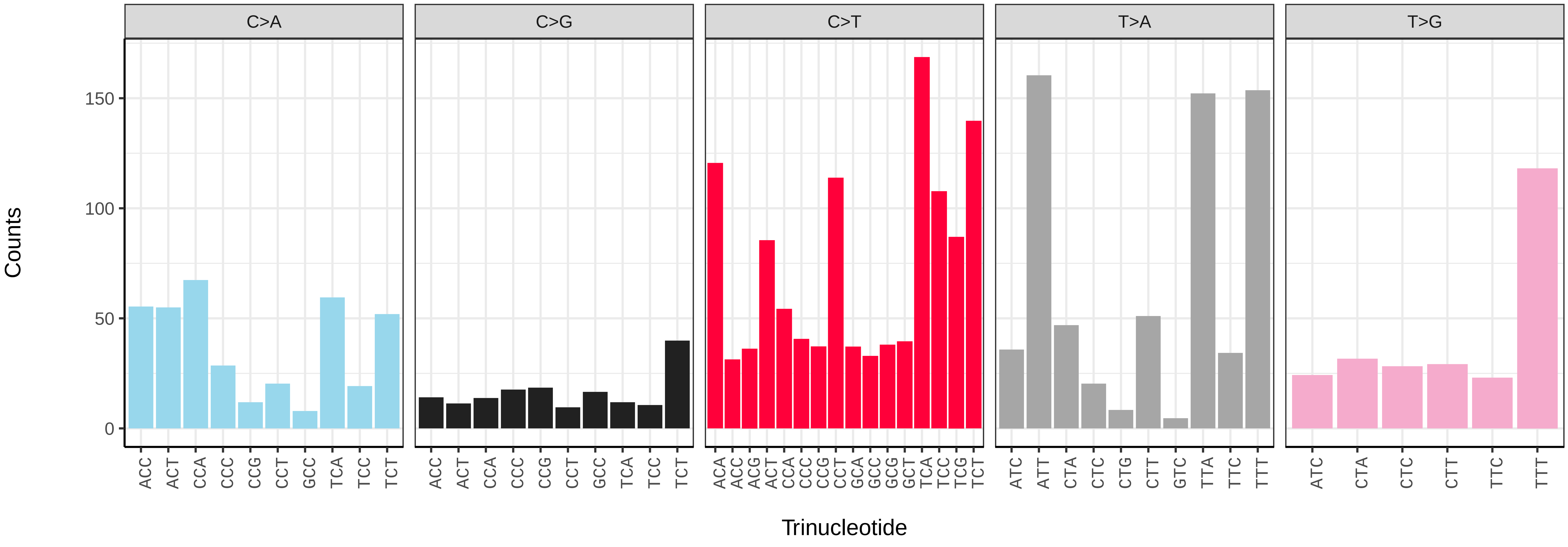

*Podabrus alpinus* (icPodAlpi1) germline mutational spectrum

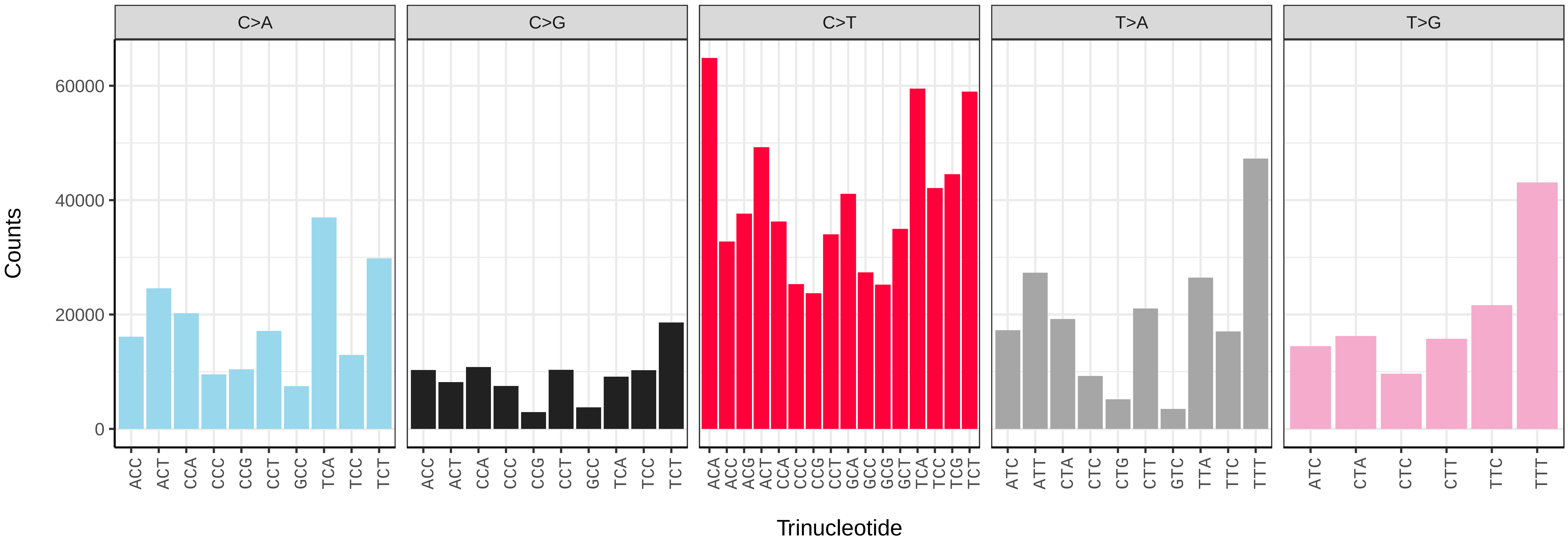


*Podabrus alpinus* (icPodAlpi1) somatic mutational spectrum (SBS96 classification)

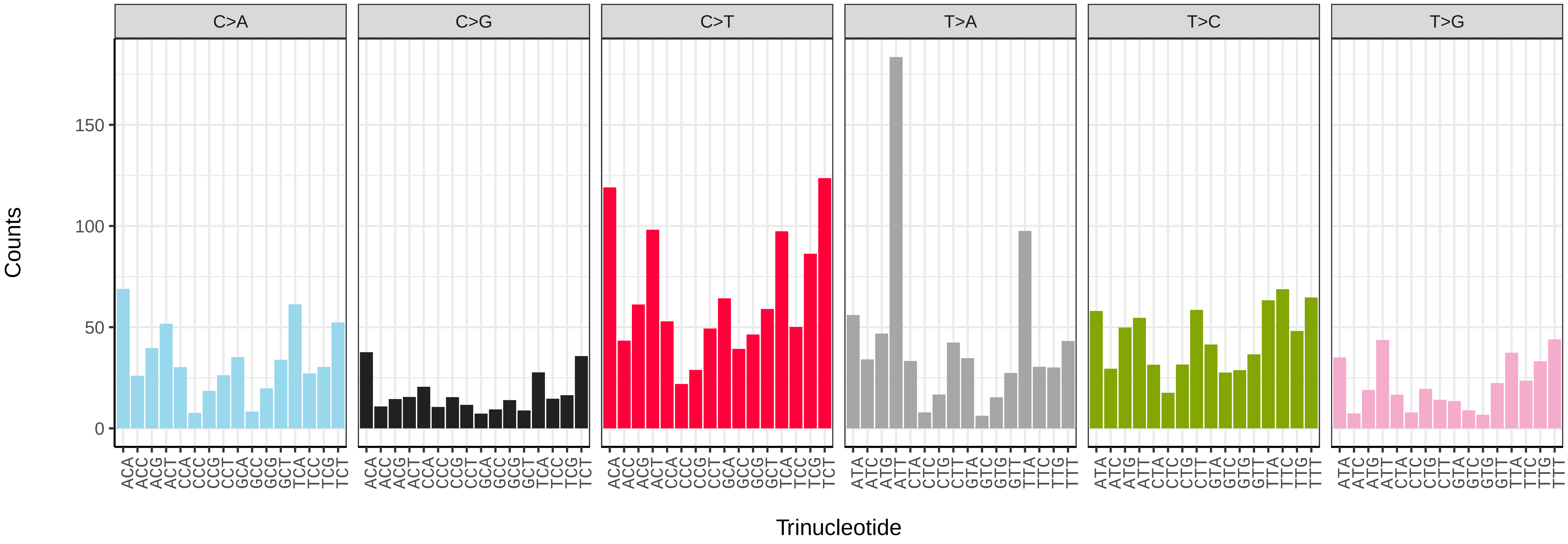


*Podabrus alpinus* (icPodAlpi1) somatic mutational spectrum (SBS52 classification)

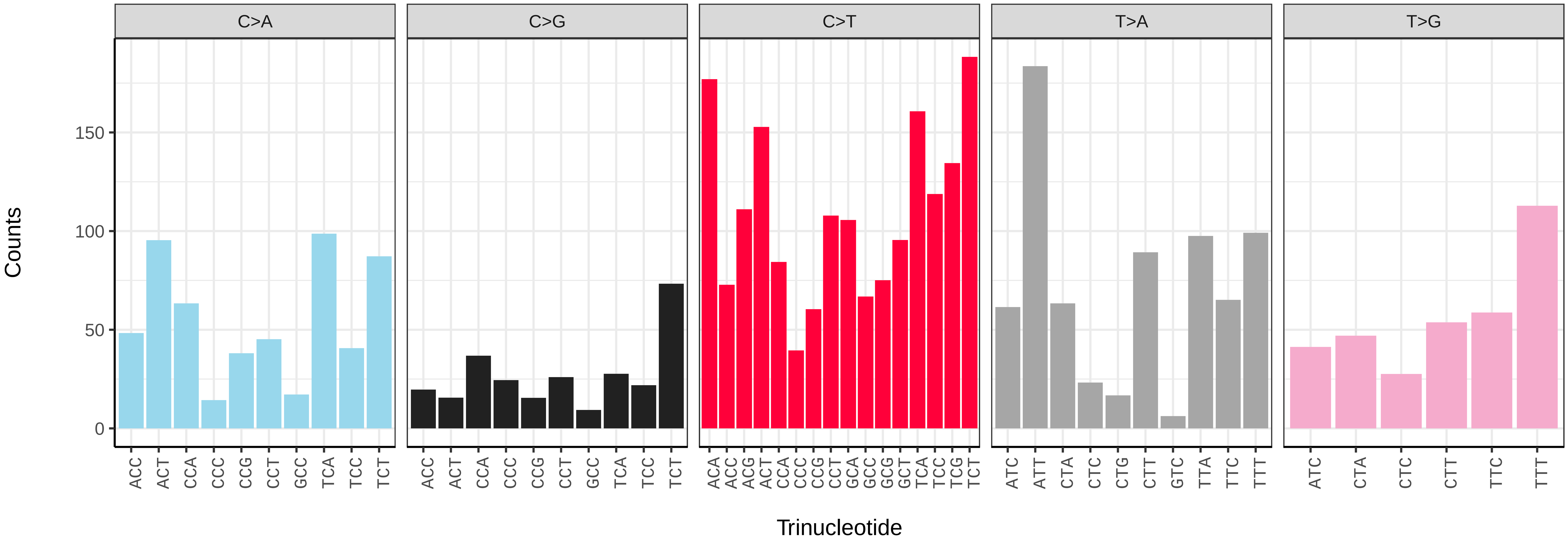

*Polydrusus cervinus* (icPolCerv1) germline mutational spectrum

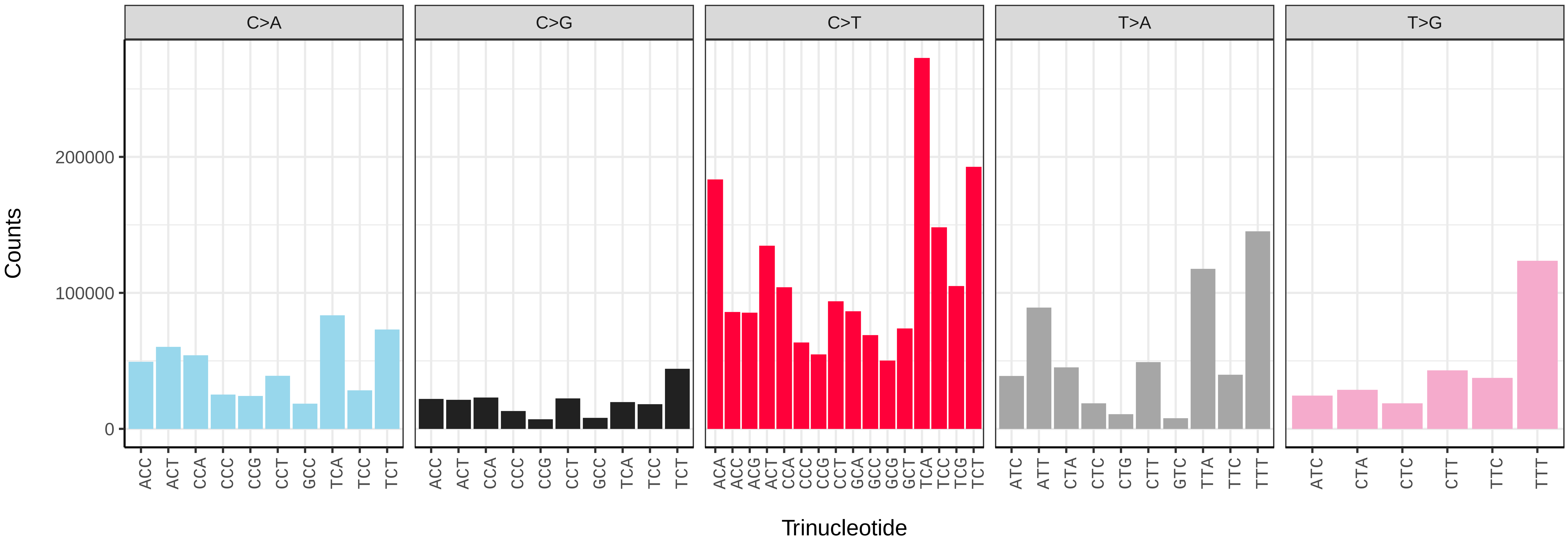


*Polydrusus cervinus* (icPolCerv1) somatic mutational spectrum (SBS96 classification)

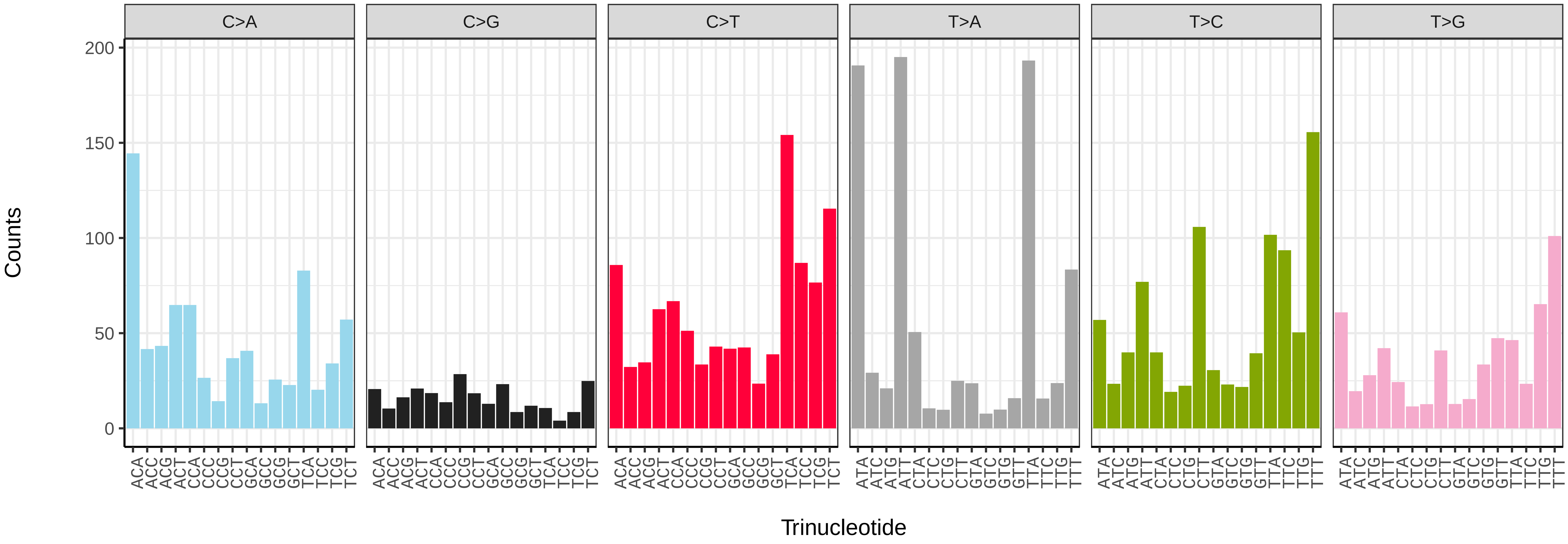


*Polydrusus cervinus* (icPolCerv1) somatic mutational spectrum (SBS52 classification)

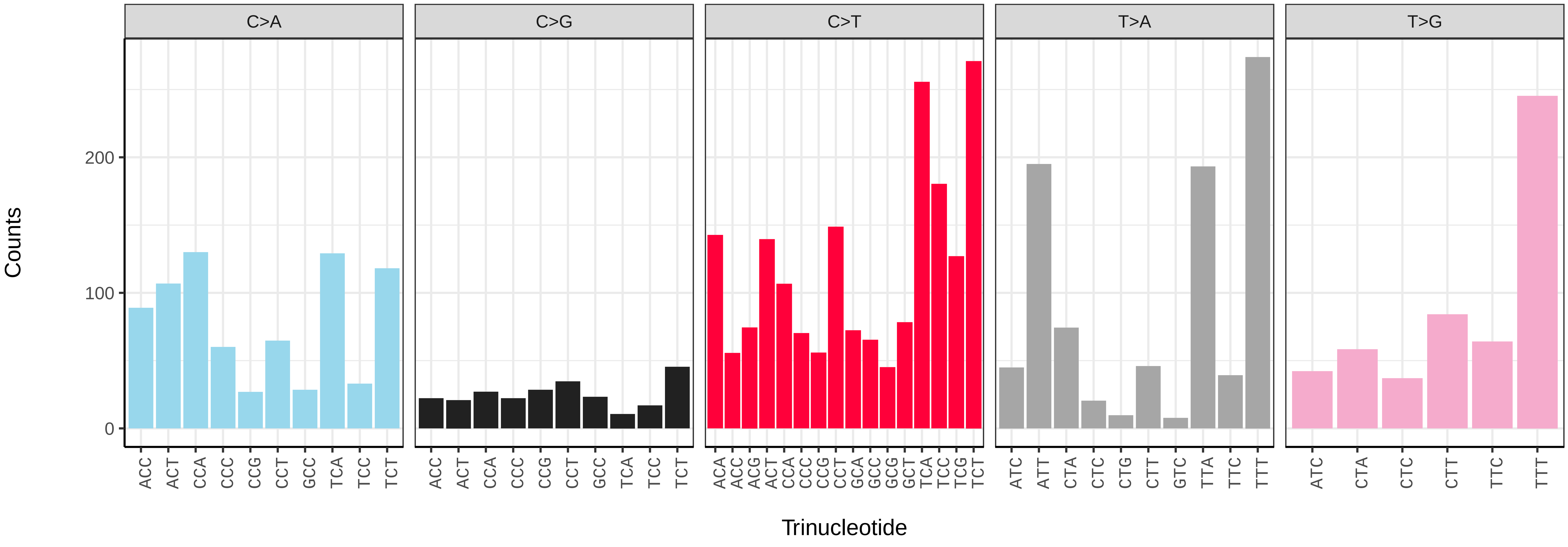

*Pterostichus madidus* (icPteMadi1) germline mutational spectrum

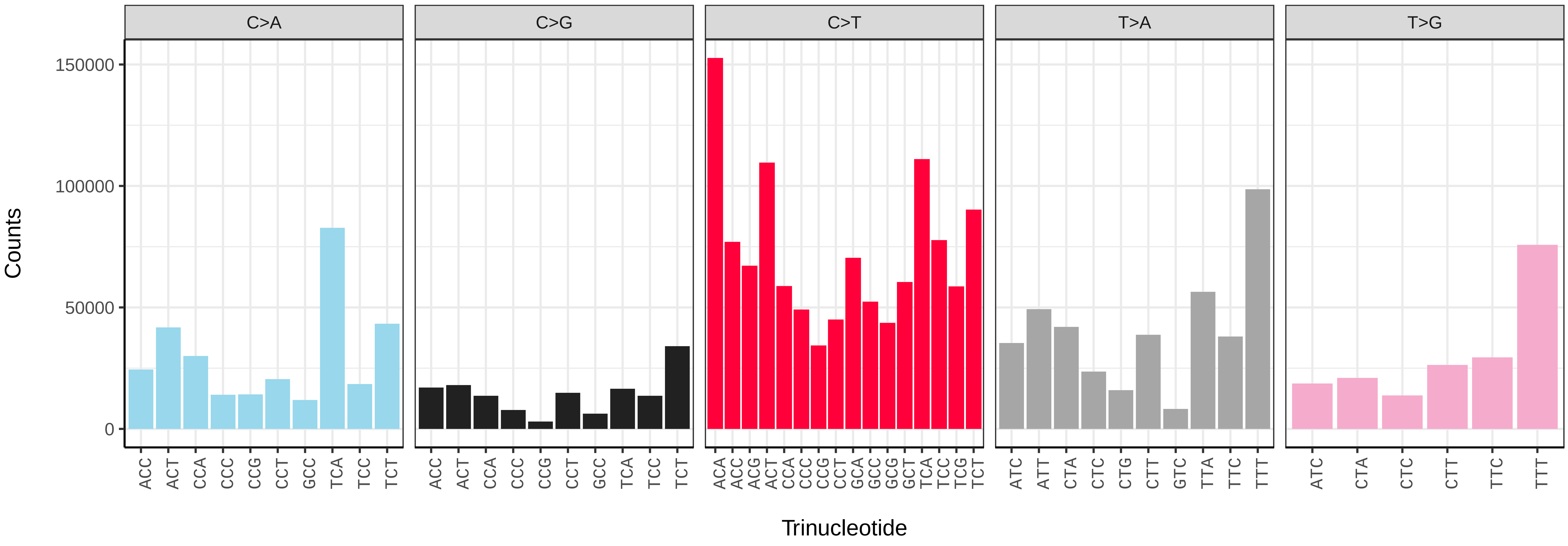


*Pterostichus madidus* (icPteMadi1) somatic mutational spectrum (SBS96 classification)

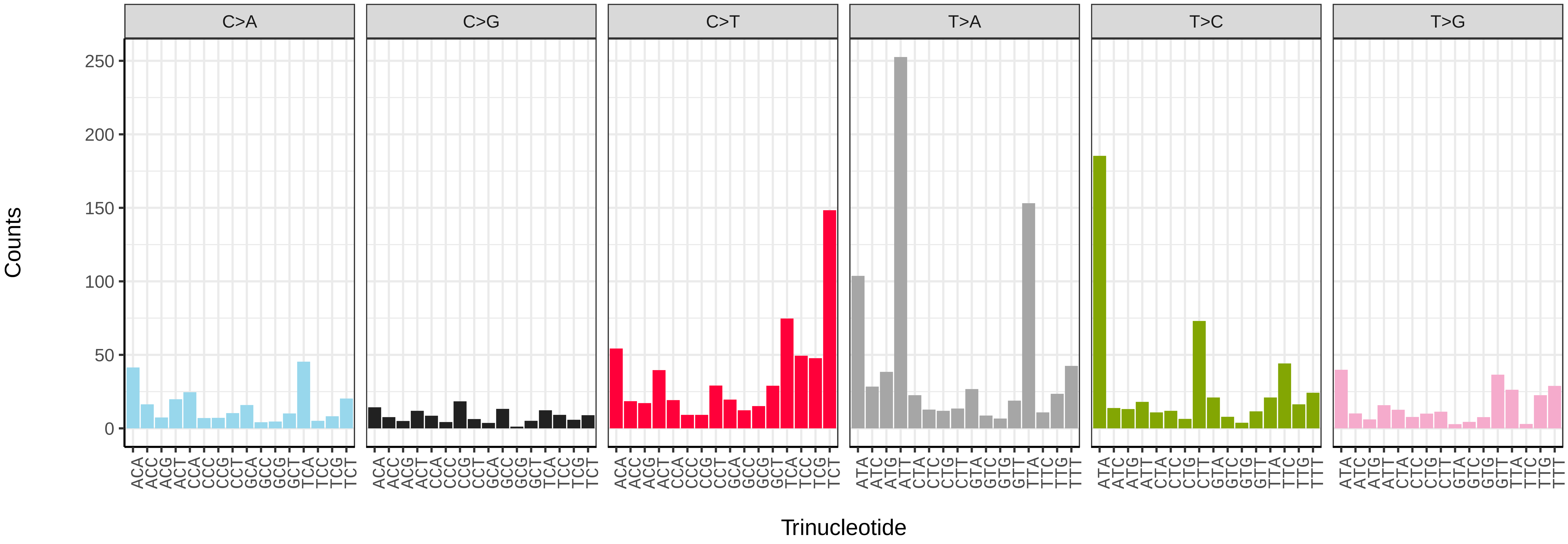


*Pterostichus madidus* (icPteMadi1) somatic mutational spectrum (SBS52 classification)

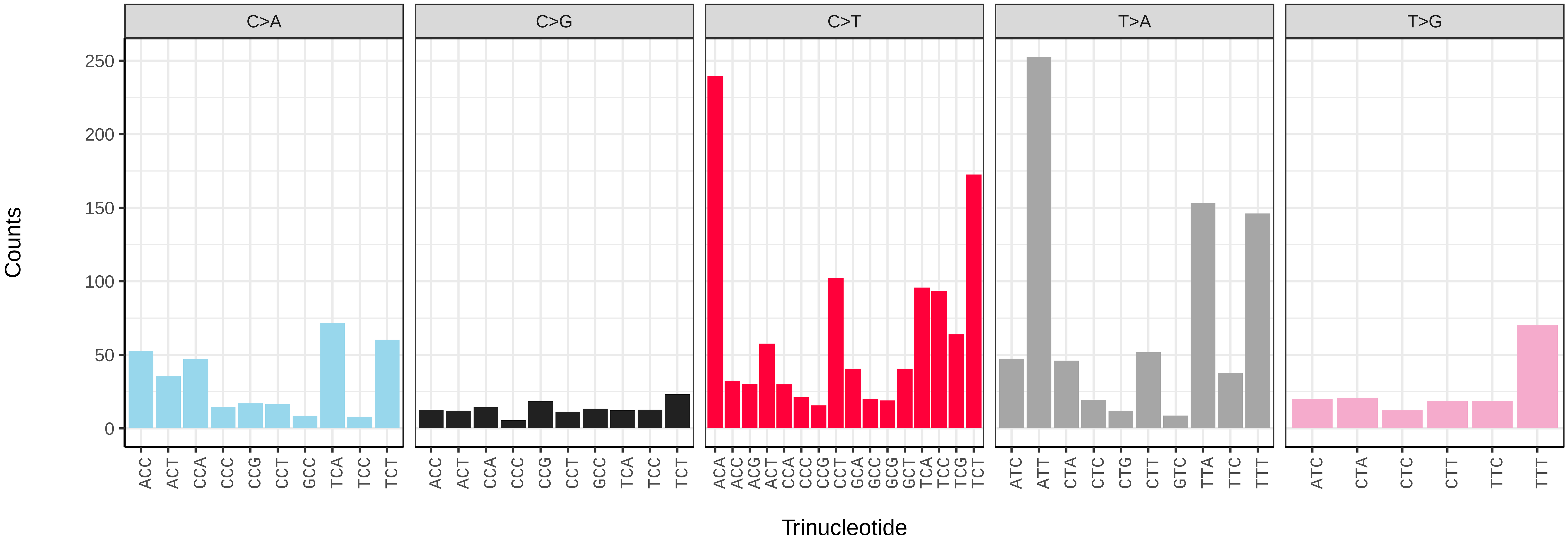

*Pterostichus madidus* (icPteMadi1.icPteMadi3) germline mutational spectrum

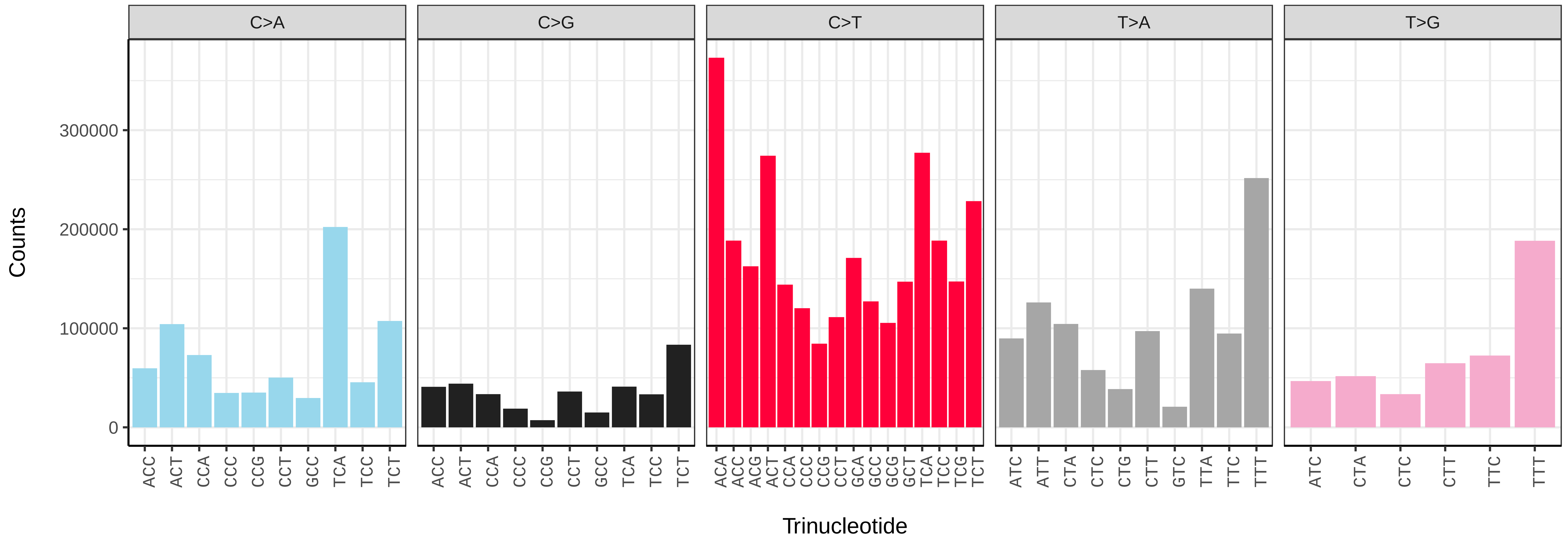


*Pterostichus madidus* (icPteMadi1.icPteMadi3) somatic mutational spectrum (SBS96 classification)

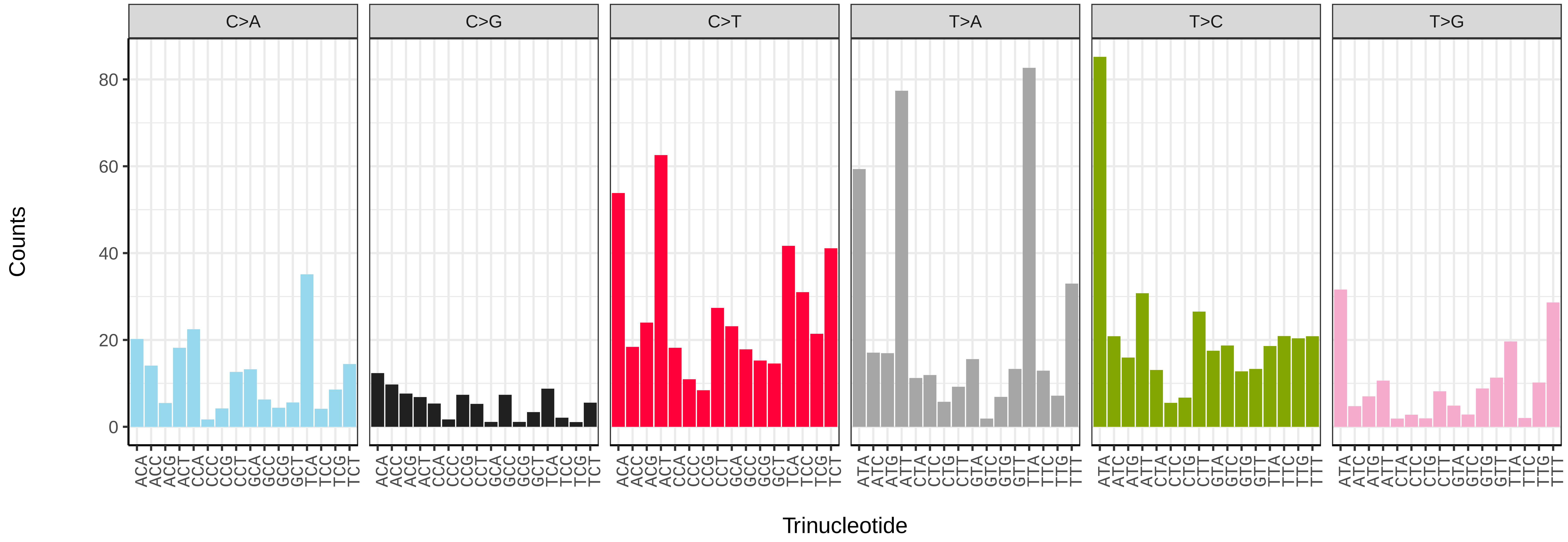


*Pterostichus madidus* (icPteMadi1.icPteMadi3) somatic mutational spectrum (SBS52 classification)

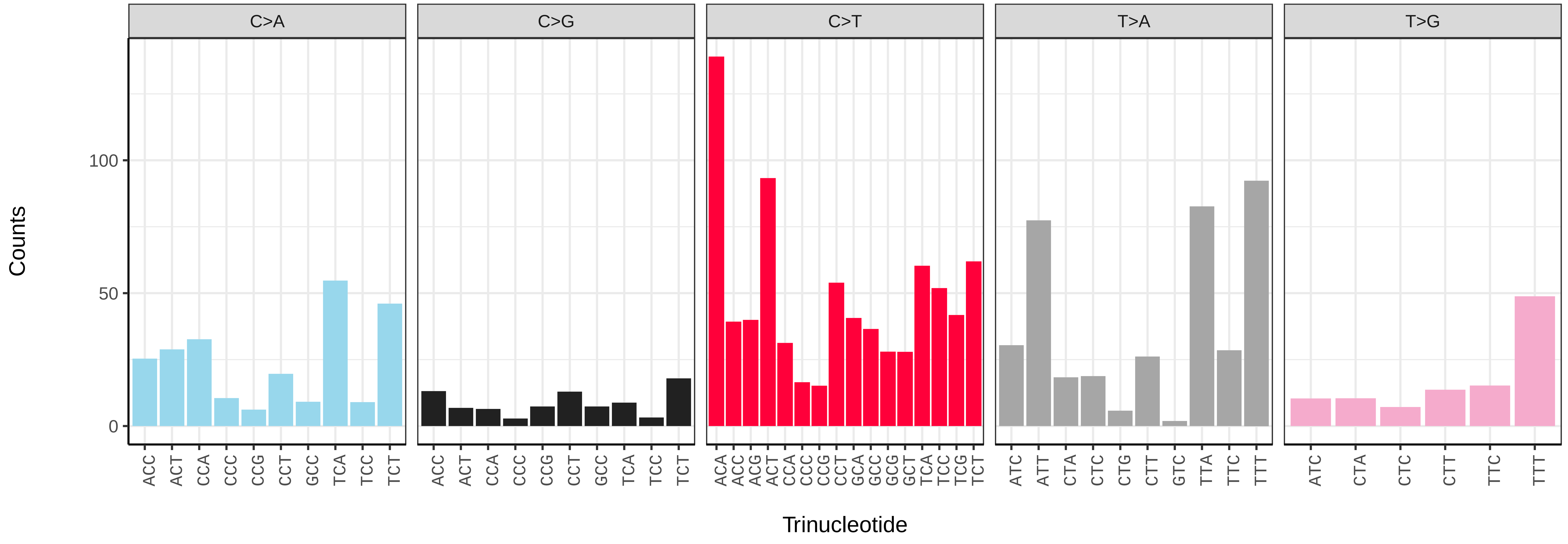

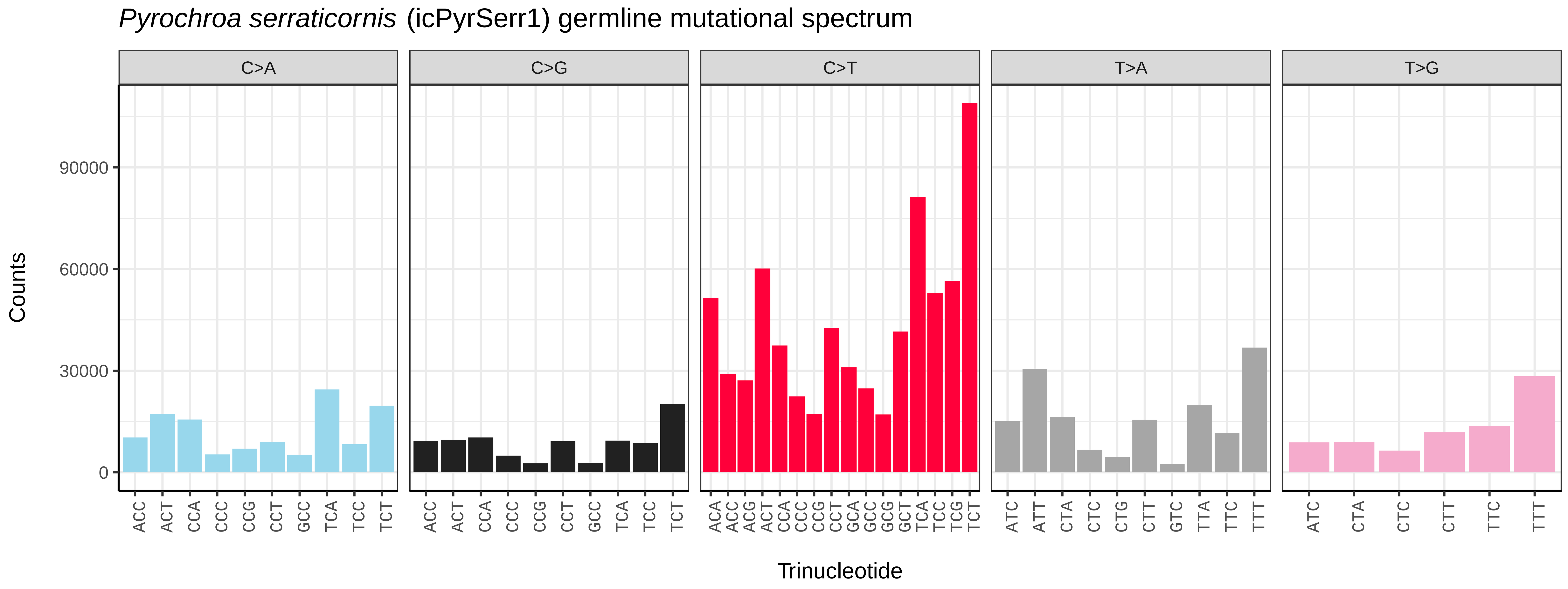

*Pyrochroa serraticornis* (icPyrSerr1) germline mutational spectrum
C>A
C>G
C>T
T>A
T>G
Counts
90000
60000
30000
0
ACC ACT CCA CCC CCG CCT GCC TCA TCC TCT
ACA ACC ACG ACT CCA CCC CCG CCT GCA GCC GCG GCT TCA TCC TCG TCT
ATC ATT CTA CTC CTG CTT GTC TTA TTC TTT
ATC CTA CTC CTT TTC TTT
Trinucleotide


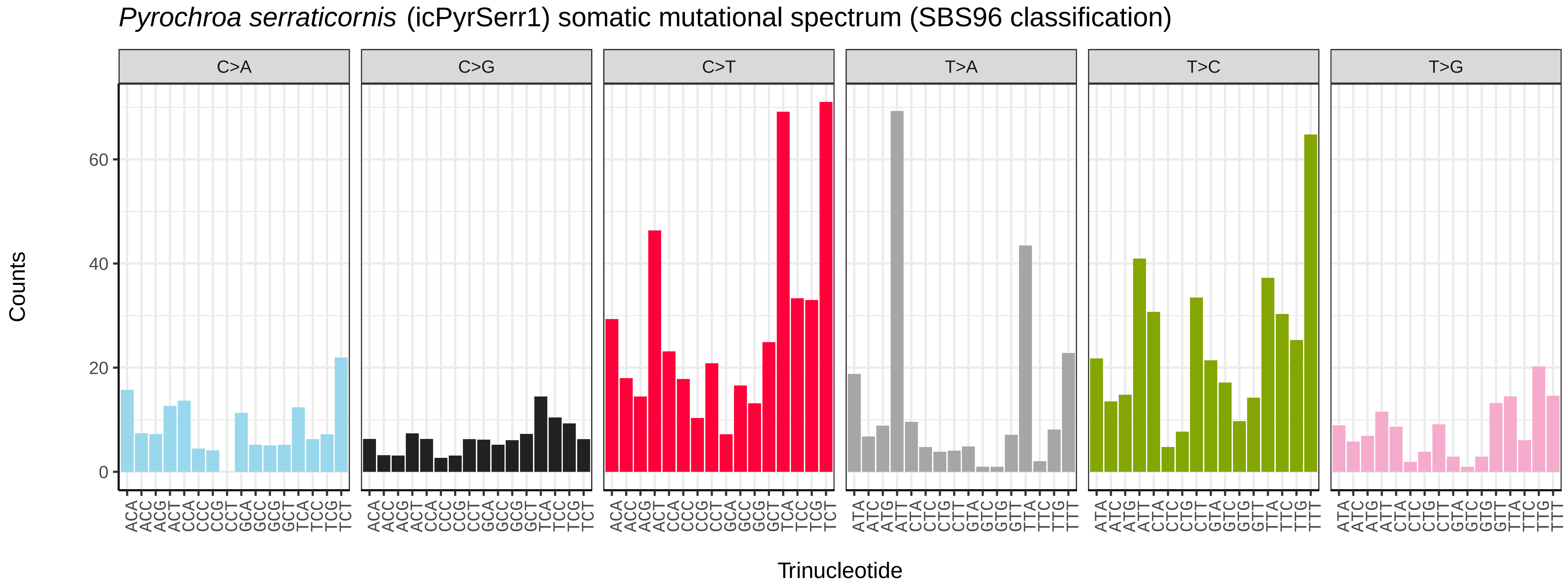

*Pyrochroa serraticornis* (icPyrSerr1) somatic mutational spectrum (SBS96 classification)
C>A
C>G
C>T
T>A
T>C
T>G
Counts
60
40
20
0
ACA ACC ACG ACT CCA CCC CCG CCT GCA GCC GCG GCT TCA TCC TCG TCT
ATA ATC ATG ATT CTA CTC CTG CTT GTA GTC GTG GTT TTA TTC TTG TTT
Trinucleotide


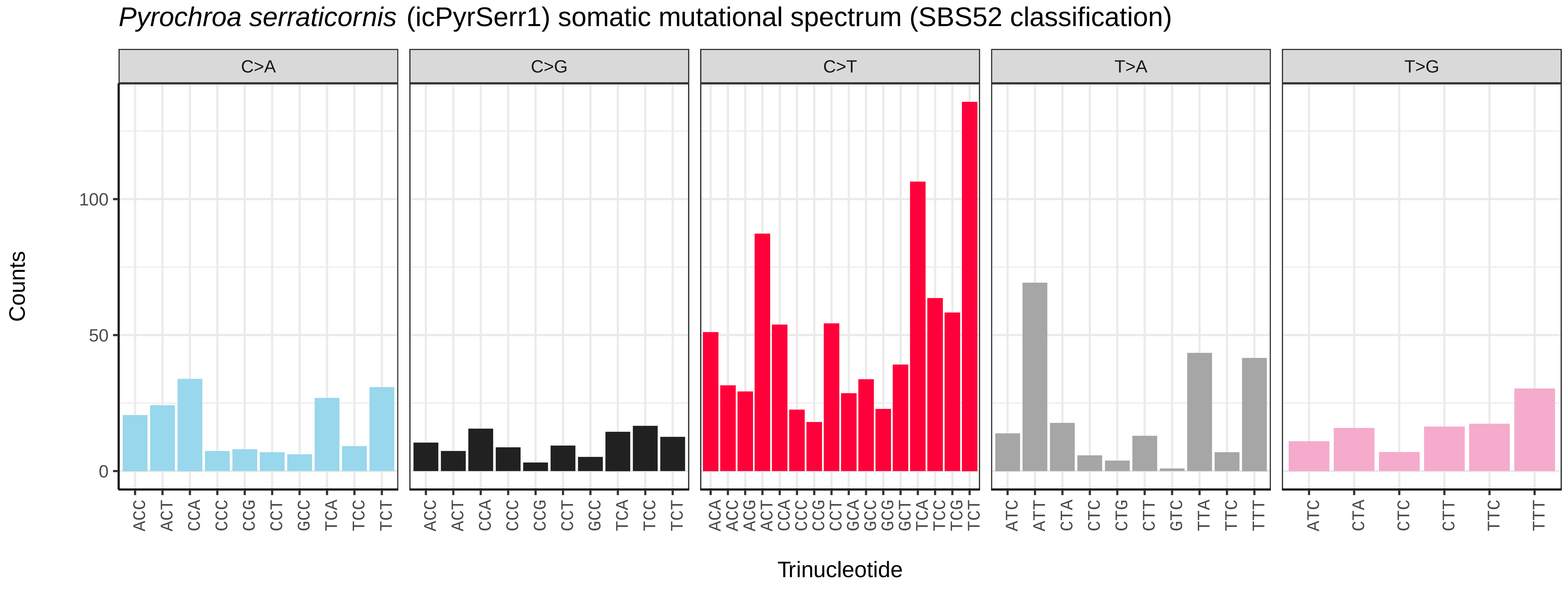

*Pyrochroa serraticornis* (icPyrSerr1) somatic mutational spectrum (SBS52 classification)
C>A
C>G
C>T
T>A
T>G
Counts
100
50
0
ACC ACT CCA CCC CCG CCT GCC TCA TCC TCT
ACA ACC ACG ACT CCA CCC CCG CCT GCA GCC GCG GCT TCA TCC TCG TCT
ATC ATT CTA CTC CTG CTT GTC TTA TTC TTT
ATC CTA CTC CTT TTC TTT
Trinucleotide

*Rhagonycha fulva* (icRhaFulv1) germline mutational spectrum

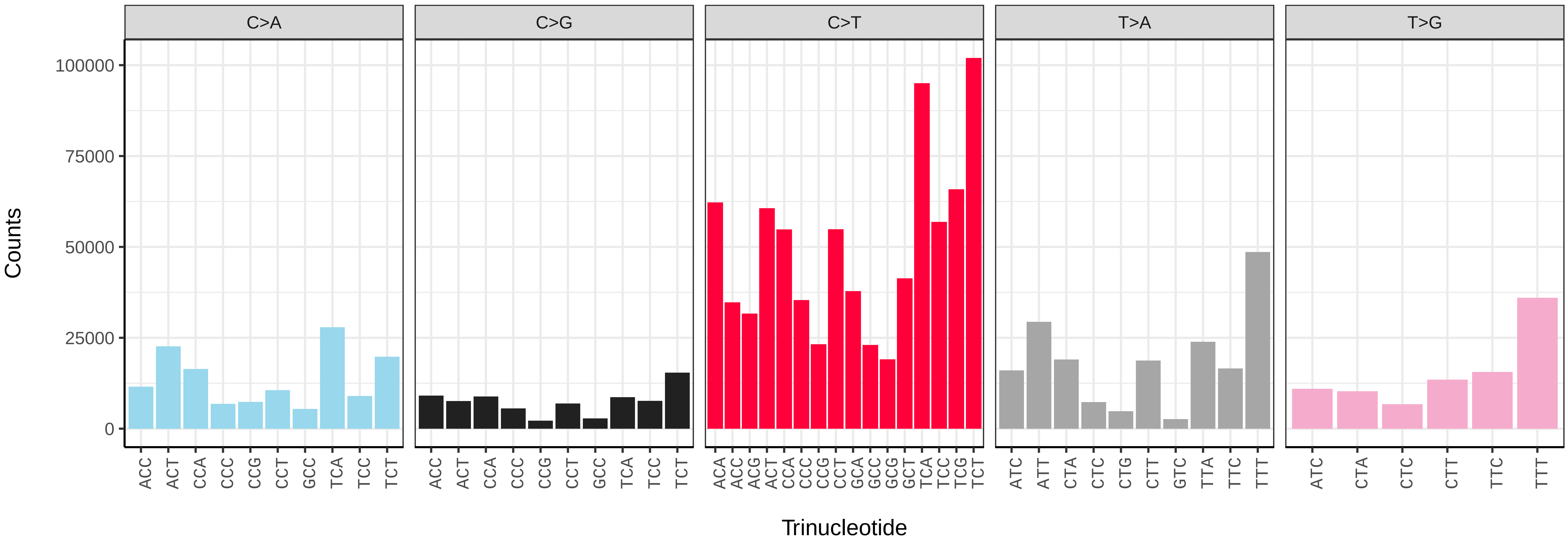


*Rhagonycha fulva* (icRhaFulv1) somatic mutational spectrum (SBS96 classification)

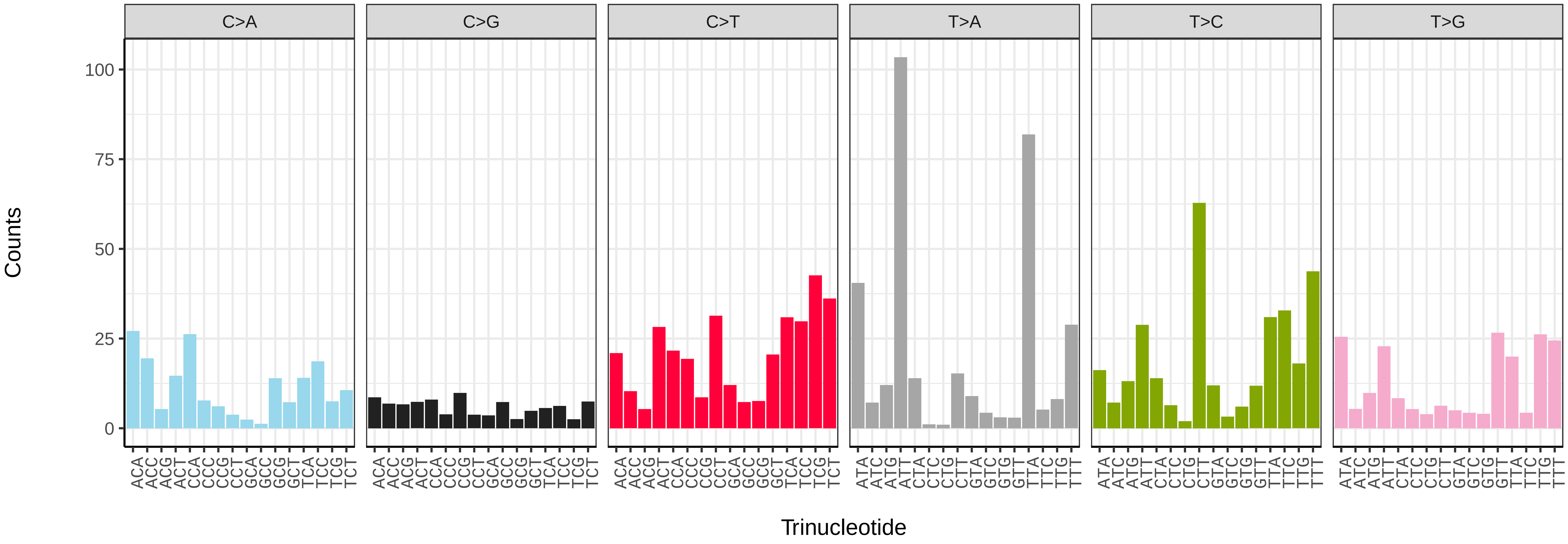


*Rhagonycha fulva* (icRhaFulv1) somatic mutational spectrum (SBS52 classification)

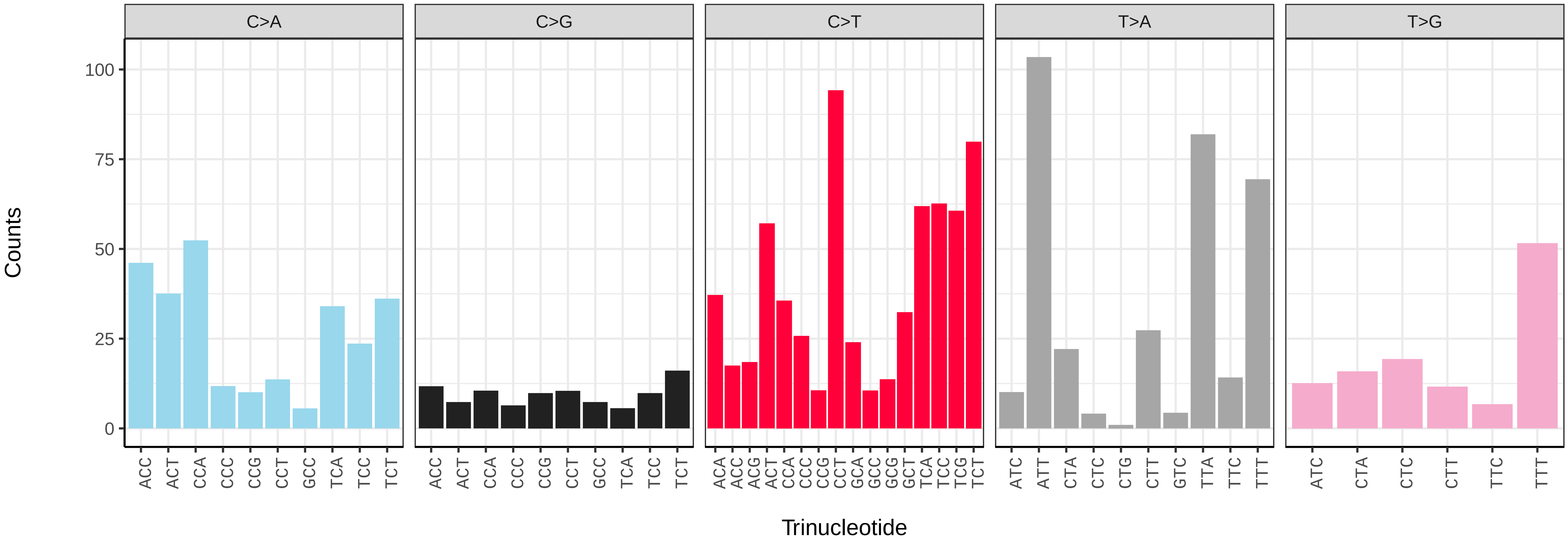

*Silpha atrata* (icSilAtra2) germline mutational spectrum

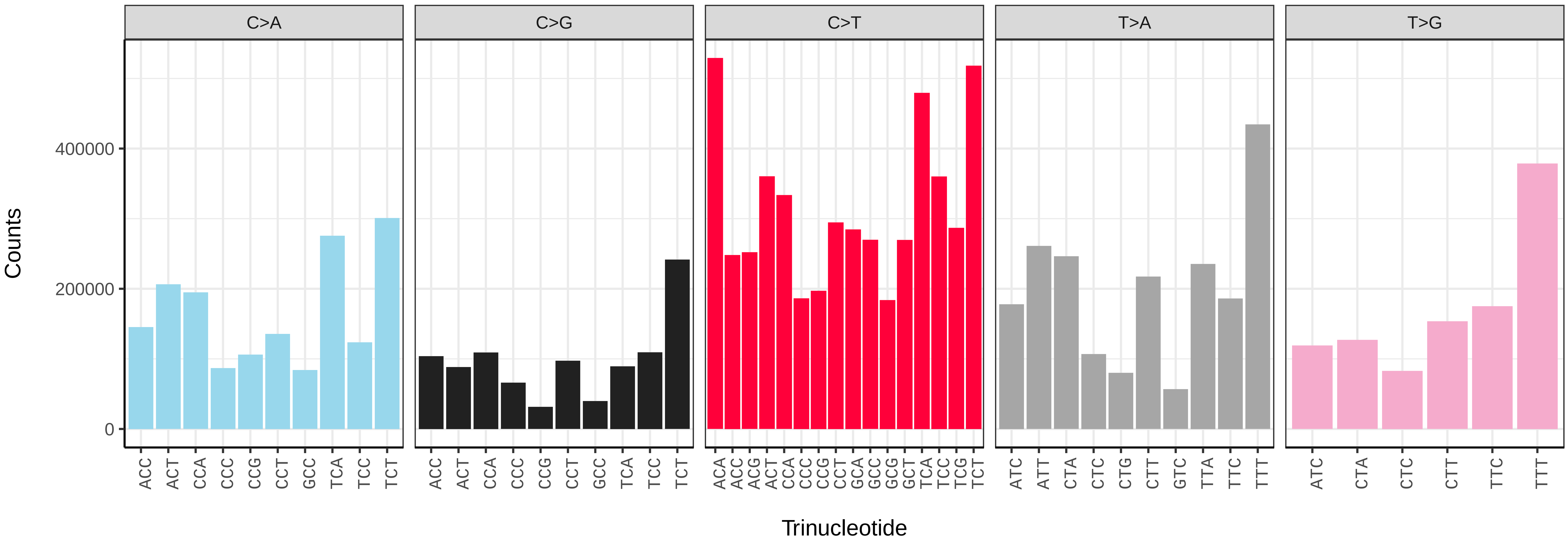


*Silpha atrata* (icSilAtra2) somatic mutational spectrum (SBS96 classification)

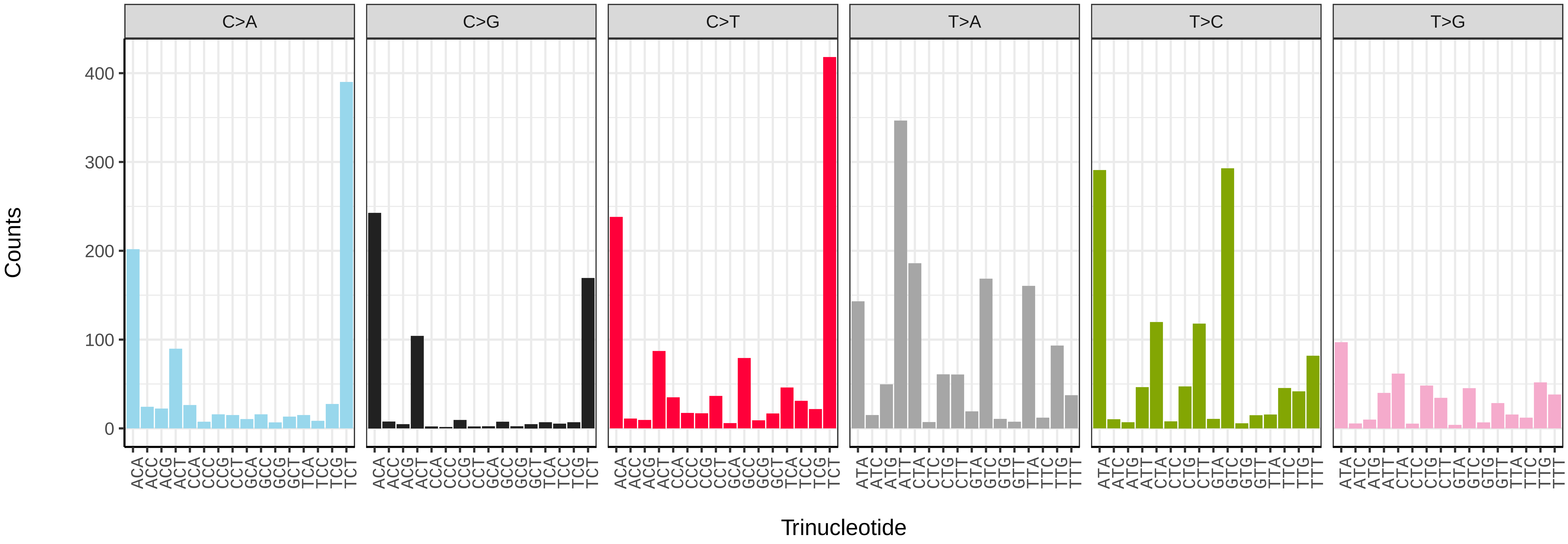


*Silpha atrata* (icSilAtra2) somatic mutational spectrum (SBS52 classification)

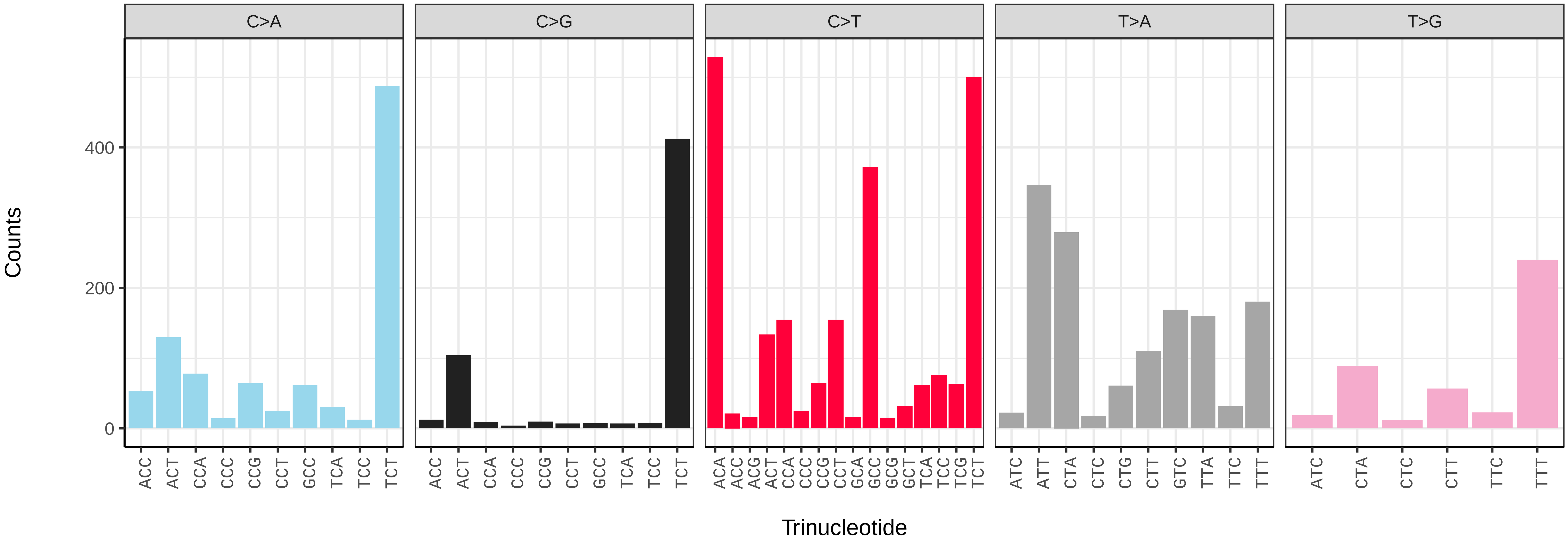

*Acrocera orbiculus* (idAcrOrbc1) germline mutational spectrum

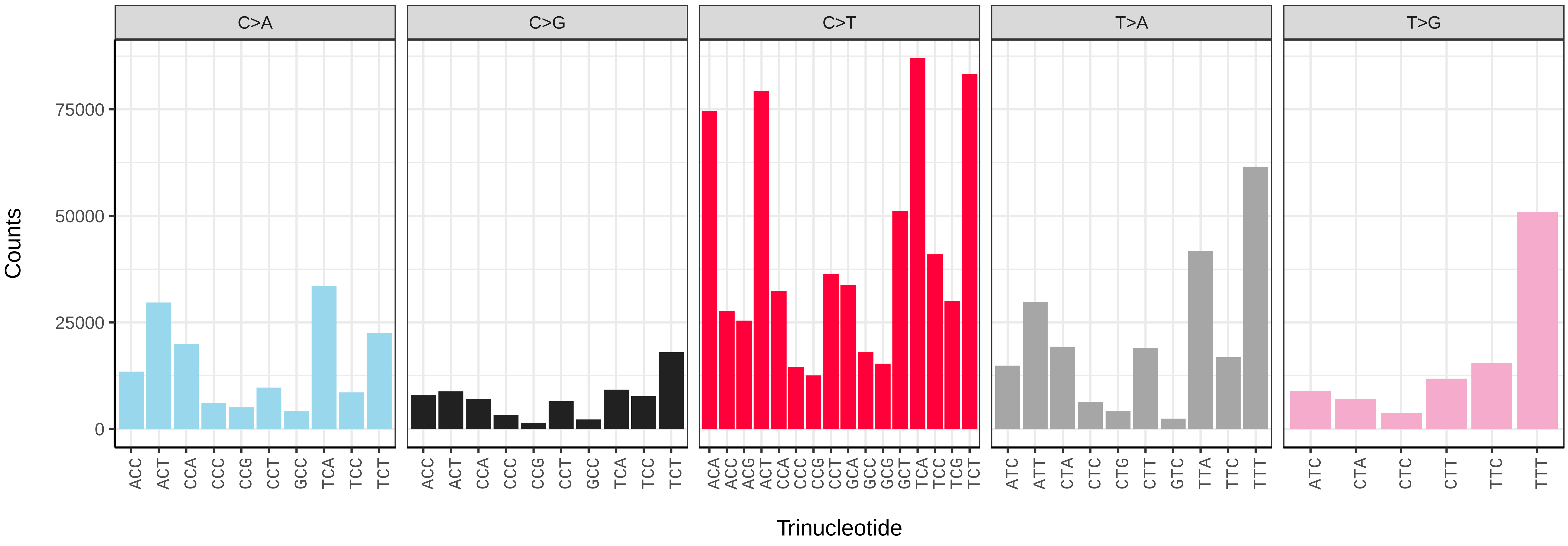


*Acrocera orbiculus* (idAcrOrbc1) somatic mutational spectrum (SBS96 classification)

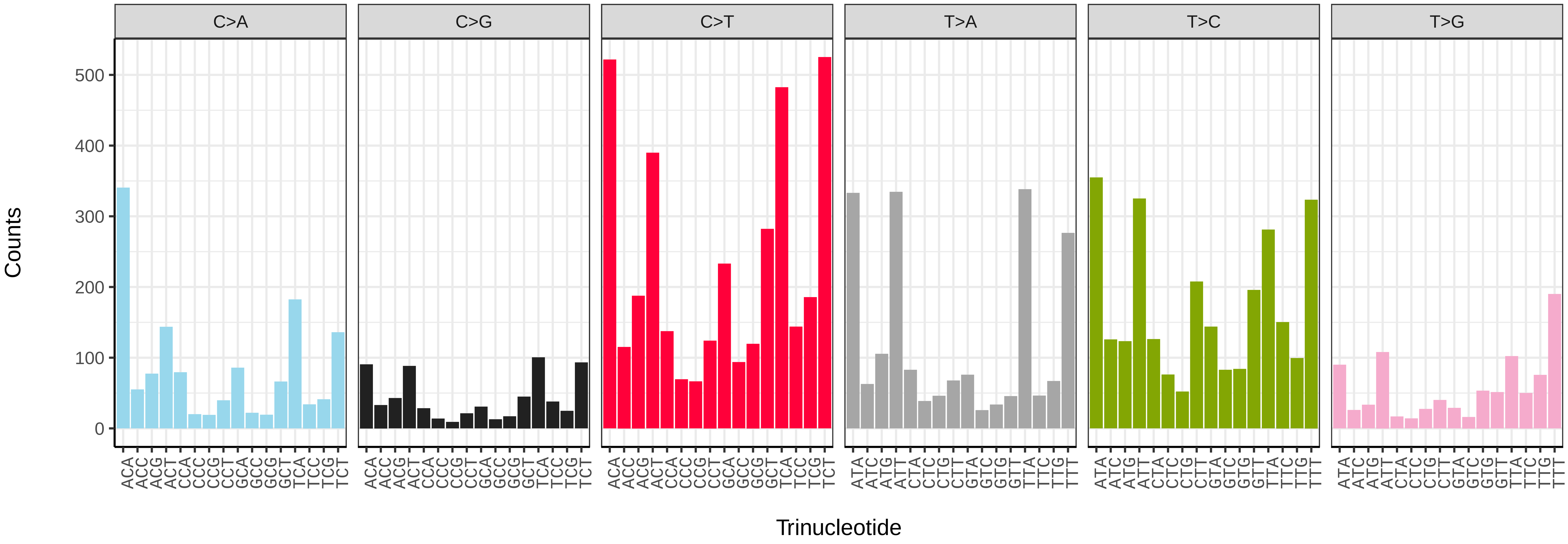


*Acrocera orbiculus* (idAcrOrbc1) somatic mutational spectrum (SBS52 classification)

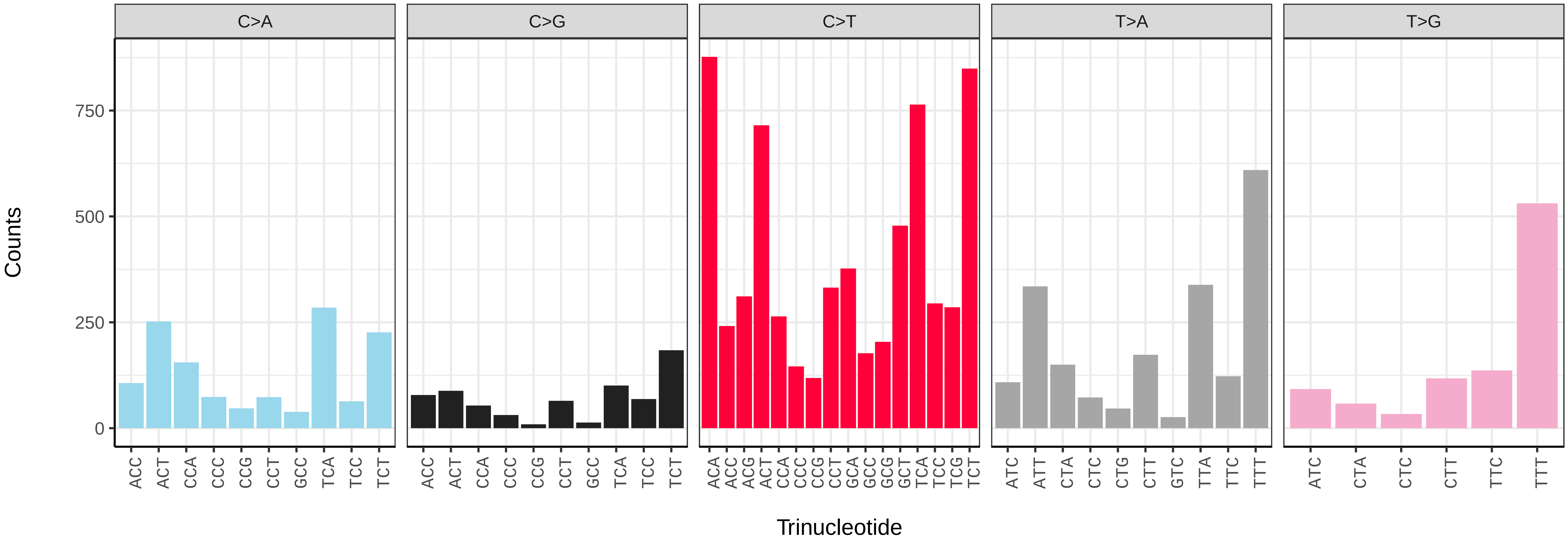

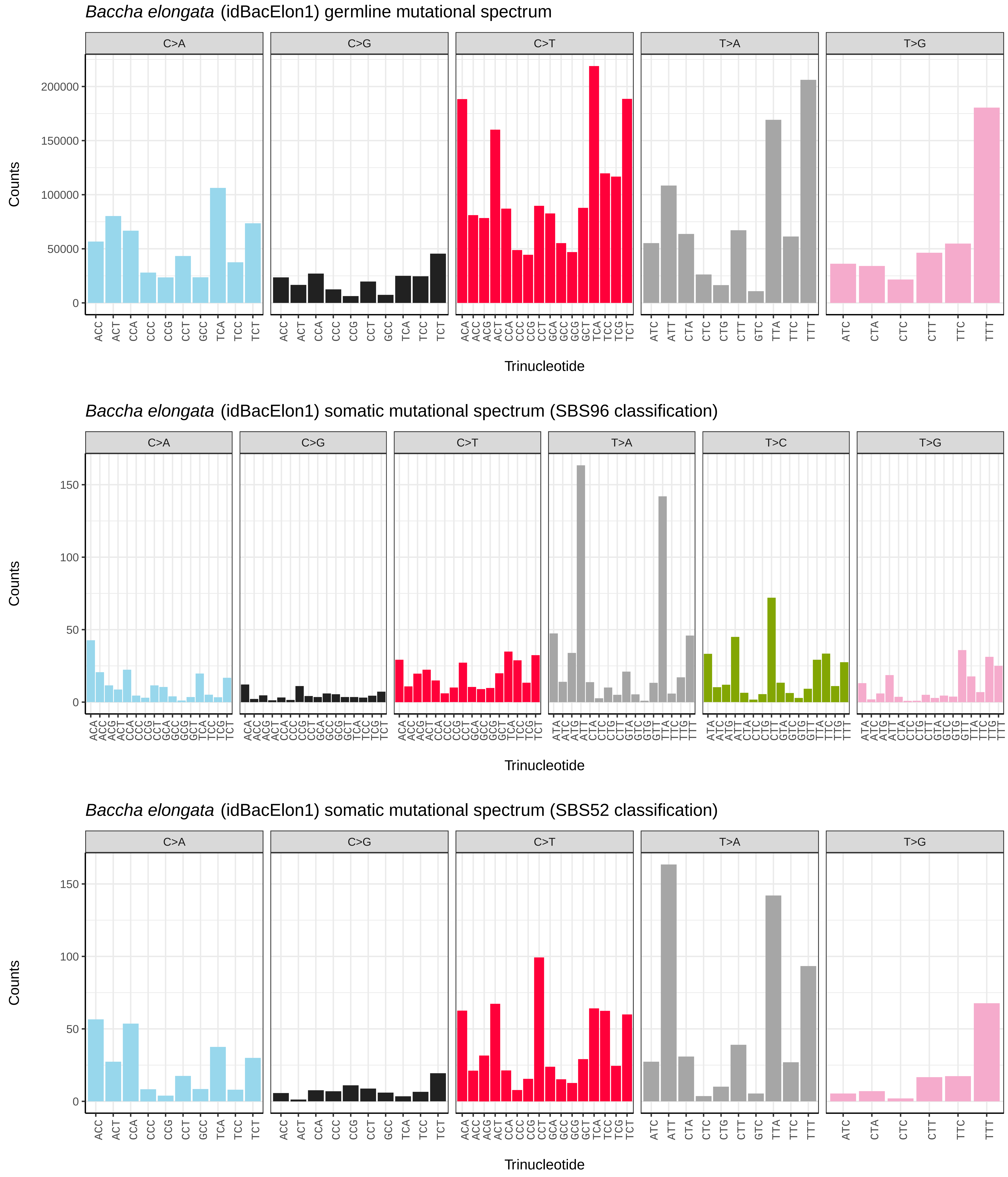

*Baccha elongata* (idBacElon1) germline mutational spectrum
C>A
C>G
C>T
T>A
T>G
Counts
0
50000
100000
150000
200000
Trinucleotide
*Baccha elongata* (idBacElon1) somatic mutational spectrum (SBS96 classification)
C>A
C>G
C>T
T>A
T>C
T>G
Counts
0
50
100
150
Trinucleotide
*Baccha elongata* (idBacElon1) somatic mutational spectrum (SBS52 classification)
C>A
C>G
C>T
T>A
T>G
Counts
0
50
100
150
Trinucleotide

*Bellardia pandia* (idBelPand1) germline mutational spectrum

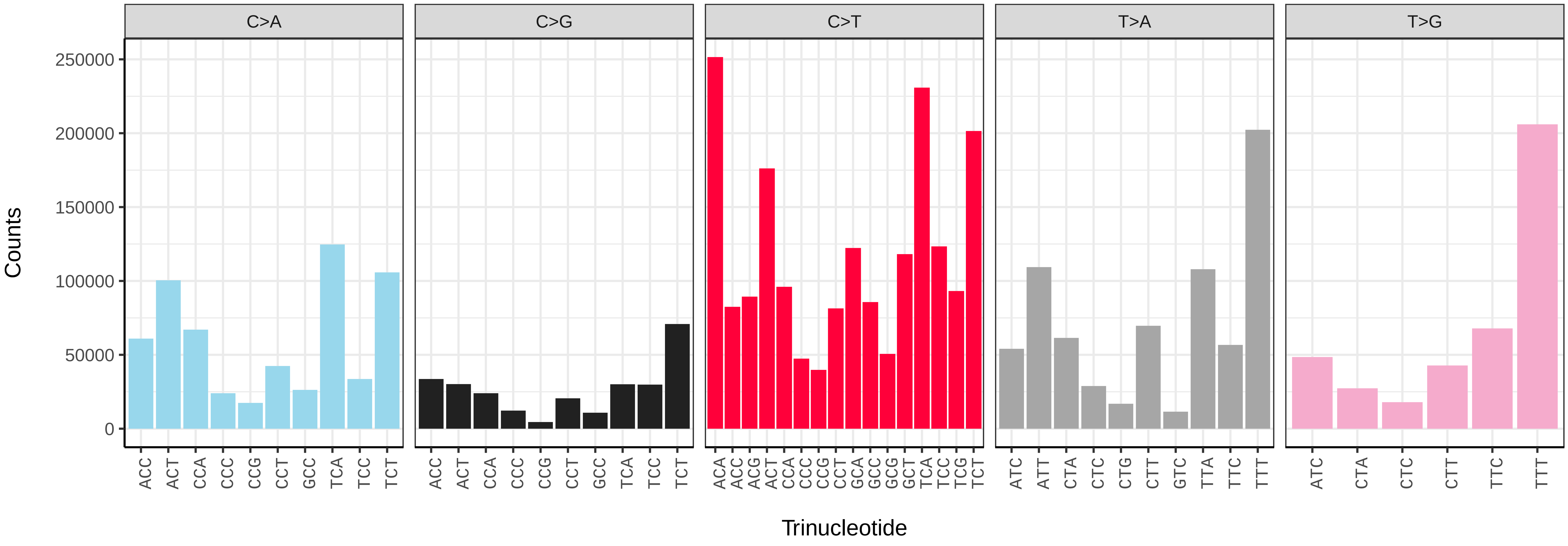


*Bellardia pandia* (idBelPand1) somatic mutational spectrum (SBS96 classification)

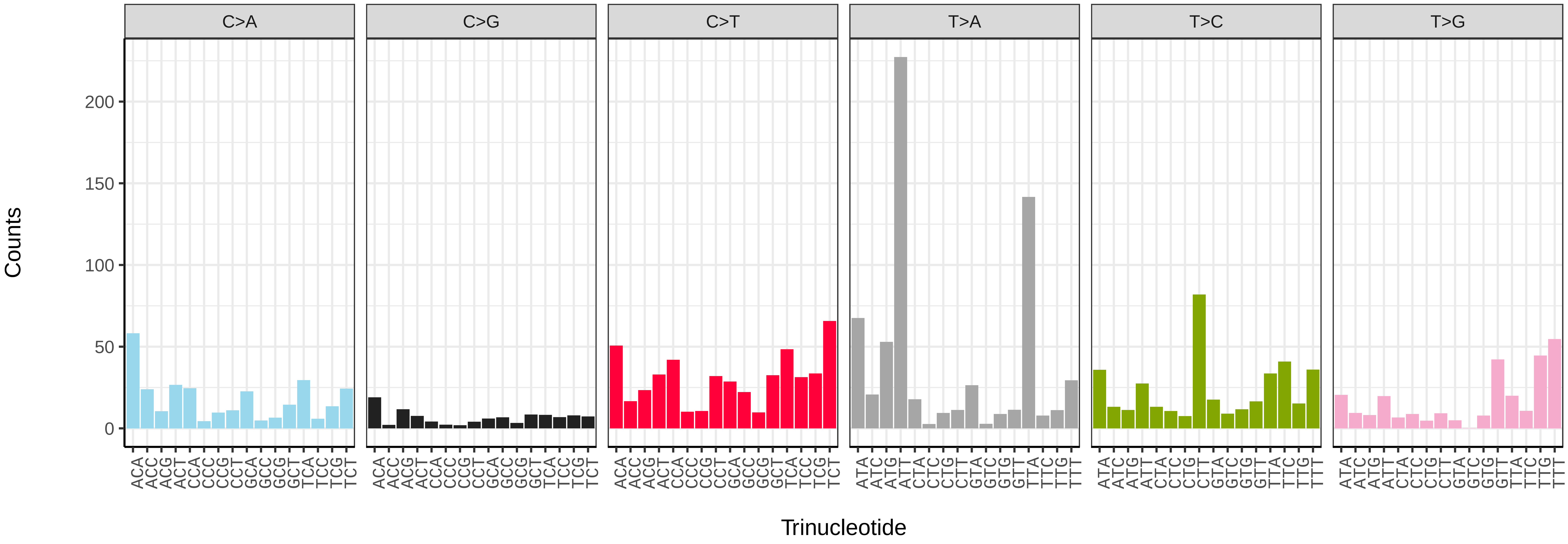


*Bellardia pandia* (idBelPand1) somatic mutational spectrum (SBS52 classification)

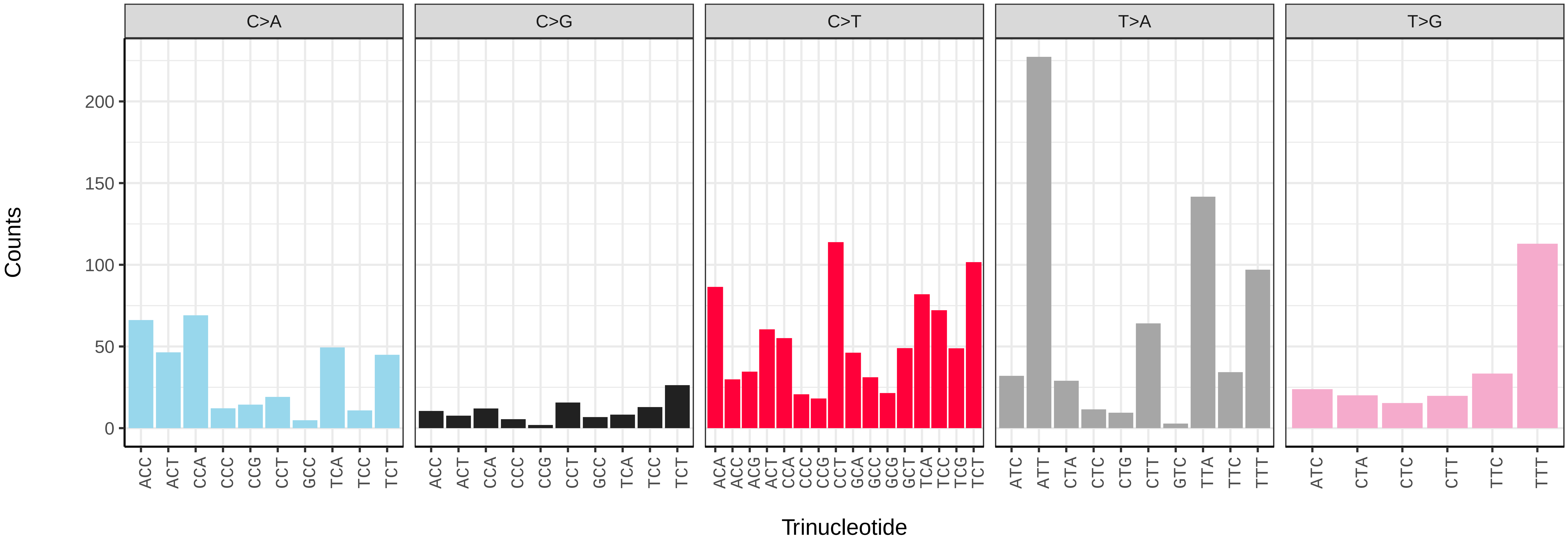

*Beris chalybata* (idBerChal2) germline mutational spectrum

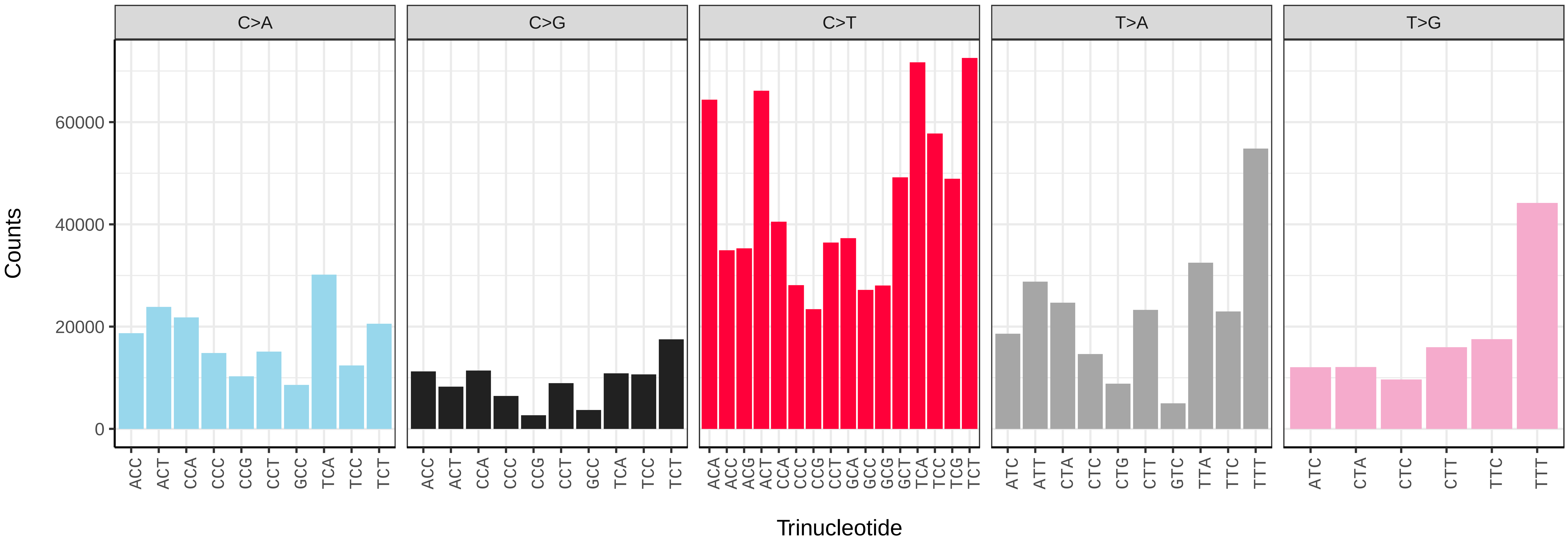


*Beris chalybata* (idBerChal2) somatic mutational spectrum (SBS96 classification)

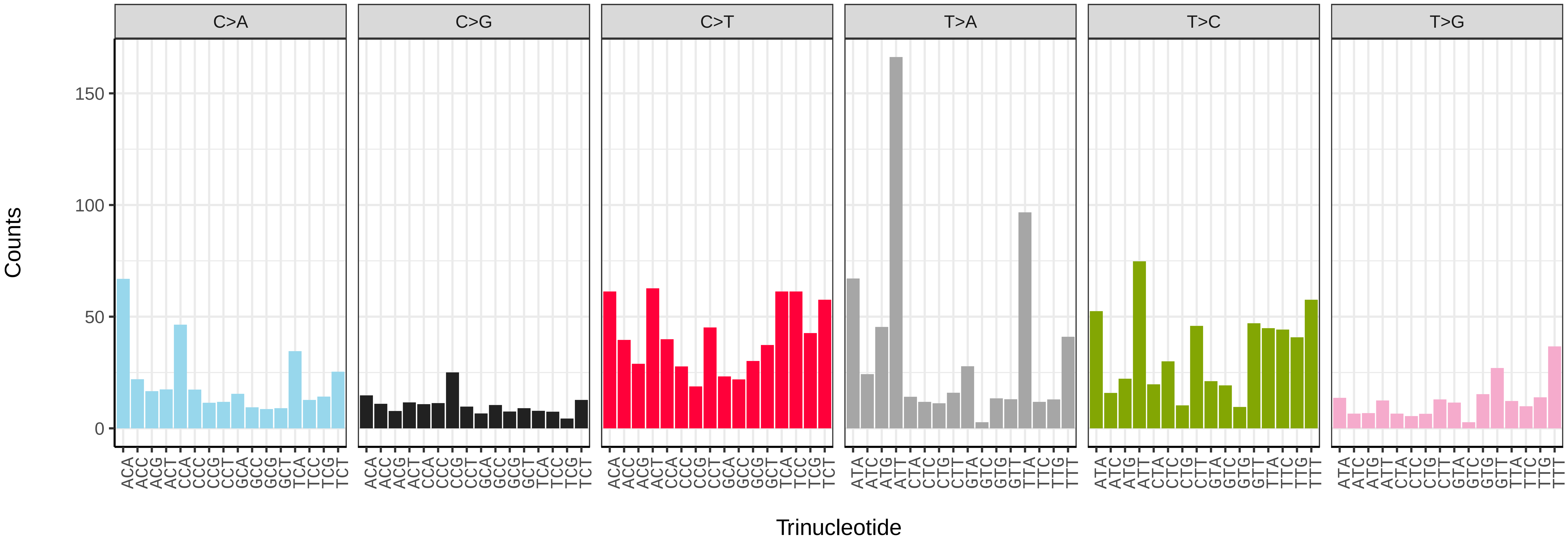


*Beris chalybata* (idBerChal2) somatic mutational spectrum (SBS52 classification)

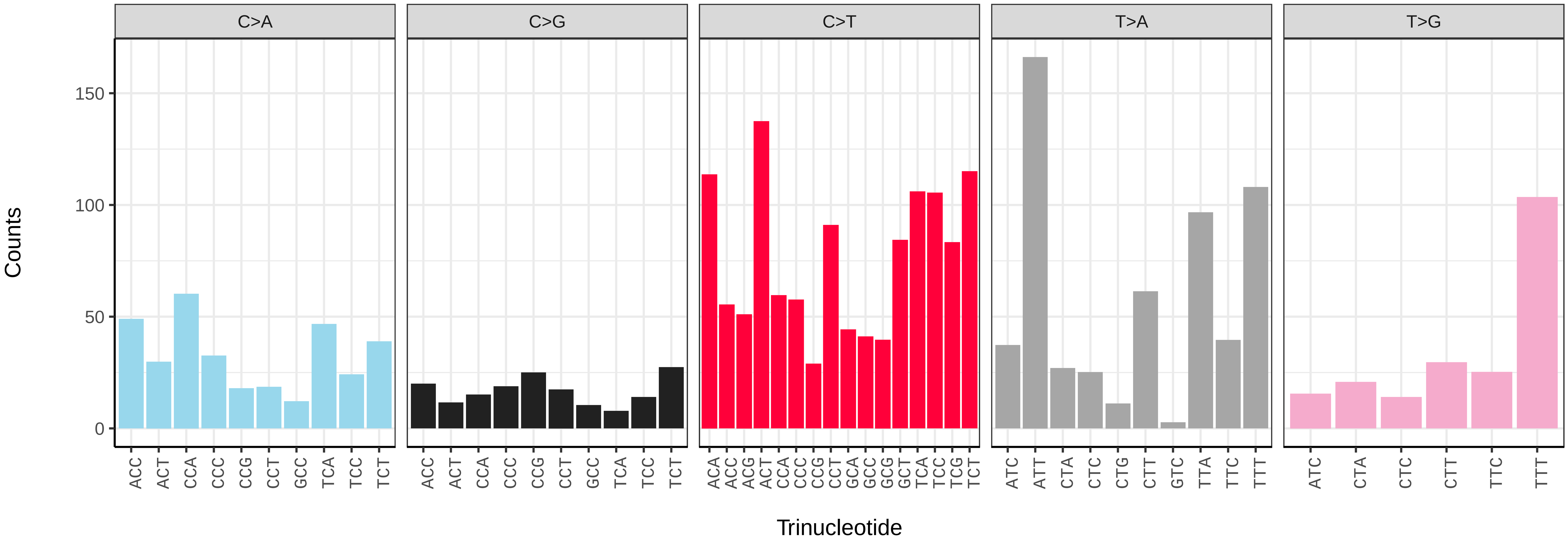

*Bibio marci* (idBibMarc1) germline mutational spectrum

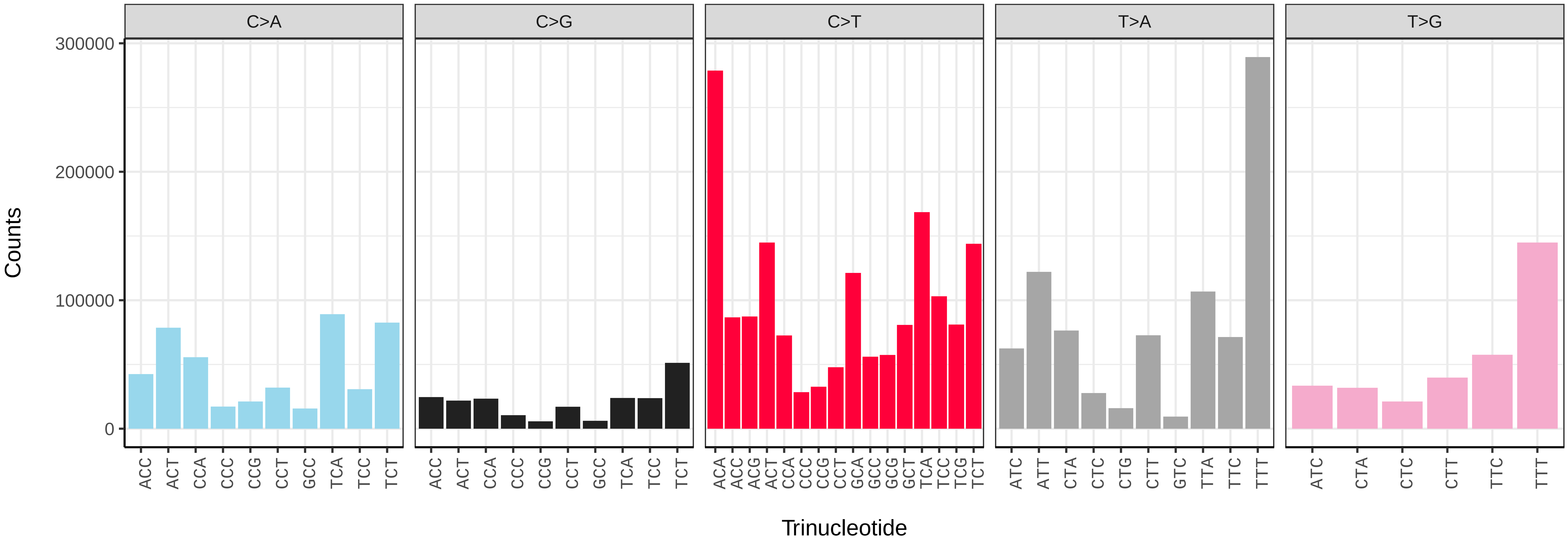


*Bibio marci* (idBibMarc1) somatic mutational spectrum (SBS96 classification)

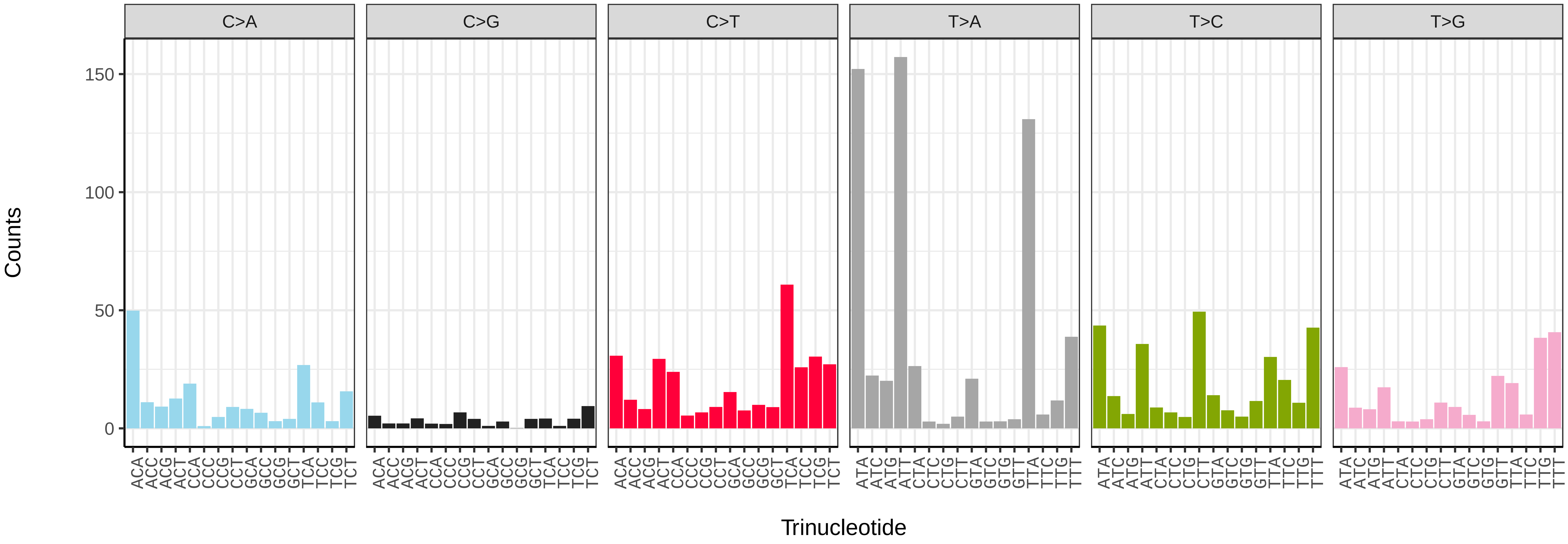


*Bibio marci* (idBibMarc1) somatic mutational spectrum (SBS52 classification)

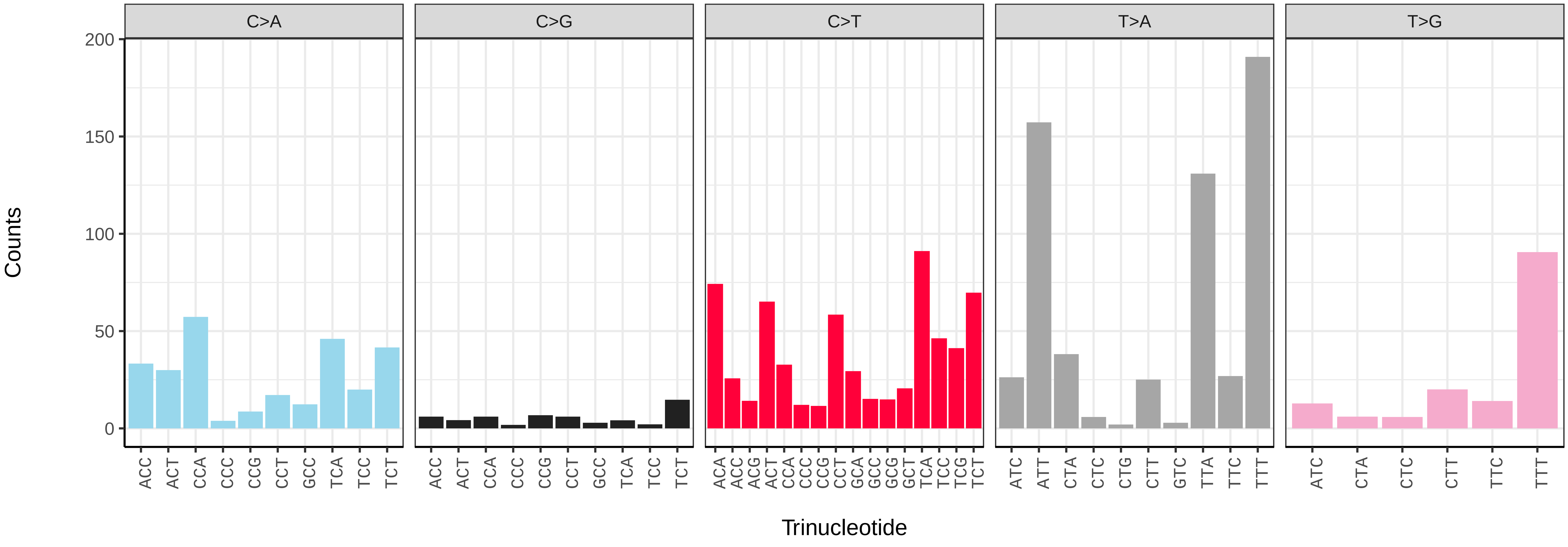

*Blera fallax* (idBleFall4) germline mutational spectrum

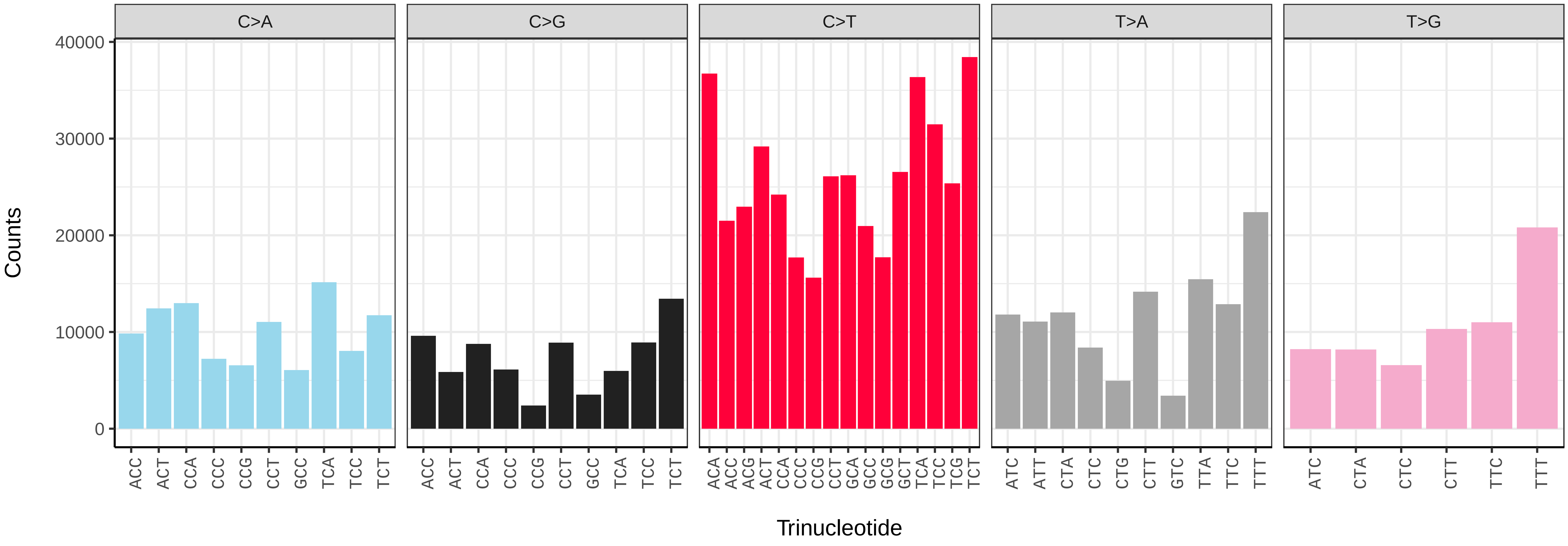


*Blera fallax* (idBleFall4) somatic mutational spectrum (SBS96 classification)

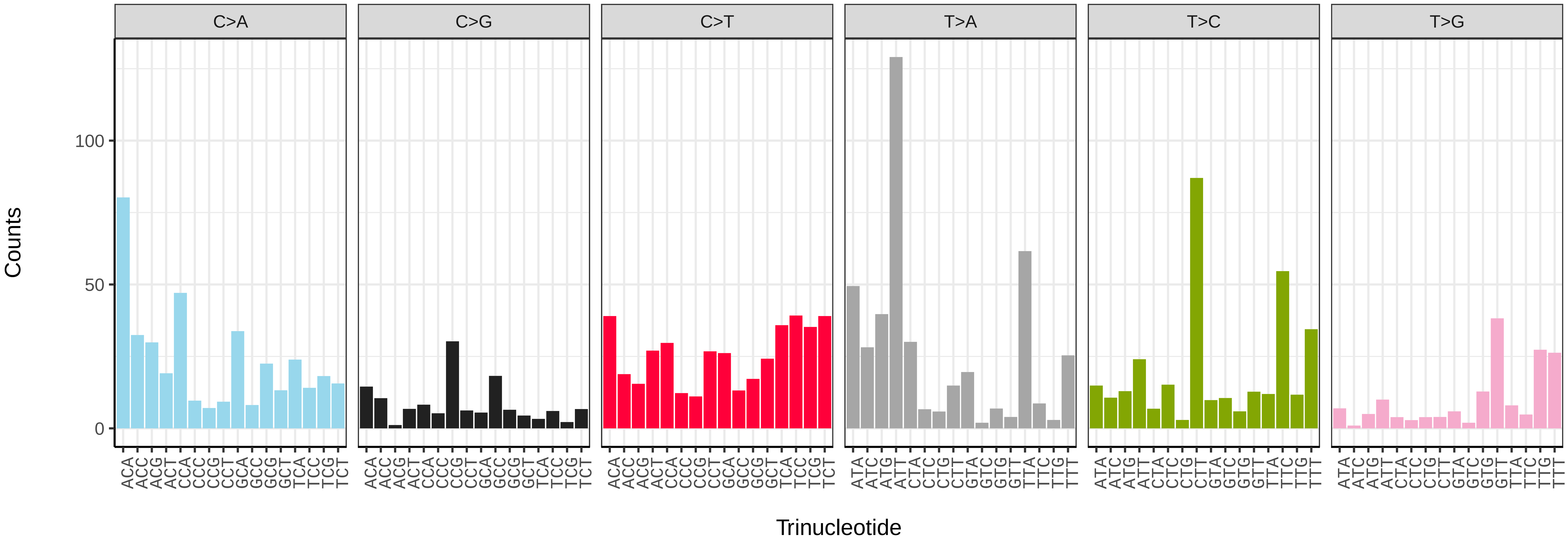


*Blera fallax* (idBleFall4) somatic mutational spectrum (SBS52 classification)

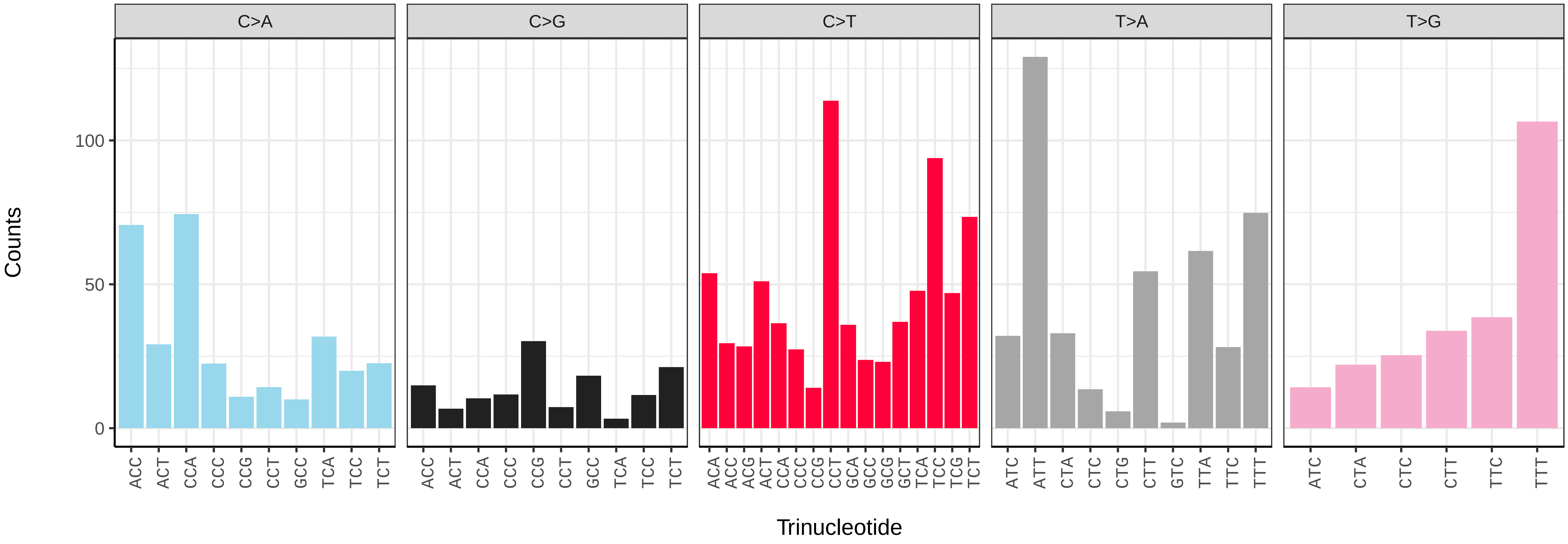

*Bombylius discolor* (idBomDisc1) germline mutational spectrum

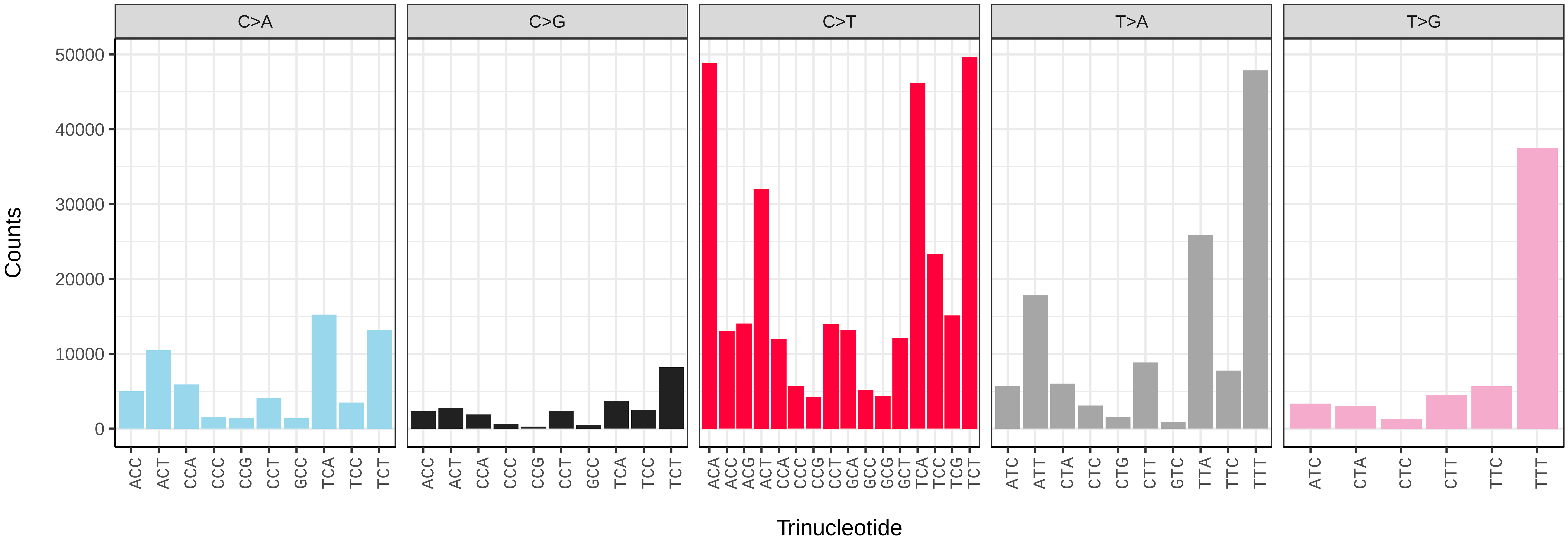


*Bombylius discolor* (idBomDisc1) somatic mutational spectrum (SBS96 classification)

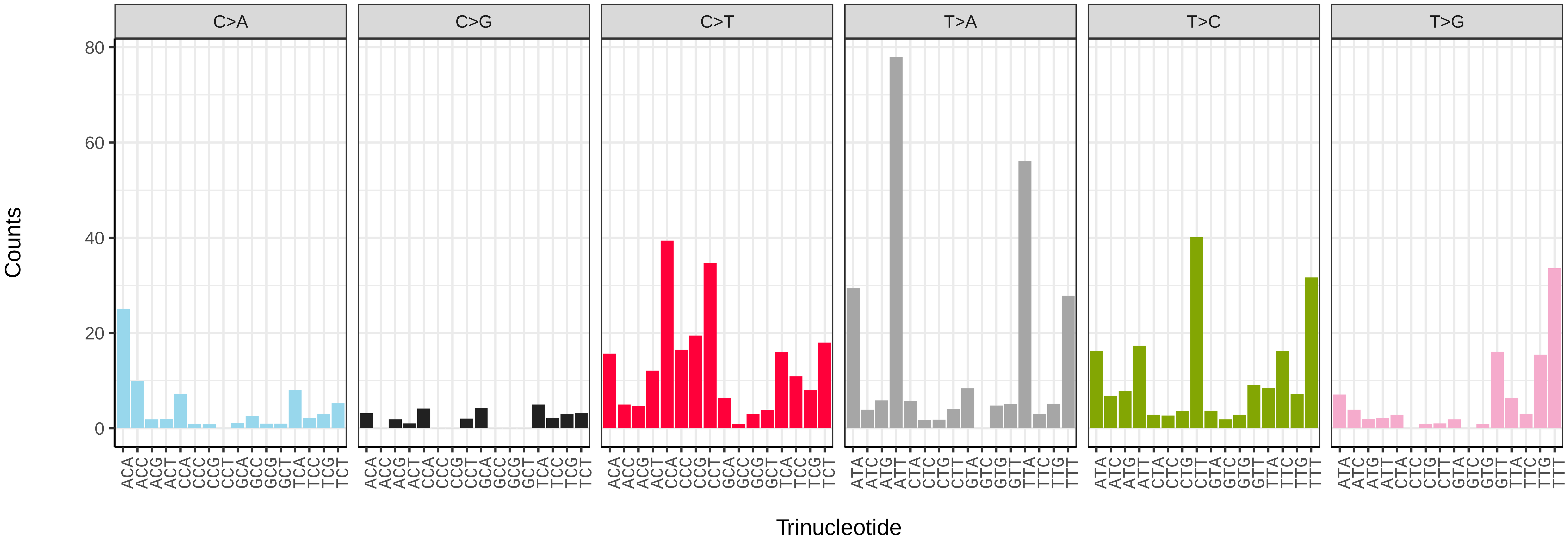


*Bombylius discolor* (idBomDisc1) somatic mutational spectrum (SBS52 classification)

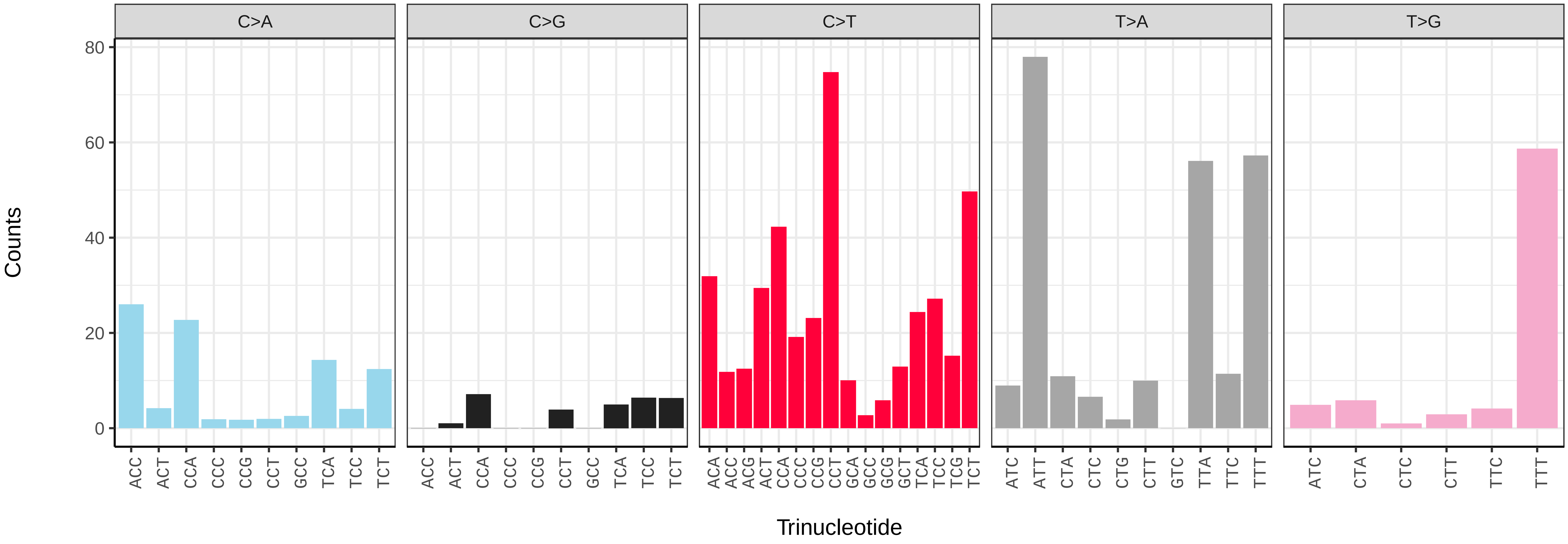

*Bombylius major* (idBomMajo1) germline mutational spectrum

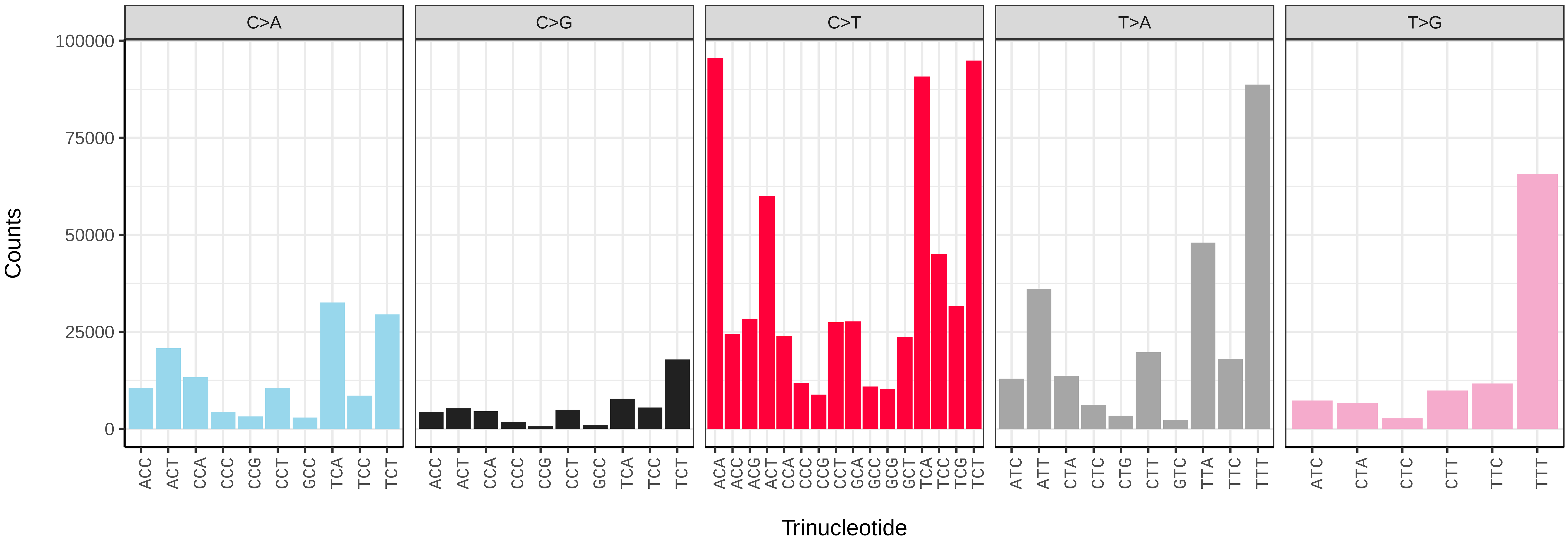


*Bombylius major* (idBomMajo1) somatic mutational spectrum (SBS96 classification)

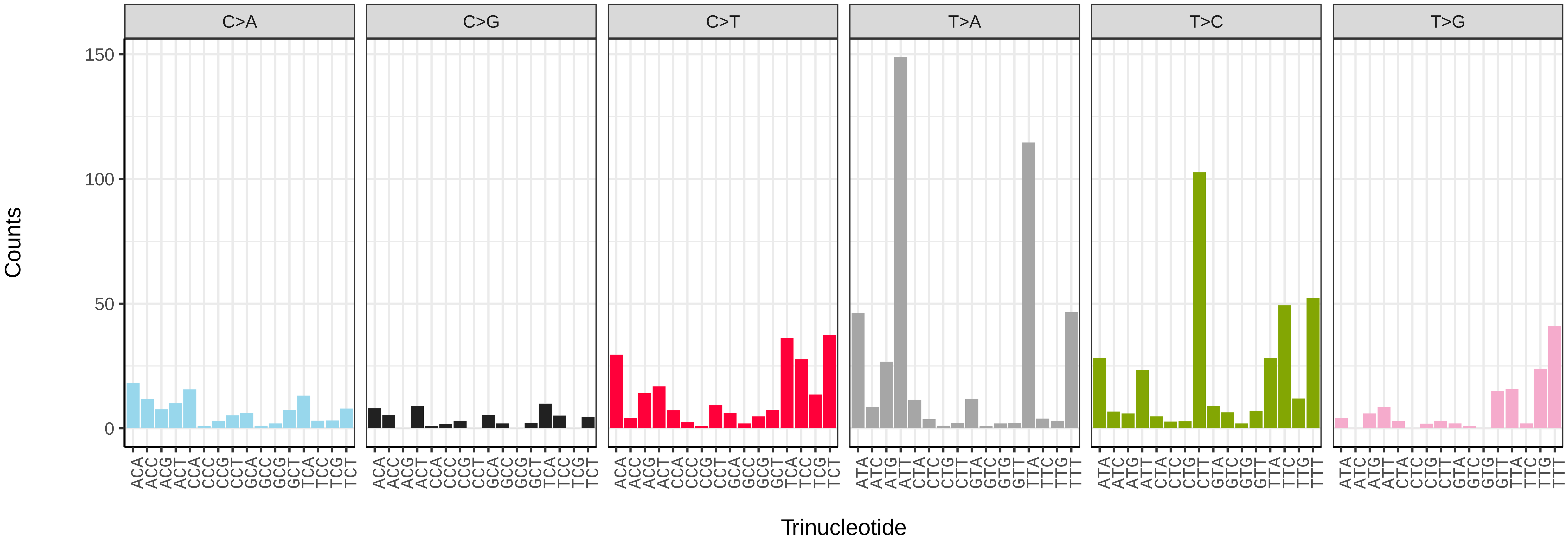


*Bombylius major* (idBomMajo1) somatic mutational spectrum (SBS52 classification)

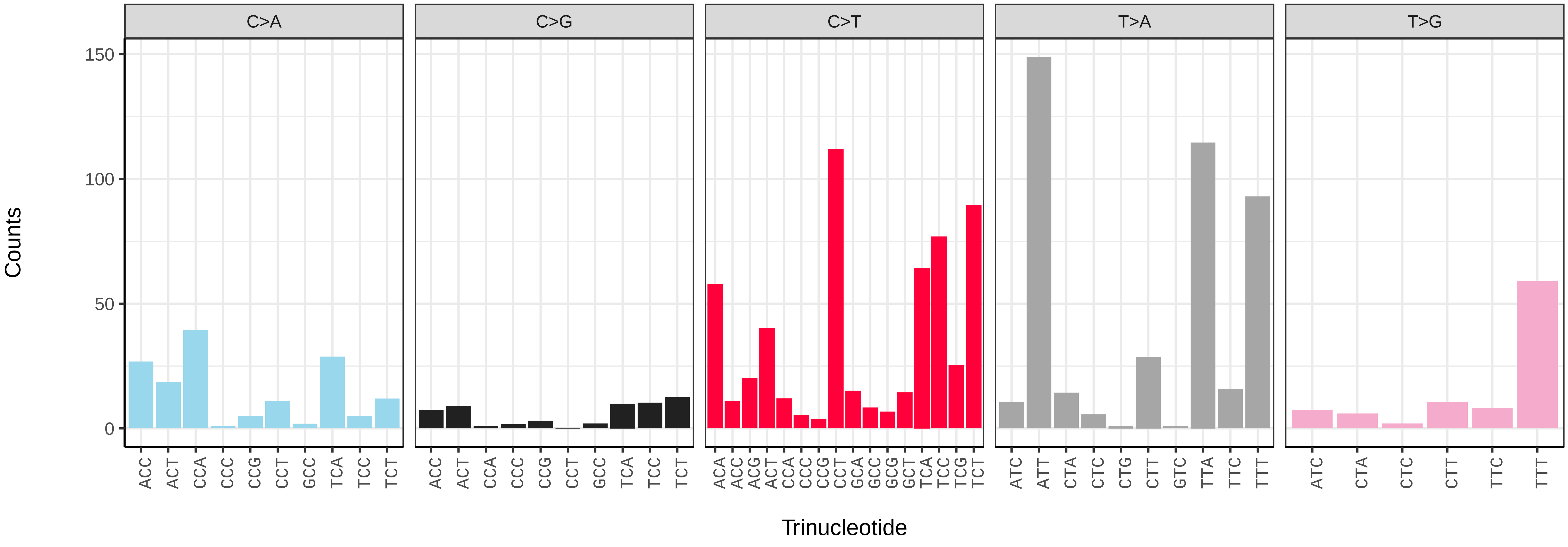

*Brachypalpus laphriformis* (idBraLaph1) germline mutational spectrum

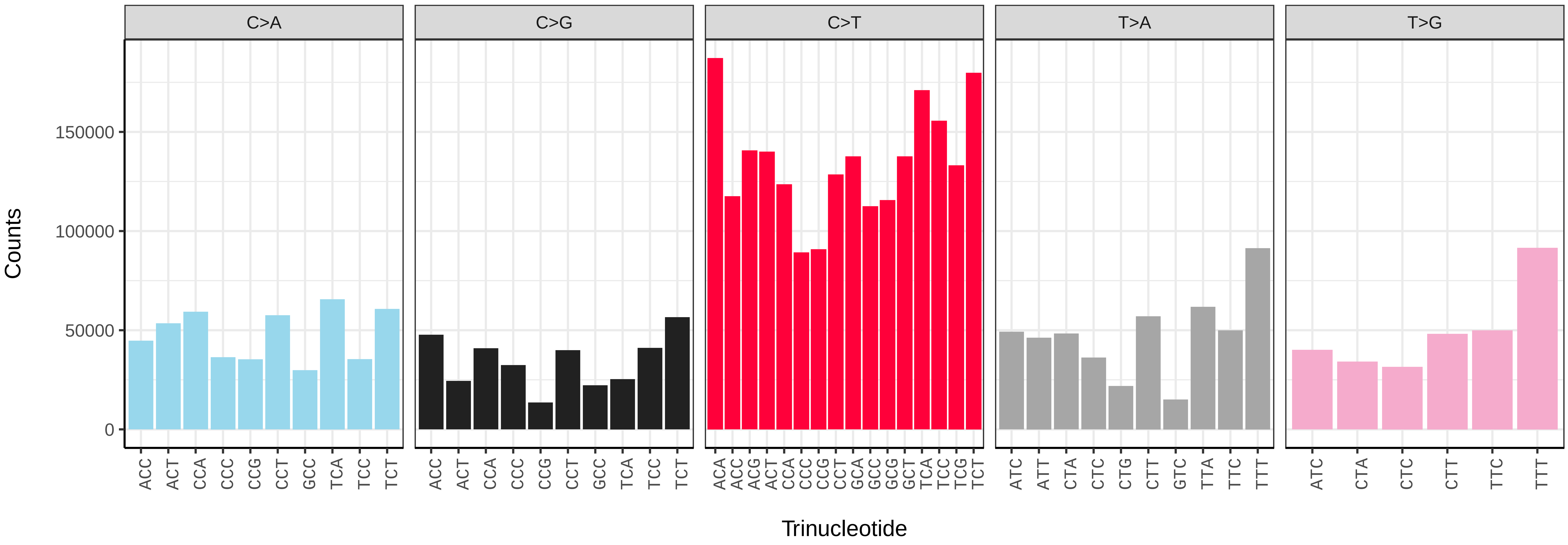


*Brachypalpus laphriformis* (idBraLaph1) somatic mutational spectrum (SBS96 classification)

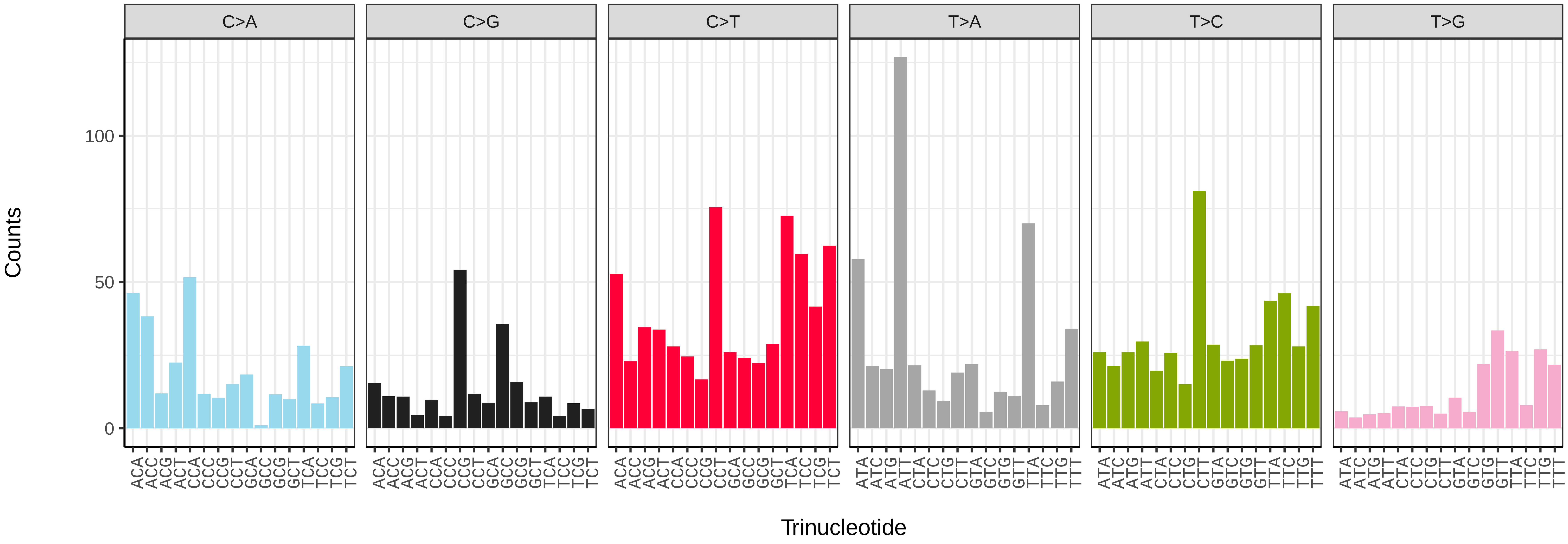


*Brachypalpus laphriformis* (idBraLaph1) somatic mutational spectrum (SBS52 classification)

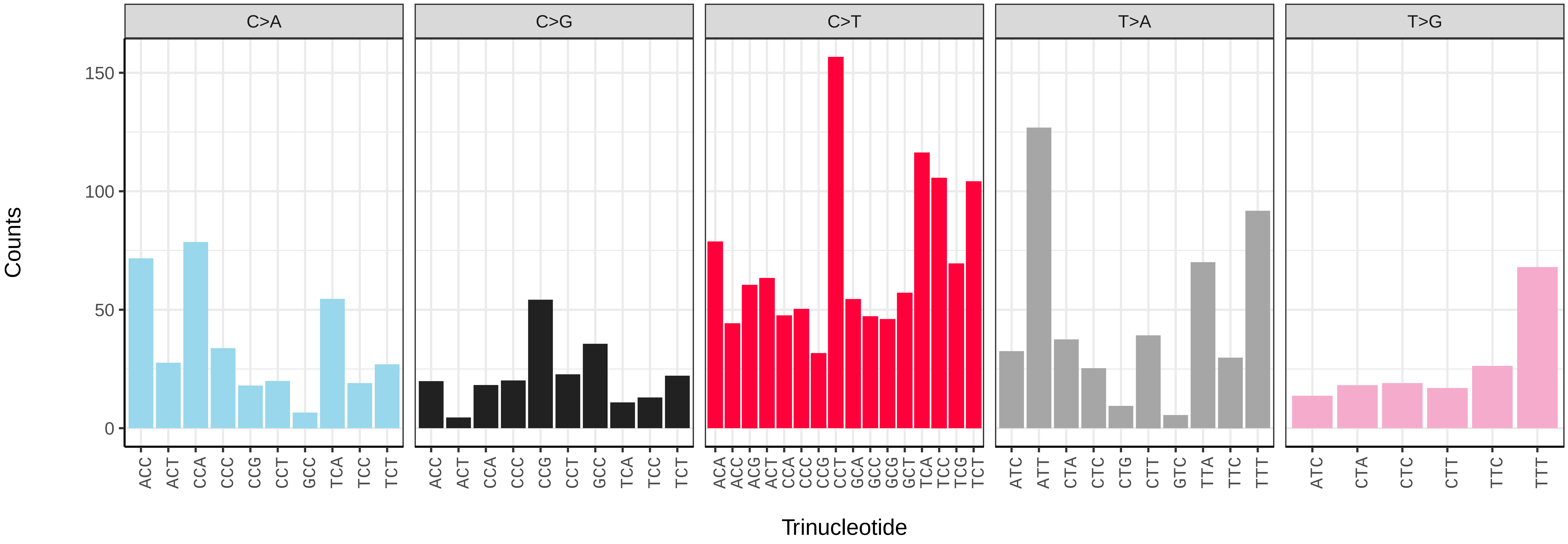

*Calliphora vomitoria* (idCalVomi1) germline mutational spectrum

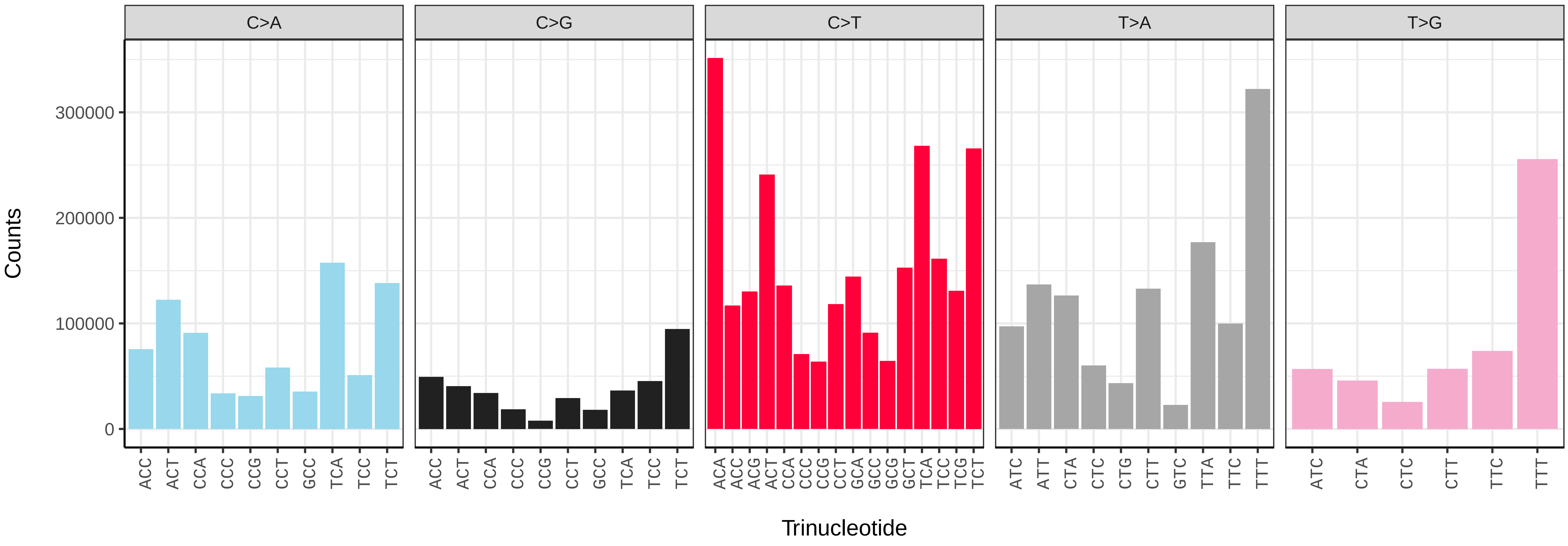


*Calliphora vomitoria* (idCalVomi1) somatic mutational spectrum (SBS96 classification)

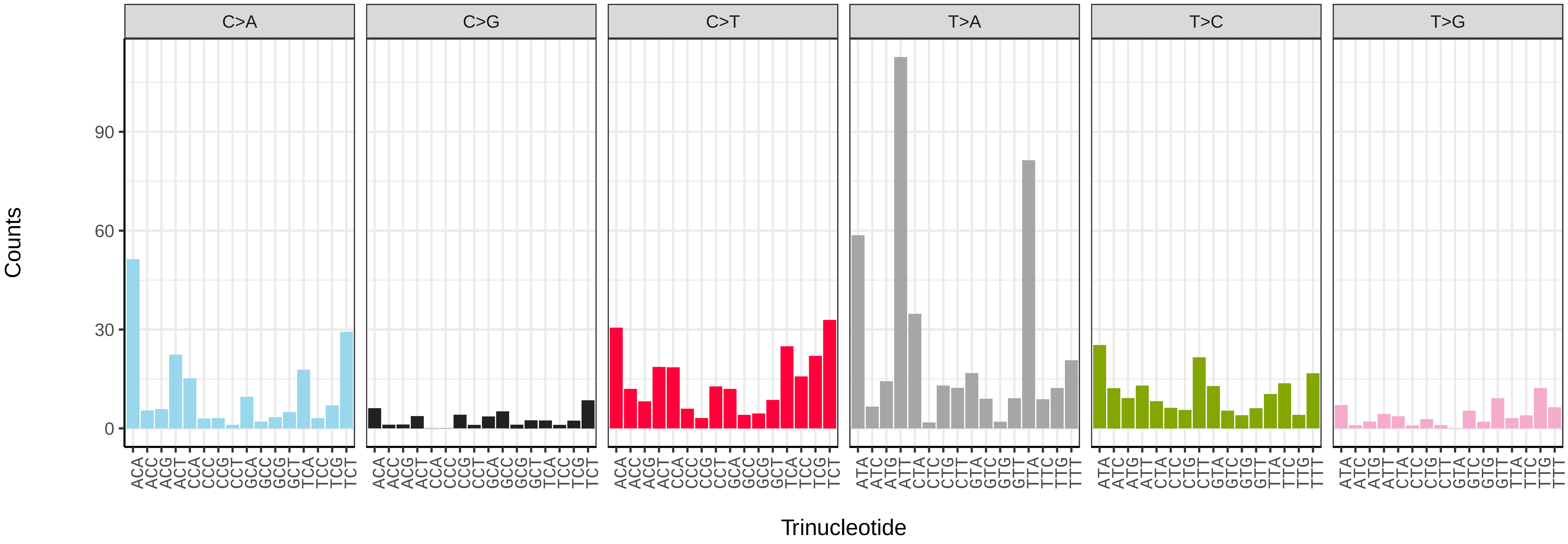


*Calliphora vomitoria* (idCalVomi1) somatic mutational spectrum (SBS52 classification)

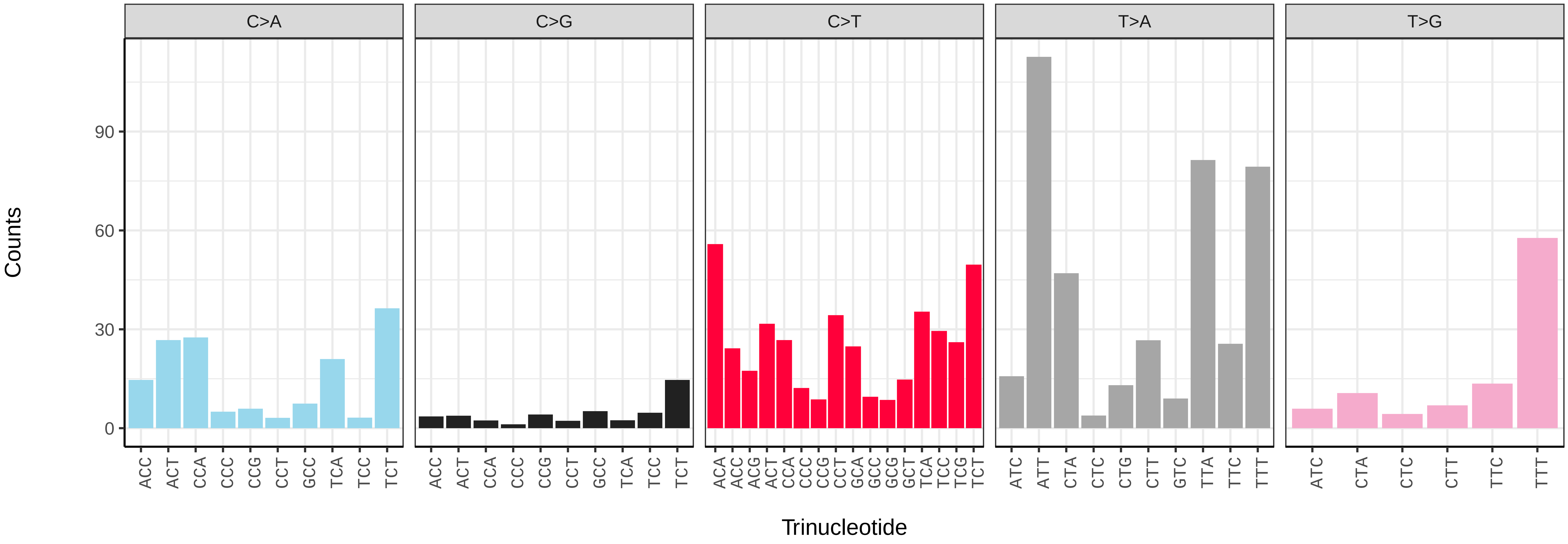

*Cheilosia impressa* (idCheImpr3) germline mutational spectrum

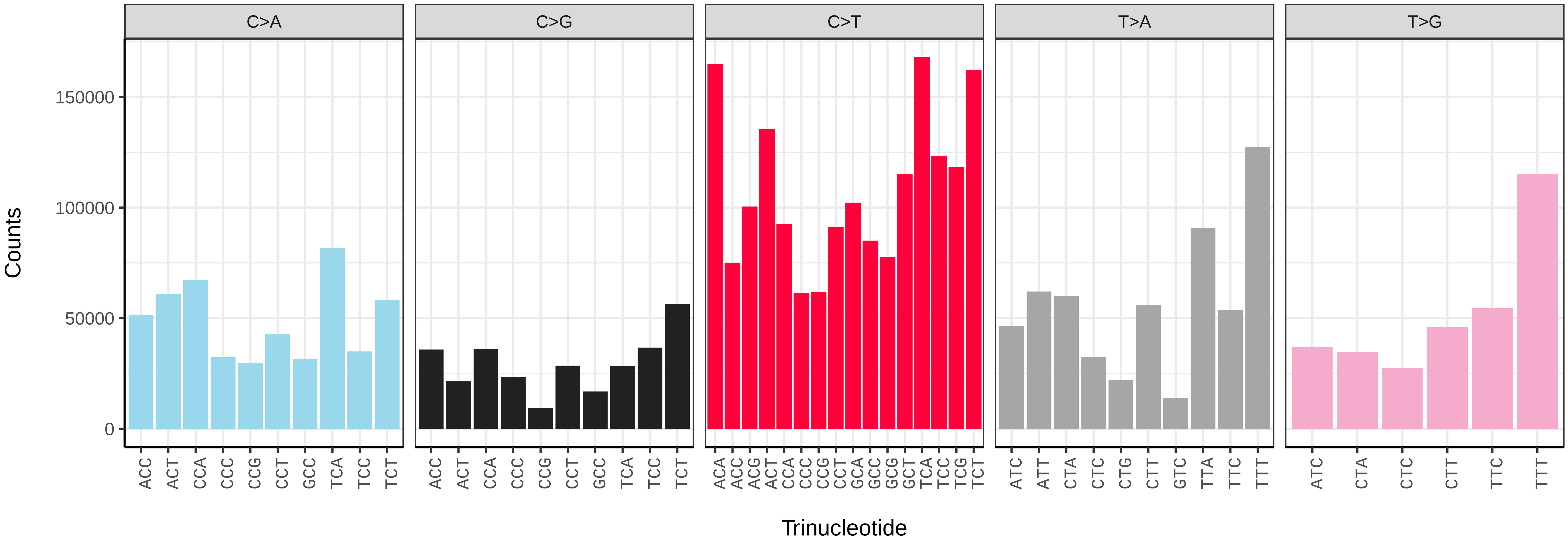


*Cheilosia impressa* (idCheImpr3) somatic mutational spectrum (SBS96 classification)

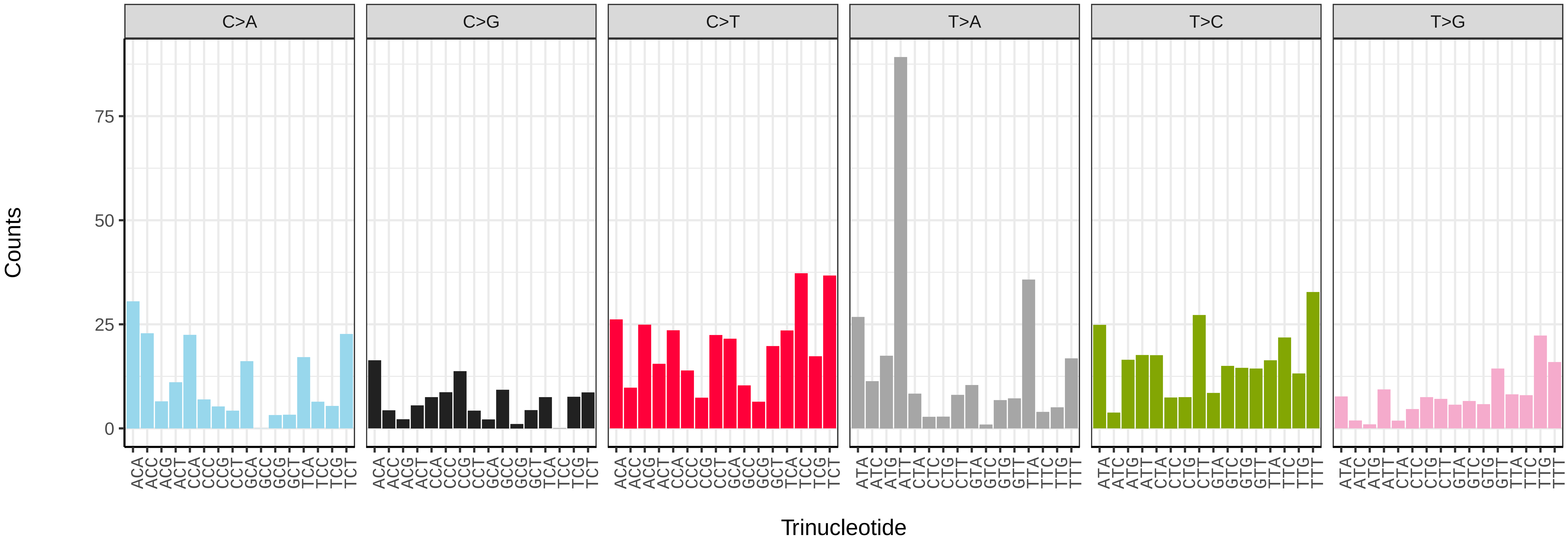


*Cheilosia impressa* (idCheImpr3) somatic mutational spectrum (SBS52 classification)

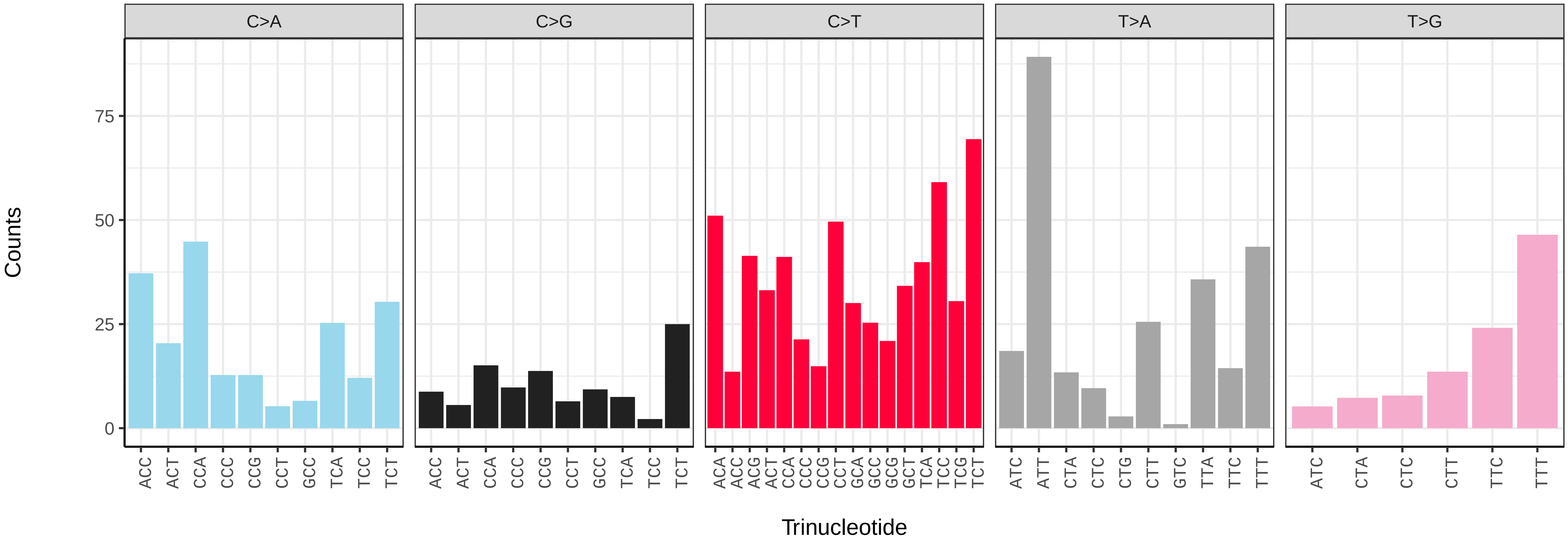

*Cheilosia pagana* (idChePaga1) germline mutational spectrum

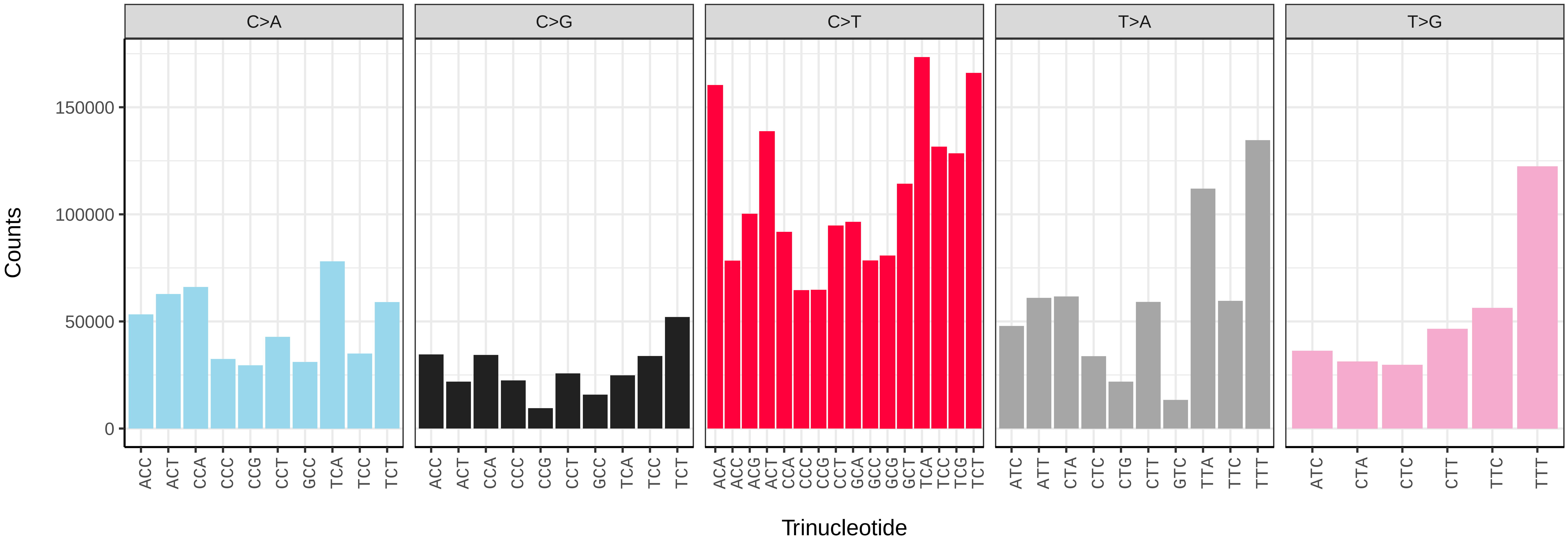


*Cheilosia pagana* (idChePaga1) somatic mutational spectrum (SBS96 classification)

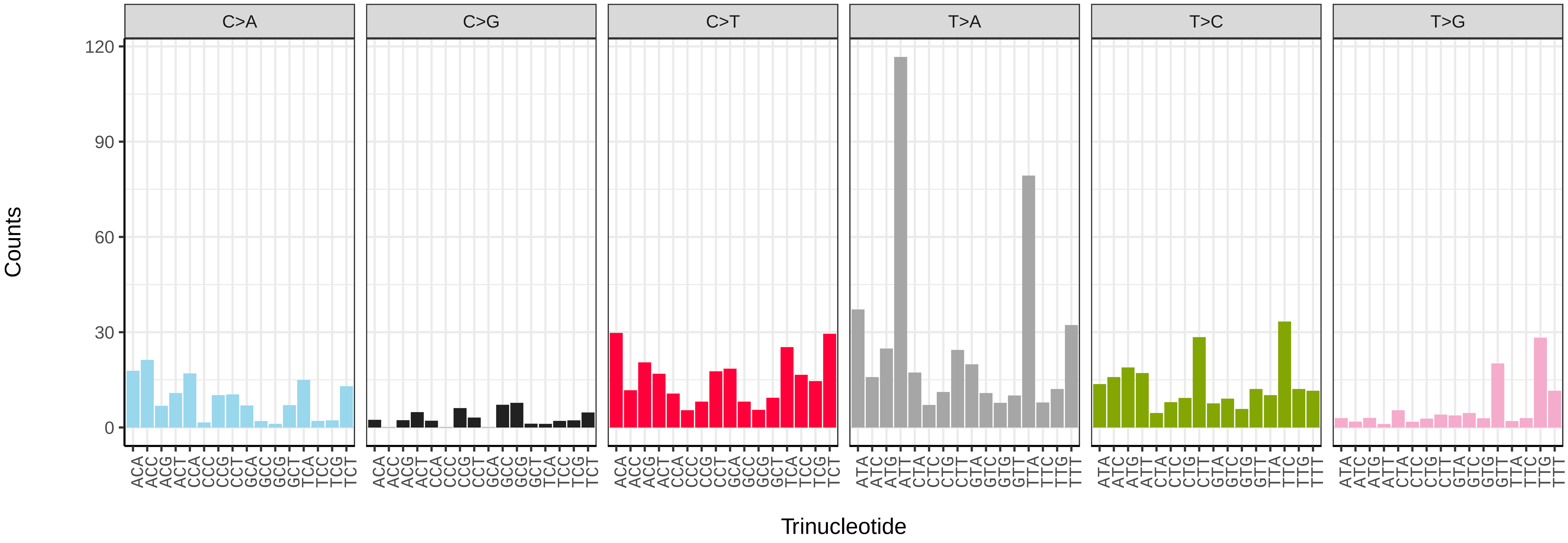


*Cheilosia pagana* (idChePaga1) somatic mutational spectrum (SBS52 classification)

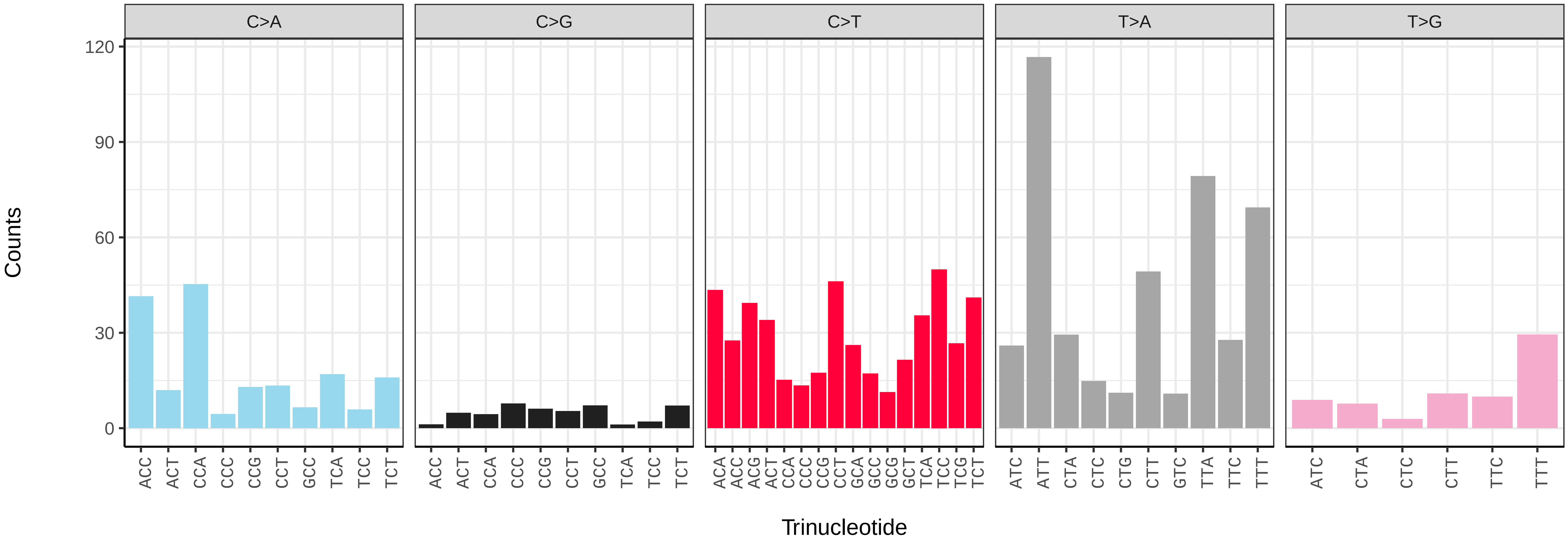

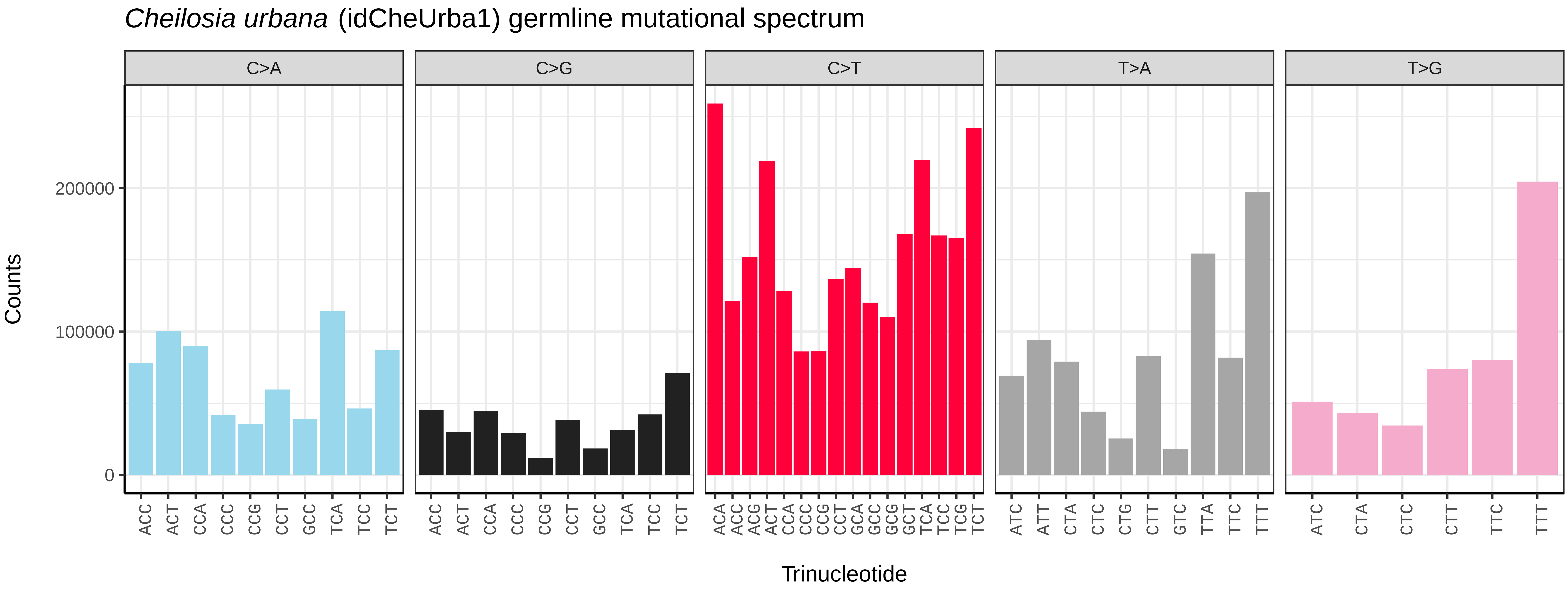
*Cheilosia urbana* (idCheUrba1) germline mutational spectrum
C>A
C>G
C>T
T>A
T>G
Counts
0
100000
200000
ACC ACT CCA CCC CCG CCT GCC TCA TCC TCT
ACA ACC ACG ACT CCA CCC CCG CCT GCA GCC GCG GCT TCA TCC TCG TCT
ATC ATT CTA CTC CTG CTT GTC TTA TTC TTT
ATC CTA CTC CTT TTC TTT
Trinucleotide

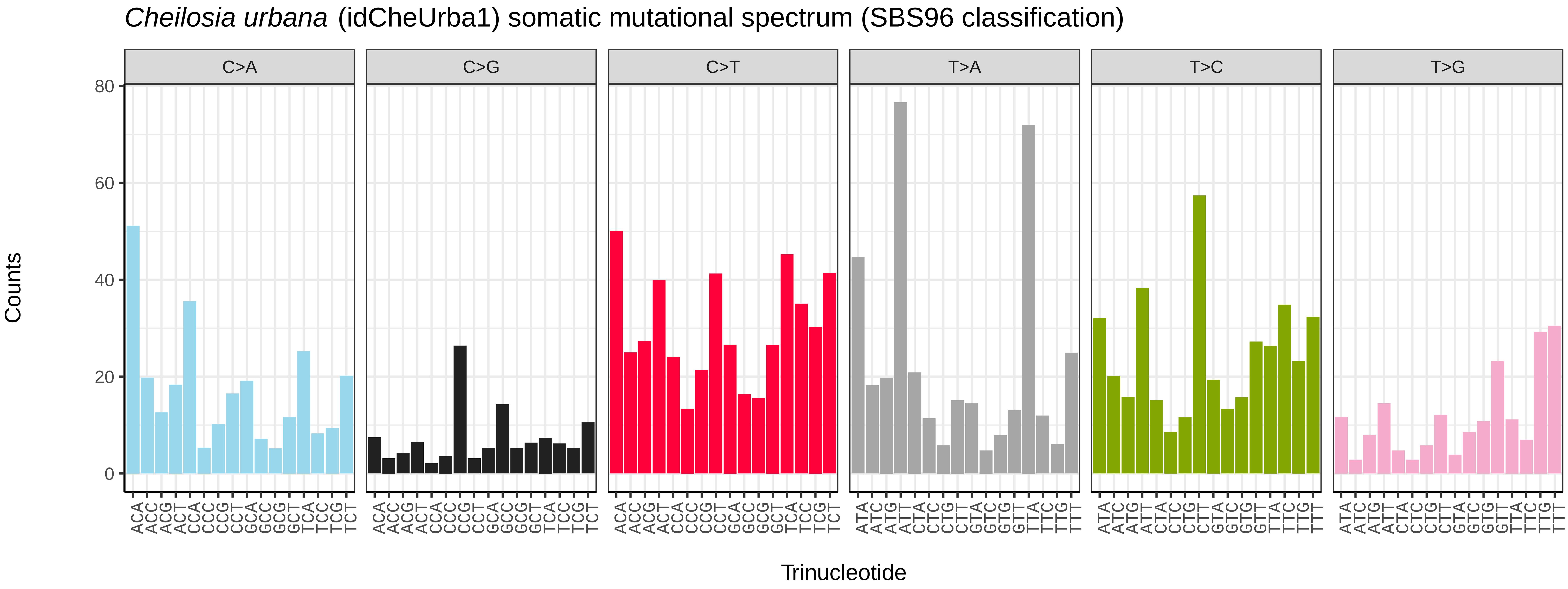
*Cheilosia urbana* (idCheUrba1) somatic mutational spectrum (SBS96 classification)
C>A
C>G
C>T
T>A
T>C
T>G
Counts
0
20
40
60
80
ACA ACC ACG ACT CCA CCC CCG CCT GCA GCC GCG GCT TCA TCC TCG TCT
ATA ATC ATG ATT CTA CTC CTG CTT GTA GTC GTG GTT TTA TTC TTG TTT
Trinucleotide

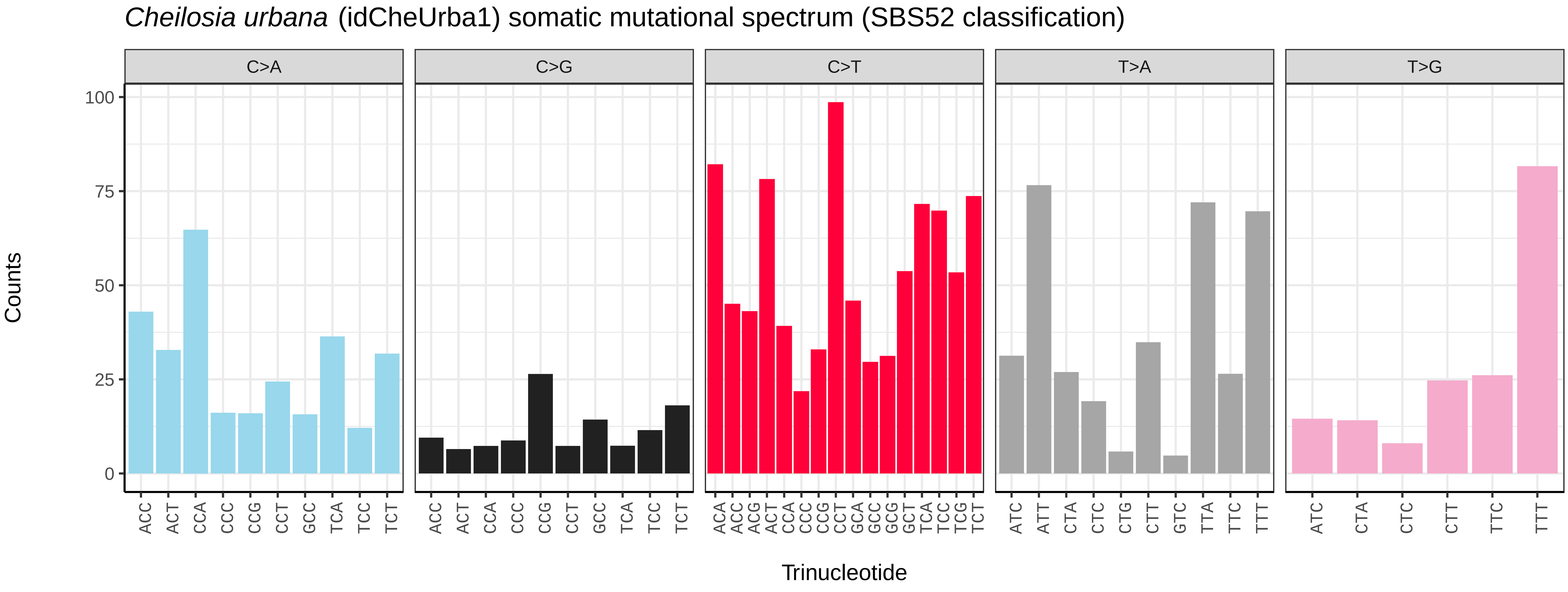
*Cheilosia urbana* (idCheUrba1) somatic mutational spectrum (SBS52 classification)
C>A
C>G
C>T
T>A
T>G
Counts
0
25
50
75
100
ACC ACT CCA CCC CCG CCT GCC TCA TCC TCT
ACA ACC ACG ACT CCA CCC CCG CCT GCA GCC GCG GCT TCA TCC TCG TCT
ATC ATT CTA CTC CTG CTT GTC TTA TTC TTT
ATC CTA CTC CTT TTC TTT
Trinucleotide

*Cheilosia vernalis* (idCheVern2) germline mutational spectrum

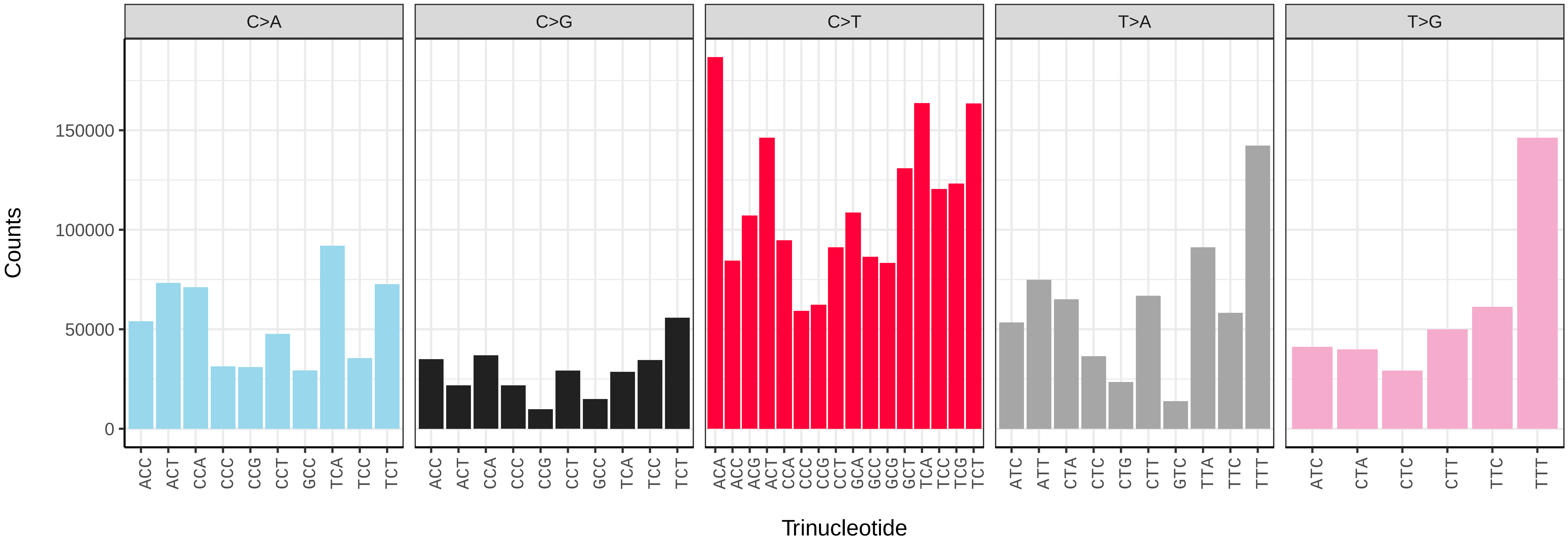


*Cheilosia vernalis* (idCheVern2) somatic mutational spectrum (SBS96 classification)

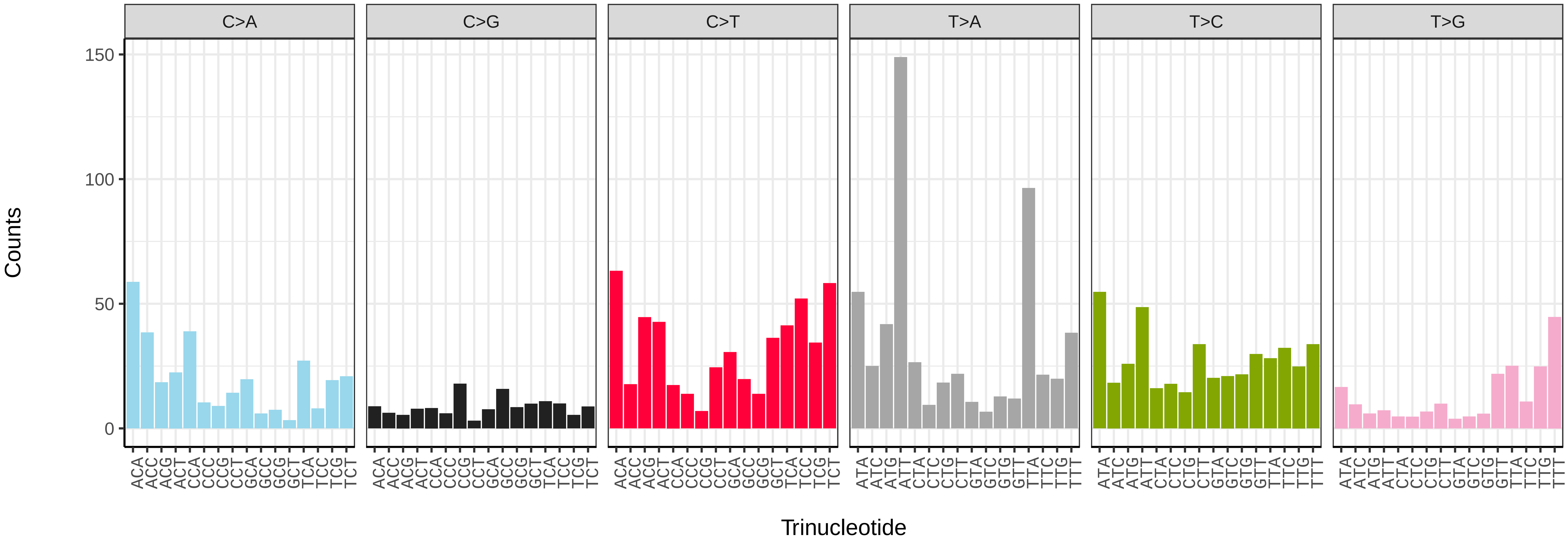


*Cheilosia vernalis* (idCheVern2) somatic mutational spectrum (SBS52 classification)

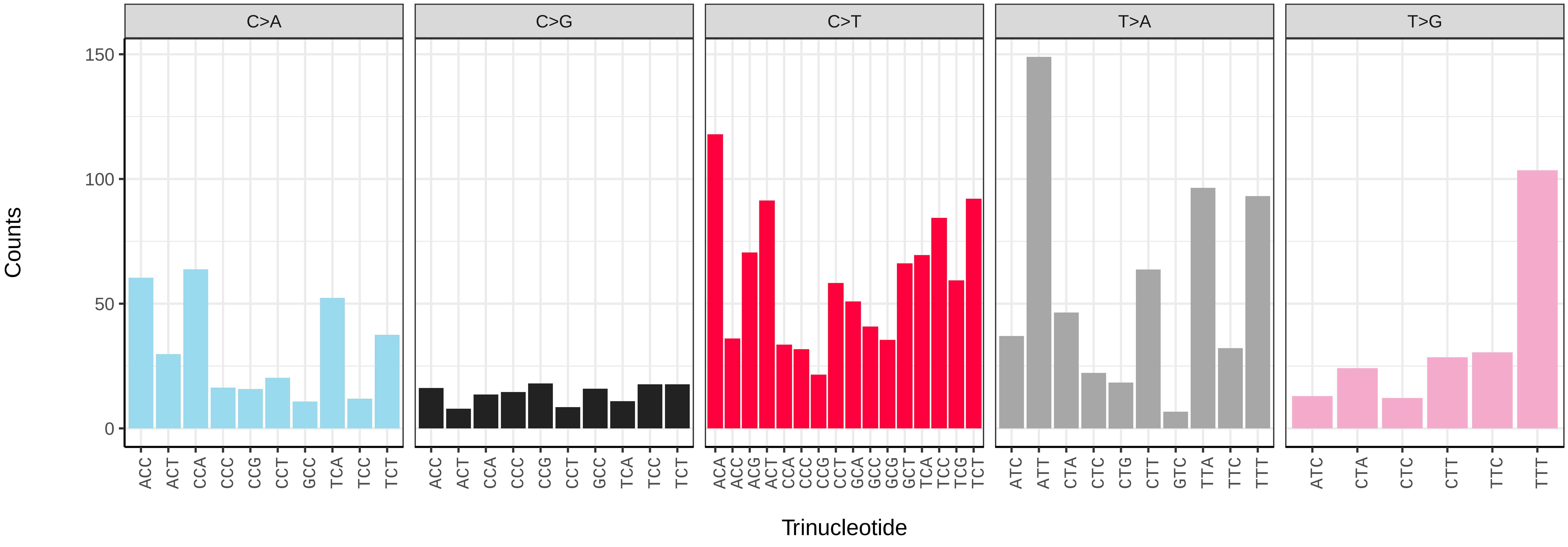

*Cheilosia vulpina* (idCheVulp2) germline mutational spectrum

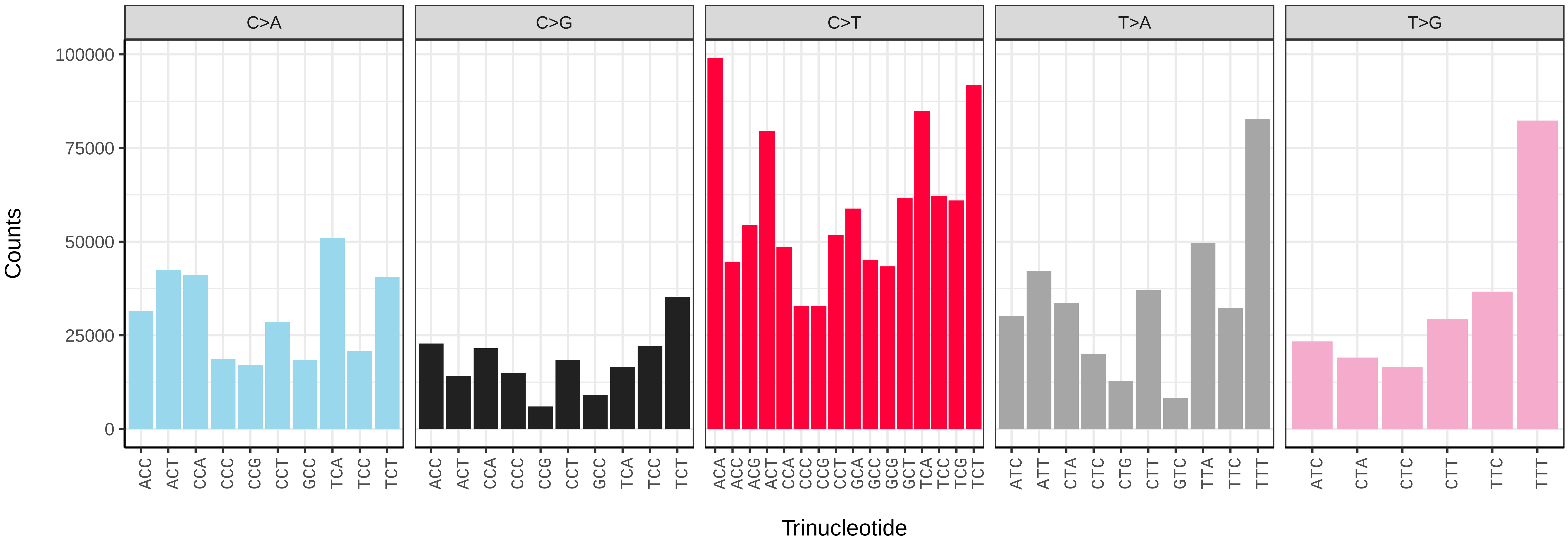


*Cheilosia vulpina* (idCheVulp2) somatic mutational spectrum (SBS96 classification)

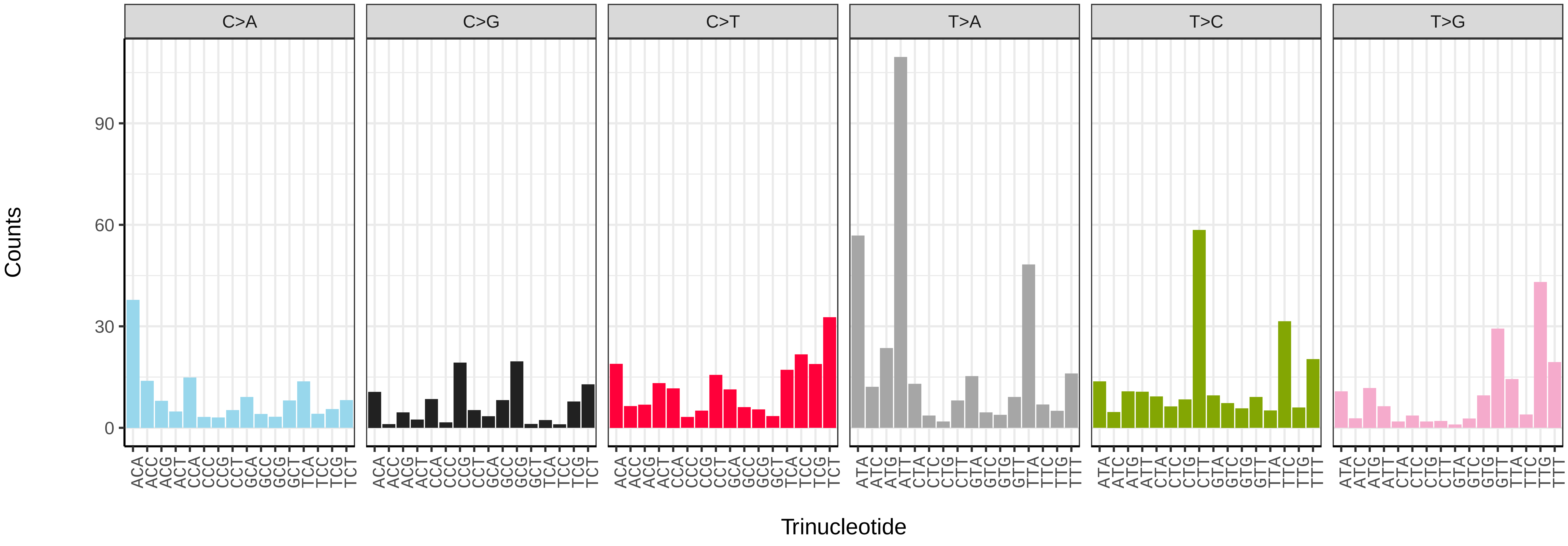


*Cheilosia vulpina* (idCheVulp2) somatic mutational spectrum (SBS52 classification)

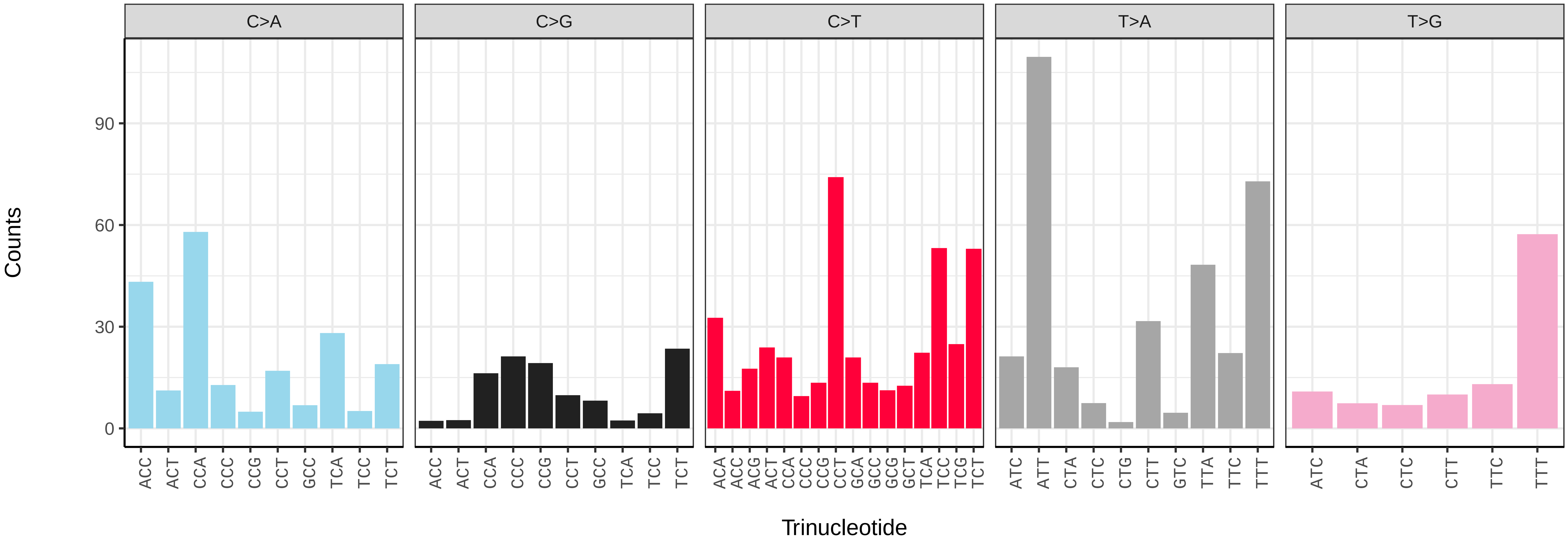

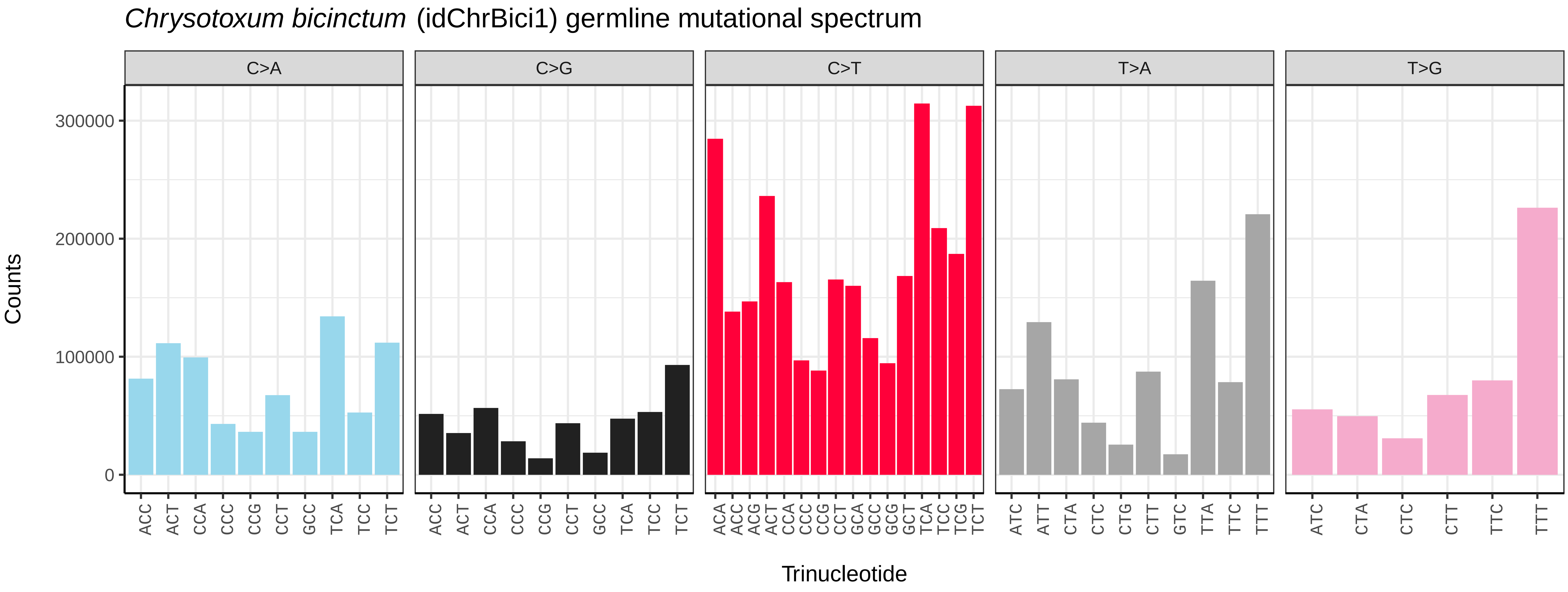
*Chrysotoxum bicinctum* (idChrBici1) germline mutational spectrum
C>A
C>G
C>T
T>A
T>G
Counts
0
100000
200000
300000
ACC ACT CCA CCC CCG CCT GCC TCA TCC TCT
ACC ACT CCA CCC CCG CCT GCC TCA TCC TCT
ACA ACC ACG ACT CCA CCC CCG CCT GCA GCC GCG GCT TCA TCC TCG TCT
ATC ATT CTA CTC CTG CTT GTC TTA TTC TTT
ATC CTA CTC CTT TTC TTT
Trinucleotide

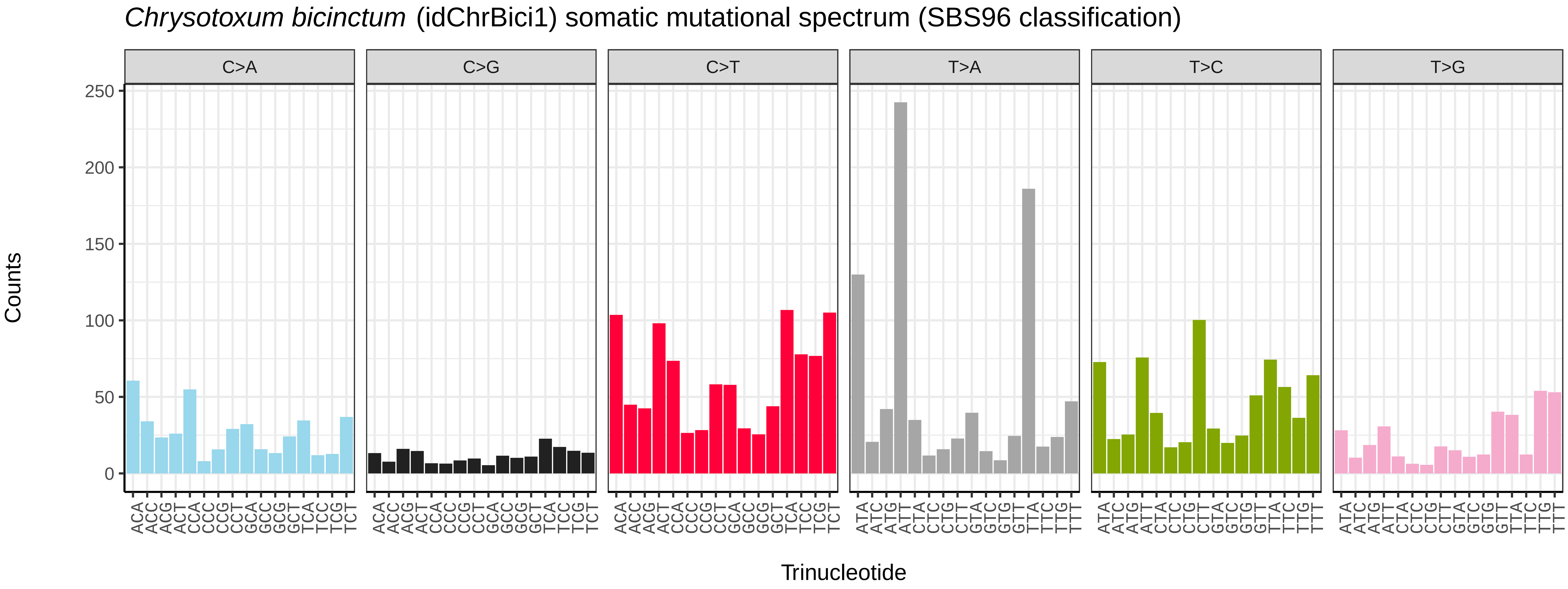
*Chrysotoxum bicinctum* (idChrBici1) somatic mutational spectrum (SBS96 classification)
C>A
C>G
C>T
T>A
T>C
T>G
Counts
0
50
100
150
200
250
ACA ACC ACG ACT CCA CCC CCG CCT GCA GCC GCG GCT TCA TCC TCG TCT
ATA ATC ATG ATT CTA CTC CTG CTT GTA GTC GTG GTT TTA TTC TTG TTT
Trinucleotide

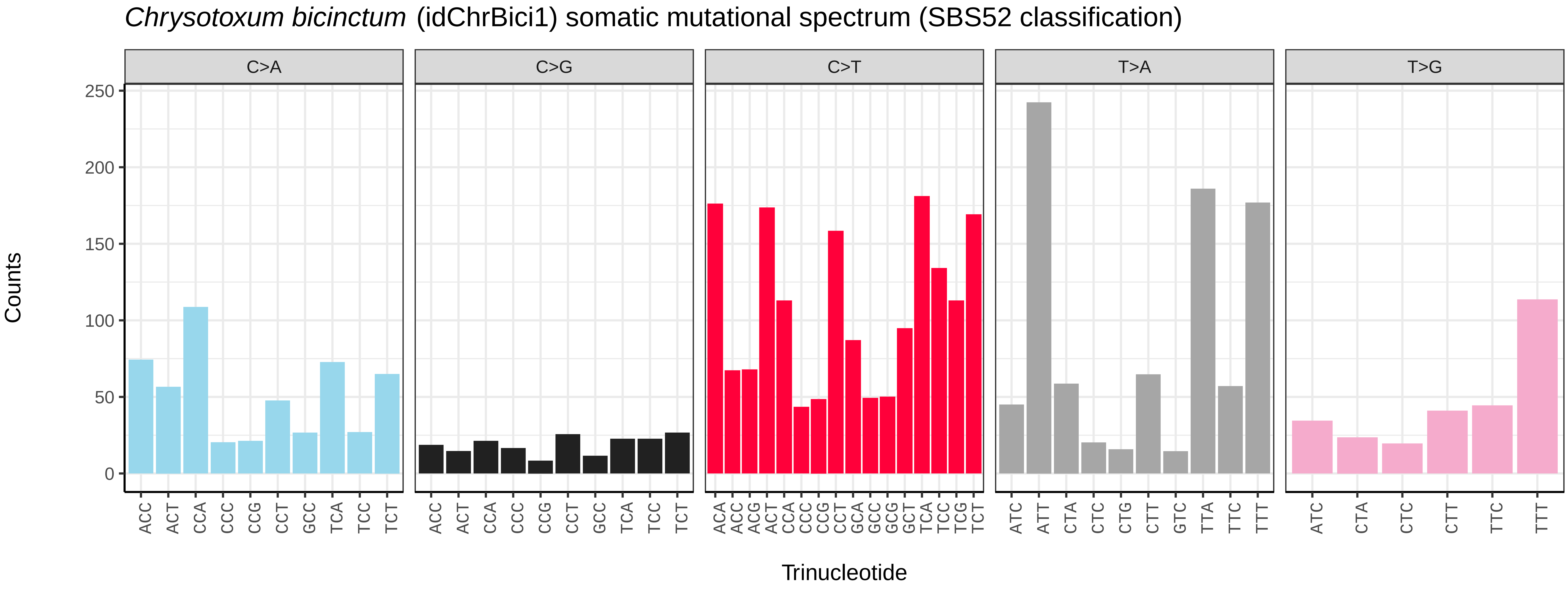
*Chrysotoxum bicinctum* (idChrBici1) somatic mutational spectrum (SBS52 classification)
C>A
C>G
C>T
T>A
T>G
Counts
0
50
100
150
200
250
ACC ACT CCA CCC CCG CCT GCC TCA TCC TCT
ACC ACT CCA CCC CCG CCT GCC TCA TCC TCT
ACA ACC ACG ACT CCA CCC CCG CCT GCA GCC GCG GCT TCA TCC TCG TCT
ATC ATT CTA CTC CTG CTT GTC TTA TTC TTT
ATC CTA CTC CTT TTC TTT
Trinucleotide

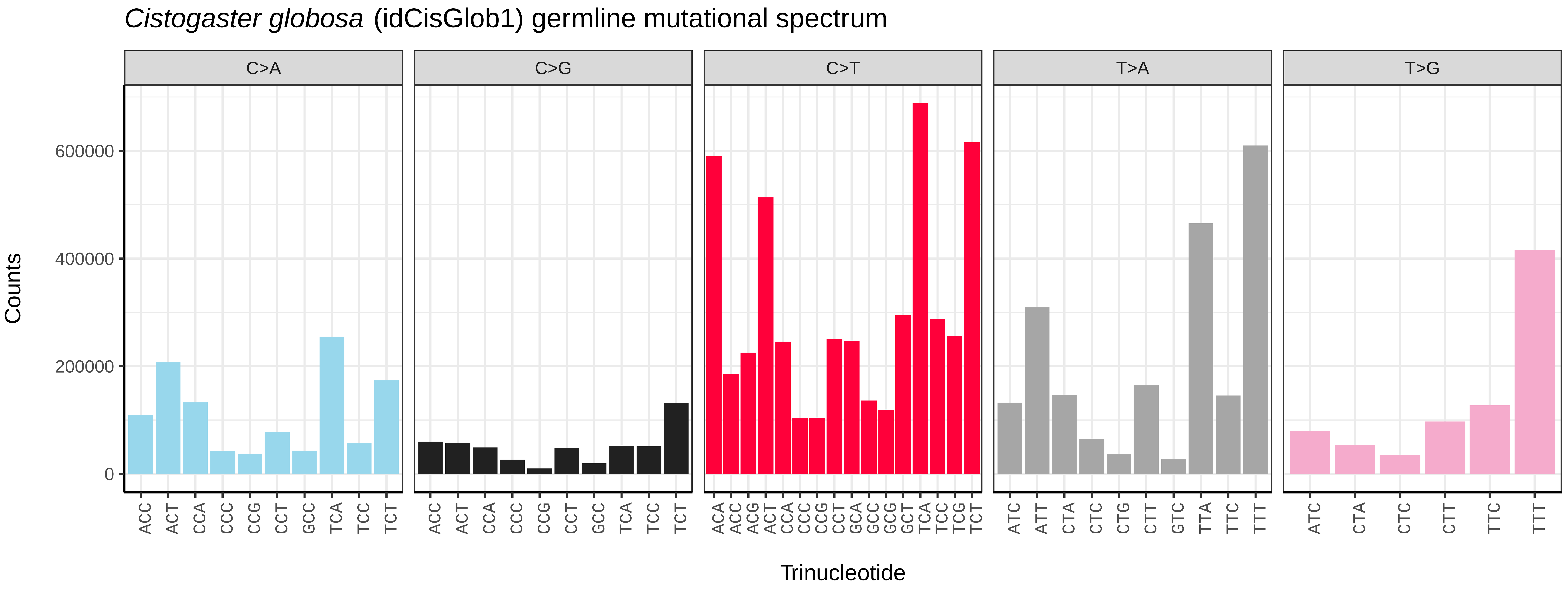

Cistogaster globosa (idCisGlob1) germline mutational spectrum
C>A
C>G
C>T
T>A
T>G
Counts
0
200000
400000
600000
Trinucleotide
ACC ACT CCA CCC CCG CCT GCC TCA TCC TCT
ACA ACC ACG ACT CCA CCC CCG CCT GCA GCC GCG GCT TCA TCC TCG TCT
ATC ATT CTA CTC CTG CTT GTC TTA TTC TTT
ATC CTA CTC CTT TTC TTT


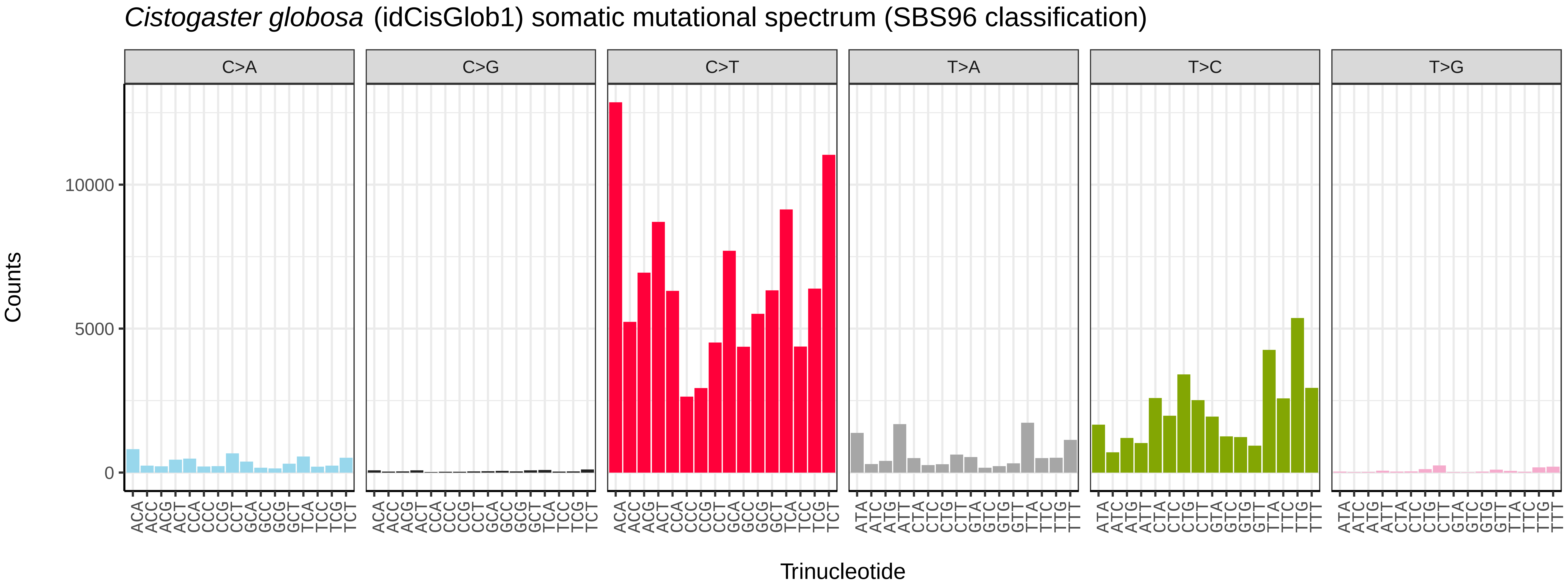

Cistogaster globosa (idCisGlob1) somatic mutational spectrum (SBS96 classification)
C>A
C>G
C>T
T>A
T>C
T>G
Counts
0
5000
10000
Trinucleotide
ACA ACC ACG ACT CCA CCC CCG CCT GCA GCC GCG GCT TCA TCC TCG TCT
ATA ATC ATG ATT CTA CTC CTG CTT GTA GTC GTG GTT TTA TTC TTG TTT


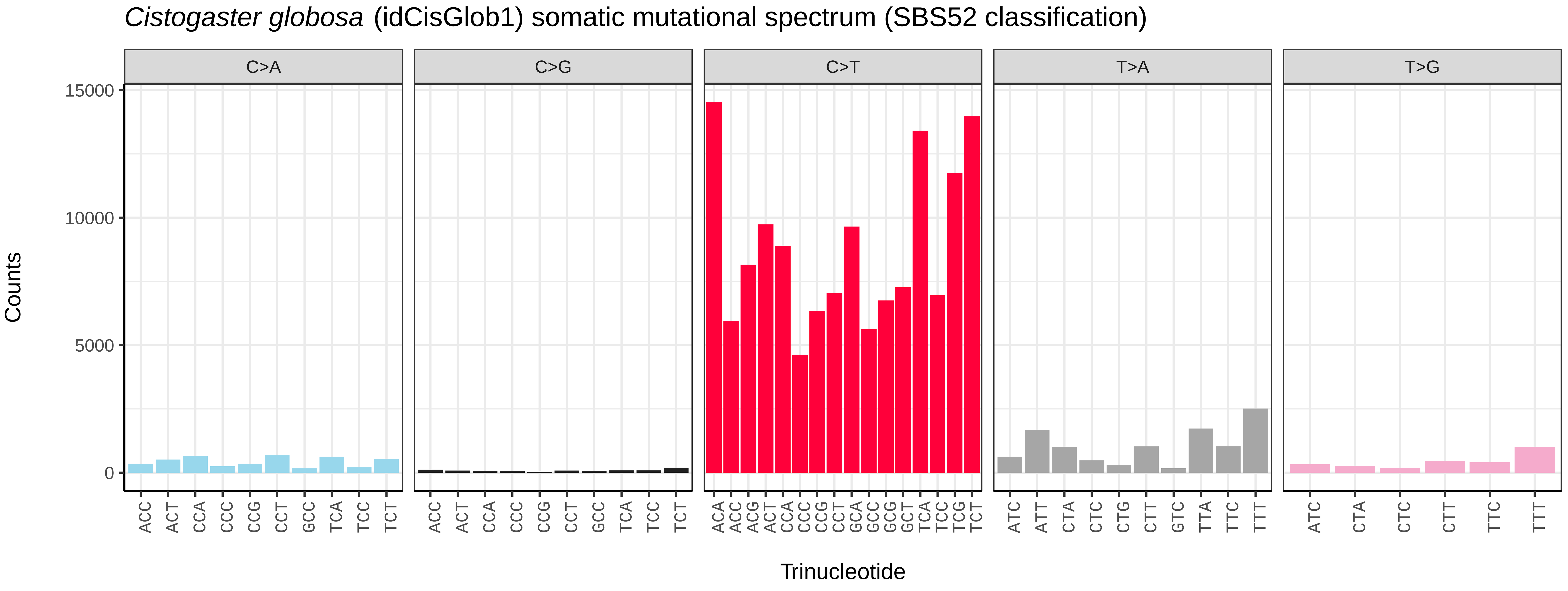

Cistogaster globosa (idCisGlob1) somatic mutational spectrum (SBS52 classification)
C>A
C>G
C>T
T>A
T>G
Counts
0
5000
10000
15000
Trinucleotide
ACC ACT CCA CCC CCG CCT GCC TCA TCC TCT
ACA ACC ACG ACT CCA CCC CCG CCT GCA GCC GCG GCT TCA TCC TCG TCT
ATC ATT CTA CTC CTG CTT GTC TTA TTC TTT
ATC CTA CTC CTT TTC TTT

*Clusia tigrina* (idCluTigr1) germline mutational spectrum

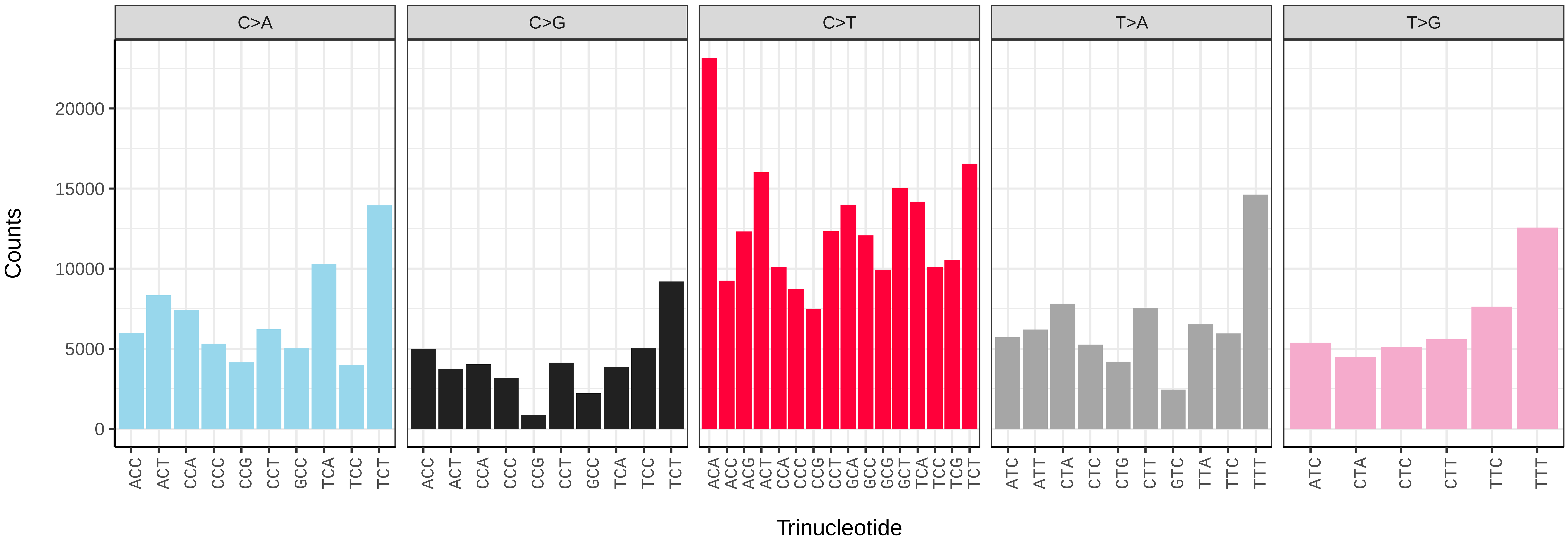


*Clusia tigrina* (idCluTigr1) somatic mutational spectrum (SBS96 classification)

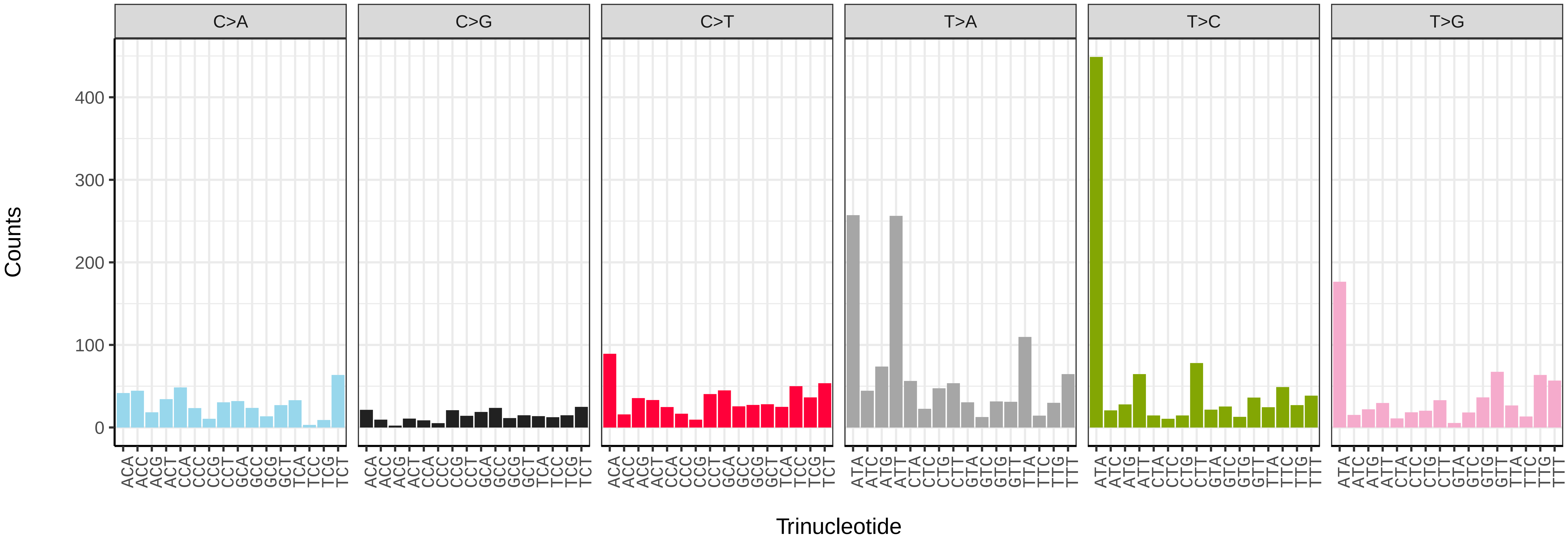


*Clusia tigrina* (idCluTigr1) somatic mutational spectrum (SBS52 classification)

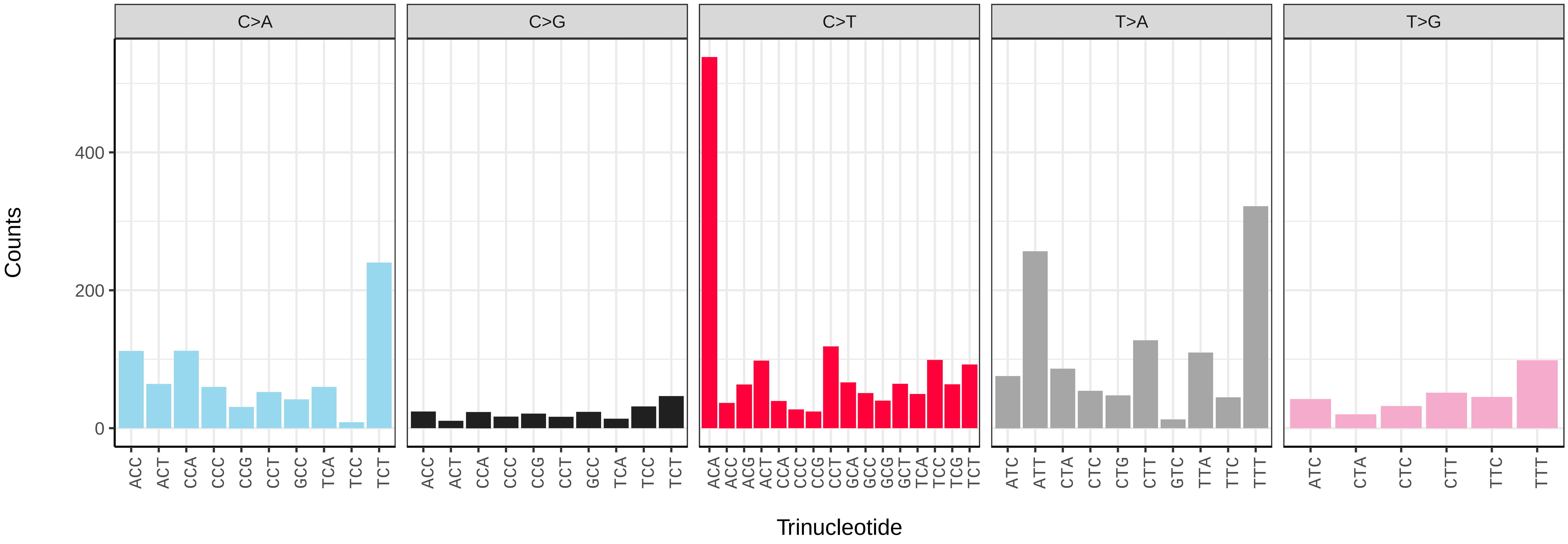

*Coelopa pilipes* (idCoePili4) germline mutational spectrum

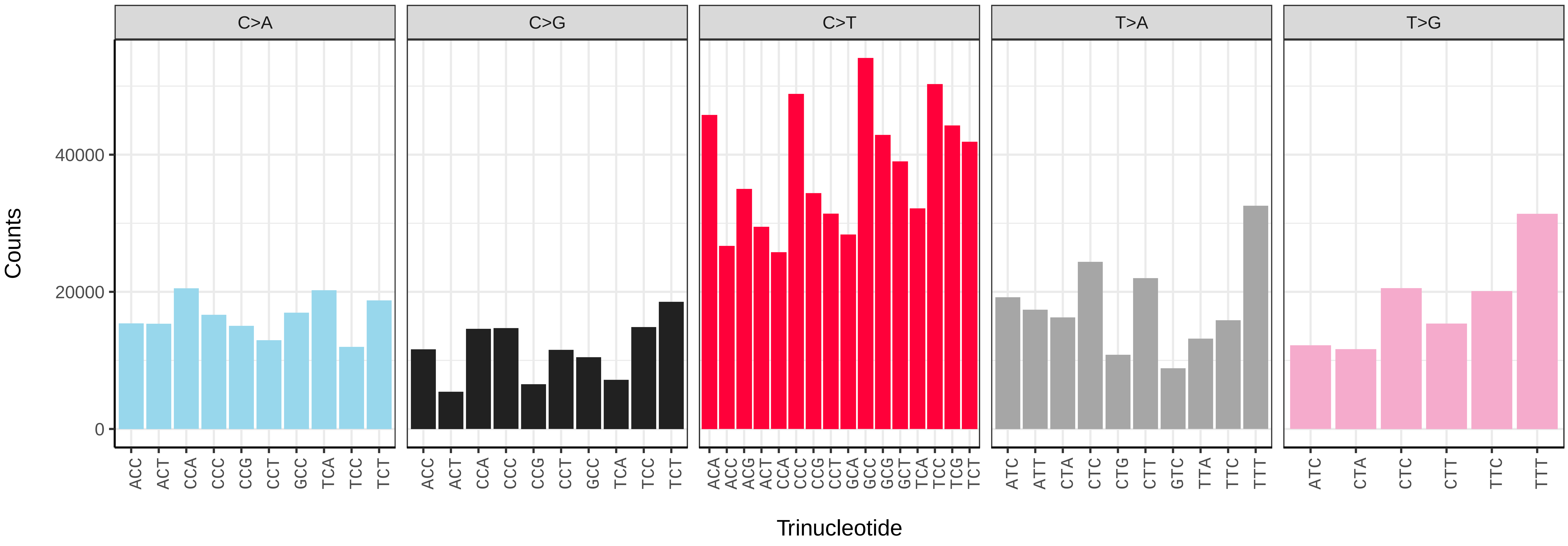


*Coelopa pilipes* (idCoePili4) somatic mutational spectrum (SBS96 classification)

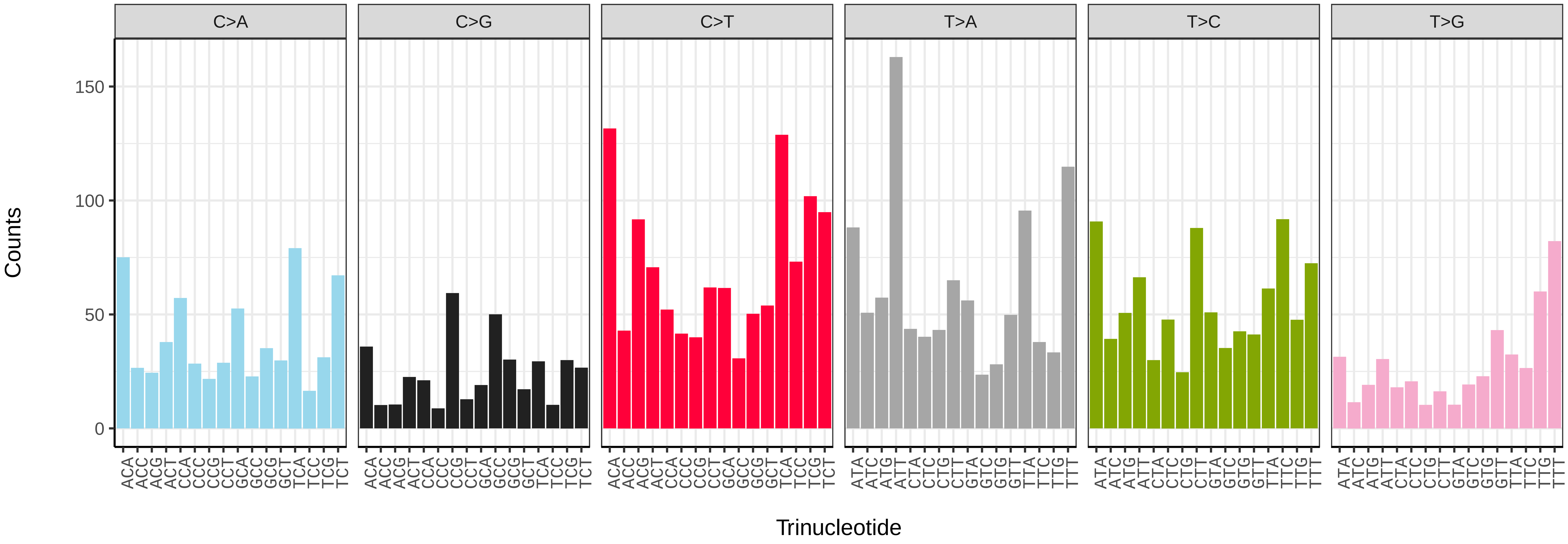


*Coelopa pilipes* (idCoePili4) somatic mutational spectrum (SBS52 classification)

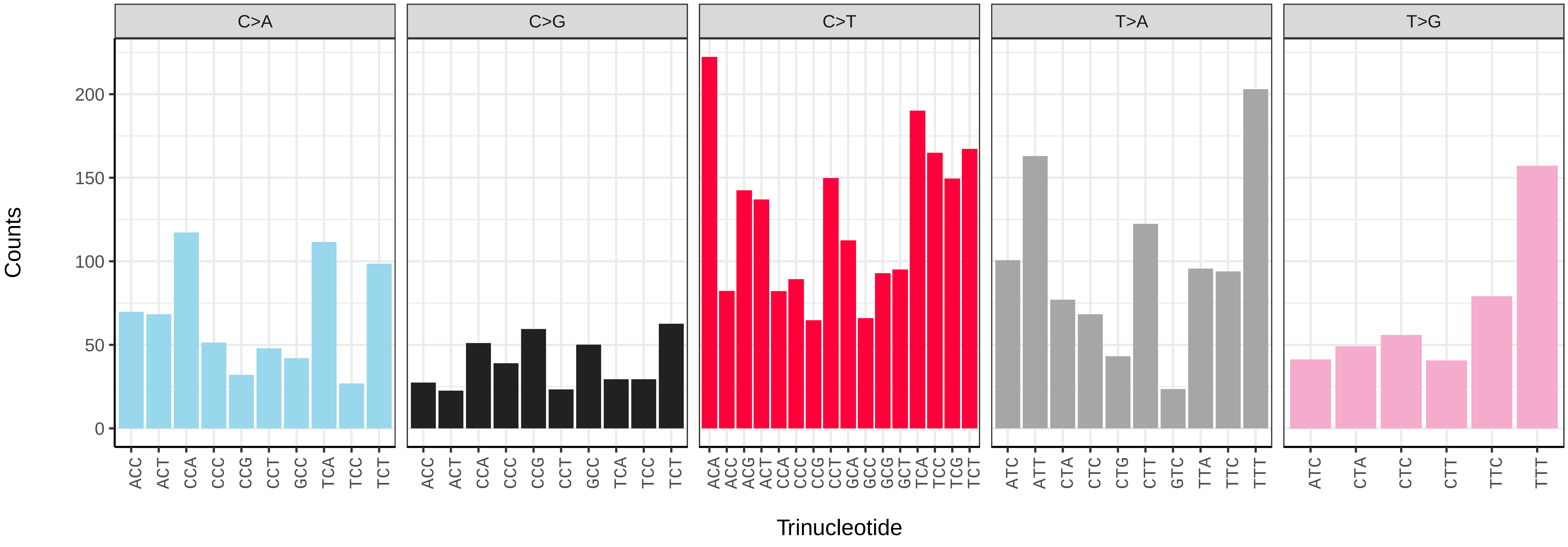

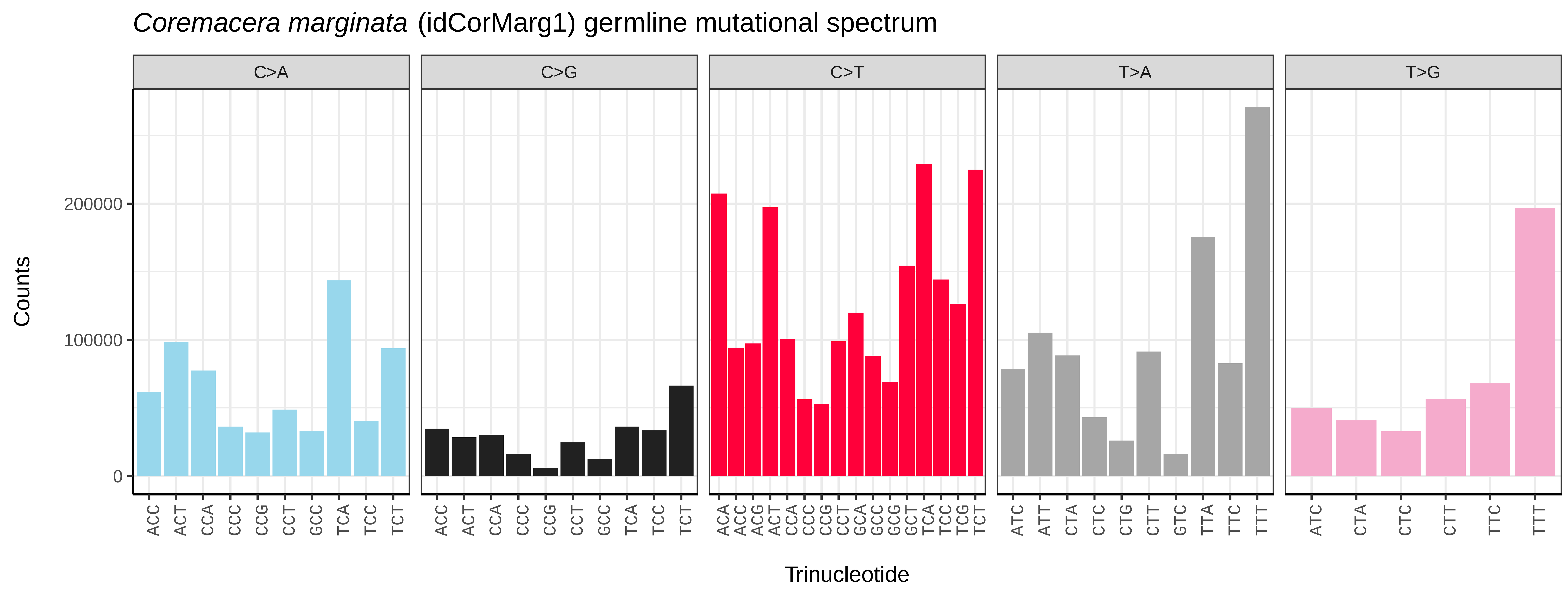
*Coremacera marginata* (idCorMarg1) germline mutational spectrum
C>A
C>G
C>T
T>A
T>G
Counts
200000
100000
0
Trinucleotide

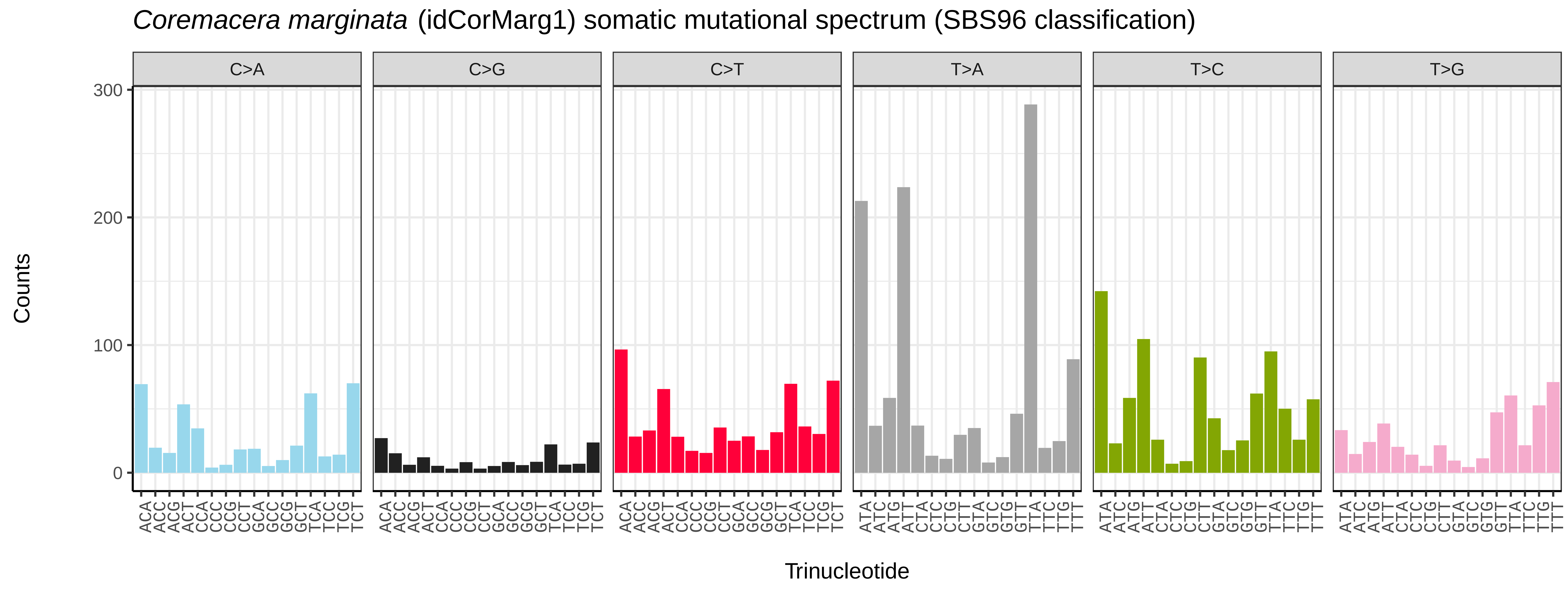
*Coremacera marginata* (idCorMarg1) somatic mutational spectrum (SBS96 classification)
C>A
C>G
C>T
T>A
T>C
T>G
Counts
300
200
100
0
Trinucleotide

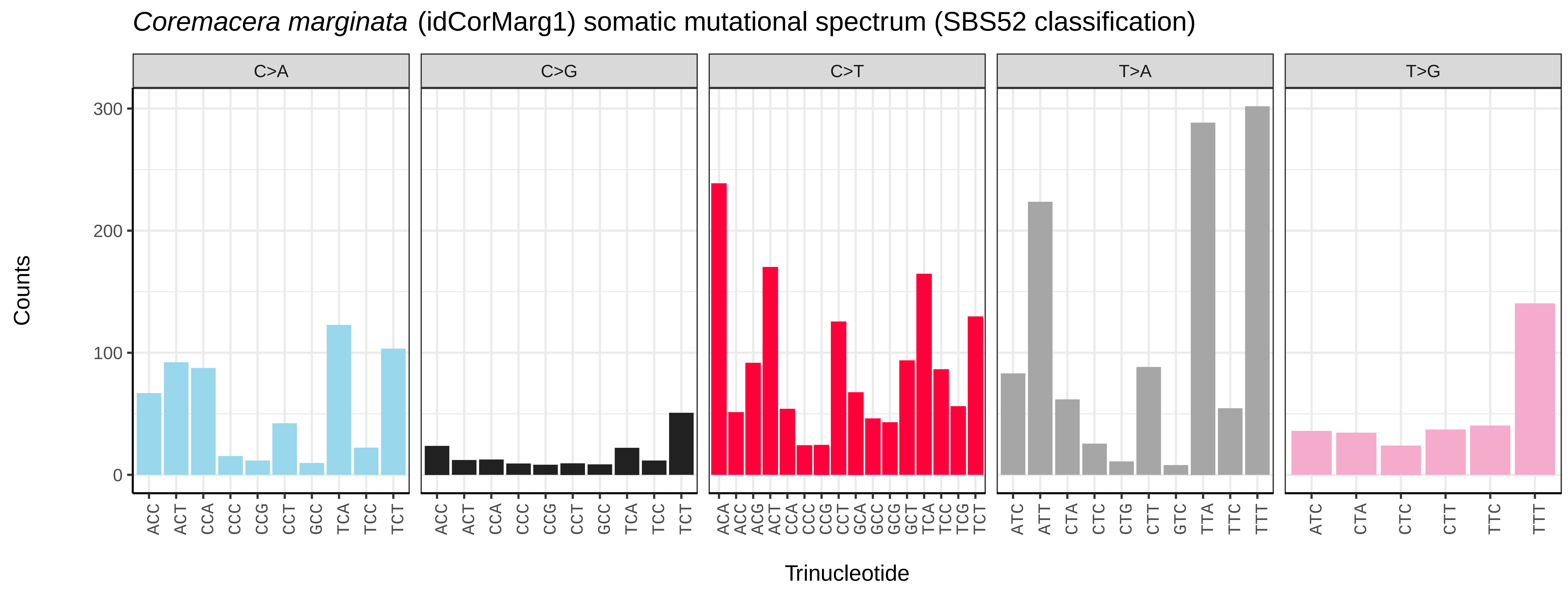
*Coremacera marginata* (idCorMarg1) somatic mutational spectrum (SBS52 classification)
C>A
C>G
C>T
T>A
T>G
Counts
300
200
100
0
Trinucleotide

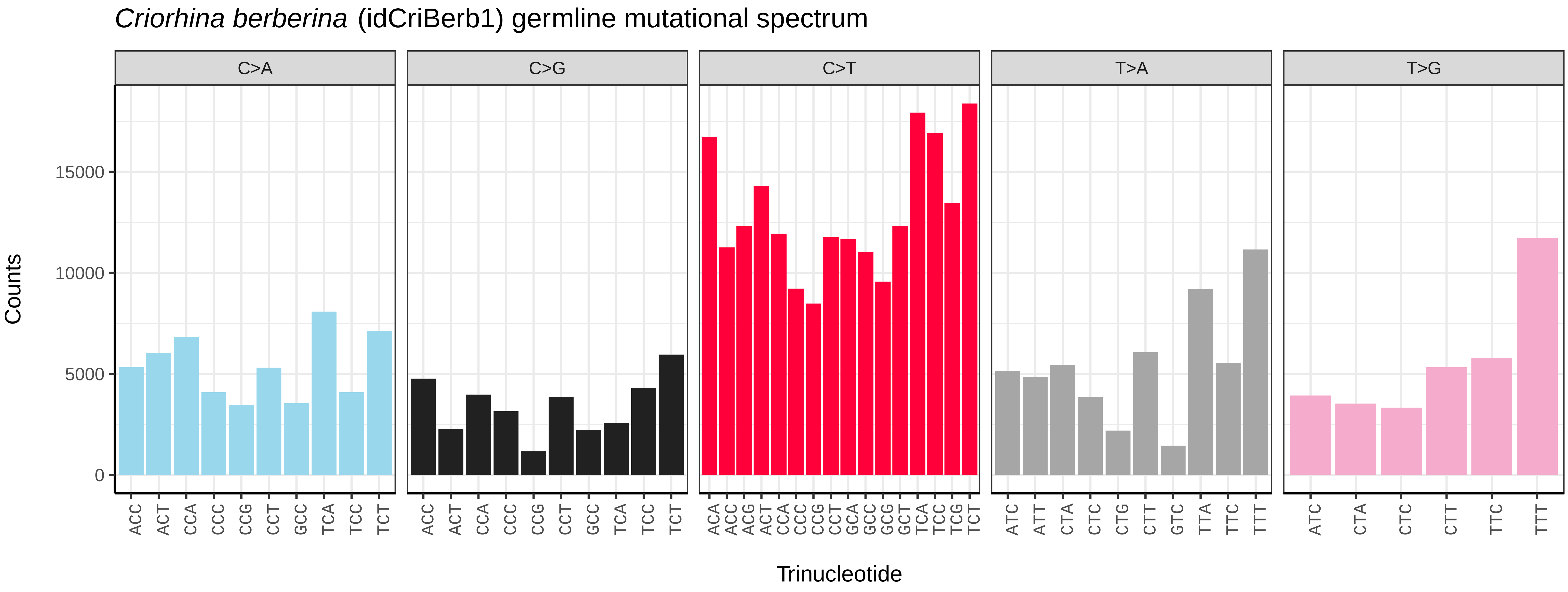
Criorhina berberina (idCriBerb1) germline mutational spectrum
C>A
C>G
C>T
T>A
T>G
Counts
Trinucleotide
0
5000
10000
15000

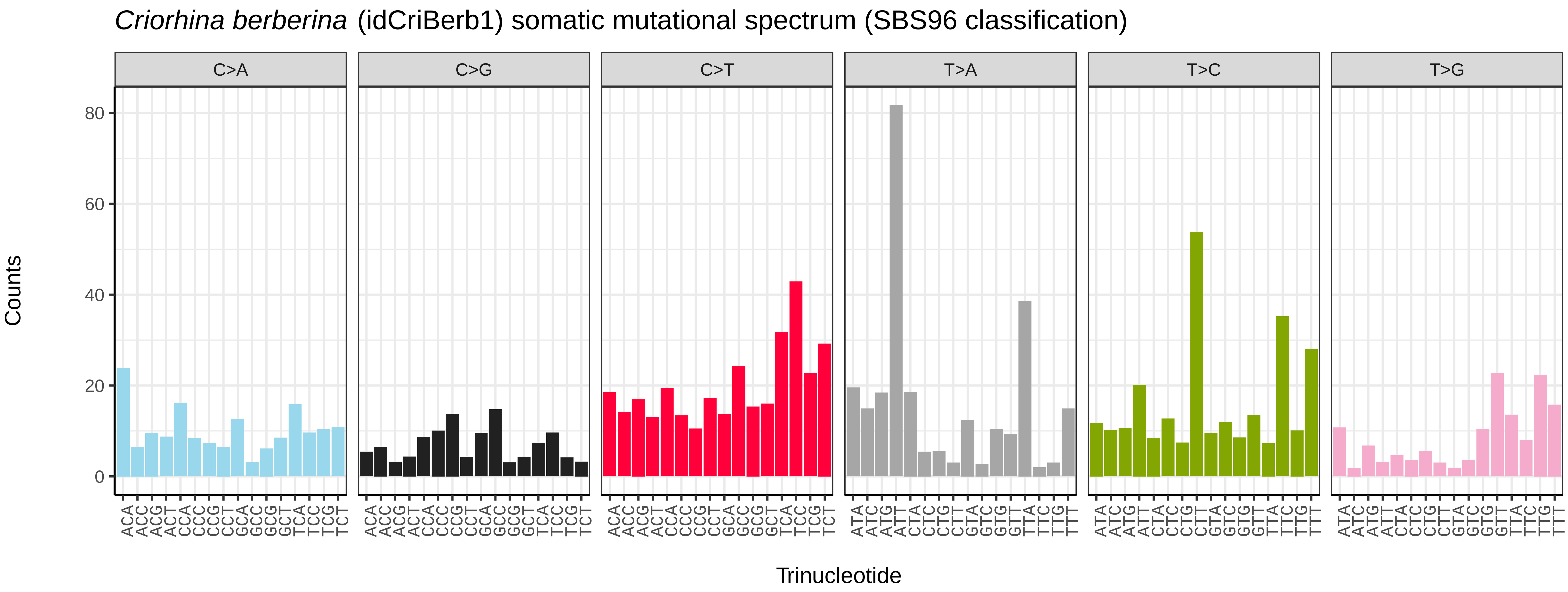
Criorhina berberina (idCriBerb1) somatic mutational spectrum (SBS96 classification)
C>A
C>G
C>T
T>A
T>C
T>G
Counts
Trinucleotide
0
20
40
60
80

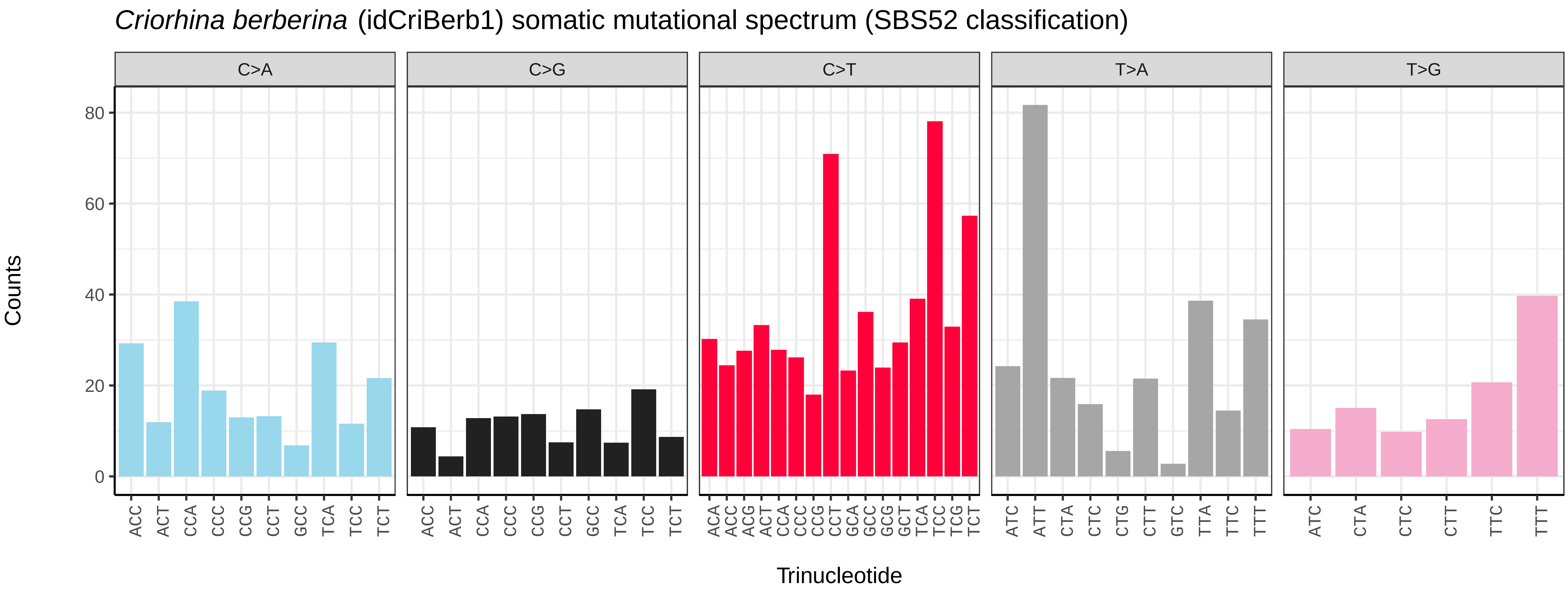
Criorhina berberina (idCriBerb1) somatic mutational spectrum (SBS52 classification)
C>A
C>G
C>T
T>A
T>G
Counts
Trinucleotide
0
20
40
60
80

*Dasysyrphus albostriatus* (idDasAlbo1) germline mutational spectrum

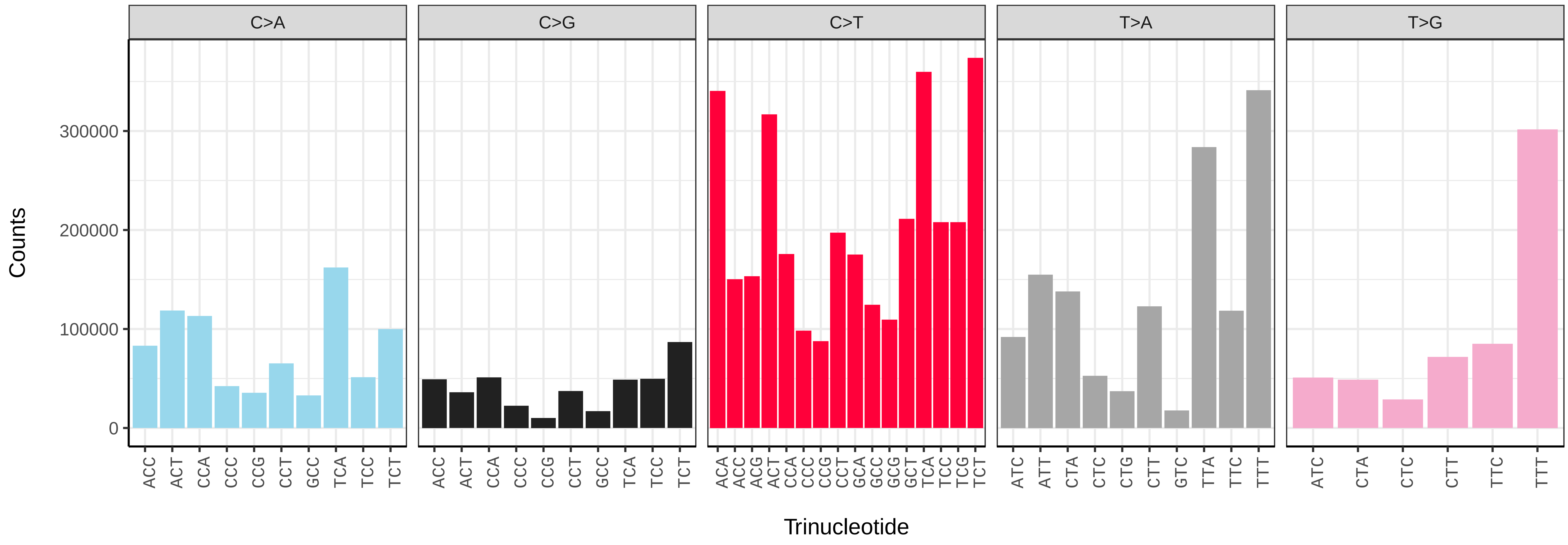


*Dasysyrphus albostriatus* (idDasAlbo1) somatic mutational spectrum (SBS96 classification)

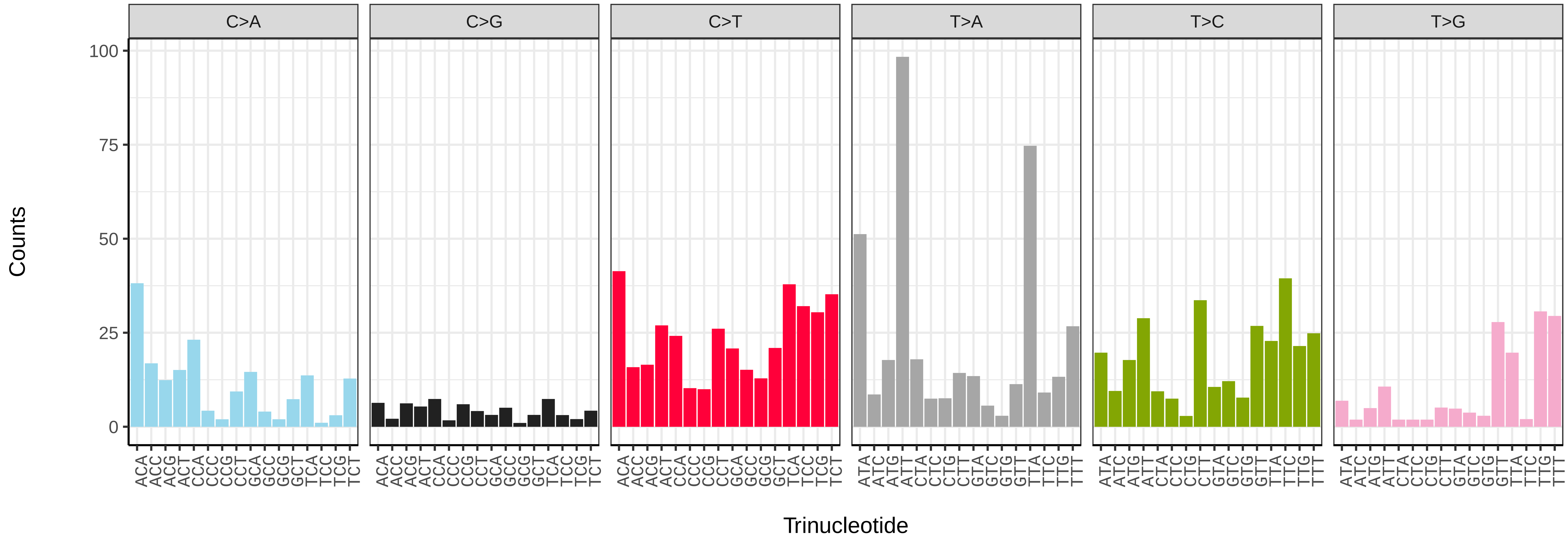


*Dasysyrphus albostriatus* (idDasAlbo1) somatic mutational spectrum (SBS52 classification)

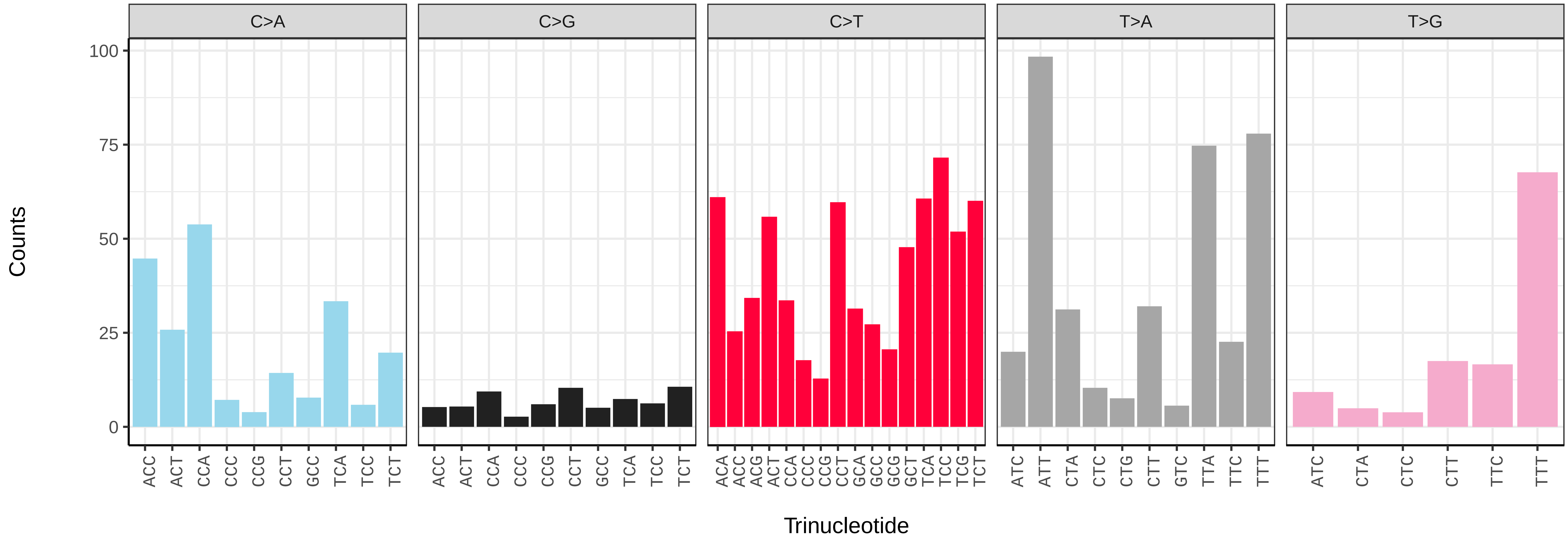

*Episyrphus balteatus* (idEpiBalt1) germline mutational spectrum

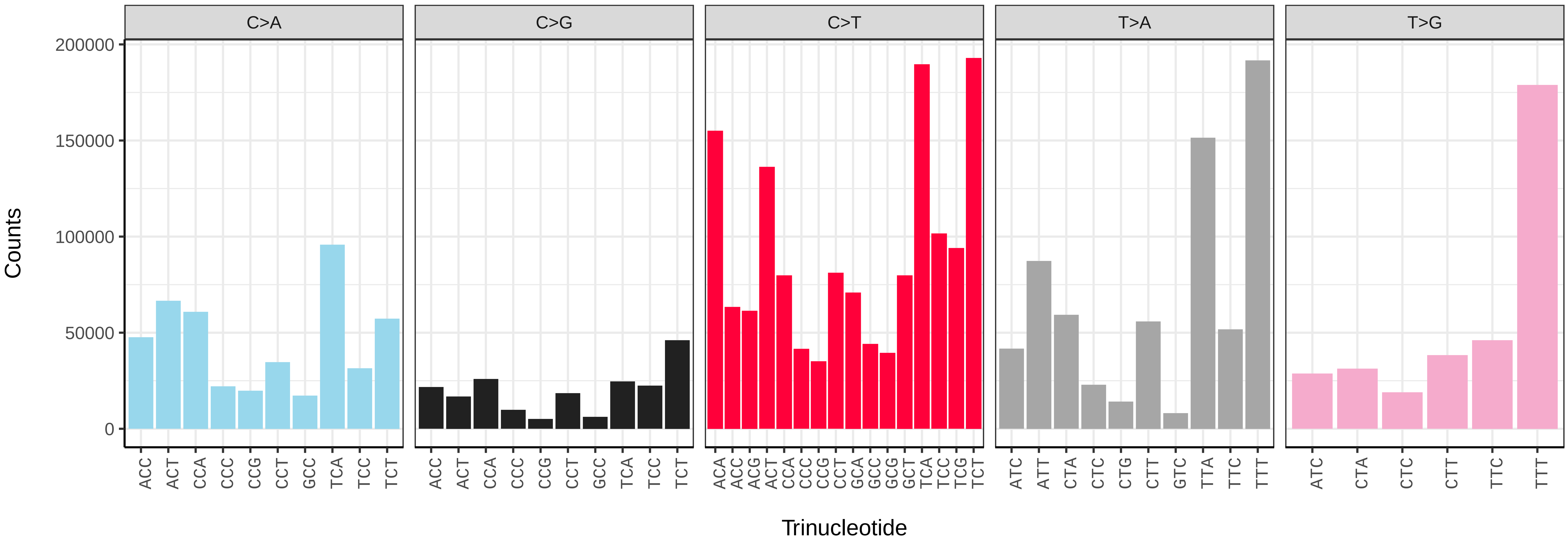


*Episyrphus balteatus* (idEpiBalt1) somatic mutational spectrum (SBS96 classification)

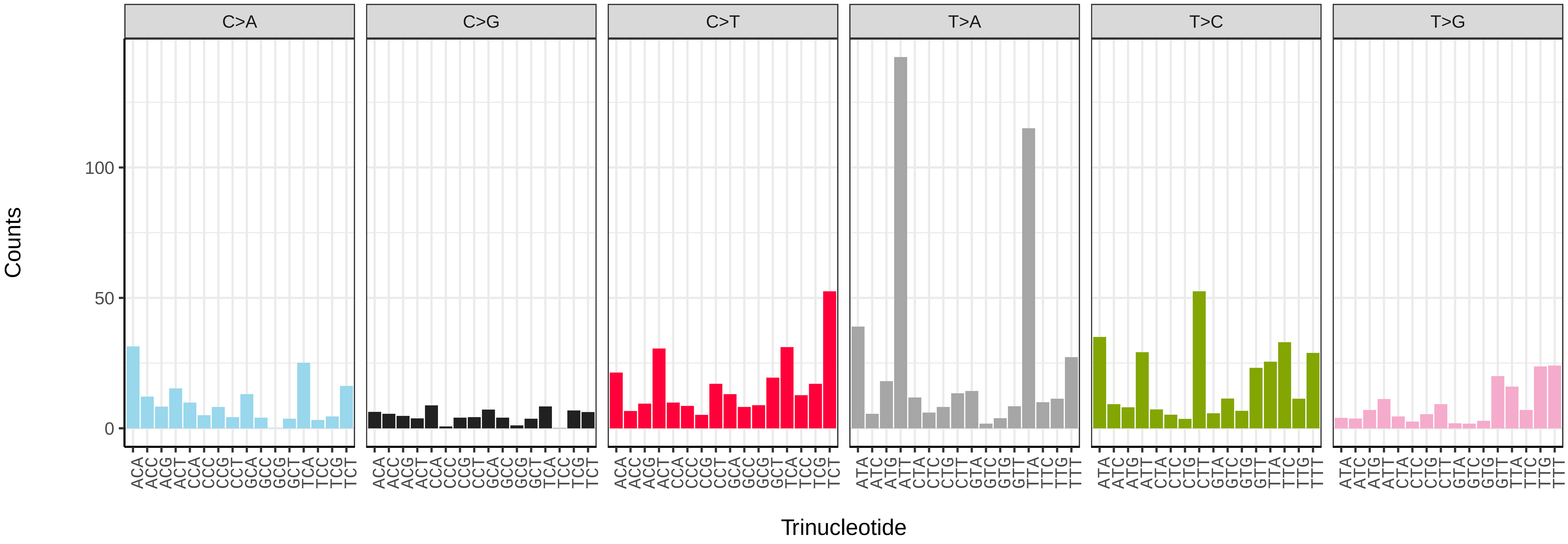


*Episyrphus balteatus* (idEpiBalt1) somatic mutational spectrum (SBS52 classification)

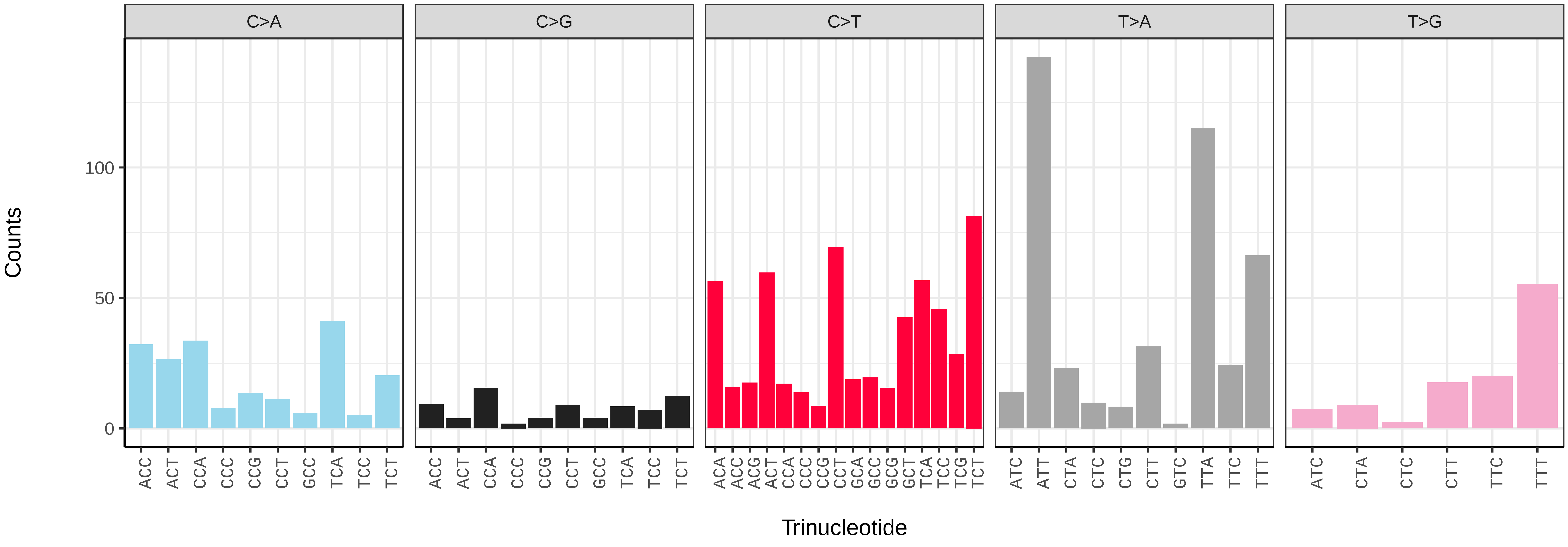

*Epistrophella euchroma* (idEpiEuco1) germline mutational spectrum

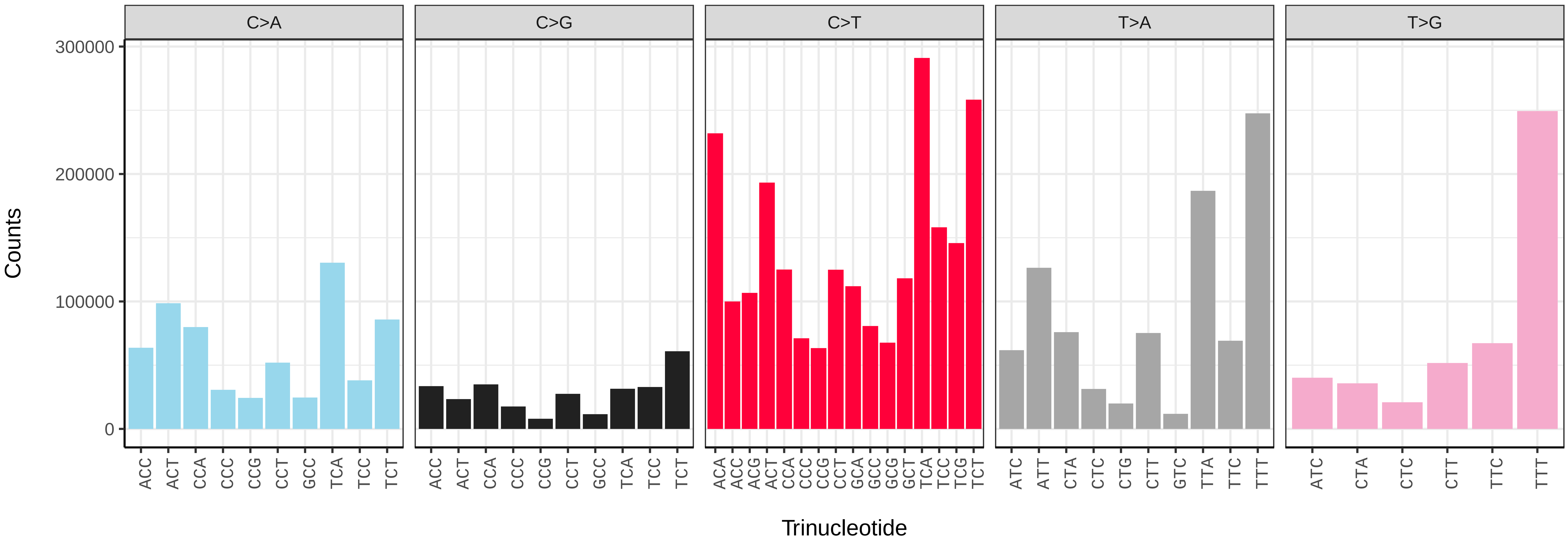


*Epistrophella euchroma* (idEpiEuco1) somatic mutational spectrum (SBS96 classification)

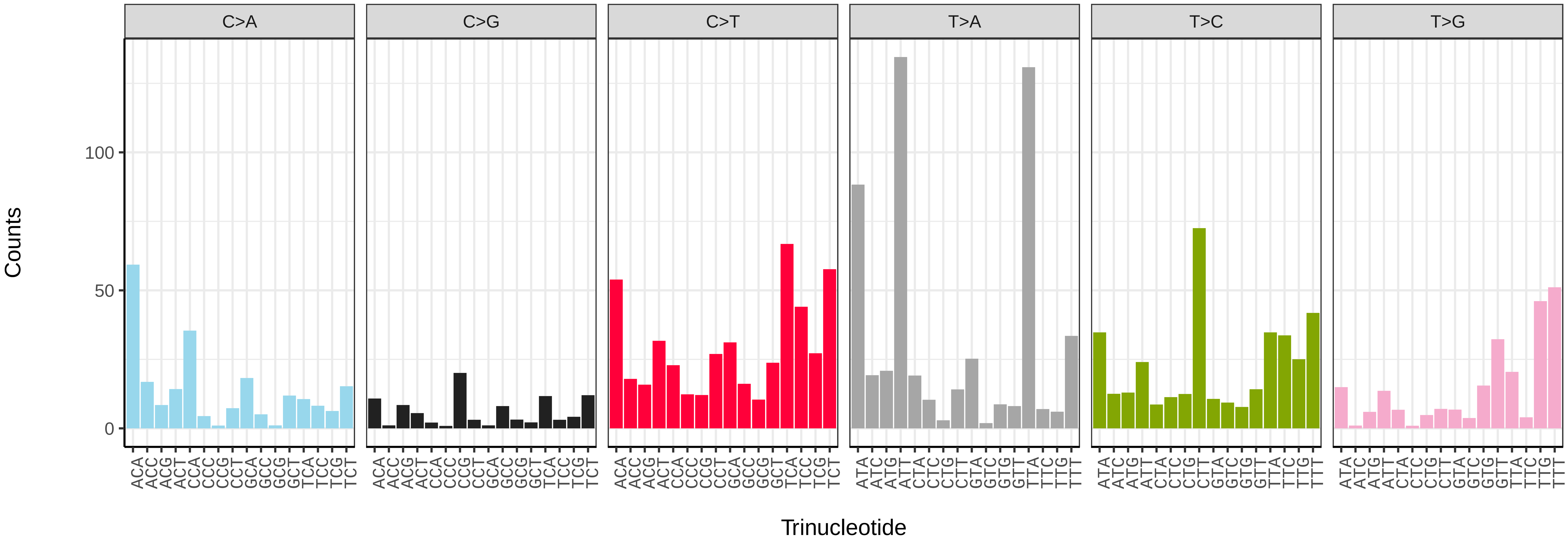


*Epistrophella euchroma* (idEpiEuco1) somatic mutational spectrum (SBS52 classification)

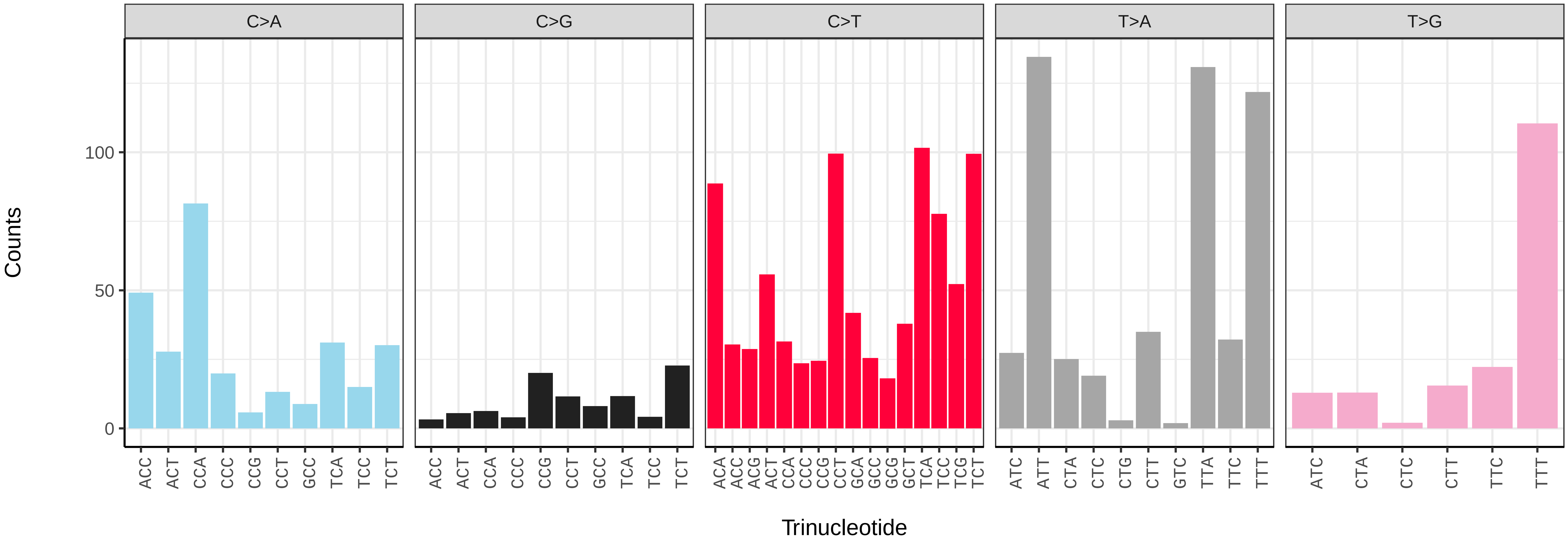

*Epistrophe grossulariae* (idEpiGros1) germline mutational spectrum

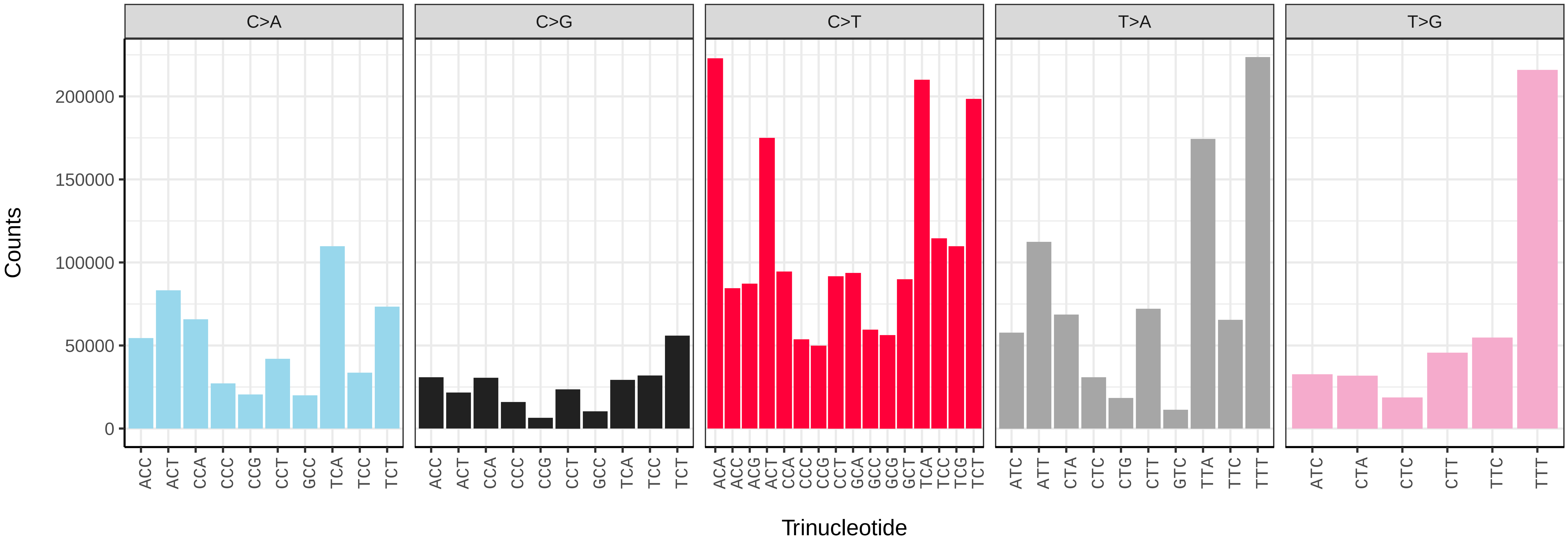


*Epistrophe grossulariae* (idEpiGros1) somatic mutational spectrum (SBS96 classification)

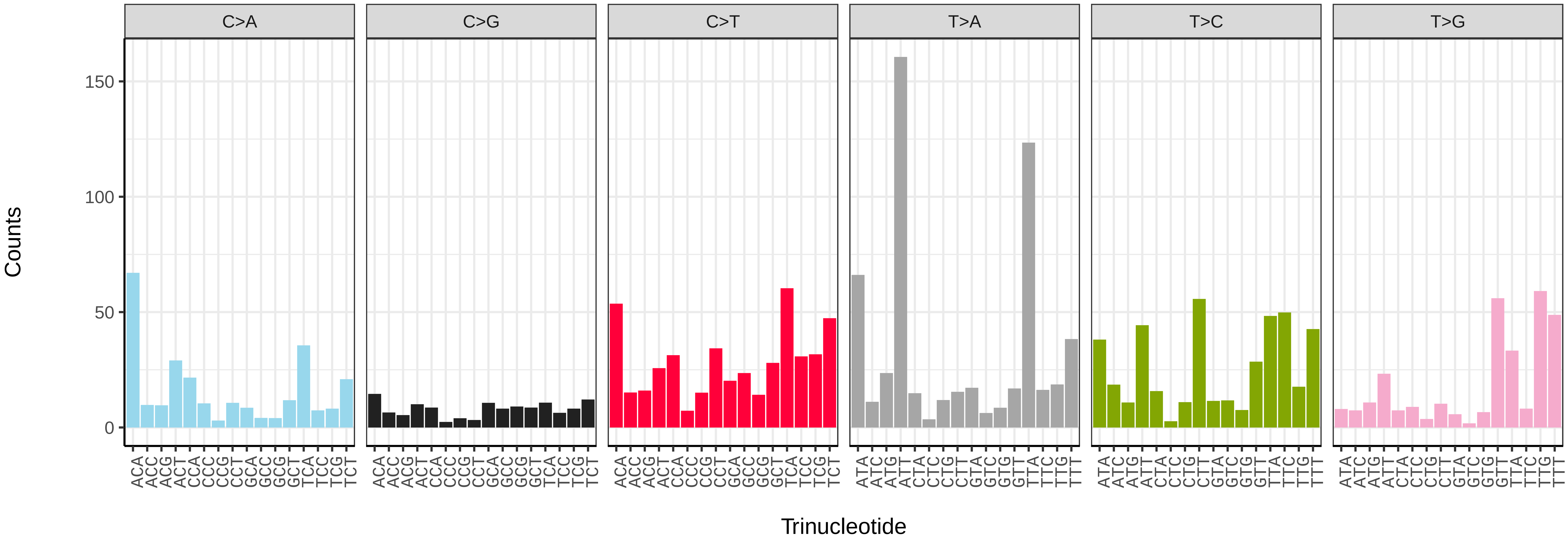


*Epistrophe grossulariae* (idEpiGros1) somatic mutational spectrum (SBS52 classification)

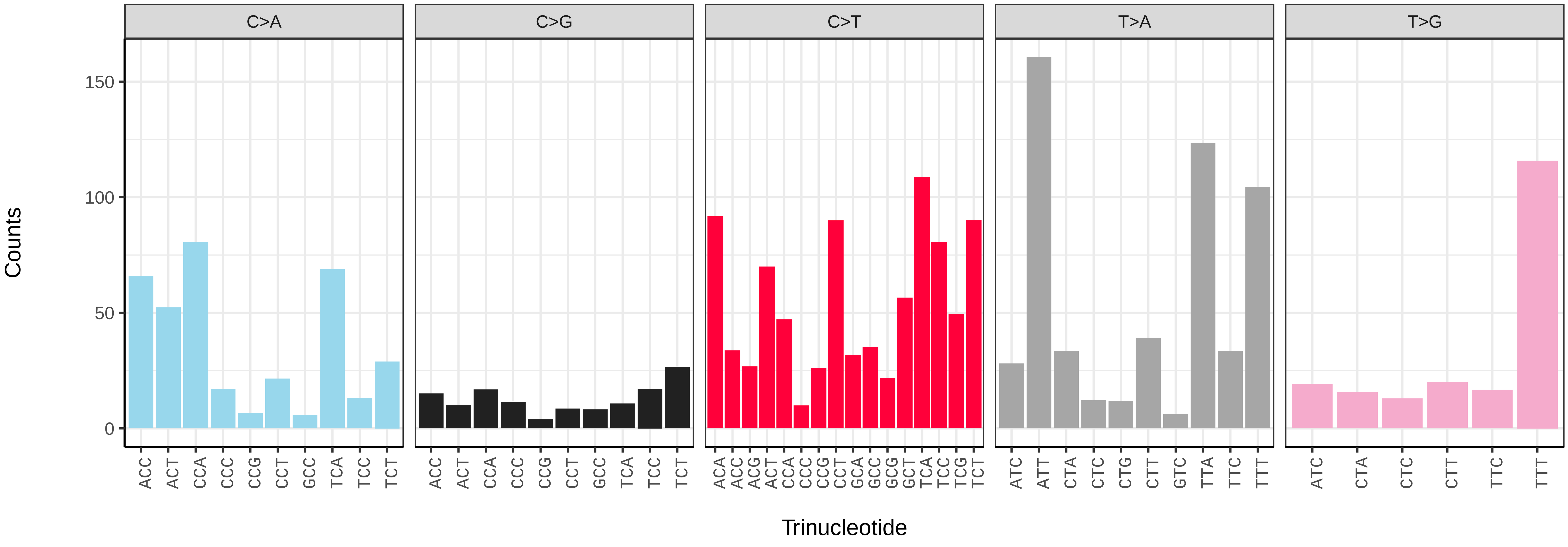

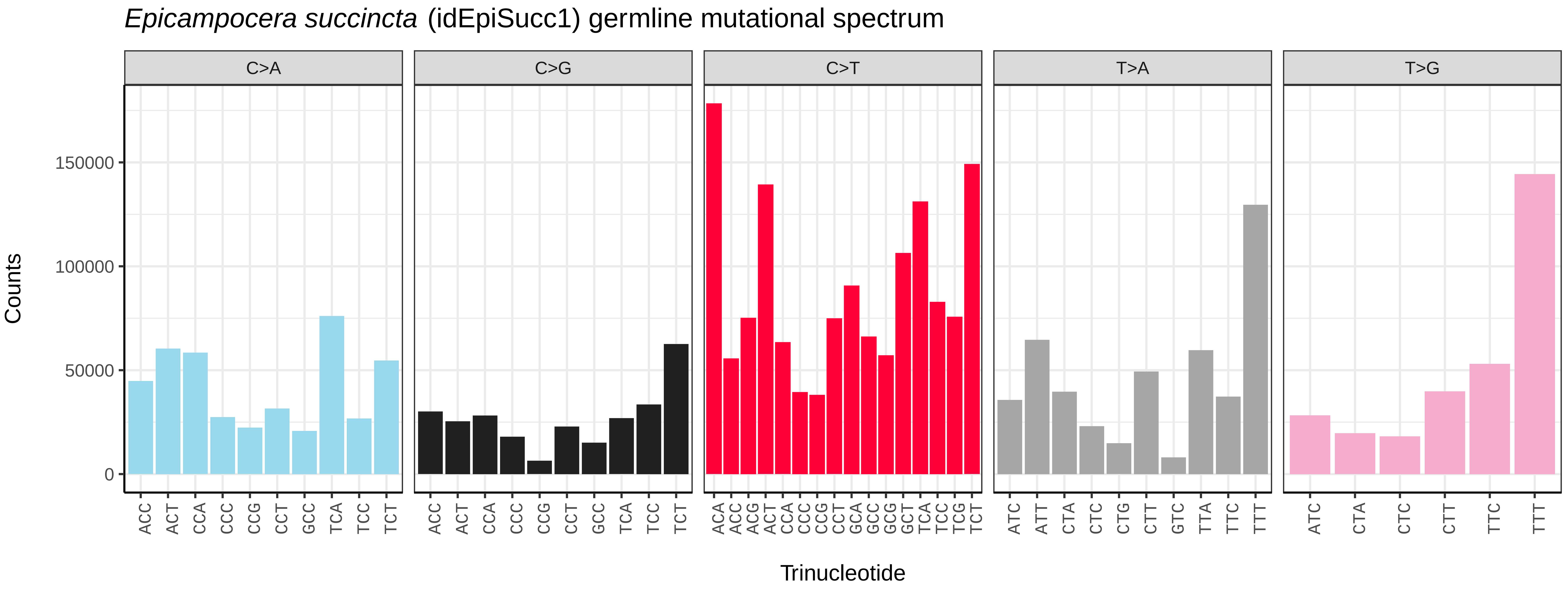
*Epicampocera succincta* (idEpiSucc1) germline mutational spectrum
C>A
C>G
C>T
T>A
T>G
Counts
150000
100000
50000
0
Trinucleotide

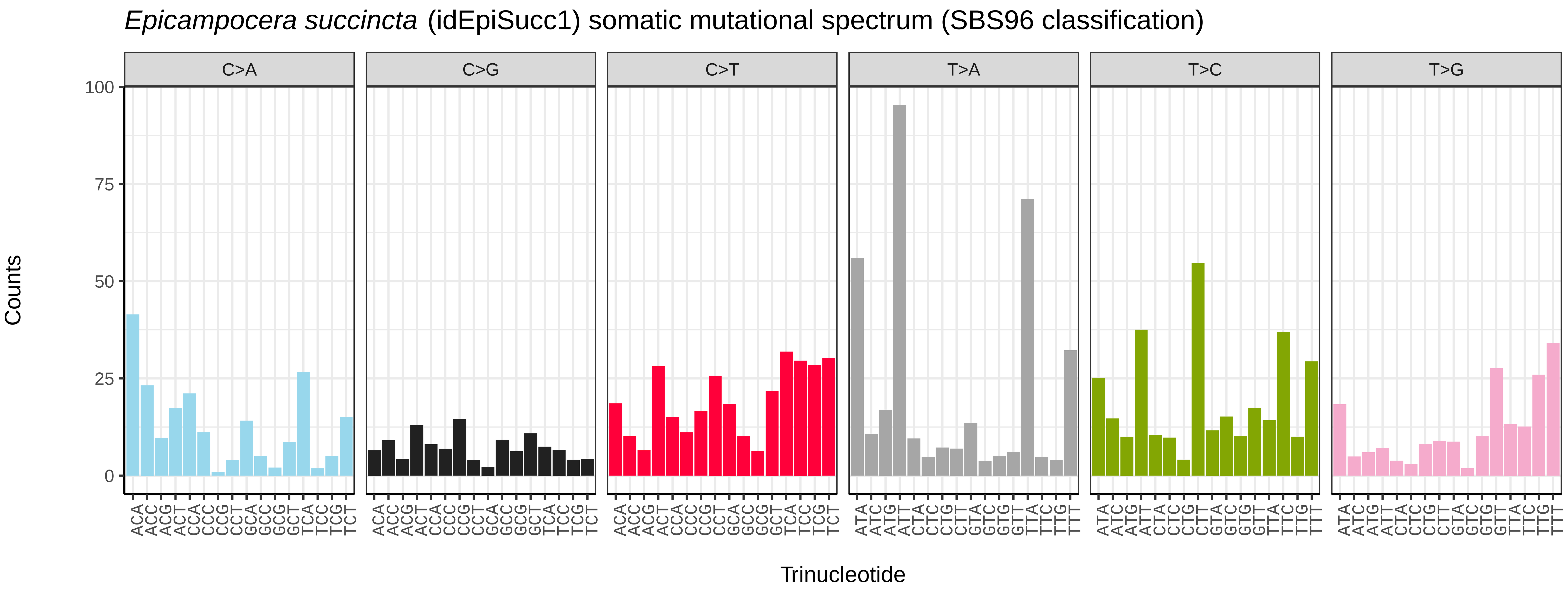
*Epicampocera succincta* (idEpiSucc1) somatic mutational spectrum (SBS96 classification)
C>A
C>G
C>T
T>A
T>C
T>G
Counts
100
75
50
25
0
Trinucleotide

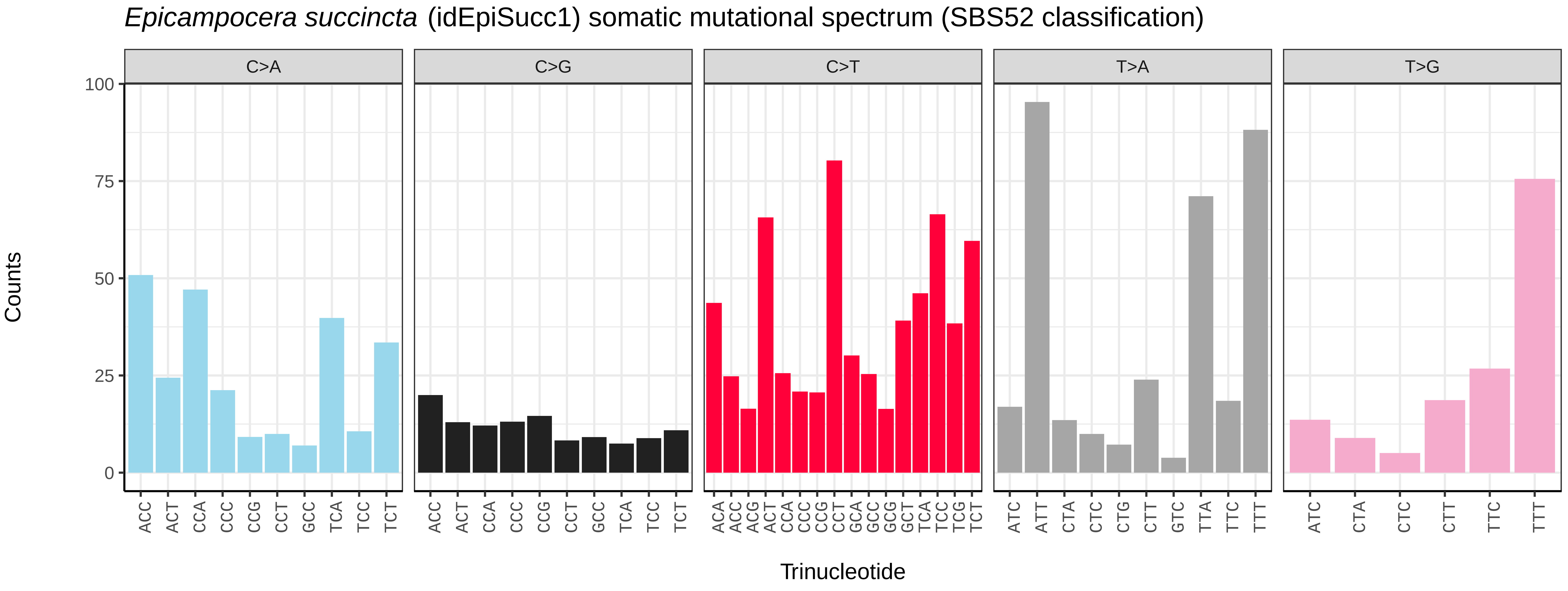
*Epicampocera succincta* (idEpiSucc1) somatic mutational spectrum (SBS52 classification)
C>A
C>G
C>T
T>A
T>G
Counts
100
75
50
25
0
Trinucleotide

*Eristalis arbustorum* (idEriArbu1) germline mutational spectrum

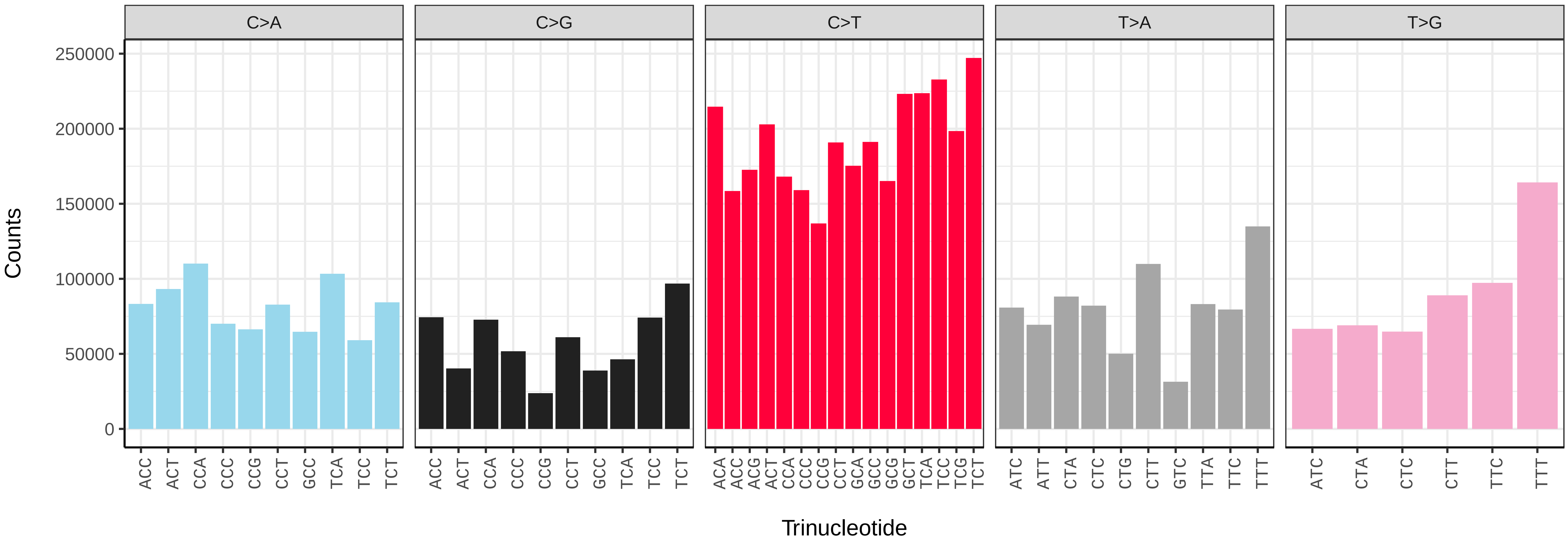


*Eristalis arbustorum* (idEriArbu1) somatic mutational spectrum (SBS96 classification)

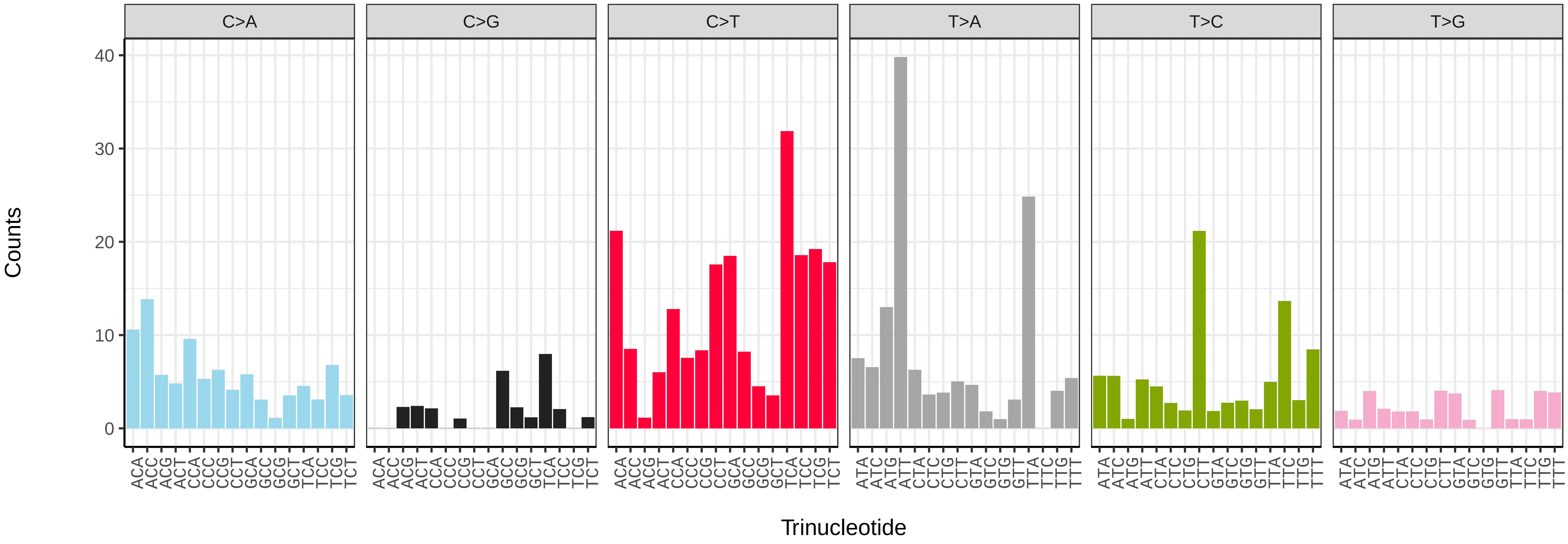


*Eristalis arbustorum* (idEriArbu1) somatic mutational spectrum (SBS52 classification)

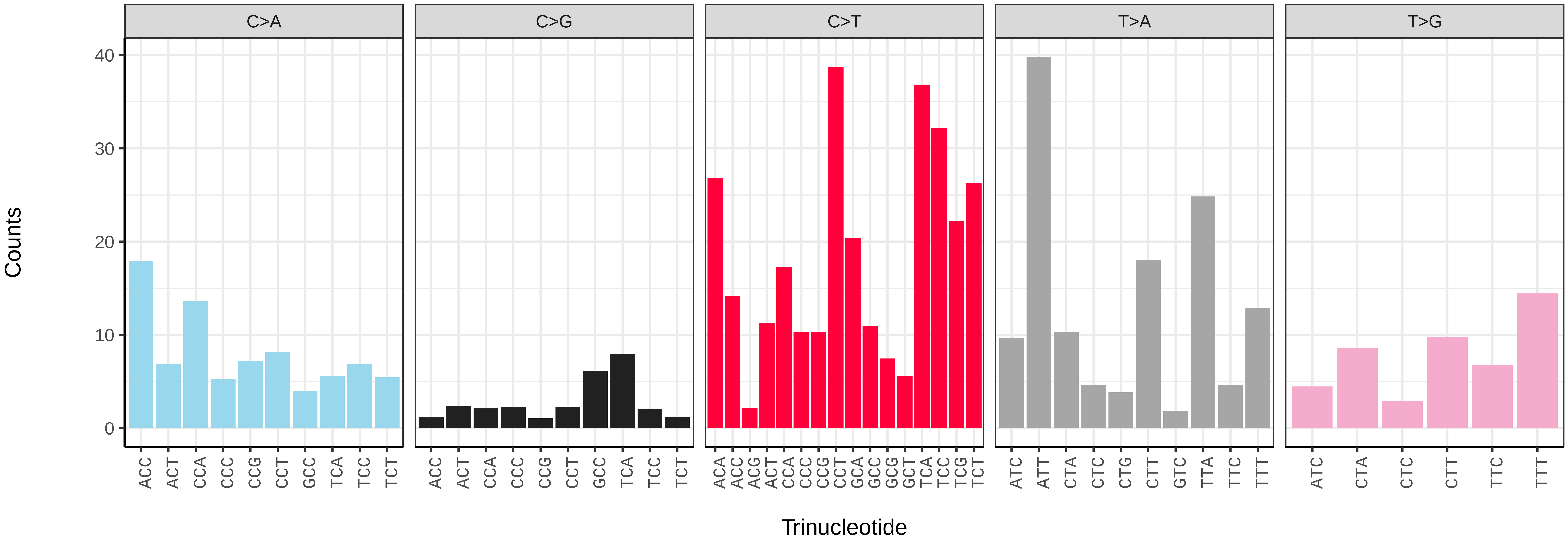

*Eristalis pertinax* (idEriPert2) germline mutational spectrum

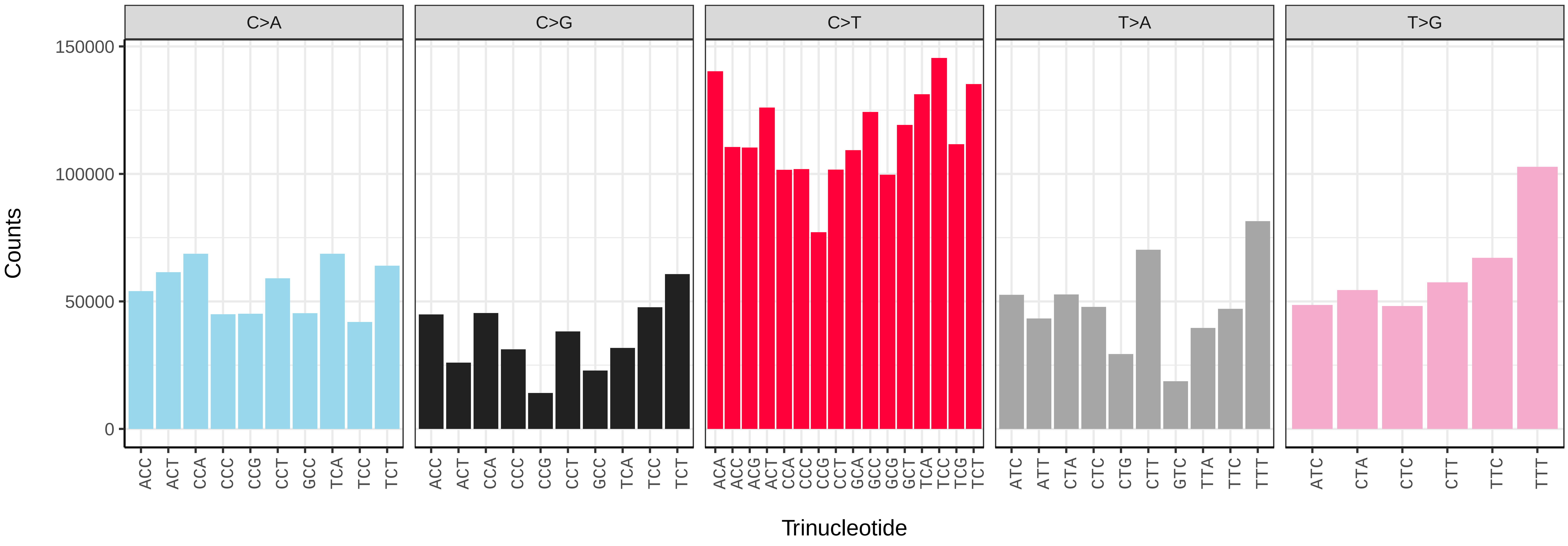


*Eristalis pertinax* (idEriPert2) somatic mutational spectrum (SBS96 classification)

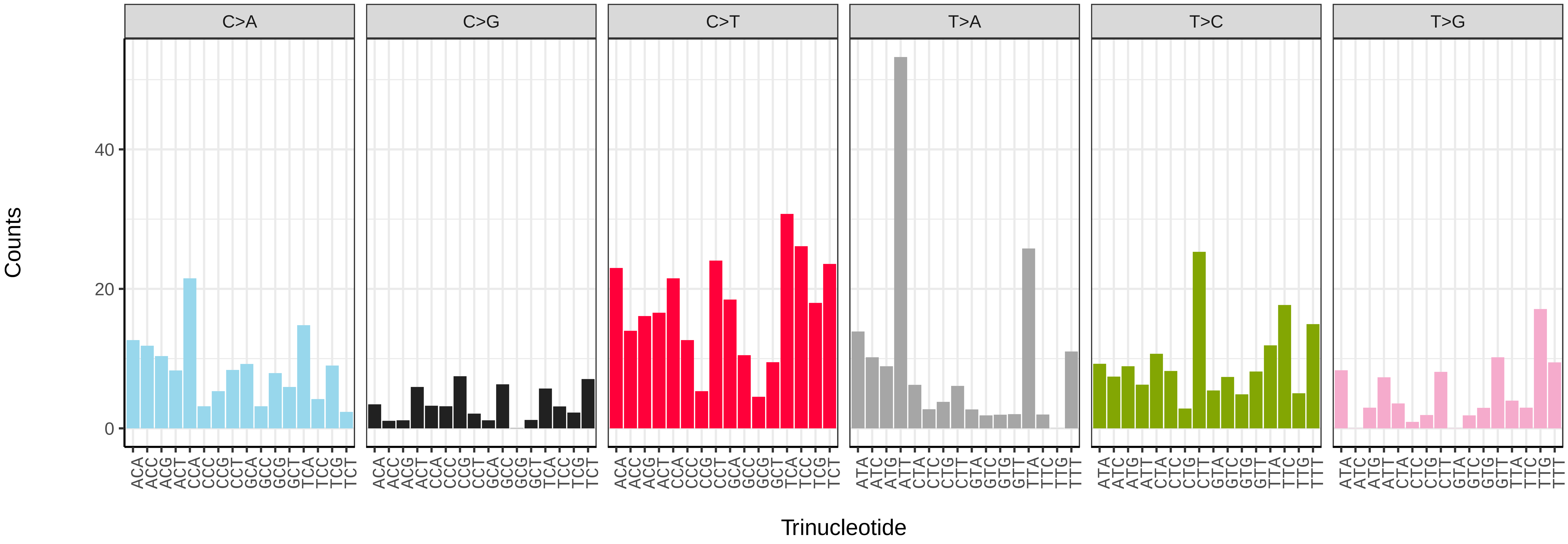


*Eristalis pertinax* (idEriPert2) somatic mutational spectrum (SBS52 classification)

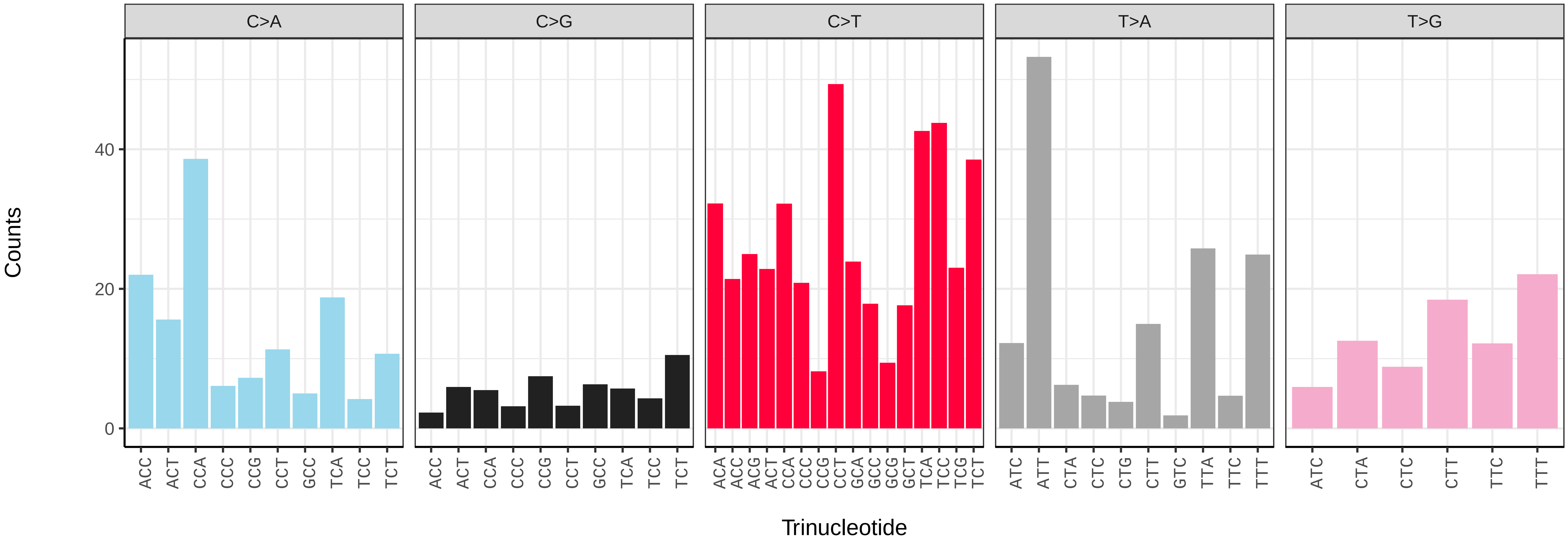

*Eristalinus sepulchralis* (idEriSepu1) germline mutational spectrum

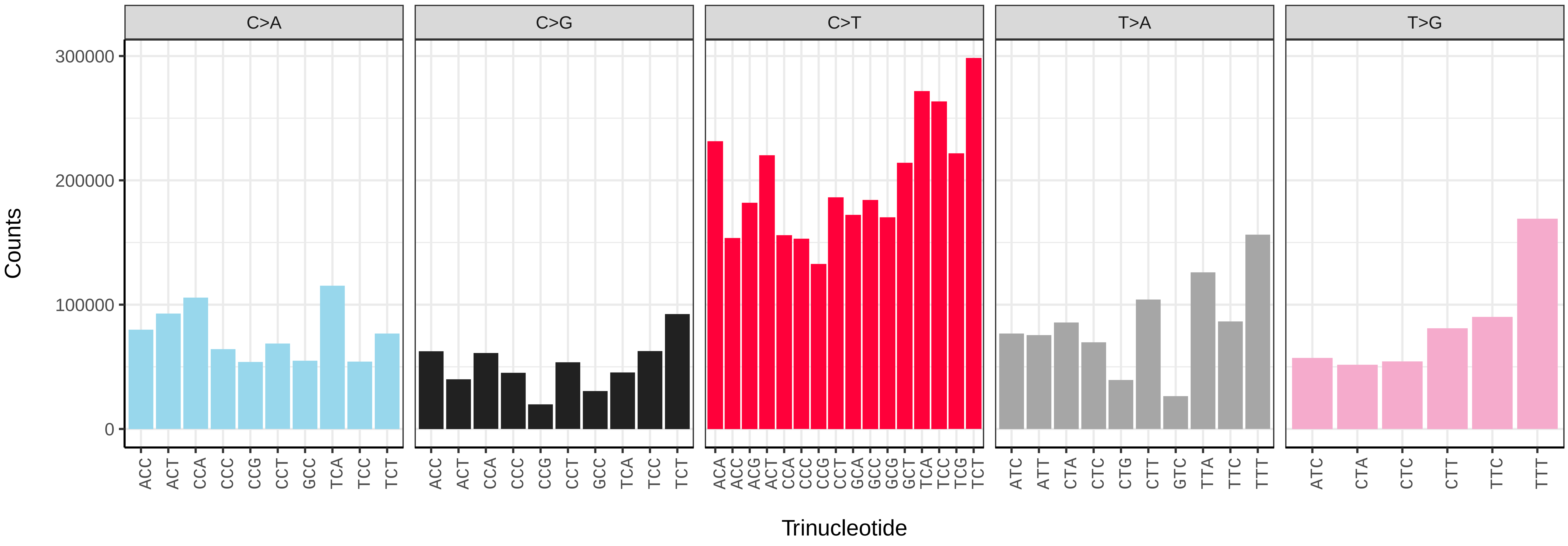


*Eristalinus sepulchralis* (idEriSepu1) somatic mutational spectrum (SBS96 classification)

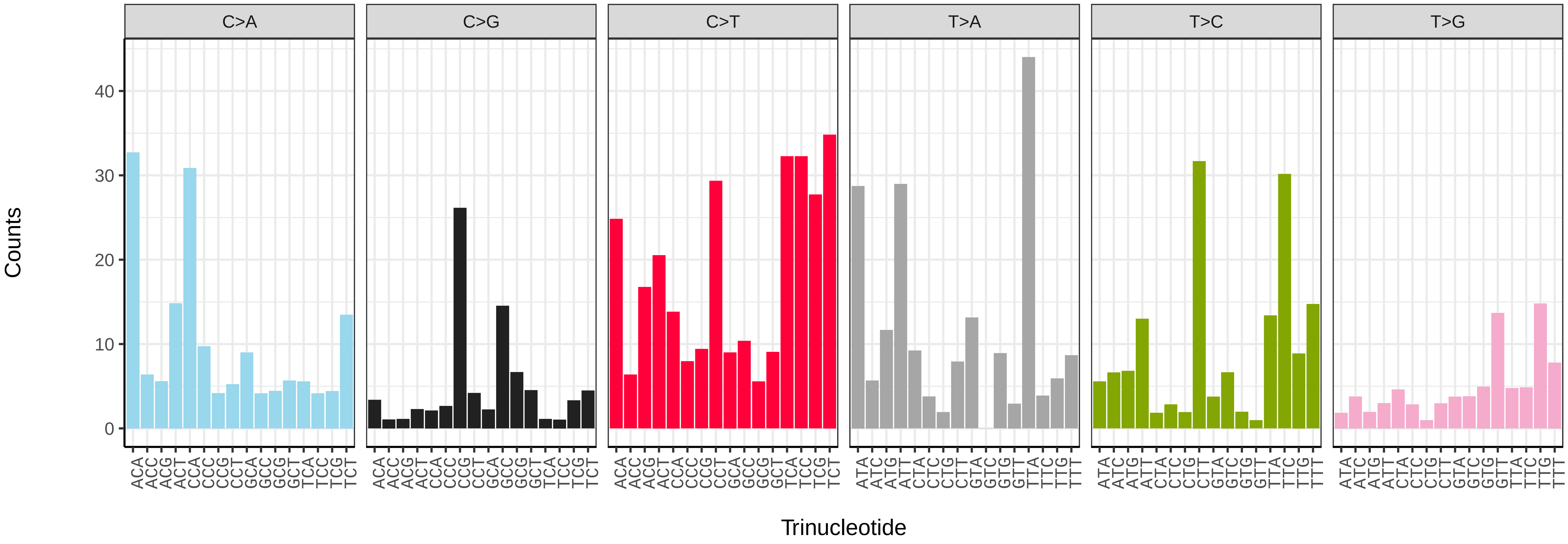


*Eristalinus sepulchralis* (idEriSepu1) somatic mutational spectrum (SBS52 classification)

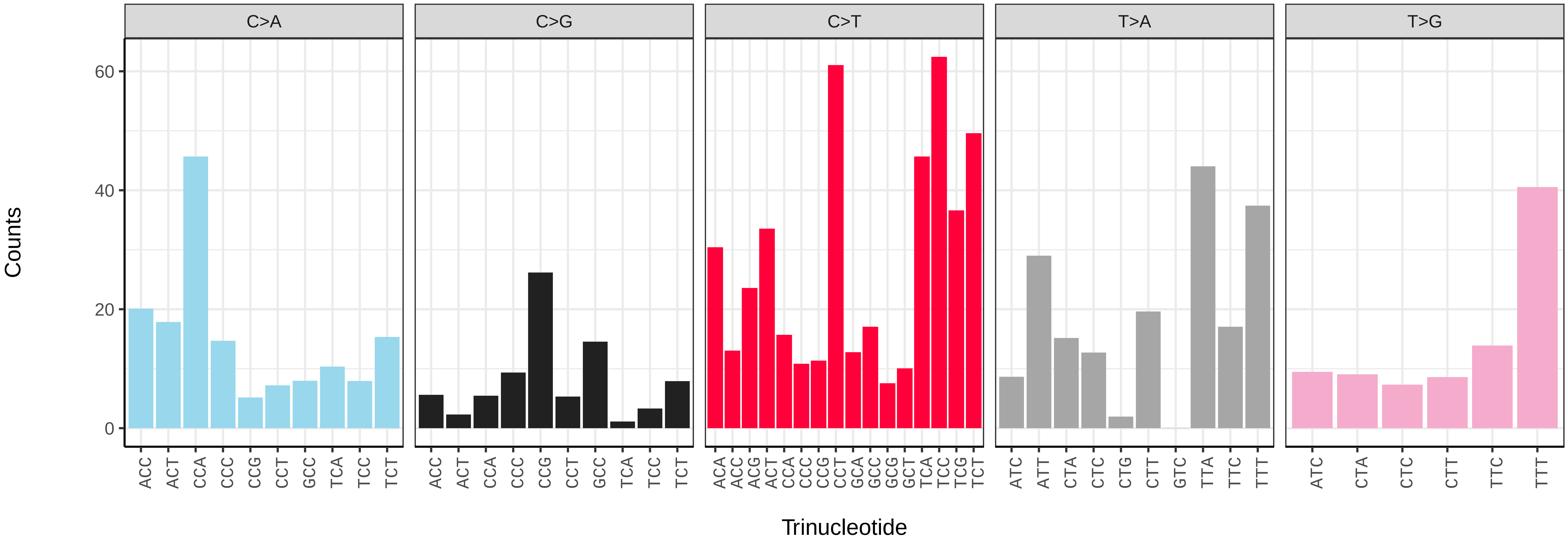

*Eristalis tenax* (idEriTena2) germline mutational spectrum

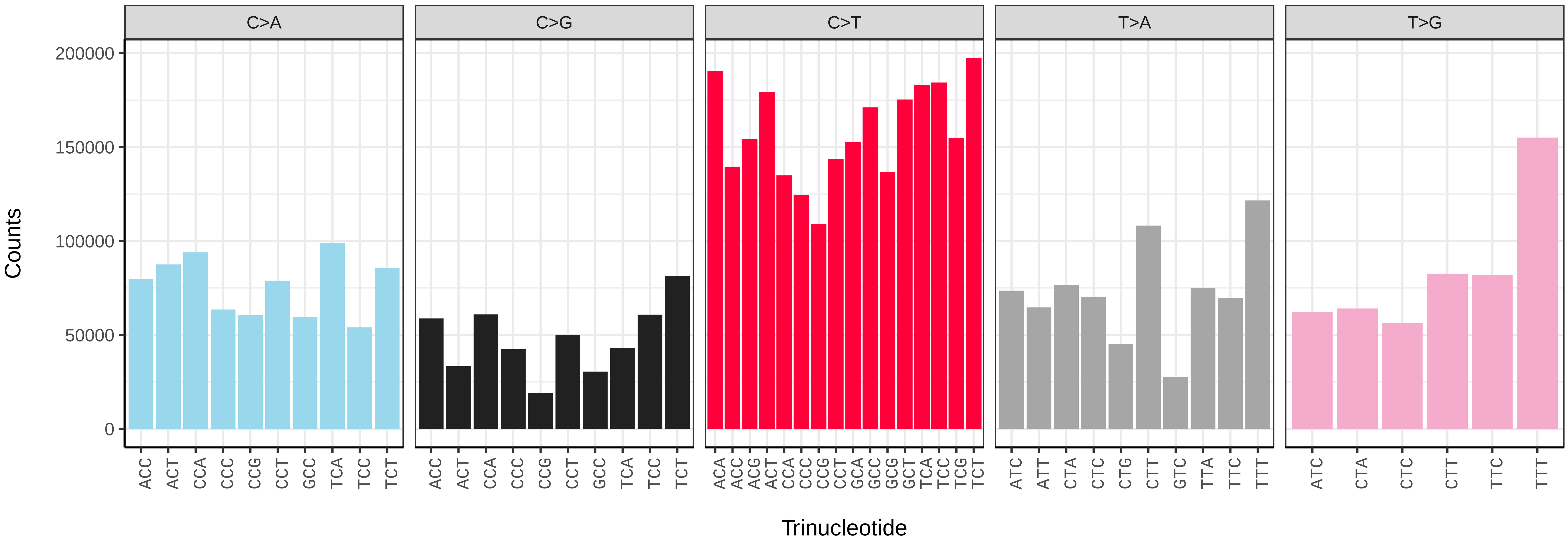


*Eristalis tenax* (idEriTena2) somatic mutational spectrum (SBS96 classification)

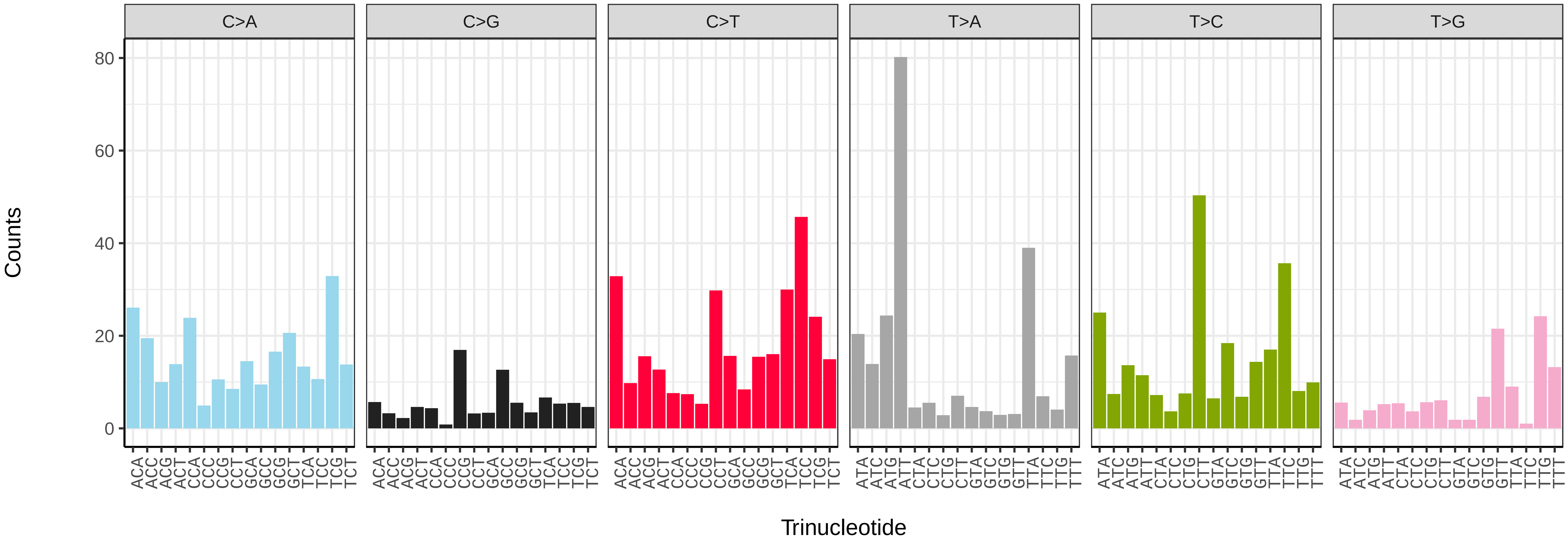


*Eristalis tenax* (idEriTena2) somatic mutational spectrum (SBS52 classification)

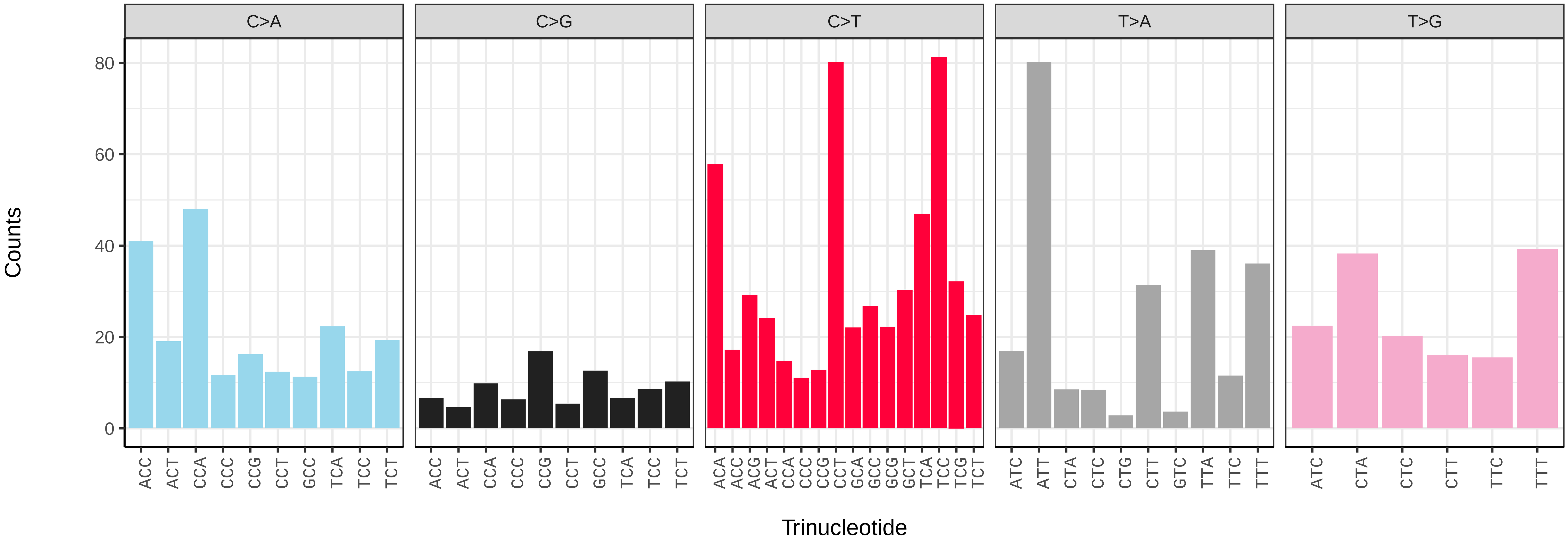

*Eupeodes corollae* (idEupCoro1) germline mutational spectrum

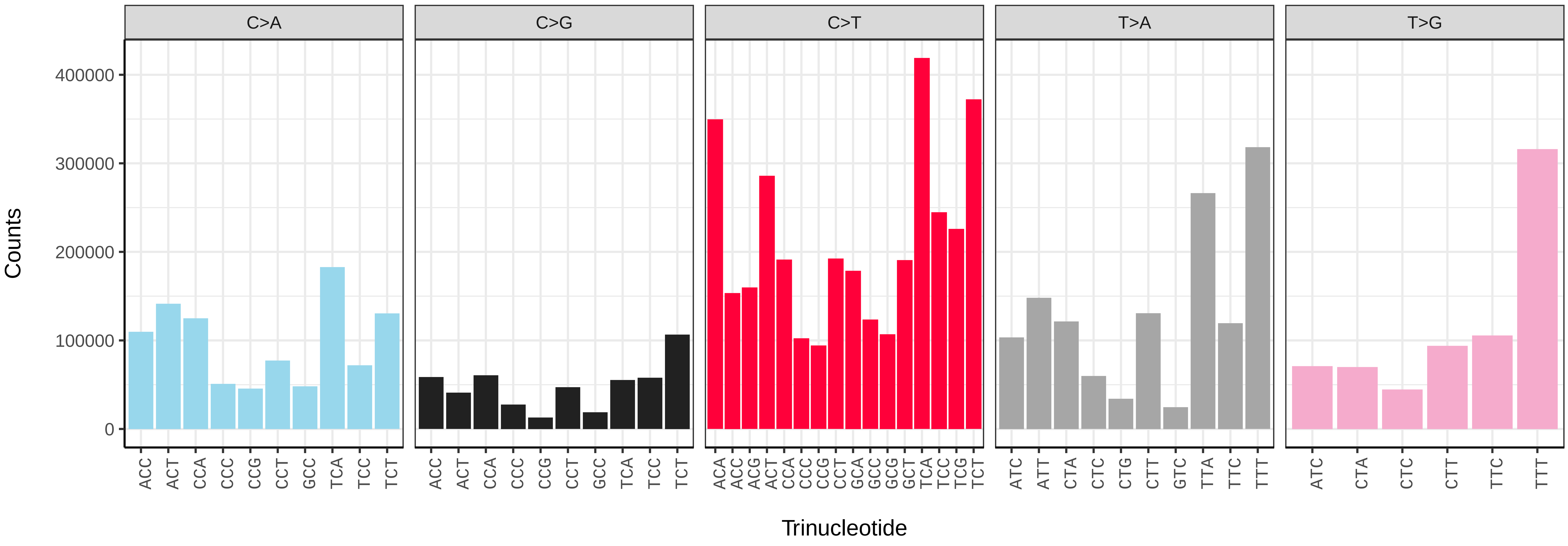


*Eupeodes corollae* (idEupCoro1) somatic mutational spectrum (SBS96 classification)

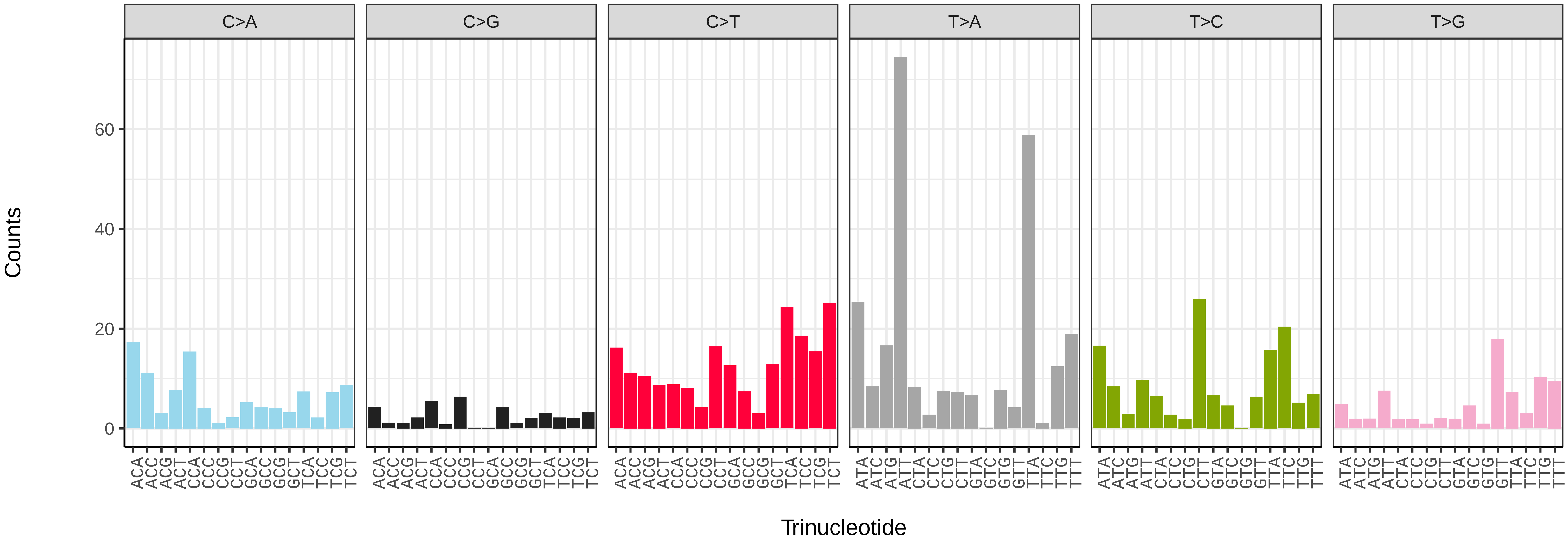


*Eupeodes corollae* (idEupCoro1) somatic mutational spectrum (SBS52 classification)

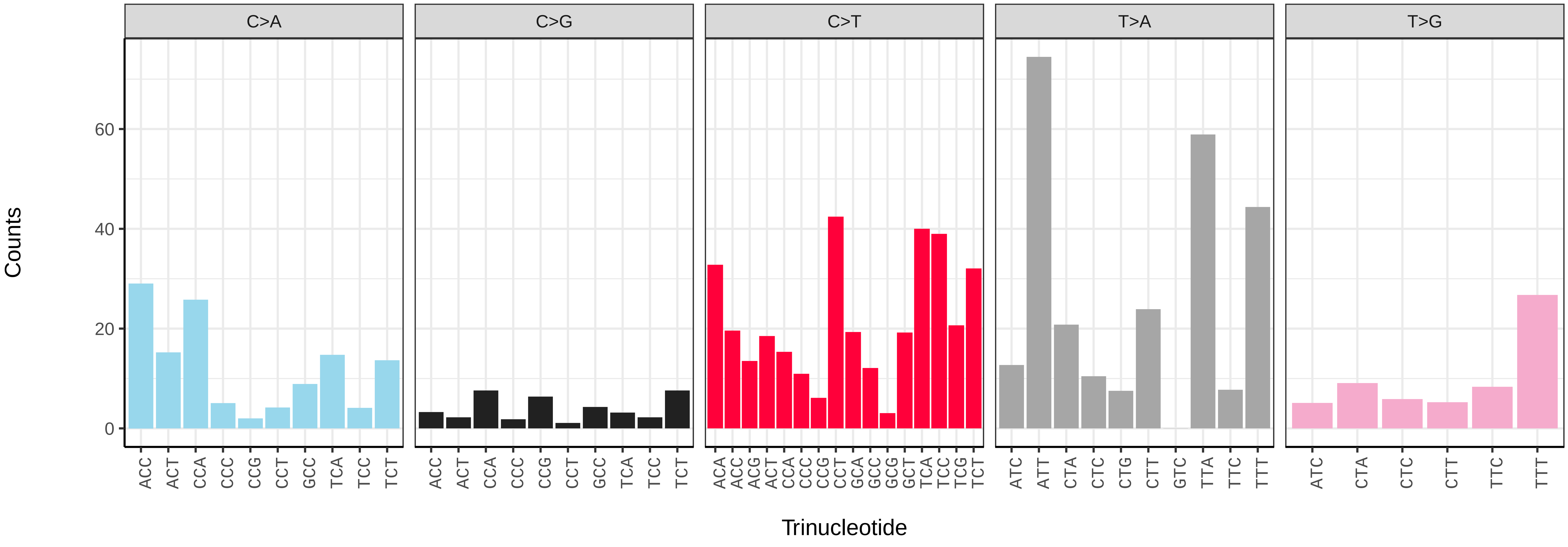

*Eupeodes latifasciatus* (idEupLati1) germline mutational spectrum

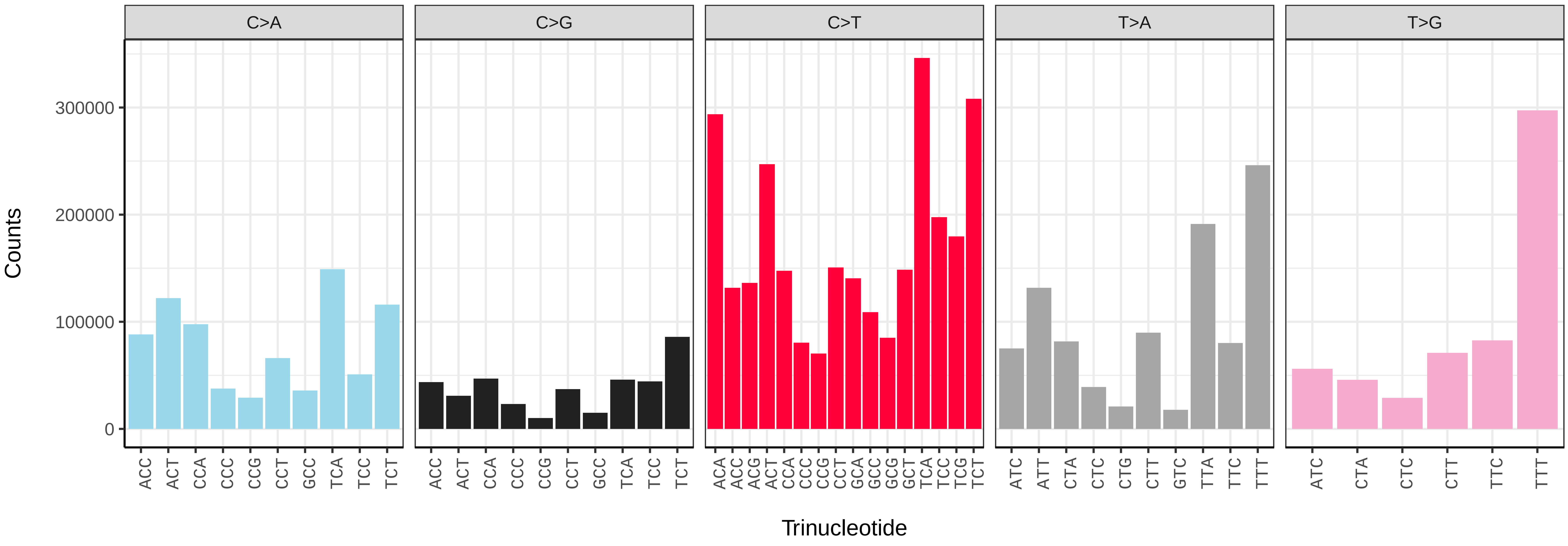


*Eupeodes latifasciatus* (idEupLati1) somatic mutational spectrum (SBS96 classification)

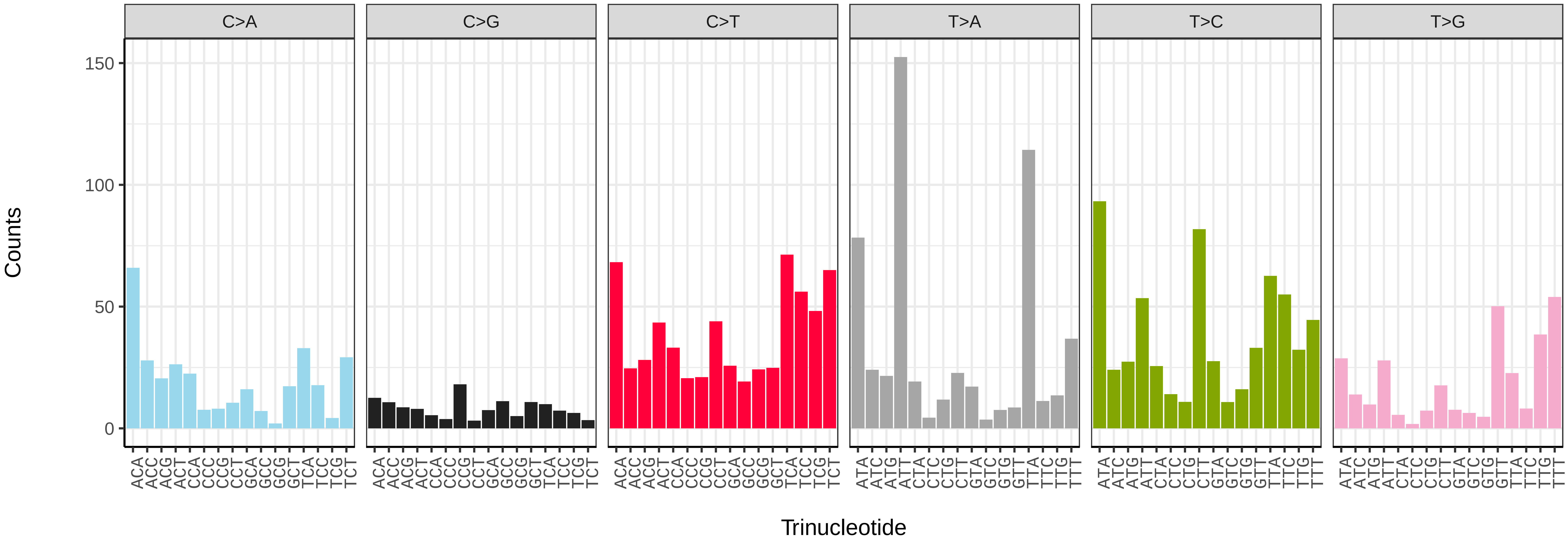


*Eupeodes latifasciatus* (idEupLati1) somatic mutational spectrum (SBS52 classification)

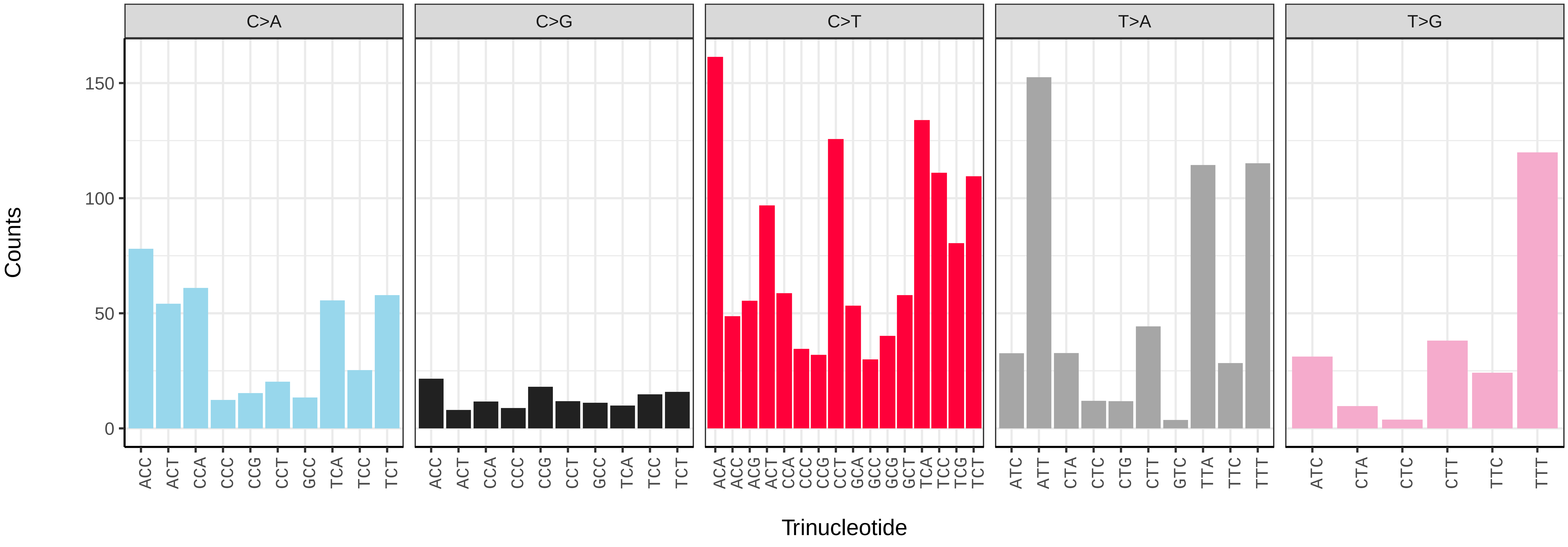

*Gymnosoma rotundatum* (idGymRotn1) germline mutational spectrum

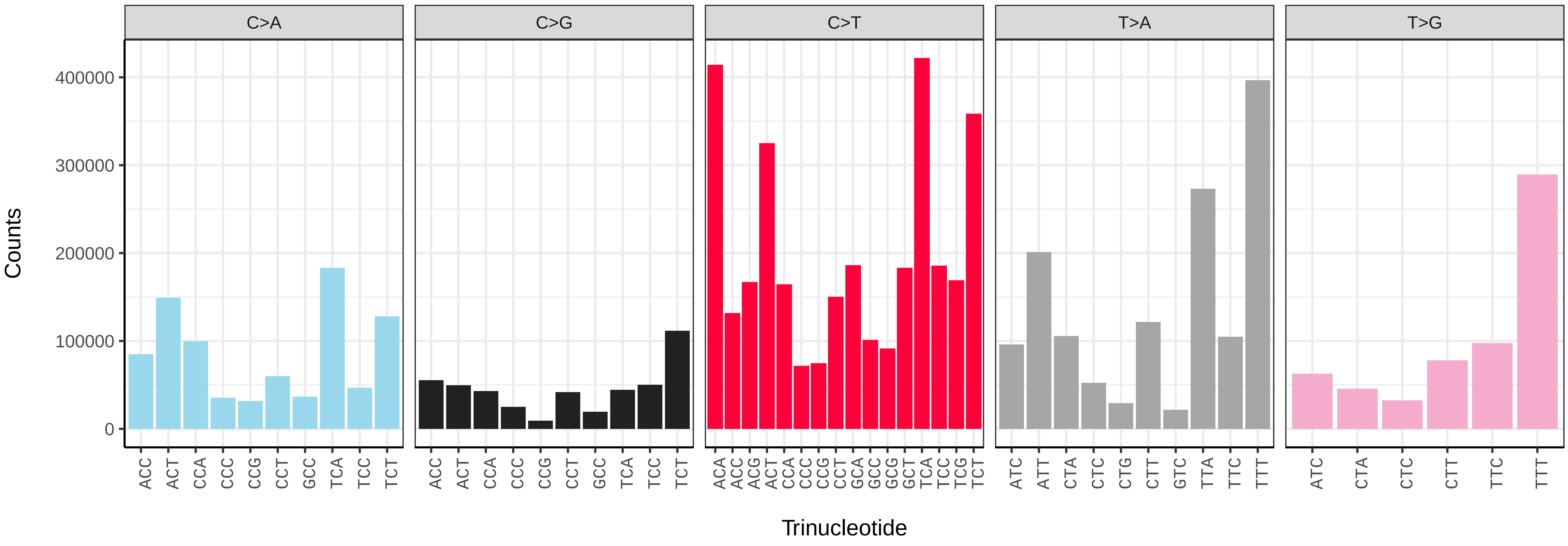


*Gymnosoma rotundatum* (idGymRotn1) somatic mutational spectrum (SBS96 classification)

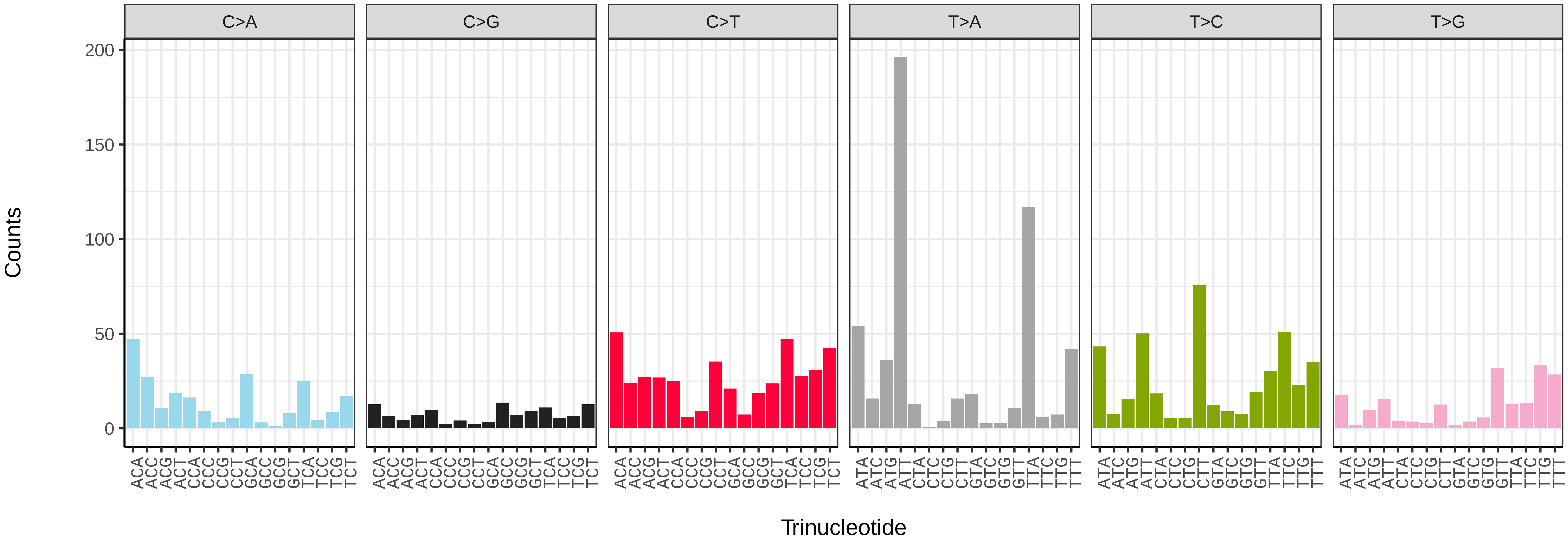


*Gymnosoma rotundatum* (idGymRotn1) somatic mutational spectrum (SBS52 classification)

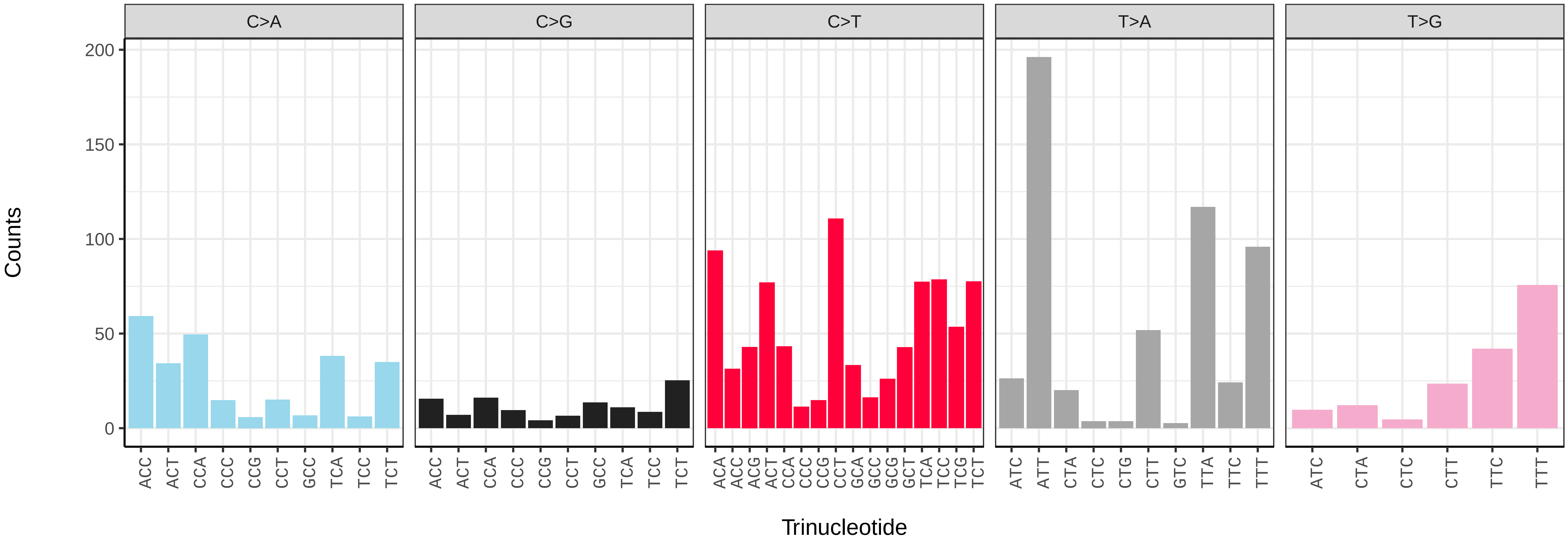

*Leucozona laternaria* (idLeuLate1) germline mutational spectrum

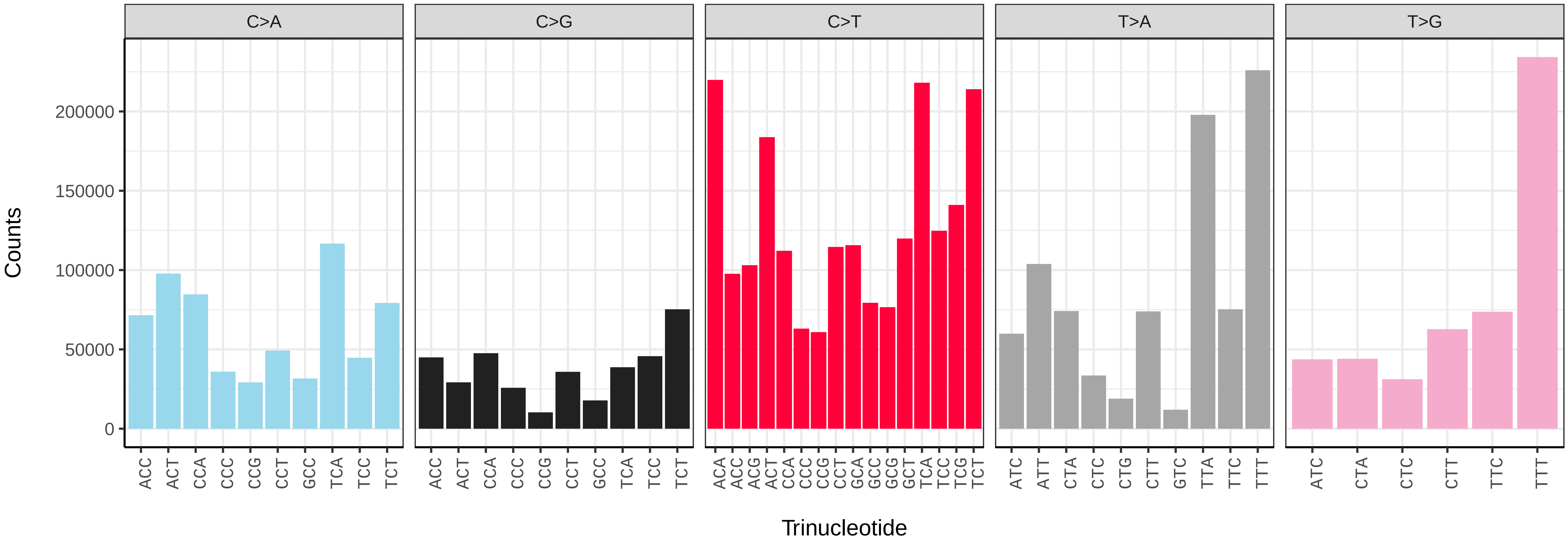


*Leucozona laternaria* (idLeuLate1) somatic mutational spectrum (SBS96 classification)

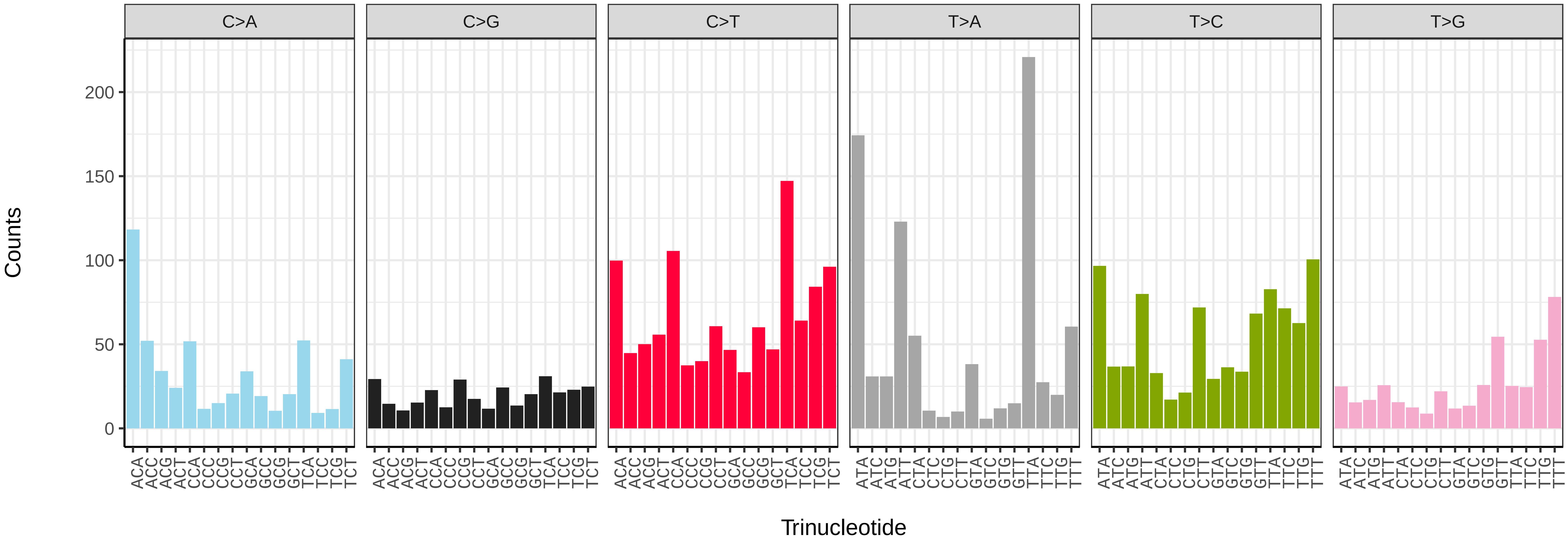


*Leucozona laternaria* (idLeuLate1) somatic mutational spectrum (SBS52 classification)

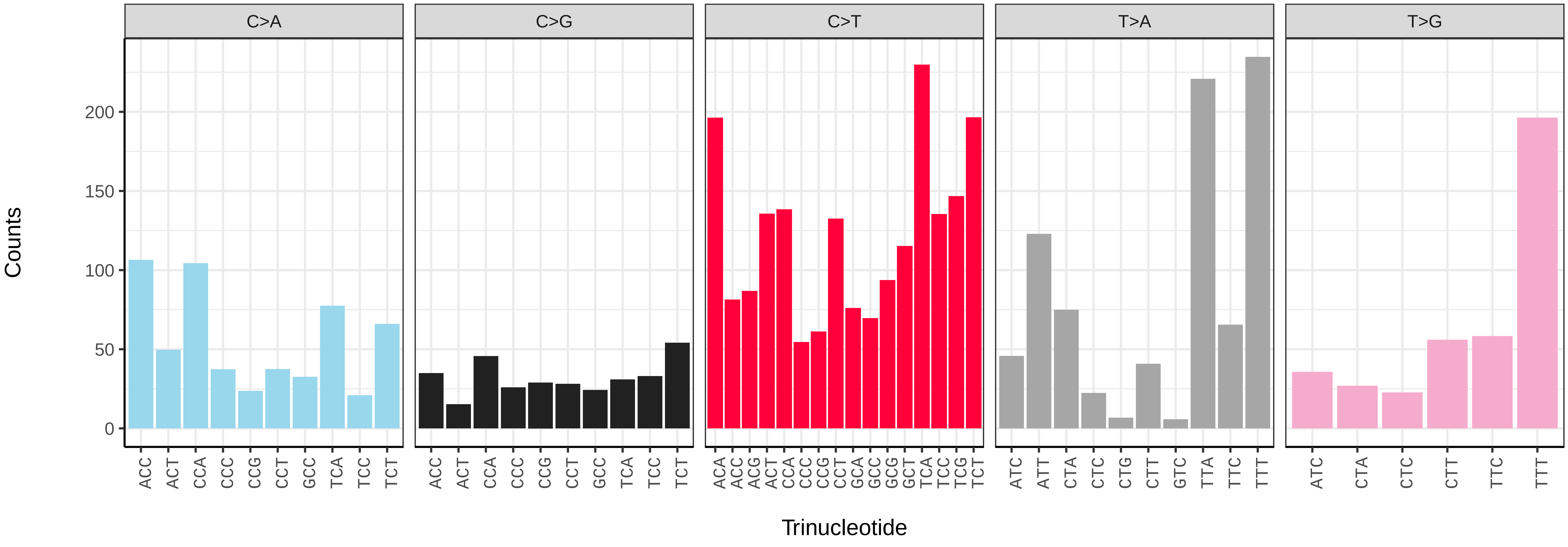

*Lypha dubia* (idLypDubi1) germline mutational spectrum

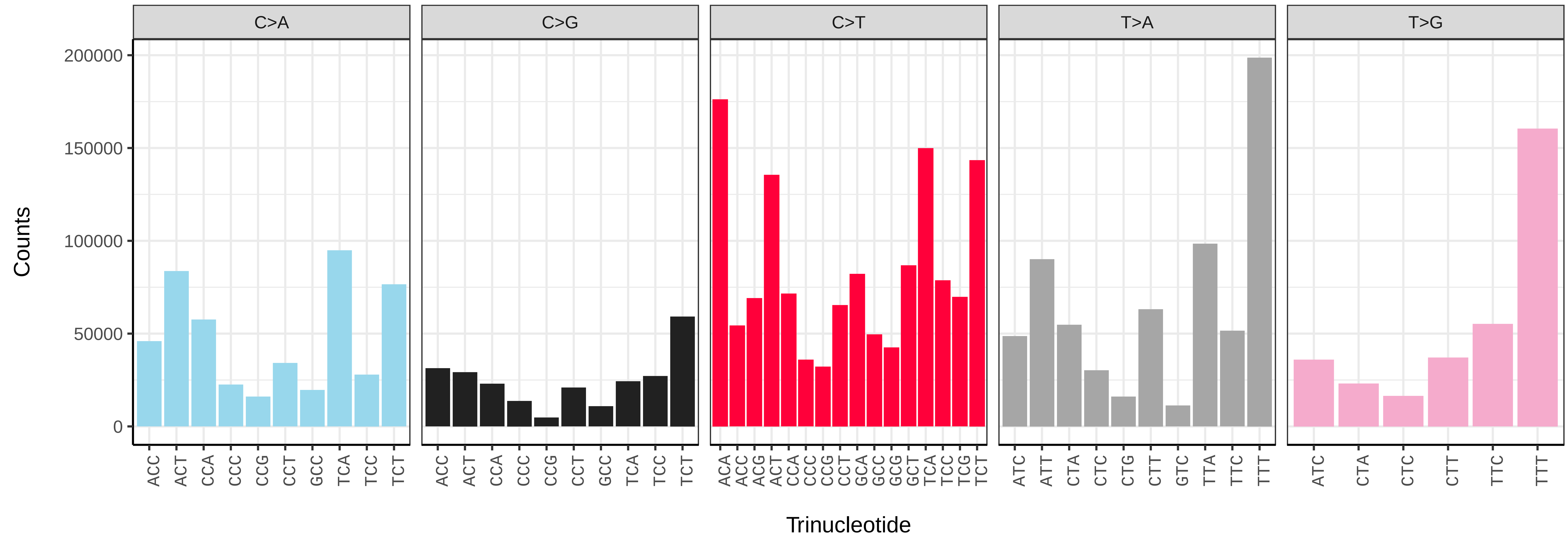


*Lypha dubia* (idLypDubi1) somatic mutational spectrum (SBS96 classification)

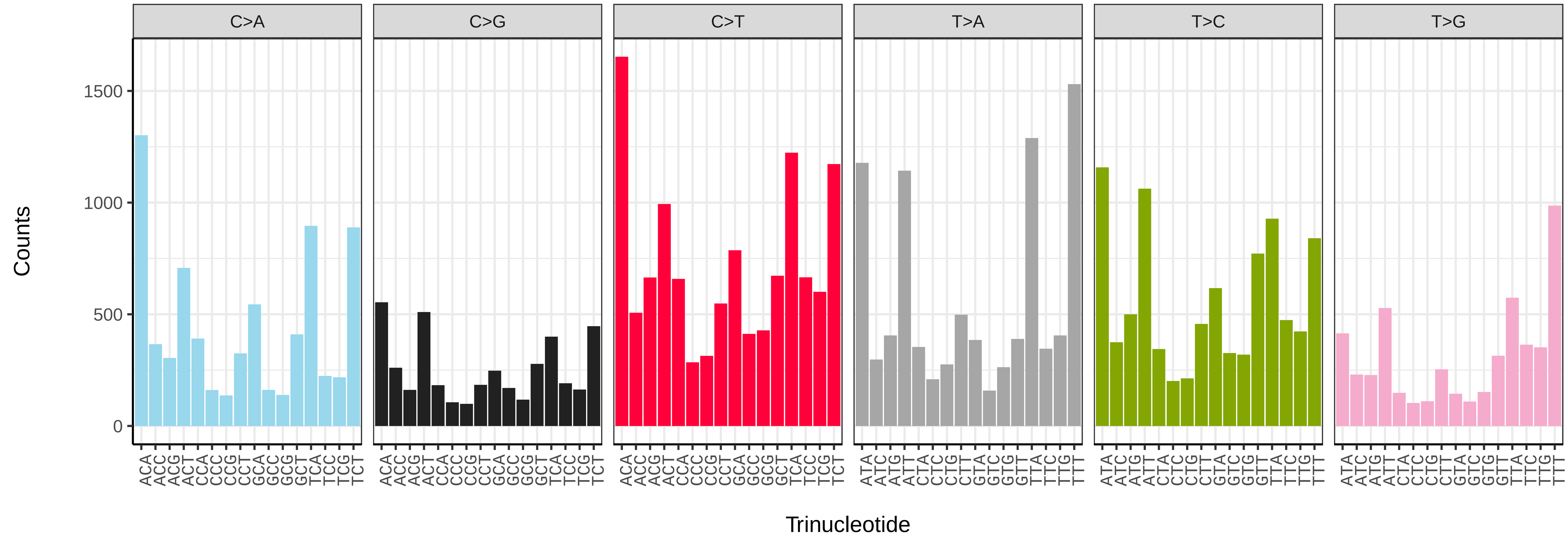


*Lypha dubia* (idLypDubi1) somatic mutational spectrum (SBS52 classification)

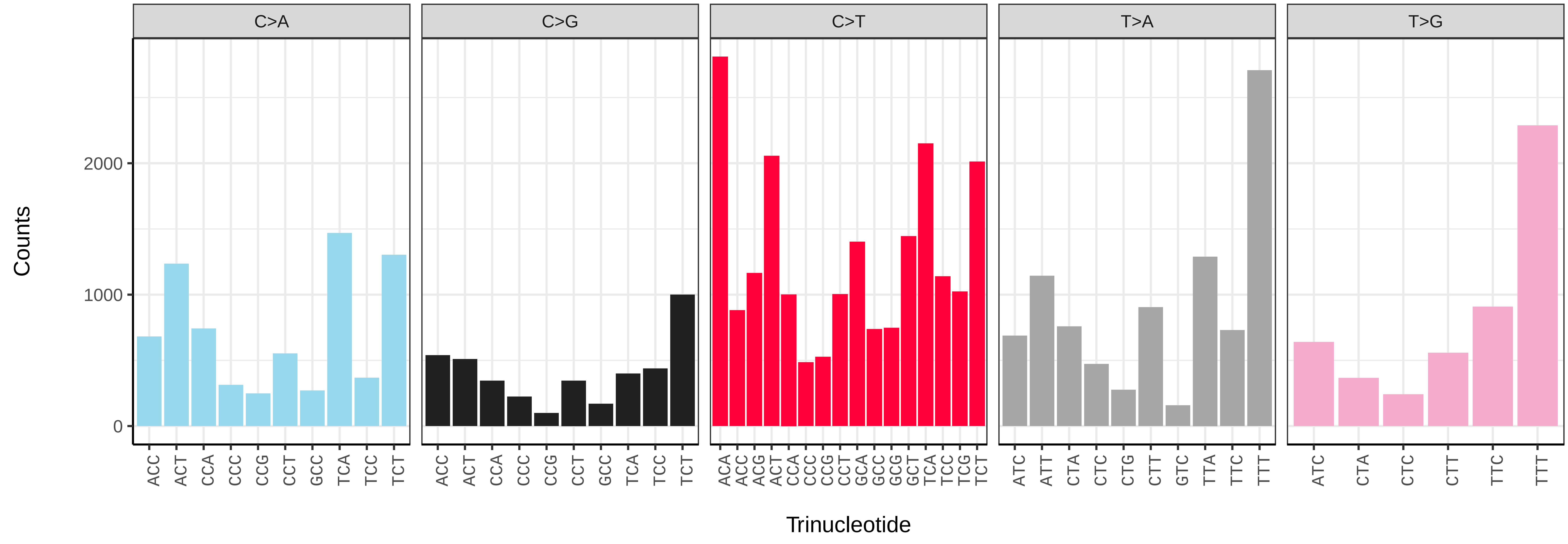

*Machimus atricapillus* (idMacAtri3) germline mutational spectrum

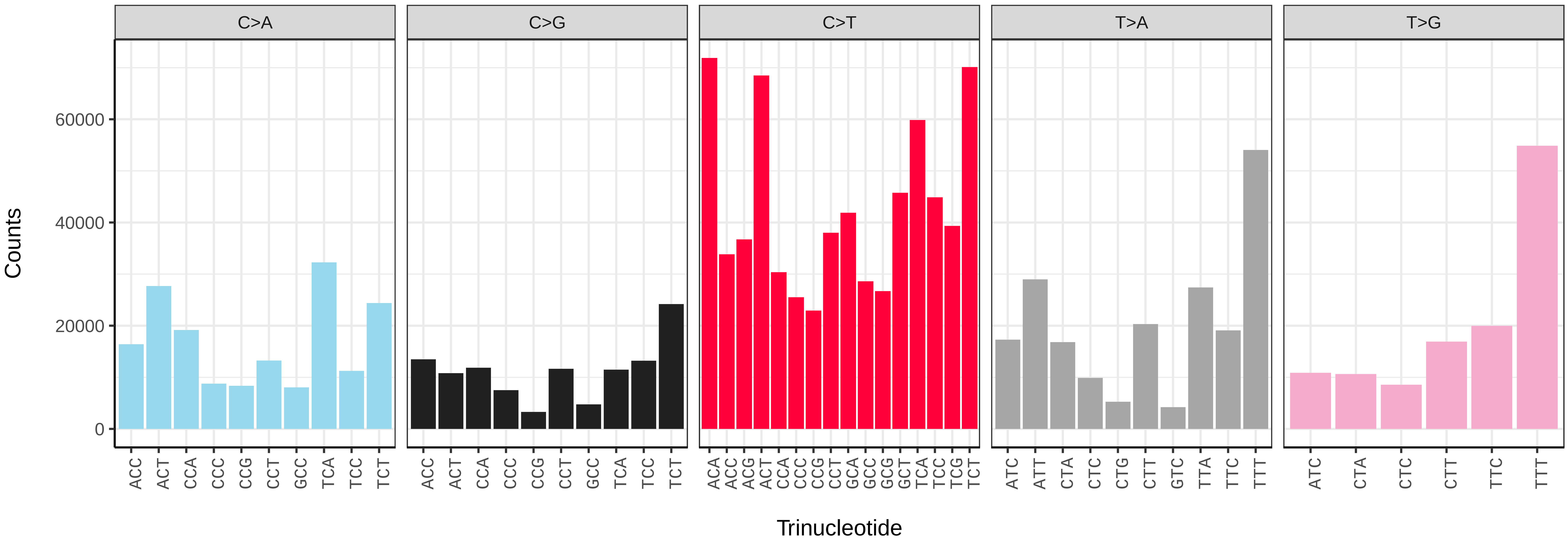


*Machimus atricapillus* (idMacAtri3) somatic mutational spectrum (SBS96 classification)

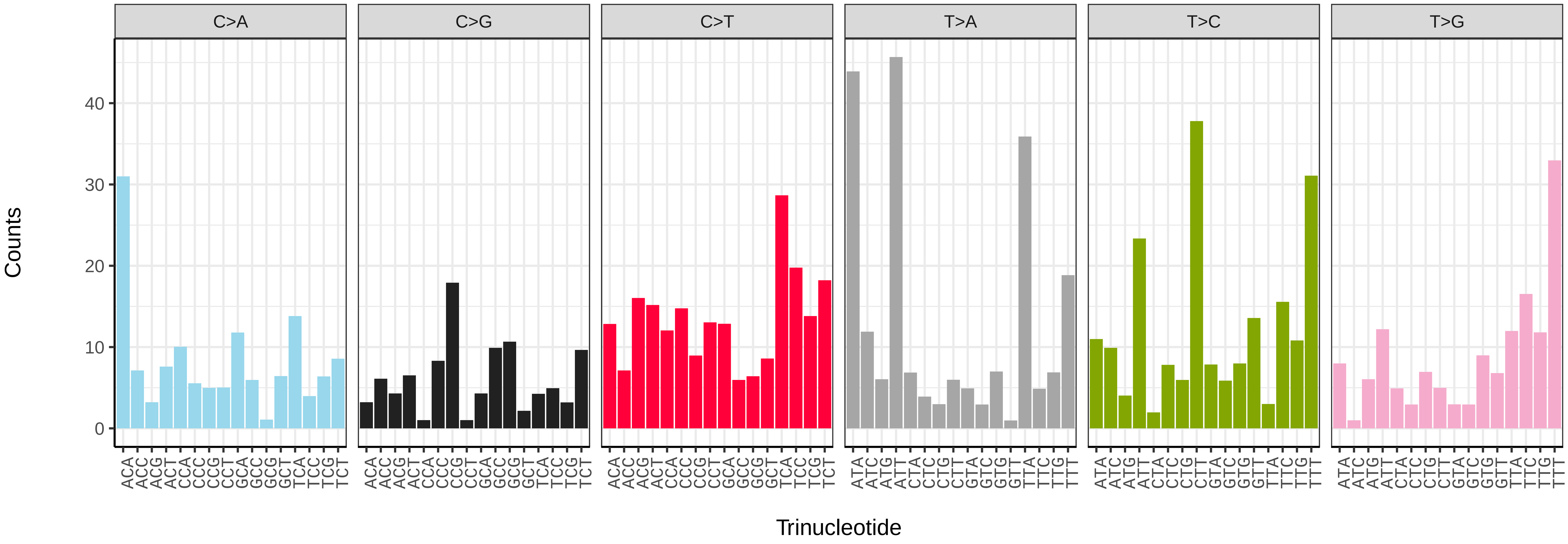


*Machimus atricapillus* (idMacAtri3) somatic mutational spectrum (SBS52 classification)

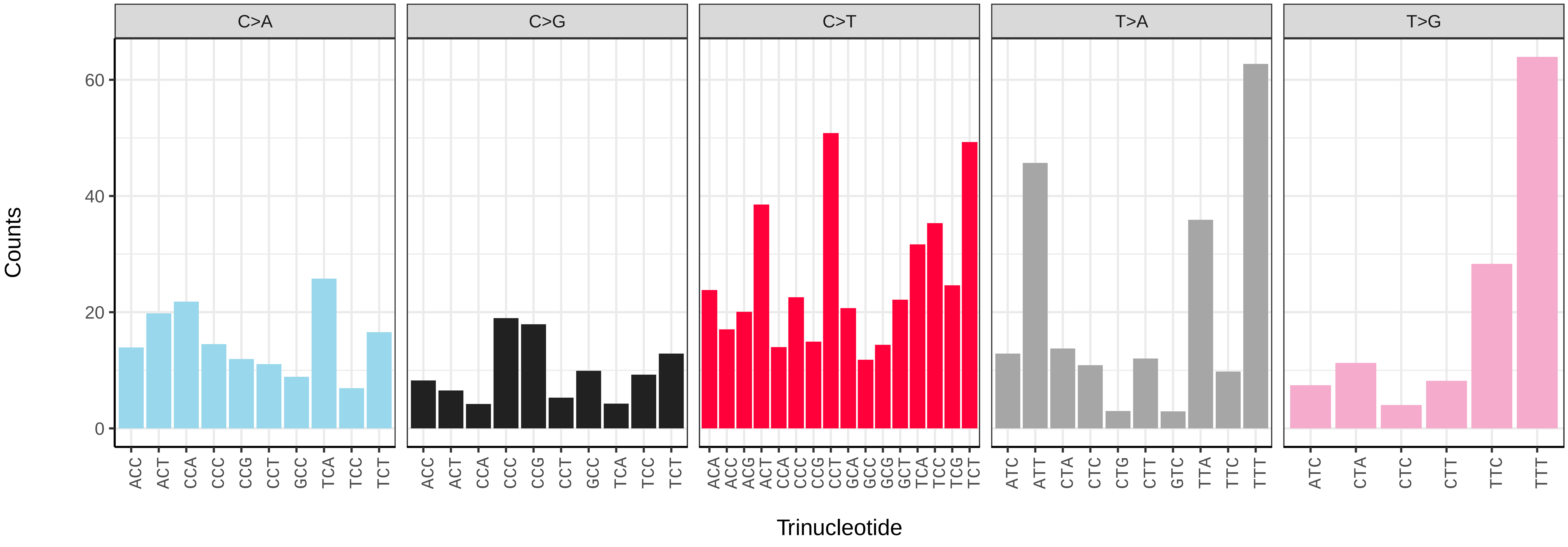

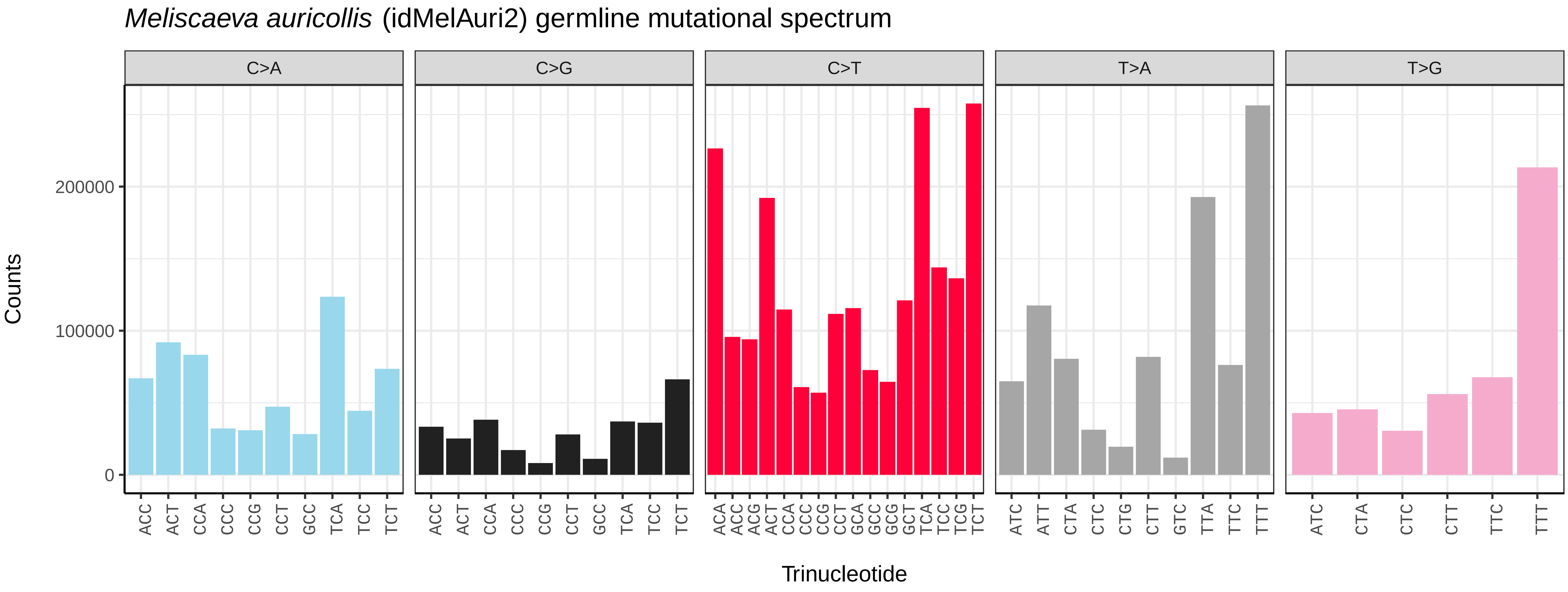
Meliscaeva auricollis (idMelAuri2) germline mutational spectrum
C>A
C>G
C>T
T>A
T>G
Counts
0
100000
200000
Trinucleotide
ACC ACT CCA CCC CCG CCT GCC TCA TCC TCT
ACA ACC ACG ACT CCA CCC CCG CCT GCA GCC GCG GCT TCA TCC TCG TCT
ATC ATT CTA CTC CTG CTT GTC TTA TTC TTT
ATC CTA CTC CTT TTC TTT

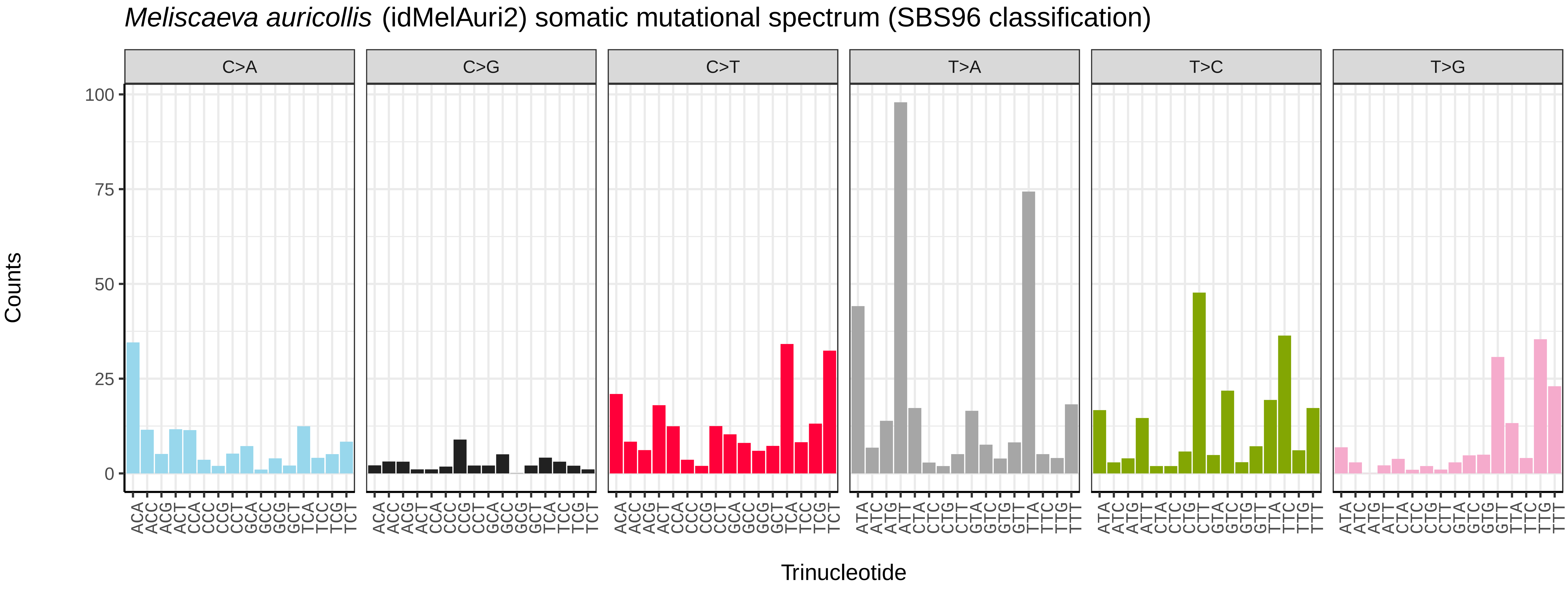
Meliscaeva auricollis (idMelAuri2) somatic mutational spectrum (SBS96 classification)
C>A
C>G
C>T
T>A
T>C
T>G
Counts
0
25
50
75
100
Trinucleotide
ACA ACC ACG ACT CCA CCC CCG CCT GCA GCC GCG GCT TCA TCC TCG TCT
ATA ATC ATG ATT CTA CTC CTG CTT GTA GTC GTG GTT TTA TTC TTG TTT

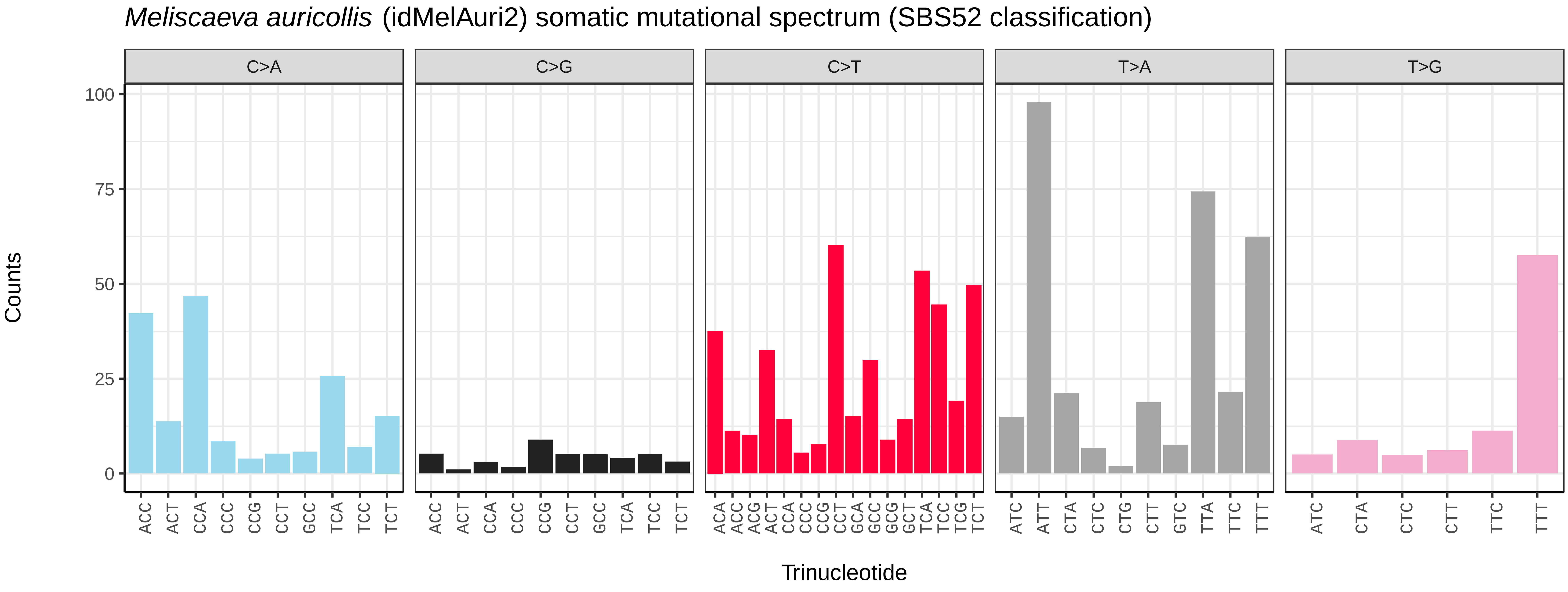
Meliscaeva auricollis (idMelAuri2) somatic mutational spectrum (SBS52 classification)
C>A
C>G
C>T
T>A
T>G
Counts
0
25
50
75
100
Trinucleotide
ACC ACT CCA CCC CCG CCT GCC TCA TCC TCT
ACA ACC ACG ACT CCA CCC CCG CCT GCA GCC GCG GCT TCA TCC TCG TCT
ATC ATT CTA CTC CTG CTT GTC TTA TTC TTT
ATC CTA CTC CTT TTC TTT

*Melanostoma mellinum* (idMelMell2) germline mutational spectrum

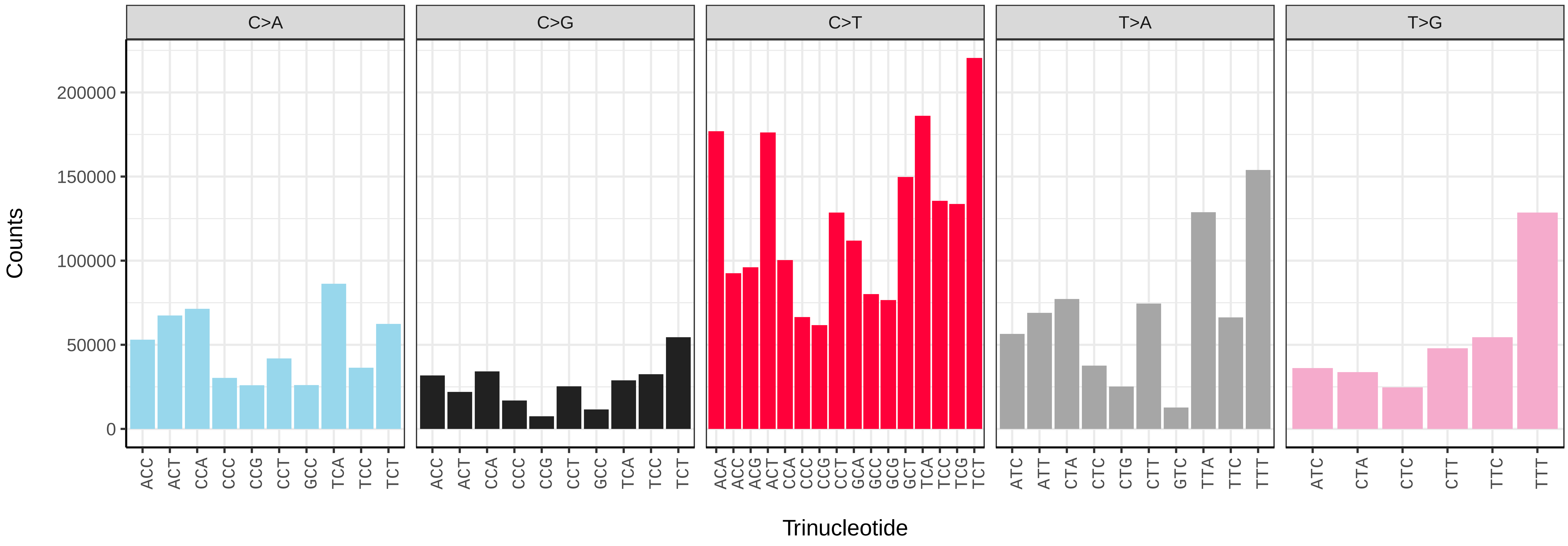


*Melanostoma mellinum* (idMelMell2) somatic mutational spectrum (SBS96 classification)

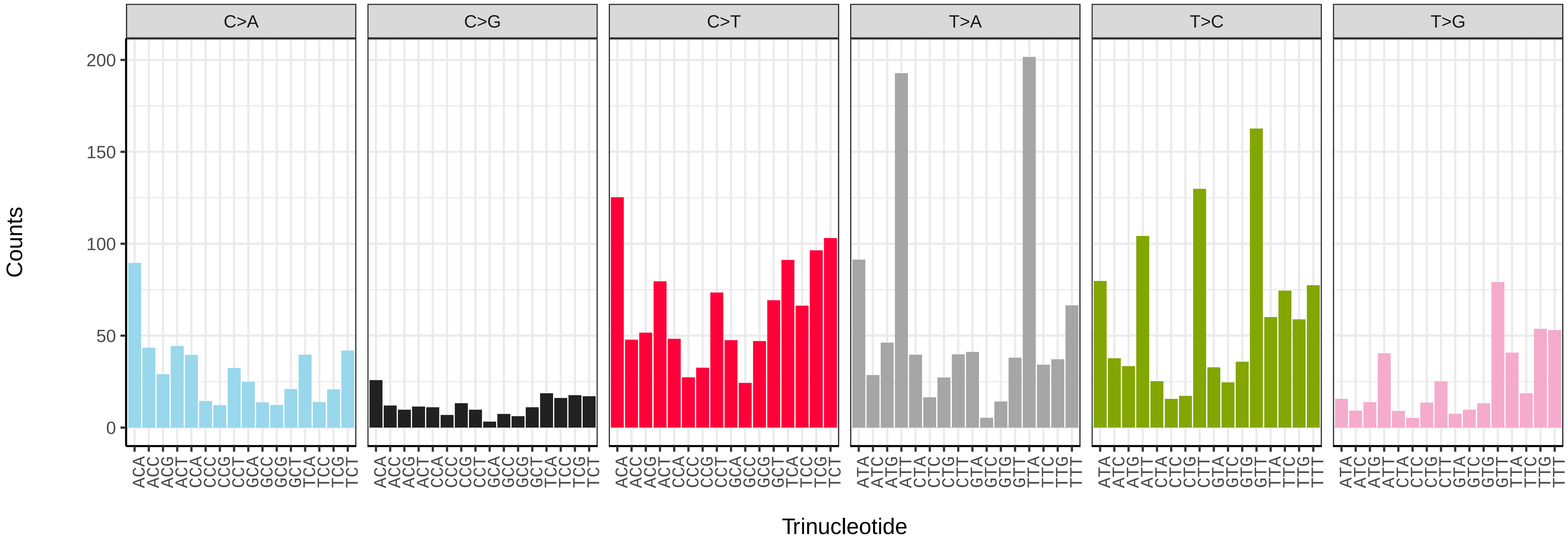


*Melanostoma mellinum* (idMelMell2) somatic mutational spectrum (SBS52 classification)

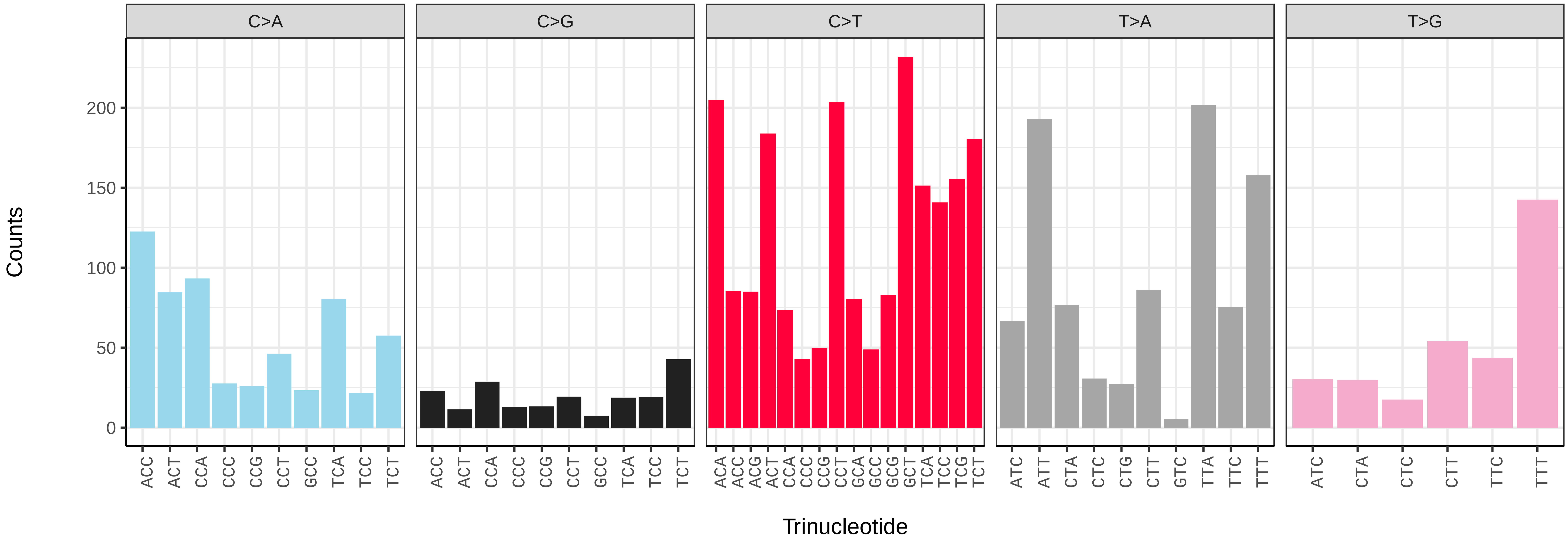

*Myathropa florea* (idMyaFlor2) germline mutational spectrum

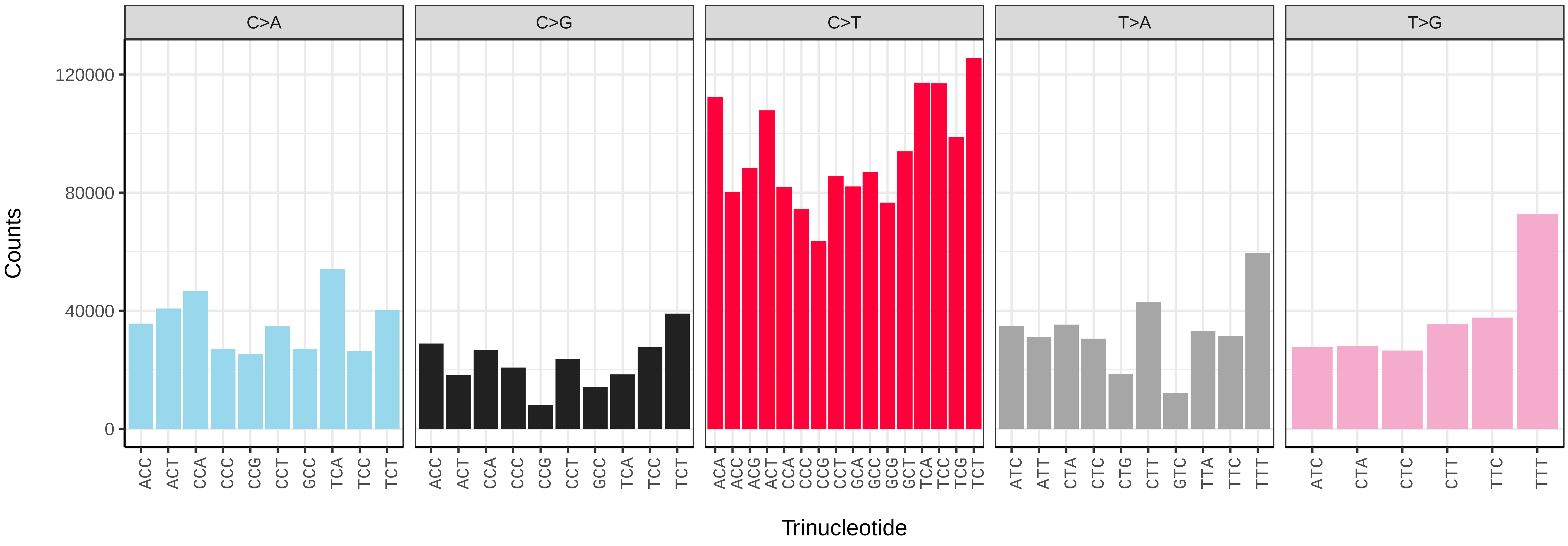


*Myathropa florea* (idMyaFlor2) somatic mutational spectrum (SBS96 classification)

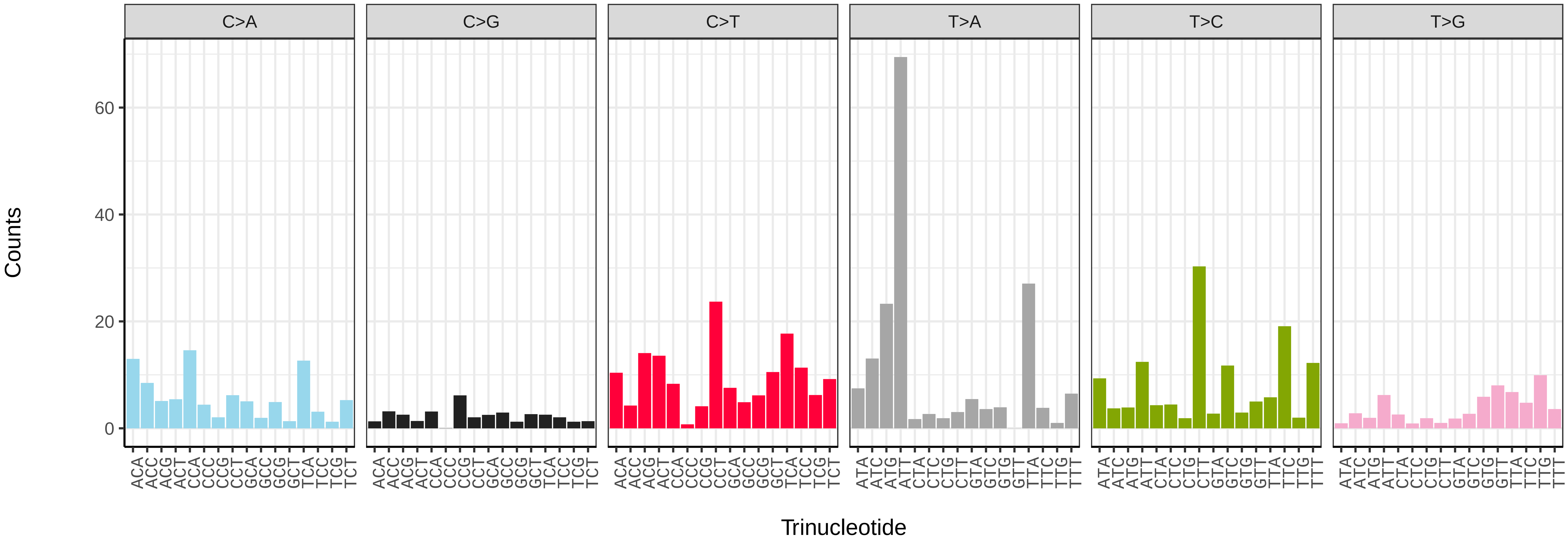


*Myathropa florea* (idMyaFlor2) somatic mutational spectrum (SBS52 classification)

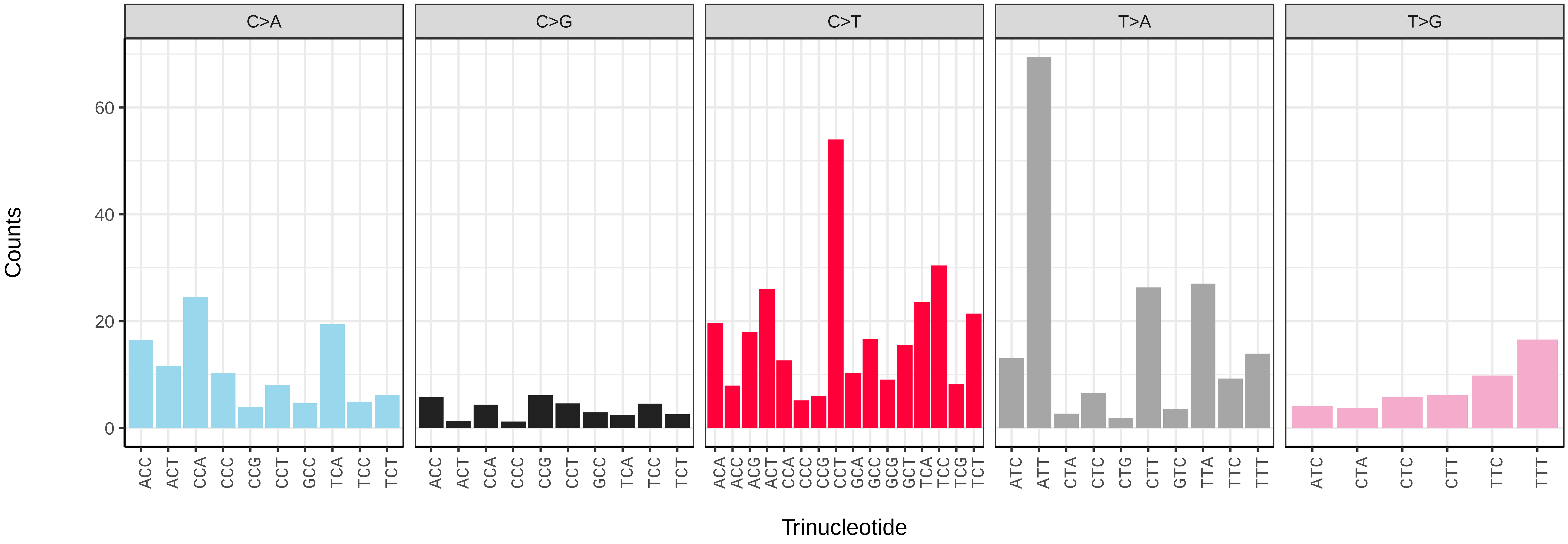

*Myopa tessellatipennis* (idMyoTess1) germline mutational spectrum

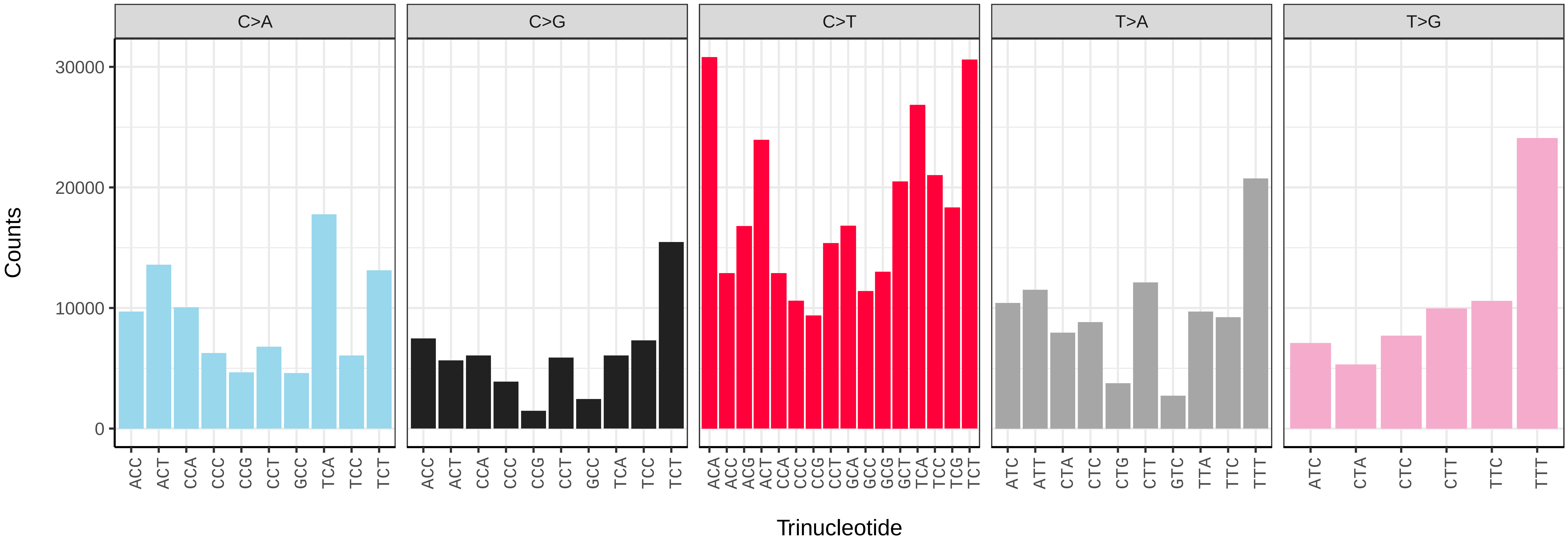


*Myopa tessellatipennis* (idMyoTess1) somatic mutational spectrum (SBS96 classification)

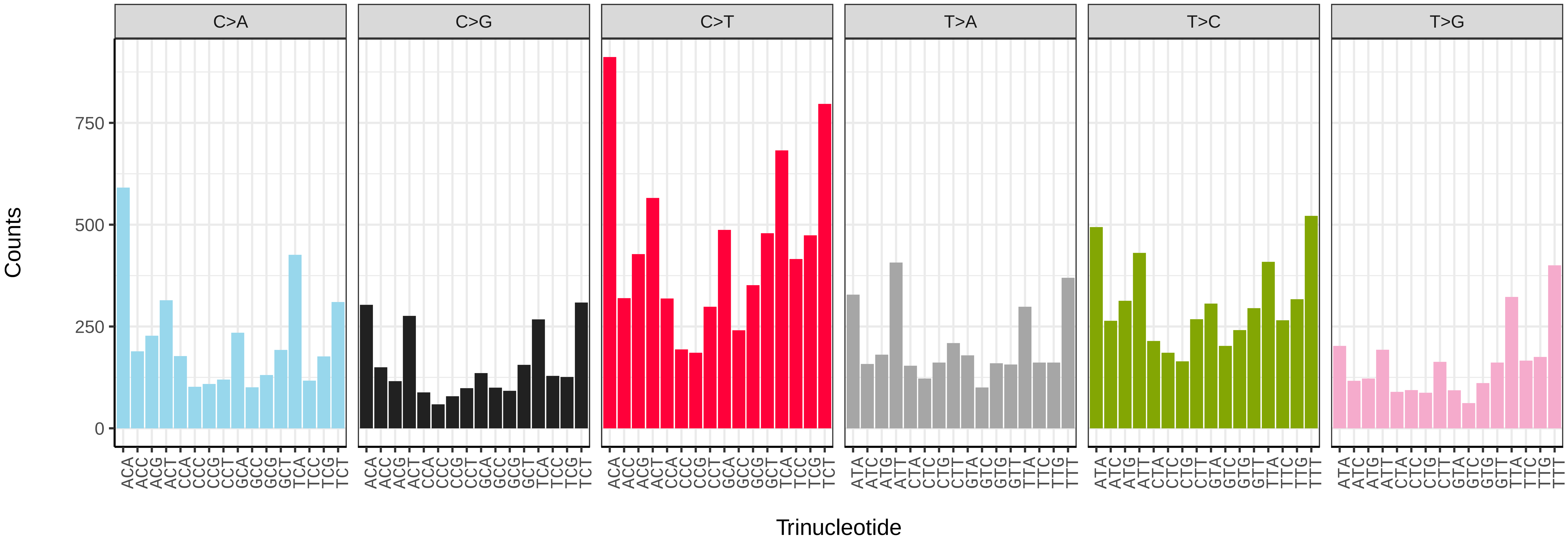


*Myopa tessellatipennis* (idMyoTess1) somatic mutational spectrum (SBS52 classification)

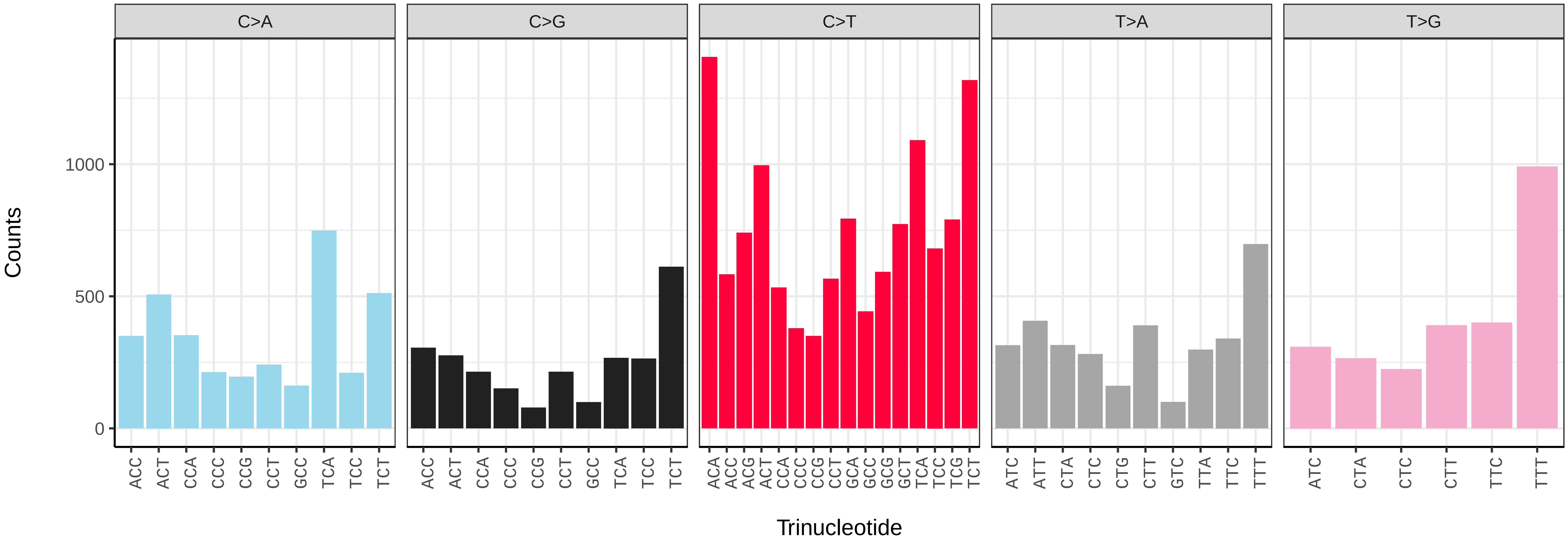

*Nemotelus nigrinus* (idNemNigr1) germline mutational spectrum

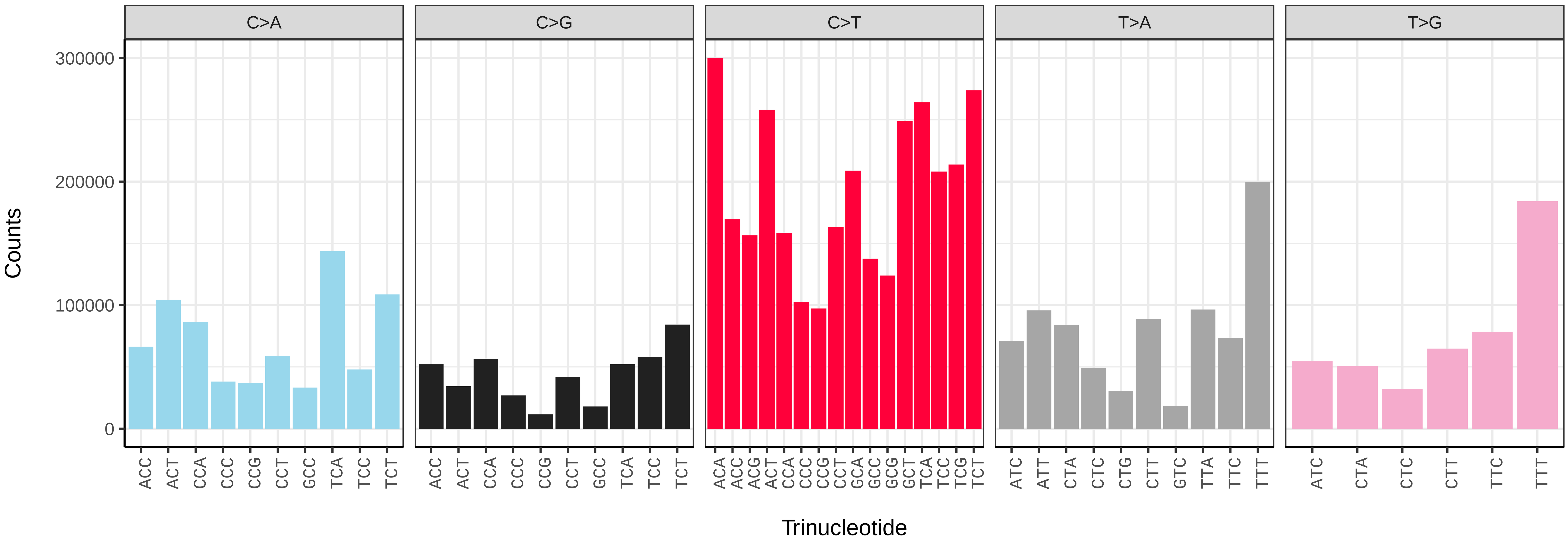


*Nemotelus nigrinus* (idNemNigr1) somatic mutational spectrum (SBS96 classification)

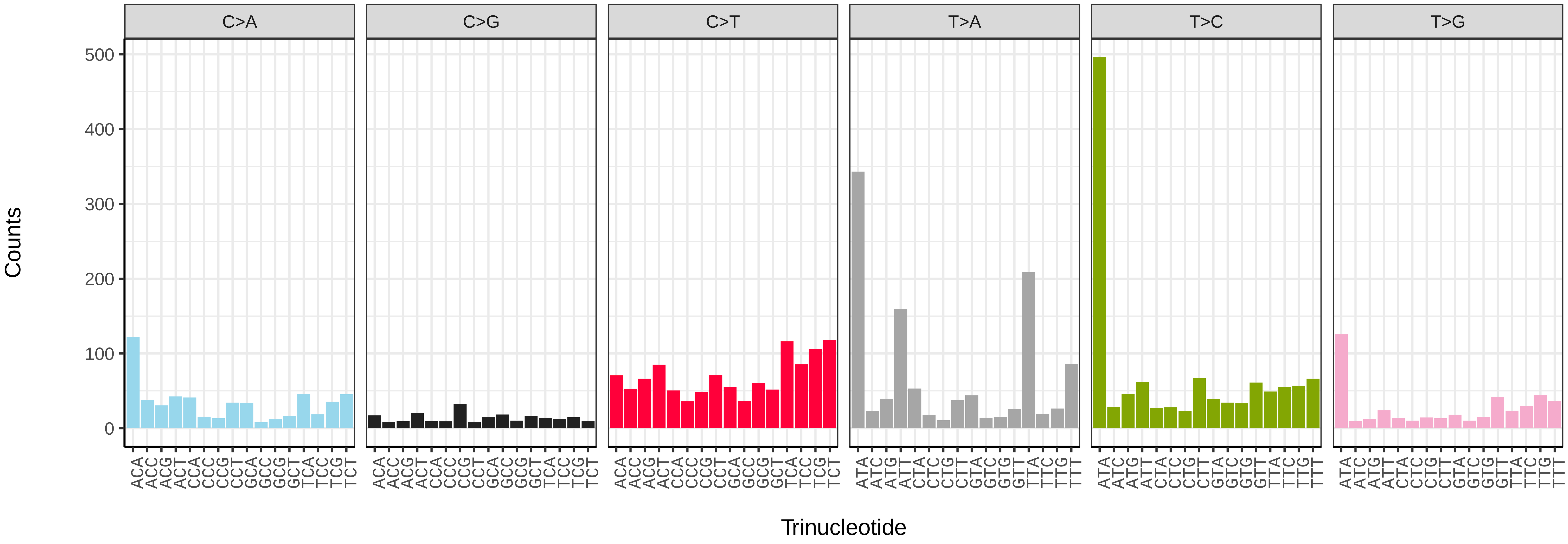


*Nemotelus nigrinus* (idNemNigr1) somatic mutational spectrum (SBS52 classification)

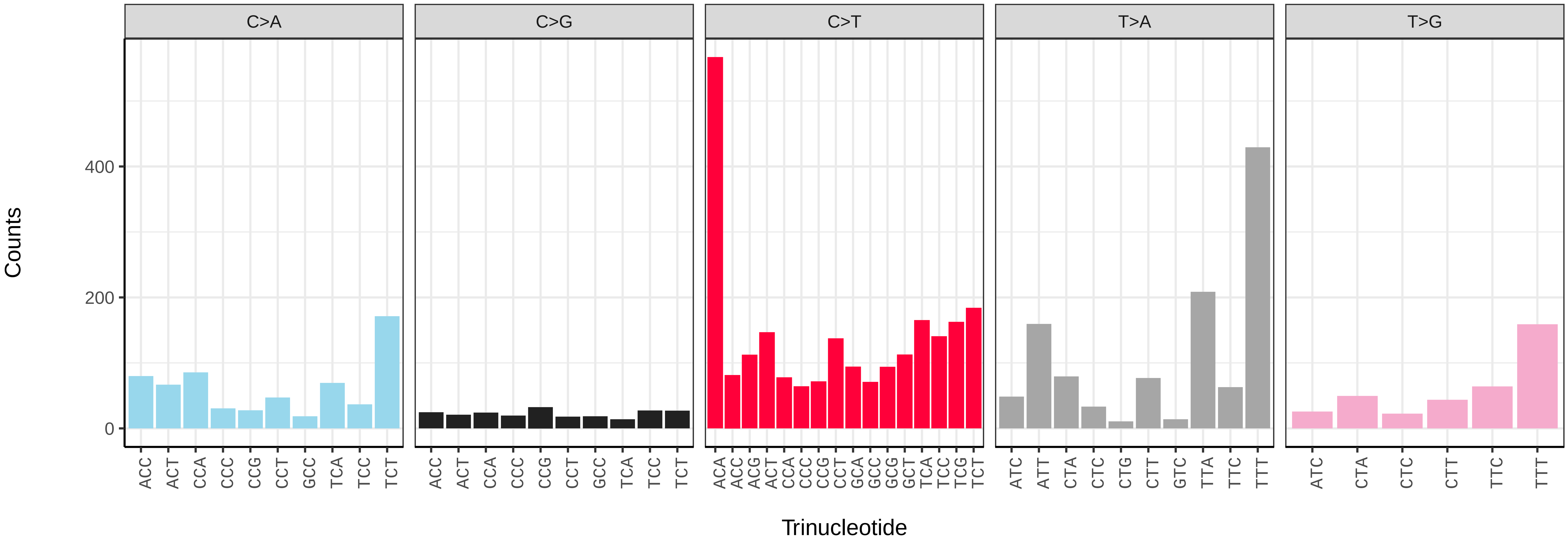

*Neoitamus cyanurus* (idNeoCyan1) germline mutational spectrum

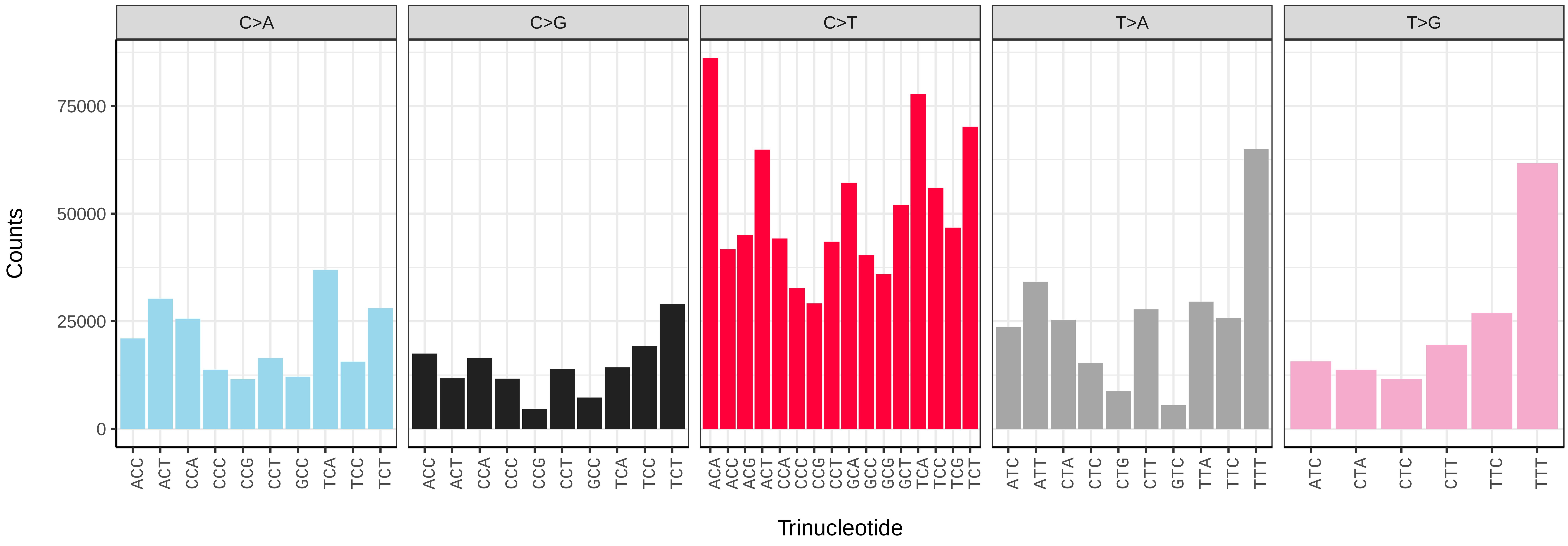


*Neoitamus cyanurus* (idNeoCyan1) somatic mutational spectrum (SBS96 classification)

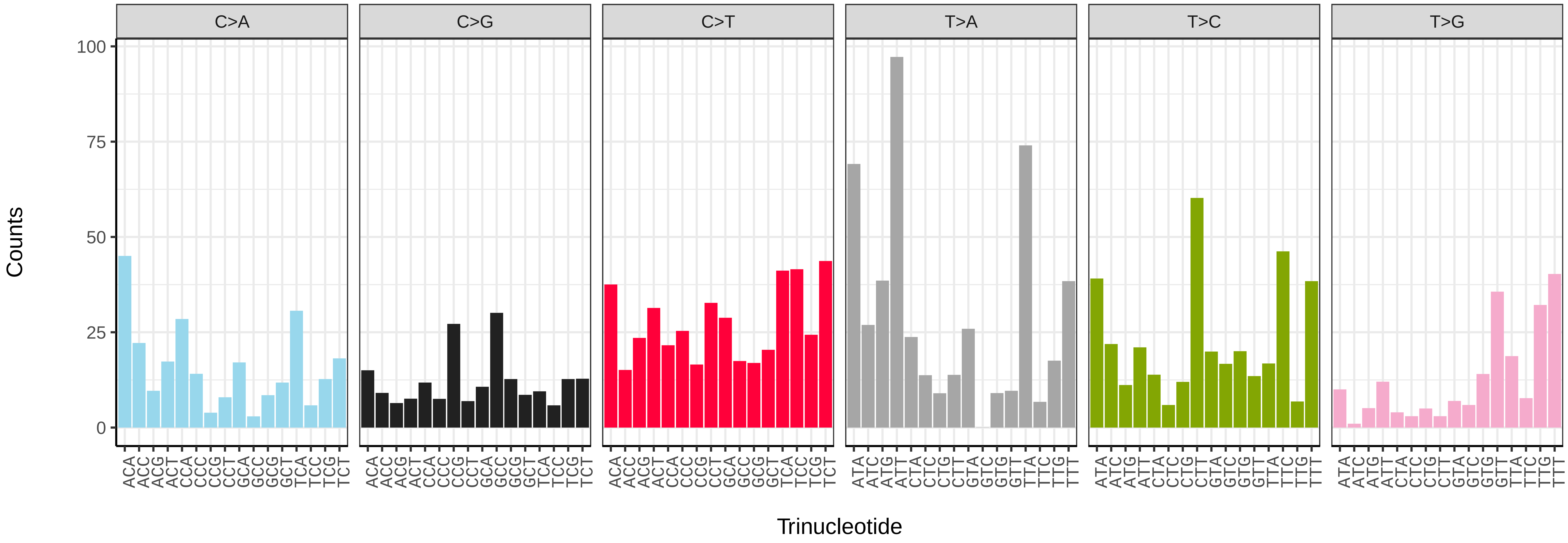


*Neoitamus cyanurus* (idNeoCyan1) somatic mutational spectrum (SBS52 classification)

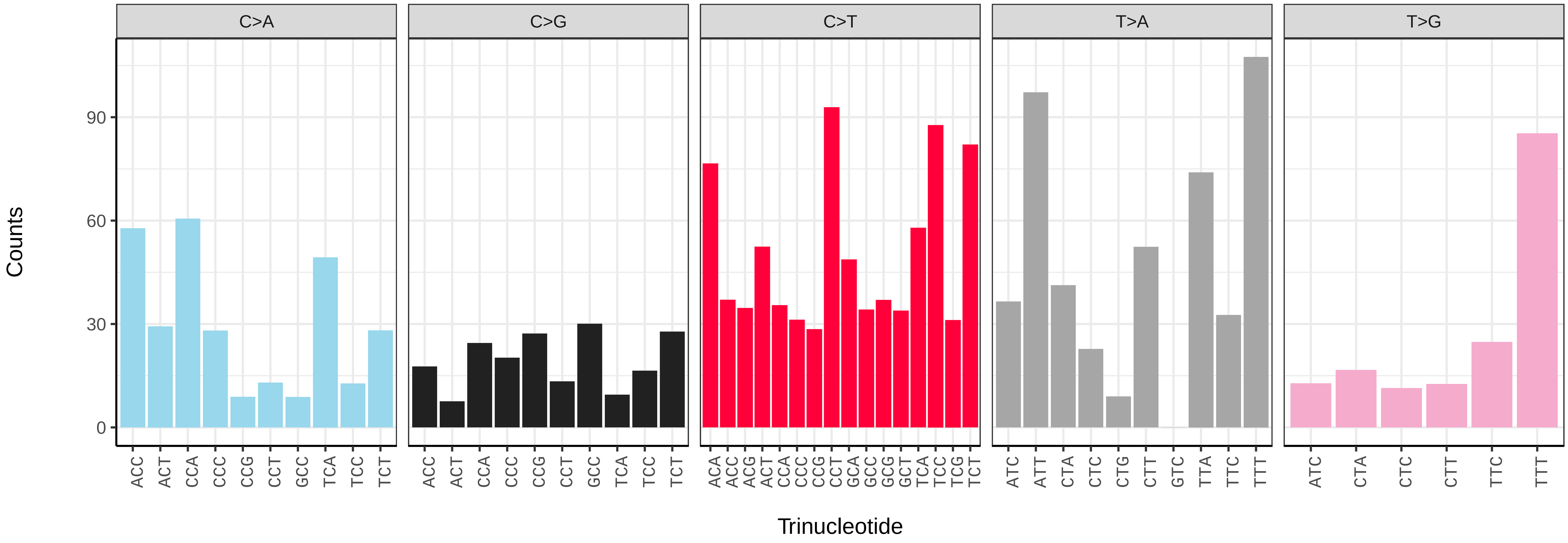

*Neoascia interrupta* (idNeoInte1) germline mutational spectrum

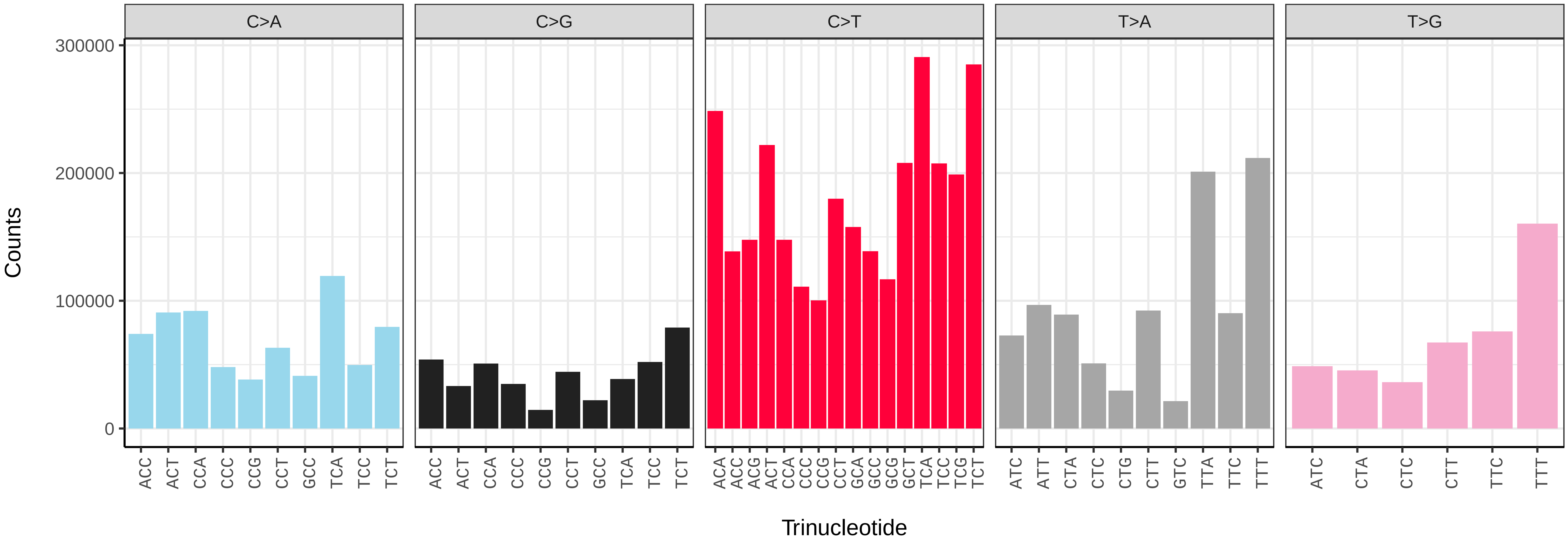


*Neoascia interrupta* (idNeoInte1) somatic mutational spectrum (SBS96 classification)

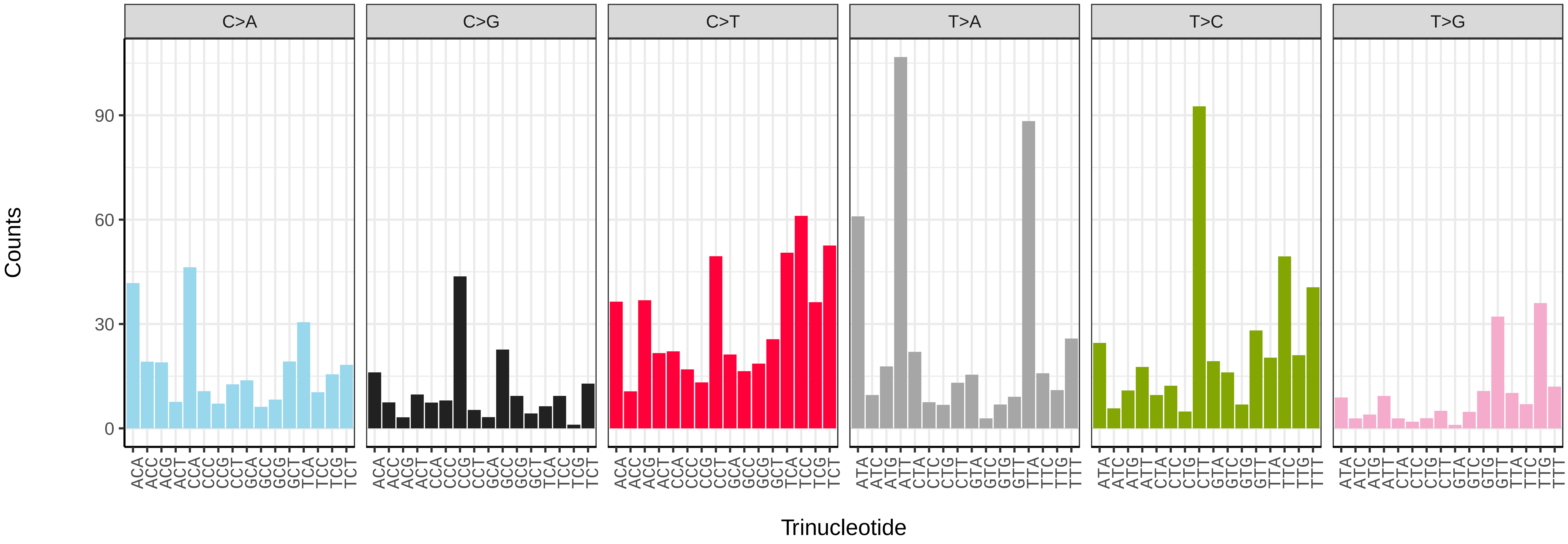


*Neoascia interrupta* (idNeoInte1) somatic mutational spectrum (SBS52 classification)

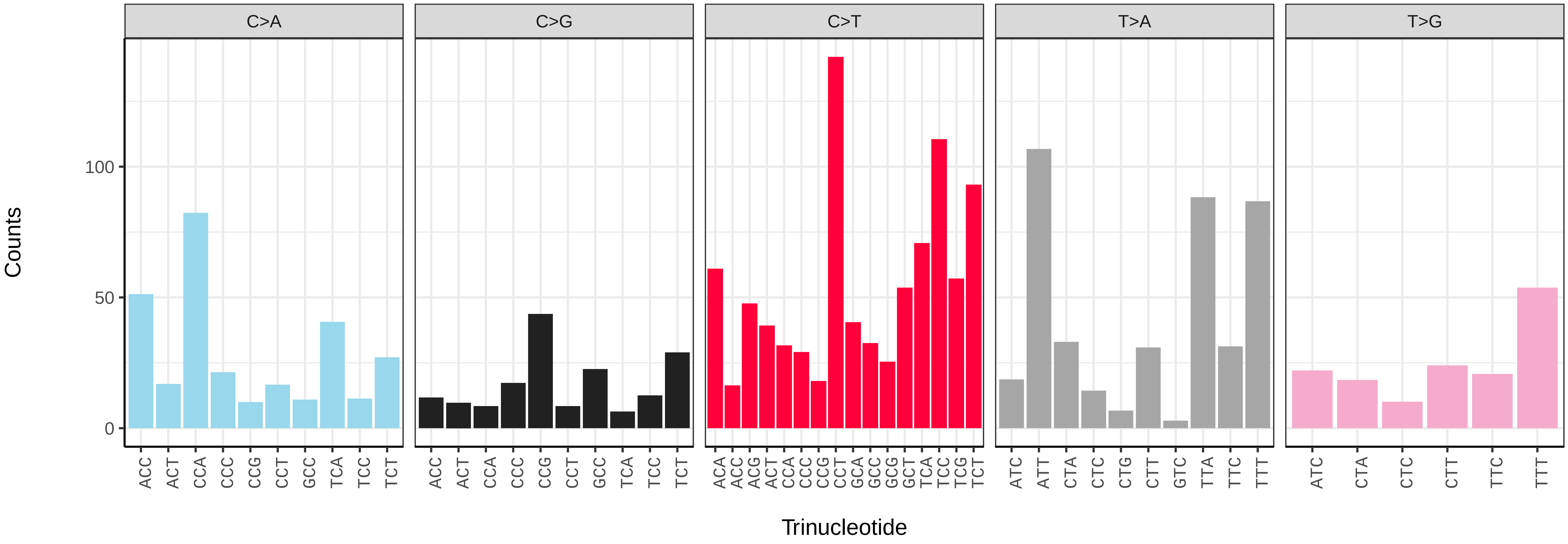

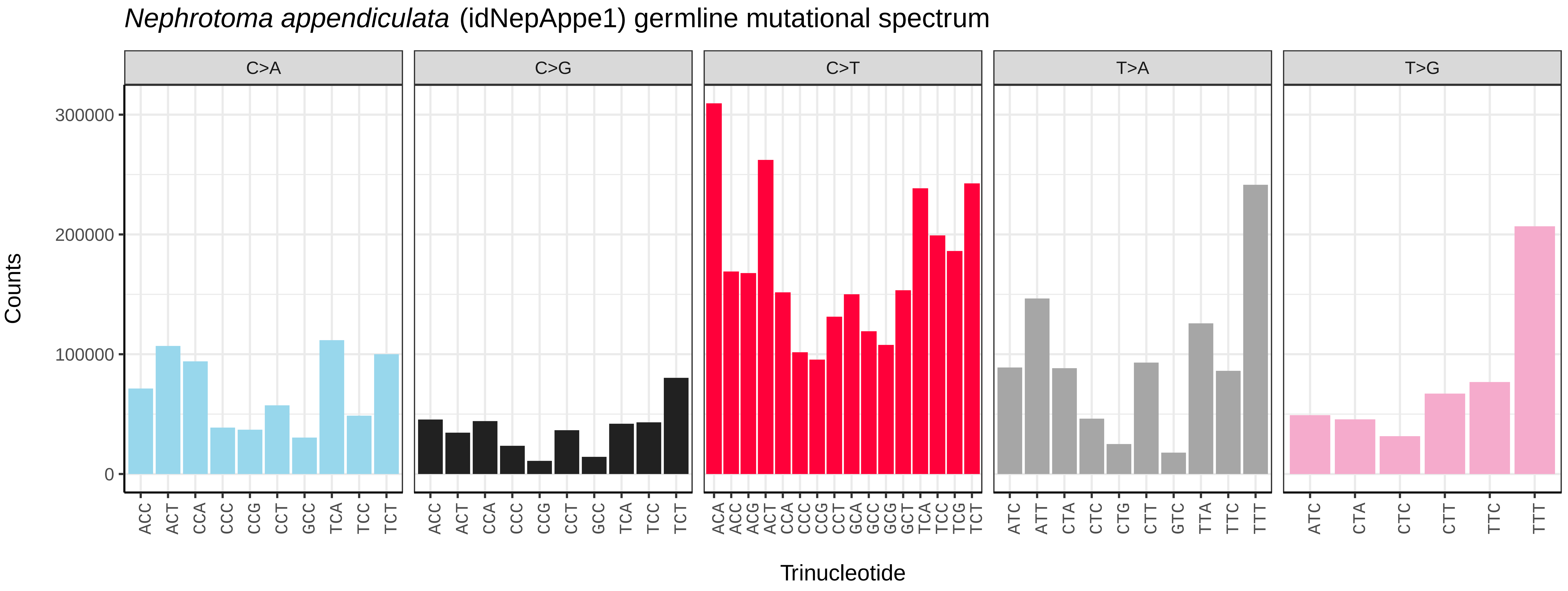
*Nephrotoma appendiculata* (idNepAppe1) germline mutational spectrum
C>A
C>G
C>T
T>A
T>G
Counts
Trinucleotide
300000
200000
100000
0

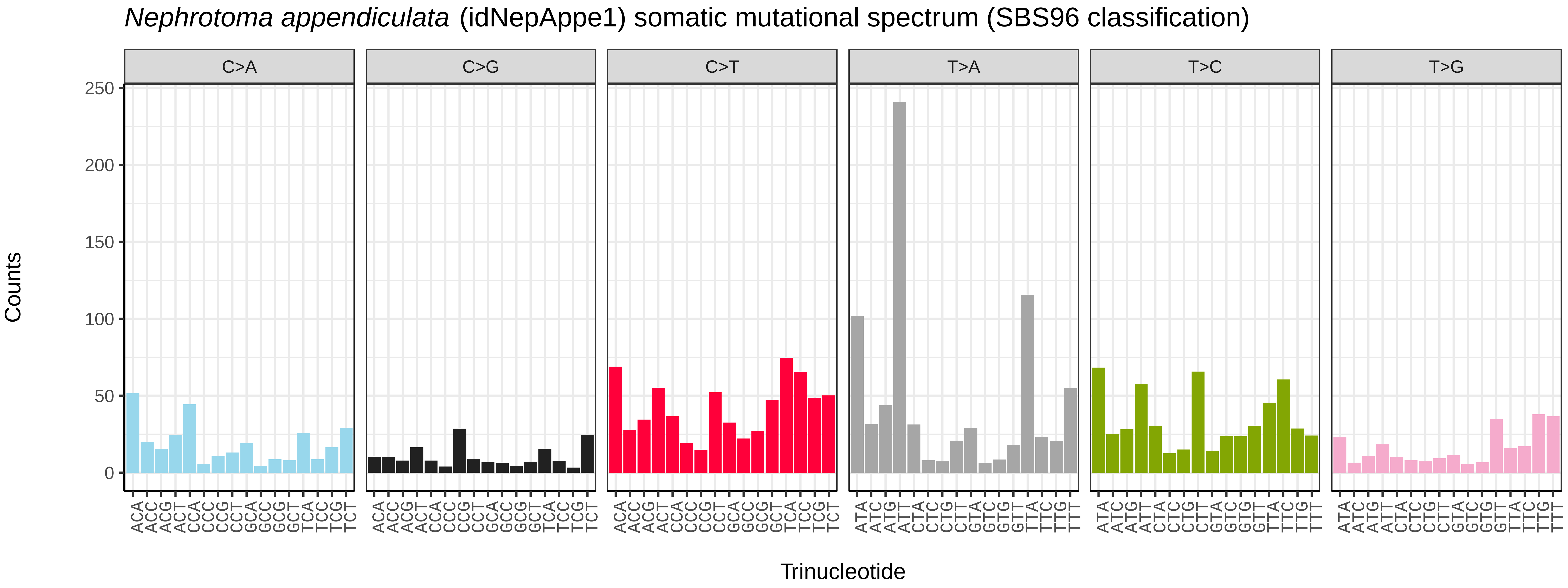
*Nephrotoma appendiculata* (idNepAppe1) somatic mutational spectrum (SBS96 classification)
C>A
C>G
C>T
T>A
T>C
T>G
Counts
Trinucleotide
250
200
150
100
50
0

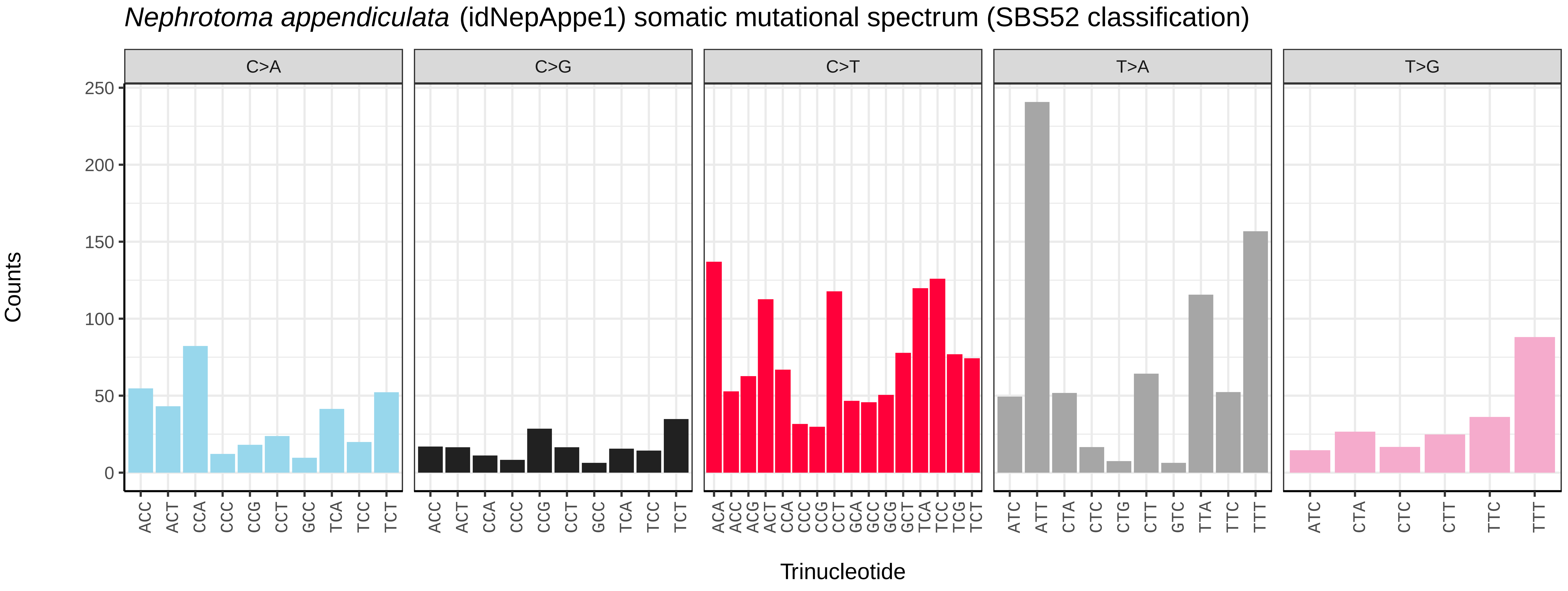
*Nephrotoma appendiculata* (idNepAppe1) somatic mutational spectrum (SBS52 classification)
C>A
C>G
C>T
T>A
T>G
Counts
Trinucleotide
250
200
150
100
50
0

*Nephrotoma flavescens* (idNepFlae1) germline mutational spectrum

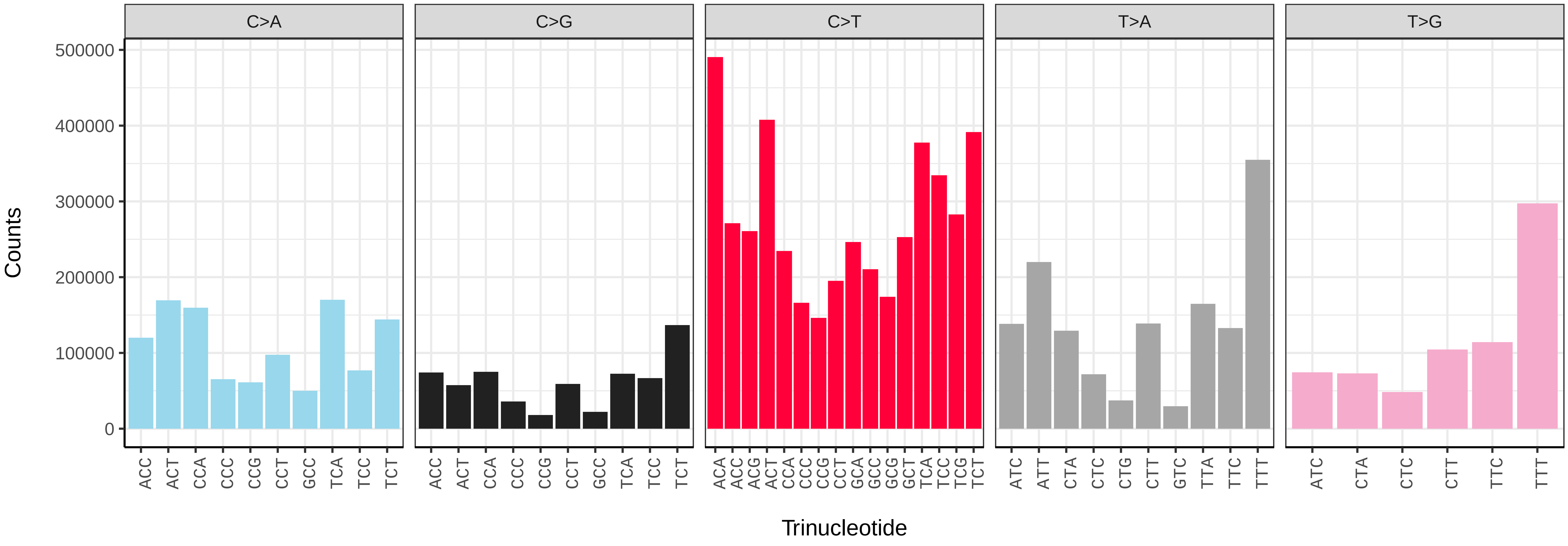


*Nephrotoma flavescens* (idNepFlae1) somatic mutational spectrum (SBS96 classification)

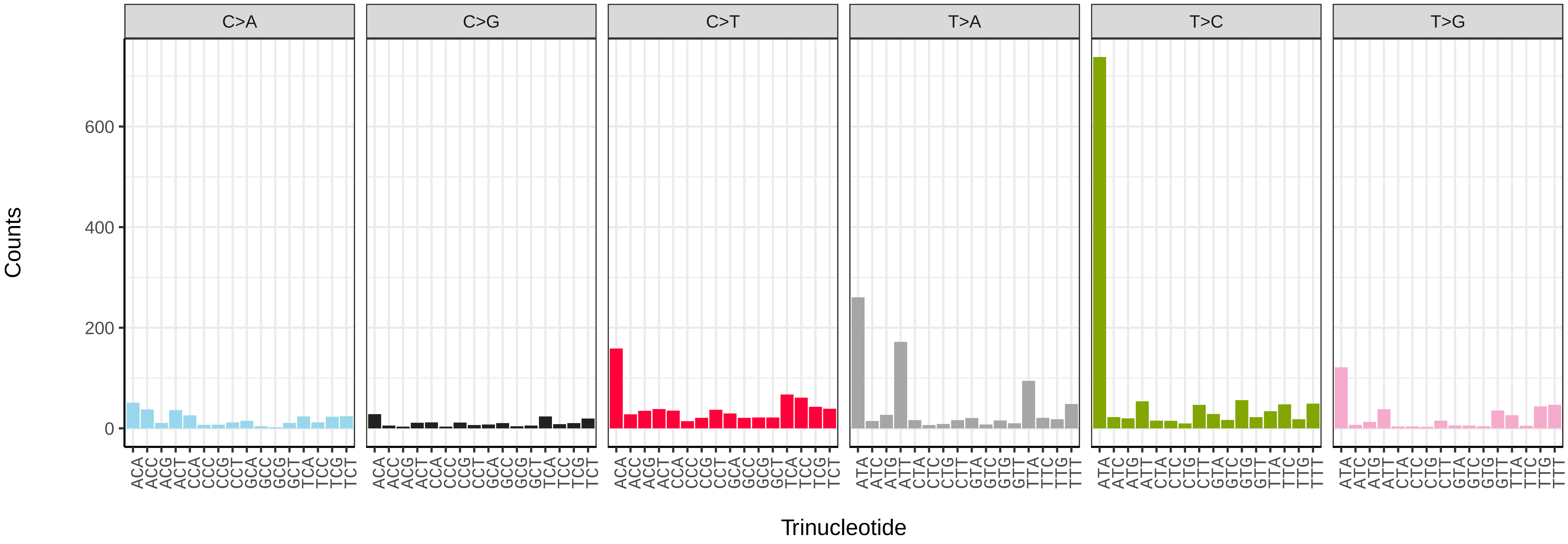


*Nephrotoma flavescens* (idNepFlae1) somatic mutational spectrum (SBS52 classification)

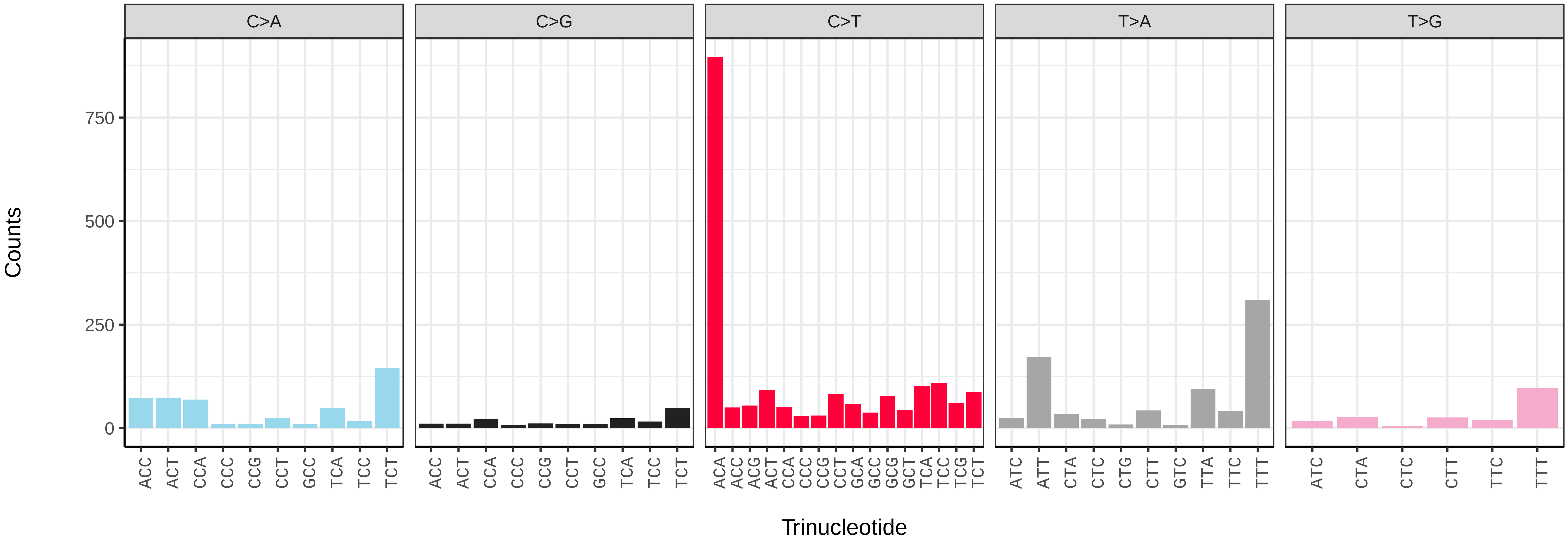

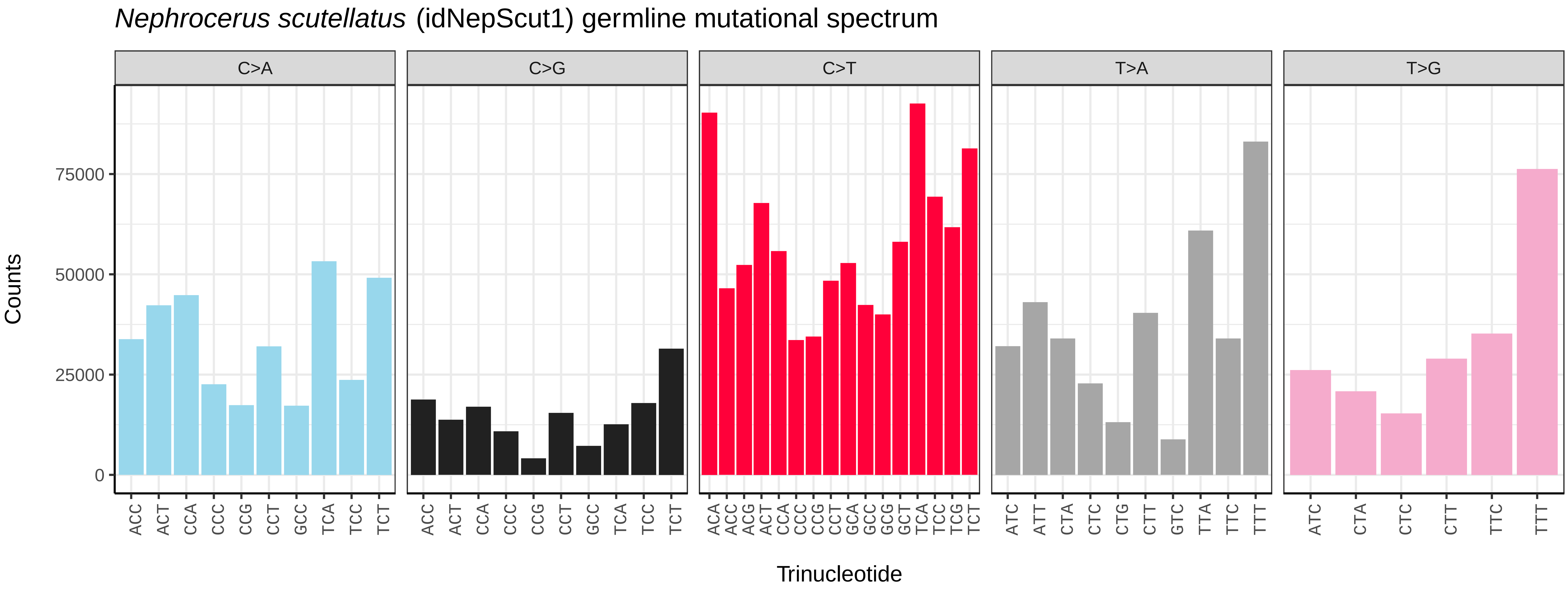
*Nephrocerus scutellatus* (idNepScut1) germline mutational spectrum
C>A
C>G
C>T
T>A
T>G
Counts
0
25000
50000
75000
Trinucleotide

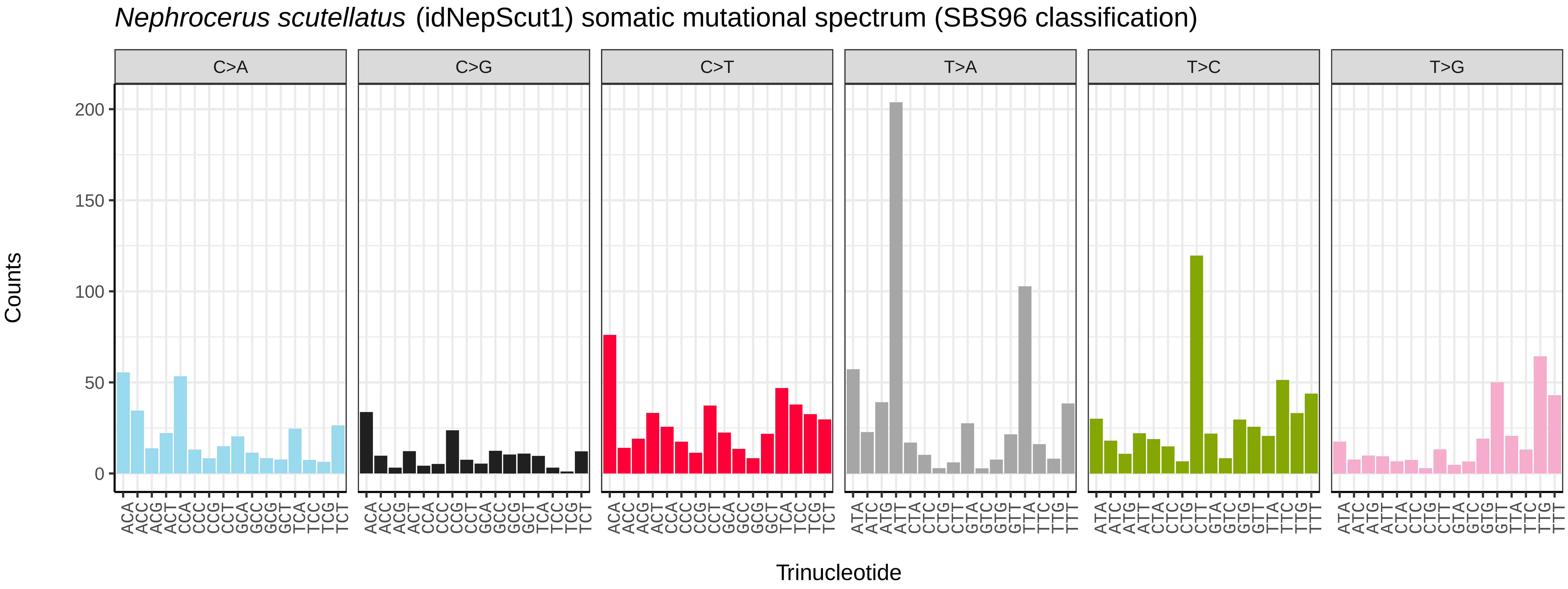
*Nephrocerus scutellatus* (idNepScut1) somatic mutational spectrum (SBS96 classification)
C>A
C>G
C>T
T>A
T>C
T>G
Counts
0
50
100
150
200
Trinucleotide

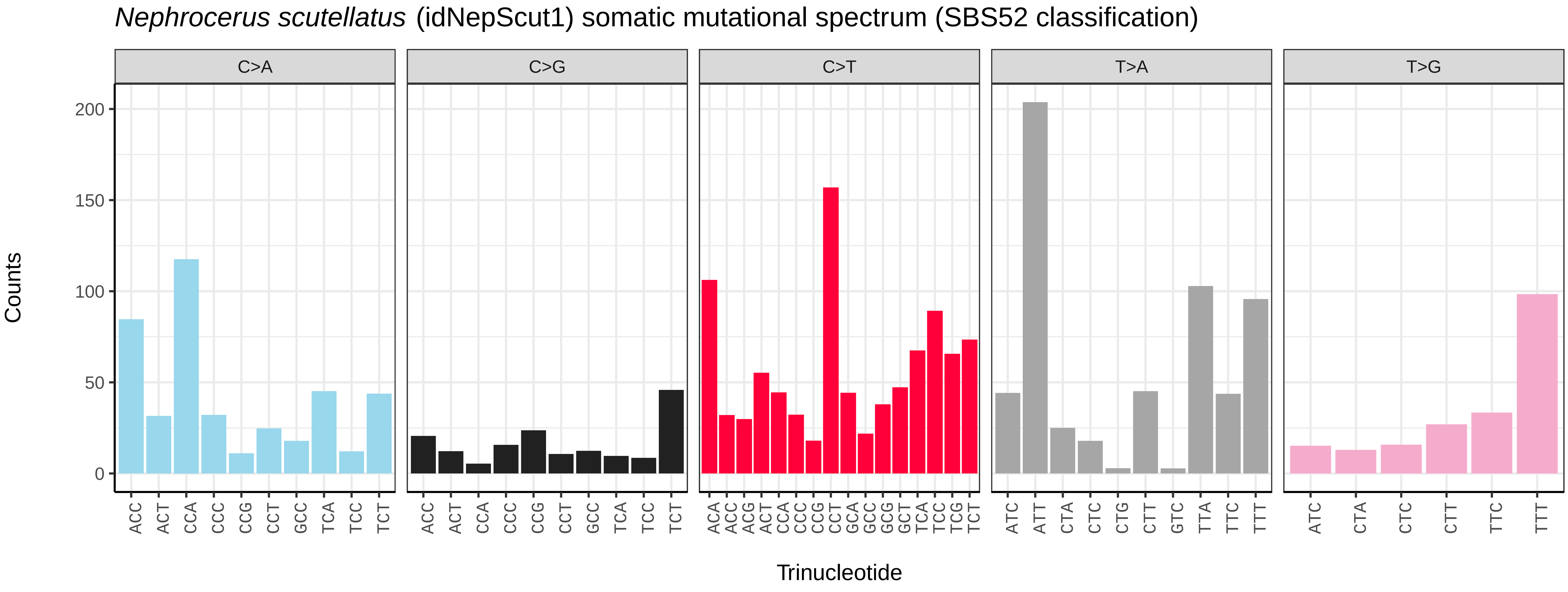
*Nephrocerus scutellatus* (idNepScut1) somatic mutational spectrum (SBS52 classification)
C>A
C>G
C>T
T>A
T>G
Counts
0
50
100
150
200
Trinucleotide

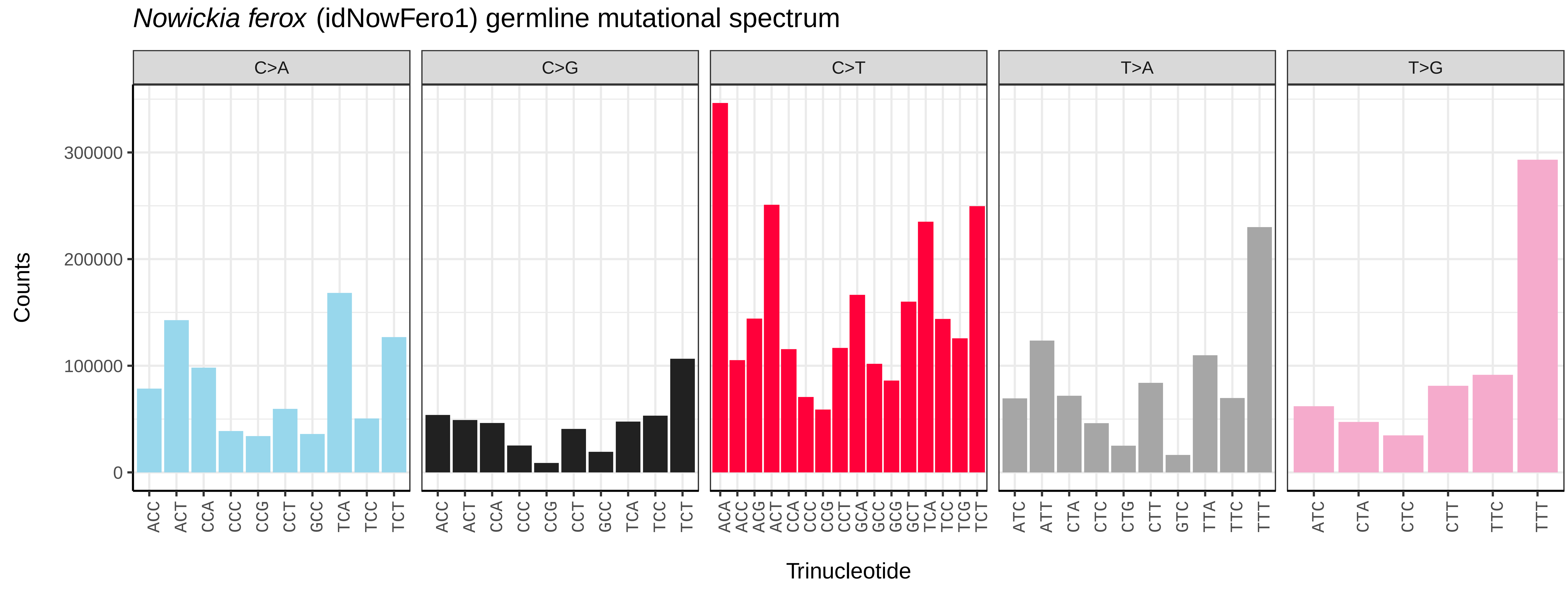
*Nowickia ferox* (idNowFero1) germline mutational spectrum
C>A
C>G
C>T
T>A
T>G
Counts
0
100000
200000
300000
Trinucleotide

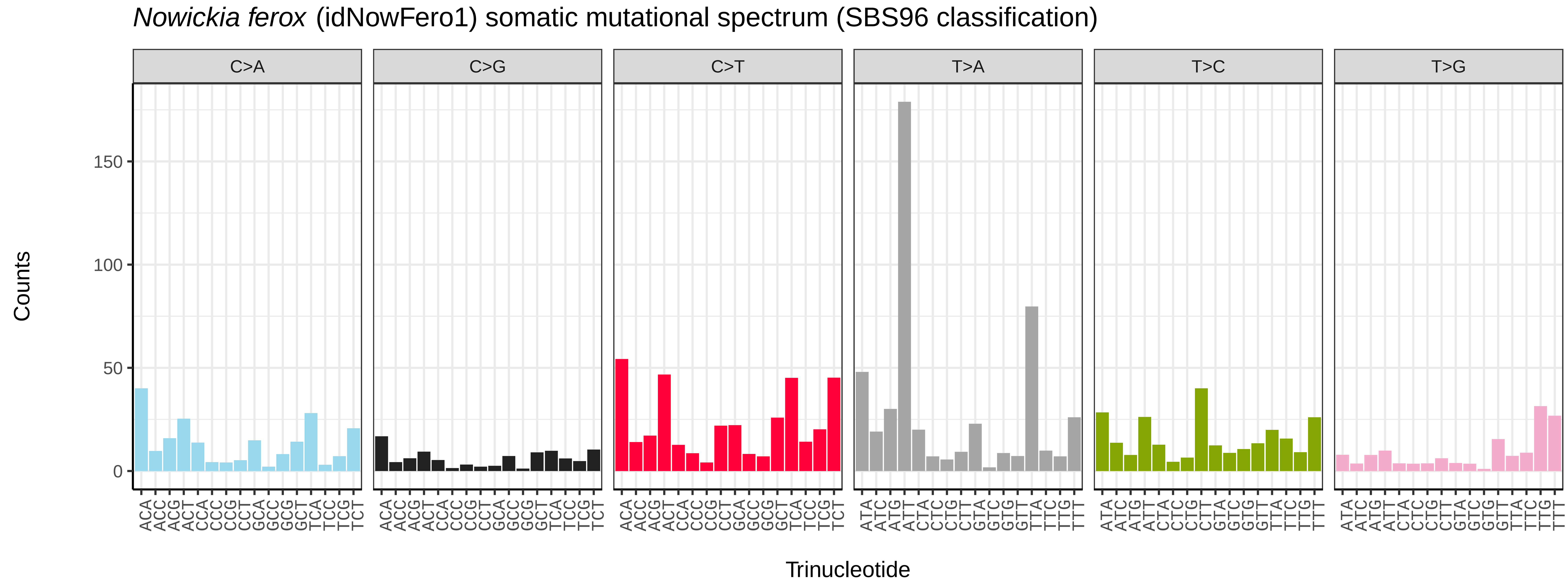
*Nowickia ferox* (idNowFero1) somatic mutational spectrum (SBS96 classification)
C>A
C>G
C>T
T>A
T>C
T>G
Counts
0
50
100
150
Trinucleotide

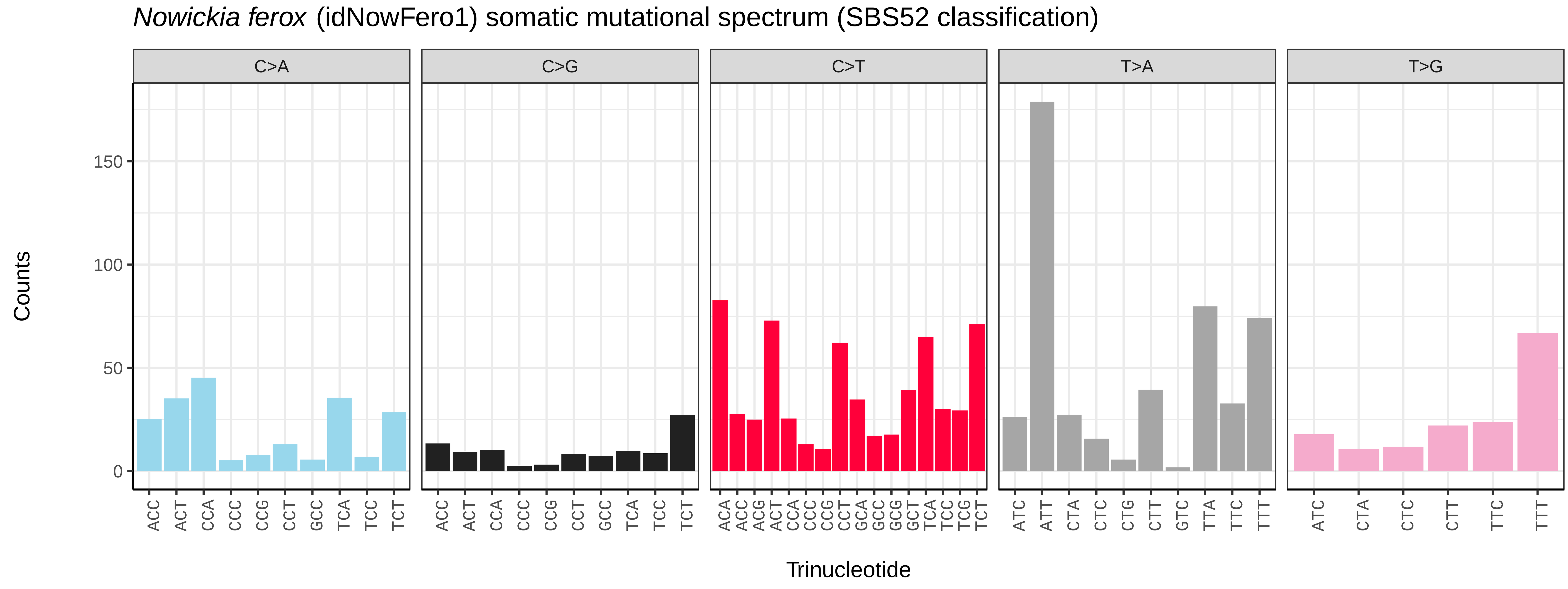
*Nowickia ferox* (idNowFero1) somatic mutational spectrum (SBS52 classification)
C>A
C>G
C>T
T>A
T>G
Counts
0
50
100
150
Trinucleotide

*Pherbina coryleti* (idPheCory1) germline mutational spectrum

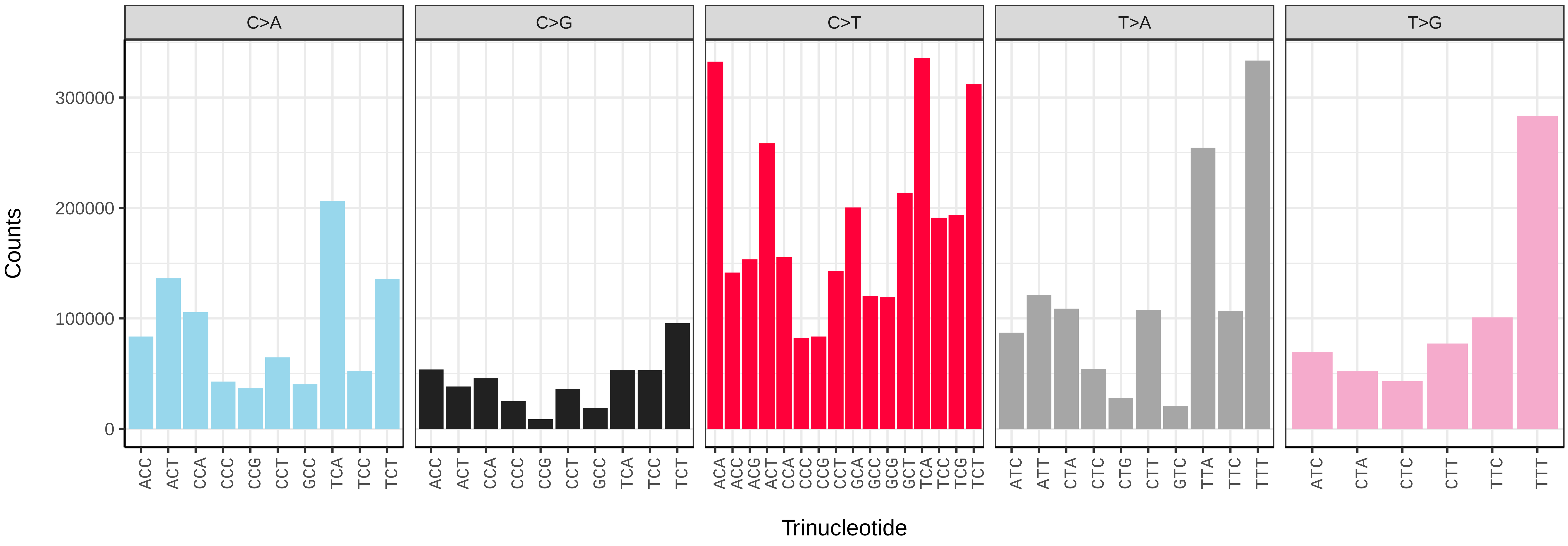


*Pherbina coryleti* (idPheCory1) somatic mutational spectrum (SBS96 classification)

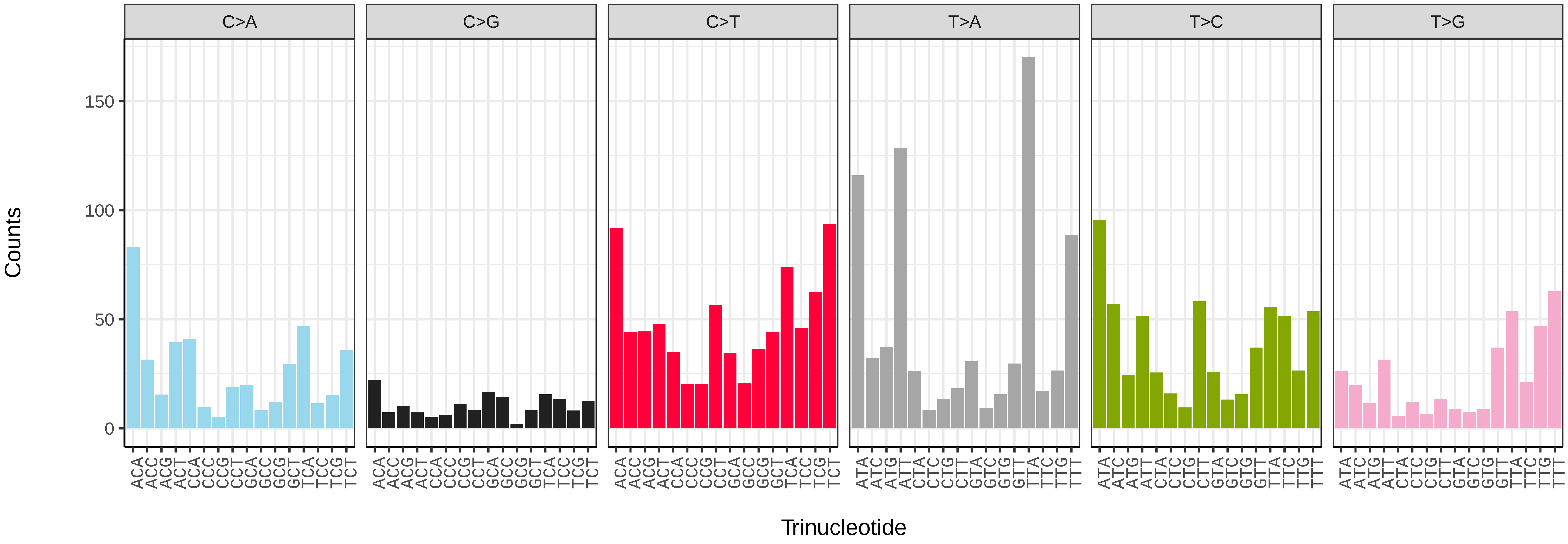


*Pherbina coryleti* (idPheCory1) somatic mutational spectrum (SBS52 classification)

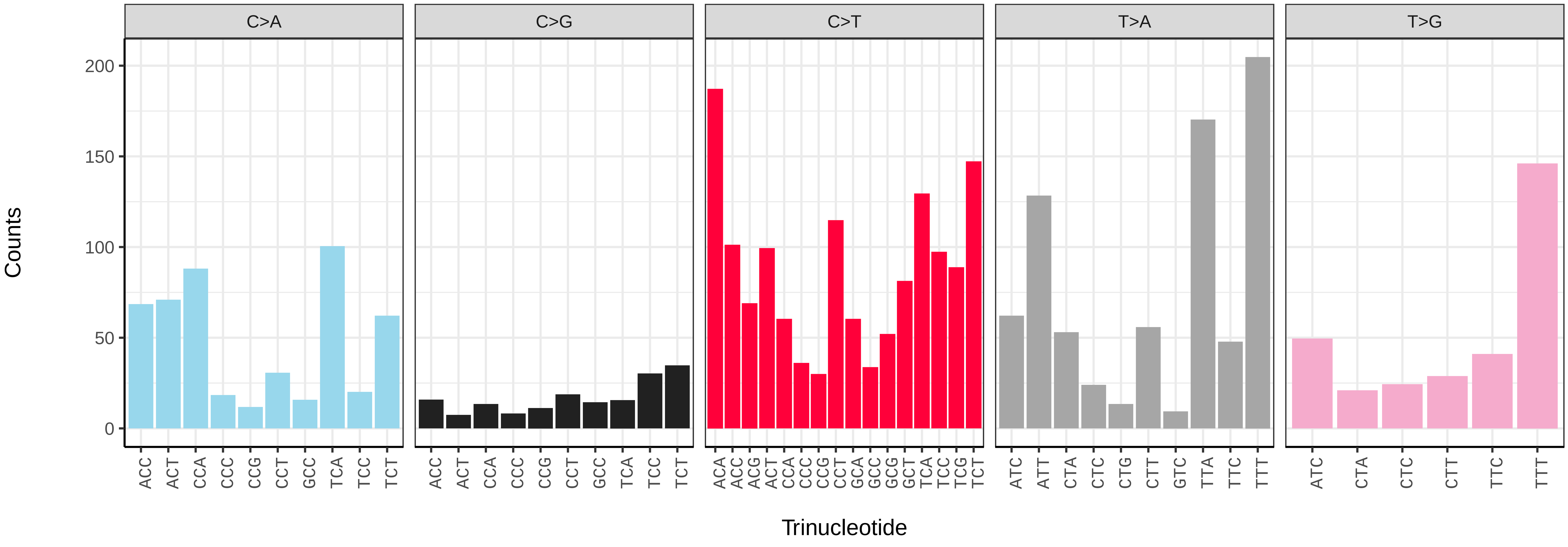

*Phyto melanocephala* (idPhyMeln1) germline mutational spectrum

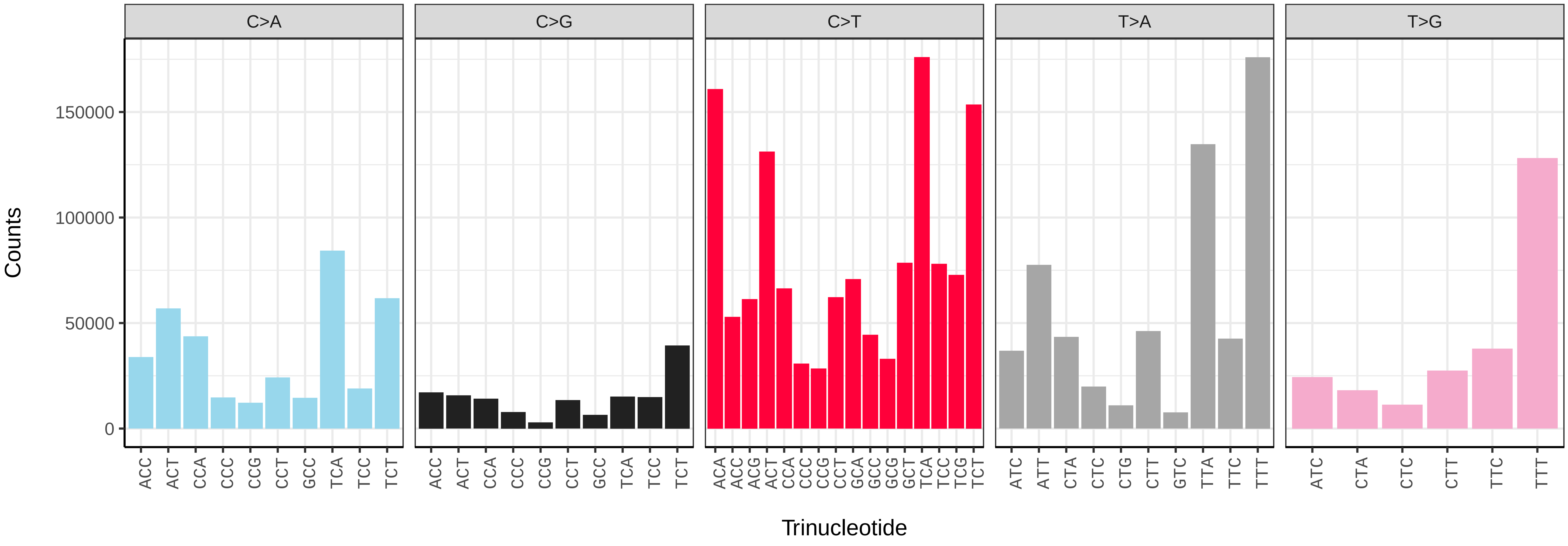


*Phyto melanocephala* (idPhyMeln1) somatic mutational spectrum (SBS96 classification)

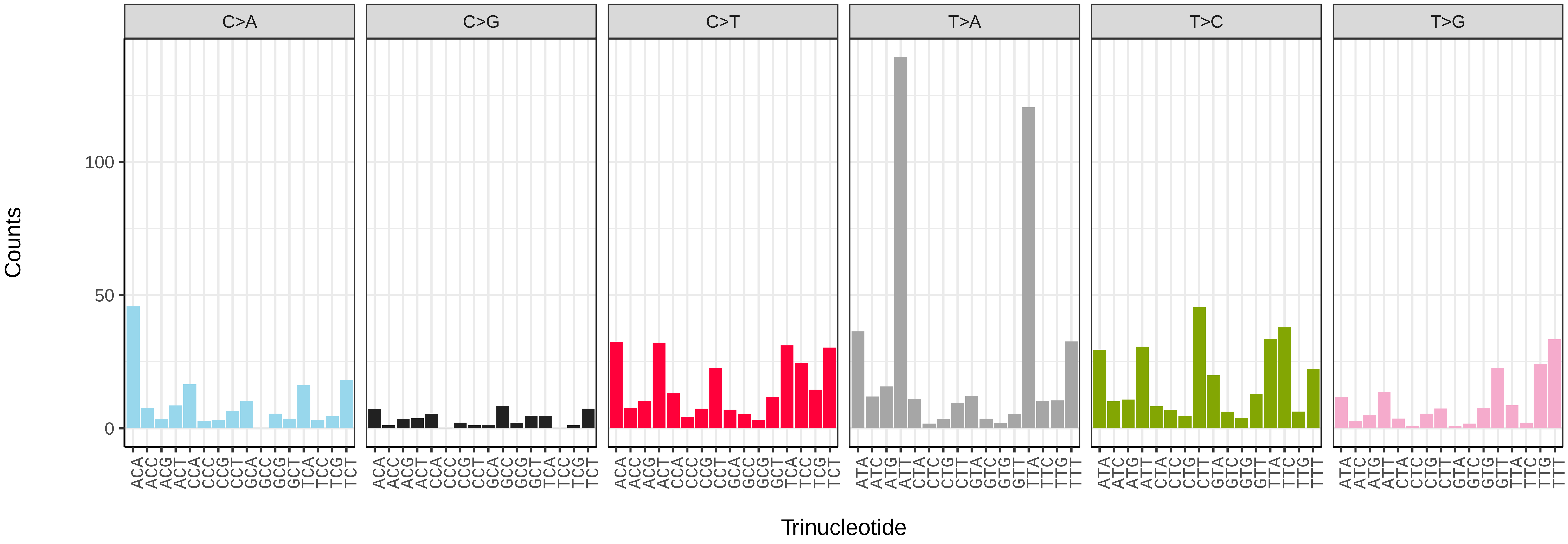


*Phyto melanocephala* (idPhyMeln1) somatic mutational spectrum (SBS52 classification)

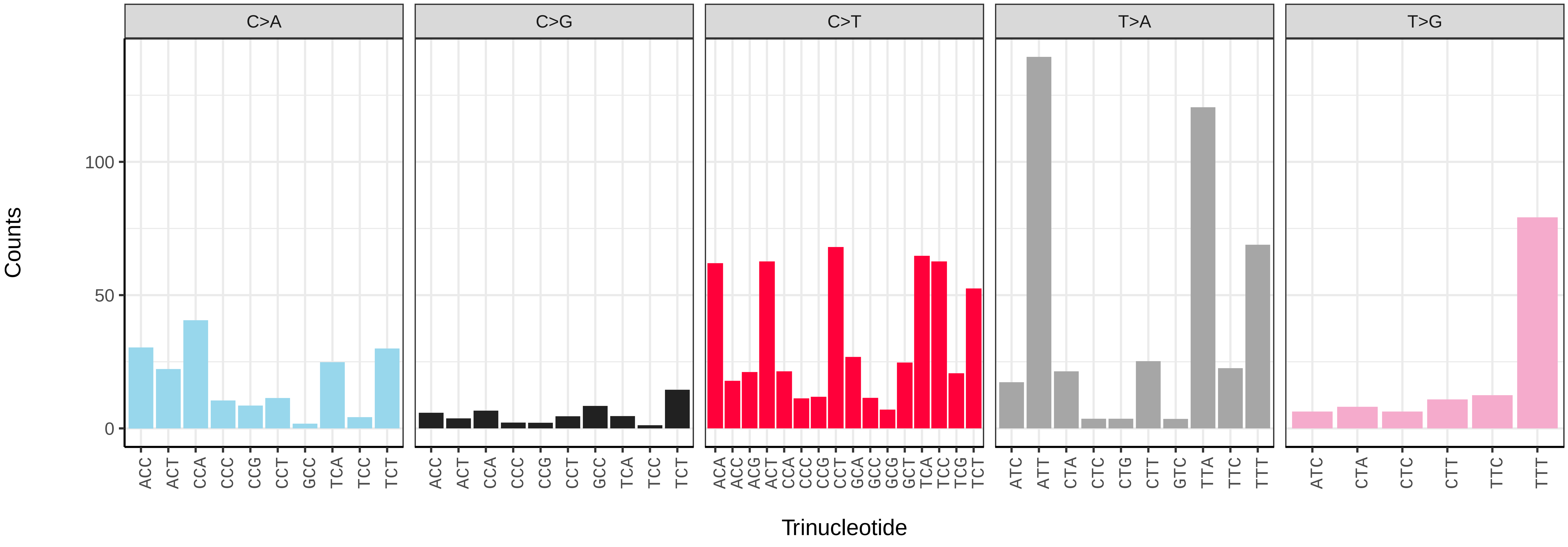

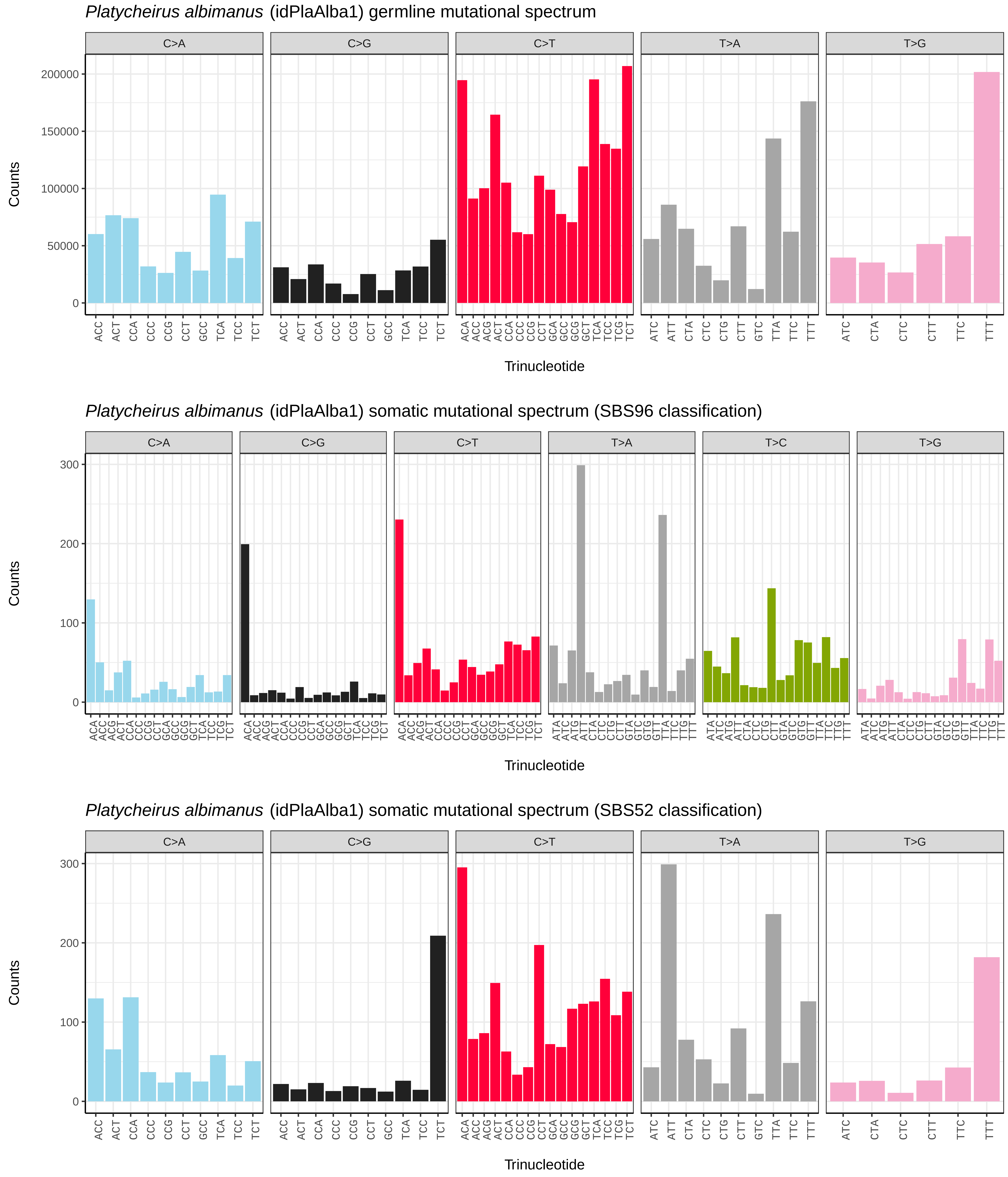
Platycheirus albimanus (idPlaAlba1) germline mutational spectrum
C>A
C>G
C>T
T>A
T>G
Counts
0
50000
100000
150000
200000
Trinucleotide
Platycheirus albimanus (idPlaAlba1) somatic mutational spectrum (SBS96 classification)
C>A
C>G
C>T
T>A
T>C
T>G
Counts
0
100
200
300
Trinucleotide
Platycheirus albimanus (idPlaAlba1) somatic mutational spectrum (SBS52 classification)
C>A
C>G
C>T
T>A
T>G
Counts
0
100
200
300
Trinucleotide

*Platycheirus albimanus* (idPlaAlba1.idPlaAlba3) germline mutational spectrum

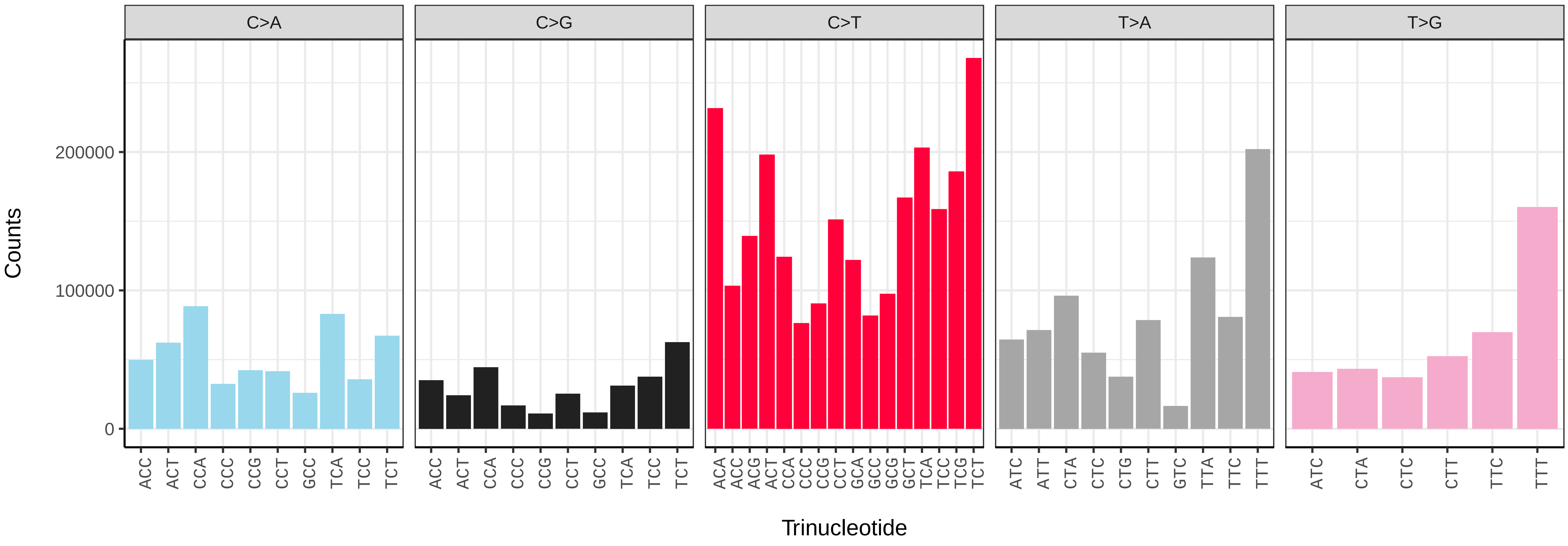


*Platycheirus albimanus* (idPlaAlba1.idPlaAlba3) somatic mutational spectrum (SBS96 classification)

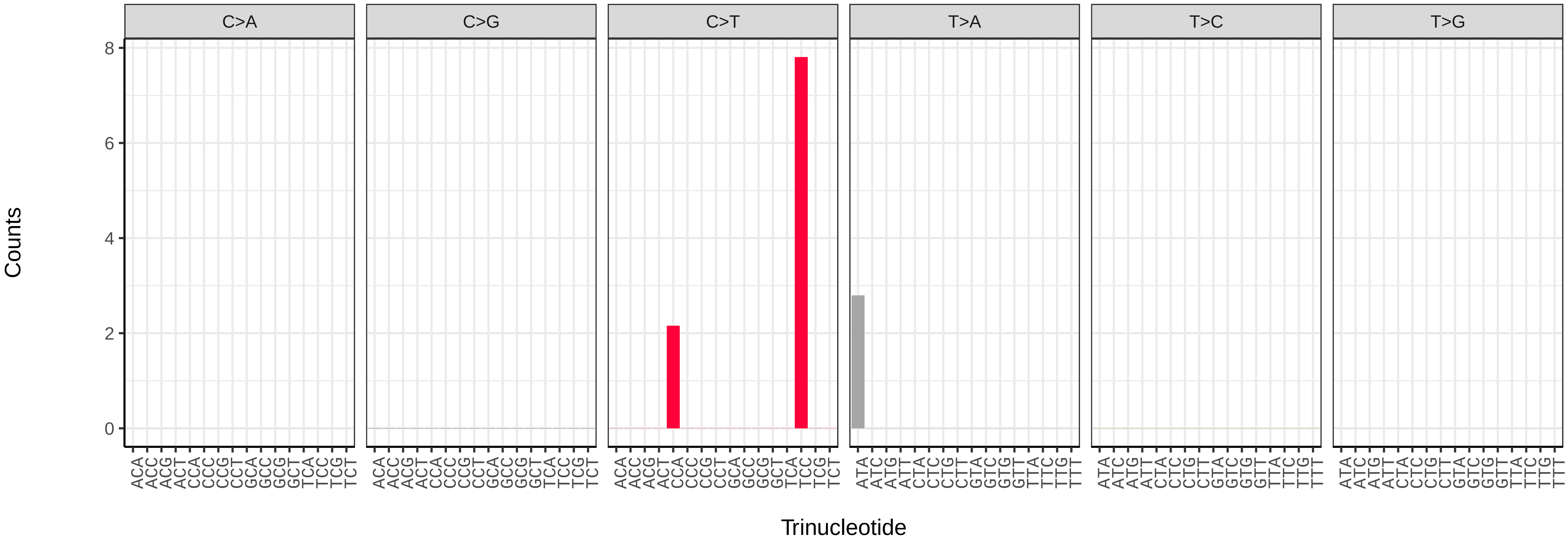


*Platycheirus albimanus* (idPlaAlba1.idPlaAlba3) somatic mutational spectrum (SBS52 classification)

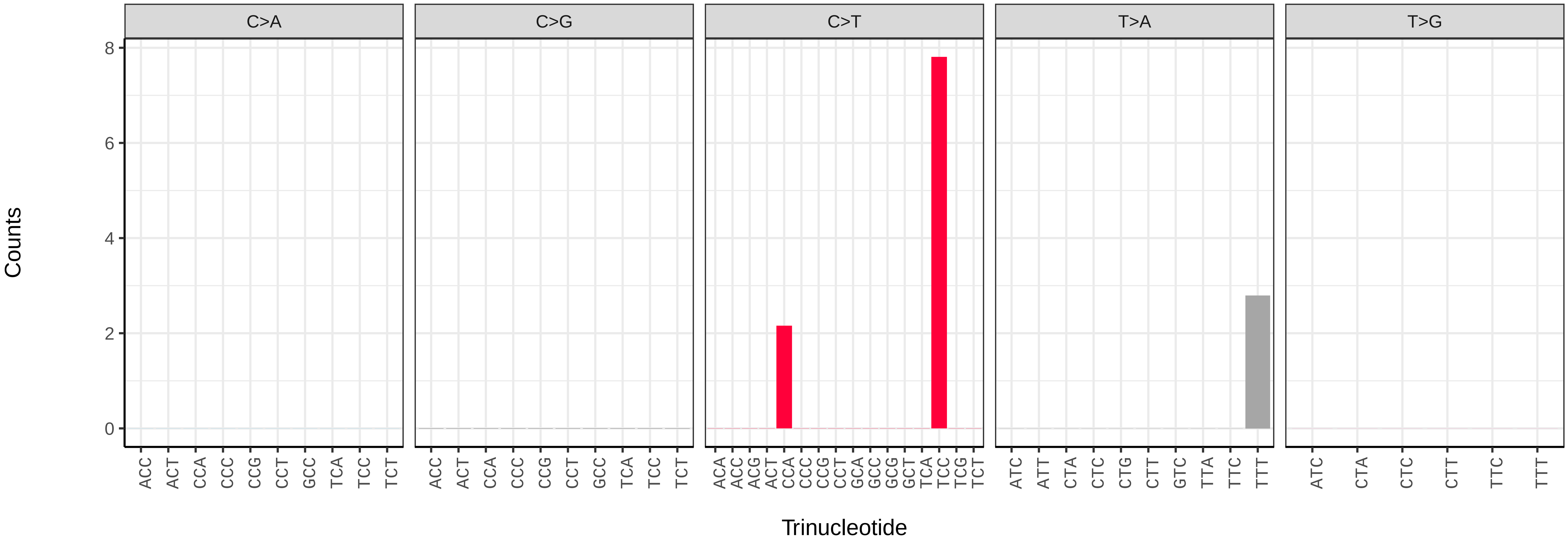

*Platycheirus albimanus* (idPlaAlba1.idPlaAlba4) germline mutational spectrum

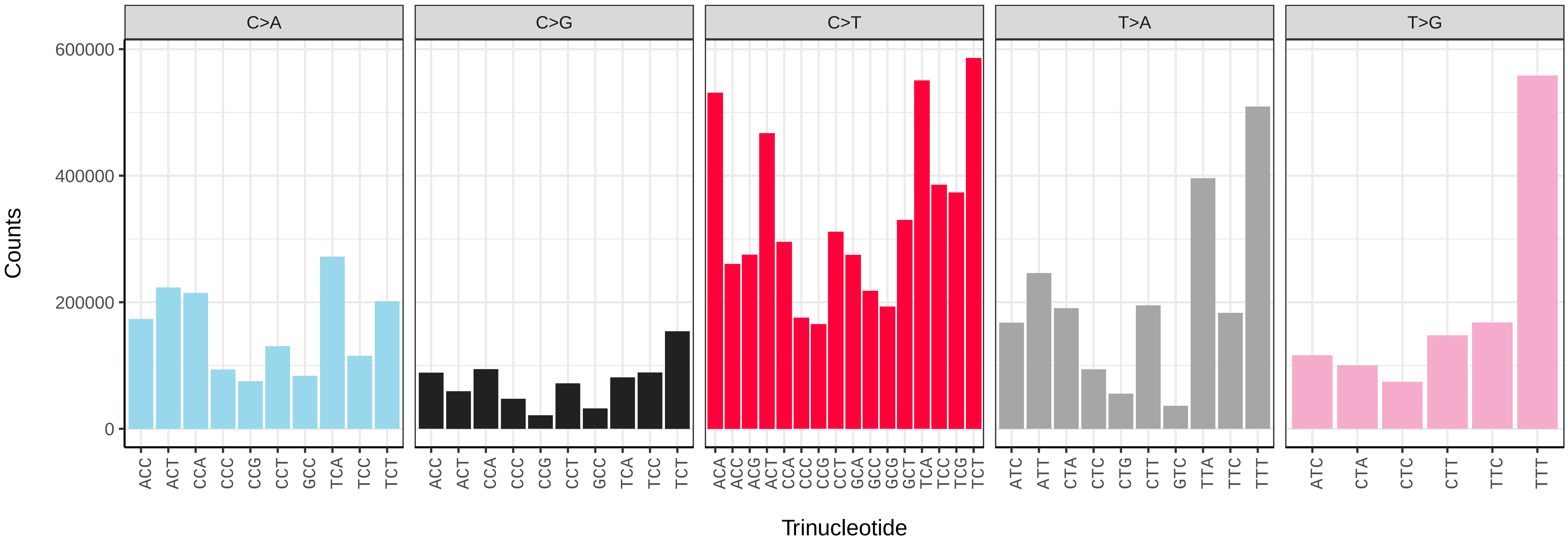


*Platycheirus albimanus* (idPlaAlba1.idPlaAlba4) somatic mutational spectrum (SBS96 classification)

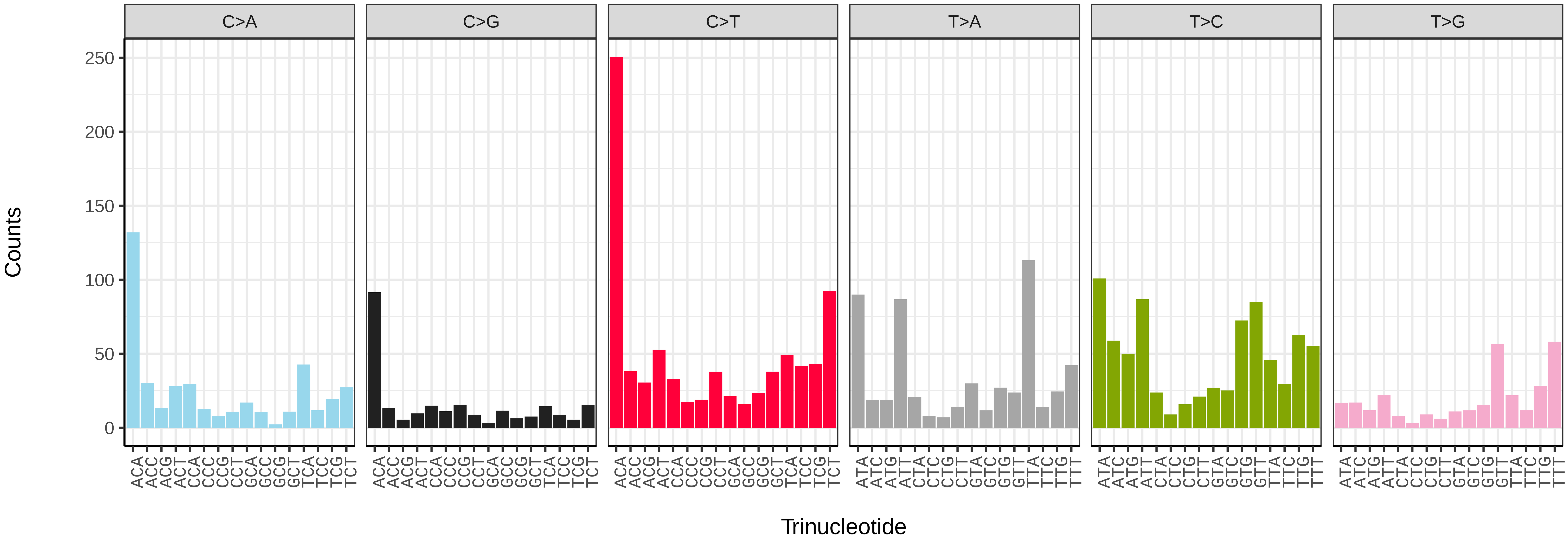


*Platycheirus albimanus* (idPlaAlba1.idPlaAlba4) somatic mutational spectrum (SBS52 classification)

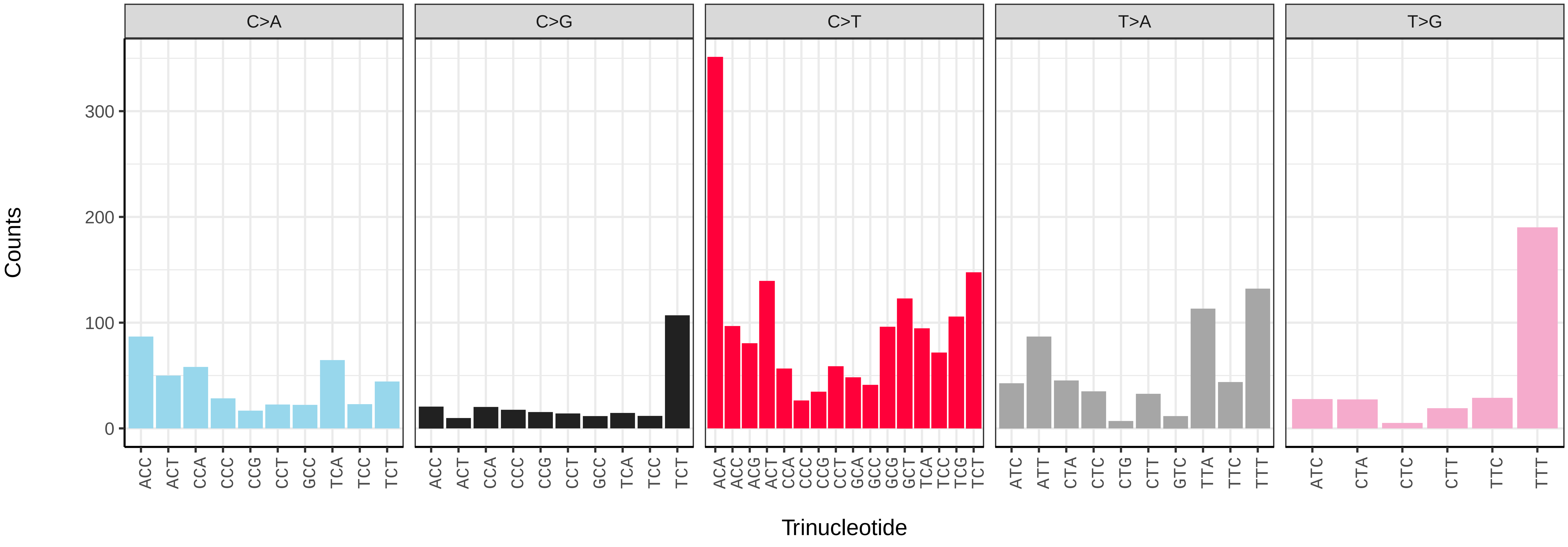

*Poecilobothrus nobilitatus* (idPoeNobi2) germline mutational spectrum

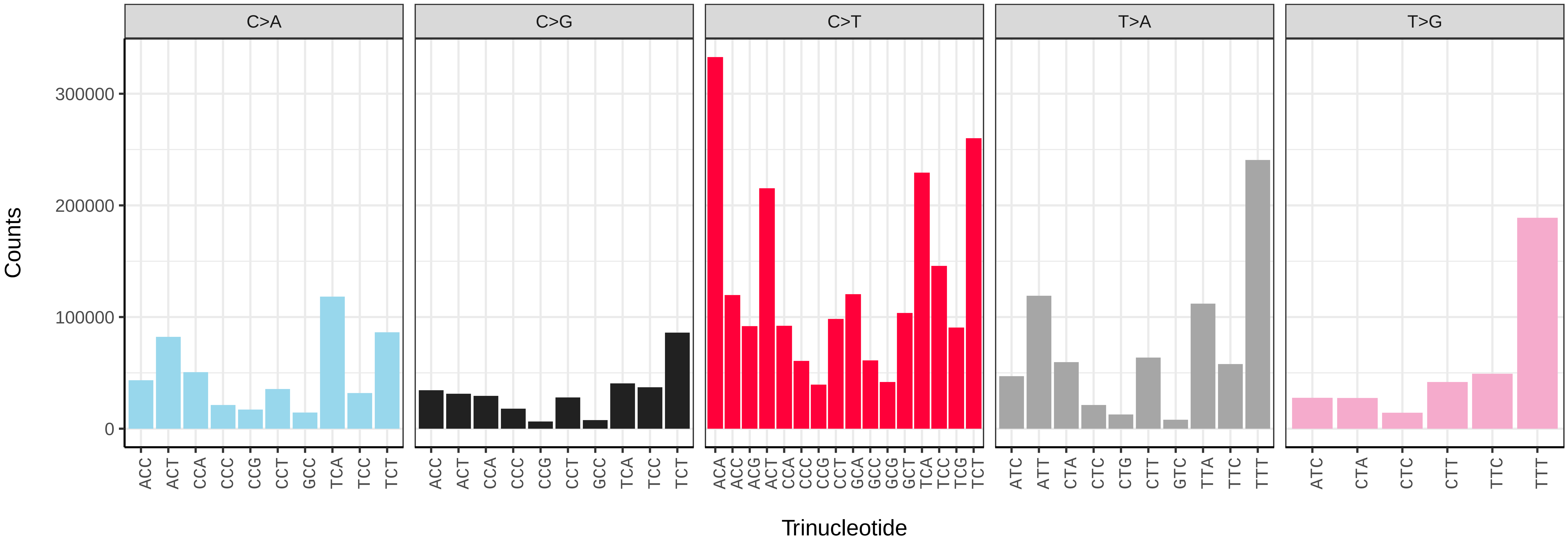


*Poecilobothrus nobilitatus* (idPoeNobi2) somatic mutational spectrum (SBS96 classification)

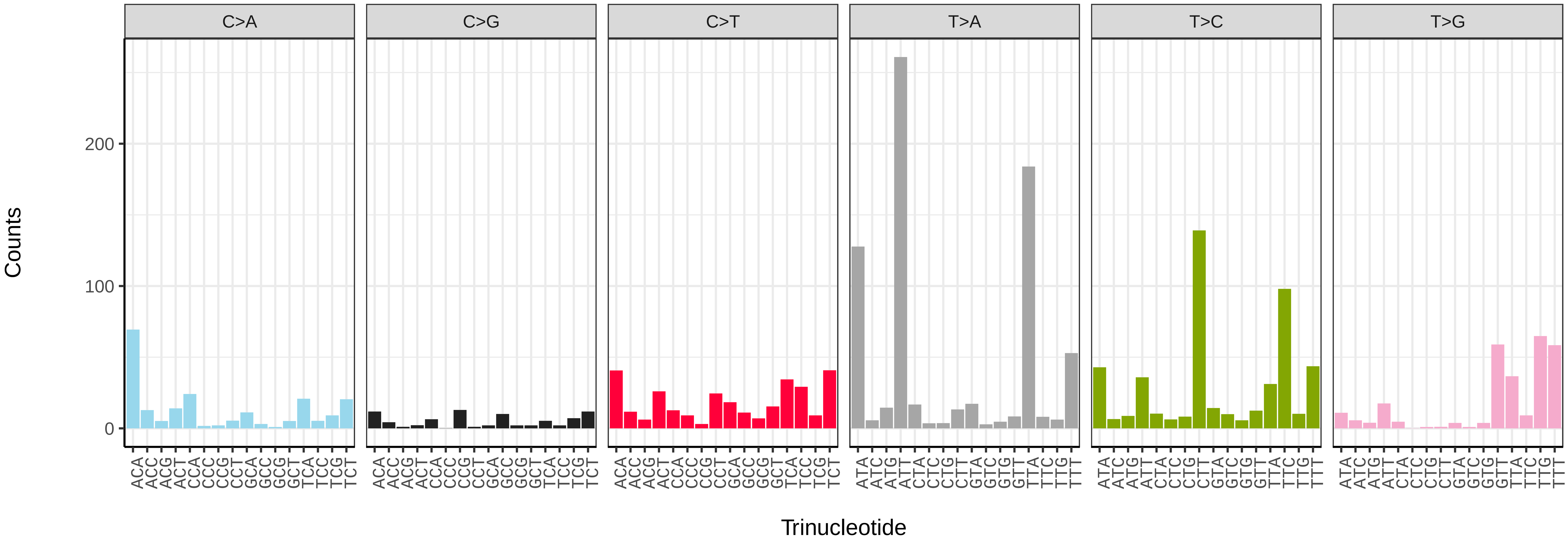


*Poecilobothrus nobilitatus* (idPoeNobi2) somatic mutational spectrum (SBS52 classification)

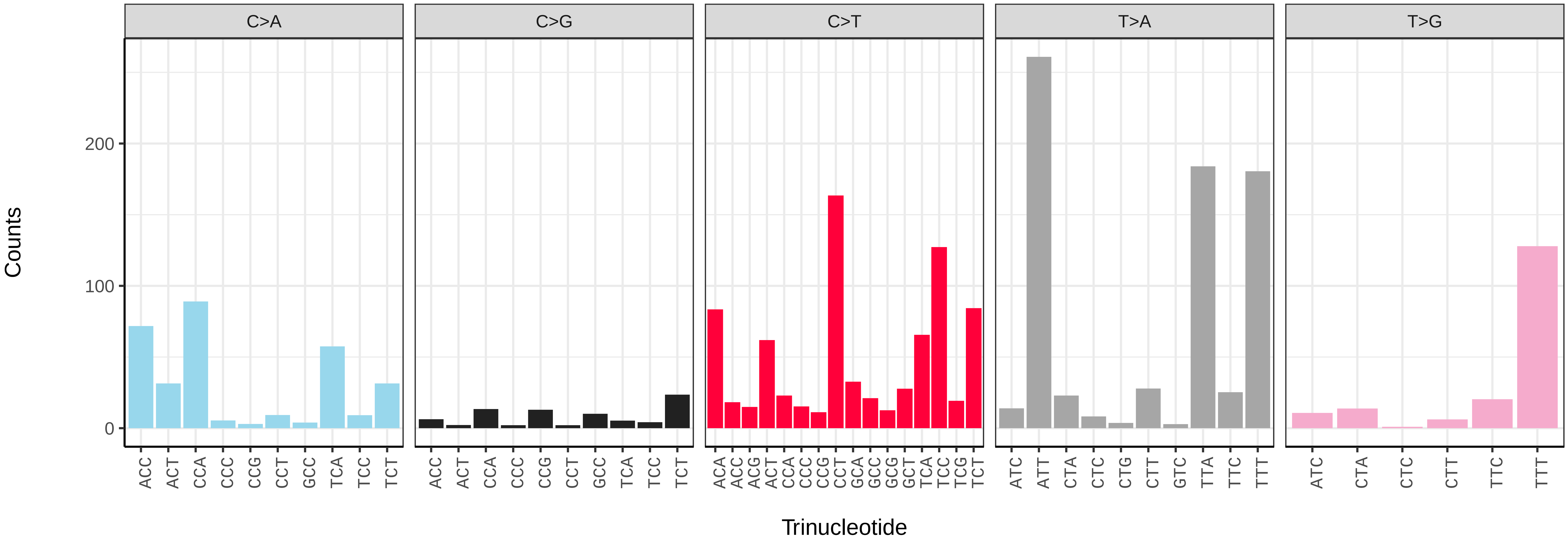

*Pollenia amentaria* (idPolAmen1) germline mutational spectrum

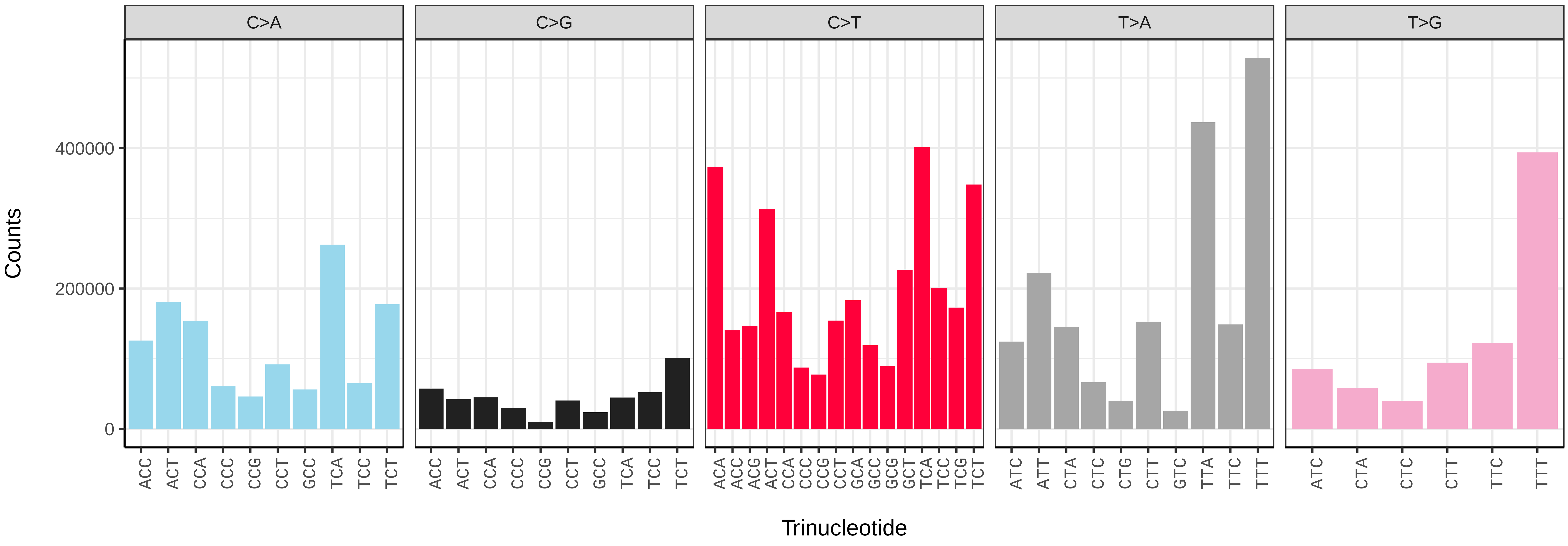


*Pollenia amentaria* (idPolAmen1) somatic mutational spectrum (SBS96 classification)

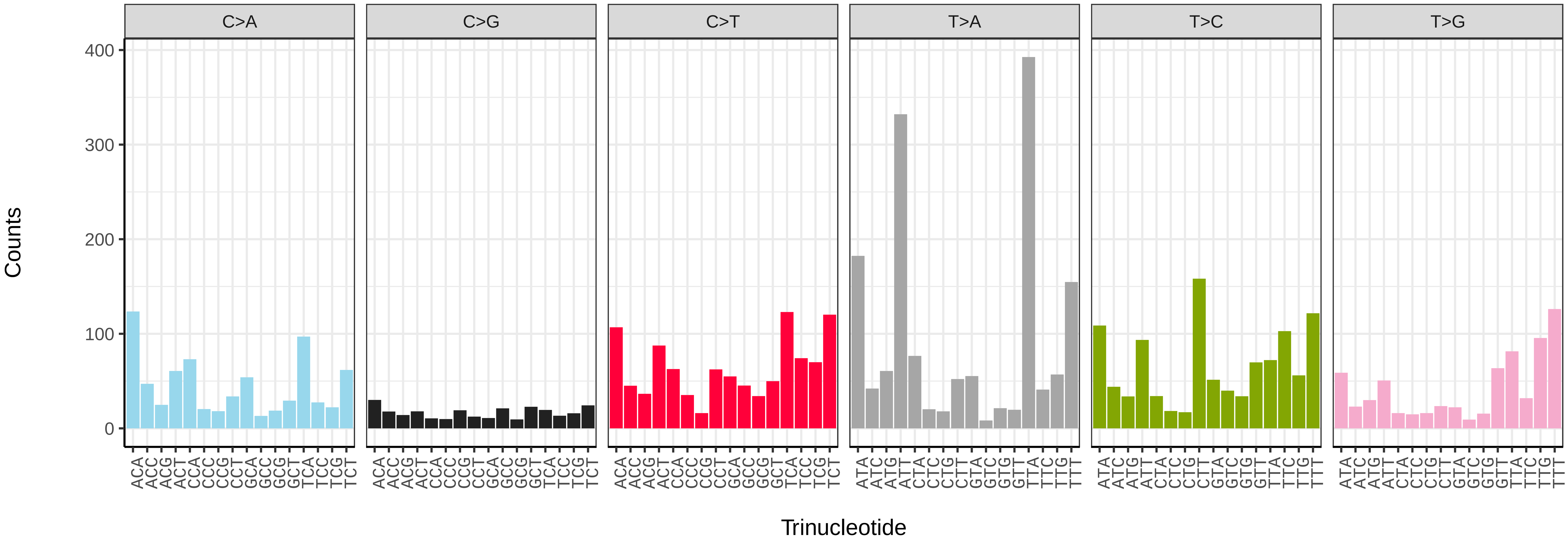


*Pollenia amentaria* (idPolAmen1) somatic mutational spectrum (SBS52 classification)

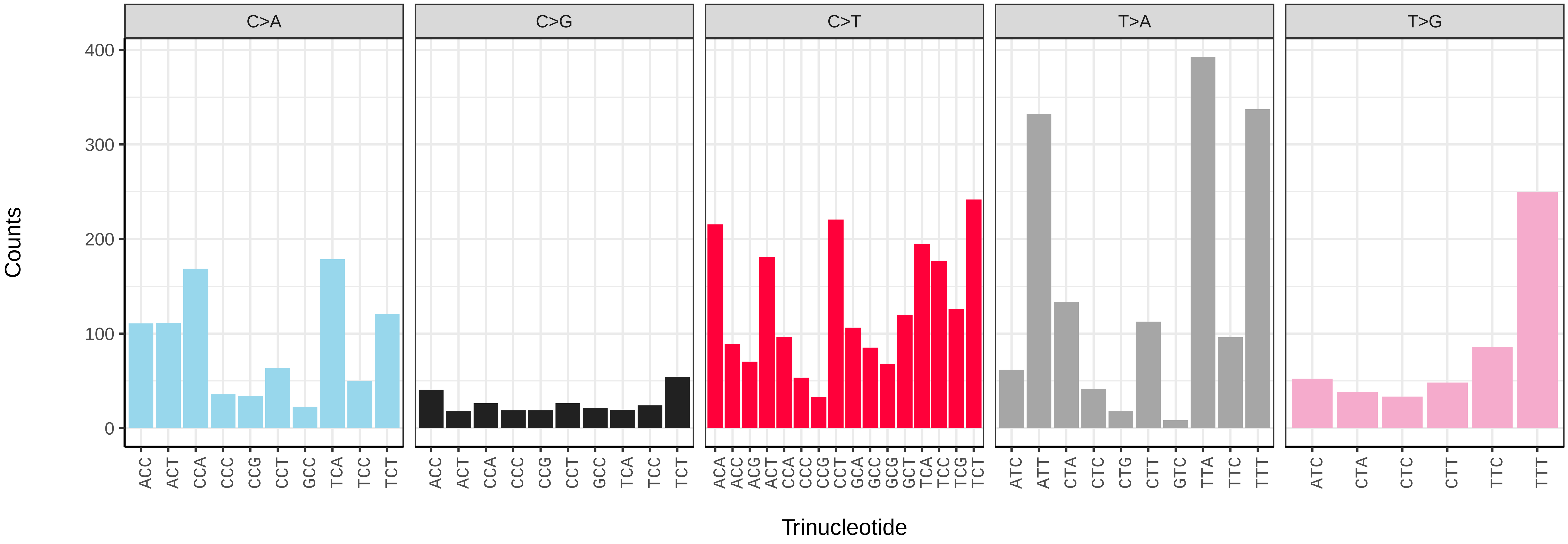

*Pollenia angustigena* (idPolAngu1) germline mutational spectrum

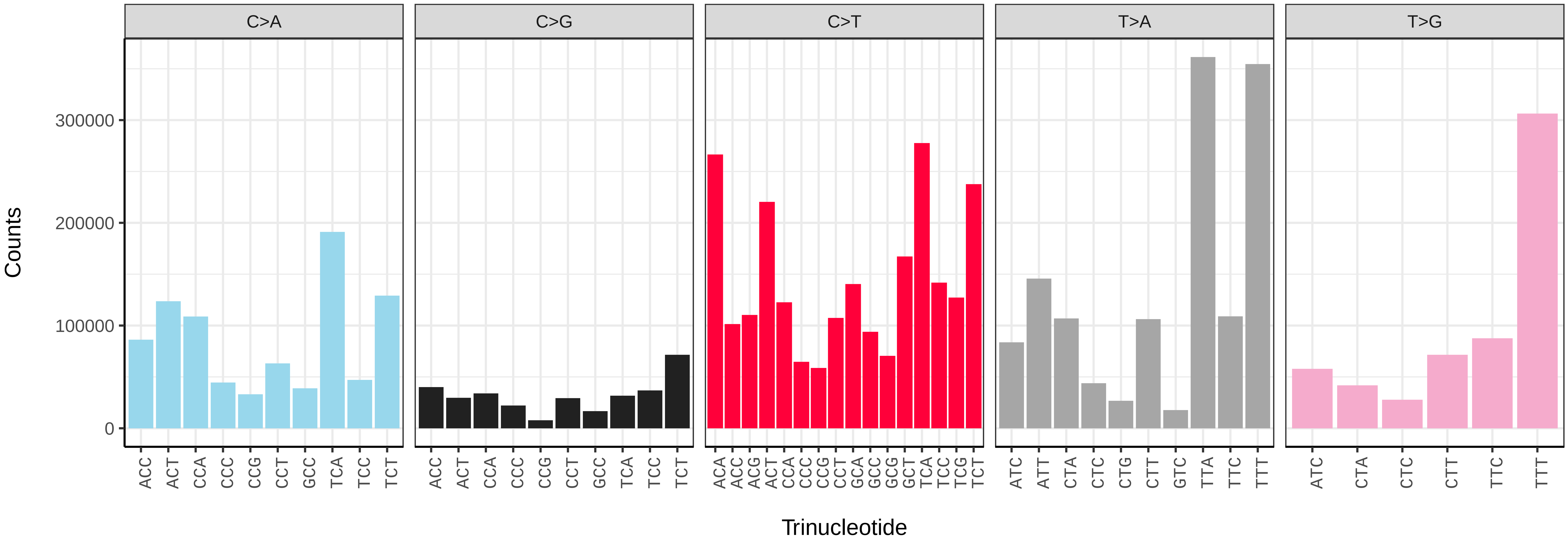


*Pollenia angustigena* (idPolAngu1) somatic mutational spectrum (SBS96 classification)

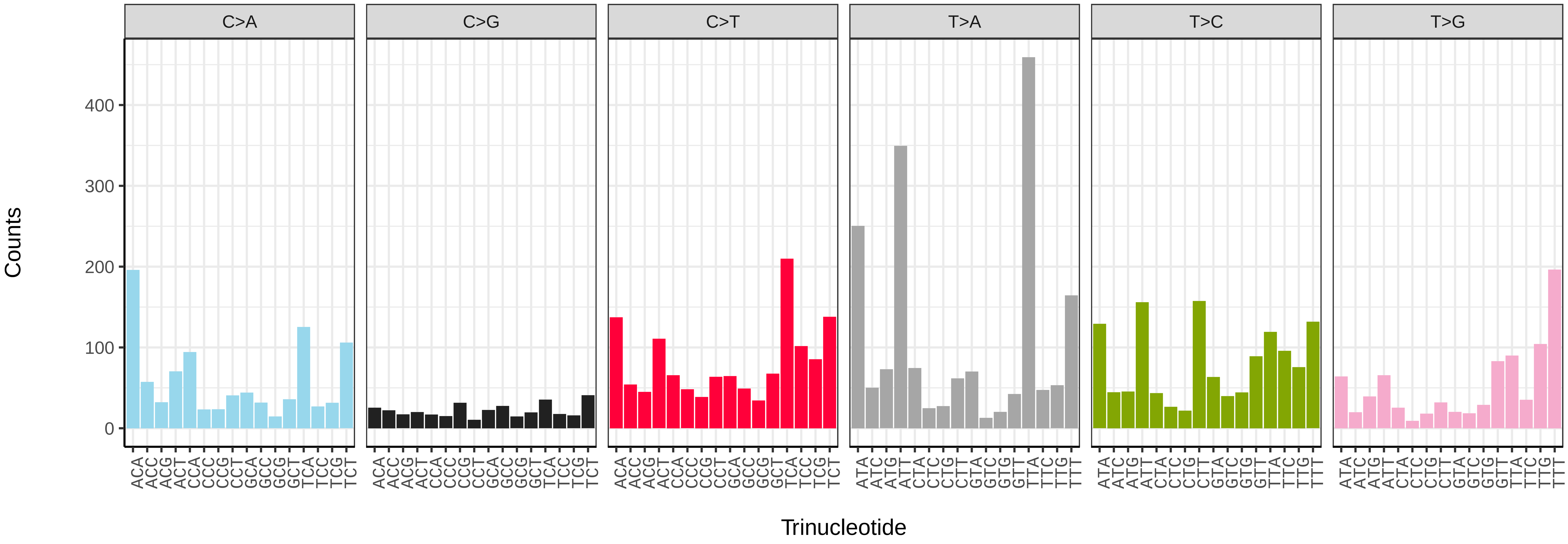


*Pollenia angustigena* (idPolAngu1) somatic mutational spectrum (SBS52 classification)

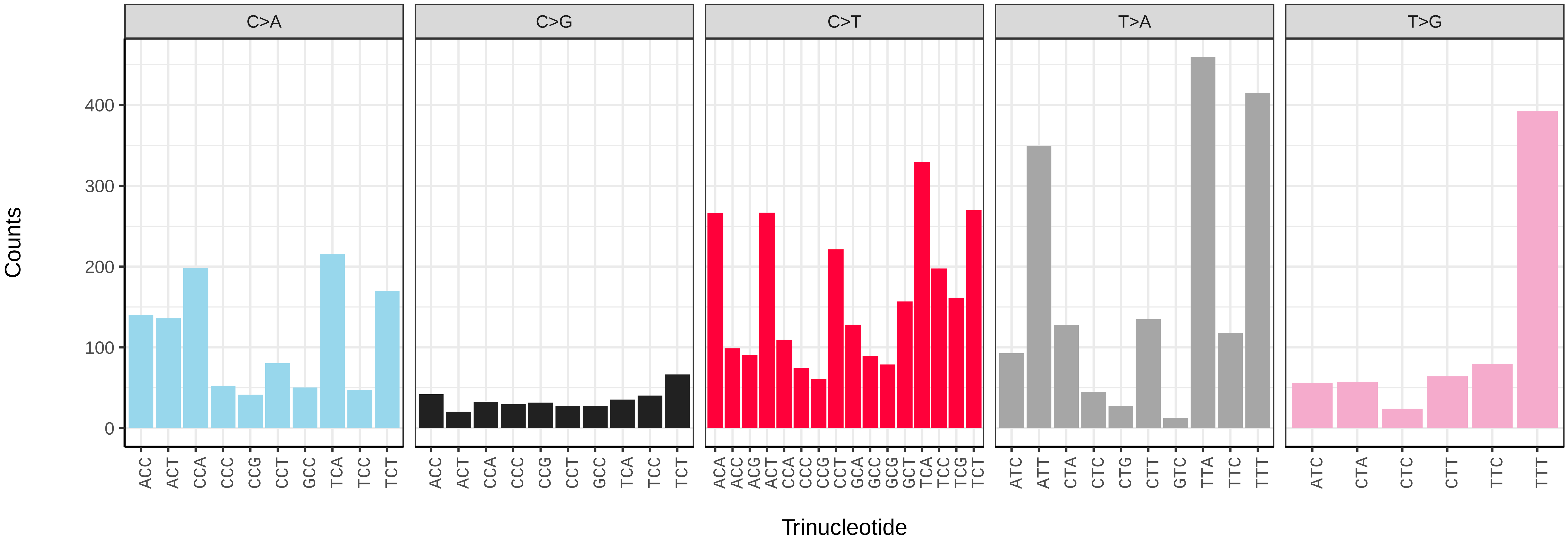

*Polietes domitor* (idPolDomi1) germline mutational spectrum

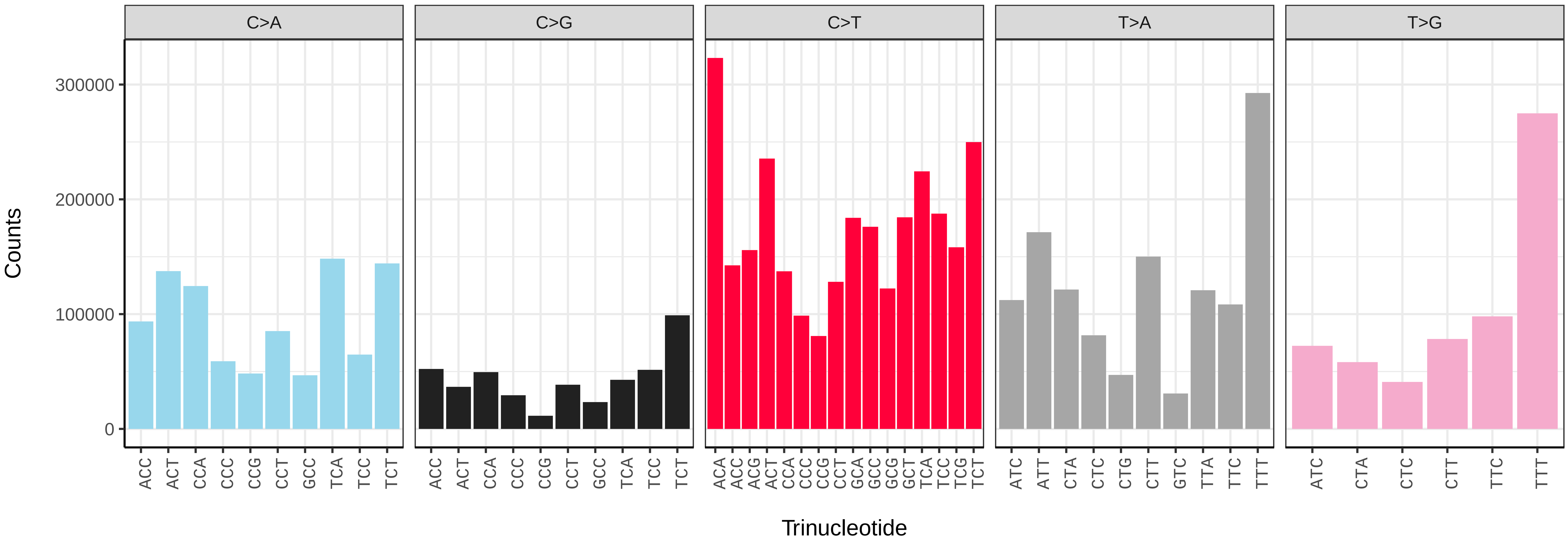


*Polietes domitor* (idPolDomi1) somatic mutational spectrum (SBS96 classification)

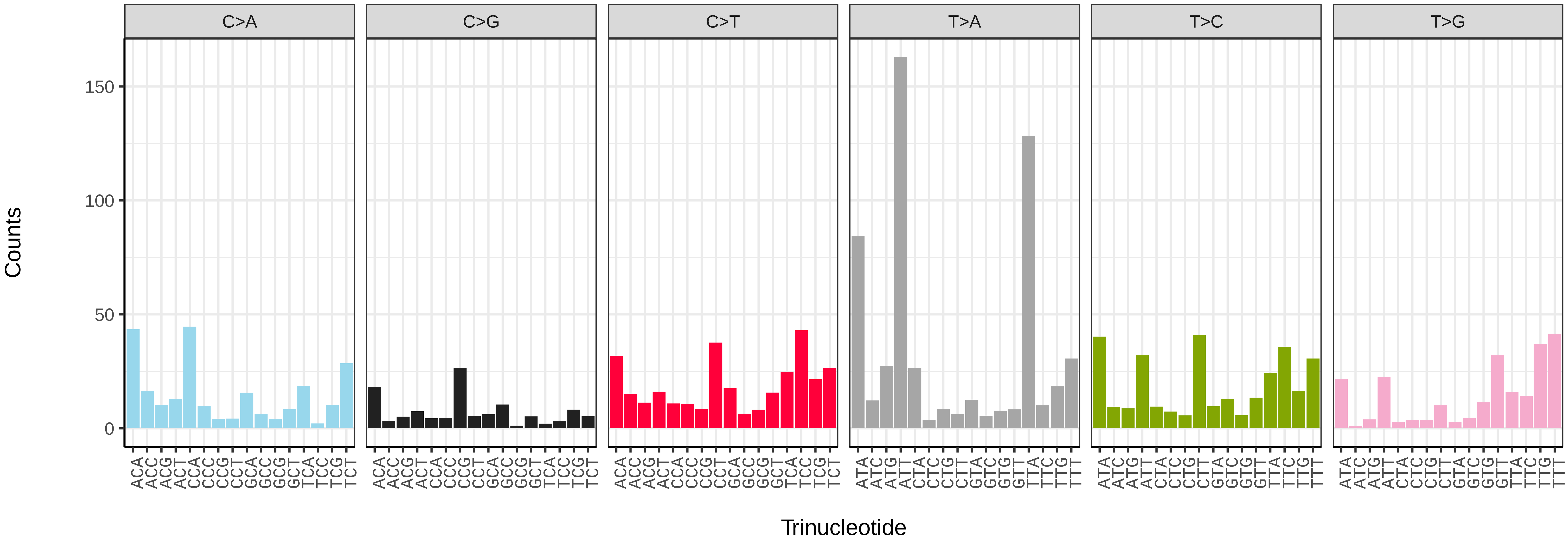


*Polietes domitor* (idPolDomi1) somatic mutational spectrum (SBS52 classification)

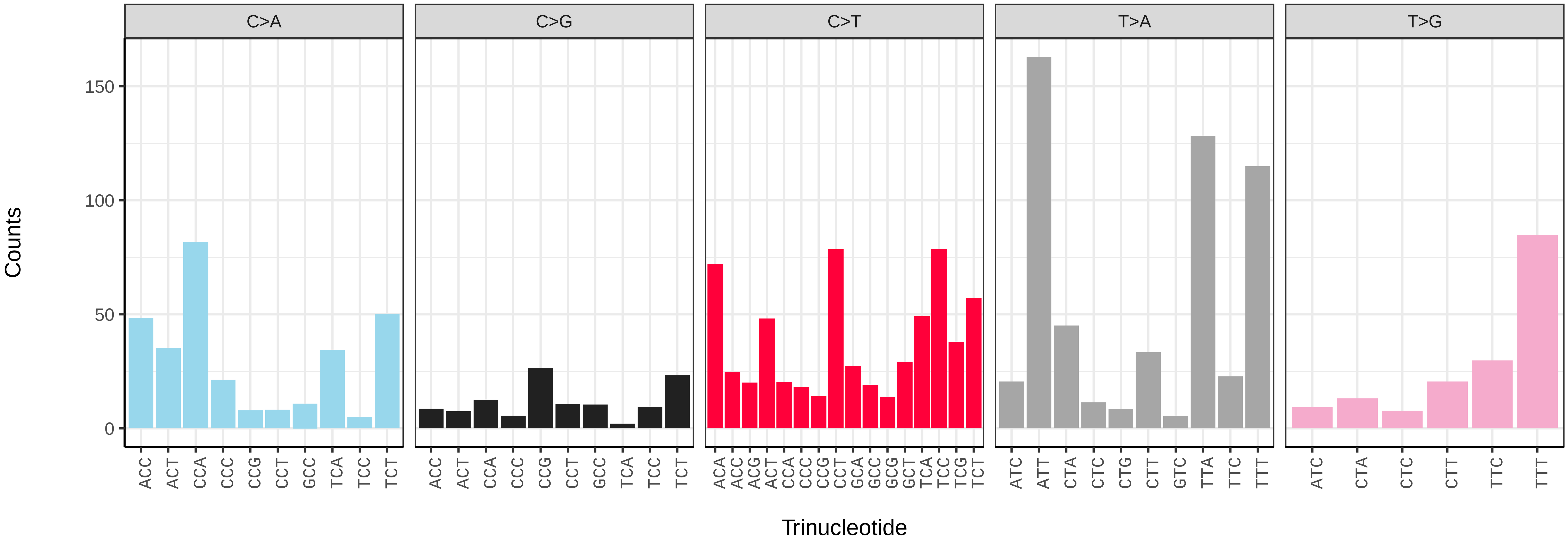

*Protocalliphora azurea* (idProAzur1) germline mutational spectrum

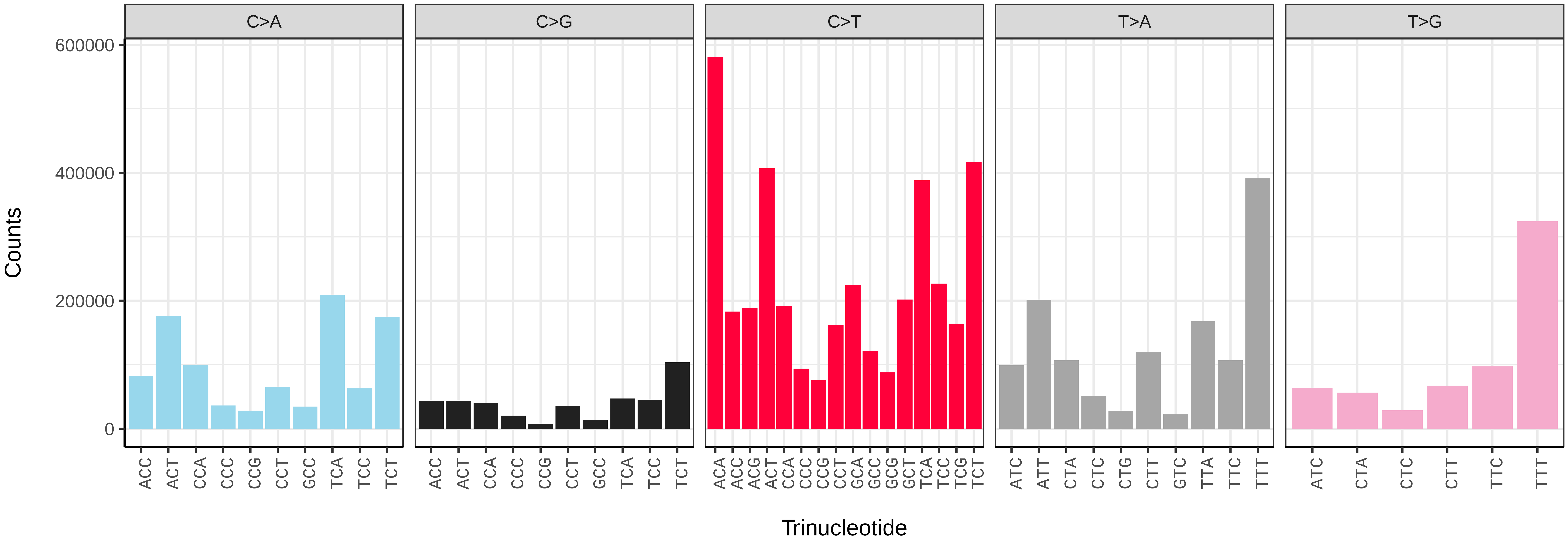


*Protocalliphora azurea* (idProAzur1) somatic mutational spectrum (SBS96 classification)

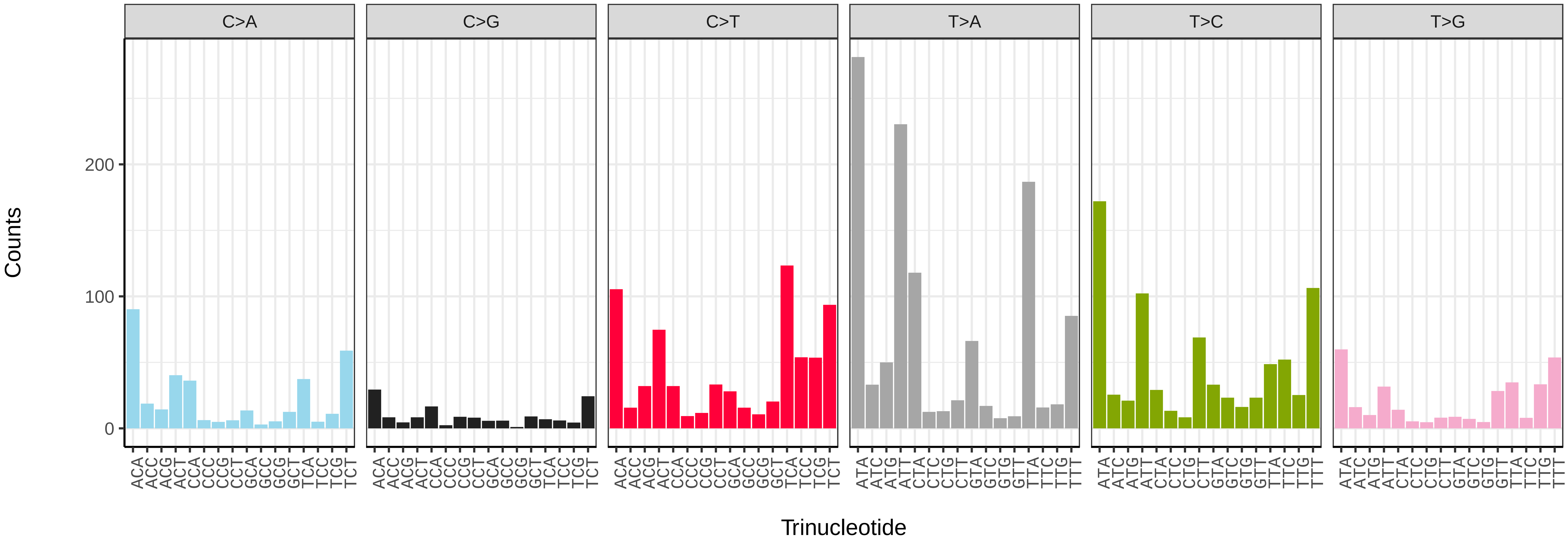


*Protocalliphora azurea* (idProAzur1) somatic mutational spectrum (SBS52 classification)

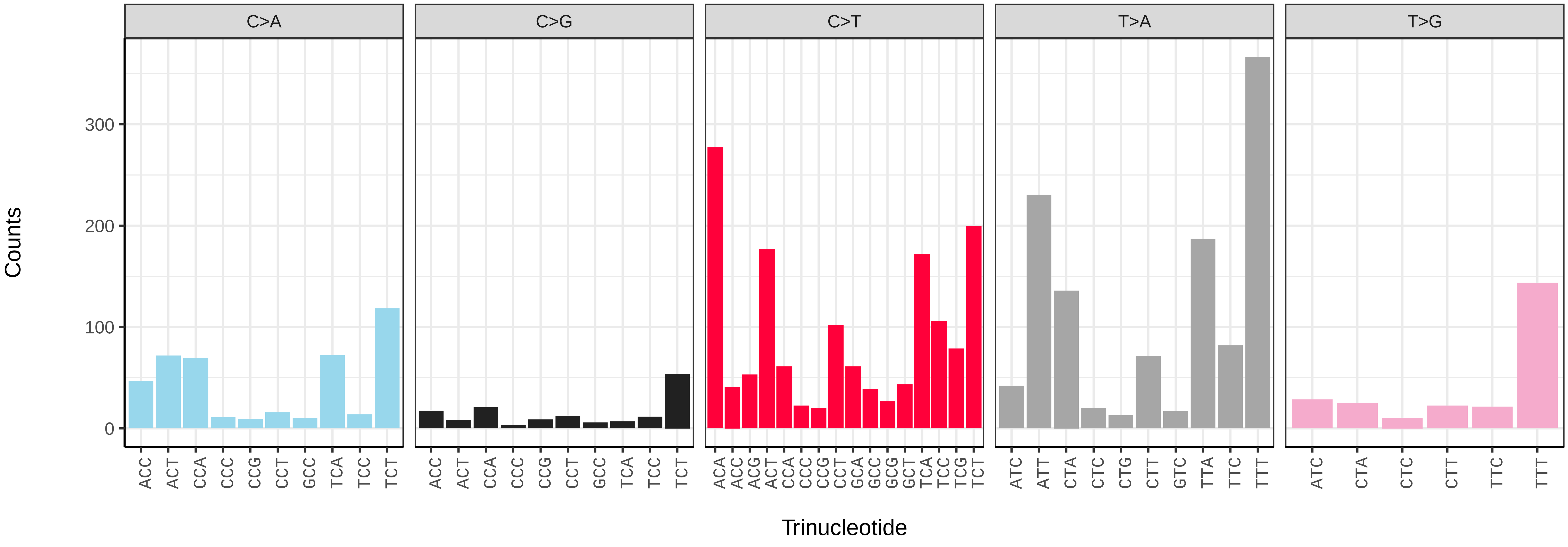

*Rhingia campestris* (idRhiCamp1) germline mutational spectrum

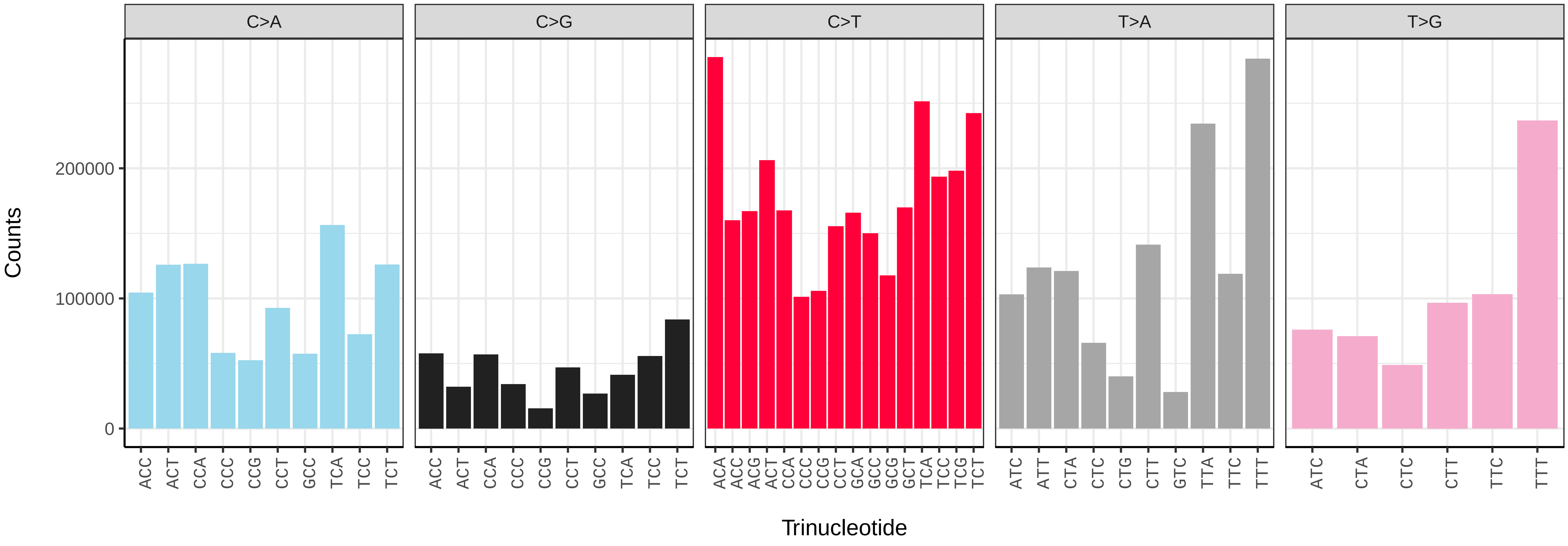


*Rhingia campestris* (idRhiCamp1) somatic mutational spectrum (SBS96 classification)

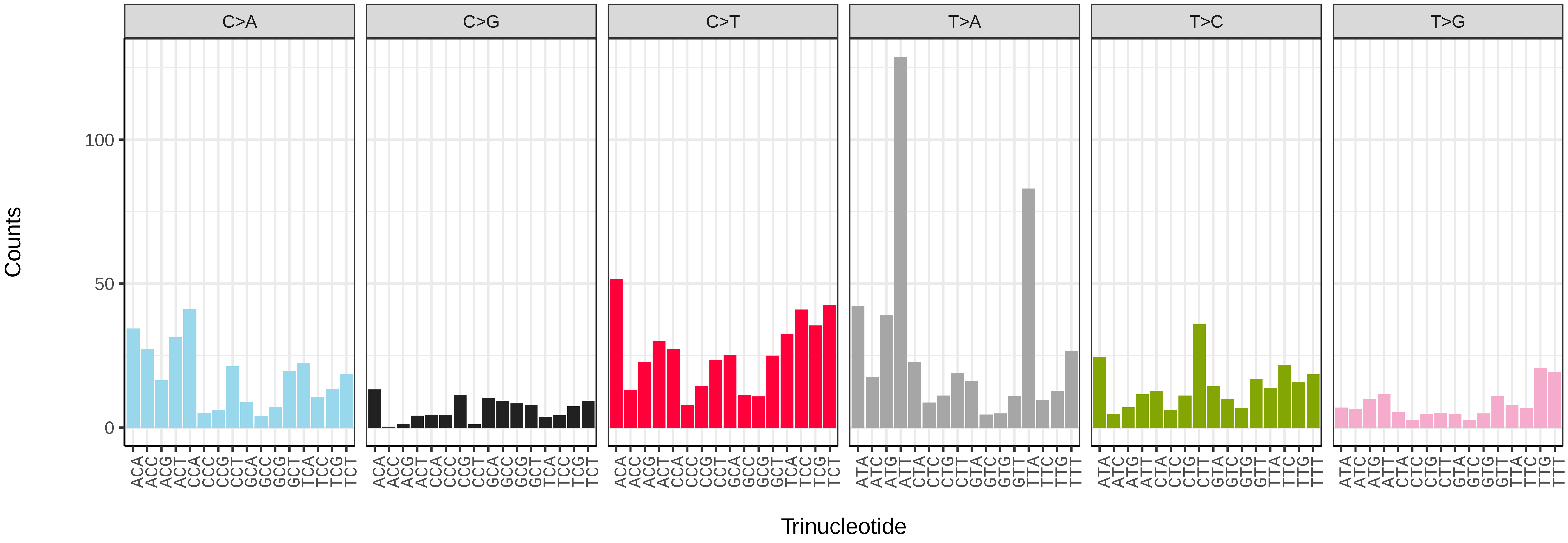


*Rhingia campestris* (idRhiCamp1) somatic mutational spectrum (SBS52 classification)

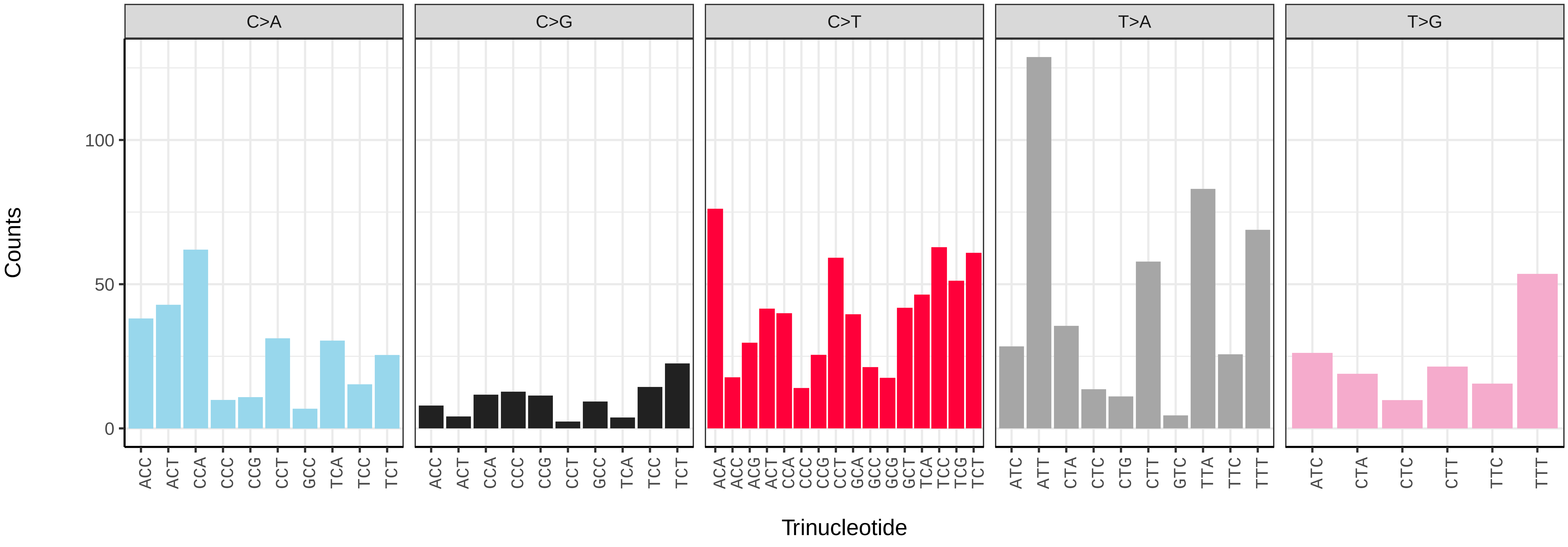

*Sarcophaga caerulescens* (idSarCaer1) germline mutational spectrum

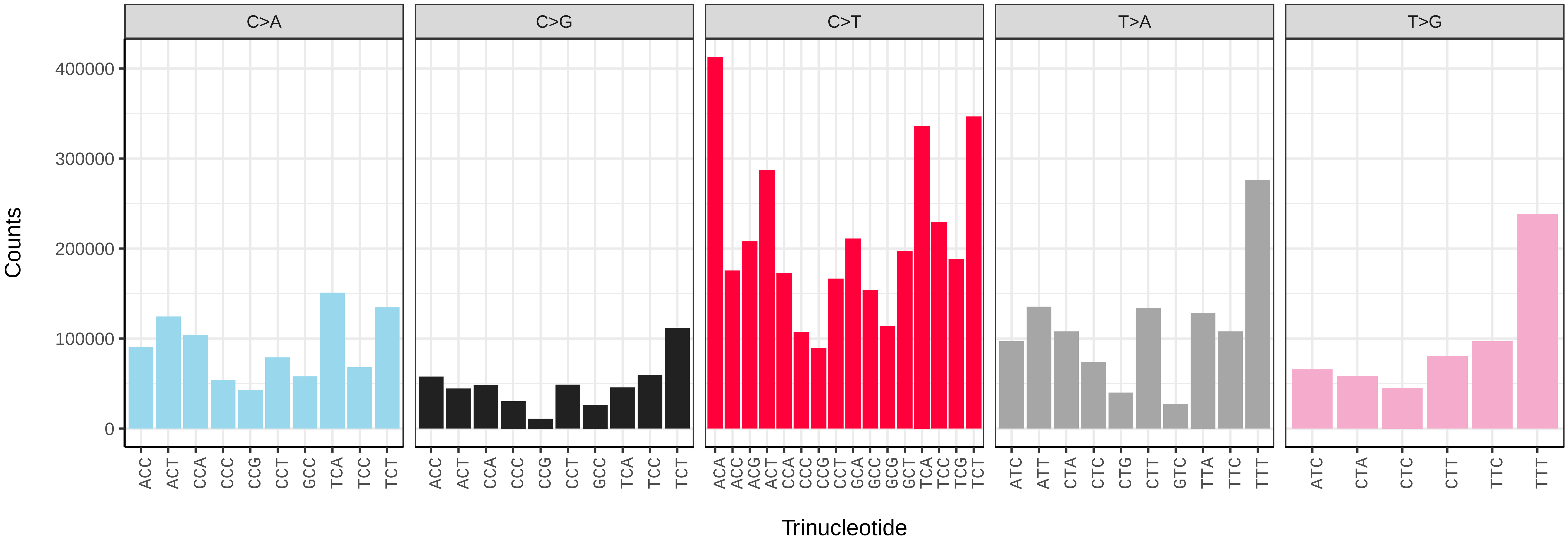


*Sarcophaga caerulescens* (idSarCaer1) somatic mutational spectrum (SBS96 classification)

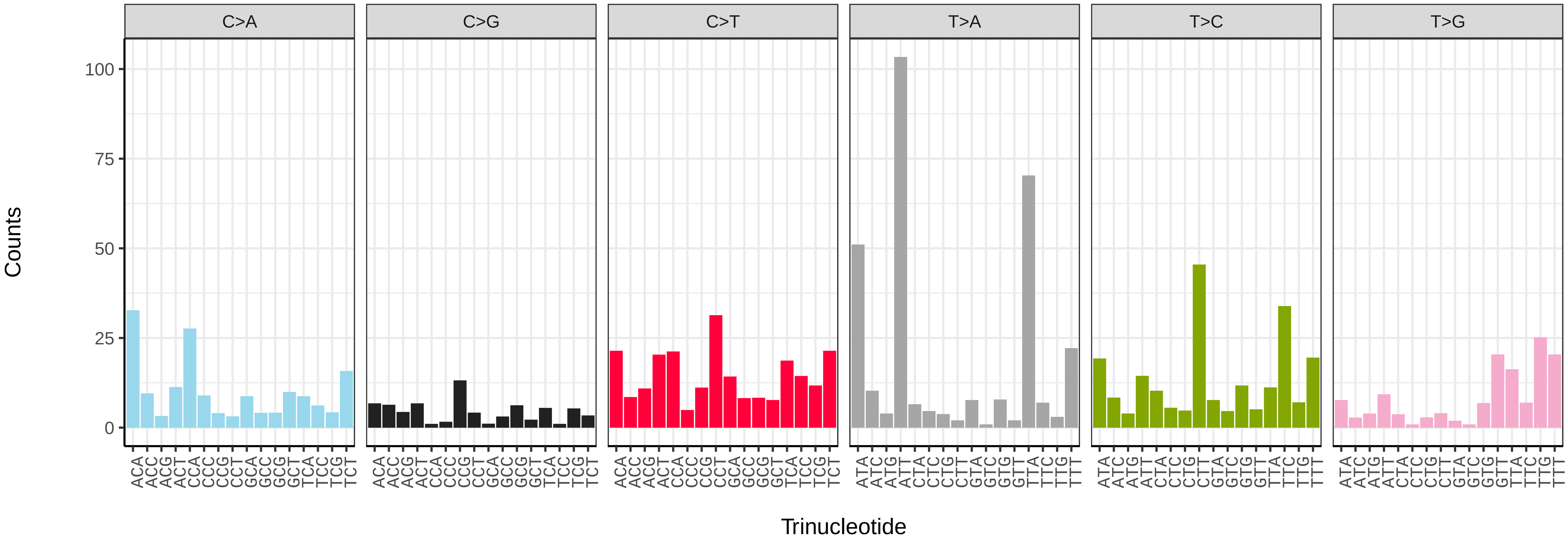


*Sarcophaga caerulescens* (idSarCaer1) somatic mutational spectrum (SBS52 classification)

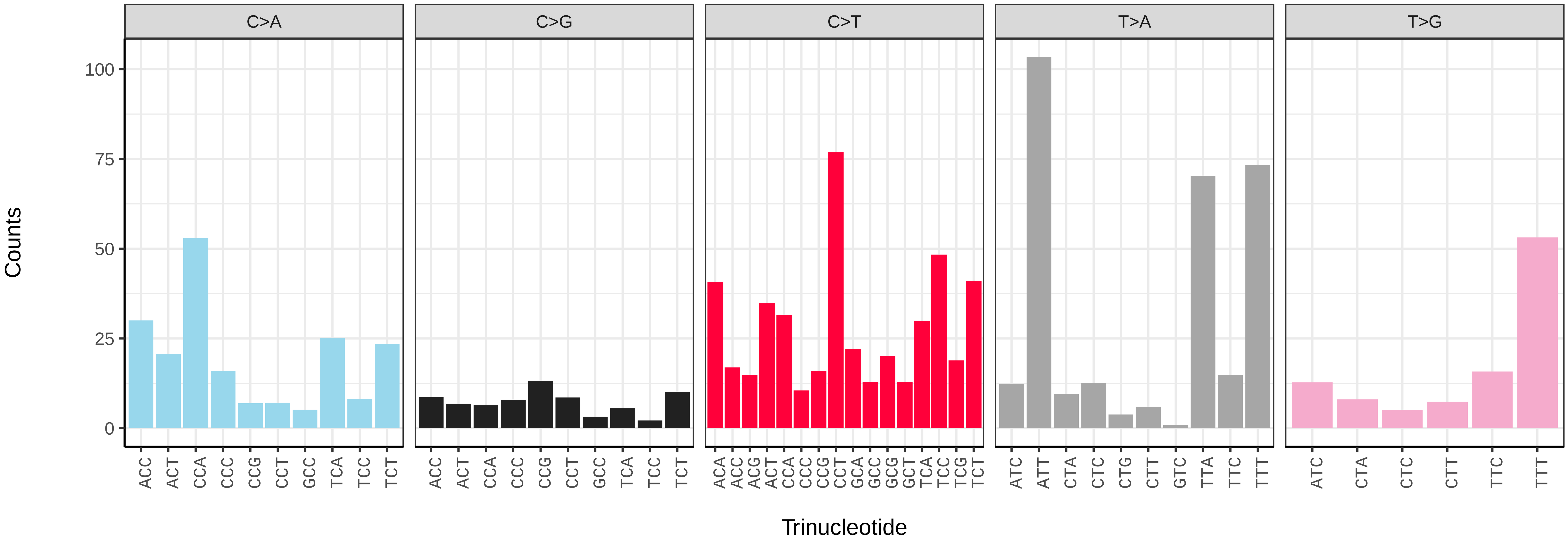

*Sarcophaga rosellei* (idSarRose1) germline mutational spectrum

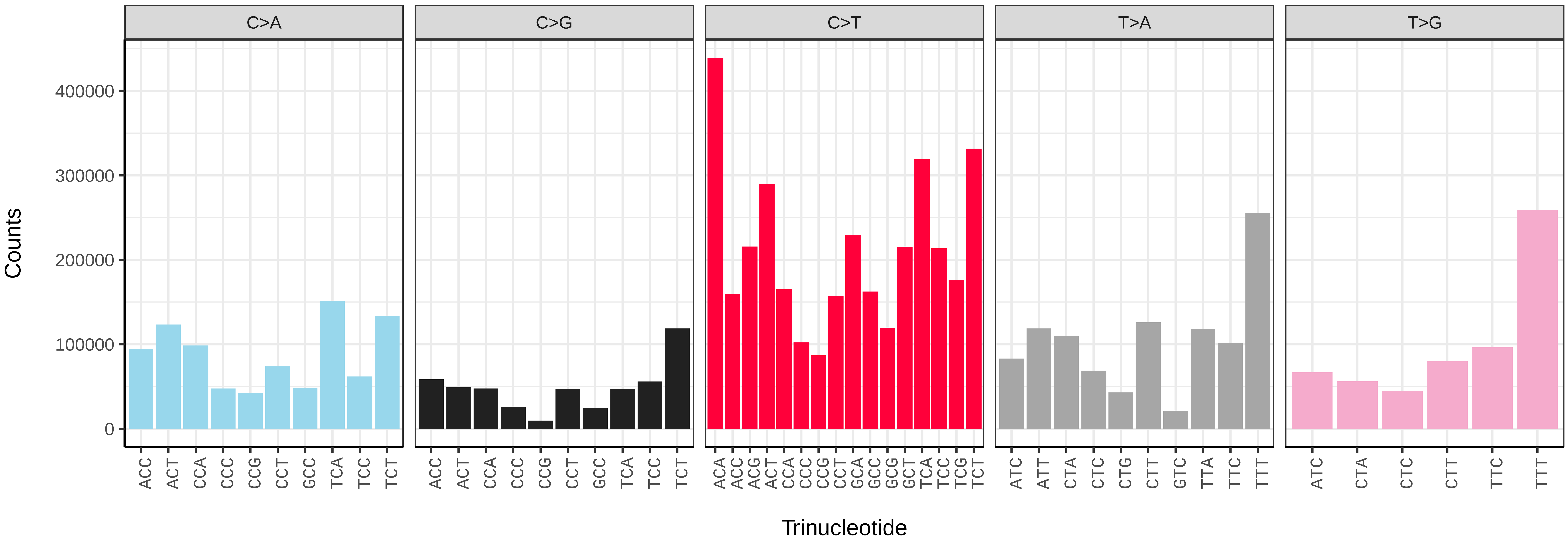


*Sarcophaga rosellei* (idSarRose1) somatic mutational spectrum (SBS96 classification)

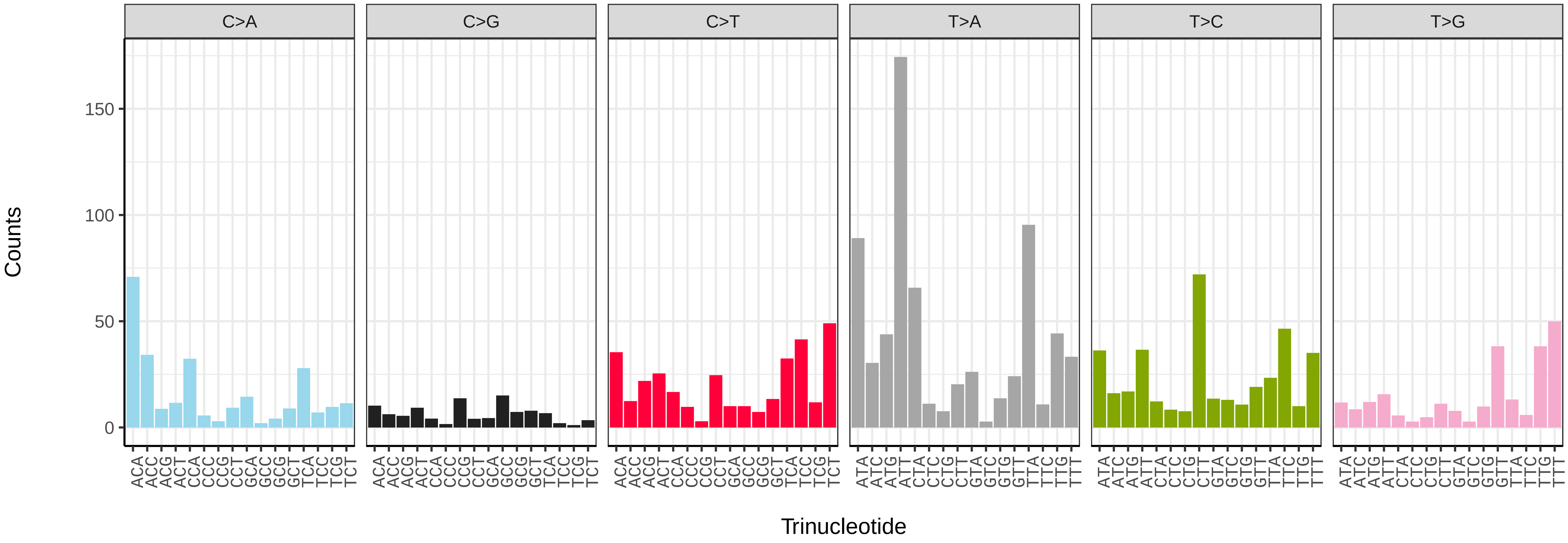


*Sarcophaga rosellei* (idSarRose1) somatic mutational spectrum (SBS52 classification)

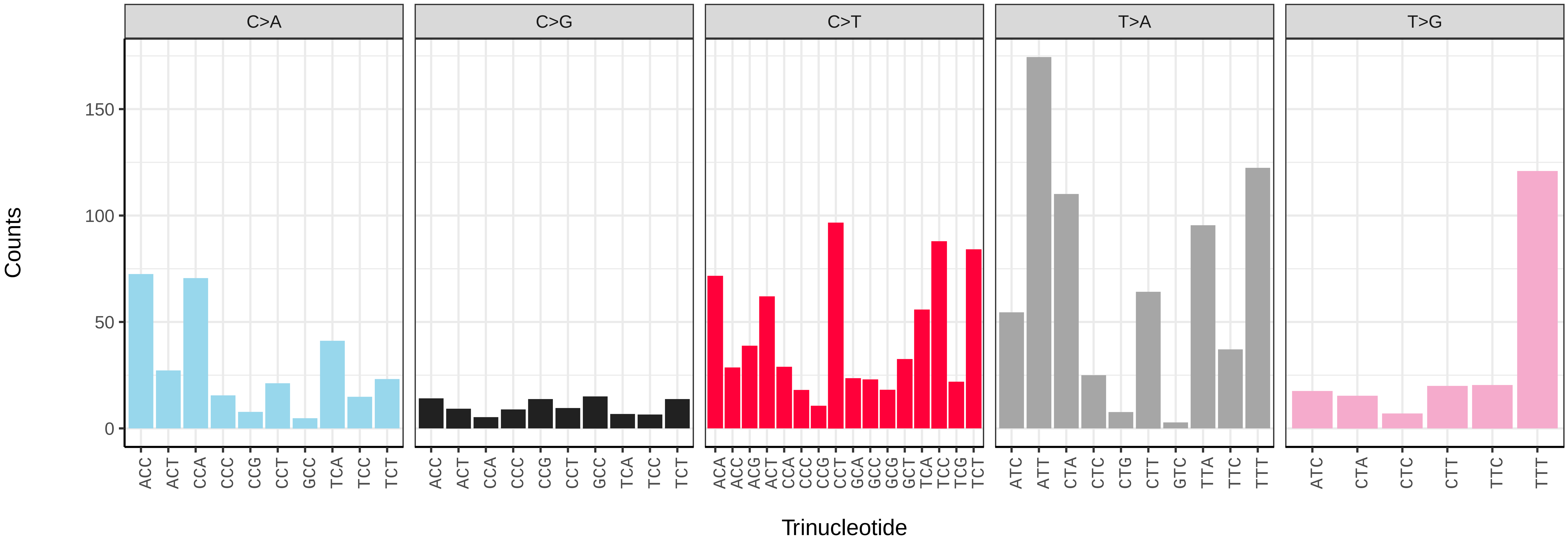

*Sarcophaga subvicina* (idSarSubv1) germline mutational spectrum

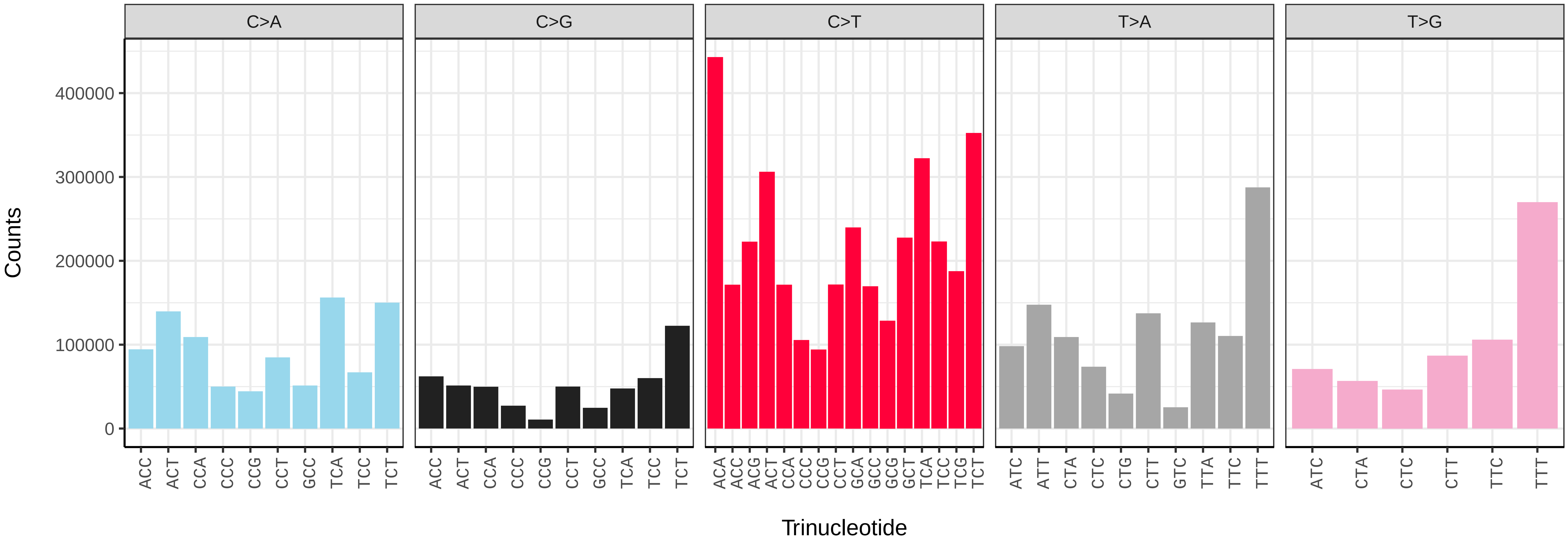


*Sarcophaga subvicina* (idSarSubv1) somatic mutational spectrum (SBS96 classification)

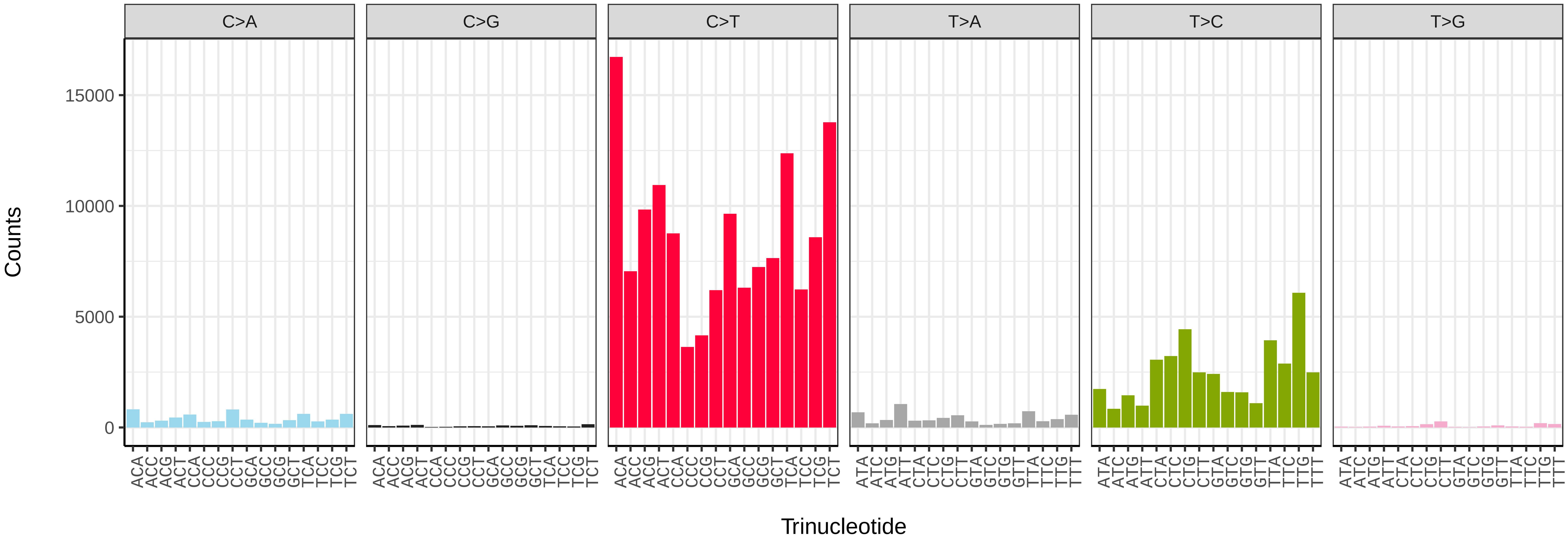


*Sarcophaga subvicina* (idSarSubv1) somatic mutational spectrum (SBS52 classification)

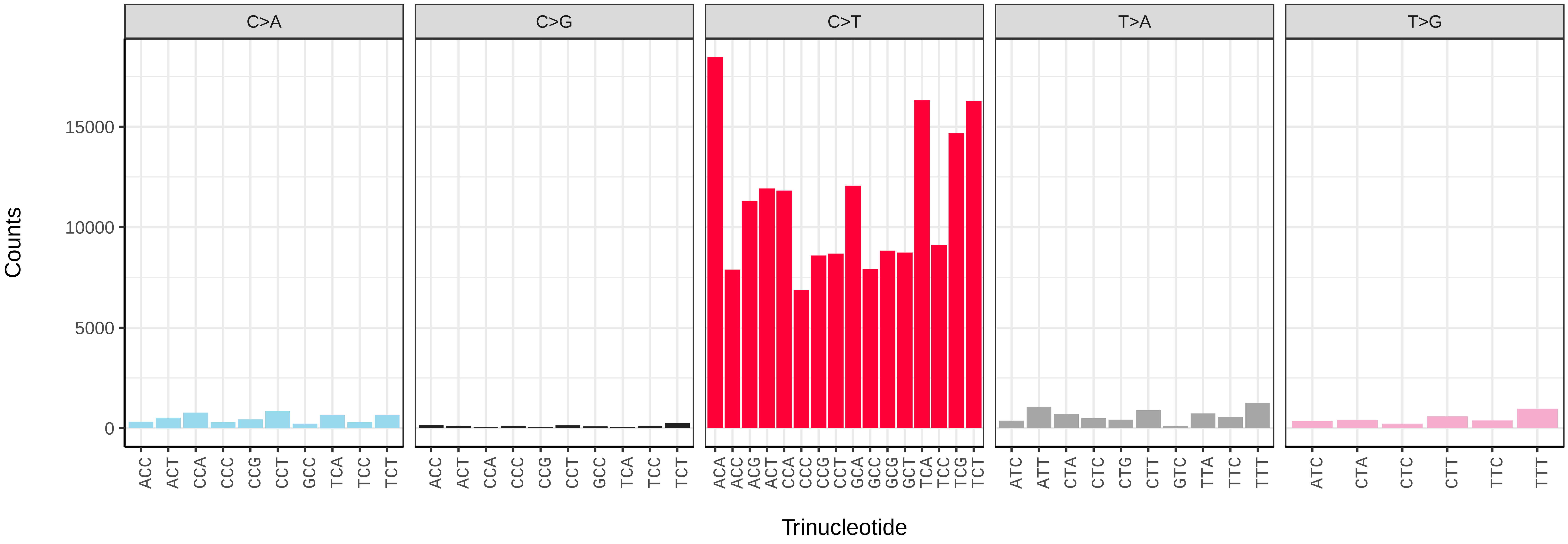

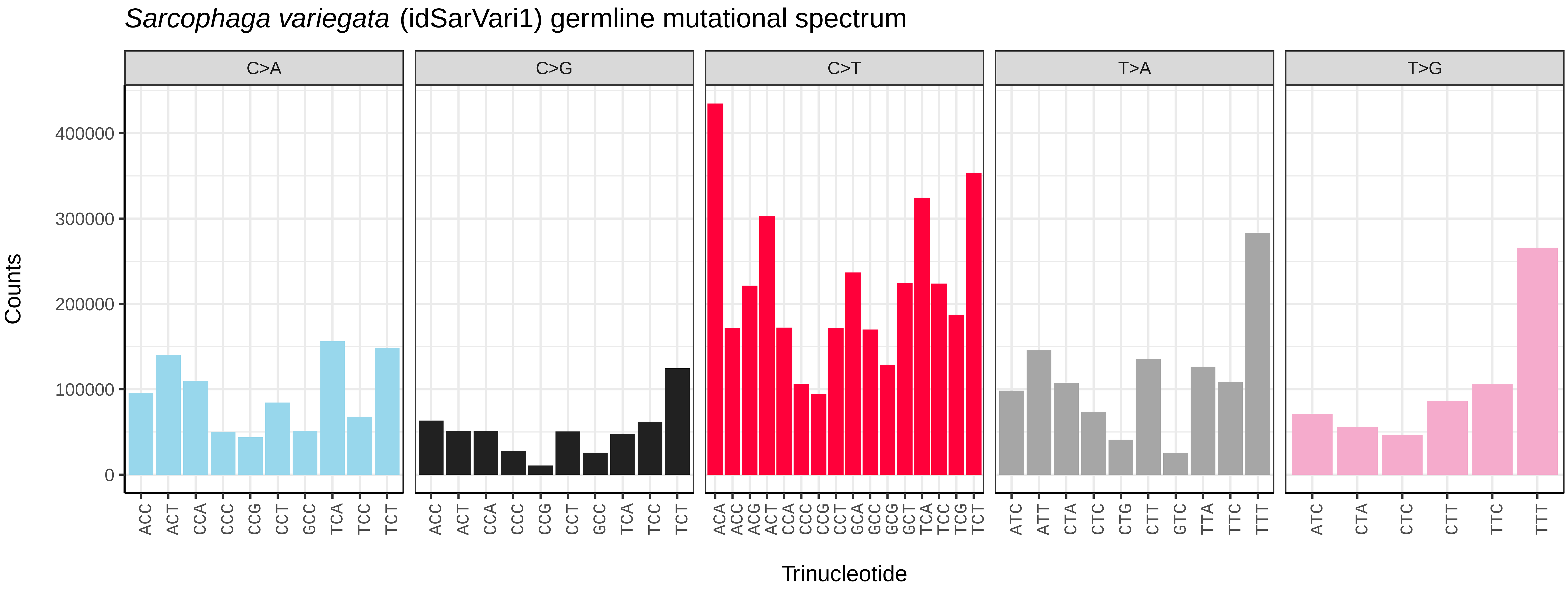
Sarcophaga variegata (idSarVari1) germline mutational spectrum
C>A
C>G
C>T
T>A
T>G
Counts
Trinucleotide

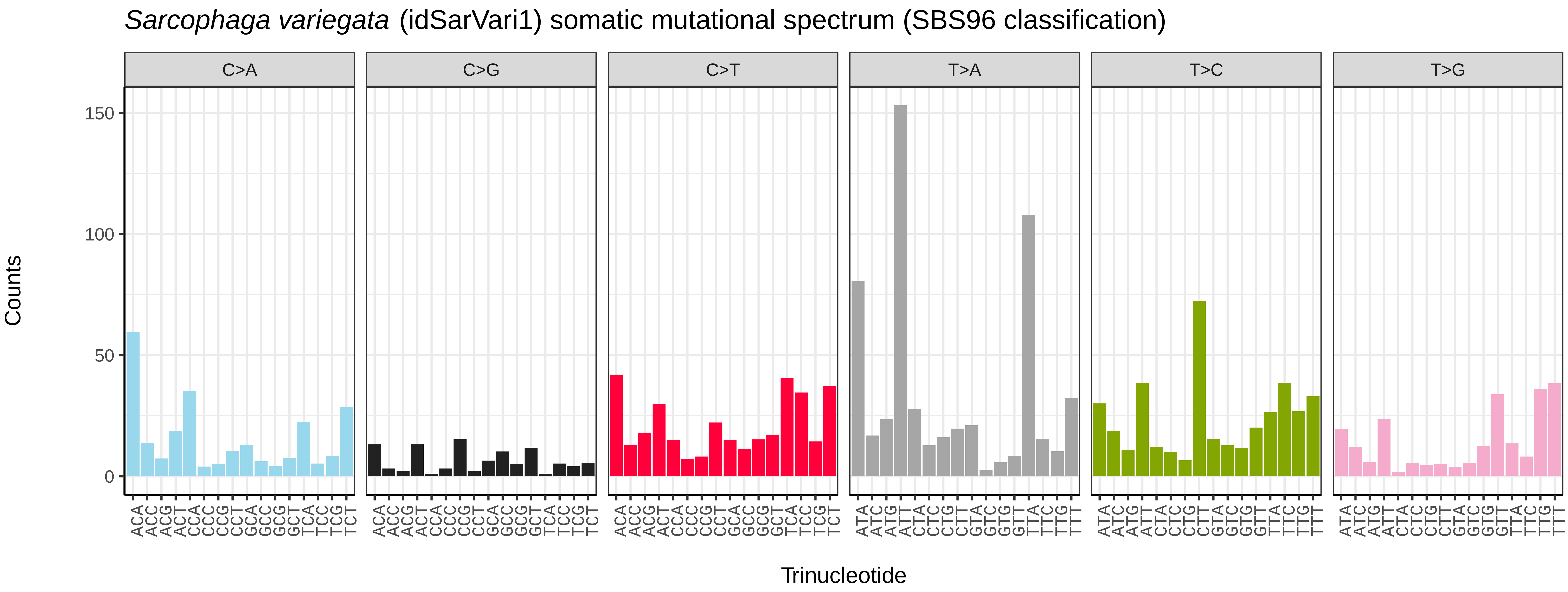
Sarcophaga variegata (idSarVari1) somatic mutational spectrum (SBS96 classification)
C>A
C>G
C>T
T>A
T>C
T>G
Counts
Trinucleotide

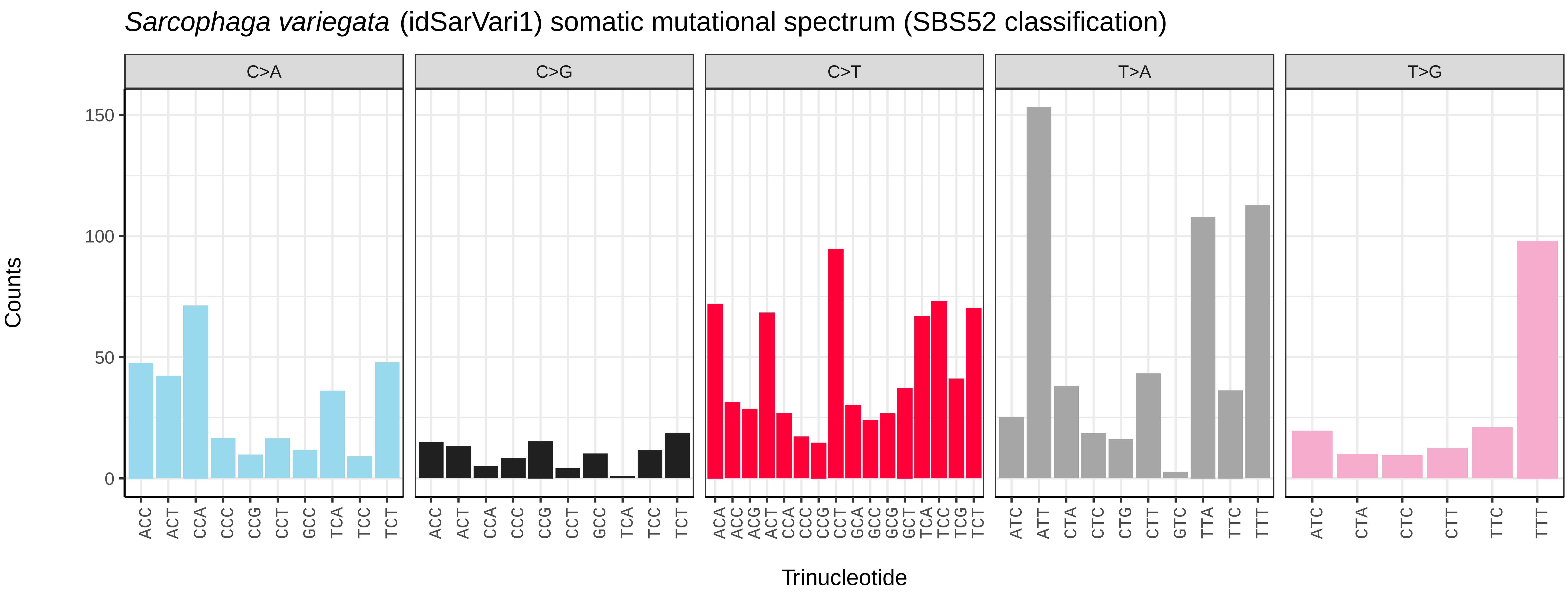
Sarcophaga variegata (idSarVari1) somatic mutational spectrum (SBS52 classification)
C>A
C>G
C>T
T>A
T>G
Counts
Trinucleotide

*Scaeva pyrastri* (idScaPyra1) germline mutational spectrum

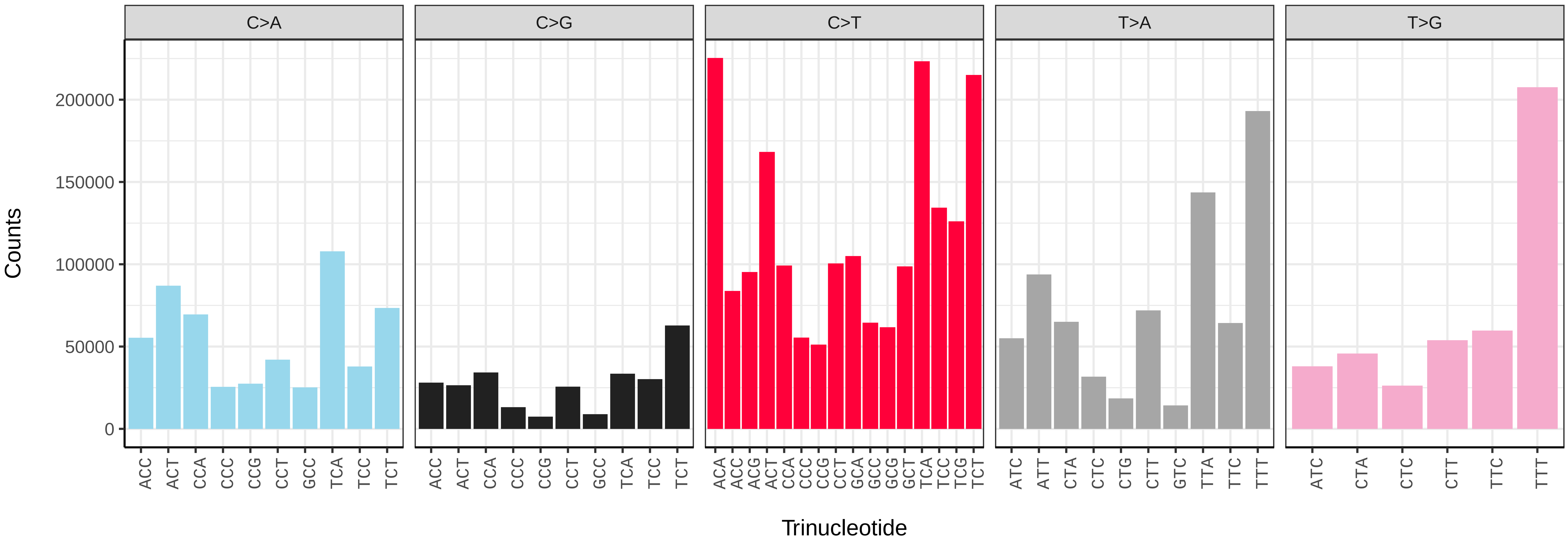


*Scaeva pyrastri* (idScaPyra1) somatic mutational spectrum (SBS96 classification)

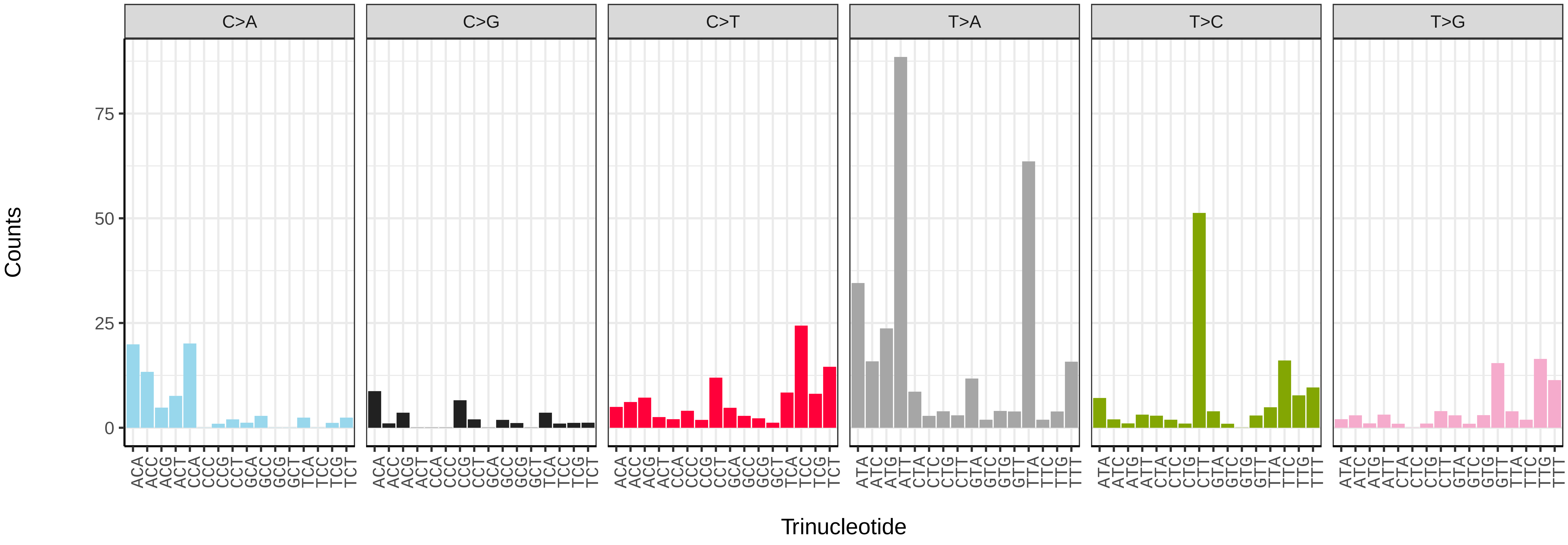


*Scaeva pyrastri* (idScaPyra1) somatic mutational spectrum (SBS52 classification)

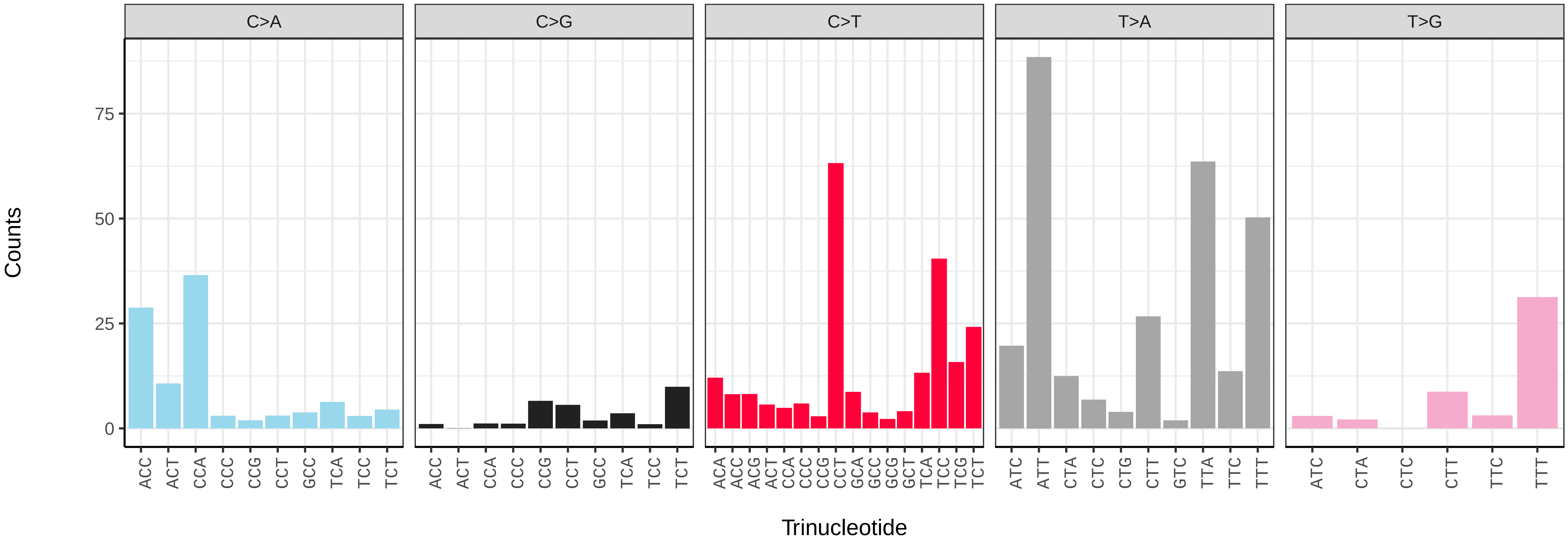

*Sicus ferrugineus* (idSicFerr1) germline mutational spectrum

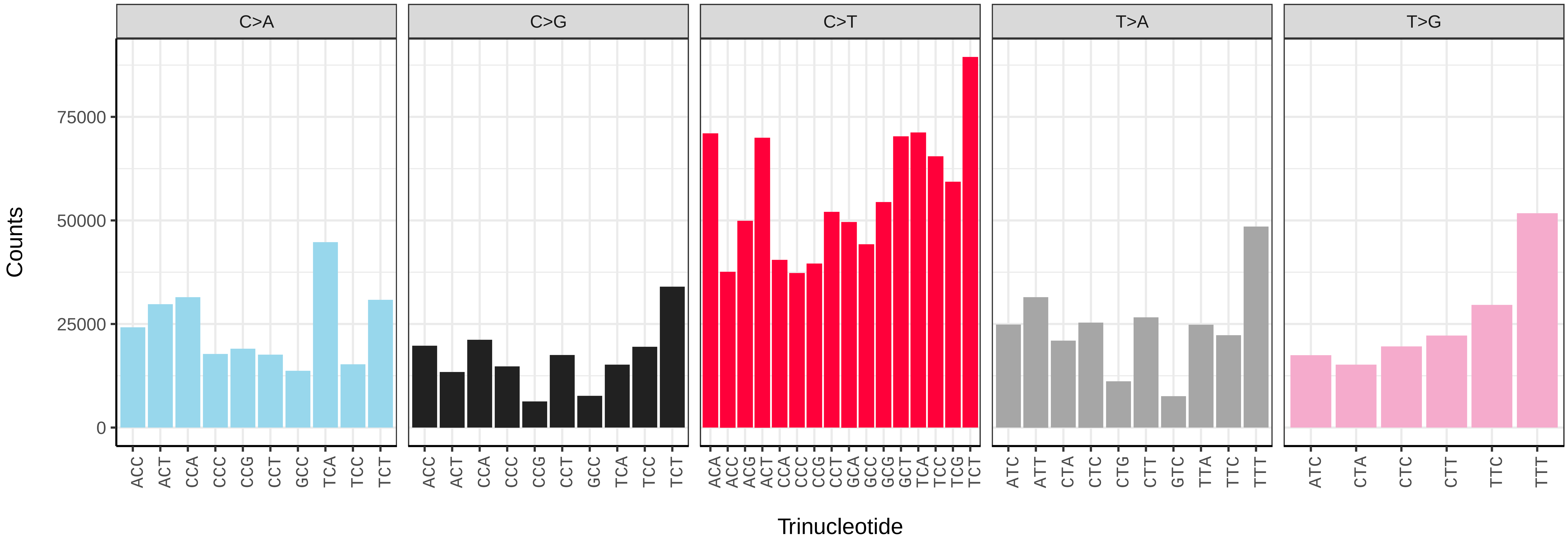


*Sicus ferrugineus* (idSicFerr1) somatic mutational spectrum (SBS96 classification)

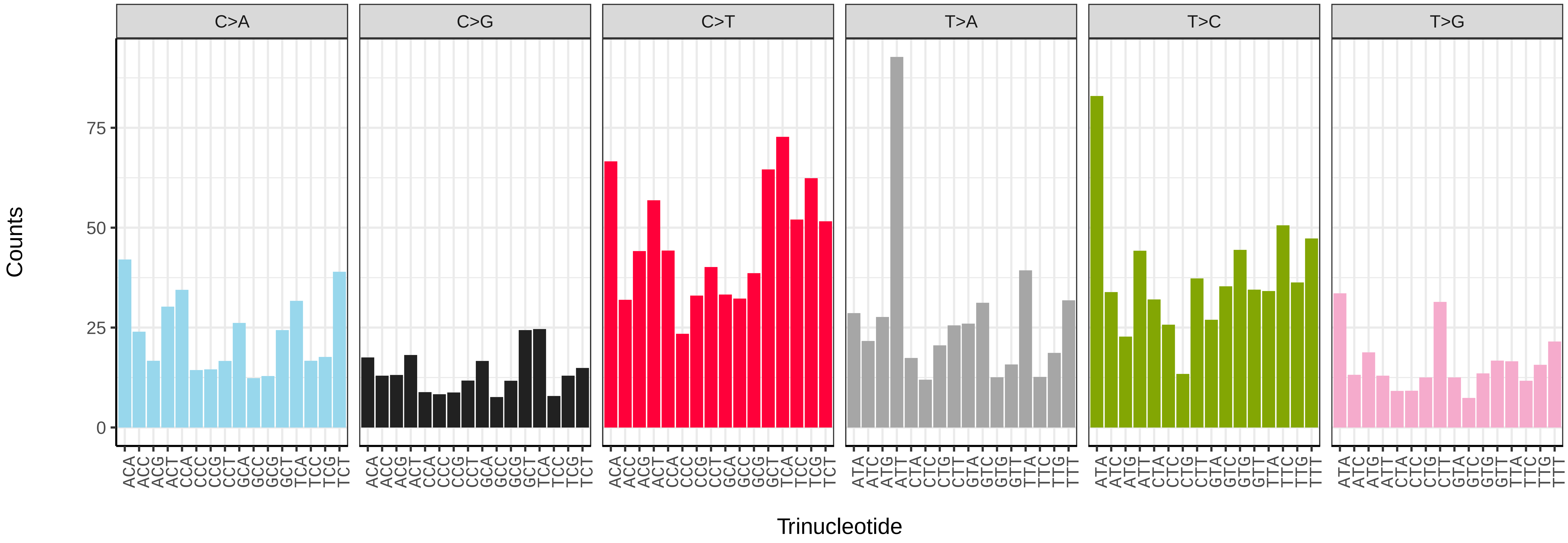


*Sicus ferrugineus* (idSicFerr1) somatic mutational spectrum (SBS52 classification)

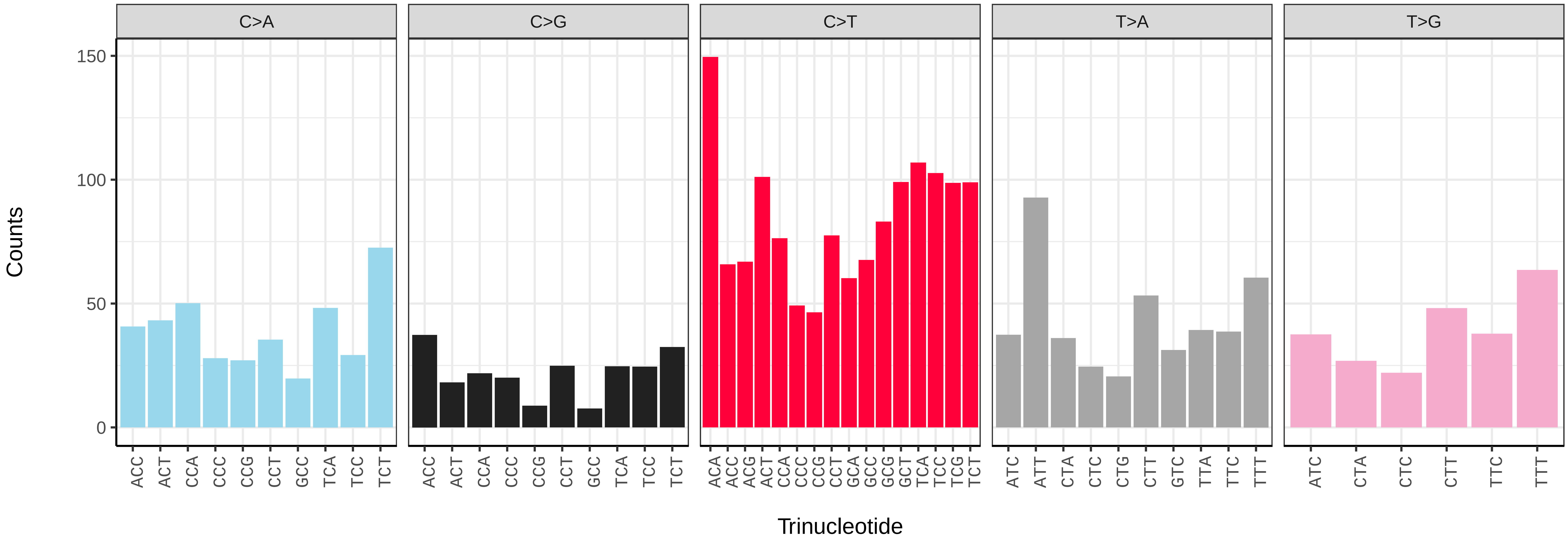

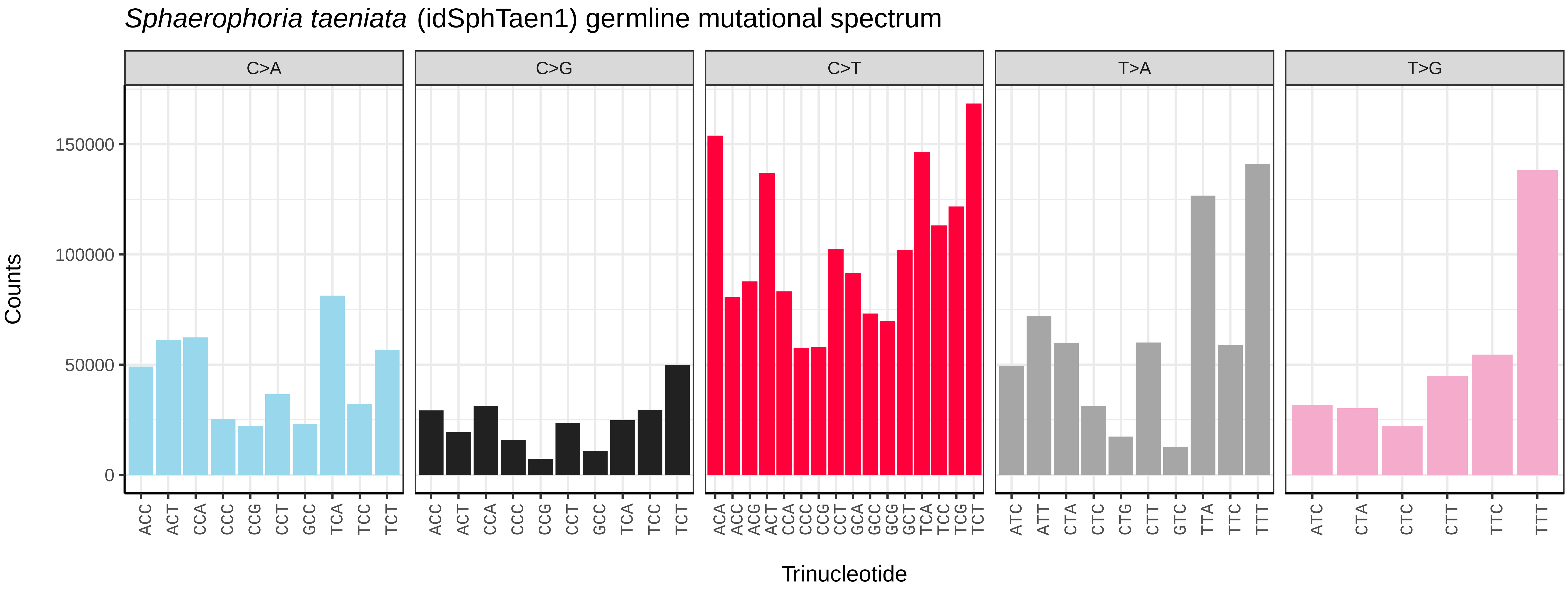
*Sphaerophoria taeniata* (idSphTaen1) germline mutational spectrum
C>A
C>G
C>T
T>A
T>G
Counts
150000
100000
50000
0
Trinucleotide

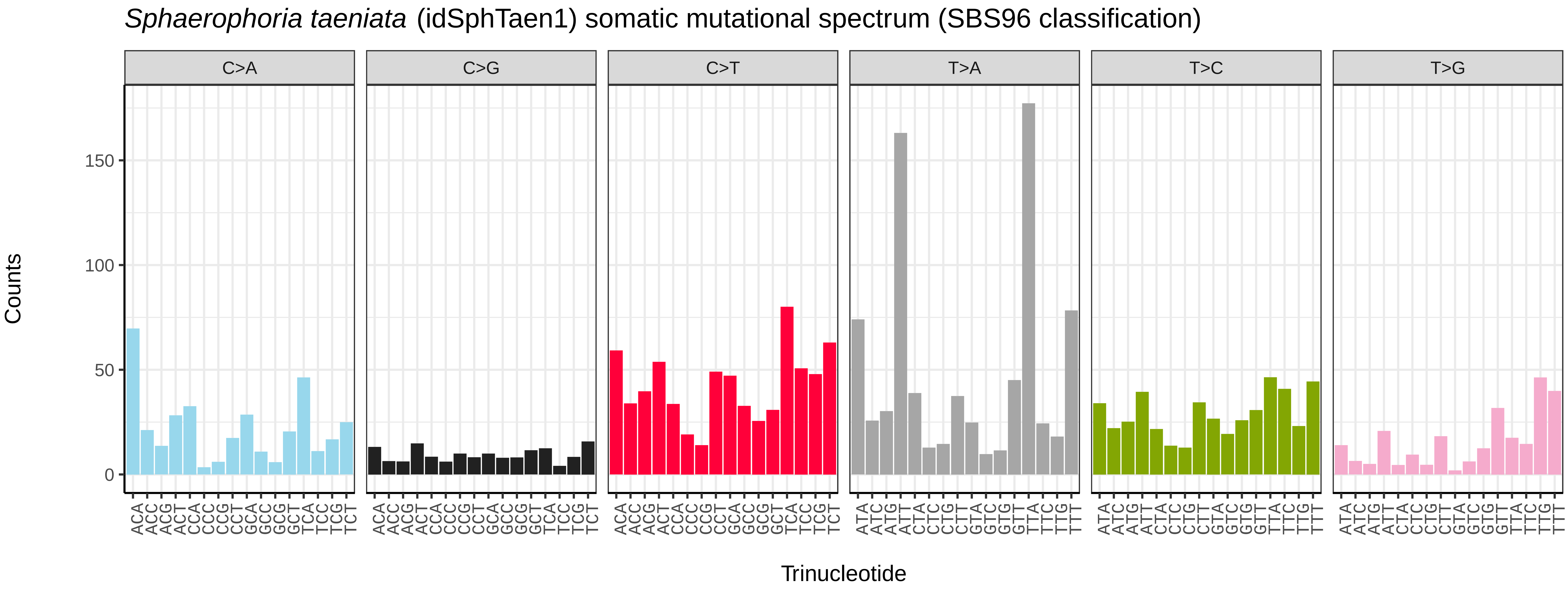
*Sphaerophoria taeniata* (idSphTaen1) somatic mutational spectrum (SBS96 classification)
C>A
C>G
C>T
T>A
T>C
T>G
Counts
150
100
50
0
Trinucleotide

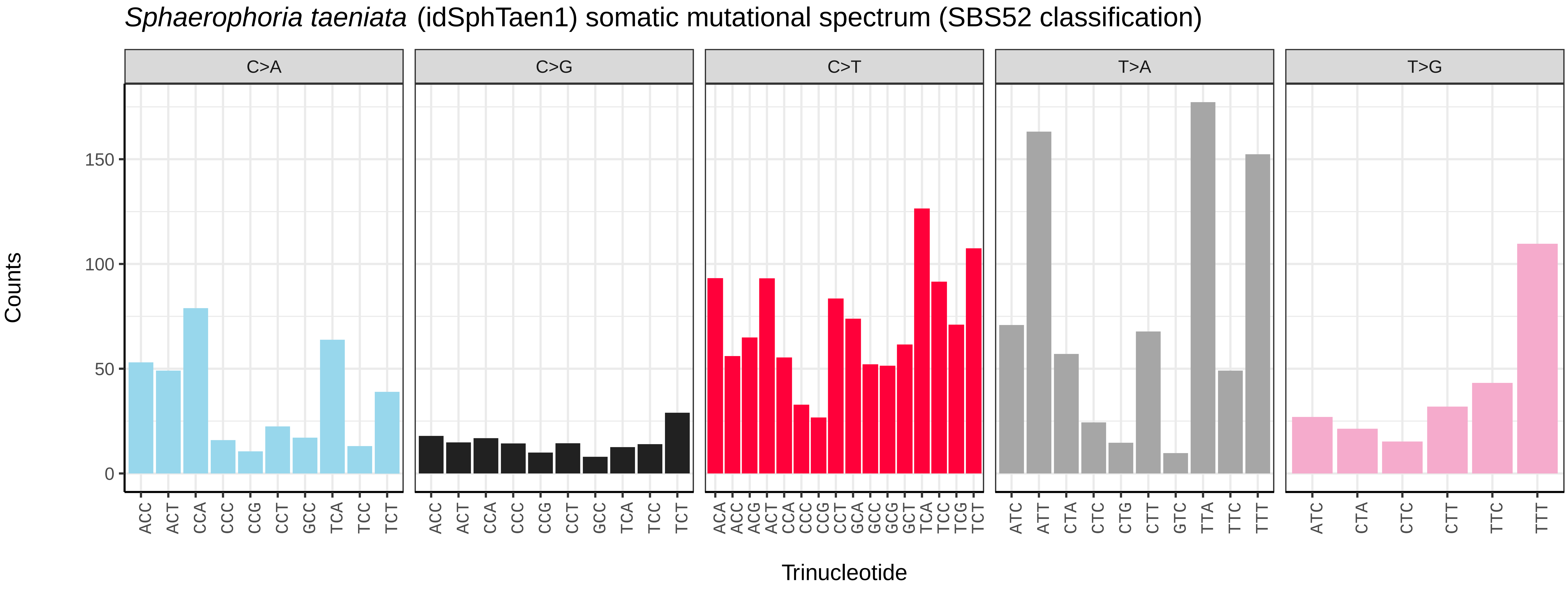
*Sphaerophoria taeniata* (idSphTaen1) somatic mutational spectrum (SBS52 classification)
C>A
C>G
C>T
T>A
T>G
Counts
150
100
50
0
Trinucleotide

*Stomorhina lunata* (idStoLuna1) germline mutational spectrum

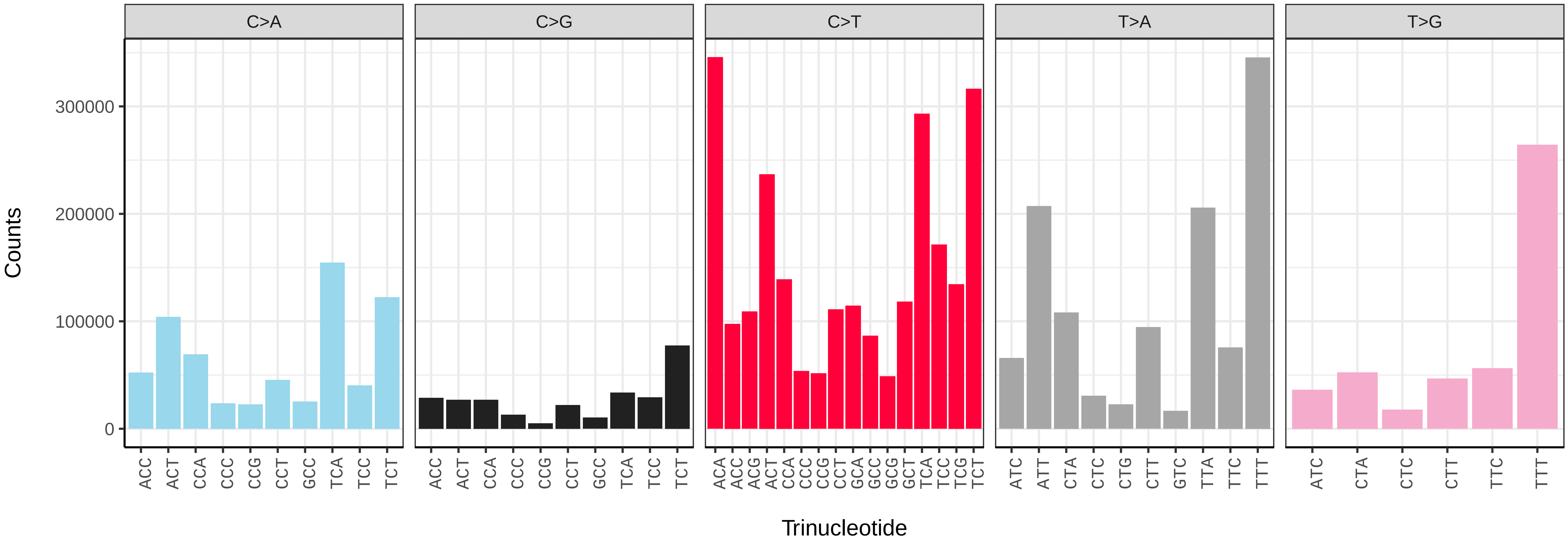


*Stomorhina lunata* (idStoLuna1) somatic mutational spectrum (SBS96 classification)

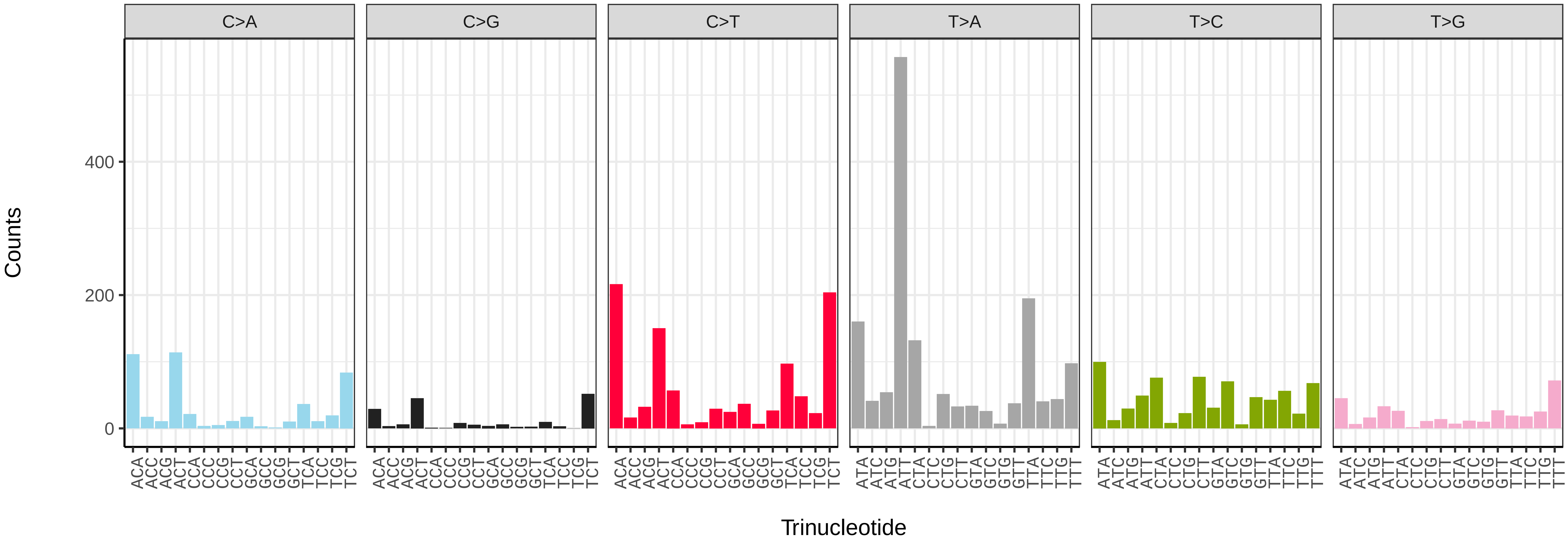


*Stomorhina lunata* (idStoLuna1) somatic mutational spectrum (SBS52 classification)

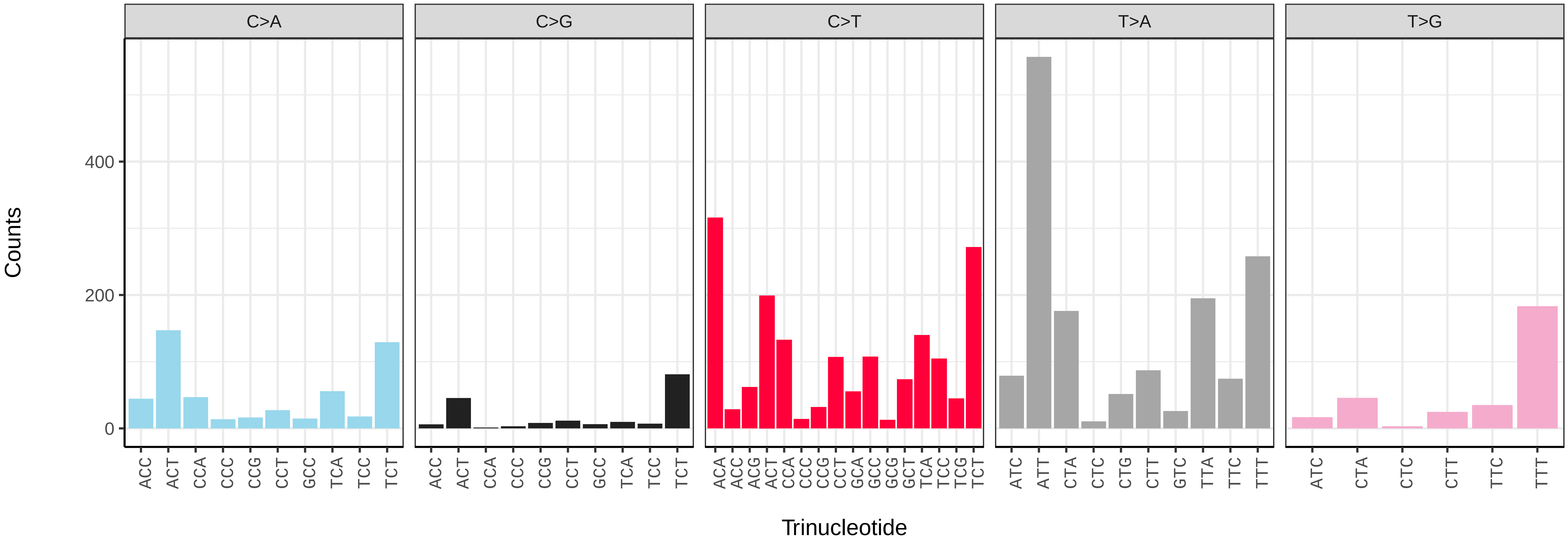

*Suillia variegata* (idSuiVari3) germline mutational spectrum

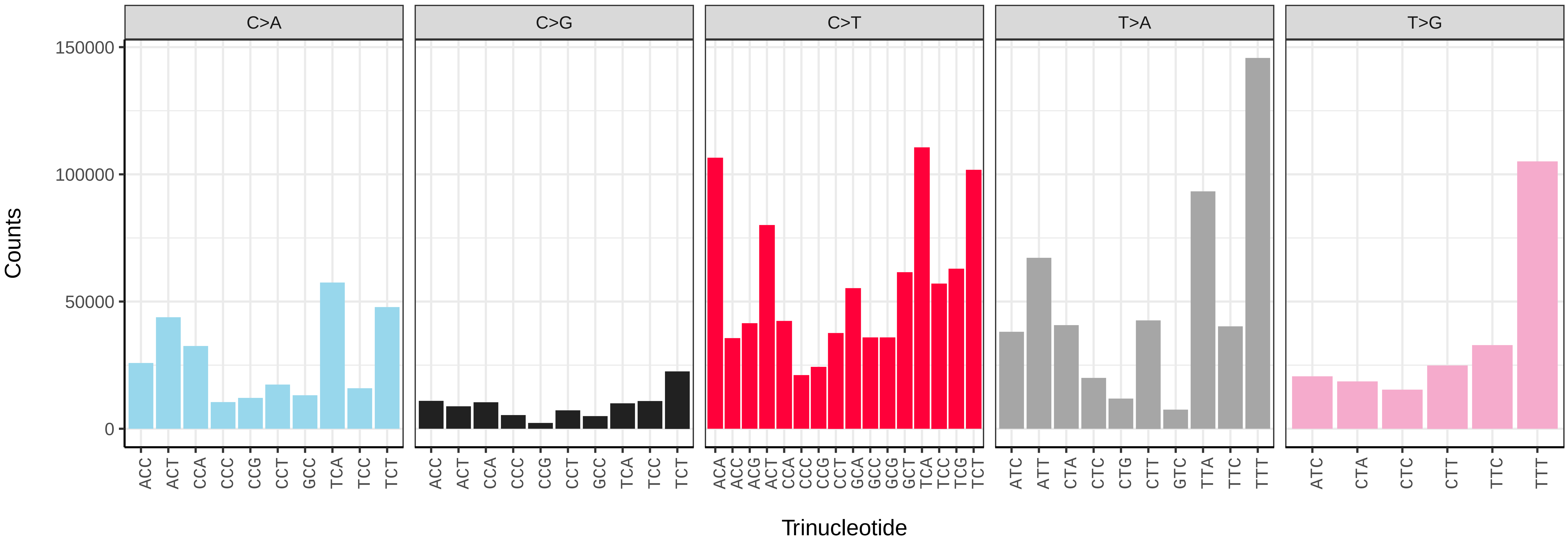


*Suillia variegata* (idSuiVari3) somatic mutational spectrum (SBS96 classification)

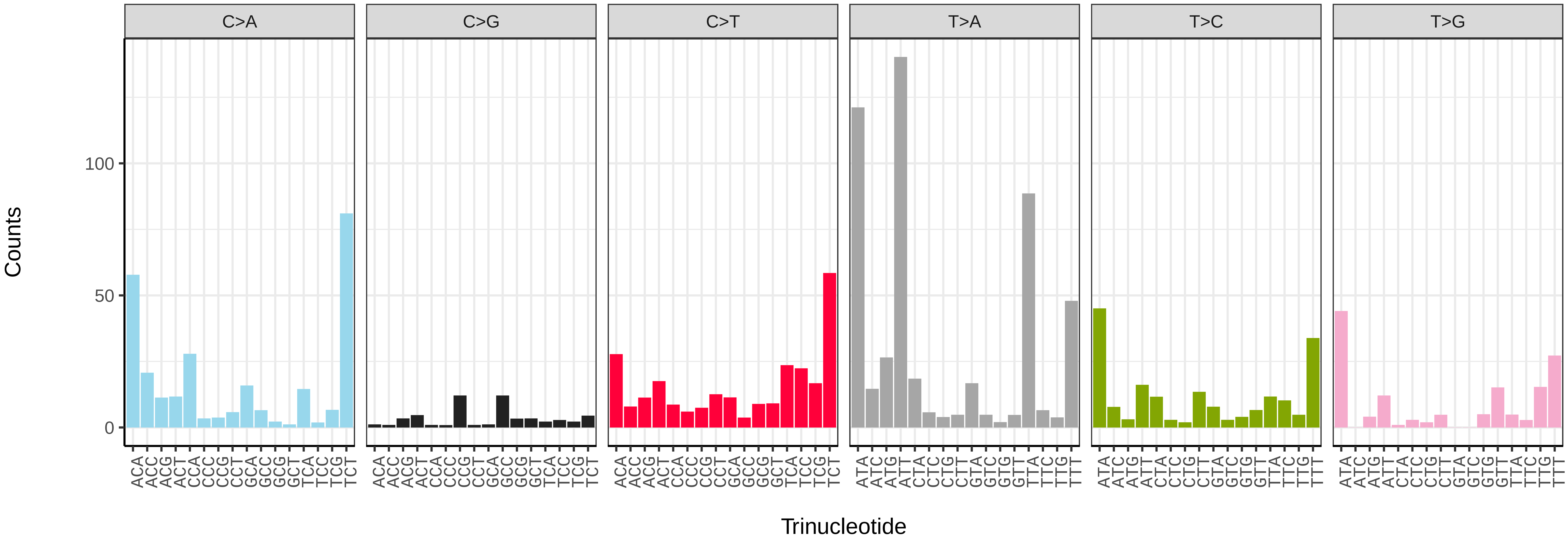


*Suillia variegata* (idSuiVari3) somatic mutational spectrum (SBS52 classification)

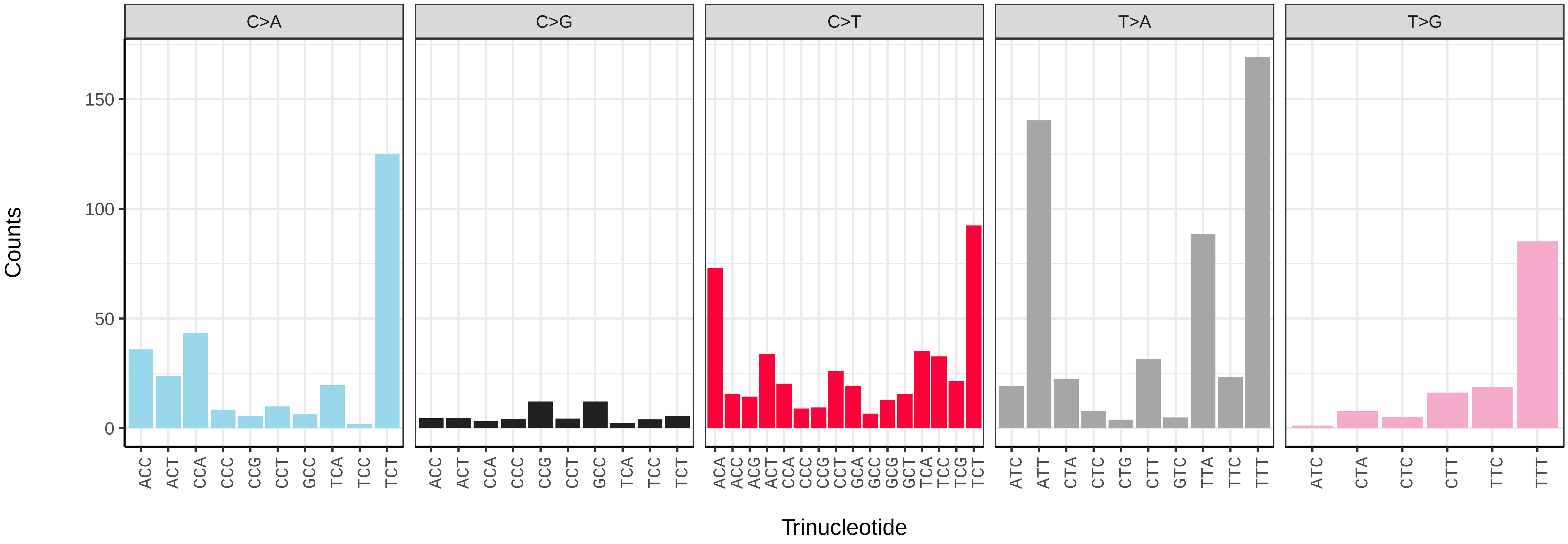

*Syritta pipiens* (idSyrPipi1) germline mutational spectrum

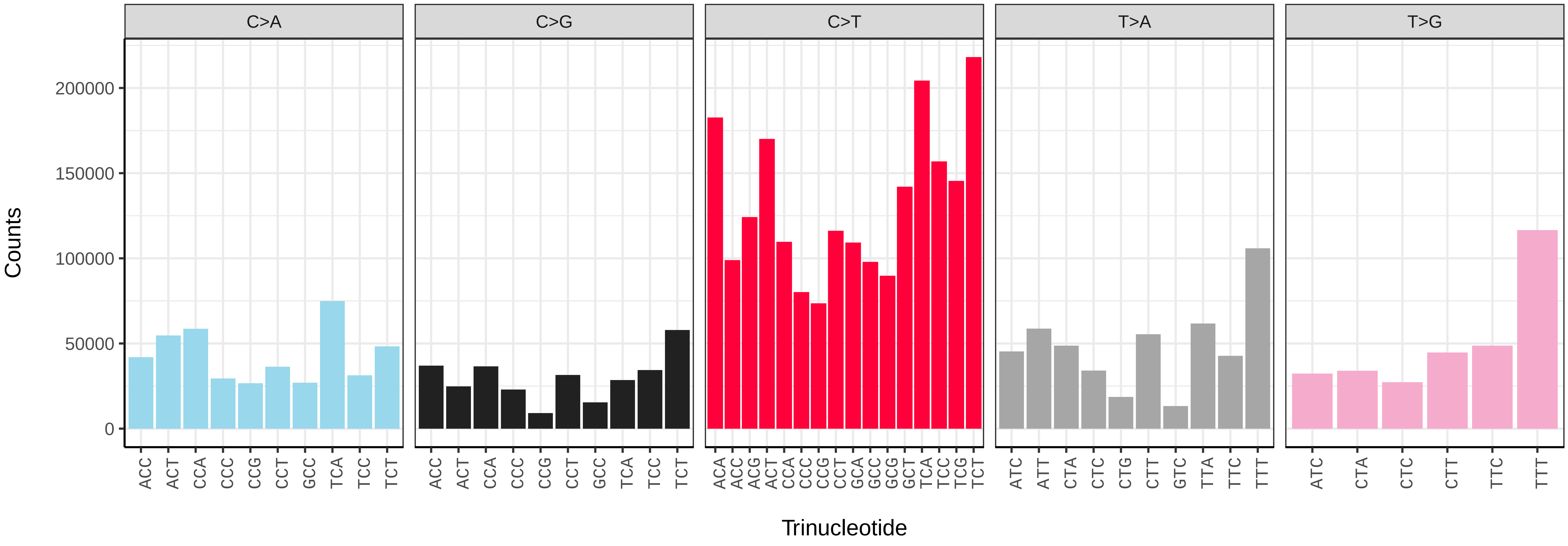


*Syritta pipiens* (idSyrPipi1) somatic mutational spectrum (SBS96 classification)

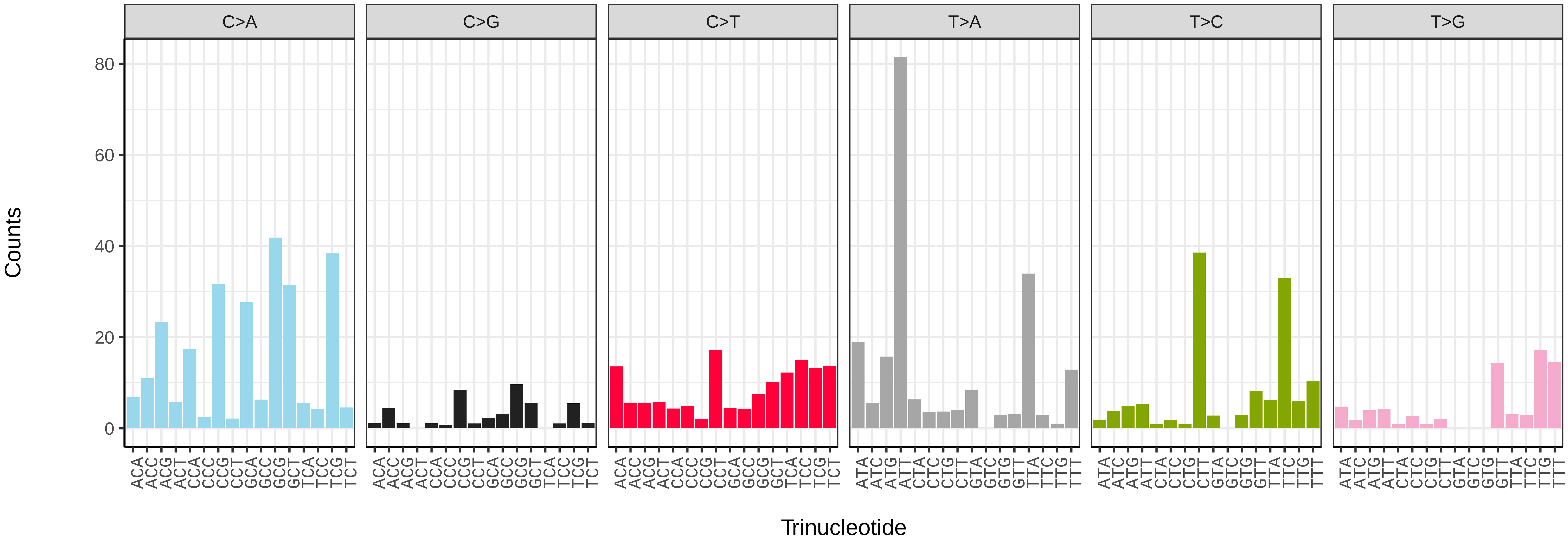


*Syritta pipiens* (idSyrPipi1) somatic mutational spectrum (SBS52 classification)

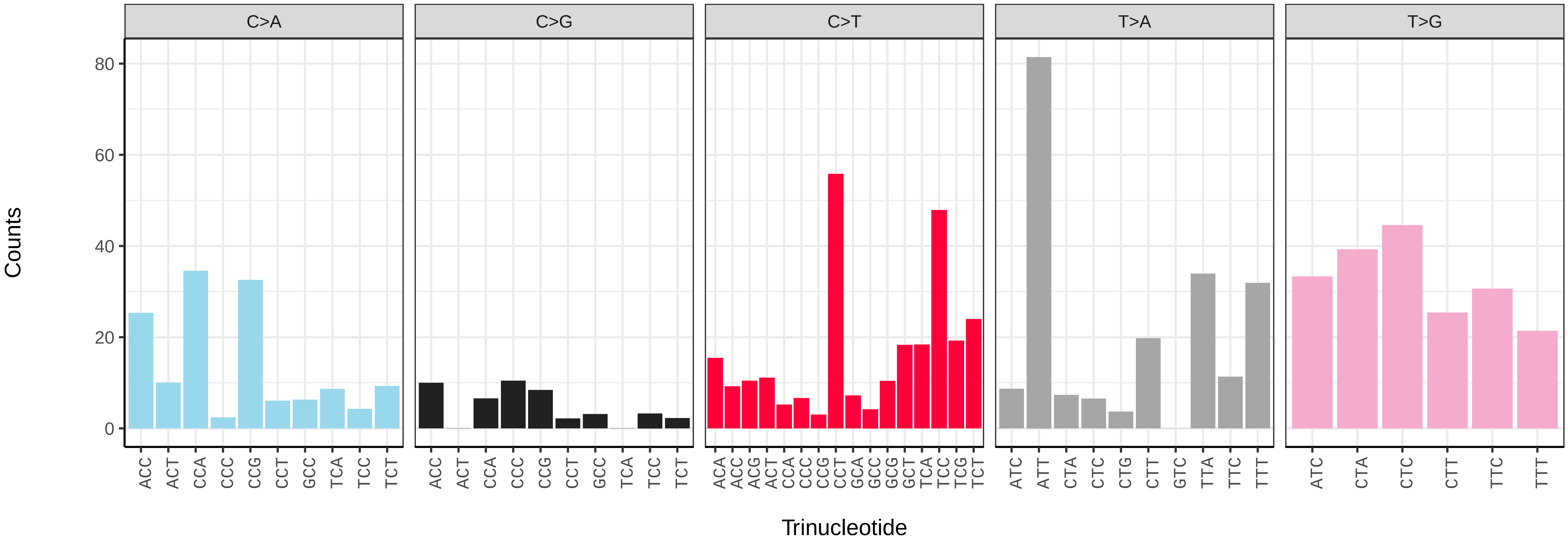

*Syritta pipiens* (idSyrPipi1.idSyrPipi10) germline mutational spectrum

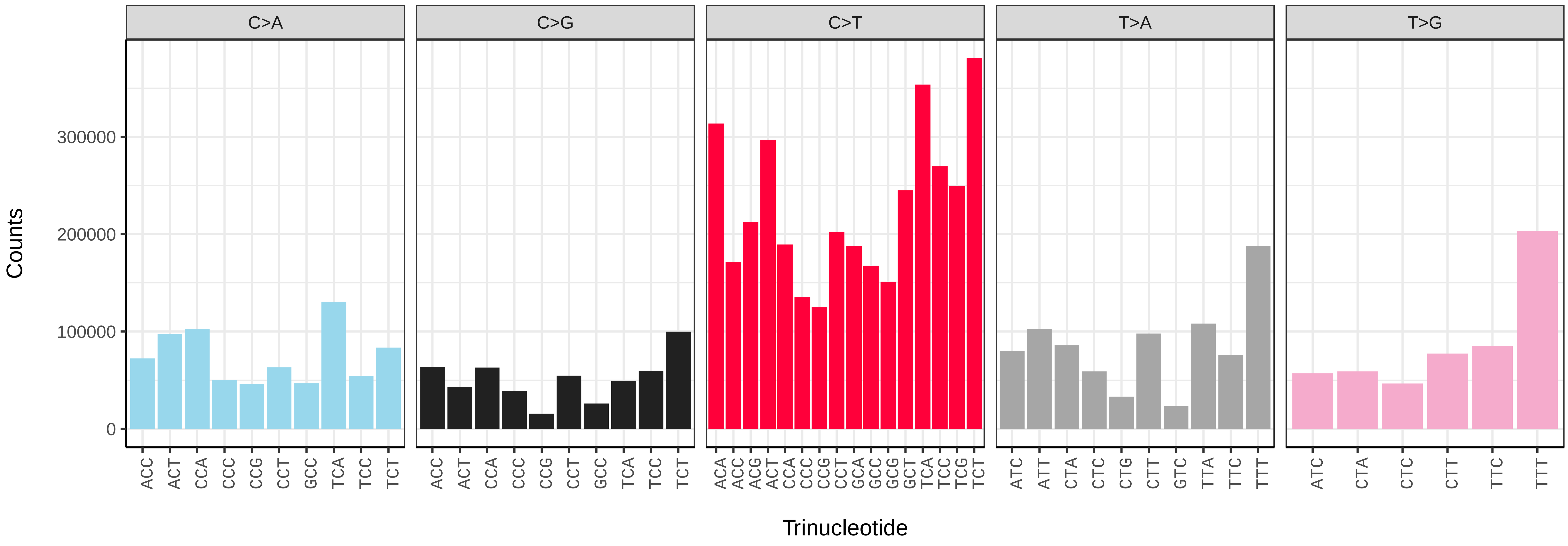


*Syritta pipiens* (idSyrPipi1.idSyrPipi10) somatic mutational spectrum (SBS96 classification)

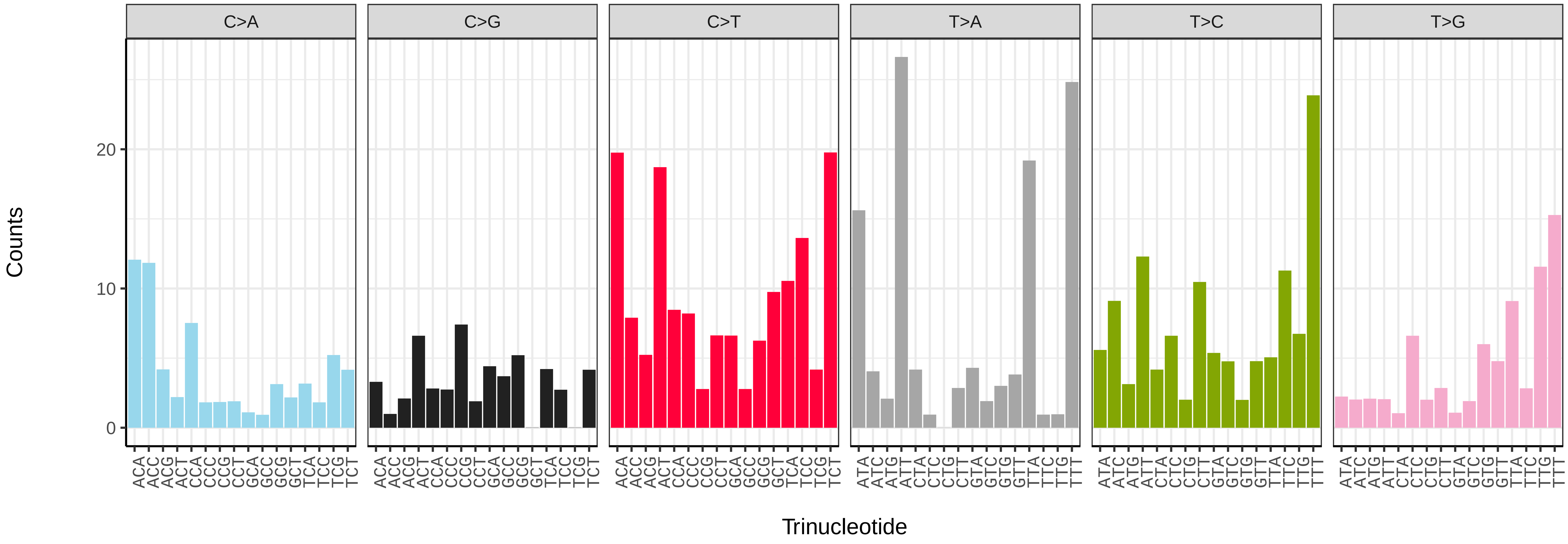


*Syritta pipiens* (idSyrPipi1.idSyrPipi10) somatic mutational spectrum (SBS52 classification)

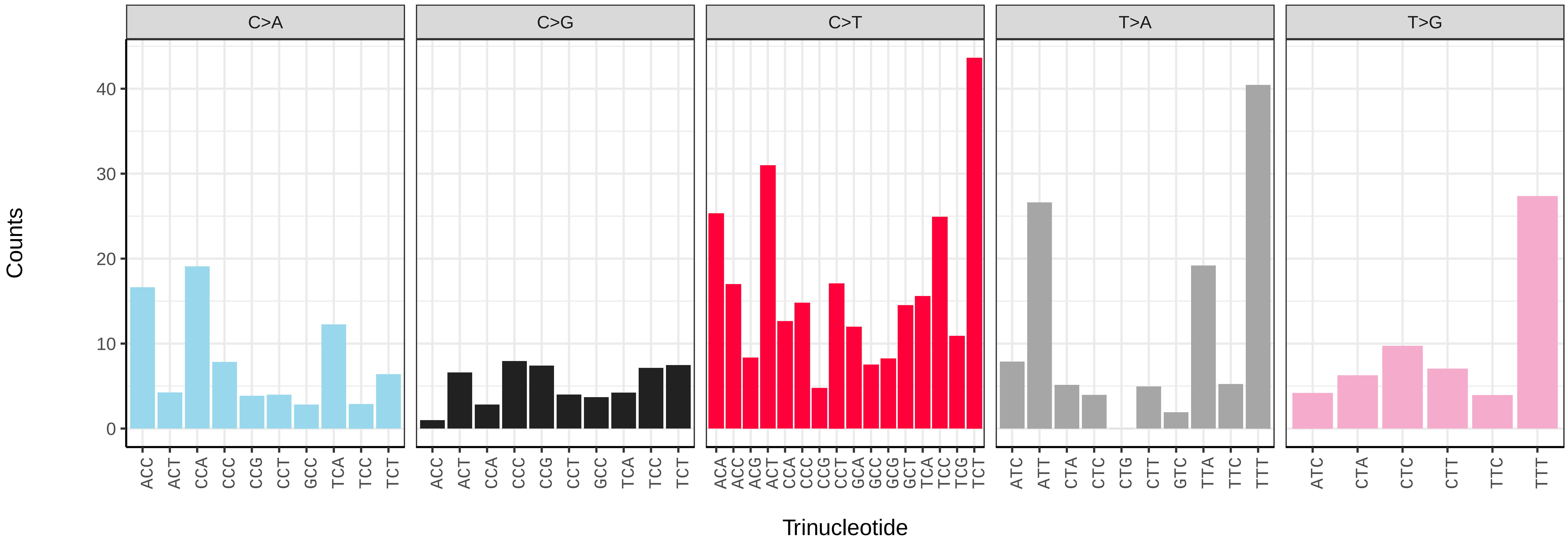

*Syritta pipiens* (idSyrPipi1.idSyrPipi11) germline mutational spectrum

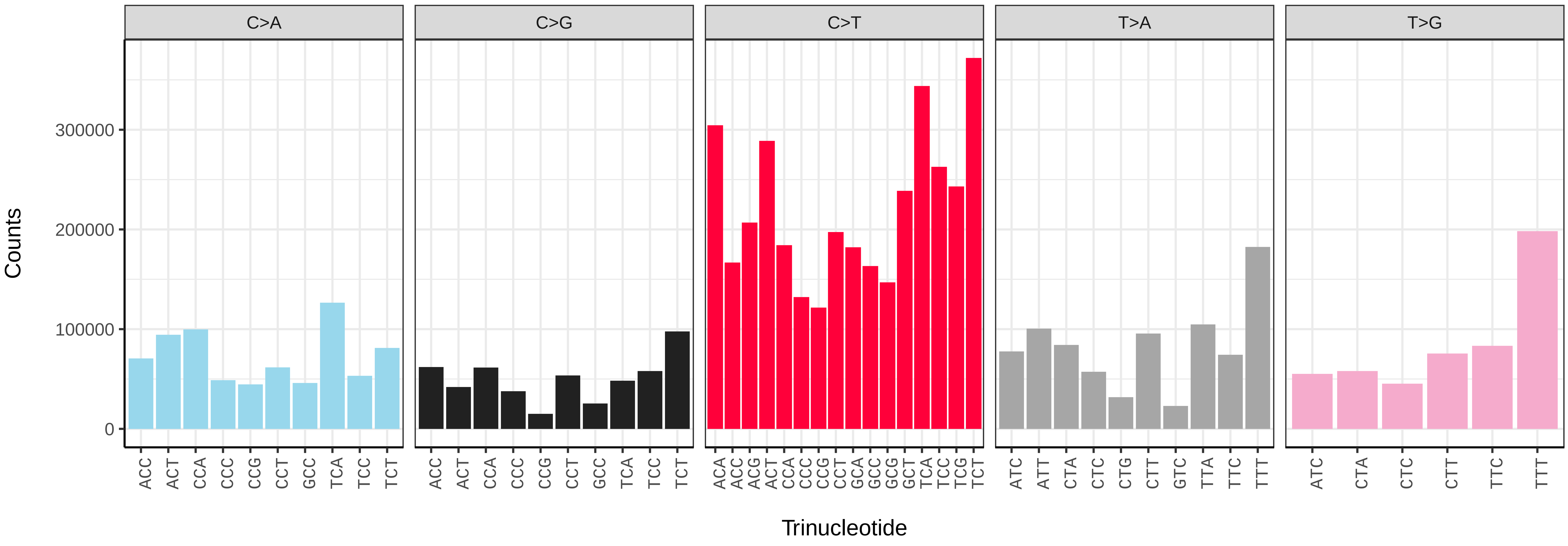


*Syritta pipiens* (idSyrPipi1.idSyrPipi11) somatic mutational spectrum (SBS96 classification)

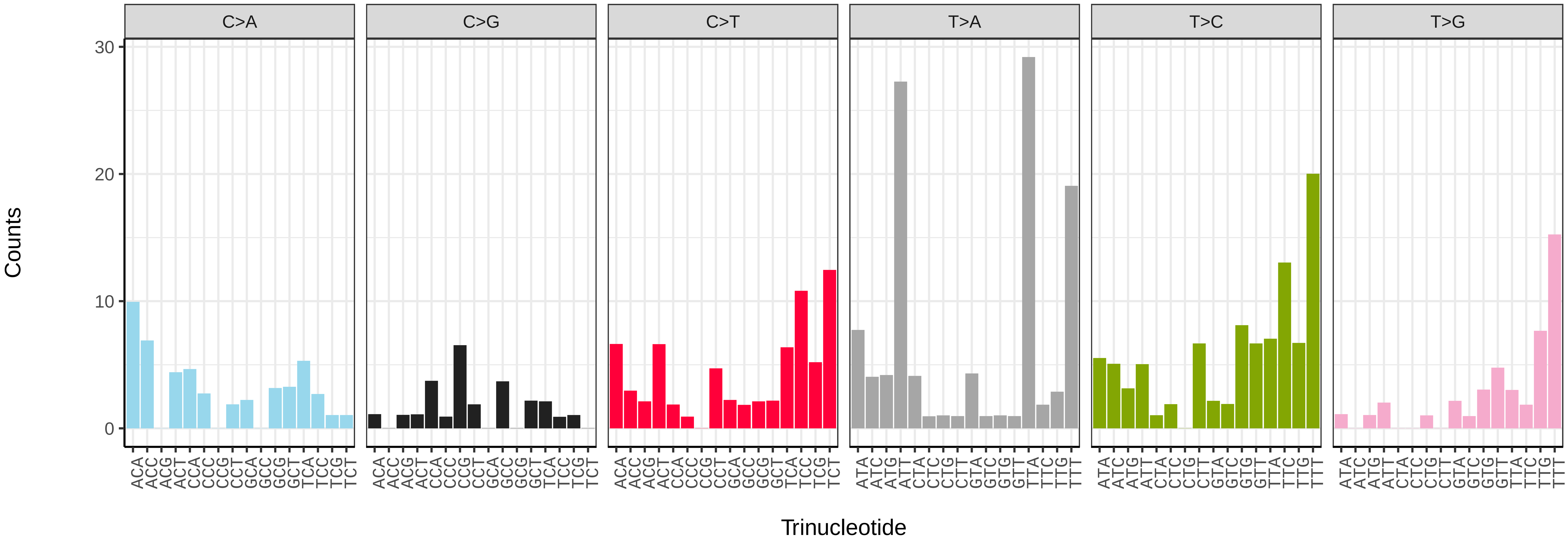


*Syritta pipiens* (idSyrPipi1.idSyrPipi11) somatic mutational spectrum (SBS52 classification)

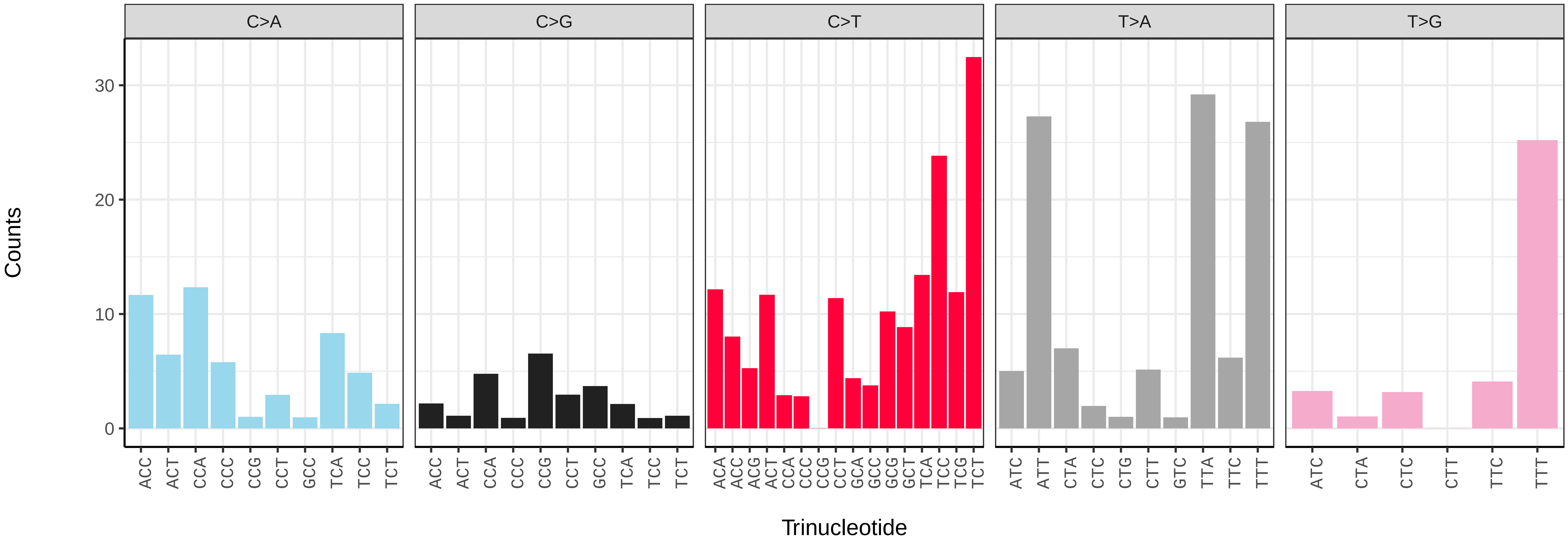

*Syritta pipiens* (idSyrPipi1.idSyrPipi2) germline mutational spectrum

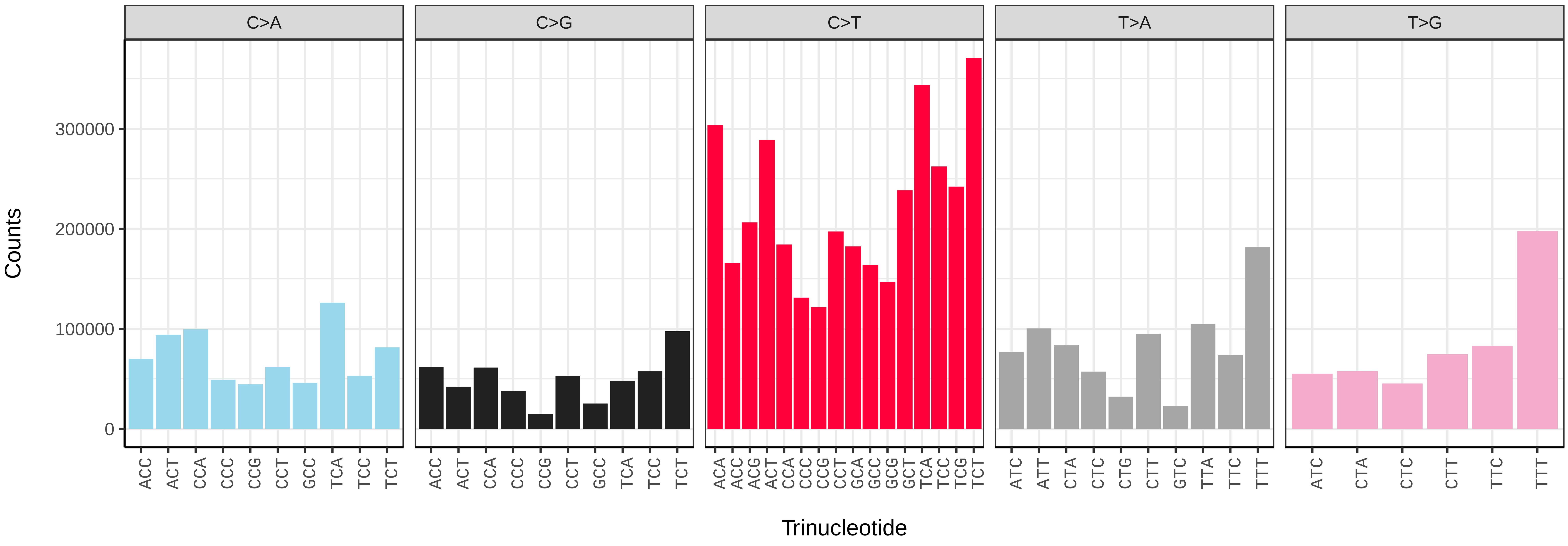


*Syritta pipiens* (idSyrPipi1.idSyrPipi2) somatic mutational spectrum (SBS96 classification)

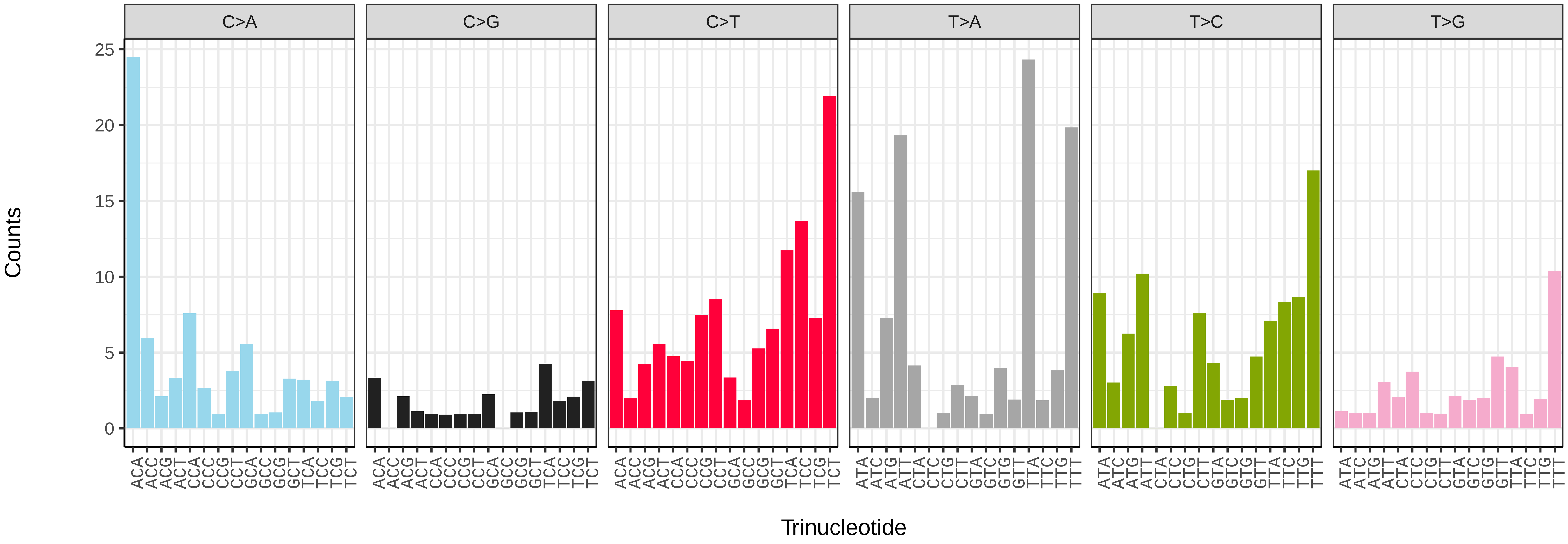


*Syritta pipiens* (idSyrPipi1.idSyrPipi2) somatic mutational spectrum (SBS52 classification)

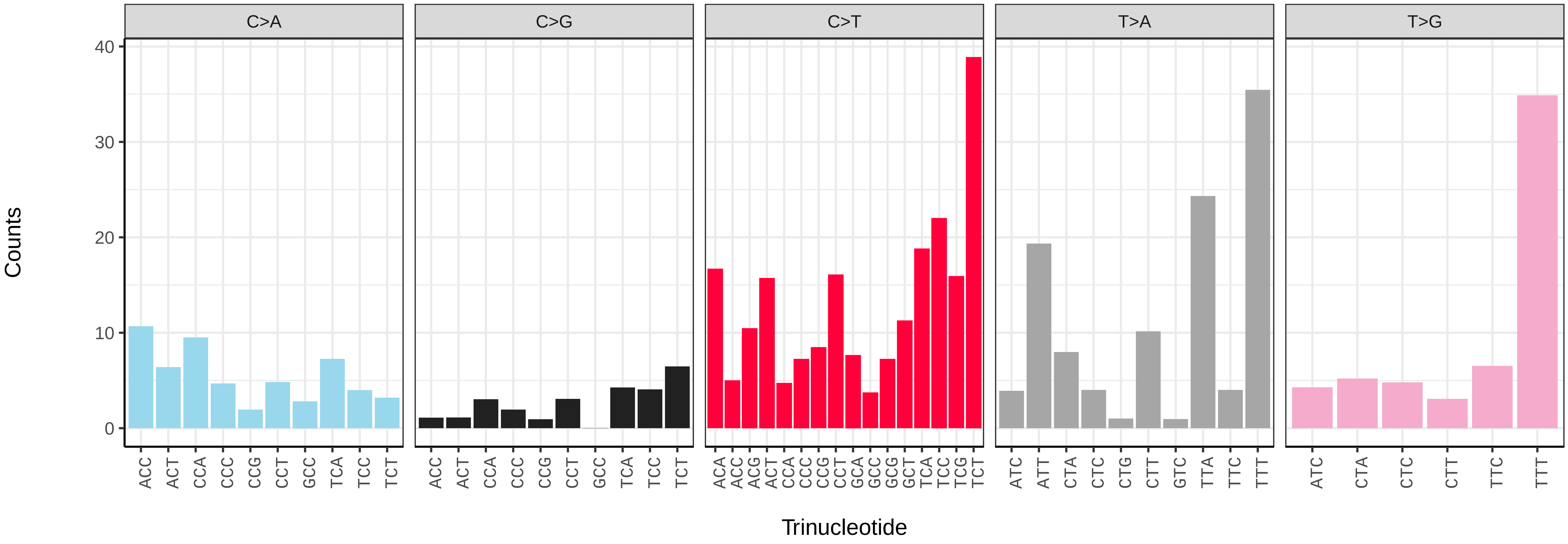

*Syritta pipiens* (idSyrPipi1.idSyrPipi5) germline mutational spectrum

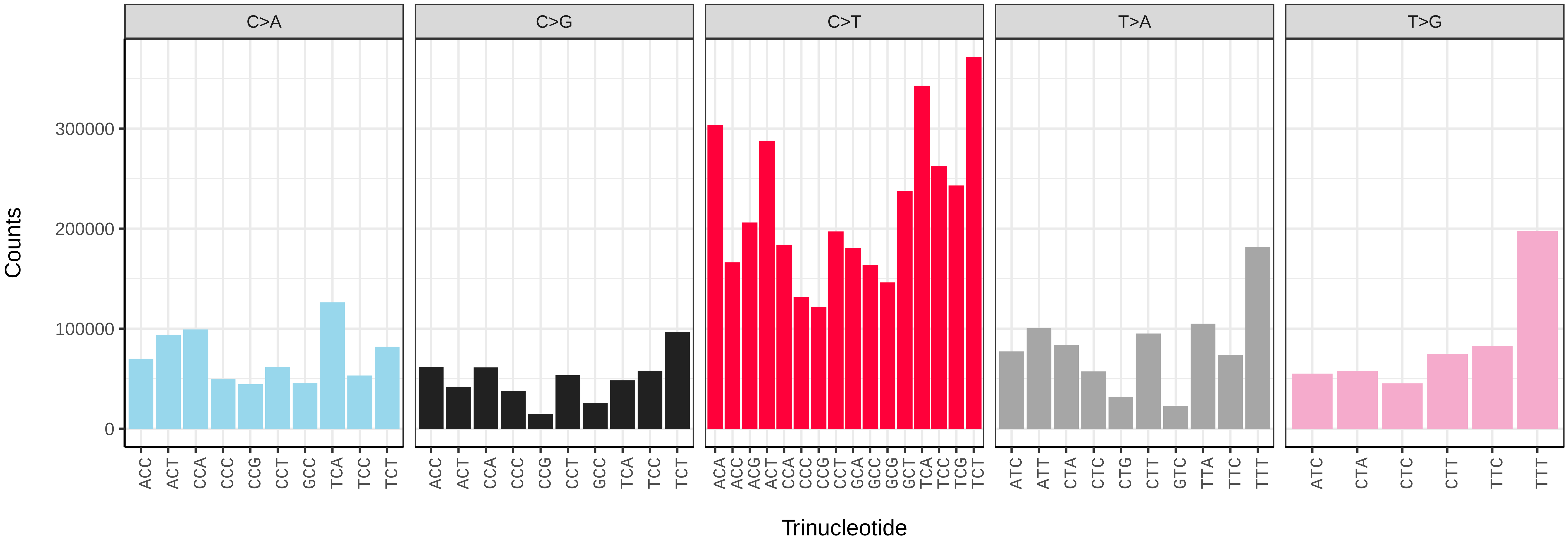


*Syritta pipiens* (idSyrPipi1.idSyrPipi5) somatic mutational spectrum (SBS96 classification)

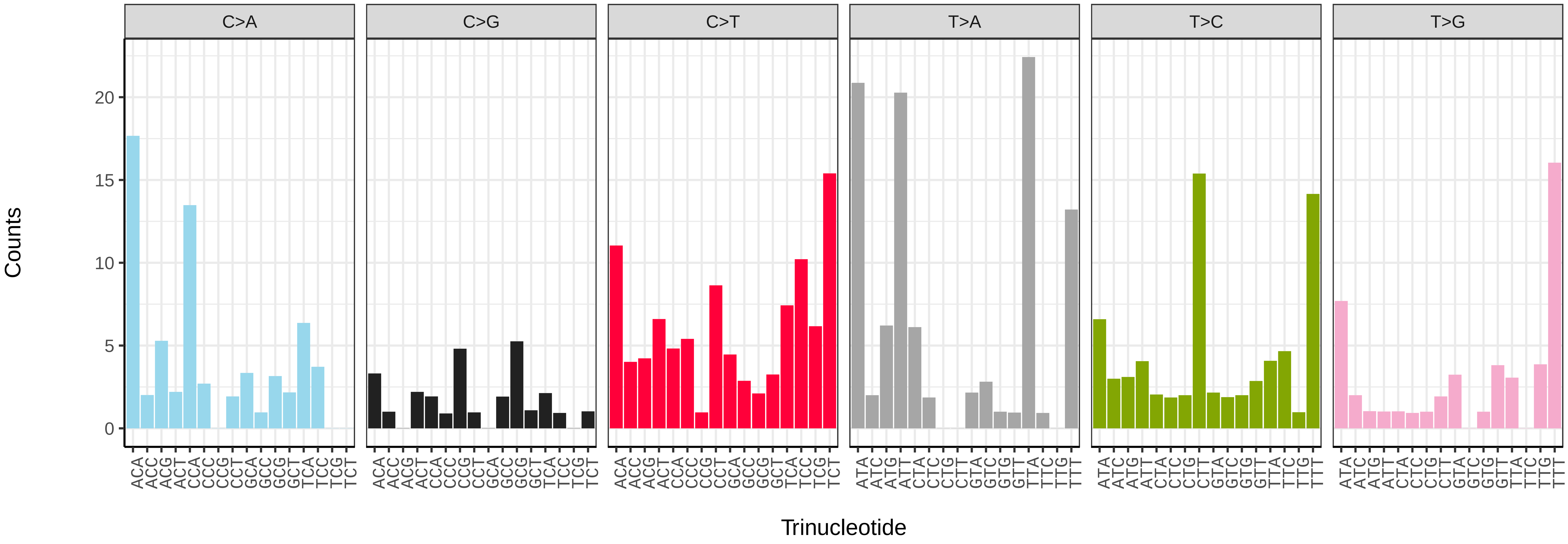


*Syritta pipiens* (idSyrPipi1.idSyrPipi5) somatic mutational spectrum (SBS52 classification)

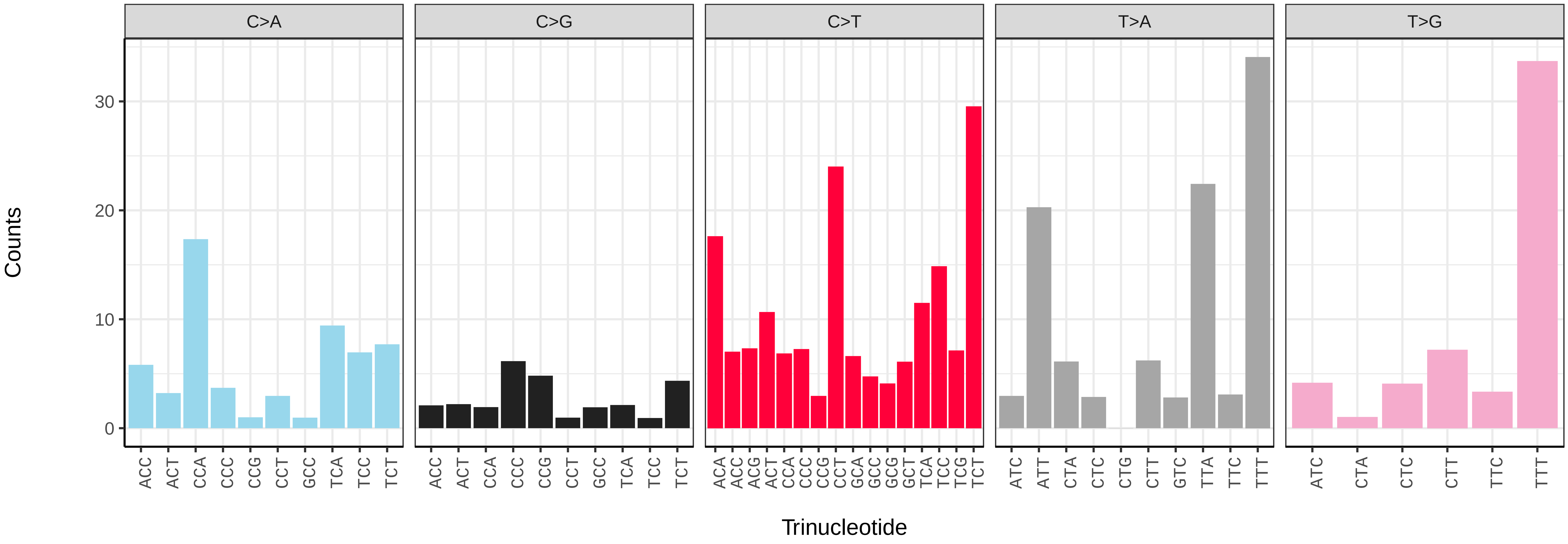

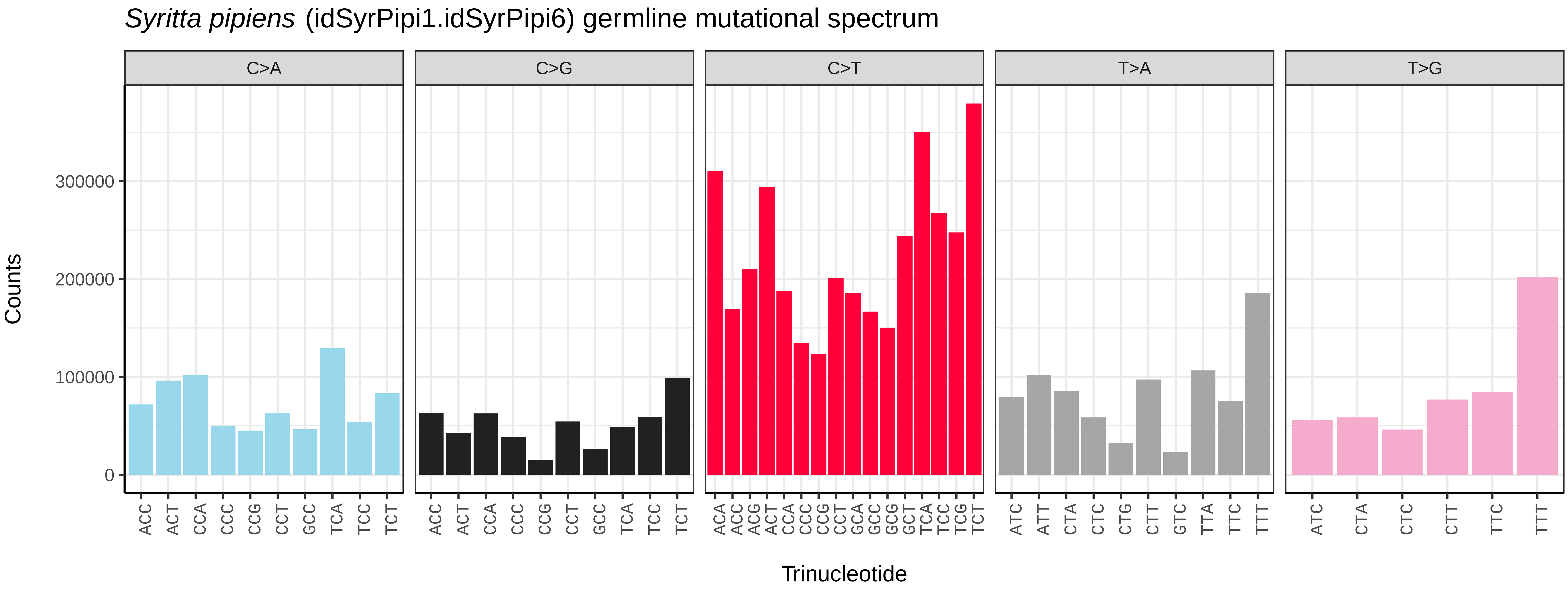
*Syritta pipiens* (idSyrPipi1.idSyrPipi6) germline mutational spectrum
C>A
C>G
C>T
T>A
T>G
Counts
Trinucleotide

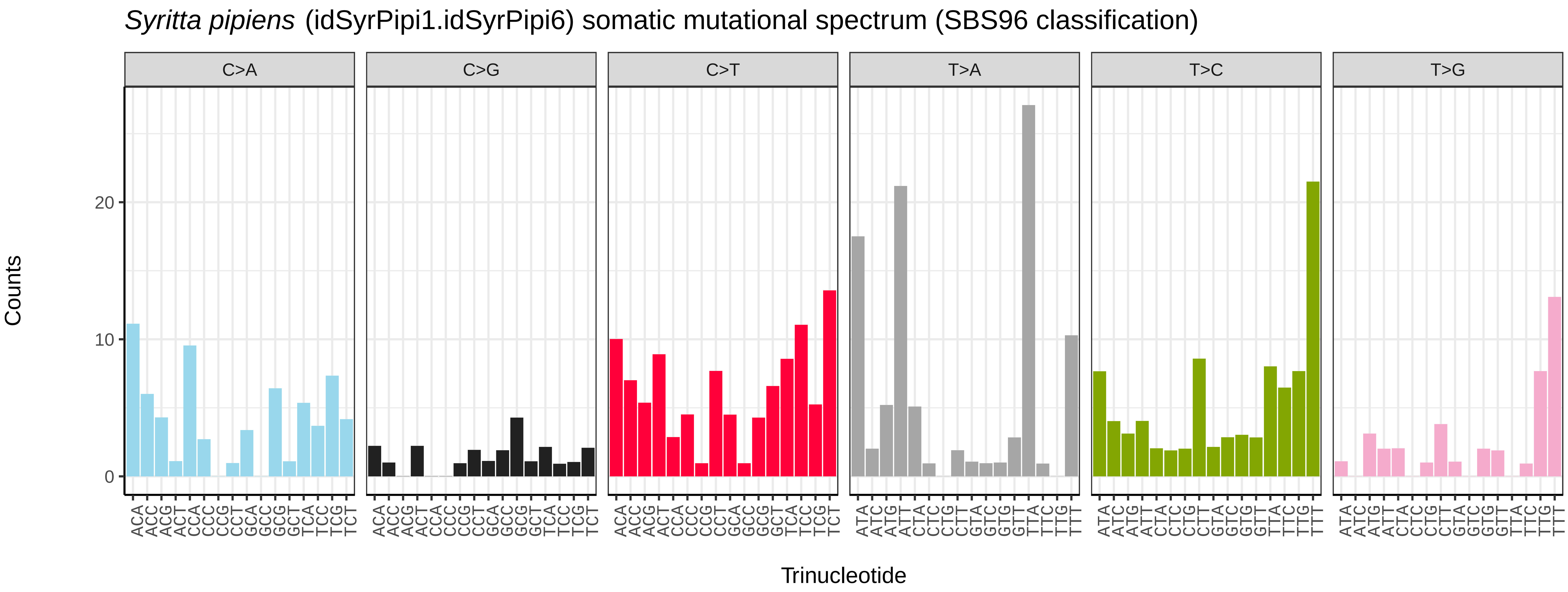
*Syritta pipiens* (idSyrPipi1.idSyrPipi6) somatic mutational spectrum (SBS96 classification)
C>A
C>G
C>T
T>A
T>C
T>G
Counts
Trinucleotide

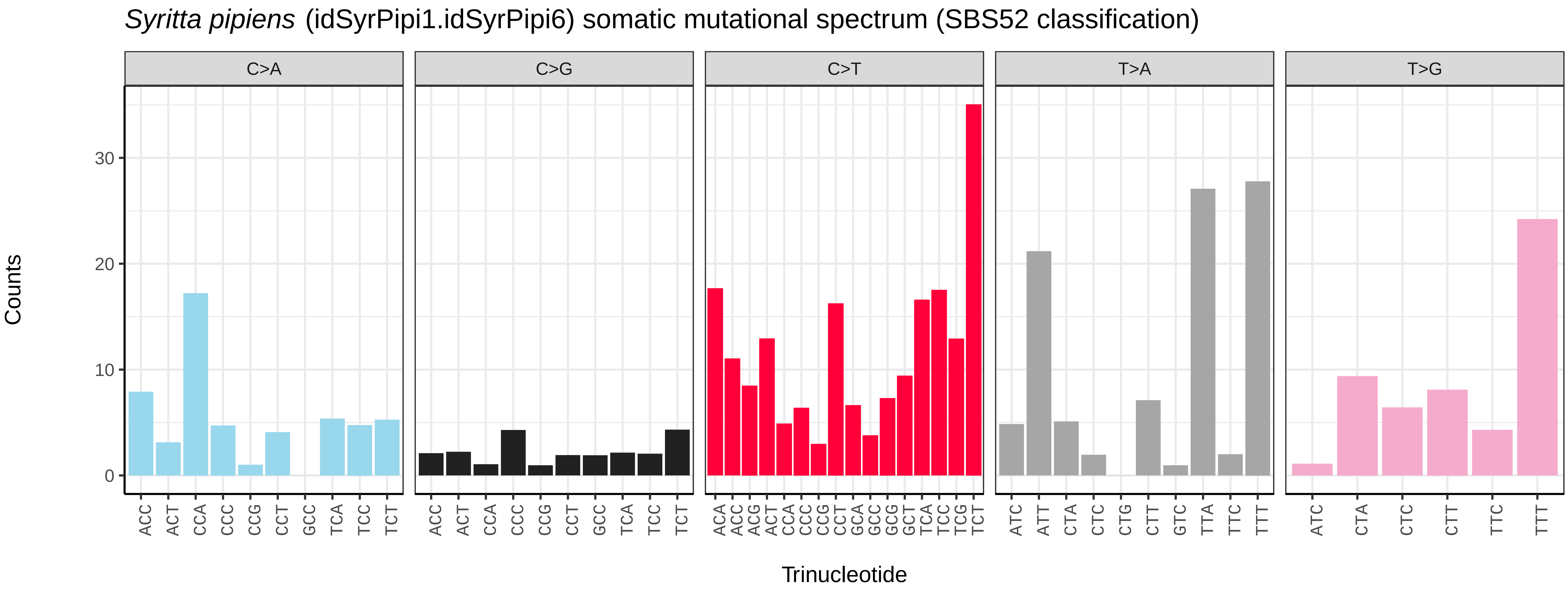
*Syritta pipiens* (idSyrPipi1.idSyrPipi6) somatic mutational spectrum (SBS52 classification)
C>A
C>G
C>T
T>A
T>G
Counts
Trinucleotide

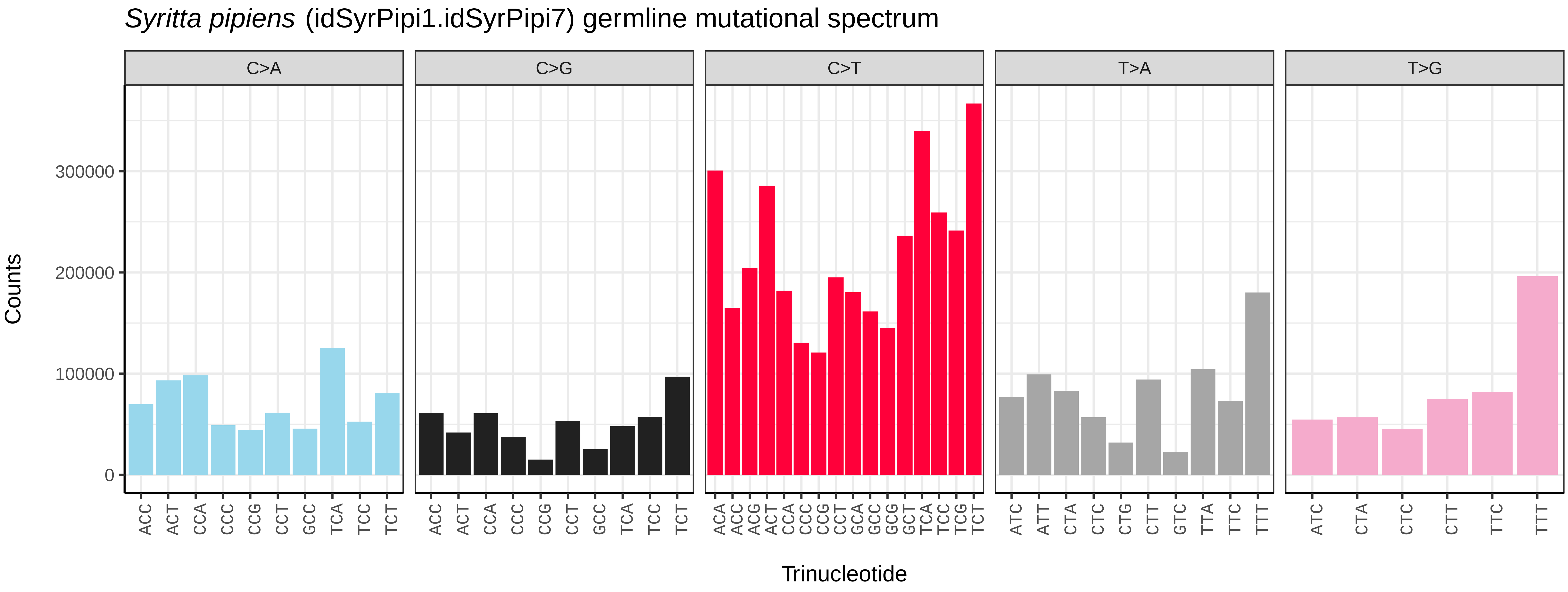
Syritta pipiens (idSyrPipi1.idSyrPipi7) germline mutational spectrum
C>A
C>G
C>T
T>A
T>G
Counts
0
100000
200000
300000
Trinucleotide
ACC ACT CCA CCC CCG CCT GCC TCA TCC TCT
ACA ACC ACG ACT CCA CCC CCG CCT GCA GCC GCG GCT TCA TCC TCG TCT
ATC ATT CTA CTC CTG CTT GTC TTA TTC TTT
ATC CTA CTC CTT TTC TTT

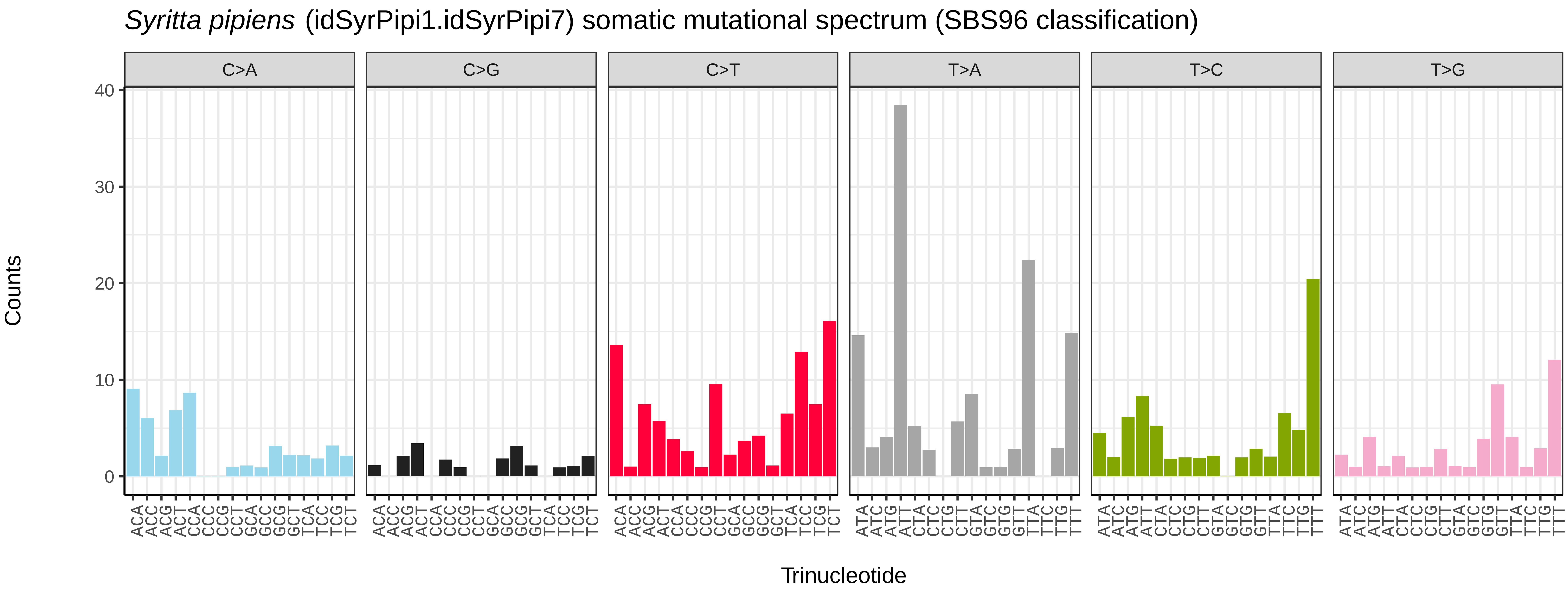
Syritta pipiens (idSyrPipi1.idSyrPipi7) somatic mutational spectrum (SBS96 classification)
C>A
C>G
C>T
T>A
T>C
T>G
Counts
0
10
20
30
40
Trinucleotide

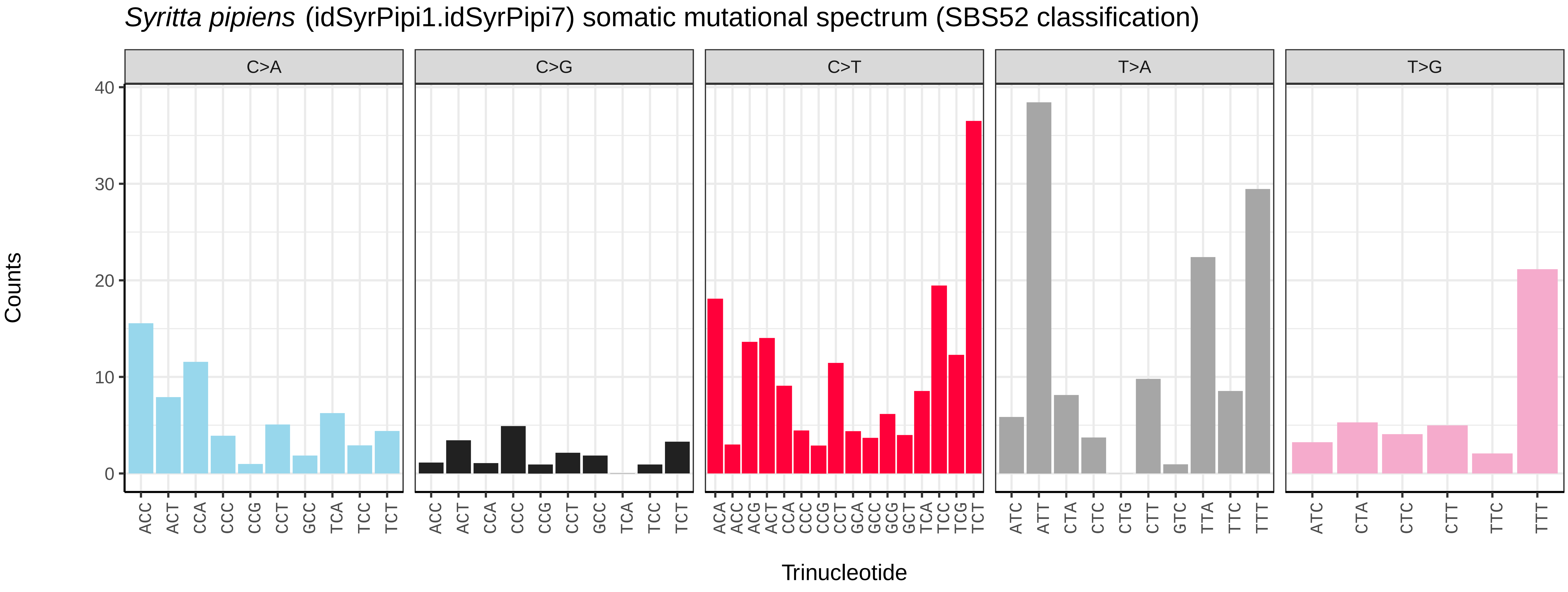
Syritta pipiens (idSyrPipi1.idSyrPipi7) somatic mutational spectrum (SBS52 classification)
C>A
C>G
C>T
T>A
T>G
Counts
0
10
20
30
40
Trinucleotide
ACC ACT CCA CCC CCG CCT GCC TCA TCC TCT
ACA ACC ACG ACT CCA CCC CCG CCT GCA GCC GCG GCT TCA TCC TCG TCT
ATC ATT CTA CTC CTG CTT GTC TTA TTC TTT
ATC CTA CTC CTT TTC TTT

*Syritta pipiens* (idSyrPipi1.idSyrPipi8) germline mutational spectrum

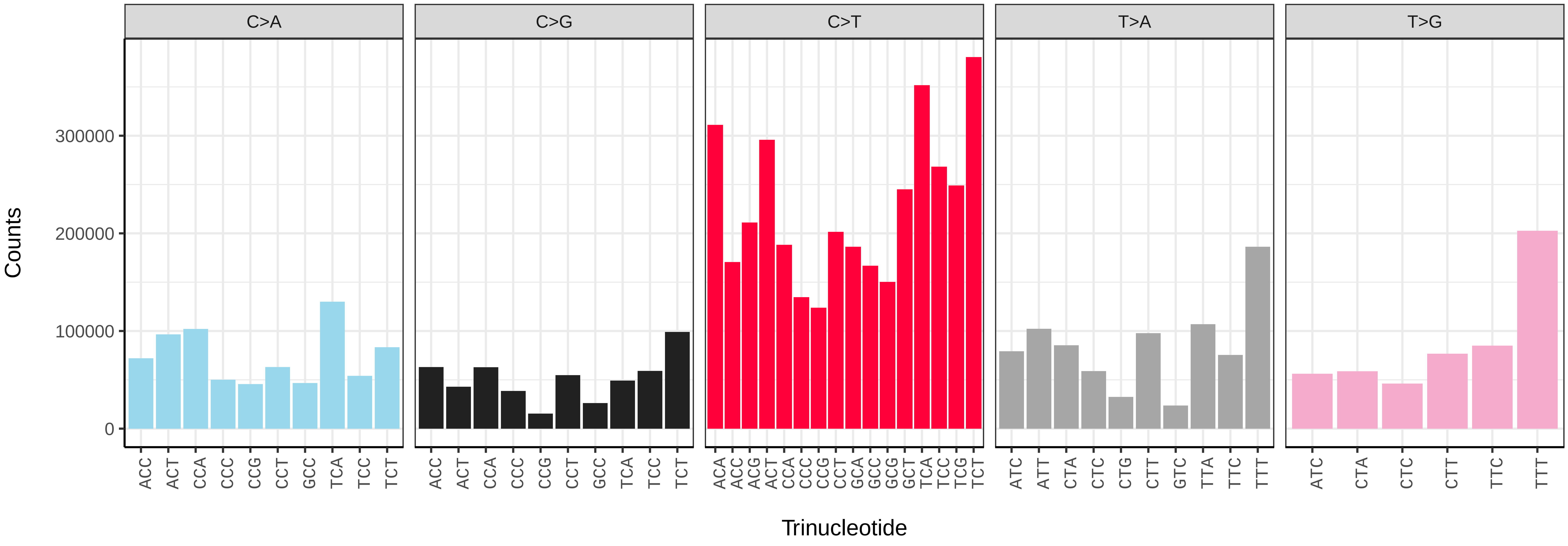


*Syritta pipiens* (idSyrPipi1.idSyrPipi8) somatic mutational spectrum (SBS96 classification)

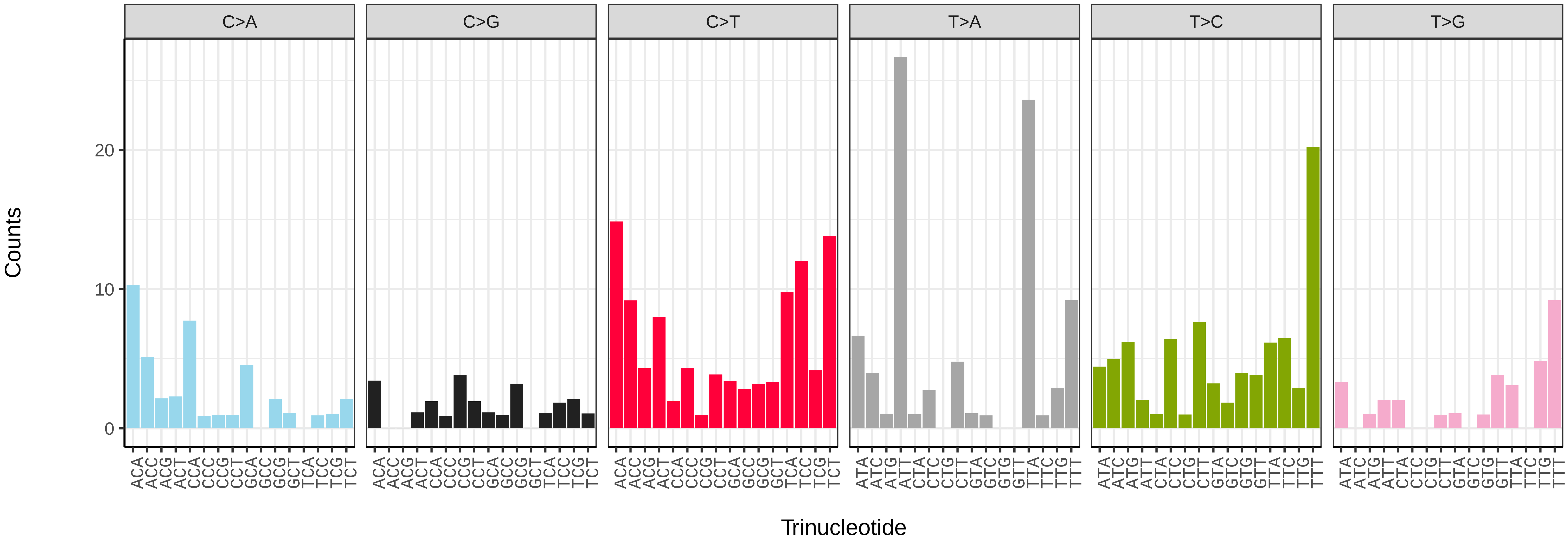


*Syritta pipiens* (idSyrPipi1.idSyrPipi8) somatic mutational spectrum (SBS52 classification)

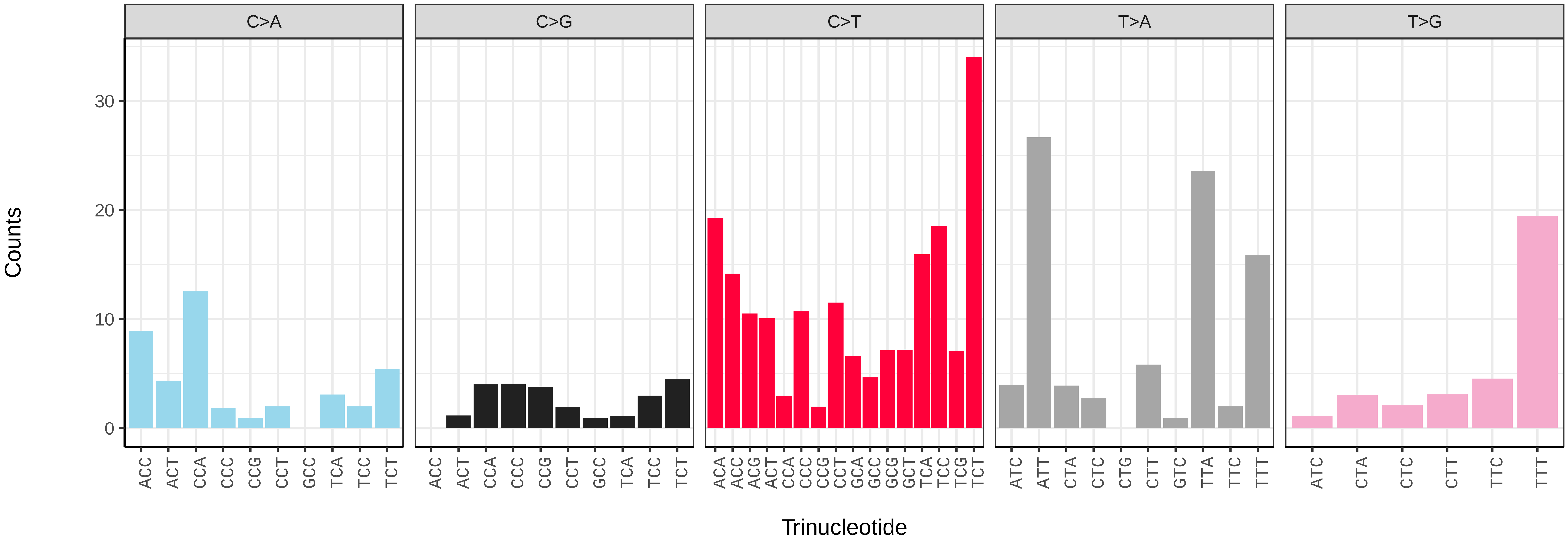

*Syritta pipiens* (idSyrPipi1.idSyrPipi9) germline mutational spectrum

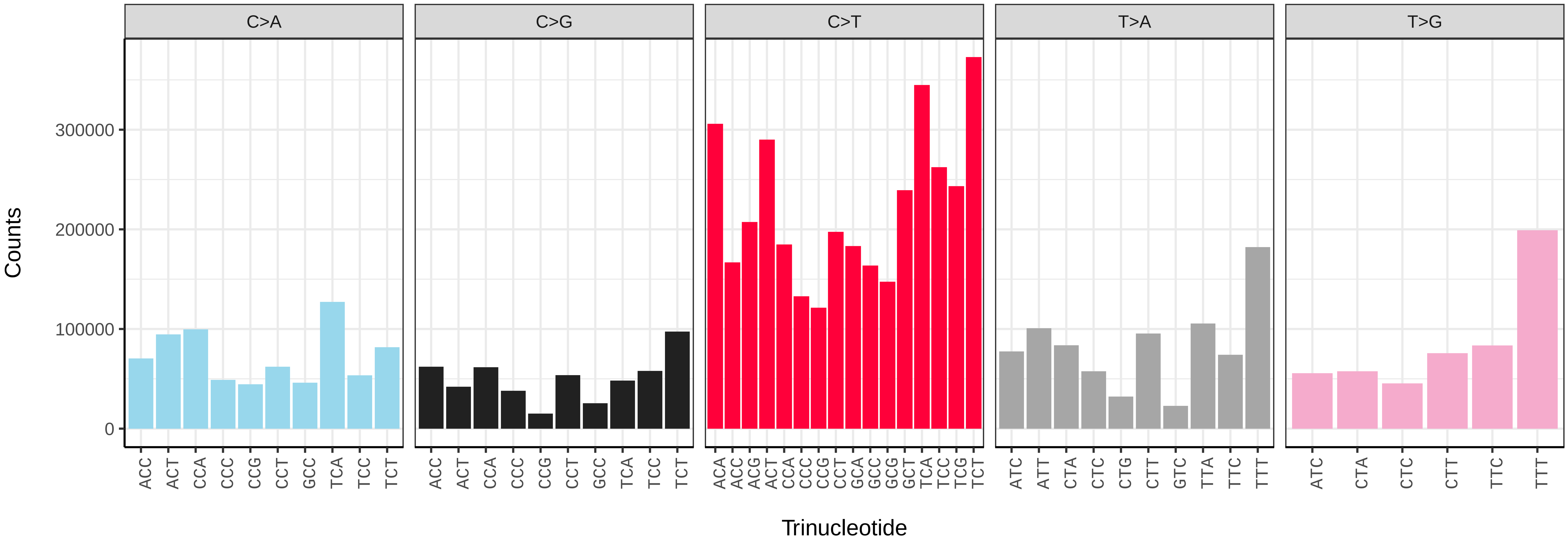


*Syritta pipiens* (idSyrPipi1.idSyrPipi9) somatic mutational spectrum (SBS96 classification)

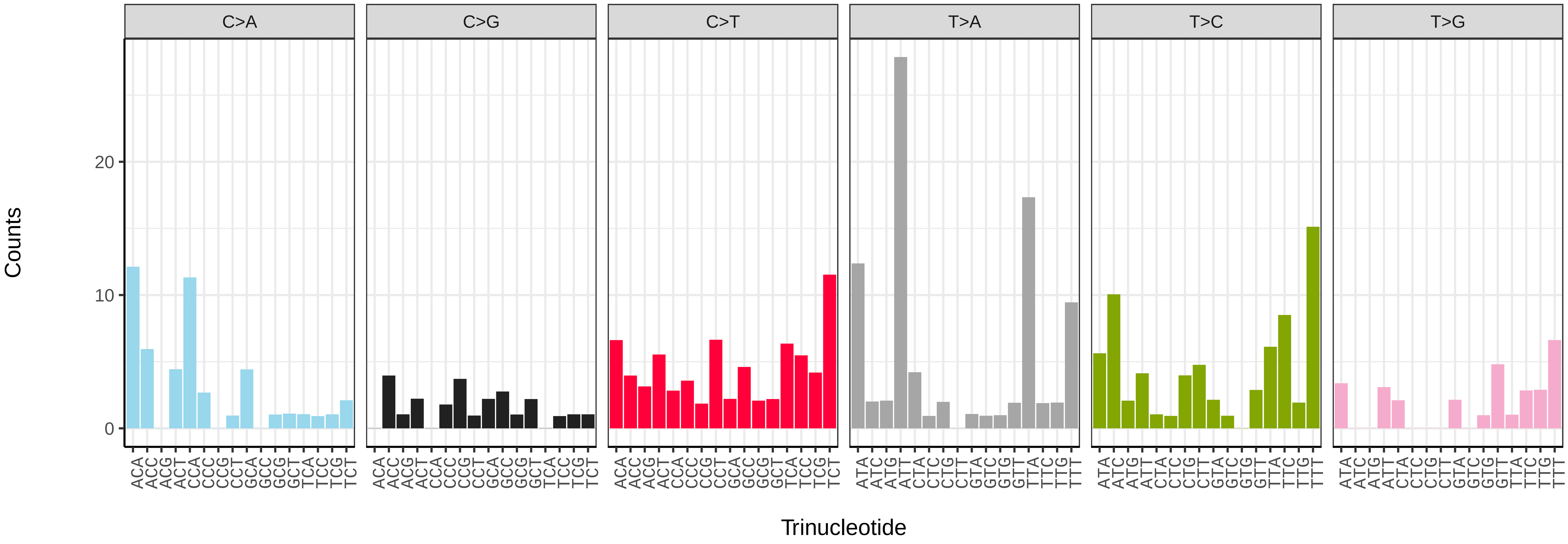


*Syritta pipiens* (idSyrPipi1.idSyrPipi9) somatic mutational spectrum (SBS52 classification)

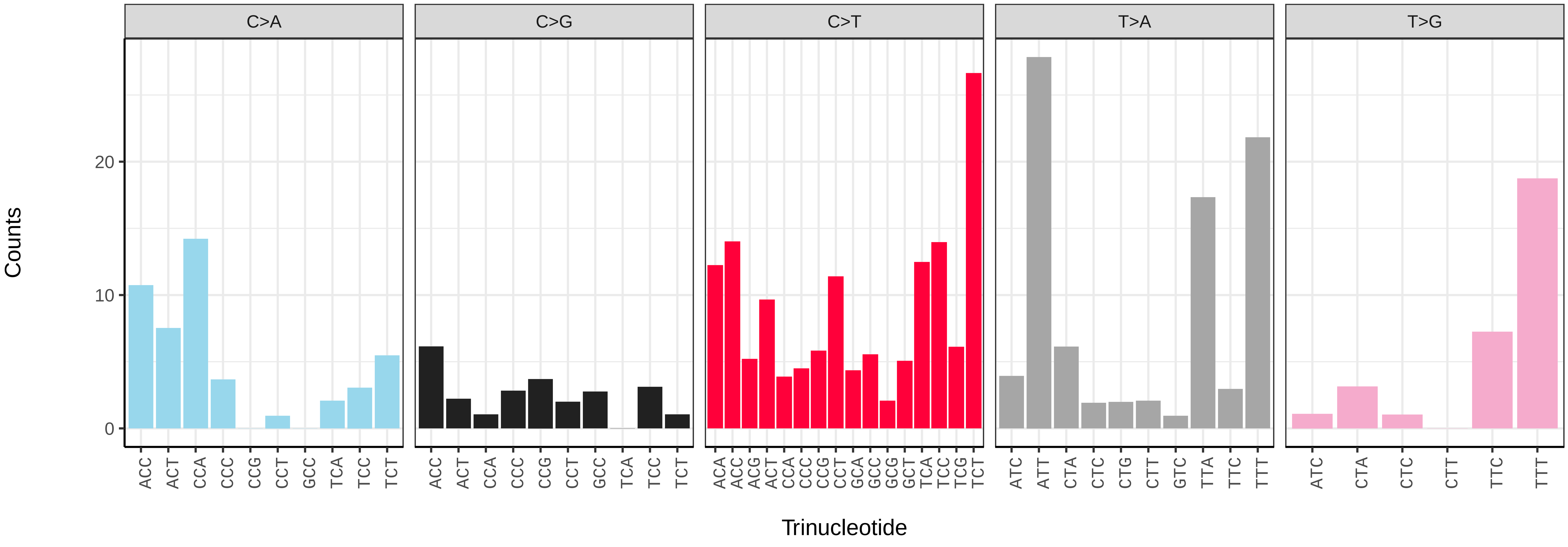

*Tachina fera* (idTacFera2) germline mutational spectrum

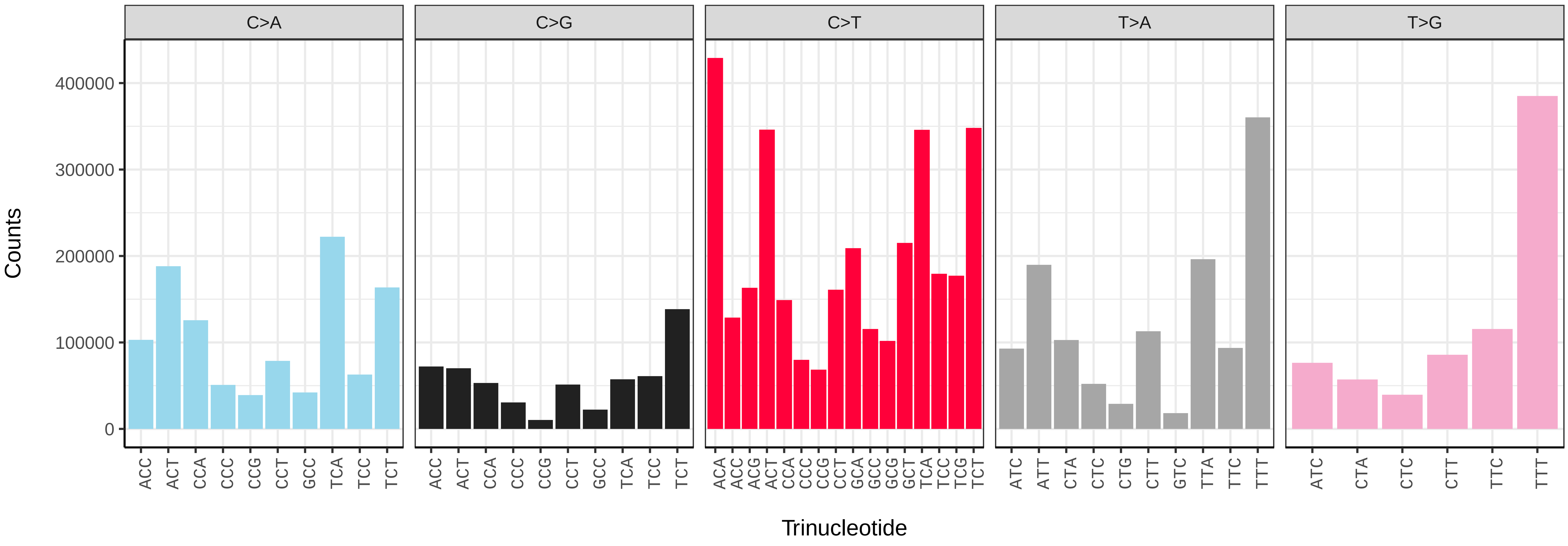


*Tachina fera* (idTacFera2) somatic mutational spectrum (SBS96 classification)

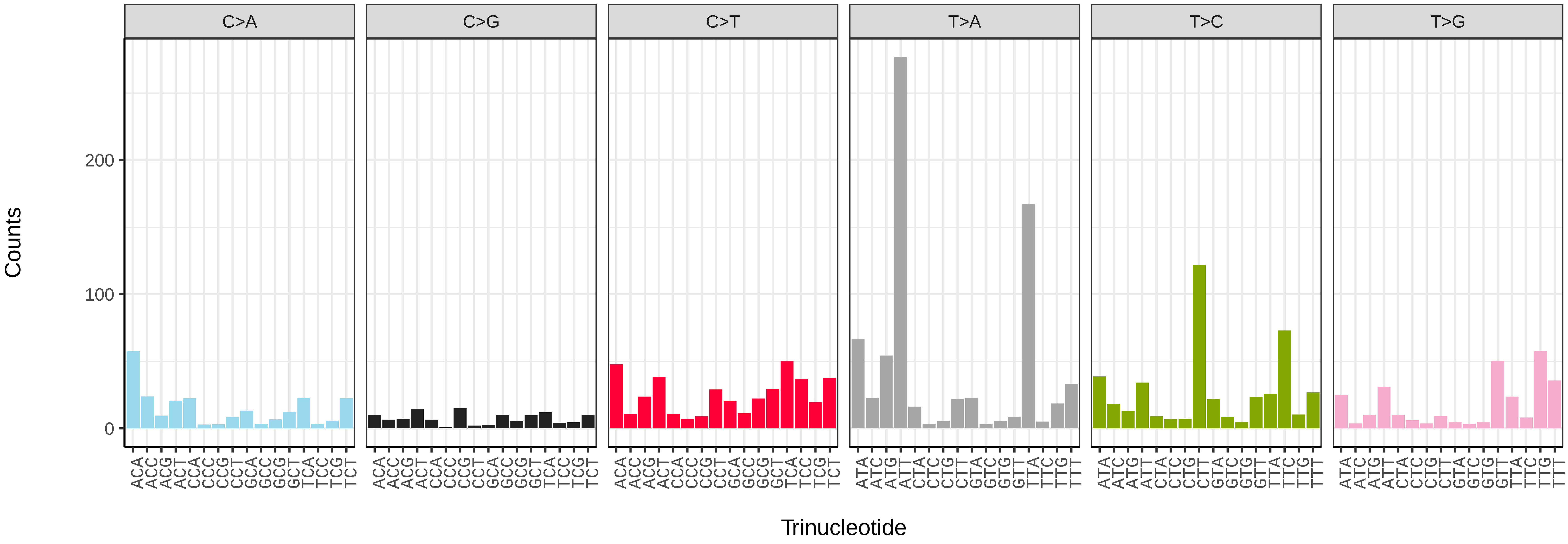


*Tachina fera* (idTacFera2) somatic mutational spectrum (SBS52 classification)

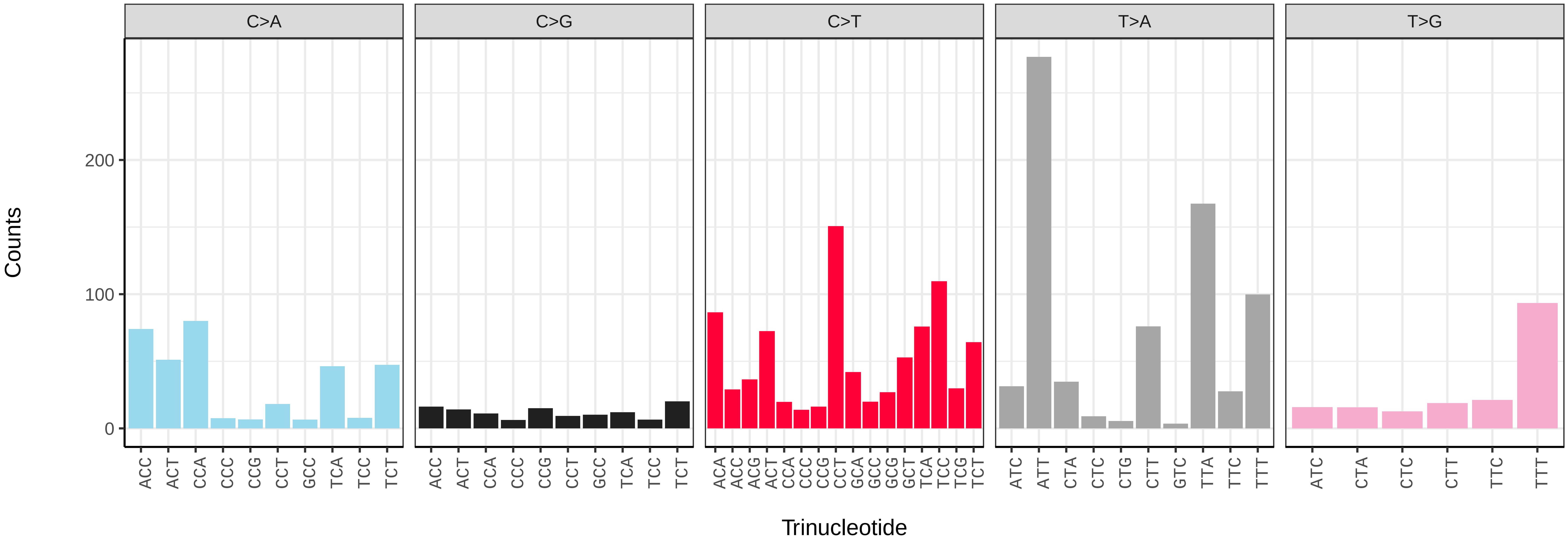

*Tachina lurida* (idTacLuri1) germline mutational spectrum

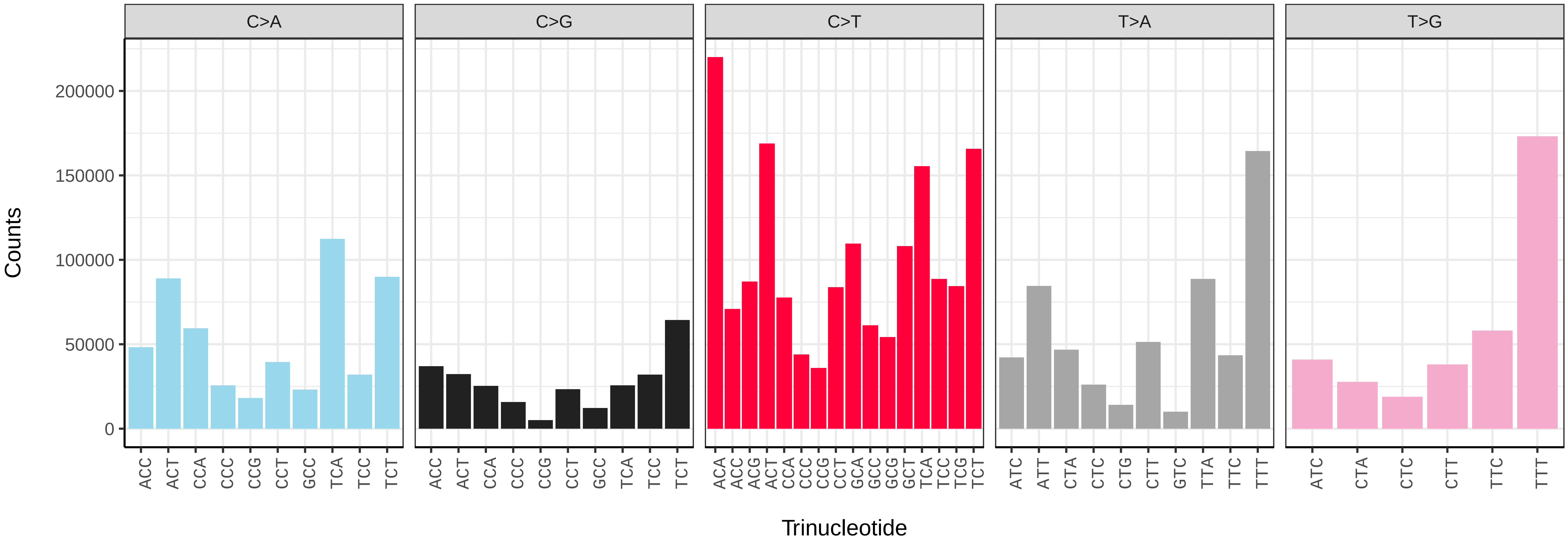


*Tachina lurida* (idTacLuri1) somatic mutational spectrum (SBS96 classification)

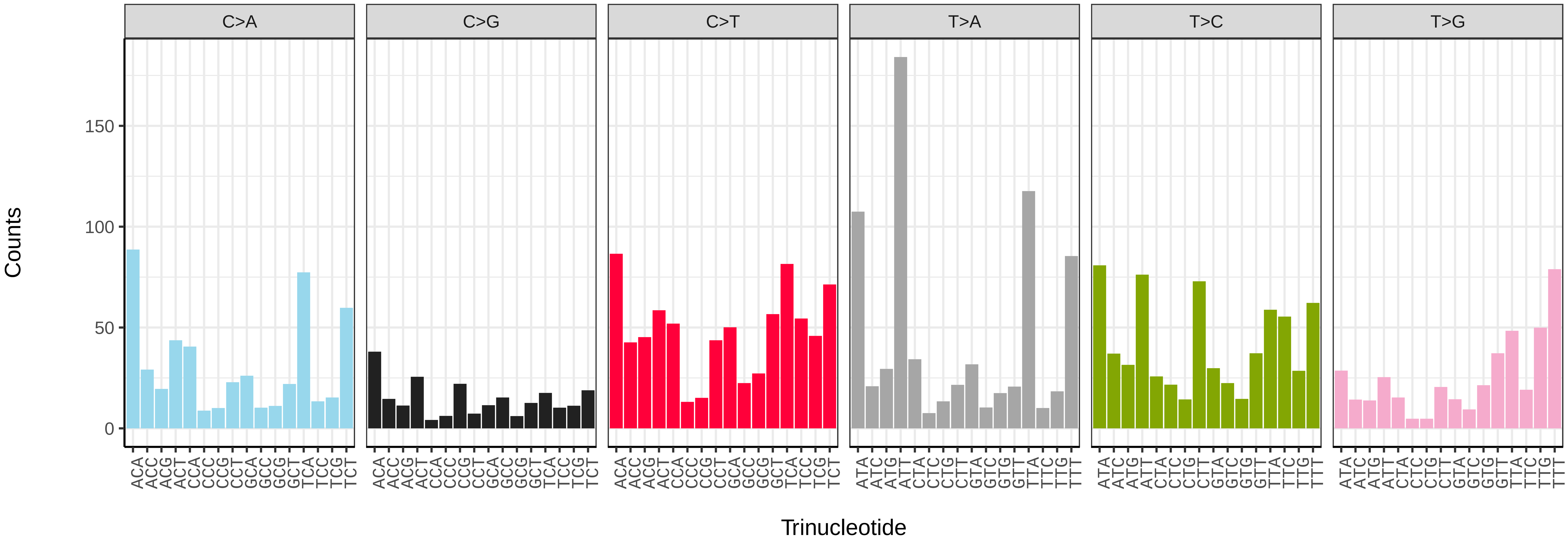


*Tachina lurida* (idTacLuri1) somatic mutational spectrum (SBS52 classification)

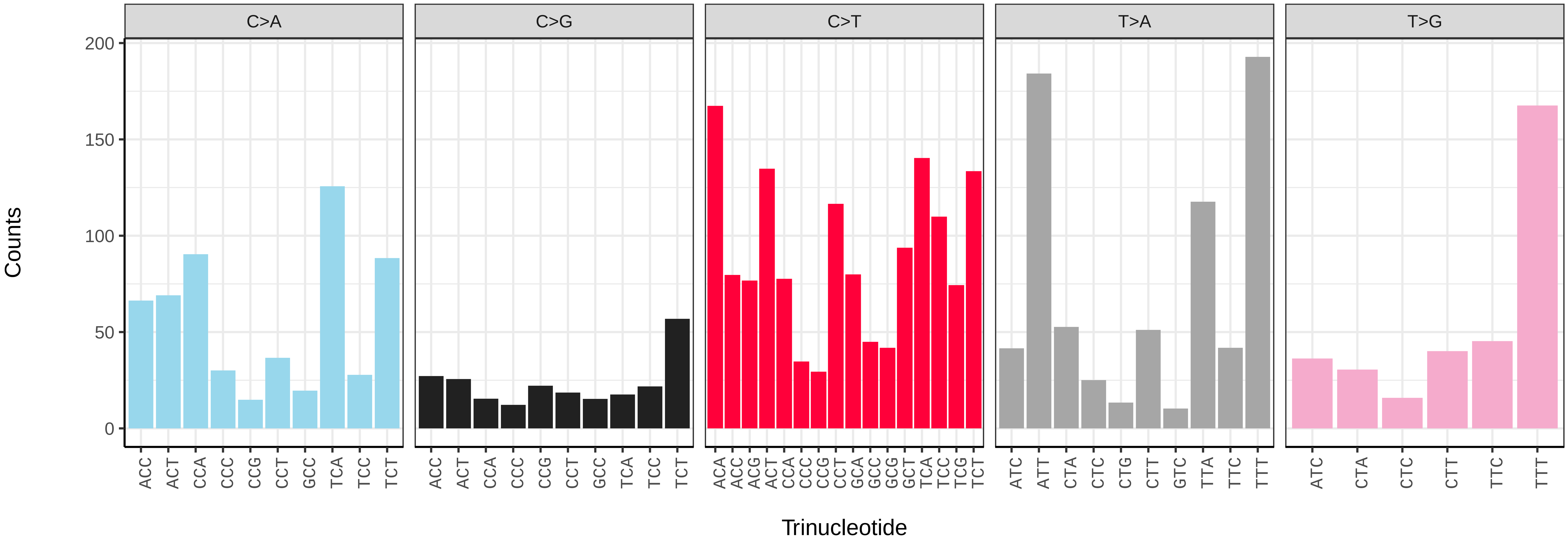

*Thecocarcelia acutangulata* (idTheAcut1) germline mutational spectrum

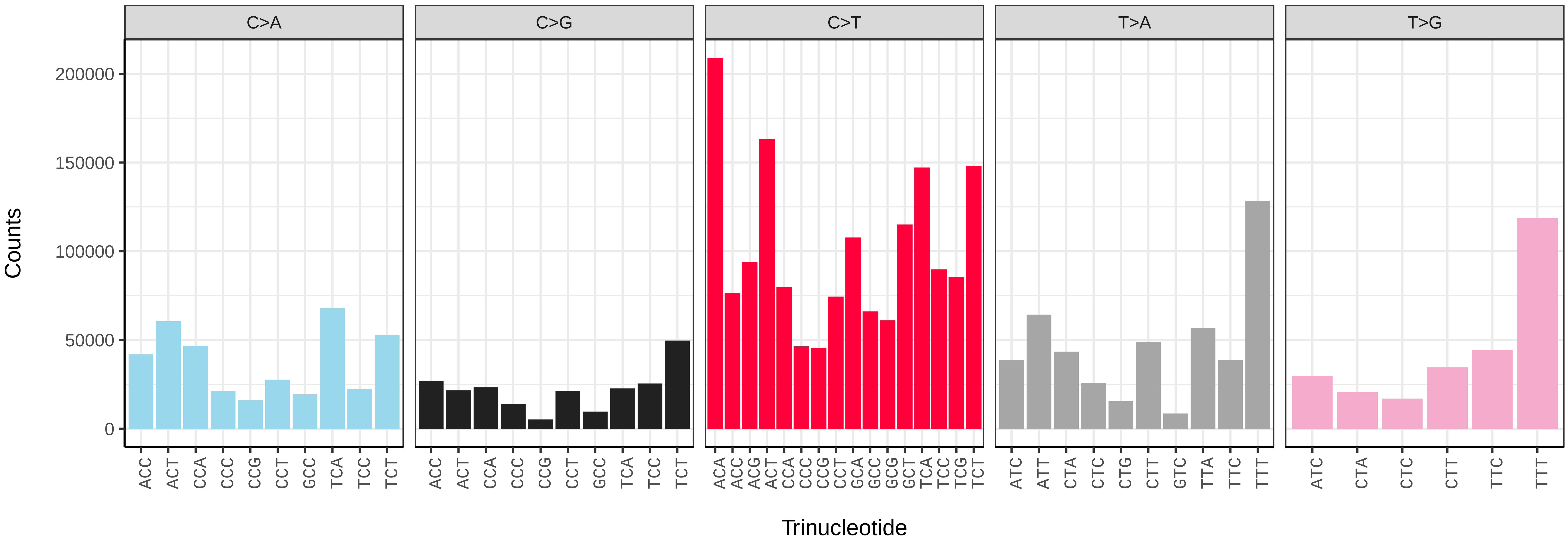


*Thecocarcelia acutangulata* (idTheAcut1) somatic mutational spectrum (SBS96 classification)

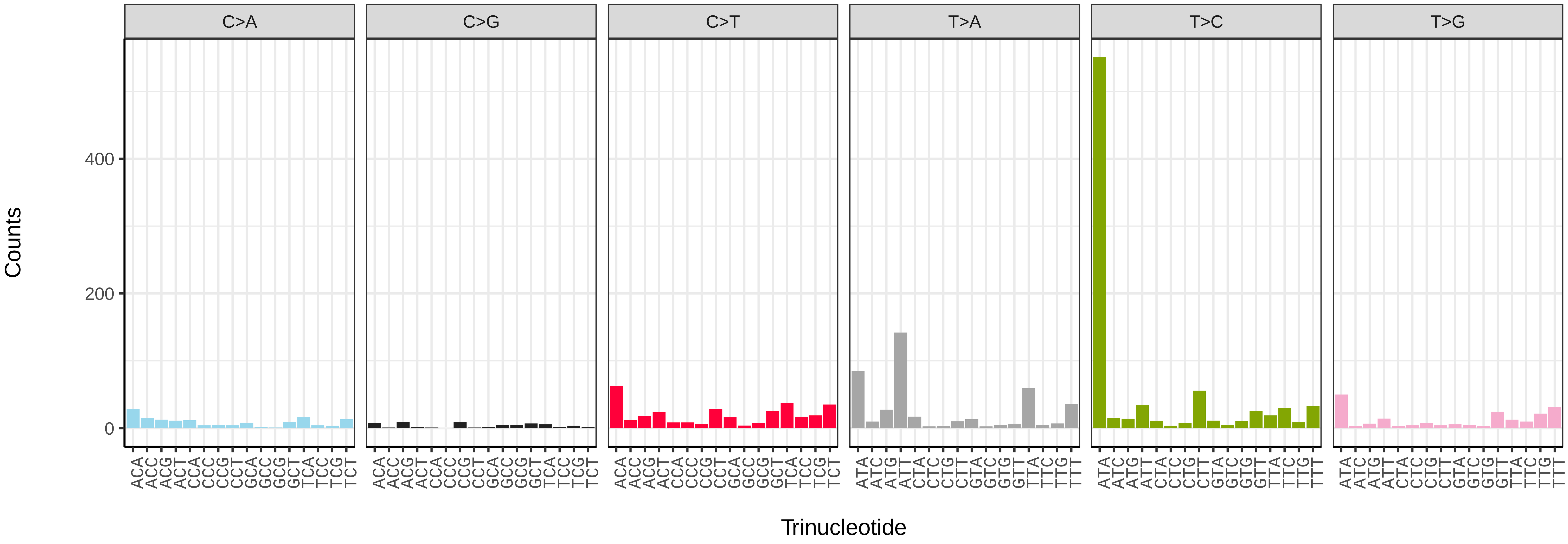


*Thecocarcelia acutangulata* (idTheAcut1) somatic mutational spectrum (SBS52 classification)

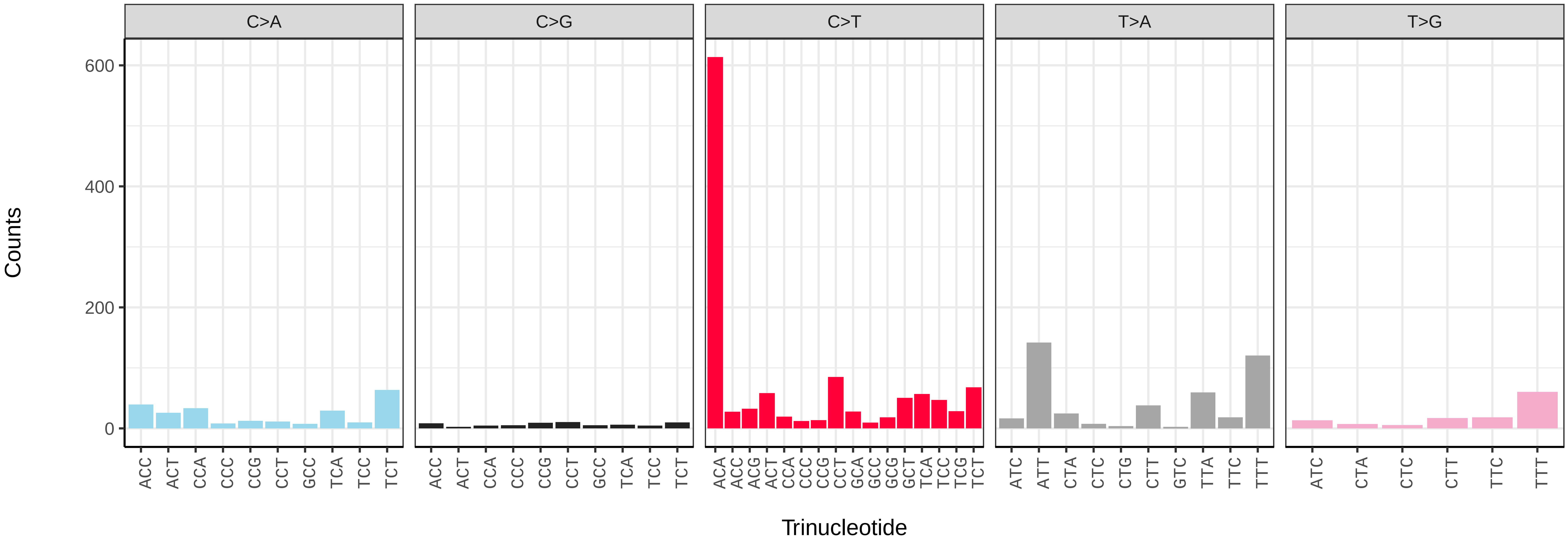

*Thecophora atra* (idTheAtra2) germline mutational spectrum

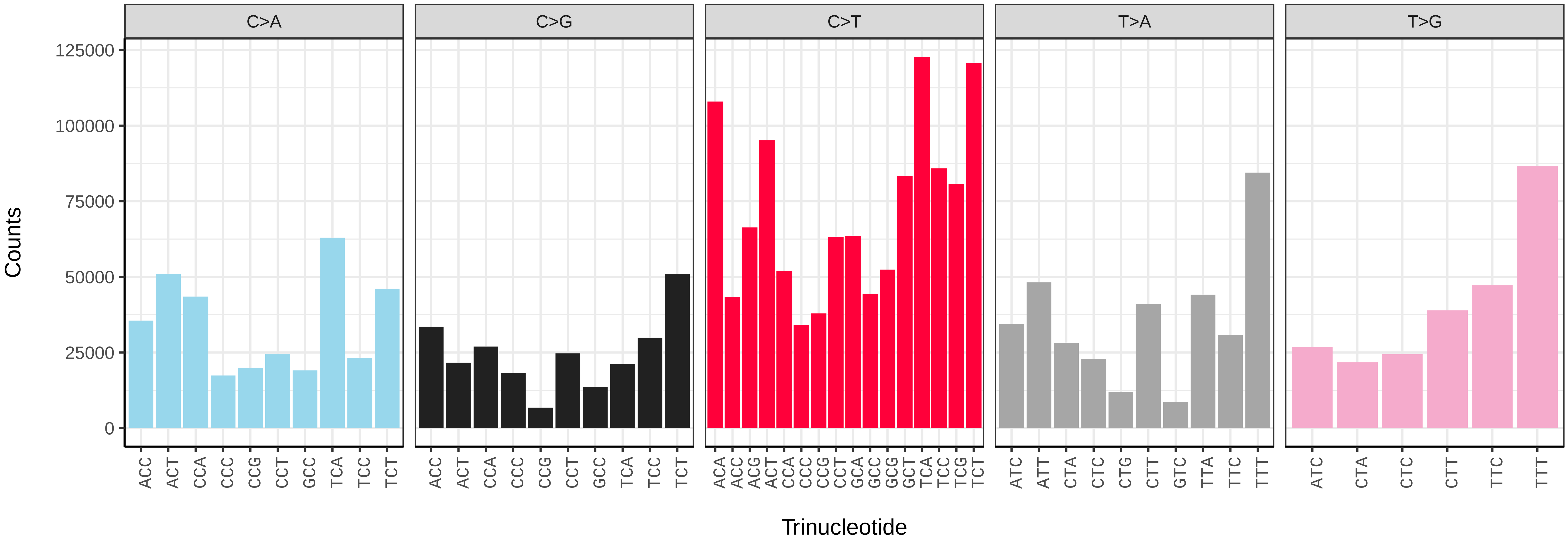


*Thecophora atra* (idTheAtra2) somatic mutational spectrum (SBS96 classification)

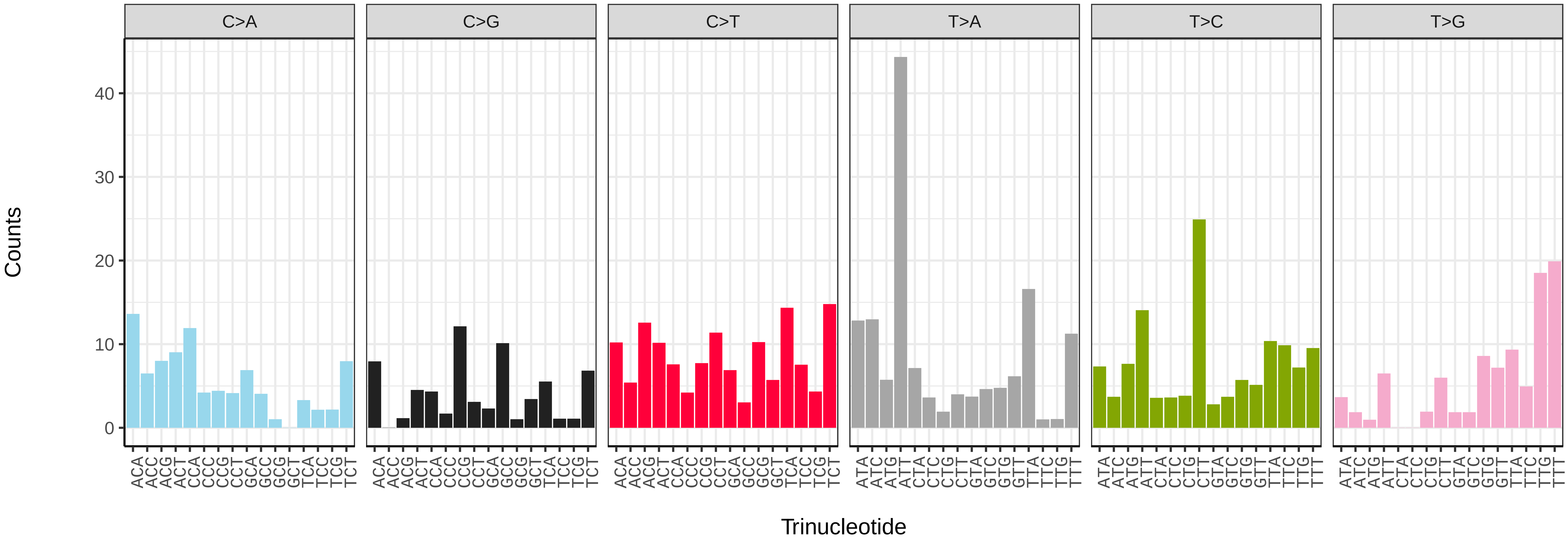


*Thecophora atra* (idTheAtra2) somatic mutational spectrum (SBS52 classification)

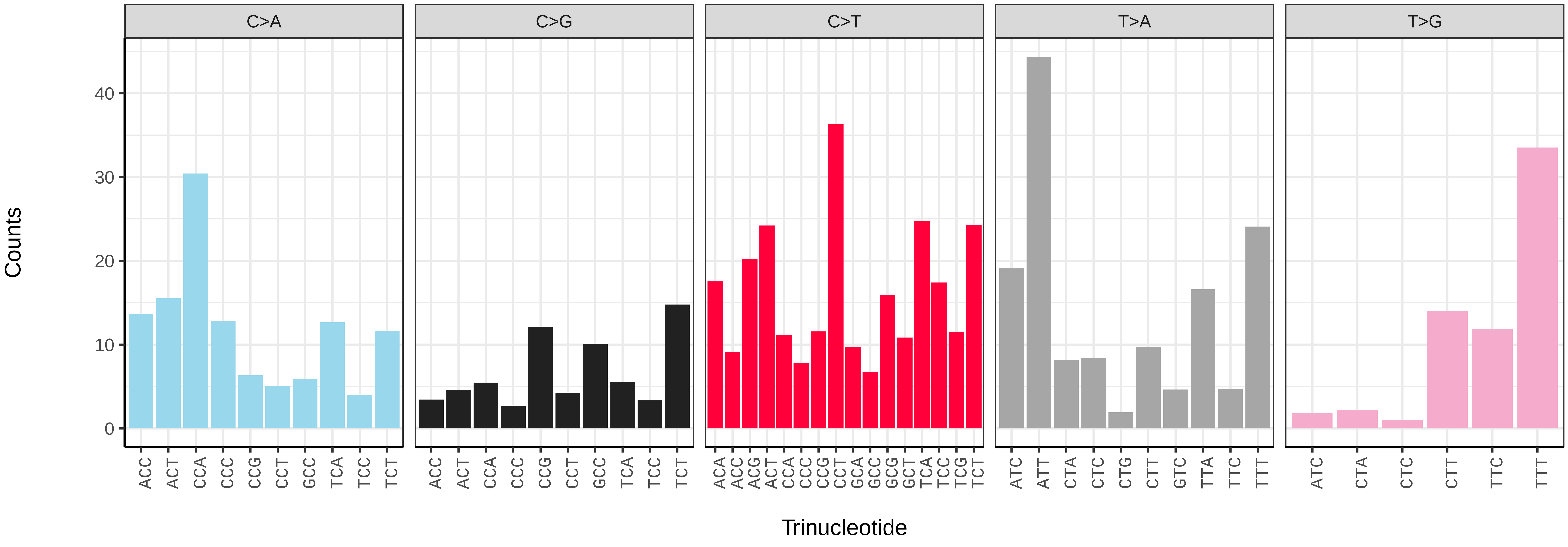

*Thelaira solivaga* (idTheSoli1) germline mutational spectrum

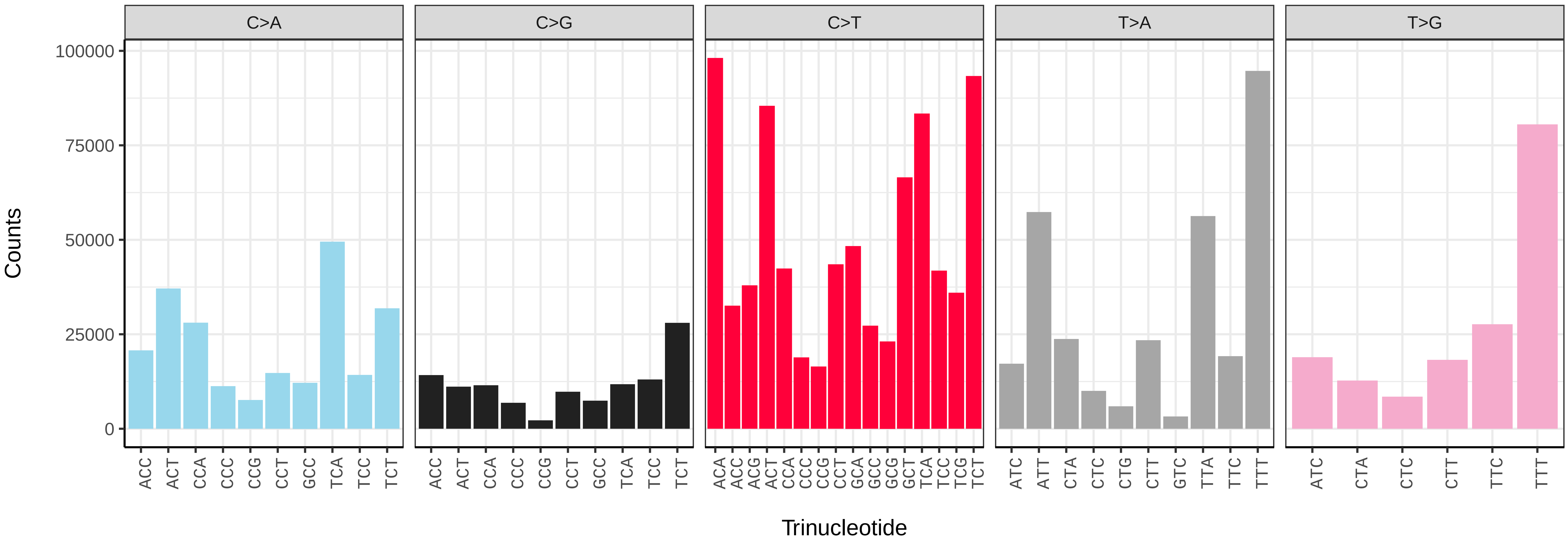


*Thelaira solivaga* (idTheSoli1) somatic mutational spectrum (SBS96 classification)

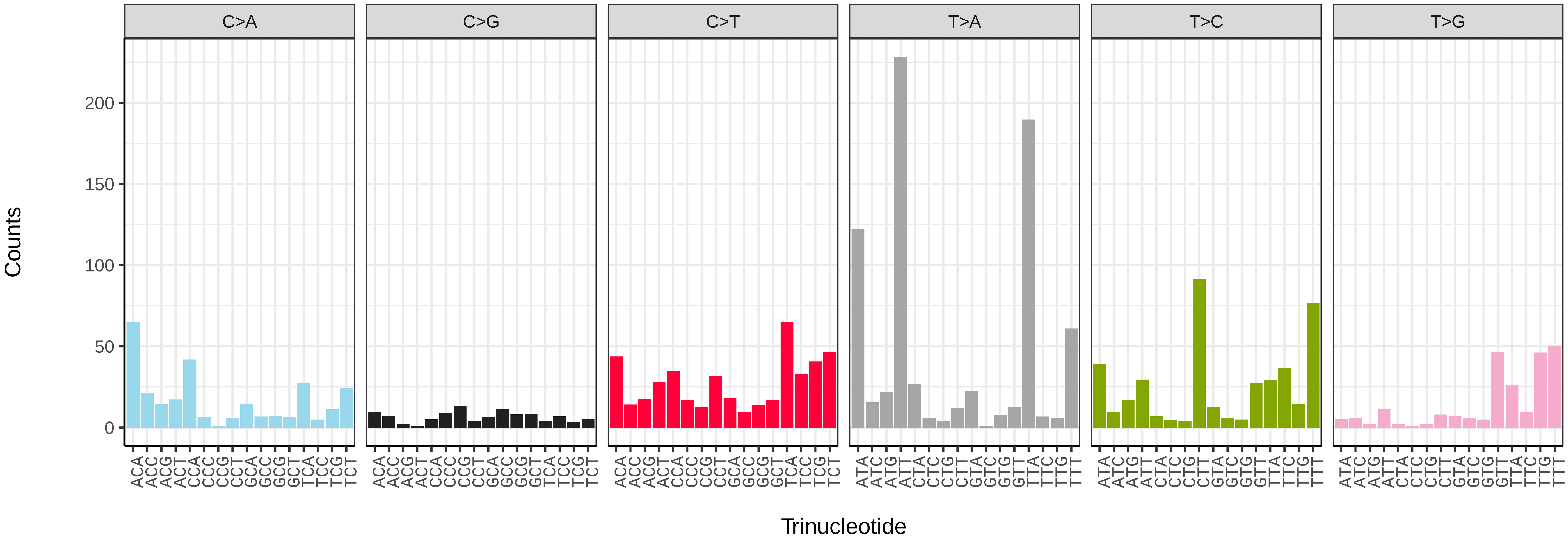


*Thelaira solivaga* (idTheSoli1) somatic mutational spectrum (SBS52 classification)

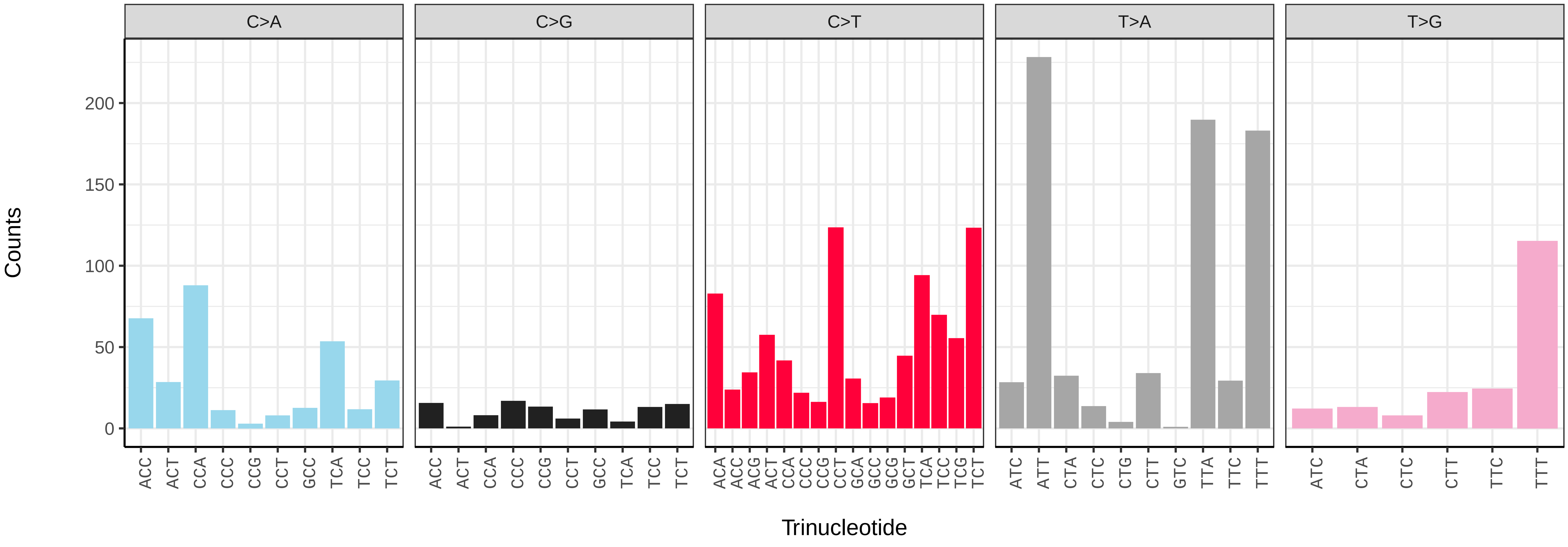

*Volucella inanis* (idVolInan1) germline mutational spectrum

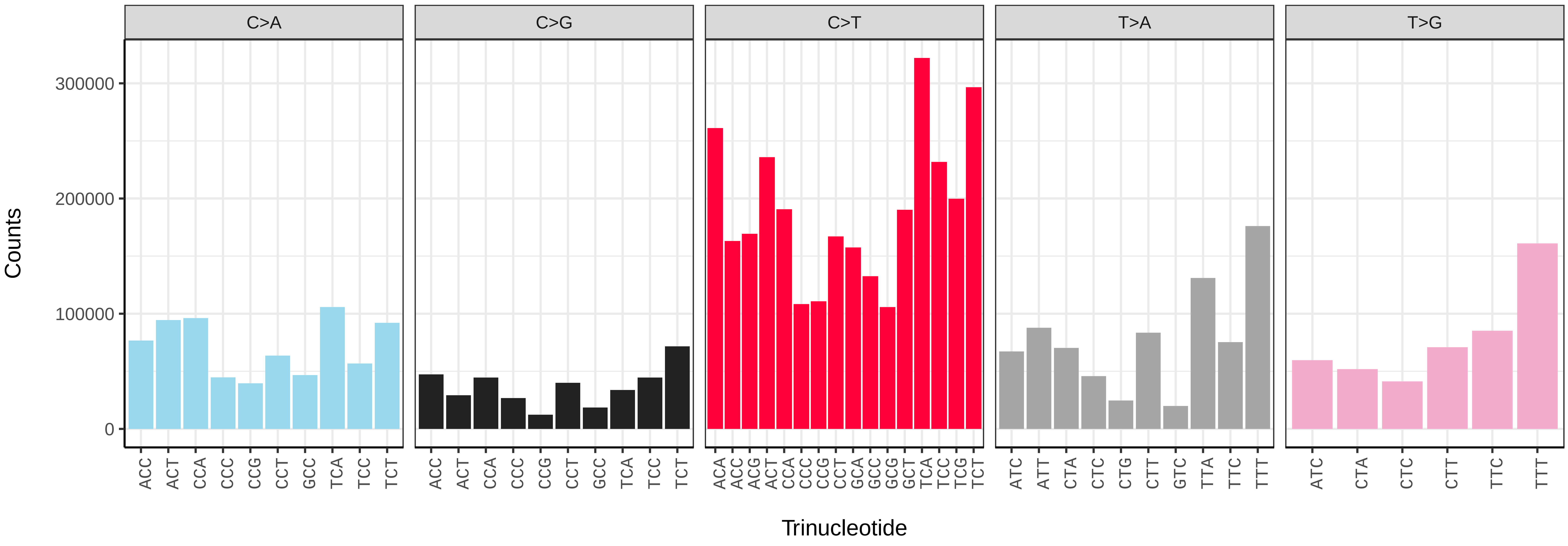


*Volucella inanis* (idVolInan1) somatic mutational spectrum (SBS96 classification)

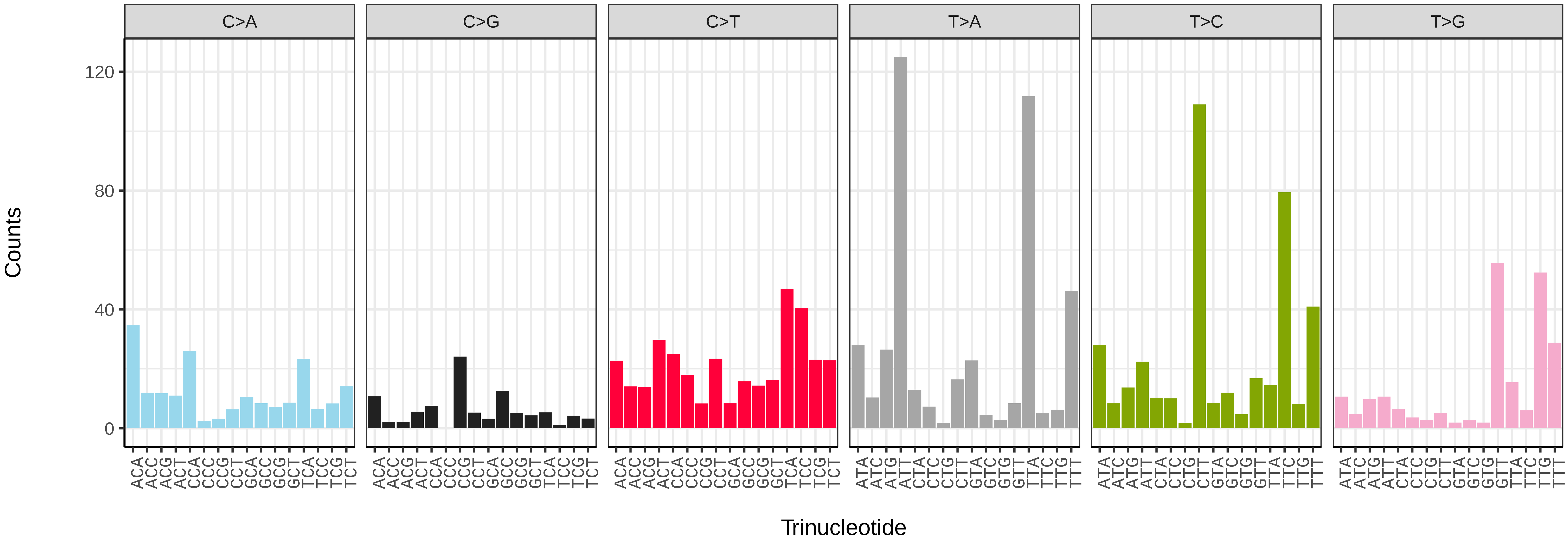


*Volucella inanis* (idVolInan1) somatic mutational spectrum (SBS52 classification)

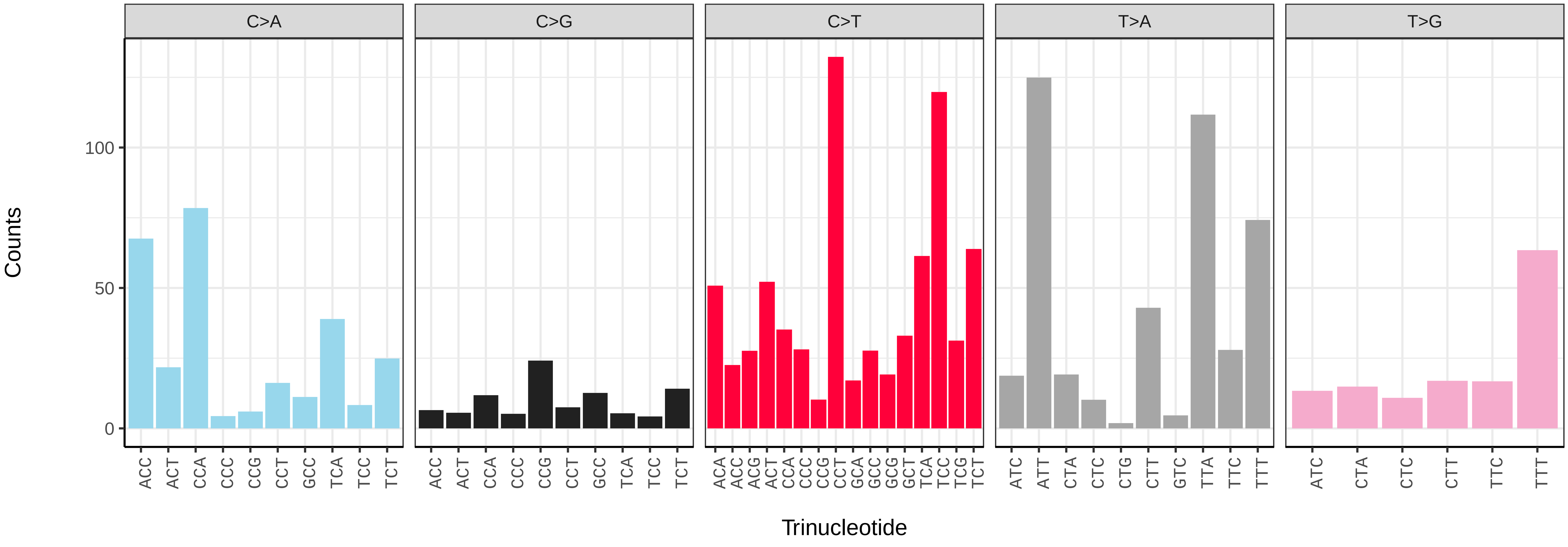

*Volucella inflata* (idVolInfl1) germline mutational spectrum

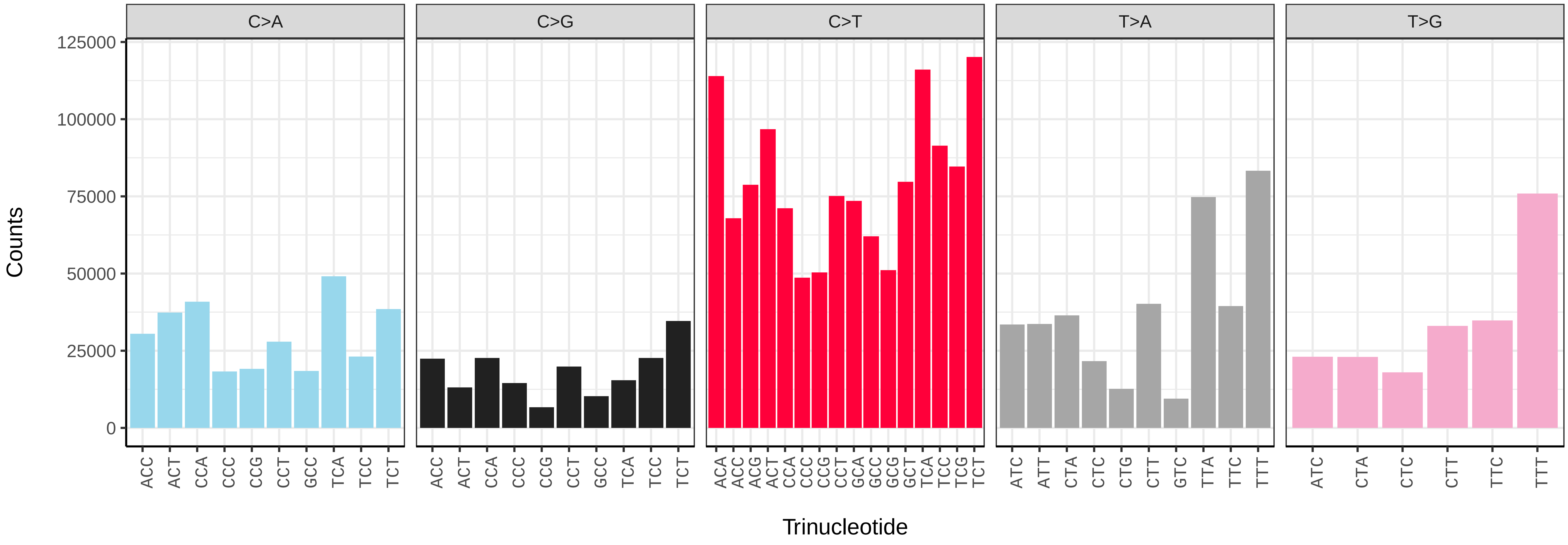


*Volucella inflata* (idVolInfl1) somatic mutational spectrum (SBS96 classification)

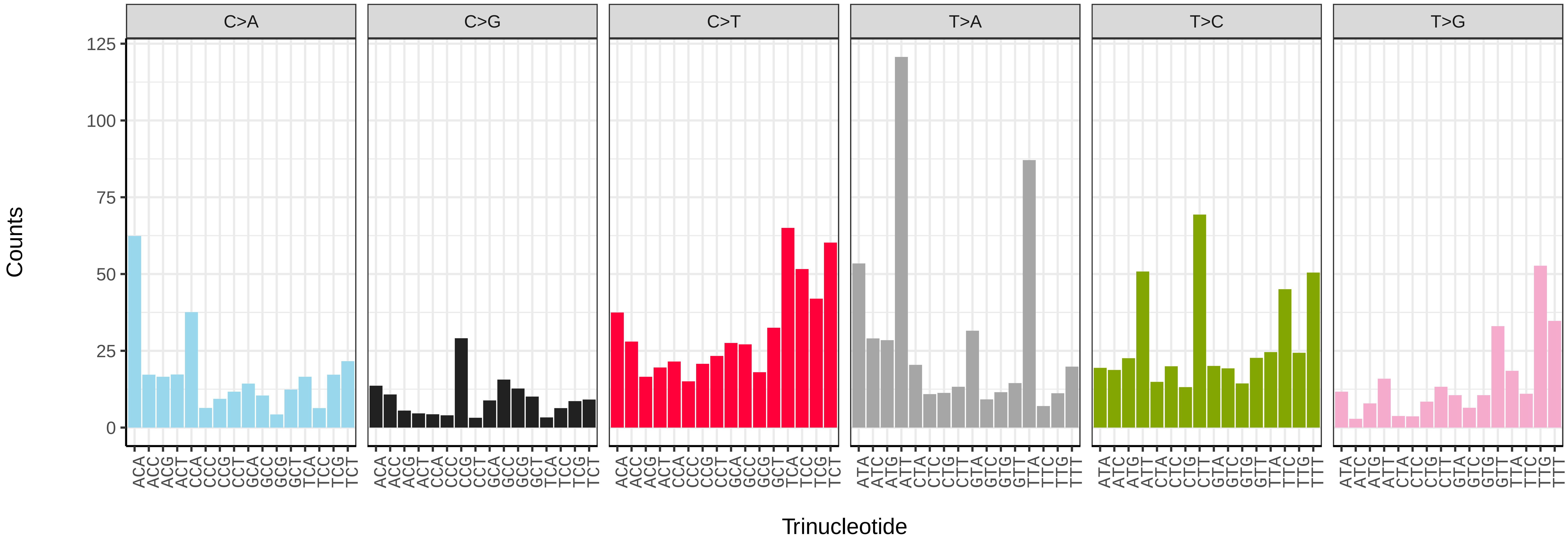


*Volucella inflata* (idVolInfl1) somatic mutational spectrum (SBS52 classification)

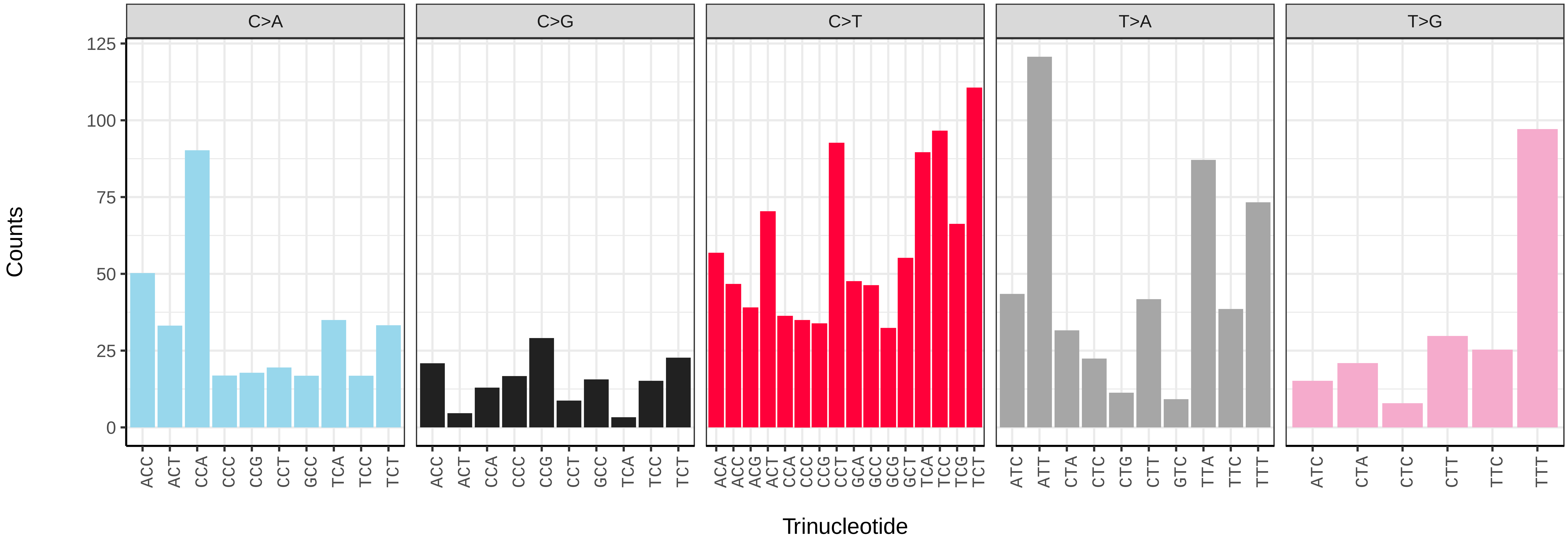

*Xanthogramma pedissequum* (idXanPedi1) germline mutational spectrum

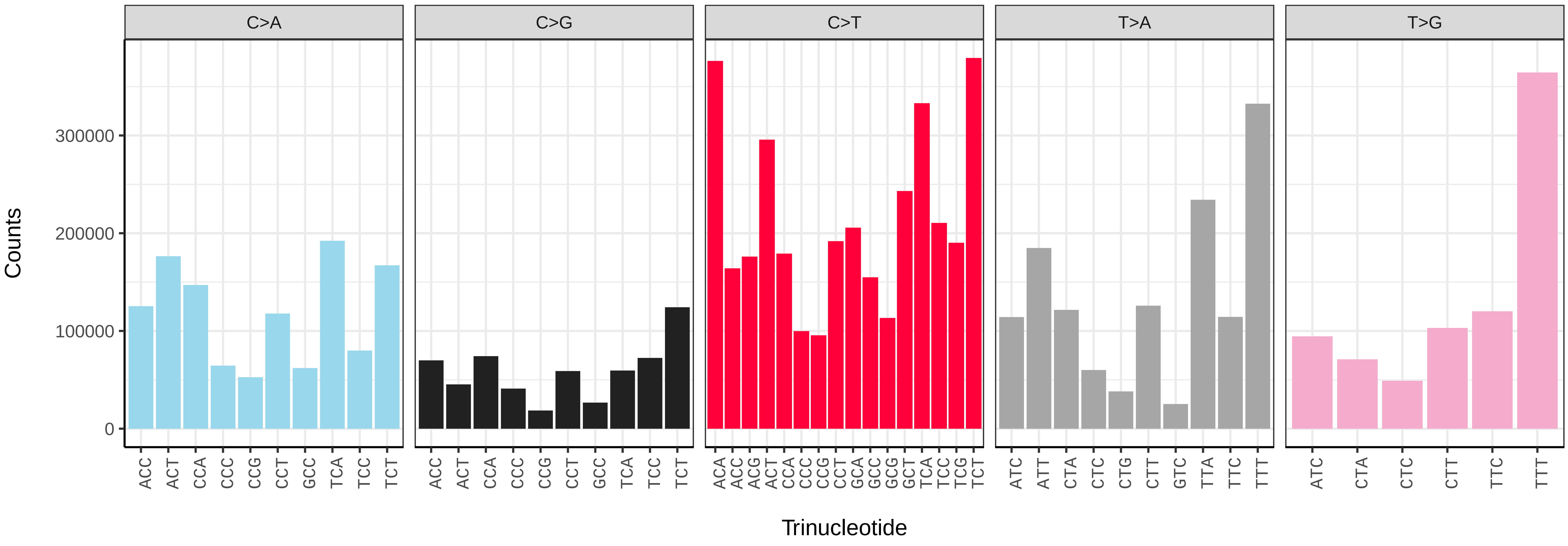


*Xanthogramma pedissequum* (idXanPedi1) somatic mutational spectrum (SBS96 classification)

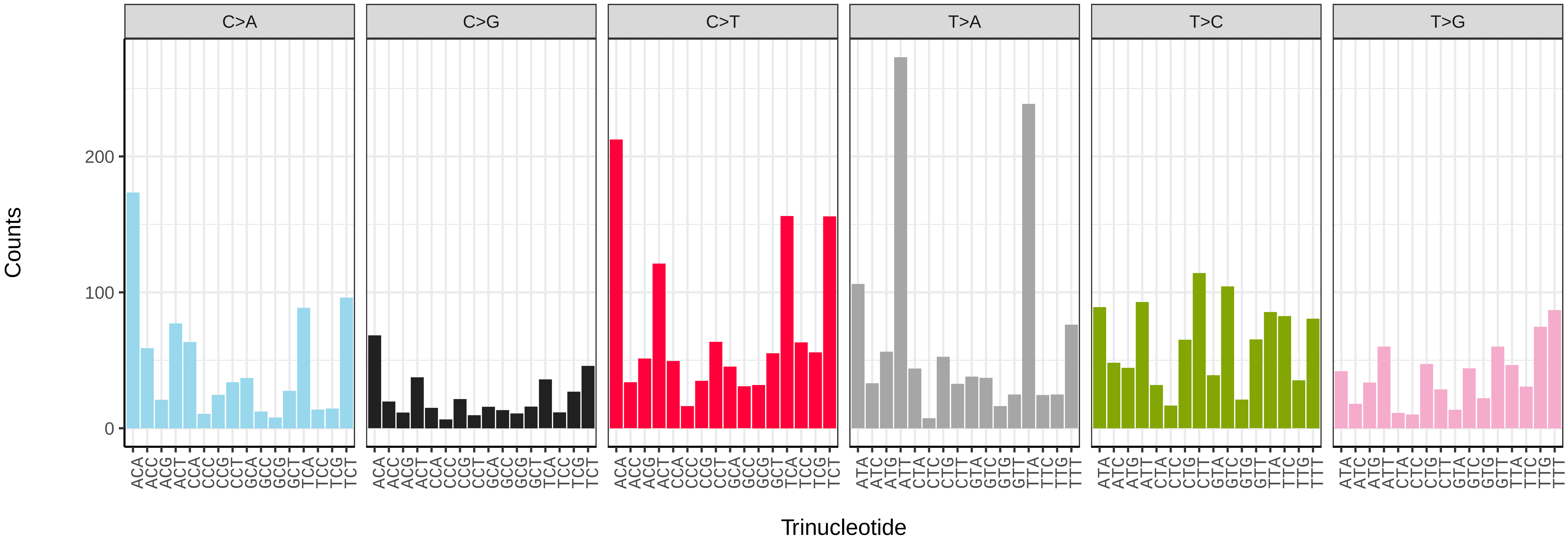


*Xanthogramma pedissequum* (idXanPedi1) somatic mutational spectrum (SBS52 classification)

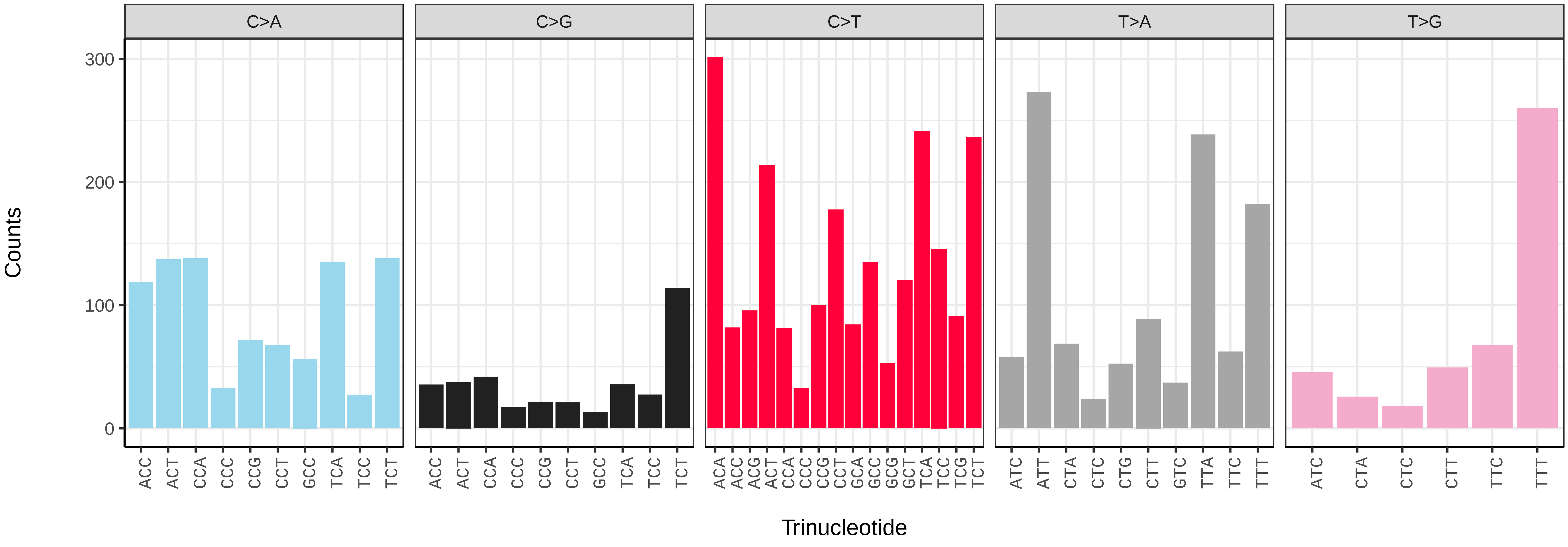

*Xylota sylvarum* (idXylSylv2) germline mutational spectrum

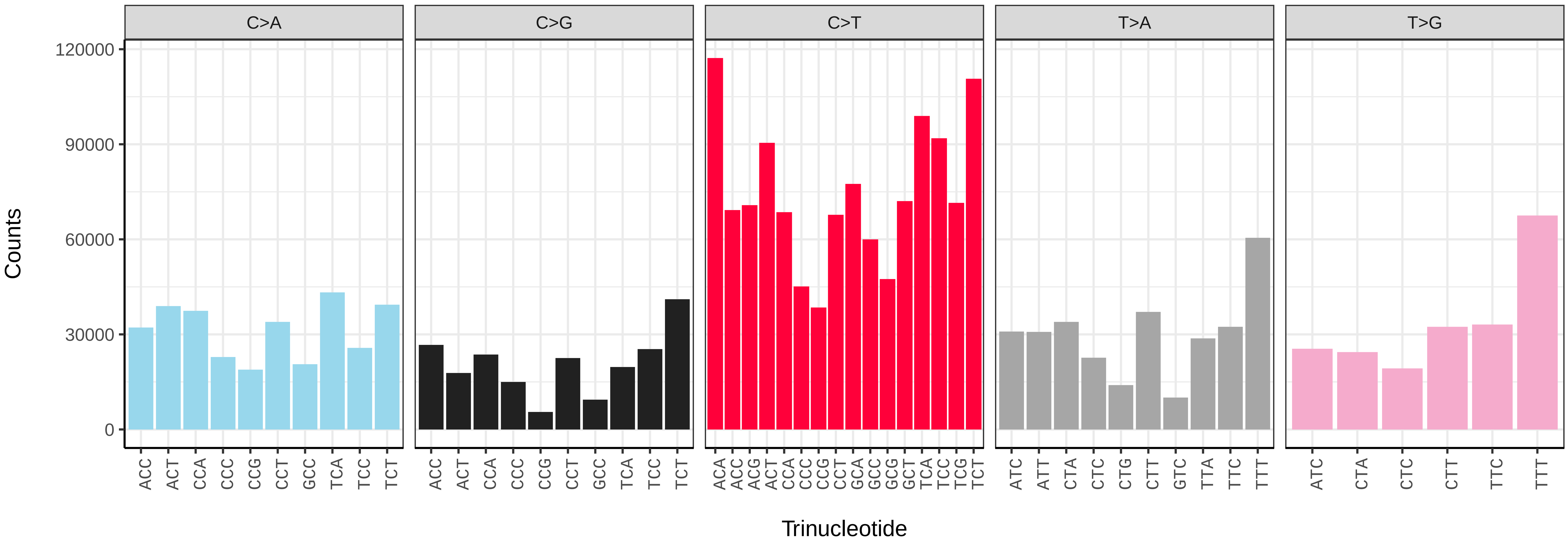


*Xylota sylvarum* (idXylSylv2) somatic mutational spectrum (SBS96 classification)

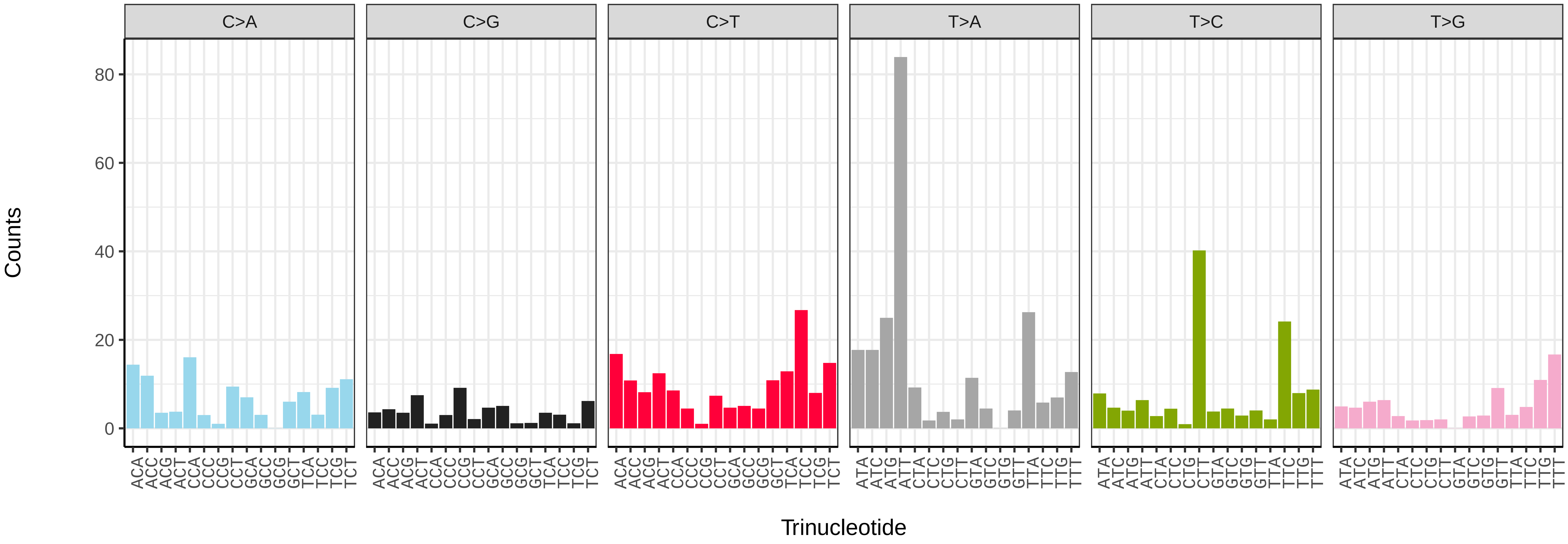


*Xylota sylvarum* (idXylSylv2) somatic mutational spectrum (SBS52 classification)

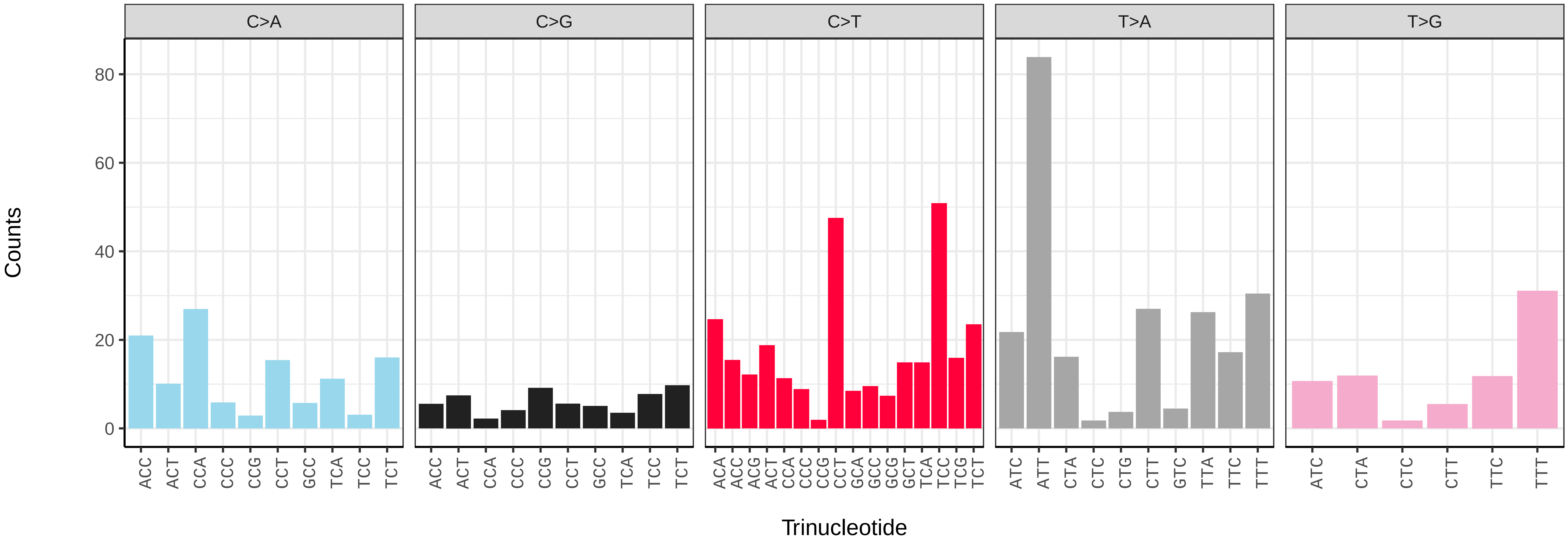

*Acanthosoma haemorrhoidale* (ihAcaHaem1) germline mutational spectrum

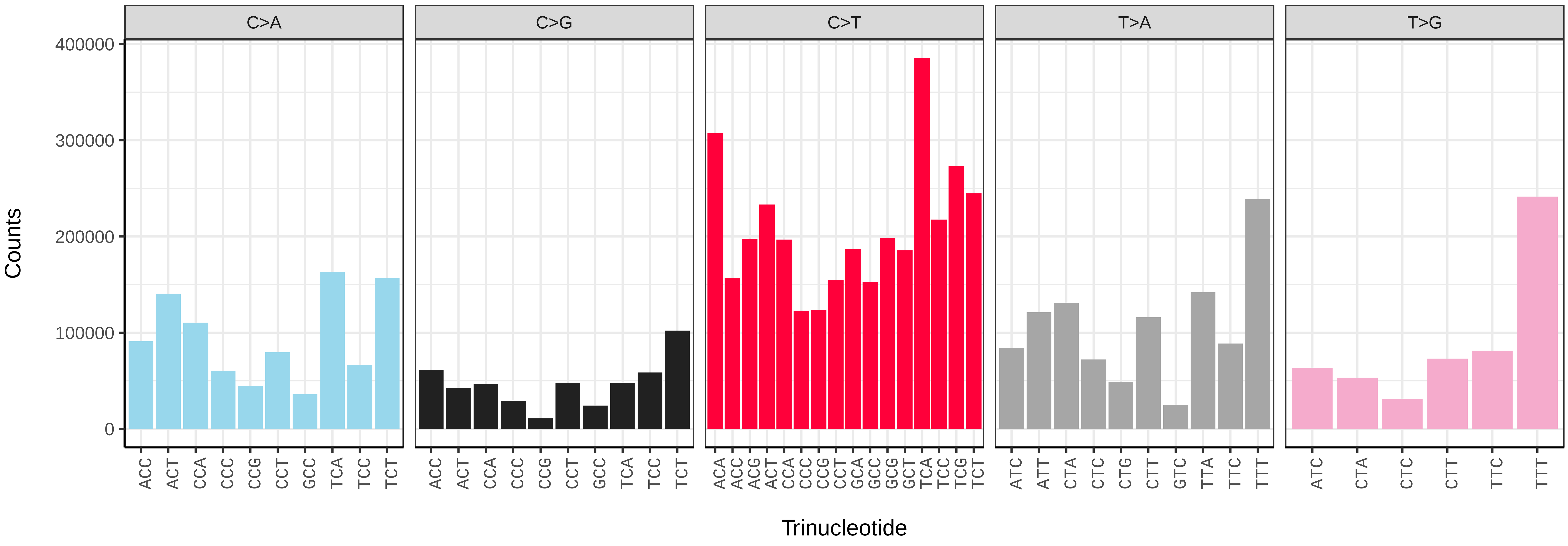


*Acanthosoma haemorrhoidale* (ihAcaHaem1) somatic mutational spectrum (SBS96 classification)

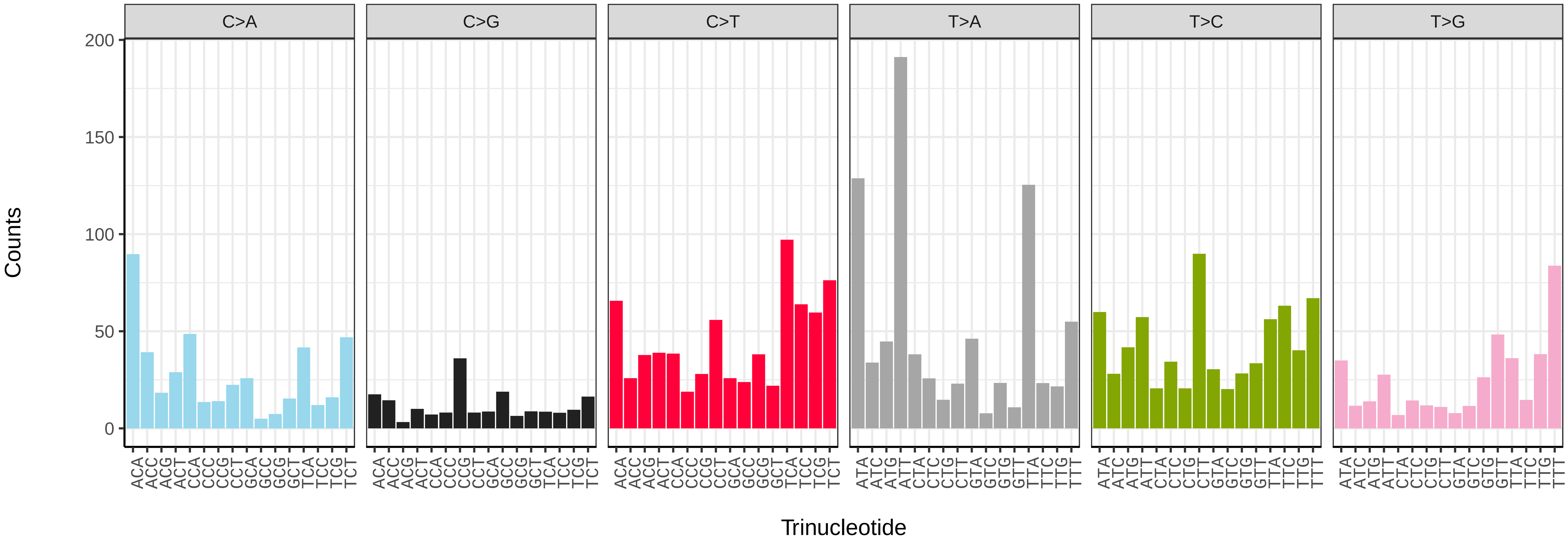


*Acanthosoma haemorrhoidale* (ihAcaHaem1) somatic mutational spectrum (SBS52 classification)

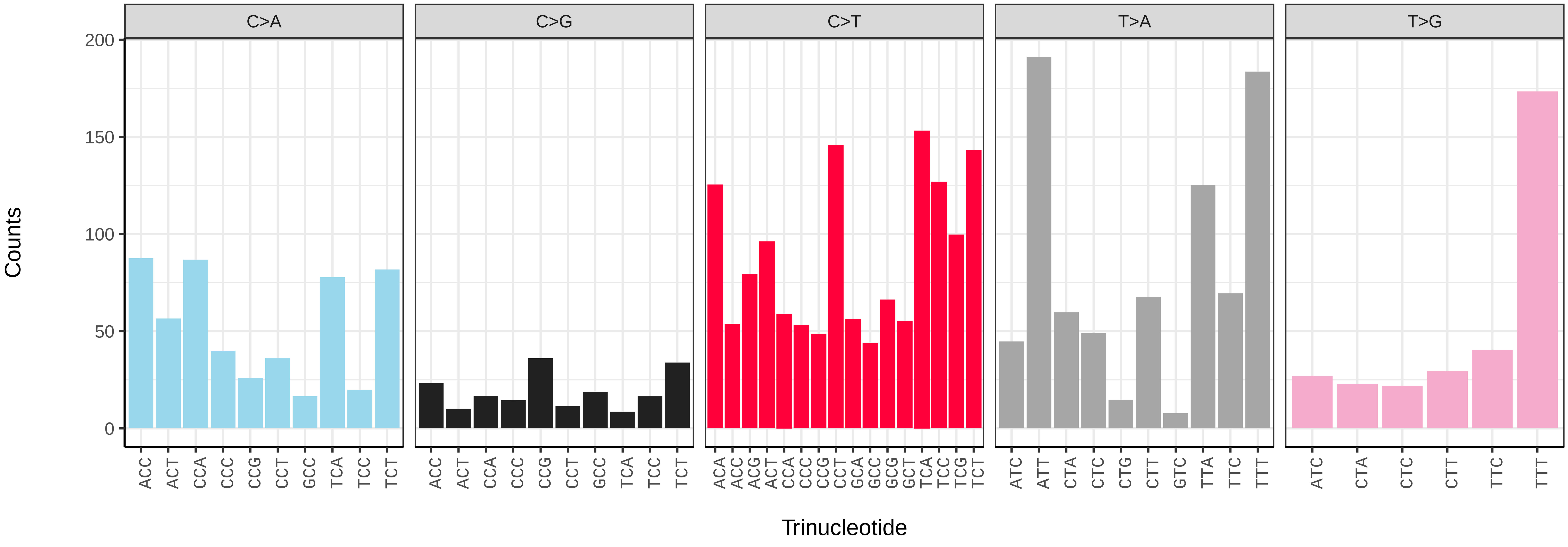

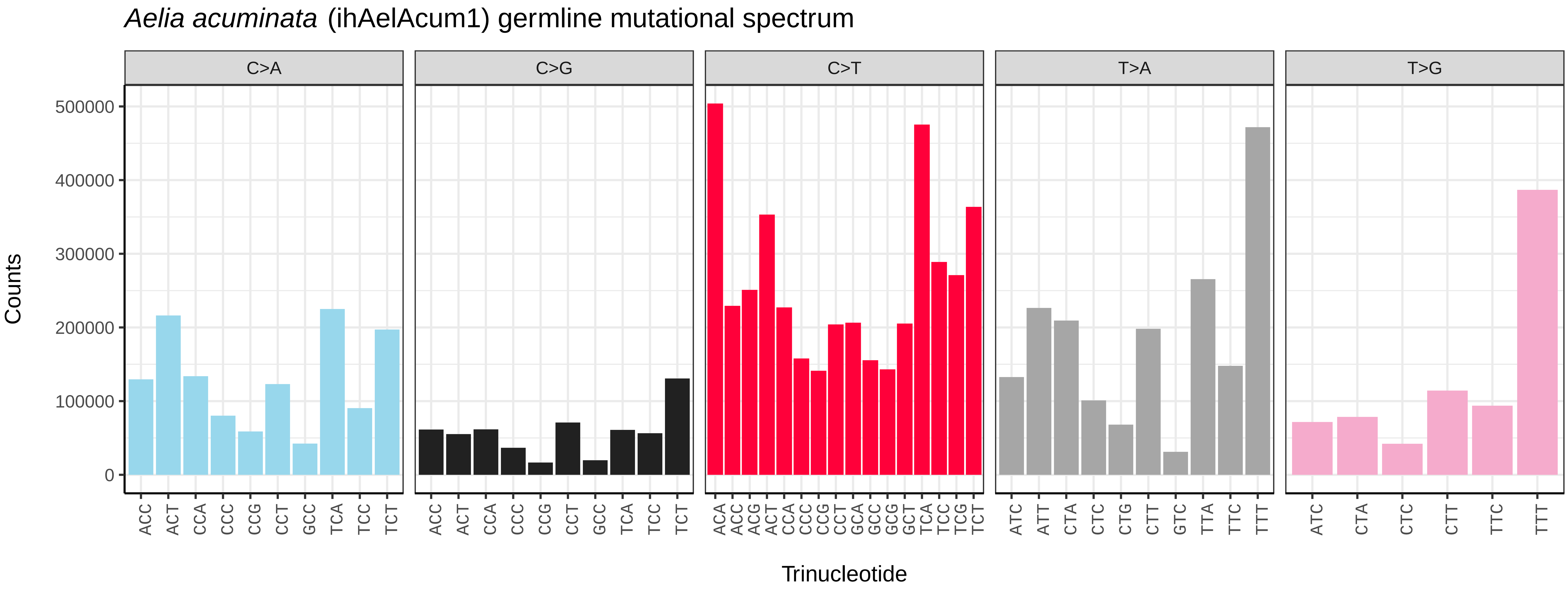
Aelia acuminata (ihAelAcum1) germline mutational spectrum
C>A
C>G
C>T
T>A
T>G
Counts
500000
400000
300000
200000
100000
0
ACC ACT CCA CCC CCG CCT GCC TCA TCC TCT
ACC ACT CCA CCC CCG CCT GCC TCA TCC TCT
ACA ACC ACG ACT CCA CCC CCG CCT GCA GCC GCG GCT TCA TCC TCG TCT
ATC ATT CTA CTC CTG CTT GTC TTA TTC TTT
ATC CTA CTC CTT TTC TTT
Trinucleotide

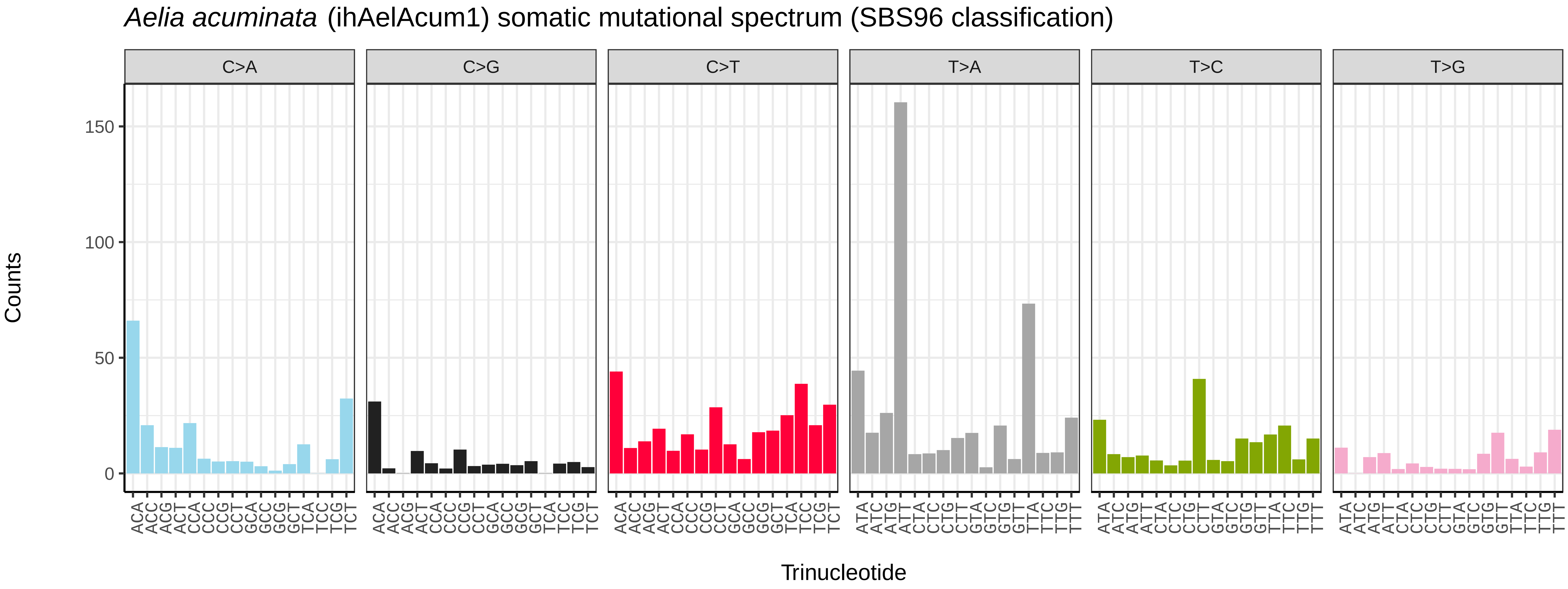
Aelia acuminata (ihAelAcum1) somatic mutational spectrum (SBS96 classification)
C>A
C>G
C>T
T>A
T>C
T>G
Counts
150
100
50
0
ACA ACC ACG ACT CCA CCC CCG CCT GCA GCC GCG GCT TCA TCC TCG TCT
ATA ATC ATG ATT CTA CTC CTG CTT GTA GTC GTG GTT TTA TTC TTG TTT
Trinucleotide

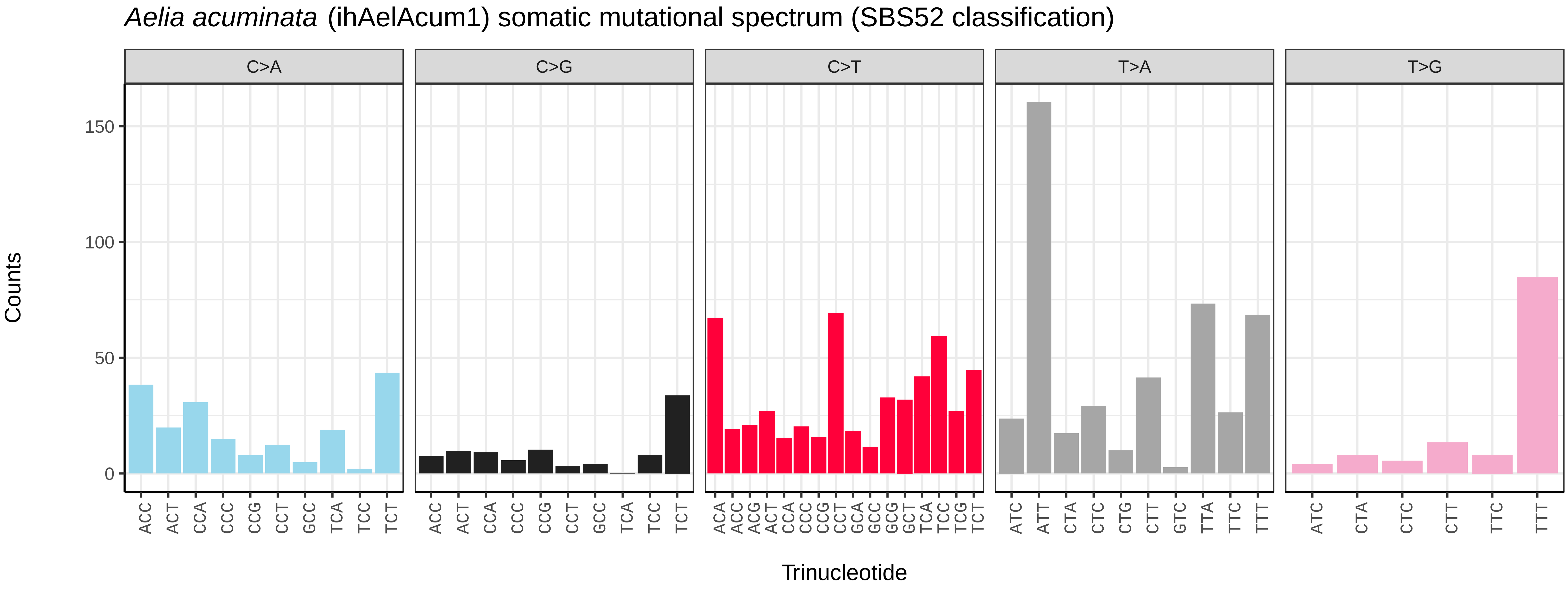
Aelia acuminata (ihAelAcum1) somatic mutational spectrum (SBS52 classification)
C>A
C>G
C>T
T>A
T>G
Counts
150
100
50
0
ACC ACT CCA CCC CCG CCT GCC TCA TCC TCT
ACC ACT CCA CCC CCG CCT GCC TCA TCC TCT
ACA ACC ACG ACT CCA CCC CCG CCT GCA GCC GCG GCT TCA TCC TCG TCT
ATC ATT CTA CTC CTG CTT GTC TTA TTC TTT
ATC CTA CTC CTT TTC TTT
Trinucleotide

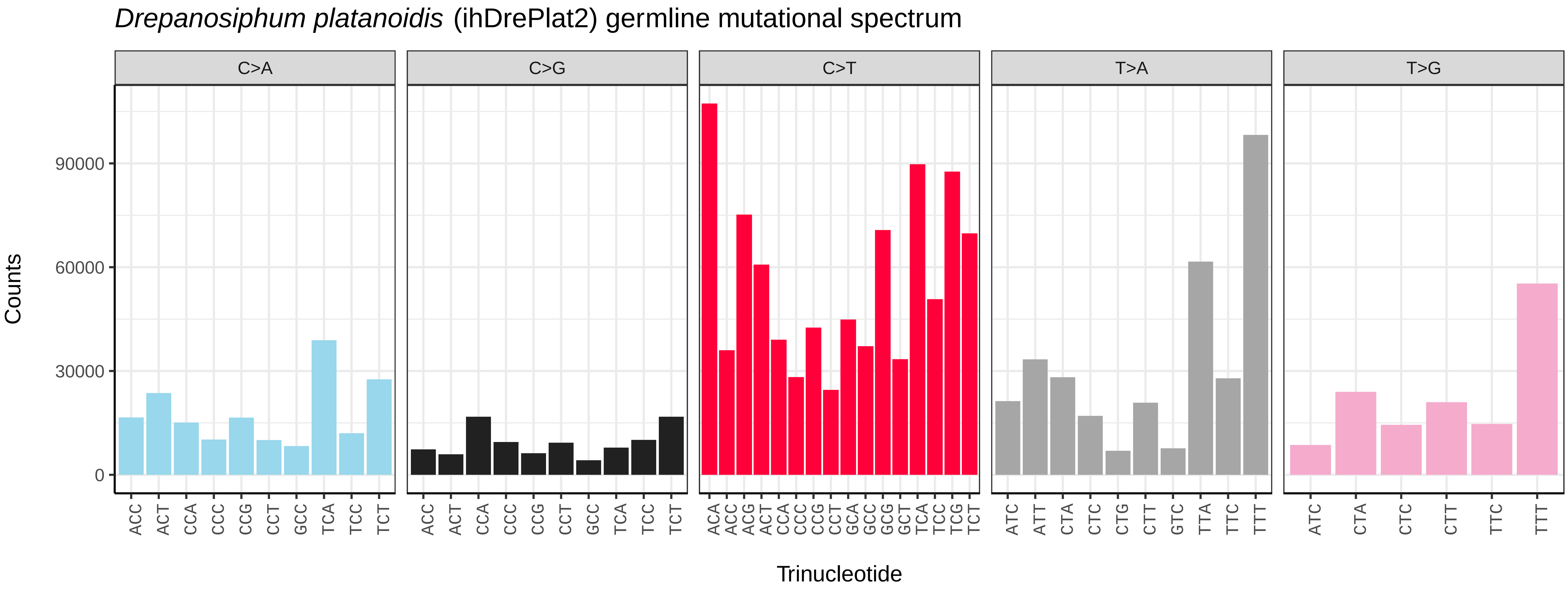
*Drepanosiphum platanoidis* (ihDrePlat2) germline mutational spectrum
C>A
C>G
C>T
T>A
T>G
Counts
90000
60000
30000
0
Trinucleotide

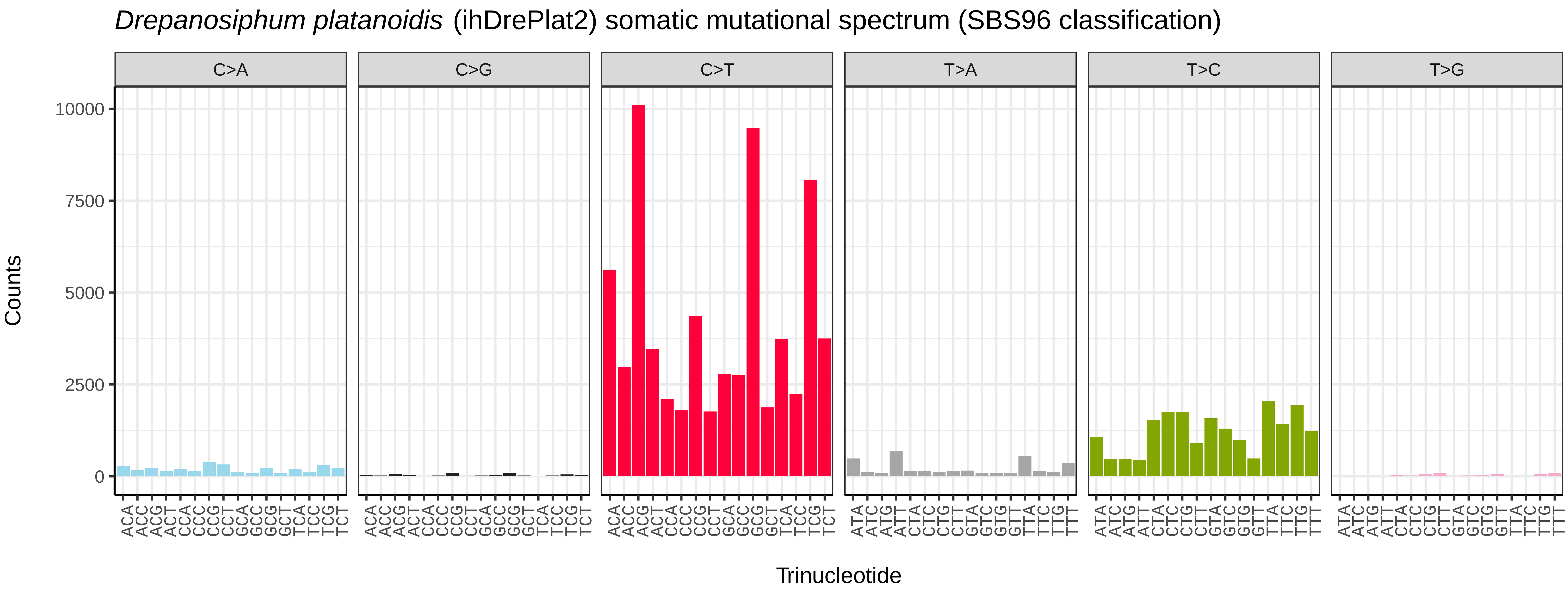
*Drepanosiphum platanoidis* (ihDrePlat2) somatic mutational spectrum (SBS96 classification)
C>A
C>G
C>T
T>A
T>C
T>G
Counts
10000
7500
5000
2500
0
Trinucleotide

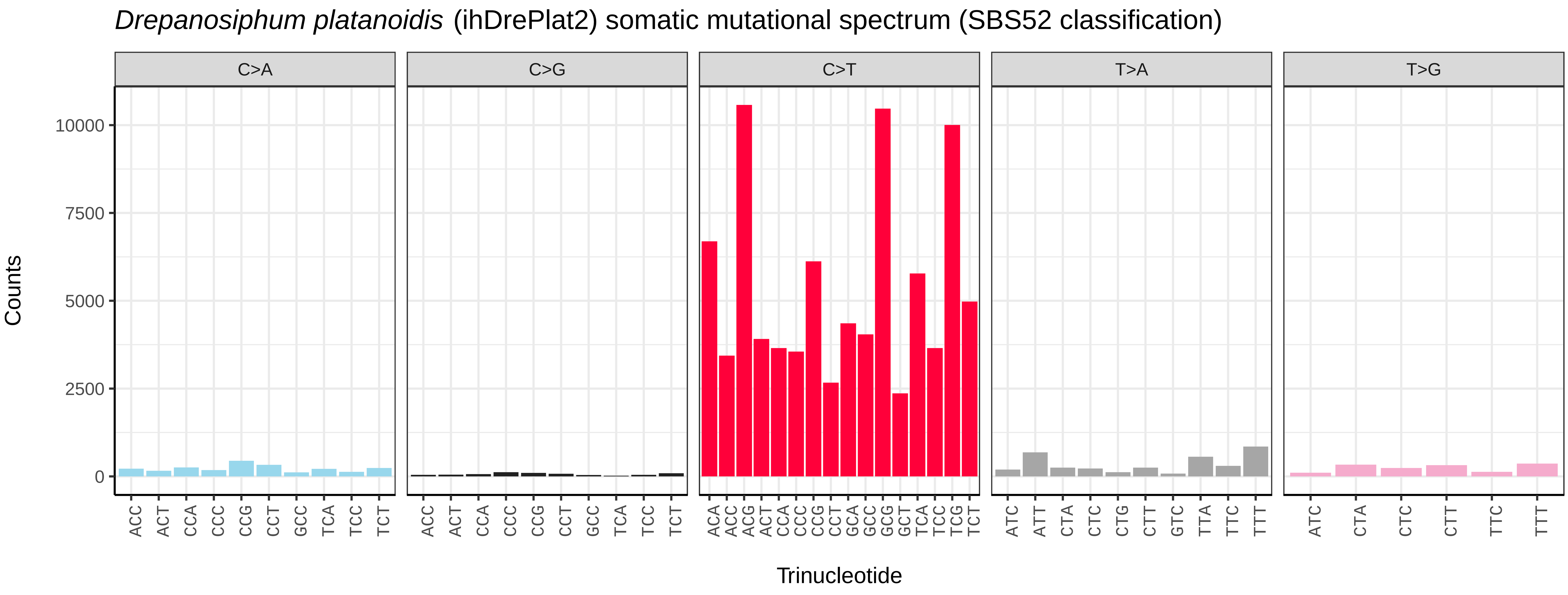
*Drepanosiphum platanoidis* (ihDrePlat2) somatic mutational spectrum (SBS52 classification)
C>A
C>G
C>T
T>A
T>G
Counts
10000
7500
5000
2500
0
Trinucleotide

*Hemaris fuciformis* (iHemFuc1.iHemFuc2) germline mutational spectrum

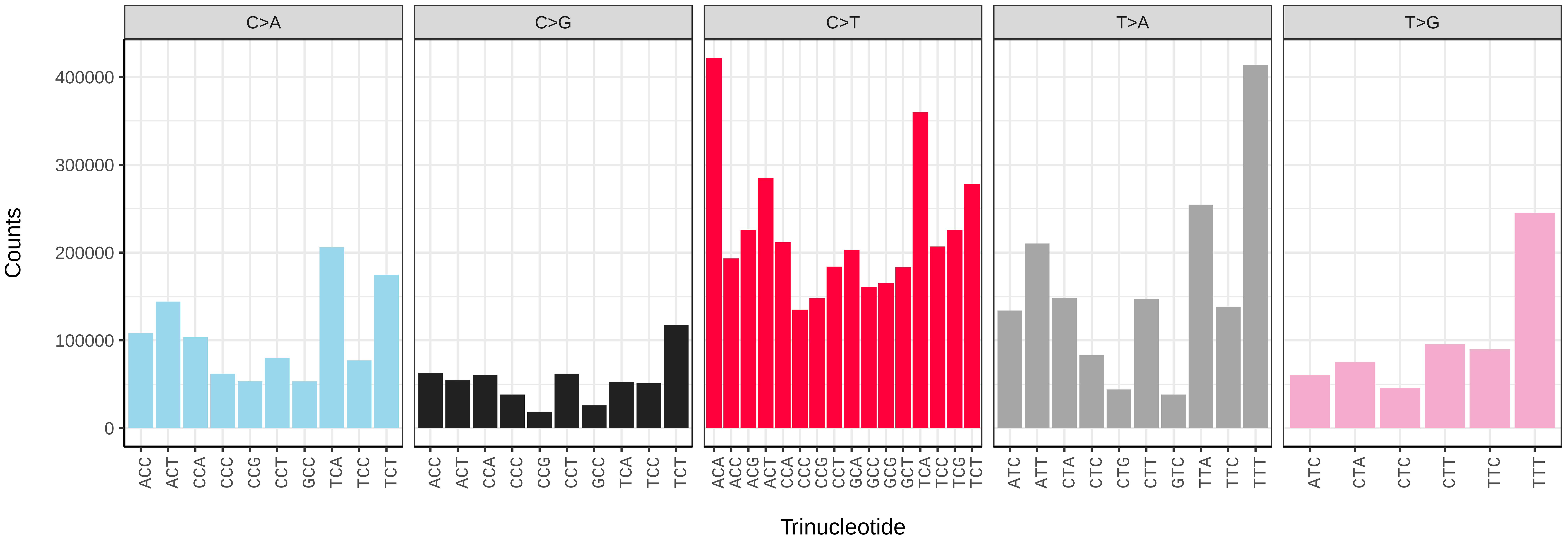


*Hemaris fuciformis* (iHemFuc1.iHemFuc2) somatic mutational spectrum (SBS96 classification)

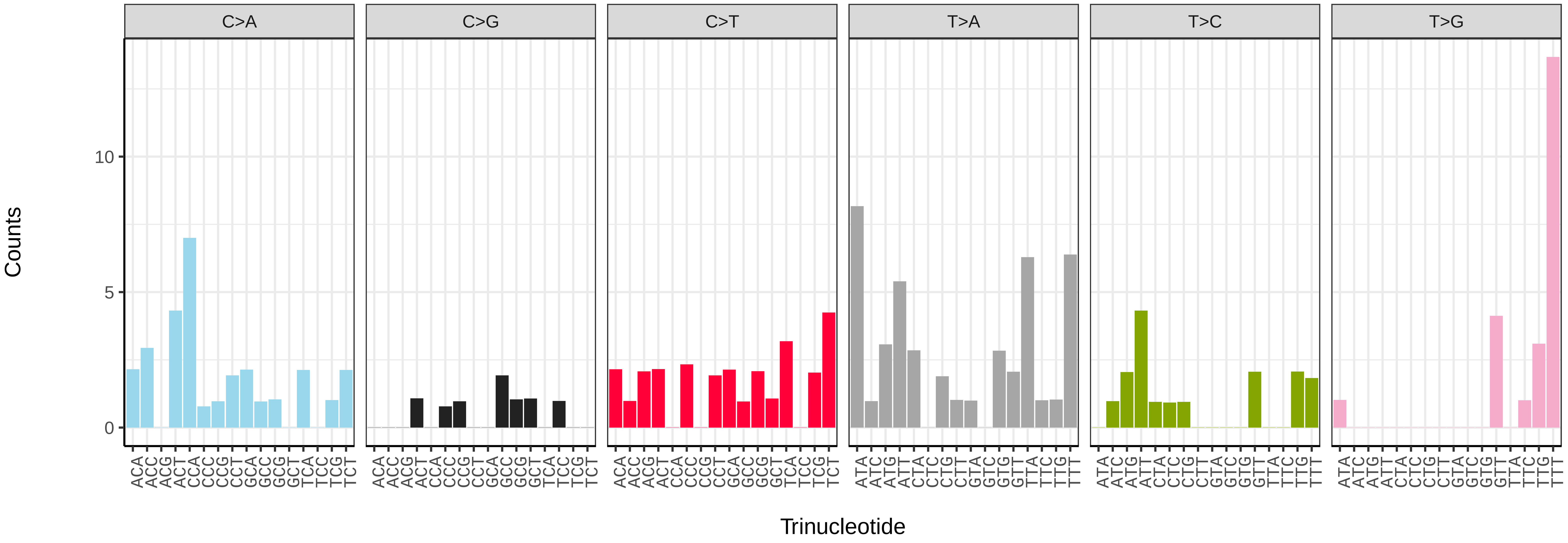


*Hemaris fuciformis* (iHemFuc1.iHemFuc2) somatic mutational spectrum (SBS52 classification)

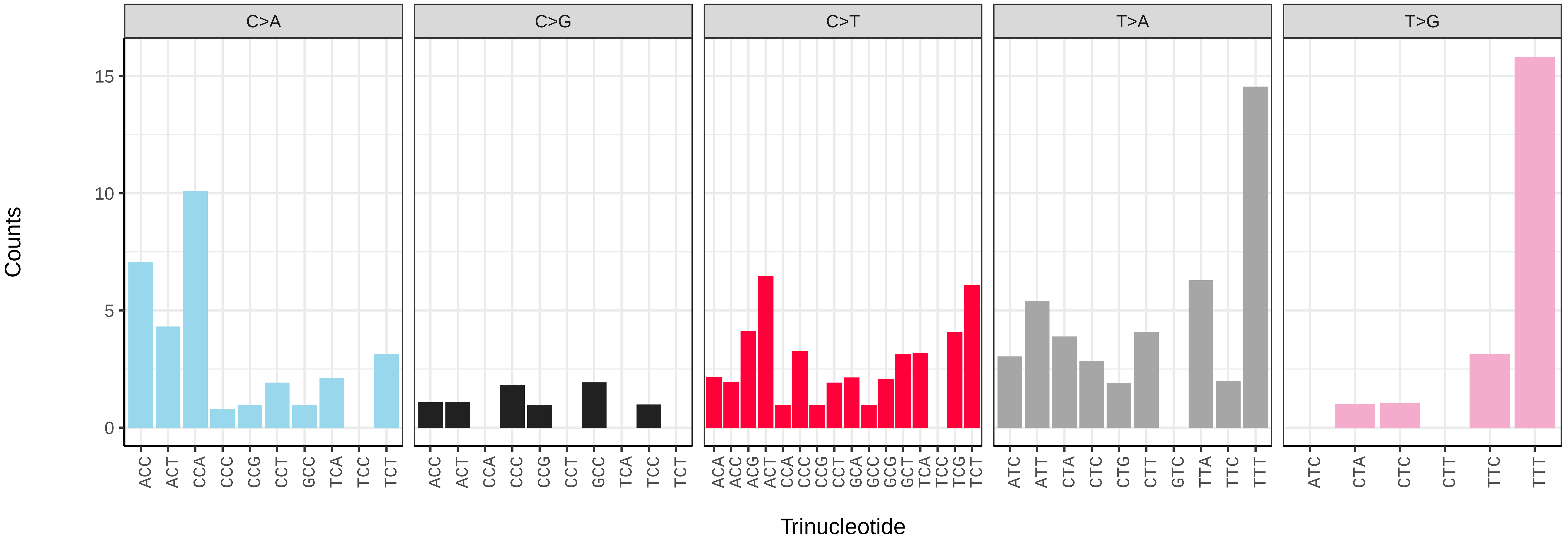

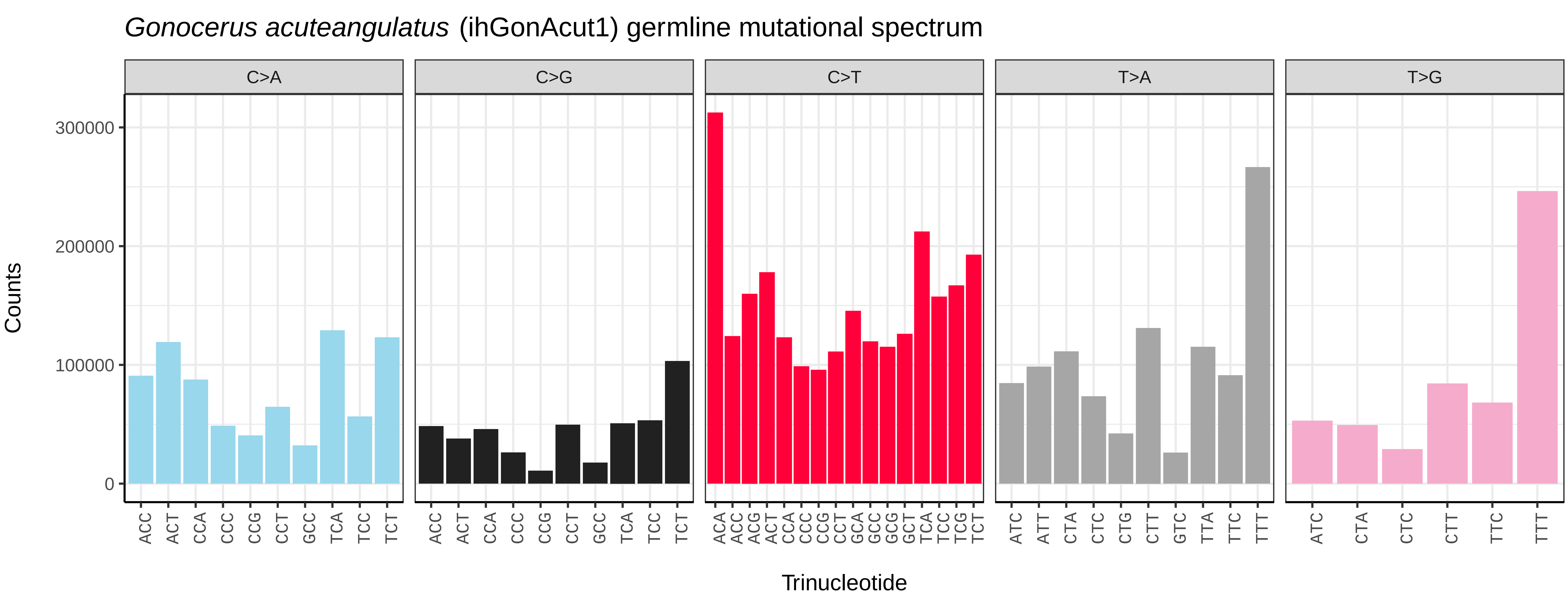
Gonocerus acuteangulatus (ihGonAcut1) germline mutational spectrum
C>A
C>G
C>T
T>A
T>G
Counts
0
100000
200000
300000
ACC ACT CCA CCC CCG CCT GCC TCA TCC TCT
ACA ACC ACG ACT CCA CCC CCG CCT GCA GCC GCG GCT TCA TCC TCG TCT
ATC ATT CTA CTC CTG CTT GTC TTA TTC TTT
ATC CTA CTC CTT TTC TTT
Trinucleotide

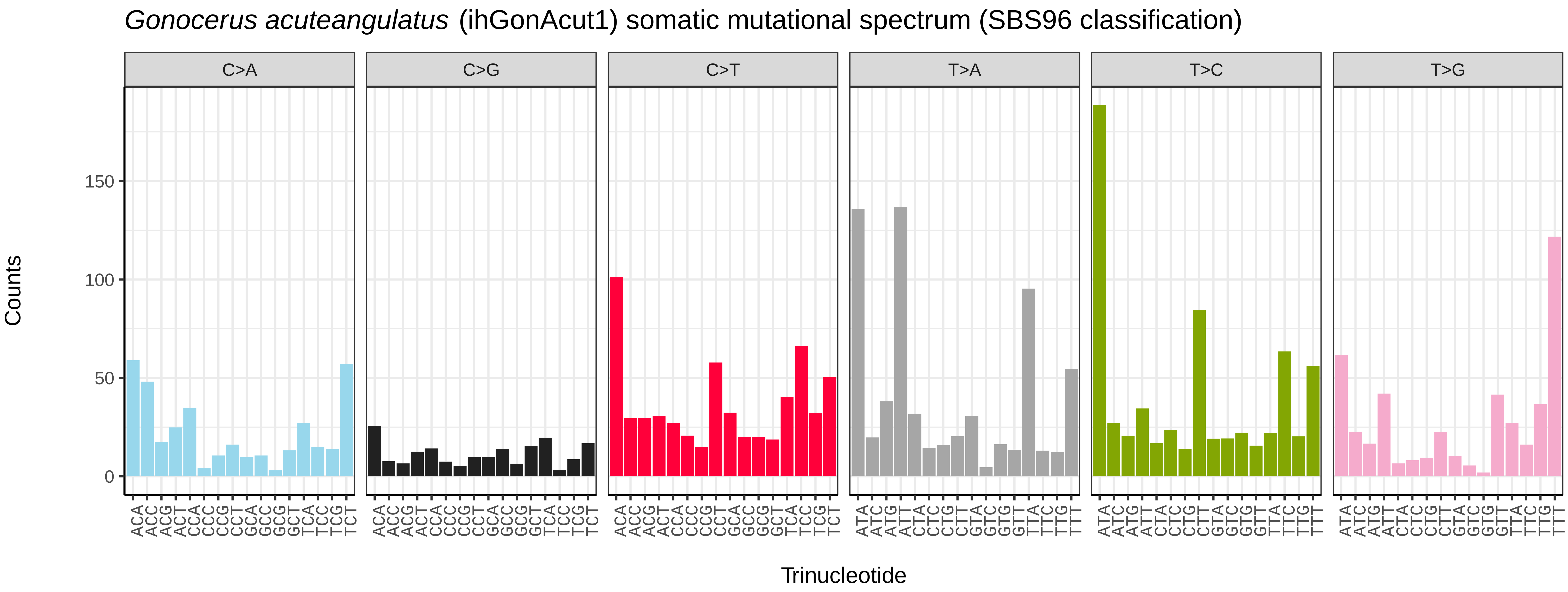
Gonocerus acuteangulatus (ihGonAcut1) somatic mutational spectrum (SBS96 classification)
C>A
C>G
C>T
T>A
T>C
T>G
Counts
0
50
100
150
ACA ACC ACG ACT CCA CCC CCG CCT GCA GCC GCG GCT TCA TCC TCG TCT
ATA ATC ATG ATT CTA CTC CTG CTT GTA GTC GTG GTT TTA TTC TTG TTT
Trinucleotide

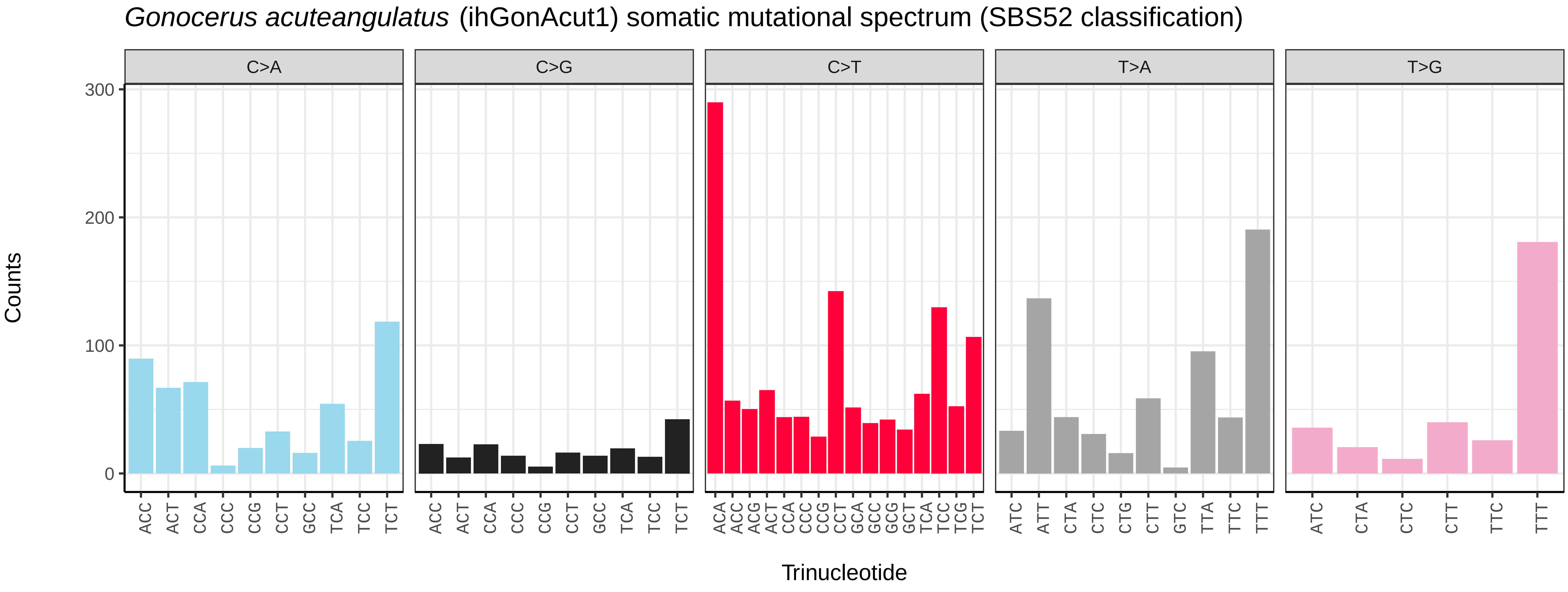
Gonocerus acuteangulatus (ihGonAcut1) somatic mutational spectrum (SBS52 classification)
C>A
C>G
C>T
T>A
T>G
Counts
0
100
200
300
ACC ACT CCA CCC CCG CCT GCC TCA TCC TCT
ACA ACC ACG ACT CCA CCC CCG CCT GCA GCC GCG GCT TCA TCC TCG TCT
ATC ATT CTA CTC CTG CTT GTC TTA TTC TTT
ATC CTA CTC CTT TTC TTT
Trinucleotide

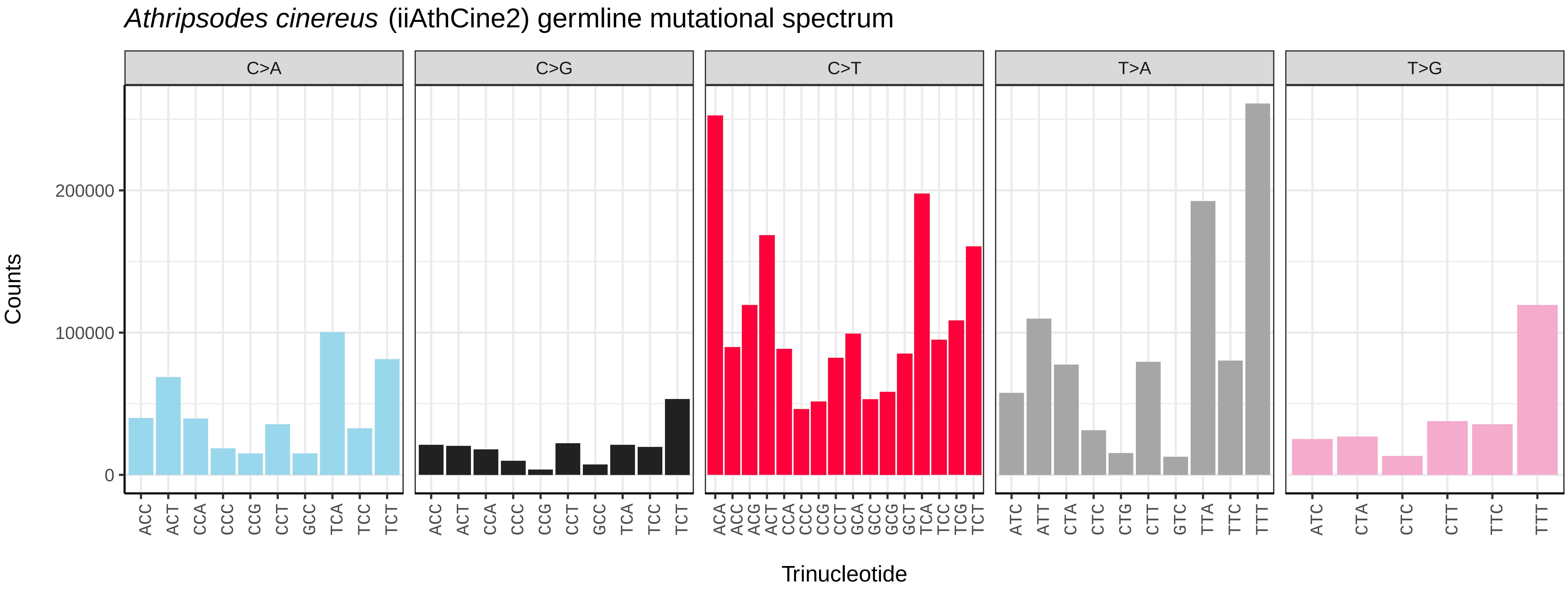
*Athripsodes cinereus* (iiAthCine2) germline mutational spectrum
C>A
C>G
C>T
T>A
T>G
Counts
0
100000
200000
ACC ACT CCA CCC CCG CCT GCC TCA TCC TCT
ACC ACT CCA CCC CCG CCT GCC TCA TCC TCT
ACA ACC ACG ACT CCA CCC CCG CCT GCA GCC GCG GCT TCA TCC TCG TCT
ATC ATT CTA CTC CTG CTT GTC TTA TTC TTT
ATC CTA CTC CTT TTC TTT
Trinucleotide

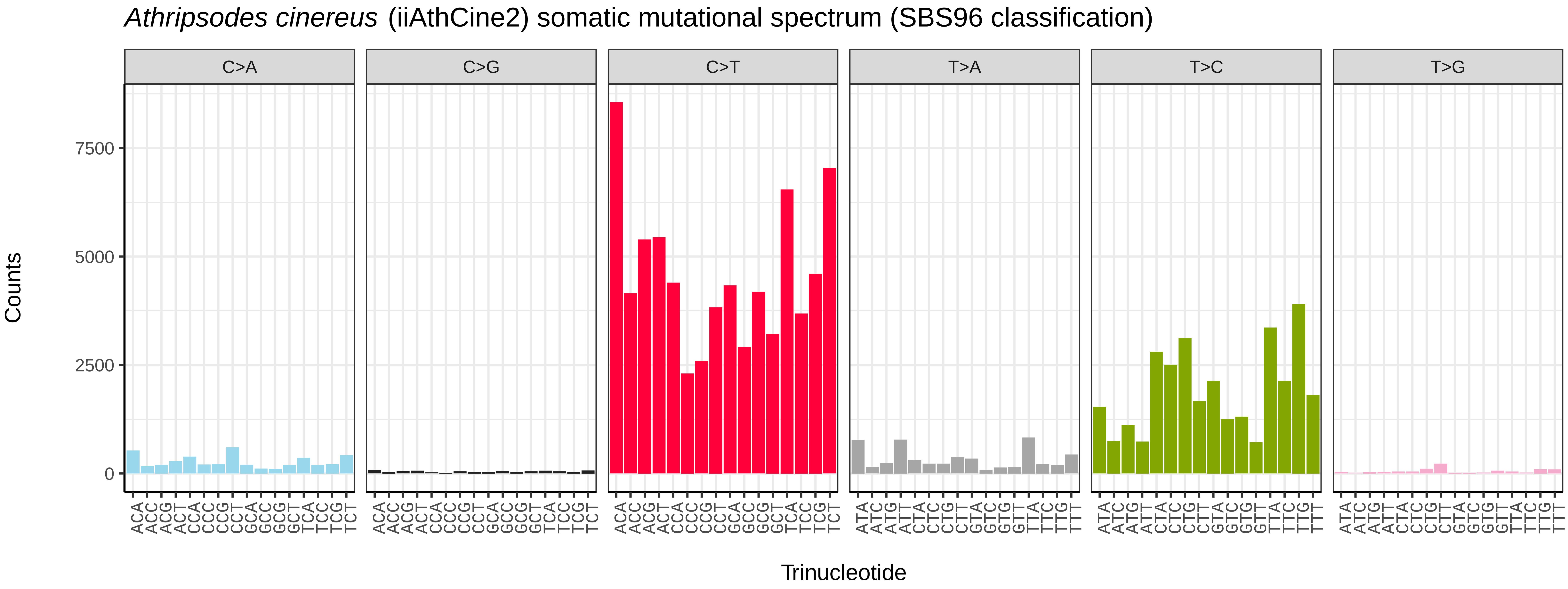
*Athripsodes cinereus* (iiAthCine2) somatic mutational spectrum (SBS96 classification)
C>A
C>G
C>T
T>A
T>C
T>G
Counts
0
2500
5000
7500
ACA ACC ACG ACT CCA CCC CCG CCT GCA GCC GCG GCT TCA TCC TCG TCT
ATA ATC ATG ATT CTA CTC CTG CTT GTA GTC GTG GTT TTA TTC TTG TTT
Trinucleotide

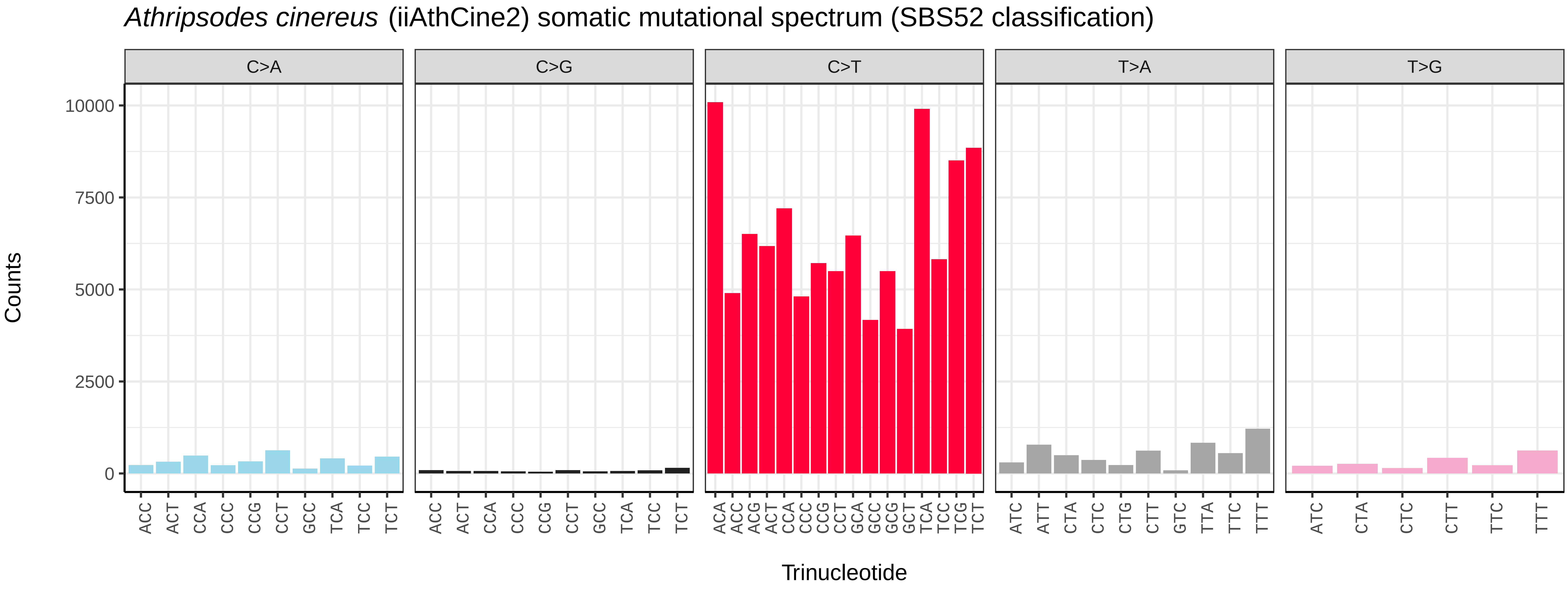
*Athripsodes cinereus* (iiAthCine2) somatic mutational spectrum (SBS52 classification)
C>A
C>G
C>T
T>A
T>G
Counts
0
2500
5000
7500
10000
ACC ACT CCA CCC CCG CCT GCC TCA TCC TCT
ACC ACT CCA CCC CCG CCT GCC TCA TCC TCT
ACA ACC ACG ACT CCA CCC CCG CCT GCA GCC GCG GCT TCA TCC TCG TCT
ATC ATT CTA CTC CTG CTT GTC TTA TTC TTT
ATC CTA CTC CTT TTC TTT
Trinucleotide

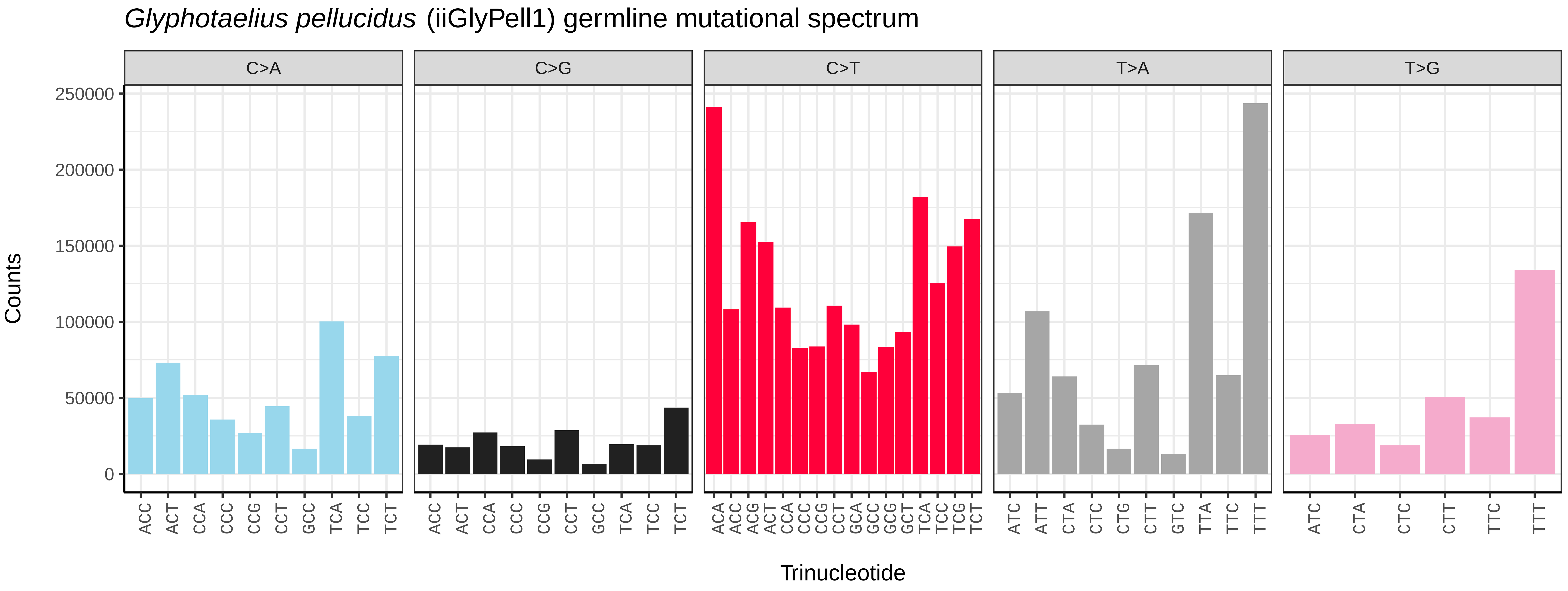
*Glyphotaelius pellucidus* (iiGlyPell1) germline mutational spectrum
C>A
C>G
C>T
T>A
T>G
Counts
0
50000
100000
150000
200000
250000
Trinucleotide

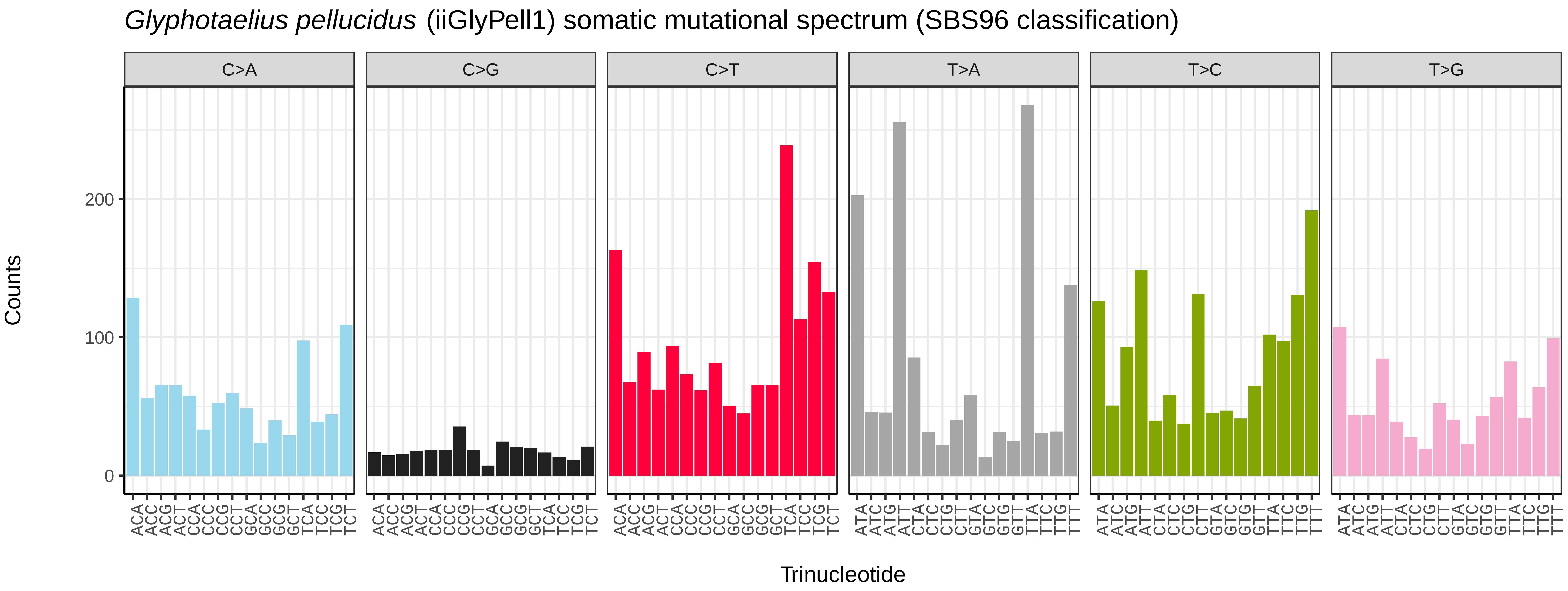
*Glyphotaelius pellucidus* (iiGlyPell1) somatic mutational spectrum (SBS96 classification)
C>A
C>G
C>T
T>A
T>C
T>G
Counts
0
100
200
Trinucleotide

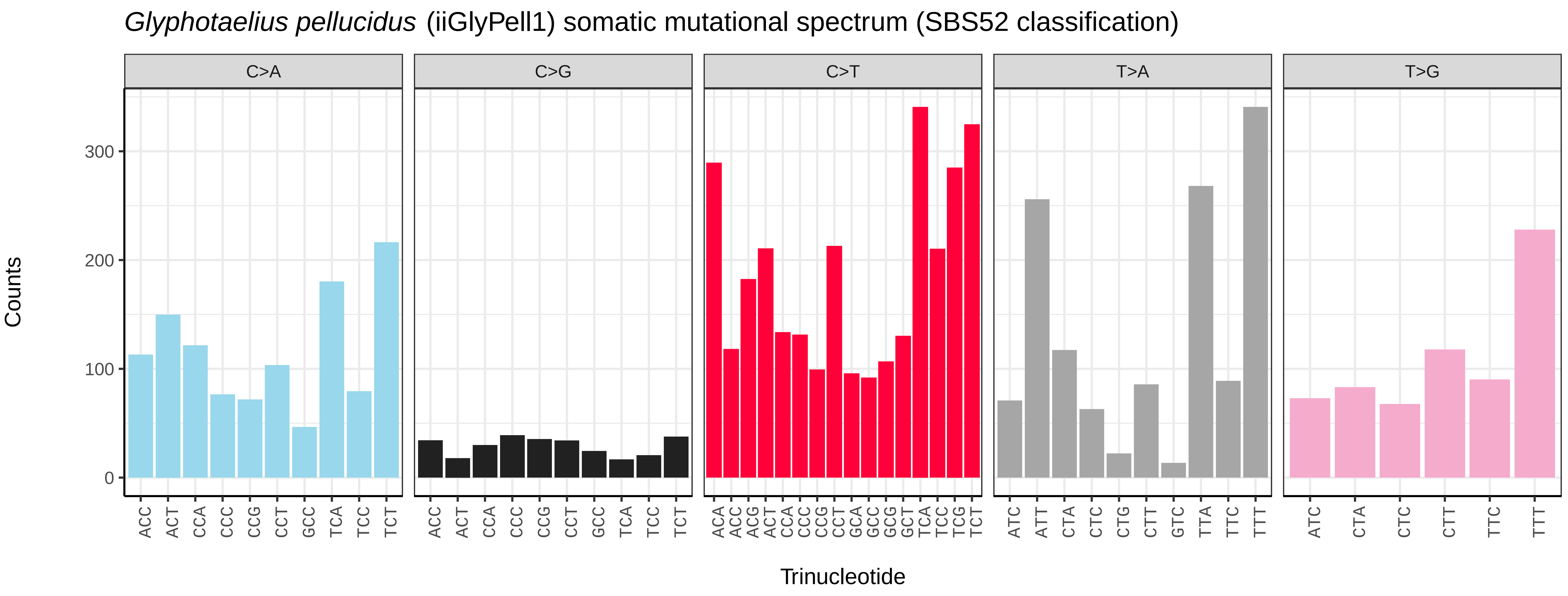
*Glyphotaelius pellucidus* (iiGlyPell1) somatic mutational spectrum (SBS52 classification)
C>A
C>G
C>T
T>A
T>G
Counts
0
100
200
300
Trinucleotide

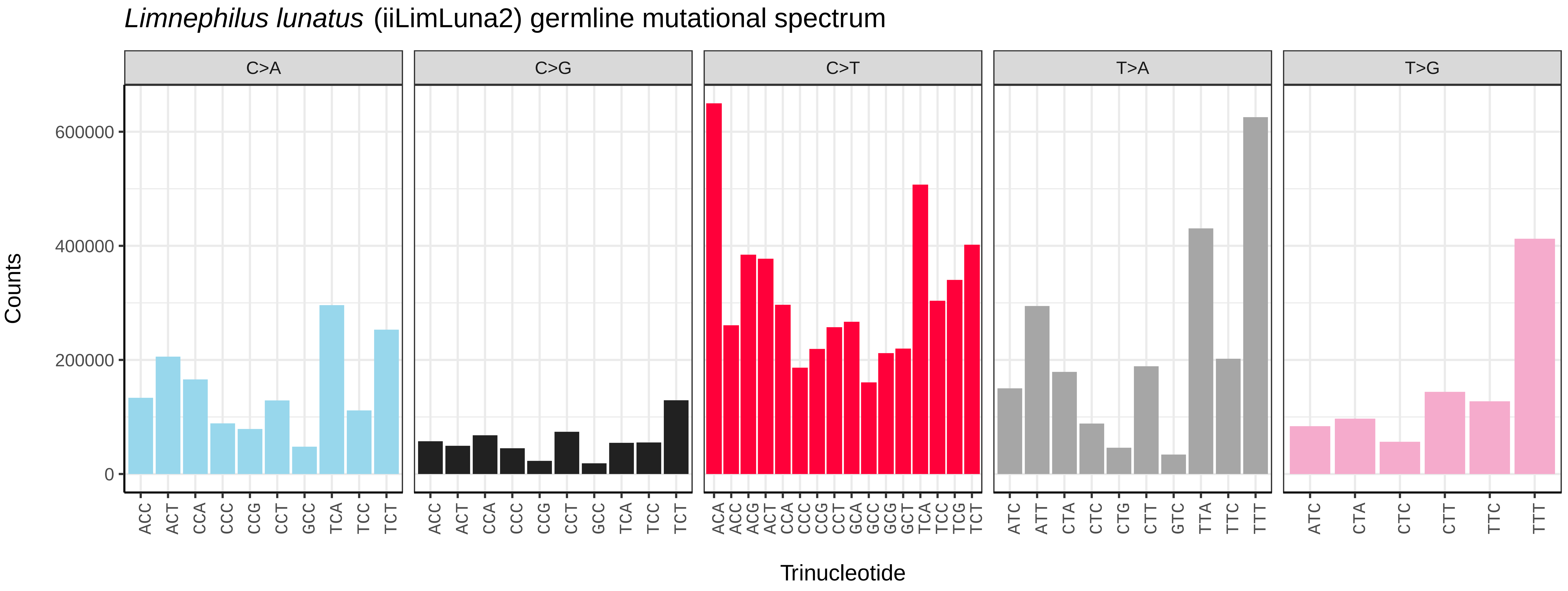
*Limnephilus lunatus* (iiLimLuna2) germline mutational spectrum
C>A
C>G
C>T
T>A
T>G
Counts
Trinucleotide
0
200000
400000
600000

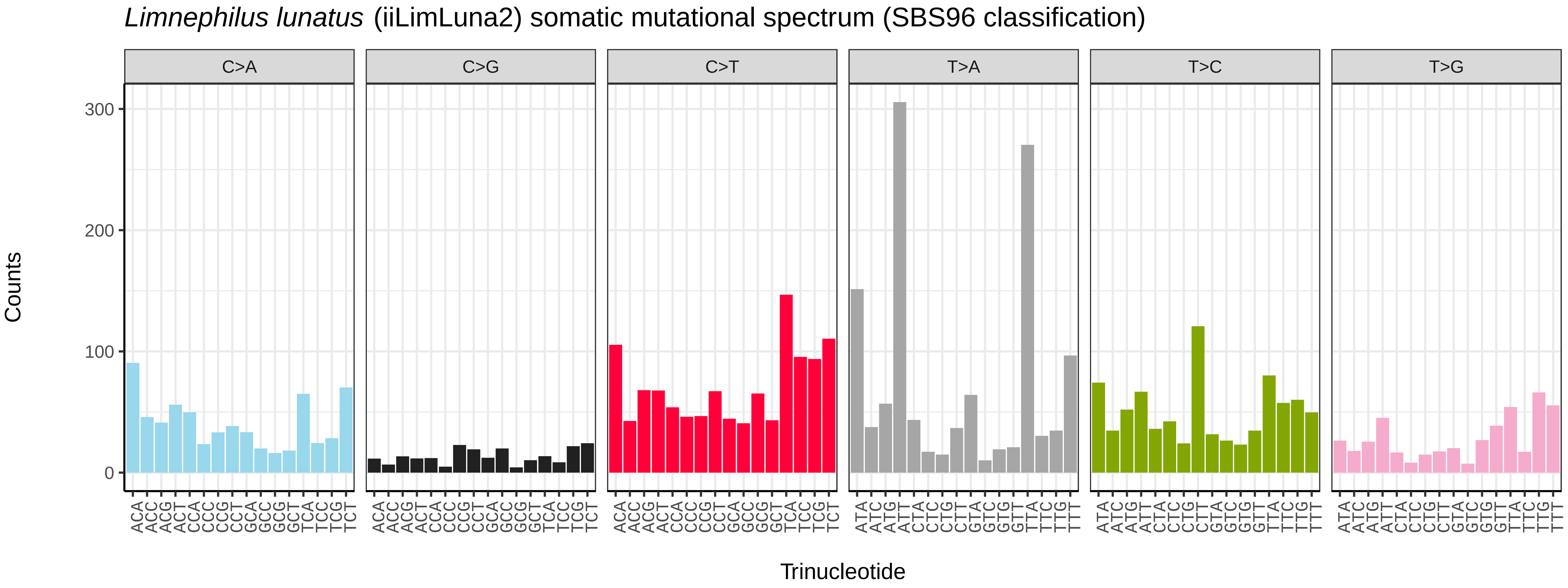
*Limnephilus lunatus* (iiLimLuna2) somatic mutational spectrum (SBS96 classification)
C>A
C>G
C>T
T>A
T>C
T>G
Counts
Trinucleotide
0
100
200
300

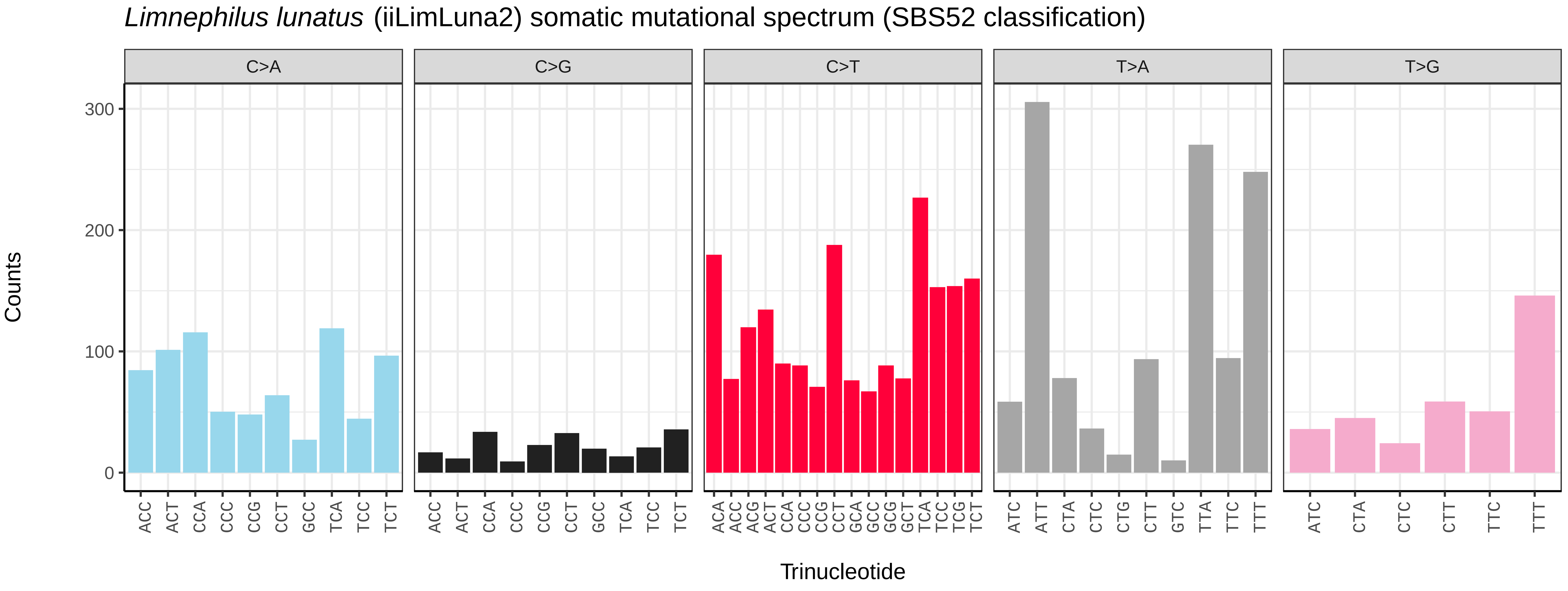
*Limnephilus lunatus* (iiLimLuna2) somatic mutational spectrum (SBS52 classification)
C>A
C>G
C>T
T>A
T>G
Counts
Trinucleotide
0
100
200
300

*Limnephilus marmoratus* (iiLimMarm1) germline mutational spectrum

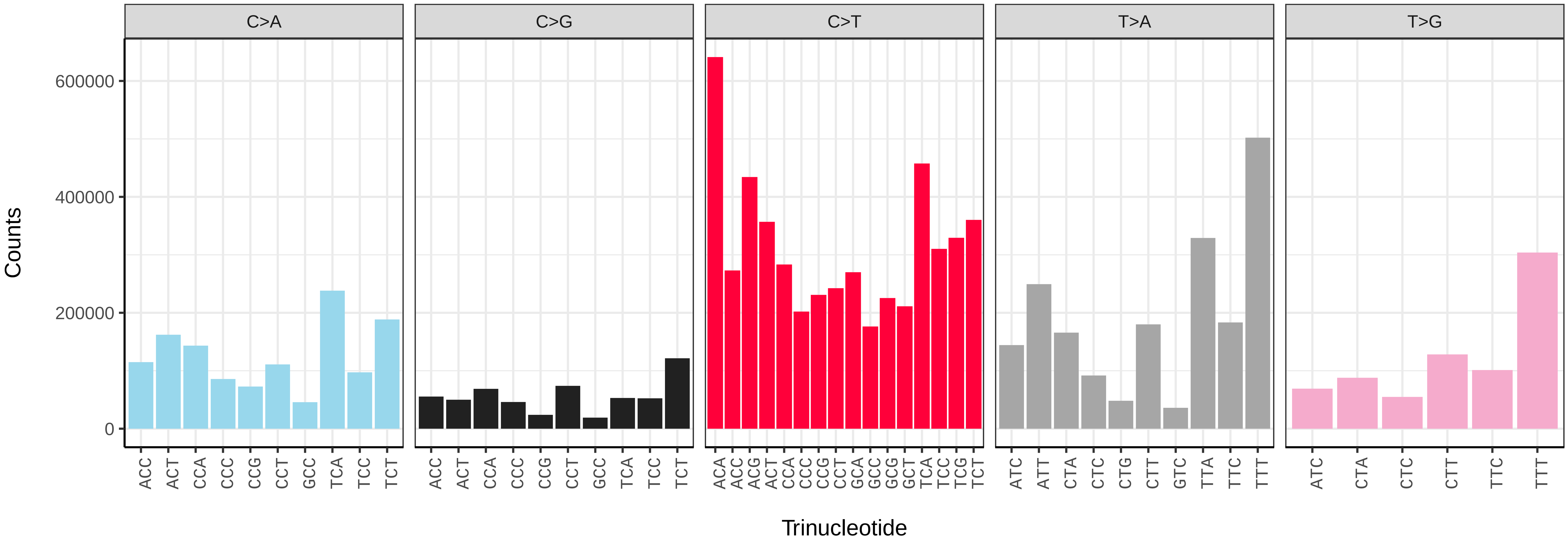


*Limnephilus marmoratus* (iiLimMarm1) somatic mutational spectrum (SBS96 classification)

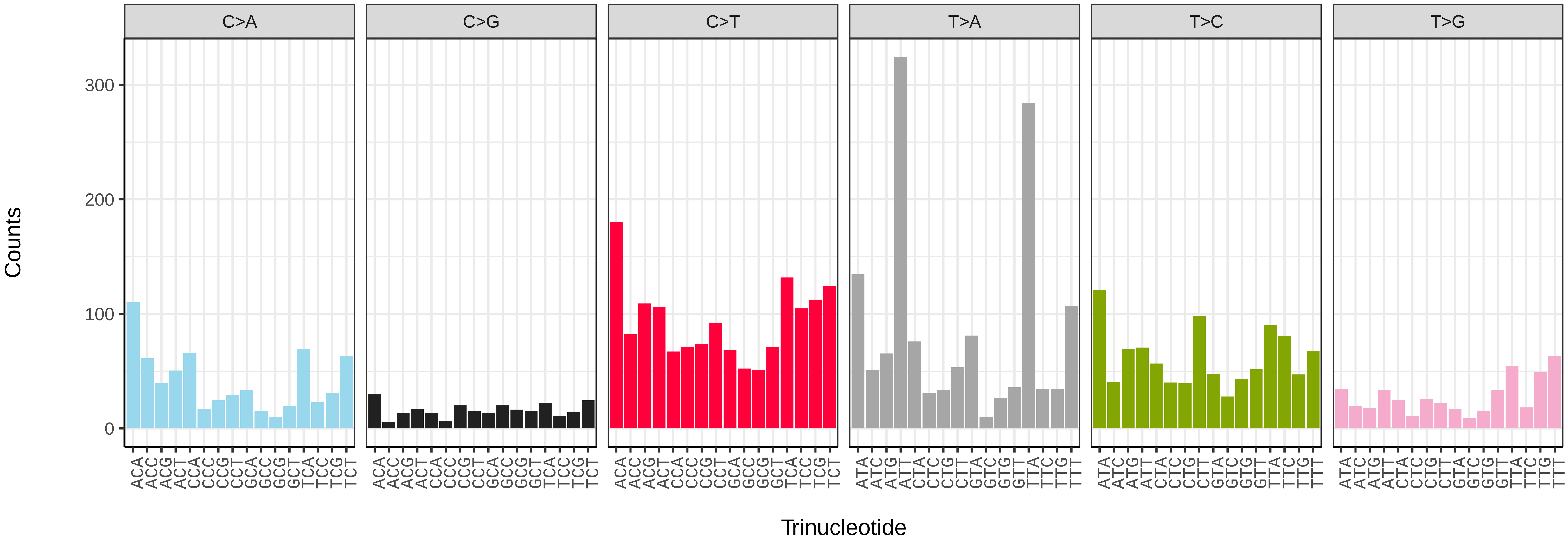


*Limnephilus marmoratus* (iiLimMarm1) somatic mutational spectrum (SBS52 classification)

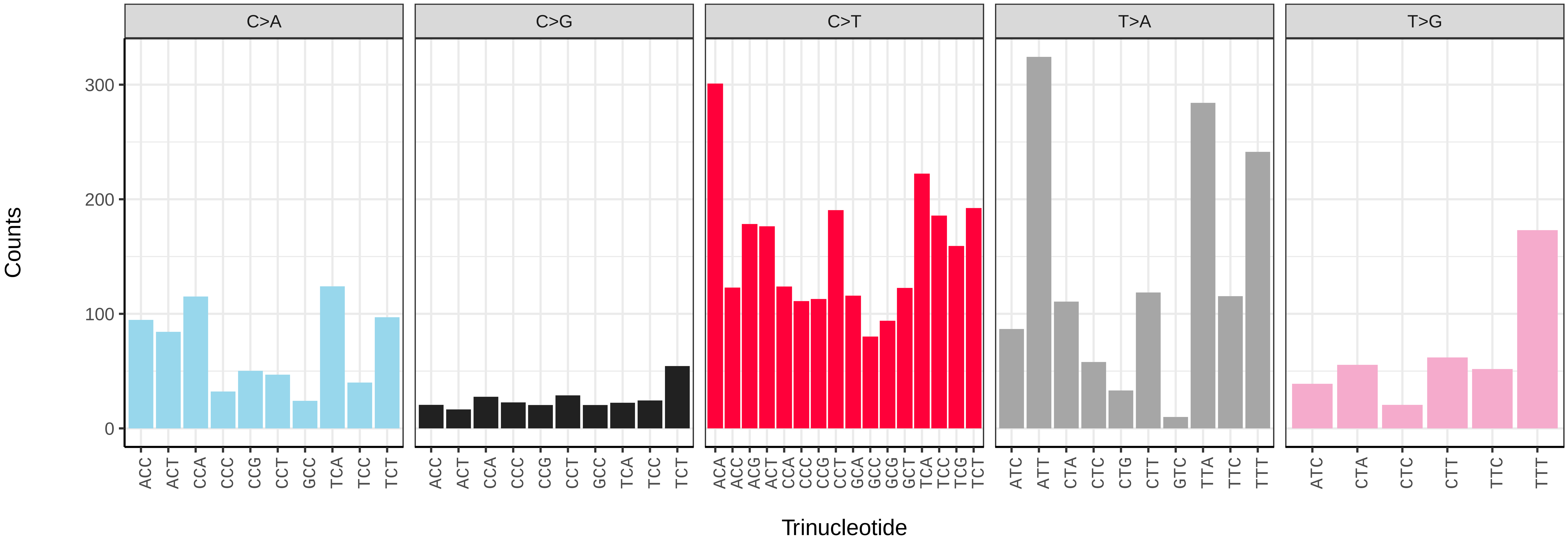

*Limnephilus rhombicus* (iiLimRhom1) germline mutational spectrum

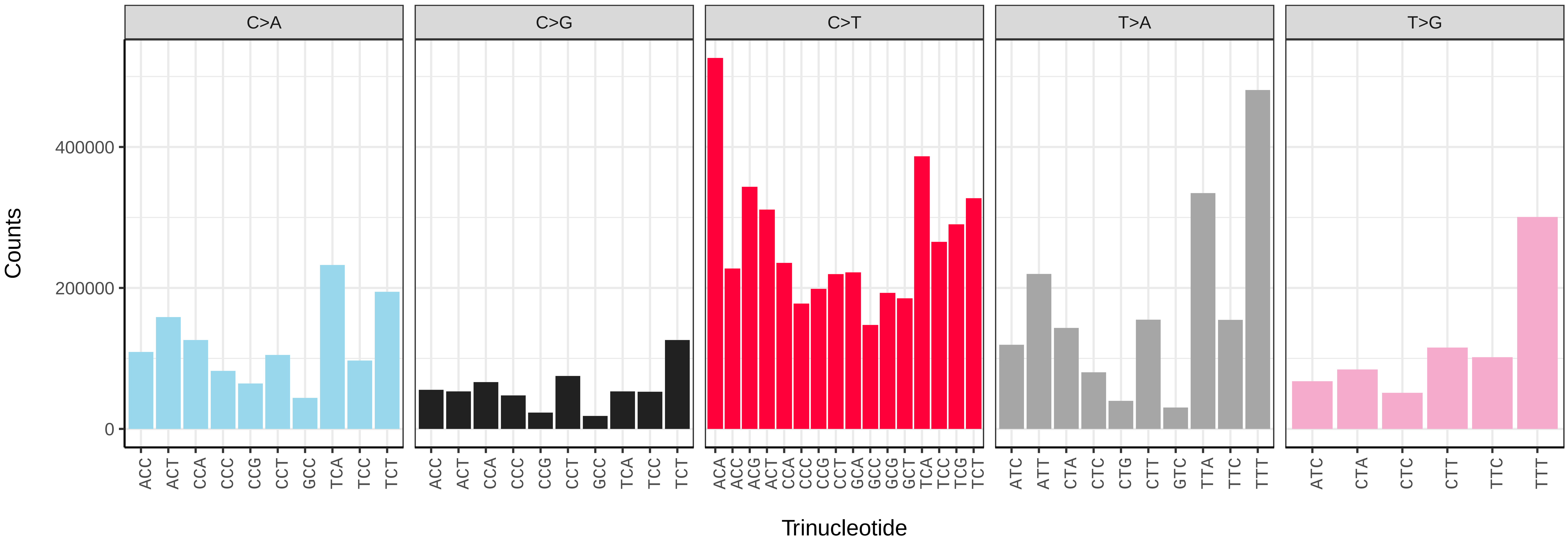


*Limnephilus rhombicus* (iiLimRhom1) somatic mutational spectrum (SBS96 classification)

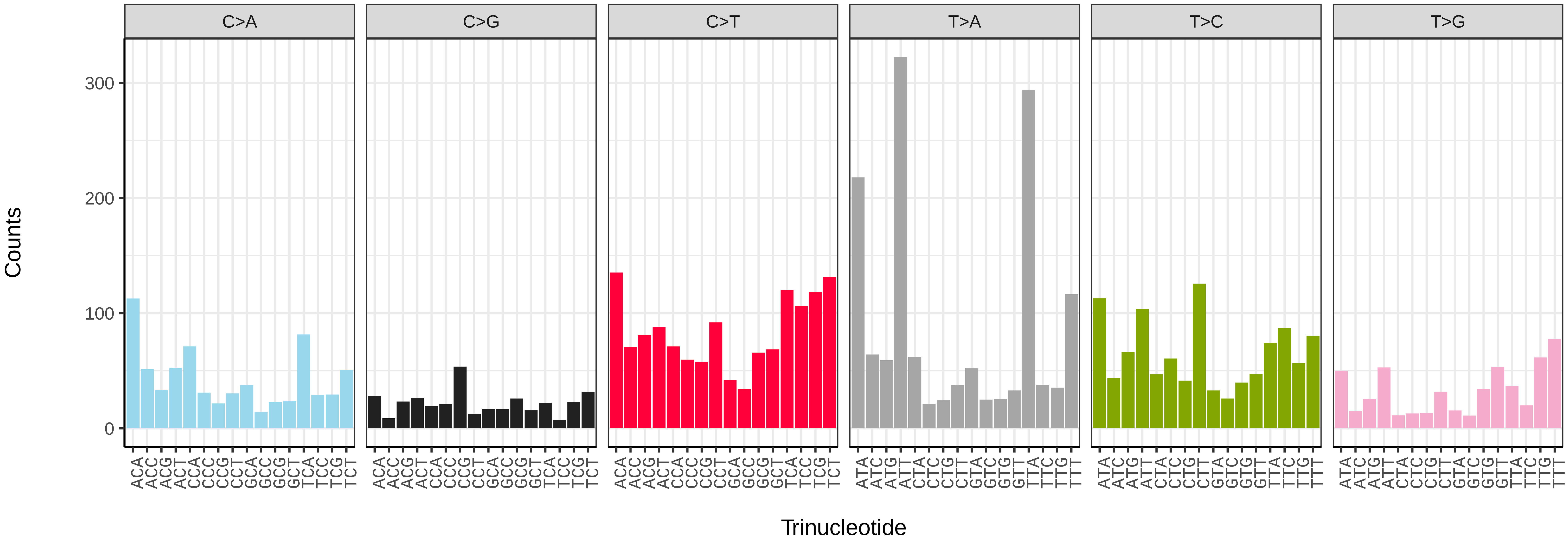


*Limnephilus rhombicus* (iiLimRhom1) somatic mutational spectrum (SBS52 classification)

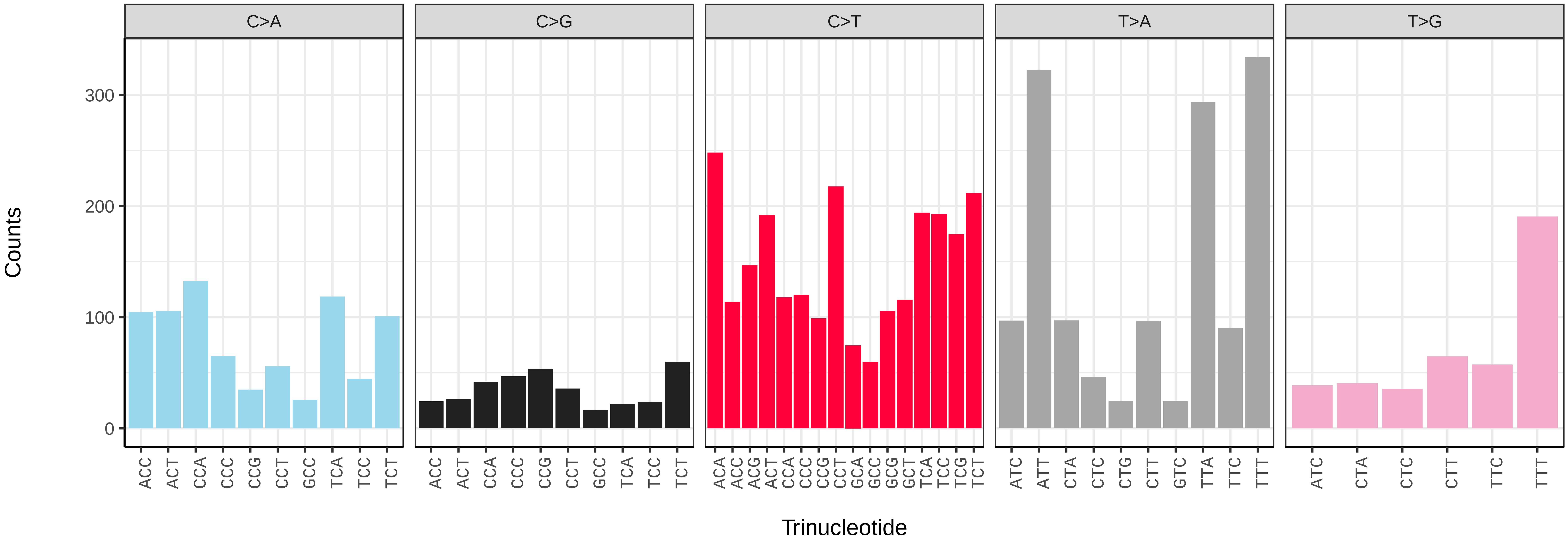

*Abrostola triplasia* (ilAbrTril1) germline mutational spectrum

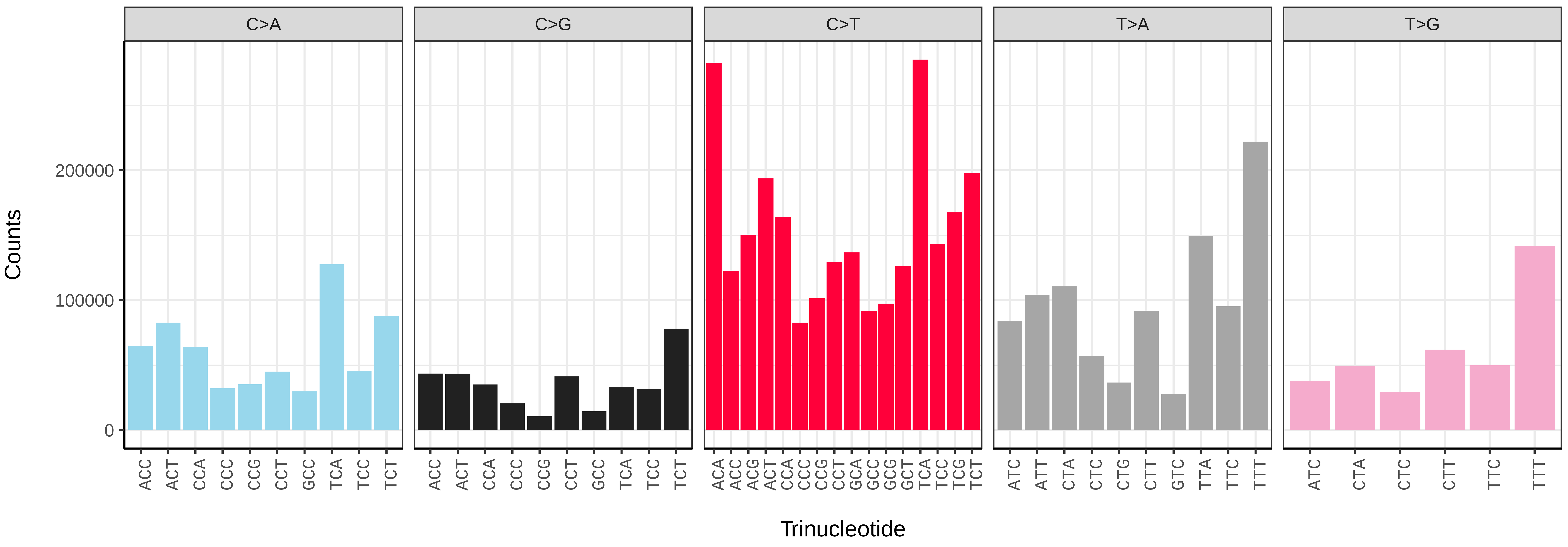


*Abrostola triplasia* (ilAbrTril1) somatic mutational spectrum (SBS96 classification)

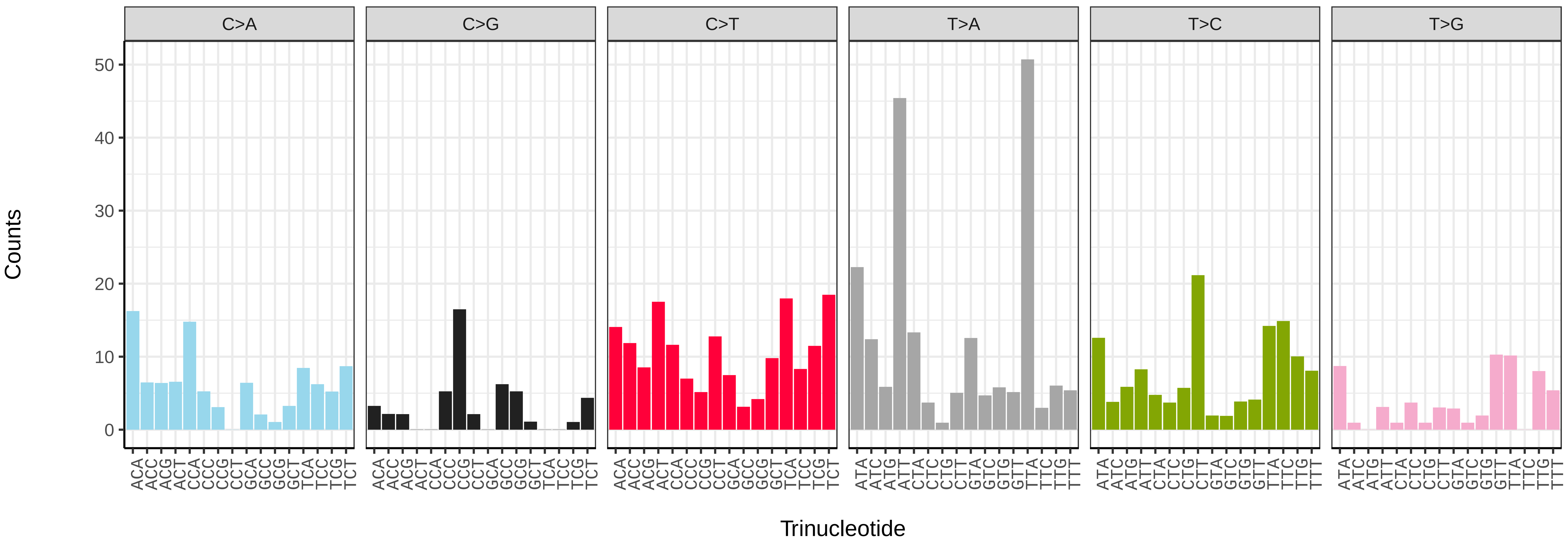


*Abrostola triplasia* (ilAbrTril1) somatic mutational spectrum (SBS52 classification)

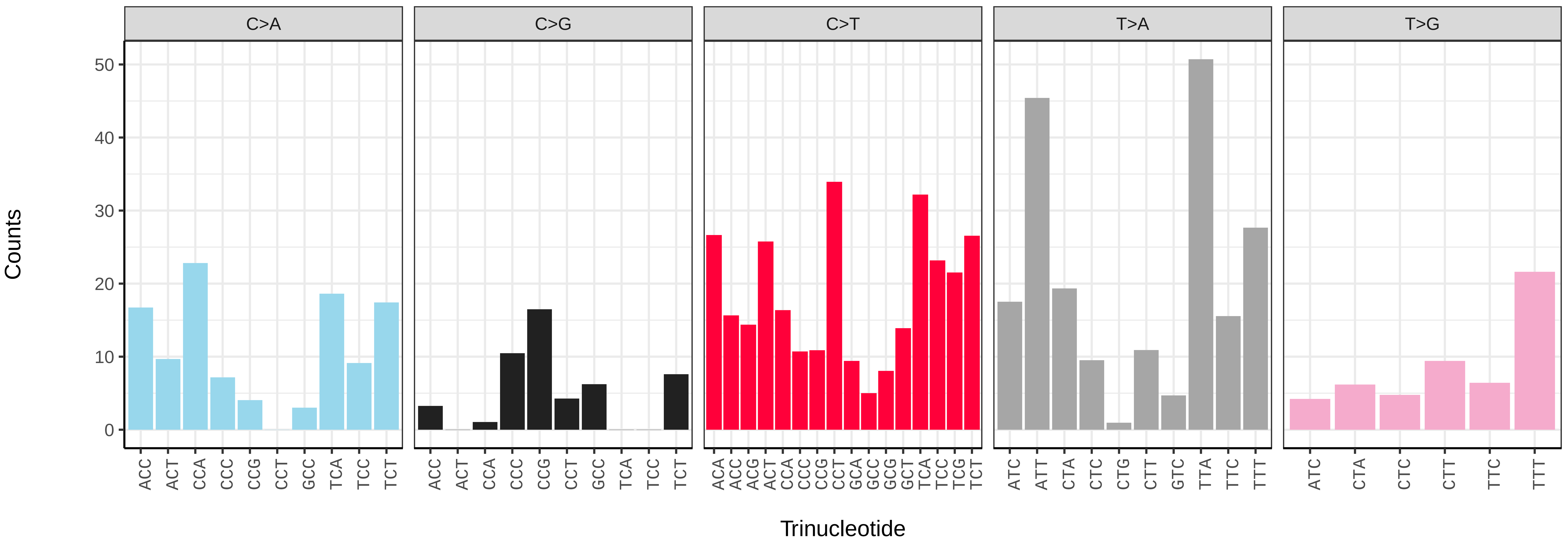

*Abrostola tripartita* (ilAbrTrip1) germline mutational spectrum

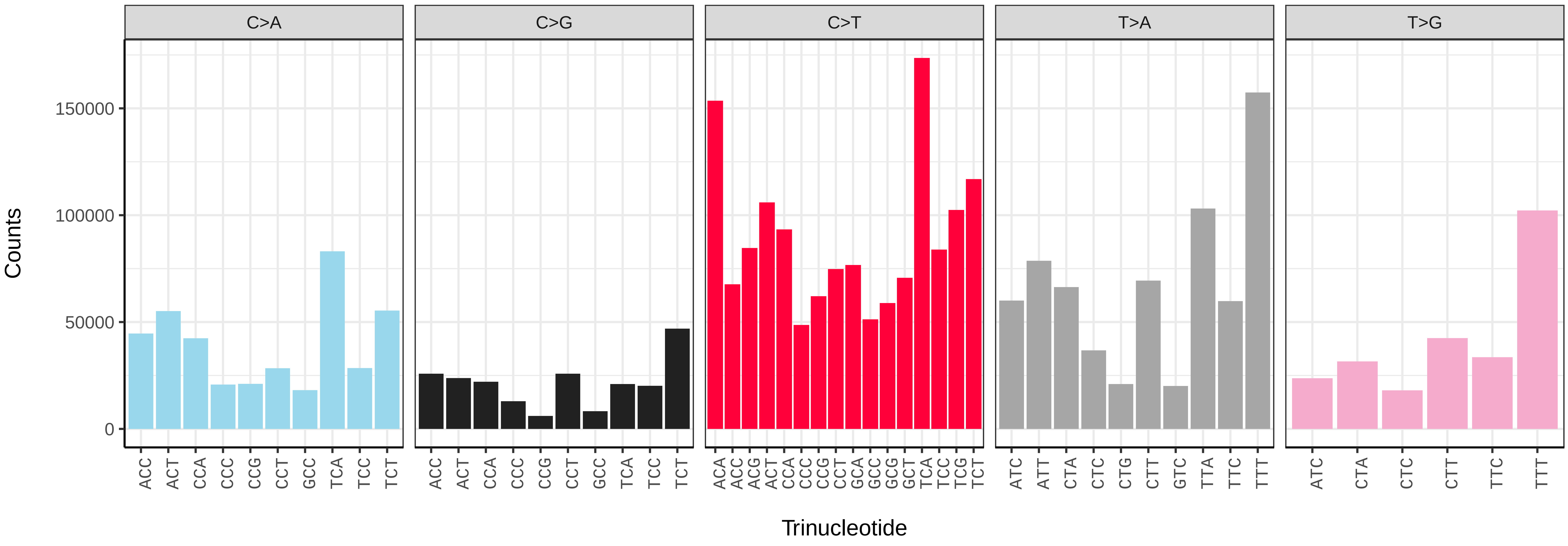


*Abrostola tripartita* (ilAbrTrip1) somatic mutational spectrum (SBS96 classification)

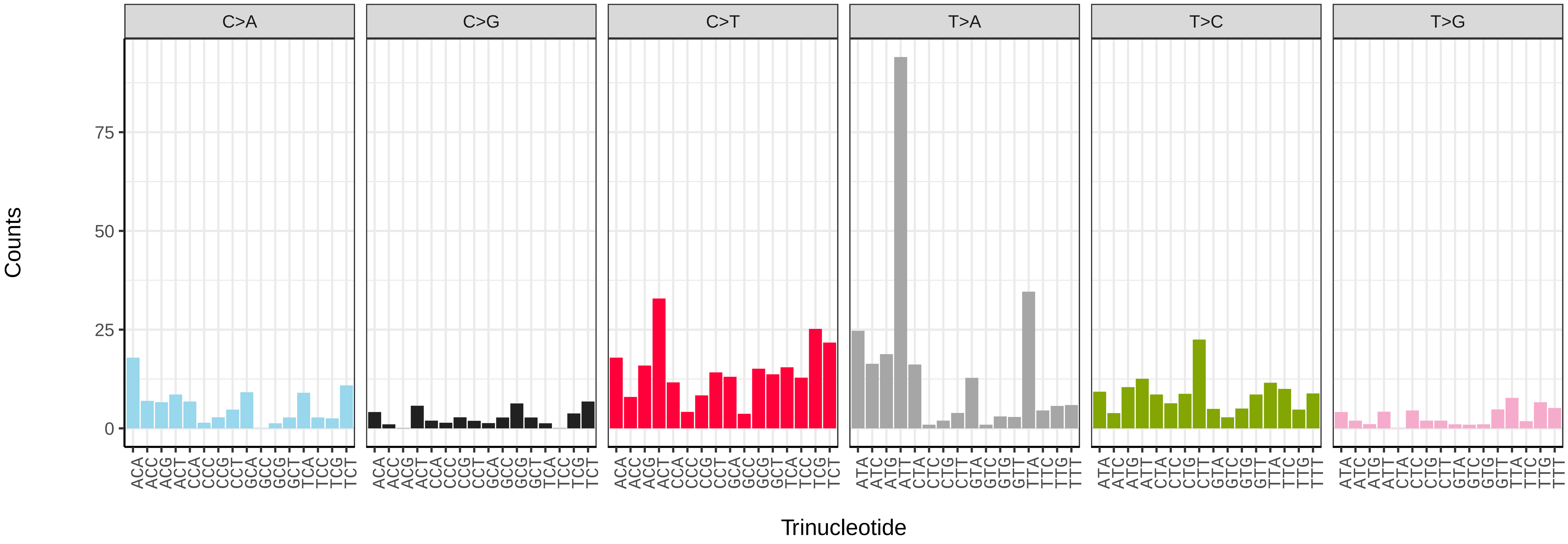


*Abrostola tripartita* (ilAbrTrip1) somatic mutational spectrum (SBS52 classification)

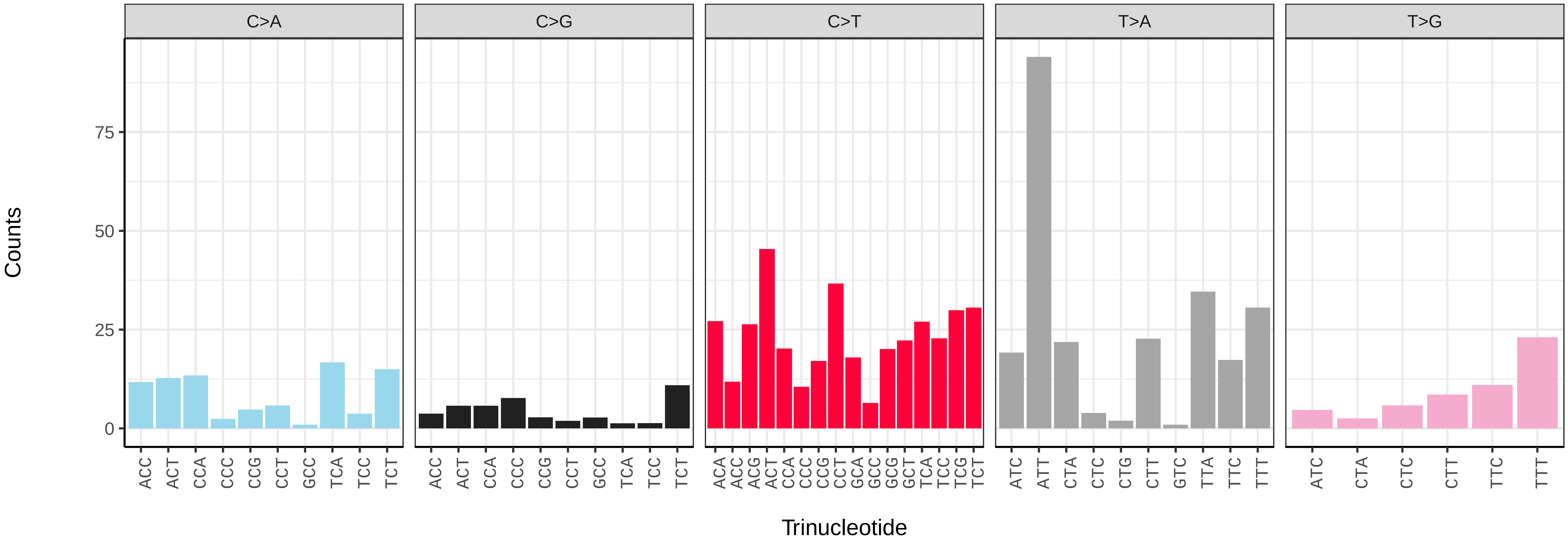

*Acentria ephemerella* (ilAceEphe1) germline mutational spectrum

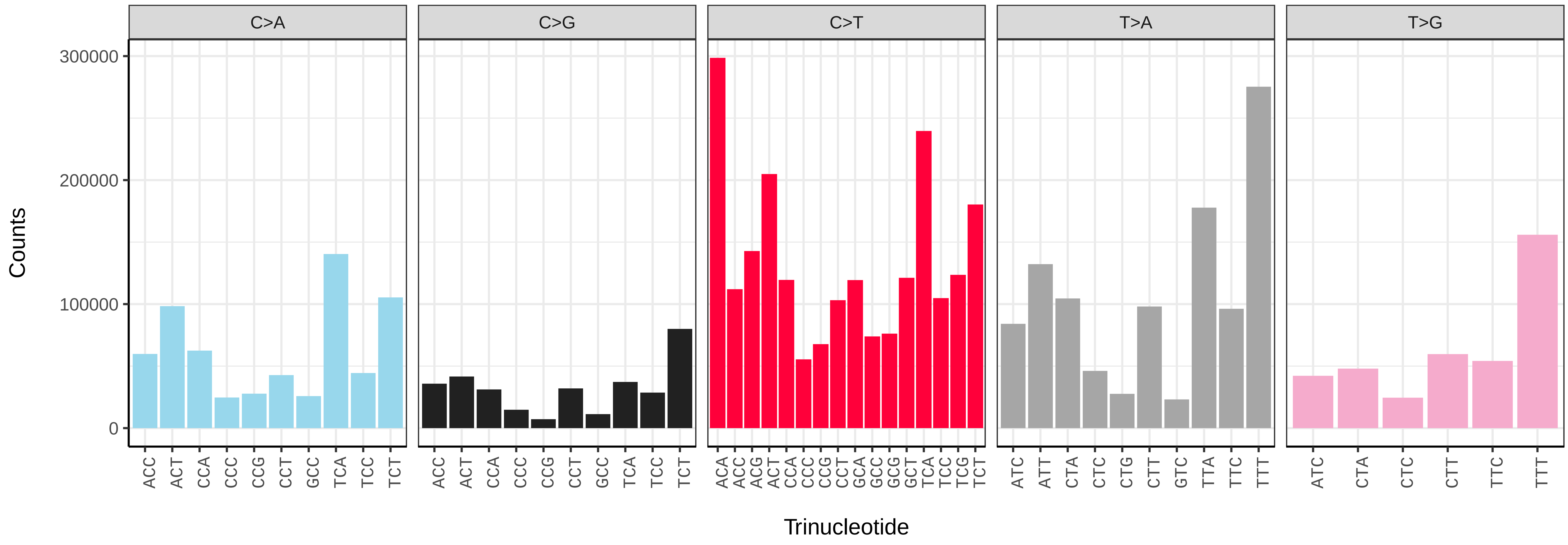


*Acentria ephemerella* (ilAceEphe1) somatic mutational spectrum (SBS96 classification)

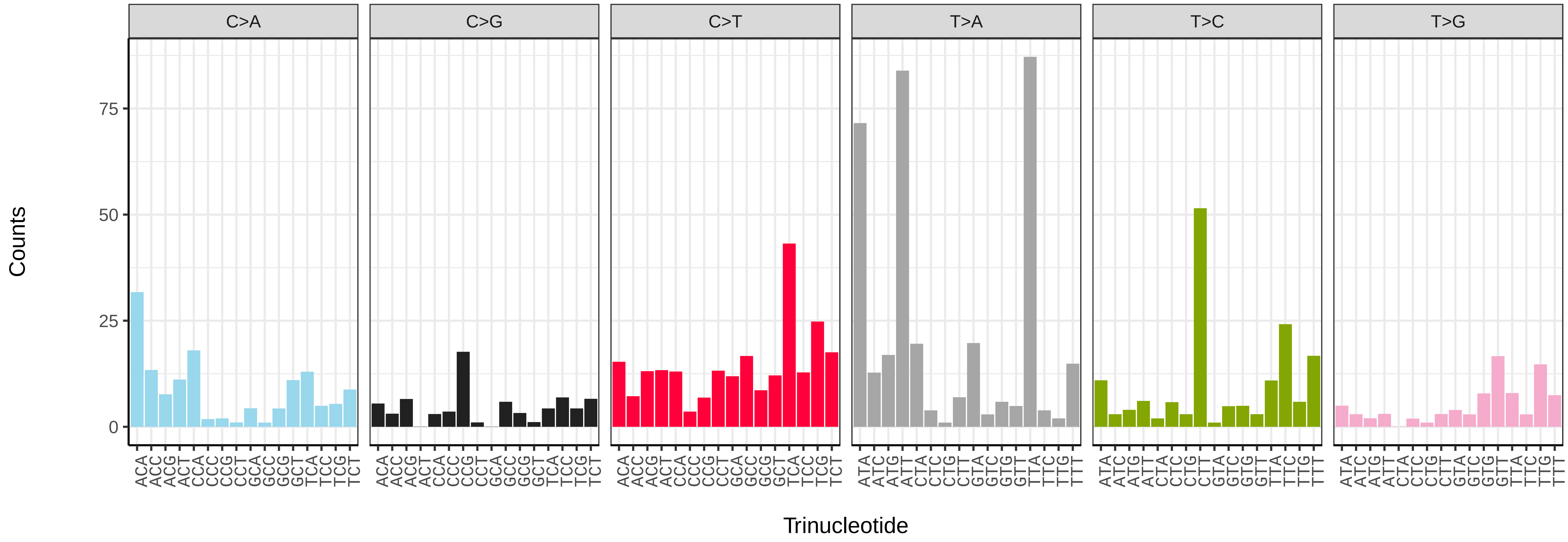


*Acentria ephemerella* (ilAceEphe1) somatic mutational spectrum (SBS52 classification)

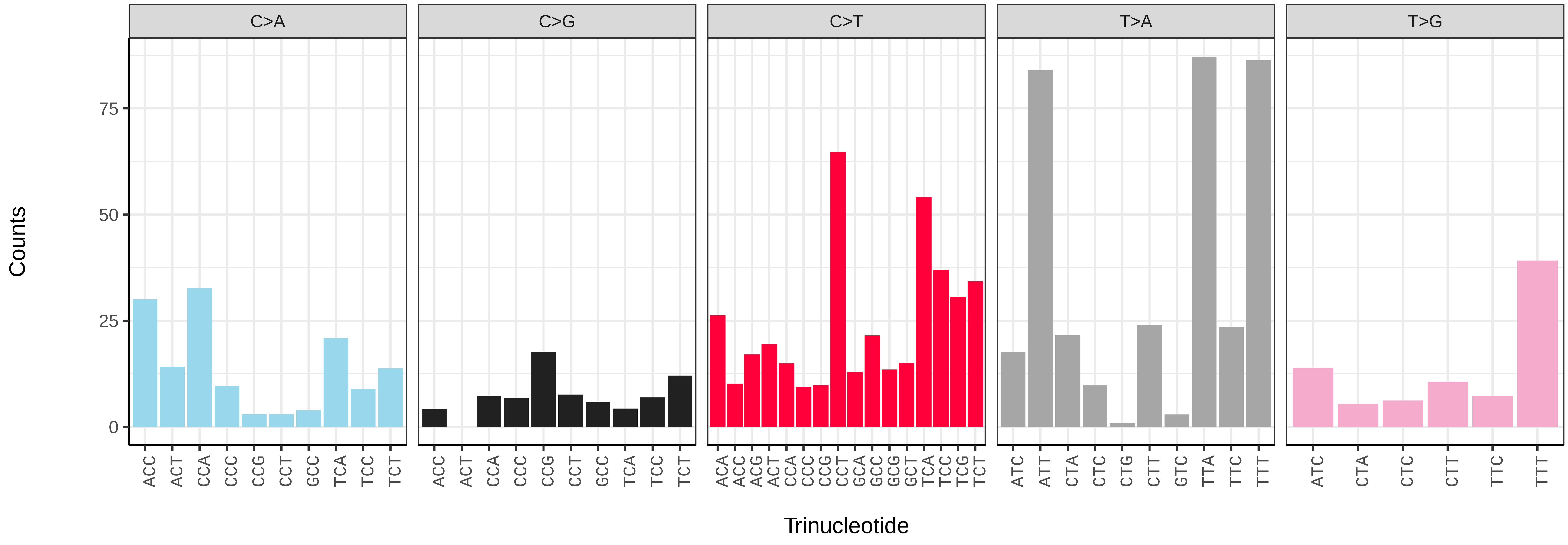

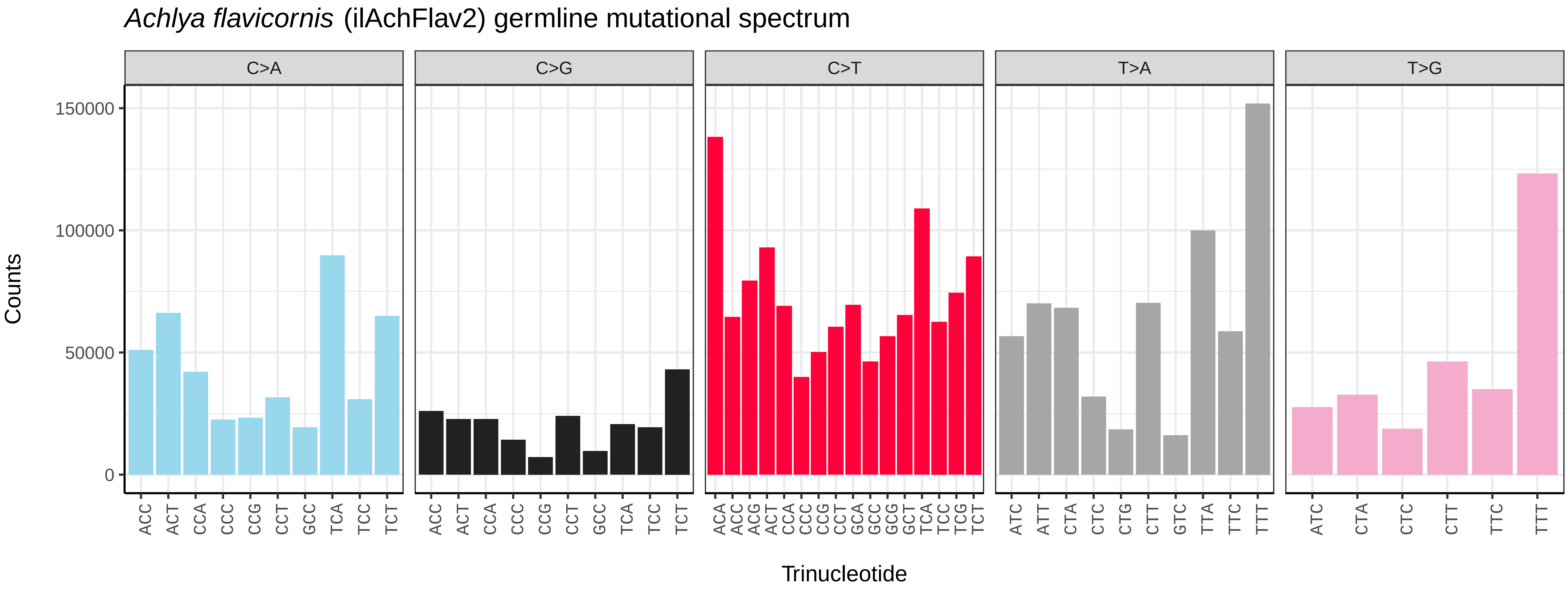
Achlya flavicornis (ilAchFlav2) germline mutational spectrum
C>A
C>G
C>T
T>A
T>G
Counts
150000
100000
50000
0
ACC ACT CCA CCC CCG CCT GCC TCA TCC TCT
ACA ACC ACG ACT CCA CCC CCG CCT GCA GCC GCG GCT TCA TCC TCG TCT
ATC ATT CTA CTC CTG CTT GTC TTA TTC TTT
ATC CTA CTC CTT TTC TTT
Trinucleotide

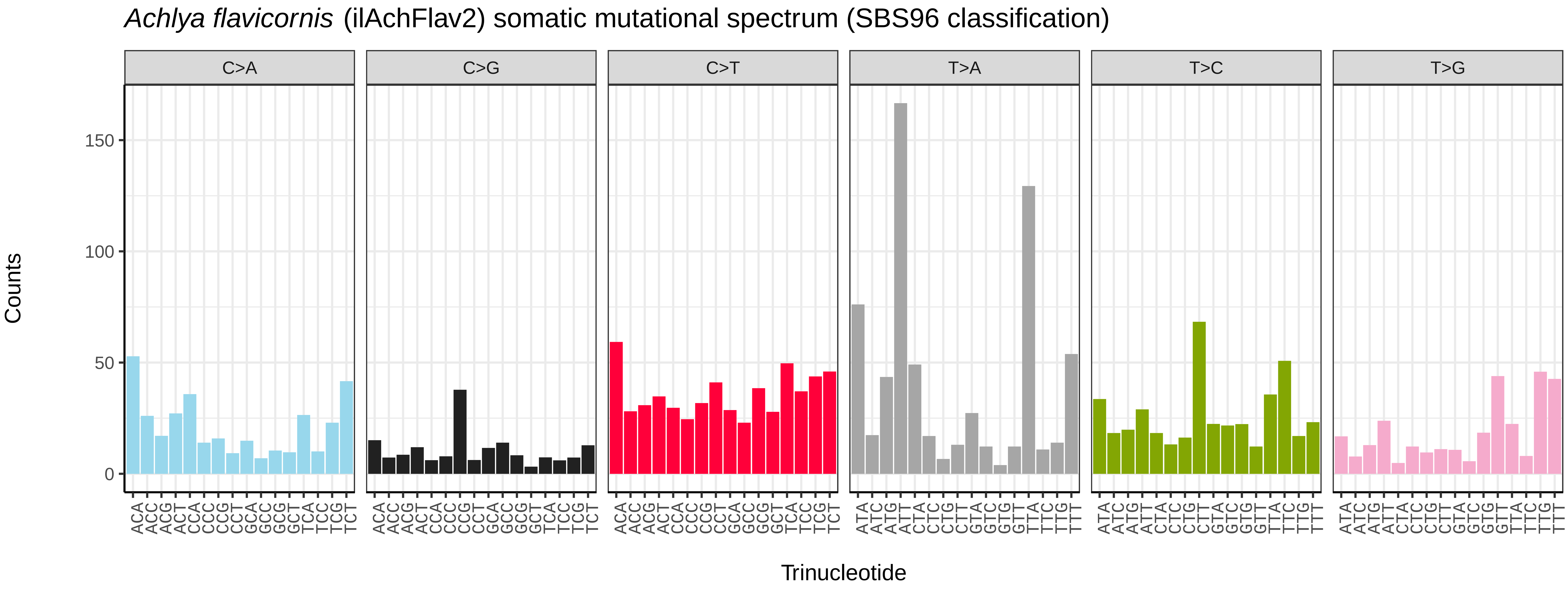
Achlya flavicornis (ilAchFlav2) somatic mutational spectrum (SBS96 classification)
C>A
C>G
C>T
T>A
T>C
T>G
Counts
150
100
50
0
ACA ACC ACG ACT CCA CCC CCG CCT GCA GCC GCG GCT TCA TCC TCG TCT
ATA ATC ATG ATT CTA CTC CTG CTT GTA GTC GTG GTT TTA TTC TTG TTT
Trinucleotide

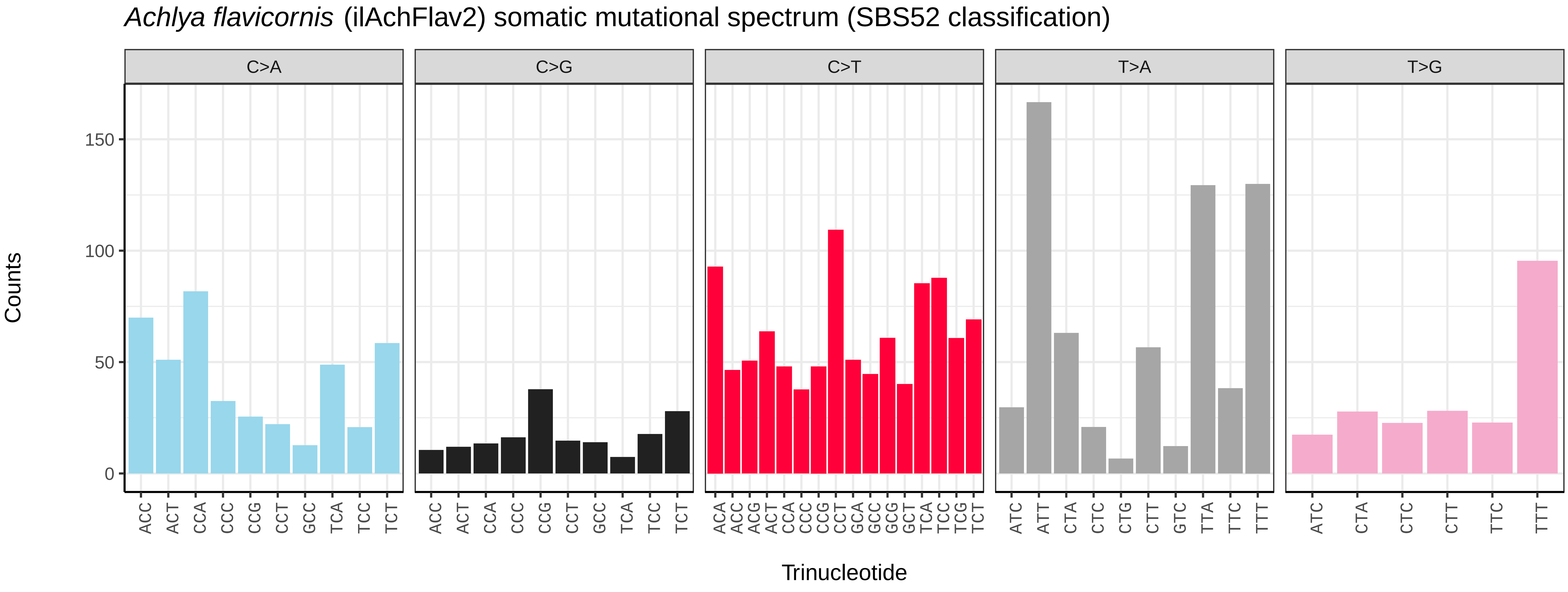
Achlya flavicornis (ilAchFlav2) somatic mutational spectrum (SBS52 classification)
C>A
C>G
C>T
T>A
T>G
Counts
150
100
50
0
ACC ACT CCA CCC CCG CCT GCC TCA TCC TCT
ACA ACC ACG ACT CCA CCC CCG CCT GCA GCC GCG GCT TCA TCC TCG TCT
ATC ATT CTA CTC CTG CTT GTC TTA TTC TTT
ATC CTA CTC CTT TTC TTT
Trinucleotide

*Acleris cristana* (ilAclCris2) germline mutational spectrum

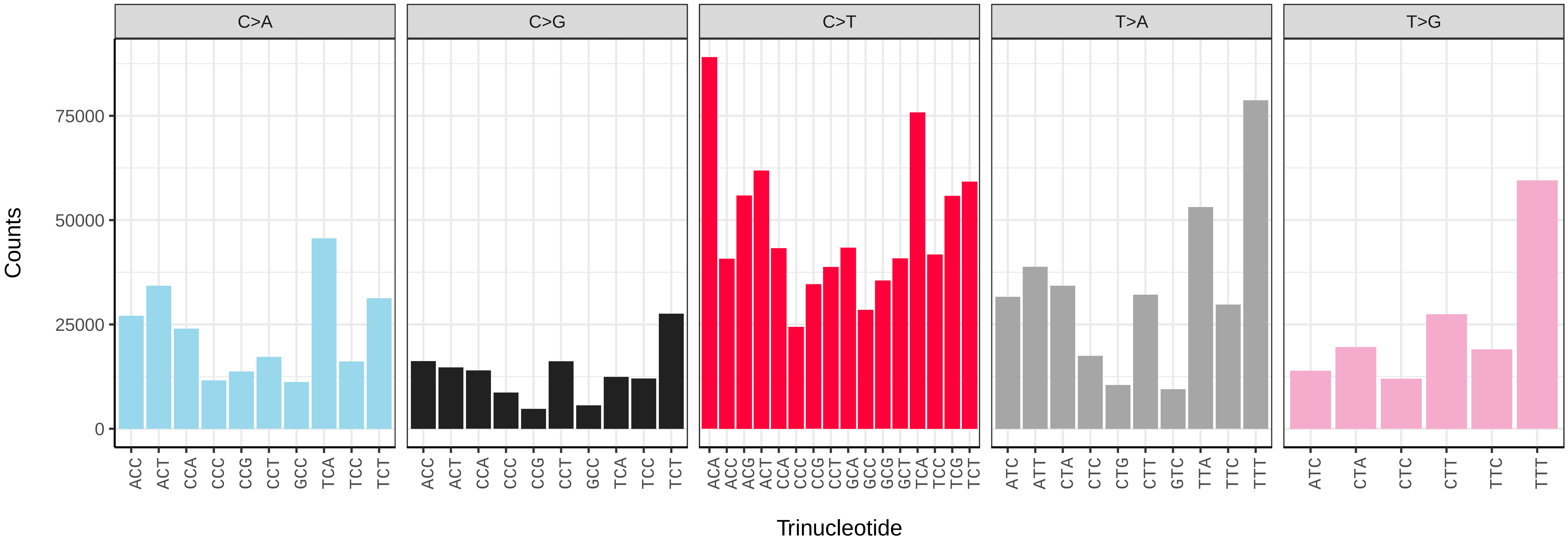


*Acleris cristana* (ilAclCris2) somatic mutational spectrum (SBS96 classification)

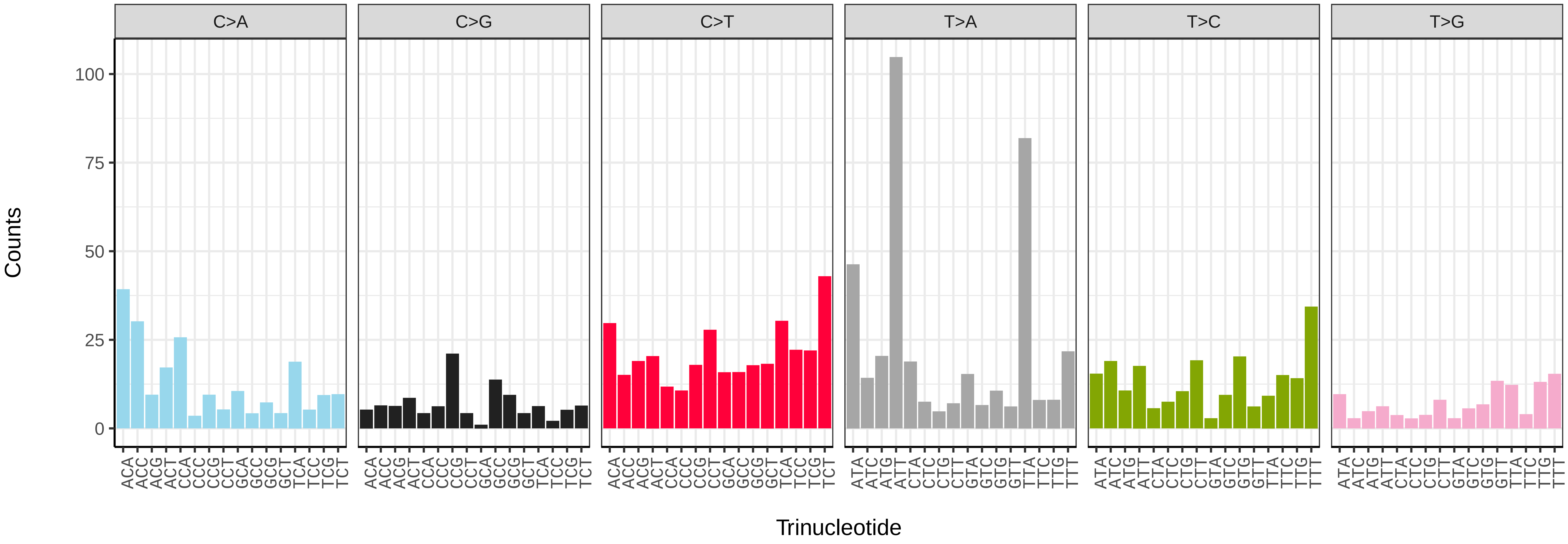


*Acleris cristana* (ilAclCris2) somatic mutational spectrum (SBS52 classification)

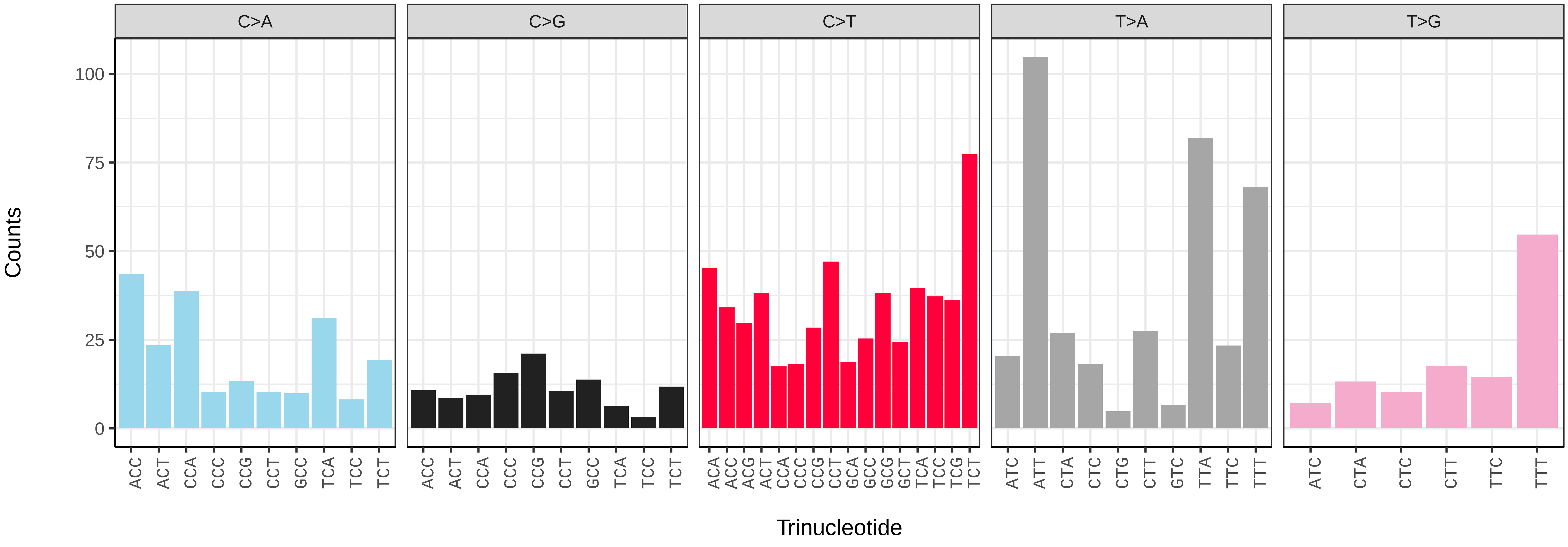

*Acleris emargana* (ilAclEmar1) germline mutational spectrum

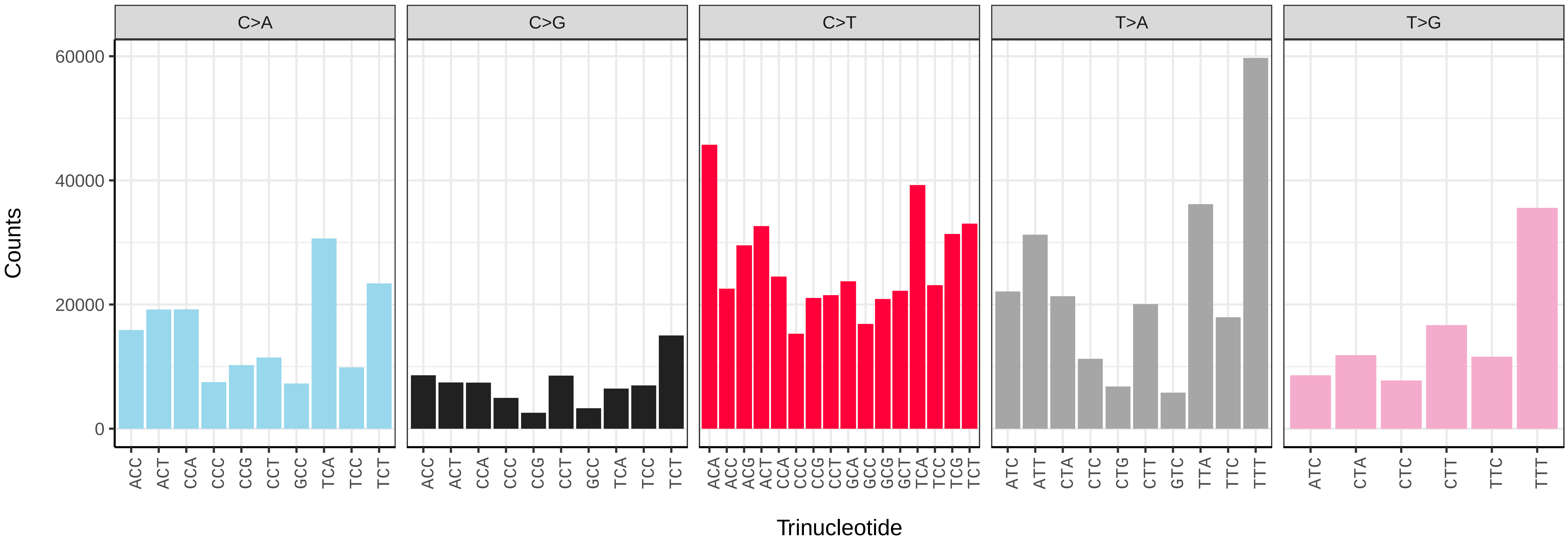


*Acleris emargana* (ilAclEmar1) somatic mutational spectrum (SBS96 classification)

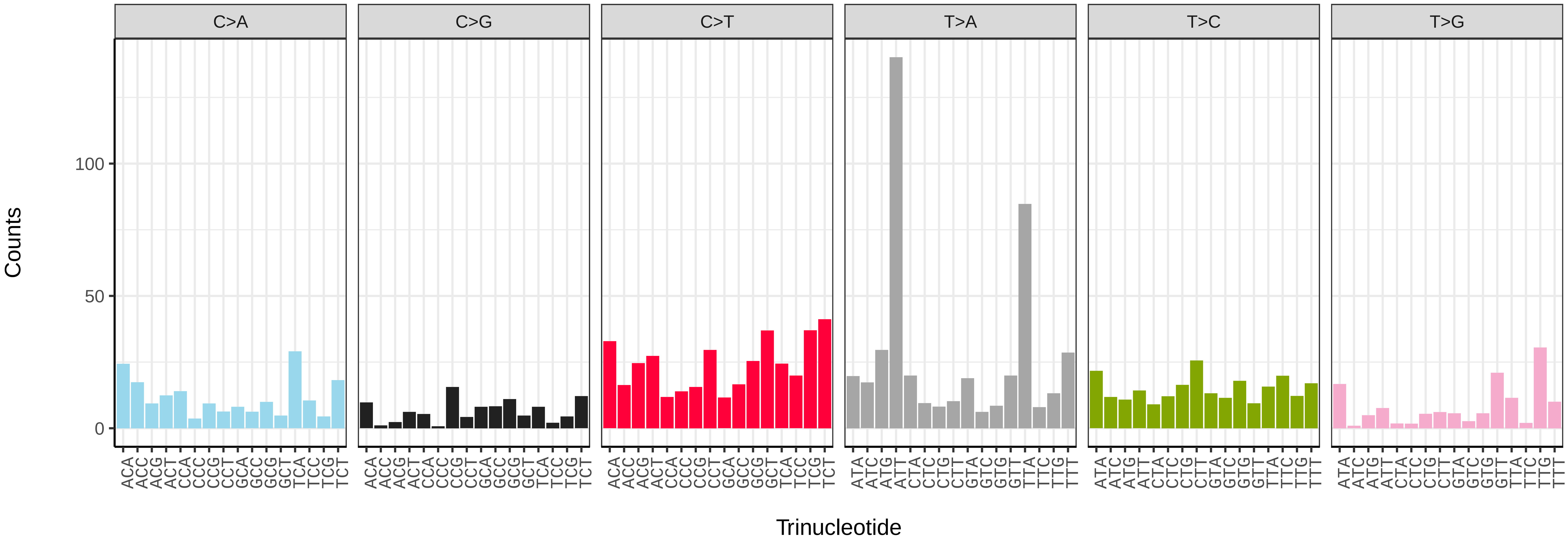


*Acleris emargana* (ilAclEmar1) somatic mutational spectrum (SBS52 classification)

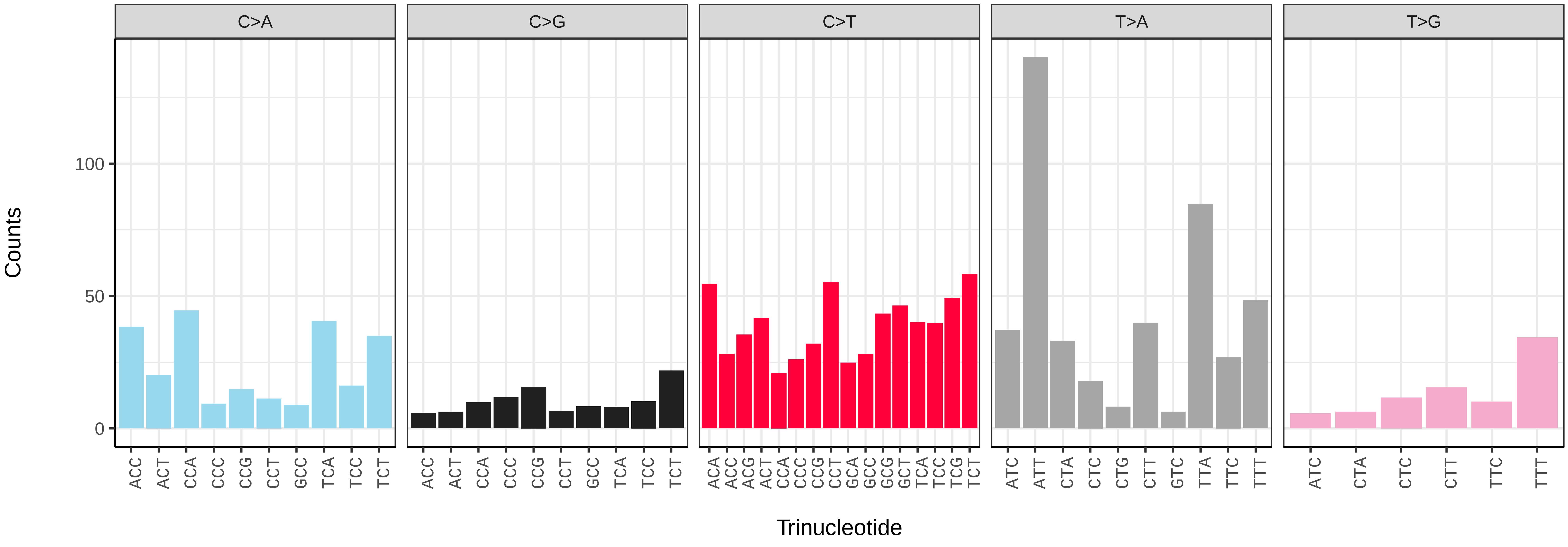

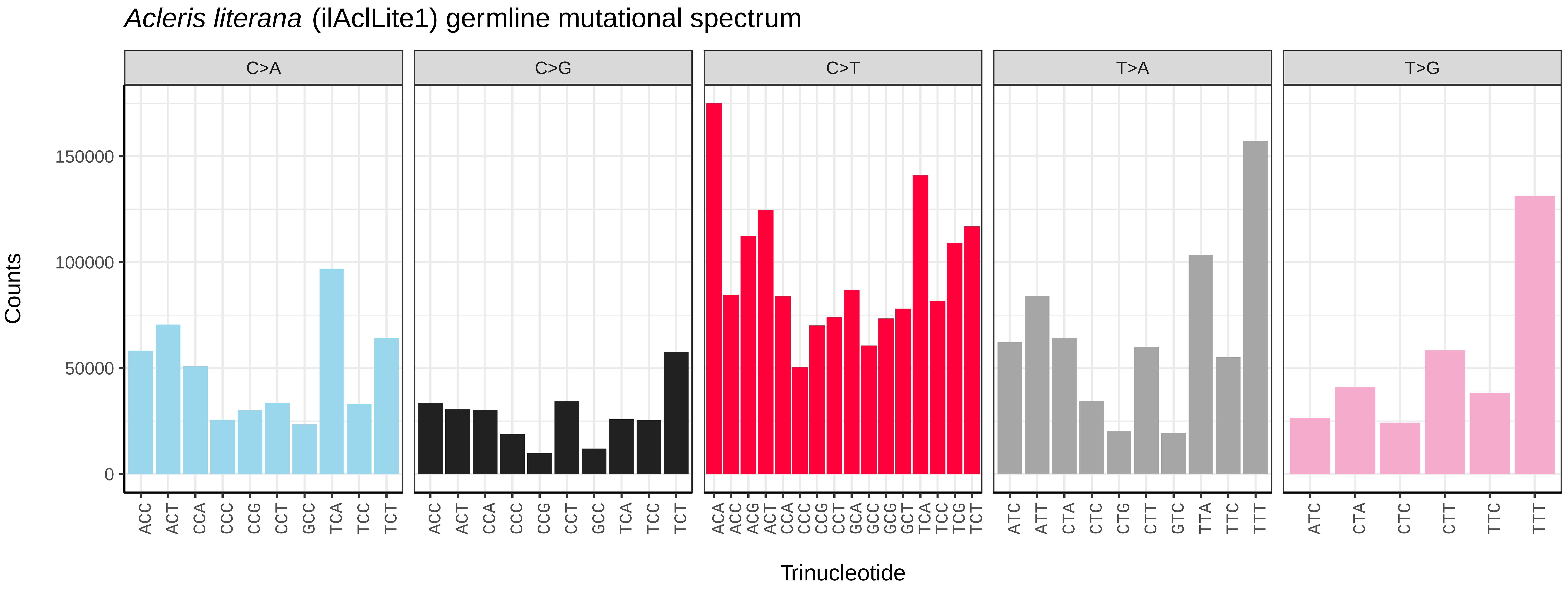
*Acleris literana* (ilAclLite1) germline mutational spectrum
C>A
C>G
C>T
T>A
T>G
Counts
0
50000
100000
150000
Trinucleotide

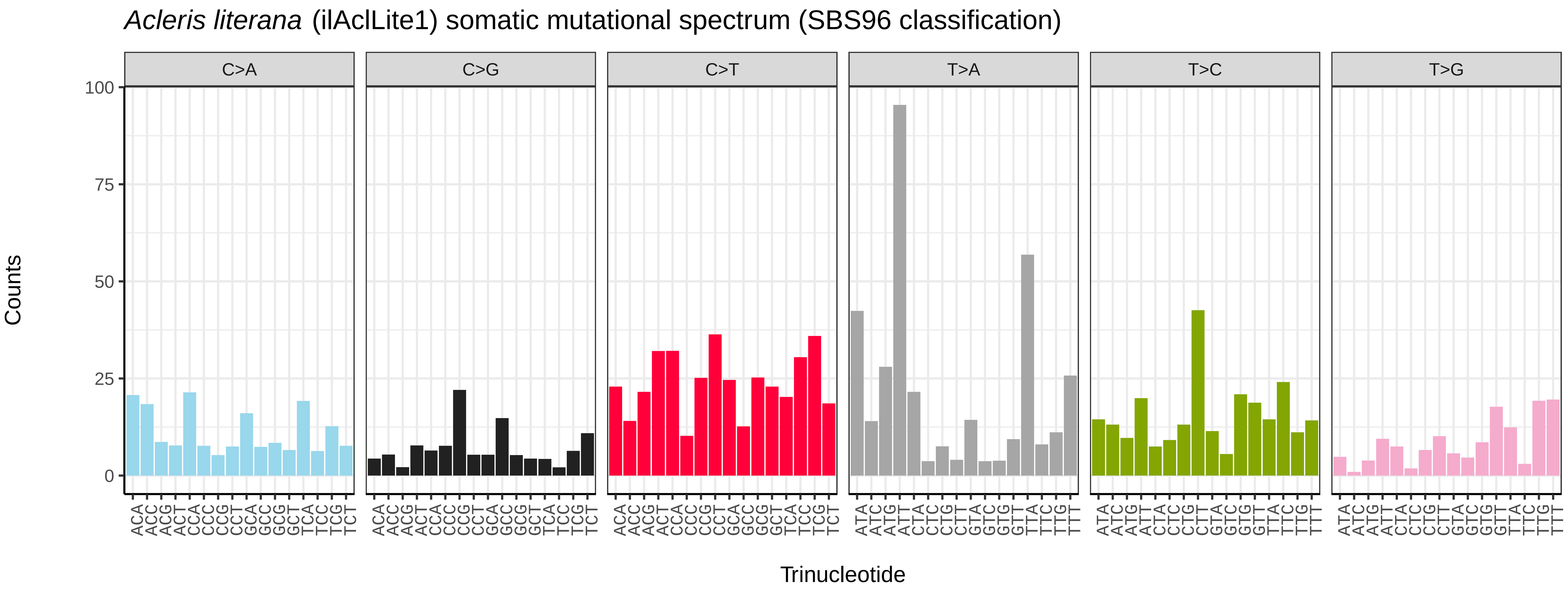
*Acleris literana* (ilAclLite1) somatic mutational spectrum (SBS96 classification)
C>A
C>G
C>T
T>A
T>C
T>G
Counts
0
25
50
75
100
Trinucleotide

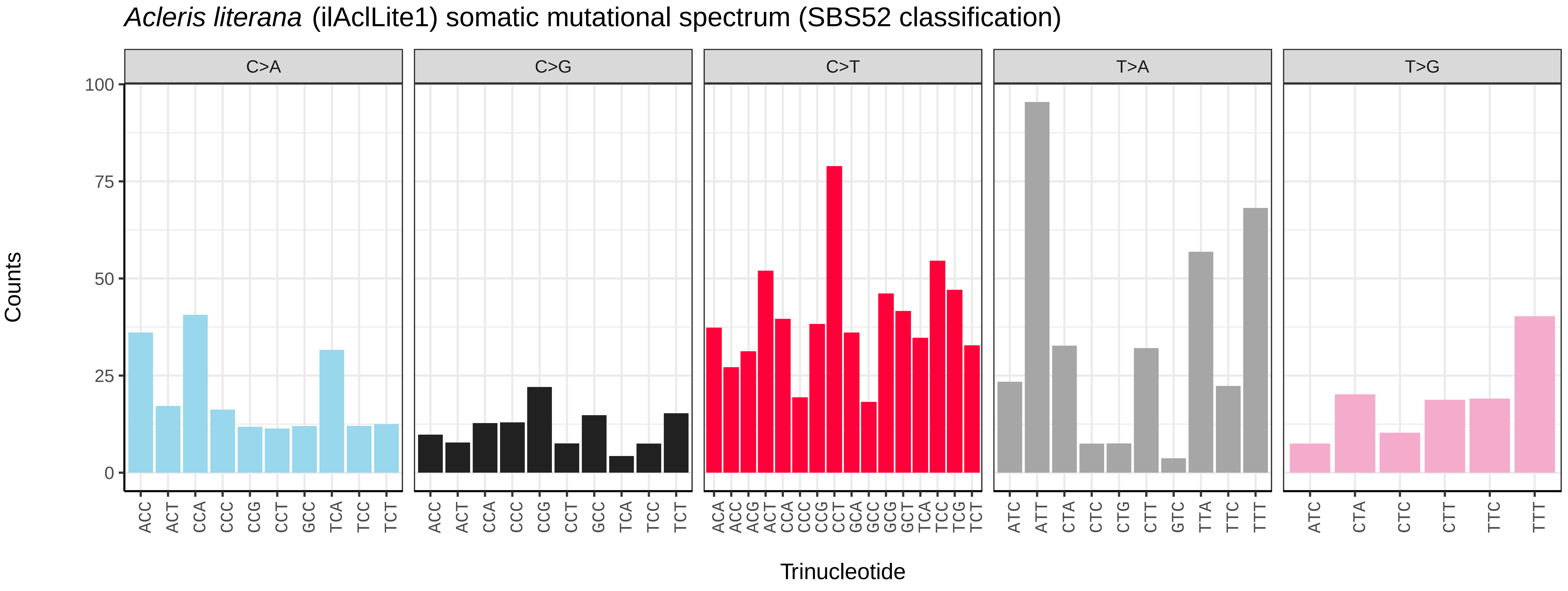
*Acleris literana* (ilAclLite1) somatic mutational spectrum (SBS52 classification)
C>A
C>G
C>T
T>A
T>G
Counts
0
25
50
75
100
Trinucleotide

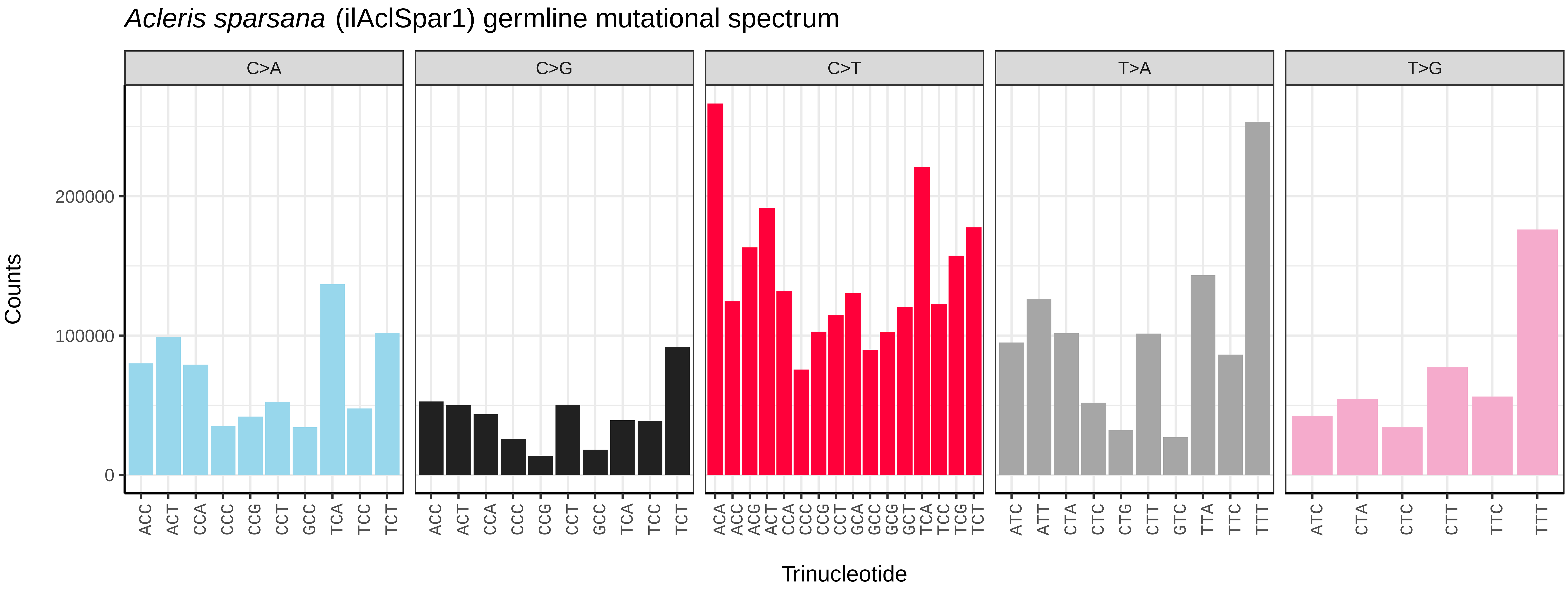
Acleris sparsana (ilAclSpar1) germline mutational spectrum
C>A
C>G
C>T
T>A
T>G
Counts
0
100000
200000
Trinucleotide

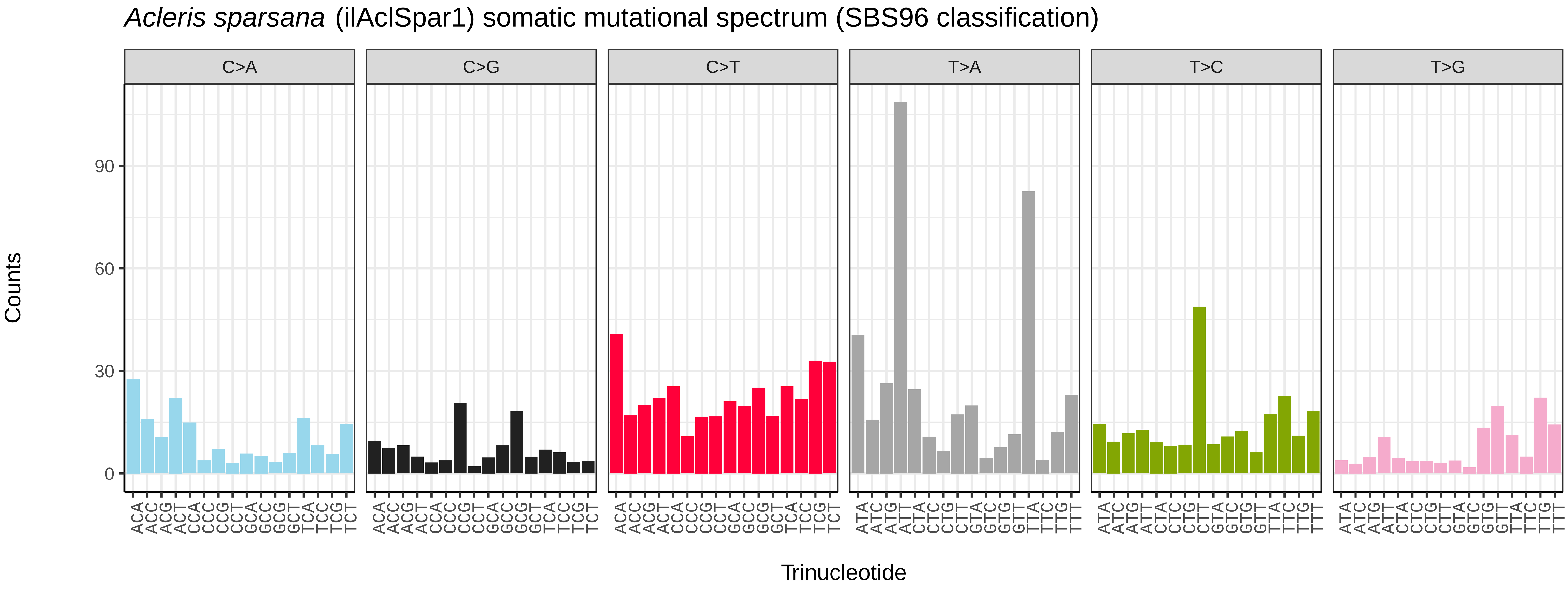
Acleris sparsana (ilAclSpar1) somatic mutational spectrum (SBS96 classification)
C>A
C>G
C>T
T>A
T>C
T>G
Counts
0
30
60
90
Trinucleotide

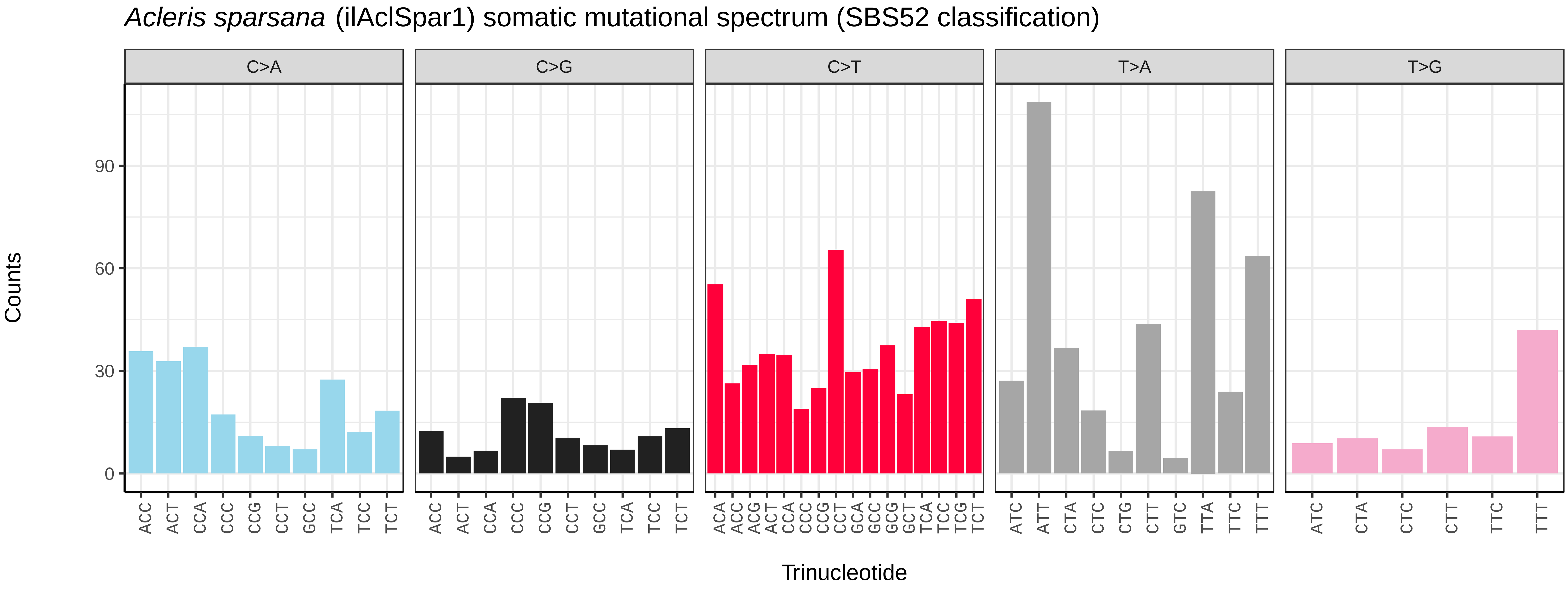
Acleris sparsana (ilAclSpar1) somatic mutational spectrum (SBS52 classification)
C>A
C>G
C>T
T>A
T>G
Counts
0
30
60
90
Trinucleotide

*Acronicta aceris* (ilAcrAcer1) germline mutational spectrum

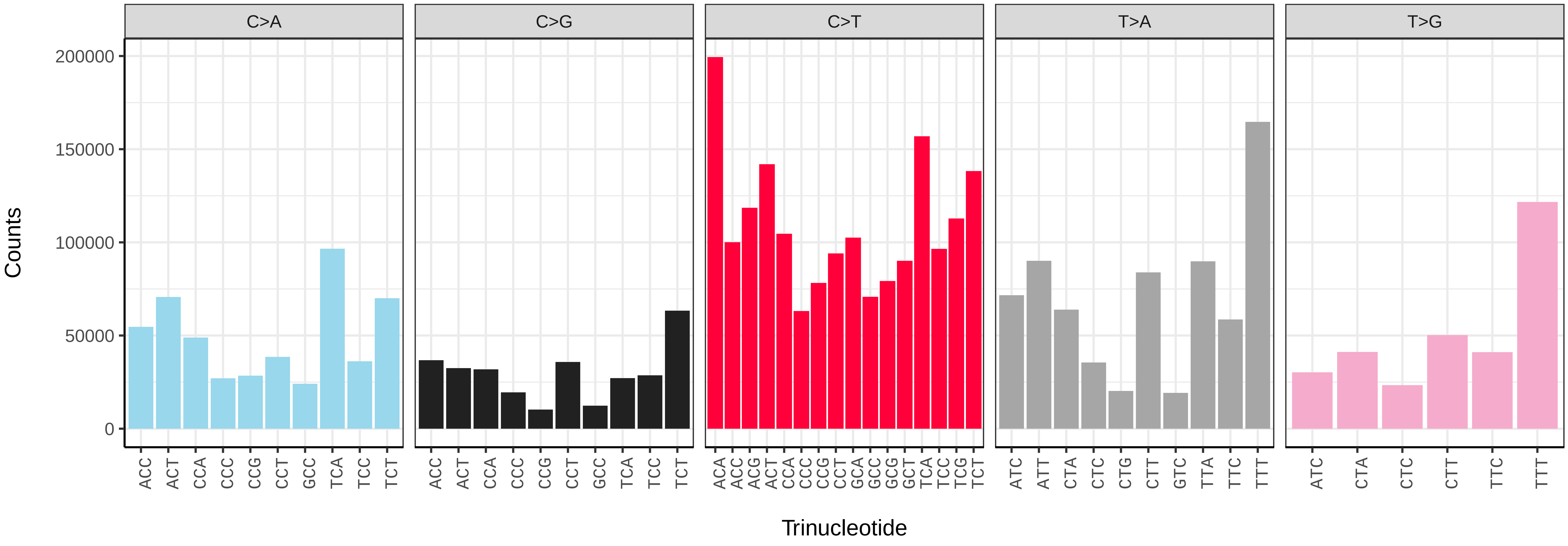


*Acronicta aceris* (ilAcrAcer1) somatic mutational spectrum (SBS96 classification)

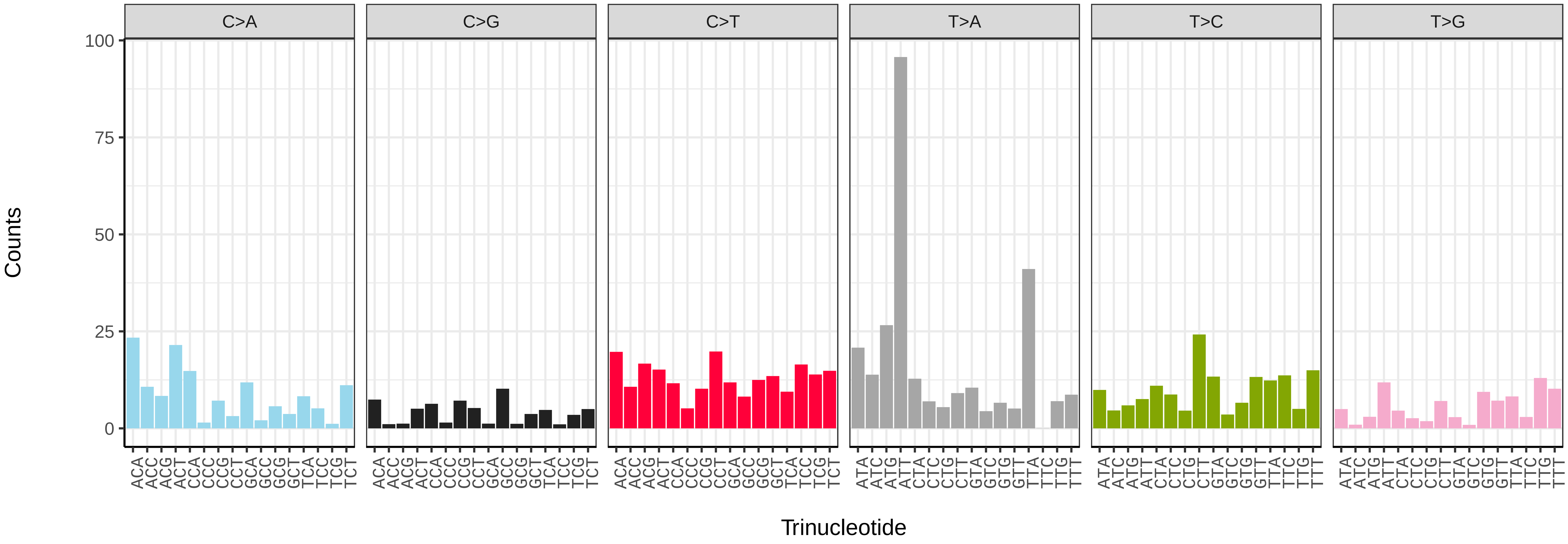


*Acronicta aceris* (ilAcrAcer1) somatic mutational spectrum (SBS52 classification)

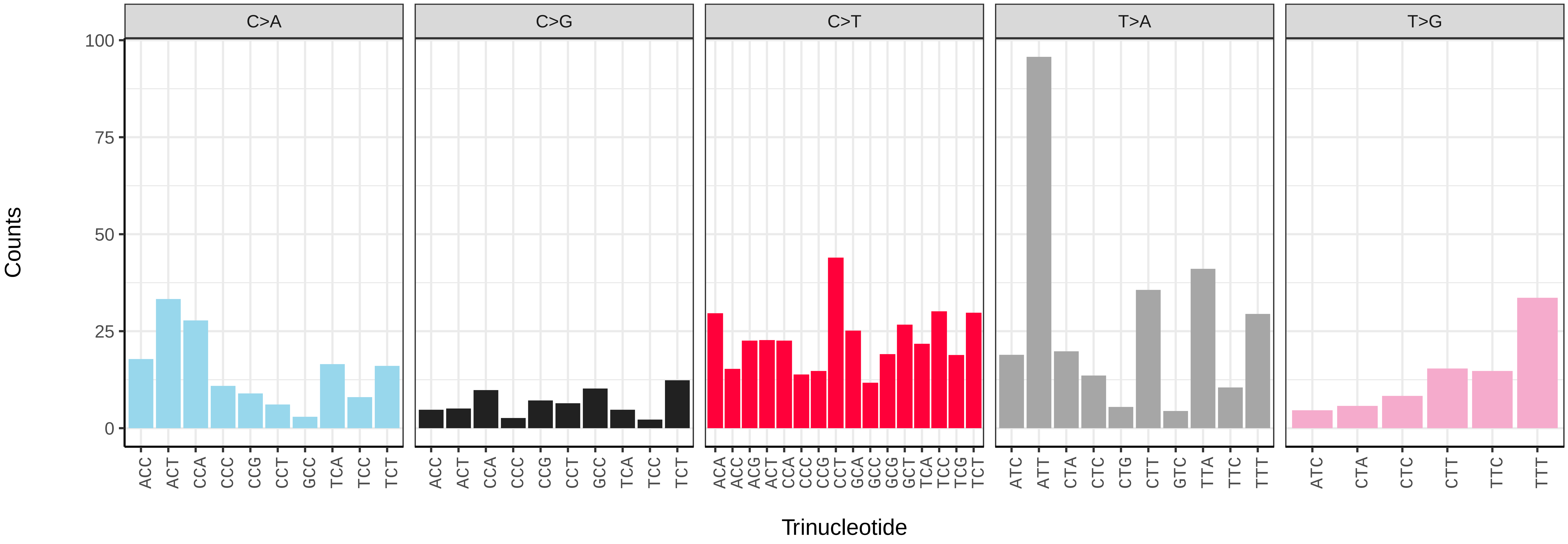

*Acronicta leporina* (ilAcrLepo1) germline mutational spectrum

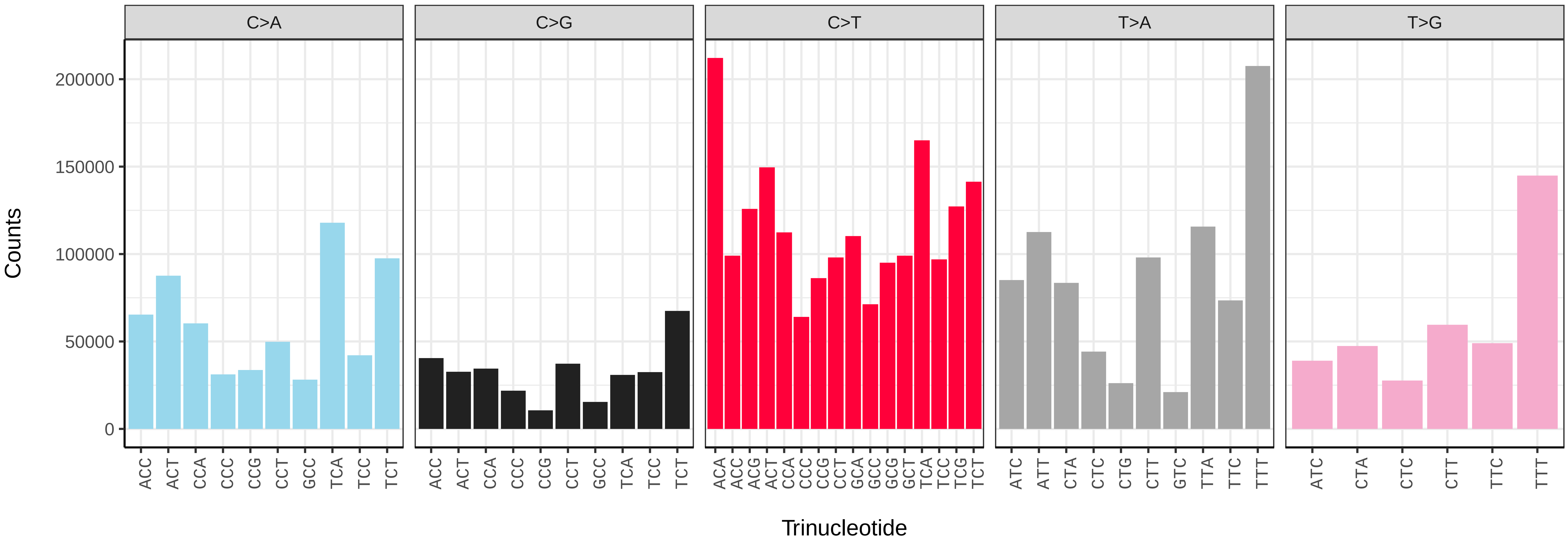


*Acronicta leporina* (ilAcrLepo1) somatic mutational spectrum (SBS96 classification)

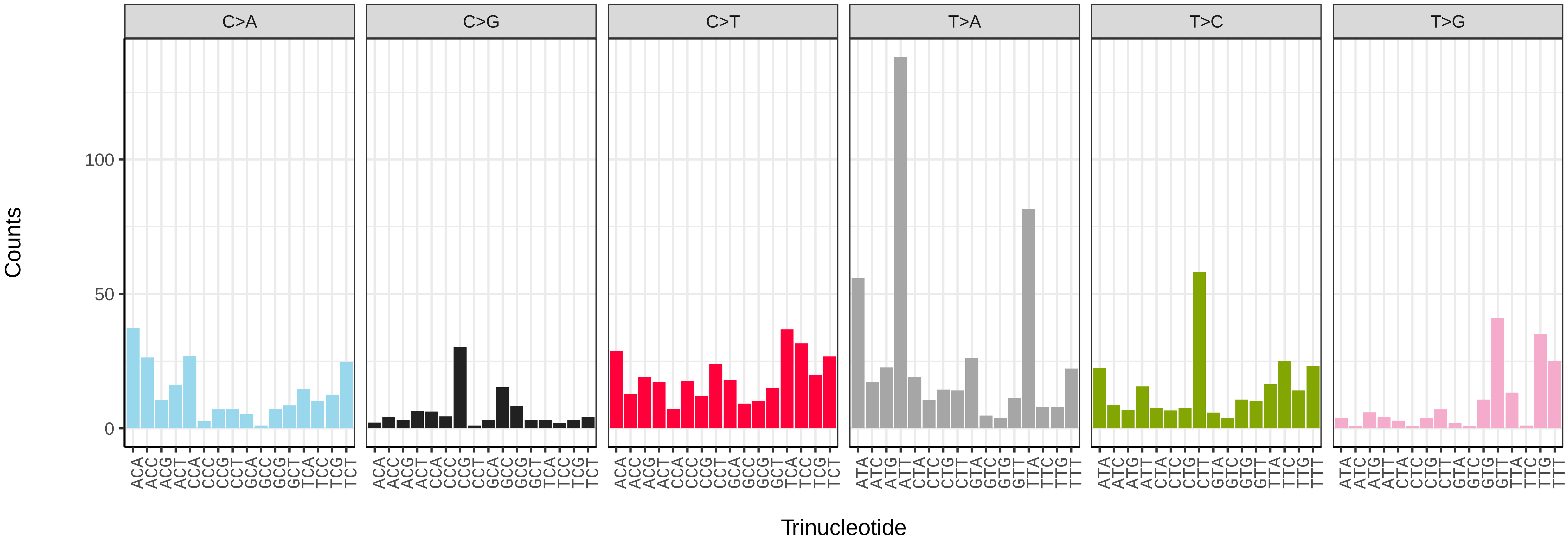


*Acronicta leporina* (ilAcrLepo1) somatic mutational spectrum (SBS52 classification)

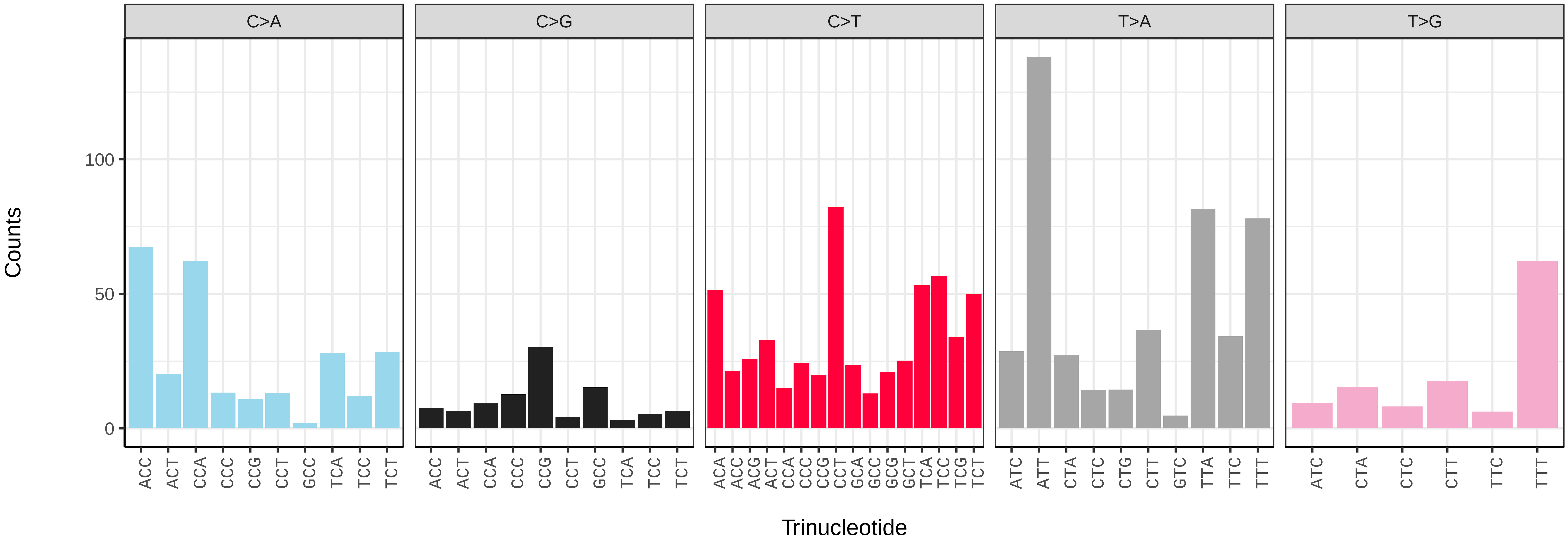

*Acronicta psi* (ilAcrPsix1) germline mutational spectrum

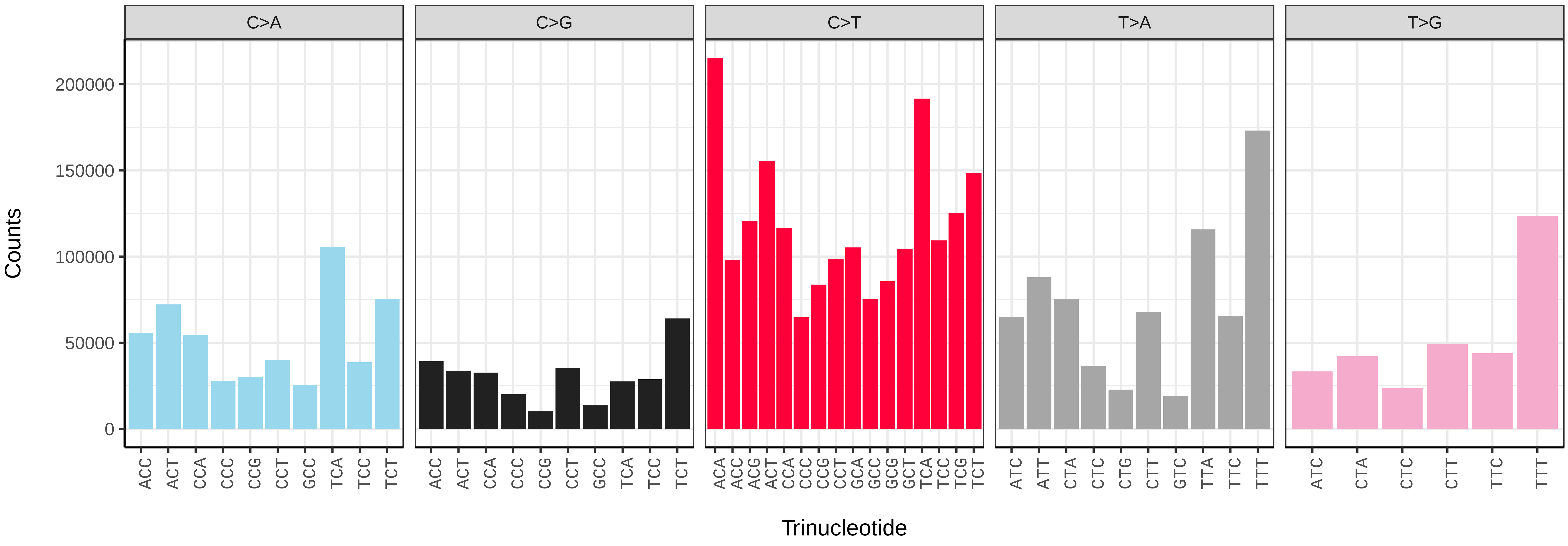


*Acronicta psi* (ilAcrPsix1) somatic mutational spectrum (SBS96 classification)

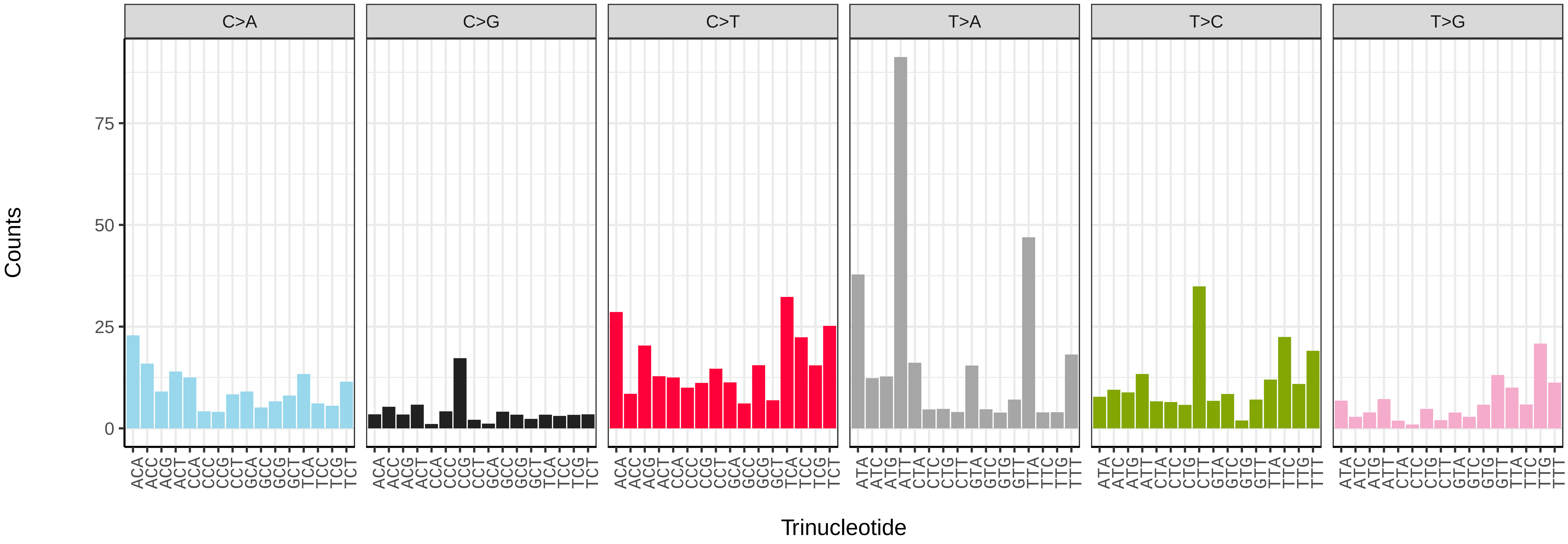


*Acronicta psi* (ilAcrPsix1) somatic mutational spectrum (SBS52 classification)

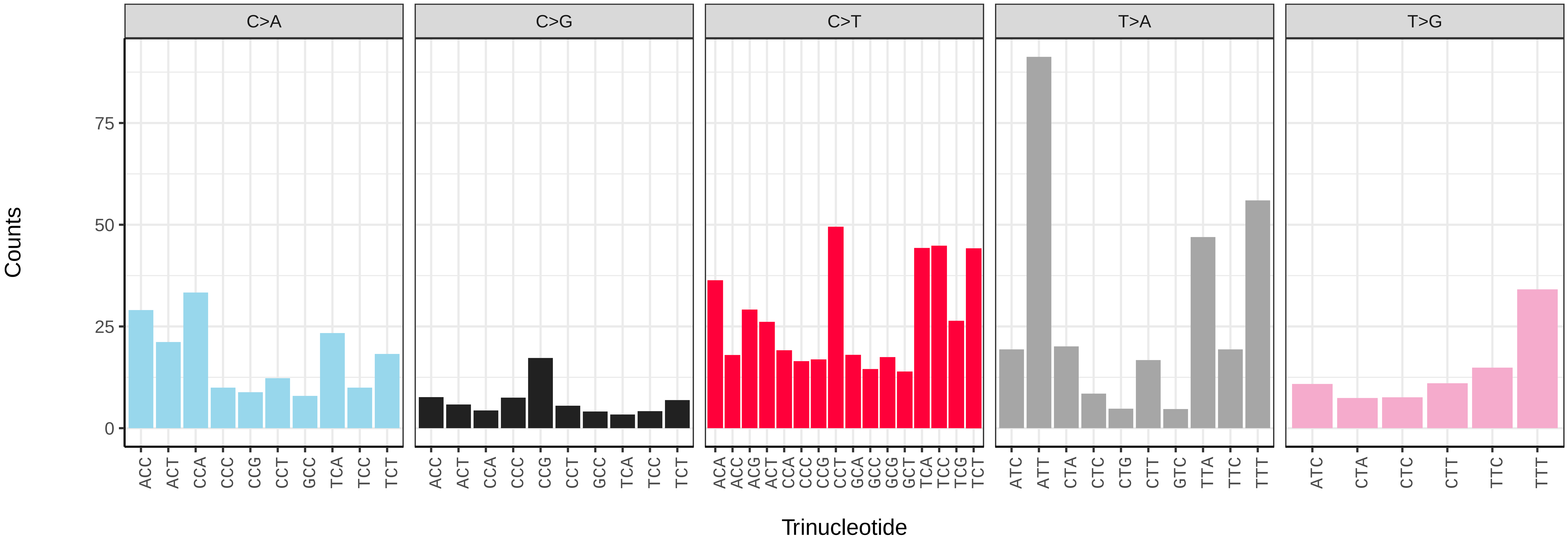

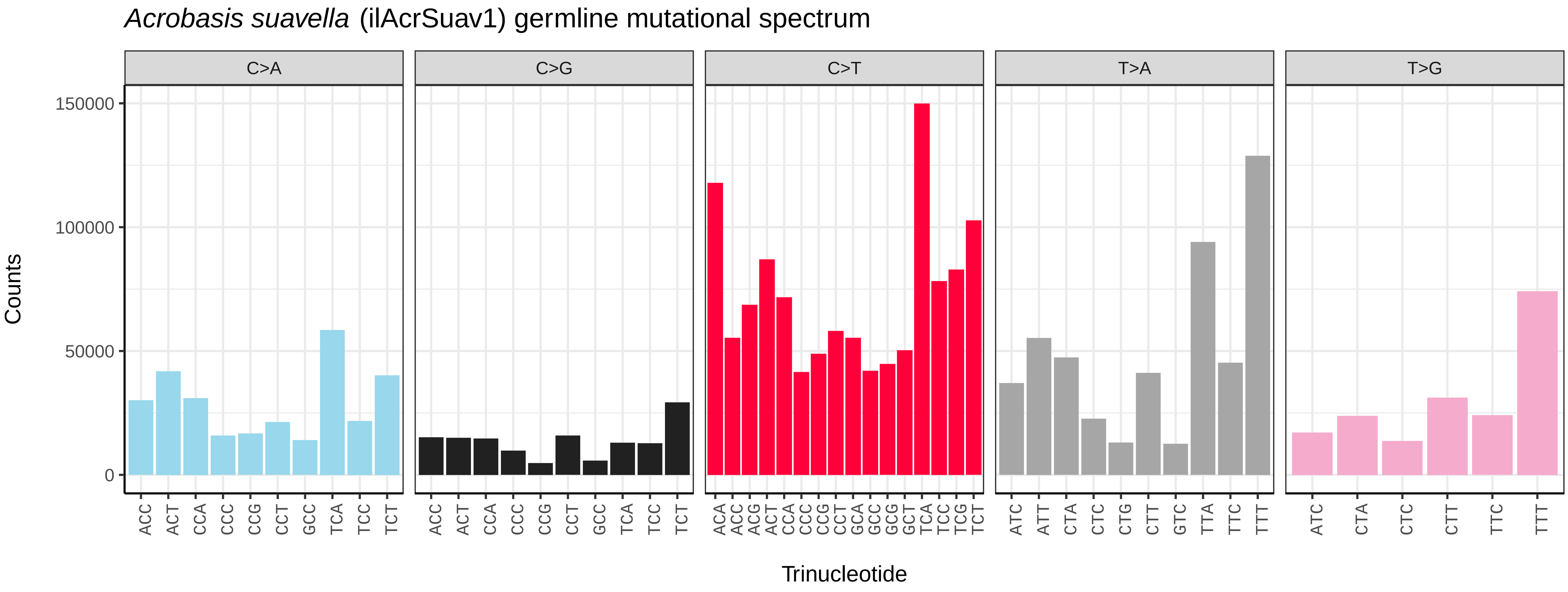

Acrobasis suavella (ilAcrSuav1) germline mutational spectrum
C>A
C>G
C>T
T>A
T>G
Counts
150000
100000
50000
0
Trinucleotide


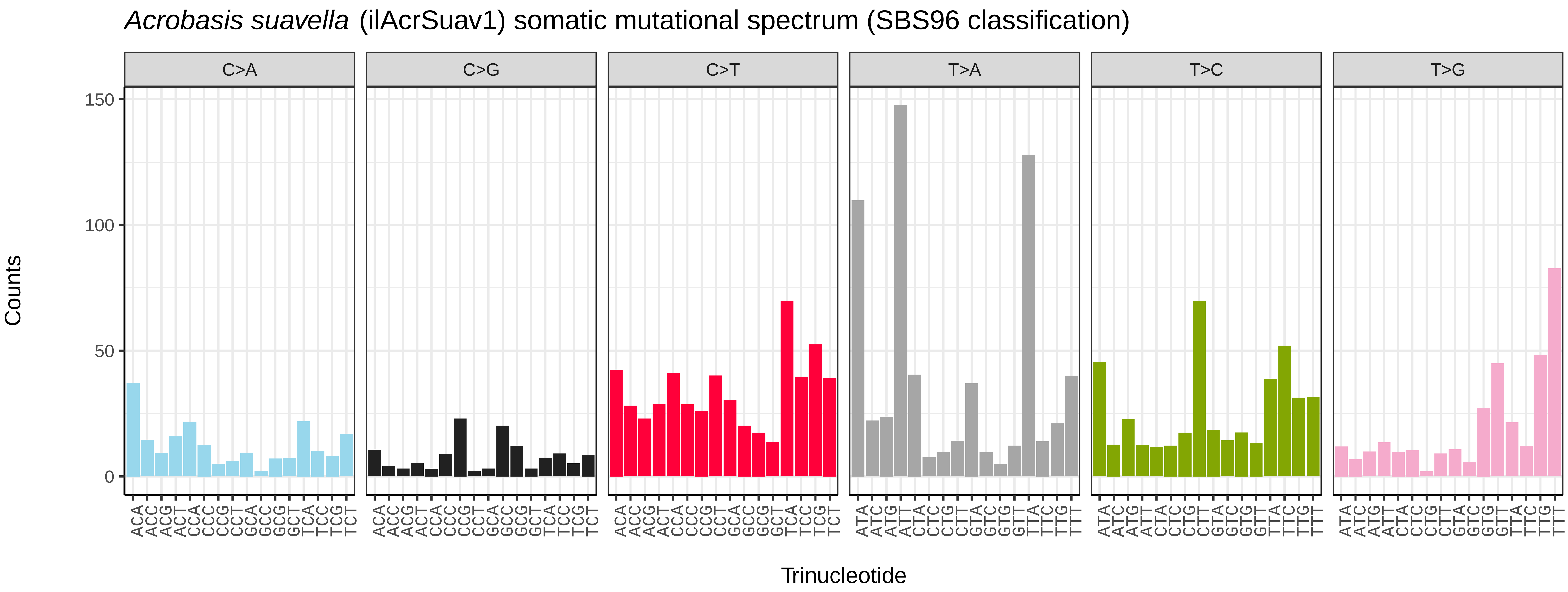

Acrobasis suavella (ilAcrSuav1) somatic mutational spectrum (SBS96 classification)
C>A
C>G
C>T
T>A
T>C
T>G
Counts
150
100
50
0
Trinucleotide


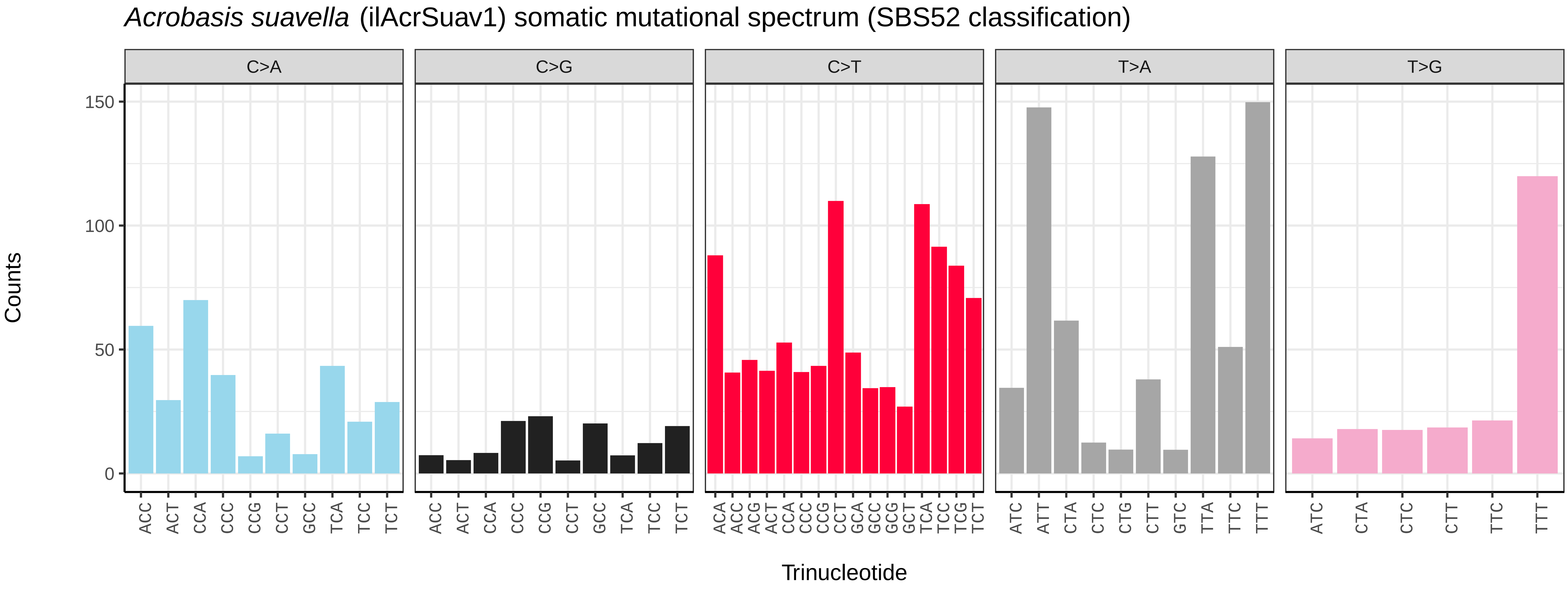

Acrobasis suavella (ilAcrSuav1) somatic mutational spectrum (SBS52 classification)
C>A
C>G
C>T
T>A
T>G
Counts
150
100
50
0
Trinucleotide

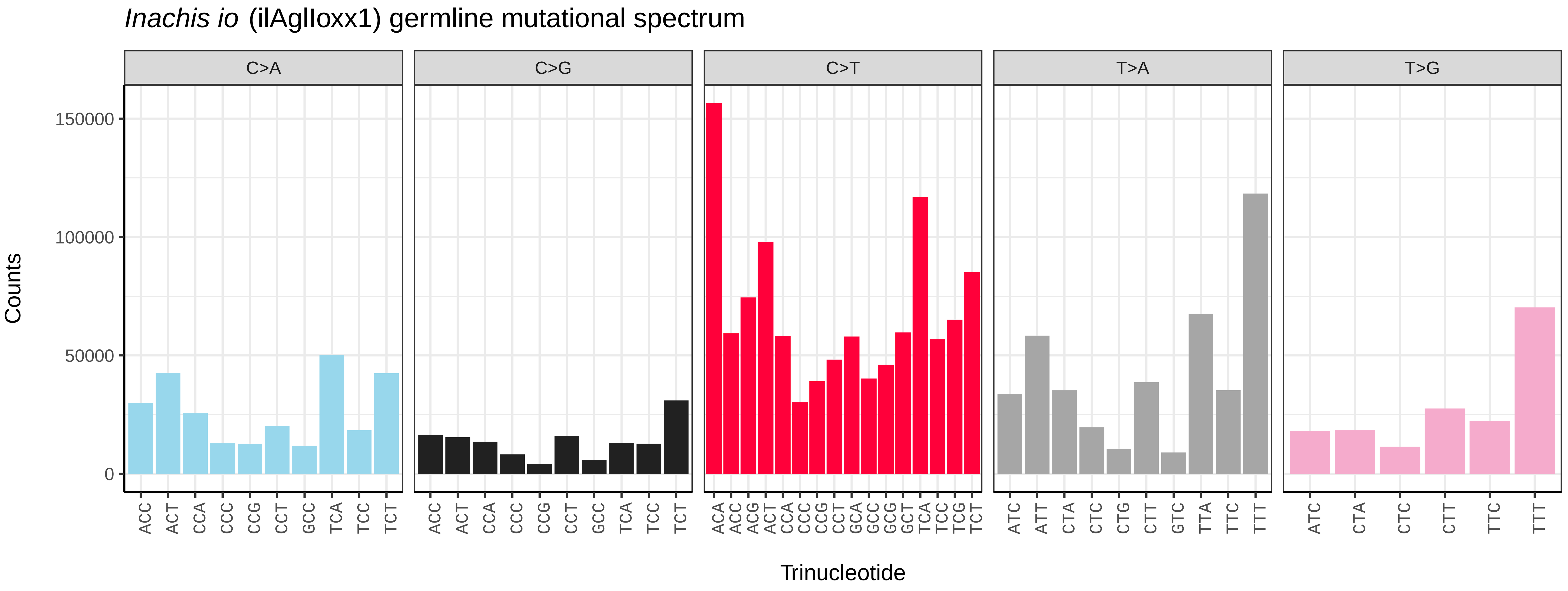
*Inachis io* (ilAglIoxx1) germline mutational spectrum
C>A
C>G
C>T
T>A
T>G
150000
100000
50000
0
Counts
Trinucleotide

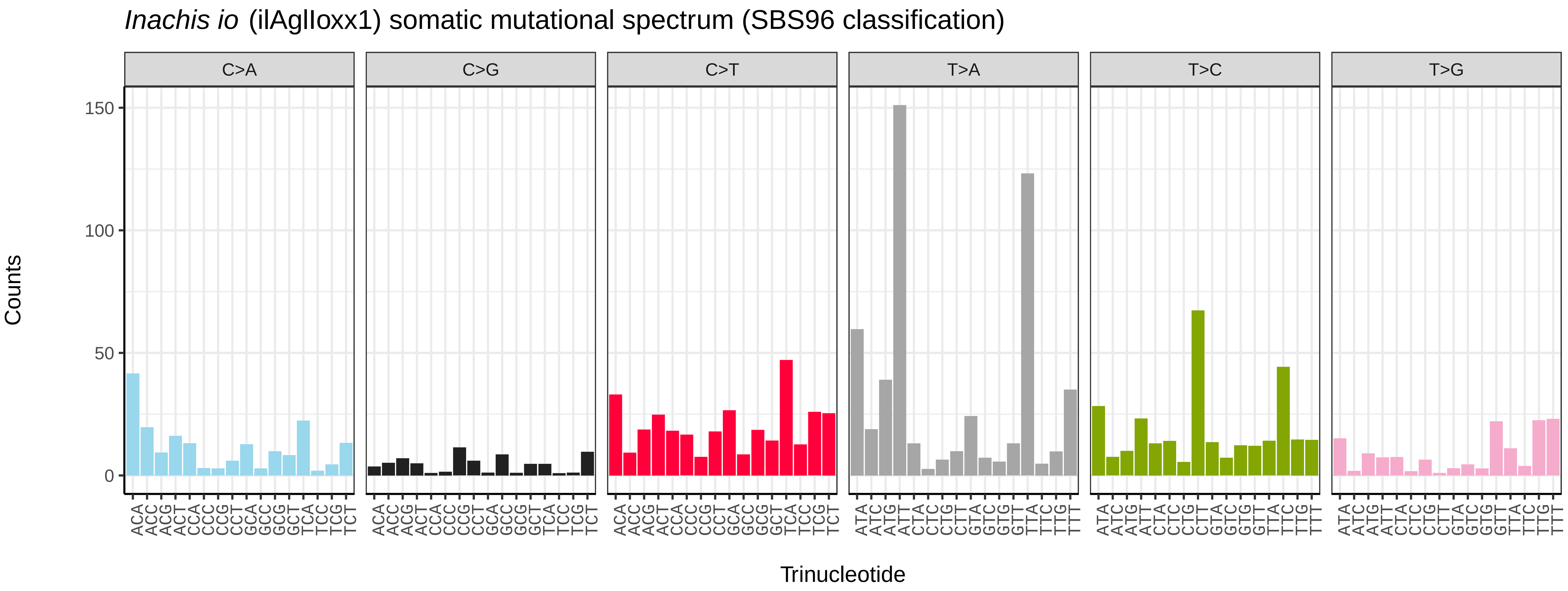
*Inachis io* (ilAglIoxx1) somatic mutational spectrum (SBS96 classification)
C>A
C>G
C>T
T>A
T>C
T>G
150
100
50
0
Counts
Trinucleotide

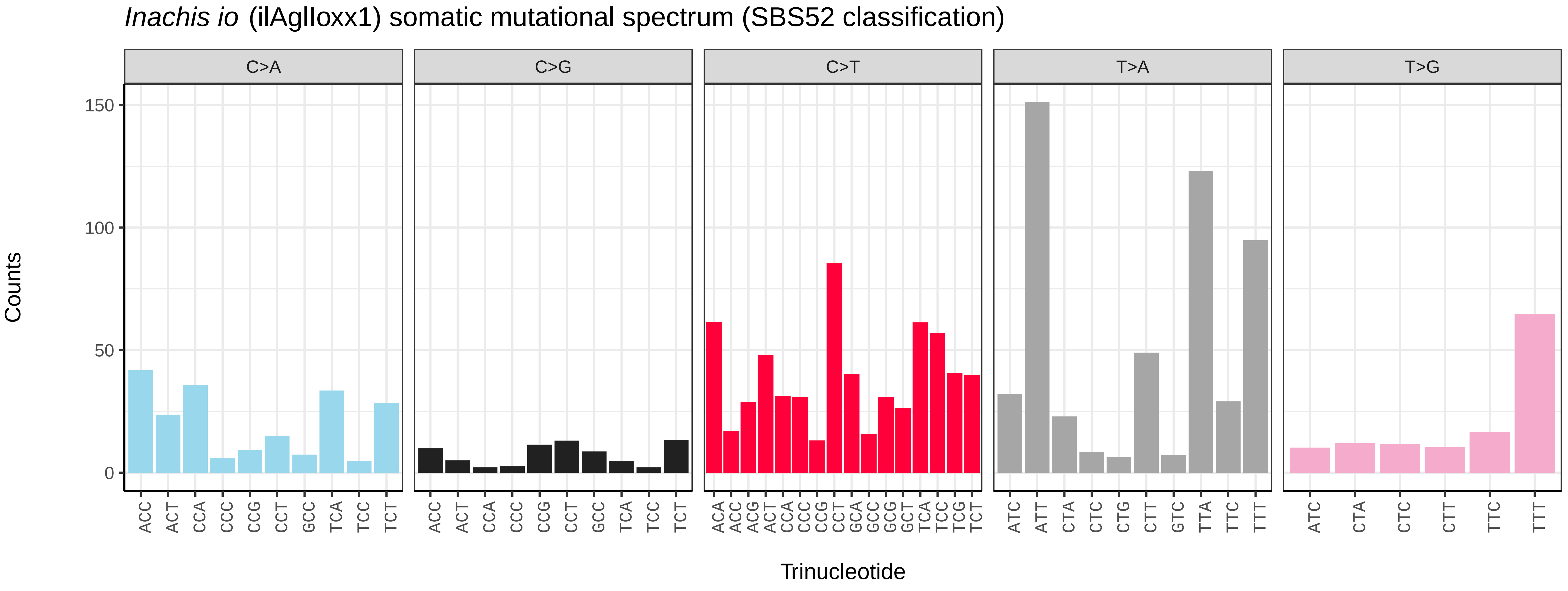
*Inachis io* (ilAglIoxx1) somatic mutational spectrum (SBS52 classification)
C>A
C>G
C>T
T>A
T>G
150
100
50
0
Counts
Trinucleotide

*Aglais urticae* (ilAglUrti1) germline mutational spectrum

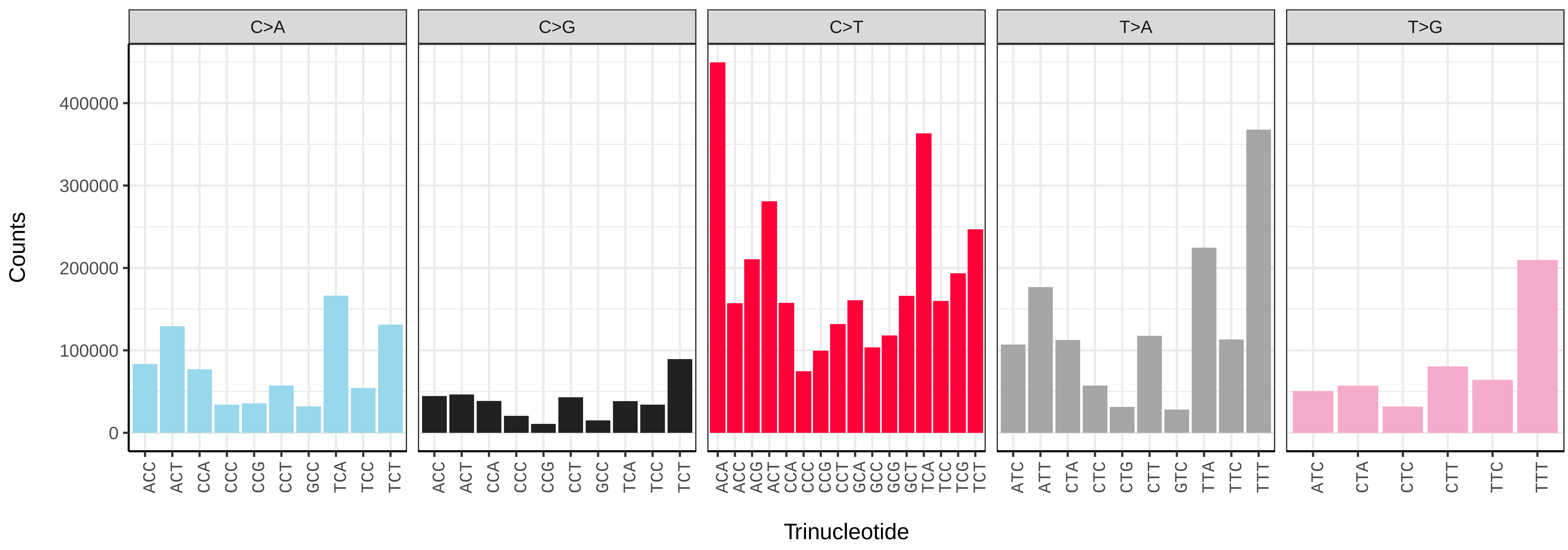


*Aglais urticae* (ilAglUrti1) somatic mutational spectrum (SBS96 classification)

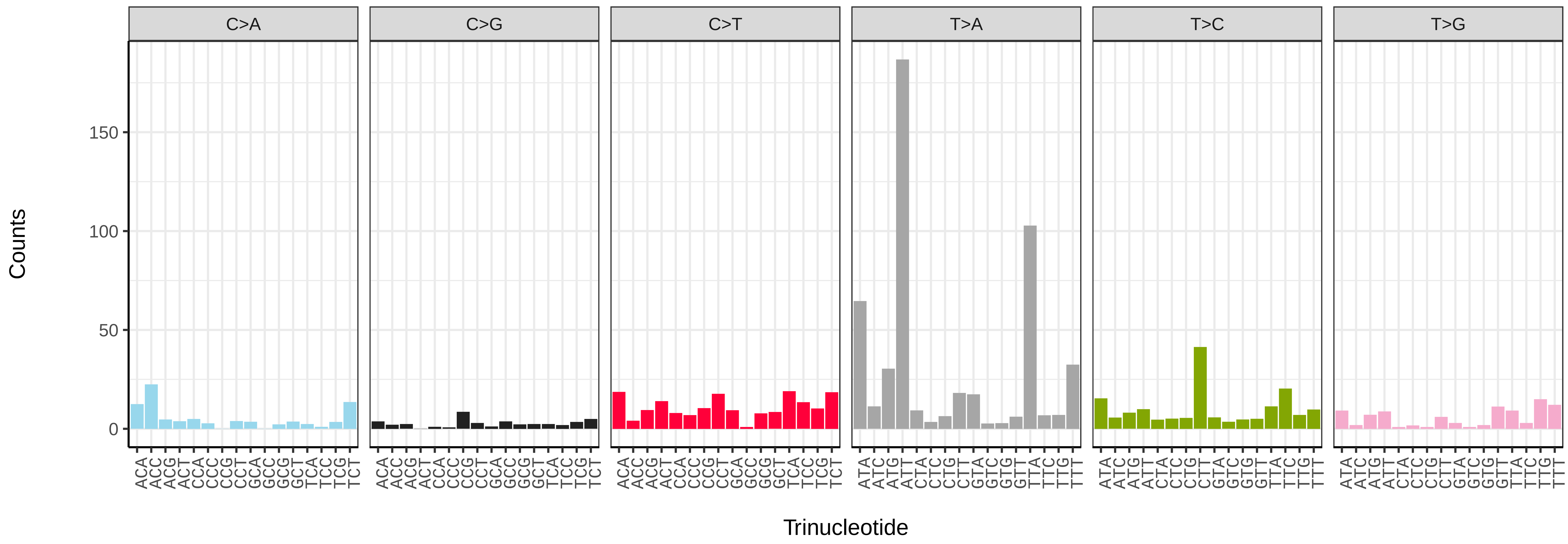


*Aglais urticae* (ilAglUrti1) somatic mutational spectrum (SBS52 classification)

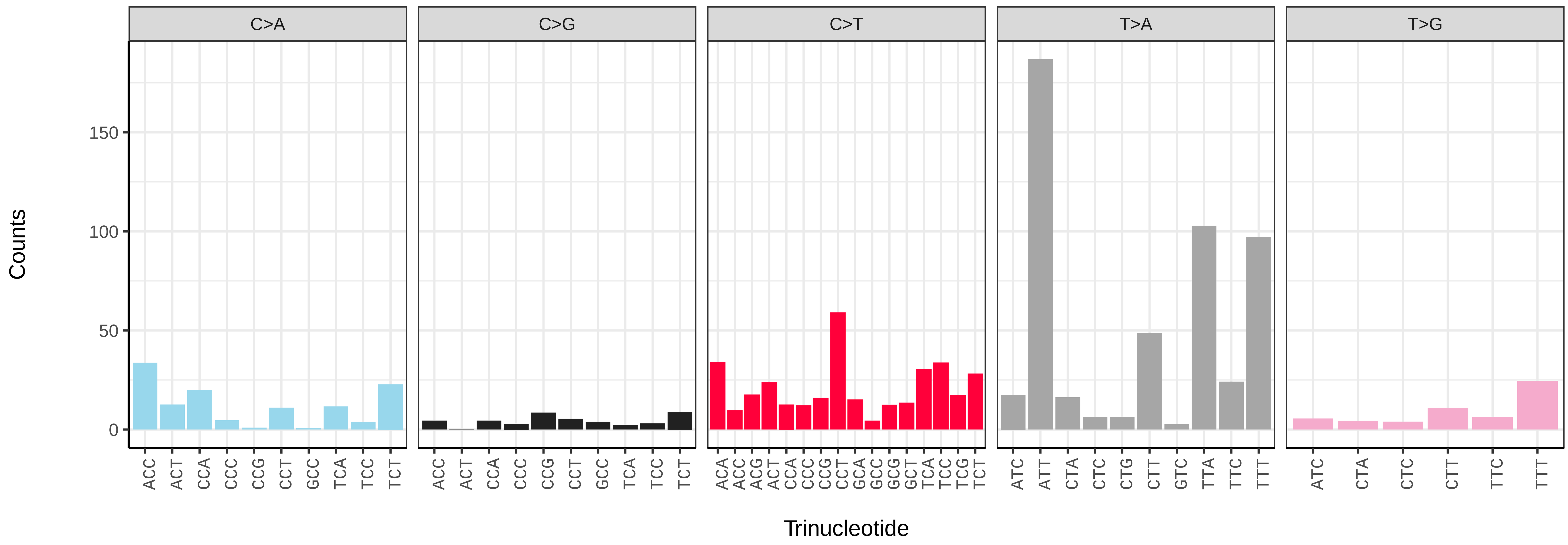

*Agonopterix arenella* (ilAgoAren1) germline mutational spectrum

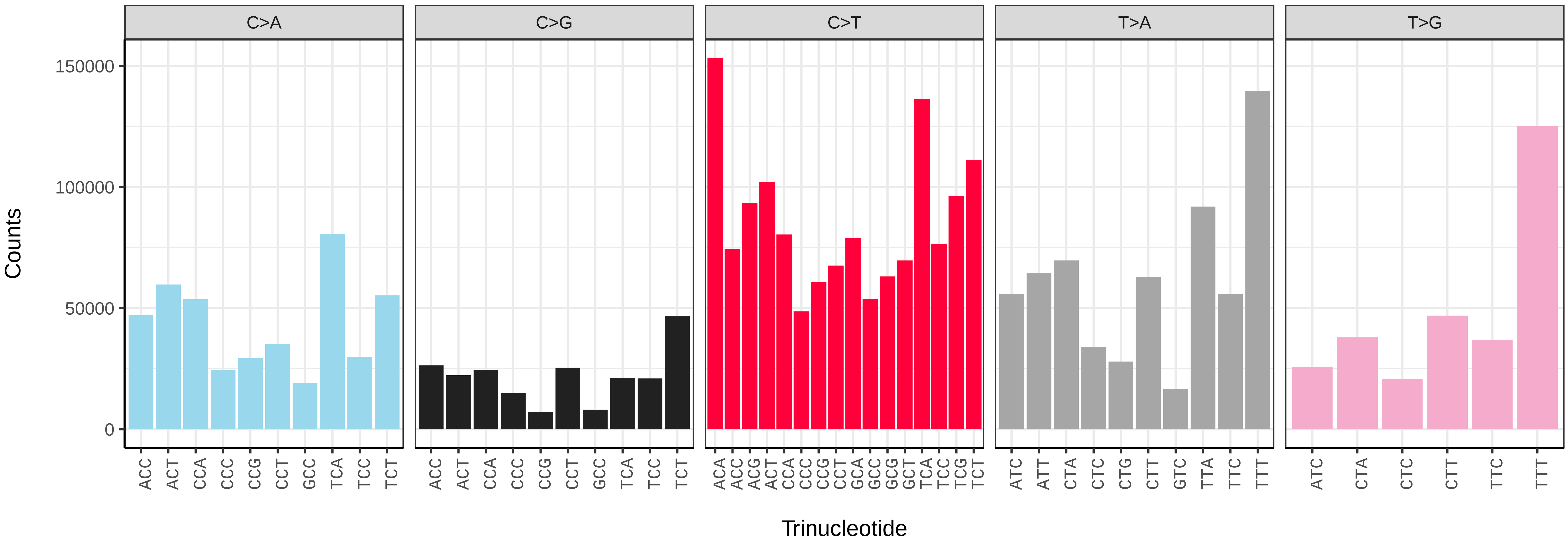


*Agonopterix arenella* (ilAgoAren1) somatic mutational spectrum (SBS96 classification)

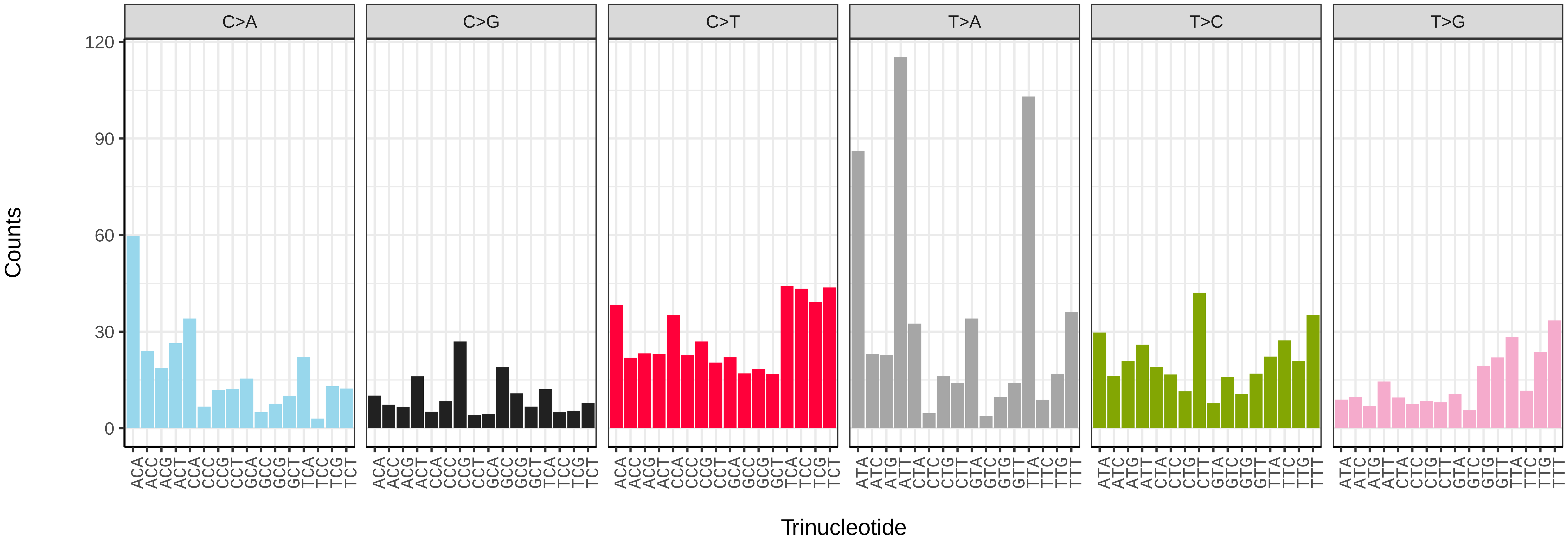


*Agonopterix arenella* (ilAgoAren1) somatic mutational spectrum (SBS52 classification)

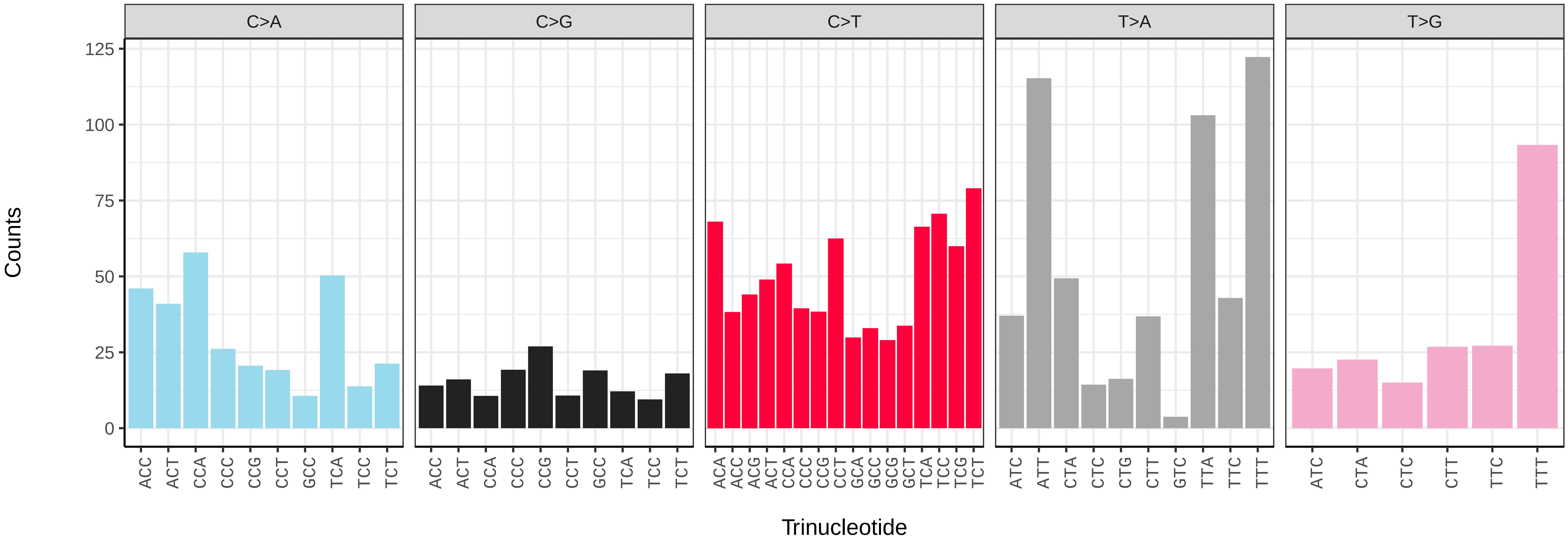

*Agonopterix subpropinquella* (ilAgoSubp1) germline mutational spectrum

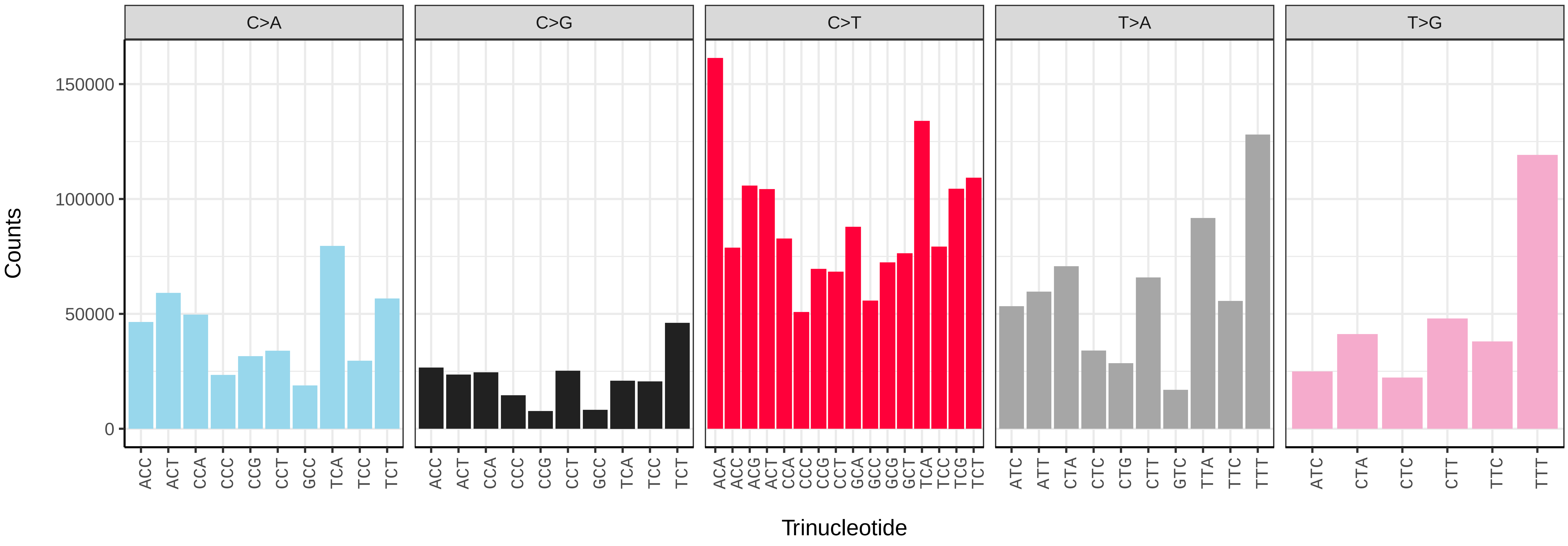


*Agonopterix subpropinquella* (ilAgoSubp1) somatic mutational spectrum (SBS96 classification)

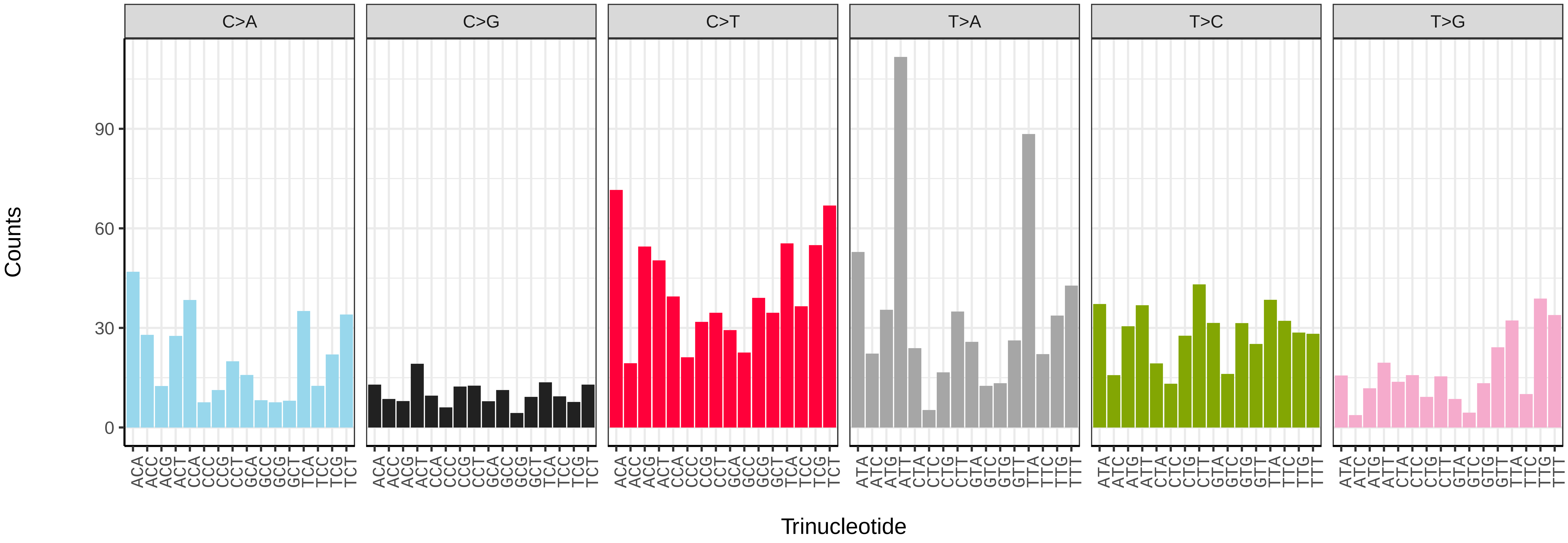


*Agonopterix subpropinquella* (ilAgoSubp1) somatic mutational spectrum (SBS52 classification)

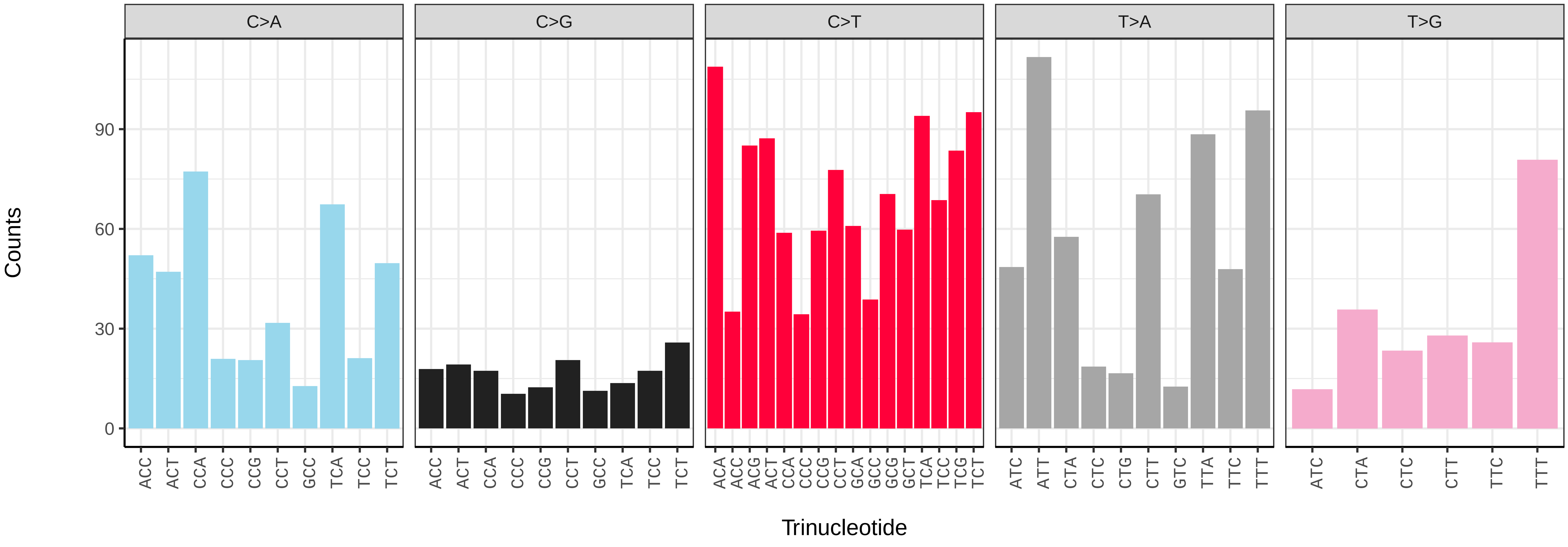

*Agriopis aurantiaria* (ilAgrAura1) germline mutational spectrum

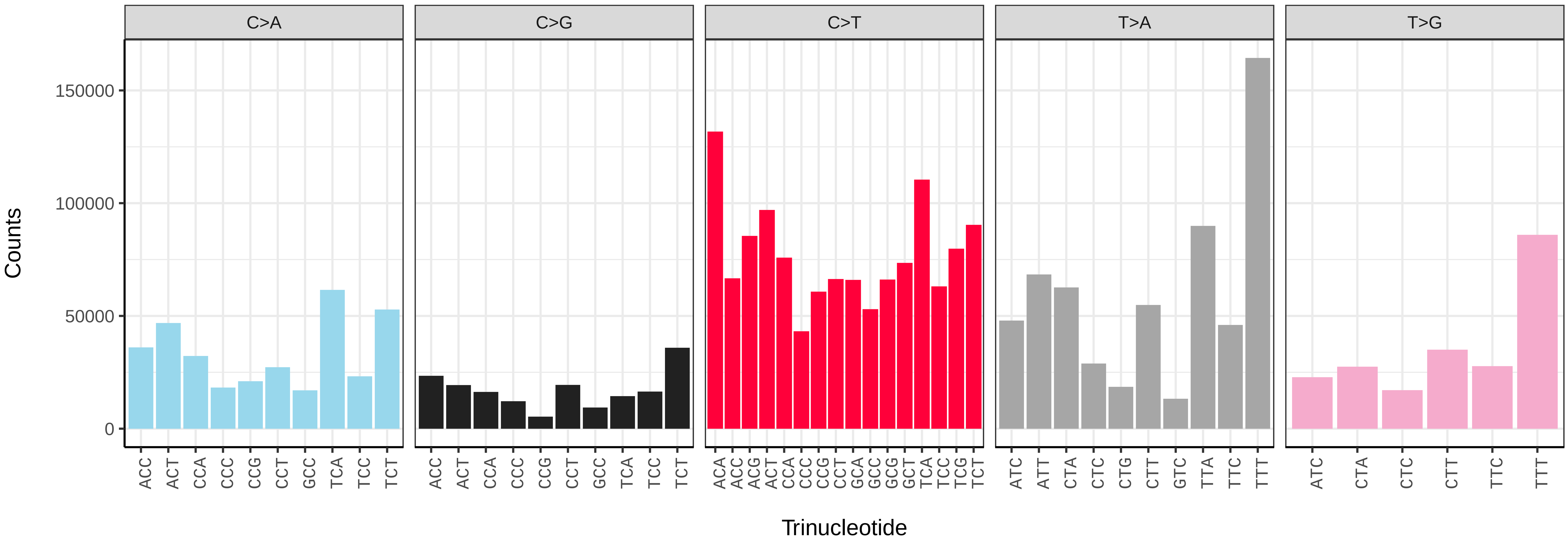


*Agriopis aurantiaria* (ilAgrAura1) somatic mutational spectrum (SBS96 classification)

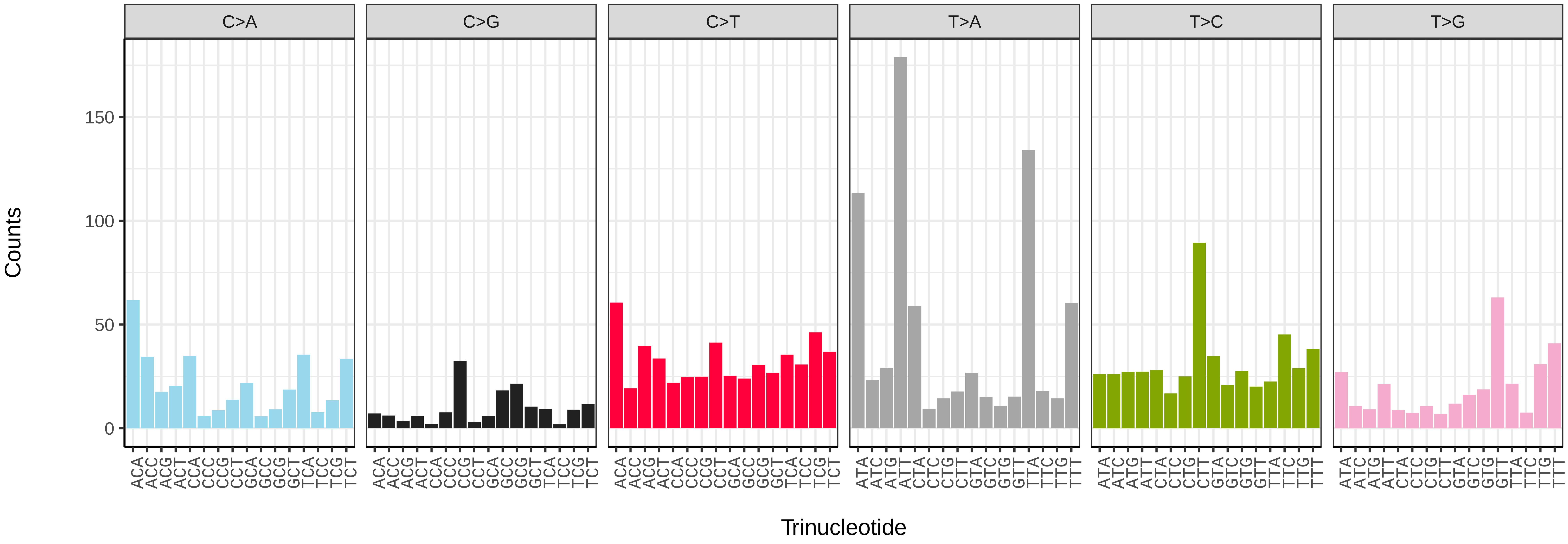


*Agriopis aurantiaria* (ilAgrAura1) somatic mutational spectrum (SBS52 classification)

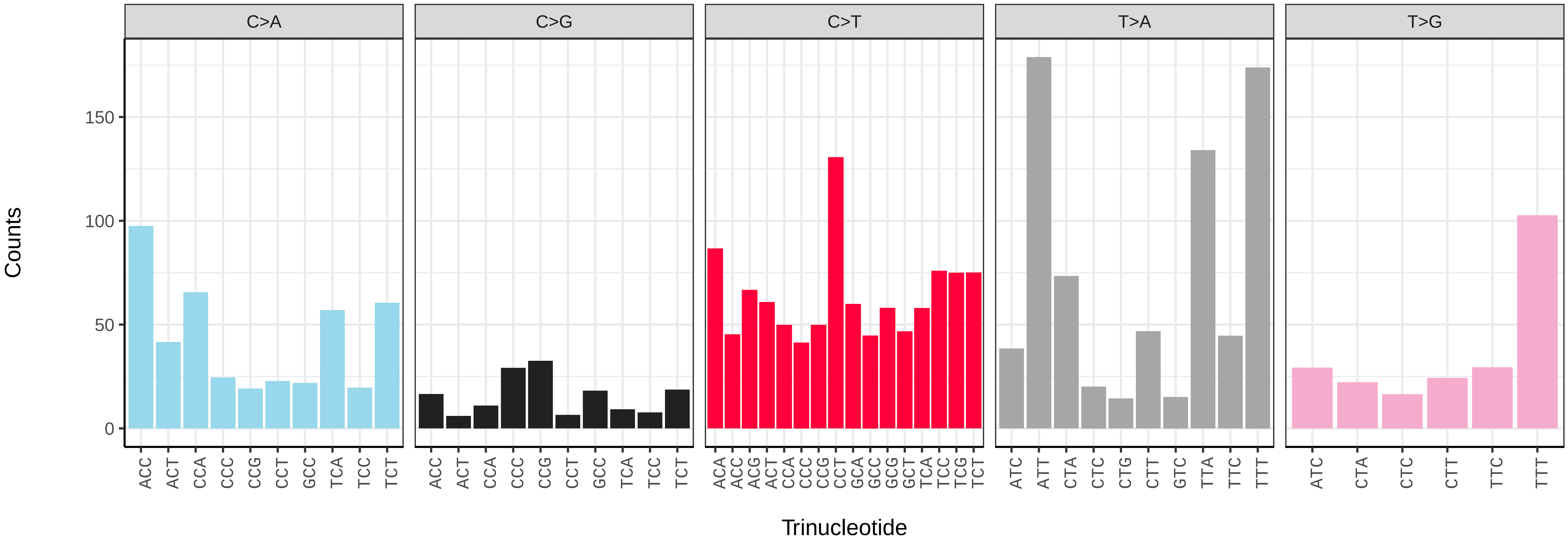

*Agrochola circellaris* (ilAgrCirc1) germline mutational spectrum

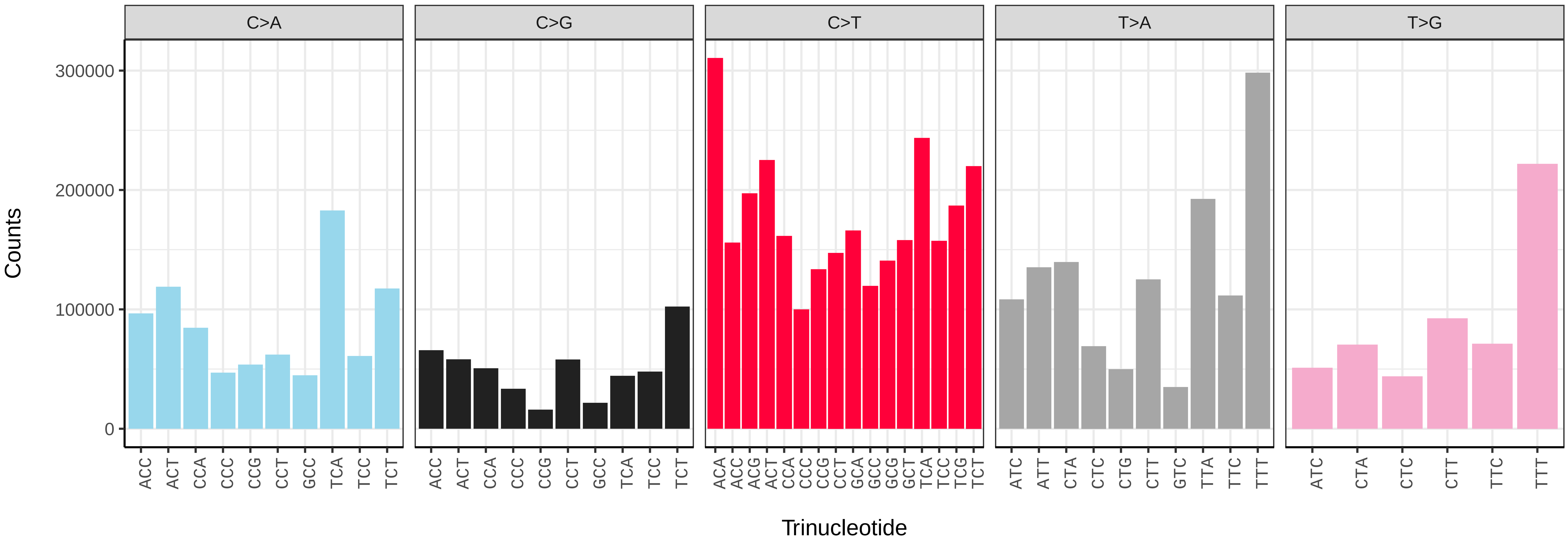


*Agrochola circellaris* (ilAgrCirc1) somatic mutational spectrum (SBS96 classification)

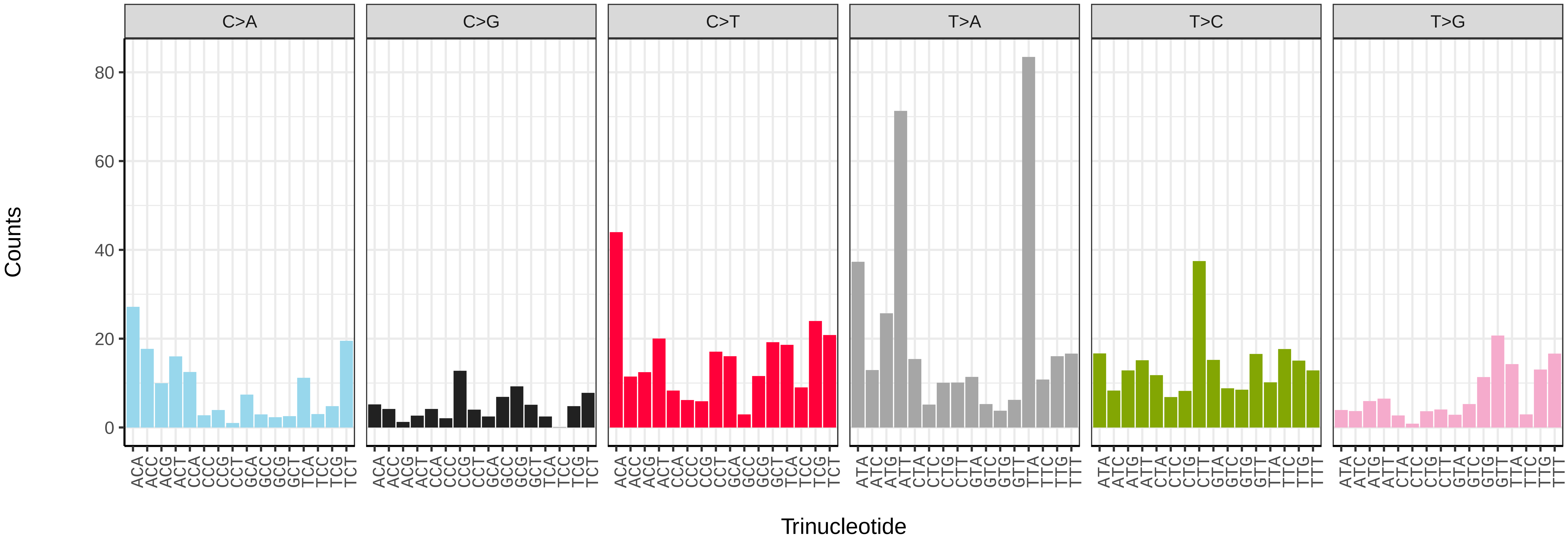


*Agrochola circellaris* (ilAgrCirc1) somatic mutational spectrum (SBS52 classification)

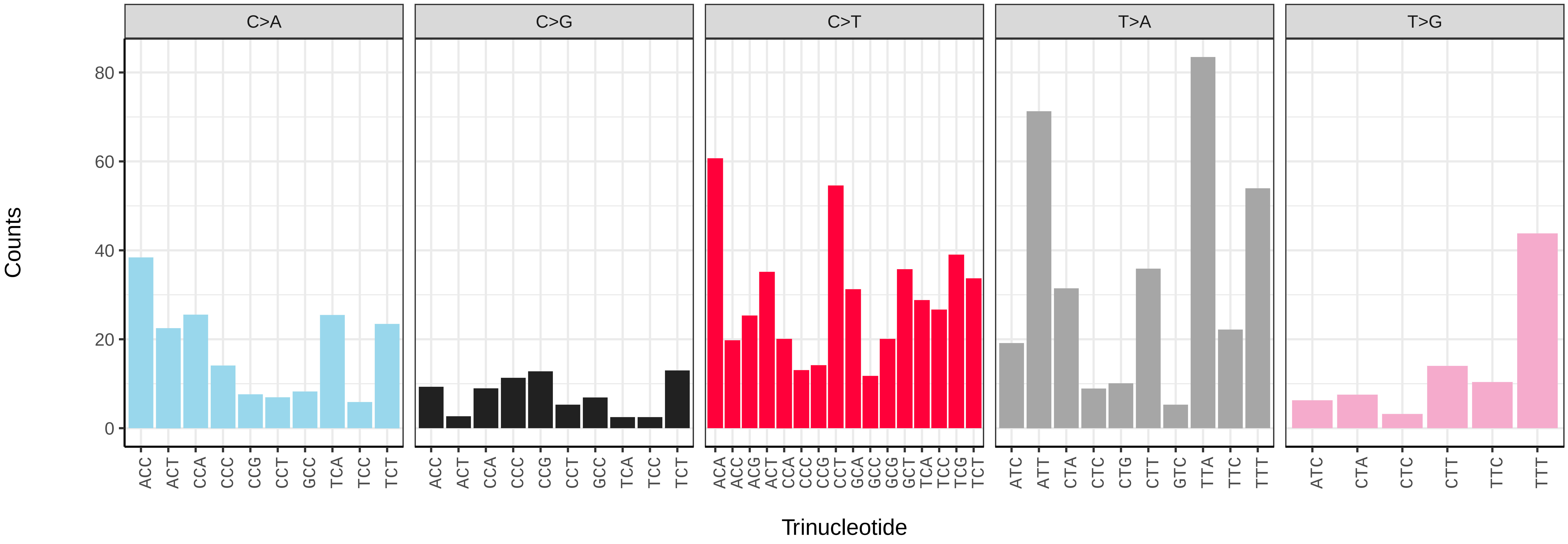

*Agriopis leucophaearia* (ilAgrLeuc1) germline mutational spectrum

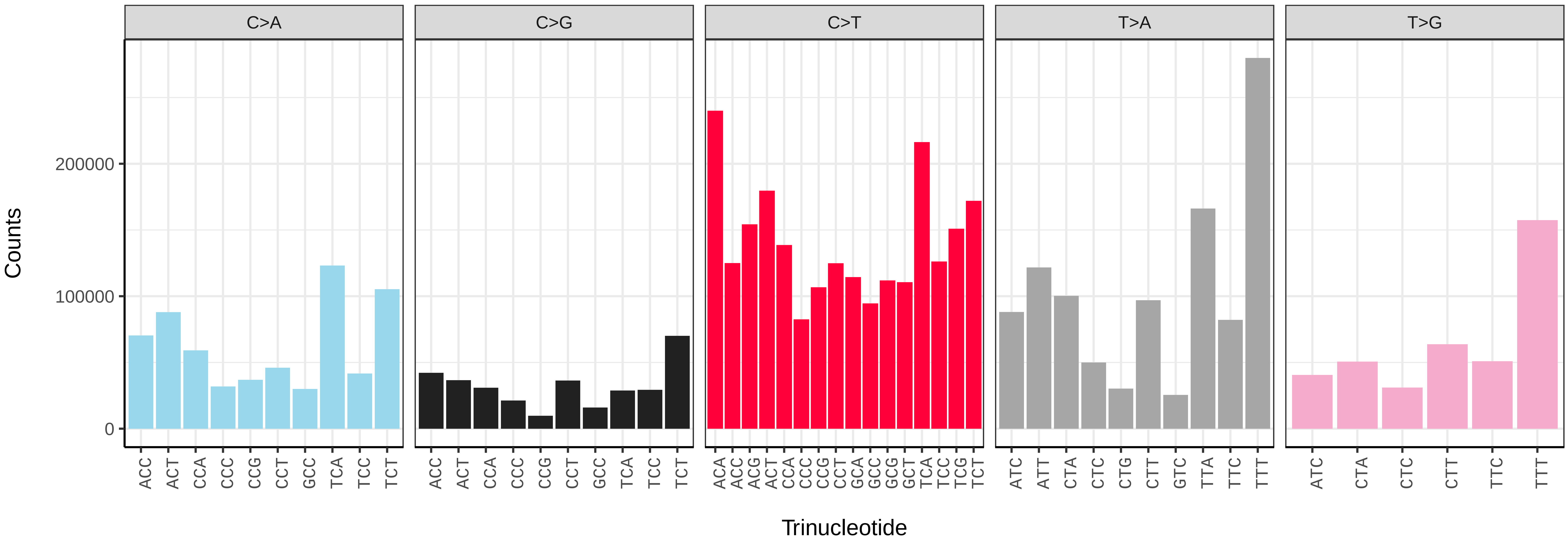


*Agriopis leucophaearia* (ilAgrLeuc1) somatic mutational spectrum (SBS96 classification)

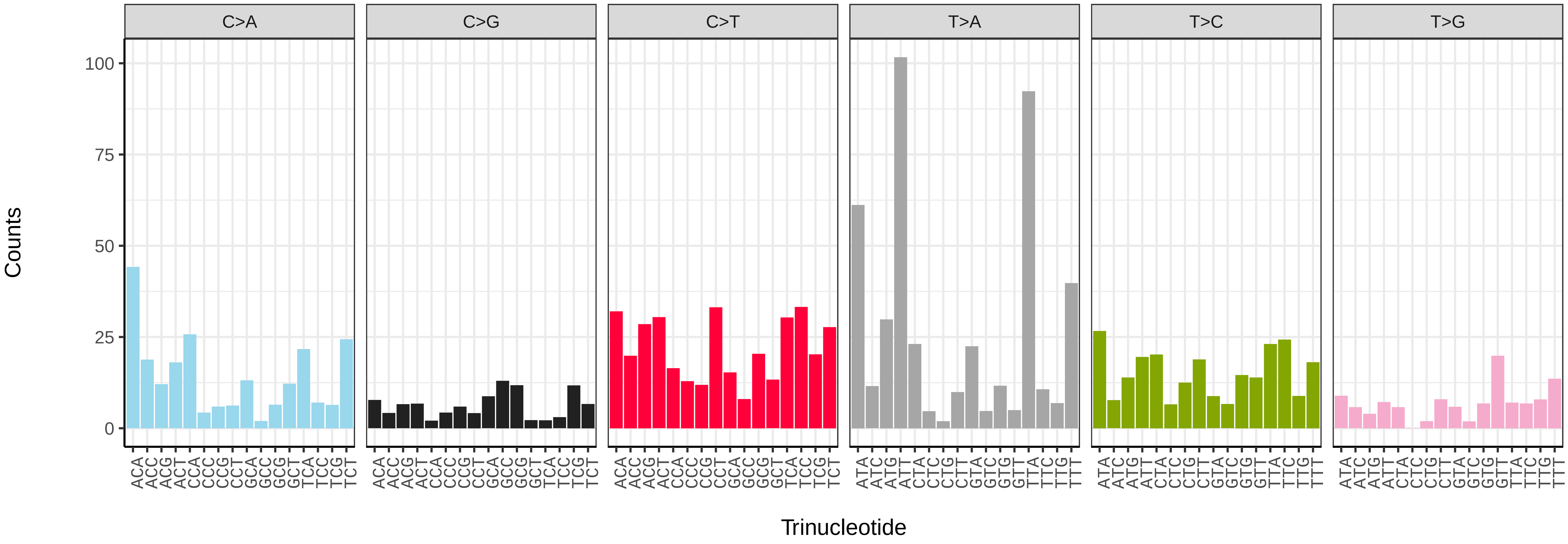


*Agriopis leucophaearia* (ilAgrLeuc1) somatic mutational spectrum (SBS52 classification)

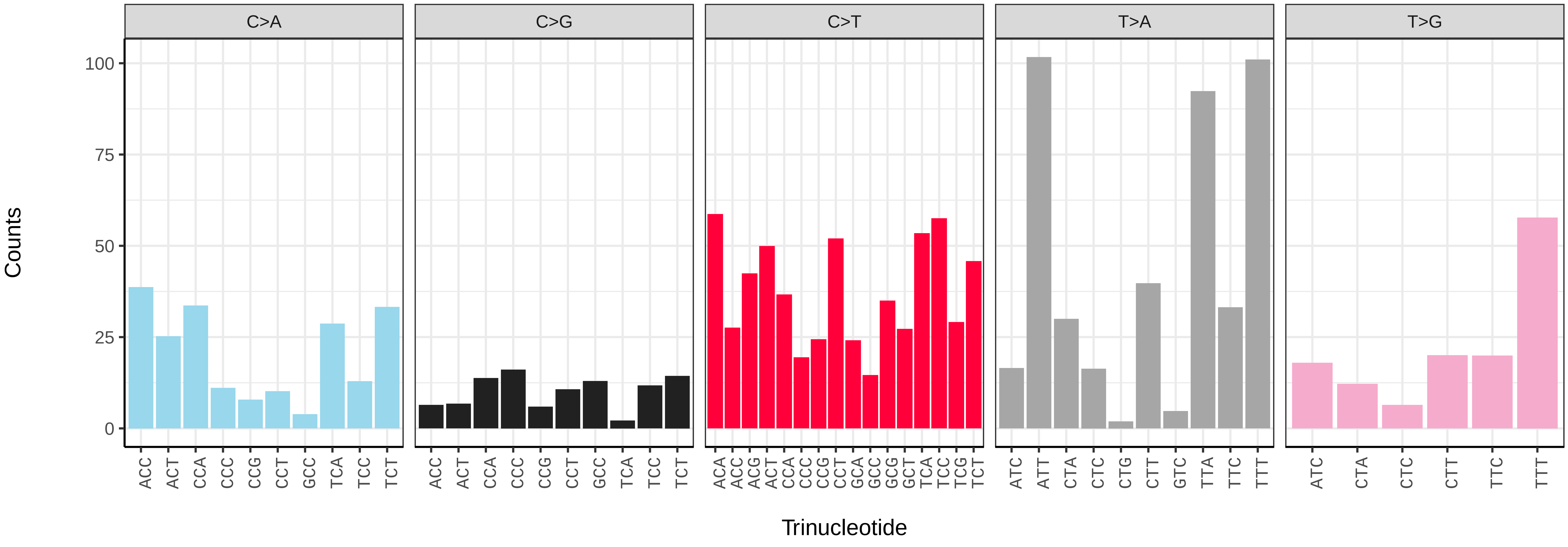

*Agrochola lota* (ilAgrLota1) germline mutational spectrum

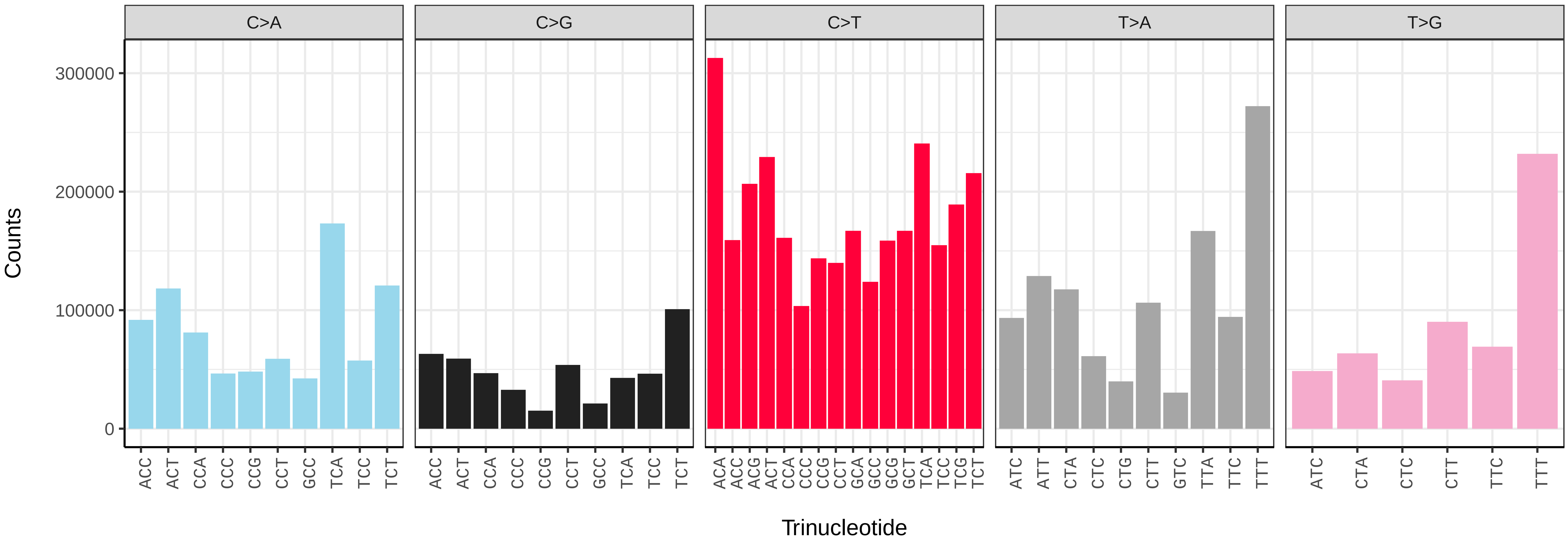


*Agrochola lota* (ilAgrLota1) somatic mutational spectrum (SBS96 classification)

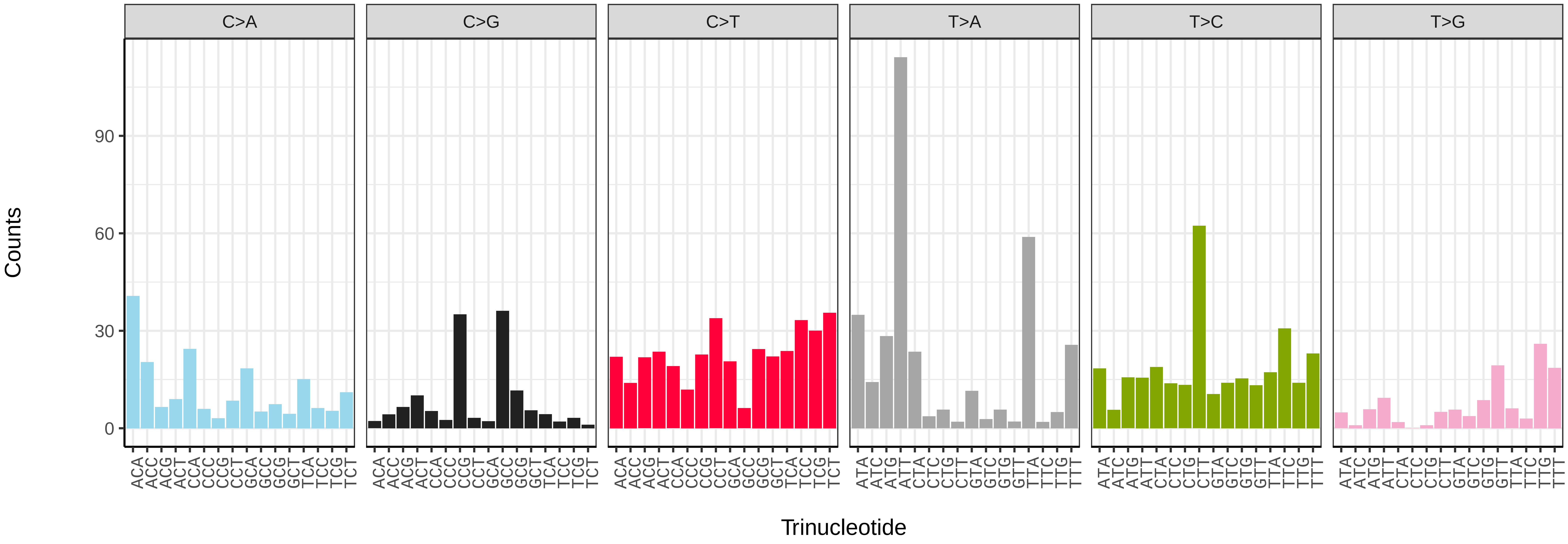


*Agrochola lota* (ilAgrLota1) somatic mutational spectrum (SBS52 classification)

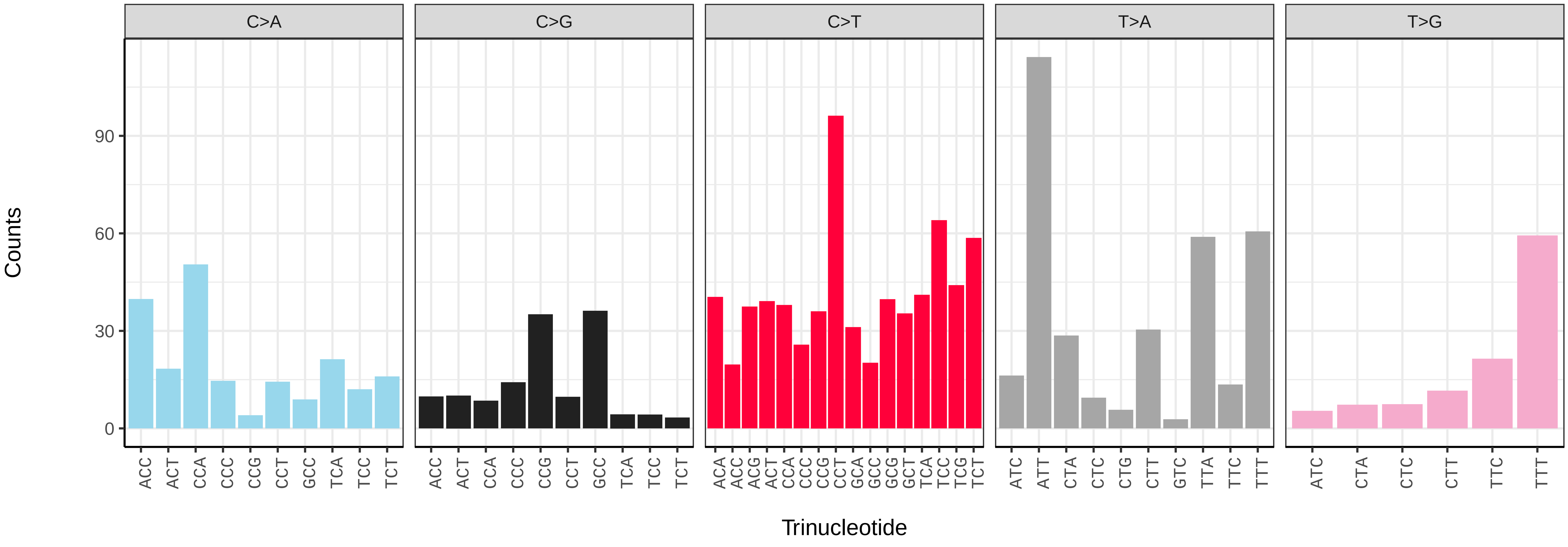

*Agrochola macilenta* (ilAgrMaci1) germline mutational spectrum

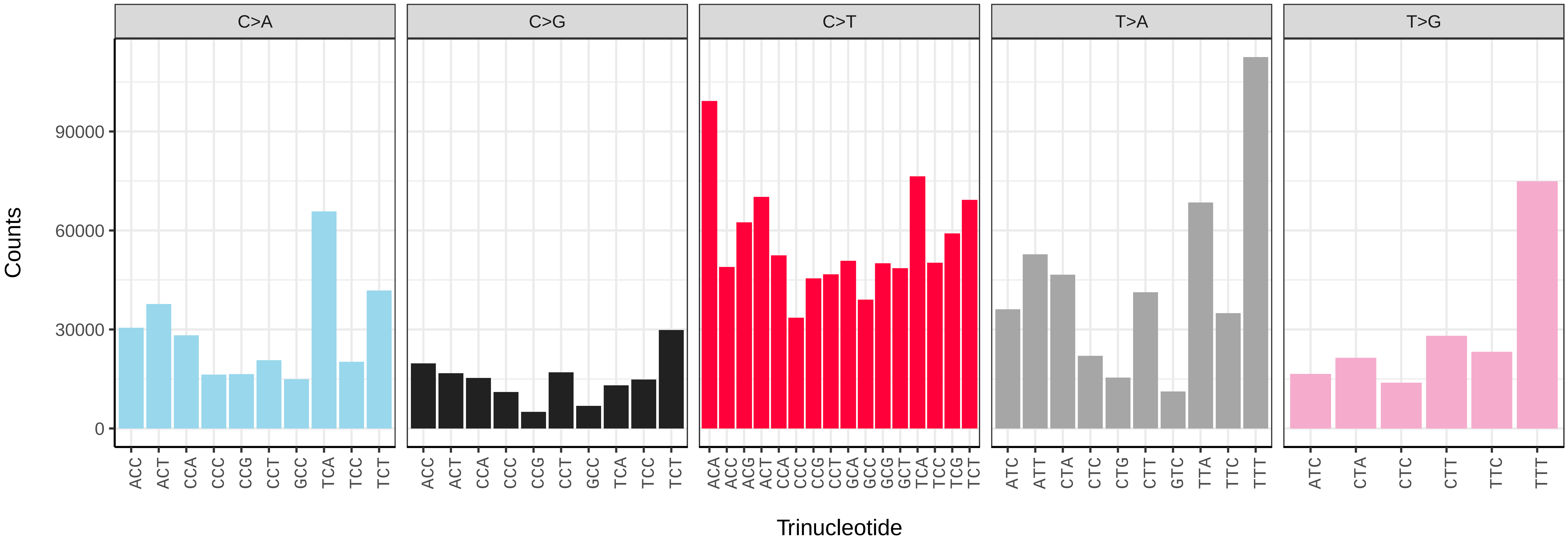


*Agrochola macilenta* (ilAgrMaci1) somatic mutational spectrum (SBS96 classification)

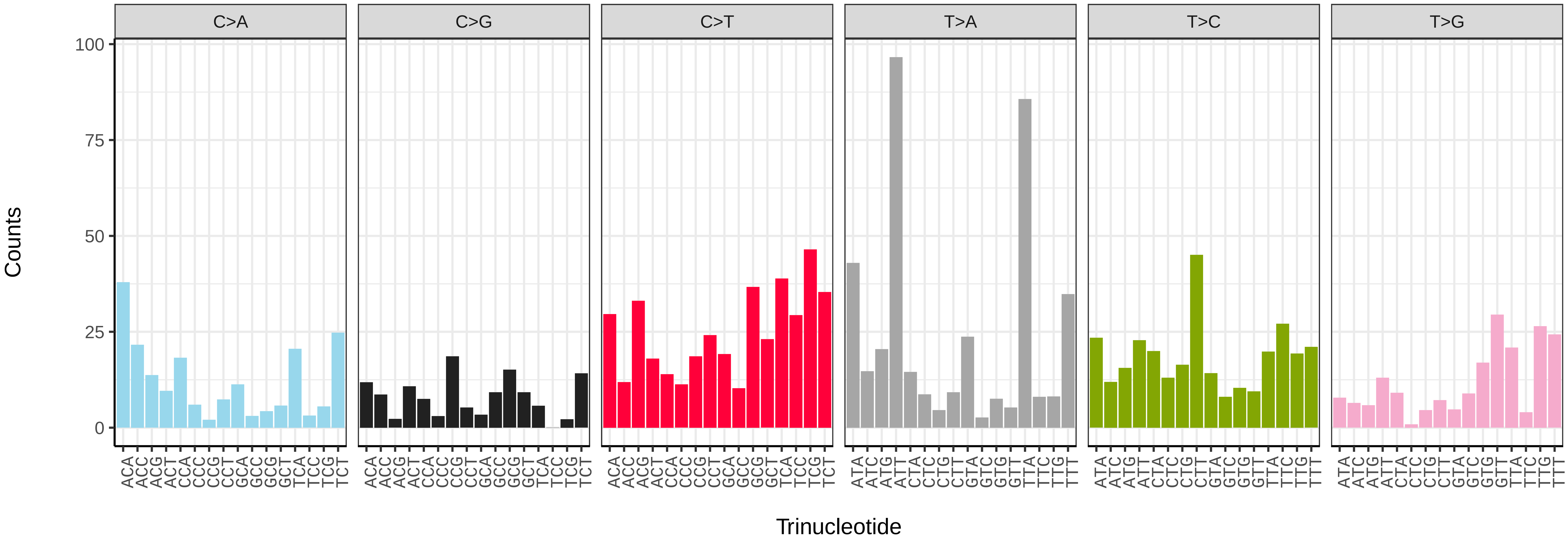


*Agrochola macilenta* (ilAgrMaci1) somatic mutational spectrum (SBS52 classification)

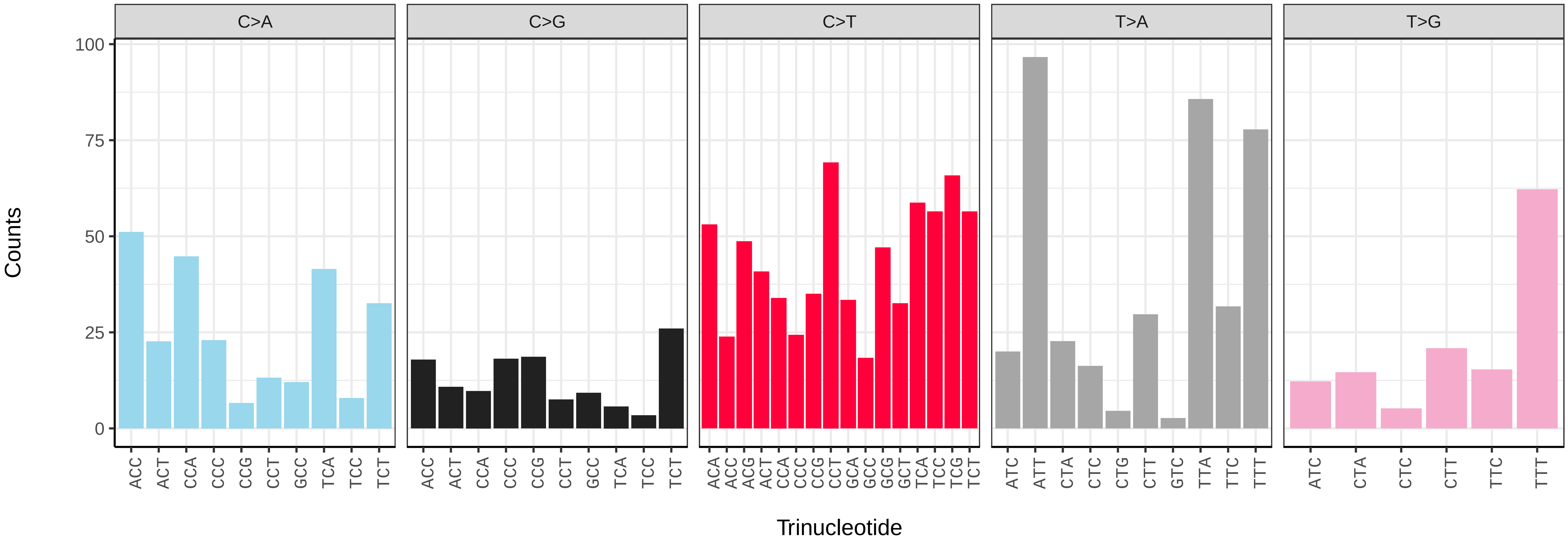

*Agriopis marginaria* (ilAgrMarg1) germline mutational spectrum

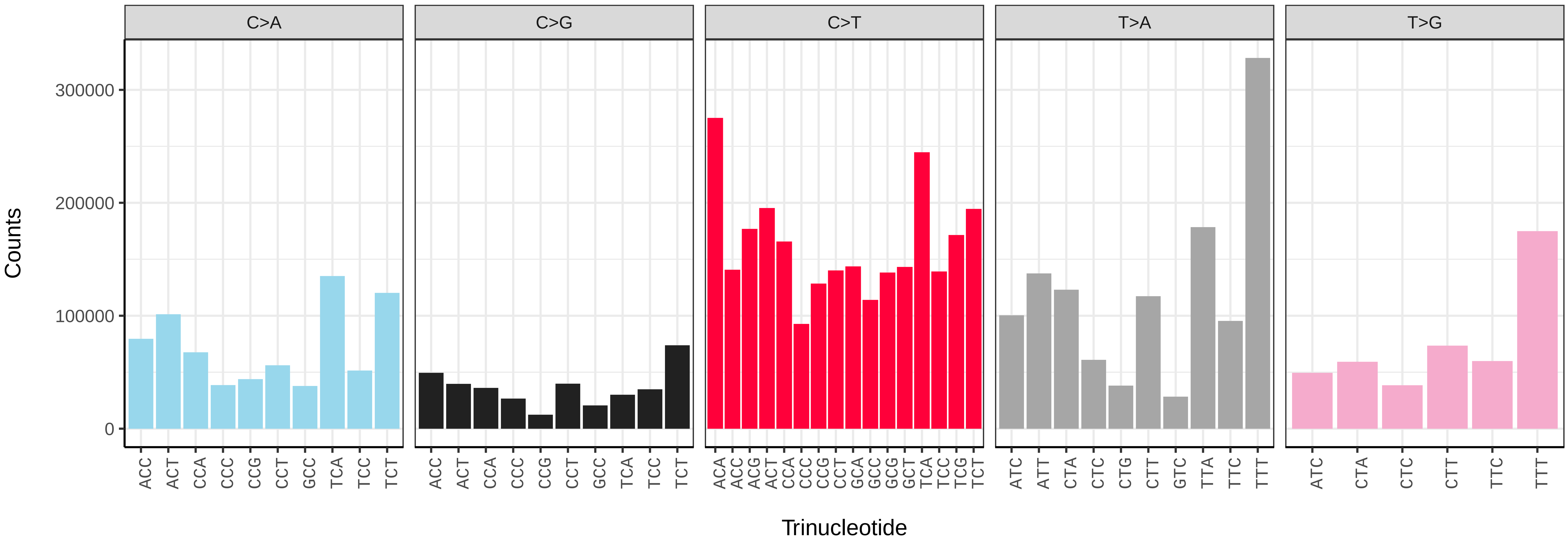


*Agriopis marginaria* (ilAgrMarg1) somatic mutational spectrum (SBS96 classification)

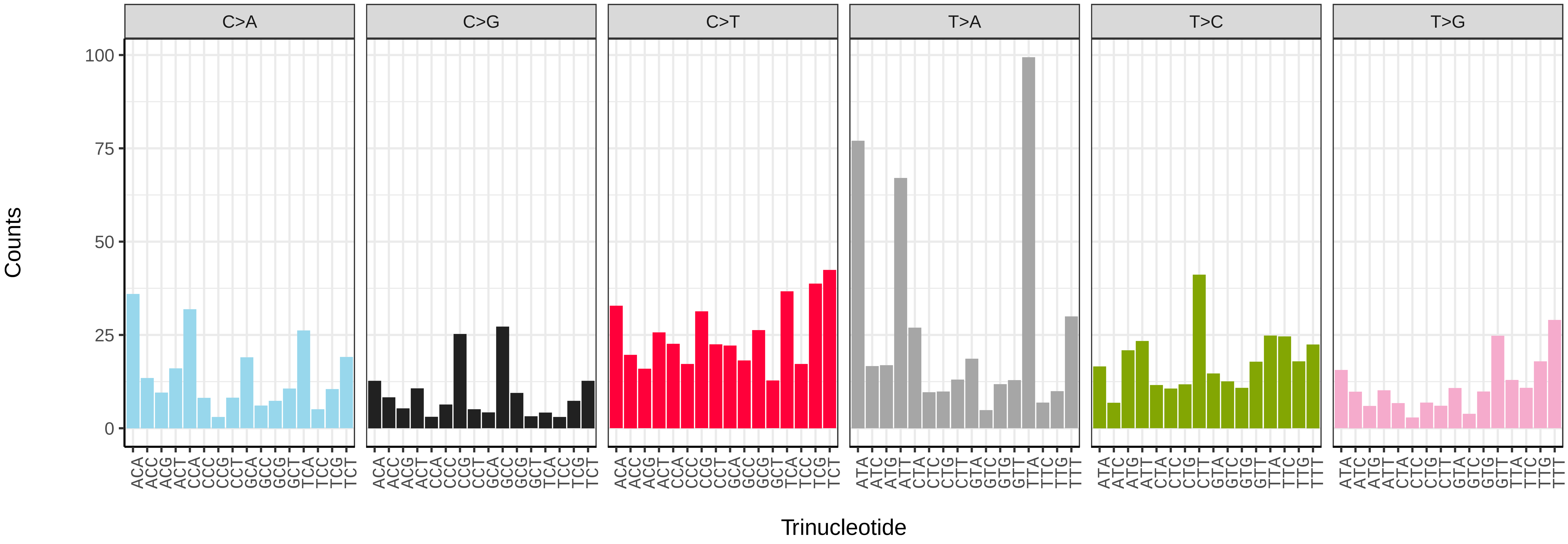


*Agriopis marginaria* (ilAgrMarg1) somatic mutational spectrum (SBS52 classification)

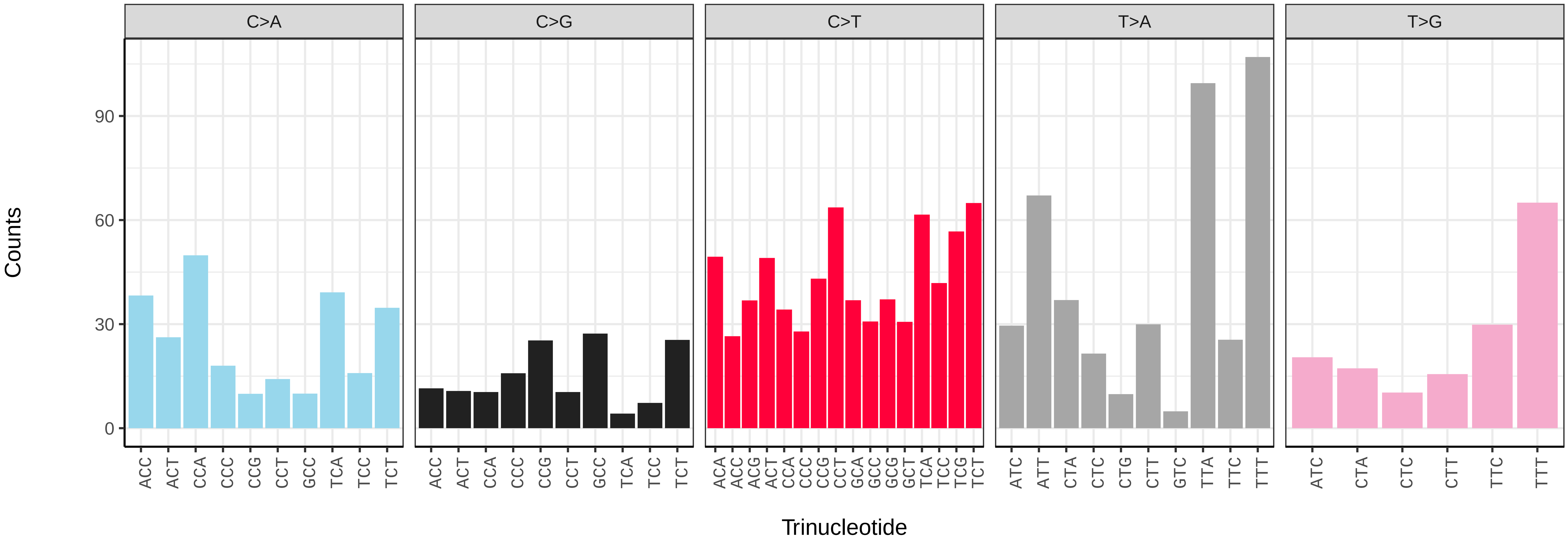

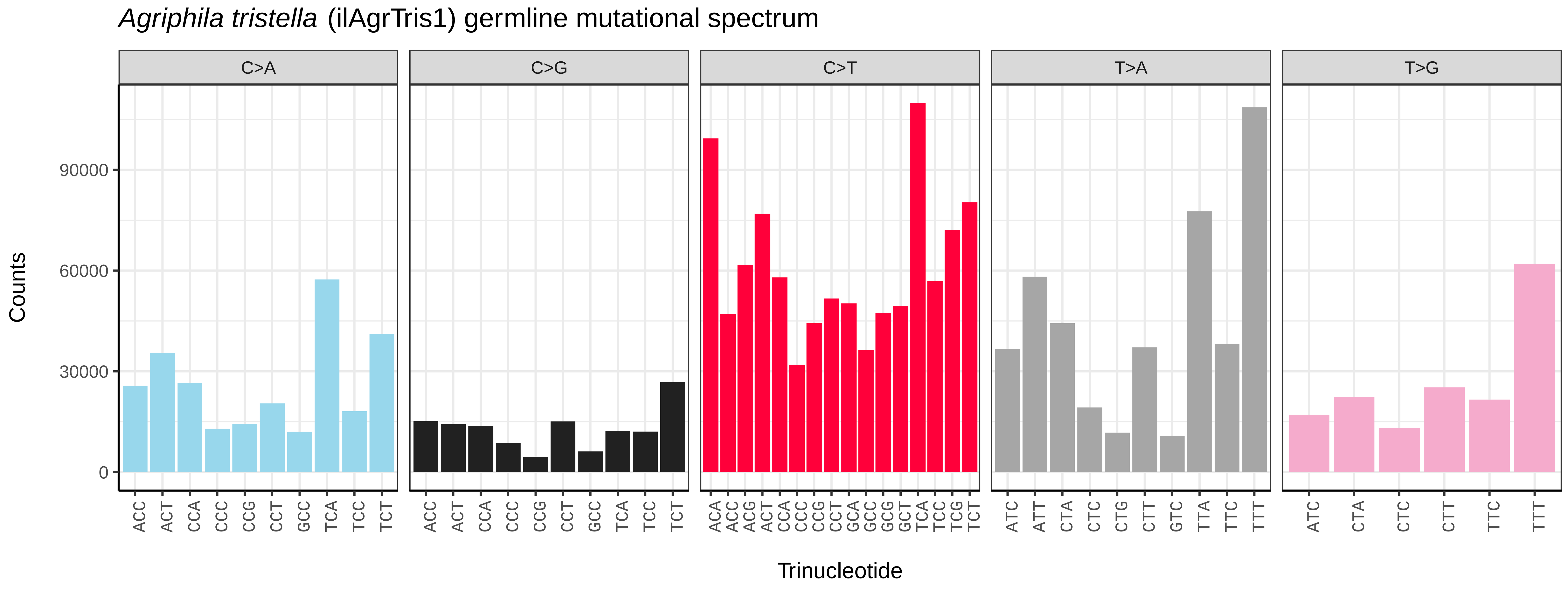
Agriphila tristella (ilAgrTris1) germline mutational spectrum
C>A
C>G
C>T
T>A
T>G
Counts
0
30000
60000
90000
ACC ACT CCA CCC CCG CCT GCC TCA TCC TCT
ACC ACT CCA CCC CCG CCT GCC TCA TCC TCT
ACA ACC ACG ACT CCA CCC CCG CCT GCA GCC GCG GCT TCA TCC TCG TCT
ATC ATT CTA CTC CTG CTT GTC TTA TTC TTT
ATC CTA CTC CTT TTC TTT
Trinucleotide

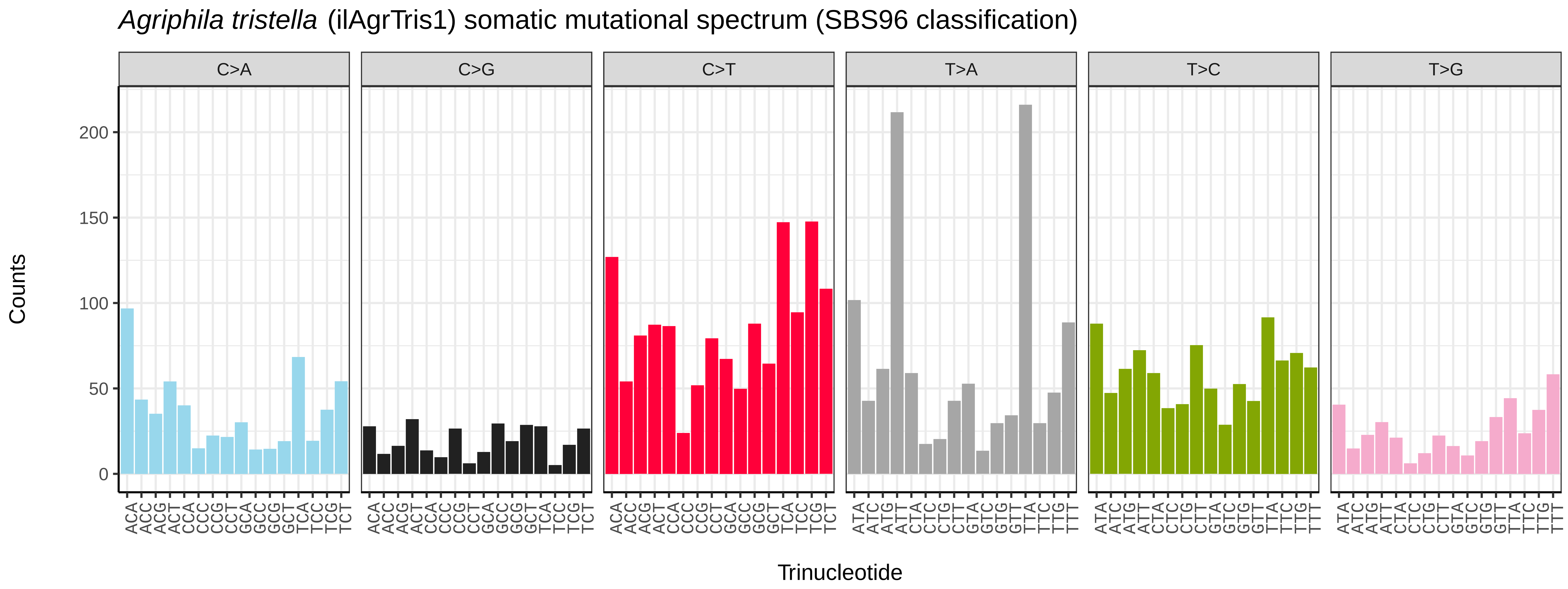
Agriphila tristella (ilAgrTris1) somatic mutational spectrum (SBS96 classification)
C>A
C>G
C>T
T>A
T>C
T>G
Counts
0
50
100
150
200
ACA ACC ACG ACT CCA CCC CCG CCT GCA GCC GCG GCT TCA TCC TCG TCT
ATA ATC ATG ATT CTA CTC CTG CTT GTA GTC GTG GTT TTA TTC TTG TTT
Trinucleotide

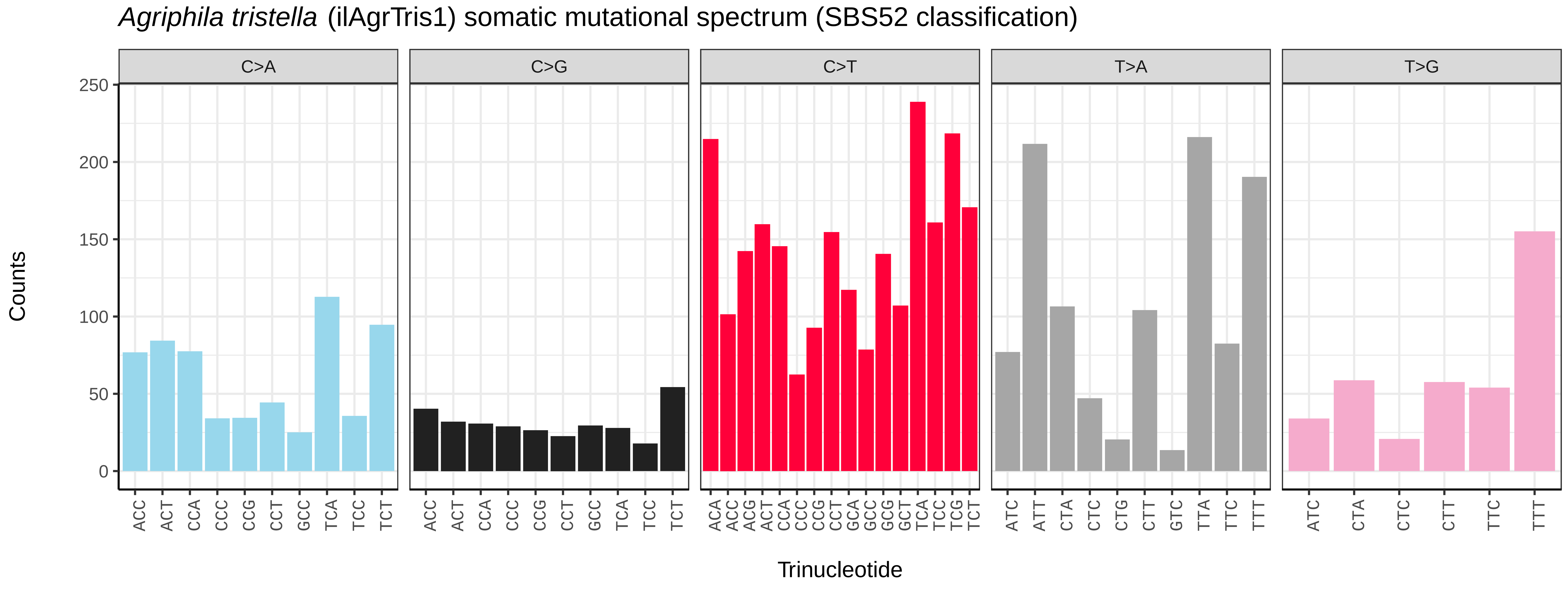
Agriphila tristella (ilAgrTris1) somatic mutational spectrum (SBS52 classification)
C>A
C>G
C>T
T>A
T>G
Counts
0
50
100
150
200
250
ACC ACT CCA CCC CCG CCT GCC TCA TCC TCT
ACA ACC ACG ACT CCA CCC CCG CCT GCA GCC GCG GCT TCA TCC TCG TCT
ATC ATT CTA CTC CTG CTT GTC TTA TTC TTT
ATC CTA CTC CTT TTC TTT
Trinucleotide

*Allophyes oxyacanthae* (ilAllOxya1) germline mutational spectrum

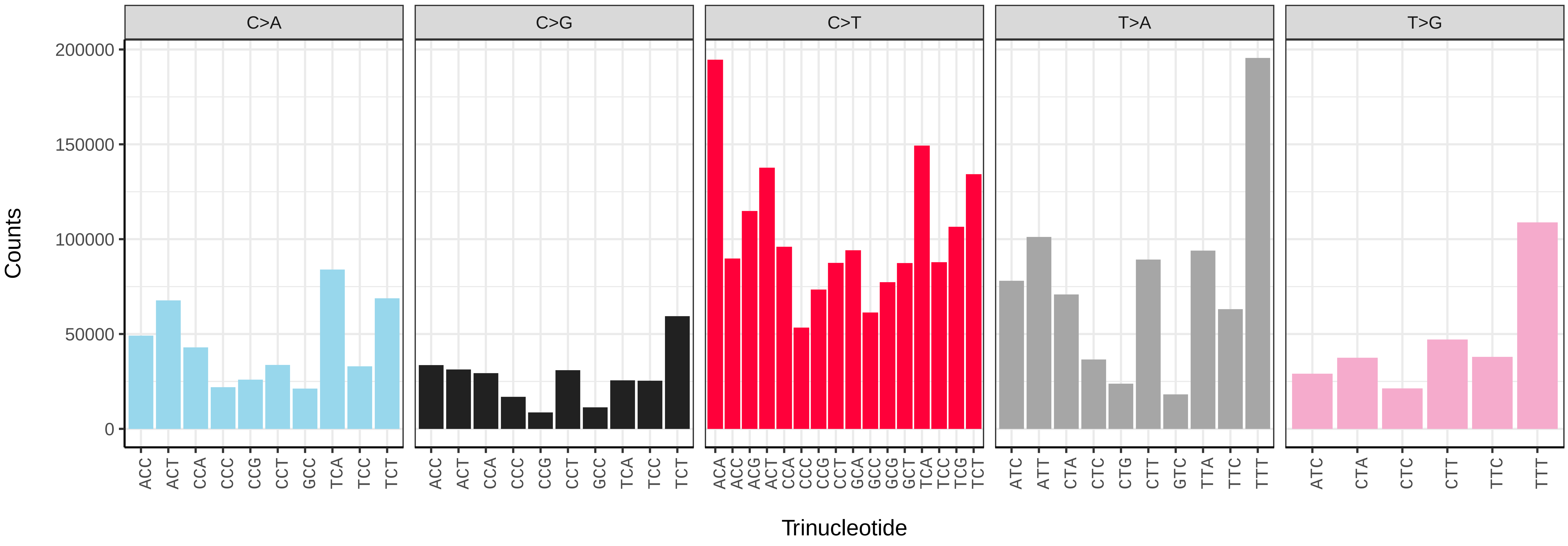


*Allophyes oxyacanthae* (ilAllOxya1) somatic mutational spectrum (SBS96 classification)

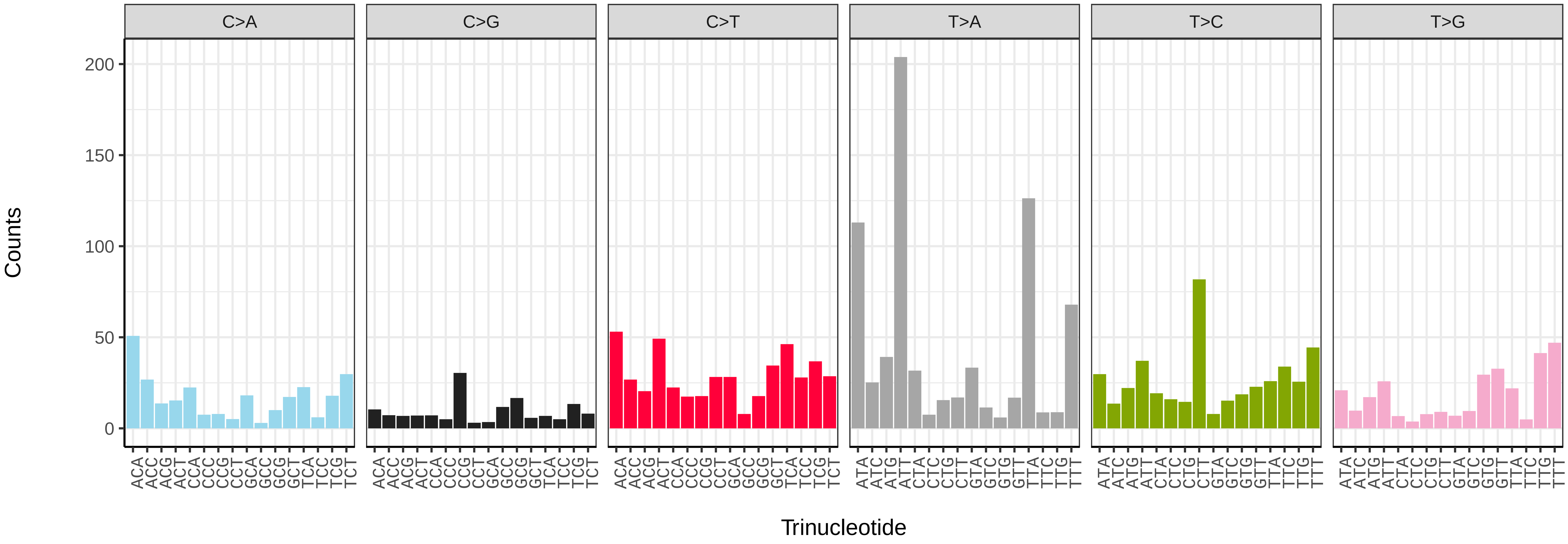


*Allophyes oxyacanthae* (ilAllOxya1) somatic mutational spectrum (SBS52 classification)

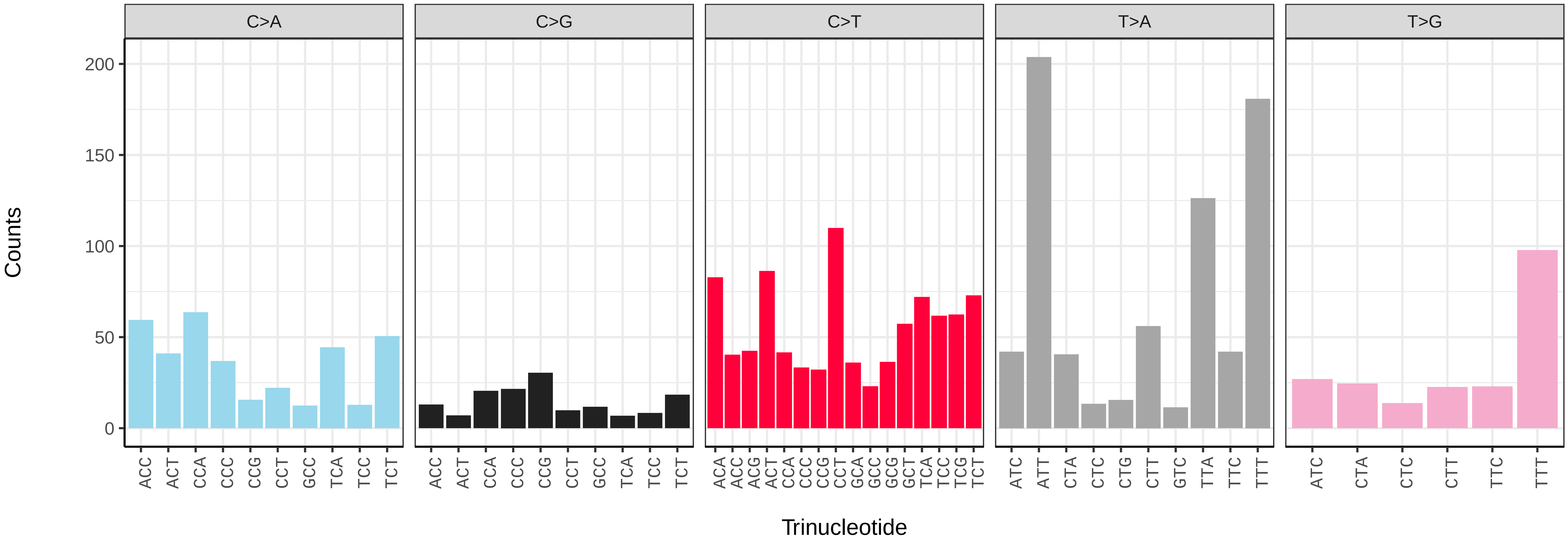

*Alsophila aescularia* (ilAlsAesc1) germline mutational spectrum

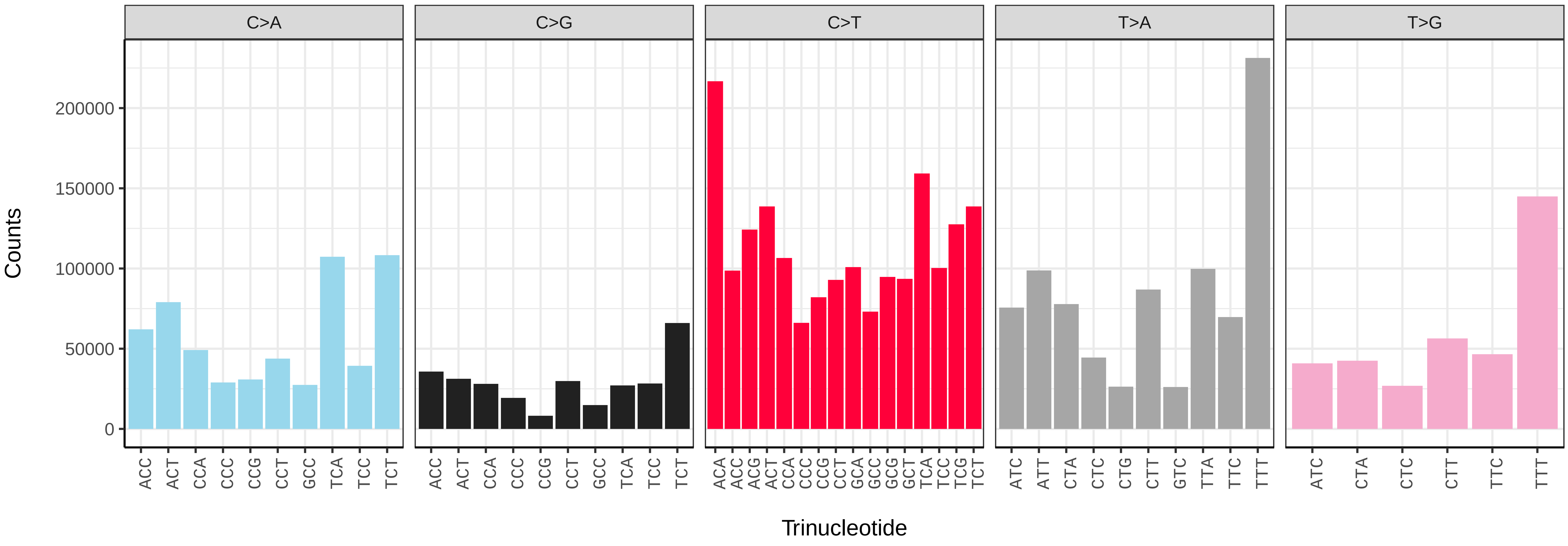


*Alsophila aescularia* (ilAlsAesc1) somatic mutational spectrum (SBS96 classification)

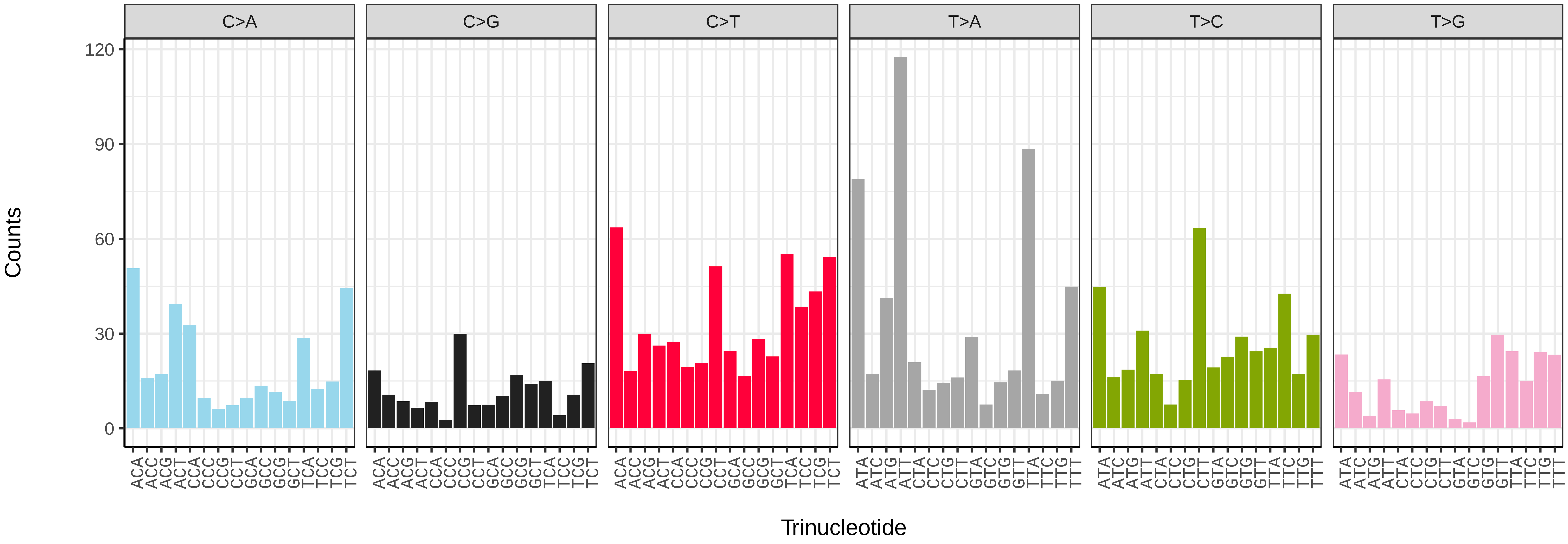


*Alsophila aescularia* (ilAlsAesc1) somatic mutational spectrum (SBS52 classification)

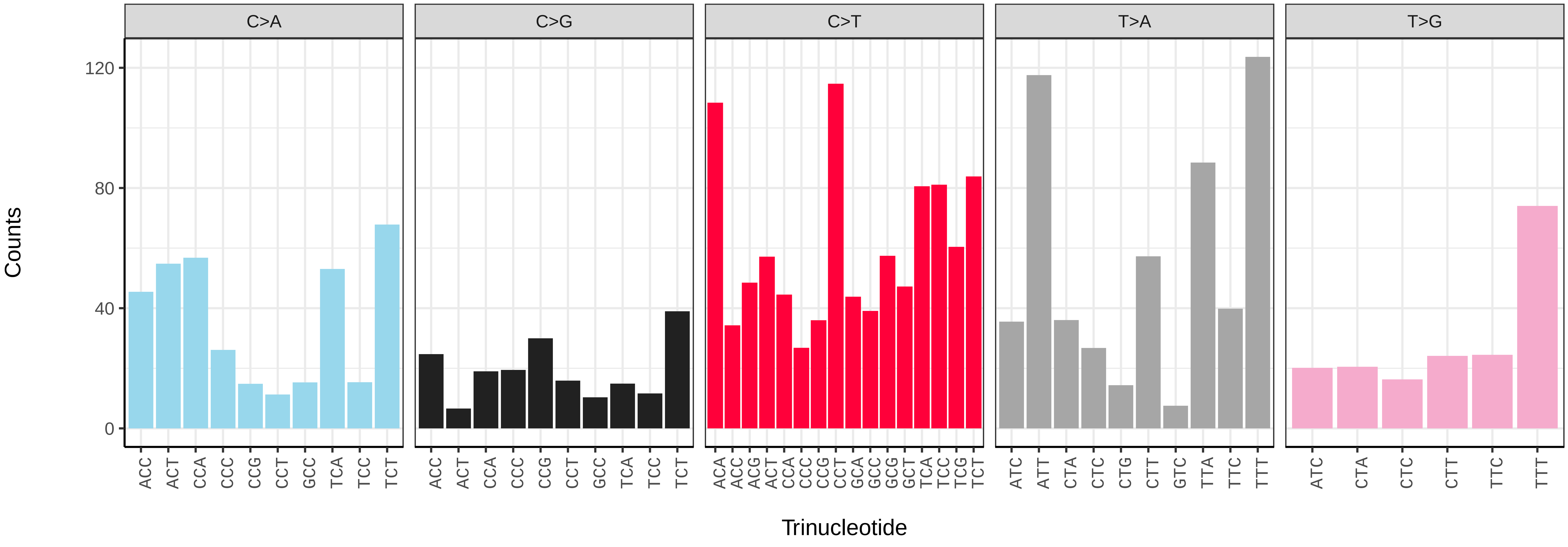

*Amphipyra berbera* (ilAmpBerb1) germline mutational spectrum

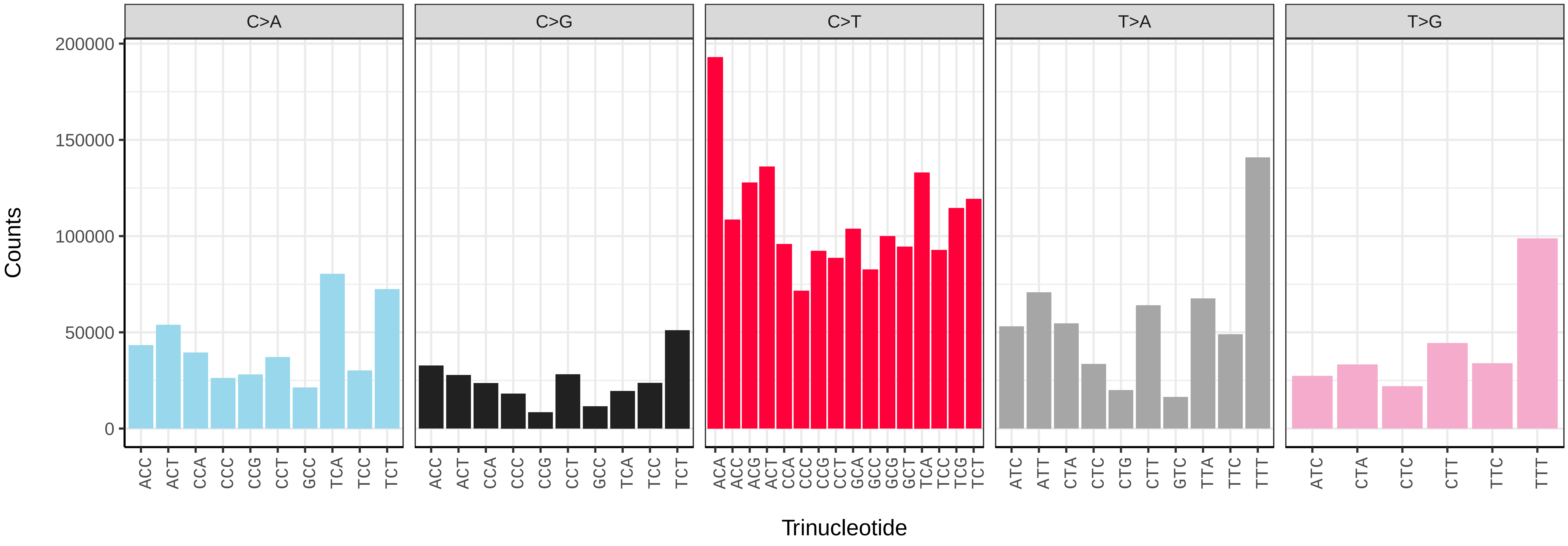


*Amphipyra berbera* (ilAmpBerb1) somatic mutational spectrum (SBS96 classification)

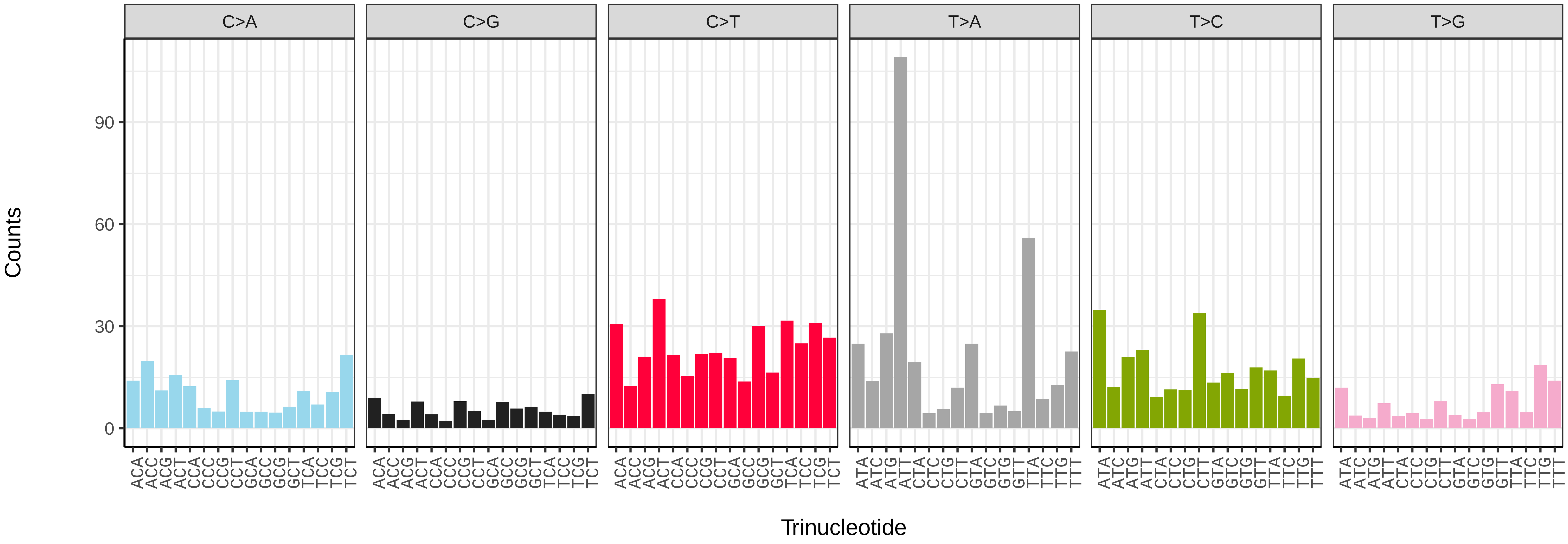


*Amphipyra berbera* (ilAmpBerb1) somatic mutational spectrum (SBS52 classification)

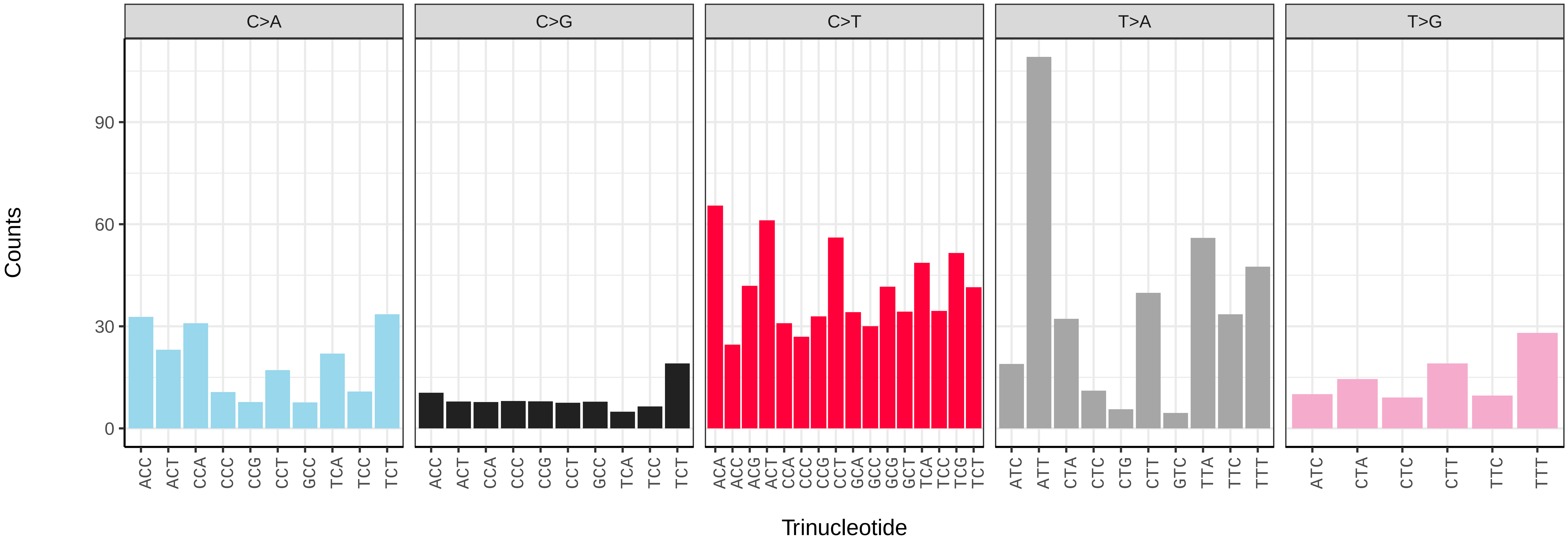

*Amphipoea lucens* (ilAmpLuce6) germline mutational spectrum

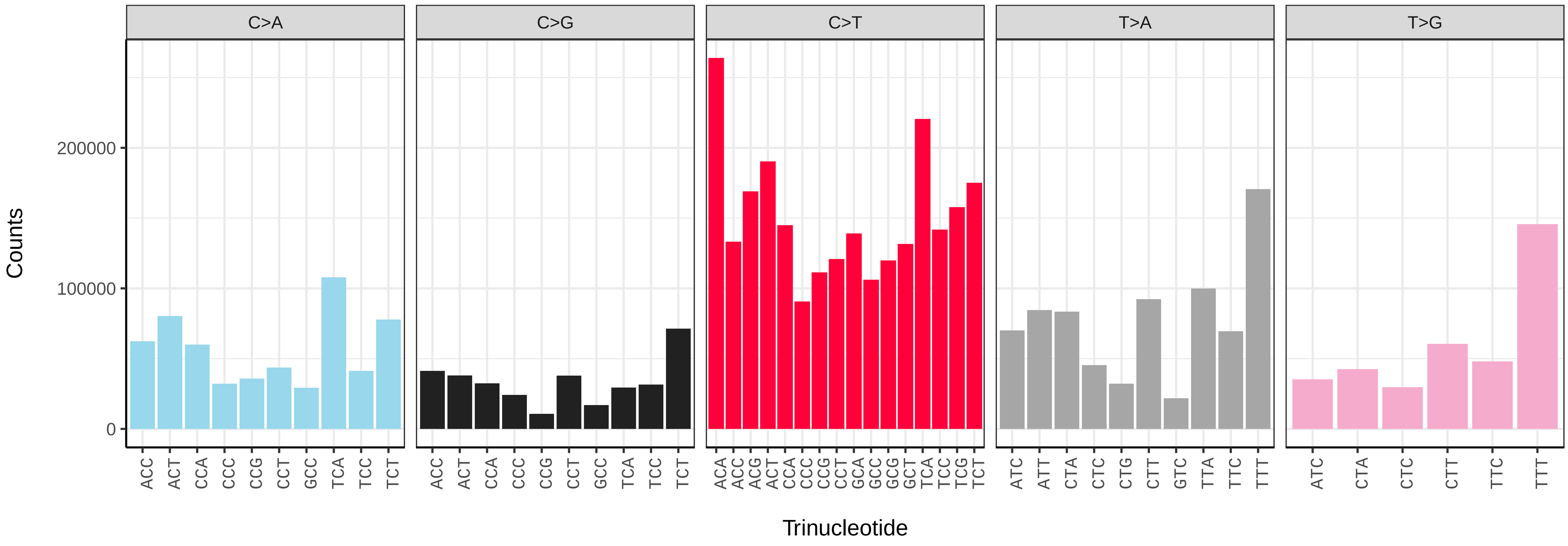


*Amphipoea lucens* (ilAmpLuce6) somatic mutational spectrum (SBS96 classification)

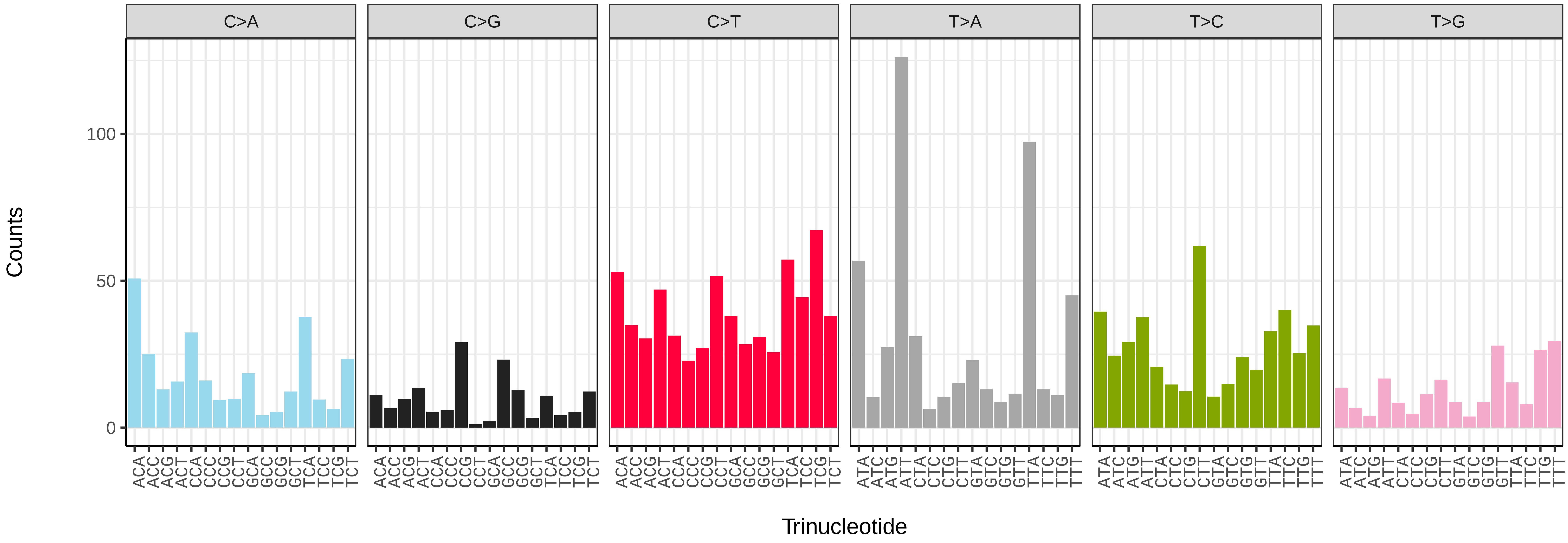


*Amphipoea lucens* (ilAmpLuce6) somatic mutational spectrum (SBS52 classification)

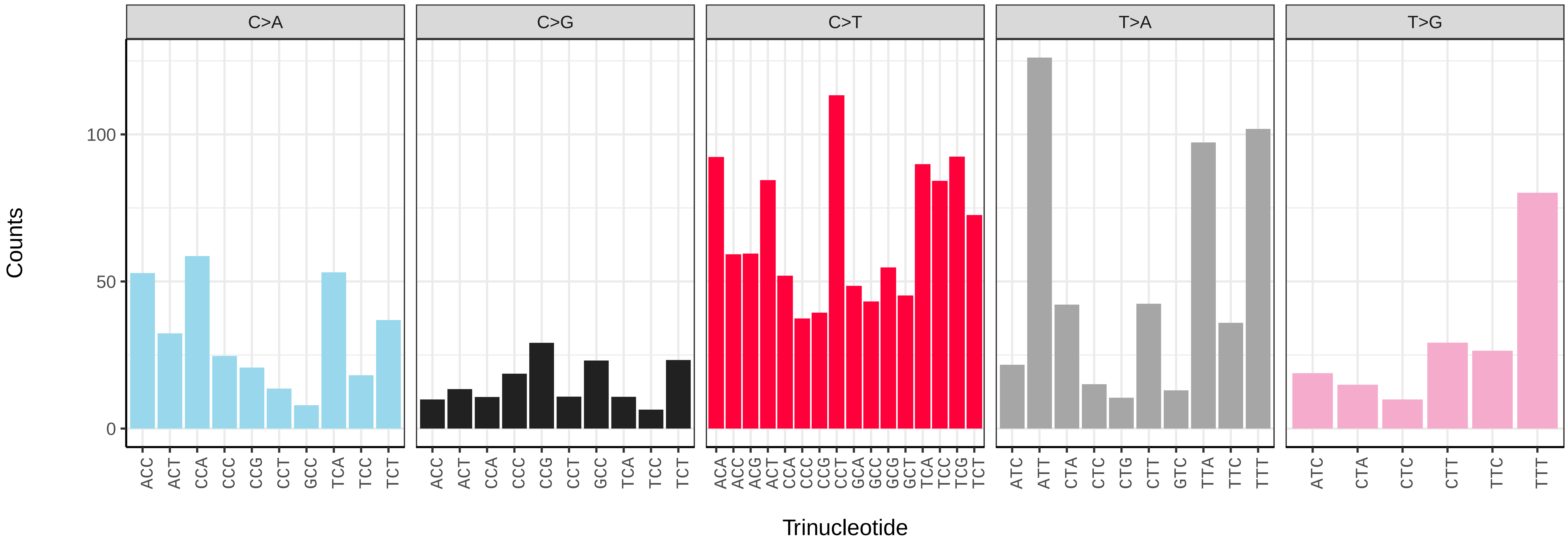

*Amphipoea oculea* (ilAmpOcul1) germline mutational spectrum

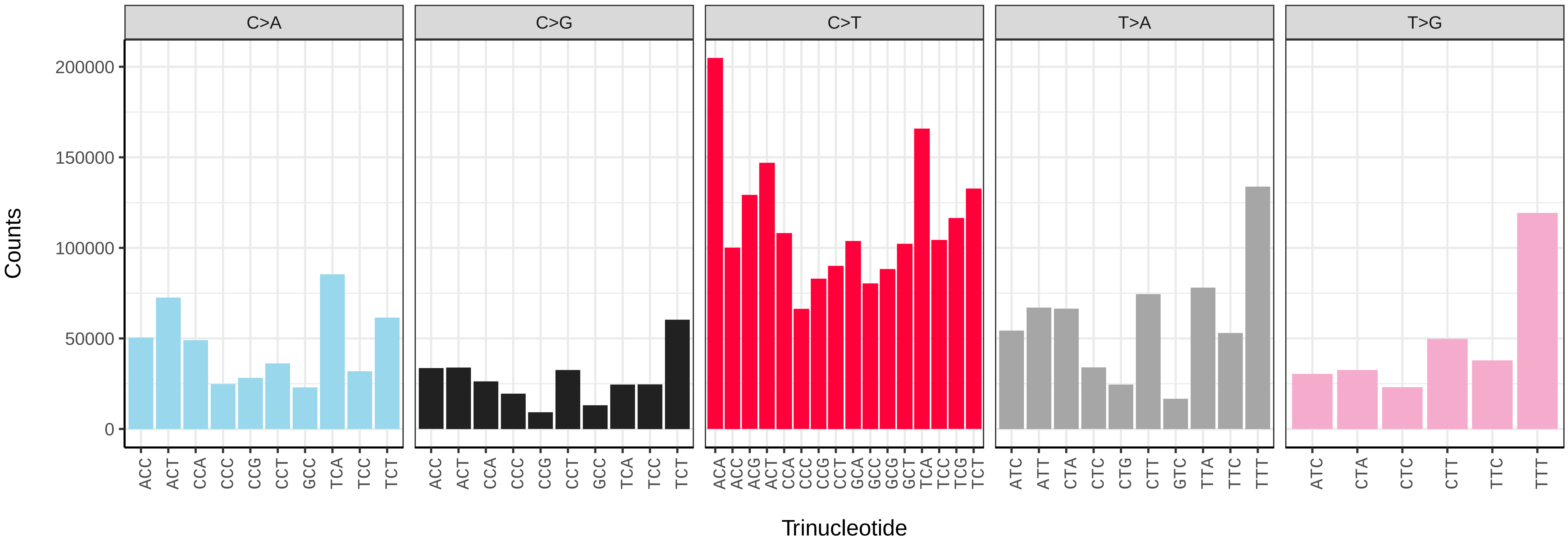


*Amphipoea oculea* (ilAmpOcul1) somatic mutational spectrum (SBS96 classification)

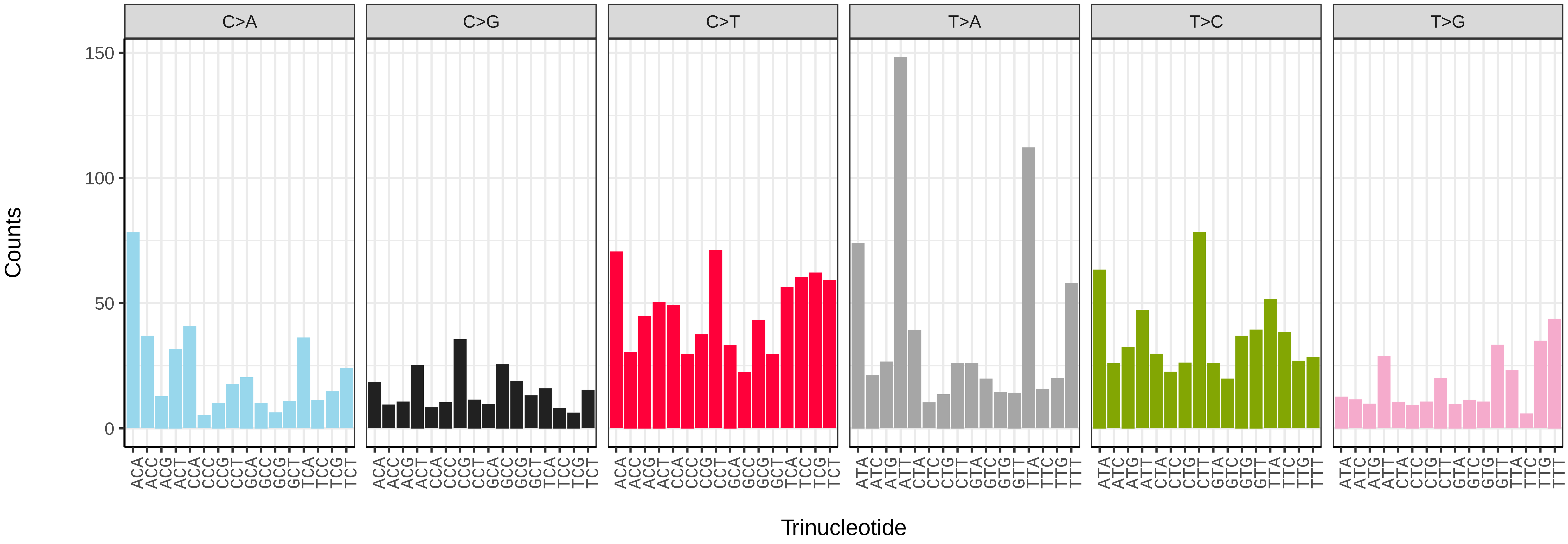


*Amphipoea oculea* (ilAmpOcul1) somatic mutational spectrum (SBS52 classification)

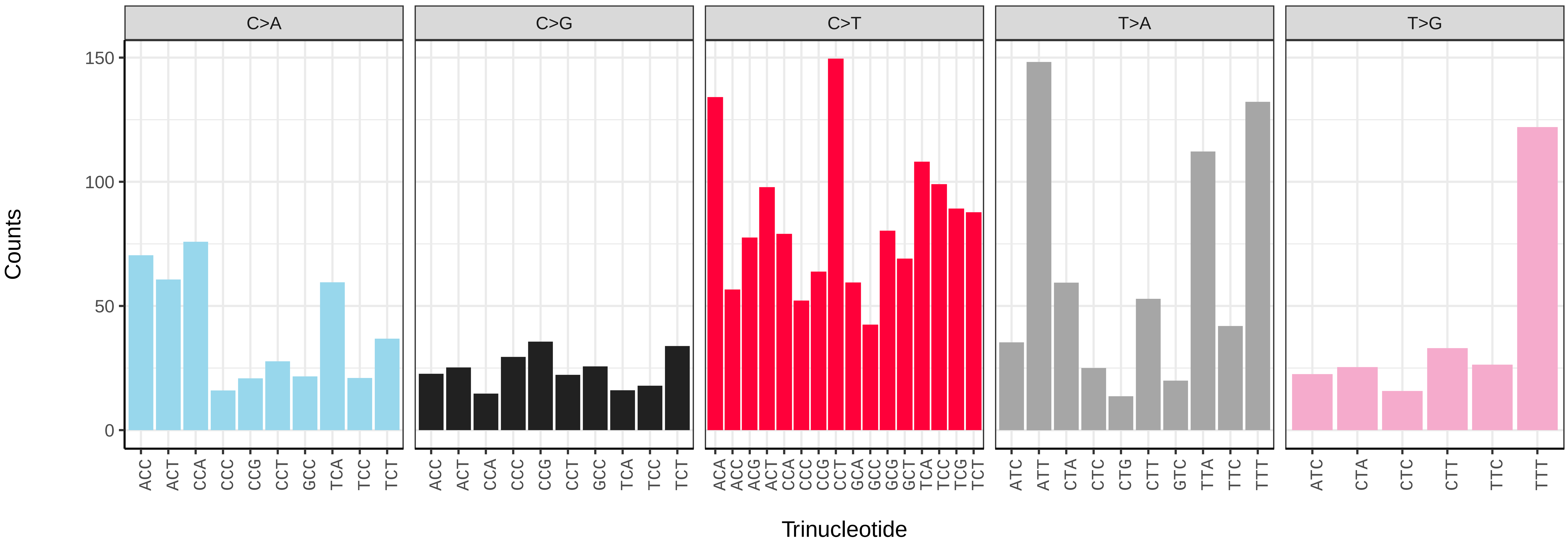

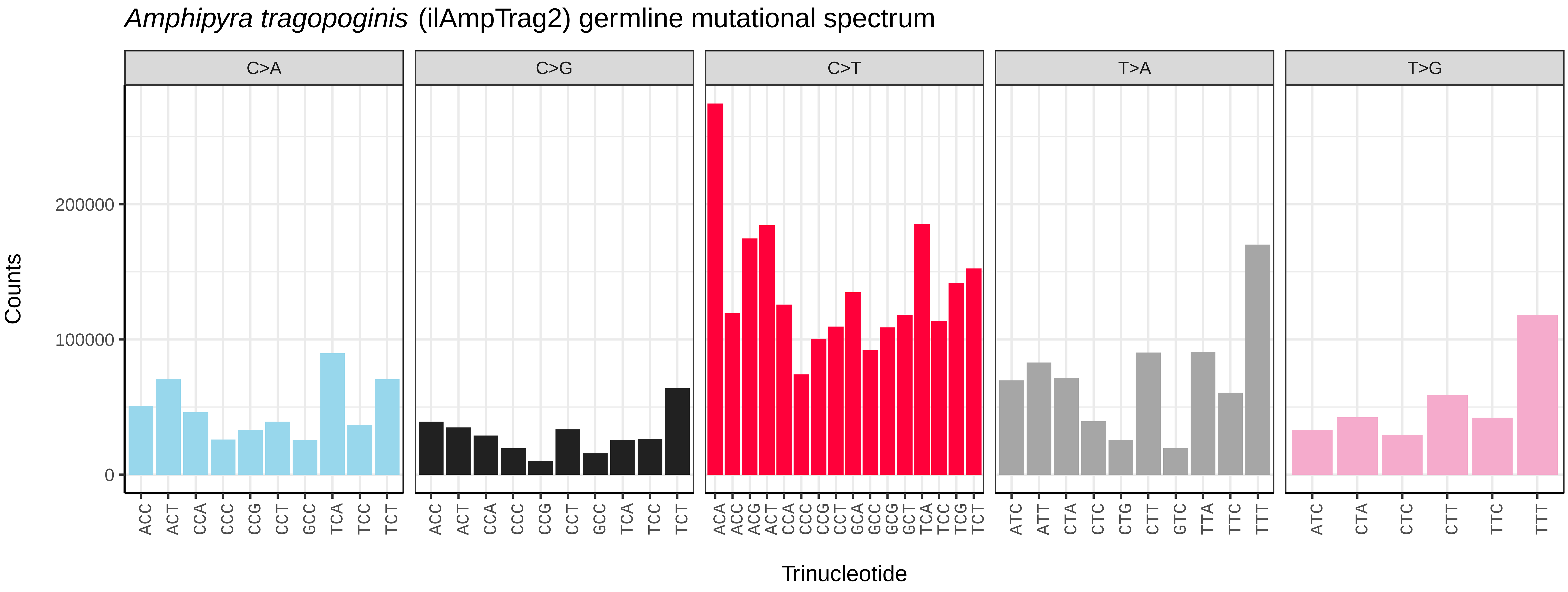
*Amphipyra tragopoginis* (ilAmpTrag2) germline mutational spectrum
C>A
C>G
C>T
T>A
T>G
Counts
Trinucleotide
0
100000
200000

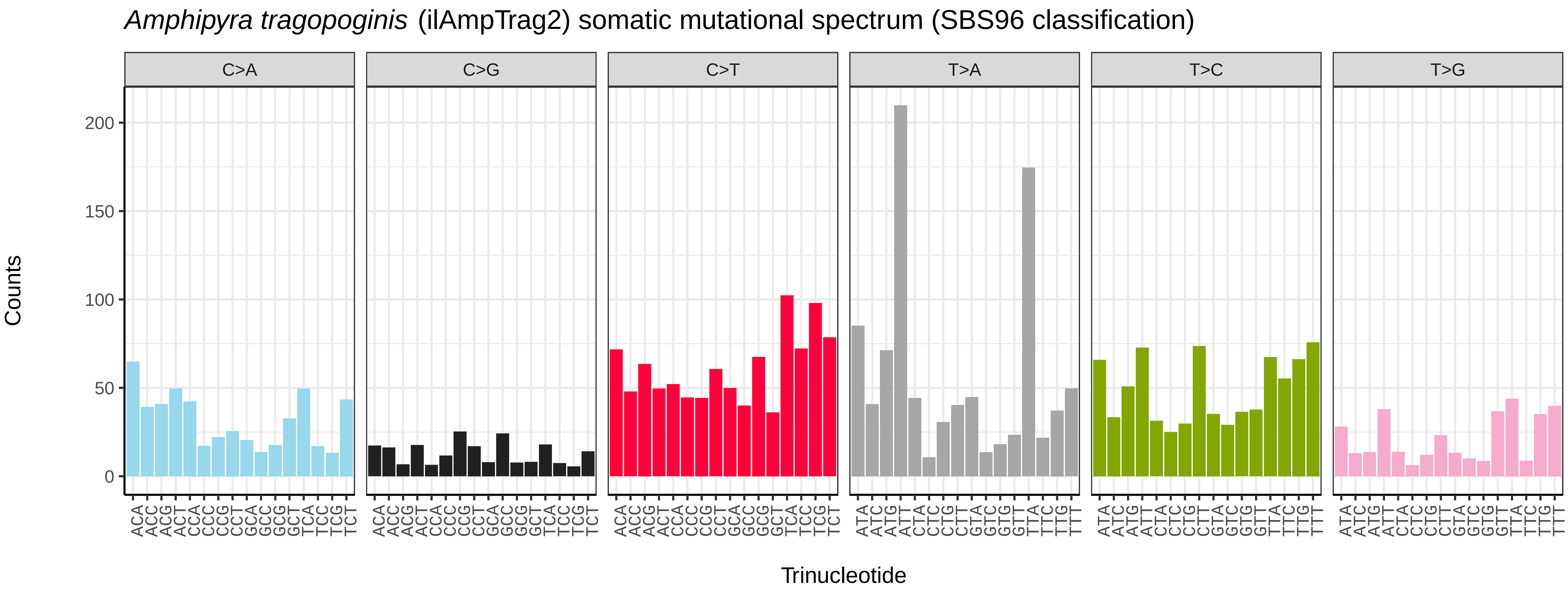
*Amphipyra tragopoginis* (ilAmpTrag2) somatic mutational spectrum (SBS96 classification)
C>A
C>G
C>T
T>A
T>C
T>G
Counts
Trinucleotide
0
50
100
150
200

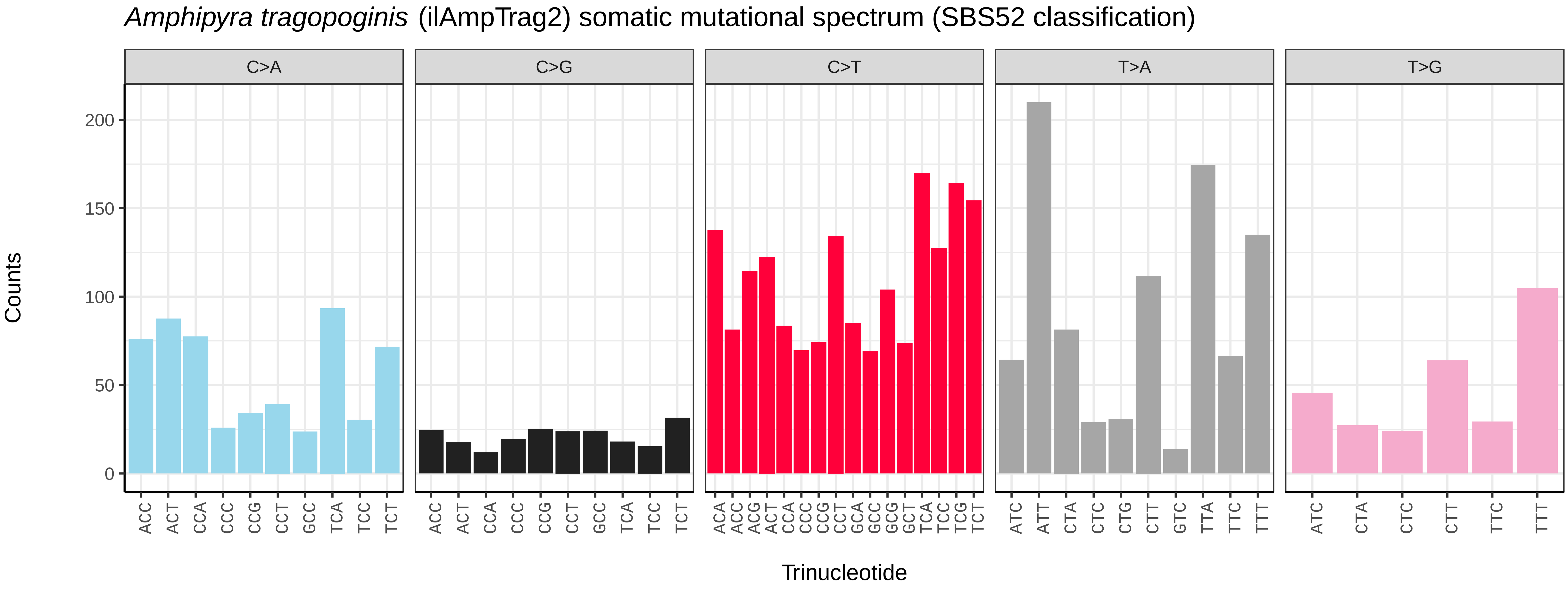
*Amphipyra tragopoginis* (ilAmpTrag2) somatic mutational spectrum (SBS52 classification)
C>A
C>G
C>T
T>A
T>G
Counts
Trinucleotide
0
50
100
150
200

*Anarsia innoxiella* (ilAnaInnx2) germline mutational spectrum

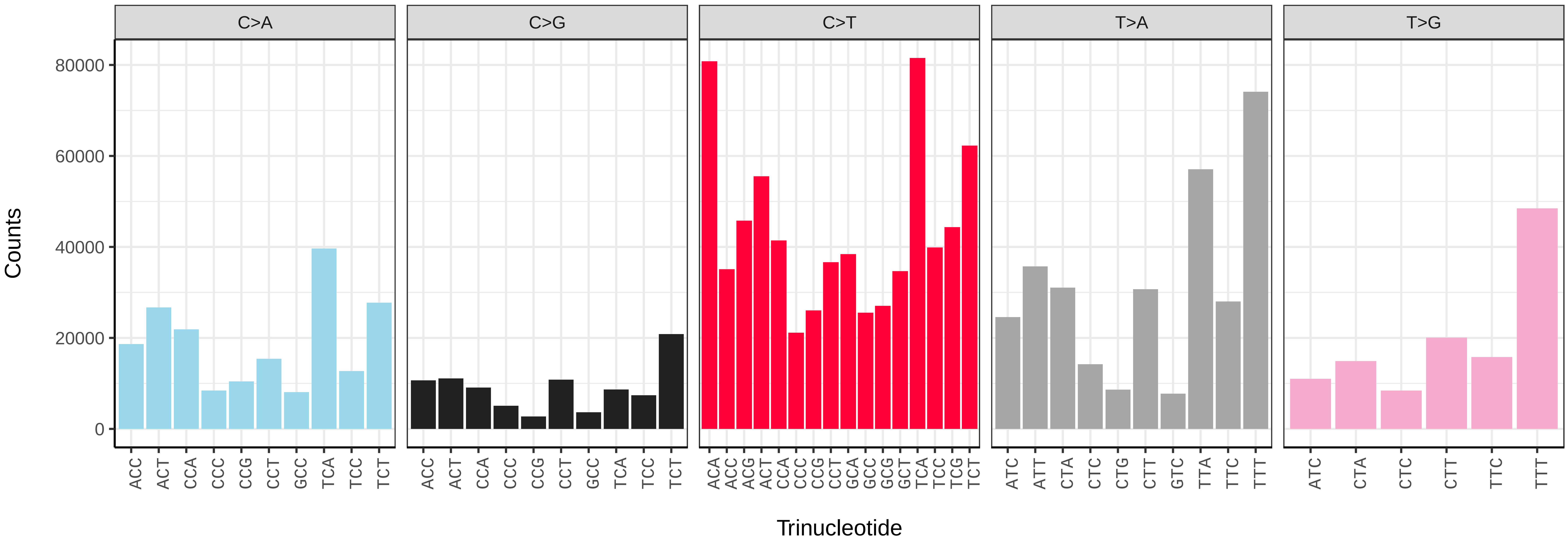


*Anarsia innoxiella* (ilAnaInnx2) somatic mutational spectrum (SBS96 classification)

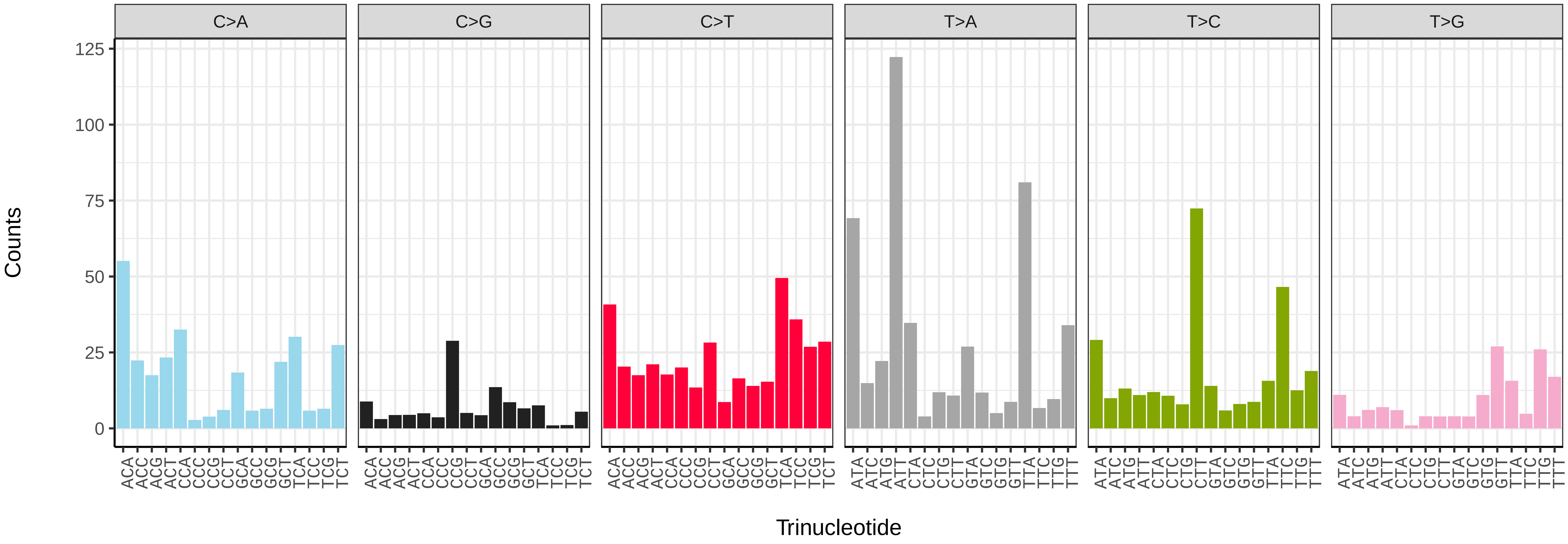


*Anarsia innoxiella* (ilAnaInnx2) somatic mutational spectrum (SBS52 classification)

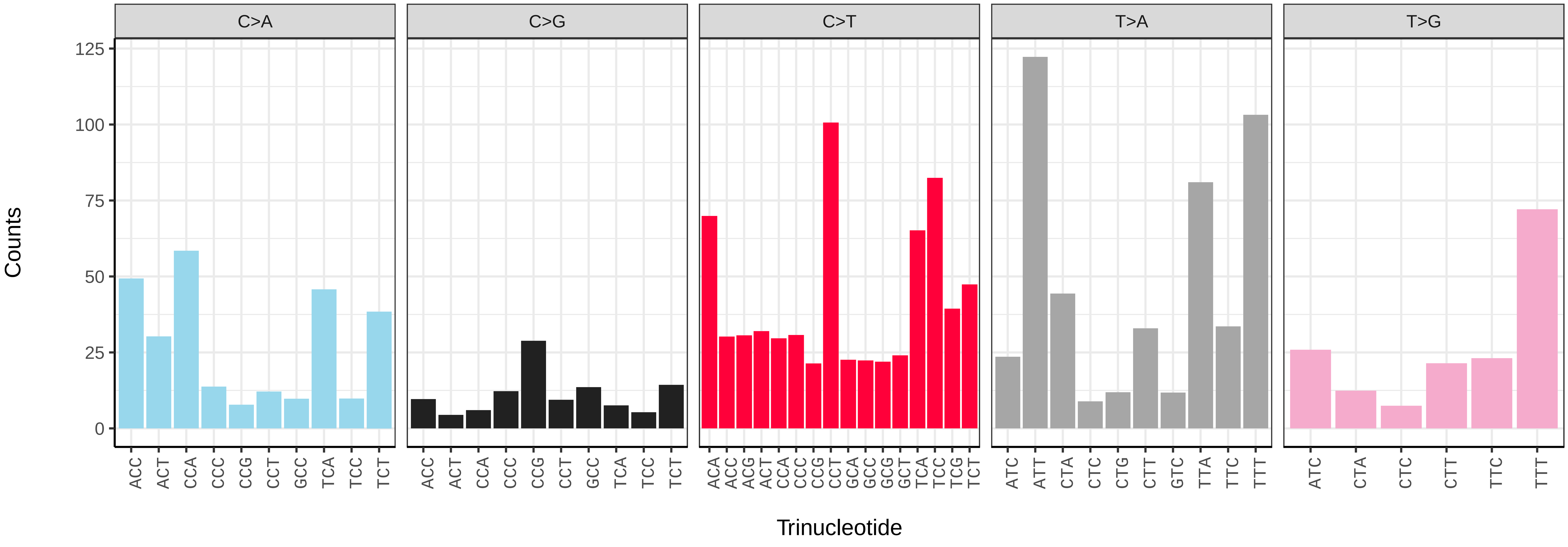

*Anorthoa munda* (ilAnoMund1) germline mutational spectrum

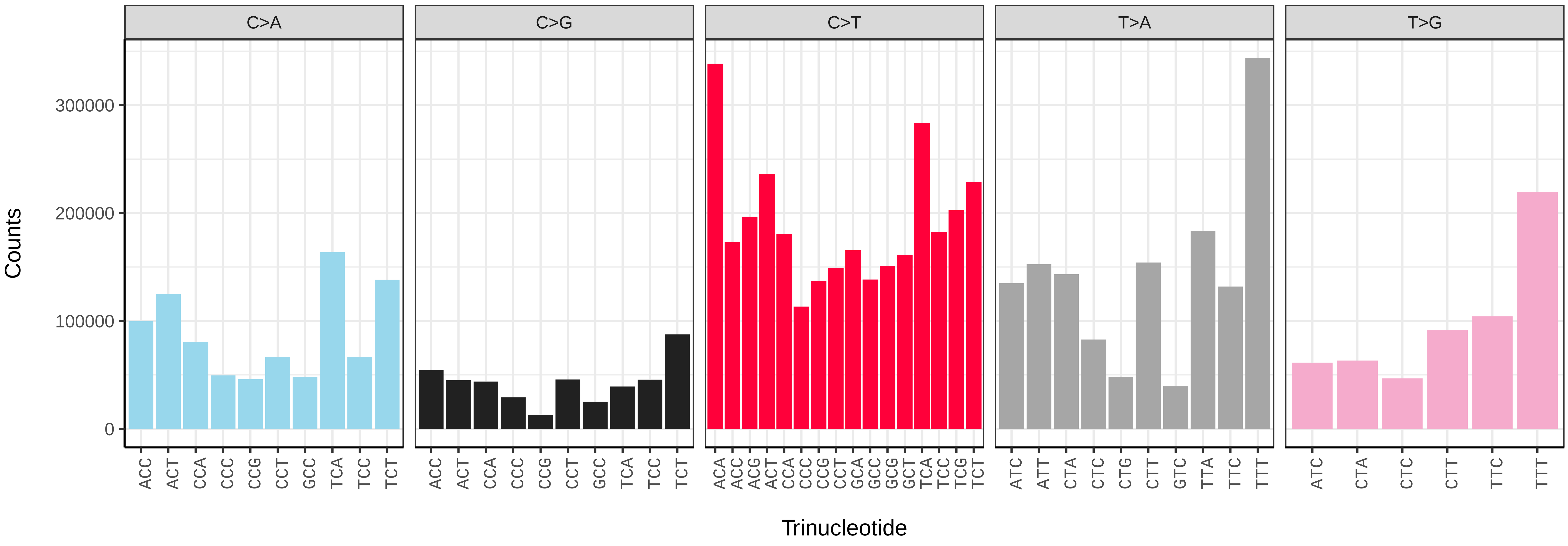


*Anorthoa munda* (ilAnoMund1) somatic mutational spectrum (SBS96 classification)

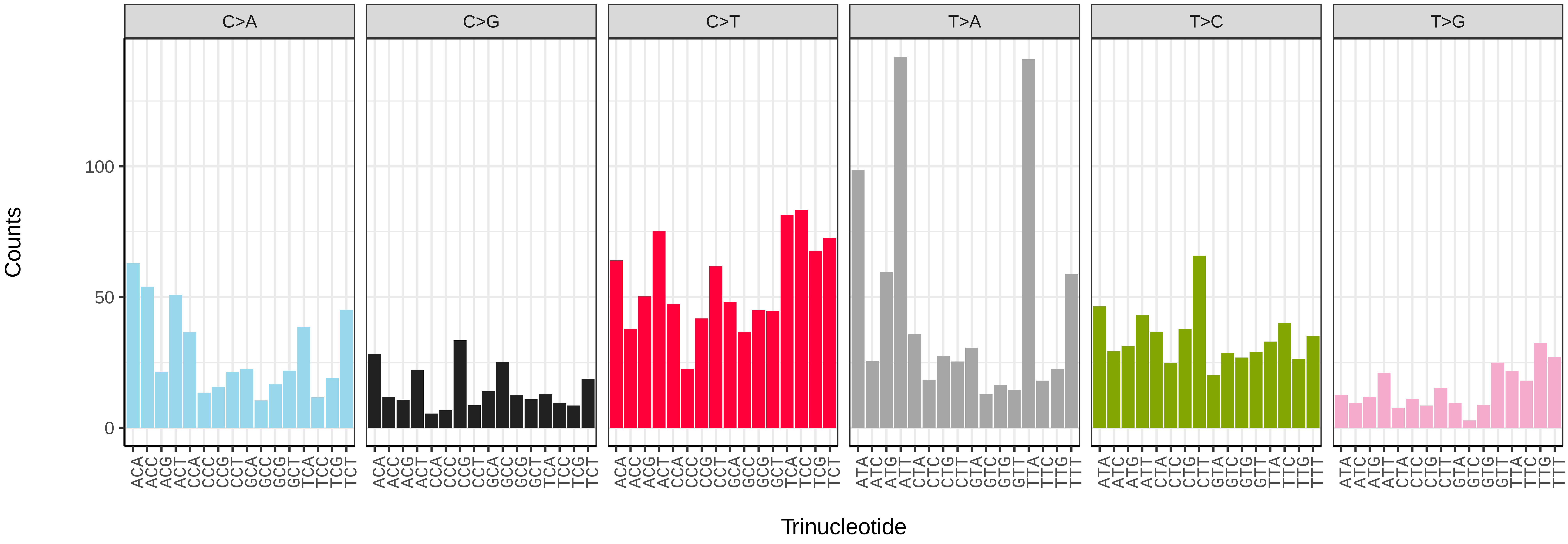


*Anorthoa munda* (ilAnoMund1) somatic mutational spectrum (SBS52 classification)

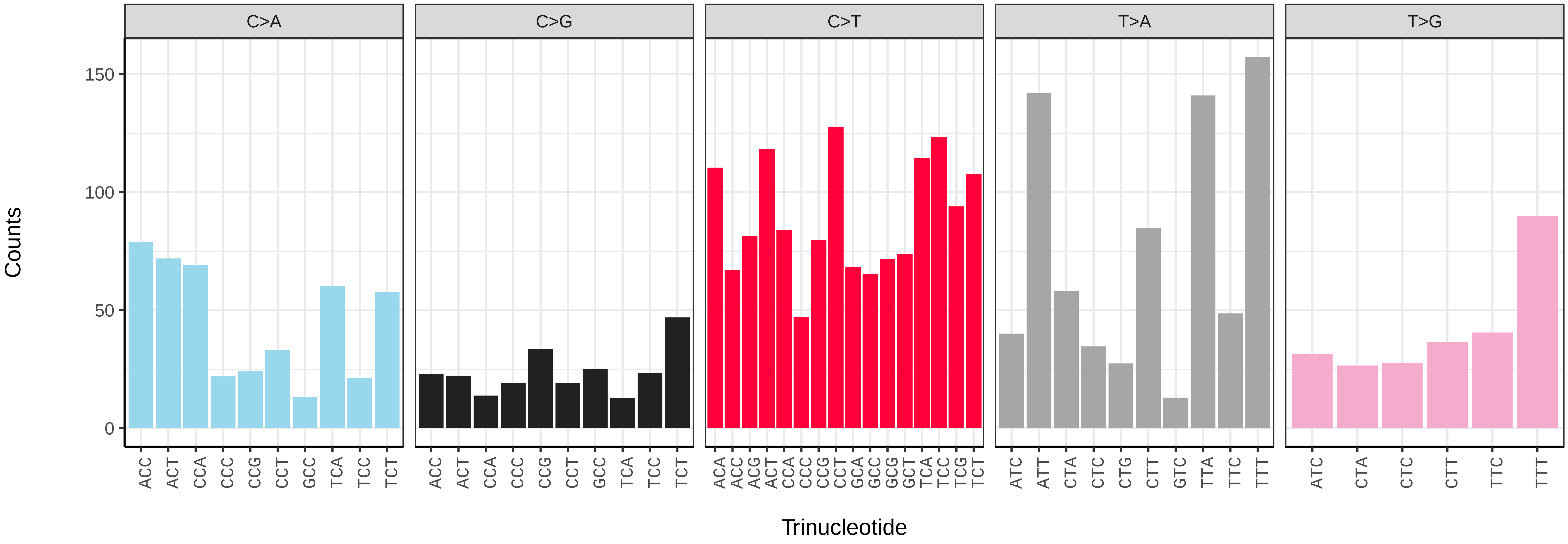

*Anthocharis cardamines* (ilAntCard3) germline mutational spectrum

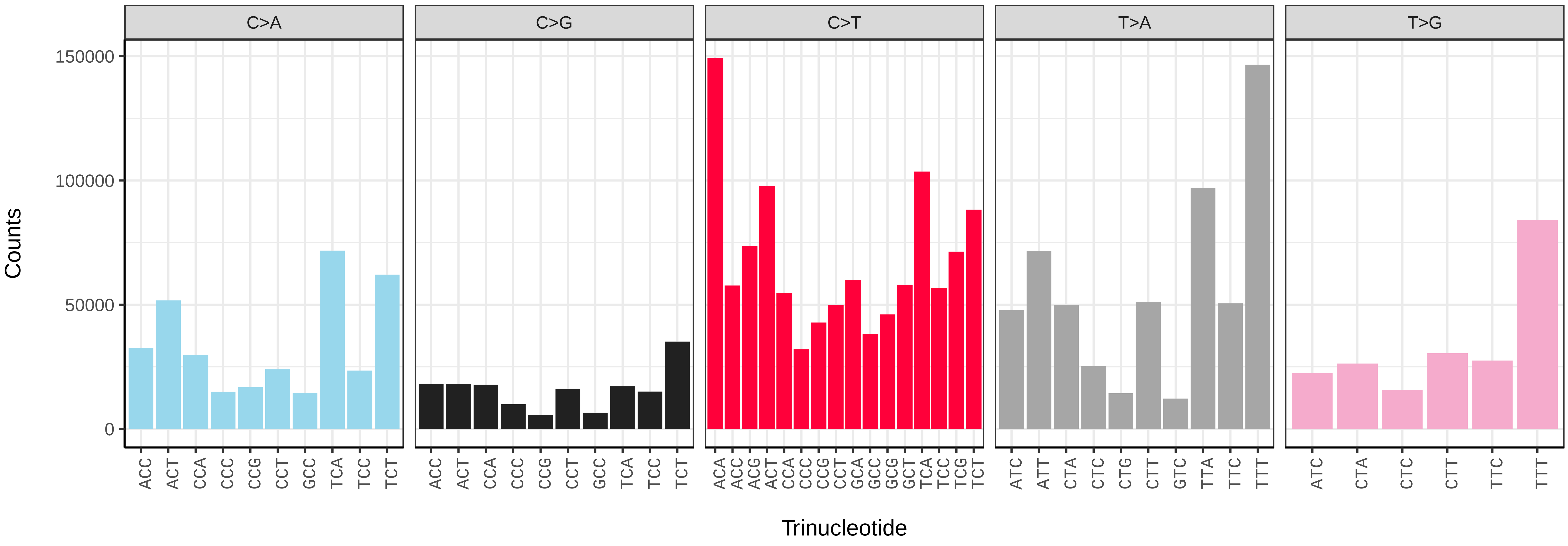


*Anthocharis cardamines* (ilAntCard3) somatic mutational spectrum (SBS96 classification)

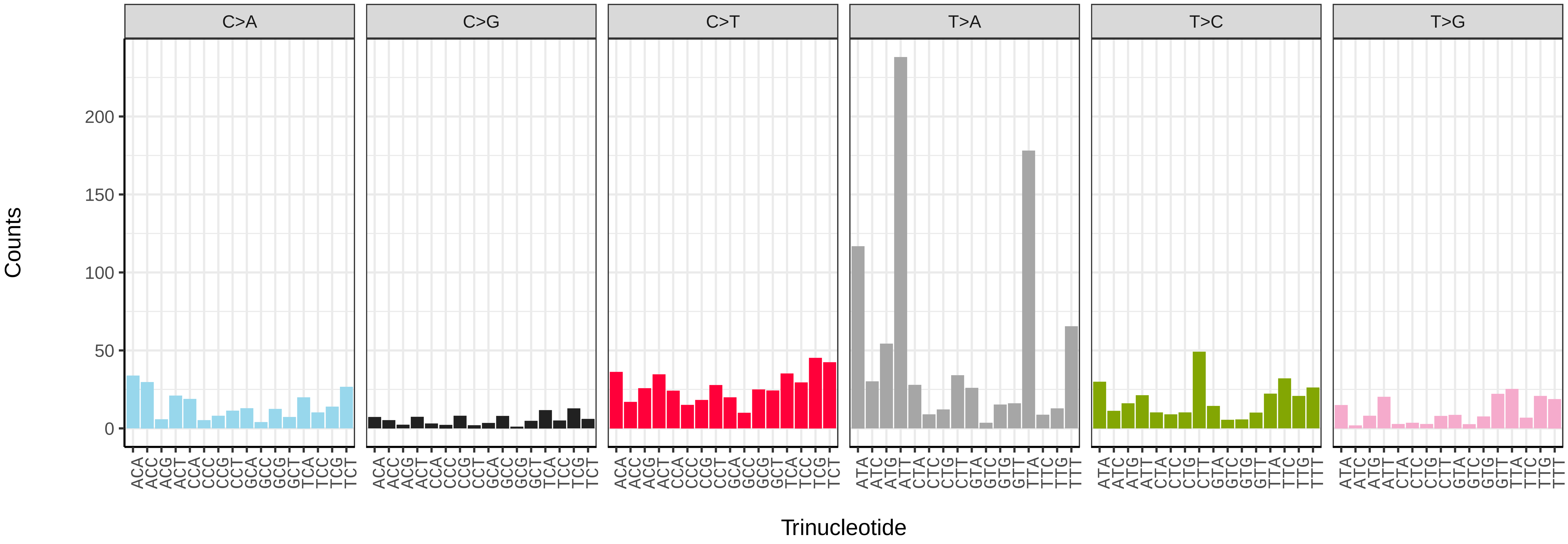


*Anthocharis cardamines* (ilAntCard3) somatic mutational spectrum (SBS52 classification)

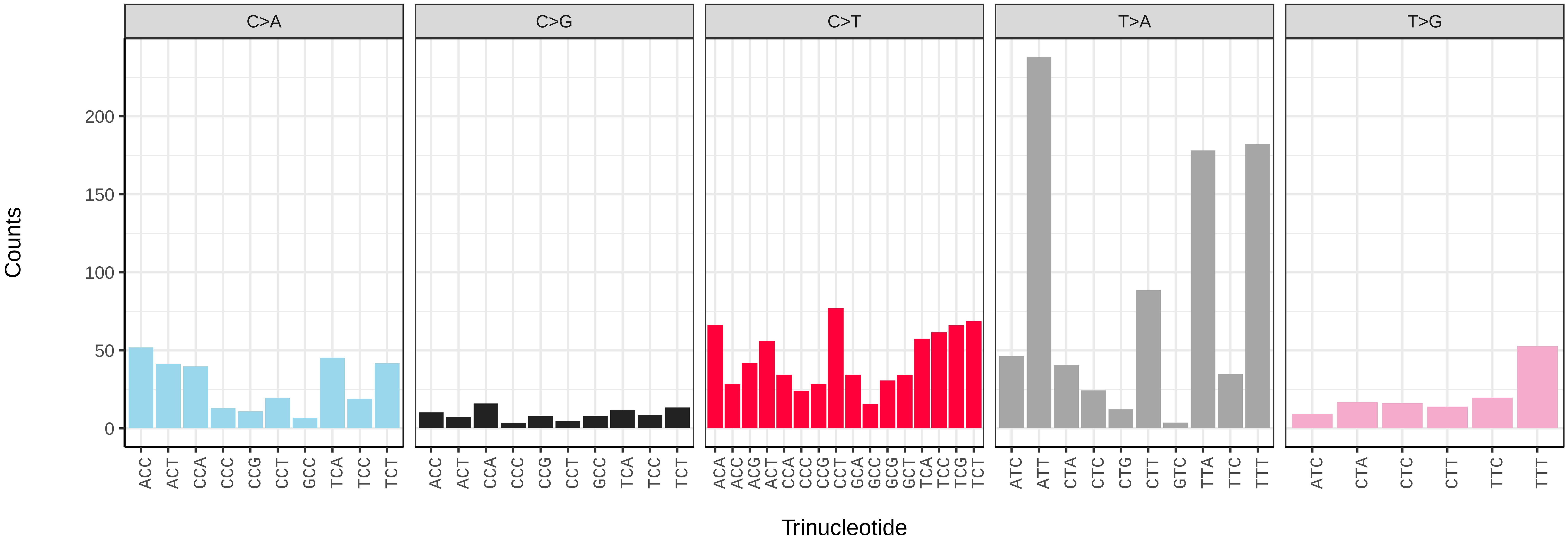

*Antitype chi* (ilAntChix2) germline mutational spectrum

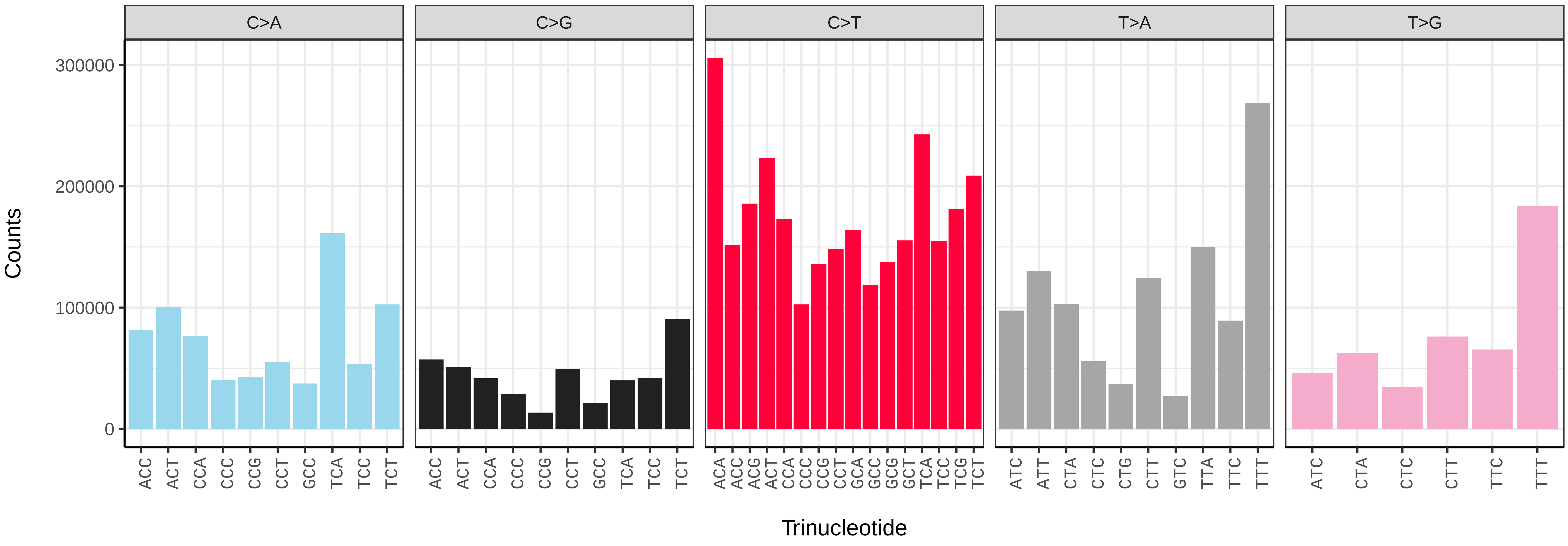


*Antitype chi* (ilAntChix2) somatic mutational spectrum (SBS96 classification)

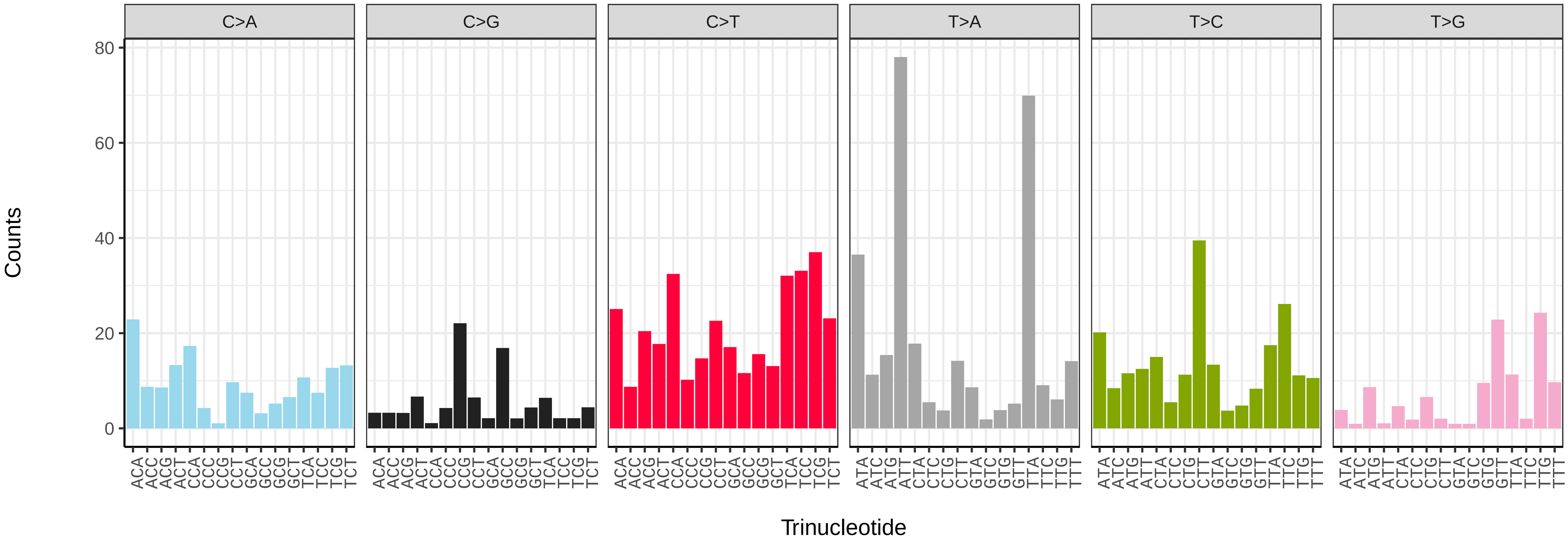


*Antitype chi* (ilAntChix2) somatic mutational spectrum (SBS52 classification)

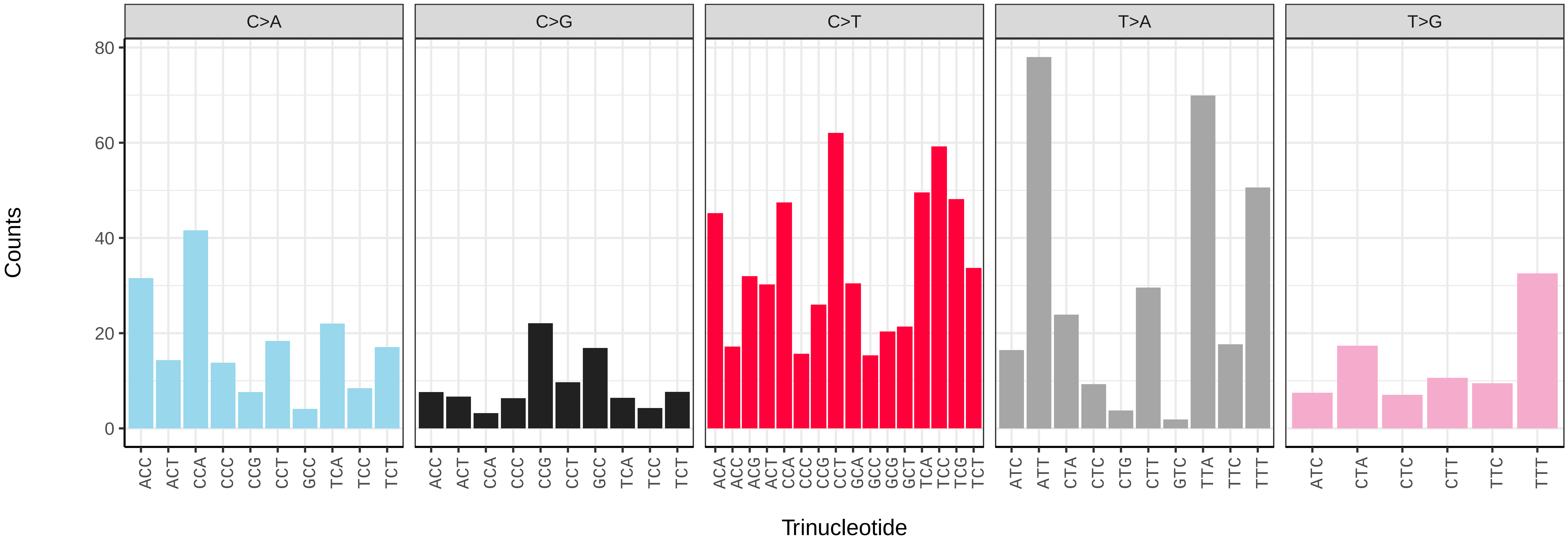

*Anticlea derivata* (ilAntDeri1) germline mutational spectrum

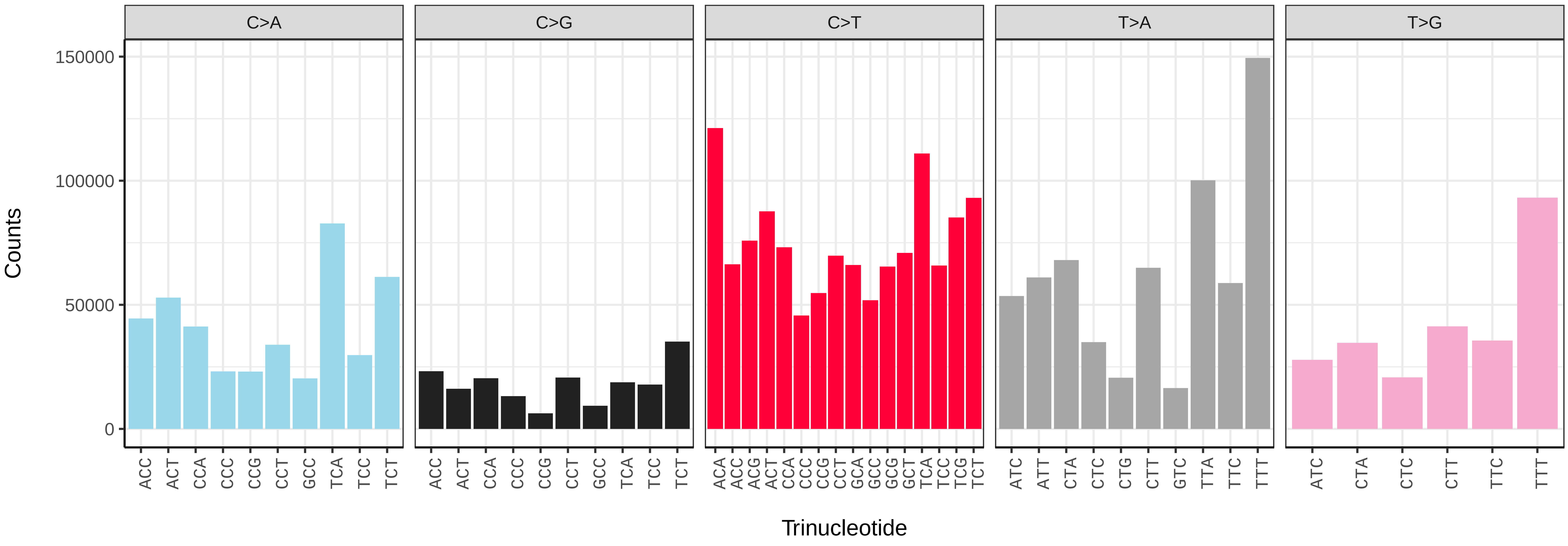


*Anticlea derivata* (ilAntDeri1) somatic mutational spectrum (SBS96 classification)

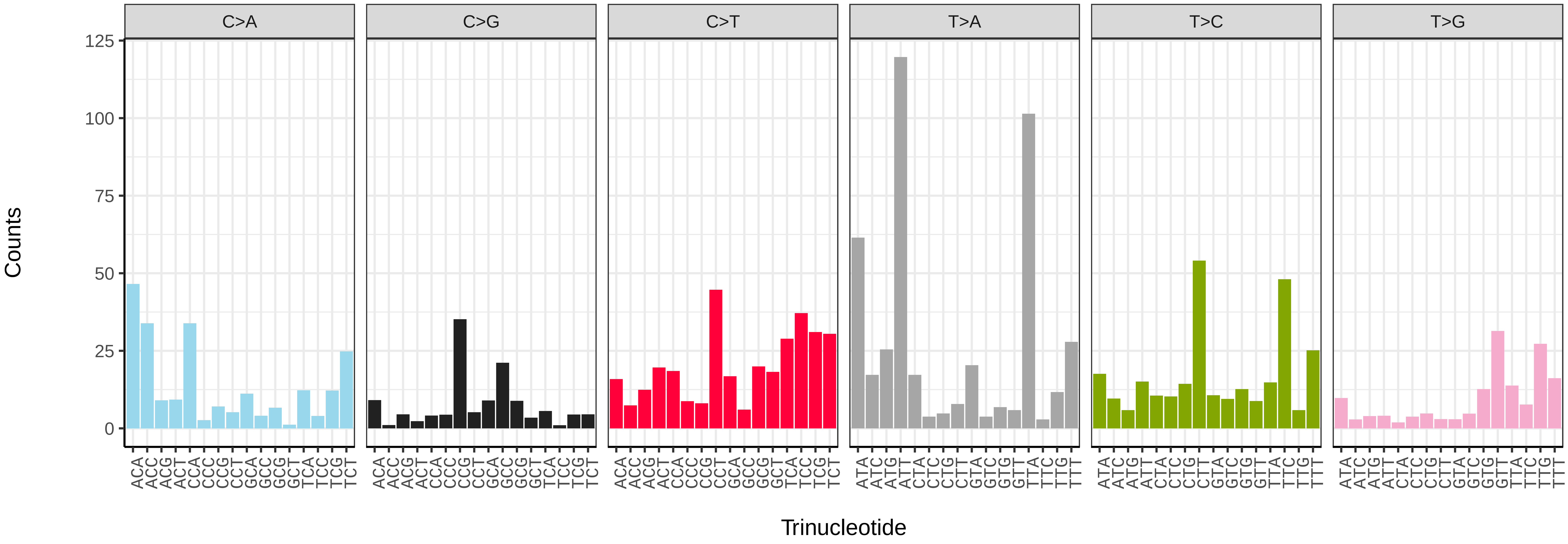


*Anticlea derivata* (ilAntDeri1) somatic mutational spectrum (SBS52 classification)

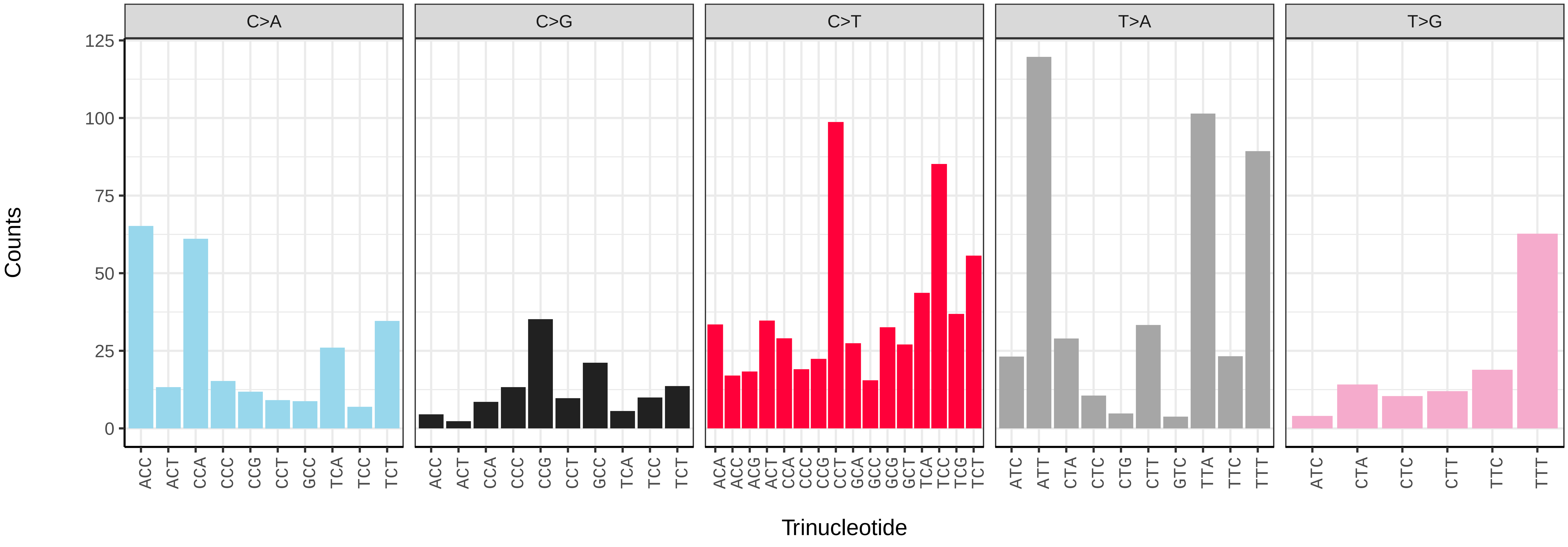

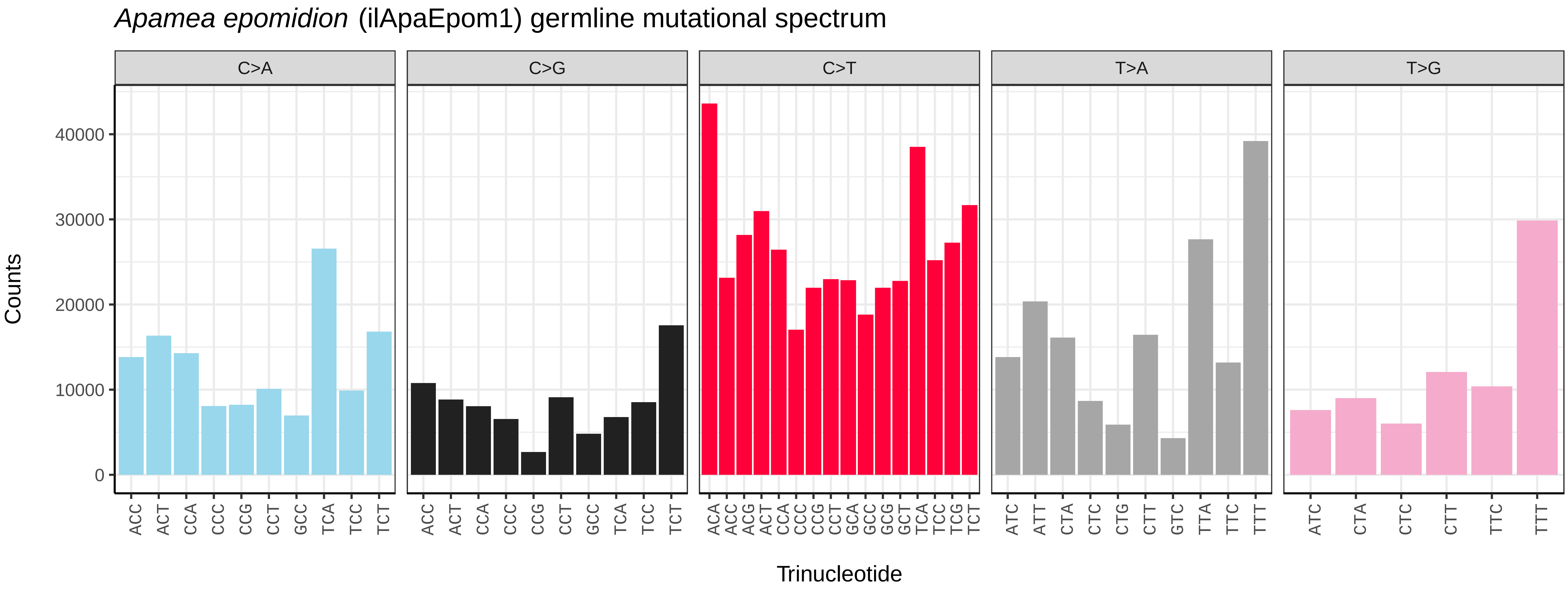
*Apamea epomidion* (ilApaEpom1) germline mutational spectrum
C>A
C>G
C>T
T>A
T>G
Counts
0
10000
20000
30000
40000
Trinucleotide
ACC ACT CCA CCC CCG CCT GCC TCA TCC TCT
ACA ACC ACG ACT CCA CCC CCG CCT GCA GCC GCG GCT TCA TCC TCG TCT
ATC ATT CTA CTC CTG CTT GTC TTA TTC TTT
ATC CTA CTC CTT TTC TTT

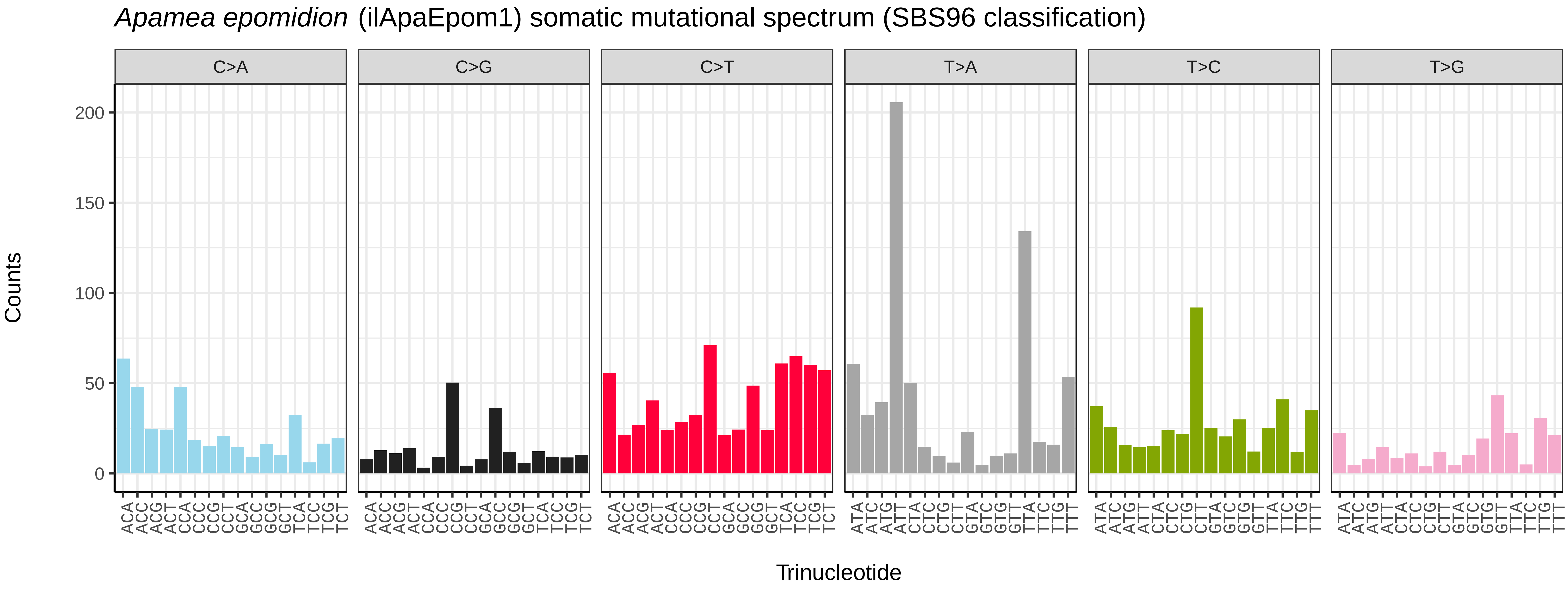
*Apamea epomidion* (ilApaEpom1) somatic mutational spectrum (SBS96 classification)
C>A
C>G
C>T
T>A
T>C
T>G
Counts
0
50
100
150
200
Trinucleotide
ACA ACC ACG ACT CCA CCC CCG CCT GCA GCC GCG GCT TCA TCC TCG TCT
ATA ATC ATG ATT CTA CTC CTG CTT GTA GTC GTG GTT TTA TTC TTG TTT

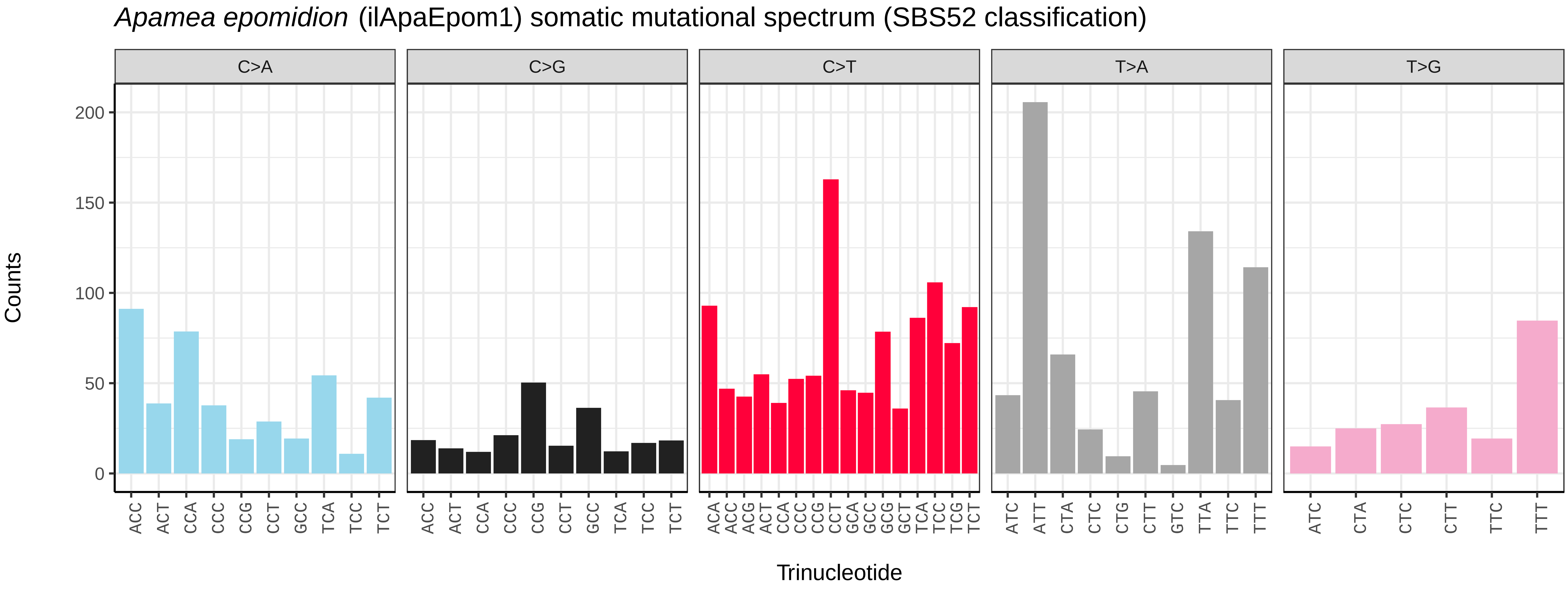
*Apamea epomidion* (ilApaEpom1) somatic mutational spectrum (SBS52 classification)
C>A
C>G
C>T
T>A
T>G
Counts
0
50
100
150
200
Trinucleotide
ACC ACT CCA CCC CCG CCT GCC TCA TCC TCT
ACA ACC ACG ACT CCA CCC CCG CCT GCA GCC GCG GCT TCA TCC TCG TCT
ATC ATT CTA CTC CTG CTT GTC TTA TTC TTT
ATC CTA CTC CTT TTC TTT

*Apamea monoglypha* (ilApaMono1) germline mutational spectrum

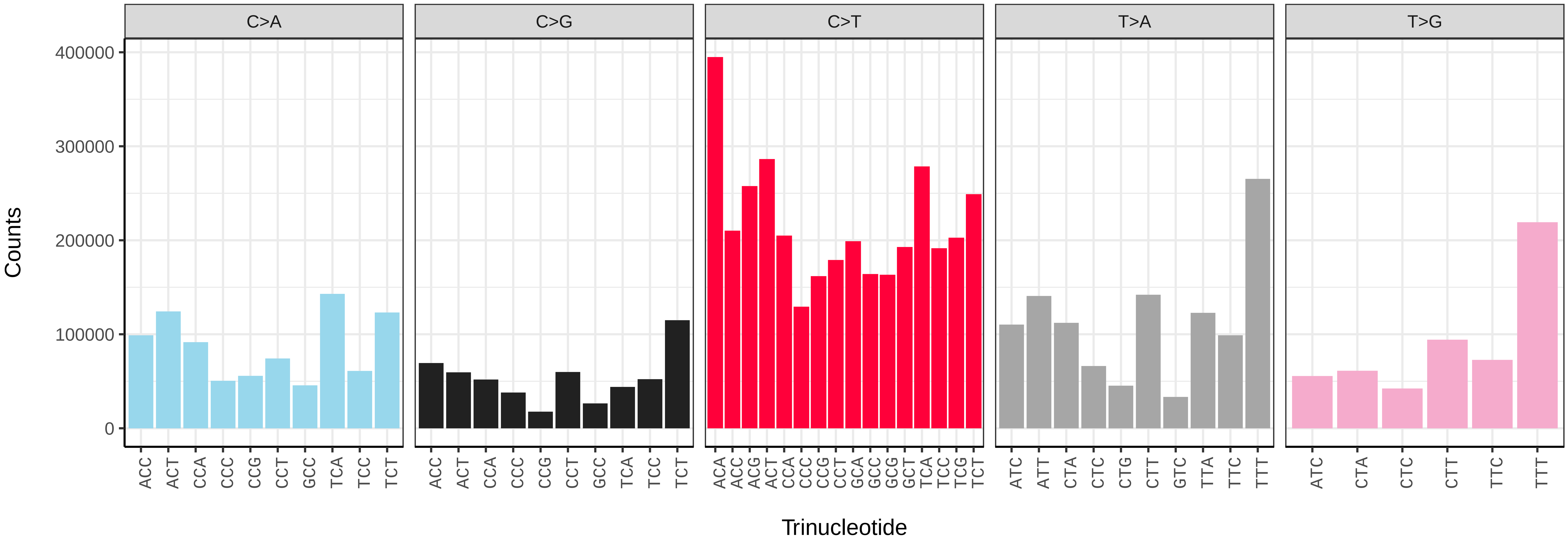


*Apamea monoglypha* (ilApaMono1) somatic mutational spectrum (SBS96 classification)

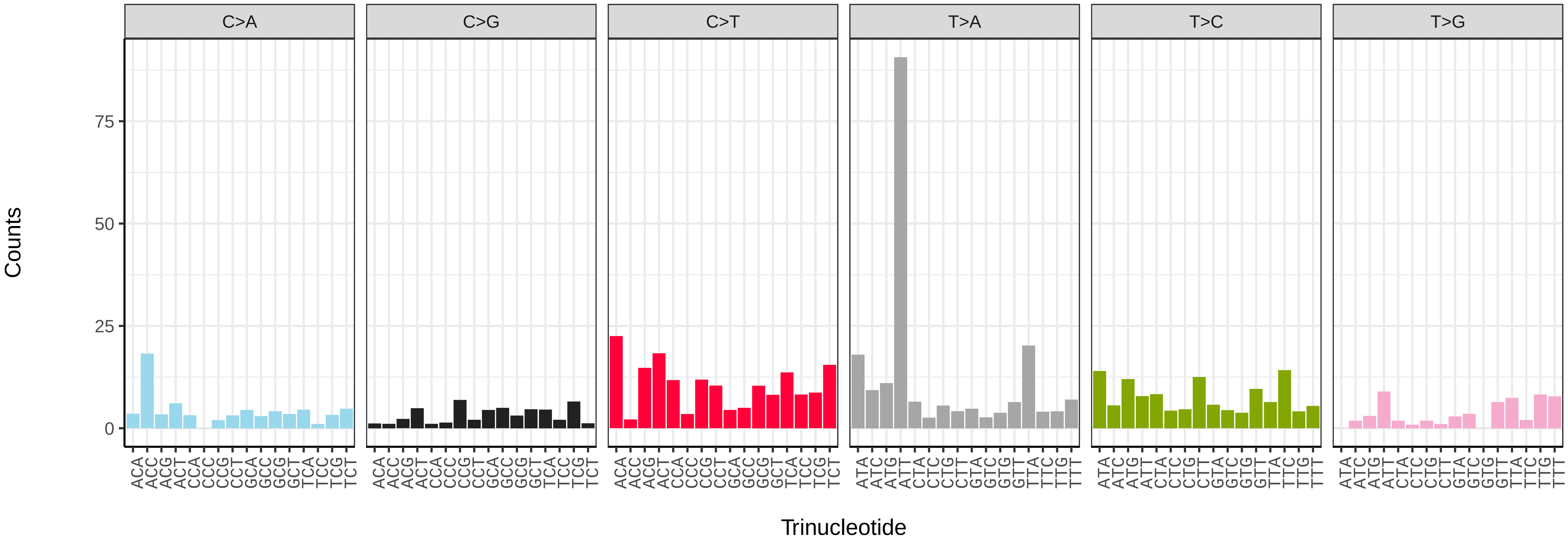


*Apamea monoglypha* (ilApaMono1) somatic mutational spectrum (SBS52 classification)

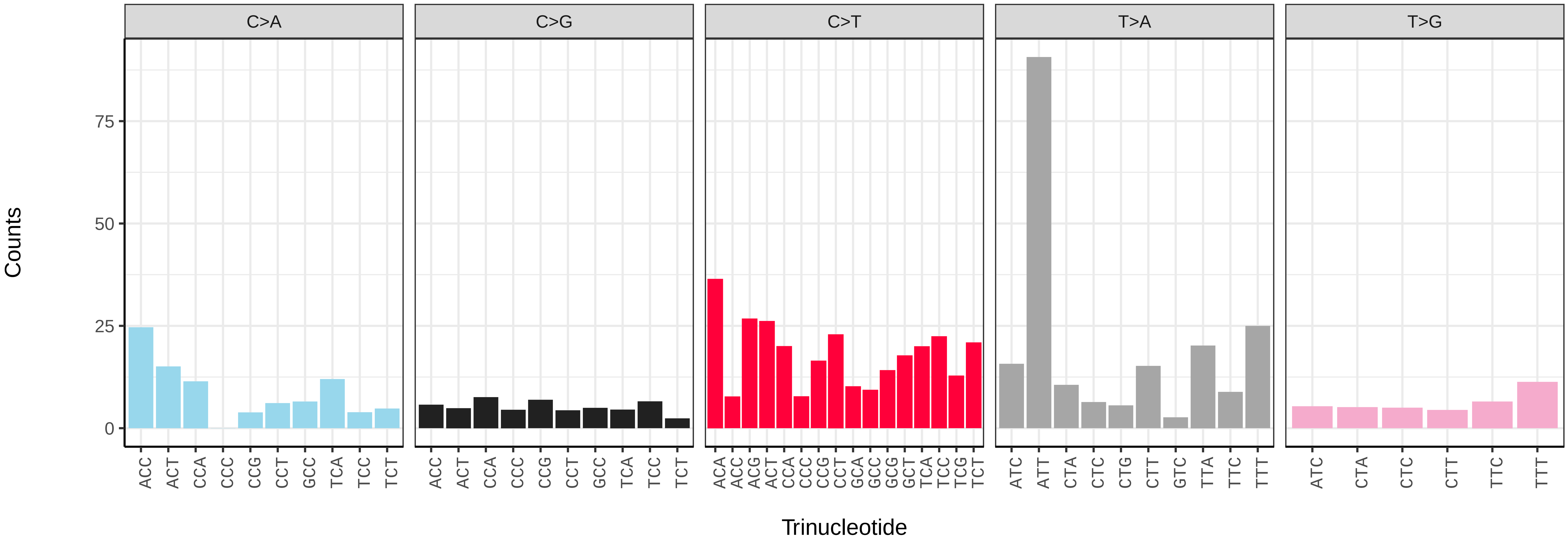

*Apamea sordens* (ilApaSord1) germline mutational spectrum

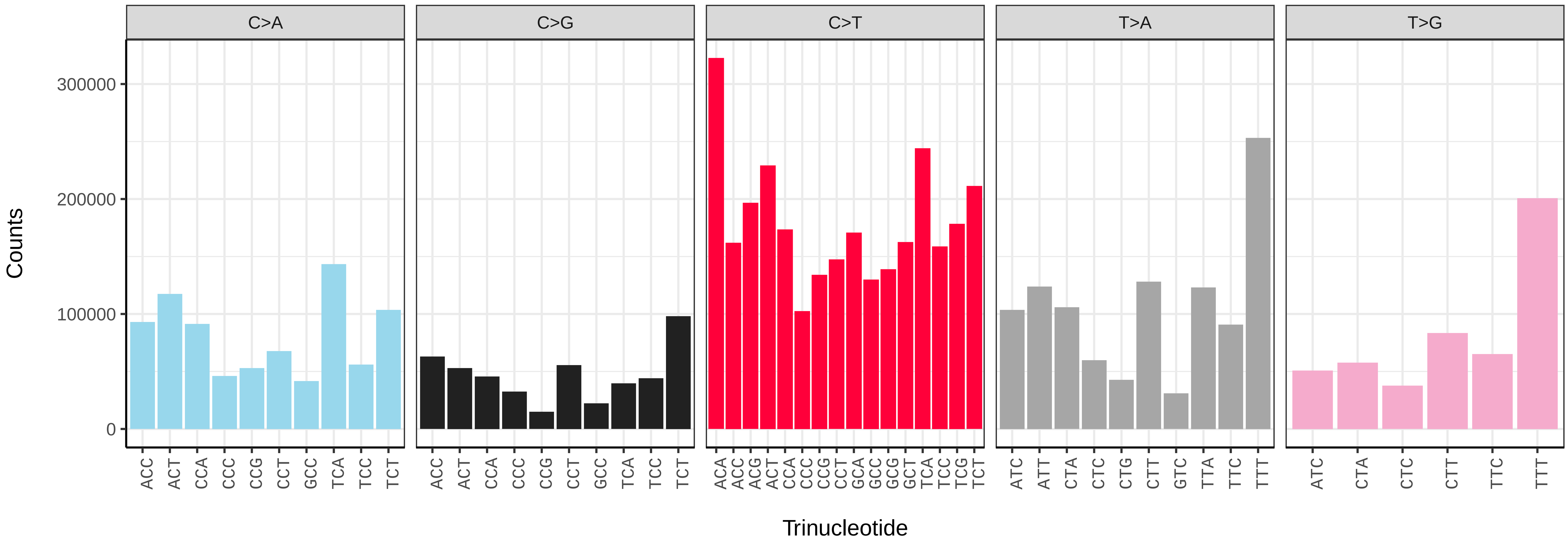


*Apamea sordens* (ilApaSord1) somatic mutational spectrum (SBS96 classification)

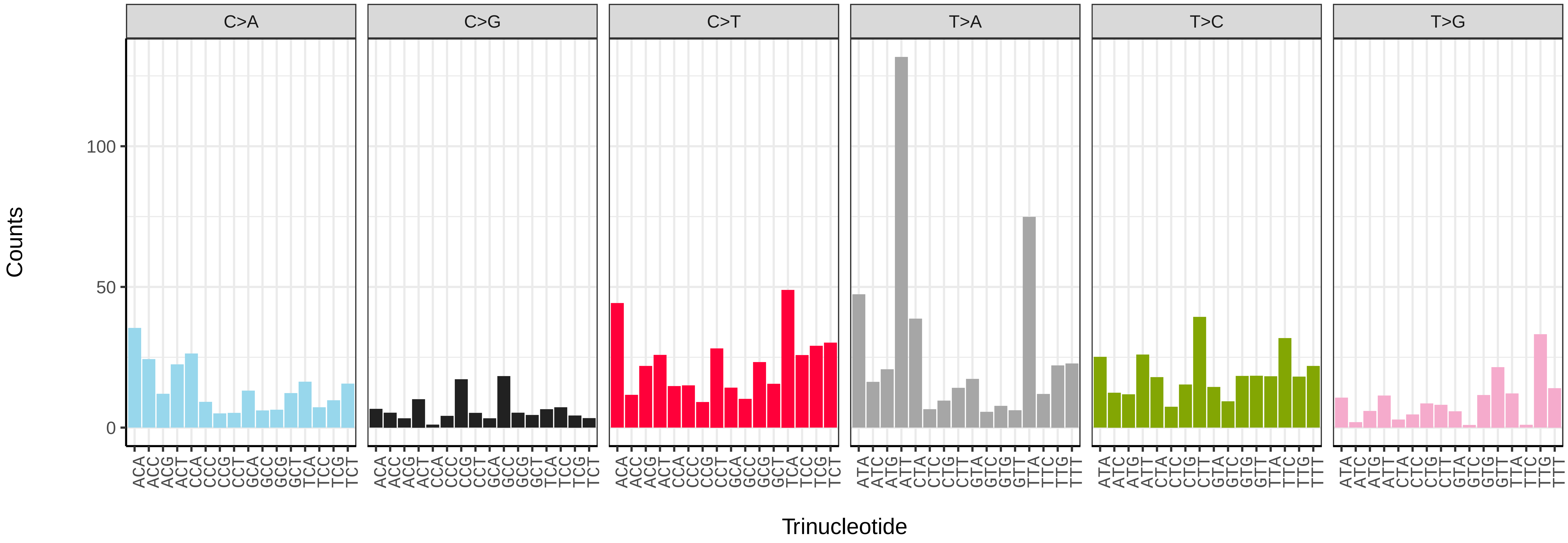


*Apamea sordens* (ilApaSord1) somatic mutational spectrum (SBS52 classification)

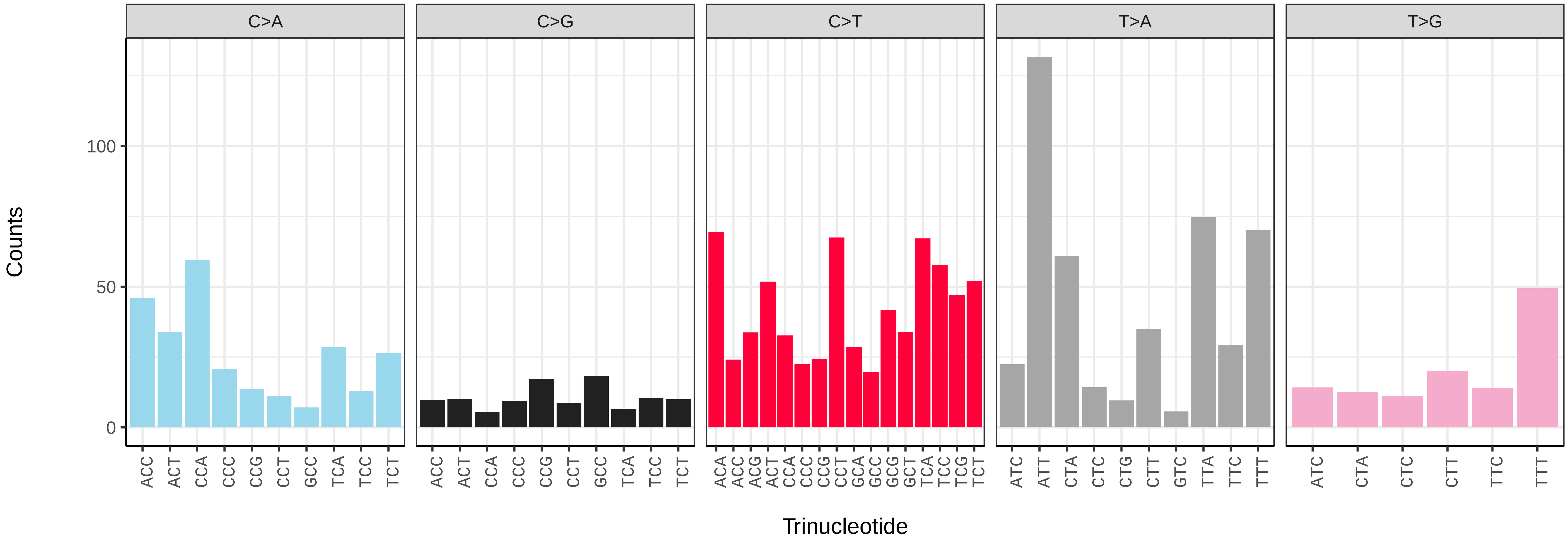

*Apeira syringaria* (ilApeSyri1) germline mutational spectrum

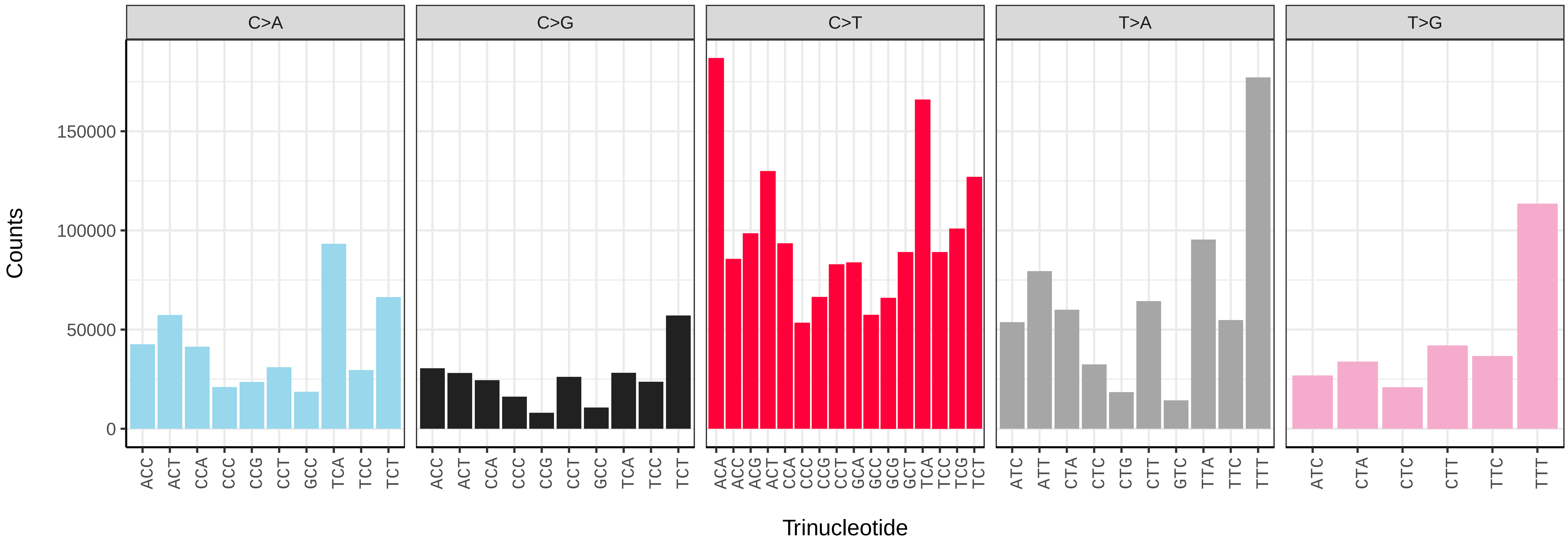


*Apeira syringaria* (ilApeSyri1) somatic mutational spectrum (SBS96 classification)

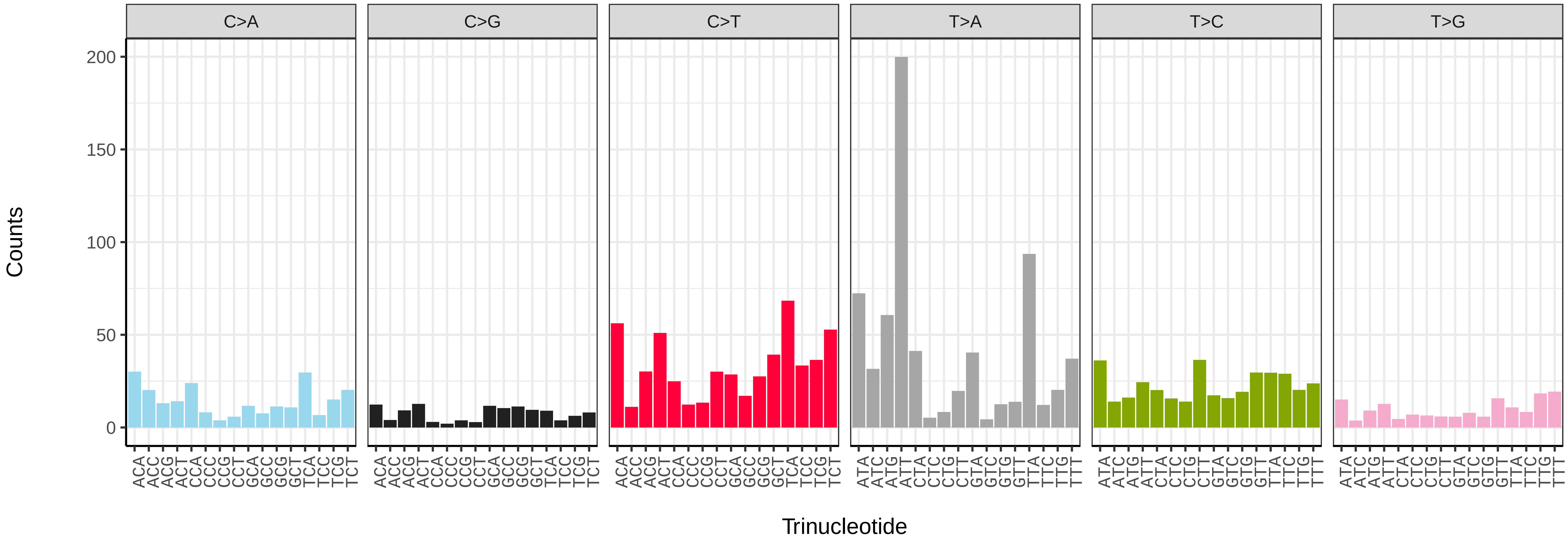


*Apeira syringaria* (ilApeSyri1) somatic mutational spectrum (SBS52 classification)

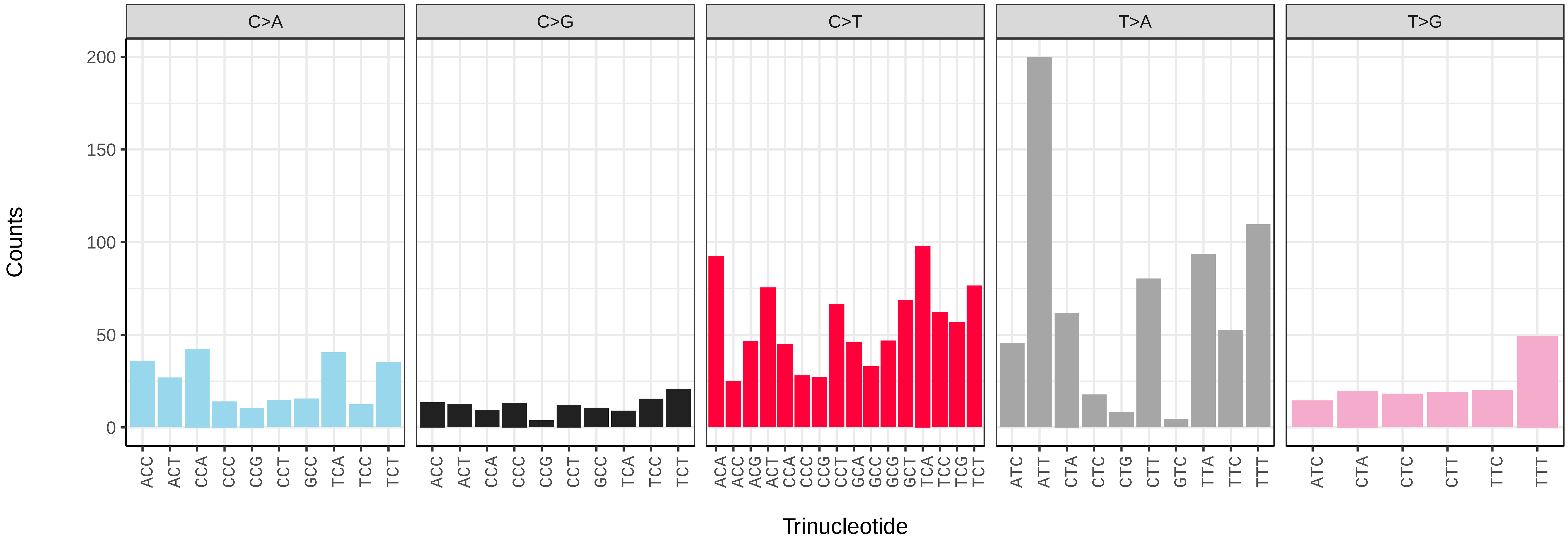

*Aplocera efformata* (ilAplEffo1) germline mutational spectrum

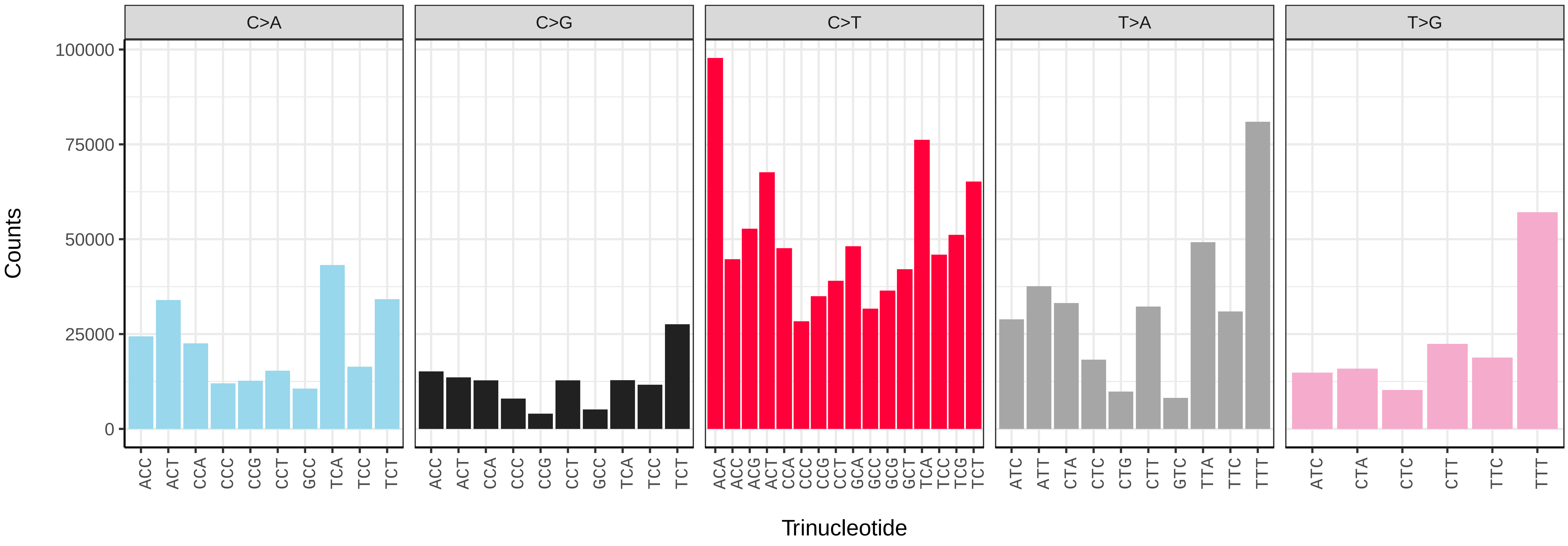


*Aplocera efformata* (ilAplEffo1) somatic mutational spectrum (SBS96 classification)

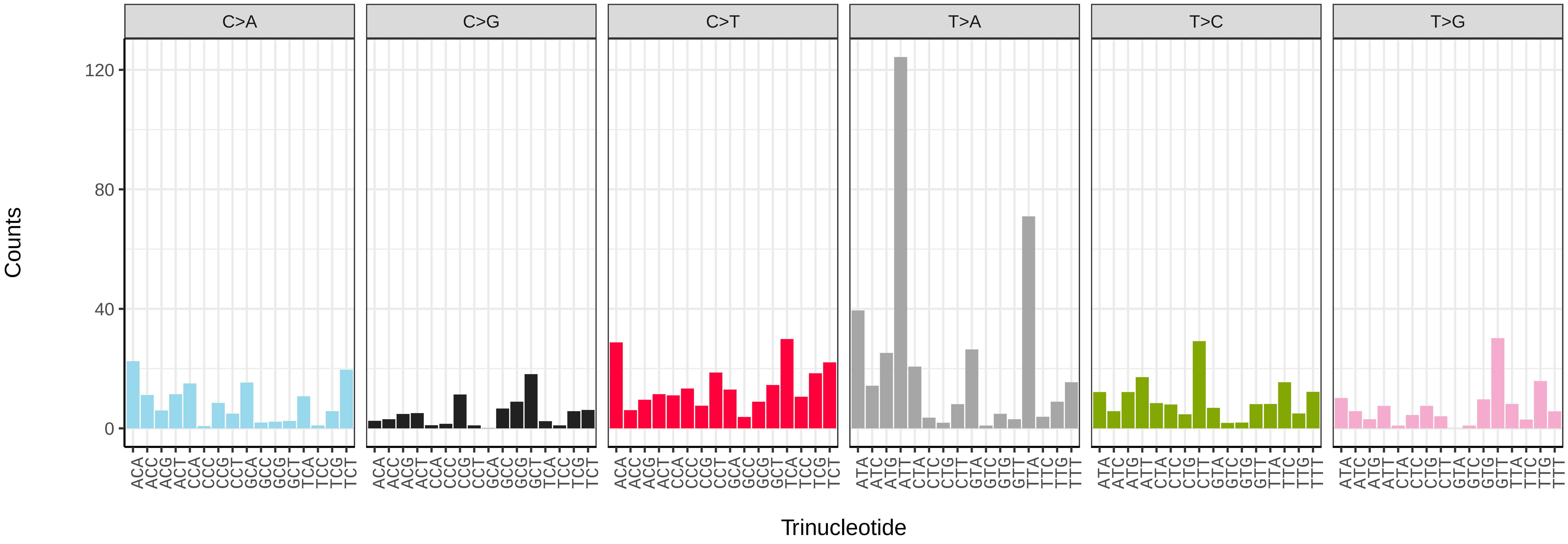


*Aplocera efformata* (ilAplEffo1) somatic mutational spectrum (SBS52 classification)

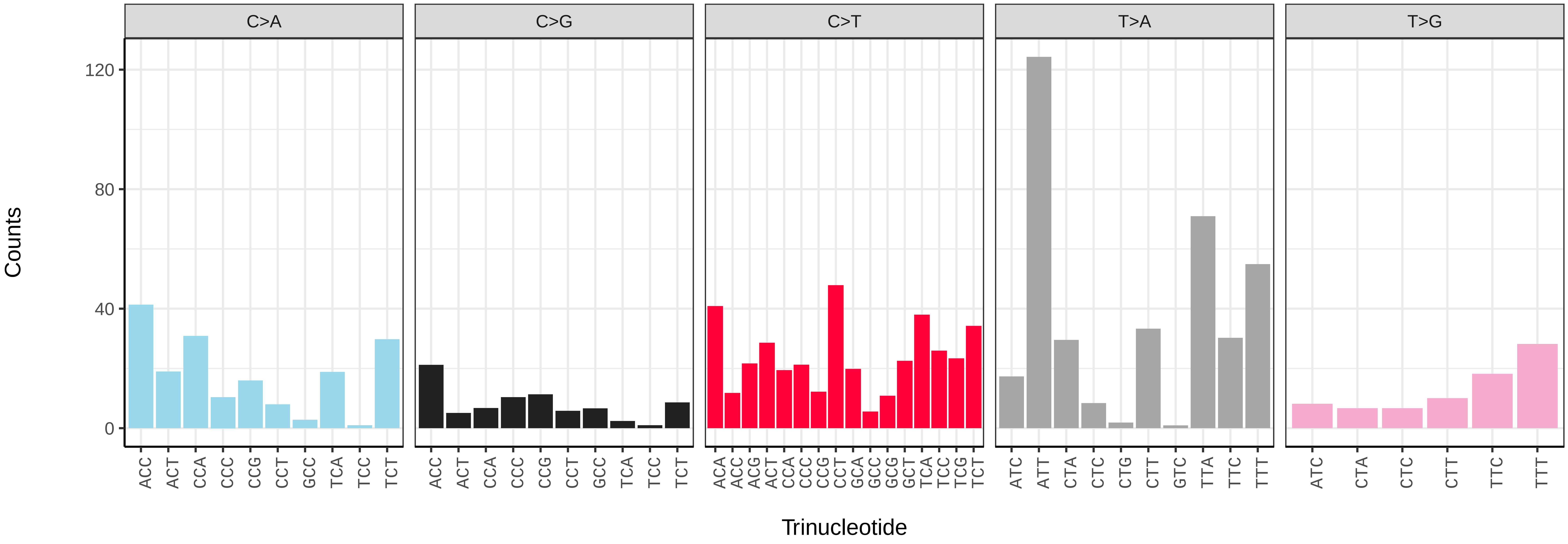

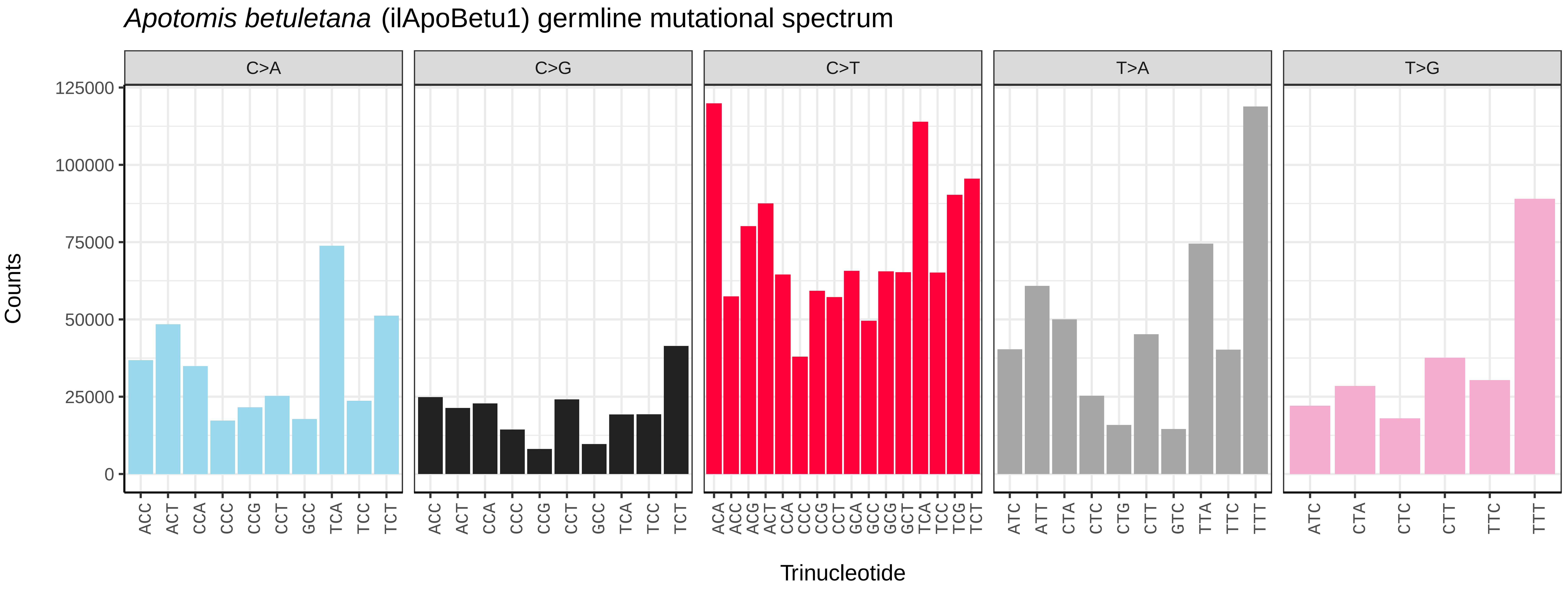
*Apotomis betuletana* (ilApoBetu1) germline mutational spectrum
C>A
C>G
C>T
T>A
T>G
Counts
125000
100000
75000
50000
25000
0
Trinucleotide

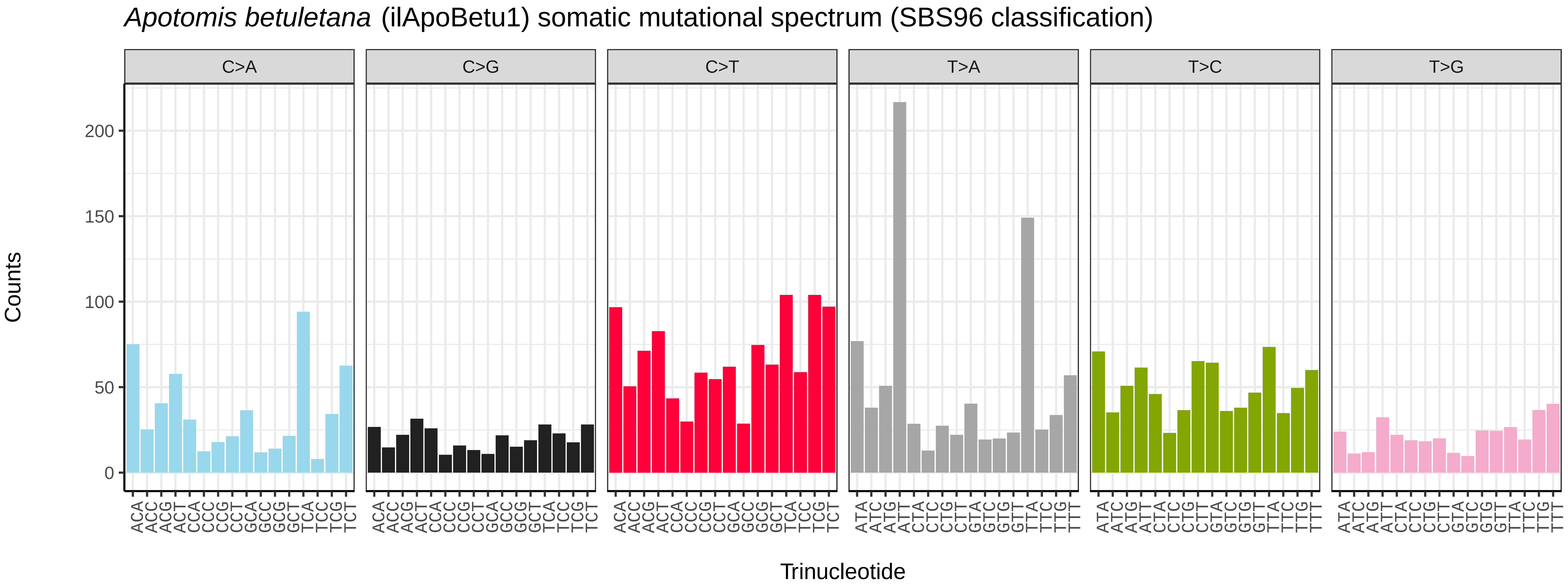
*Apotomis betuletana* (ilApoBetu1) somatic mutational spectrum (SBS96 classification)
C>A
C>G
C>T
T>A
T>C
T>G
Counts
200
150
100
50
0
Trinucleotide

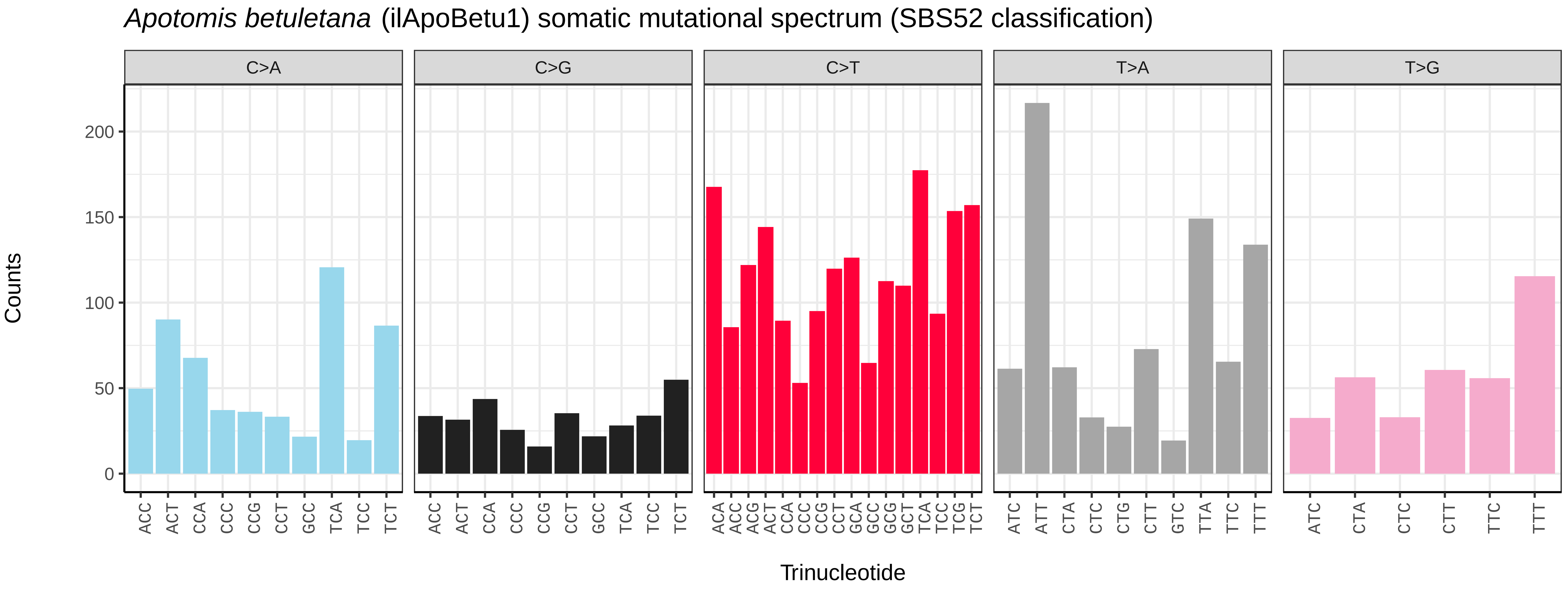
*Apotomis betuletana* (ilApoBetu1) somatic mutational spectrum (SBS52 classification)
C>A
C>G
C>T
T>A
T>G
Counts
200
150
100
50
0
Trinucleotide

*Apomyelois bistriatella* (ilApoBist1) germline mutational spectrum

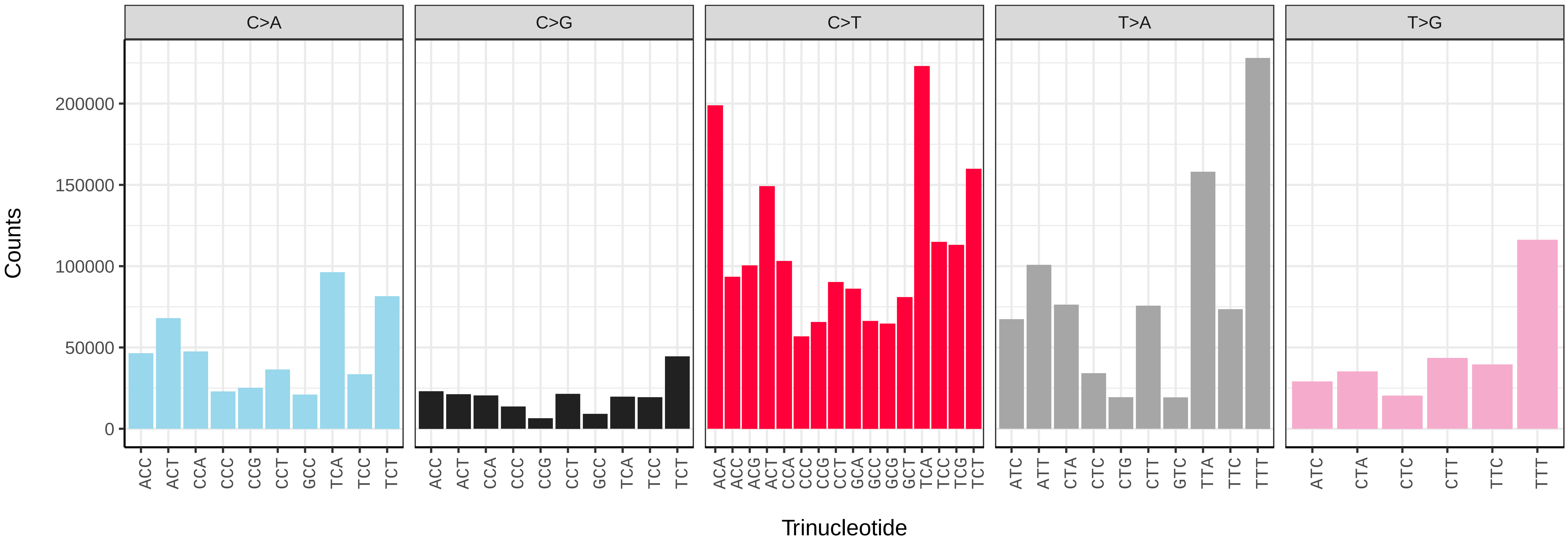


*Apomyelois bistriatella* (ilApoBist1) somatic mutational spectrum (SBS96 classification)

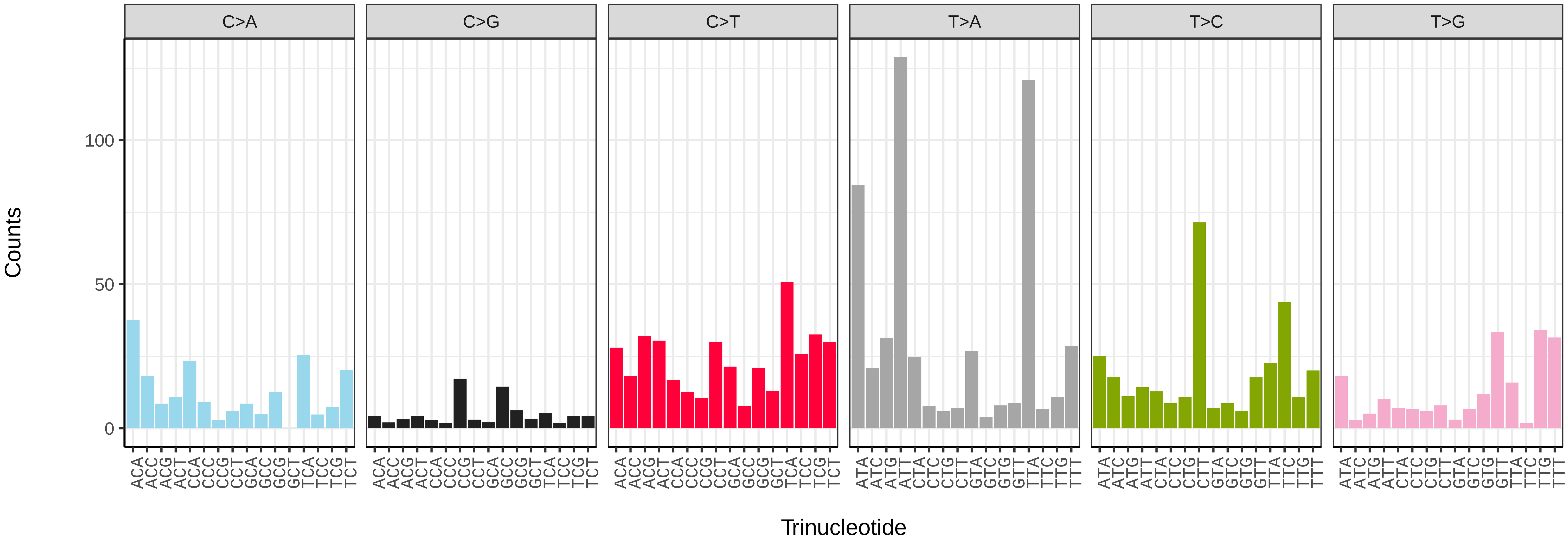


*Apomyelois bistriatella* (ilApoBist1) somatic mutational spectrum (SBS52 classification)

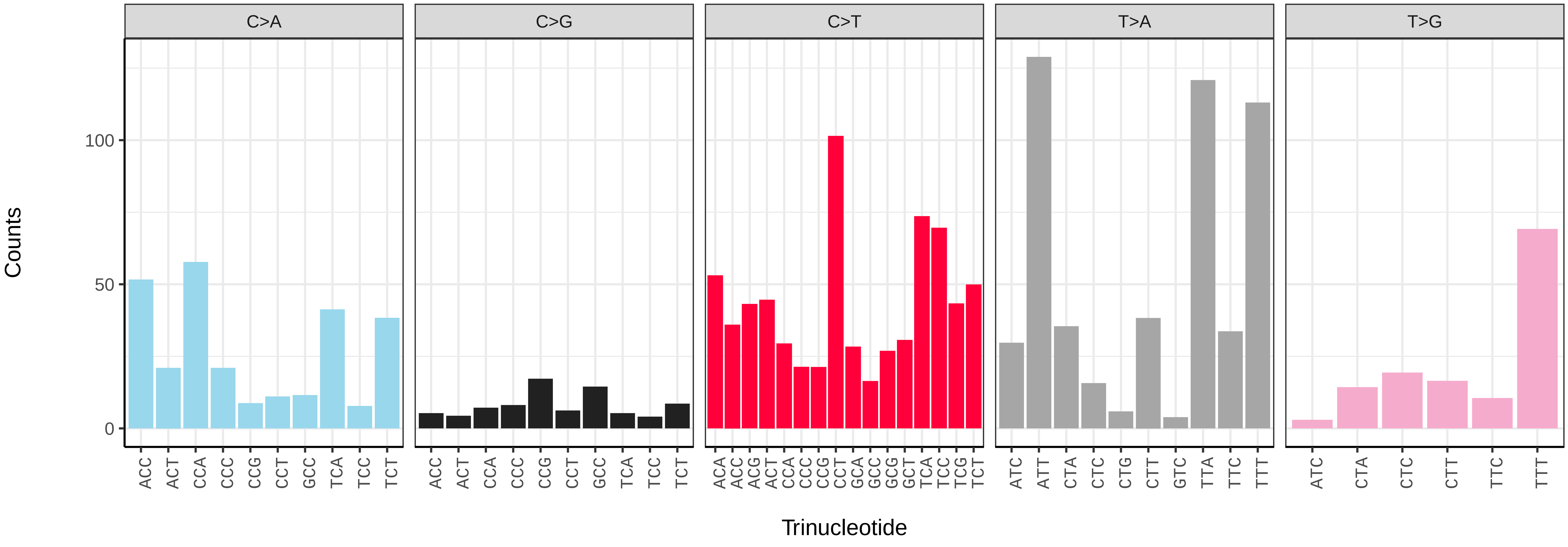

*Apotomis capreana* (ilApoCapr1) germline mutational spectrum

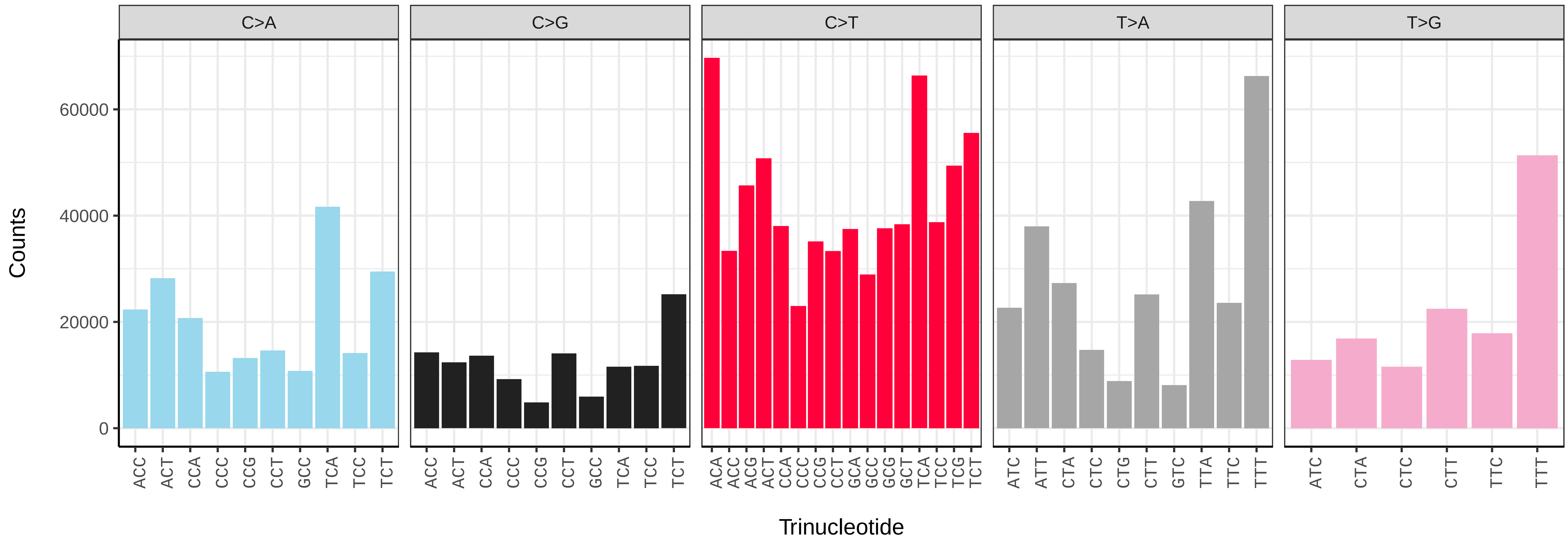


*Apotomis capreana* (ilApoCapr1) somatic mutational spectrum (SBS96 classification)

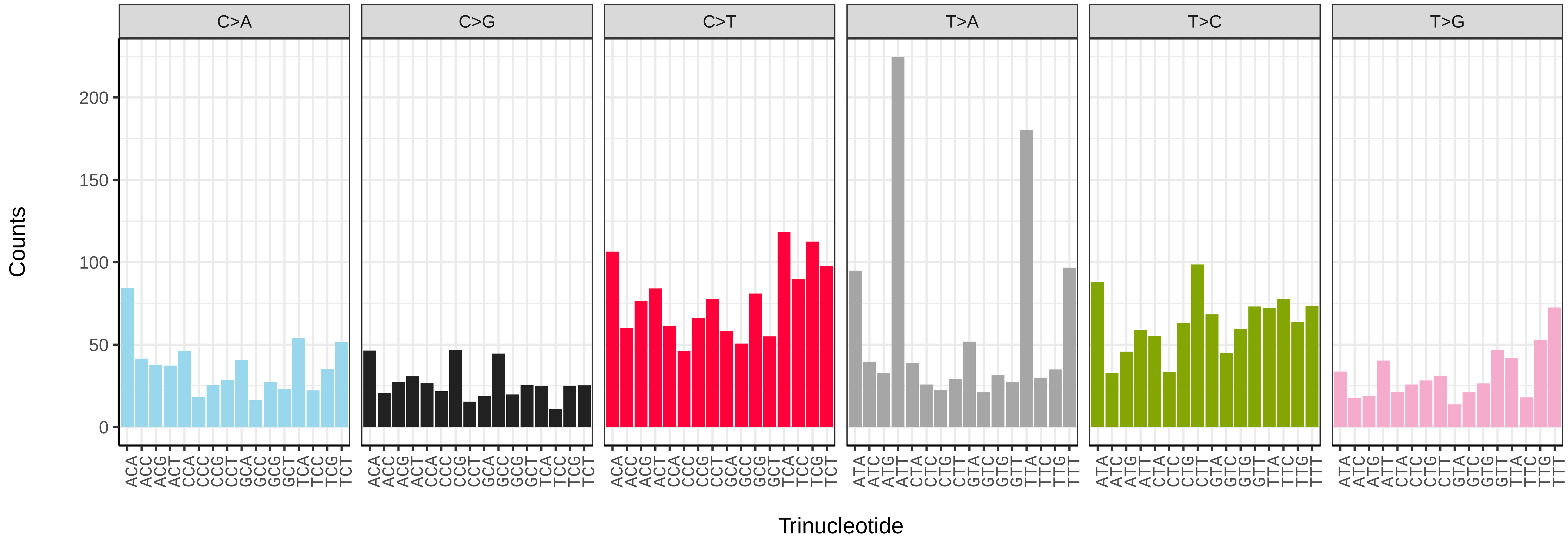


*Apotomis capreana* (ilApoCapr1) somatic mutational spectrum (SBS52 classification)

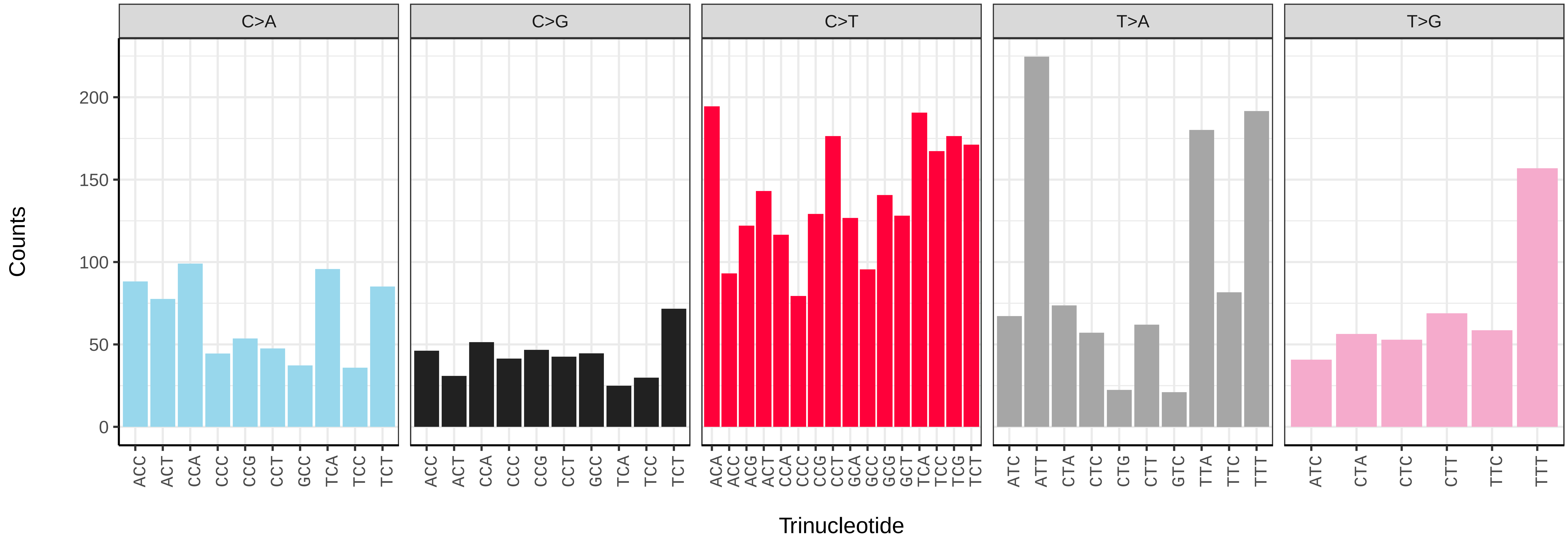

*Aporia crataegi* (ilApoCrat1) germline mutational spectrum

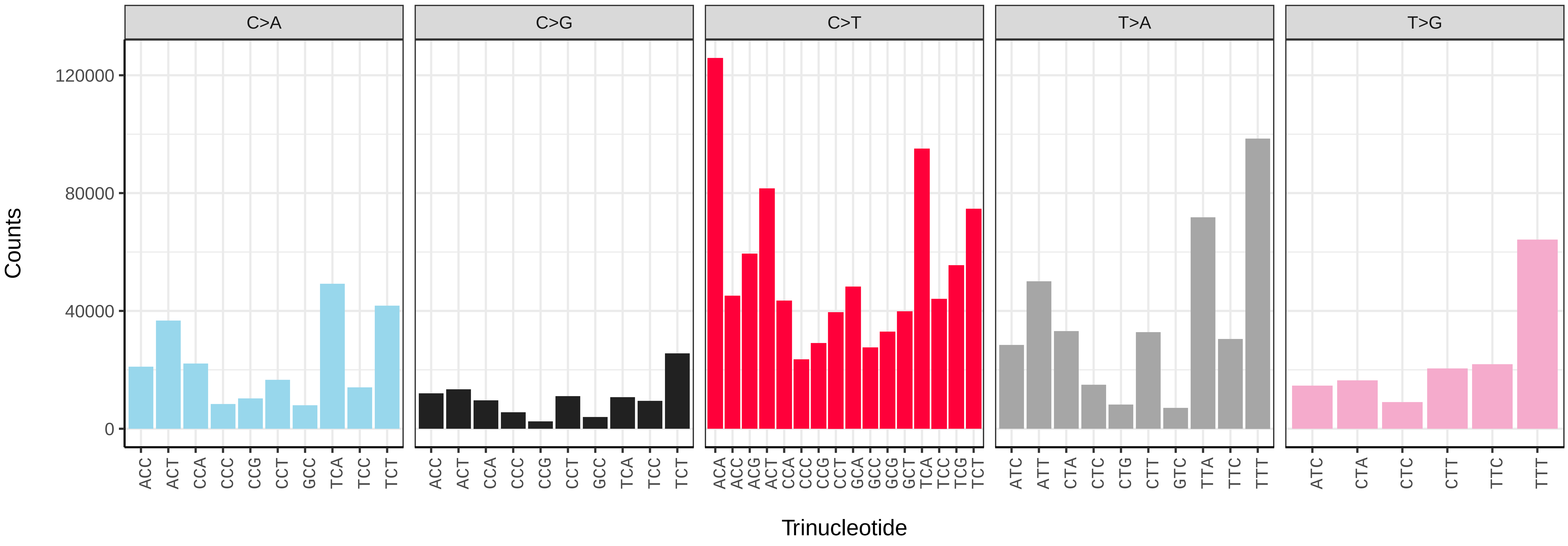


*Aporia crataegi* (ilApoCrat1) somatic mutational spectrum (SBS96 classification)

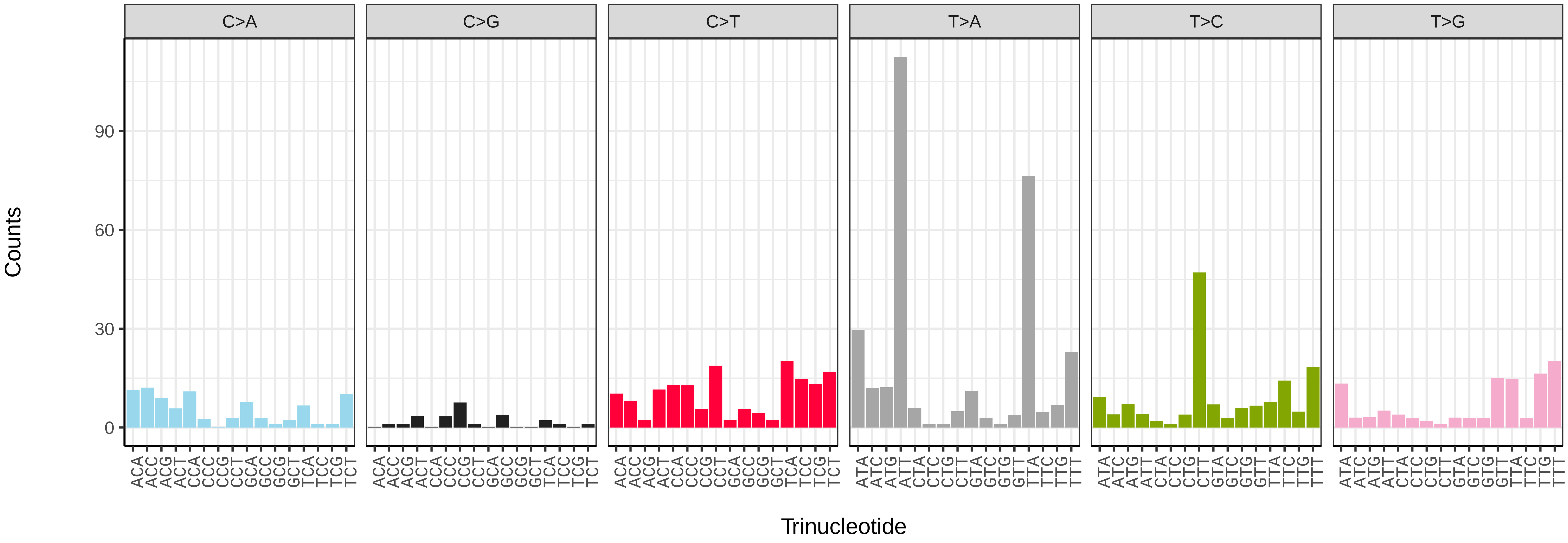


*Aporia crataegi* (ilApoCrat1) somatic mutational spectrum (SBS52 classification)

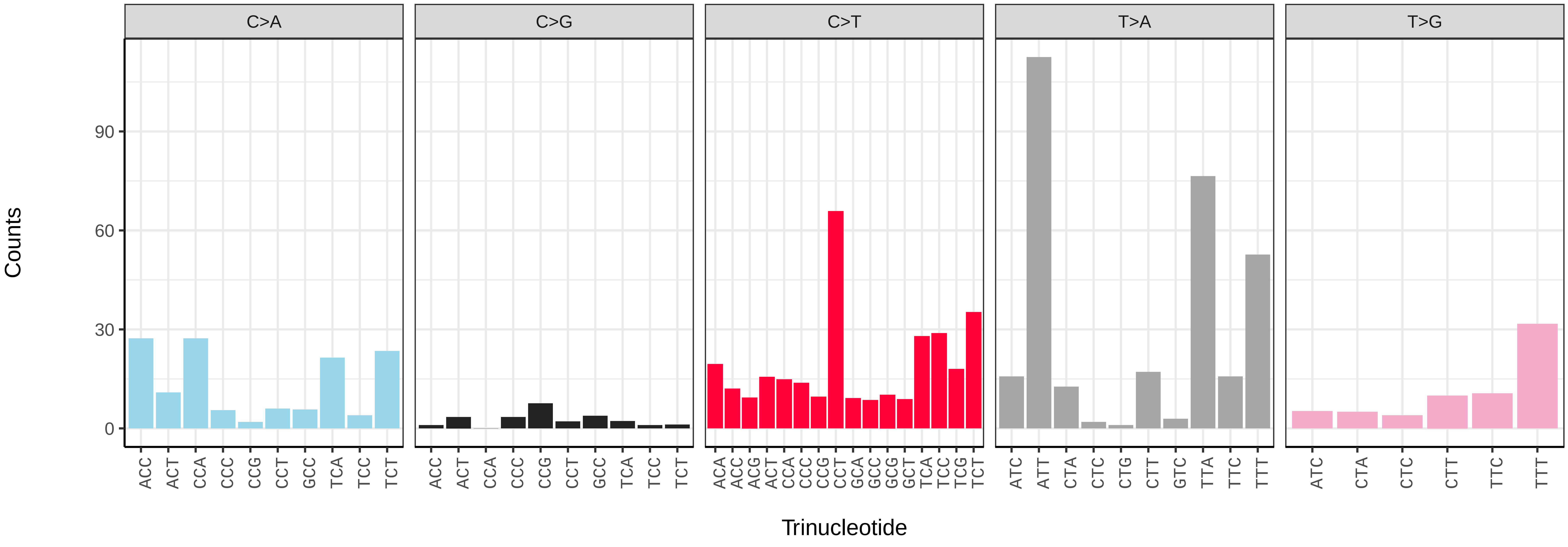

*Apocheima hispidaria* (ilApoHisp1) germline mutational spectrum

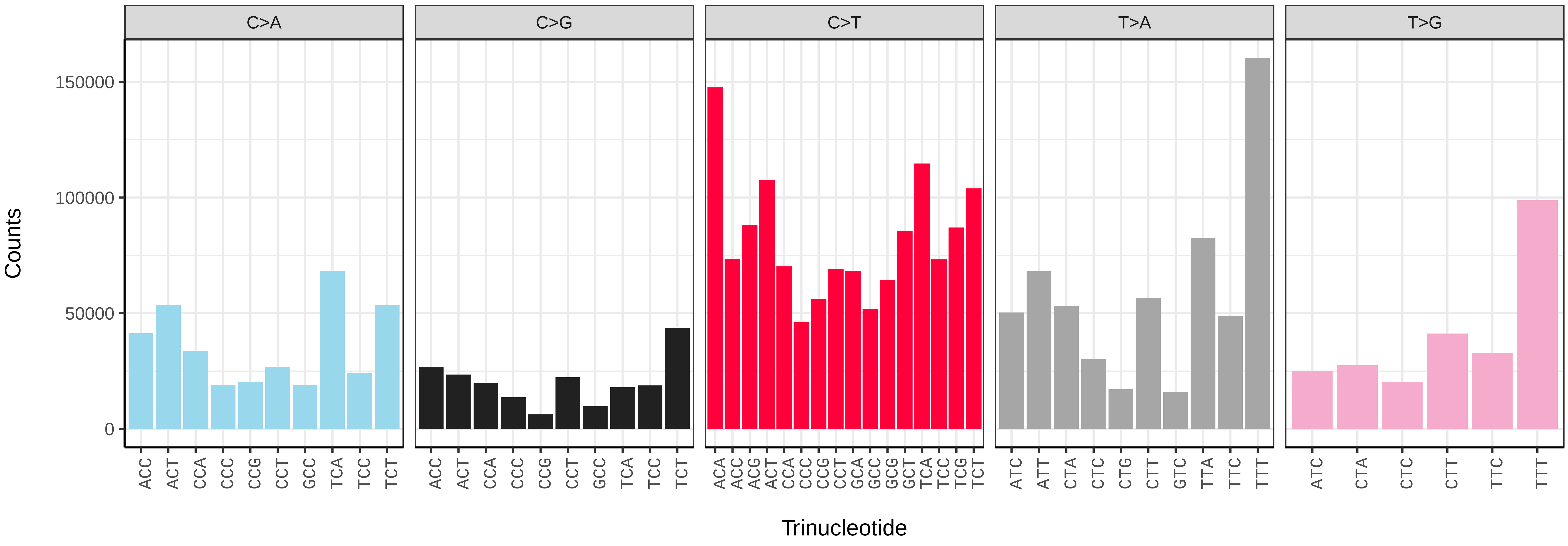


*Apocheima hispidaria* (ilApoHisp1) somatic mutational spectrum (SBS96 classification)

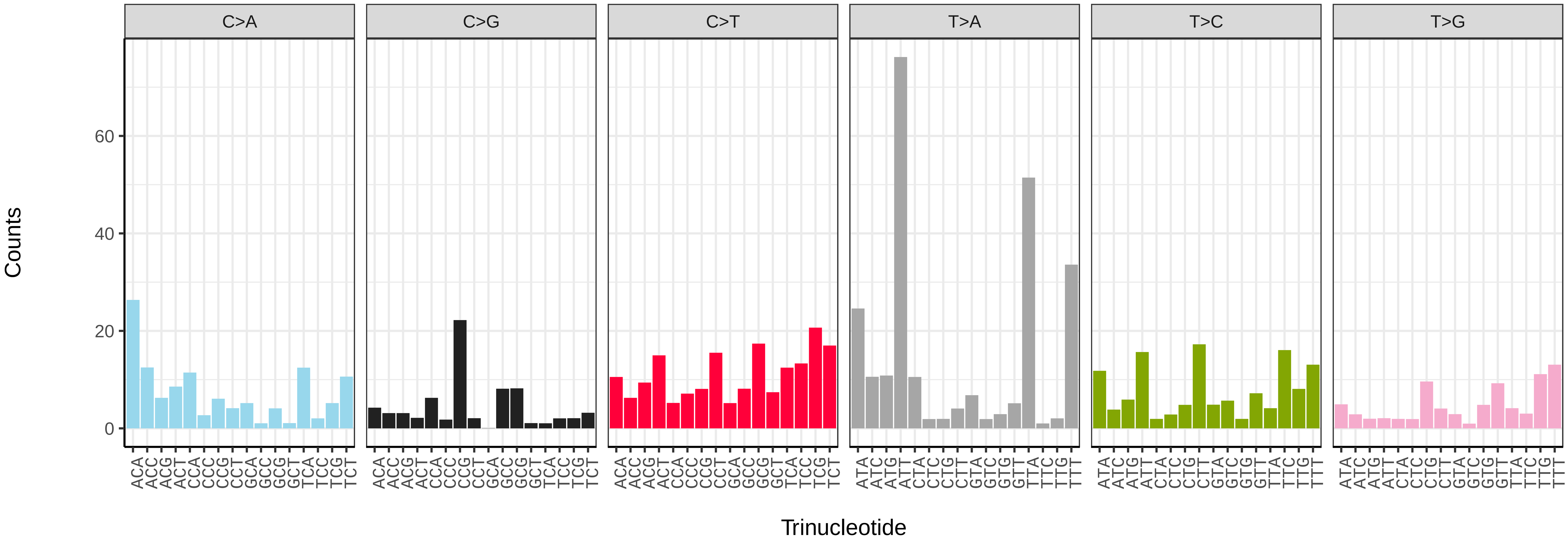


*Apocheima hispidaria* (ilApoHisp1) somatic mutational spectrum (SBS52 classification)

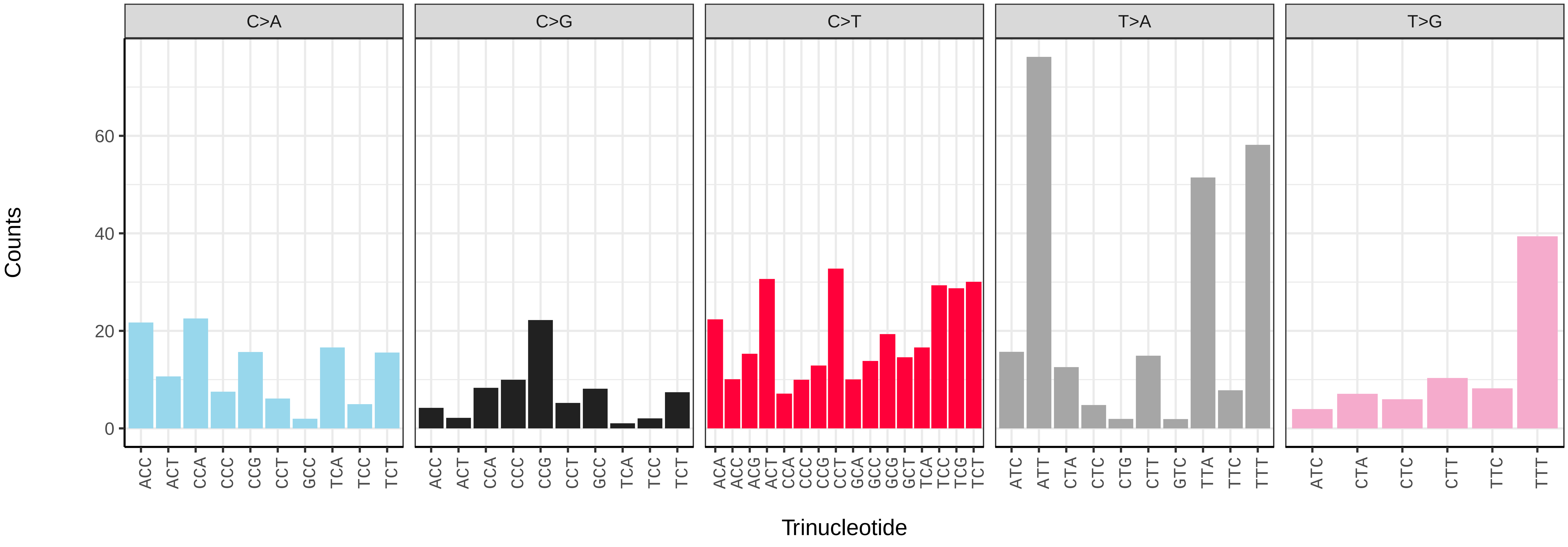

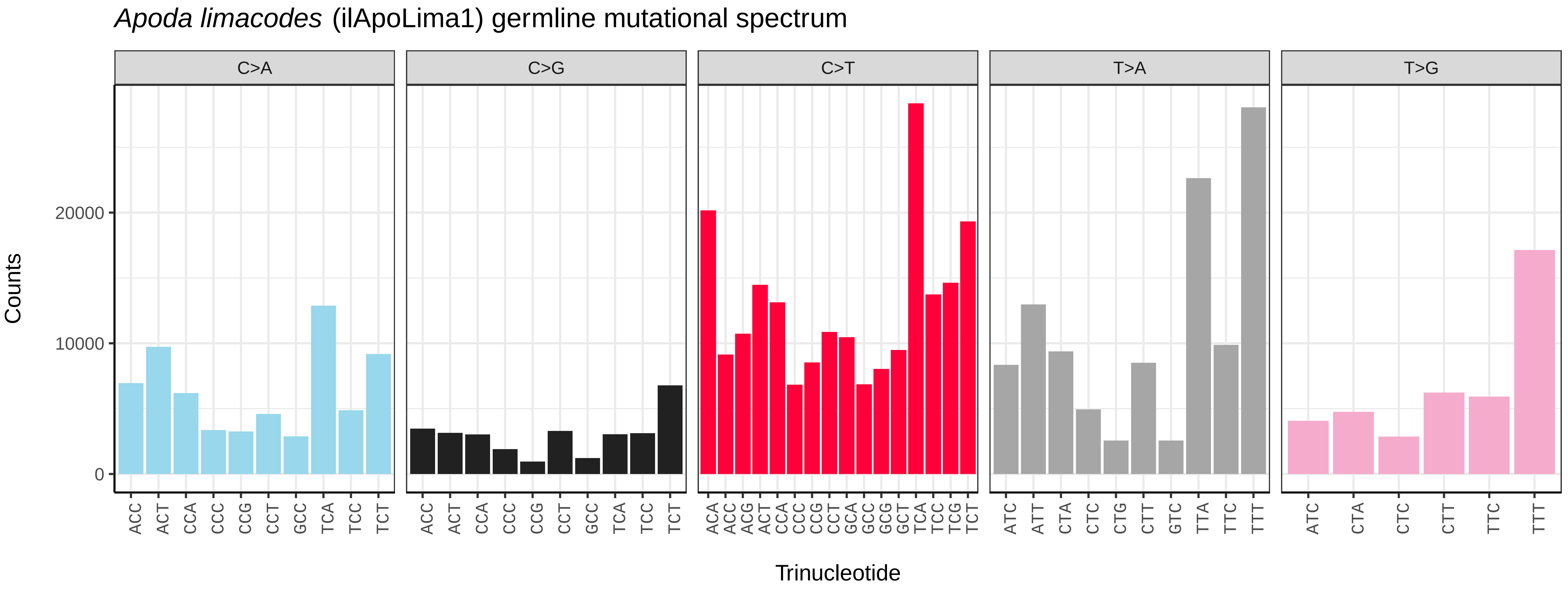
Apoda limacodes (ilApoLima1) germline mutational spectrum
C>A
C>G
C>T
T>A
T>G
Counts
Trinucleotide

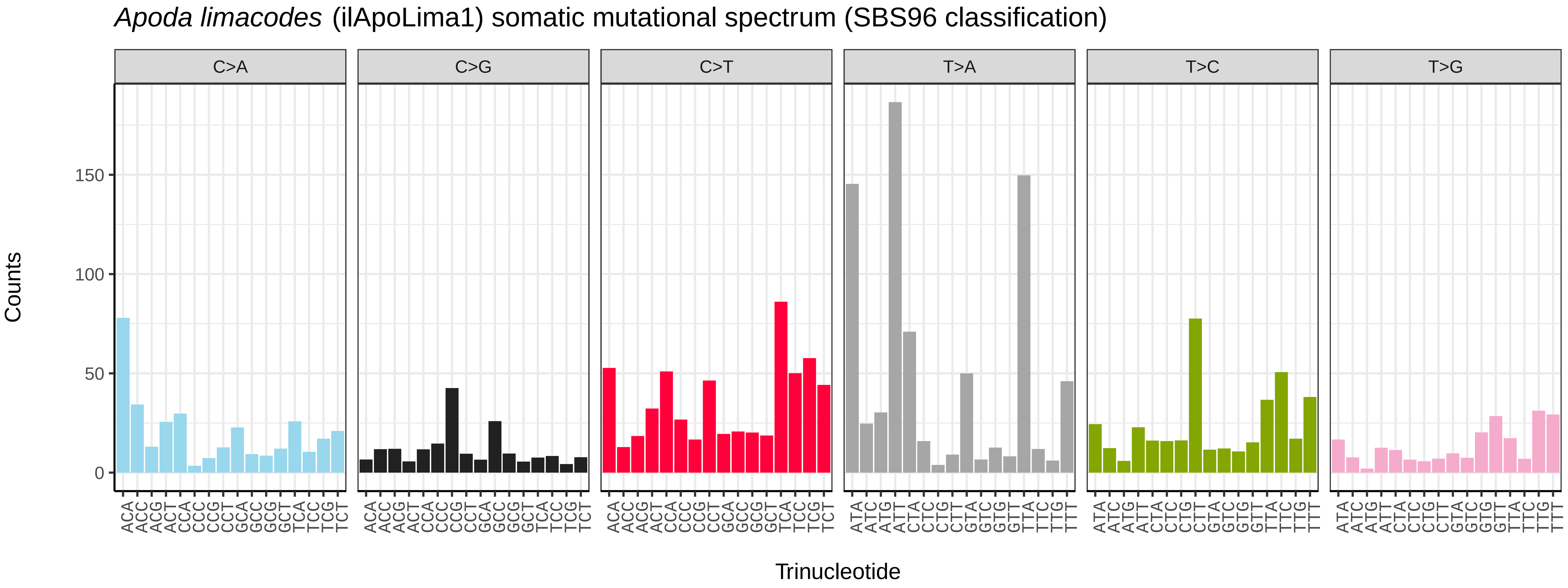
Apoda limacodes (ilApoLima1) somatic mutational spectrum (SBS96 classification)
C>A
C>G
C>T
T>A
T>C
T>G
Counts
Trinucleotide

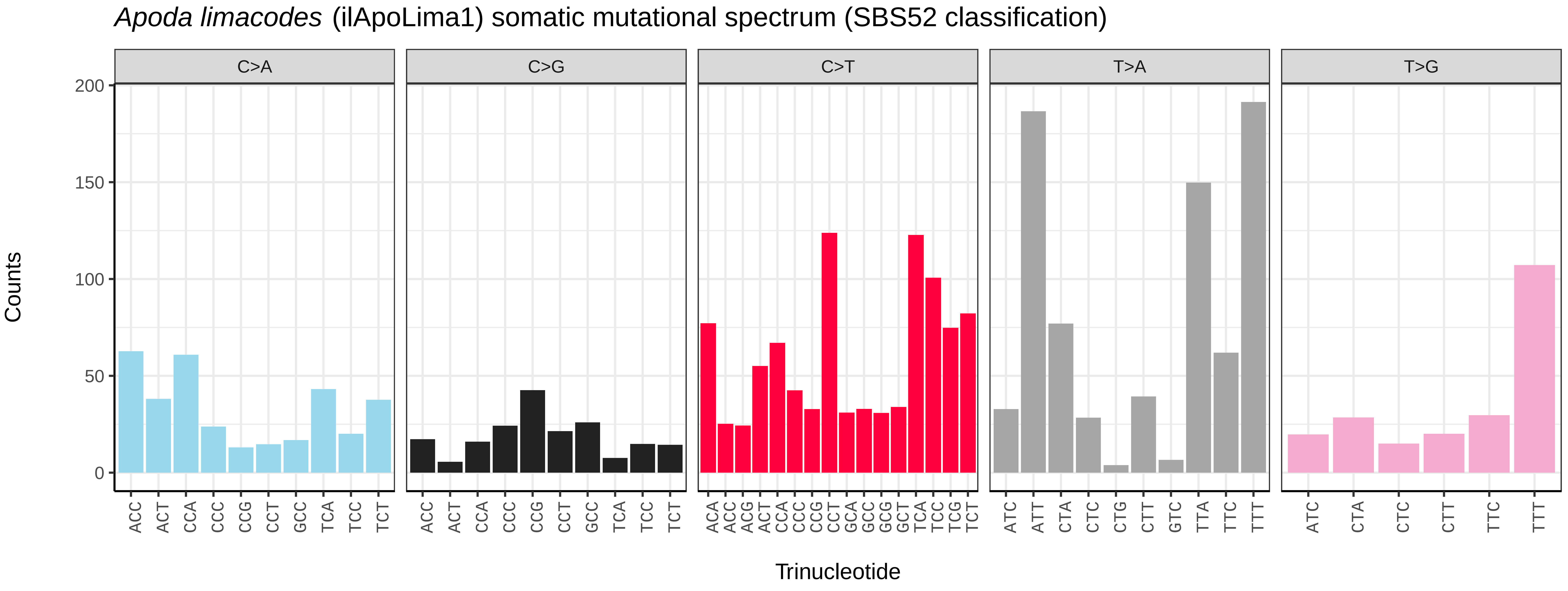
Apoda limacodes (ilApoLima1) somatic mutational spectrum (SBS52 classification)
C>A
C>G
C>T
T>A
T>G
Counts
Trinucleotide

*Aporophyla lueneburgensis* (ilApoLuen1) germline mutational spectrum

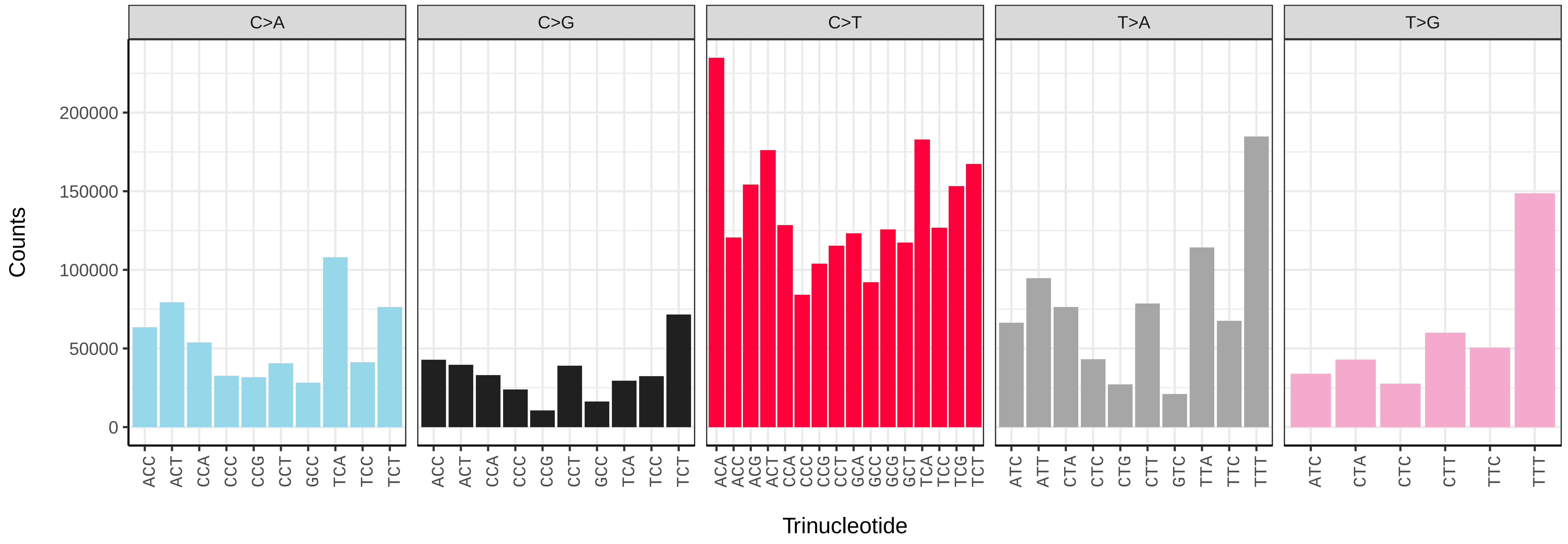


*Aporophyla lueneburgensis* (ilApoLuen1) somatic mutational spectrum (SBS96 classification)

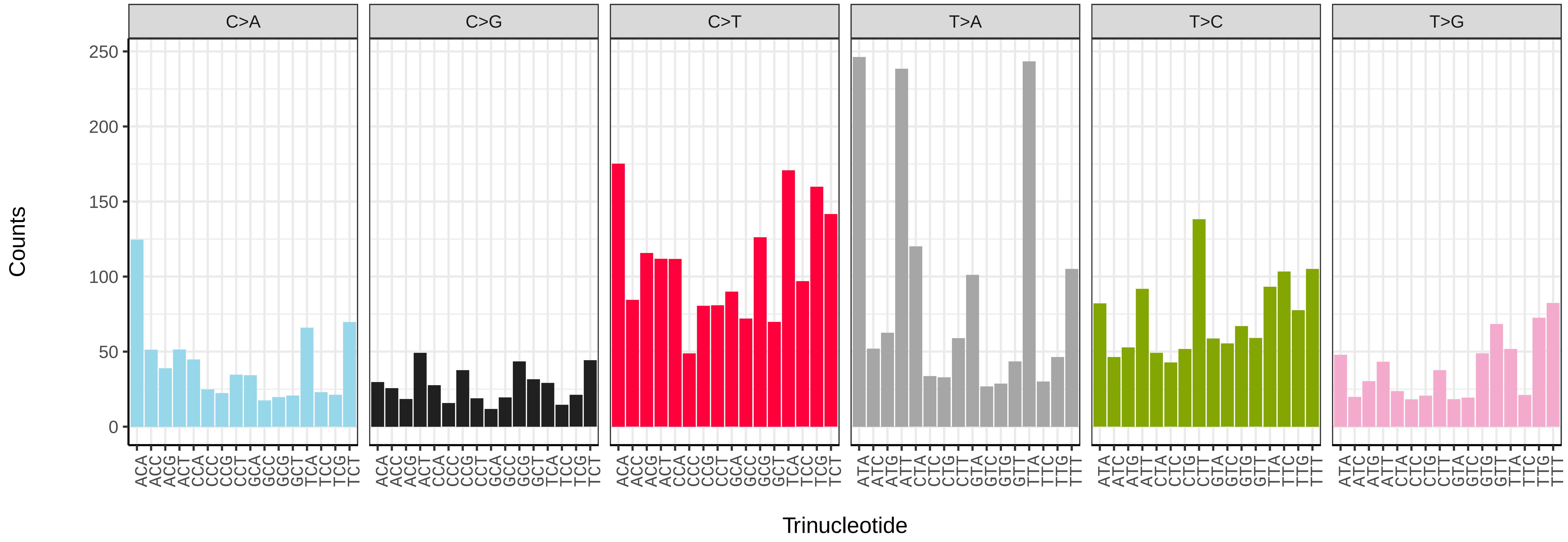


*Aporophyla lueneburgensis* (ilApoLuen1) somatic mutational spectrum (SBS52 classification)

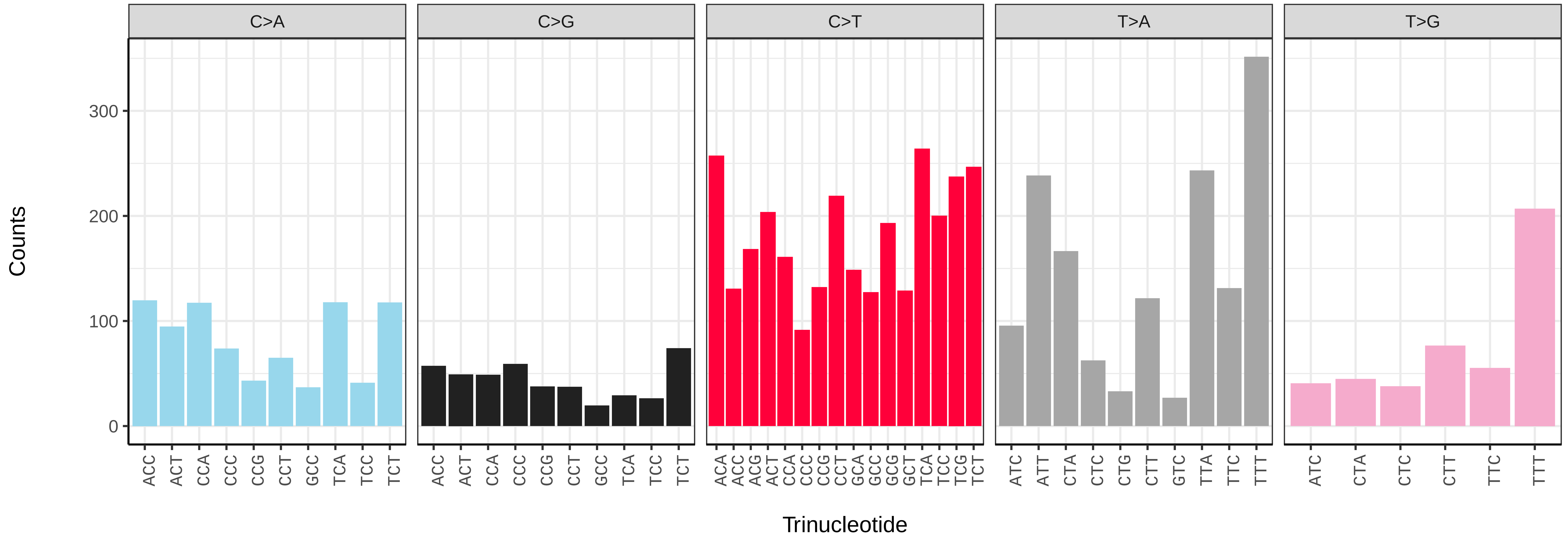

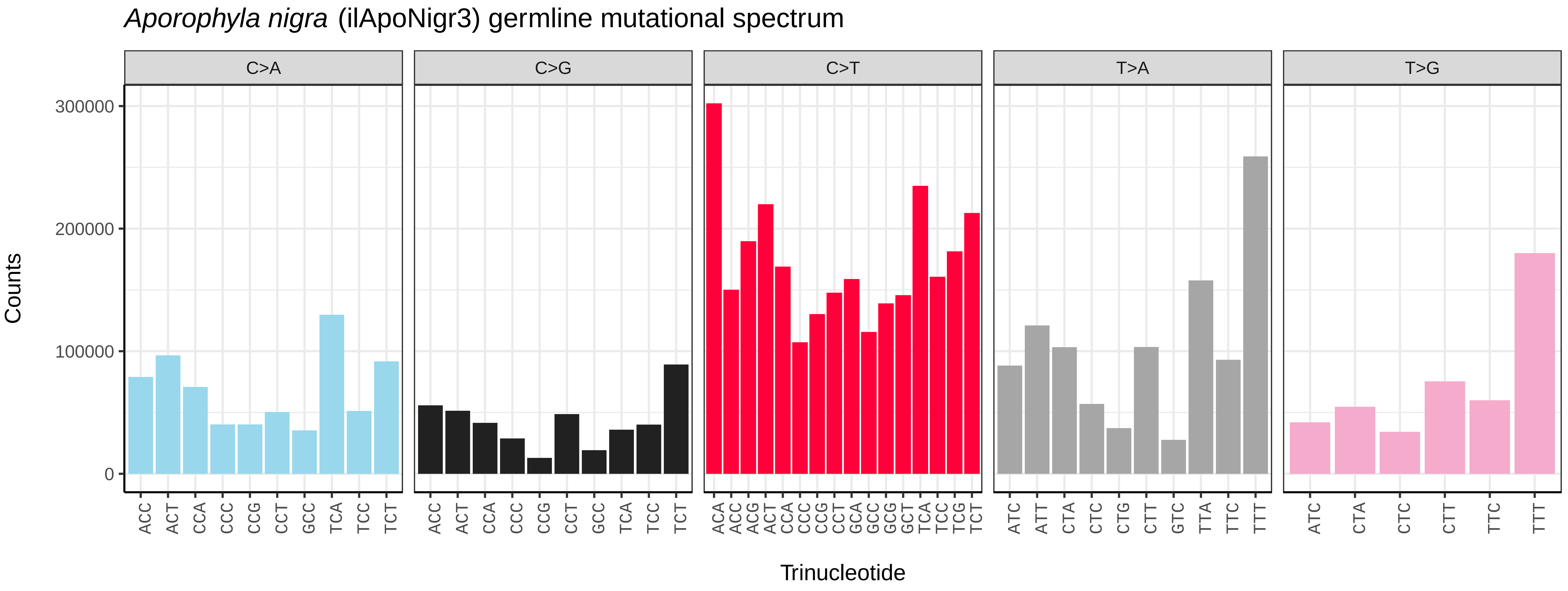
Aporophyla nigra (ilApoNigr3) germline mutational spectrum
C>A
C>G
C>T
T>A
T>G
Counts
300000
200000
100000
0
ACC ACT CCA CCC CCG CCT GCC TCA TCC TCT
ACA ACC ACG ACT CCA CCC CCG CCT GCA GCC GCG GCT TCA TCC TCG TCT
ATC ATT CTA CTC CTG CTT GTC TTA TTC TTT
ATC CTA CTC CTT TTC TTT
Trinucleotide

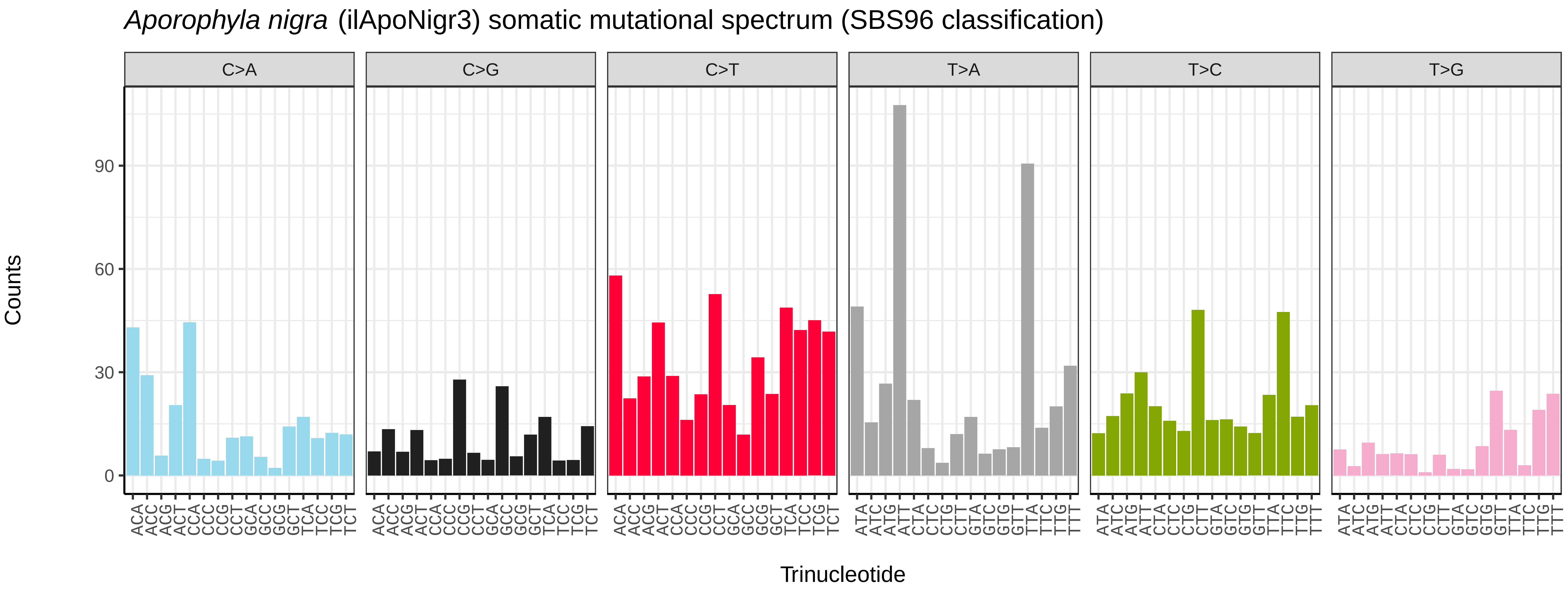
Aporophyla nigra (ilApoNigr3) somatic mutational spectrum (SBS96 classification)
C>A
C>G
C>T
T>A
T>C
T>G
Counts
90
60
30
0
ACA ACC ACG ACT CCA CCC CCG CCT GCA GCC GCG GCT TCA TCC TCG TCT
ATA ATC ATG ATT CTA CTC CTG CTT GTA GTC GTG GTT TTA TTC TTG TTT
Trinucleotide

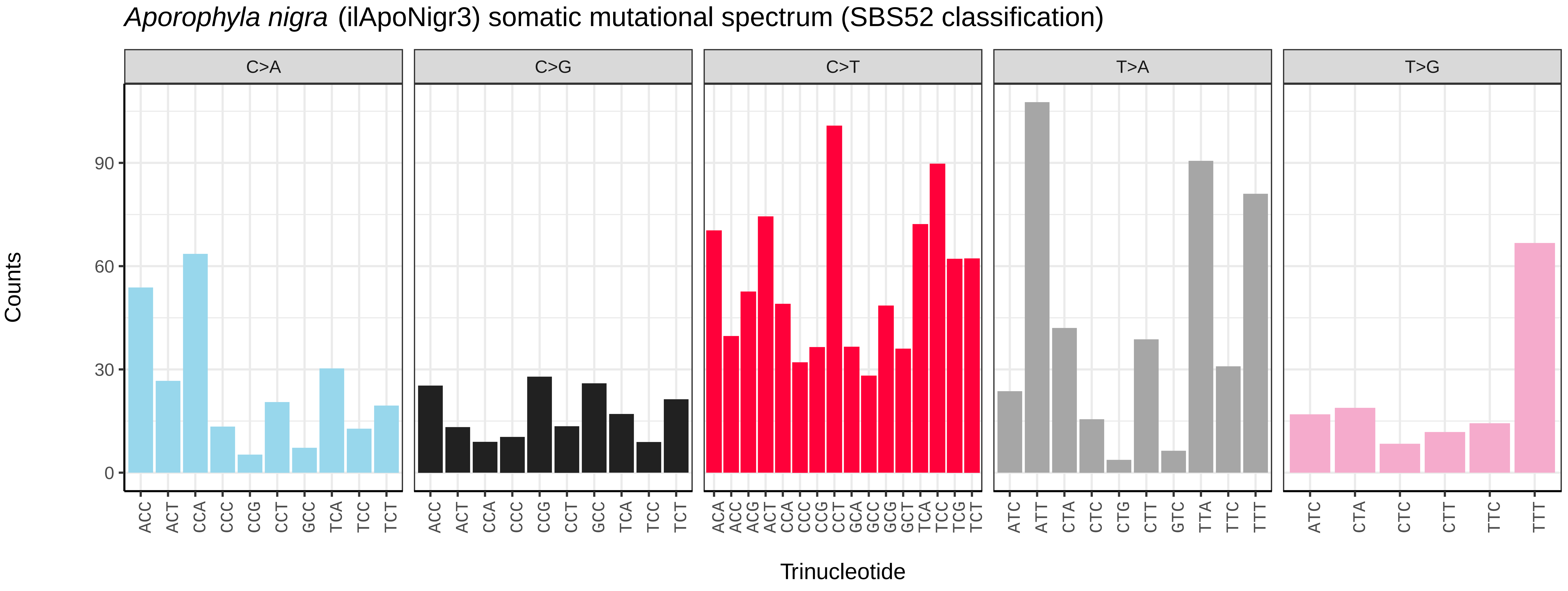
Aporophyla nigra (ilApoNigr3) somatic mutational spectrum (SBS52 classification)
C>A
C>G
C>T
T>A
T>G
Counts
90
60
30
0
ACC ACT CCA CCC CCG CCT GCC TCA TCC TCT
ACA ACC ACG ACT CCA CCC CCG CCT GCA GCC GCG GCT TCA TCC TCG TCT
ATC ATT CTA CTC CTG CTT GTC TTA TTC TTT
ATC CTA CTC CTT TTC TTT
Trinucleotide

*Apotomis turbidana* (ilApoTurb1) germline mutational spectrum

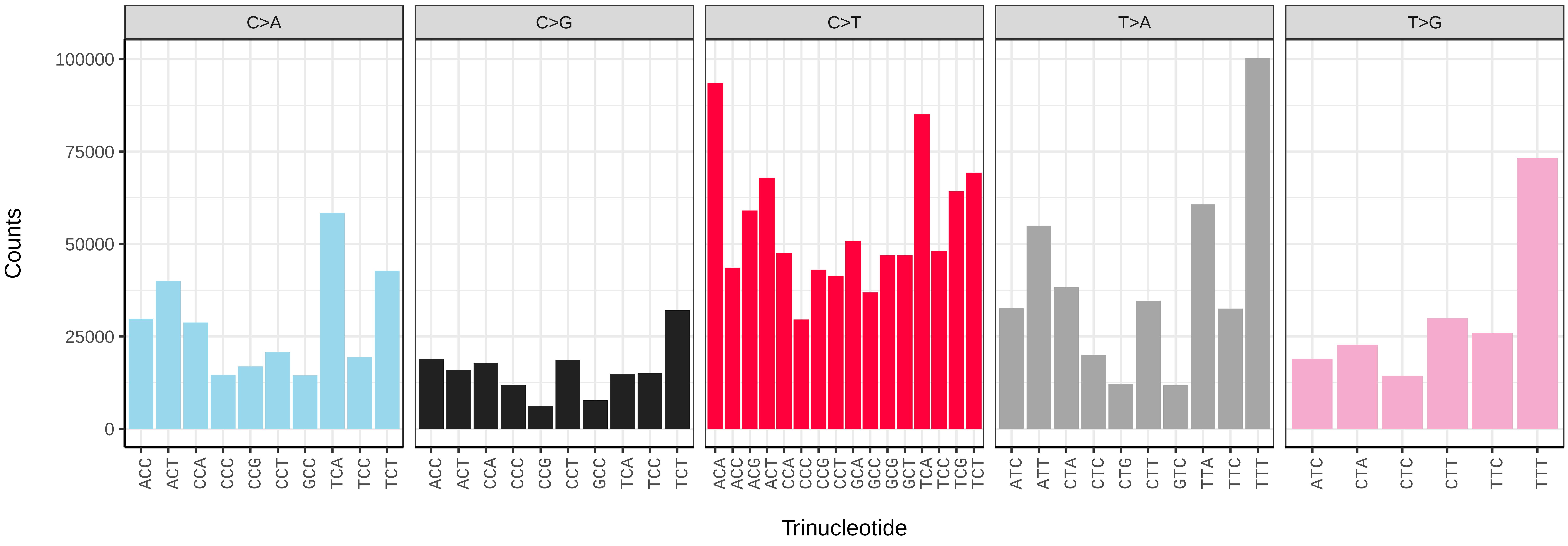


*Apotomis turbidana* (ilApoTurb1) somatic mutational spectrum (SBS96 classification)

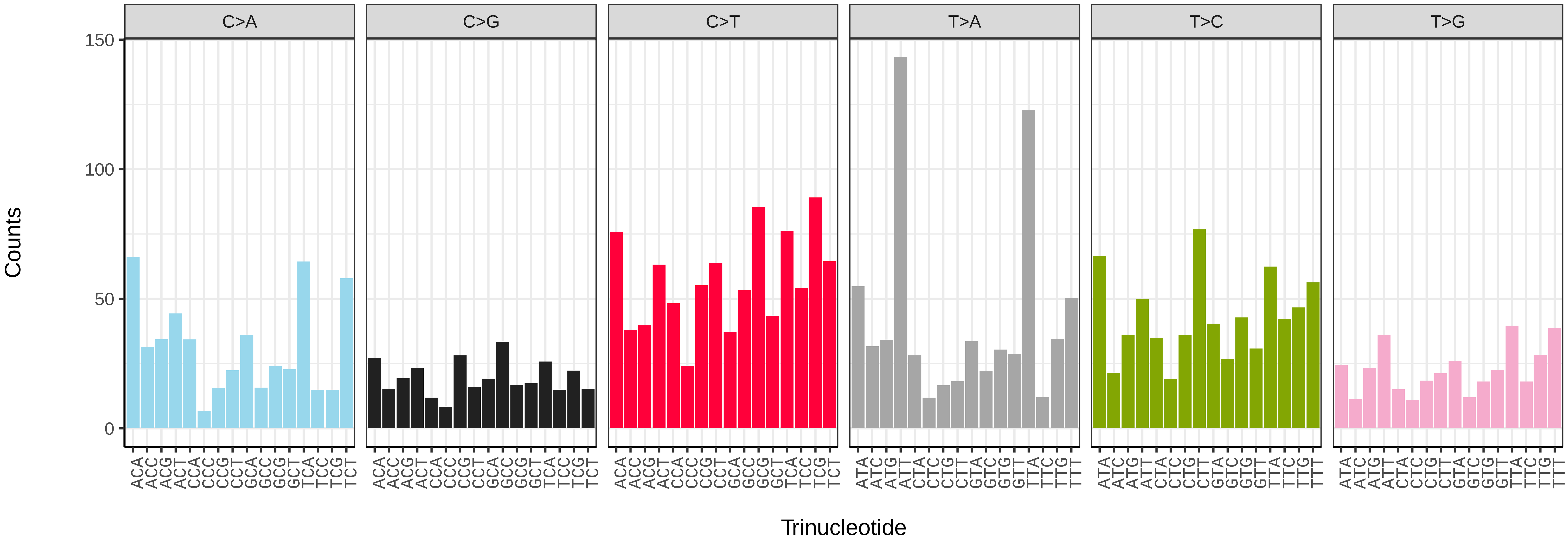


*Apotomis turbidana* (ilApoTurb1) somatic mutational spectrum (SBS52 classification)

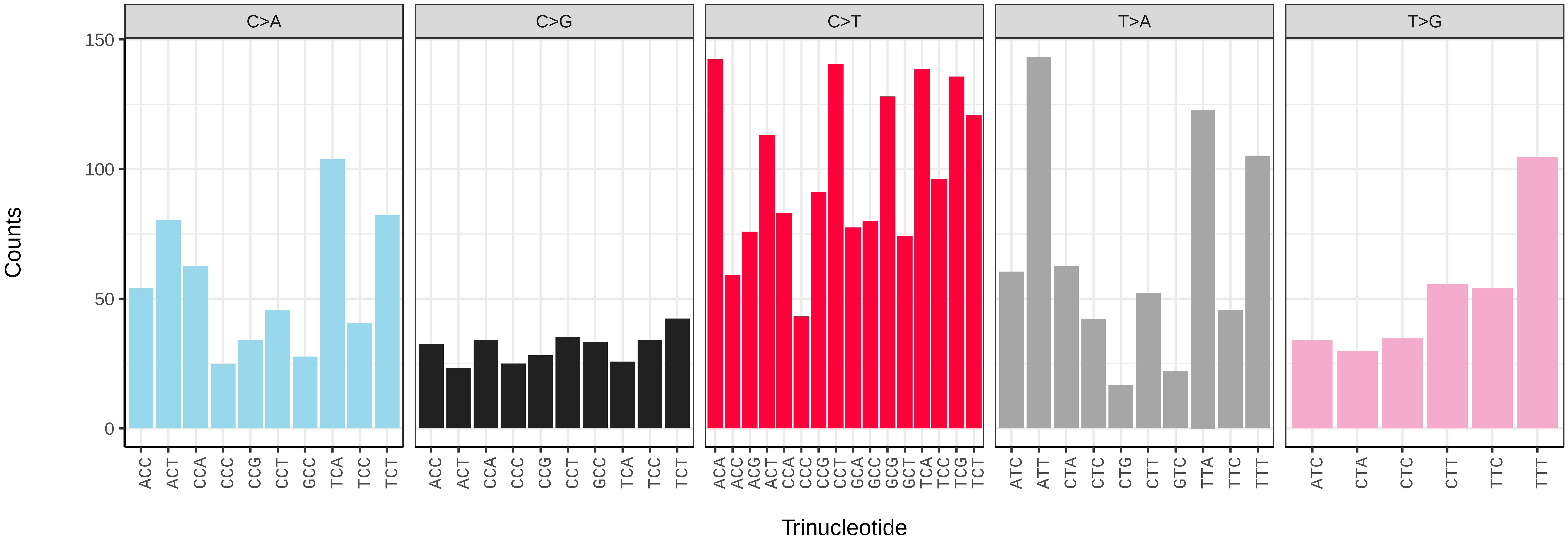

*Archips crataeganus* (ilArcCraa1) germline mutational spectrum

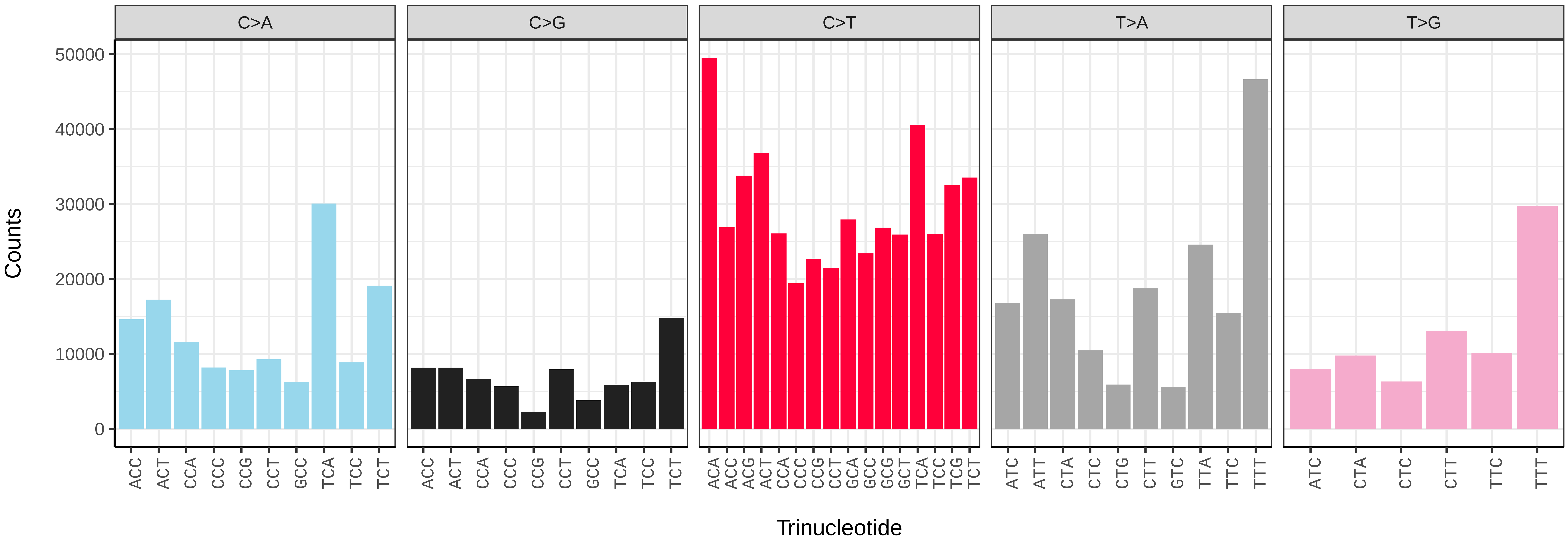


*Archips crataeganus* (ilArcCraa1) somatic mutational spectrum (SBS96 classification)

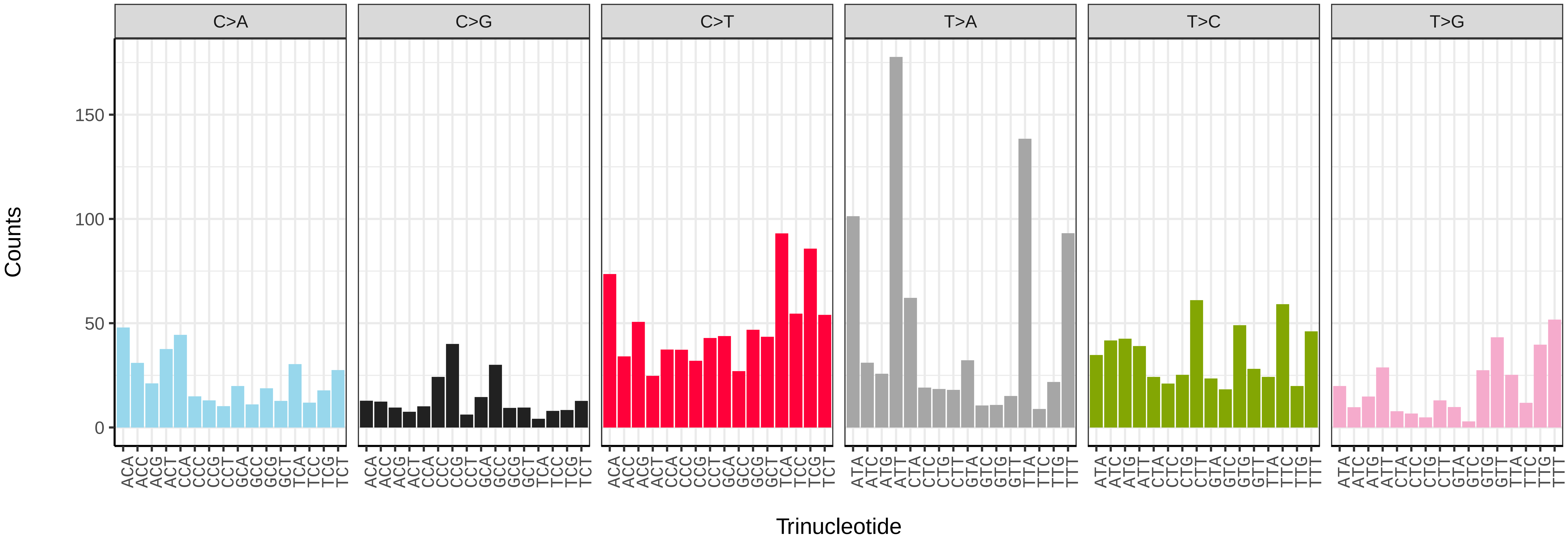


*Archips crataeganus* (ilArcCraa1) somatic mutational spectrum (SBS52 classification)

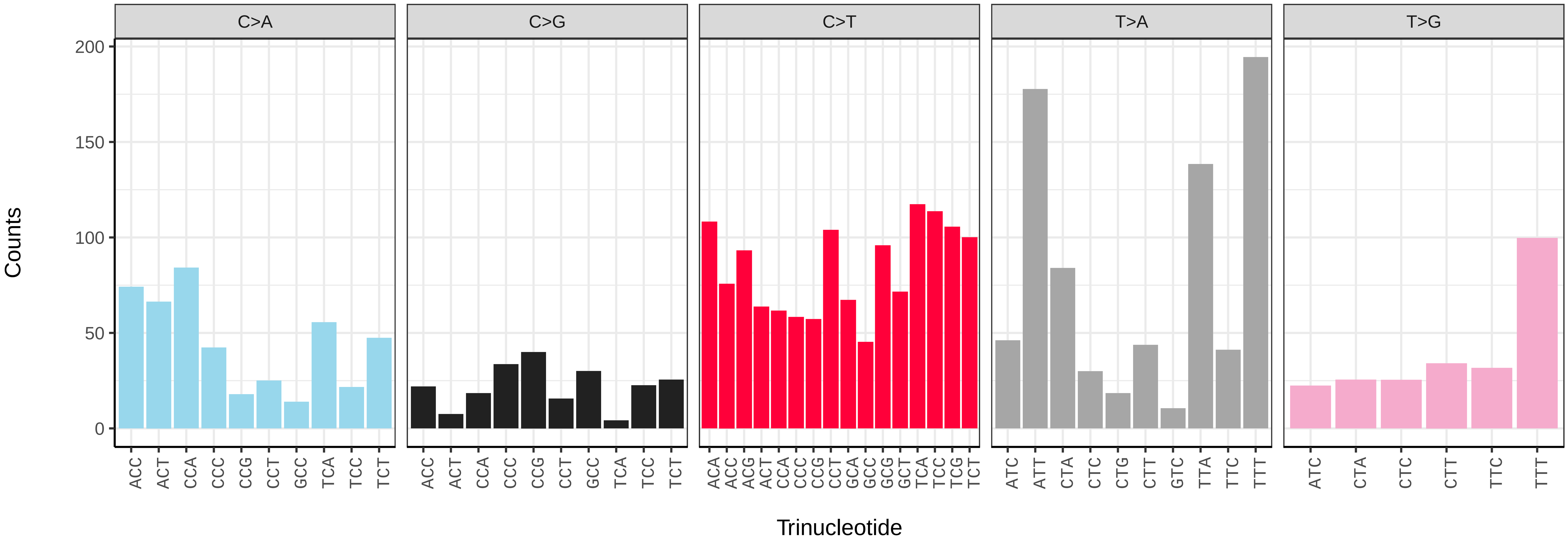

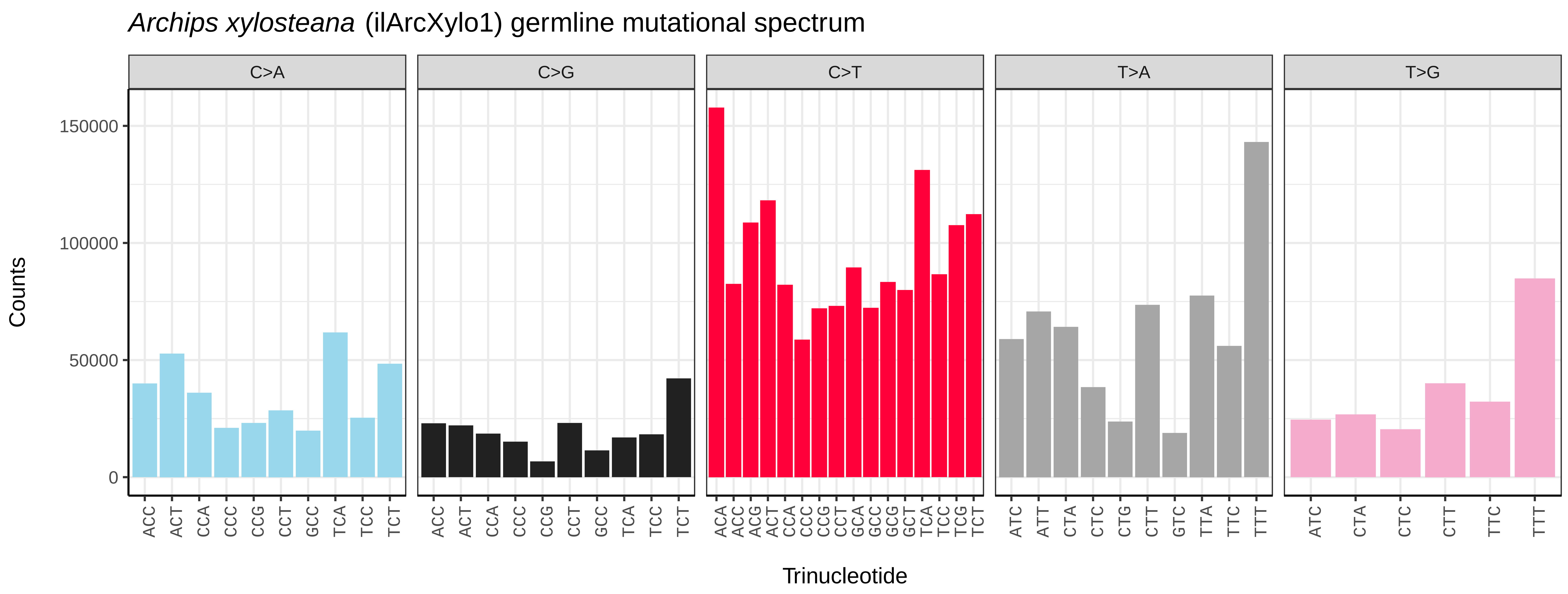
Archips xylosteana (ilArcXylo1) germline mutational spectrum
C>A
C>G
C>T
T>A
T>G
Counts
0
50000
100000
150000
ACC ACT CCA CCC CCG CCT GCC TCA TCC TCT
ACC ACT CCA CCC CCG CCT GCC TCA TCC TCT
ACA ACC ACG ACT CCA CCC CCG CCT GCA GCC GCG GCT TCA TCC TCG TCT
ATC ATT CTA CTC CTG CTT GTC TTA TTC TTT
ATC CTA CTC CTT TTC TTT
Trinucleotide

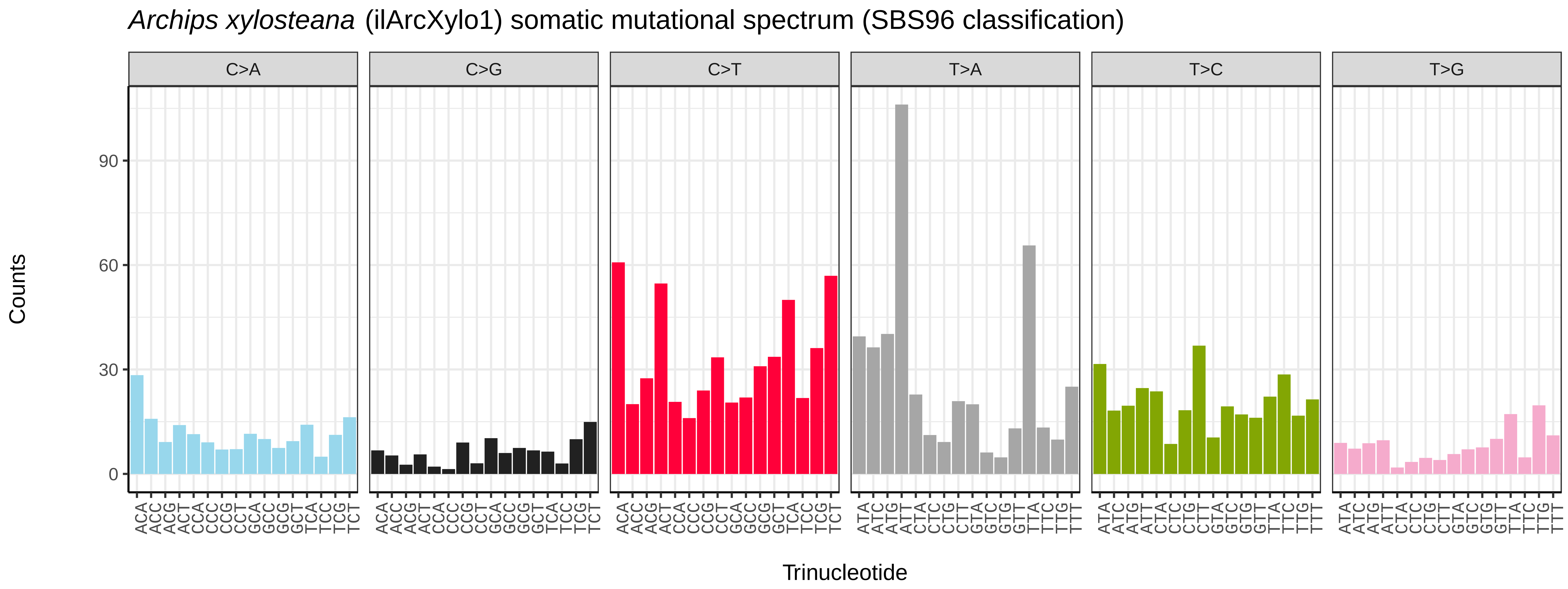
Archips xylosteana (ilArcXylo1) somatic mutational spectrum (SBS96 classification)
C>A
C>G
C>T
T>A
T>C
T>G
Counts
0
30
60
90
ACA ACC ACG ACT CCA CCC CCG CCT GCA GCC GCG GCT TCA TCC TCG TCT
ATA ATC ATG ATT CTA CTC CTG CTT GTA GTC GTG GTT TTA TTC TTG TTT
Trinucleotide

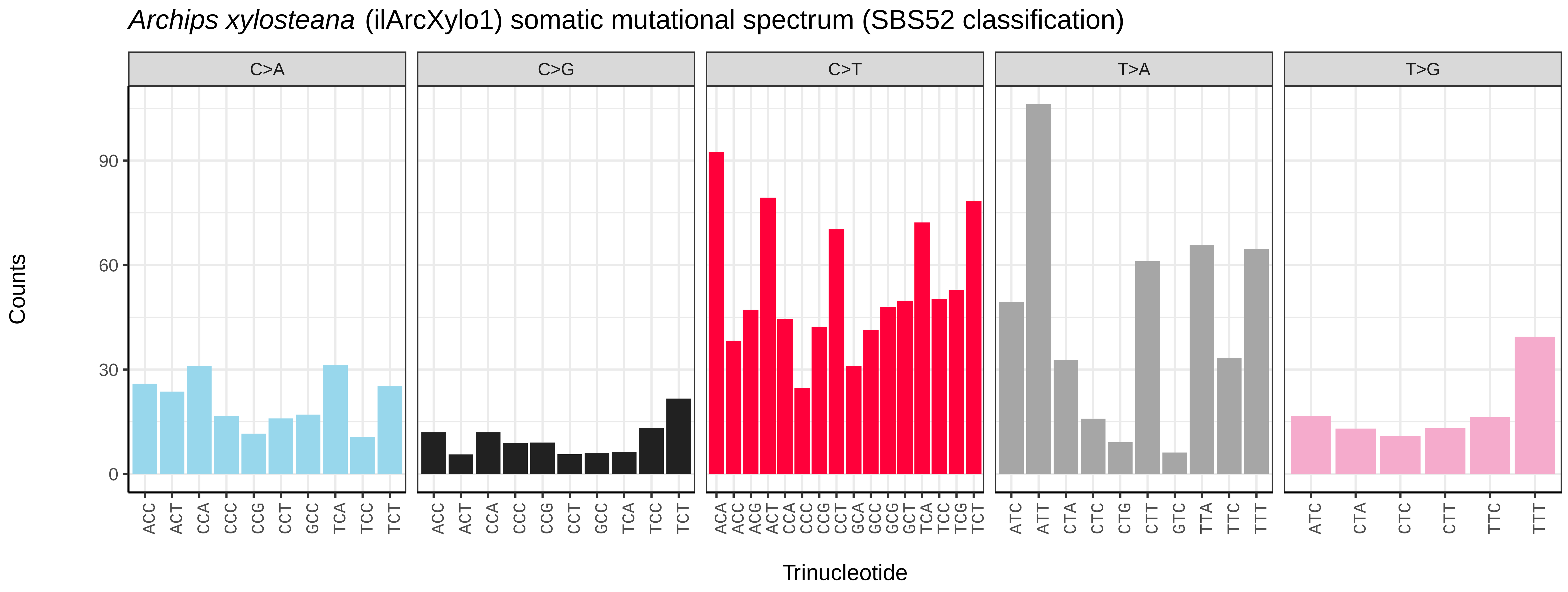
Archips xylosteana (ilArcXylo1) somatic mutational spectrum (SBS52 classification)
C>A
C>G
C>T
T>A
T>G
Counts
0
30
60
90
ACC ACT CCA CCC CCG CCT GCC TCA TCC TCT
ACC ACT CCA CCC CCG CCT GCC TCA TCC TCT
ACA ACC ACG ACT CCA CCC CCG CCT GCA GCC GCG GCT TCA TCC TCG TCT
ATC ATT CTA CTC CTG CTT GTC TTA TTC TTT
ATC CTA CTC CTT TTC TTT
Trinucleotide

*Aricia agestis* (ilAriAges1) germline mutational spectrum

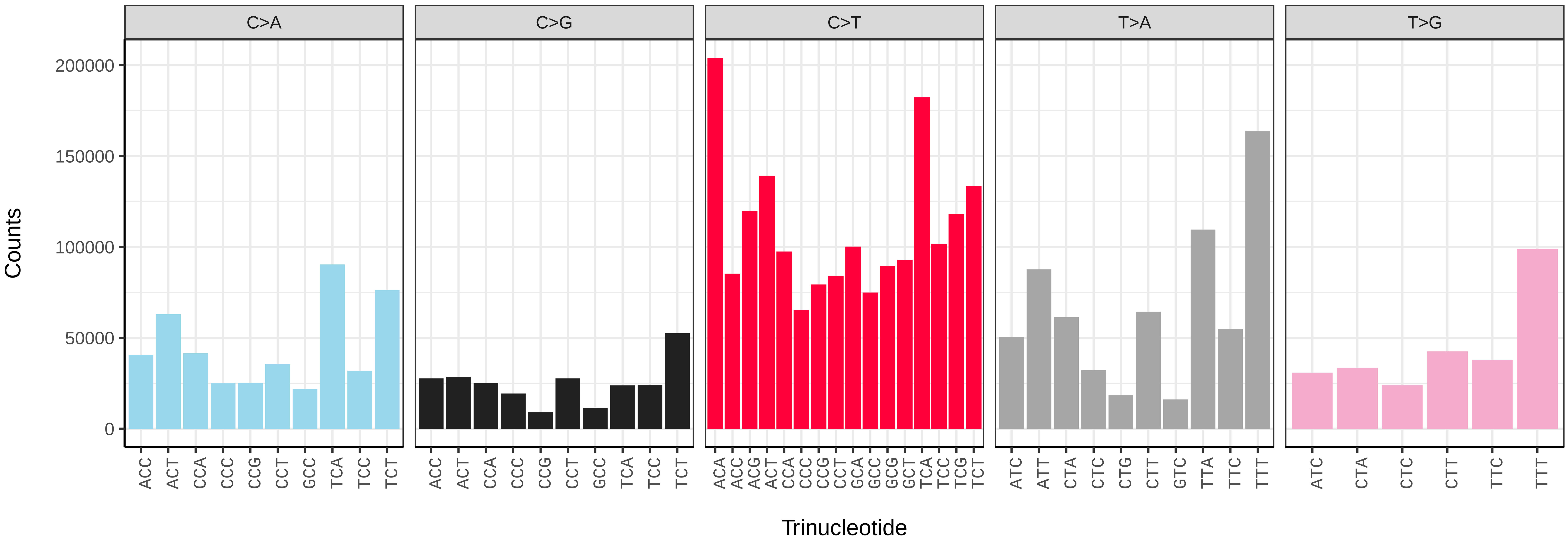


*Aricia agestis* (ilAriAges1) somatic mutational spectrum (SBS96 classification)

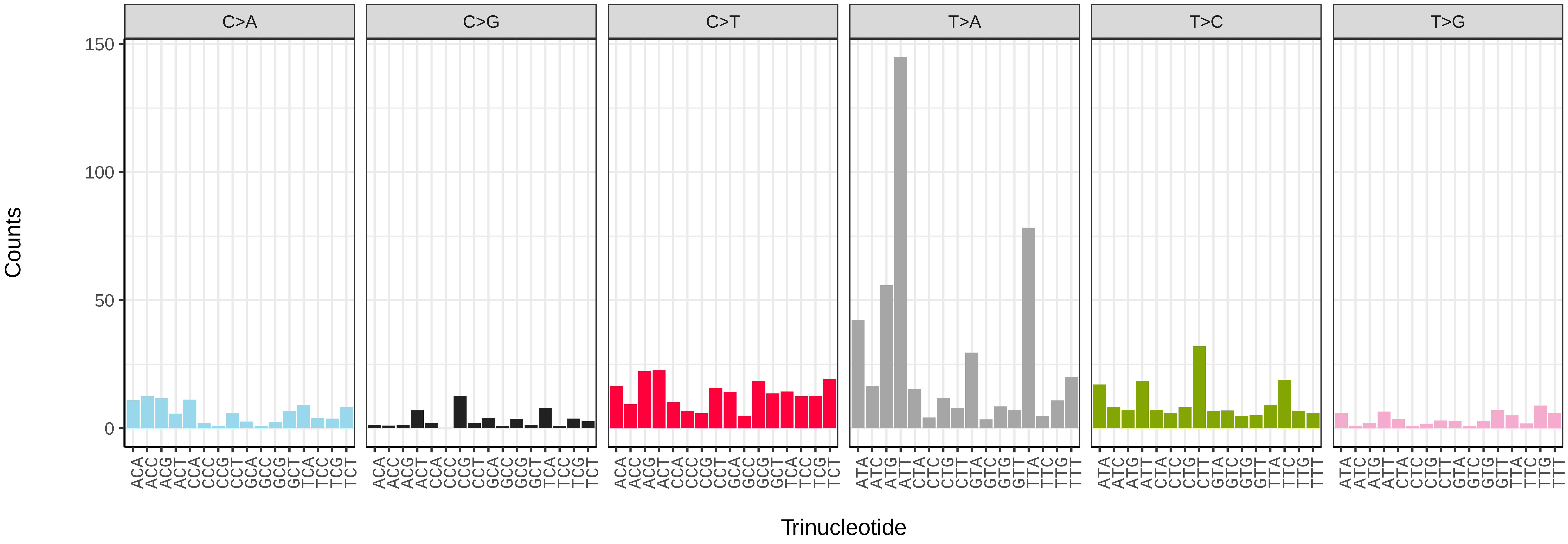


*Aricia agestis* (ilAriAges1) somatic mutational spectrum (SBS52 classification)

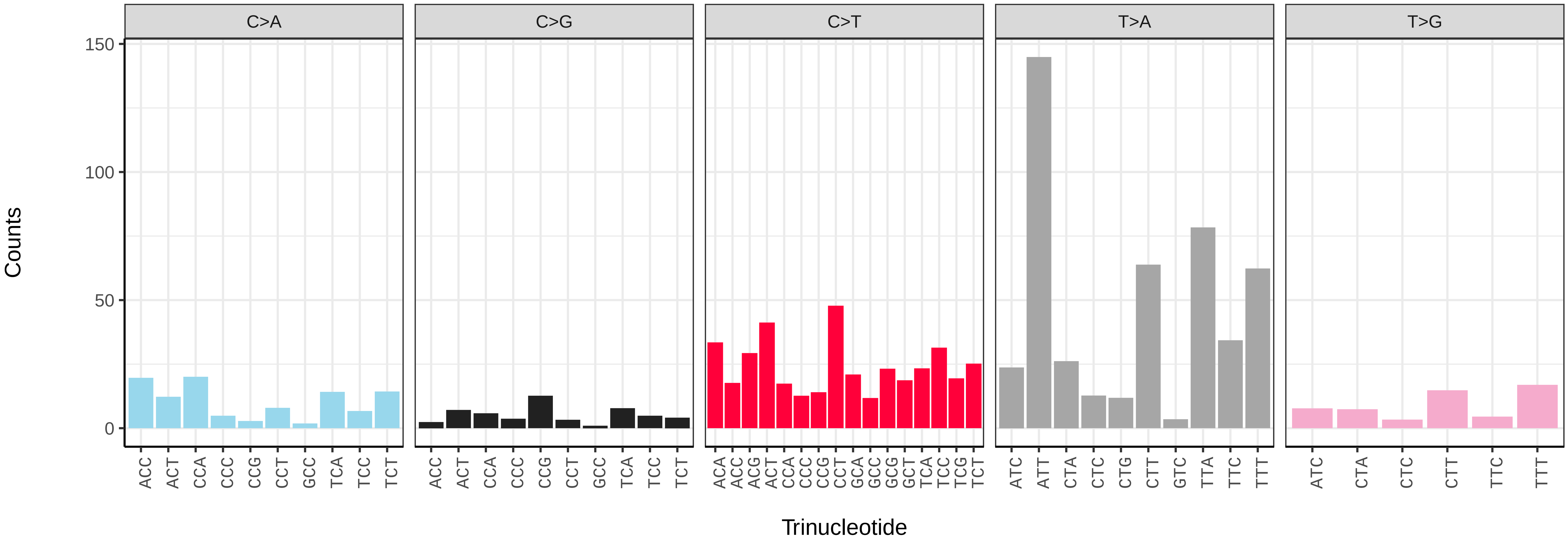

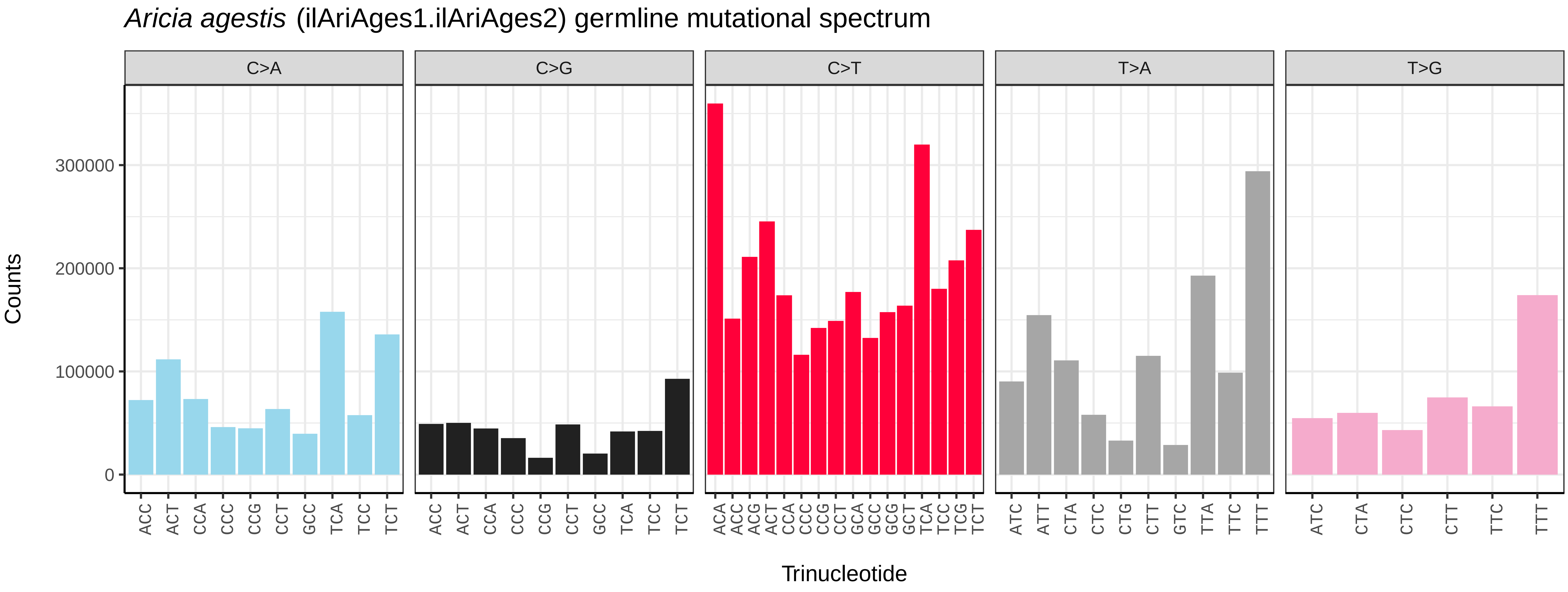
*Aricia agestis* (ilAriAges1.ilAriAges2) germline mutational spectrum
C>A
C>G
C>T
T>A
T>G
Counts
0
100000
200000
300000
Trinucleotide

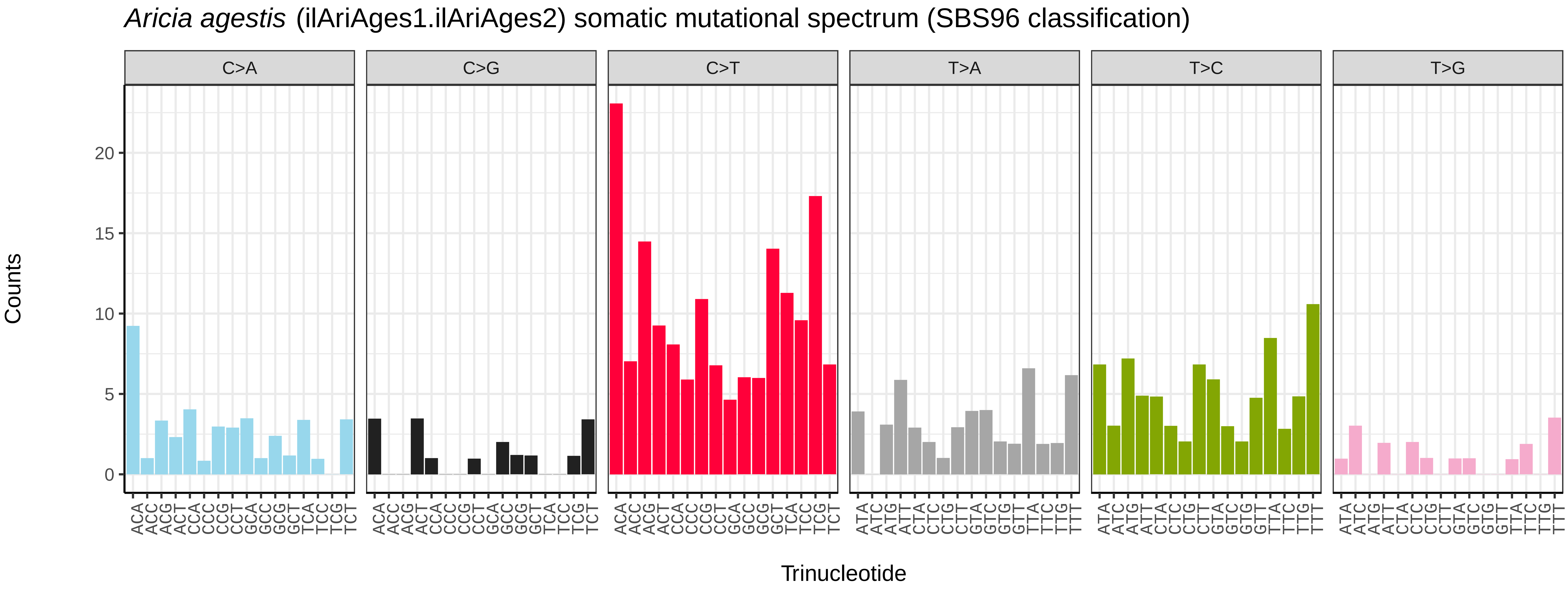
*Aricia agestis* (ilAriAges1.ilAriAges2) somatic mutational spectrum (SBS96 classification)
C>A
C>G
C>T
T>A
T>C
T>G
Counts
0
5
10
15
20
Trinucleotide

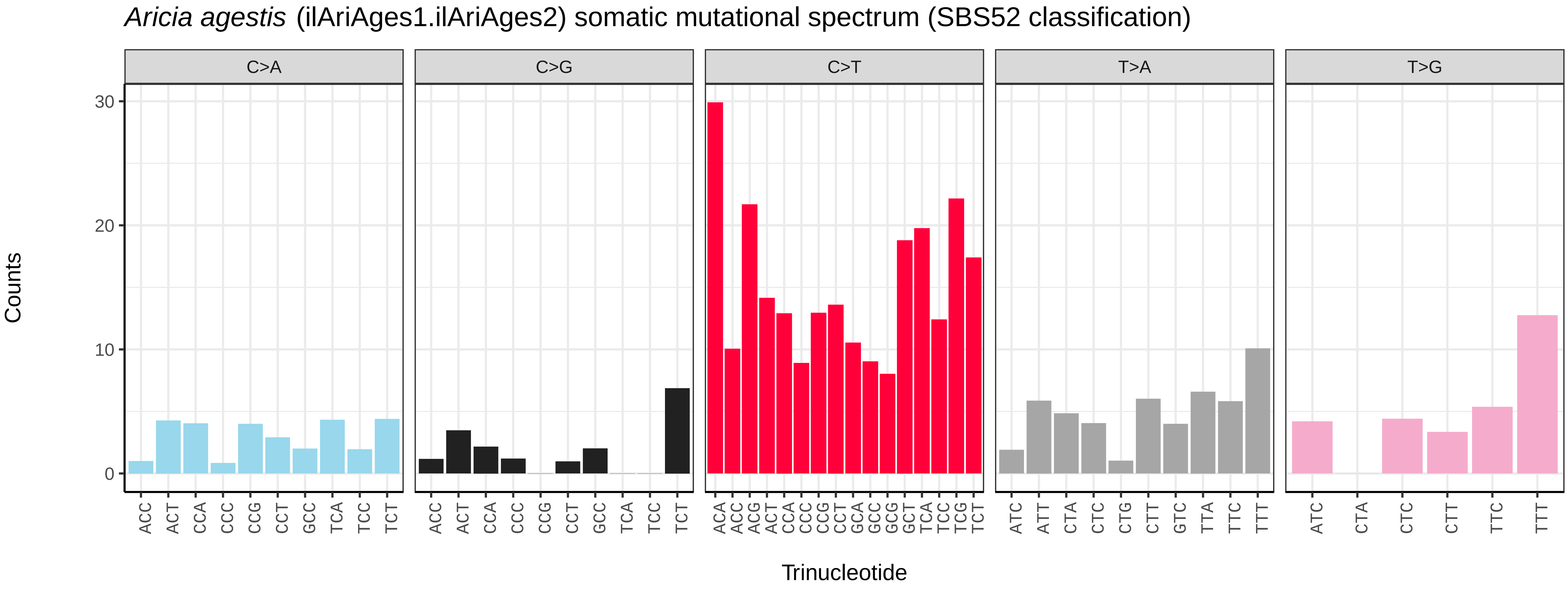
*Aricia agestis* (ilAriAges1.ilAriAges2) somatic mutational spectrum (SBS52 classification)
C>A
C>G
C>T
T>A
T>G
Counts
0
10
20
30
Trinucleotide

*Aricia artaxerxes* (ilAriArta2) germline mutational spectrum

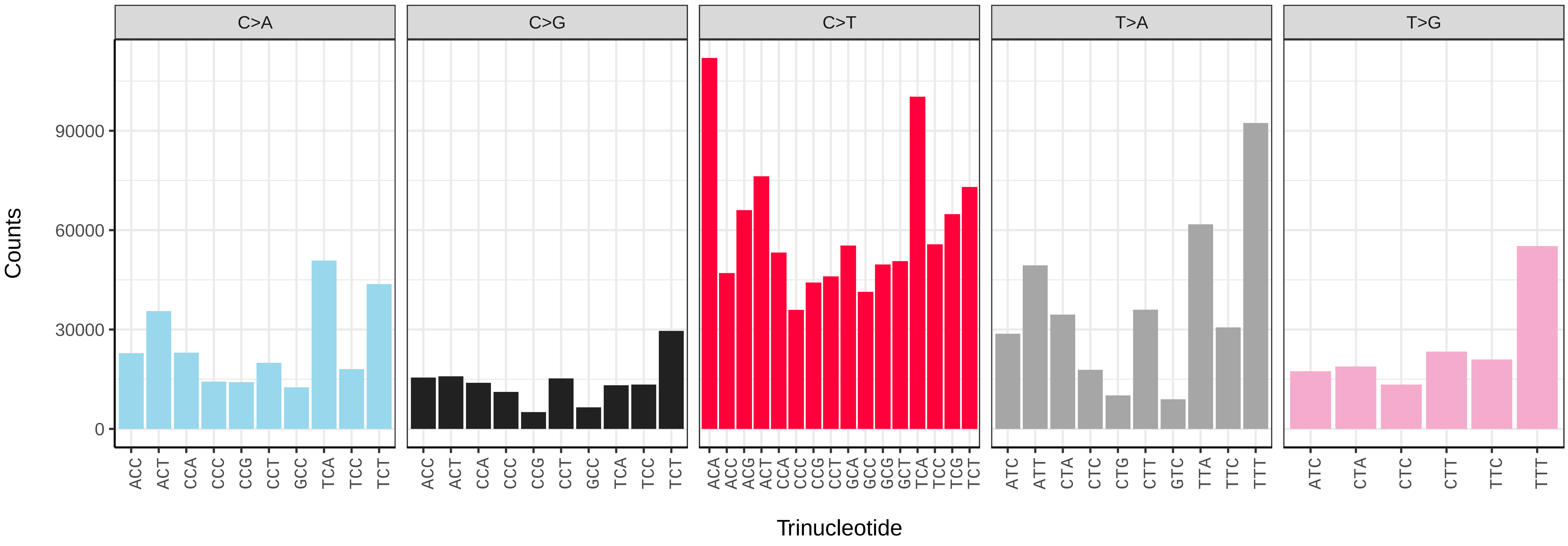


*Aricia artaxerxes* (ilAriArta2) somatic mutational spectrum (SBS96 classification)

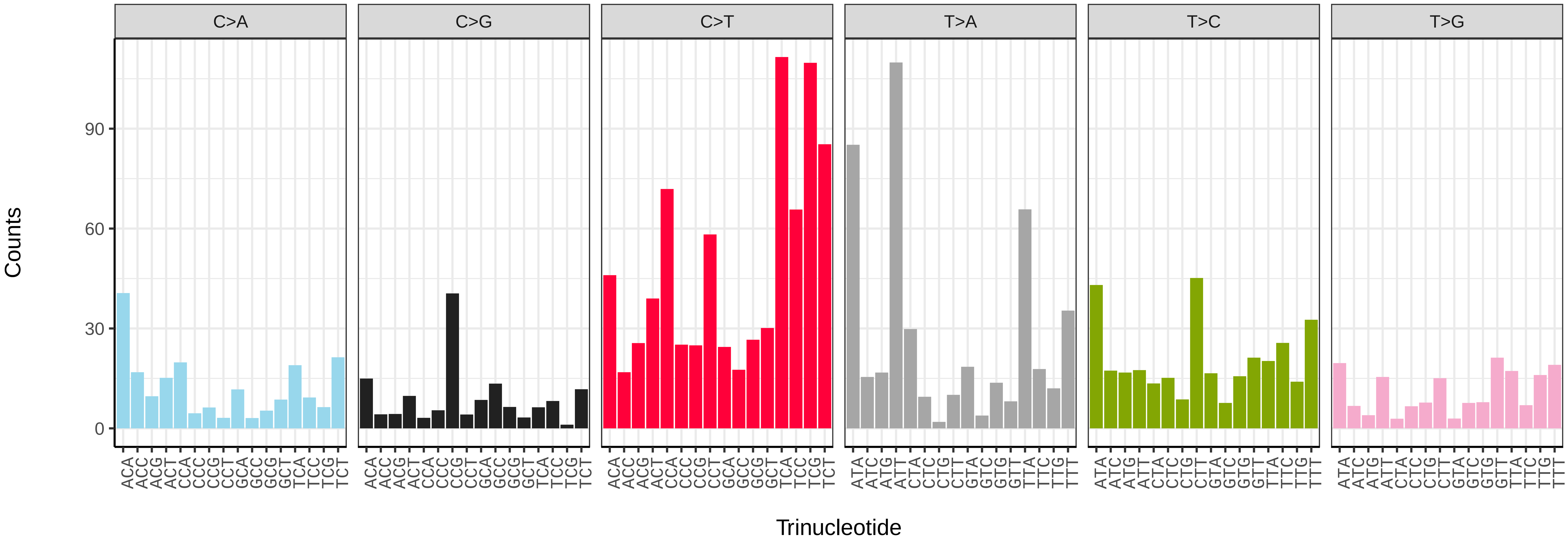


*Aricia artaxerxes* (ilAriArta2) somatic mutational spectrum (SBS52 classification)

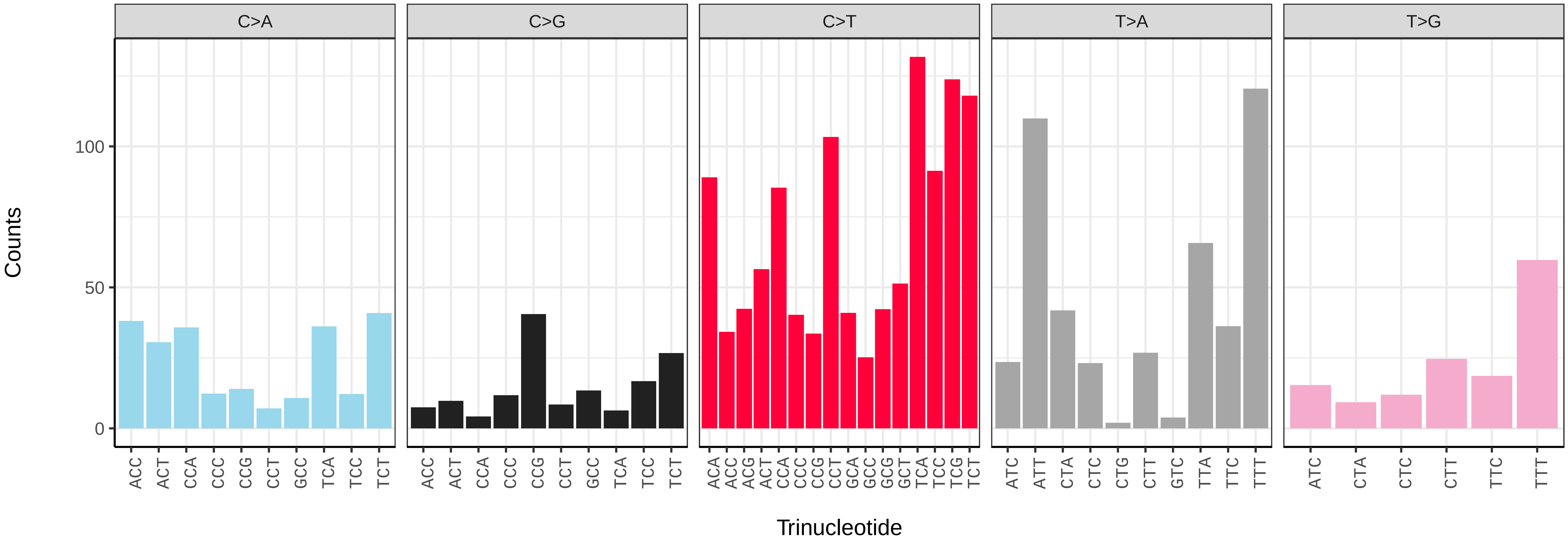

*Atethmia centrago* (ilAteCent1) germline mutational spectrum

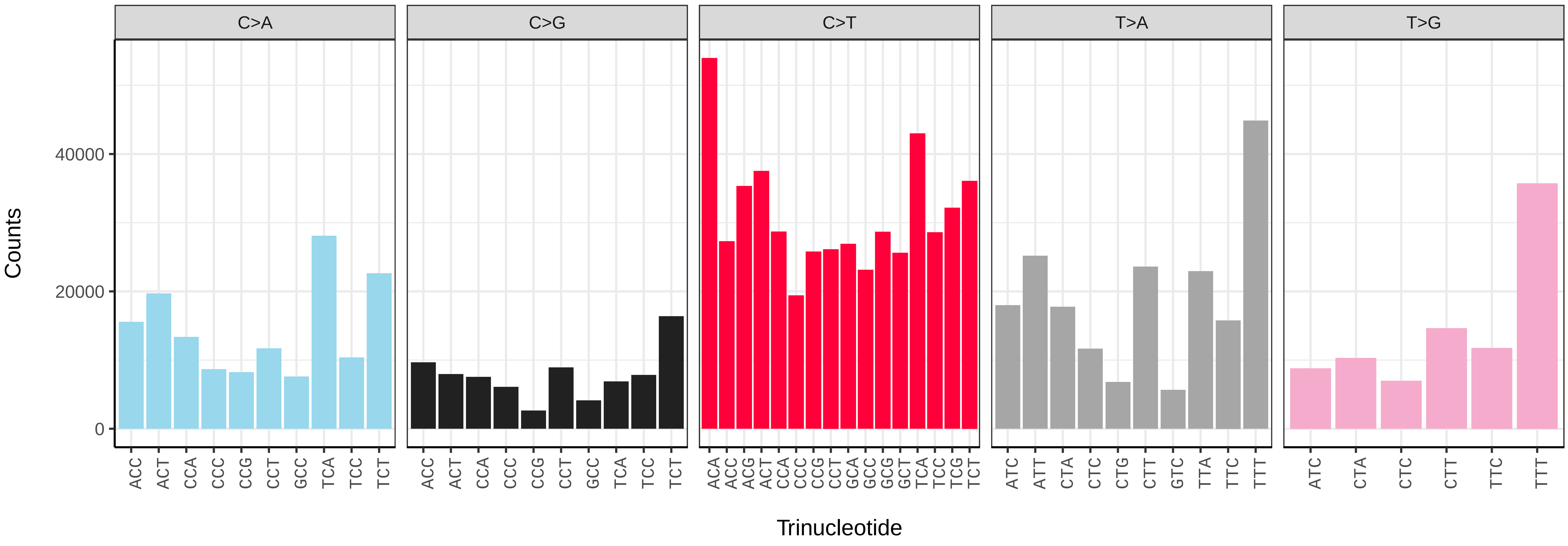


*Atethmia centrago* (ilAteCent1) somatic mutational spectrum (SBS96 classification)

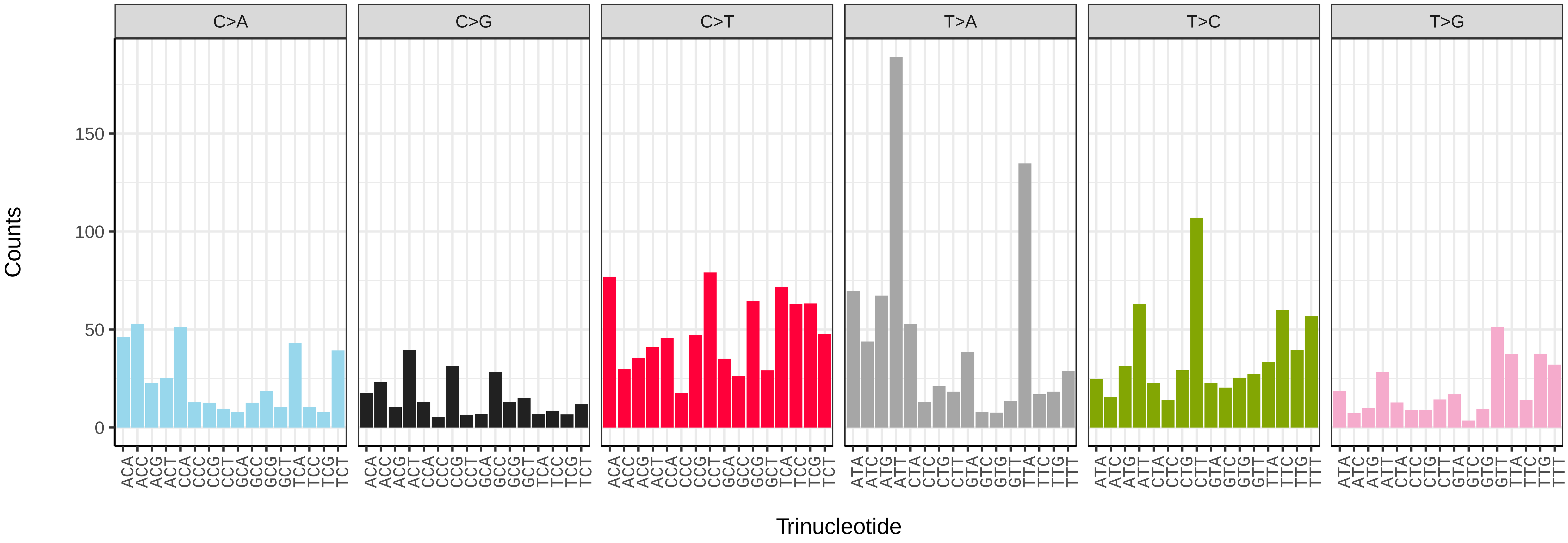


*Atethmia centrago* (ilAteCent1) somatic mutational spectrum (SBS52 classification)

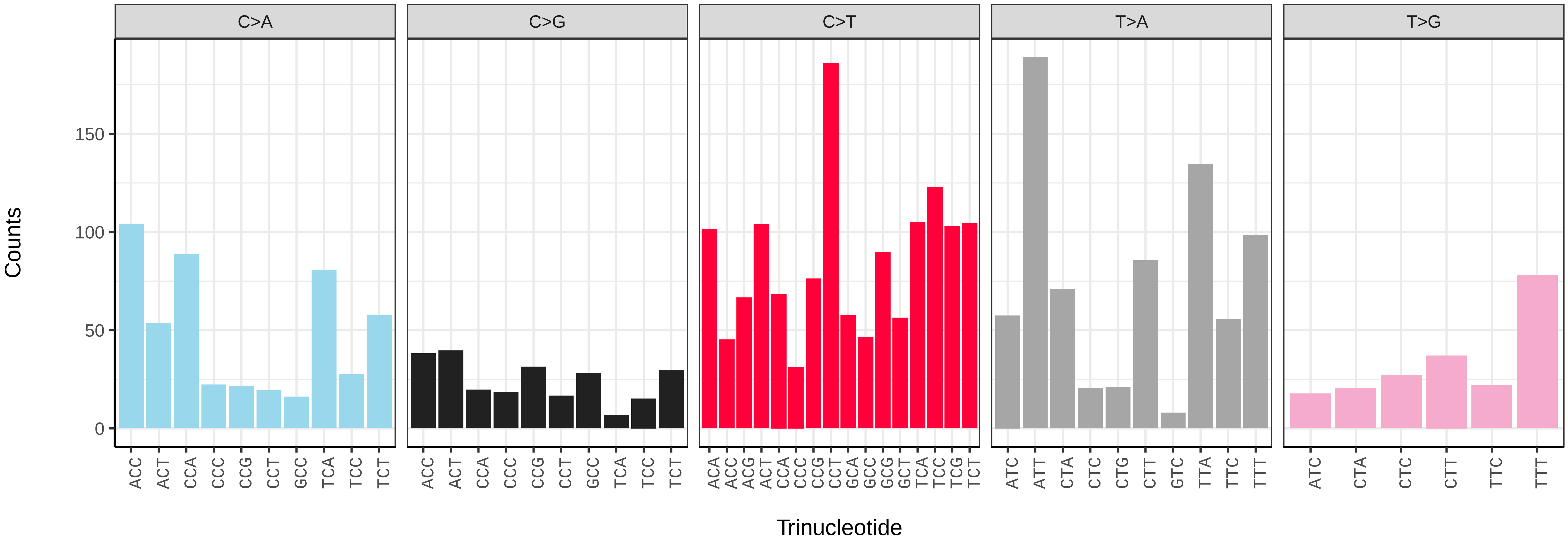

*Athrips mouffetella* (ilAthMouf1) germline mutational spectrum

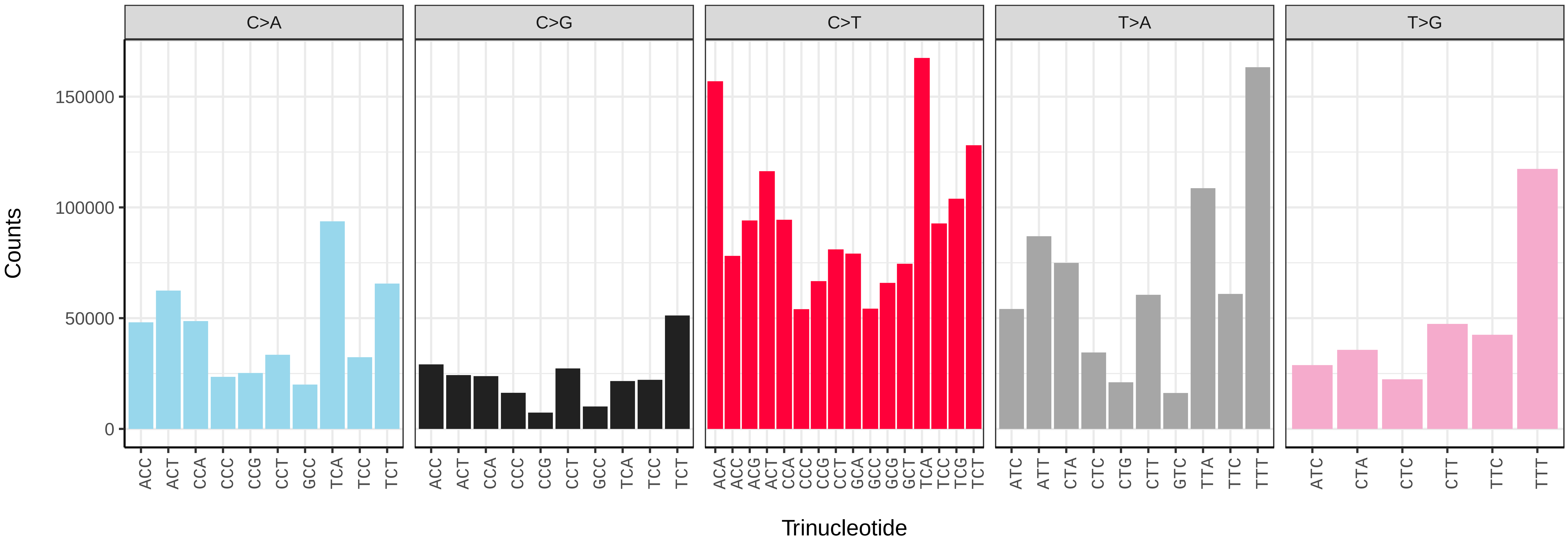


*Athrips mouffetella* (ilAthMouf1) somatic mutational spectrum (SBS96 classification)

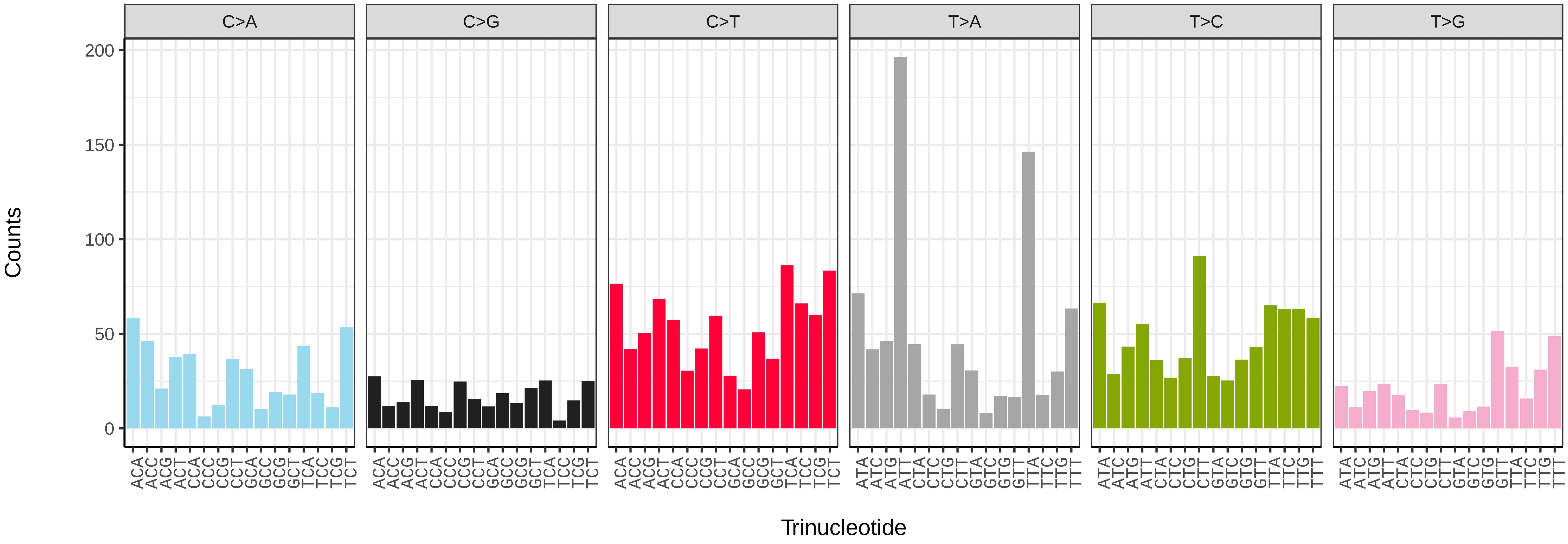


*Athrips mouffetella* (ilAthMouf1) somatic mutational spectrum (SBS52 classification)

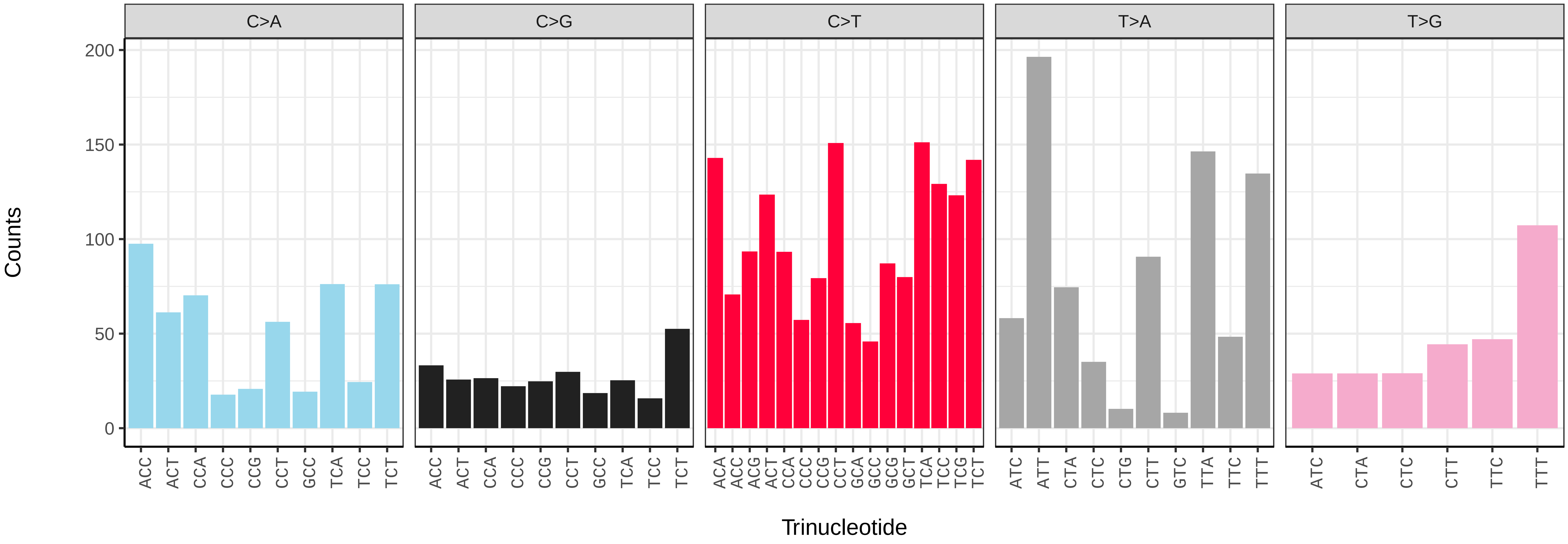

*Autographa gamma* (ilAutGamm1) germline mutational spectrum

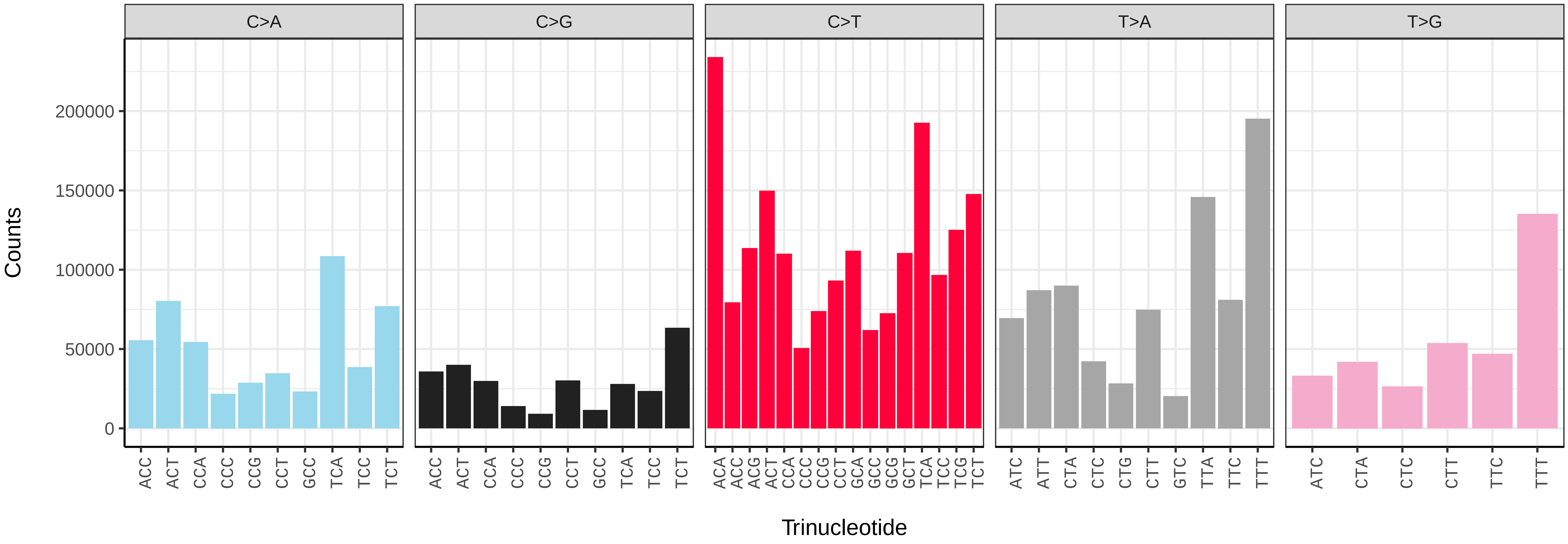


*Autographa gamma* (ilAutGamm1) somatic mutational spectrum (SBS96 classification)

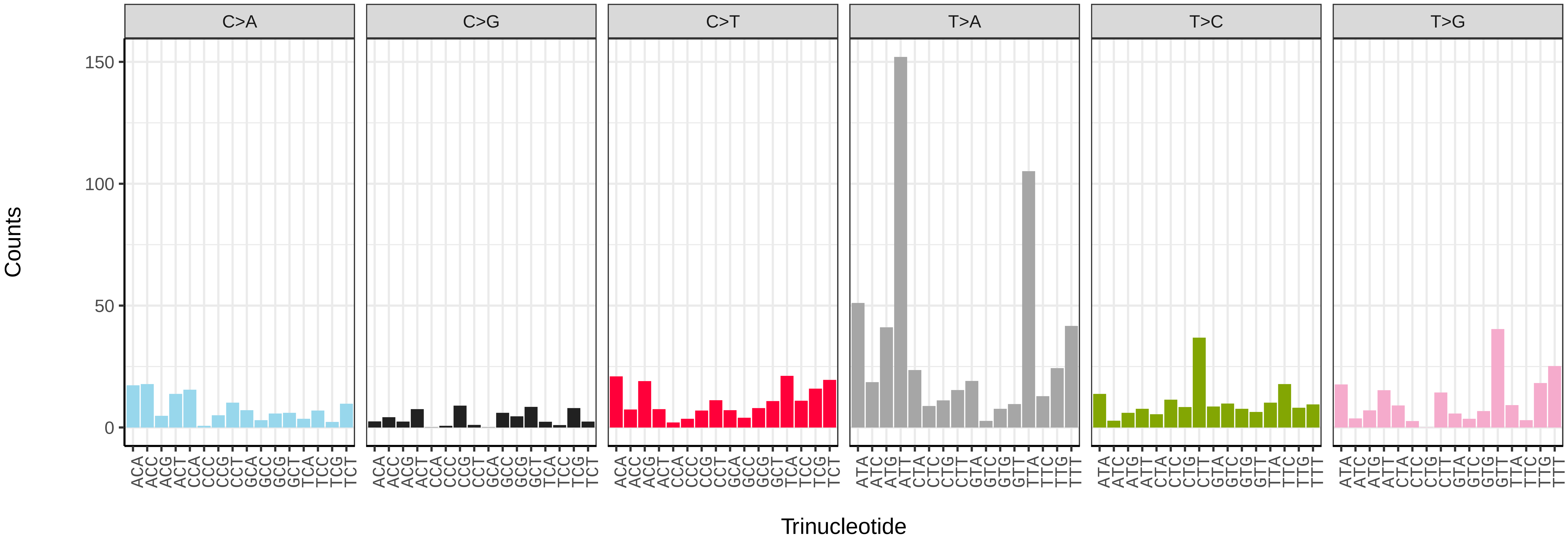


*Autographa gamma* (ilAutGamm1) somatic mutational spectrum (SBS52 classification)

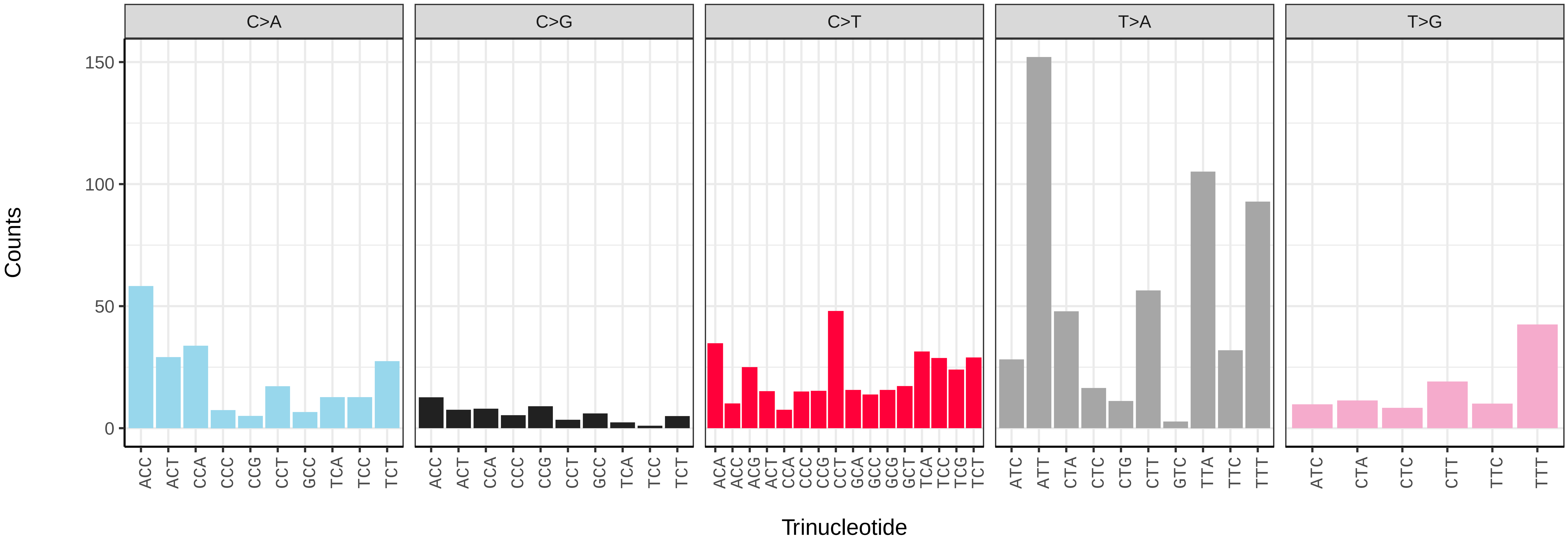

*Autographa pulchrina* (ilAutPulc1) germline mutational spectrum

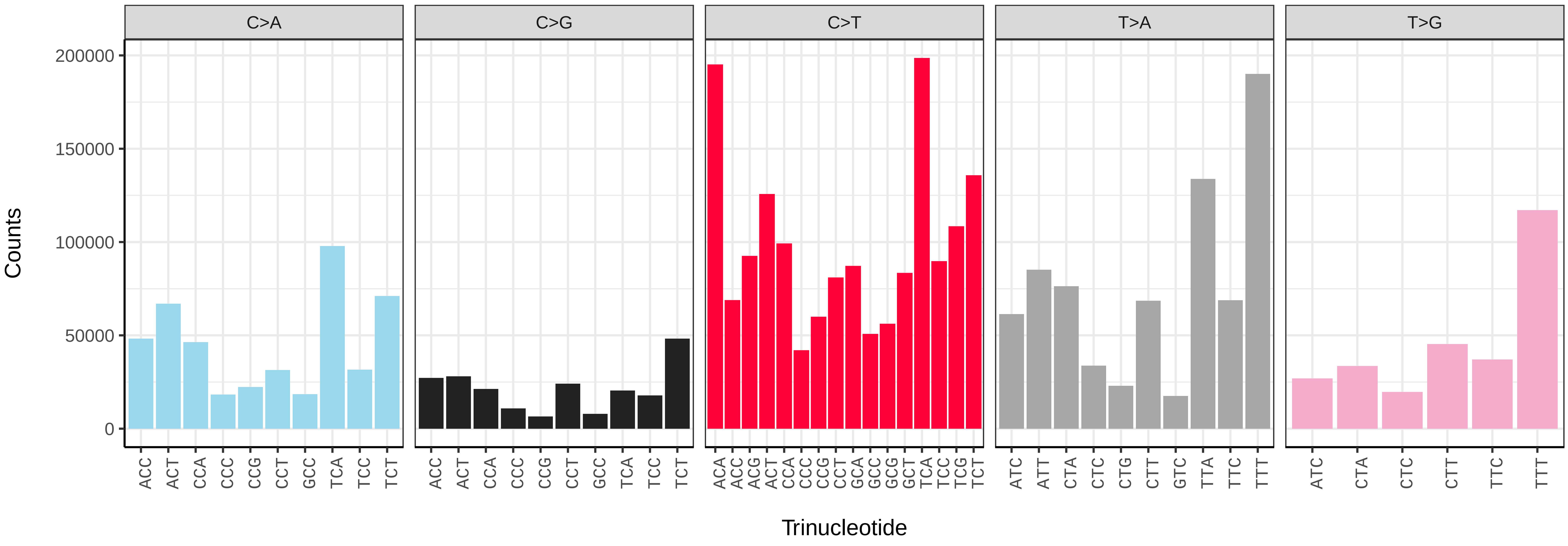


*Autographa pulchrina* (ilAutPulc1) somatic mutational spectrum (SBS96 classification)

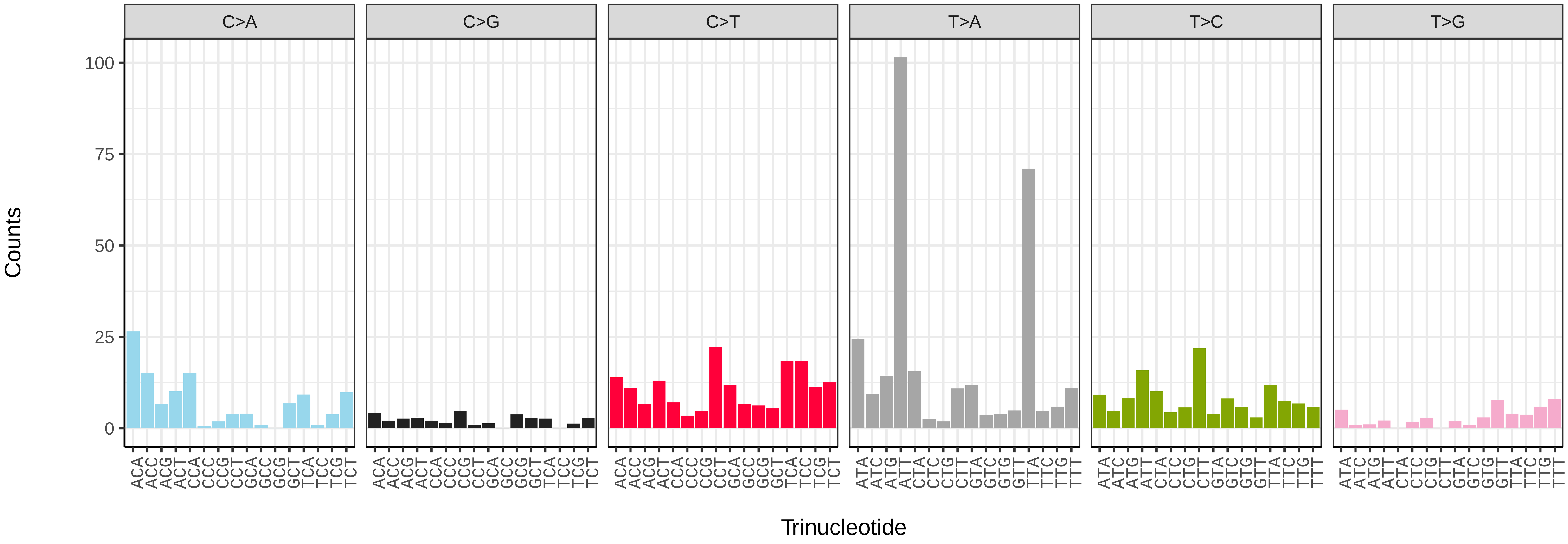


*Autographa pulchrina* (ilAutPulc1) somatic mutational spectrum (SBS52 classification)

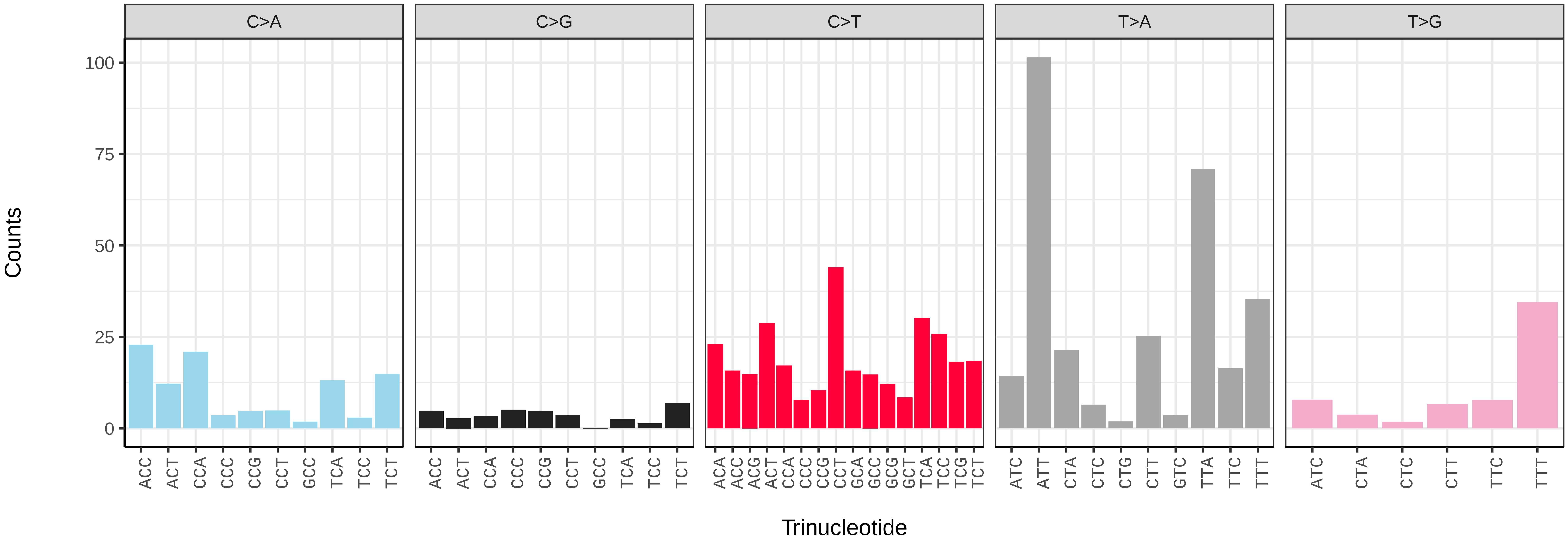

*Bembecia ichneumoniformis* (ilBemIchn1) germline mutational spectrum

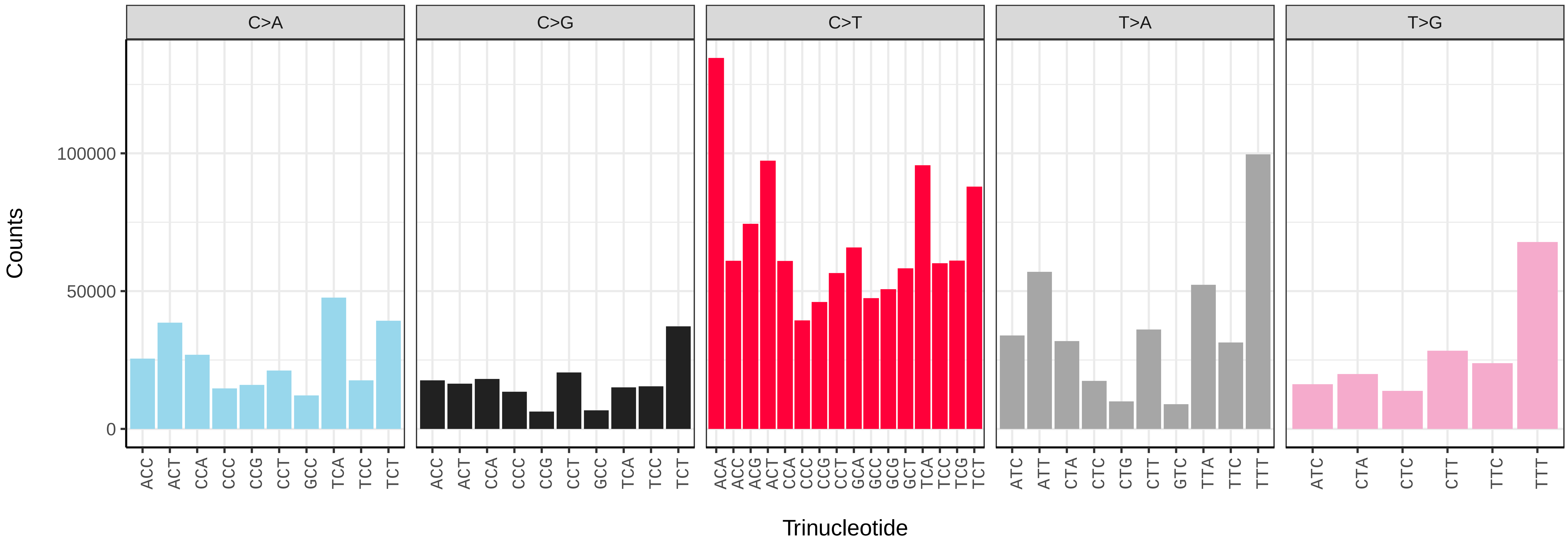


*Bembecia ichneumoniformis* (ilBemIchn1) somatic mutational spectrum (SBS96 classification)

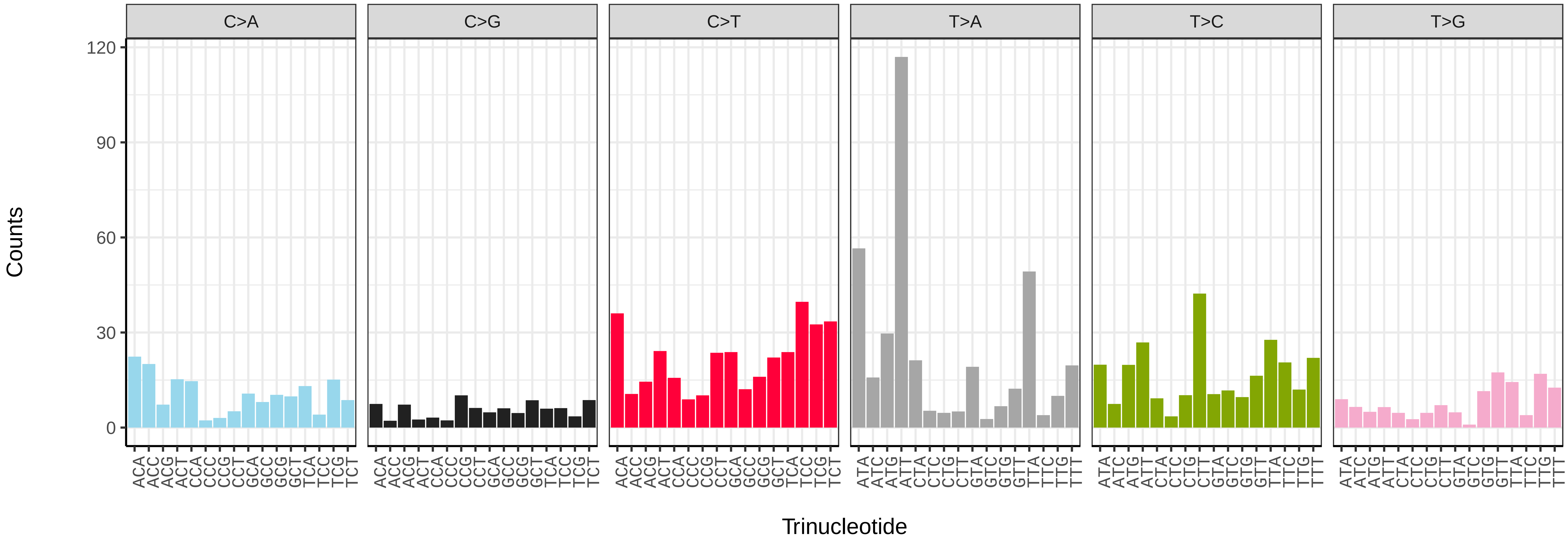


*Bembecia ichneumoniformis* (ilBemIchn1) somatic mutational spectrum (SBS52 classification)

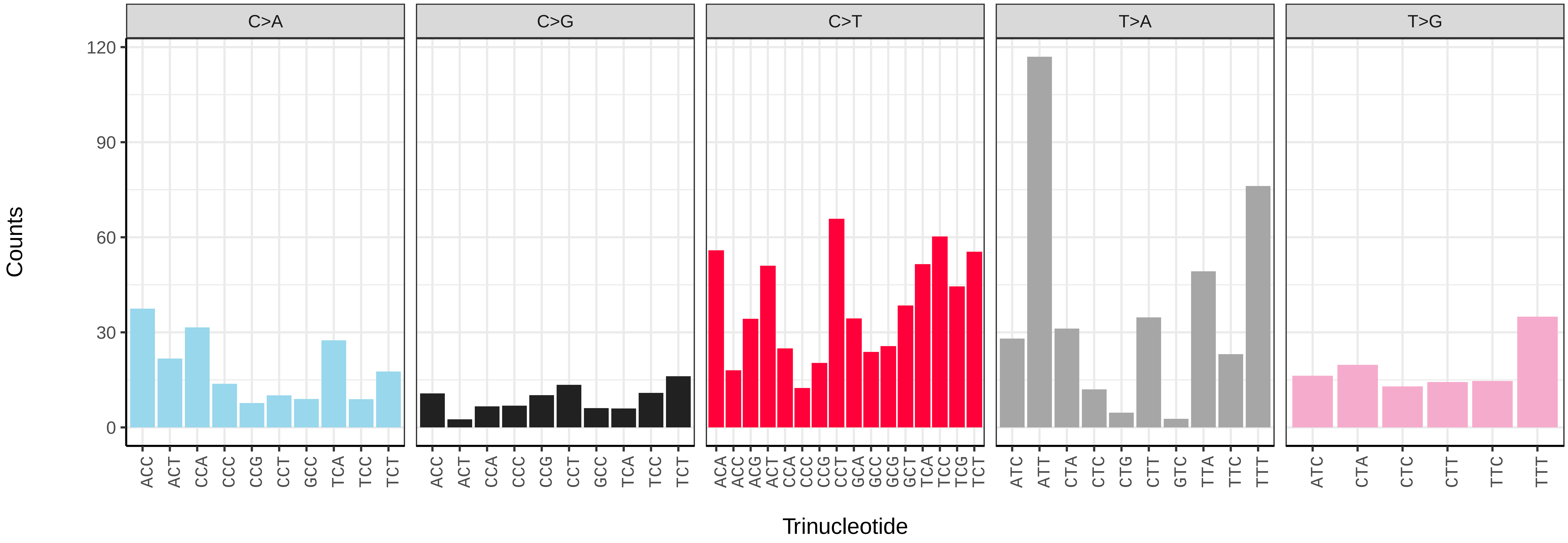

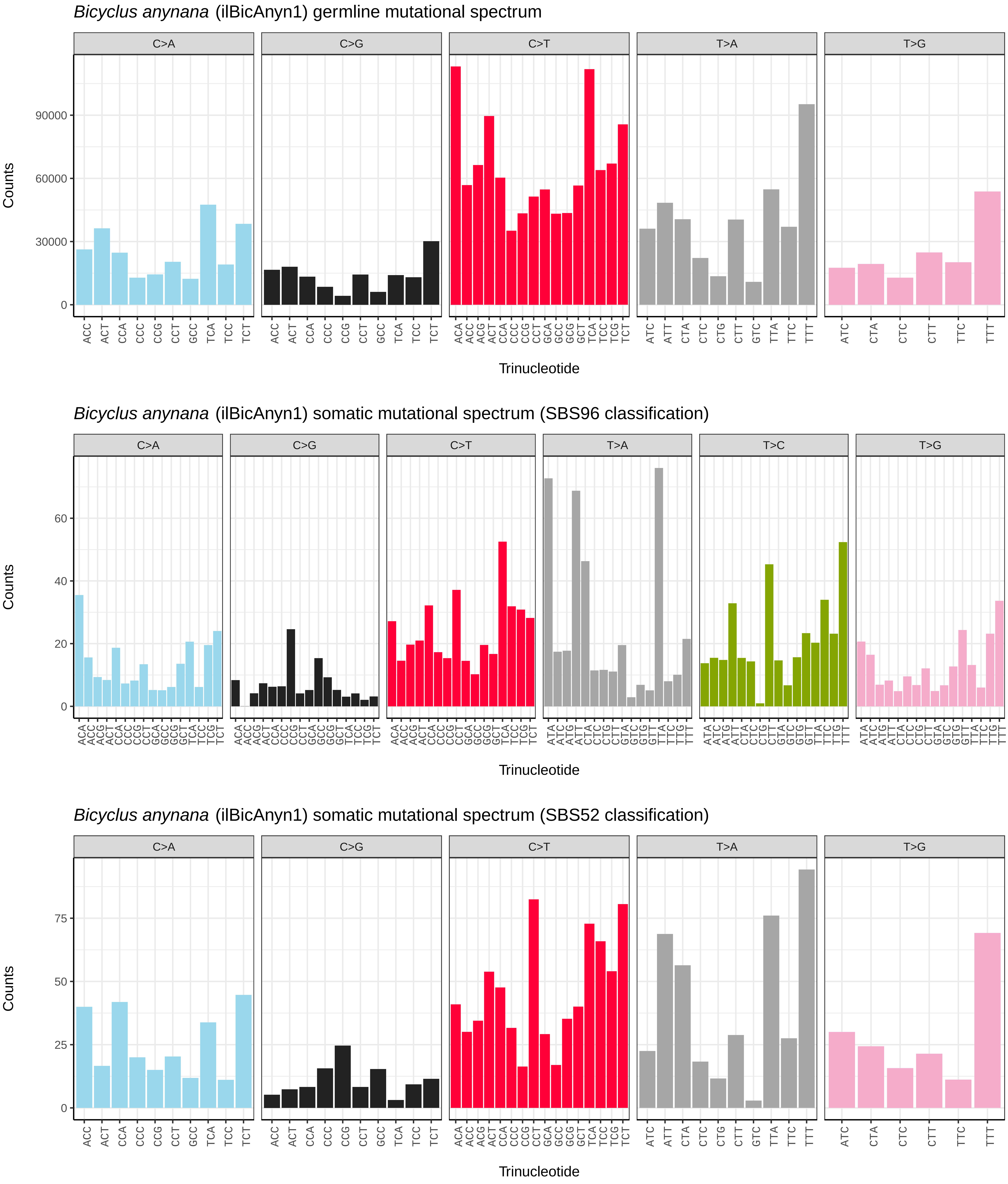
*Bicyclus anynana* (ilBicAnyn1) germline mutational spectrum
C>A
C>G
C>T
T>A
T>G
Counts
0
30000
60000
90000
Trinucleotide
*Bicyclus anynana* (ilBicAnyn1) somatic mutational spectrum (SBS96 classification)
C>A
C>G
C>T
T>A
T>C
T>G
Counts
0
20
40
60
Trinucleotide
*Bicyclus anynana* (ilBicAnyn1) somatic mutational spectrum (SBS52 classification)
C>A
C>G
C>T
T>A
T>G
Counts
0
25
50
75
Trinucleotide

*Biston betularia* (ilBisBetu1) germline mutational spectrum

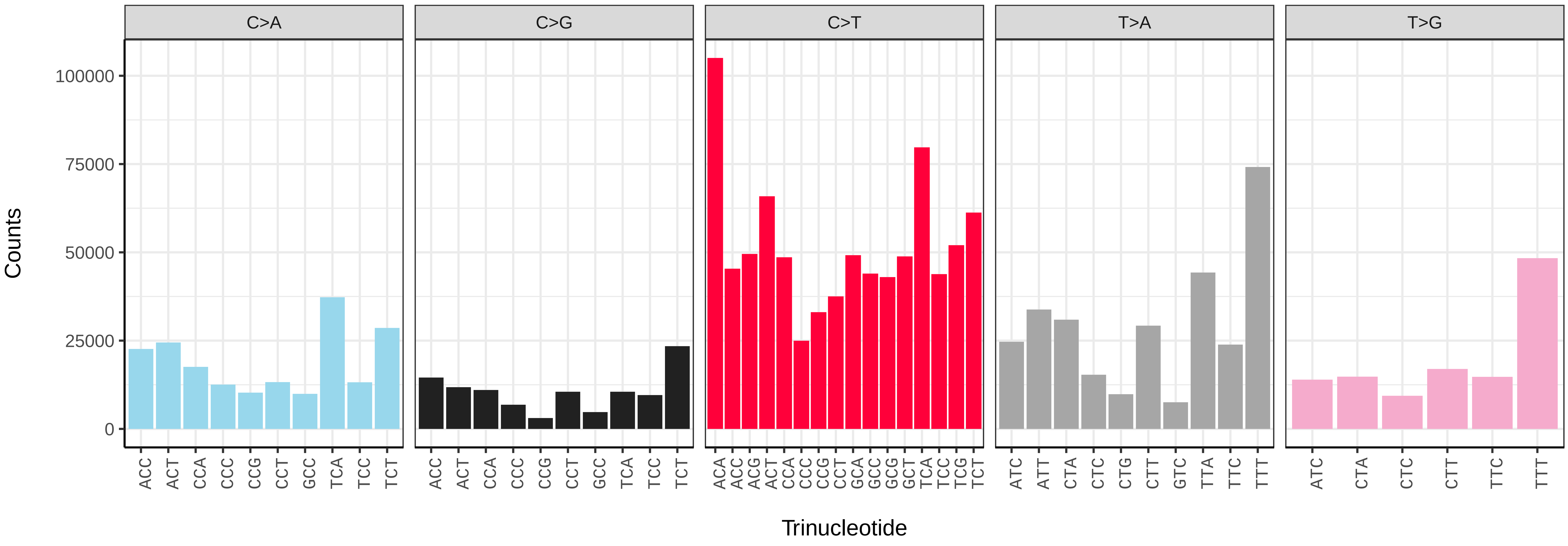


*Biston betularia* (ilBisBetu1) somatic mutational spectrum (SBS96 classification)

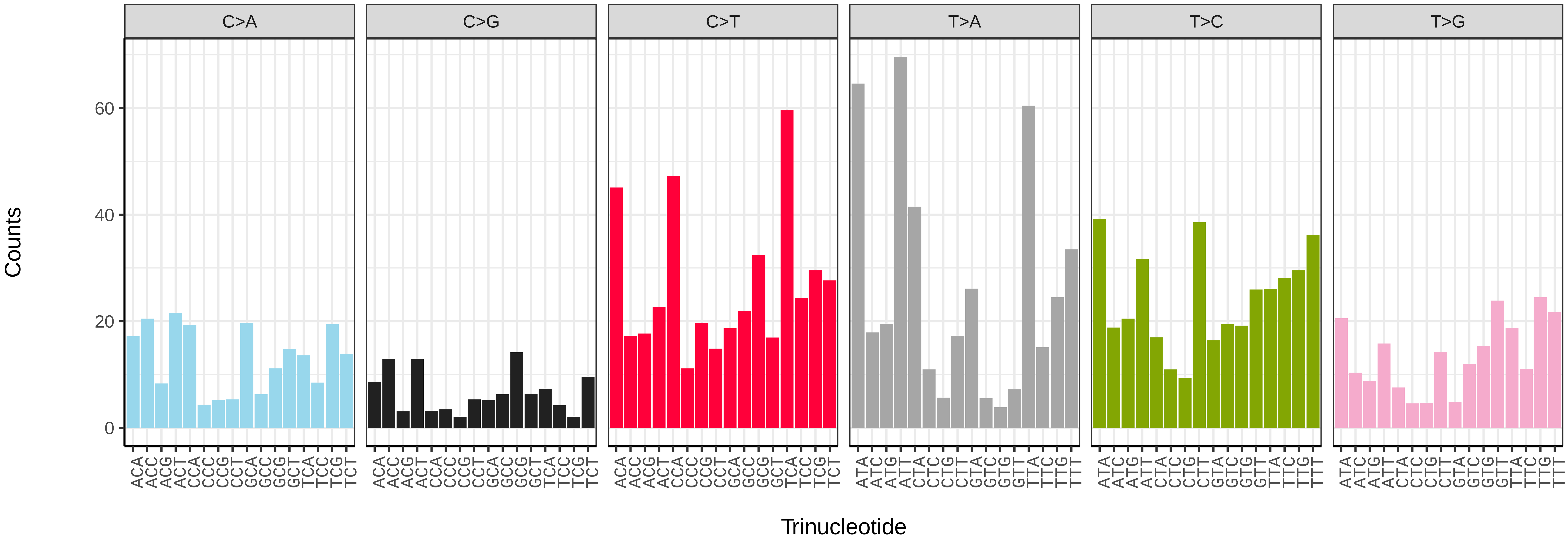


*Biston betularia* (ilBisBetu1) somatic mutational spectrum (SBS52 classification)

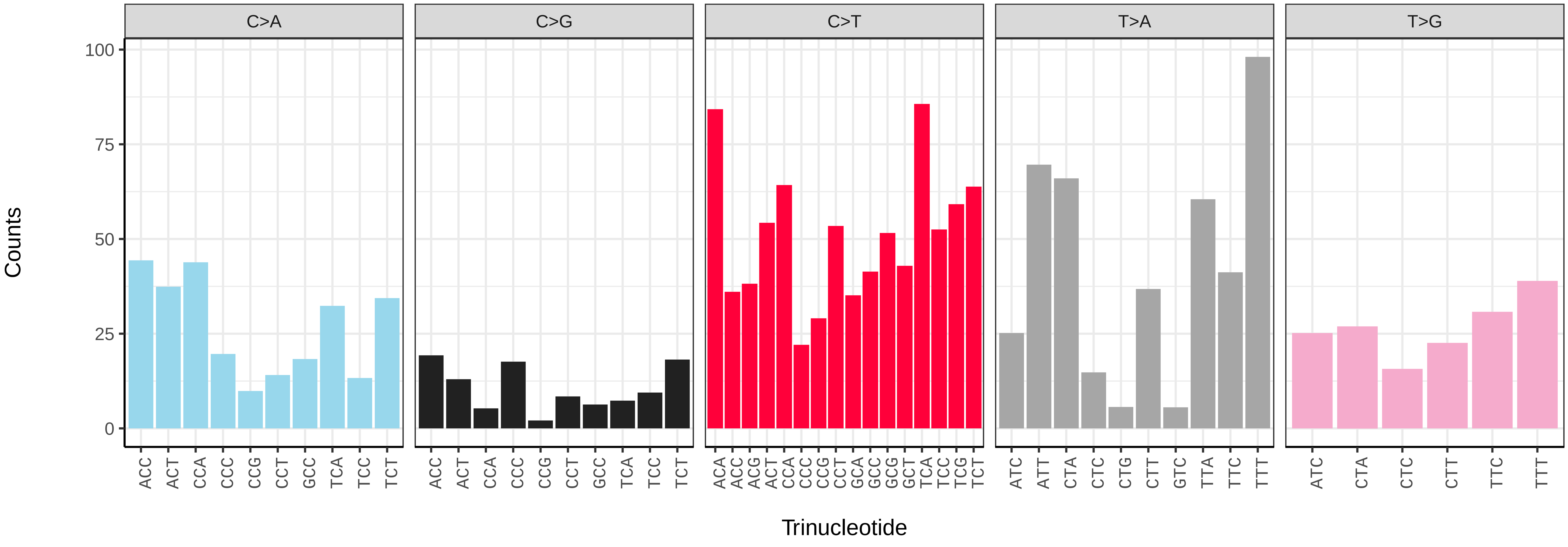

*Blastobasis adustella* (ilBlaAdus2) germline mutational spectrum

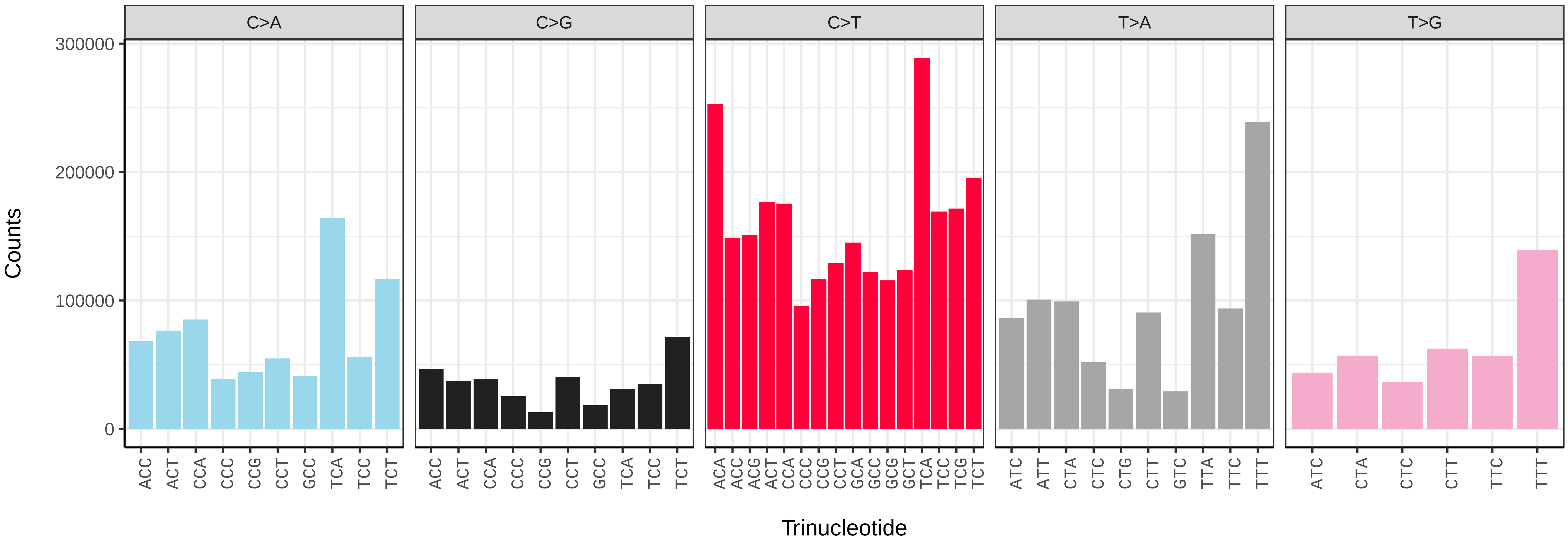


*Blastobasis adustella* (ilBlaAdus2) somatic mutational spectrum (SBS96 classification)

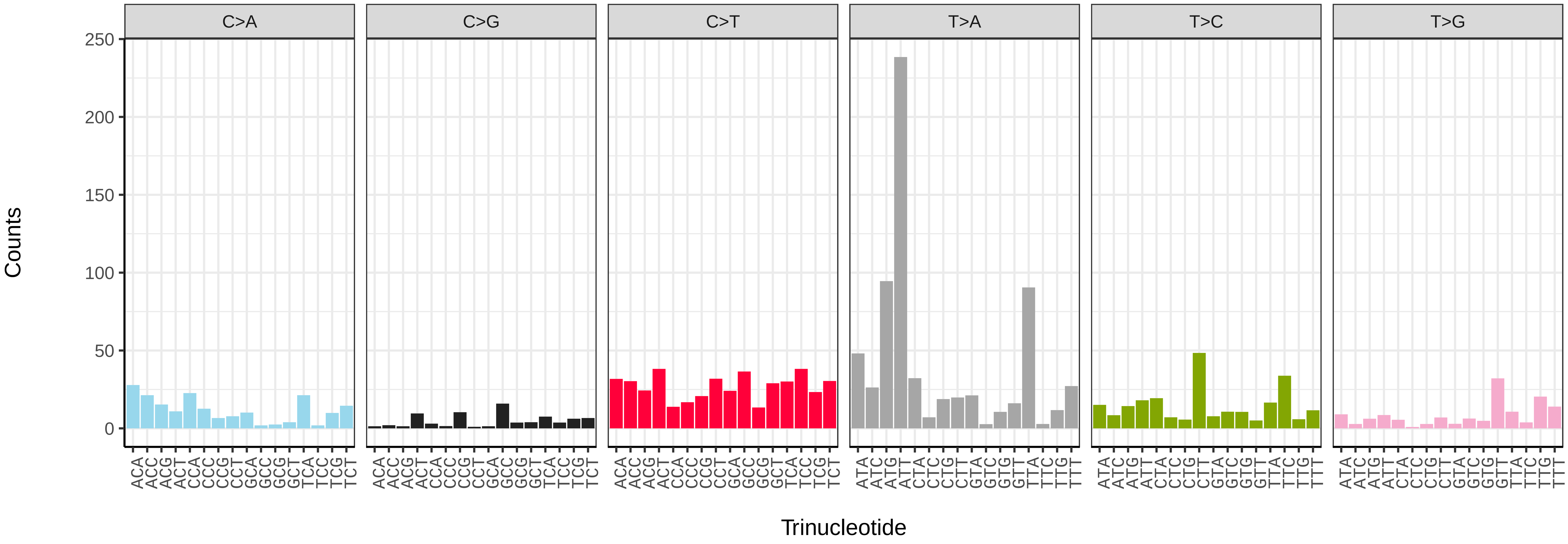


*Blastobasis adustella* (ilBlaAdus2) somatic mutational spectrum (SBS52 classification)

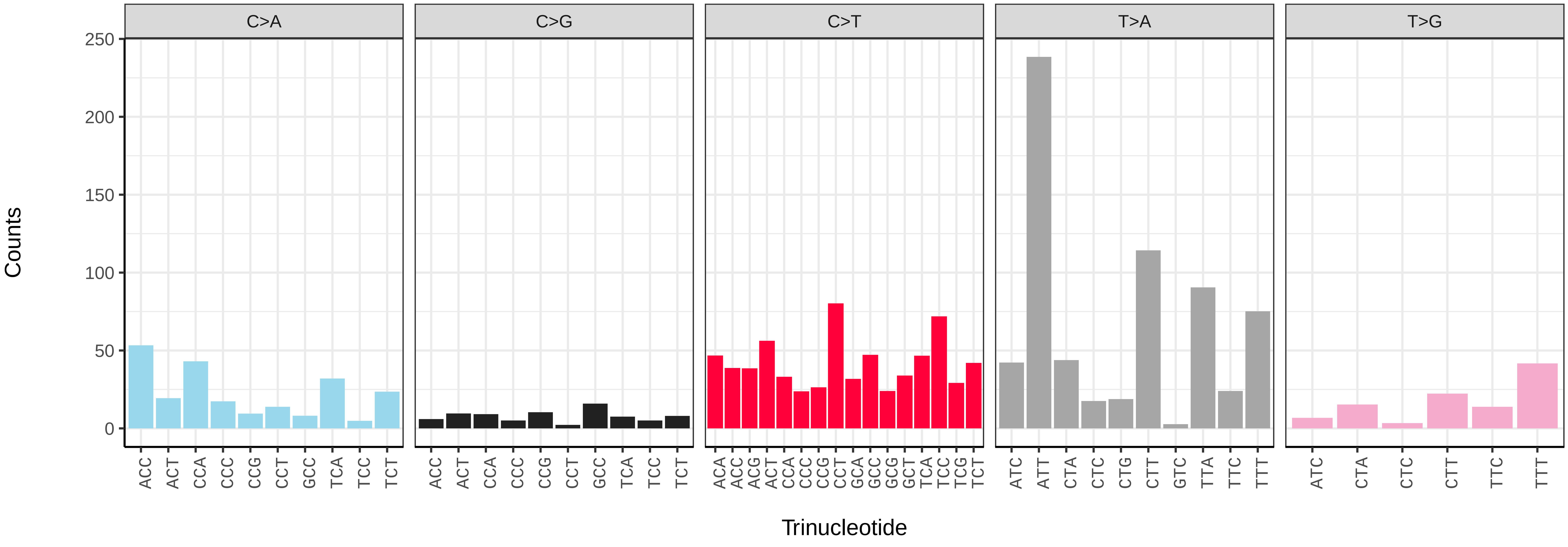

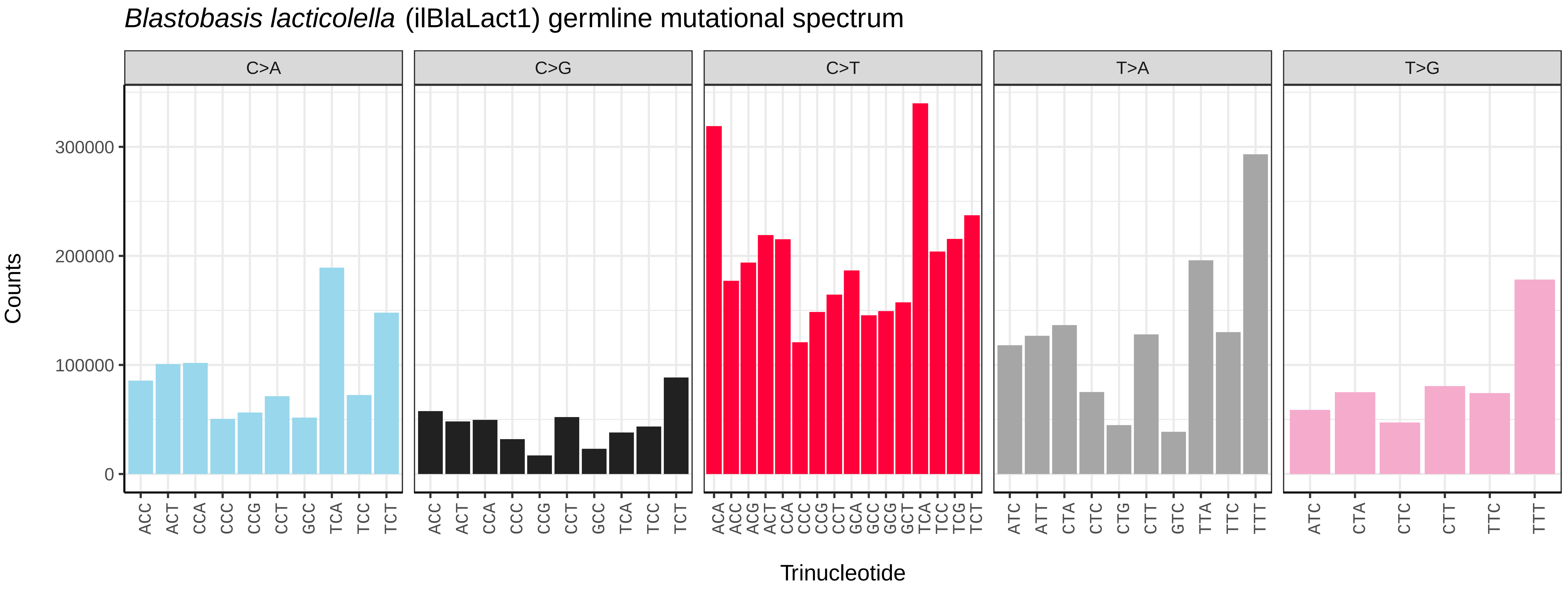
*Blastobasis lacticolella* (ilBlaLact1) germline mutational spectrum
C>A
C>G
C>T
T>A
T>G
Counts
0
100000
200000
300000
ACC ACT CCA CCC CCG CCT GCC TCA TCC TCT
ACC ACT CCA CCC CCG CCT GCC TCA TCC TCT
ACA ACC ACG ACT CCA CCC CCG CCT GCA GCC GCG GCT TCA TCC TCG TCT
ATC ATT CTA CTC CTG CTT GTC TTA TTC TTT
ATC CTA CTC CTT TTC TTT
Trinucleotide

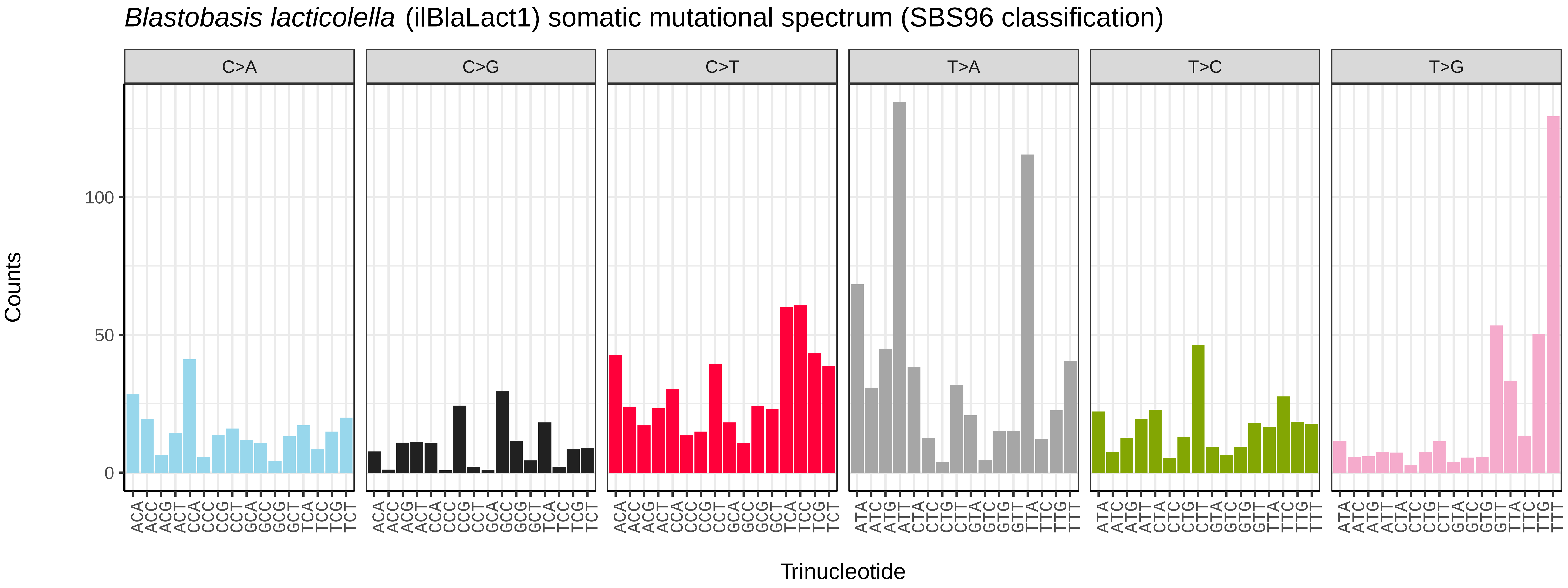
*Blastobasis lacticolella* (ilBlaLact1) somatic mutational spectrum (SBS96 classification)
C>A
C>G
C>T
T>A
T>C
T>G
Counts
0
50
100
ACA ACC ACG ACT CCA CCC CCG CCT GCA GCC GCG GCT TCA TCC TCG TCT
ACA ACC ACG ACT CCA CCC CCG CCT GCA GCC GCG GCT TCA TCC TCG TCT
ACA ACC ACG ACT CCA CCC CCG CCT GCA GCC GCG GCT TCA TCC TCG TCT
ATA ATC ATG ATT CTA CTC CTG CTT GTA GTC GTG GTT TTA TTC TTG TTT
ATA ATC ATG ATT CTA CTC CTG CTT GTA GTC GTG GTT TTA TTC TTG TTT
ATA ATC ATG ATT CTA CTC CTG CTT GTA GTC GTG GTT TTA TTC TTG TTT
Trinucleotide

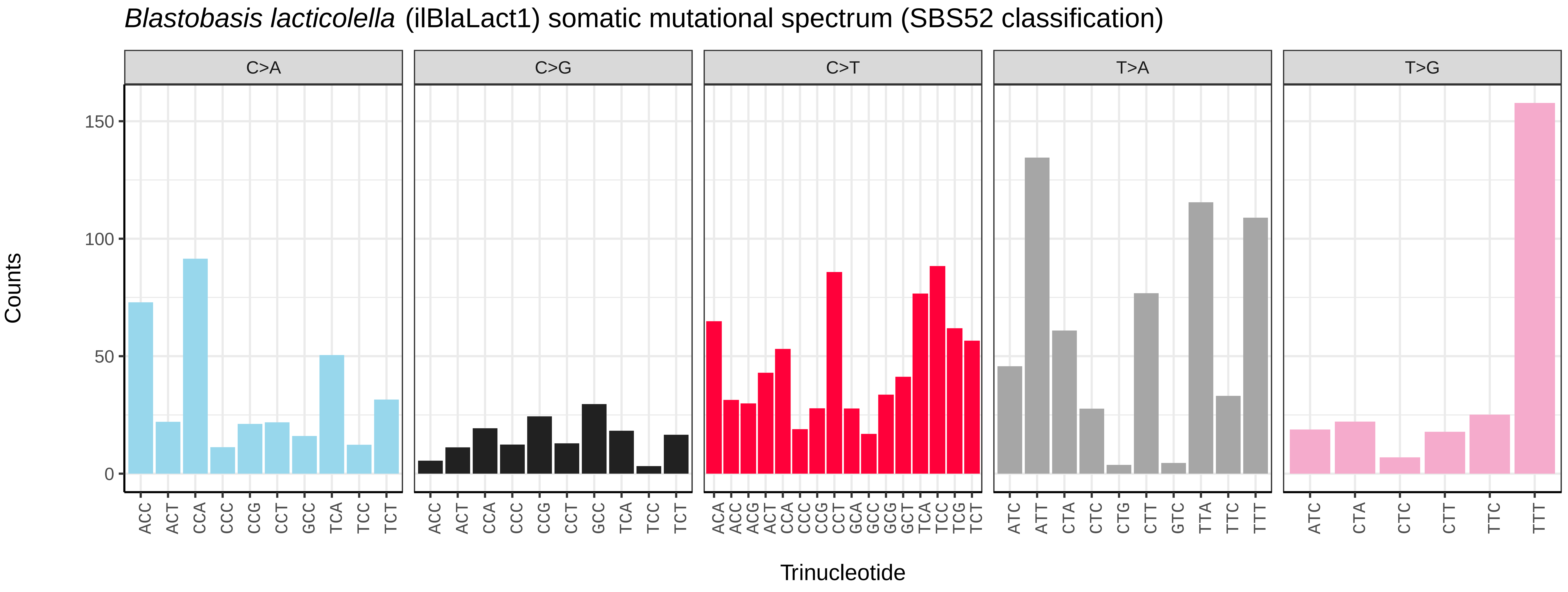
*Blastobasis lacticolella* (ilBlaLact1) somatic mutational spectrum (SBS52 classification)
C>A
C>G
C>T
T>A
T>G
Counts
0
50
100
150
ACC ACT CCA CCC CCG CCT GCC TCA TCC TCT
ACC ACT CCA CCC CCG CCT GCC TCA TCC TCT
ACA ACC ACG ACT CCA CCC CCG CCT GCA GCC GCG GCT TCA TCC TCG TCT
ATC ATT CTA CTC CTG CTT GTC TTA TTC TTT
ATC CTA CTC CTT TTC TTT
Trinucleotide

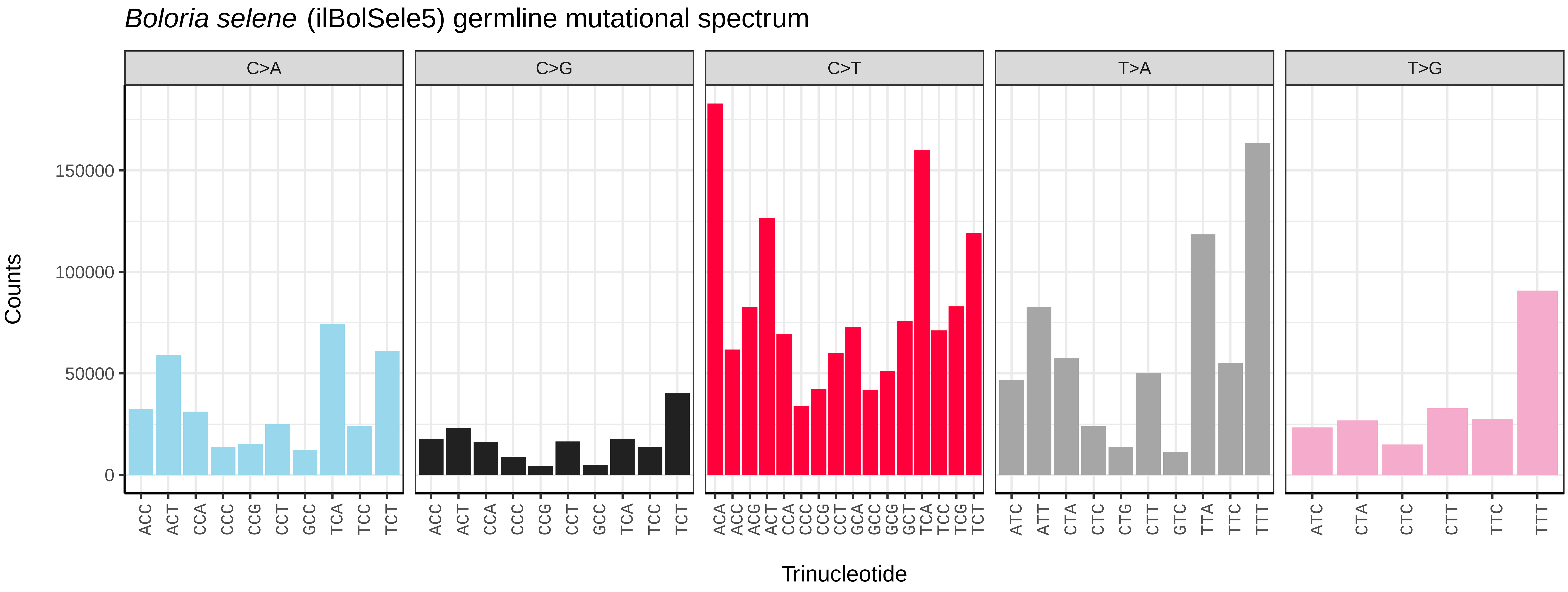
Boloria selene (ilBolSele5) germline mutational spectrum
C>A
C>G
C>T
T>A
T>G
Counts
0
50000
100000
150000
ACC ACT CCA CCC CCG CCT GCC TCA TCC TCT
ACA ACC ACG ACT CCA CCC CCG CCT GCA GCC GCG GCT TCA TCC TCG TCT
ATC ATT CTA CTC CTG CTT GTC TTA TTC TTT
ATC CTA CTC CTT TTC TTT
Trinucleotide

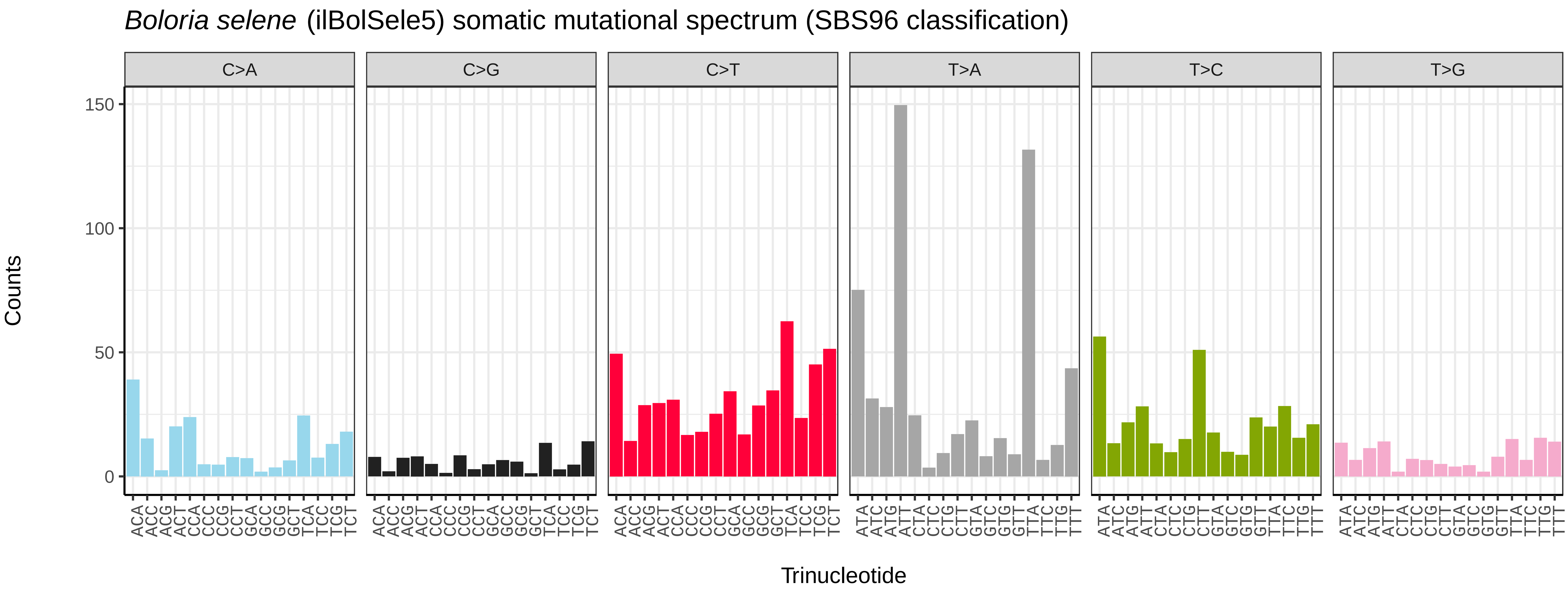
Boloria selene (ilBolSele5) somatic mutational spectrum (SBS96 classification)
C>A
C>G
C>T
T>A
T>C
T>G
Counts
0
50
100
150
ACA ACC ACG ACT CCA CCC CCG CCT GCA GCC GCG GCT TCA TCC TCG TCT
ATA ATC ATG ATT CTA CTC CTG CTT GTA GTC GTG GTT TTA TTC TTG TTT
Trinucleotide

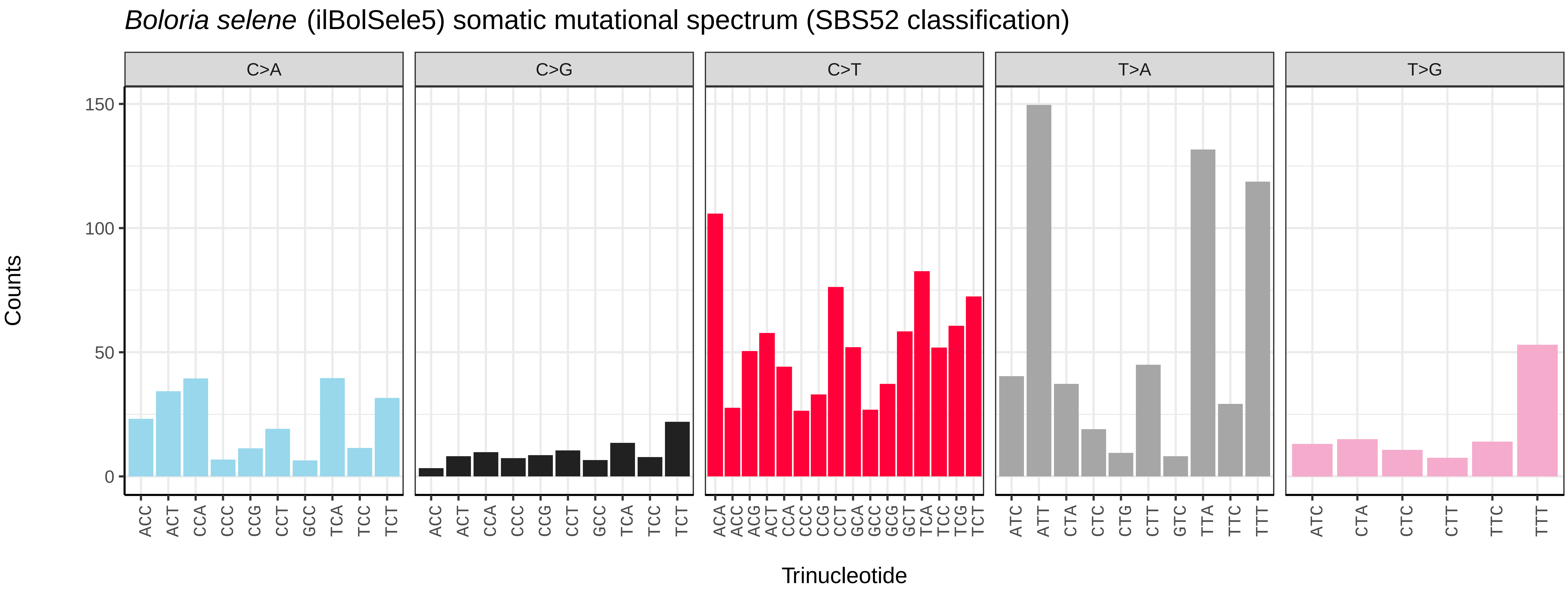
Boloria selene (ilBolSele5) somatic mutational spectrum (SBS52 classification)
C>A
C>G
C>T
T>A
T>G
Counts
0
50
100
150
ACC ACT CCA CCC CCG CCT GCC TCA TCC TCT
ACA ACC ACG ACT CCA CCC CCG CCT GCA GCC GCG GCT TCA TCC TCG TCT
ATC ATT CTA CTC CTG CTT GTC TTA TTC TTT
ATC CTA CTC CTT TTC TTT
Trinucleotide

*Brachylomia viminalis* (ilBraVimi1) germline mutational spectrum

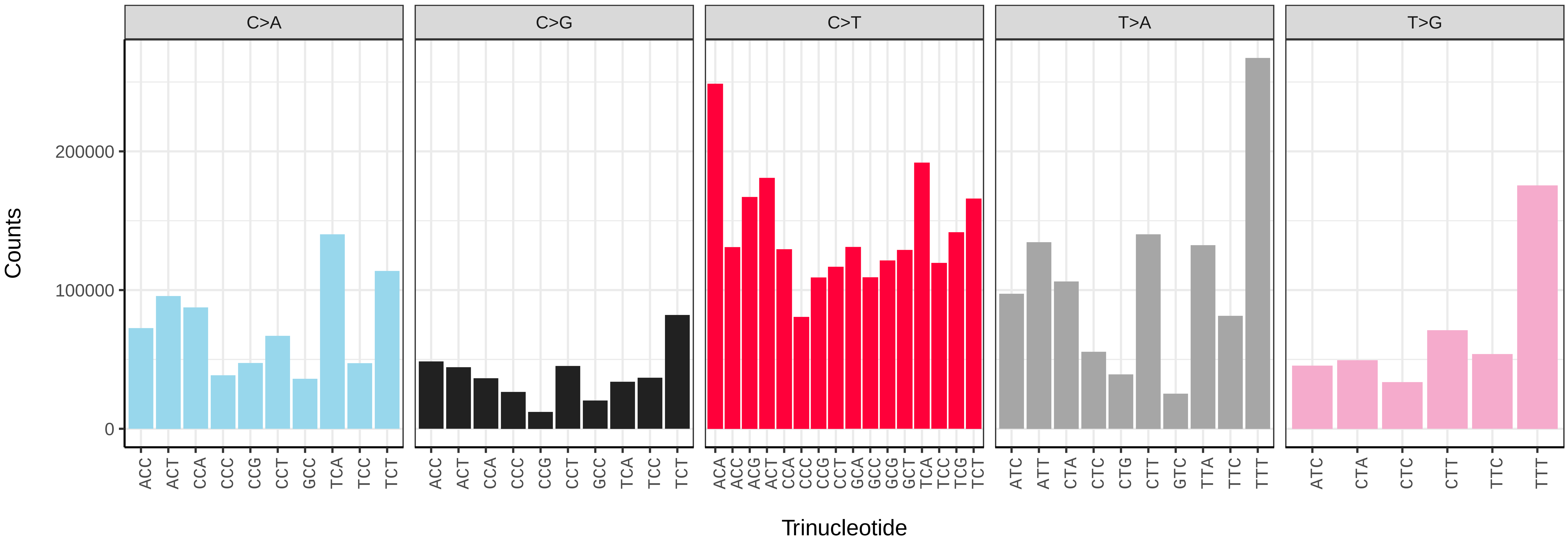


*Brachylomia viminalis* (ilBraVimi1) somatic mutational spectrum (SBS96 classification)

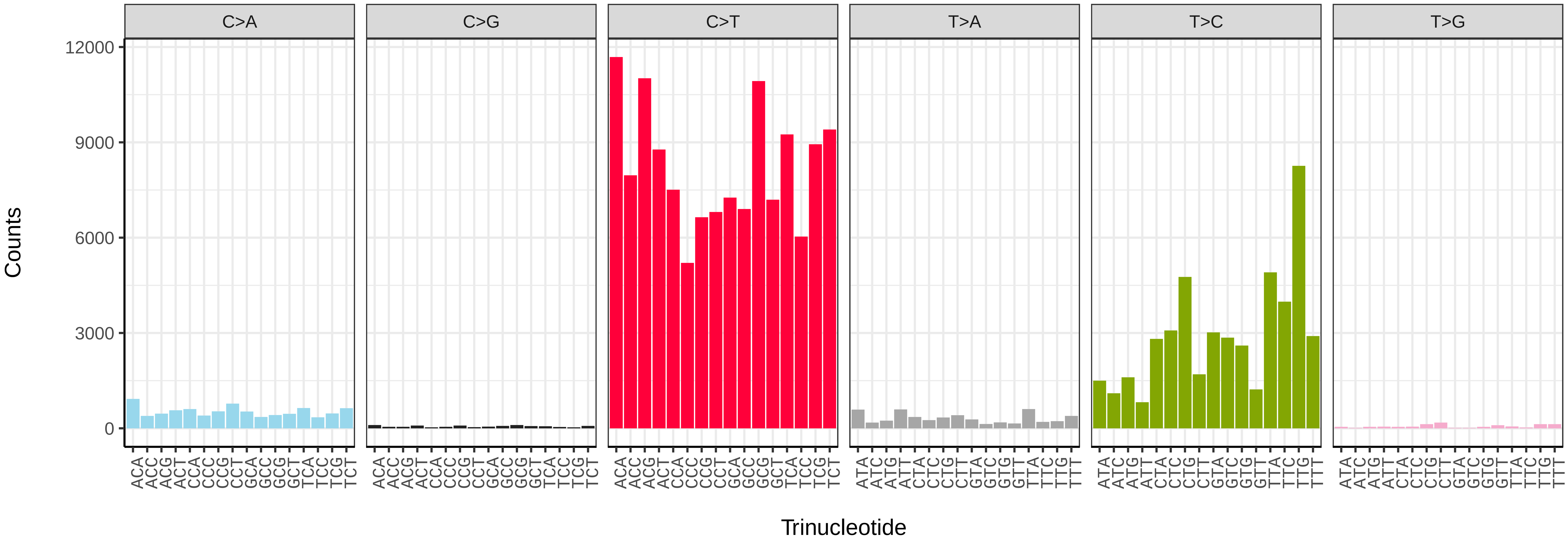


*Brachylomia viminalis* (ilBraVimi1) somatic mutational spectrum (SBS52 classification)

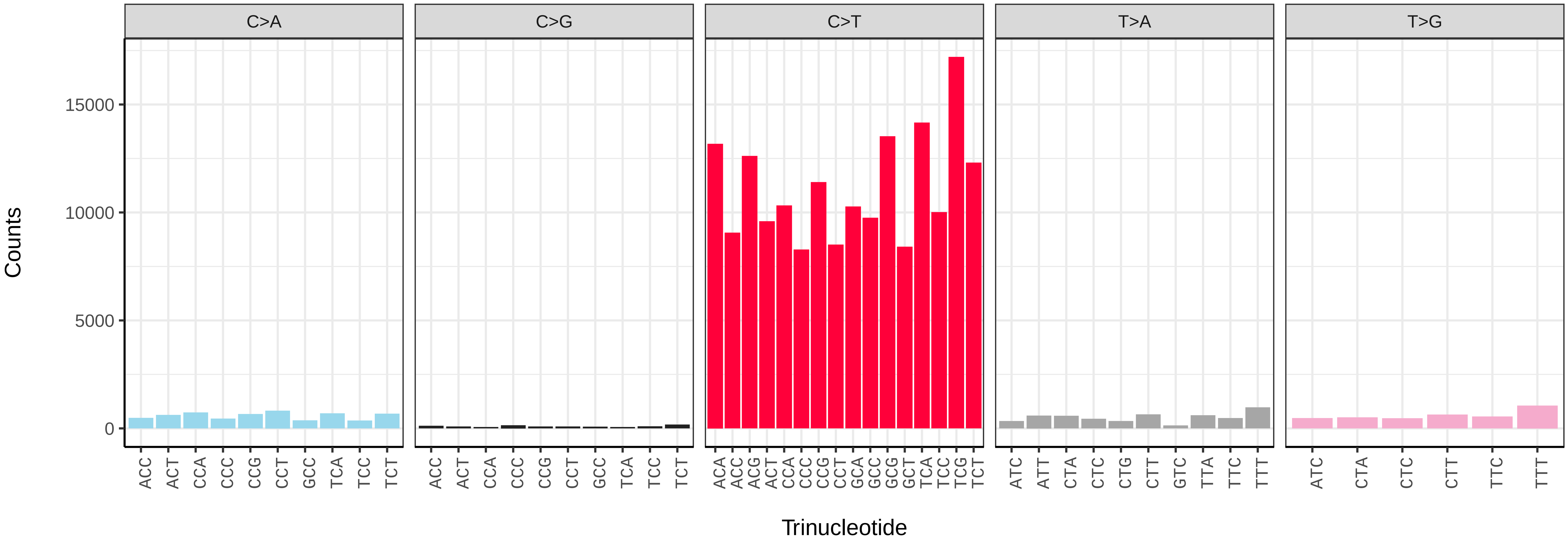

*Calamotropha paludella* (ilCalPalu1) germline mutational spectrum

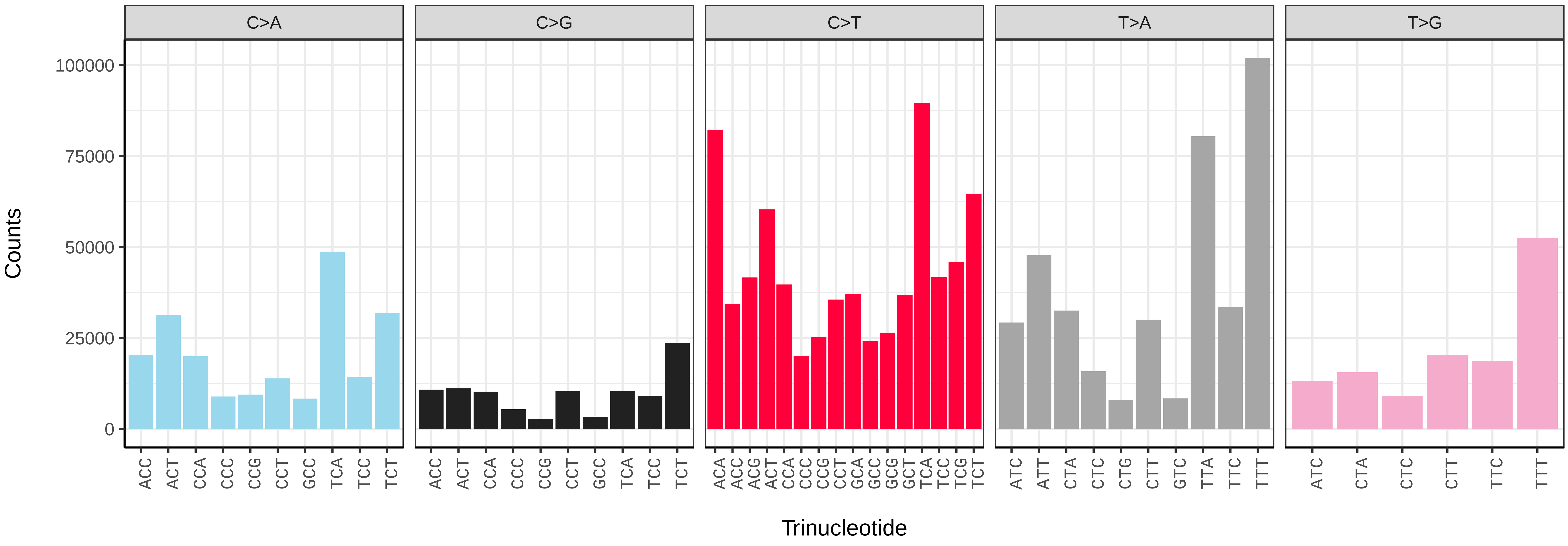


*Calamotropha paludella* (ilCalPalu1) somatic mutational spectrum (SBS96 classification)

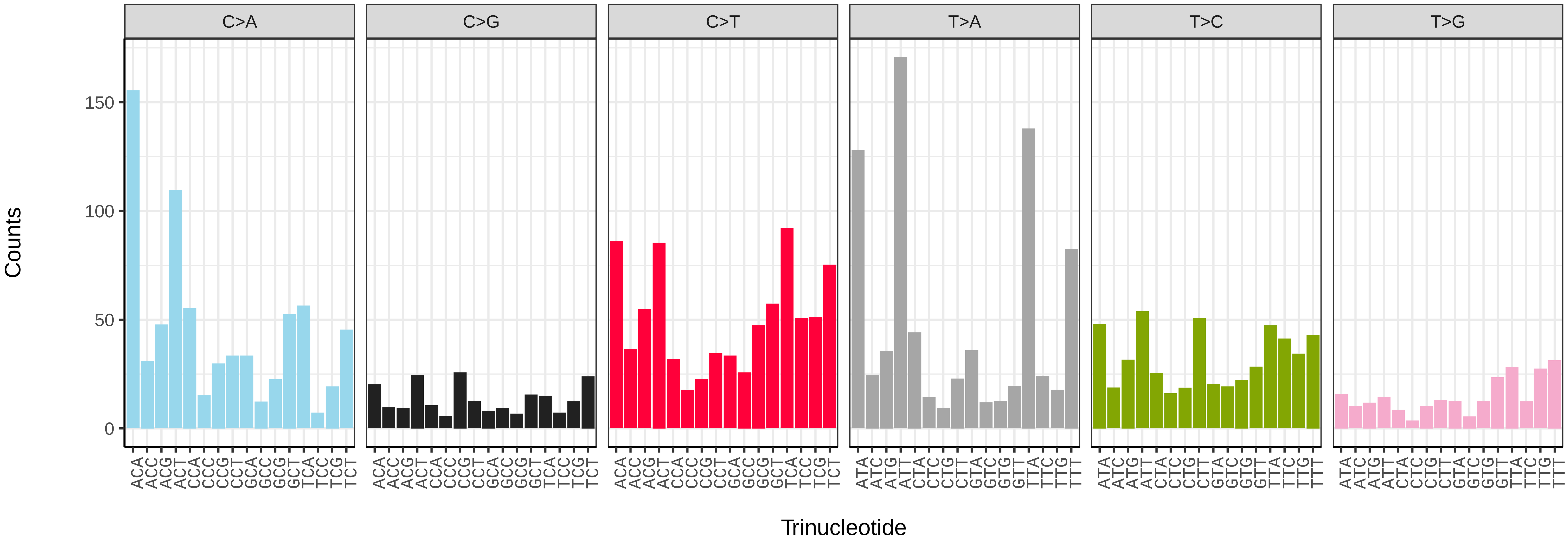


*Calamotropha paludella* (ilCalPalu1) somatic mutational spectrum (SBS52 classification)

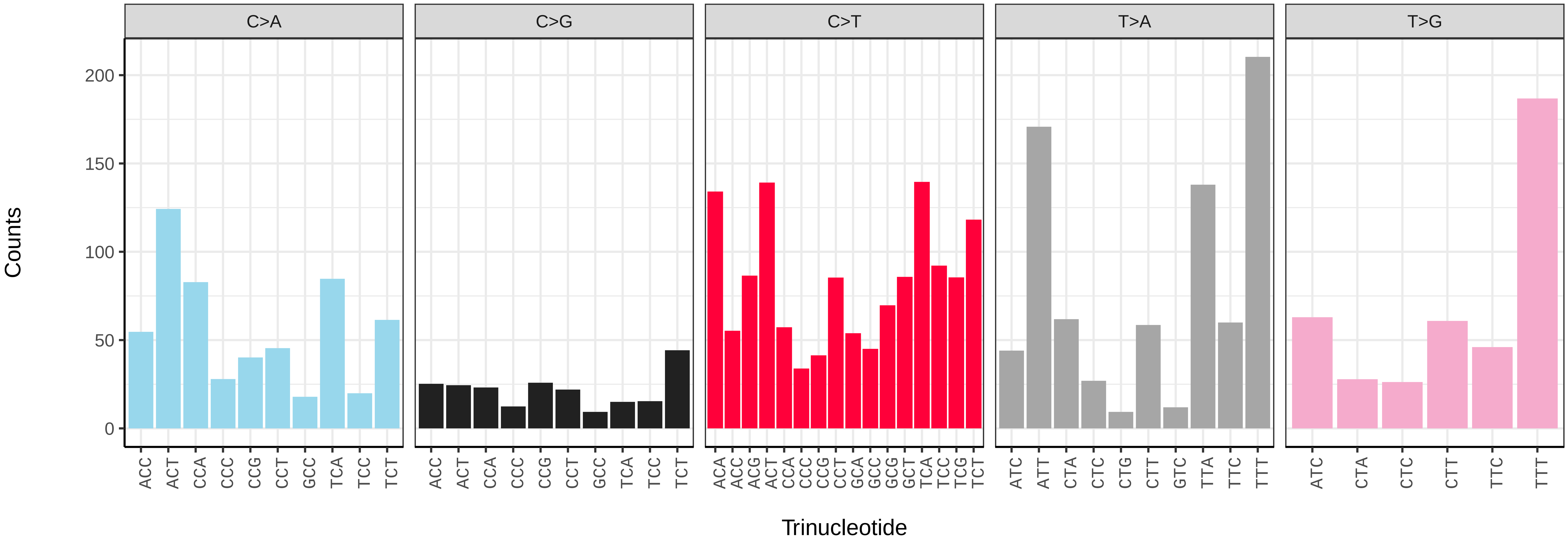

*Campaea margaritaria* (ilCamMarg1) germline mutational spectrum

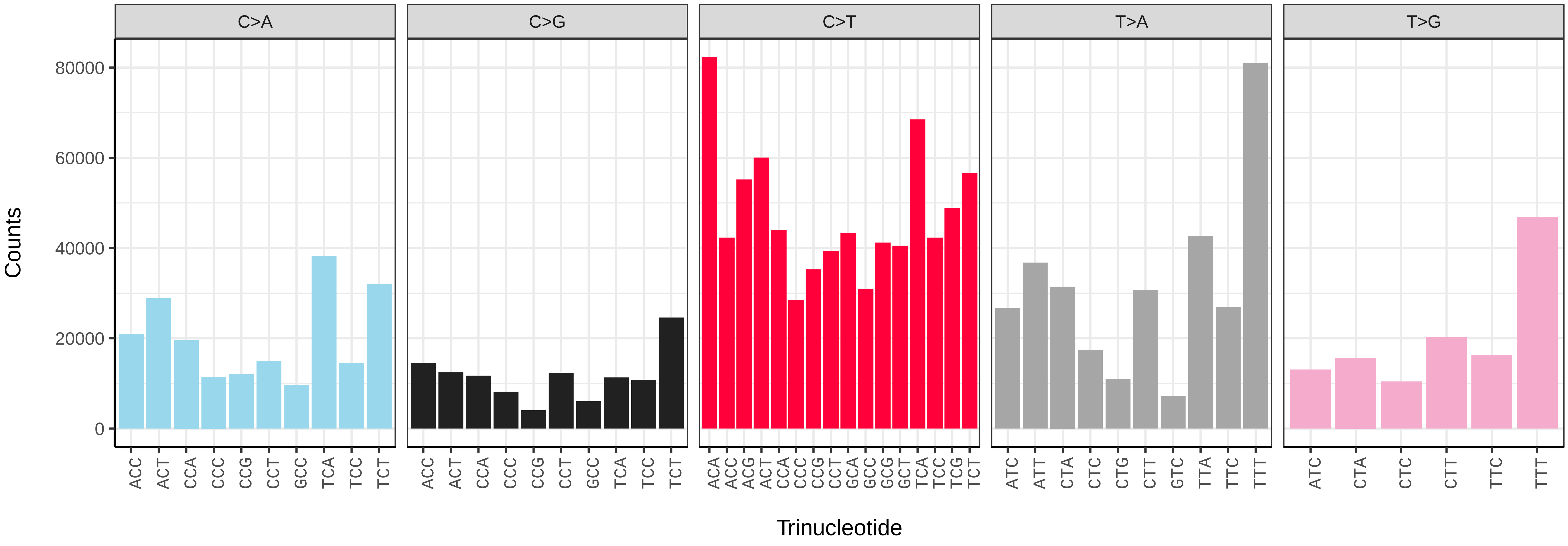


*Campaea margaritaria* (ilCamMarg1) somatic mutational spectrum (SBS96 classification)

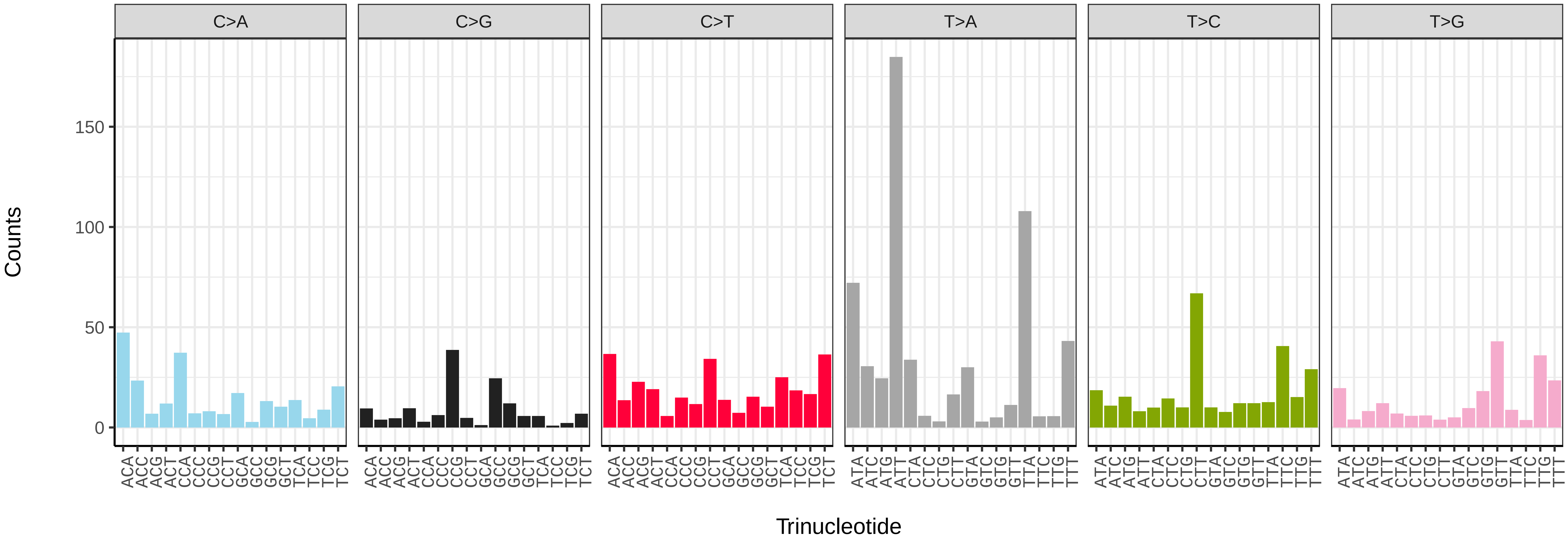


*Campaea margaritaria* (ilCamMarg1) somatic mutational spectrum (SBS52 classification)

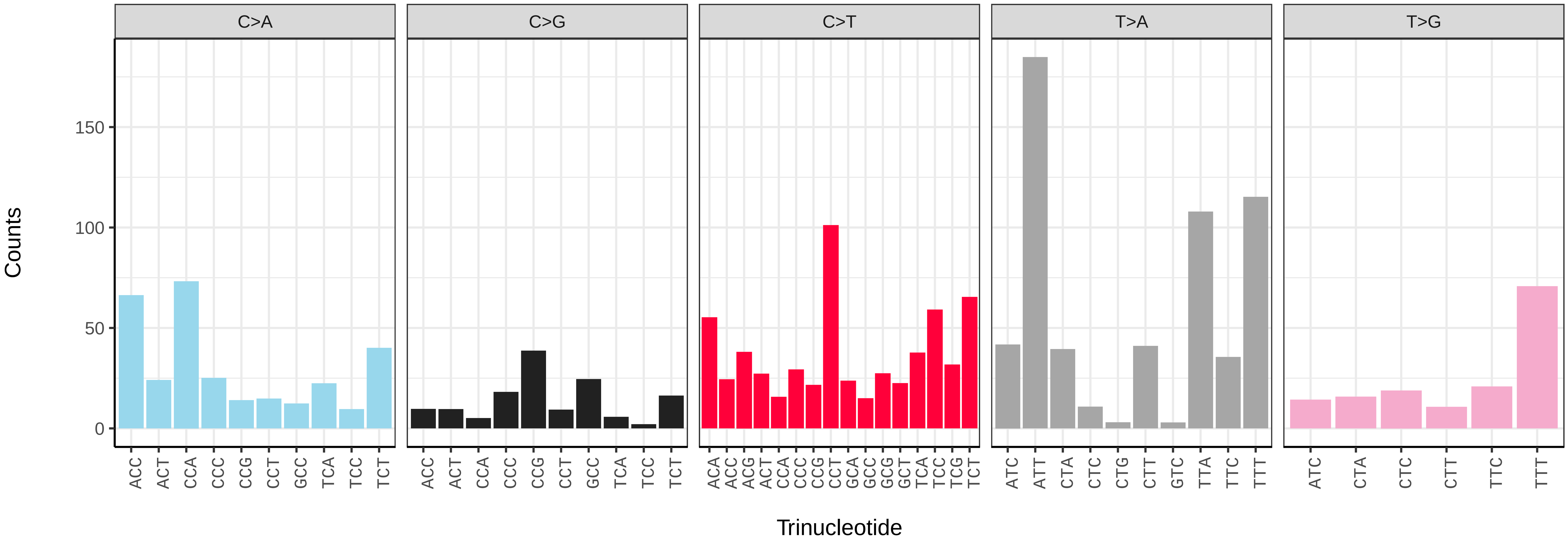

*Caradrina clavipalpis* (ilCarClav1) germline mutational spectrum

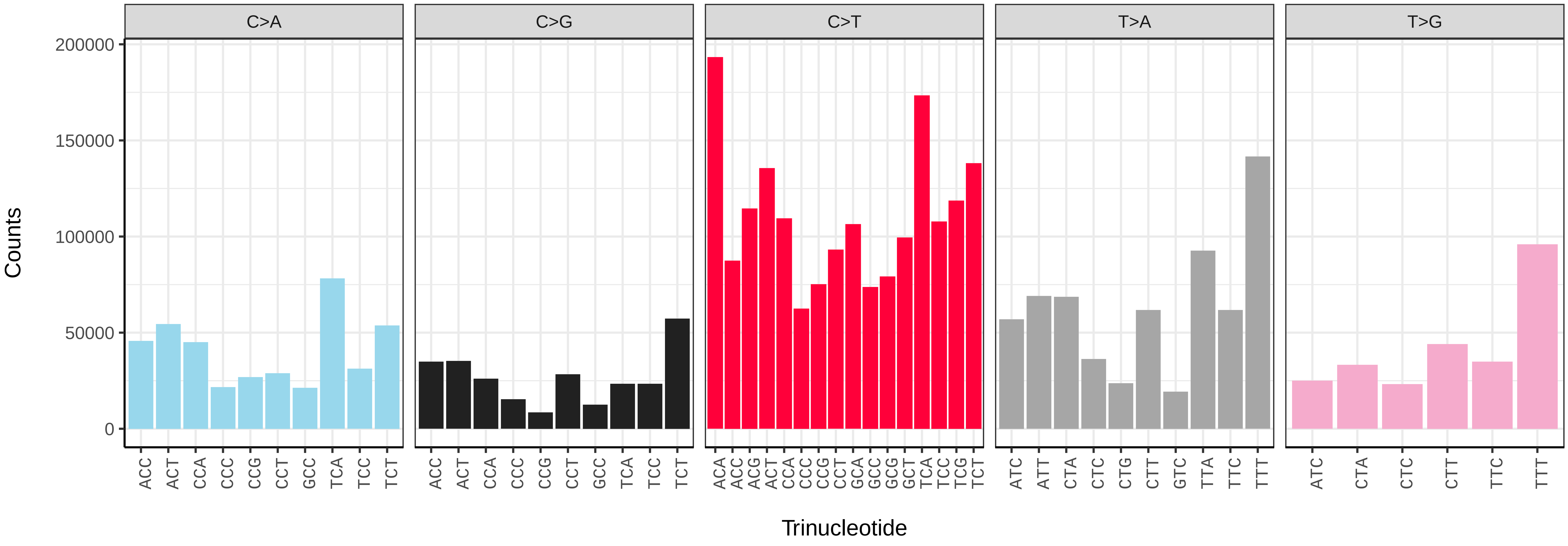


*Caradrina clavipalpis* (ilCarClav1) somatic mutational spectrum (SBS96 classification)

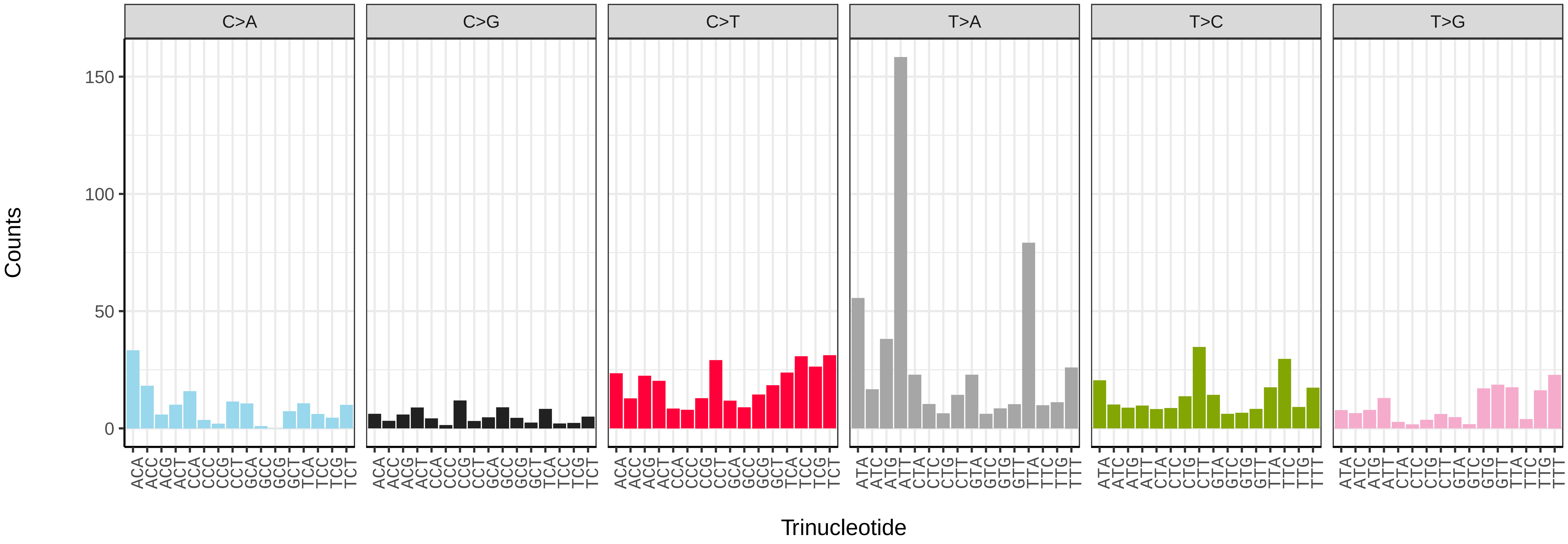


*Caradrina clavipalpis* (ilCarClav1) somatic mutational spectrum (SBS52 classification)

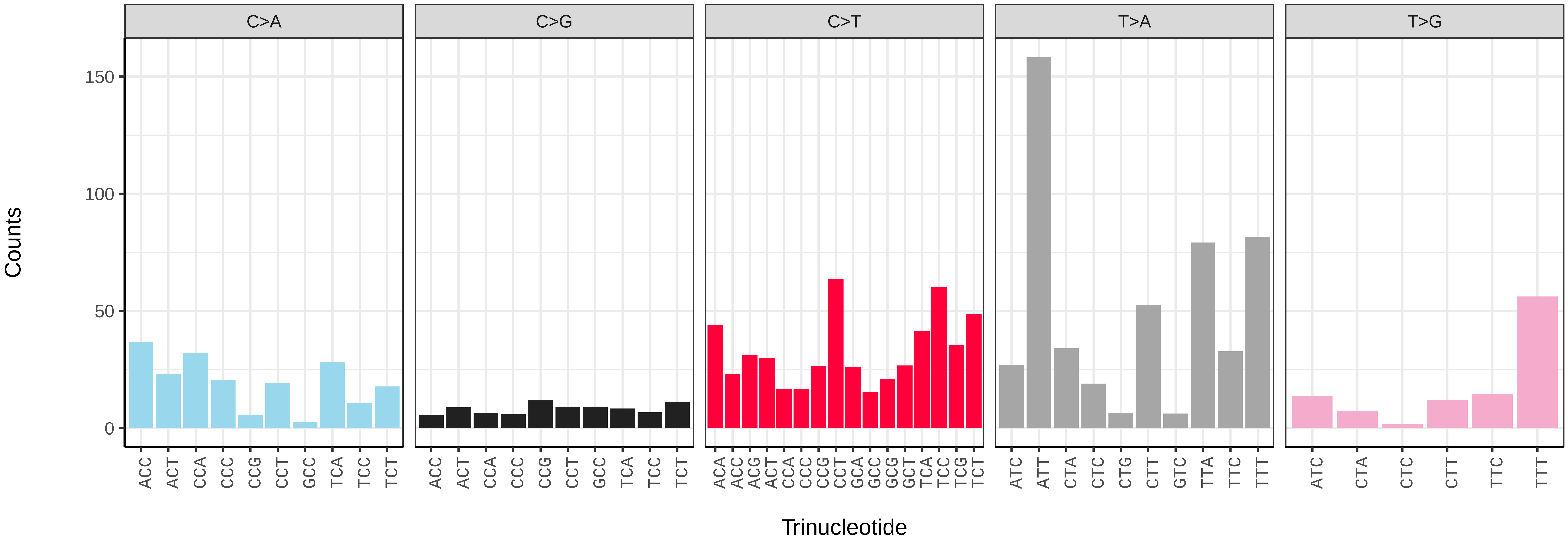

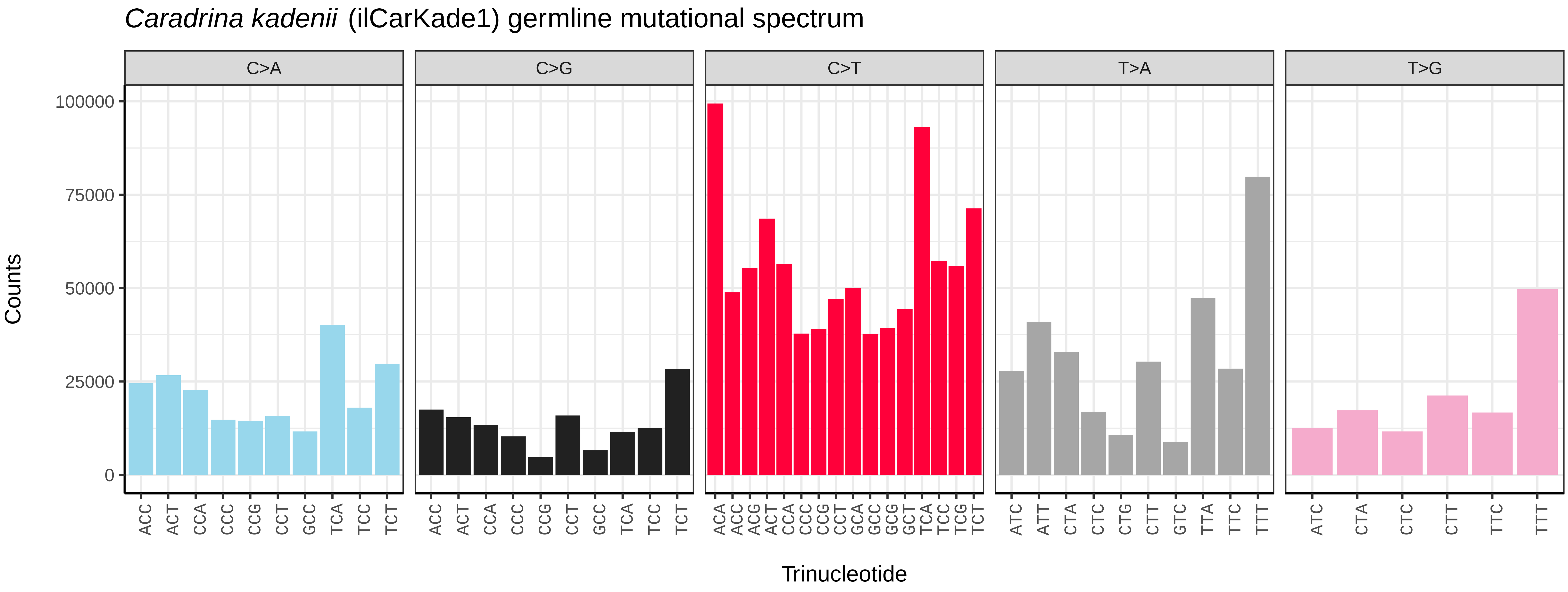
*Caradrina kadenii* (ilCarKade1) germline mutational spectrum
C>A
C>G
C>T
T>A
T>G
Counts
100000
75000
50000
25000
0
ACC ACT CCA CCC CCG CCT GCC TCA TCC TCT
ACC ACT CCA CCC CCG CCT GCC TCA TCC TCT
ACA ACC ACG ACT CCA CCC CCG CCT GCA GCC GCG GCT TCA TCC TCG TCT
ATC ATT CTA CTC CTG CTT GTC TTA TTC TTT
ATC CTA CTC CTT TTC TTT
Trinucleotide

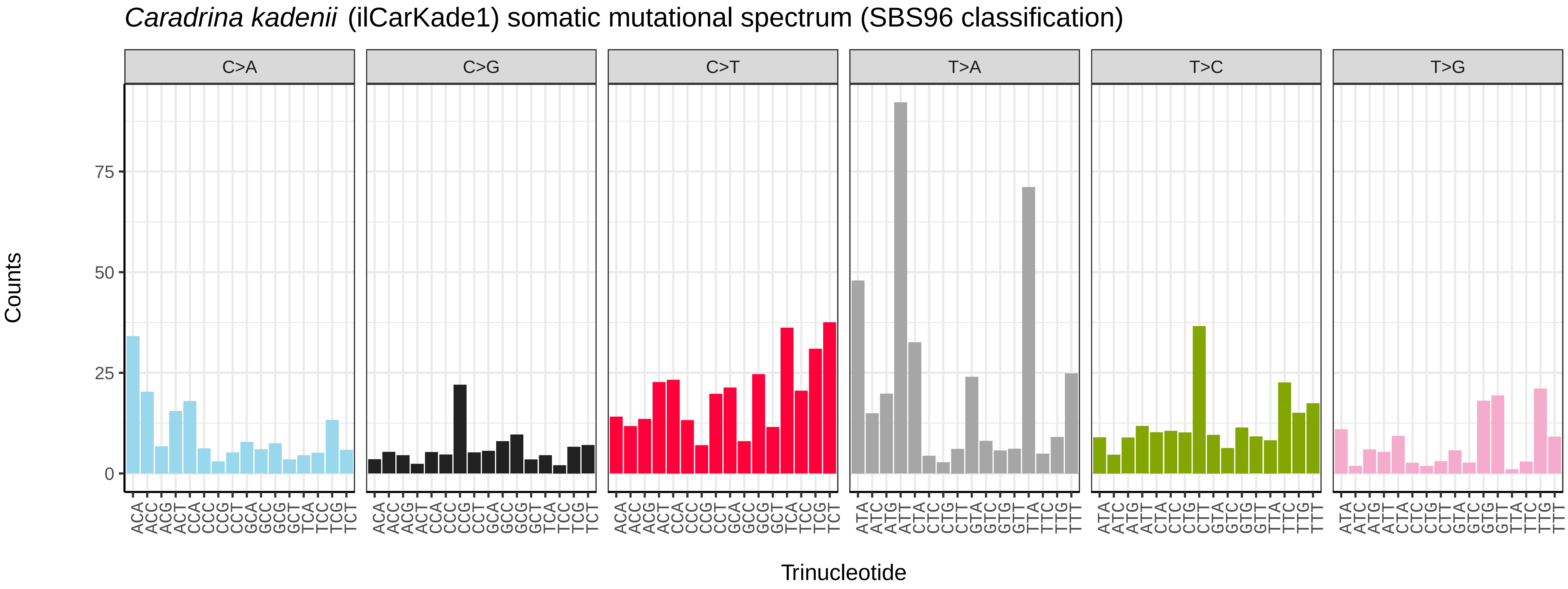
*Caradrina kadenii* (ilCarKade1) somatic mutational spectrum (SBS96 classification)
C>A
C>G
C>T
T>A
T>C
T>G
Counts
75
50
25
0
ACA ACC ACG ACT CCA CCC CCG CCT GCA GCC GCG GCT TCA TCC TCG TCT
ATA ATC ATG ATT CTA CTC CTG CTT GTA GTC GTG GTT TTA TTC TTG TTT
Trinucleotide

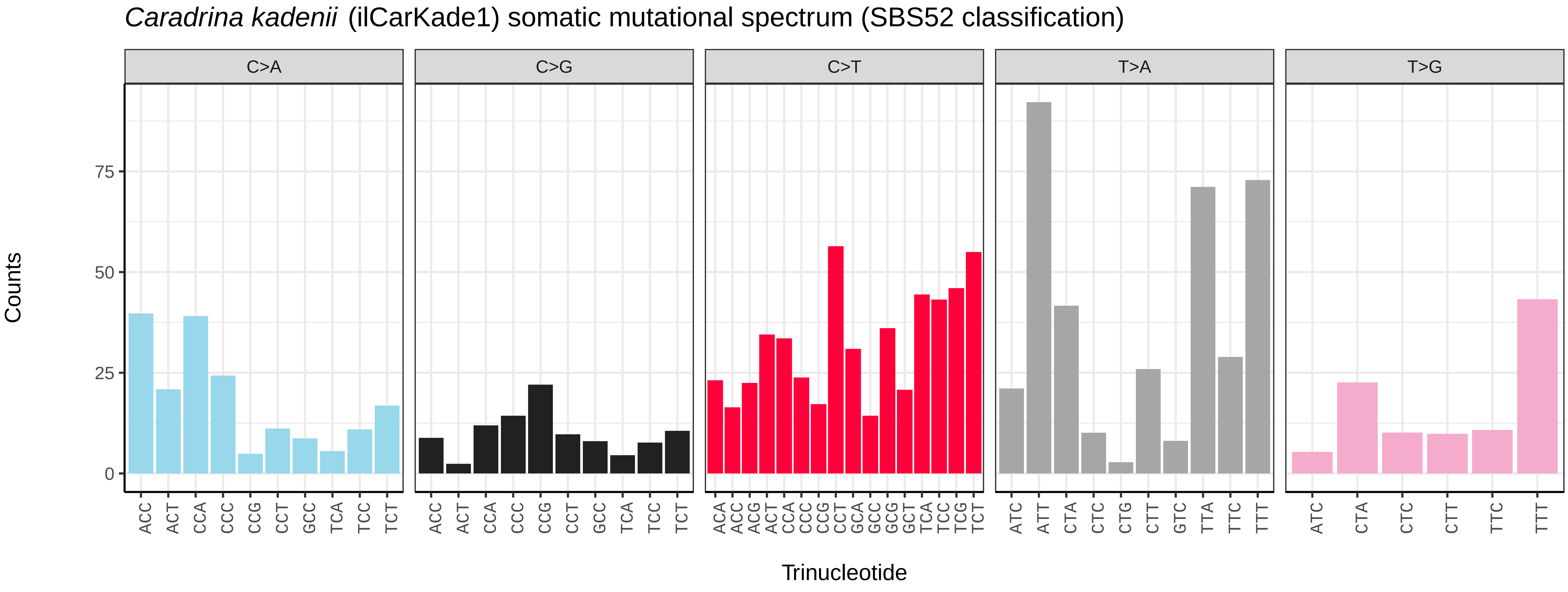
*Caradrina kadenii* (ilCarKade1) somatic mutational spectrum (SBS52 classification)
C>A
C>G
C>T
T>A
T>G
Counts
75
50
25
0
ACC ACT CCA CCC CCG CCT GCC TCA TCC TCT
ACC ACT CCA CCC CCG CCT GCC TCA TCC TCT
ACA ACC ACG ACT CCA CCC CCG CCT GCA GCC GCG GCT TCA TCC TCG TCT
ATC ATT CTA CTC CTG CTT GTC TTA TTC TTT
ATC CTA CTC CTT TTC TTT
Trinucleotide

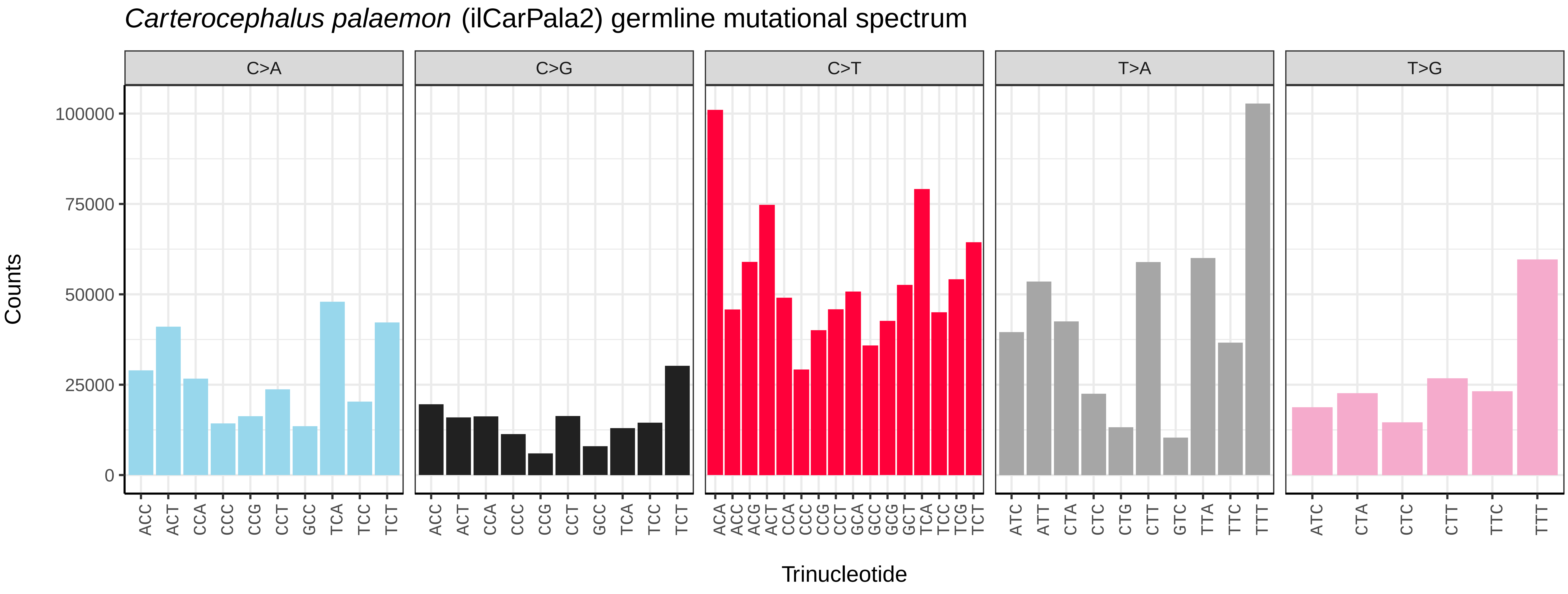
*Carterocephalus palaemon* (ilCarPala2) germline mutational spectrum
C>A
C>G
C>T
T>A
T>G
Counts
0
25000
50000
75000
100000
ACC ACT CCA CCC CCG CCT GCC TCA TCC TCT
ACC ACT CCA CCC CCG CCT GCC TCA TCC TCT
ACA ACC ACG ACT CCA CCC CCG CCT GCA GCC GCG GCT TCA TCC TCG TCT
ATC ATT CTA CTC CTG CTT GTC TTA TTC TTT
ATC CTA CTC CTT TTC TTT
Trinucleotide

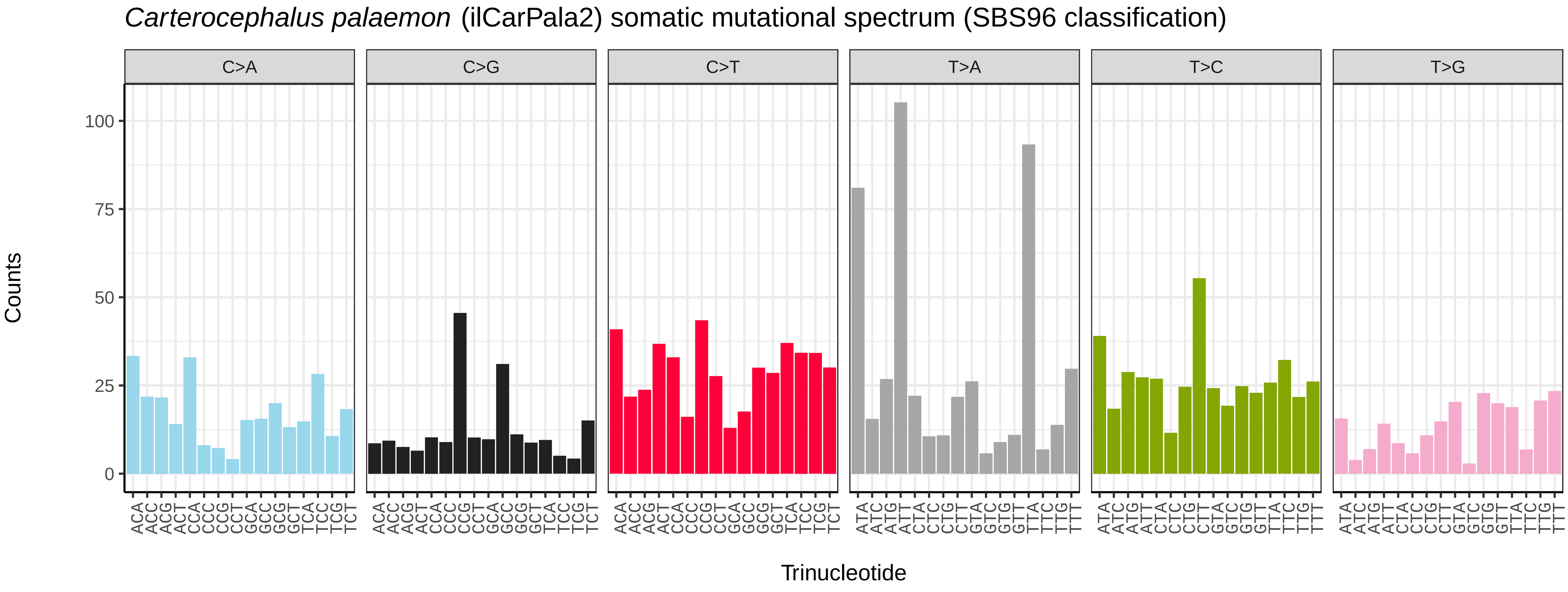
*Carterocephalus palaemon* (ilCarPala2) somatic mutational spectrum (SBS96 classification)
C>A
C>G
C>T
T>A
T>C
T>G
Counts
0
25
50
75
100
ACA ACC ACG ACT CCA CCC CCG CCT GCA GCC GCG GCT TCA TCC TCG TCT
ACA ACC ACG ACT CCA CCC CCG CCT GCA GCC GCG GCT TCA TCC TCG TCT
ACA ACC ACG ACT CCA CCC CCG CCT GCA GCC GCG GCT TCA TCC TCG TCT
ATA ATC ATG ATT CTA CTC CTG CTT GTA GTC GTG GTT TTA TTC TTG TTT
ATA ATC ATG ATT CTA CTC CTG CTT GTA GTC GTG GTT TTA TTC TTG TTT
ATA ATC ATG ATT CTA CTC CTG CTT GTA GTC GTG GTT TTA TTC TTG TTT
Trinucleotide

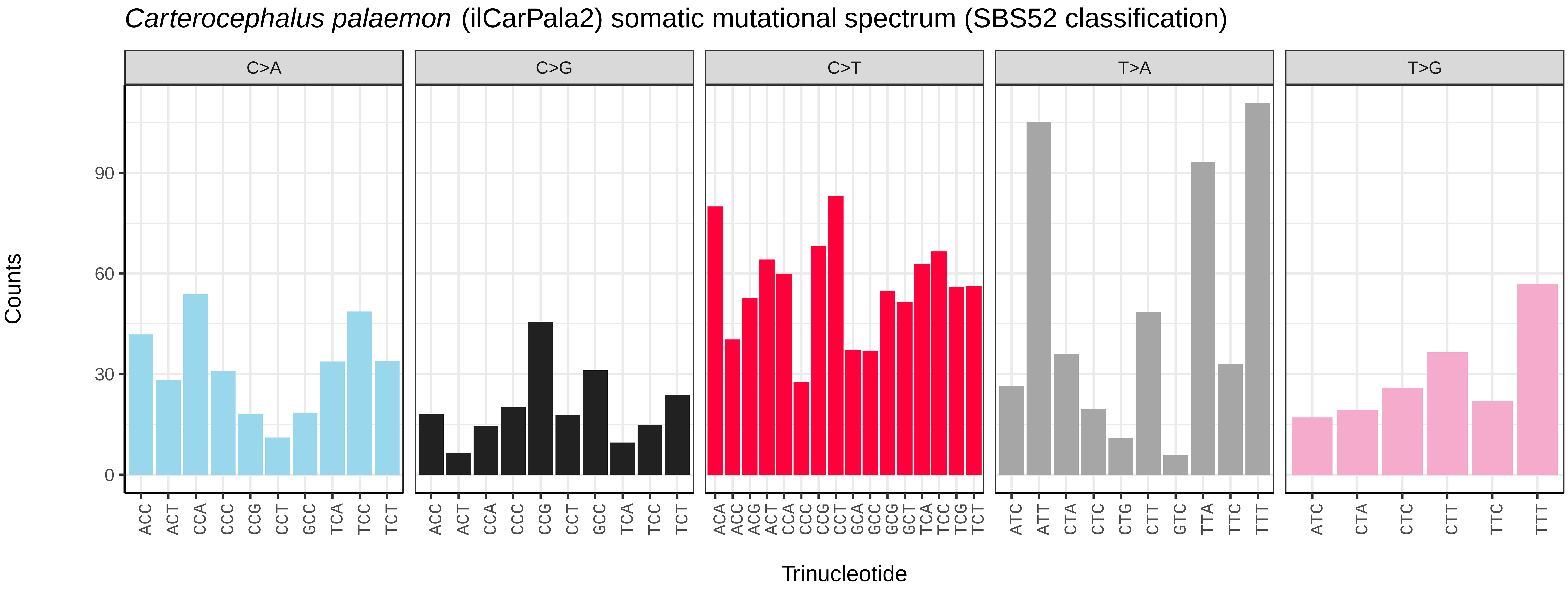
*Carterocephalus palaemon* (ilCarPala2) somatic mutational spectrum (SBS52 classification)
C>A
C>G
C>T
T>A
T>G
Counts
0
30
60
90
ACC ACT CCA CCC CCG CCT GCC TCA TCC TCT
ACC ACT CCA CCC CCG CCT GCC TCA TCC TCT
ACA ACC ACG ACT CCA CCC CCG CCT GCA GCC GCG GCT TCA TCC TCG TCT
ATC ATT CTA CTC CTG CTT GTC TTA TTC TTT
ATC CTA CTC CTT TTC TTT
Trinucleotide

*Carcina quercana* (ilCarQuer1) germline mutational spectrum

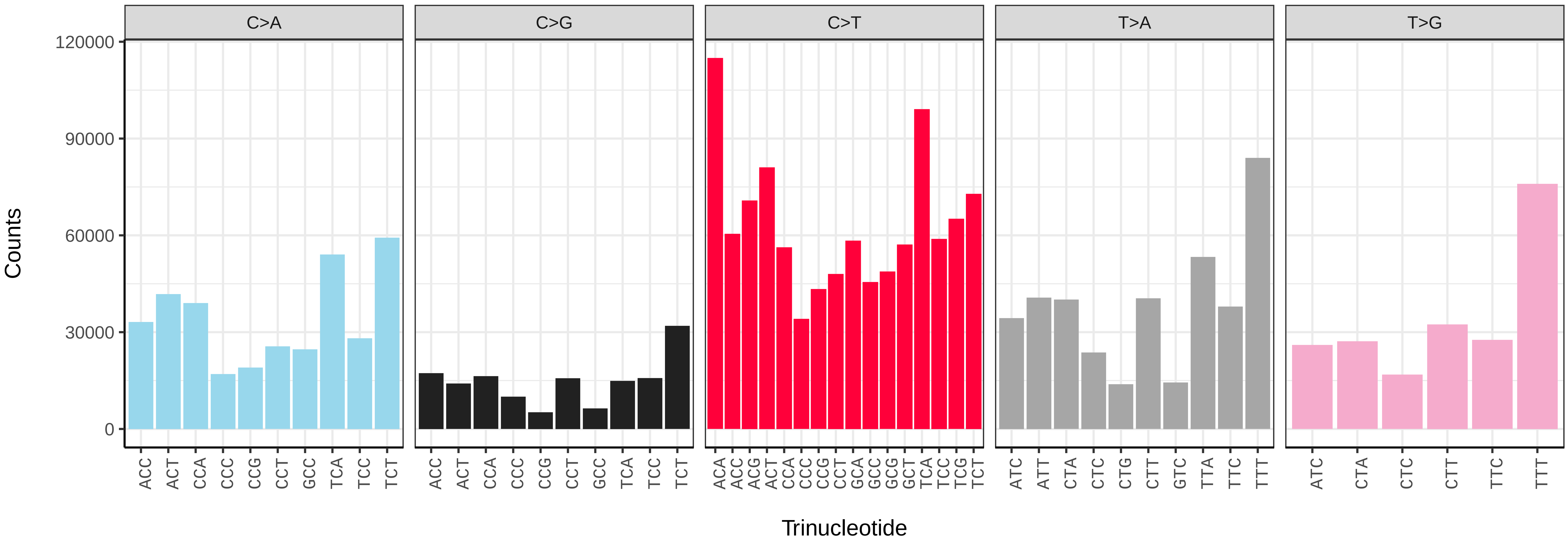


*Carcina quercana* (ilCarQuer1) somatic mutational spectrum (SBS96 classification)

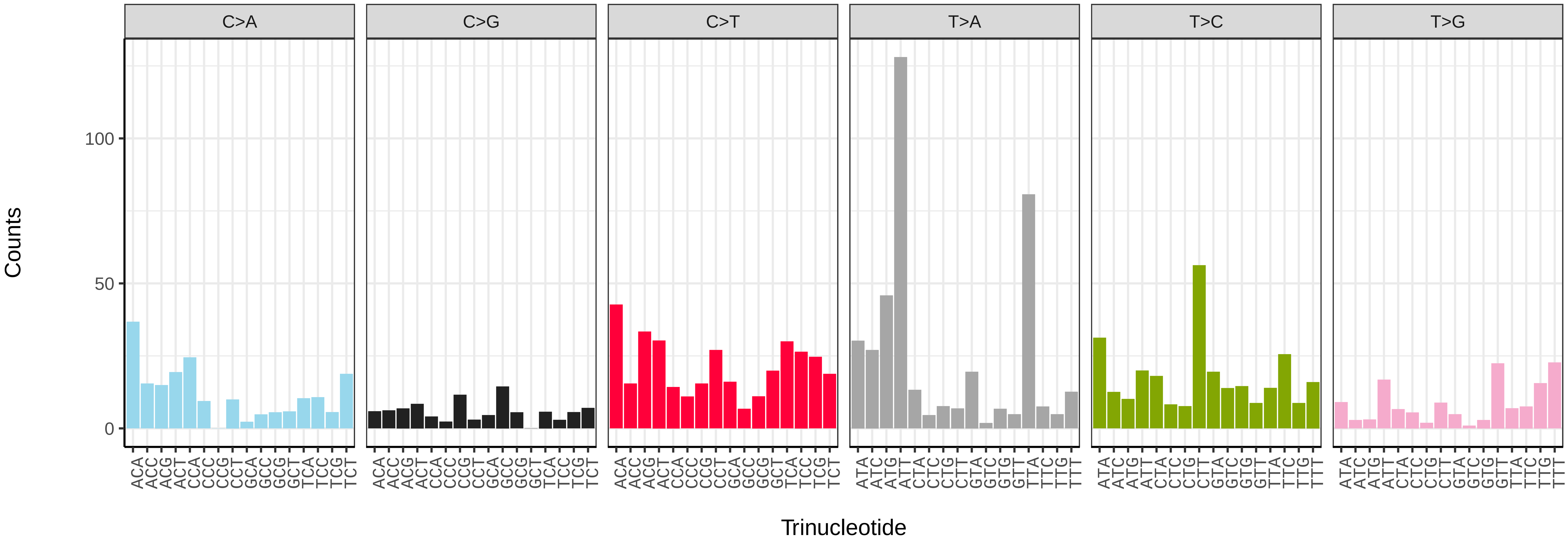


*Carcina quercana* (ilCarQuer1) somatic mutational spectrum (SBS52 classification)

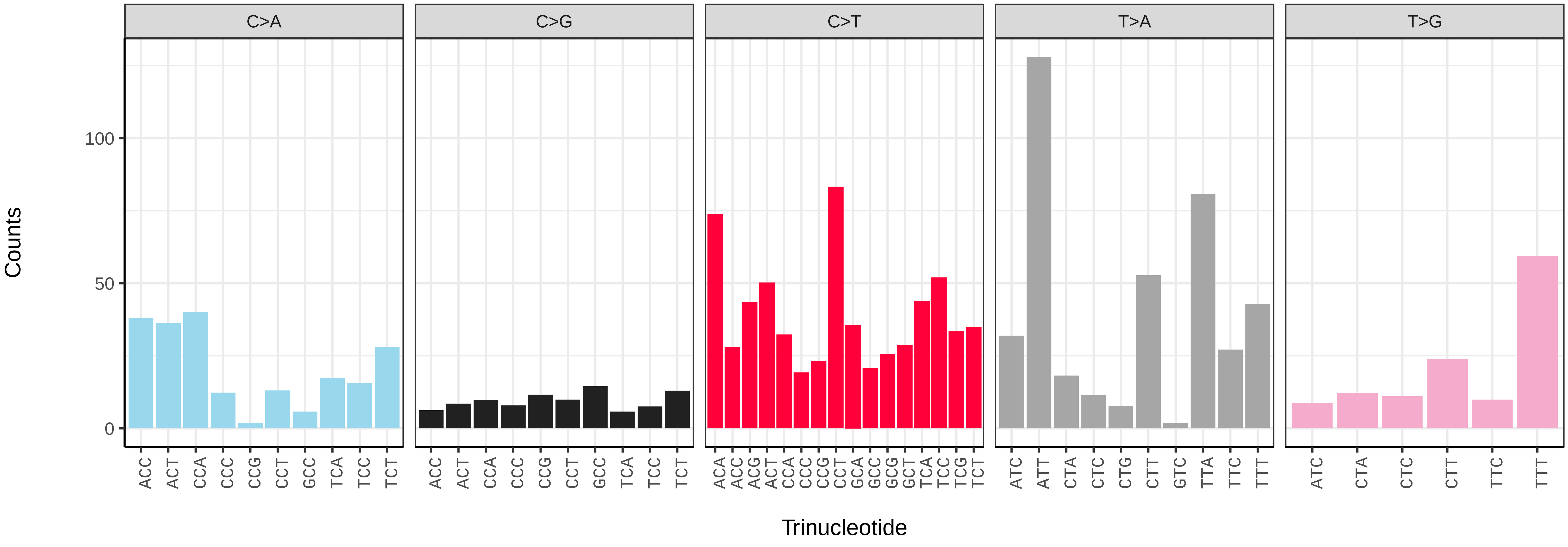

*Catocala fraxini* (ilCatFrax1) germline mutational spectrum

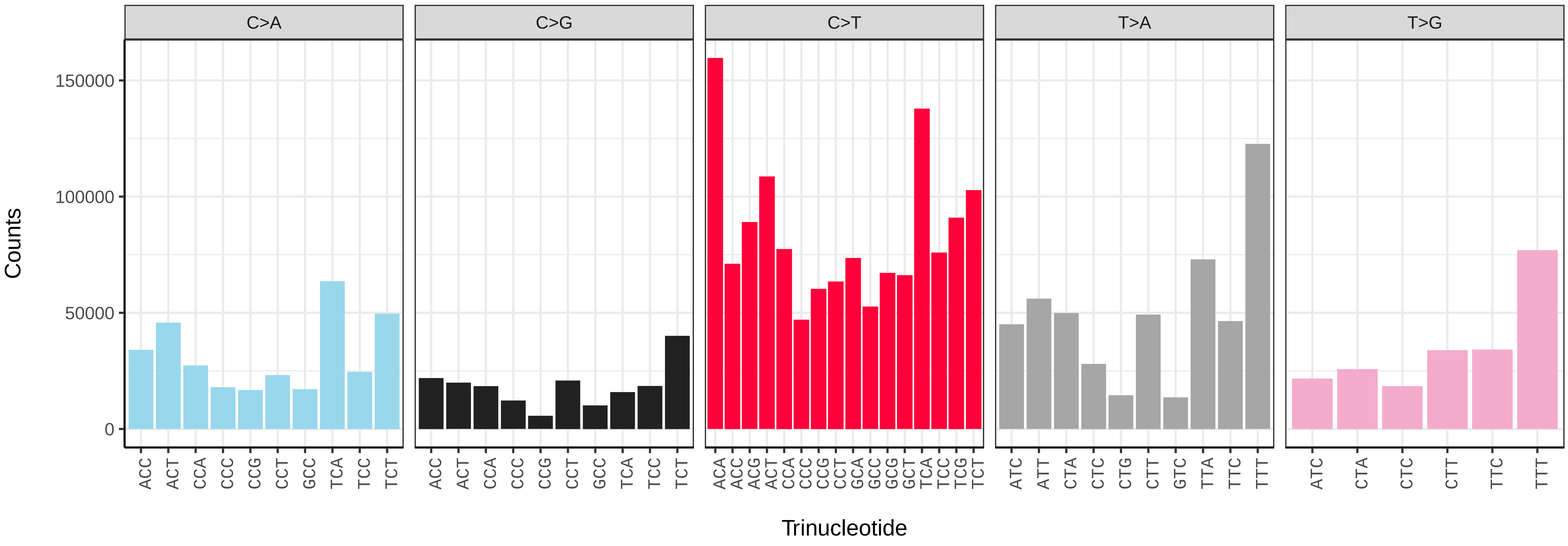


*Catocala fraxini* (ilCatFrax1) somatic mutational spectrum (SBS96 classification)

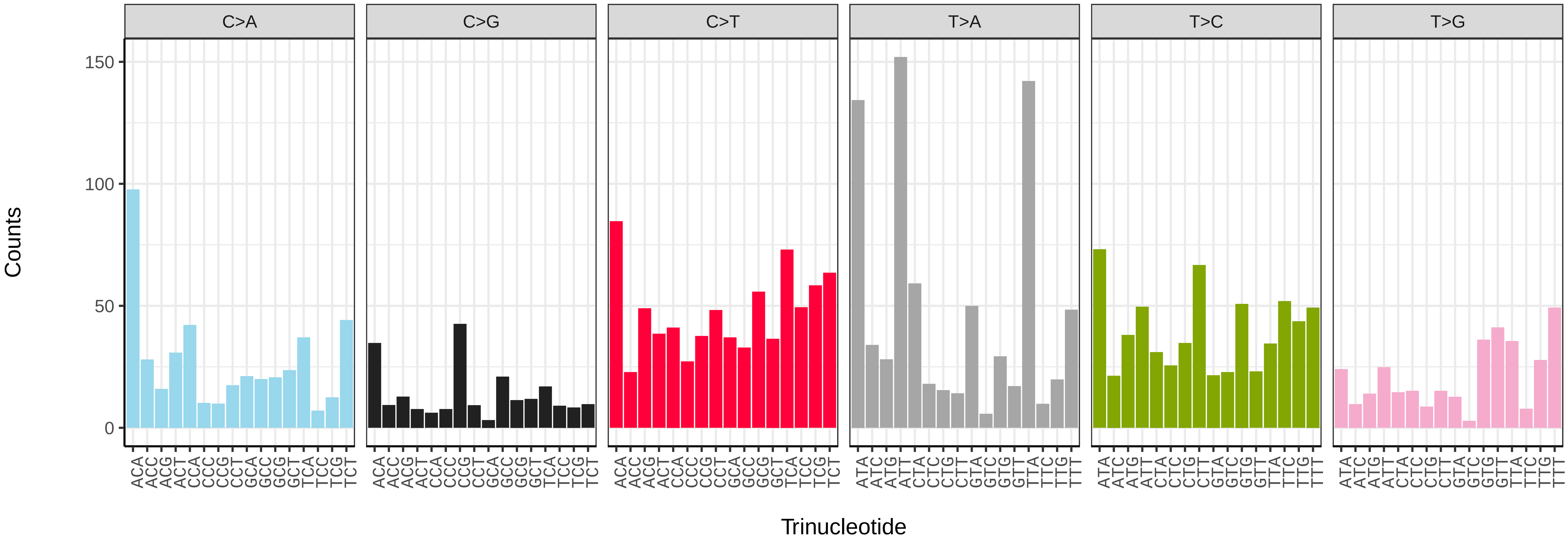


*Catocala fraxini* (ilCatFrax1) somatic mutational spectrum (SBS52 classification)

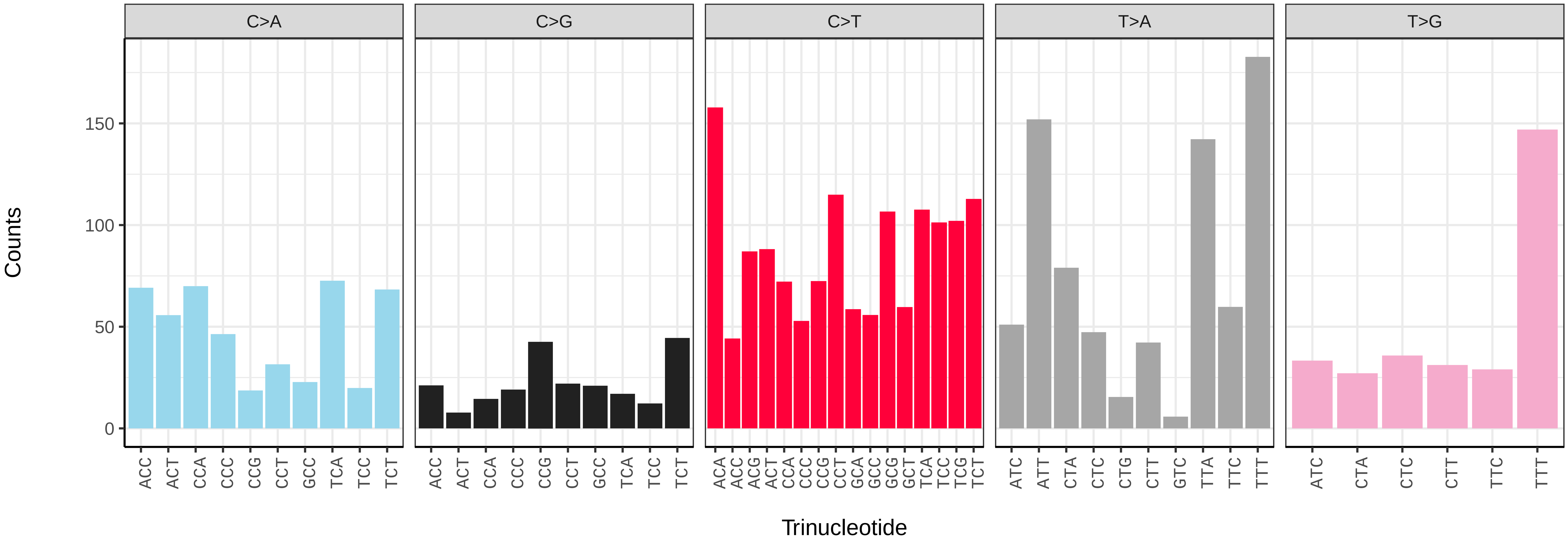

*Celastrina argiolus* (ilCelArgi3) germline mutational spectrum

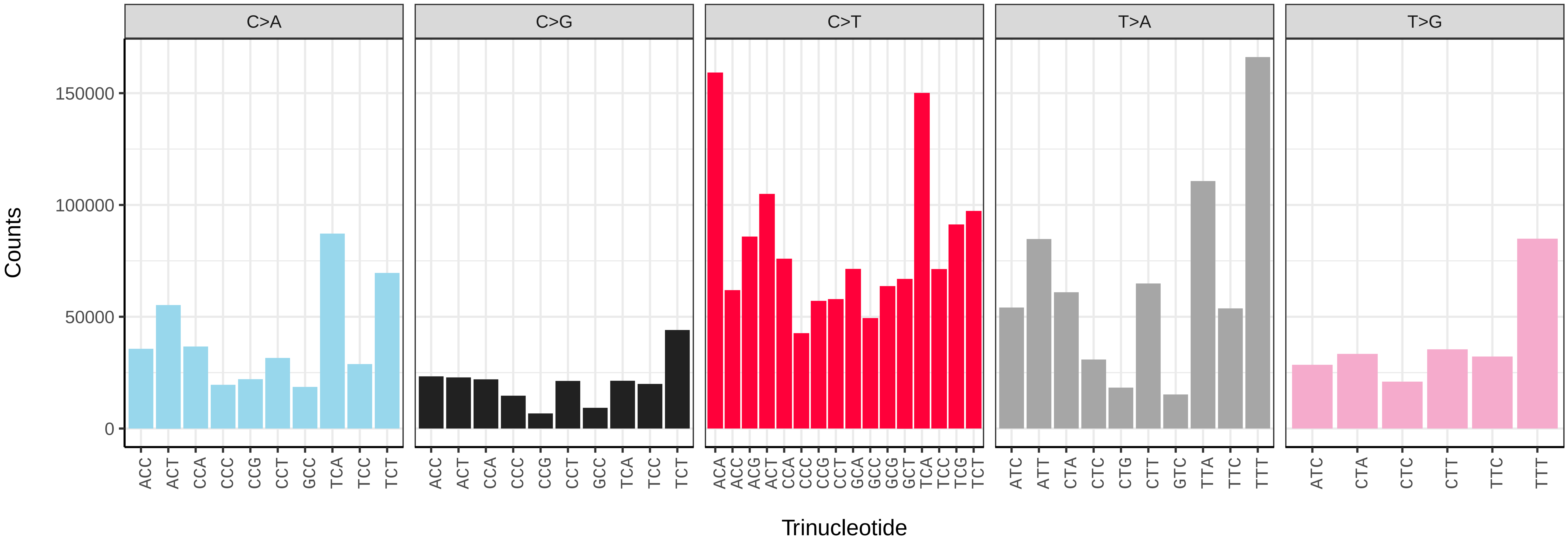


*Celastrina argiolus* (ilCelArgi3) somatic mutational spectrum (SBS96 classification)

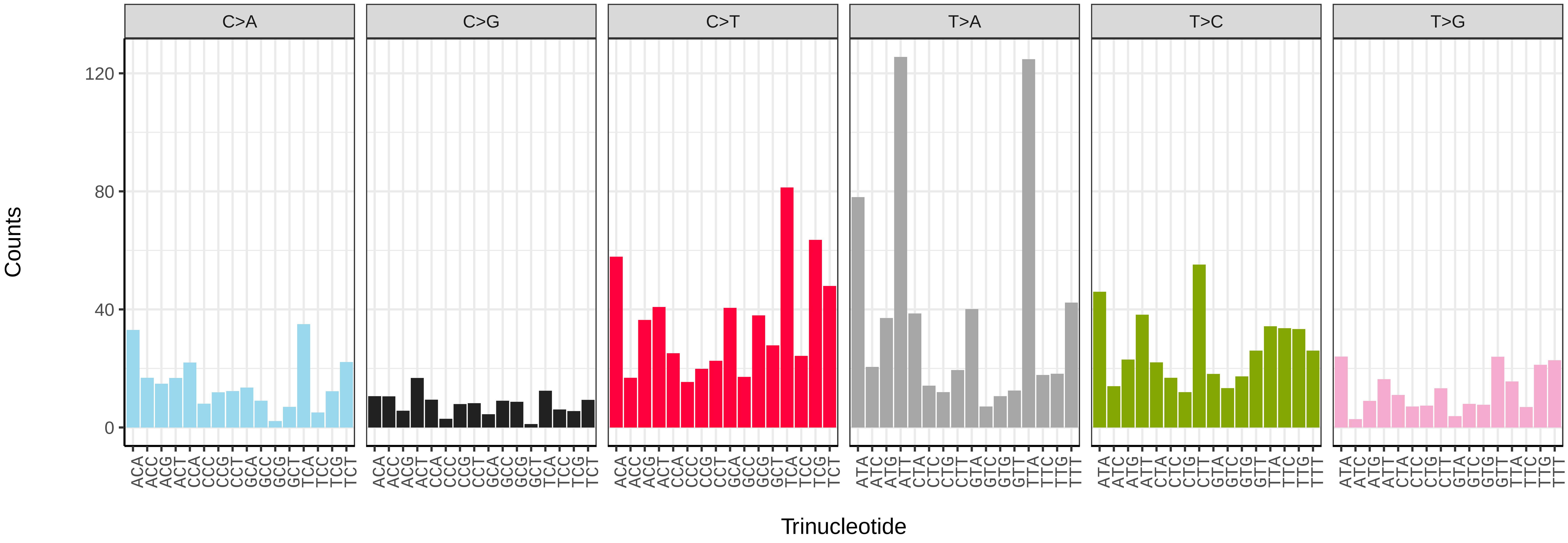


*Celastrina argiolus* (ilCelArgi3) somatic mutational spectrum (SBS52 classification)

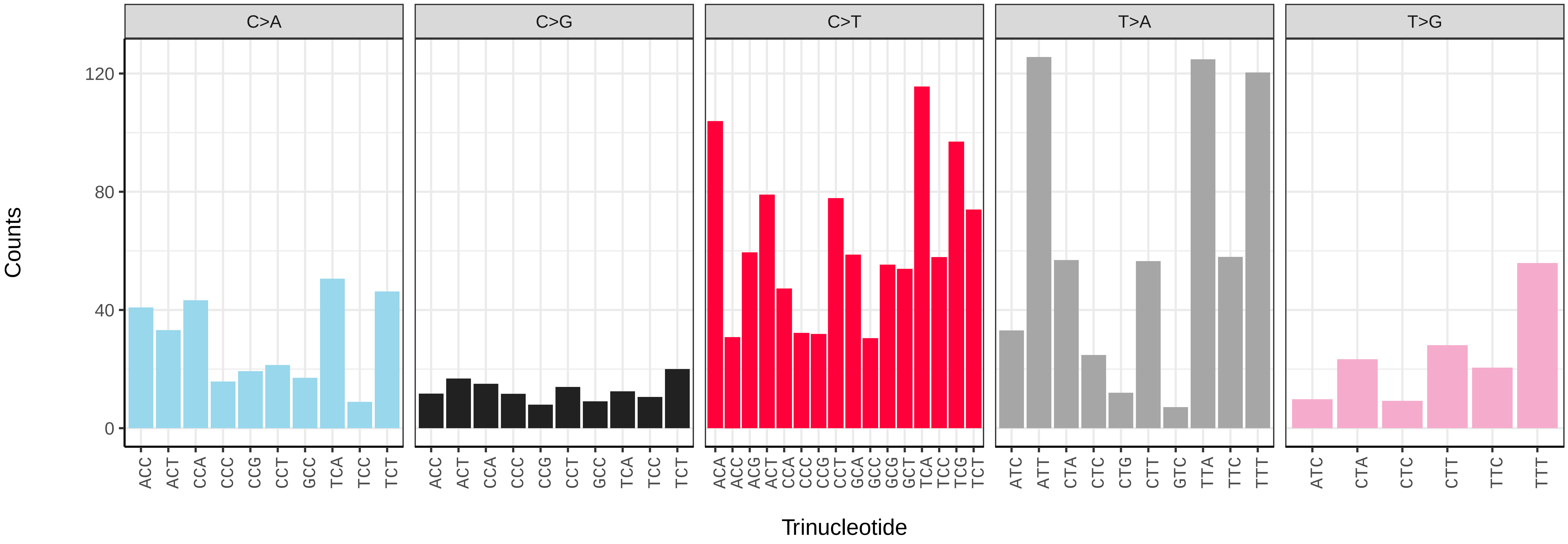

*Charanyca ferruginea* (ilChaFerr1) germline mutational spectrum

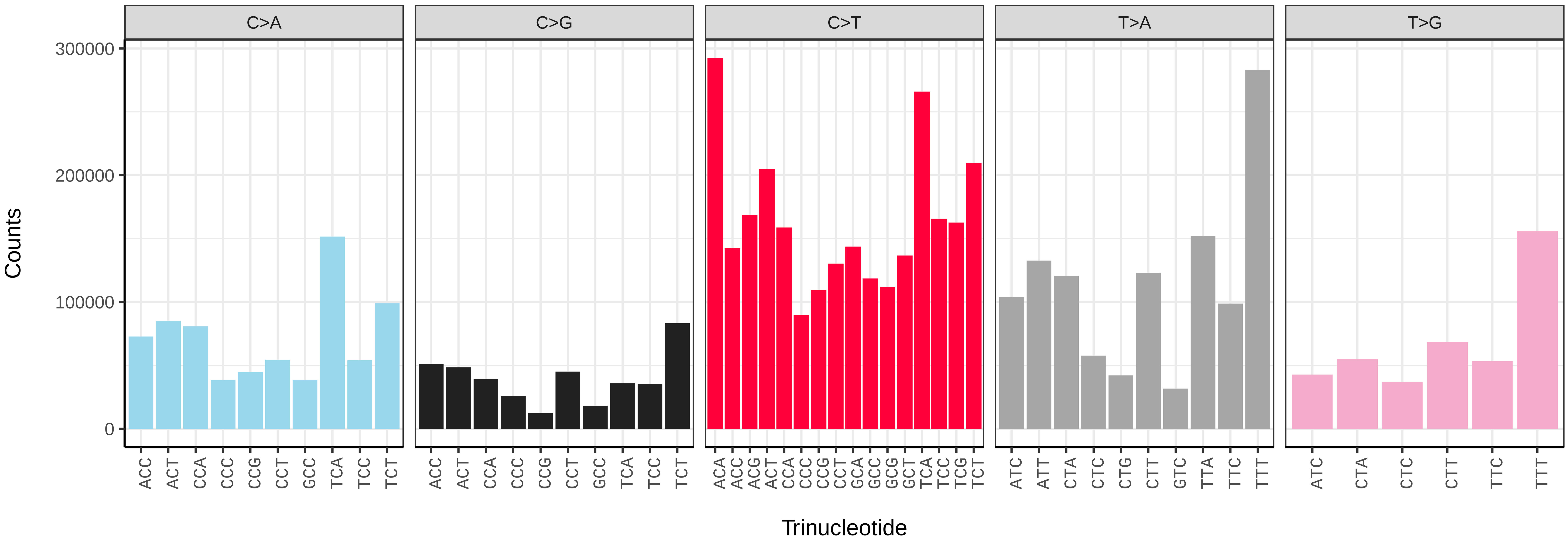


*Charanyca ferruginea* (ilChaFerr1) somatic mutational spectrum (SBS96 classification)

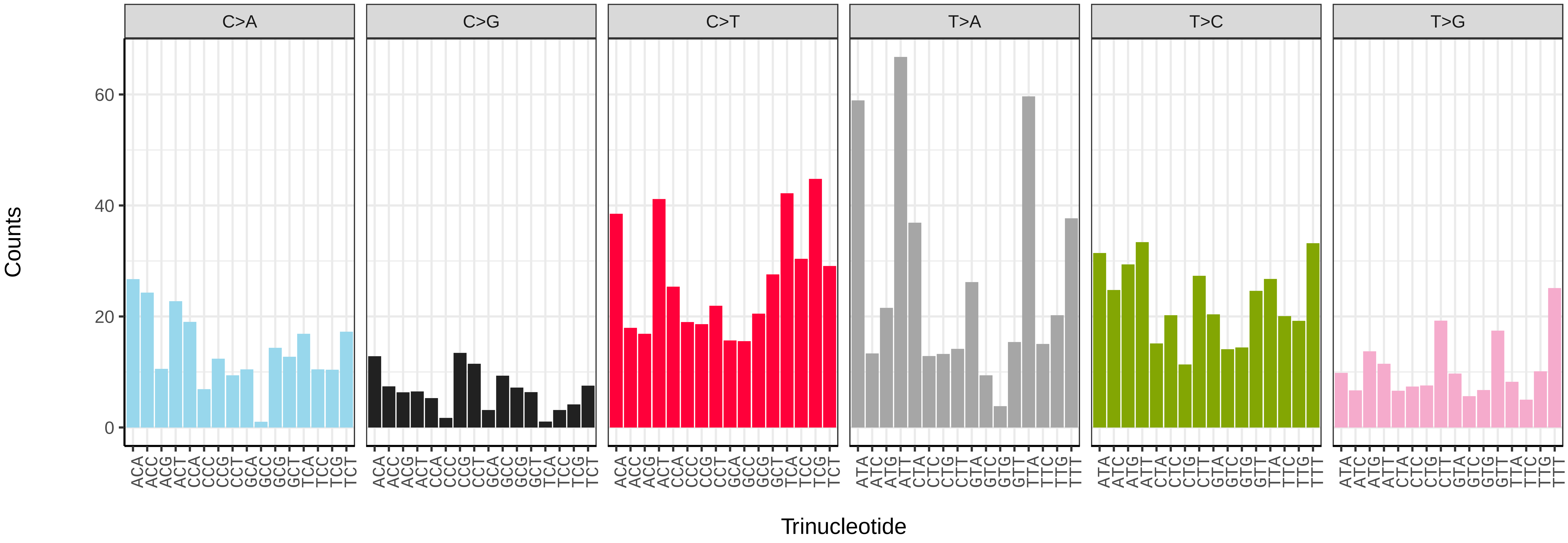


*Charanyca ferruginea* (ilChaFerr1) somatic mutational spectrum (SBS52 classification)

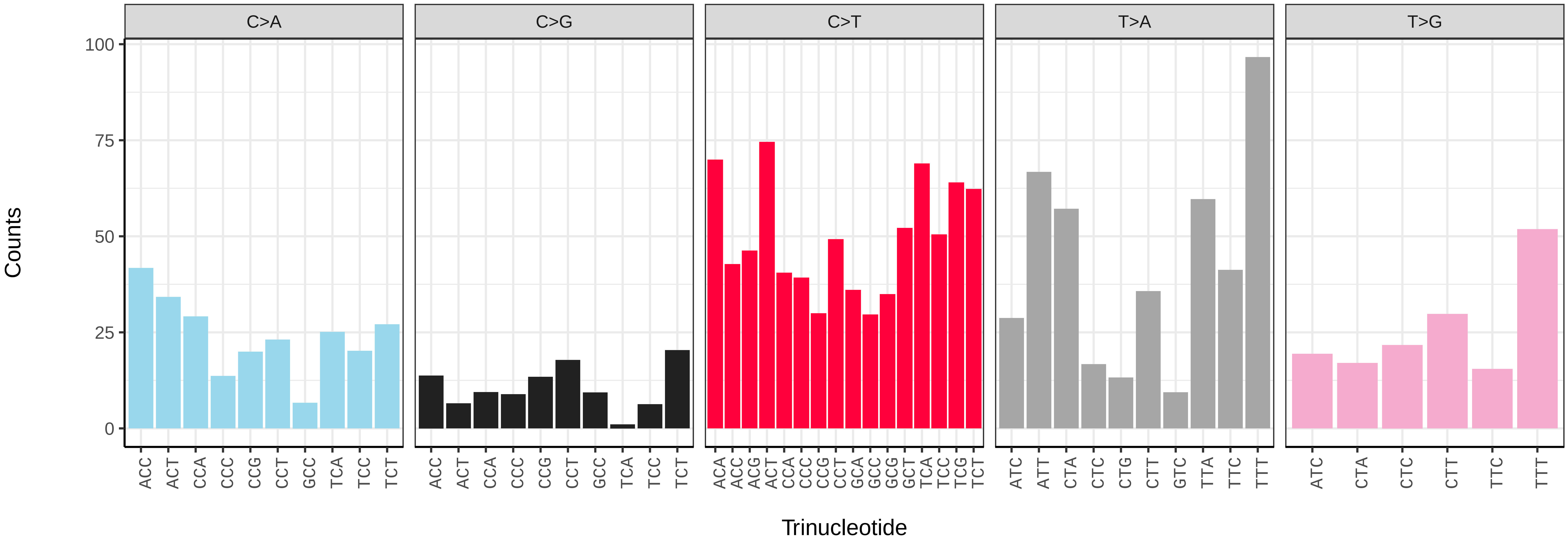

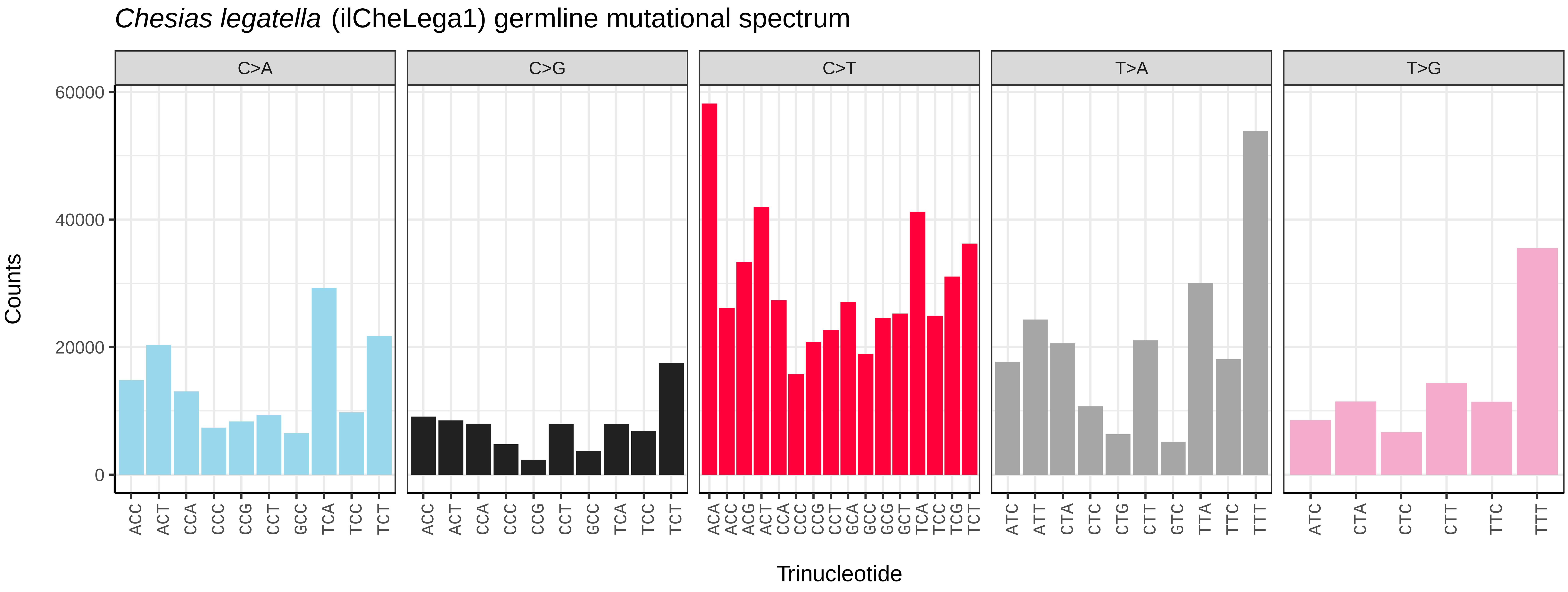
Chesias legatella (ilCheLega1) germline mutational spectrum
C>A
C>G
C>T
T>A
T>G
Counts
Trinucleotide
0
20000
40000
60000

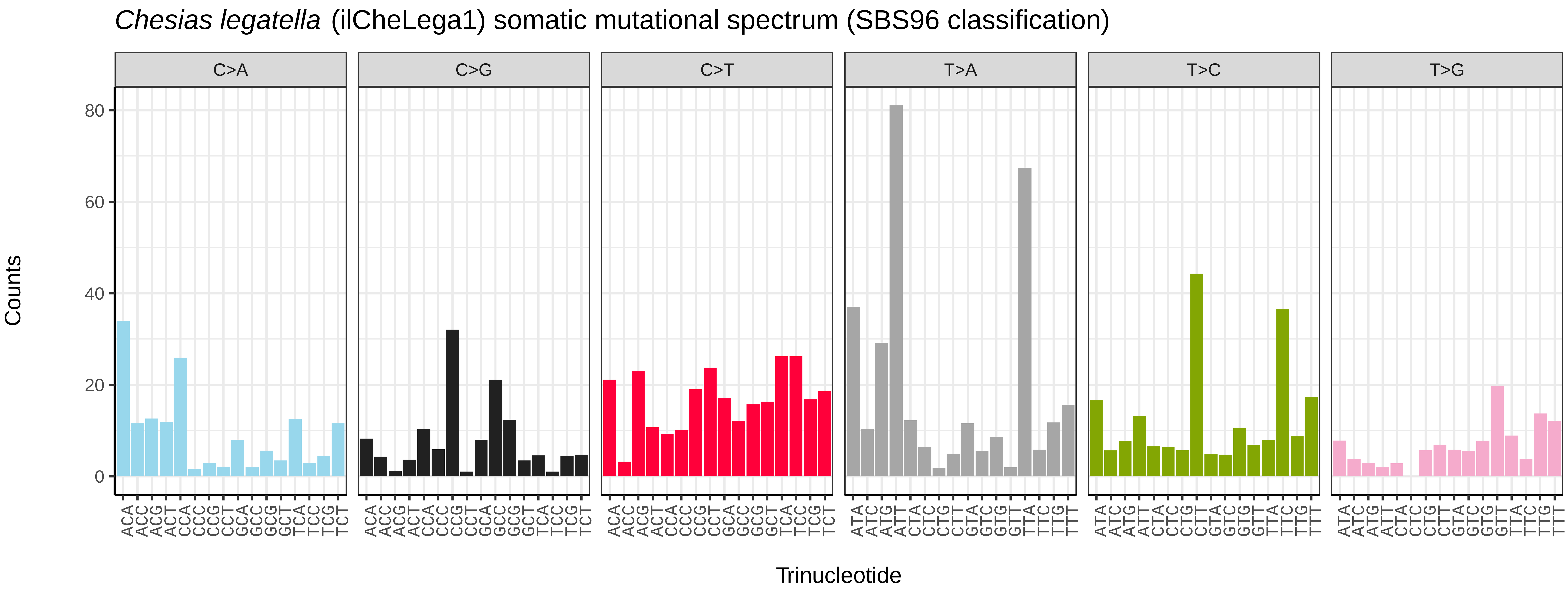
Chesias legatella (ilCheLega1) somatic mutational spectrum (SBS96 classification)
C>A
C>G
C>T
T>A
T>C
T>G
Counts
Trinucleotide
0
20
40
60
80

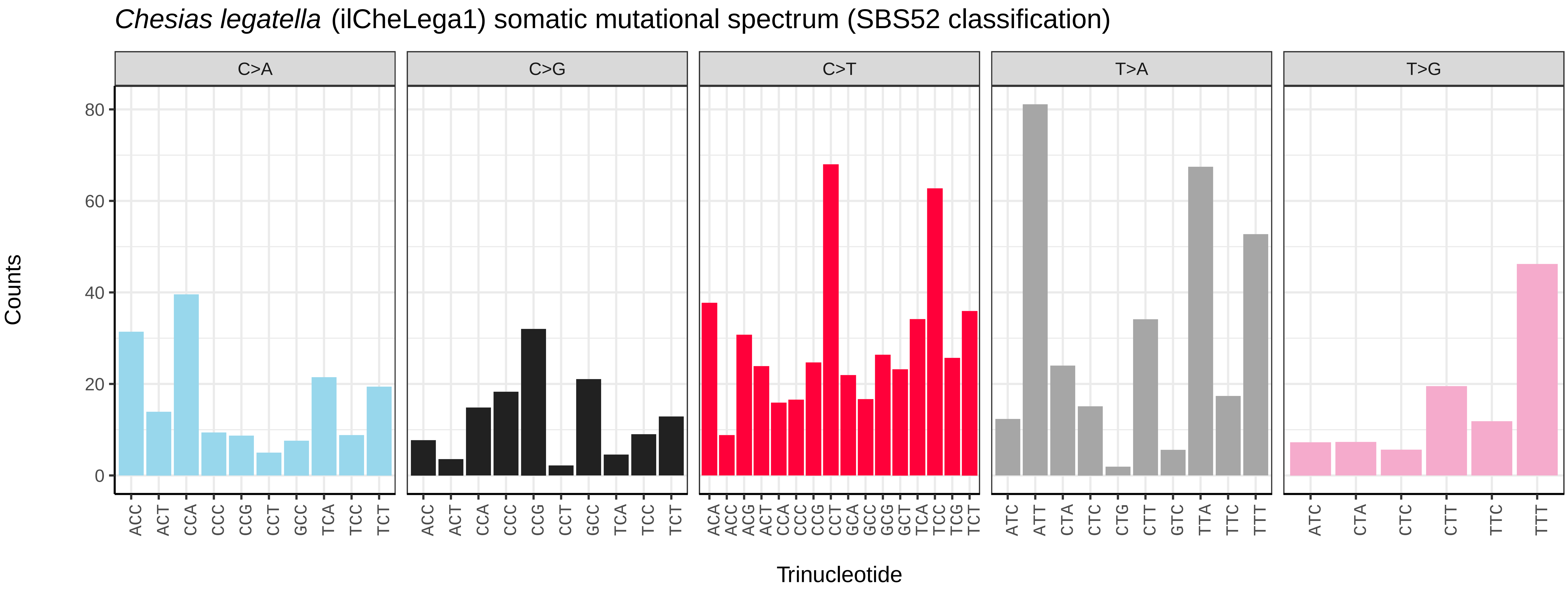
Chesias legatella (ilCheLega1) somatic mutational spectrum (SBS52 classification)
C>A
C>G
C>T
T>A
T>G
Counts
Trinucleotide
0
20
40
60
80

*Chloroclysta siterata* (ilChlSite2) germline mutational spectrum

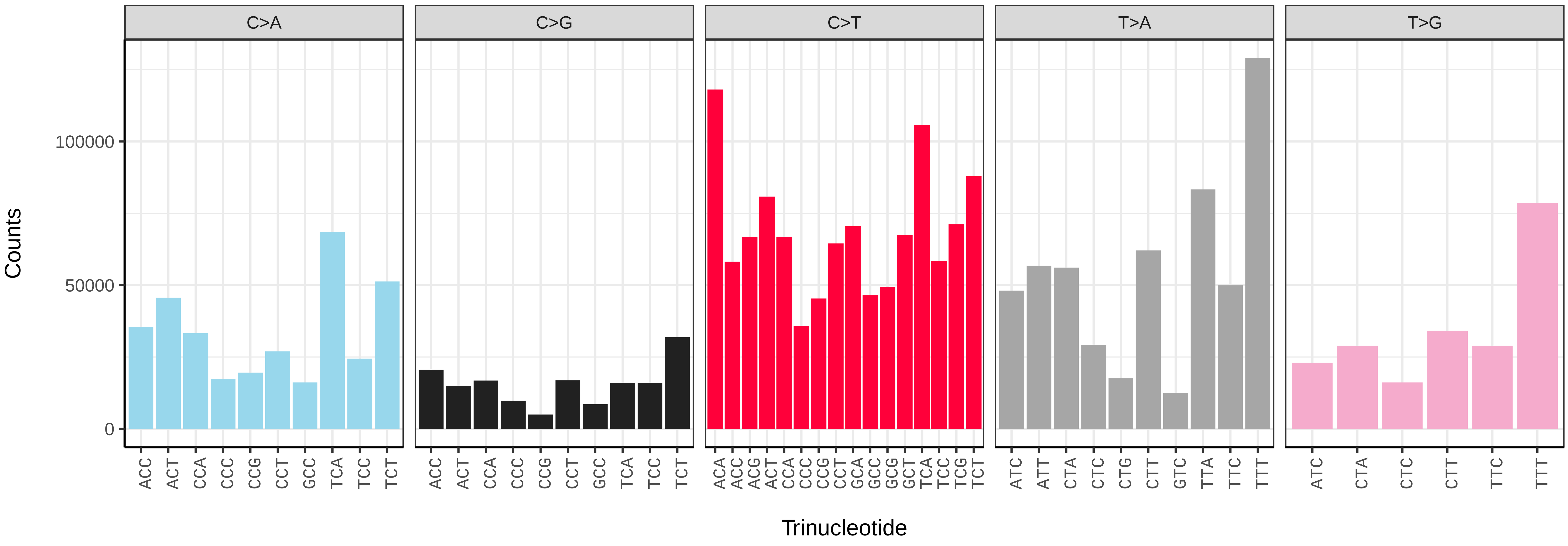


*Chloroclysta siterata* (ilChlSite2) somatic mutational spectrum (SBS96 classification)

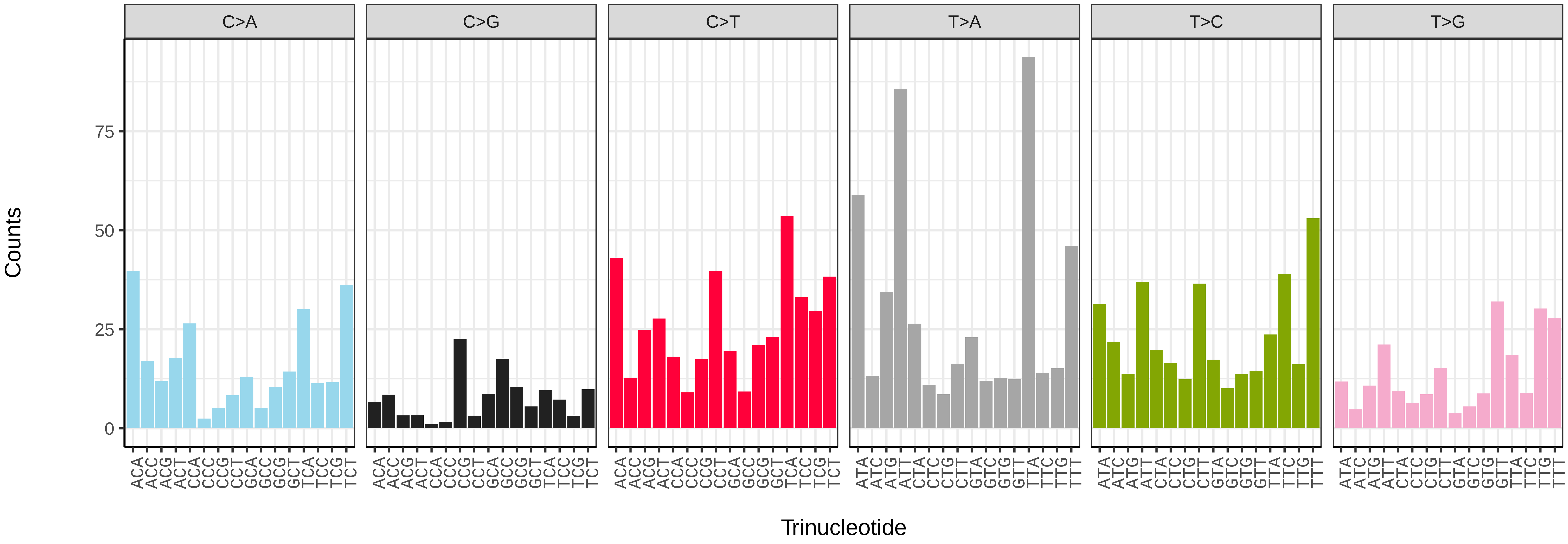


*Chloroclysta siterata* (ilChlSite2) somatic mutational spectrum (SBS52 classification)

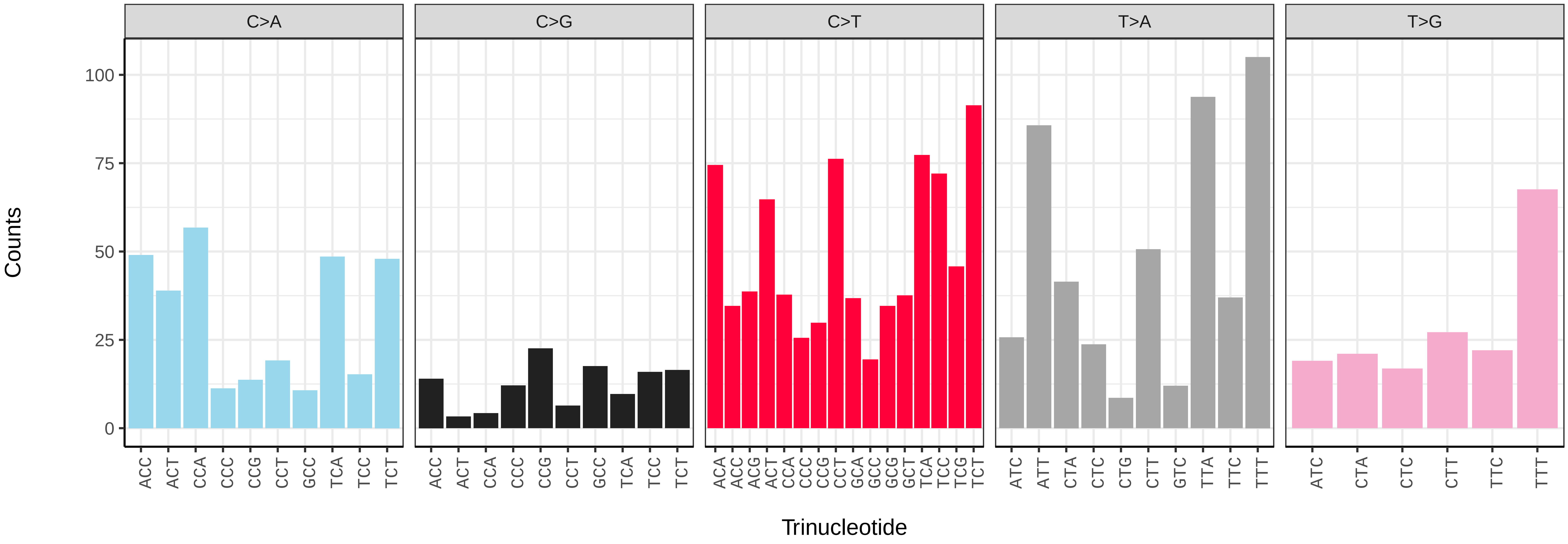

*Chrysoteuchia culmella* (ilChrCulm1) germline mutational spectrum

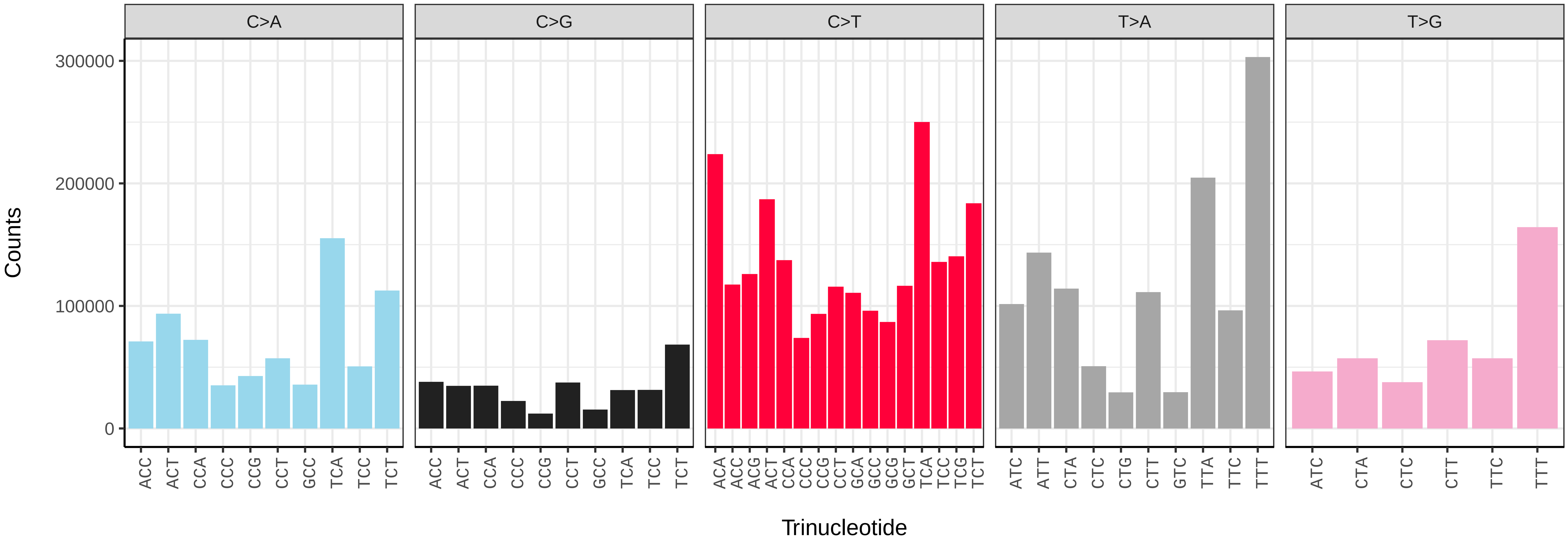


*Chrysoteuchia culmella* (ilChrCulm1) somatic mutational spectrum (SBS96 classification)

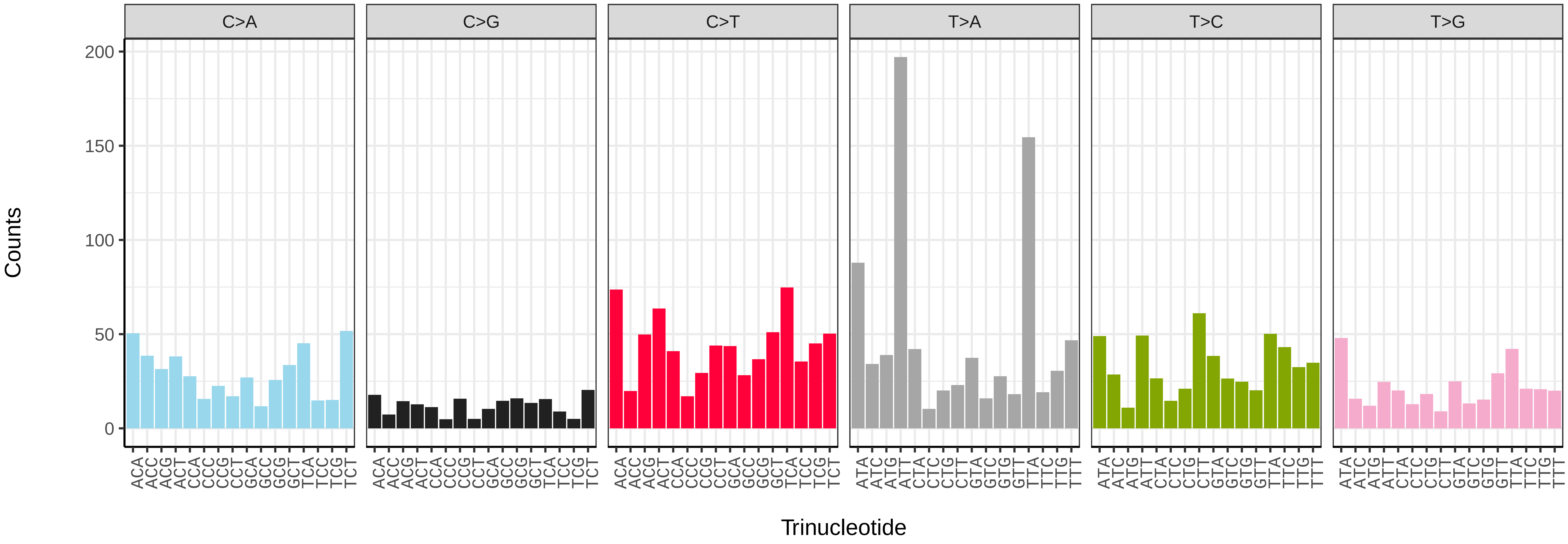


*Chrysoteuchia culmella* (ilChrCulm1) somatic mutational spectrum (SBS52 classification)

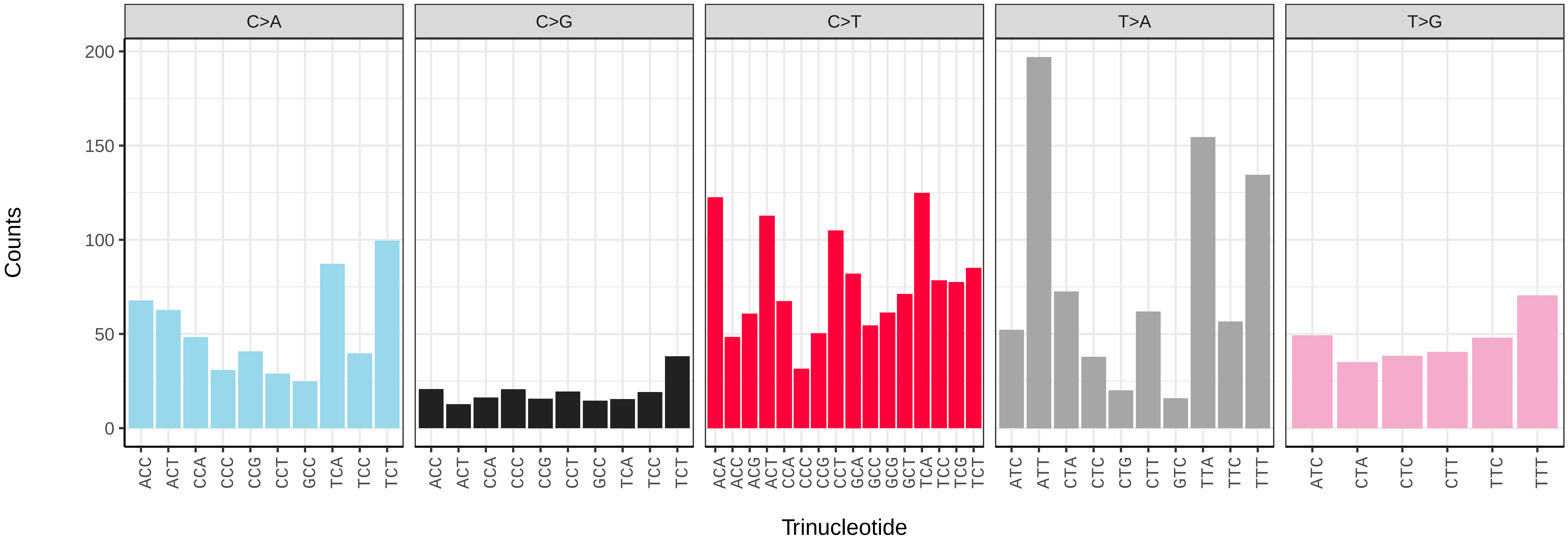

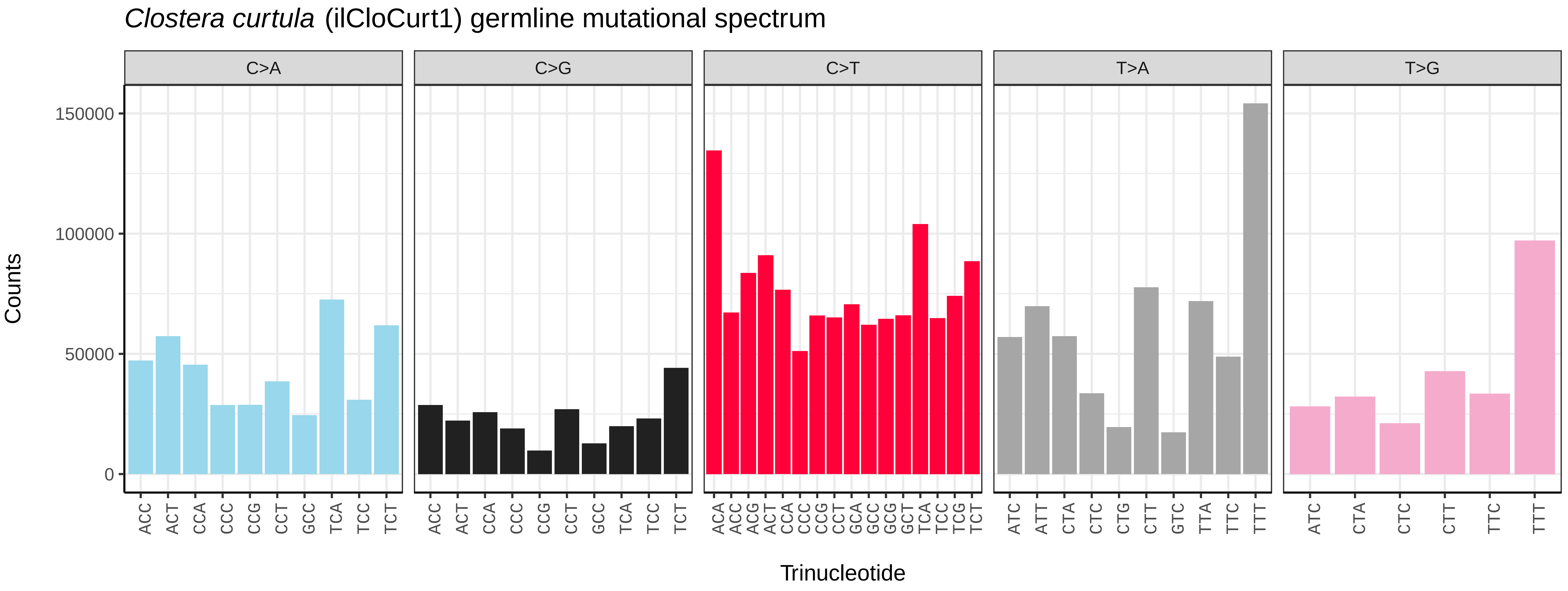
Clostera curtula (ilCloCurt1) germline mutational spectrum
C>A
C>G
C>T
T>A
T>G
Counts
0
50000
100000
150000
ACC ACT CCA CCC CCG CCT GCC TCA TCC TCT
ACC ACT CCA CCC CCG CCT GCC TCA TCC TCT
ACA ACC ACG ACT CCA CCC CCG CCT GCA GCC GCG GCT TCA TCC TCG TCT
ATC ATT CTA CTC CTG CTT GTC TTA TTC TTT
ATC CTA CTC CTT TTC TTT
Trinucleotide

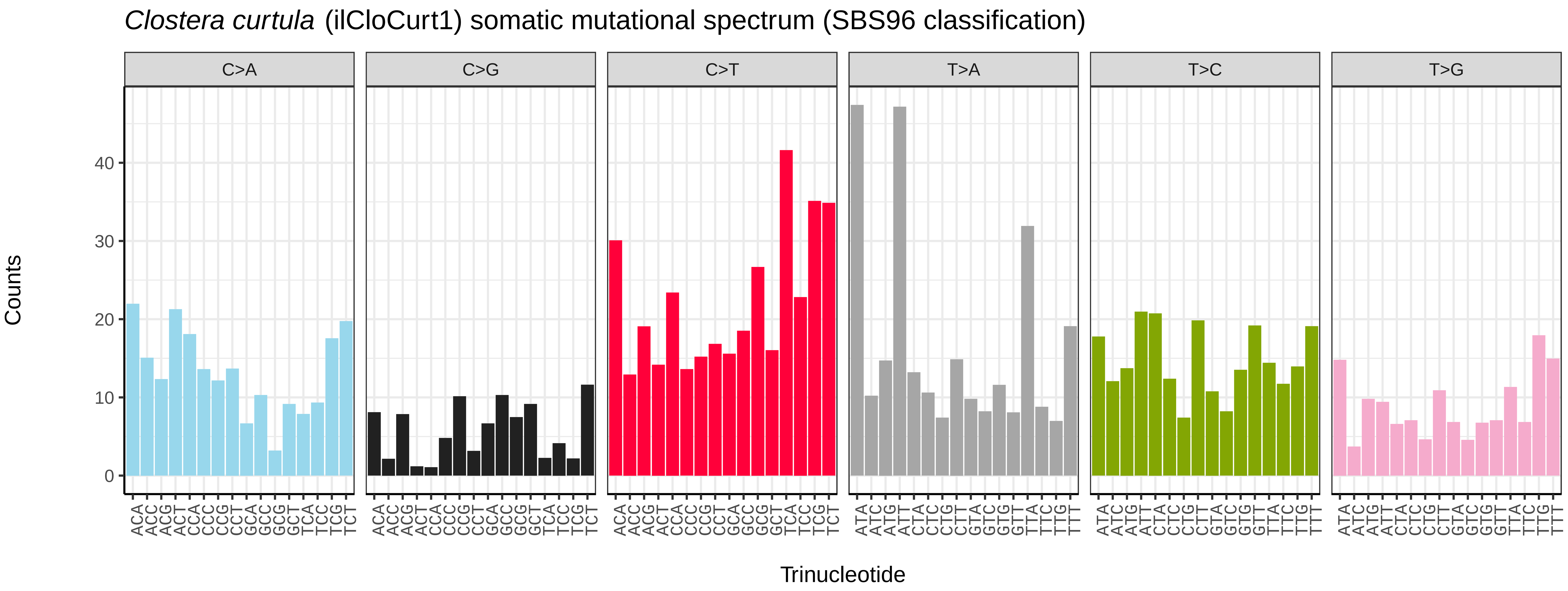
Clostera curtula (ilCloCurt1) somatic mutational spectrum (SBS96 classification)
C>A
C>G
C>T
T>A
T>C
T>G
Counts
0
10
20
30
40
ACA ACC ACG ACT CCA CCC CCG CCT GCA GCC GCG GCT TCA TCC TCG TCT
ACA ACC ACG ACT CCA CCC CCG CCT GCA GCC GCG GCT TCA TCC TCG TCT
ACA ACC ACG ACT CCA CCC CCG CCT GCA GCC GCG GCT TCA TCC TCG TCT
ATA ATC ATG ATT CTA CTC CTG CTT GTA GTC GTG GTT TTA TTC TTG TTT
ATA ATC ATG ATT CTA CTC CTG CTT GTA GTC GTG GTT TTA TTC TTG TTT
ATA ATC ATG ATT CTA CTC CTG CTT GTA GTC GTG GTT TTA TTC TTG TTT
Trinucleotide

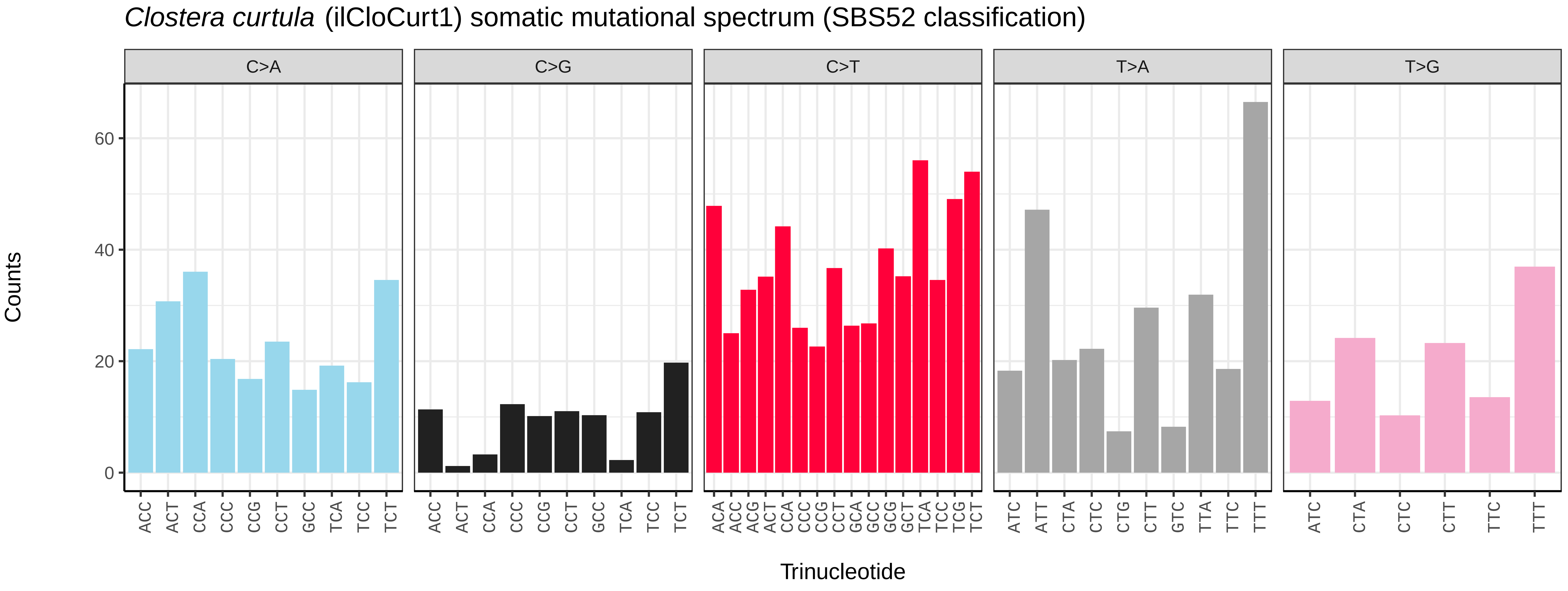
Clostera curtula (ilCloCurt1) somatic mutational spectrum (SBS52 classification)
C>A
C>G
C>T
T>A
T>G
Counts
0
20
40
60
ACC ACT CCA CCC CCG CCT GCC TCA TCC TCT
ACC ACT CCA CCC CCG CCT GCC TCA TCC TCT
ACA ACC ACG ACT CCA CCC CCG CCT GCA GCC GCG GCT TCA TCC TCG TCT
ATC ATT CTA CTC CTG CTT GTC TTA TTC TTT
ATC CTA CTC CTT TTC TTT
Trinucleotide

*Colias croceus* (ilColCroc2) germline mutational spectrum

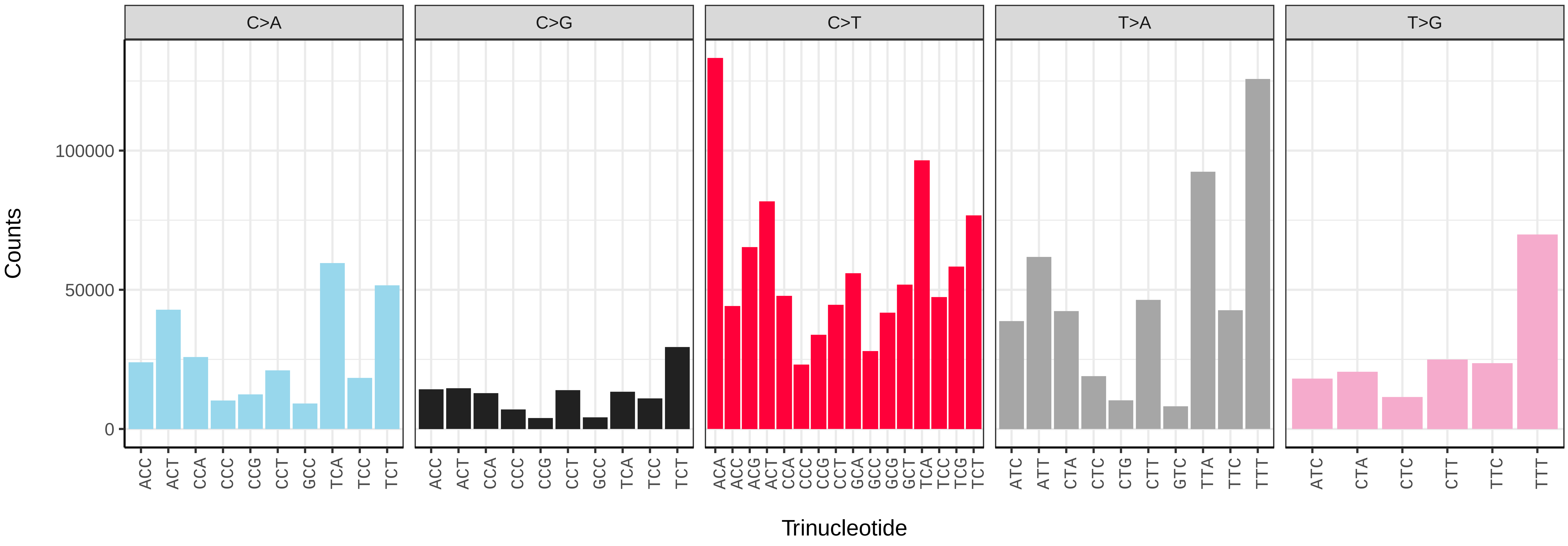


*Colias croceus* (ilColCroc2) somatic mutational spectrum (SBS96 classification)

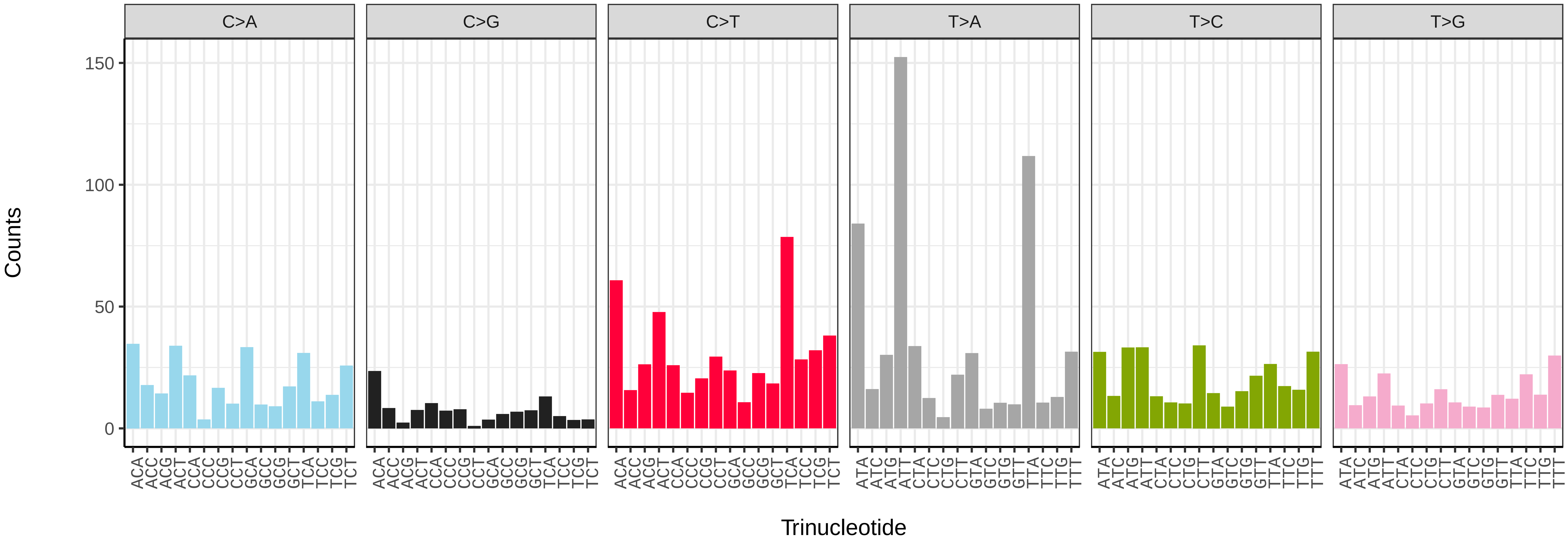


*Colias croceus* (ilColCroc2) somatic mutational spectrum (SBS52 classification)

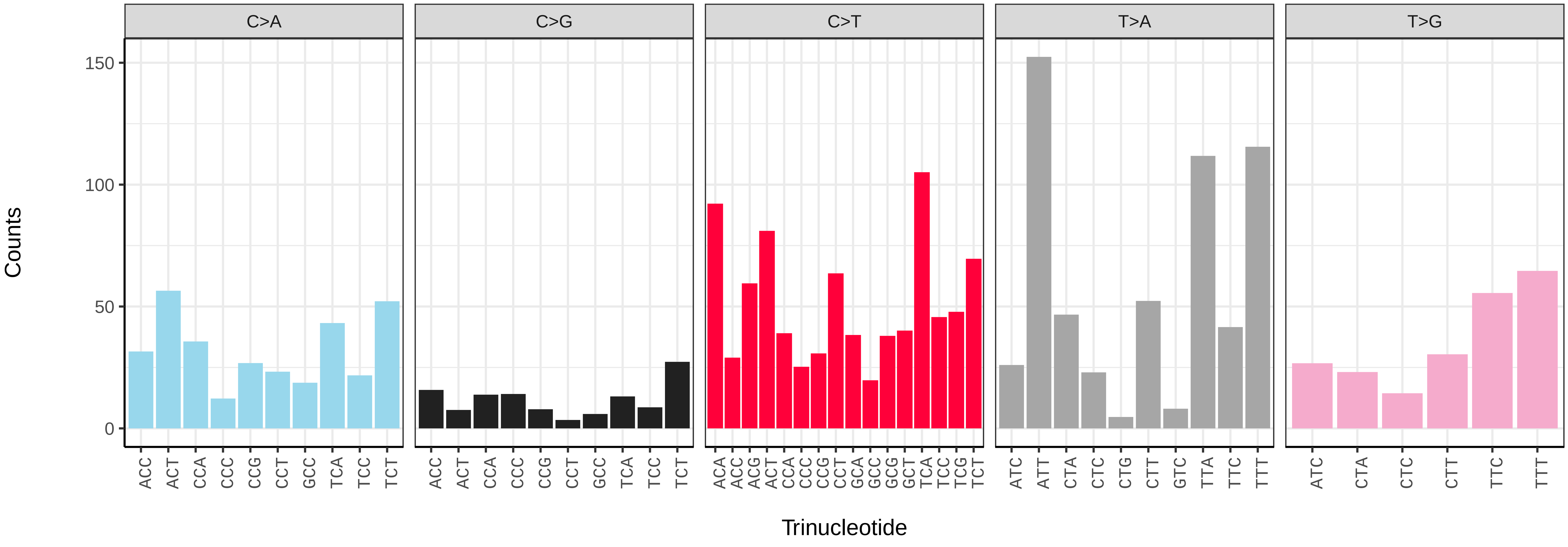

*Coleophora flavipennella* (ilColFlav1) germline mutational spectrum

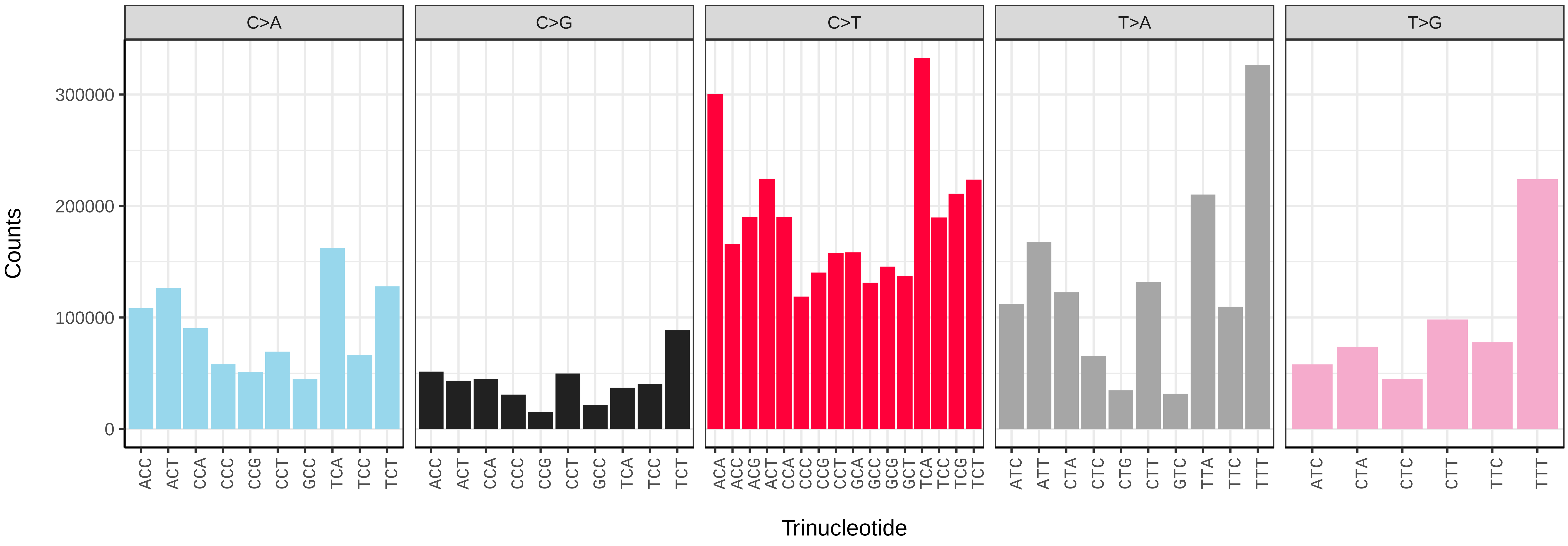


*Coleophora flavipennella* (ilColFlav1) somatic mutational spectrum (SBS96 classification)

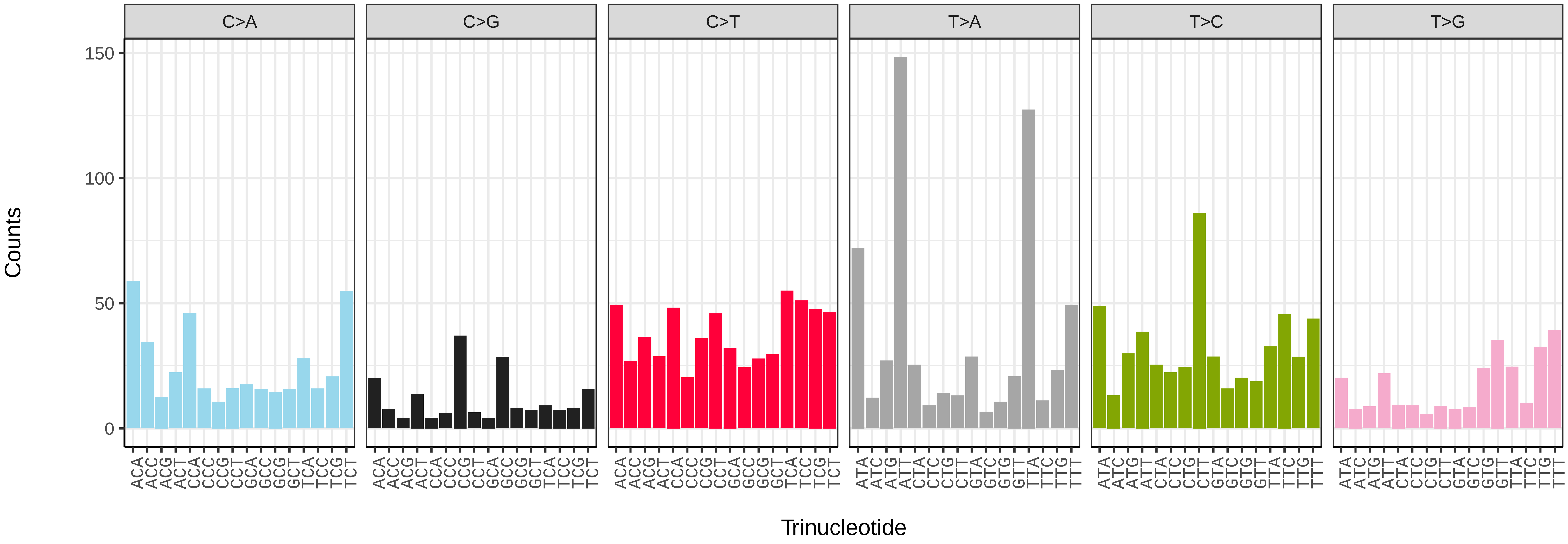


*Coleophora flavipennella* (ilColFlav1) somatic mutational spectrum (SBS52 classification)

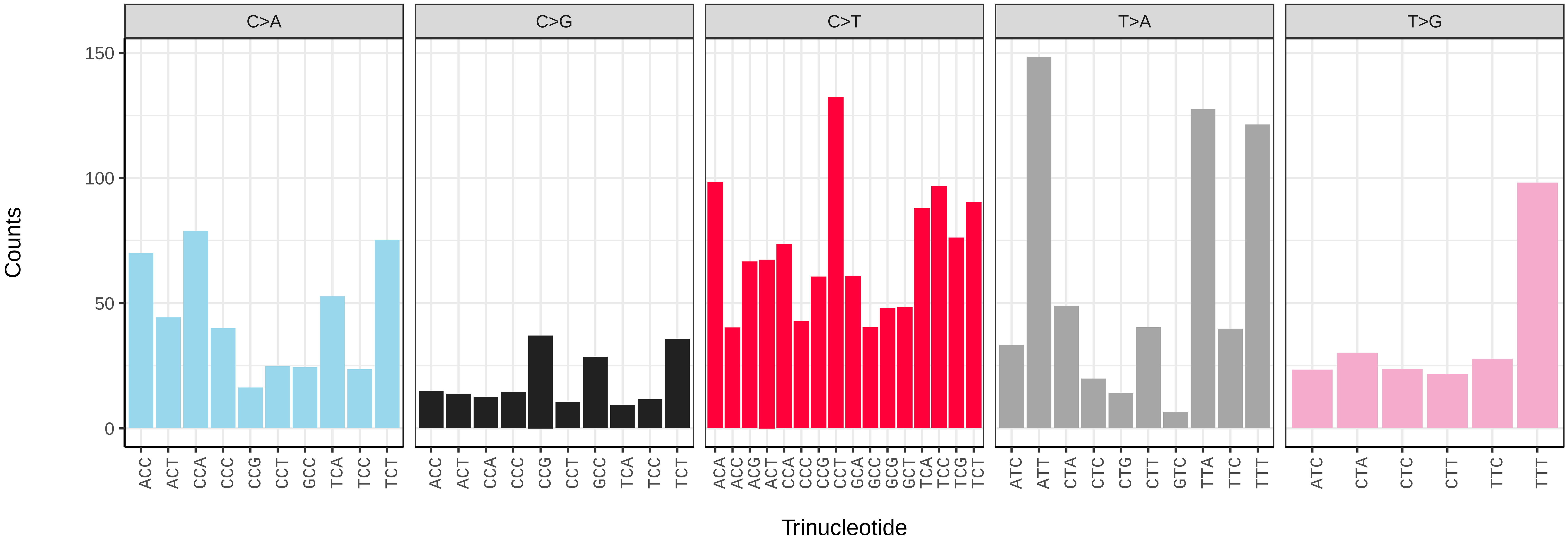

*Conistra vaccinii* (ilConVacc3) germline mutational spectrum

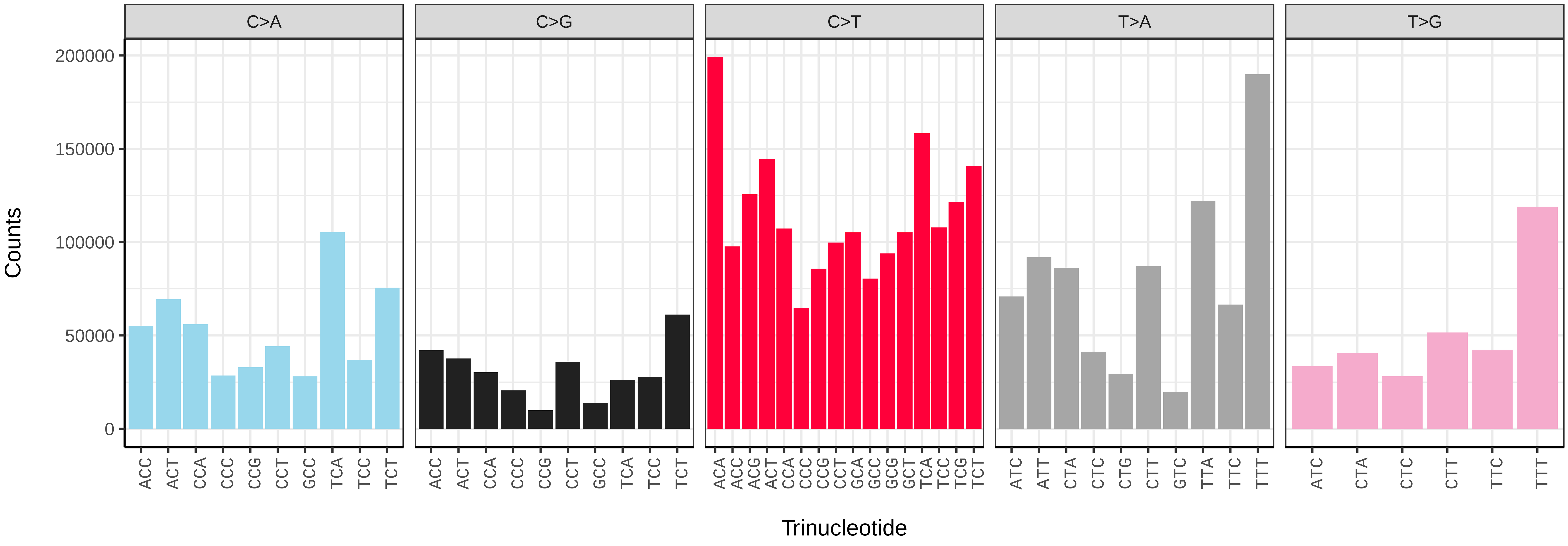


*Conistra vaccinii* (ilConVacc3) somatic mutational spectrum (SBS96 classification)

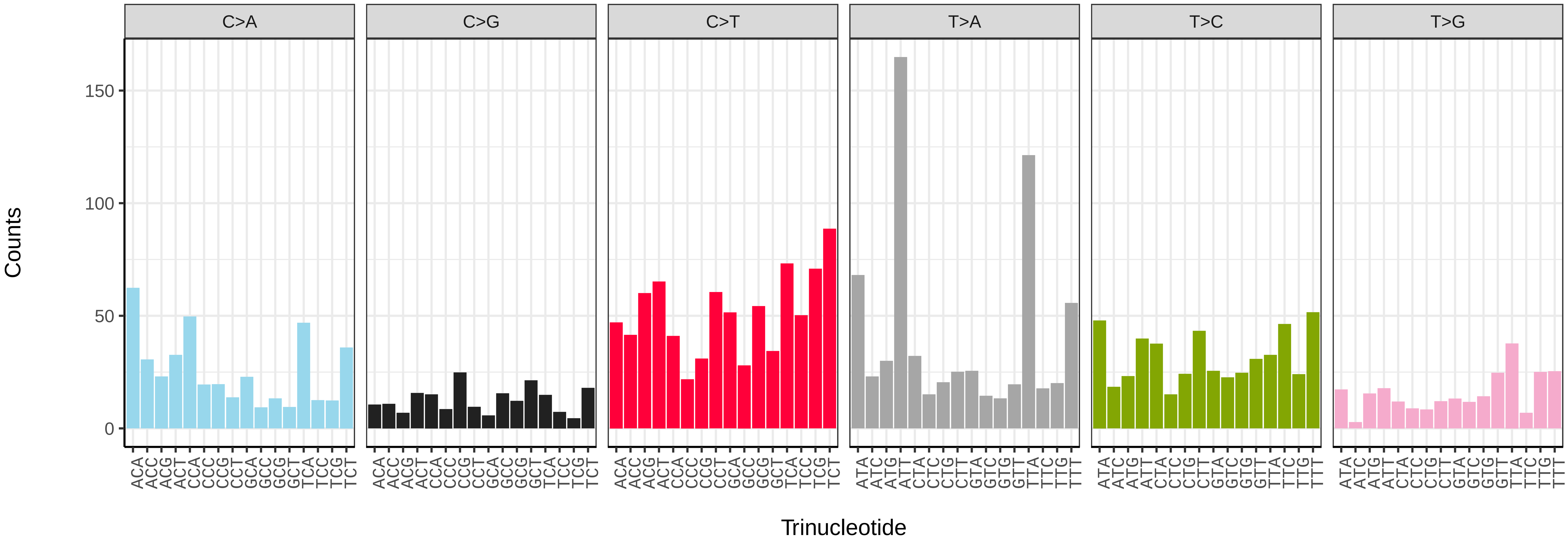


*Conistra vaccinii* (ilConVacc3) somatic mutational spectrum (SBS52 classification)

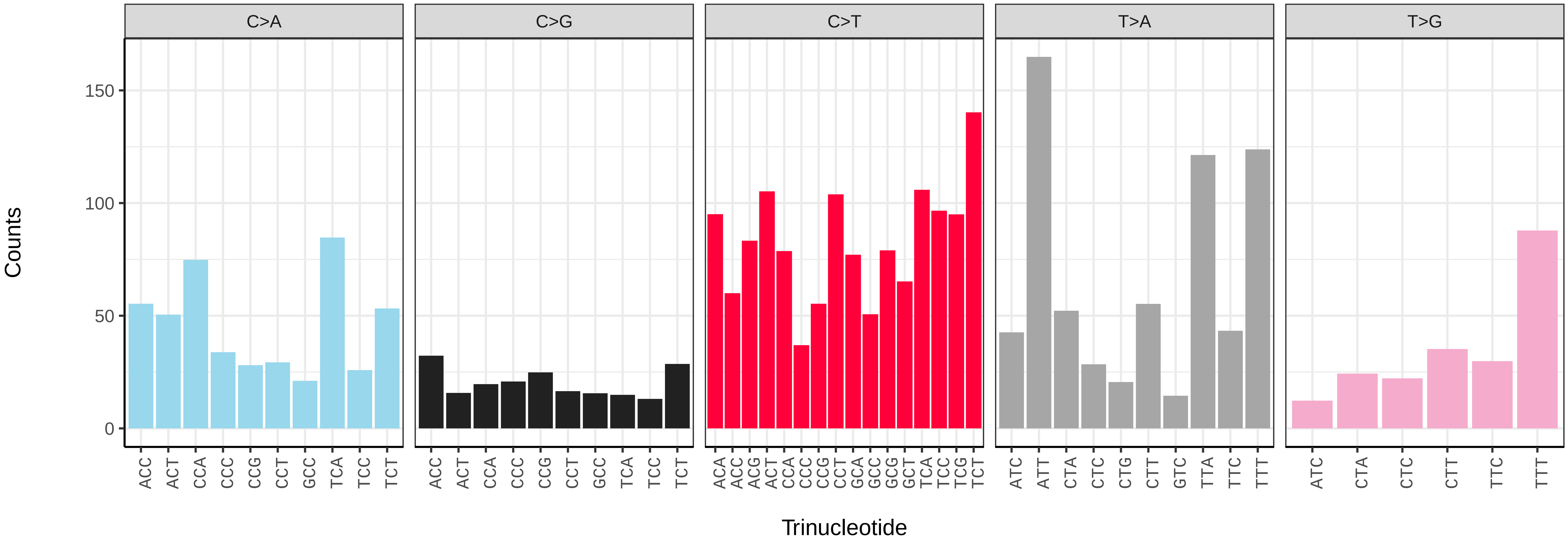

*Cosmia pyralina* (ilCosPyra2) germline mutational spectrum

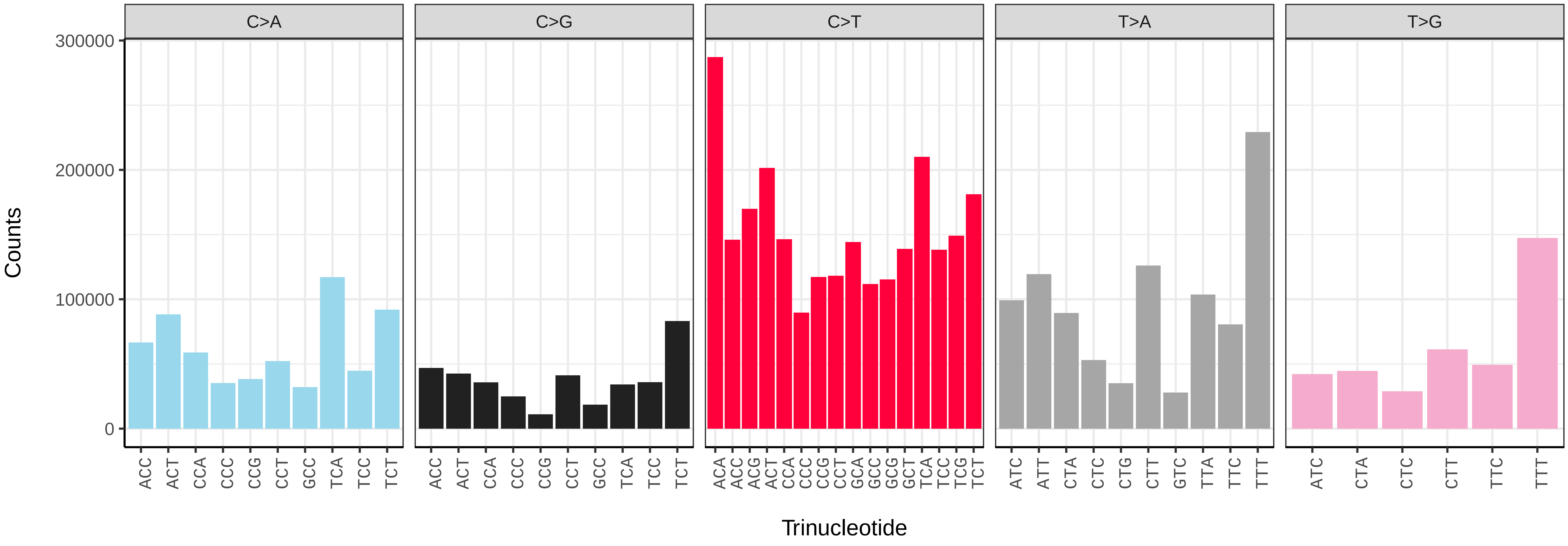


*Cosmia pyralina* (ilCosPyra2) somatic mutational spectrum (SBS96 classification)

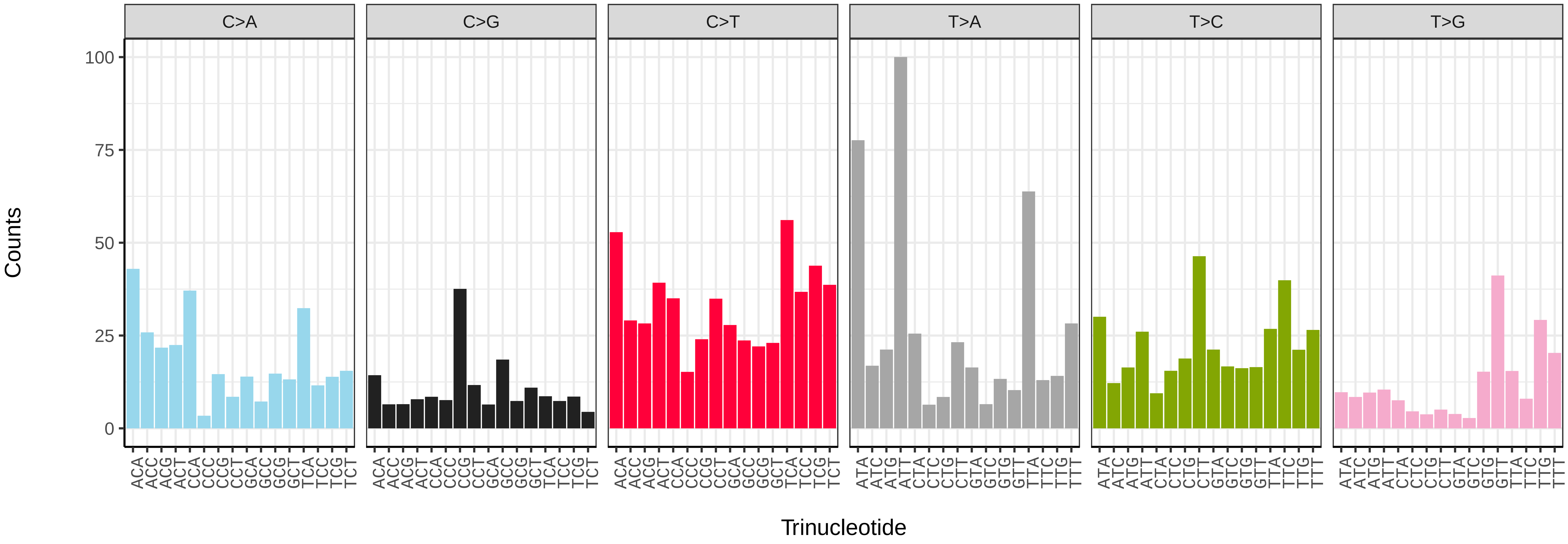


*Cosmia pyralina* (ilCosPyra2) somatic mutational spectrum (SBS52 classification)

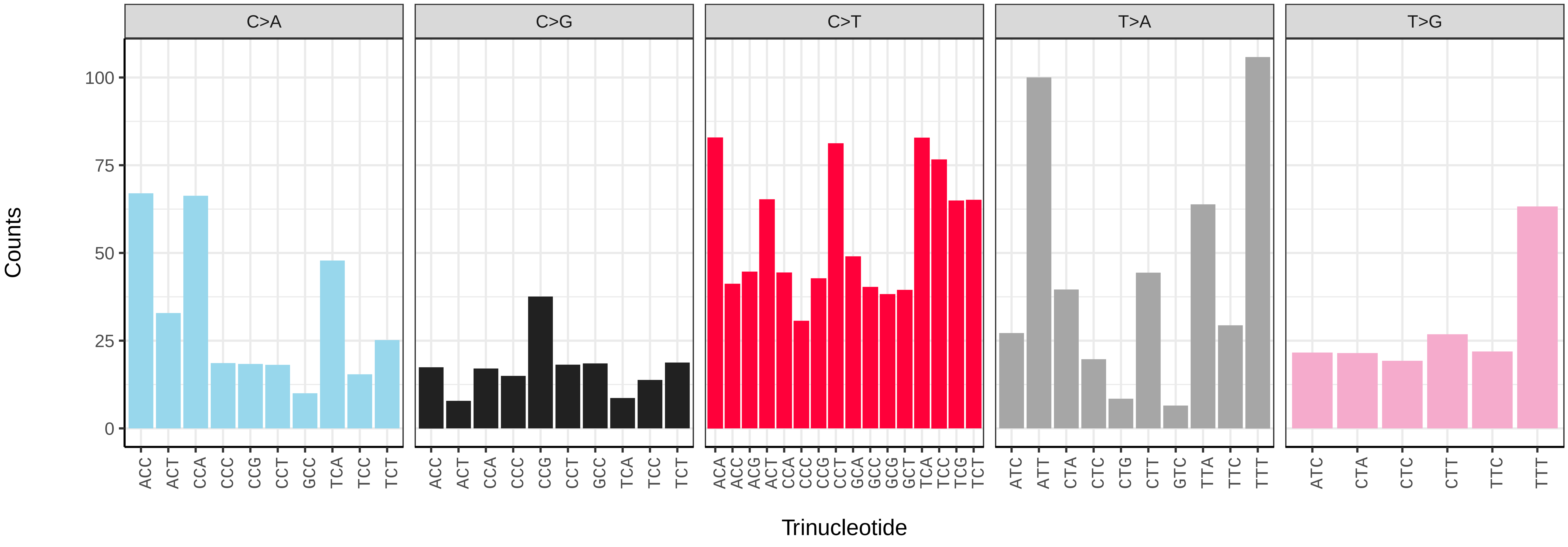

*Cosmia trapezina* (ilCosTrap1) germline mutational spectrum

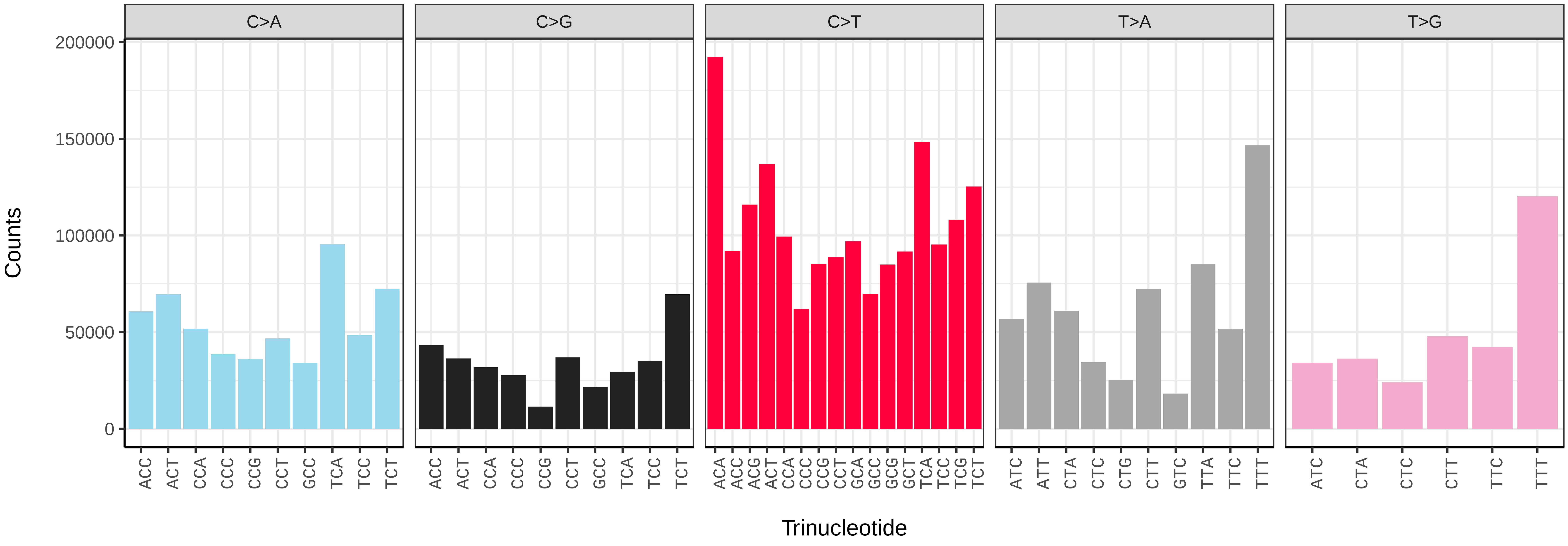


*Cosmia trapezina* (ilCosTrap1) somatic mutational spectrum (SBS96 classification)

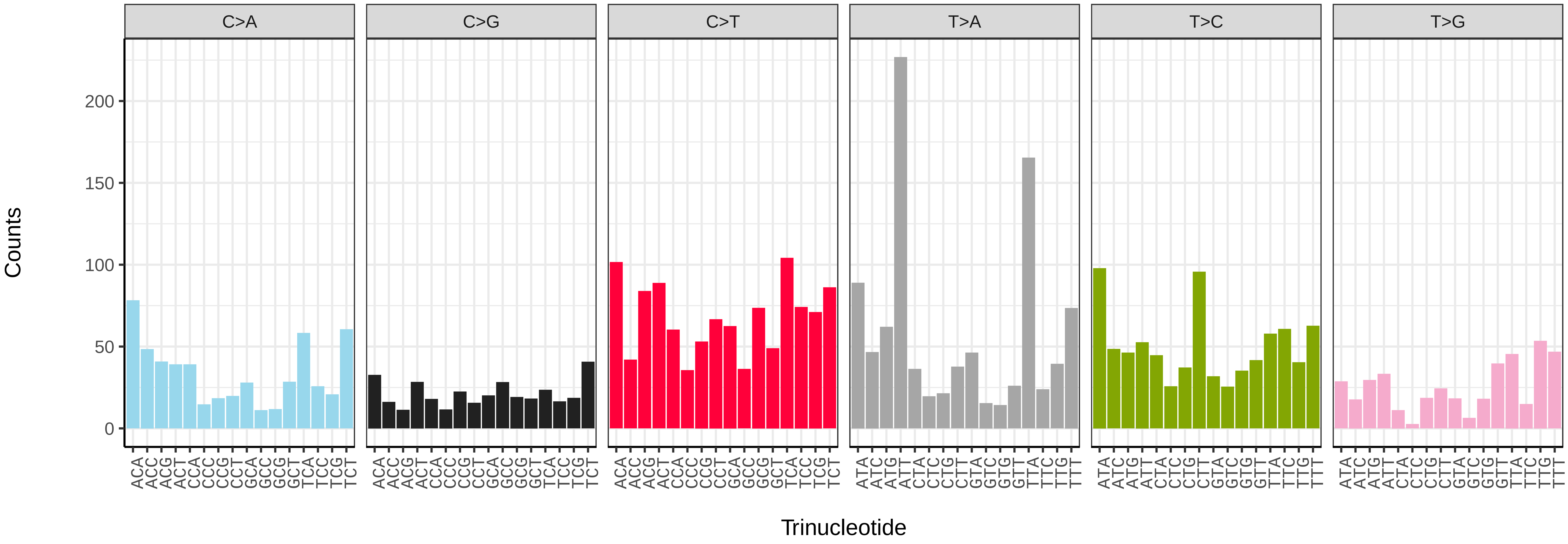


*Cosmia trapezina* (ilCosTrap1) somatic mutational spectrum (SBS52 classification)

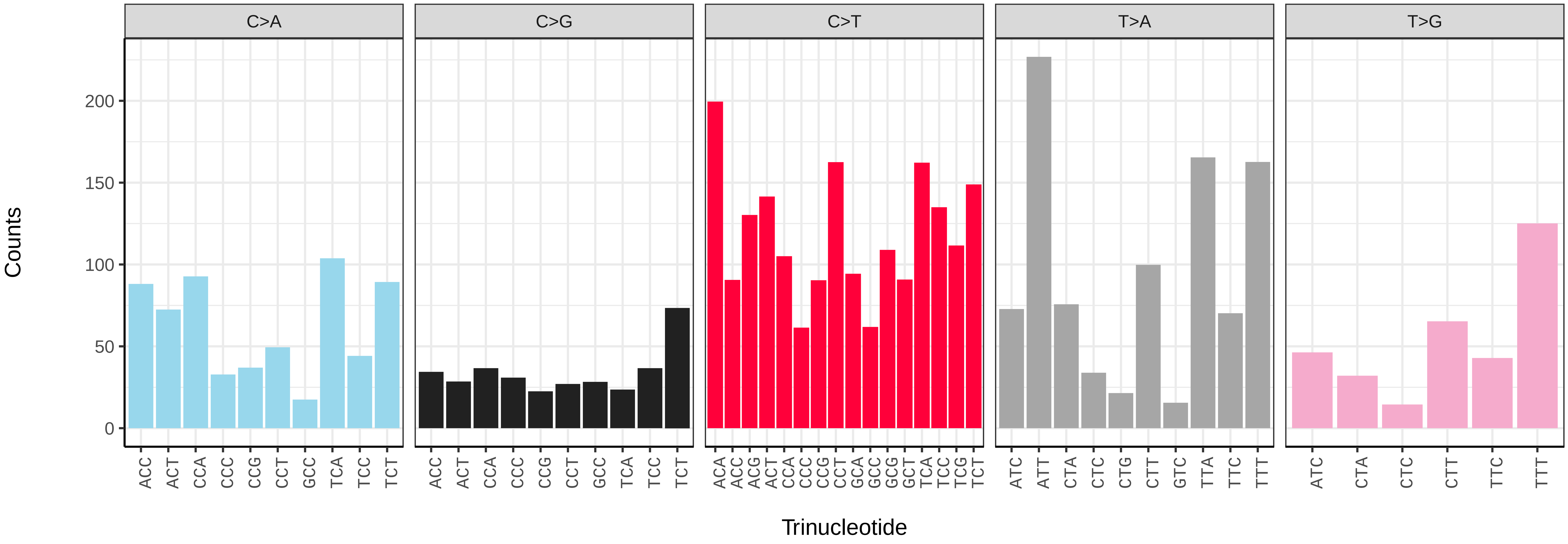

*Craniophora ligustri* (ilCraLigu1) germline mutational spectrum

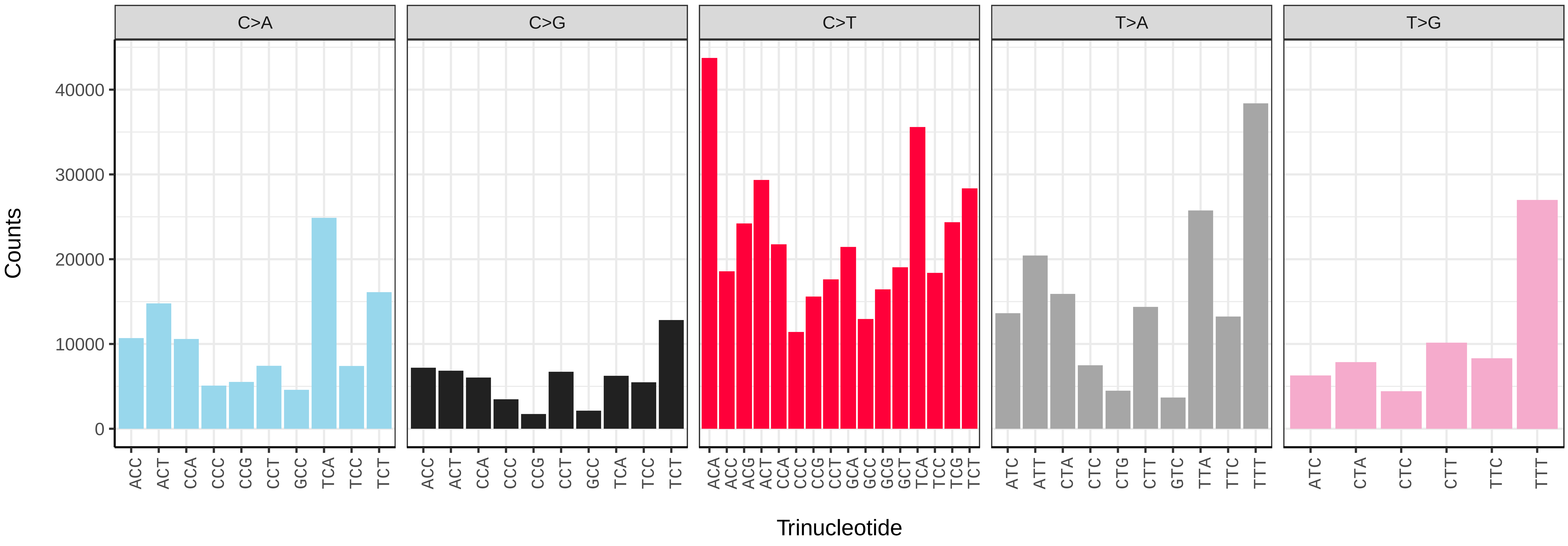


*Craniophora ligustri* (ilCraLigu1) somatic mutational spectrum (SBS96 classification)

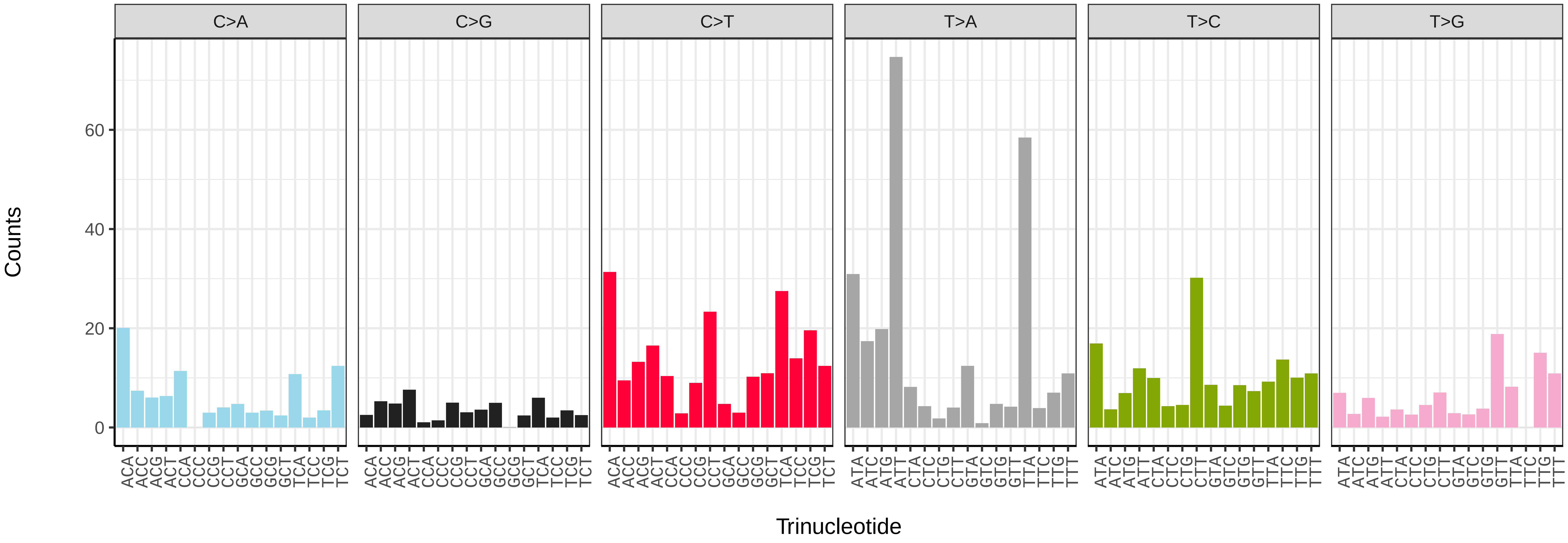


*Craniophora ligustri* (ilCraLigu1) somatic mutational spectrum (SBS52 classification)

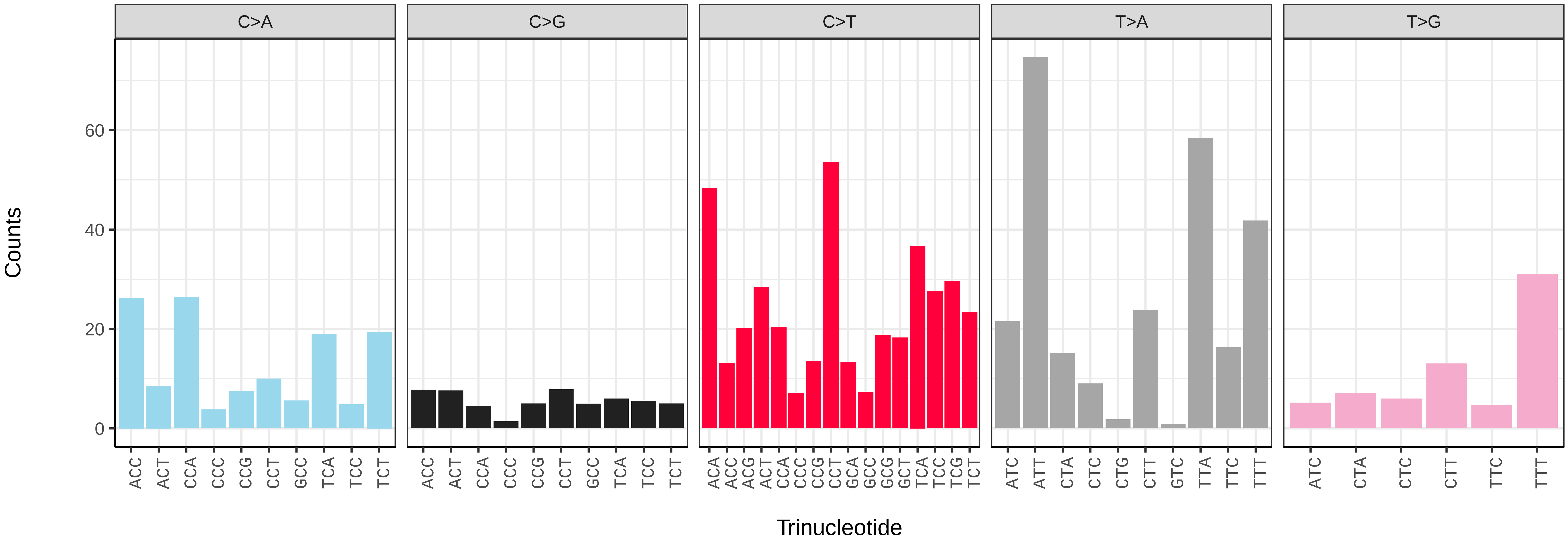

*Crocallis elinguaria* (ilCroElin1) germline mutational spectrum

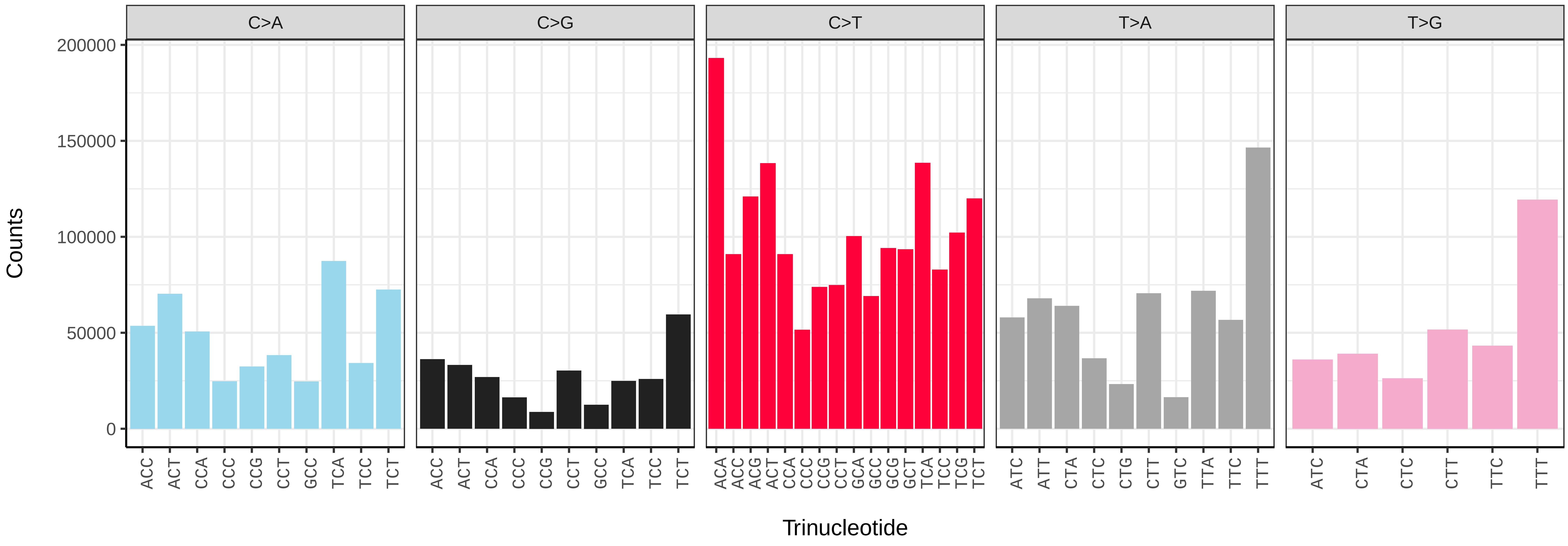


*Crocallis elinguaria* (ilCroElin1) somatic mutational spectrum (SBS96 classification)

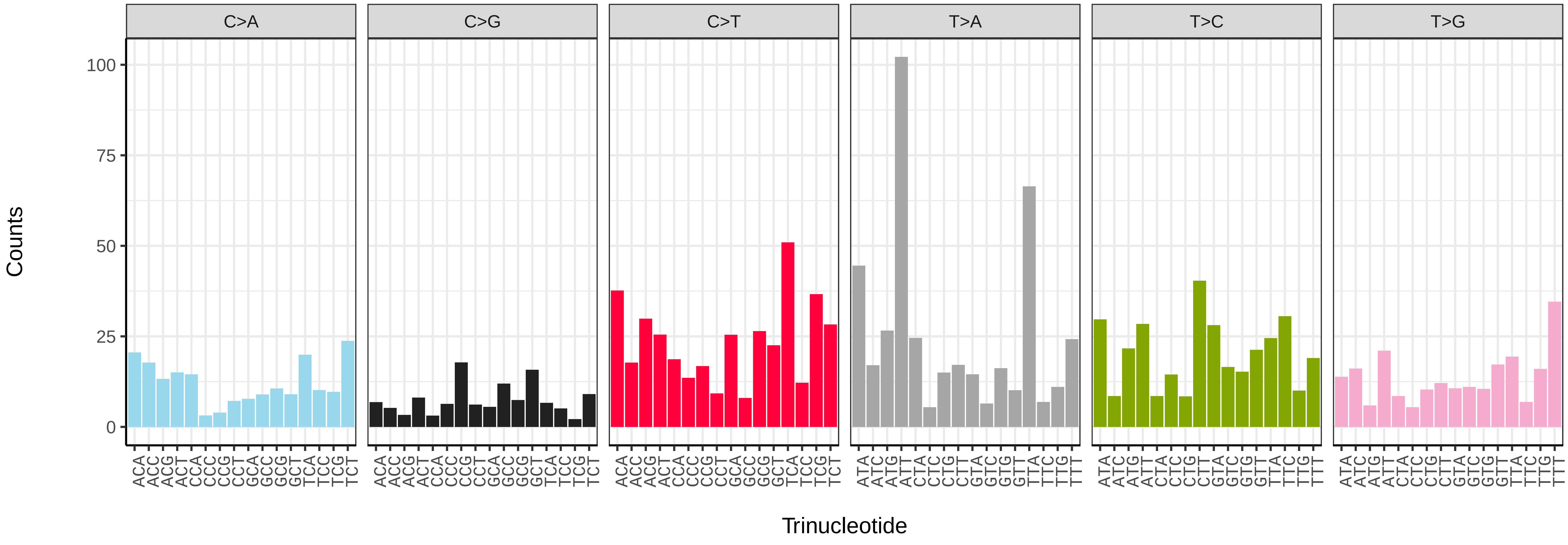


*Crocallis elinguaria* (ilCroElin1) somatic mutational spectrum (SBS52 classification)

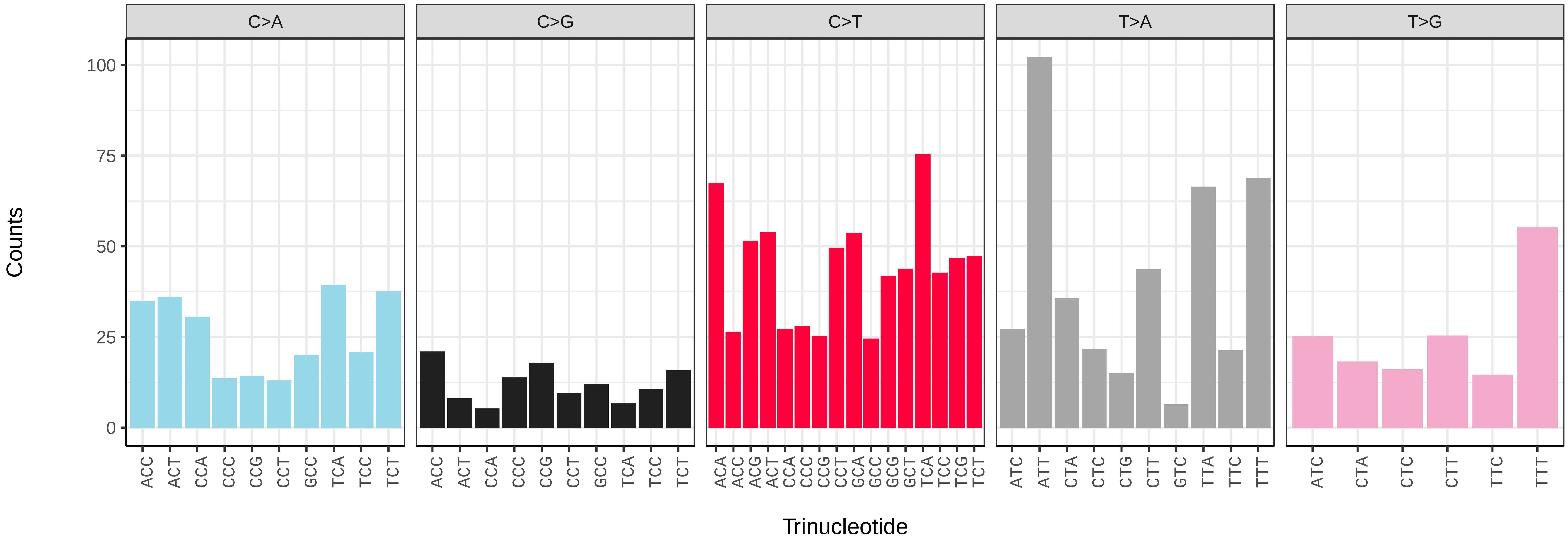

*Cyaniris semiargus* (ilCyaSemi1) germline mutational spectrum

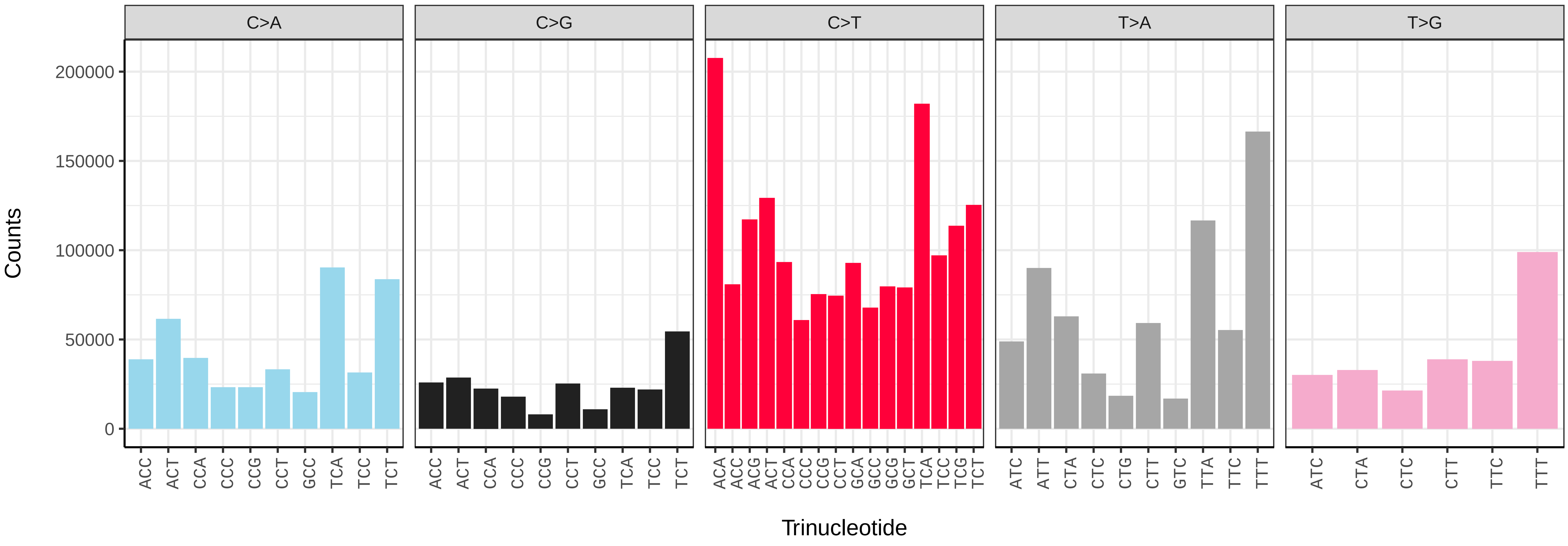


*Cyaniris semiargus* (ilCyaSemi1) somatic mutational spectrum (SBS96 classification)

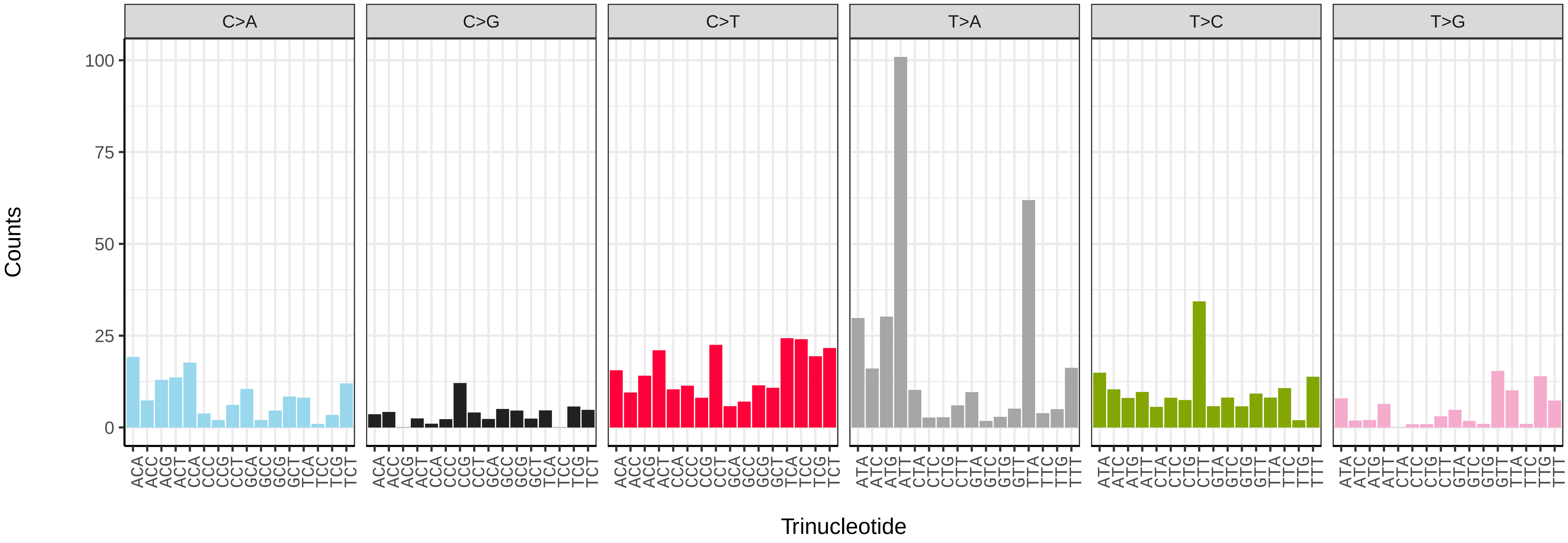


*Cyaniris semiargus* (ilCyaSemi1) somatic mutational spectrum (SBS52 classification)

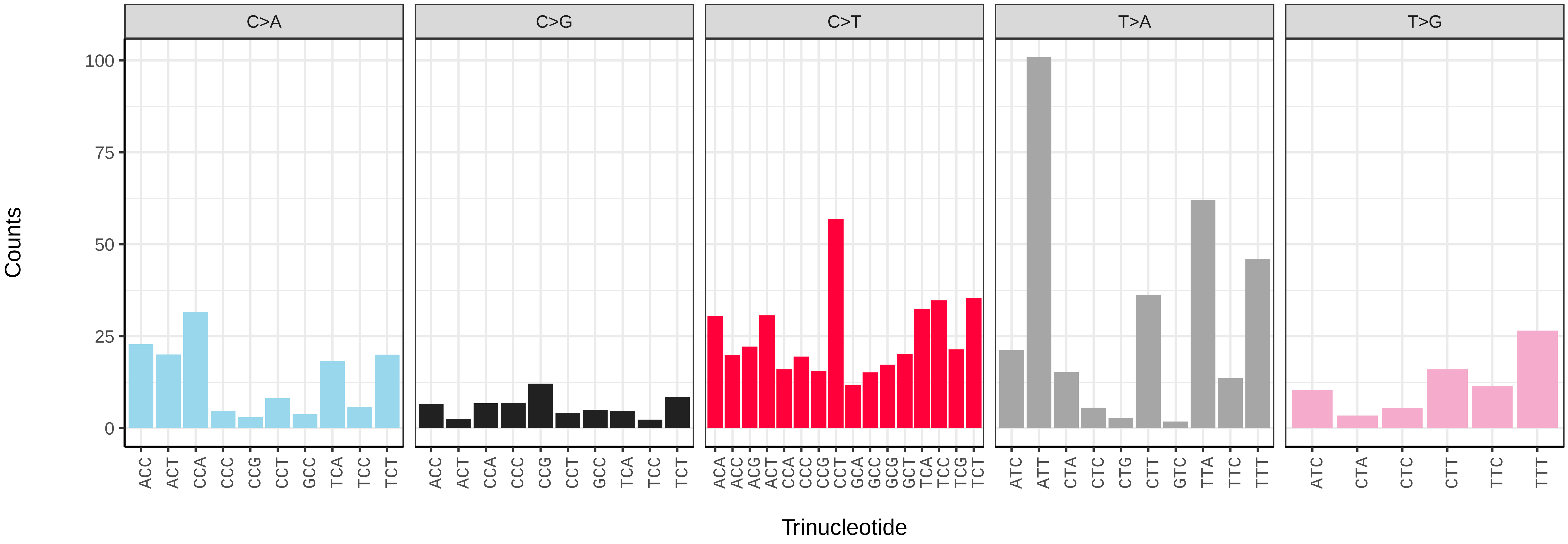

*Cybosia mesomella* (ilCybMeso1) germline mutational spectrum

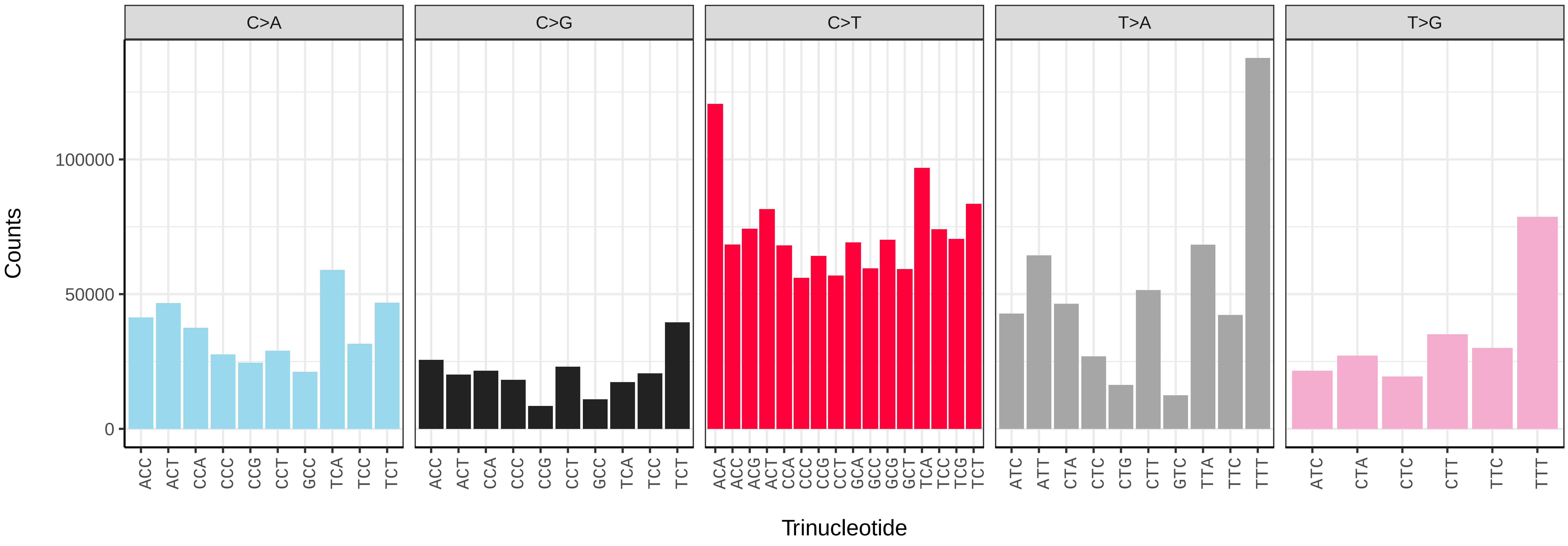


*Cybosia mesomella* (ilCybMeso1) somatic mutational spectrum (SBS96 classification)

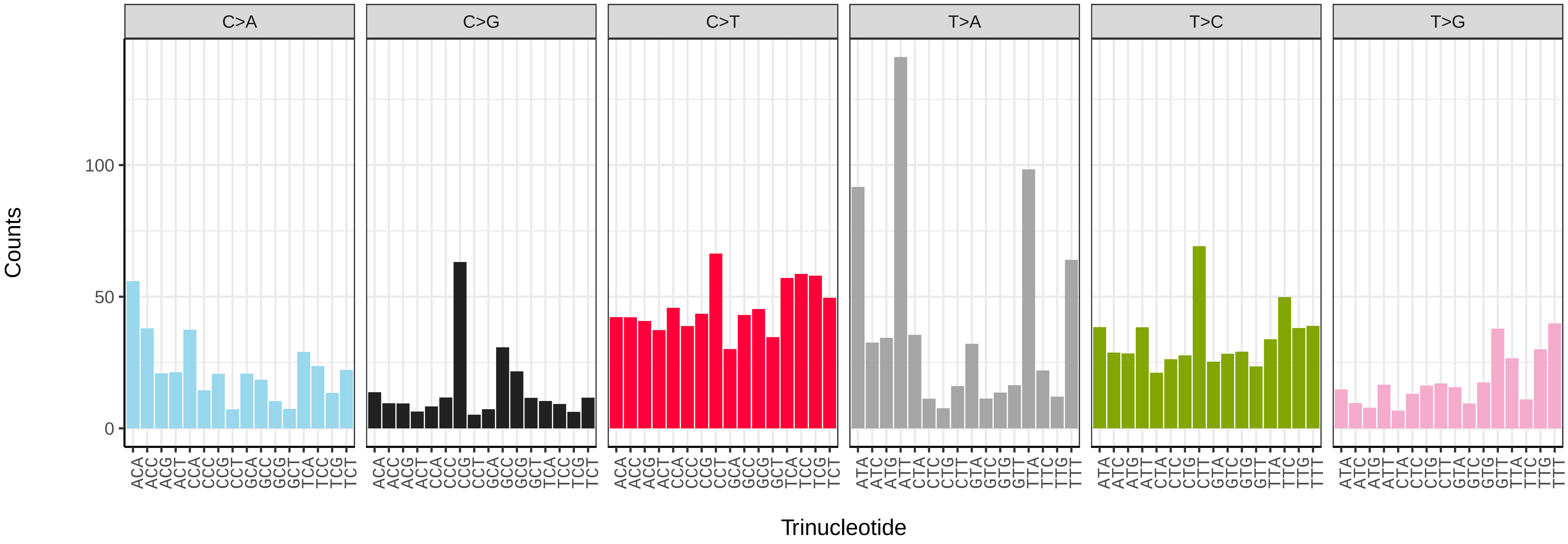


*Cybosia mesomella* (ilCybMeso1) somatic mutational spectrum (SBS52 classification)

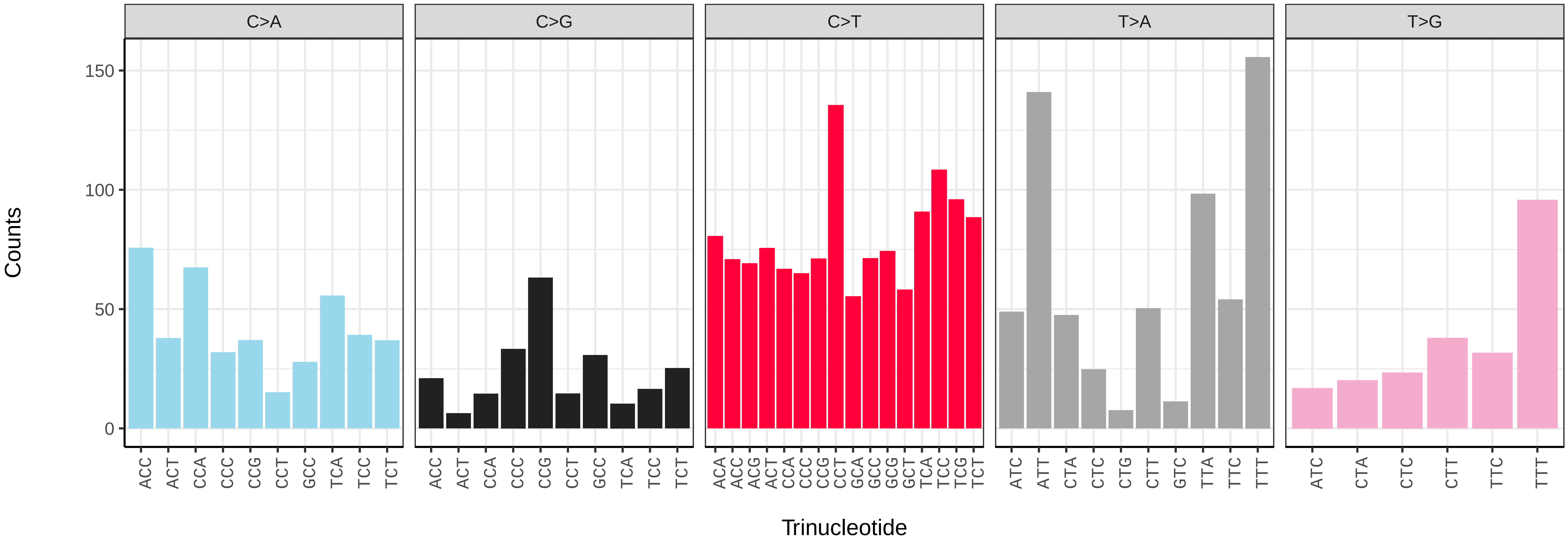

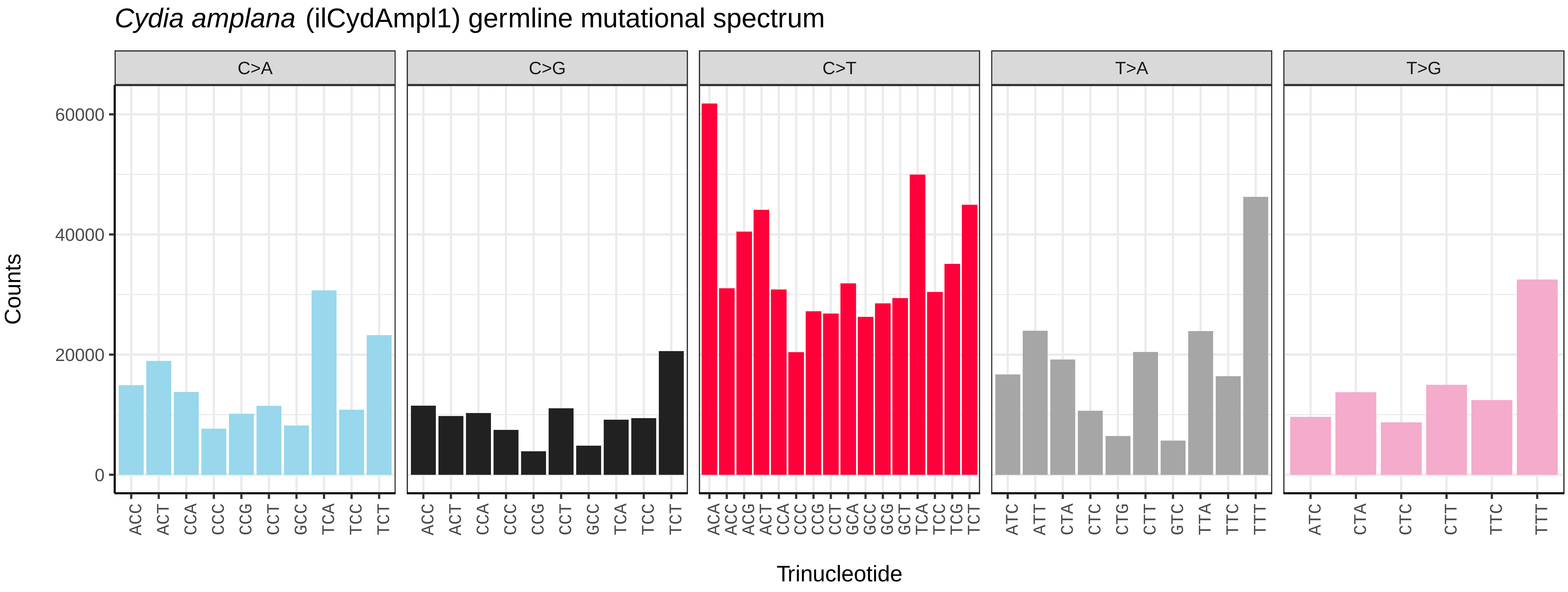

Cydia amplana (ilCydAmpl1) germline mutational spectrum
C>A
C>G
C>T
T>A
T>G
Counts
60000
40000
20000
0
Trinucleotide


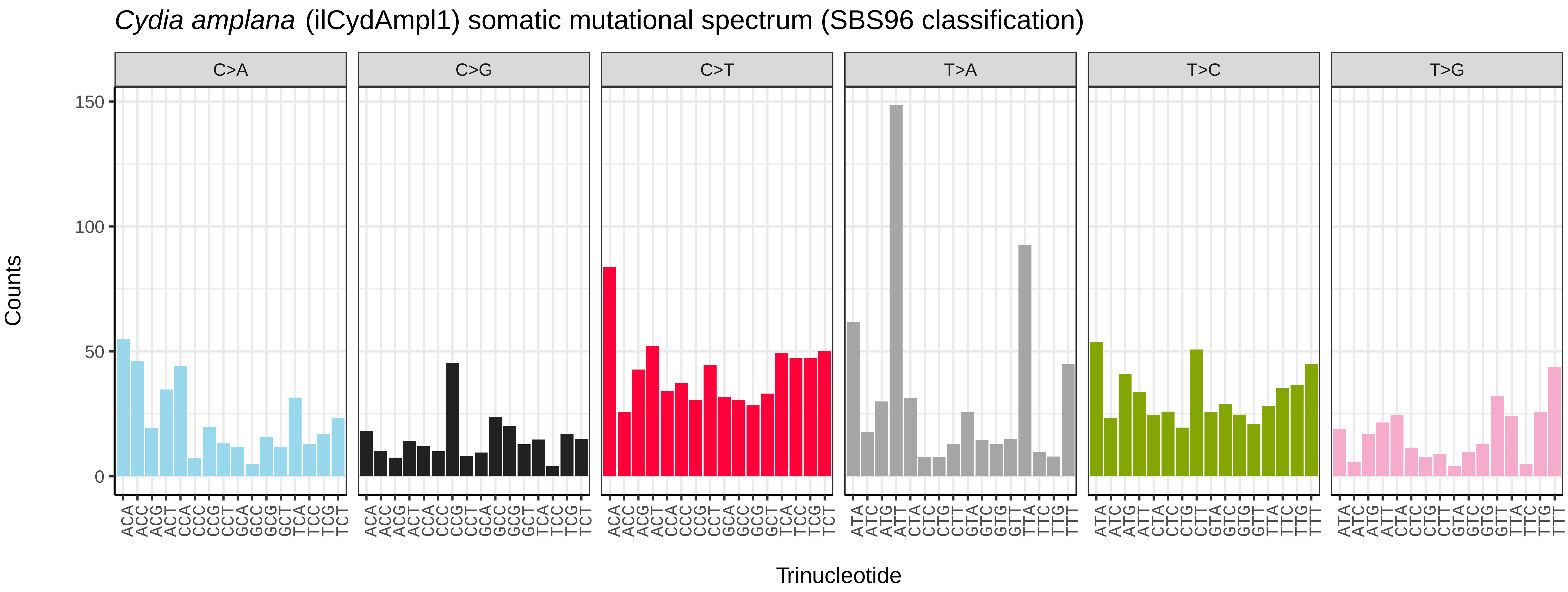

Cydia amplana (ilCydAmpl1) somatic mutational spectrum (SBS96 classification)
C>A
C>G
C>T
T>A
T>C
T>G
Counts
150
100
50
0
Trinucleotide


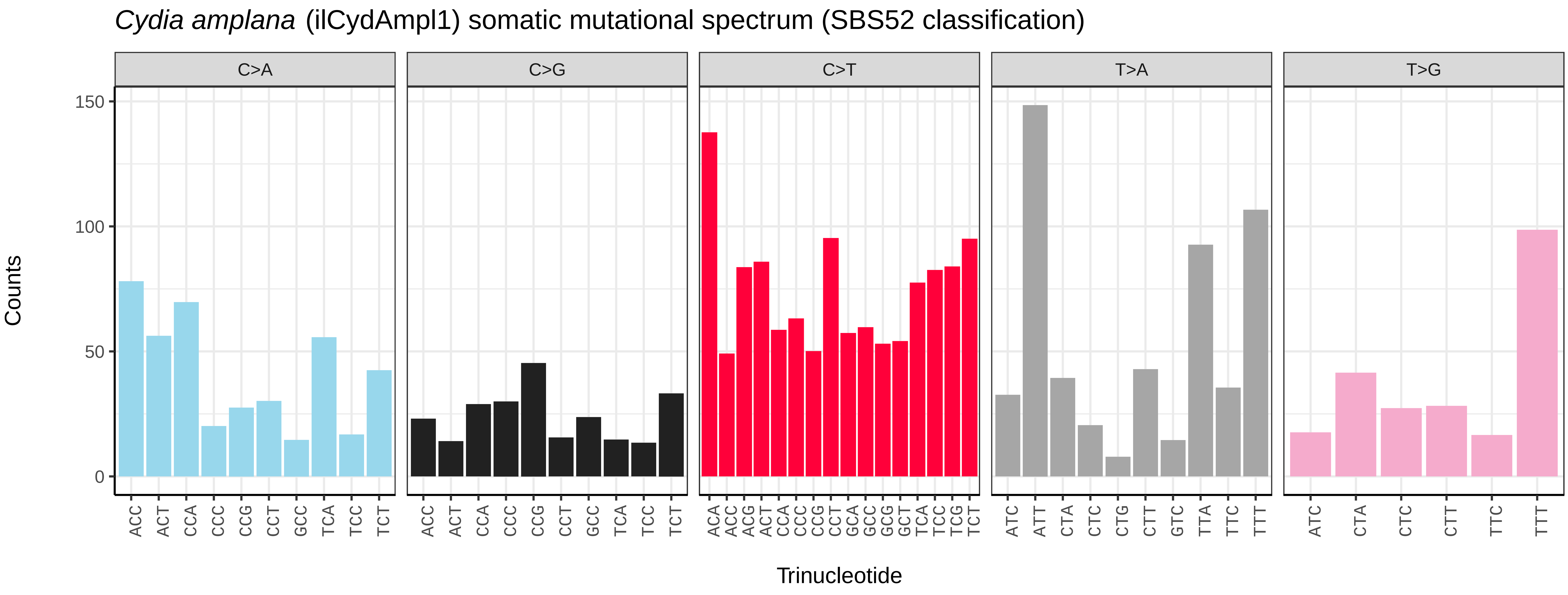

Cydia amplana (ilCydAmpl1) somatic mutational spectrum (SBS52 classification)
C>A
C>G
C>T
T>A
T>G
Counts
150
100
50
0
Trinucleotide

*Cydia splendana* (ilCydSple1) germline mutational spectrum

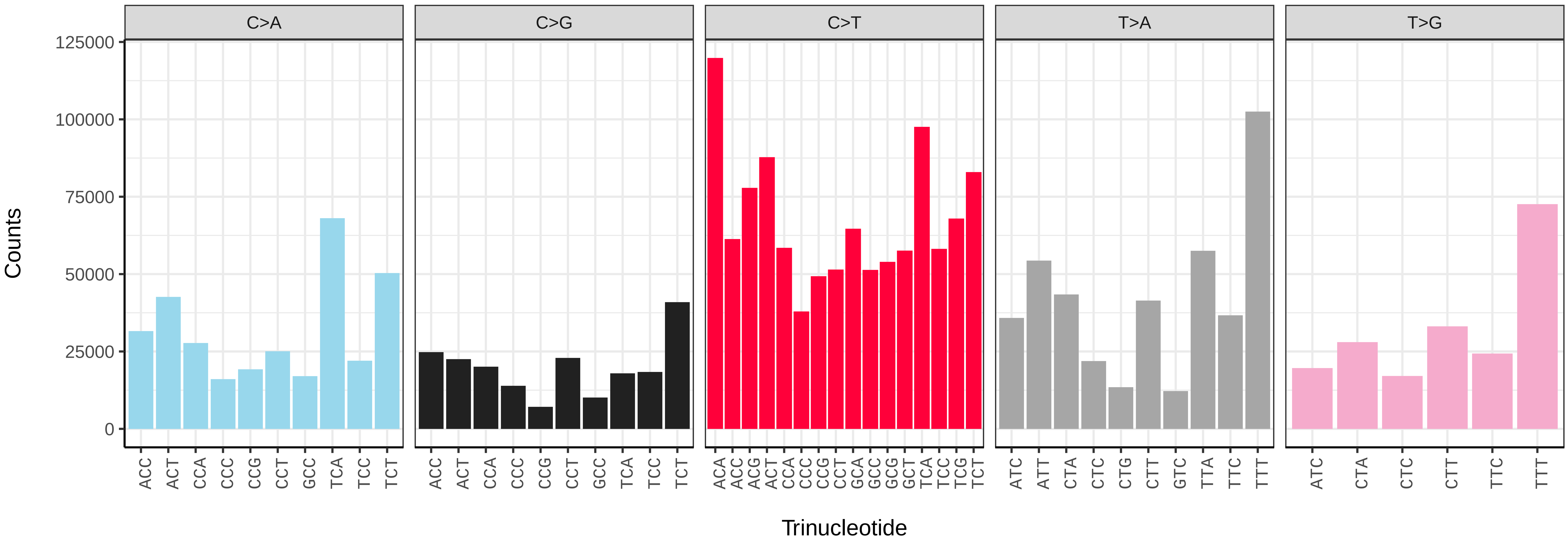


*Cydia splendana* (ilCydSple1) somatic mutational spectrum (SBS96 classification)

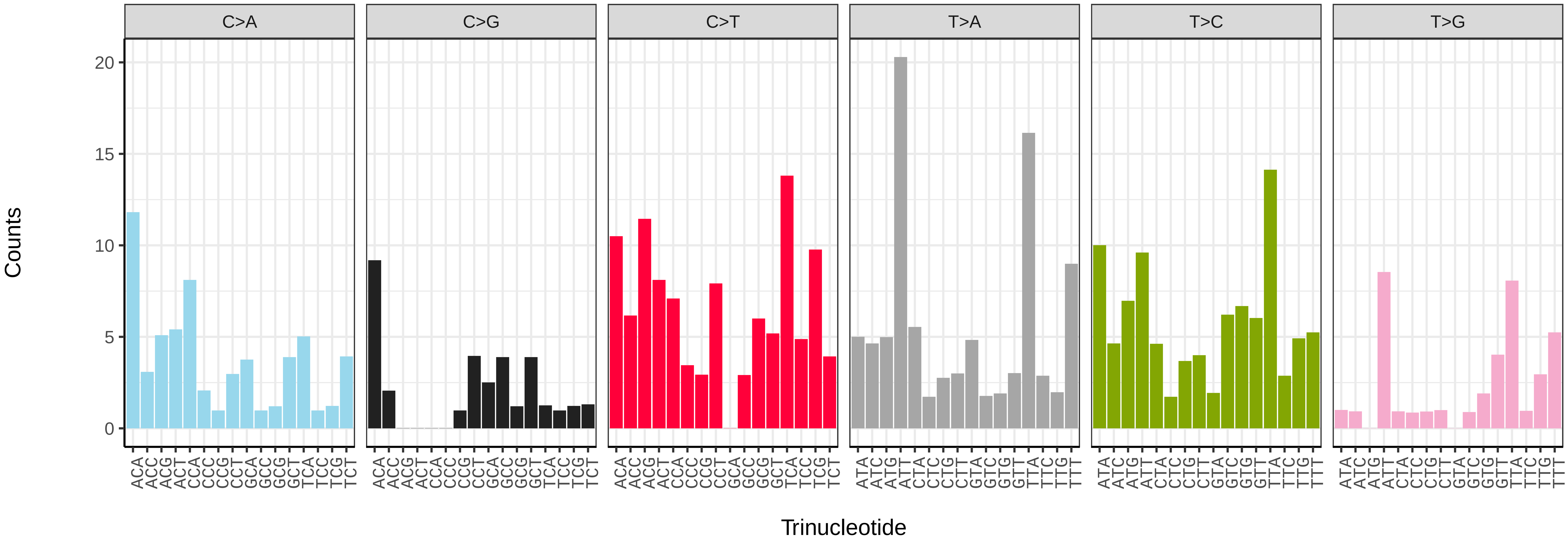


*Cydia splendana* (ilCydSple1) somatic mutational spectrum (SBS52 classification)

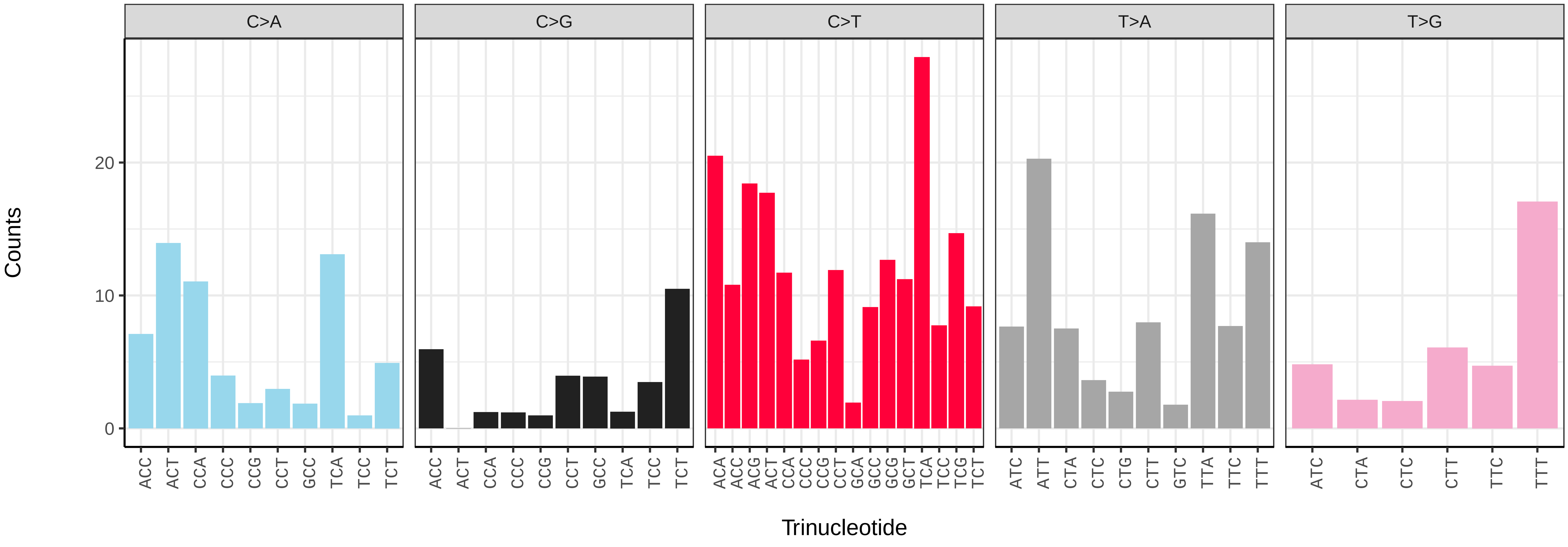

*Cydia strobilella* (ilCydStro3) germline mutational spectrum

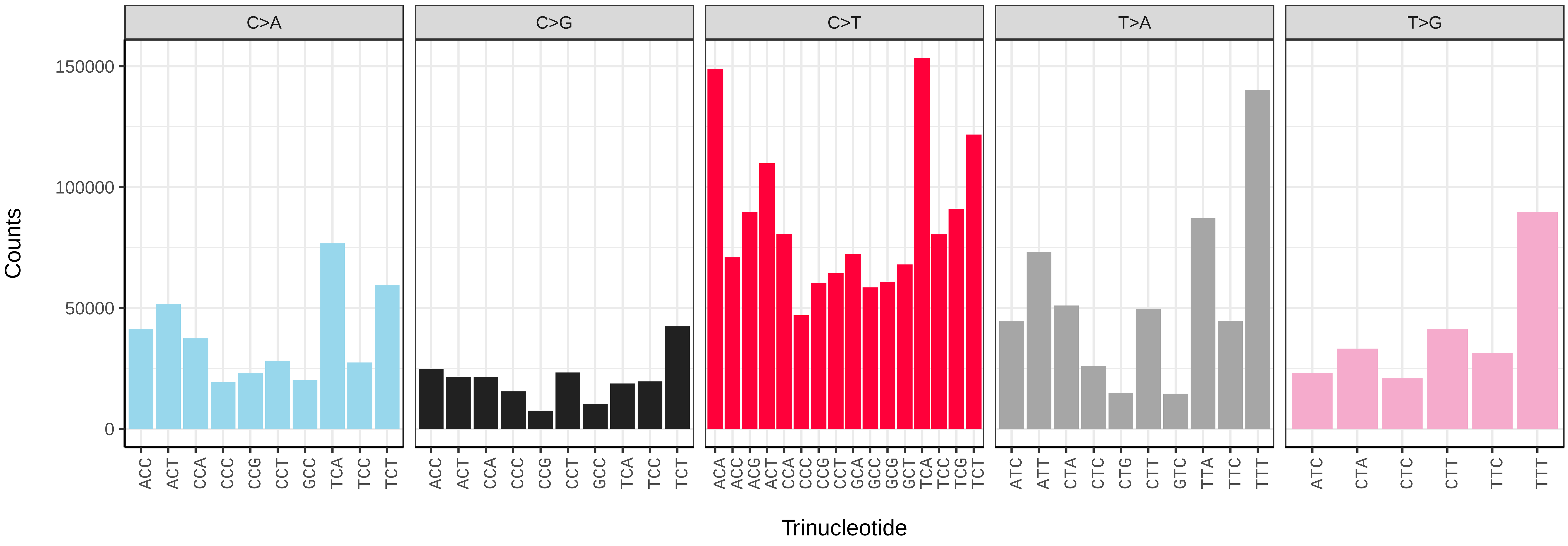


*Cydia strobilella* (ilCydStro3) somatic mutational spectrum (SBS96 classification)

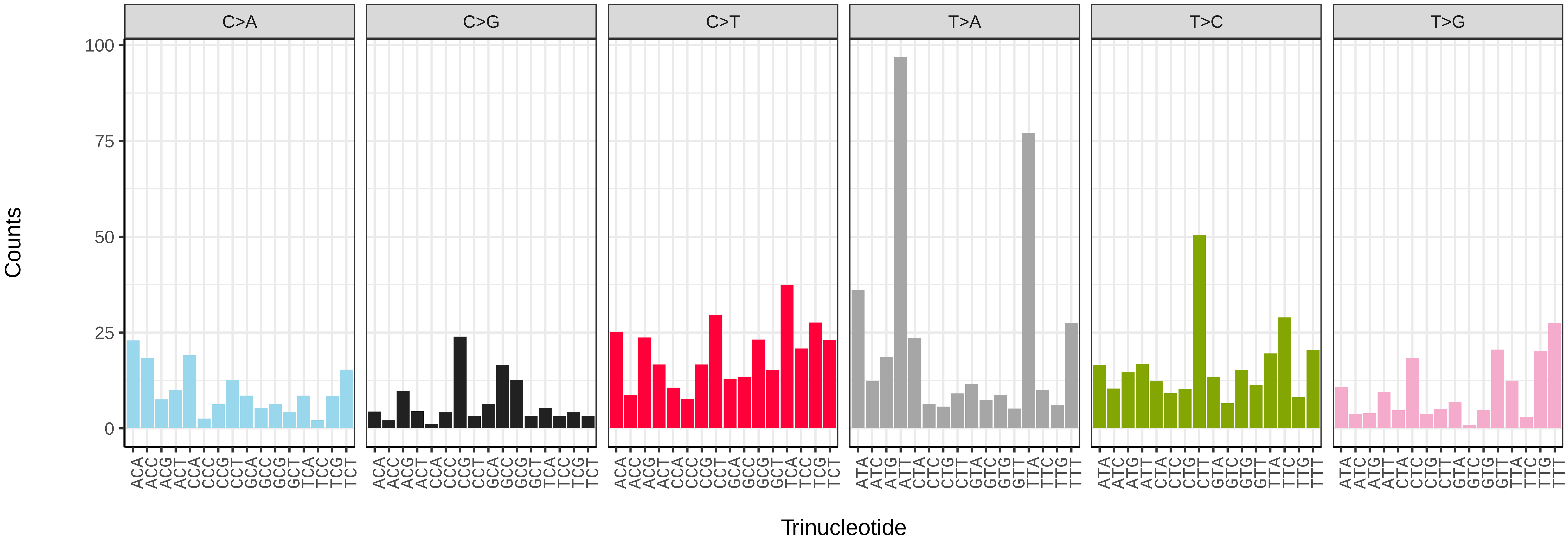


*Cydia strobilella* (ilCydStro3) somatic mutational spectrum (SBS52 classification)

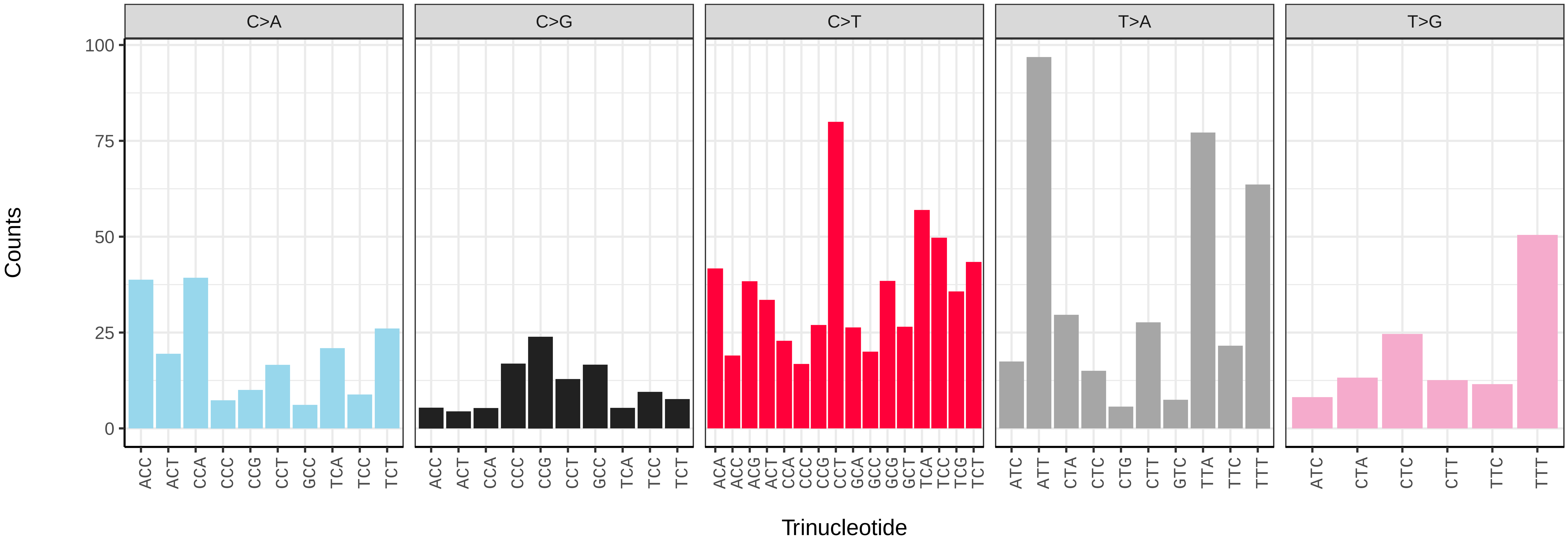

*Cydia strobilella* (ilCydStro3.ilCydStro1) germline mutational spectrum

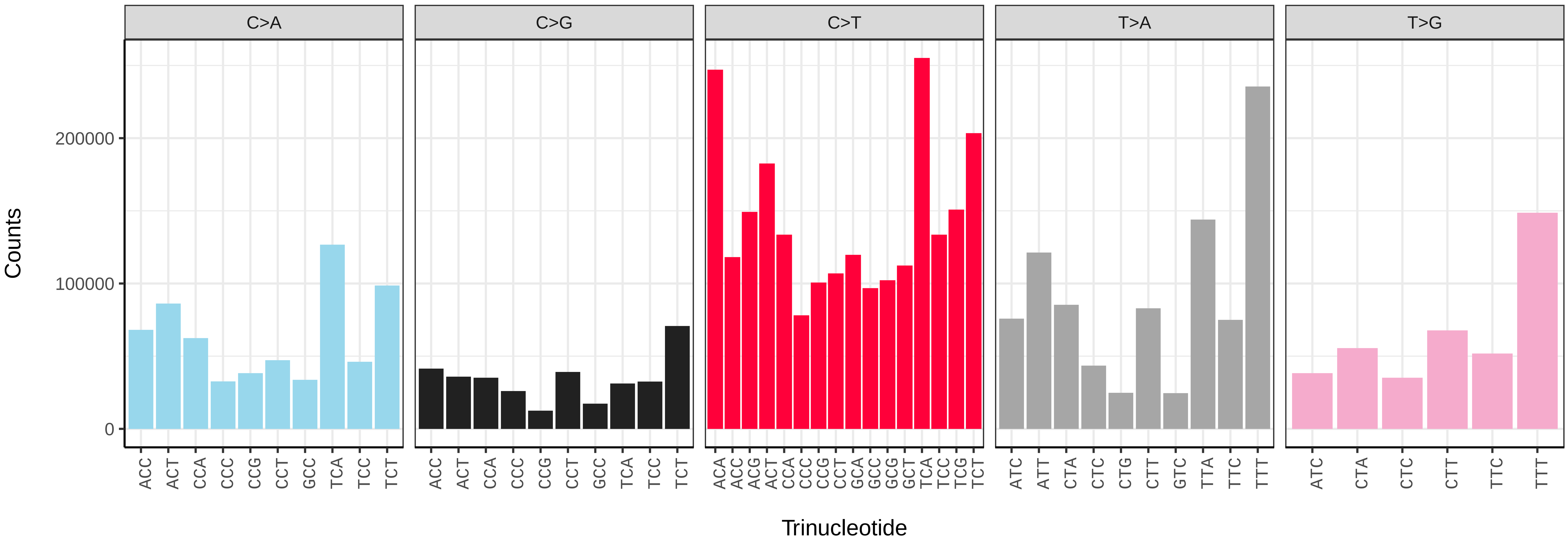


*Cydia strobilella* (ilCydStro3.ilCydStro1) somatic mutational spectrum (SBS96 classification)

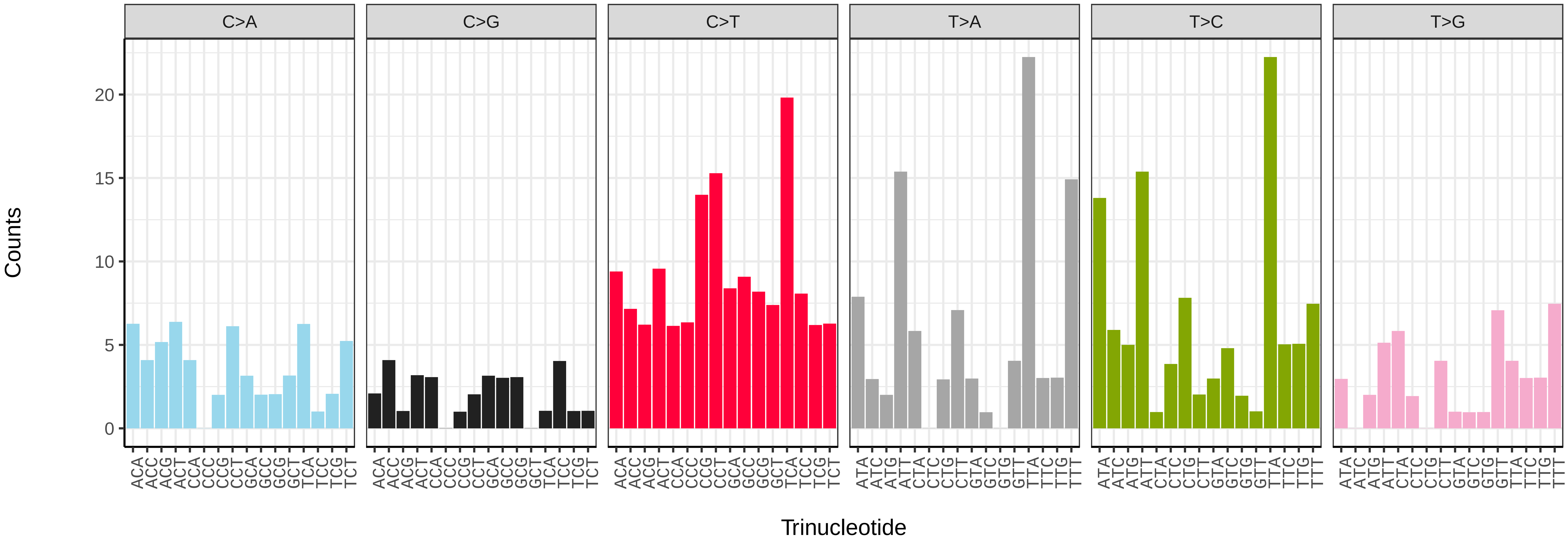


*Cydia strobilella* (ilCydStro3.ilCydStro1) somatic mutational spectrum (SBS52 classification)

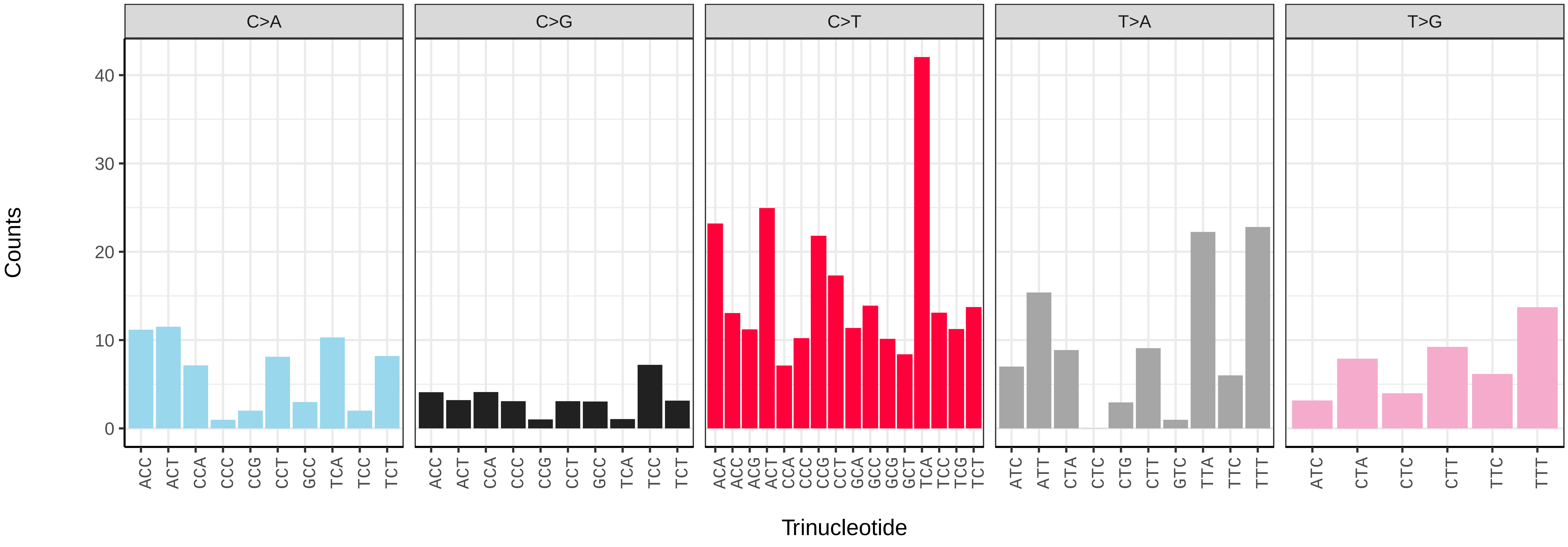

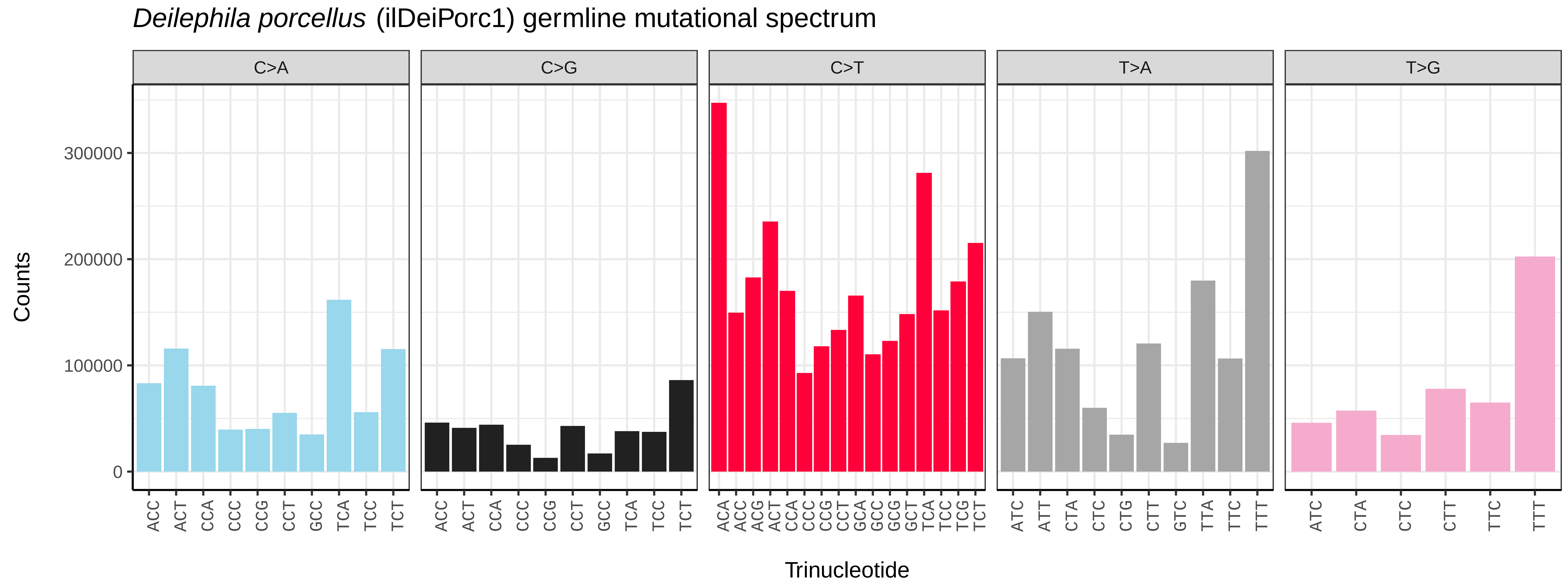
Deilephila porcellus (ilDeiPorc1) germline mutational spectrum
C>A
C>G
C>T
T>A
T>G
Counts
0
100000
200000
300000
ACC ACT CCA CCC CCG CCT GCC TCA TCC TCT
ACA ACC ACG ACT CCA CCC CCG CCT GCA GCC GCG GCT TCA TCC TCG TCT
ATC ATT CTA CTC CTG CTT GTC TTA TTC TTT
ATC CTA CTC CTT TTC TTT
Trinucleotide

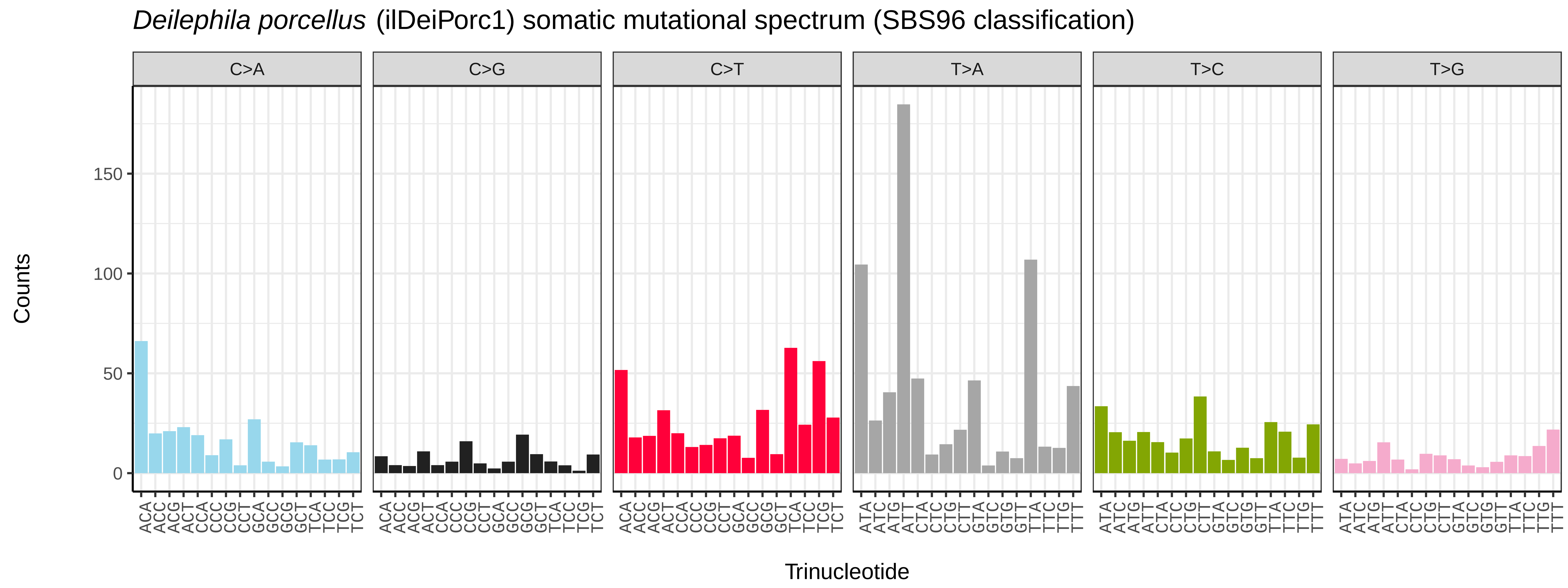
Deilephila porcellus (ilDeiPorc1) somatic mutational spectrum (SBS96 classification)
C>A
C>G
C>T
T>A
T>C
T>G
Counts
0
50
100
150
ACA ACC ACG ACT CCA CCC CCG CCT GCA GCC GCG GCT TCA TCC TCG TCT
ATA ATC ATG ATT CTA CTC CTG CTT GTA GTC GTG GTT TTA TTC TTG TTT
Trinucleotide

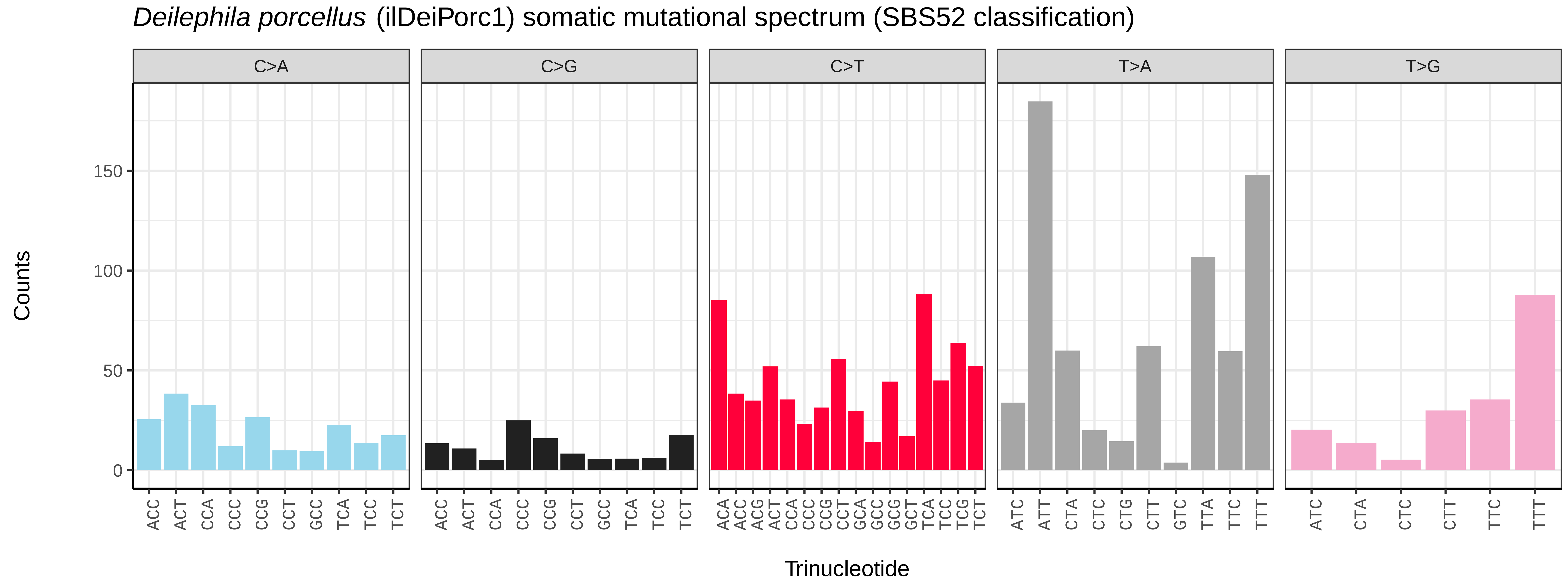
Deilephila porcellus (ilDeiPorc1) somatic mutational spectrum (SBS52 classification)
C>A
C>G
C>T
T>A
T>G
Counts
0
50
100
150
ACC ACT CCA CCC CCG CCT GCC TCA TCC TCT
ACA ACC ACG ACT CCA CCC CCG CCT GCA GCC GCG GCT TCA TCC TCG TCT
ATC ATT CTA CTC CTG CTT GTC TTA TTC TTT
ATC CTA CTC CTT TTC TTT
Trinucleotide

*Diachrysia chrysitis* (ilDiaChry1) germline mutational spectrum

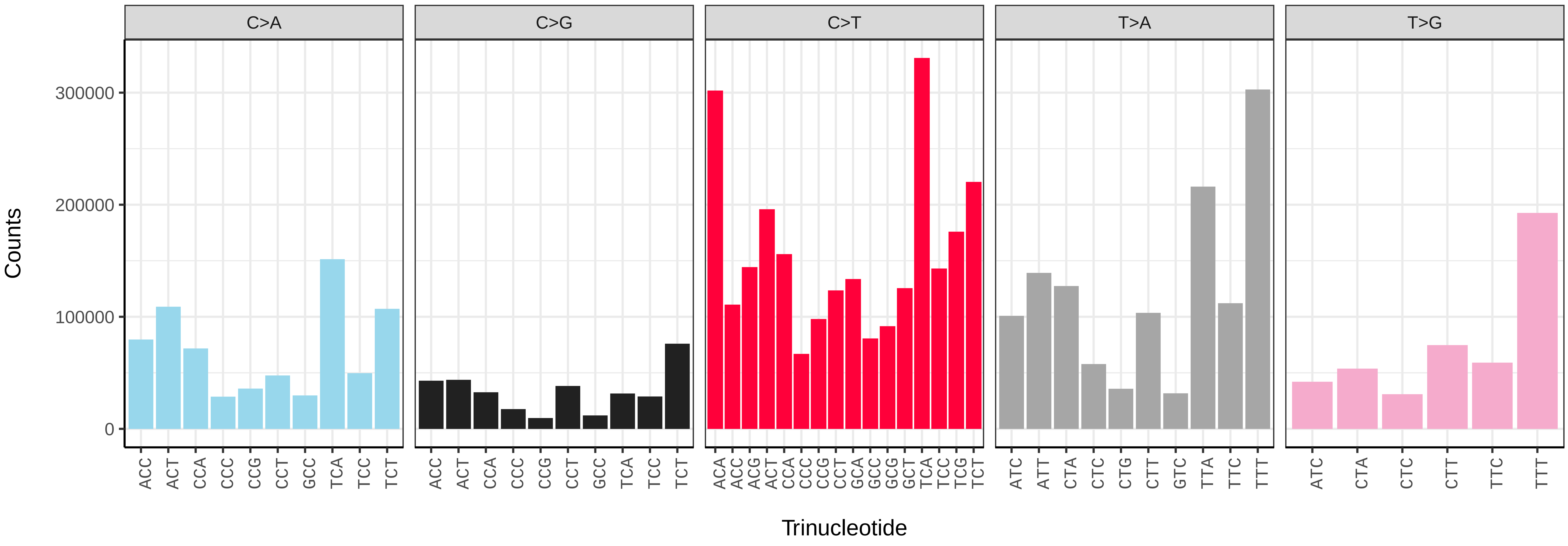


*Diachrysia chrysitis* (ilDiaChry1) somatic mutational spectrum (SBS96 classification)

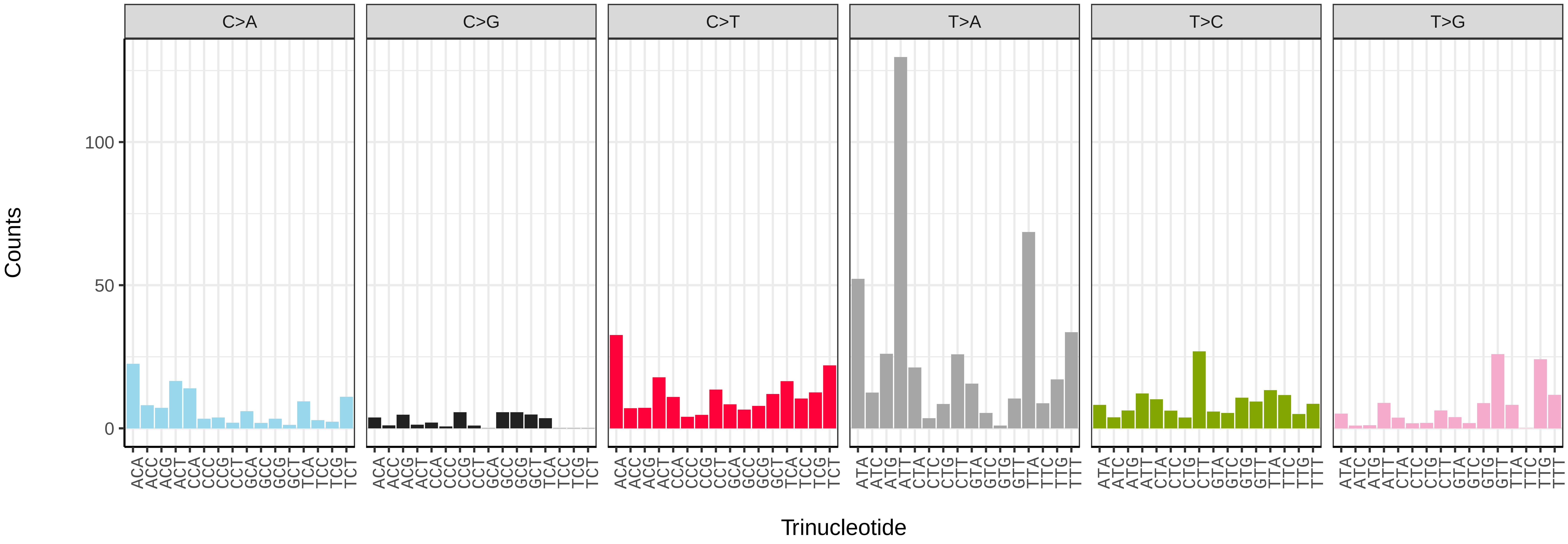


*Diachrysia chrysitis* (ilDiaChry1) somatic mutational spectrum (SBS52 classification)

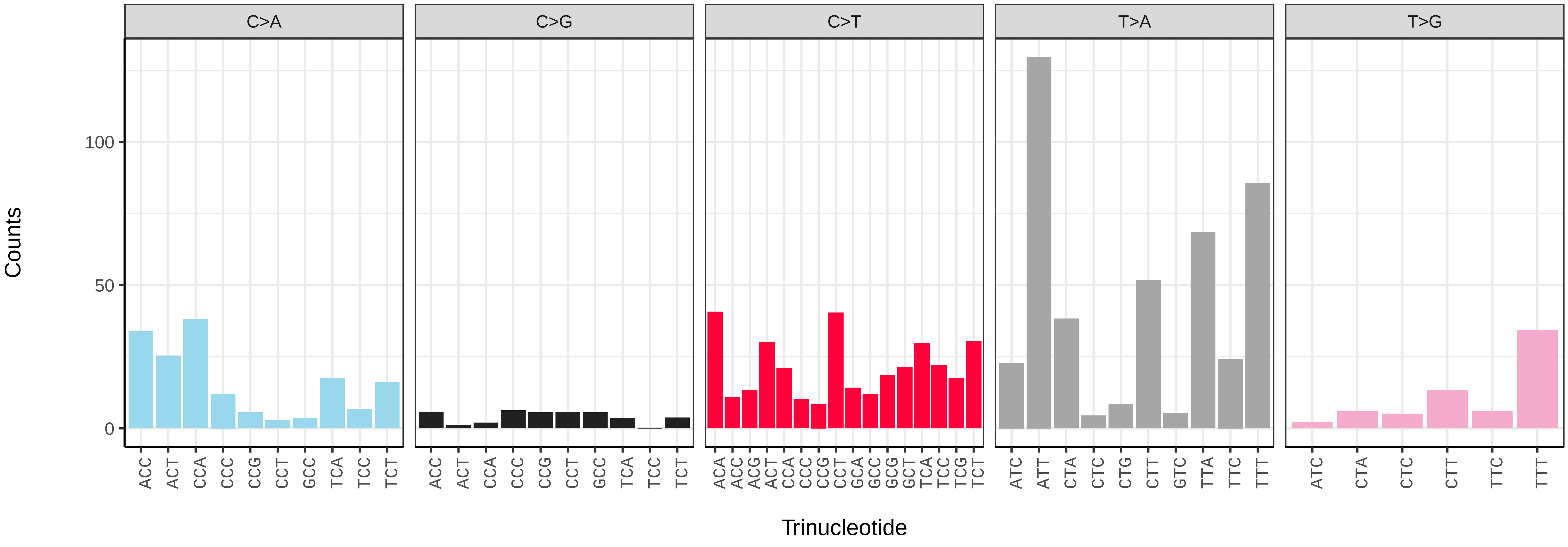

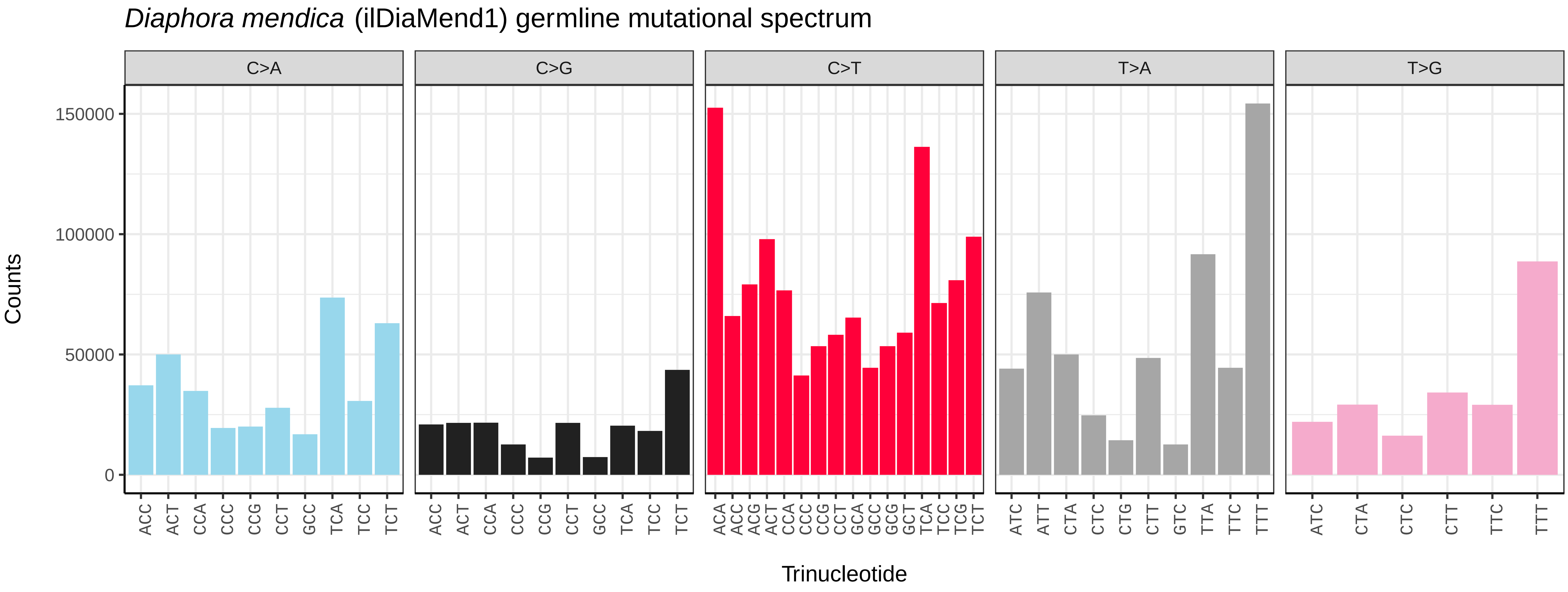
*Diaphora mendica* (ilDiaMend1) germline mutational spectrum
C>A
C>G
C>T
T>A
T>G
Counts
0
50000
100000
150000
ACC ACT CCA CCC CCG CCT GCC TCA TCC TCT
ACA ACC ACG ACT CCA CCC CCG CCT GCA GCC GCG GCT TCA TCC TCG TCT
ATC ATT CTA CTC CTG CTT GTC TTA TTC TTT
ATC CTA CTC CTT TTC TTT
Trinucleotide

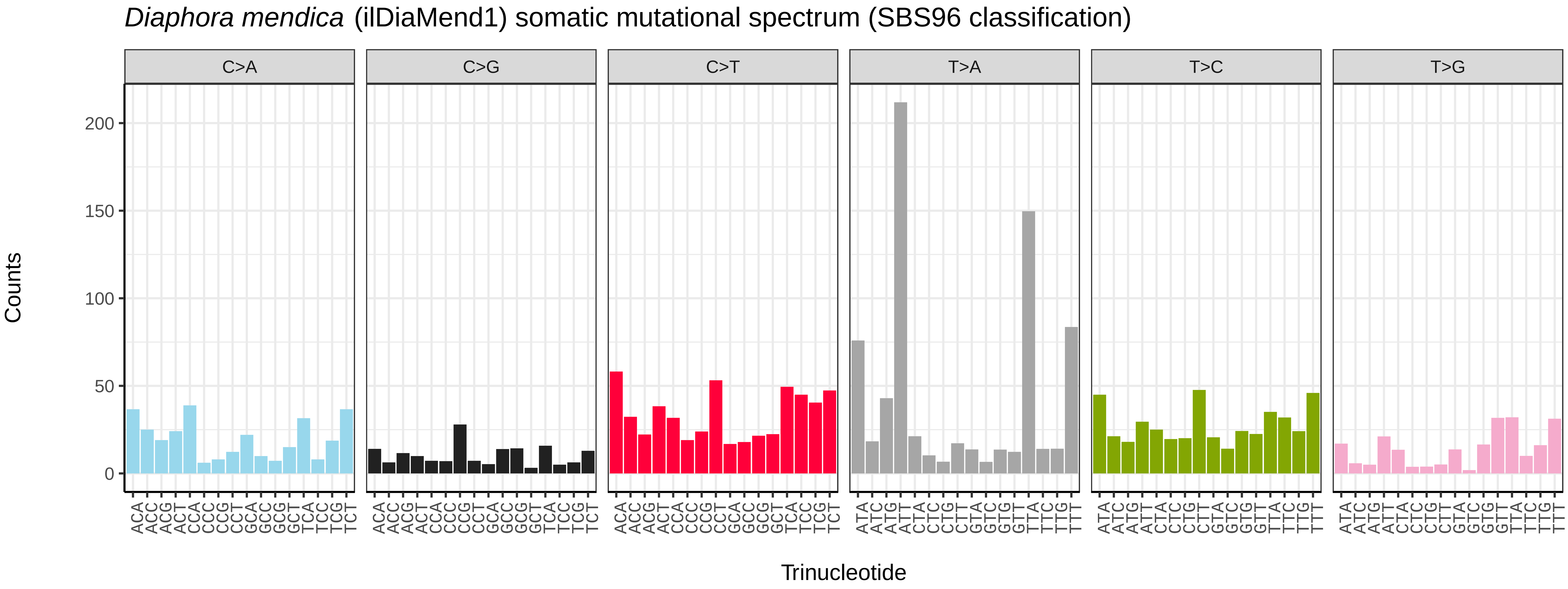
*Diaphora mendica* (ilDiaMend1) somatic mutational spectrum (SBS96 classification)
C>A
C>G
C>T
T>A
T>C
T>G
Counts
0
50
100
150
200
ACA ACC ACG ACT CCA CCC CCG CCT GCA GCC GCG GCT TCA TCC TCG TCT
ATA ATC ATG ATT CTA CTC CTG CTT GTA GTC GTG GTT TTA TTC TTG TTT
Trinucleotide

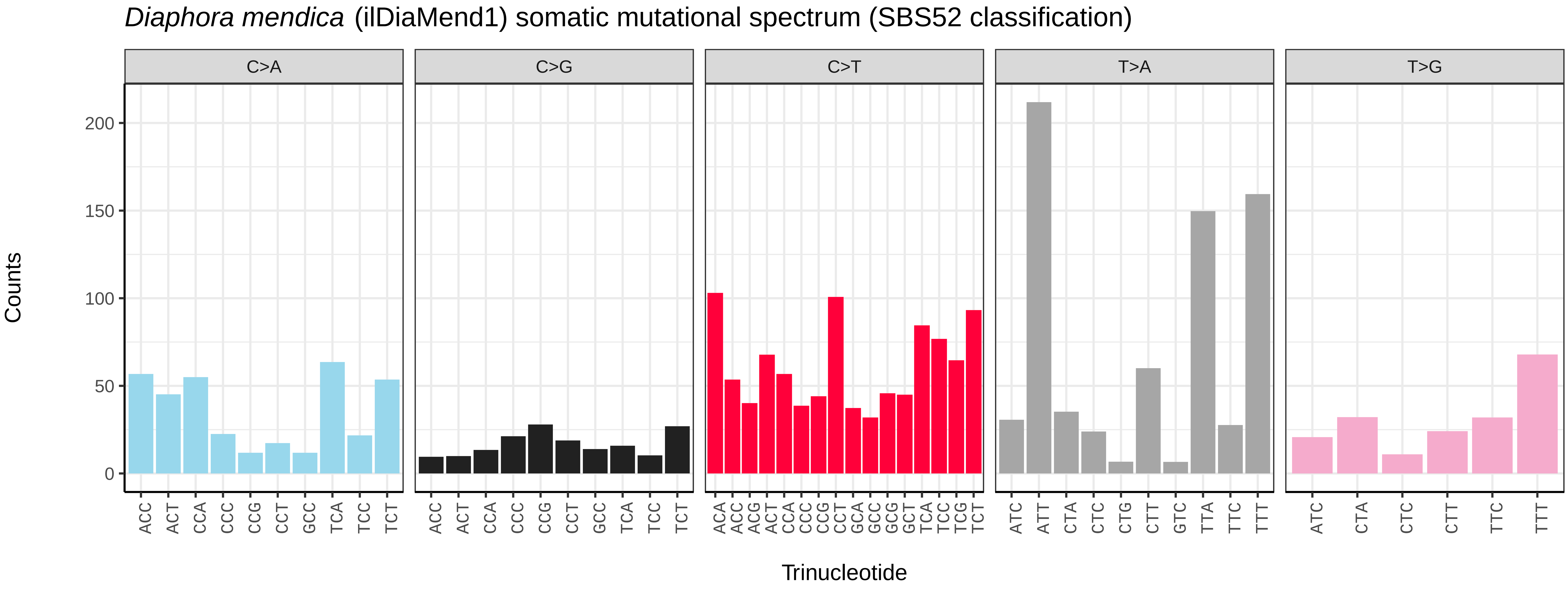
*Diaphora mendica* (ilDiaMend1) somatic mutational spectrum (SBS52 classification)
C>A
C>G
C>T
T>A
T>G
Counts
0
50
100
150
200
ACC ACT CCA CCC CCG CCT GCC TCA TCC TCT
ACA ACC ACG ACT CCA CCC CCG CCT GCA GCC GCG GCT TCA TCC TCG TCT
ATC ATT CTA CTC CTG CTT GTC TTA TTC TTT
ATC CTA CTC CTT TTC TTT
Trinucleotide

*Diarsia rubi* (ilDiaRubi1) germline mutational spectrum

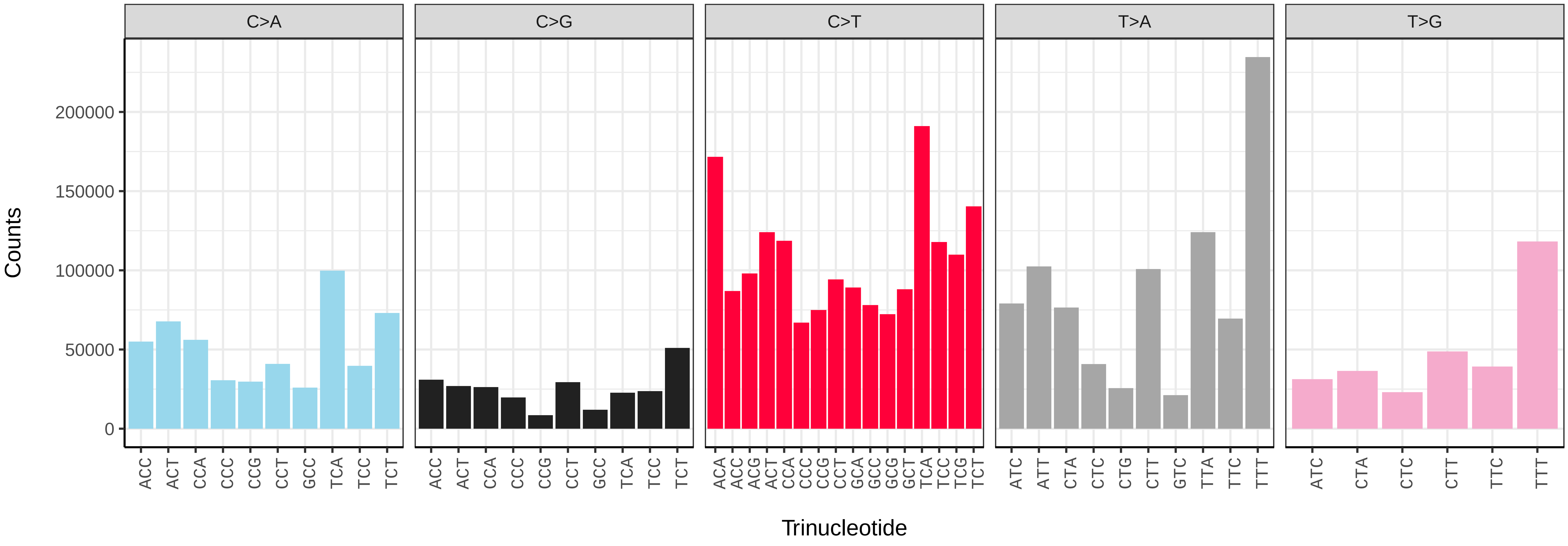


*Diarsia rubi* (ilDiaRubi1) somatic mutational spectrum (SBS96 classification)

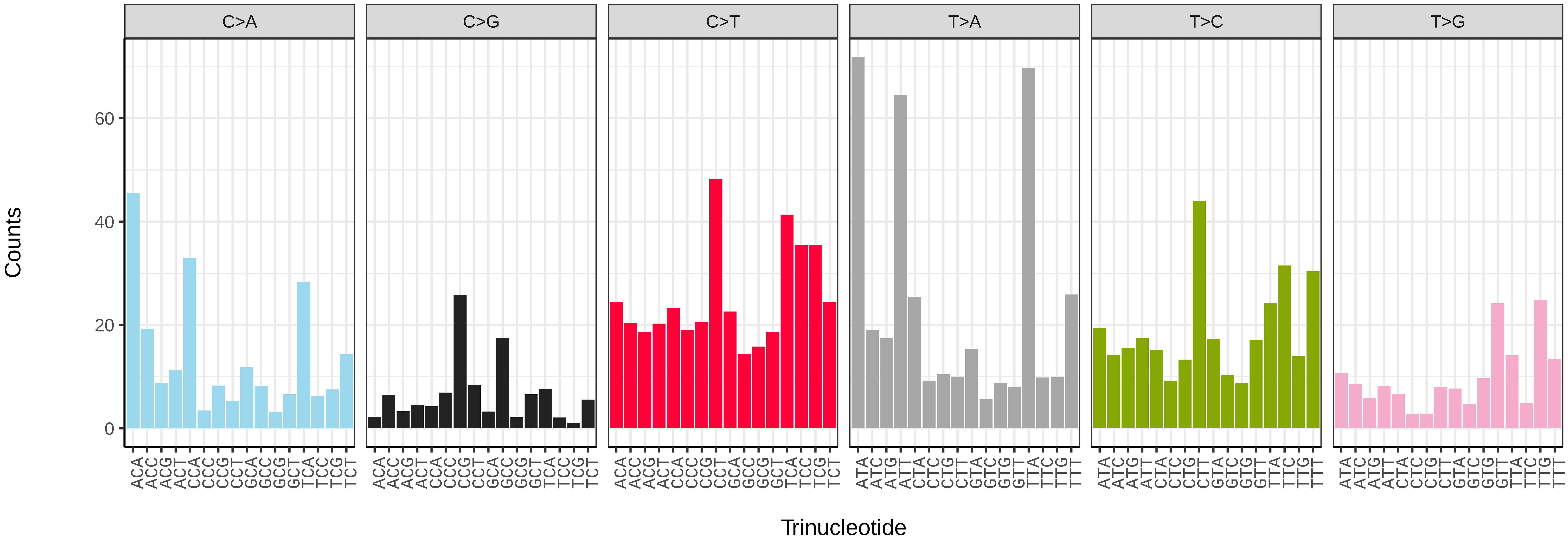


*Diarsia rubi* (ilDiaRubi1) somatic mutational spectrum (SBS52 classification)

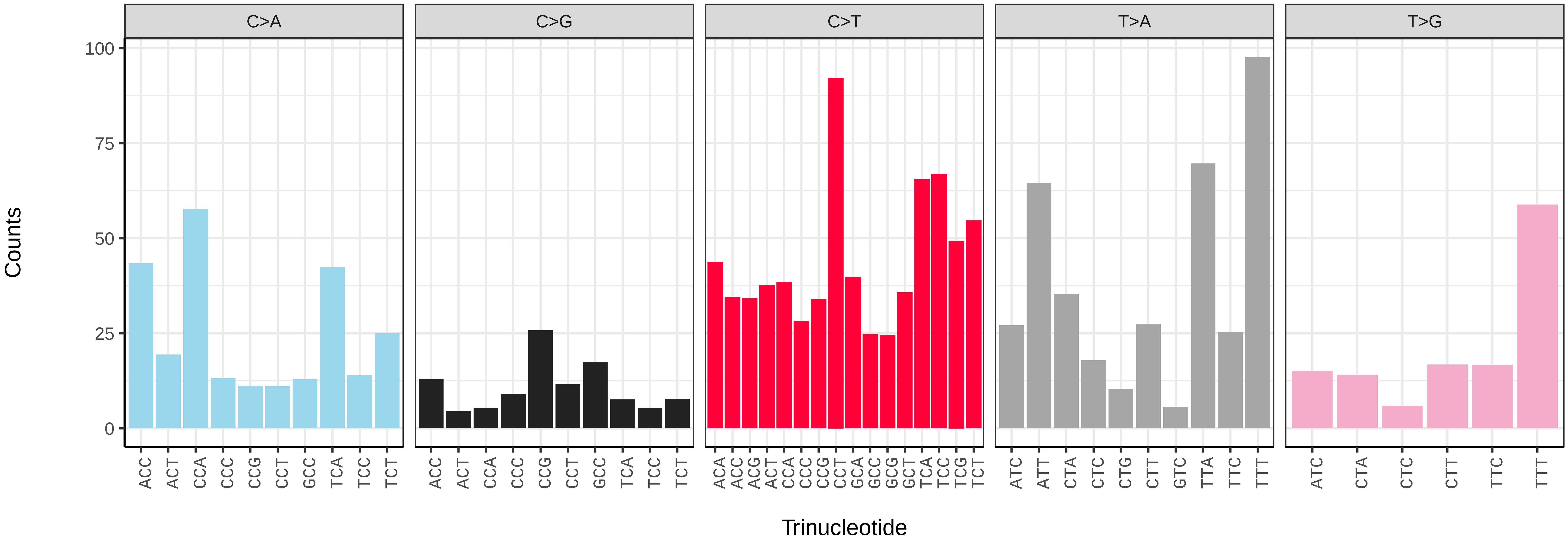

*Dicycla oo* (ilDicOoxx2) germline mutational spectrum

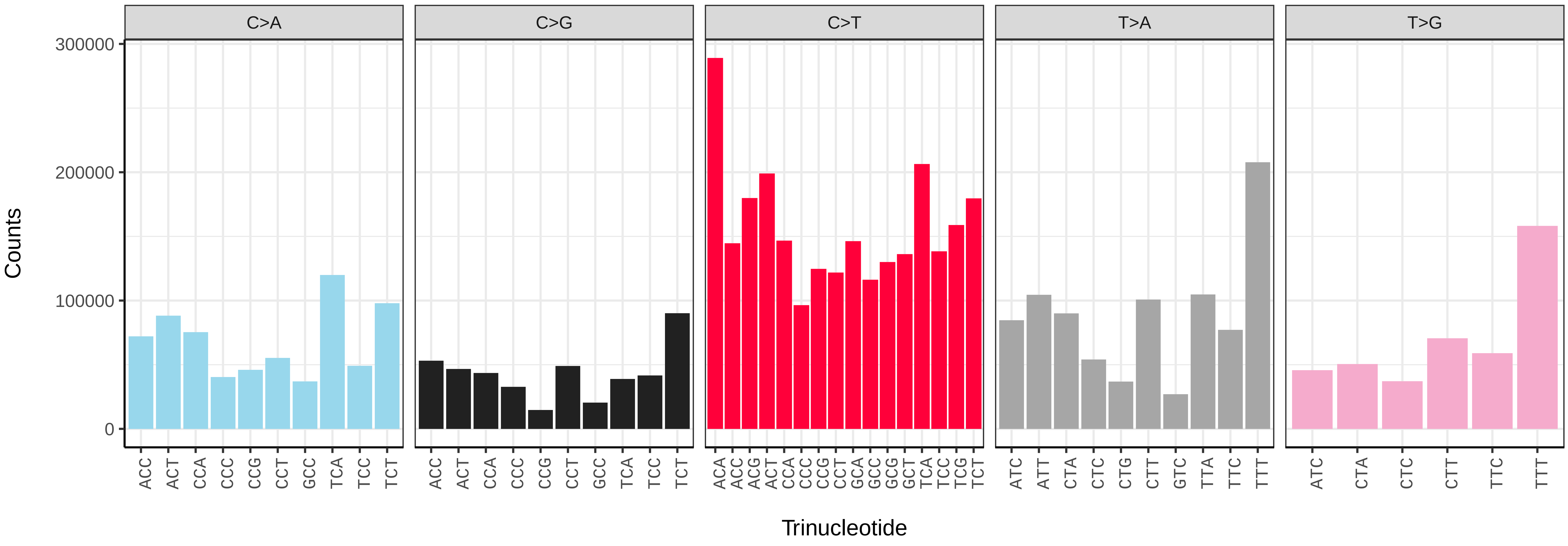


*Dicycla oo* (ilDicOoxx2) somatic mutational spectrum (SBS96 classification)

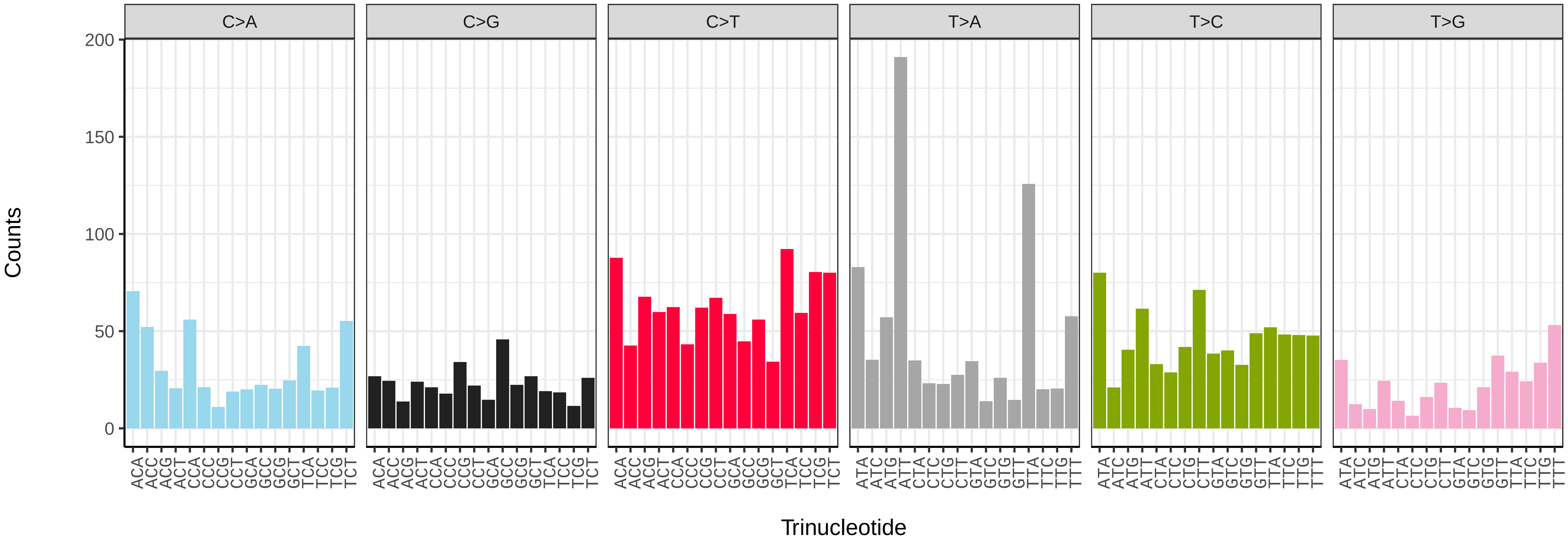


*Dicycla oo* (ilDicOoxx2) somatic mutational spectrum (SBS52 classification)

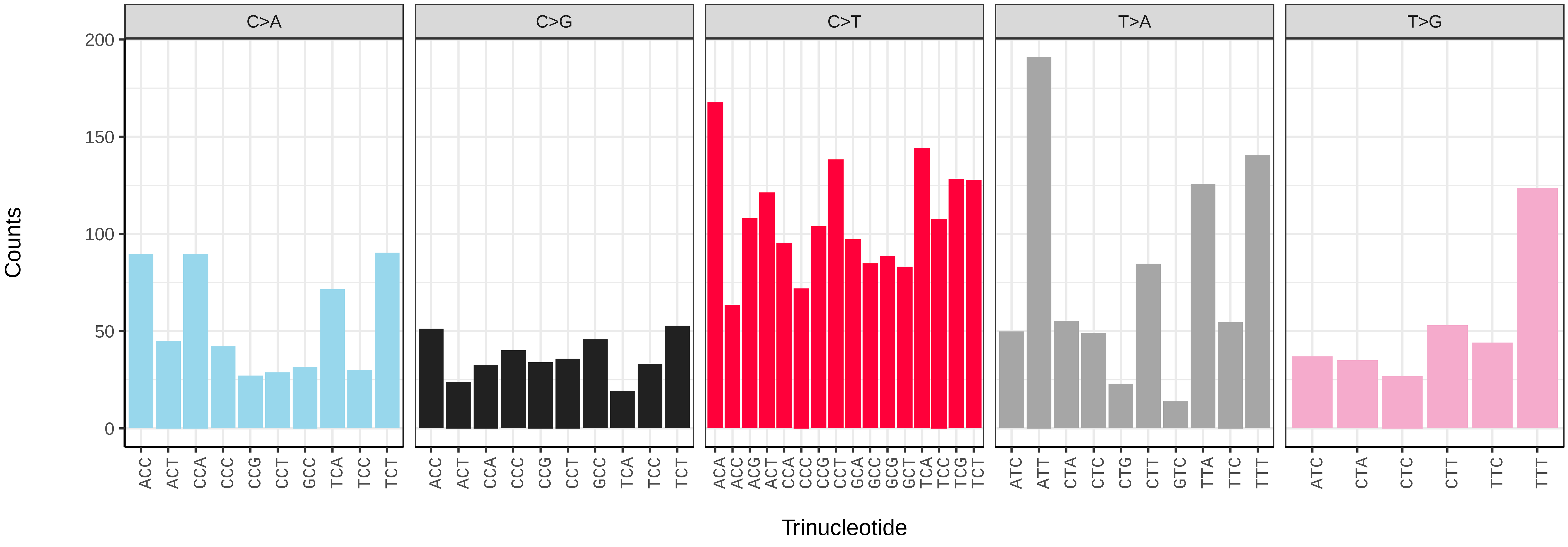

*Diloba caeruleocephala* (ilDilCaer1) germline mutational spectrum

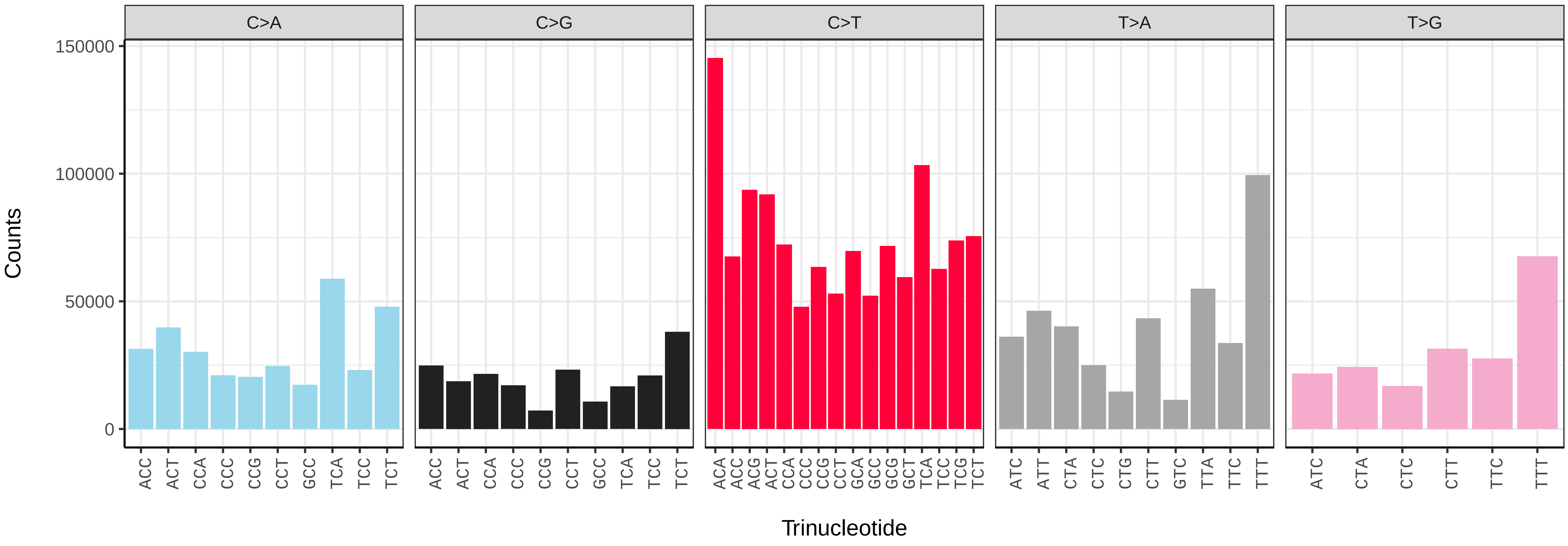


*Diloba caeruleocephala* (ilDilCaer1) somatic mutational spectrum (SBS96 classification)

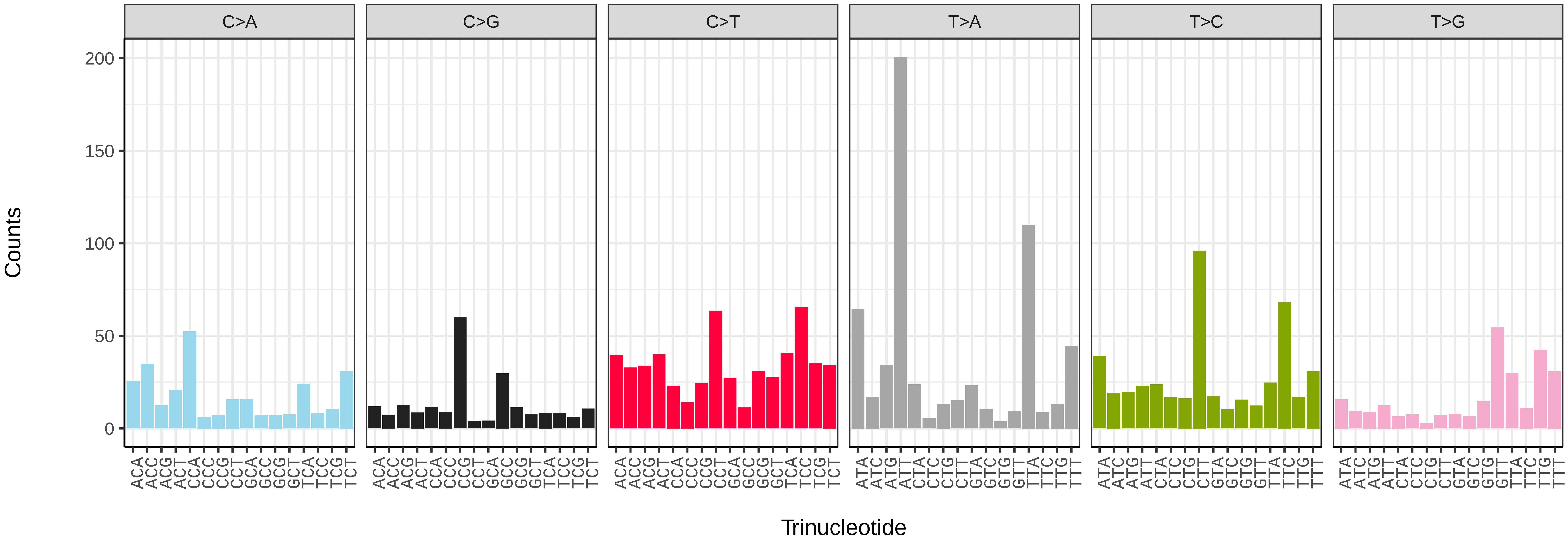


*Diloba caeruleocephala* (ilDilCaer1) somatic mutational spectrum (SBS52 classification)

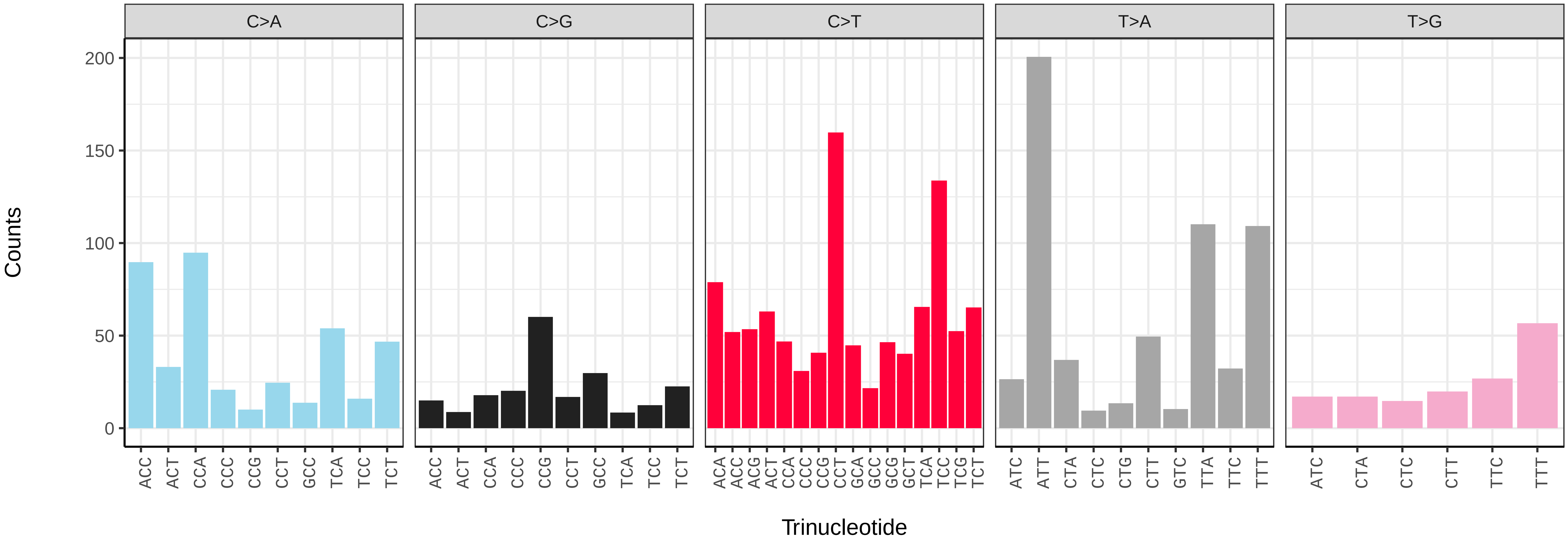

*Drepana falcataria* (ilDreFalc1) germline mutational spectrum

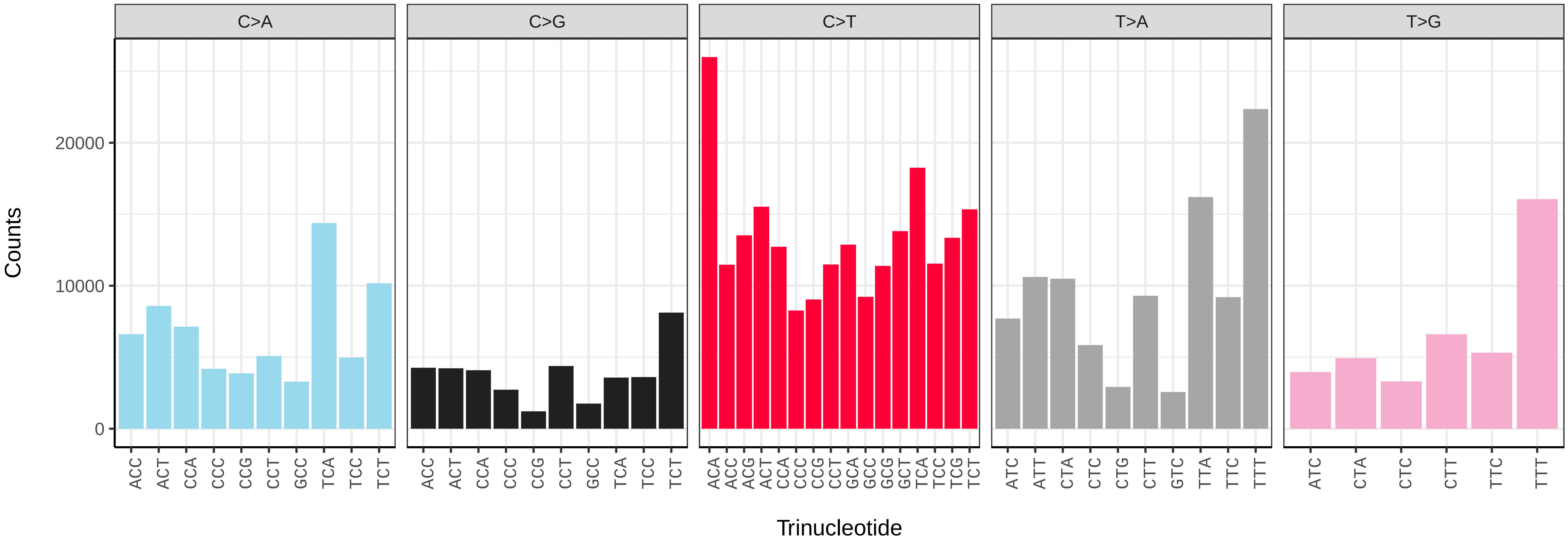


*Drepana falcataria* (ilDreFalc1) somatic mutational spectrum (SBS96 classification)

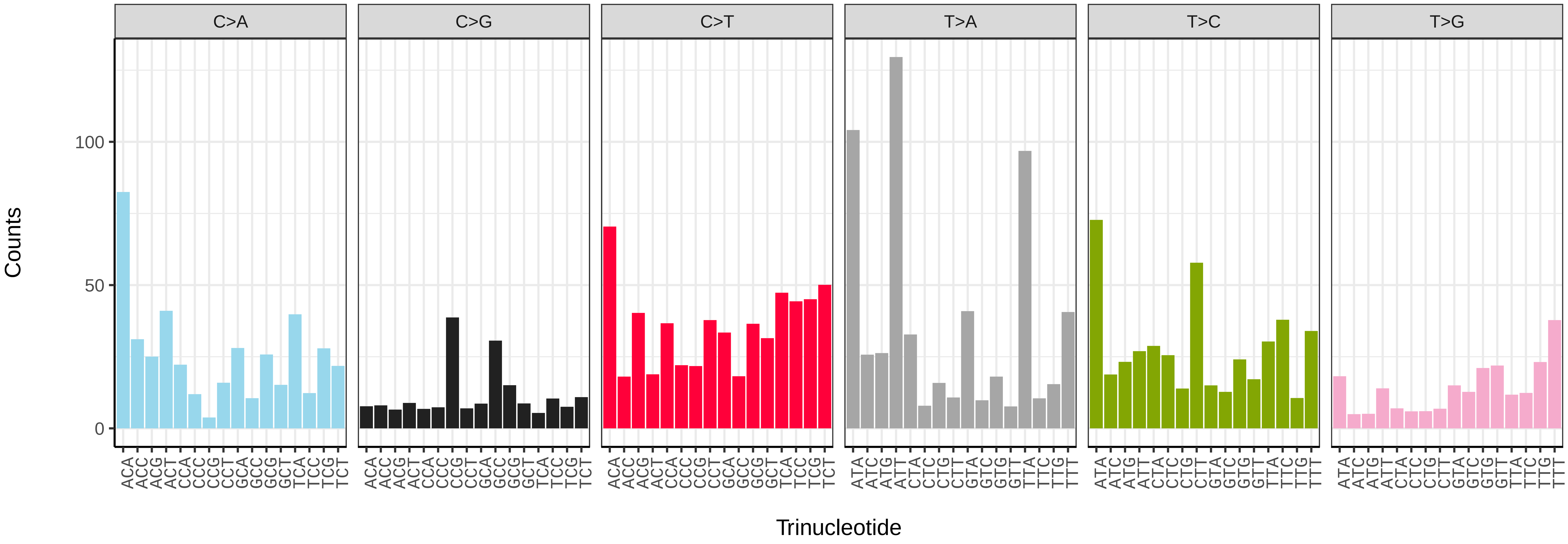


*Drepana falcataria* (ilDreFalc1) somatic mutational spectrum (SBS52 classification)

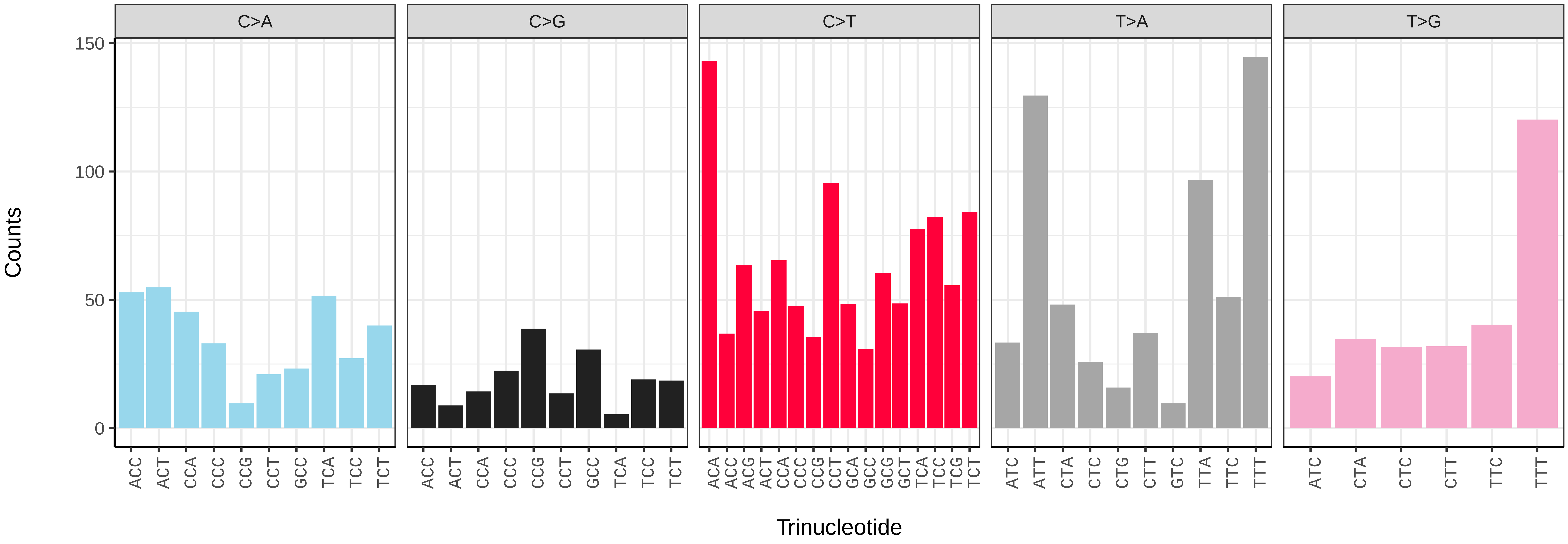

*Dryobotodes eremita* (ilDryErem1) germline mutational spectrum

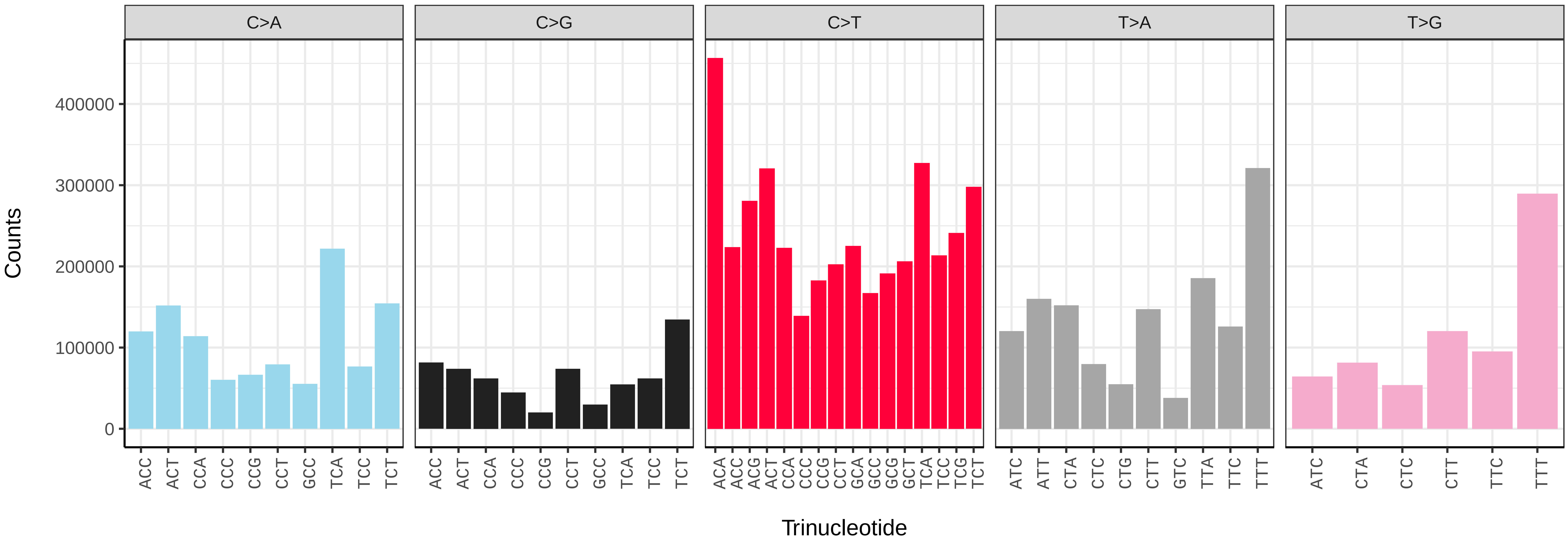


*Dryobotodes eremita* (ilDryErem1) somatic mutational spectrum (SBS96 classification)

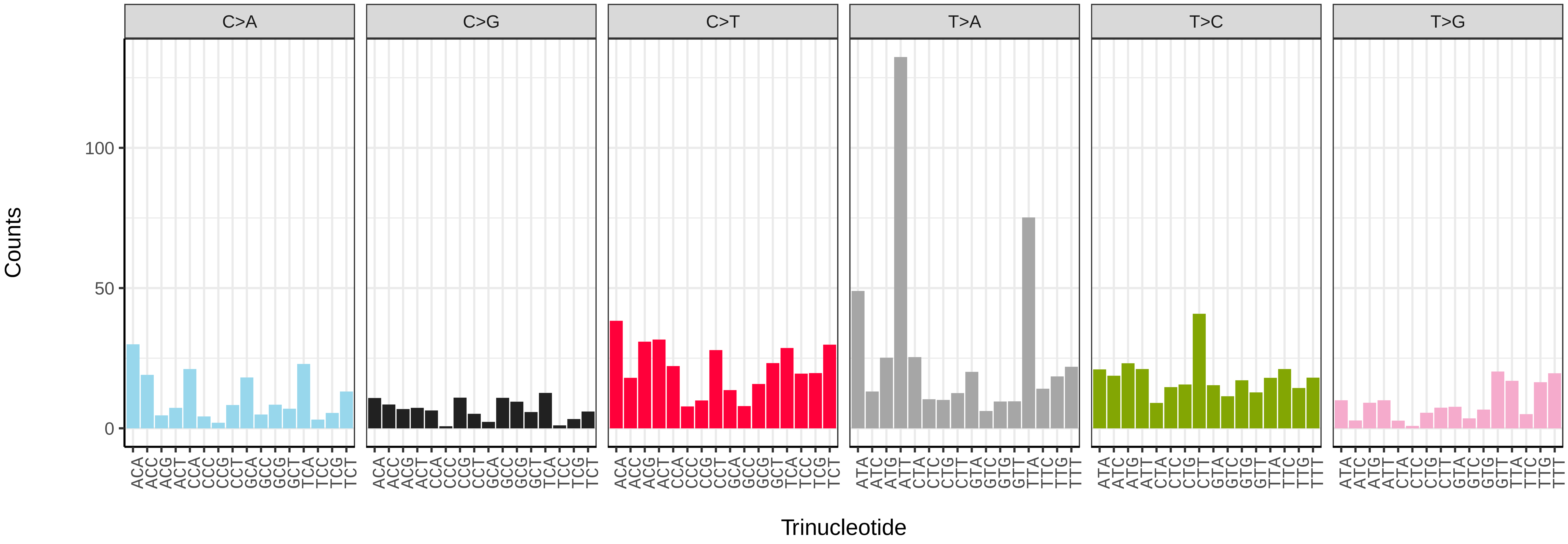


*Dryobotodes eremita* (ilDryErem1) somatic mutational spectrum (SBS52 classification)

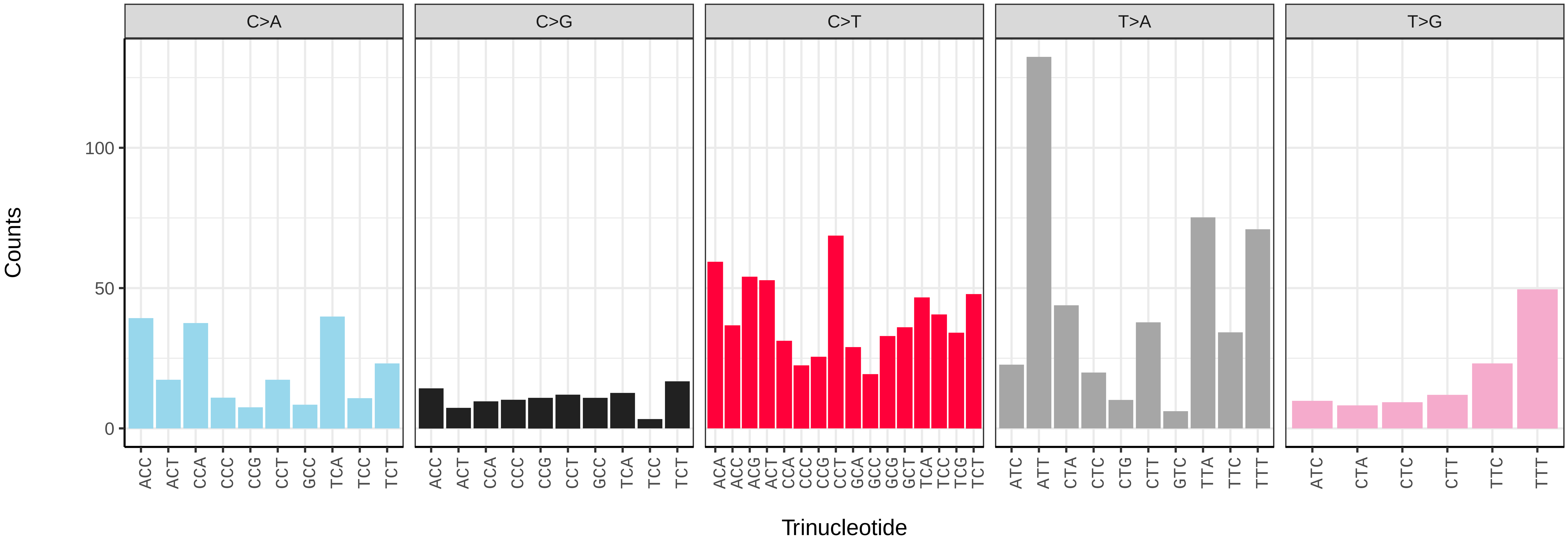

*Dryobota labecula* (ilDryLabe1) germline mutational spectrum

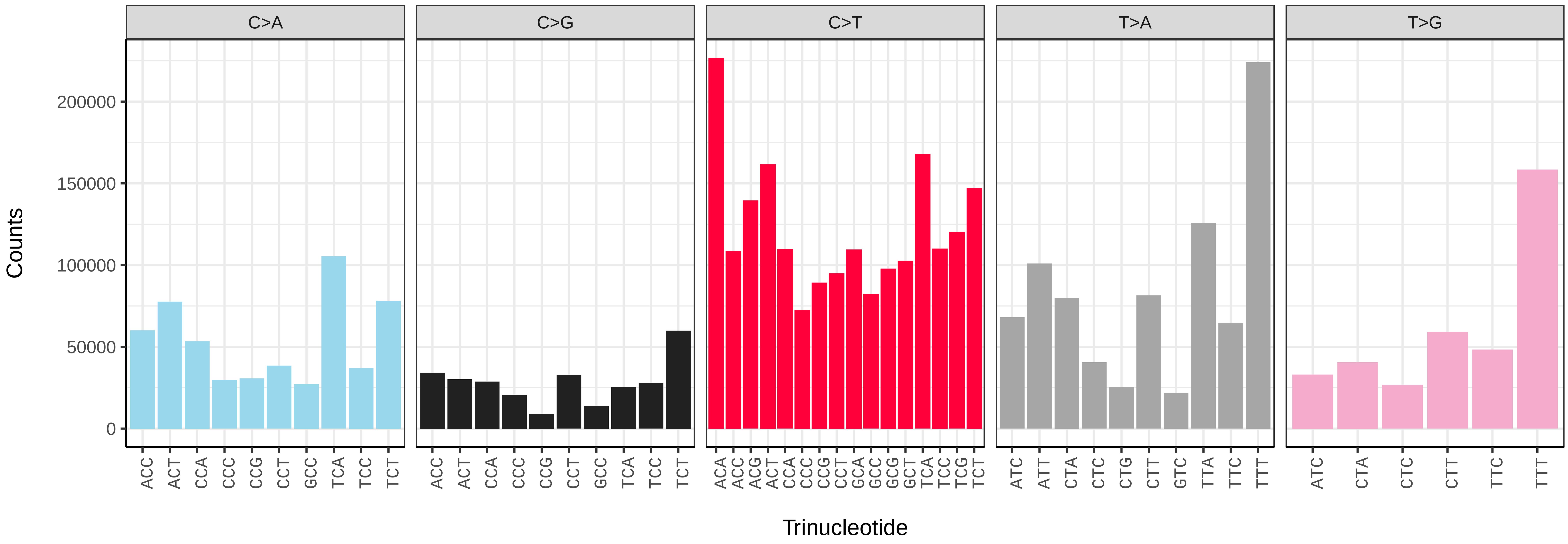


*Dryobota labecula* (ilDryLabe1) somatic mutational spectrum (SBS96 classification)

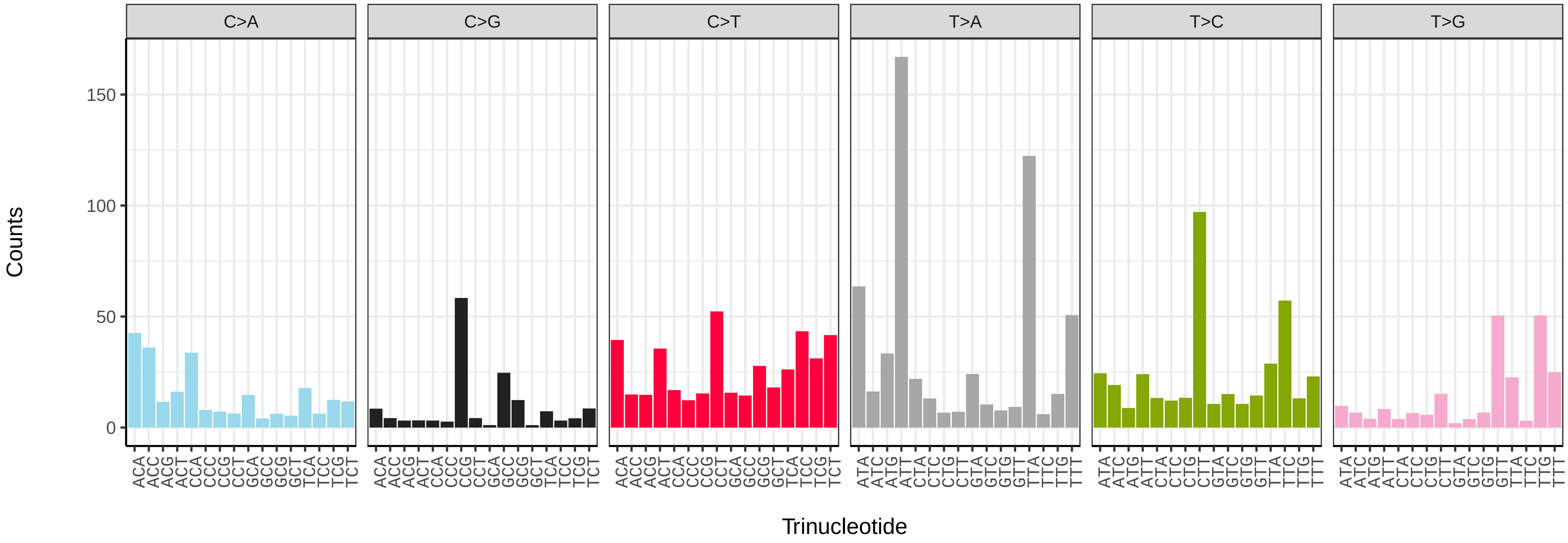


*Dryobota labecula* (ilDryLabe1) somatic mutational spectrum (SBS52 classification)

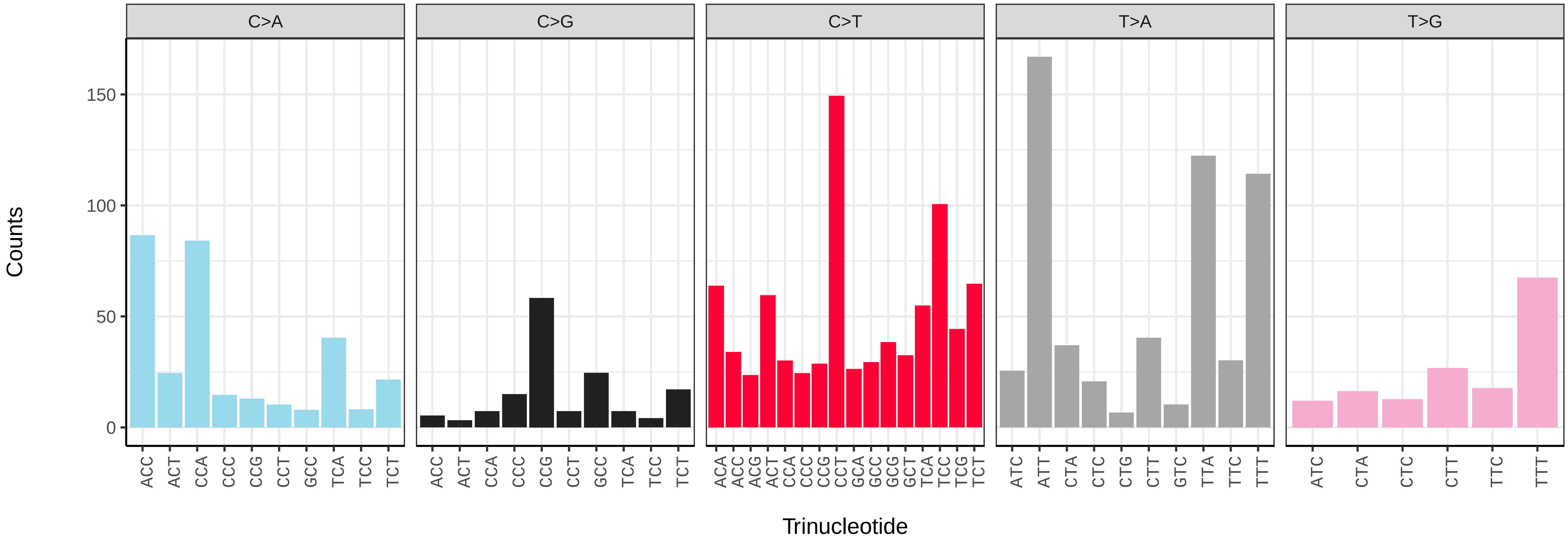

*Drymonia ruficornis* (ilDryRufi1) germline mutational spectrum

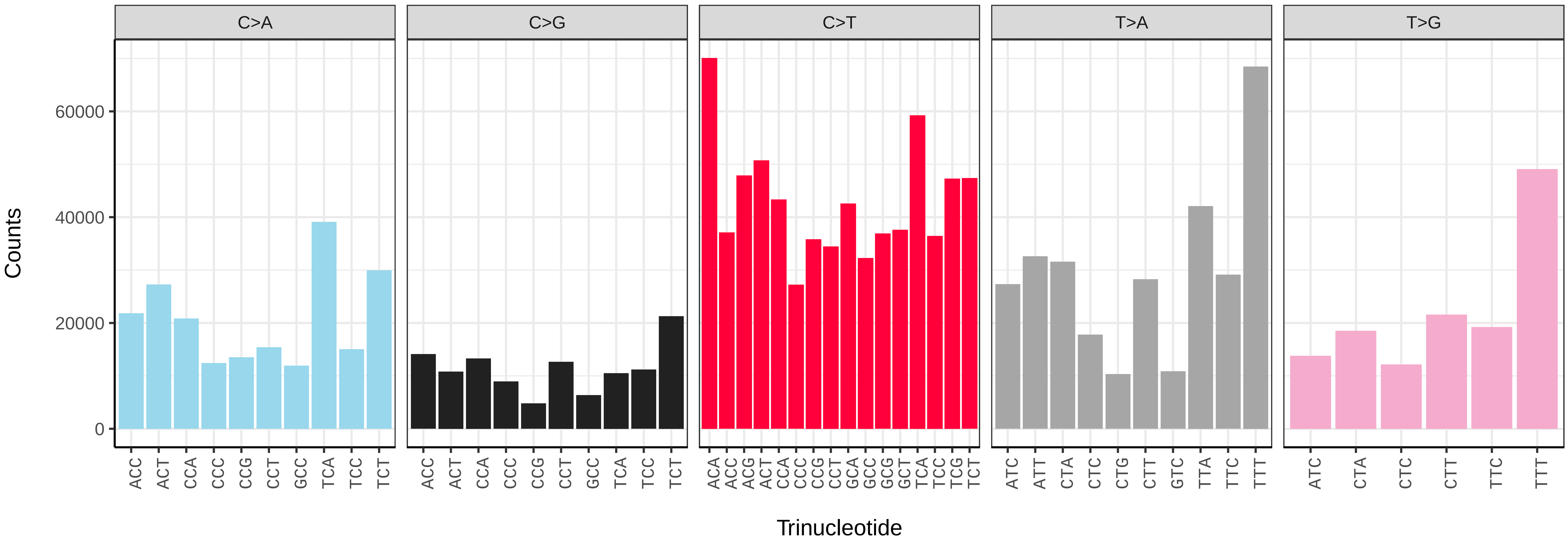


*Drymonia ruficornis* (ilDryRufi1) somatic mutational spectrum (SBS96 classification)

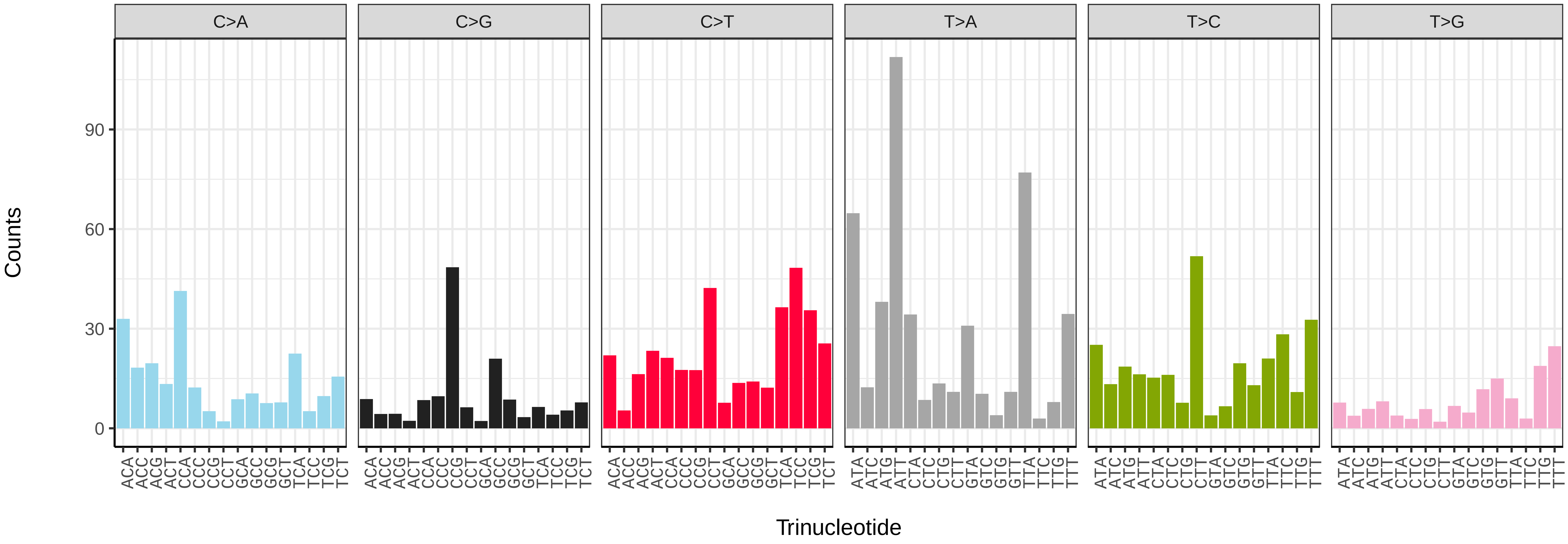


*Drymonia ruficornis* (ilDryRufi1) somatic mutational spectrum (SBS52 classification)

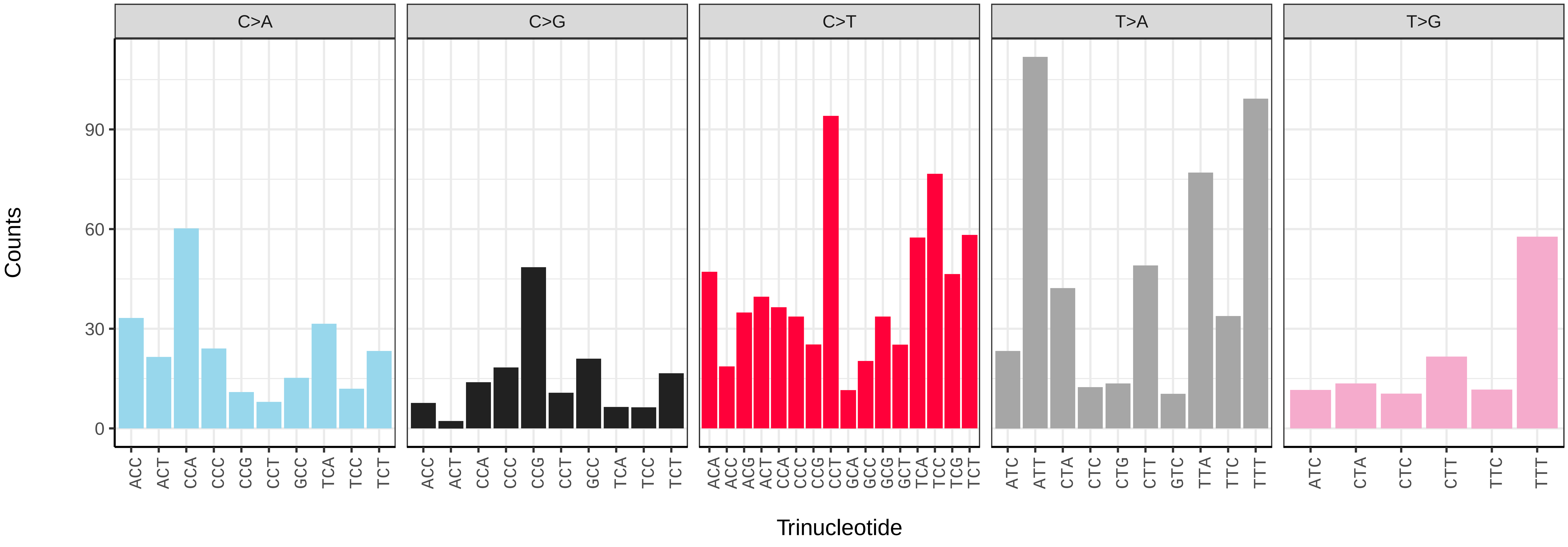

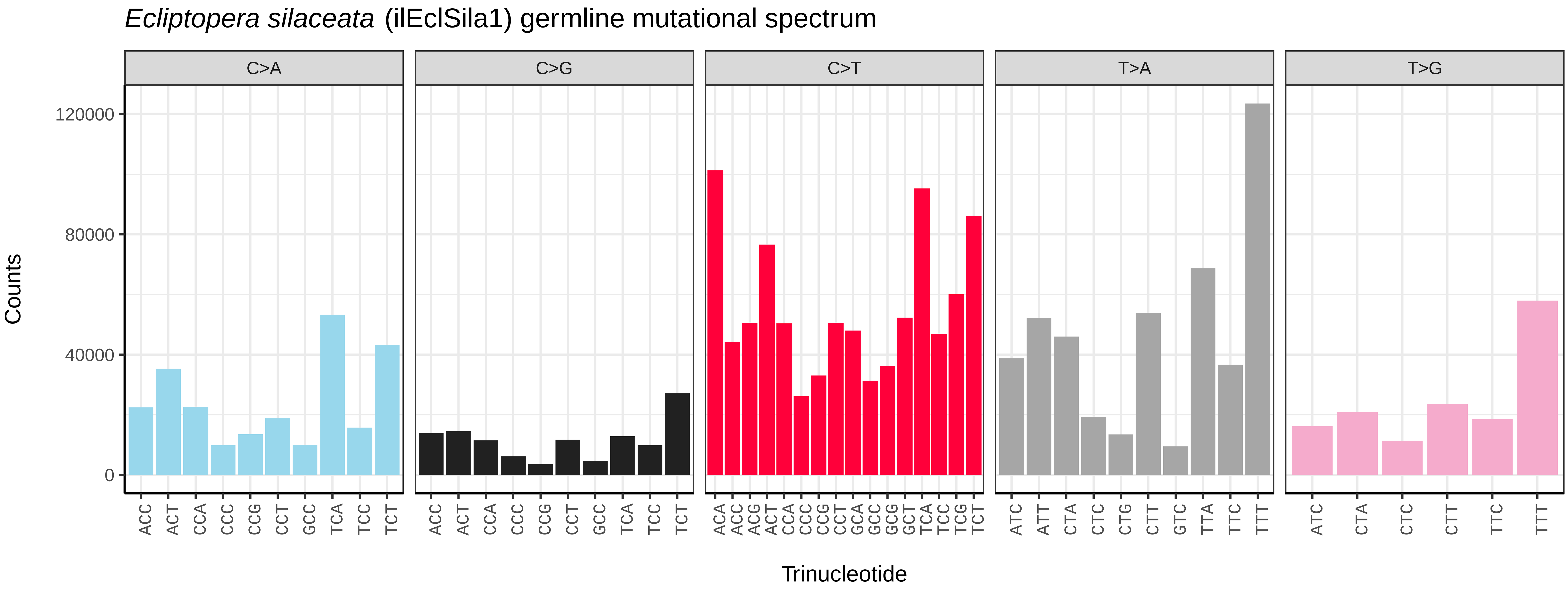
*Ecliptopera silaceata* (ilEclSila1) germline mutational spectrum
C>A
C>G
C>T
T>A
T>G
Counts
0
40000
80000
120000
Trinucleotide

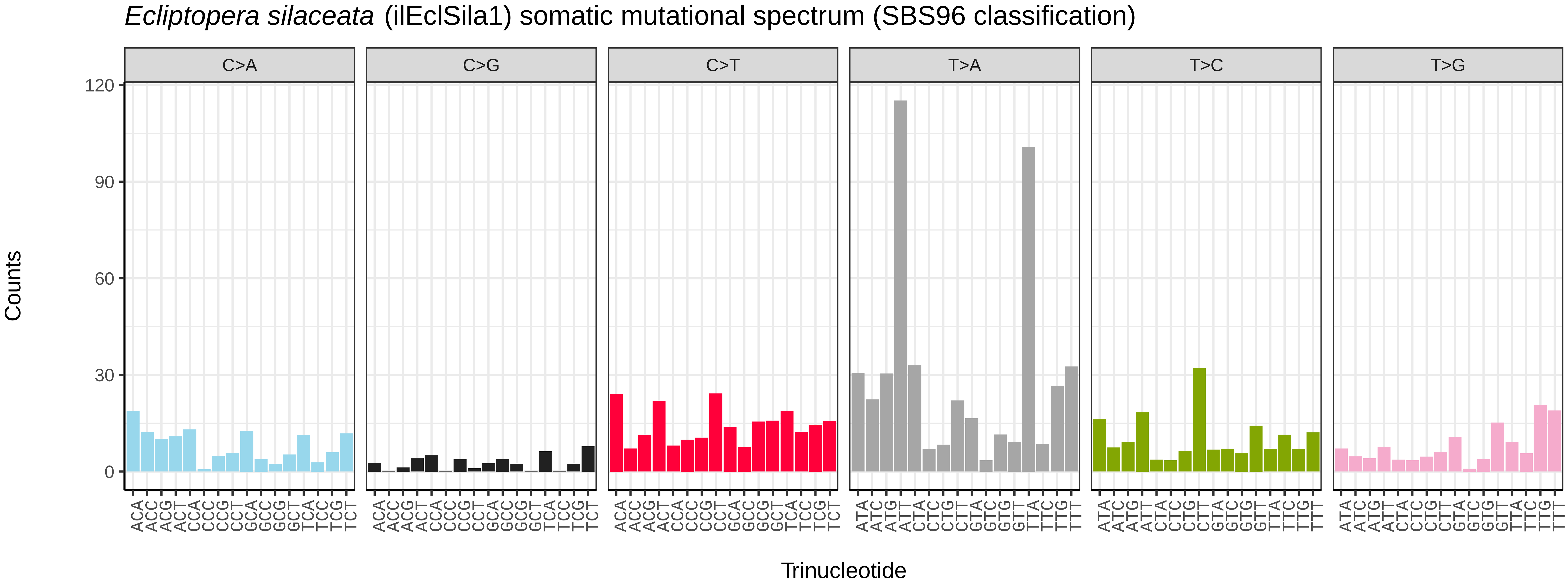
*Ecliptopera silaceata* (ilEclSila1) somatic mutational spectrum (SBS96 classification)
C>A
C>G
C>T
T>A
T>C
T>G
Counts
0
30
60
90
120
Trinucleotide

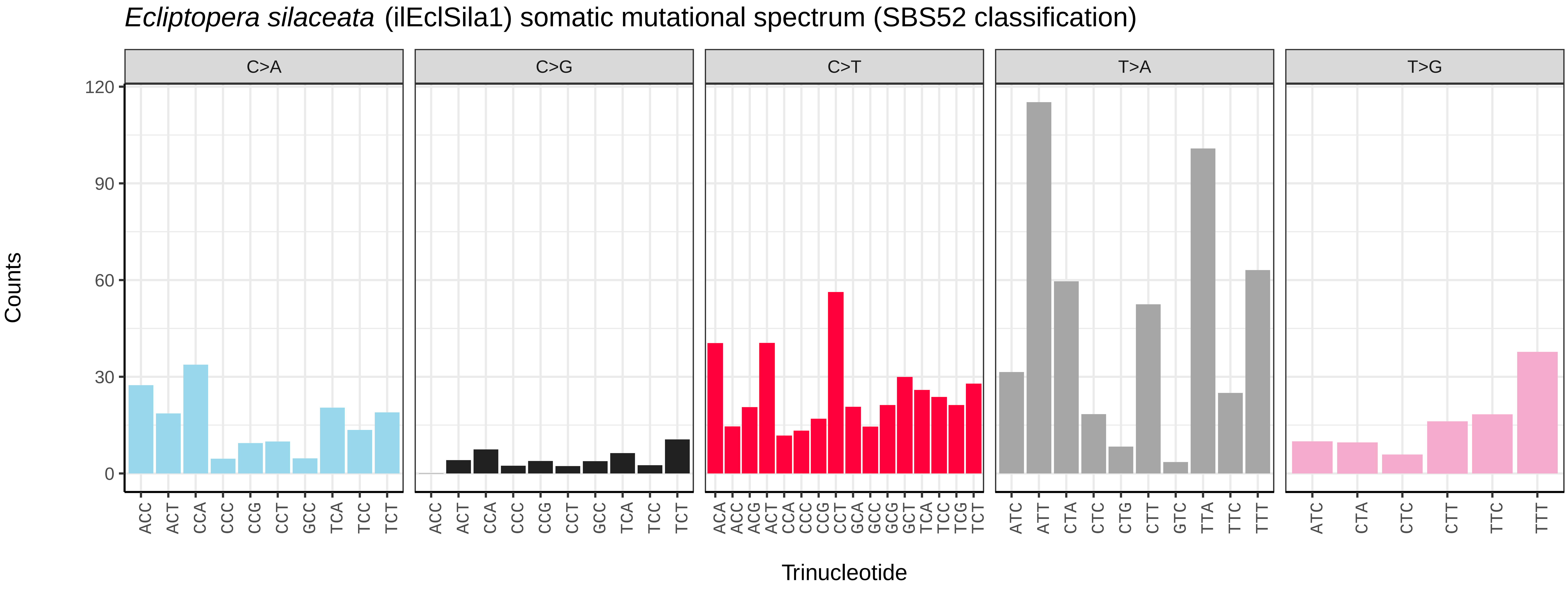
*Ecliptopera silaceata* (ilEclSila1) somatic mutational spectrum (SBS52 classification)
C>A
C>G
C>T
T>A
T>G
Counts
0
30
60
90
120
Trinucleotide

*Eilema caniola* (ilEilCani1) germline mutational spectrum

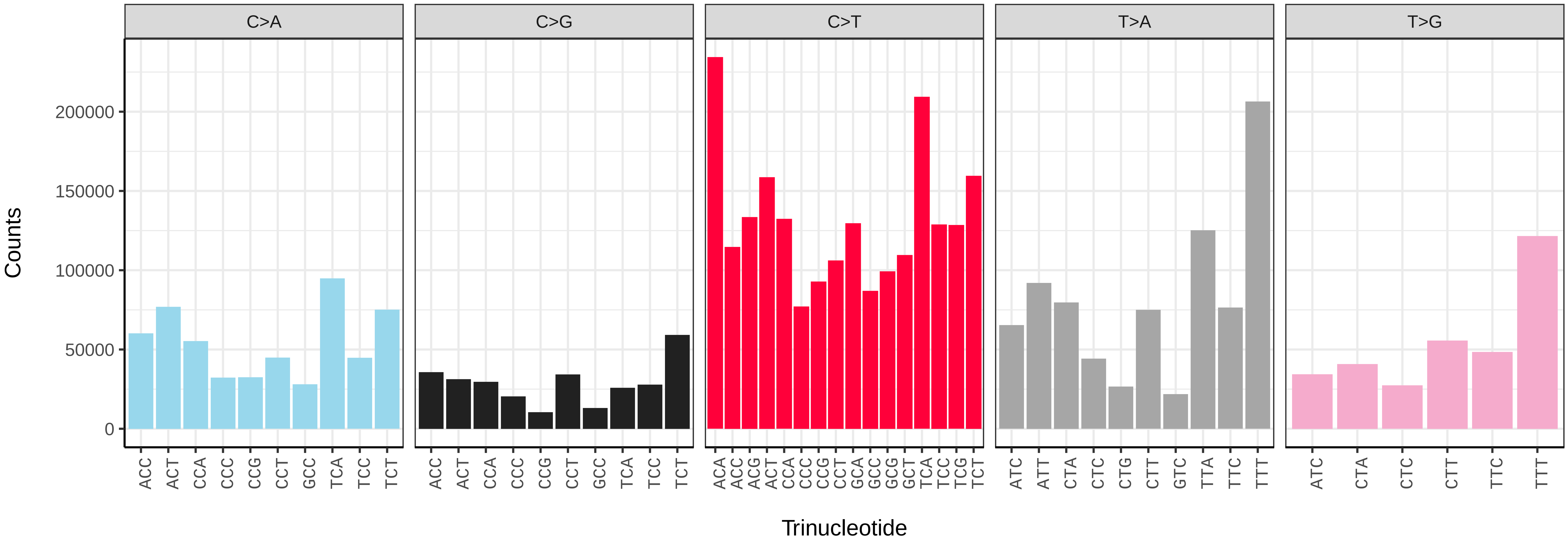


*Eilema caniola* (ilEilCani1) somatic mutational spectrum (SBS96 classification)

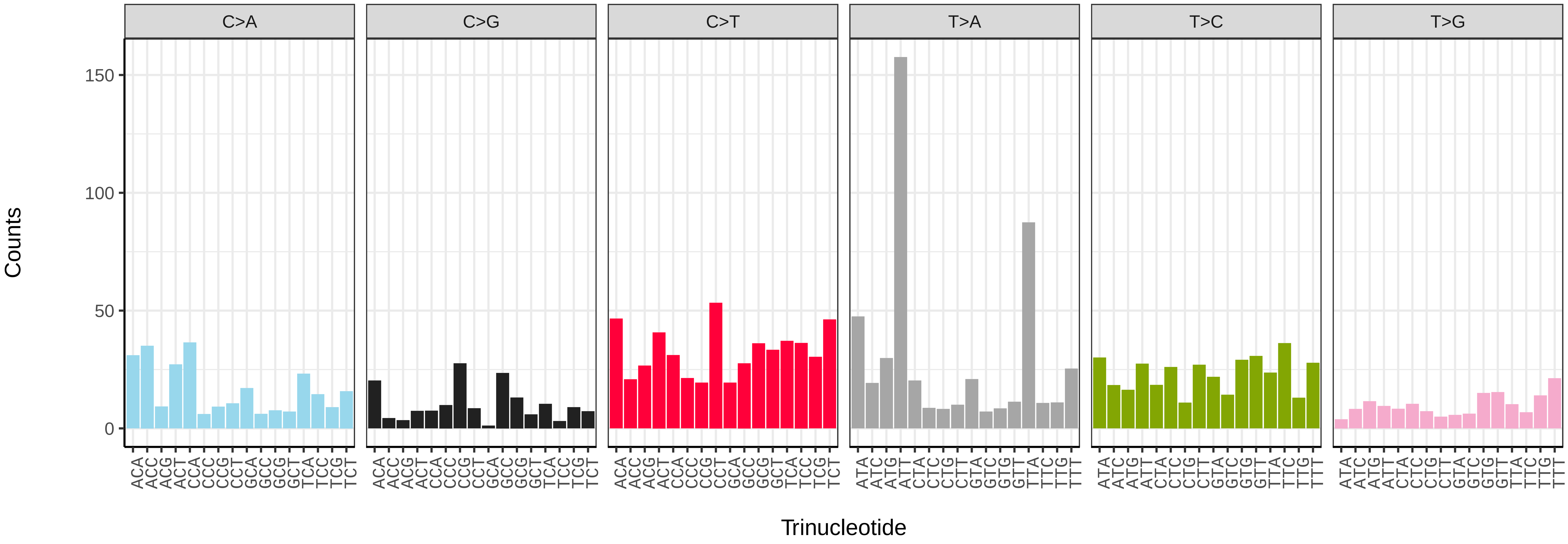


*Eilema caniola* (ilEilCani1) somatic mutational spectrum (SBS52 classification)

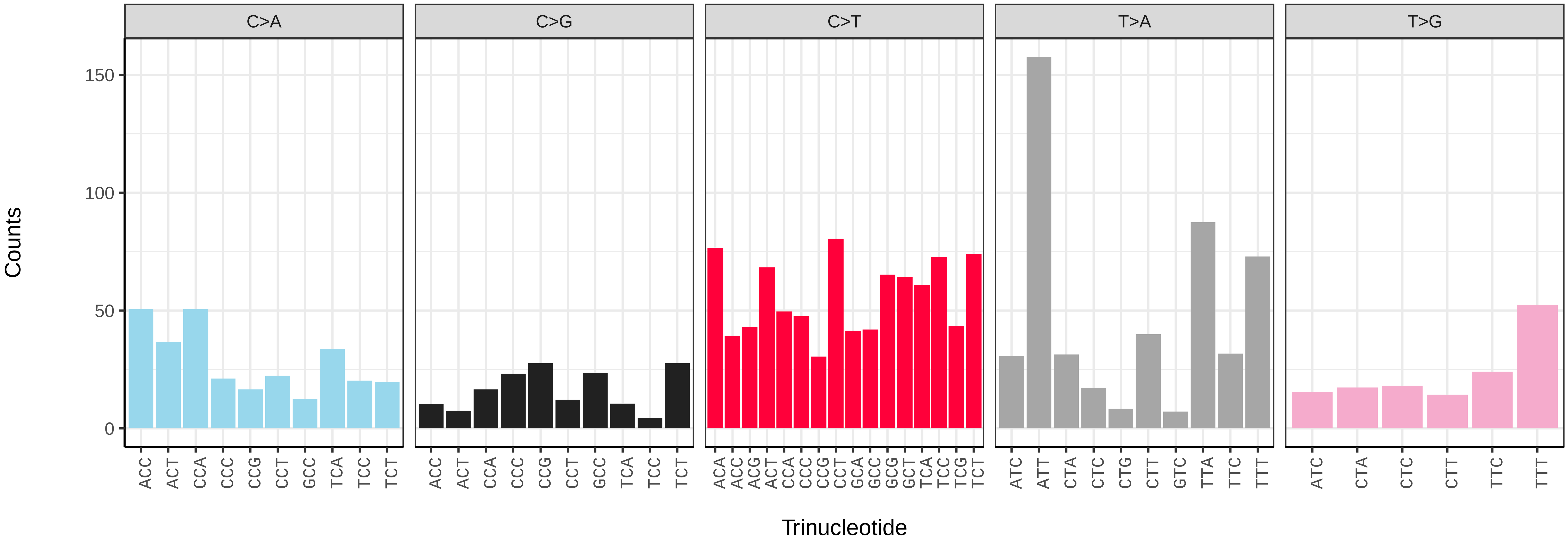

*Eilema depressum* (ilEilDepe1) germline mutational spectrum

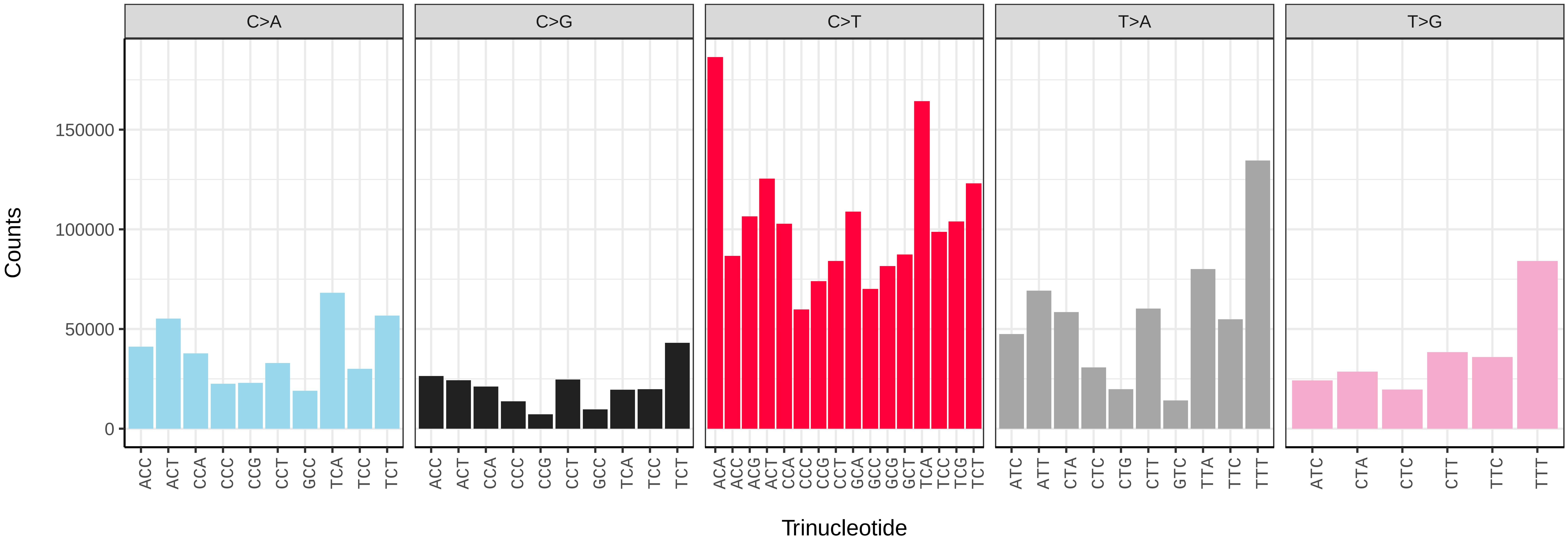


*Eilema depressum* (ilEilDepe1) somatic mutational spectrum (SBS96 classification)

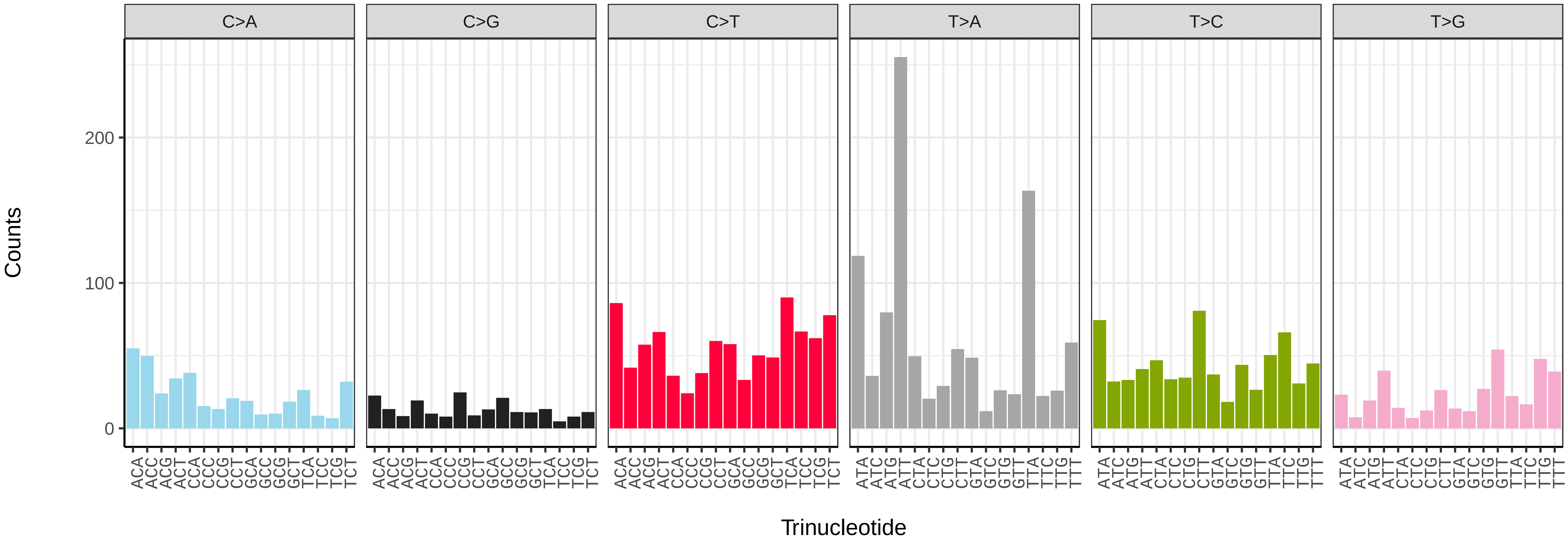


*Eilema depressum* (ilEilDepe1) somatic mutational spectrum (SBS52 classification)

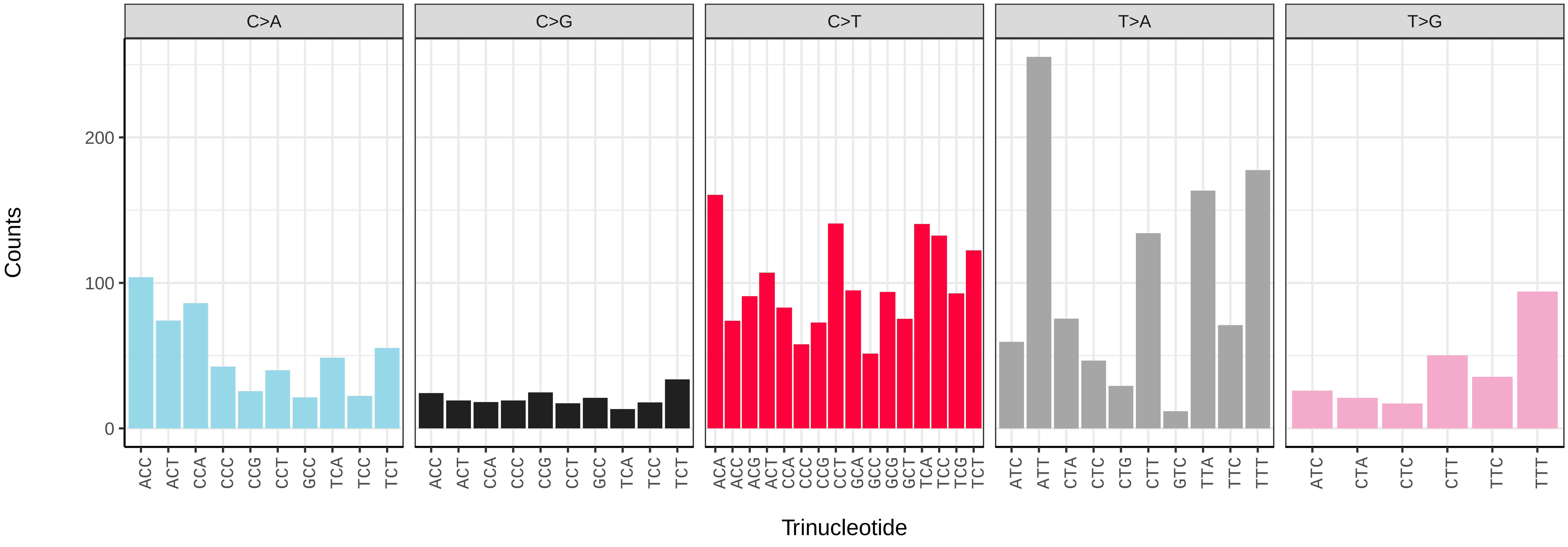

*Eilema sororcula* (ilEilSoro1) germline mutational spectrum

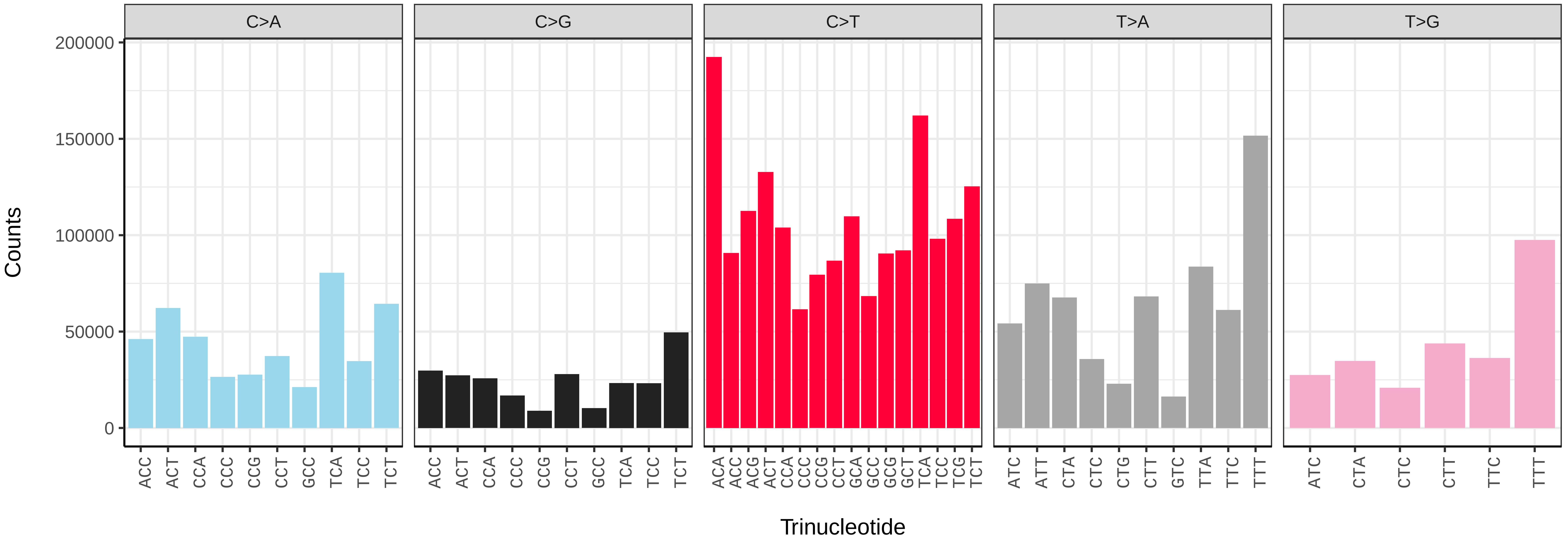


*Eilema sororcula* (ilEilSoro1) somatic mutational spectrum (SBS96 classification)

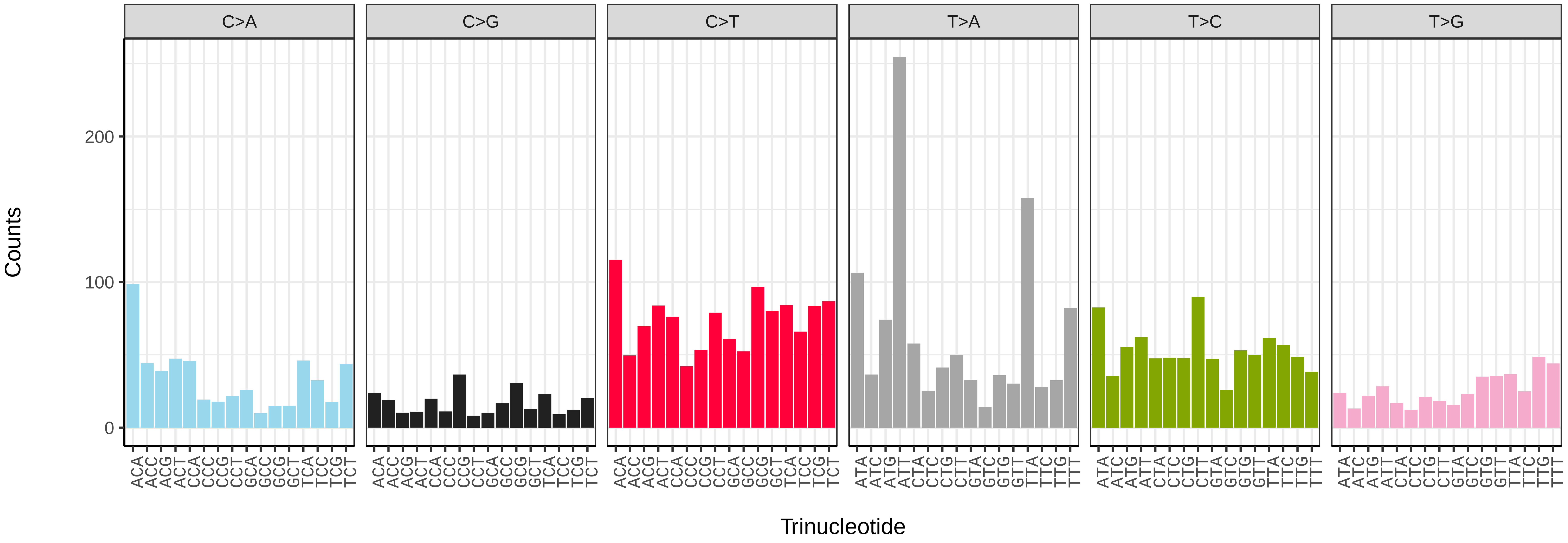


*Eilema sororcula* (ilEilSoro1) somatic mutational spectrum (SBS52 classification)

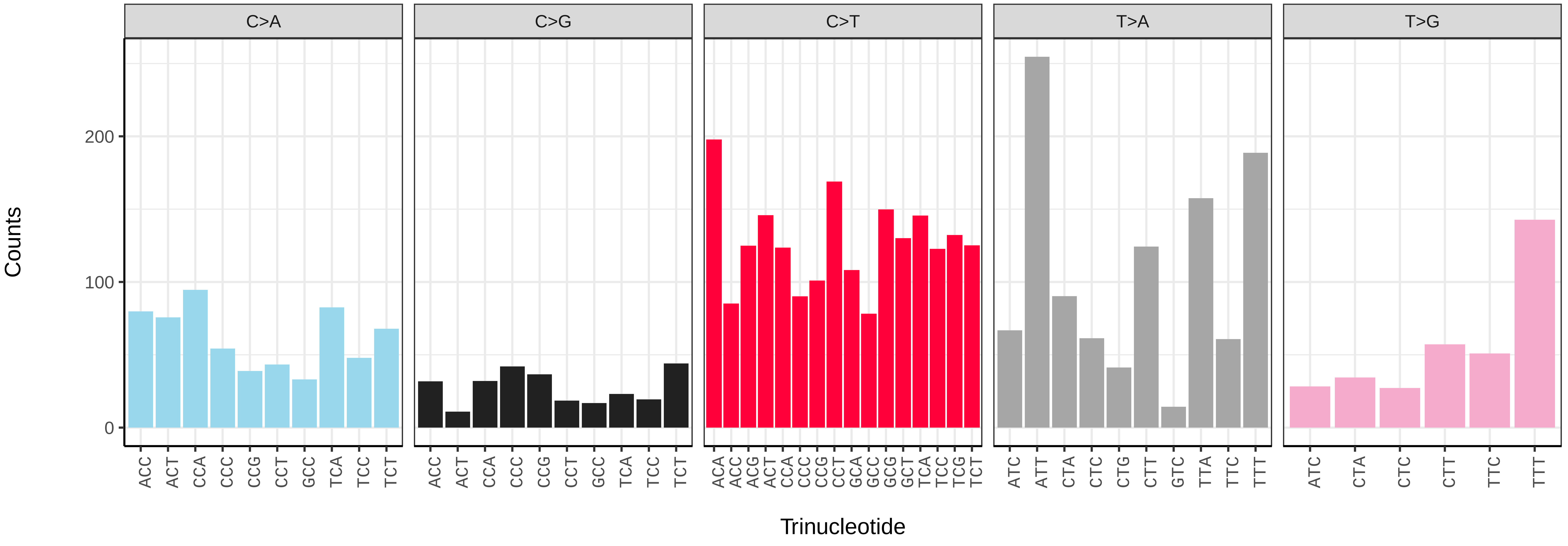

*Electrophaes corylata* (ilEleCory1) germline mutational spectrum

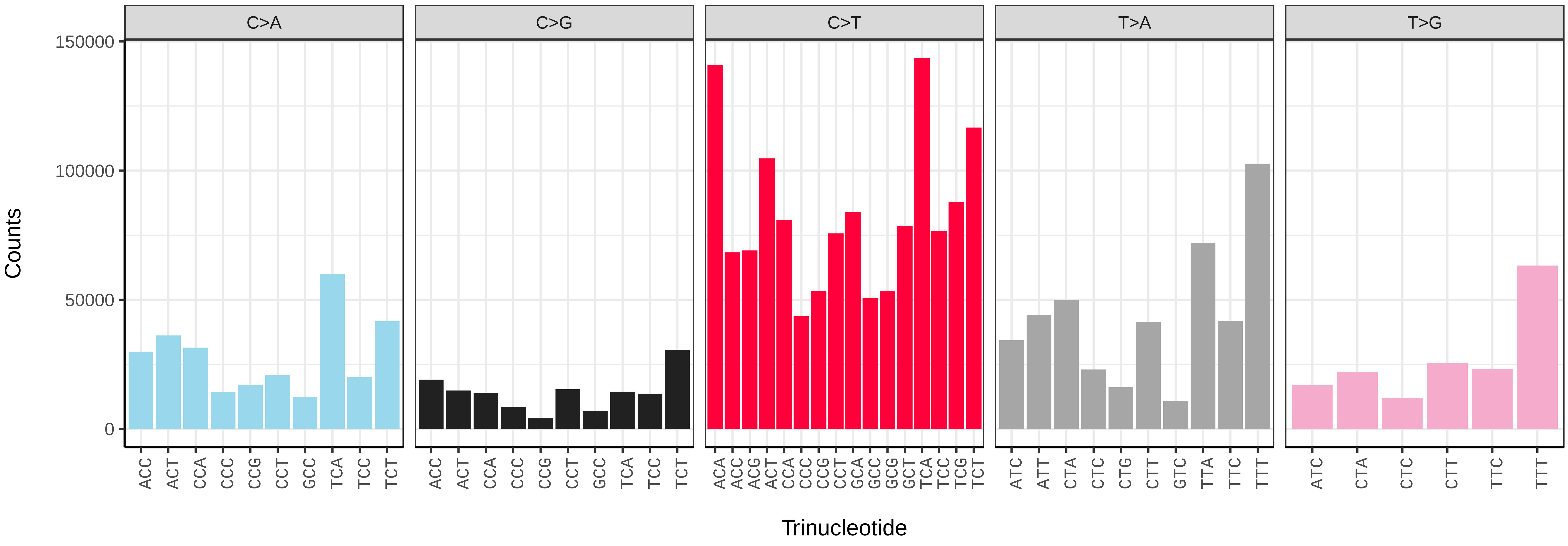


*Electrophaes corylata* (ilEleCory1) somatic mutational spectrum (SBS96 classification)

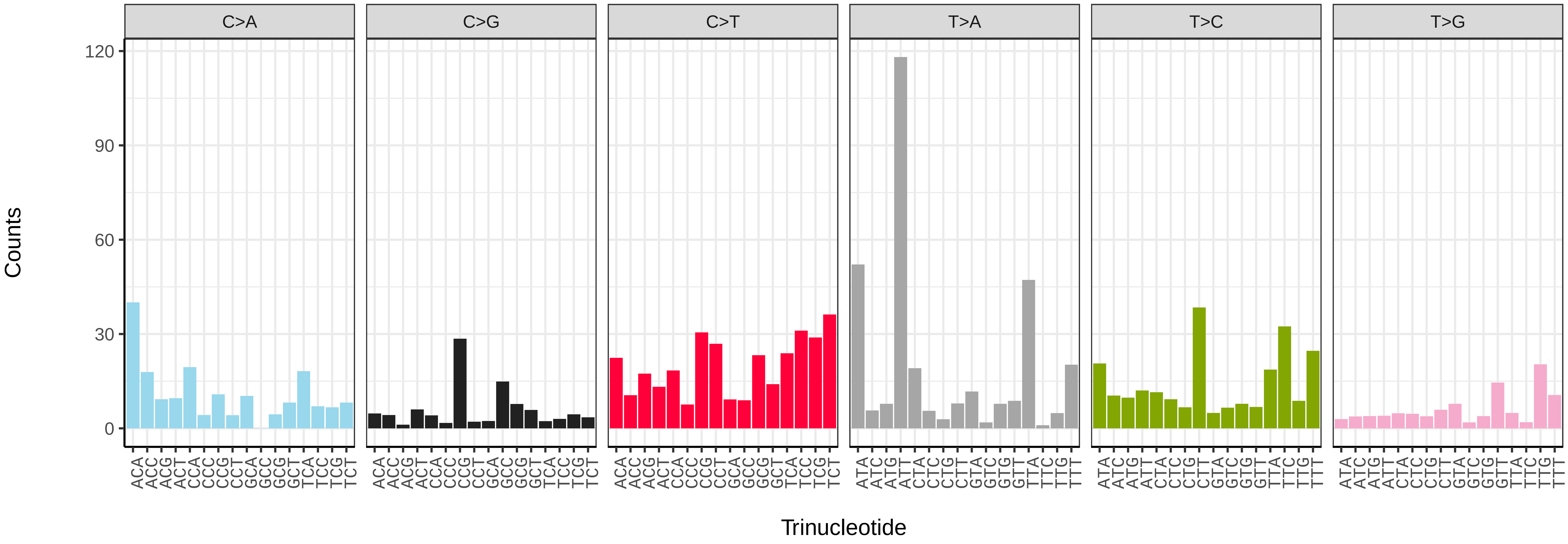


*Electrophaes corylata* (ilEleCory1) somatic mutational spectrum (SBS52 classification)

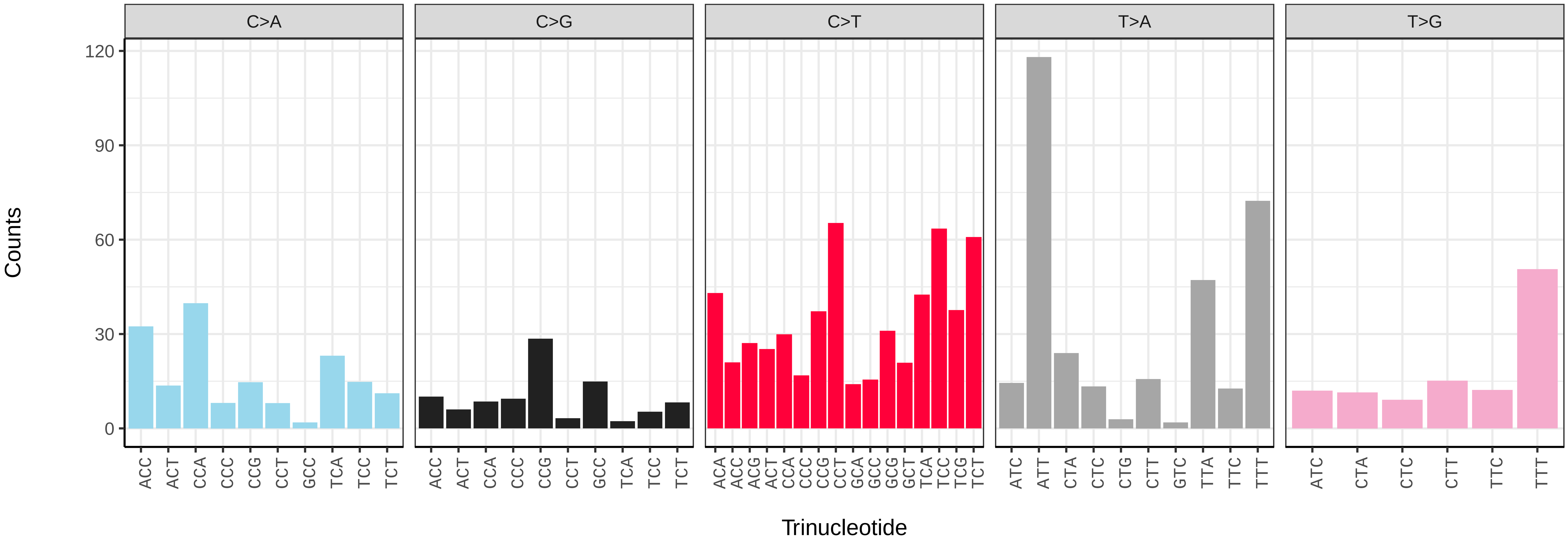

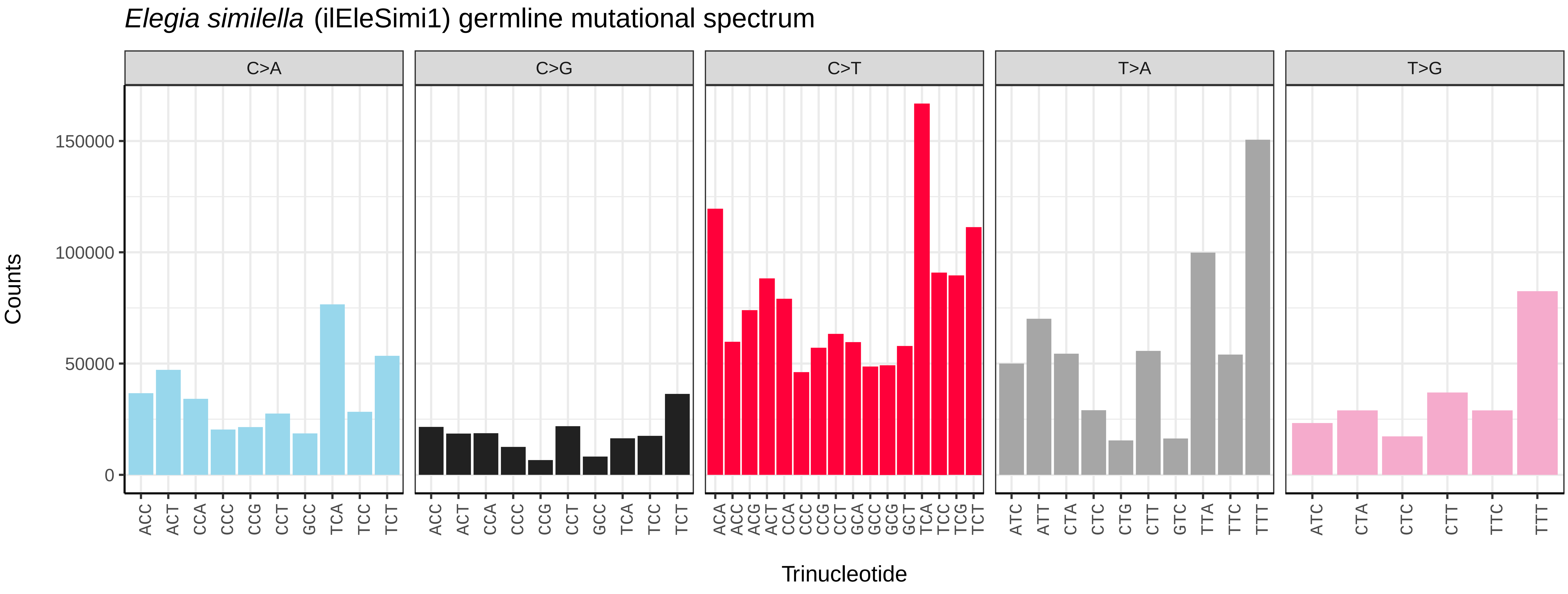
Elegia similella (ilEleSimi1) germline mutational spectrum
C>A
C>G
C>T
T>A
T>G
Counts
0
50000
100000
150000
Trinucleotide

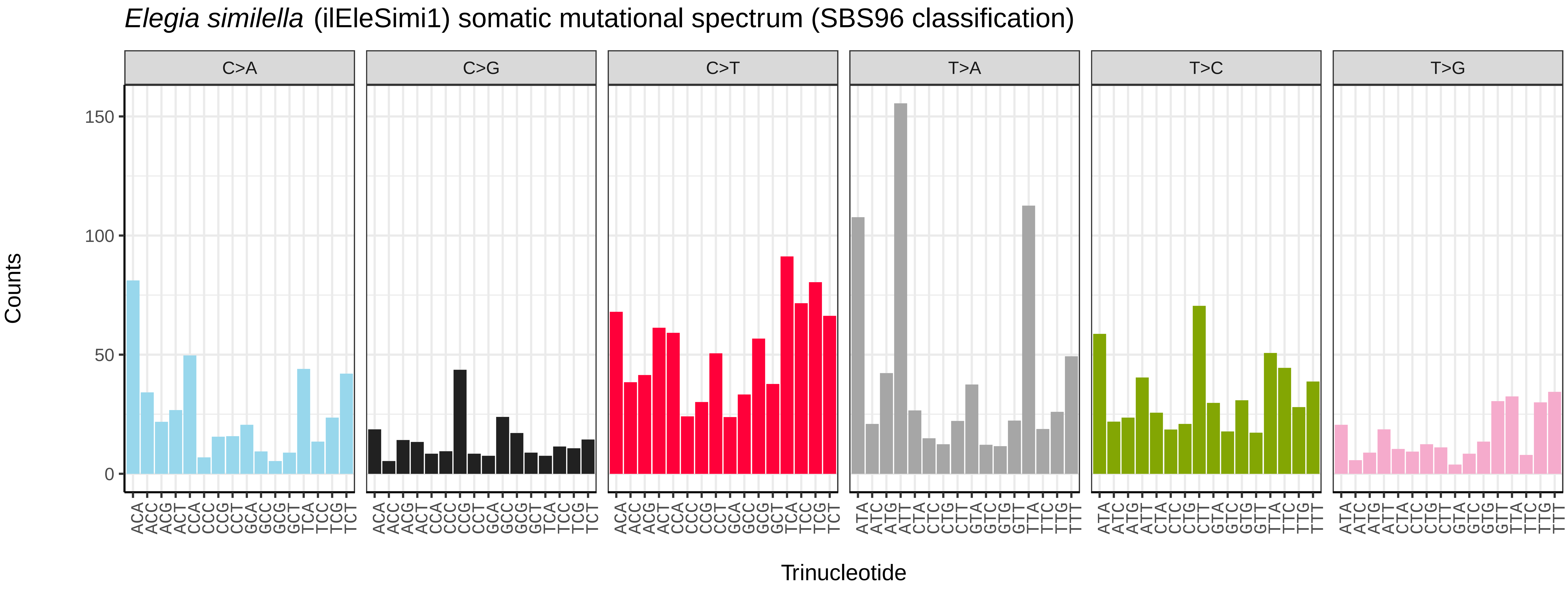
Elegia similella (ilEleSimi1) somatic mutational spectrum (SBS96 classification)
C>A
C>G
C>T
T>A
T>C
T>G
Counts
0
50
100
150
Trinucleotide

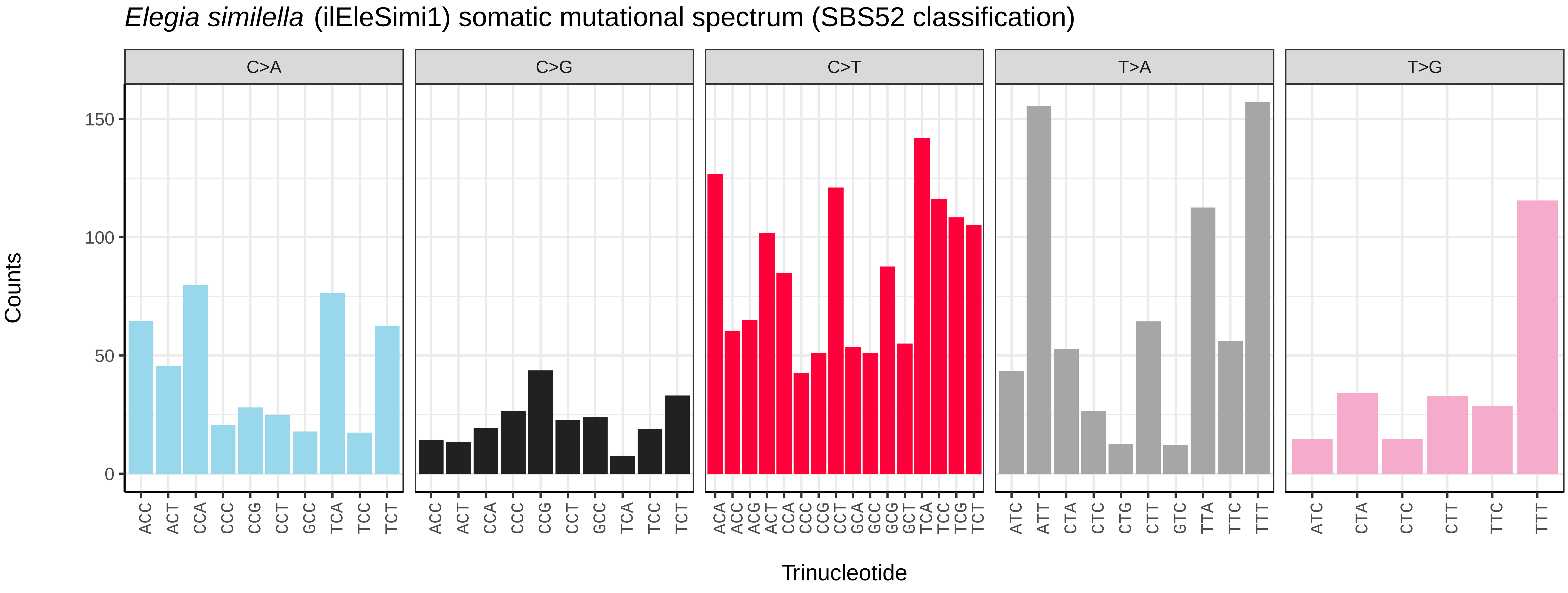
Elegia similella (ilEleSimi1) somatic mutational spectrum (SBS52 classification)
C>A
C>G
C>T
T>A
T>G
Counts
0
50
100
150
Trinucleotide

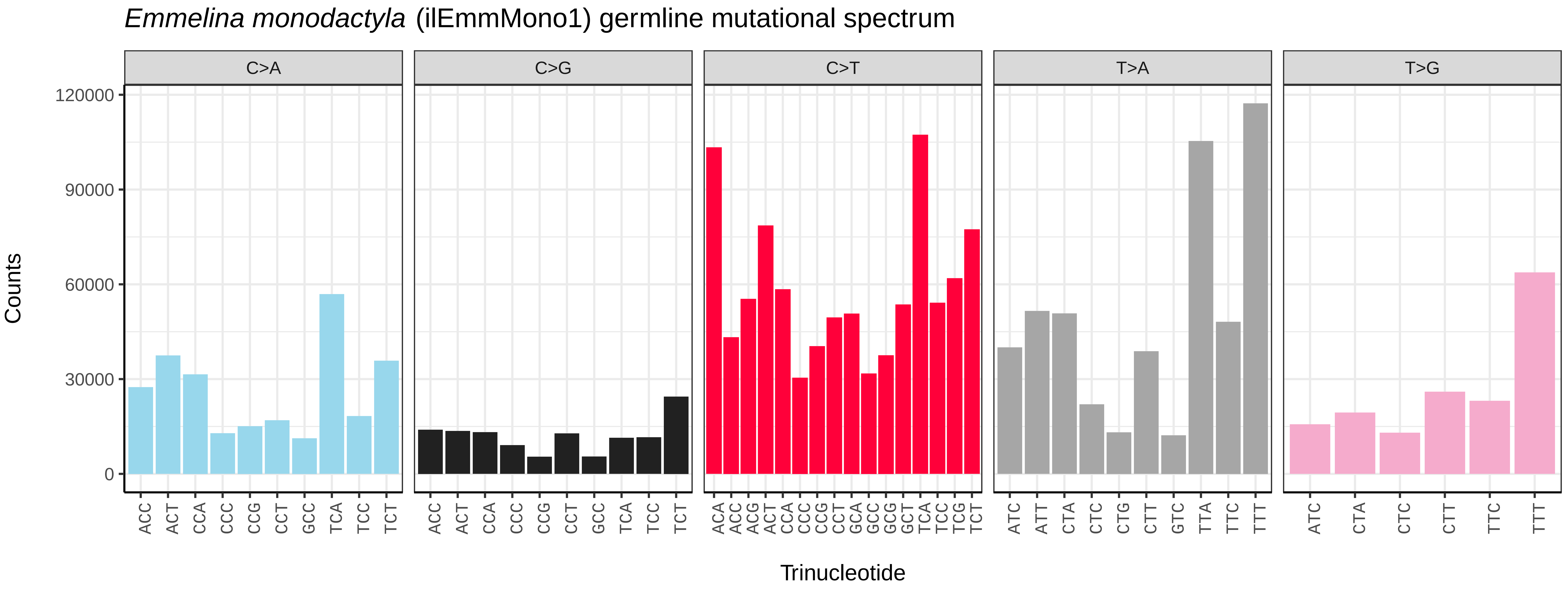
*Emmelina monodactyla* (ilEmmMono1) germline mutational spectrum
C>A
C>G
C>T
T>A
T>G
0
30000
60000
90000
120000
Counts
Trinucleotide

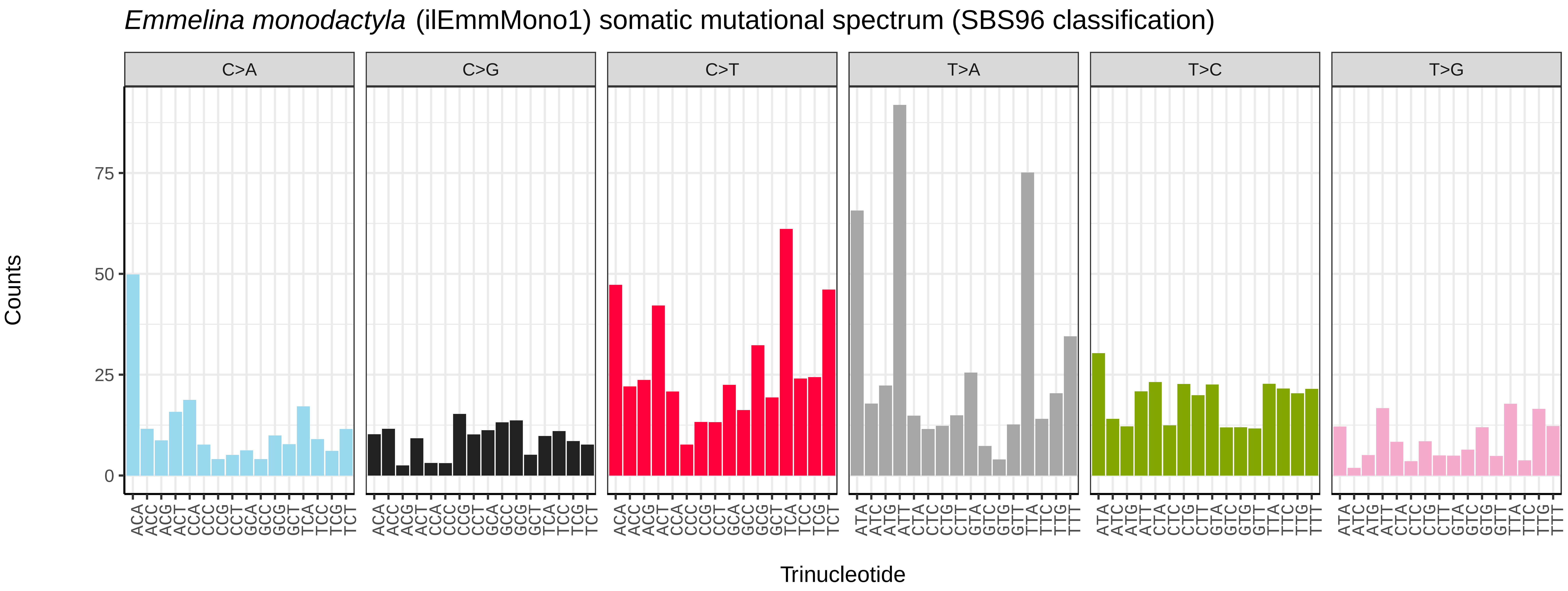
*Emmelina monodactyla* (ilEmmMono1) somatic mutational spectrum (SBS96 classification)
C>A
C>G
C>T
T>A
T>C
T>G
0
25
50
75
Counts
Trinucleotide

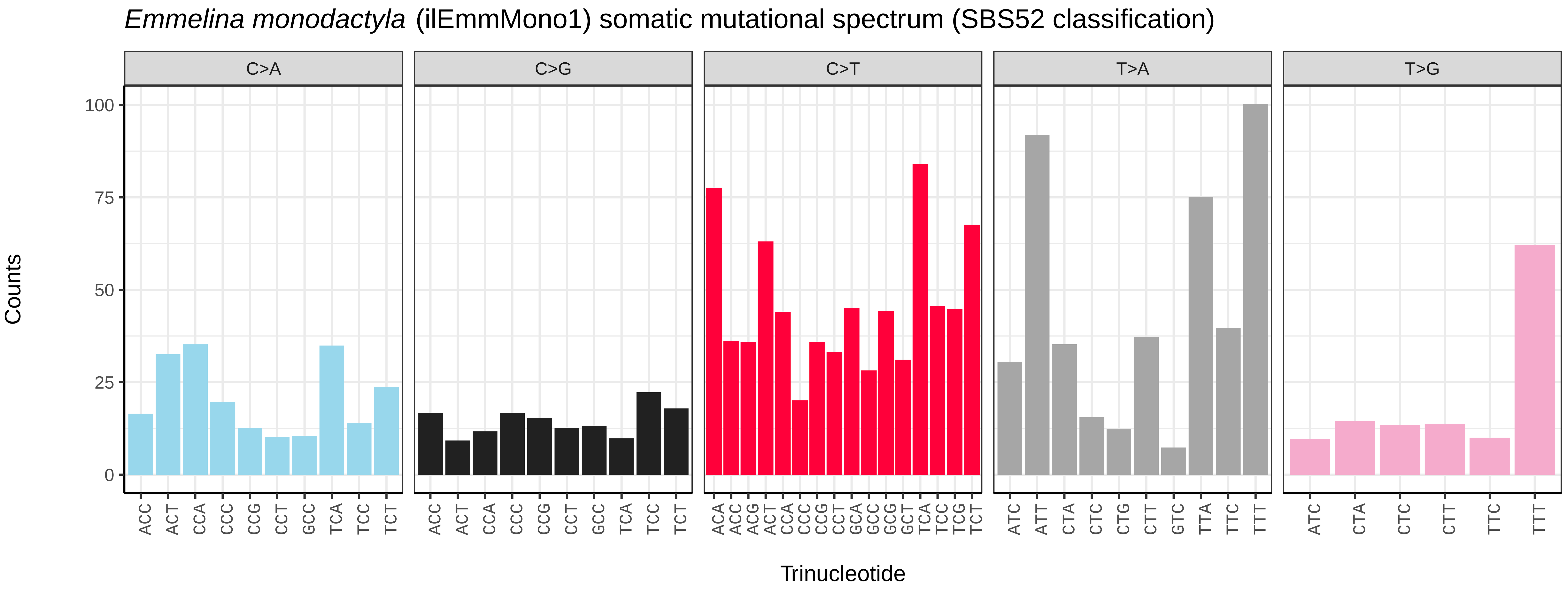
*Emmelina monodactyla* (ilEmmMono1) somatic mutational spectrum (SBS52 classification)
C>A
C>G
C>T
T>A
T>G
0
25
50
75
100
Counts
Trinucleotide

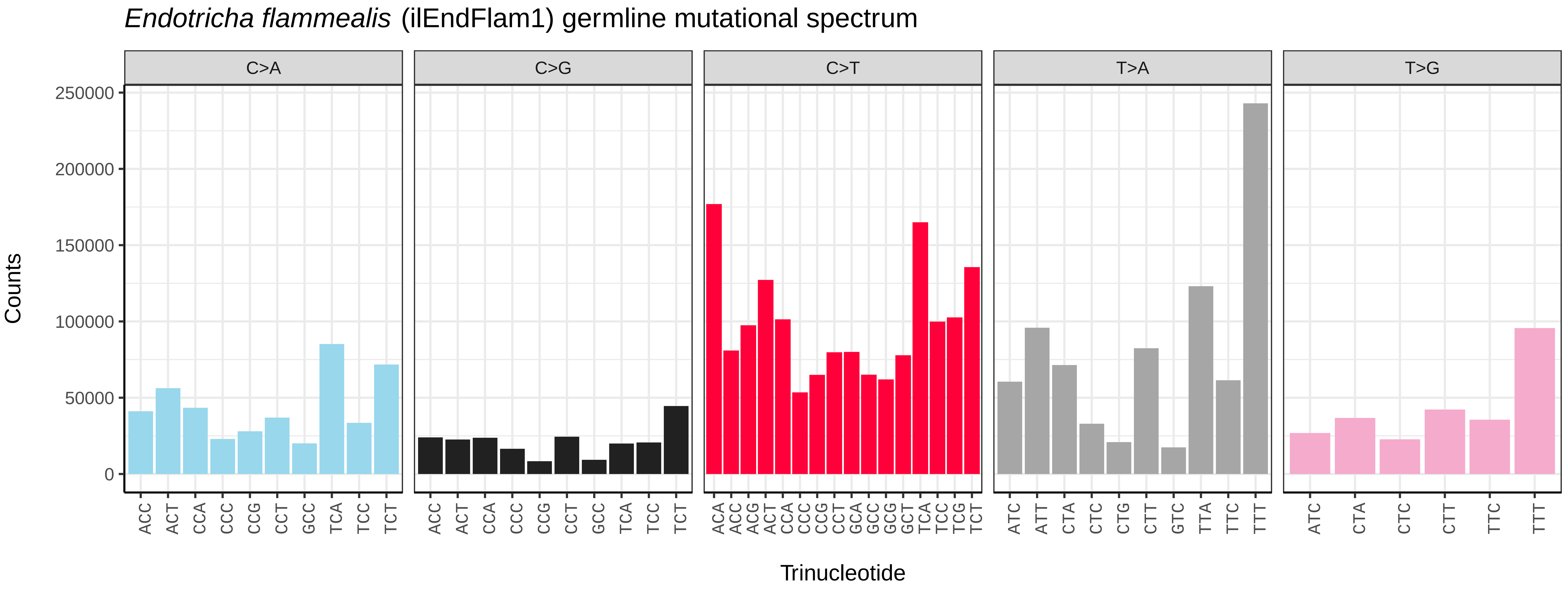
*Endotricha flammealis* (ilEndFlam1) germline mutational spectrum
C>A
C>G
C>T
T>A
T>G
Counts
0
50000
100000
150000
200000
250000
Trinucleotide

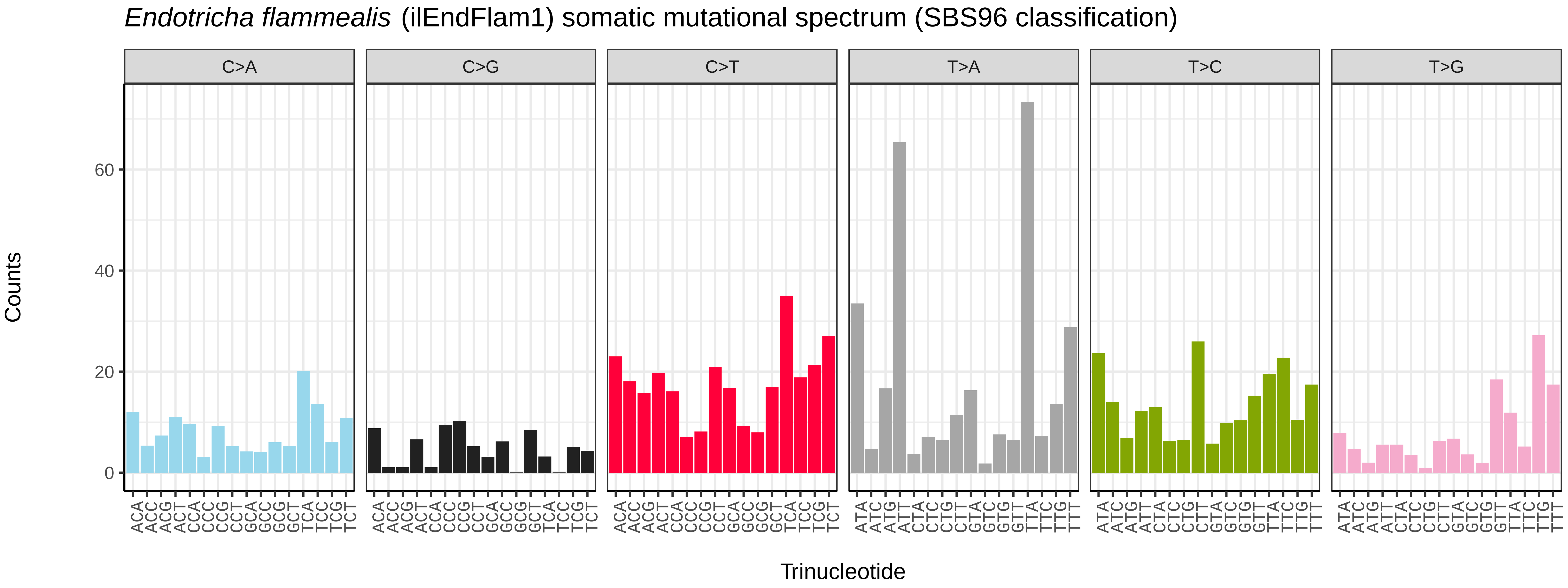
*Endotricha flammealis* (ilEndFlam1) somatic mutational spectrum (SBS96 classification)
C>A
C>G
C>T
T>A
T>C
T>G
Counts
0
20
40
60
Trinucleotide

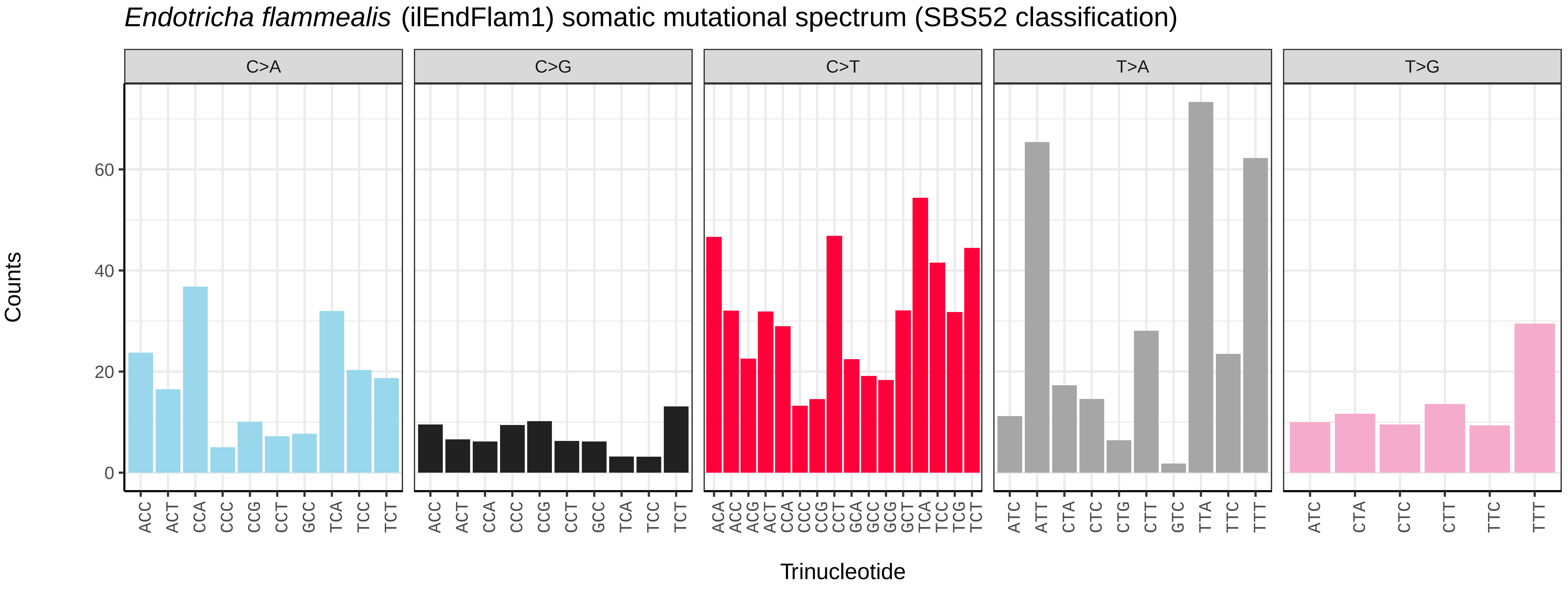
*Endotricha flammealis* (ilEndFlam1) somatic mutational spectrum (SBS52 classification)
C>A
C>G
C>T
T>A
T>G
Counts
0
20
40
60
Trinucleotide

*Ennomos fuscantarius* (ilEnnFusc2) germline mutational spectrum

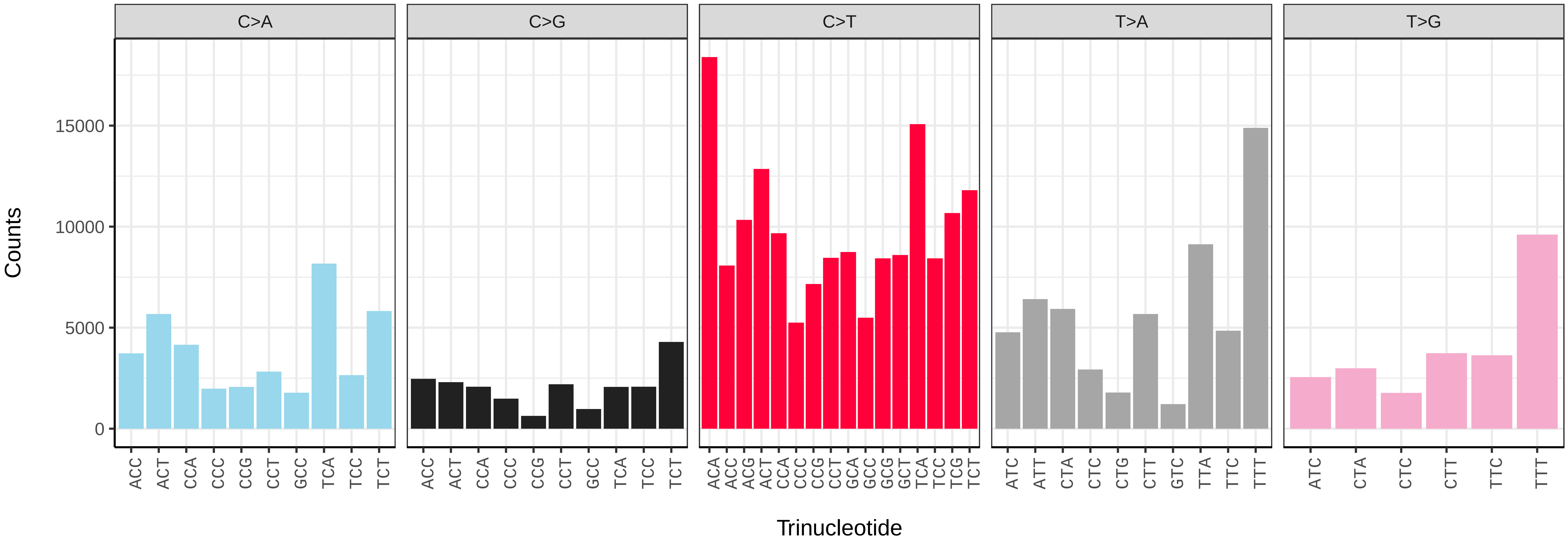


*Ennomos fuscantarius* (ilEnnFusc2) somatic mutational spectrum (SBS96 classification)

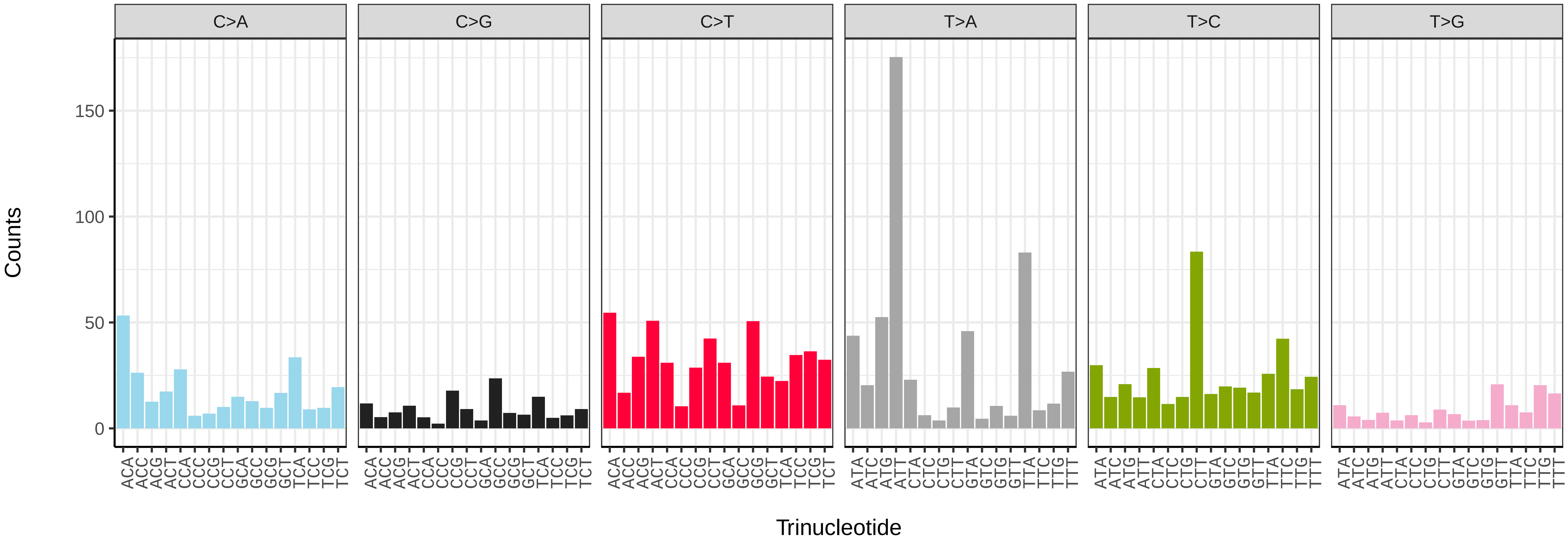


*Ennomos fuscantarius* (ilEnnFusc2) somatic mutational spectrum (SBS52 classification)

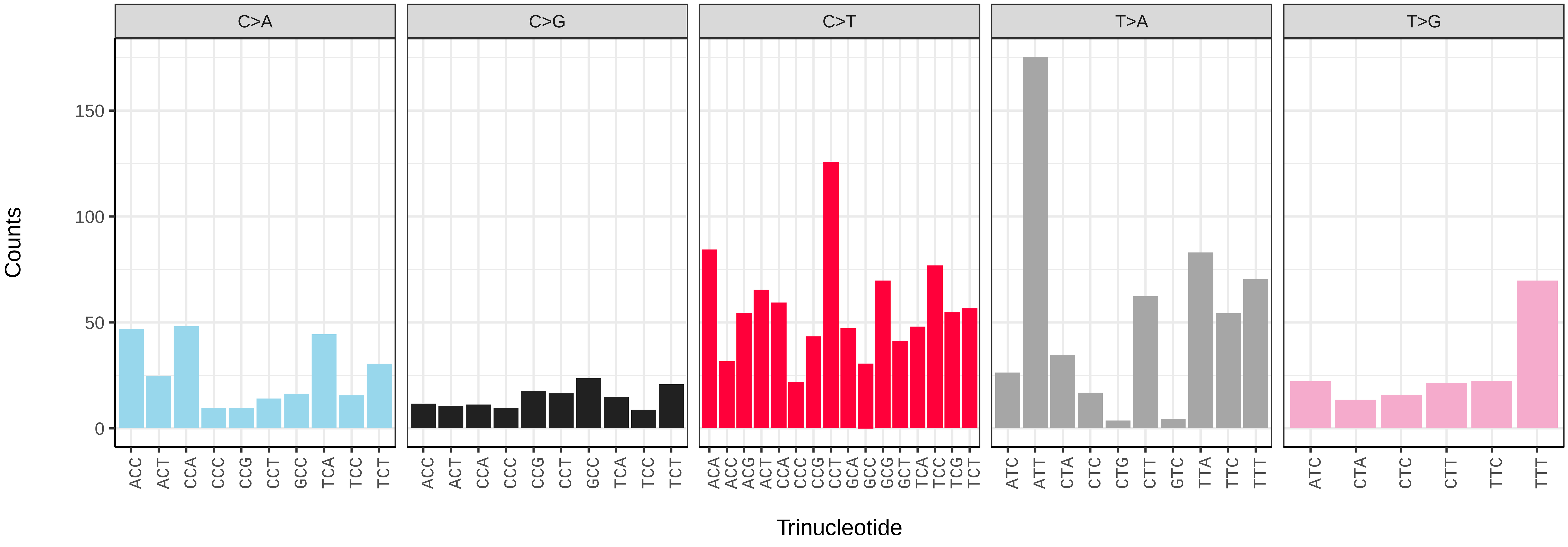

*Ennomos quercinarius* (ilEnnQuei1) germline mutational spectrum

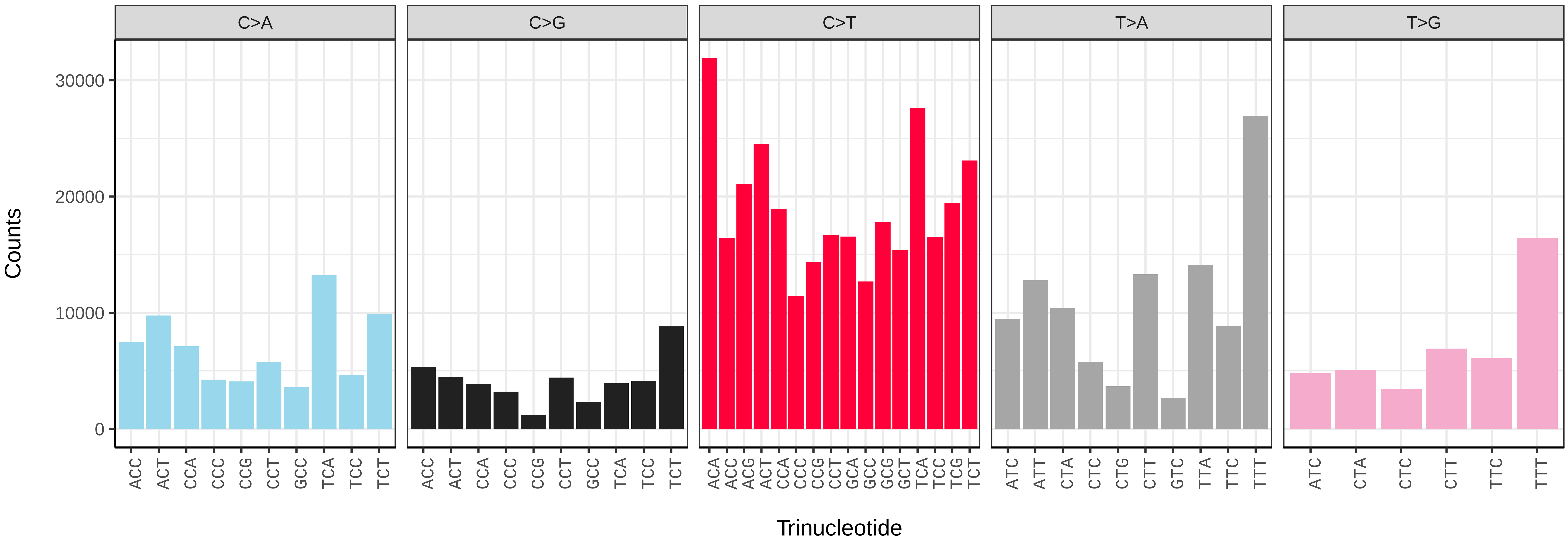


*Ennomos quercinarius* (ilEnnQuei1) somatic mutational spectrum (SBS96 classification)

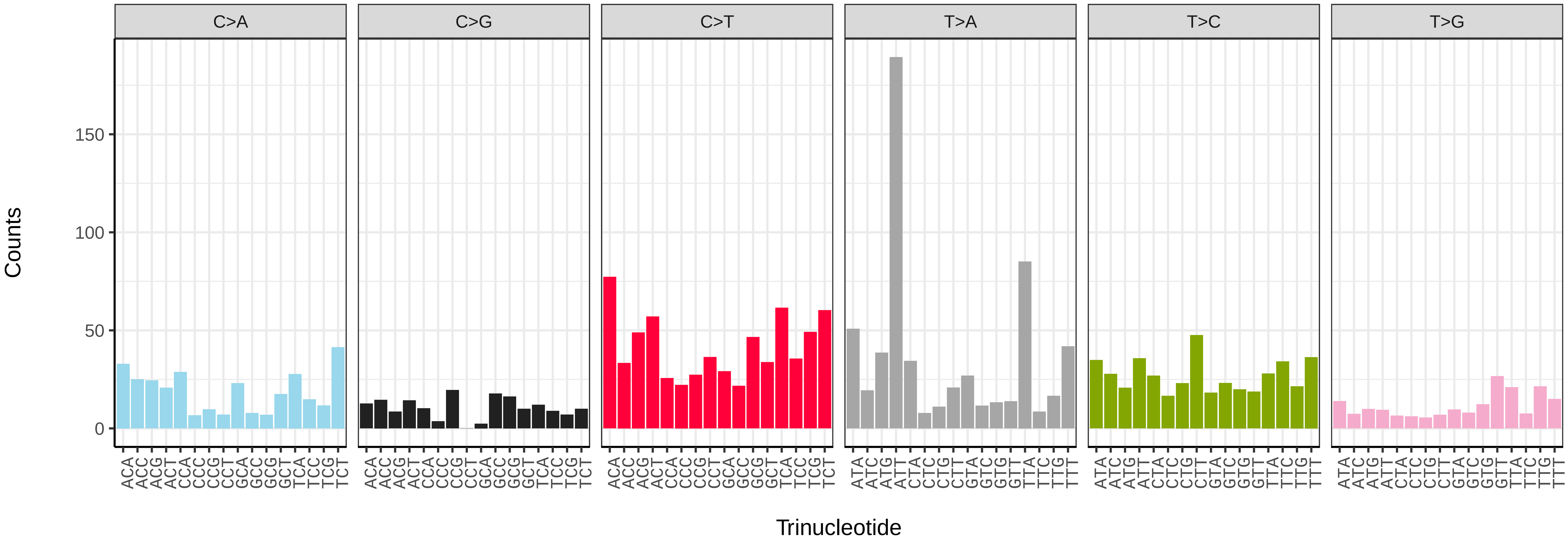


*Ennomos quercinarius* (ilEnnQuei1) somatic mutational spectrum (SBS52 classification)

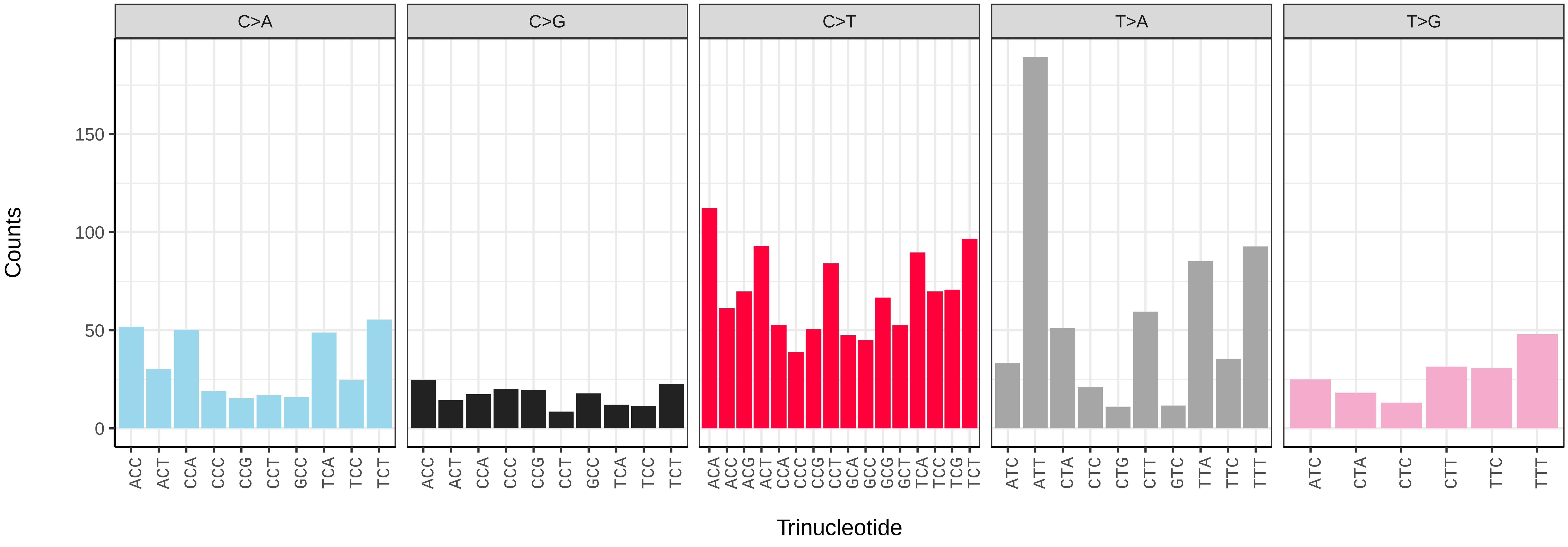

*Epinotia bilunana* (ilEpiBilu1) germline mutational spectrum

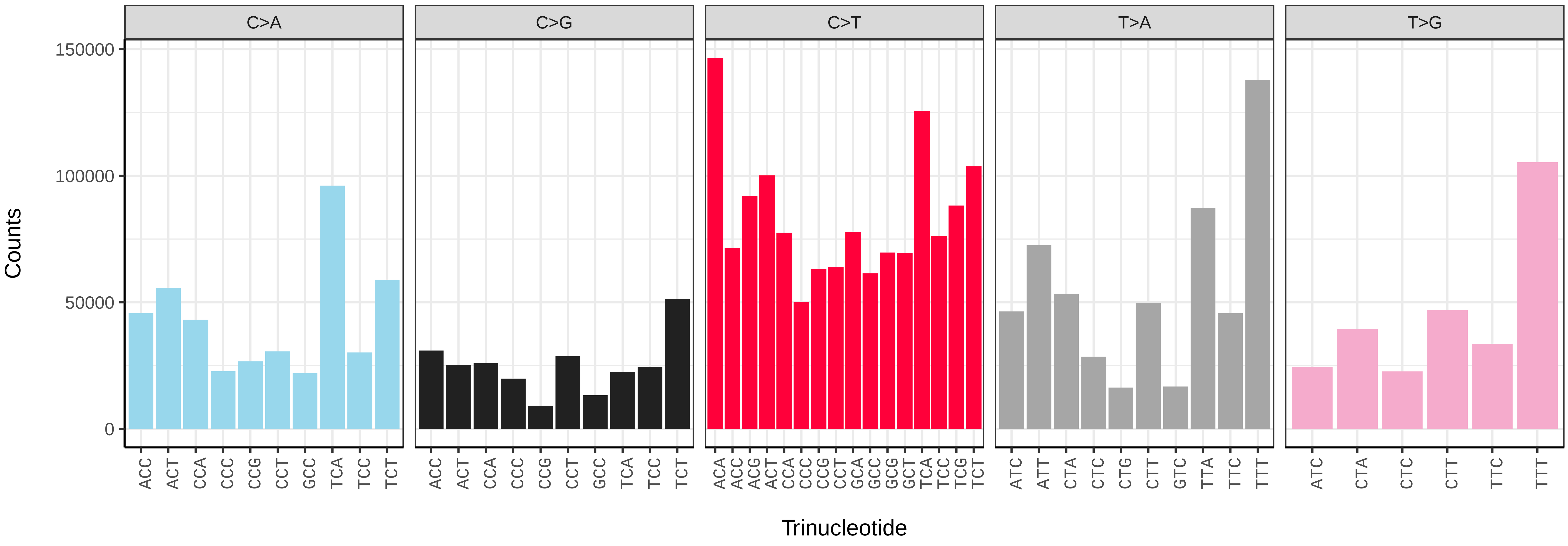


*Epinotia bilunana* (ilEpiBilu1) somatic mutational spectrum (SBS96 classification)

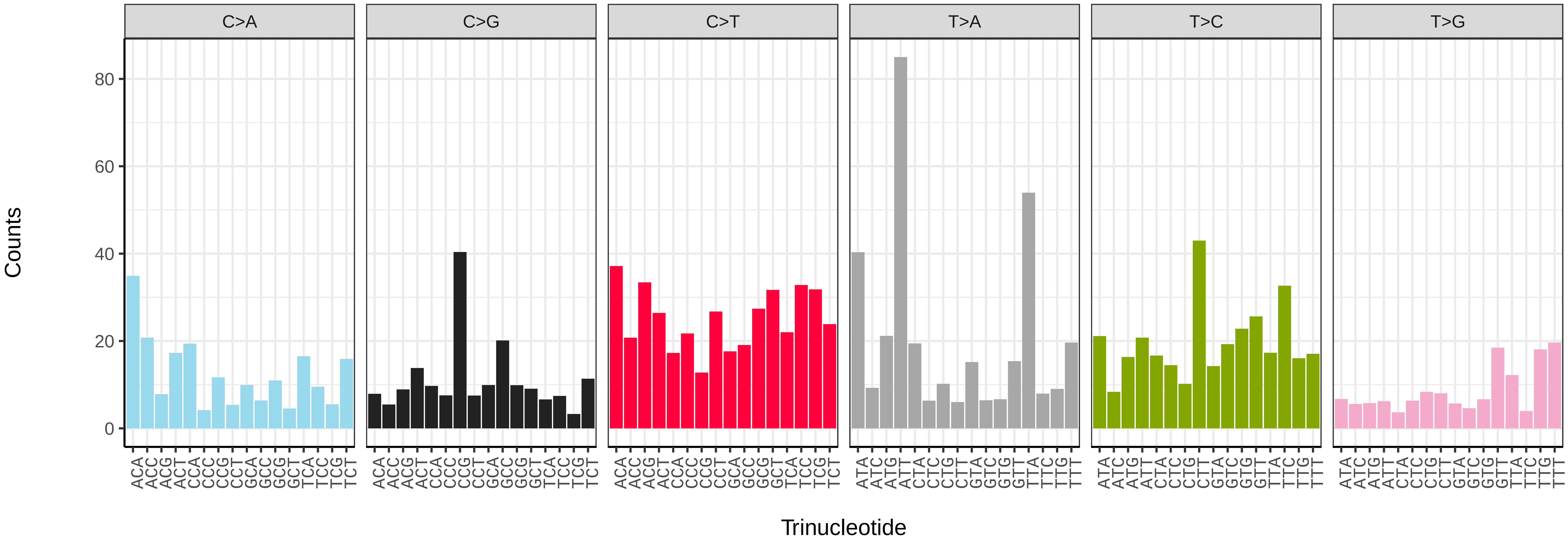


*Epinotia bilunana* (ilEpiBilu1) somatic mutational spectrum (SBS52 classification)

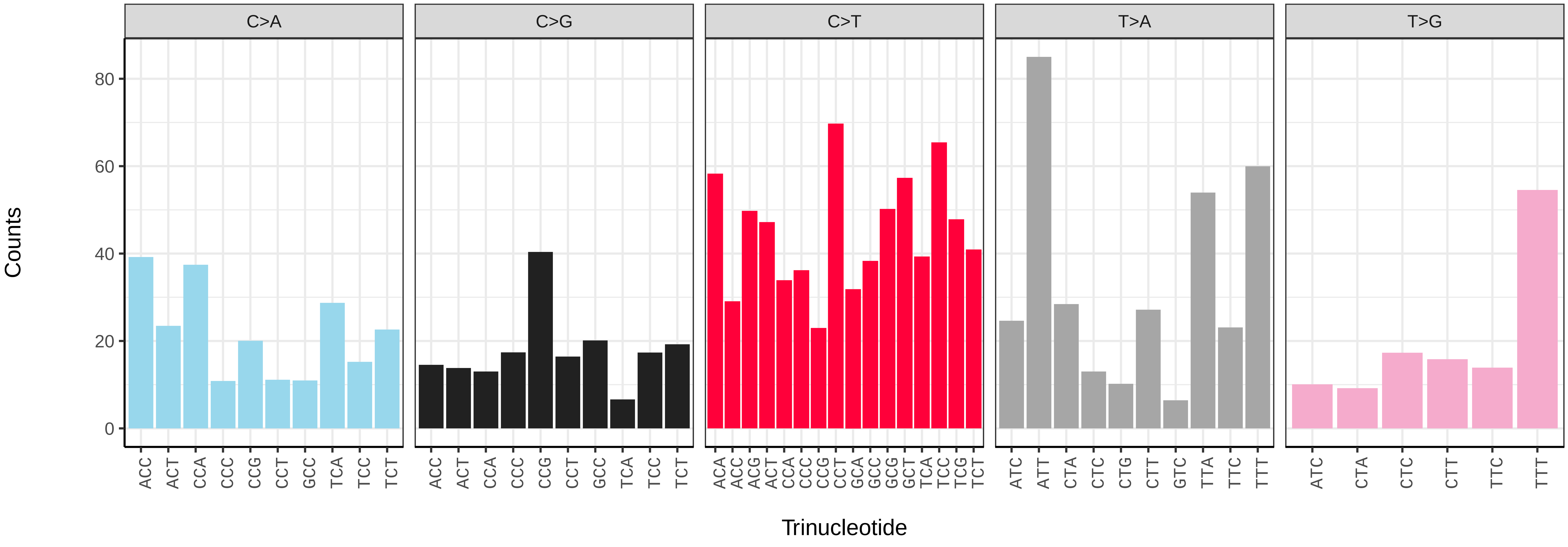

*Epinotia demarniana* (ilEpiDema1) germline mutational spectrum

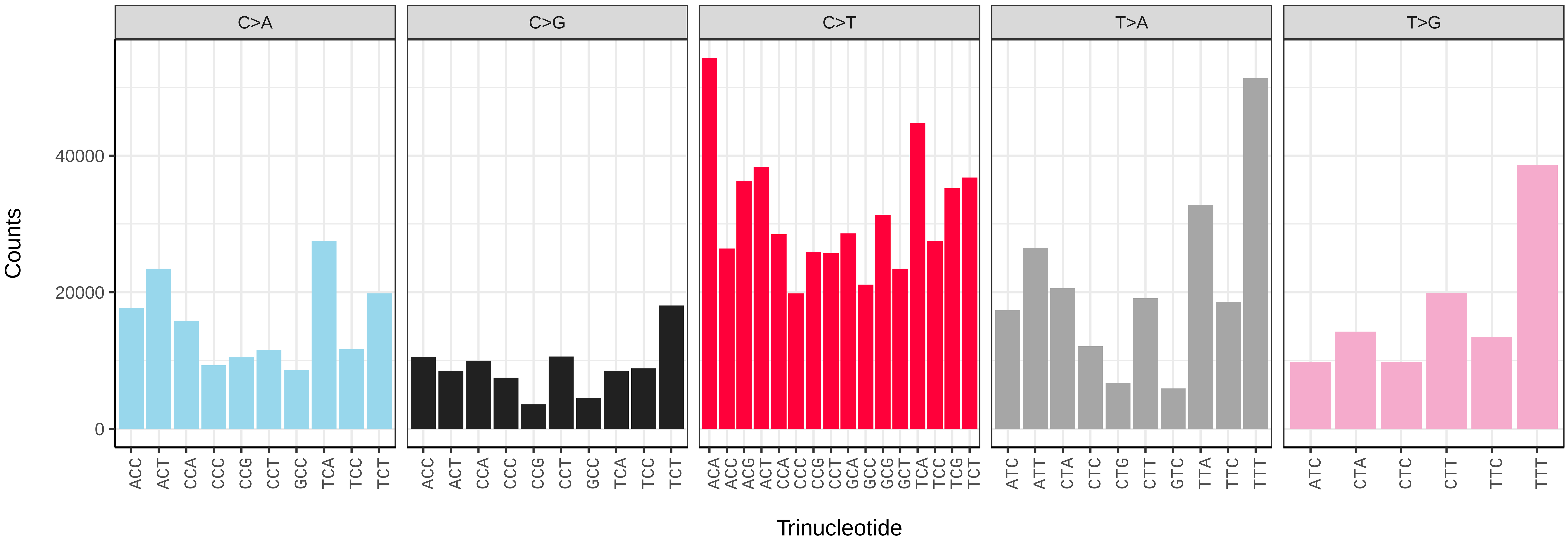


*Epinotia demarniana* (ilEpiDema1) somatic mutational spectrum (SBS96 classification)

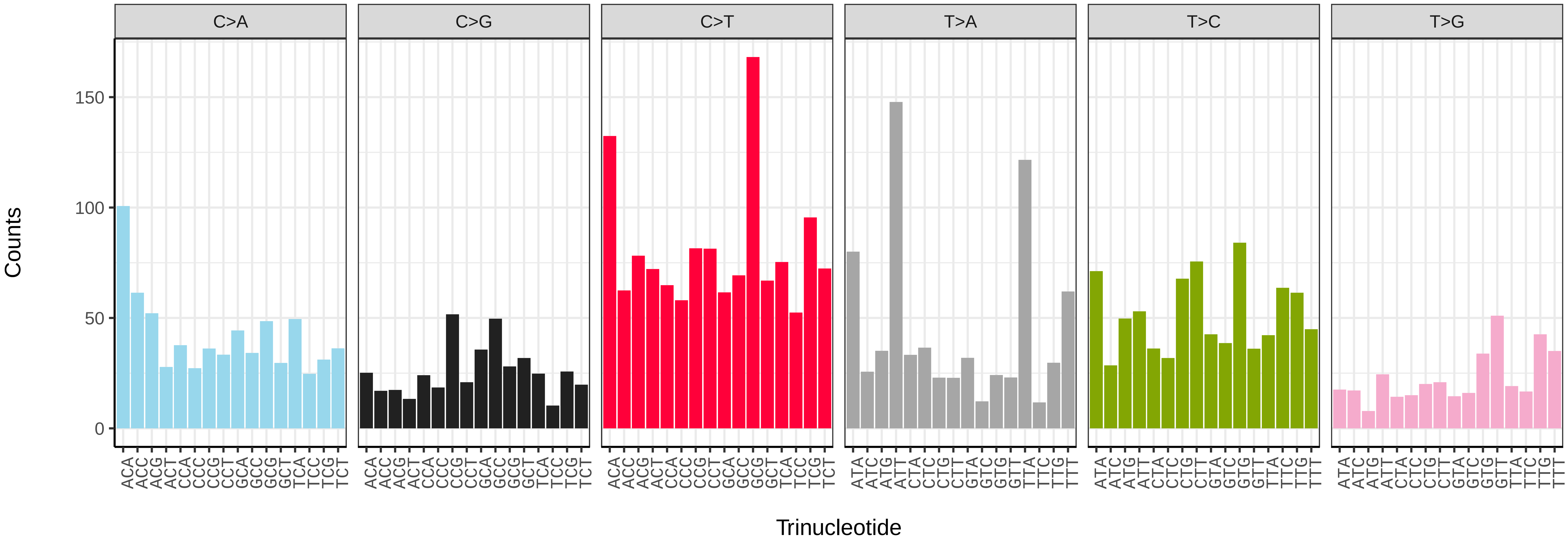


*Epinotia demarniana* (ilEpiDema1) somatic mutational spectrum (SBS52 classification)

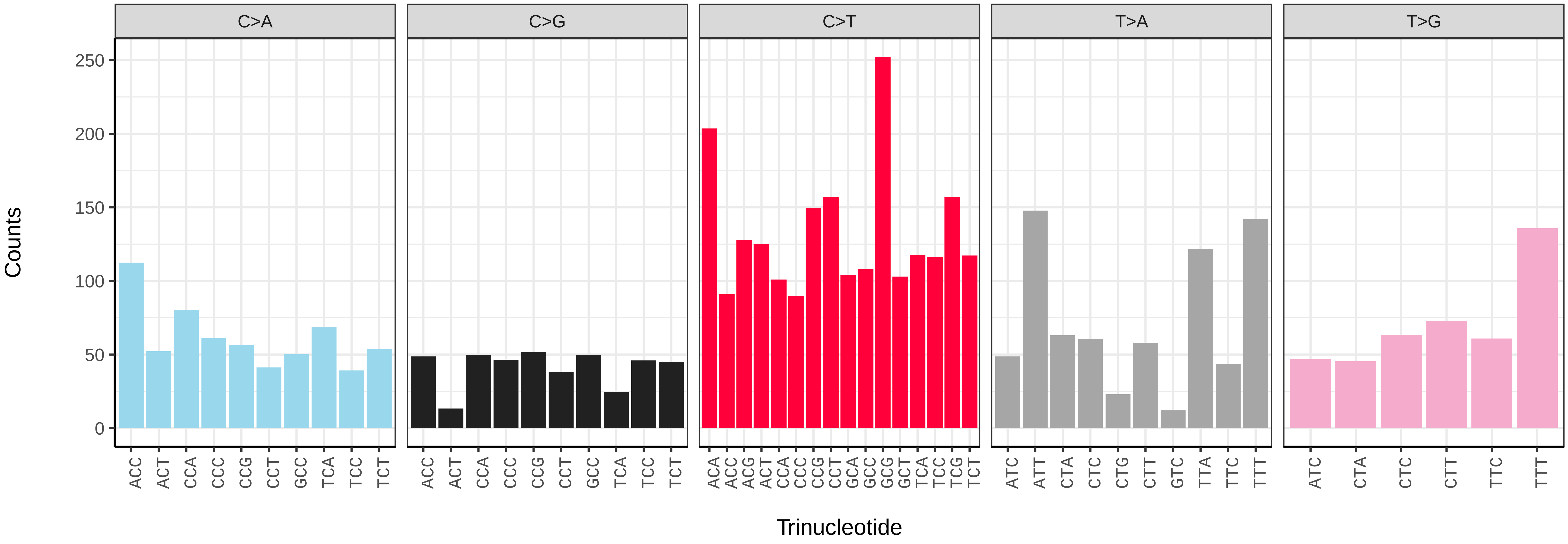

*Epinotia nisella* (ilEpiNise2) germline mutational spectrum

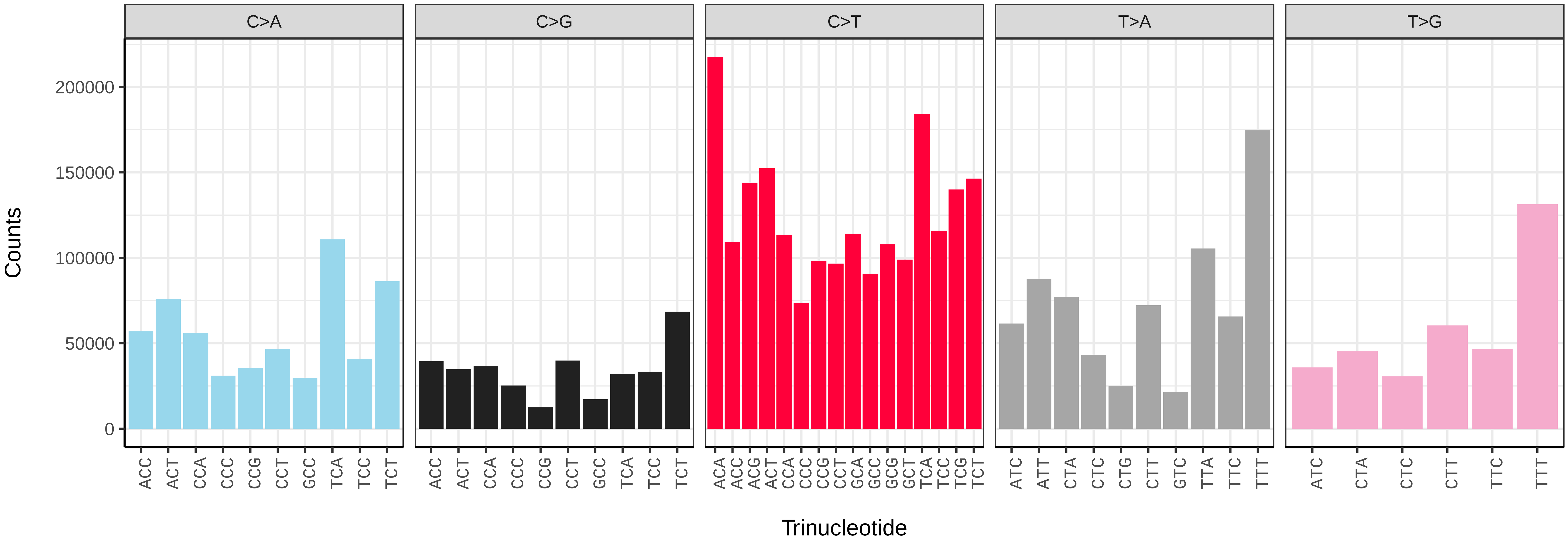


*Epinotia nisella* (ilEpiNise2) somatic mutational spectrum (SBS96 classification)

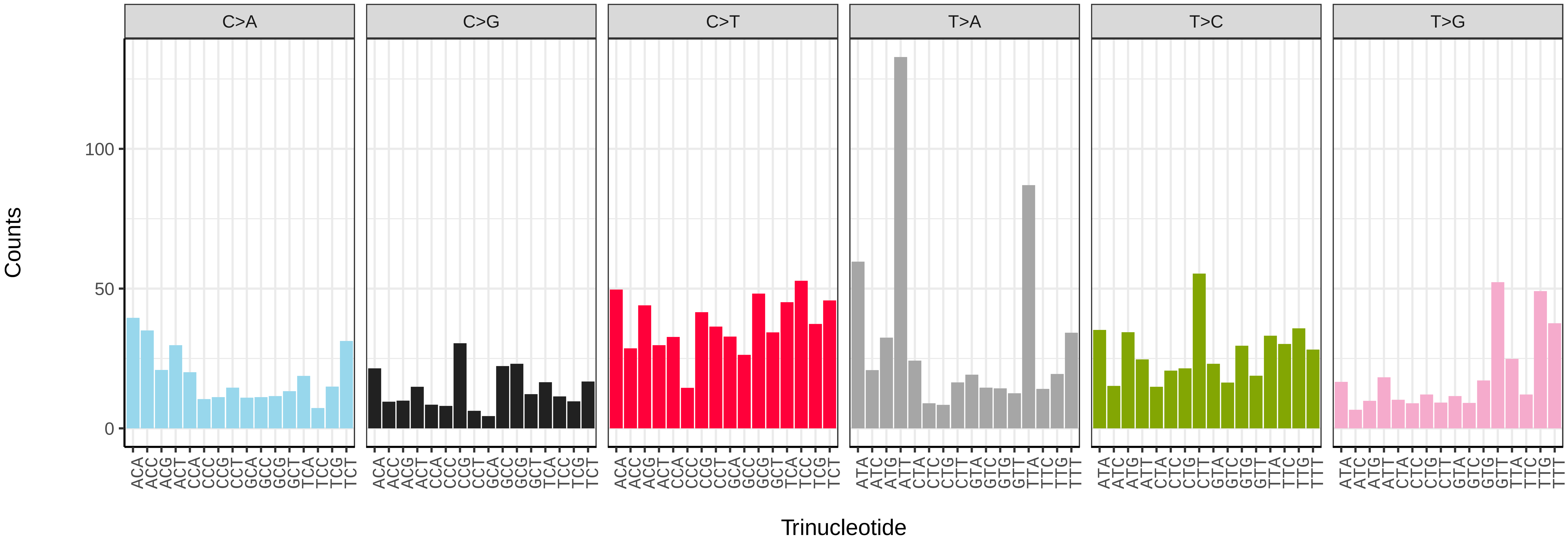


*Epinotia nisella* (ilEpiNise2) somatic mutational spectrum (SBS52 classification)

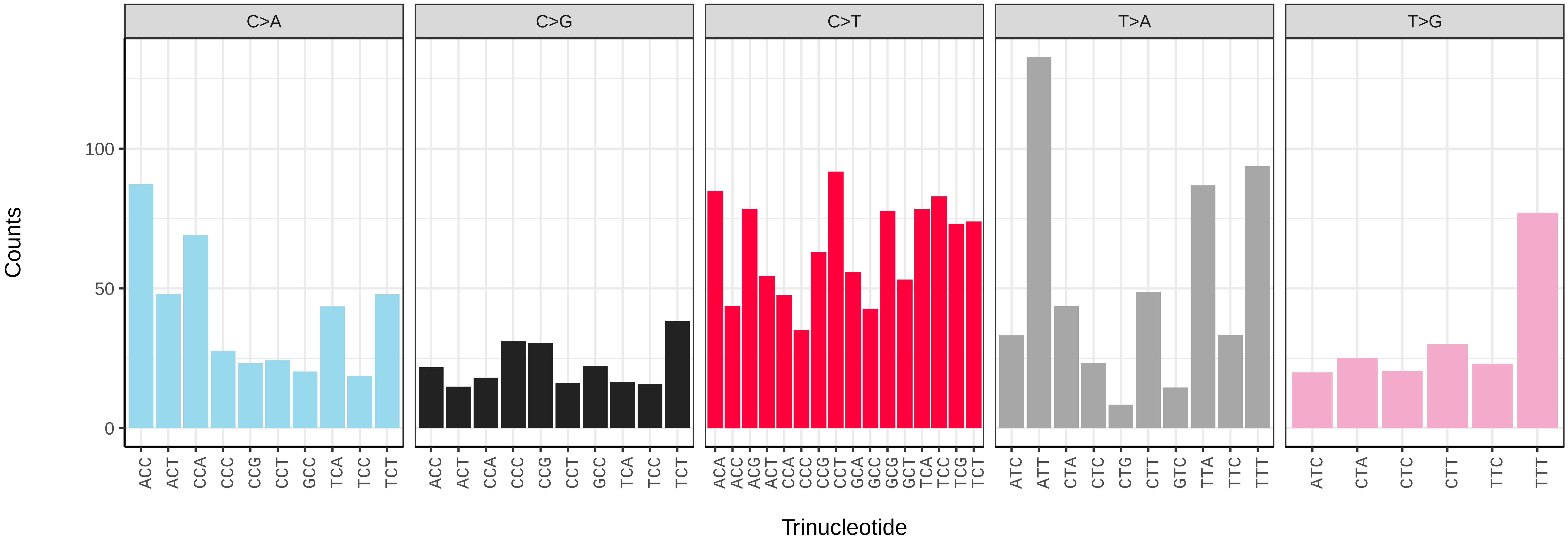

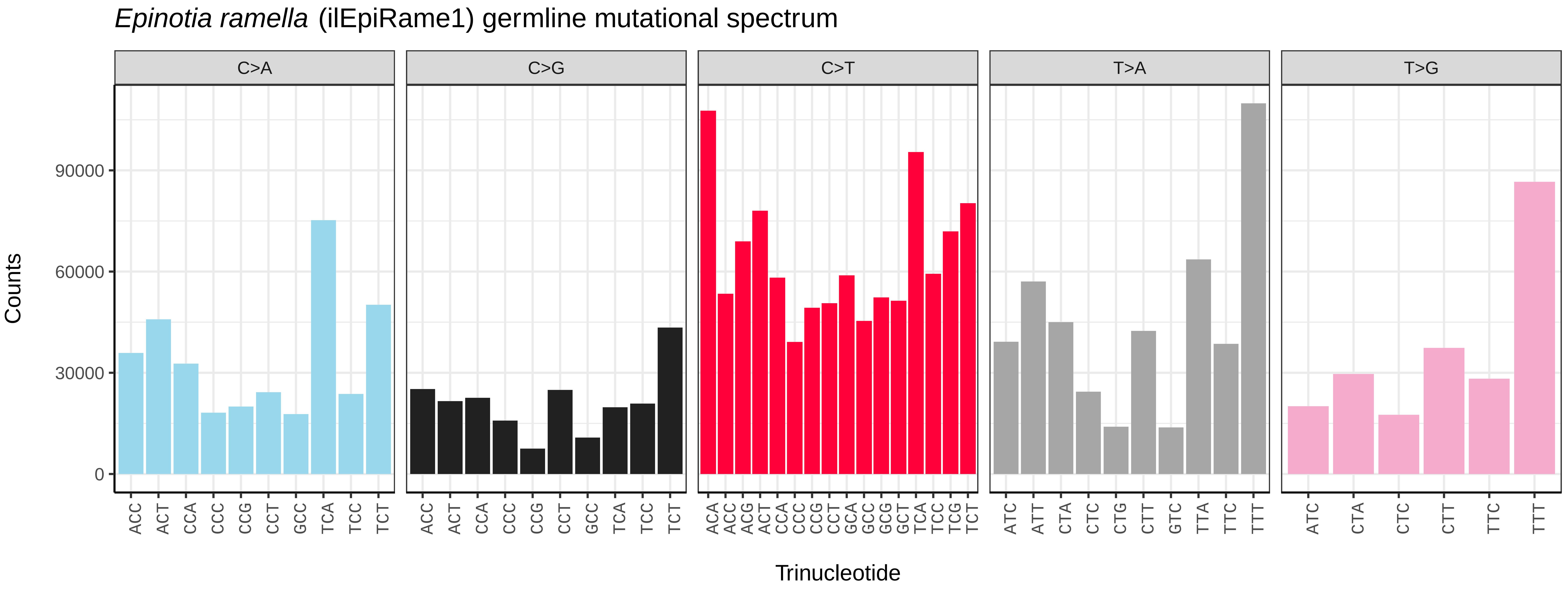
*Epinotia ramella* (ilEpiRame1) germline mutational spectrum
C>A
C>G
C>T
T>A
T>G
Counts
0
30000
60000
90000
Trinucleotide

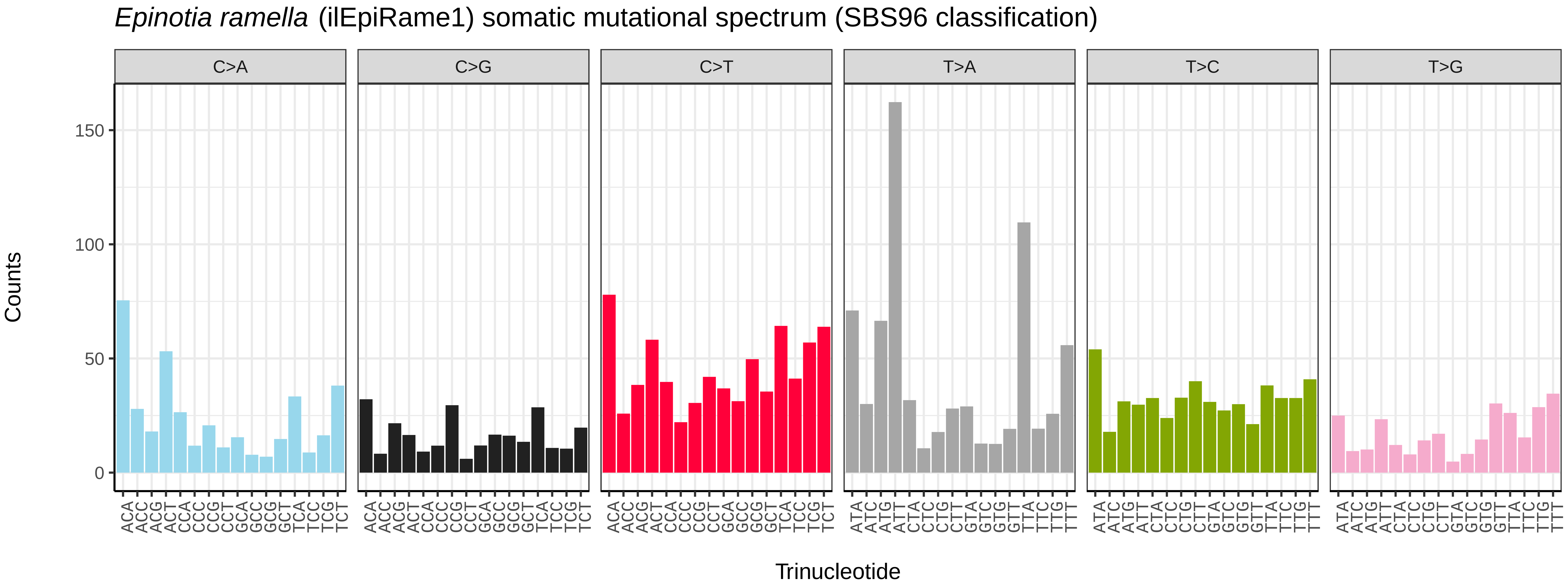
*Epinotia ramella* (ilEpiRame1) somatic mutational spectrum (SBS96 classification)
C>A
C>G
C>T
T>A
T>C
T>G
Counts
0
50
100
150
Trinucleotide

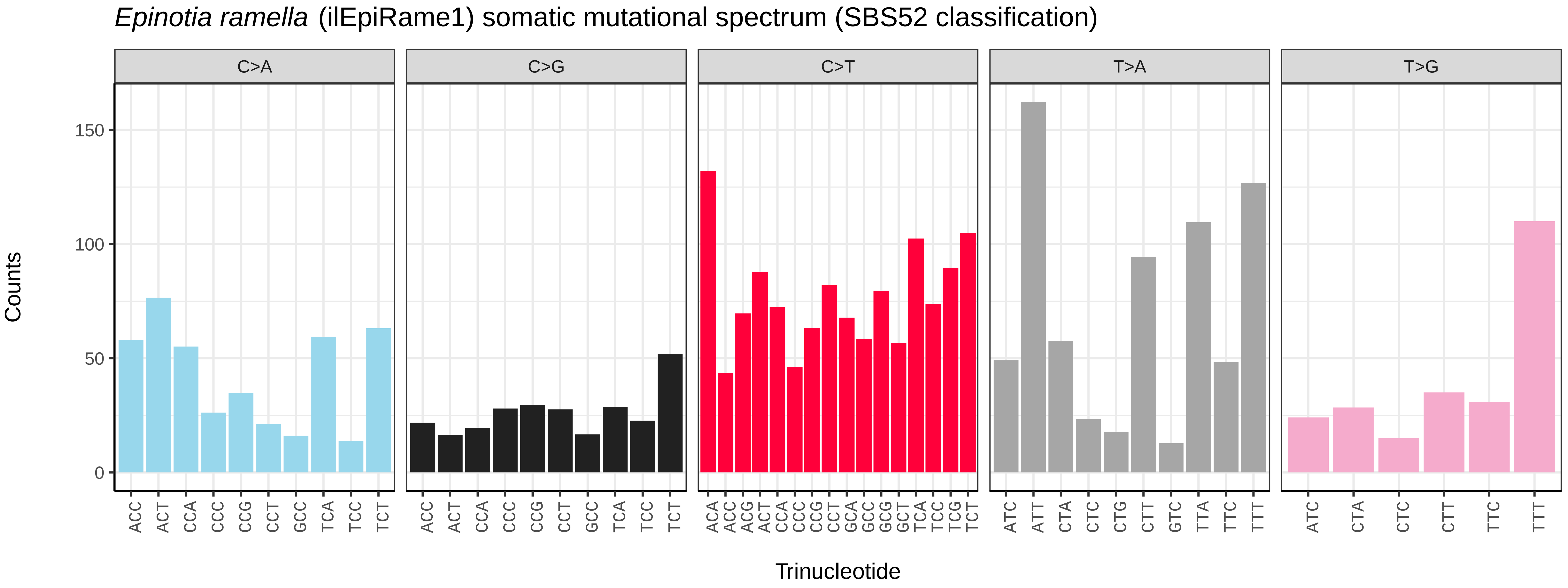
*Epinotia ramella* (ilEpiRame1) somatic mutational spectrum (SBS52 classification)
C>A
C>G
C>T
T>A
T>G
Counts
0
50
100
150
Trinucleotide

*Erannis defoliaria* (ilEraDefo1) germline mutational spectrum

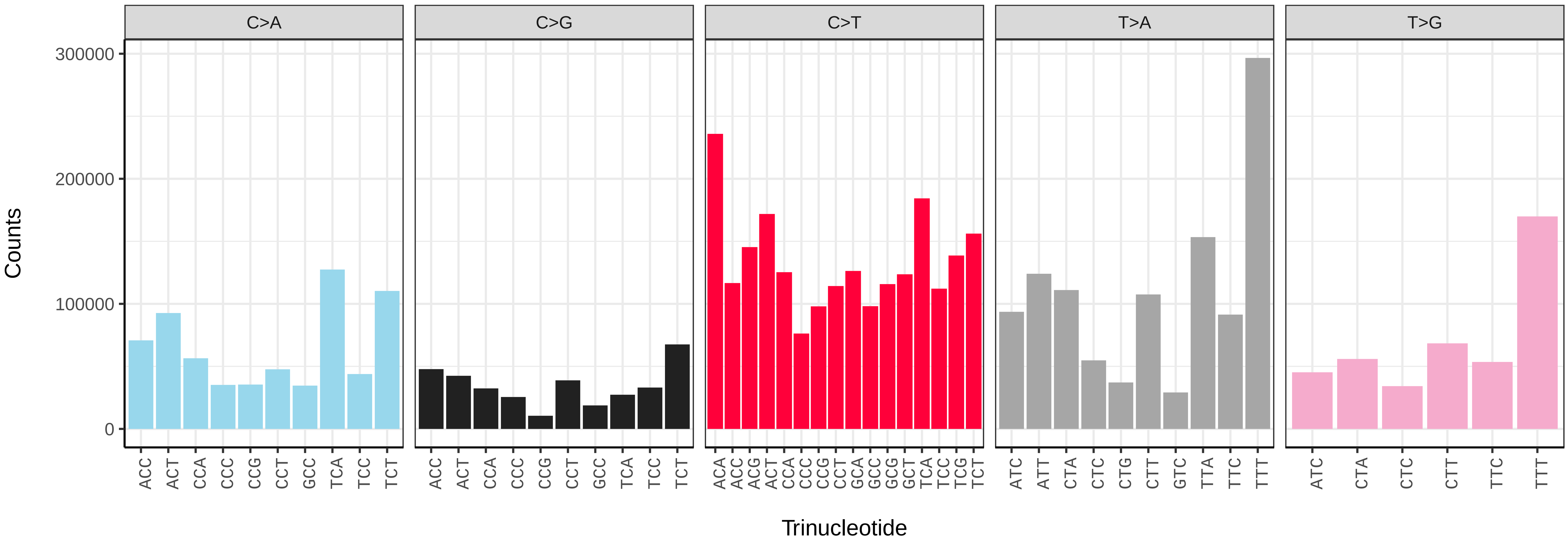


*Erannis defoliaria* (ilEraDefo1) somatic mutational spectrum (SBS96 classification)

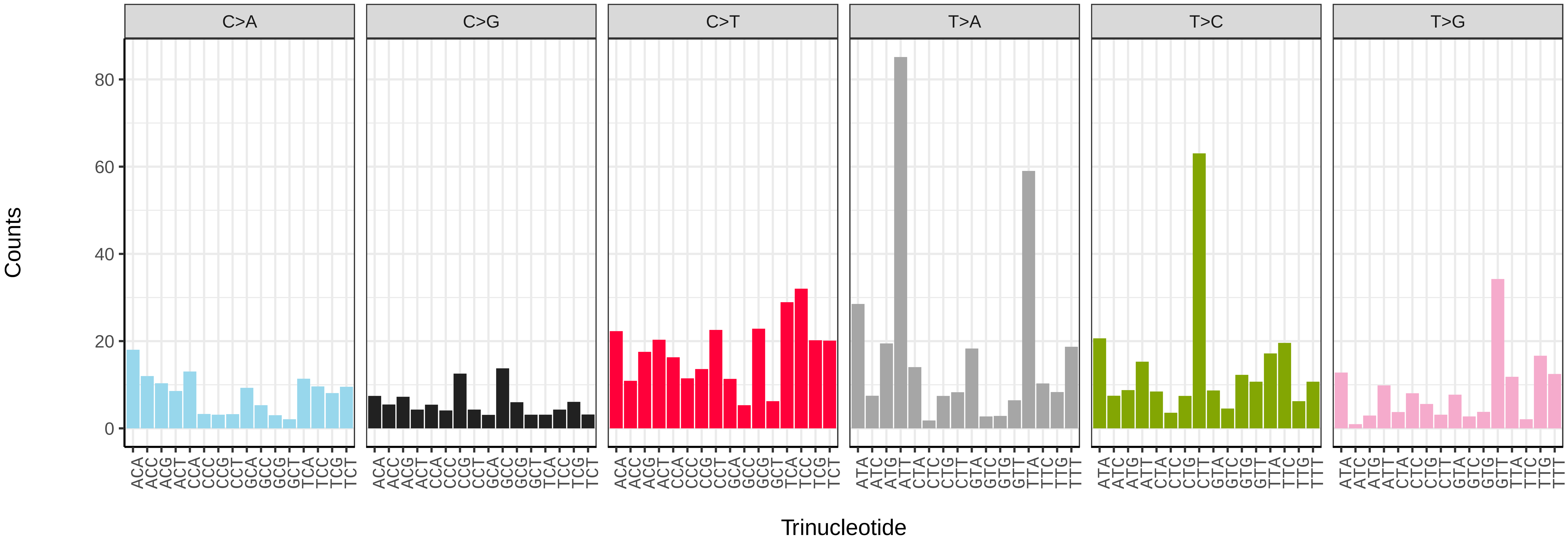


*Erannis defoliaria* (ilEraDefo1) somatic mutational spectrum (SBS52 classification)

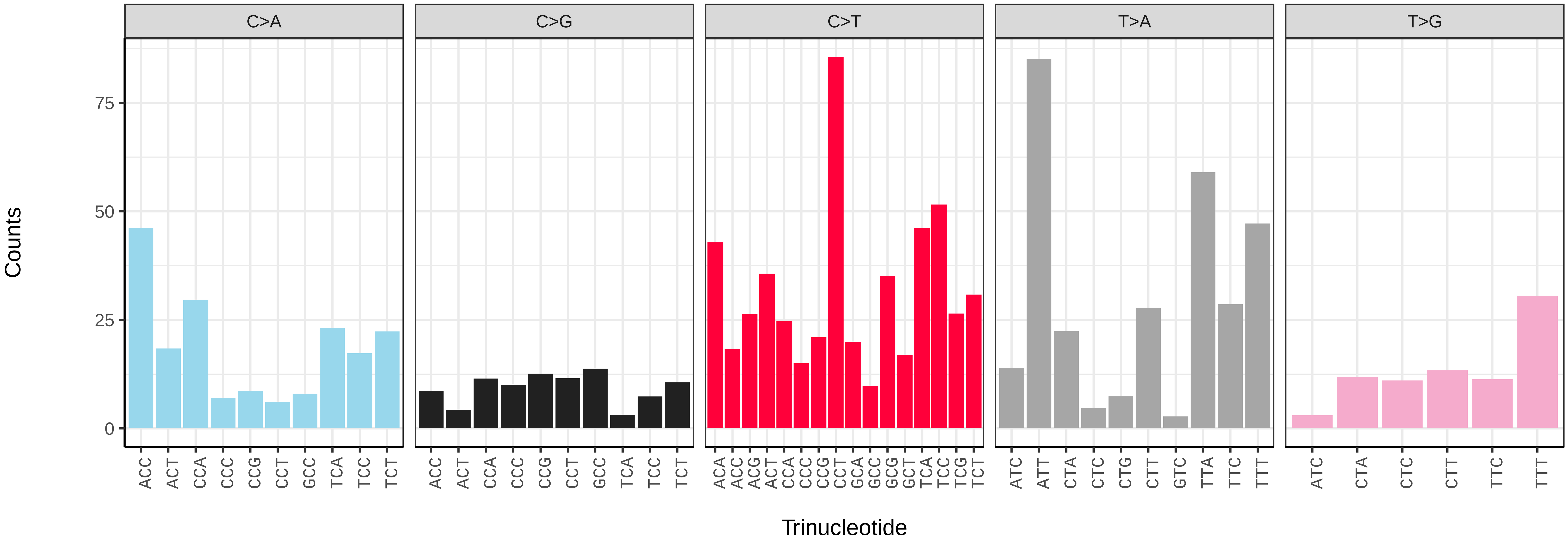

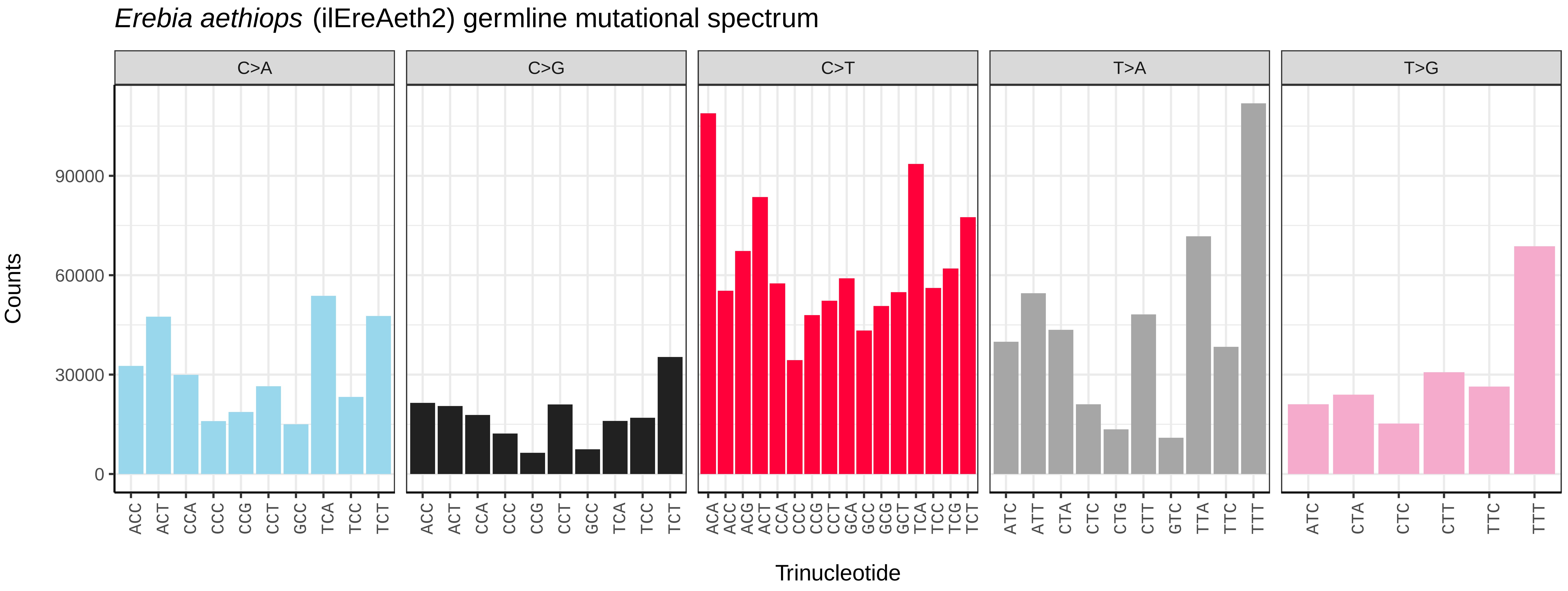
Erebia aethiops (ilEreAeth2) germline mutational spectrum
C>A
C>G
C>T
T>A
T>G
Counts
0
30000
60000
90000
Trinucleotide

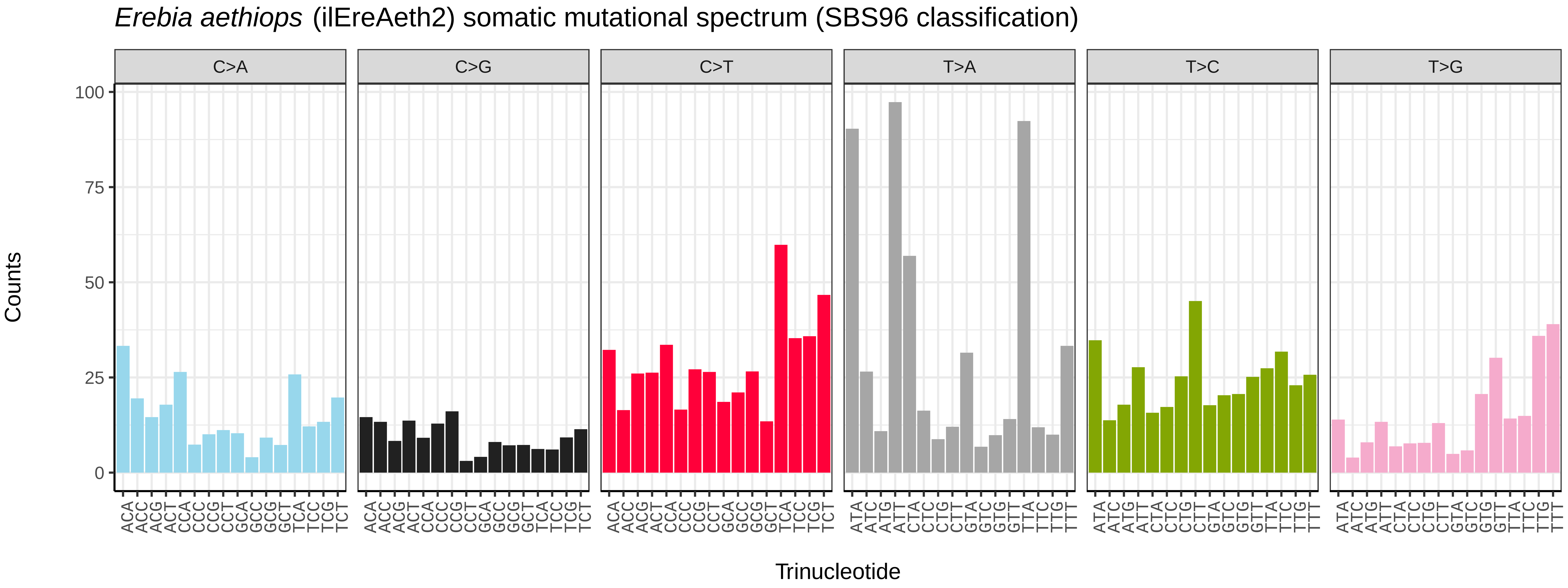
Erebia aethiops (ilEreAeth2) somatic mutational spectrum (SBS96 classification)
C>A
C>G
C>T
T>A
T>C
T>G
Counts
0
25
50
75
100
Trinucleotide

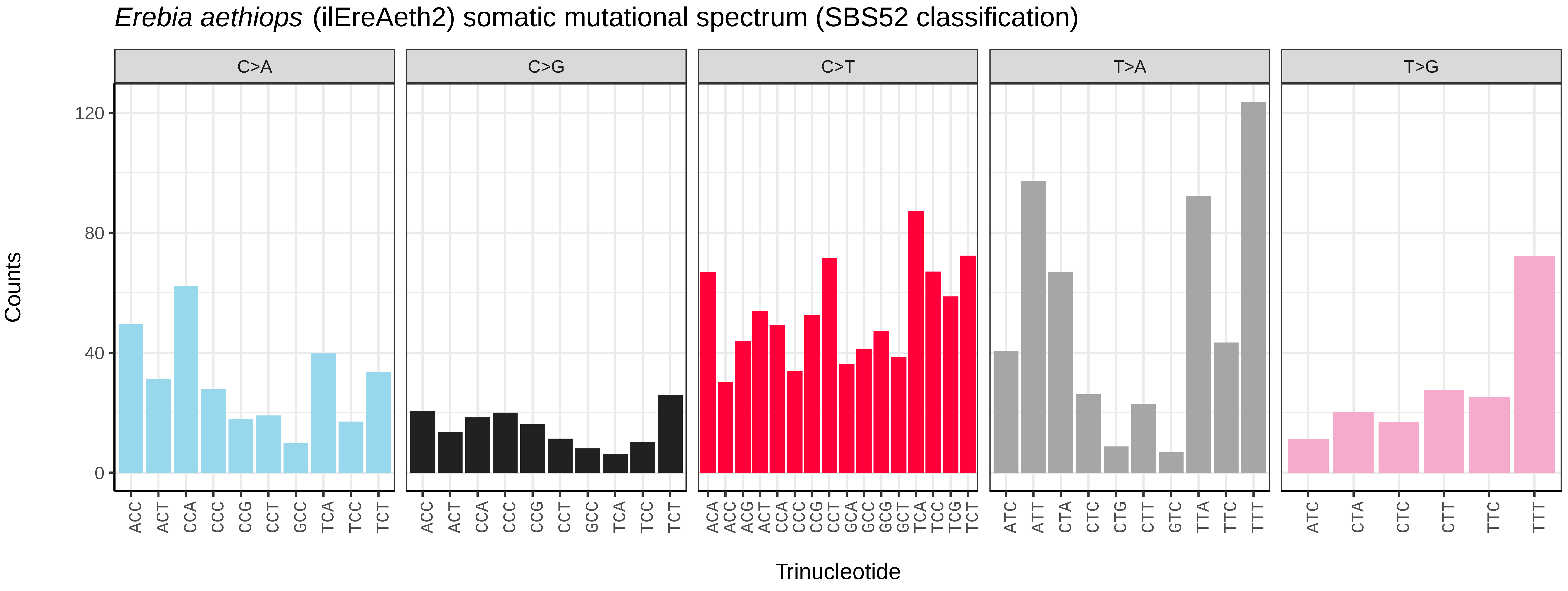
Erebia aethiops (ilEreAeth2) somatic mutational spectrum (SBS52 classification)
C>A
C>G
C>T
T>A
T>G
Counts
0
40
80
120
Trinucleotide

*Erebia ligea* (ilEreLige1) germline mutational spectrum

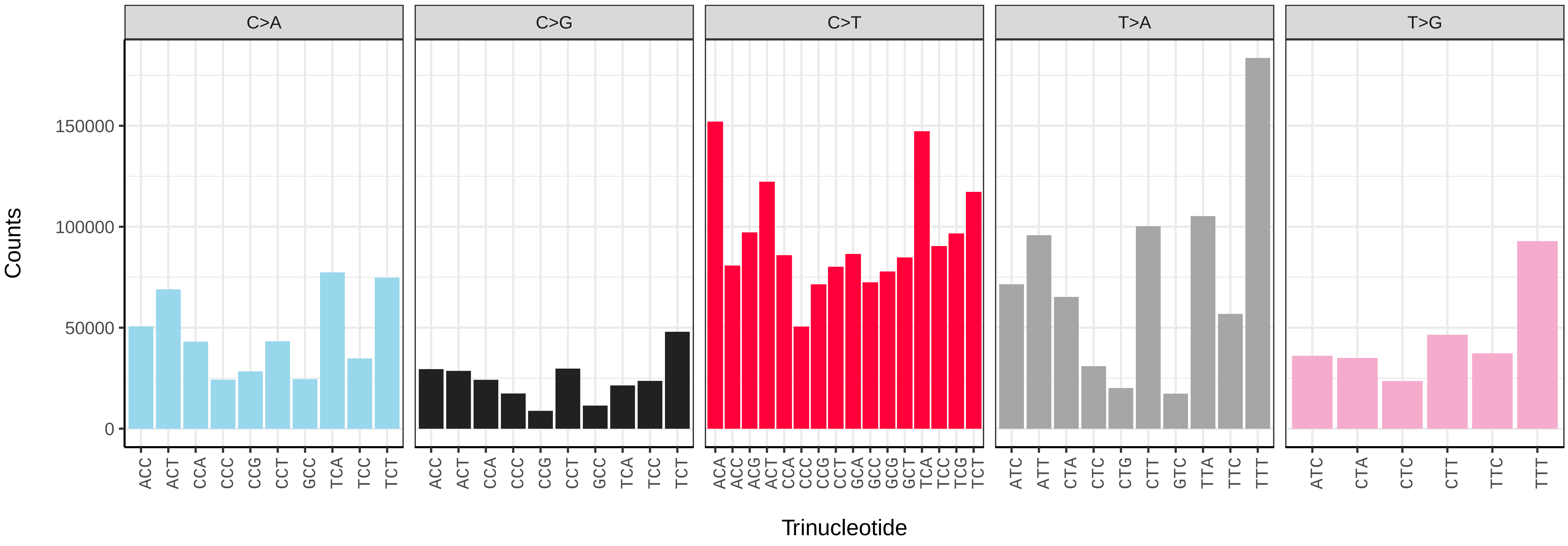


*Erebia ligea* (ilEreLige1) somatic mutational spectrum (SBS96 classification)

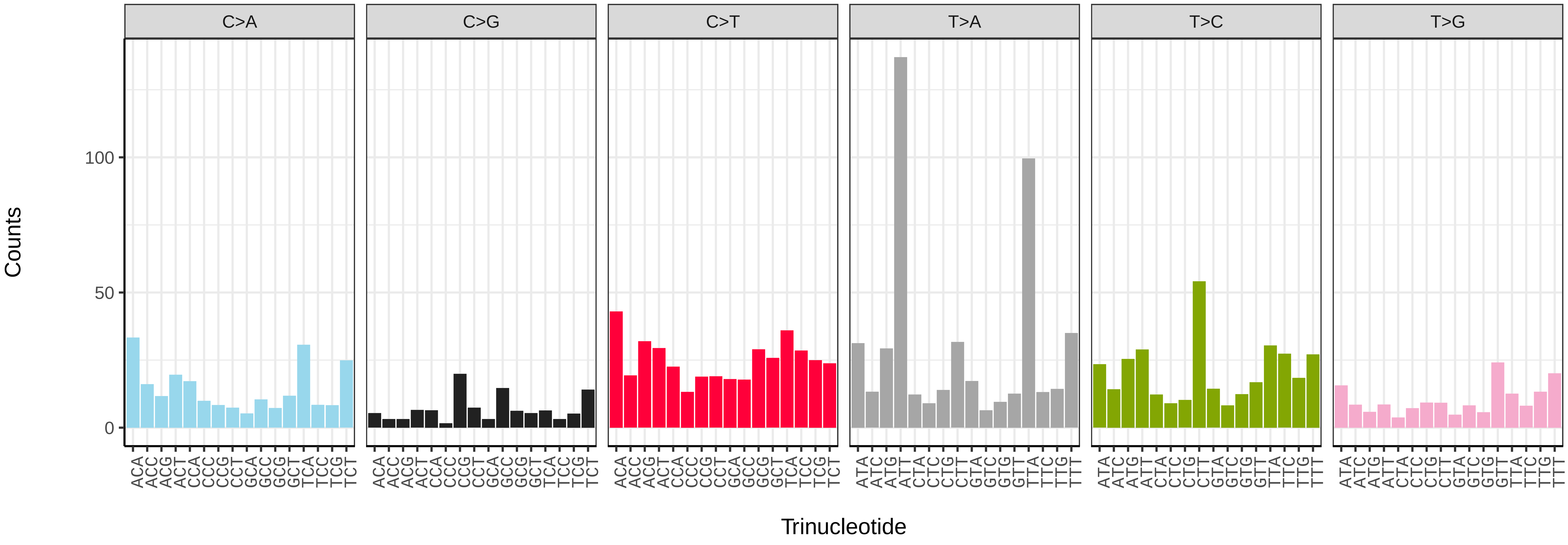


*Erebia ligea* (ilEreLige1) somatic mutational spectrum (SBS52 classification)

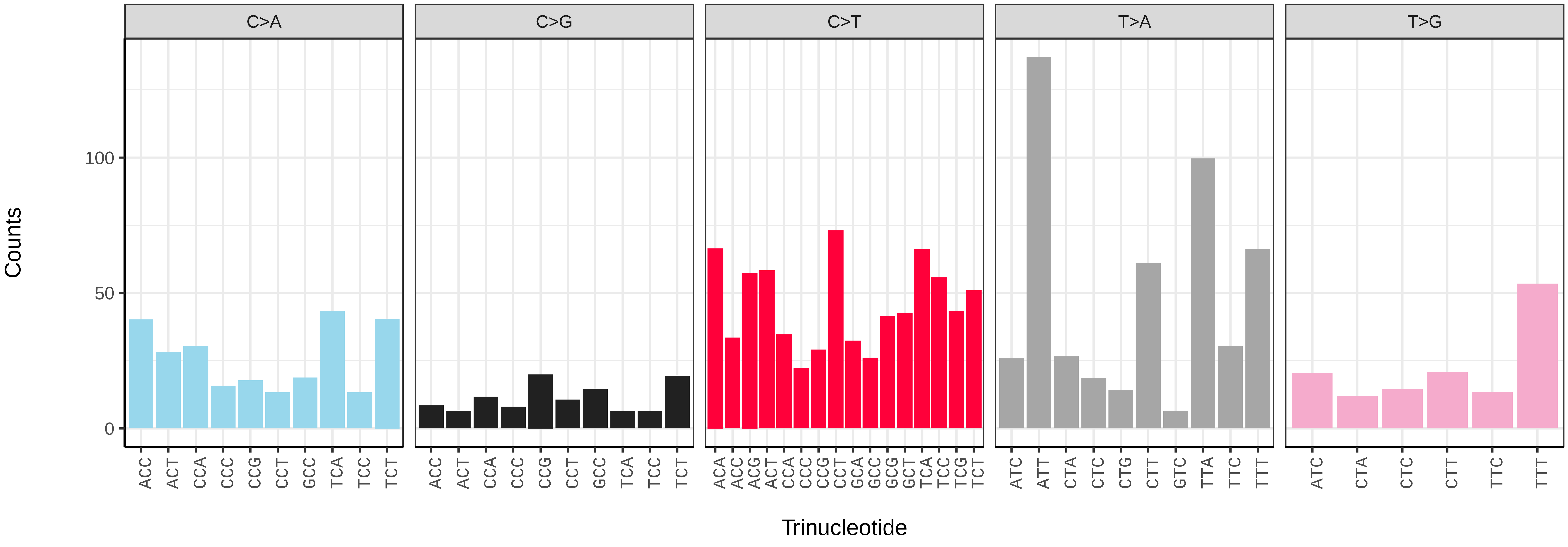

*Erynnis tages* (ilEryTage1) germline mutational spectrum

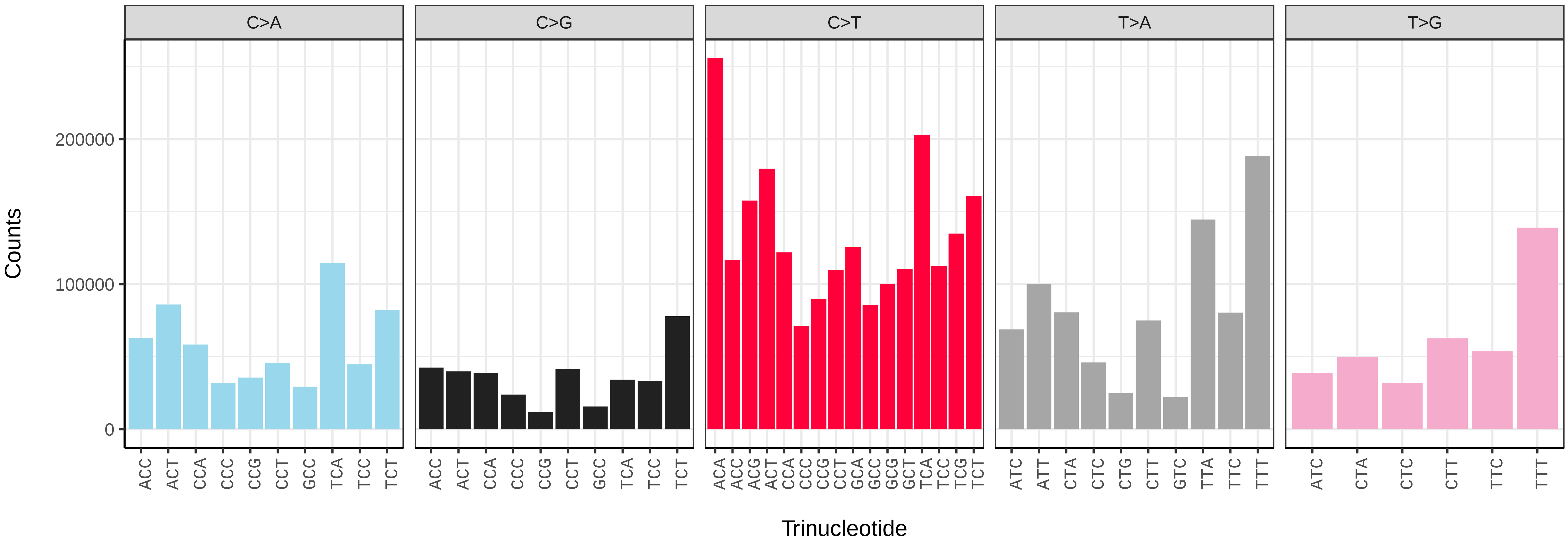


*Erynnis tages* (ilEryTage1) somatic mutational spectrum (SBS96 classification)

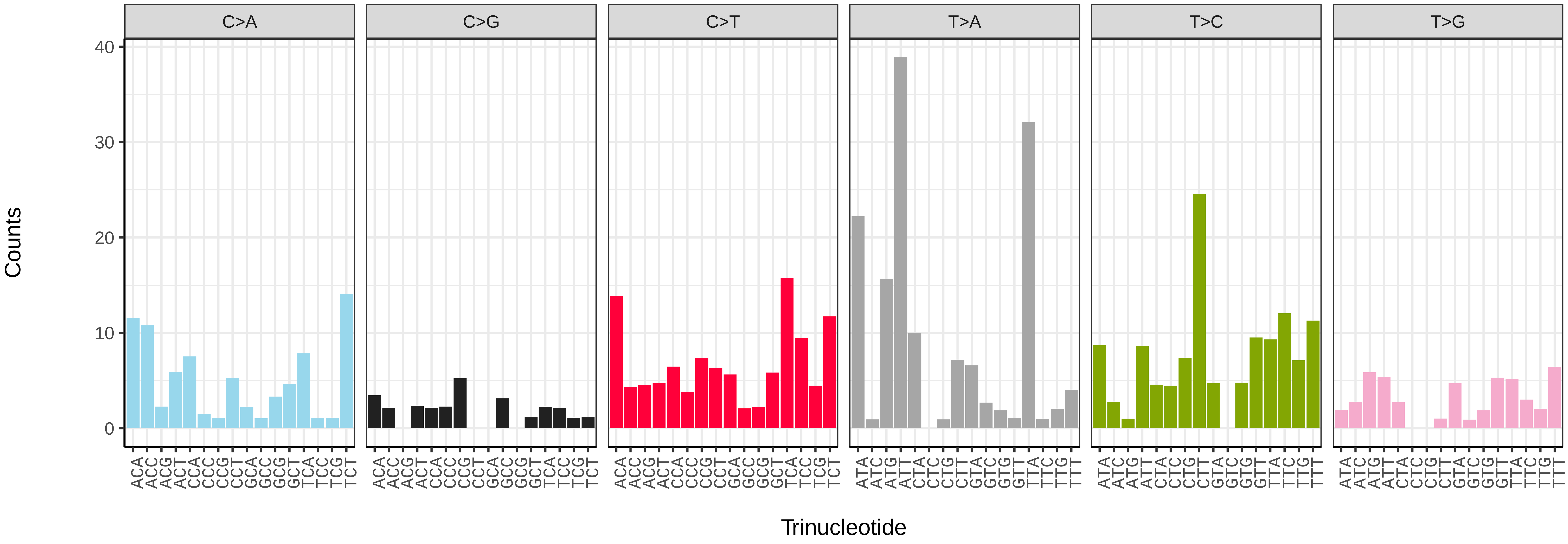


*Erynnis tages* (ilEryTage1) somatic mutational spectrum (SBS52 classification)

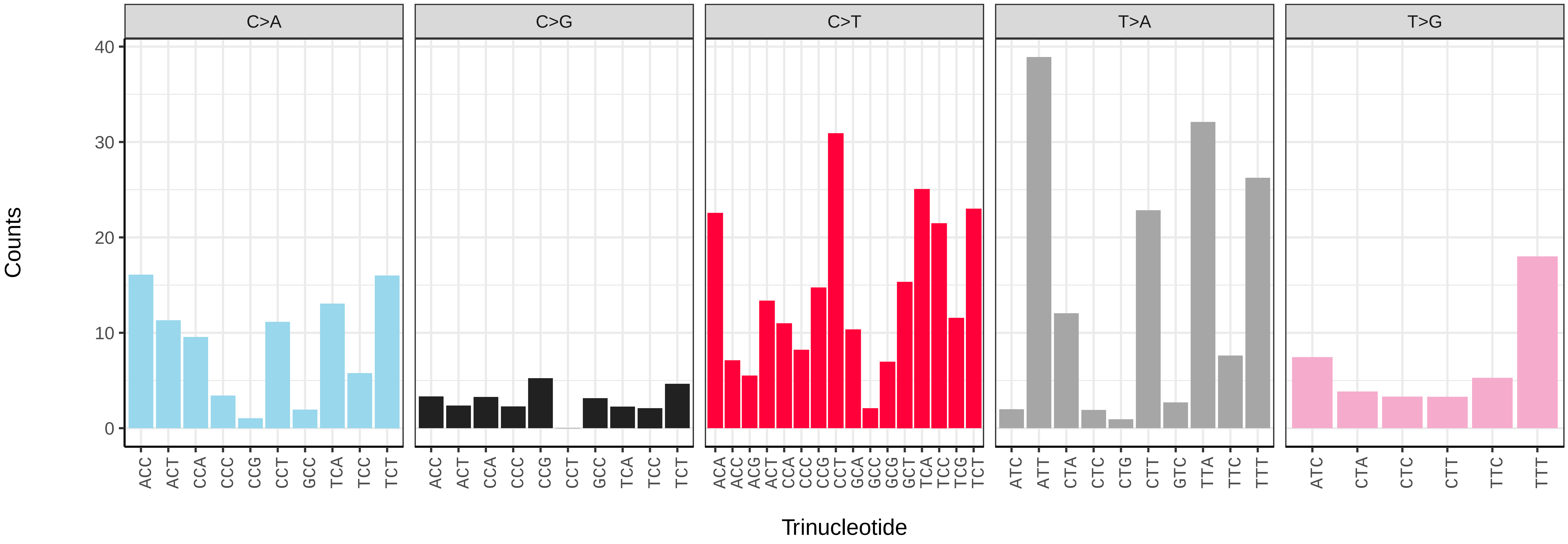

*Esperia sulphurella* (ilEspSulp1) germline mutational spectrum

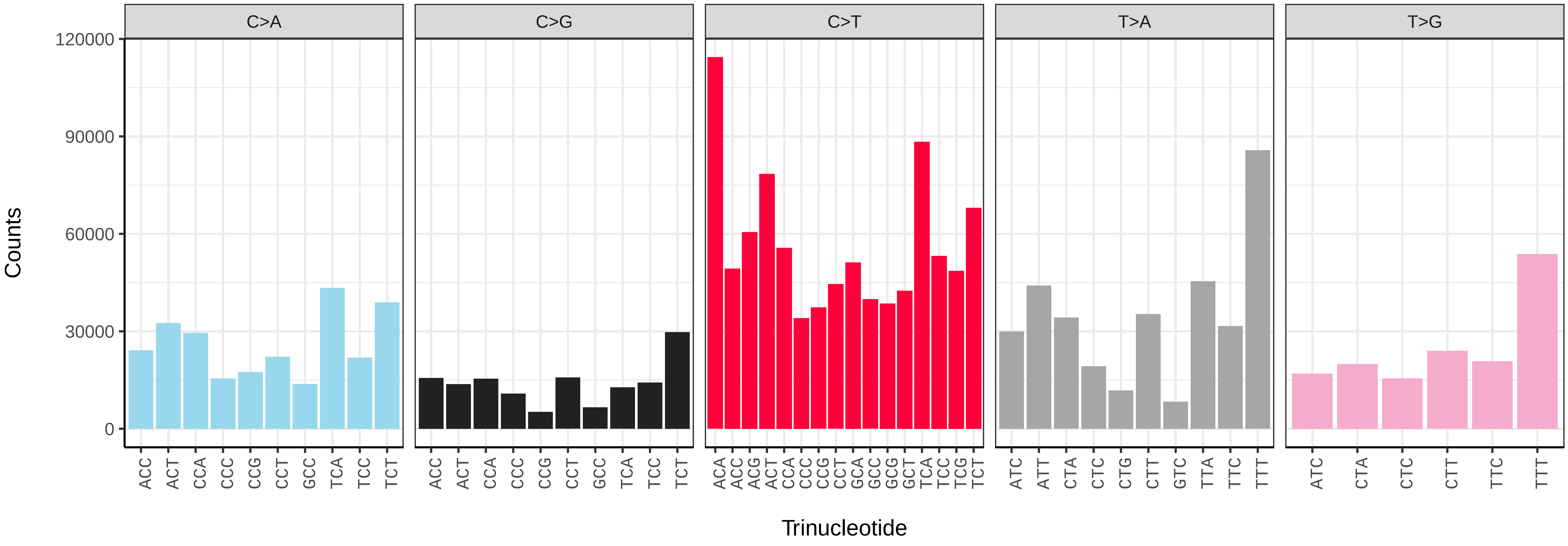


*Esperia sulphurella* (ilEspSulp1) somatic mutational spectrum (SBS96 classification)

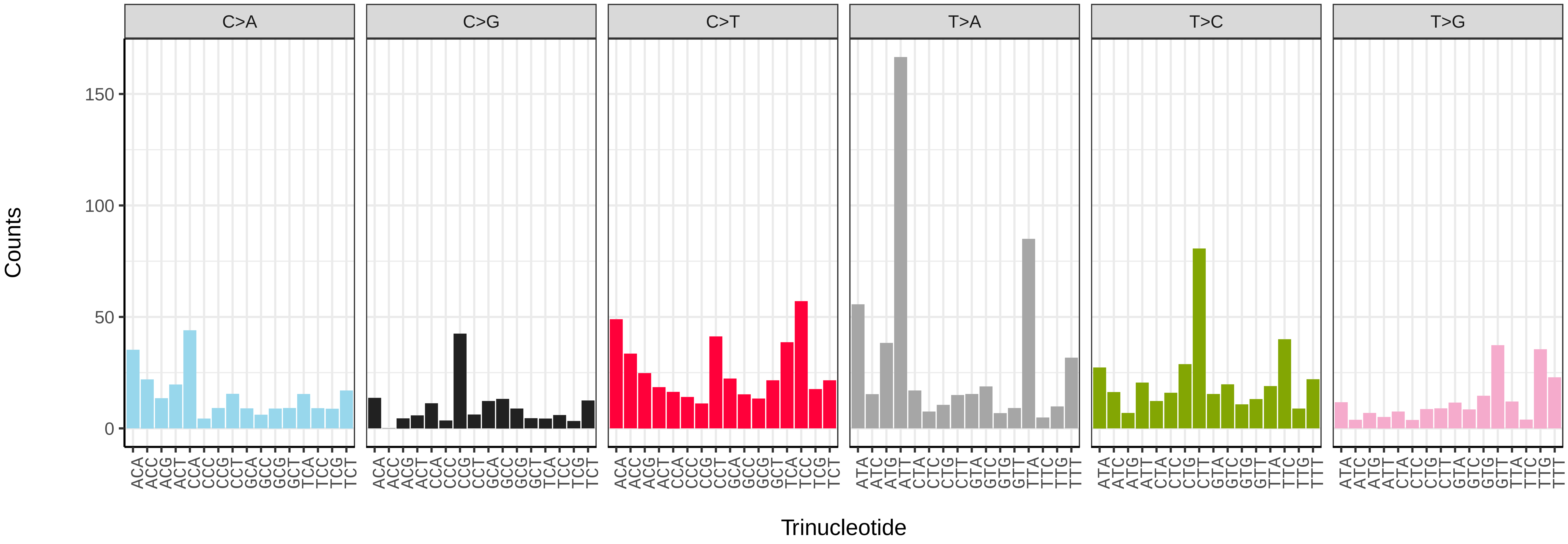


*Esperia sulphurella* (ilEspSulp1) somatic mutational spectrum (SBS52 classification)

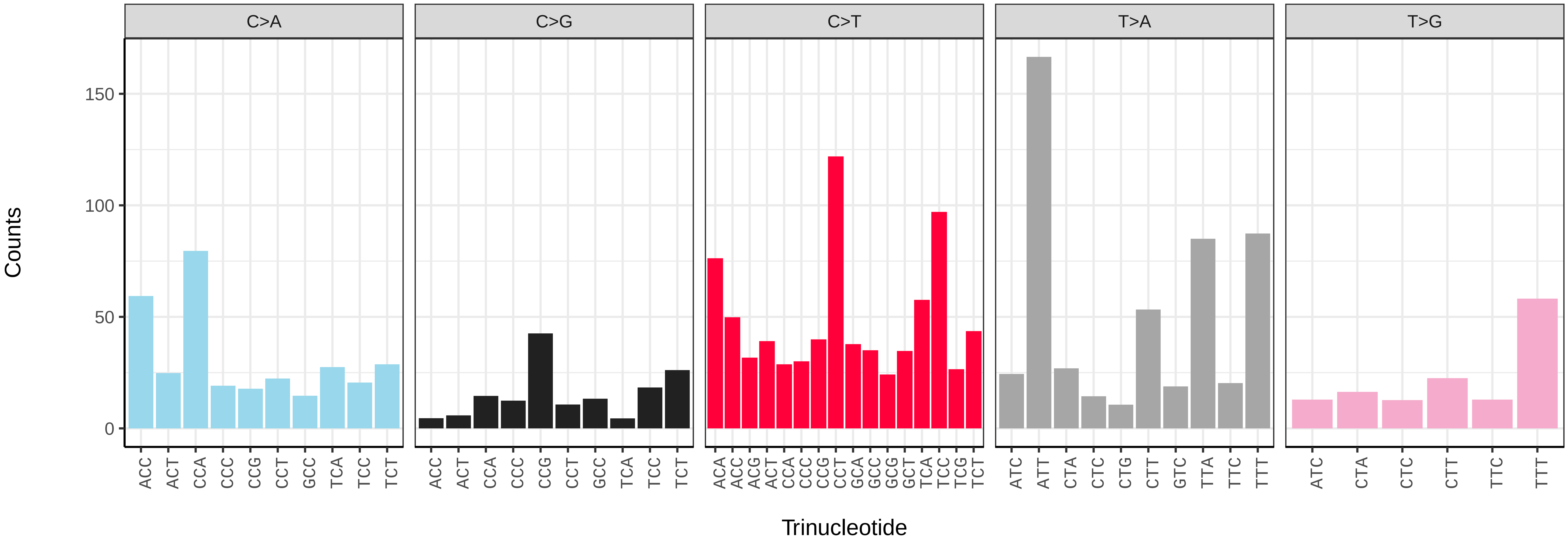

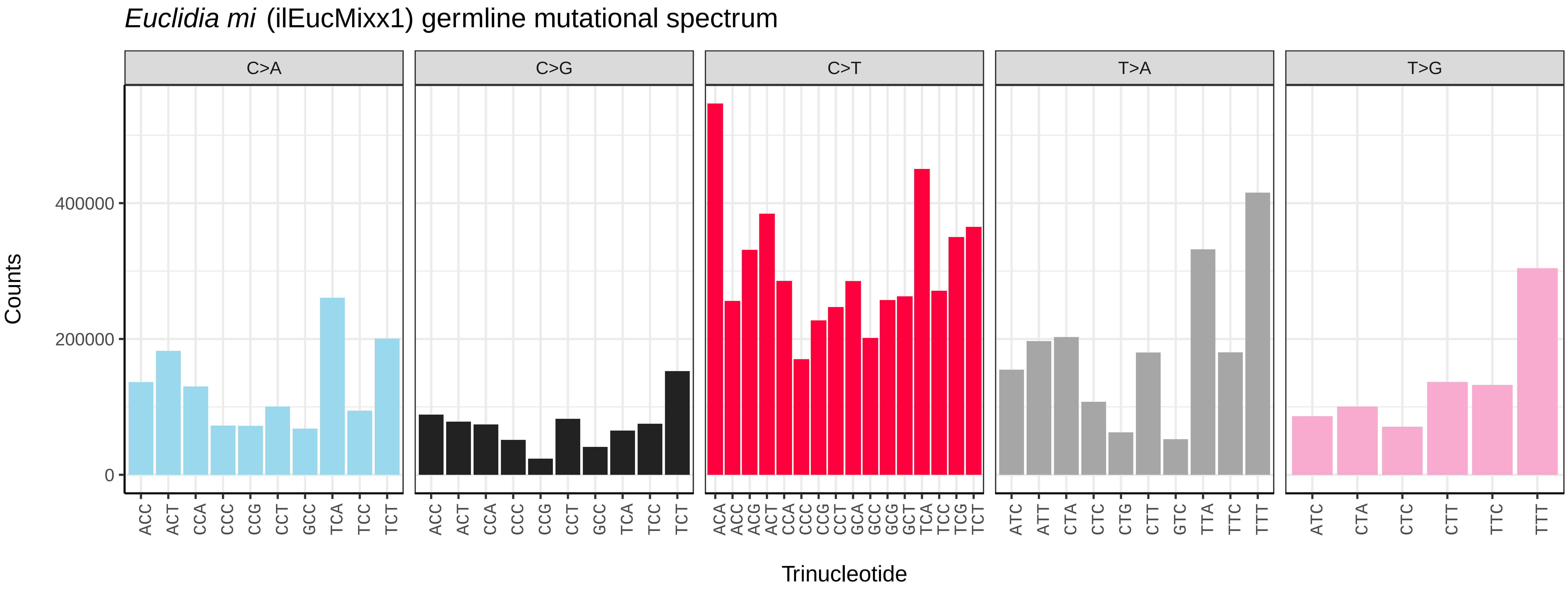
Euclidia mi (ilEucMixx1) germline mutational spectrum
C>A
C>G
C>T
T>A
T>G
Counts
0
200000
400000
Trinucleotide
ACC ACT CCA CCC CCG CCT GCC TCA TCC TCT
ACA ACC ACG ACT CCA CCC CCG CCT GCA GCC GCG GCT TCA TCC TCG TCT
ATC ATT CTA CTC CTG CTT GTC TTA TTC TTT
ATC CTA CTC CTT TTC TTT

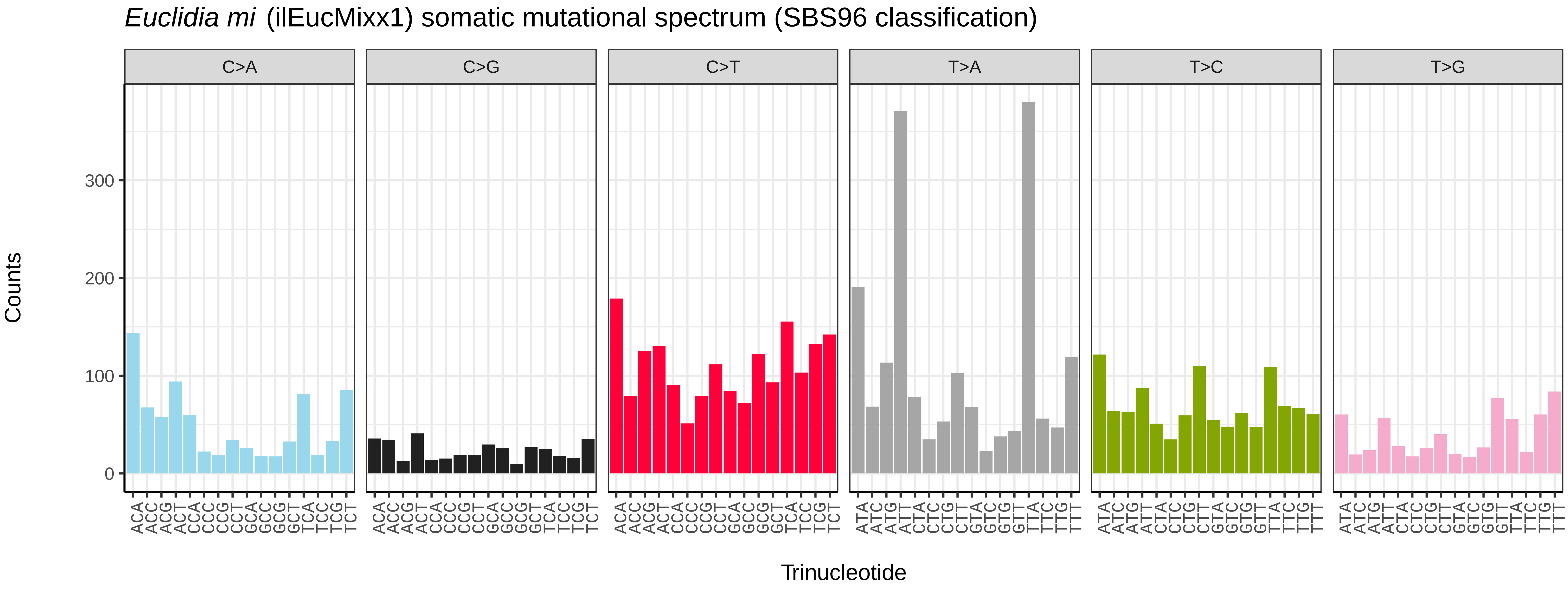
Euclidia mi (ilEucMixx1) somatic mutational spectrum (SBS96 classification)
C>A
C>G
C>T
T>A
T>C
T>G
Counts
0
100
200
300
Trinucleotide
ACA ACC ACG ACT CCA CCC CCG CCT GCA GCC GCG GCT TCA TCC TCG TCT
ATA ATC ATG ATT CTA CTC CTG CTT GTA GTC GTG GTT TTA TTC TTG TTT

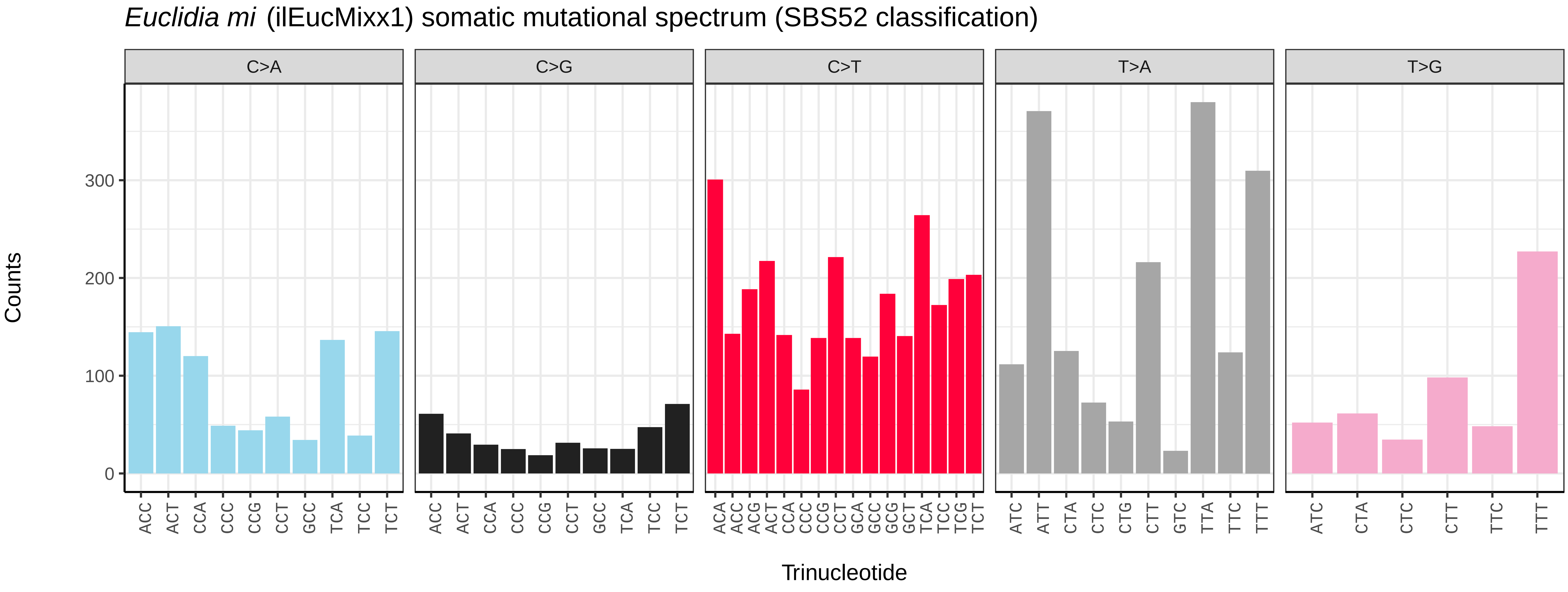
Euclidia mi (ilEucMixx1) somatic mutational spectrum (SBS52 classification)
C>A
C>G
C>T
T>A
T>G
Counts
0
100
200
300
Trinucleotide
ACC ACT CCA CCC CCG CCT GCC TCA TCC TCT
ACA ACC ACG ACT CCA CCC CCG CCT GCA GCC GCG GCT TCA TCC TCG TCT
ATC ATT CTA CTC CTG CTT GTC TTA TTC TTT
ATC CTA CTC CTT TTC TTT

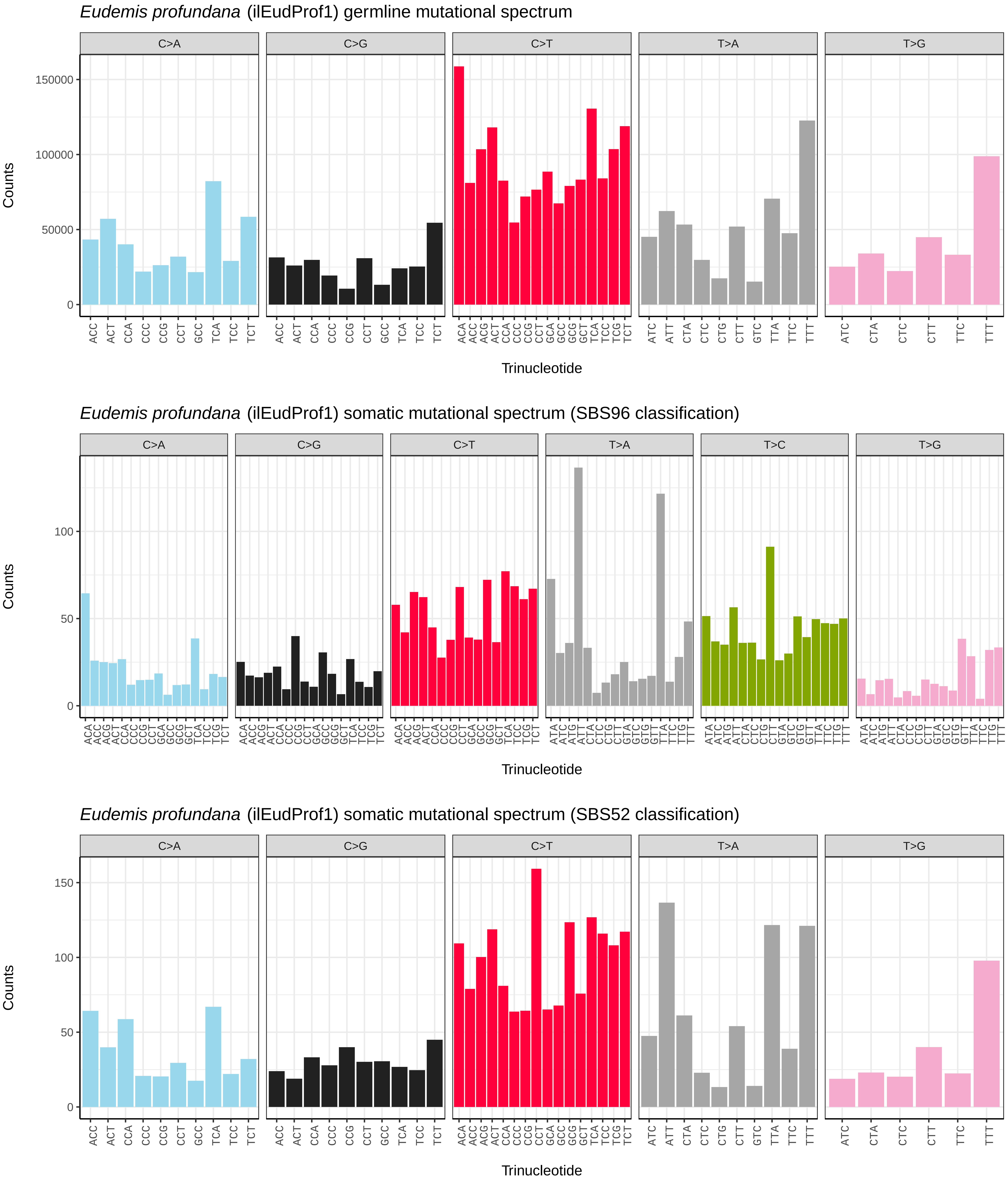

*Eudemis profundana* (ilEudProf1) germline mutational spectrum
C>A
C>G
C>T
T>A
T>G
Counts
150000
100000
50000
0
ACC ACT CCA CCC CCG CCT GCC TCA TCC TCT
ACA ACC ACG ACT CCA CCC CCG CCT GCA GCC GCG GCT TCA TCC TCG TCT
ATC ATT CTA CTC CTG CTT GTC TTA TTC TTT
ATC CTA CTC CTT TTC TTT
Trinucleotide
*Eudemis profundana* (ilEudProf1) somatic mutational spectrum (SBS96 classification)
C>A
C>G
C>T
T>A
T>C
T>G
Counts
100
50
0
ACA ACC ACG ACT CCA CCC CCG CCT GCA GCC GCG GCT TCA TCC TCG TCT
ATA ATC ATG ATT CTA CTC CTG CTT GTA GTC GTG GTT TTA TTC TTG TTT
Trinucleotide
*Eudemis profundana* (ilEudProf1) somatic mutational spectrum (SBS52 classification)
C>A
C>G
C>T
T>A
T>G
Counts
150
100
50
0
ACC ACT CCA CCC CCG CCT GCC TCA TCC TCT
ACA ACC ACG ACT CCA CCC CCG CCT GCA GCC GCG GCT TCA TCC TCG TCT
ATC ATT CTA CTC CTG CTT GTC TTA TTC TTT
ATC CTA CTC CTT TTC TTT
Trinucleotide

*Eugnorisma glareosa* (ilEugGlar6) germline mutational spectrum

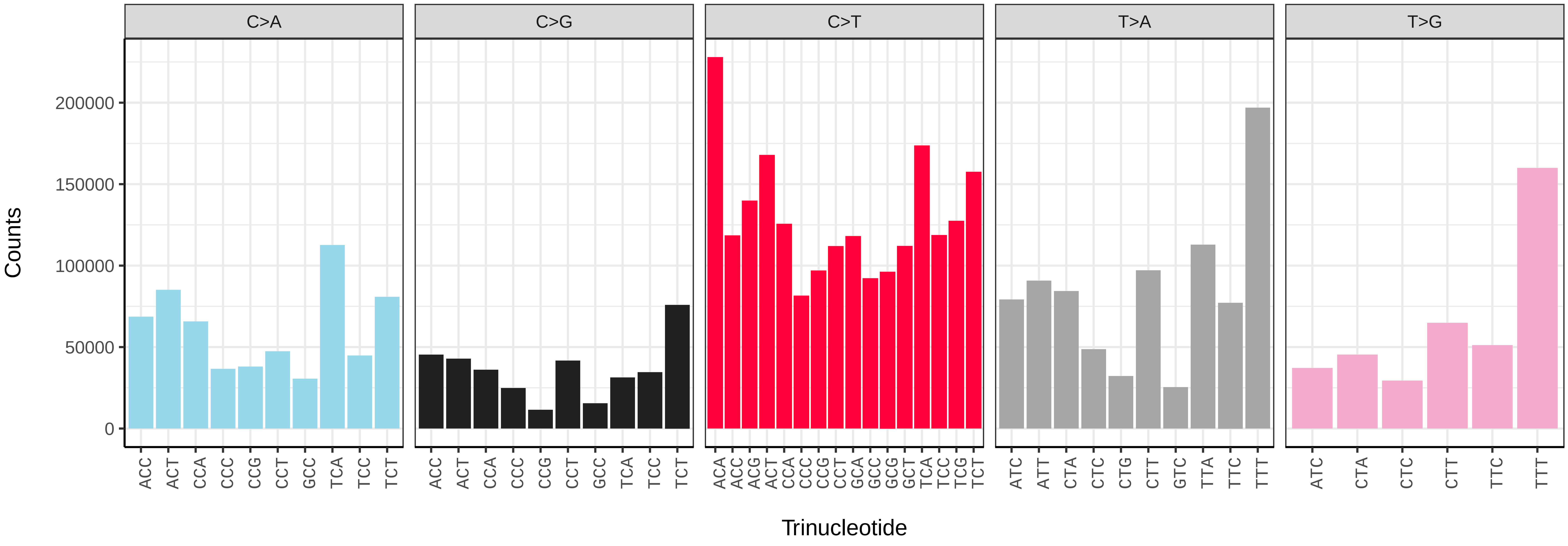


*Eugnorisma glareosa* (ilEugGlar6) somatic mutational spectrum (SBS96 classification)

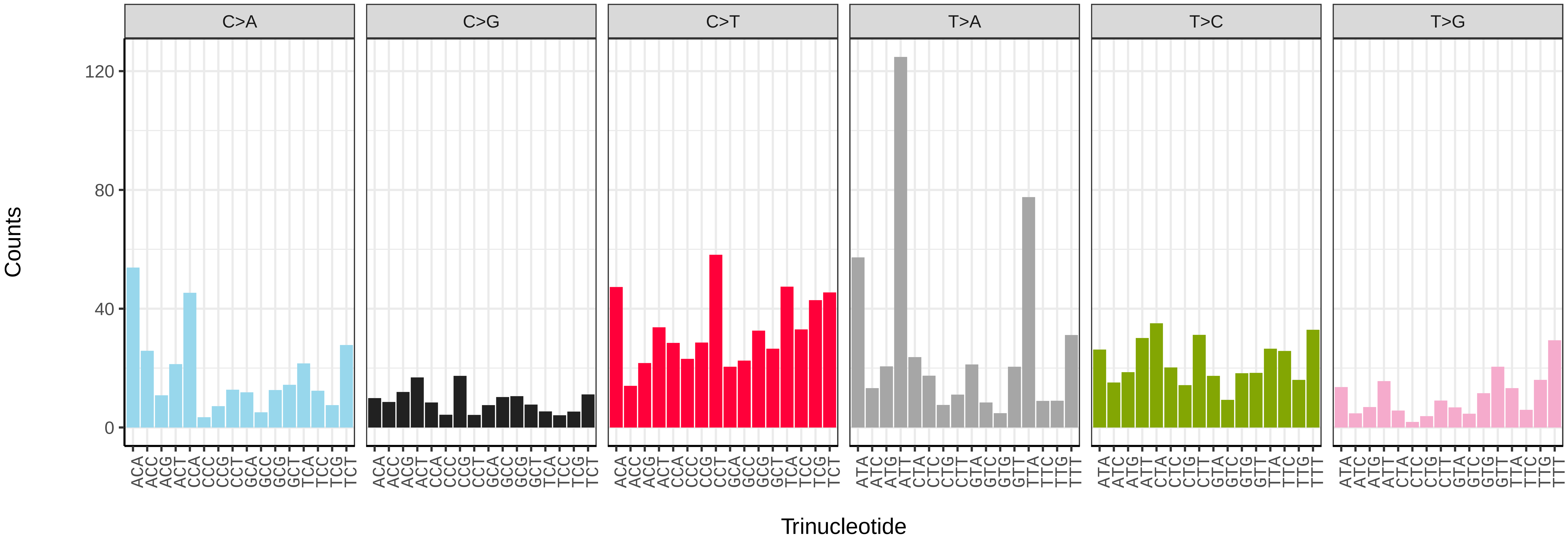


*Eugnorisma glareosa* (ilEugGlar6) somatic mutational spectrum (SBS52 classification)

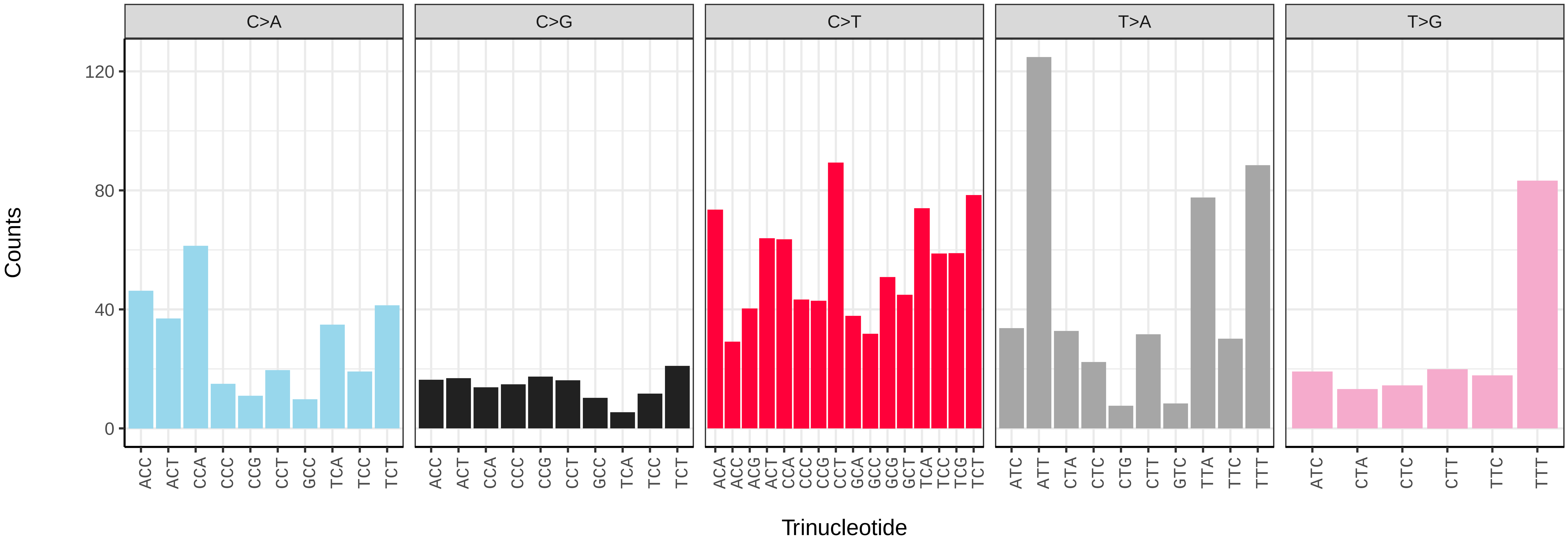

*Eulithis prunata* (ilEulPrun1) germline mutational spectrum

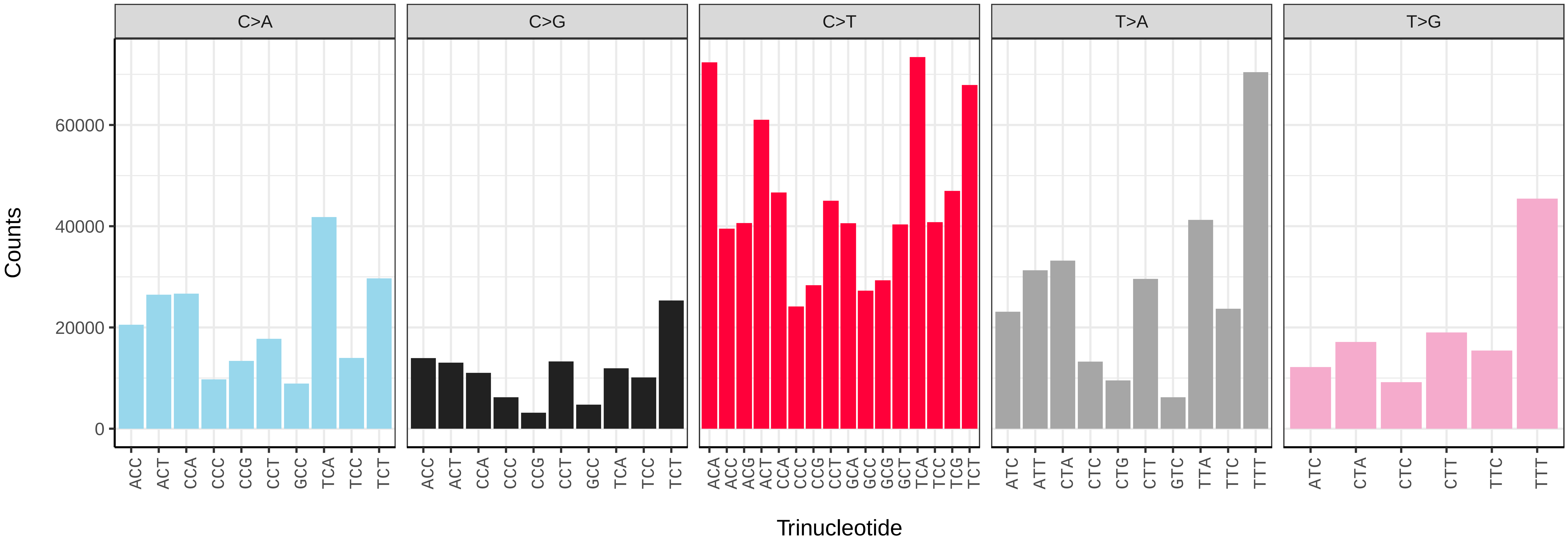


*Eulithis prunata* (ilEulPrun1) somatic mutational spectrum (SBS96 classification)

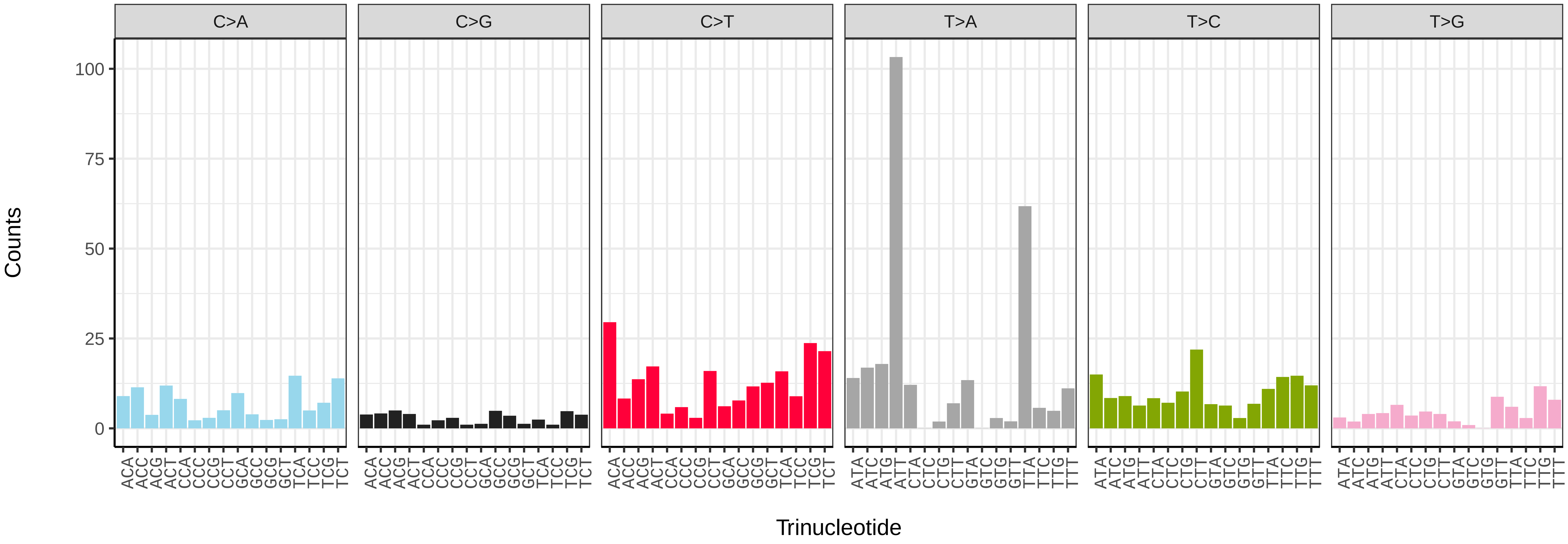


*Eulithis prunata* (ilEulPrun1) somatic mutational spectrum (SBS52 classification)

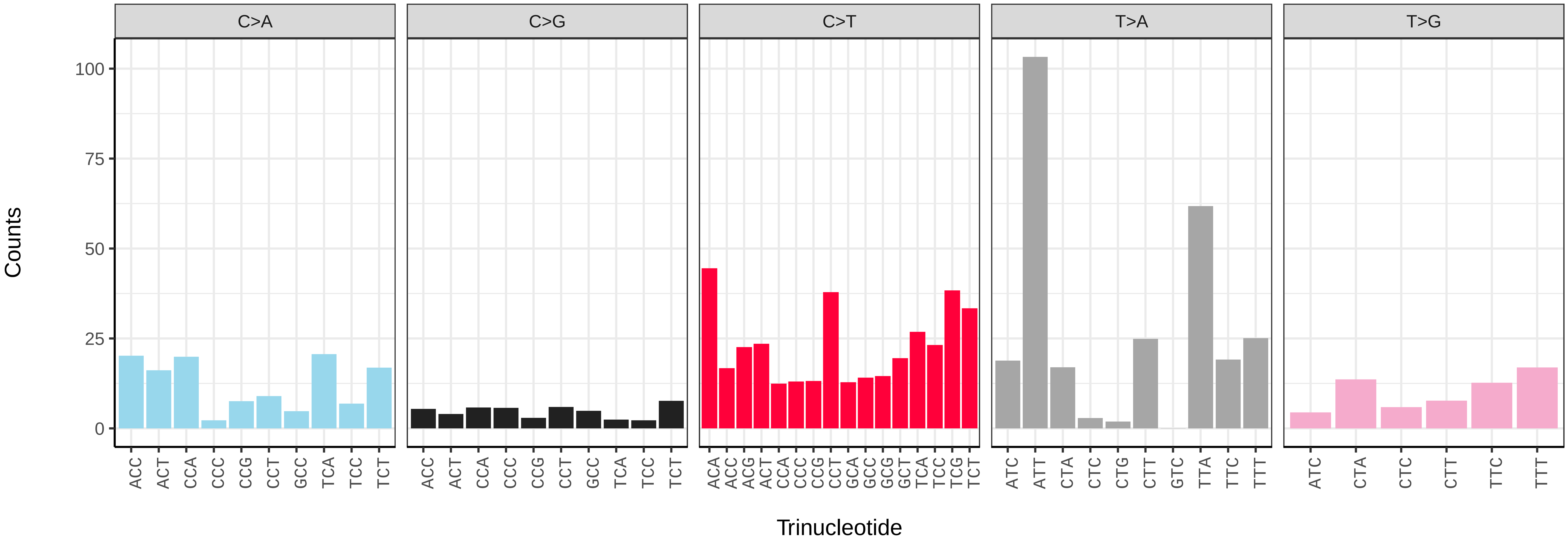

*Eulithis testata* (ilEulTest2) germline mutational spectrum

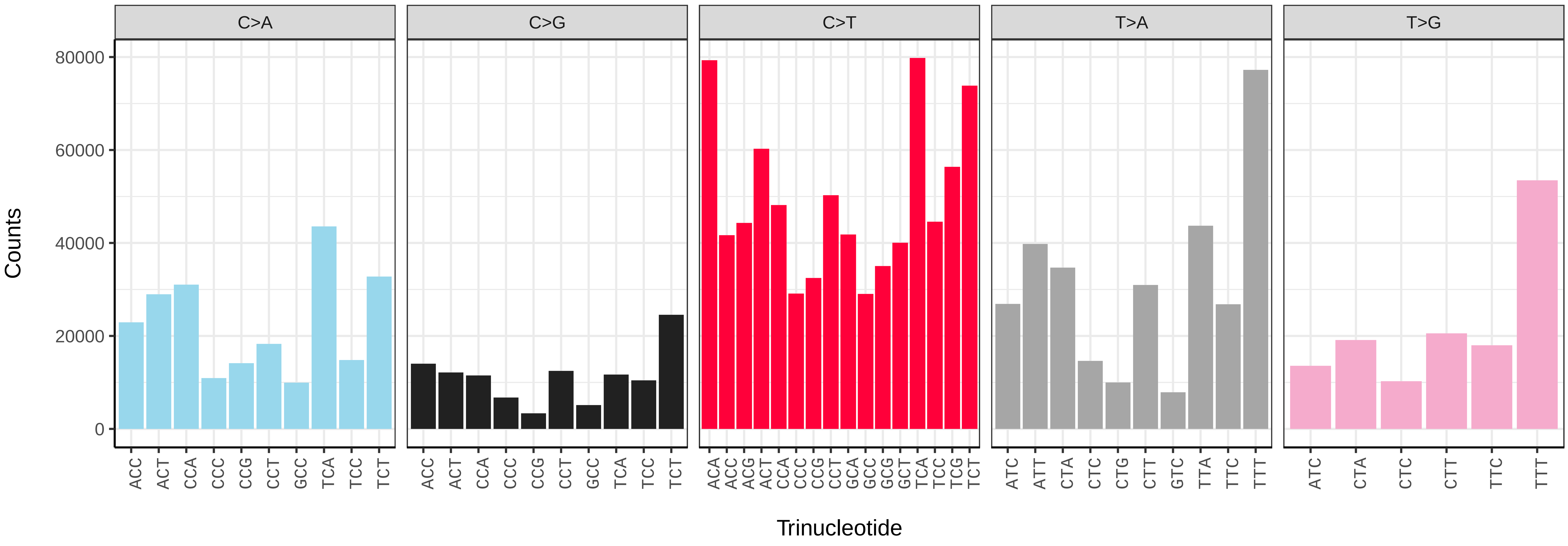


*Eulithis testata* (ilEulTest2) somatic mutational spectrum (SBS96 classification)

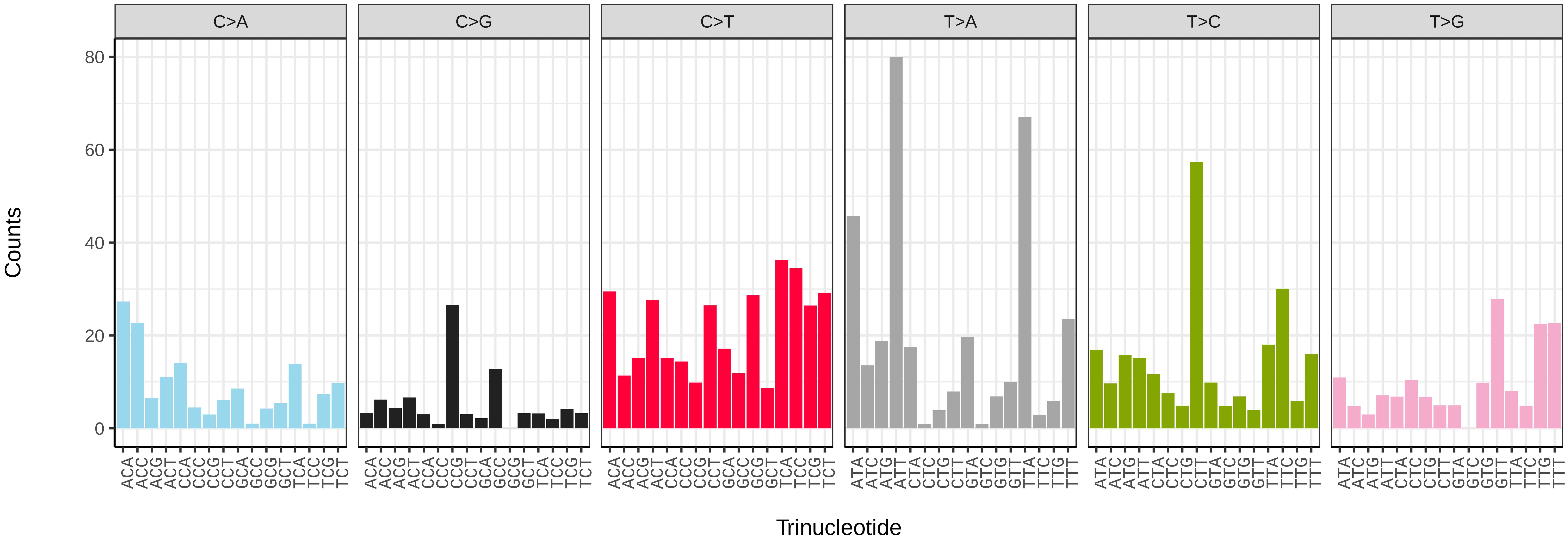


*Eulithis testata* (ilEulTest2) somatic mutational spectrum (SBS52 classification)

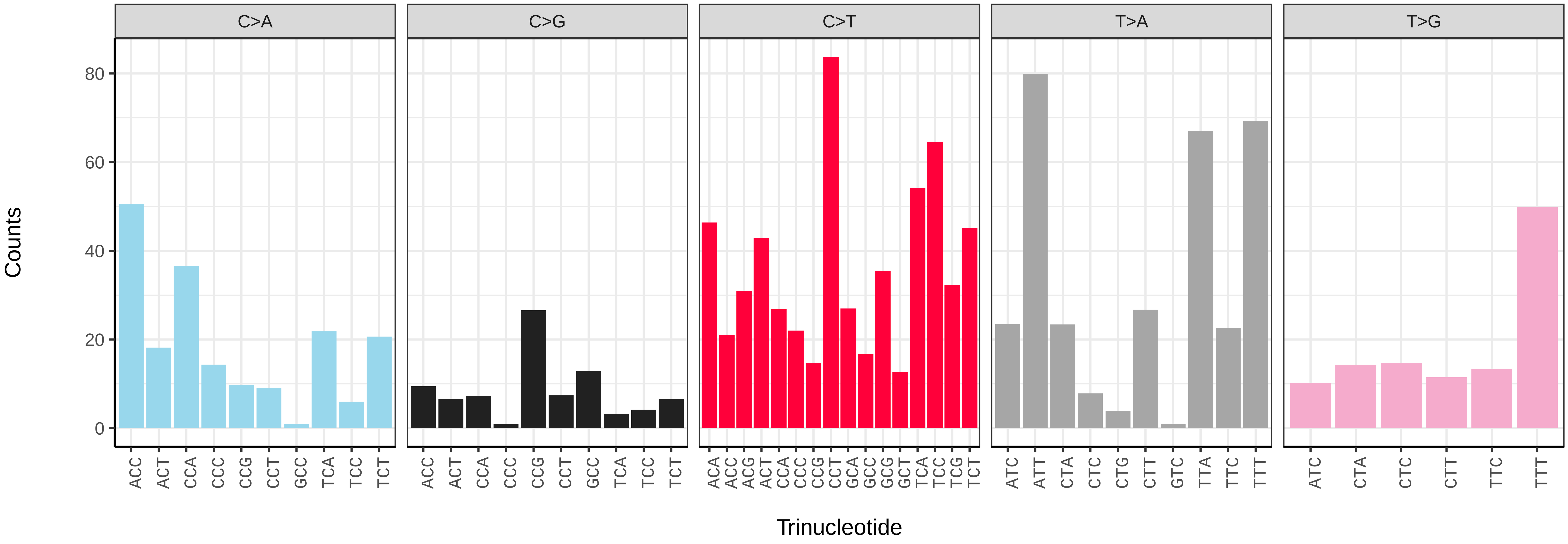

*Eupithecia abbreviata* (ilEupAbbr2) germline mutational spectrum

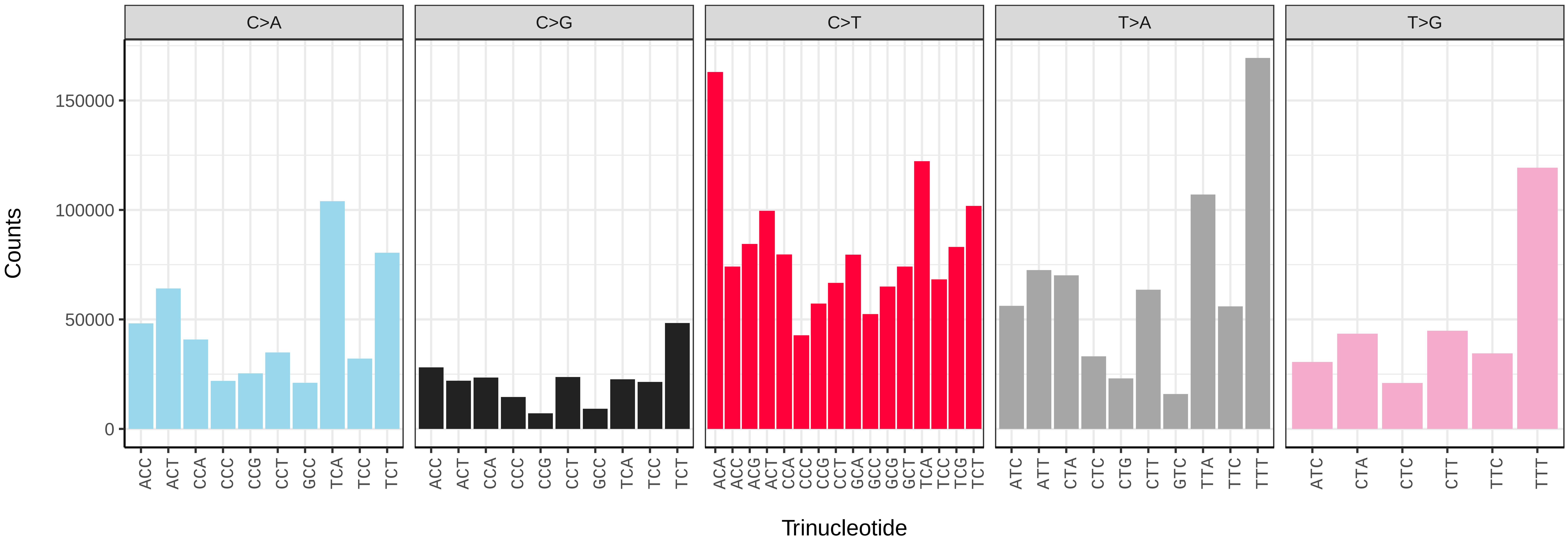


*Eupithecia abbreviata* (ilEupAbbr2) somatic mutational spectrum (SBS96 classification)

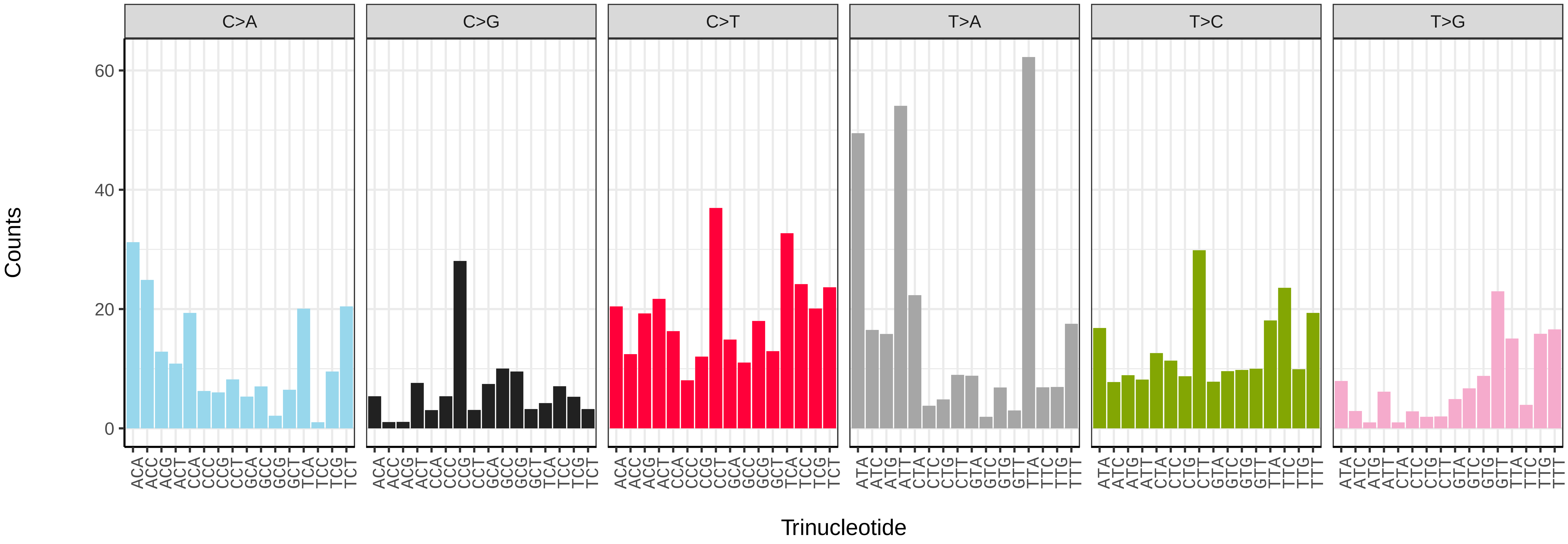


*Eupithecia abbreviata* (ilEupAbbr2) somatic mutational spectrum (SBS52 classification)

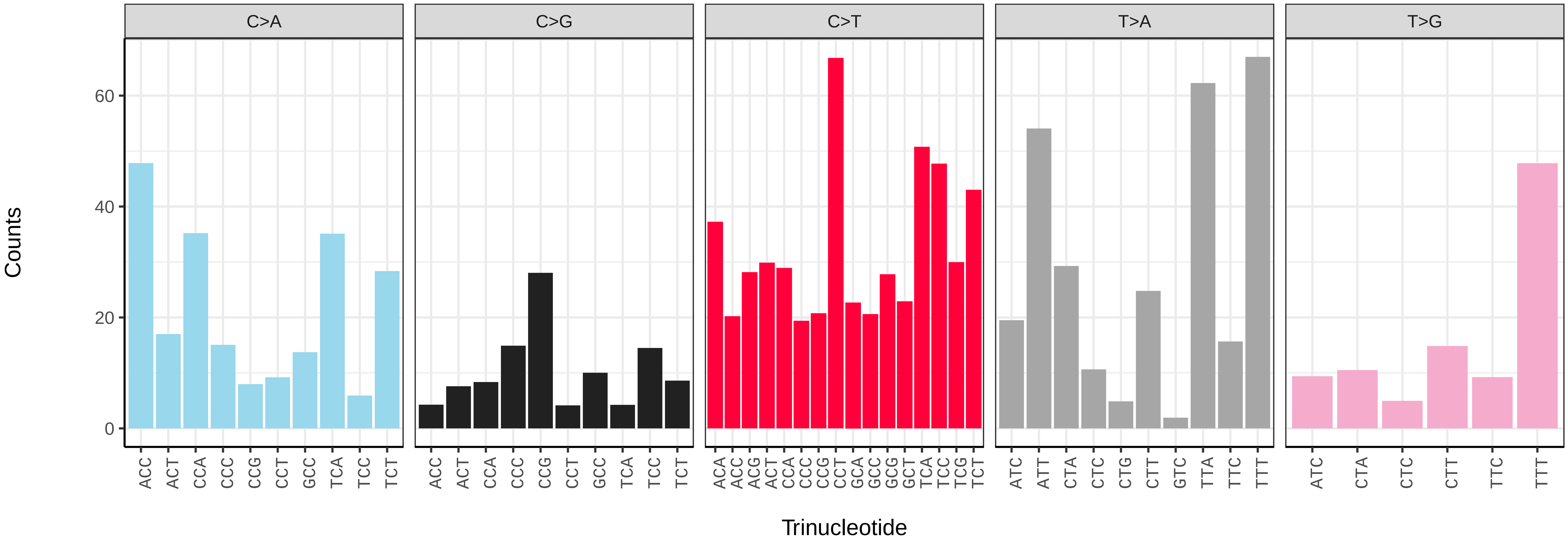

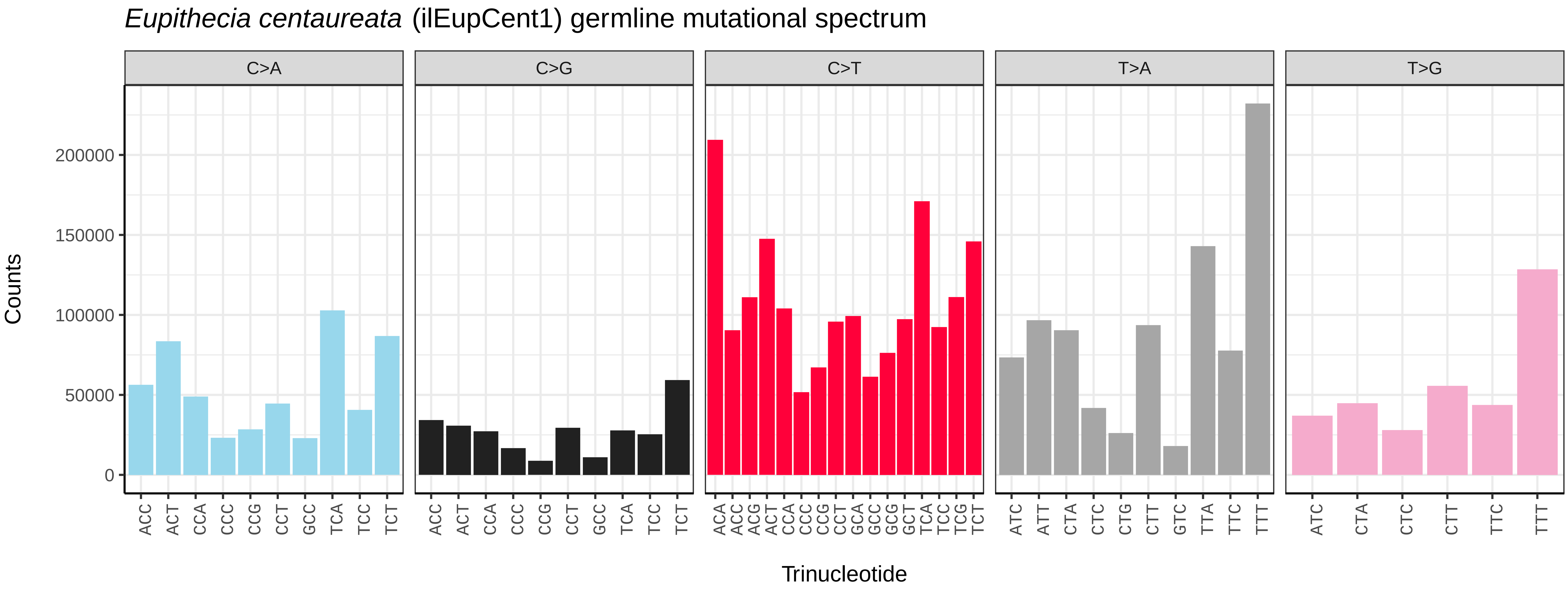
*Eupithecia centaureata* (ilEupCent1) germline mutational spectrum
C>A
C>G
C>T
T>A
T>G
Counts
0
50000
100000
150000
200000
Trinucleotide

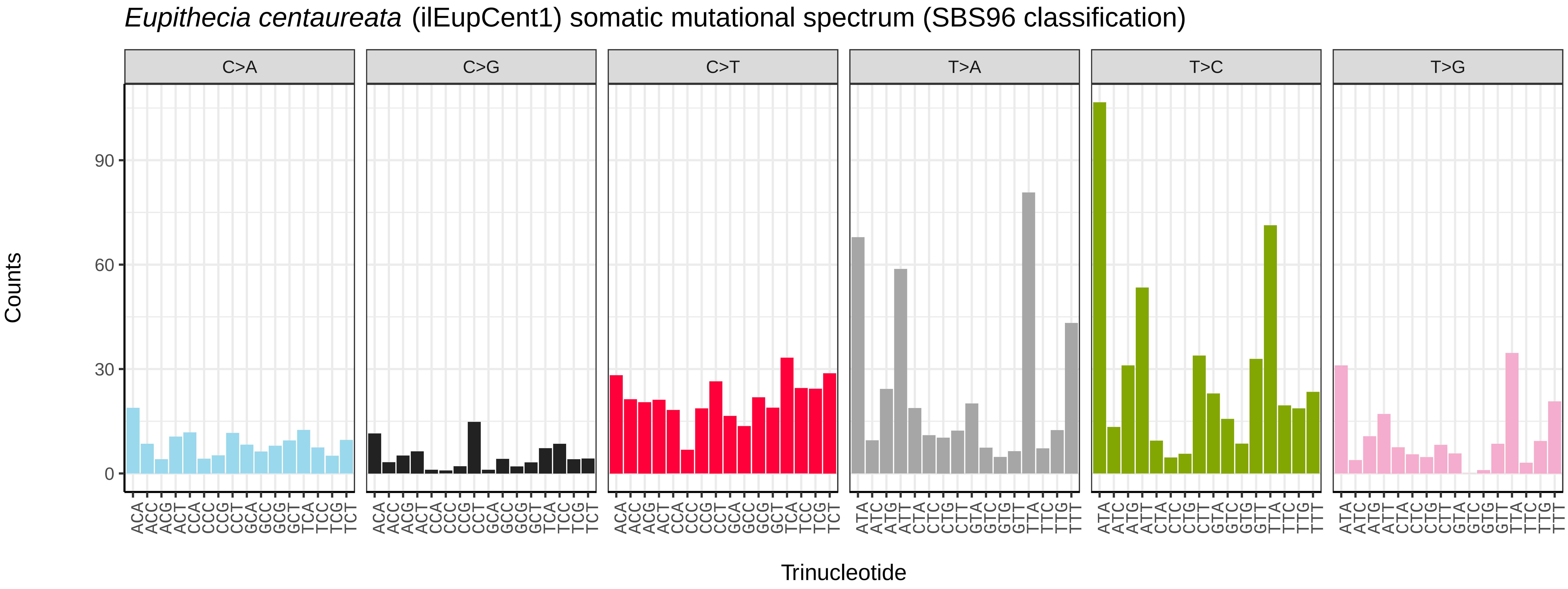
*Eupithecia centaureata* (ilEupCent1) somatic mutational spectrum (SBS96 classification)
C>A
C>G
C>T
T>A
T>C
T>G
Counts
0
30
60
90
Trinucleotide

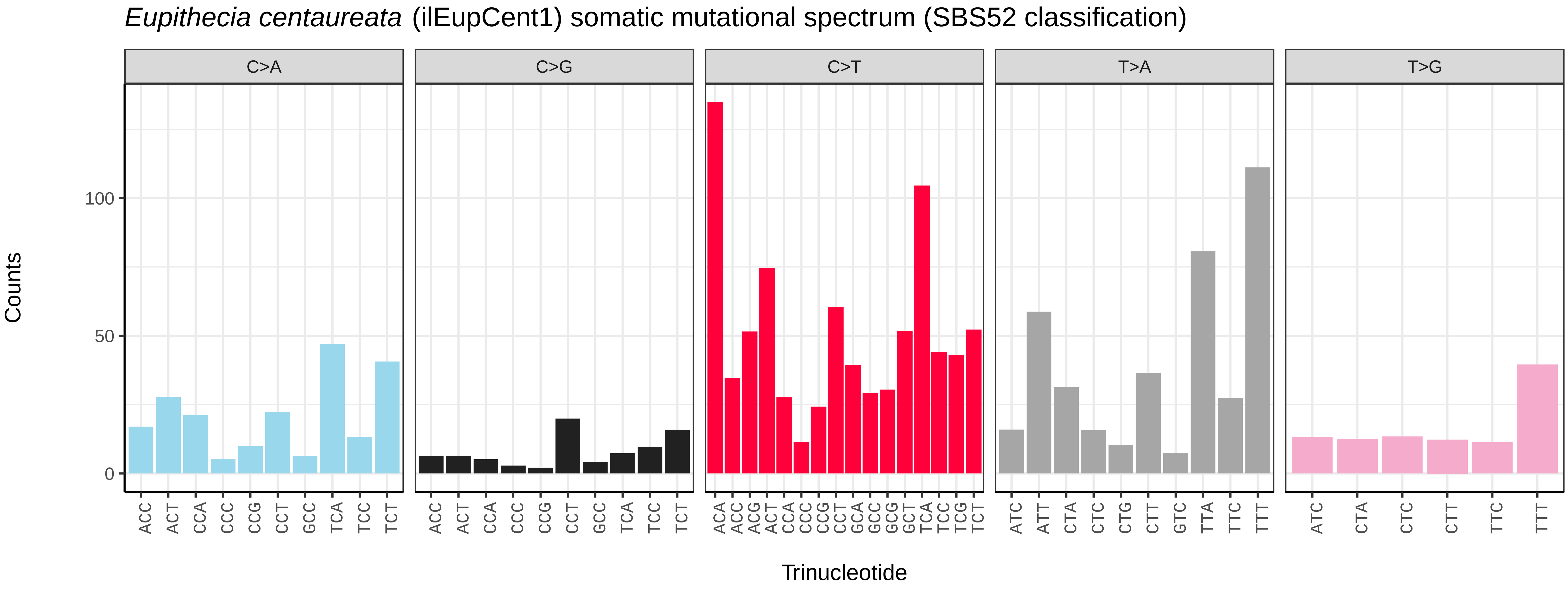
*Eupithecia centaureata* (ilEupCent1) somatic mutational spectrum (SBS52 classification)
C>A
C>G
C>T
T>A
T>G
Counts
0
50
100
Trinucleotide

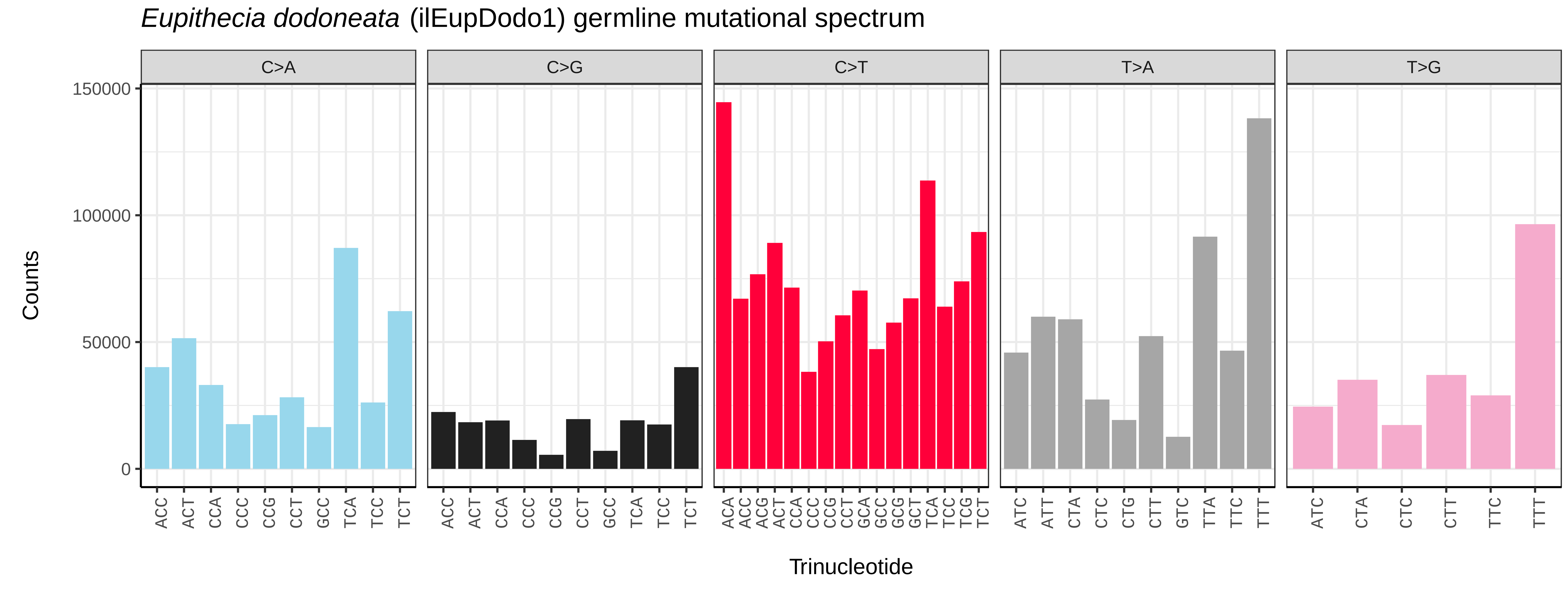
Eupithecia dodoneata (ilEupDodo1) germline mutational spectrum
C>A
C>G
C>T
T>A
T>G
Counts
0
50000
100000
150000
Trinucleotide

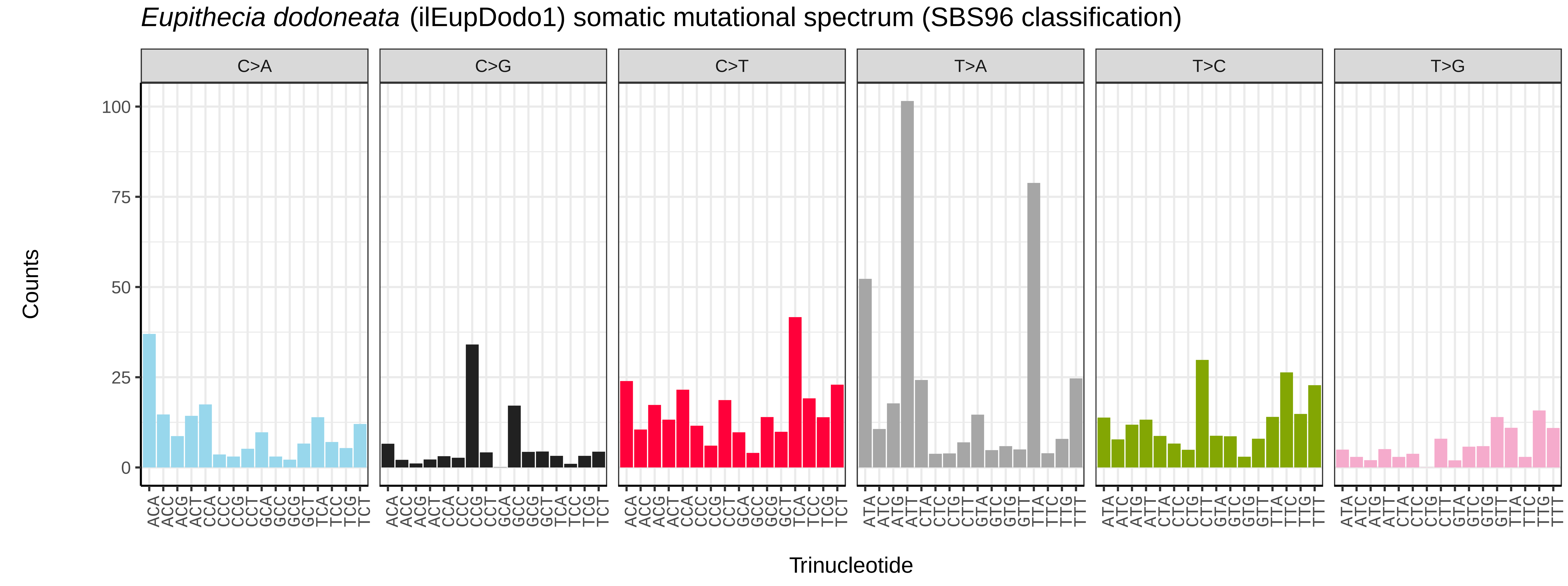
Eupithecia dodoneata (ilEupDodo1) somatic mutational spectrum (SBS96 classification)
C>A
C>G
C>T
T>A
T>C
T>G
Counts
0
25
50
75
100
Trinucleotide

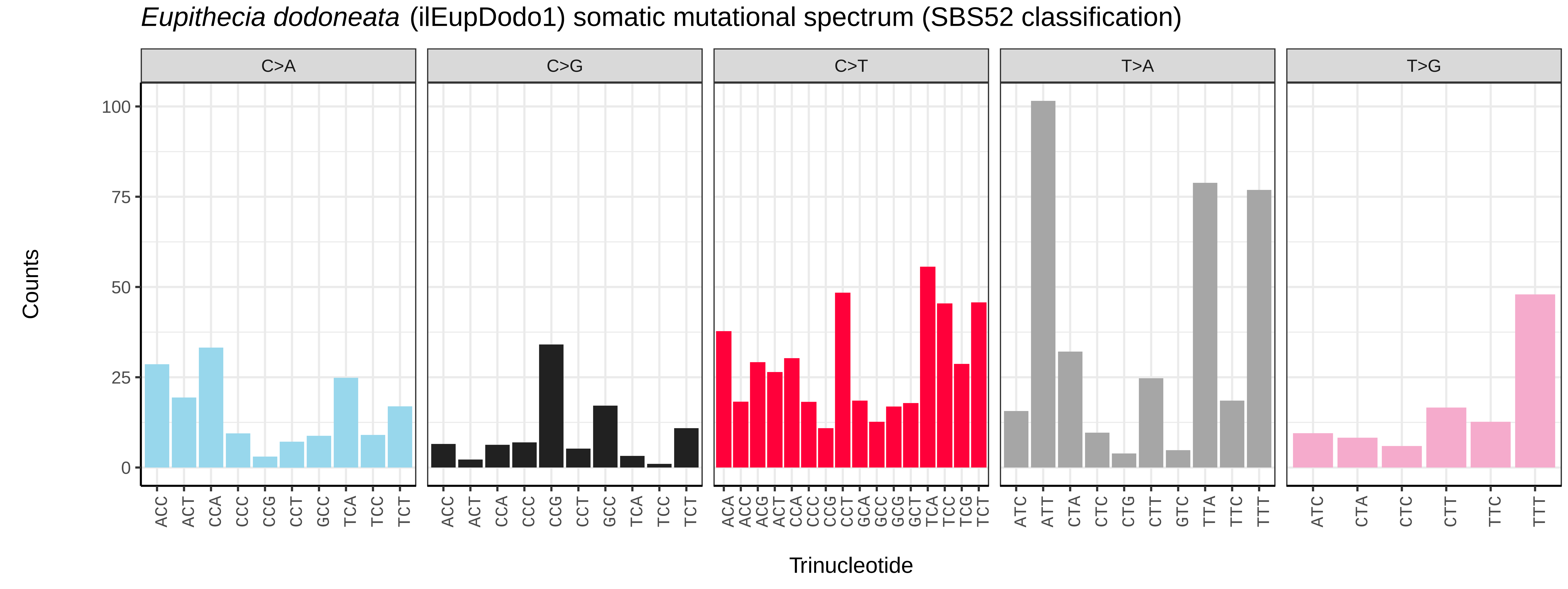
Eupithecia dodoneata (ilEupDodo1) somatic mutational spectrum (SBS52 classification)
C>A
C>G
C>T
T>A
T>G
Counts
0
25
50
75
100
Trinucleotide

*Eupithecia exiguata* (ilEupExig1) germline mutational spectrum

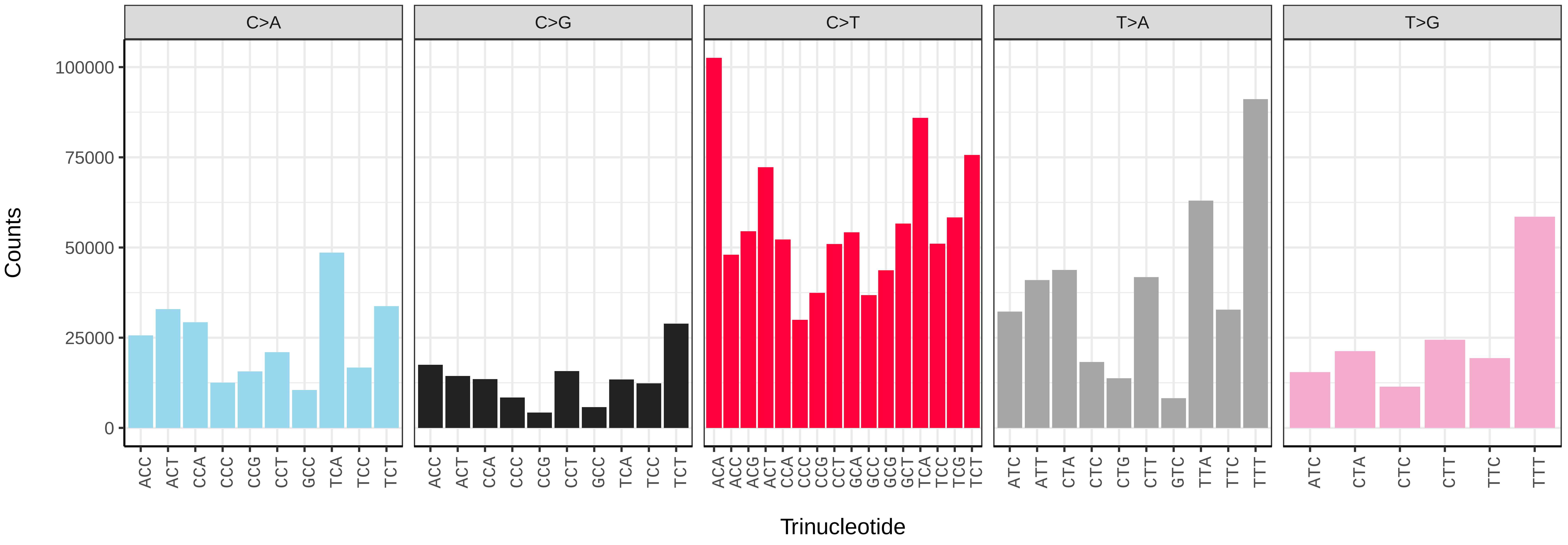


*Eupithecia exiguata* (ilEupExig1) somatic mutational spectrum (SBS96 classification)

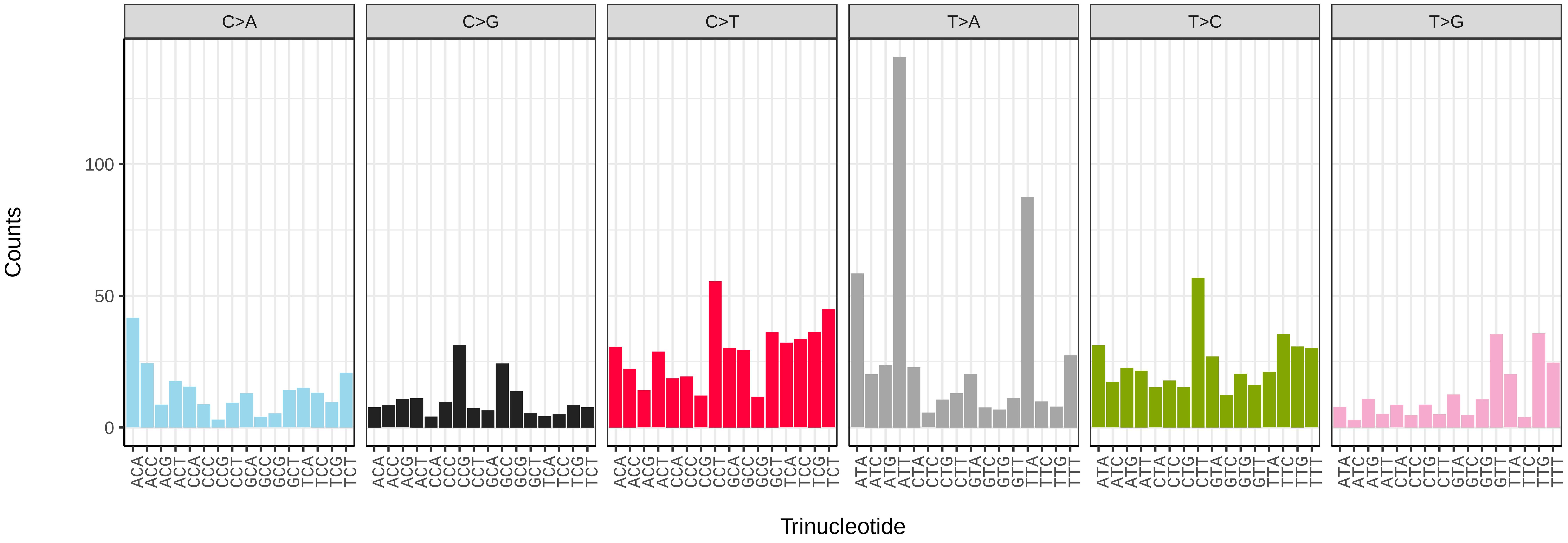


*Eupithecia exiguata* (ilEupExig1) somatic mutational spectrum (SBS52 classification)

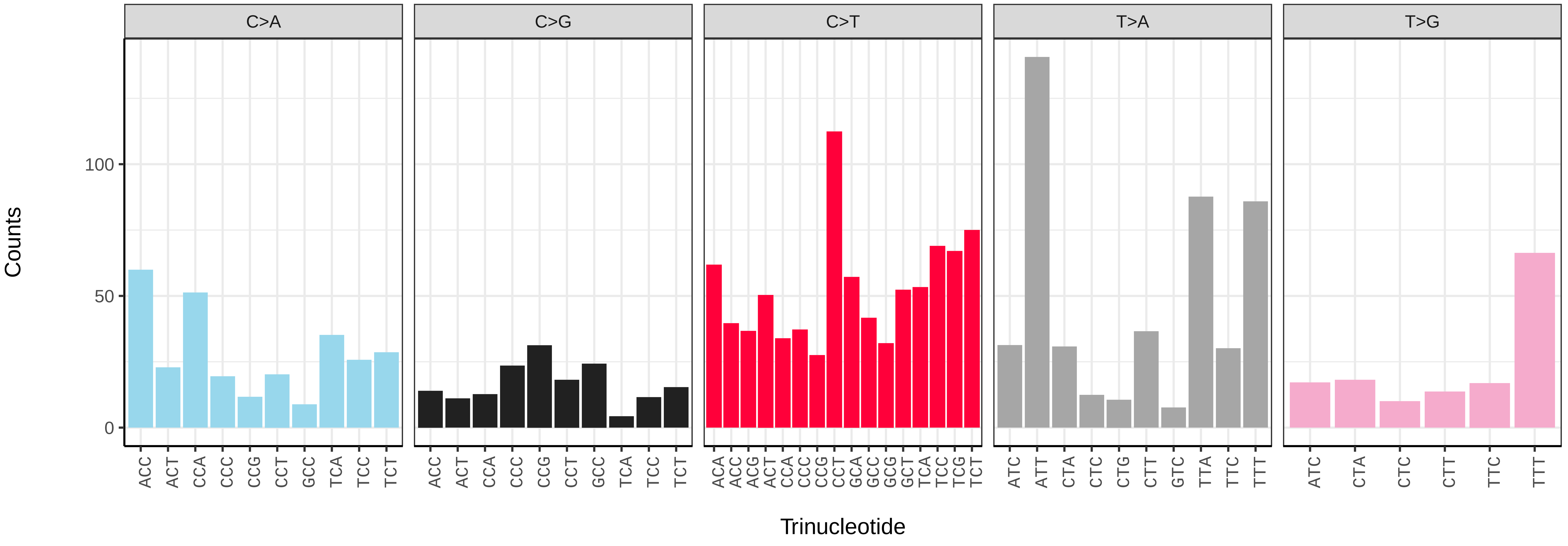

*Eupithecia insigniata* (ilEupInsi1) germline mutational spectrum

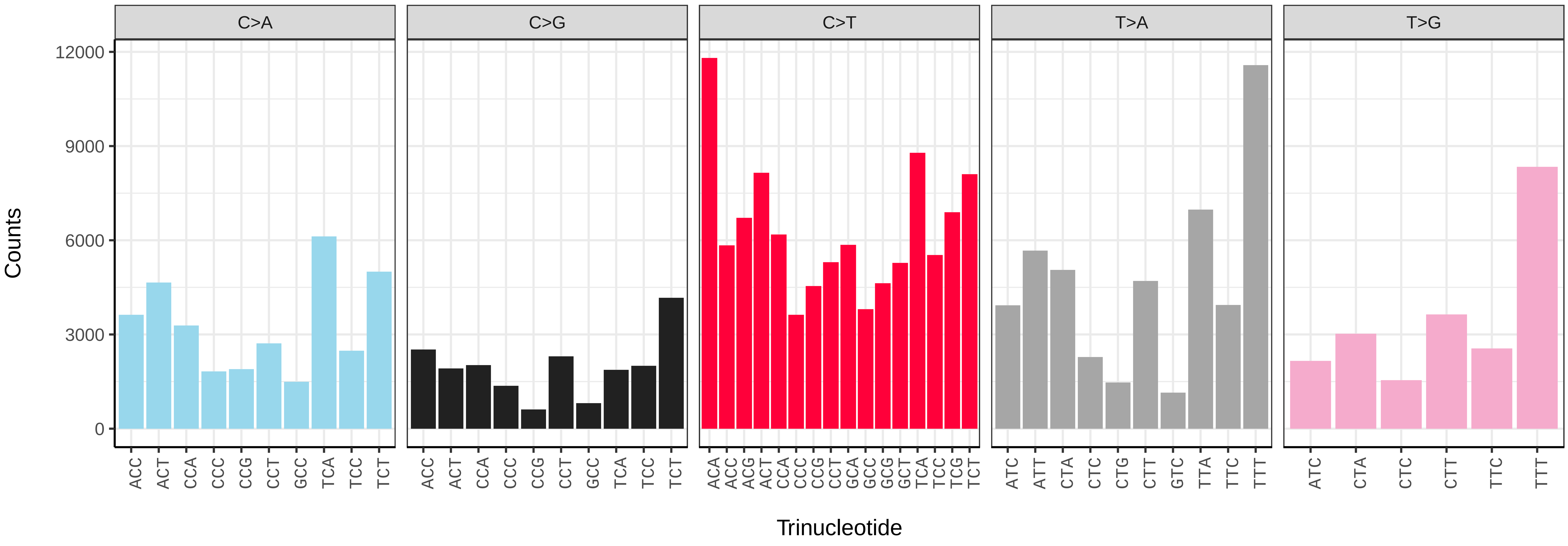


*Eupithecia insigniata* (ilEupInsi1) somatic mutational spectrum (SBS96 classification)

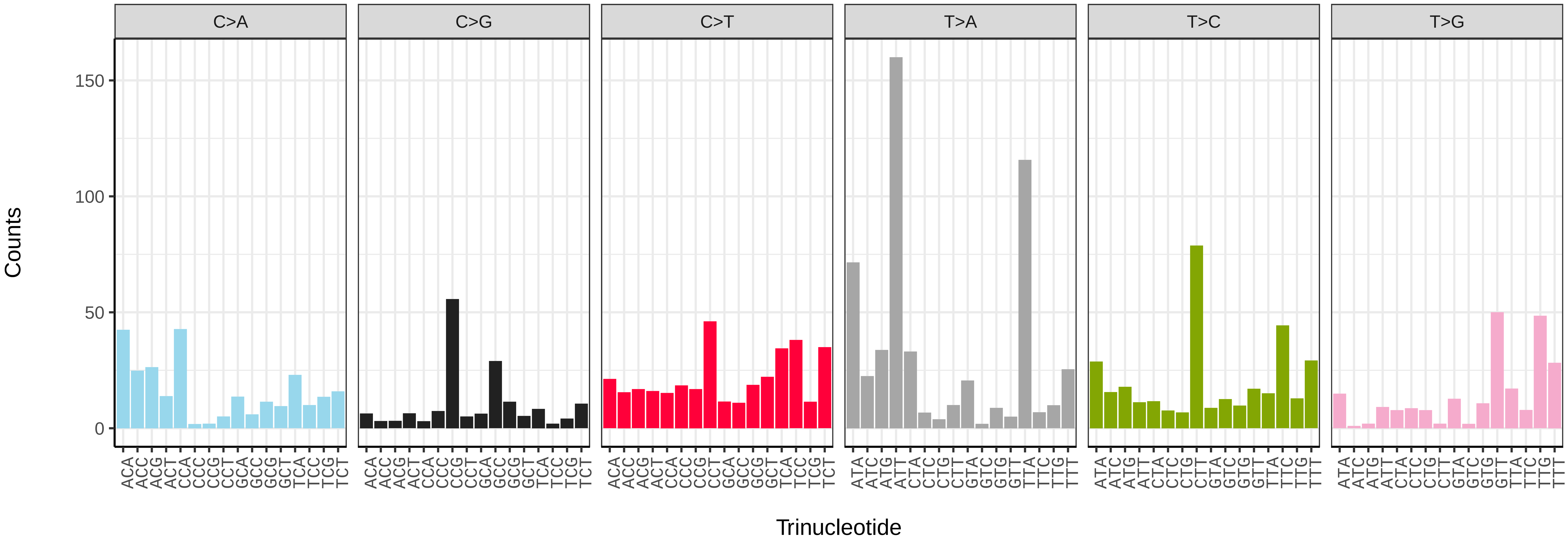


*Eupithecia insigniata* (ilEupInsi1) somatic mutational spectrum (SBS52 classification)

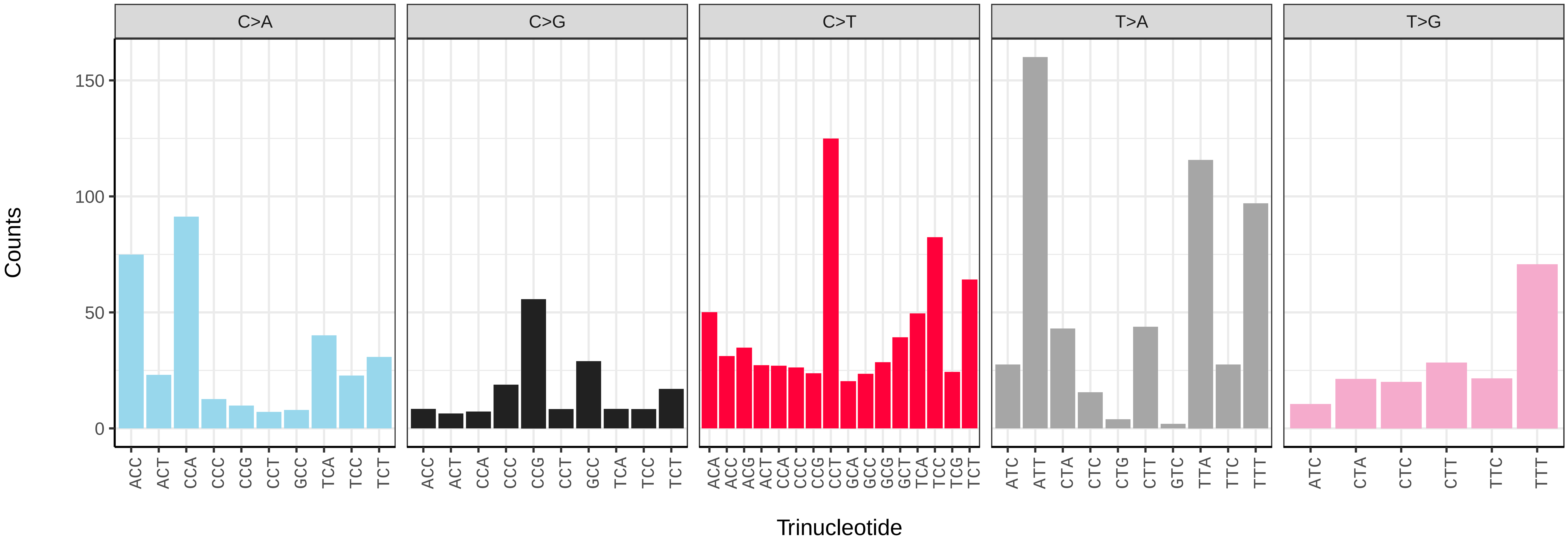

*Euplexia lucipara* (ilEupLuci1) germline mutational spectrum

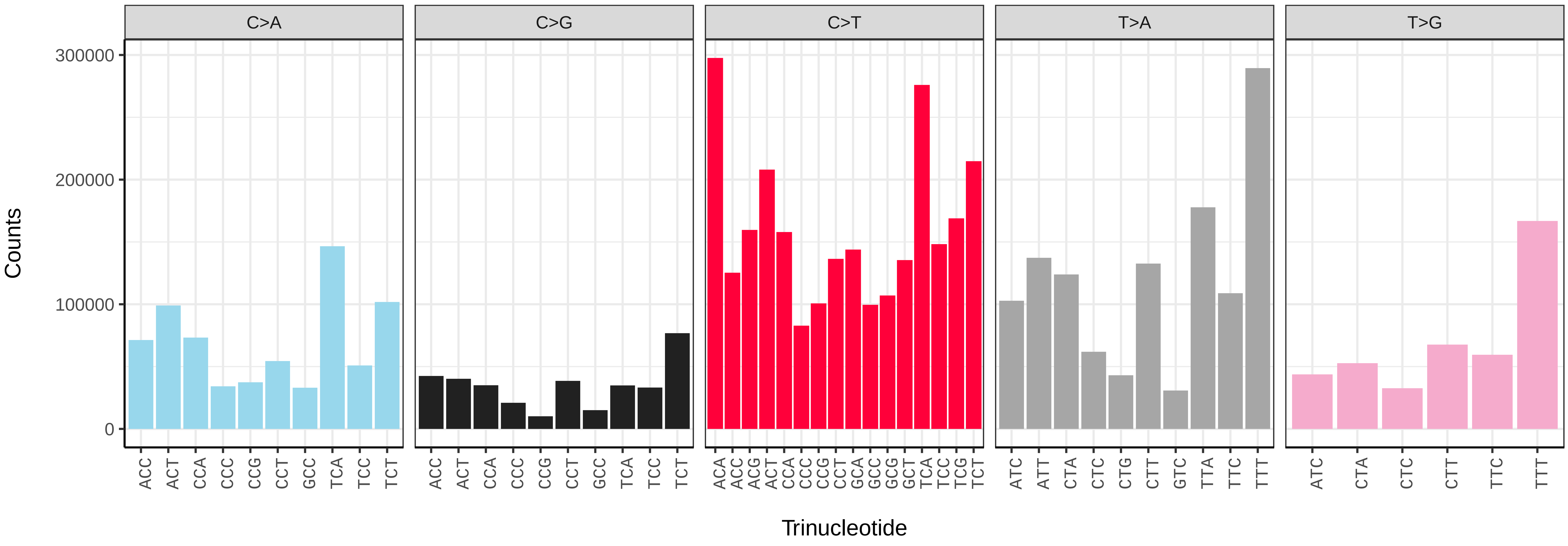


*Euplexia lucipara* (ilEupLuci1) somatic mutational spectrum (SBS96 classification)

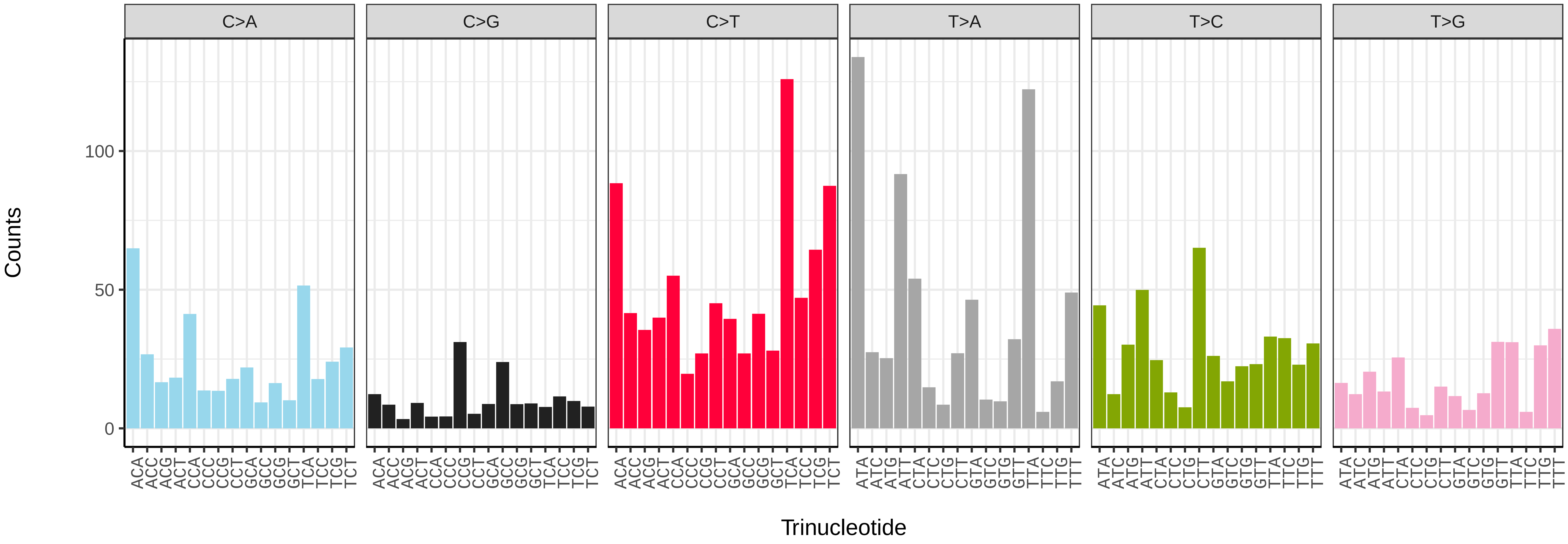


*Euplexia lucipara* (ilEupLuci1) somatic mutational spectrum (SBS52 classification)

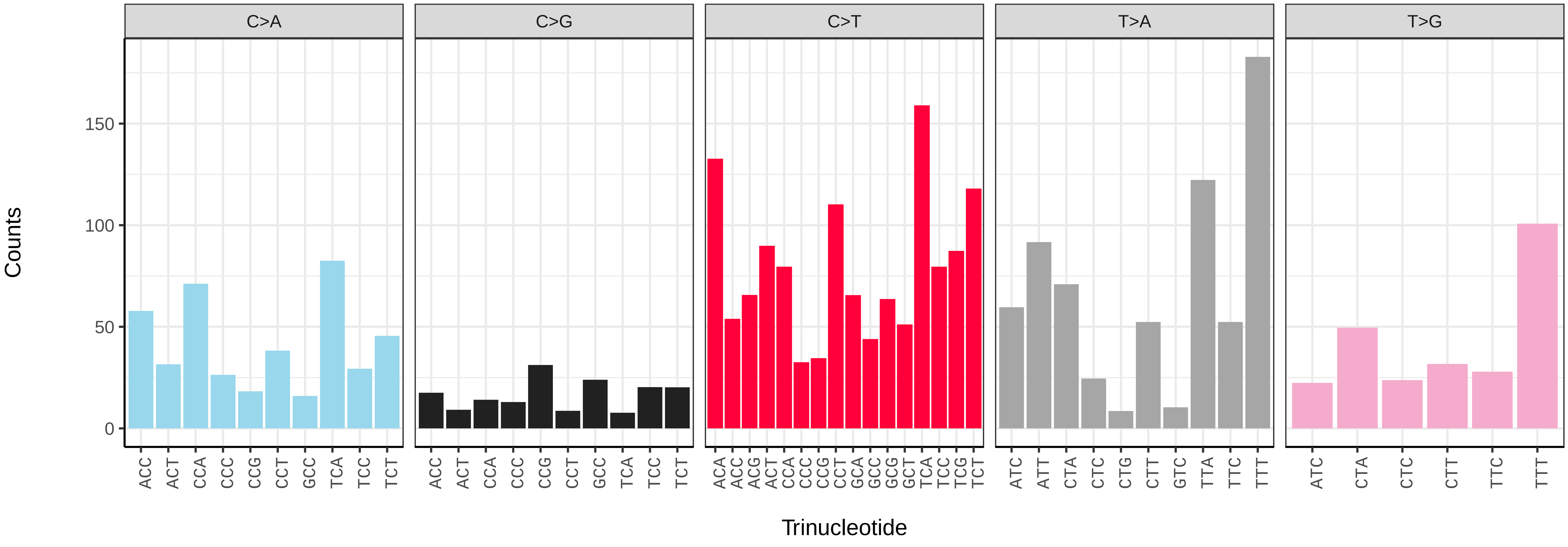

*Euproctis similis* (ilEupSimi1) germline mutational spectrum

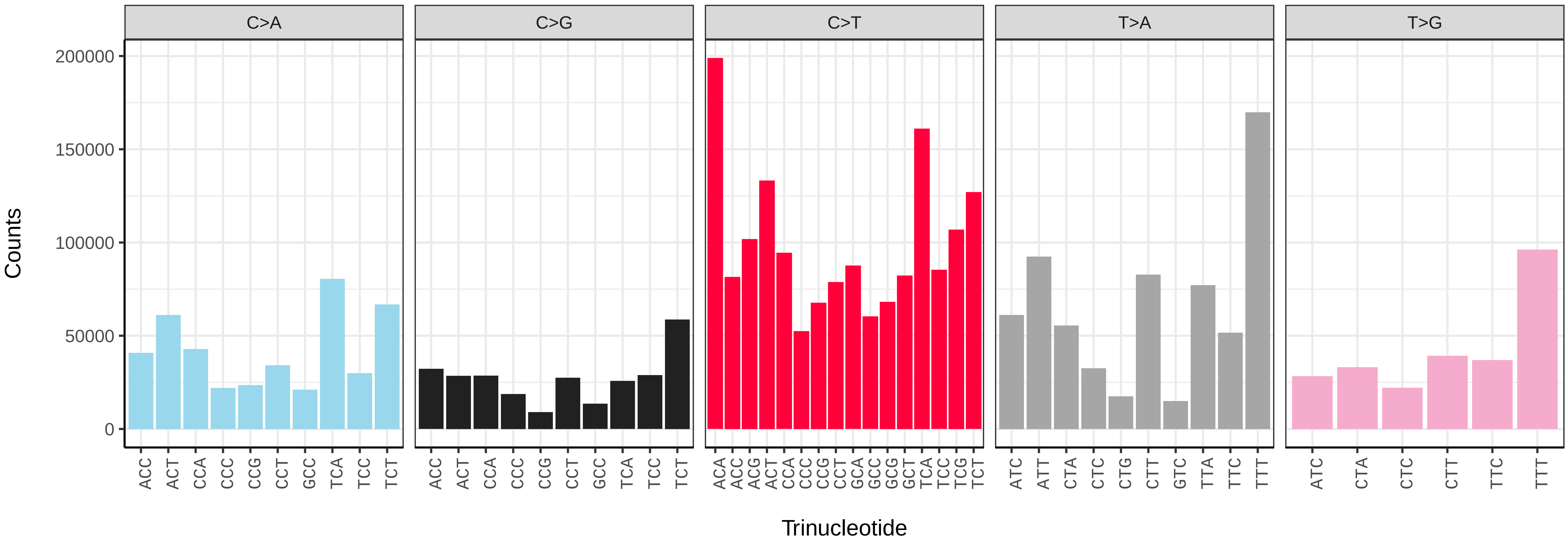


*Euproctis similis* (ilEupSimi1) somatic mutational spectrum (SBS96 classification)

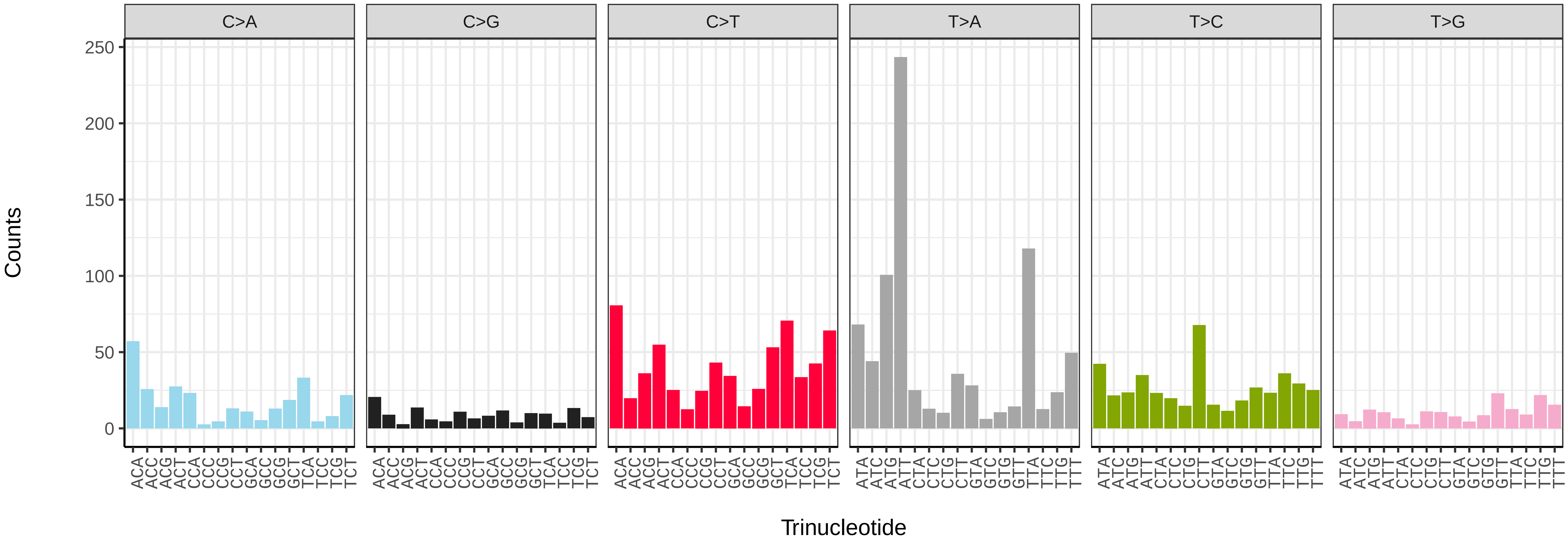


*Euproctis similis* (ilEupSimi1) somatic mutational spectrum (SBS52 classification)

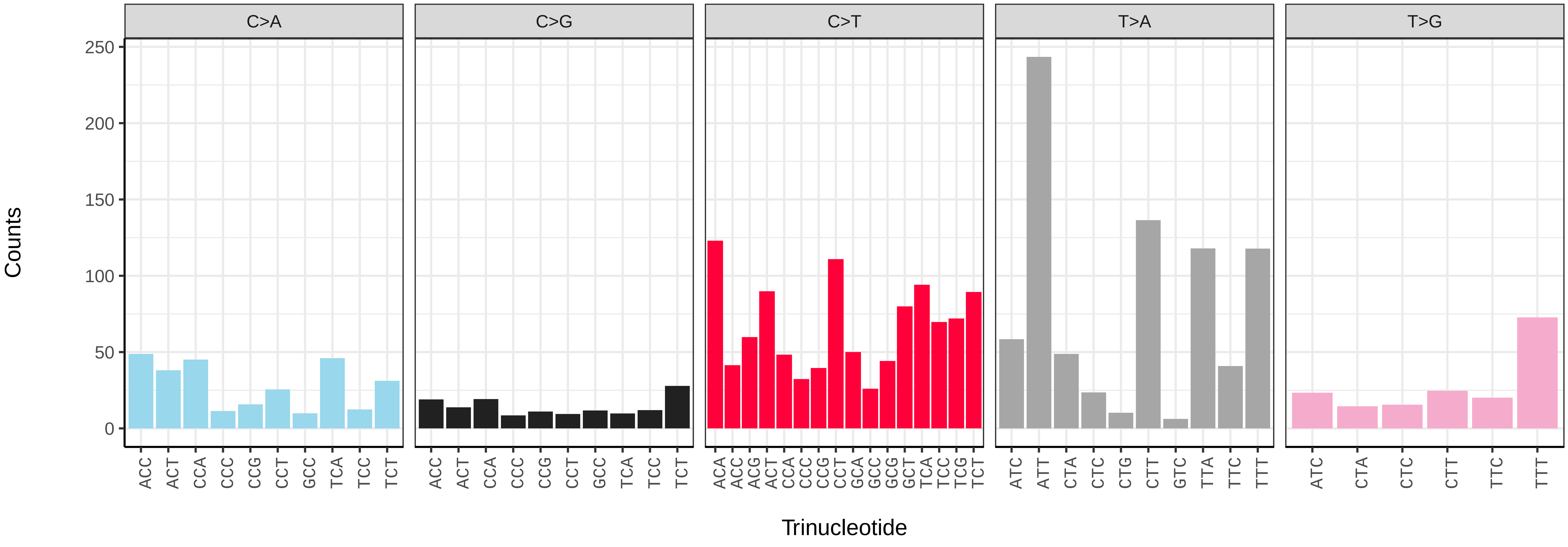

*Eupsilia transversa* (ilEupTran1) germline mutational spectrum

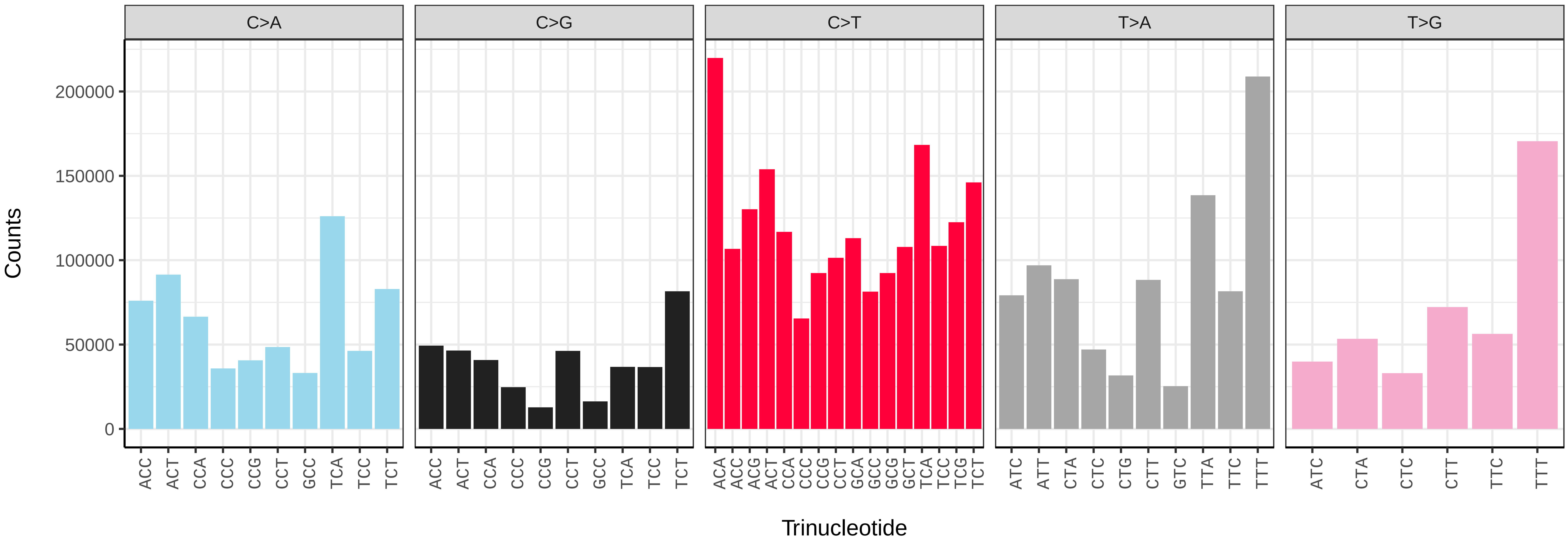


*Eupsilia transversa* (ilEupTran1) somatic mutational spectrum (SBS96 classification)

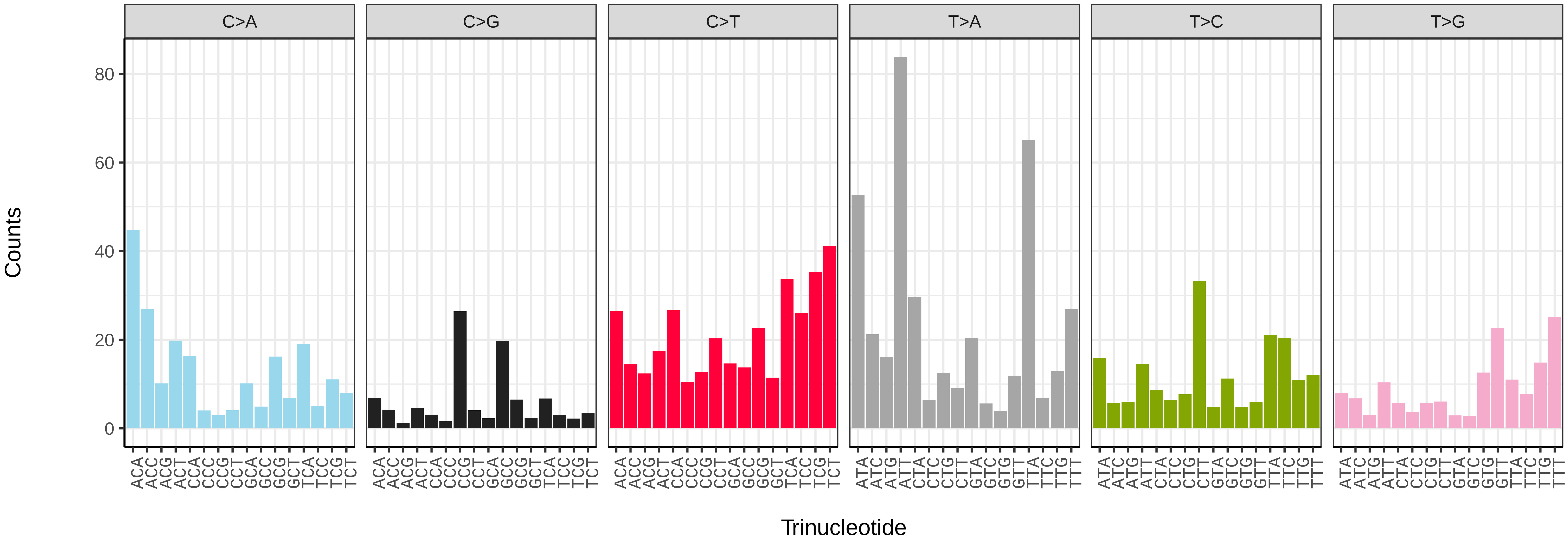


*Eupsilia transversa* (ilEupTran1) somatic mutational spectrum (SBS52 classification)

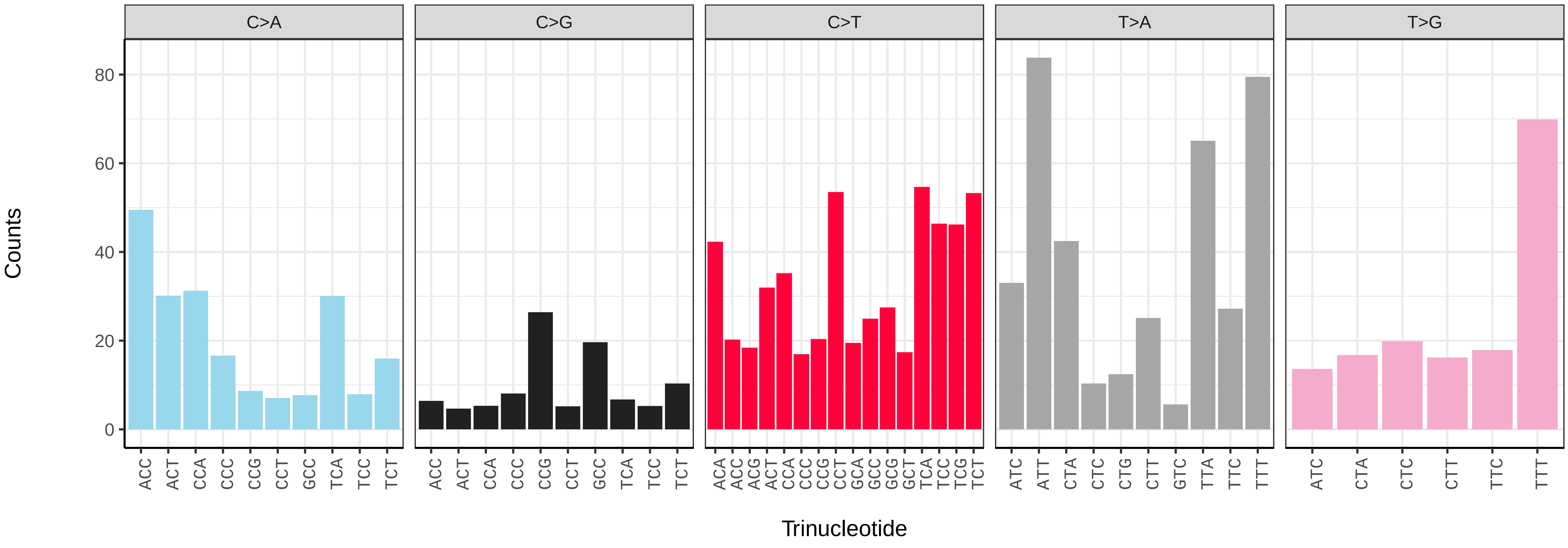

*Eupithecia vulgata* (ilEupVulg1) germline mutational spectrum

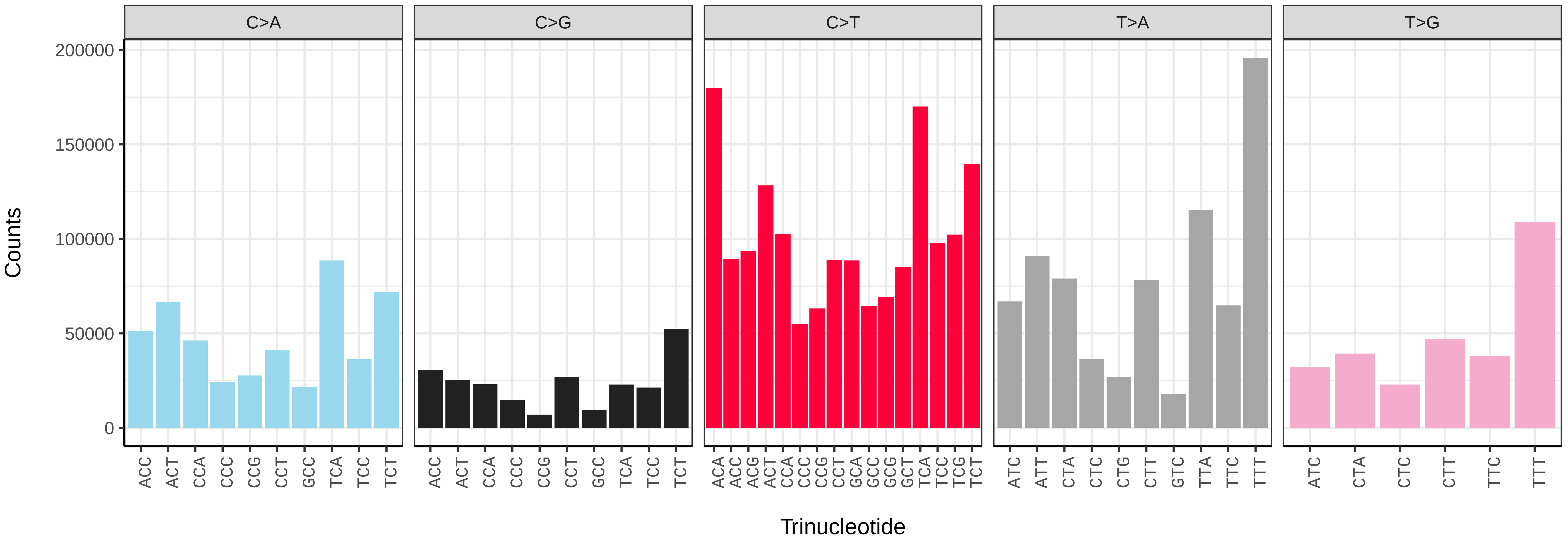


*Eupithecia vulgata* (ilEupVulg1) somatic mutational spectrum (SBS96 classification)

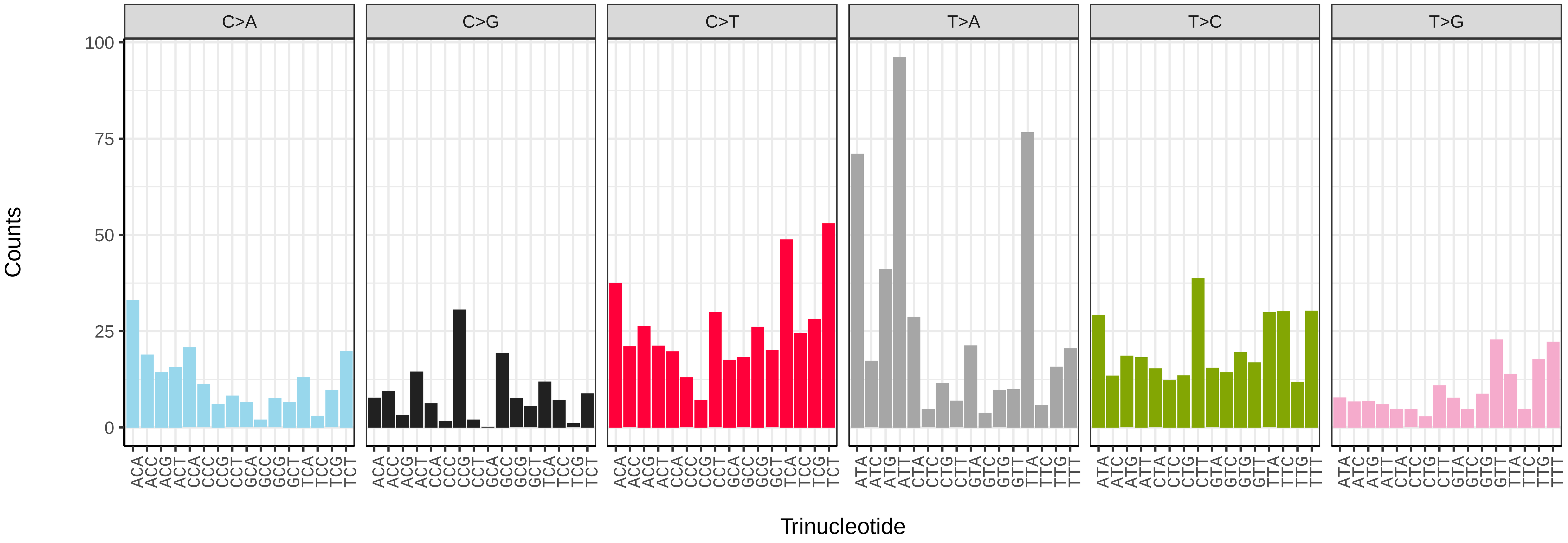


*Eupithecia vulgata* (ilEupVulg1) somatic mutational spectrum (SBS52 classification)

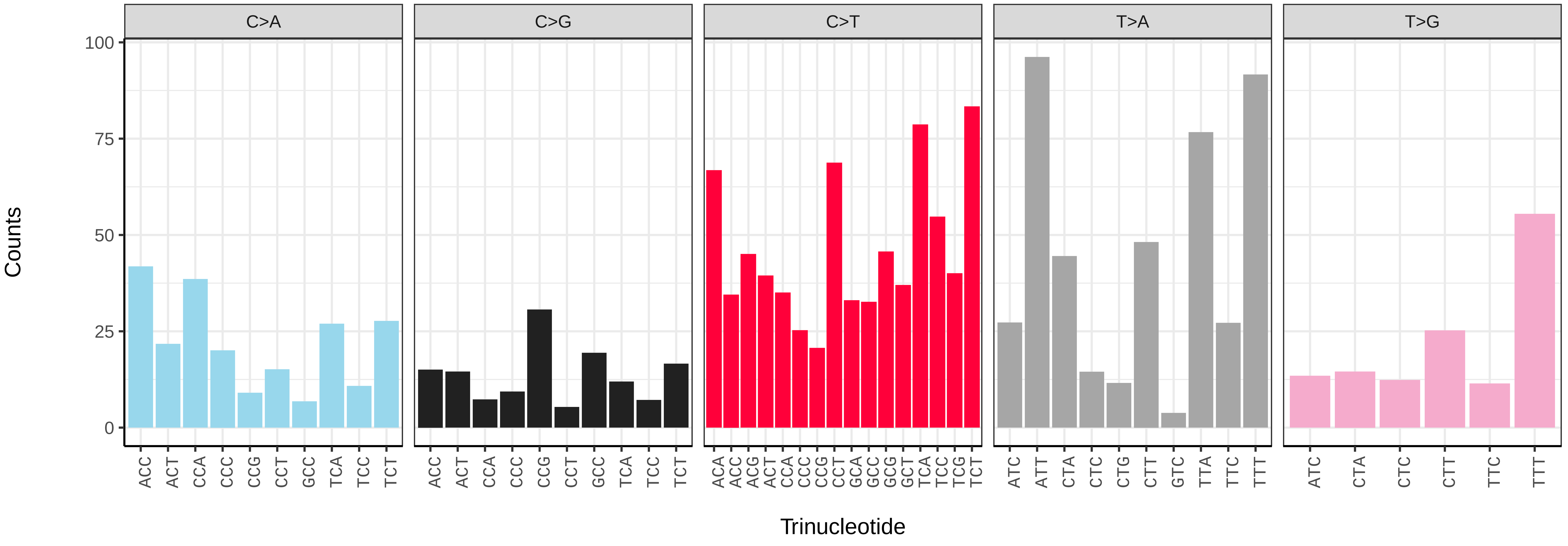

*Euzophera pinguis* (ilEuzPing2) germline mutational spectrum

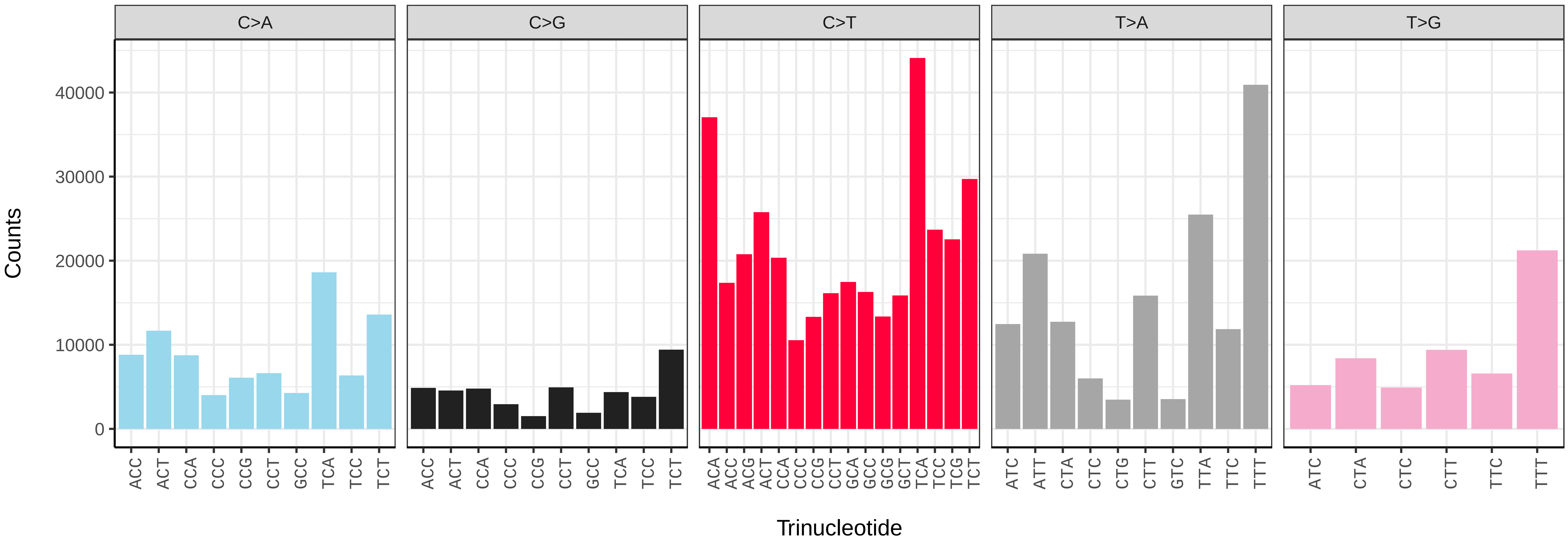


*Euzophera pinguis* (ilEuzPing2) somatic mutational spectrum (SBS96 classification)

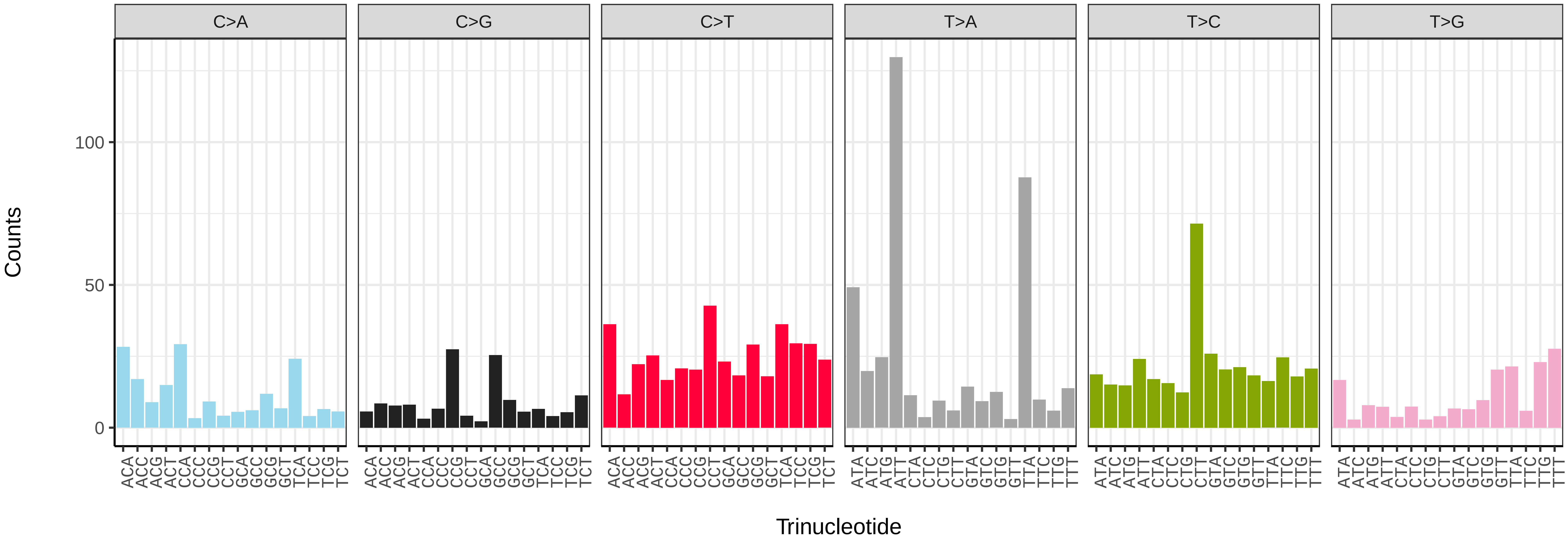


*Euzophera pinguis* (ilEuzPing2) somatic mutational spectrum (SBS52 classification)

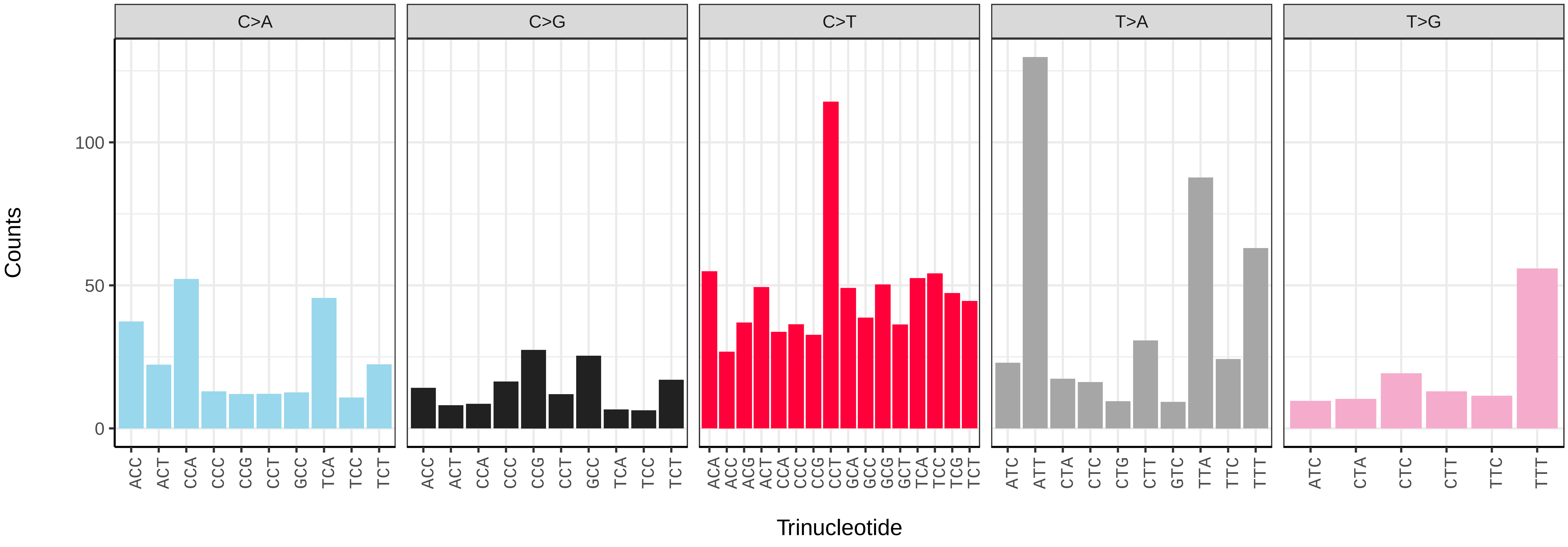

*Fabriciana adippe* (ilFabAdip1) germline mutational spectrum

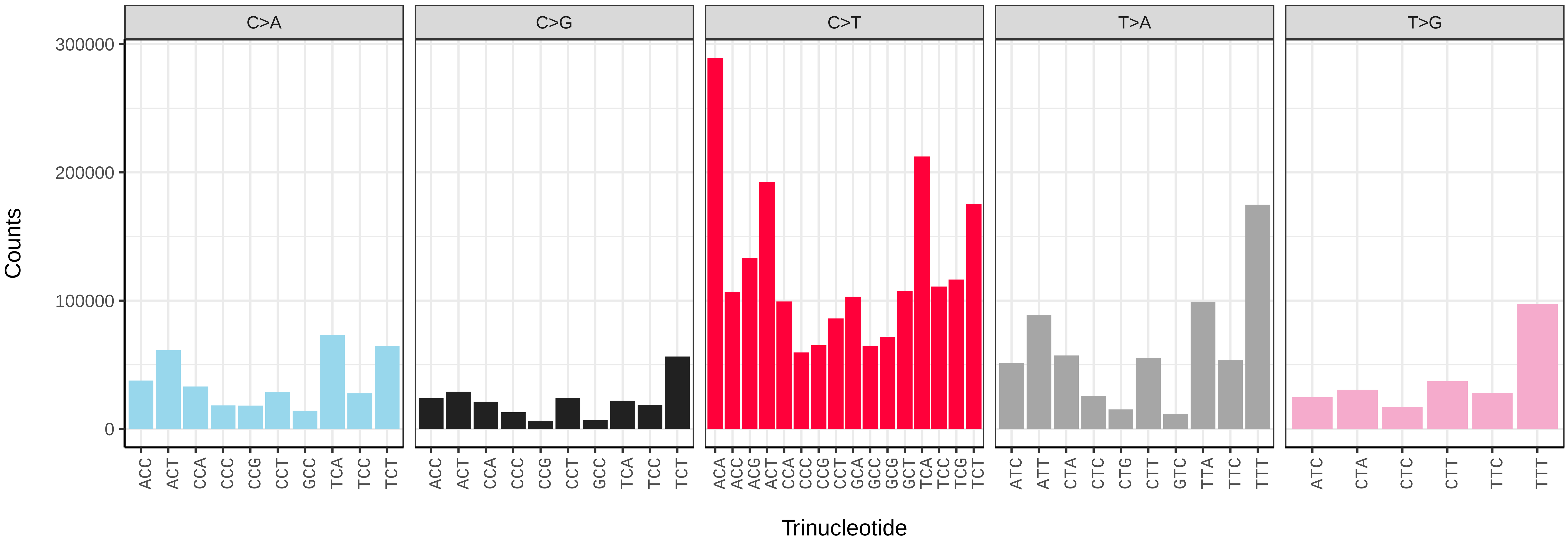


*Fabriciana adippe* (ilFabAdip1) somatic mutational spectrum (SBS96 classification)

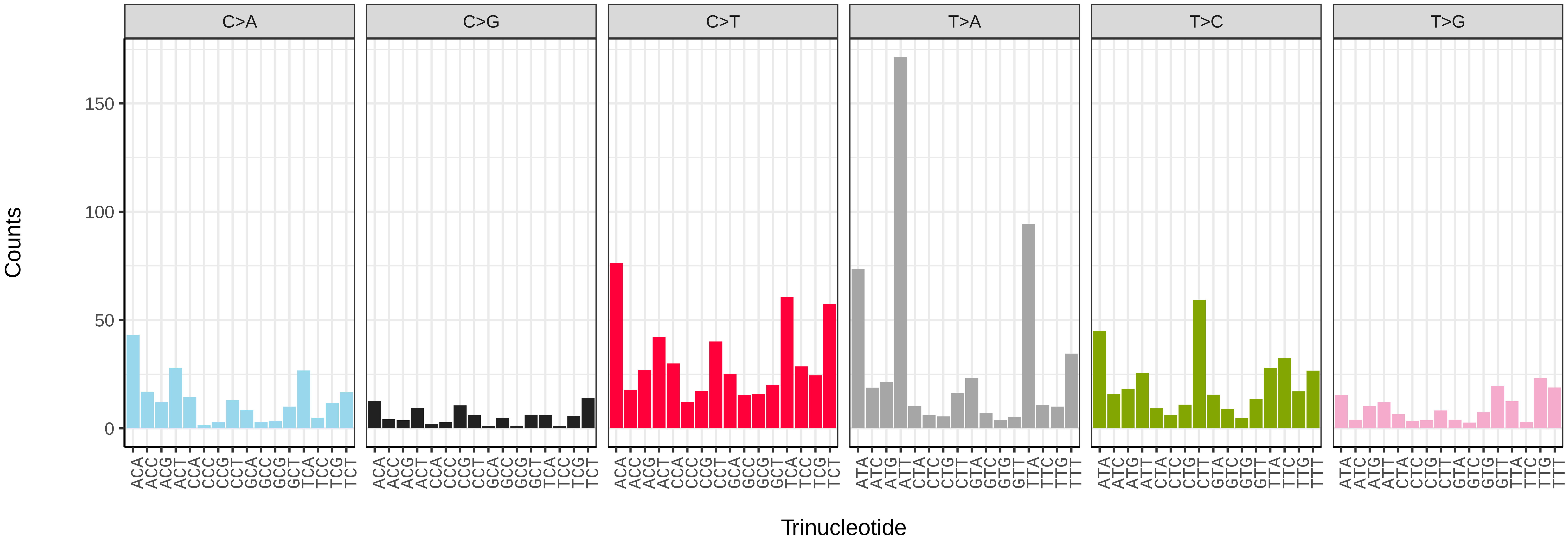


*Fabriciana adippe* (ilFabAdip1) somatic mutational spectrum (SBS52 classification)

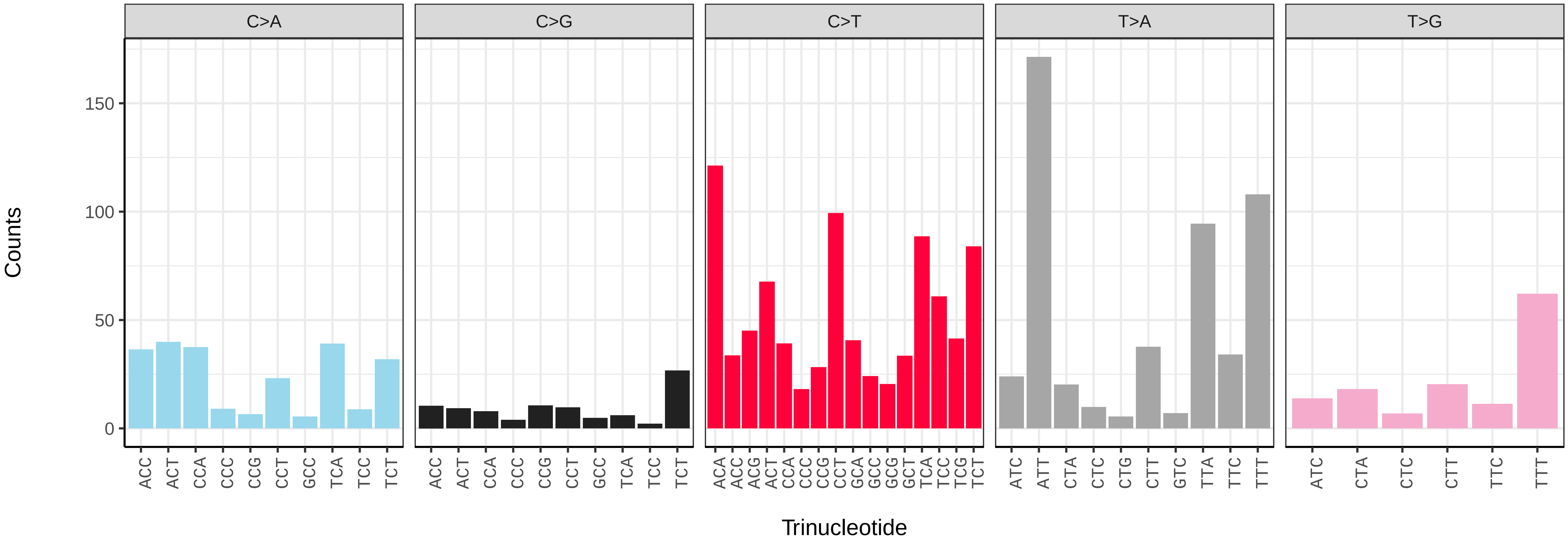

*Furcula furcula* (ilFurFurc1) germline mutational spectrum

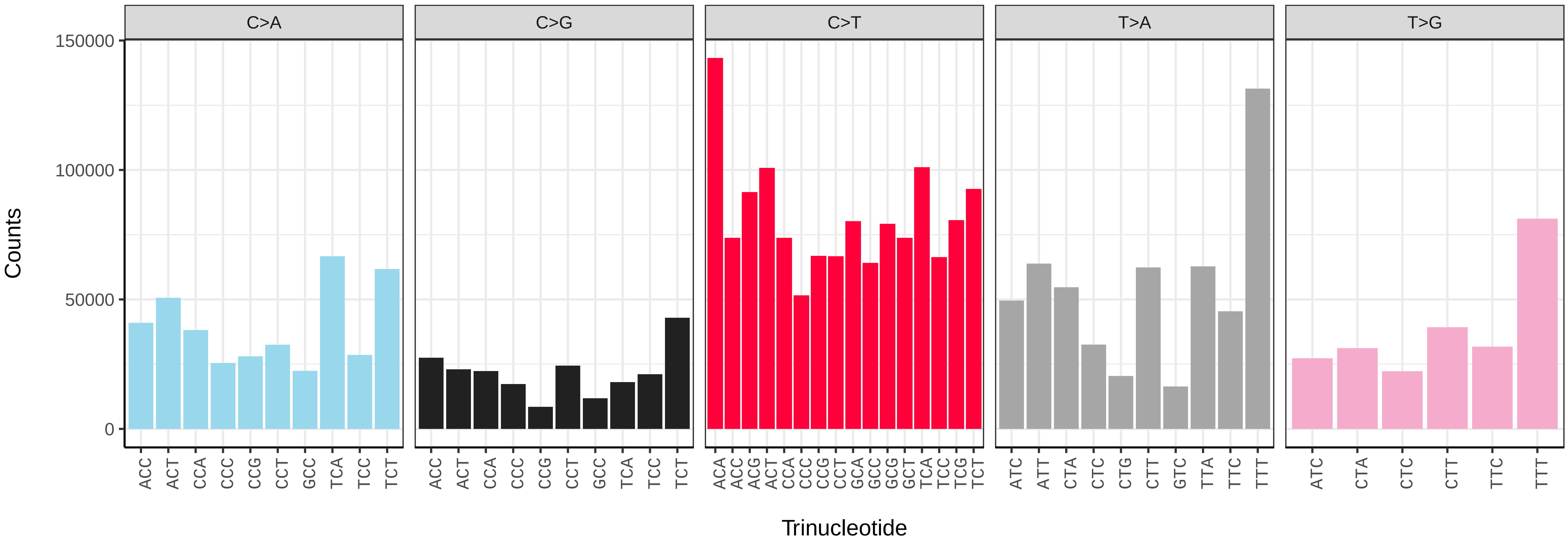


*Furcula furcula* (ilFurFurc1) somatic mutational spectrum (SBS96 classification)

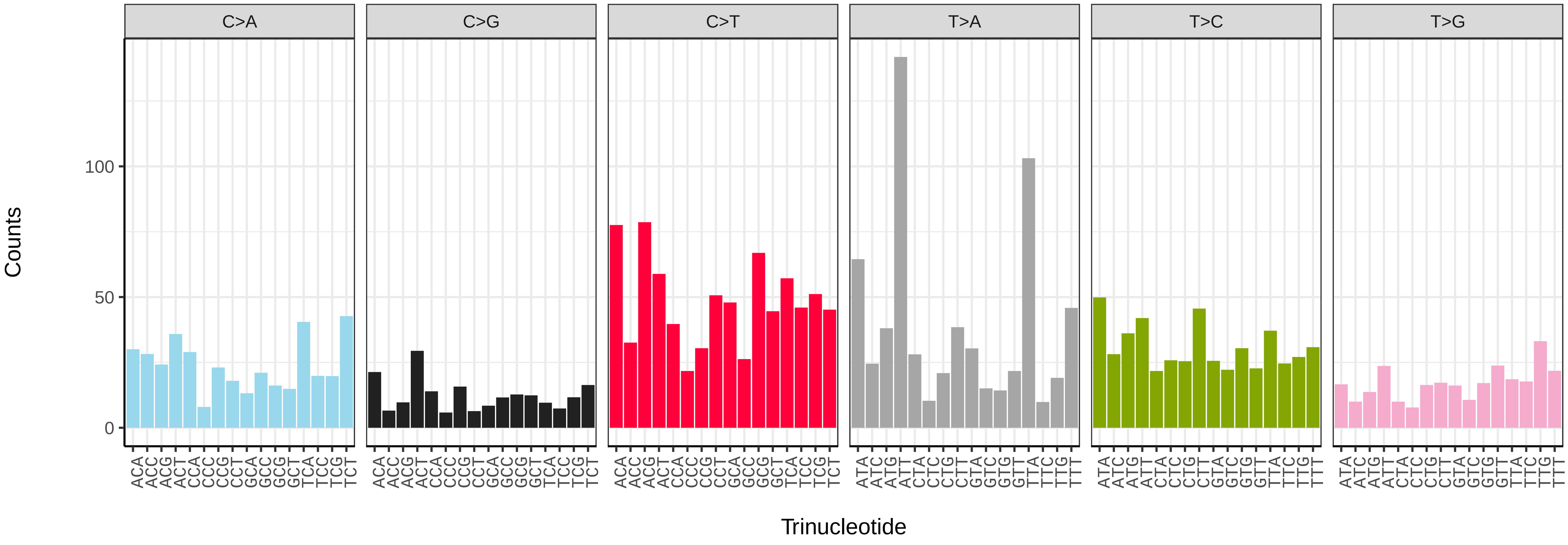


*Furcula furcula* (ilFurFurc1) somatic mutational spectrum (SBS52 classification)

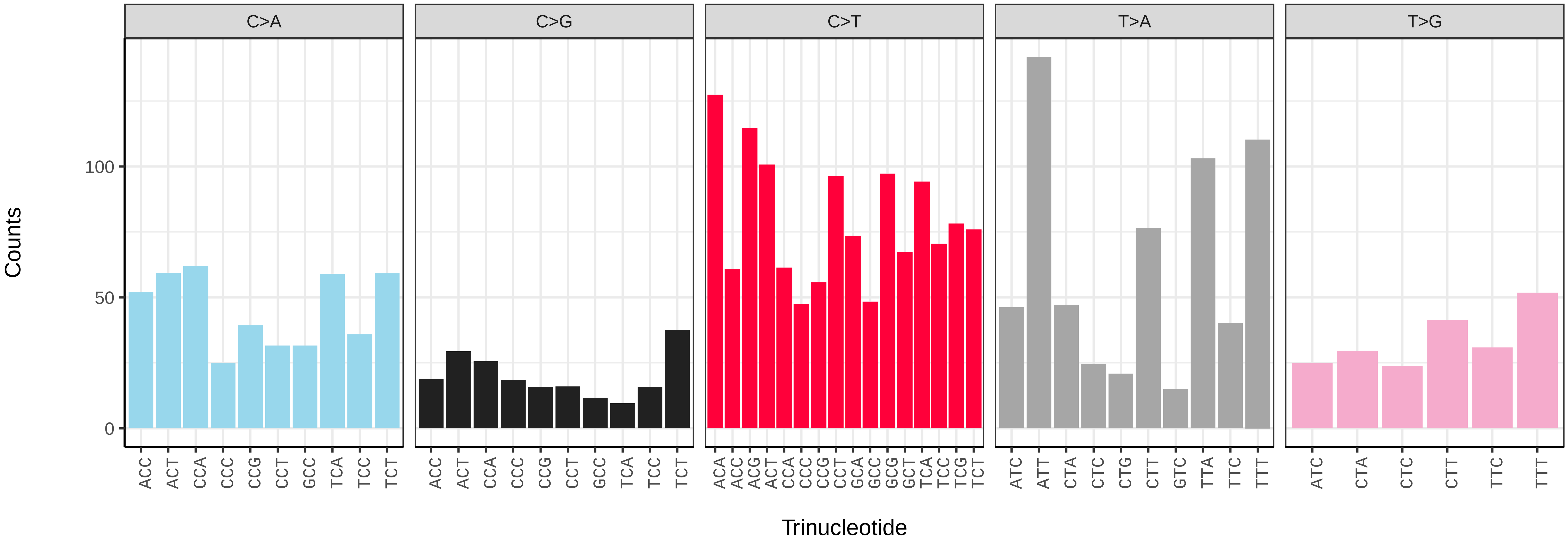

*Gandaritis pyraliata* (ilGanPyra1) germline mutational spectrum

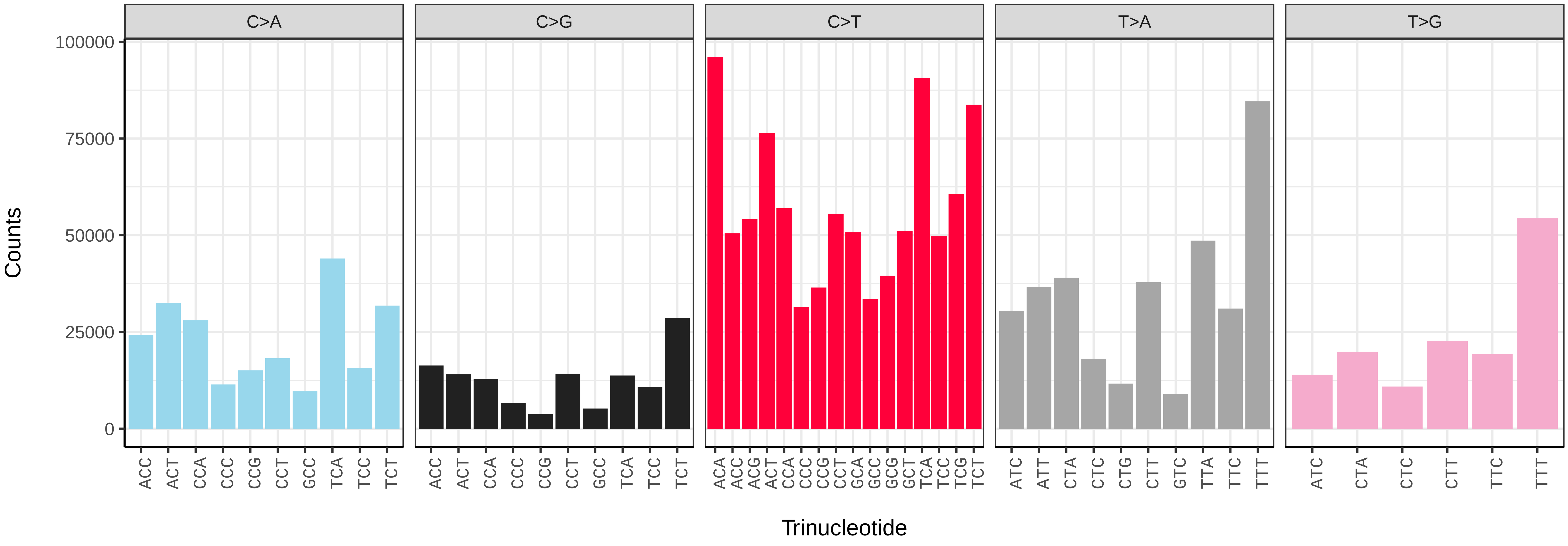


*Gandaritis pyraliata* (ilGanPyra1) somatic mutational spectrum (SBS96 classification)

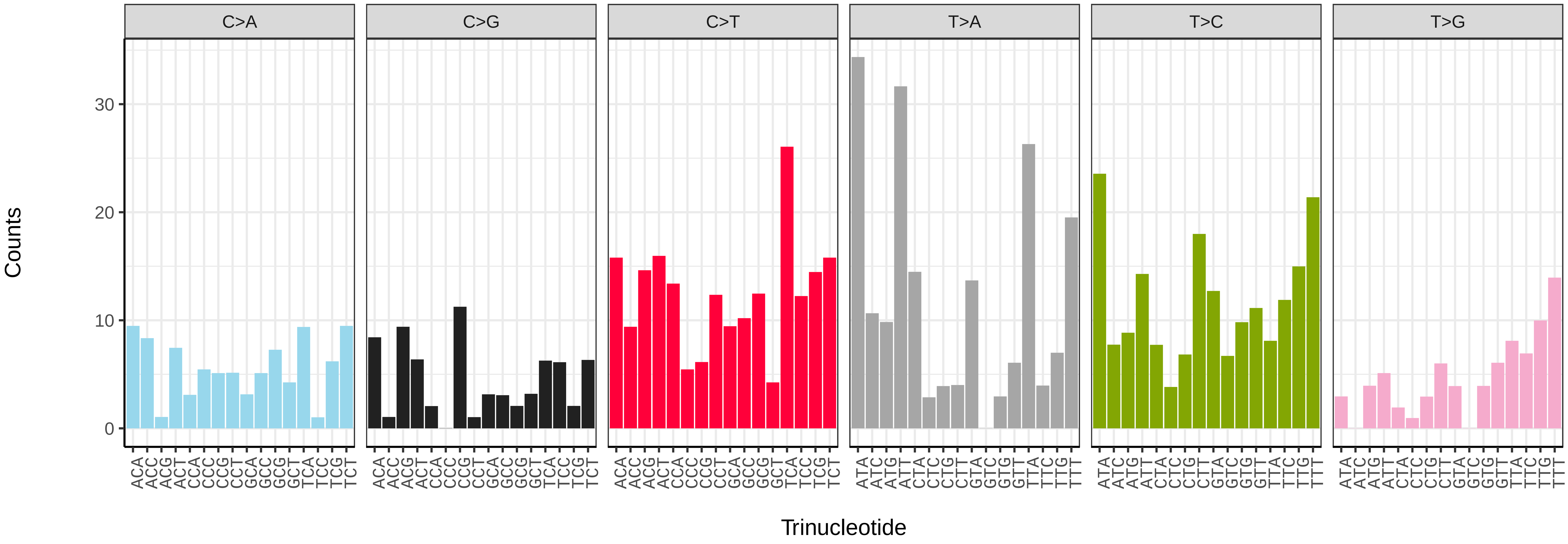


*Gandaritis pyraliata* (ilGanPyra1) somatic mutational spectrum (SBS52 classification)

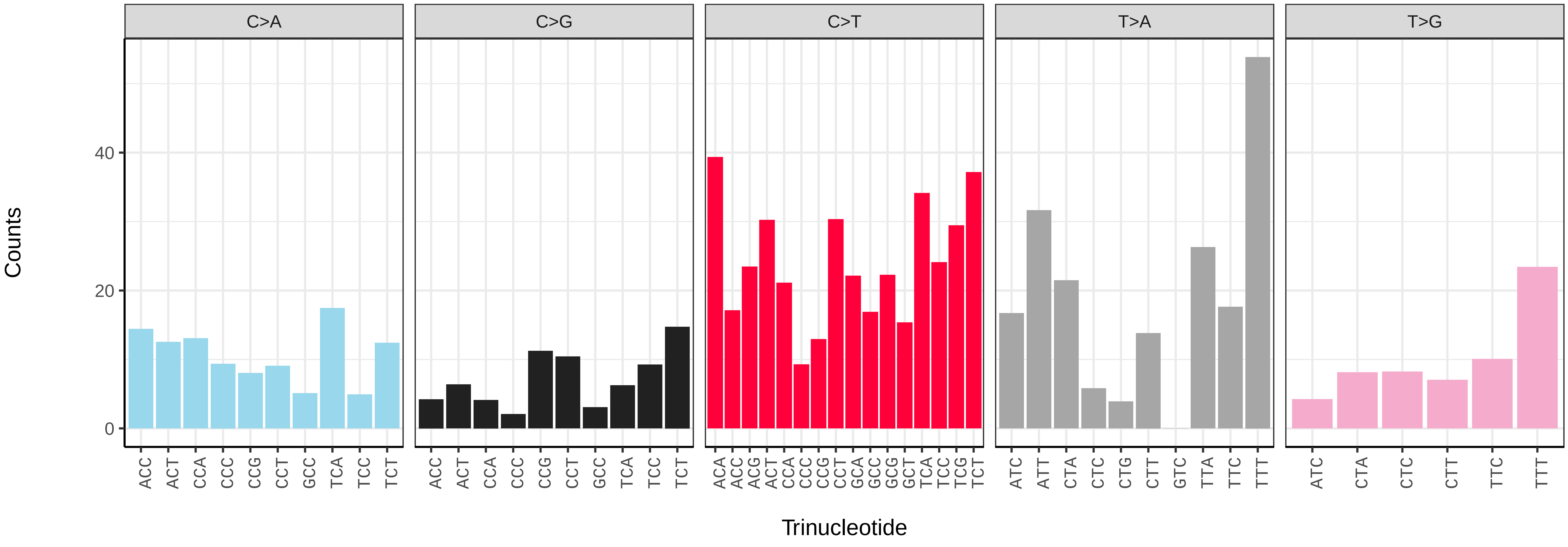

*Glaucopsyche alexis* (ilGlaAlex1) germline mutational spectrum

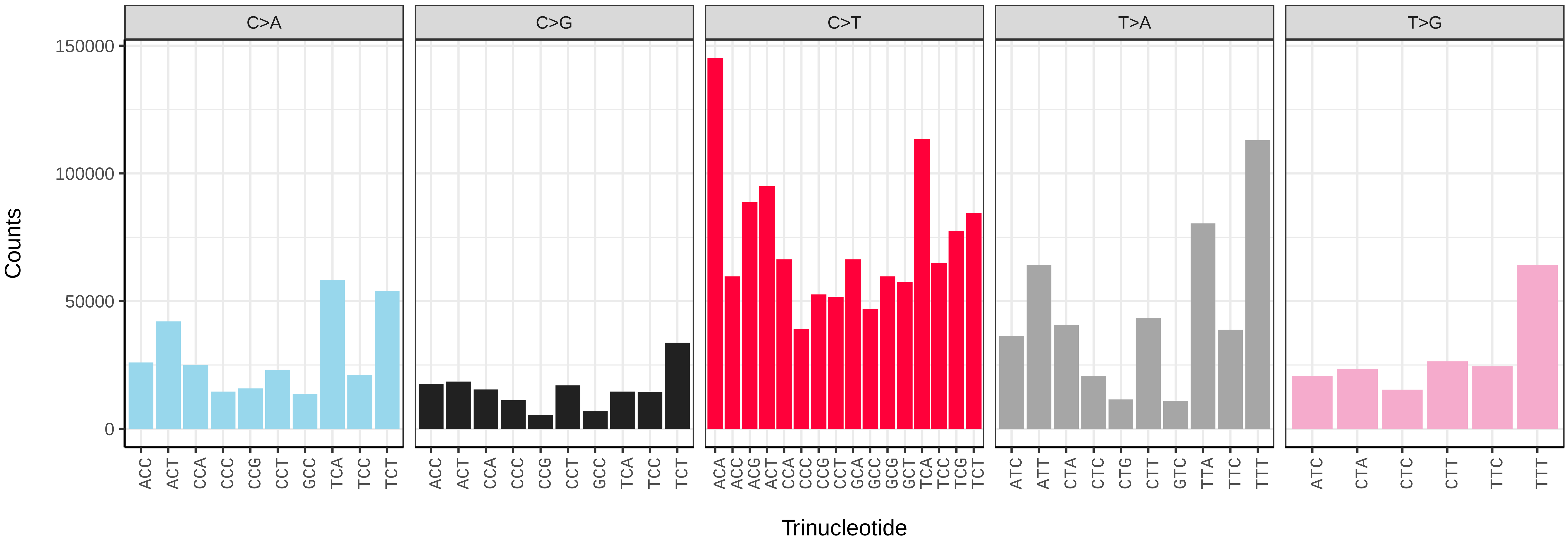


*Glaucopsyche alexis* (ilGlaAlex1) somatic mutational spectrum (SBS96 classification)

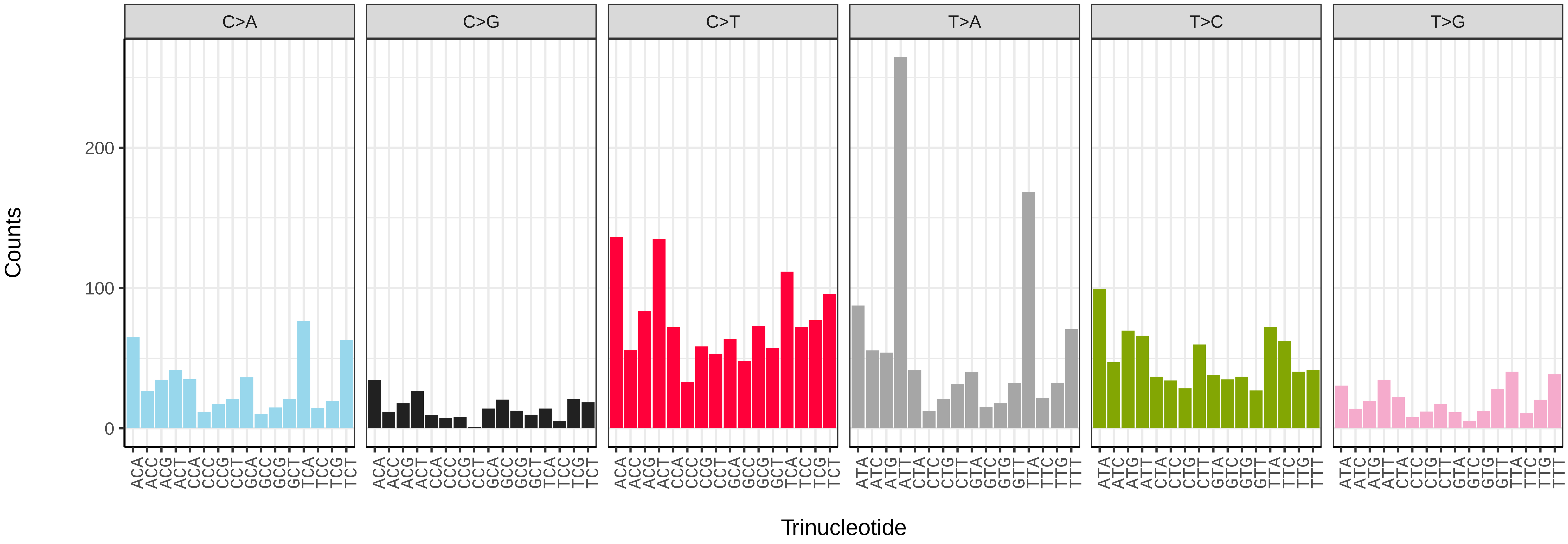


*Glaucopsyche alexis* (ilGlaAlex1) somatic mutational spectrum (SBS52 classification)

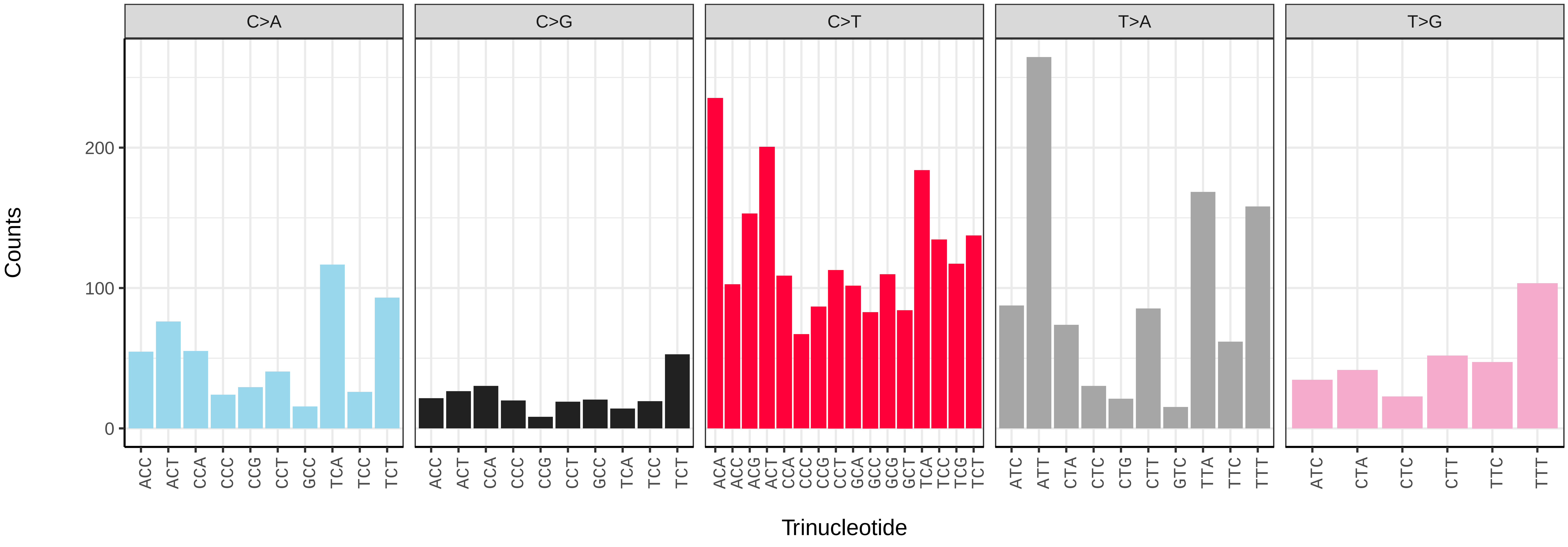

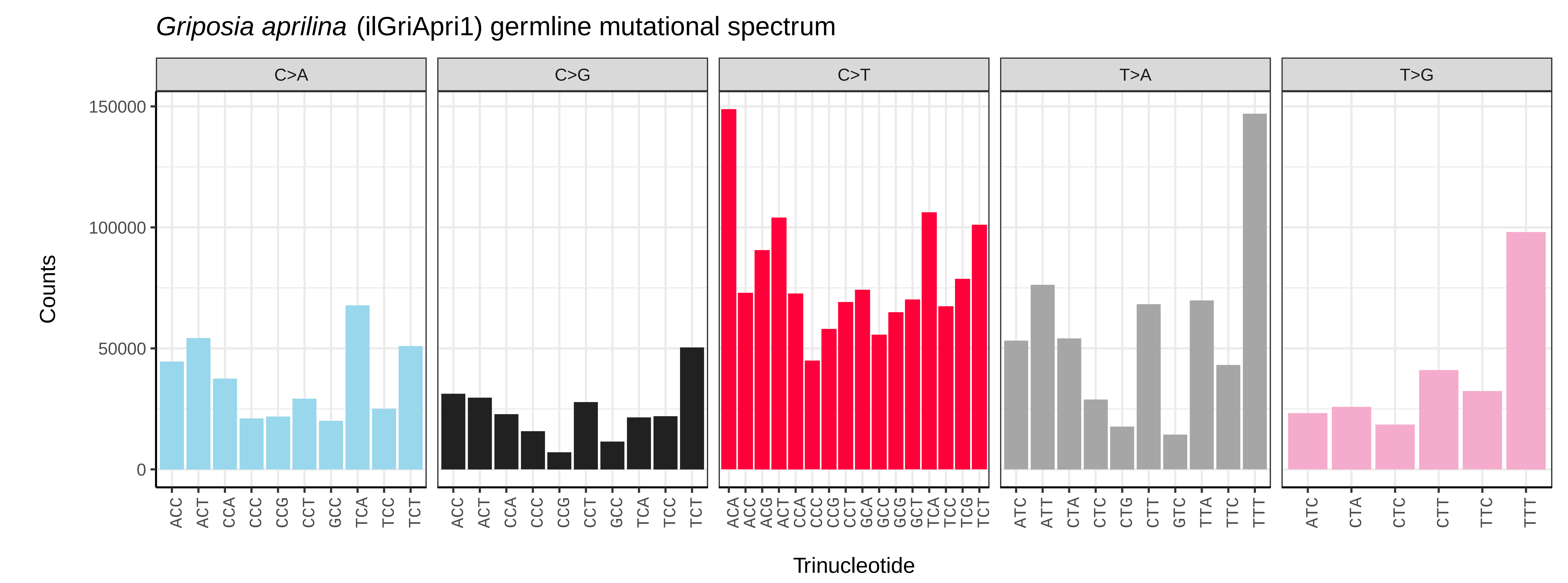
*Griposia aprilina* (ilGriApri1) germline mutational spectrum
C>A
C>G
C>T
T>A
T>G
Counts
150000
100000
50000
0
Trinucleotide

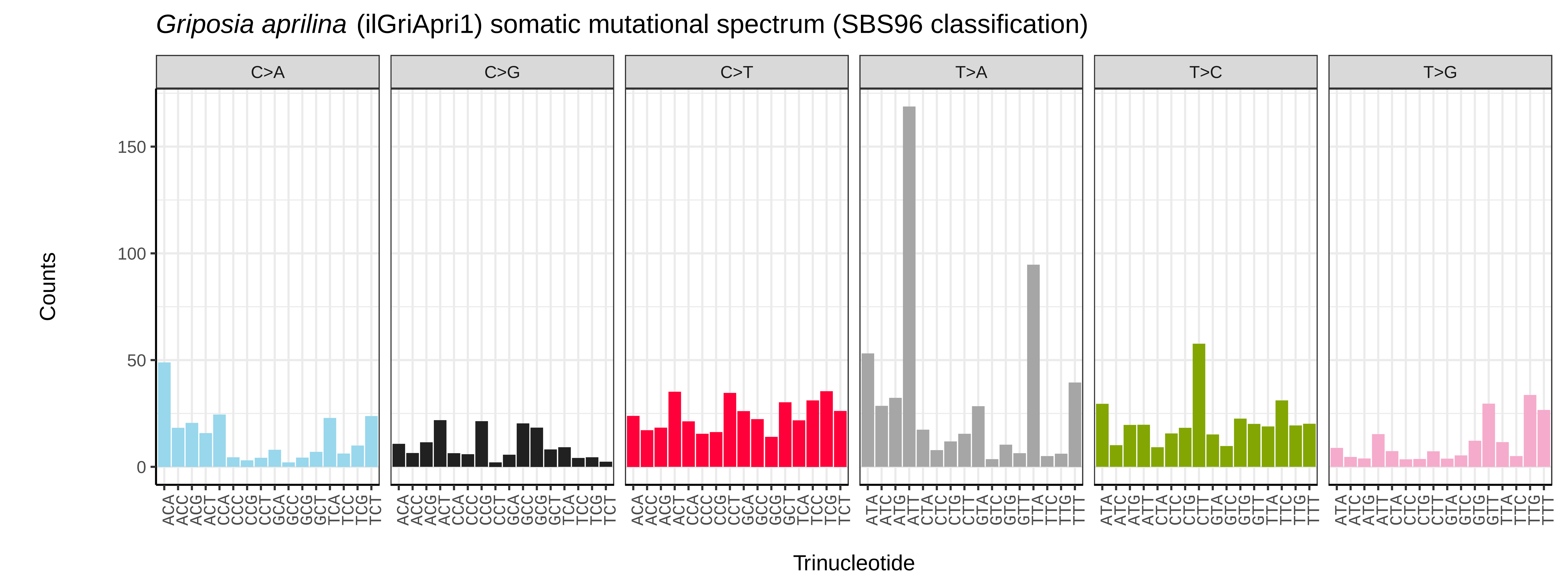
*Griposia aprilina* (ilGriApri1) somatic mutational spectrum (SBS96 classification)
C>A
C>G
C>T
T>A
T>C
T>G
Counts
150
100
50
0
Trinucleotide

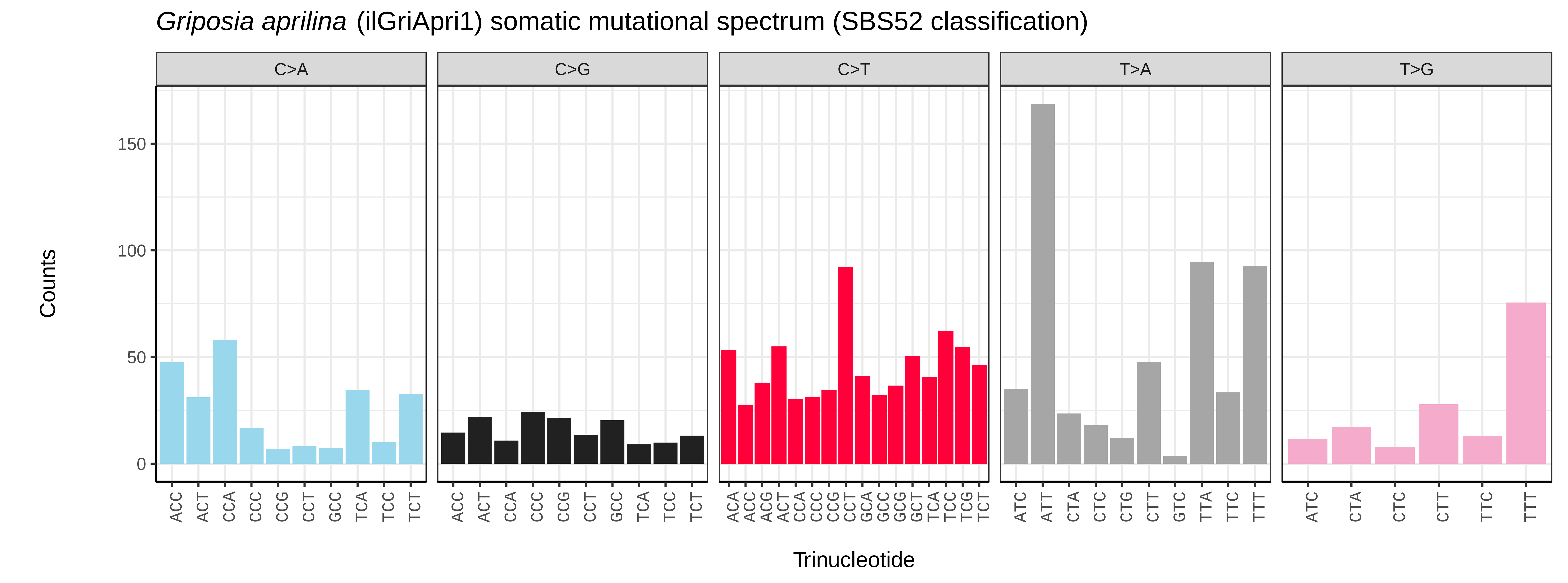
*Griposia aprilina* (ilGriApri1) somatic mutational spectrum (SBS52 classification)
C>A
C>G
C>T
T>A
T>G
Counts
150
100
50
0
Trinucleotide

*Gymnoscelis rufifasciata* (ilGymRufi1) germline mutational spectrum

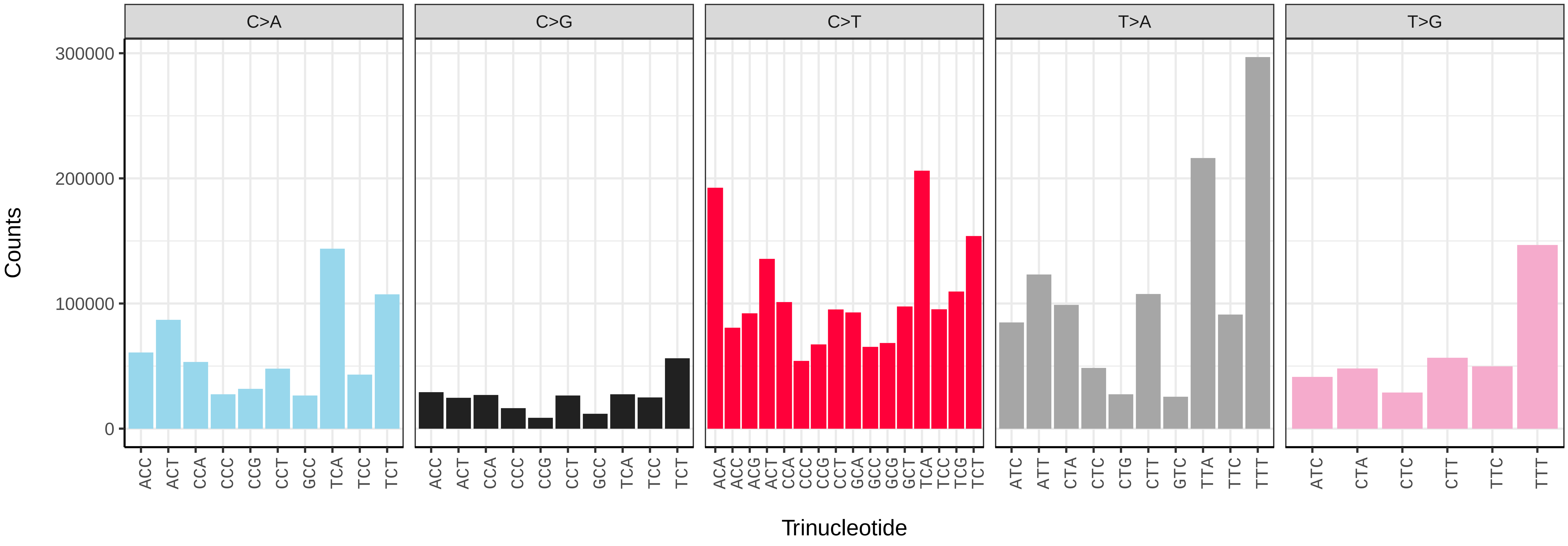


*Gymnoscelis rufifasciata* (ilGymRufi1) somatic mutational spectrum (SBS96 classification)

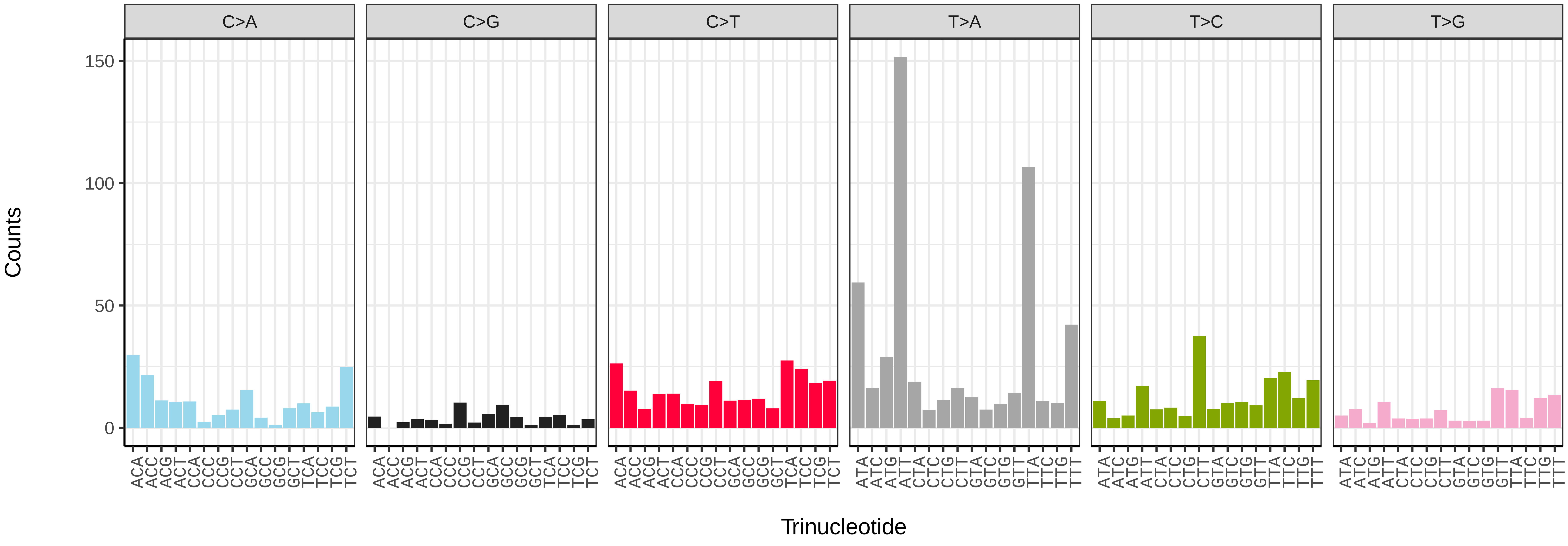


*Gymnoscelis rufifasciata* (ilGymRufi1) somatic mutational spectrum (SBS52 classification)

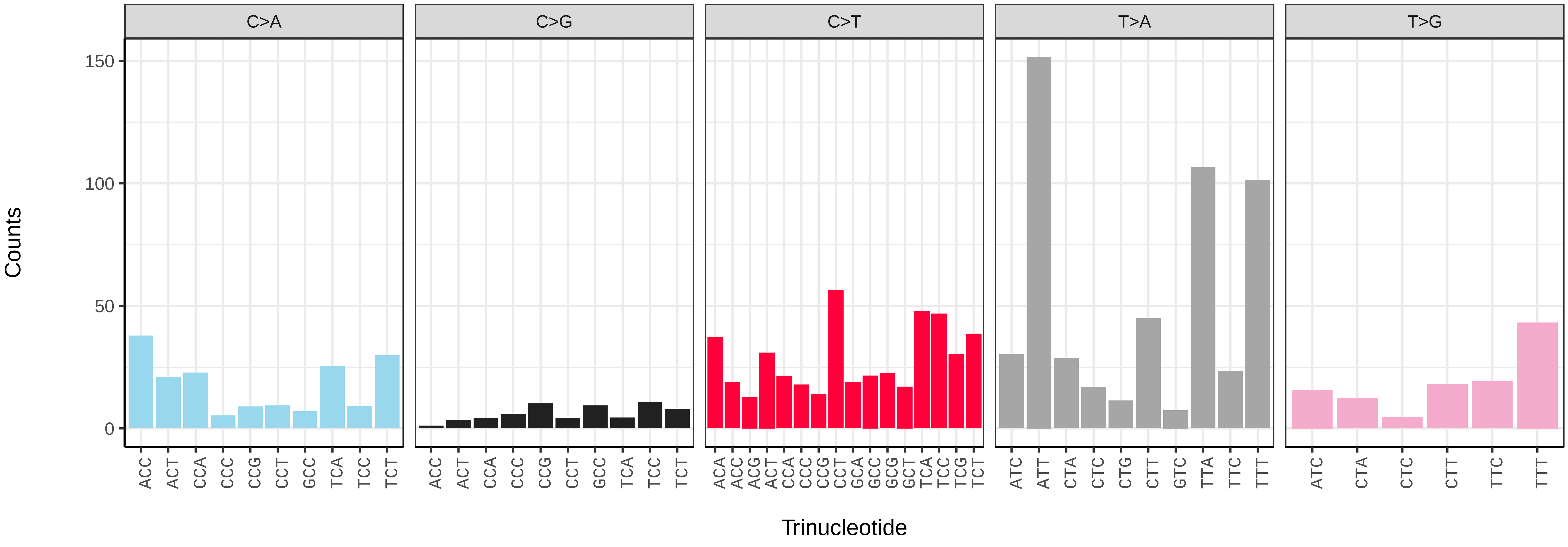

*Habrosyne pyritoides* (ilHabPyri1) germline mutational spectrum

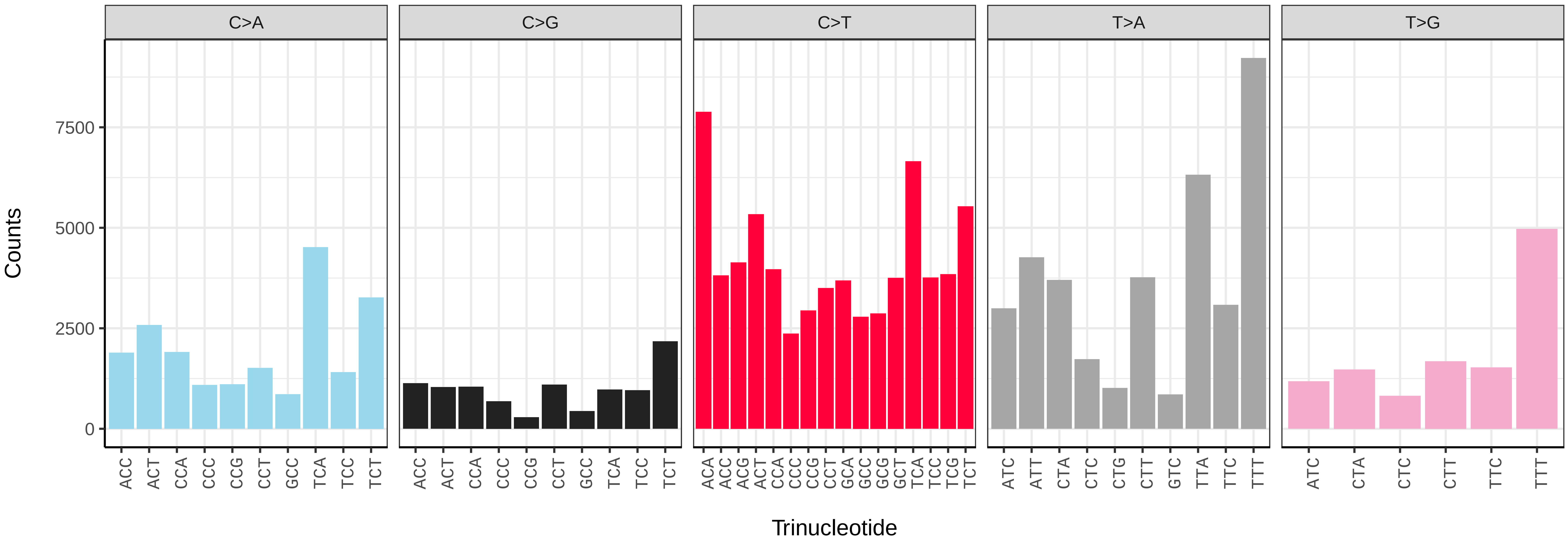


*Habrosyne pyritoides* (ilHabPyri1) somatic mutational spectrum (SBS96 classification)

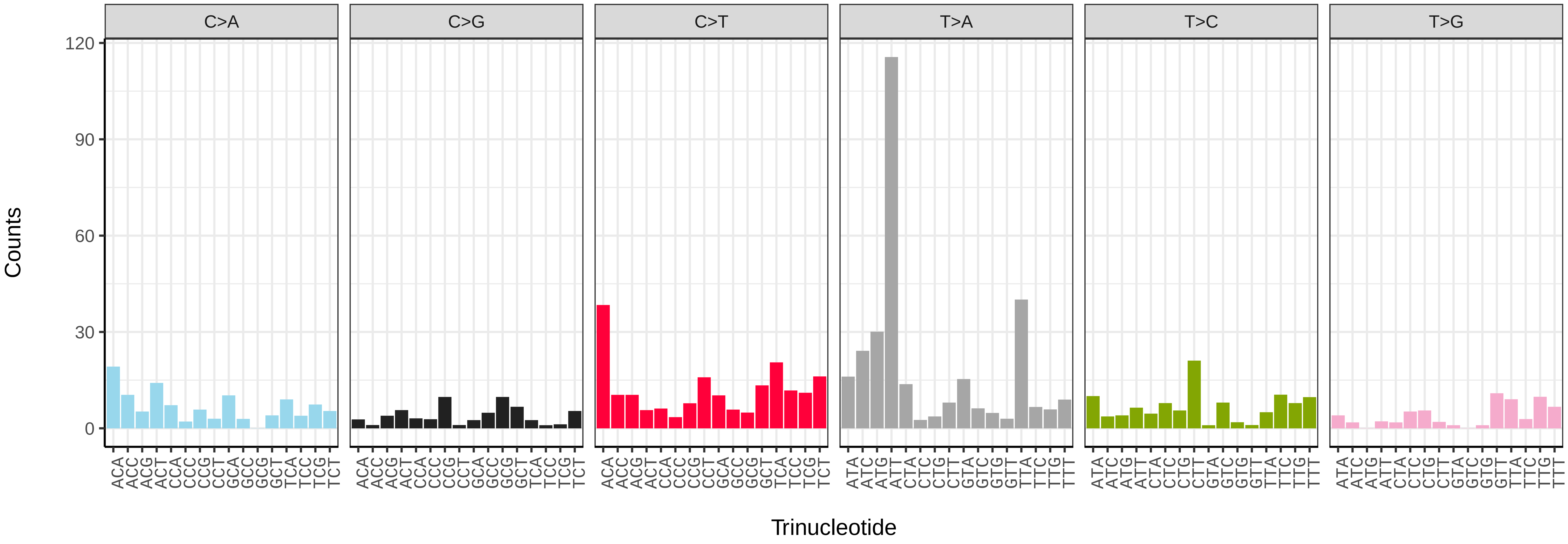


*Habrosyne pyritoides* (ilHabPyri1) somatic mutational spectrum (SBS52 classification)

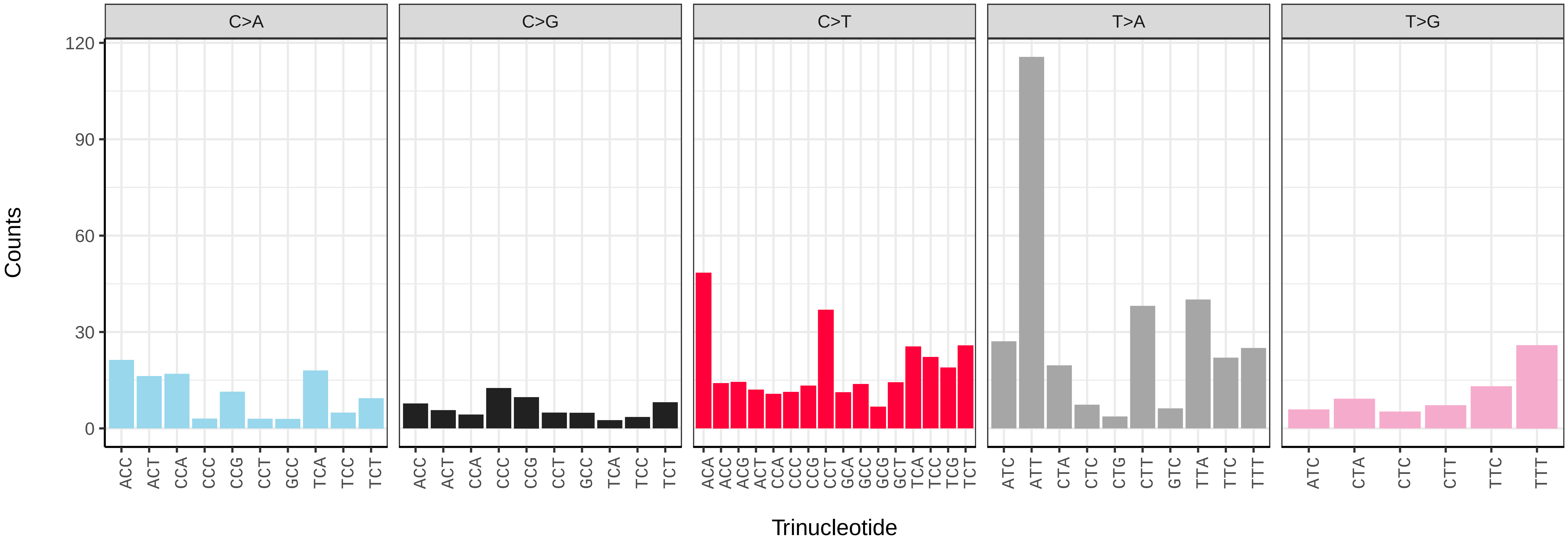

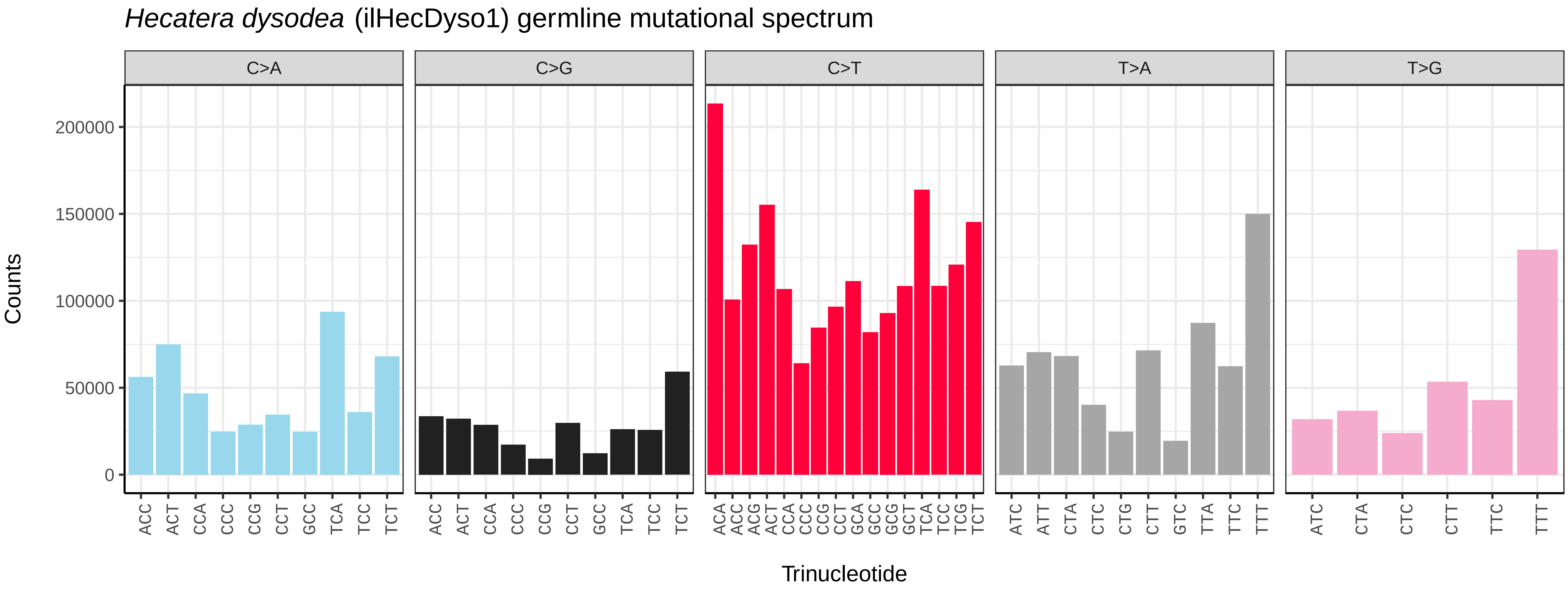
*Hecatera dysodea* (ilHecDyso1) germline mutational spectrum
C>A
C>G
C>T
T>A
T>G
Counts
200000
150000
100000
50000
0
ACC ACT CCA CCC CCG CCT GCC TCA TCC TCT
ACC ACT CCA CCC CCG CCT GCC TCA TCC TCT
ACA ACC ACG ACT CCA CCC CCG CCT GCA GCC GCG GCT TCA TCC TCG TCT
ATC ATT CTA CTC CTG CTT GTC TTA TTC TTT
ATC CTA CTC CTT TTC TTT
Trinucleotide

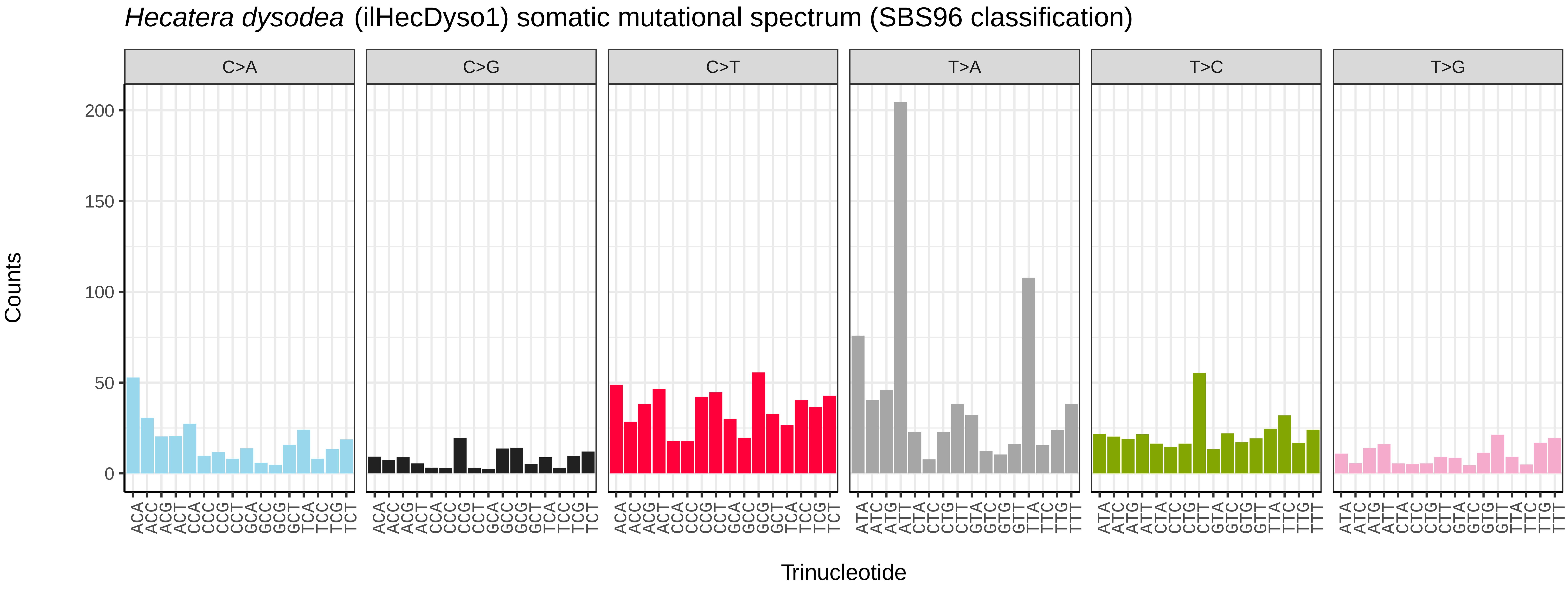
*Hecatera dysodea* (ilHecDyso1) somatic mutational spectrum (SBS96 classification)
C>A
C>G
C>T
T>A
T>C
T>G
Counts
200
150
100
50
0
ACA ACC ACG ACT CCA CCC CCG CCT GCA GCC GCG GCT TCA TCC TCG TCT
ACA ACC ACG ACT CCA CCC CCG CCT GCA GCC GCG GCT TCA TCC TCG TCT
ACA ACC ACG ACT CCA CCC CCG CCT GCA GCC GCG GCT TCA TCC TCG TCT
ATA ATC ATG ATT CTA CTC CTG CTT GTA GTC GTG GTT TTA TTC TTG TTT
ATA ATC ATG ATT CTA CTC CTG CTT GTA GTC GTG GTT TTA TTC TTG TTT
ATA ATC ATG ATT CTA CTC CTG CTT GTA GTC GTG GTT TTA TTC TTG TTT
Trinucleotide

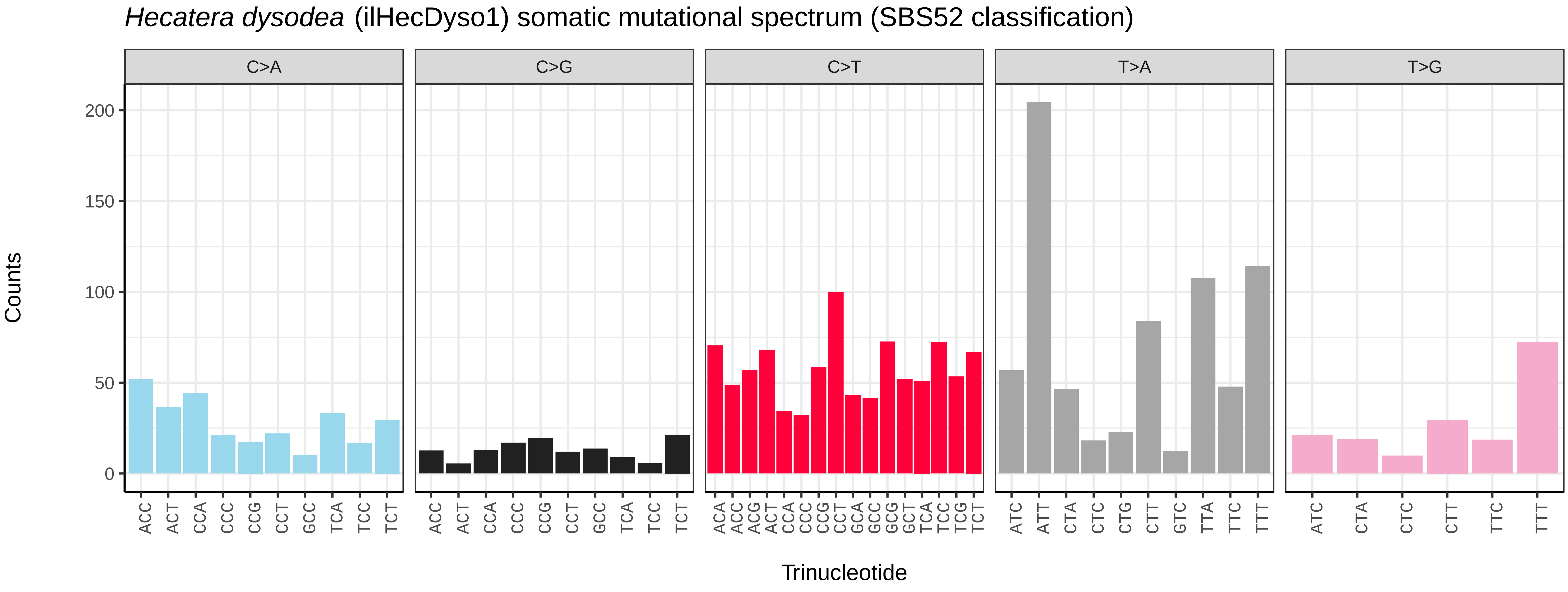
*Hecatera dysodea* (ilHecDyso1) somatic mutational spectrum (SBS52 classification)
C>A
C>G
C>T
T>A
T>G
Counts
200
150
100
50
0
ACC ACT CCA CCC CCG CCT GCC TCA TCC TCT
ACC ACT CCA CCC CCG CCT GCC TCA TCC TCT
ACA ACC ACG ACT CCA CCC CCG CCT GCA GCC GCG GCT TCA TCC TCG TCT
ATC ATT CTA CTC CTG CTT GTC TTA TTC TTT
ATC CTA CTC CTT TTC TTT
Trinucleotide

*Hedya salicella* (ilHedSali1) germline mutational spectrum

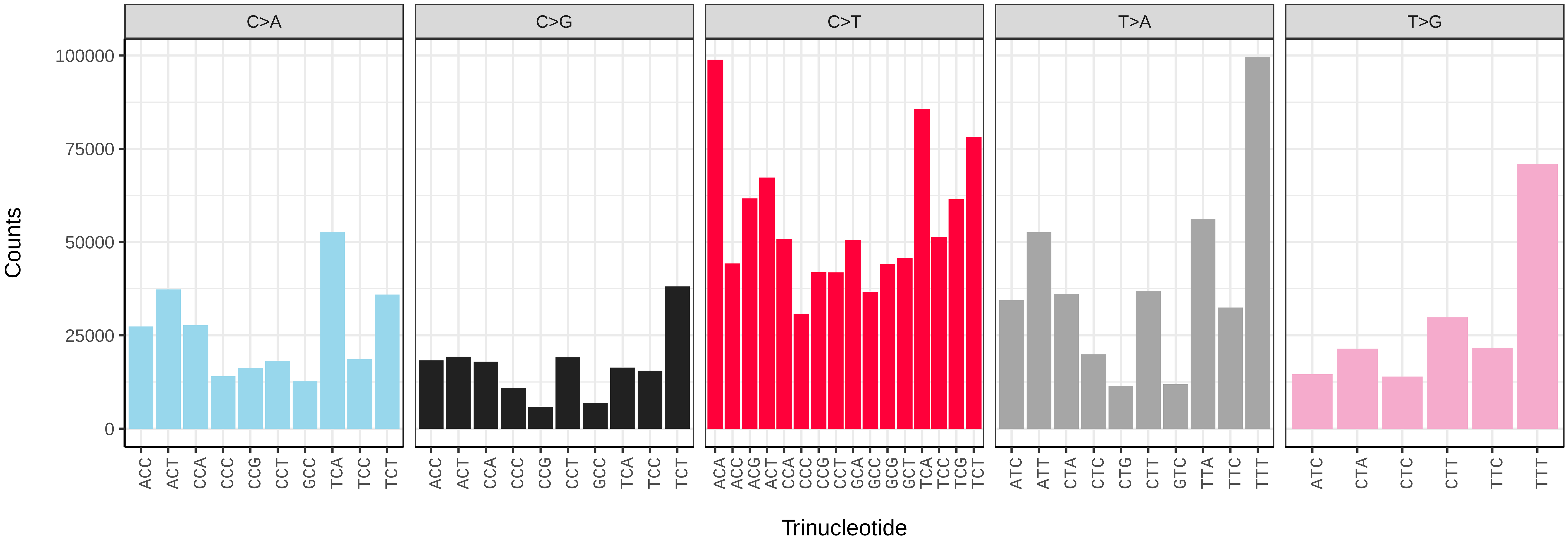


*Hedya salicella* (ilHedSali1) somatic mutational spectrum (SBS96 classification)

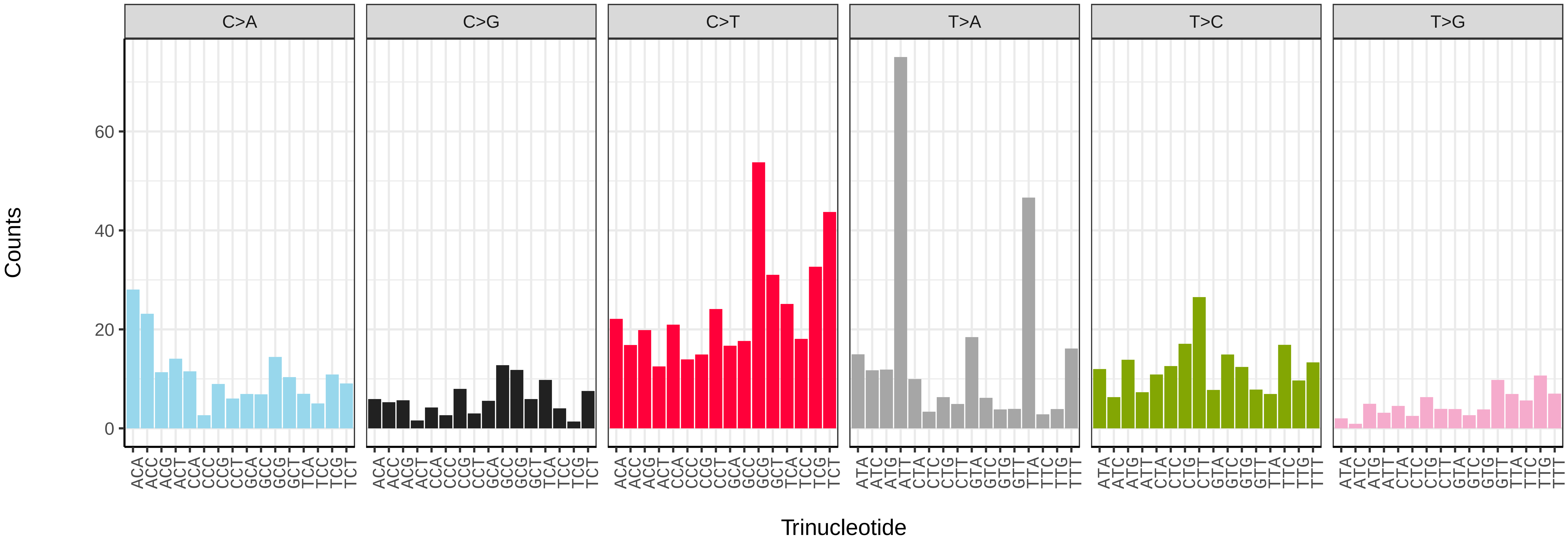


*Hedya salicella* (ilHedSali1) somatic mutational spectrum (SBS52 classification)

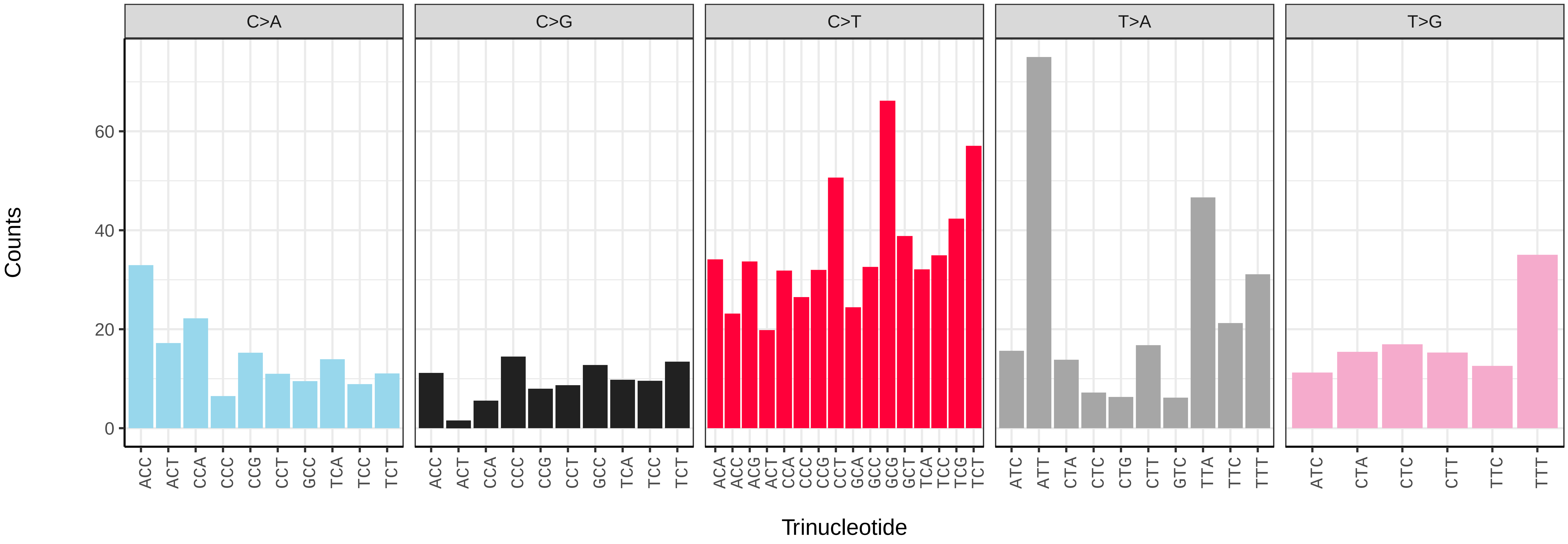

*Hemithea aestivaria* (ilHemAest2) germline mutational spectrum

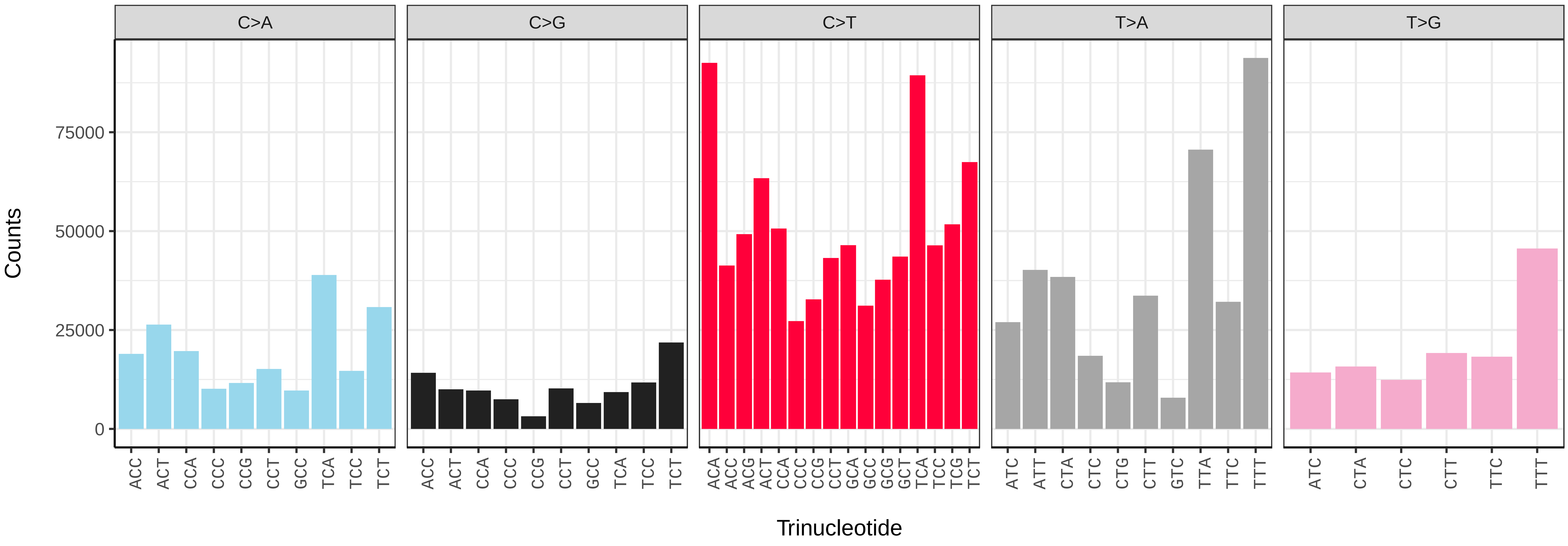


*Hemithea aestivaria* (ilHemAest2) somatic mutational spectrum (SBS96 classification)

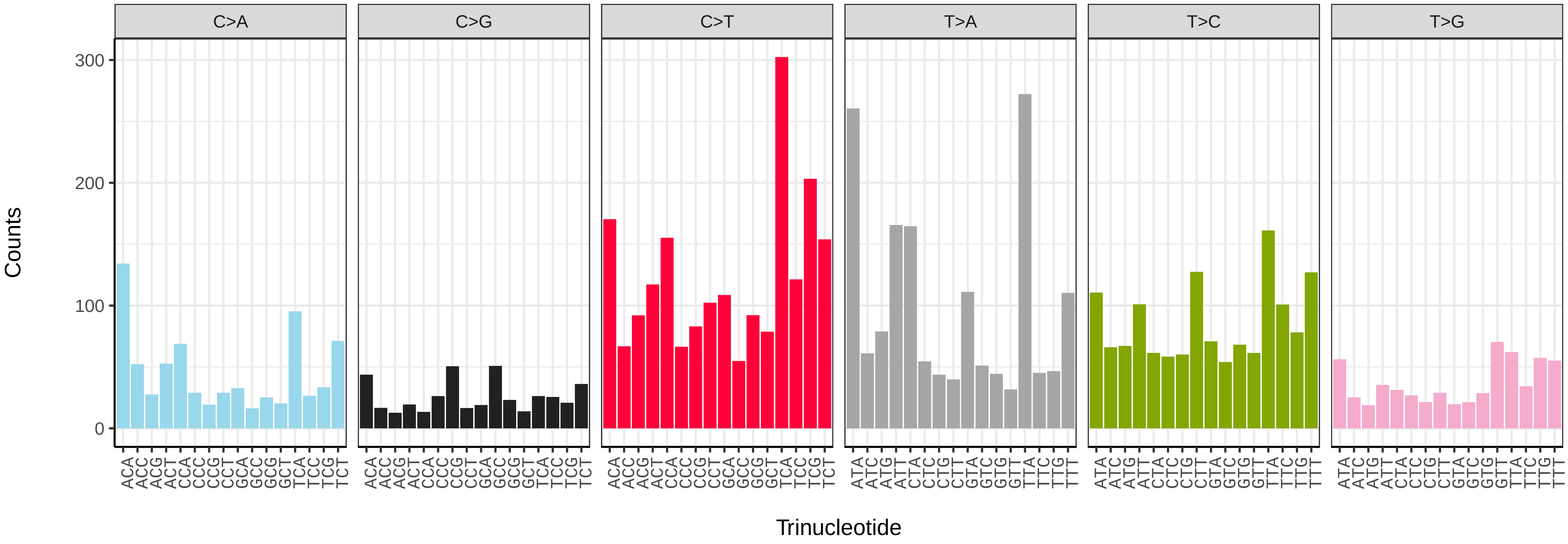


*Hemithea aestivaria* (ilHemAest2) somatic mutational spectrum (SBS52 classification)

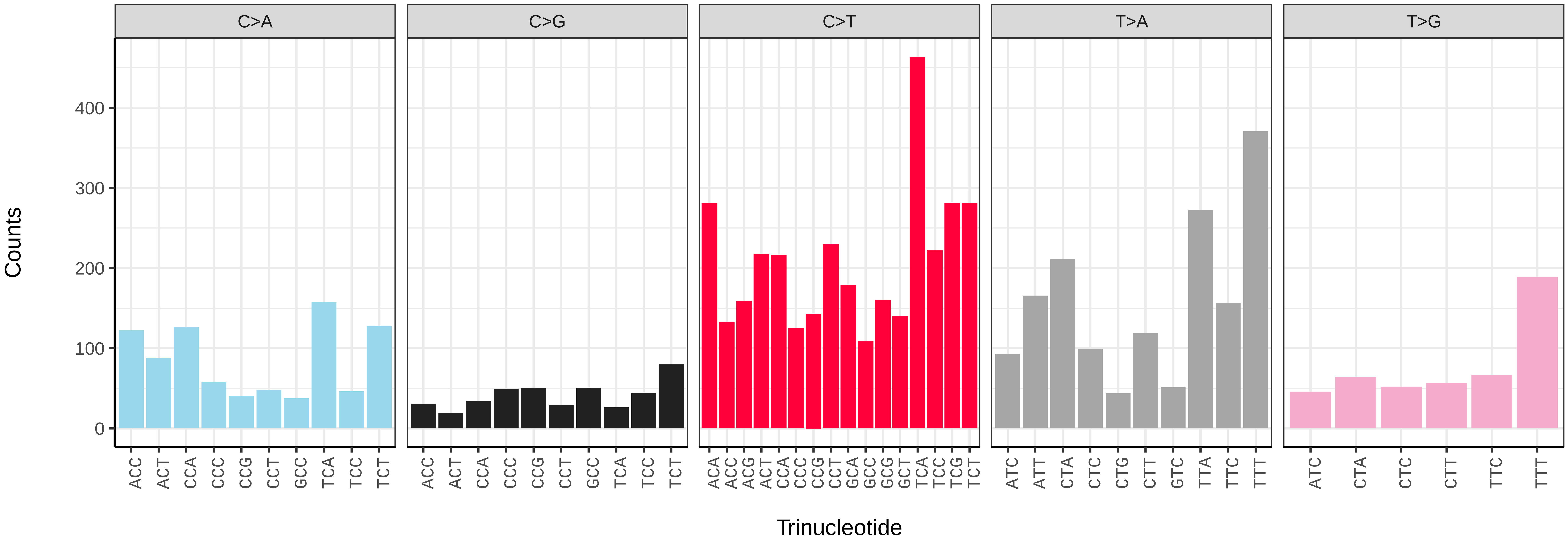

*Hemistola chrysoprasaria* (ilHemChry1) germline mutational spectrum

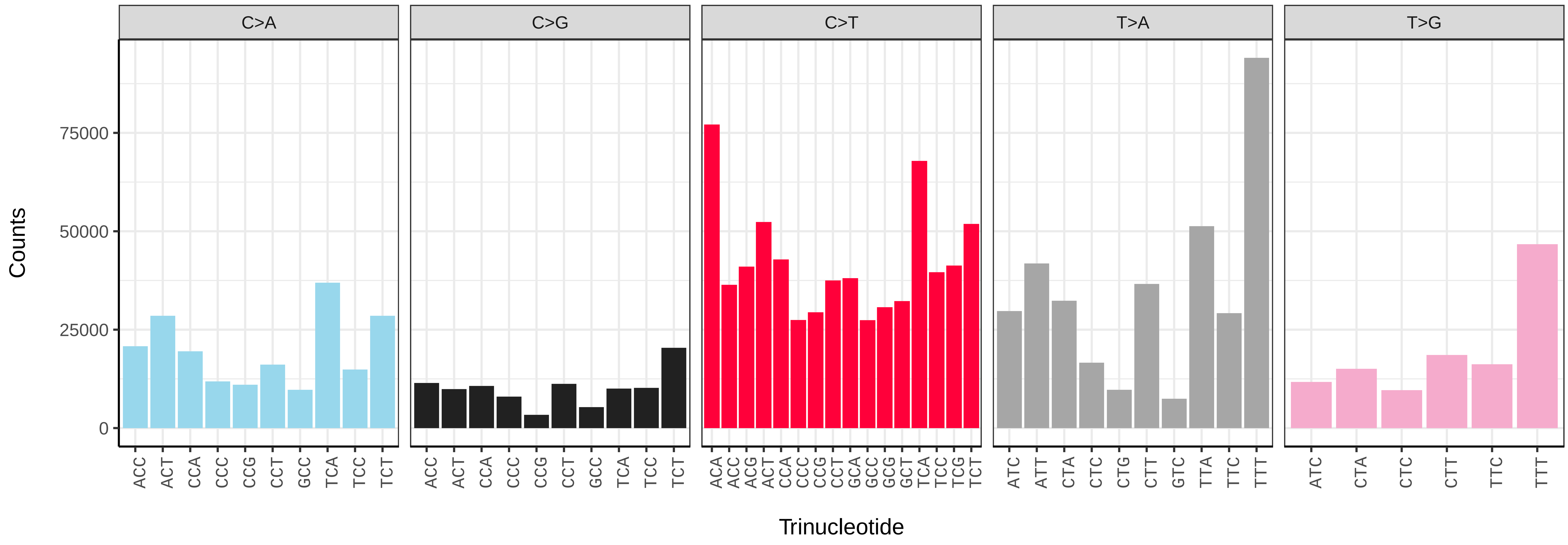


*Hemistola chrysoprasaria* (ilHemChry1) somatic mutational spectrum (SBS96 classification)

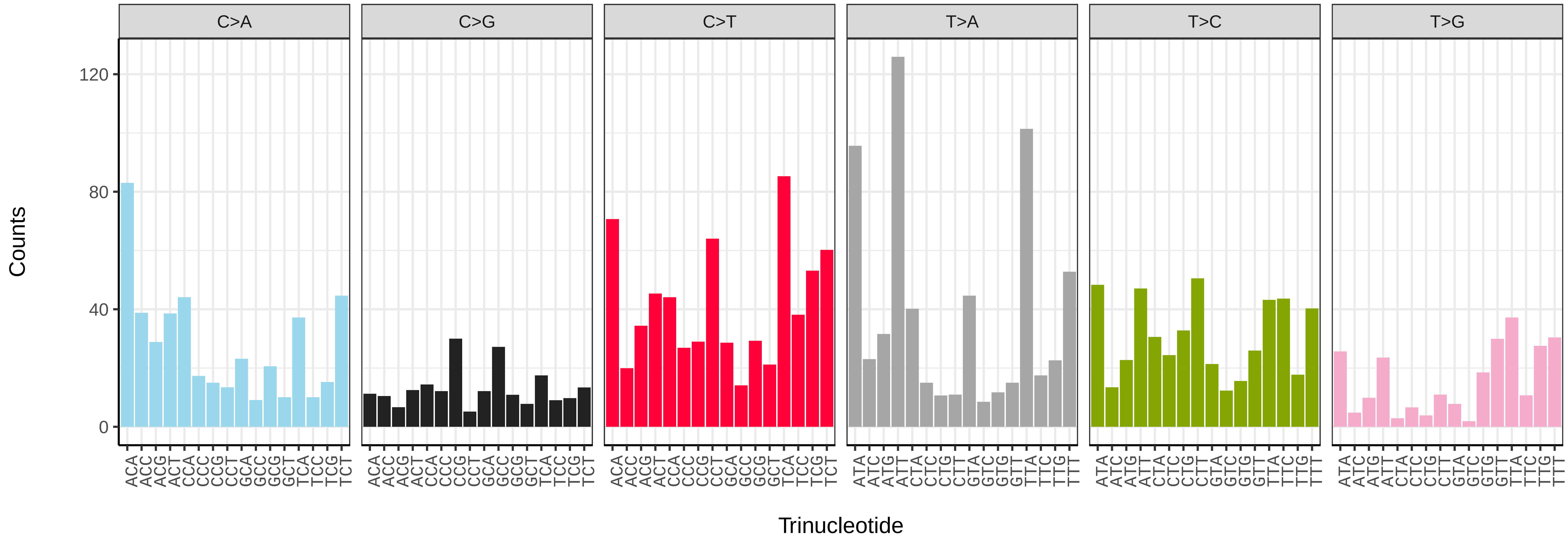


*Hemistola chrysoprasaria* (ilHemChry1) somatic mutational spectrum (SBS52 classification)

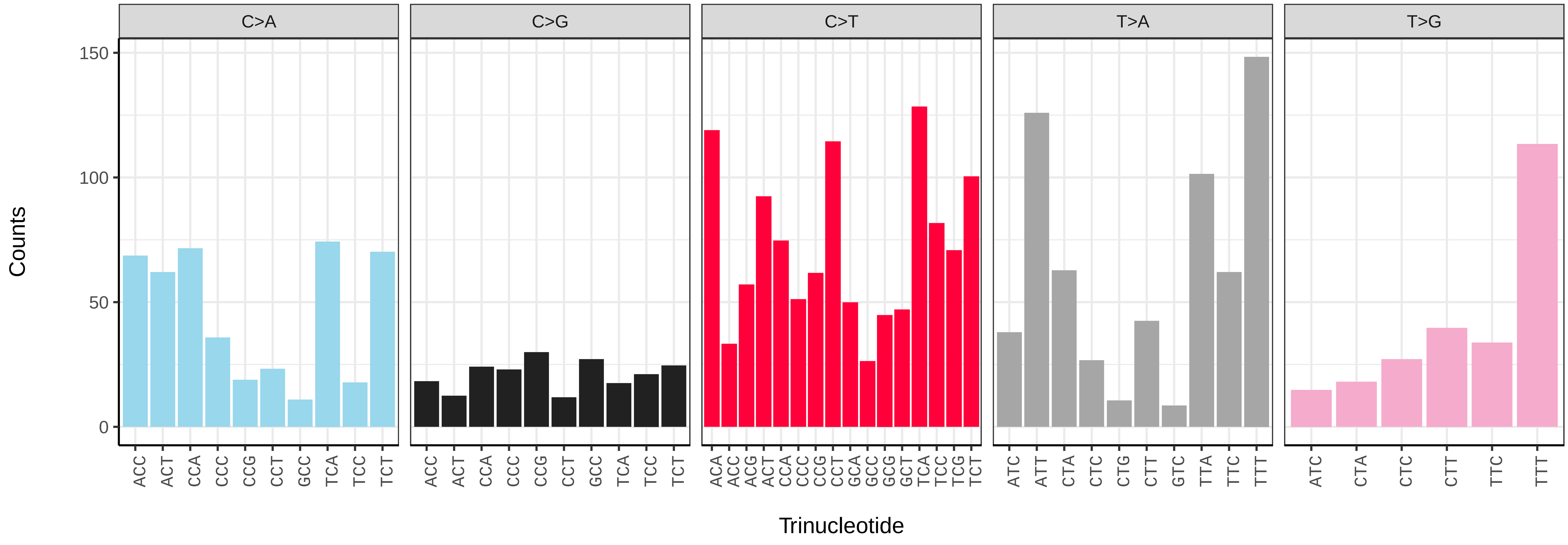

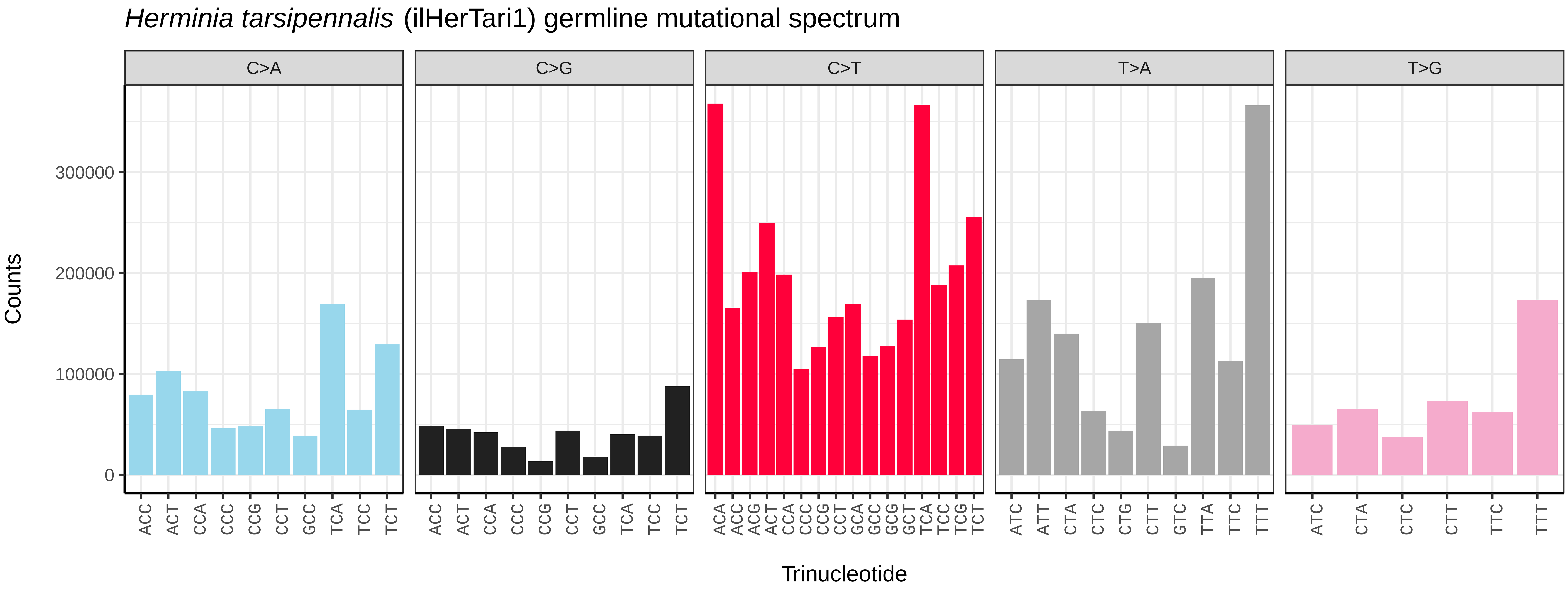
*Herminia tarsipennalis* (ilHerTari1) germline mutational spectrum
C>A
C>G
C>T
T>A
T>G
Counts
0
100000
200000
300000
ACC ACT CCA CCC CCG CCT GCC TCA TCC TCT
ACA ACC ACG ACT CCA CCC CCG CCT GCA GCC GCG GCT TCA TCC TCG TCT
ATC ATT CTA CTC CTG CTT GTC TTA TTC TTT
ATC CTA CTC CTT TTC TTT
Trinucleotide

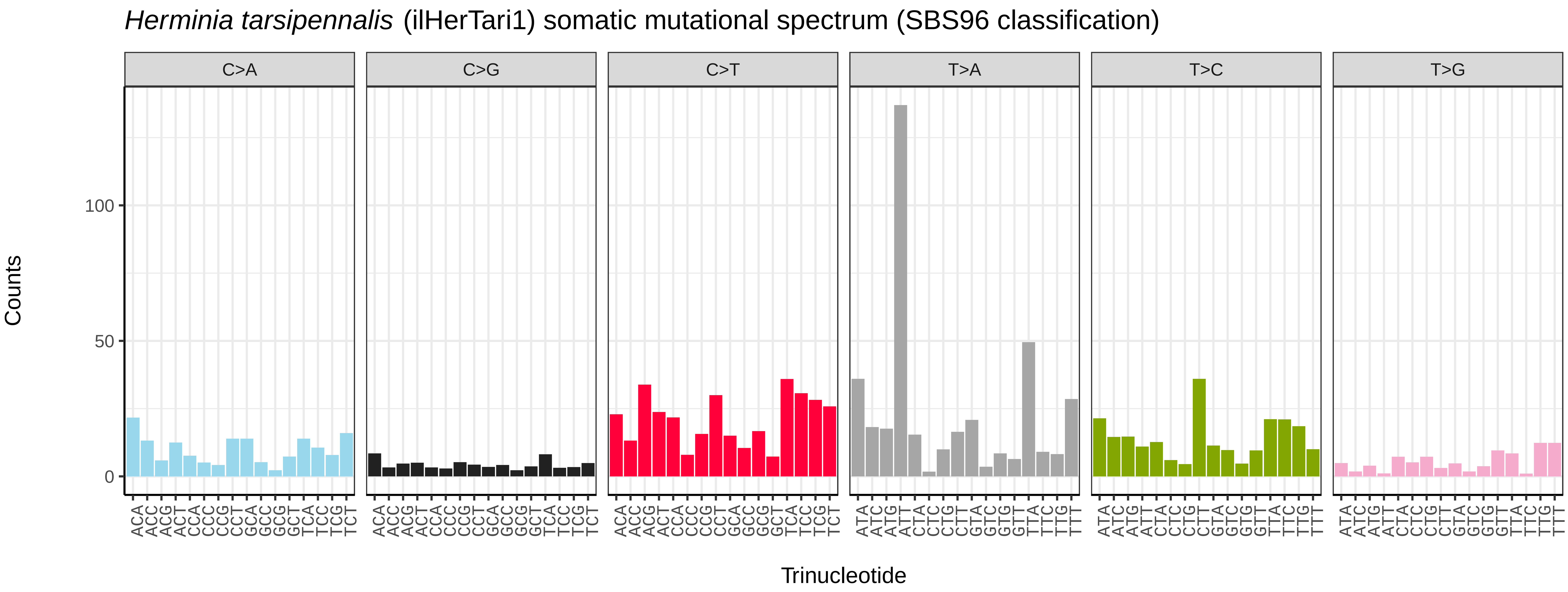
*Herminia tarsipennalis* (ilHerTari1) somatic mutational spectrum (SBS96 classification)
C>A
C>G
C>T
T>A
T>C
T>G
Counts
0
50
100
ACA ACC ACG ACT CCA CCC CCG CCT GCA GCC GCG GCT TCA TCC TCG TCT
ATA ATC ATG ATT CTA CTC CTG CTT GTA GTC GTG GTT TTA TTC TTG TTT
Trinucleotide

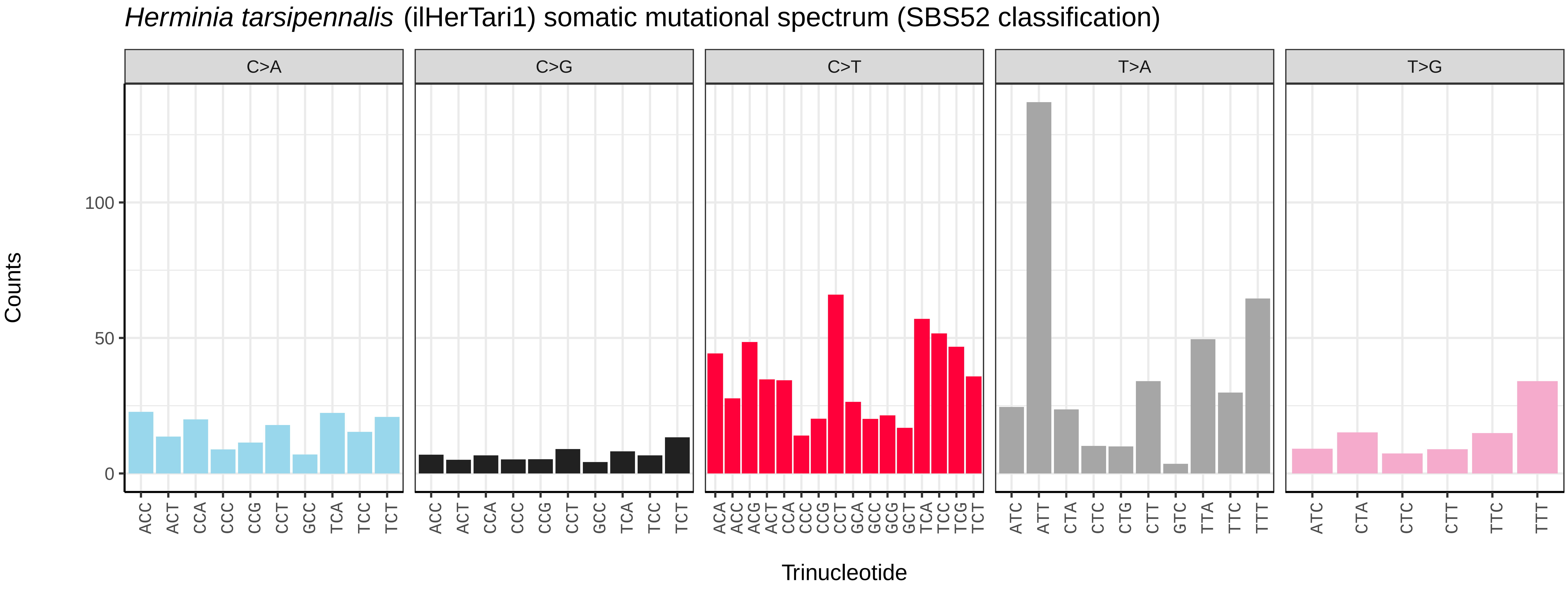
*Herminia tarsipennalis* (ilHerTari1) somatic mutational spectrum (SBS52 classification)
C>A
C>G
C>T
T>A
T>G
Counts
0
50
100
ACC ACT CCA CCC CCG CCT GCC TCA TCC TCT
ACA ACC ACG ACT CCA CCC CCG CCT GCA GCC GCG GCT TCA TCC TCG TCT
ATC ATT CTA CTC CTG CTT GTC TTA TTC TTT
ATC CTA CTC CTT TTC TTT
Trinucleotide

*Hesperia comma* (ilHesComm1) germline mutational spectrum

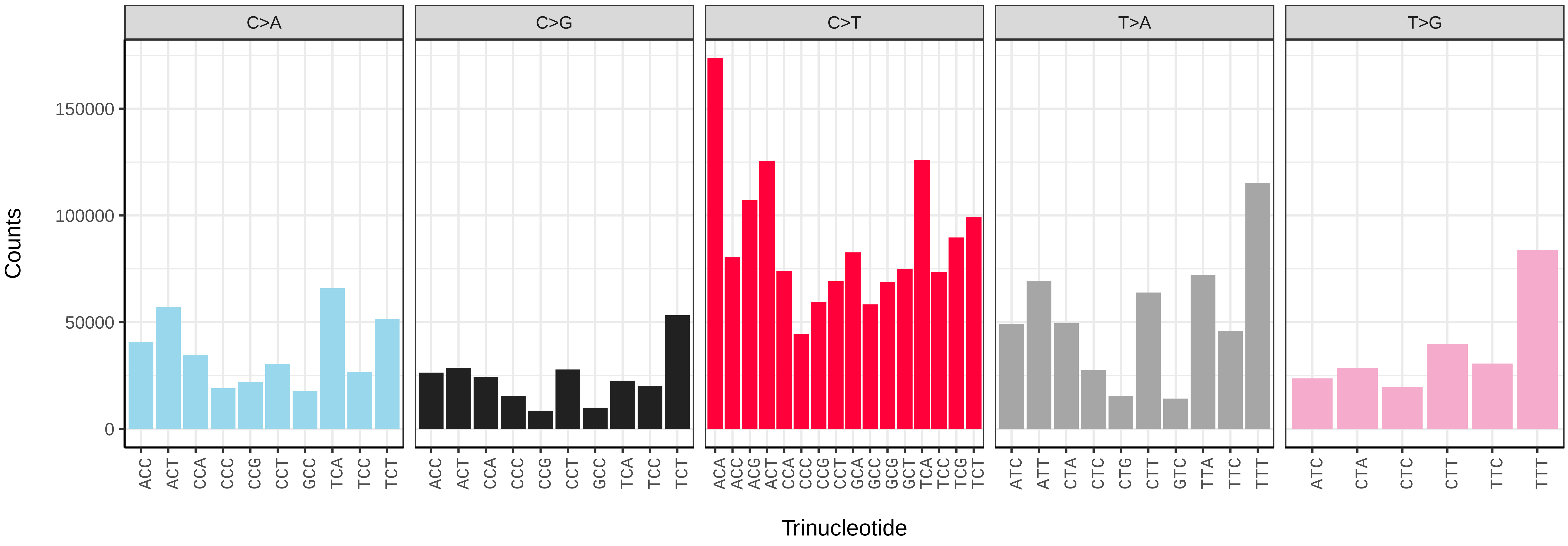


*Hesperia comma* (ilHesComm1) somatic mutational spectrum (SBS96 classification)

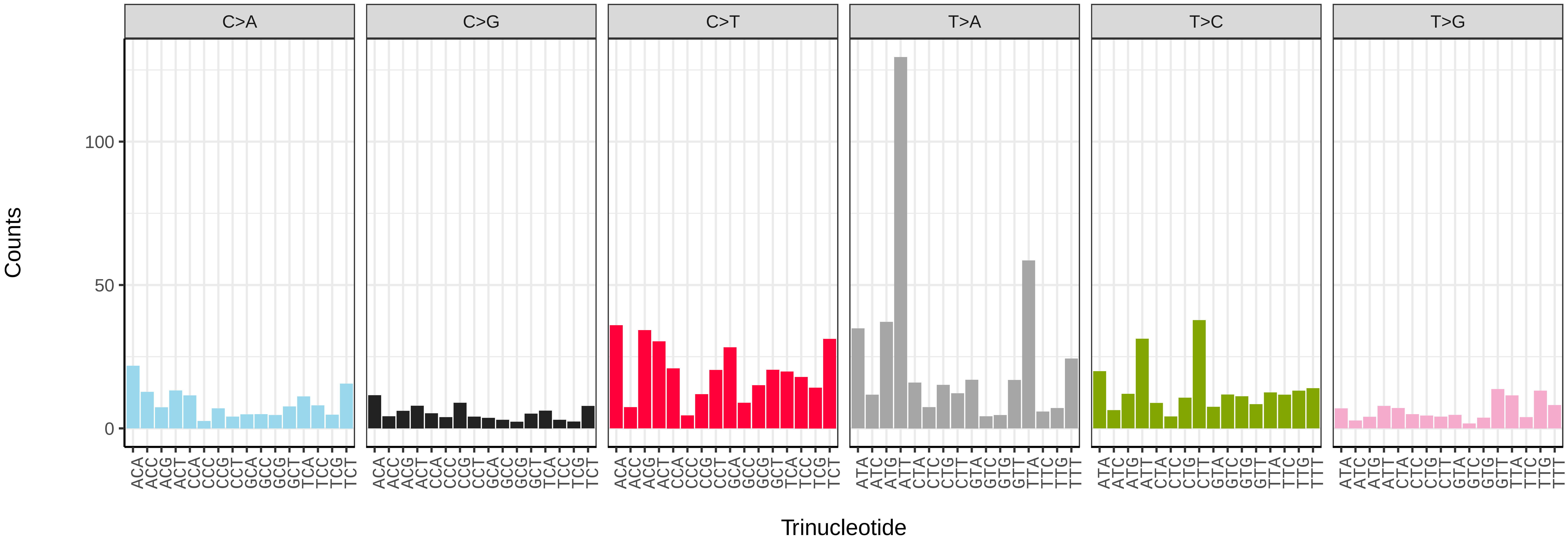


*Hesperia comma* (ilHesComm1) somatic mutational spectrum (SBS52 classification)

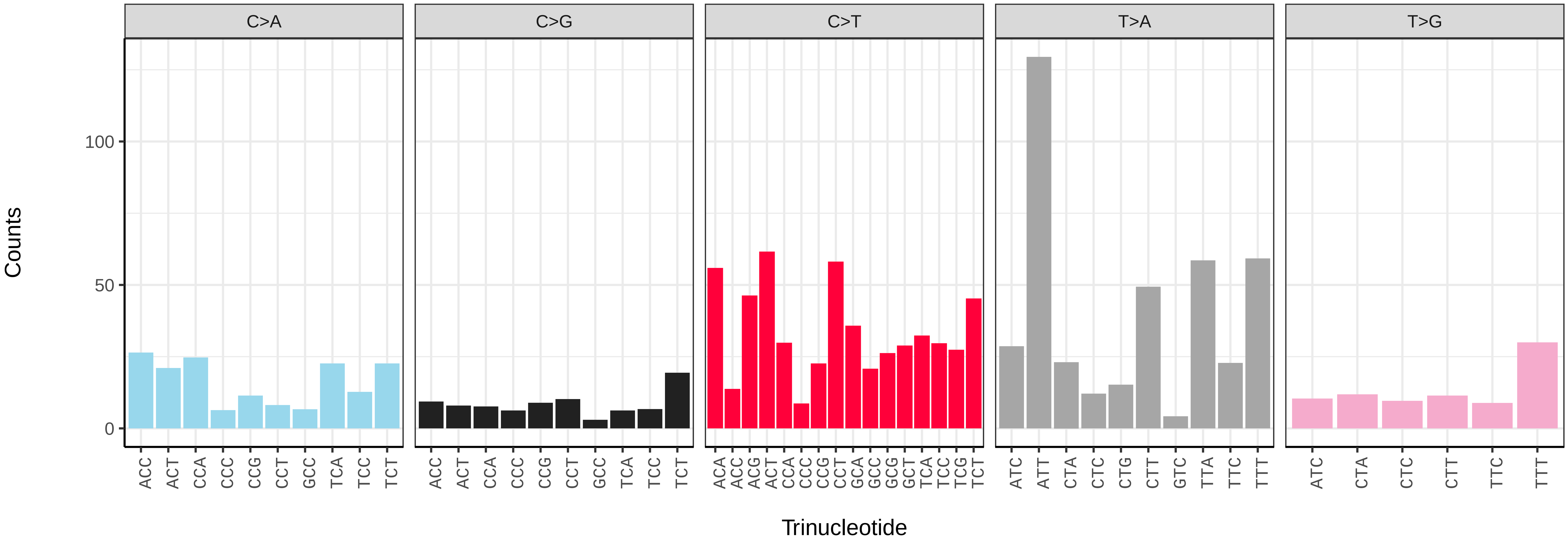

*Hipparchia semele* (ilHipSeme1) germline mutational spectrum

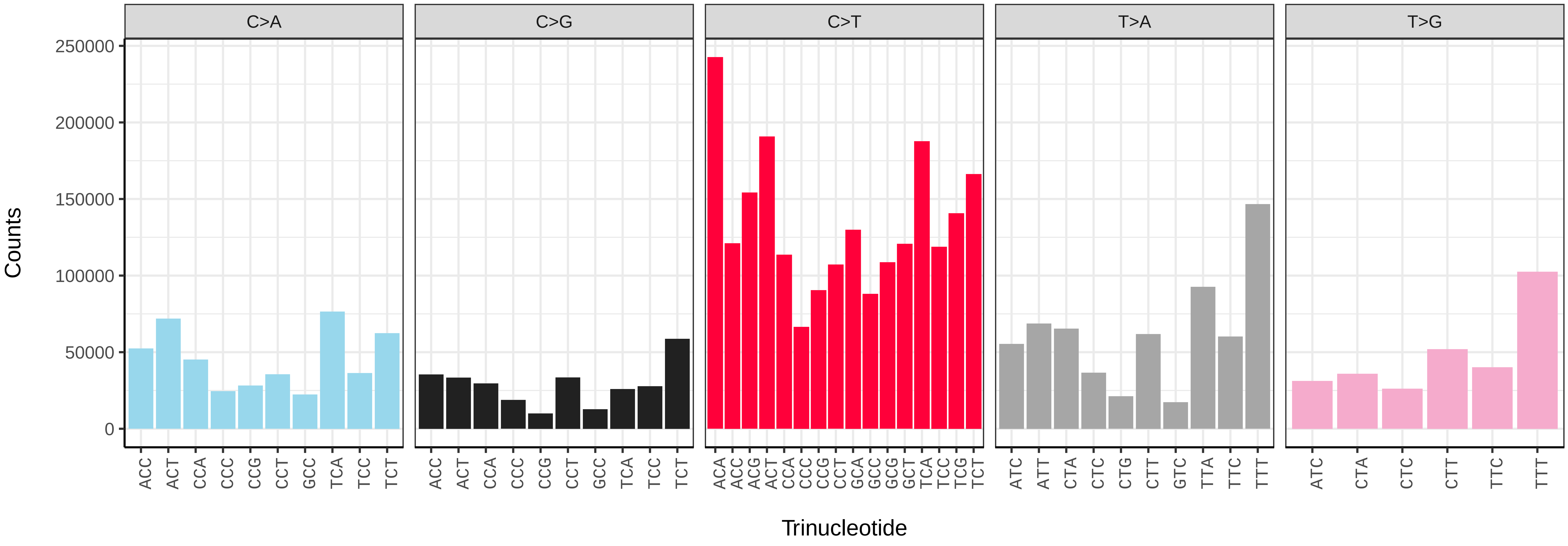


*Hipparchia semele* (ilHipSeme1) somatic mutational spectrum (SBS96 classification)

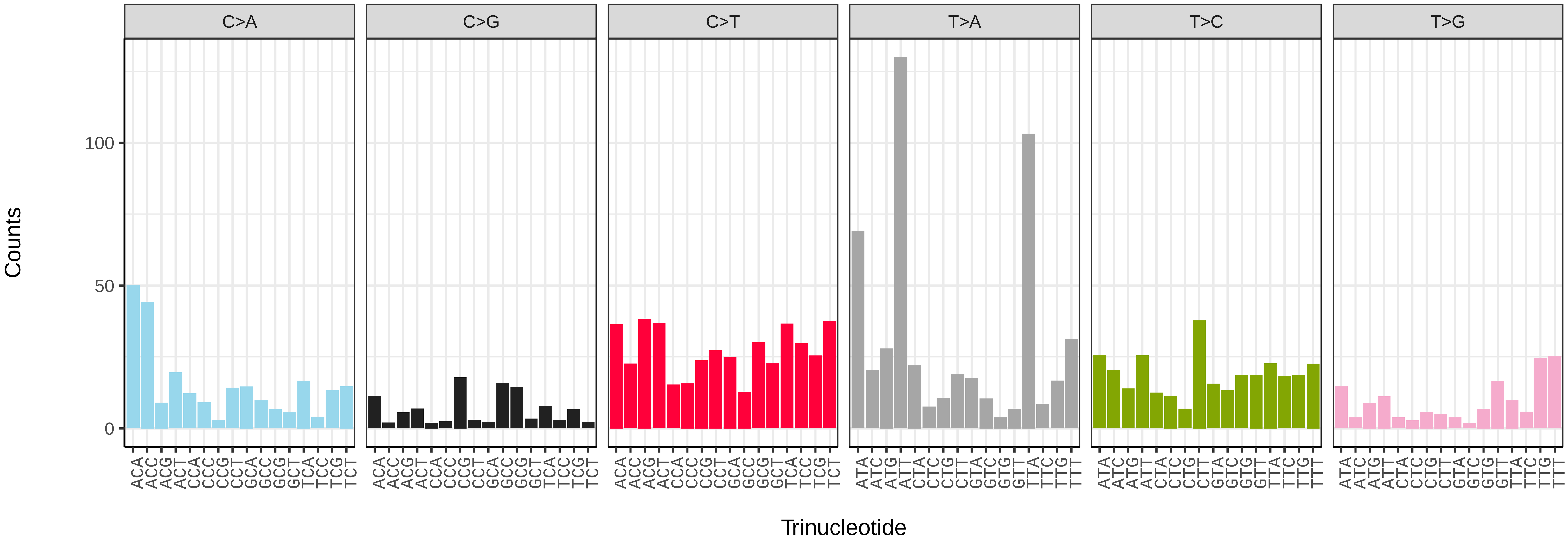


*Hipparchia semele* (ilHipSeme1) somatic mutational spectrum (SBS52 classification)

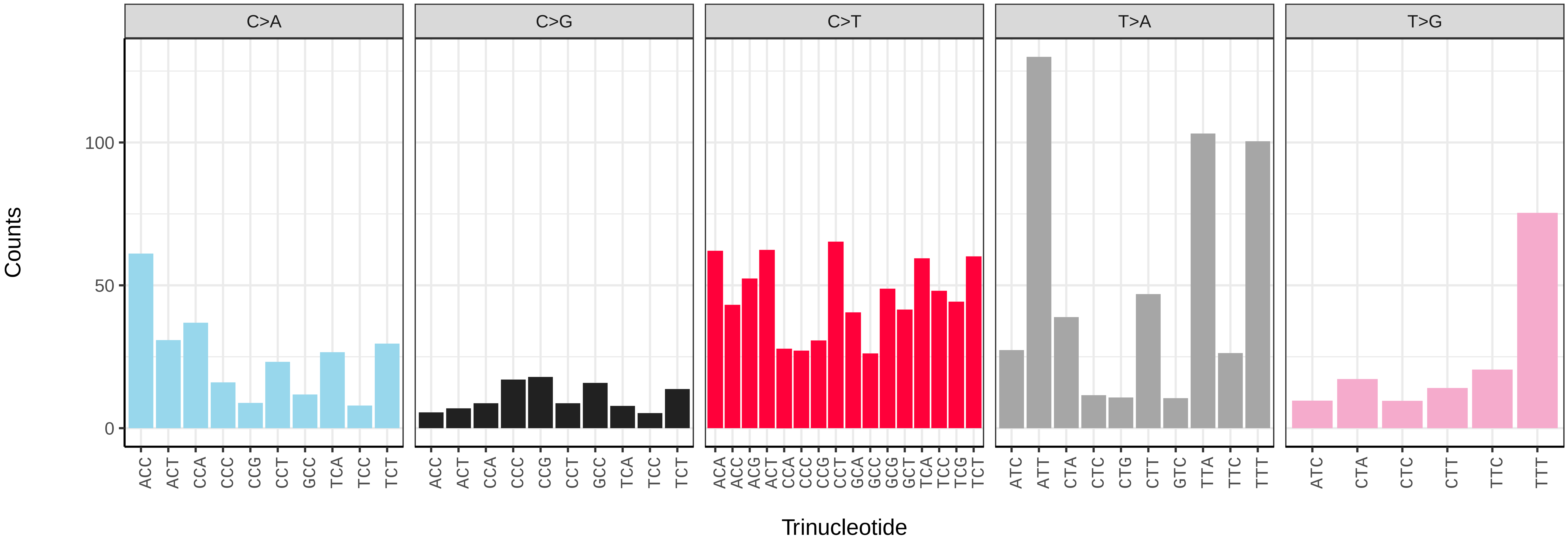

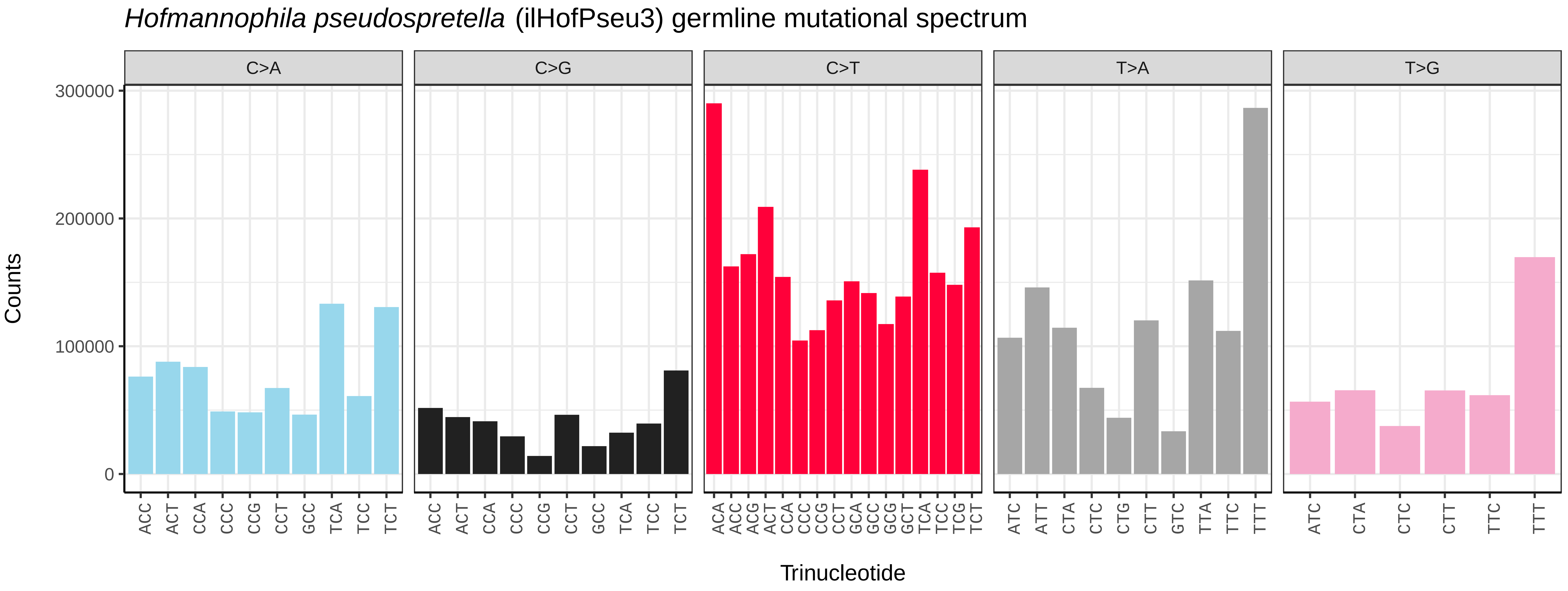
*Hofmannophila pseudospretella* (ilHofPseu3) germline mutational spectrum
C>A
C>G
C>T
T>A
T>G
Counts
0
100000
200000
300000
ACC ACT CCA CCC CCG CCT GCC TCA TCC TCT
ACC ACT CCA CCC CCG CCT GCC TCA TCC TCT
ACA ACC ACG ACT CCA CCC CCG CCT GCA GCC GCG GCT TCA TCC TCG TCT
ATC ATT CTA CTC CTG CTT GTC TTA TTC TTT
ATC CTA CTC CTT TTC TTT
Trinucleotide

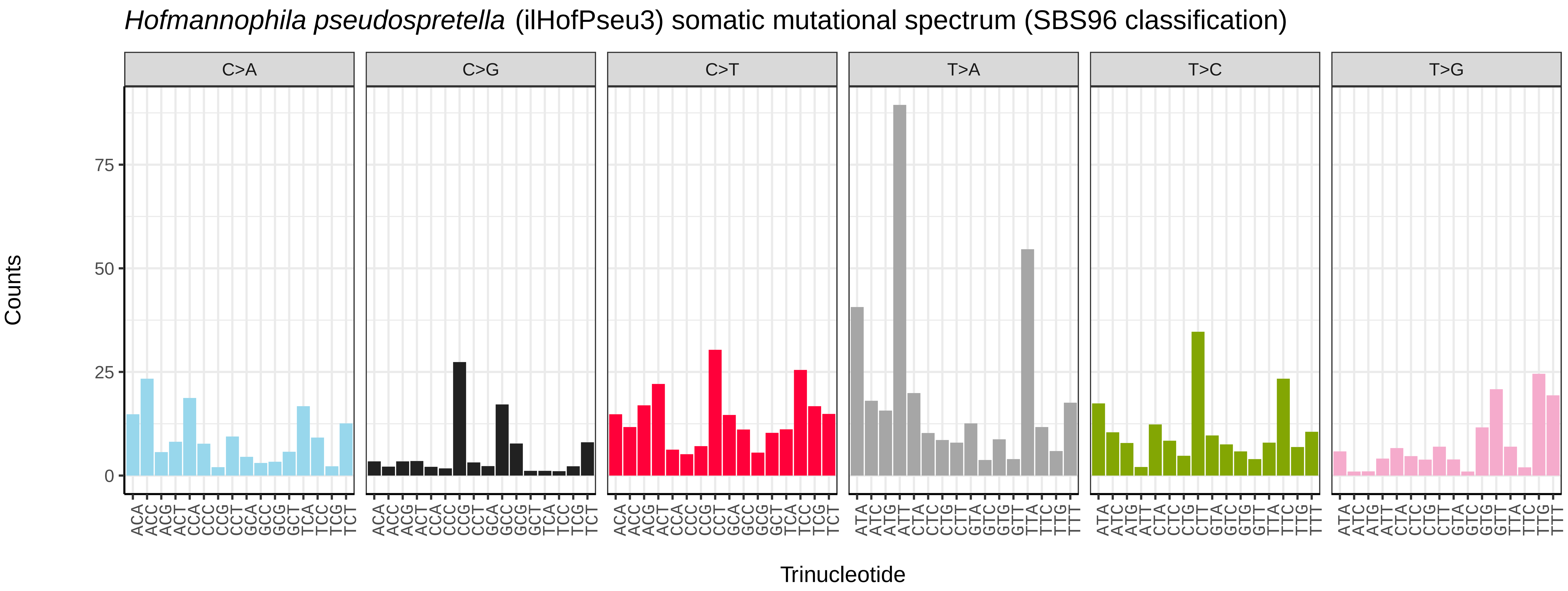
*Hofmannophila pseudospretella* (ilHofPseu3) somatic mutational spectrum (SBS96 classification)
C>A
C>G
C>T
T>A
T>C
T>G
Counts
0
25
50
75
ACA ACC ACG ACT CCA CCC CCG CCT GCA GCC GCG GCT TCA TCC TCG TCT
ACA ACC ACG ACT CCA CCC CCG CCT GCA GCC GCG GCT TCA TCC TCG TCT
ACA ACC ACG ACT CCA CCC CCG CCT GCA GCC GCG GCT TCA TCC TCG TCT
ATA ATC ATG ATT CTA CTC CTG CTT GTA GTC GTG GTT TTA TTC TTG TTT
ATA ATC ATG ATT CTA CTC CTG CTT GTA GTC GTG GTT TTA TTC TTG TTT
ATA ATC ATG ATT CTA CTC CTG CTT GTA GTC GTG GTT TTA TTC TTG TTT
Trinucleotide

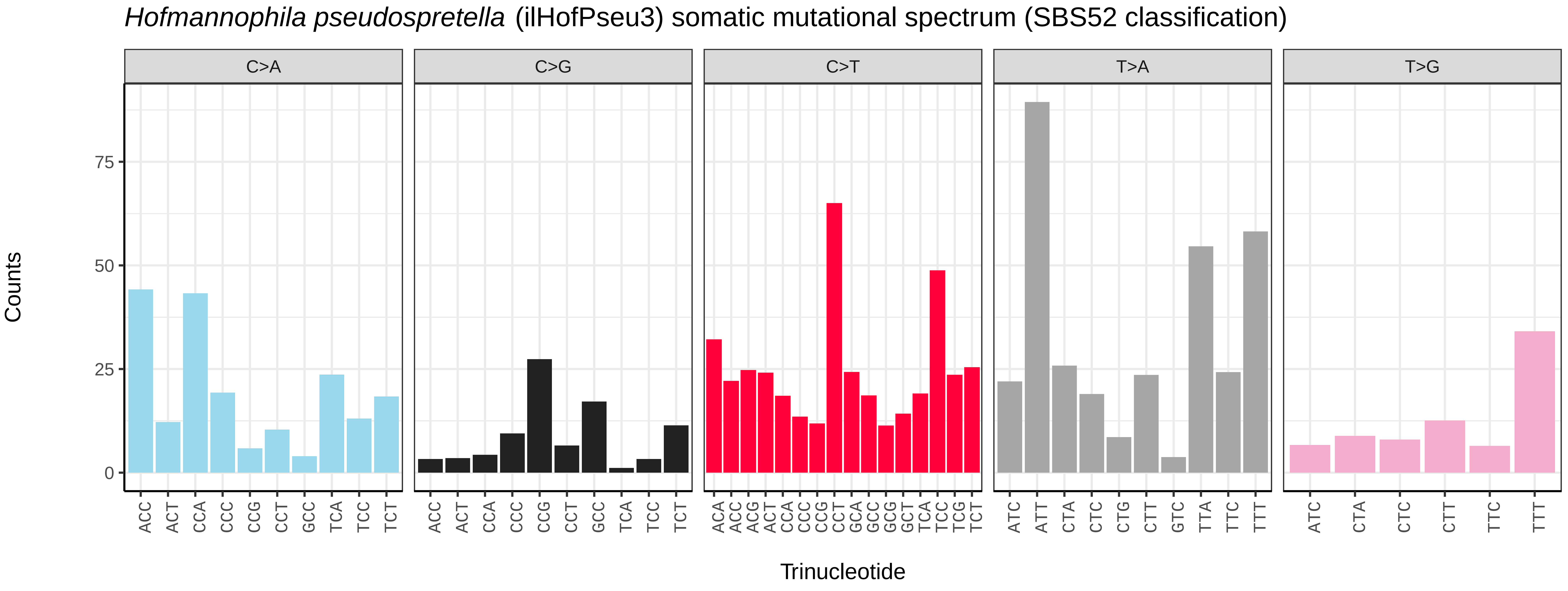
*Hofmannophila pseudospretella* (ilHofPseu3) somatic mutational spectrum (SBS52 classification)
C>A
C>G
C>T
T>A
T>G
Counts
0
25
50
75
ACC ACT CCA CCC CCG CCT GCC TCA TCC TCT
ACC ACT CCA CCC CCG CCT GCC TCA TCC TCT
ACA ACC ACG ACT CCA CCC CCG CCT GCA GCC GCG GCT TCA TCC TCG TCT
ATC ATT CTA CTC CTG CTT GTC TTA TTC TTT
ATC CTA CTC CTT TTC TTT
Trinucleotide

*Hydriomena furcata* (ilHydFurc1) germline mutational spectrum

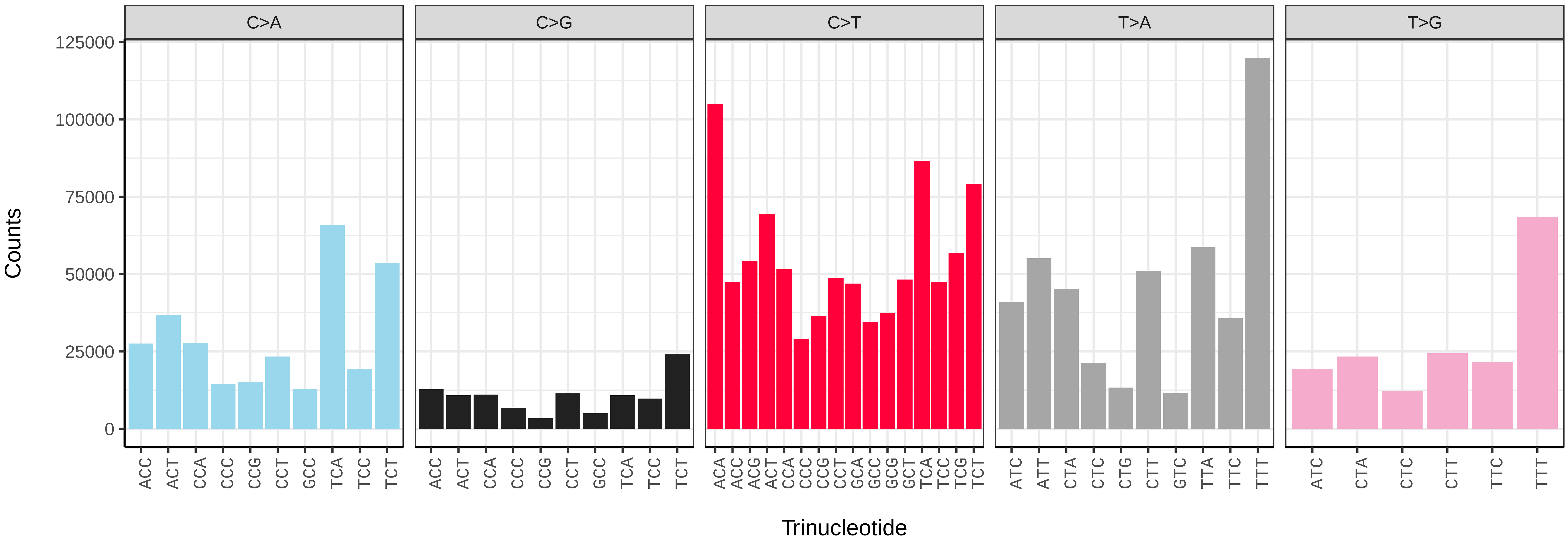


*Hydriomena furcata* (ilHydFurc1) somatic mutational spectrum (SBS96 classification)

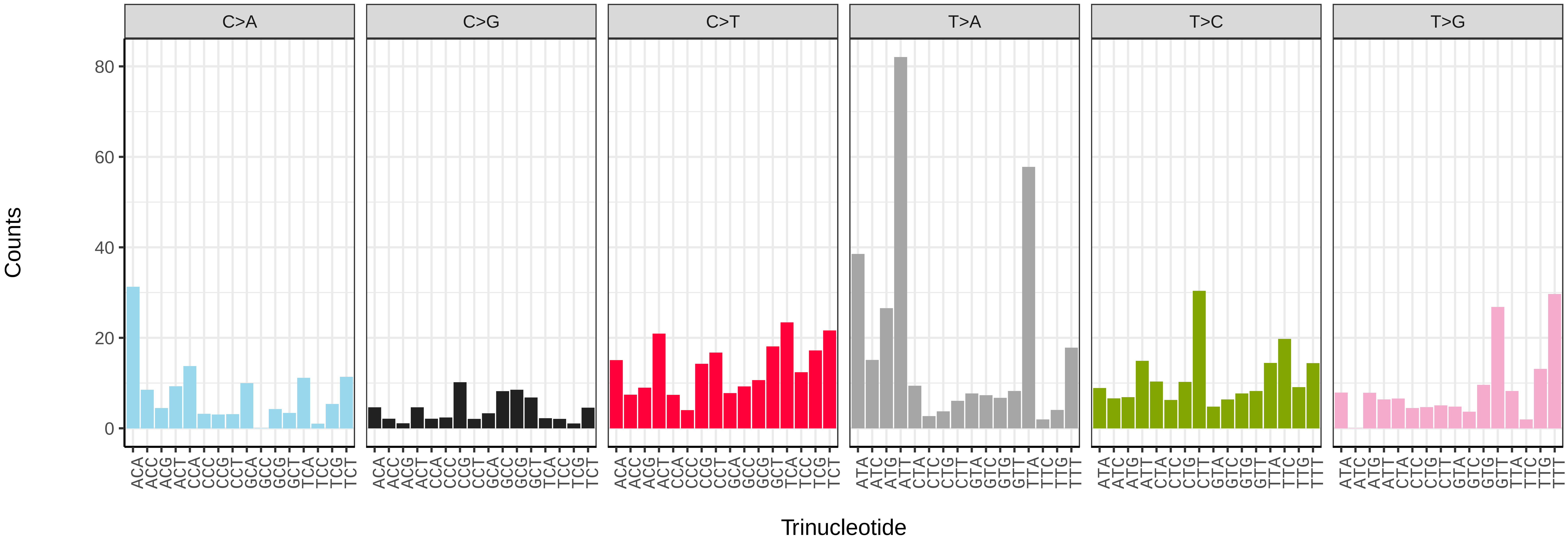


*Hydriomena furcata* (ilHydFurc1) somatic mutational spectrum (SBS52 classification)

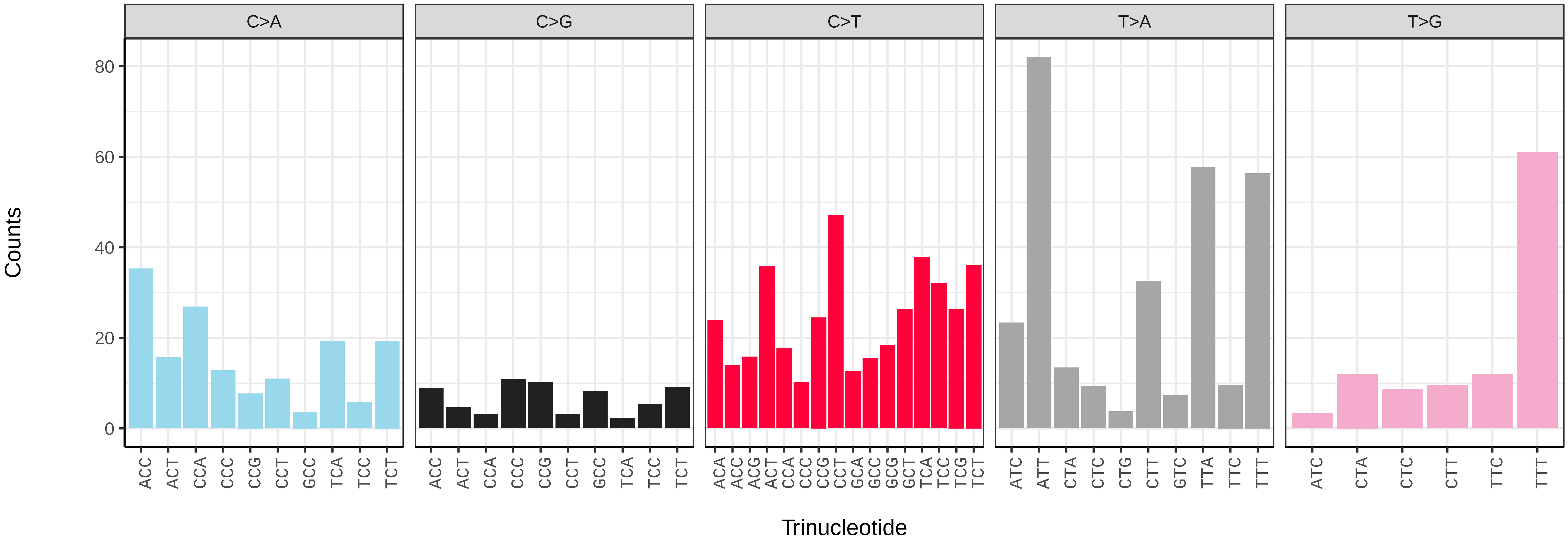

*Hydraecia micacea* (ilHydMica1) germline mutational spectrum

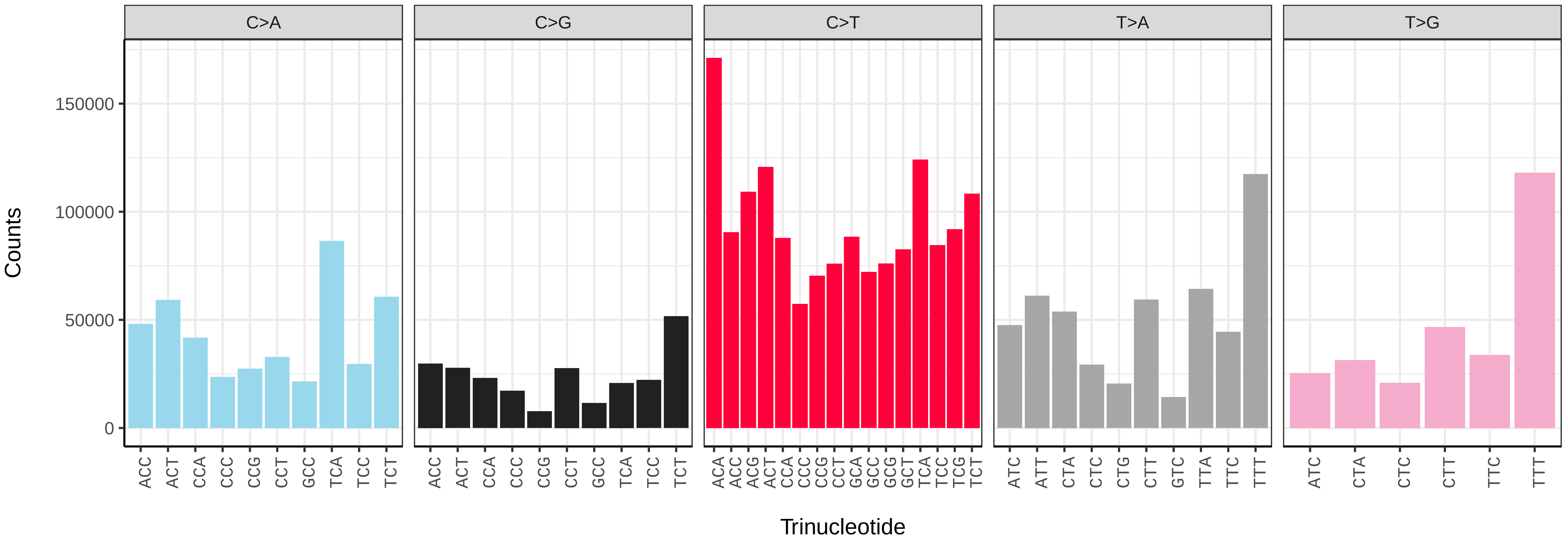


*Hydraecia micacea* (ilHydMica1) somatic mutational spectrum (SBS96 classification)

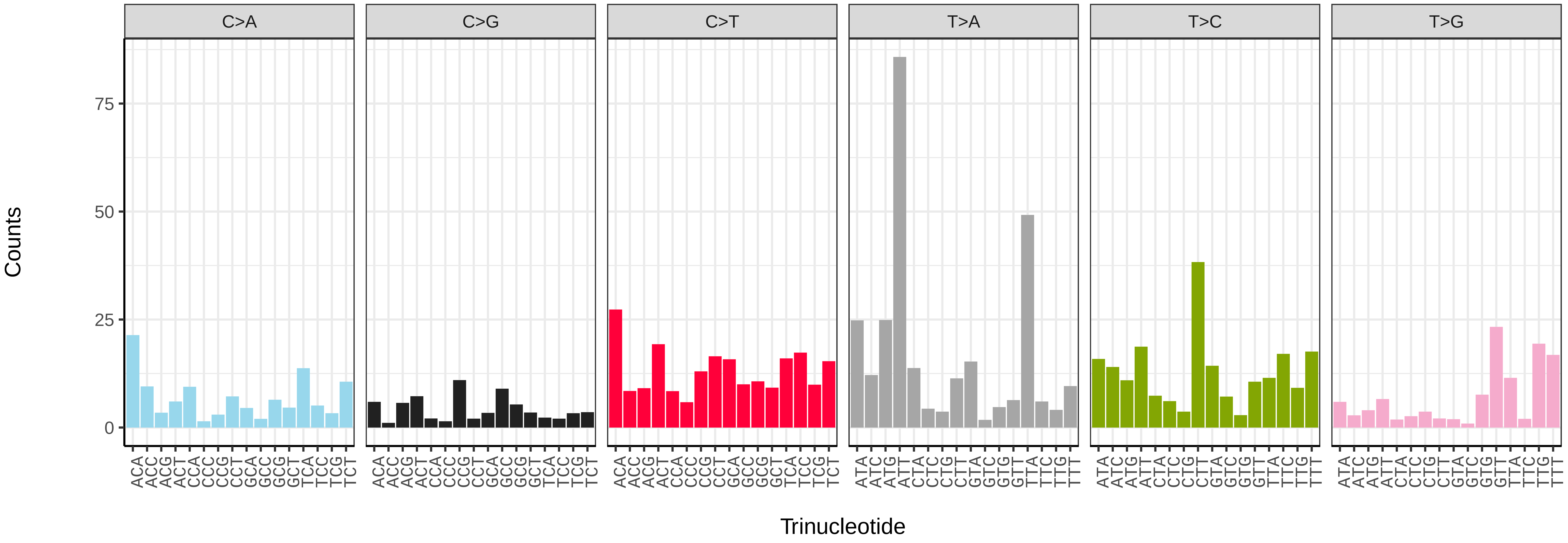


*Hydraecia micacea* (ilHydMica1) somatic mutational spectrum (SBS52 classification)

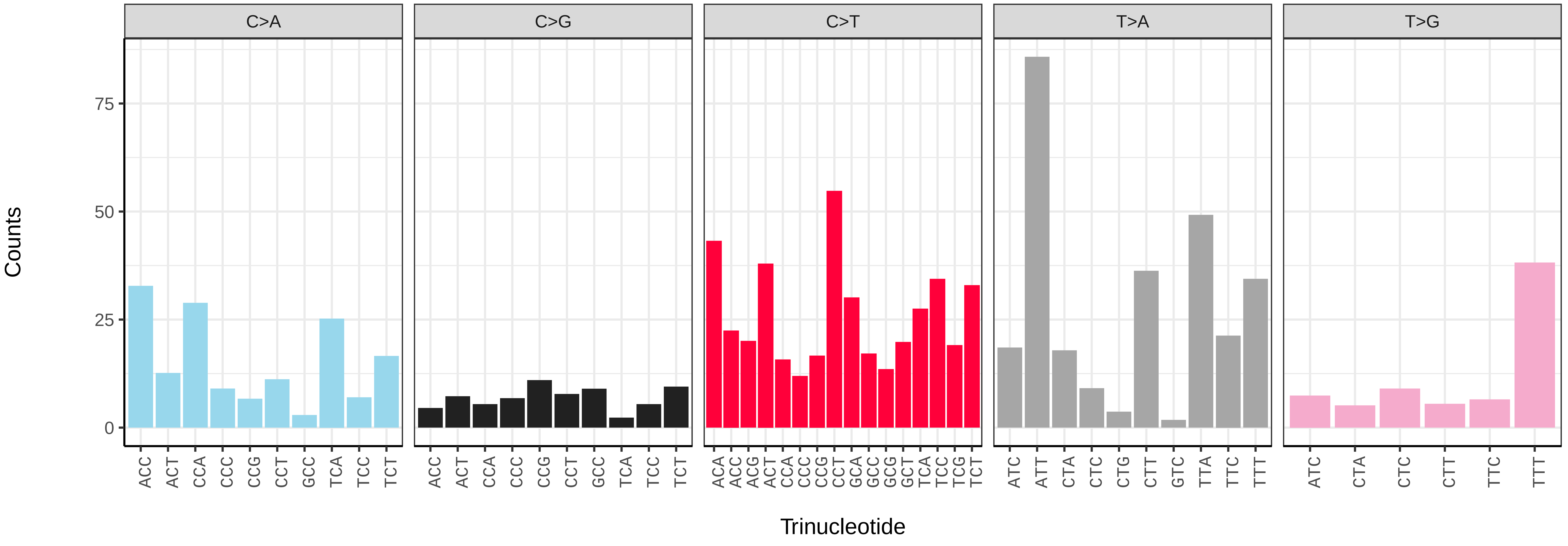

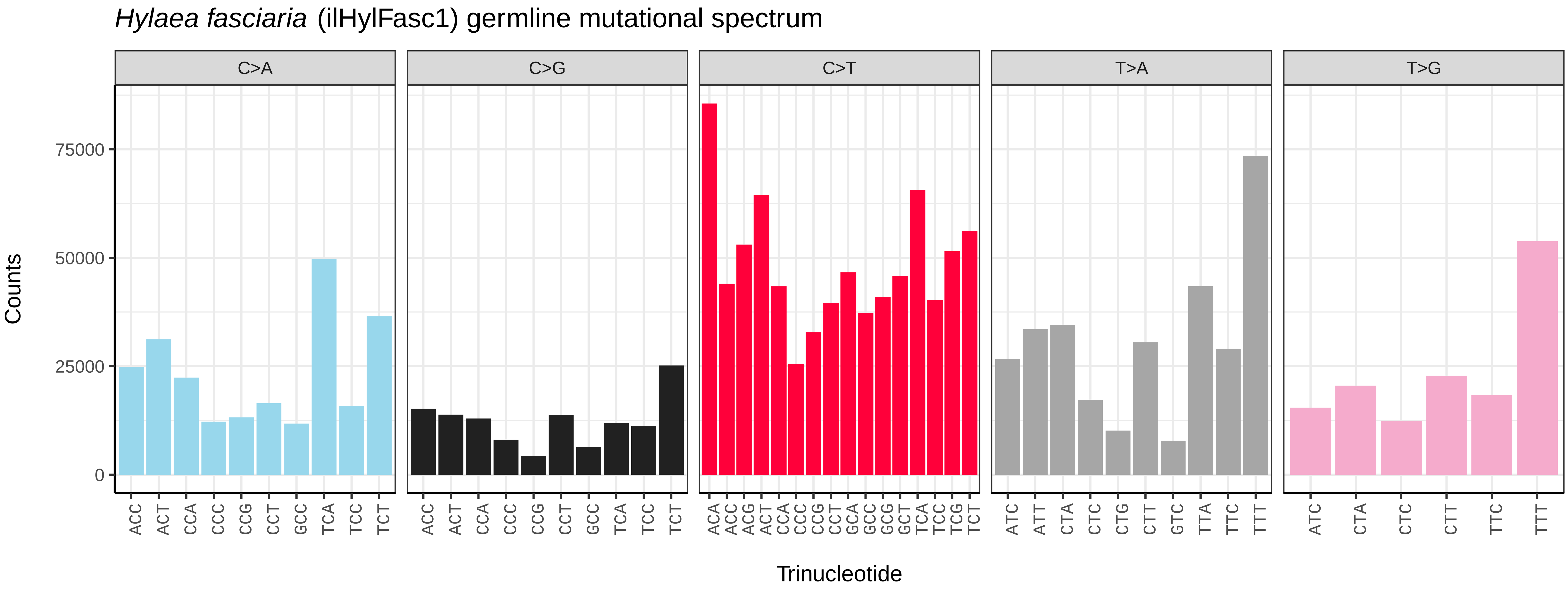
*Hylaea fasciaria* (ilHylFasc1) germline mutational spectrum
C>A
C>G
C>T
T>A
T>G
Counts
0
25000
50000
75000
Trinucleotide

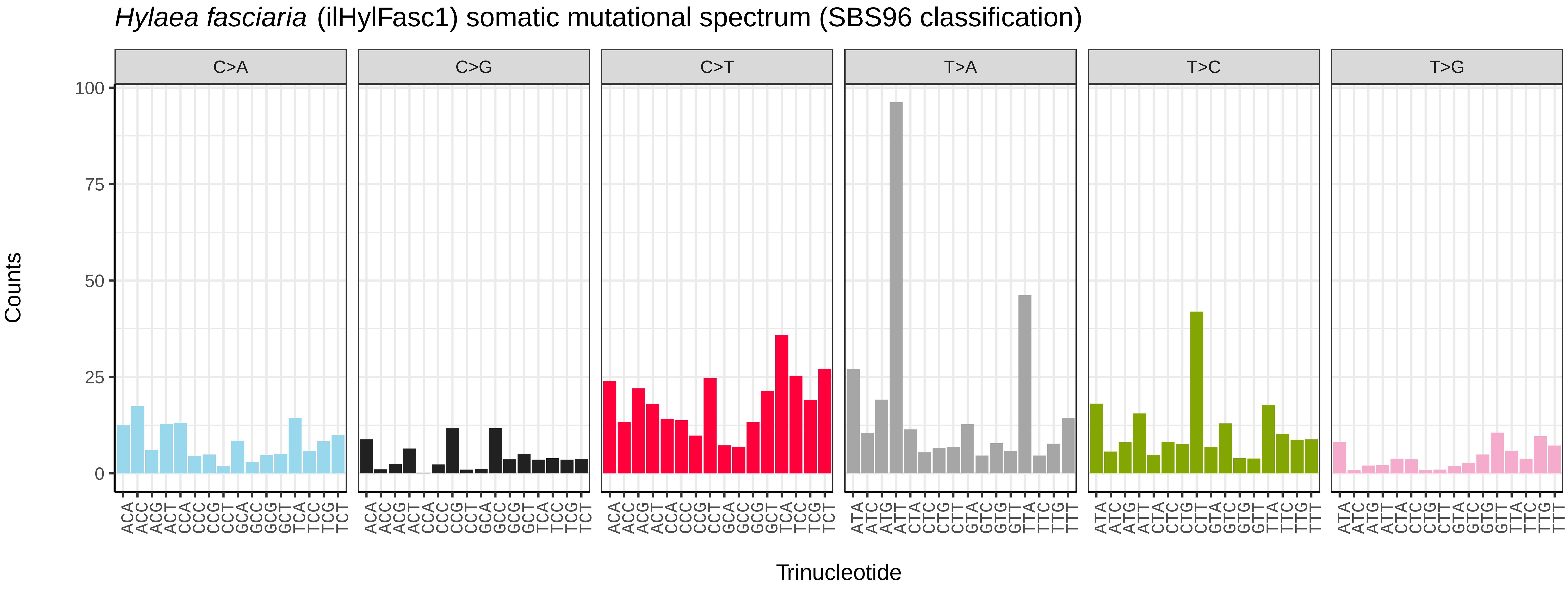
*Hylaea fasciaria* (ilHylFasc1) somatic mutational spectrum (SBS96 classification)
C>A
C>G
C>T
T>A
T>C
T>G
Counts
0
25
50
75
100
Trinucleotide

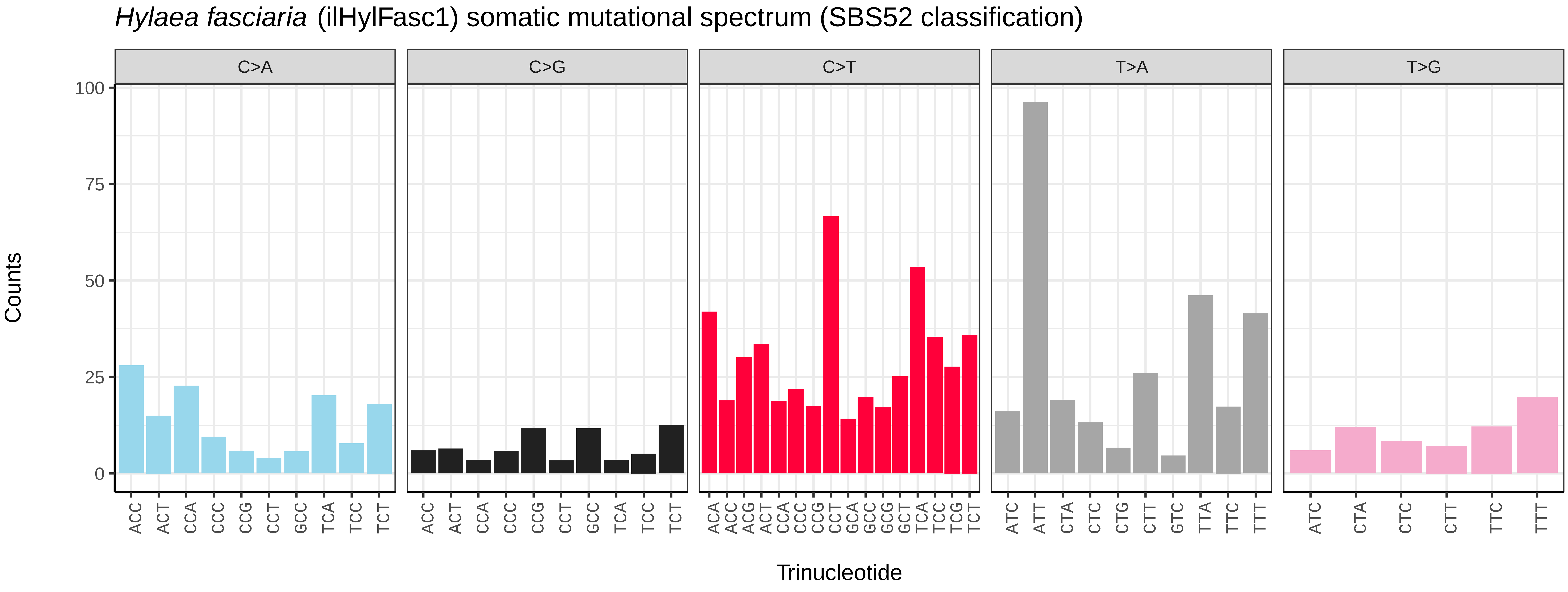
*Hylaea fasciaria* (ilHylFasc1) somatic mutational spectrum (SBS52 classification)
C>A
C>G
C>T
T>A
T>G
Counts
0
25
50
75
100
Trinucleotide

*Hypsopygia costalis* (ilHypCost1) germline mutational spectrum

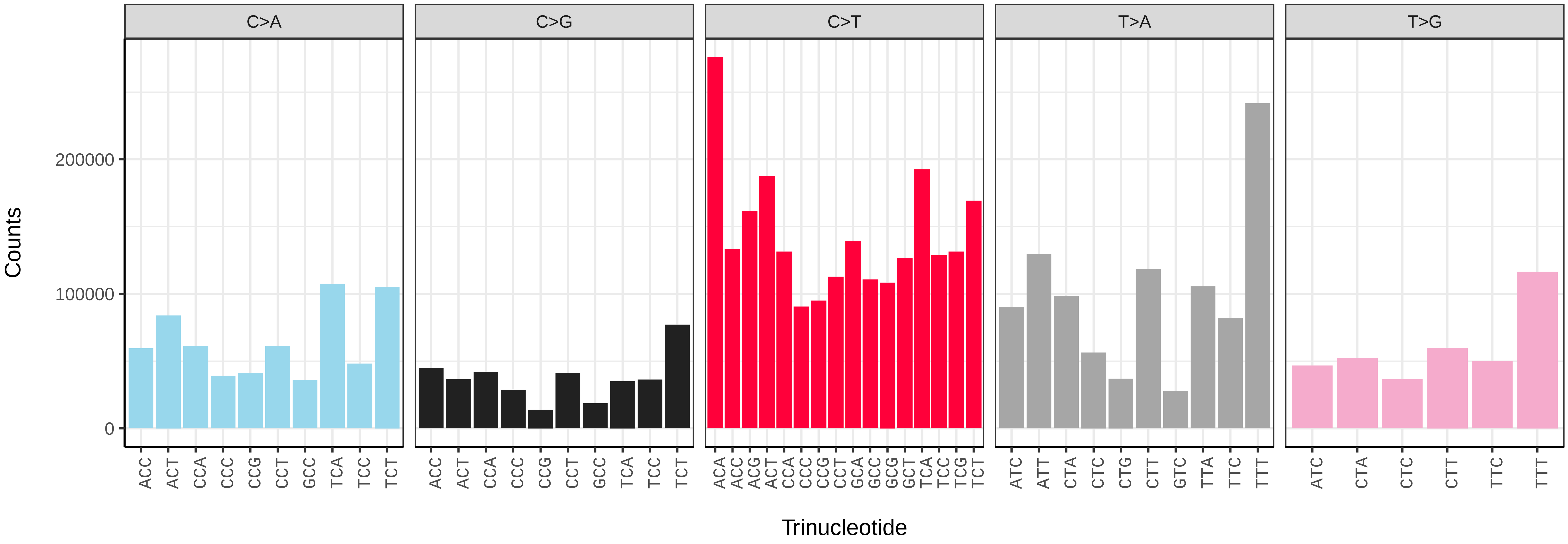


*Hypsopygia costalis* (ilHypCost1) somatic mutational spectrum (SBS96 classification)

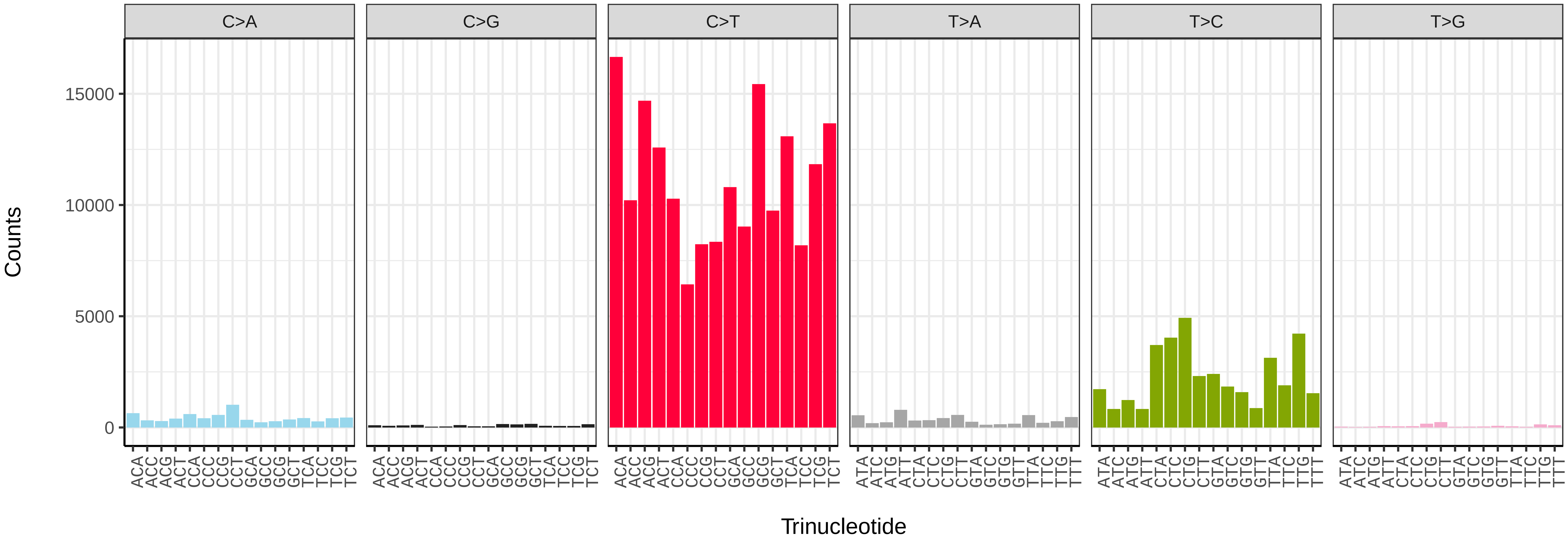


*Hypsopygia costalis* (ilHypCost1) somatic mutational spectrum (SBS52 classification)

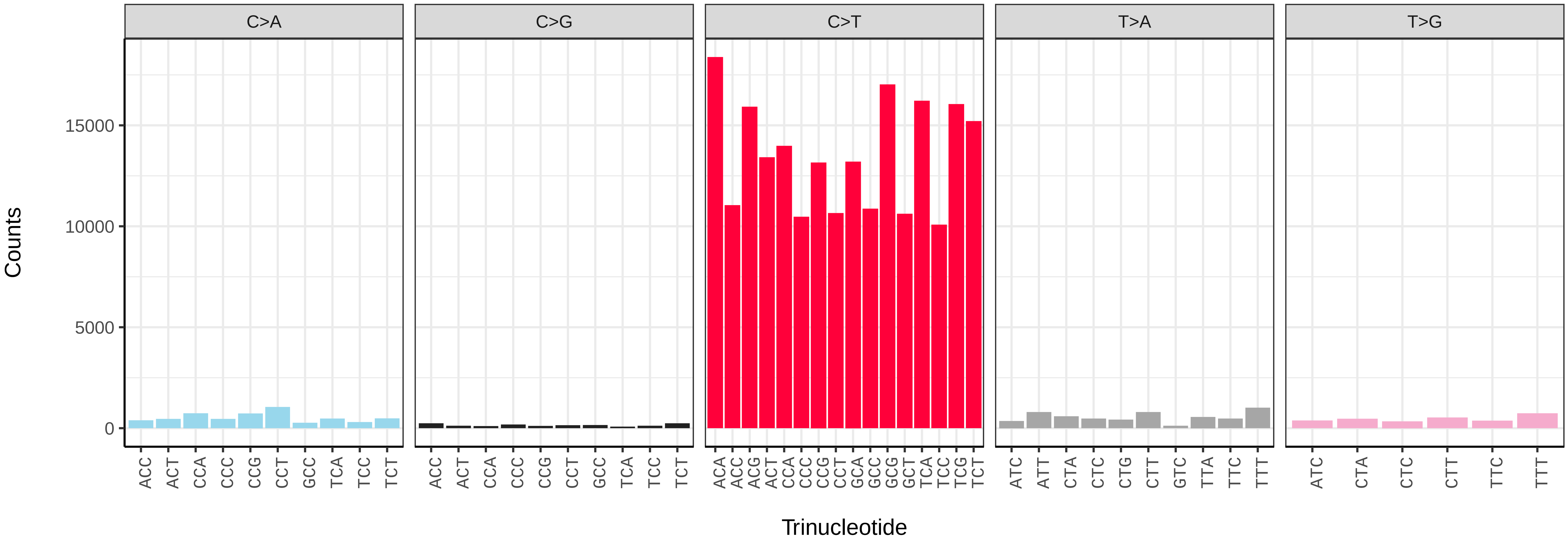

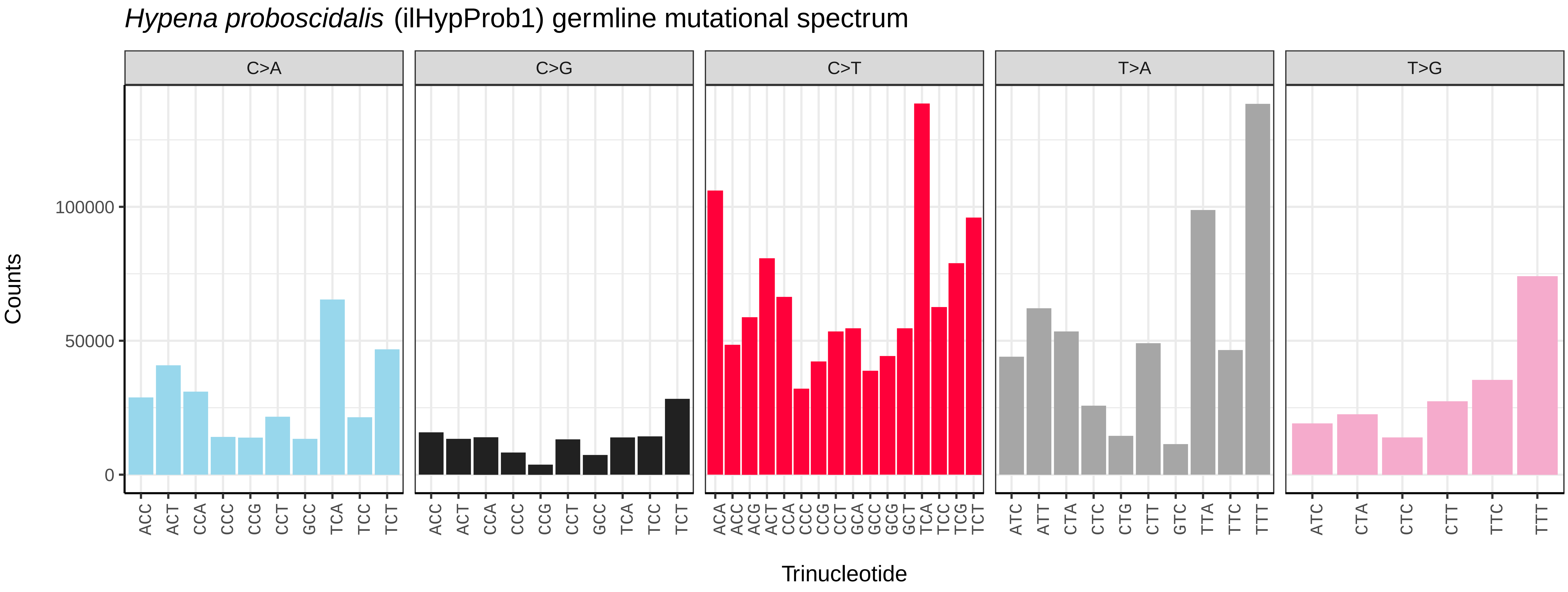
*Hypena proboscidalis* (ilHypProb1) germline mutational spectrum
C>A
C>G
C>T
T>A
T>G
Counts
100000
50000
0
ACC ACT CCA CCC CCG CCT GCC TCA TCC TCT
ACC ACT CCA CCC CCG CCT GCC TCA TCC TCT
ACA ACC ACG ACT CCA CCC CCG CCT GCA GCC GCG GCT TCA TCC TCG TCT
ATC ATT CTA CTC CTG CTT GTC TTA TTC TTT
ATC CTA CTC CTT TTC TTT
Trinucleotide

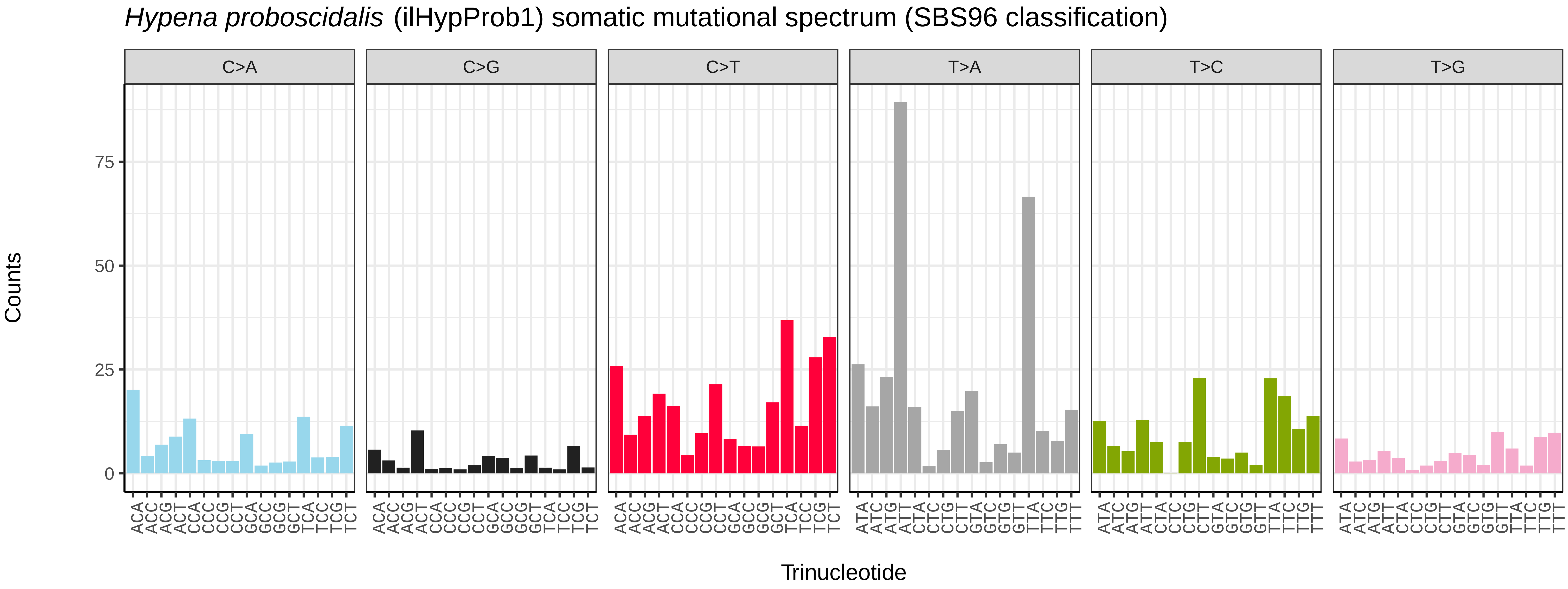
*Hypena proboscidalis* (ilHypProb1) somatic mutational spectrum (SBS96 classification)
C>A
C>G
C>T
T>A
T>C
T>G
Counts
75
50
25
0
ACA ACC ACG ACT CCA CCC CCG CCT GCA GCC GCG GCT TCA TCC TCG TCT
ATA ATC ATG ATT CTA CTC CTG CTT GTA GTC GTG GTT TTA TTC TTG TTT
Trinucleotide

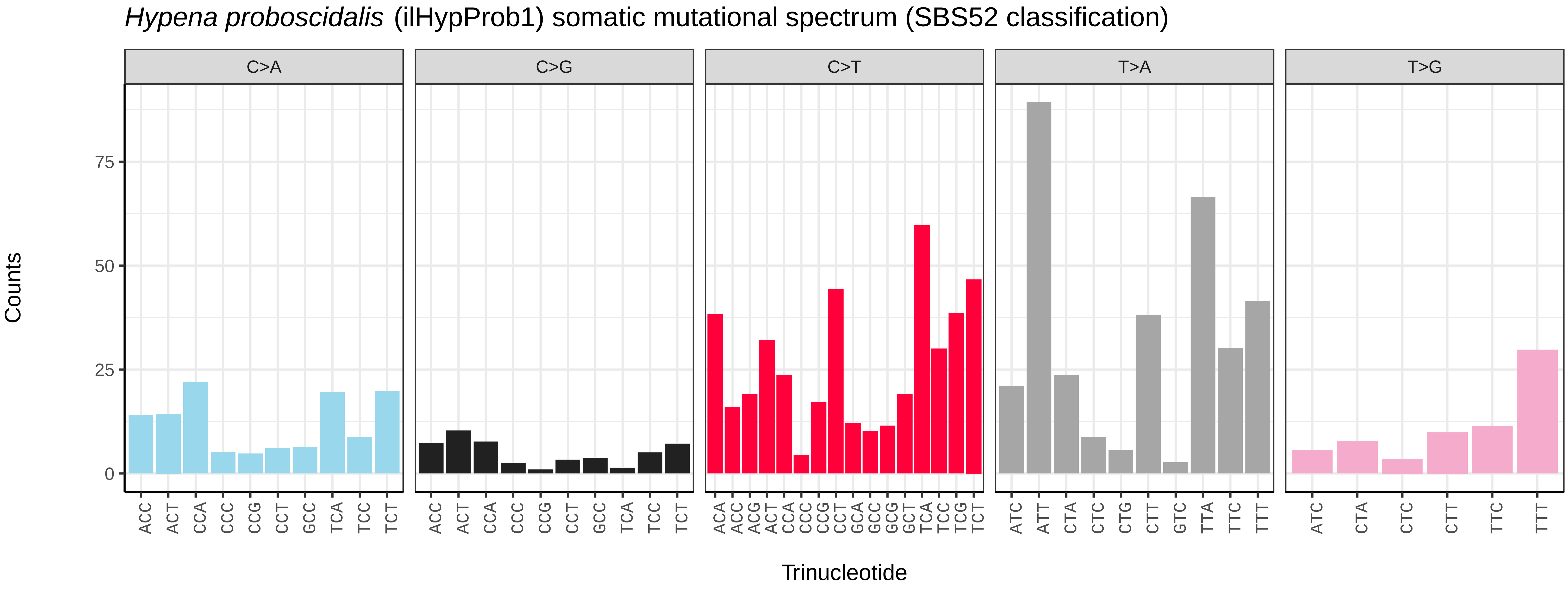
*Hypena proboscidalis* (ilHypProb1) somatic mutational spectrum (SBS52 classification)
C>A
C>G
C>T
T>A
T>G
Counts
75
50
25
0
ACC ACT CCA CCC CCG CCT GCC TCA TCC TCT
ACC ACT CCA CCC CCG CCT GCC TCA TCC TCT
ACA ACC ACG ACT CCA CCC CCG CCT GCA GCC GCG GCT TCA TCC TCG TCT
ATC ATT CTA CTC CTG CTT GTC TTA TTC TTT
ATC CTA CTC CTT TTC TTT
Trinucleotide

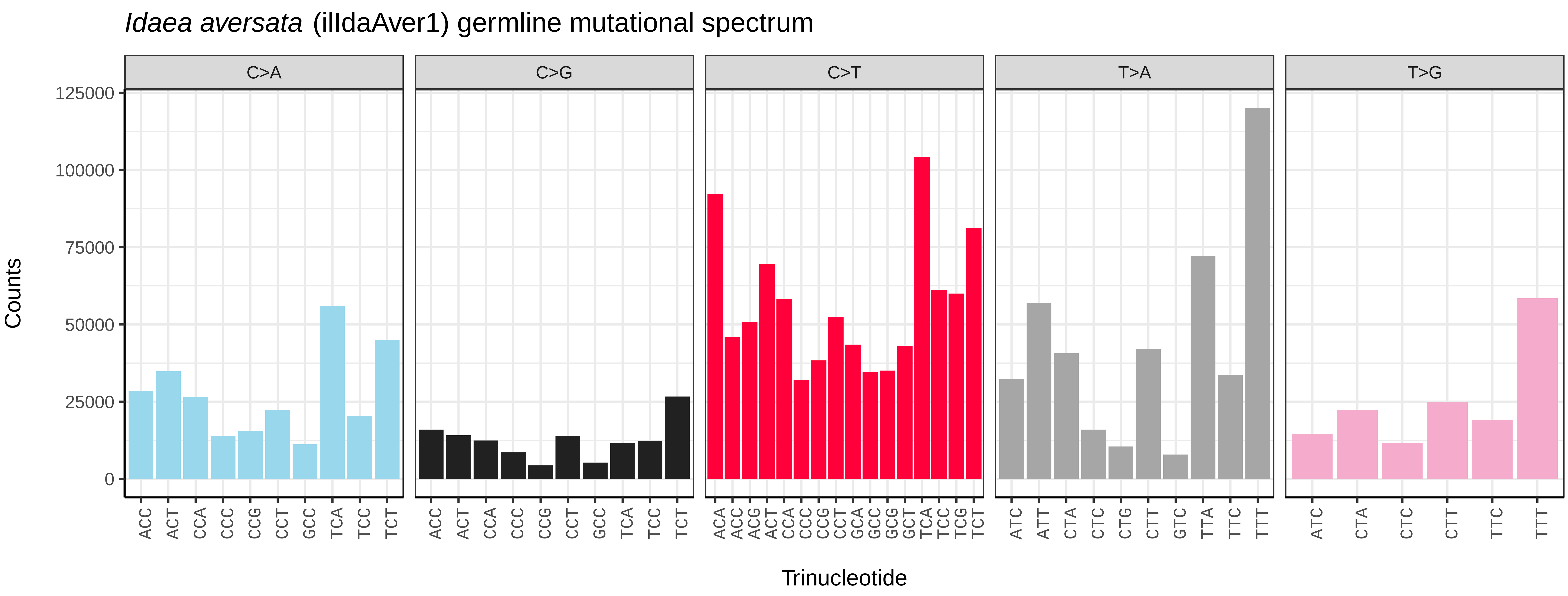
*Idaea aversata* (ilIdaAver1) germline mutational spectrum
C>A
C>G
C>T
T>A
T>G
Counts
125000
100000
75000
50000
25000
0
Trinucleotide

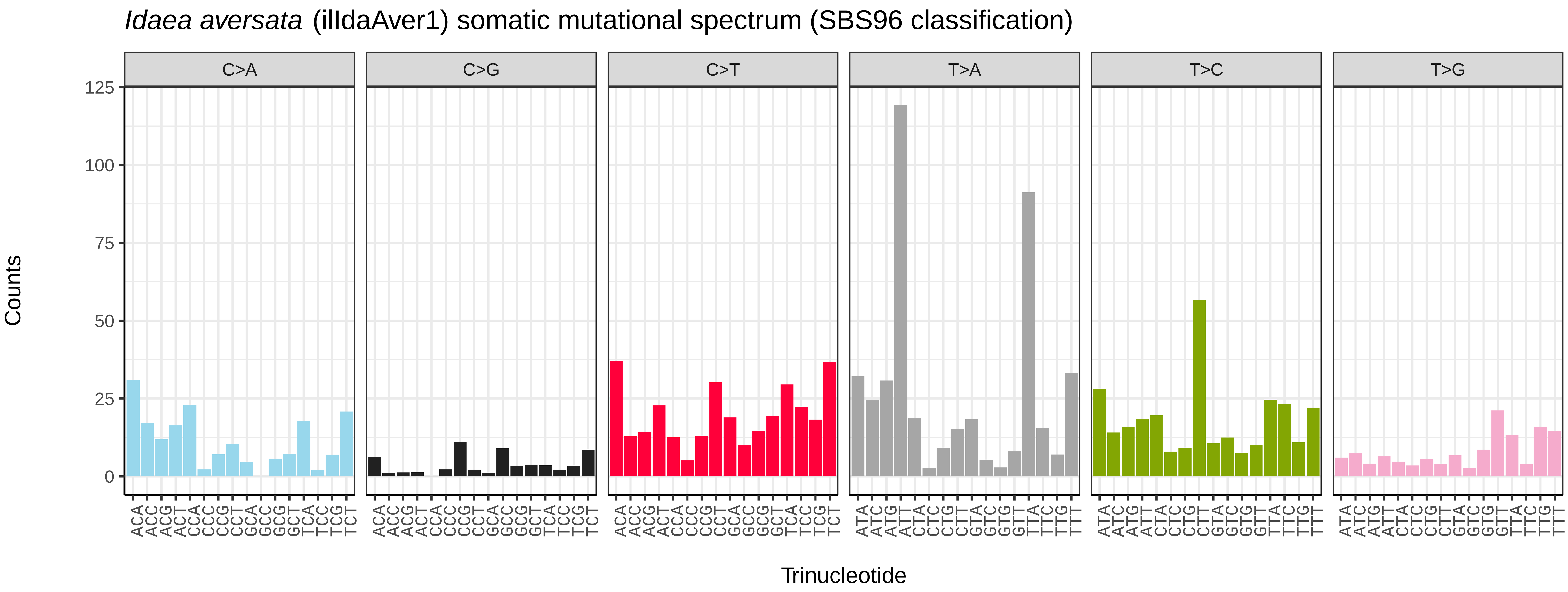
*Idaea aversata* (ilIdaAver1) somatic mutational spectrum (SBS96 classification)
C>A
C>G
C>T
T>A
T>C
T>G
Counts
125
100
75
50
25
0
Trinucleotide

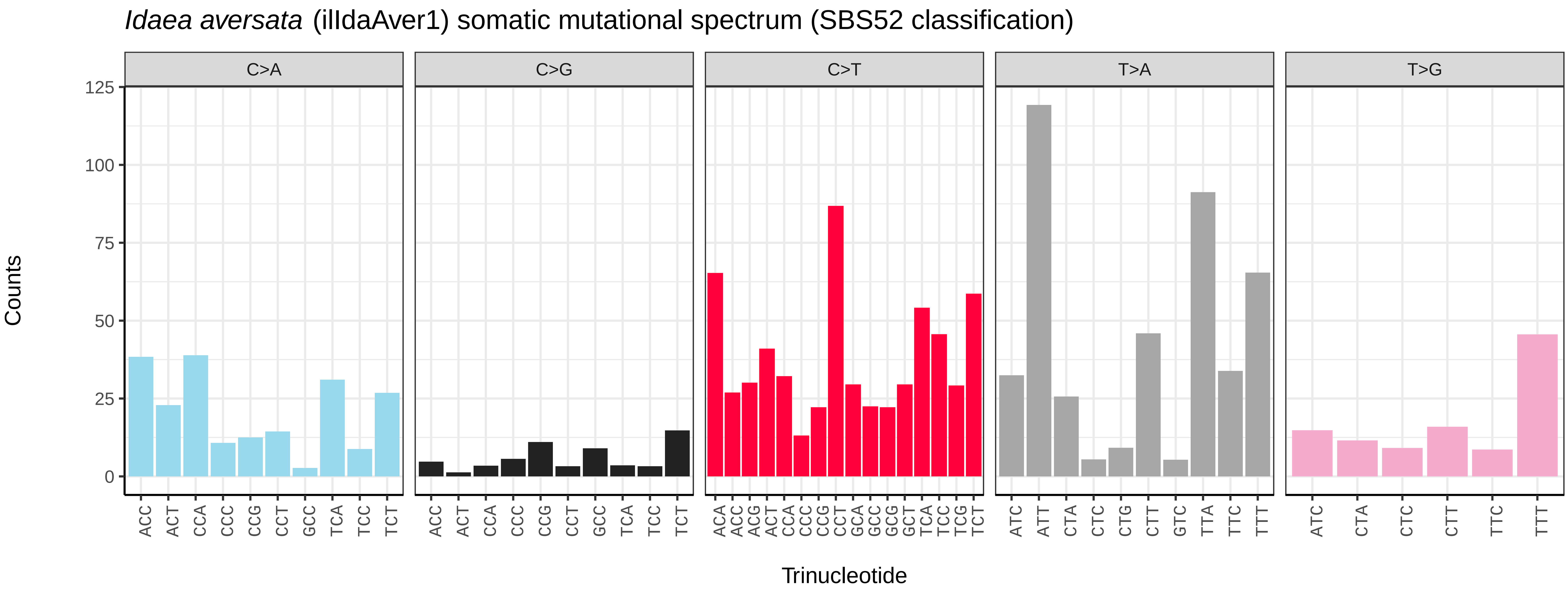
*Idaea aversata* (ilIdaAver1) somatic mutational spectrum (SBS52 classification)
C>A
C>G
C>T
T>A
T>G
Counts
125
100
75
50
25
0
Trinucleotide

*Incurvaria masculella* (ilIncMasc1) germline mutational spectrum

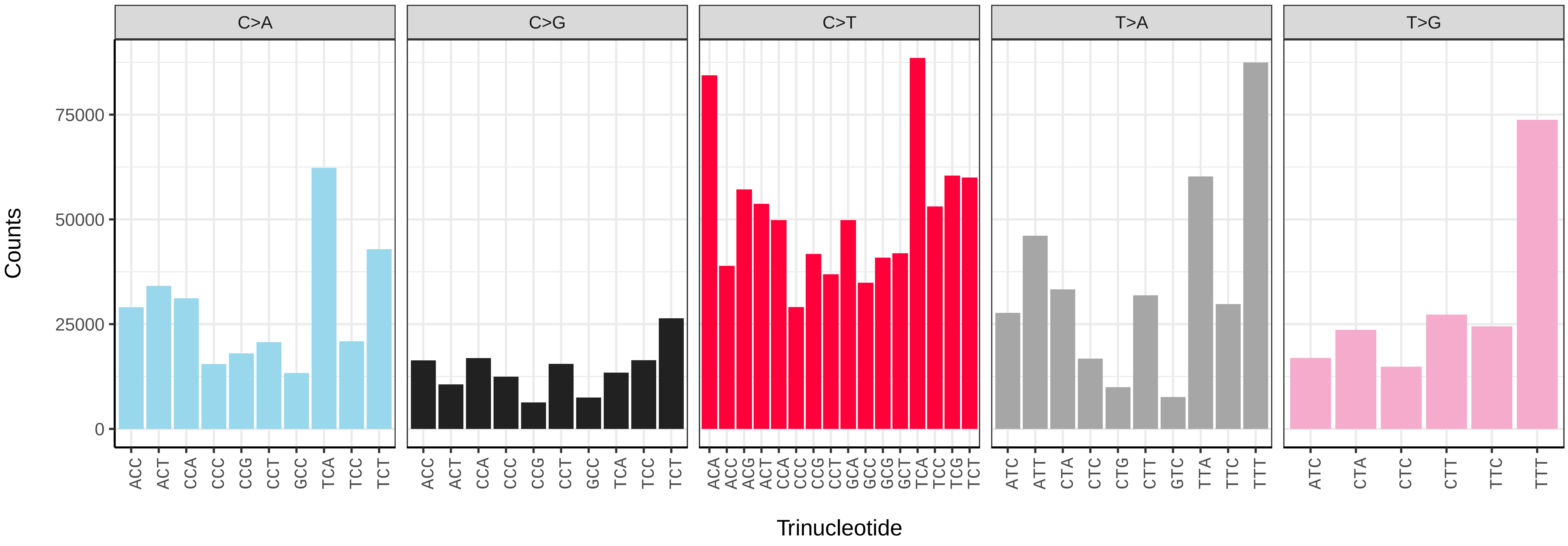


*Incurvaria masculella* (ilIncMasc1) somatic mutational spectrum (SBS96 classification)

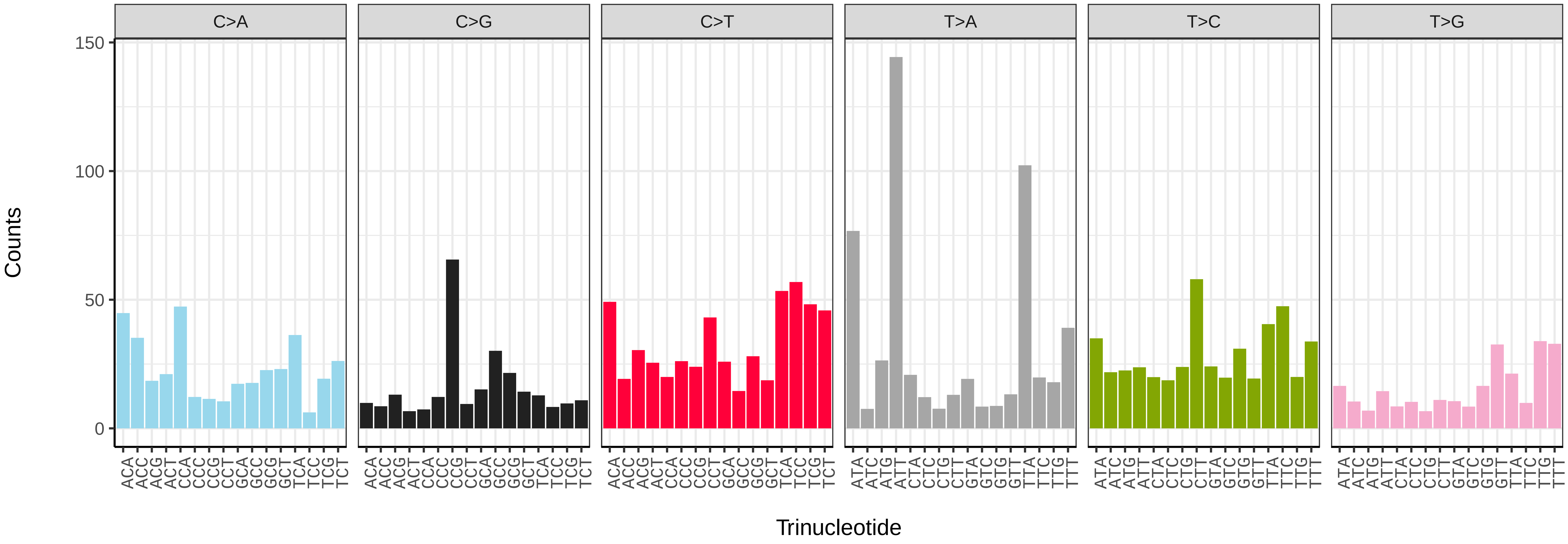


*Incurvaria masculella* (ilIncMasc1) somatic mutational spectrum (SBS52 classification)

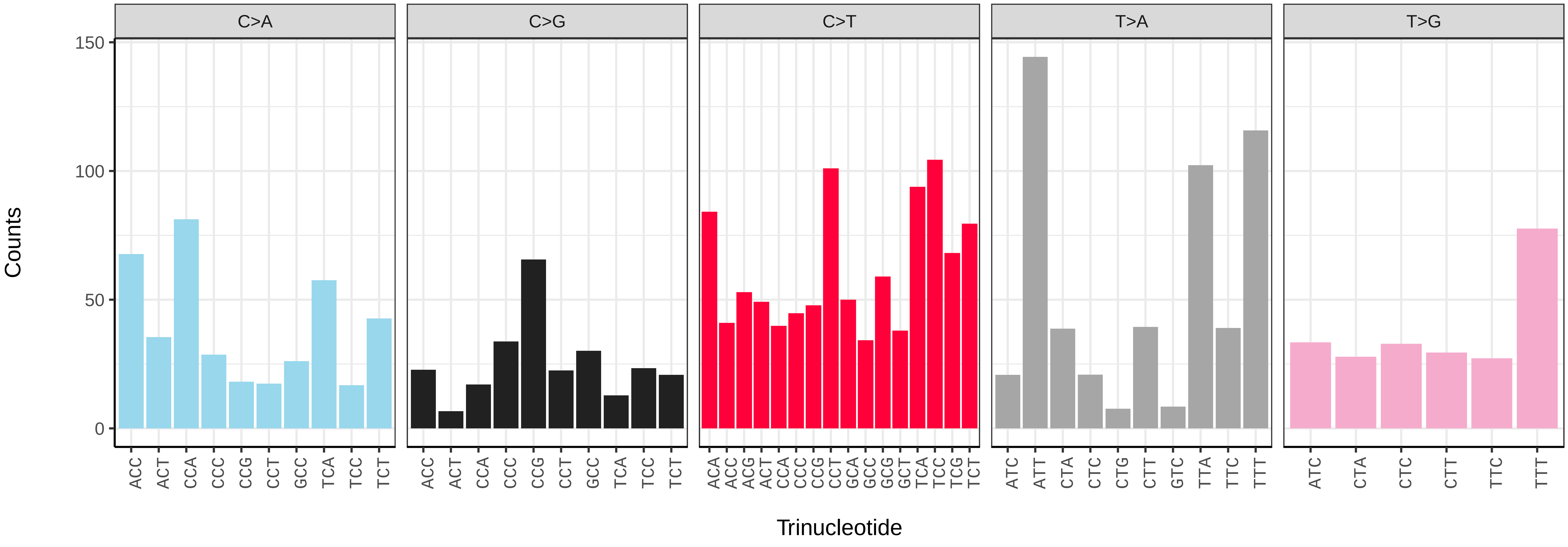

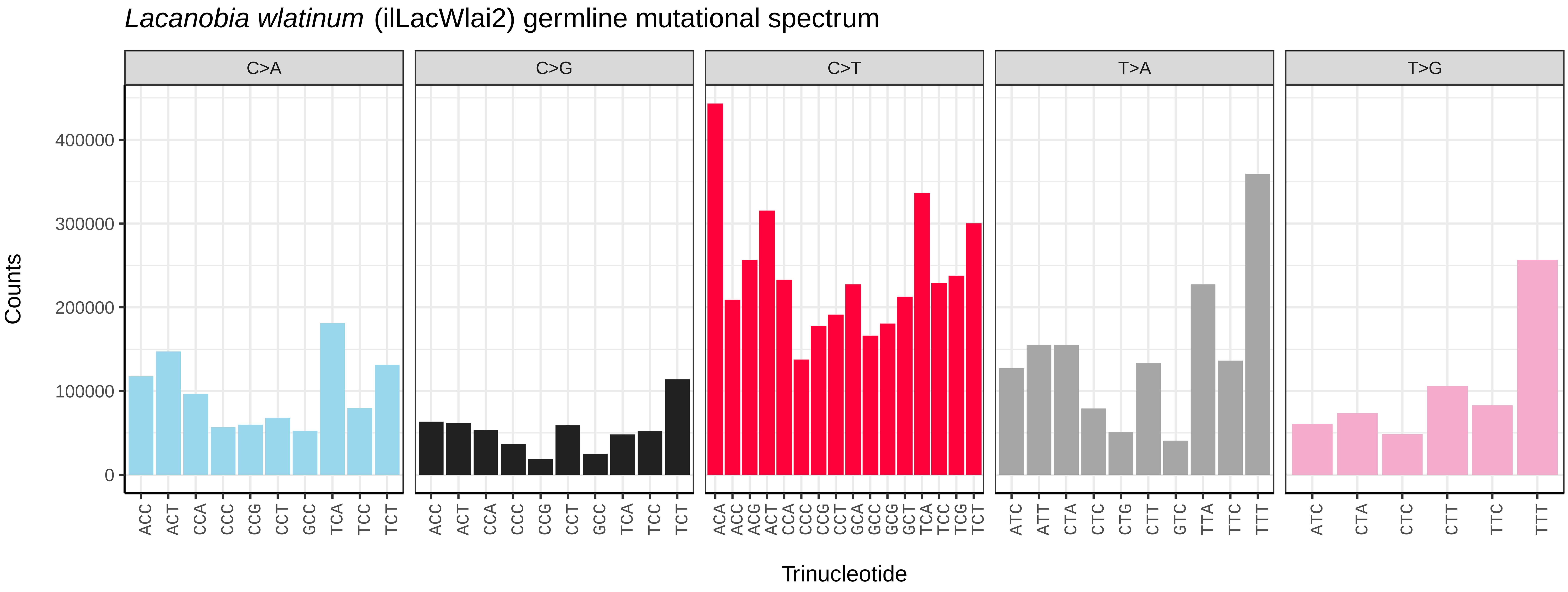
*Lacanobia wlatinum* (ilLacWlai2) germline mutational spectrum
C>A
C>G
C>T
T>A
T>G
Counts
400000
300000
200000
100000
0
ACC ACT CCA CCC CCG CCT GCC TCA TCC TCT
ACC ACT CCA CCC CCG CCT GCC TCA TCC TCT
ACA ACC ACG ACT CCA CCC CCG CCT GCA GCC GCG GCT TCA TCC TCG TCT
ATC ATT CTA CTC CTG CTT GTC TTA TTC TTT
ATC CTA CTC CTT TTC TTT
Trinucleotide

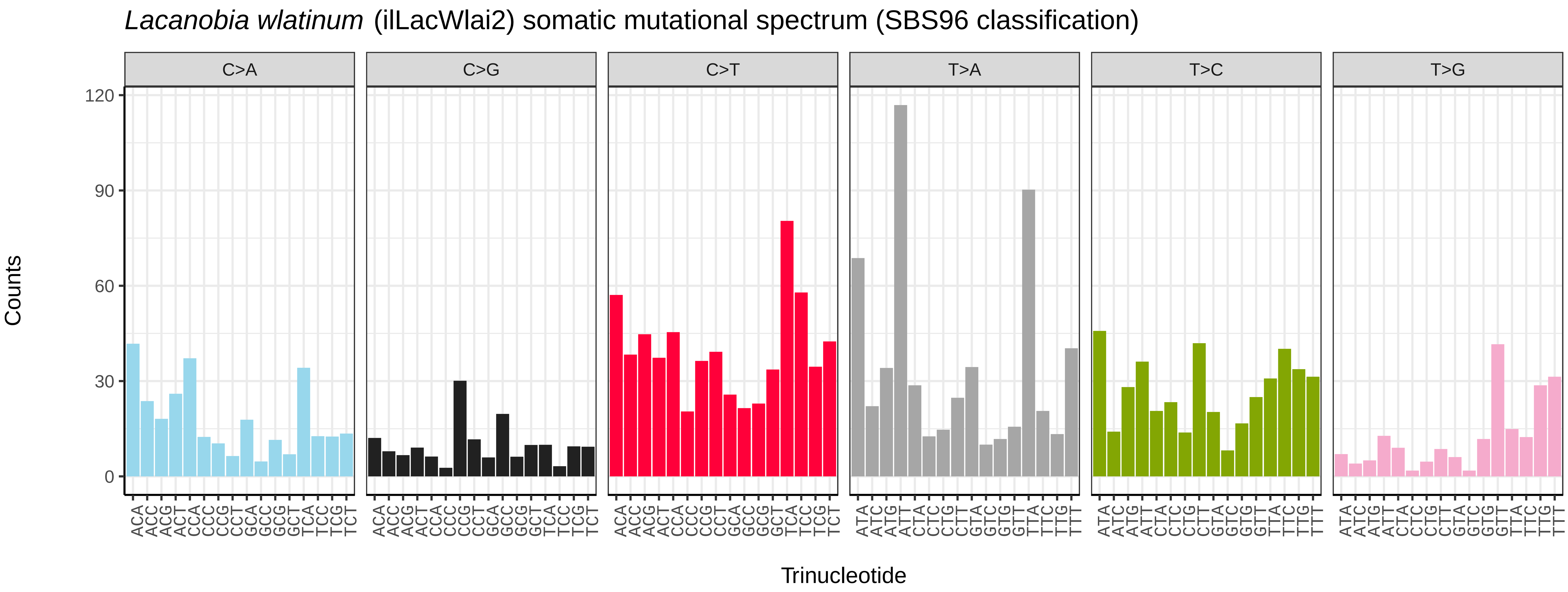
*Lacanobia wlatinum* (ilLacWlai2) somatic mutational spectrum (SBS96 classification)
C>A
C>G
C>T
T>A
T>C
T>G
Counts
120
90
60
30
0
ACA ACC ACG ACT CCA CCC CCG CCT GCA GCC GCG GCT TCA TCC TCG TCT
ATA ATC ATG ATT CTA CTC CTG CTT GTA GTC GTG GTT TTA TTC TTG TTT
Trinucleotide

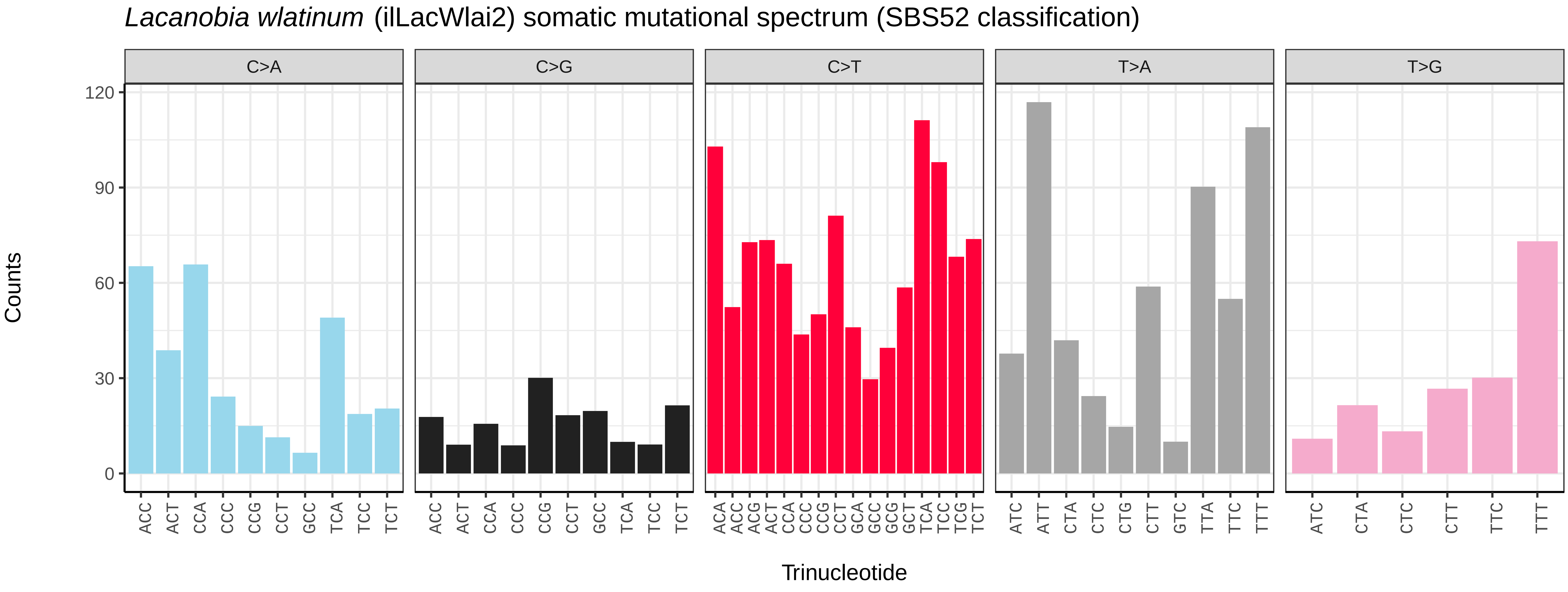
*Lacanobia wlatinum* (ilLacWlai2) somatic mutational spectrum (SBS52 classification)
C>A
C>G
C>T
T>A
T>G
Counts
120
90
60
30
0
ACC ACT CCA CCC CCG CCT GCC TCA TCC TCT
ACC ACT CCA CCC CCG CCT GCC TCA TCC TCT
ACA ACC ACG ACT CCA CCC CCG CCT GCA GCC GCG GCT TCA TCC TCG TCT
ATC ATT CTA CTC CTG CTT GTC TTA TTC TTT
ATC CTA CTC CTT TTC TTT
Trinucleotide

*Lampropteryx suffumata* (ilLamSuff1) germline mutational spectrum

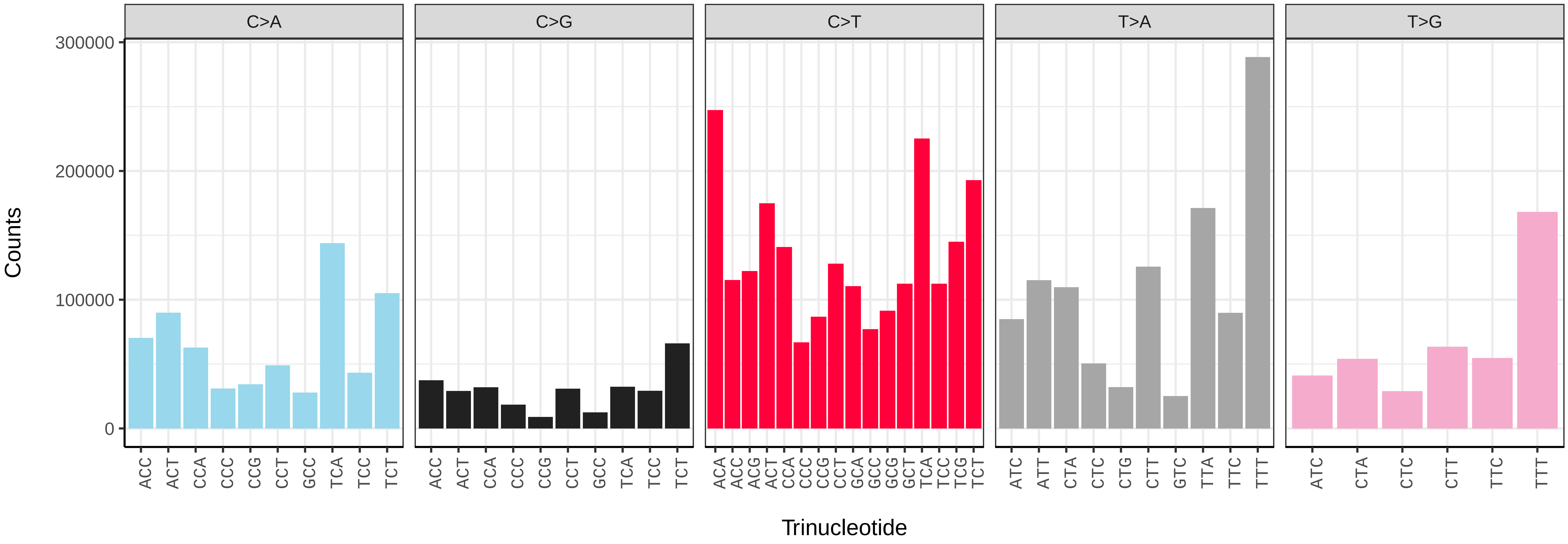


*Lampropteryx suffumata* (ilLamSuff1) somatic mutational spectrum (SBS96 classification)

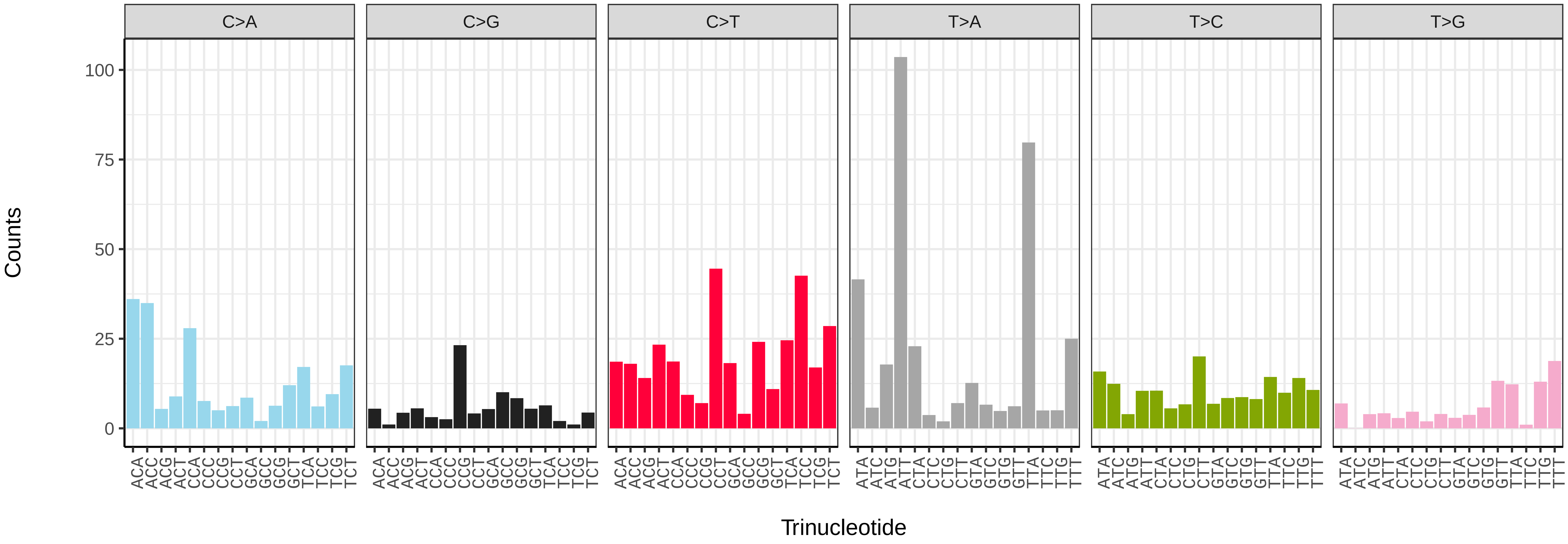


*Lampropteryx suffumata* (ilLamSuff1) somatic mutational spectrum (SBS52 classification)

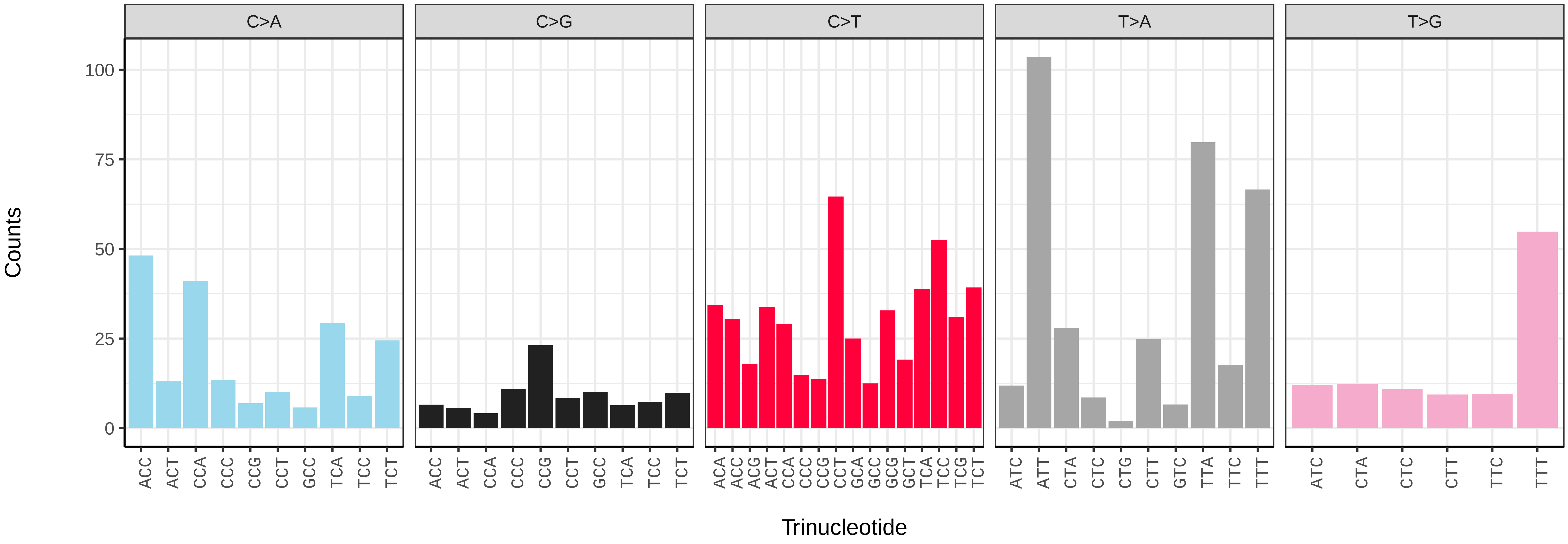

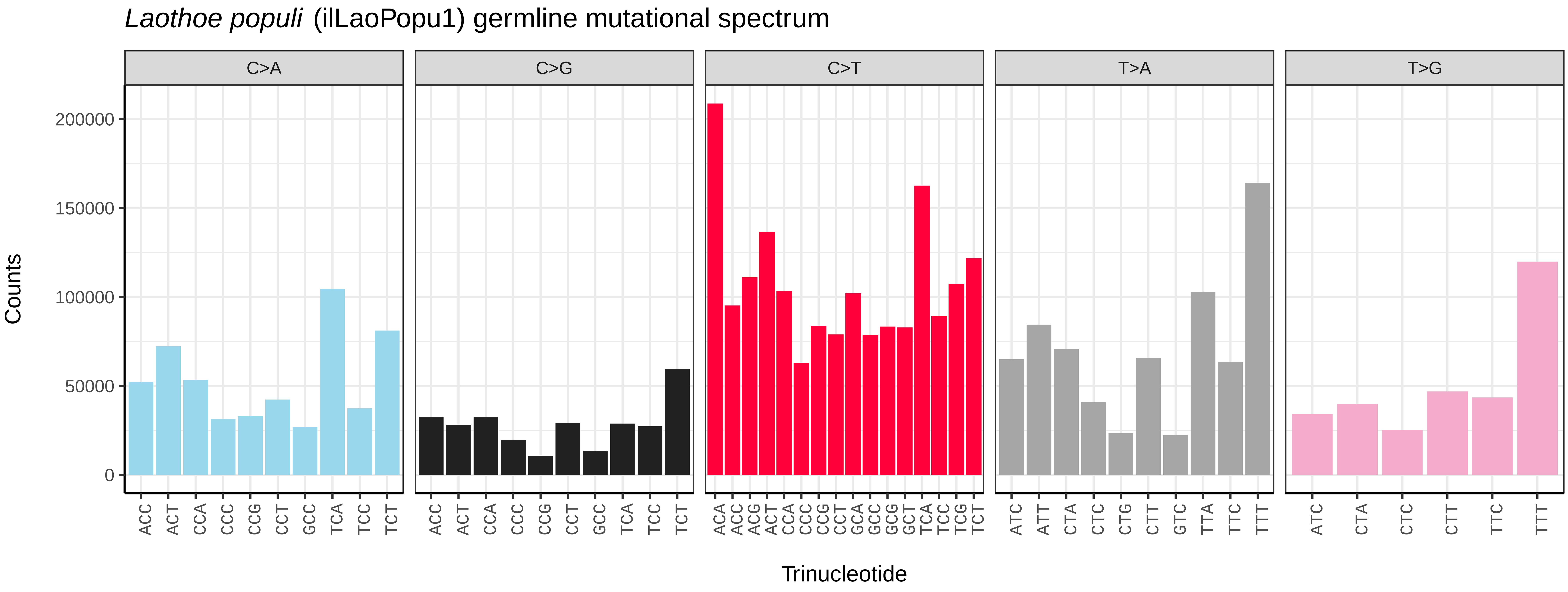
Laothoe populi (ilLaoPopu1) germline mutational spectrum
C>A
C>G
C>T
T>A
T>G
Counts
Trinucleotide

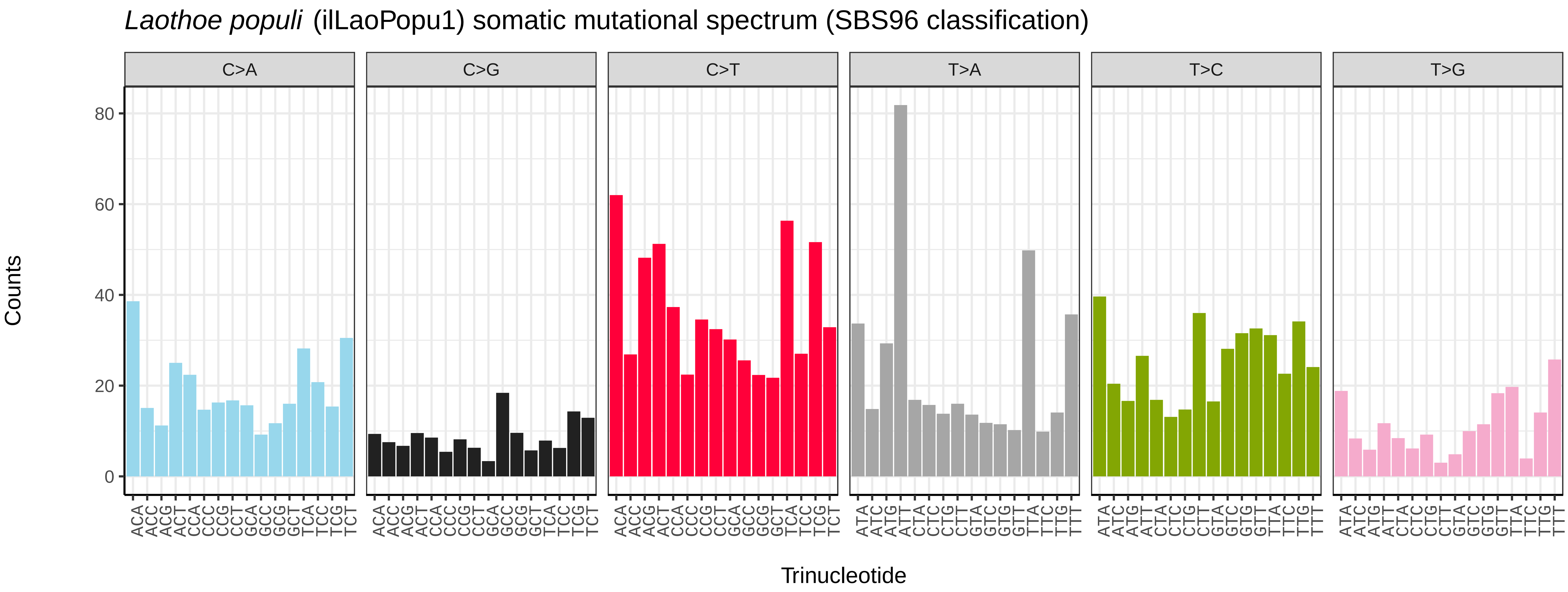
Laothoe populi (ilLaoPopu1) somatic mutational spectrum (SBS96 classification)
C>A
C>G
C>T
T>A
T>C
T>G
Counts
Trinucleotide

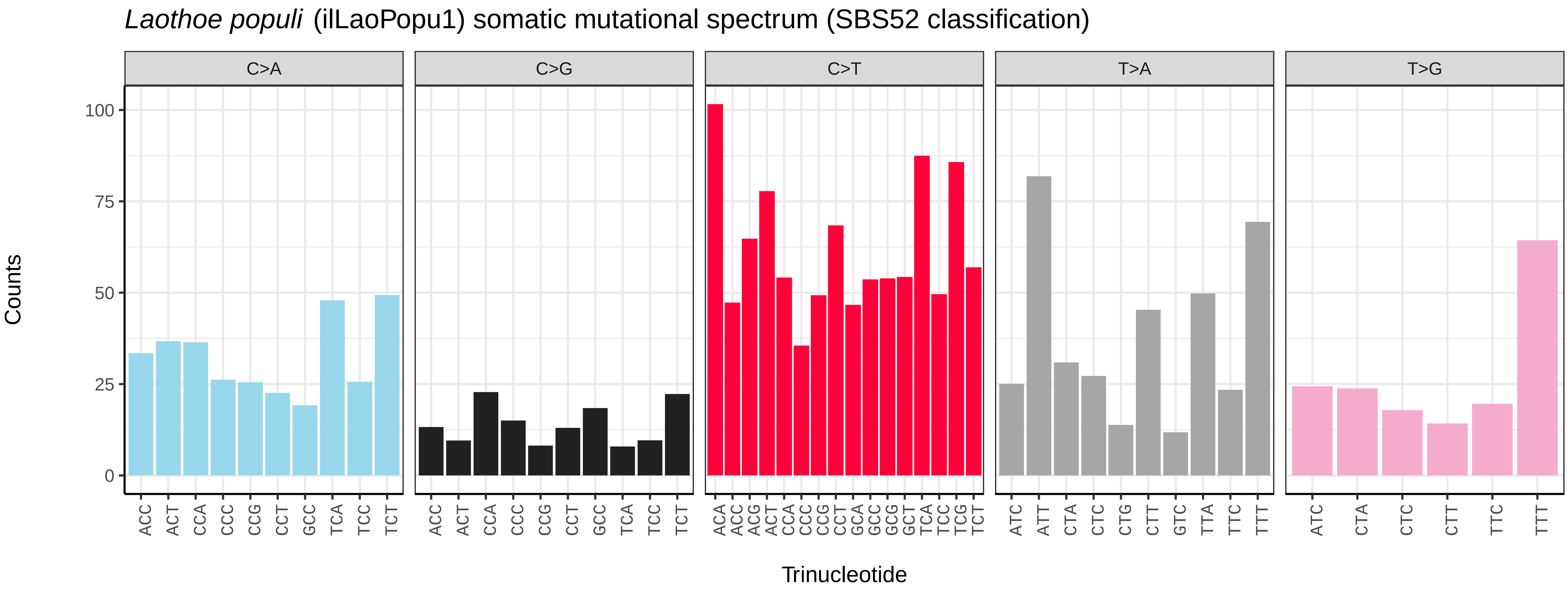
Laothoe populi (ilLaoPopu1) somatic mutational spectrum (SBS52 classification)
C>A
C>G
C>T
T>A
T>G
Counts
Trinucleotide

*Laspeyria flexula* (ilLasFlex1) germline mutational spectrum

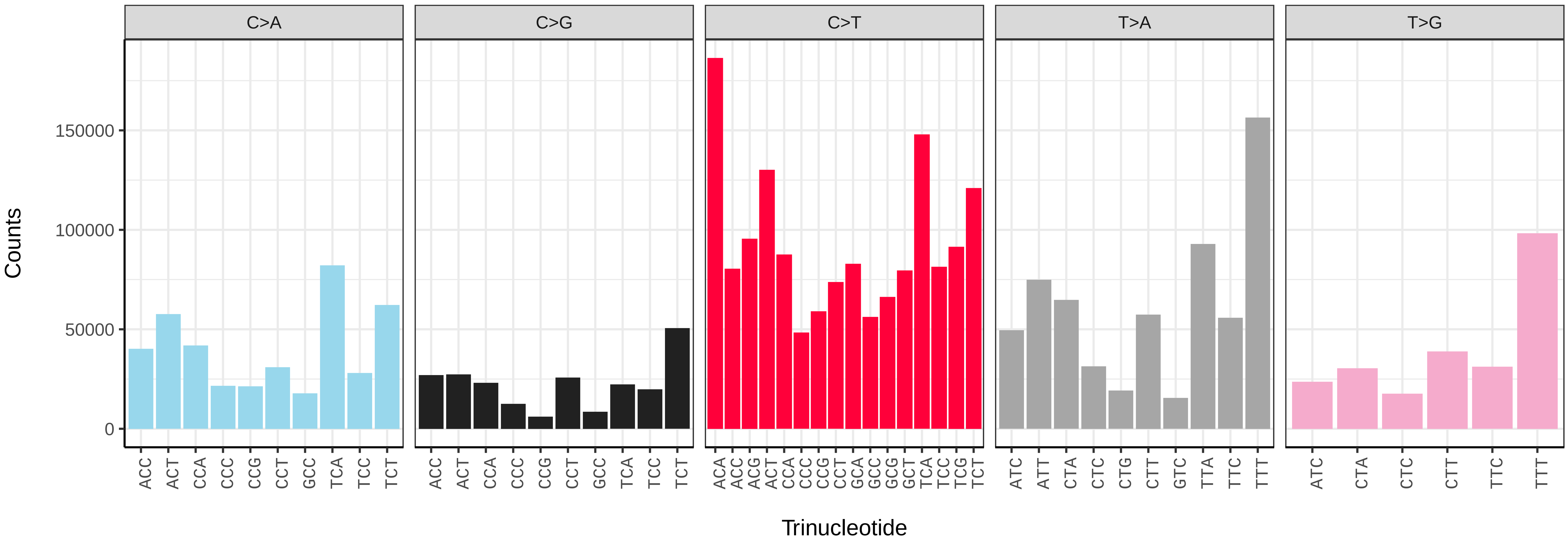


*Laspeyria flexula* (ilLasFlex1) somatic mutational spectrum (SBS96 classification)

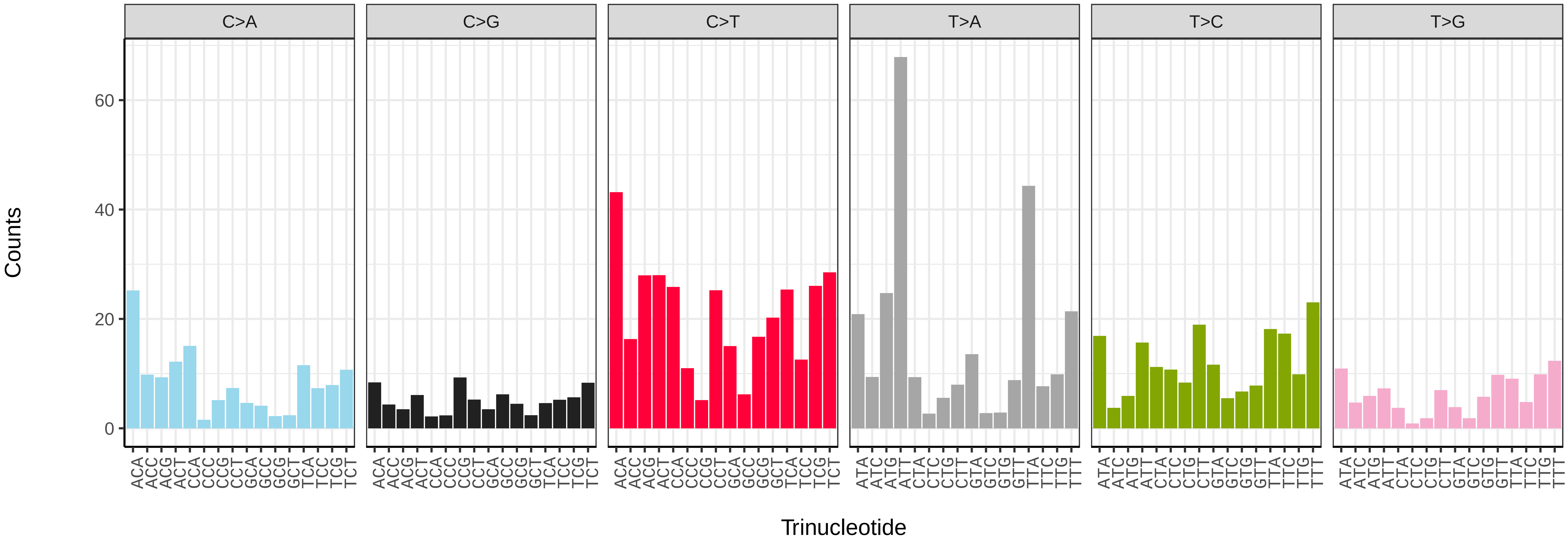


*Laspeyria flexula* (ilLasFlex1) somatic mutational spectrum (SBS52 classification)

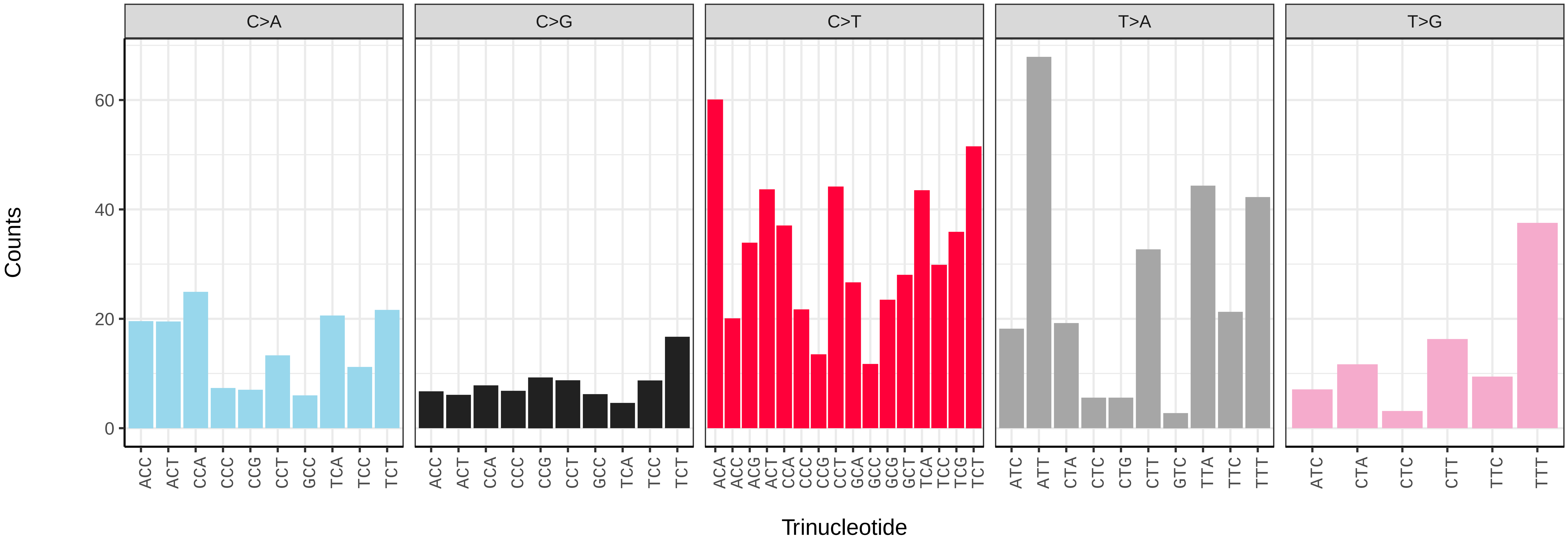

*Lasiommata megera* (ilLasMege1) germline mutational spectrum

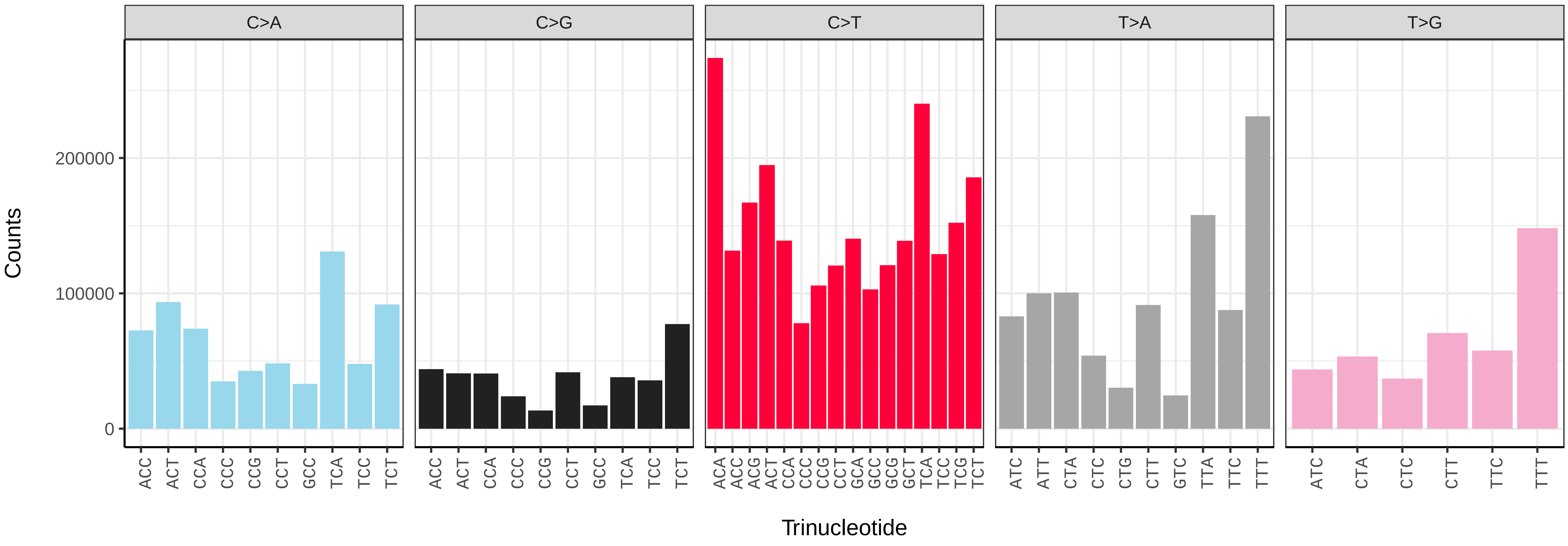


*Lasiommata megera* (ilLasMege1) somatic mutational spectrum (SBS96 classification)

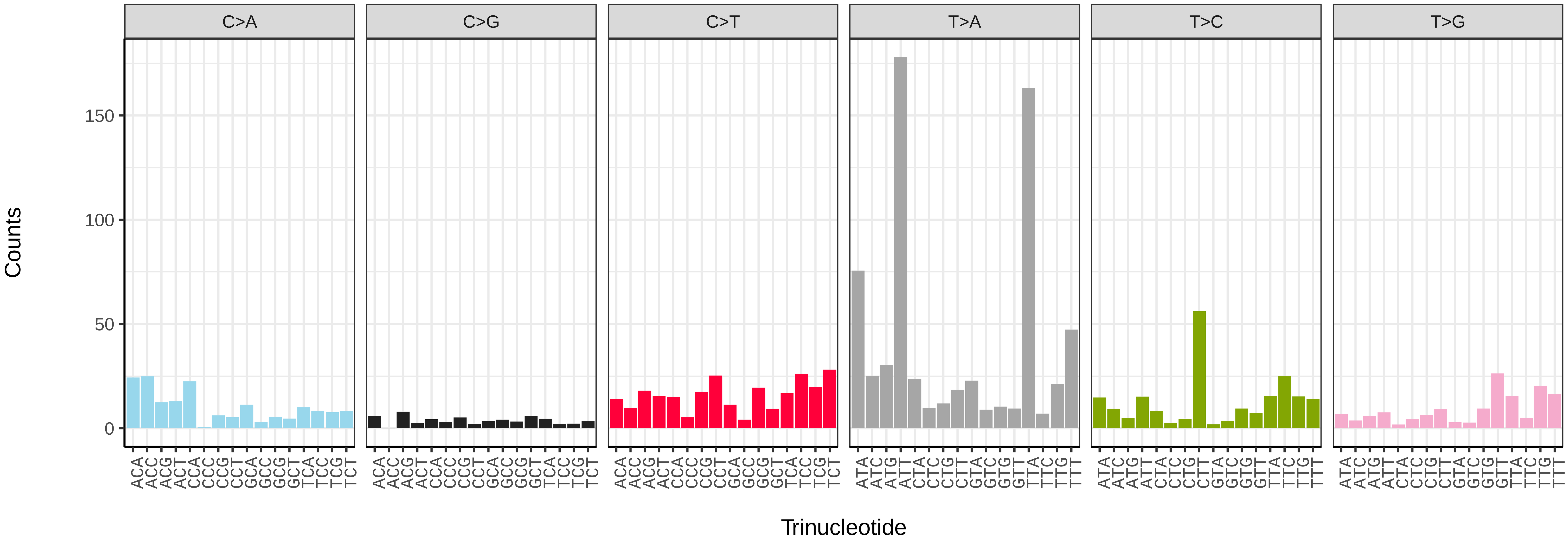


*Lasiommata megera* (ilLasMege1) somatic mutational spectrum (SBS52 classification)

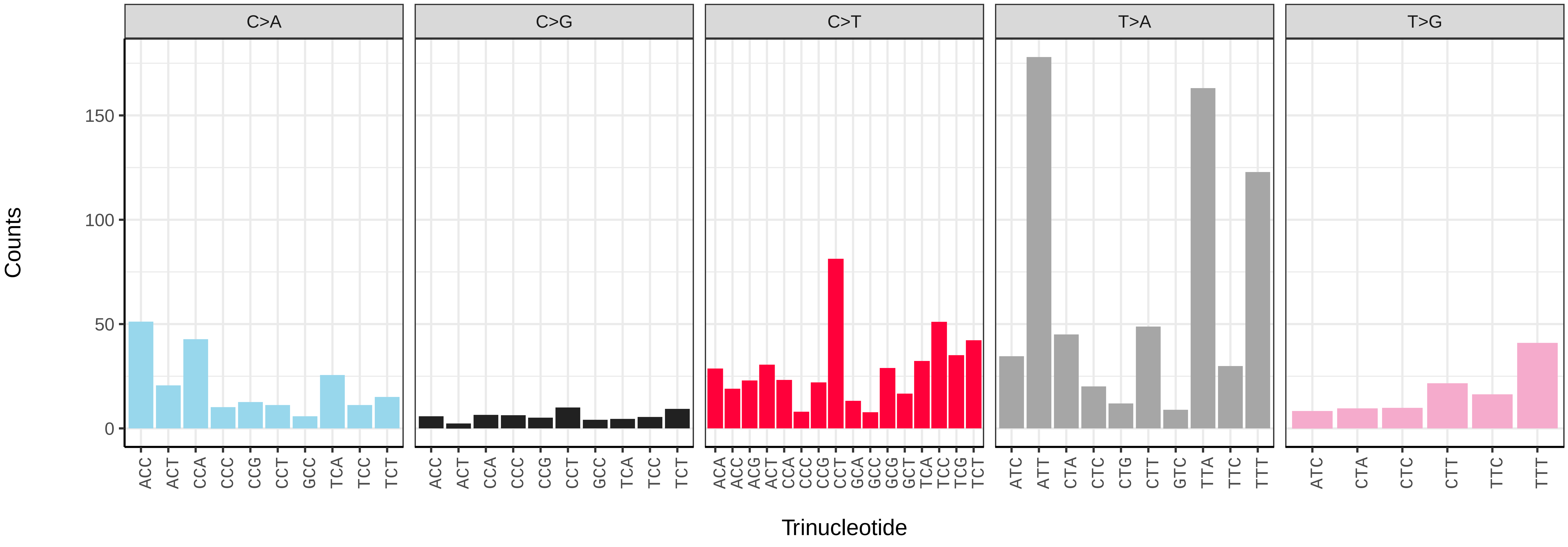

*Lathronympha strigana* (ilLatStri6) germline mutational spectrum

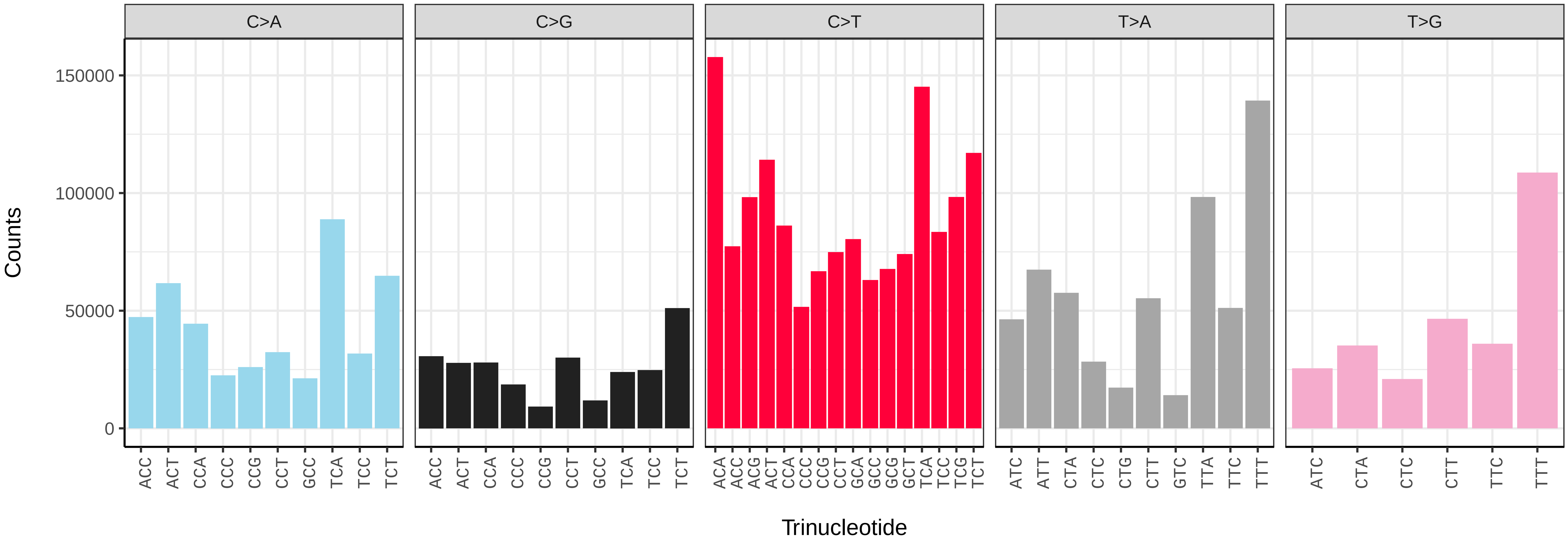


*Lathronympha strigana* (ilLatStri6) somatic mutational spectrum (SBS96 classification)

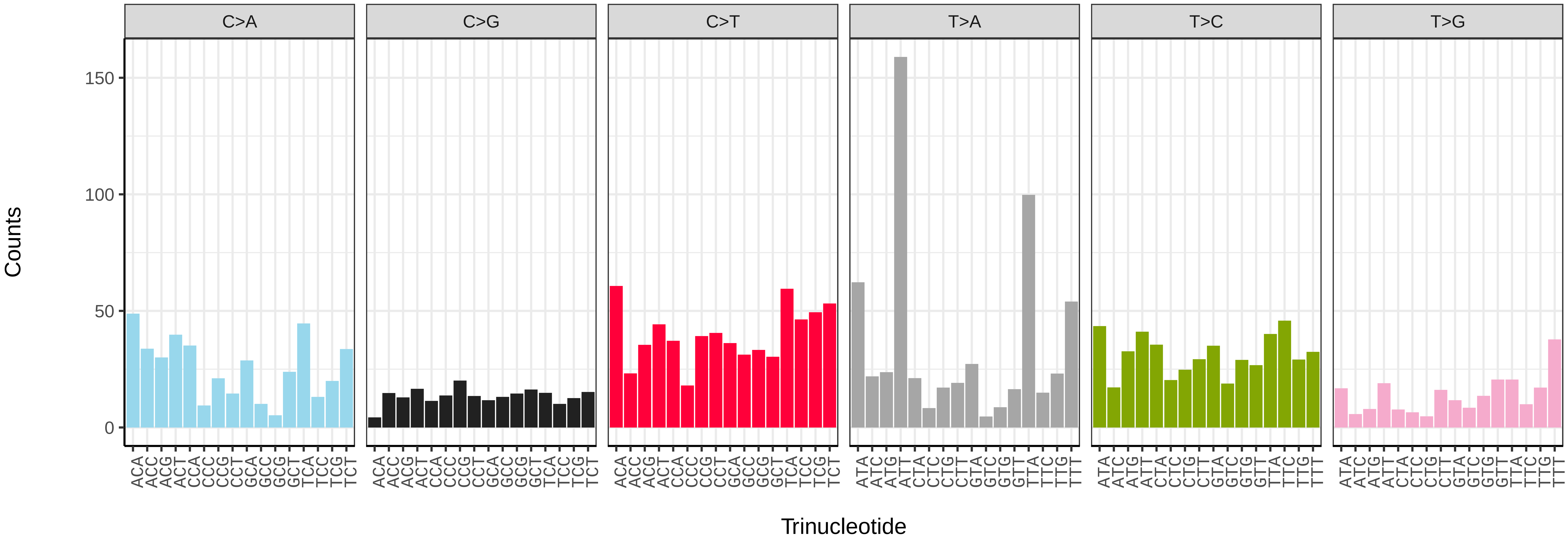


*Lathronympha strigana* (ilLatStri6) somatic mutational spectrum (SBS52 classification)

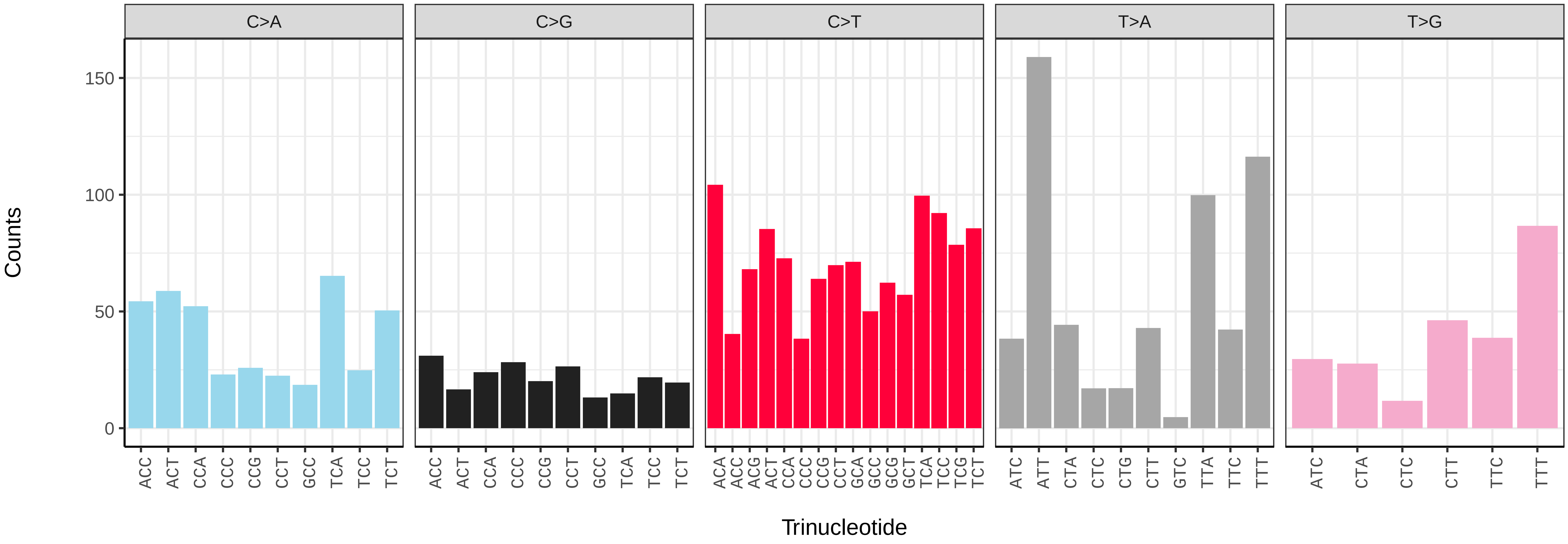

*Leptidea sinapis* (ilLepSina1) germline mutational spectrum

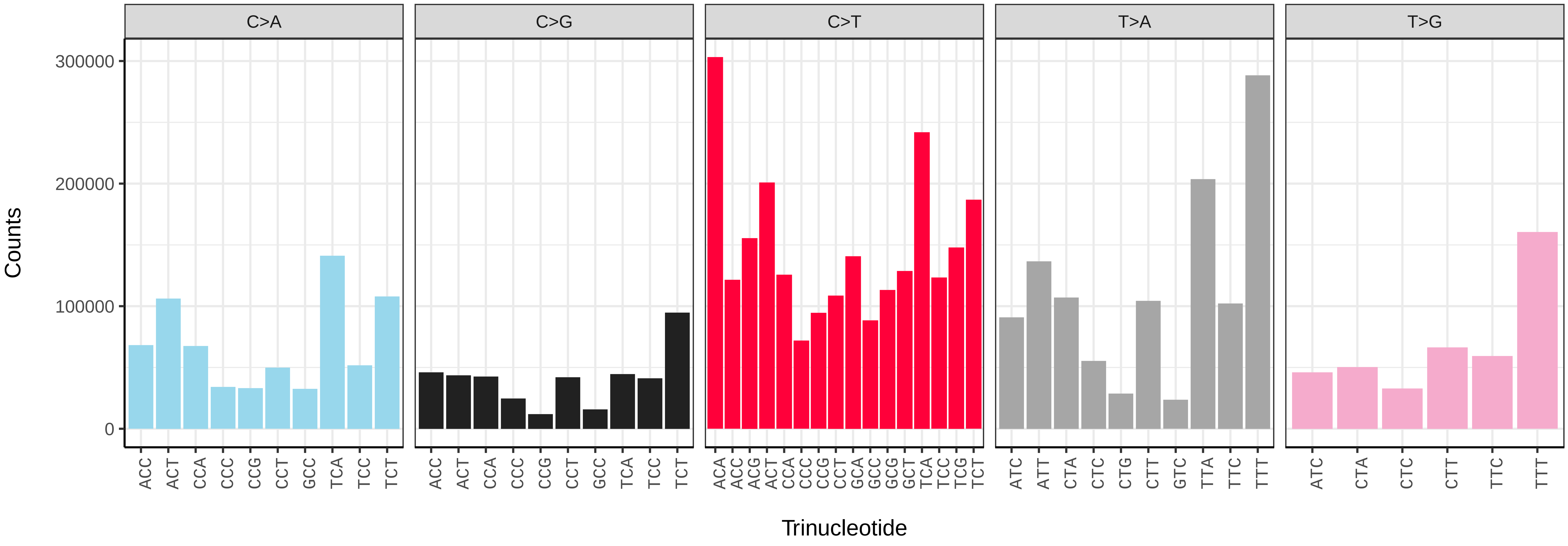


*Leptidea sinapis* (ilLepSina1) somatic mutational spectrum (SBS96 classification)

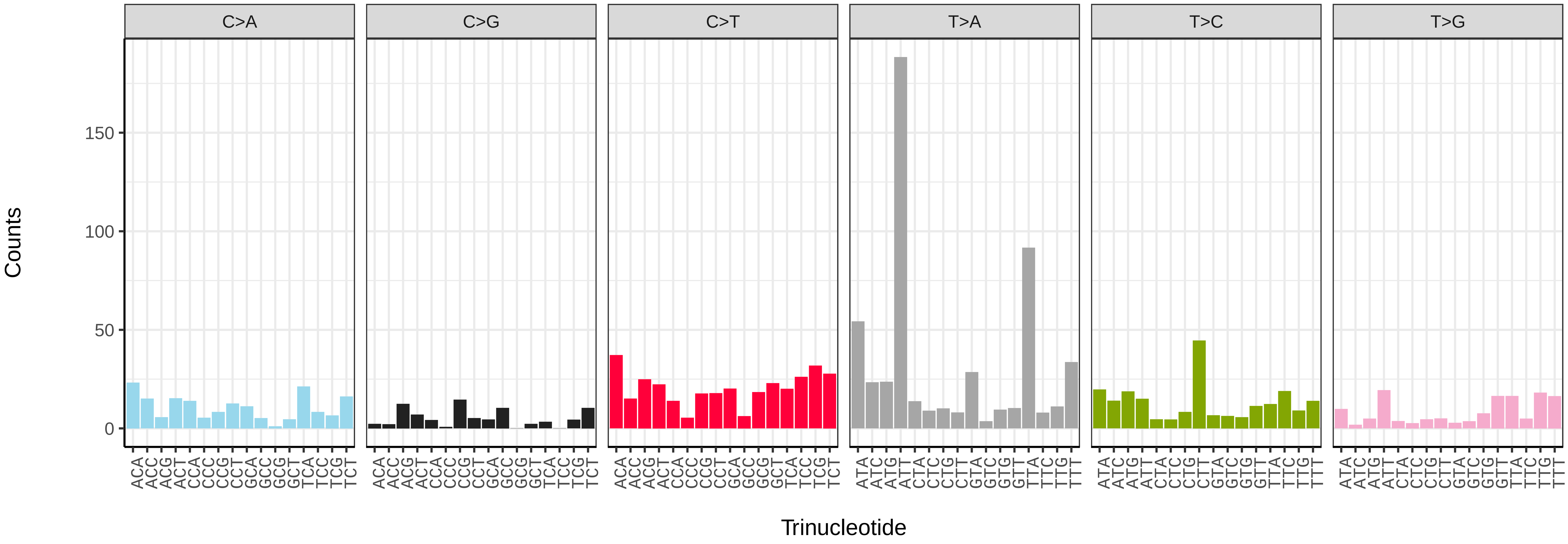


*Leptidea sinapis* (ilLepSina1) somatic mutational spectrum (SBS52 classification)

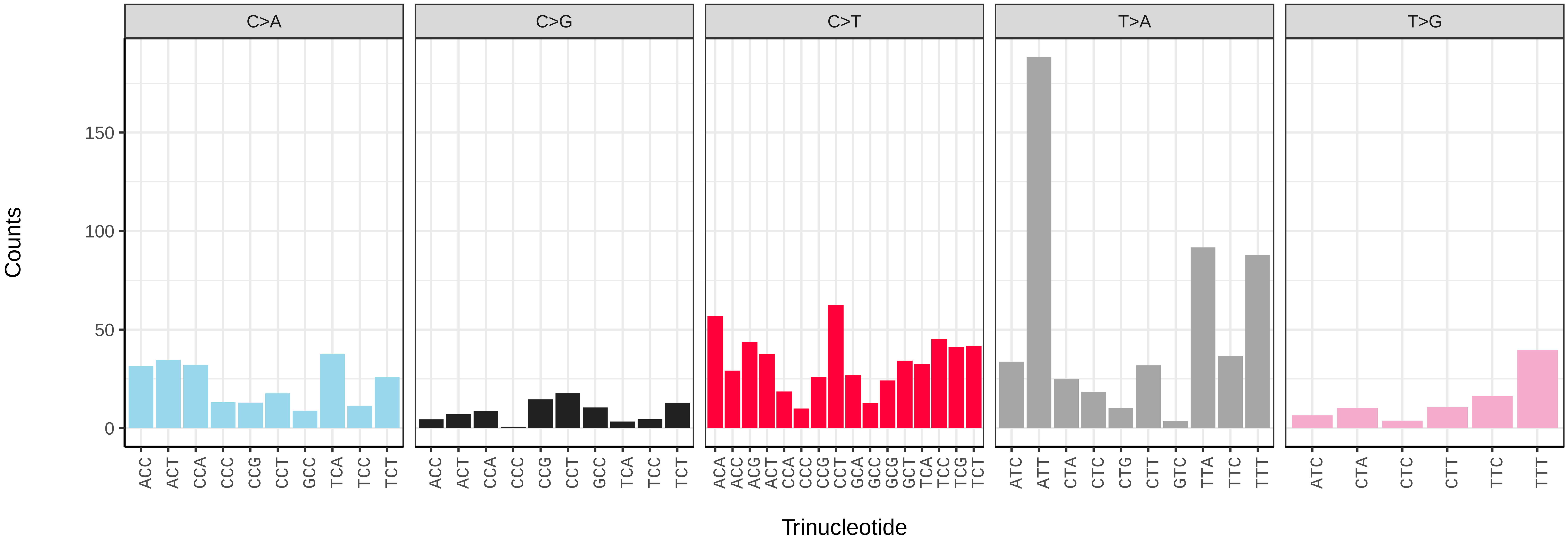

*Leucoma salicis* (ilLeuSali1) germline mutational spectrum

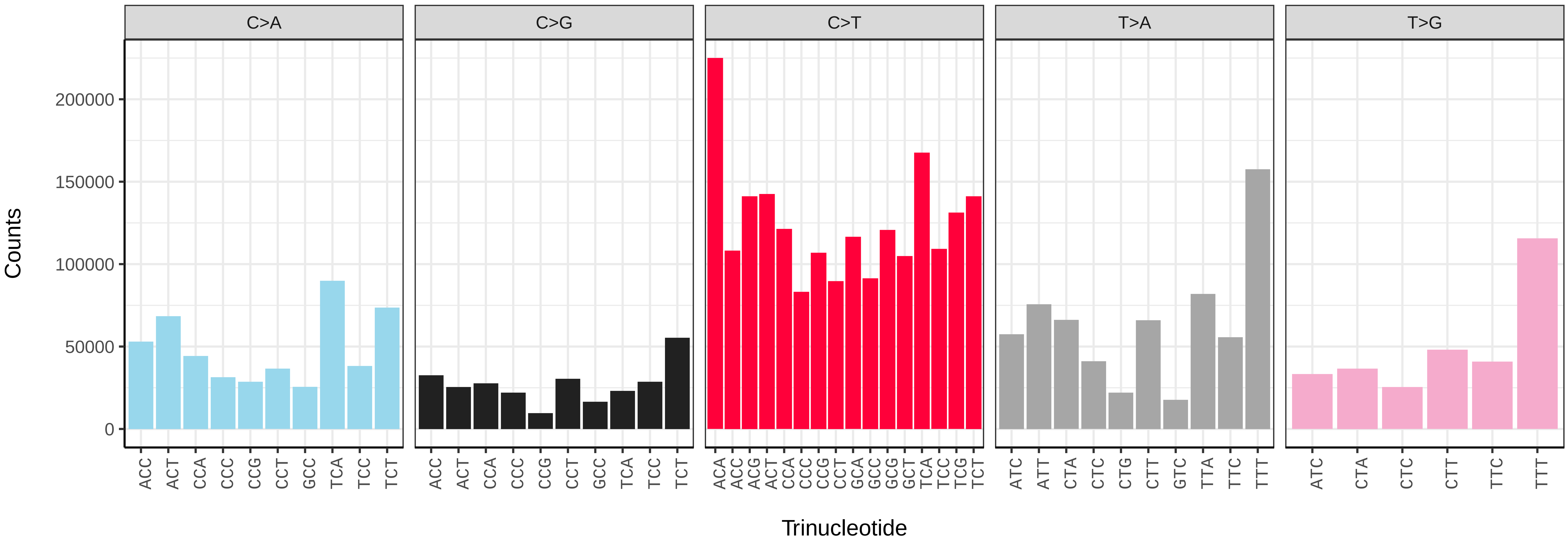


*Leucoma salicis* (ilLeuSali1) somatic mutational spectrum (SBS96 classification)

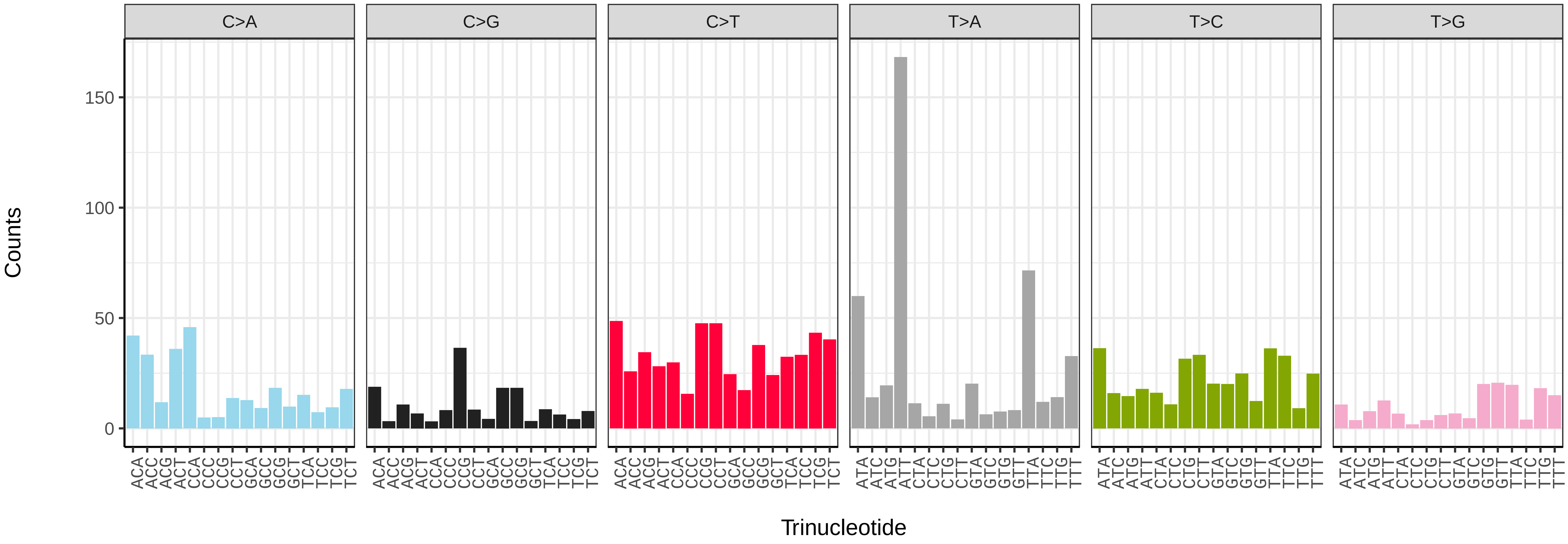


*Leucoma salicis* (ilLeuSali1) somatic mutational spectrum (SBS52 classification)

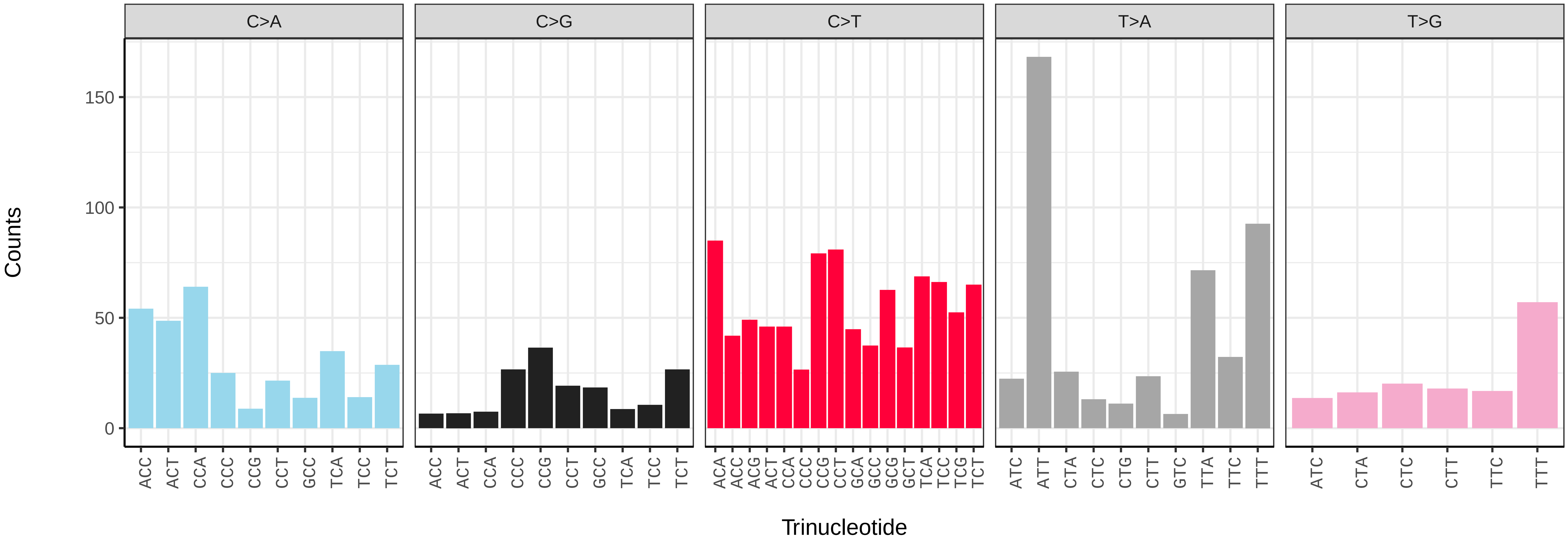

*Ligdia adustata* (ilLigAdus1) germline mutational spectrum

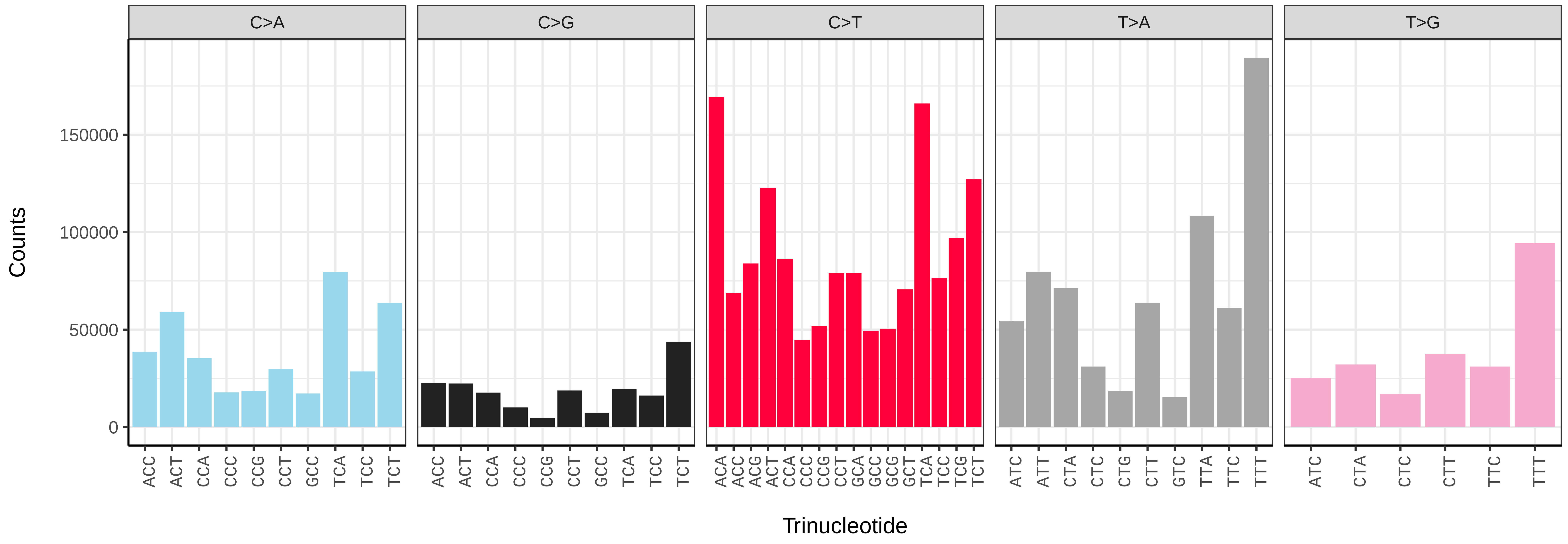


*Ligdia adustata* (ilLigAdus1) somatic mutational spectrum (SBS96 classification)

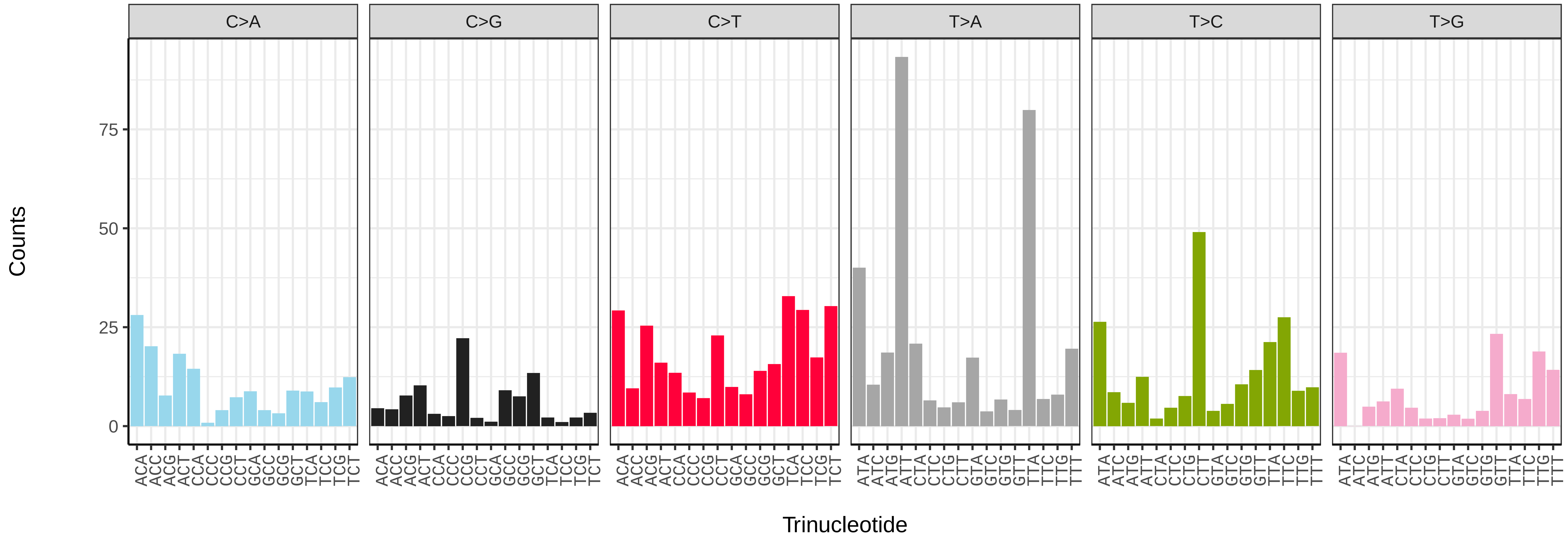


*Ligdia adustata* (ilLigAdus1) somatic mutational spectrum (SBS52 classification)

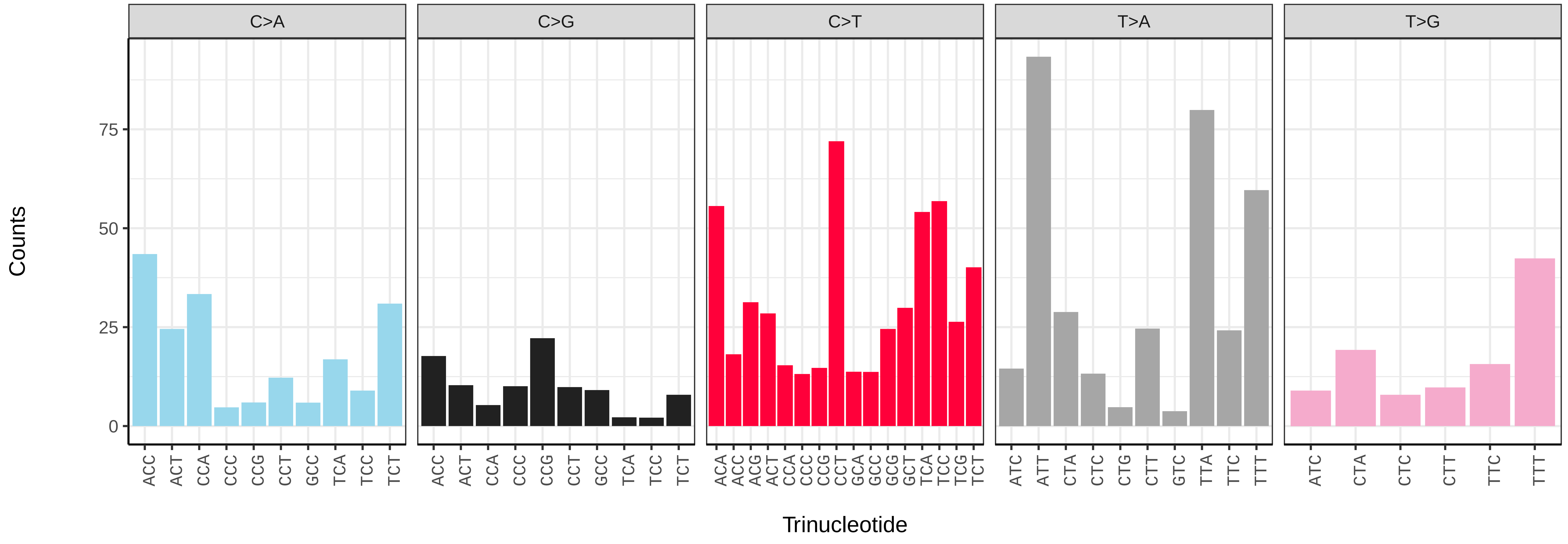

### *Limenitis camilla* (ilLimCami1) germline mutational spectrum

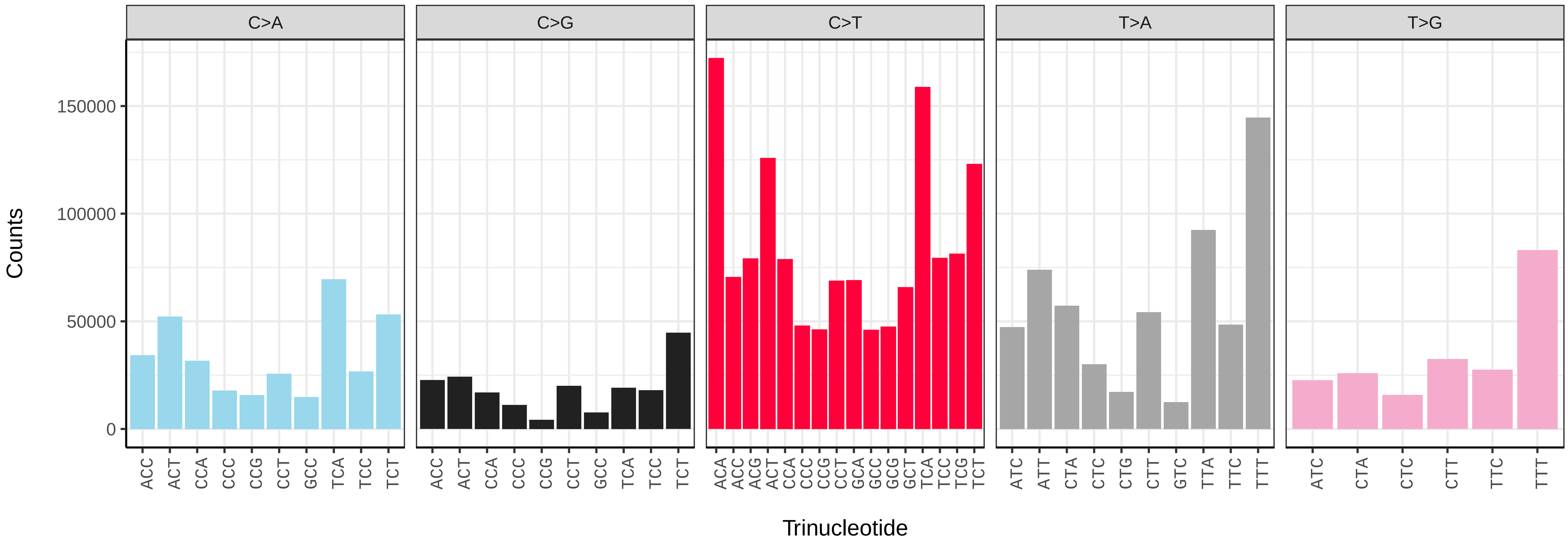


### *Limenitis camilla* (ilLimCami1) somatic mutational spectrum (SBS96 classification)

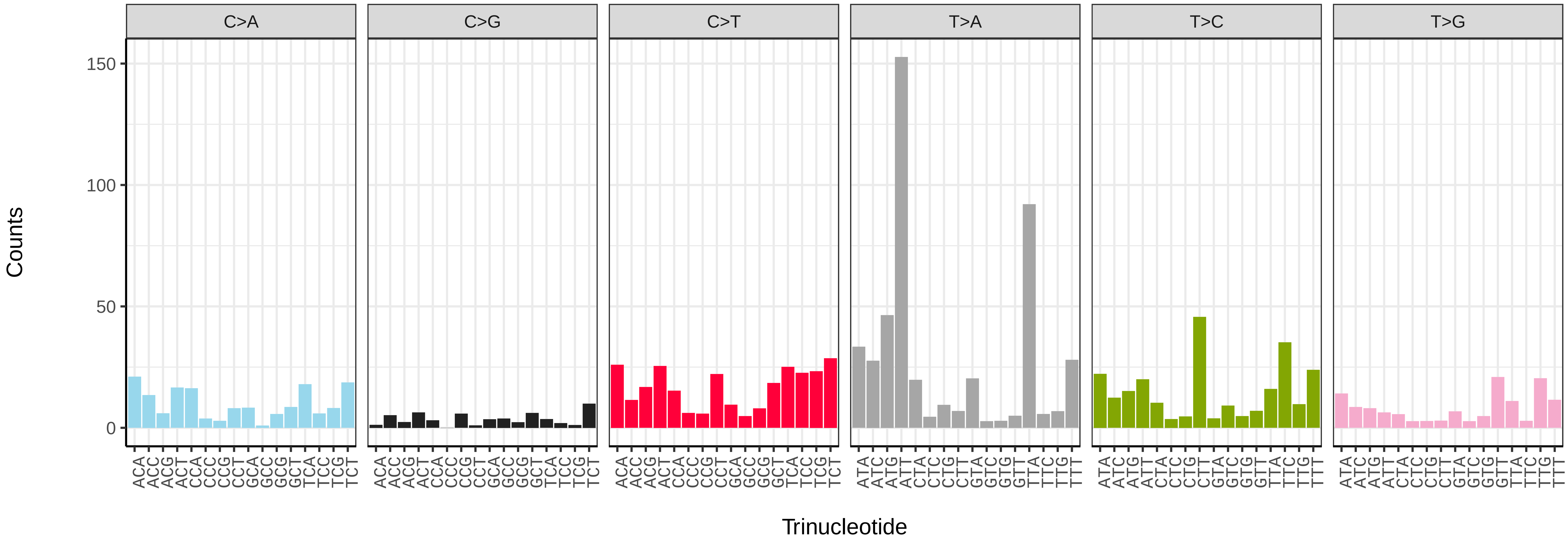


### *Limenitis camilla* (ilLimCami1) somatic mutational spectrum (SBS52 classification)

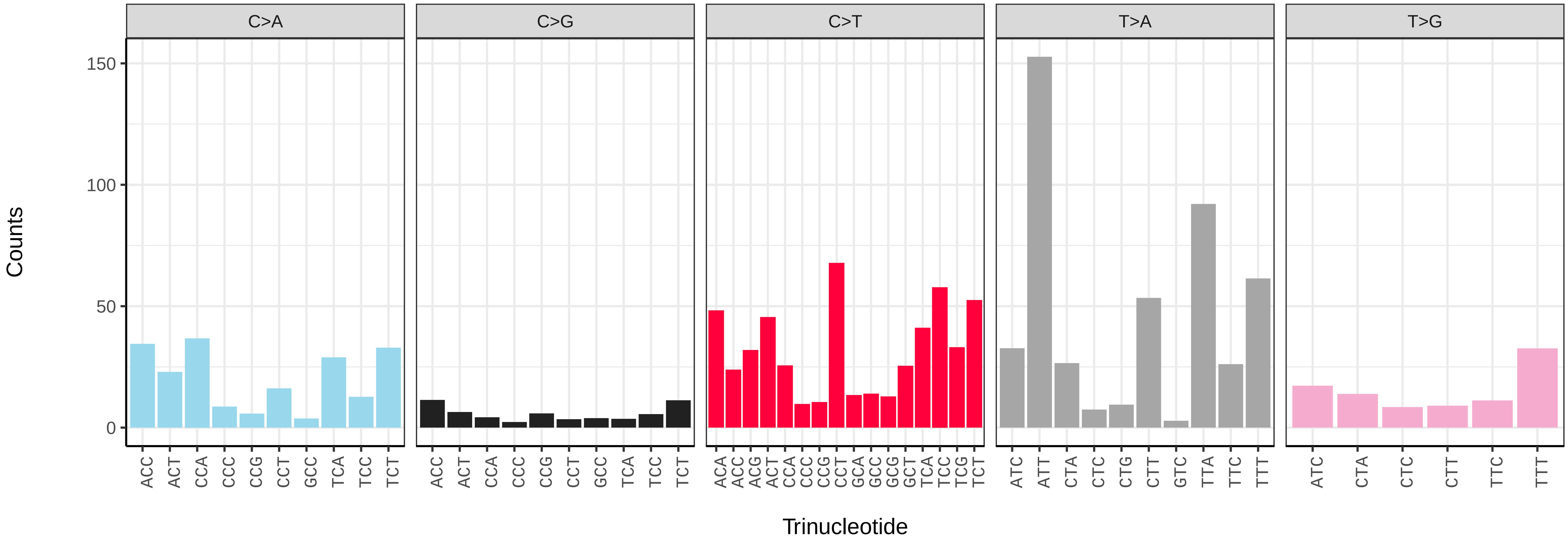

*Lithophane ornitopus* (ilLitOrni1) germline mutational spectrum

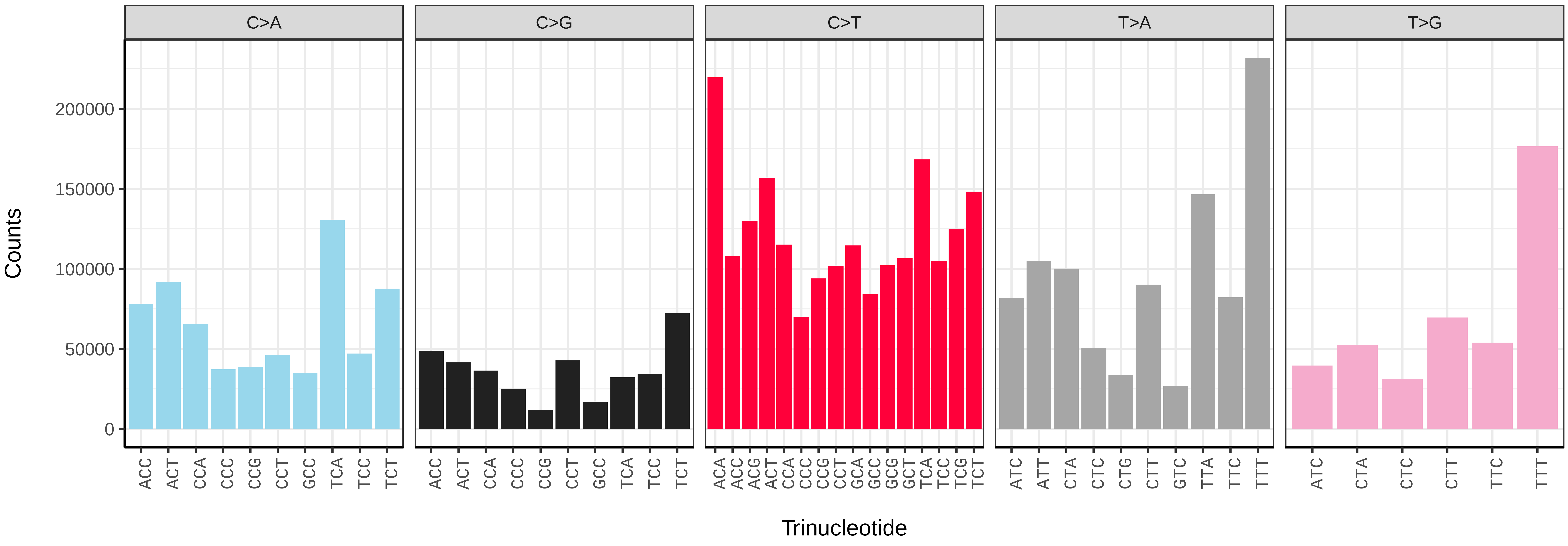


*Lithophane ornitopus* (ilLitOrni1) somatic mutational spectrum (SBS96 classification)

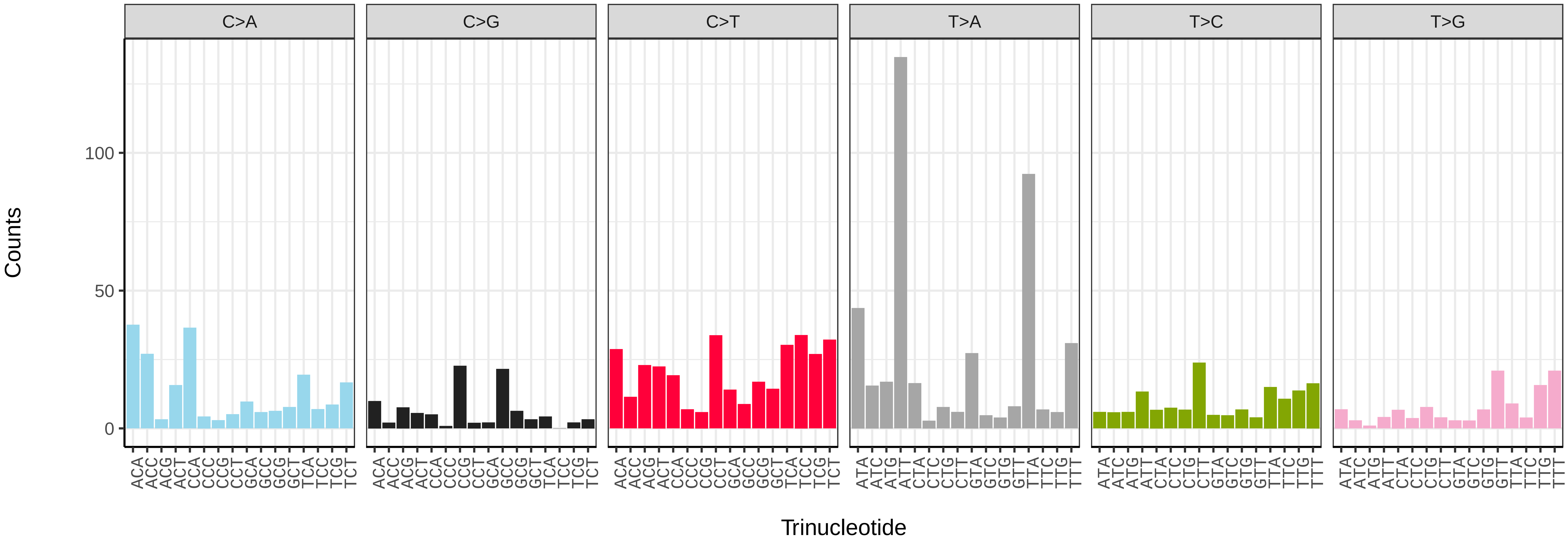


*Lithophane ornitopus* (ilLitOrni1) somatic mutational spectrum (SBS52 classification)

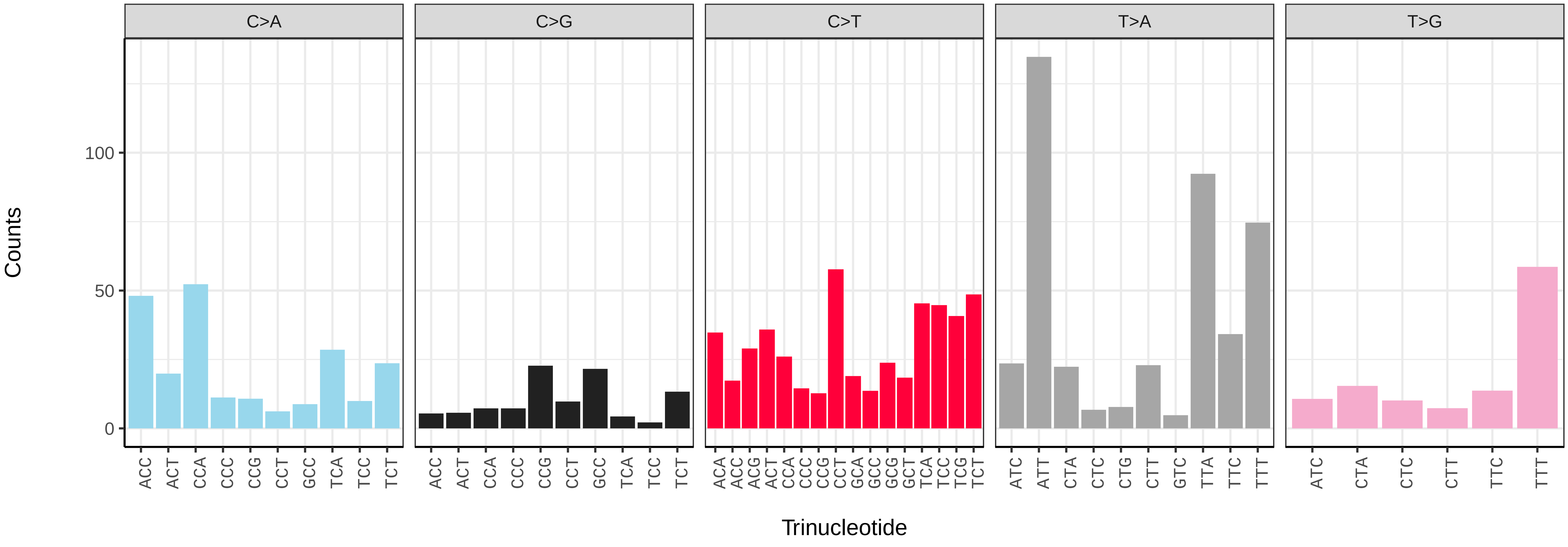

*Lithophane semibrunnea* (ilLitSemi2) germline mutational spectrum

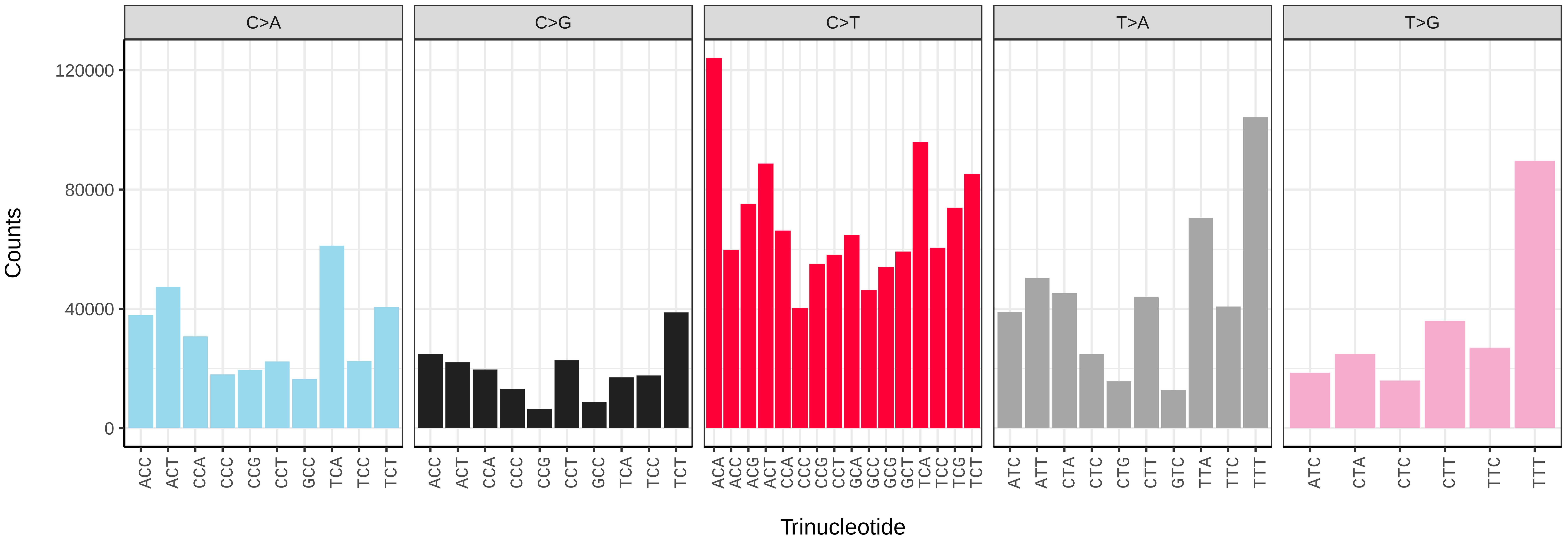


*Lithophane semibrunnea* (ilLitSemi2) somatic mutational spectrum (SBS96 classification)

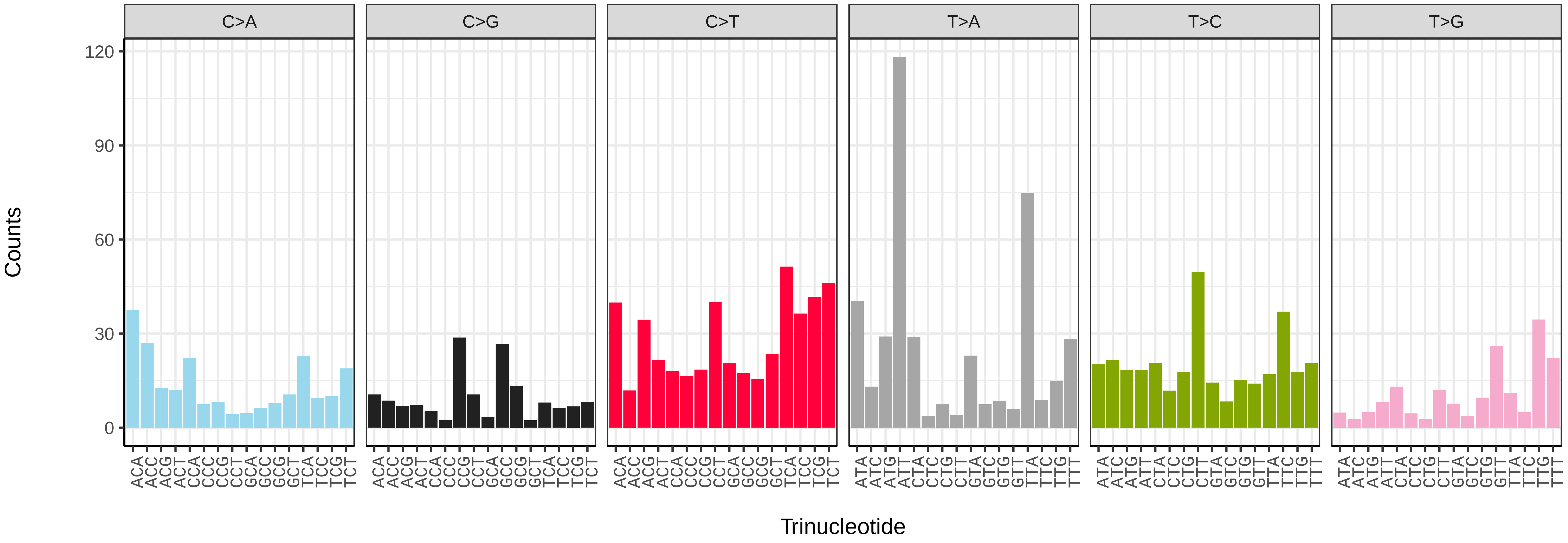


*Lithophane semibrunnea* (ilLitSemi2) somatic mutational spectrum (SBS52 classification)

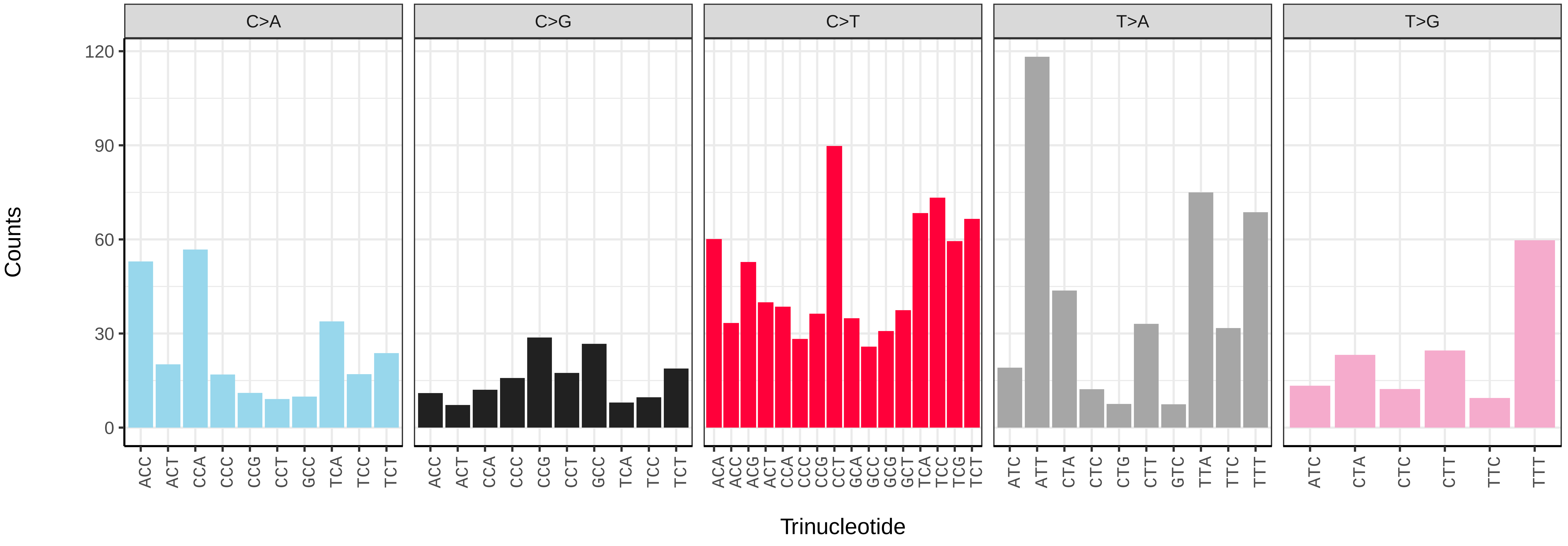

*Lithophane socia* (ilLitSoci1) germline mutational spectrum

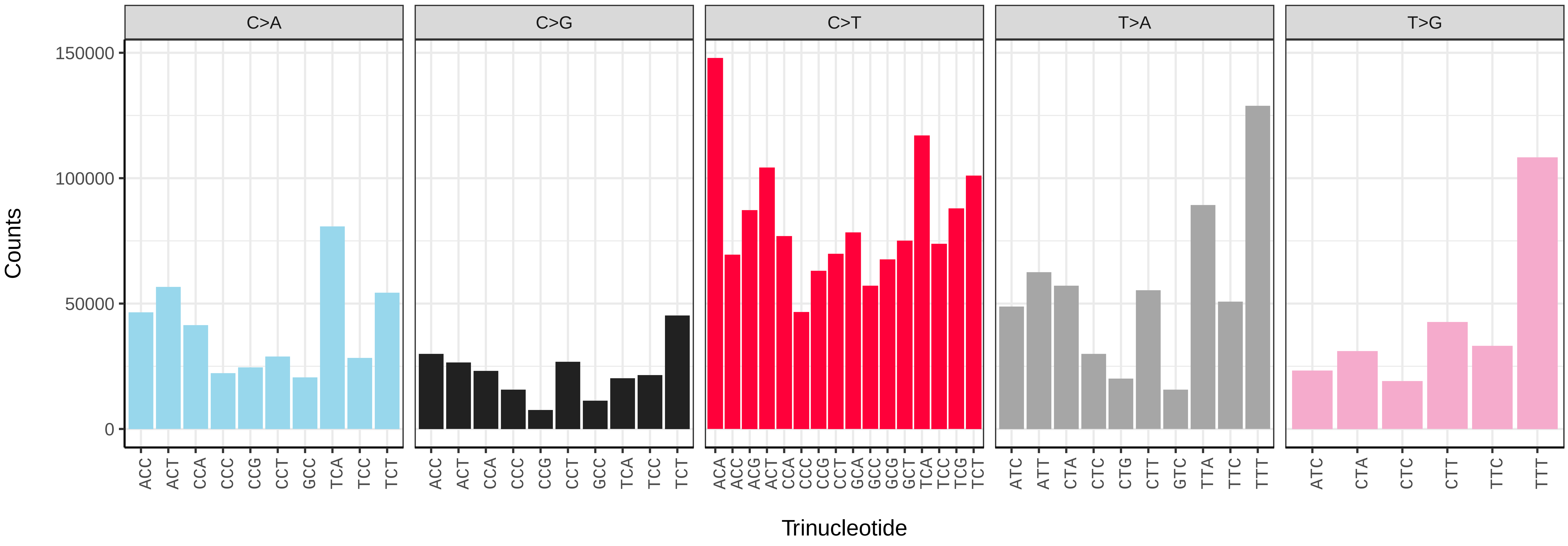


*Lithophane socia* (ilLitSoci1) somatic mutational spectrum (SBS96 classification)

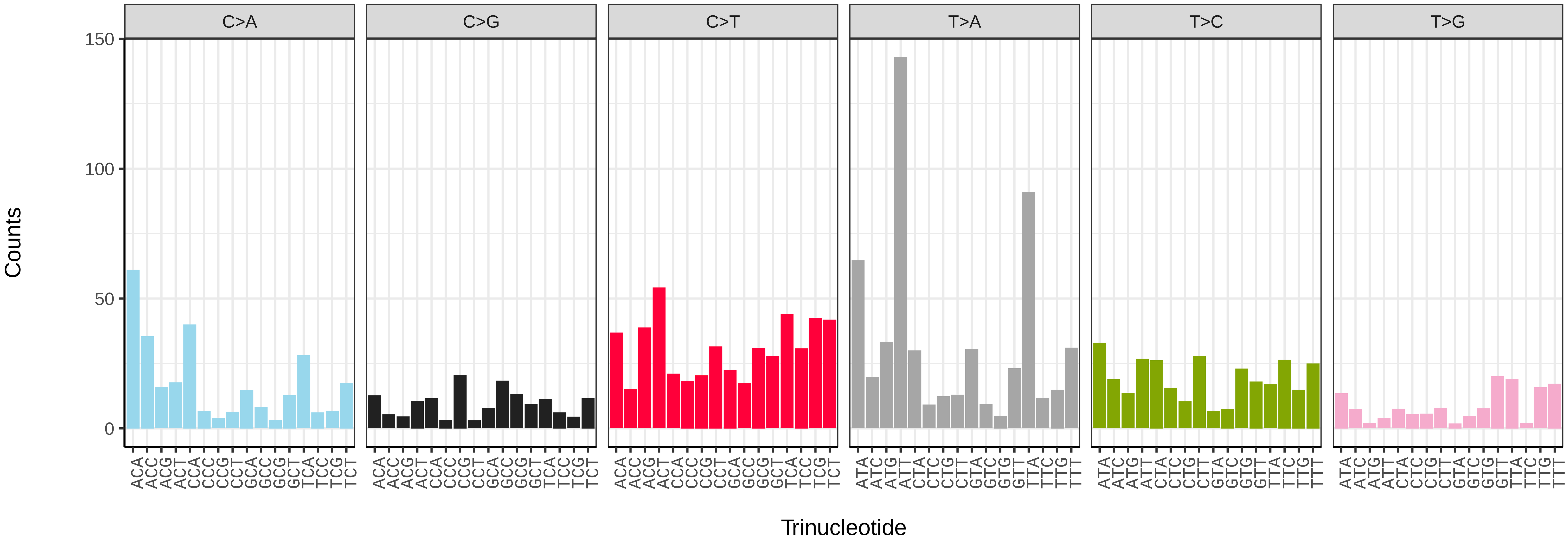


*Lithophane socia* (ilLitSoci1) somatic mutational spectrum (SBS52 classification)

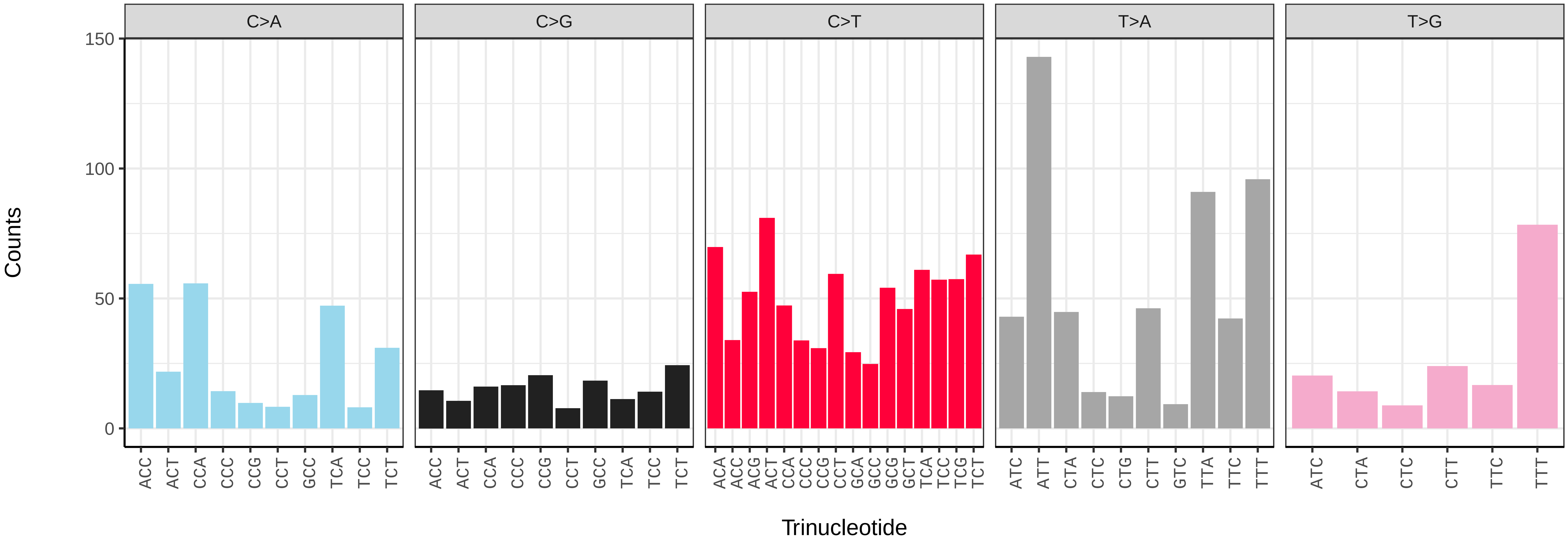

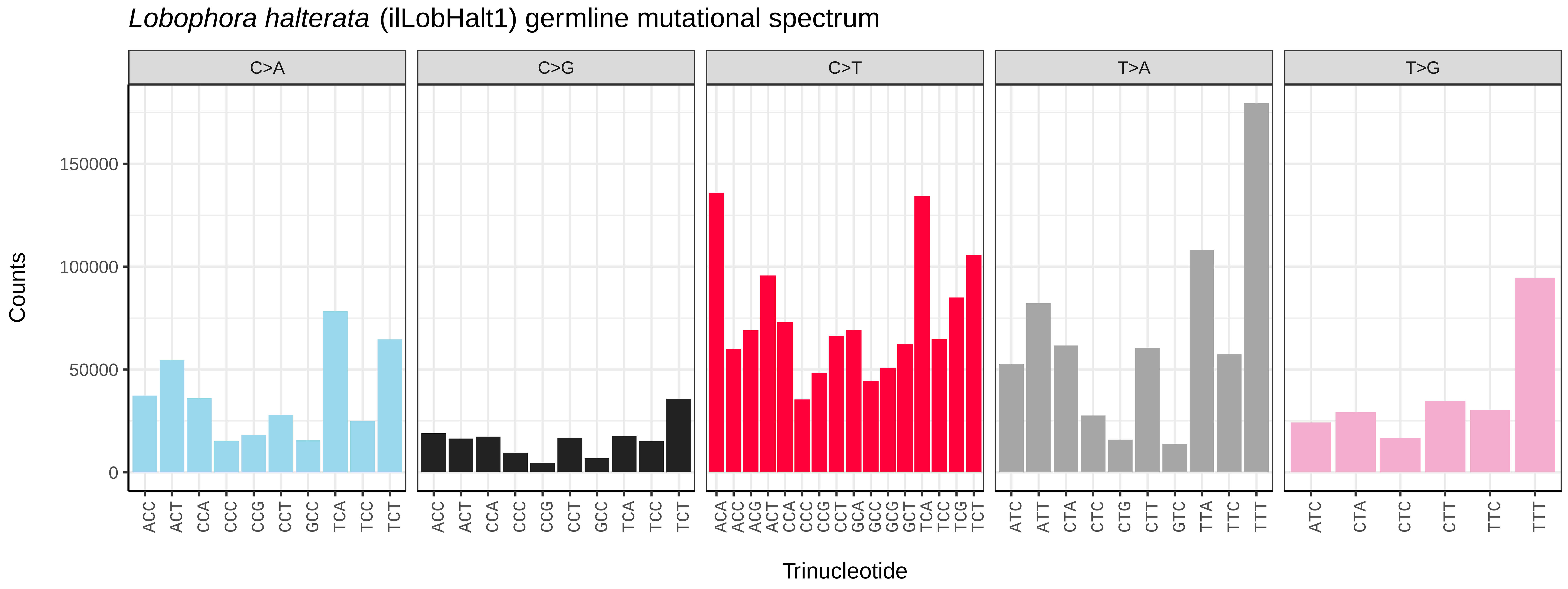
Lobophora halterata (ilLobHalt1) germline mutational spectrum
C>A
C>G
C>T
T>A
T>G
Counts
0
50000
100000
150000
Trinucleotide
ACC ACT CCA CCC CCG CCT GCC TCA TCC TCT
ACA ACC ACG ACT CCA CCC CCG CCT GCA GCC GCG GCT TCA TCC TCG TCT
ATC ATT CTA CTC CTG CTT GTC TTA TTC TTT
ATC CTA CTC CTT TTC TTT

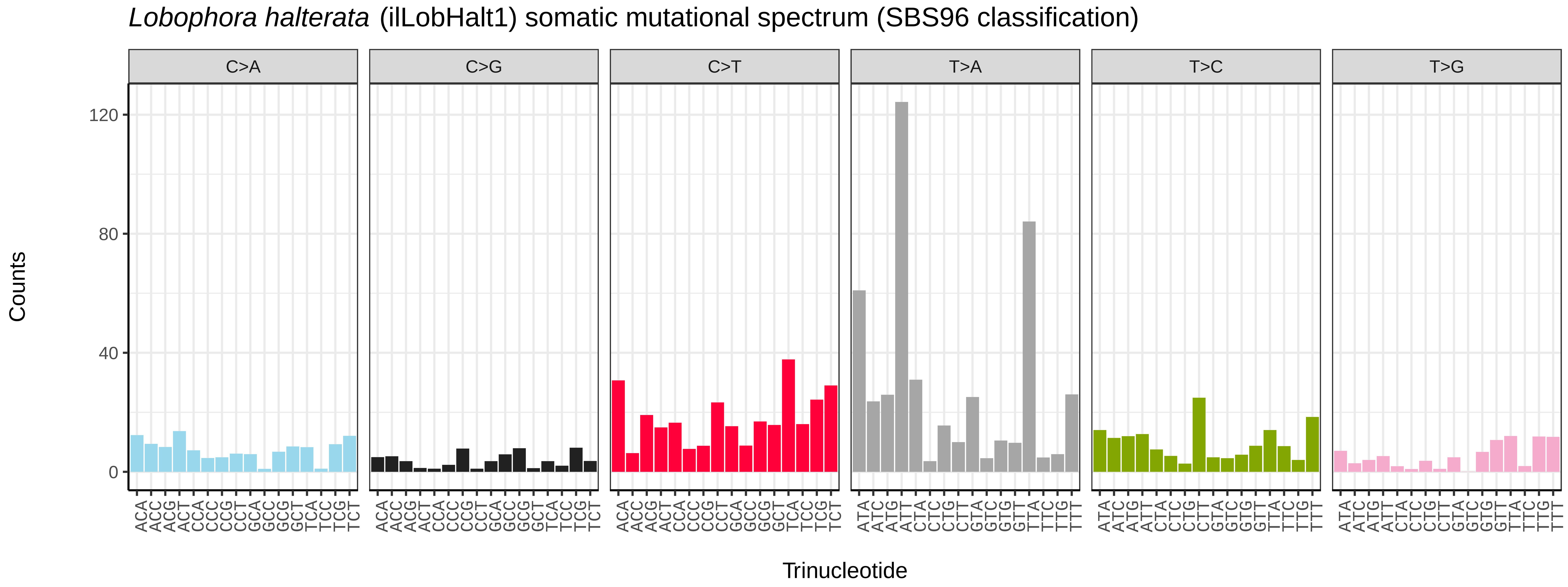
Lobophora halterata (ilLobHalt1) somatic mutational spectrum (SBS96 classification)
C>A
C>G
C>T
T>A
T>C
T>G
Counts
0
40
80
120
Trinucleotide
ACA ACC ACG ACT CCA CCC CCG CCT GCA GCC GCG GCT TCA TCC TCG TCT
ATA ATC ATG ATT CTA CTC CTG CTT GTA GTC GTG GTT TTA TTC TTG TTT

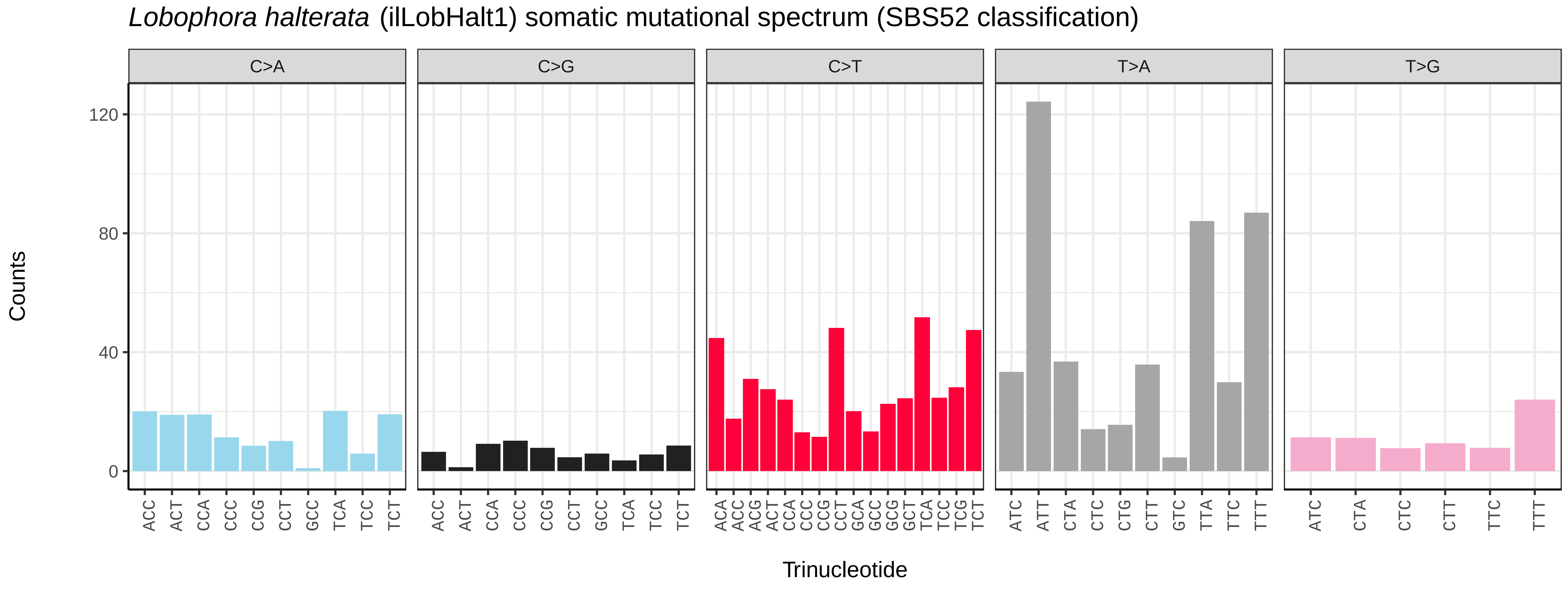
Lobophora halterata (ilLobHalt1) somatic mutational spectrum (SBS52 classification)
C>A
C>G
C>T
T>A
T>G
Counts
0
40
80
120
Trinucleotide
ACC ACT CCA CCC CCG CCT GCC TCA TCC TCT
ACA ACC ACG ACT CCA CCC CCG CCT GCA GCC GCG GCT TCA TCC TCG TCT
ATC ATT CTA CTC CTG CTT GTC TTA TTC TTT
ATC CTA CTC CTT TTC TTT

*Lomographa bimaculata* (ilLomBima1) germline mutational spectrum

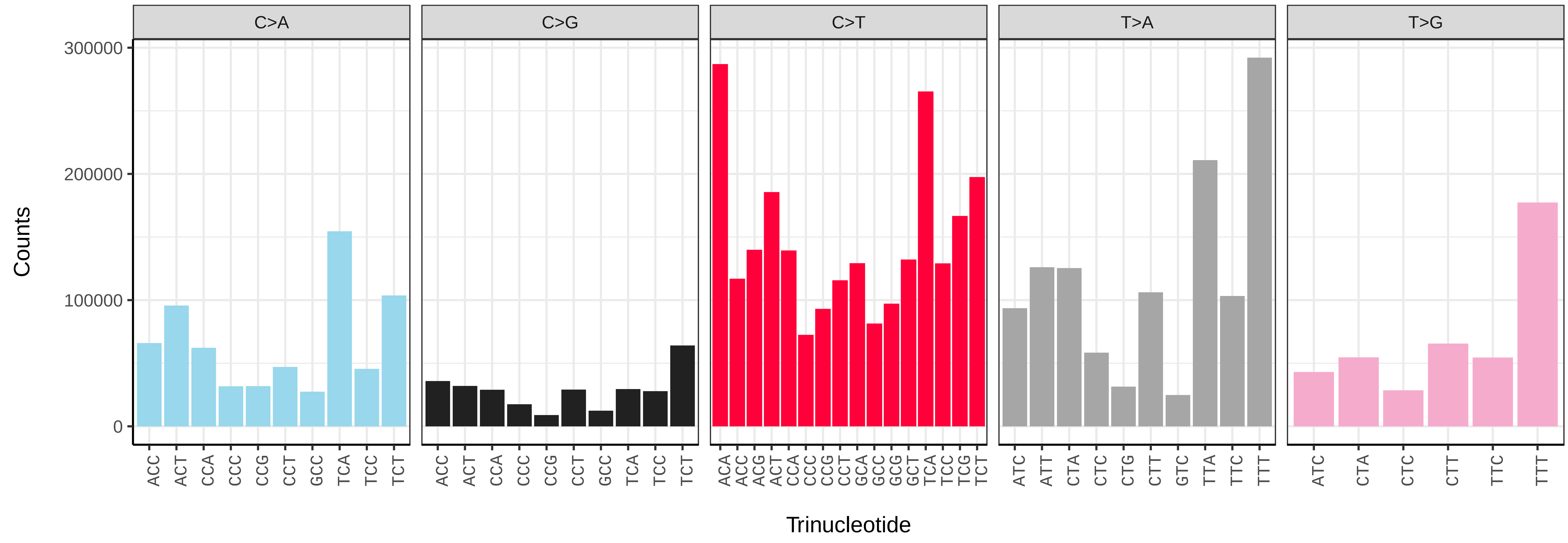


*Lomographa bimaculata* (ilLomBima1) somatic mutational spectrum (SBS96 classification)

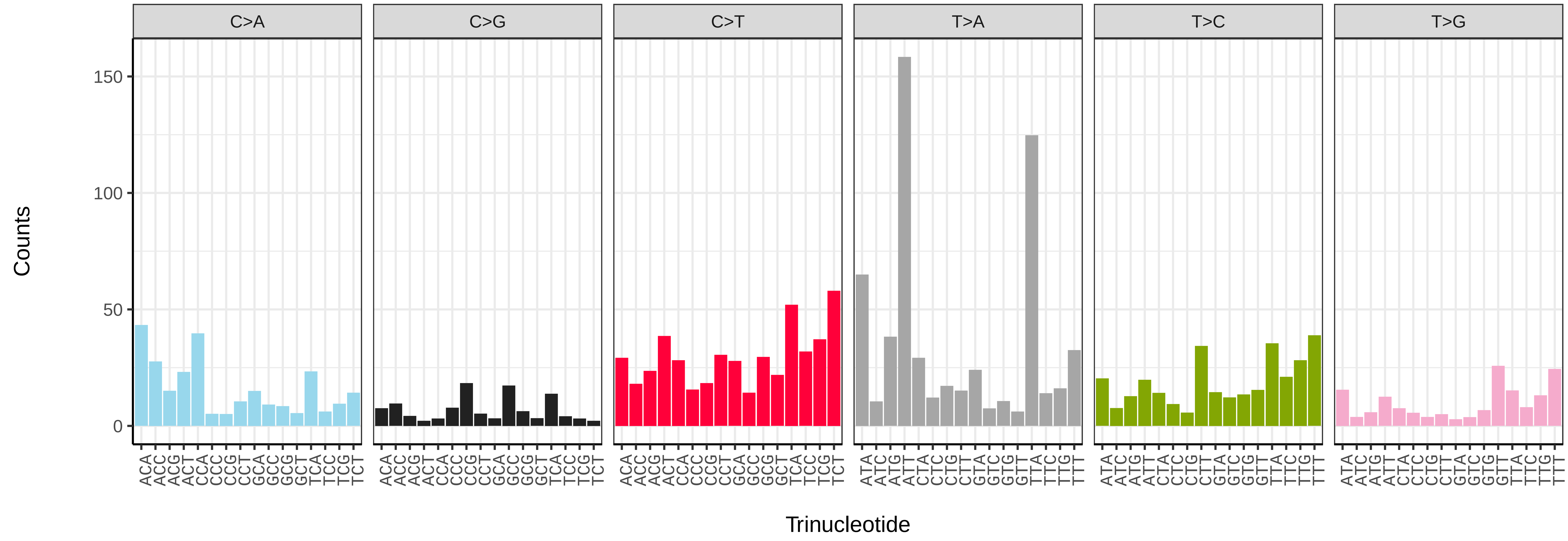


*Lomographa bimaculata* (ilLomBima1) somatic mutational spectrum (SBS52 classification)

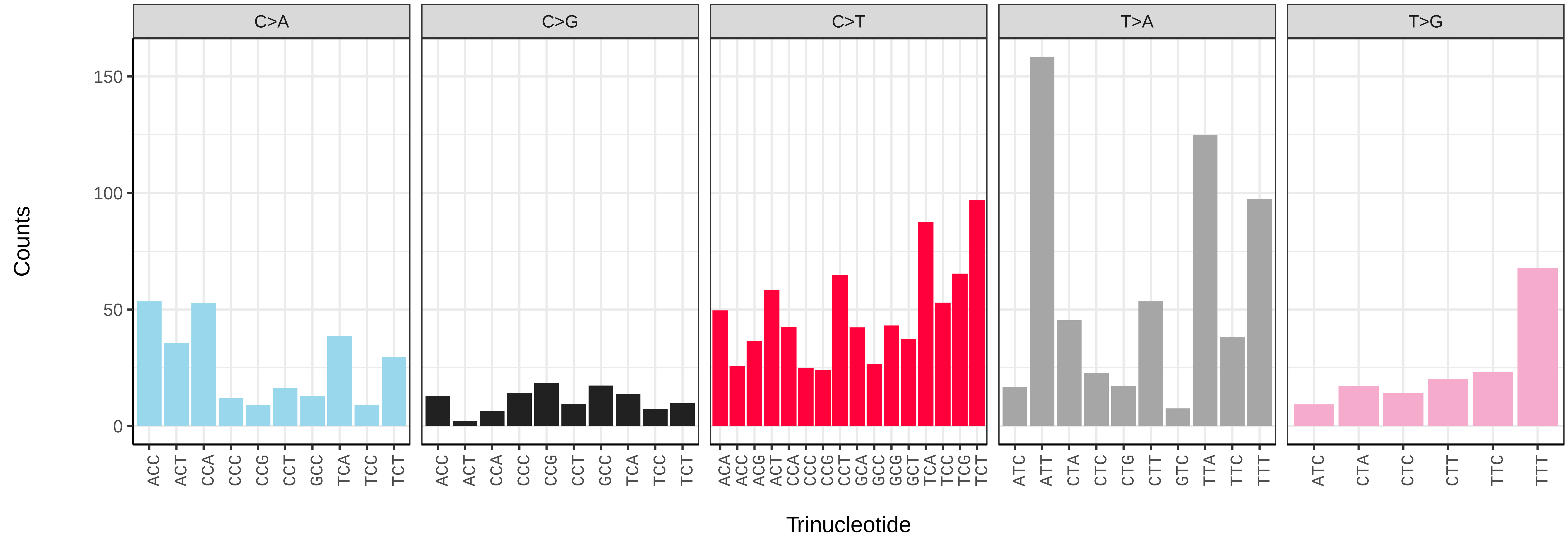

*Luperina testacea* (ilLupTest1) germline mutational spectrum

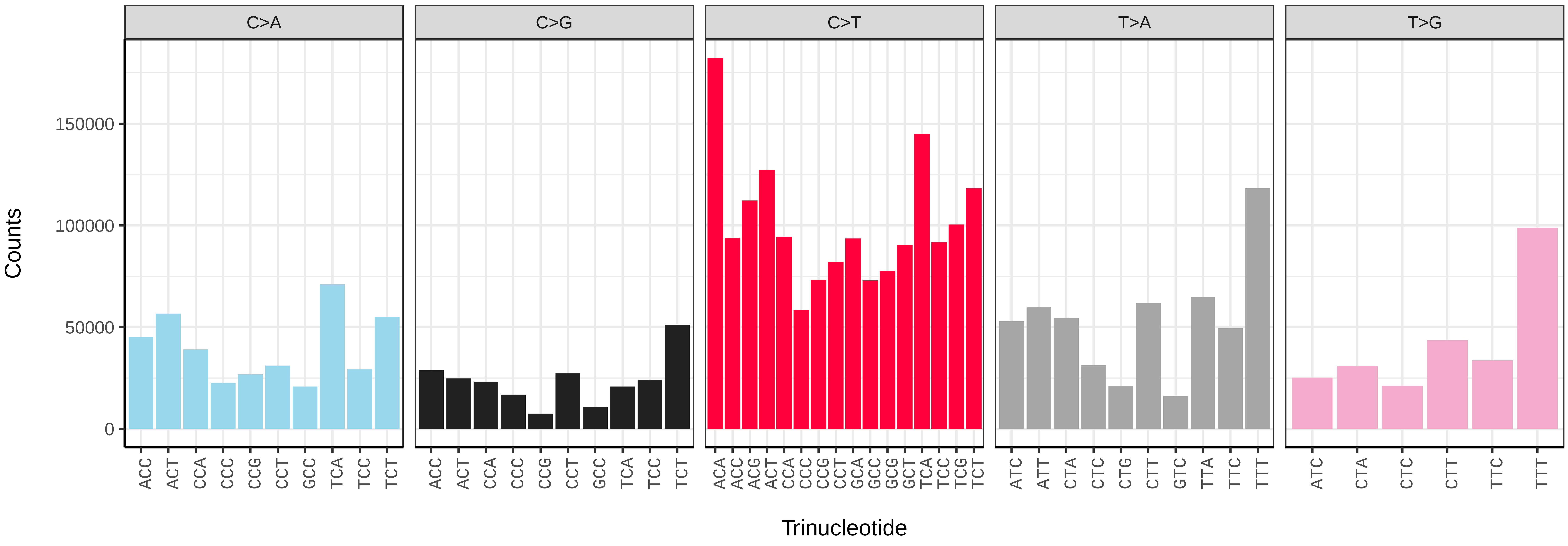


*Luperina testacea* (ilLupTest1) somatic mutational spectrum (SBS96 classification)

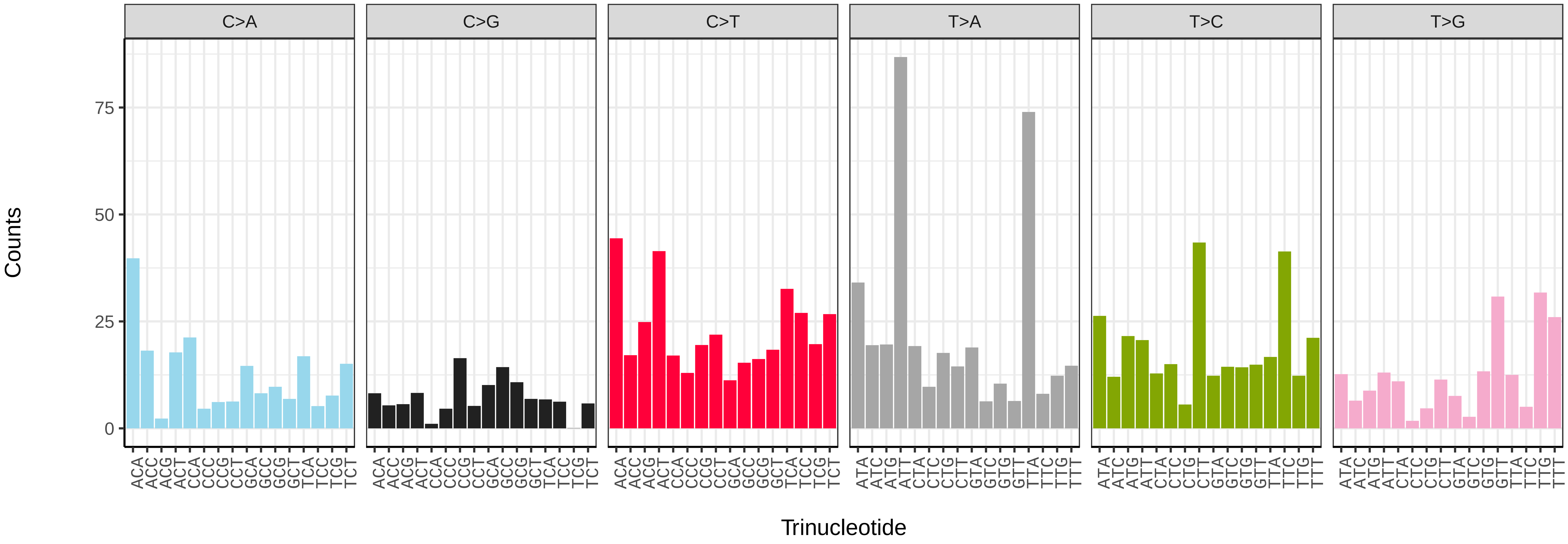


*Luperina testacea* (ilLupTest1) somatic mutational spectrum (SBS52 classification)

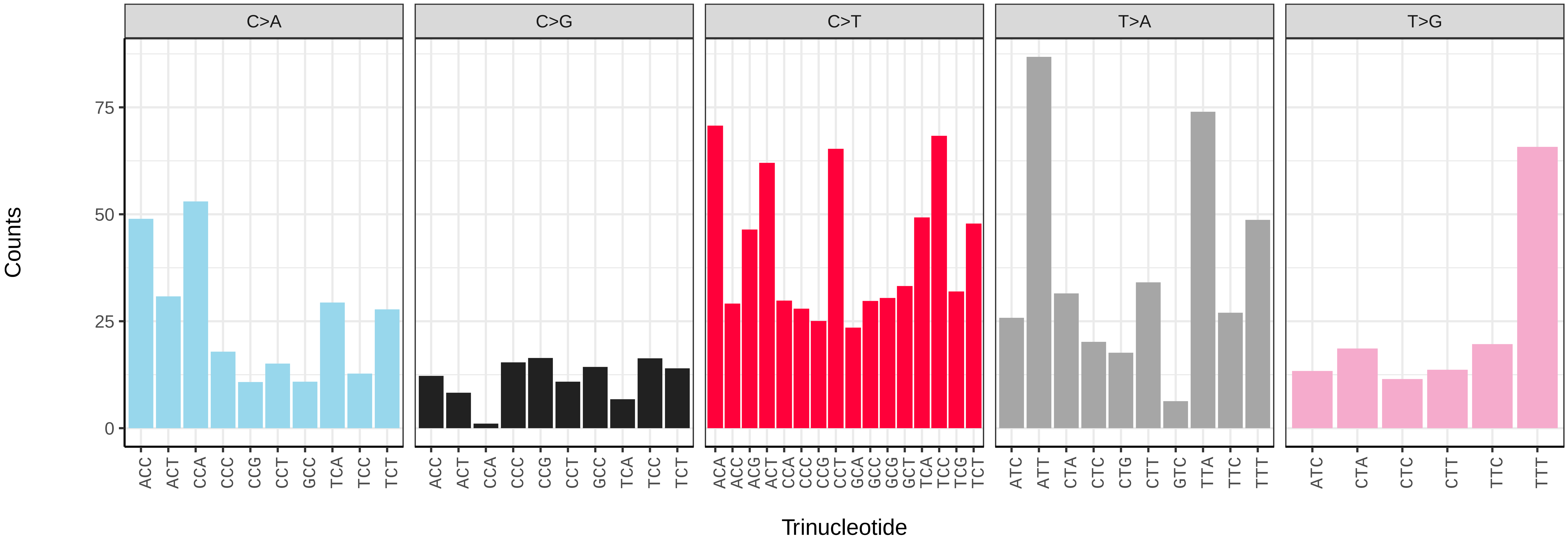

*Lycia hirtaria* (ilLycHirt1) germline mutational spectrum

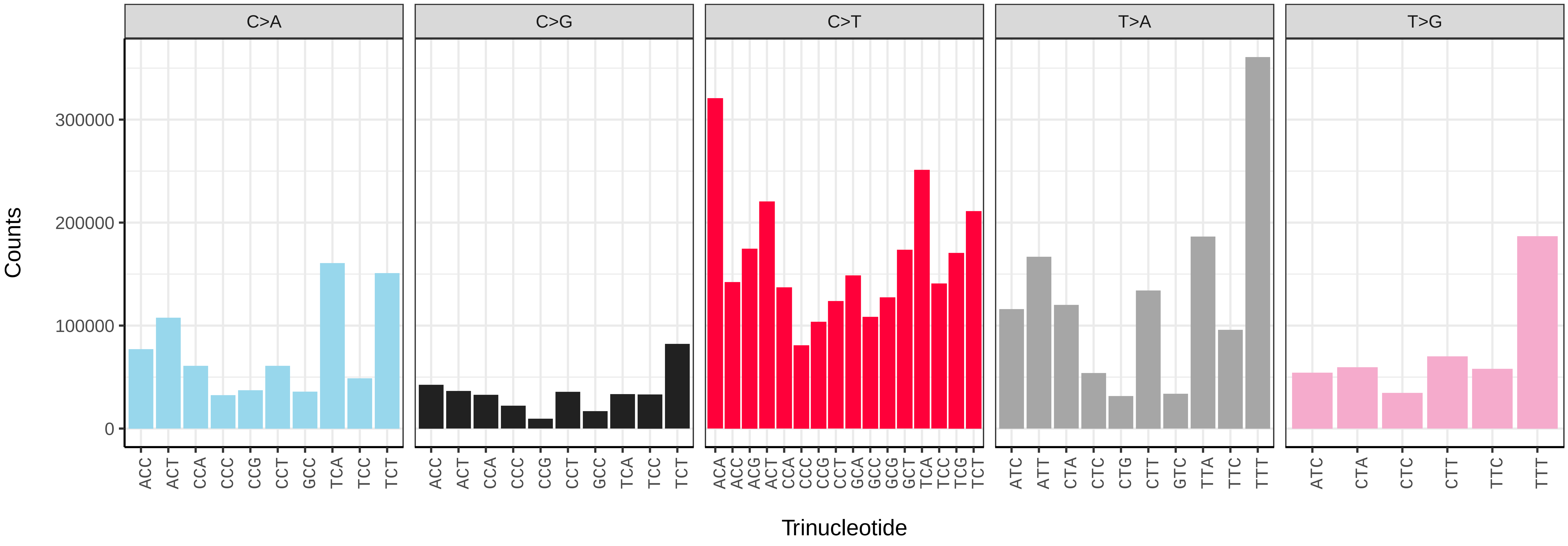


*Lycia hirtaria* (ilLycHirt1) somatic mutational spectrum (SBS96 classification)

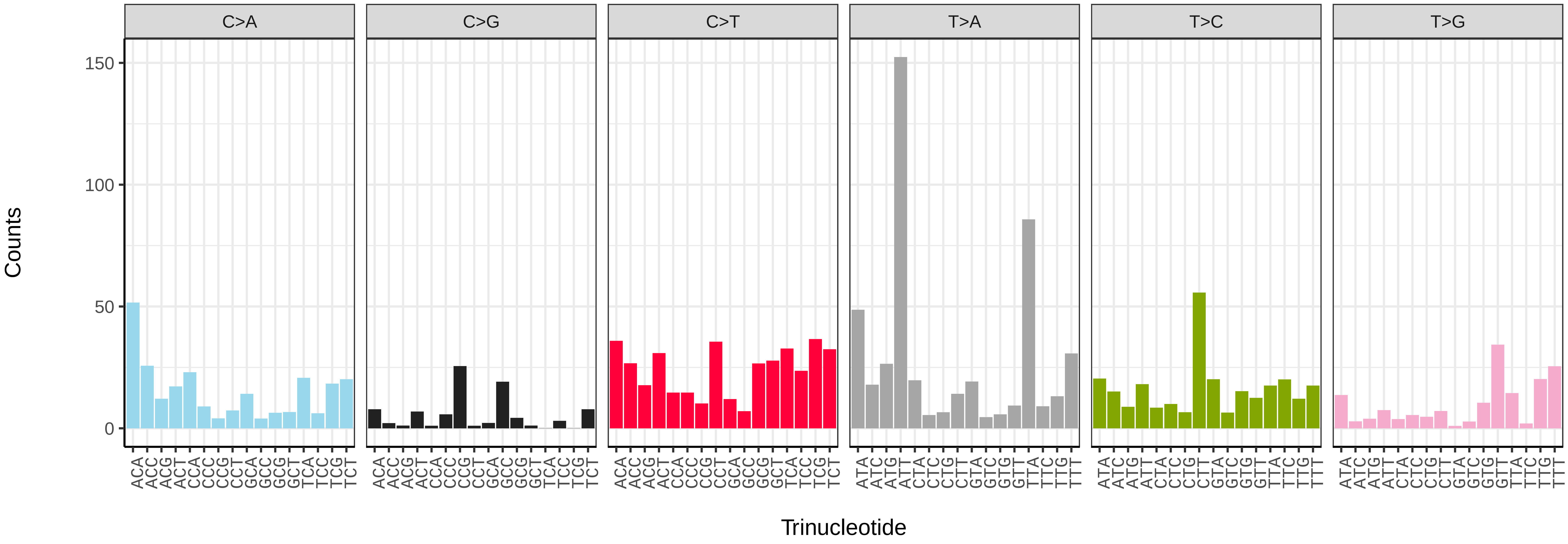


*Lycia hirtaria* (ilLycHirt1) somatic mutational spectrum (SBS52 classification)

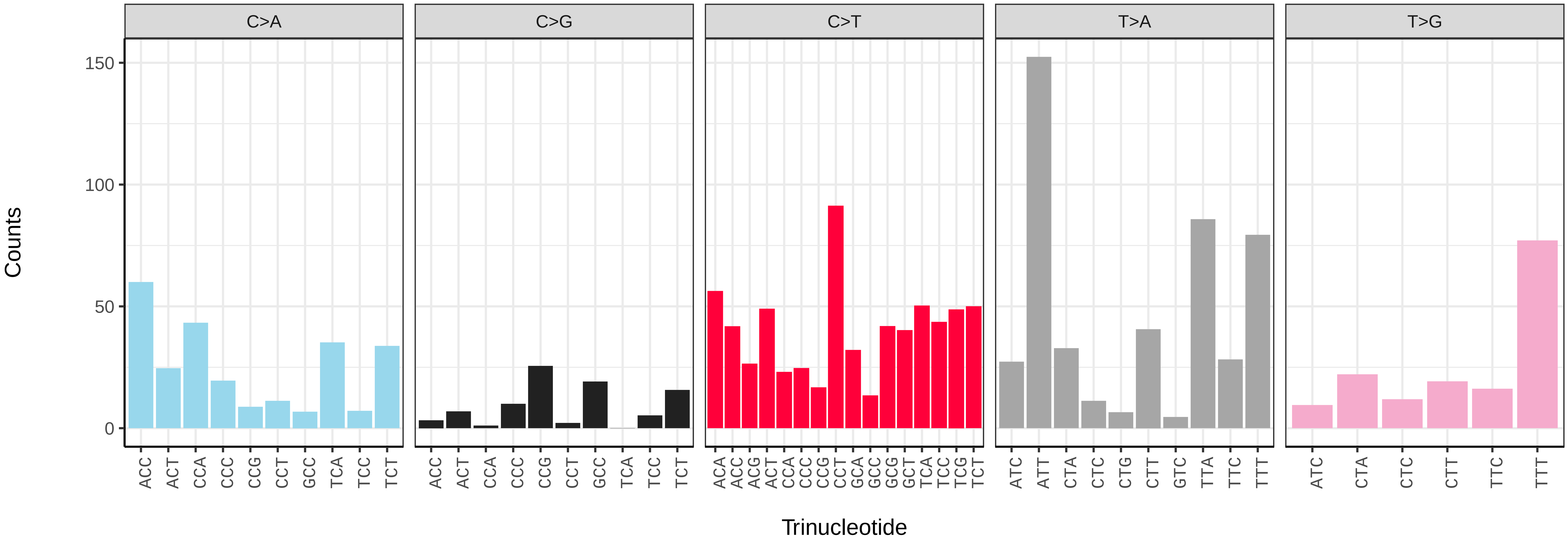

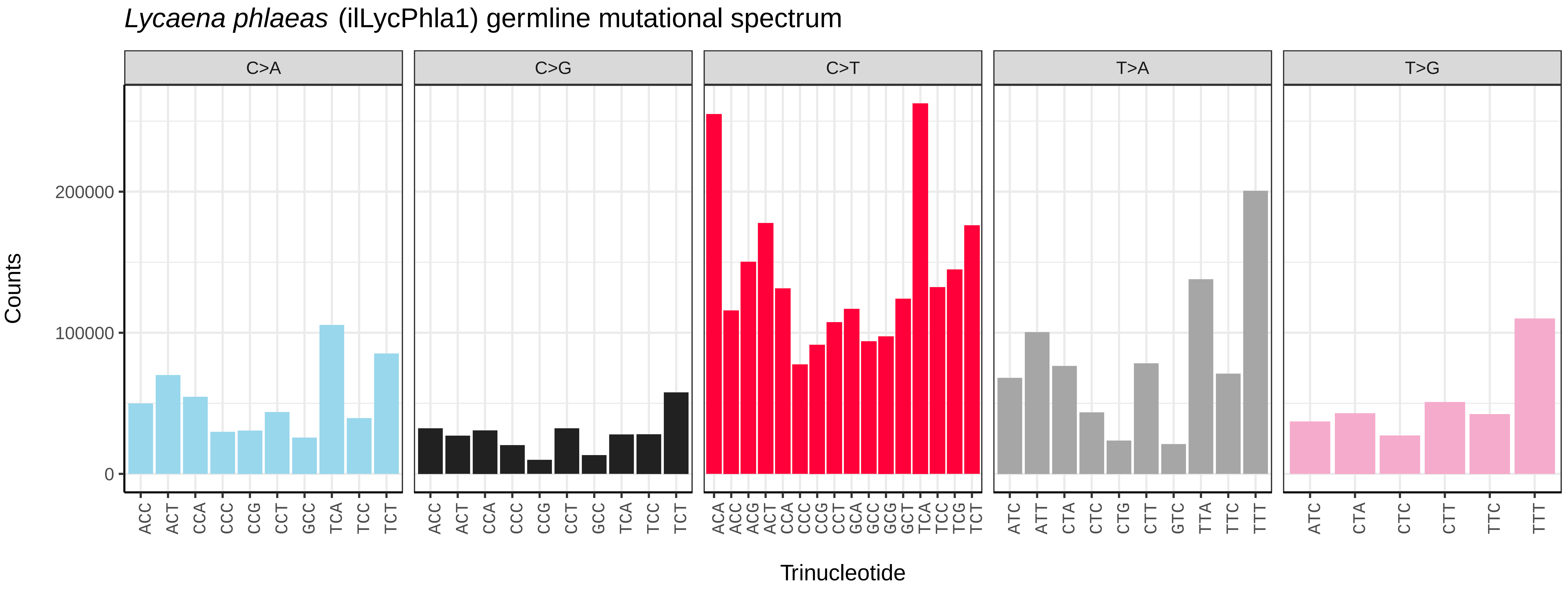
*Lycaena phlaeas* (ilLycPhla1) germline mutational spectrum
C>A
C>G
C>T
T>A
T>G
Counts
0
100000
200000
ACC ACT CCA CCC CCG CCT GCC TCA TCC TCT
ACA ACC ACG ACT CCA CCC CCG CCT GCA GCC GCG GCT TCA TCC TCG TCT
ATC ATT CTA CTC CTG CTT GTC TTA TTC TTT
ATC CTA CTC CTT TTC TTT
Trinucleotide

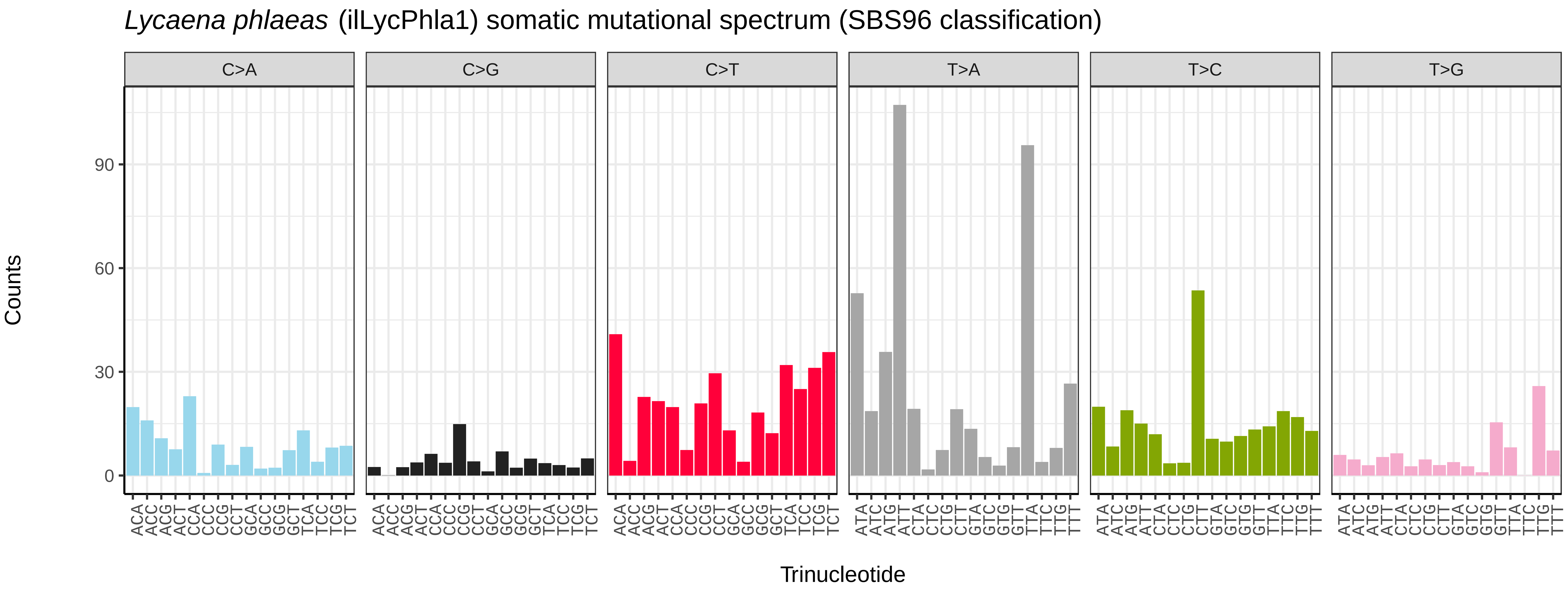
*Lycaena phlaeas* (ilLycPhla1) somatic mutational spectrum (SBS96 classification)
C>A
C>G
C>T
T>A
T>C
T>G
Counts
0
30
60
90
ACA ACC ACG ACT CCA CCC CCG CCT GCA GCC GCG GCT TCA TCC TCG TCT
ATA ATC ATG ATT CTA CTC CTG CTT GTA GTC GTG GTT TTA TTC TTG TTT
Trinucleotide

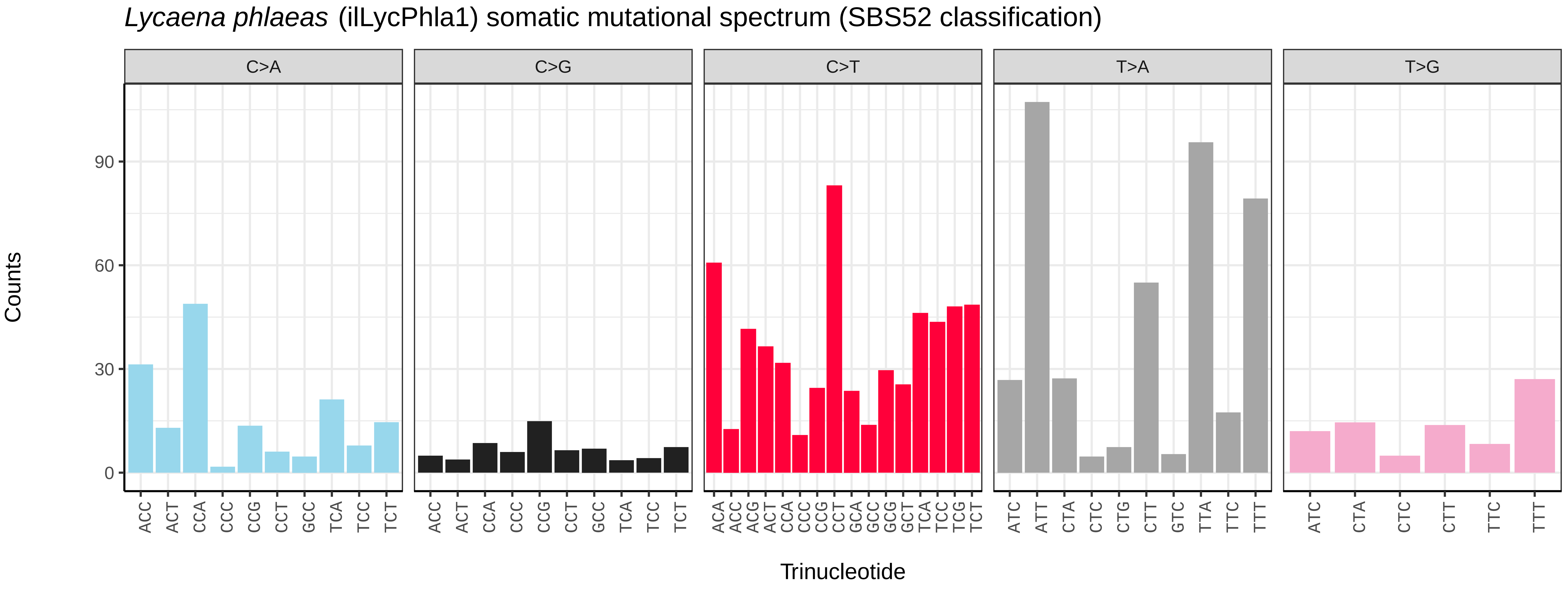
*Lycaena phlaeas* (ilLycPhla1) somatic mutational spectrum (SBS52 classification)
C>A
C>G
C>T
T>A
T>G
Counts
0
30
60
90
ACC ACT CCA CCC CCG CCT GCC TCA TCC TCT
ACA ACC ACG ACT CCA CCC CCG CCT GCA GCC GCG GCT TCA TCC TCG TCT
ATC ATT CTA CTC CTG CTT GTC TTA TTC TTT
ATC CTA CTC CTT TTC TTT
Trinucleotide

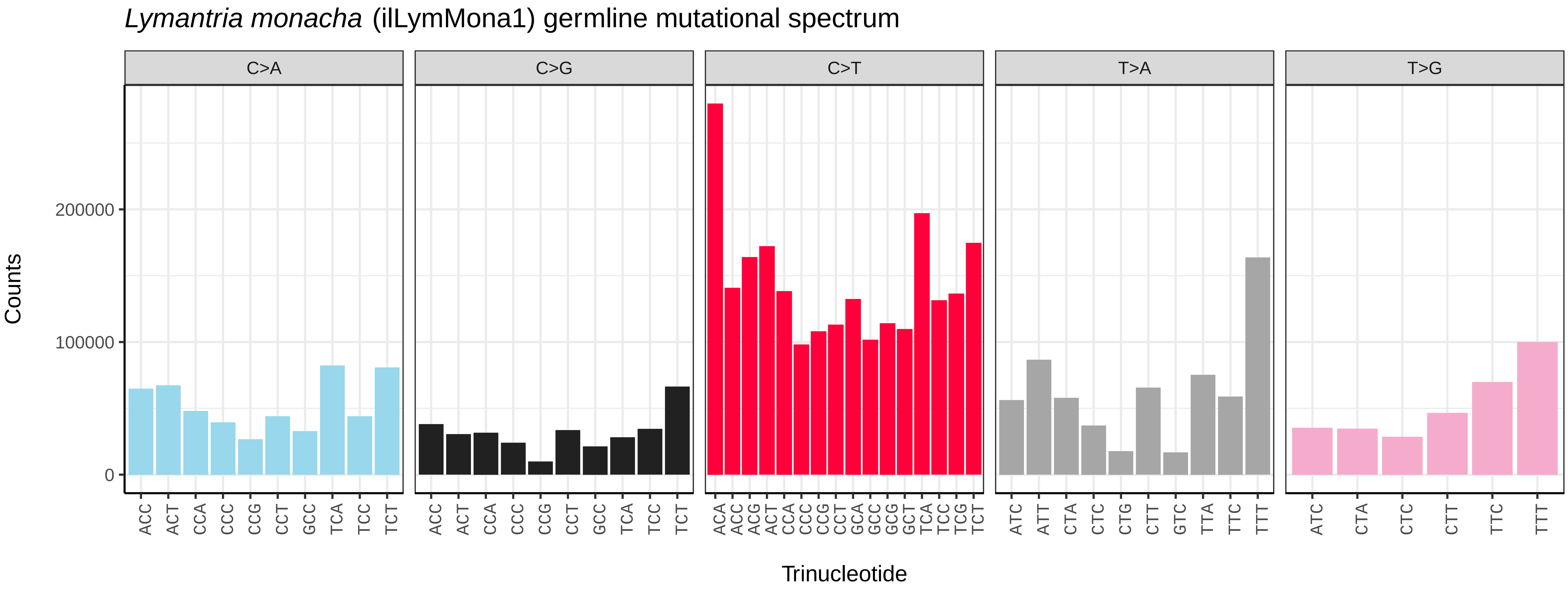
*Lymantria monacha* (ilLymMona1) germline mutational spectrum
C>A
C>G
C>T
T>A
T>G
Counts
200000
100000
0
ACC ACT CCA CCC CCG CCT GCC TCA TCC TCT
ACA ACC ACG ACT CCA CCC CCG CCT GCA GCC GCG GCT TCA TCC TCG TCT
ATC ATT CTA CTC CTG CTT GTC TTA TTC TTT
ATC CTA CTC CTT TTC TTT
Trinucleotide

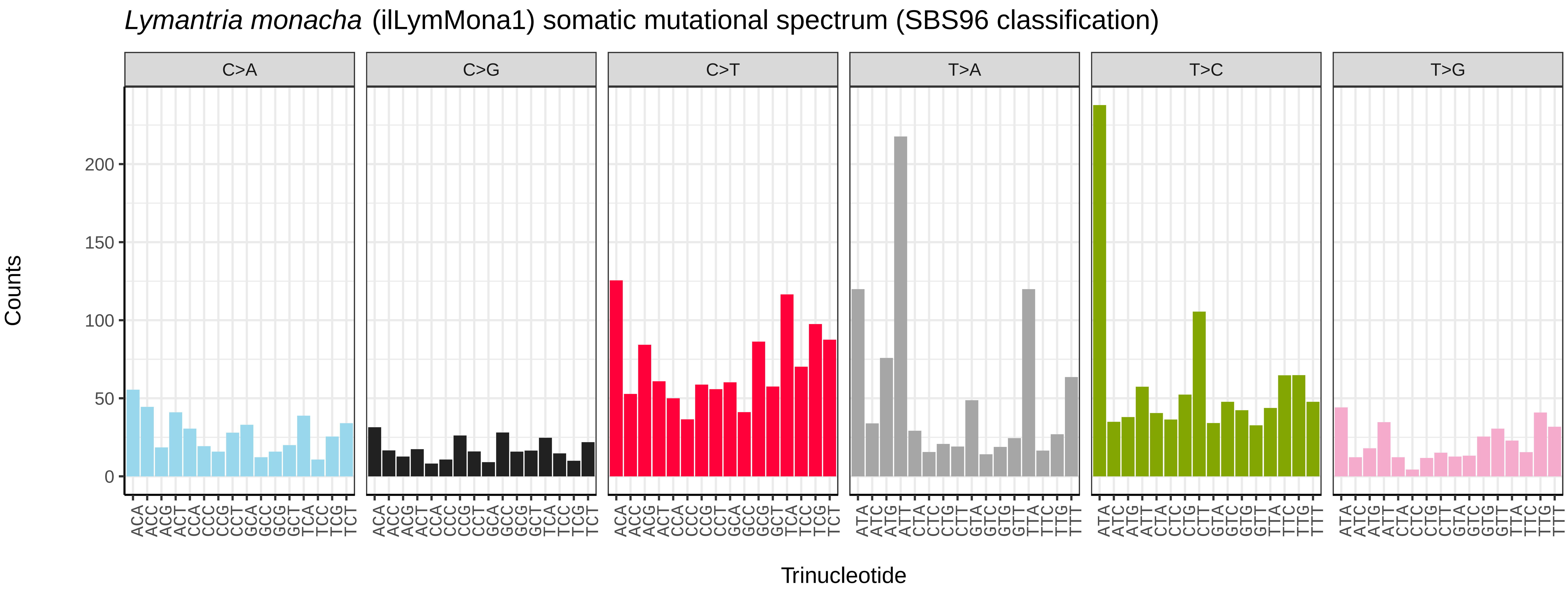
*Lymantria monacha* (ilLymMona1) somatic mutational spectrum (SBS96 classification)
C>A
C>G
C>T
T>A
T>C
T>G
Counts
200
150
100
50
0
ACA ACC ACG ACT CCA CCC CCG CCT GCA GCC GCG GCT TCA TCC TCG TCT
ATA ATC ATG ATT CTA CTC CTG CTT GTA GTC GTG GTT TTA TTC TTG TTT
Trinucleotide

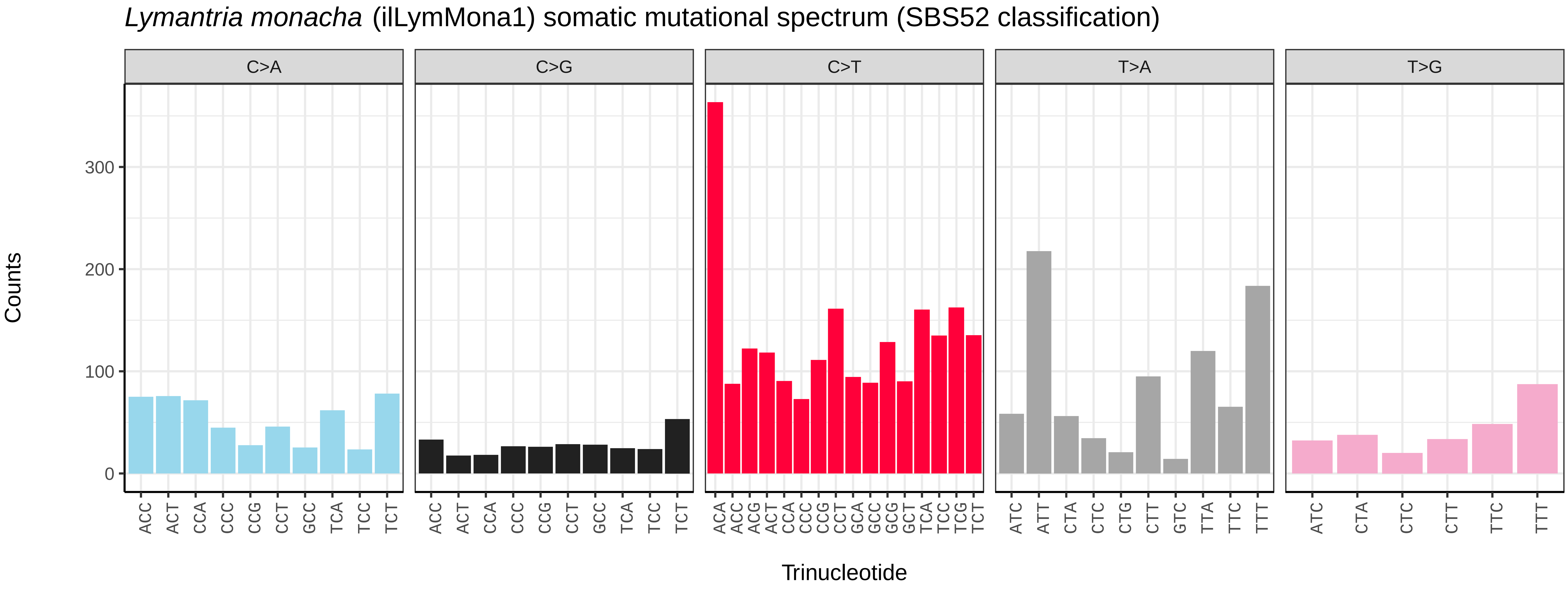
*Lymantria monacha* (ilLymMona1) somatic mutational spectrum (SBS52 classification)
C>A
C>G
C>T
T>A
T>G
Counts
300
200
100
0
ACC ACT CCA CCC CCG CCT GCC TCA TCC TCT
ACA ACC ACG ACT CCA CCC CCG CCT GCA GCC GCG GCT TCA TCC TCG TCT
ATC ATT CTA CTC CTG CTT GTC TTA TTC TTT
ATC CTA CTC CTT TTC TTT
Trinucleotide

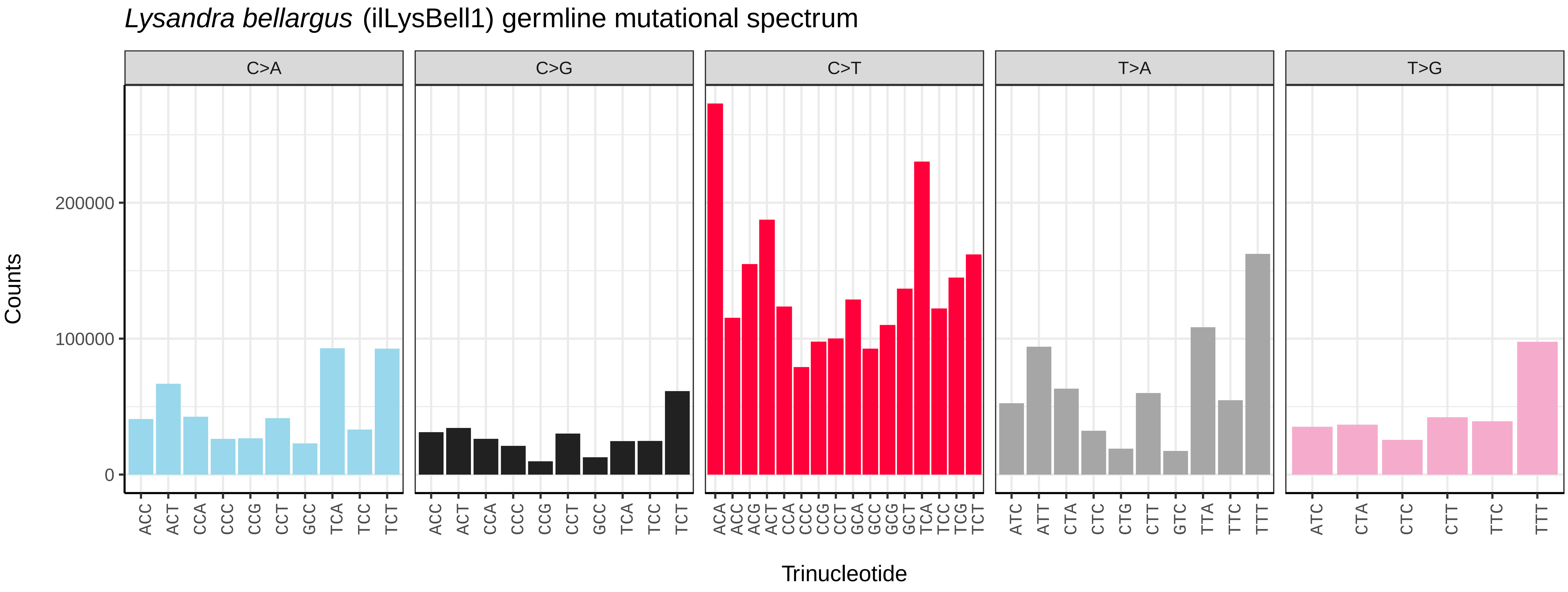
*Lysandra bellargus* (ilLysBell1) germline mutational spectrum
C>A
C>G
C>T
T>A
T>G
Counts
0
100000
200000
Trinucleotide

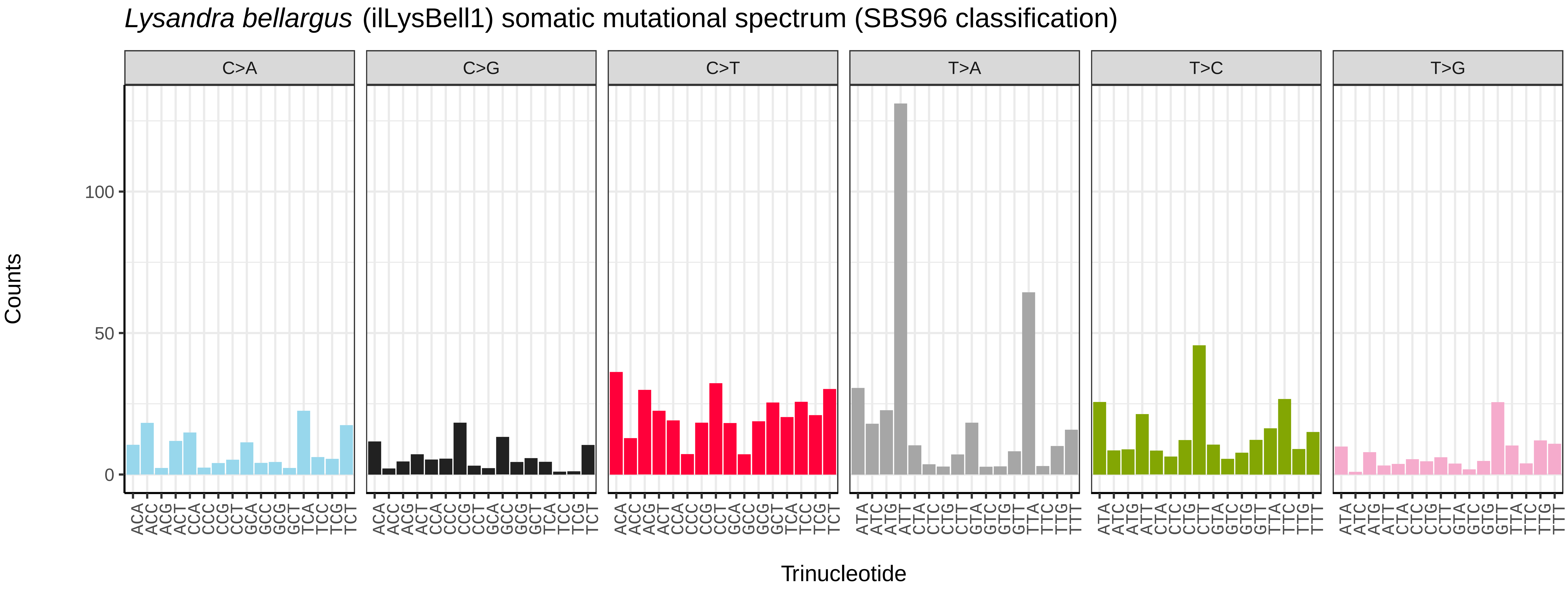
*Lysandra bellargus* (ilLysBell1) somatic mutational spectrum (SBS96 classification)
C>A
C>G
C>T
T>A
T>C
T>G
Counts
0
50
100
Trinucleotide

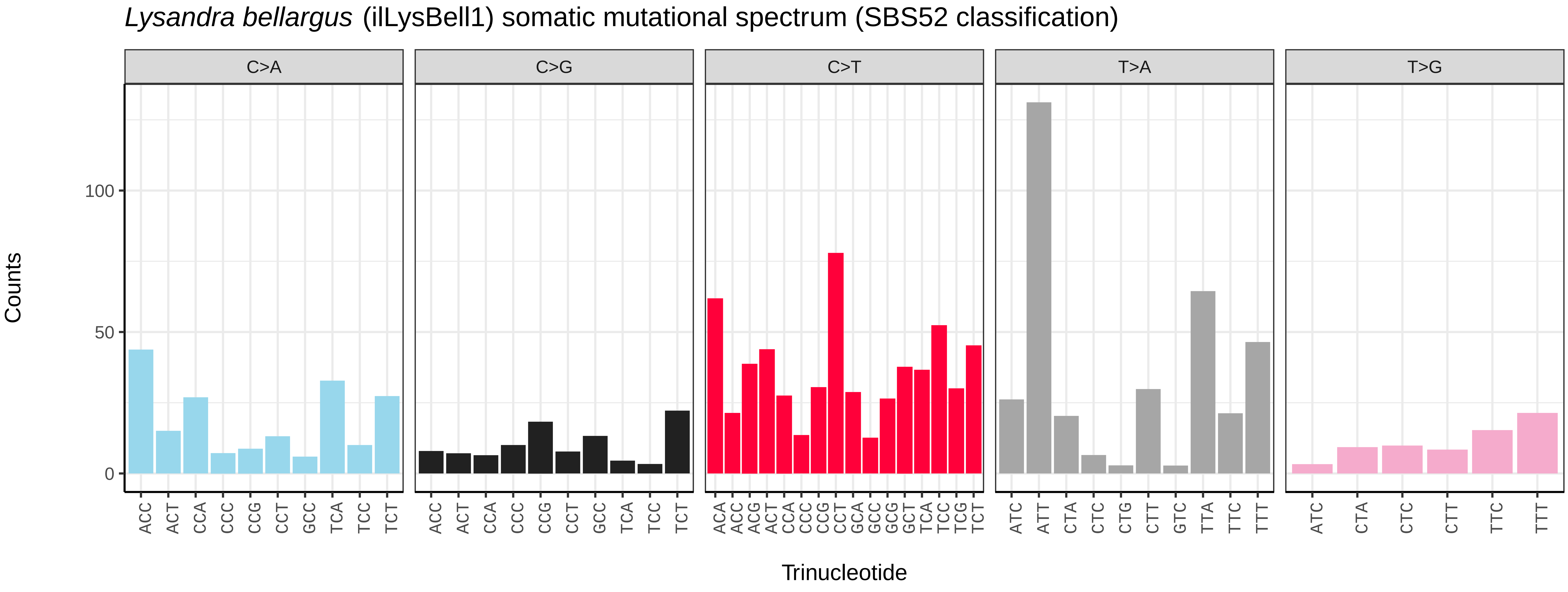
*Lysandra bellargus* (ilLysBell1) somatic mutational spectrum (SBS52 classification)
C>A
C>G
C>T
T>A
T>G
Counts
0
50
100
Trinucleotide

*Lysandra coridon* (ilLysCori1) germline mutational spectrum

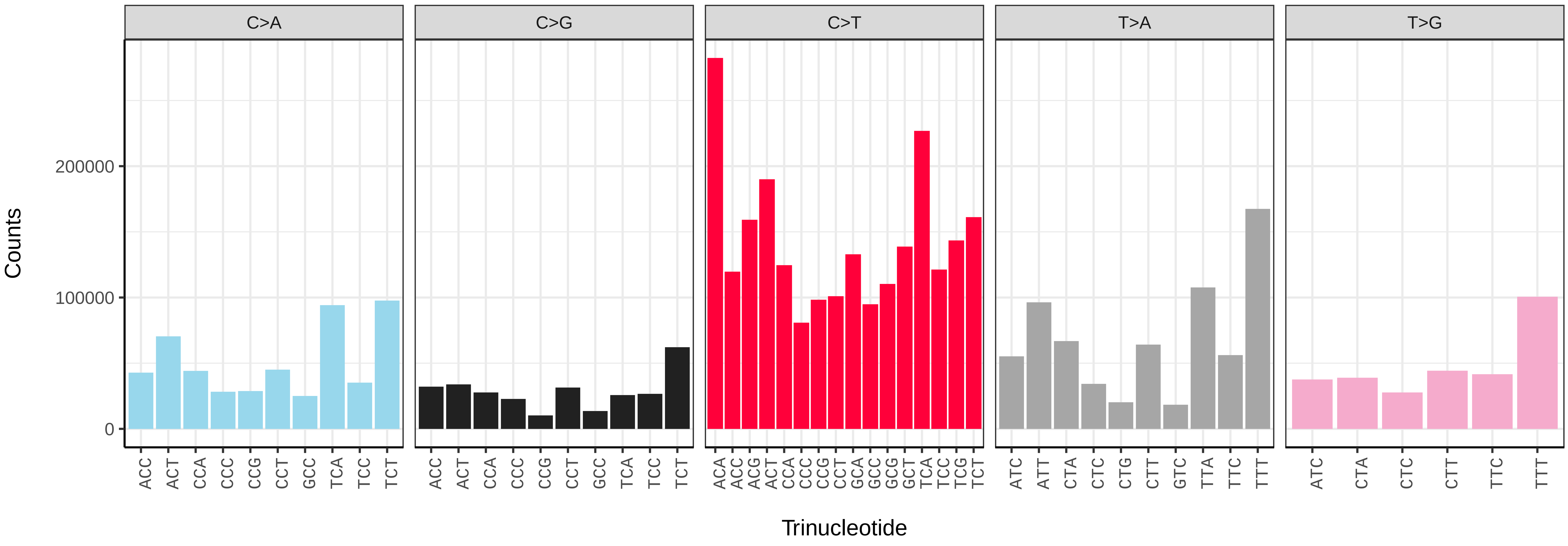


*Lysandra coridon* (ilLysCori1) somatic mutational spectrum (SBS96 classification)

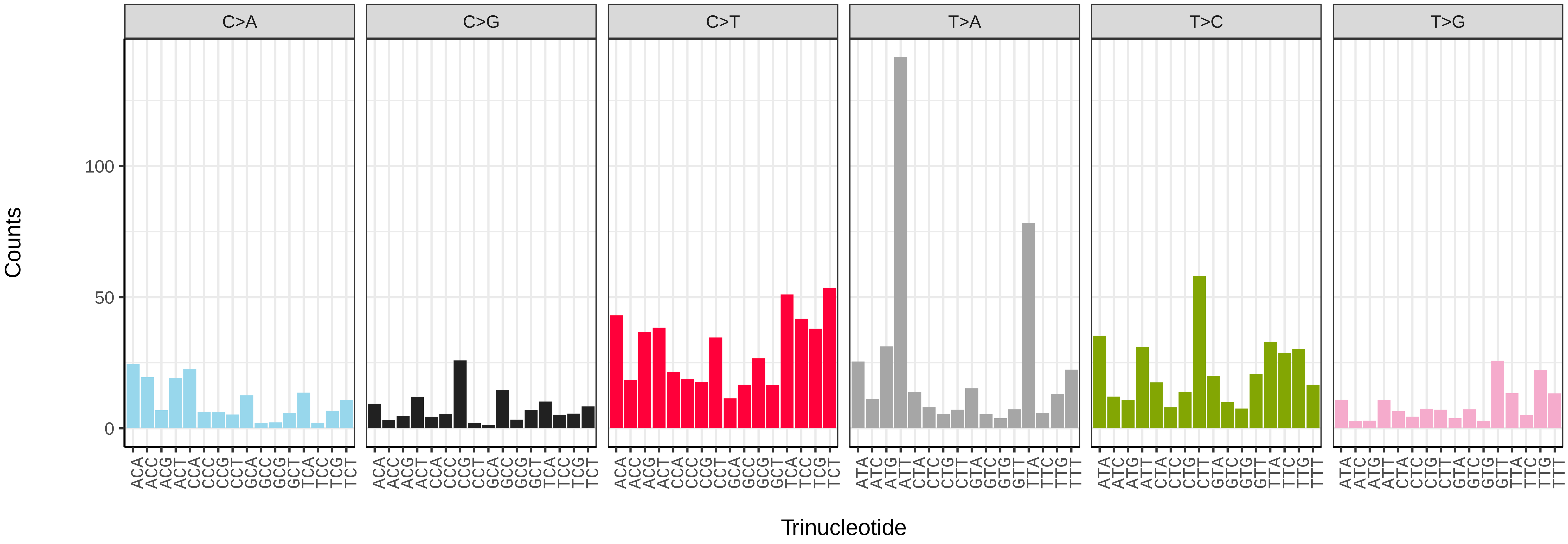


*Lysandra coridon* (ilLysCori1) somatic mutational spectrum (SBS52 classification)

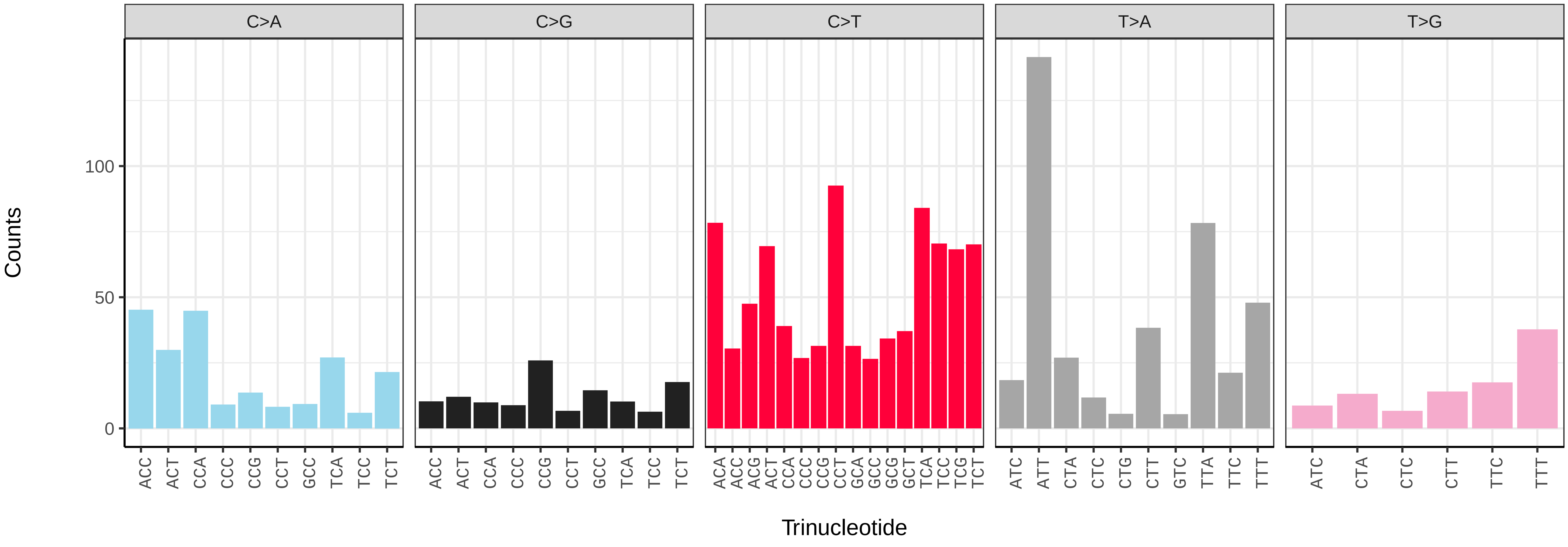

*Macaria notata* (ilMacNota1) germline mutational spectrum

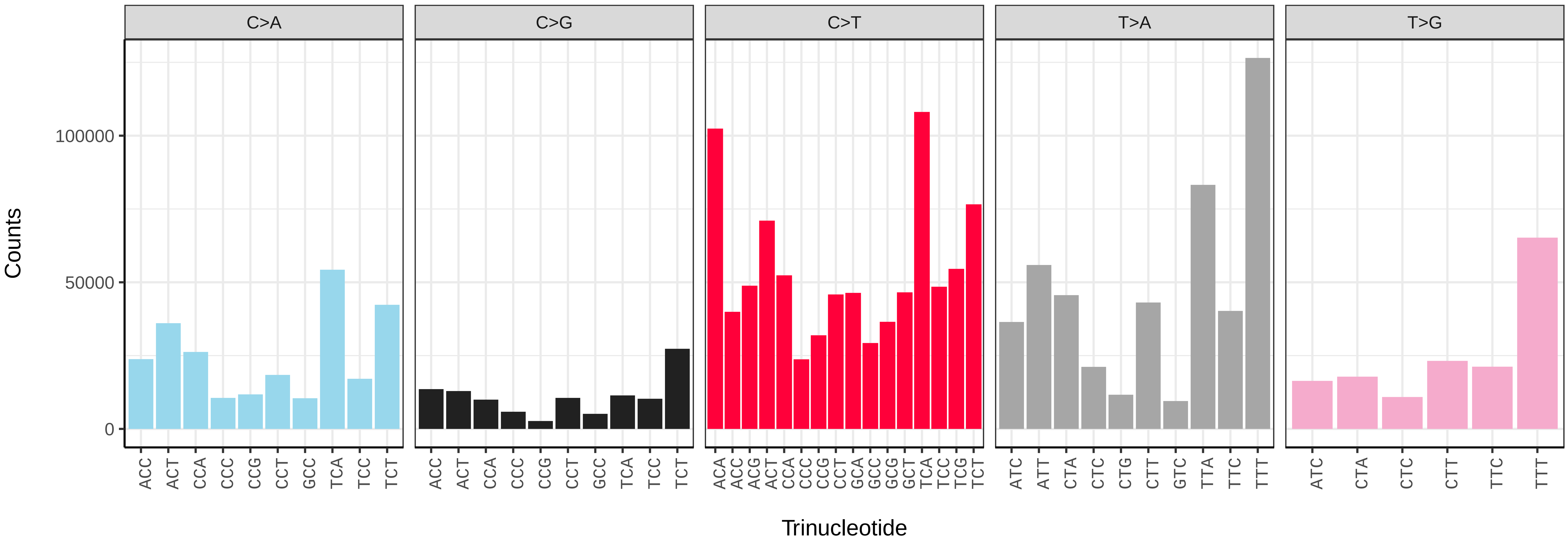


*Macaria notata* (ilMacNota1) somatic mutational spectrum (SBS96 classification)

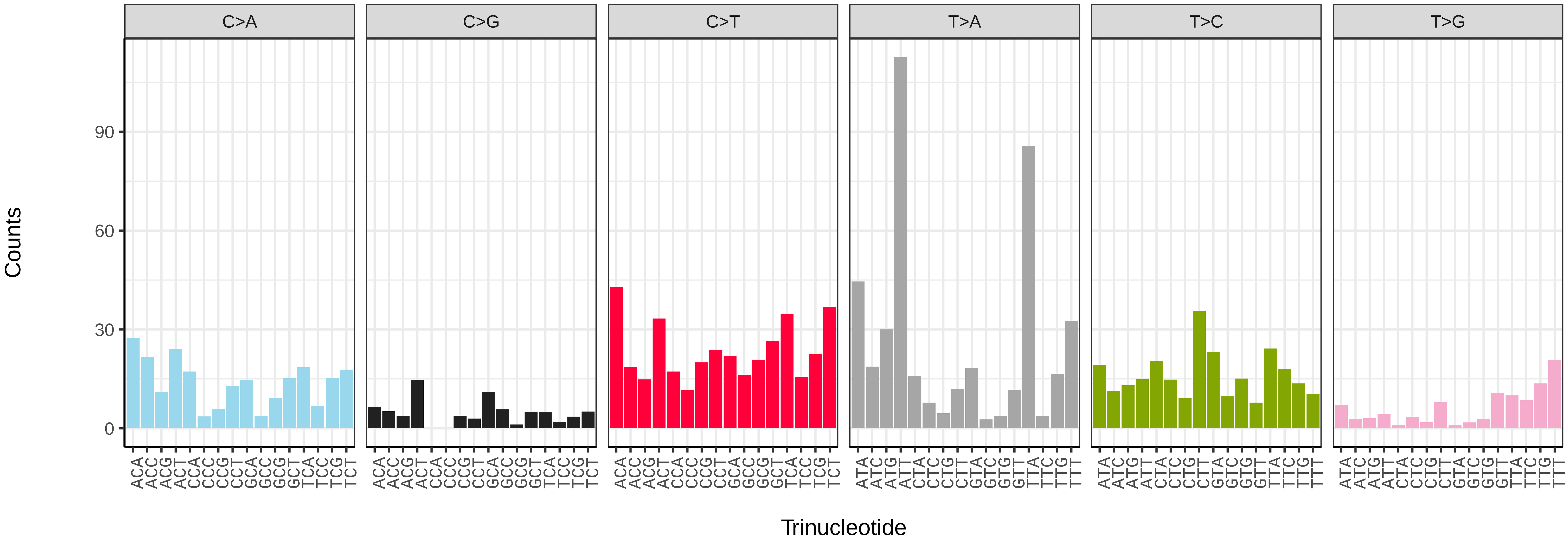


*Macaria notata* (ilMacNota1) somatic mutational spectrum (SBS52 classification)

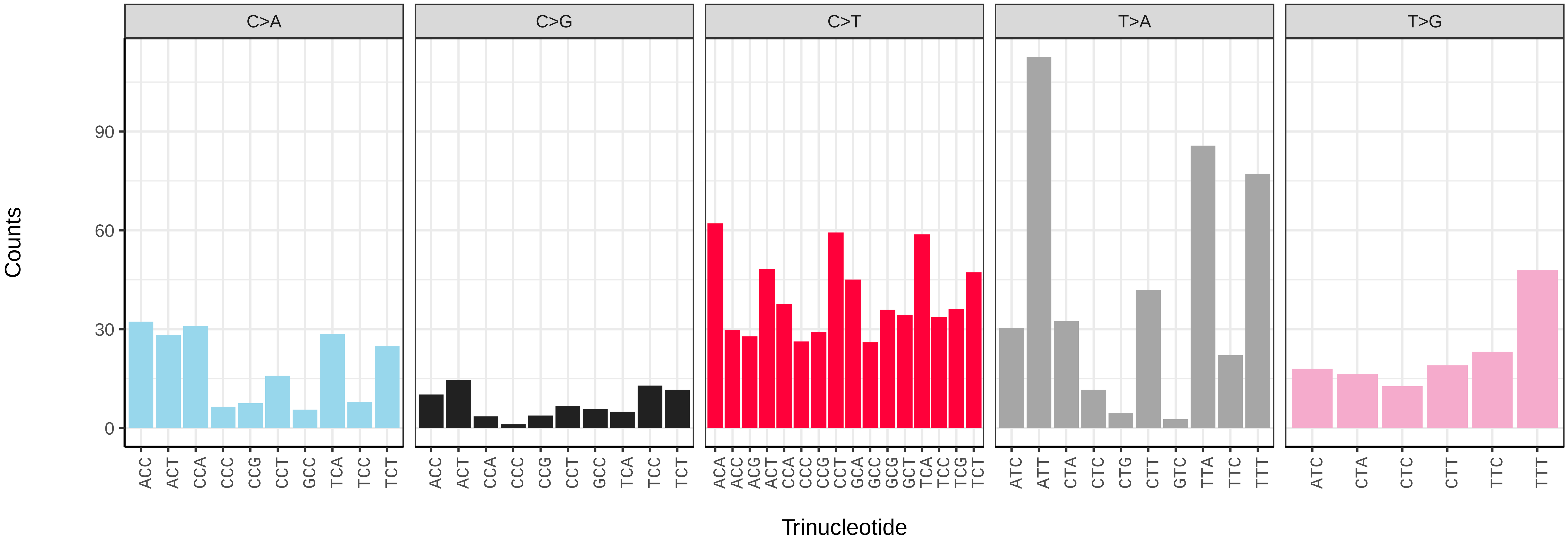

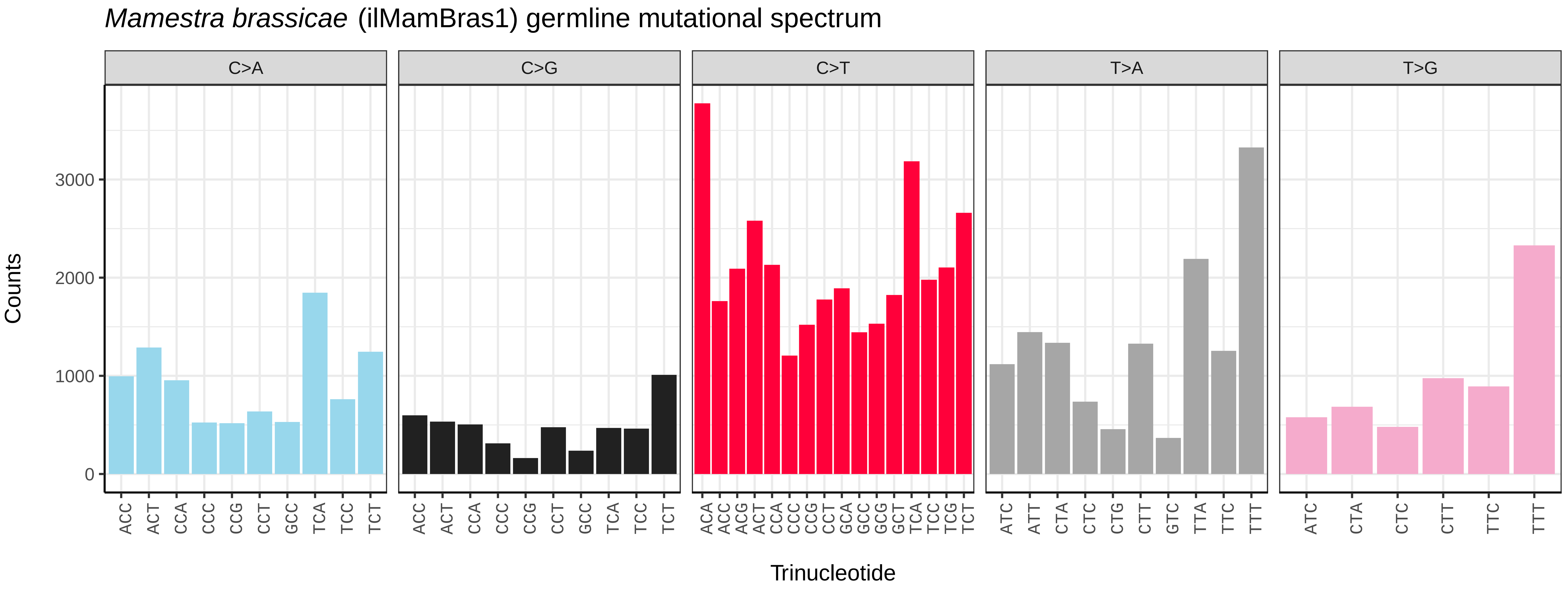
*Mamestra brassicae* (ilMamBras1) germline mutational spectrum
C>A
C>G
C>T
T>A
T>G
Counts
0
1000
2000
3000
ACC ACT CCA CCC CCG CCT GCC TCA TCC TCT
ACC ACT CCA CCC CCG CCT GCC TCA TCC TCT
ACA ACC ACG ACT CCA CCC CCG CCT GCA GCC GCG GCT TCA TCC TCG TCT
ATC ATT CTA CTC CTG CTT GTC TTA TTC TTT
ATC CTA CTC CTT TTC TTT
Trinucleotide

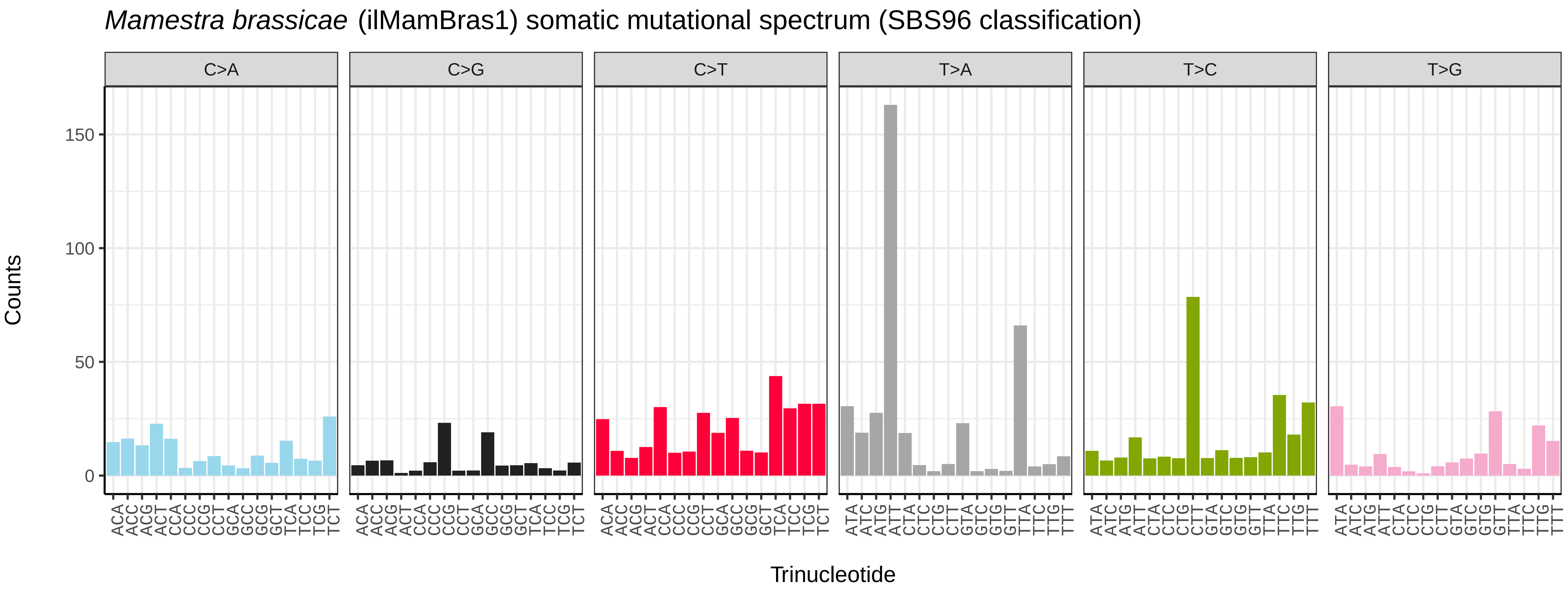
*Mamestra brassicae* (ilMamBras1) somatic mutational spectrum (SBS96 classification)
C>A
C>G
C>T
T>A
T>C
T>G
Counts
0
50
100
150
ACA ACC ACG ACT CCA CCC CCG CCT GCA GCC GCG GCT TCA TCC TCG TCT
ACA ACC ACG ACT CCA CCC CCG CCT GCA GCC GCG GCT TCA TCC TCG TCT
ACA ACC ACG ACT CCA CCC CCG CCT GCA GCC GCG GCT TCA TCC TCG TCT
ATA ATC ATG ATT CTA CTC CTG CTT GTA GTC GTG GTT TTA TTC TTG TTT
ATA ATC ATG ATT CTA CTC CTG CTT GTA GTC GTG GTT TTA TTC TTG TTT
ATA ATC ATG ATT CTA CTC CTG CTT GTA GTC GTG GTT TTA TTC TTG TTT
Trinucleotide

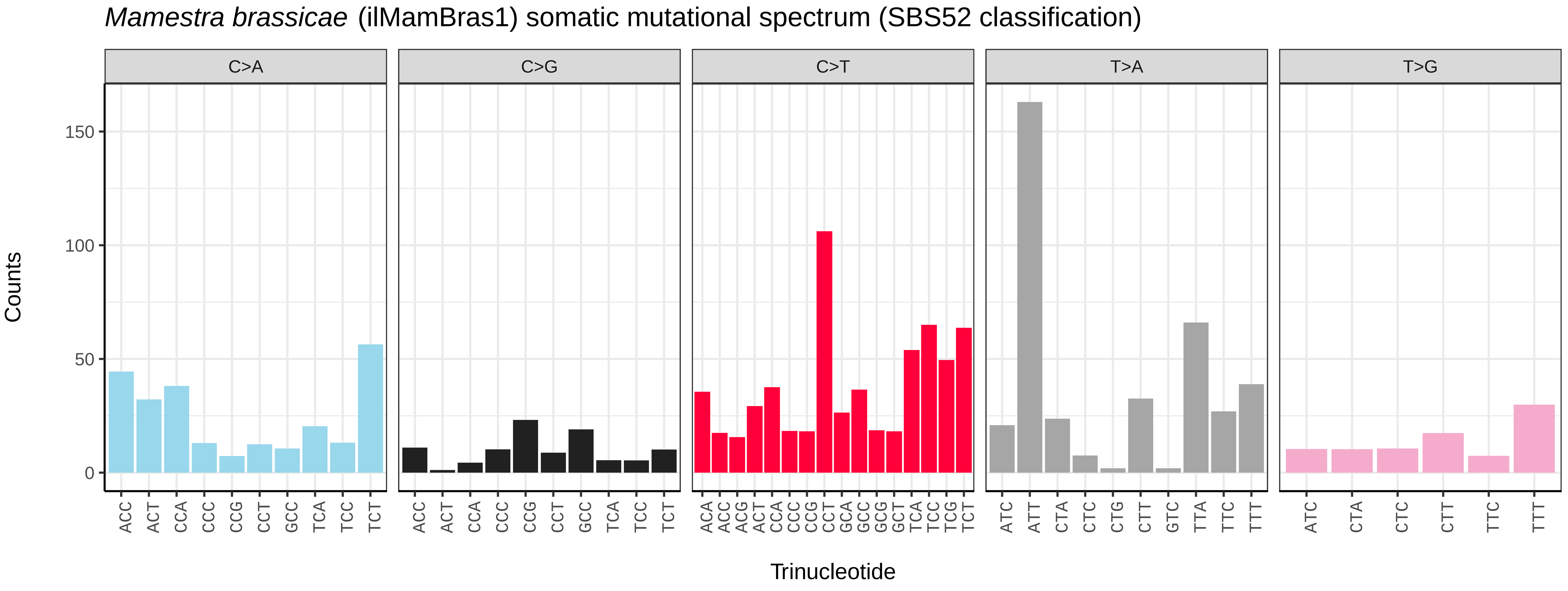
*Mamestra brassicae* (ilMamBras1) somatic mutational spectrum (SBS52 classification)
C>A
C>G
C>T
T>A
T>G
Counts
0
50
100
150
ACC ACT CCA CCC CCG CCT GCC TCA TCC TCT
ACC ACT CCA CCC CCG CCT GCC TCA TCC TCT
ACA ACC ACG ACT CCA CCC CCG CCT GCA GCC GCG GCT TCA TCC TCG TCT
ATC ATT CTA CTC CTG CTT GTC TTA TTC TTT
ATC CTA CTC CTT TTC TTT
Trinucleotide

*Maniola jurtina* (ilManJurt1) germline mutational spectrum

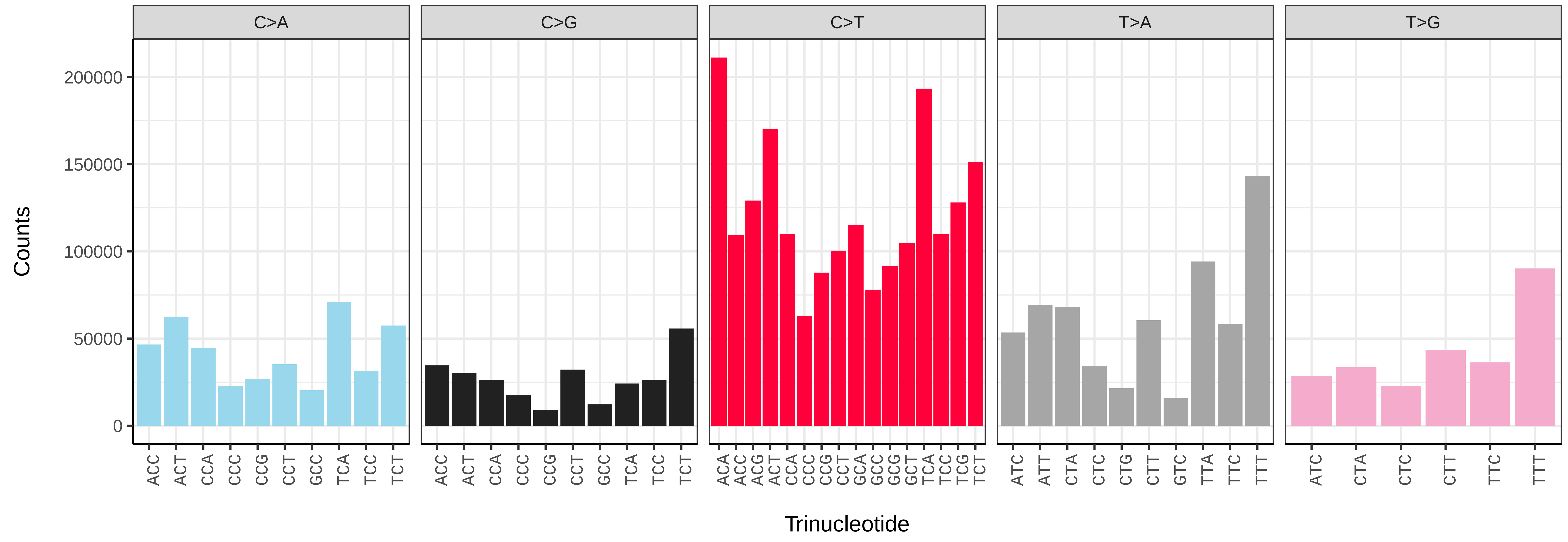


*Maniola jurtina* (ilManJurt1) somatic mutational spectrum (SBS96 classification)

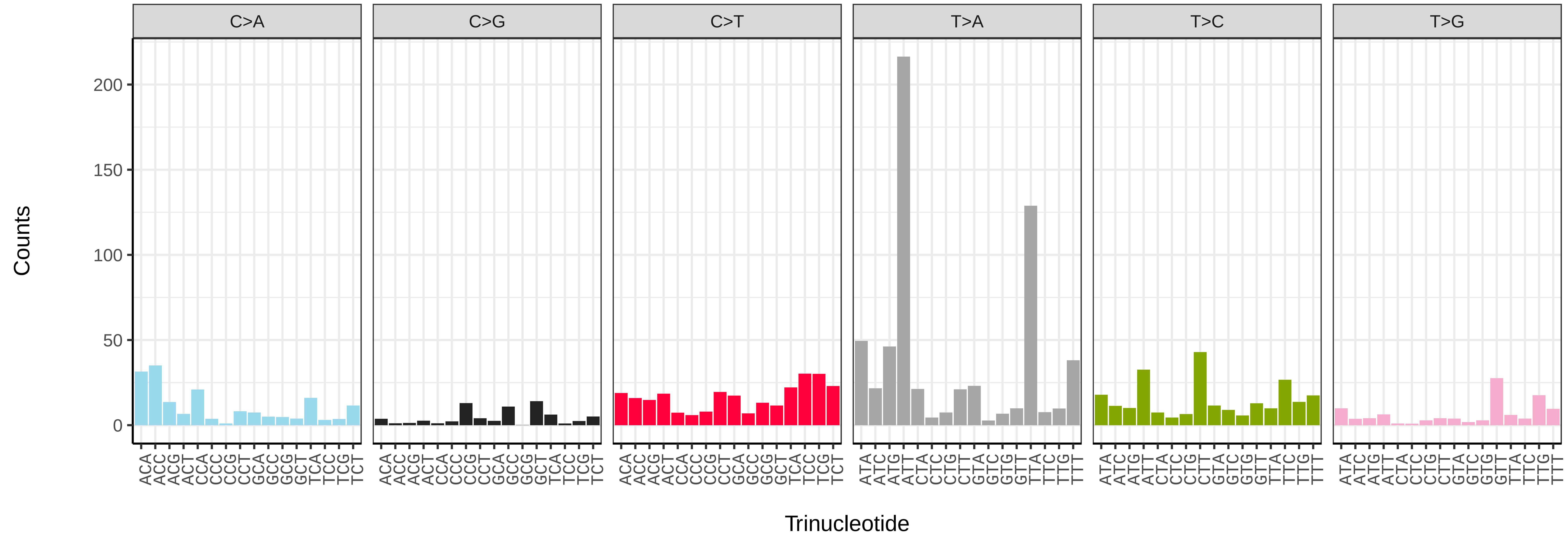


*Maniola jurtina* (ilManJurt1) somatic mutational spectrum (SBS52 classification)

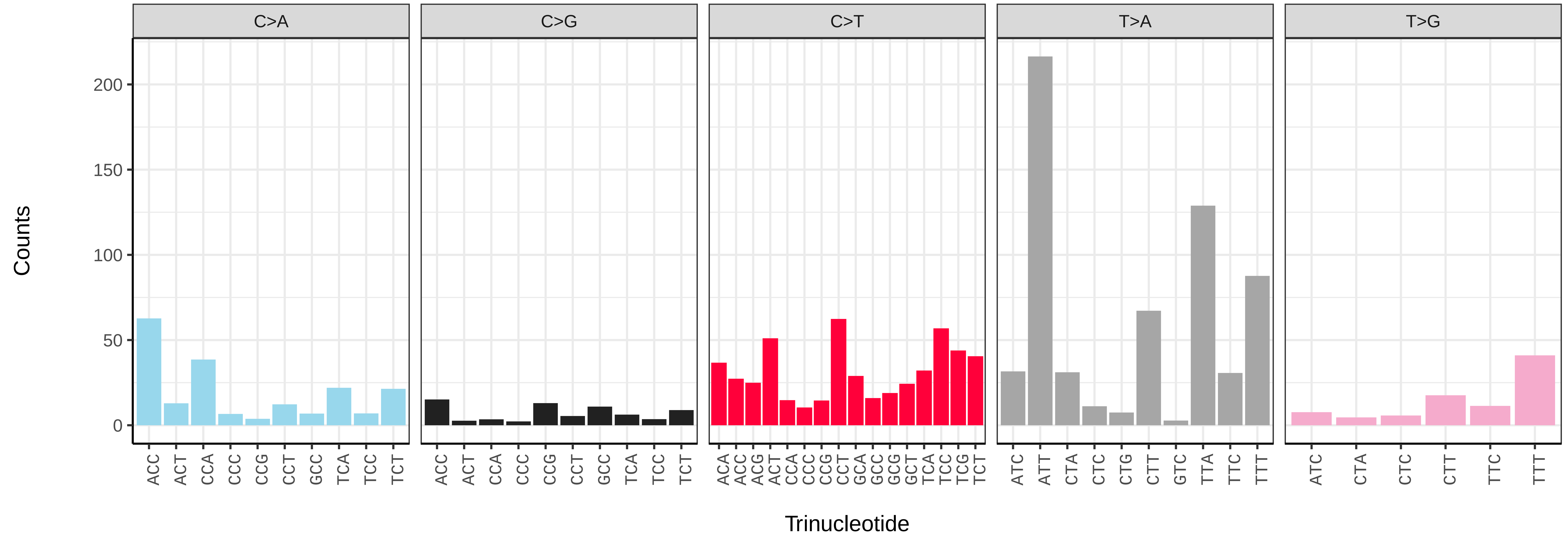

*Maniola jurtina* (ilManJurt1.ilManJurt4) germline mutational spectrum

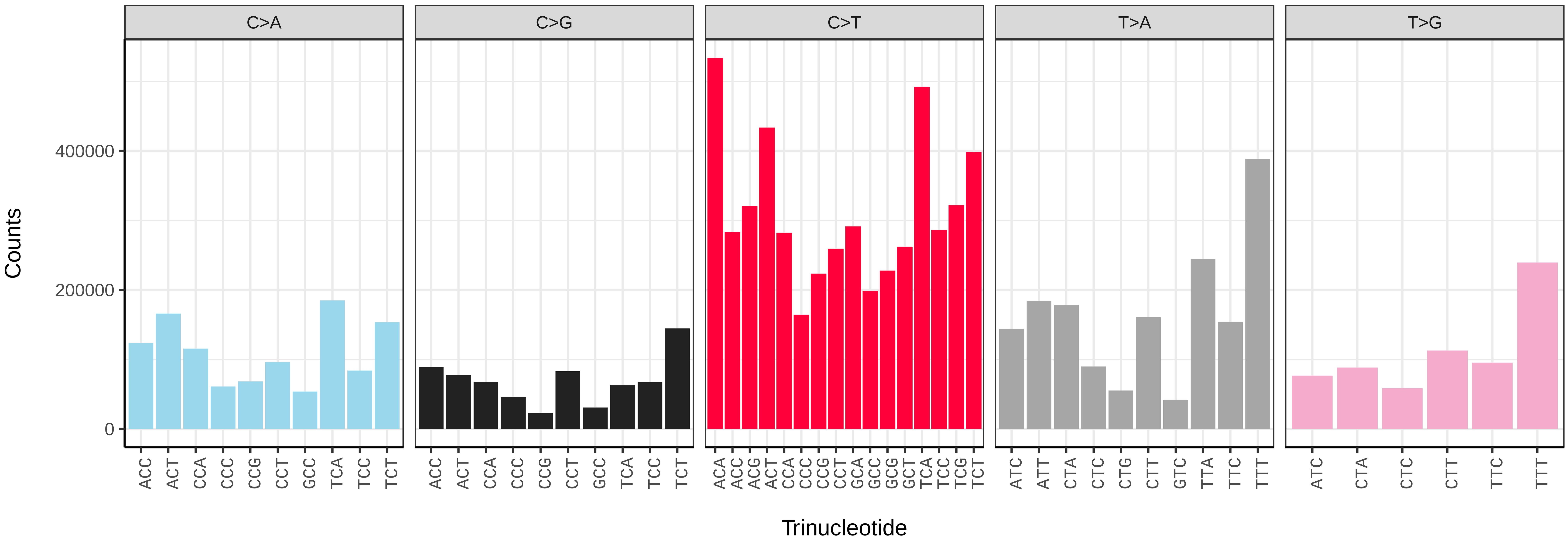


*Maniola jurtina* (ilManJurt1.ilManJurt4) somatic mutational spectrum (SBS96 classification)

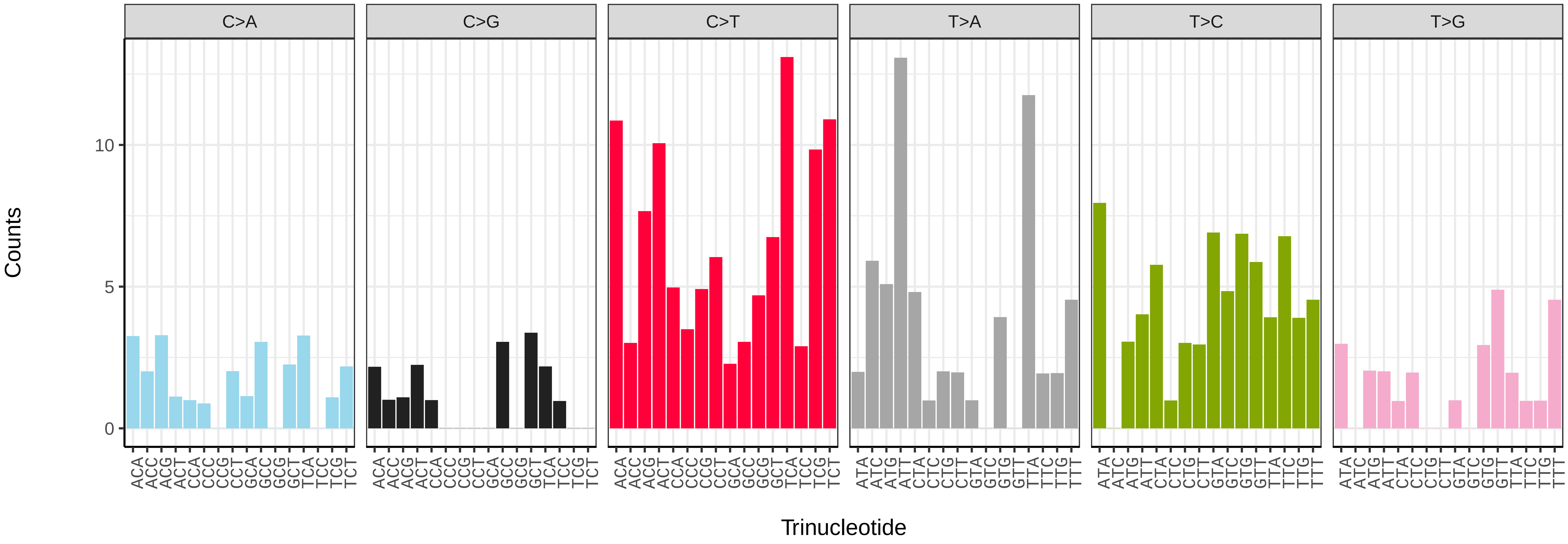


*Maniola jurtina* (ilManJurt1.ilManJurt4) somatic mutational spectrum (SBS52 classification)

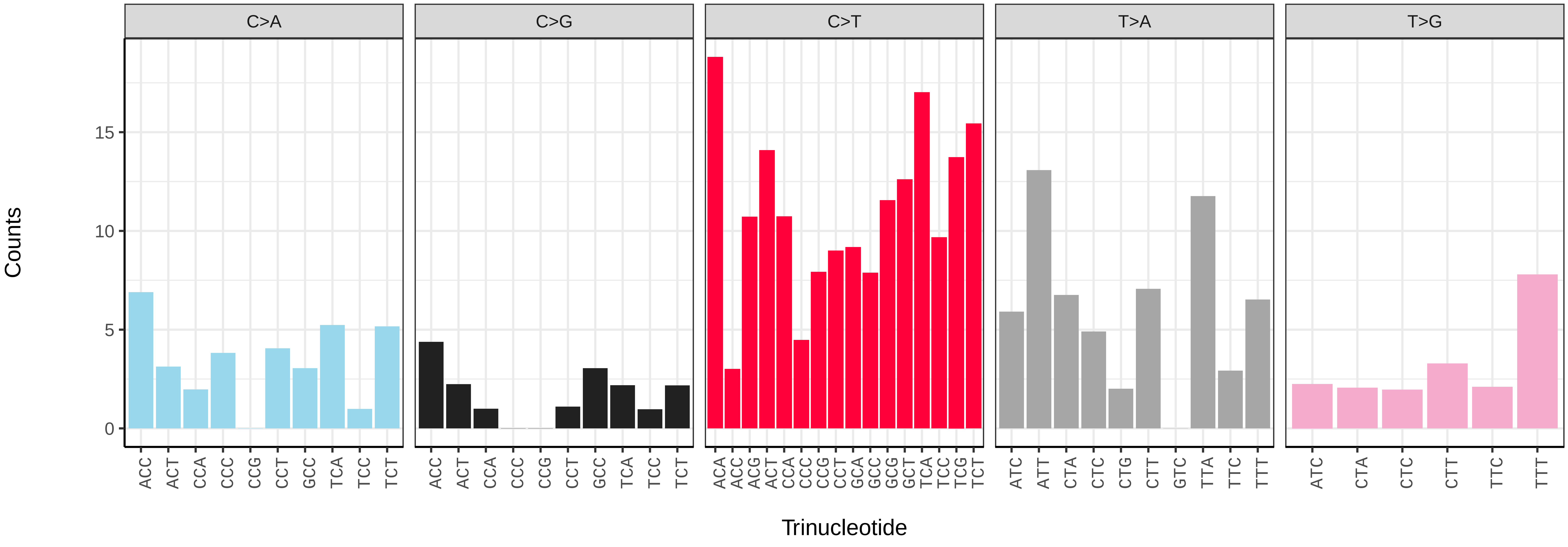

*Marasmarcha lunaedactyla* (ilMarLuna1) germline mutational spectrum

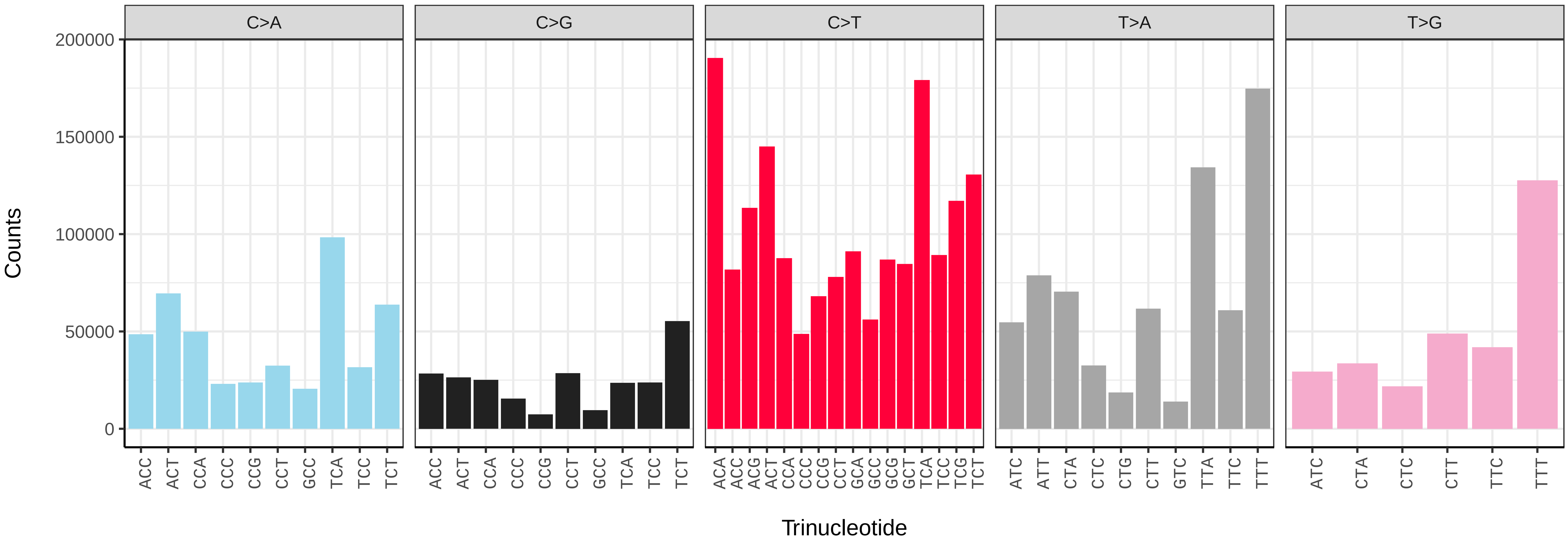


*Marasmarcha lunaedactyla* (ilMarLuna1) somatic mutational spectrum (SBS96 classification)

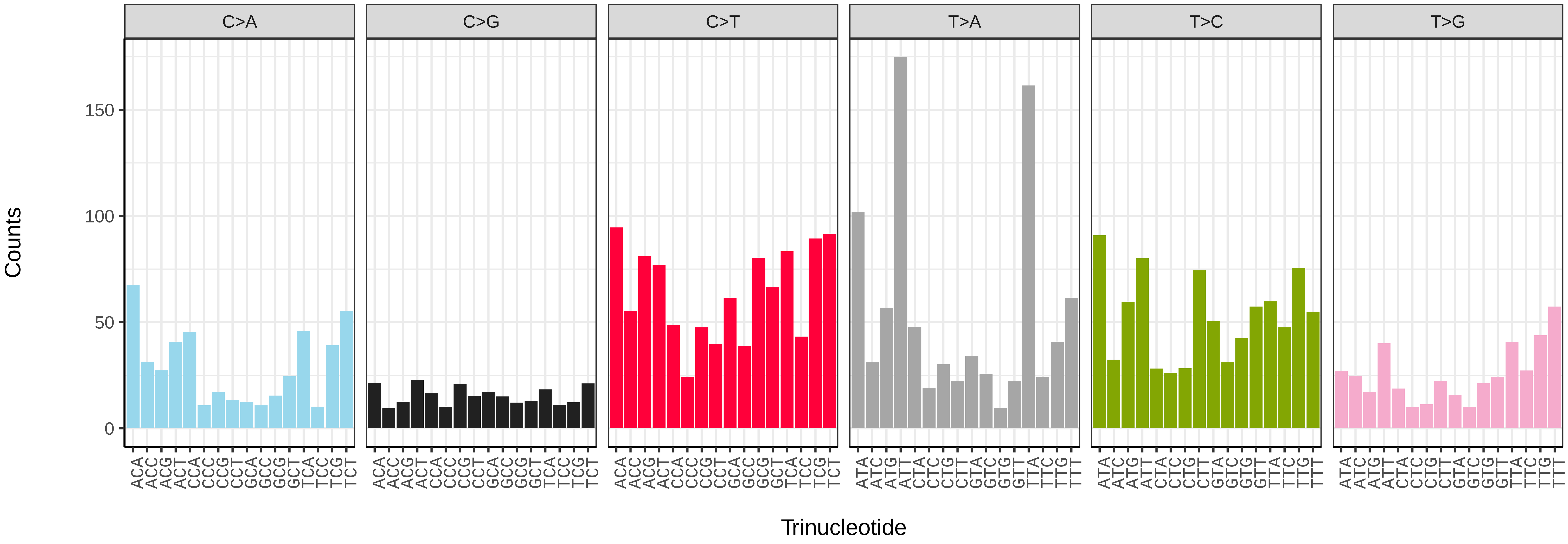


*Marasmarcha lunaedactyla* (ilMarLuna1) somatic mutational spectrum (SBS52 classification)

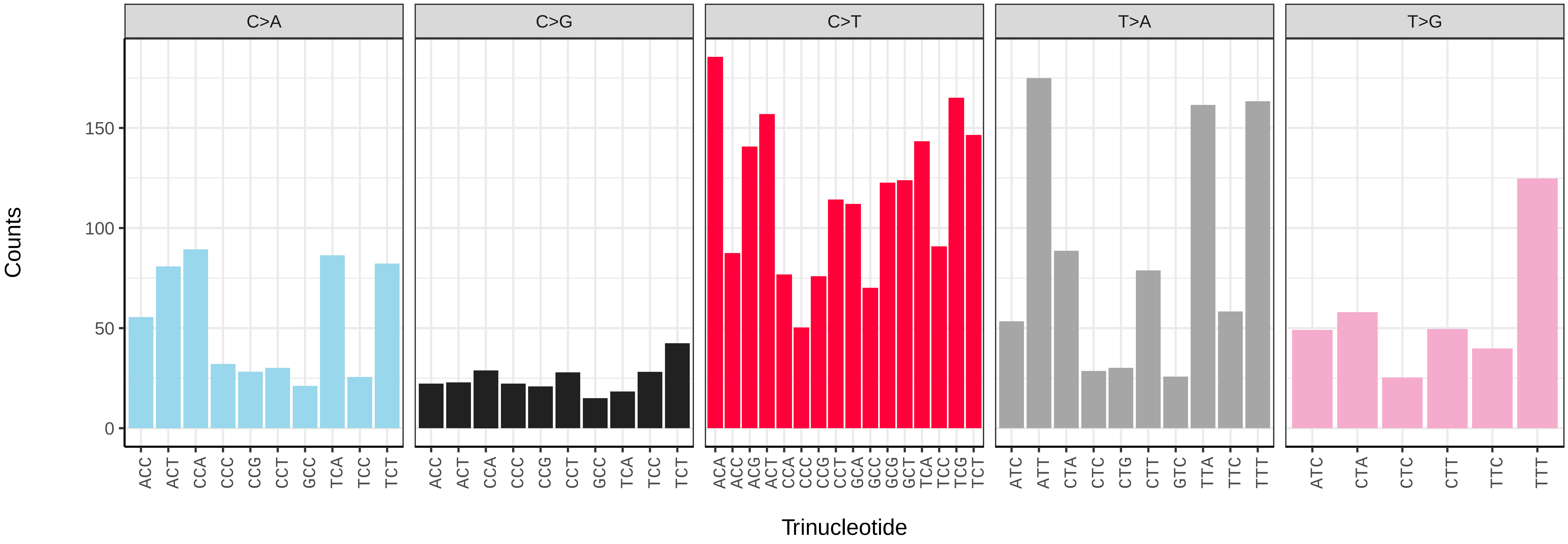

*Meganola albula* (ilMegAlbu1) germline mutational spectrum

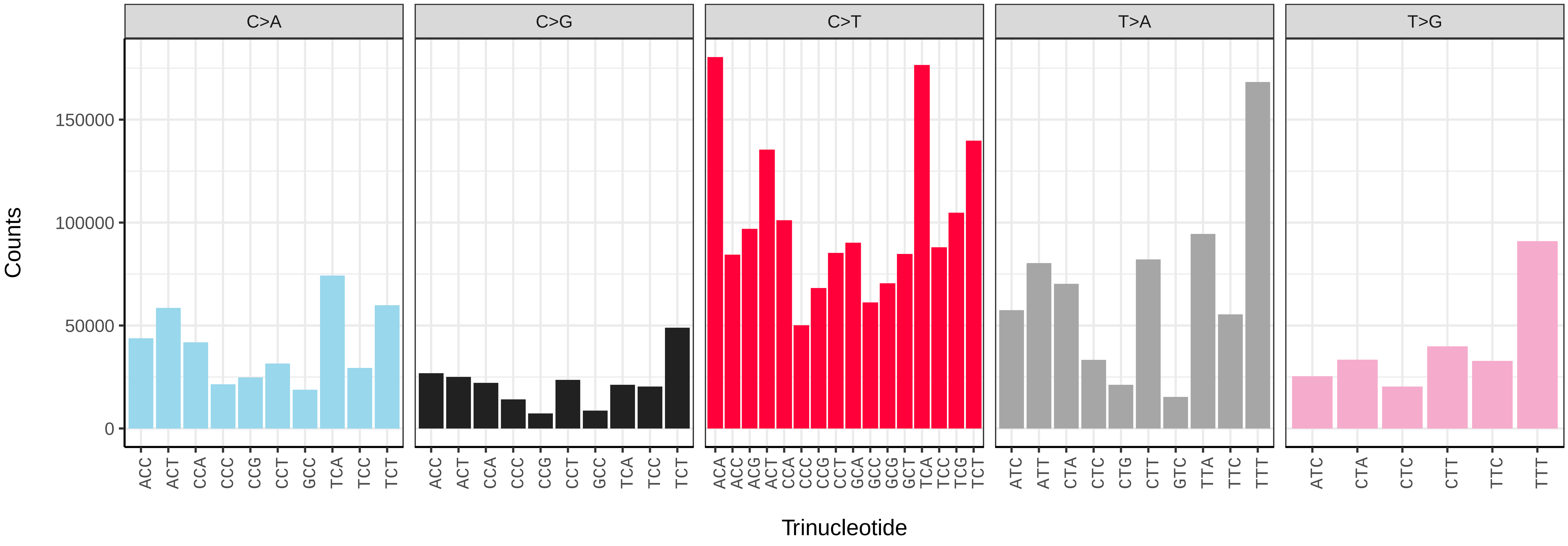


*Meganola albula* (ilMegAlbu1) somatic mutational spectrum (SBS96 classification)

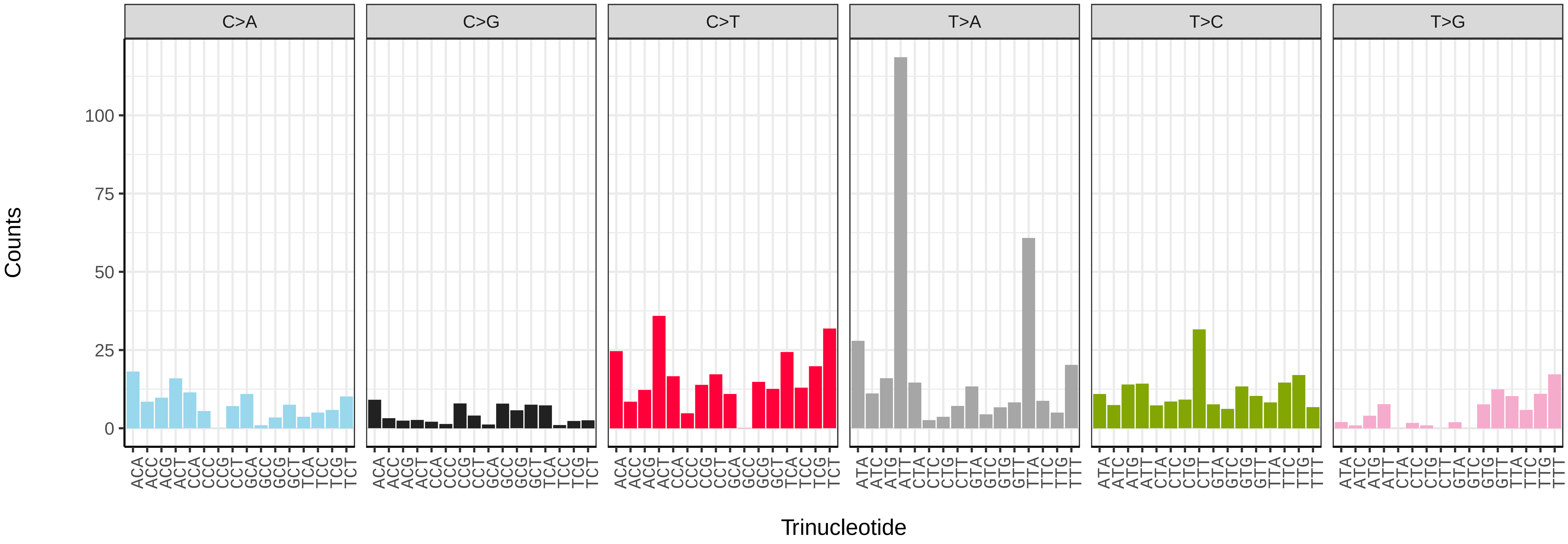


*Meganola albula* (ilMegAlbu1) somatic mutational spectrum (SBS52 classification)

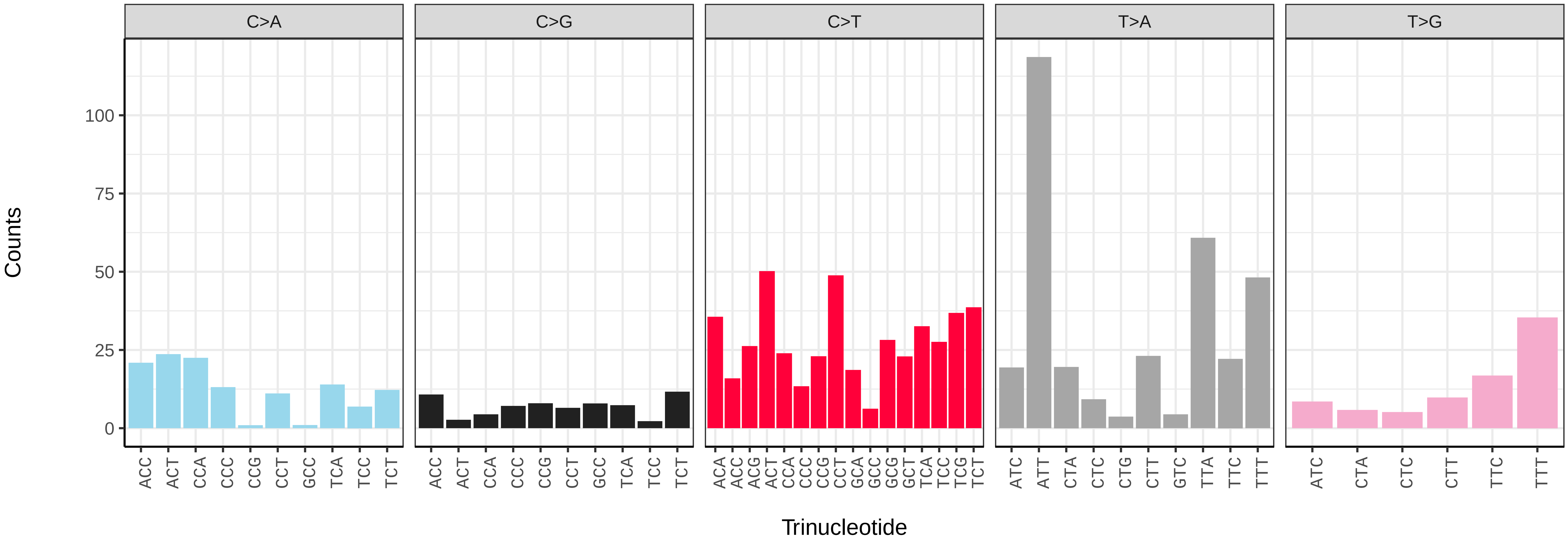

*Mellicta athalia* (ilMelAtha1) germline mutational spectrum

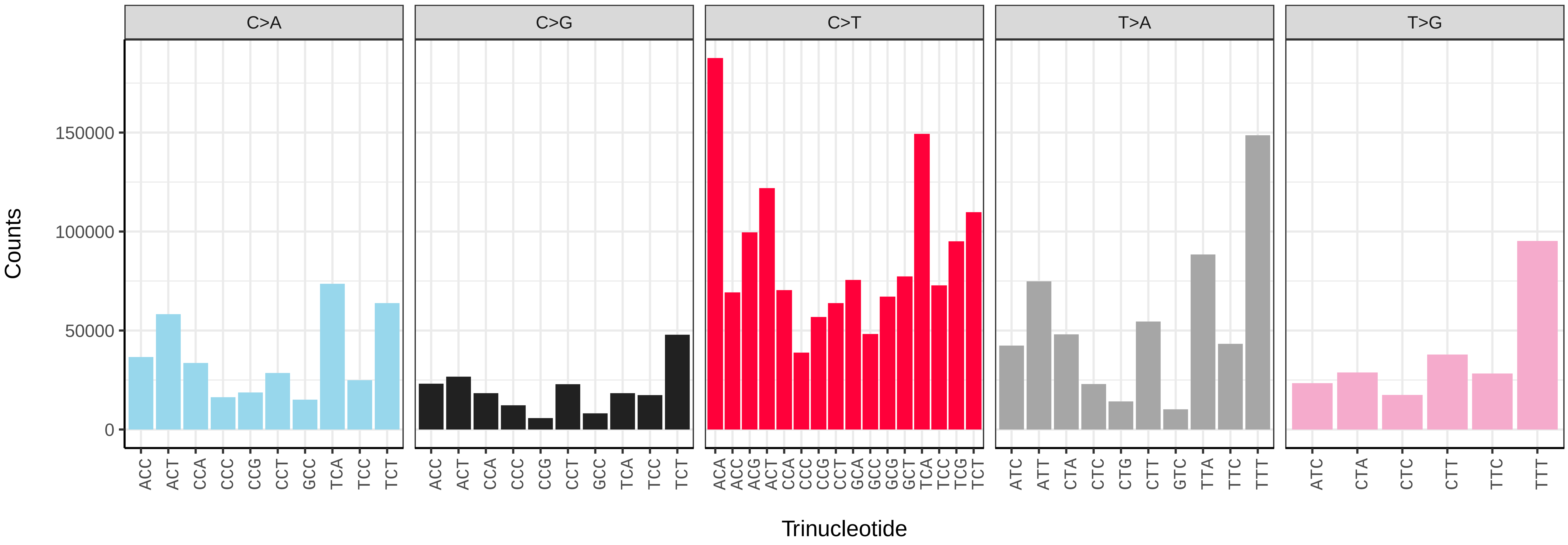


*Mellicta athalia* (ilMelAtha1) somatic mutational spectrum (SBS96 classification)

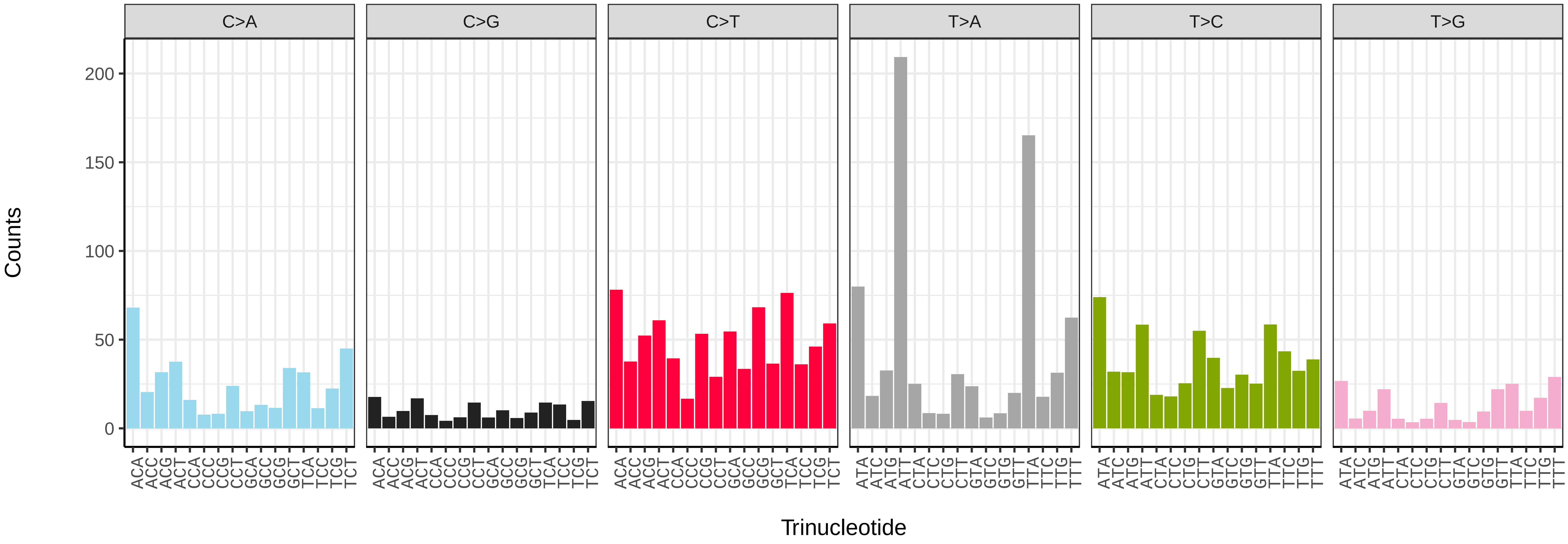


*Mellicta athalia* (ilMelAtha1) somatic mutational spectrum (SBS52 classification)

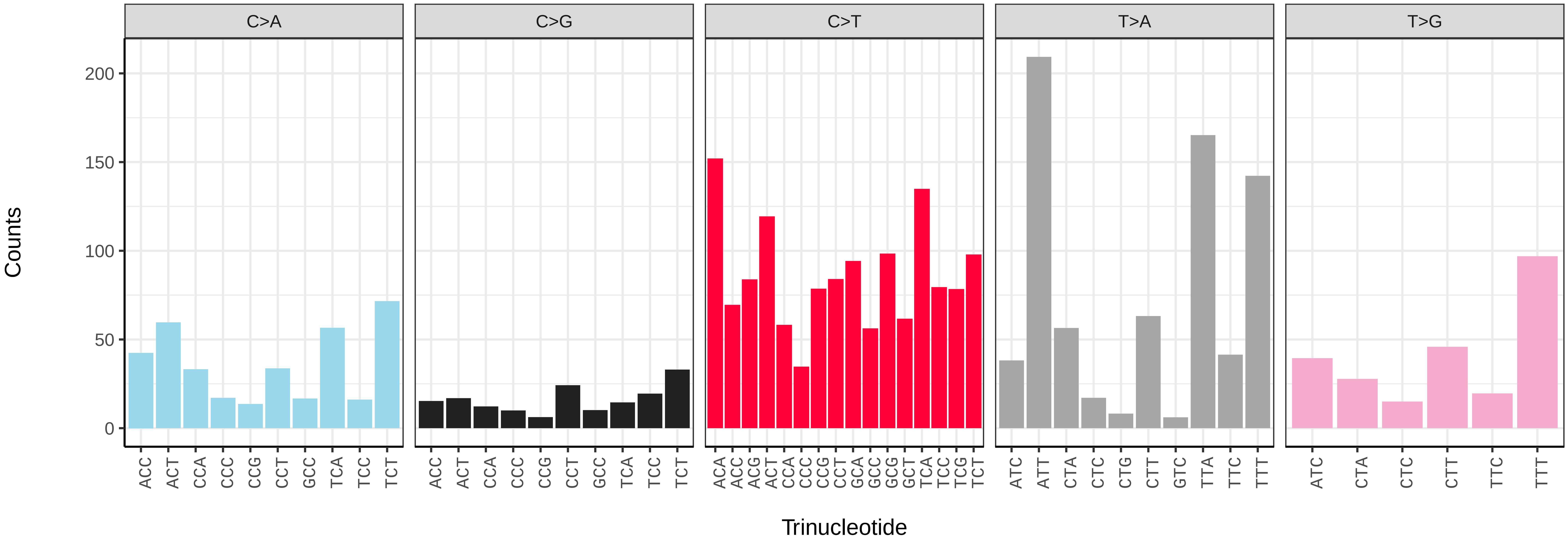

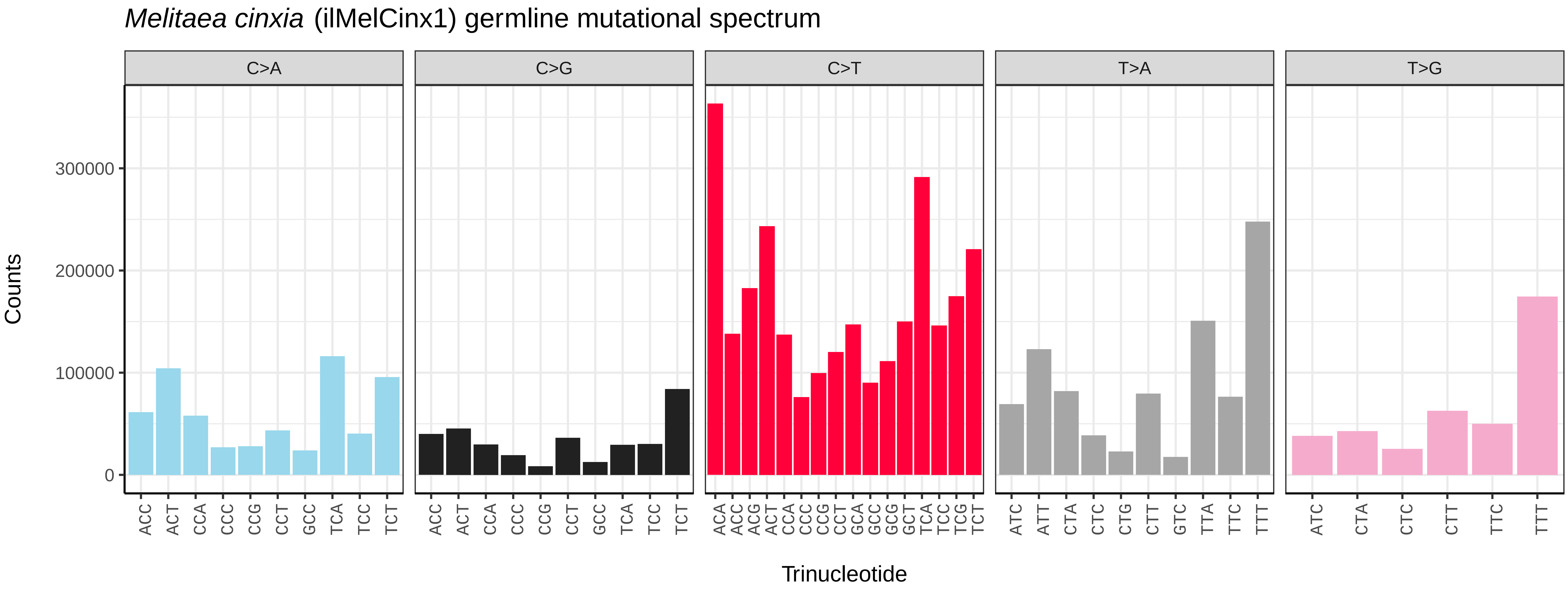
*Melitaea cinxia* (ilMelCinx1) germline mutational spectrum
C>A
C>G
C>T
T>A
T>G
Counts
0
100000
200000
300000
Trinucleotide

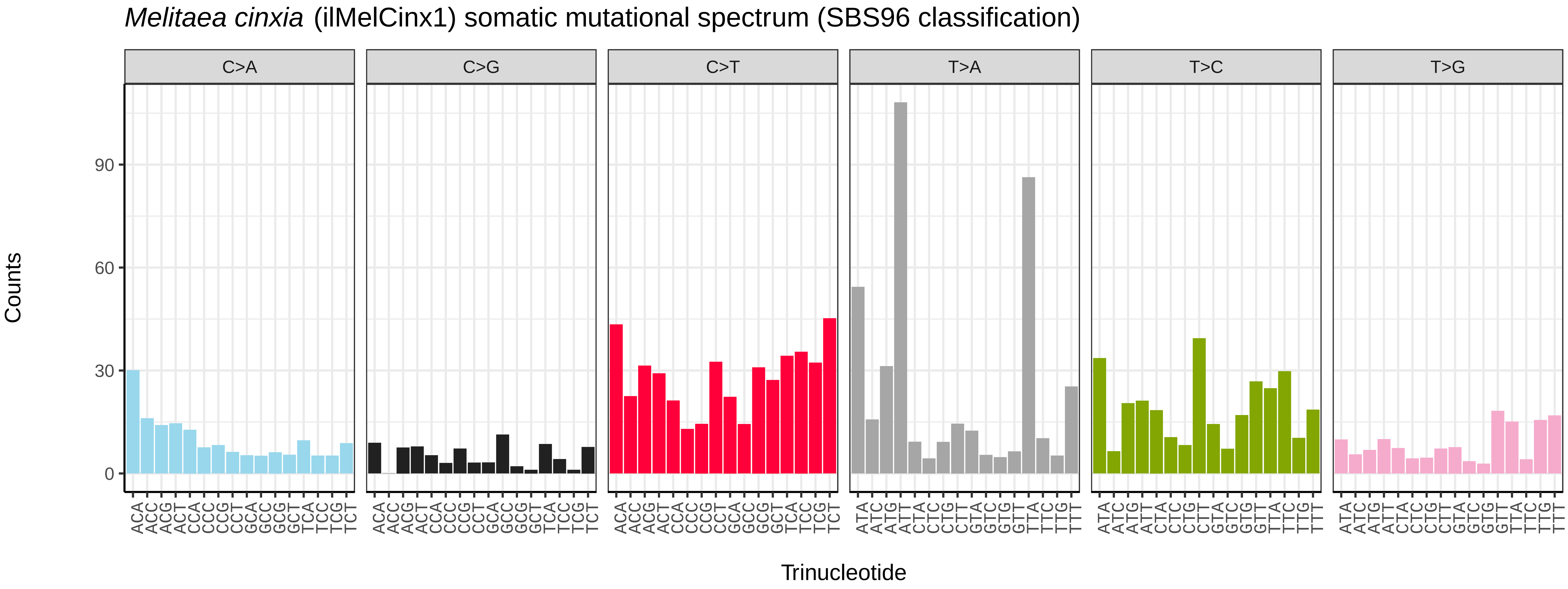
*Melitaea cinxia* (ilMelCinx1) somatic mutational spectrum (SBS96 classification)
C>A
C>G
C>T
T>A
T>C
T>G
Counts
0
30
60
90
Trinucleotide

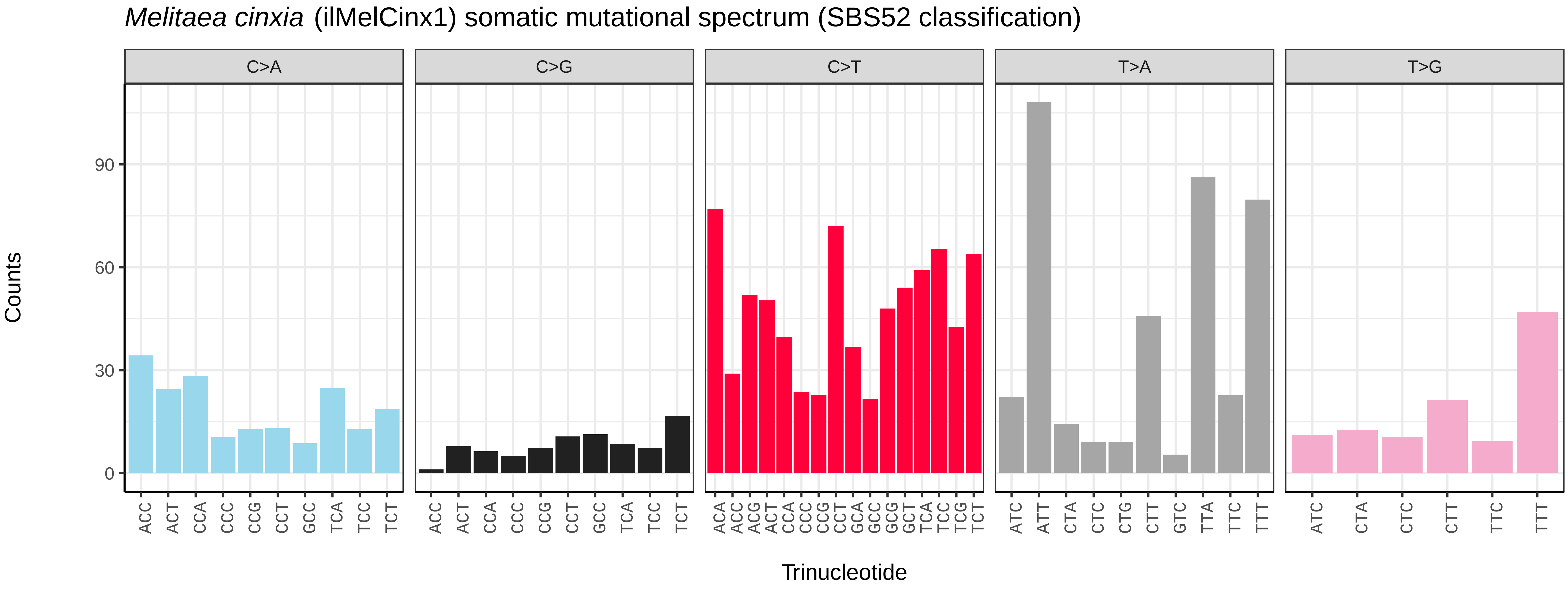
*Melitaea cinxia* (ilMelCinx1) somatic mutational spectrum (SBS52 classification)
C>A
C>G
C>T
T>A
T>G
Counts
0
30
60
90
Trinucleotide

*Melanargia galathea* (ilMelGala2) germline mutational spectrum

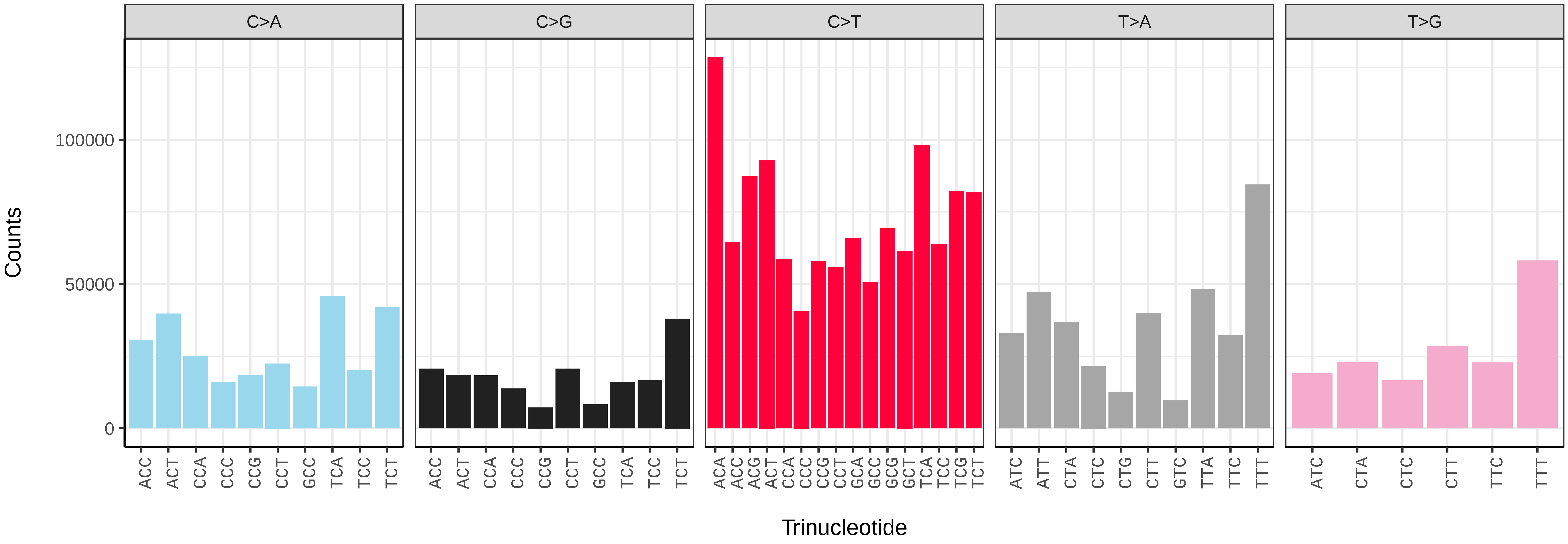


*Melanargia galathea* (ilMelGala2) somatic mutational spectrum (SBS96 classification)

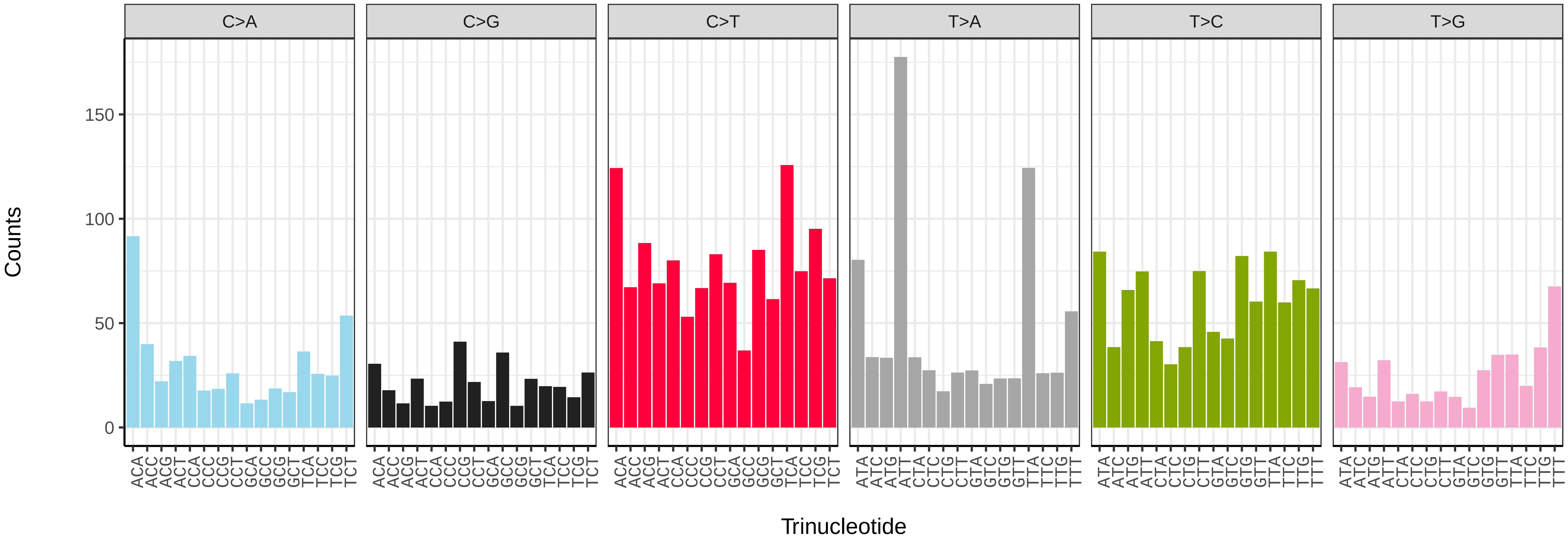


*Melanargia galathea* (ilMelGala2) somatic mutational spectrum (SBS52 classification)

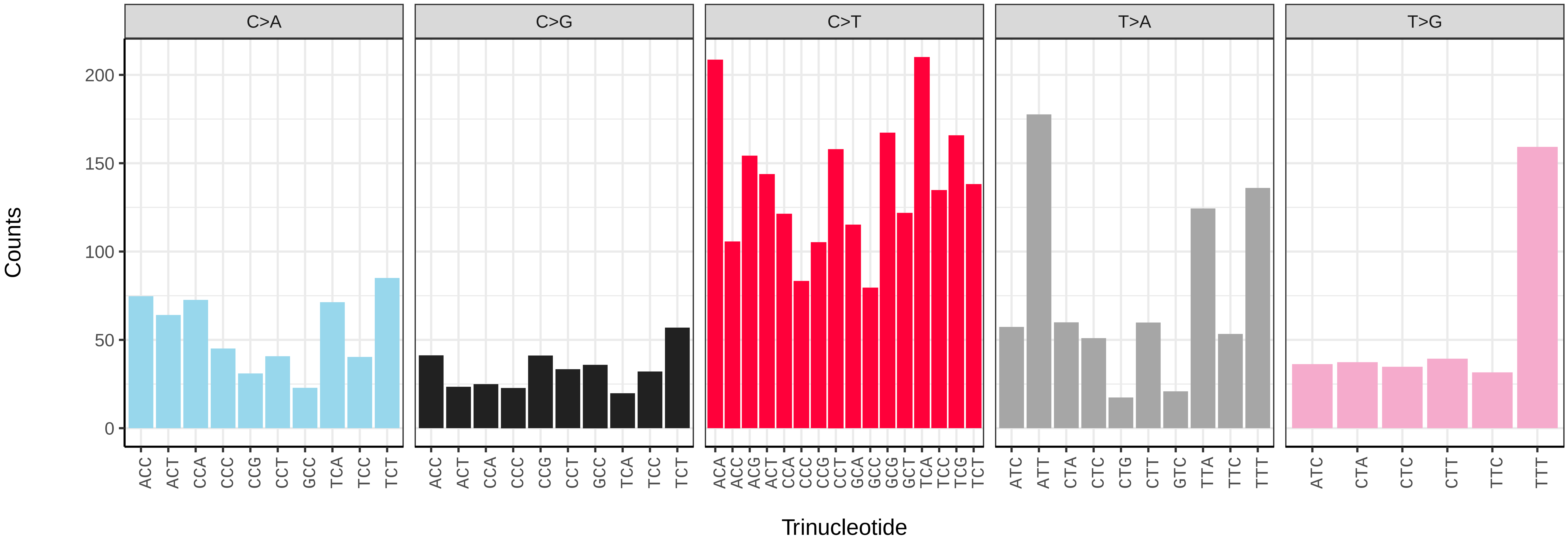

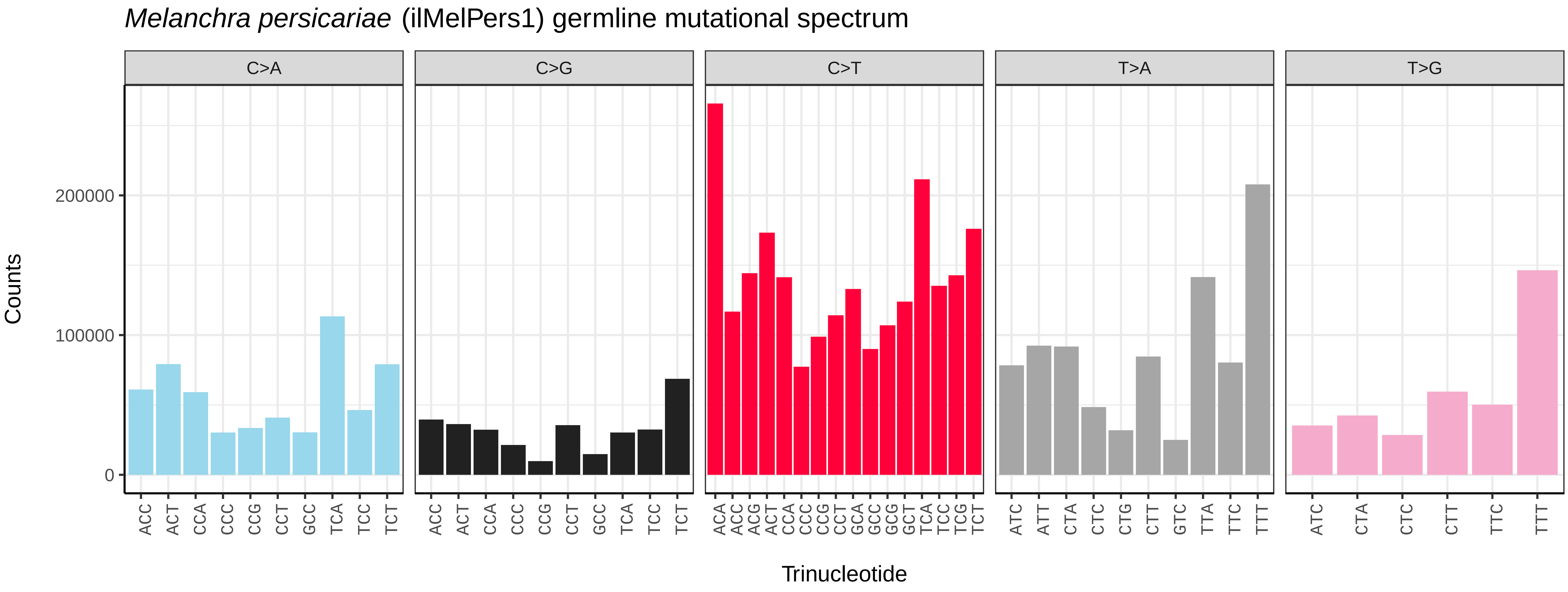
Melanchra persicariae (ilMelPers1) germline mutational spectrum
C>A
C>G
C>T
T>A
T>G
Counts
200000
100000
0
ACC ACT CCA CCC CCG CCT GCC TCA TCC TCT
ACC ACT CCA CCC CCG CCT GCC TCA TCC TCT
ACA ACC ACG ACT CCA CCC CCG CCT GCA GCC GCG GCT TCA TCC TCG TCT
ATC ATT CTA CTC CTG CTT GTC TTA TTC TTT
ATC CTA CTC CTT TTC TTT
Trinucleotide

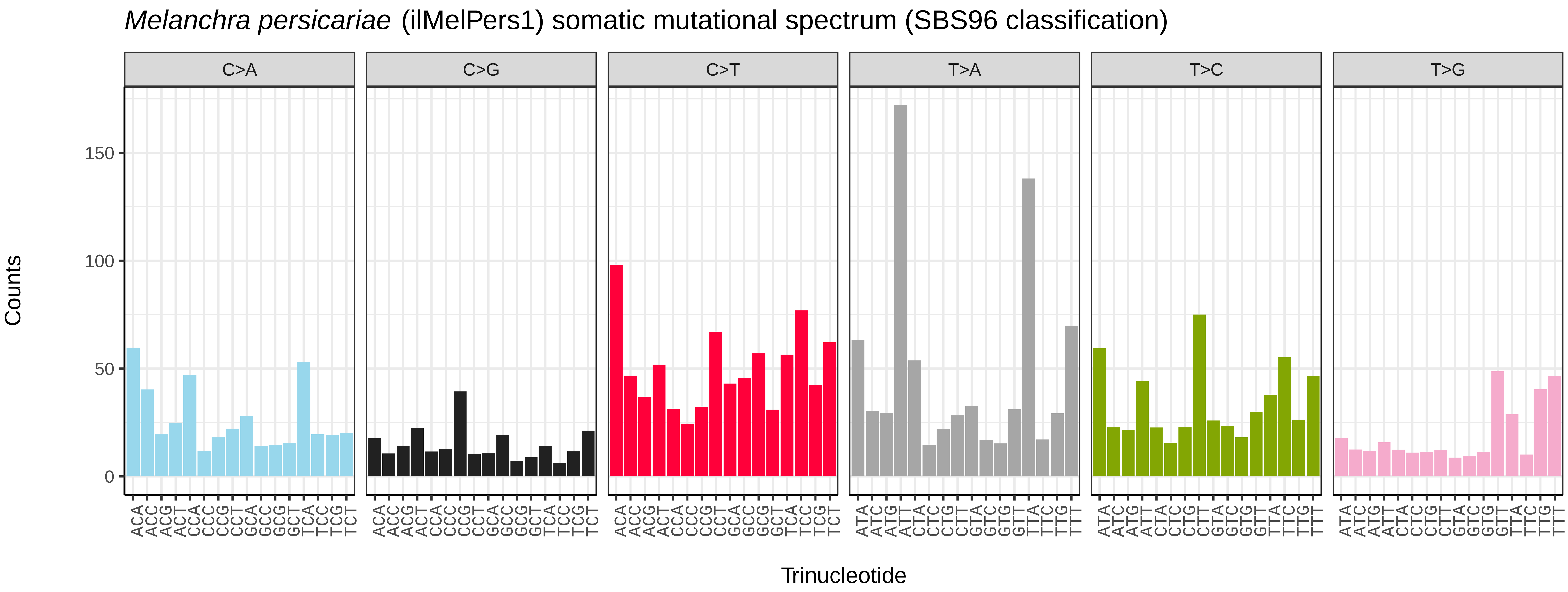
Melanchra persicariae (ilMelPers1) somatic mutational spectrum (SBS96 classification)
C>A
C>G
C>T
T>A
T>C
T>G
Counts
150
100
50
0
ACA ACC ACG ACT CCA CCC CCG CCT GCA GCC GCG GCT TCA TCC TCG TCT
ACA ACC ACG ACT CCA CCC CCG CCT GCA GCC GCG GCT TCA TCC TCG TCT
ACA ACC ACG ACT CCA CCC CCG CCT GCA GCC GCG GCT TCA TCC TCG TCT
ATA ATC ATG ATT CTA CTC CTG CTT GTA GTC GTG GTT TTA TTC TTG TTT
ATA ATC ATG ATT CTA CTC CTG CTT GTA GTC GTG GTT TTA TTC TTG TTT
ATA ATC ATG ATT CTA CTC CTG CTT GTA GTC GTG GTT TTA TTC TTG TTT
Trinucleotide

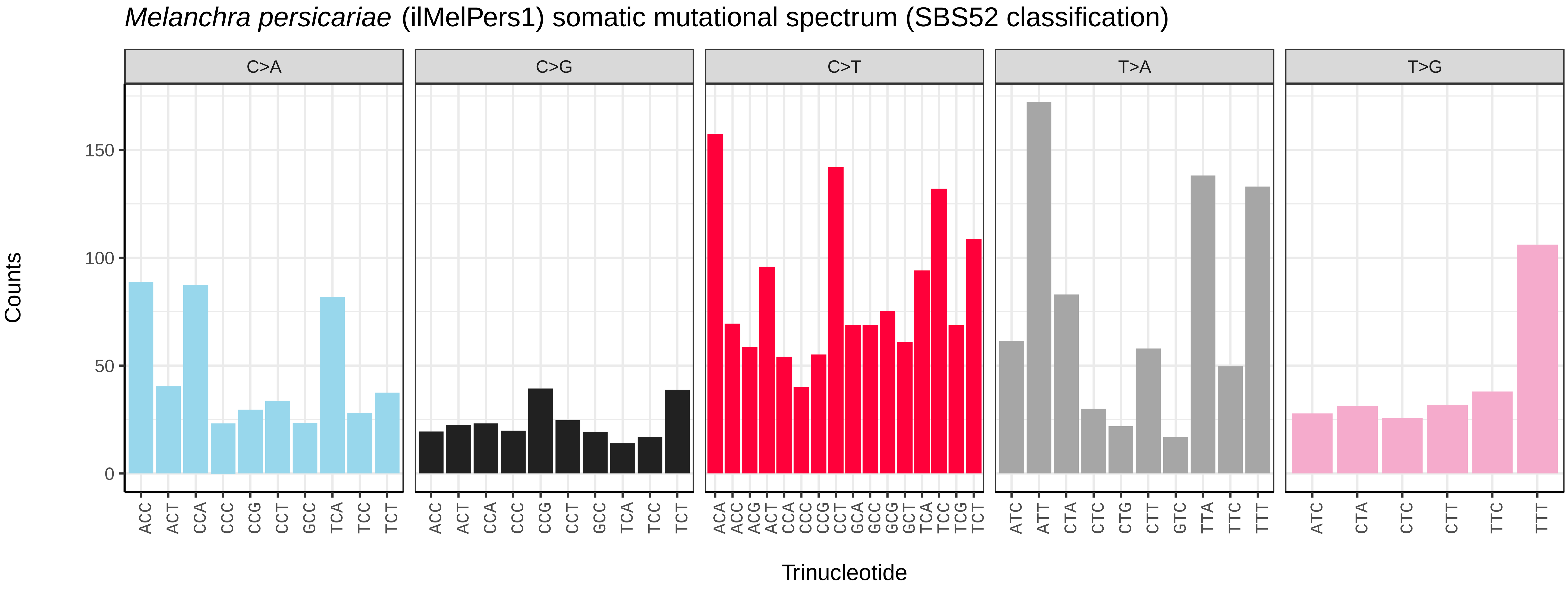
Melanchra persicariae (ilMelPers1) somatic mutational spectrum (SBS52 classification)
C>A
C>G
C>T
T>A
T>G
Counts
150
100
50
0
ACC ACT CCA CCC CCG CCT GCC TCA TCC TCT
ACC ACT CCA CCC CCG CCT GCC TCA TCC TCT
ACA ACC ACG ACT CCA CCC CCG CCT GCA GCC GCG GCT TCA TCC TCG TCT
ATC ATT CTA CTC CTG CTT GTC TTA TTC TTT
ATC CTA CTC CTT TTC TTT
Trinucleotide

*Mesoligia furuncula* (ilMesFuru1) germline mutational spectrum

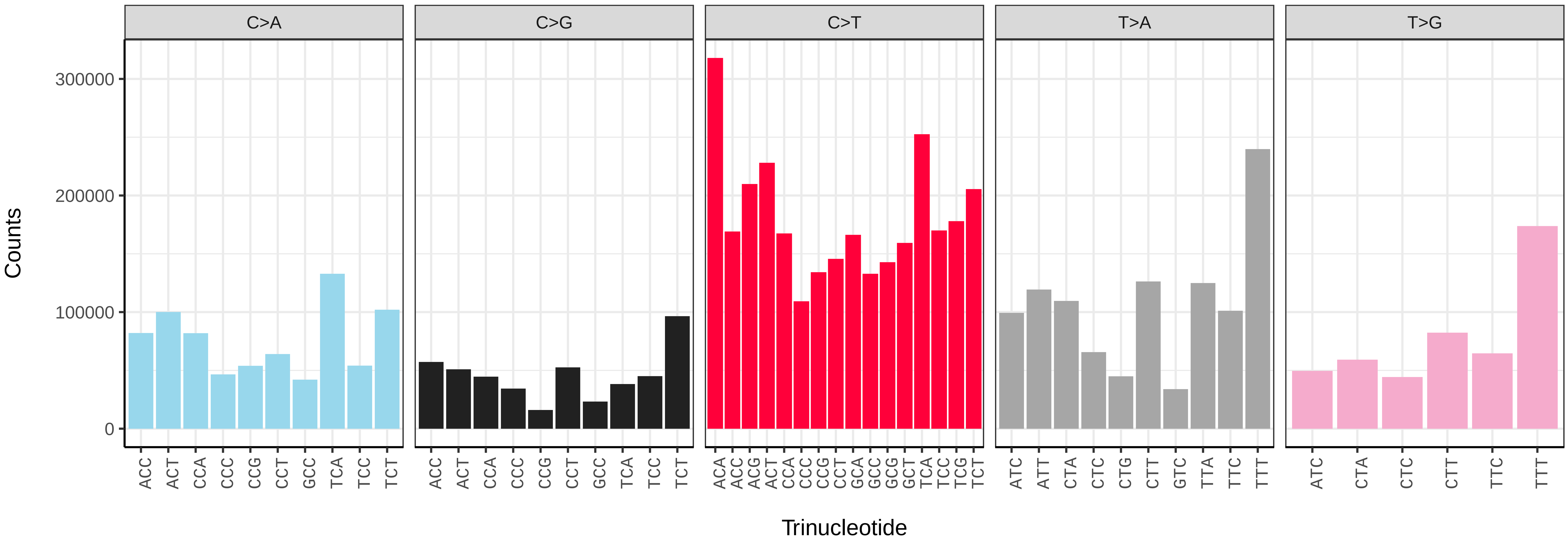


*Mesoligia furuncula* (ilMesFuru1) somatic mutational spectrum (SBS96 classification)

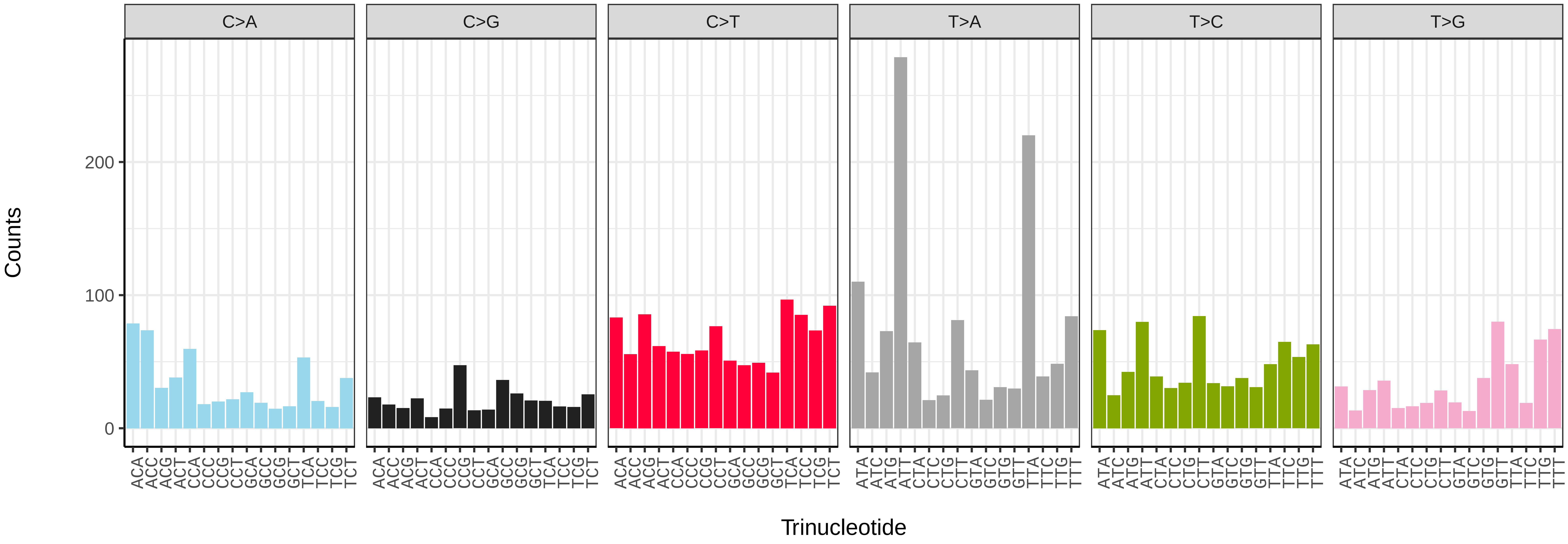


*Mesoligia furuncula* (ilMesFuru1) somatic mutational spectrum (SBS52 classification)

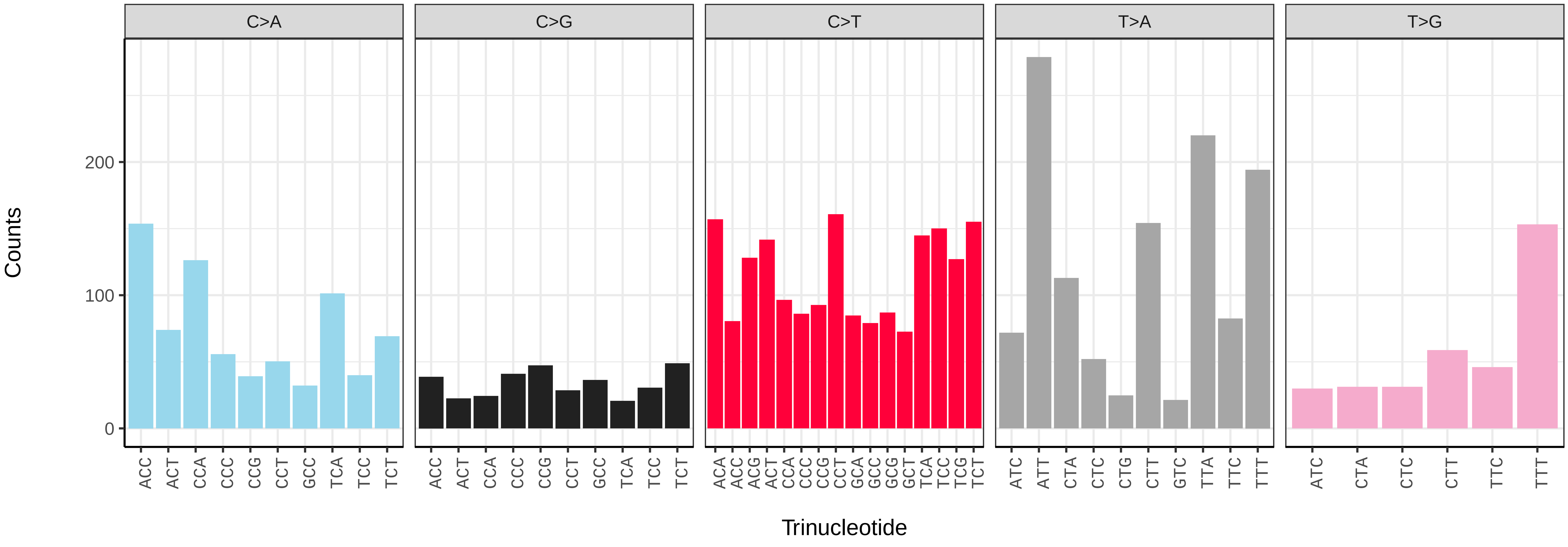

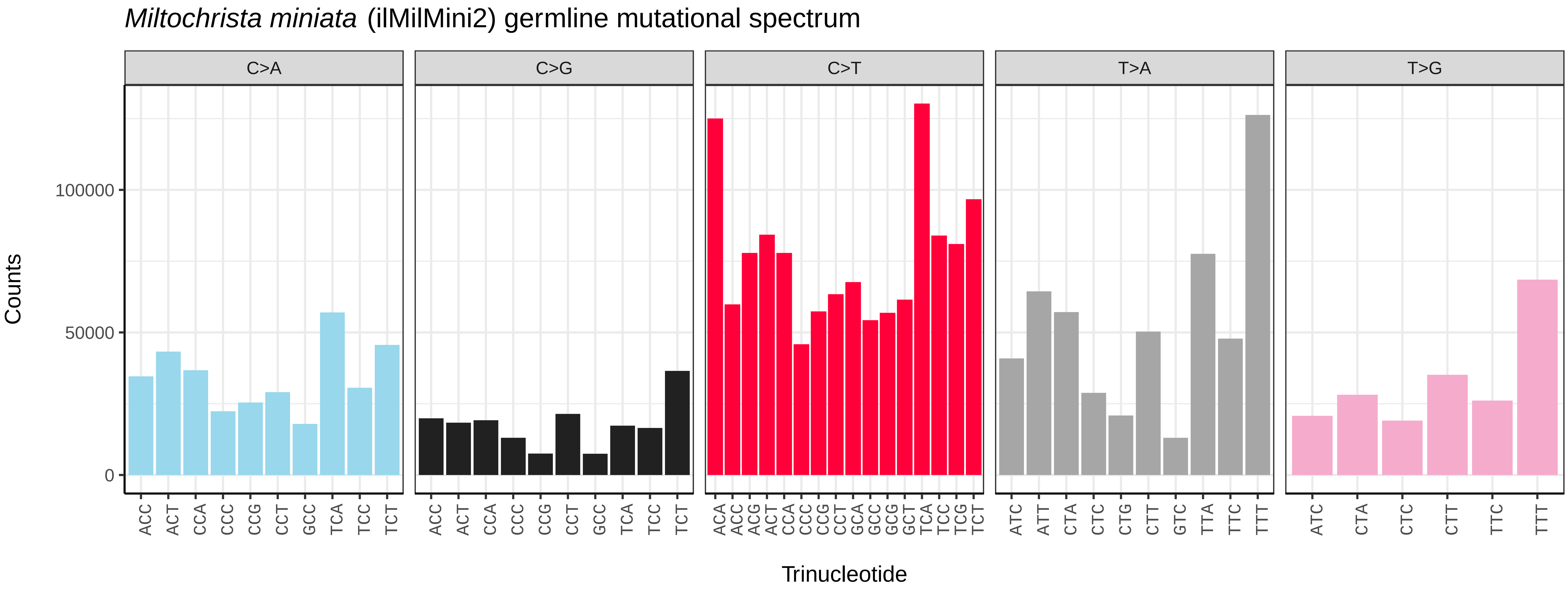
*Miltochrista miniata* (ilMilMini2) germline mutational spectrum
C>A
C>G
C>T
T>A
T>G
Counts
0
50000
100000
ACC ACT CCA CCC CCG CCT GCC TCA TCC TCT
ACC ACT CCA CCC CCG CCT GCC TCA TCC TCT
ACA ACC ACG ACT CCA CCC CCG CCT GCA GCC GCG GCT TCA TCC TCG TCT
ATC ATT CTA CTC CTG CTT GTC TTA TTC TTT
ATC CTA CTC CTT TTC TTT
Trinucleotide

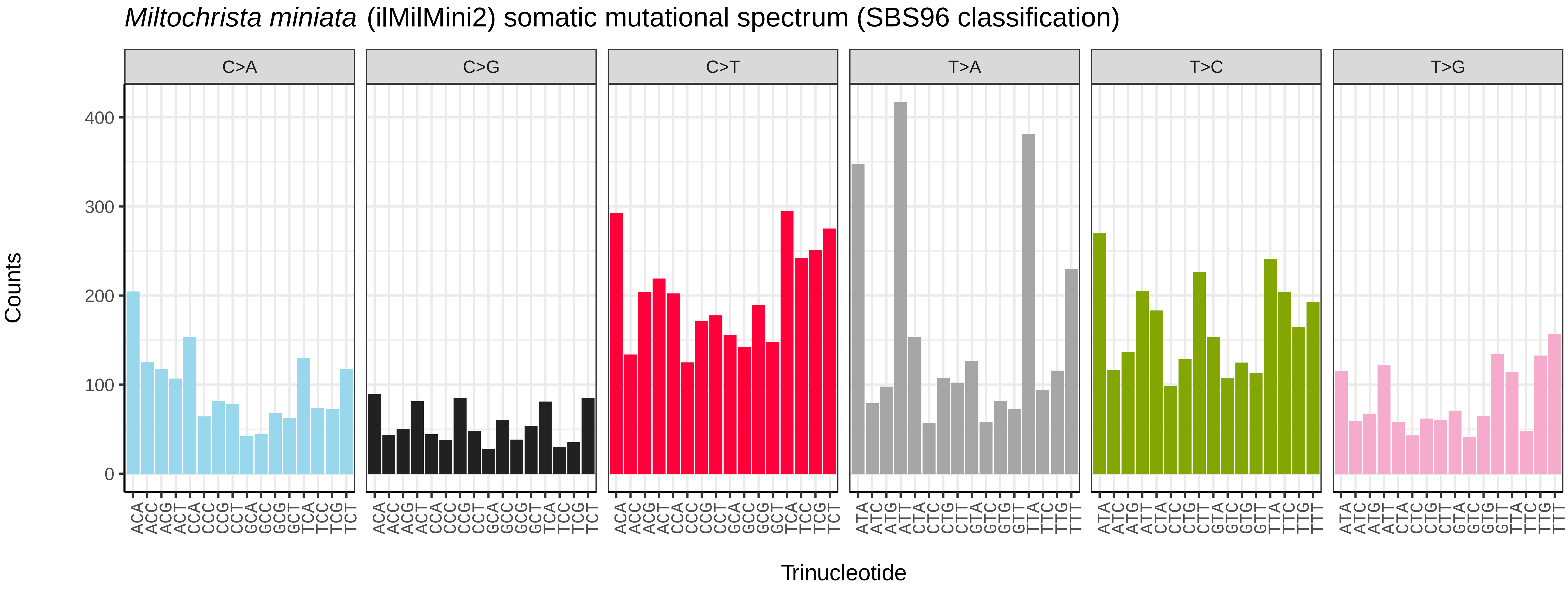
*Miltochrista miniata* (ilMilMini2) somatic mutational spectrum (SBS96 classification)
C>A
C>G
C>T
T>A
T>C
T>G
Counts
0
100
200
300
400
ACA ACC ACG ACT CCA CCC CCG CCT GCA GCC GCG GCT TCA TCC TCG TCT
ACA ACC ACG ACT CCA CCC CCG CCT GCA GCC GCG GCT TCA TCC TCG TCT
ACA ACC ACG ACT CCA CCC CCG CCT GCA GCC GCG GCT TCA TCC TCG TCT
ATA ATC ATG ATT CTA CTC CTG CTT GTA GTC GTG GTT TTA TTC TTG TTT
ATA ATC ATG ATT CTA CTC CTG CTT GTA GTC GTG GTT TTA TTC TTG TTT
ATA ATC ATG ATT CTA CTC CTG CTT GTA GTC GTG GTT TTA TTC TTG TTT
Trinucleotide

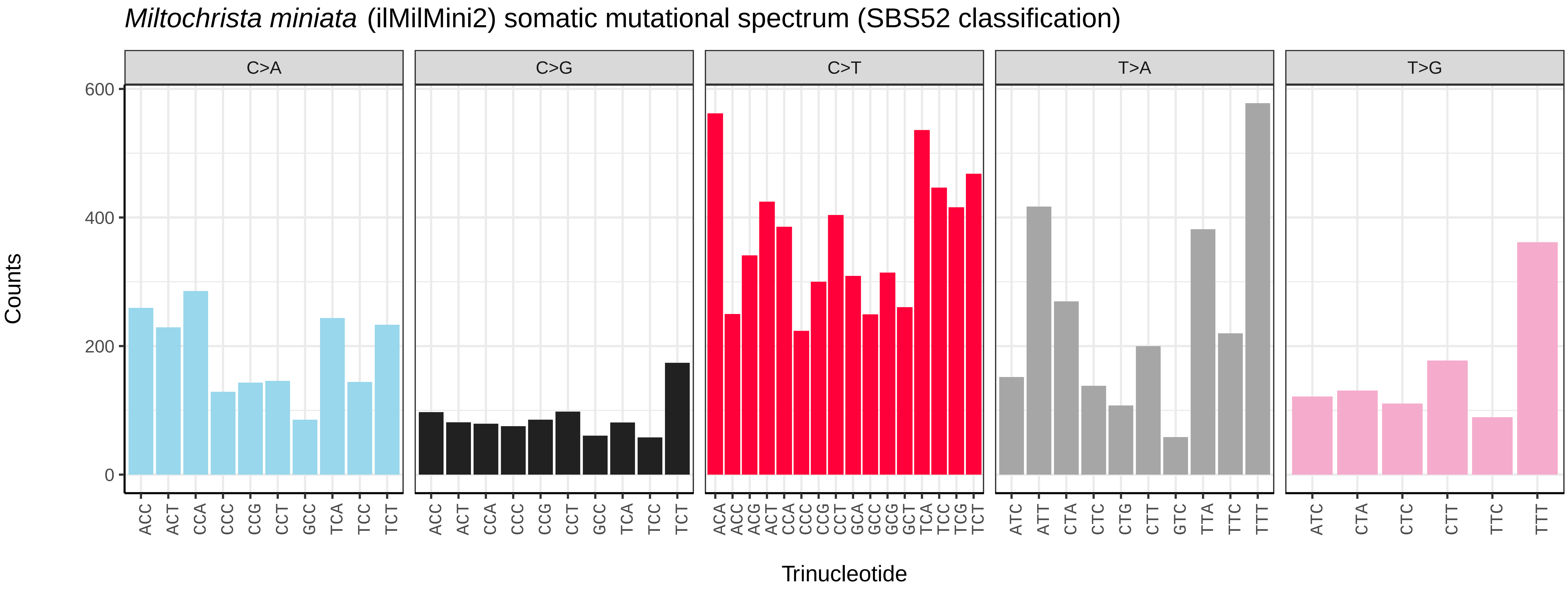
*Miltochrista miniata* (ilMilMini2) somatic mutational spectrum (SBS52 classification)
C>A
C>G
C>T
T>A
T>G
Counts
0
200
400
600
ACC ACT CCA CCC CCG CCT GCC TCA TCC TCT
ACC ACT CCA CCC CCG CCT GCC TCA TCC TCT
ACA ACC ACG ACT CCA CCC CCG CCT GCA GCC GCG GCT TCA TCC TCG TCT
ATC ATT CTA CTC CTG CTT GTC TTA TTC TTT
ATC CTA CTC CTT TTC TTT
Trinucleotide

*Mimas tiliae* (ilMimTili1) germline mutational spectrum

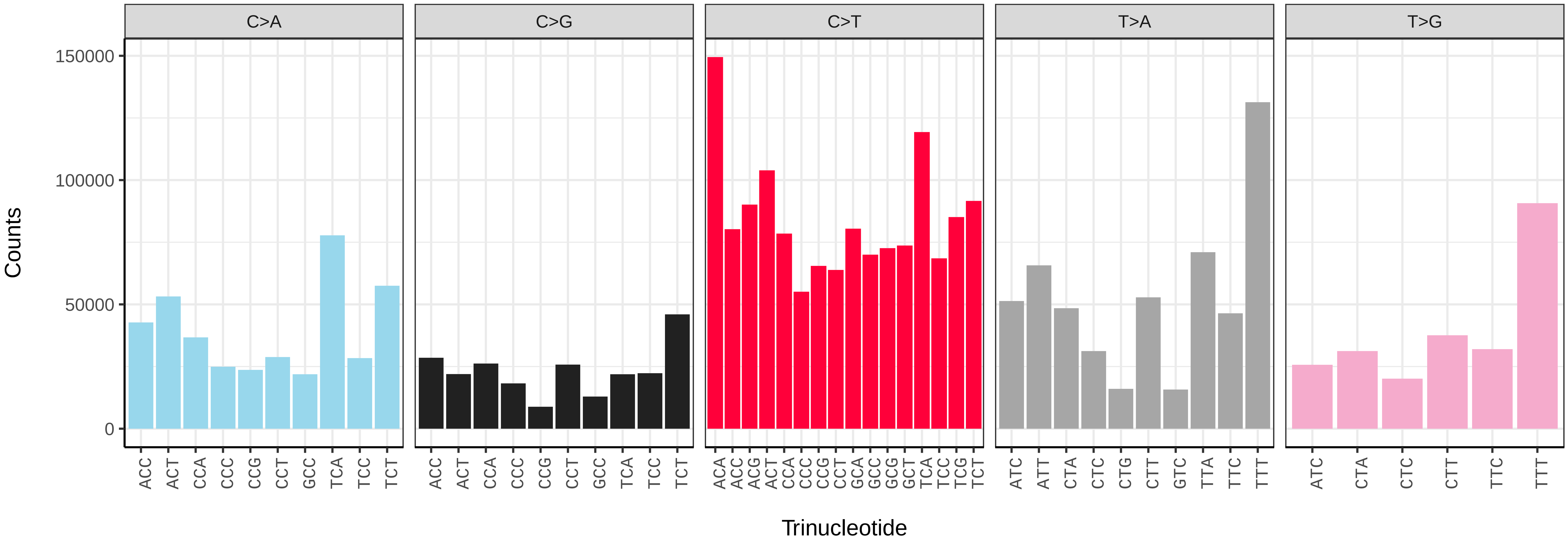


*Mimas tiliae* (ilMimTili1) somatic mutational spectrum (SBS96 classification)

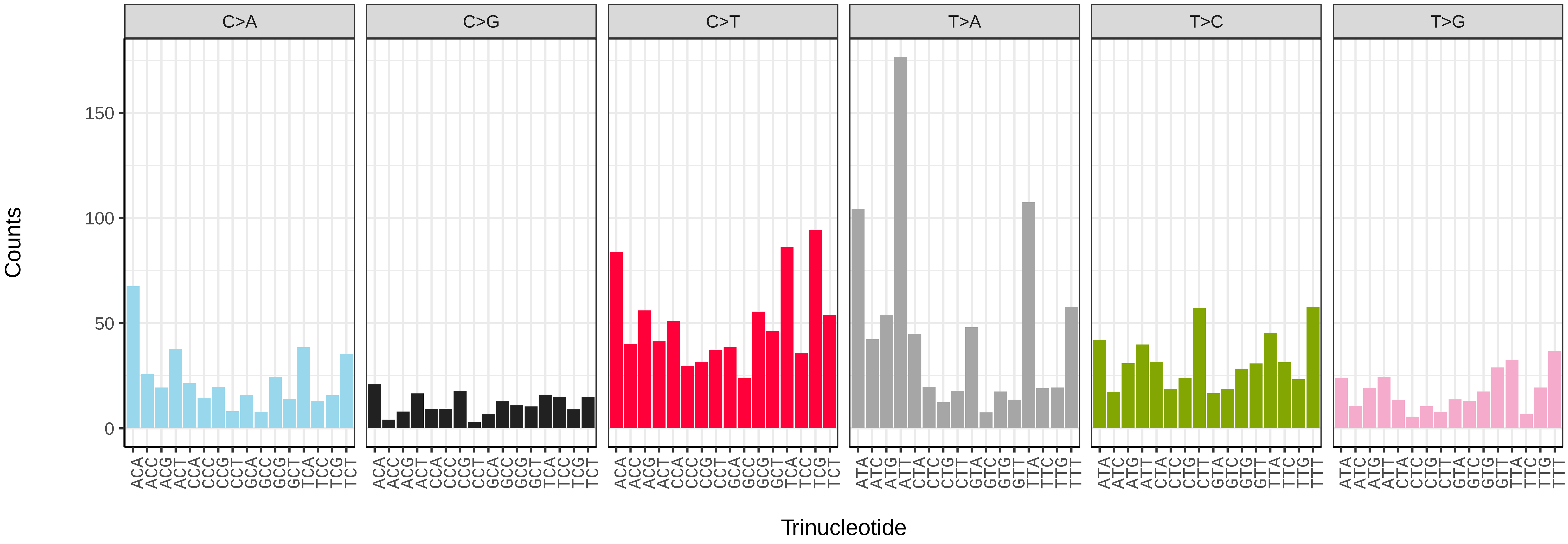


*Mimas tiliae* (ilMimTili1) somatic mutational spectrum (SBS52 classification)

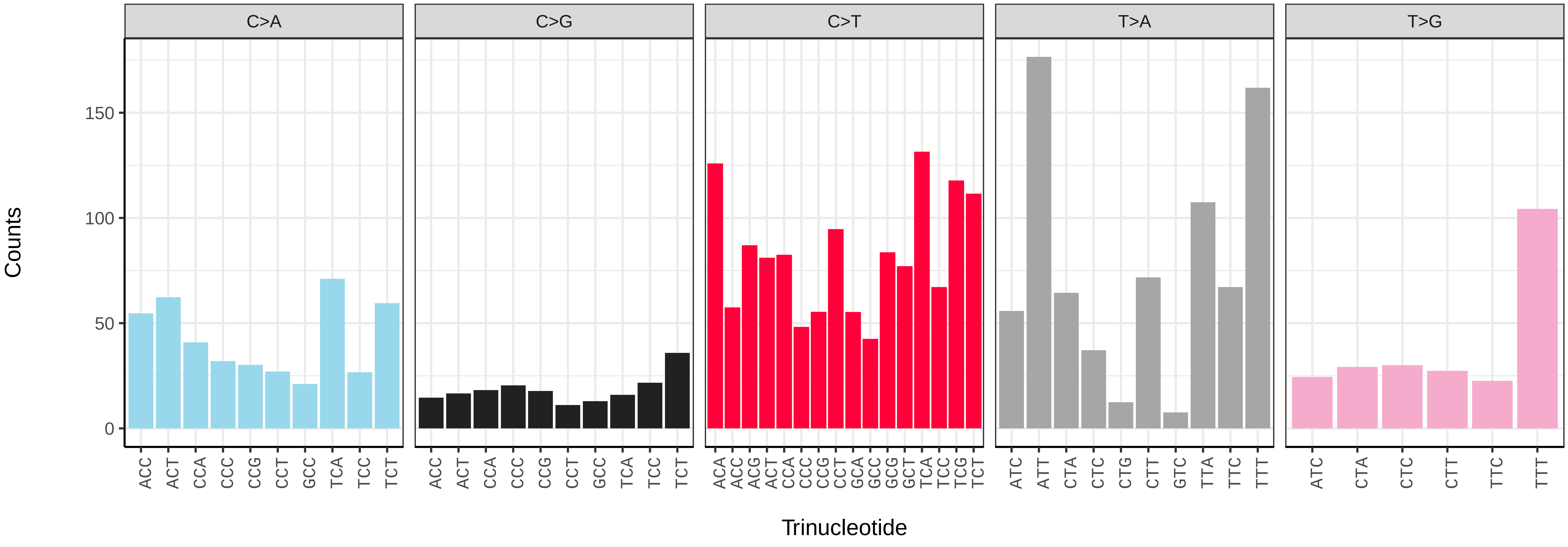

*Monopis laevigella* (ilMonLaev2) germline mutational spectrum

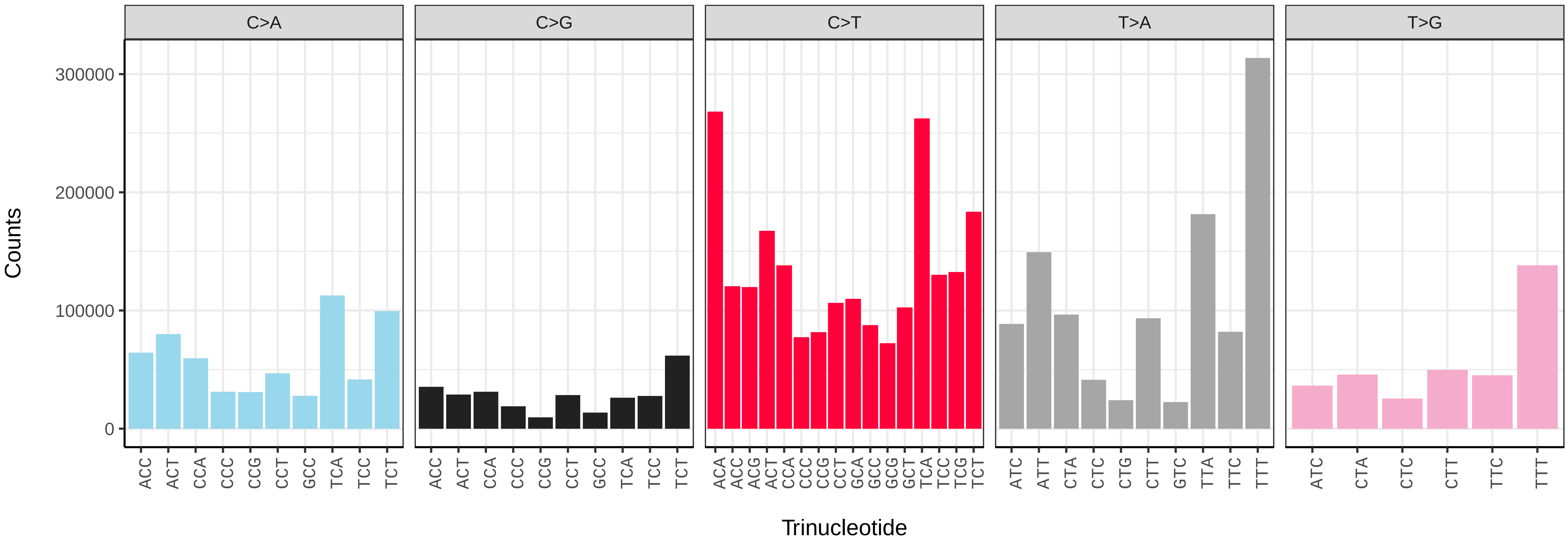


*Monopis laevigella* (ilMonLaev2) somatic mutational spectrum (SBS96 classification)

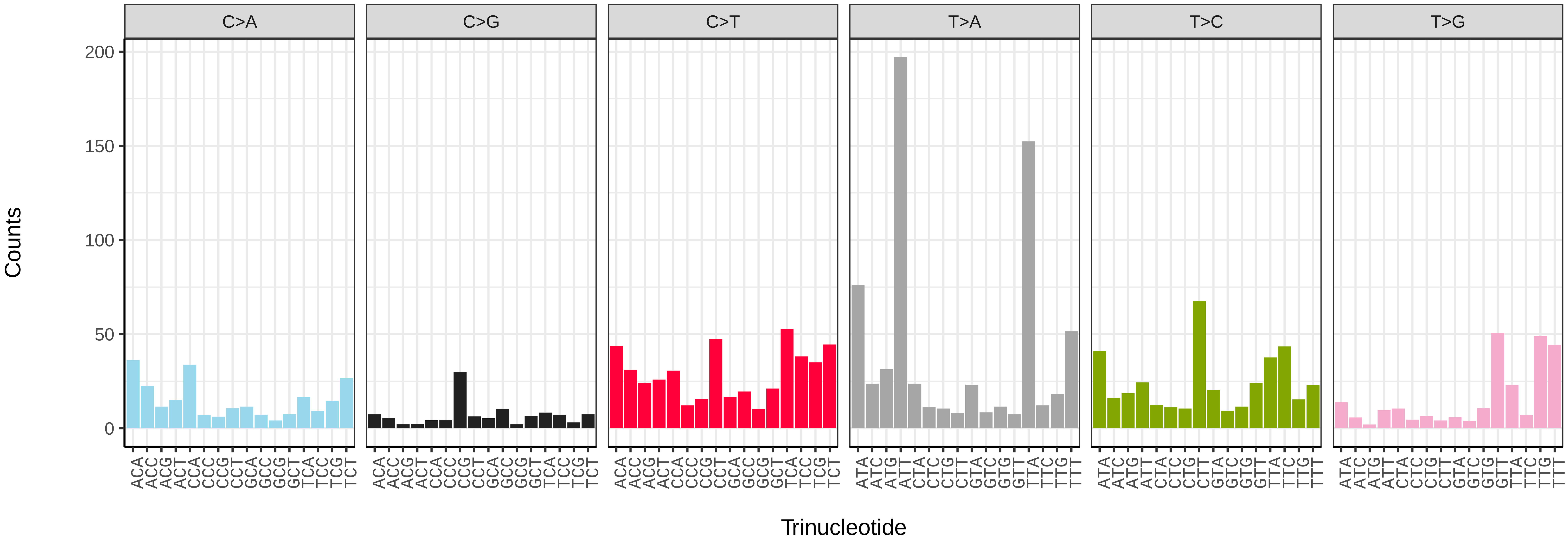


*Monopis laevigella* (ilMonLaev2) somatic mutational spectrum (SBS52 classification)

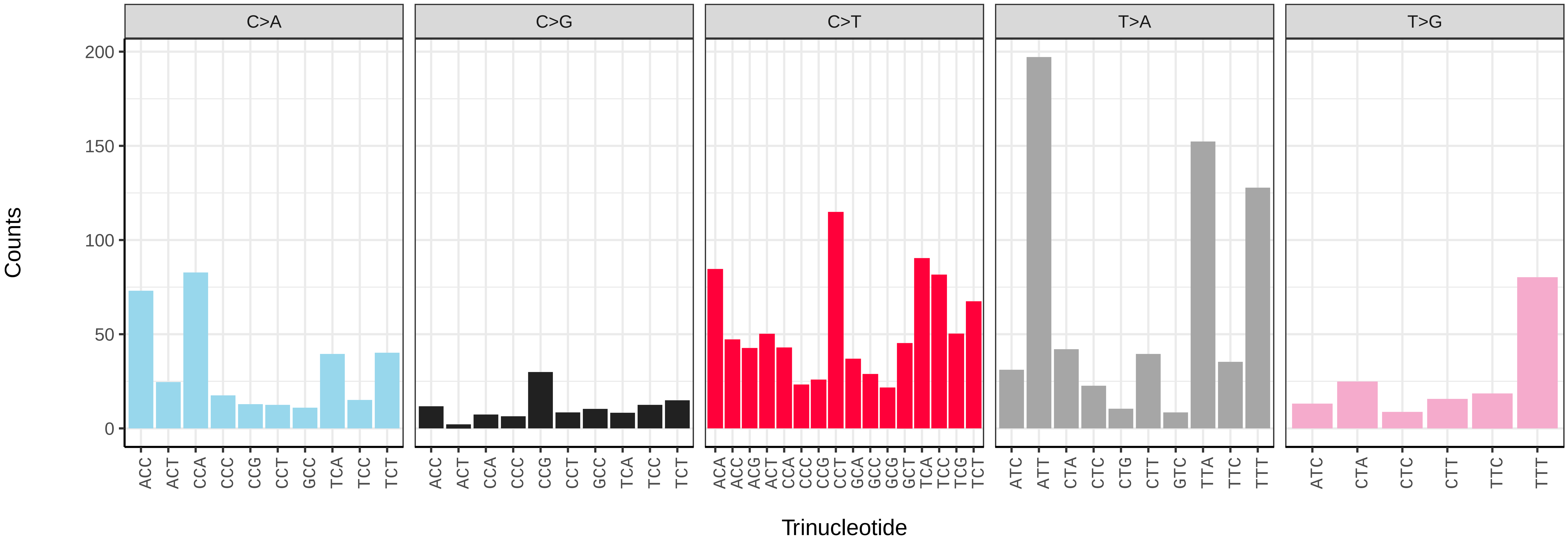

*Monopis laevigella* (ilMonLaev2.ilMonLaev1) germline mutational spectrum

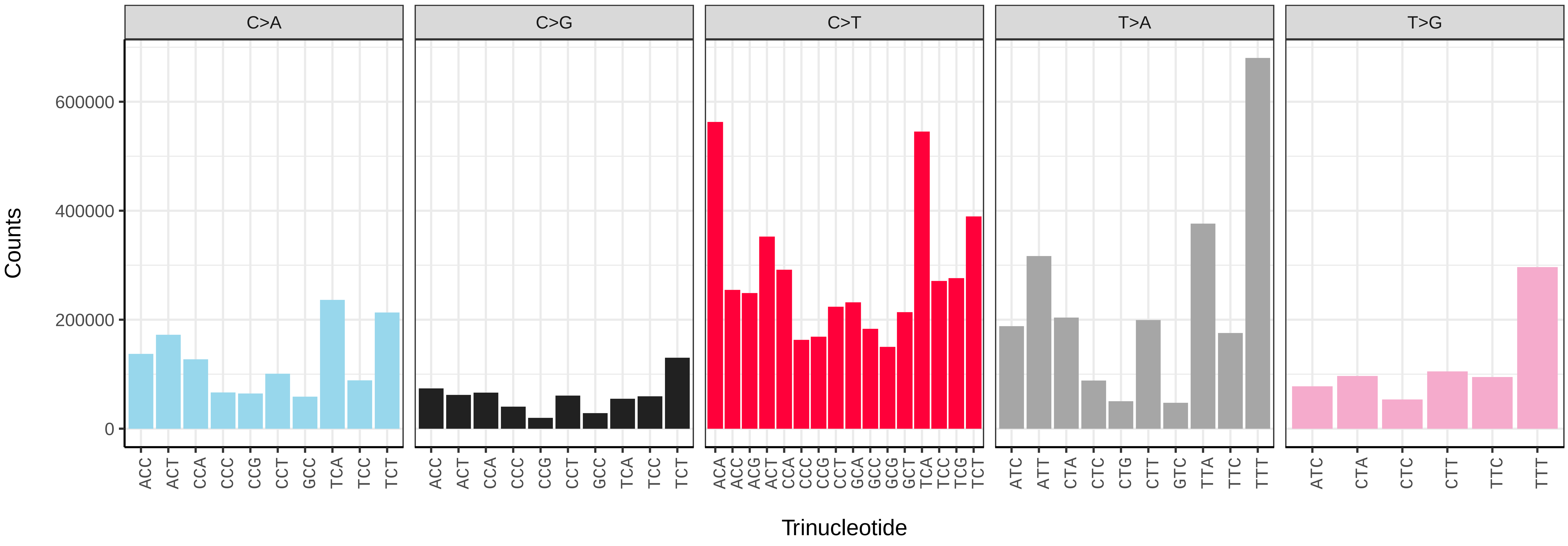


*Monopis laevigella* (ilMonLaev2.ilMonLaev1) somatic mutational spectrum (SBS96 classification)

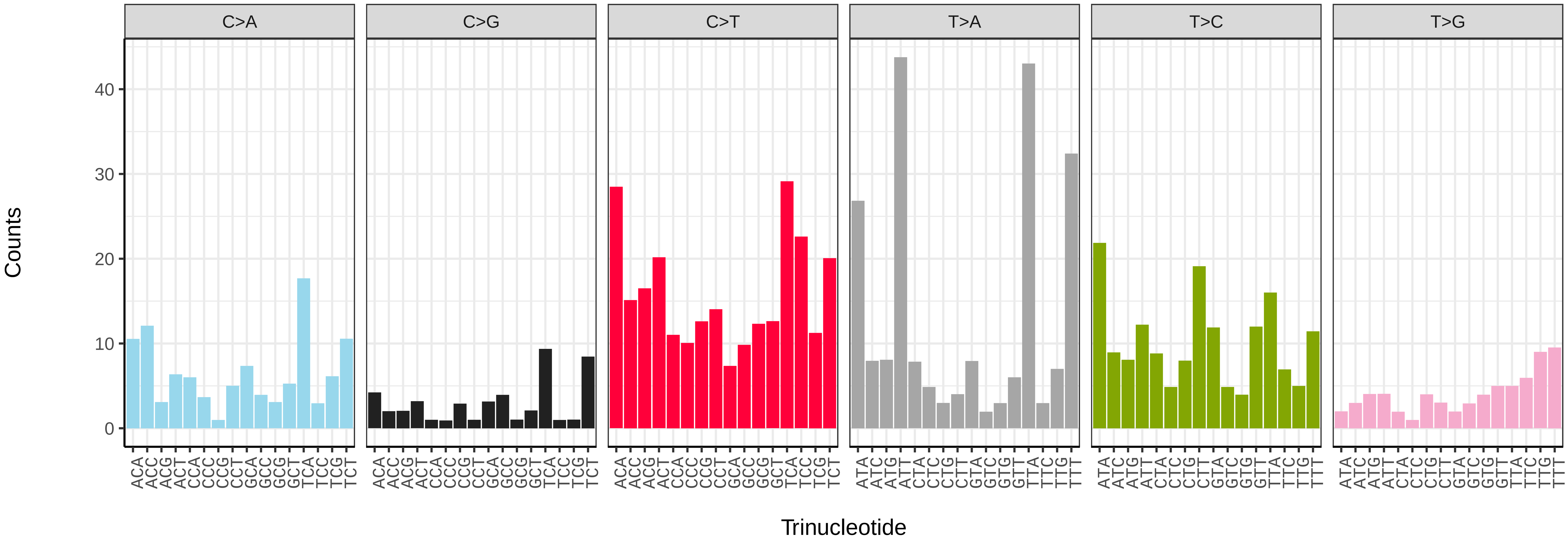


*Monopis laevigella* (ilMonLaev2.ilMonLaev1) somatic mutational spectrum (SBS52 classification)

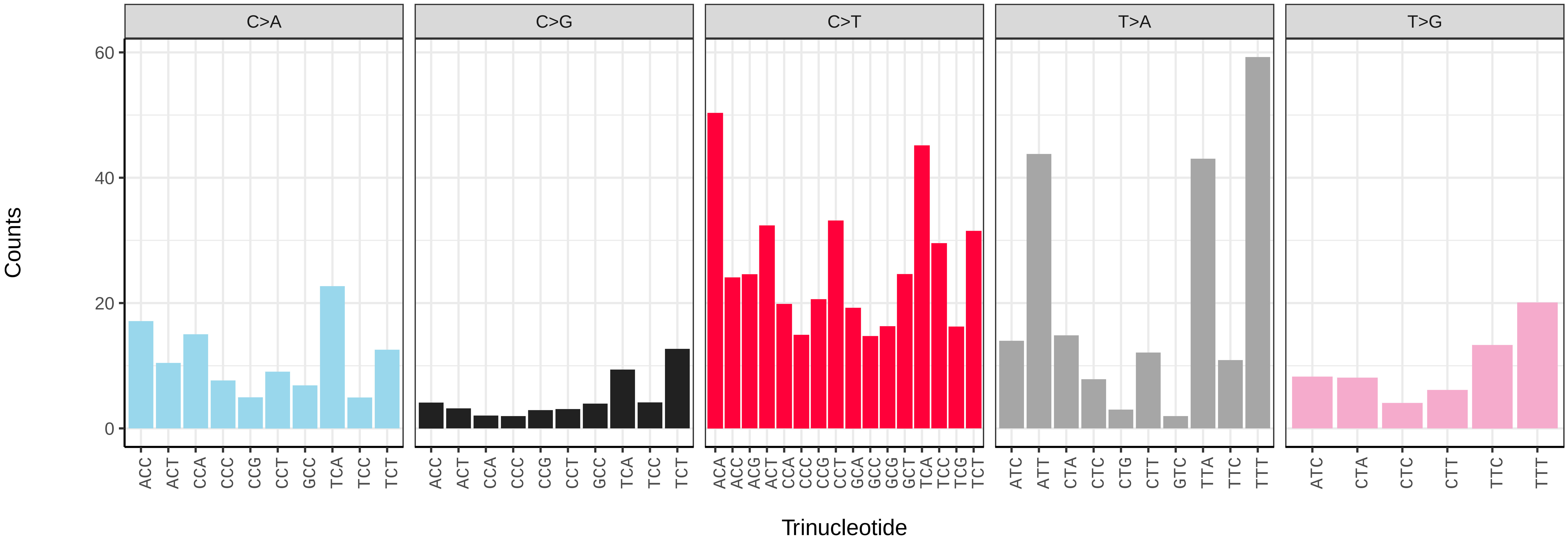

*Musotima nitidalis* (ilMusNiti1) germline mutational spectrum

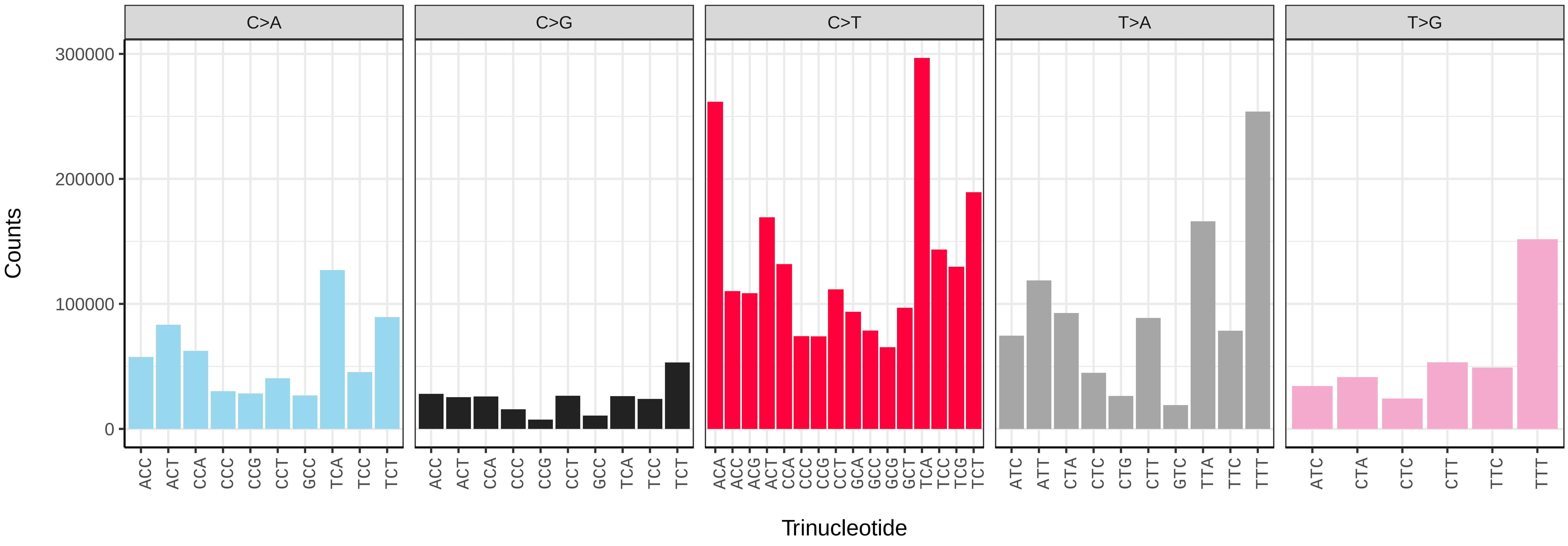


*Musotima nitidalis* (ilMusNiti1) somatic mutational spectrum (SBS96 classification)

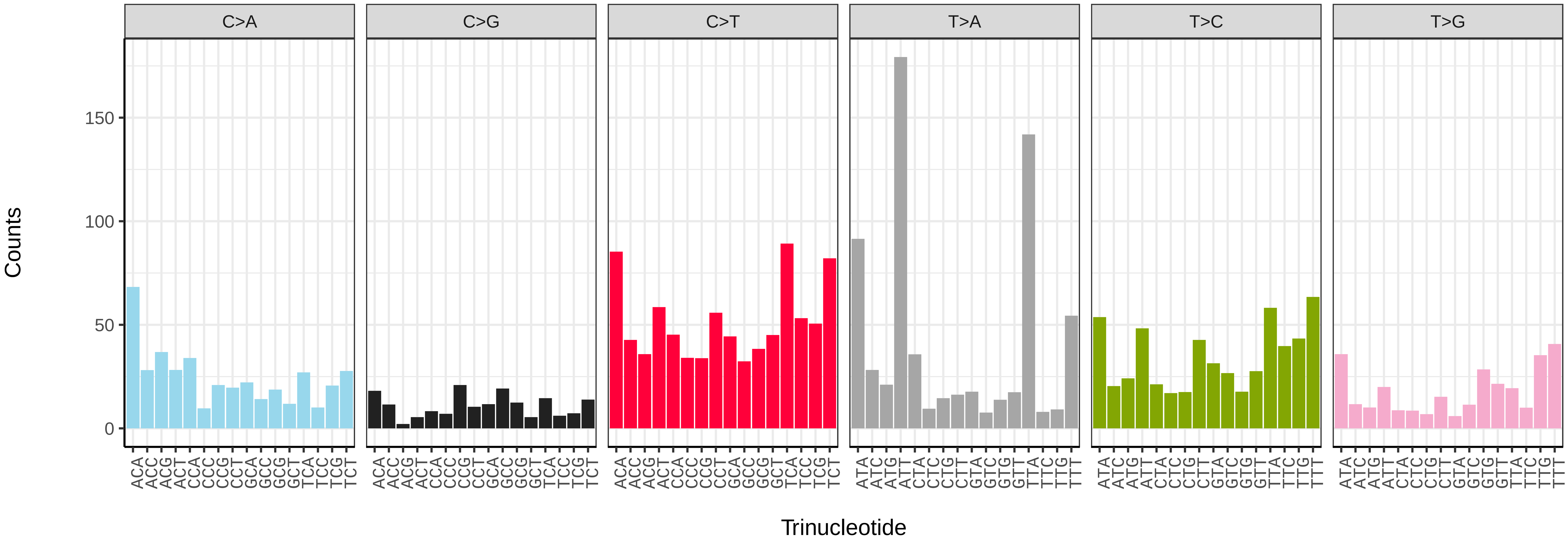


*Musotima nitidalis* (ilMusNiti1) somatic mutational spectrum (SBS52 classification)

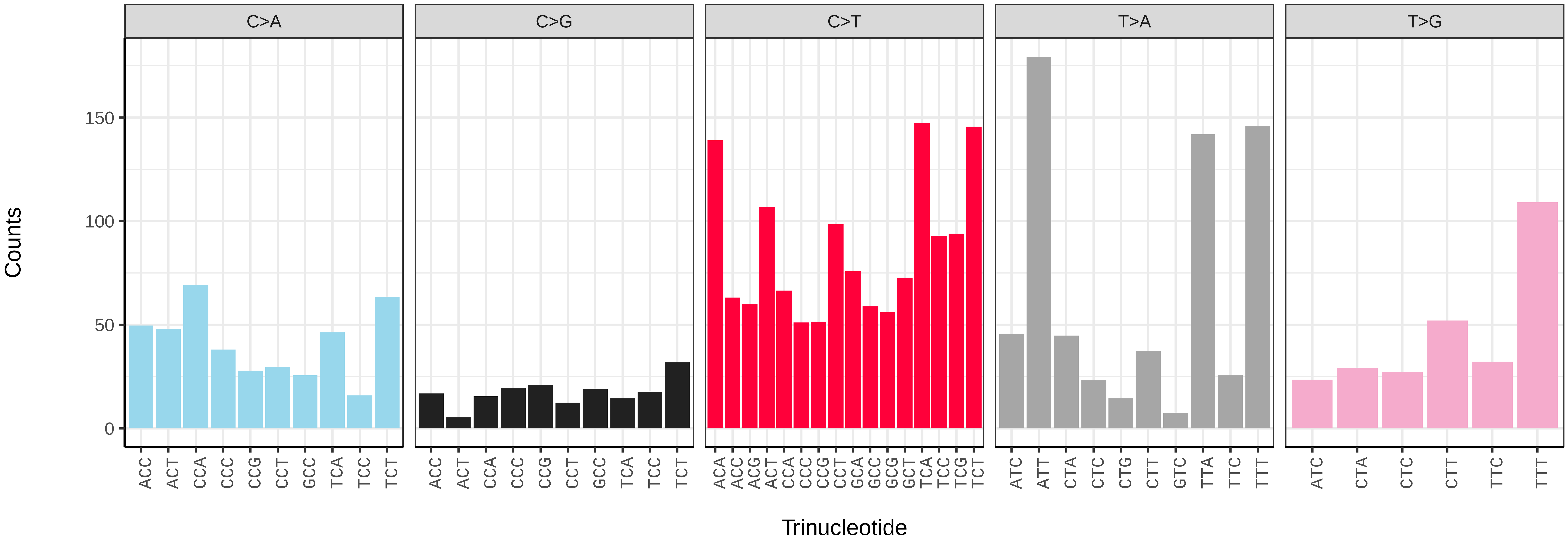

*Mythimna albipuncta* (ilMytAlbi1) germline mutational spectrum

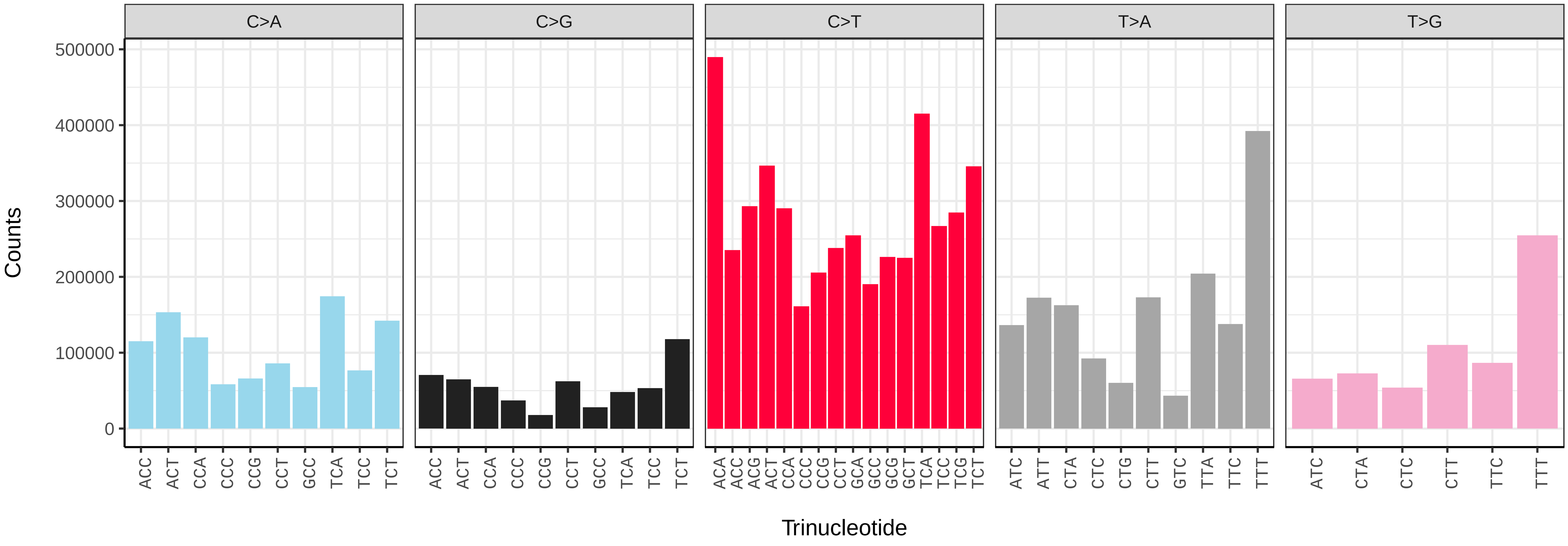


*Mythimna albipuncta* (ilMytAlbi1) somatic mutational spectrum (SBS96 classification)

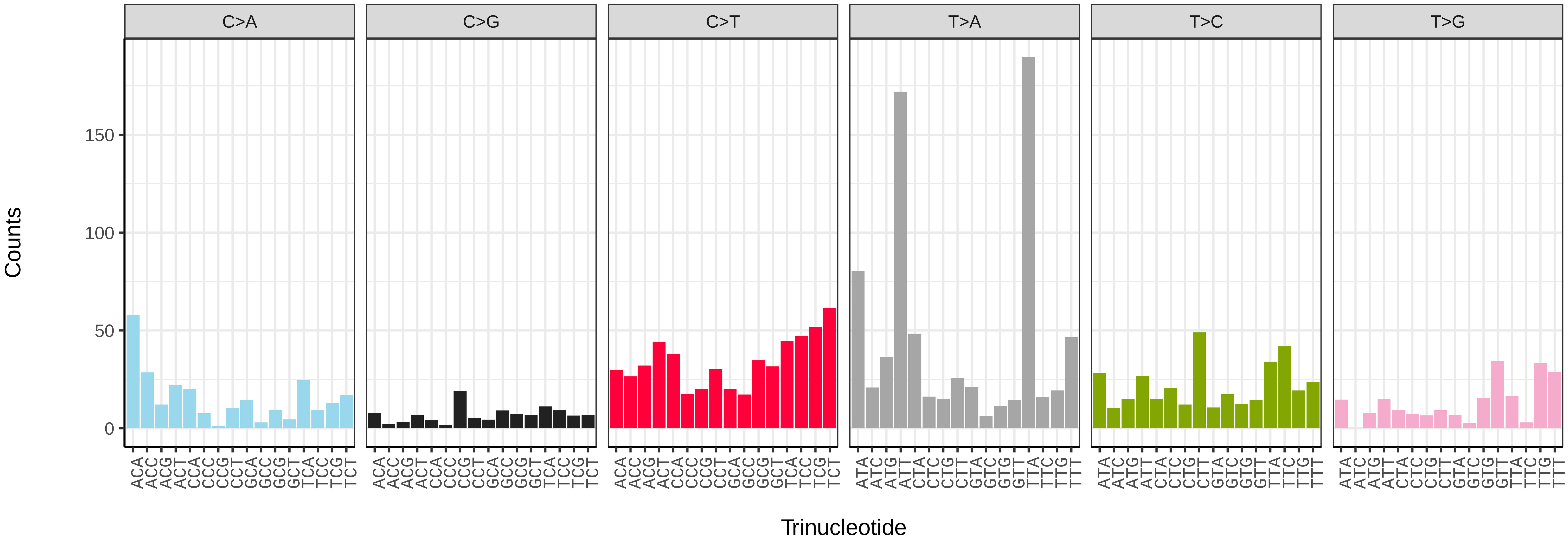


*Mythimna albipuncta* (ilMytAlbi1) somatic mutational spectrum (SBS52 classification)

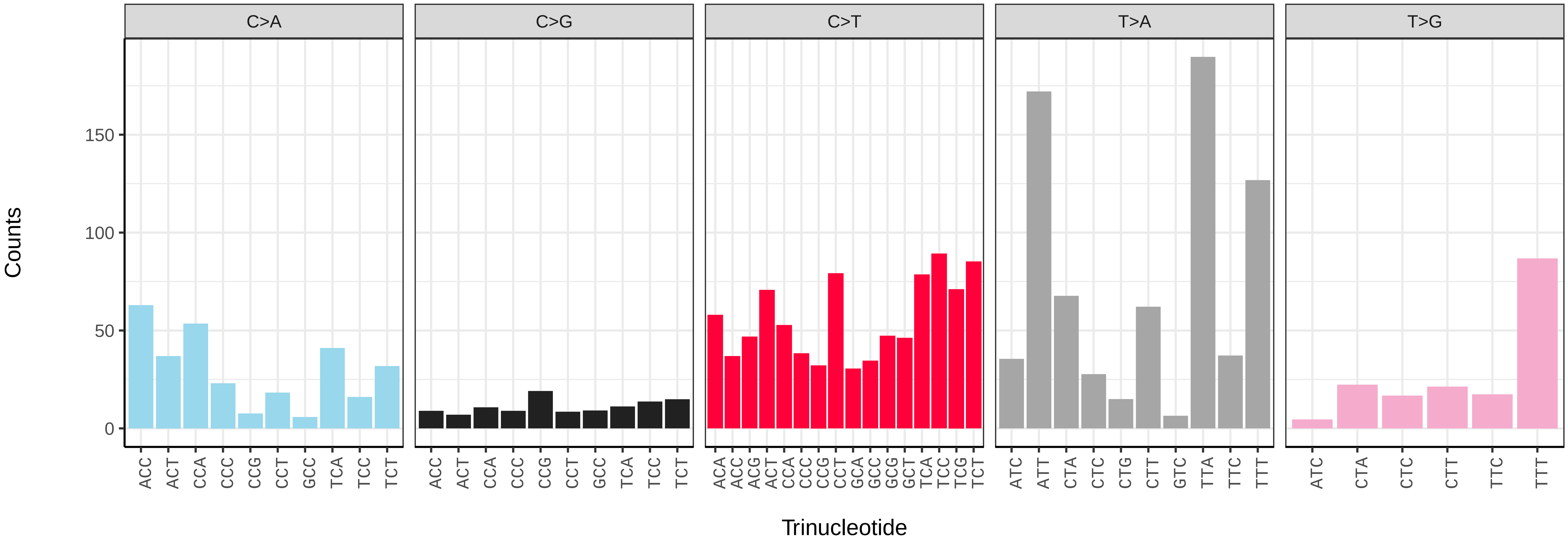

*Mythimna ferrago* (ilMytFerr1) germline mutational spectrum

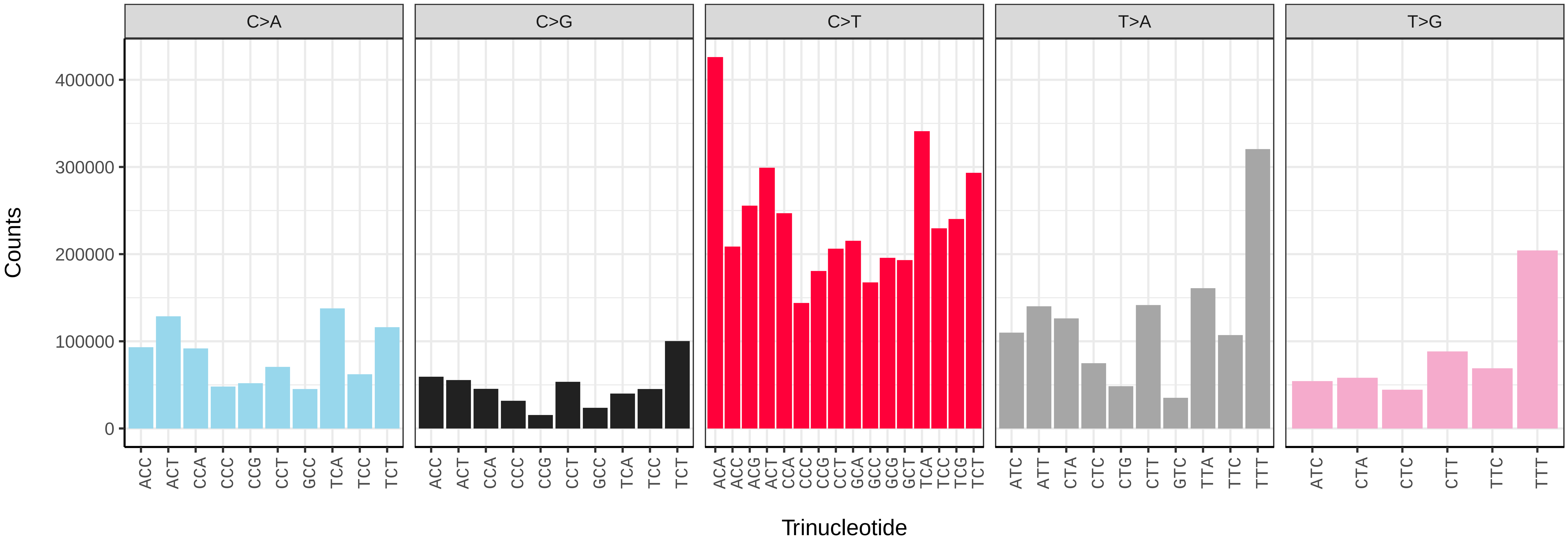


*Mythimna ferrago* (ilMytFerr1) somatic mutational spectrum (SBS96 classification)

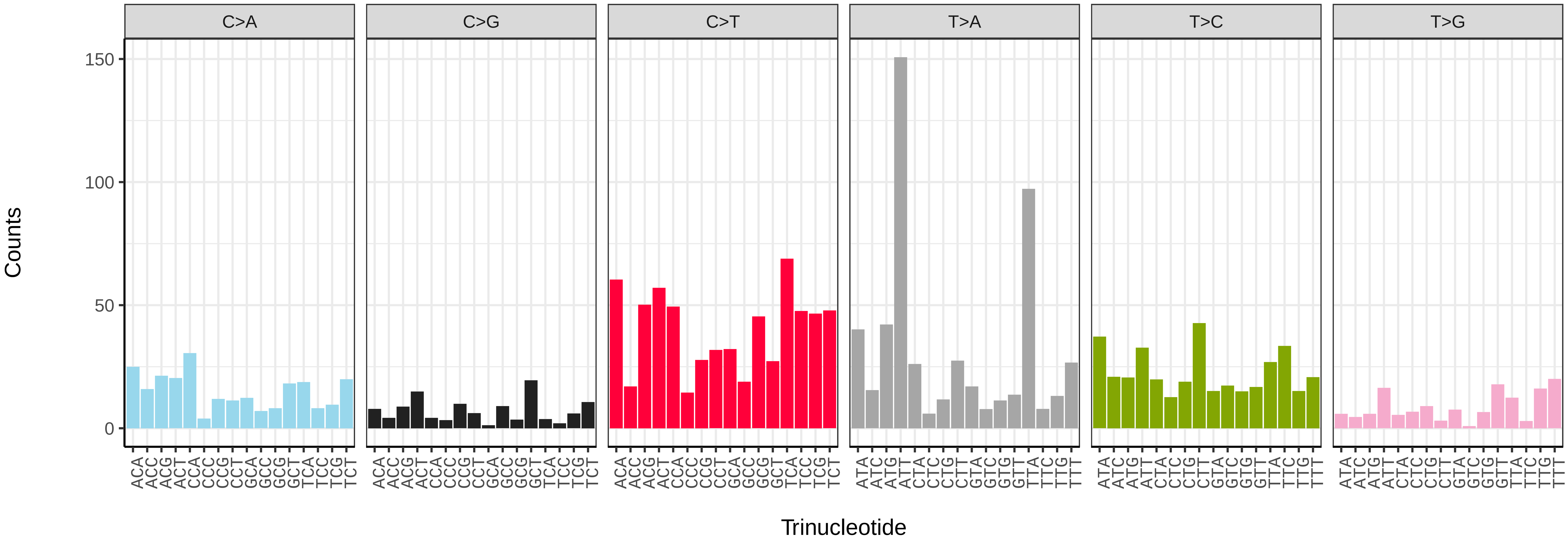


*Mythimna ferrago* (ilMytFerr1) somatic mutational spectrum (SBS52 classification)

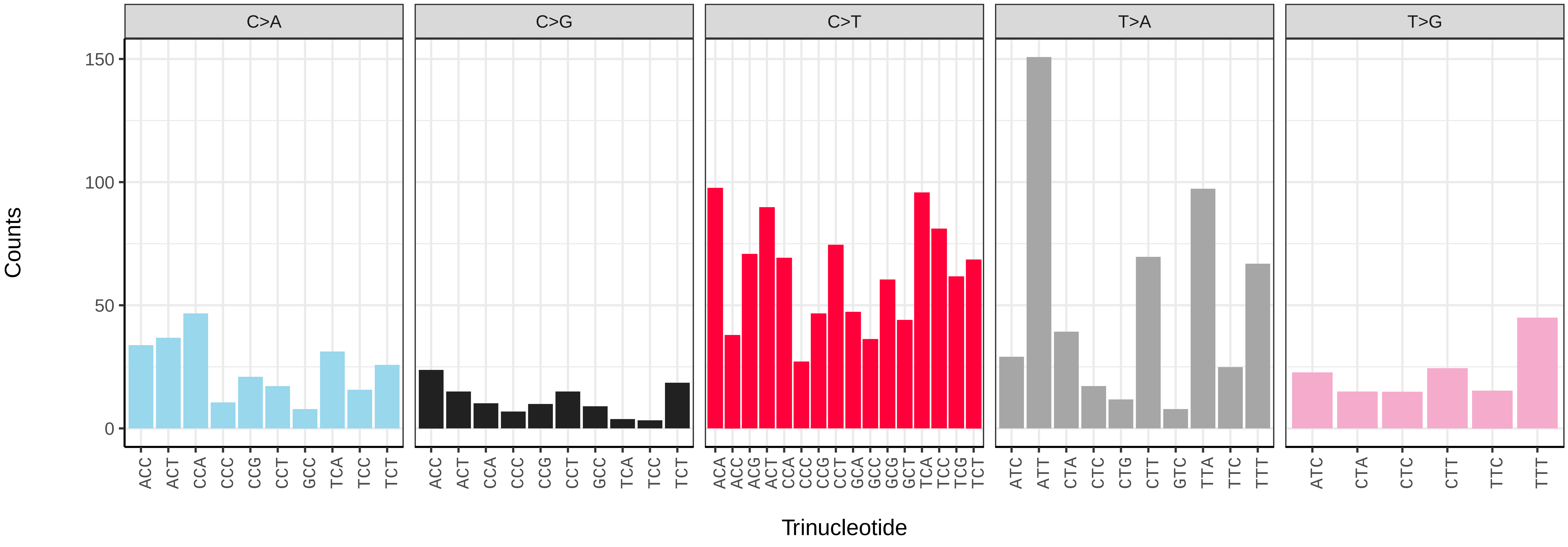

*Mythimna impura* (ilMytImpu1) germline mutational spectrum

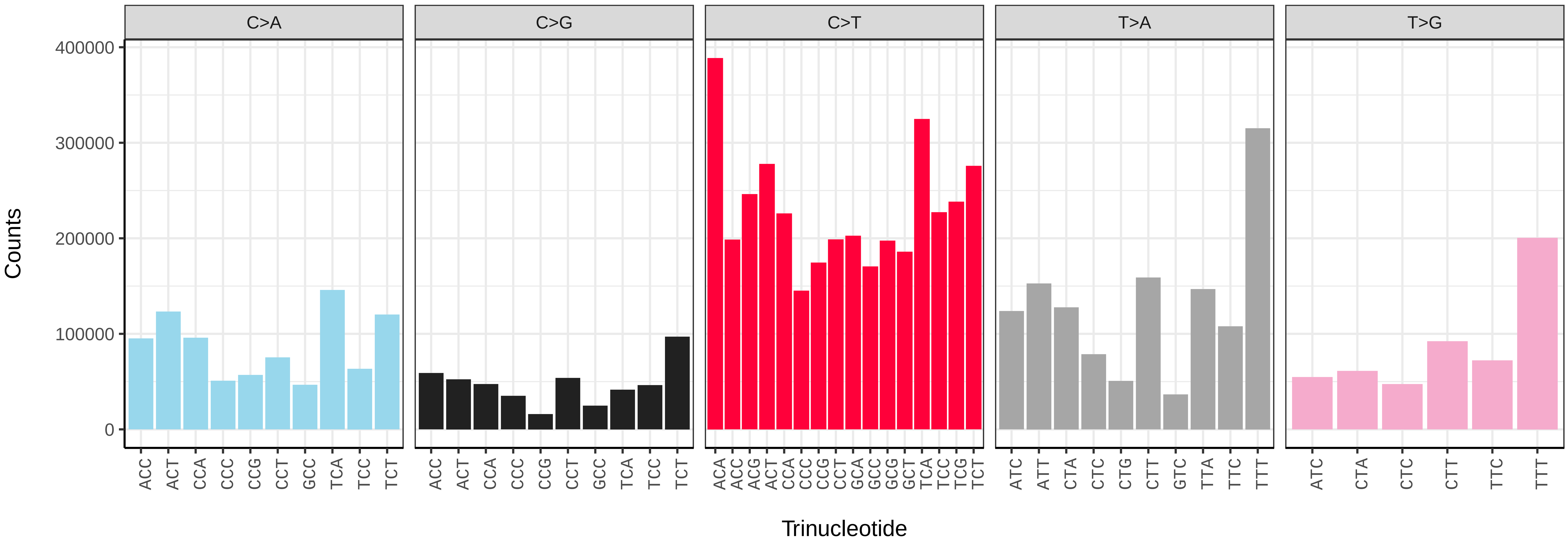


*Mythimna impura* (ilMytImpu1) somatic mutational spectrum (SBS96 classification)

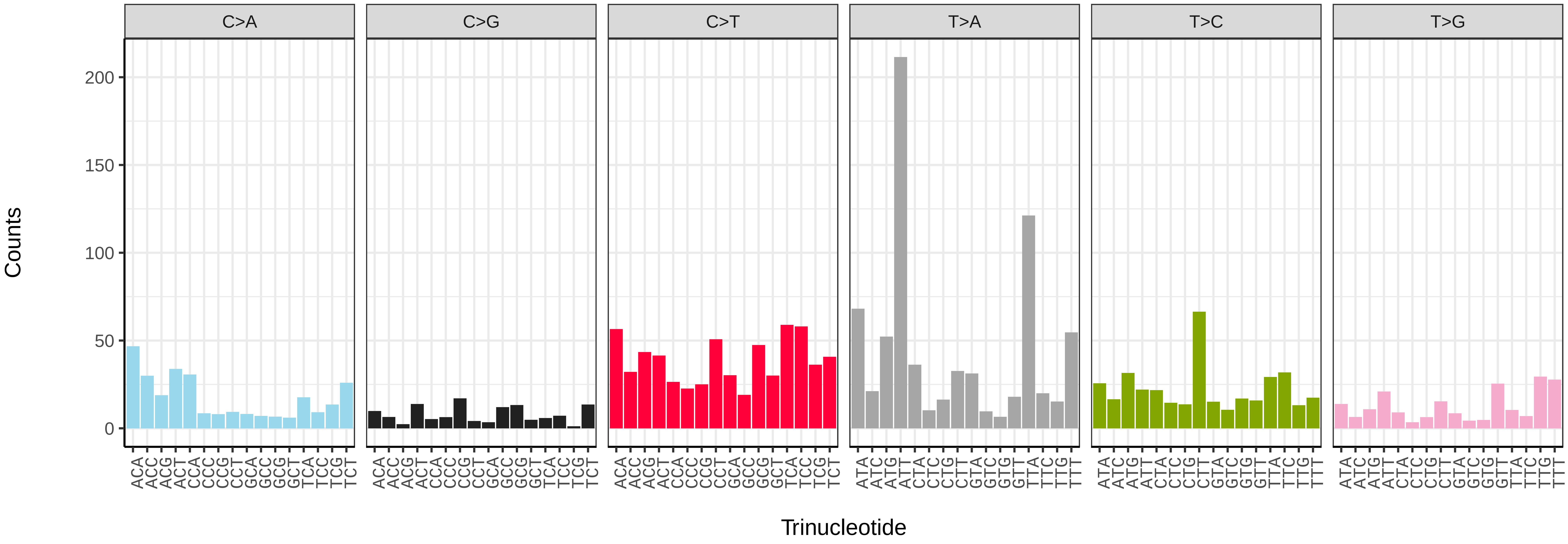


*Mythimna impura* (ilMytImpu1) somatic mutational spectrum (SBS52 classification)

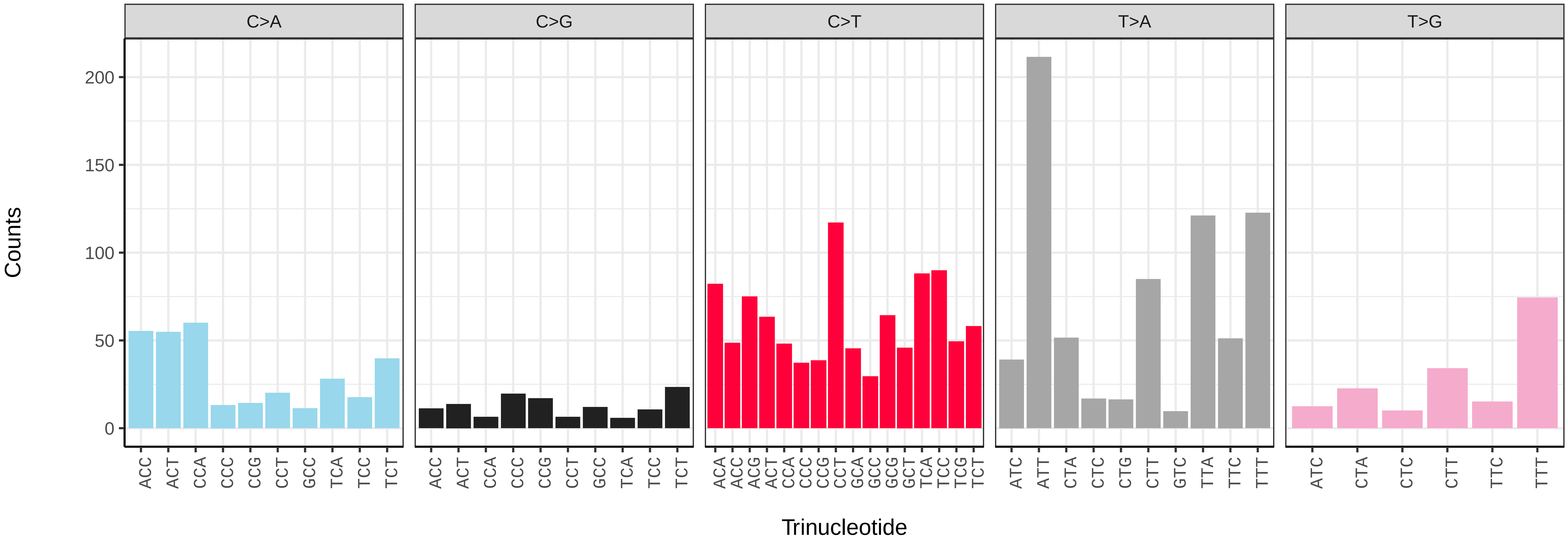

*Nematopogon swammerdamellus* (ilNemSwae1) germline mutational spectrum

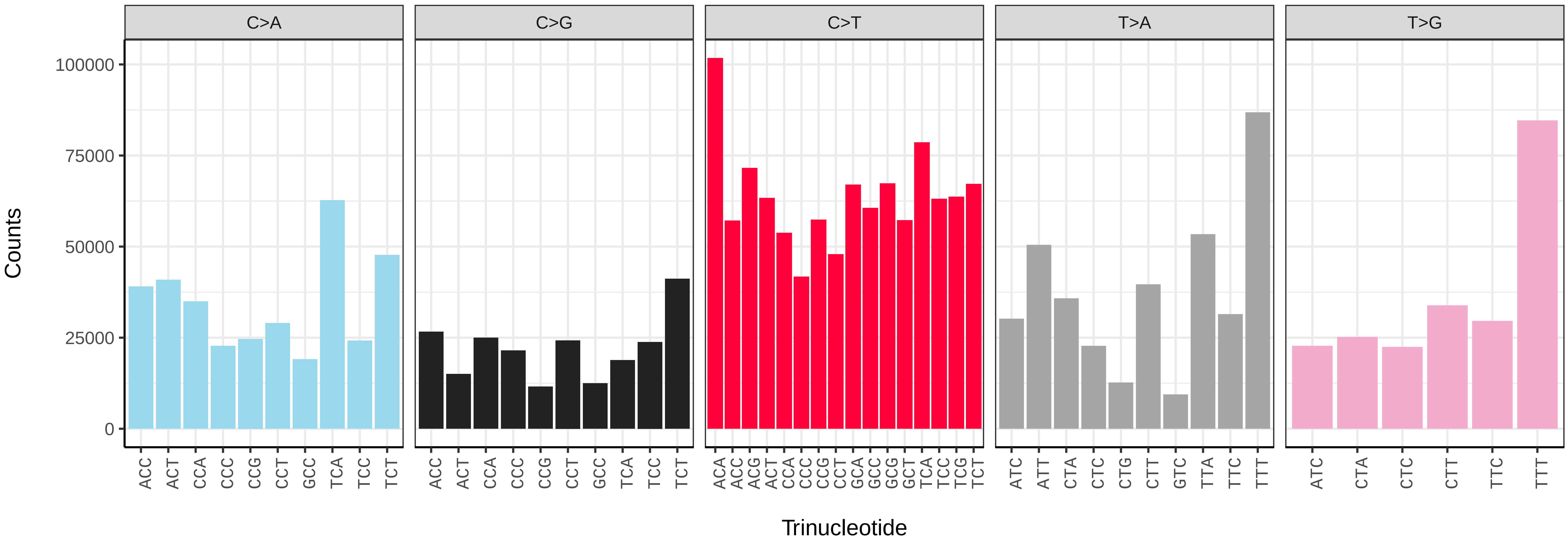


*Nematopogon swammerdamellus* (ilNemSwae1) somatic mutational spectrum (SBS96 classification)

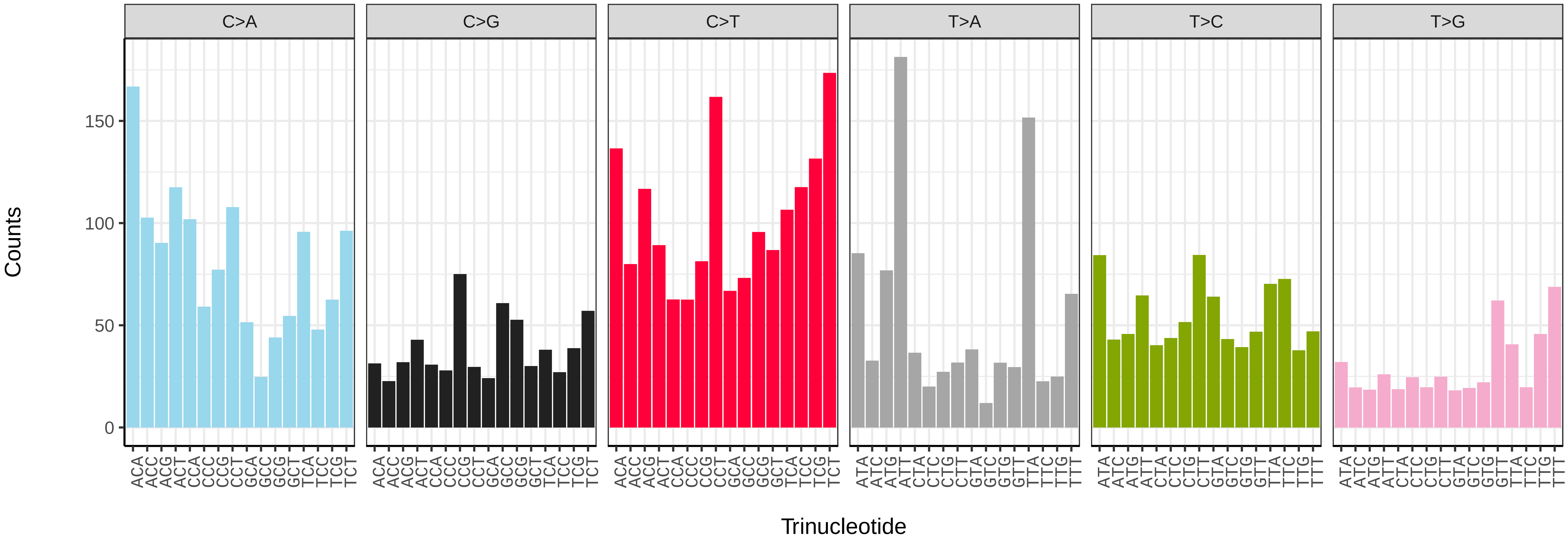


*Nematopogon swammerdamellus* (ilNemSwae1) somatic mutational spectrum (SBS52 classification)

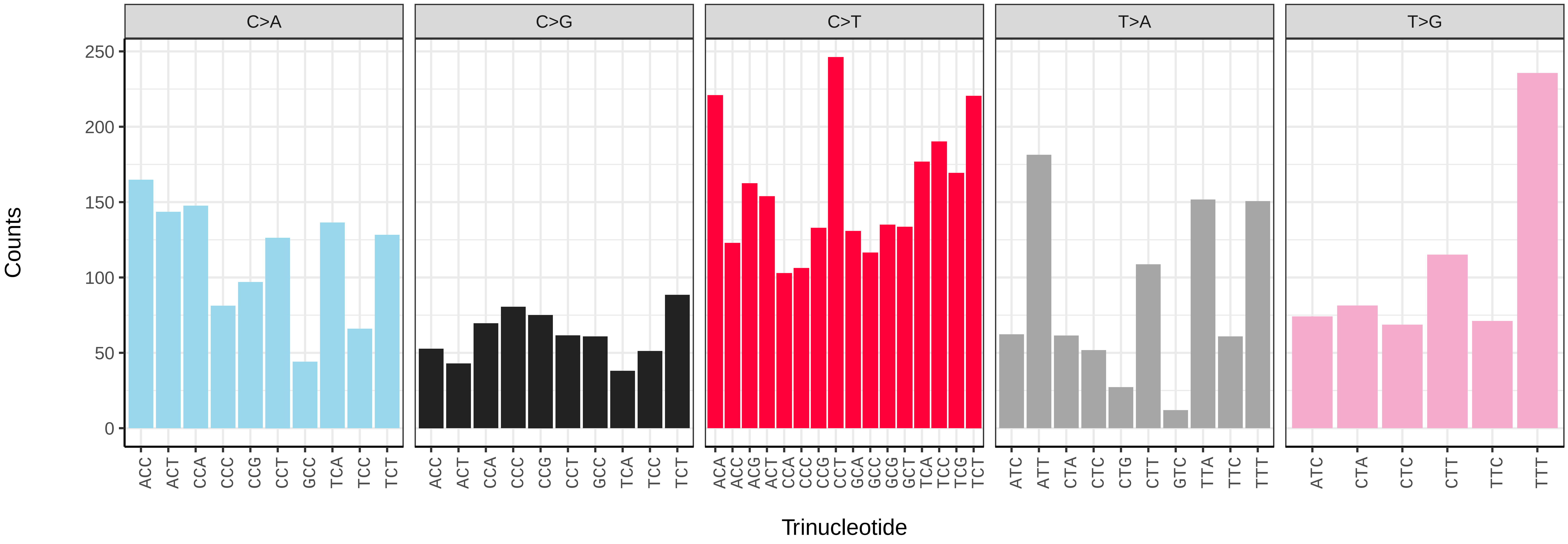

*Noctua fimbriata* (ilNocFimb1) germline mutational spectrum

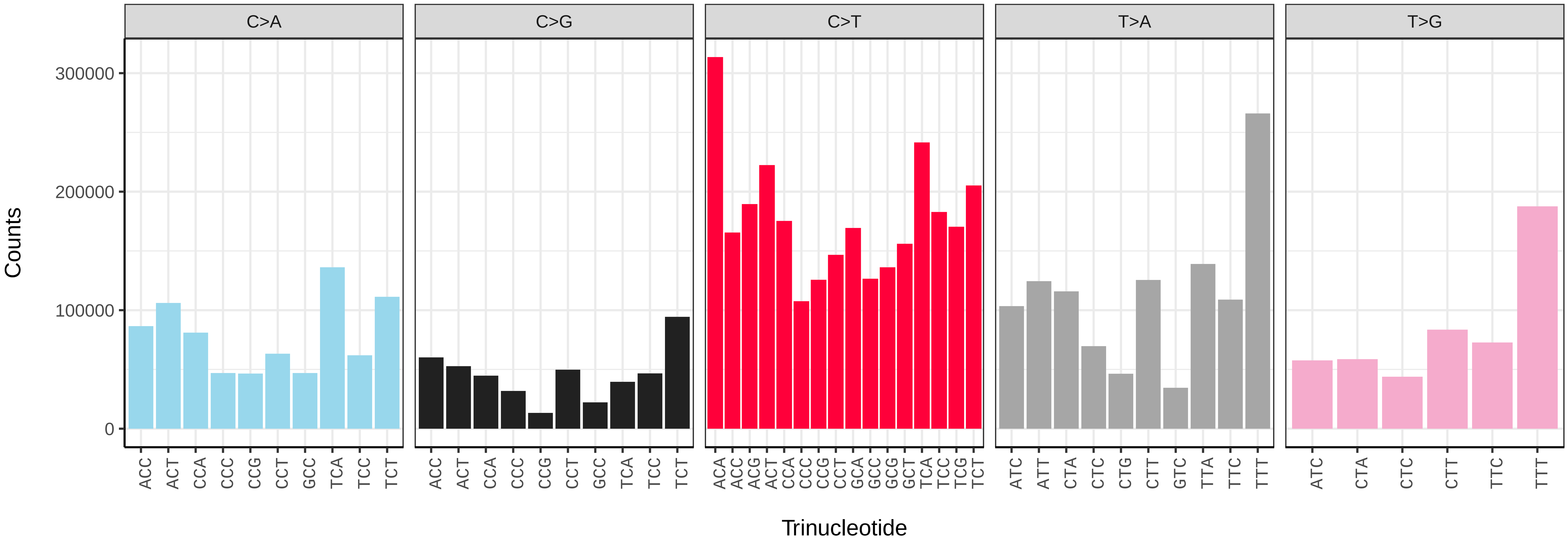


*Noctua fimbriata* (ilNocFimb1) somatic mutational spectrum (SBS96 classification)

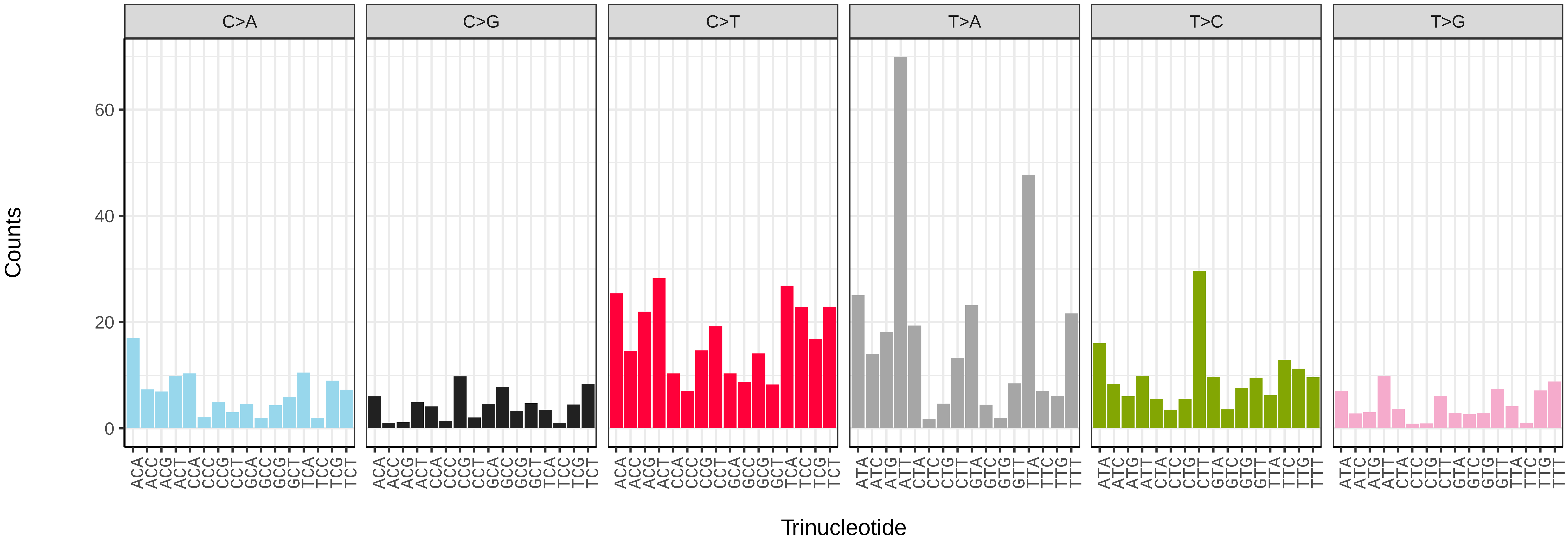


*Noctua fimbriata* (ilNocFimb1) somatic mutational spectrum (SBS52 classification)

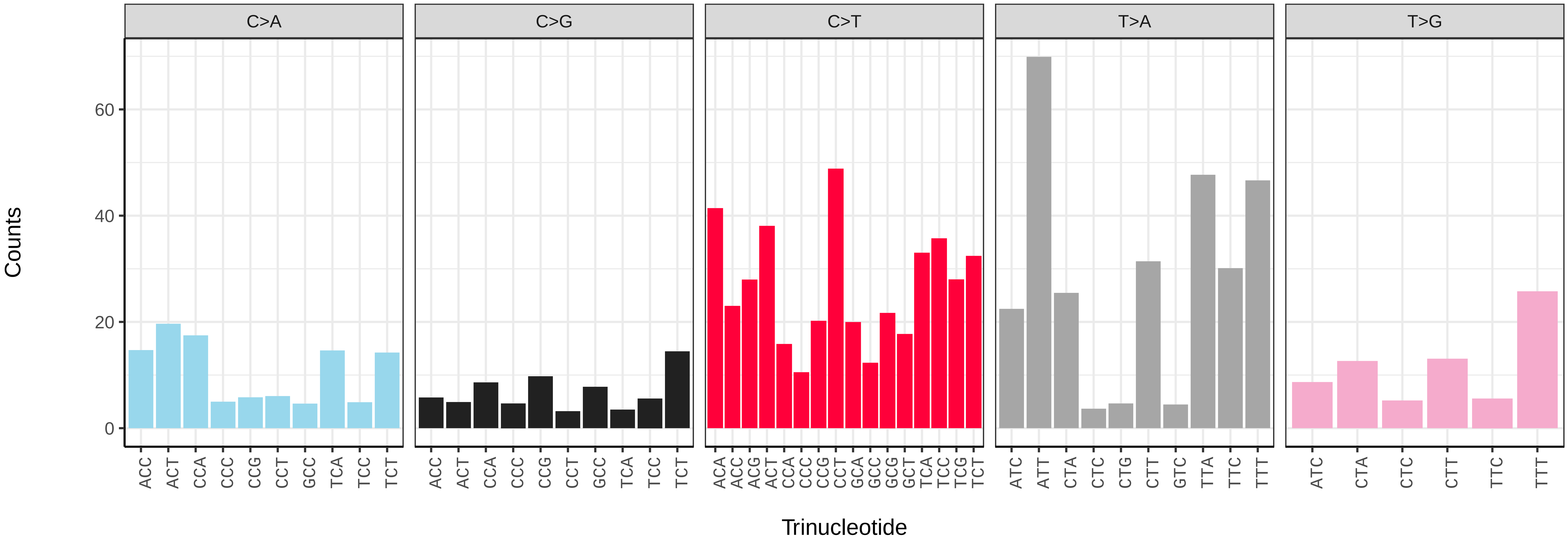

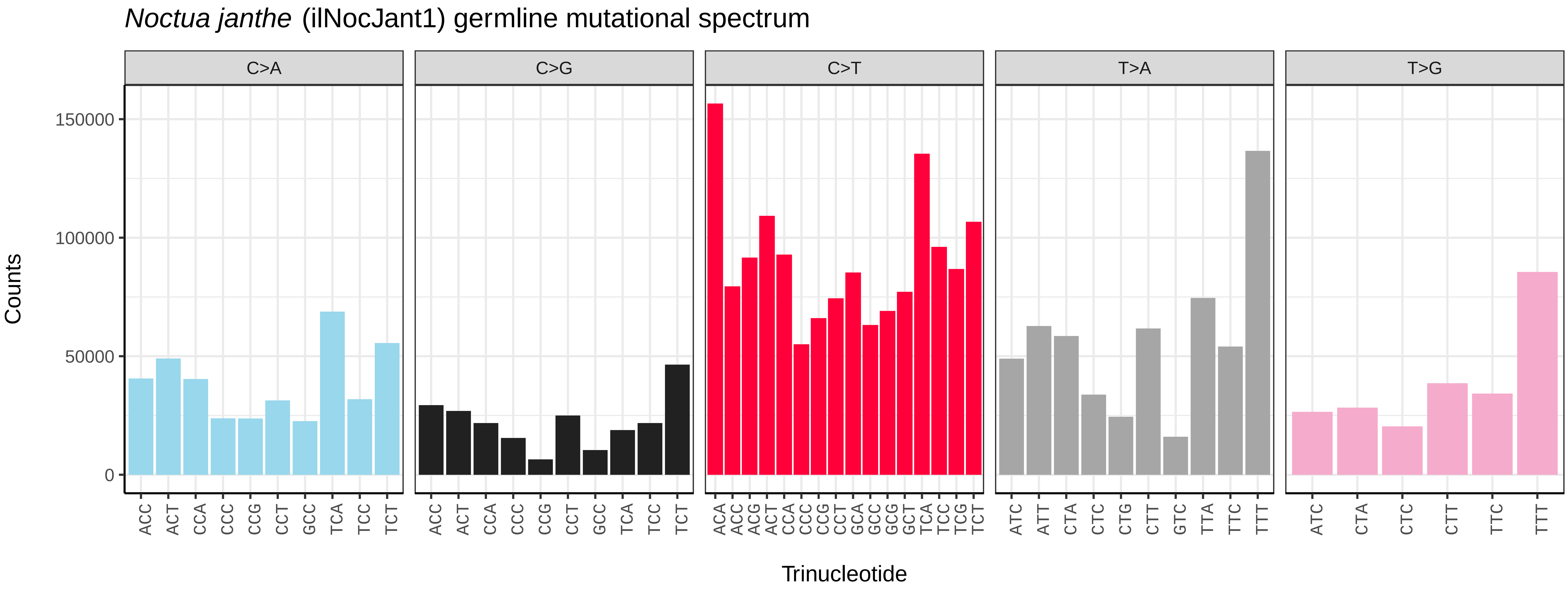
*Noctua janthe* (ilNocJant1) germline mutational spectrum
C>A
C>G
C>T
T>A
T>G
Counts
150000
100000
50000
0
ACC ACT CCA CCC CCG CCT GCC TCA TCC TCT
ACC ACT CCA CCC CCG CCT GCC TCA TCC TCT
ACA ACC ACG ACT CCA CCC CCG CCT GCA GCC GCG GCT TCA TCC TCG TCT
ATC ATT CTA CTC CTG CTT GTC TTA TTC TTT
ATC CTA CTC CTT TTC TTT
Trinucleotide

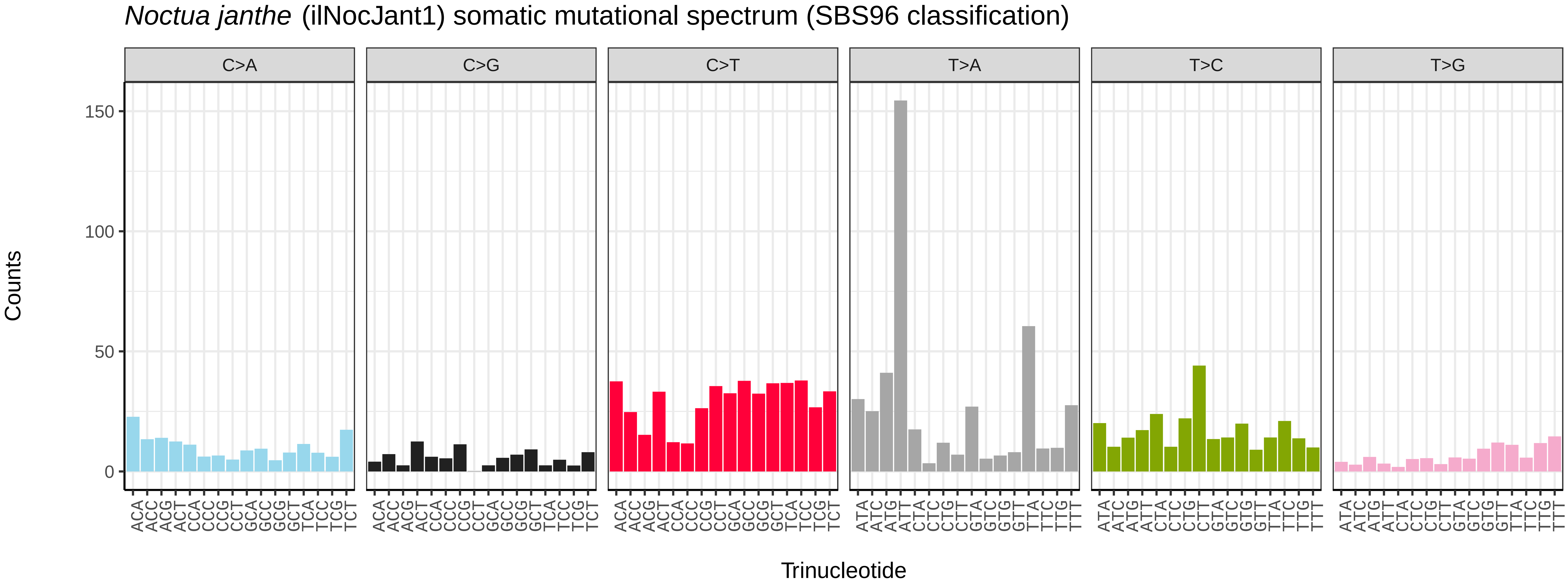
*Noctua janthe* (ilNocJant1) somatic mutational spectrum (SBS96 classification)
C>A
C>G
C>T
T>A
T>C
T>G
Counts
150
100
50
0
ACA ACC ACG ACT CCA CCC CCG CCT GCA GCC GCG GCT TCA TCC TCG TCT
ATA ATC ATG ATT CTA CTC CTG CTT GTA GTC GTG GTT TTA TTC TTG TTT
Trinucleotide

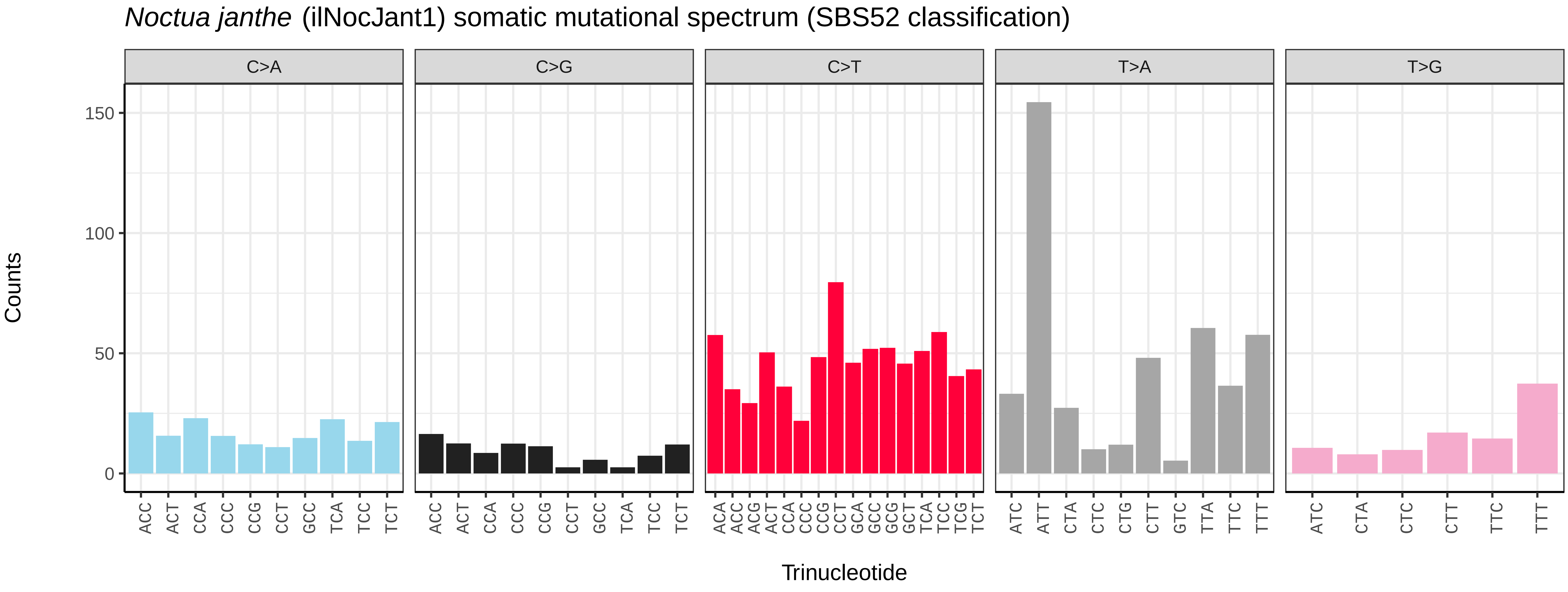
*Noctua janthe* (ilNocJant1) somatic mutational spectrum (SBS52 classification)
C>A
C>G
C>T
T>A
T>G
Counts
150
100
50
0
ACC ACT CCA CCC CCG CCT GCC TCA TCC TCT
ACC ACT CCA CCC CCG CCT GCC TCA TCC TCT
ACA ACC ACG ACT CCA CCC CCG CCT GCA GCC GCG GCT TCA TCC TCG TCT
ATC ATT CTA CTC CTG CTT GTC TTA TTC TTT
ATC CTA CTC CTT TTC TTT
Trinucleotide

*Noctua pronuba* (ilNocPron1) germline mutational spectrum

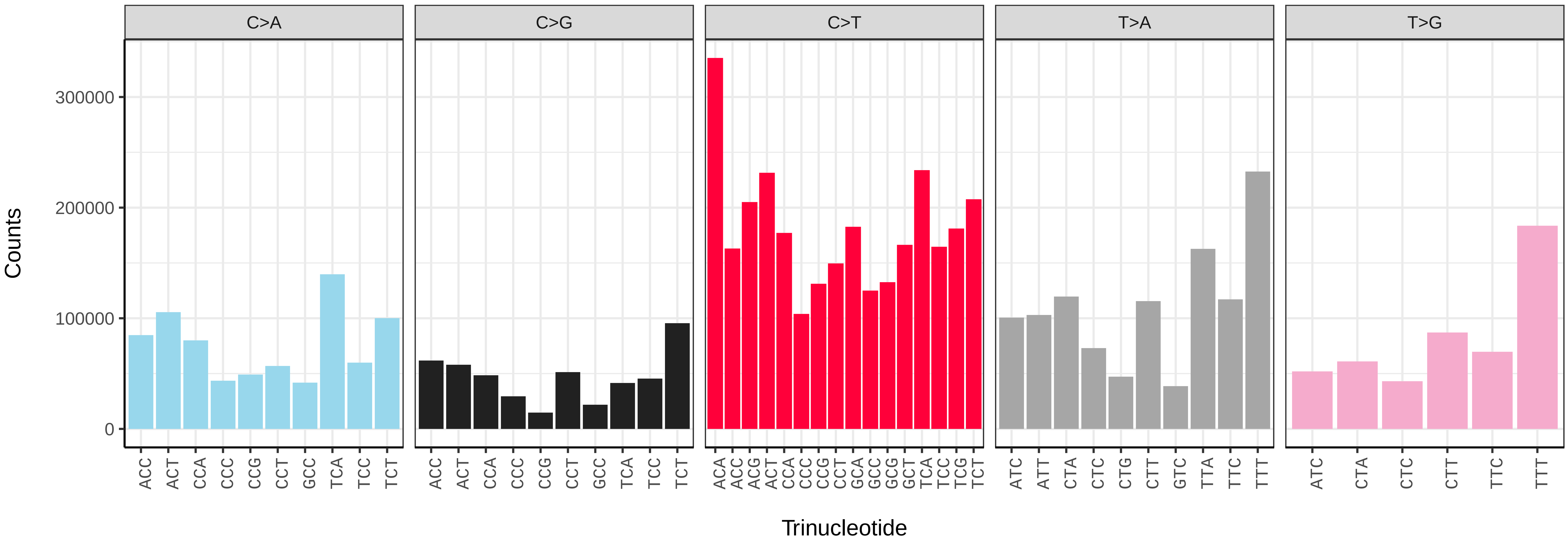


*Noctua pronuba* (ilNocPron1) somatic mutational spectrum (SBS96 classification)

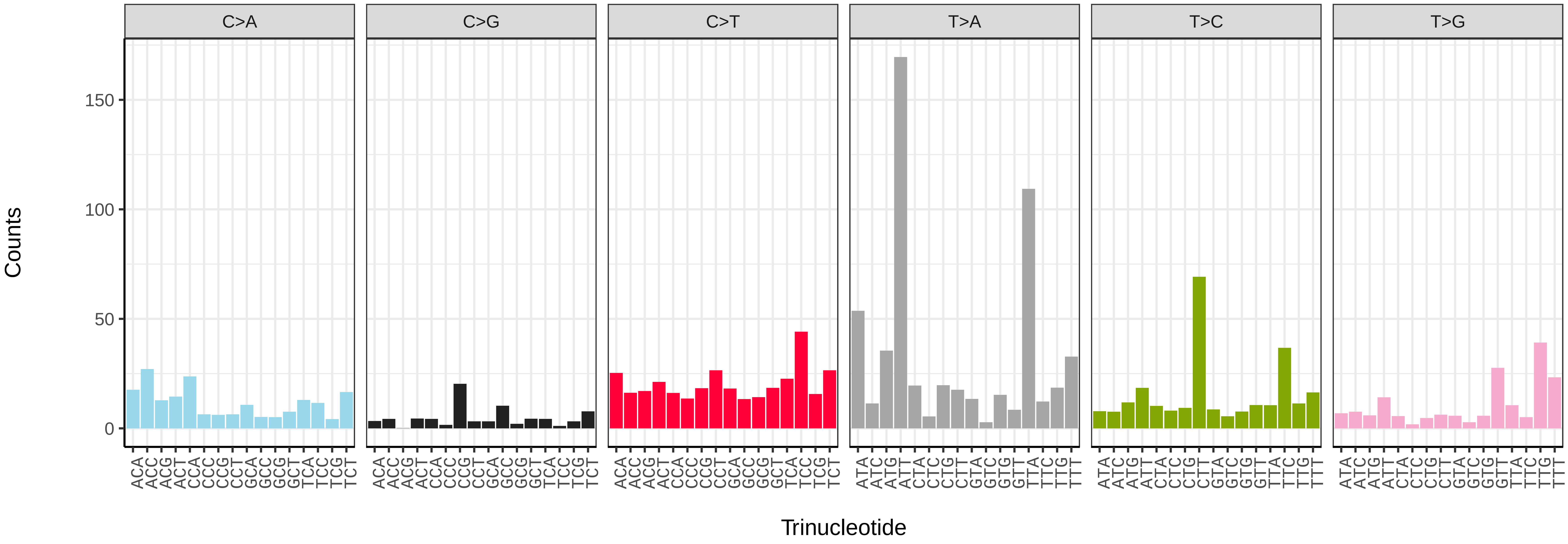


*Noctua pronuba* (ilNocPron1) somatic mutational spectrum (SBS52 classification)

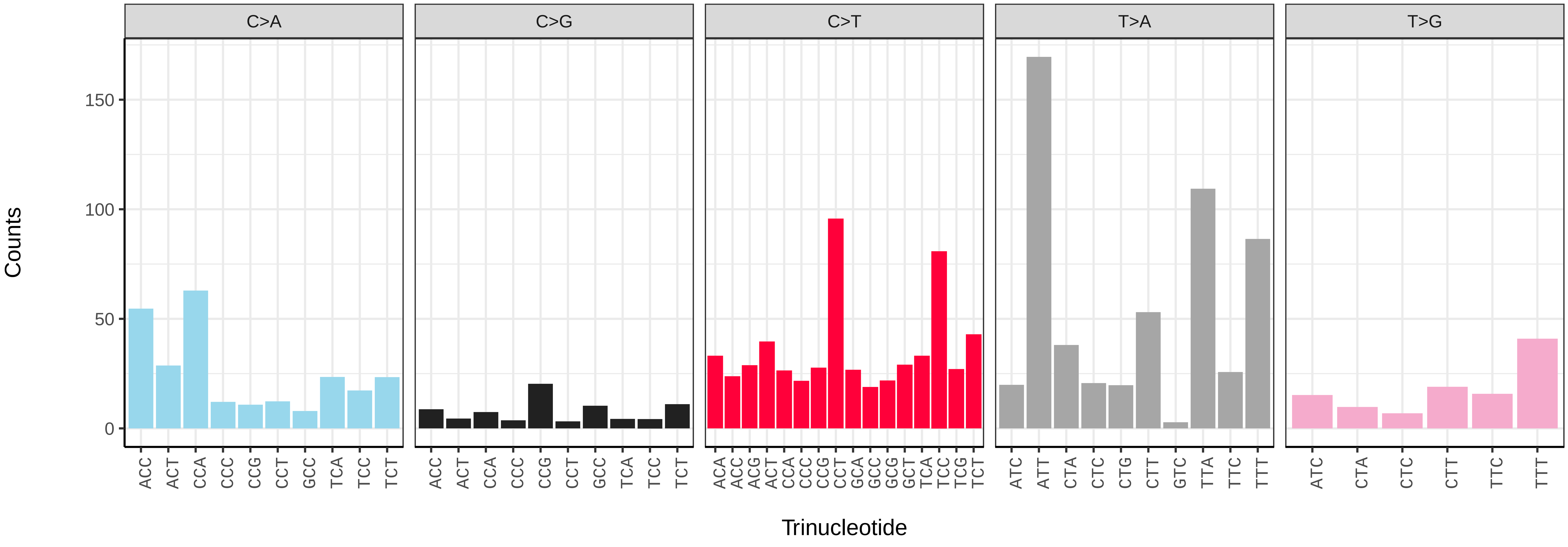

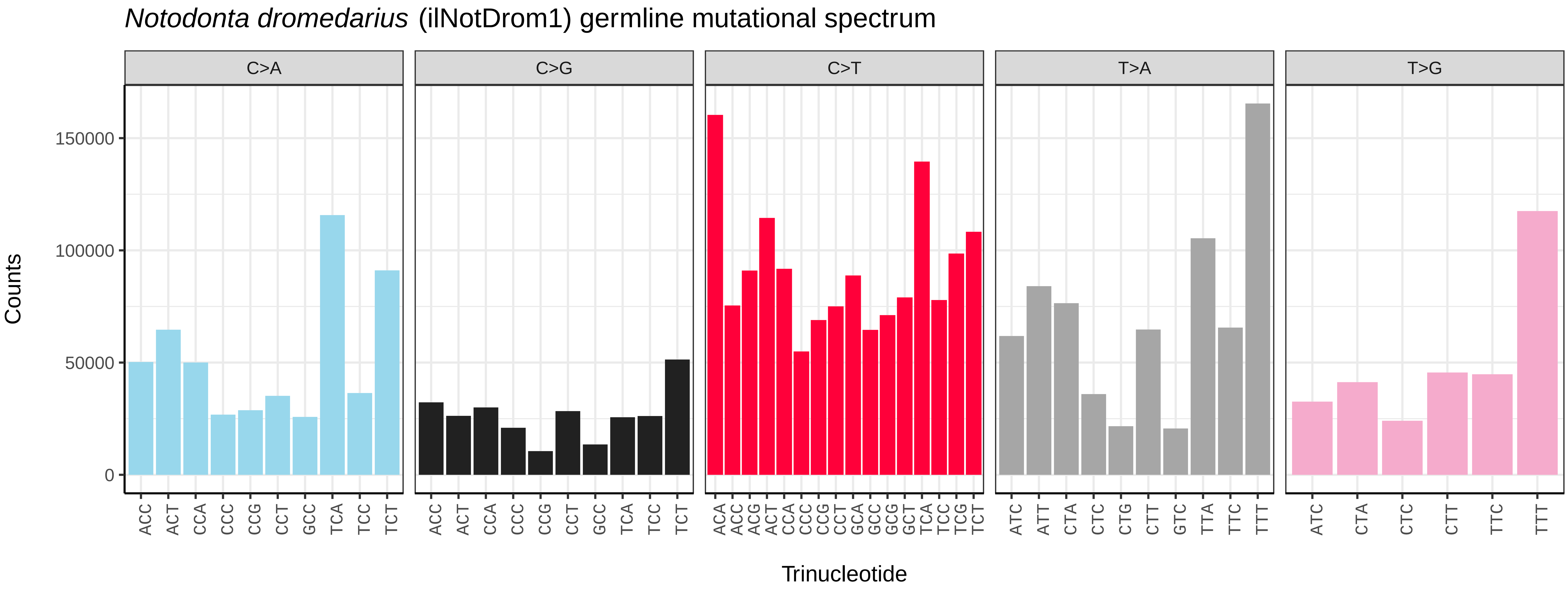
*Notodonta dromedarius* (ilNotDrom1) germline mutational spectrum
C>A
C>G
C>T
T>A
T>G
Counts
150000
100000
50000
0
Trinucleotide

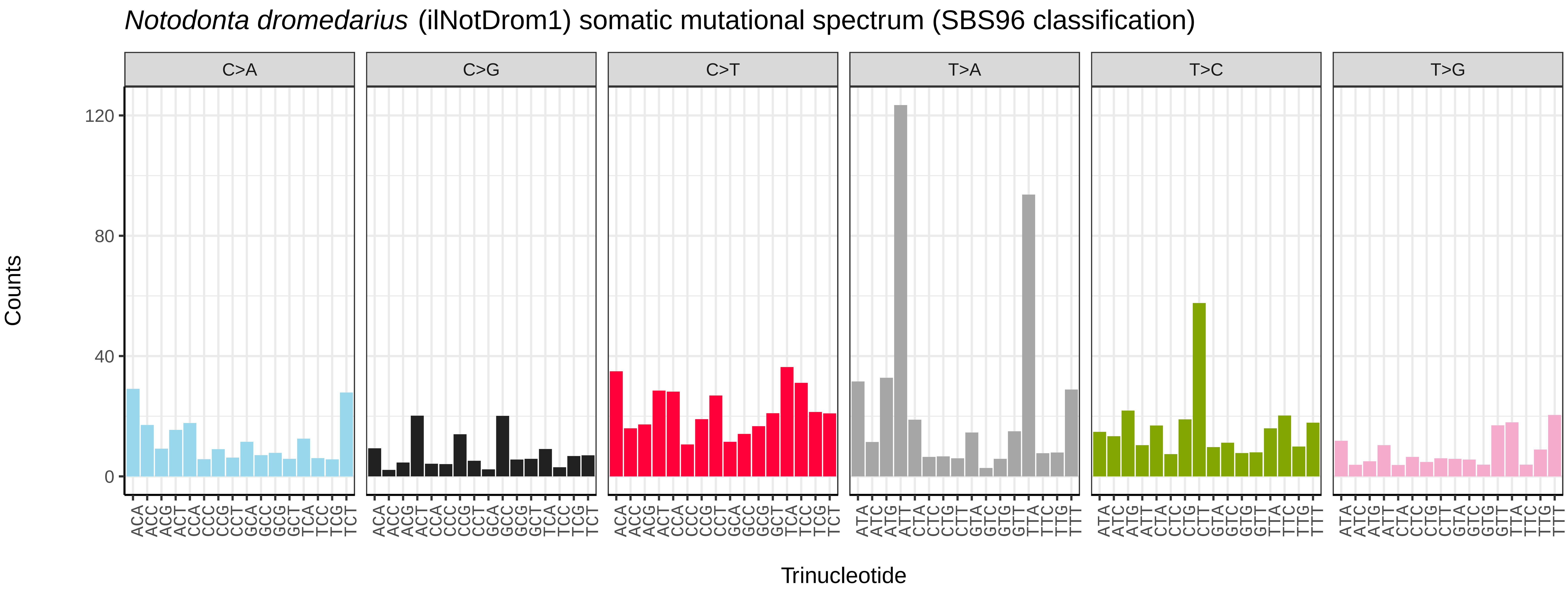
*Notodonta dromedarius* (ilNotDrom1) somatic mutational spectrum (SBS96 classification)
C>A
C>G
C>T
T>A
T>C
T>G
Counts
120
80
40
0
Trinucleotide

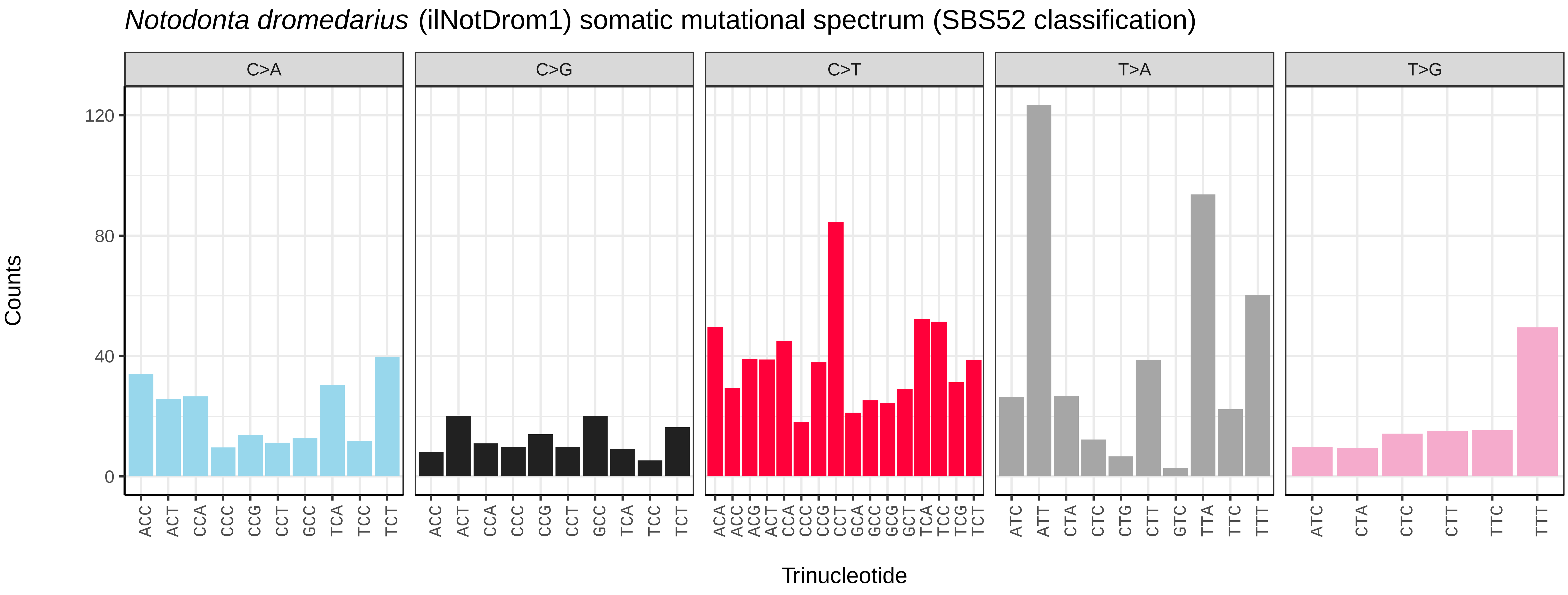
*Notodonta dromedarius* (ilNotDrom1) somatic mutational spectrum (SBS52 classification)
C>A
C>G
C>T
T>A
T>G
Counts
120
80
40
0
Trinucleotide

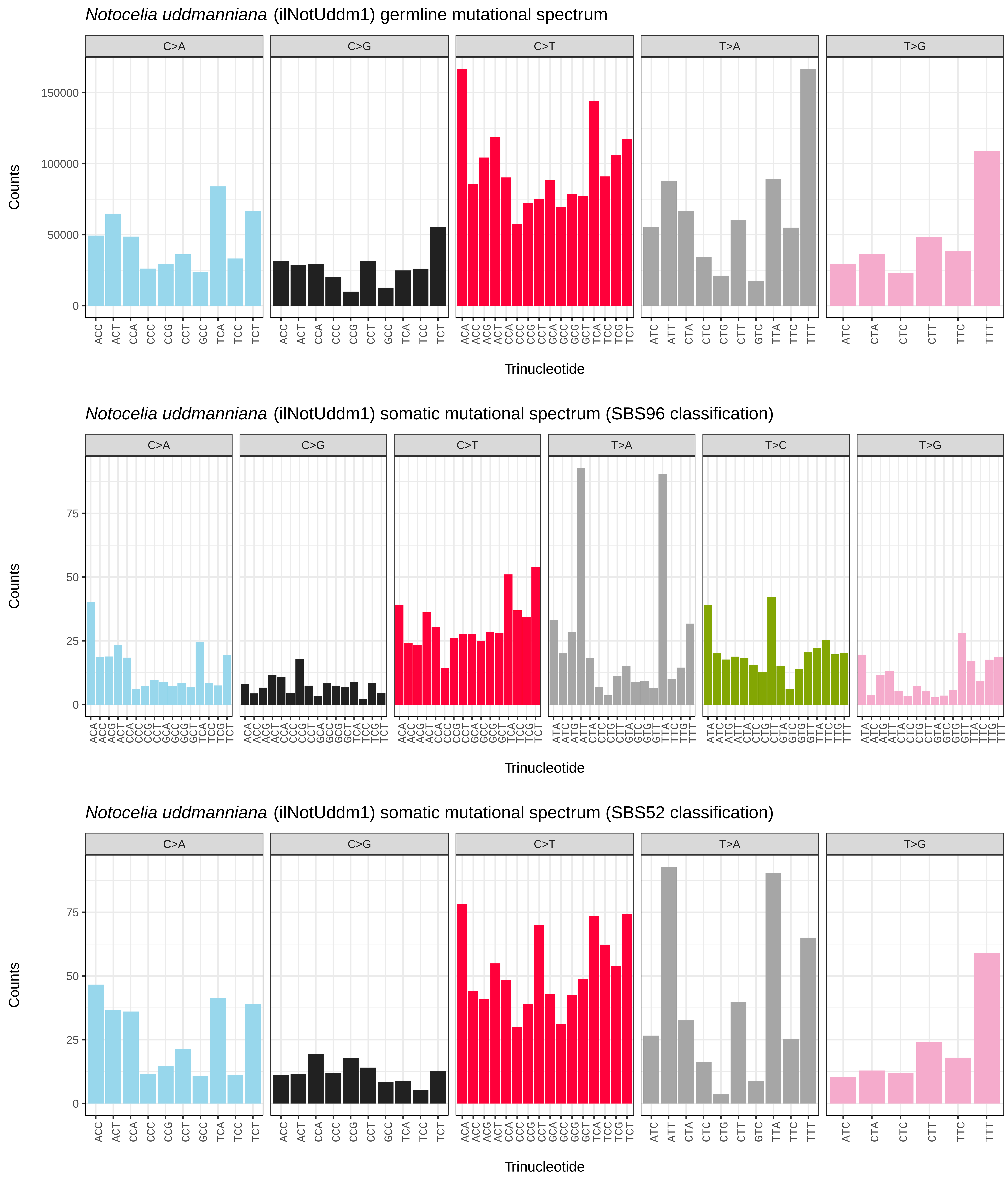

Notocelia uddmanniana (ilNotUddm1) germline mutational spectrum
C>A
C>G
C>T
T>A
T>G
Counts
0
50000
100000
150000
Trinucleotide
Notocelia uddmanniana (ilNotUddm1) somatic mutational spectrum (SBS96 classification)
C>A
C>G
C>T
T>A
T>C
T>G
Counts
0
25
50
75
Trinucleotide
Notocelia uddmanniana (ilNotUddm1) somatic mutational spectrum (SBS52 classification)
C>A
C>G
C>T
T>A
T>G
Counts
0
25
50
75
Trinucleotide

*Notodonta ziczac* (ilNotZicz1) germline mutational spectrum

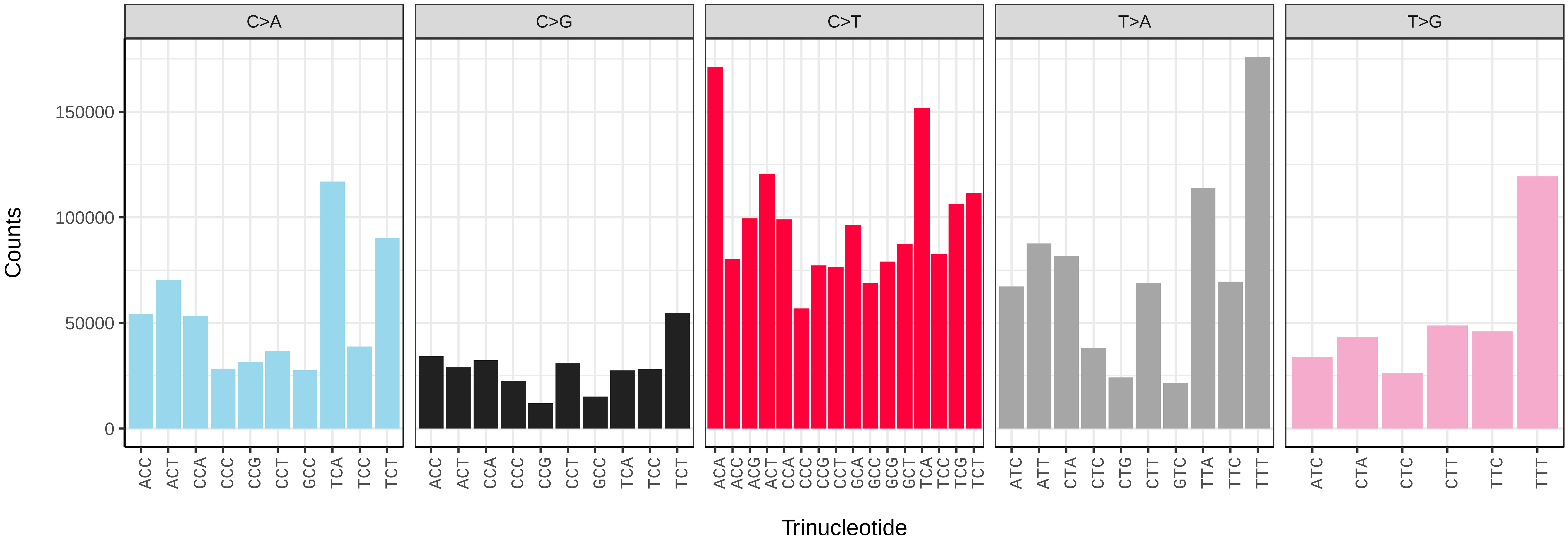


*Notodonta ziczac* (ilNotZicz1) somatic mutational spectrum (SBS96 classification)

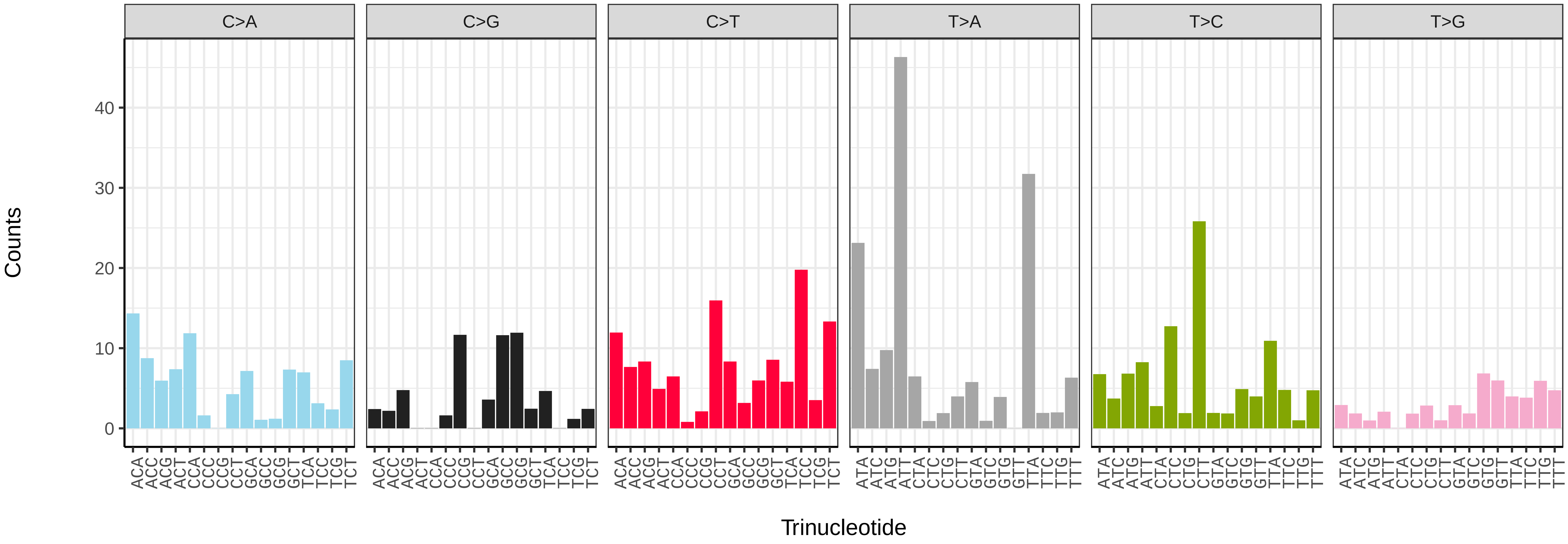


*Notodonta ziczac* (ilNotZicz1) somatic mutational spectrum (SBS52 classification)

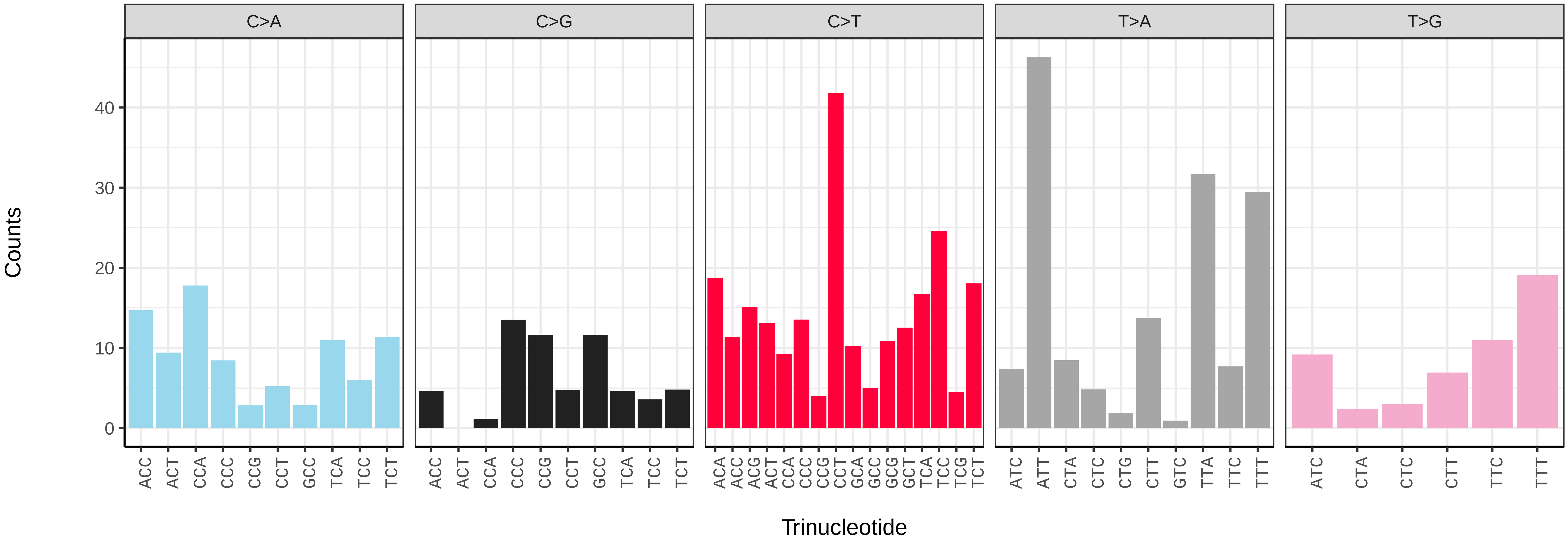

*Nycteola revayana* (ilNycReva1) germline mutational spectrum

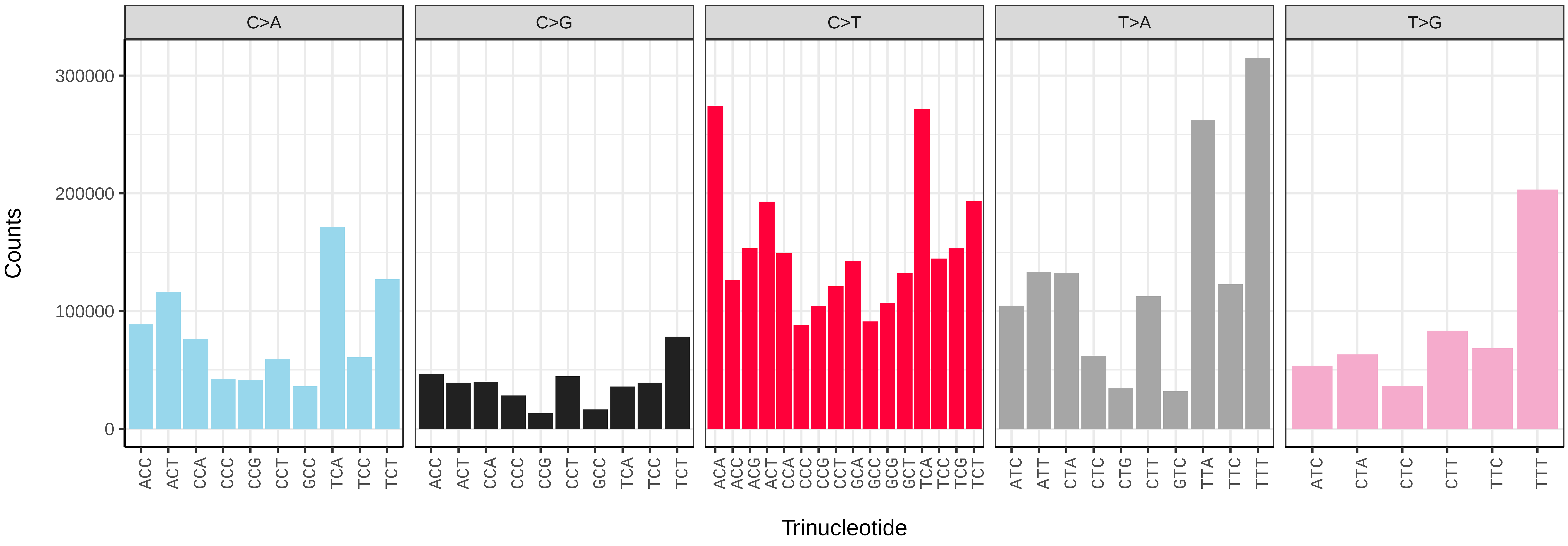


*Nycteola revayana* (ilNycReva1) somatic mutational spectrum (SBS96 classification)

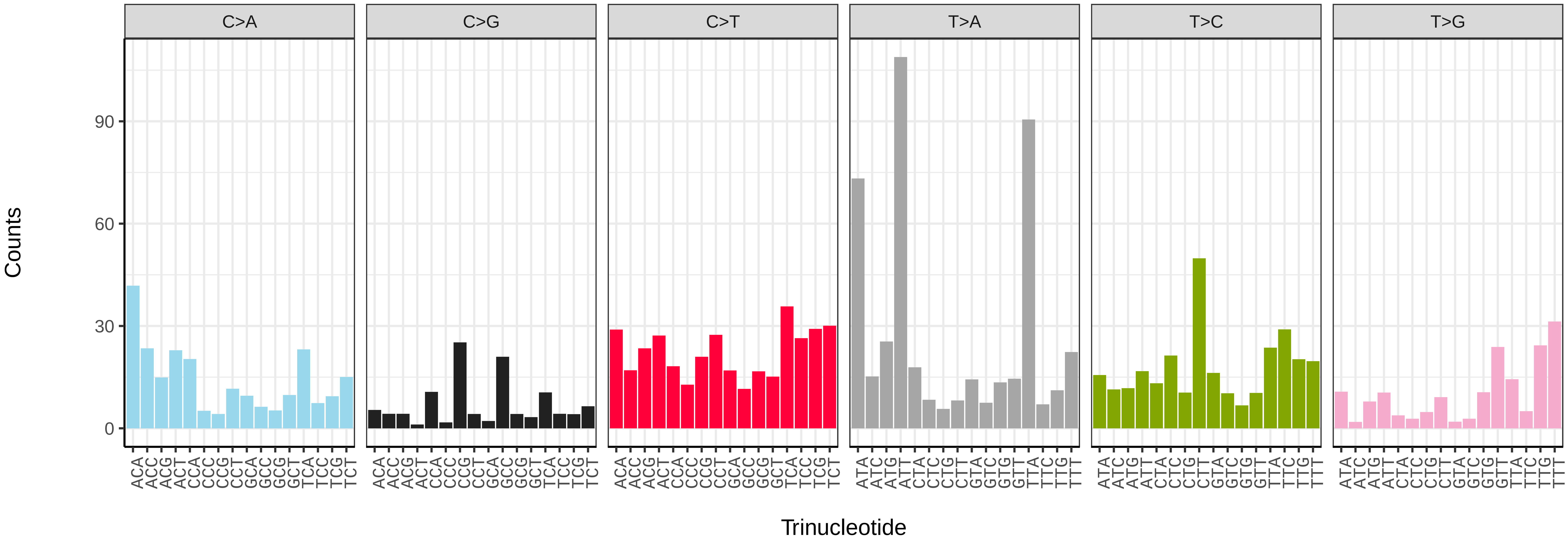


*Nycteola revayana* (ilNycReva1) somatic mutational spectrum (SBS52 classification)

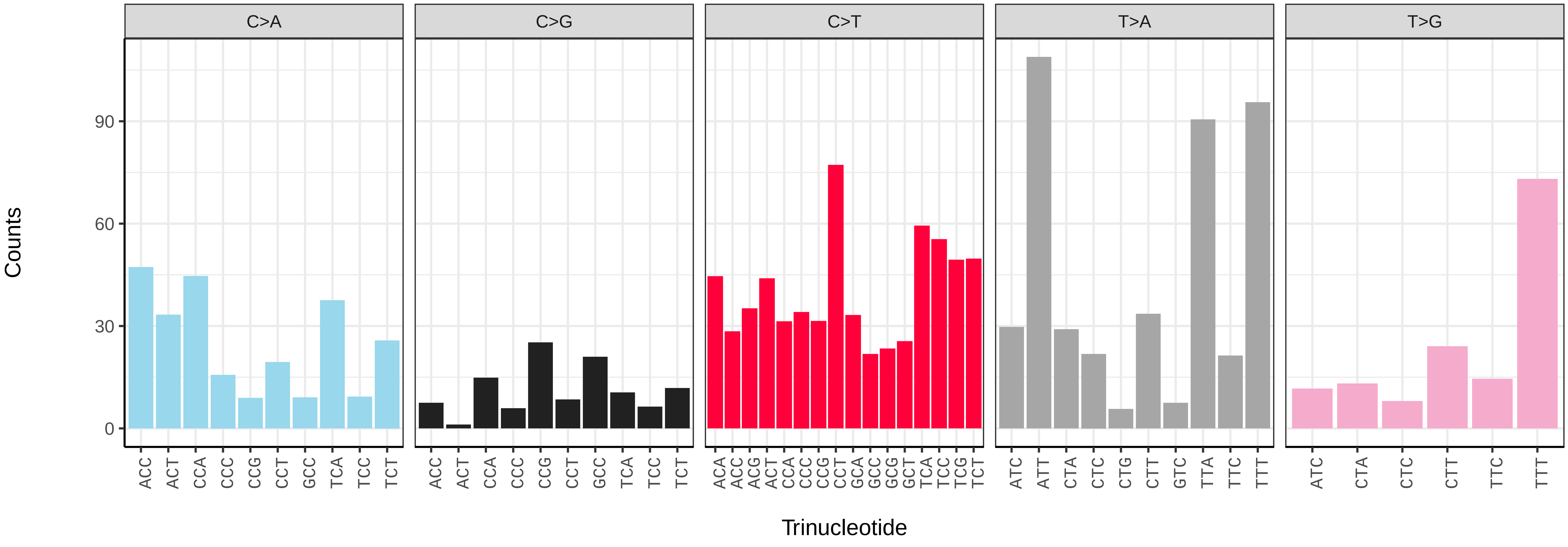

*Nymphula nitidulata* (ilNymNiti1) germline mutational spectrum

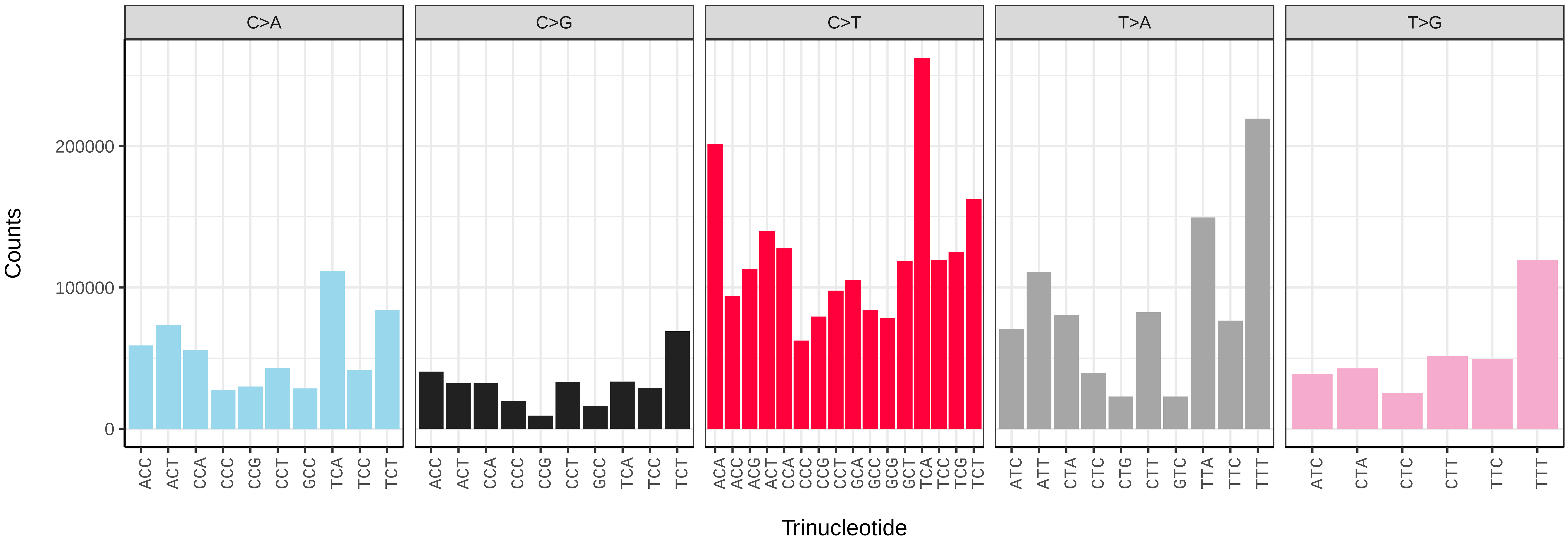


*Nymphula nitidulata* (ilNymNiti1) somatic mutational spectrum (SBS96 classification)

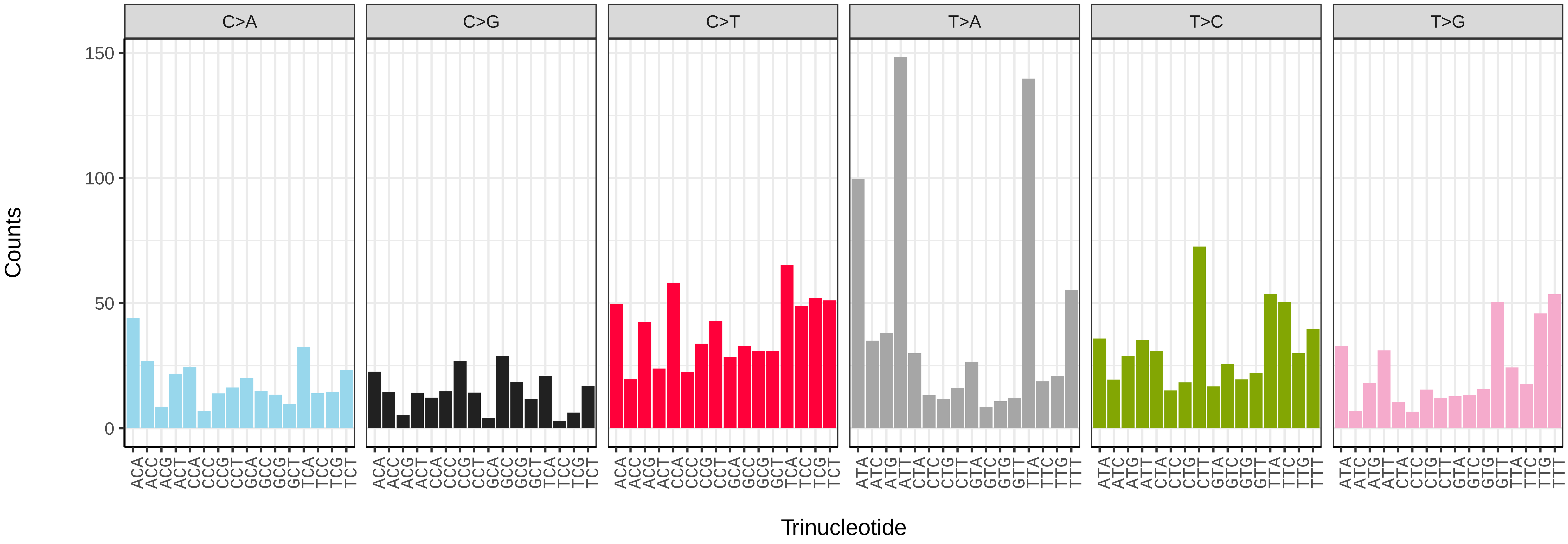


*Nymphula nitidulata* (ilNymNiti1) somatic mutational spectrum (SBS52 classification)

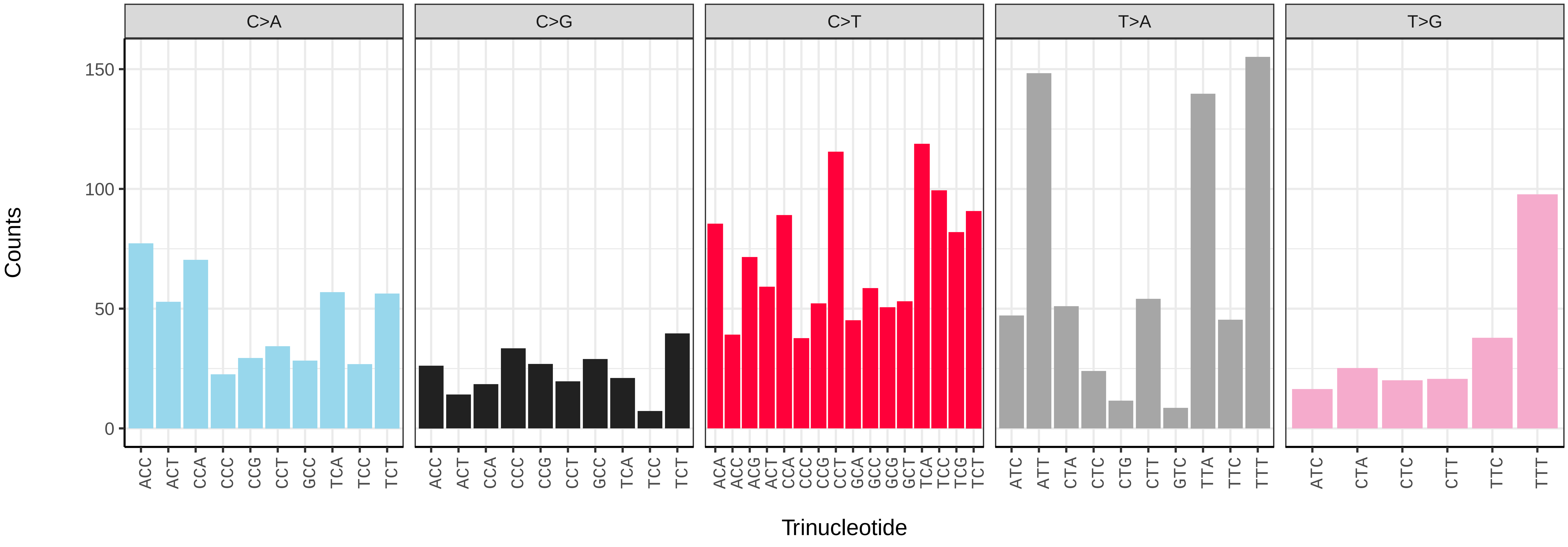

*Nymphalis polychloros* (ilNymPoly1) germline mutational spectrum

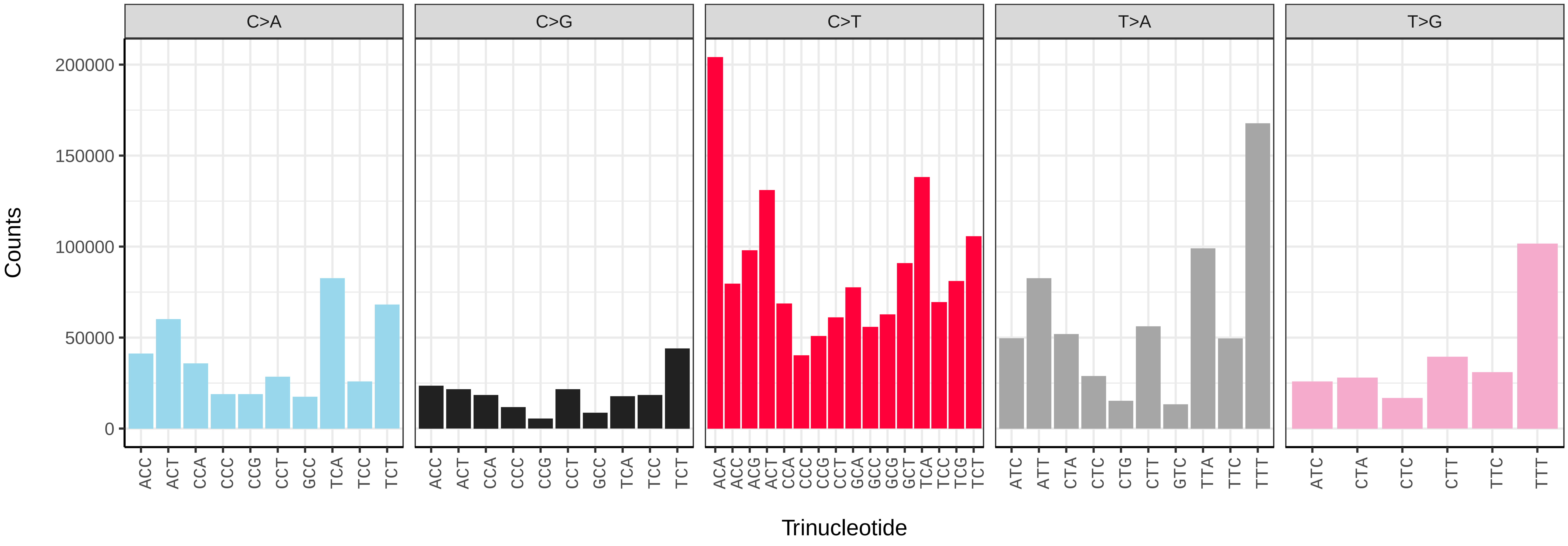


*Nymphalis polychloros* (ilNymPoly1) somatic mutational spectrum (SBS96 classification)

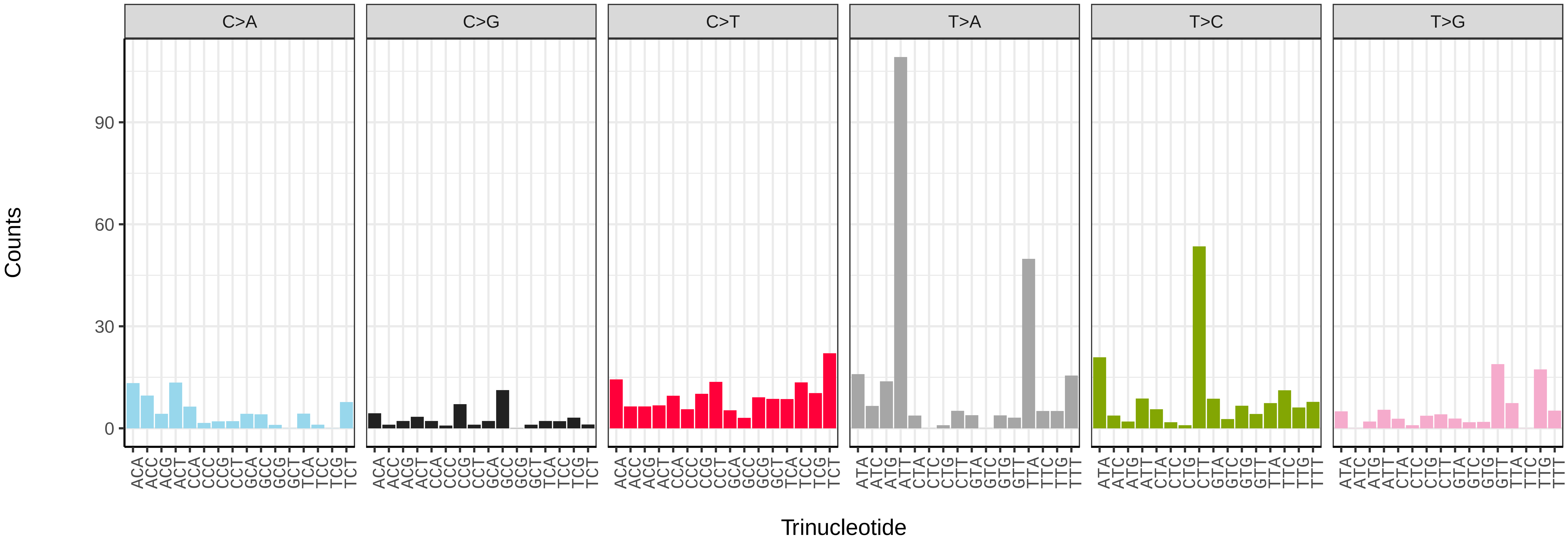


*Nymphalis polychloros* (ilNymPoly1) somatic mutational spectrum (SBS52 classification)

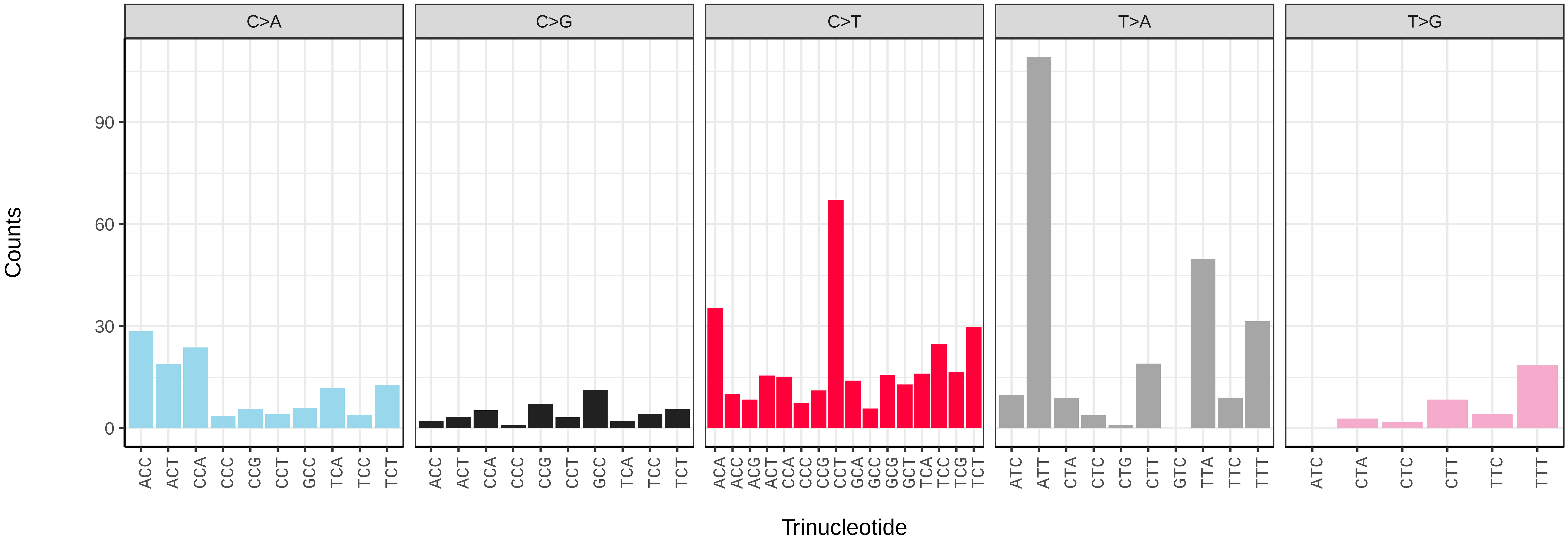

*Ochropleura plecta* (ilOchPlec1) germline mutational spectrum

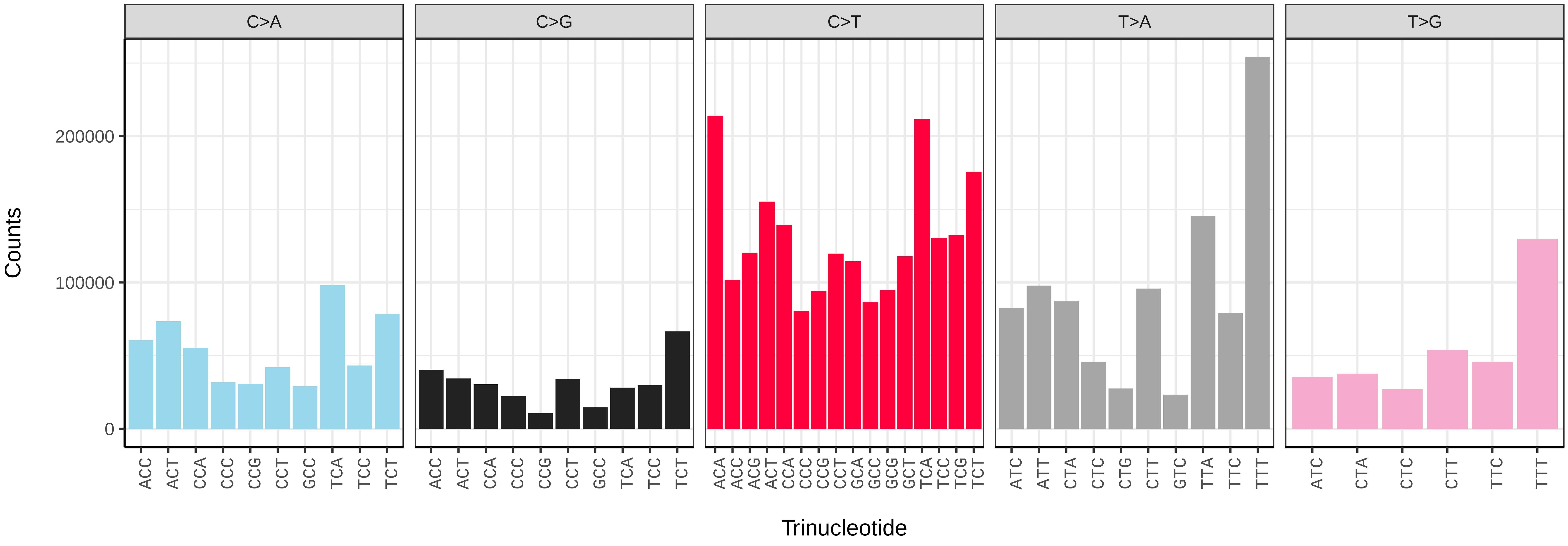


*Ochropleura plecta* (ilOchPlec1) somatic mutational spectrum (SBS96 classification)

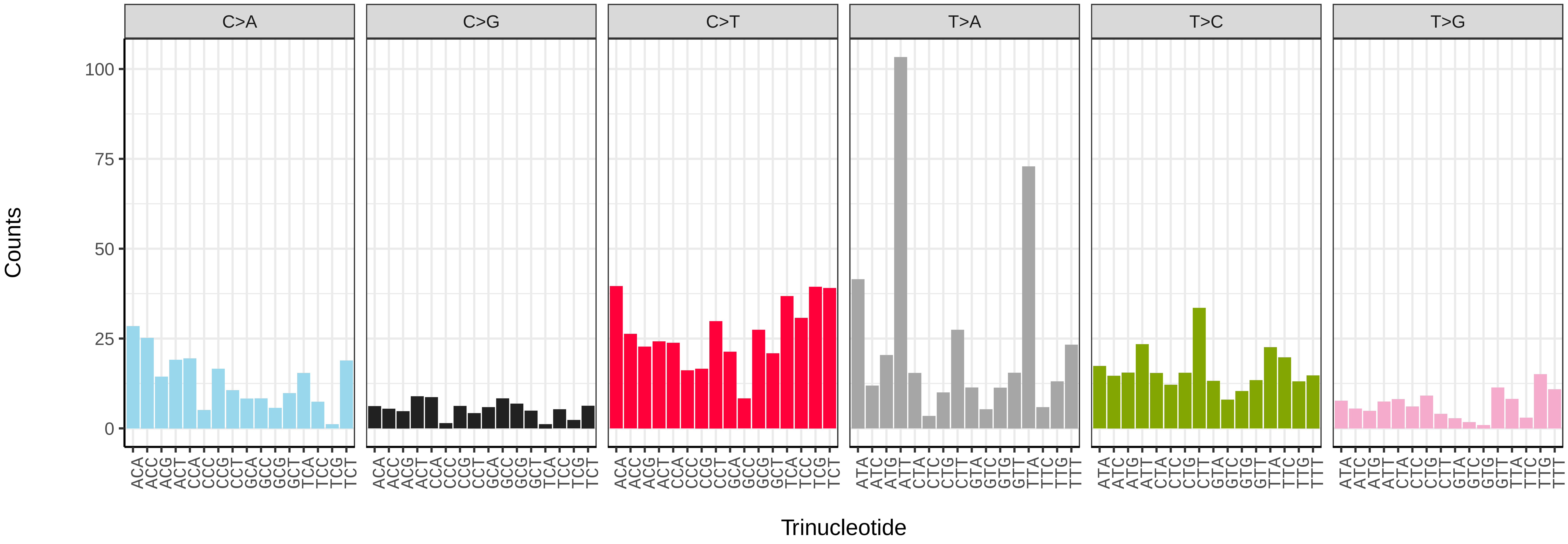


*Ochropleura plecta* (ilOchPlec1) somatic mutational spectrum (SBS52 classification)

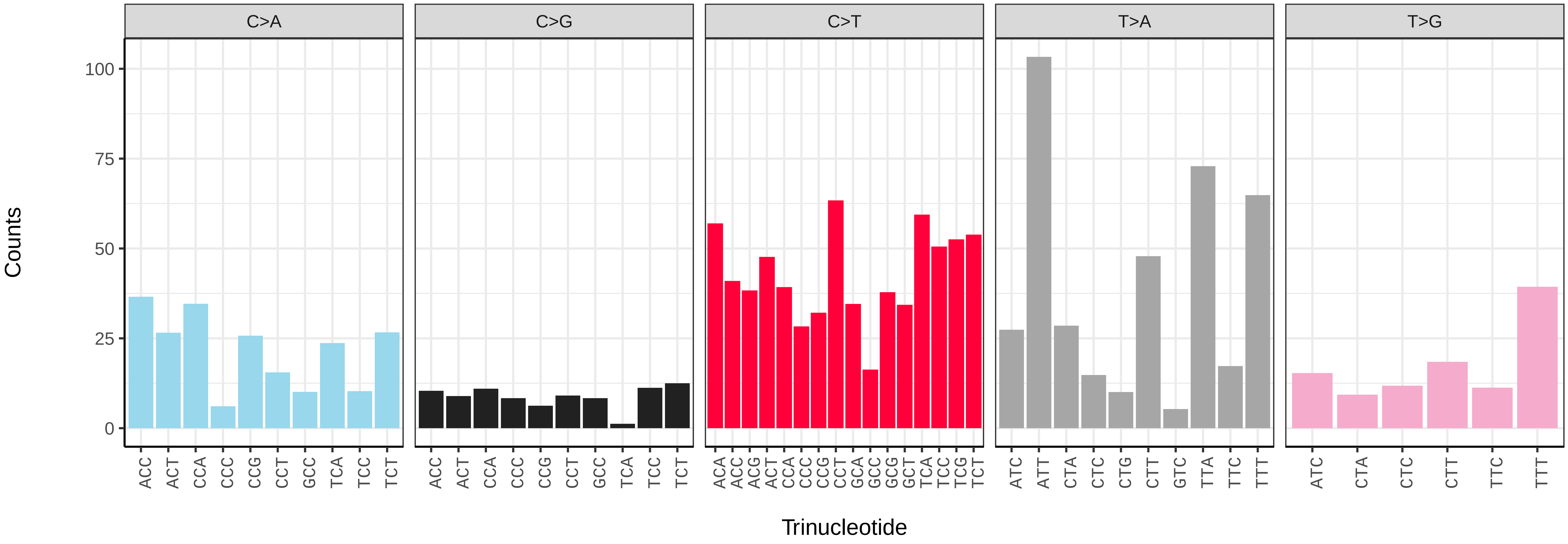

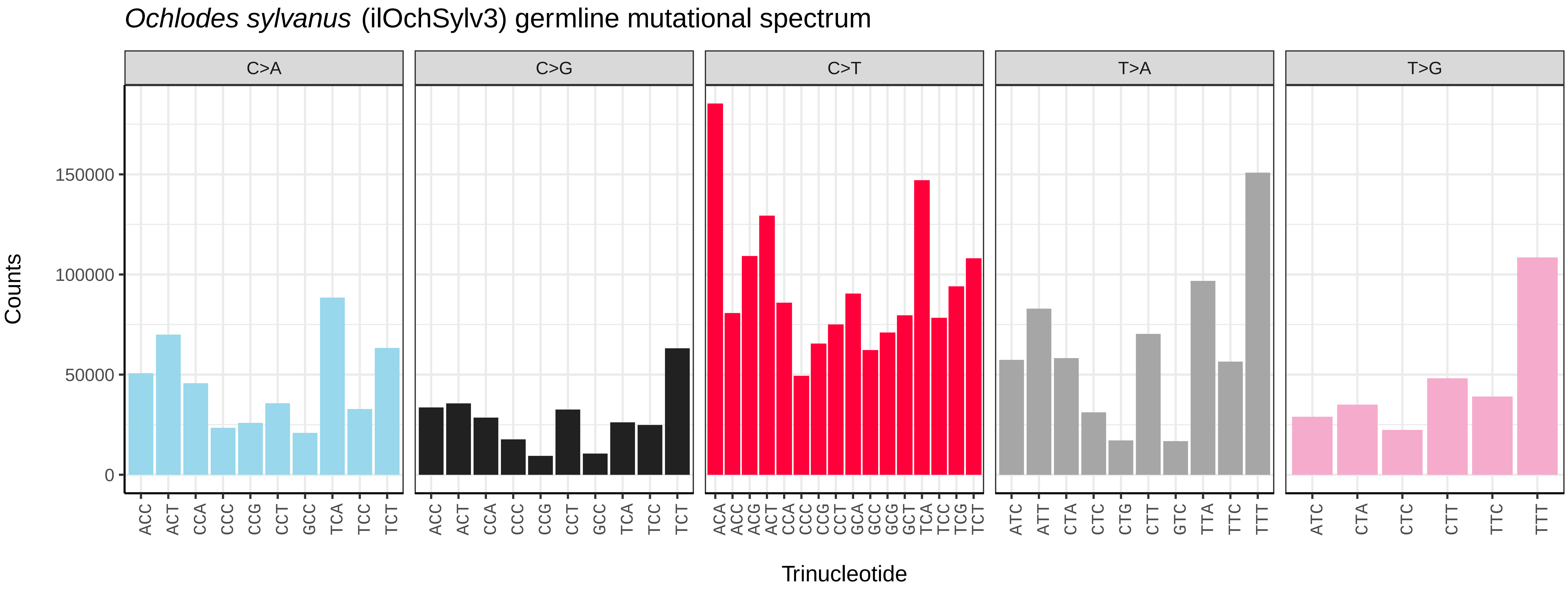
Ochlodes sylvanus (ilOchSylv3) germline mutational spectrum
C>A
C>G
C>T
T>A
T>G
Counts
150000
100000
50000
0
Trinucleotide

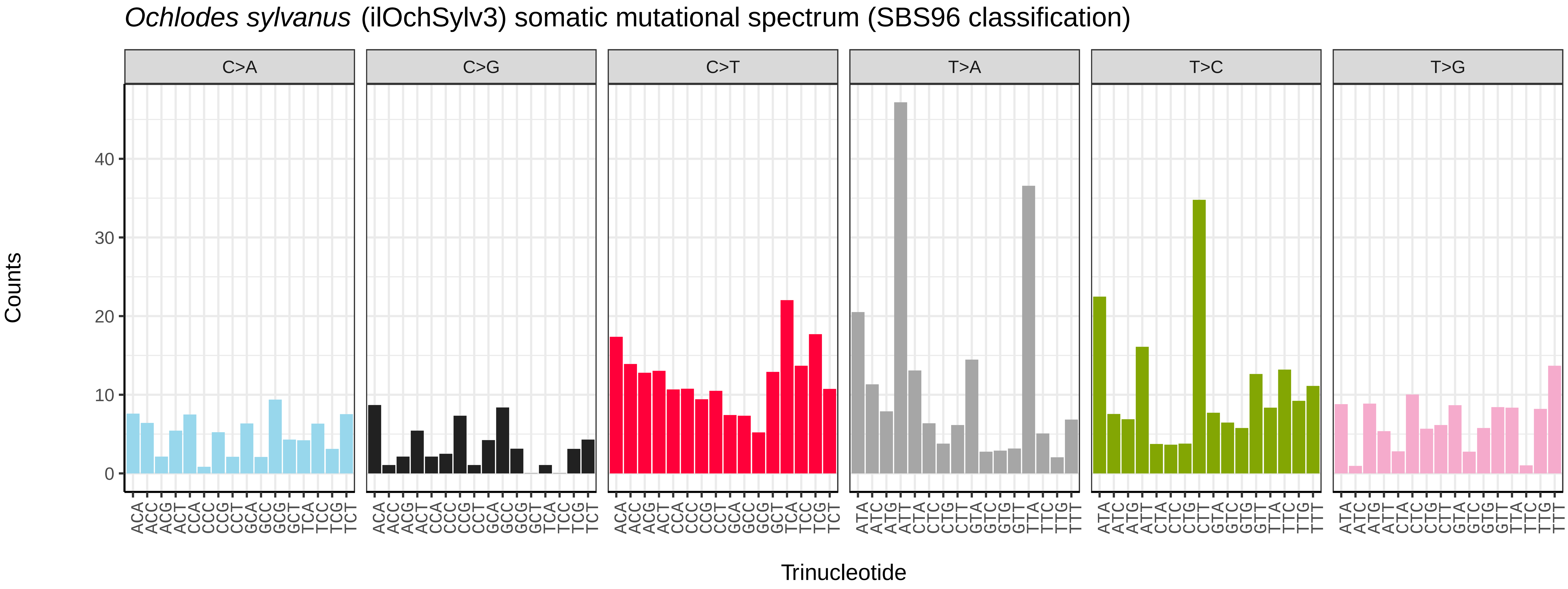
Ochlodes sylvanus (ilOchSylv3) somatic mutational spectrum (SBS96 classification)
C>A
C>G
C>T
T>A
T>C
T>G
Counts
40
30
20
10
0
Trinucleotide

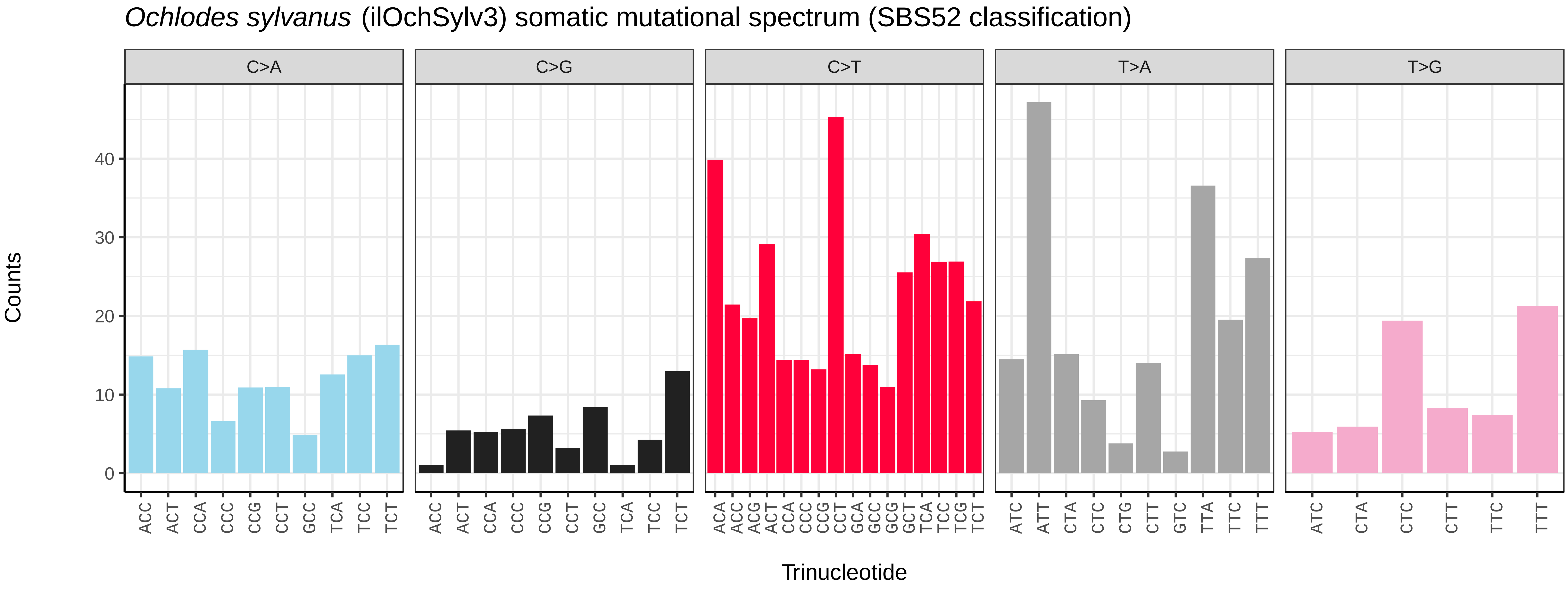
Ochlodes sylvanus (ilOchSylv3) somatic mutational spectrum (SBS52 classification)
C>A
C>G
C>T
T>A
T>G
Counts
40
30
20
10
0
Trinucleotide

*Oligia latruncula* (ilOliLatr1) germline mutational spectrum

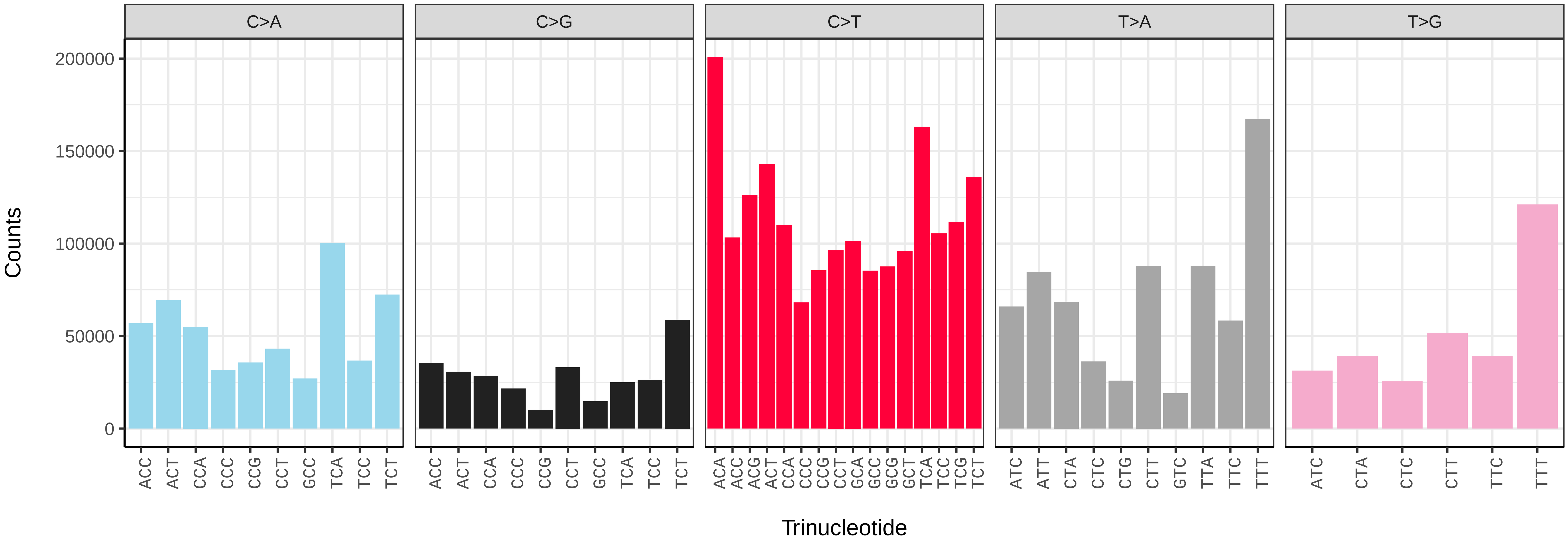


*Oligia latruncula* (ilOliLatr1) somatic mutational spectrum (SBS96 classification)

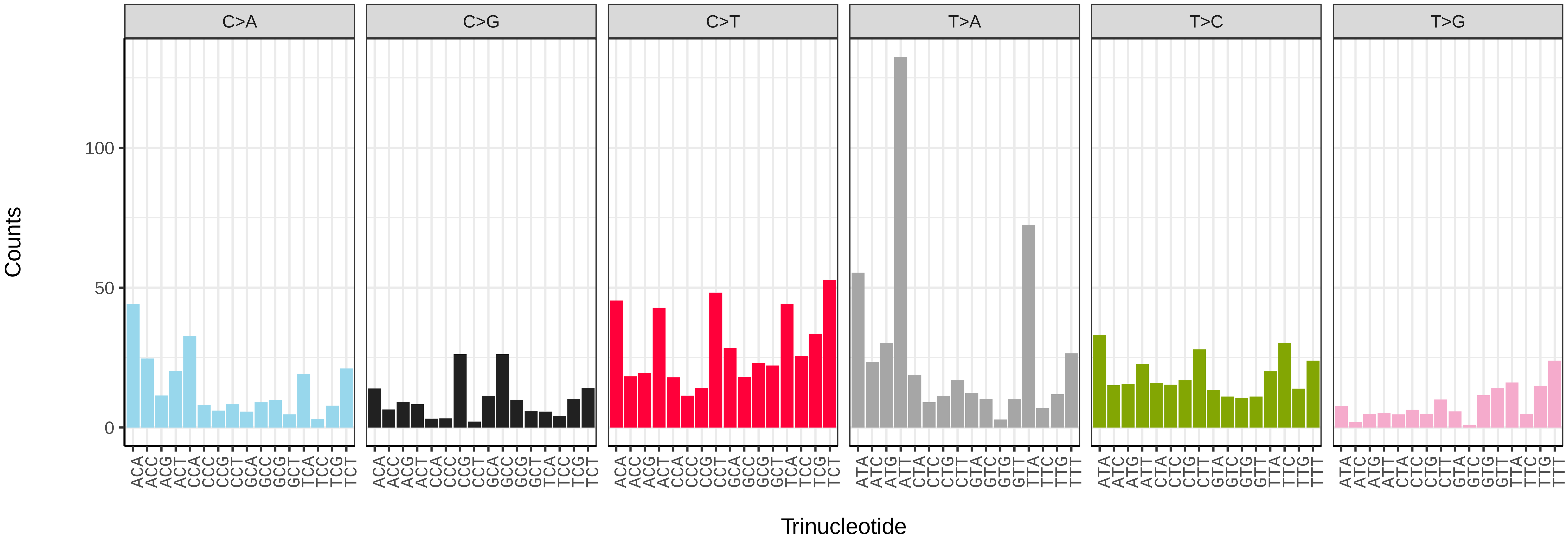


*Oligia latruncula* (ilOliLatr1) somatic mutational spectrum (SBS52 classification)

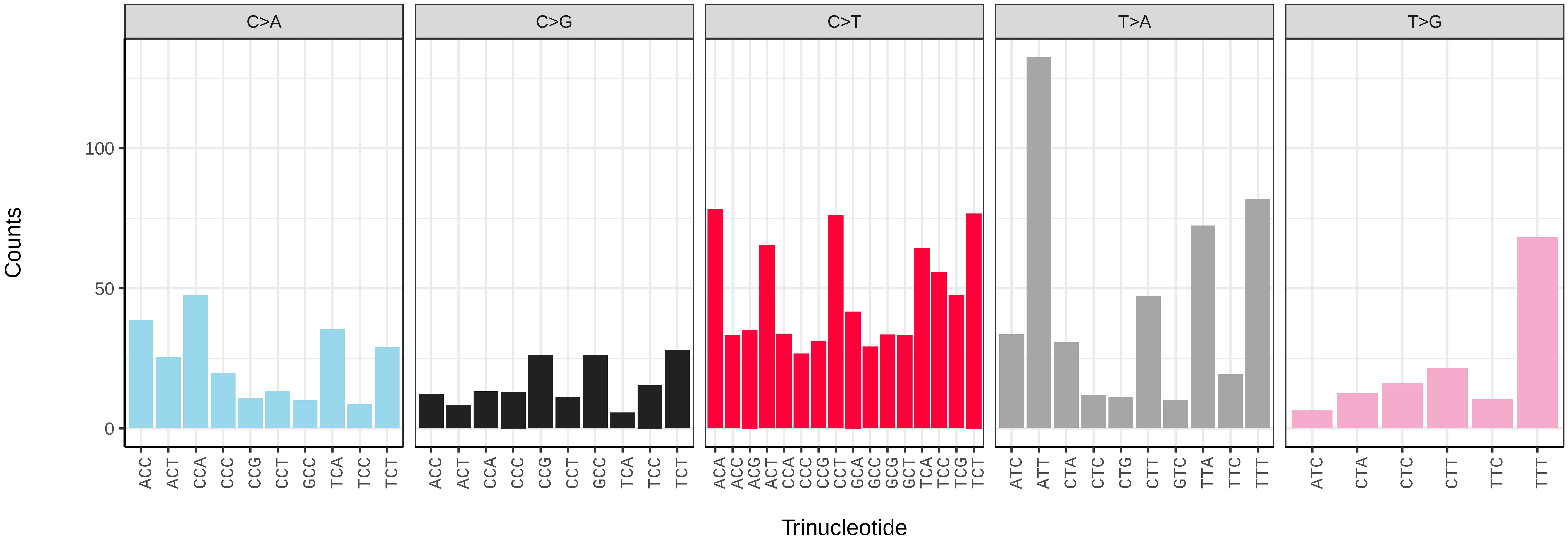

*Omphaloscelis lunosa* (ilOmpLuno1) germline mutational spectrum

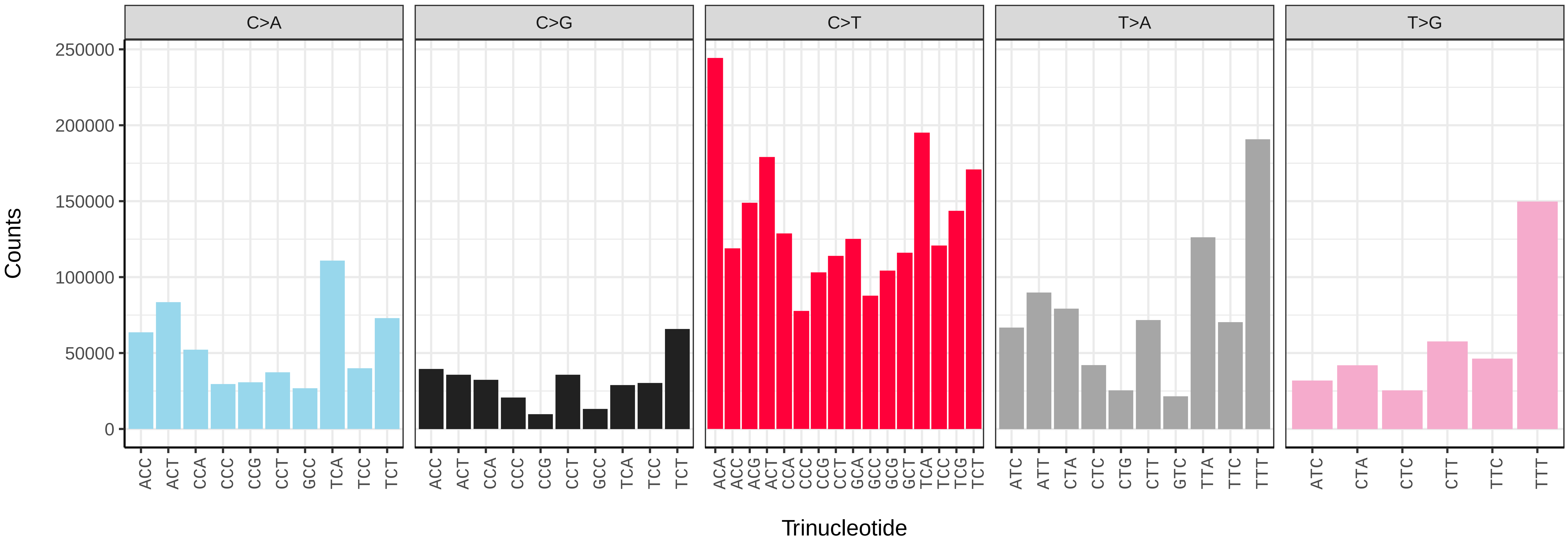


*Omphaloscelis lunosa* (ilOmpLuno1) somatic mutational spectrum (SBS96 classification)

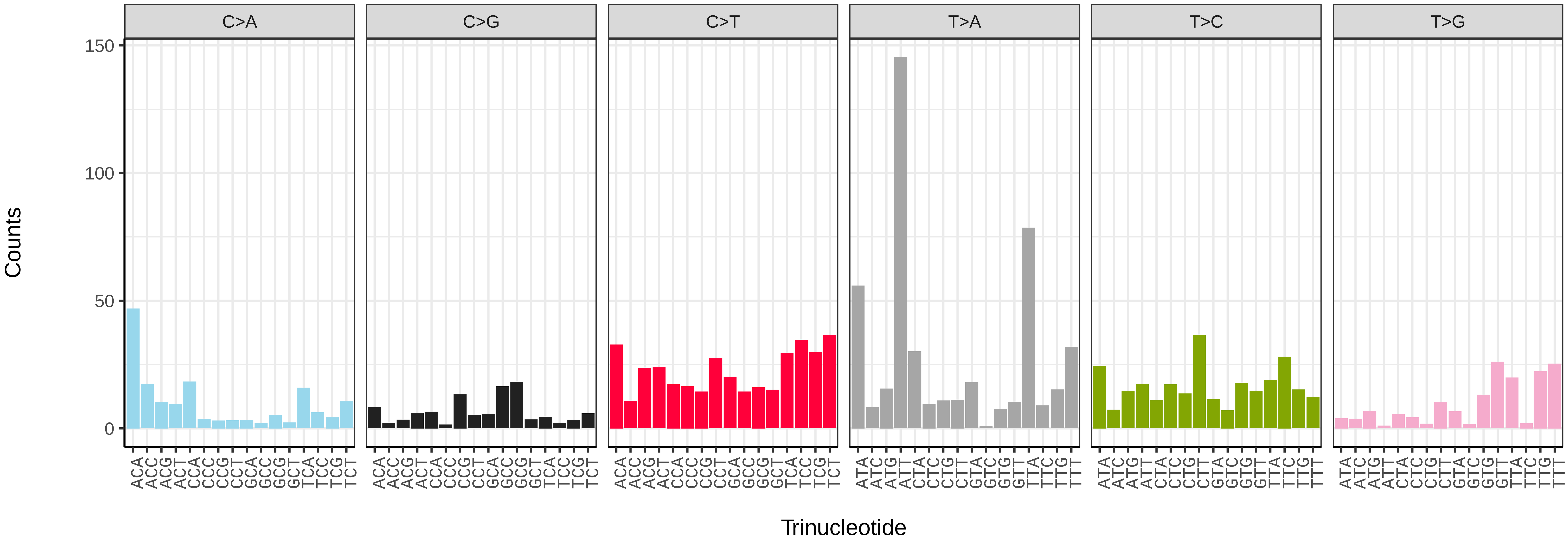


*Omphaloscelis lunosa* (ilOmpLuno1) somatic mutational spectrum (SBS52 classification)

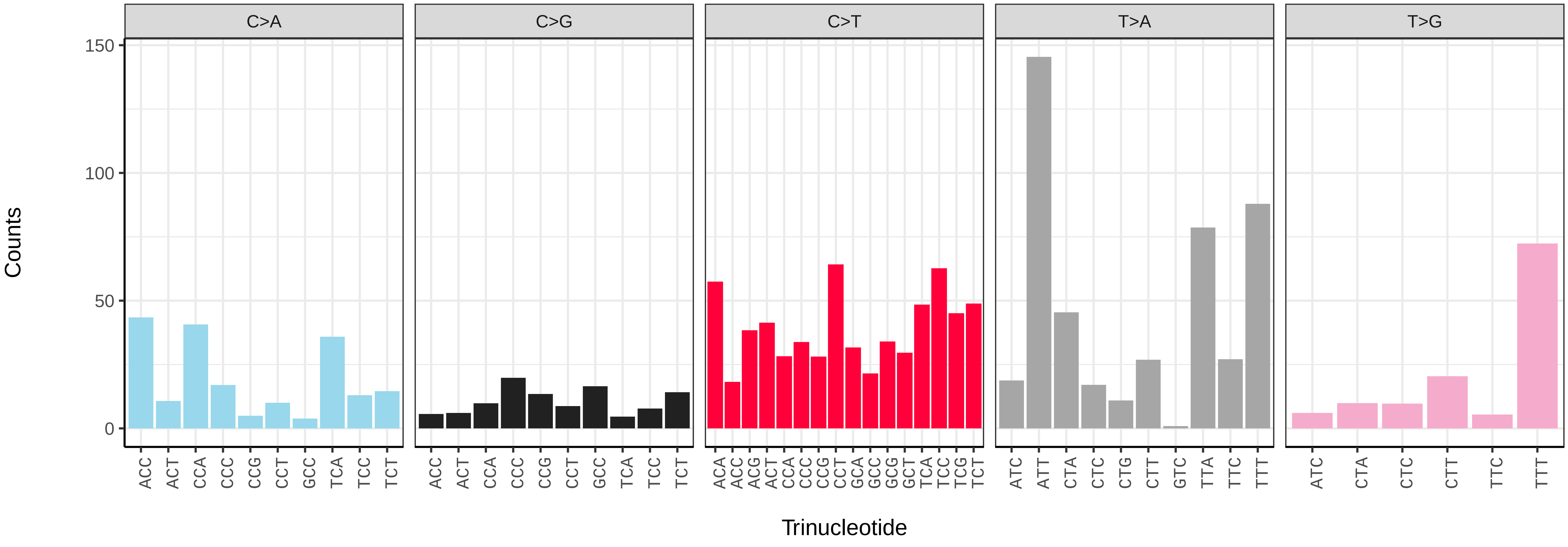

*Operophtera brumata* (ilOpeBrum1) germline mutational spectrum

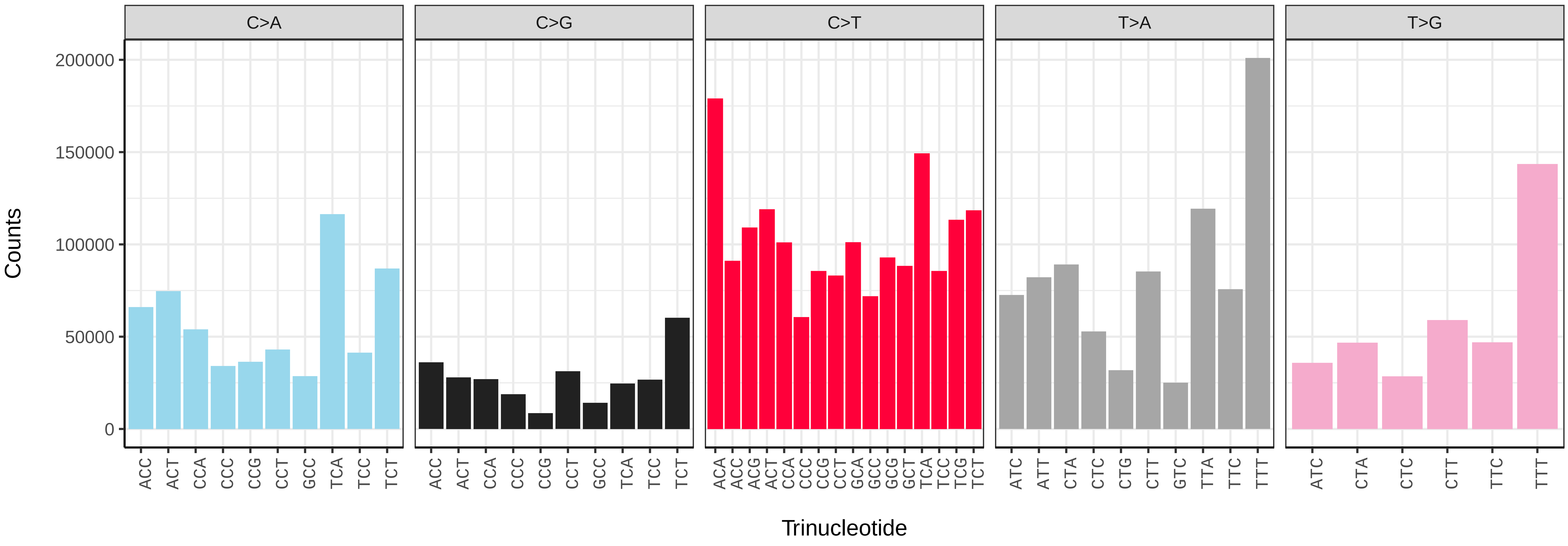


*Operophtera brumata* (ilOpeBrum1) somatic mutational spectrum (SBS96 classification)

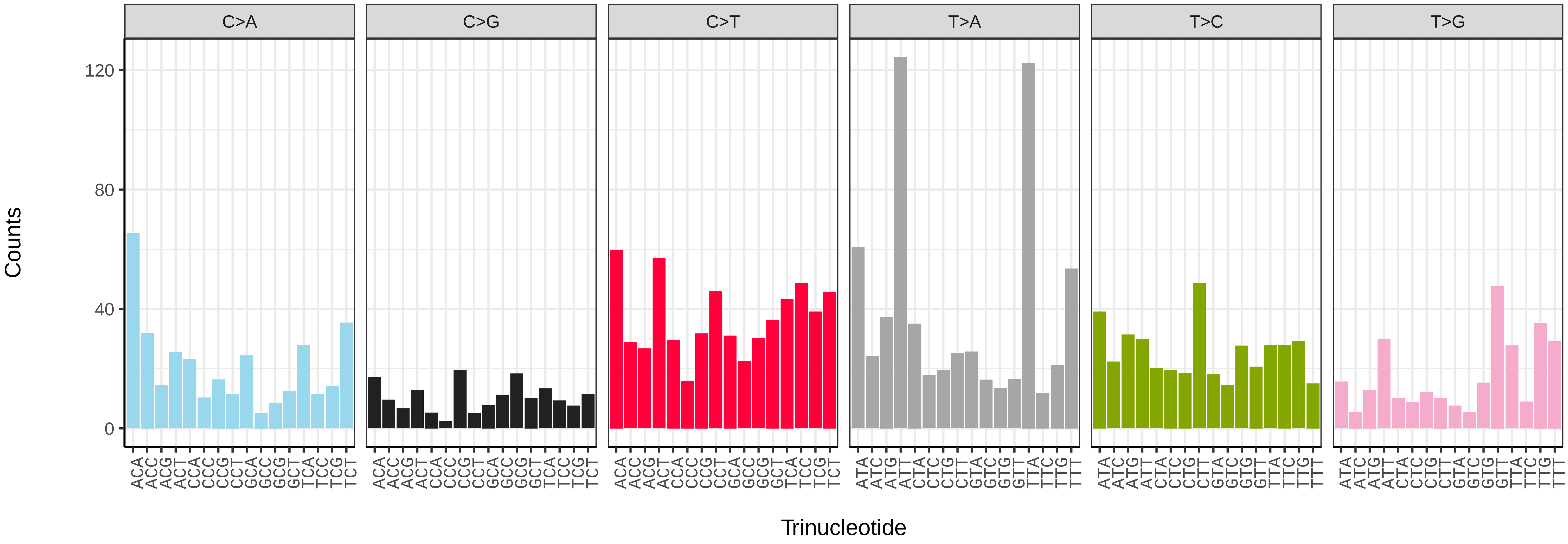


*Operophtera brumata* (ilOpeBrum1) somatic mutational spectrum (SBS52 classification)

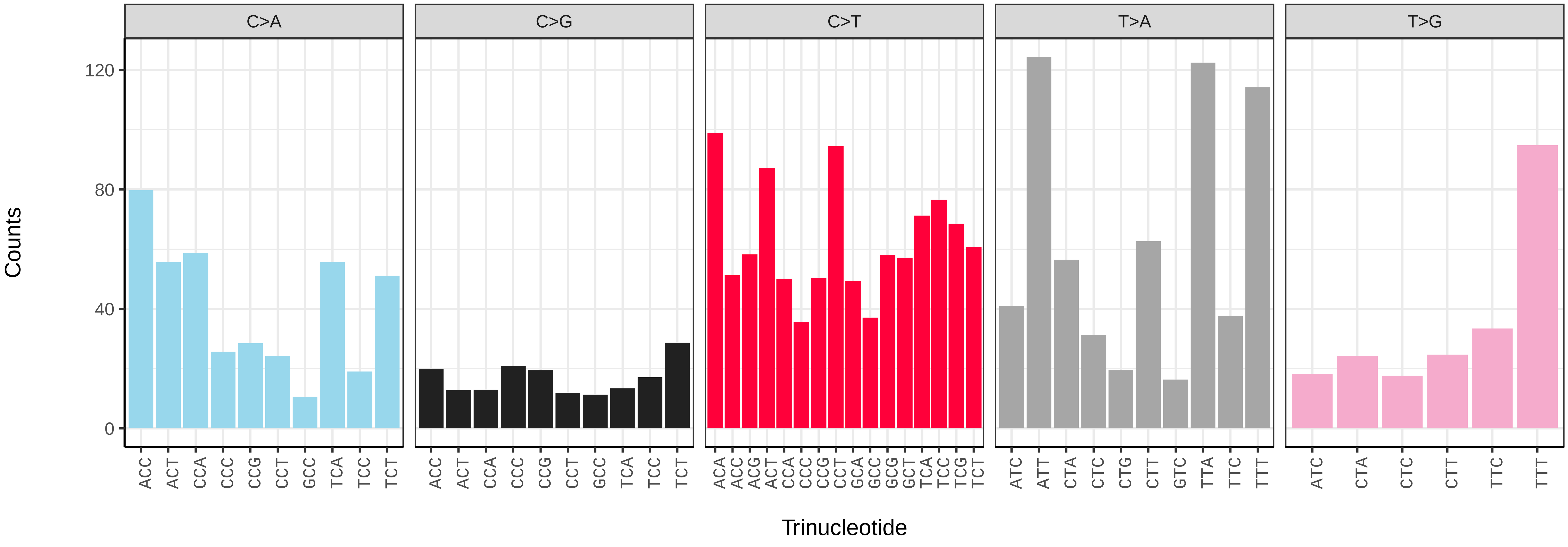

*Opisthograptis luteolata* (ilOpiLute1) germline mutational spectrum

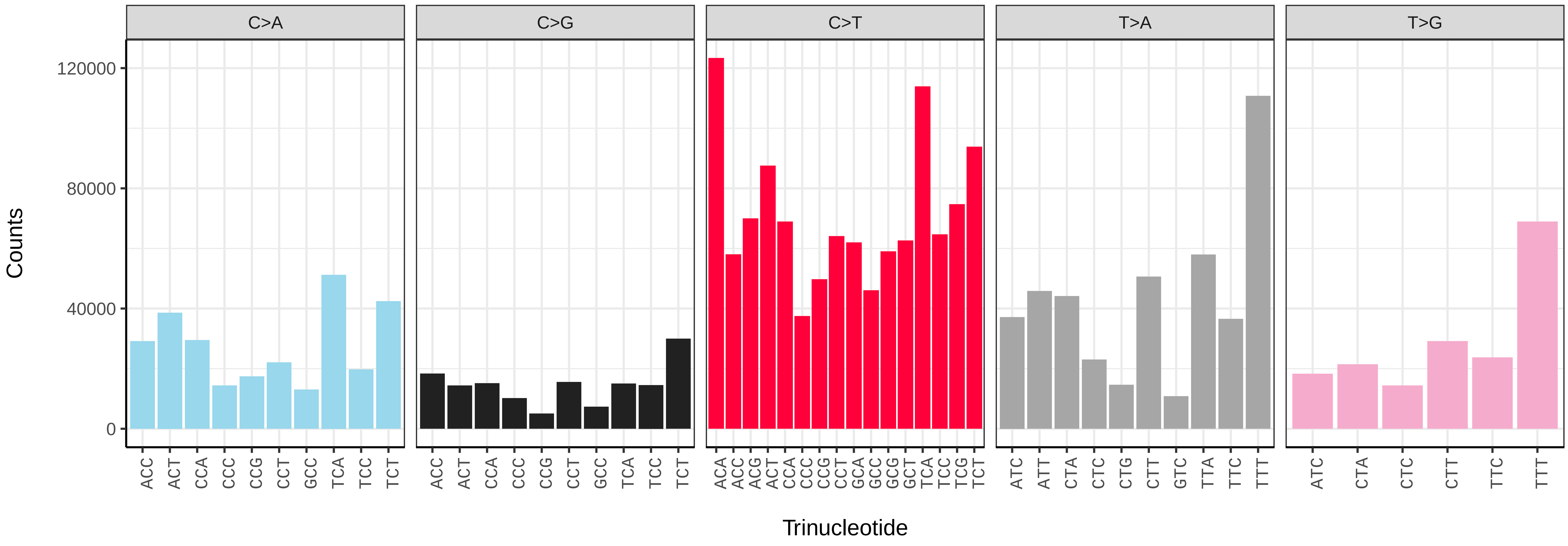


*Opisthograptis luteolata* (ilOpiLute1) somatic mutational spectrum (SBS96 classification)

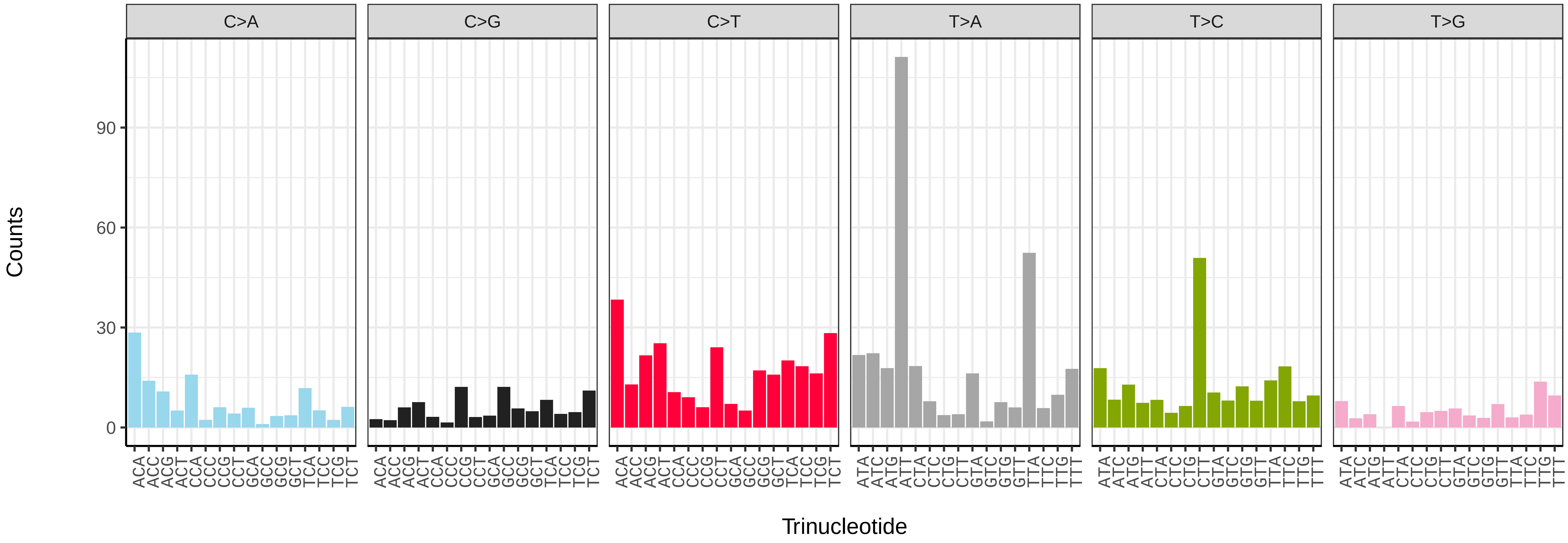


*Opisthograptis luteolata* (ilOpiLute1) somatic mutational spectrum (SBS52 classification)

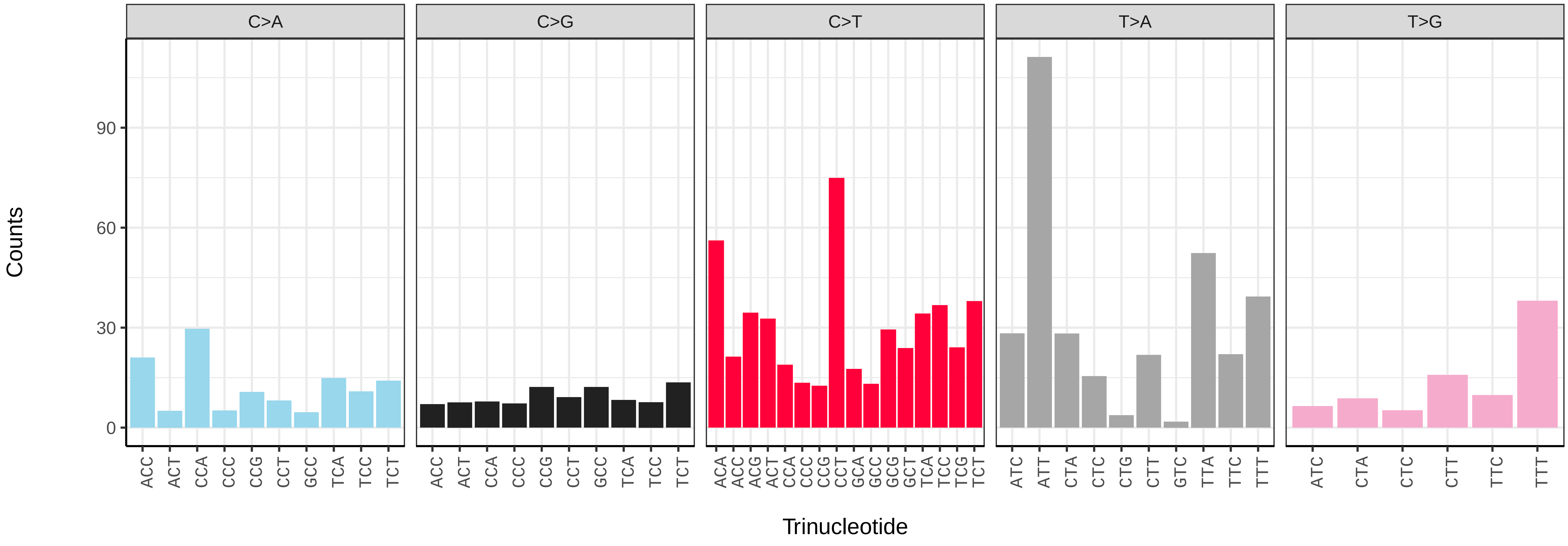

*Orgyia antiqua* (ilOrgAnti1) germline mutational spectrum

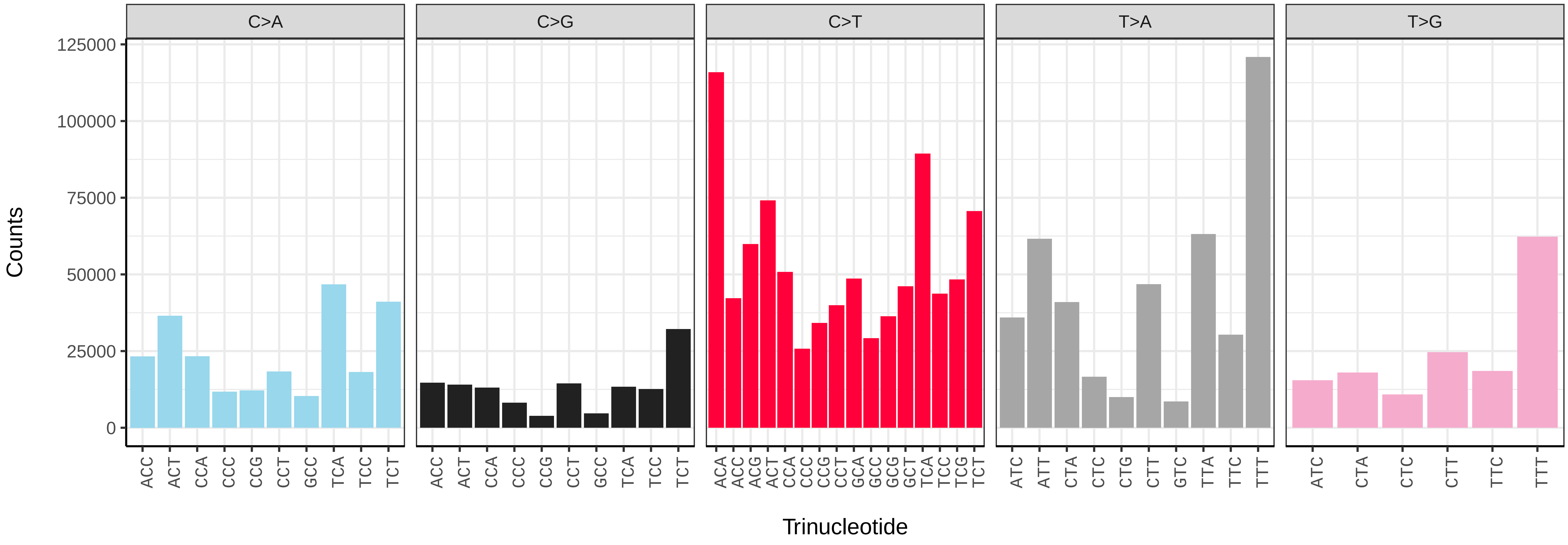


*Orgyia antiqua* (ilOrgAnti1) somatic mutational spectrum (SBS96 classification)

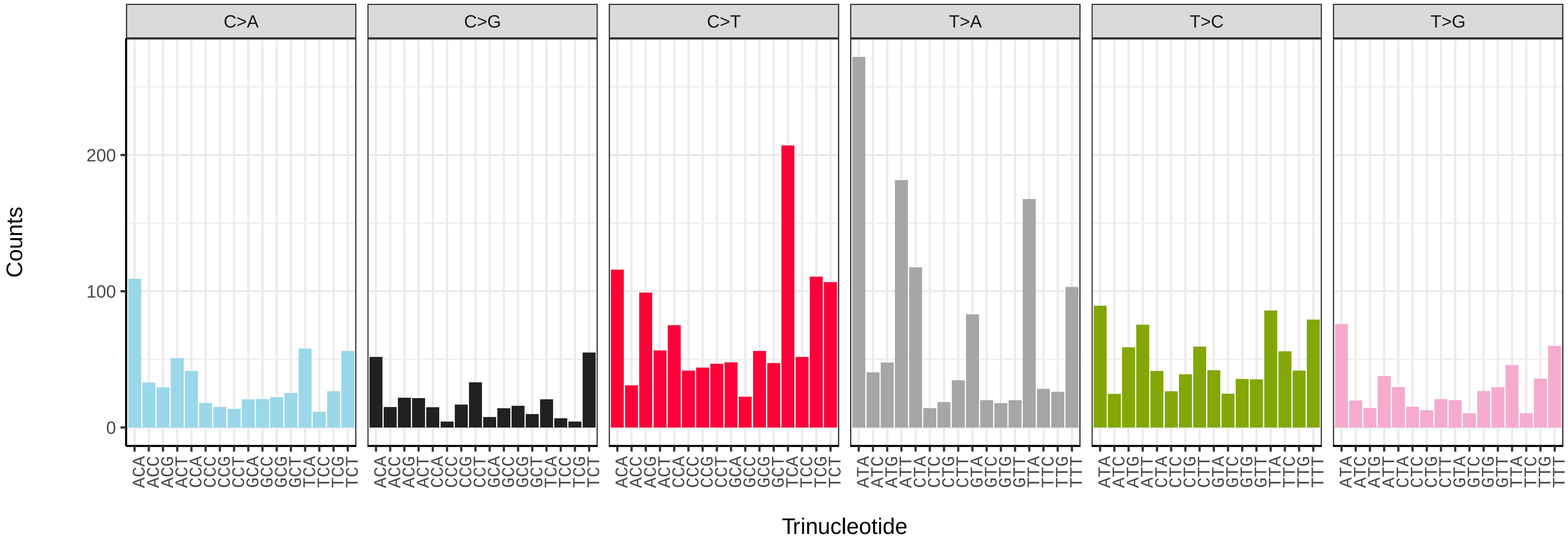


*Orgyia antiqua* (ilOrgAnti1) somatic mutational spectrum (SBS52 classification)

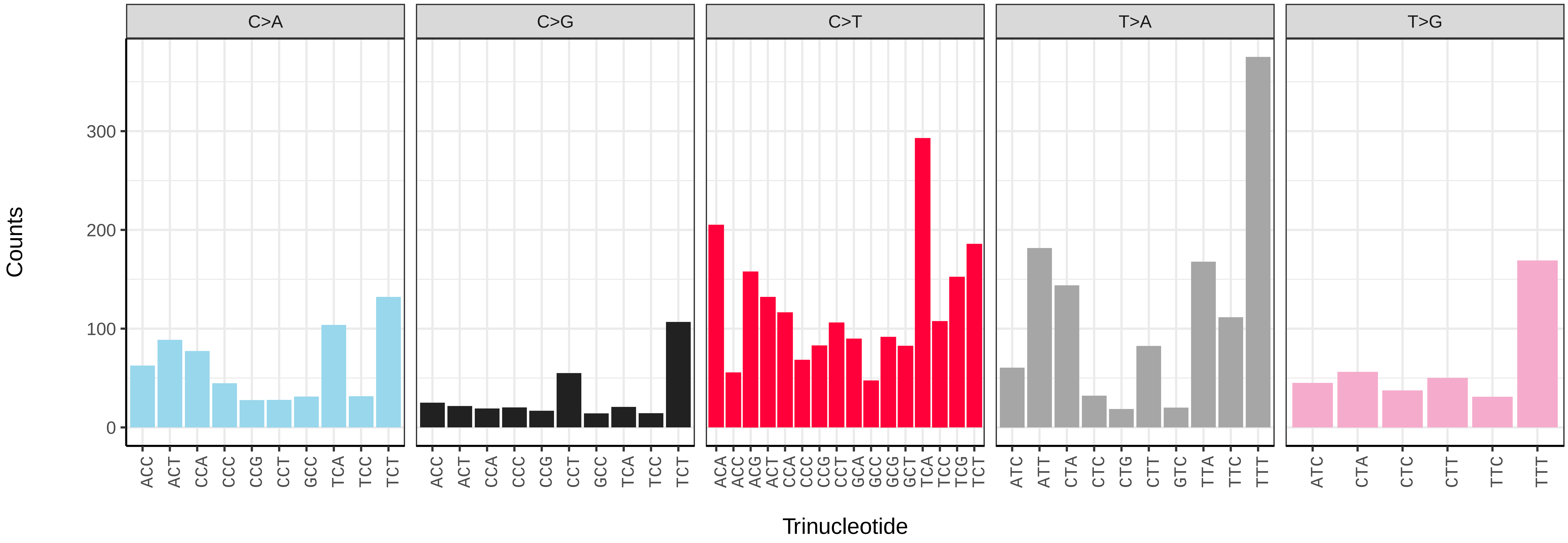

*Orgyia antiqua* (ilOrgAnti1.ilOrgAnti2) germline mutational spectrum

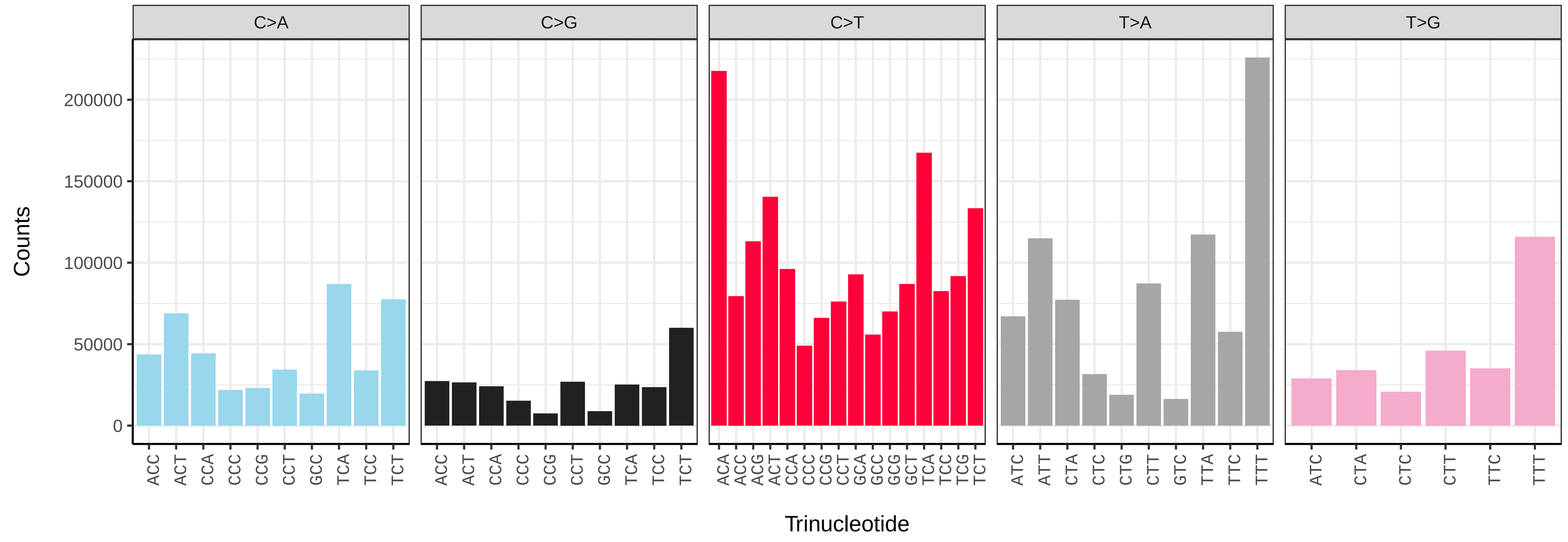


*Orgyia antiqua* (ilOrgAnti1.ilOrgAnti2) somatic mutational spectrum (SBS96 classification)

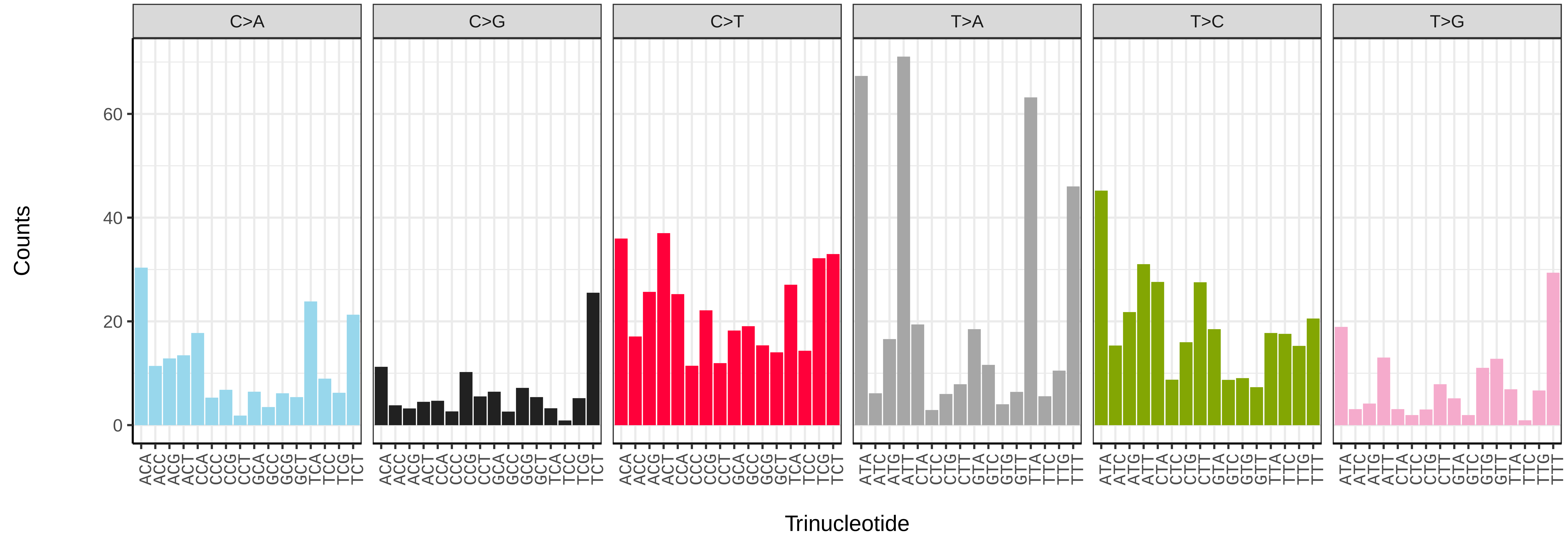


*Orgyia antiqua* (ilOrgAnti1.ilOrgAnti2) somatic mutational spectrum (SBS52 classification)

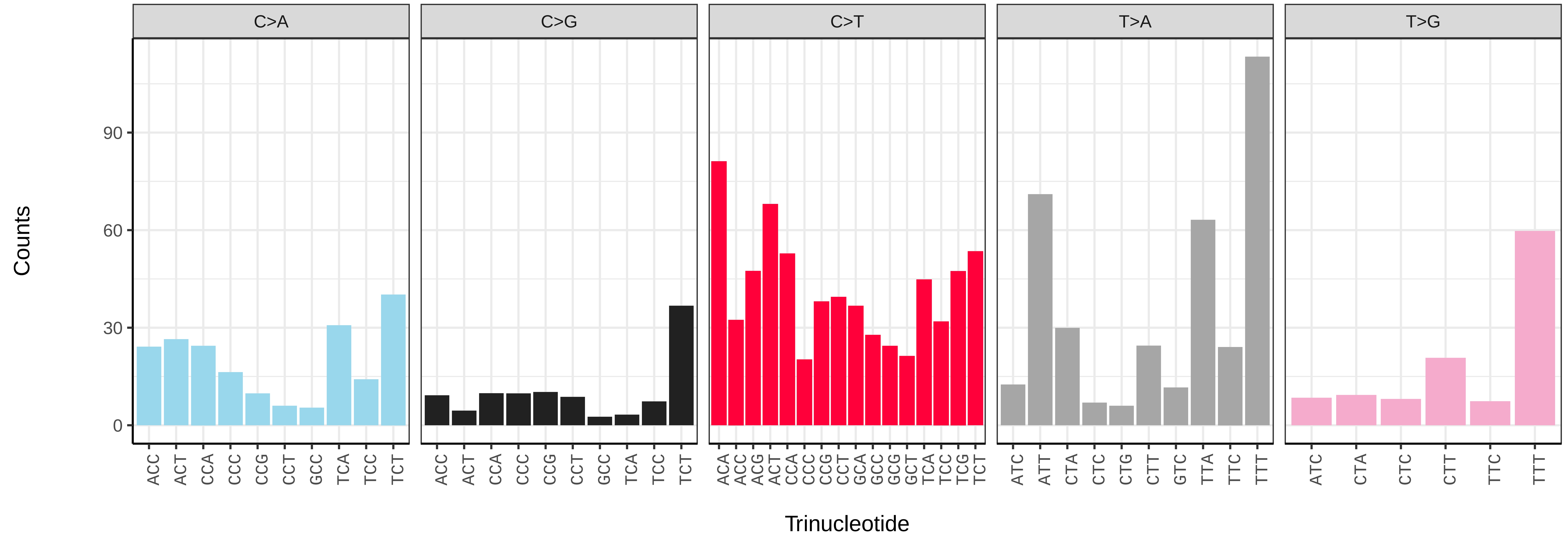

*Orthosia gracilis* (ilOrtGrac1) germline mutational spectrum

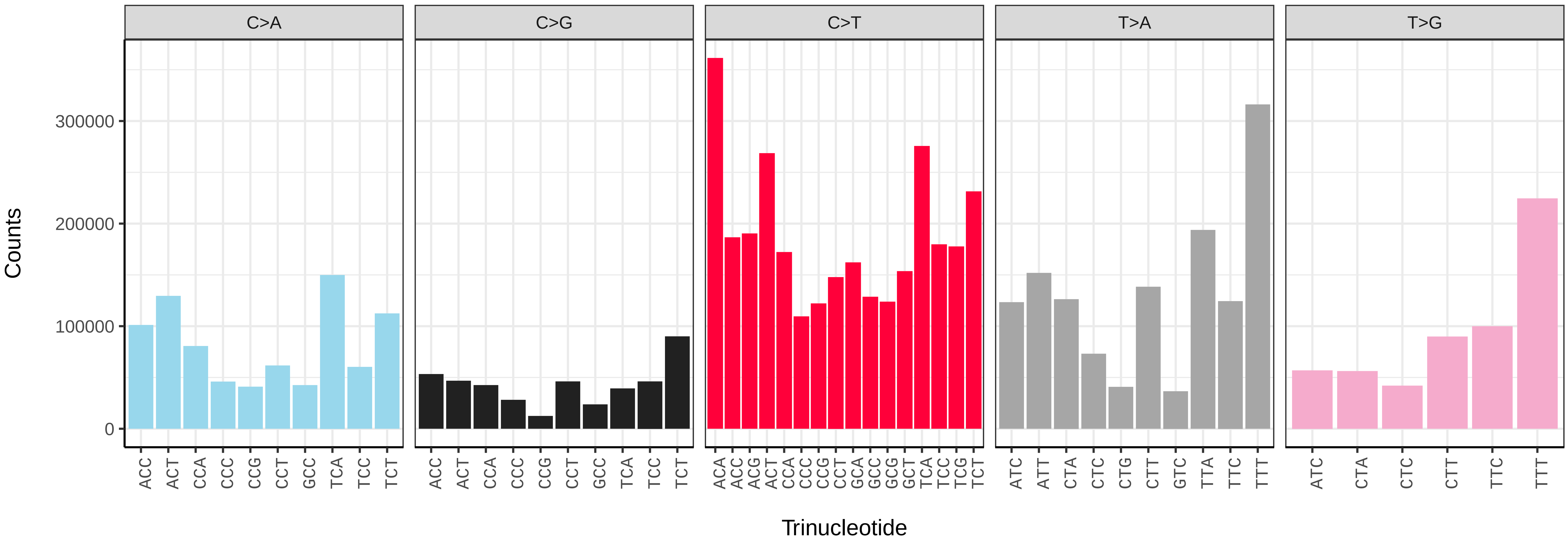


*Orthosia gracilis* (ilOrtGrac1) somatic mutational spectrum (SBS96 classification)

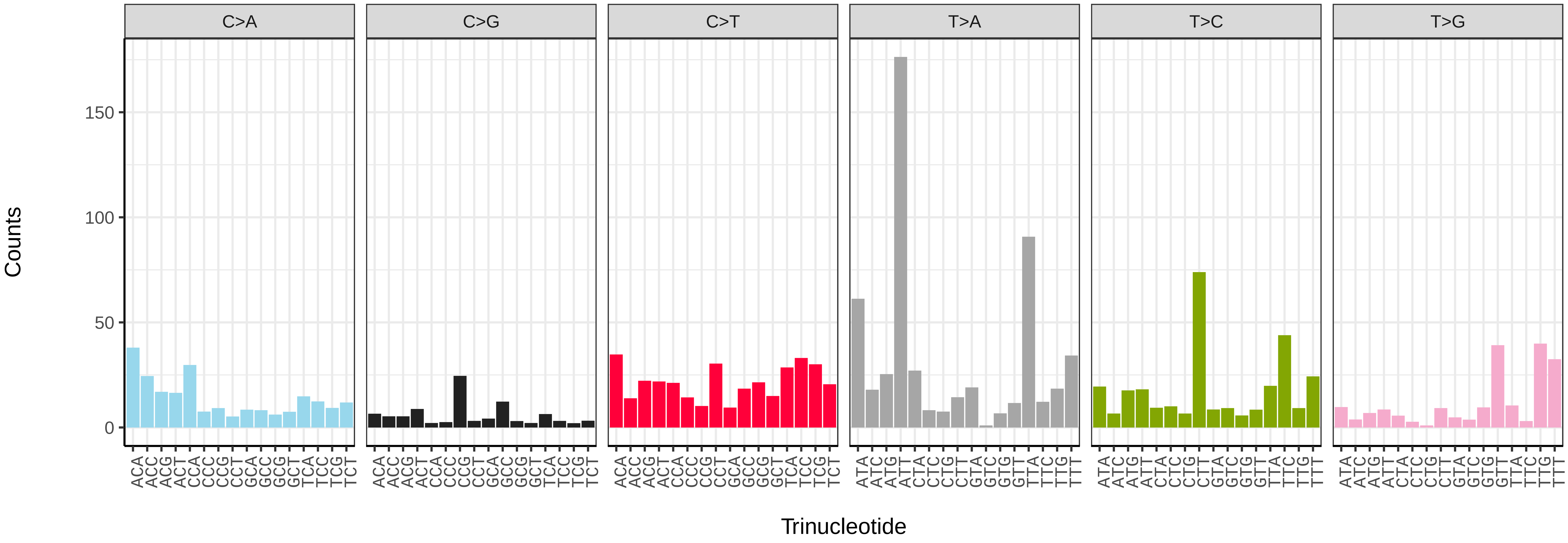


*Orthosia gracilis* (ilOrtGrac1) somatic mutational spectrum (SBS52 classification)

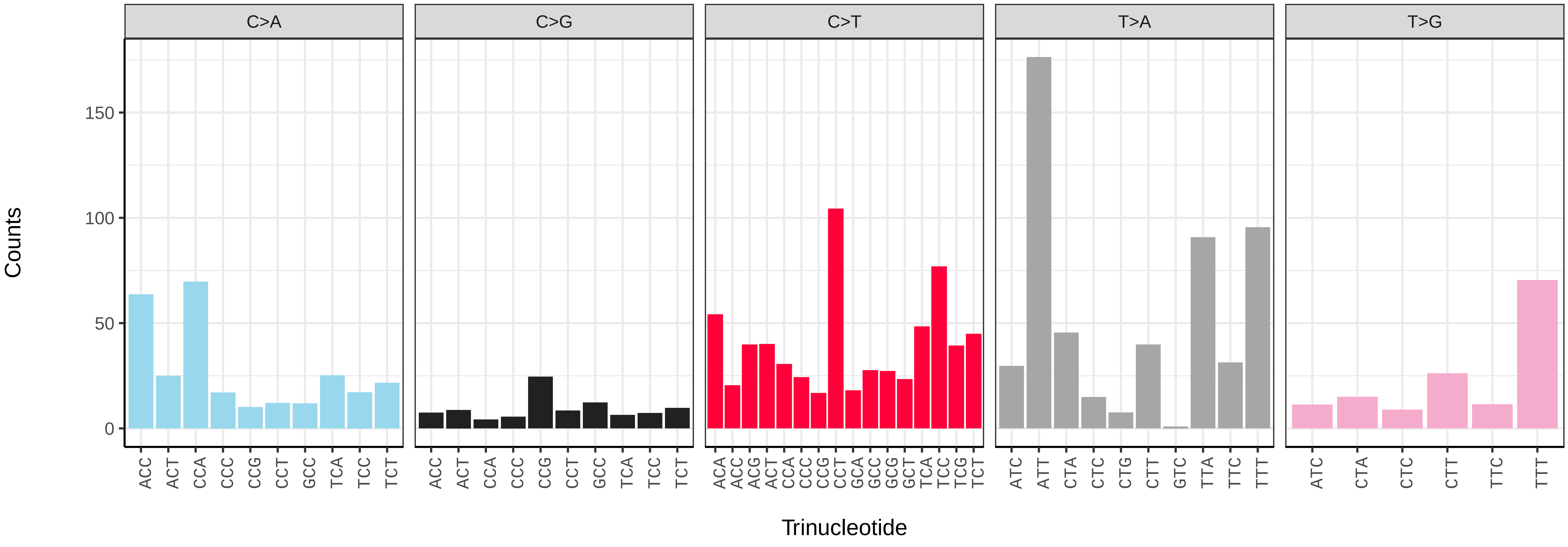

*Orthosia incerta* (ilOrtInce1) germline mutational spectrum

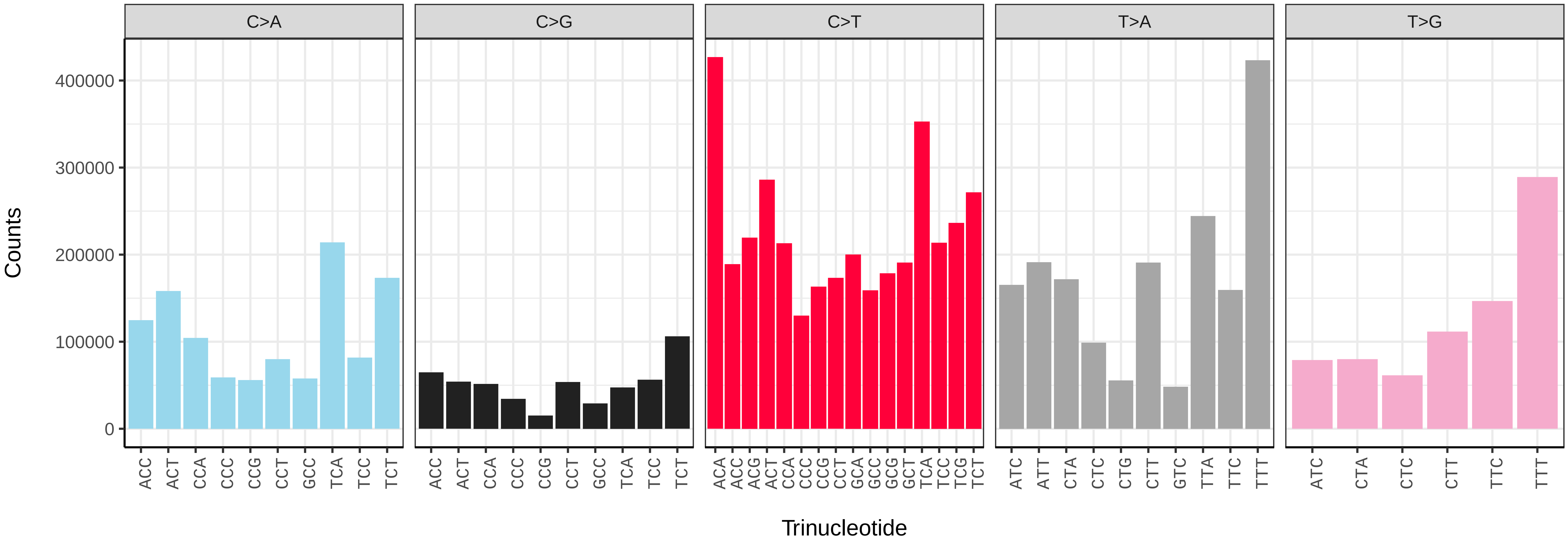


*Orthosia incerta* (ilOrtInce1) somatic mutational spectrum (SBS96 classification)

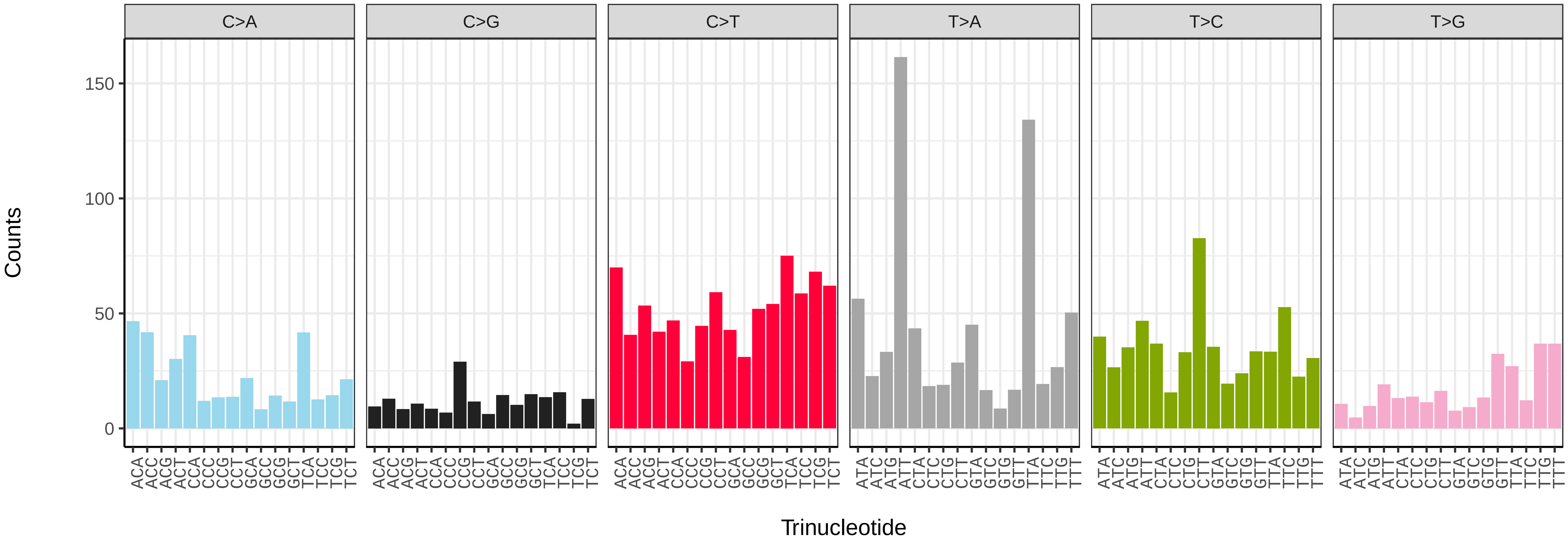


*Orthosia incerta* (ilOrtInce1) somatic mutational spectrum (SBS52 classification)

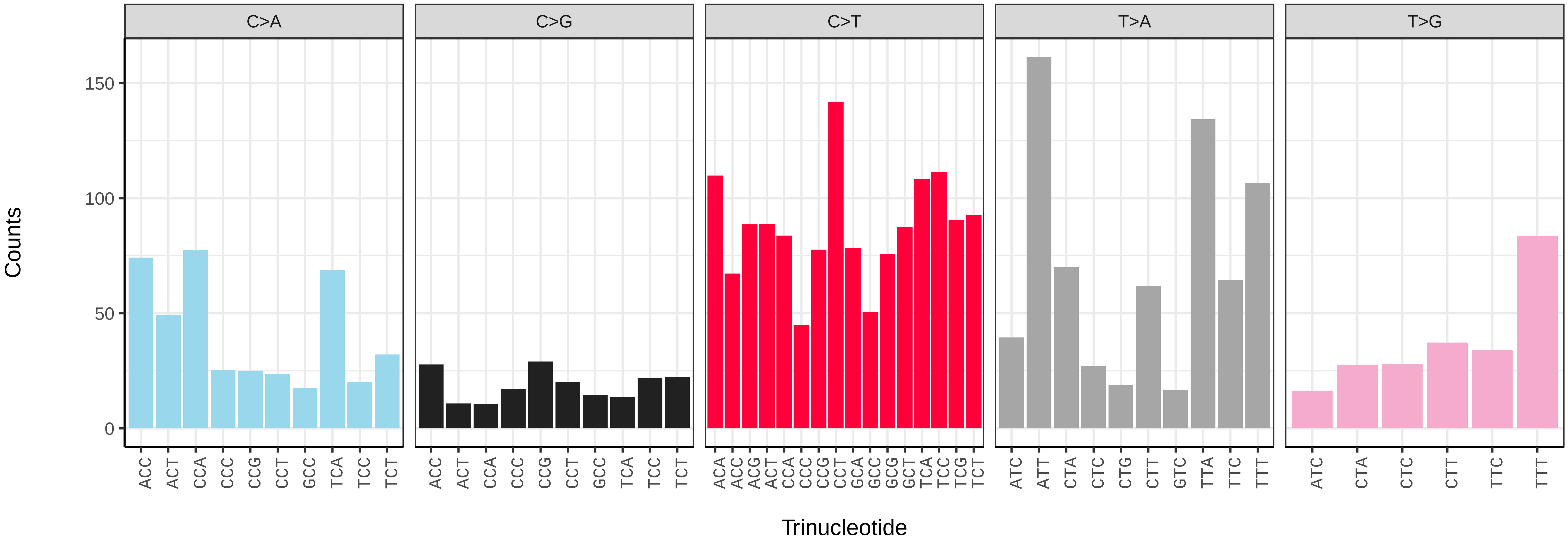

*Pammene aurita* (ilPamAuri1) germline mutational spectrum

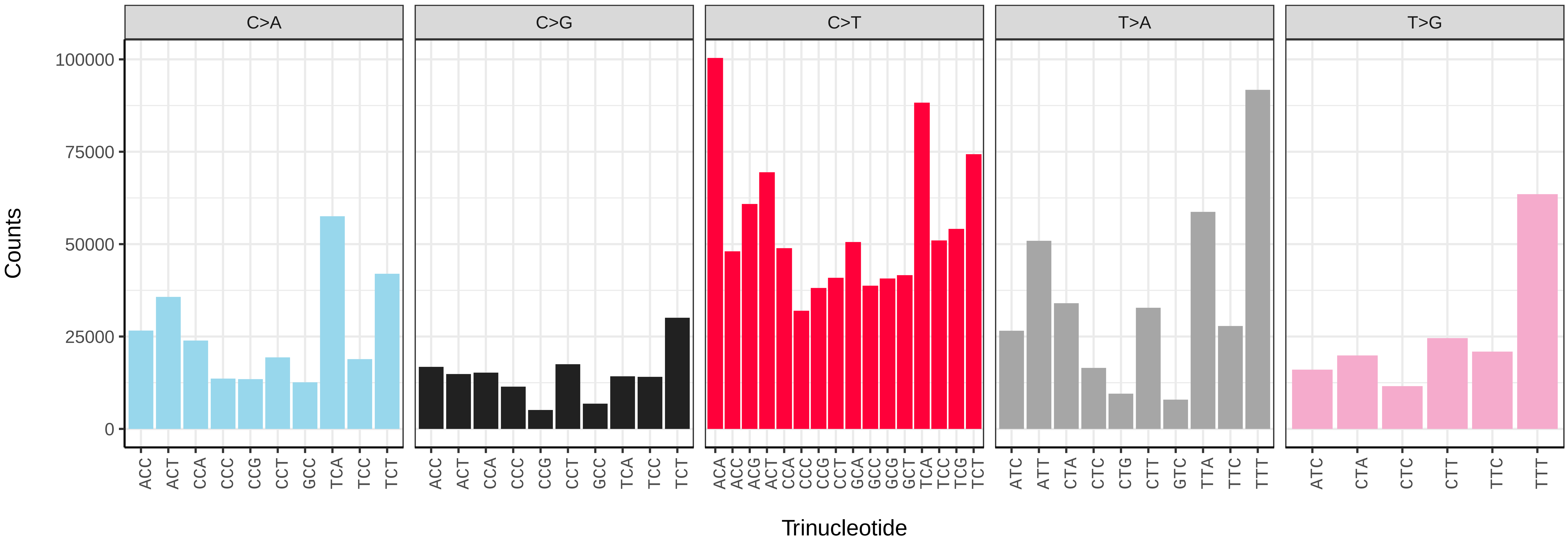


*Pammene aurita* (ilPamAuri1) somatic mutational spectrum (SBS96 classification)

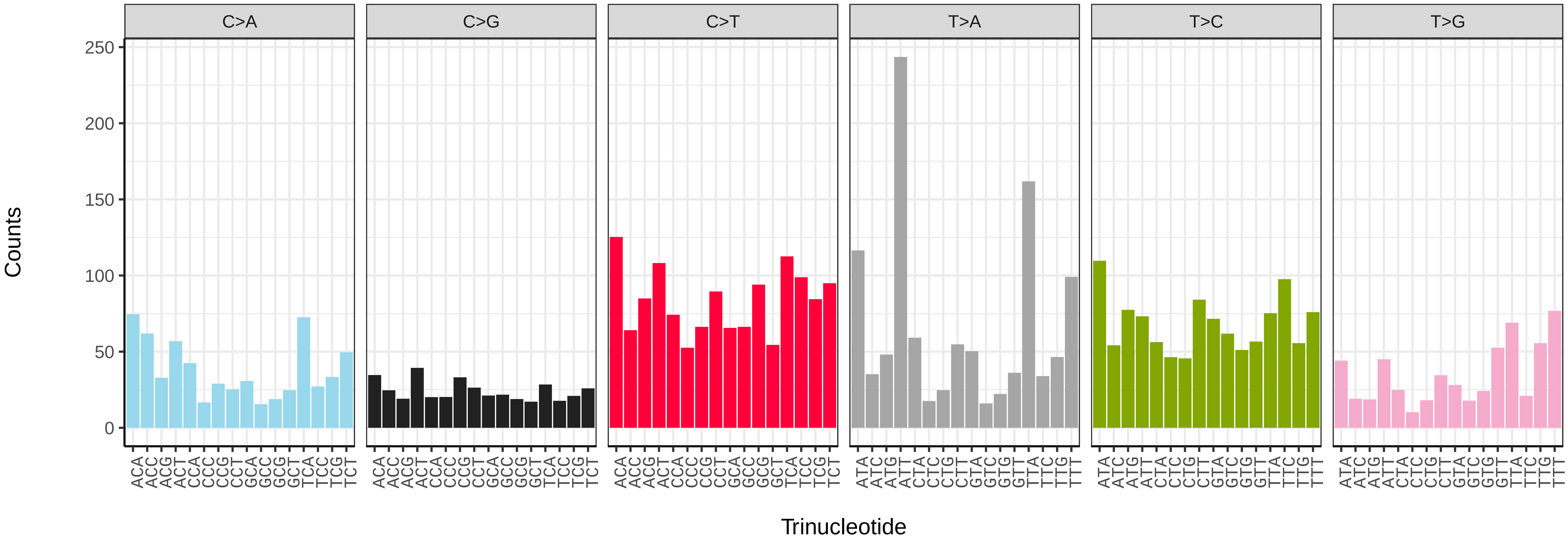


*Pammene aurita* (ilPamAuri1) somatic mutational spectrum (SBS52 classification)

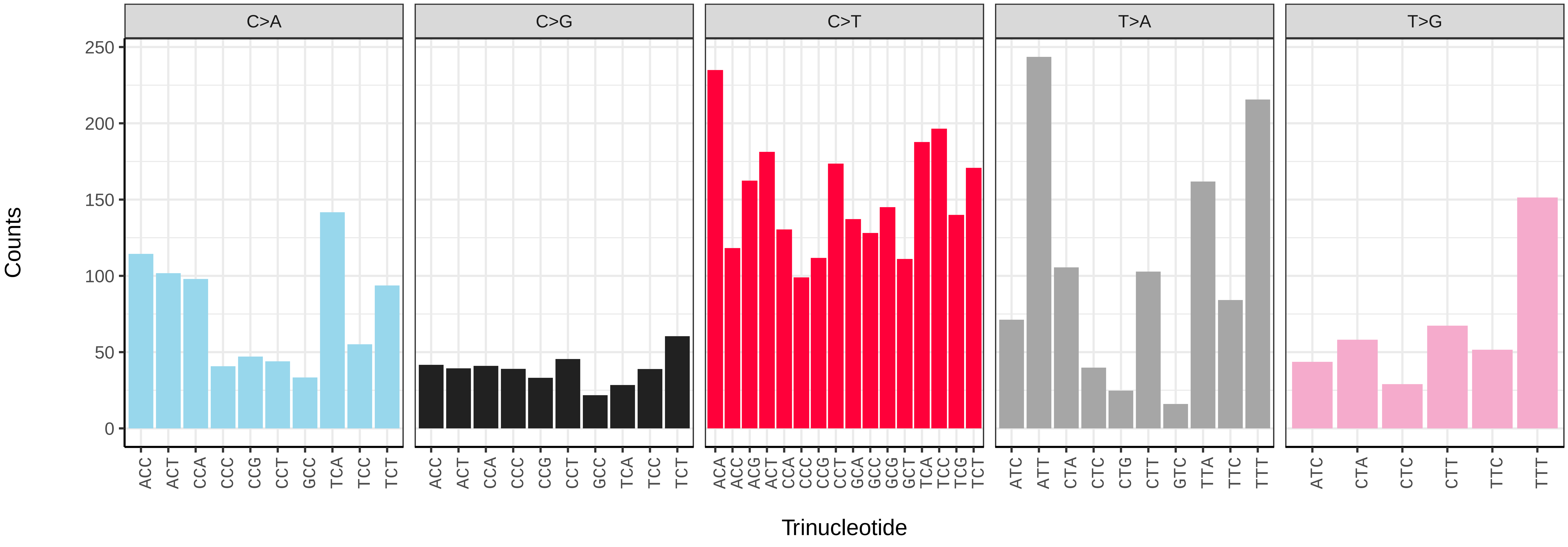

*Pammene fasciana* (ilPamFasc1) germline mutational spectrum

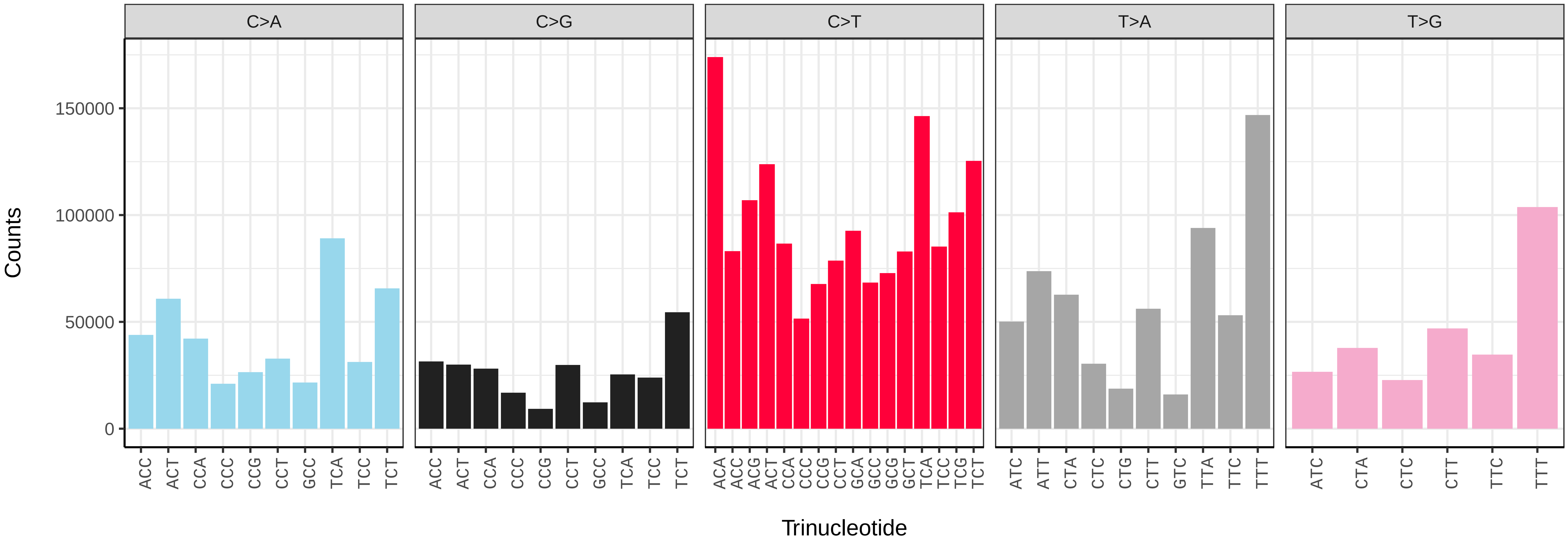


*Pammene fasciana* (ilPamFasc1) somatic mutational spectrum (SBS96 classification)

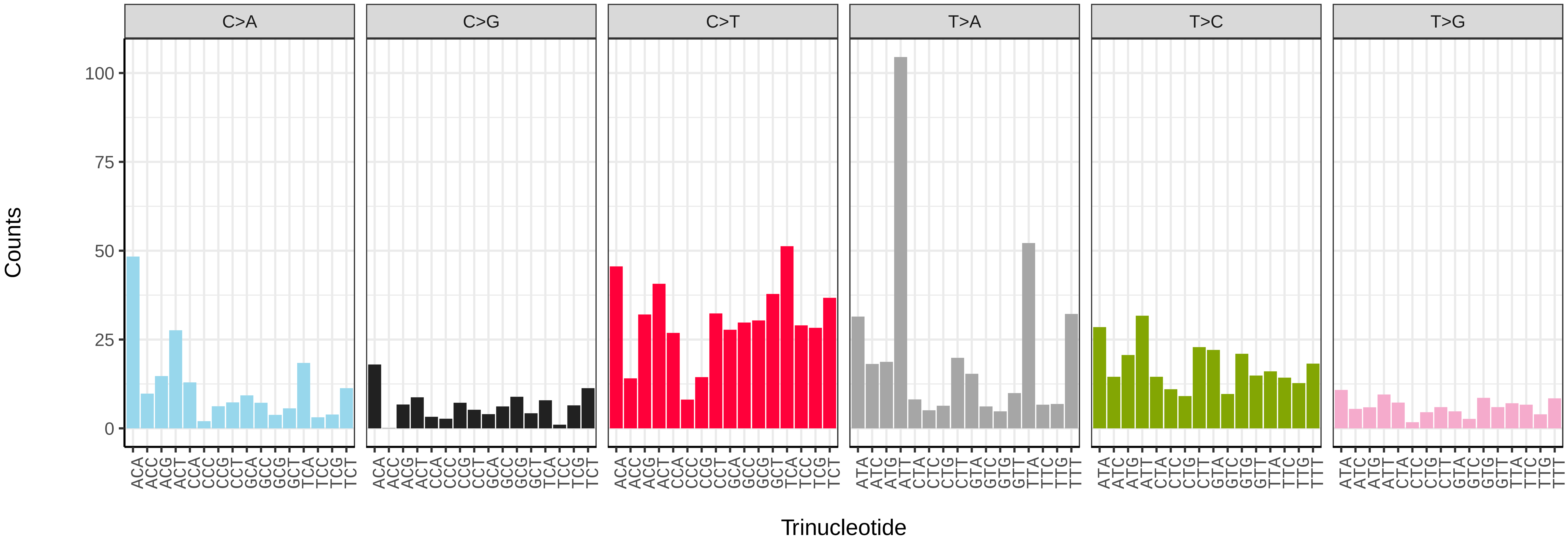


*Pammene fasciana* (ilPamFasc1) somatic mutational spectrum (SBS52 classification)

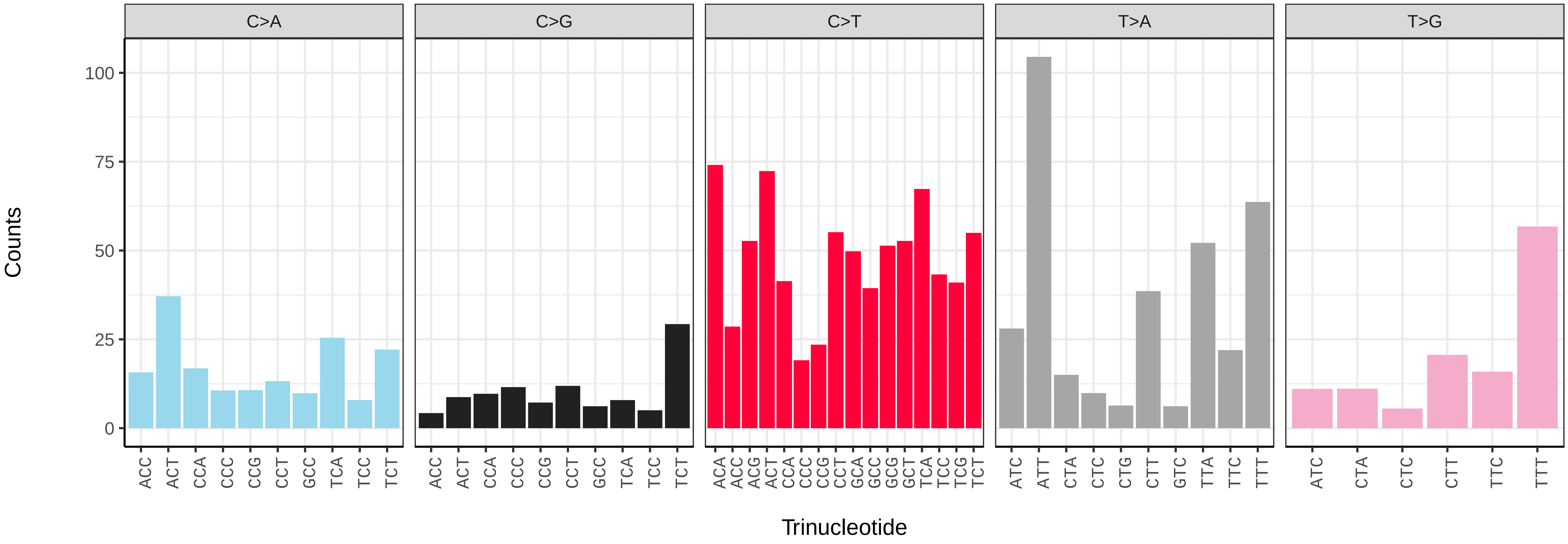

*Pandemis cinnamomeana* (ilPanCinn1) germline mutational spectrum

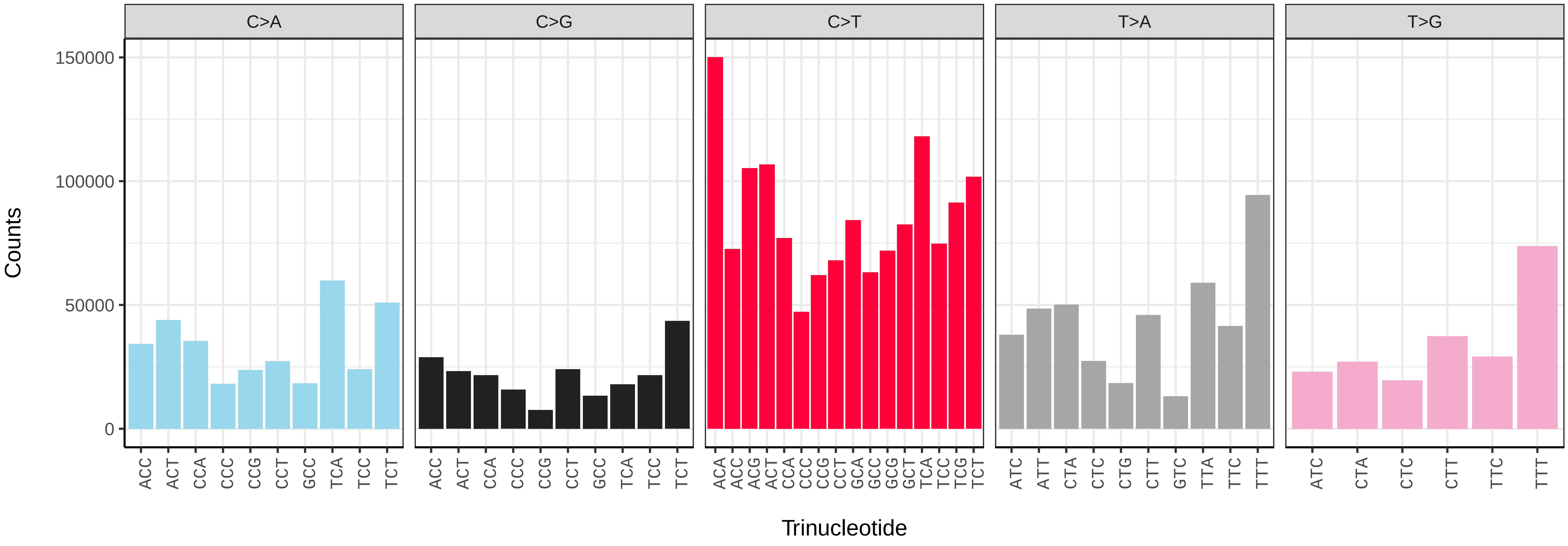


*Pandemis cinnamomeana* (ilPanCinn1) somatic mutational spectrum (SBS96 classification)

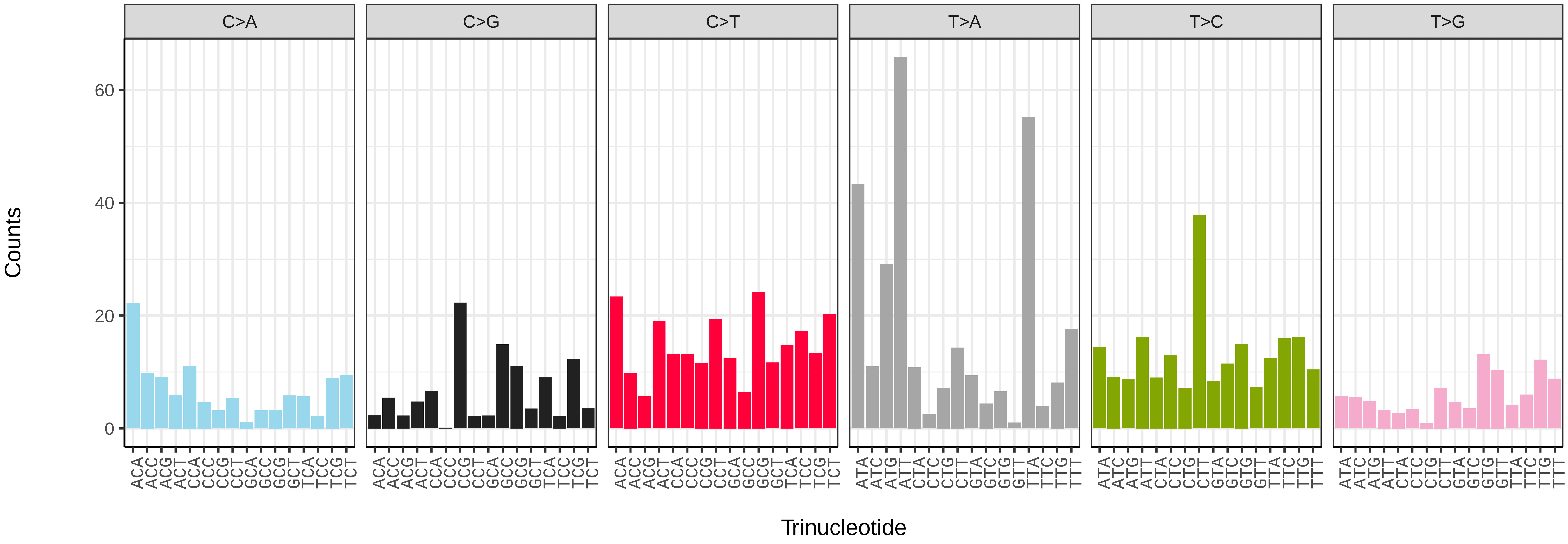


*Pandemis cinnamomeana* (ilPanCinn1) somatic mutational spectrum (SBS52 classification)

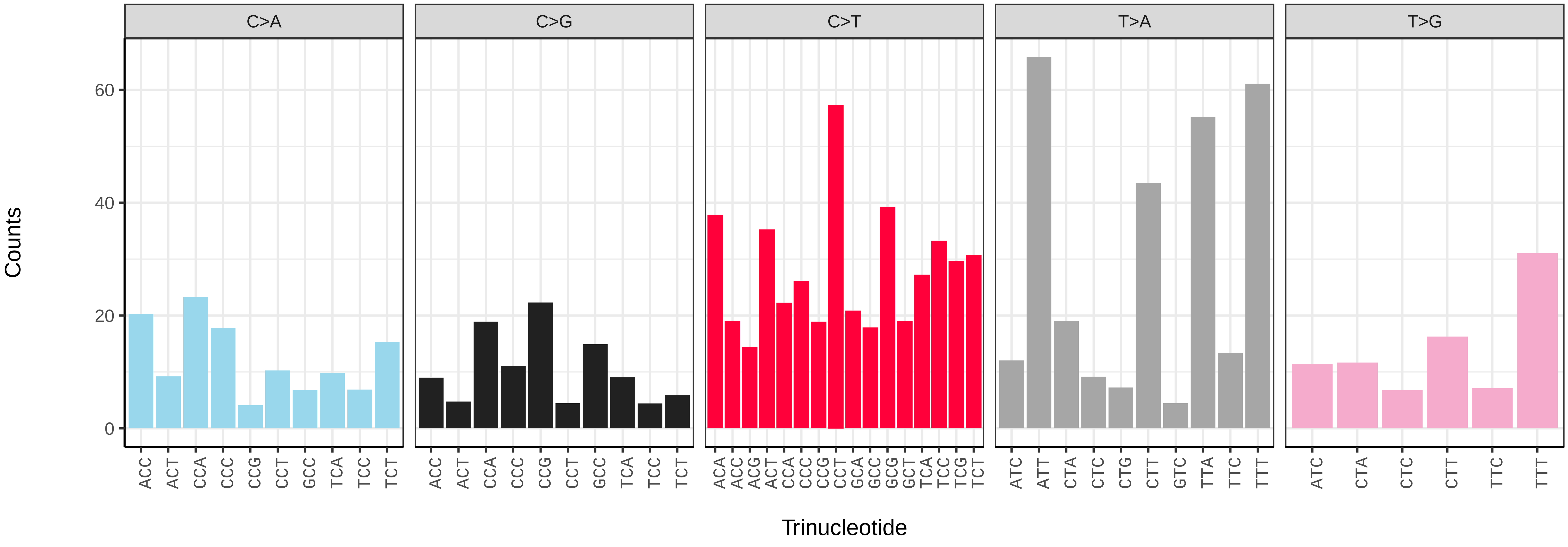

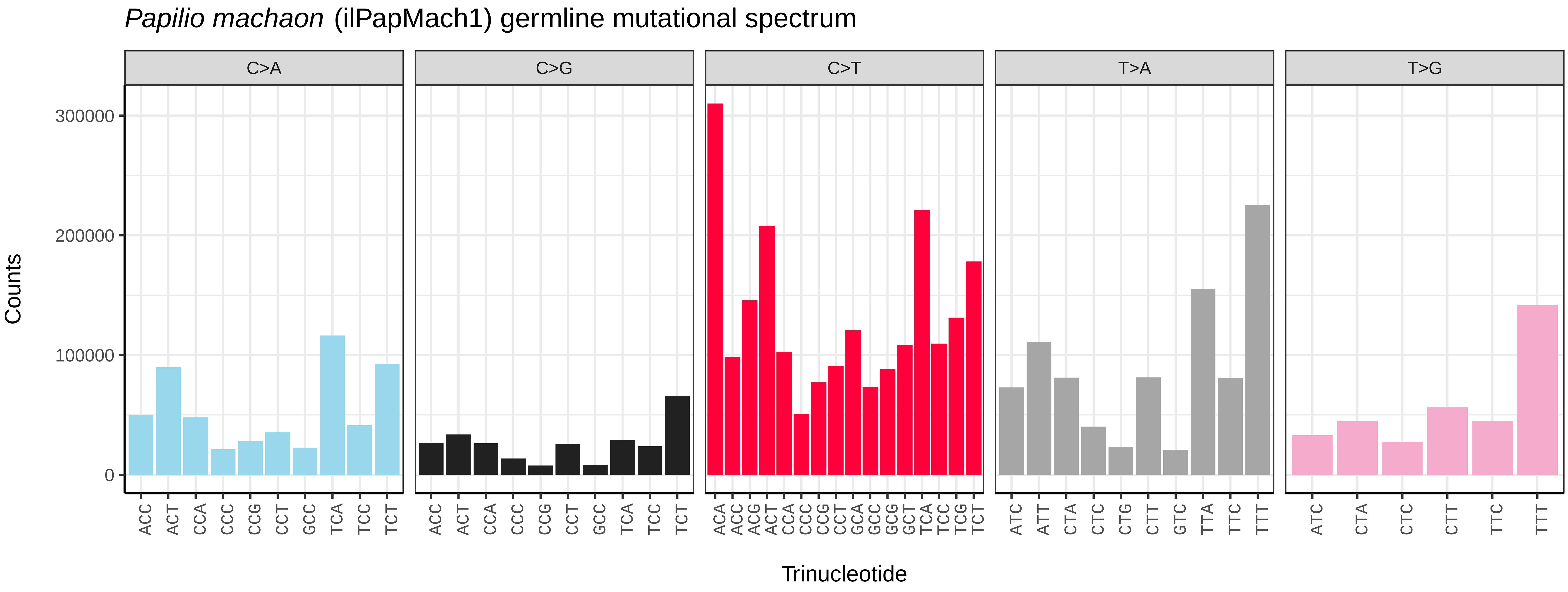
*Papilio machaon* (ilPapMach1) germline mutational spectrum
C>A
C>G
C>T
T>A
T>G
Counts
0
100000
200000
300000
ACC ACT CCA CCC CCG CCT GCC TCA TCC TCT
ACC ACT CCA CCC CCG CCT GCC TCA TCC TCT
ACA ACC ACG ACT CCA CCC CCG CCT GCA GCC GCG GCT TCA TCC TCG TCT
ATC ATT CTA CTC CTG CTT GTC TTA TTC TTT
ATC CTA CTC CTT TTC TTT
Trinucleotide

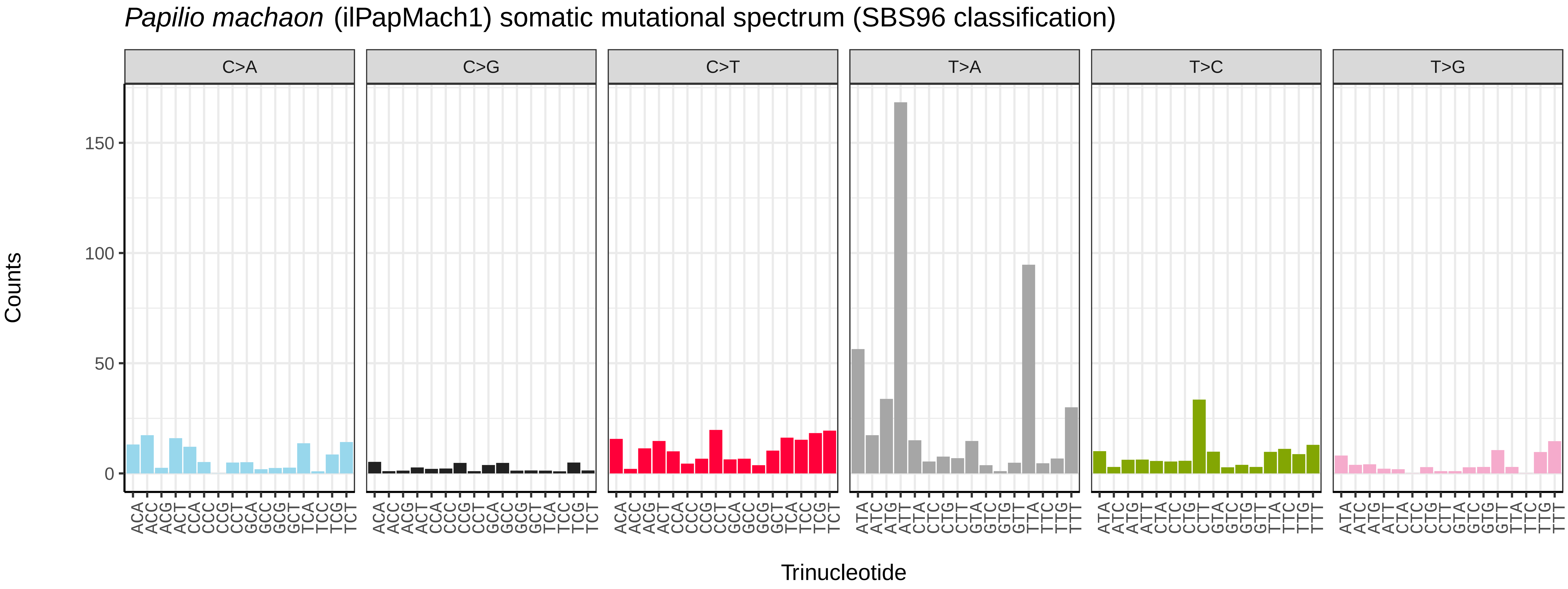
*Papilio machaon* (ilPapMach1) somatic mutational spectrum (SBS96 classification)
C>A
C>G
C>T
T>A
T>C
T>G
Counts
0
50
100
150
ACA ACC ACG ACT CCA CCC CCG CCT GCA GCC GCG GCT TCA TCC TCG TCT
ATA ATC ATG ATT CTA CTC CTG CTT GTA GTC GTG GTT TTA TTC TTG TTT
Trinucleotide

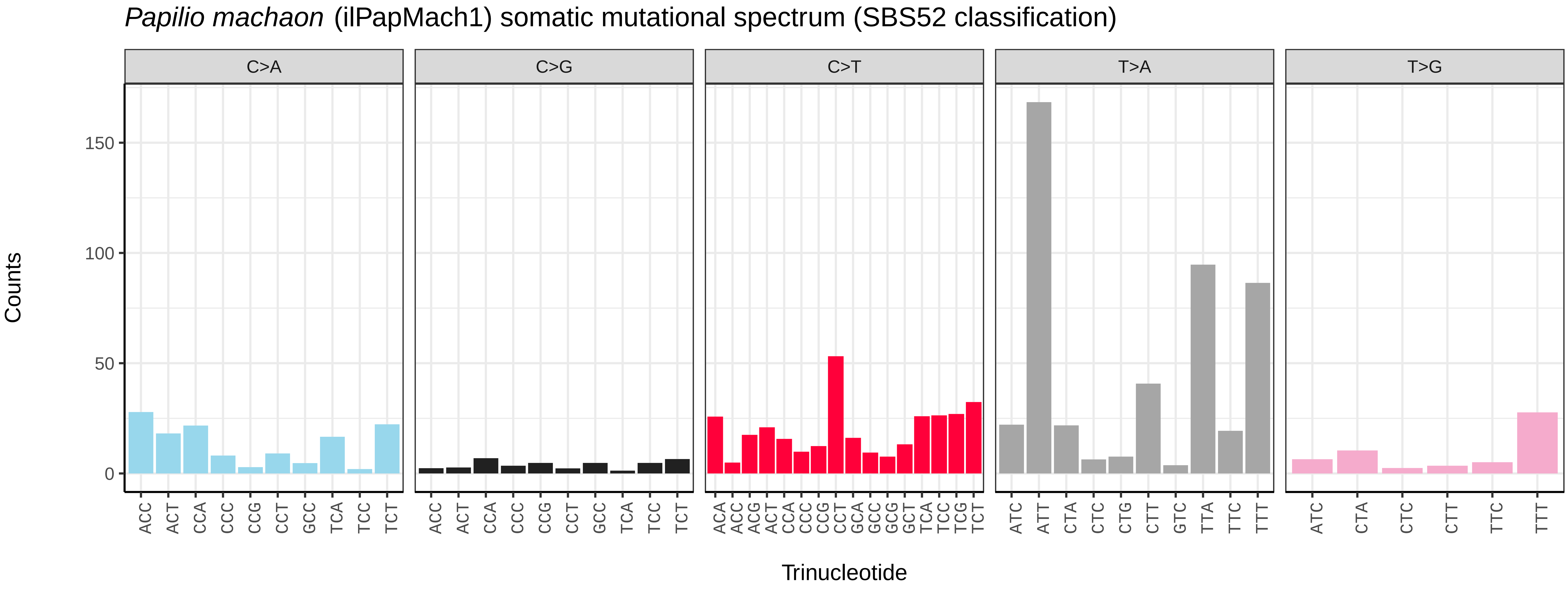
*Papilio machaon* (ilPapMach1) somatic mutational spectrum (SBS52 classification)
C>A
C>G
C>T
T>A
T>G
Counts
0
50
100
150
ACC ACT CCA CCC CCG CCT GCC TCA TCC TCT
ACC ACT CCA CCC CCG CCT GCC TCA TCC TCT
ACA ACC ACG ACT CCA CCC CCG CCT GCA GCC GCG GCT TCA TCC TCG TCT
ATC ATT CTA CTC CTG CTT GTC TTA TTC TTT
ATC CTA CTC CTT TTC TTT
Trinucleotide

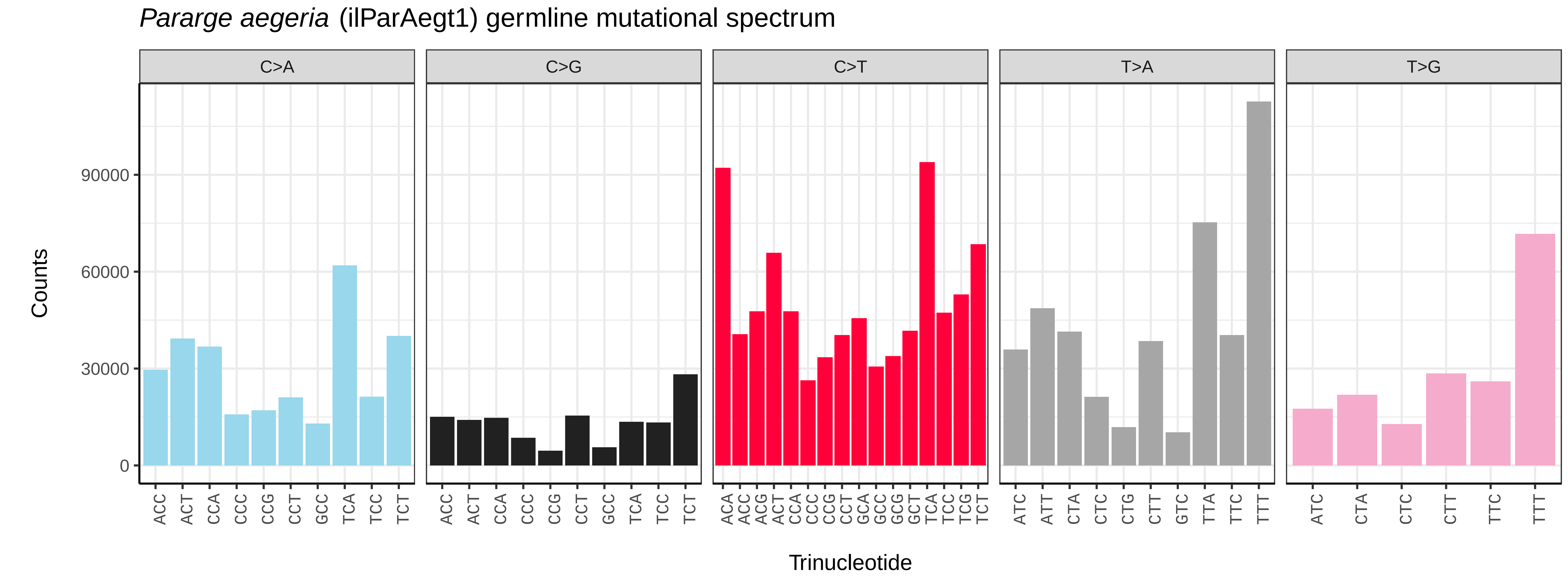
*Pararge aegeria* (ilParAegt1) germline mutational spectrum
C>A
C>G
C>T
T>A
T>G
Counts
0
30000
60000
90000
ACC ACT CCA CCC CCG CCT GCC TCA TCC TCT
ACA ACC ACG ACT CCA CCC CCG CCT GCA GCC GCG GCT TCA TCC TCG TCT
ATC ATT CTA CTC CTG CTT GTC TTA TTC TTT
ATC CTA CTC CTT TTC TTT
Trinucleotide

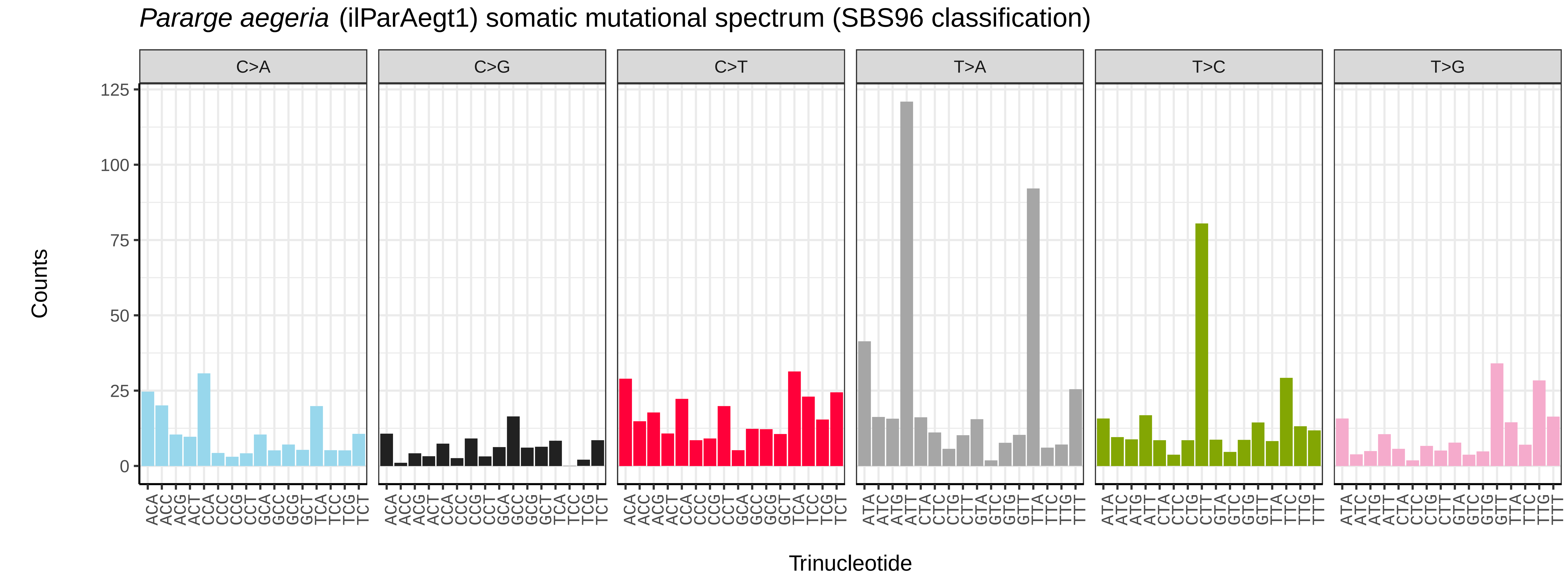
*Pararge aegeria* (ilParAegt1) somatic mutational spectrum (SBS96 classification)
C>A
C>G
C>T
T>A
T>C
T>G
Counts
0
25
50
75
100
125
ACA ACC ACG ACT CCA CCC CCG CCT GCA GCC GCG GCT TCA TCC TCG TCT
ATA ATC ATG ATT CTA CTC CTG CTT GTA GTC GTG GTT TTA TTC TTG TTT
Trinucleotide

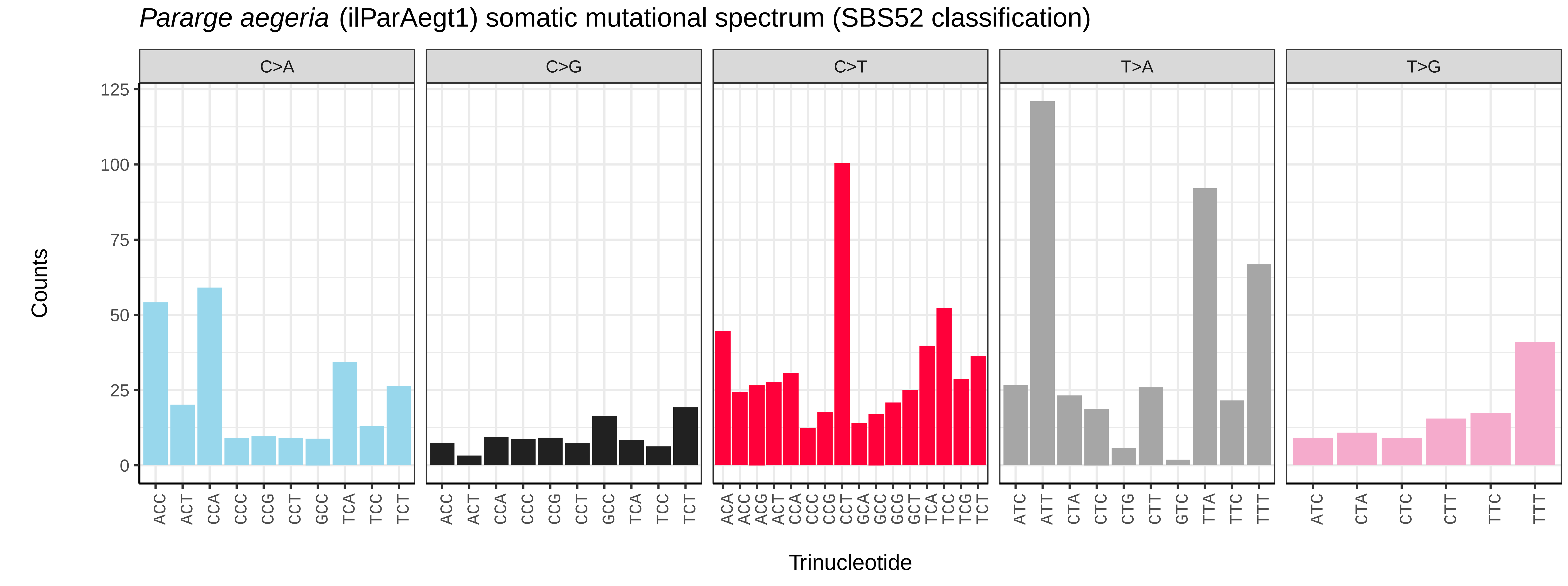
*Pararge aegeria* (ilParAegt1) somatic mutational spectrum (SBS52 classification)
C>A
C>G
C>T
T>A
T>G
Counts
0
25
50
75
100
125
ACC ACT CCA CCC CCG CCT GCC TCA TCC TCT
ACA ACC ACG ACT CCA CCC CCG CCT GCA GCC GCG GCT TCA TCC TCG TCT
ATC ATT CTA CTC CTG CTT GTC TTA TTC TTT
ATC CTA CTC CTT TTC TTT
Trinucleotide

*Parapoynx stratiotata* (ilParStra1) germline mutational spectrum

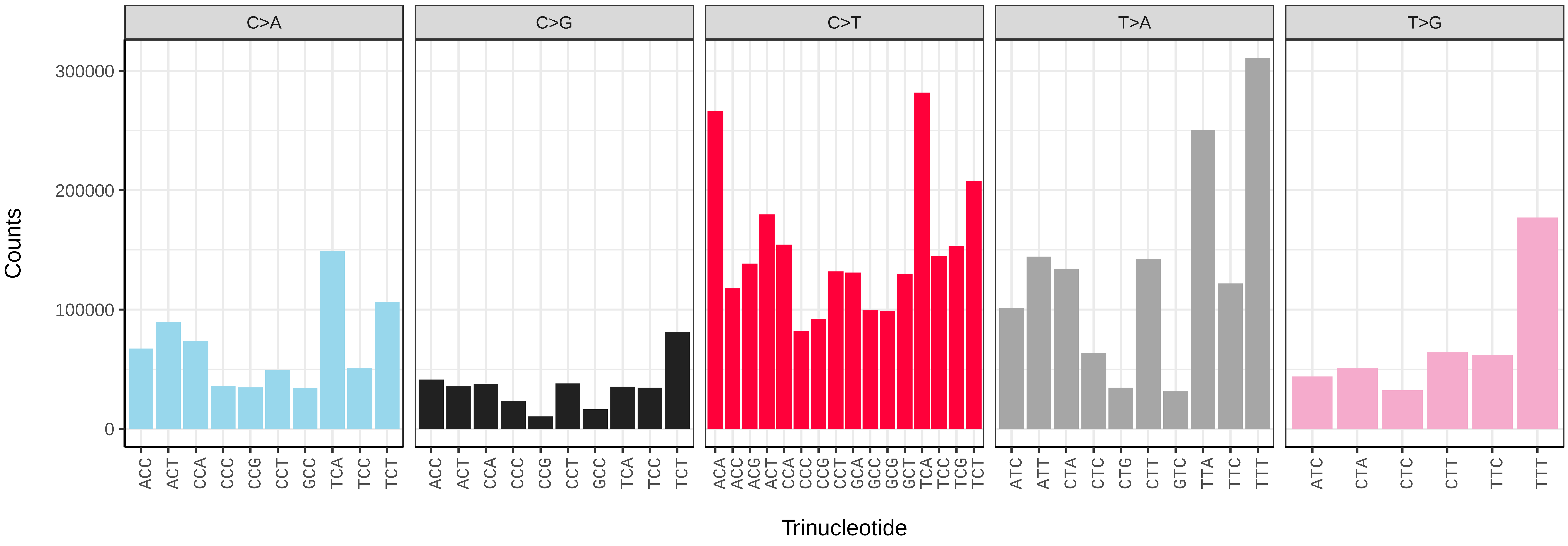


*Parapoynx stratiotata* (ilParStra1) somatic mutational spectrum (SBS96 classification)

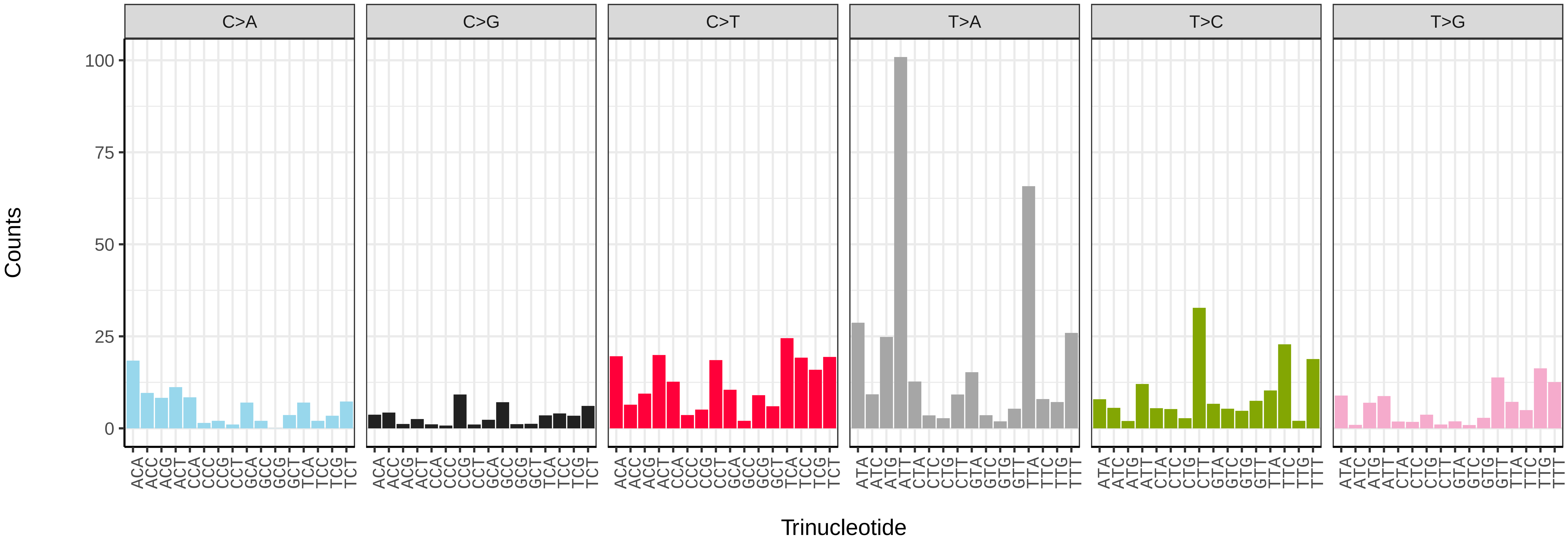


*Parapoynx stratiotata* (ilParStra1) somatic mutational spectrum (SBS52 classification)

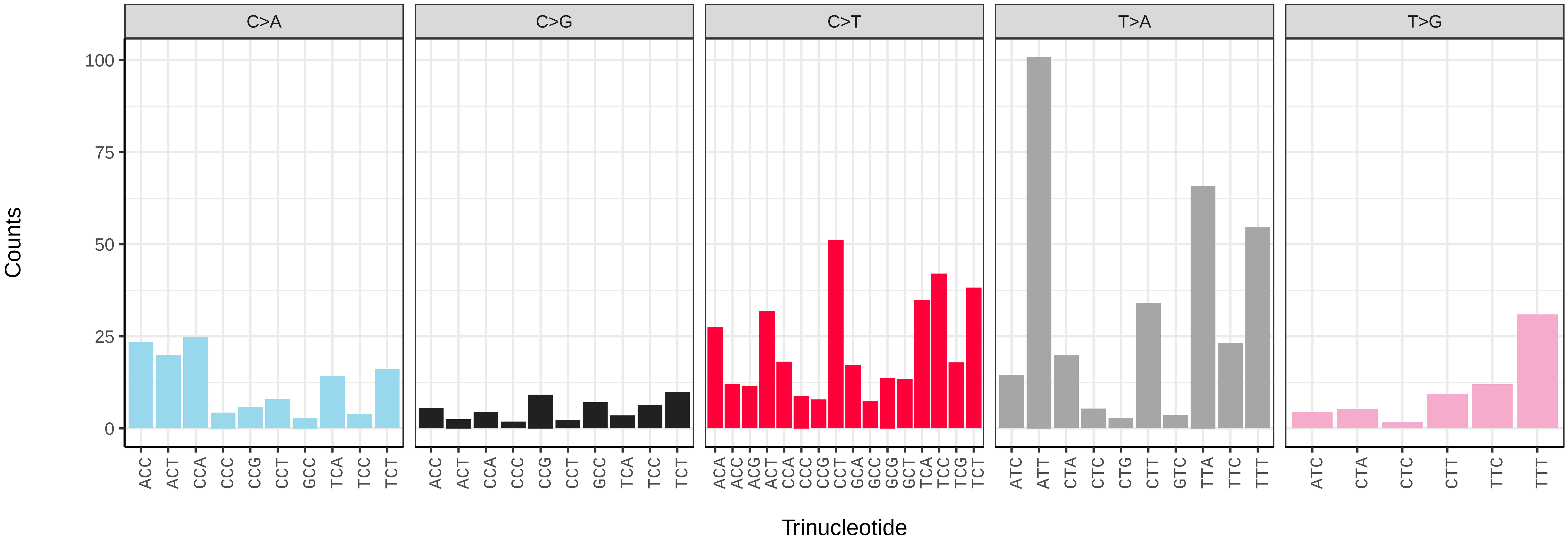

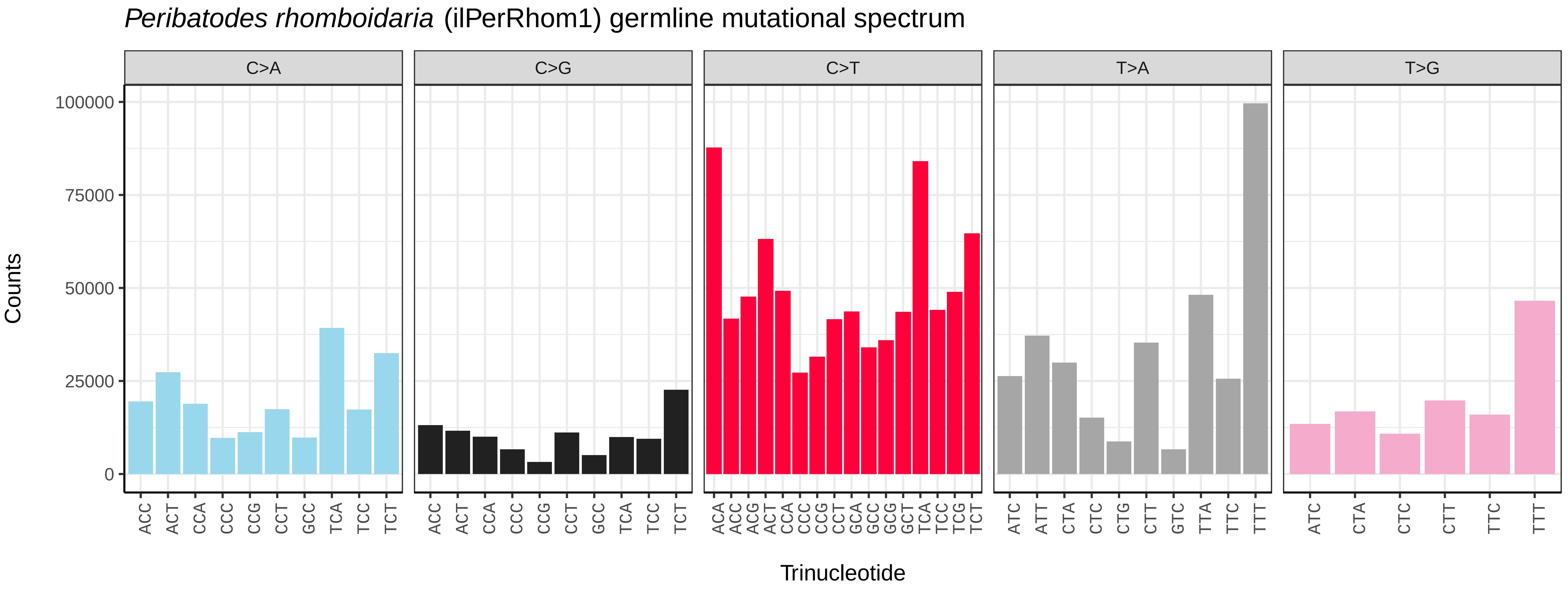
*Peribatodes rhomboidaria* (ilPerRhom1) germline mutational spectrum
C>A
C>G
C>T
T>A
T>G
Counts
0
25000
50000
75000
100000
ACC ACT CCA CCC CCG CCT GCC TCA TCC TCT
ACC ACT CCA CCC CCG CCT GCC TCA TCC TCT
ACA ACC ACG ACT CCA CCC CCG CCT GCA GCC GCG GCT TCA TCC TCG TCT
ATC ATT CTA CTC CTG CTT GTC TTA TTC TTT
ATC CTA CTC CTT TTC TTT
Trinucleotide

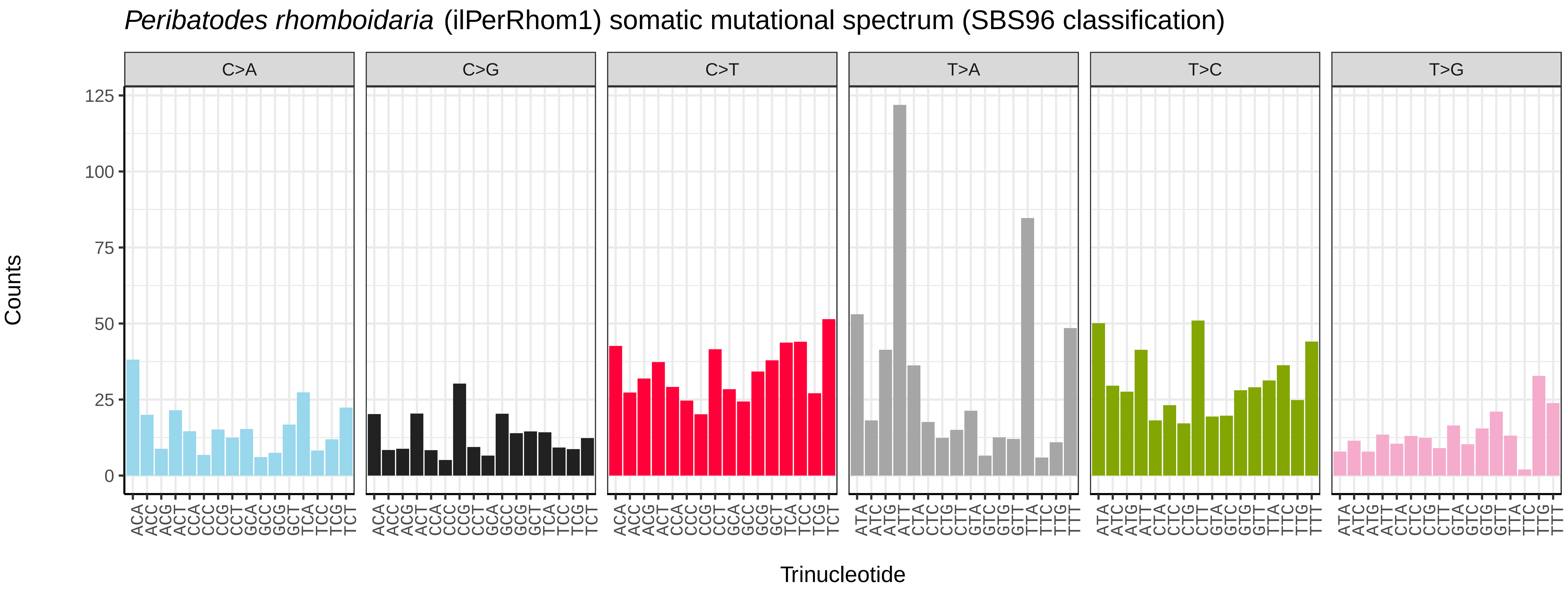
*Peribatodes rhomboidaria* (ilPerRhom1) somatic mutational spectrum (SBS96 classification)
C>A
C>G
C>T
T>A
T>C
T>G
Counts
0
25
50
75
100
125
ACA ACC ACG ACT CCA CCC CCG CCT GCA GCC GCG GCT TCA TCC TCG TCT
ACA ACC ACG ACT CCA CCC CCG CCT GCA GCC GCG GCT TCA TCC TCG TCT
ACA ACC ACG ACT CCA CCC CCG CCT GCA GCC GCG GCT TCA TCC TCG TCT
ATA ATC ATG ATT CTA CTC CTG CTT GTA GTC GTG GTT TTA TTC TTG TTT
ATA ATC ATG ATT CTA CTC CTG CTT GTA GTC GTG GTT TTA TTC TTG TTT
ATA ATC ATG ATT CTA CTC CTG CTT GTA GTC GTG GTT TTA TTC TTG TTT
Trinucleotide

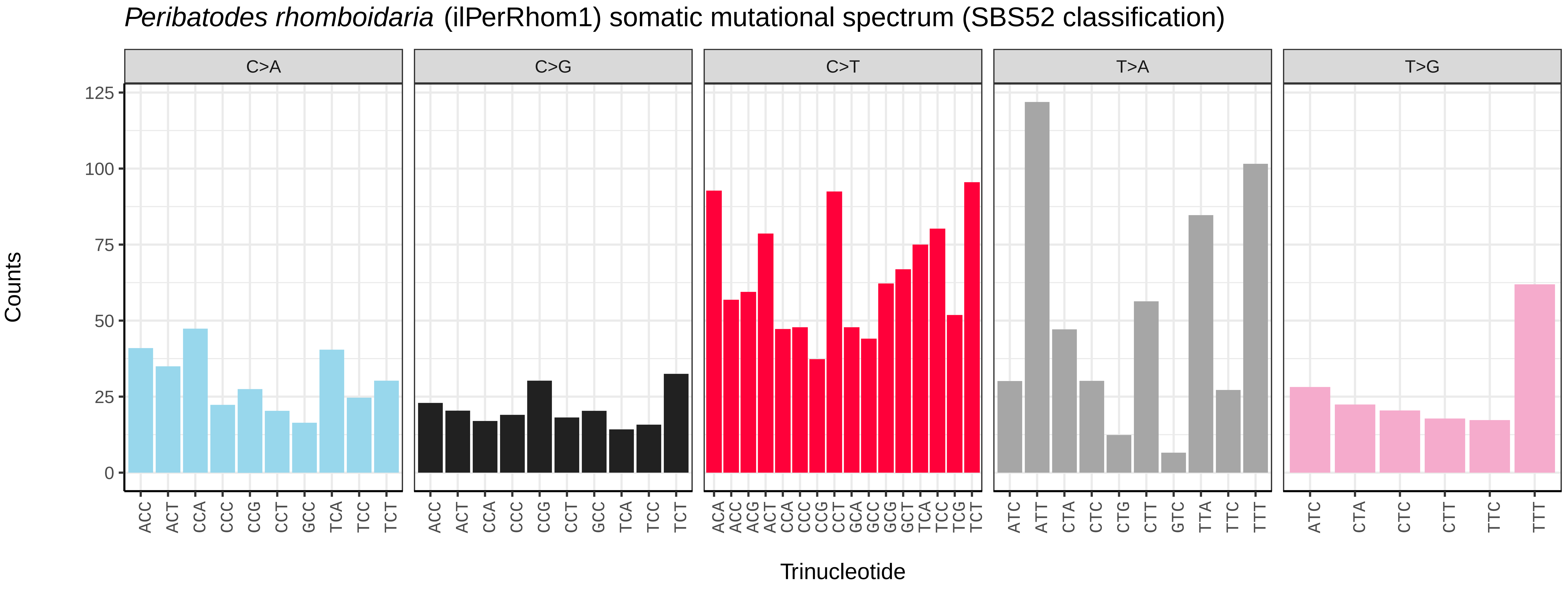
*Peribatodes rhomboidaria* (ilPerRhom1) somatic mutational spectrum (SBS52 classification)
C>A
C>G
C>T
T>A
T>G
Counts
0
25
50
75
100
125
ACC ACT CCA CCC CCG CCT GCC TCA TCC TCT
ACC ACT CCA CCC CCG CCT GCC TCA TCC TCT
ACA ACC ACG ACT CCA CCC CCG CCT GCA GCC GCG GCT TCA TCC TCG TCT
ATC ATT CTA CTC CTG CTT GTC TTA TTC TTT
ATC CTA CTC CTT TTC TTT
Trinucleotide

*Phalera bucephala* (ilPhaBuce1) germline mutational spectrum

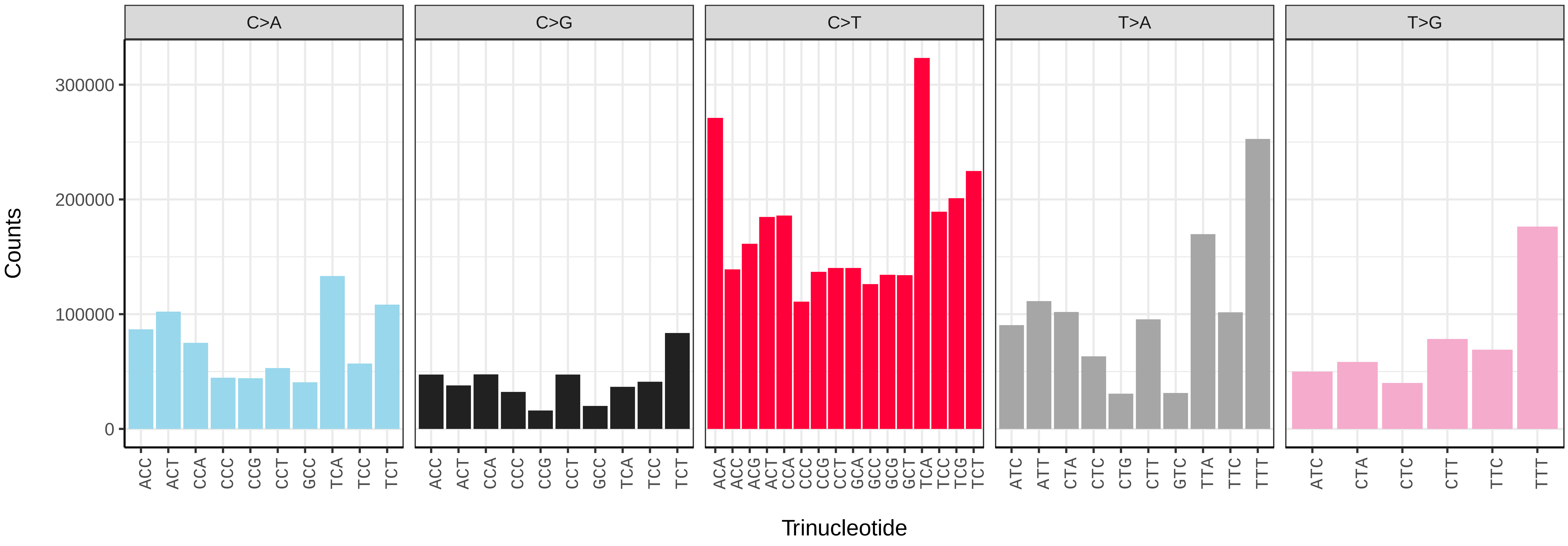


*Phalera bucephala* (ilPhaBuce1) somatic mutational spectrum (SBS96 classification)

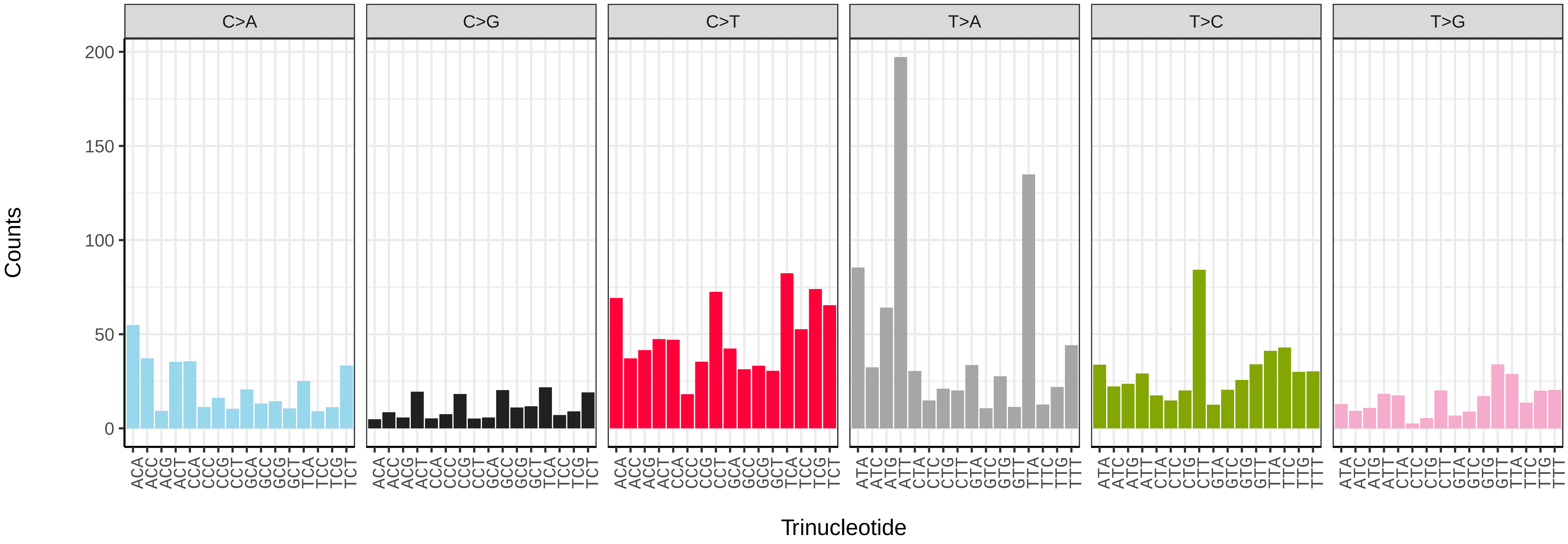


*Phalera bucephala* (ilPhaBuce1) somatic mutational spectrum (SBS52 classification)

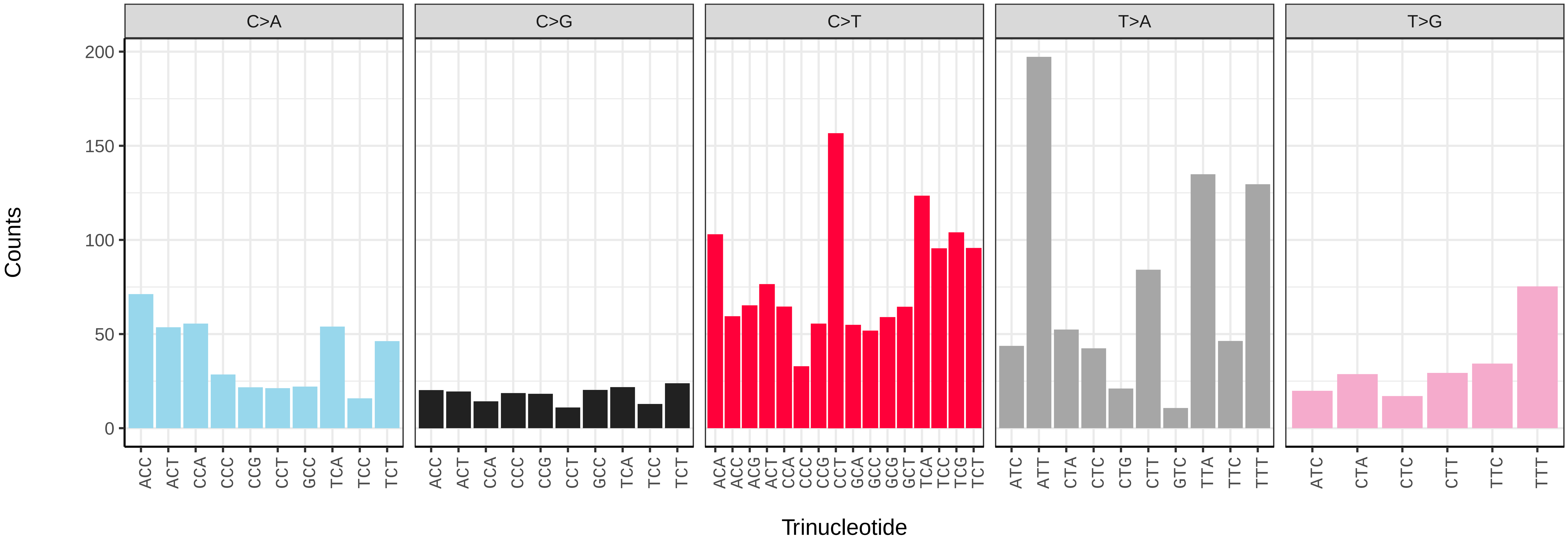

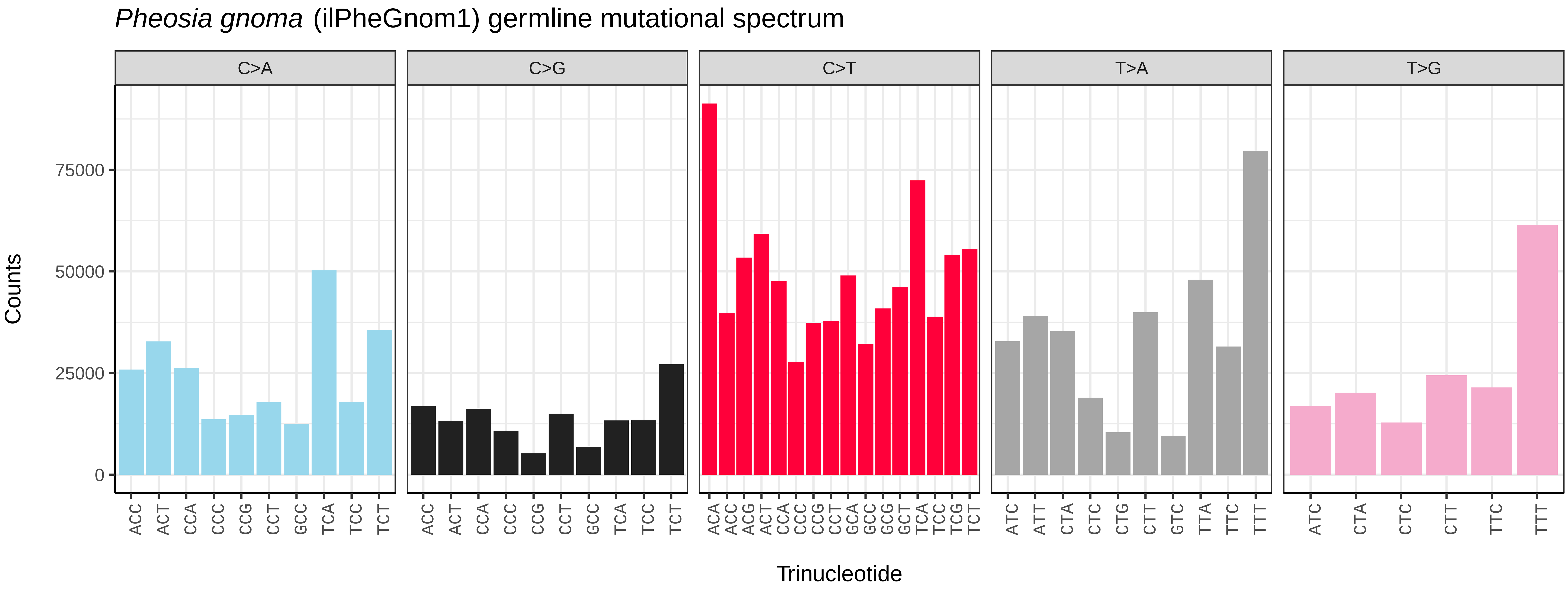
*Pheosia gnoma* (ilPheGnom1) germline mutational spectrum
C>A
C>G
C>T
T>A
T>G
Counts
0
25000
50000
75000
ACC ACT CCA CCC CCG CCT GCC TCA TCC TCT
ACC ACT CCA CCC CCG CCT GCC TCA TCC TCT
ACA ACC ACG ACT CCA CCC CCG CCT GCA GCC GCG GCT TCA TCC TCG TCT
ATC ATT CTA CTC CTG CTT GTC TTA TTC TTT
ATC CTA CTC CTT TTC TTT
Trinucleotide

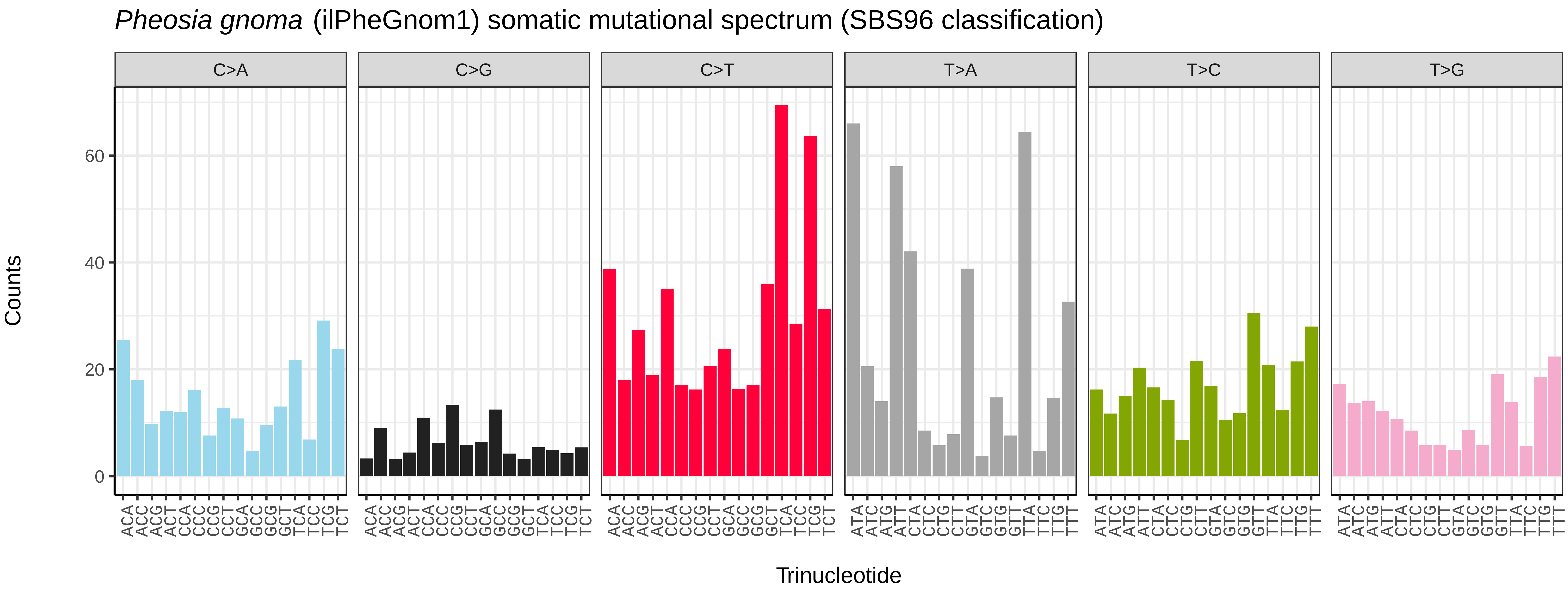
*Pheosia gnoma* (ilPheGnom1) somatic mutational spectrum (SBS96 classification)
C>A
C>G
C>T
T>A
T>C
T>G
Counts
0
20
40
60
ACA ACC ACG ACT CCA CCC CCG CCT GCA GCC GCG GCT TCA TCC TCG TCT
ACA ACC ACG ACT CCA CCC CCG CCT GCA GCC GCG GCT TCA TCC TCG TCT
ACA ACC ACG ACT CCA CCC CCG CCT GCA GCC GCG GCT TCA TCC TCG TCT
ATA ATC ATG ATT CTA CTC CTG CTT GTA GTC GTG GTT TTA TTC TTG TTT
ATA ATC ATG ATT CTA CTC CTG CTT GTA GTC GTG GTT TTA TTC TTG TTT
ATA ATC ATG ATT CTA CTC CTG CTT GTA GTC GTG GTT TTA TTC TTG TTT
Trinucleotide

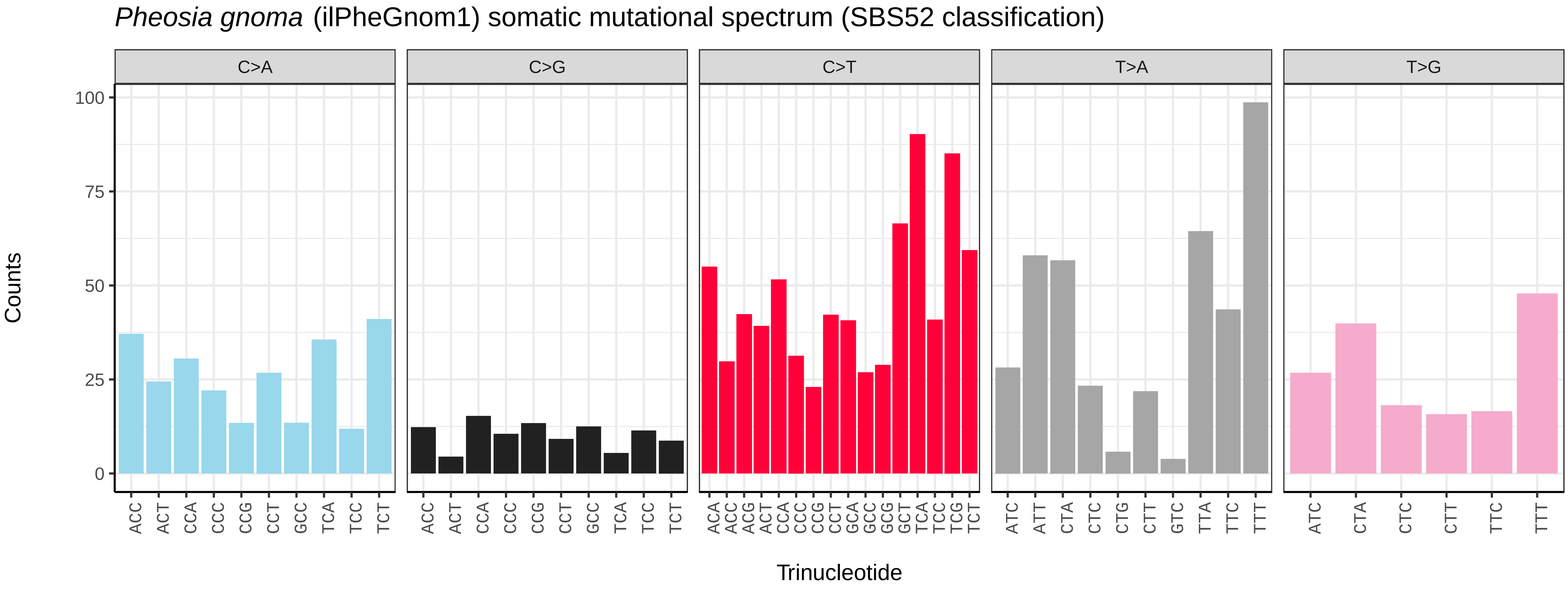
*Pheosia gnoma* (ilPheGnom1) somatic mutational spectrum (SBS52 classification)
C>A
C>G
C>T
T>A
T>G
Counts
0
25
50
75
100
ACC ACT CCA CCC CCG CCT GCC TCA TCC TCT
ACC ACT CCA CCC CCG CCT GCC TCA TCC TCT
ACA ACC ACG ACT CCA CCC CCG CCT GCA GCC GCG GCT TCA TCC TCG TCT
ATC ATT CTA CTC CTG CTT GTC TTA TTC TTT
ATC CTA CTC CTT TTC TTT
Trinucleotide

*Pheosia tremula* (ilPheTrem1) germline mutational spectrum

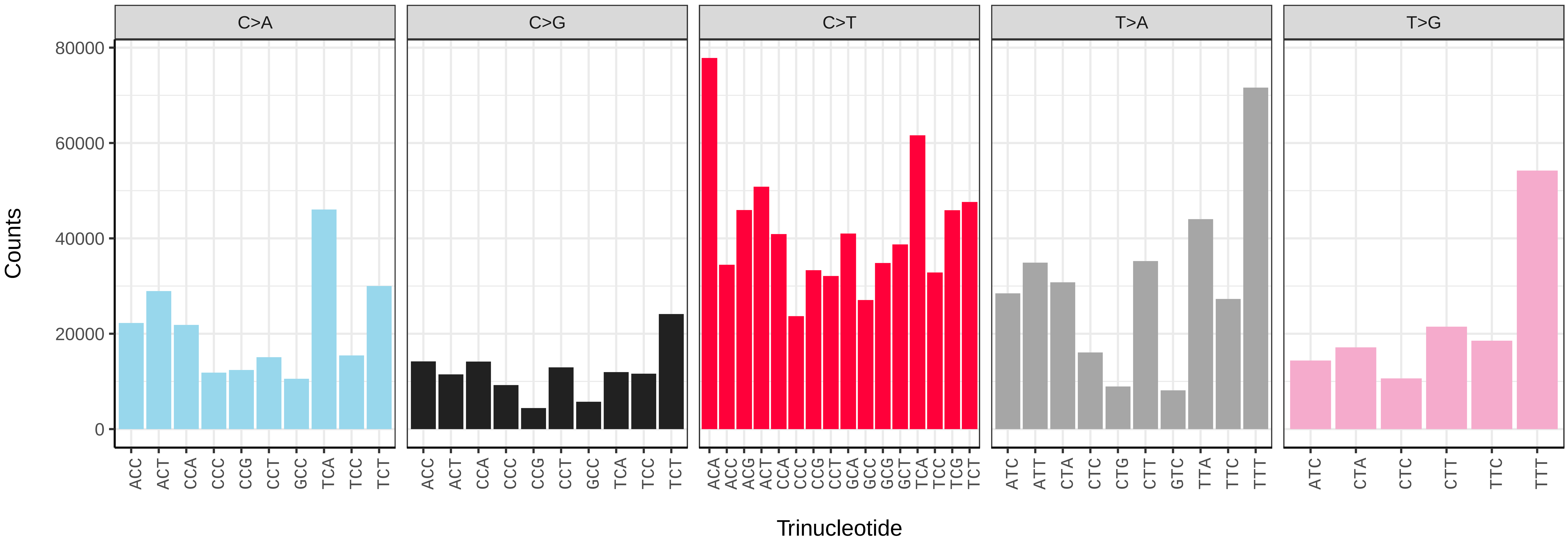


*Pheosia tremula* (ilPheTrem1) somatic mutational spectrum (SBS96 classification)

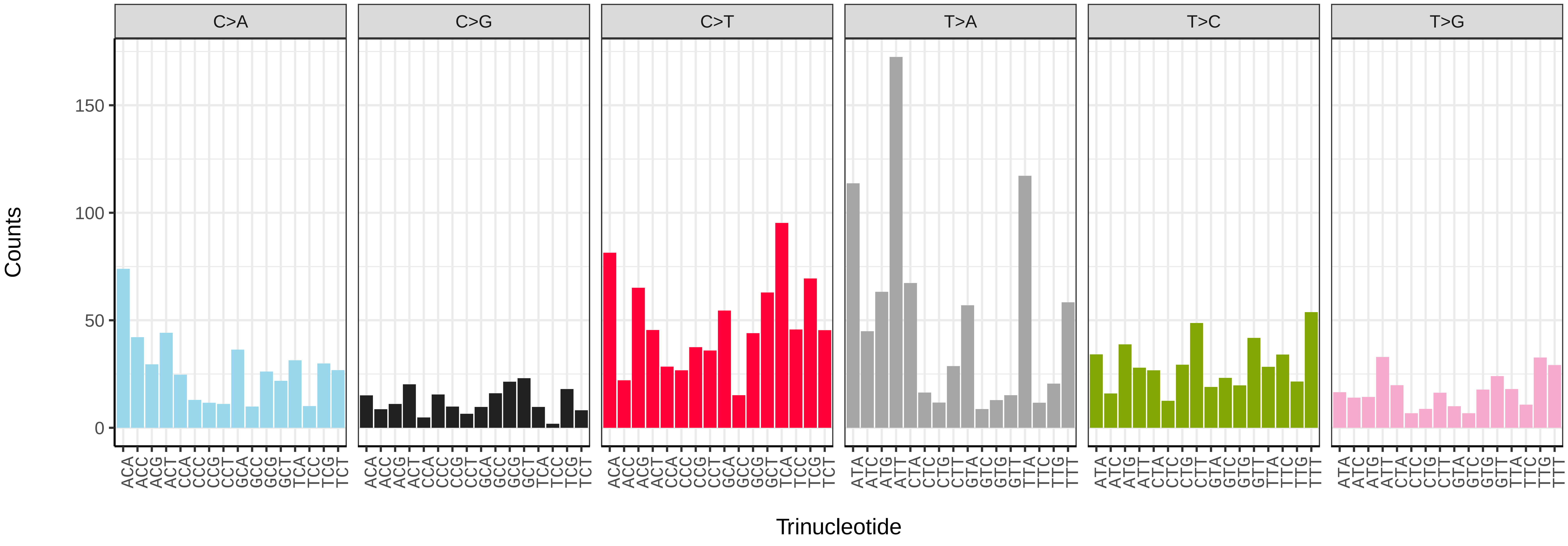


*Pheosia tremula* (ilPheTrem1) somatic mutational spectrum (SBS52 classification)

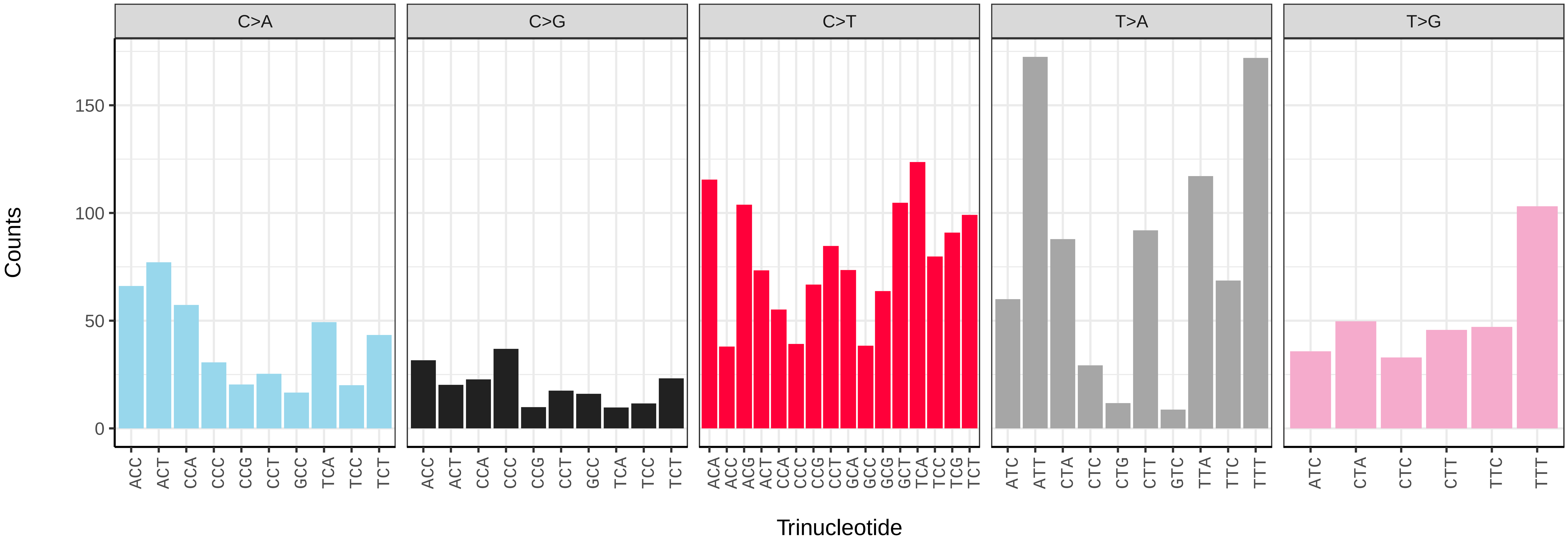

*Philereme vetulata* (ilPhiVetu1) germline mutational spectrum

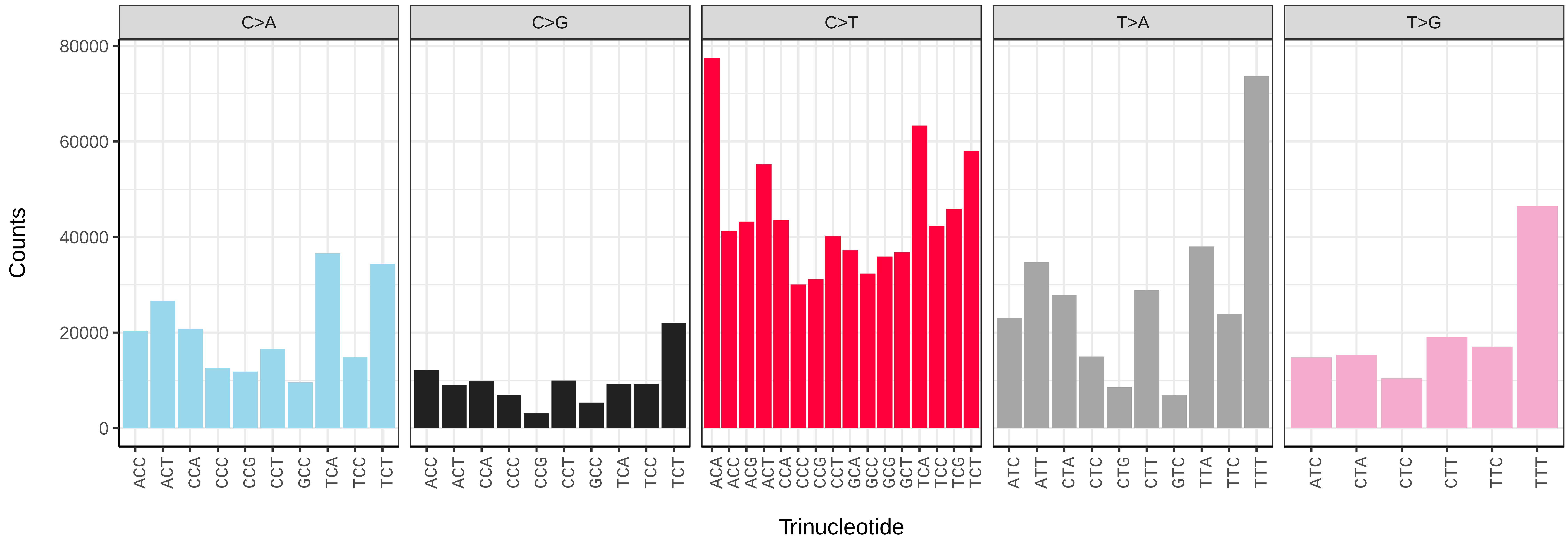


*Philereme vetulata* (ilPhiVetu1) somatic mutational spectrum (SBS96 classification)

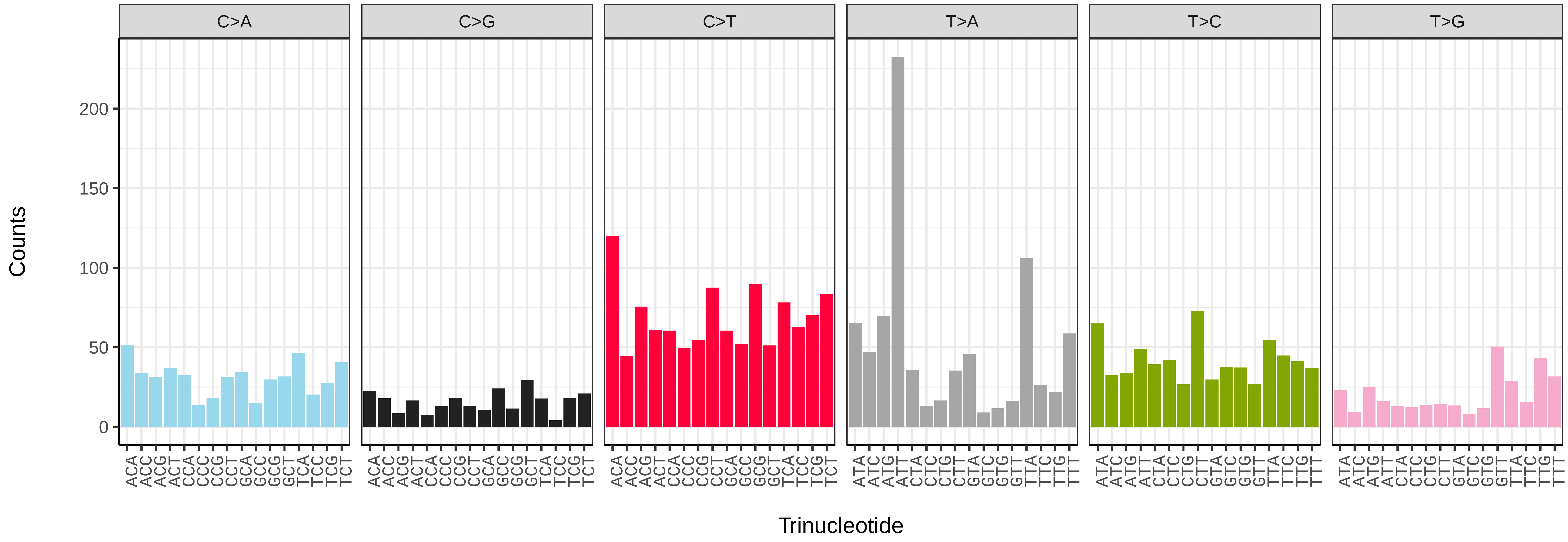


*Philereme vetulata* (ilPhiVetu1) somatic mutational spectrum (SBS52 classification)

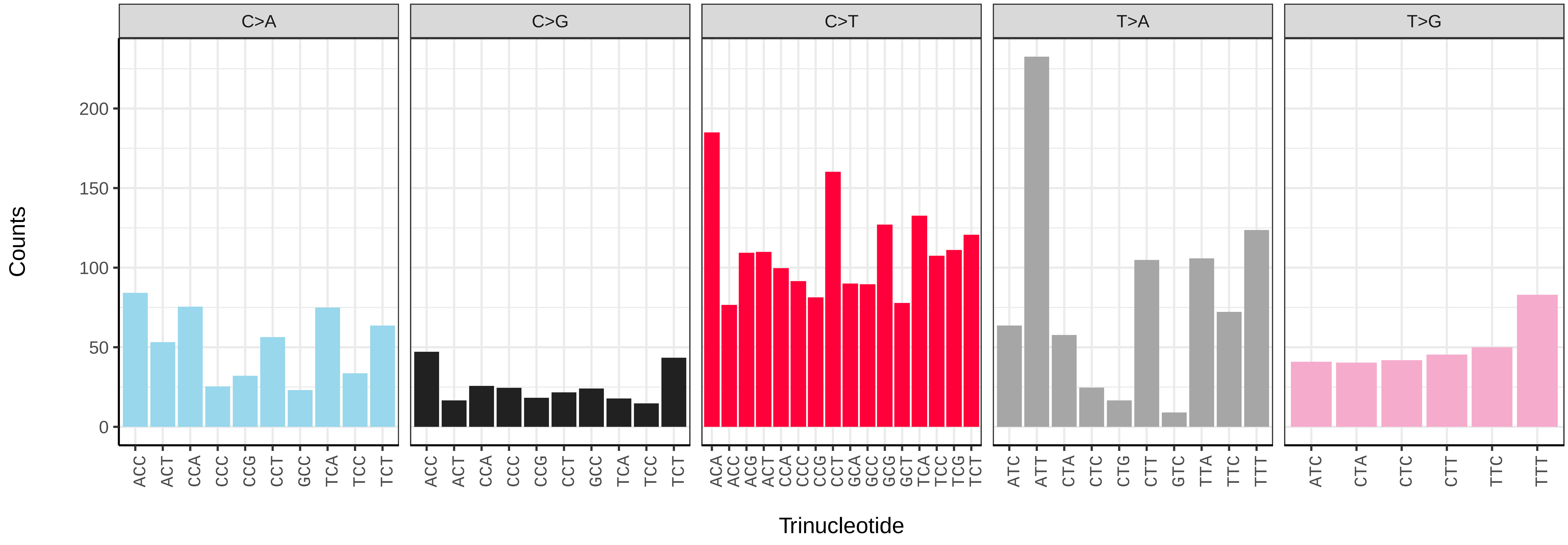

*Phlogophora meticulosa* (ilPhlMeti2) germline mutational spectrum

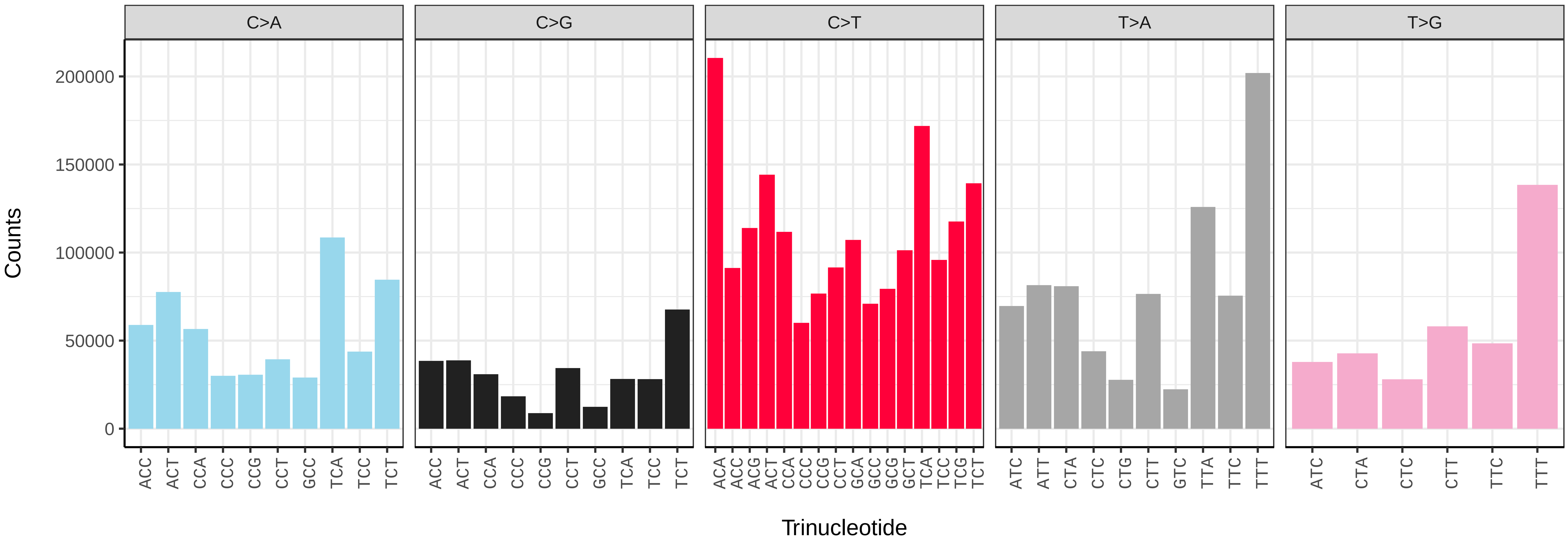


*Phlogophora meticulosa* (ilPhlMeti2) somatic mutational spectrum (SBS96 classification)

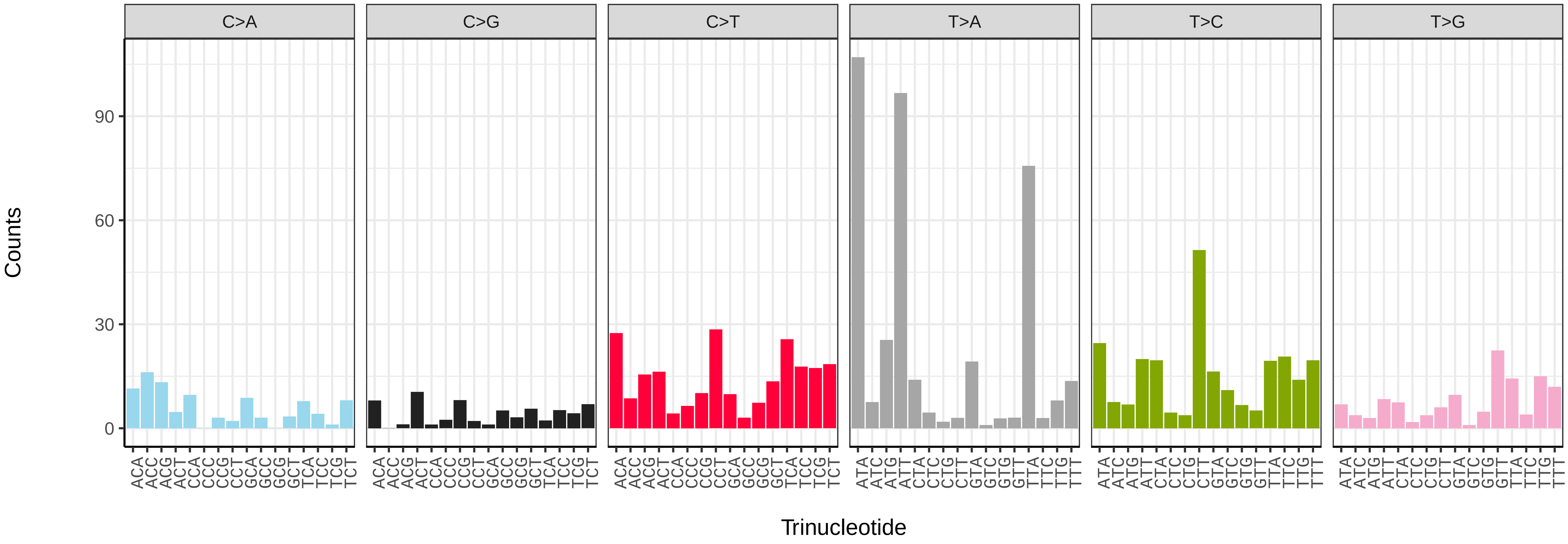


*Phlogophora meticulosa* (ilPhlMeti2) somatic mutational spectrum (SBS52 classification)

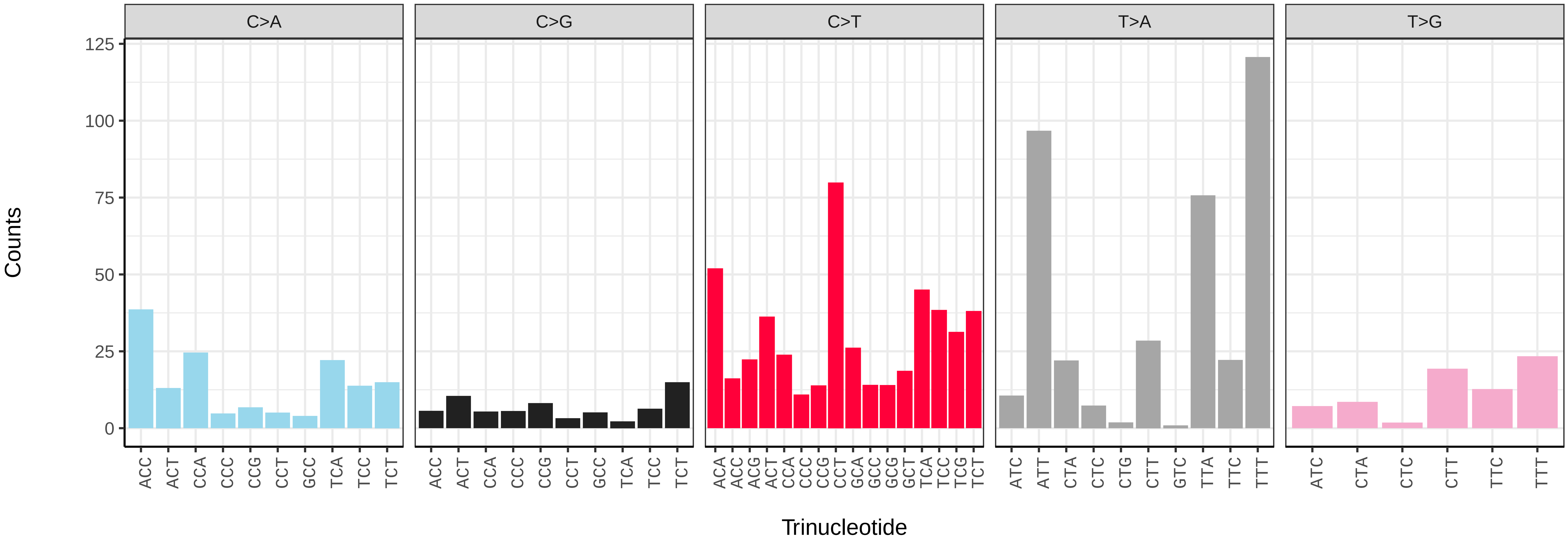

*Phragmatobia fuliginosa* (ilPhrFuli1) germline mutational spectrum

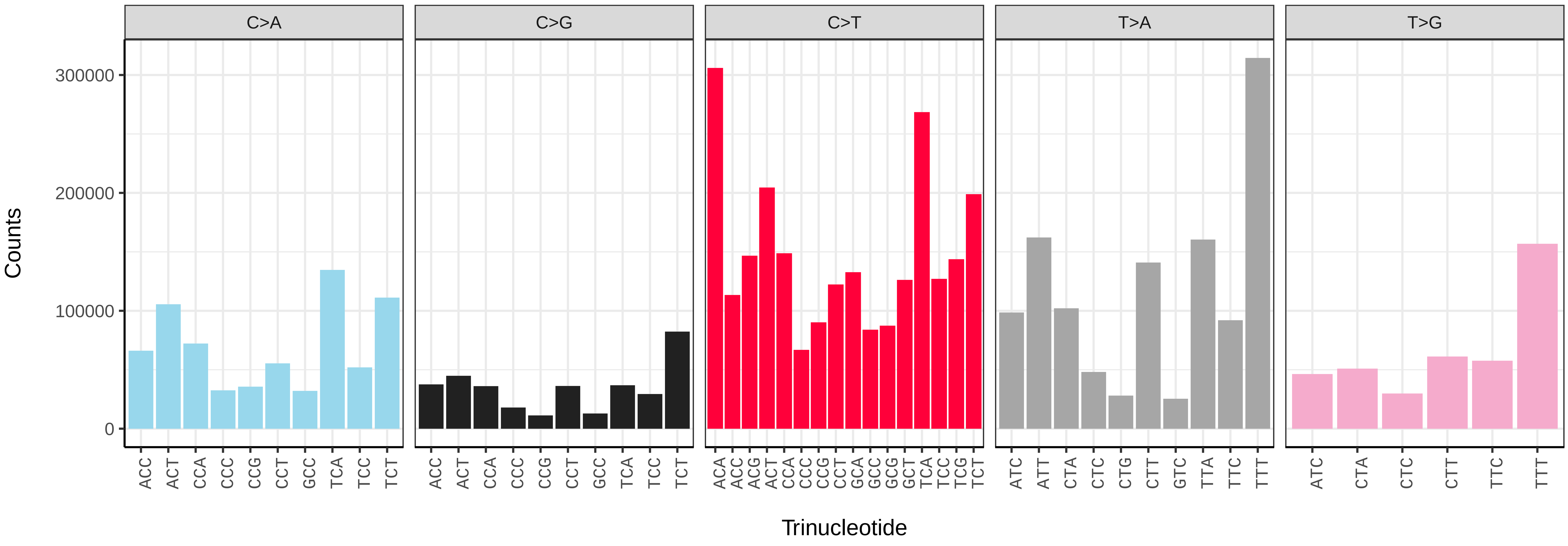


*Phragmatobia fuliginosa* (ilPhrFuli1) somatic mutational spectrum (SBS96 classification)

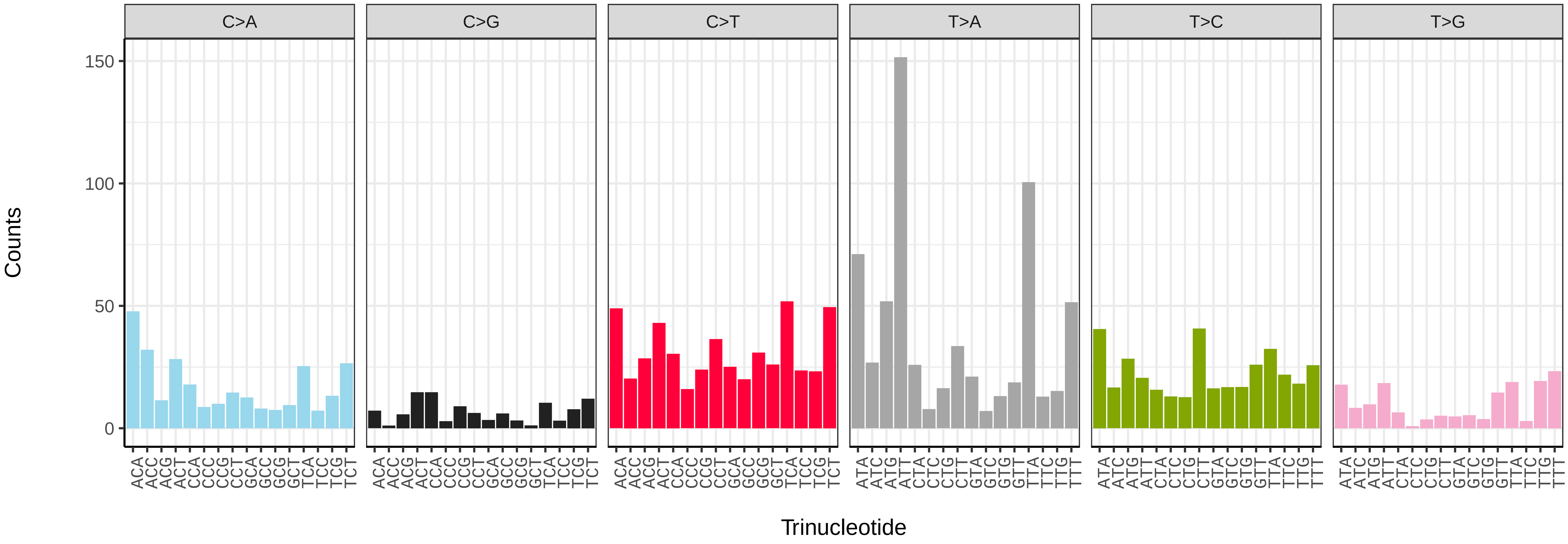


*Phragmatobia fuliginosa* (ilPhrFuli1) somatic mutational spectrum (SBS52 classification)

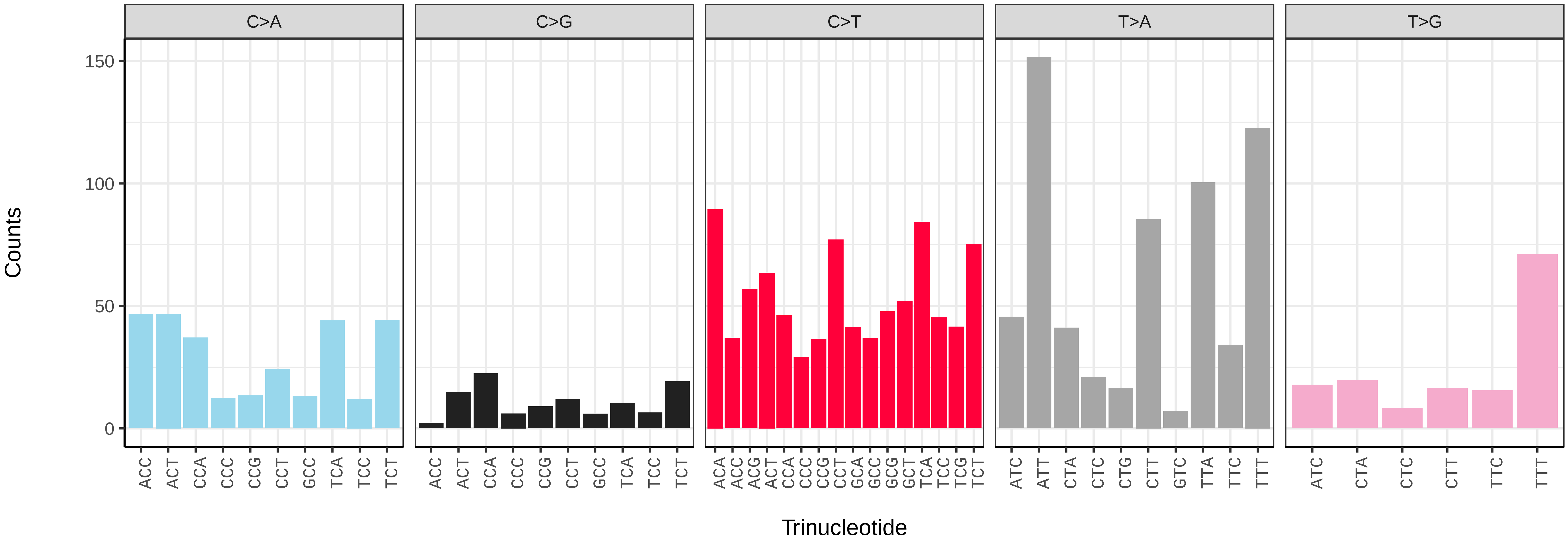

*Pieris brassicae* (ilPieBrab1) germline mutational spectrum

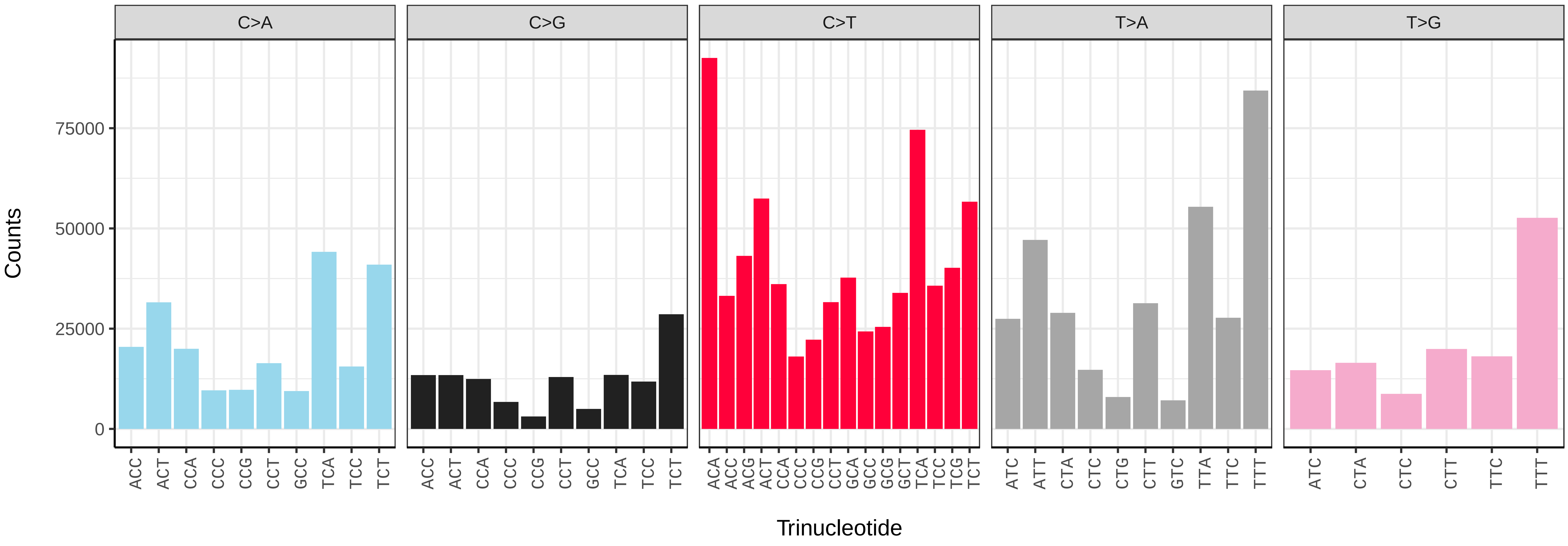


*Pieris brassicae* (ilPieBrab1) somatic mutational spectrum (SBS96 classification)

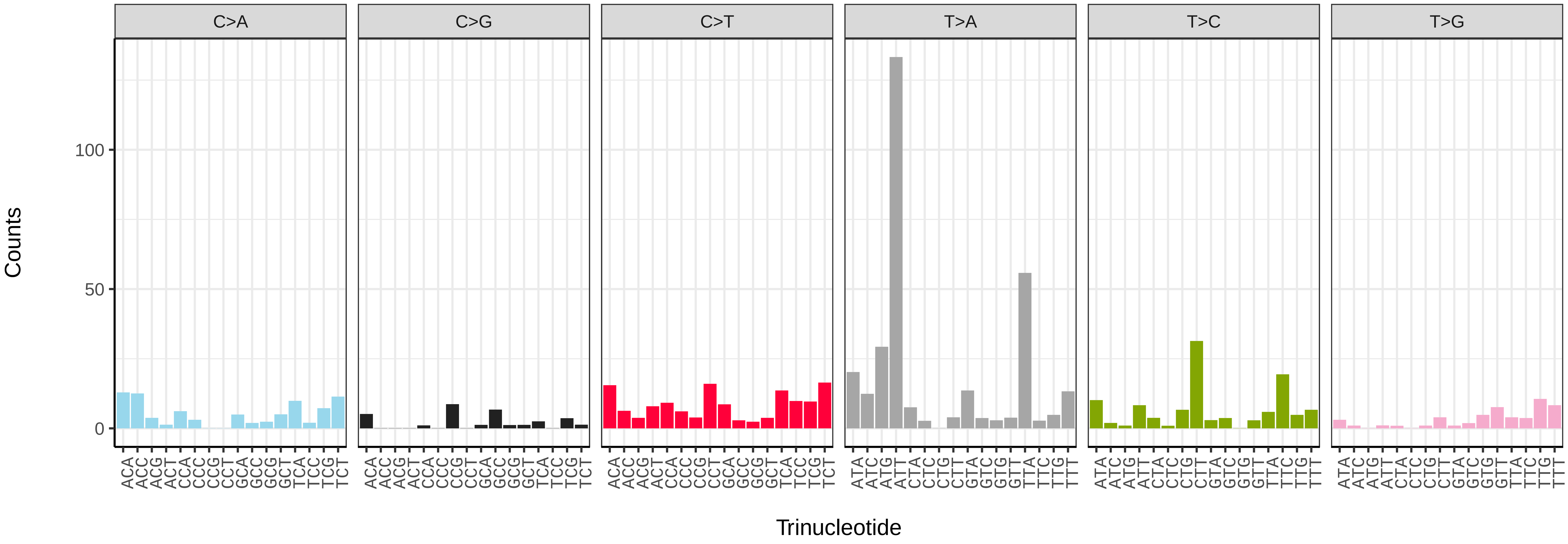


*Pieris brassicae* (ilPieBrab1) somatic mutational spectrum (SBS52 classification)

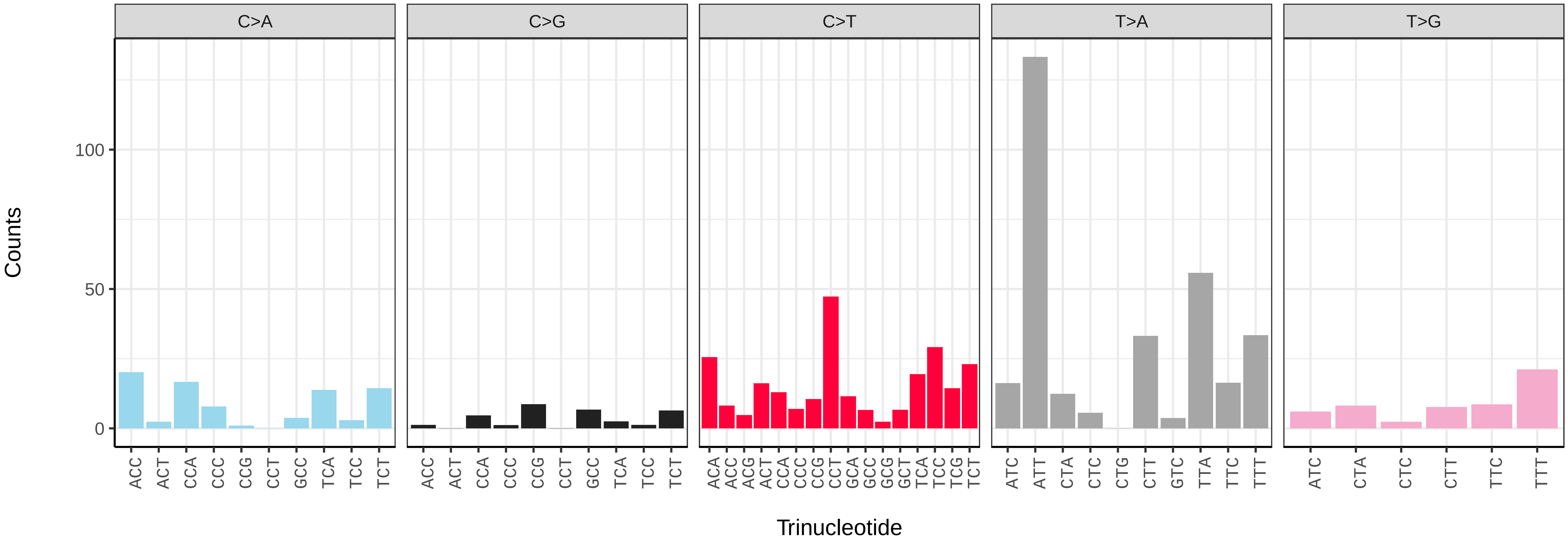

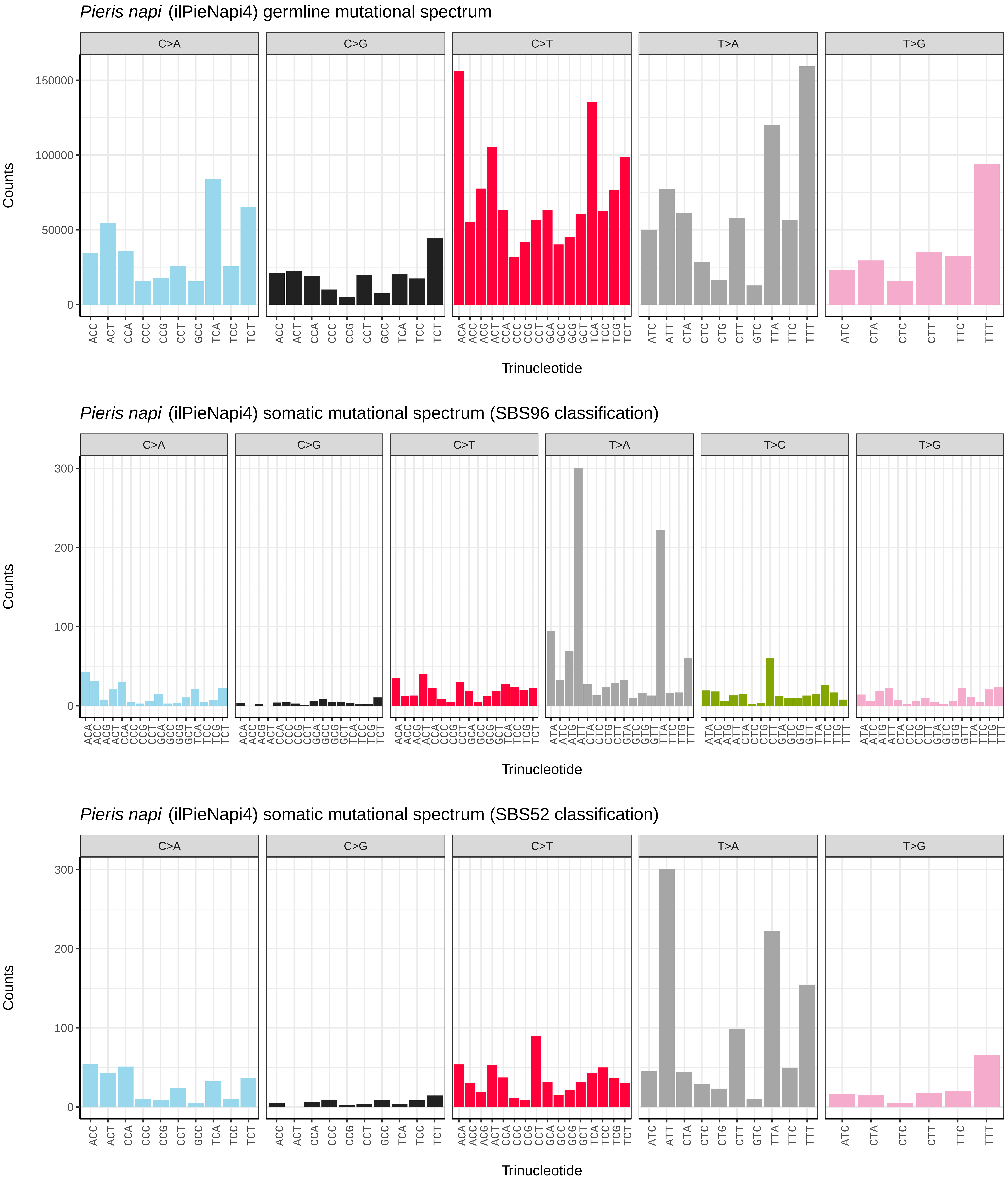
Pieris napi (ilPieNapi4) germline mutational spectrum
C>A
C>G
C>T
T>A
T>G
Counts
150000
100000
50000
0
Trinucleotide
Pieris napi (ilPieNapi4) somatic mutational spectrum (SBS96 classification)
C>A
C>G
C>T
T>A
T>C
T>G
Counts
300
200
100
0
Trinucleotide
Pieris napi (ilPieNapi4) somatic mutational spectrum (SBS52 classification)
C>A
C>G
C>T
T>A
T>G
Counts
300
200
100
0
Trinucleotide

*Pieris napi* (ilPieNapi4.ilPieNapi1) germline mutational spectrum

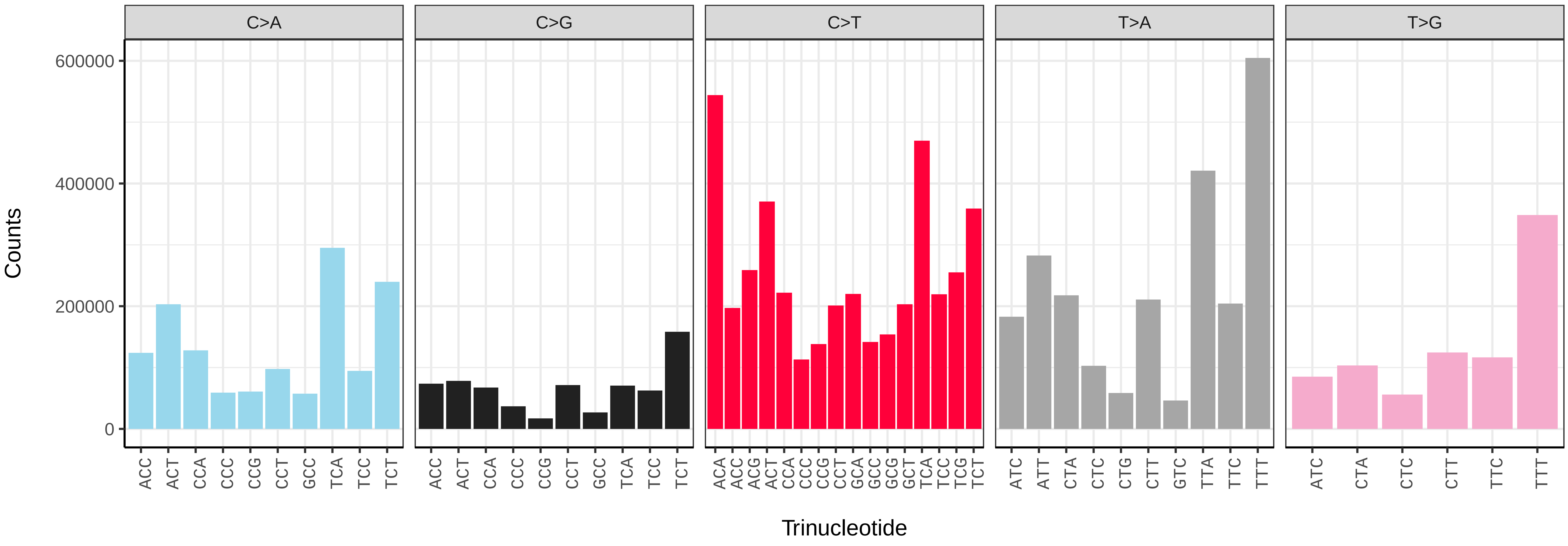


*Pieris napi* (ilPieNapi4.ilPieNapi1) somatic mutational spectrum (SBS96 classification)

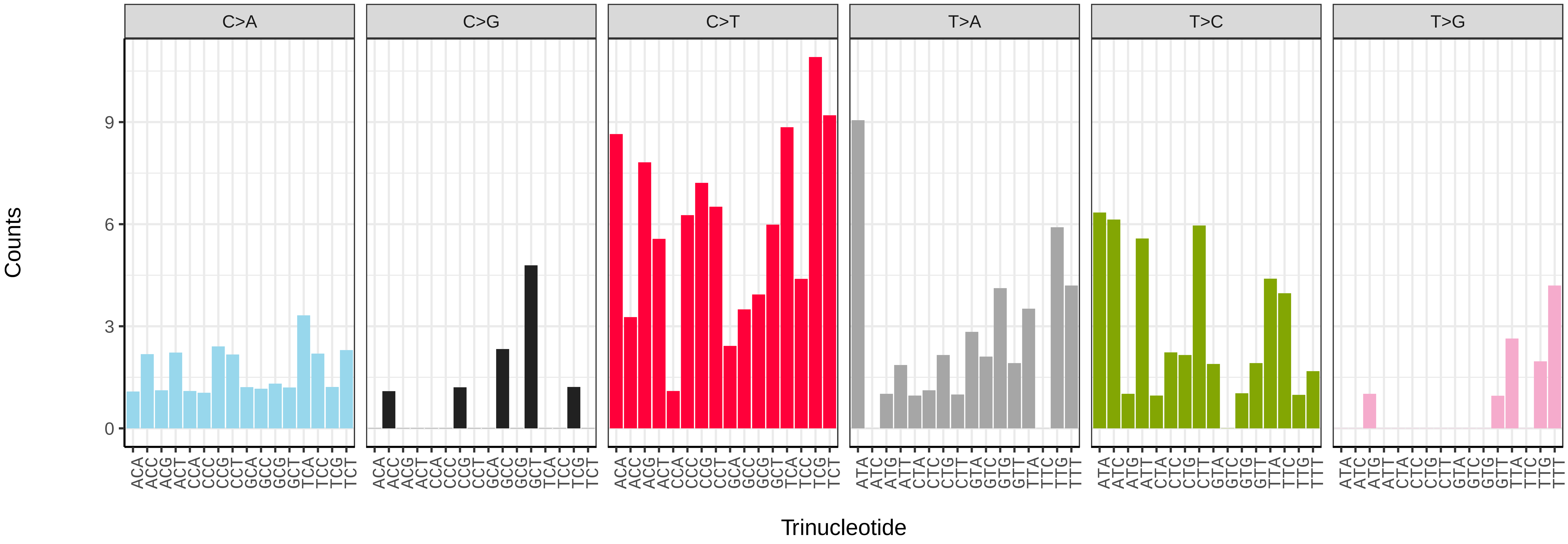


*Pieris napi* (ilPieNapi4.ilPieNapi1) somatic mutational spectrum (SBS52 classification)

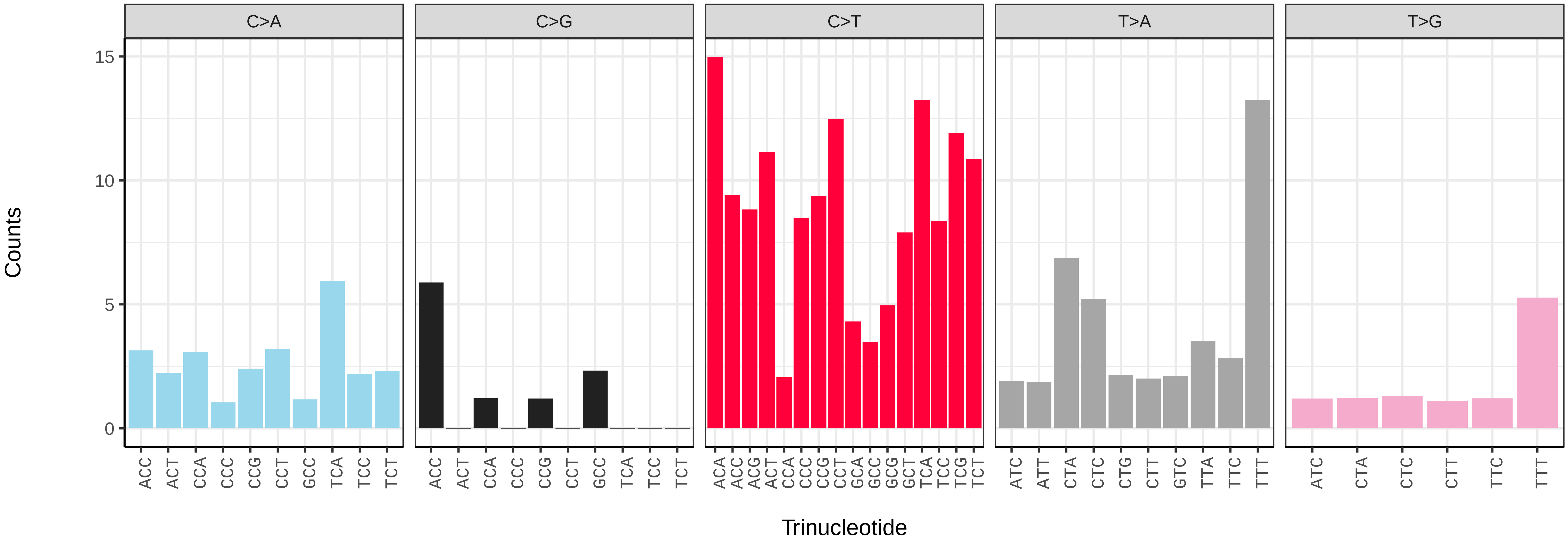

*Pieris rapae* (ilPieRapa1) germline mutational spectrum

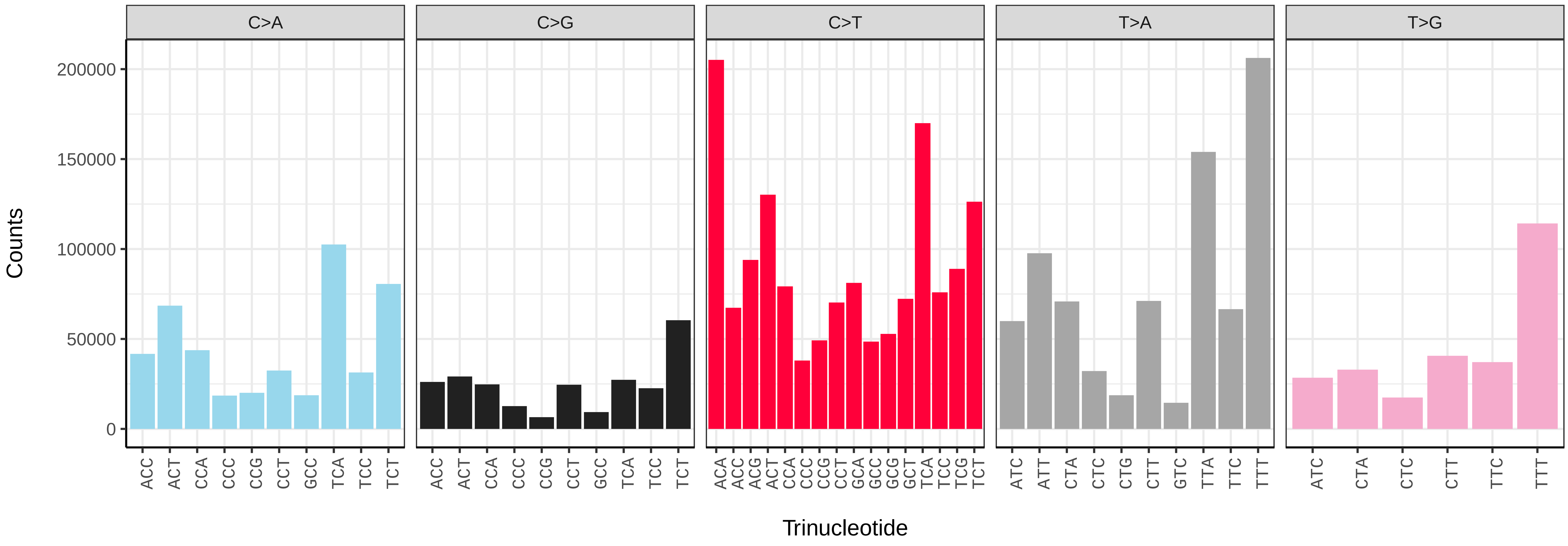


*Pieris rapae* (ilPieRapa1) somatic mutational spectrum (SBS96 classification)

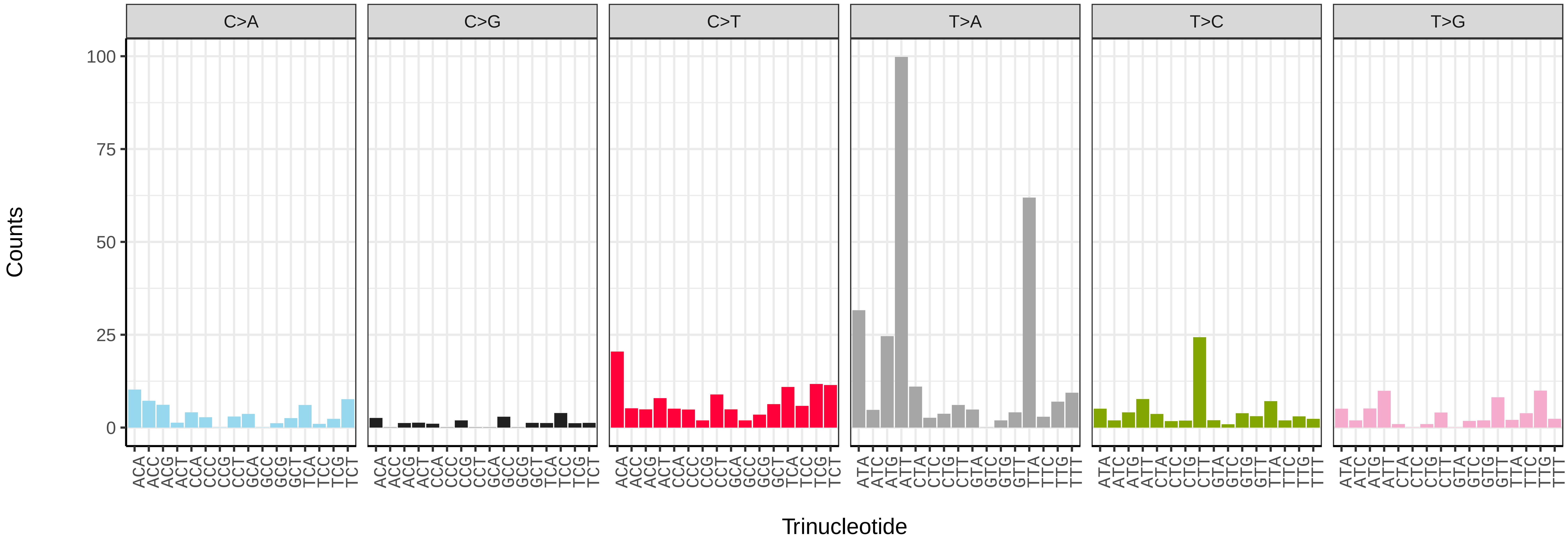


*Pieris rapae* (ilPieRapa1) somatic mutational spectrum (SBS52 classification)

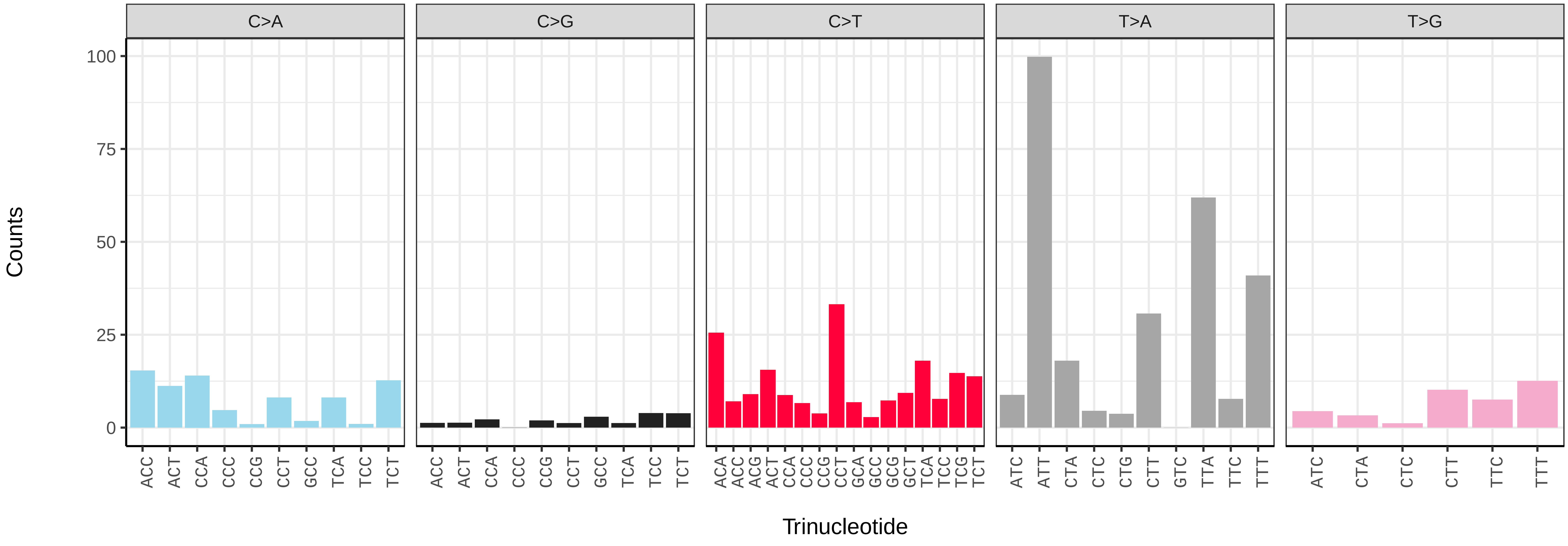

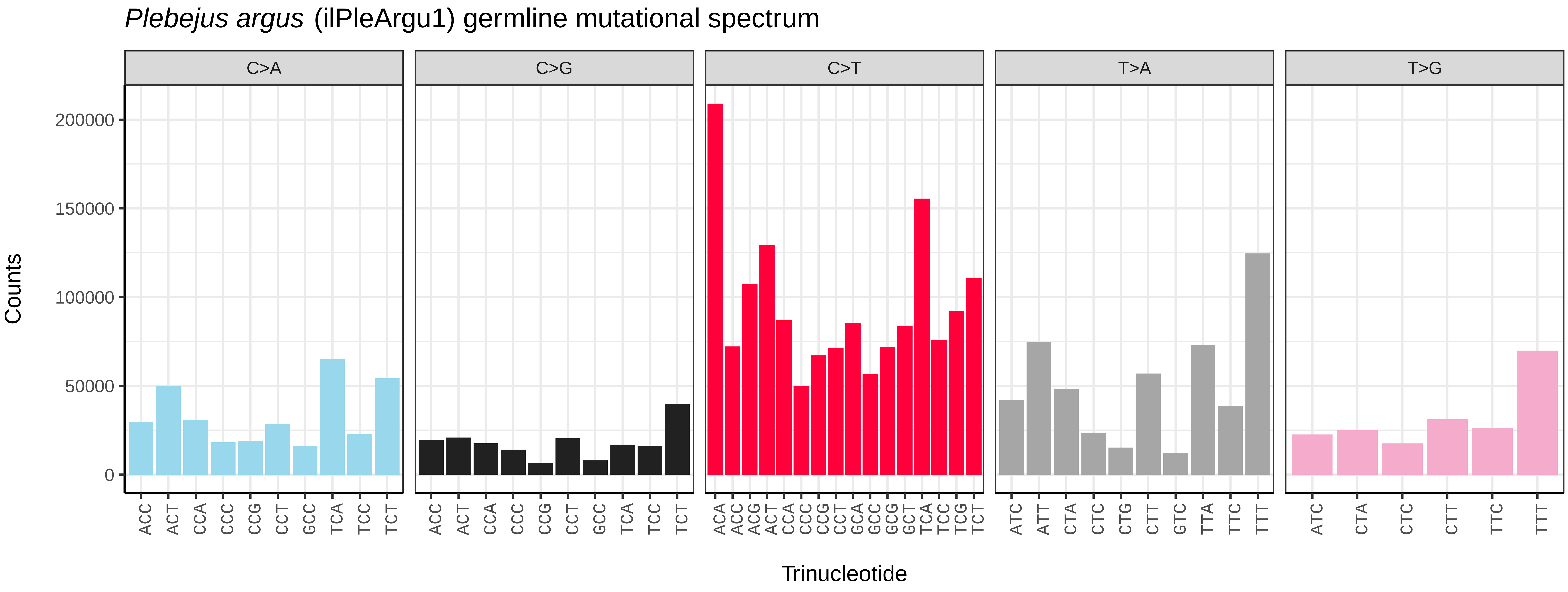
*Plebejus argus* (ilPleArgu1) germline mutational spectrum
C>A
C>G
C>T
T>A
T>G
Counts
0
50000
100000
150000
200000
ACC ACT CCA CCC CCG CCT GCC TCA TCC TCT
ACC ACT CCA CCC CCG CCT GCC TCA TCC TCT
ACA ACC ACG ACT CCA CCC CCG CCT GCA GCC GCG GCT TCA TCC TCG TCT
ATC ATT CTA CTC CTG CTT GTC TTA TTC TTT
ATC CTA CTC CTT TTC TTT
Trinucleotide

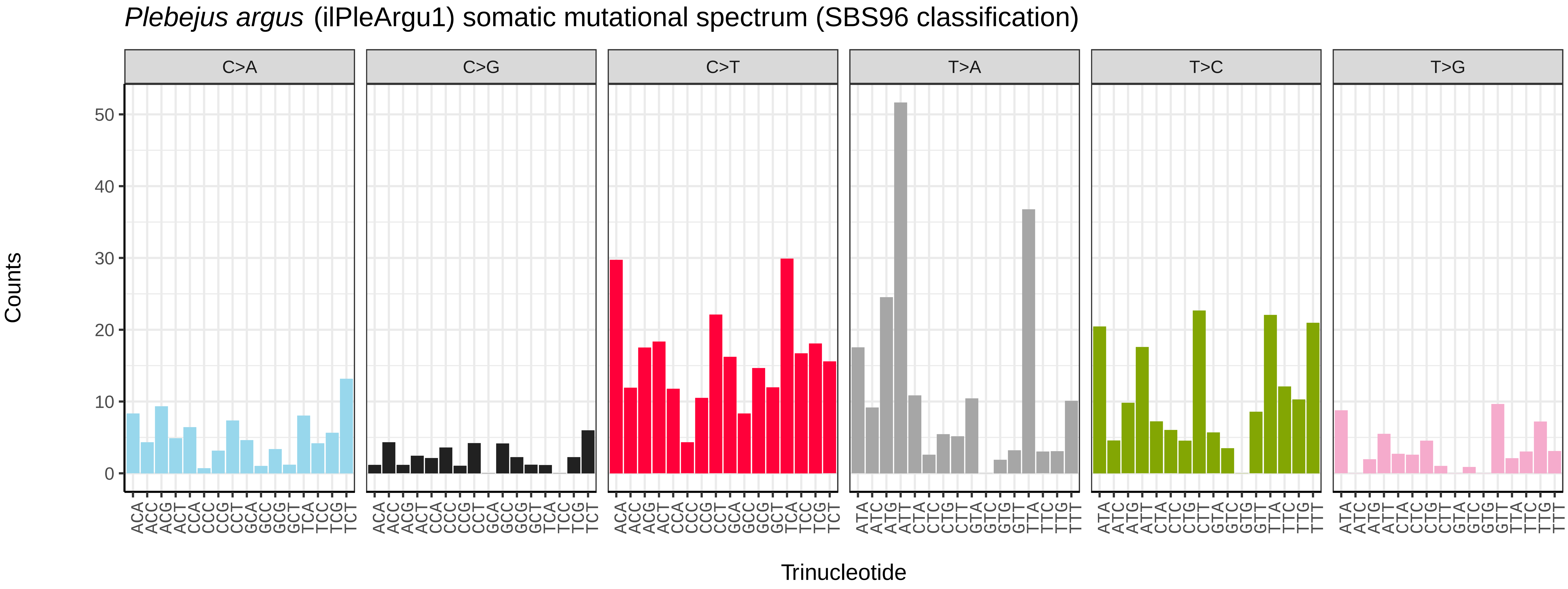
*Plebejus argus* (ilPleArgu1) somatic mutational spectrum (SBS96 classification)
C>A
C>G
C>T
T>A
T>C
T>G
Counts
0
10
20
30
40
50
ACA ACC ACG ACT CCA CCC CCG CCT GCA GCC GCG GCT TCA TCC TCG TCT
ATA ATC ATG ATT CTA CTC CTG CTT GTA GTC GTG GTT TTA TTC TTG TTT
Trinucleotide

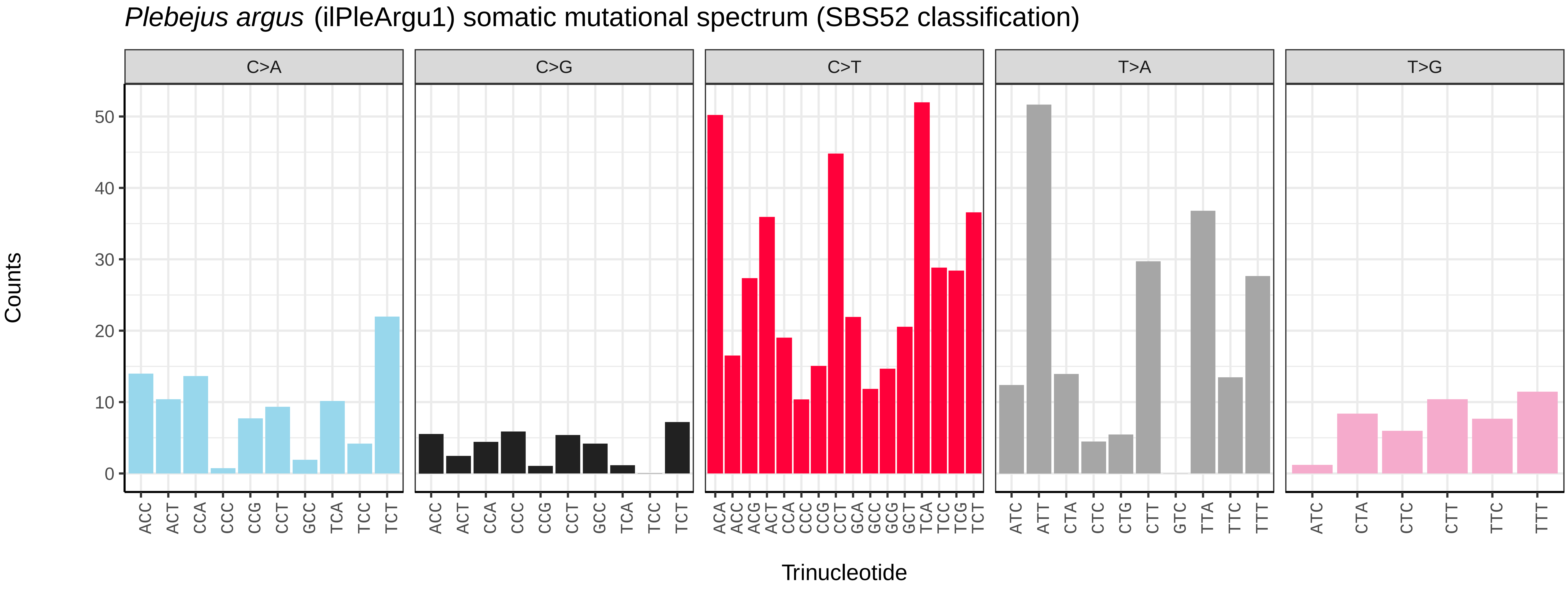
*Plebejus argus* (ilPleArgu1) somatic mutational spectrum (SBS52 classification)
C>A
C>G
C>T
T>A
T>G
Counts
0
10
20
30
40
50
ACC ACT CCA CCC CCG CCT GCC TCA TCC TCT
ACC ACT CCA CCC CCG CCT GCC TCA TCC TCT
ACA ACC ACG ACT CCA CCC CCG CCT GCA GCC GCG GCT TCA TCC TCG TCT
ATC ATT CTA CTC CTG CTT GTC TTA TTC TTT
ATC CTA CTC CTT TTC TTT
Trinucleotide

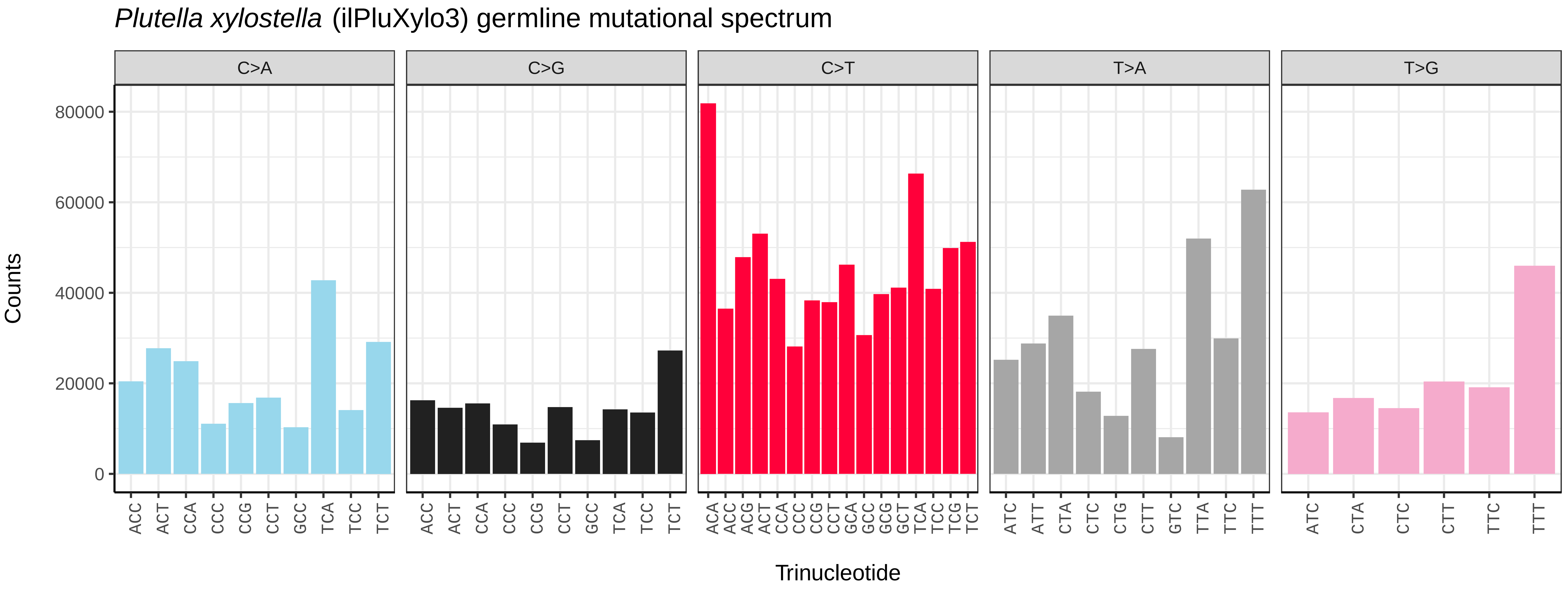
Plutella xylostella (ilPluXylo3) germline mutational spectrum
C>A
C>G
C>T
T>A
T>G
Counts
0
20000
40000
60000
80000
ACC ACT CCA CCC CCG CCT GCC TCA TCC TCT
ACA ACC ACG ACT CCA CCC CCG CCT GCA GCC GCG GCT TCA TCC TCG TCT
ATC ATT CTA CTC CTG CTT GTC TTA TTC TTT
ATC CTA CTC CTT TTC TTT
Trinucleotide

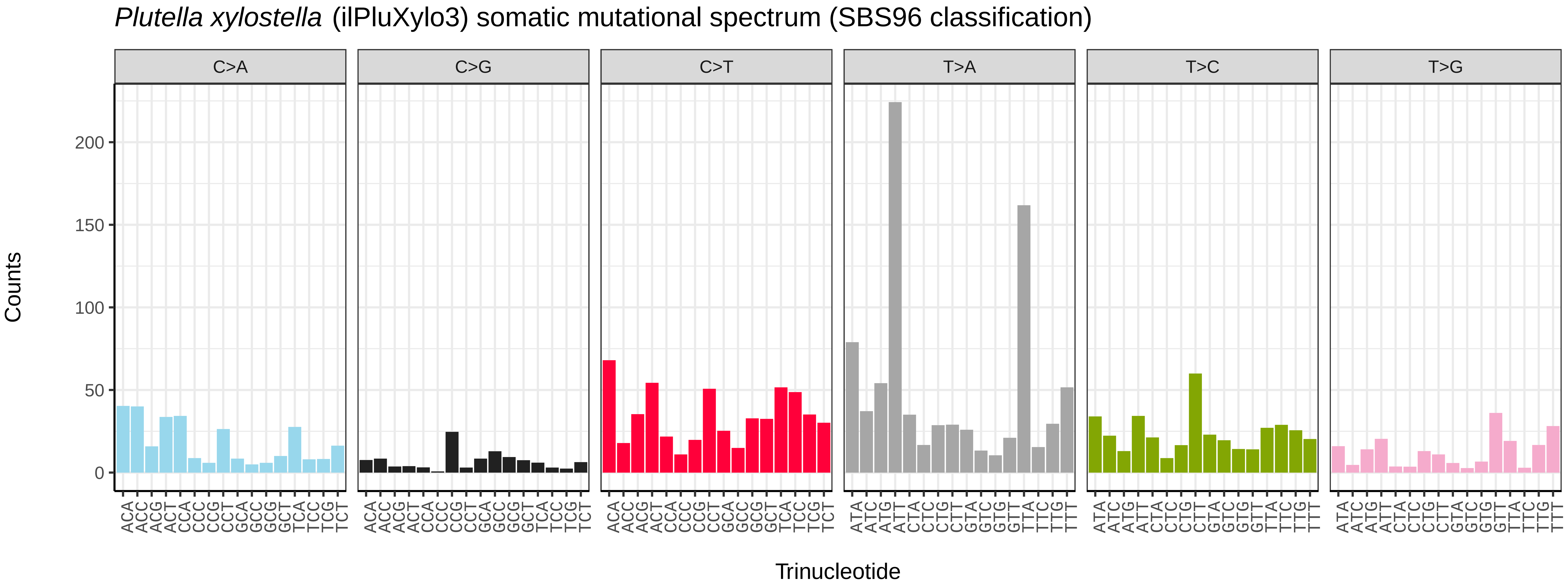
Plutella xylostella (ilPluXylo3) somatic mutational spectrum (SBS96 classification)
C>A
C>G
C>T
T>A
T>C
T>G
Counts
0
50
100
150
200
Trinucleotide

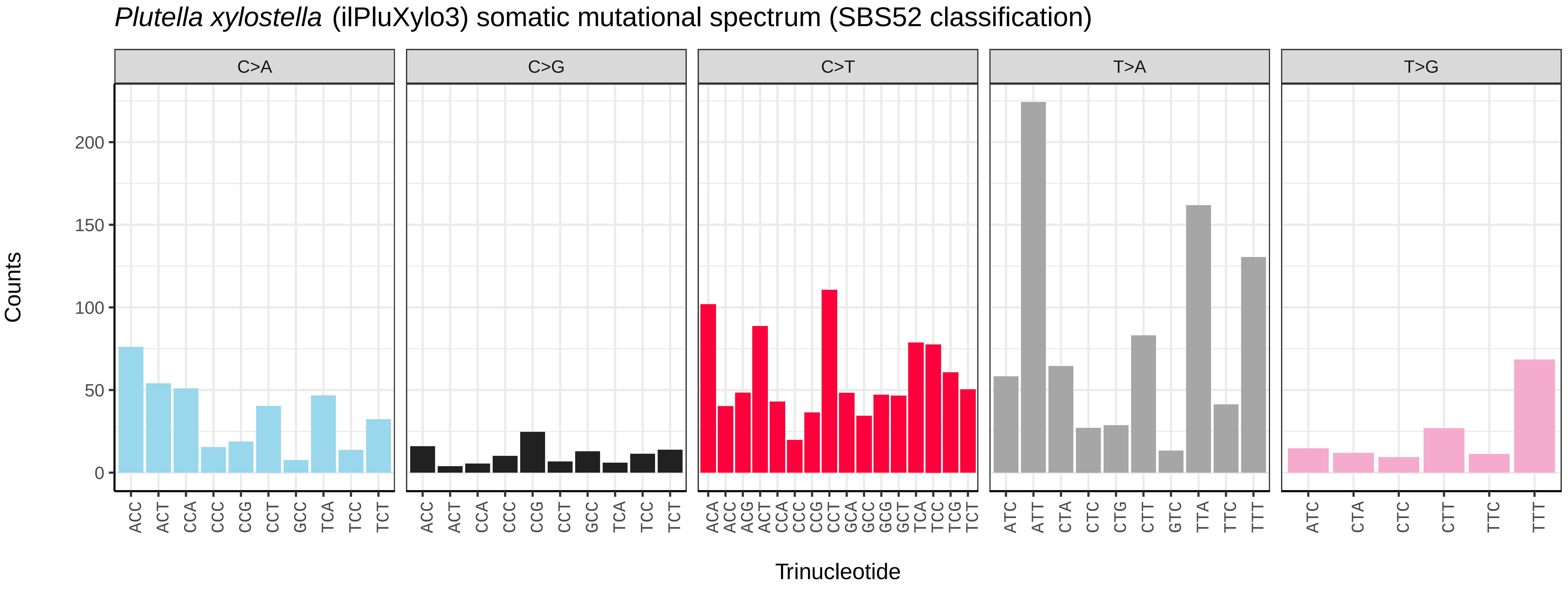
Plutella xylostella (ilPluXylo3) somatic mutational spectrum (SBS52 classification)
C>A
C>G
C>T
T>A
T>G
Counts
0
50
100
150
200
ACC ACT CCA CCC CCG CCT GCC TCA TCC TCT
ACA ACC ACG ACT CCA CCC CCG CCT GCA GCC GCG GCT TCA TCC TCG TCT
ATC ATT CTA CTC CTG CTT GTC TTA TTC TTT
ATC CTA CTC CTT TTC TTT
Trinucleotide

*Polyommatus icarus* (ilPolIcar1) germline mutational spectrum

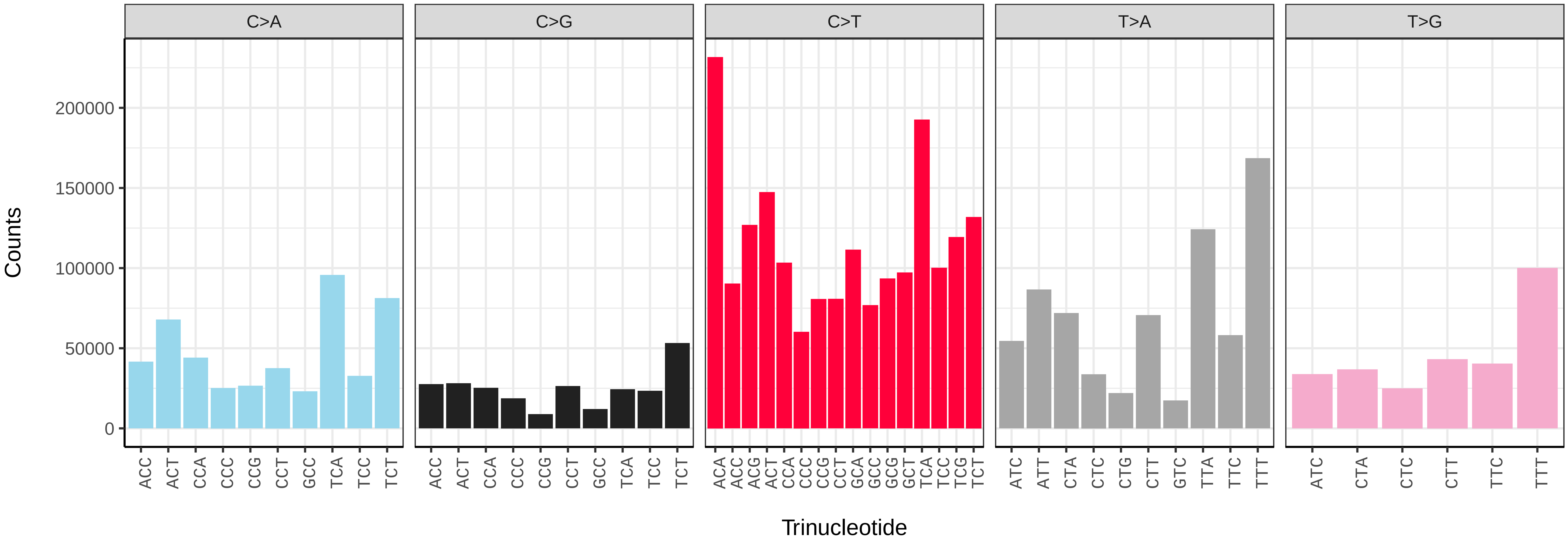


*Polyommatus icarus* (ilPolIcar1) somatic mutational spectrum (SBS96 classification)

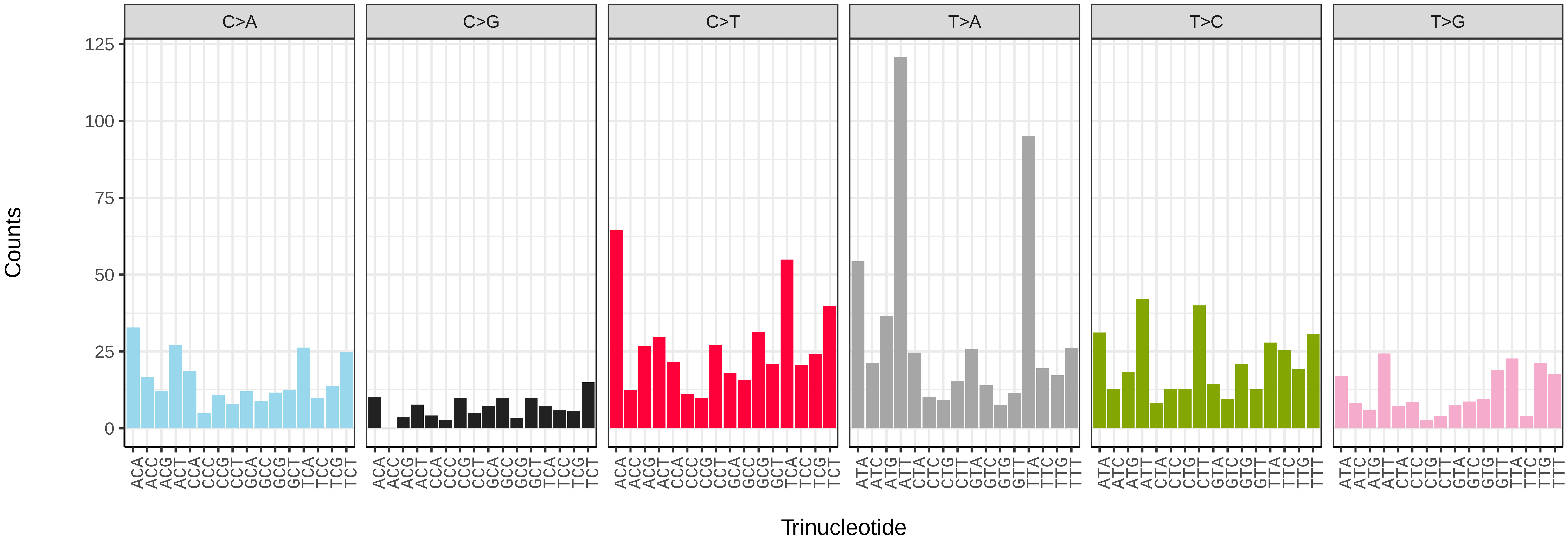


*Polyommatus icarus* (ilPolIcar1) somatic mutational spectrum (SBS52 classification)

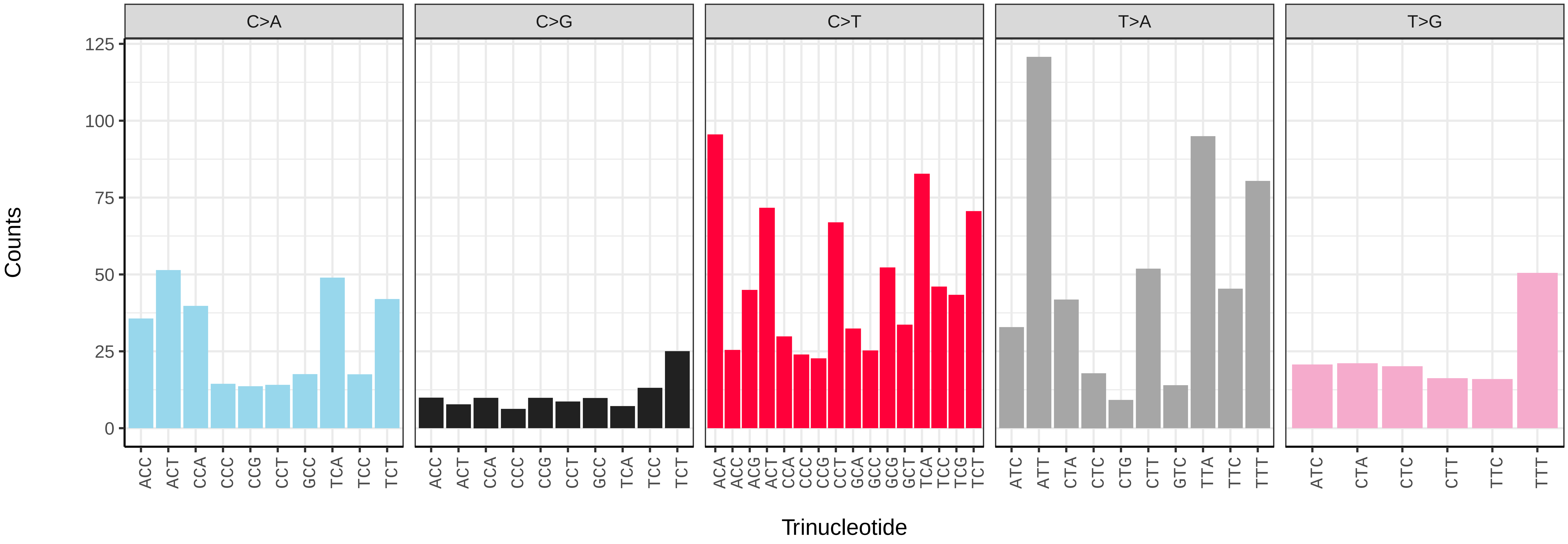

*Polymixis lichenea* (ilPolLich1) germline mutational spectrum

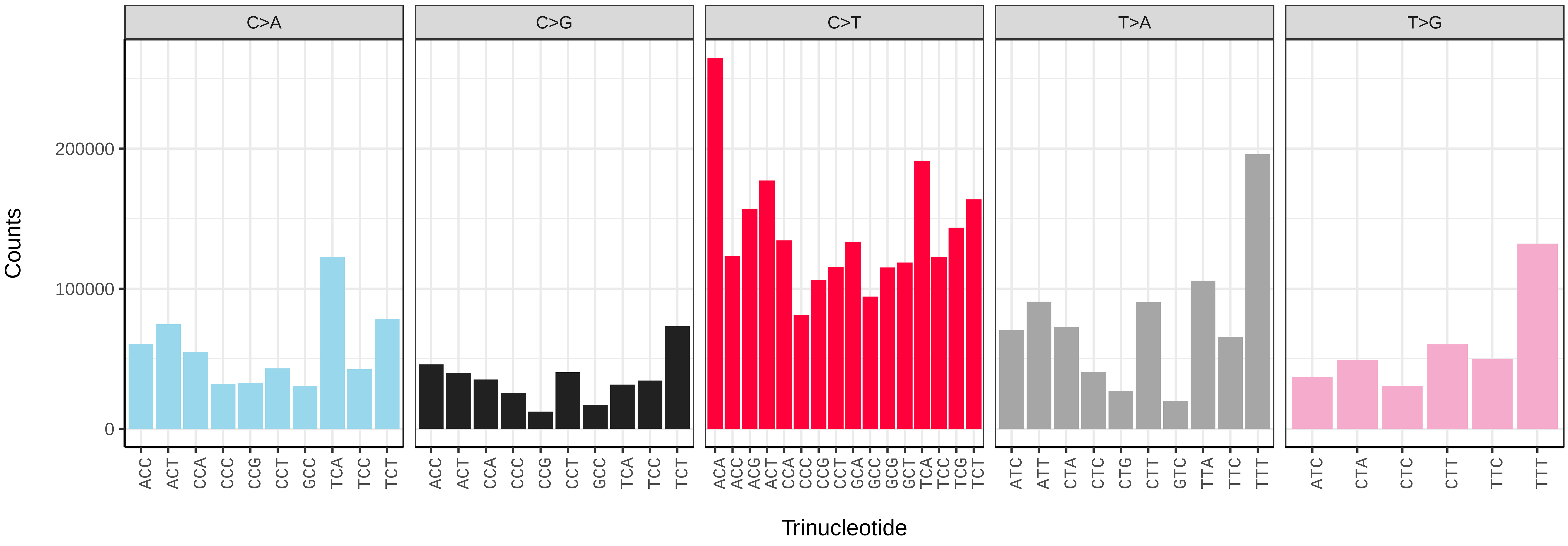


*Polymixis lichenea* (ilPolLich1) somatic mutational spectrum (SBS96 classification)

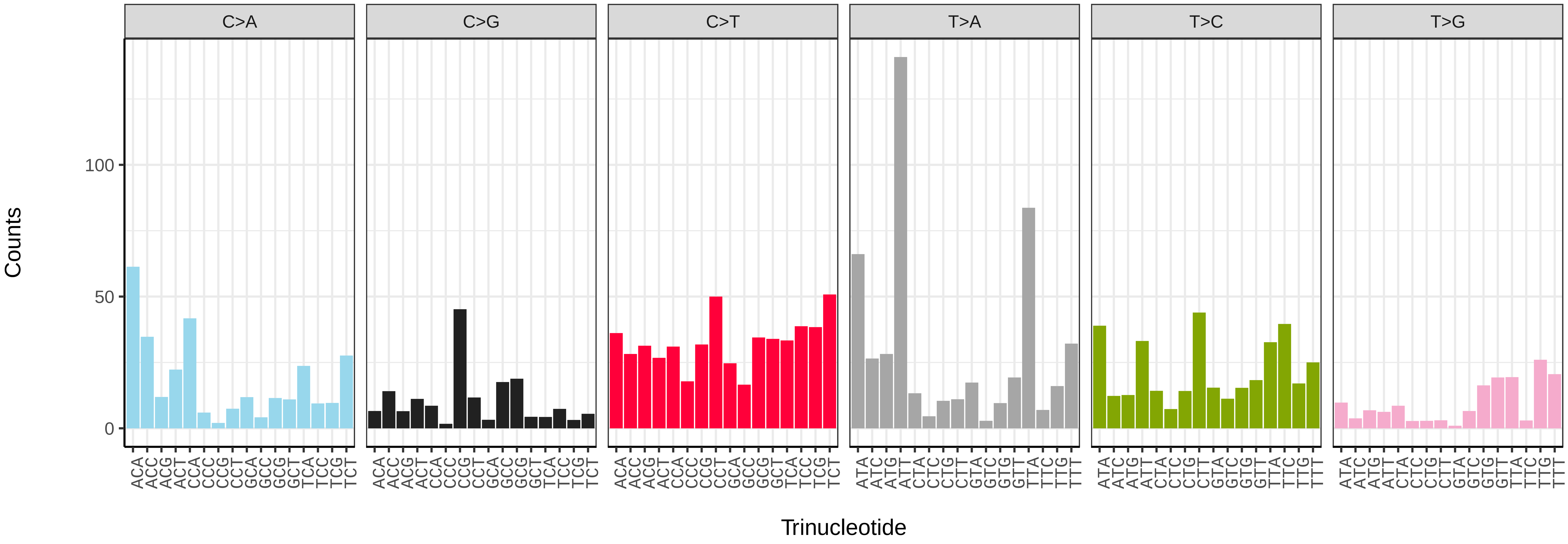


*Polymixis lichenea* (ilPolLich1) somatic mutational spectrum (SBS52 classification)

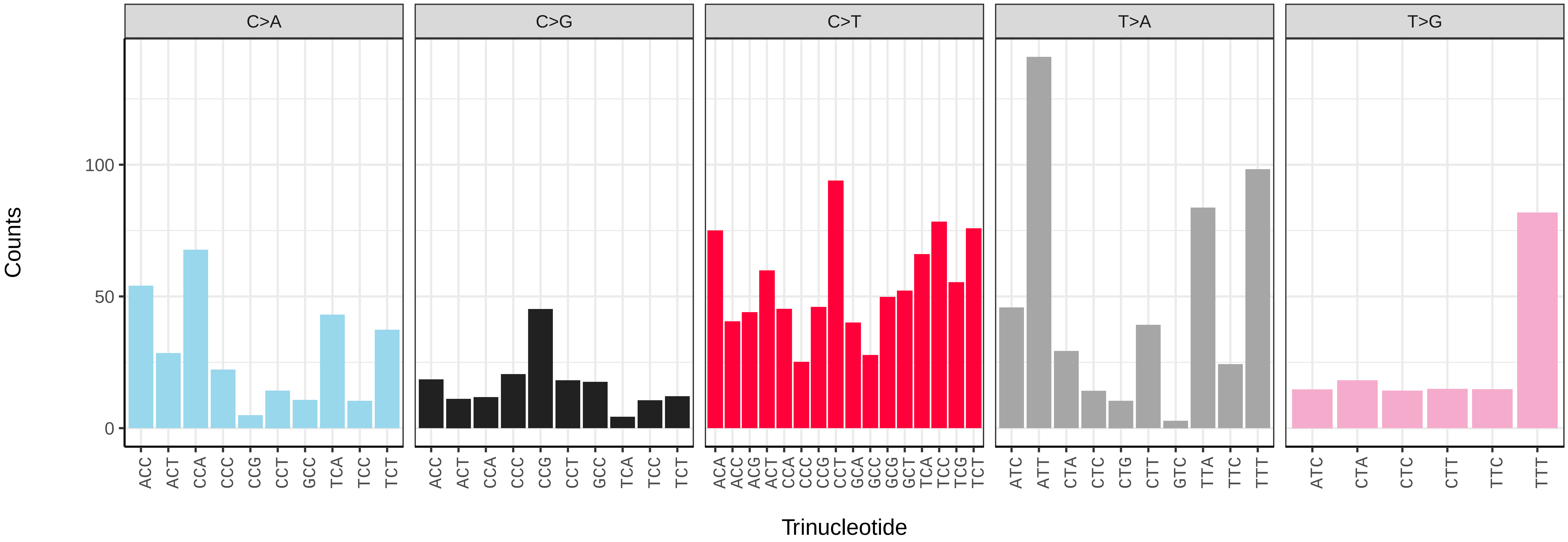

*Protodeltote pygarga* (ilProPyga1) germline mutational spectrum

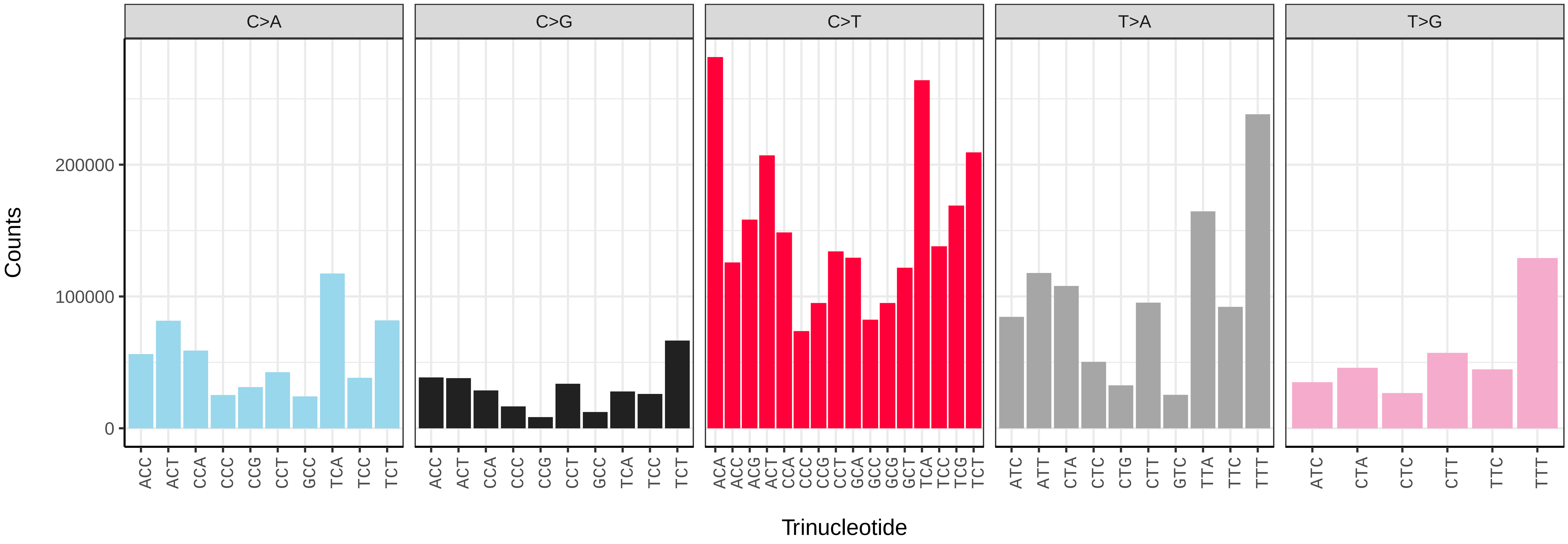


*Protodeltote pygarga* (ilProPyga1) somatic mutational spectrum (SBS96 classification)

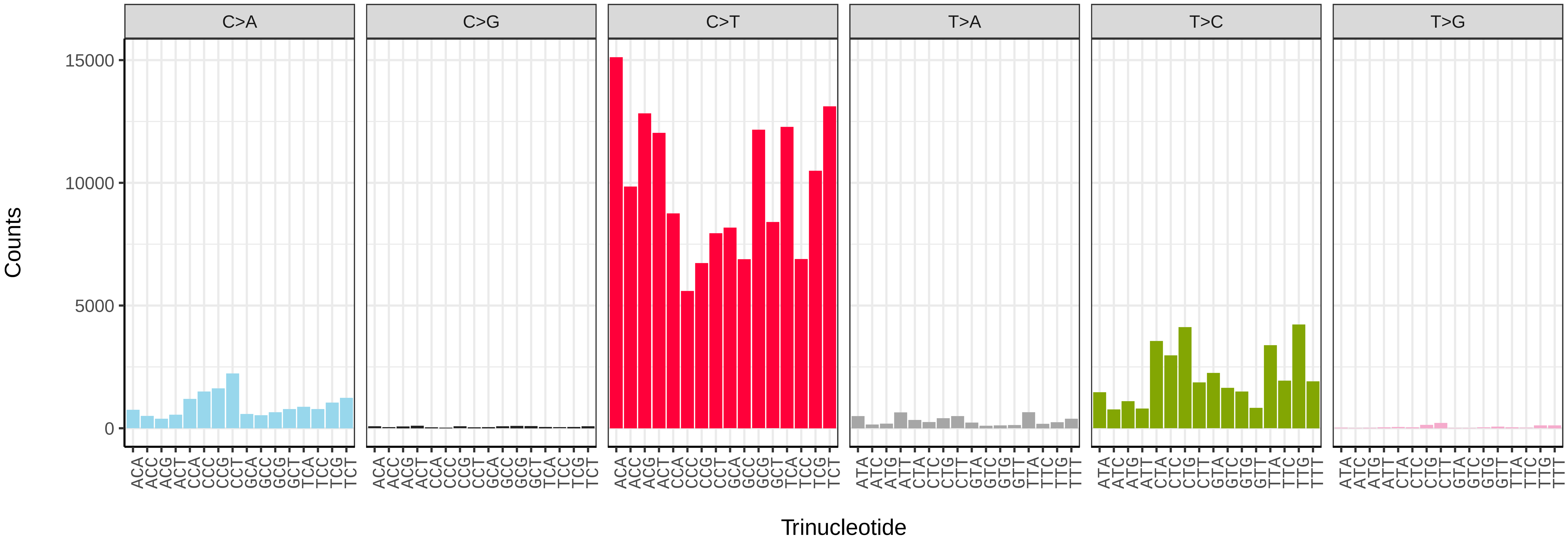


*Protodeltote pygarga* (ilProPyga1) somatic mutational spectrum (SBS52 classification)

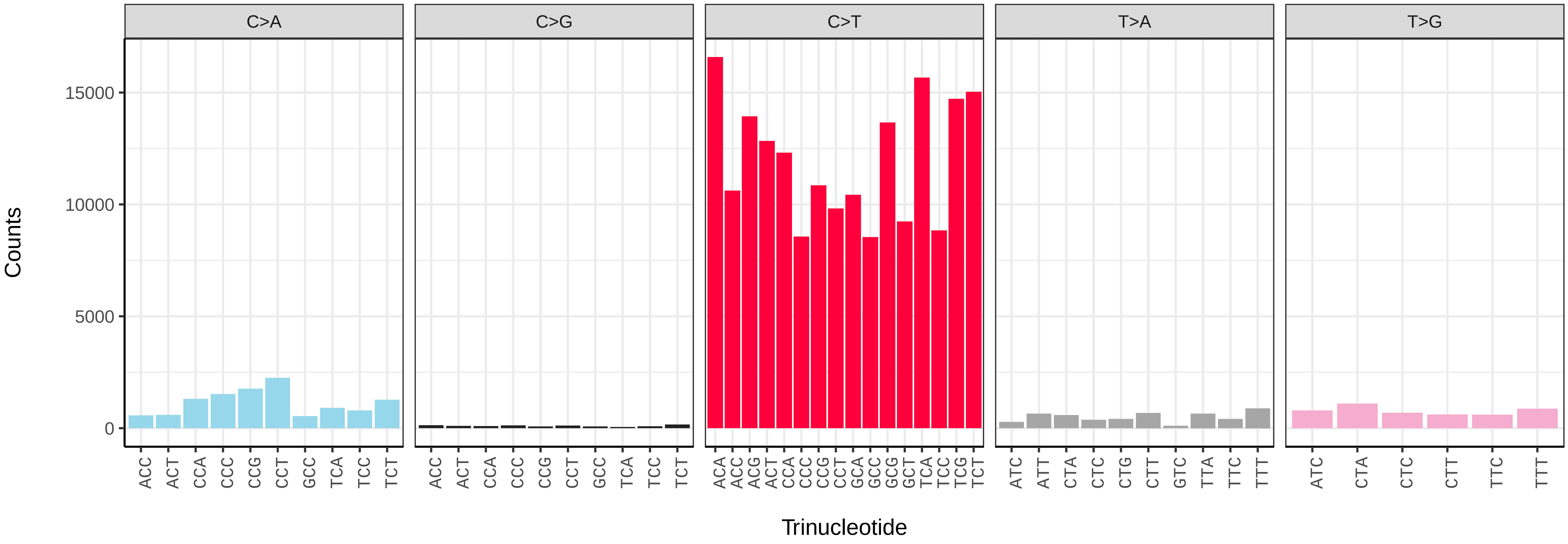

*Ptilodon capucinus* (ilPtiCapc1) germline mutational spectrum

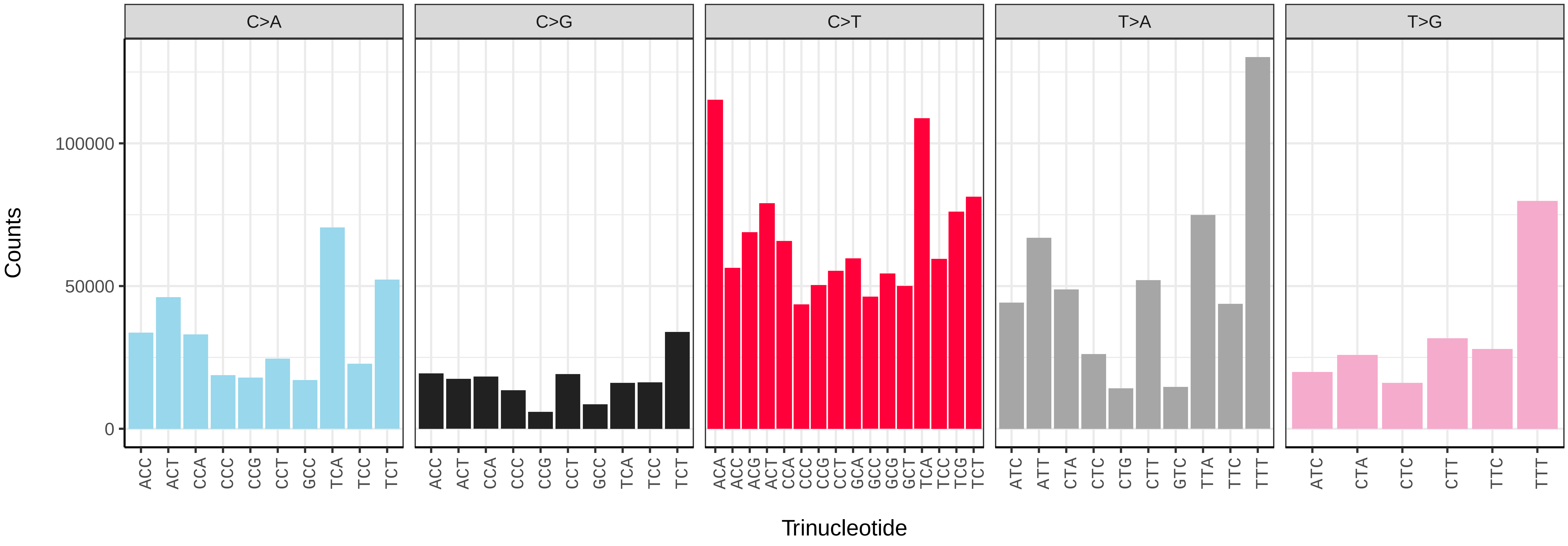


*Ptilodon capucinus* (ilPtiCapc1) somatic mutational spectrum (SBS96 classification)

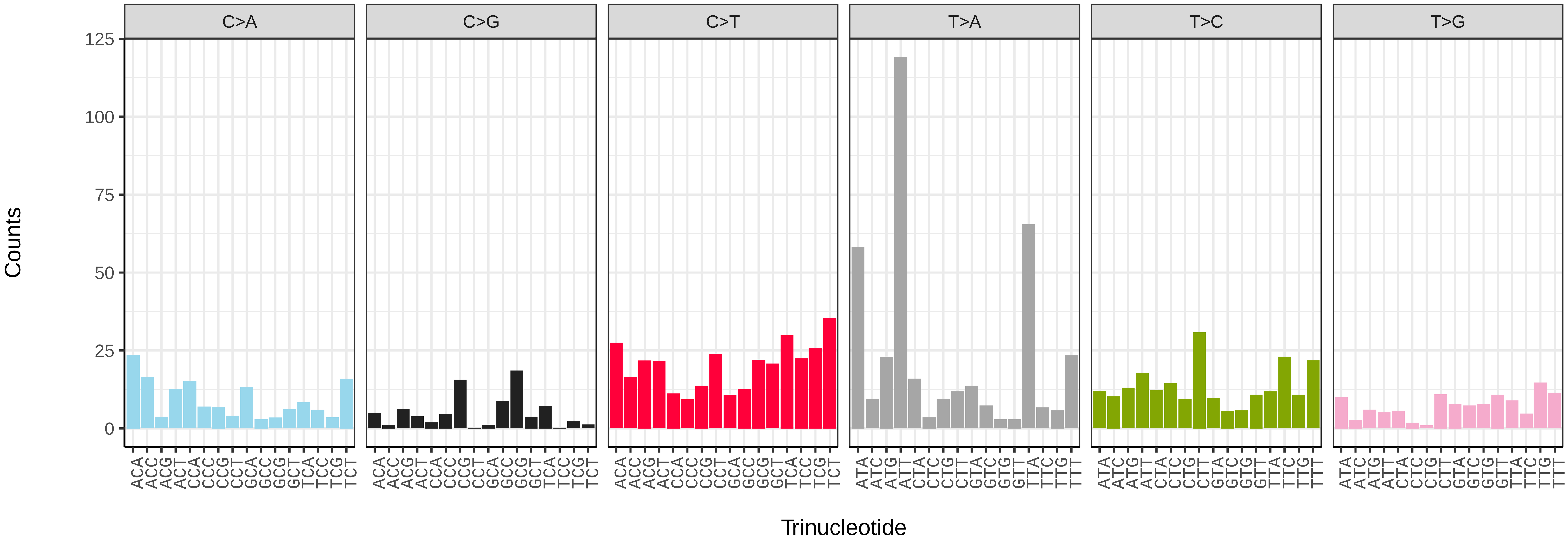


*Ptilodon capucinus* (ilPtiCapc1) somatic mutational spectrum (SBS52 classification)

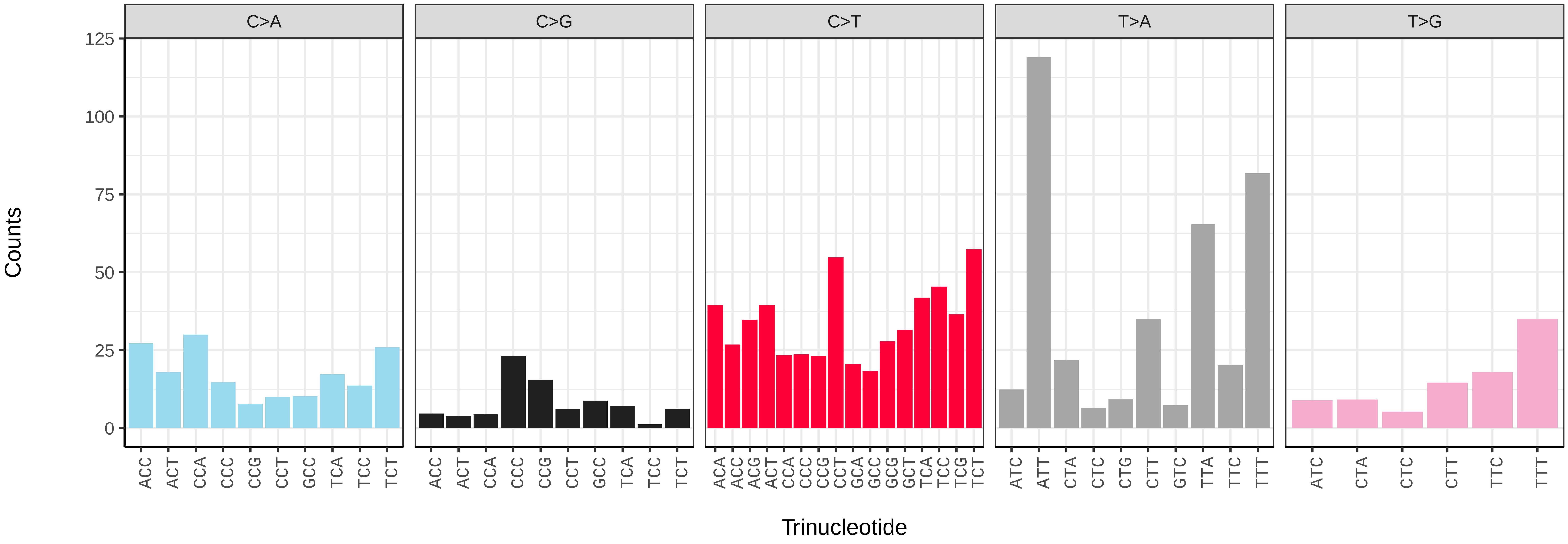

*Pyralis farinalis* (ilPyrFari1) germline mutational spectrum

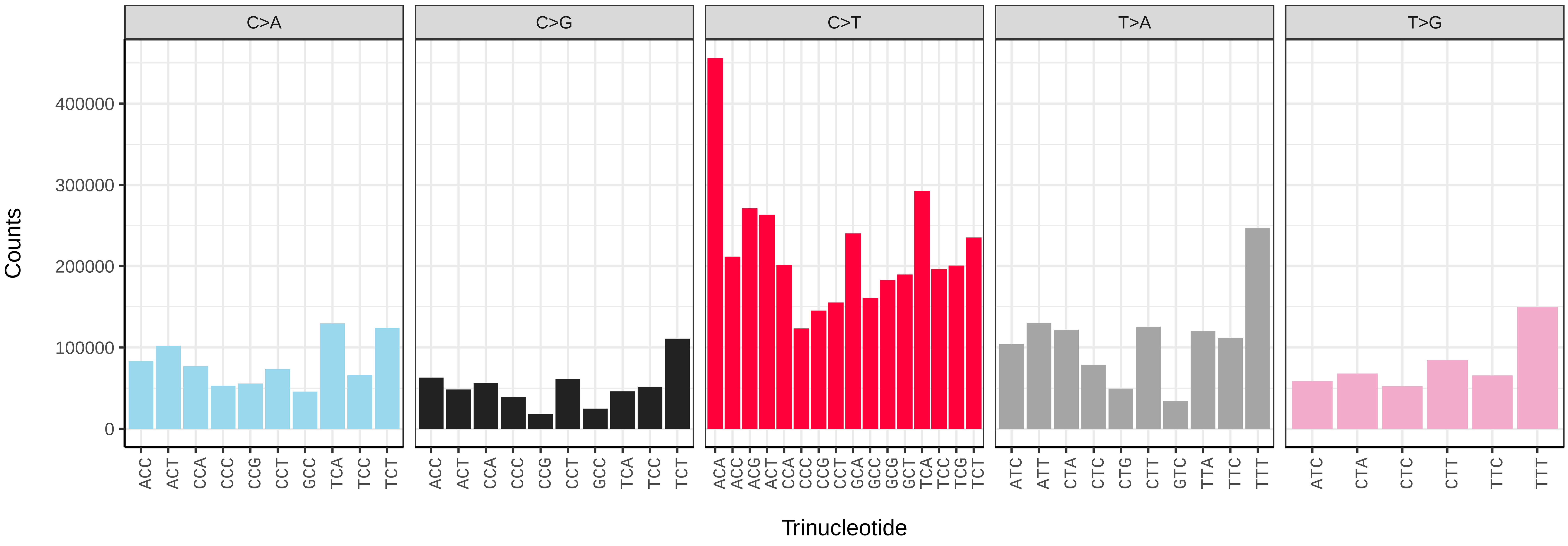


*Pyralis farinalis* (ilPyrFari1) somatic mutational spectrum (SBS96 classification)

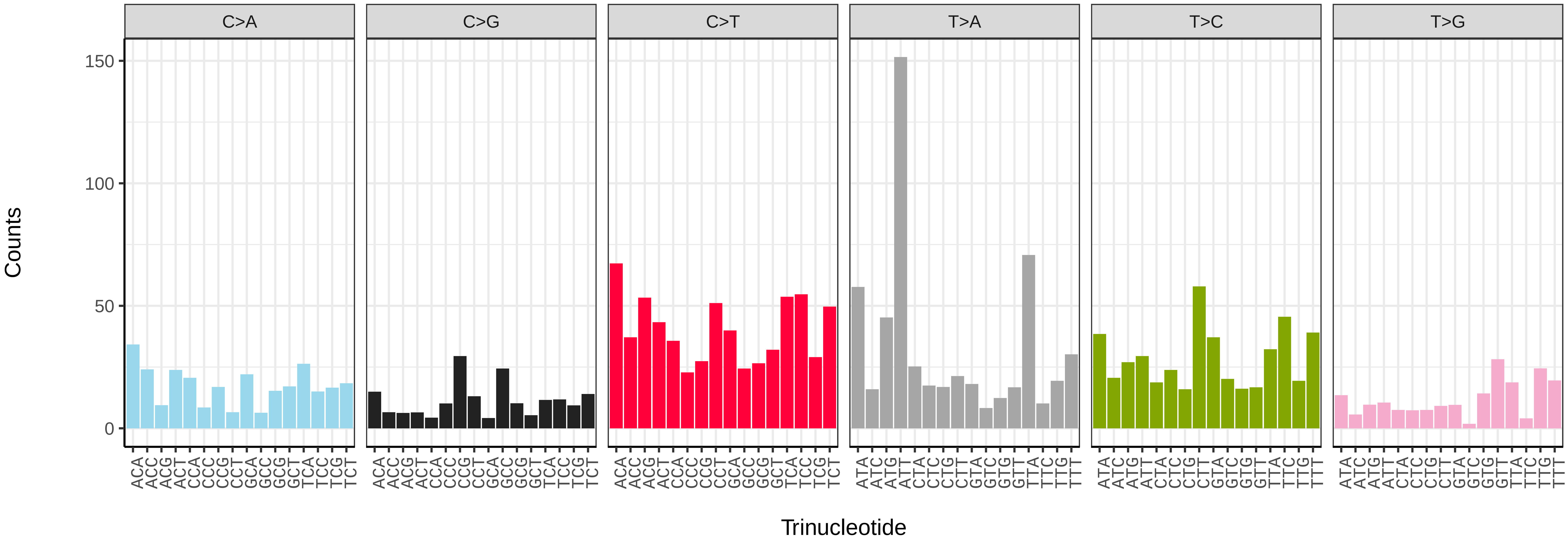


*Pyralis farinalis* (ilPyrFari1) somatic mutational spectrum (SBS52 classification)

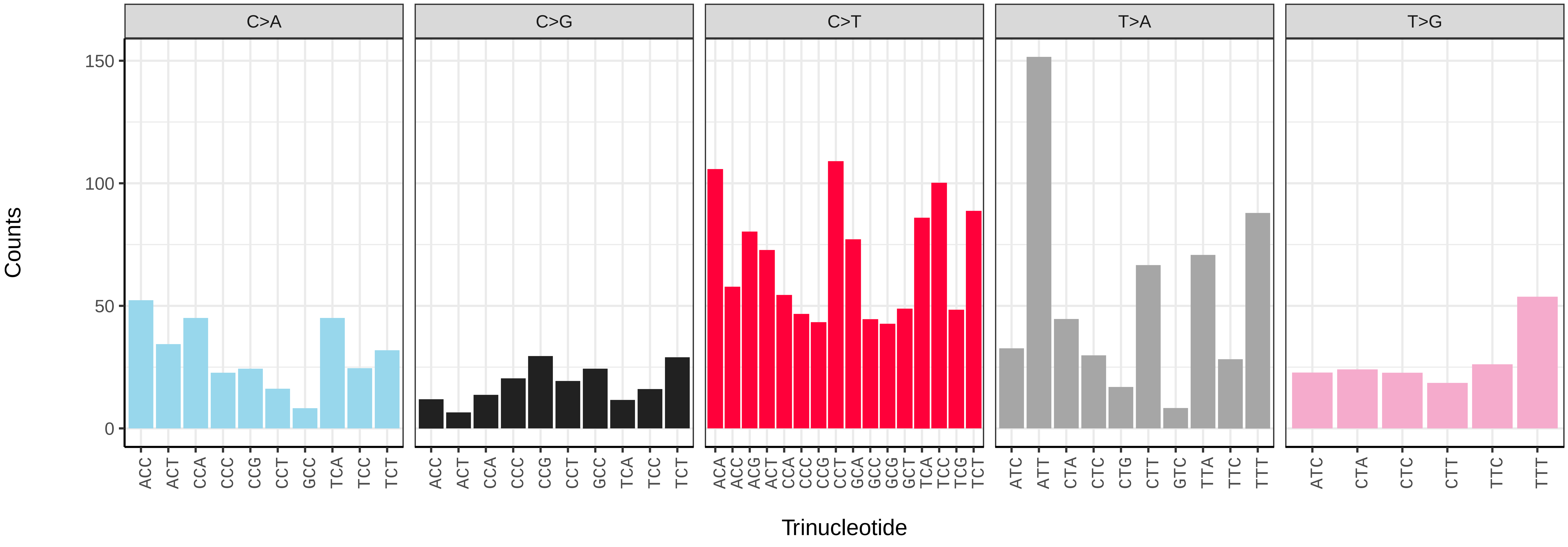

*Pyrgus malvae* (ilPyrMalv3) germline mutational spectrum

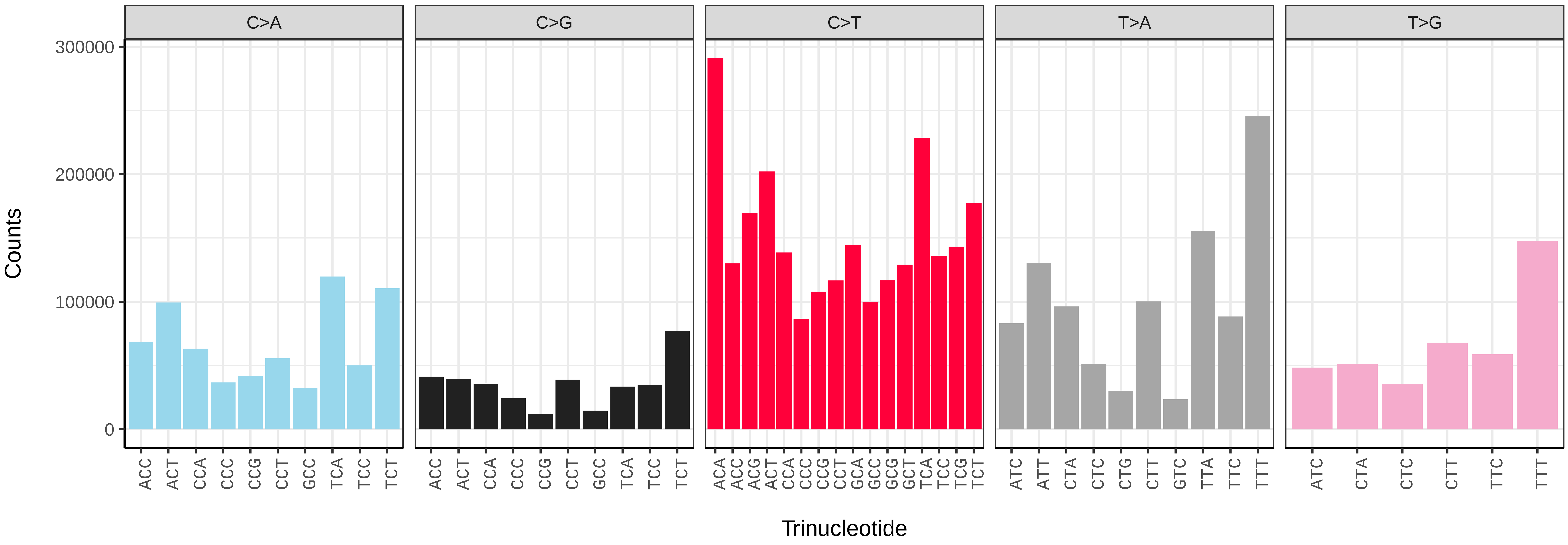


*Pyrgus malvae* (ilPyrMalv3) somatic mutational spectrum (SBS96 classification)

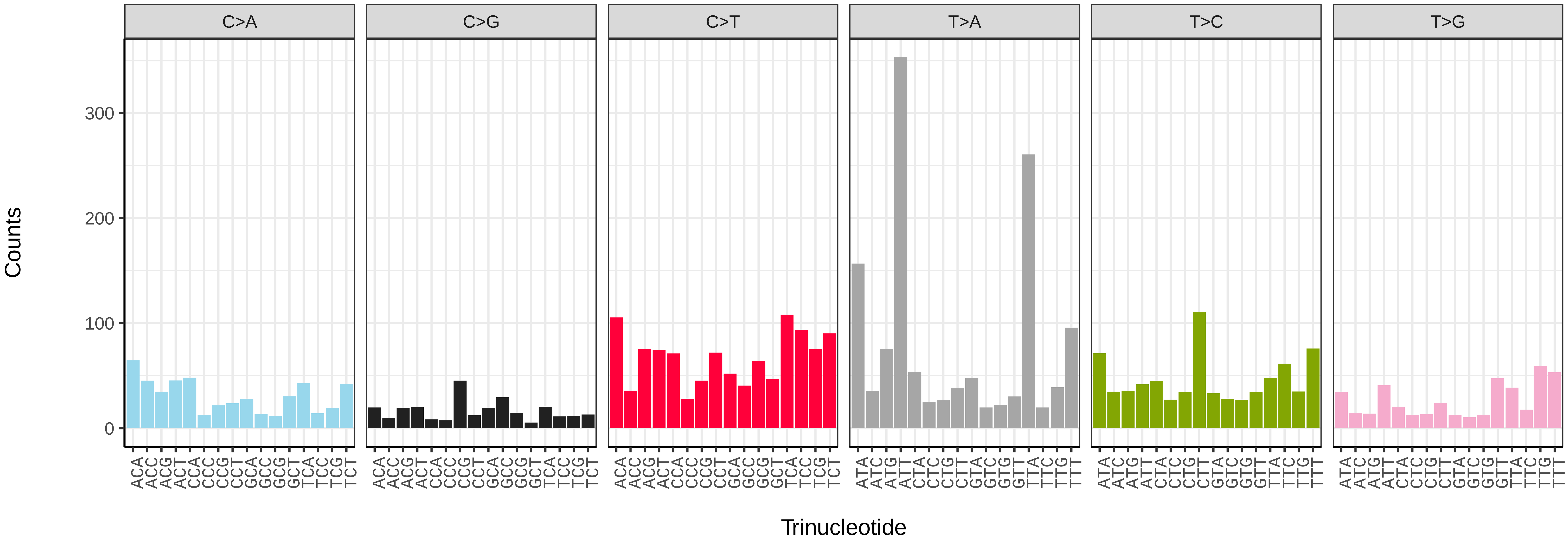


*Pyrgus malvae* (ilPyrMalv3) somatic mutational spectrum (SBS52 classification)

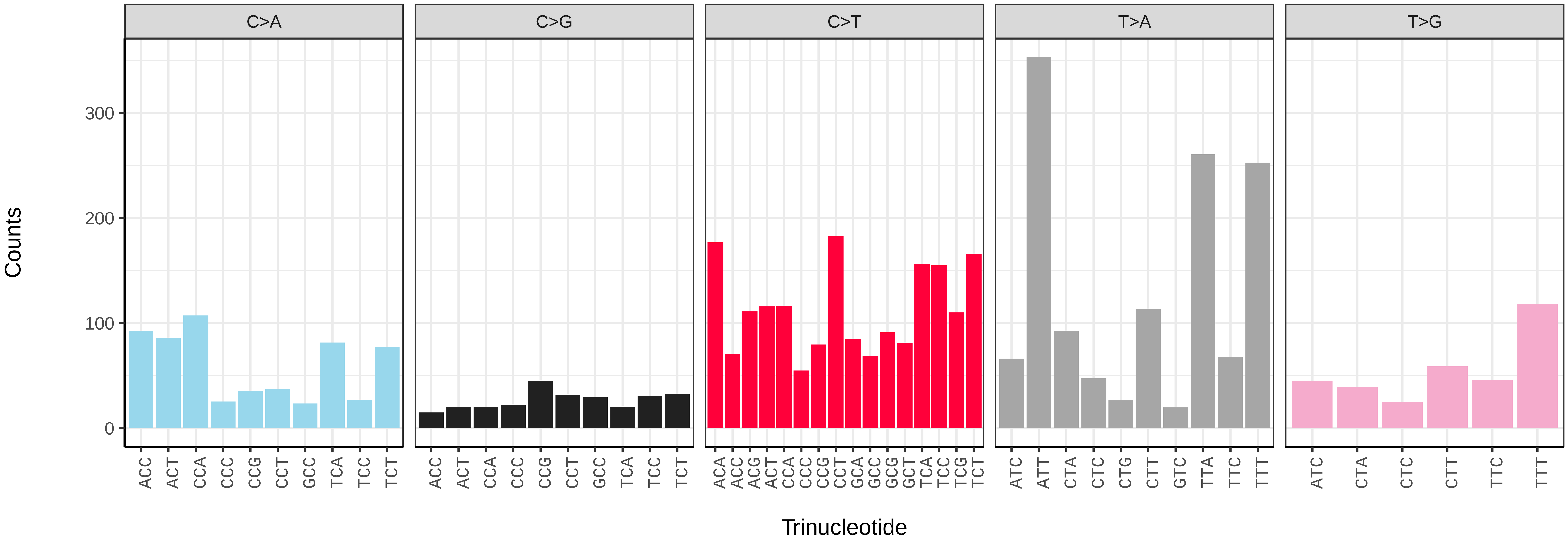

*Saturnia pavonia* (ilSatPavo1) germline mutational spectrum

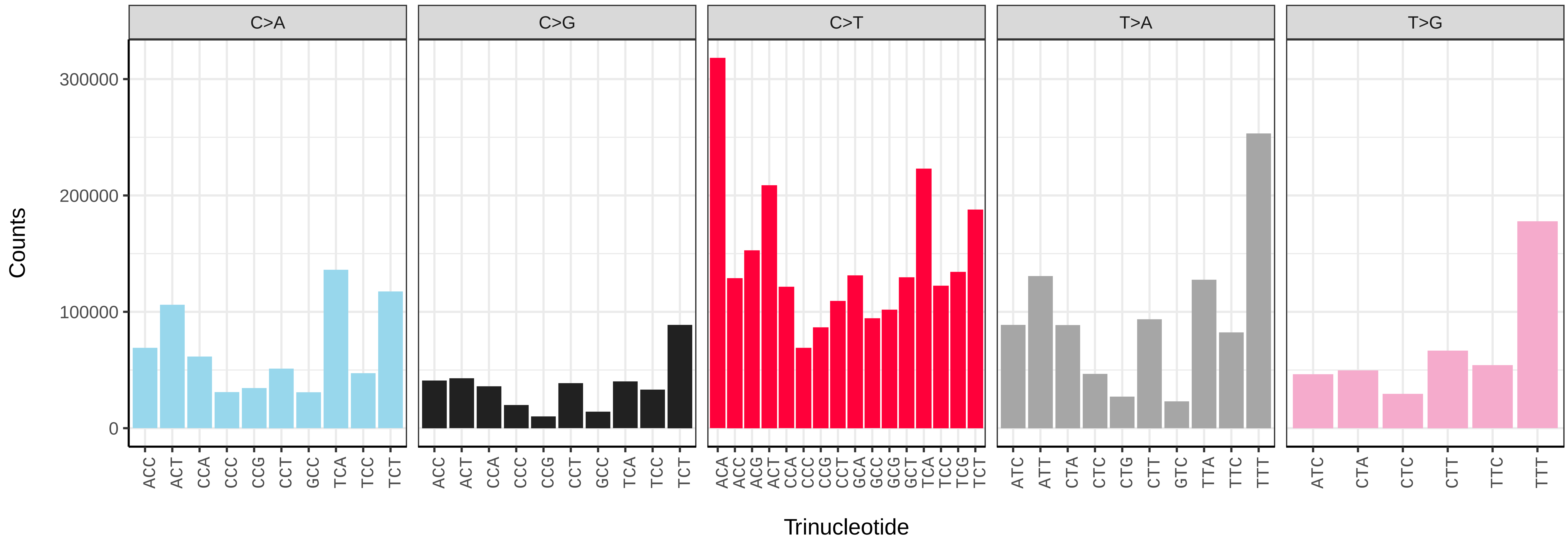


*Saturnia pavonia* (ilSatPavo1) somatic mutational spectrum (SBS96 classification)

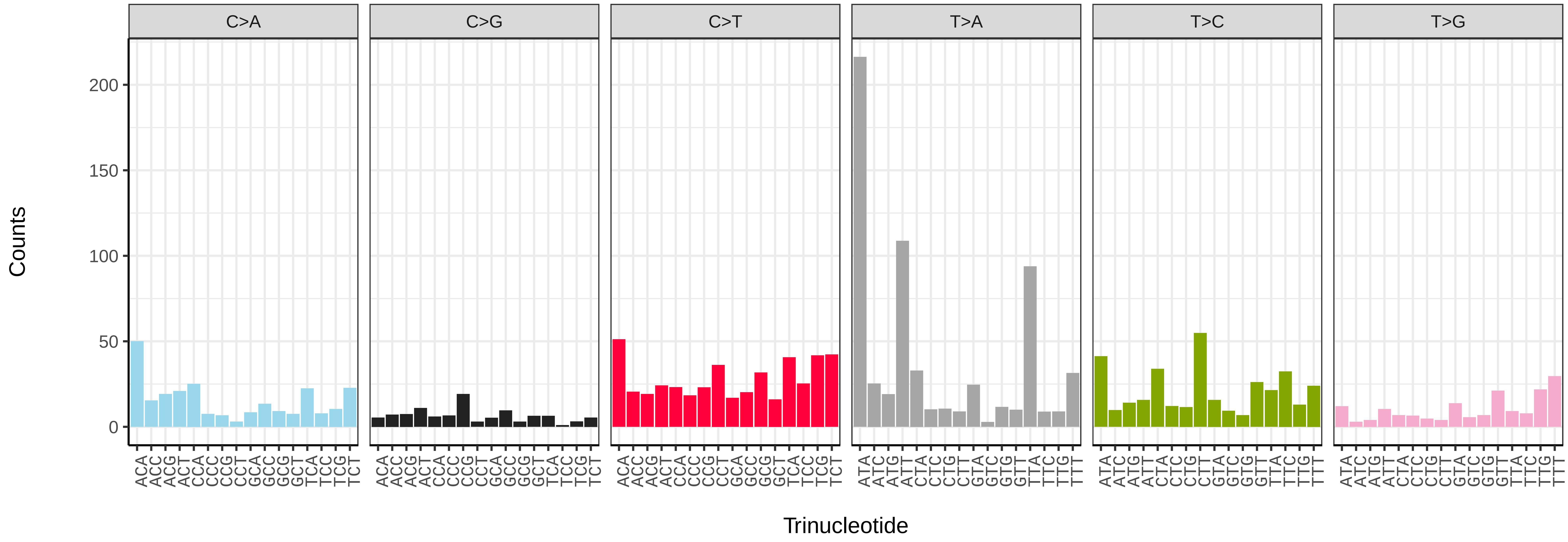


*Saturnia pavonia* (ilSatPavo1) somatic mutational spectrum (SBS52 classification)

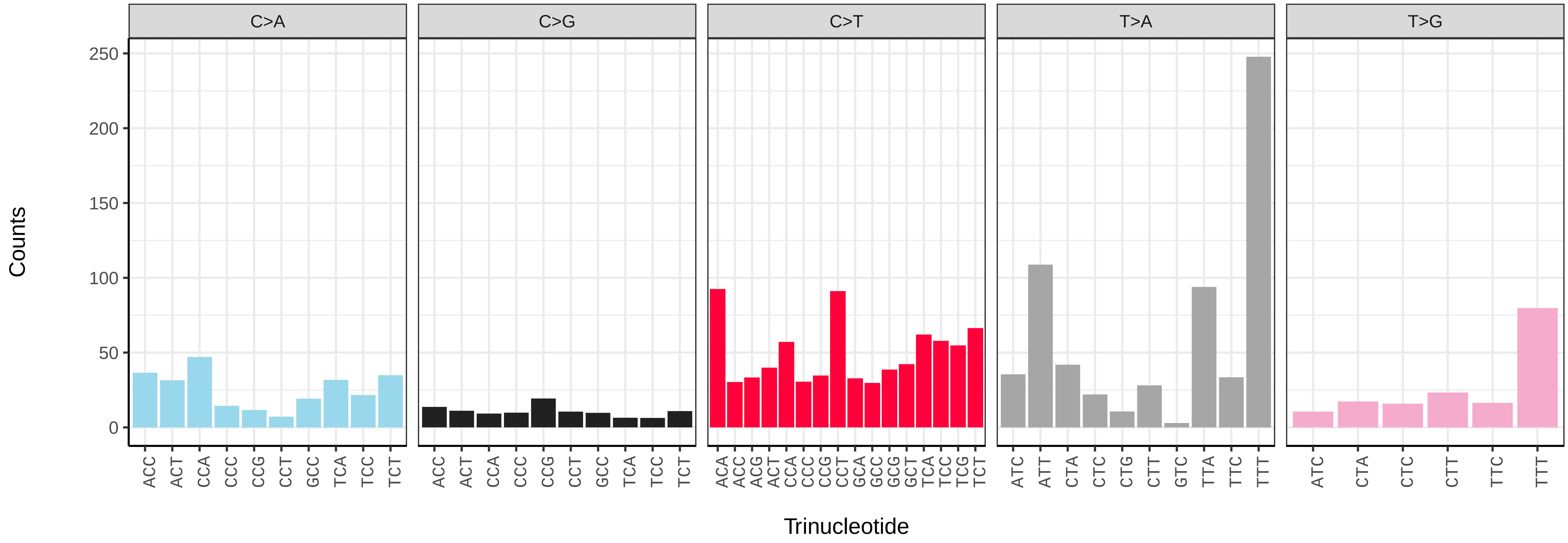

*Schrankia costaestrigalis* (ilSchCost1) germline mutational spectrum

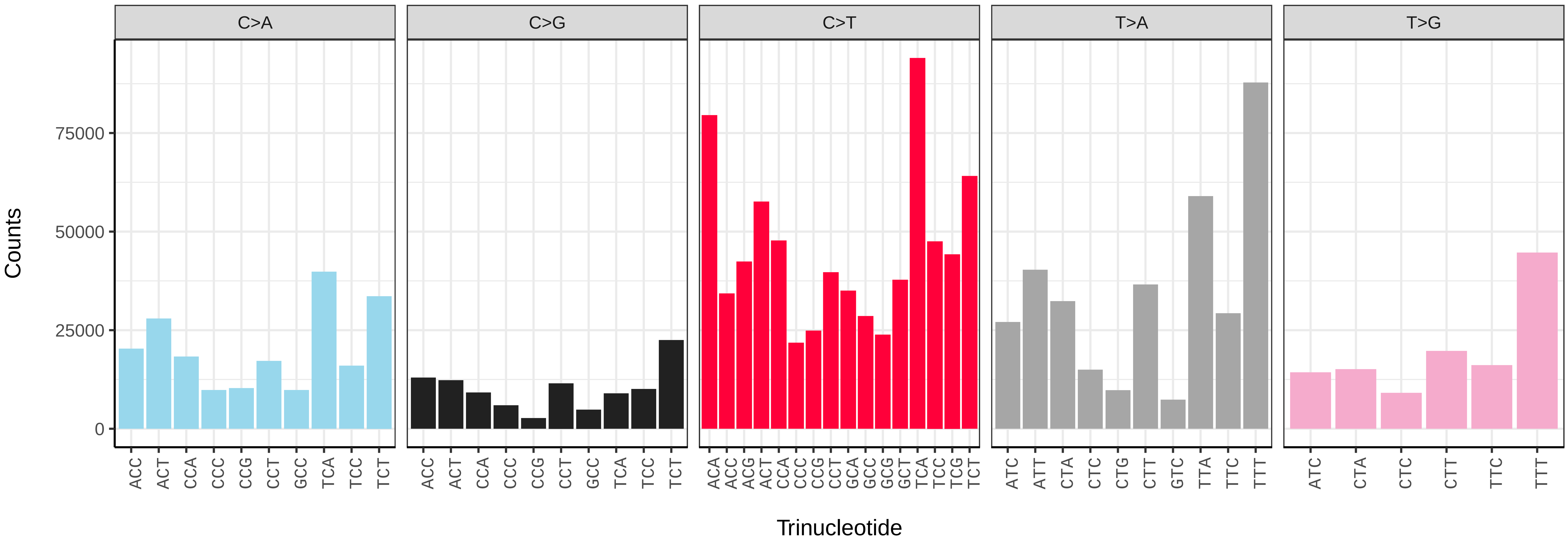


*Schrankia costaestrigalis* (ilSchCost1) somatic mutational spectrum (SBS96 classification)

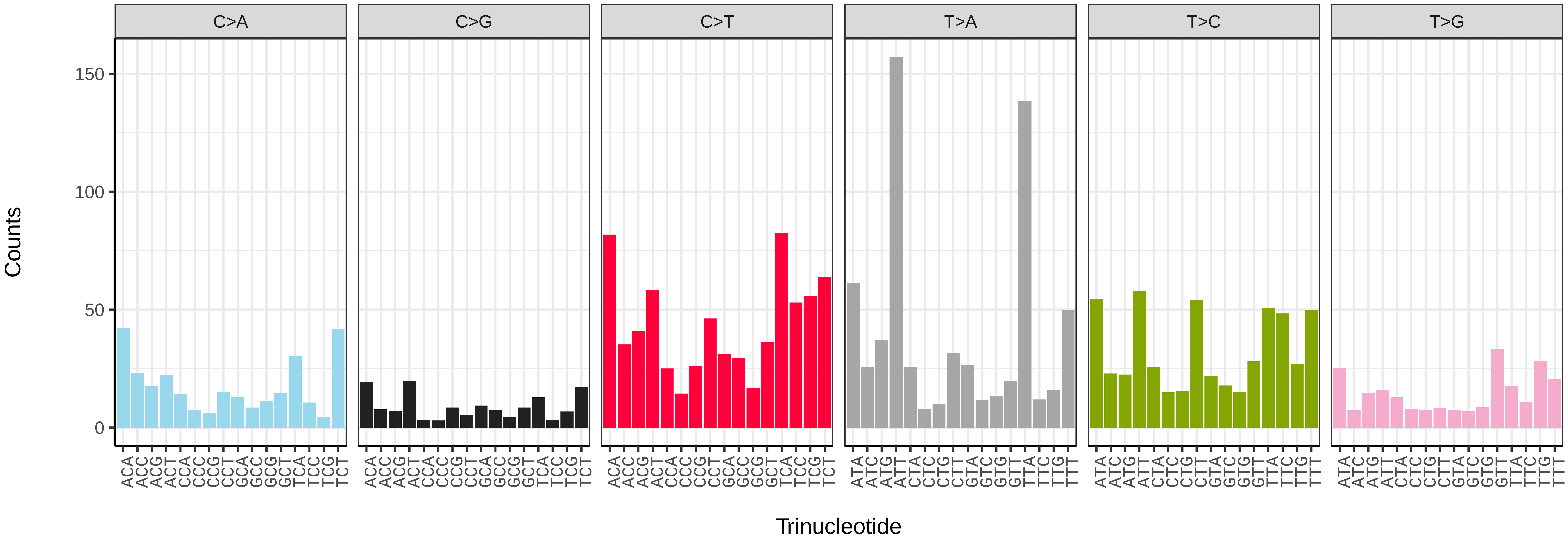


*Schrankia costaestrigalis* (ilSchCost1) somatic mutational spectrum (SBS52 classification)

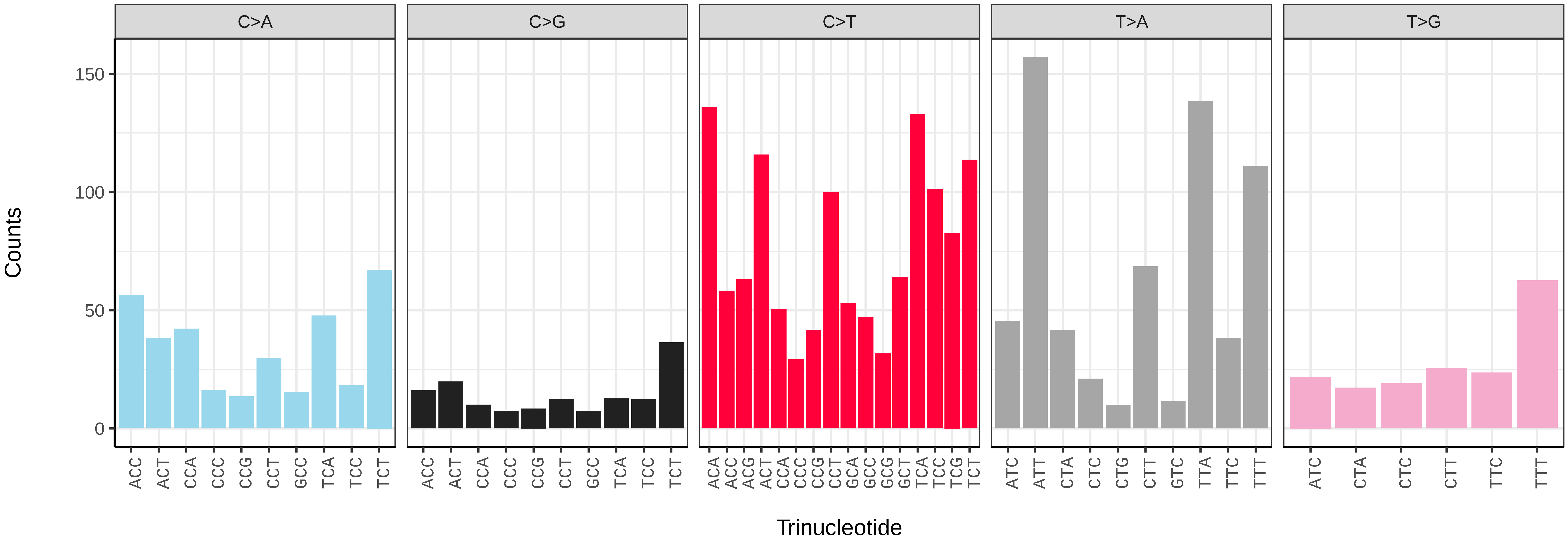

*Selenia dentaria* (ilSelDent1) germline mutational spectrum

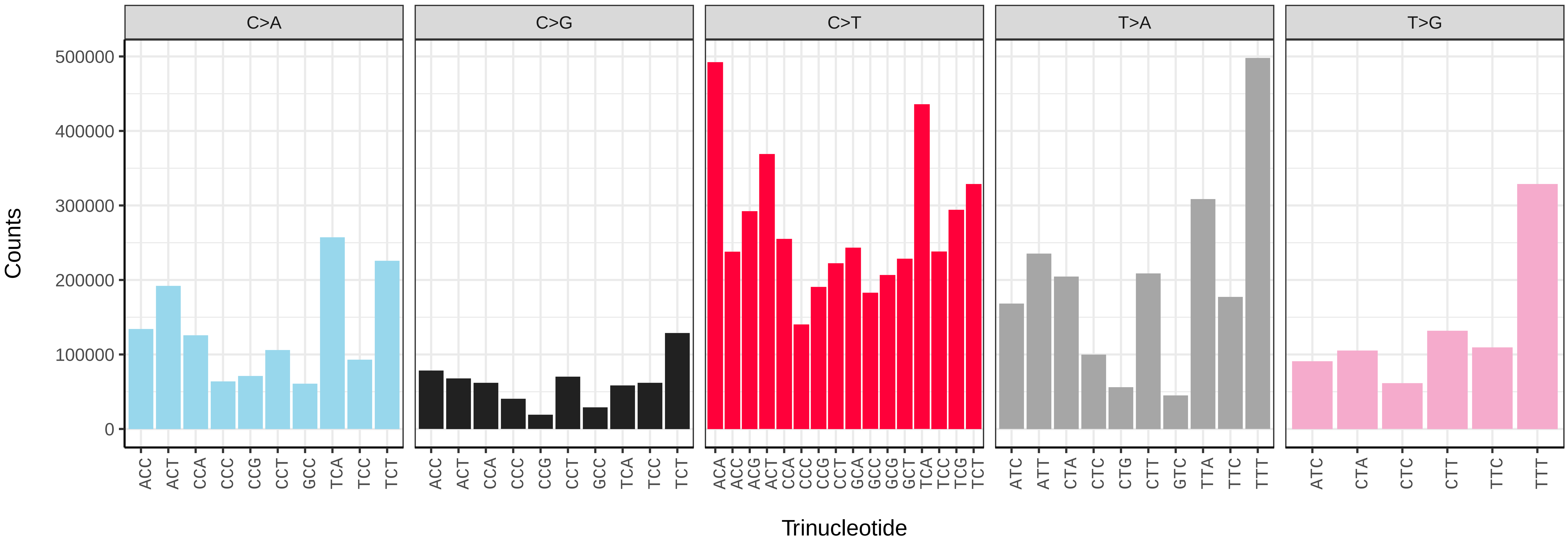


*Selenia dentaria* (ilSelDent1) somatic mutational spectrum (SBS96 classification)

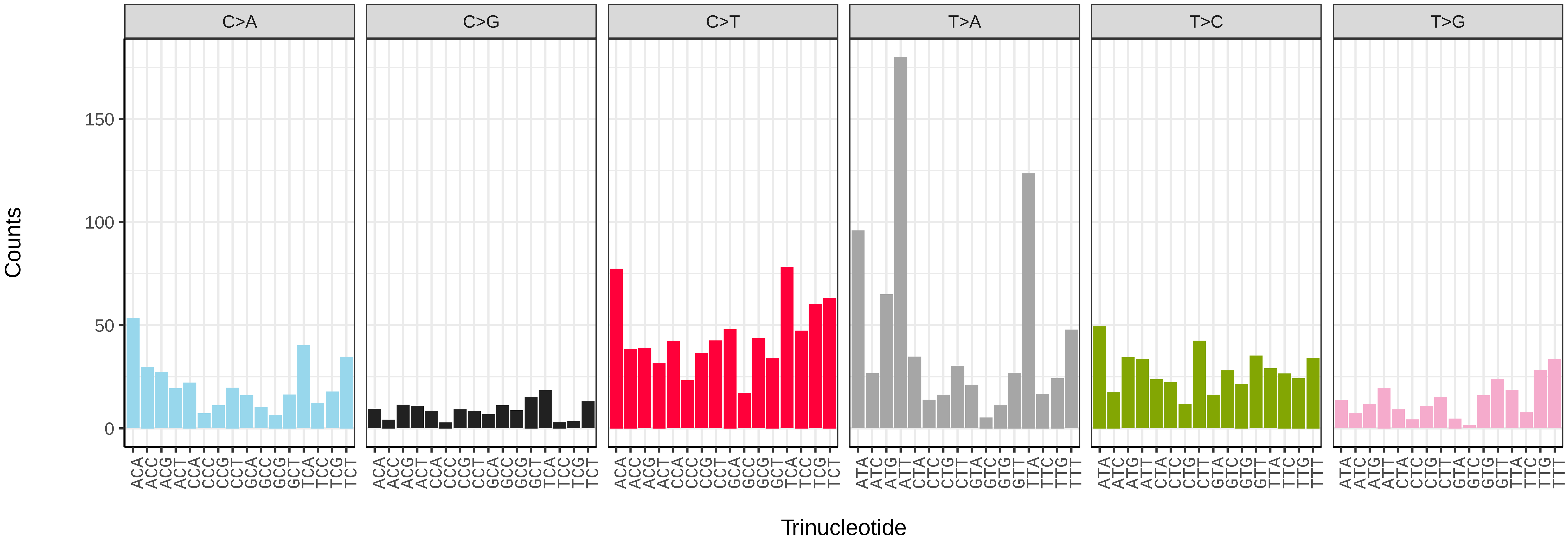


*Selenia dentaria* (ilSelDent1) somatic mutational spectrum (SBS52 classification)

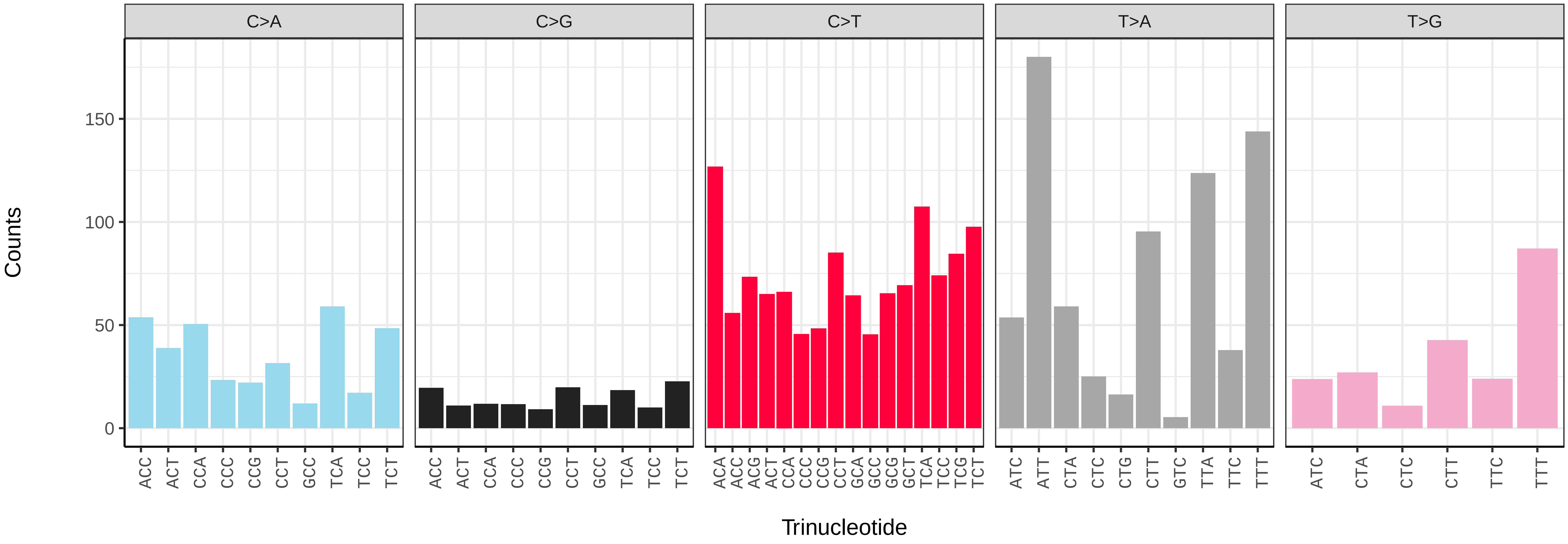

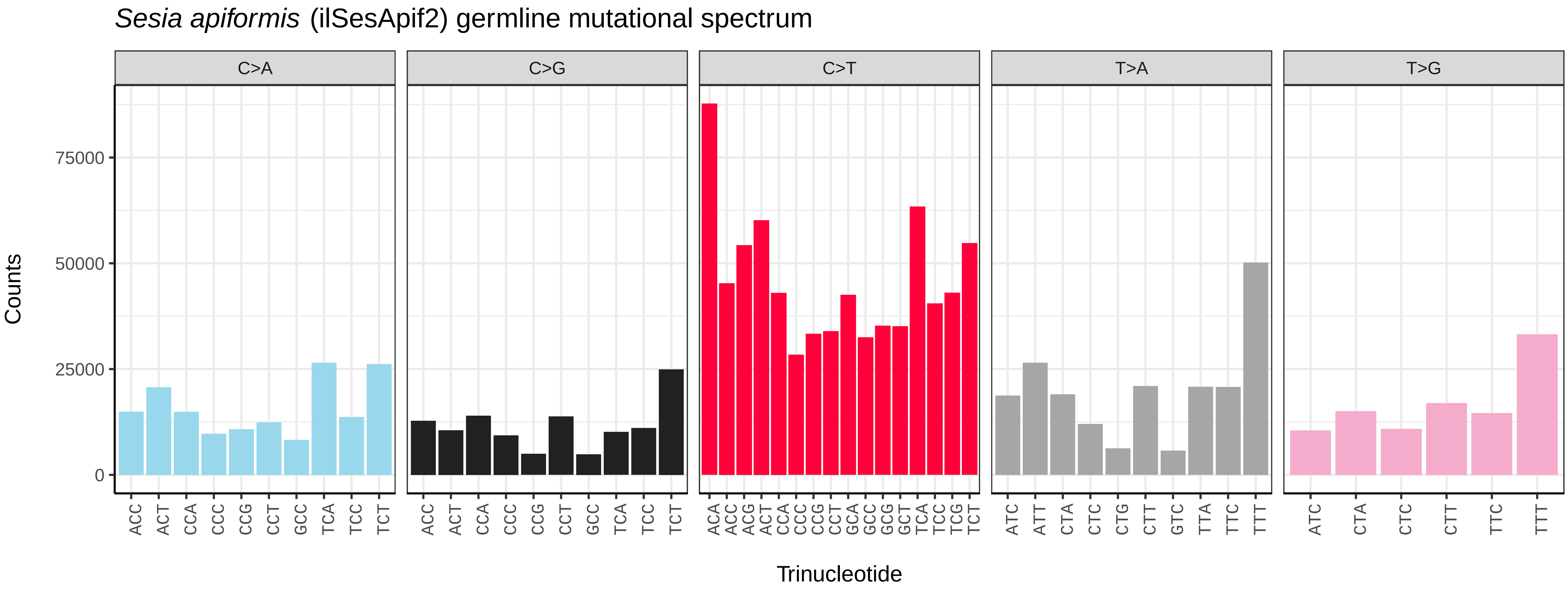
Sesia apiformis (ilSesApif2) germline mutational spectrum
C>A
C>G
C>T
T>A
T>G
Counts
0
25000
50000
75000
ACC ACT CCA CCC CCG CCT GCC TCA TCC TCT
ACA ACC ACG ACT CCA CCC CCG CCT GCA GCC GCG GCT TCA TCC TCG TCT
ATC ATT CTA CTC CTG CTT GTC TTA TTC TTT
ATC CTA CTC CTT TTC TTT
Trinucleotide

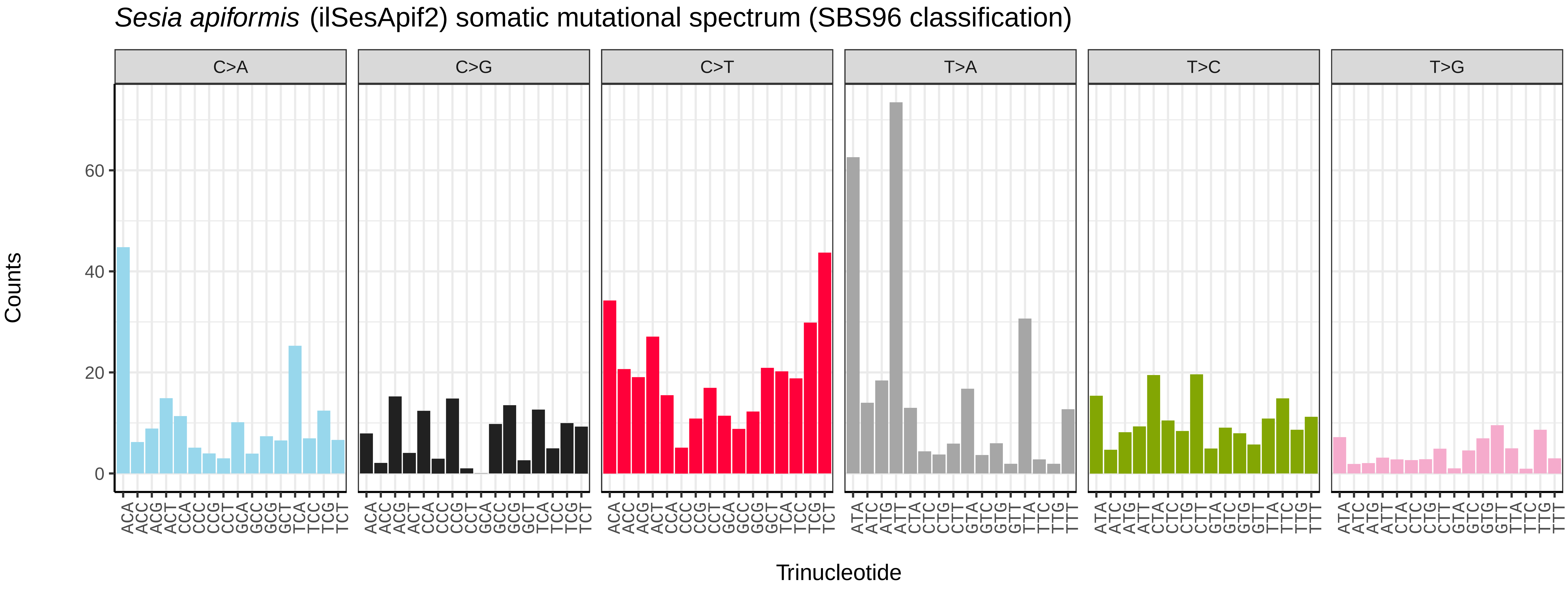
Sesia apiformis (ilSesApif2) somatic mutational spectrum (SBS96 classification)
C>A
C>G
C>T
T>A
T>C
T>G
Counts
0
20
40
60
ACA ACC ACG ACT CCA CCC CCG CCT GCA GCC GCG GCT TCA TCC TCG TCT
ATA ATC ATG ATT CTA CTC CTG CTT GTA GTC GTG GTT TTA TTC TTG TTT
Trinucleotide

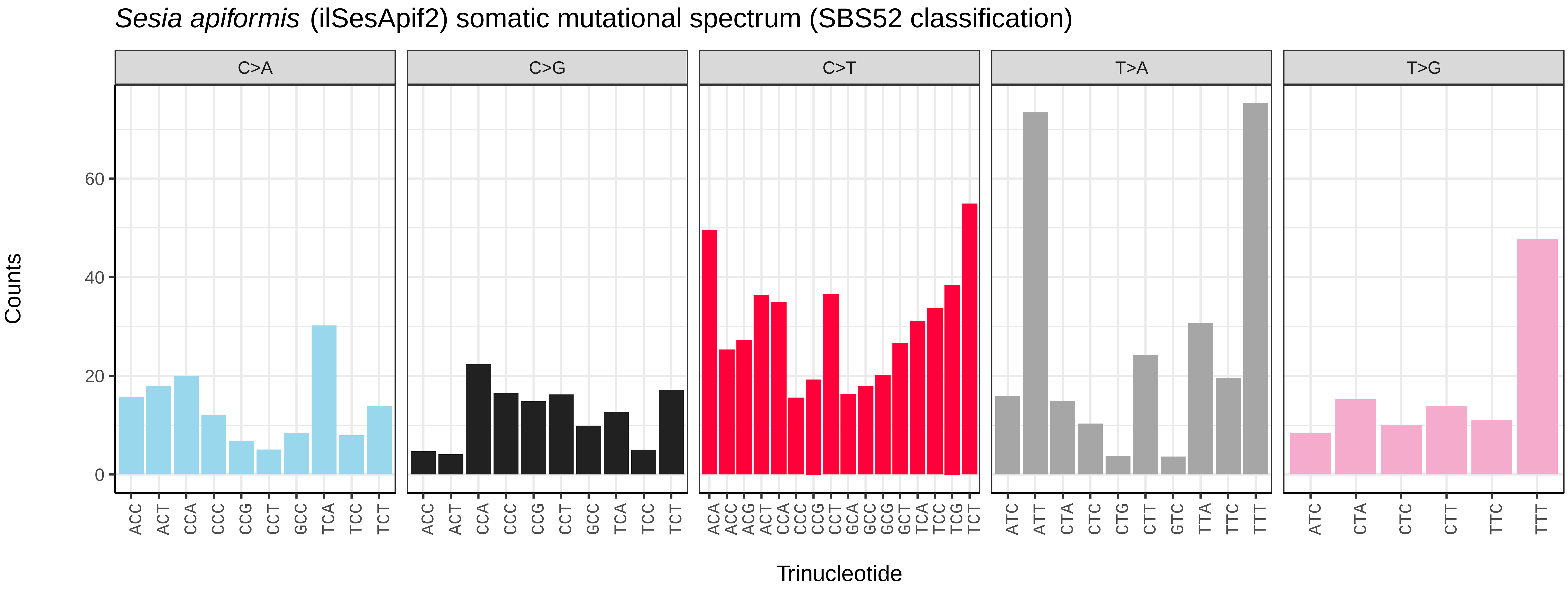
Sesia apiformis (ilSesApif2) somatic mutational spectrum (SBS52 classification)
C>A
C>G
C>T
T>A
T>G
Counts
0
20
40
60
ACC ACT CCA CCC CCG CCT GCC TCA TCC TCT
ACA ACC ACG ACT CCA CCC CCG CCT GCA GCC GCG GCT TCA TCC TCG TCT
ATC ATT CTA CTC CTG CTT GTC TTA TTC TTT
ATC CTA CTC CTT TTC TTT
Trinucleotide

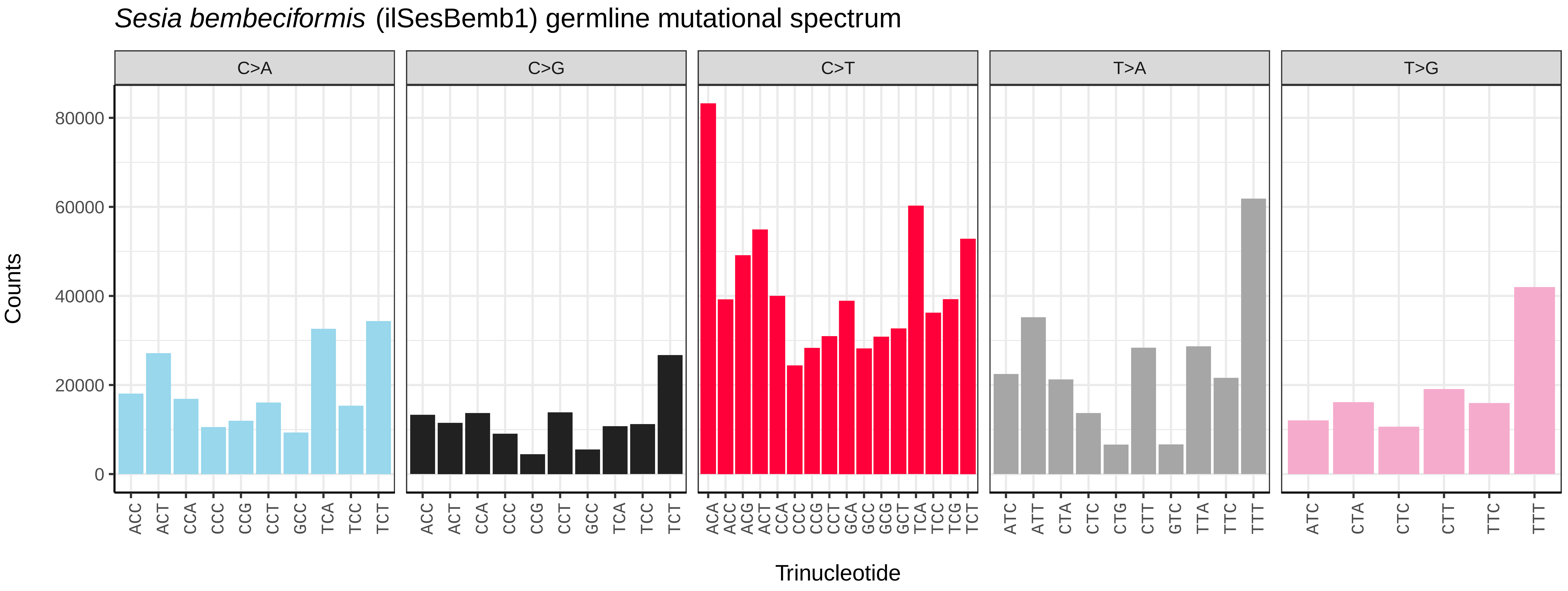
*Sesia bembeciformis* (ilSesBemb1) germline mutational spectrum
C>A
C>G
C>T
T>A
T>G
Counts
0
20000
40000
60000
80000
ACC ACT CCA CCC CCG CCT GCC TCA TCC TCT
ACA ACC ACG ACT CCA CCC CCG CCT GCA GCC GCG GCT TCA TCC TCG TCT
ATC ATT CTA CTC CTG CTT GTC TTA TTC TTT
ATC CTA CTC CTT TTC TTT
Trinucleotide

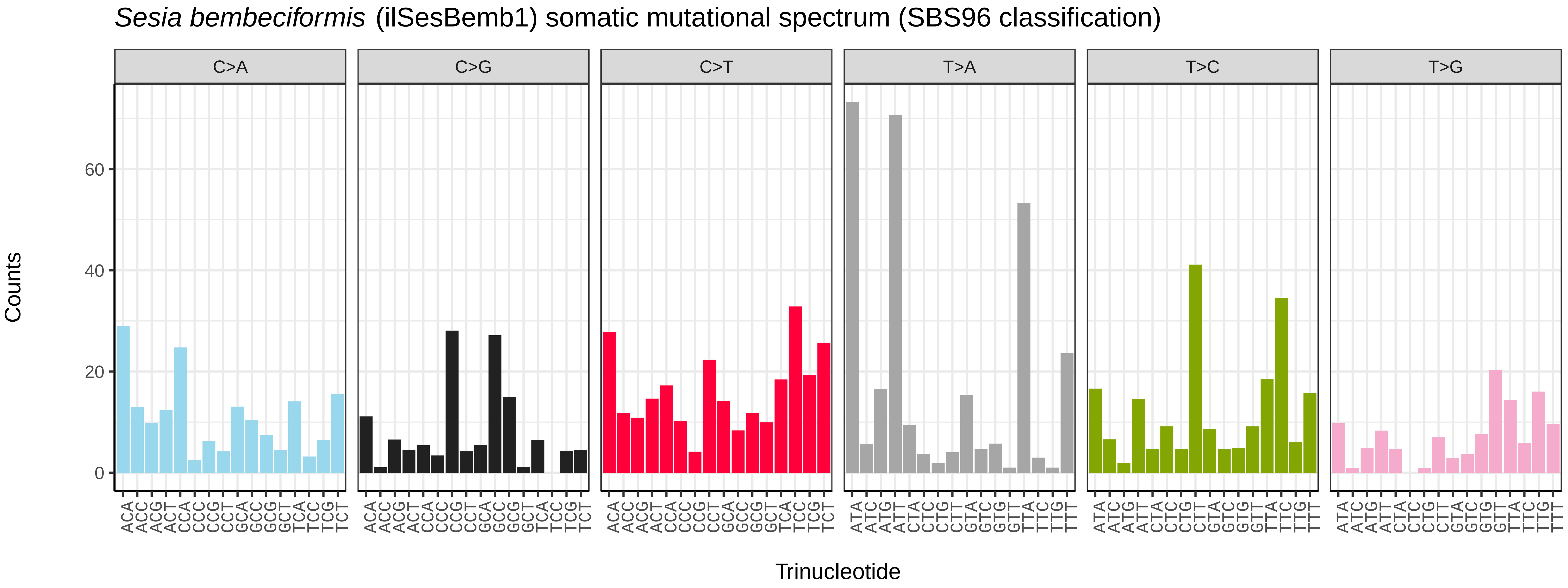
*Sesia bembeciformis* (ilSesBemb1) somatic mutational spectrum (SBS96 classification)
C>A
C>G
C>T
T>A
T>C
T>G
Counts
0
20
40
60
ACA ACC ACG ACT CCA CCC CCG CCT GCA GCC GCG GCT TCA TCC TCG TCT
ATA ATC ATG ATT CTA CTC CTG CTT GTA GTC GTG GTT TTA TTC TTG TTT
Trinucleotide

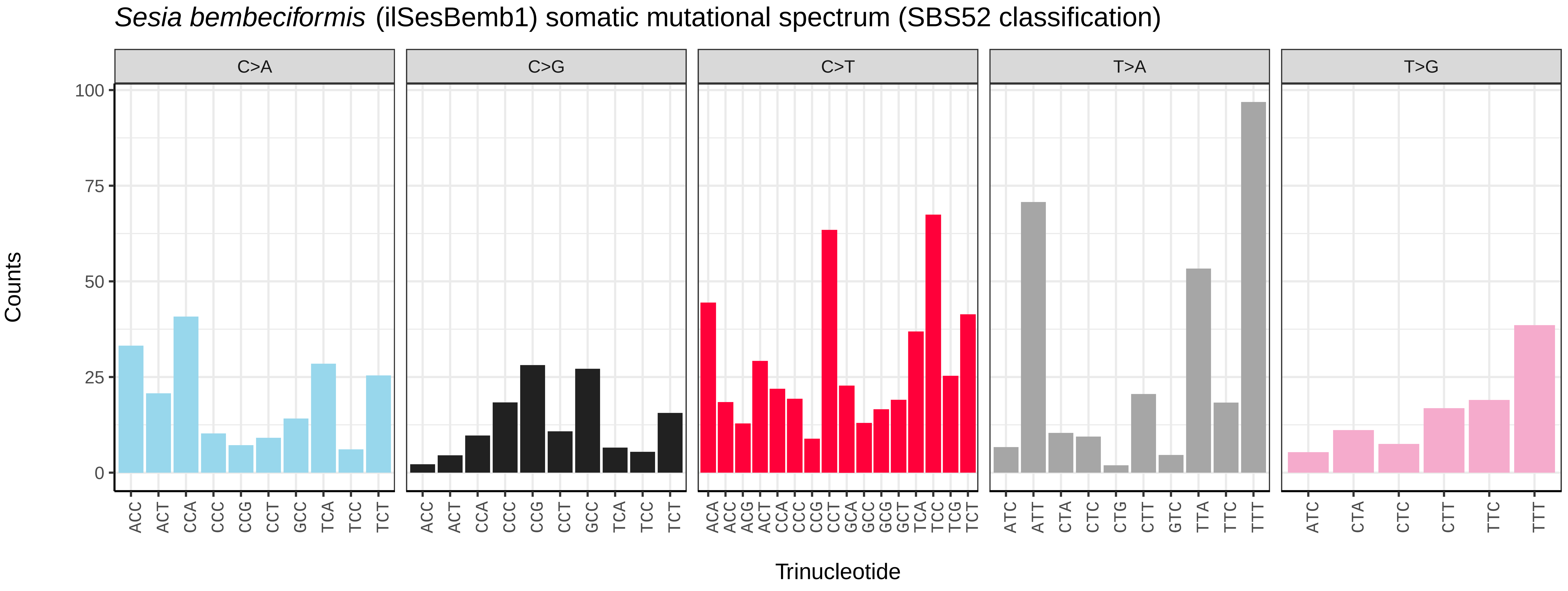
*Sesia bembeciformis* (ilSesBemb1) somatic mutational spectrum (SBS52 classification)
C>A
C>G
C>T
T>A
T>G
Counts
0
25
50
75
100
ACC ACT CCA CCC CCG CCT GCC TCA TCC TCT
ACA ACC ACG ACT CCA CCC CCG CCT GCA GCC GCG GCT TCA TCC TCG TCT
ATC ATT CTA CTC CTG CTT GTC TTA TTC TTT
ATC CTA CTC CTT TTC TTT
Trinucleotide

*Shargacucullia verbasci* (ilShaVerb5) germline mutational spectrum

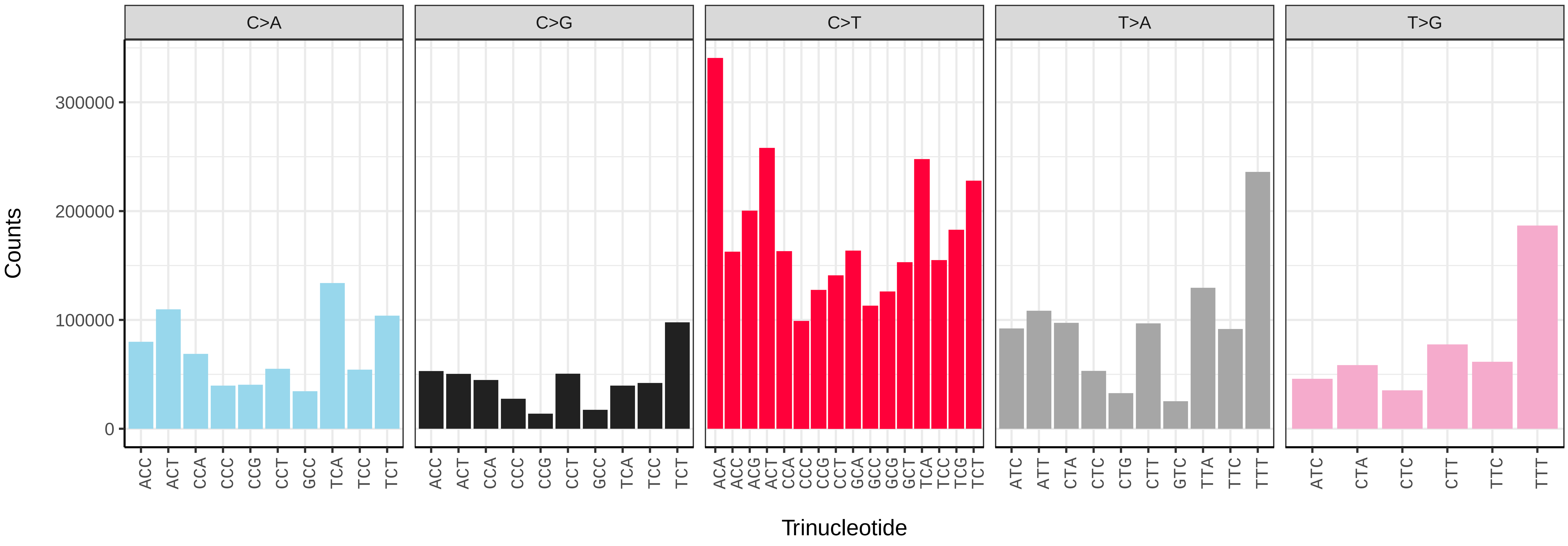


*Shargacucullia verbasci* (ilShaVerb5) somatic mutational spectrum (SBS96 classification)

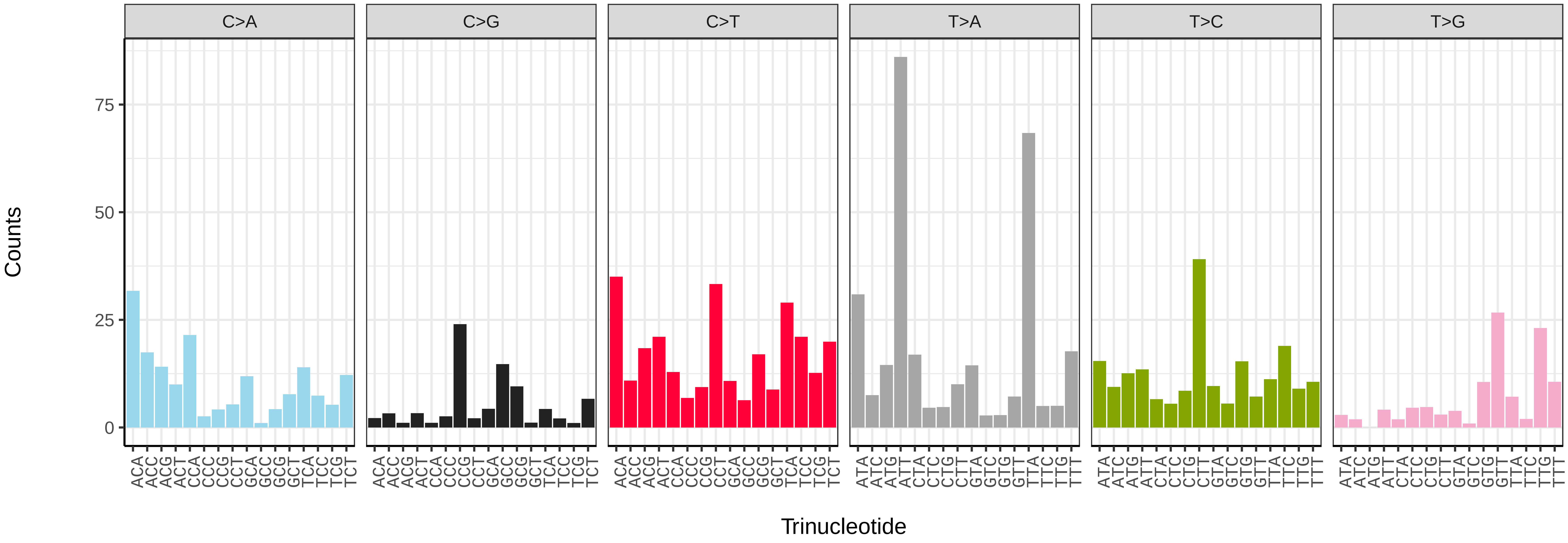


*Shargacucullia verbasci* (ilShaVerb5) somatic mutational spectrum (SBS52 classification)

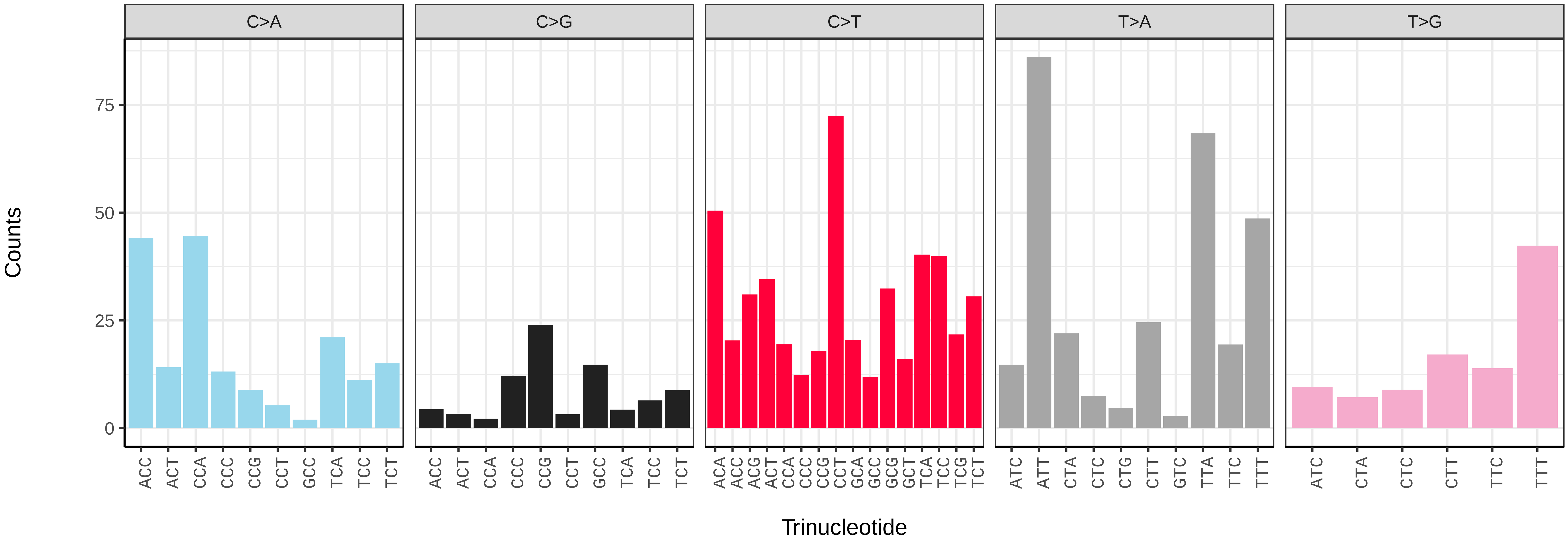

*Sphinx pinastri* (ilSphPina1) germline mutational spectrum

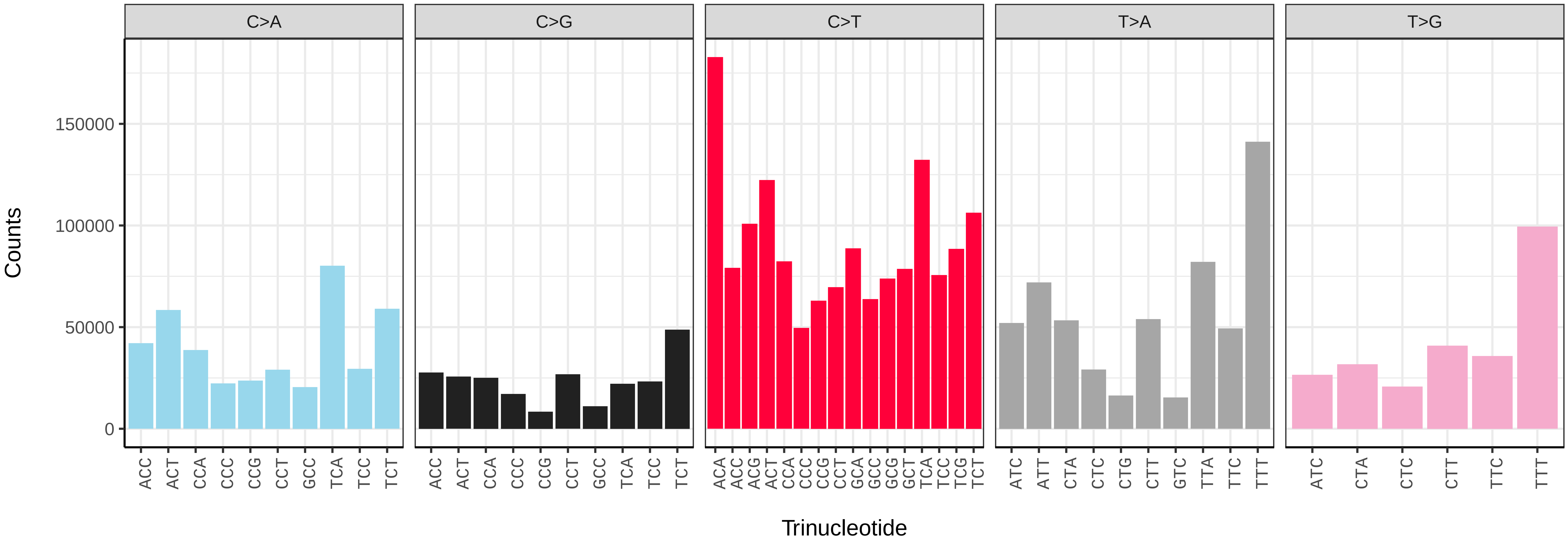


*Sphinx pinastri* (ilSphPina1) somatic mutational spectrum (SBS96 classification)

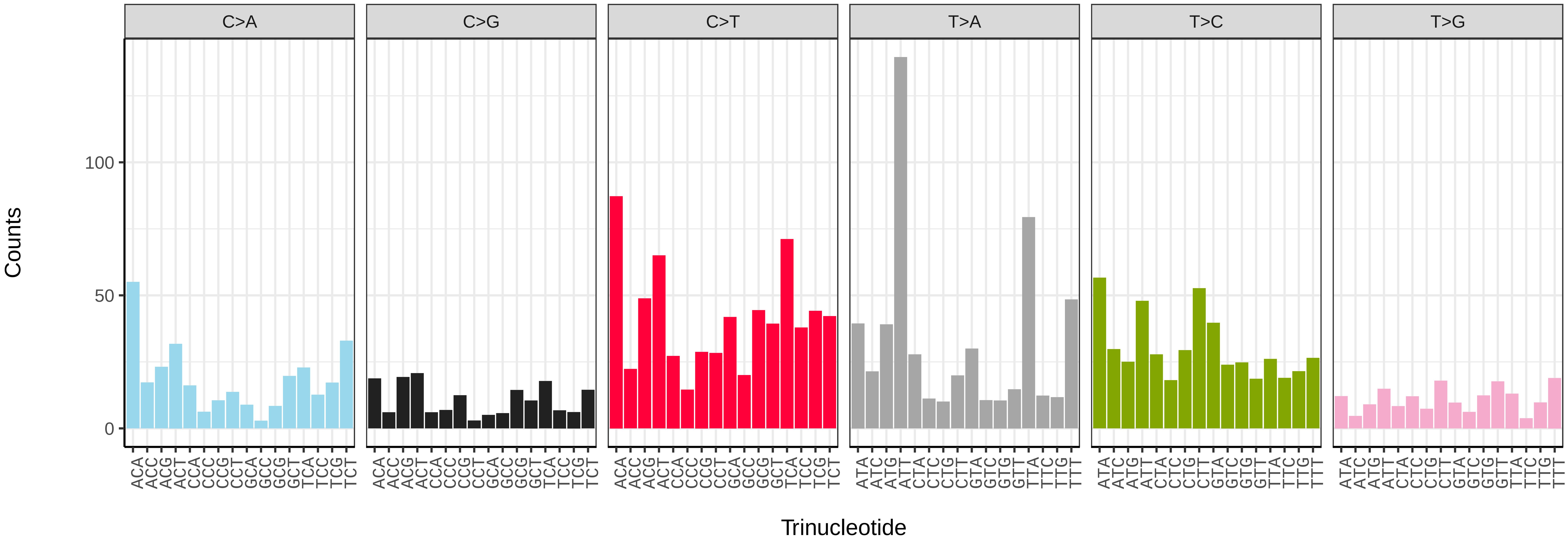


*Sphinx pinastri* (ilSphPina1) somatic mutational spectrum (SBS52 classification)

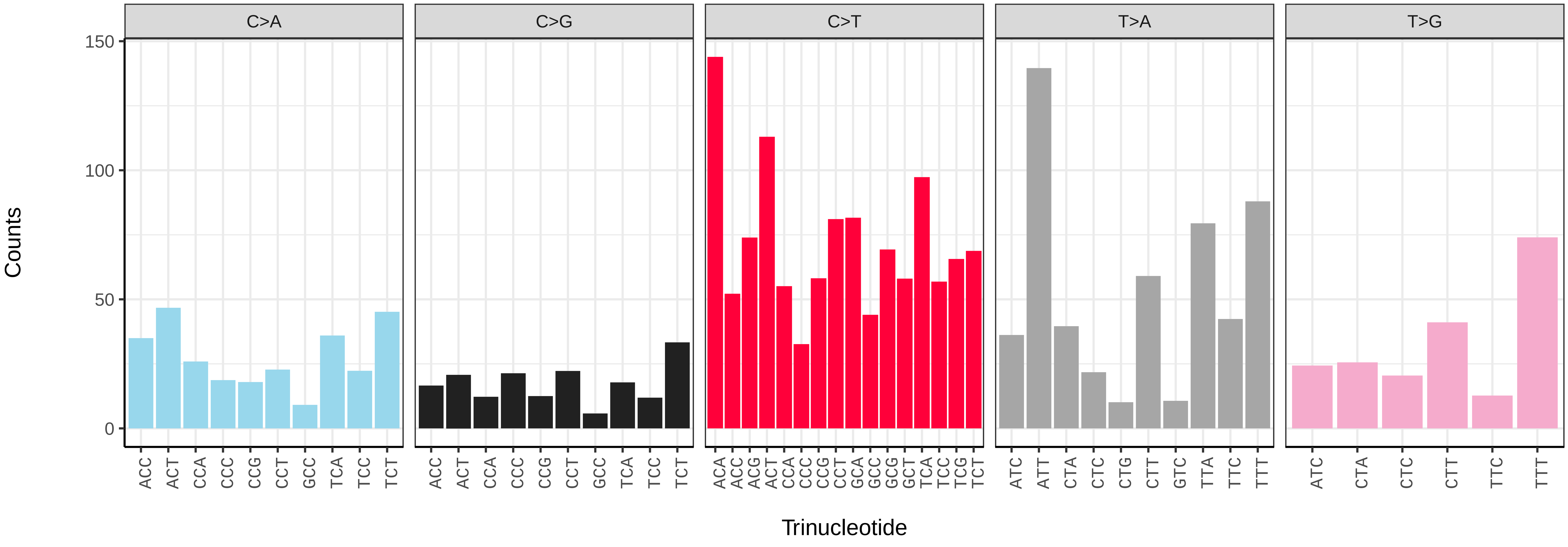

*Spilosoma lubricipeda* (ilSpiLubr1) germline mutational spectrum

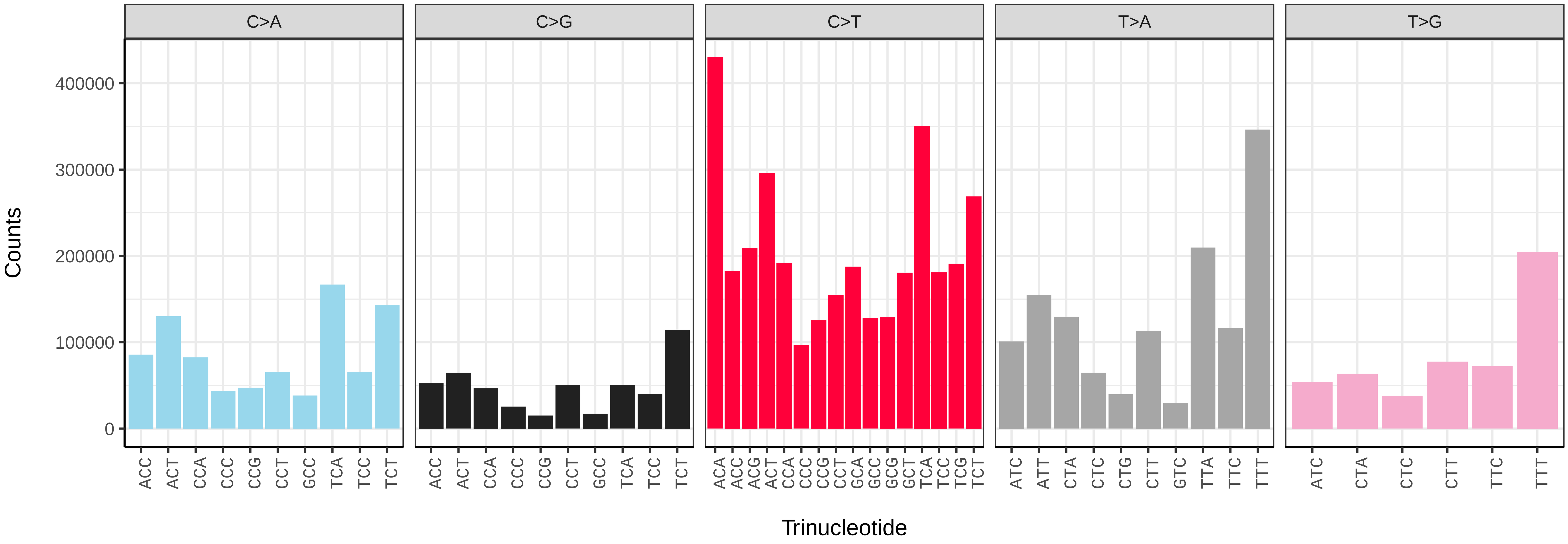


*Spilosoma lubricipeda* (ilSpiLubr1) somatic mutational spectrum (SBS96 classification)

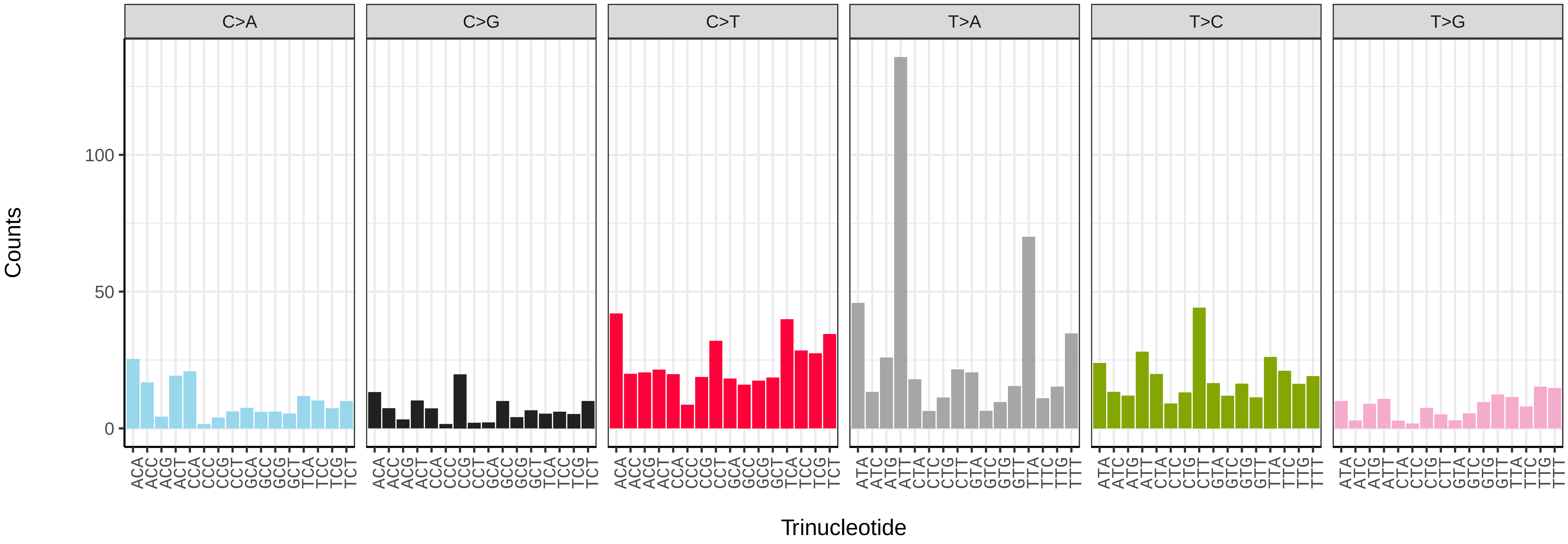


*Spilosoma lubricipeda* (ilSpiLubr1) somatic mutational spectrum (SBS52 classification)

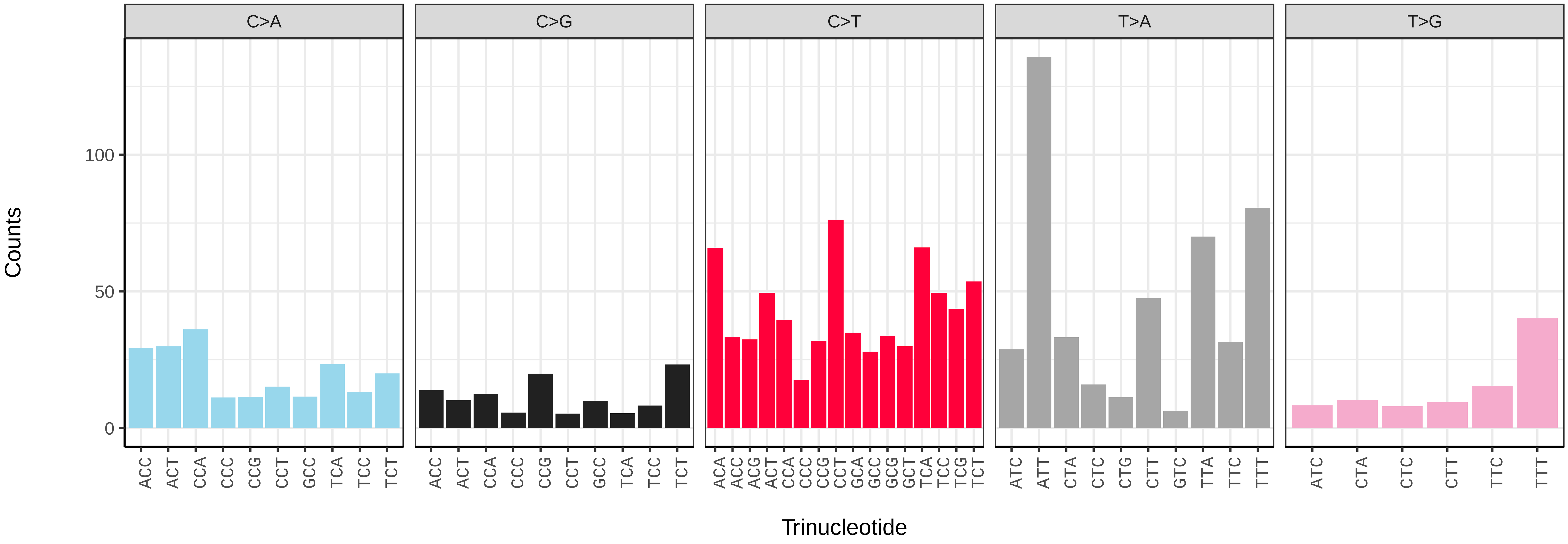

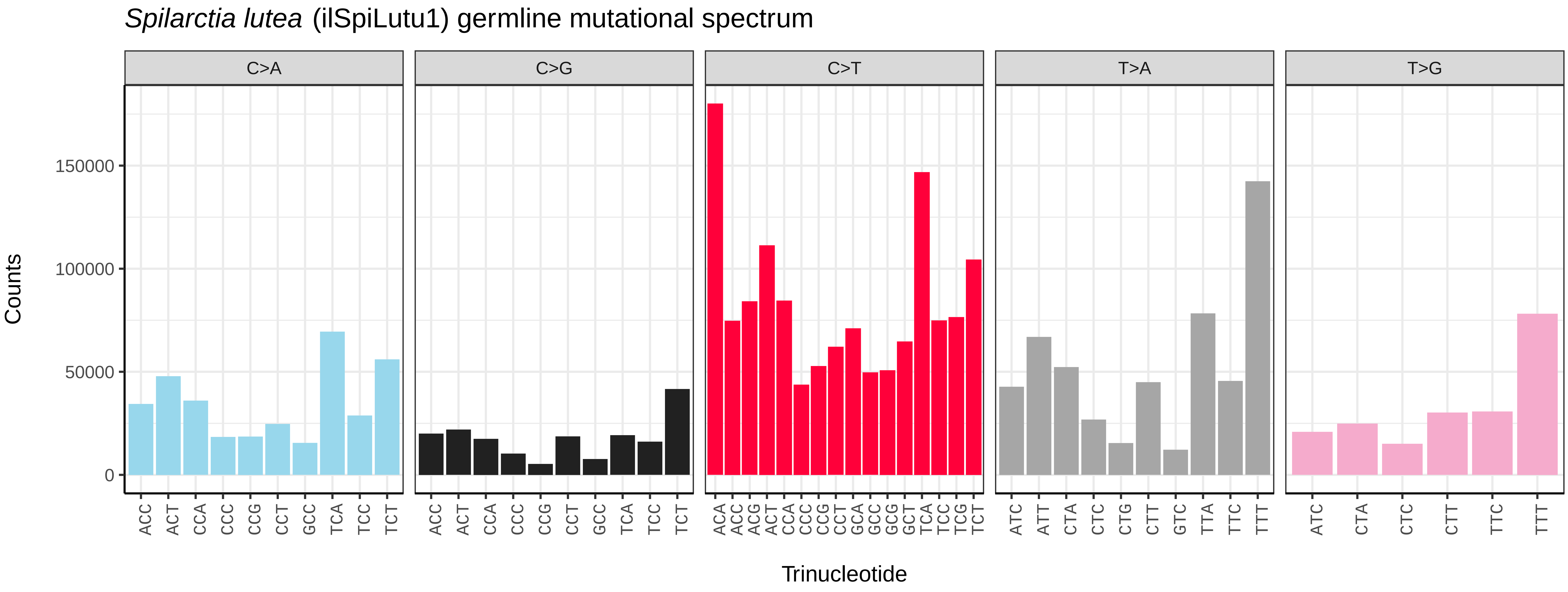
Spilarctia lutea (ilSpiLutu1) germline mutational spectrum
C>A
C>G
C>T
T>A
T>G
Counts
150000
100000
50000
0
ACC ACT CCA CCC CCG CCT GCC TCA TCC TCT
ACC ACT CCA CCC CCG CCT GCC TCA TCC TCT
ACA ACC ACG ACT CCA CCC CCG CCT GCA GCC GCG GCT TCA TCC TCG TCT
ATC ATT CTA CTC CTG CTT GTC TTA TTC TTT
ATC CTA CTC CTT TTC TTT
Trinucleotide

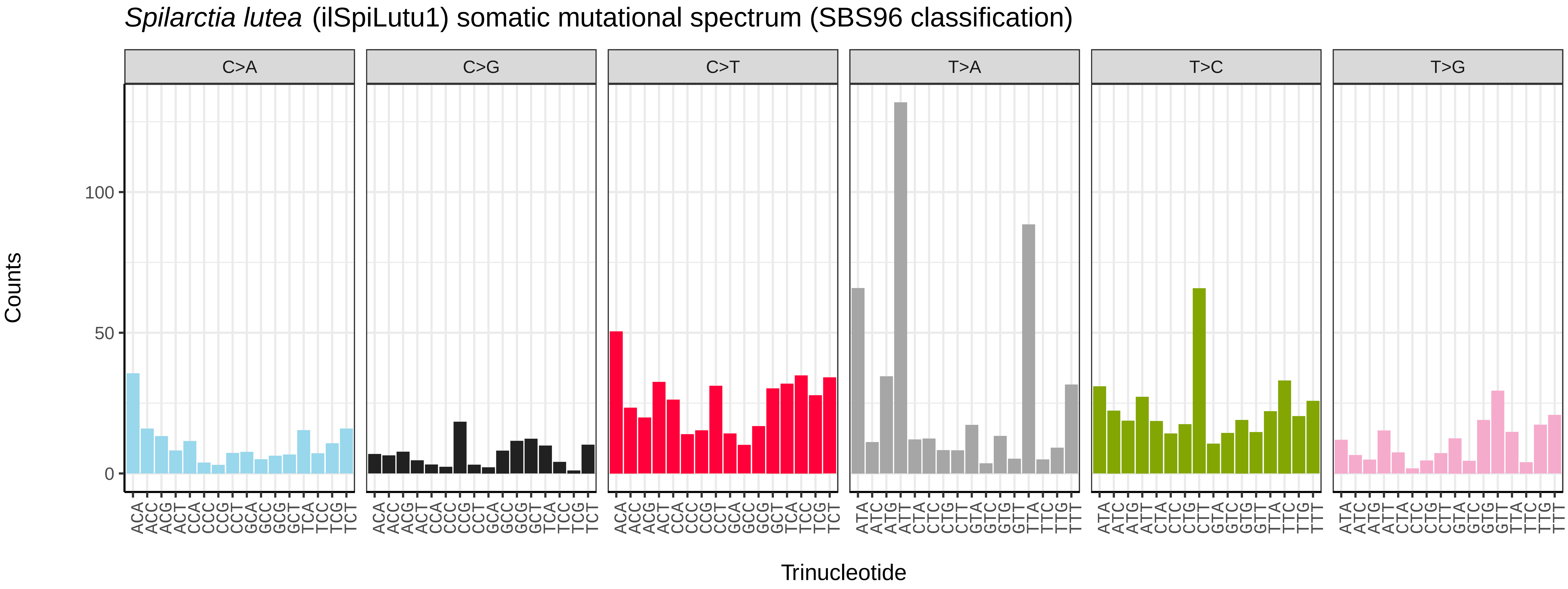
Spilarctia lutea (ilSpiLutu1) somatic mutational spectrum (SBS96 classification)
C>A
C>G
C>T
T>A
T>C
T>G
Counts
100
50
0
ACA ACC ACG ACT CCA CCC CCG CCT GCA GCC GCG GCT TCA TCC TCG TCT
ACA ACC ACG ACT CCA CCC CCG CCT GCA GCC GCG GCT TCA TCC TCG TCT
ACA ACC ACG ACT CCA CCC CCG CCT GCA GCC GCG GCT TCA TCC TCG TCT
ATA ATC ATG ATT CTA CTC CTG CTT GTA GTC GTG GTT TTA TTC TTG TTT
ATA ATC ATG ATT CTA CTC CTG CTT GTA GTC GTG GTT TTA TTC TTG TTT
ATA ATC ATG ATT CTA CTC CTG CTT GTA GTC GTG GTT TTA TTC TTG TTT
Trinucleotide

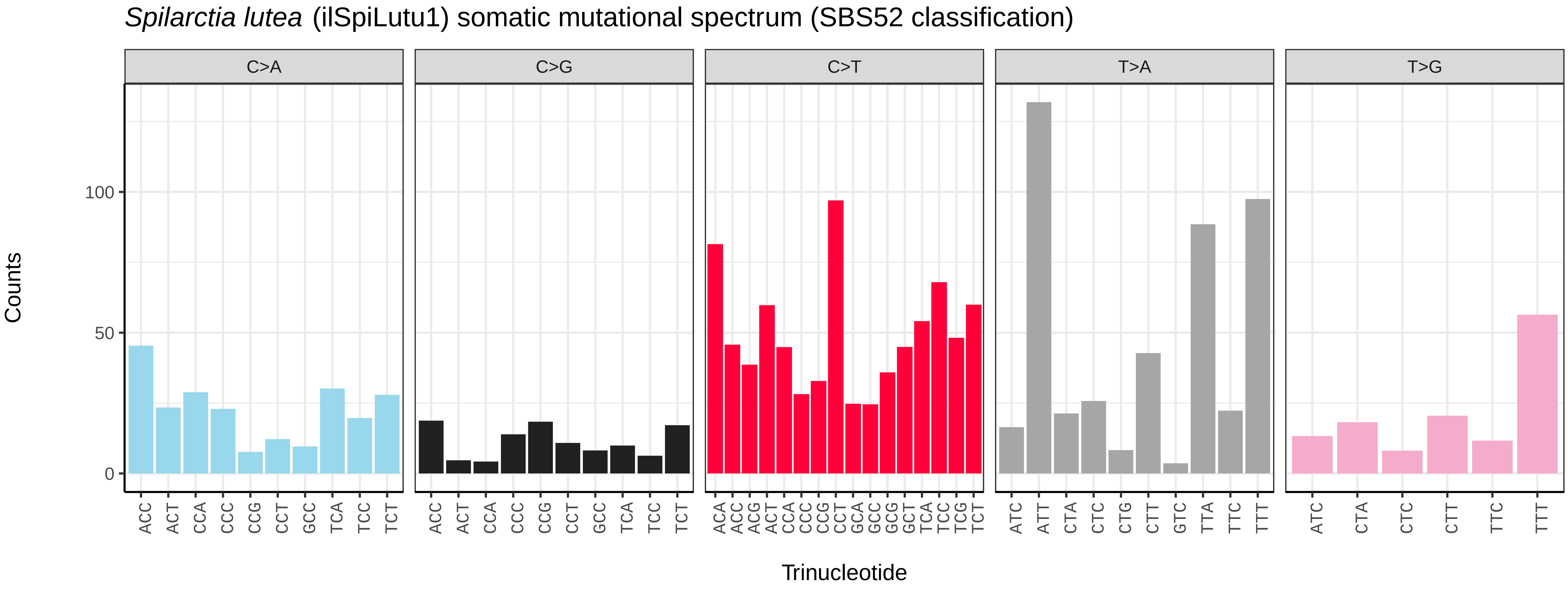
Spilarctia lutea (ilSpiLutu1) somatic mutational spectrum (SBS52 classification)
C>A
C>G
C>T
T>A
T>G
Counts
100
50
0
ACC ACT CCA CCC CCG CCT GCC TCA TCC TCT
ACC ACT CCA CCC CCG CCT GCC TCA TCC TCT
ACA ACC ACG ACT CCA CCC CCG CCT GCA GCC GCG GCT TCA TCC TCG TCT
ATC ATT CTA CTC CTG CTT GTC TTA TTC TTT
ATC CTA CTC CTT TTC TTT
Trinucleotide

*Stenoptilia bipunctidactyla* (ilSteBipu1) germline mutational spectrum

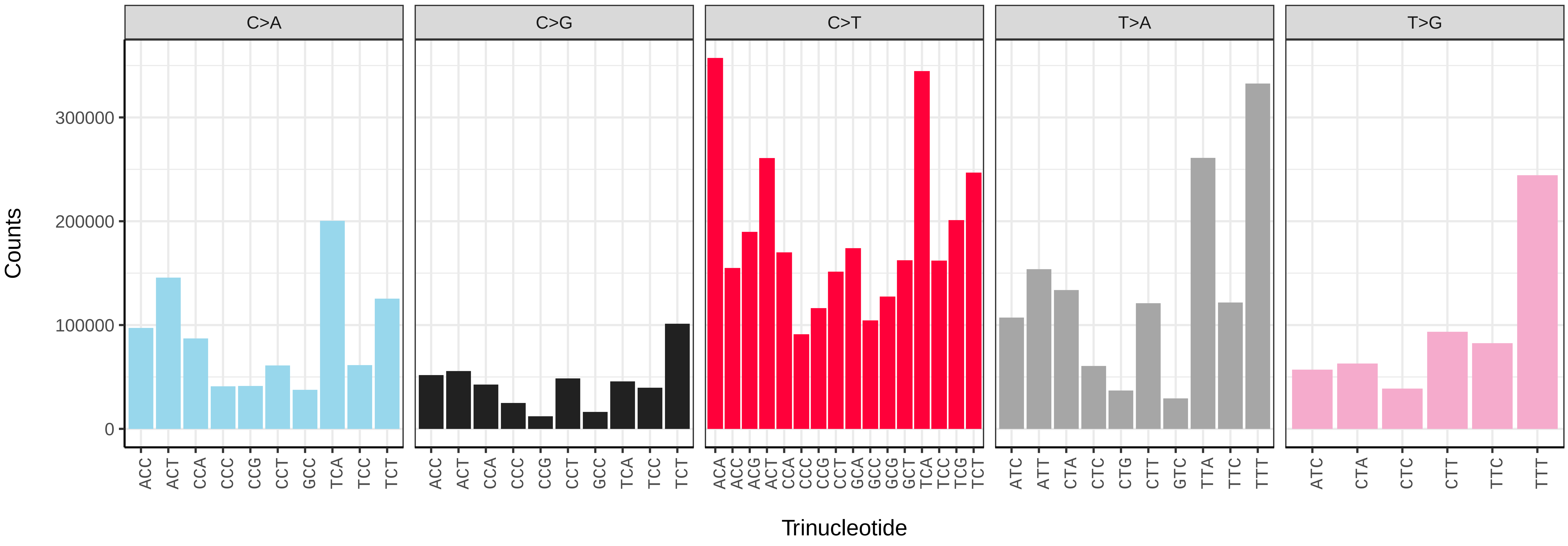


*Stenoptilia bipunctidactyla* (ilSteBipu1) somatic mutational spectrum (SBS96 classification)

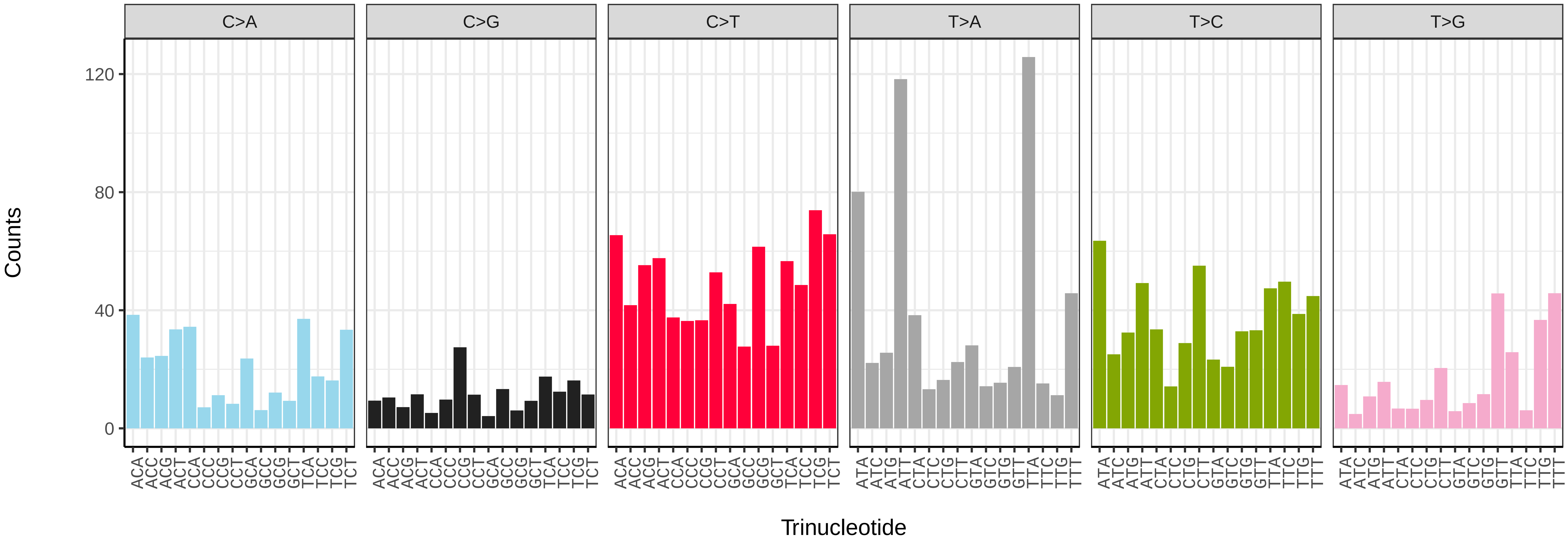


*Stenoptilia bipunctidactyla* (ilSteBipu1) somatic mutational spectrum (SBS52 classification)

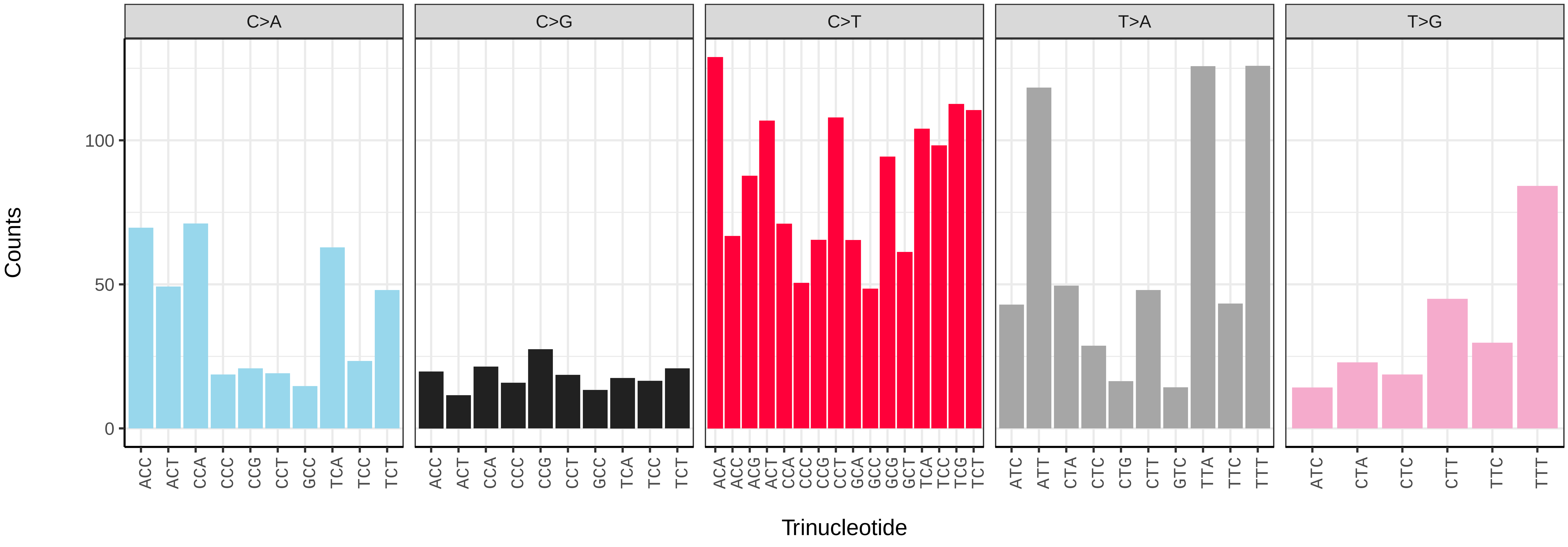

*Synanthedon andrenaeformis* (ilSynAndr1) germline mutational spectrum

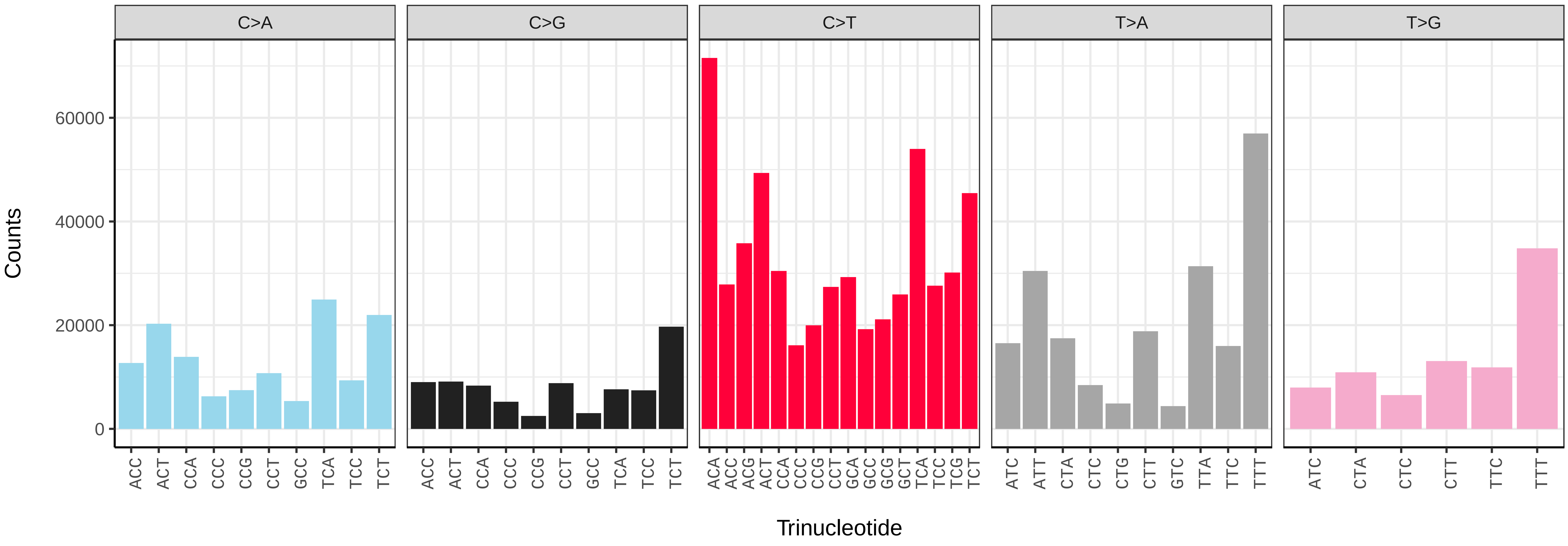


*Synanthedon andrenaeformis* (ilSynAndr1) somatic mutational spectrum (SBS96 classification)

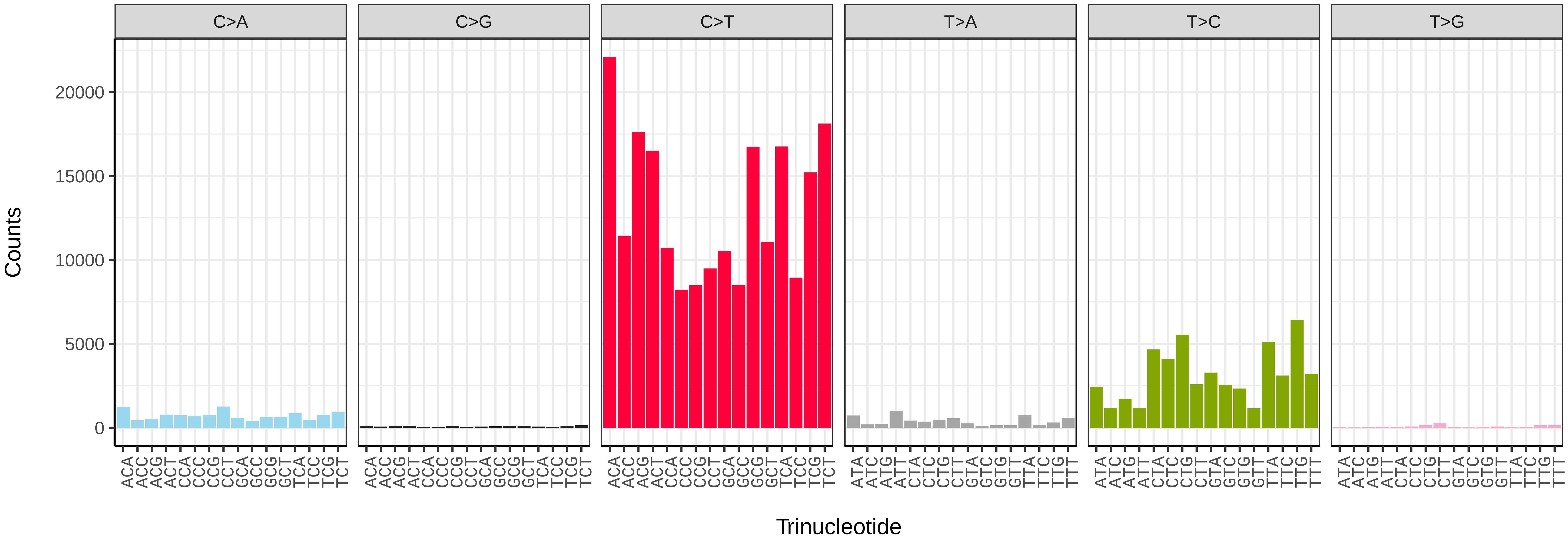


*Synanthedon andrenaeformis* (ilSynAndr1) somatic mutational spectrum (SBS52 classification)

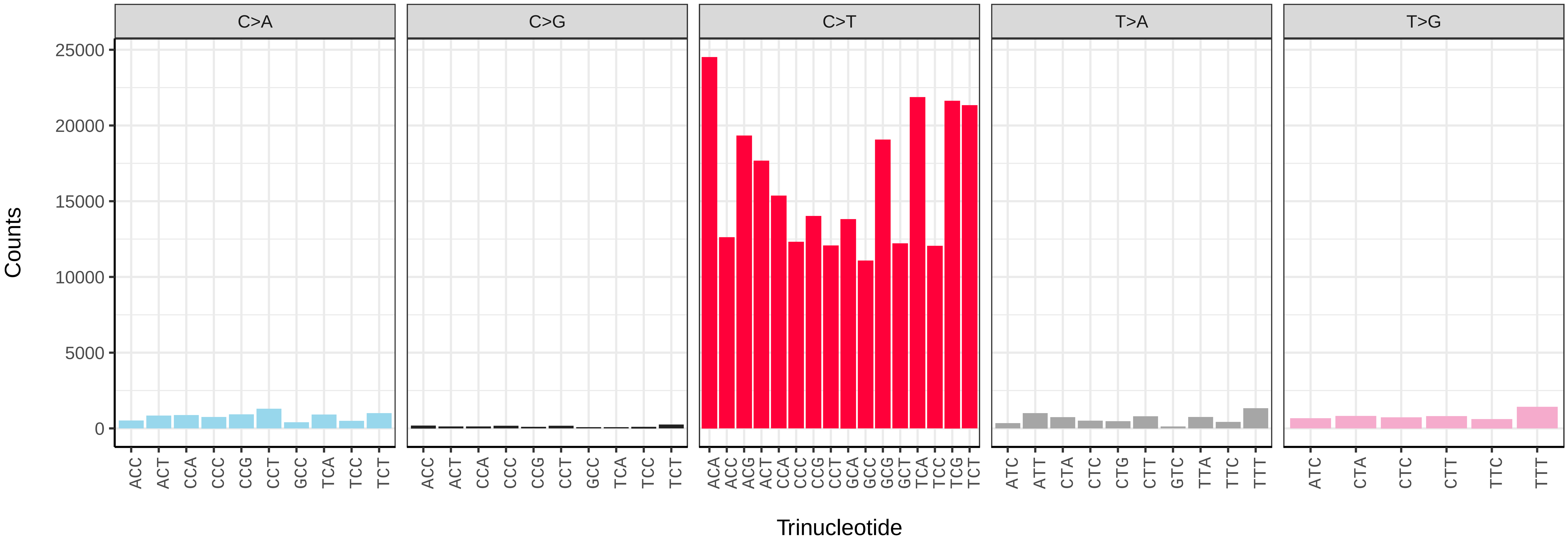

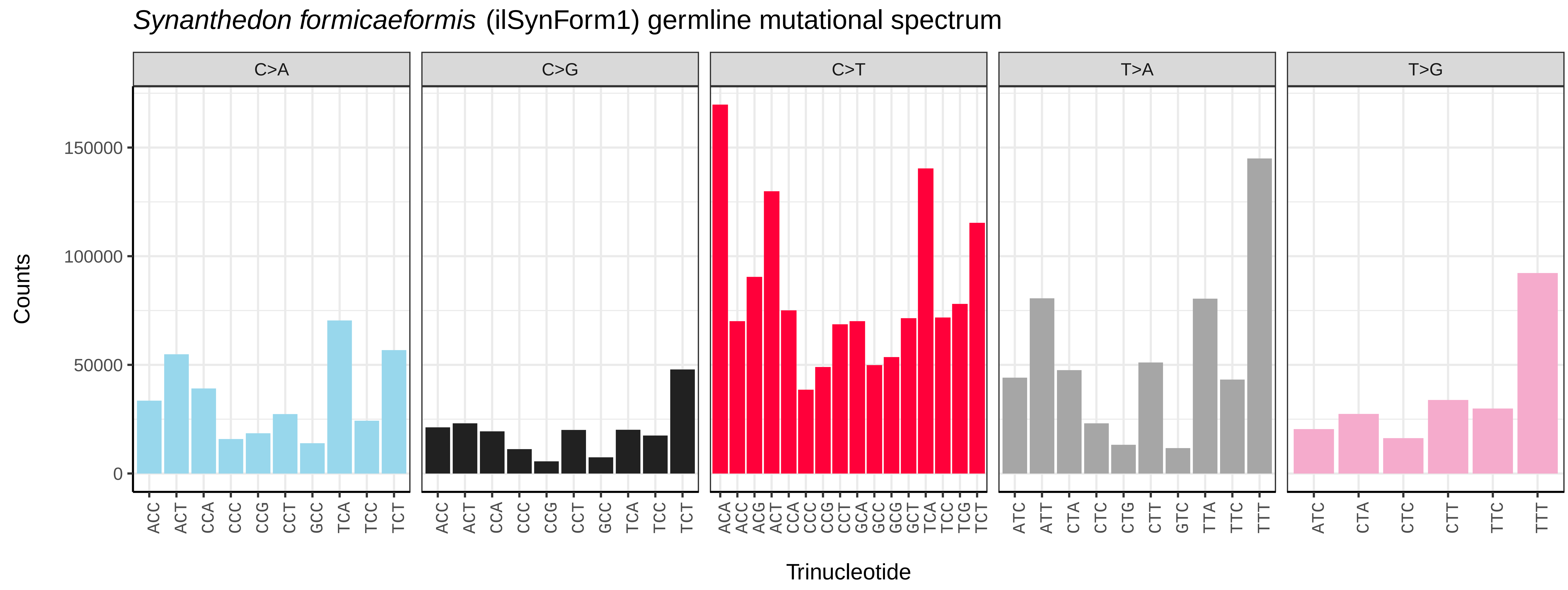
Synanthedon formicaeformis (ilSynForm1) germline mutational spectrum
C>A
C>G
C>T
T>A
T>G
Counts
0
50000
100000
150000
Trinucleotide

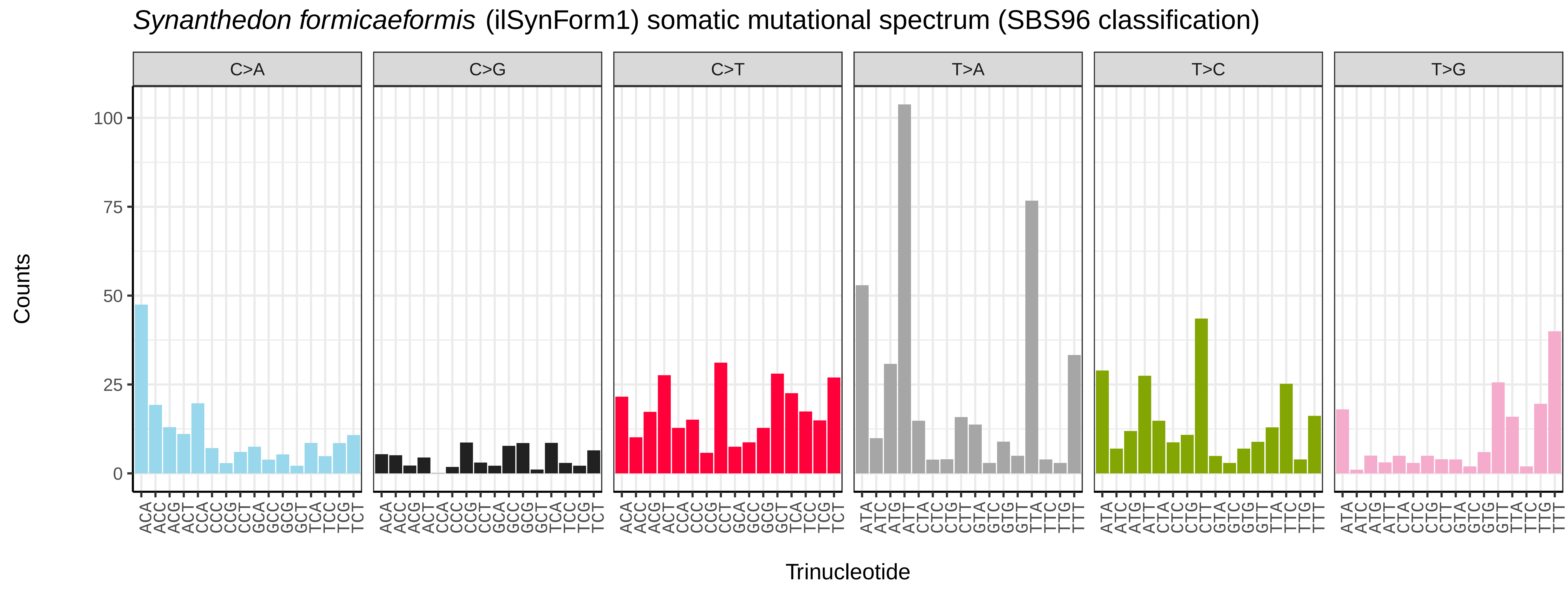
Synanthedon formicaeformis (ilSynForm1) somatic mutational spectrum (SBS96 classification)
C>A
C>G
C>T
T>A
T>C
T>G
Counts
0
25
50
75
100
Trinucleotide

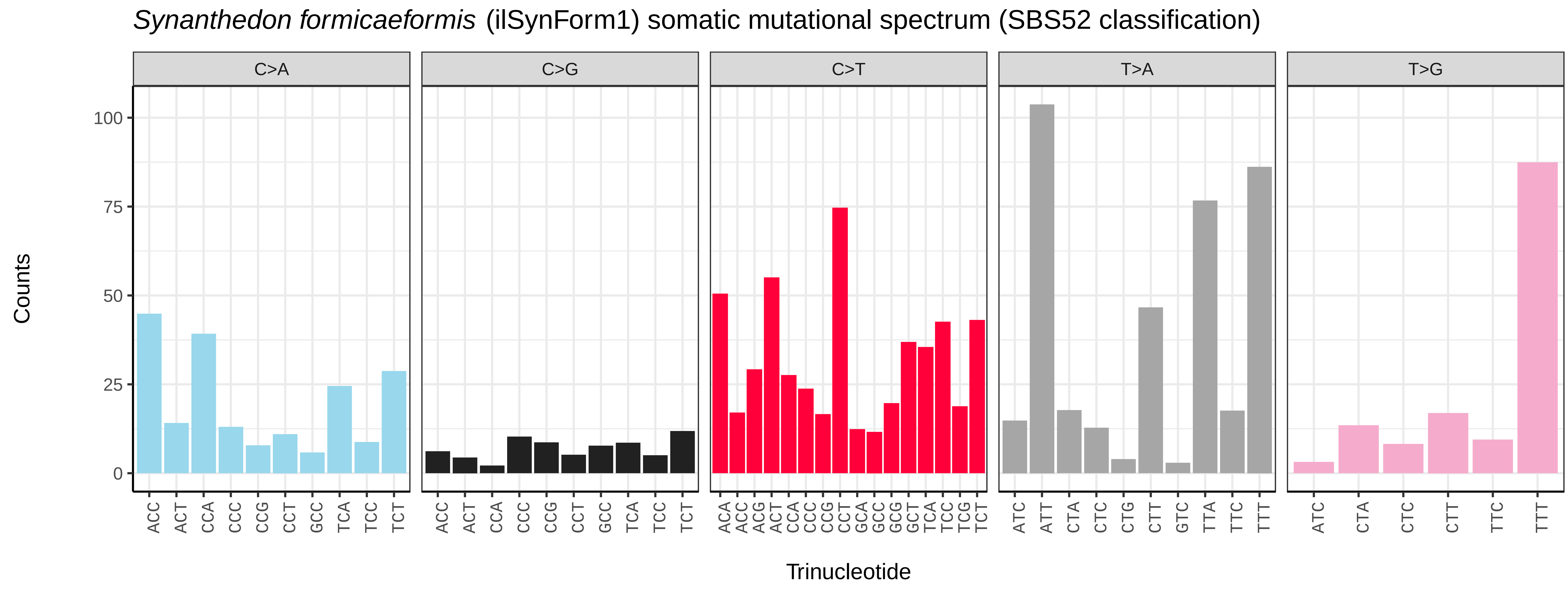
Synanthedon formicaeformis (ilSynForm1) somatic mutational spectrum (SBS52 classification)
C>A
C>G
C>T
T>A
T>G
Counts
0
25
50
75
100
Trinucleotide

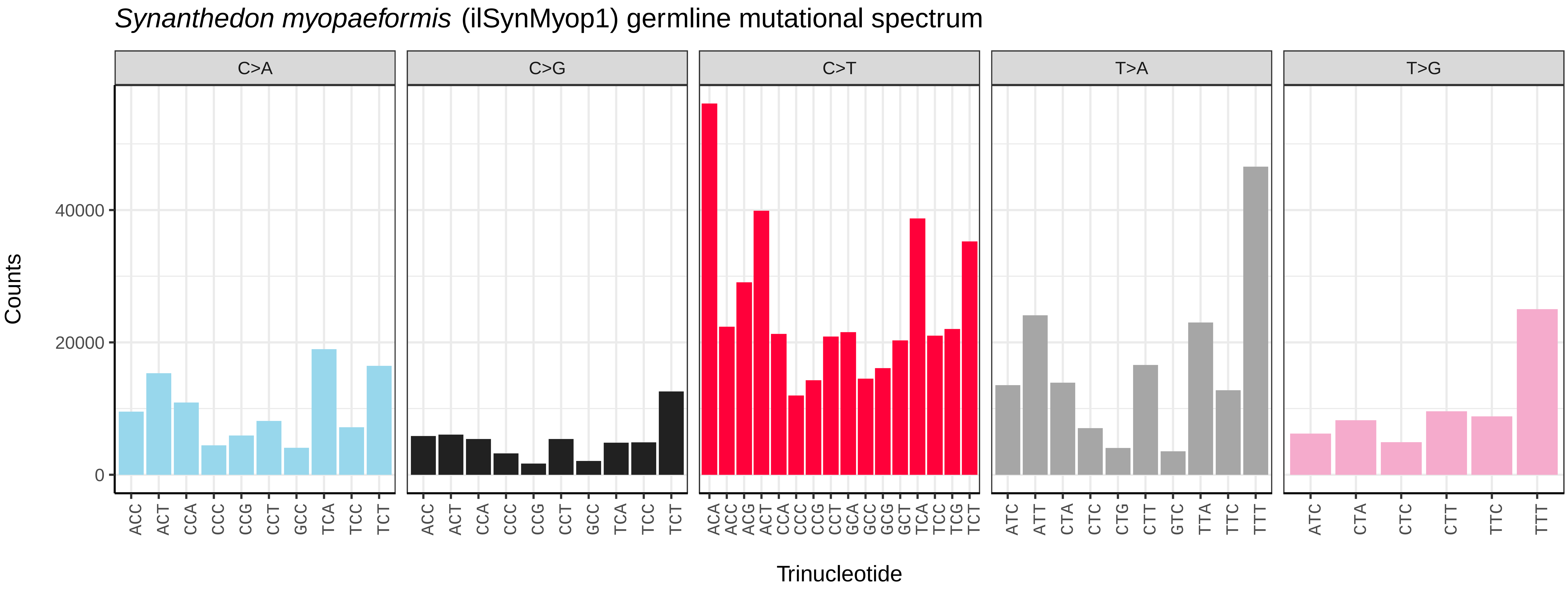
*Synanthedon myopaeformis* (ilSynMyop1) germline mutational spectrum
C>A
C>G
C>T
T>A
T>G
Counts
0
20000
40000
ACC ACT CCA CCC CCG CCT GCC TCA TCC TCT
ACA ACC ACG ACT CCA CCC CCG CCT GCA GCC GCG GCT TCA TCC TCG TCT
ATC ATT CTA CTC CTG CTT GTC TTA TTC TTT
ATC CTA CTC CTT TTC TTT
Trinucleotide

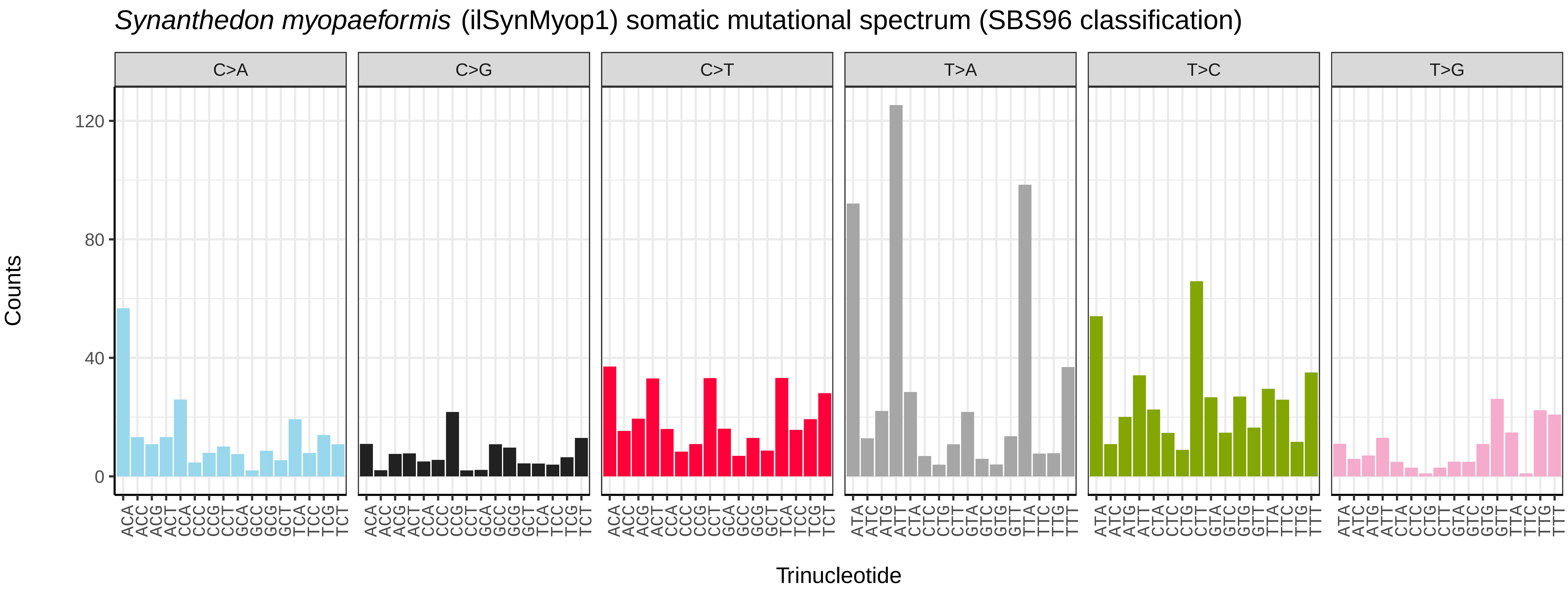
*Synanthedon myopaeformis* (ilSynMyop1) somatic mutational spectrum (SBS96 classification)
C>A
C>G
C>T
T>A
T>C
T>G
Counts
0
40
80
120
ACA ACC ACG ACT CCA CCC CCG CCT GCA GCC GCG GCT TCA TCC TCG TCT
ATA ATC ATG ATT CTA CTC CTG CTT GTA GTC GTG GTT TTA TTC TTG TTT
Trinucleotide

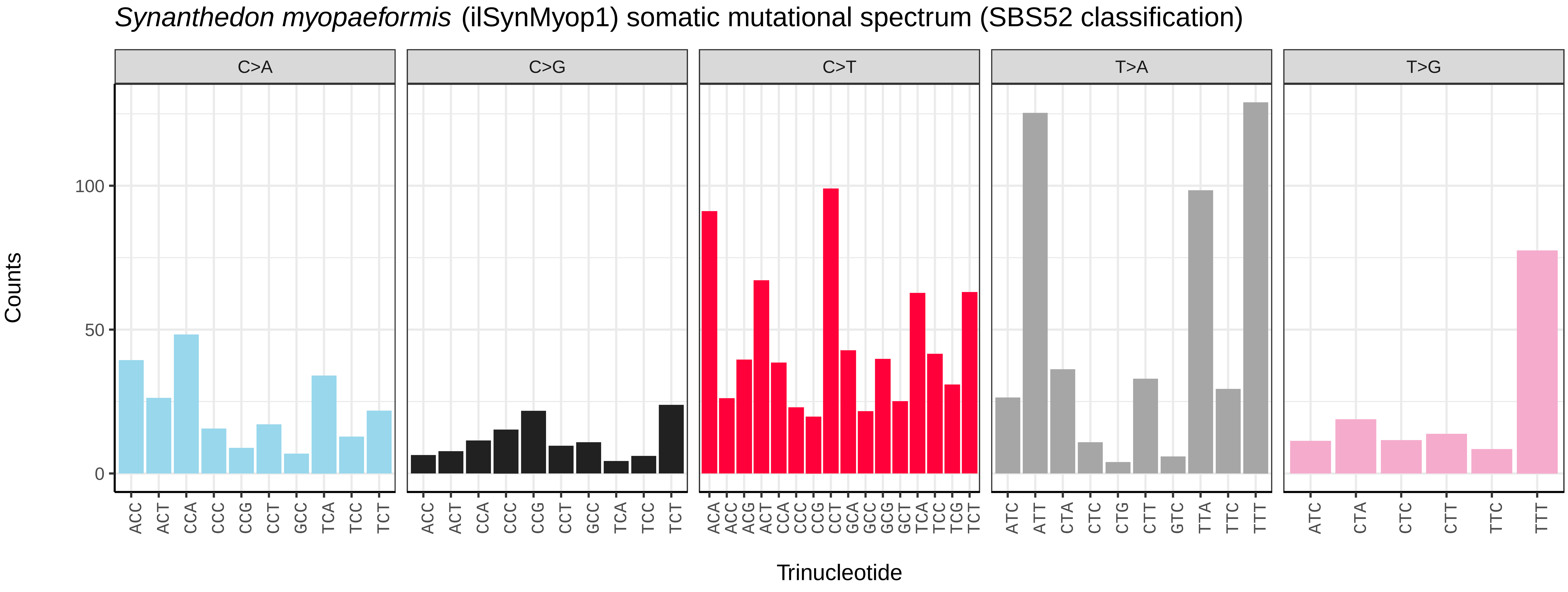
*Synanthedon myopaeformis* (ilSynMyop1) somatic mutational spectrum (SBS52 classification)
C>A
C>G
C>T
T>A
T>G
Counts
0
50
100
ACC ACT CCA CCC CCG CCT GCC TCA TCC TCT
ACA ACC ACG ACT CCA CCC CCG CCT GCA GCC GCG GCT TCA TCC TCG TCT
ATC ATT CTA CTC CTG CTT GTC TTA TTC TTT
ATC CTA CTC CTT TTC TTT
Trinucleotide

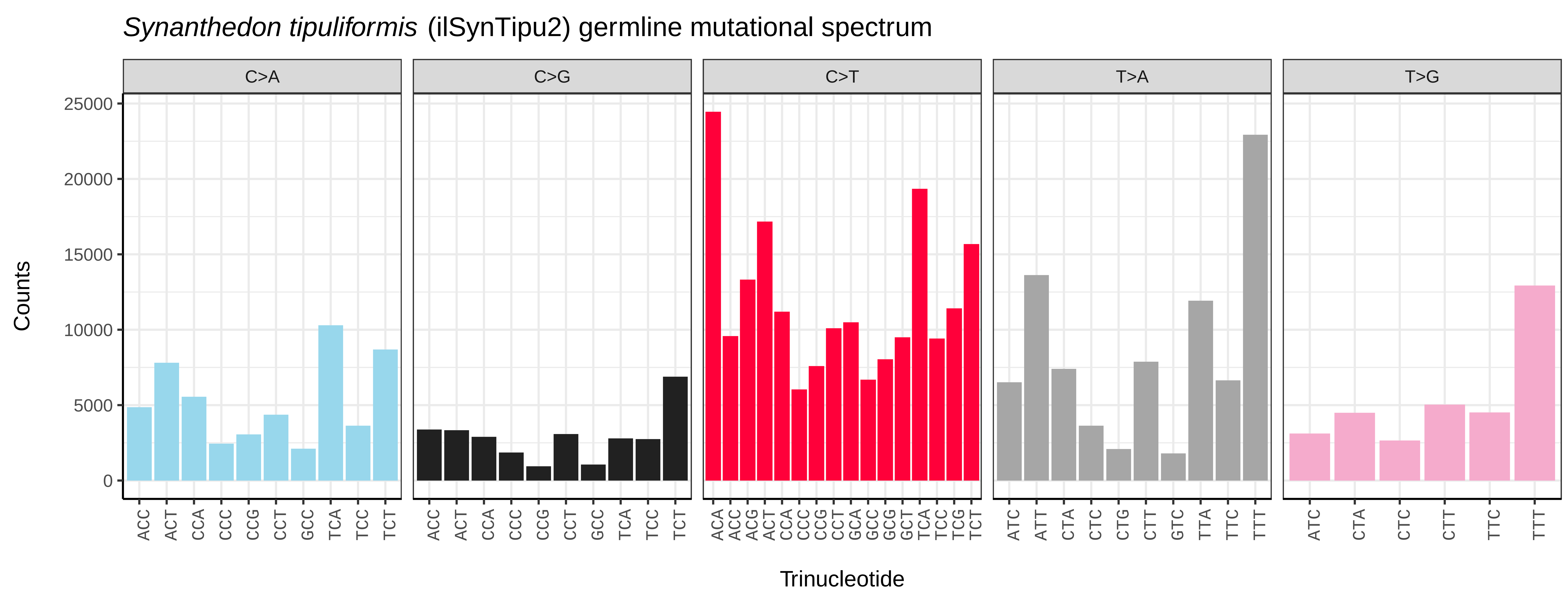
*Synanthedon tipuliformis* (ilSynTipu2) germline mutational spectrum
C>A
C>G
C>T
T>A
T>G
Counts
25000
20000
15000
10000
5000
0
ACC ACT CCA CCC CCG CCT GCC TCA TCC TCT
ACA ACC ACG ACT CCA CCC CCG CCT GCA GCC GCG GCT TCA TCC TCG TCT
ATC ATT CTA CTC CTG CTT GTC TTA TTC TTT
ATC CTA CTC CTT TTC TTT
Trinucleotide

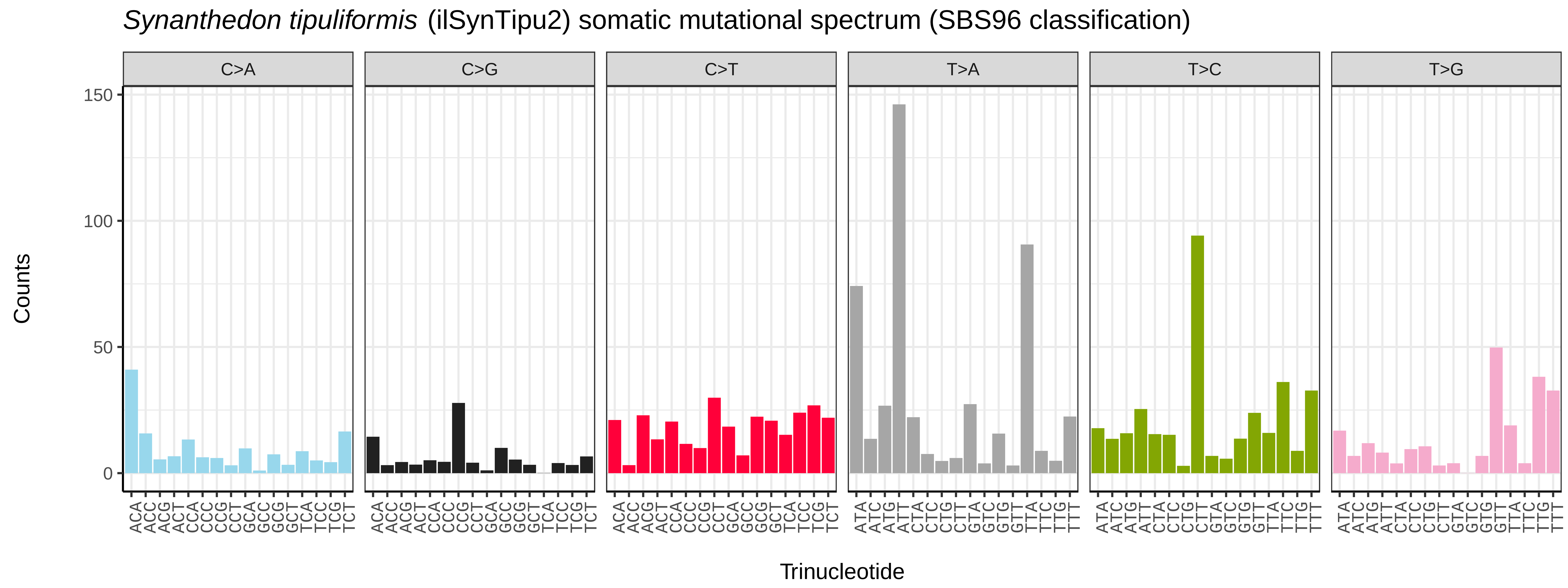
*Synanthedon tipuliformis* (ilSynTipu2) somatic mutational spectrum (SBS96 classification)
C>A
C>G
C>T
T>A
T>C
T>G
Counts
150
100
50
0
ACA ACC ACG ACT CCA CCC CCG CCT GCA GCC GCG GCT TCA TCC TCG TCT
ATA ATC ATG ATT CTA CTC CTG CTT GTA GTC GTG GTT TTA TTC TTG TTT
Trinucleotide

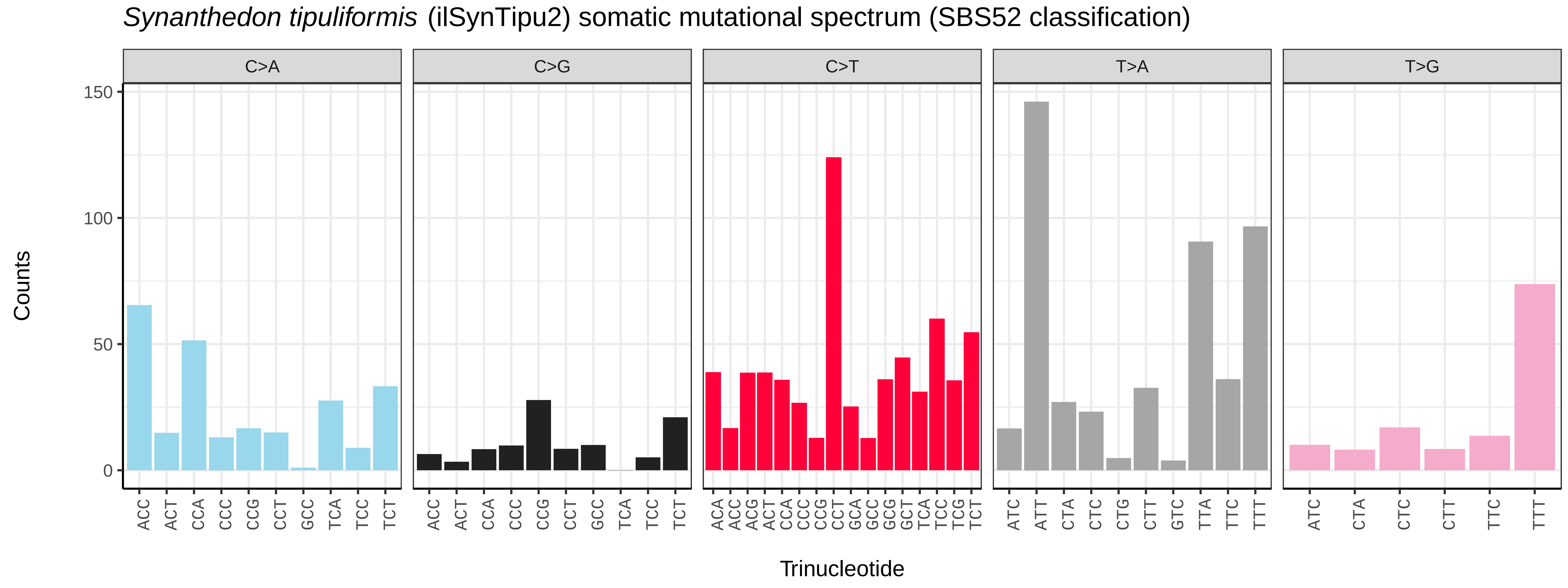
*Synanthedon tipuliformis* (ilSynTipu2) somatic mutational spectrum (SBS52 classification)
C>A
C>G
C>T
T>A
T>G
Counts
150
100
50
0
ACC ACT CCA CCC CCG CCT GCC TCA TCC TCT
ACA ACC ACG ACT CCA CCC CCG CCT GCA GCC GCG GCT TCA TCC TCG TCT
ATC ATT CTA CTC CTG CTT GTC TTA TTC TTT
ATC CTA CTC CTT TTC TTT
Trinucleotide

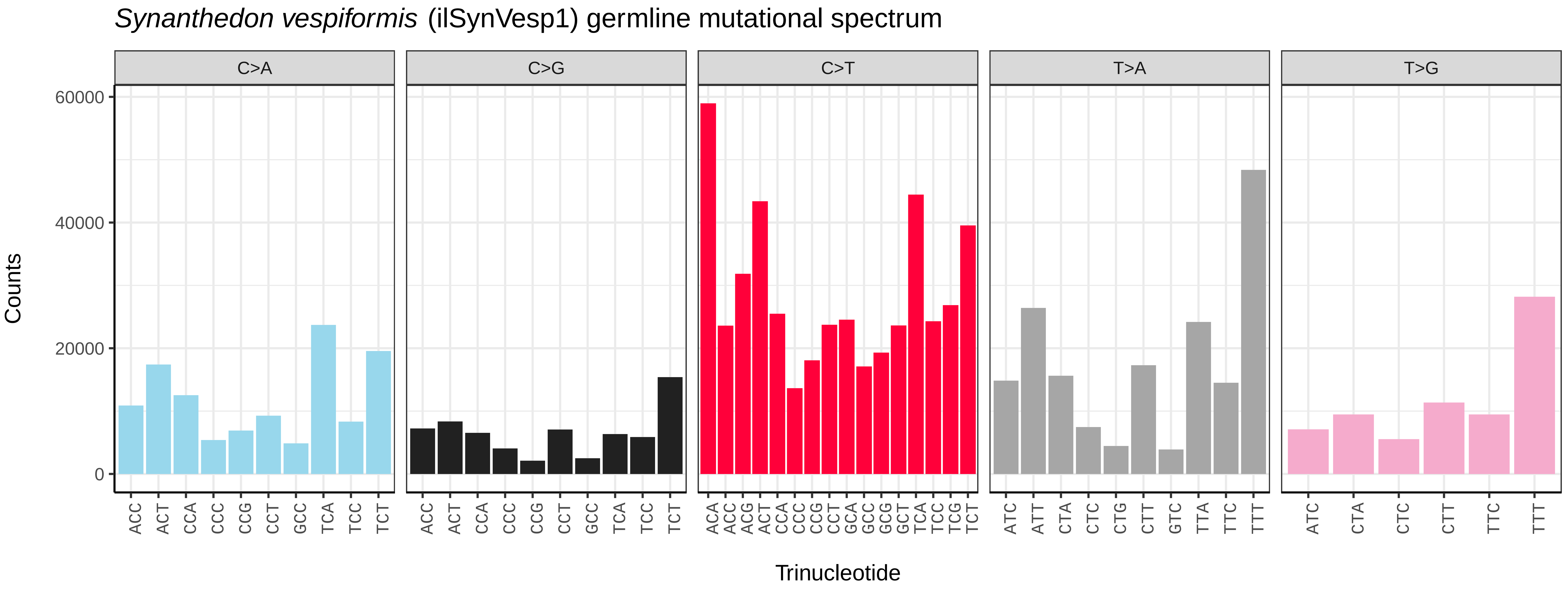
Synanthedon vespiformis (ilSynVesp1) germline mutational spectrum
C>A
C>G
C>T
T>A
T>G
Counts
0
20000
40000
60000
Trinucleotide

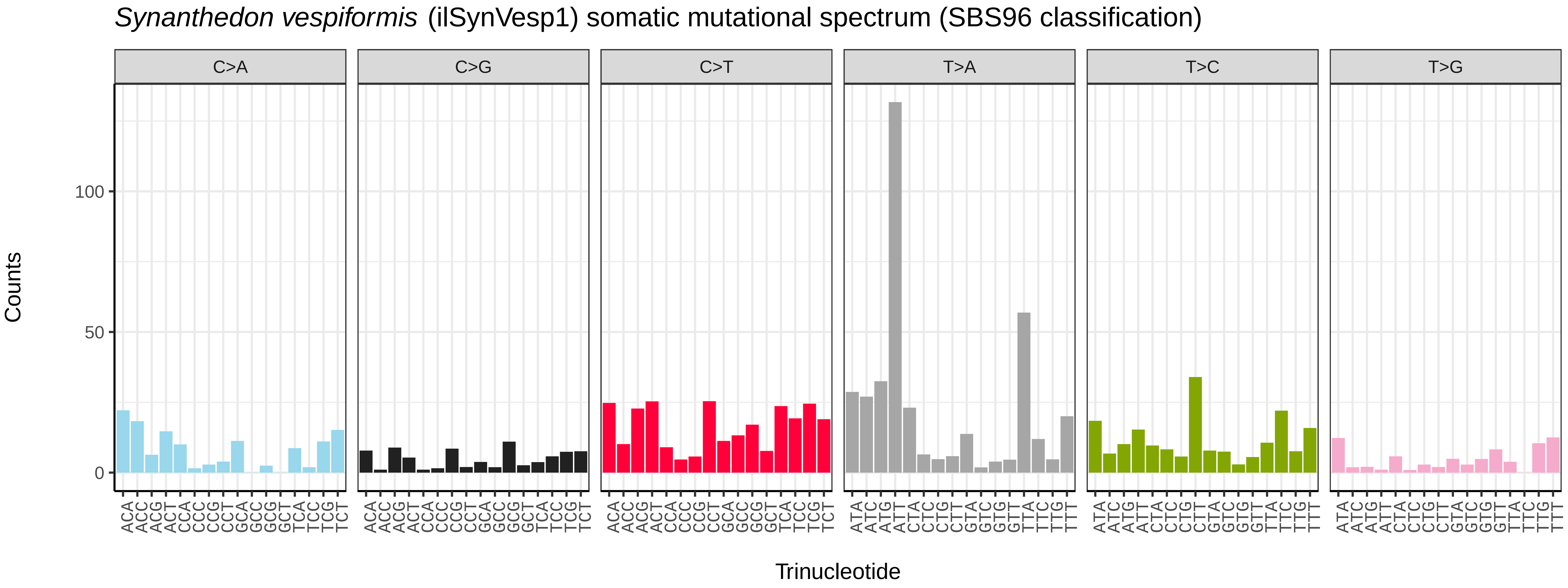
Synanthedon vespiformis (ilSynVesp1) somatic mutational spectrum (SBS96 classification)
C>A
C>G
C>T
T>A
T>C
T>G
Counts
0
50
100
Trinucleotide

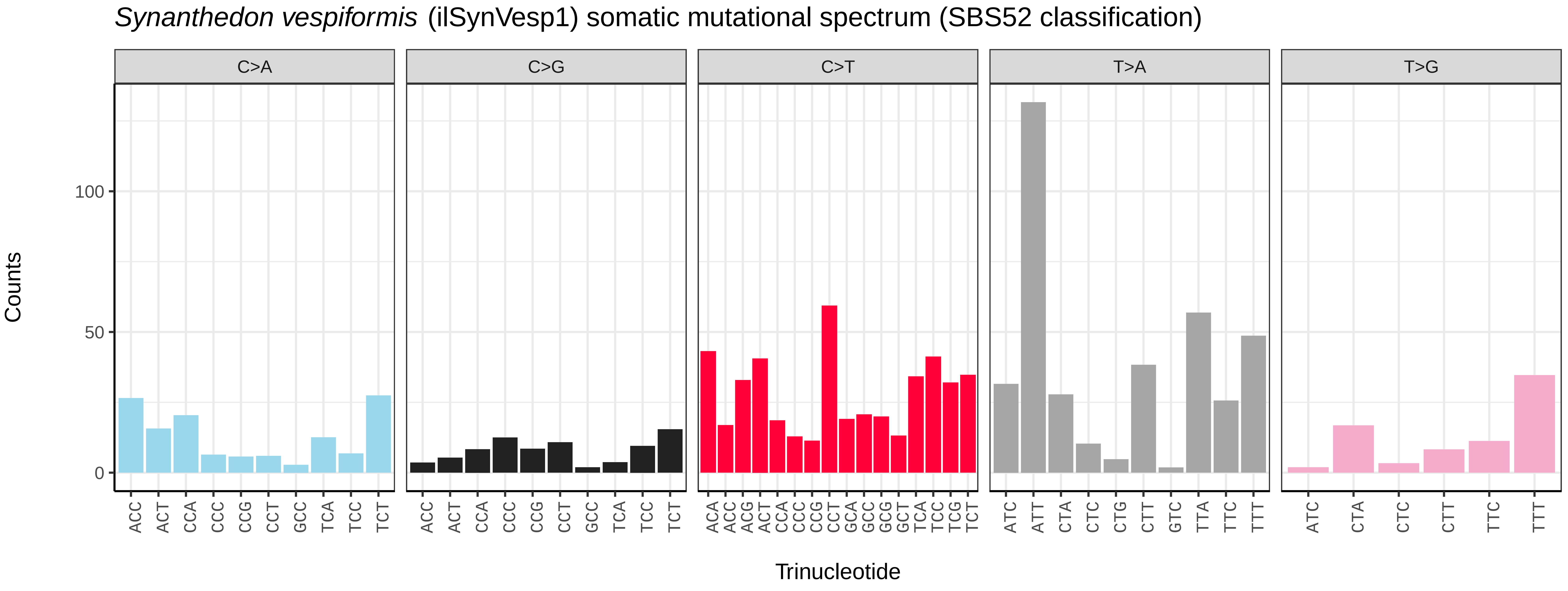
Synanthedon vespiformis (ilSynVesp1) somatic mutational spectrum (SBS52 classification)
C>A
C>G
C>T
T>A
T>G
Counts
0
50
100
Trinucleotide

*Teleiodes luculella* (ilTelLucu1) germline mutational spectrum

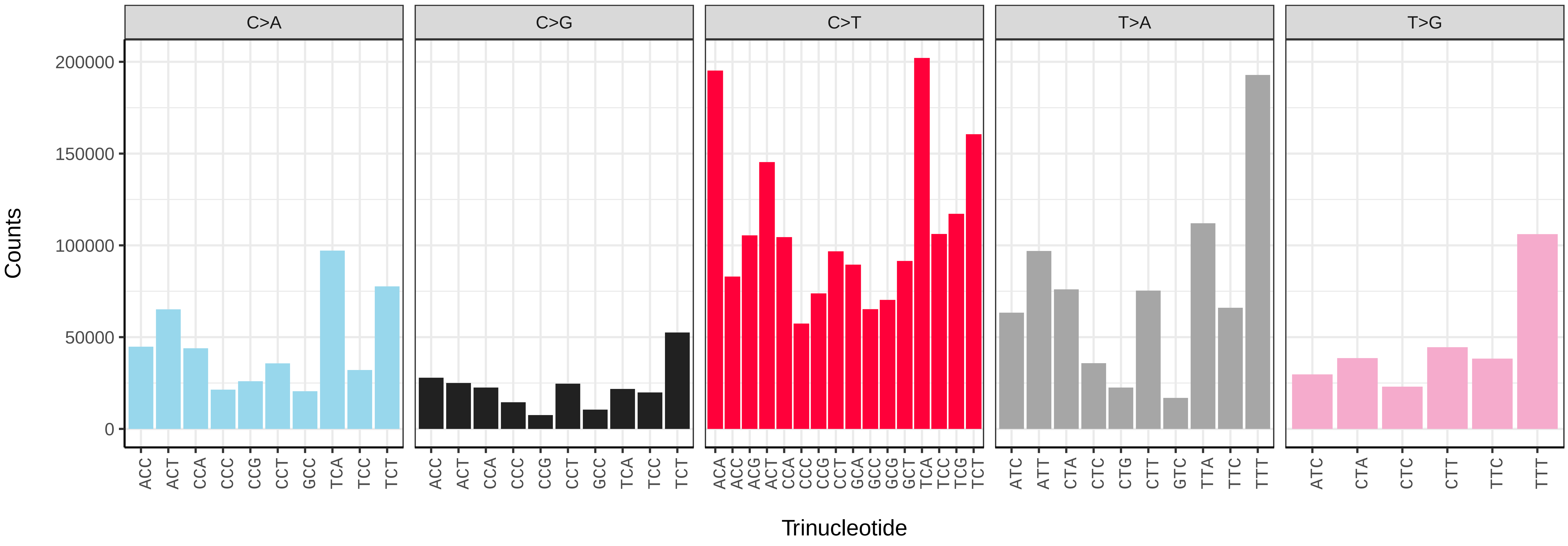


*Teleiodes luculella* (ilTelLucu1) somatic mutational spectrum (SBS96 classification)

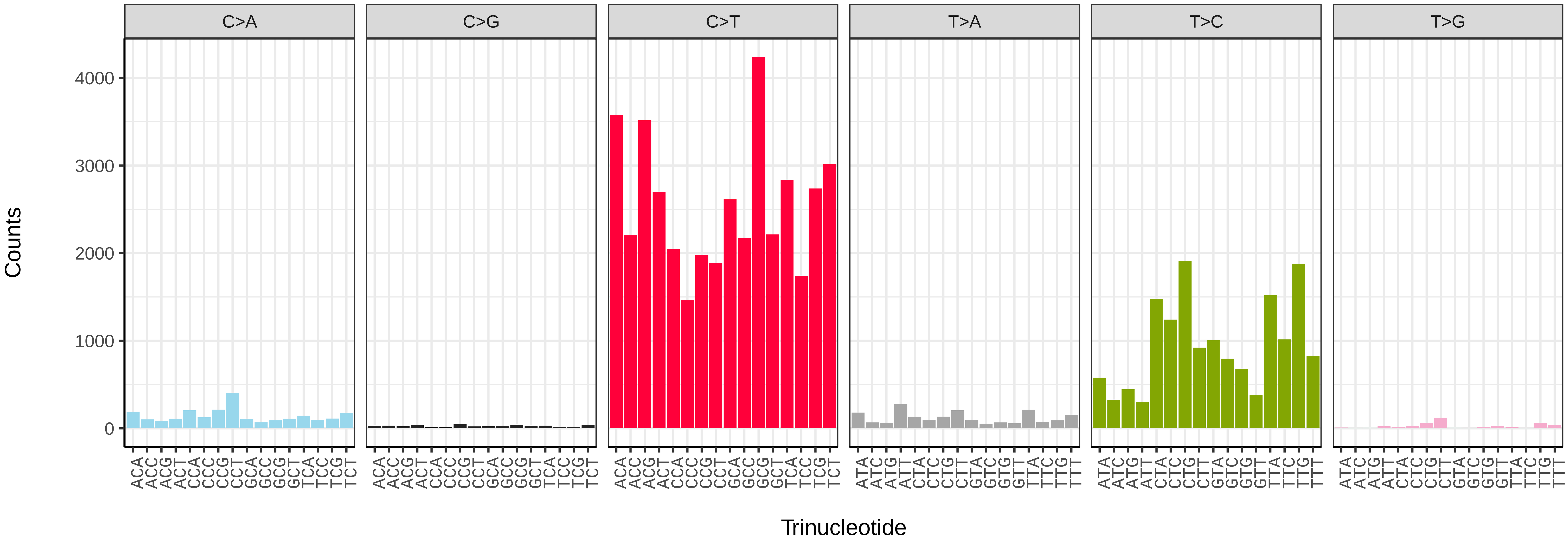


*Teleiodes luculella* (ilTelLucu1) somatic mutational spectrum (SBS52 classification)

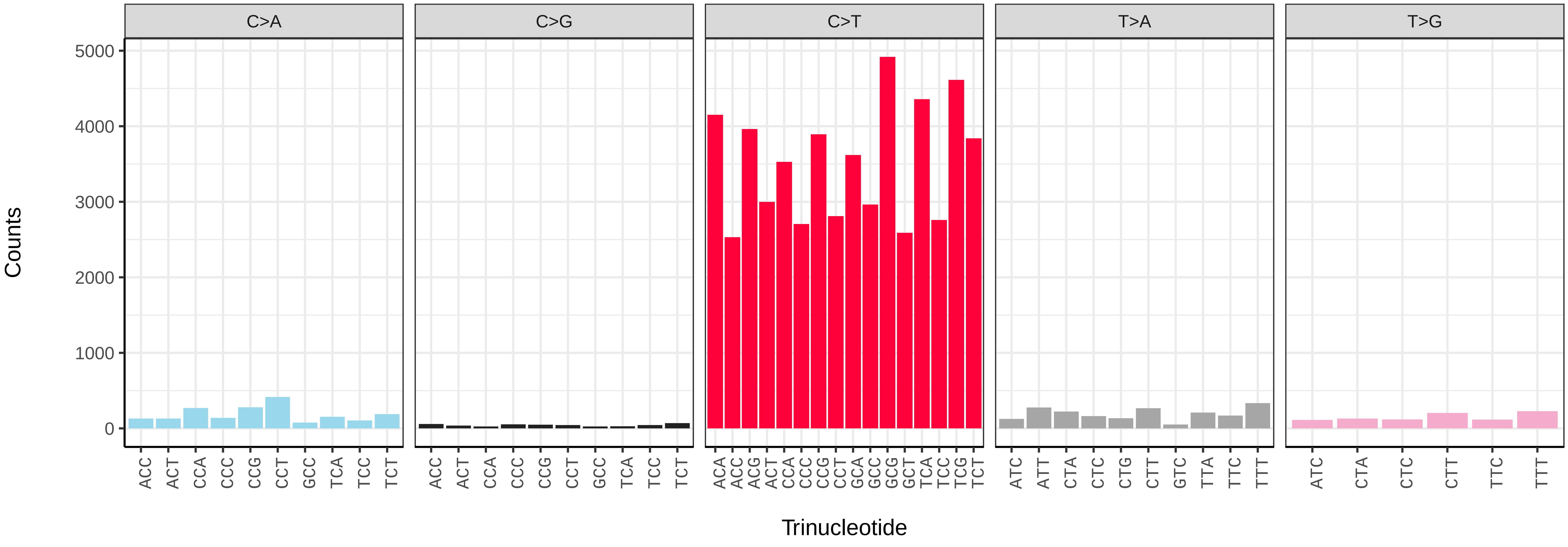

*Thalpophila matura* (ilThaMatu1) germline mutational spectrum

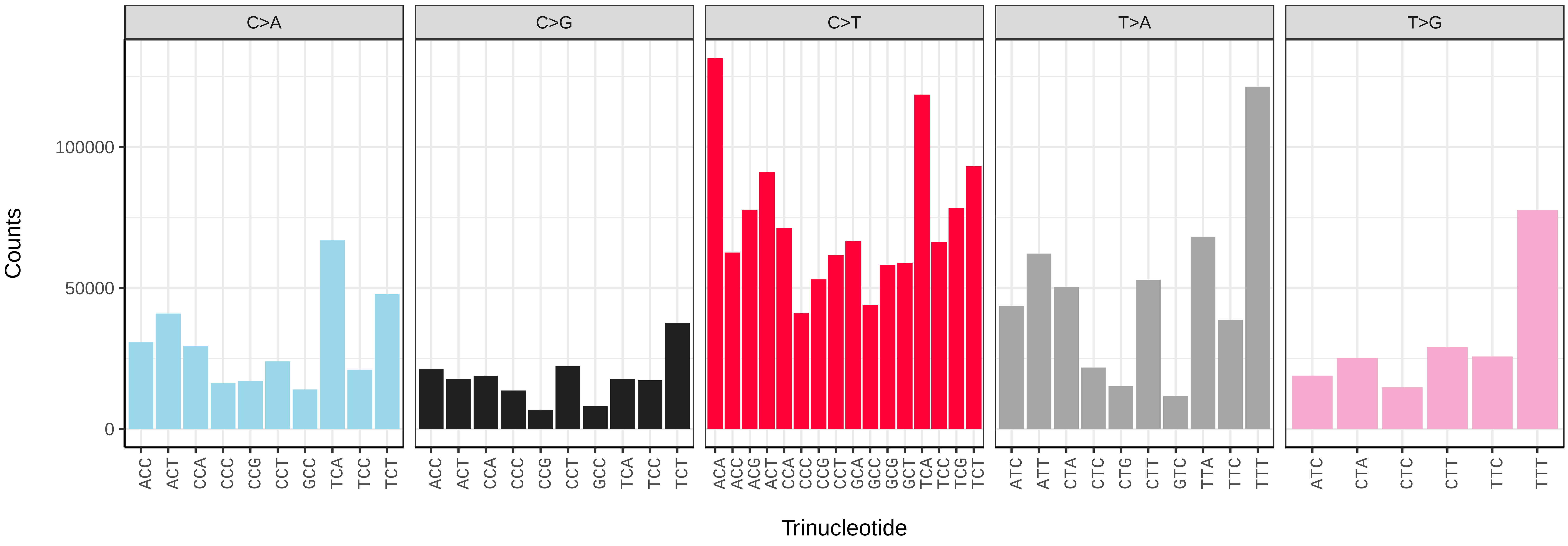


*Thalpophila matura* (ilThaMatu1) somatic mutational spectrum (SBS96 classification)

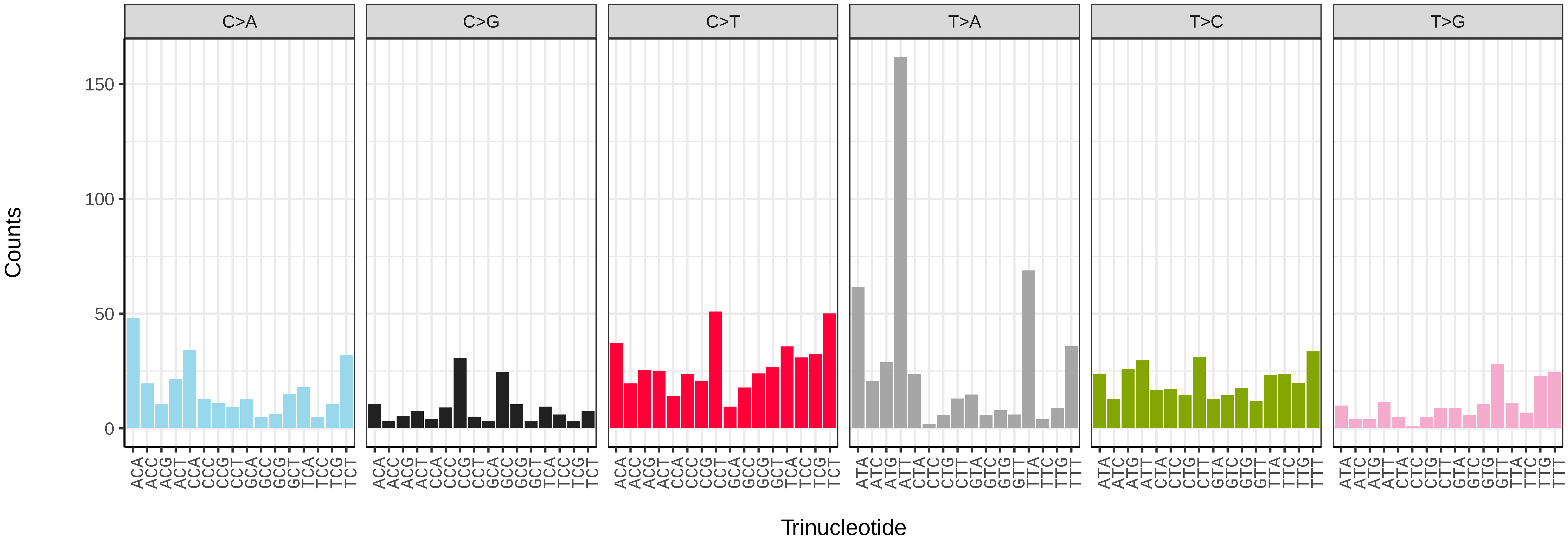


*Thalpophila matura* (ilThaMatu1) somatic mutational spectrum (SBS52 classification)

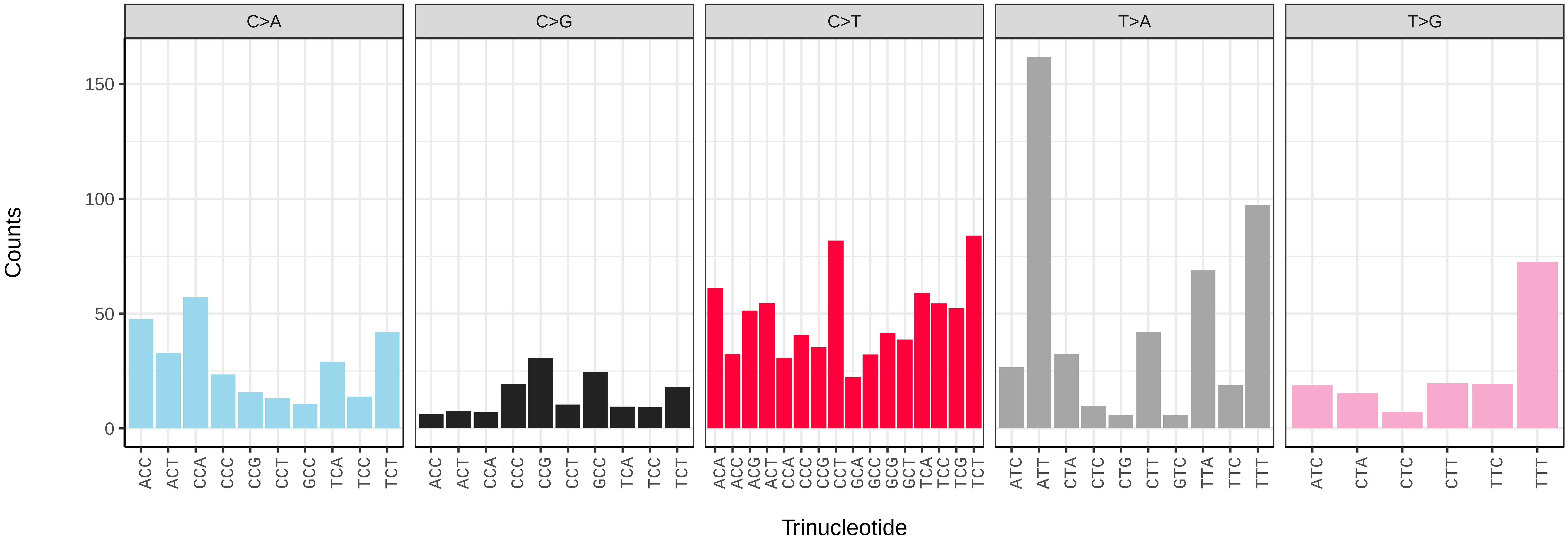

*Thera britannica* (ilTheBrit1) germline mutational spectrum

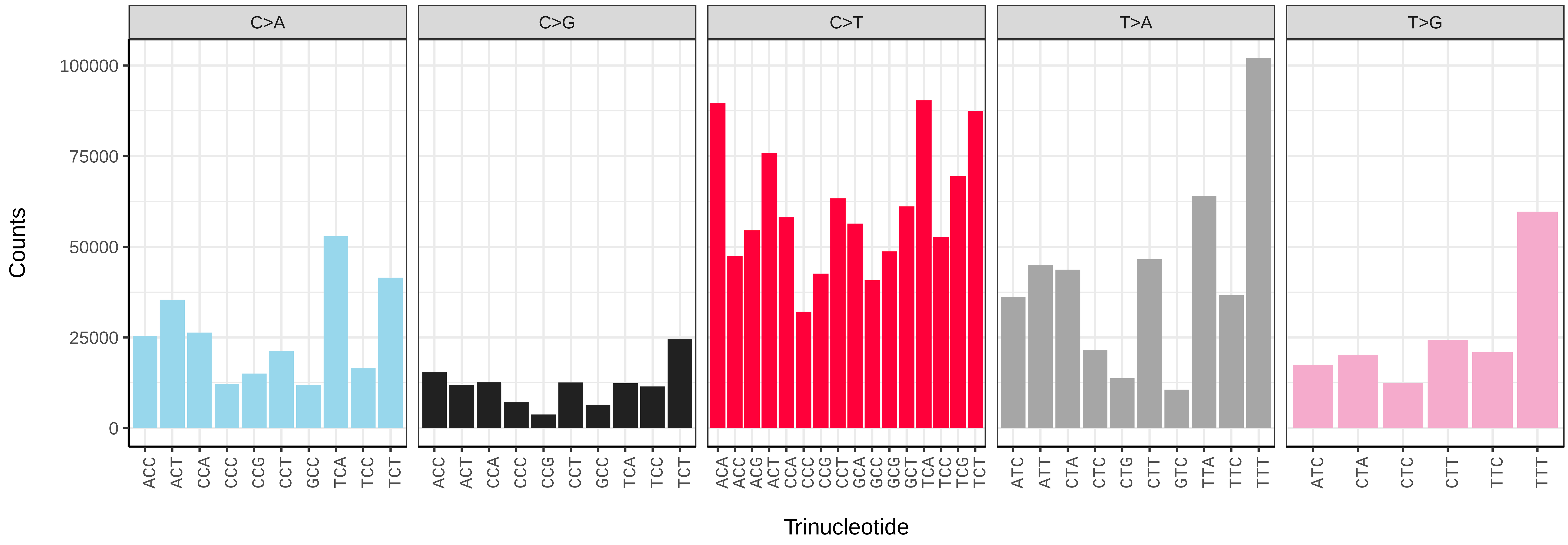


*Thera britannica* (ilTheBrit1) somatic mutational spectrum (SBS96 classification)

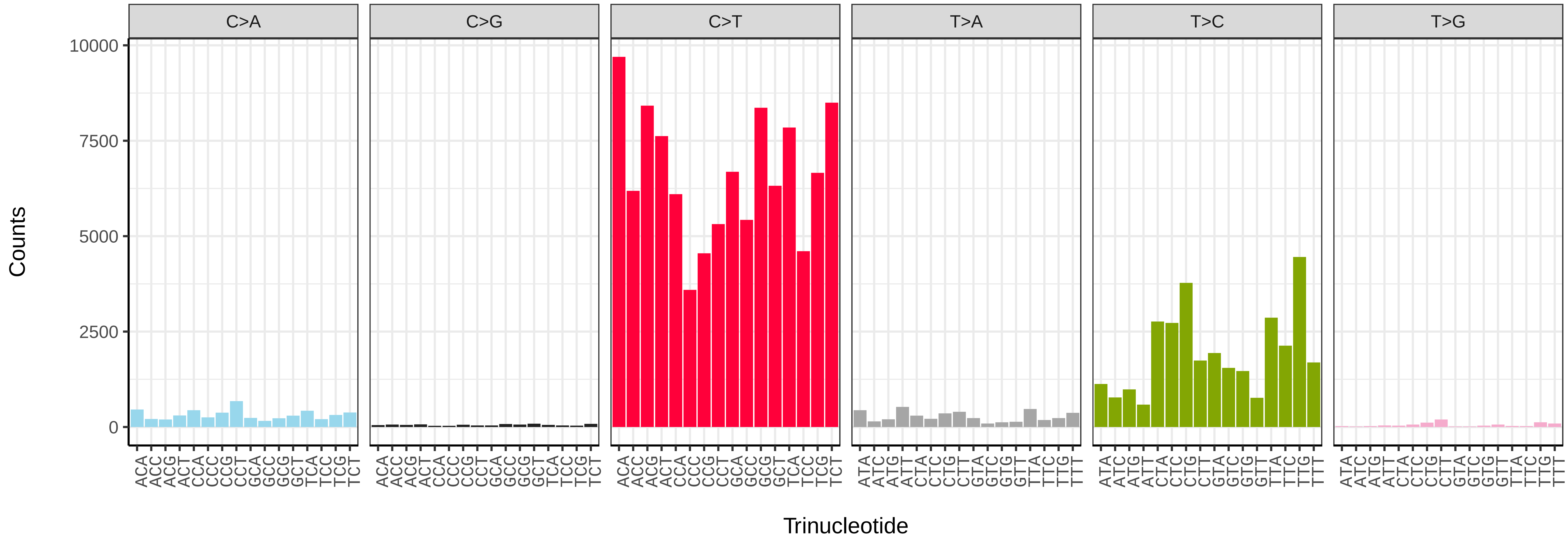


*Thera britannica* (ilTheBrit1) somatic mutational spectrum (SBS52 classification)

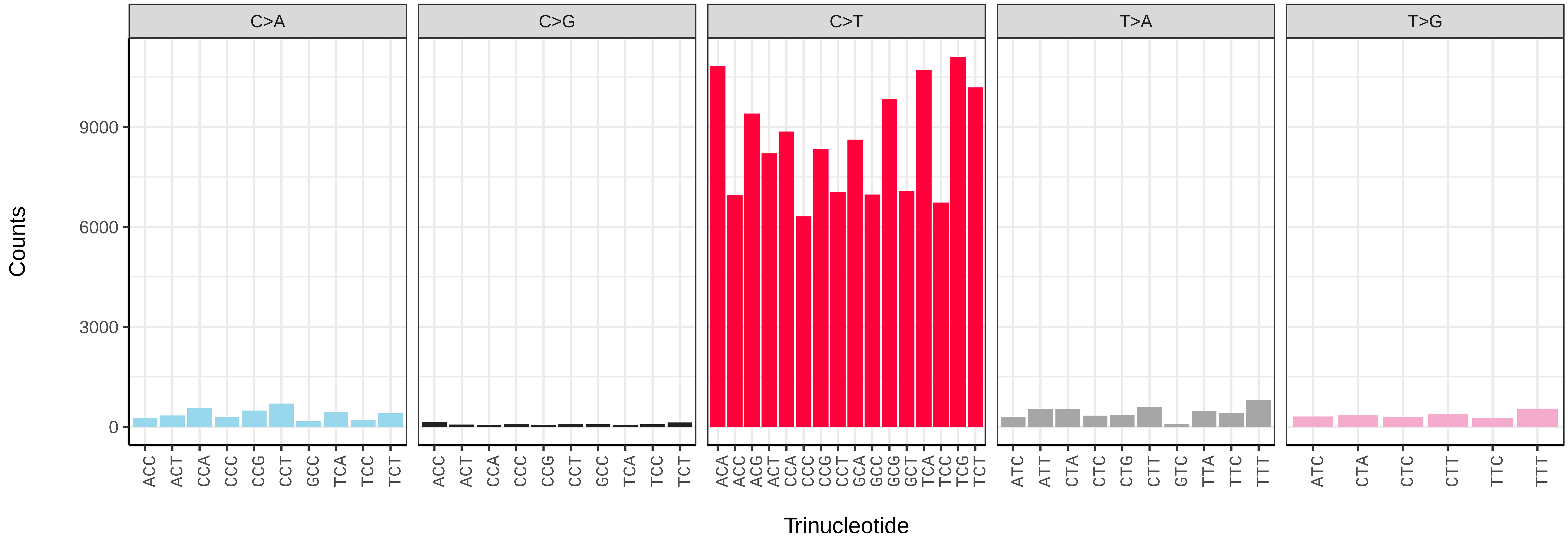

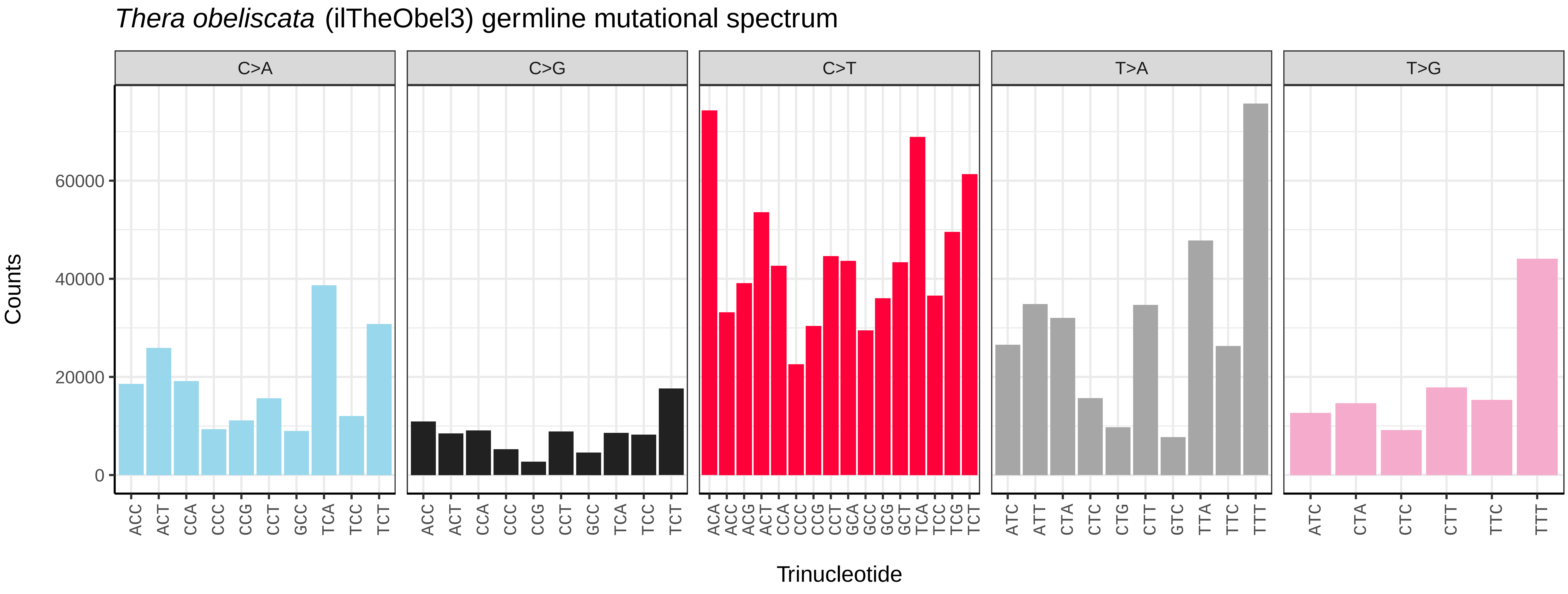
*Thera obeliscata* (ilTheObel3) germline mutational spectrum
C>A
C>G
C>T
T>A
T>G
Counts
0
20000
40000
60000
ACC ACT CCA CCC CCG CCT GCC TCA TCC TCT
ACC ACT CCA CCC CCG CCT GCC TCA TCC TCT
ACA ACC ACG ACT CCA CCC CCG CCT GCA GCC GCG GCT TCA TCC TCG TCT
ATC ATT CTA CTC CTG CTT GTC TTA TTC TTT
ATC CTA CTC CTT TTC TTT
Trinucleotide

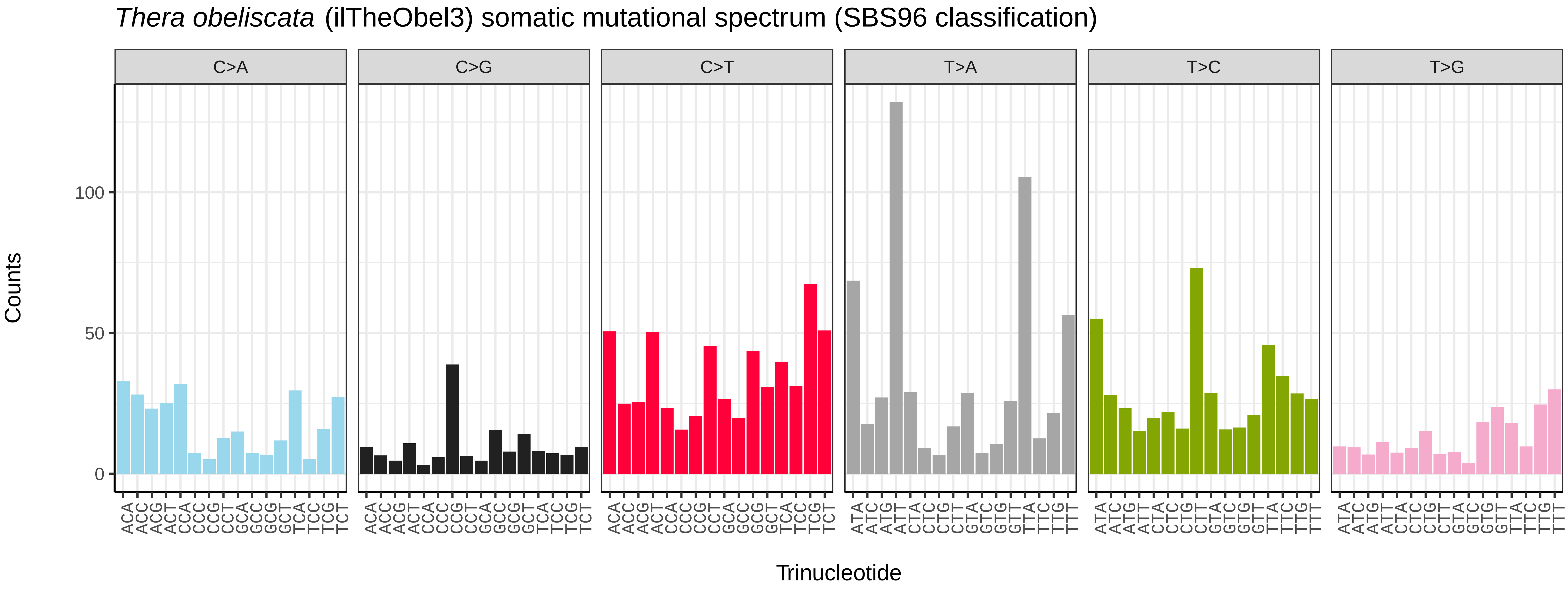
*Thera obeliscata* (ilTheObel3) somatic mutational spectrum (SBS96 classification)
C>A
C>G
C>T
T>A
T>C
T>G
Counts
0
50
100
ACA ACC ACG ACT CCA CCC CCG CCT GCA GCC GCG GCT TCA TCC TCG TCT
ATA ATC ATG ATT CTA CTC CTG CTT GTA GTC GTG GTT TTA TTC TTG TTT
Trinucleotide

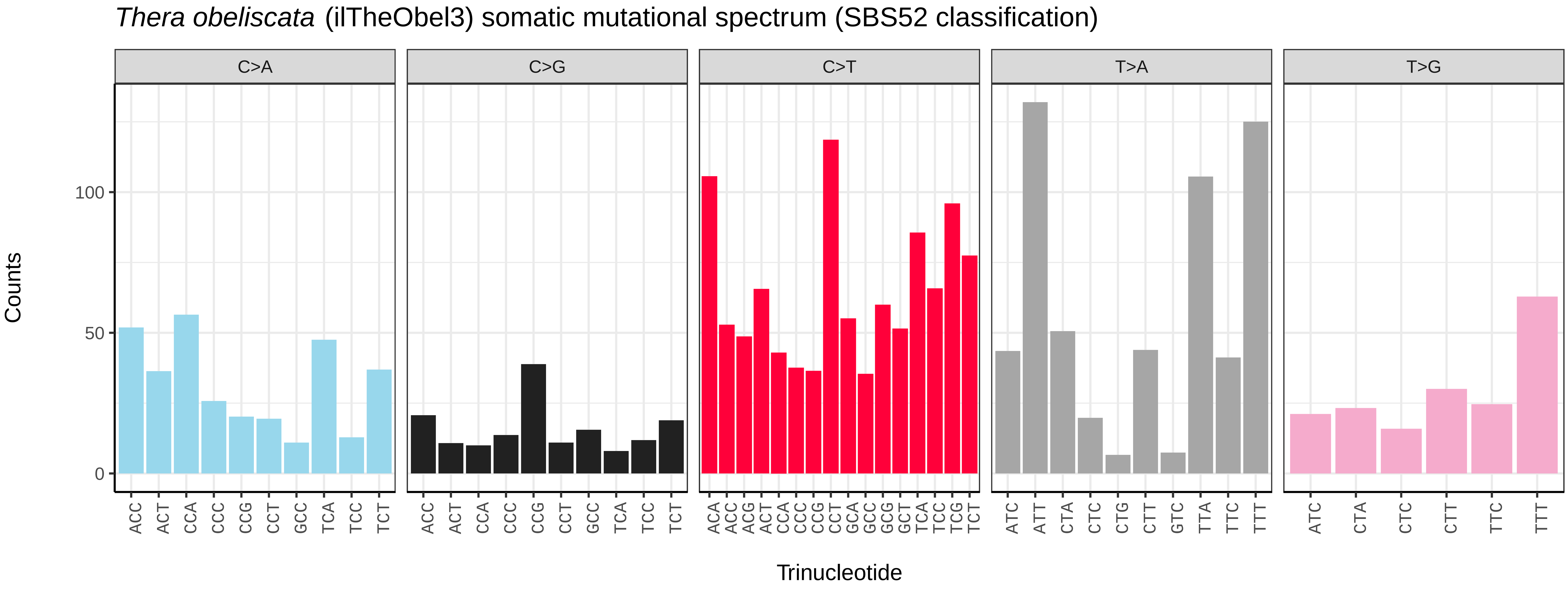
*Thera obeliscata* (ilTheObel3) somatic mutational spectrum (SBS52 classification)
C>A
C>G
C>T
T>A
T>G
Counts
0
50
100
ACC ACT CCA CCC CCG CCT GCC TCA TCC TCT
ACC ACT CCA CCC CCG CCT GCC TCA TCC TCT
ACA ACC ACG ACT CCA CCC CCG CCT GCA GCC GCG GCT TCA TCC TCG TCT
ATC ATT CTA CTC CTG CTT GTC TTA TTC TTT
ATC CTA CTC CTT TTC TTT
Trinucleotide

*Tholera decimalis* (ilThoDeci1) germline mutational spectrum

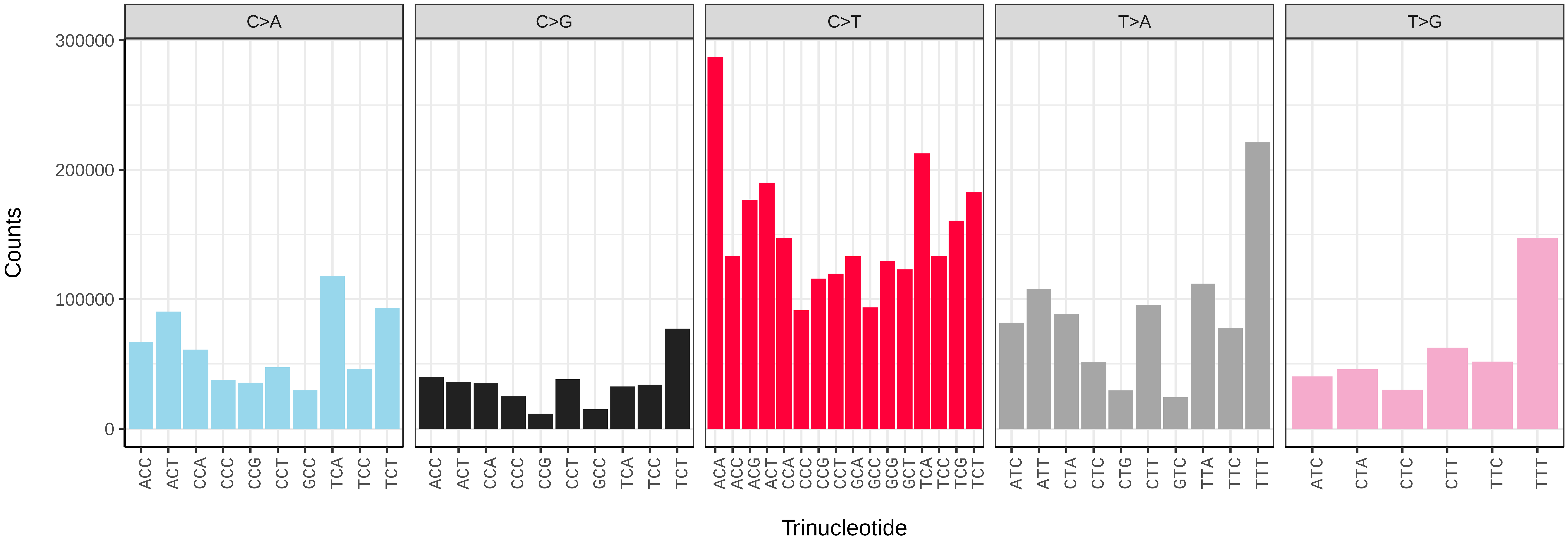


*Tholera decimalis* (ilThoDeci1) somatic mutational spectrum (SBS96 classification)

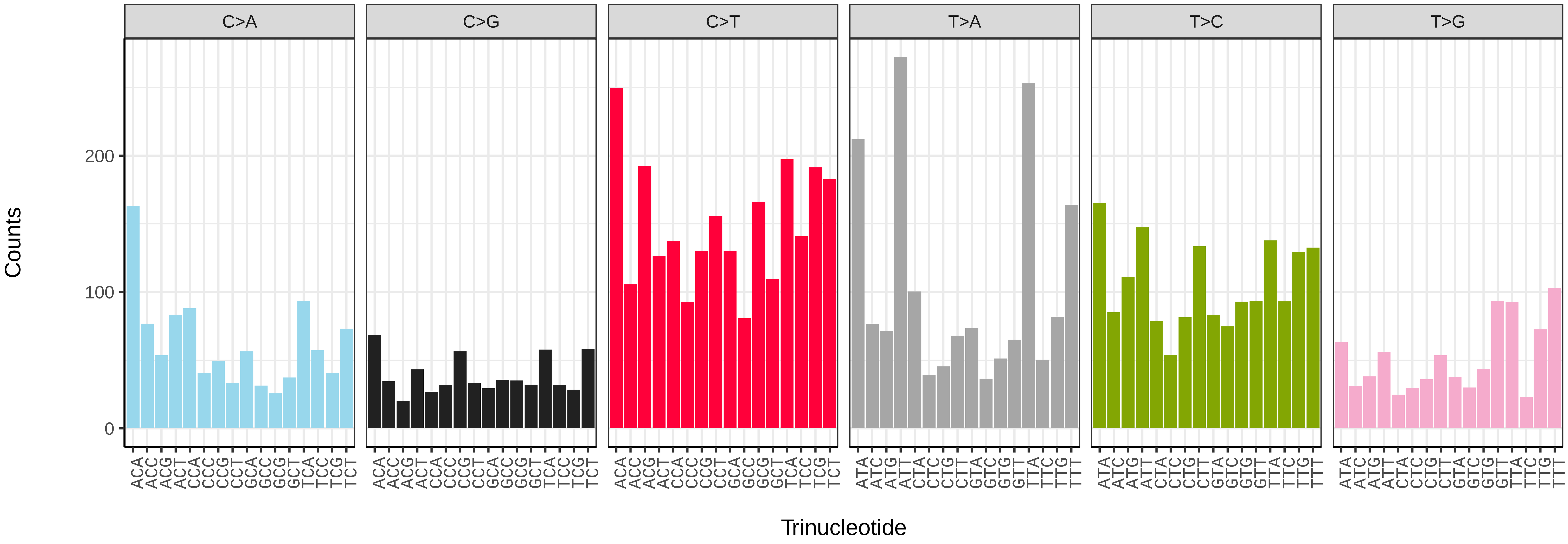


*Tholera decimalis* (ilThoDeci1) somatic mutational spectrum (SBS52 classification)

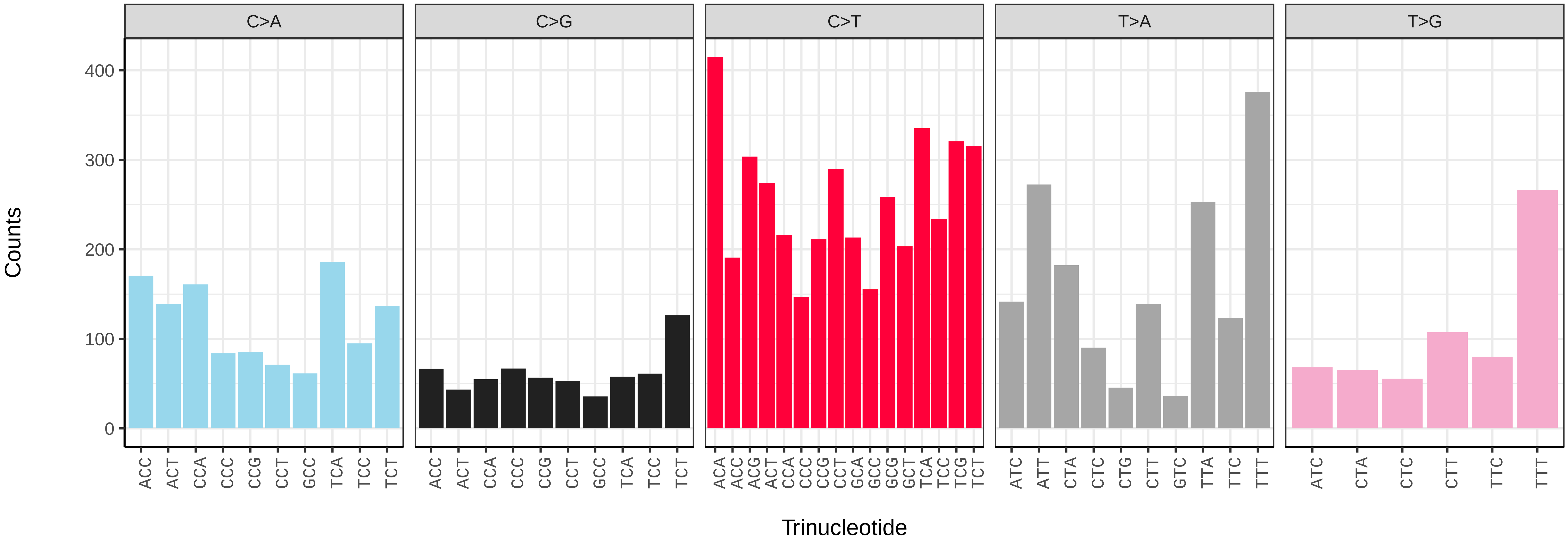

*Thumatha senex* (ilThuSene1) germline mutational spectrum

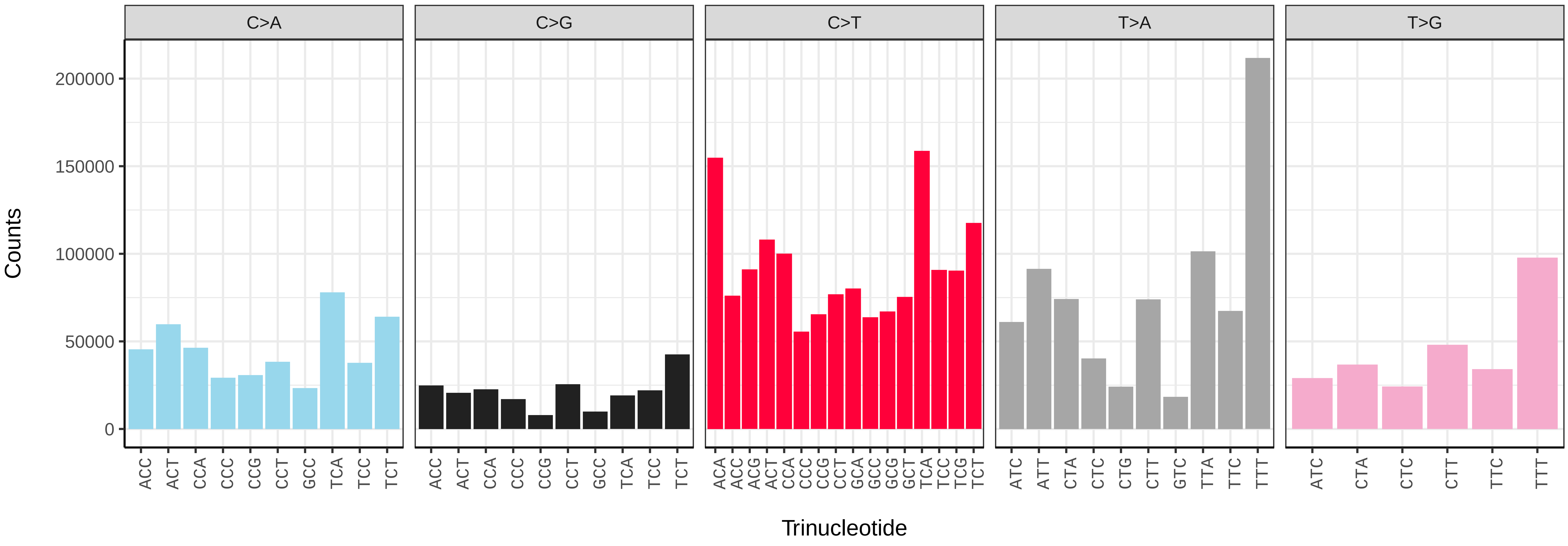


*Thumatha senex* (ilThuSene1) somatic mutational spectrum (SBS96 classification)

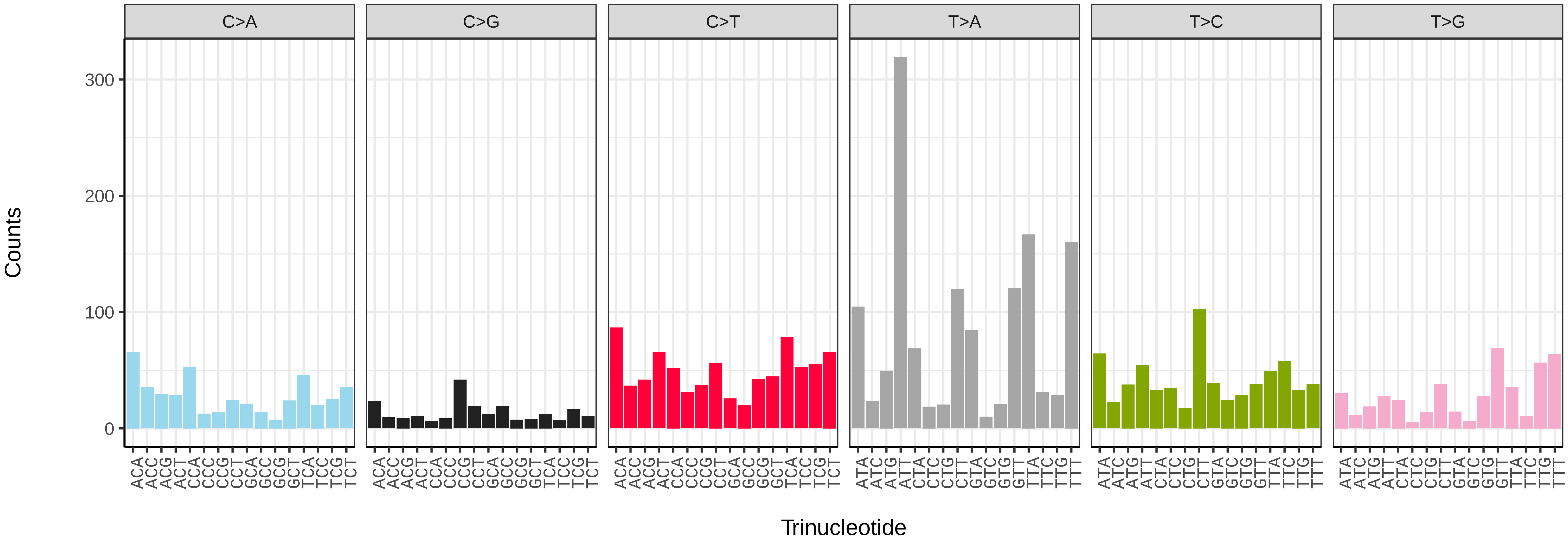


*Thumatha senex* (ilThuSene1) somatic mutational spectrum (SBS52 classification)

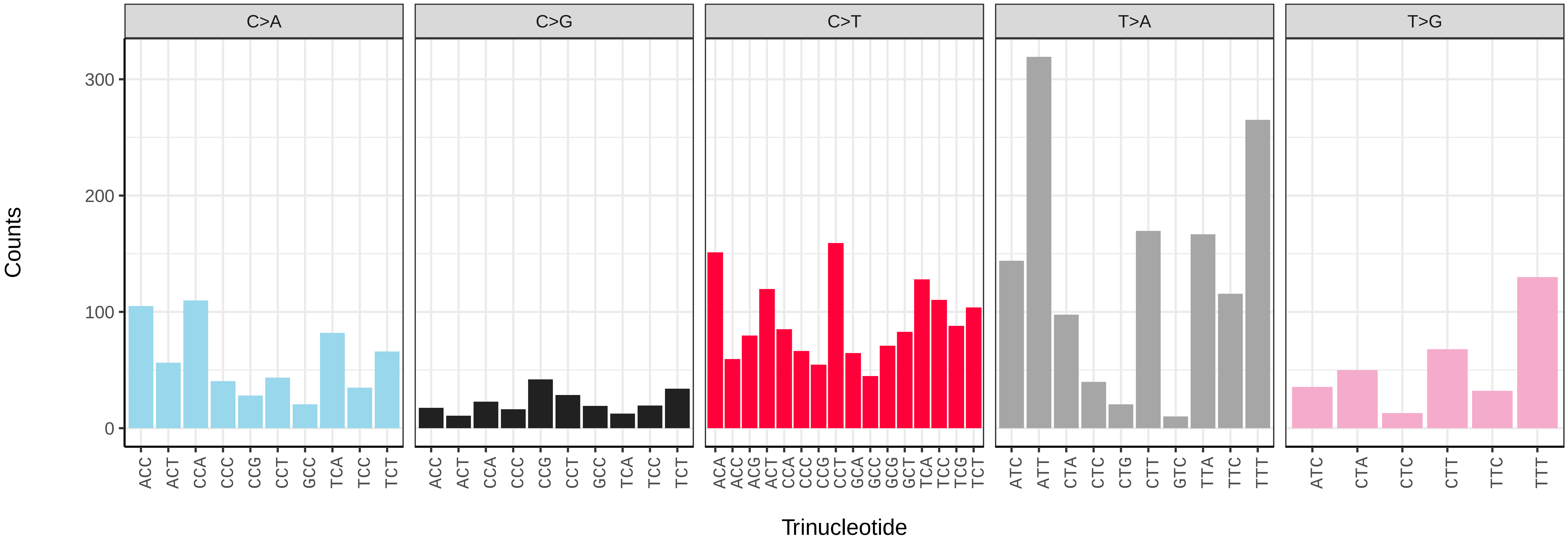

*Thyatira batis* (ilThyBati1) germline mutational spectrum

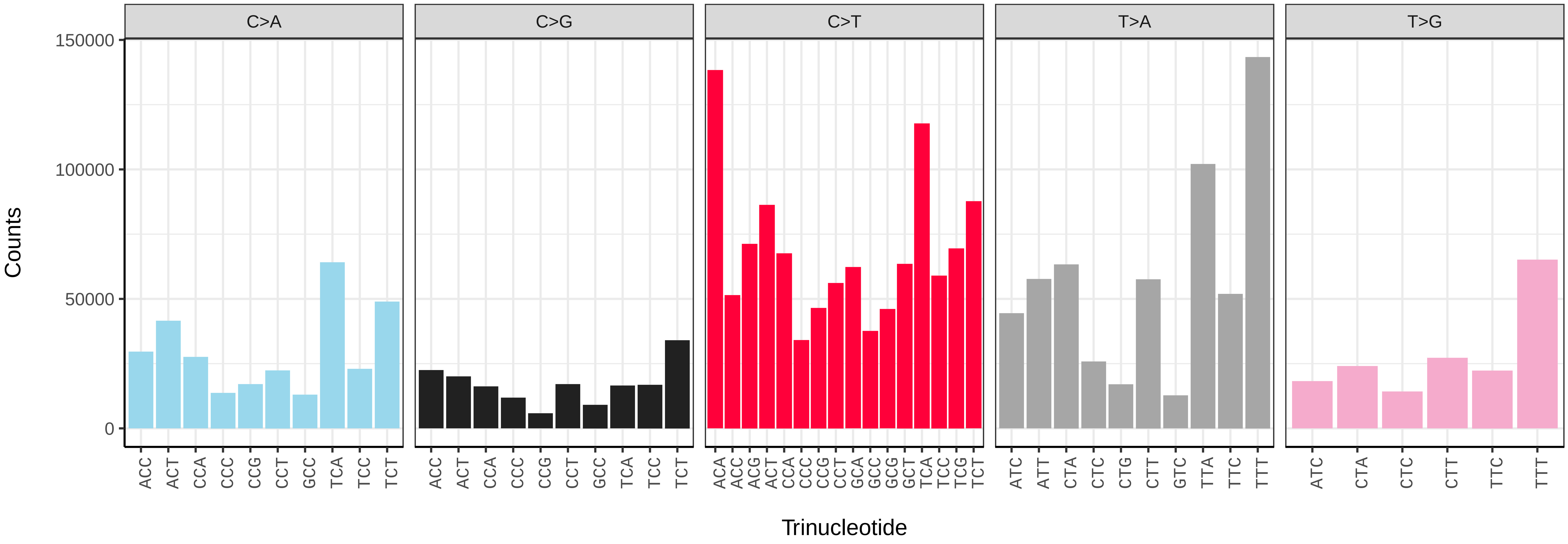


*Thyatira batis* (ilThyBati1) somatic mutational spectrum (SBS96 classification)

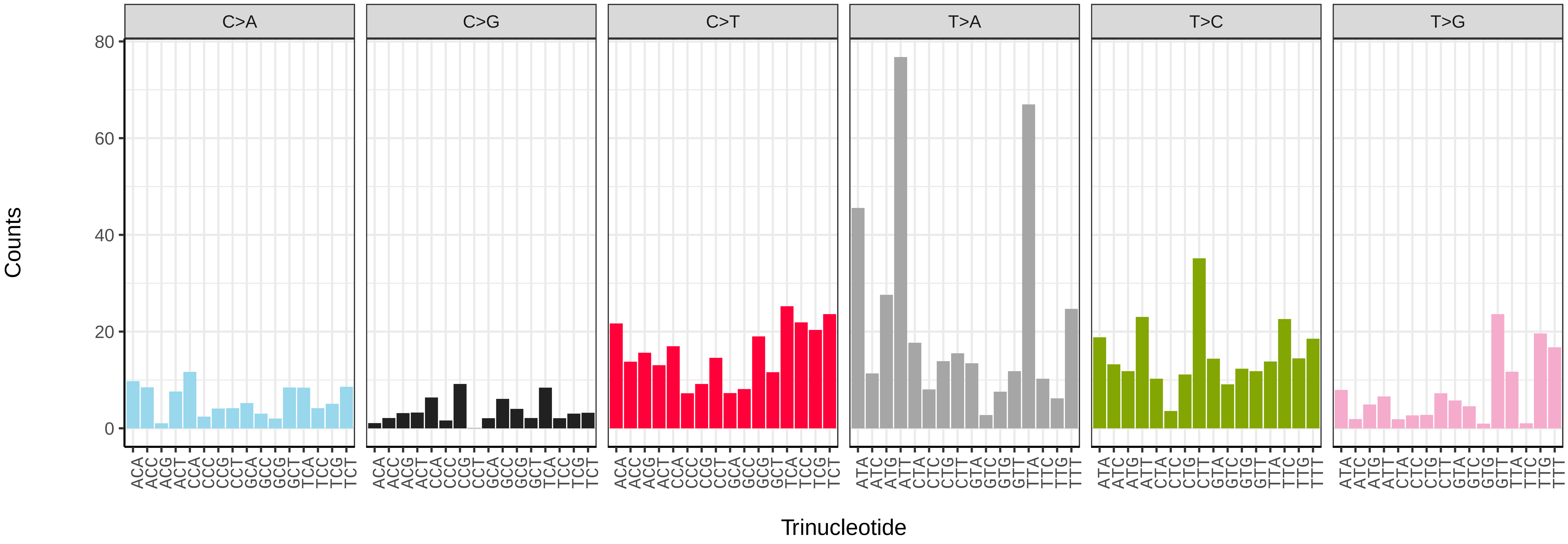


*Thyatira batis* (ilThyBati1) somatic mutational spectrum (SBS52 classification)

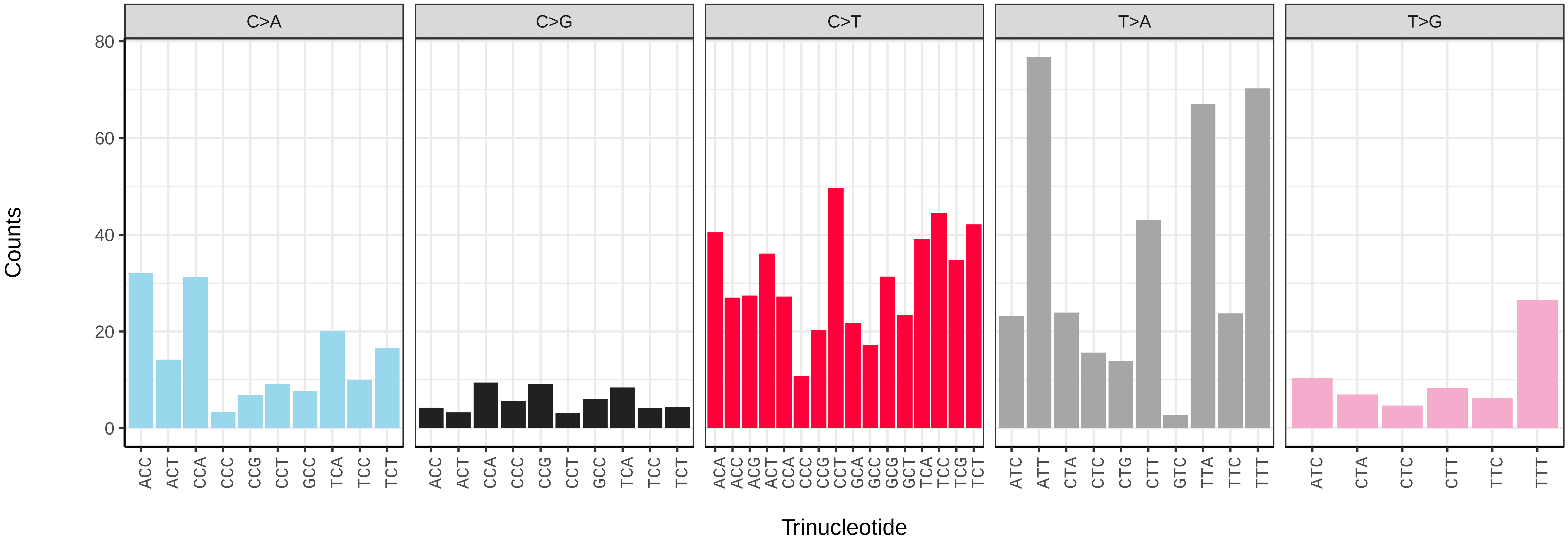

*Thymelicus sylvestris* (ilThySylv1) germline mutational spectrum

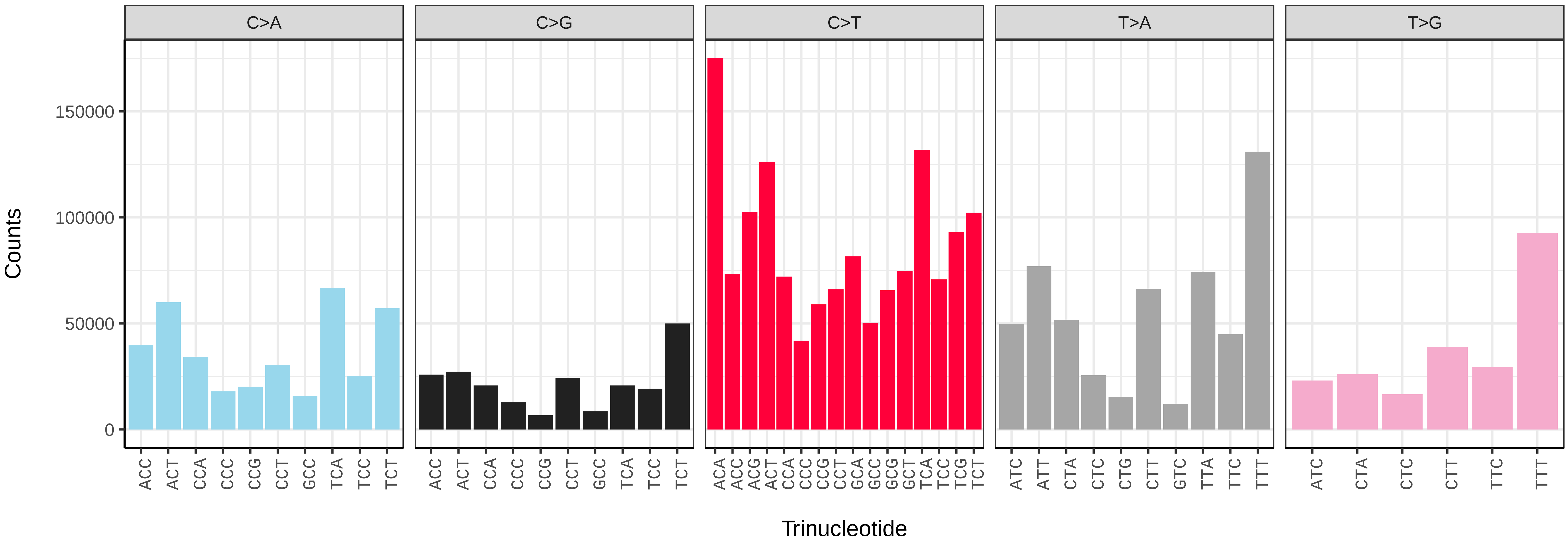


*Thymelicus sylvestris* (ilThySylv1) somatic mutational spectrum (SBS96 classification)

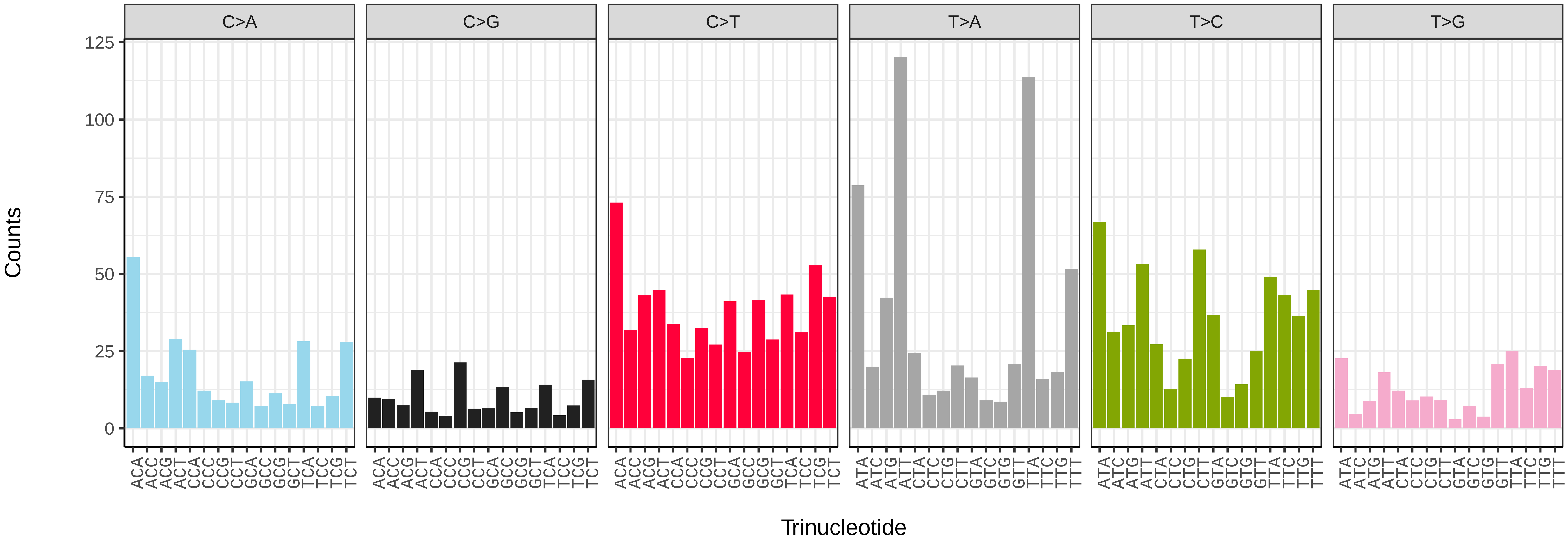


*Thymelicus sylvestris* (ilThySylv1) somatic mutational spectrum (SBS52 classification)

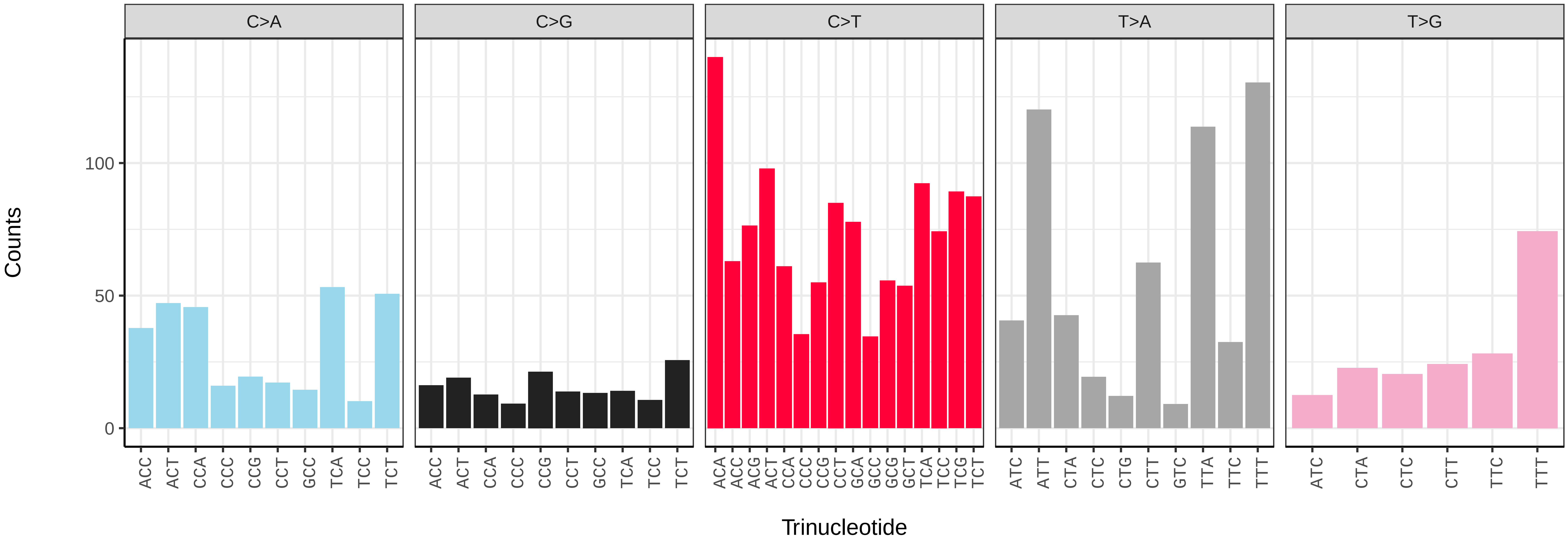

*Tiliacea aurago* (ilTilAura1) germline mutational spectrum

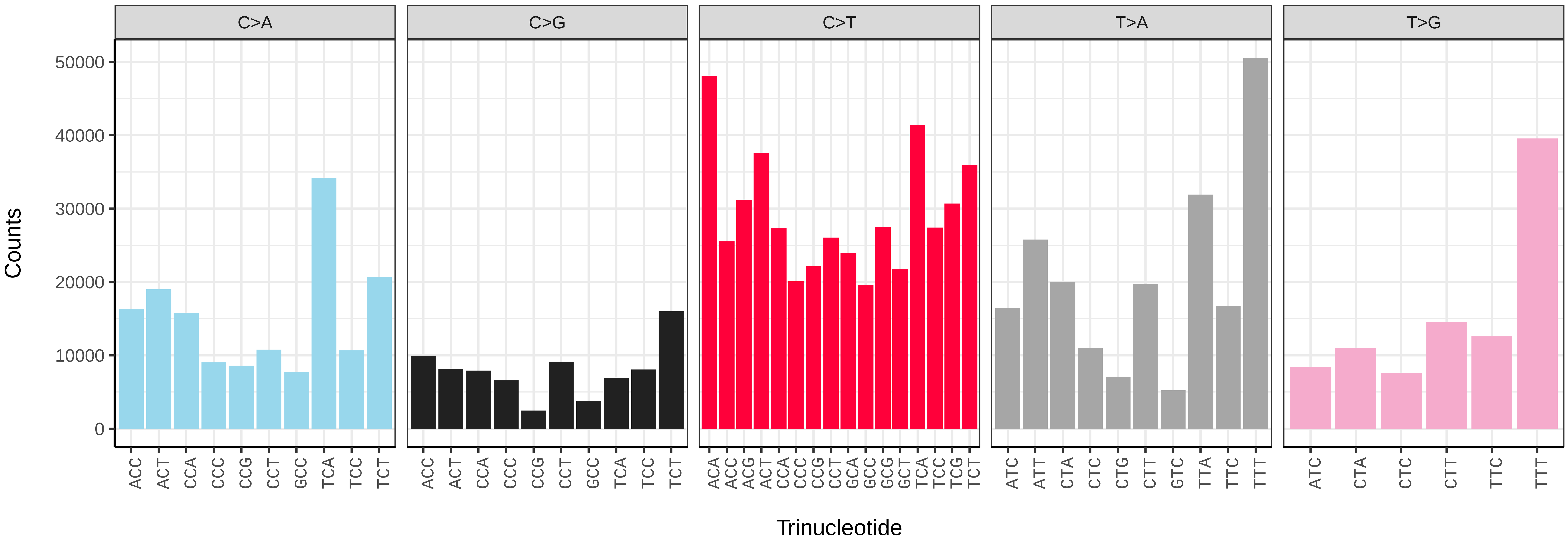


*Tiliacea aurago* (ilTilAura1) somatic mutational spectrum (SBS96 classification)

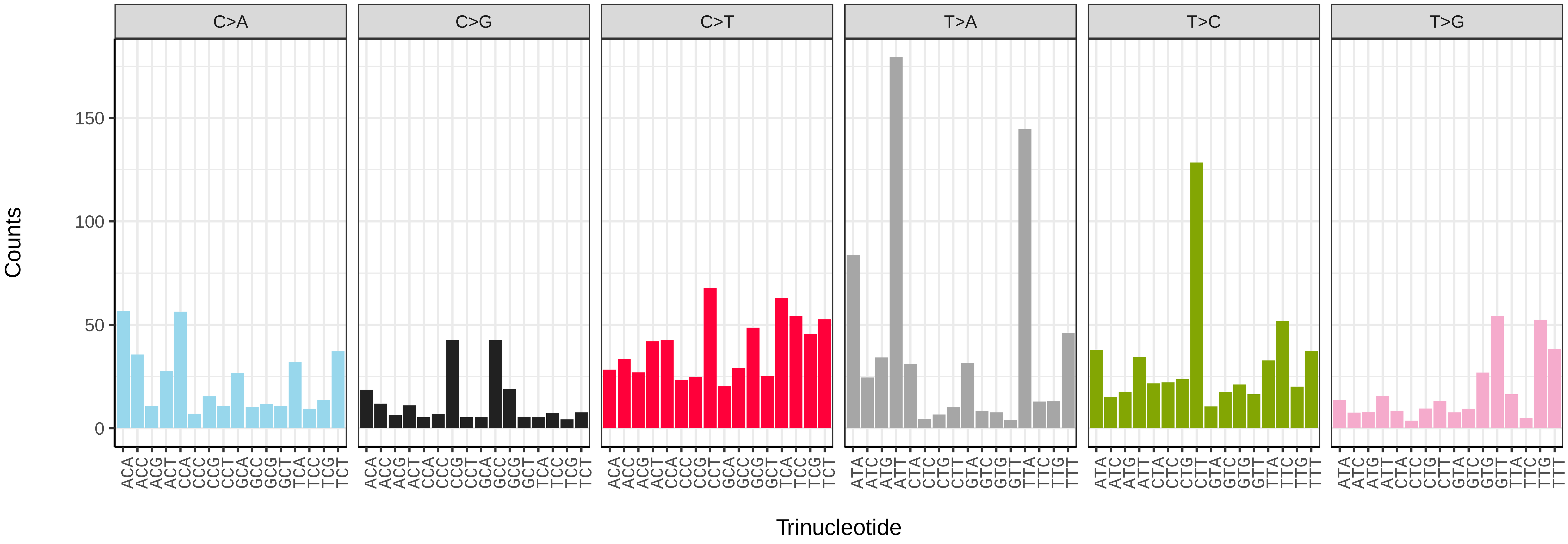


*Tiliacea aurago* (ilTilAura1) somatic mutational spectrum (SBS52 classification)

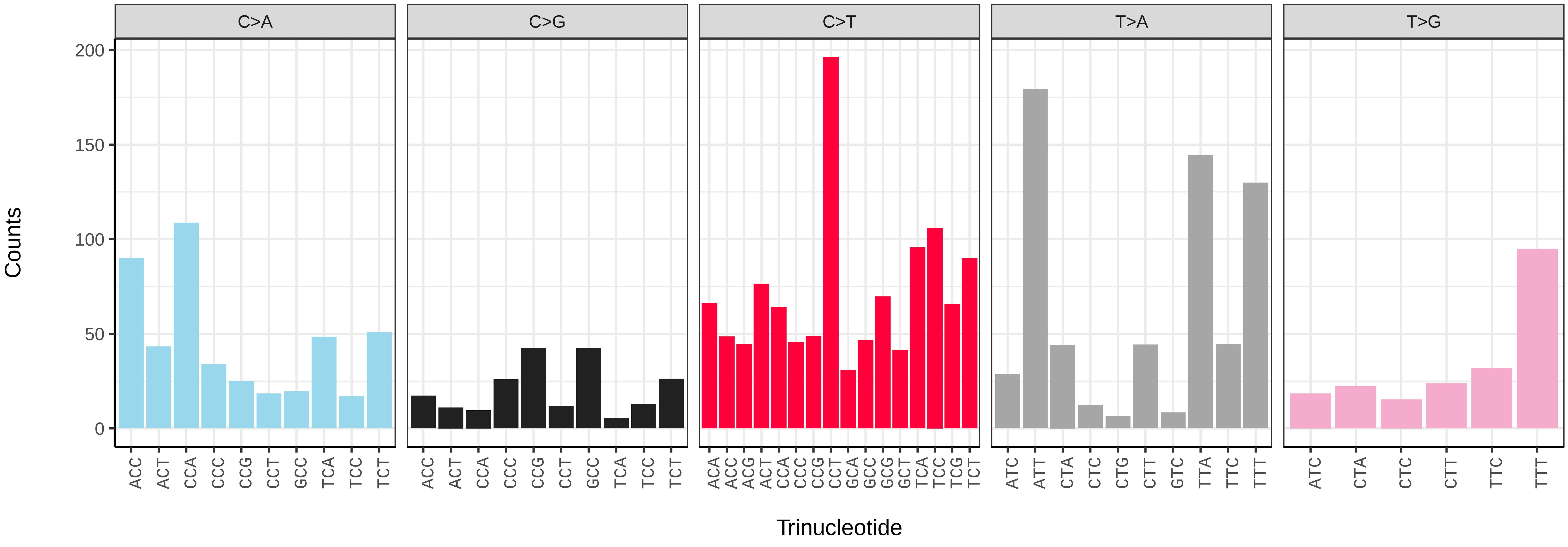

*Tinea pellionella* (ilTinPell1) germline mutational spectrum

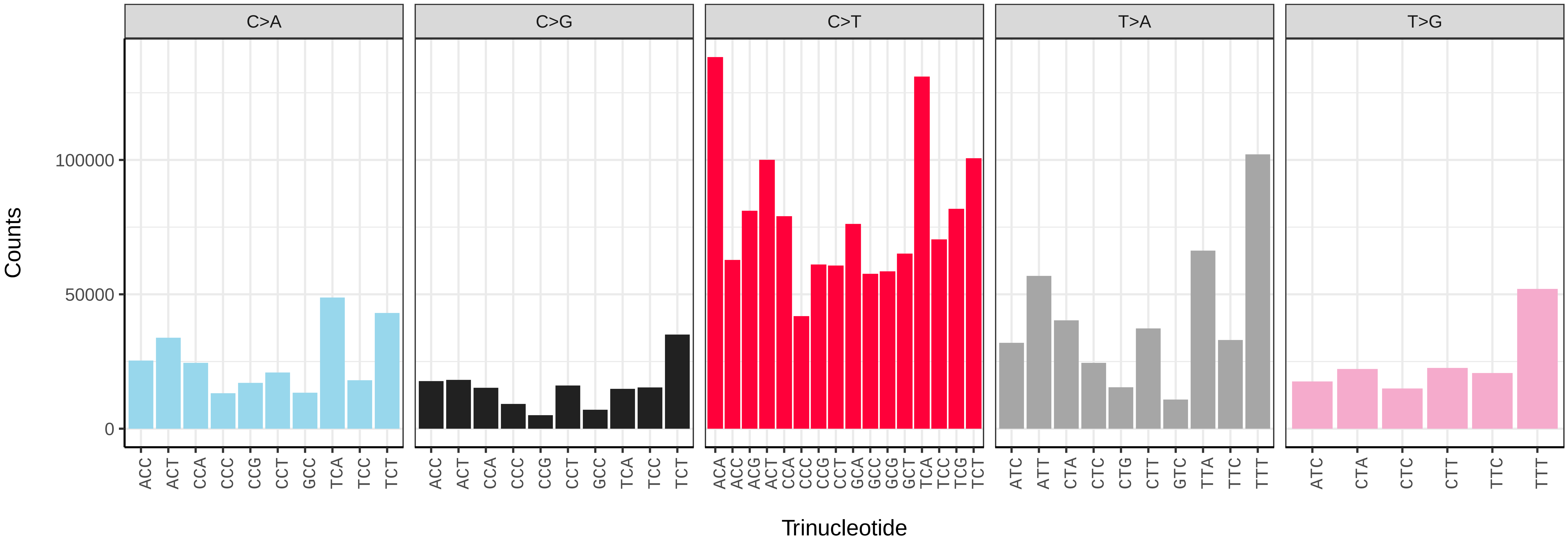


*Tinea pellionella* (ilTinPell1) somatic mutational spectrum (SBS96 classification)

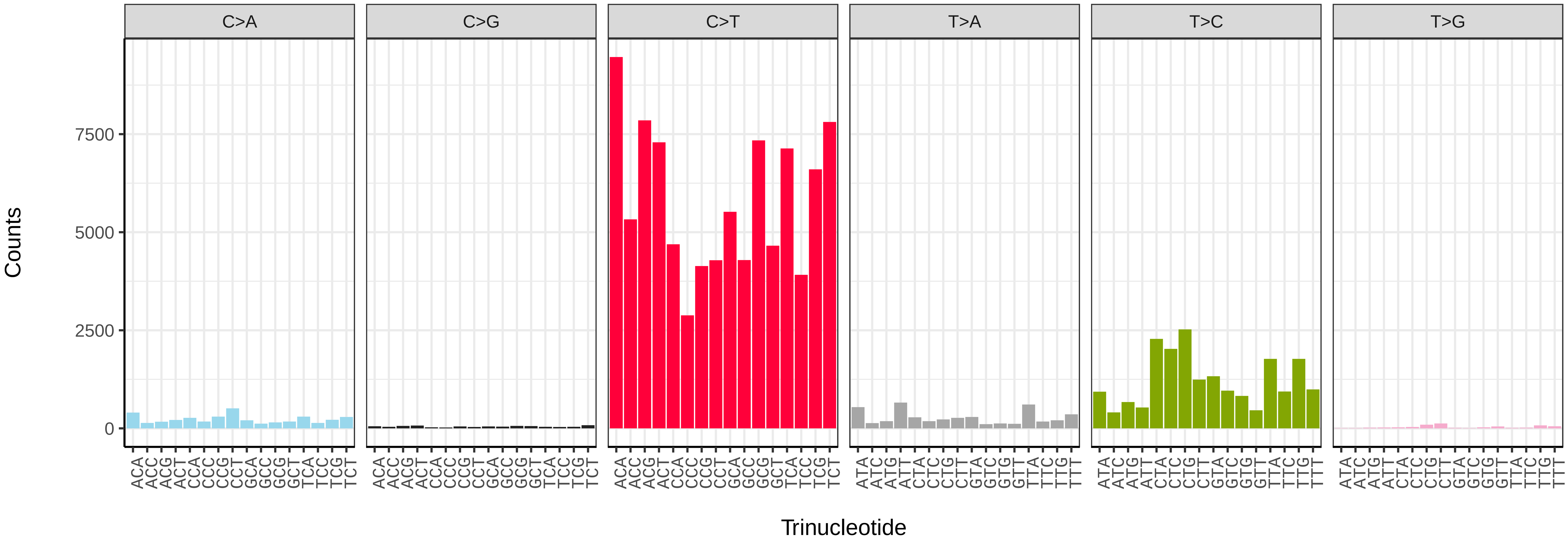


*Tinea pellionella* (ilTinPell1) somatic mutational spectrum (SBS52 classification)

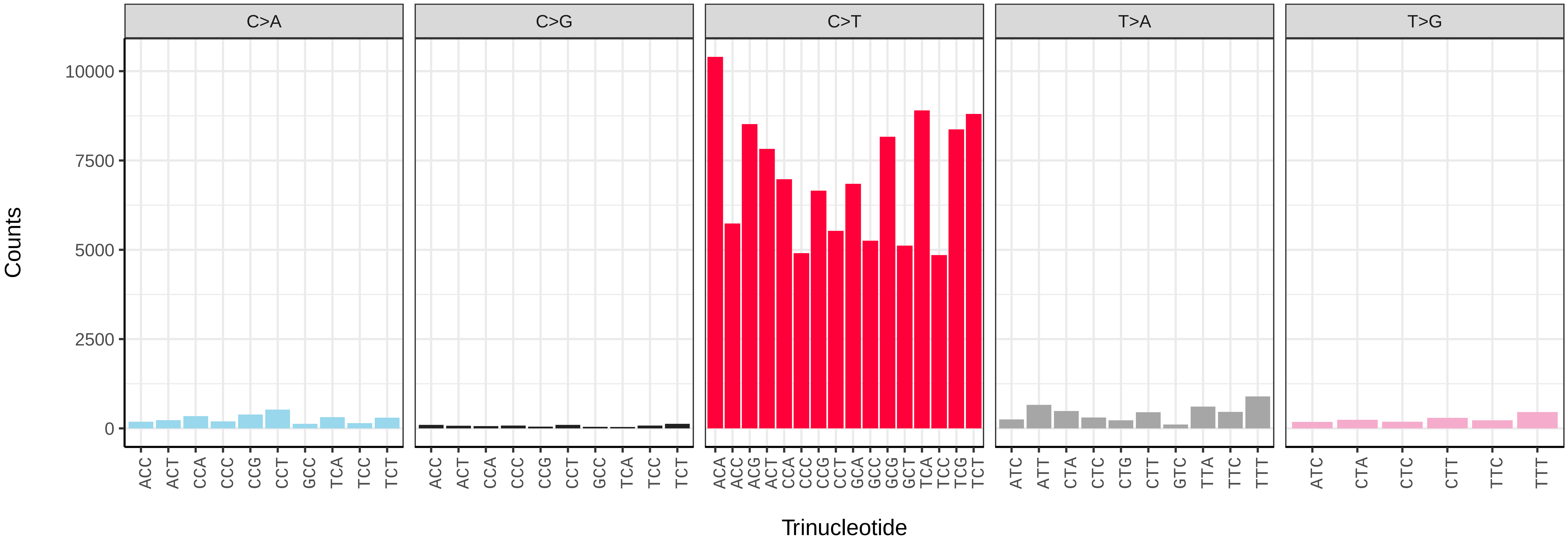

*Tinea semifulvella* (ilTinSemi1) germline mutational spectrum

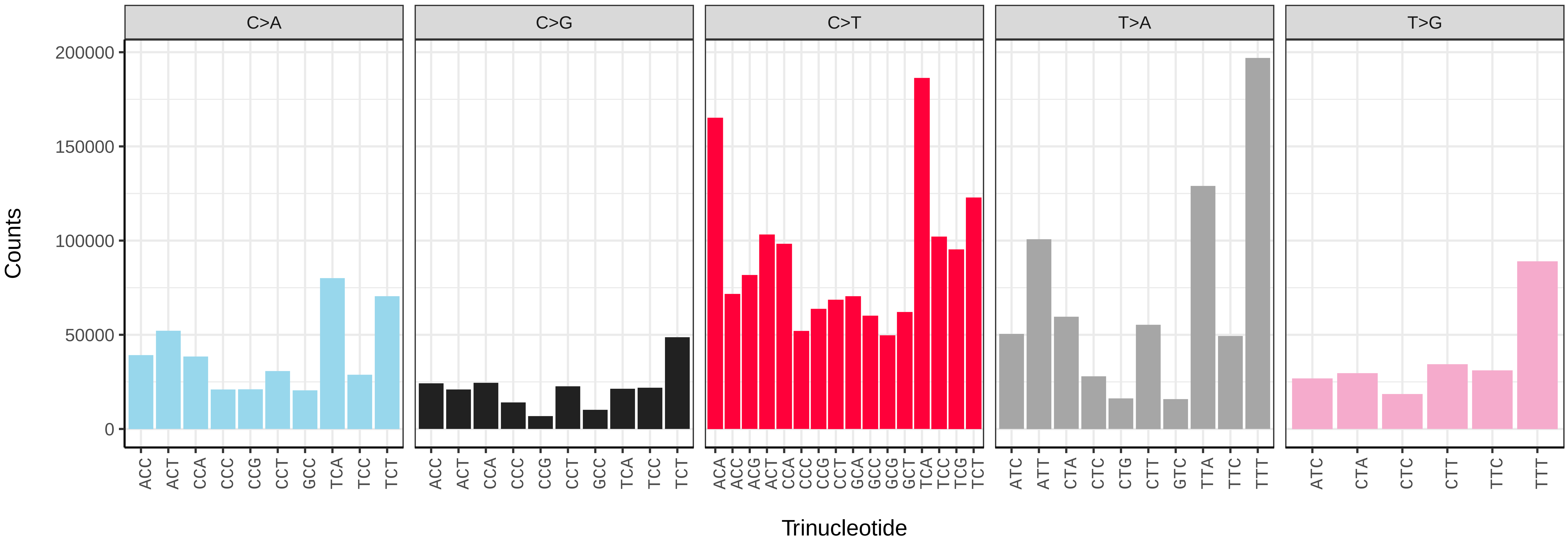


*Tinea semifulvella* (ilTinSemi1) somatic mutational spectrum (SBS96 classification)

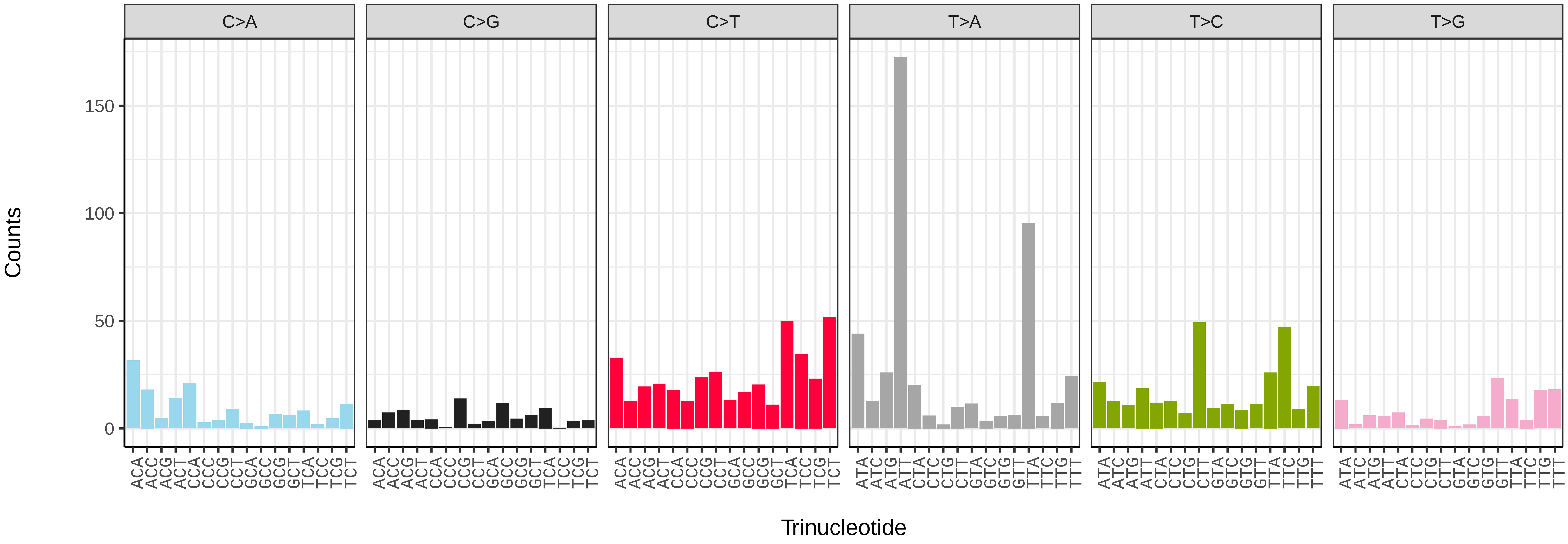


*Tinea semifulvella* (ilTinSemi1) somatic mutational spectrum (SBS52 classification)

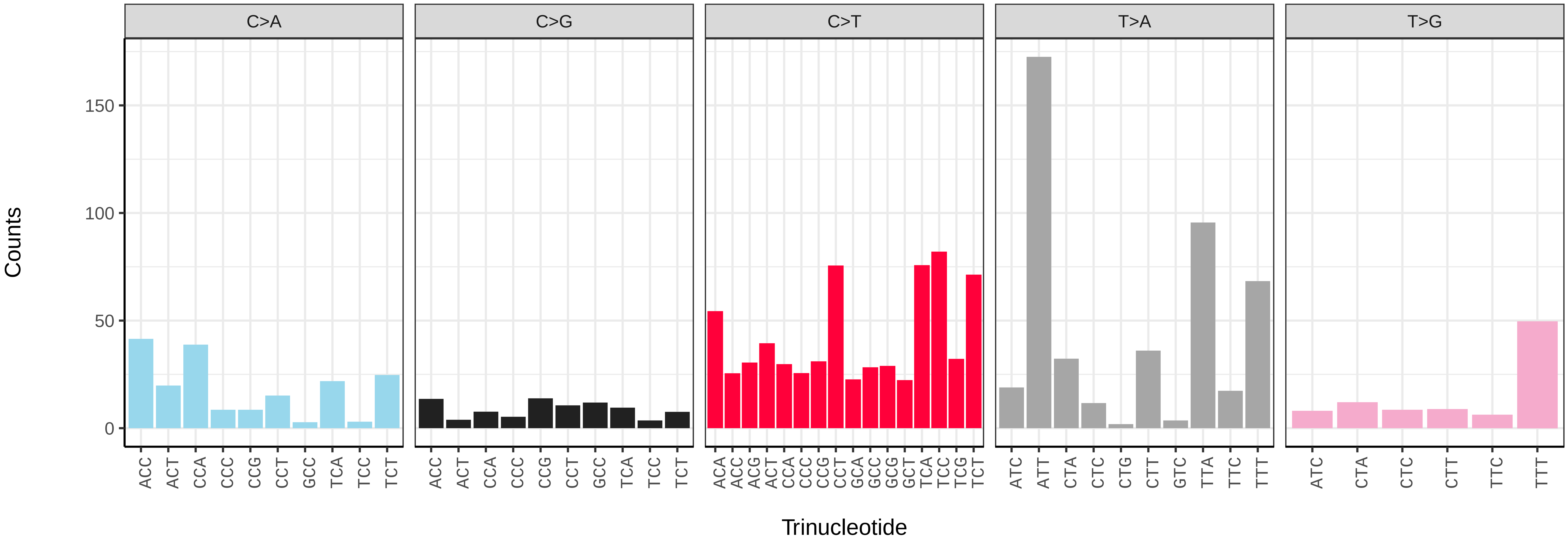

*Tinea trinotella* (ilTinTrin1) germline mutational spectrum

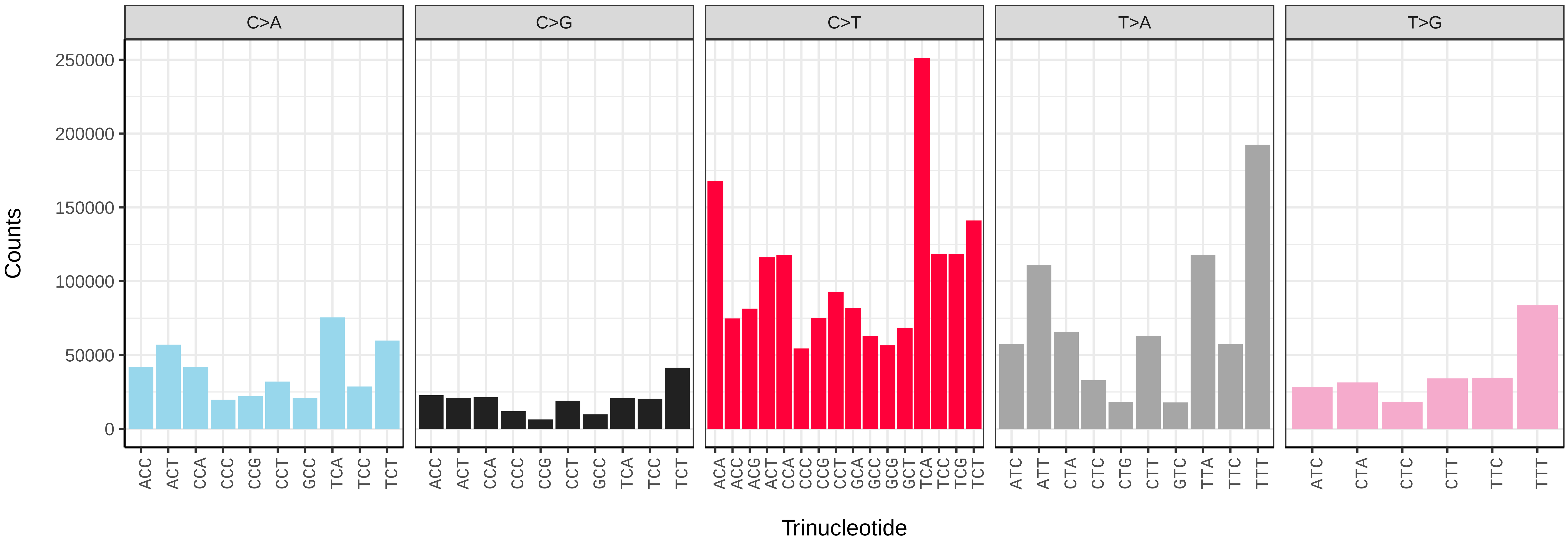


*Tinea trinotella* (ilTinTrin1) somatic mutational spectrum (SBS96 classification)

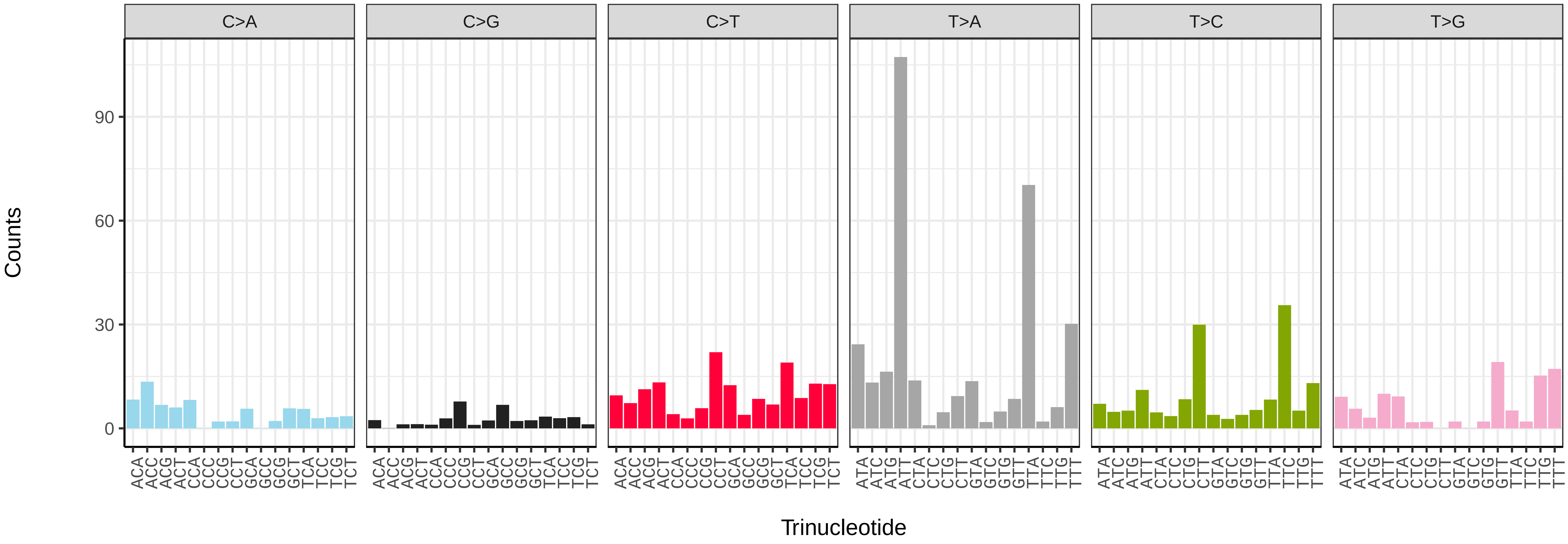


*Tinea trinotella* (ilTinTrin1) somatic mutational spectrum (SBS52 classification)

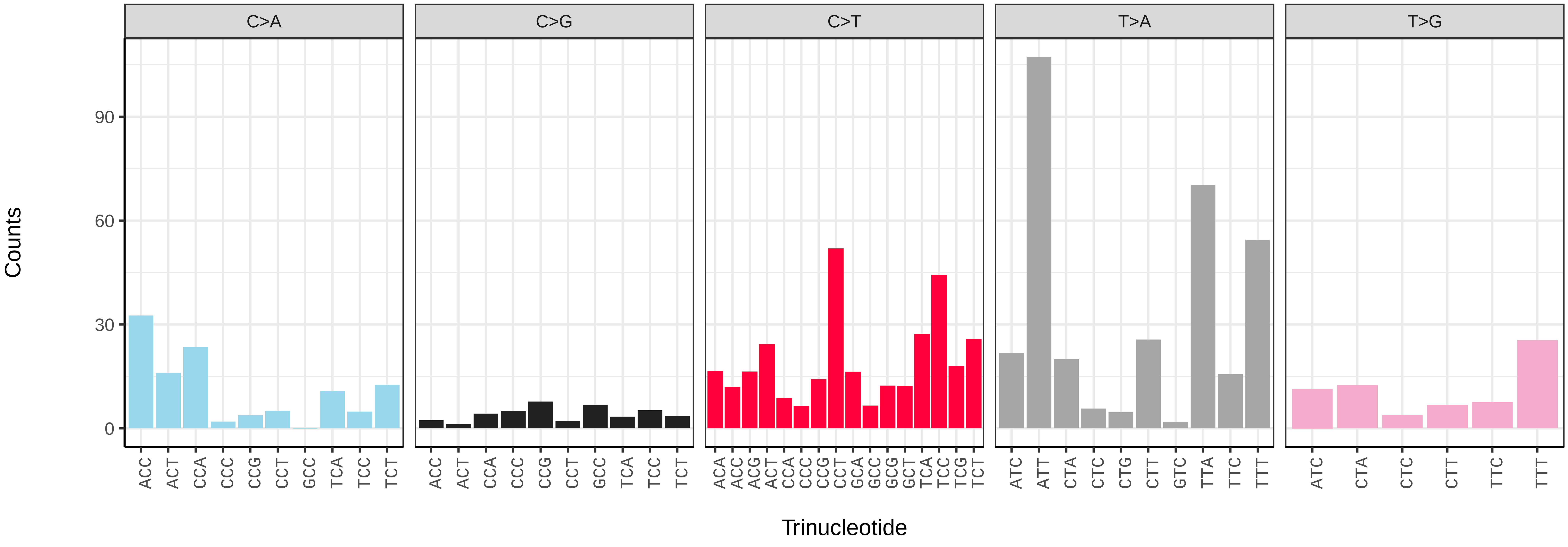

*Tortricodes alternella* (ilTorAlte1) germline mutational spectrum

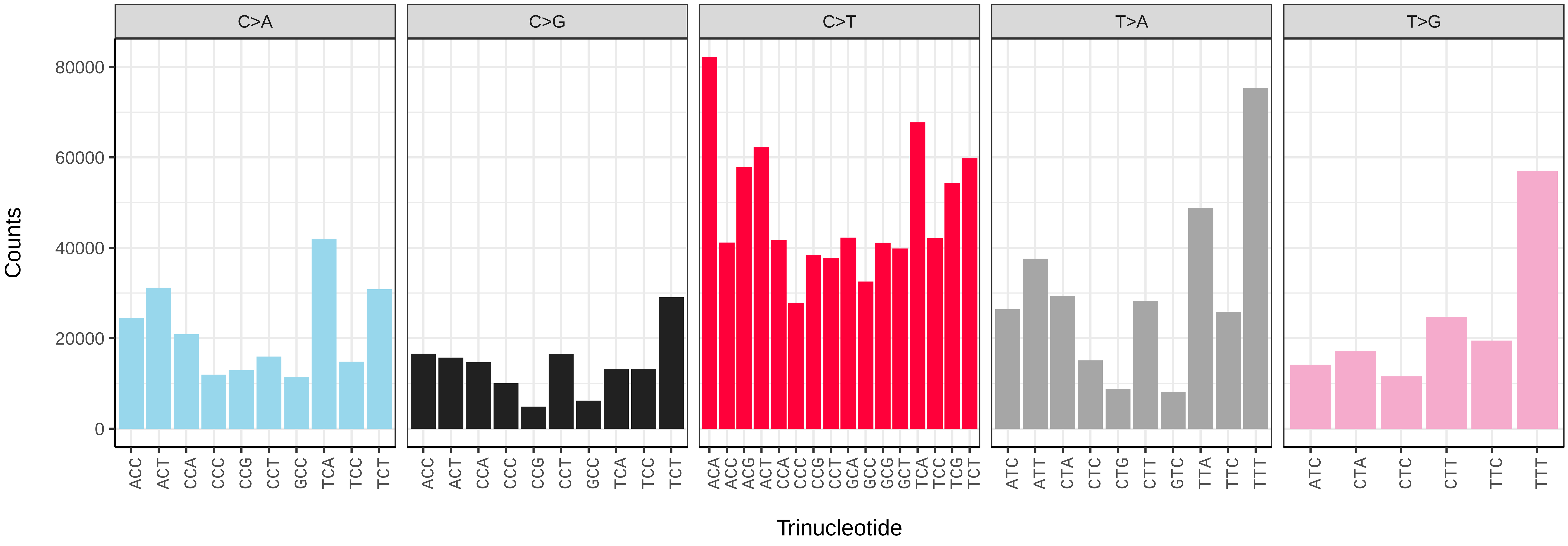


*Tortricodes alternella* (ilTorAlte1) somatic mutational spectrum (SBS96 classification)

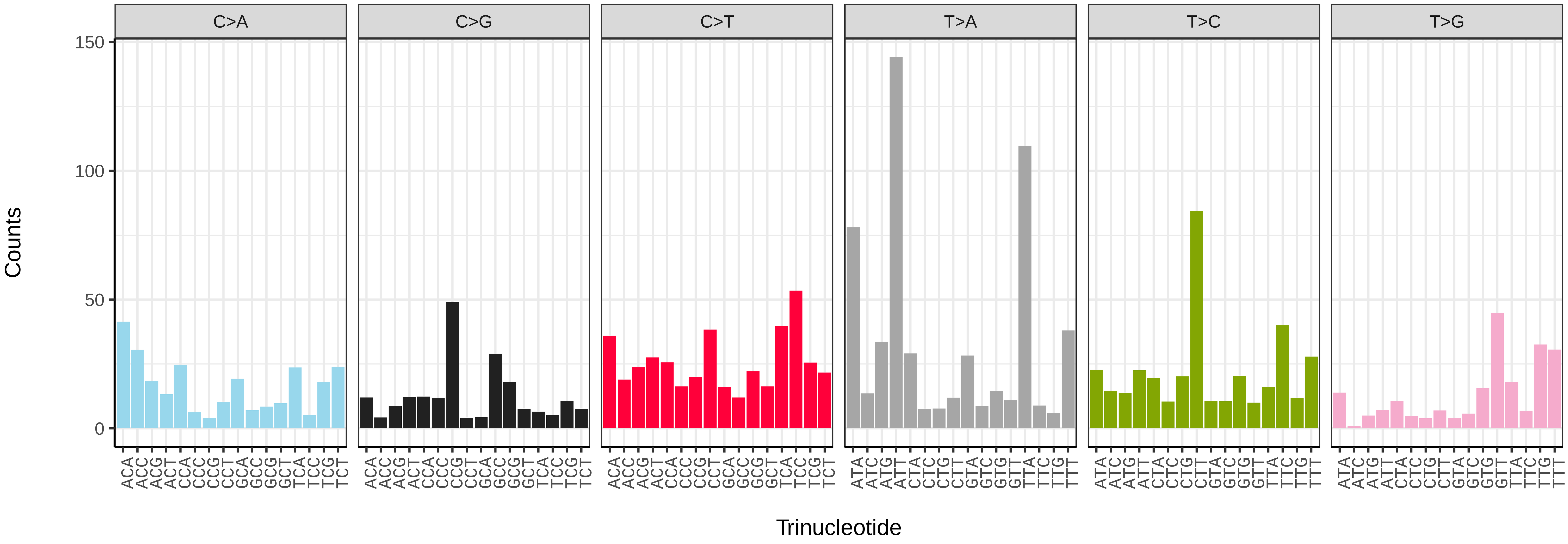


*Tortricodes alternella* (ilTorAlte1) somatic mutational spectrum (SBS52 classification)

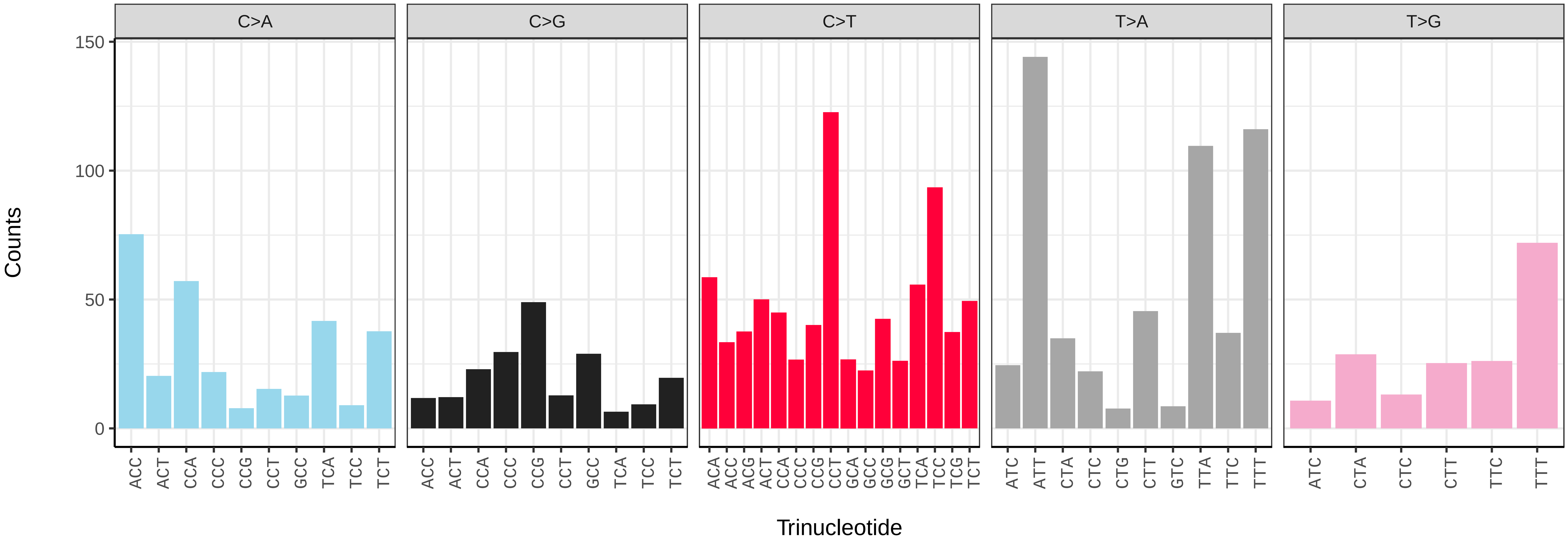

*Trisateles emortualis* (ilTriEmor1) germline mutational spectrum

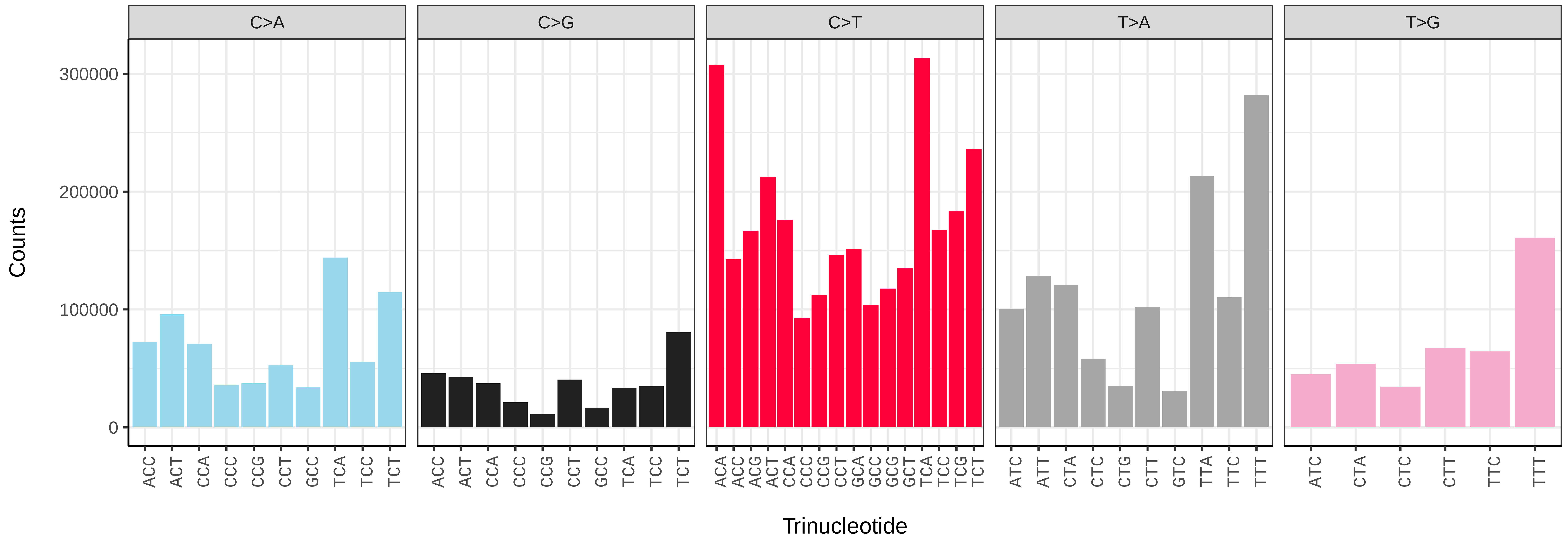


*Trisateles emortualis* (ilTriEmor1) somatic mutational spectrum (SBS96 classification)

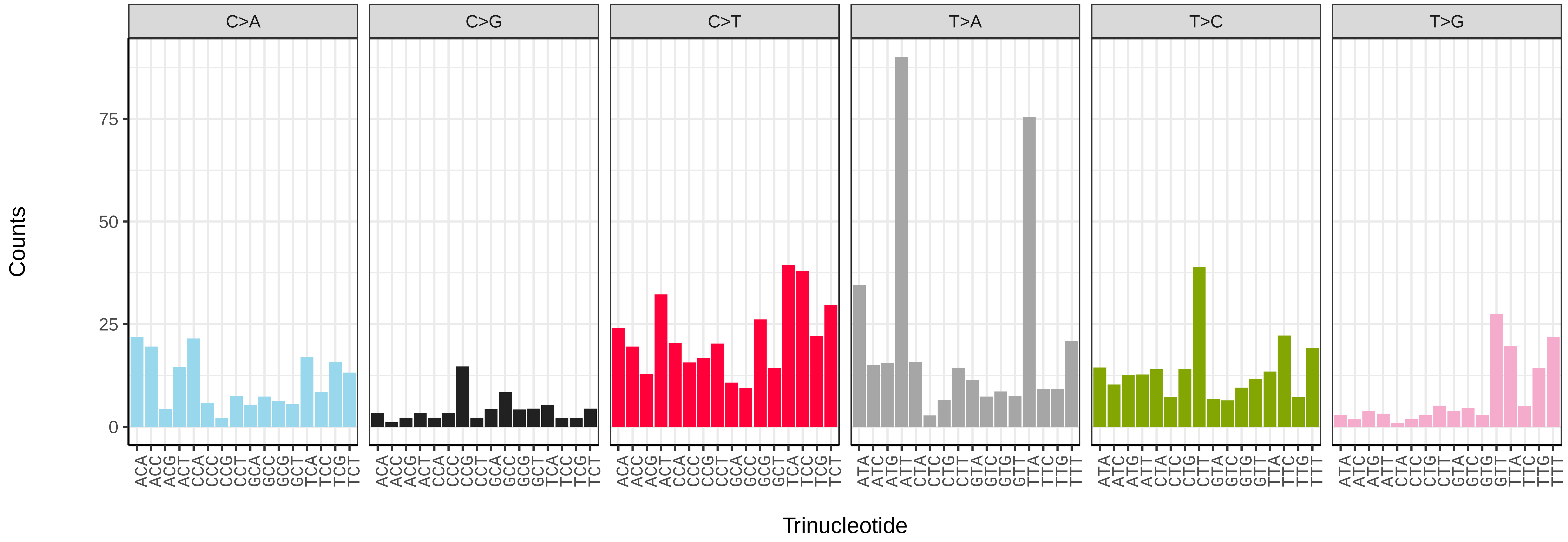


*Trisateles emortualis* (ilTriEmor1) somatic mutational spectrum (SBS52 classification)

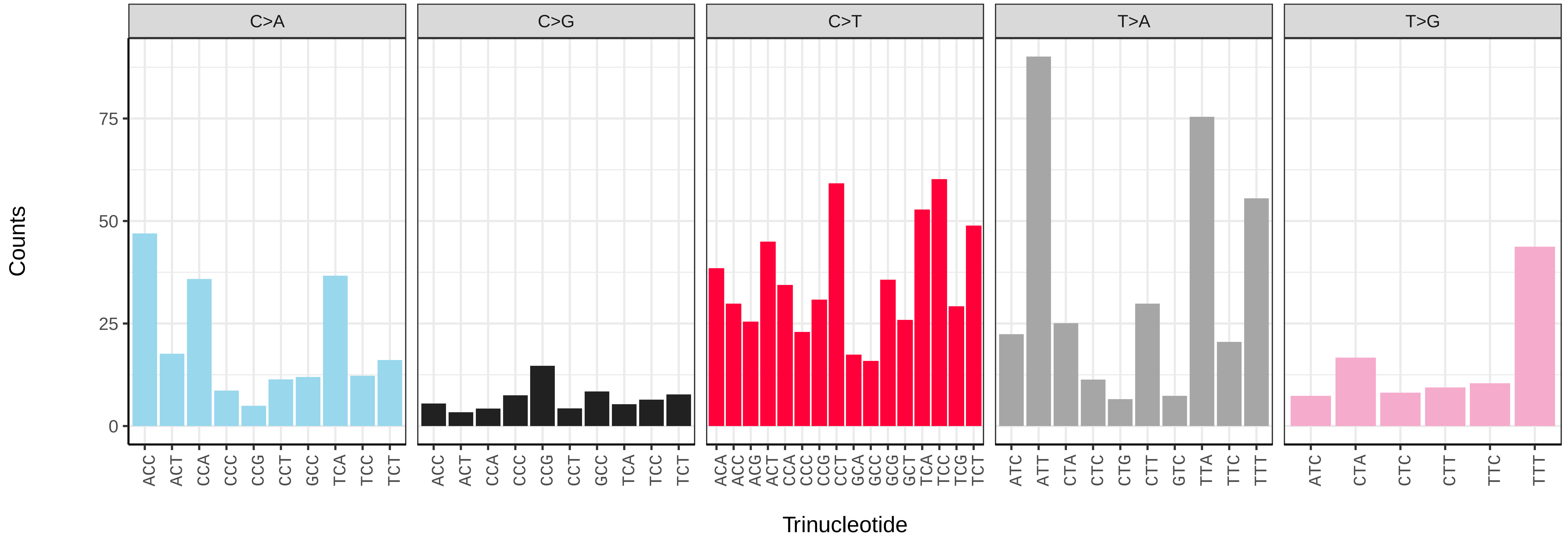

*Tyria jacobaeae* (ilTyrJaco4) germline mutational spectrum

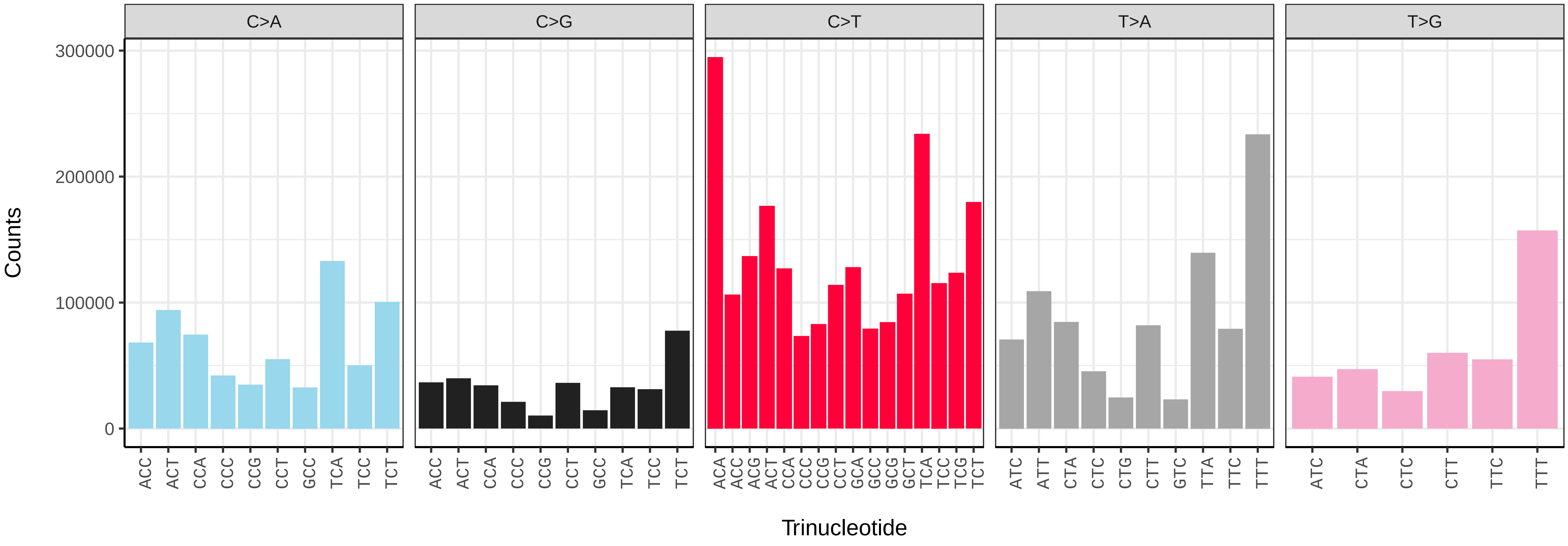


*Tyria jacobaeae* (ilTyrJaco4) somatic mutational spectrum (SBS96 classification)

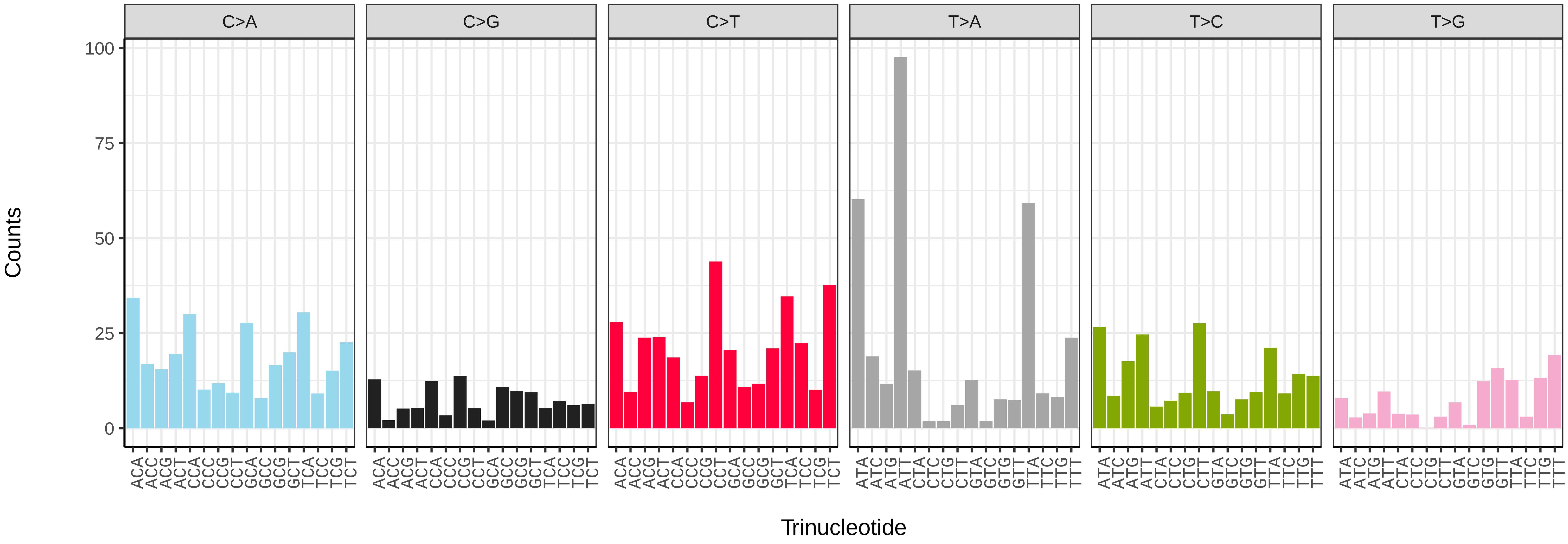


*Tyria jacobaeae* (ilTyrJaco4) somatic mutational spectrum (SBS52 classification)

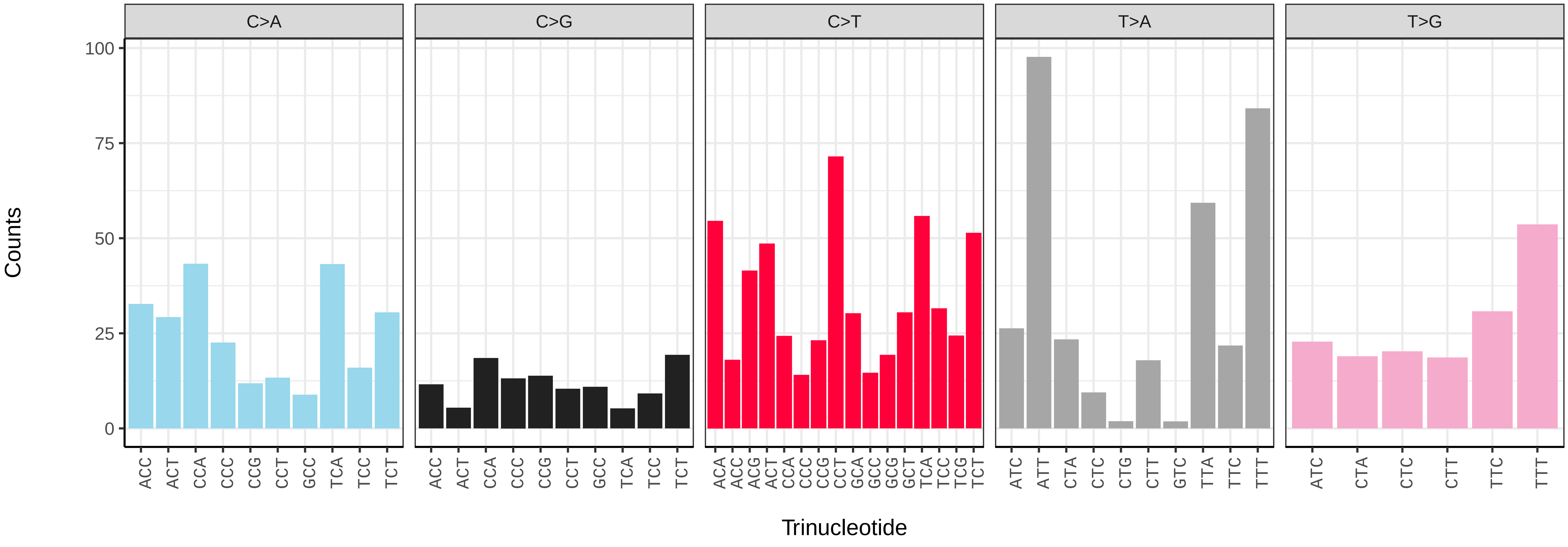

*Udea olivalis* (ilUdeOliv2) germline mutational spectrum

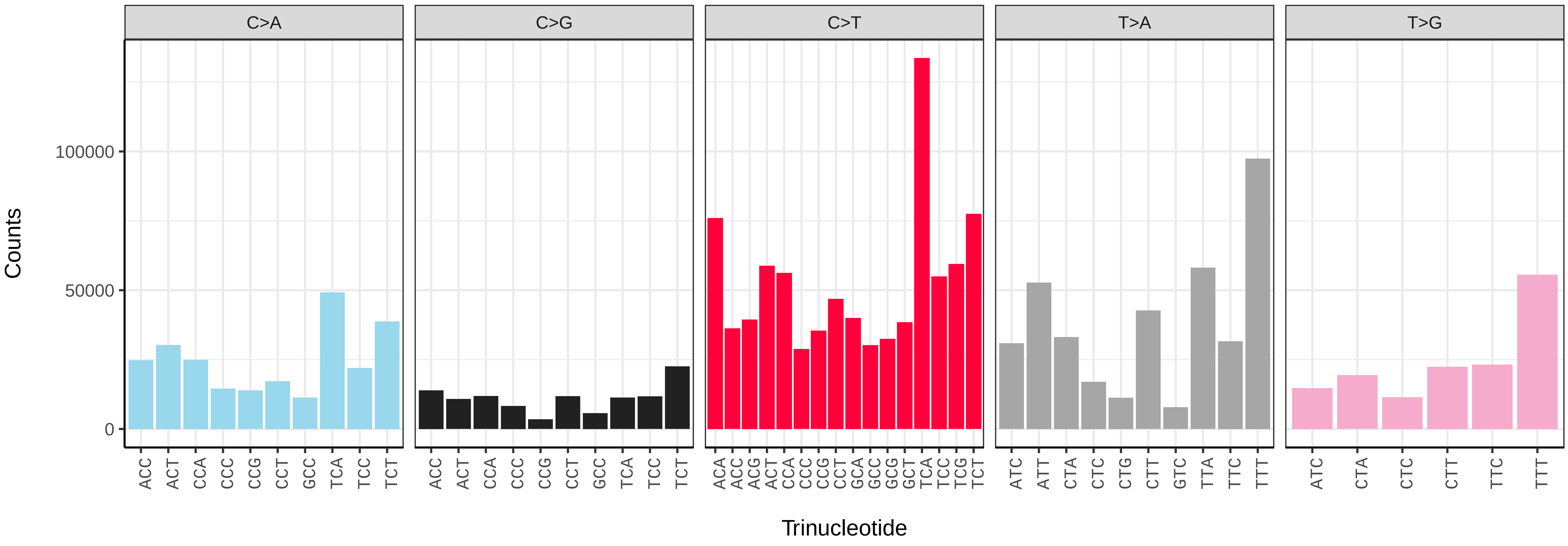


*Udea olivalis* (ilUdeOliv2) somatic mutational spectrum (SBS96 classification)

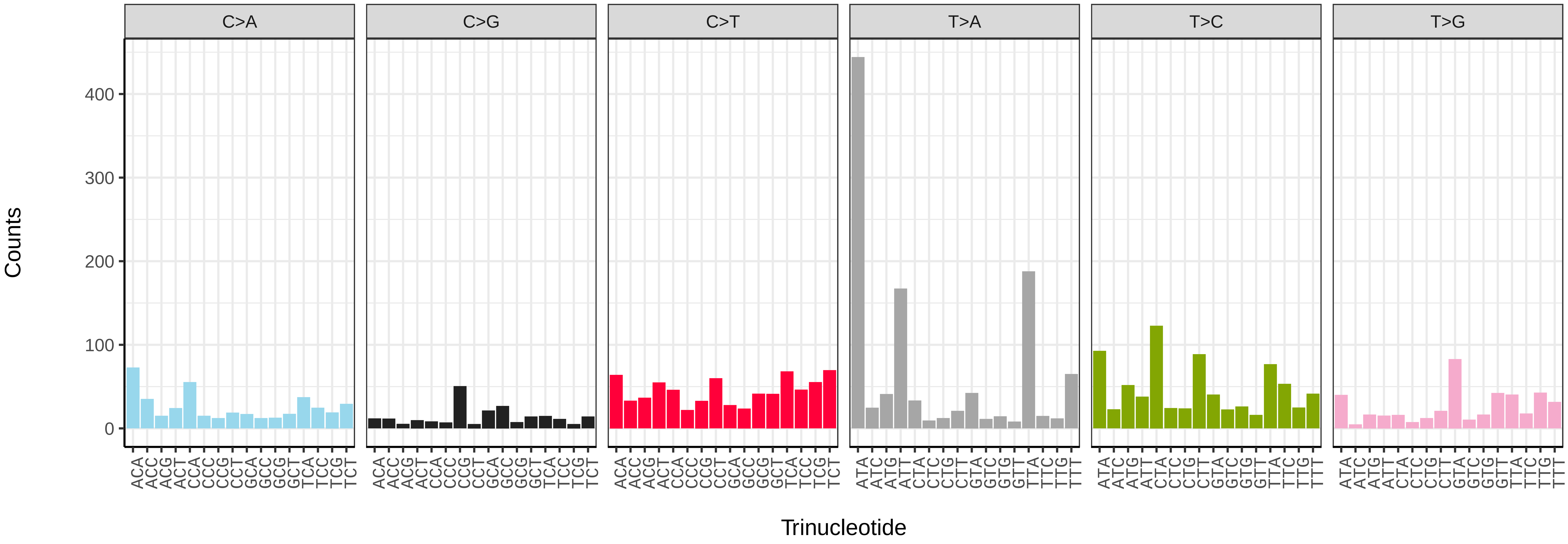


*Udea olivalis* (ilUdeOliv2) somatic mutational spectrum (SBS52 classification)

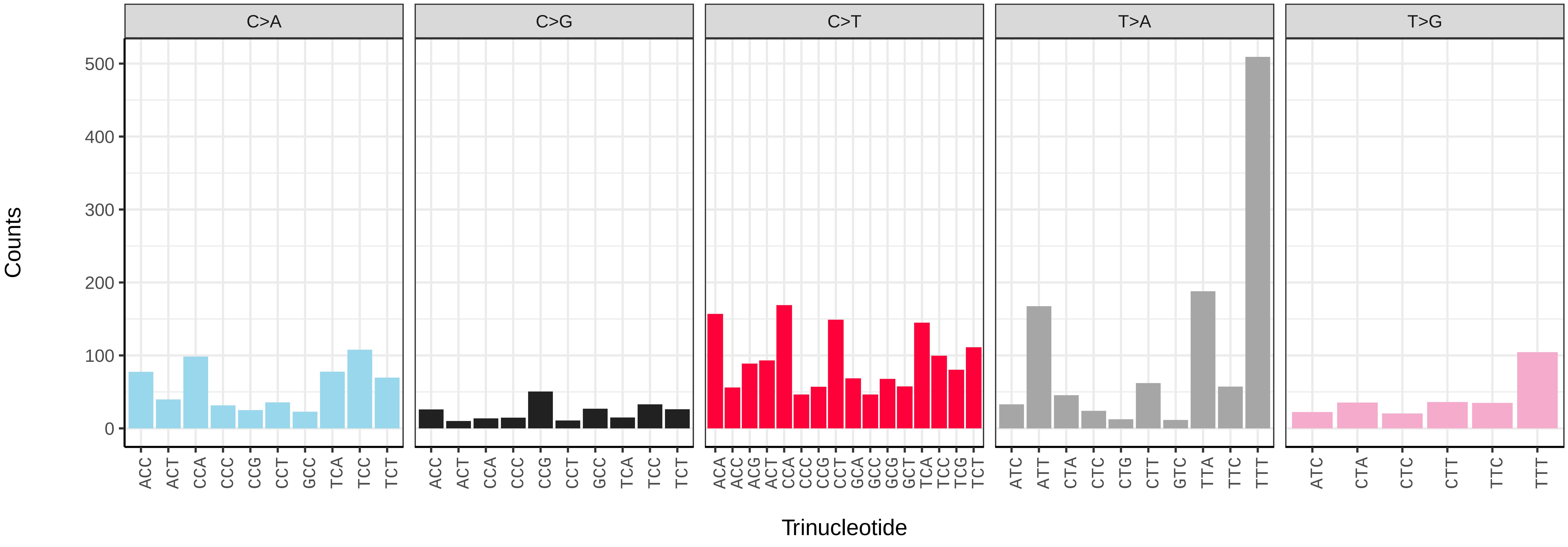

*Vanessa atalanta* (ilVanAtal1) germline mutational spectrum

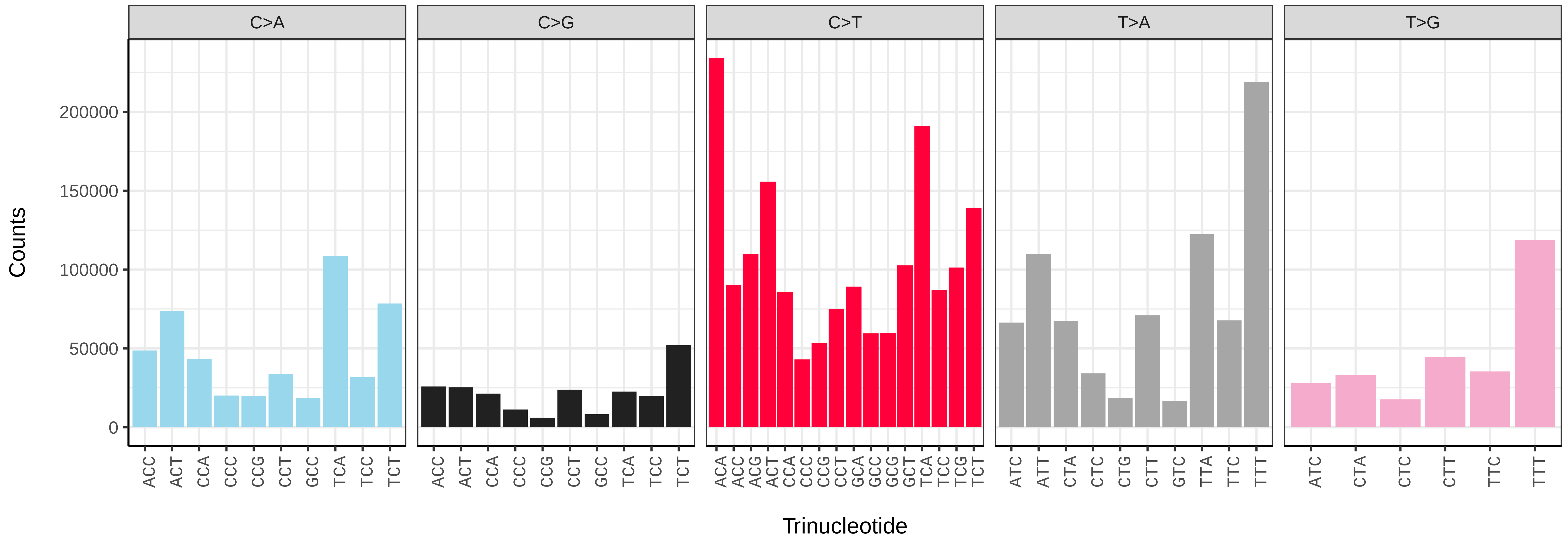


*Vanessa atalanta* (ilVanAtal1) somatic mutational spectrum (SBS96 classification)

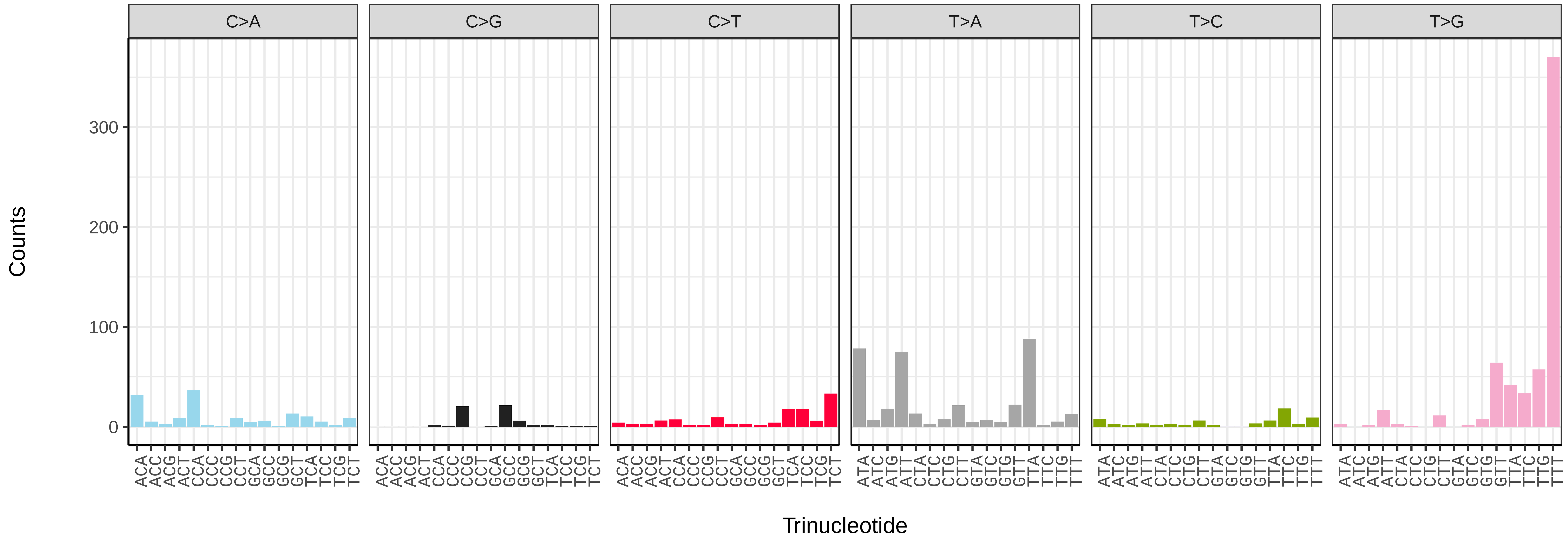


*Vanessa atalanta* (ilVanAtal1) somatic mutational spectrum (SBS52 classification)

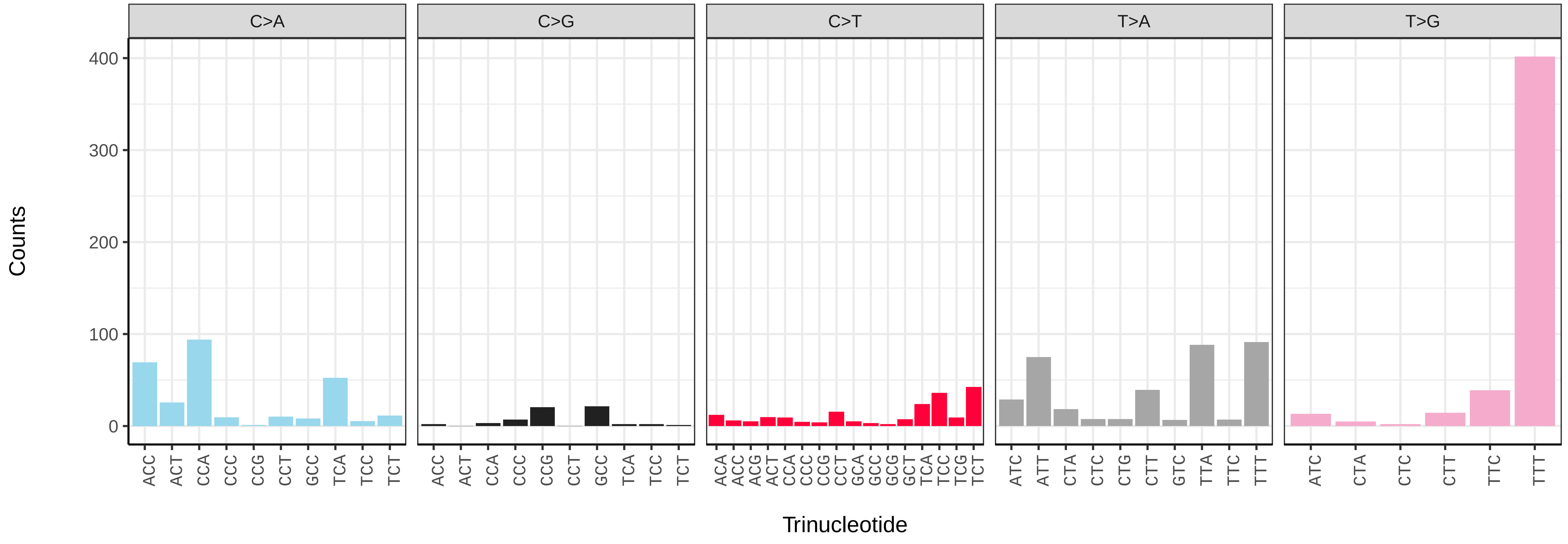

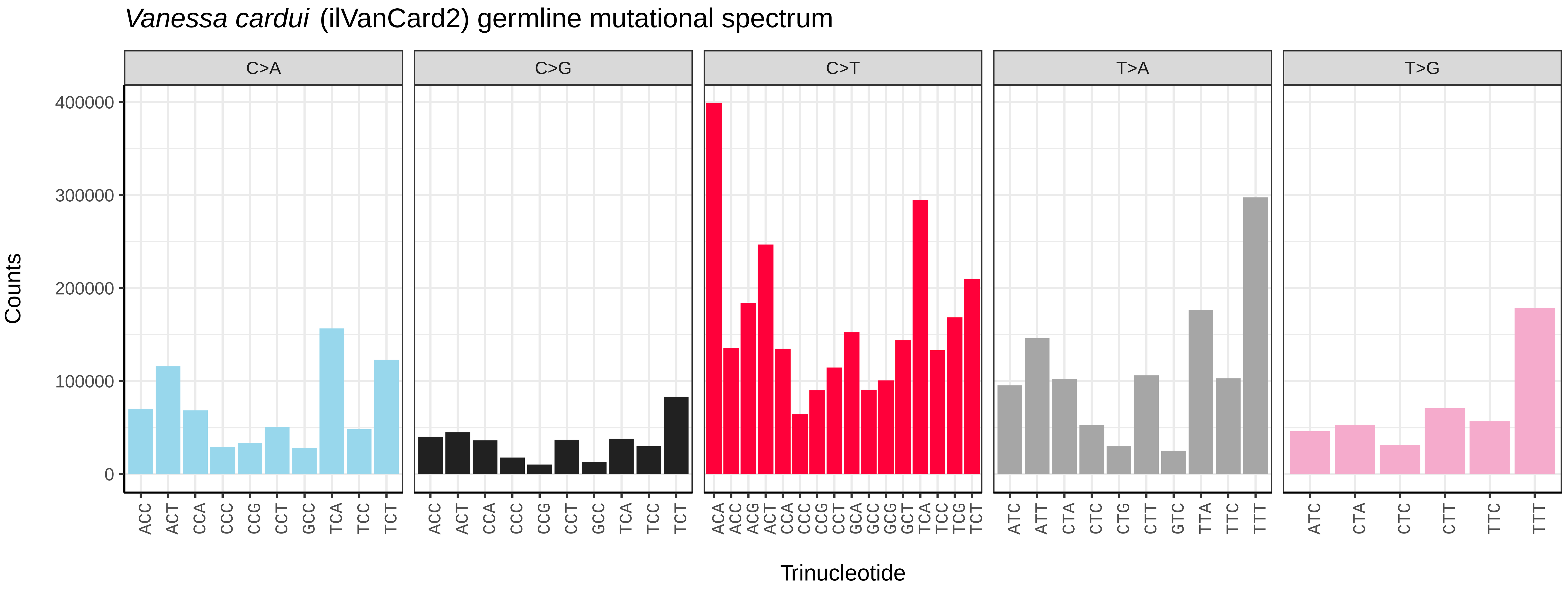
*Vanessa cardui* (ilVanCard2) germline mutational spectrum
C>A
C>G
C>T
T>A
T>G
Counts
0
100000
200000
300000
400000
ACC ACT CCA CCC CCG CCT GCC TCA TCC TCT
ACC ACT CCA CCC CCG CCT GCC TCA TCC TCT
ACA ACC ACG ACT CCA CCC CCG CCT GCA GCC GCG GCT TCA TCC TCG TCT
ATC ATT CTA CTC CTG CTT GTC TTA TTC TTT
ATC CTA CTC CTT TTC TTT
Trinucleotide

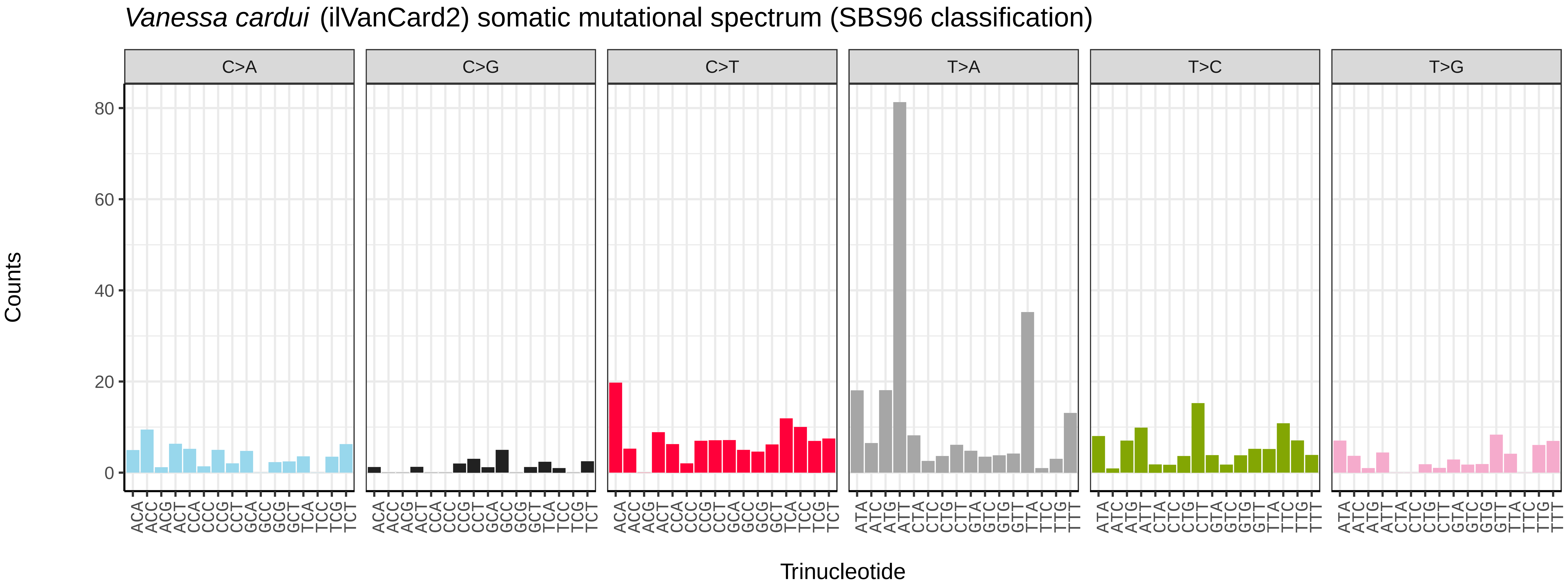
*Vanessa cardui* (ilVanCard2) somatic mutational spectrum (SBS96 classification)
C>A
C>G
C>T
T>A
T>C
T>G
Counts
0
20
40
60
80
ACA ACC ACG ACT CCA CCC CCG CCT GCA GCC GCG GCT TCA TCC TCG TCT
ACA ACC ACG ACT CCA CCC CCG CCT GCA GCC GCG GCT TCA TCC TCG TCT
ACA ACC ACG ACT CCA CCC CCG CCT GCA GCC GCG GCT TCA TCC TCG TCT
ATA ATC ATG ATT CTA CTC CTG CTT GTA GTC GTG GTT TTA TTC TTG TTT
ATA ATC ATG ATT CTA CTC CTG CTT GTA GTC GTG GTT TTA TTC TTG TTT
ATA ATC ATG ATT CTA CTC CTG CTT GTA GTC GTG GTT TTA TTC TTG TTT
Trinucleotide

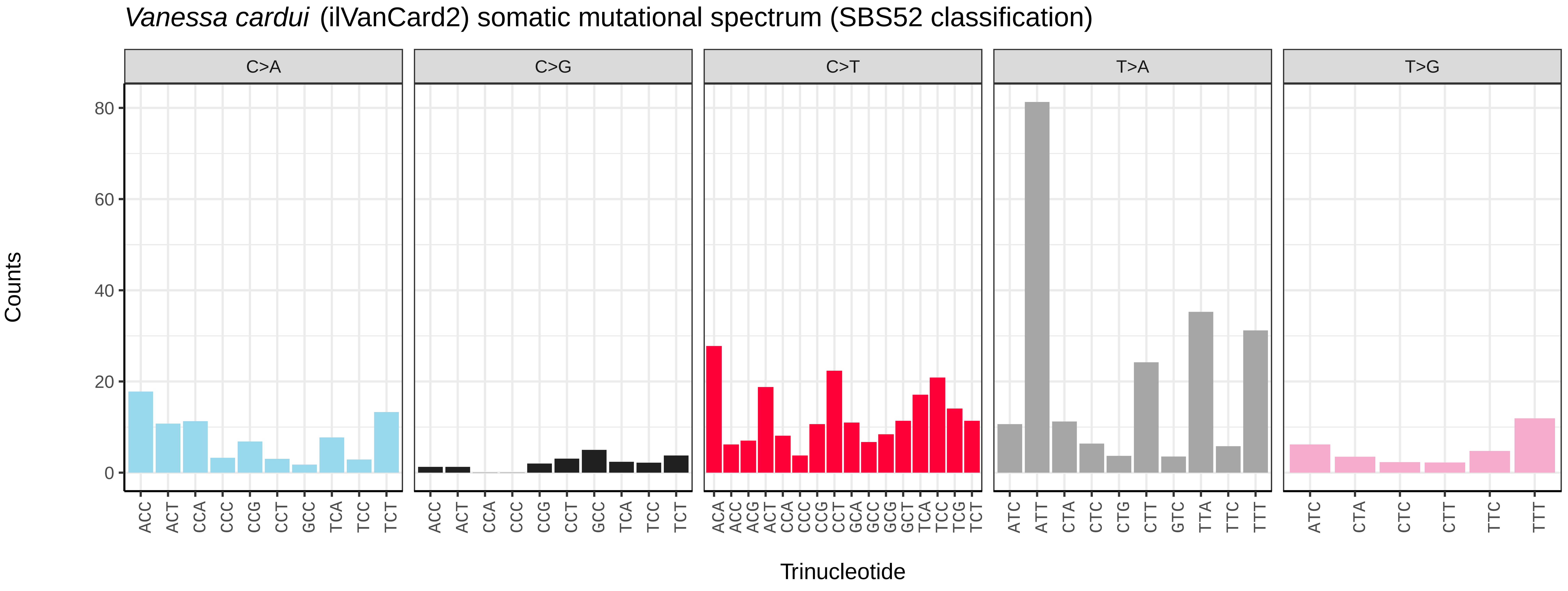
*Vanessa cardui* (ilVanCard2) somatic mutational spectrum (SBS52 classification)
C>A
C>G
C>T
T>A
T>G
Counts
0
20
40
60
80
ACC ACT CCA CCC CCG CCT GCC TCA TCC TCT
ACC ACT CCA CCC CCG CCT GCC TCA TCC TCT
ACA ACC ACG ACT CCA CCC CCG CCT GCA GCC GCG GCT TCA TCC TCG TCT
ATC ATT CTA CTC CTG CTT GTC TTA TTC TTT
ATC CTA CTC CTT TTC TTT
Trinucleotide

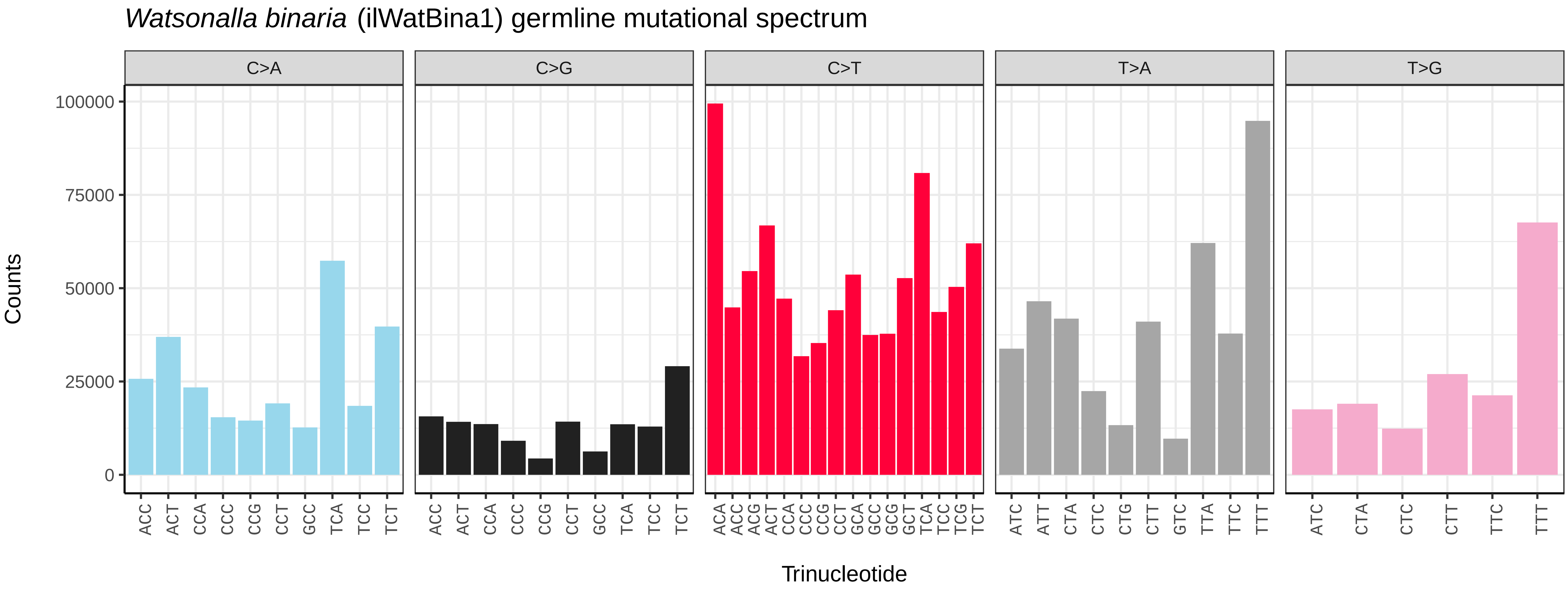
*Watsonalla binaria* (ilWatBina1) germline mutational spectrum
C>A
C>G
C>T
T>A
T>G
100000
75000
50000
25000
0
Counts
ACC ACT CCA CCC CCG CCT GCC TCA TCC TCT
ACA ACC ACG ACT CCA CCC CCG CCT GCA GCC GCG GCT TCA TCC TCG TCT
ATC ATT CTA CTC CTG CTT GTC TTA TTC TTT
ATC CTA CTC CTT TTC TTT
Trinucleotide

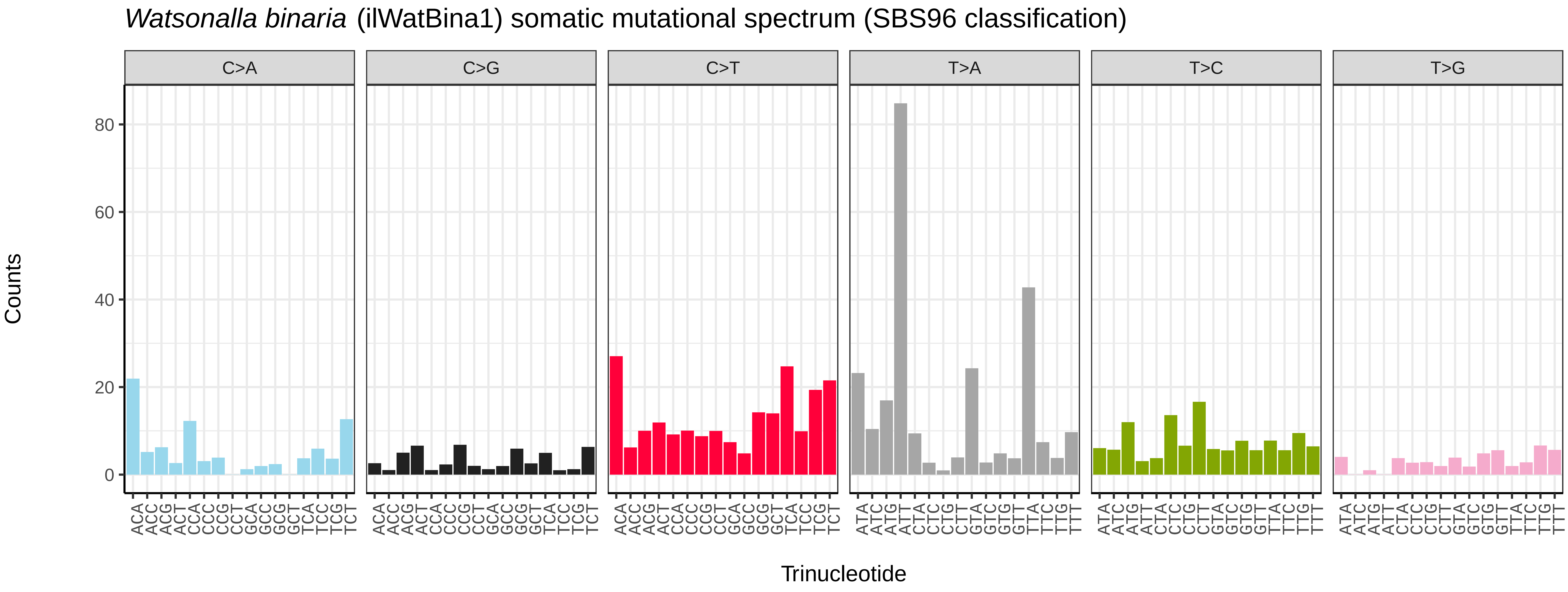
*Watsonalla binaria* (ilWatBina1) somatic mutational spectrum (SBS96 classification)
C>A
C>G
C>T
T>A
T>C
T>G
80
60
40
20
0
Counts
ACA ACC ACG ACT CCA CCC CCG CCT GCA GCC GCG GCT TCA TCC TCG TCT
ATA ATC ATG ATT CTA CTC CTG CTT GTA GTC GTG GTT TTA TTC TTG TTT
Trinucleotide

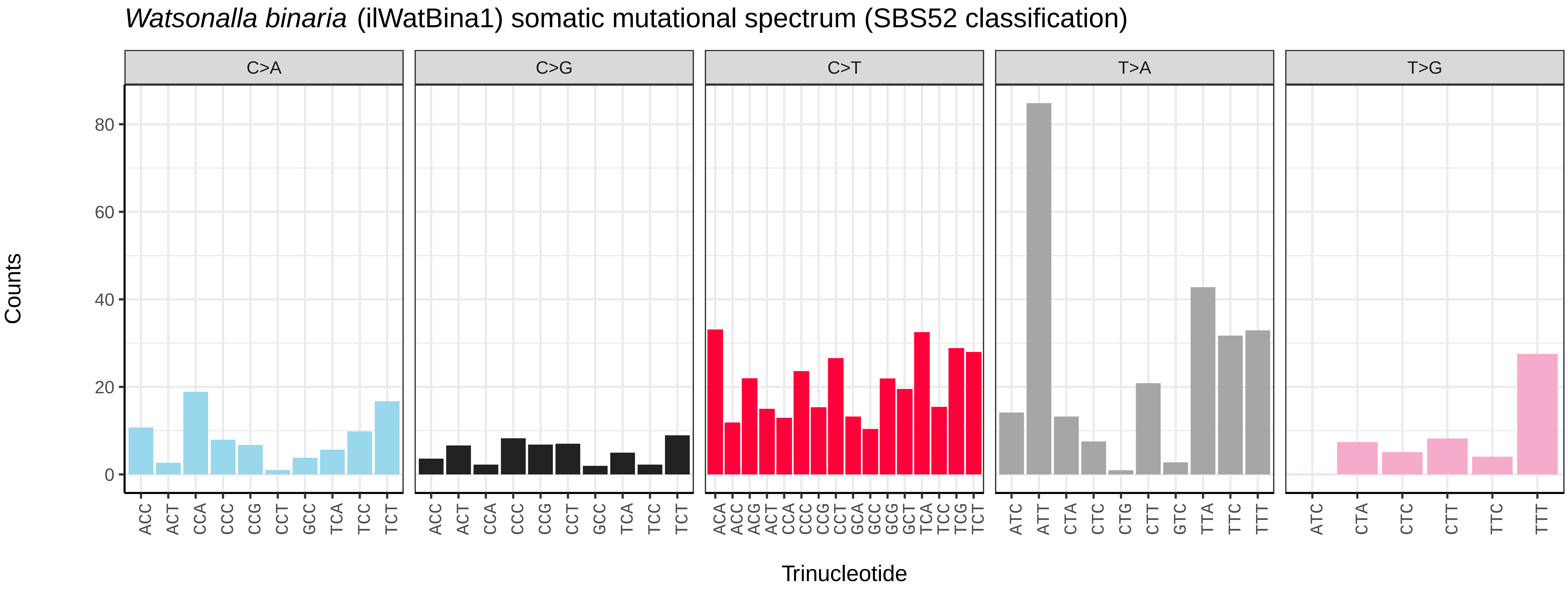
*Watsonalla binaria* (ilWatBina1) somatic mutational spectrum (SBS52 classification)
C>A
C>G
C>T
T>A
T>G
80
60
40
20
0
Counts
ACC ACT CCA CCC CCG CCT GCC TCA TCC TCT
ACA ACC ACG ACT CCA CCC CCG CCT GCA GCC GCG GCT TCA TCC TCG TCT
ATC ATT CTA CTC CTG CTT GTC TTA TTC TTT
ATC CTA CTC CTT TTC TTT
Trinucleotide

*Xanthia icteritia* (ilXanIcte2) germline mutational spectrum

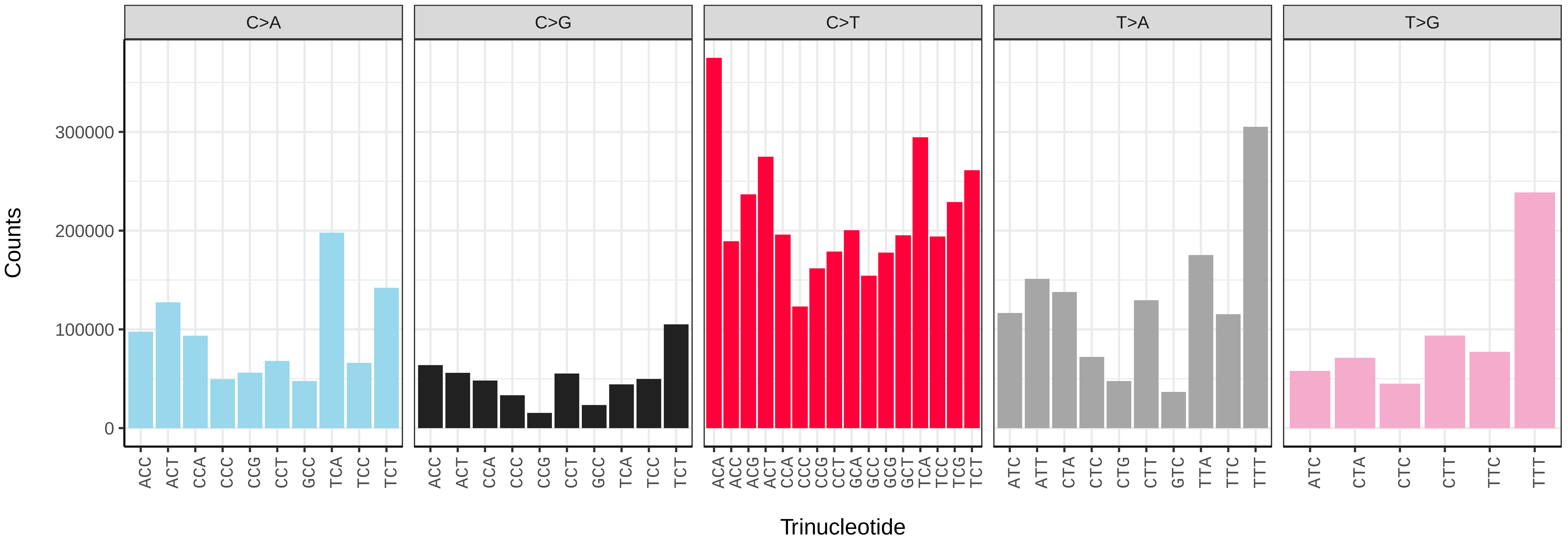


*Xanthia icteritia* (ilXanIcte2) somatic mutational spectrum (SBS96 classification)

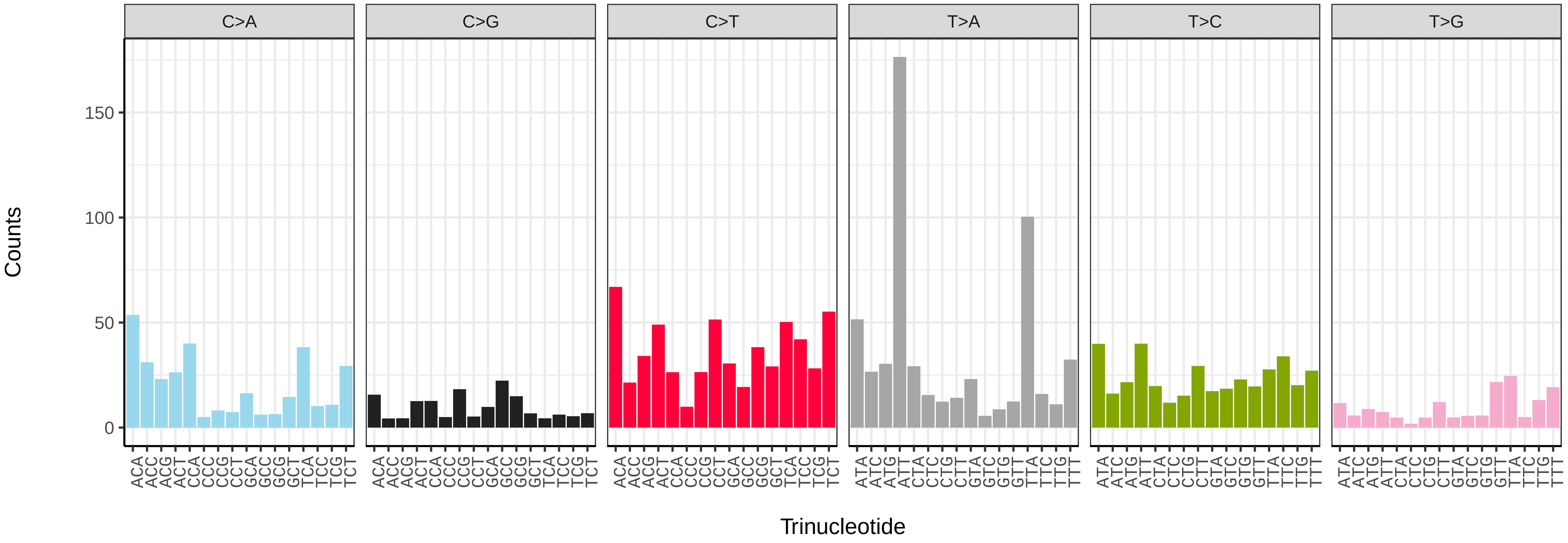


*Xanthia icteritia* (ilXanIcte2) somatic mutational spectrum (SBS52 classification)

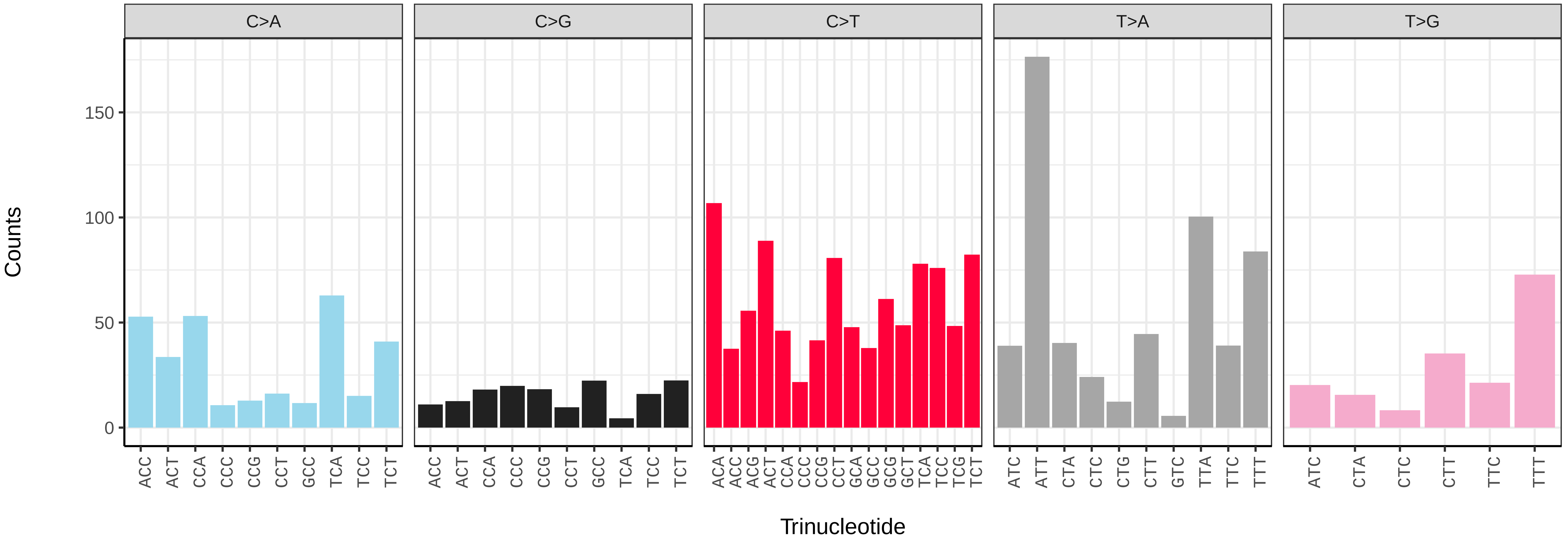

*Xanthorhoe spadicearia* (ilXanSpad1) germline mutational spectrum

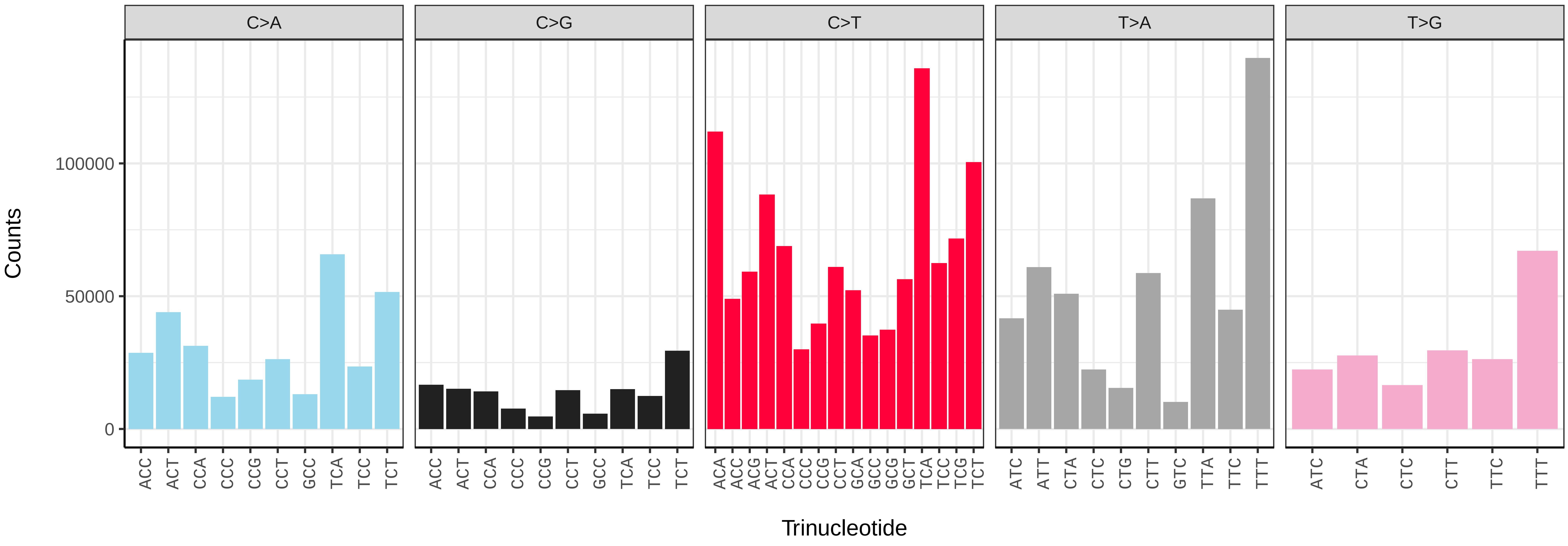


*Xanthorhoe spadicearia* (ilXanSpad1) somatic mutational spectrum (SBS96 classification)

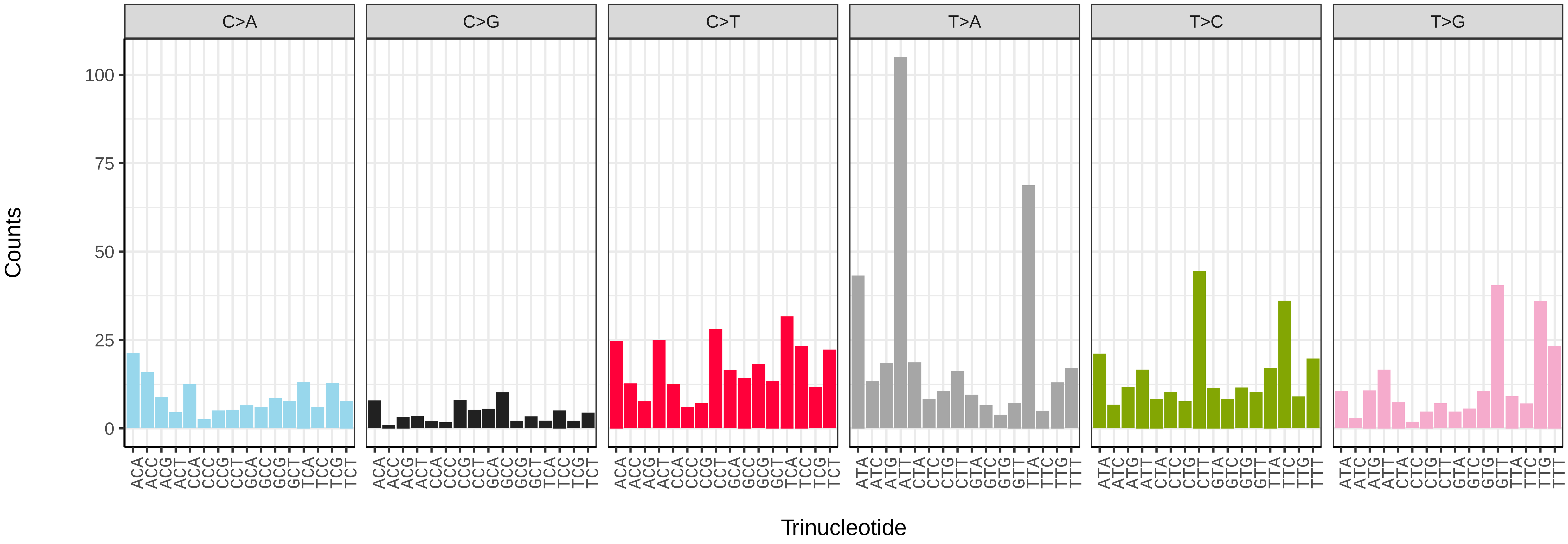


*Xanthorhoe spadicearia* (ilXanSpad1) somatic mutational spectrum (SBS52 classification)

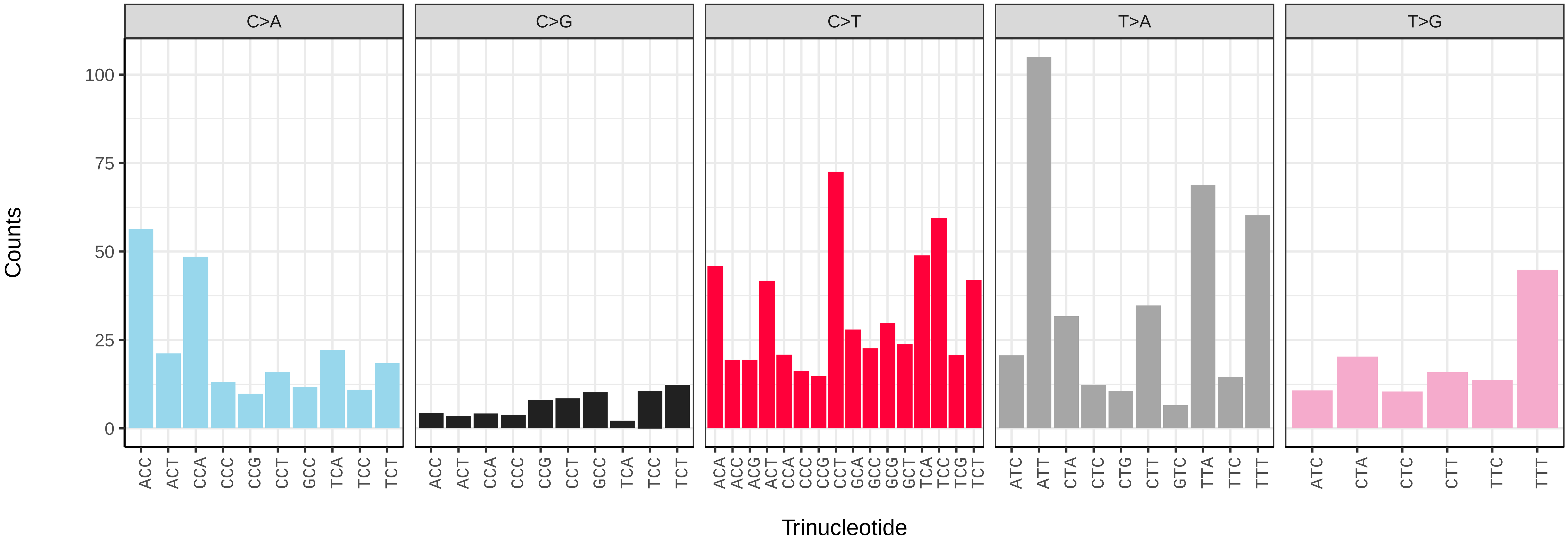

*Xestia c−nigrum* (ilXesCnig1) germline mutational spectrum

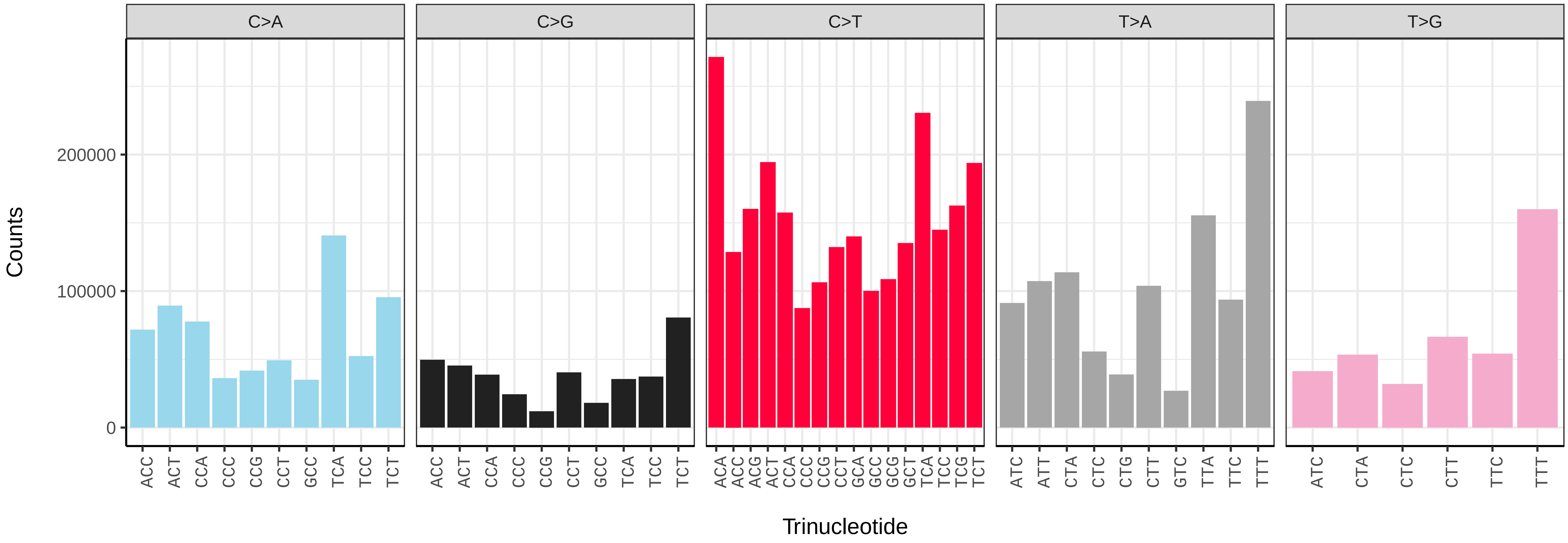


*Xestia c−nigrum* (ilXesCnig1) somatic mutational spectrum (SBS96 classification)

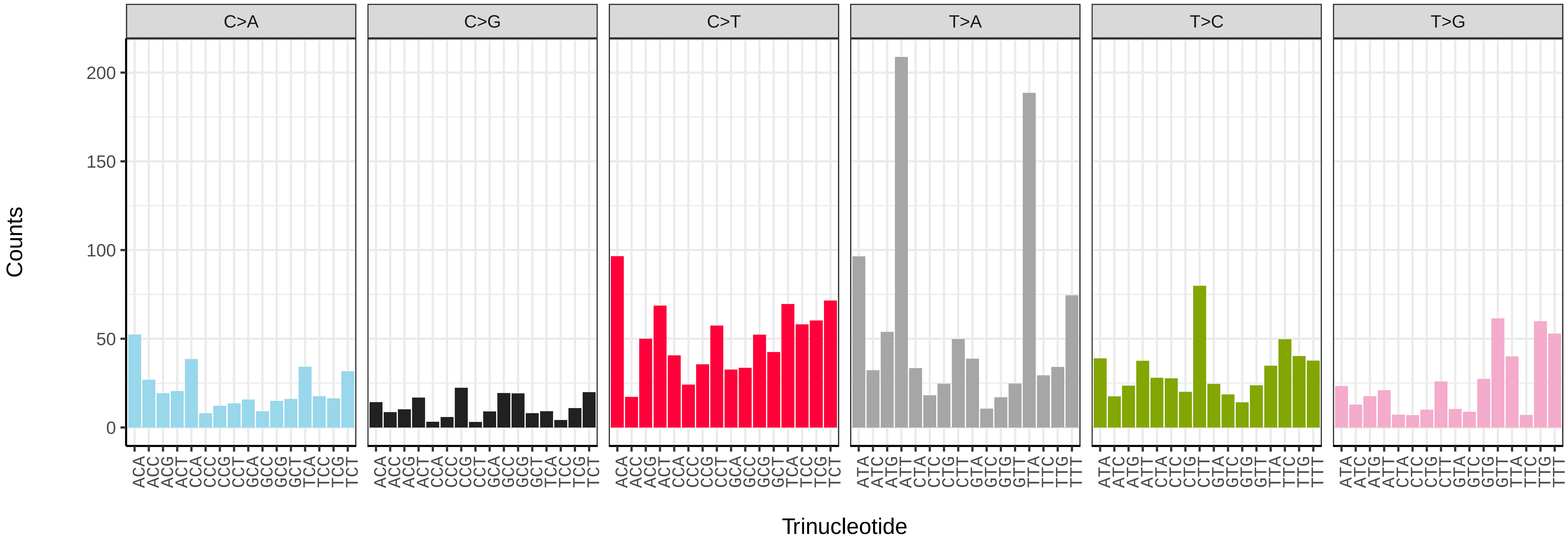


*Xestia c−nigrum* (ilXesCnig1) somatic mutational spectrum (SBS52 classification)

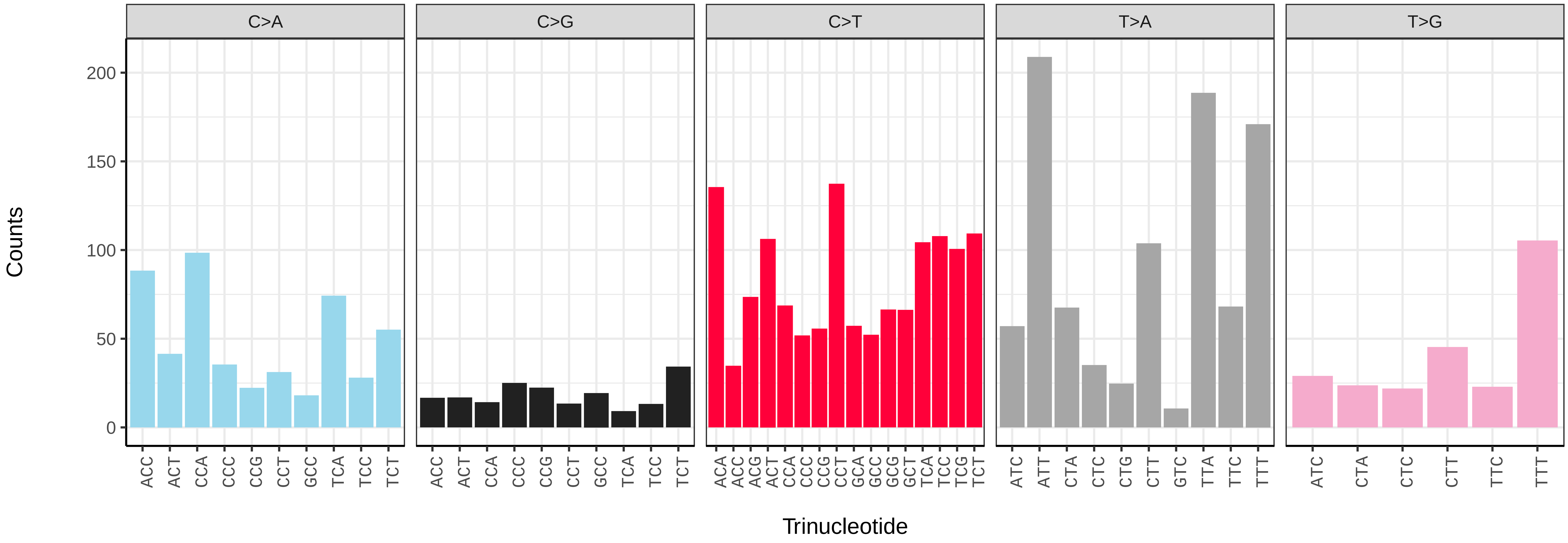

*Xestia sexstrigata* (ilXesSexs1) germline mutational spectrum

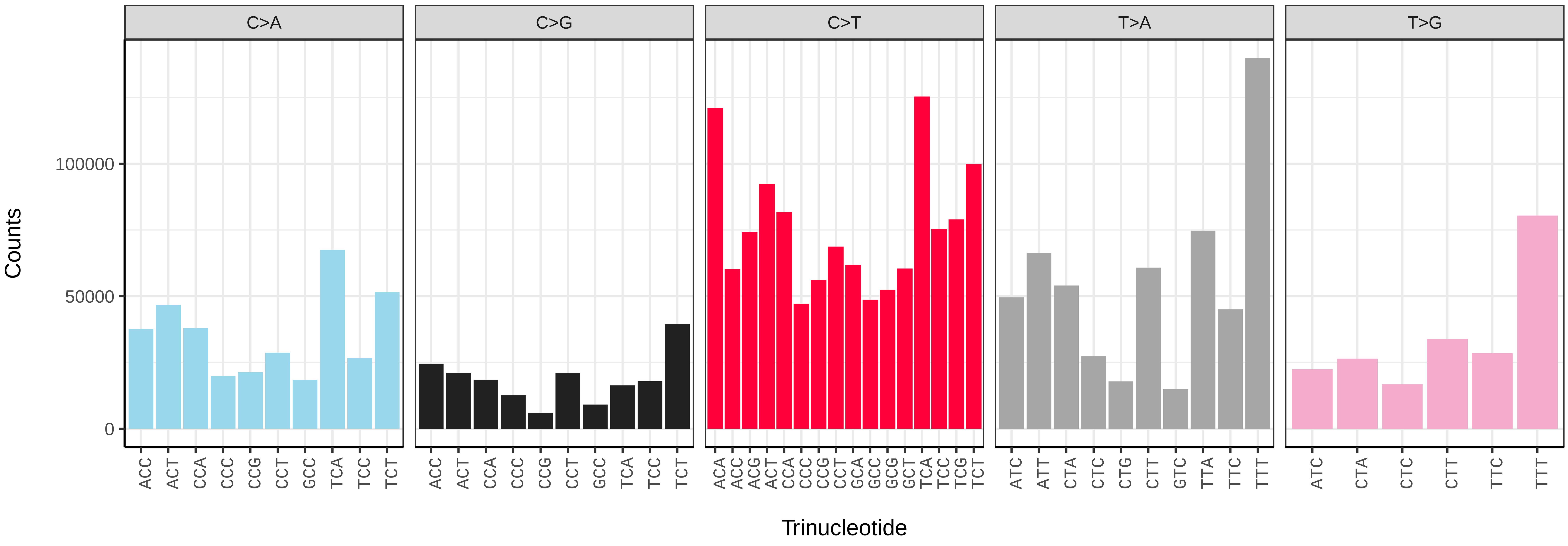


*Xestia sexstrigata* (ilXesSexs1) somatic mutational spectrum (SBS96 classification)

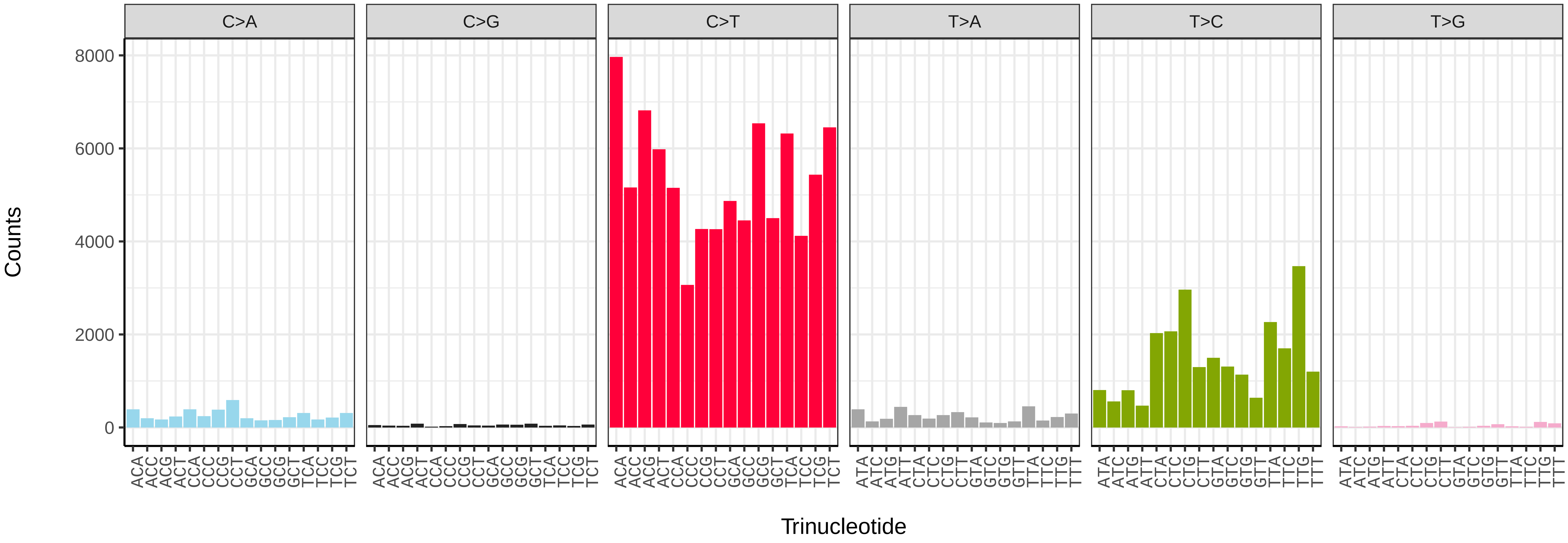


*Xestia sexstrigata* (ilXesSexs1) somatic mutational spectrum (SBS52 classification)

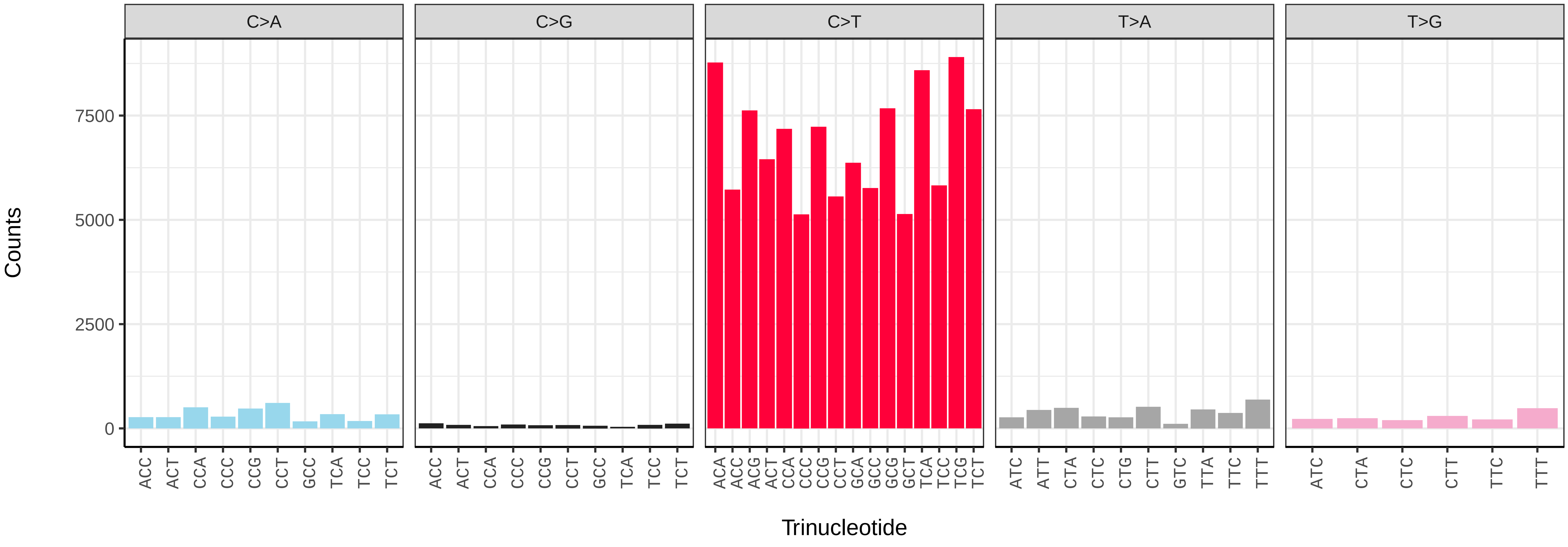

*Xestia xanthographa* (ilXesXant1) germline mutational spectrum

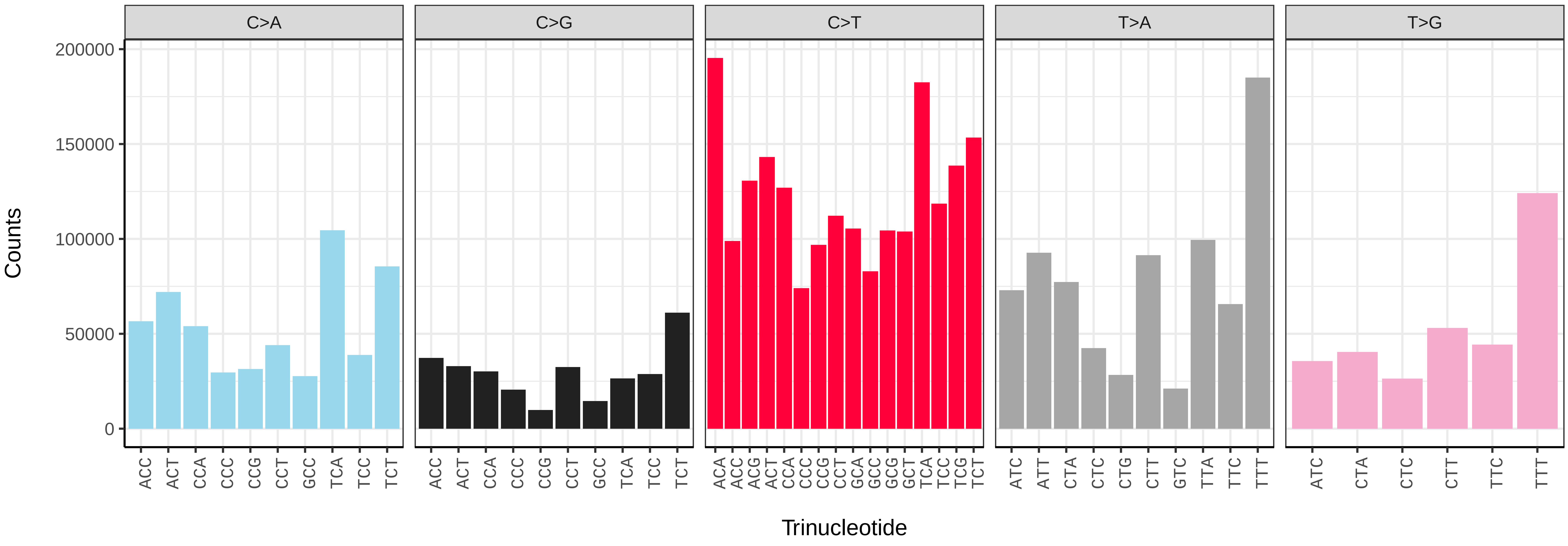


*Xestia xanthographa* (ilXesXant1) somatic mutational spectrum (SBS96 classification)

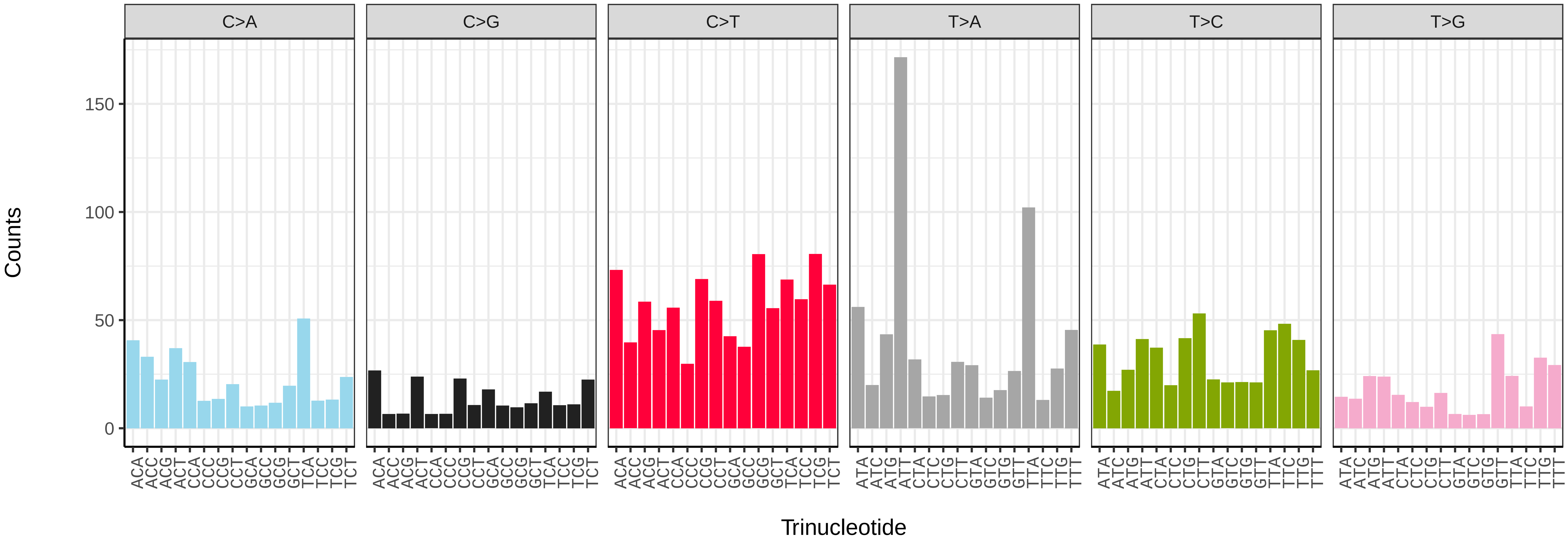


*Xestia xanthographa* (ilXesXant1) somatic mutational spectrum (SBS52 classification)

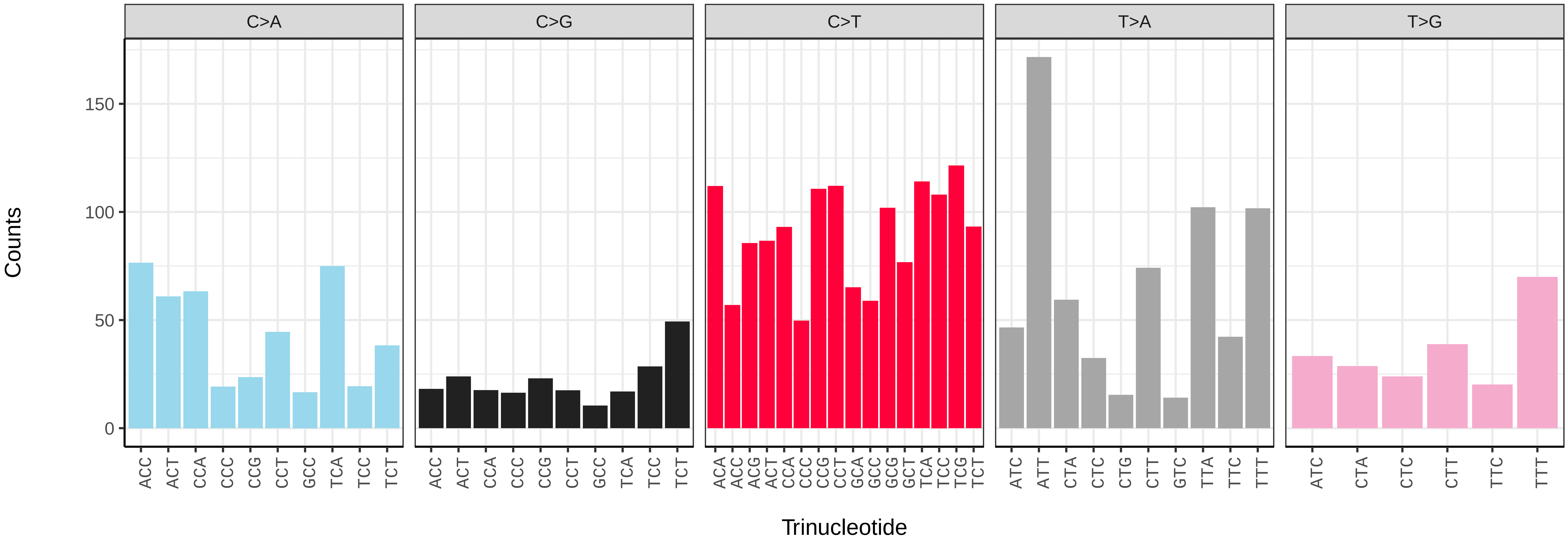

*Xylocampa areola* (ilXylAreo1) germline mutational spectrum

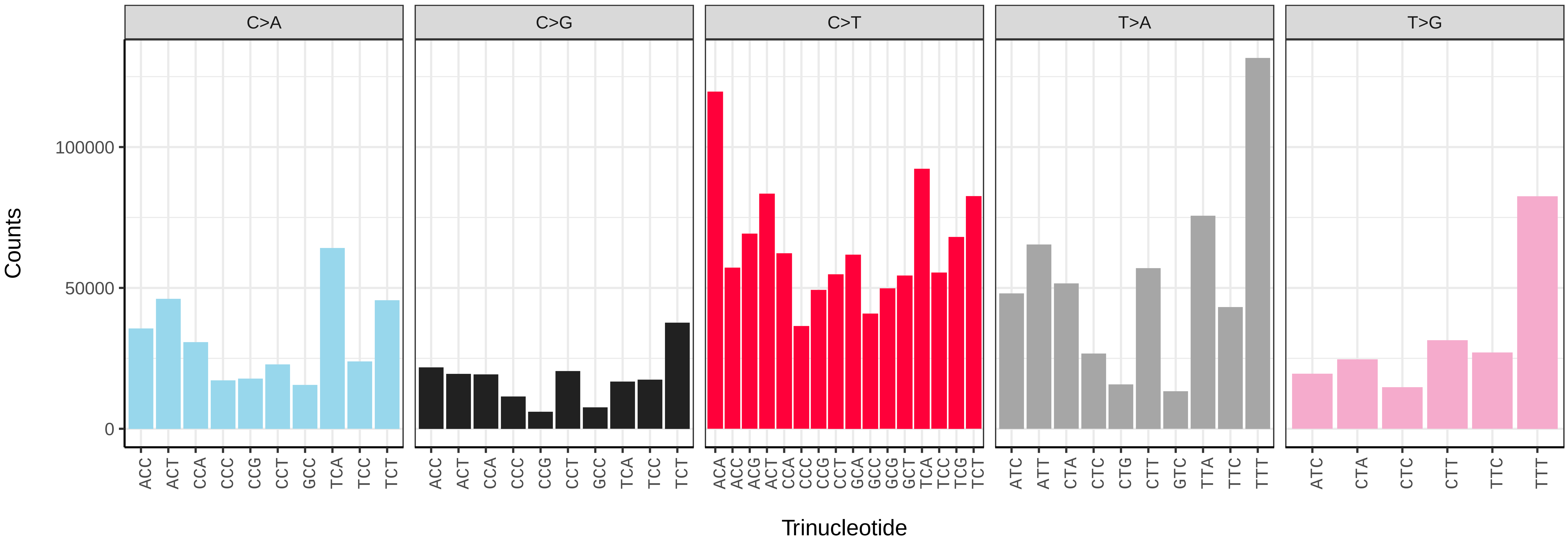


*Xylocampa areola* (ilXylAreo1) somatic mutational spectrum (SBS96 classification)

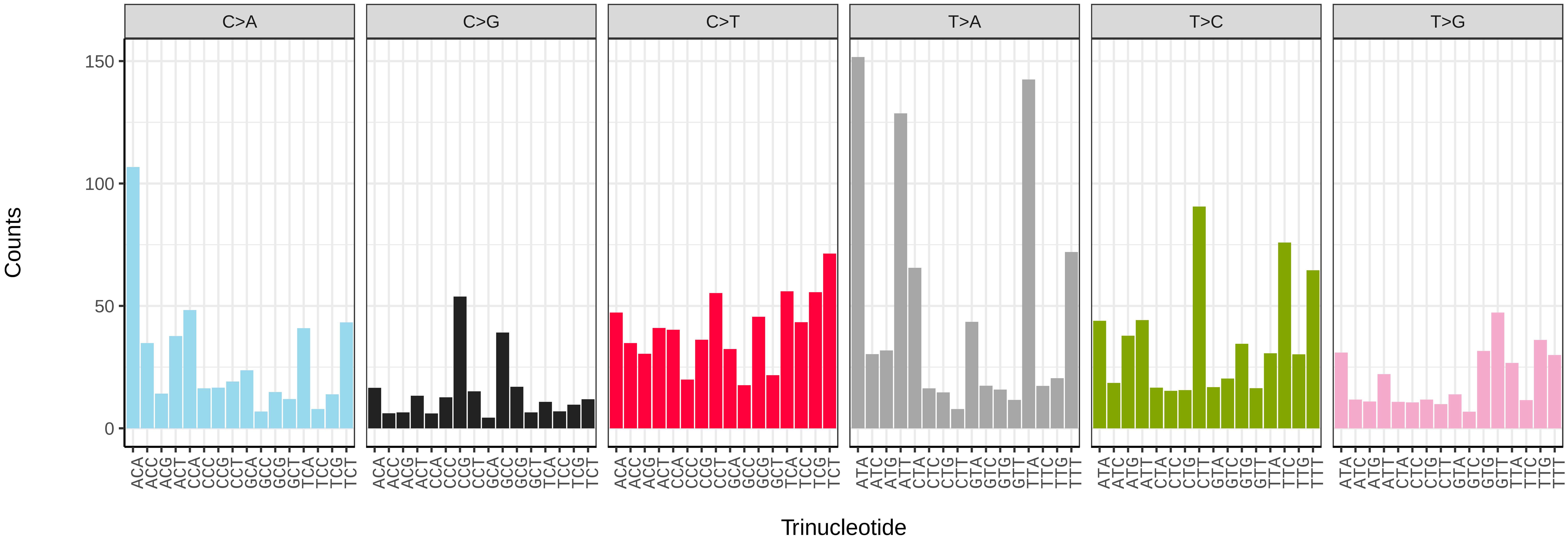


*Xylocampa areola* (ilXylAreo1) somatic mutational spectrum (SBS52 classification)

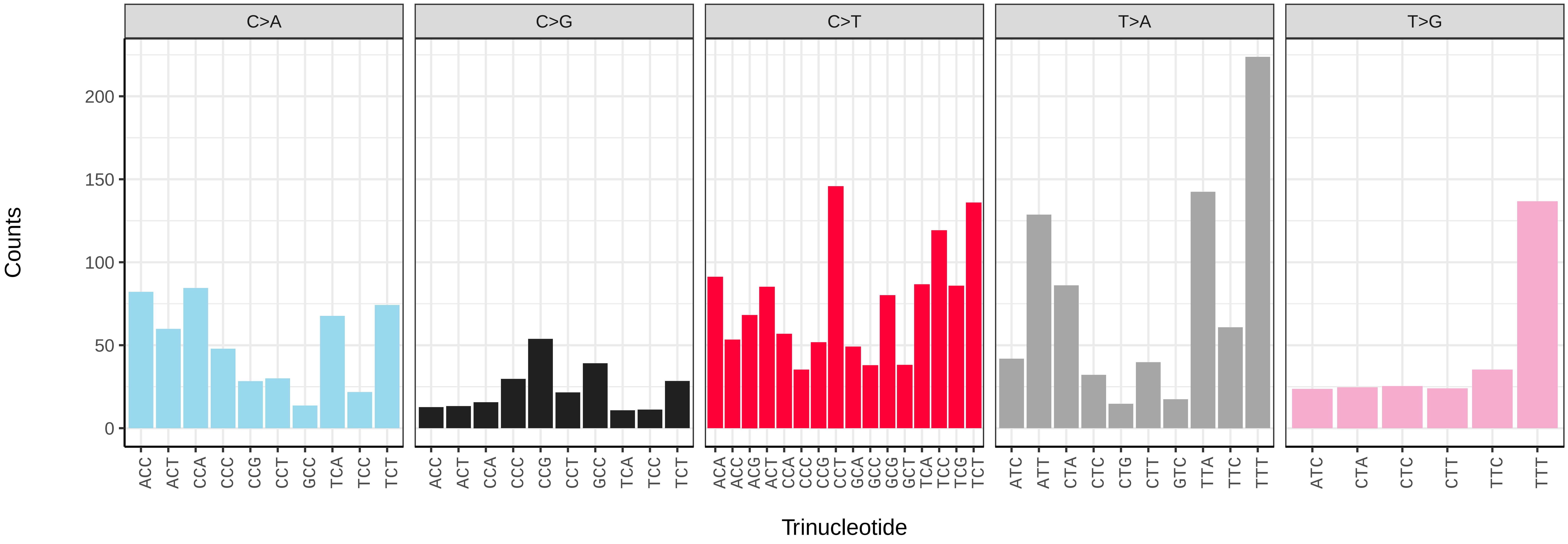

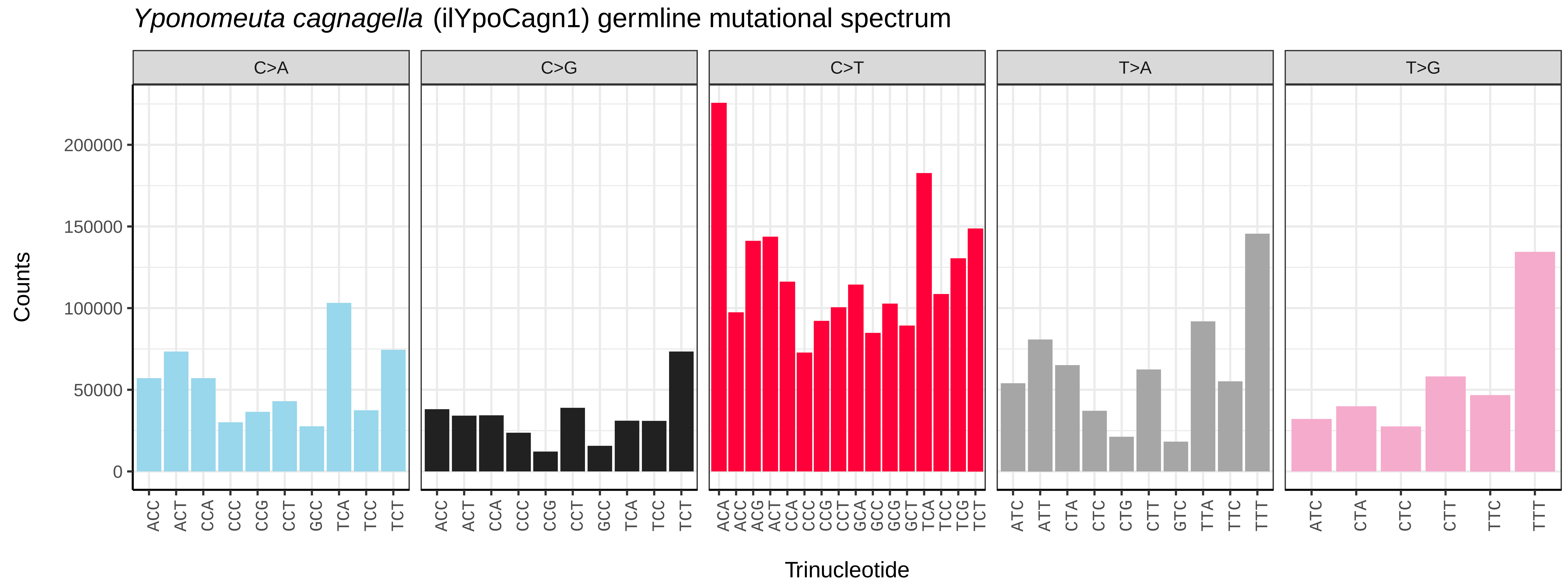
*Yponomeuta cagnagella* (ilYpoCagn1) germline mutational spectrum
C>A
C>G
C>T
T>A
T>G
Counts
0
50000
100000
150000
200000
Trinucleotide

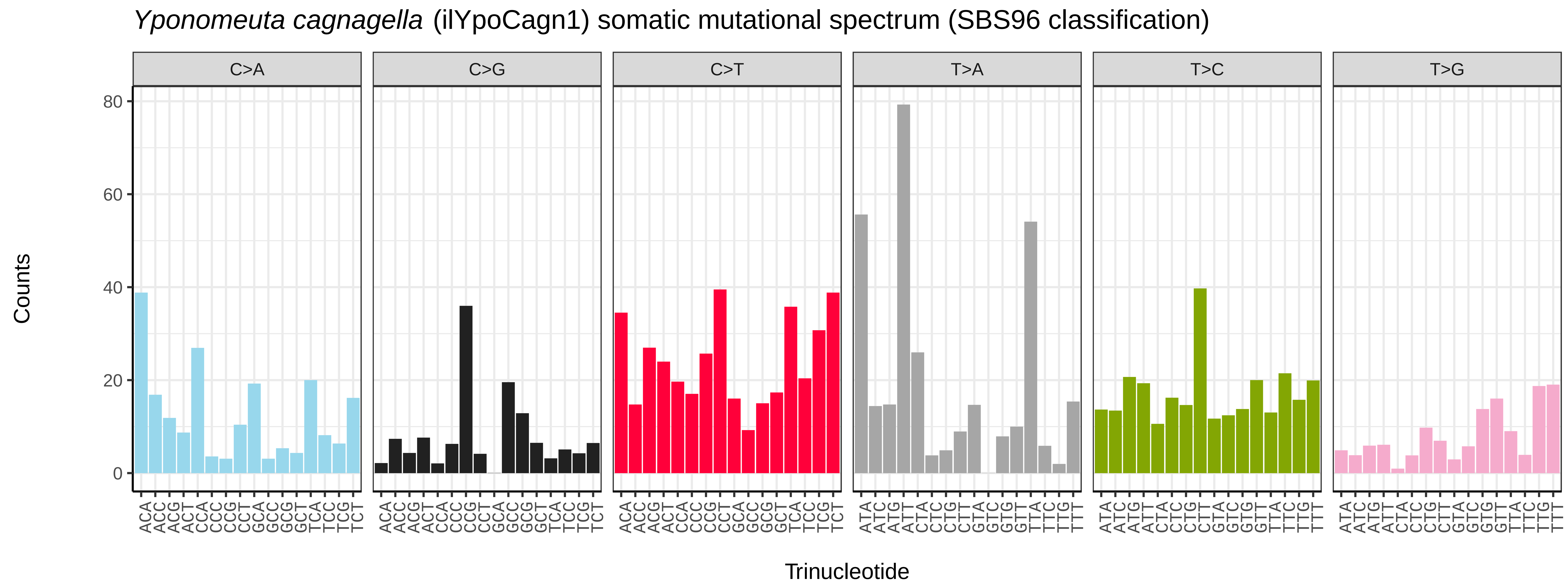
*Yponomeuta cagnagella* (ilYpoCagn1) somatic mutational spectrum (SBS96 classification)
C>A
C>G
C>T
T>A
T>C
T>G
Counts
0
20
40
60
80
Trinucleotide

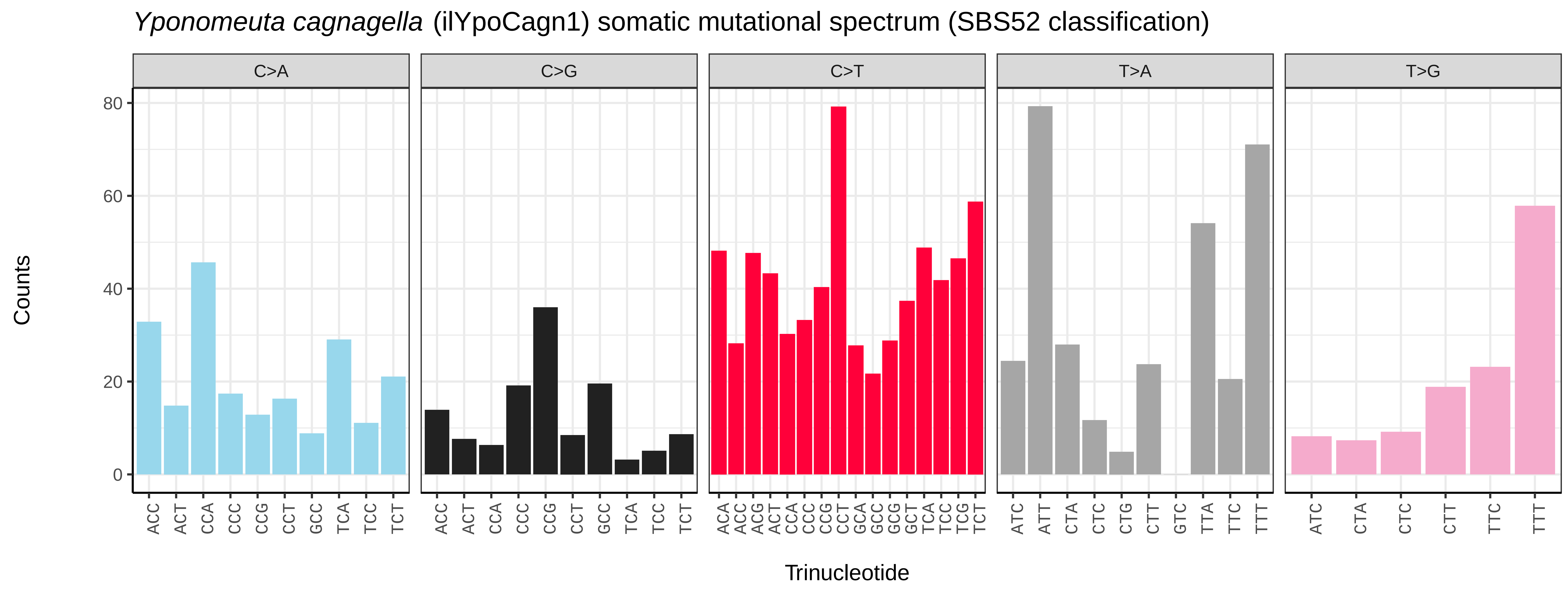
*Yponomeuta cagnagella* (ilYpoCagn1) somatic mutational spectrum (SBS52 classification)
C>A
C>G
C>T
T>A
T>G
Counts
0
20
40
60
80
Trinucleotide

*Yponomeuta cagnagella* (ilYpoCagn1.ilYpoCagn5) germline mutational spectrum

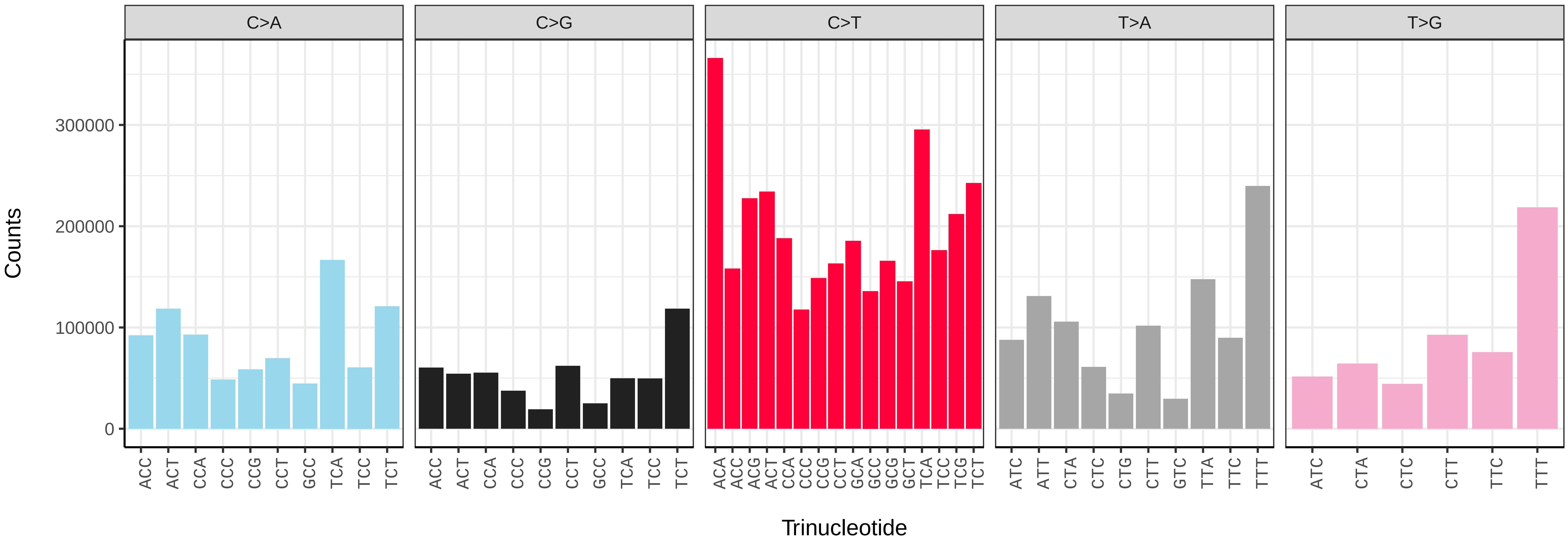


*Yponomeuta cagnagella* (ilYpoCagn1.ilYpoCagn5) somatic mutational spectrum (SBS96 classification)

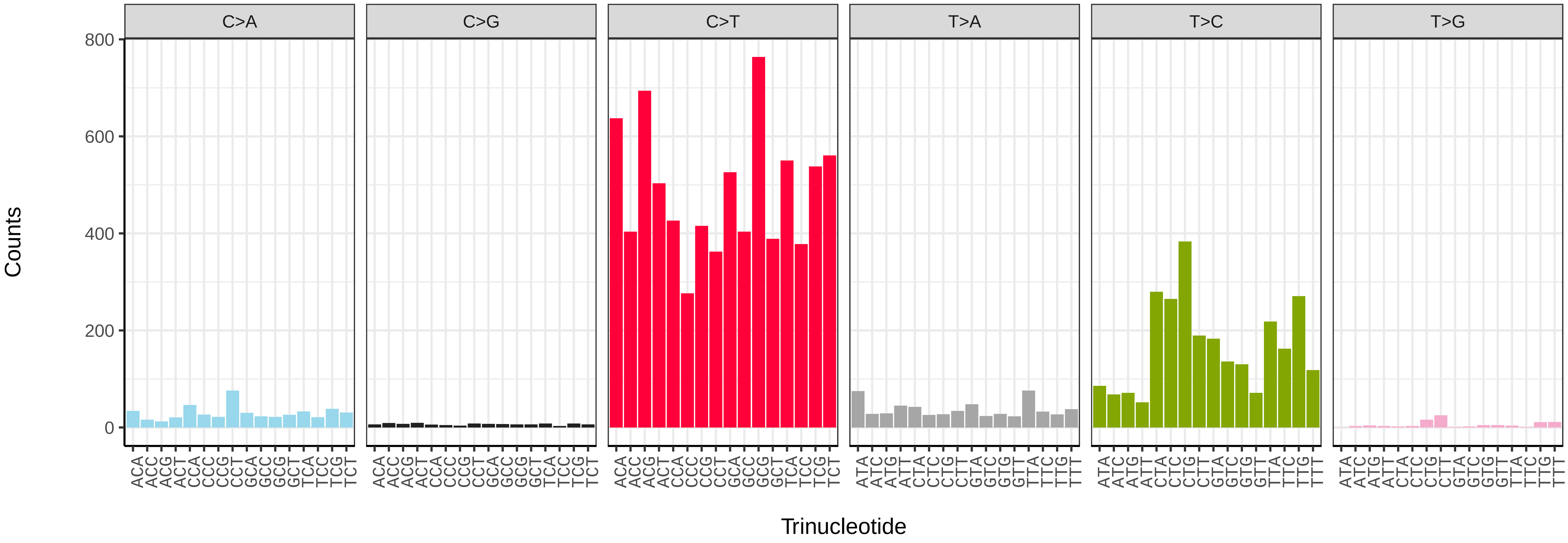


*Yponomeuta cagnagella* (ilYpoCagn1.ilYpoCagn5) somatic mutational spectrum (SBS52 classification)

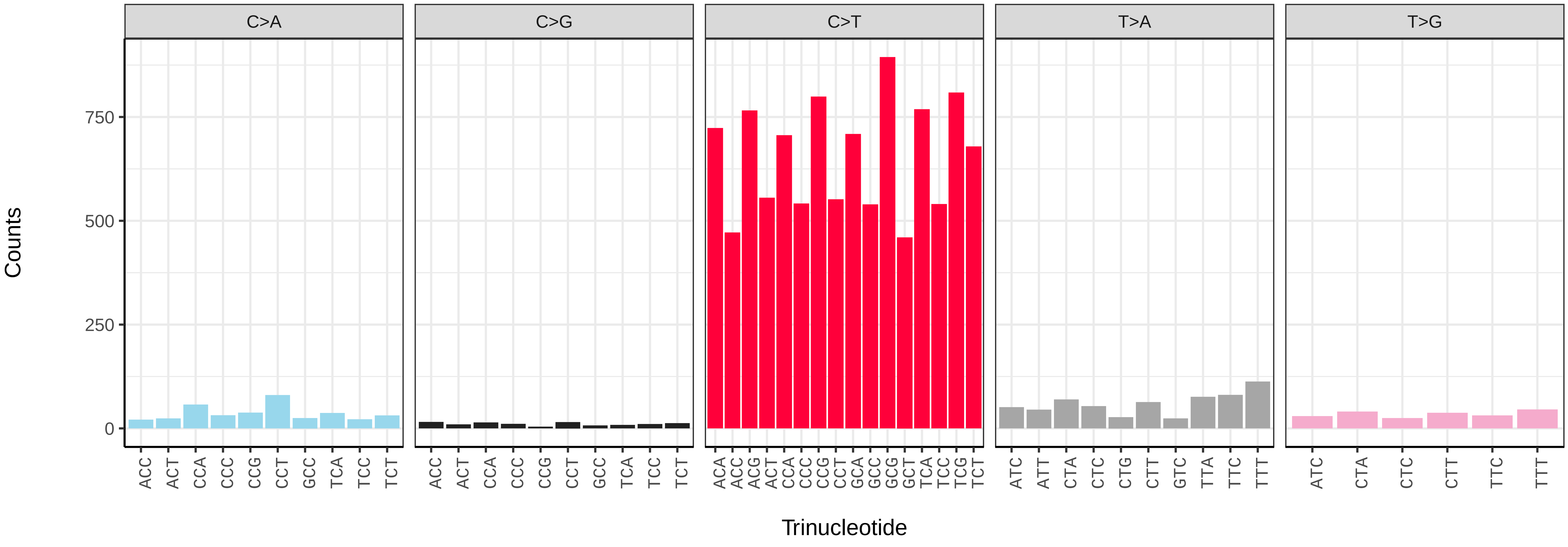

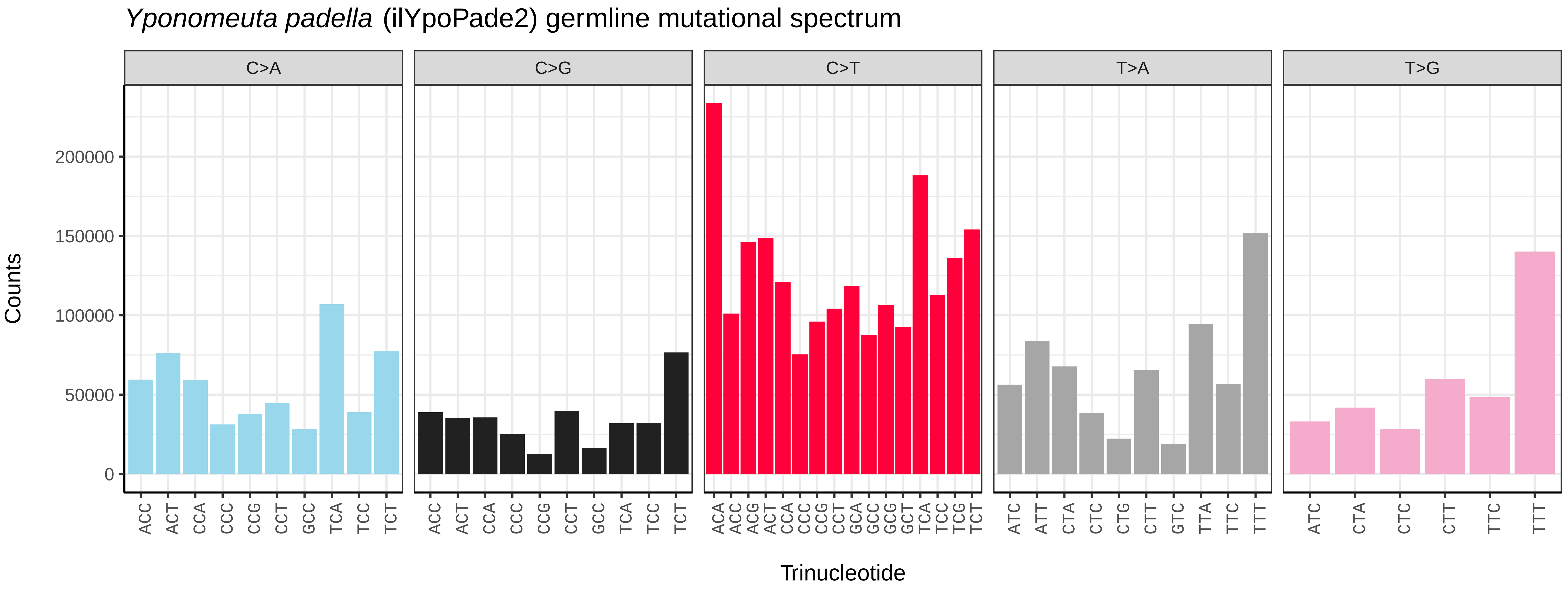
Yponomeuta padella (ilYpoPade2) germline mutational spectrum
C>A
C>G
C>T
T>A
T>G
Counts
0
50000
100000
150000
200000
ACC ACT CCA CCC CCG CCT GCC TCA TCC TCT
ACA ACC ACG ACT CCA CCC CCG CCT GCA GCC GCG GCT TCA TCC TCG TCT
ATC ATT CTA CTC CTG CTT GTC TTA TTC TTT
ATC CTA CTC CTT TTC TTT
Trinucleotide

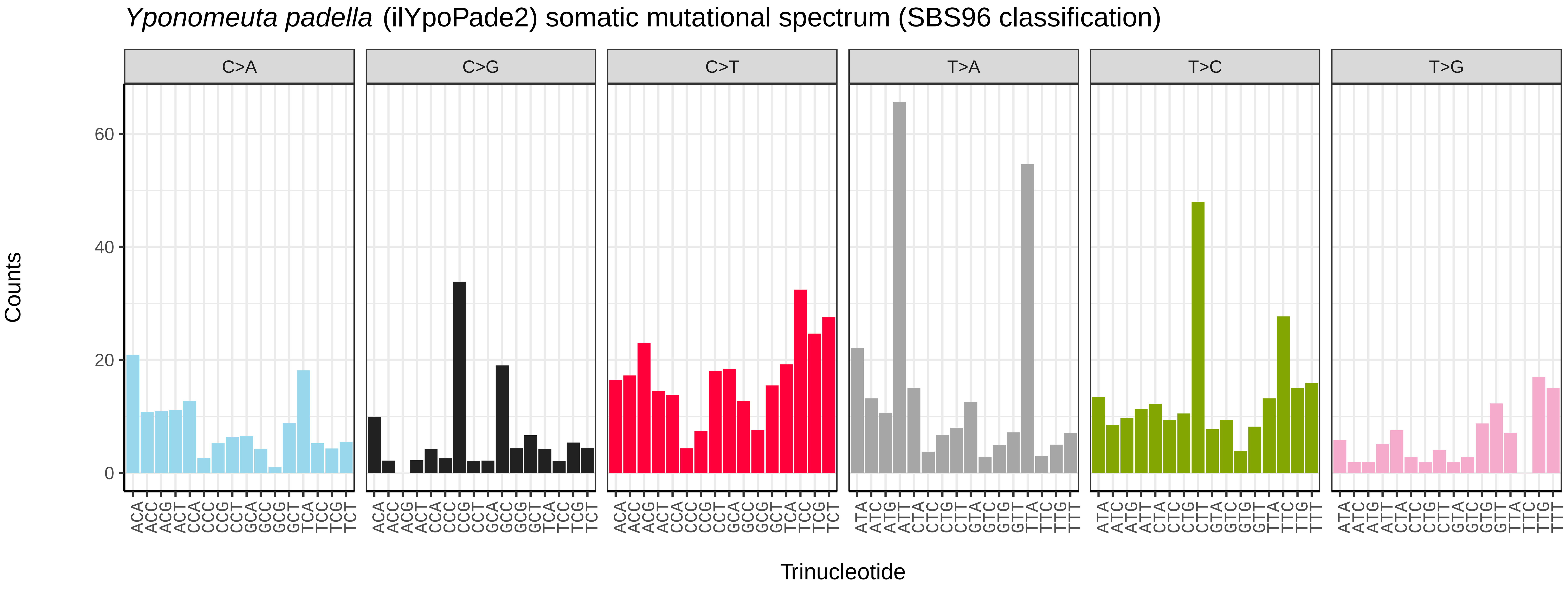
Yponomeuta padella (ilYpoPade2) somatic mutational spectrum (SBS96 classification)
C>A
C>G
C>T
T>A
T>C
T>G
Counts
0
20
40
60
ACA ACC ACG ACT CCA CCC CCG CCT GCA GCC GCG GCT TCA TCC TCG TCT
ATA ATC ATG ATT CTA CTC CTG CTT GTA GTC GTG GTT TTA TTC TTG TTT
Trinucleotide

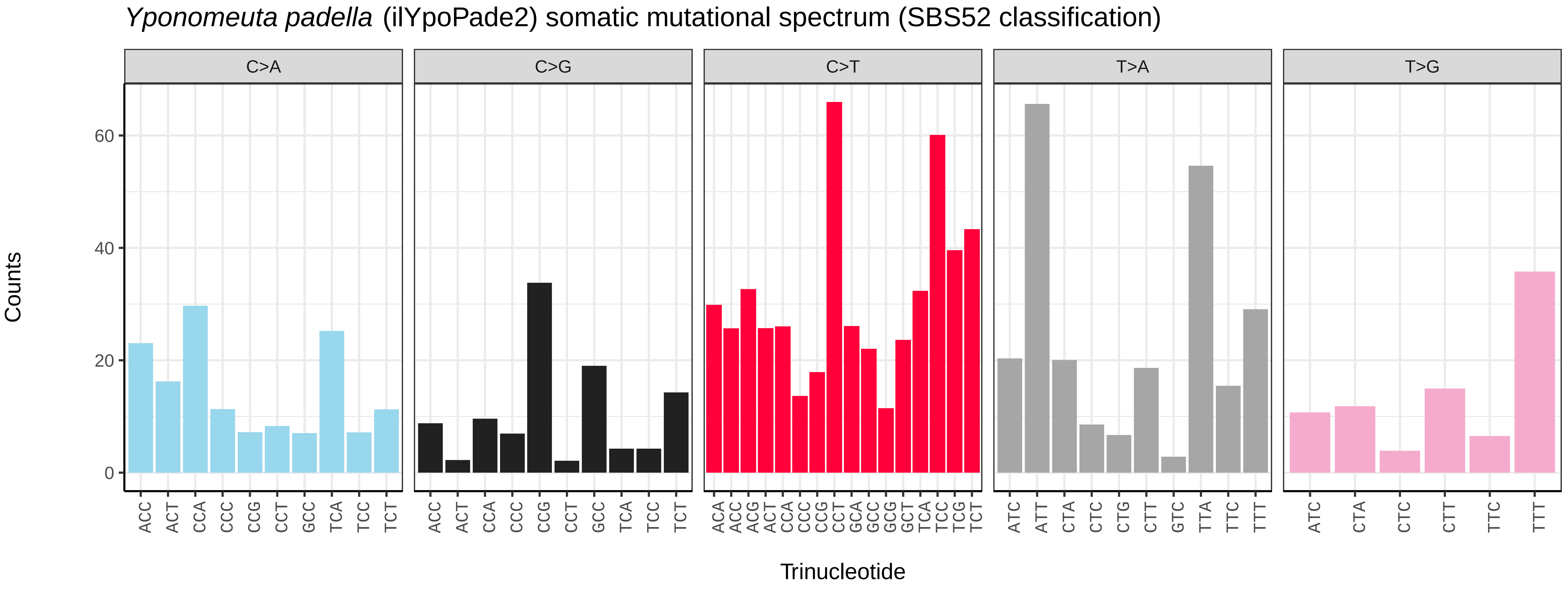
Yponomeuta padella (ilYpoPade2) somatic mutational spectrum (SBS52 classification)
C>A
C>G
C>T
T>A
T>G
Counts
0
20
40
60
ACC ACT CCA CCC CCG CCT GCC TCA TCC TCT
ACA ACC ACG ACT CCA CCC CCG CCT GCA GCC GCG GCT TCA TCC TCG TCT
ATC ATT CTA CTC CTG CTT GTC TTA TTC TTT
ATC CTA CTC CTT TTC TTT
Trinucleotide

*Yponomeuta padella* (ilYpoPade2.ilYpoPade1) germline mutational spectrum

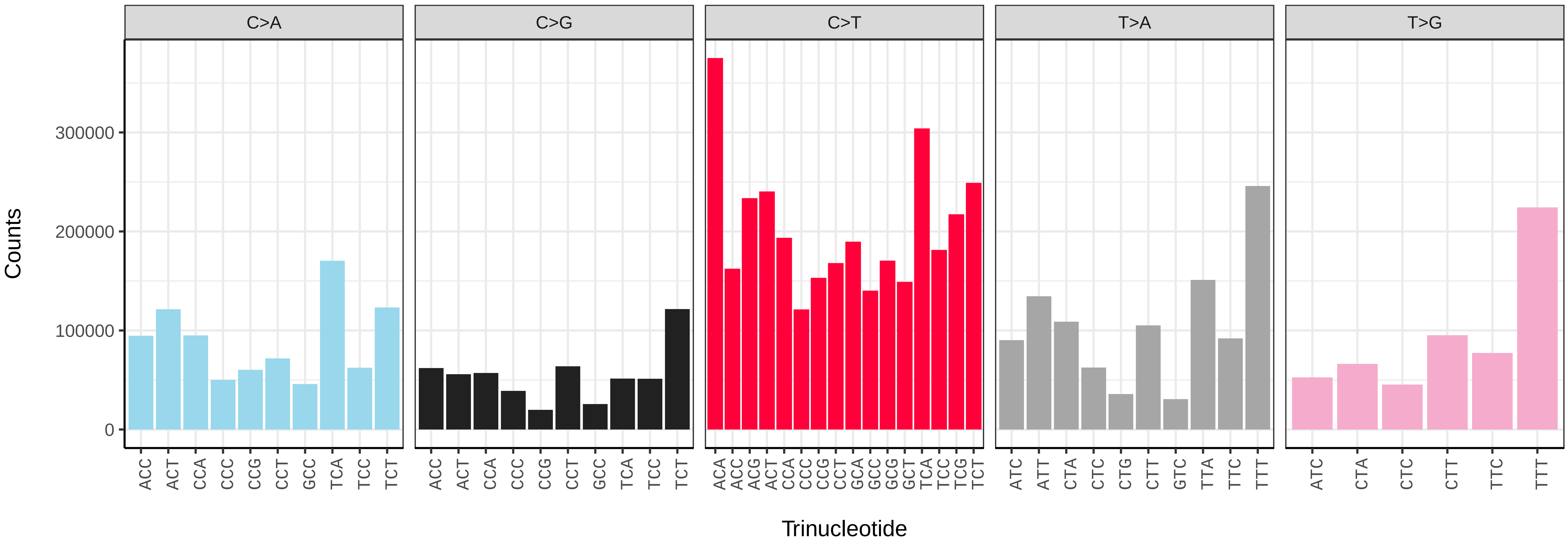


*Yponomeuta padella* (ilYpoPade2.ilYpoPade1) somatic mutational spectrum (SBS96 classification)

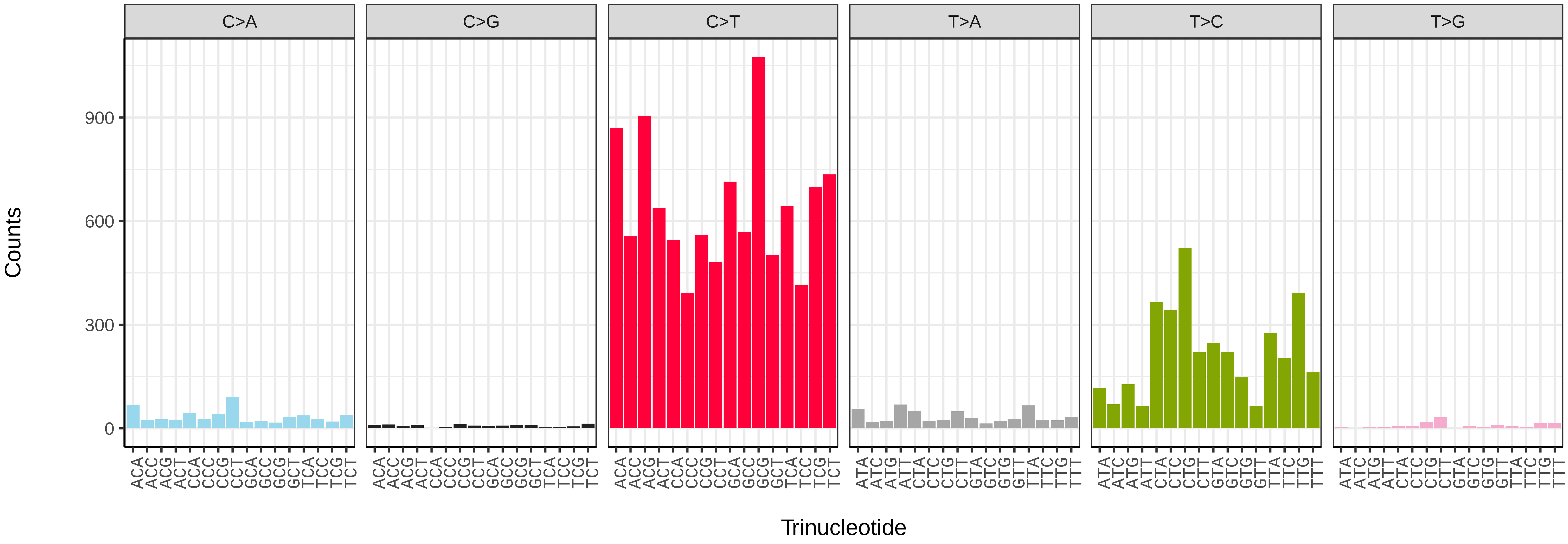


*Yponomeuta padella* (ilYpoPade2.ilYpoPade1) somatic mutational spectrum (SBS52 classification)

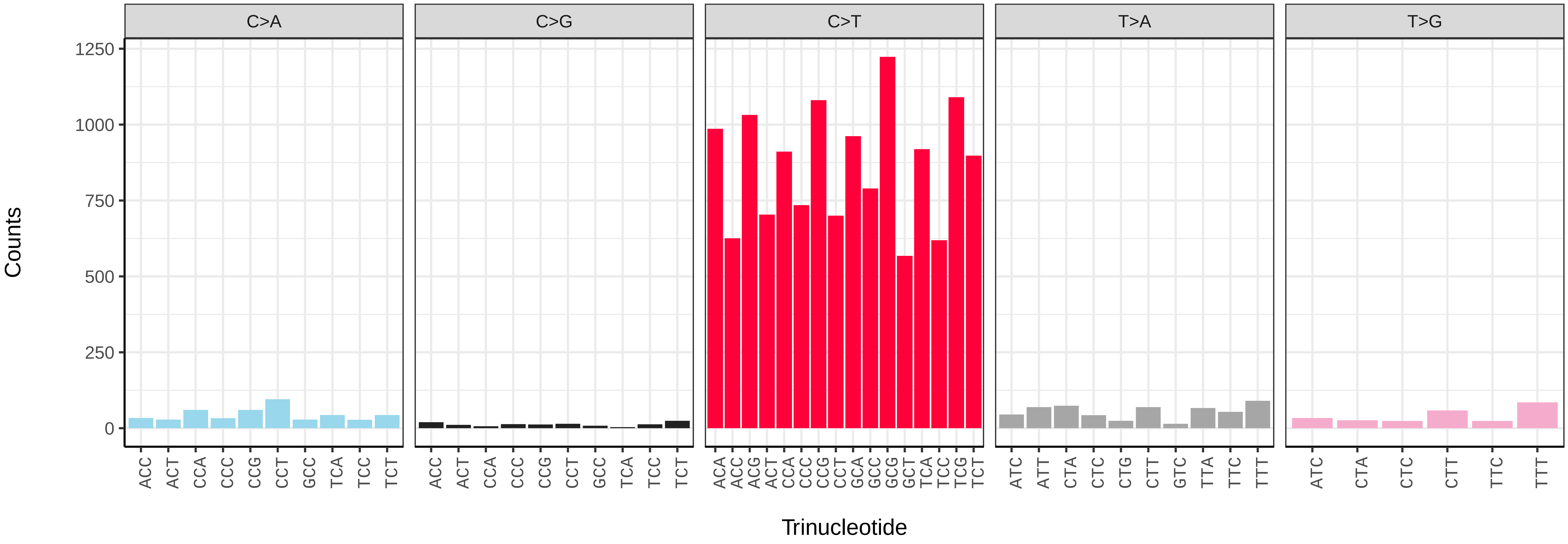

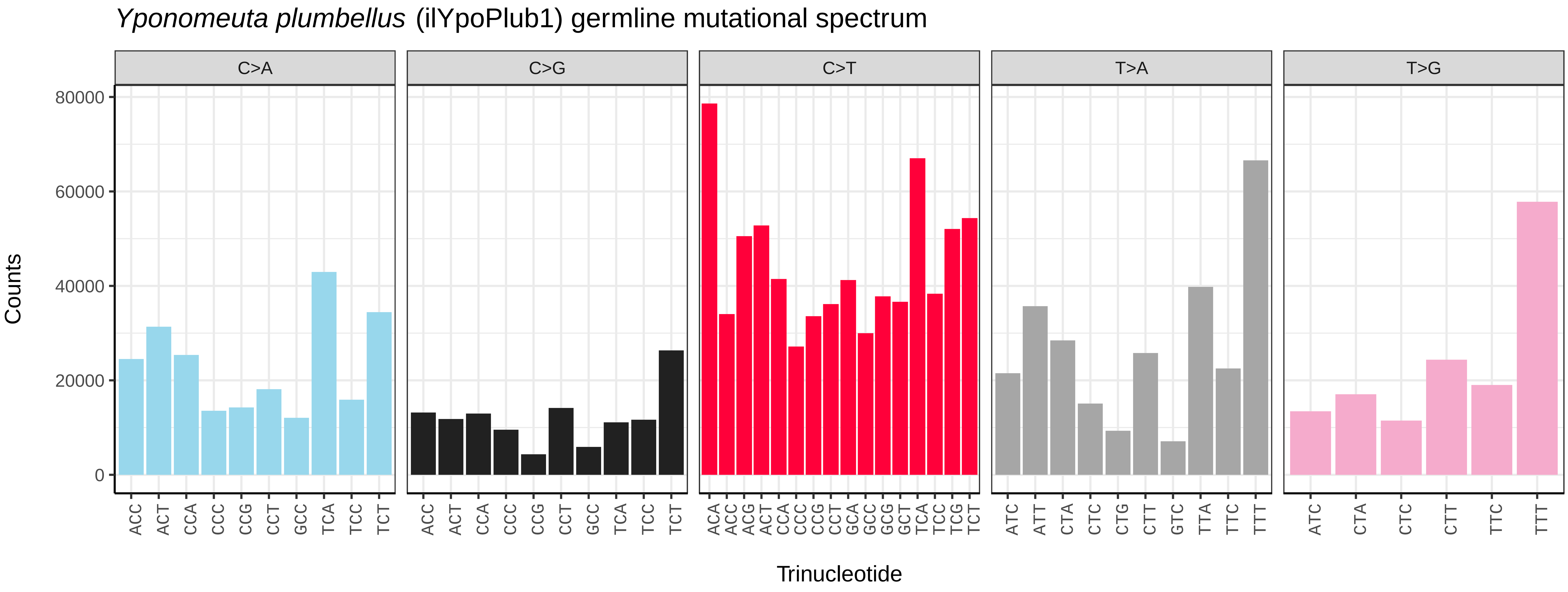
Yponomeuta plumbellus (ilYpoPlub1) germline mutational spectrum
C>A
C>G
C>T
T>A
T>G
Counts
0
20000
40000
60000
80000
Trinucleotide

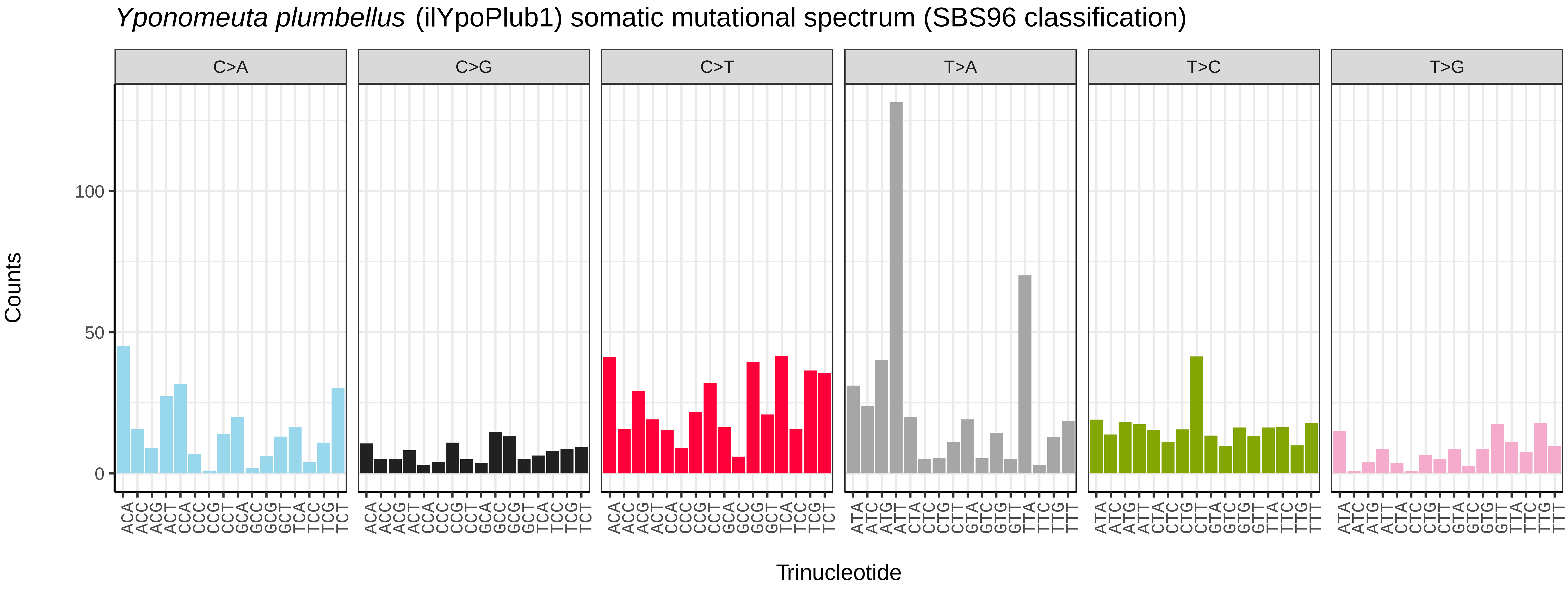
Yponomeuta plumbellus (ilYpoPlub1) somatic mutational spectrum (SBS96 classification)
C>A
C>G
C>T
T>A
T>C
T>G
Counts
0
50
100
Trinucleotide

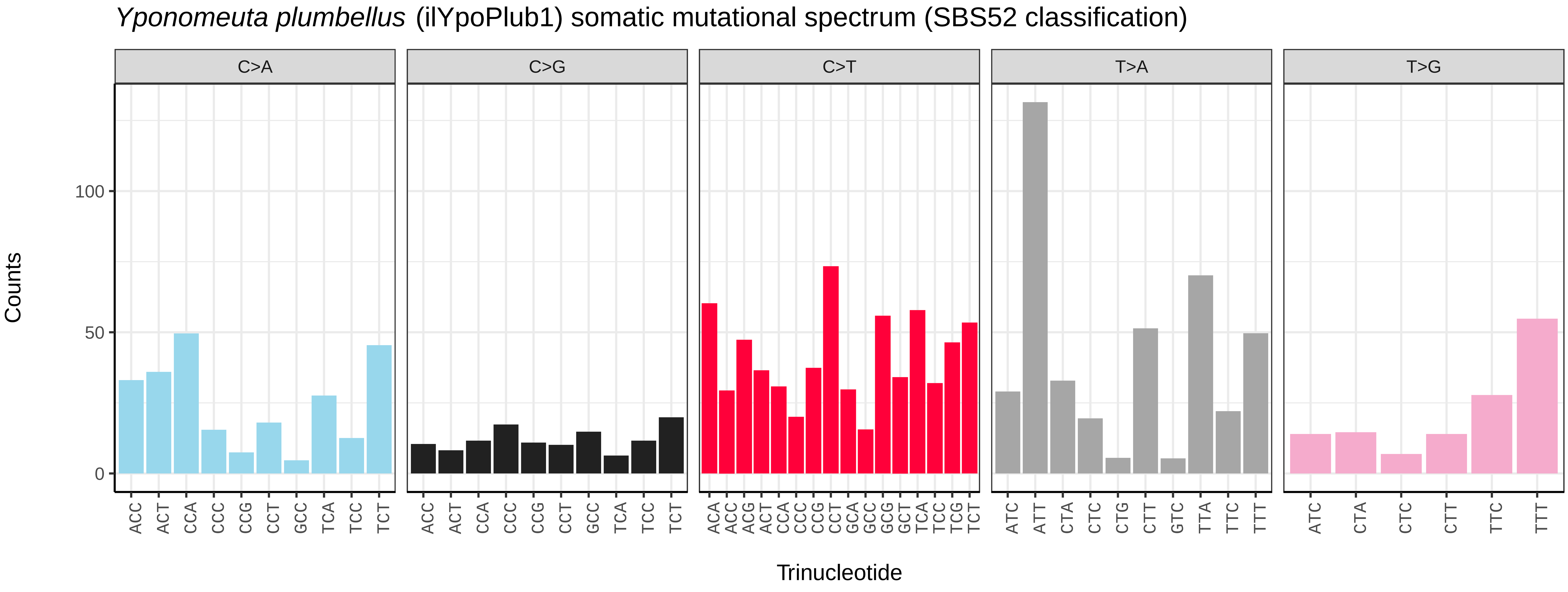
Yponomeuta plumbellus (ilYpoPlub1) somatic mutational spectrum (SBS52 classification)
C>A
C>G
C>T
T>A
T>G
Counts
0
50
100
Trinucleotide

*Yponomeuta rorrellus* (ilYpoRore1) germline mutational spectrum

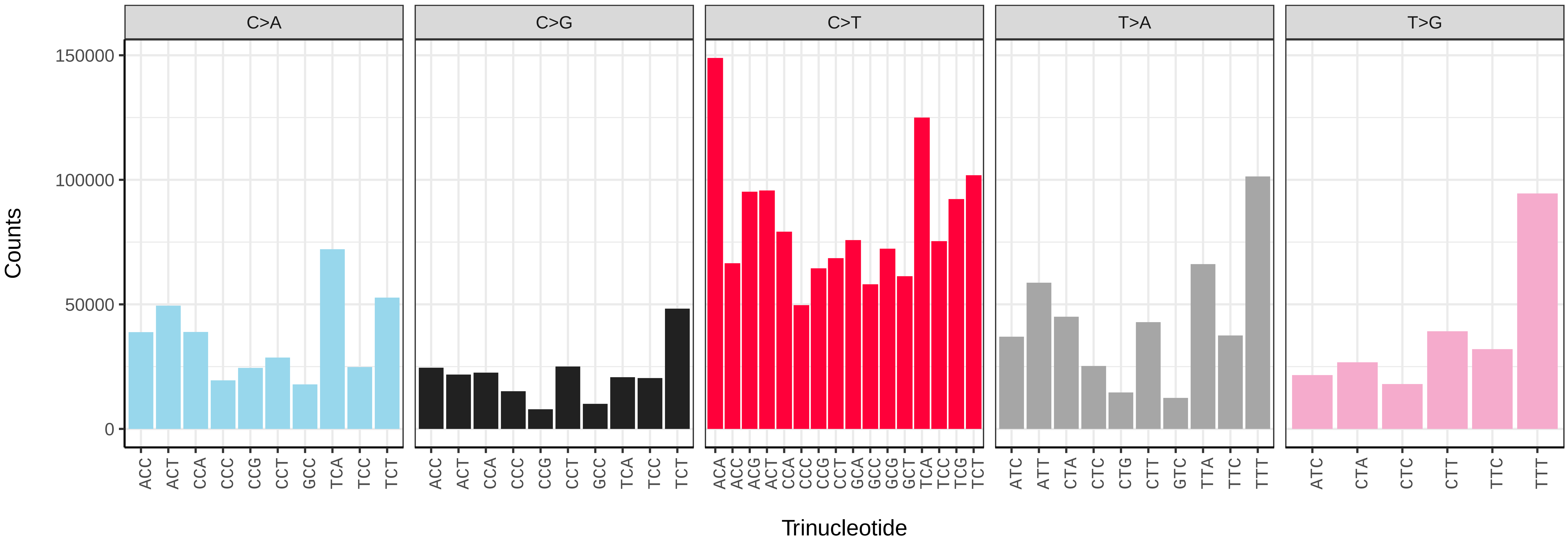


*Yponomeuta rorrellus* (ilYpoRore1) somatic mutational spectrum (SBS96 classification)

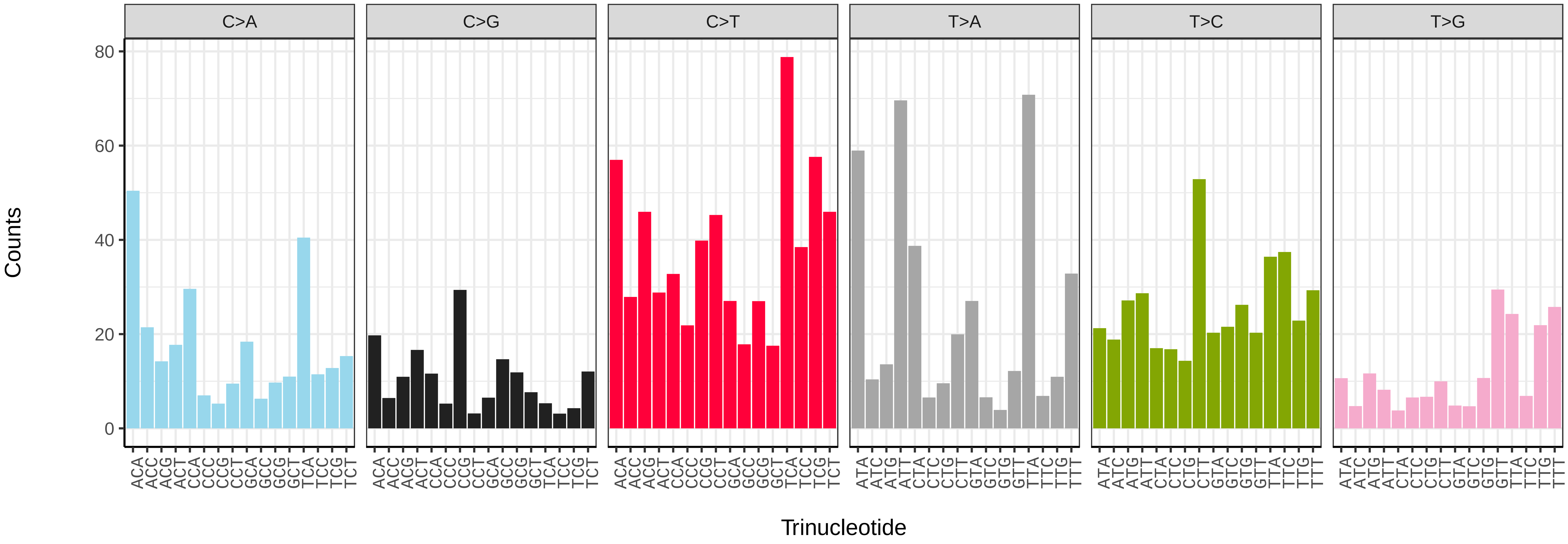


*Yponomeuta rorrellus* (ilYpoRore1) somatic mutational spectrum (SBS52 classification)

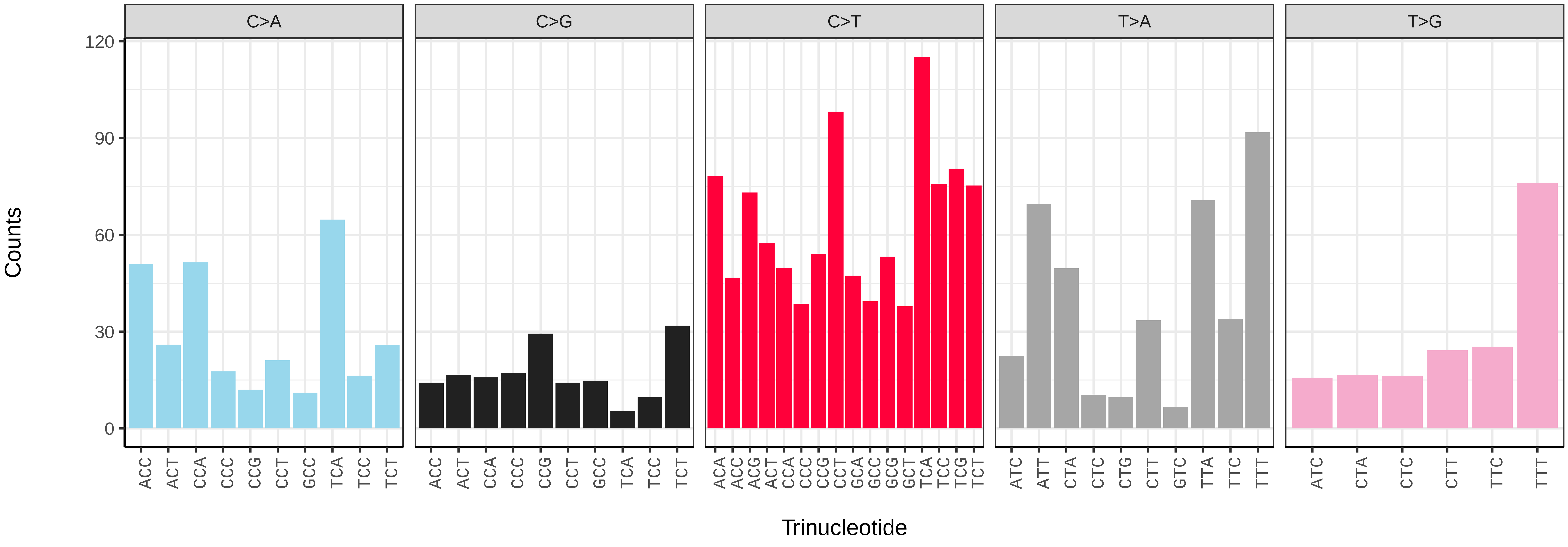

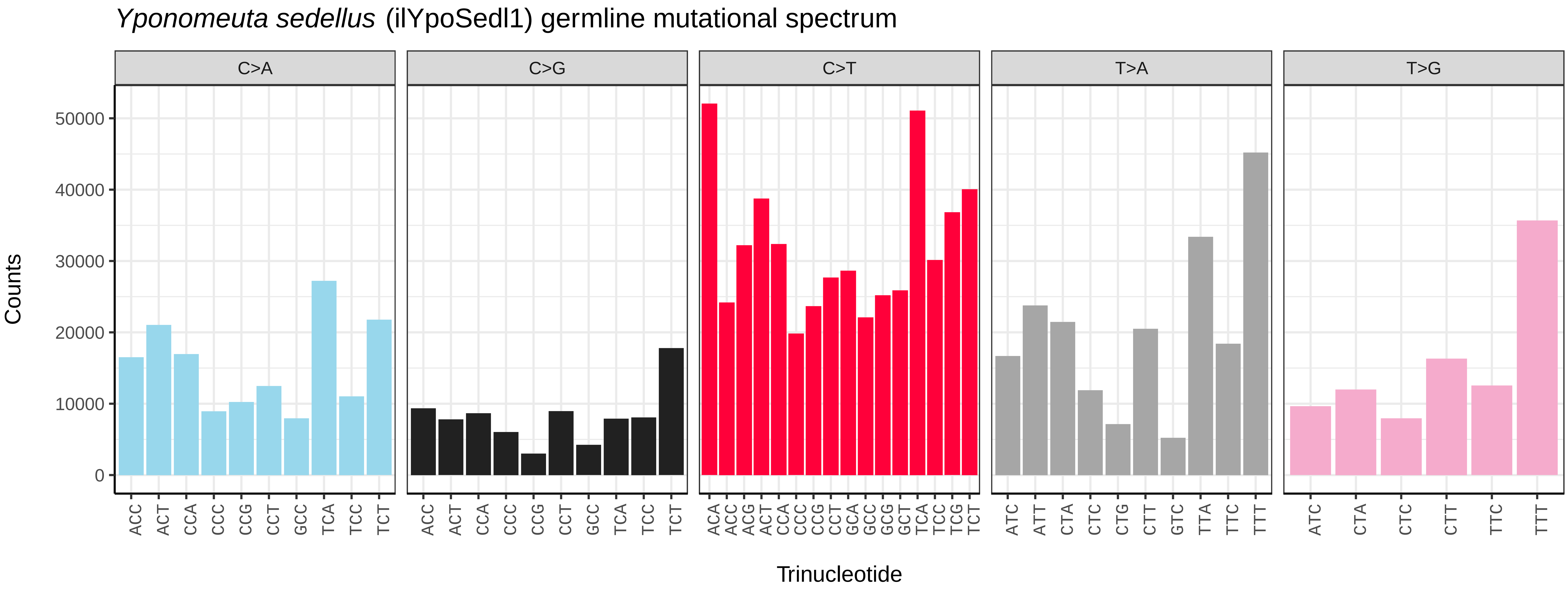
Yponomeuta sedellus (ilYpoSedl1) germline mutational spectrum
C>A
C>G
C>T
T>A
T>G
Counts
Trinucleotide

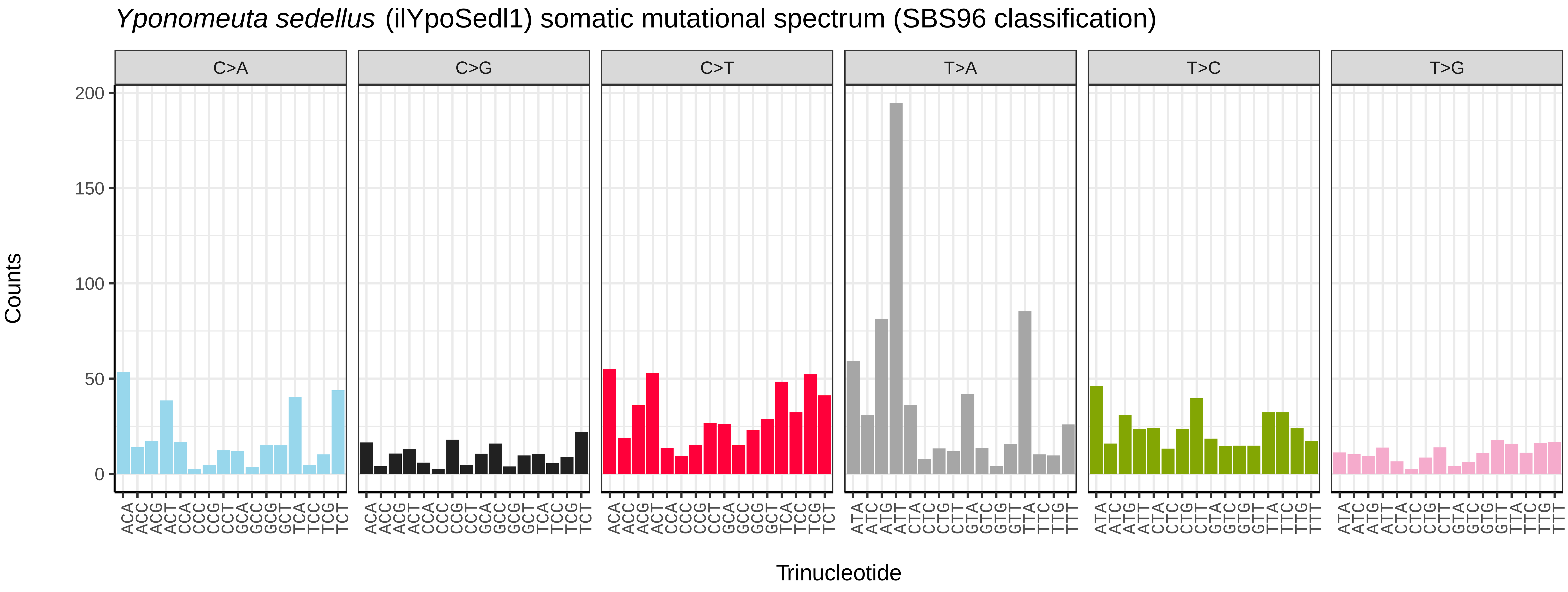
Yponomeuta sedellus (ilYpoSedl1) somatic mutational spectrum (SBS96 classification)
C>A
C>G
C>T
T>A
T>C
T>G
Counts
Trinucleotide

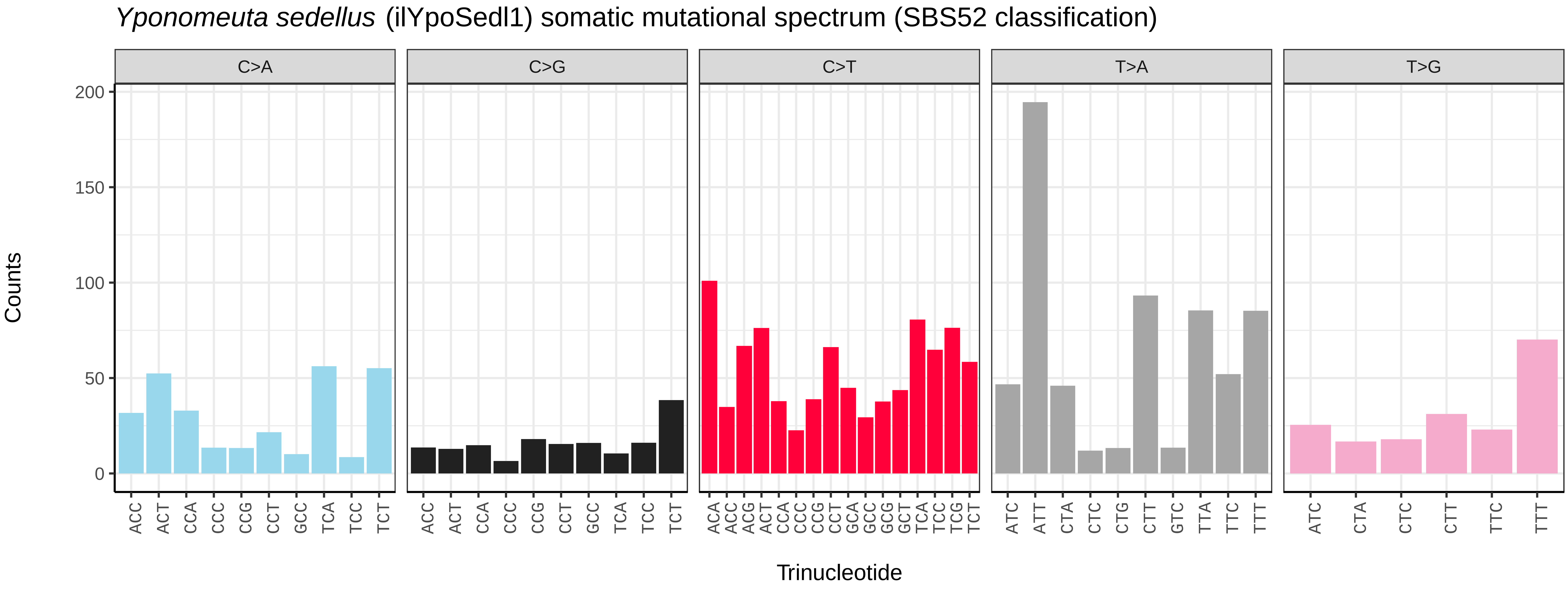
Yponomeuta sedellus (ilYpoSedl1) somatic mutational spectrum (SBS52 classification)
C>A
C>G
C>T
T>A
T>G
Counts
Trinucleotide

*Ypsolopha scabrella* (ilYpsScab1) germline mutational spectrum

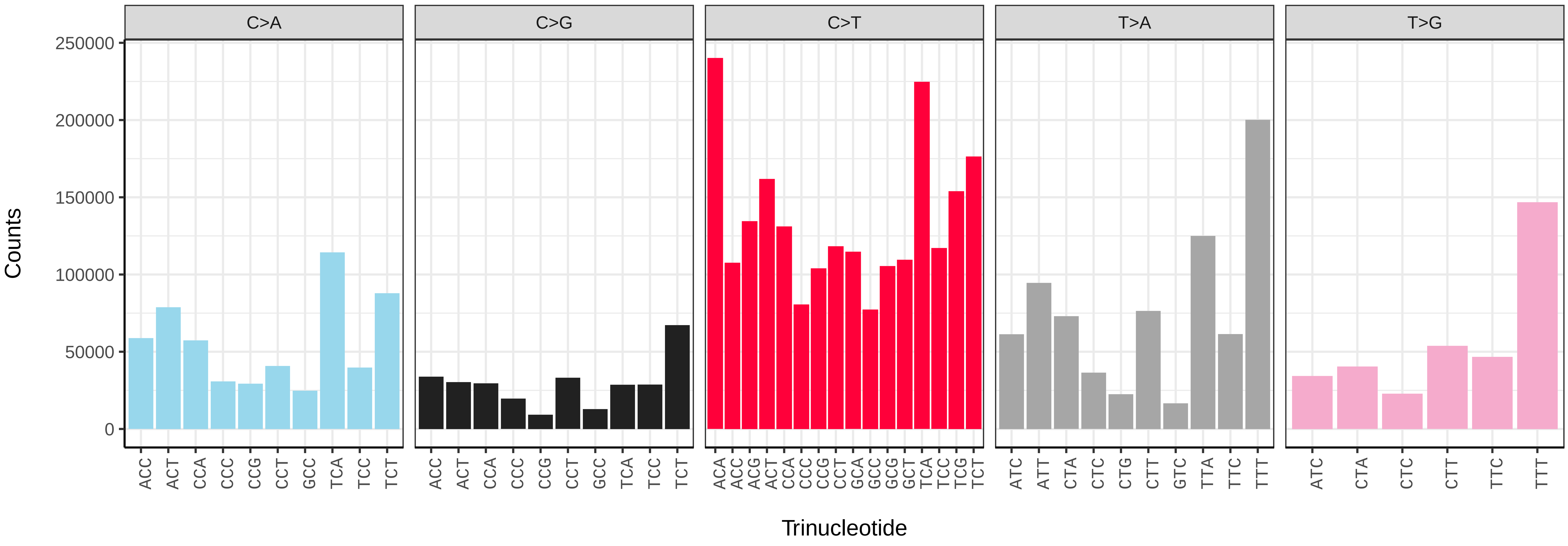


*Ypsolopha scabrella* (ilYpsScab1) somatic mutational spectrum (SBS96 classification)

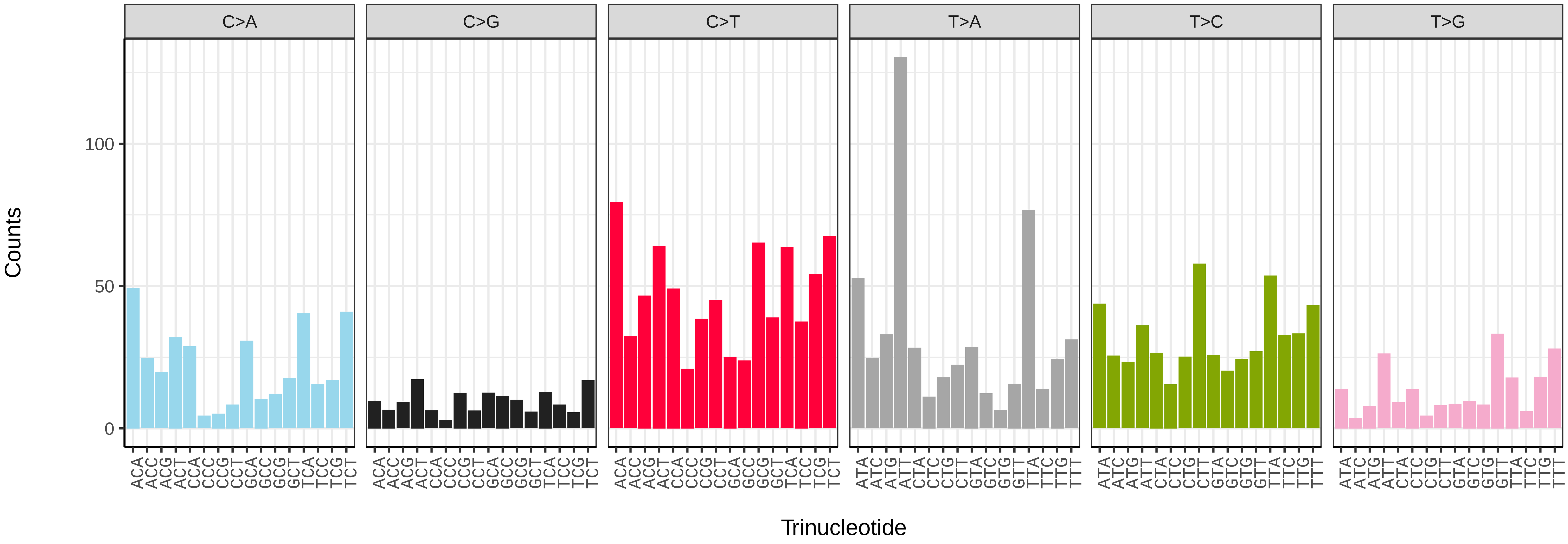


*Ypsolopha scabrella* (ilYpsScab1) somatic mutational spectrum (SBS52 classification)

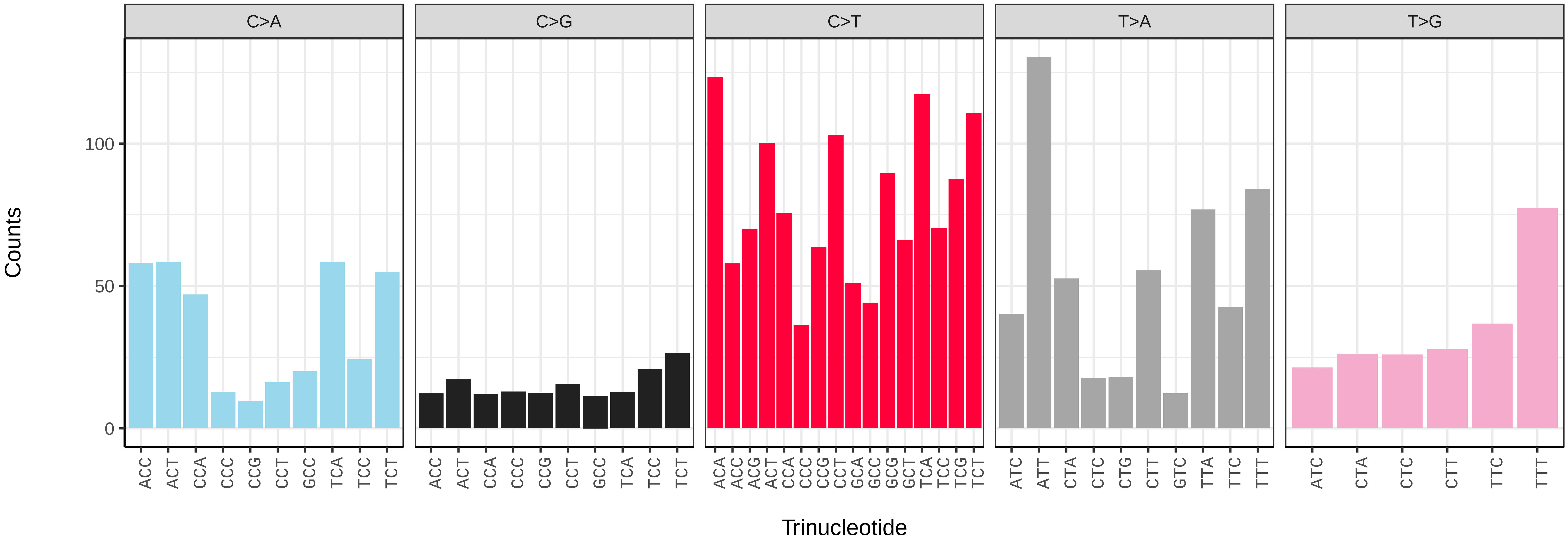

*Ypsolopha sequella* (ilYpsSequ2) germline mutational spectrum

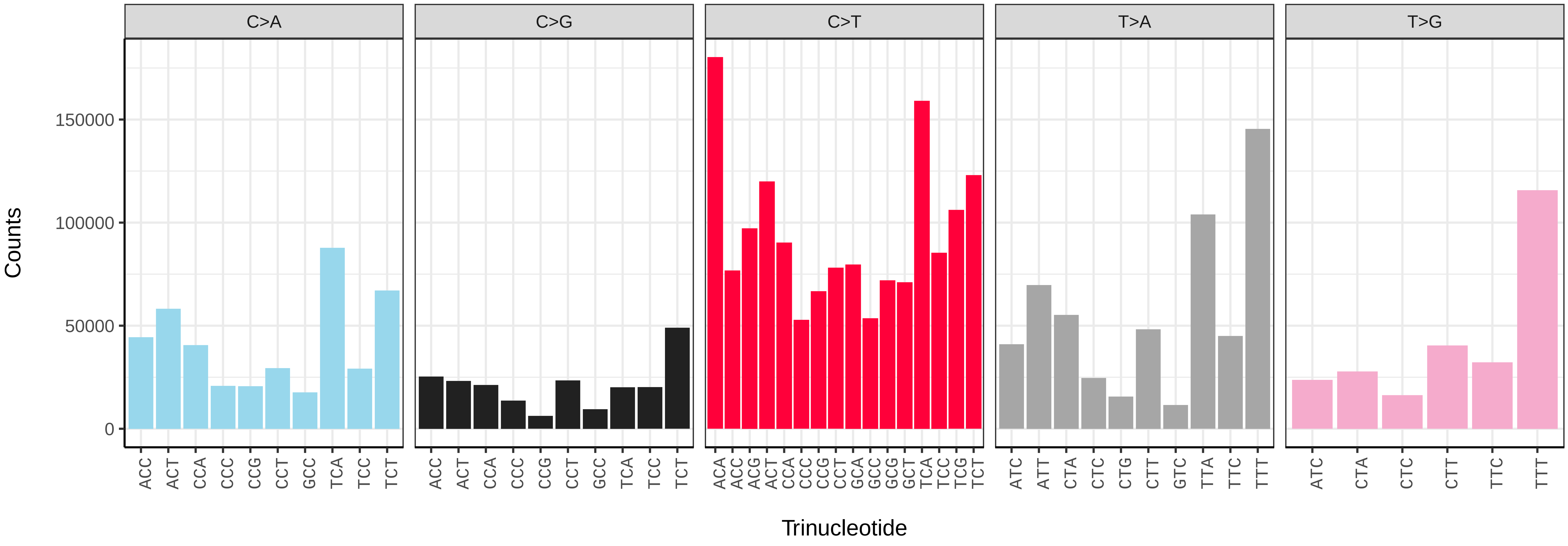


*Ypsolopha sequella* (ilYpsSequ2) somatic mutational spectrum (SBS96 classification)

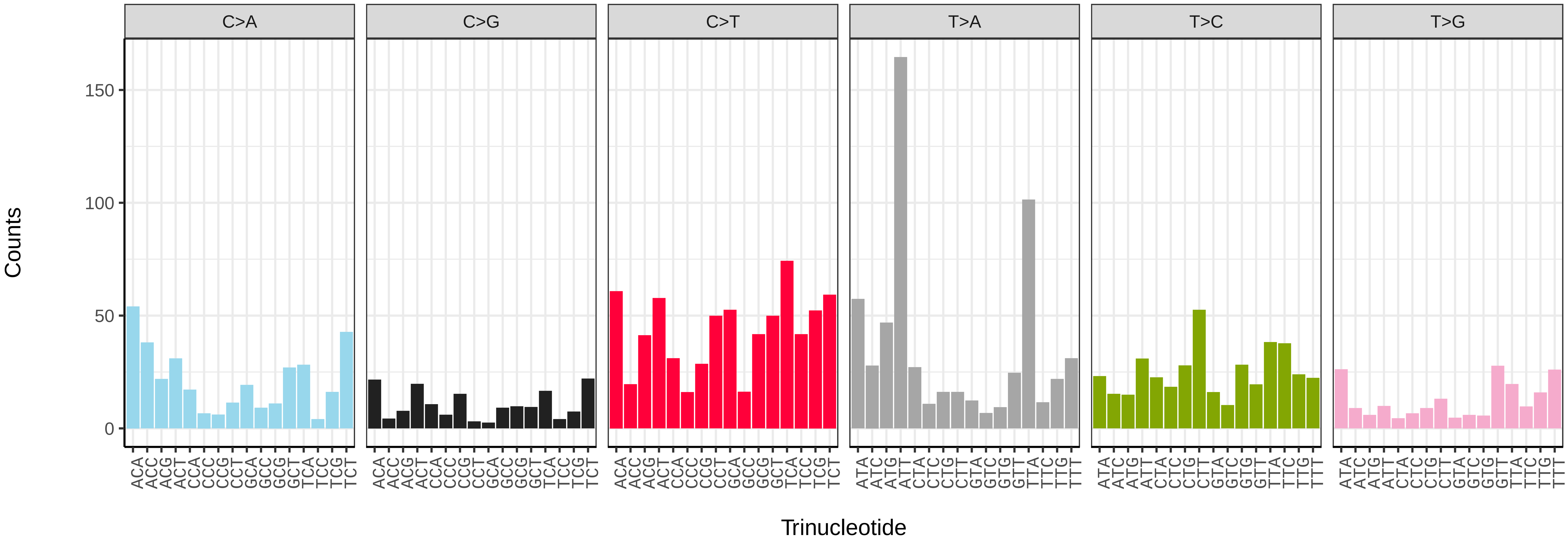


*Ypsolopha sequella* (ilYpsSequ2) somatic mutational spectrum (SBS52 classification)

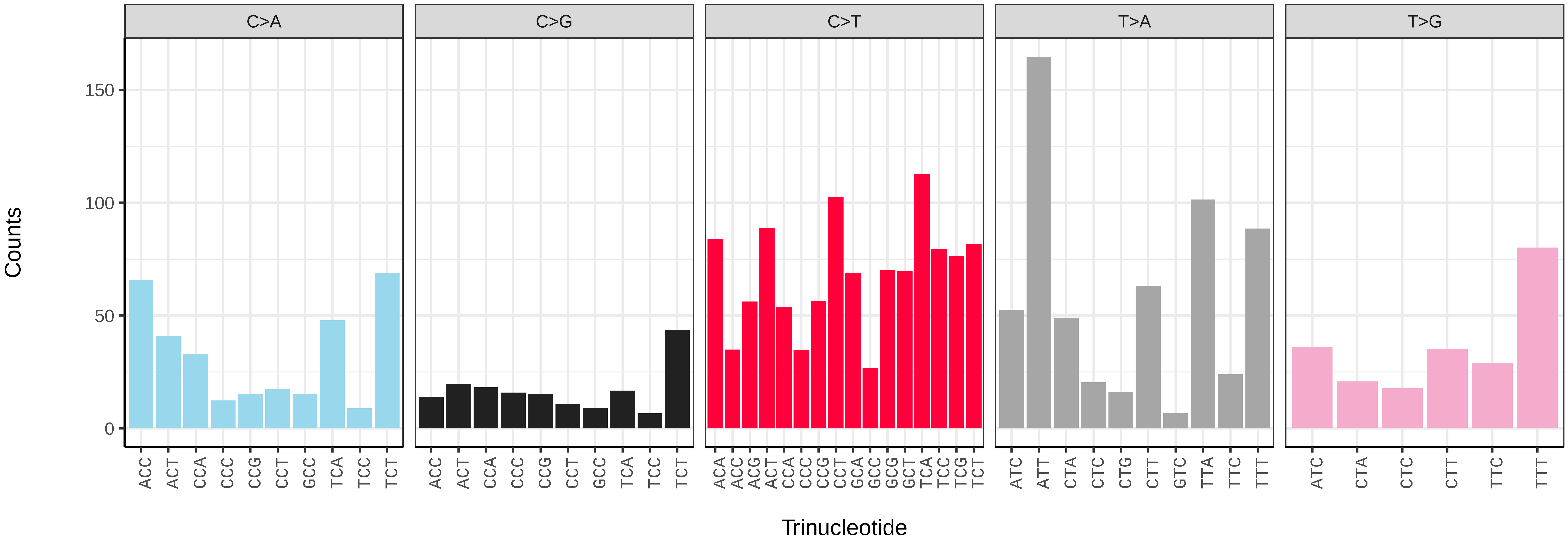

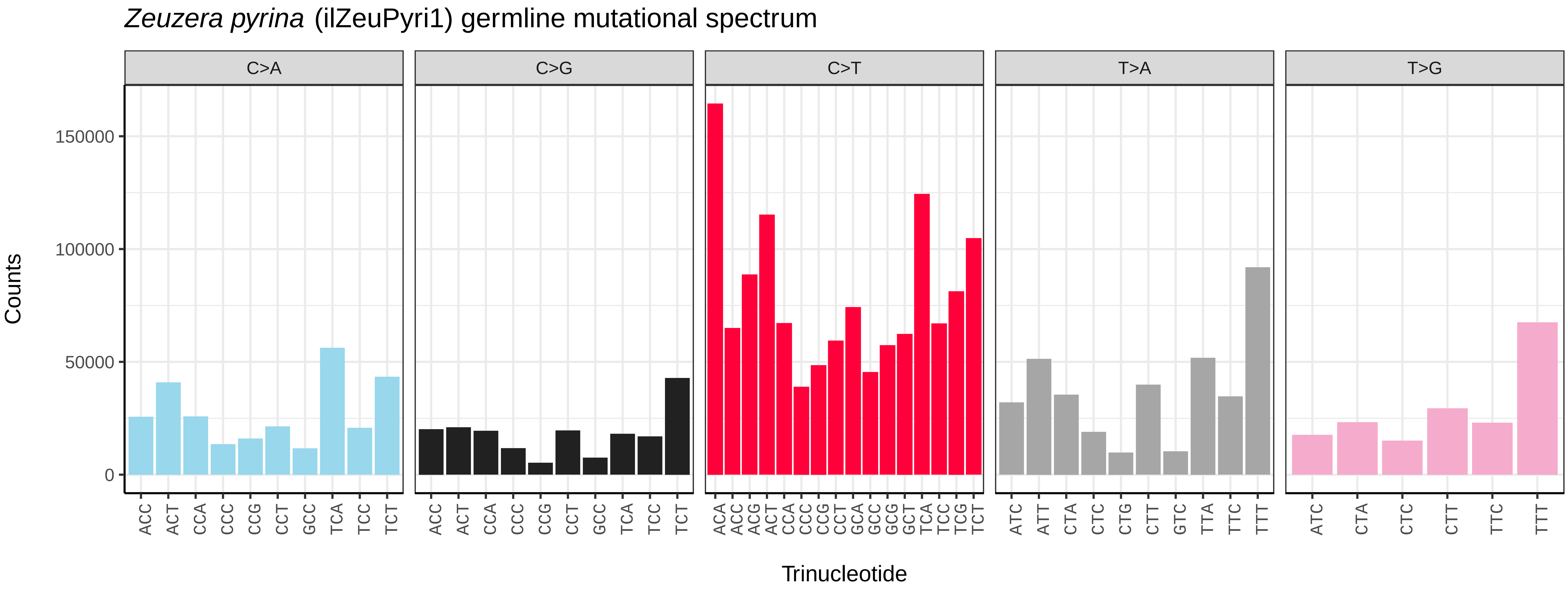
*Zeuzera pyrina* (ilZeuPyri1) germline mutational spectrum
C>A
C>G
C>T
T>A
T>G
Counts
150000
100000
50000
0
ACC ACT CCA CCC CCG CCT GCC TCA TCC TCT
ACA ACC ACG ACT CCA CCC CCG CCT GCA GCC GCG GCT TCA TCC TCG TCT
ATC ATT CTA CTC CTG CTT GTC TTA TTC TTT
ATC CTA CTC CTT TTC TTT
Trinucleotide

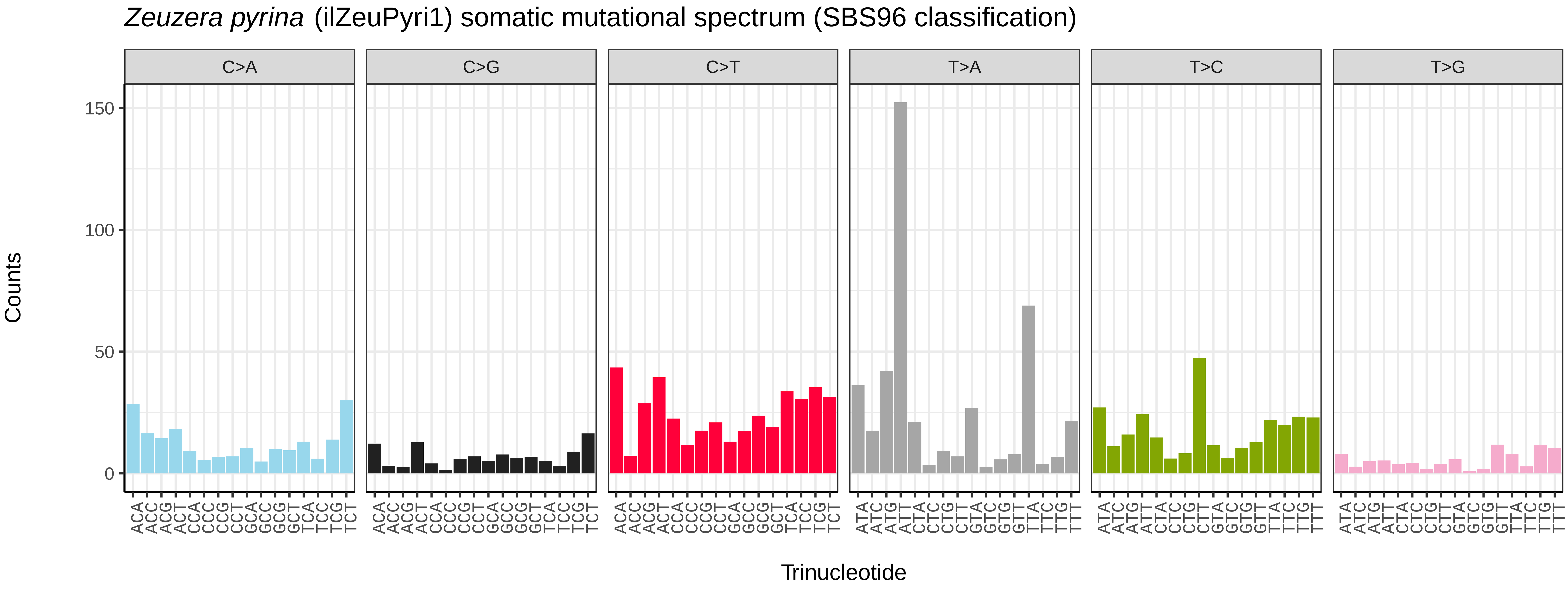
*Zeuzera pyrina* (ilZeuPyri1) somatic mutational spectrum (SBS96 classification)
C>A
C>G
C>T
T>A
T>C
T>G
Counts
150
100
50
0
Trinucleotide

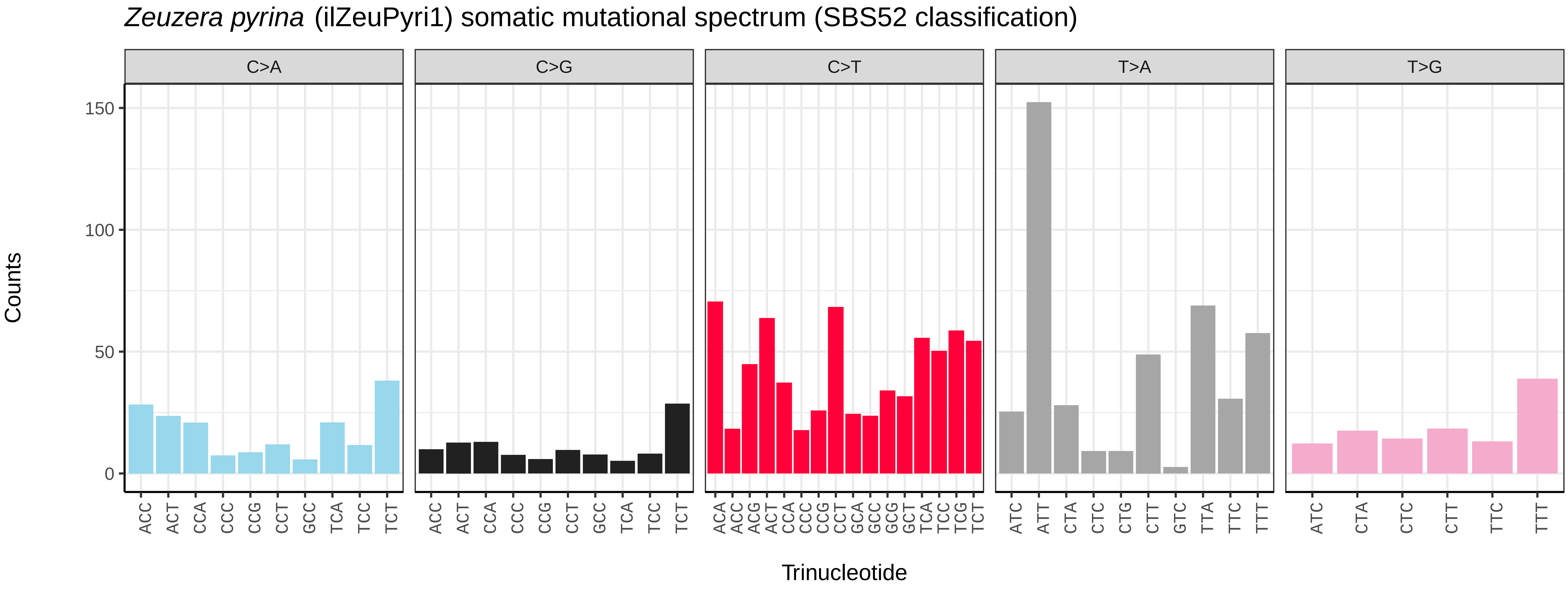
*Zeuzera pyrina* (ilZeuPyri1) somatic mutational spectrum (SBS52 classification)
C>A
C>G
C>T
T>A
T>G
Counts
150
100
50
0
ACC ACT CCA CCC CCG CCT GCC TCA TCC TCT
ACA ACC ACG ACT CCA CCC CCG CCT GCA GCC GCG GCT TCA TCC TCG TCT
ATC ATT CTA CTC CTG CTT GTC TTA TTC TTT
ATC CTA CTC CTT TTC TTT
Trinucleotide

*Zygaena filipendulae* (ilZygFili1) germline mutational spectrum

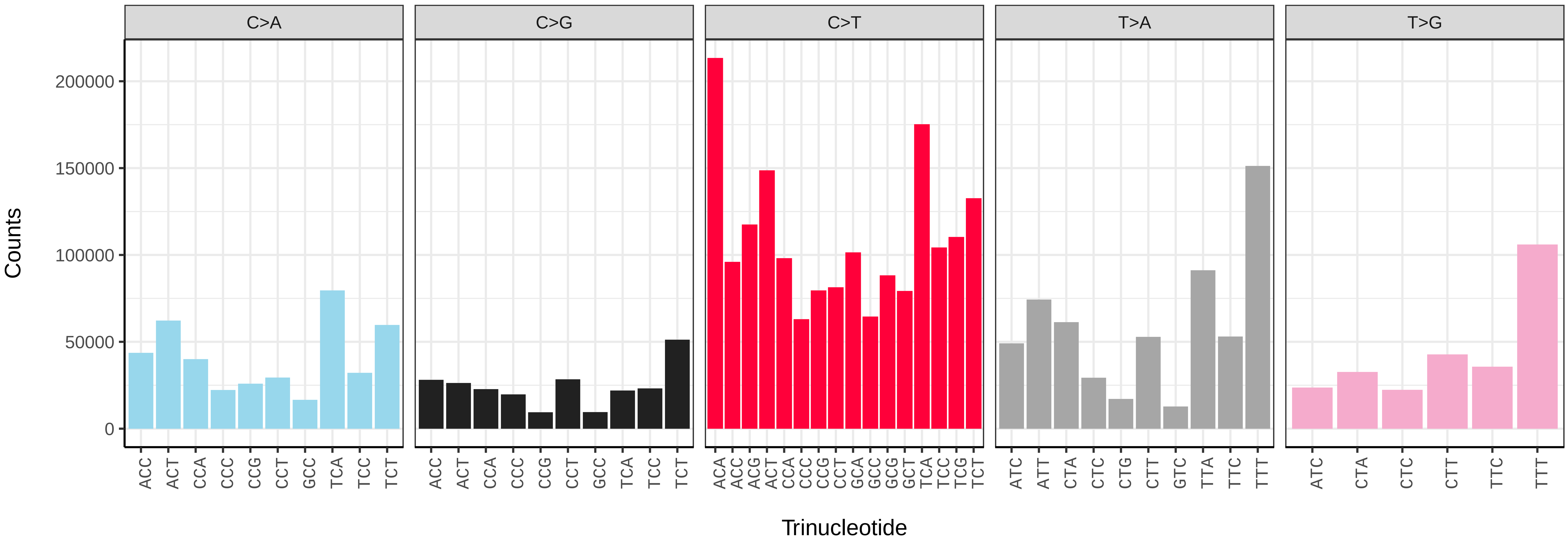


*Zygaena filipendulae* (ilZygFili1) somatic mutational spectrum (SBS96 classification)

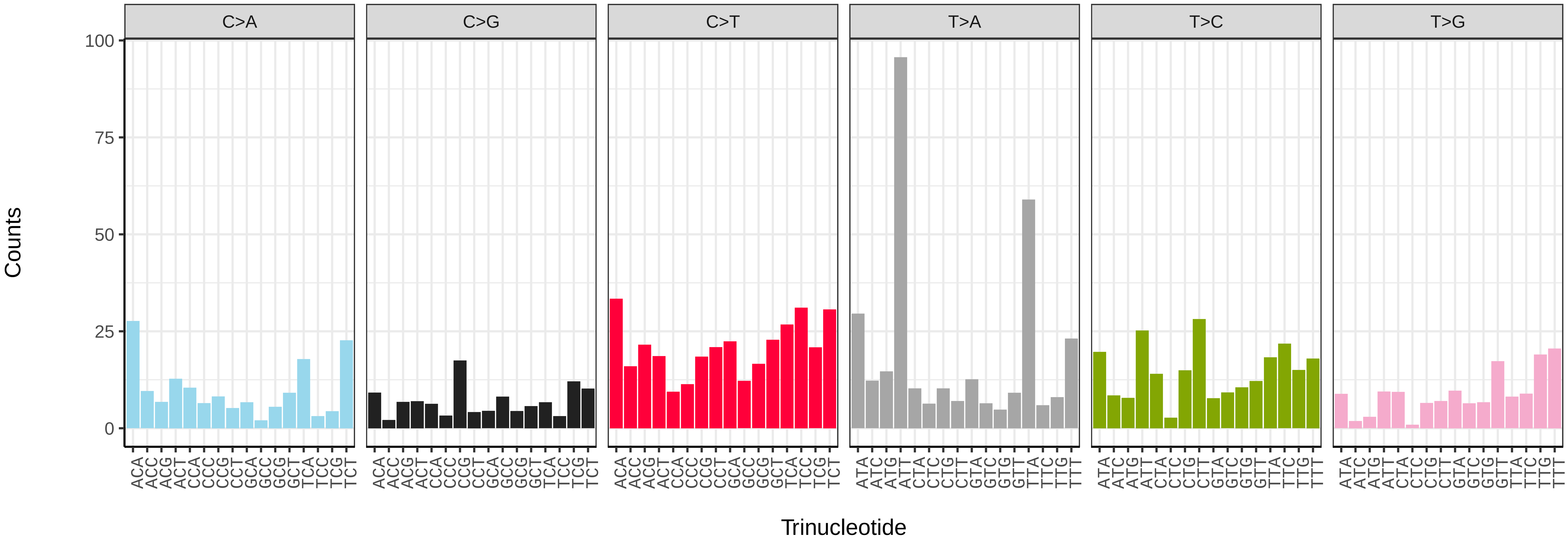


*Zygaena filipendulae* (ilZygFili1) somatic mutational spectrum (SBS52 classification)

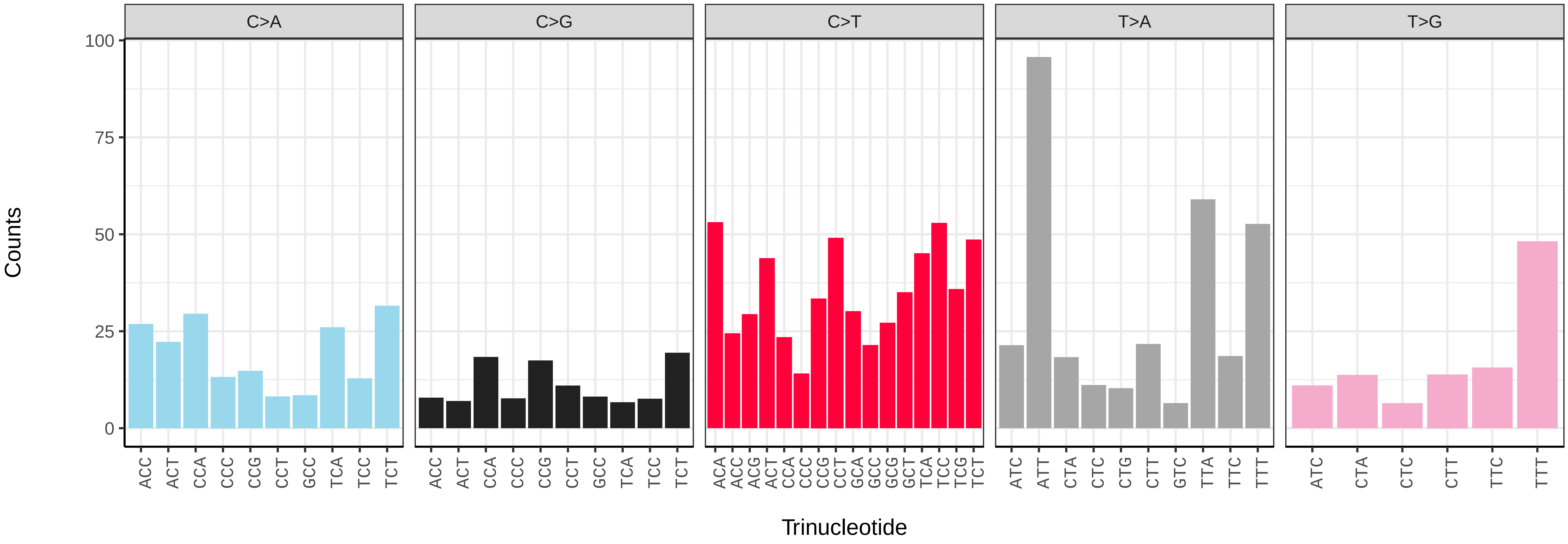

*Chrysoperla carnea* (inChrCarn1) germline mutational spectrum

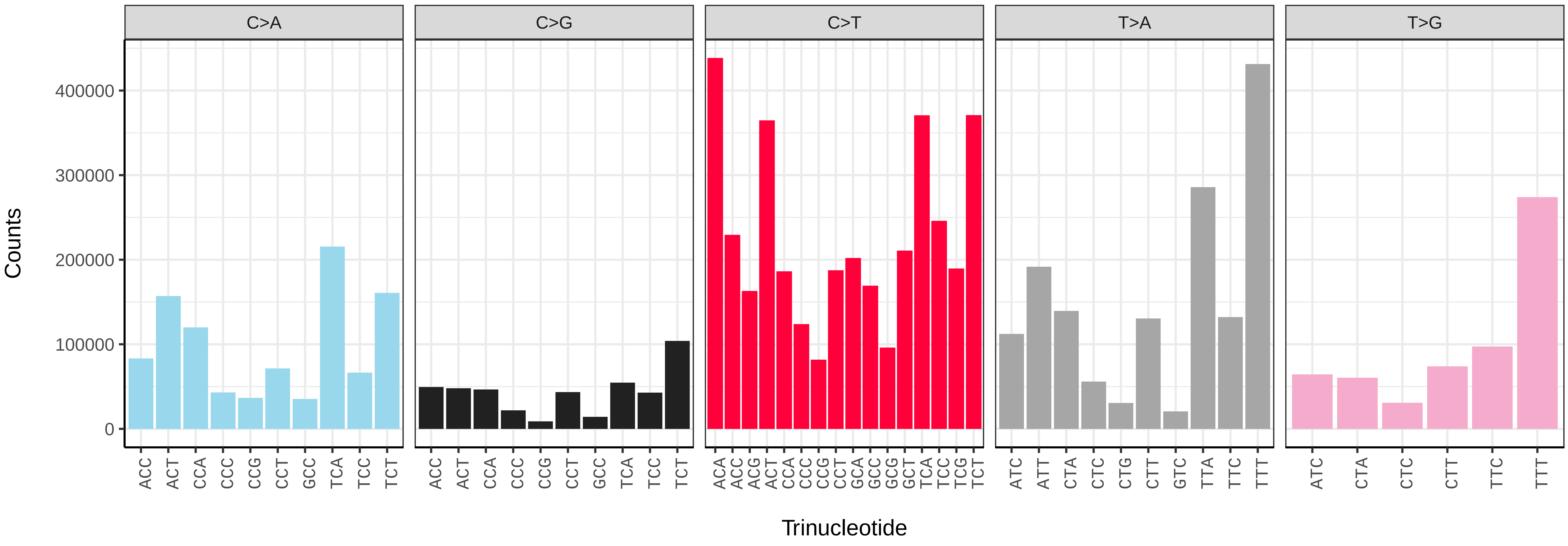


*Chrysoperla carnea* (inChrCarn1) somatic mutational spectrum (SBS96 classification)

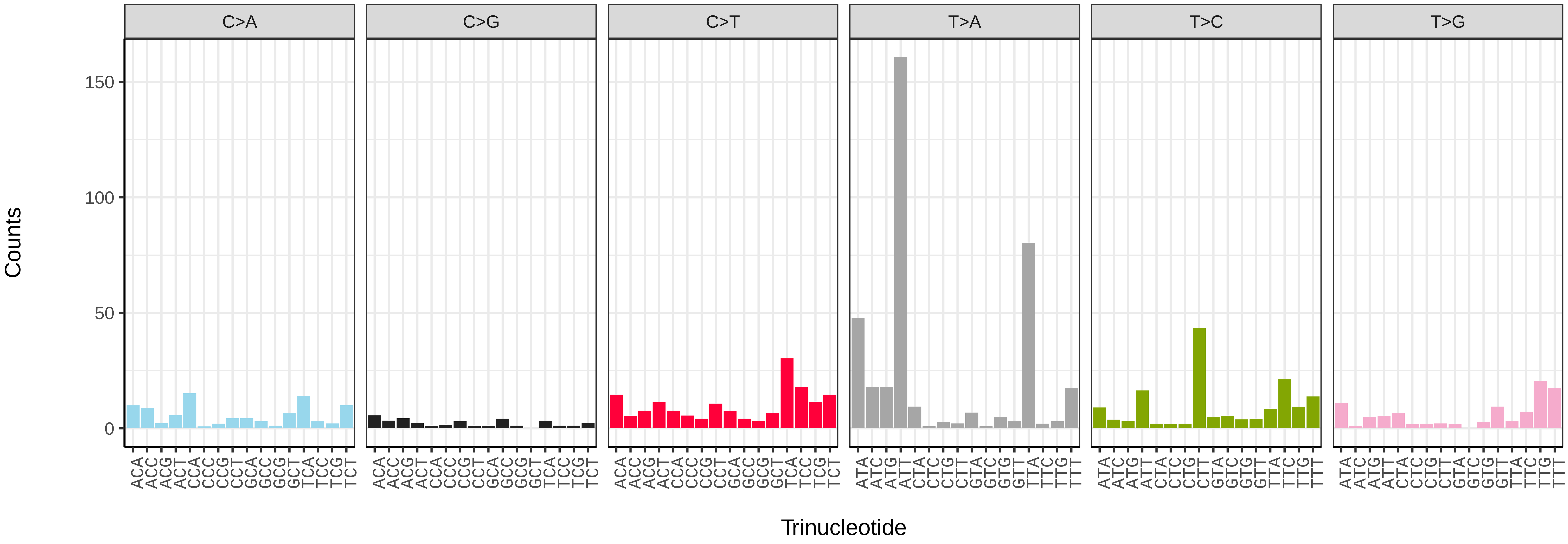


*Chrysoperla carnea* (inChrCarn1) somatic mutational spectrum (SBS52 classification)

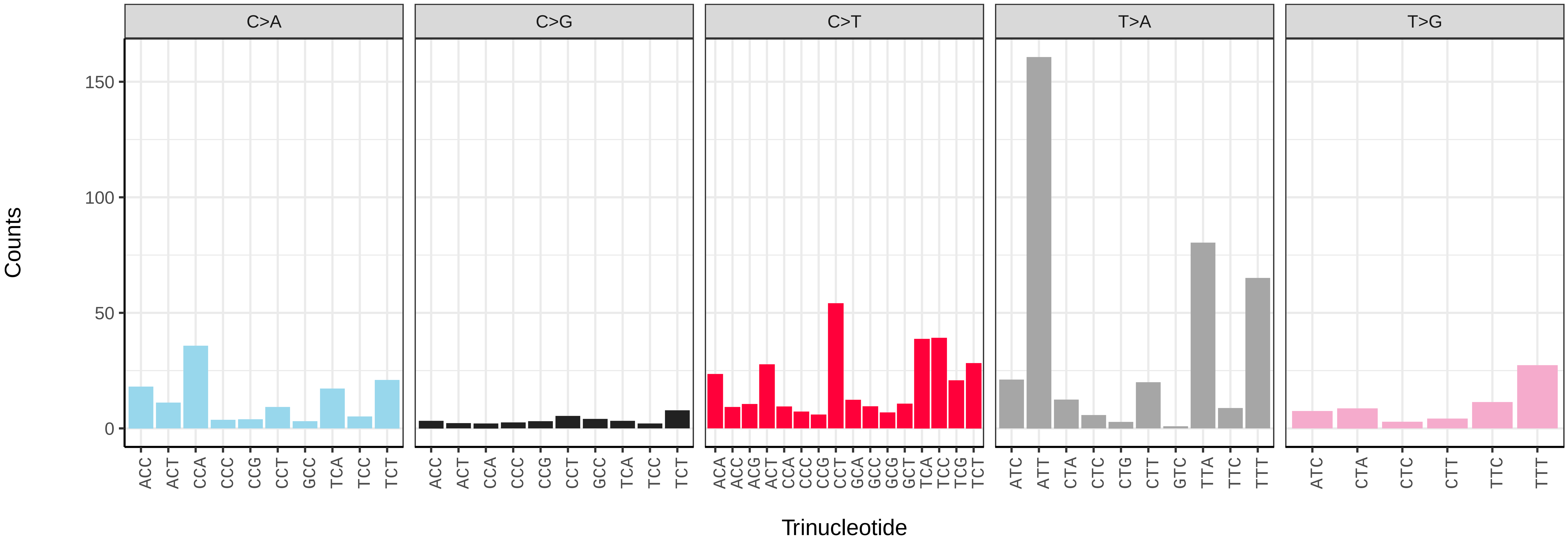

*Ischnura elegans* (ioIscEleg1) germline mutational spectrum

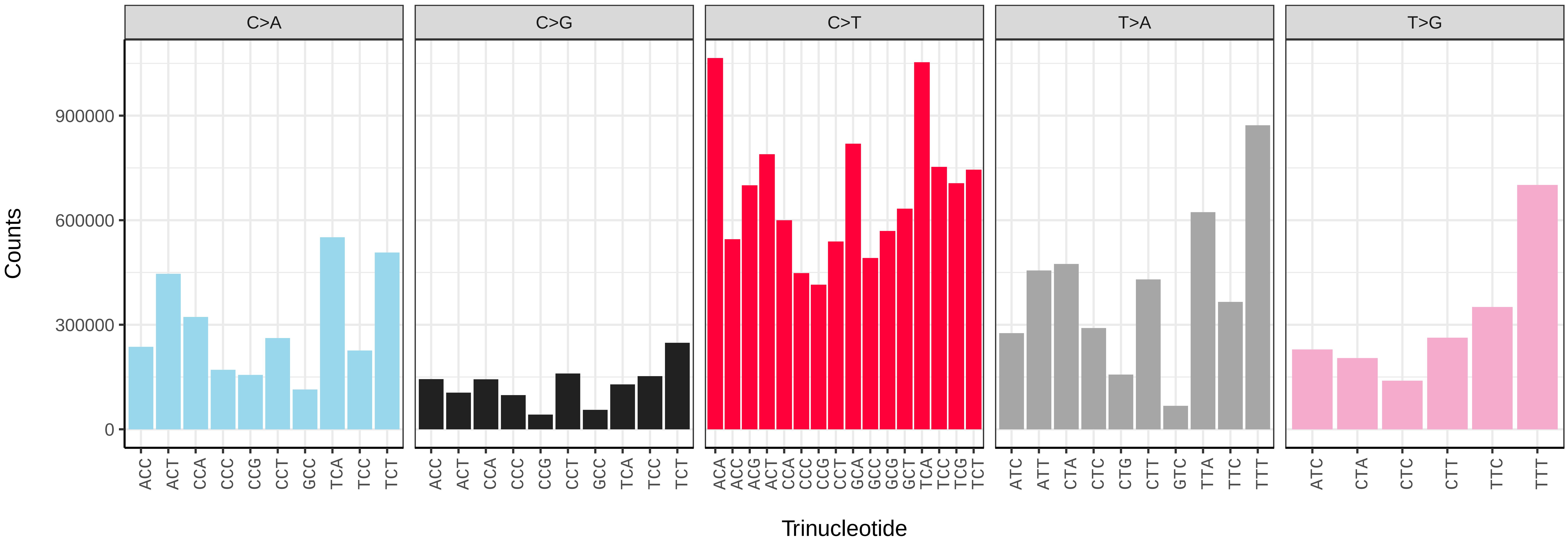


*Ischnura elegans* (ioIscEleg1) somatic mutational spectrum (SBS96 classification)

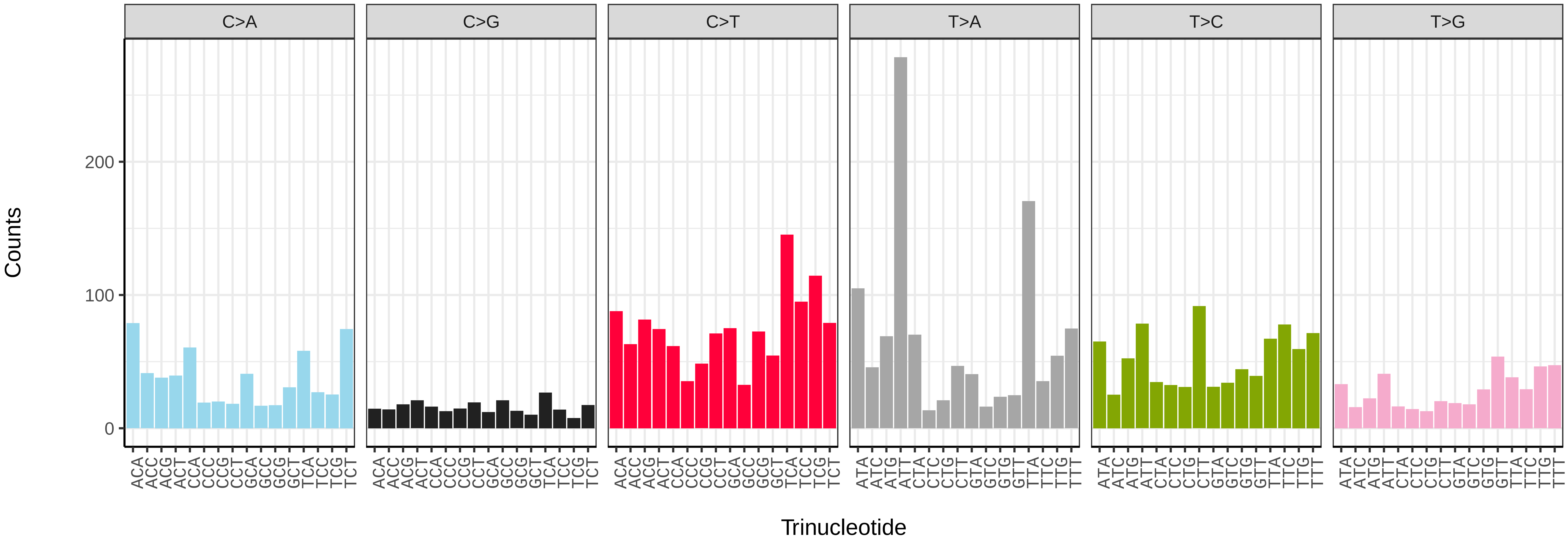


*Ischnura elegans* (ioIscEleg1) somatic mutational spectrum (SBS52 classification)

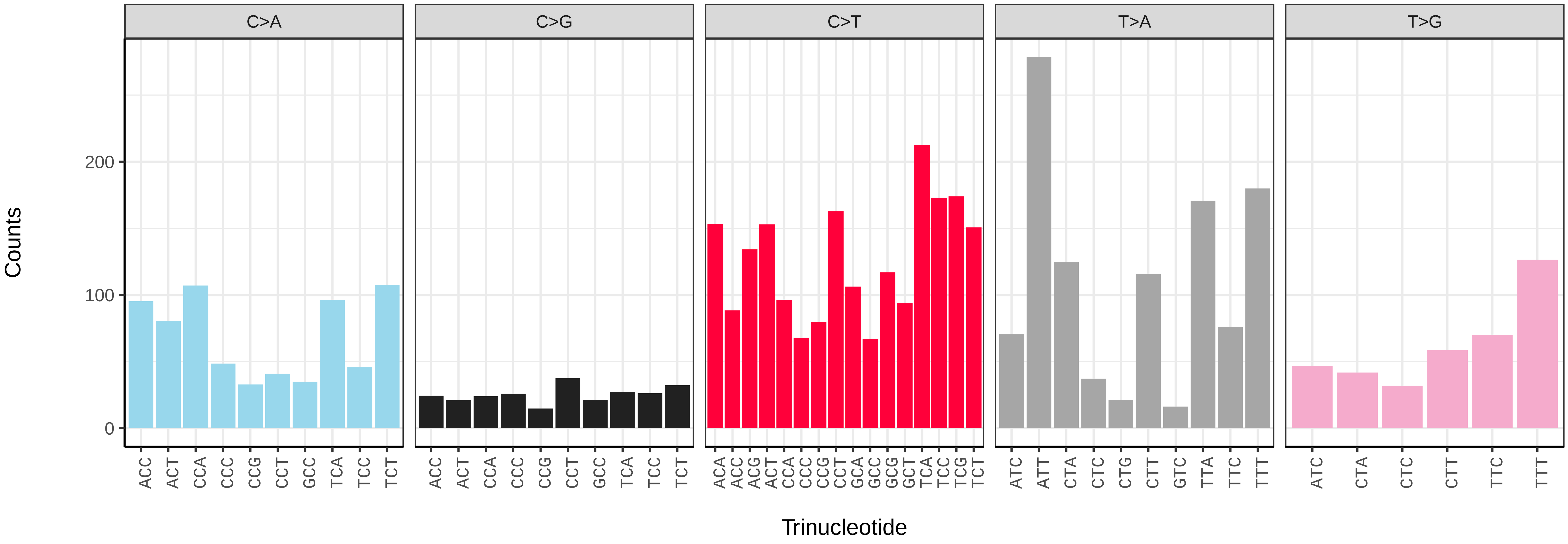

*Platycnemis pennipes* (ioPlaPenn1) germline mutational spectrum

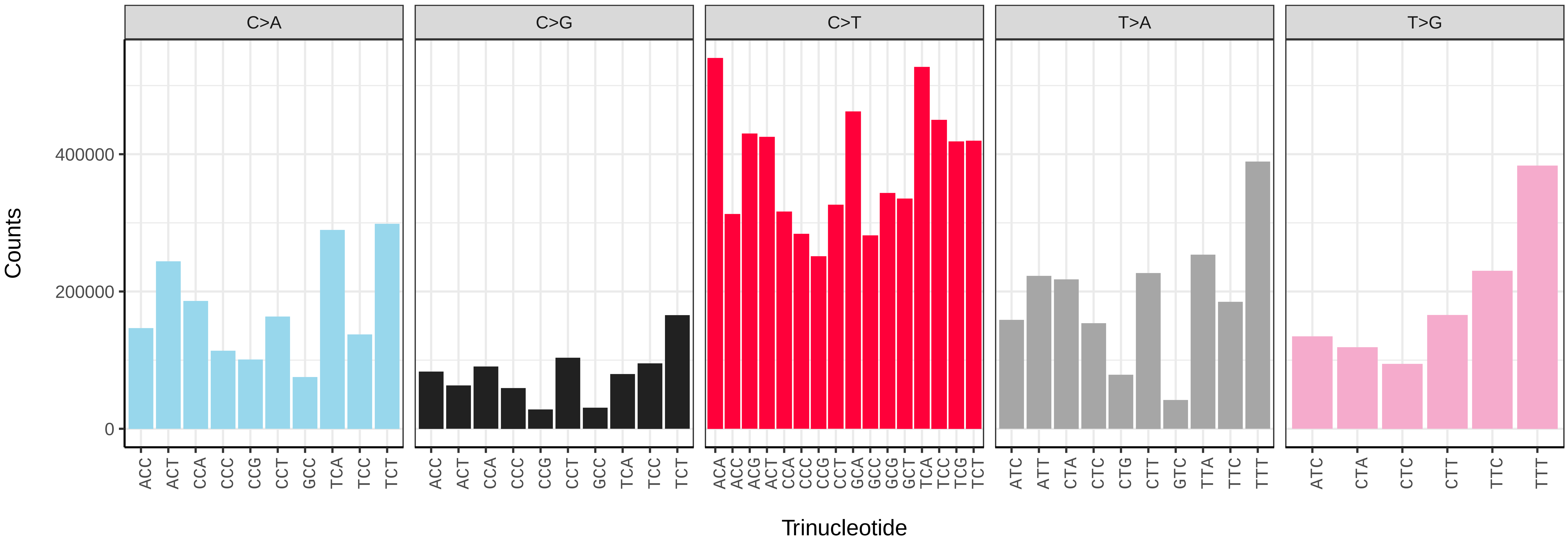


*Platycnemis pennipes* (ioPlaPenn1) somatic mutational spectrum (SBS96 classification)

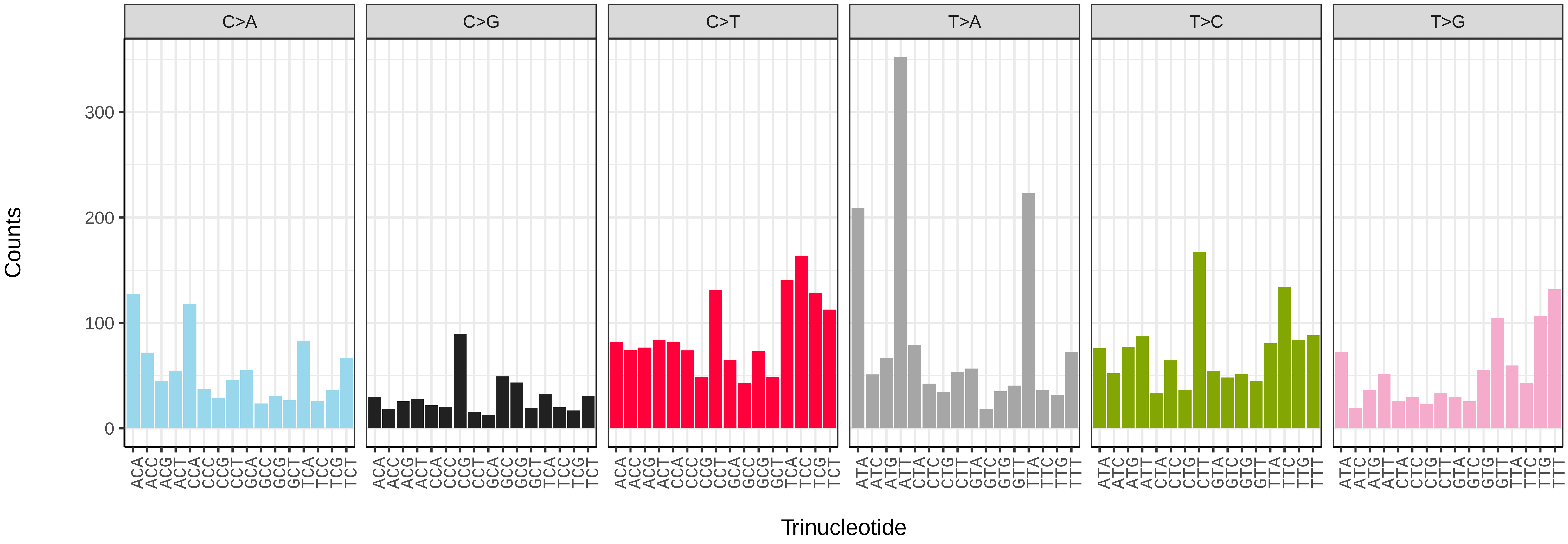


*Platycnemis pennipes* (ioPlaPenn1) somatic mutational spectrum (SBS52 classification)

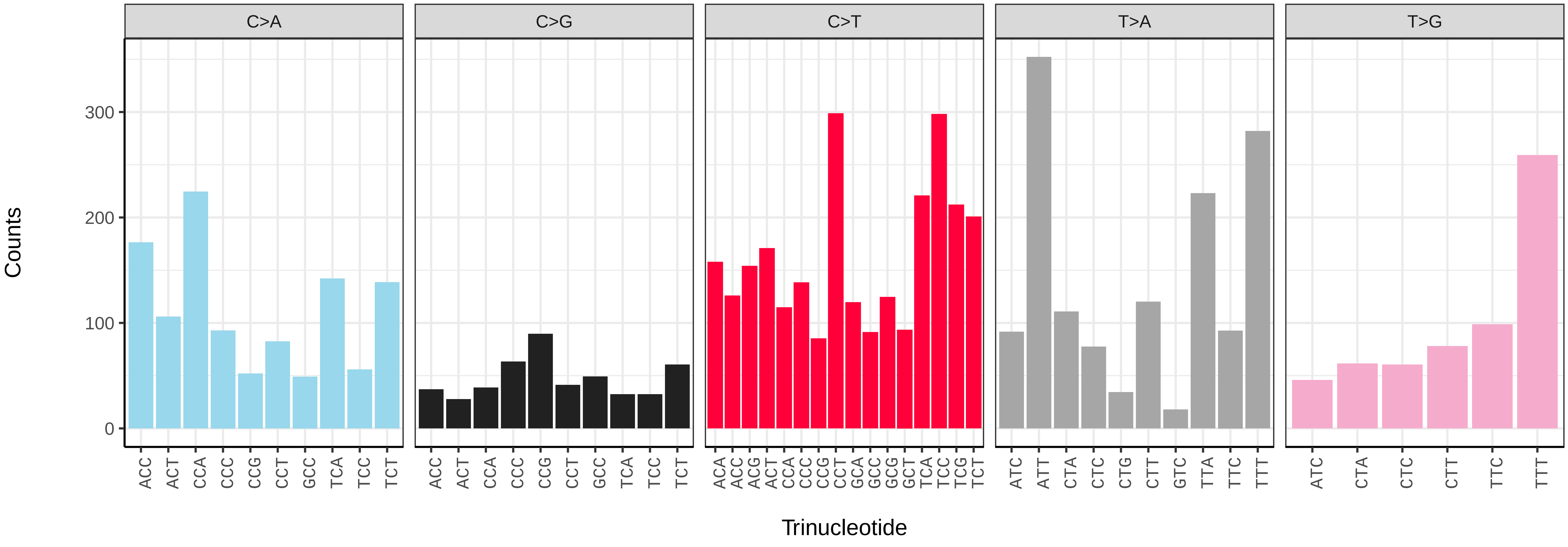

*Sympetrum striolatum* (ioSymStri1) germline mutational spectrum

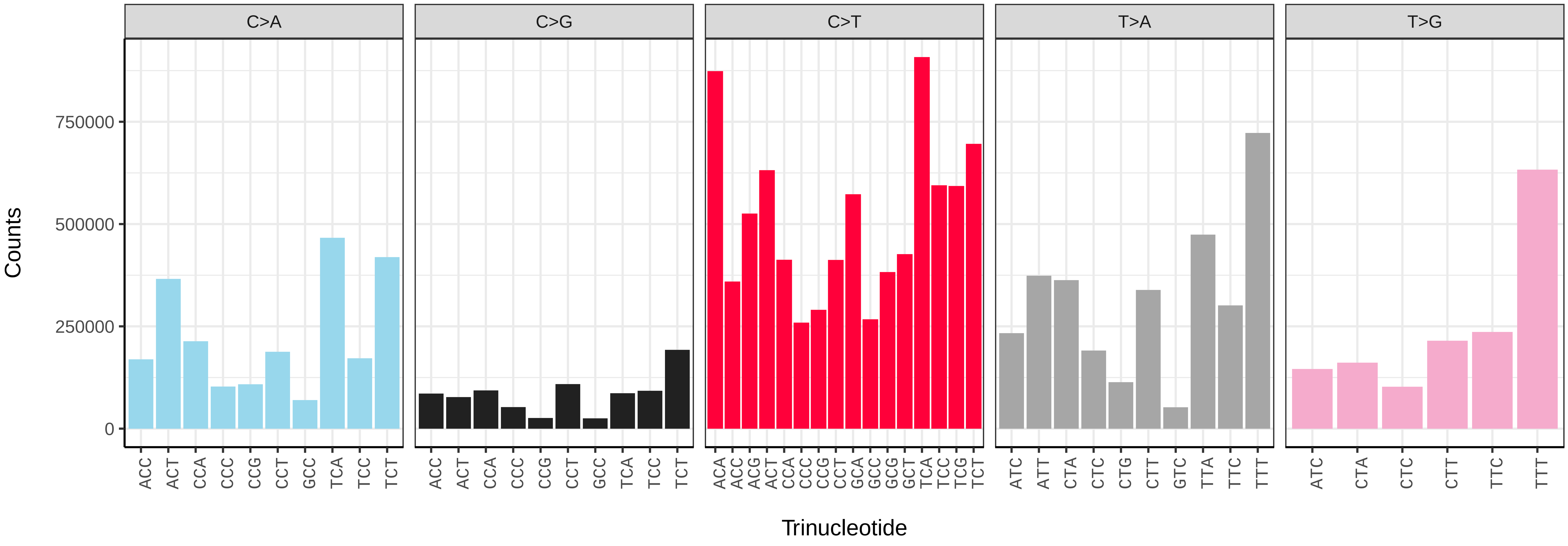


*Sympetrum striolatum* (ioSymStri1) somatic mutational spectrum (SBS96 classification)

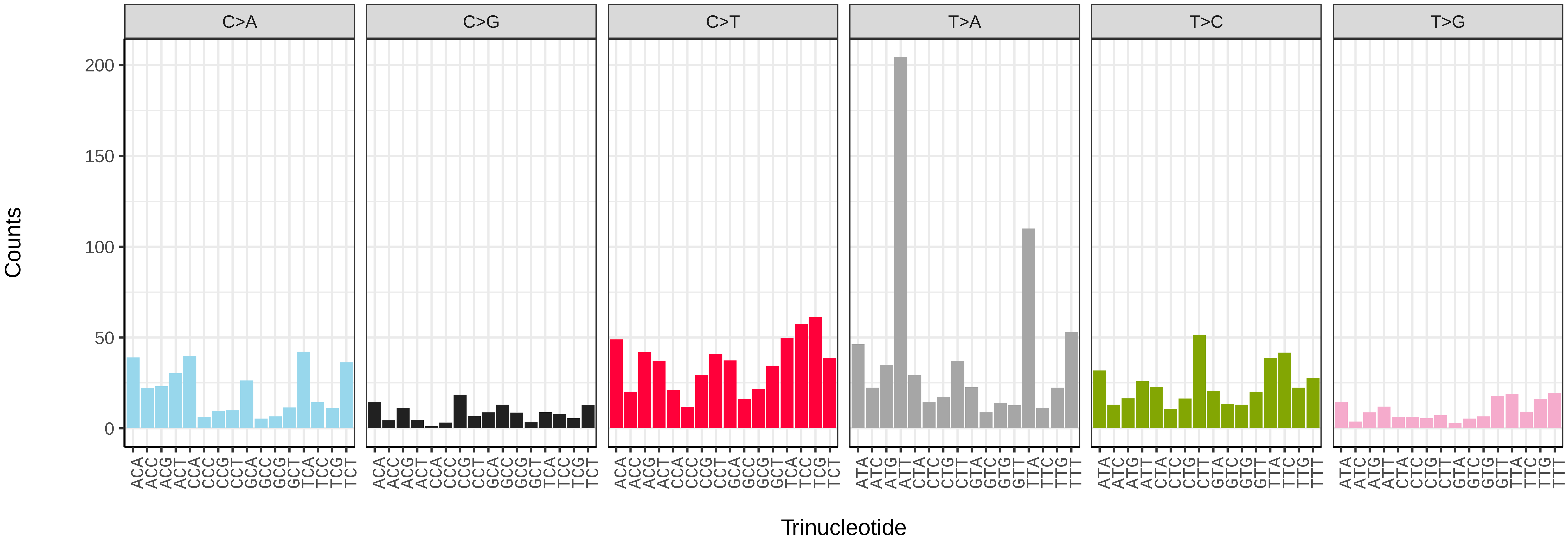


*Sympetrum striolatum* (ioSymStri1) somatic mutational spectrum (SBS52 classification)

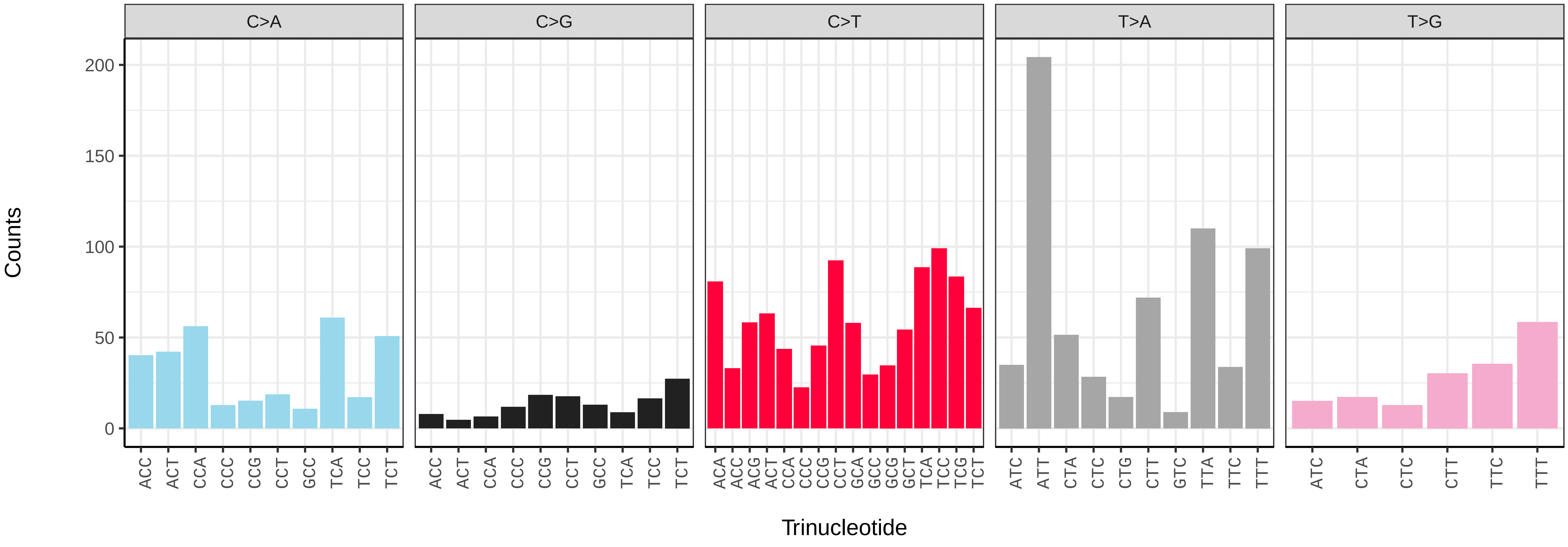

*Isoperla grammatica* (ipIsoGram3) germline mutational spectrum

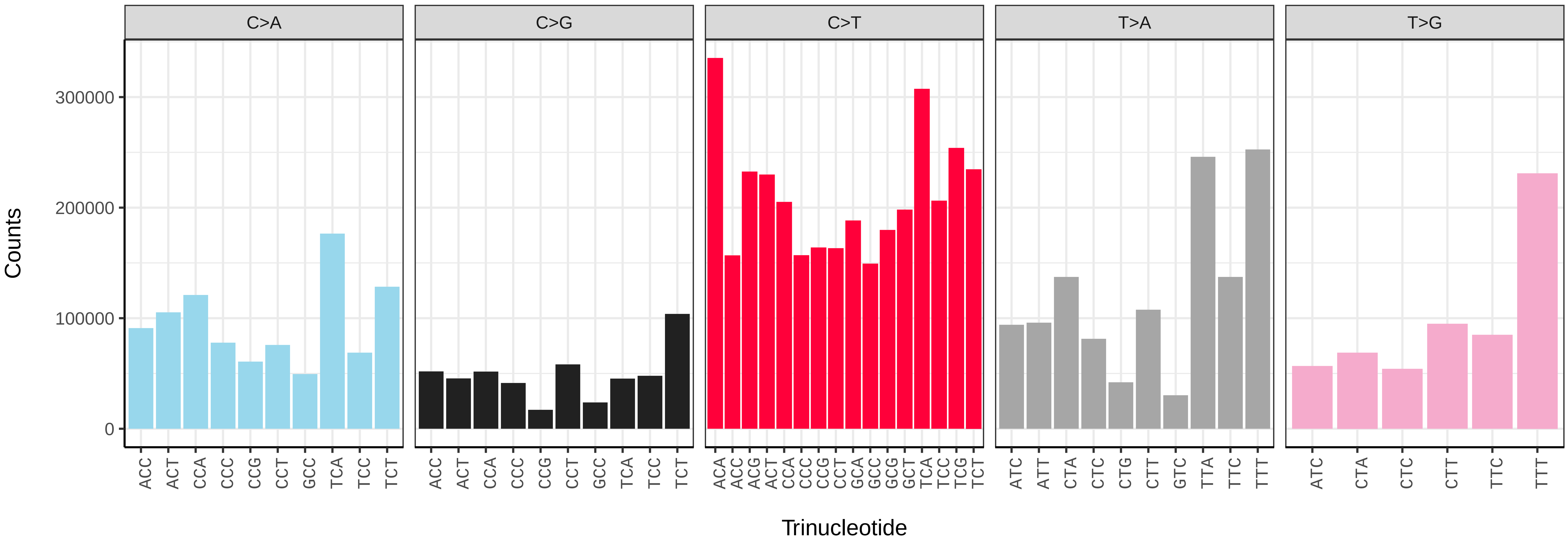


*Isoperla grammatica* (ipIsoGram3) somatic mutational spectrum (SBS96 classification)

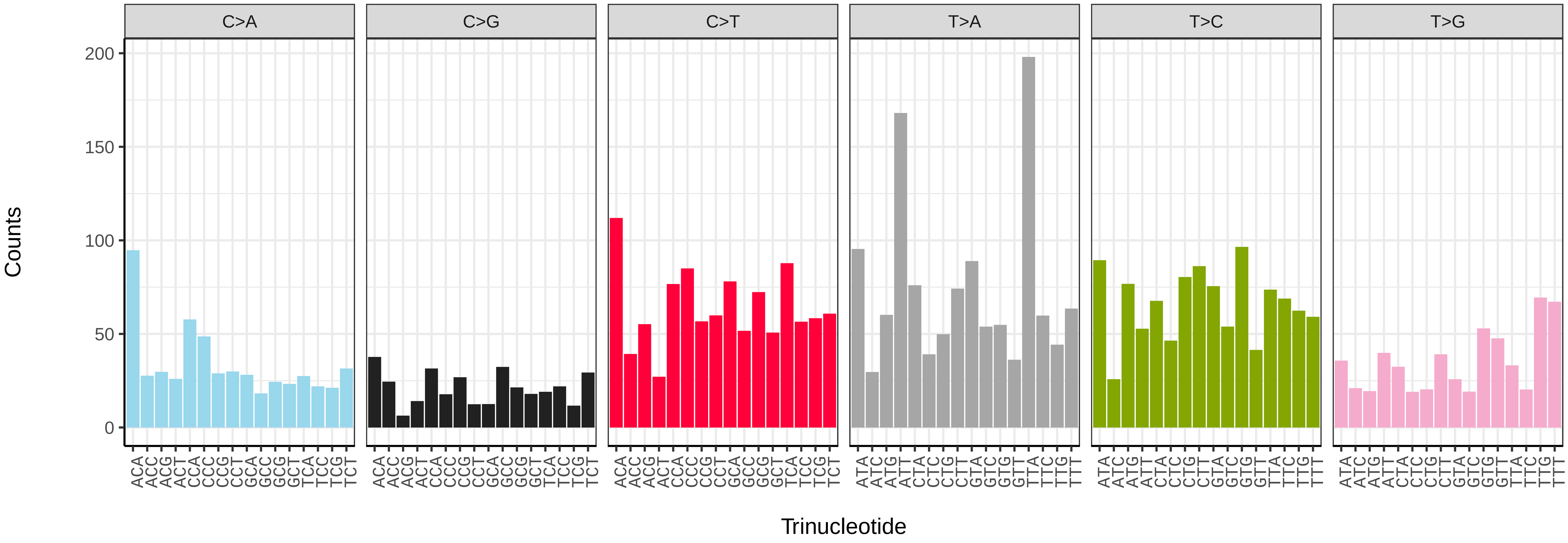


*Isoperla grammatica* (ipIsoGram3) somatic mutational spectrum (SBS52 classification)

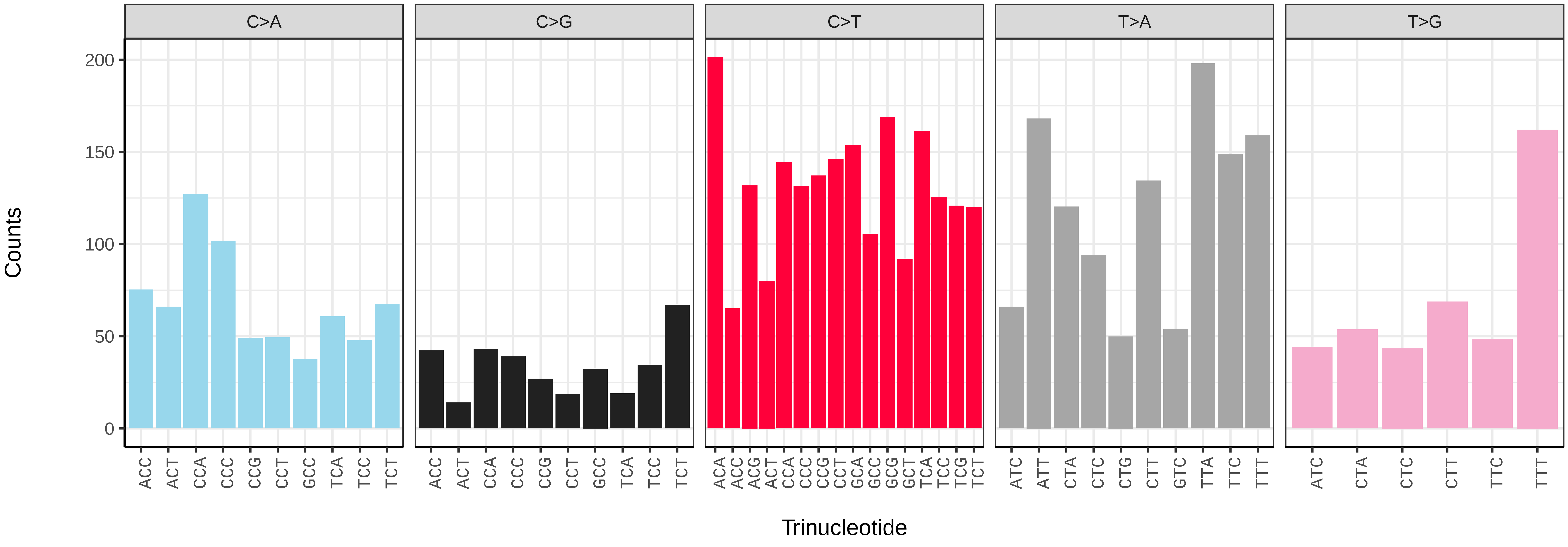

*Leuctra nigra* (ipLeuNigr2) germline mutational spectrum

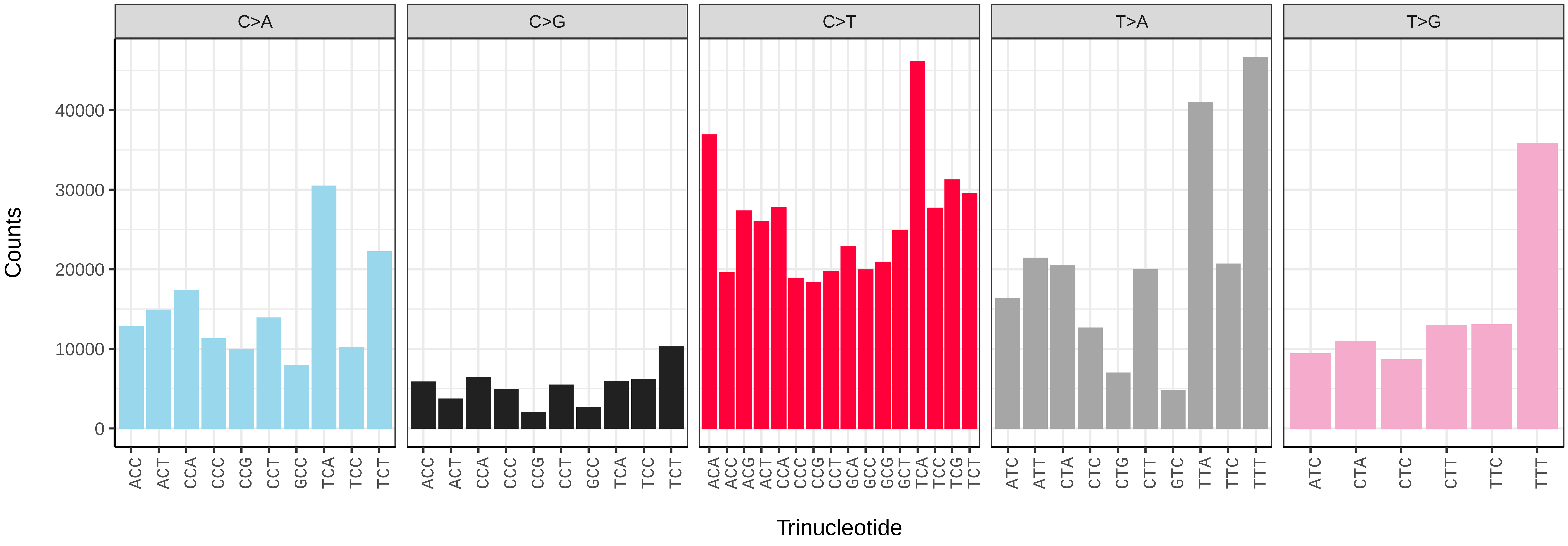


*Leuctra nigra* (ipLeuNigr2) somatic mutational spectrum (SBS96 classification)

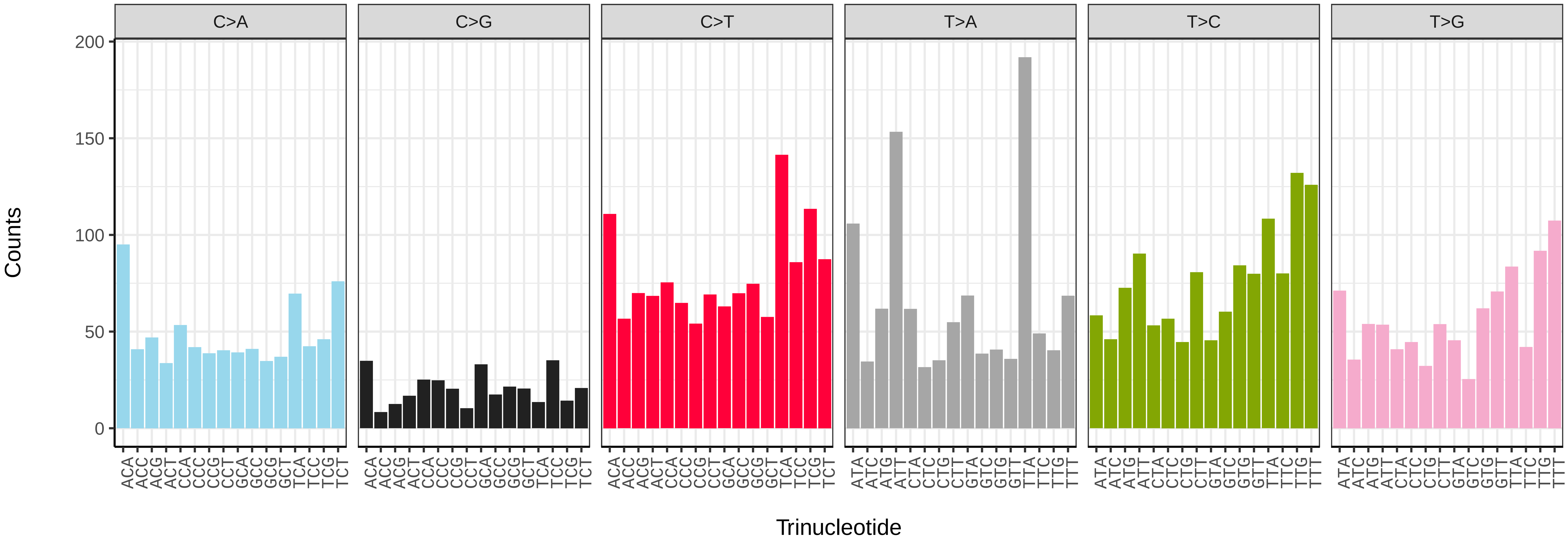


*Leuctra nigra* (ipLeuNigr2) somatic mutational spectrum (SBS52 classification)

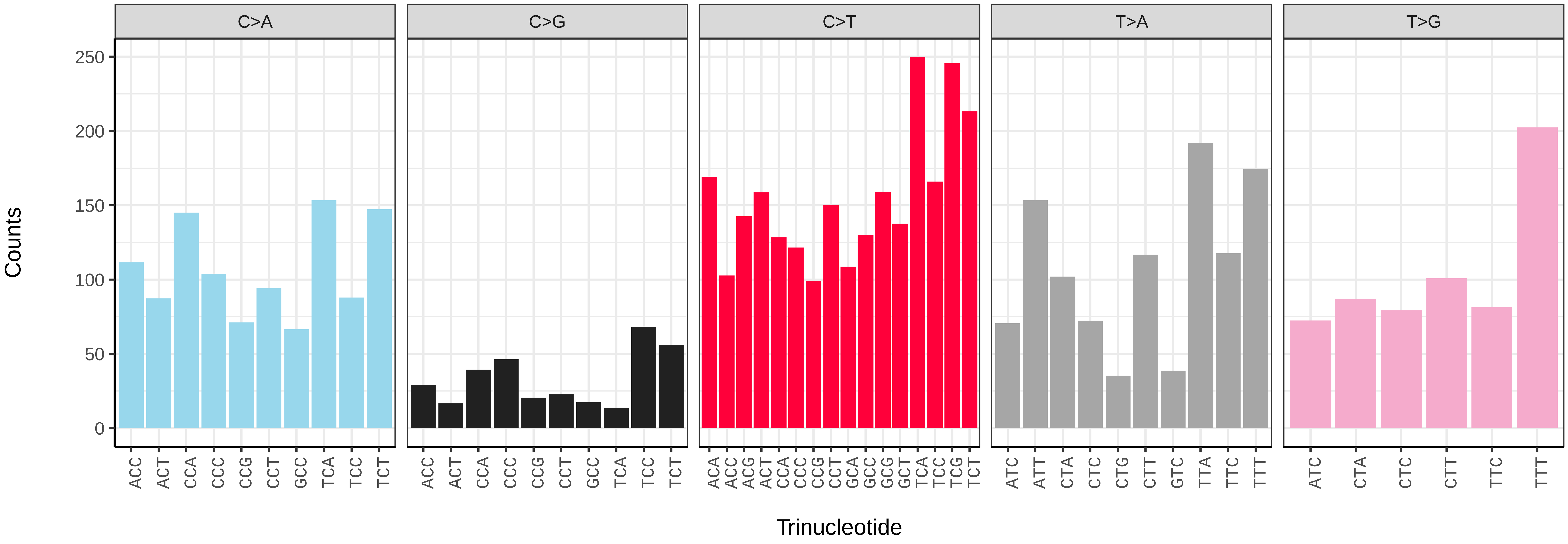

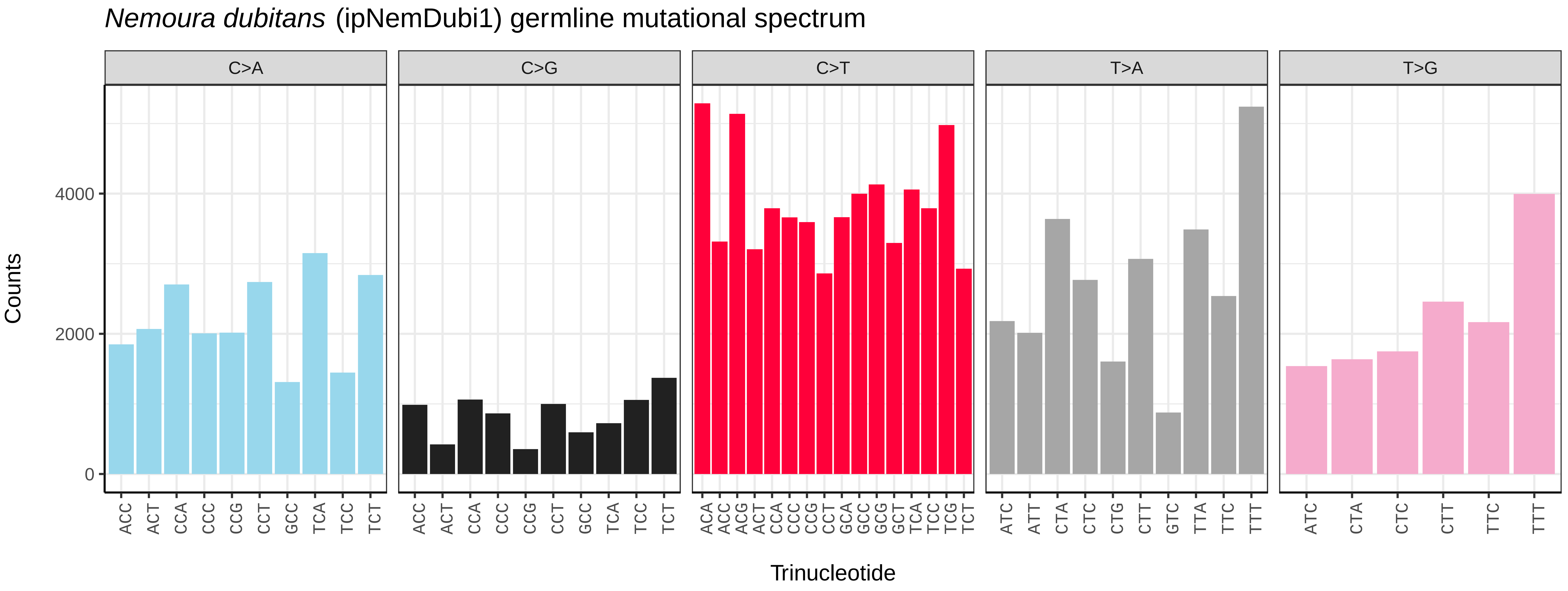
*Nemoura dubitans* (ipNemDubi1) germline mutational spectrum
C>A
C>G
C>T
T>A
T>G
Counts
Trinucleotide

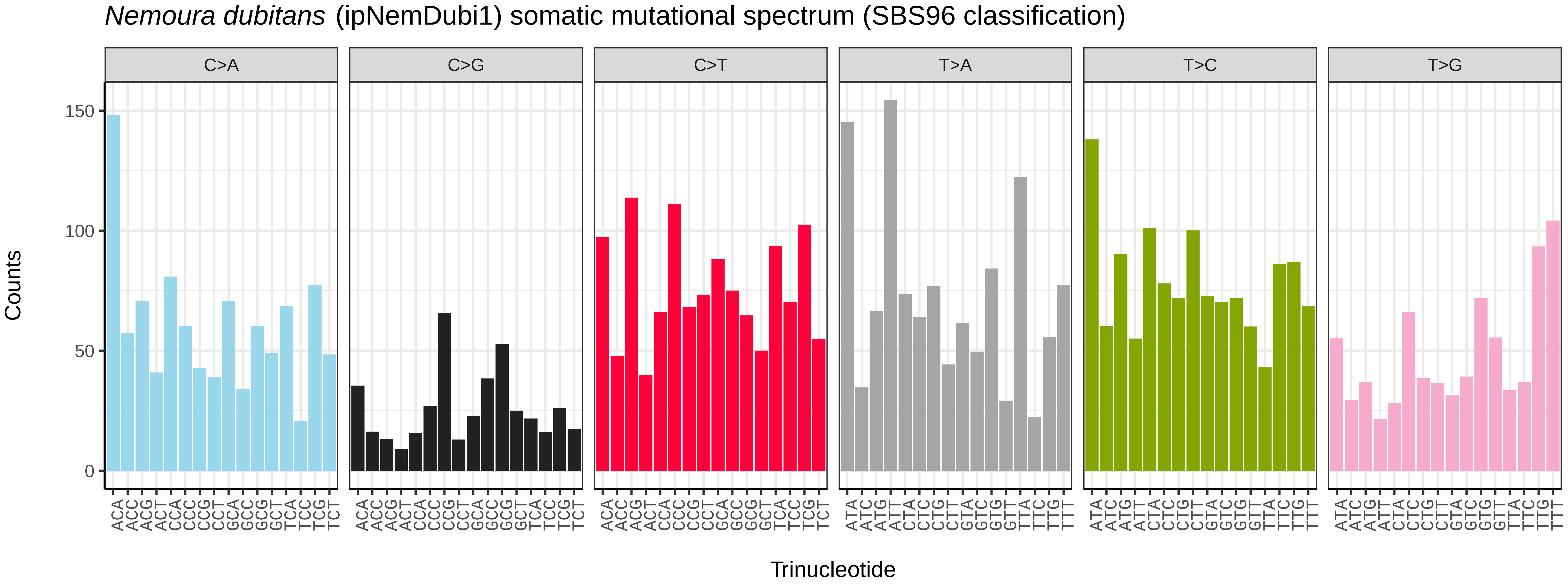
*Nemoura dubitans* (ipNemDubi1) somatic mutational spectrum (SBS96 classification)
C>A
C>G
C>T
T>A
T>C
T>G
Counts
Trinucleotide

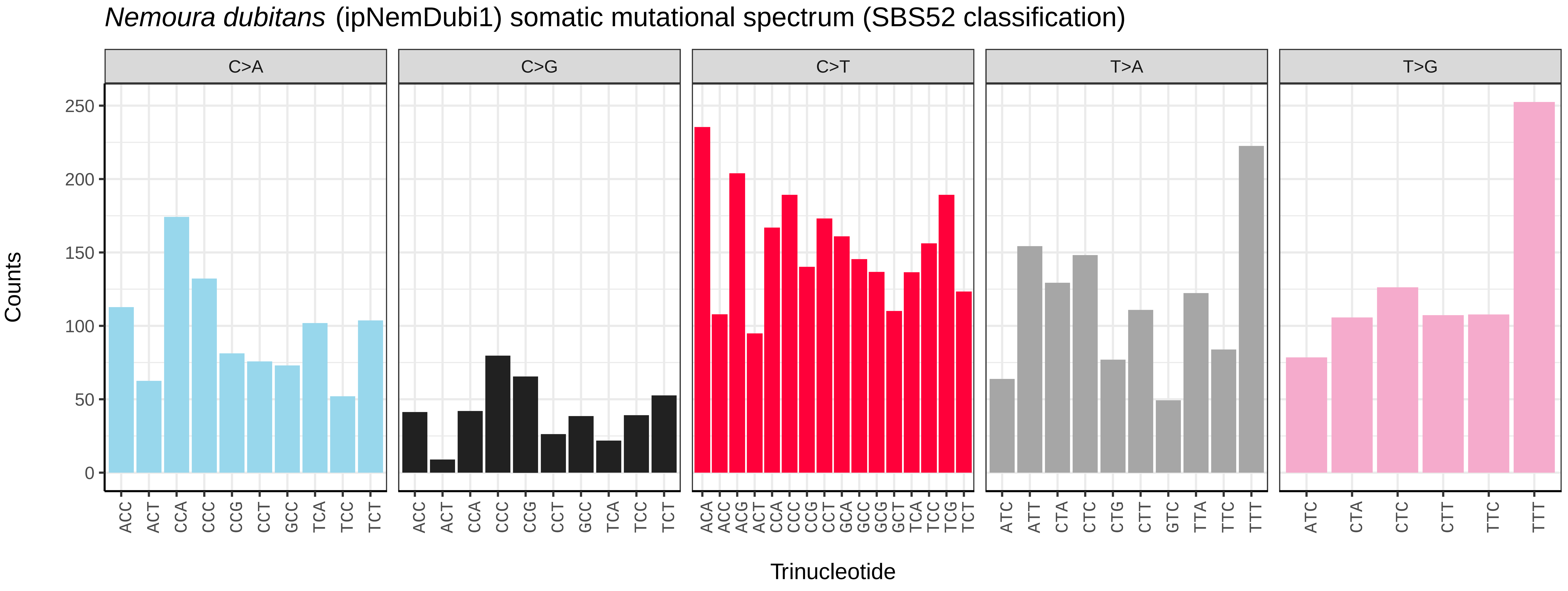
*Nemoura dubitans* (ipNemDubi1) somatic mutational spectrum (SBS52 classification)
C>A
C>G
C>T
T>A
T>G
Counts
Trinucleotide

*Nemurella pictetii* (ipNemPict2) germline mutational spectrum

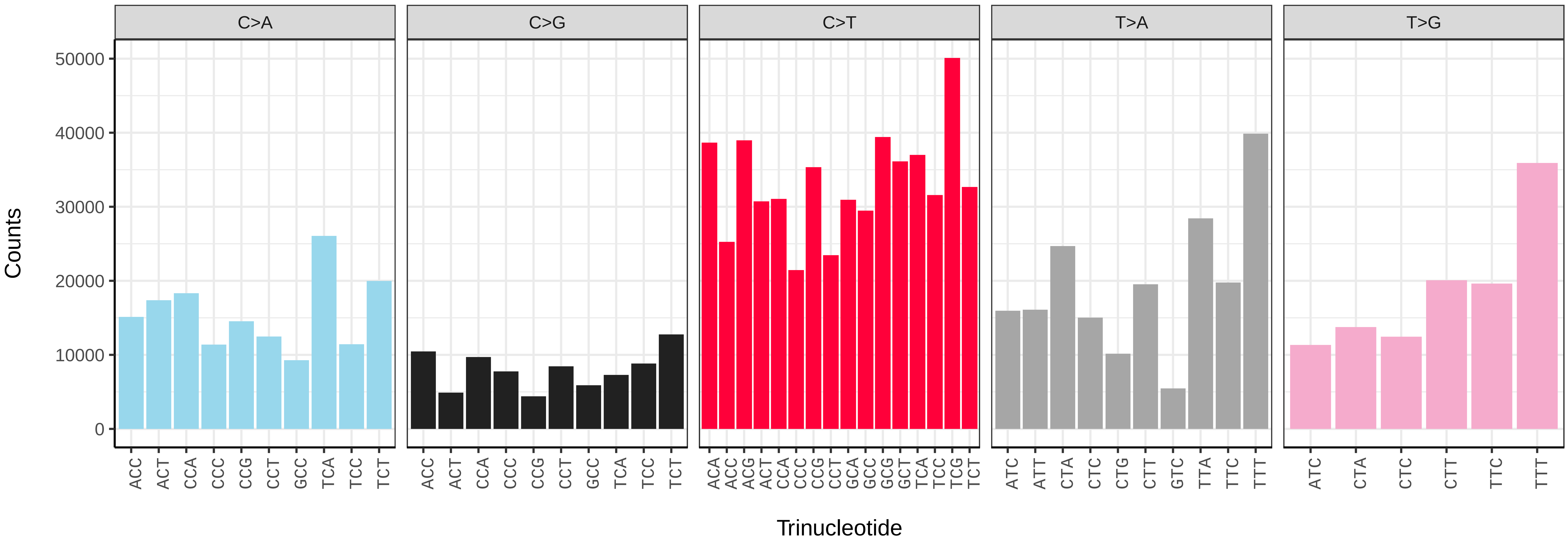


*Nemurella pictetii* (ipNemPict2) somatic mutational spectrum (SBS96 classification)

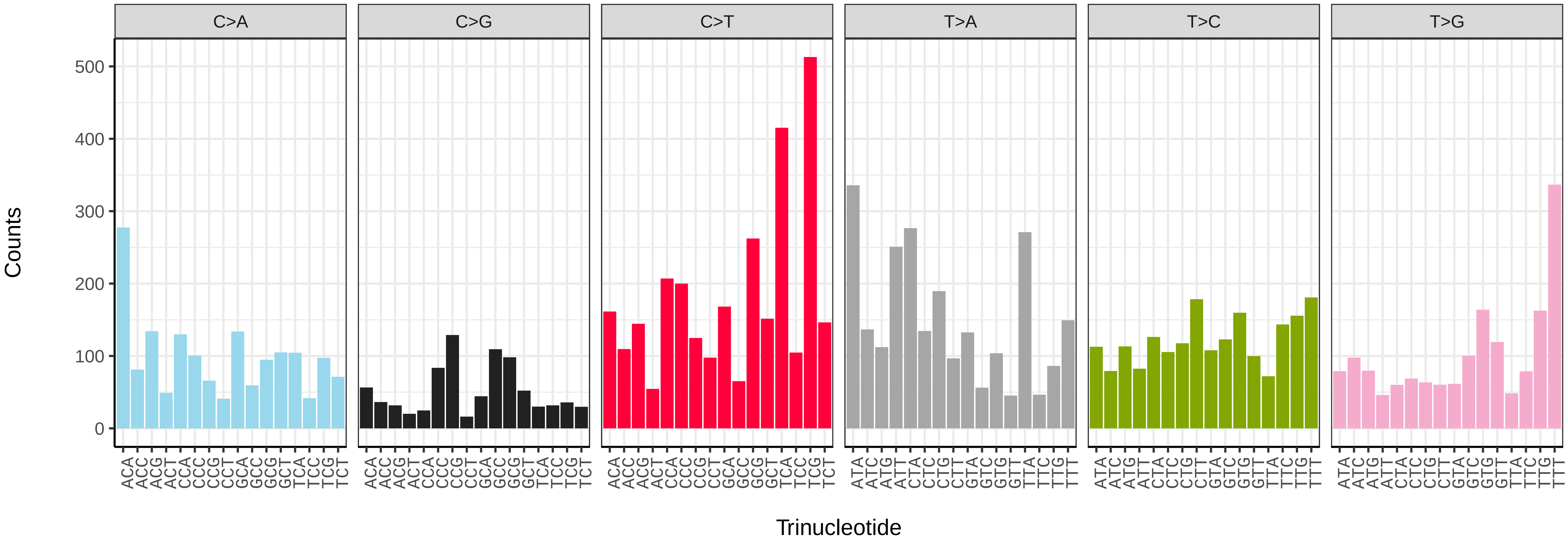


*Nemurella pictetii* (ipNemPict2) somatic mutational spectrum (SBS52 classification)

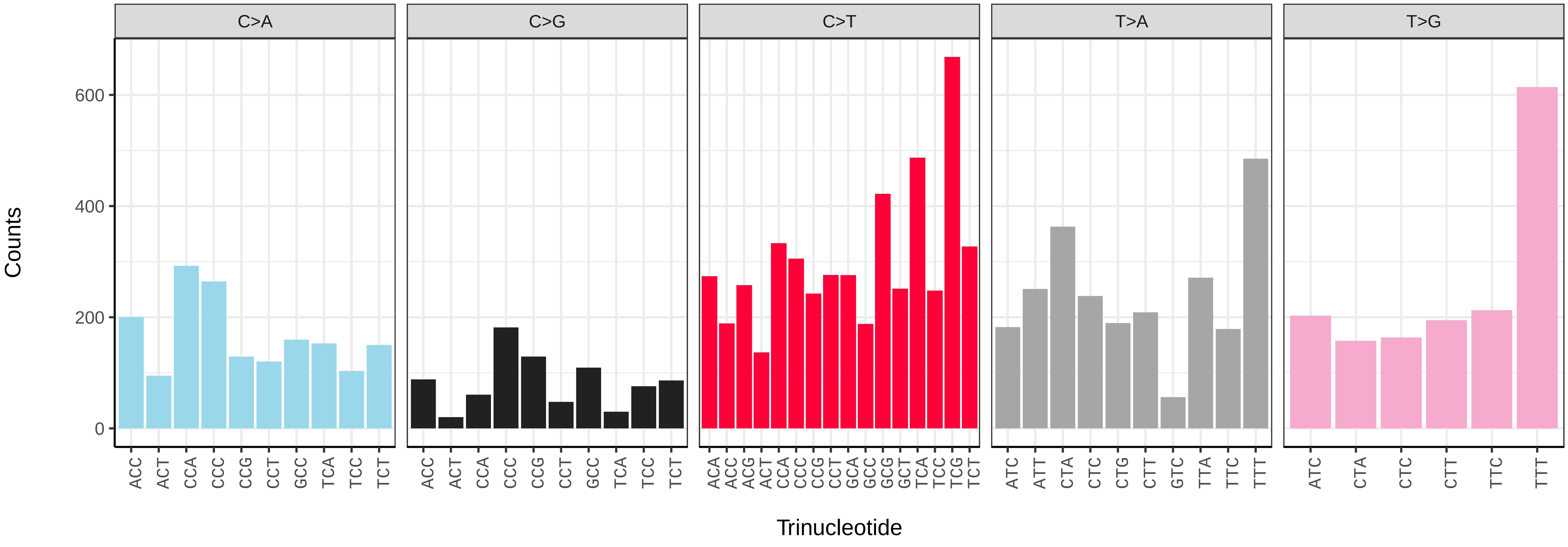

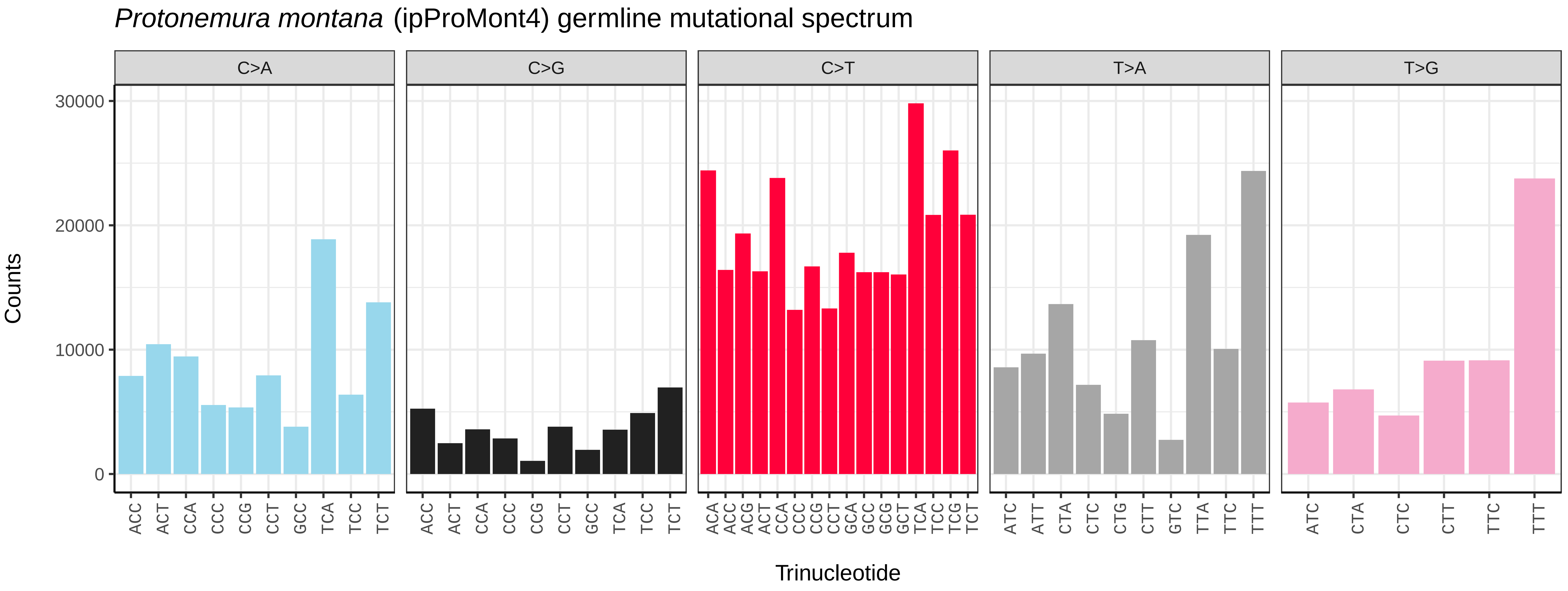
*Protonemura montana* (ipProMont4) germline mutational spectrum
C>A
C>G
C>T
T>A
T>G
Counts
30000
20000
10000
0
ACC ACT CCA CCC CCG CCT GCC TCA TCC TCT
ACA ACC ACG ACT CCA CCC CCG CCT GCA GCC GCG GCT TCA TCC TCG TCT
ATC ATT CTA CTC CTG CTT GTC TTA TTC TTT
ATC CTA CTC CTT TTC TTT
Trinucleotide

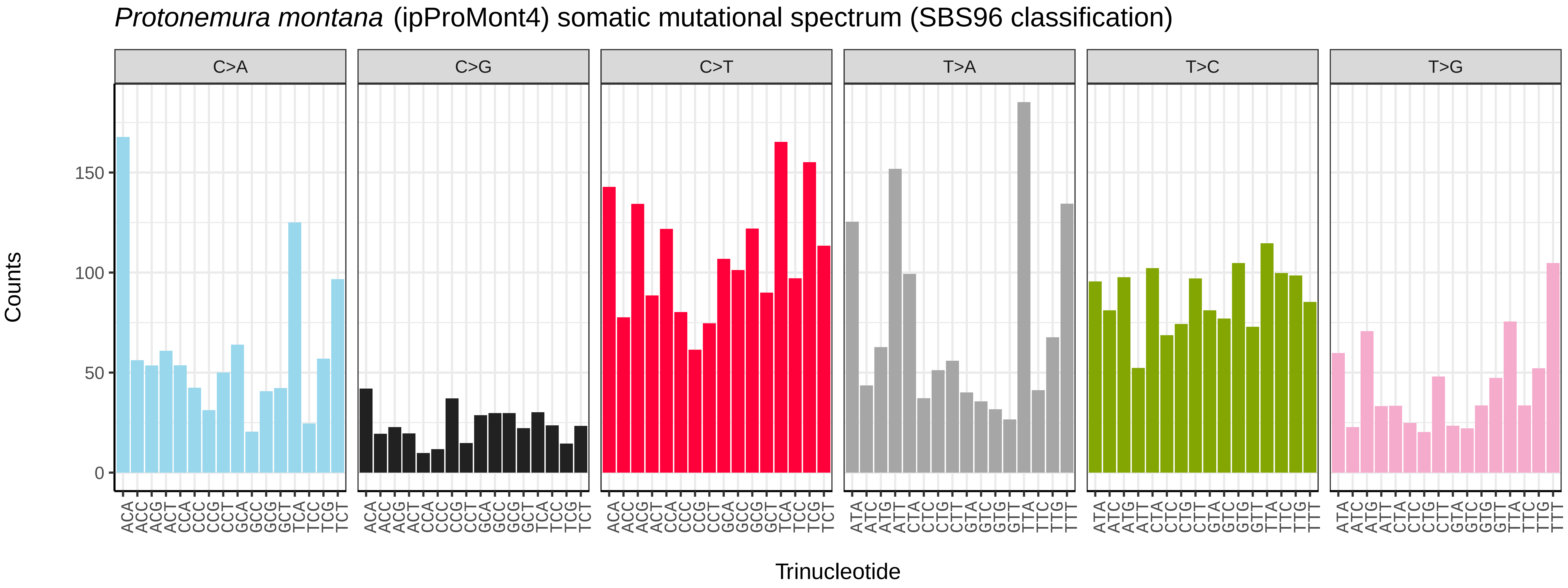
*Protonemura montana* (ipProMont4) somatic mutational spectrum (SBS96 classification)
C>A
C>G
C>T
T>A
T>C
T>G
Counts
150
100
50
0
ACA ACC ACG ACT CCA CCC CCG CCT GCA GCC GCG GCT TCA TCC TCG TCT
ATA ATC ATG ATT CTA CTC CTG CTT GTA GTC GTG GTT TTA TTC TTG TTT
Trinucleotide

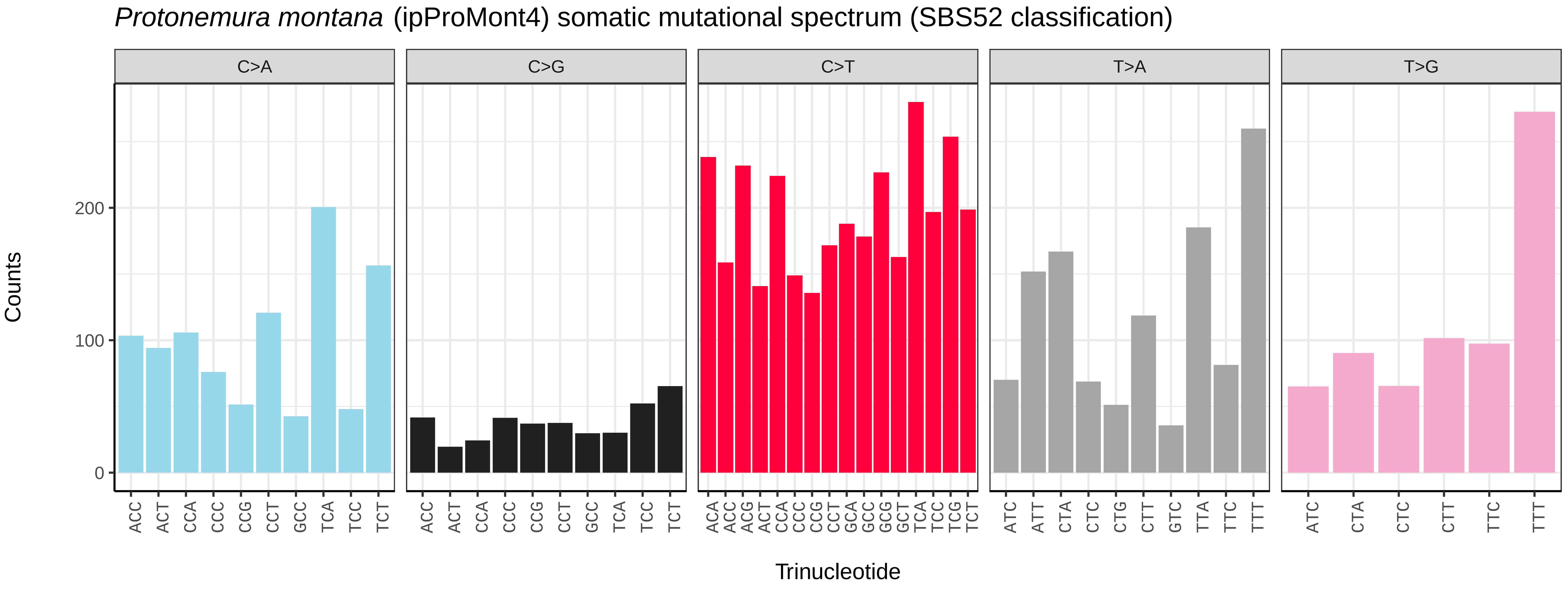
*Protonemura montana* (ipProMont4) somatic mutational spectrum (SBS52 classification)
C>A
C>G
C>T
T>A
T>G
Counts
200
100
0
ACC ACT CCA CCC CCG CCT GCC TCA TCC TCT
ACA ACC ACG ACT CCA CCC CCG CCT GCA GCC GCG GCT TCA TCC TCG TCT
ATC ATT CTA CTC CTG CTT GTC TTA TTC TTT
ATC CTA CTC CTT TTC TTT
Trinucleotide

*Ancistrocerus nigricornis* (iyAncNigr1) germline mutational spectrum

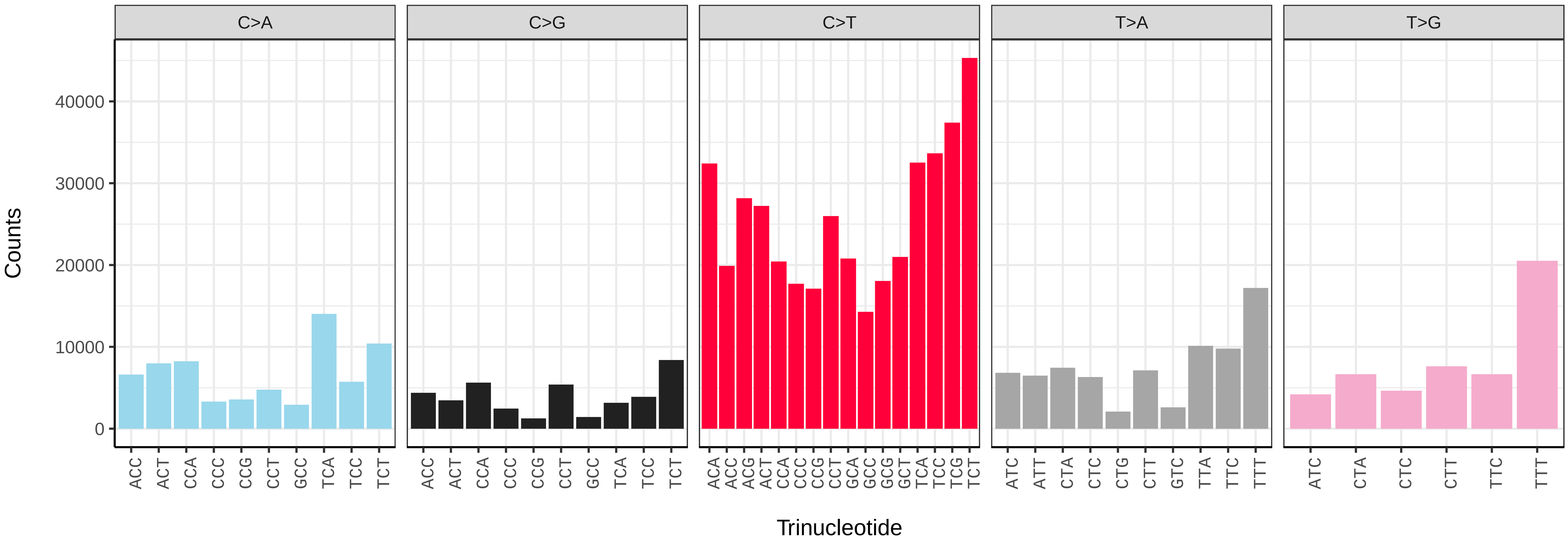


*Ancistrocerus nigricornis* (iyAncNigr1) somatic mutational spectrum (SBS96 classification)

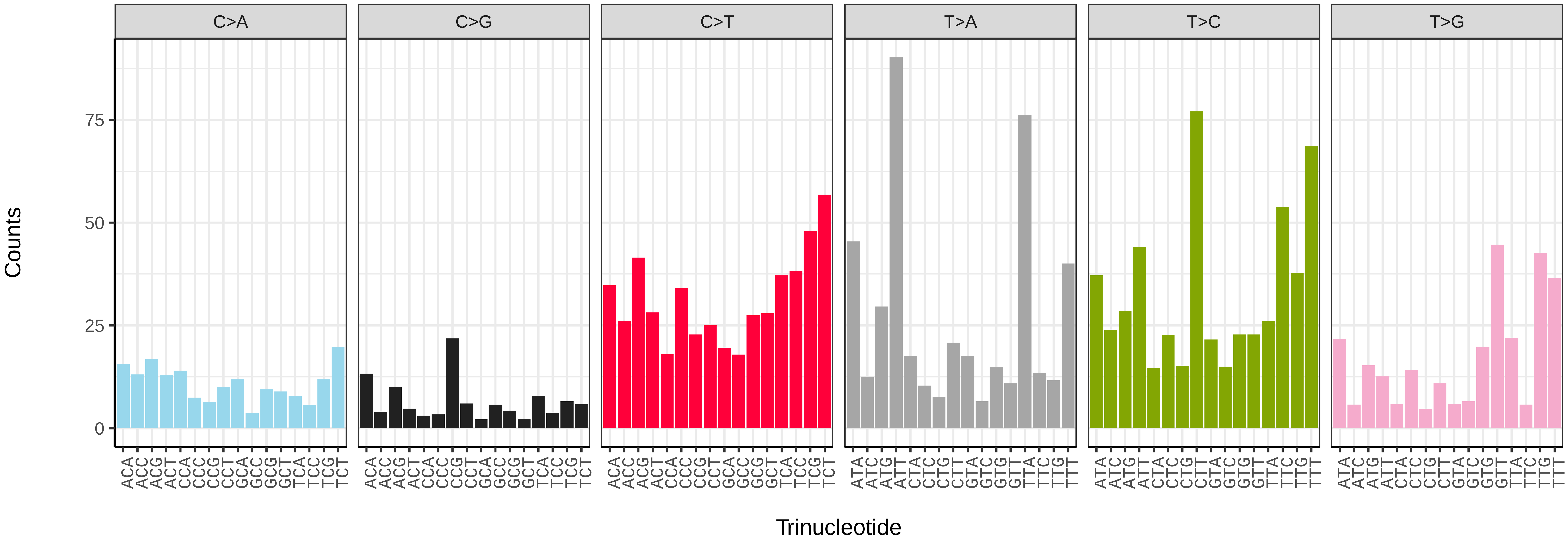


*Ancistrocerus nigricornis* (iyAncNigr1) somatic mutational spectrum (SBS52 classification)

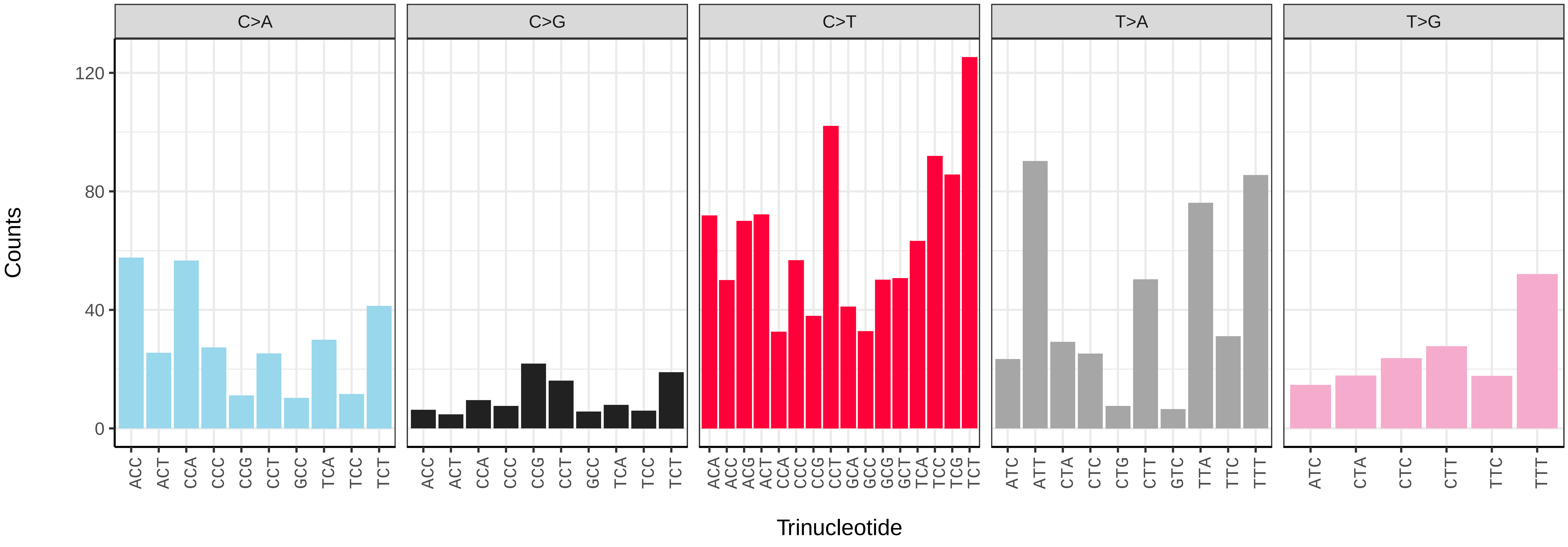

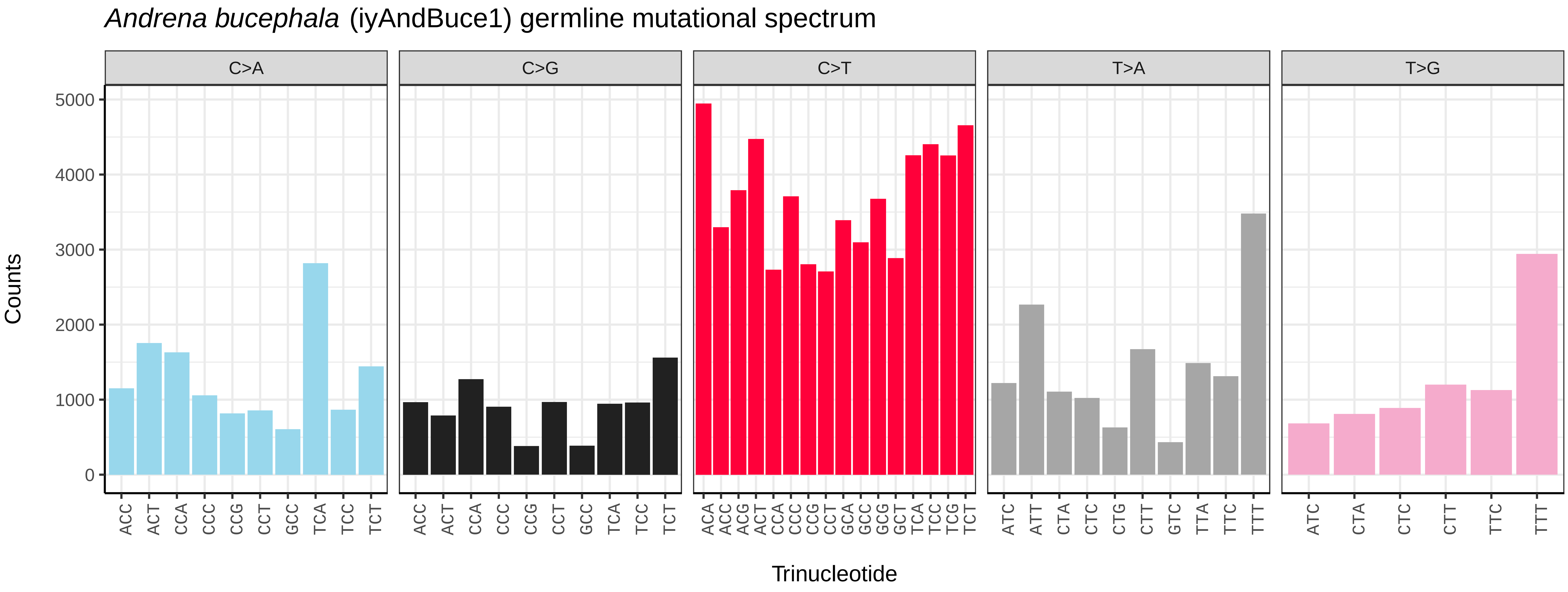
Andrena bucephala (iyAndBuce1) germline mutational spectrum
C>A
C>G
C>T
T>A
T>G
Counts
0
1000
2000
3000
4000
5000
ACC ACT CCA CCC CCG CCT GCC TCA TCC TCT
ACA ACC ACG ACT CCA CCC CCG CCT GCA GCC GCG GCT TCA TCC TCG TCT
ATC ATT CTA CTC CTG CTT GTC TTA TTC TTT
ATC CTA CTC CTT TTC TTT
Trinucleotide

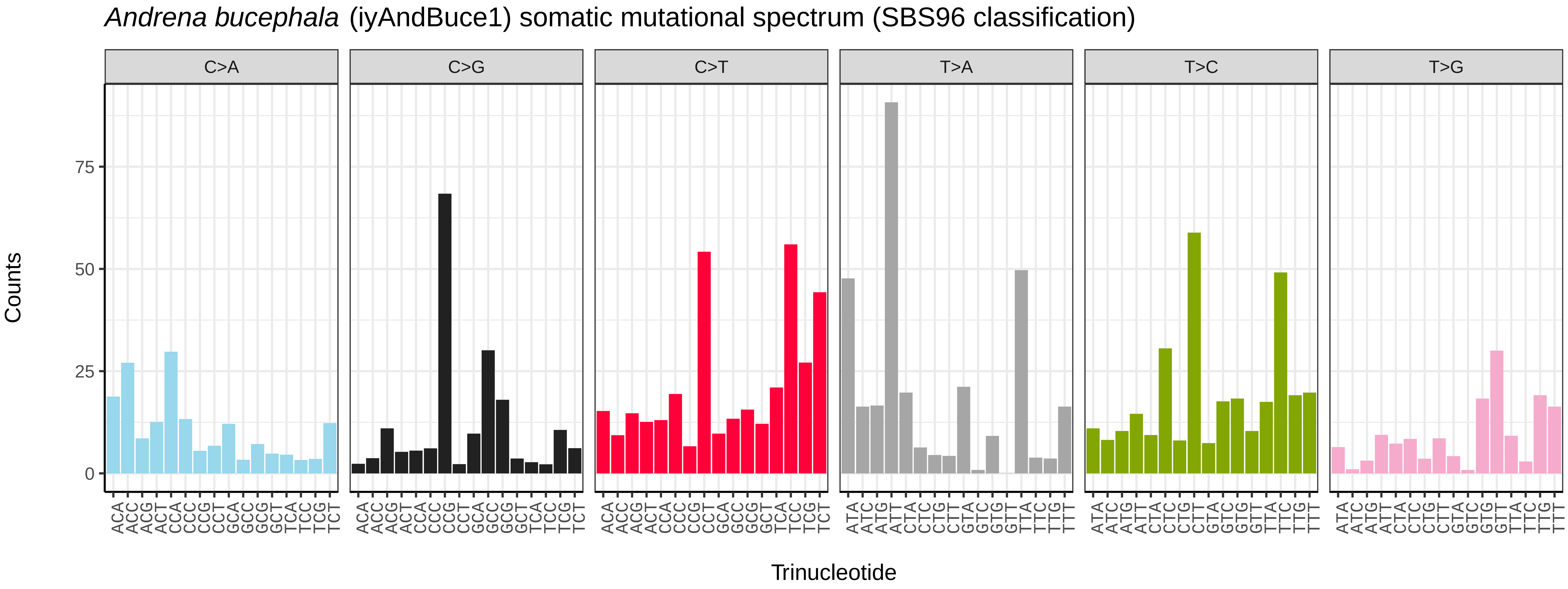
Andrena bucephala (iyAndBuce1) somatic mutational spectrum (SBS96 classification)
C>A
C>G
C>T
T>A
T>C
T>G
Counts
0
25
50
75
ACA ACC ACG ACT CCA CCC CCG CCT GCA GCC GCG GCT TCA TCC TCG TCT
ATA ATC ATG ATT CTA CTC CTG CTT GTA GTC GTG GTT TTA TTC TTG TTT
Trinucleotide

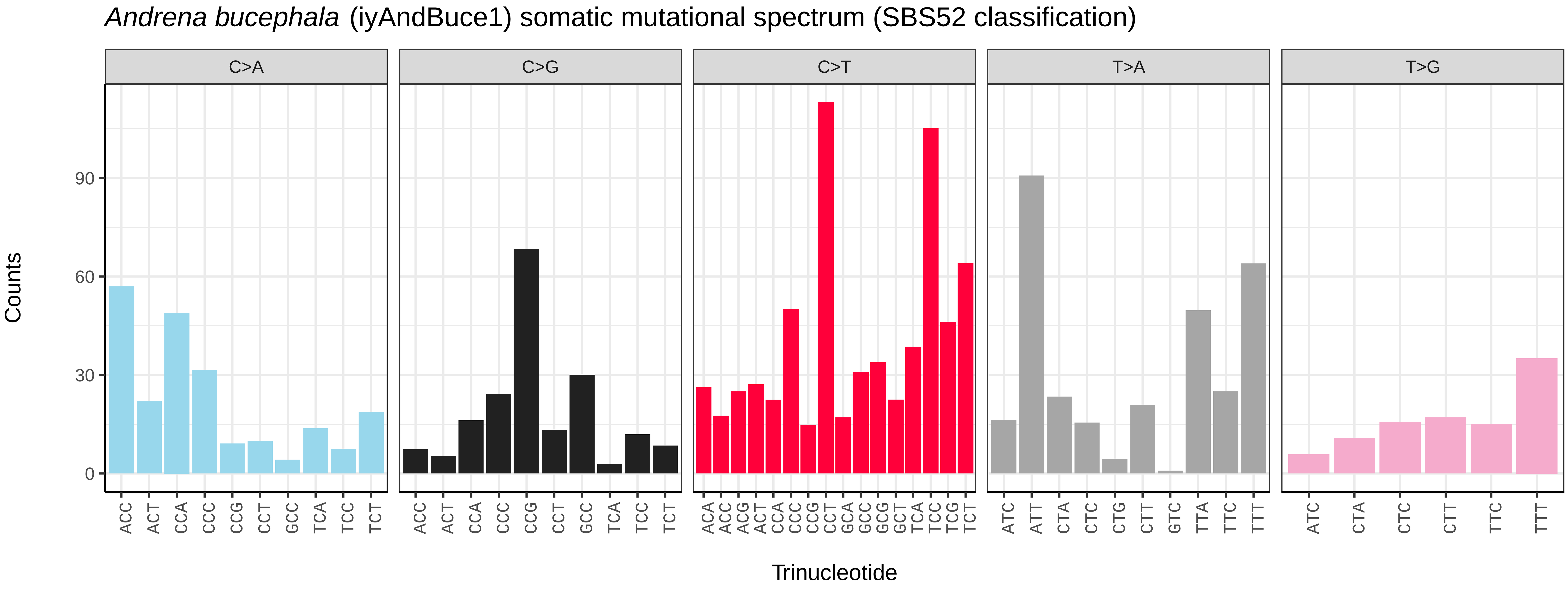
Andrena bucephala (iyAndBuce1) somatic mutational spectrum (SBS52 classification)
C>A
C>G
C>T
T>A
T>G
Counts
0
30
60
90
ACC ACT CCA CCC CCG CCT GCC TCA TCC TCT
ACA ACC ACG ACT CCA CCC CCG CCT GCA GCC GCG GCT TCA TCC TCG TCT
ATC ATT CTA CTC CTG CTT GTC TTA TTC TTT
ATC CTA CTC CTT TTC TTT
Trinucleotide

*Andrena dorsata* (iyAndDors1) germline mutational spectrum

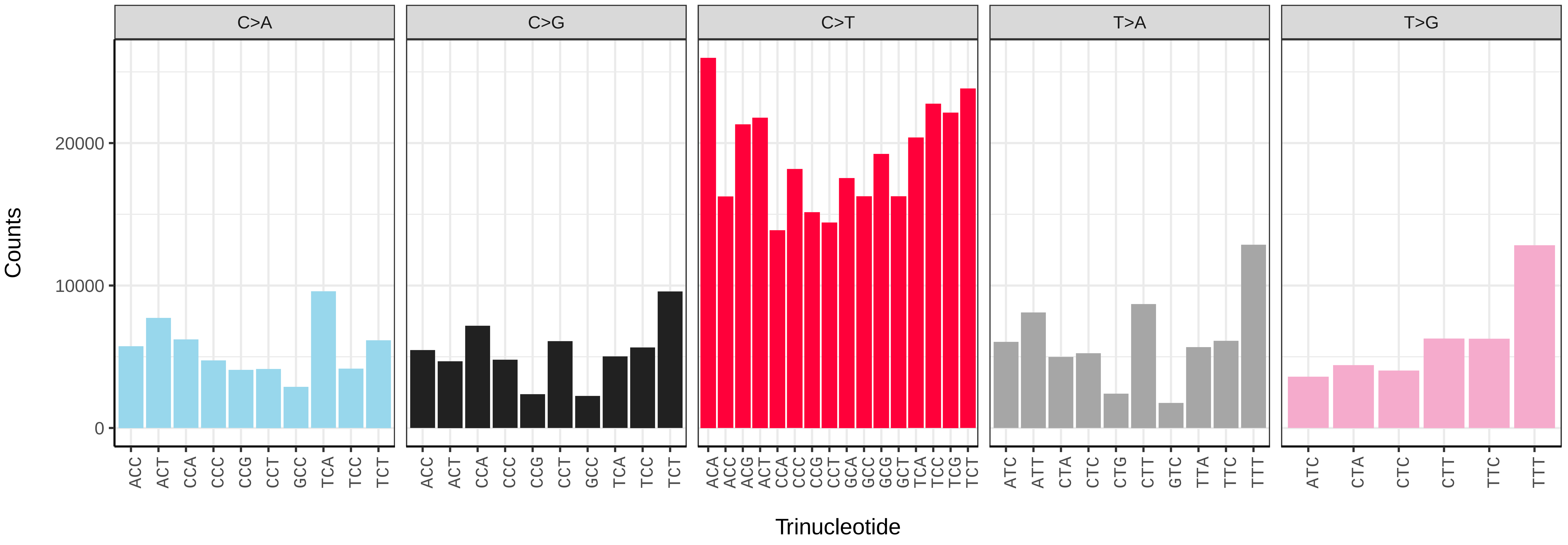


*Andrena dorsata* (iyAndDors1) somatic mutational spectrum (SBS96 classification)

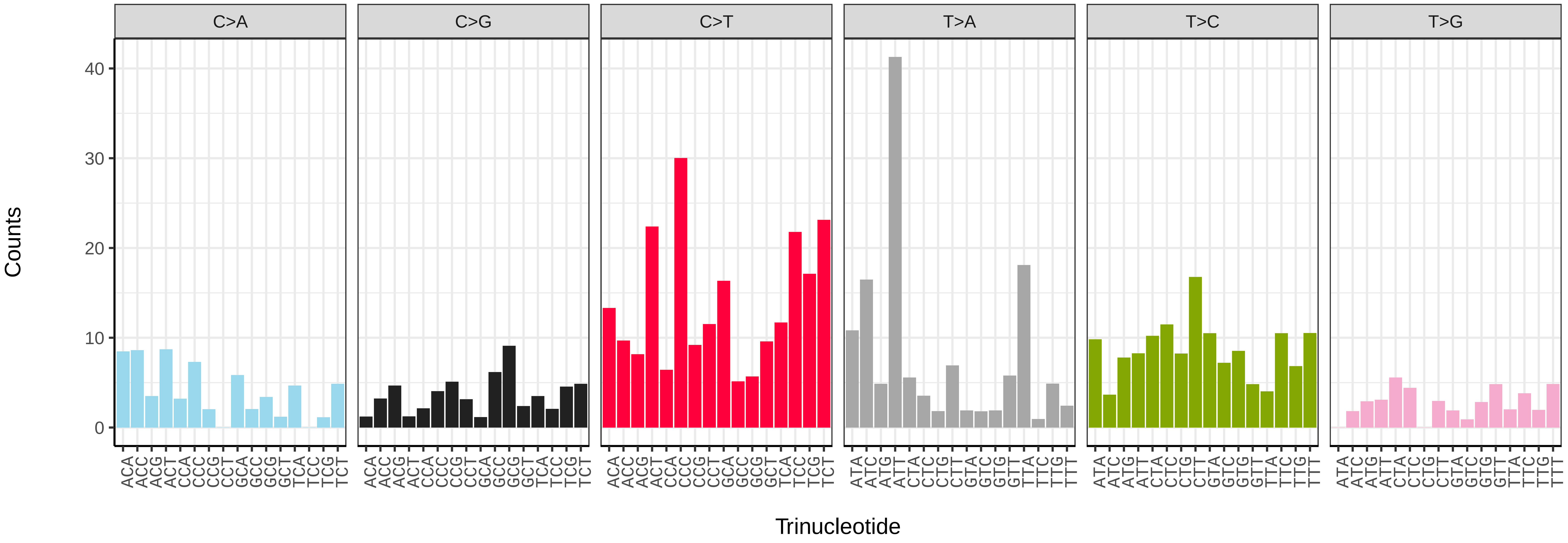


*Andrena dorsata* (iyAndDors1) somatic mutational spectrum (SBS52 classification)

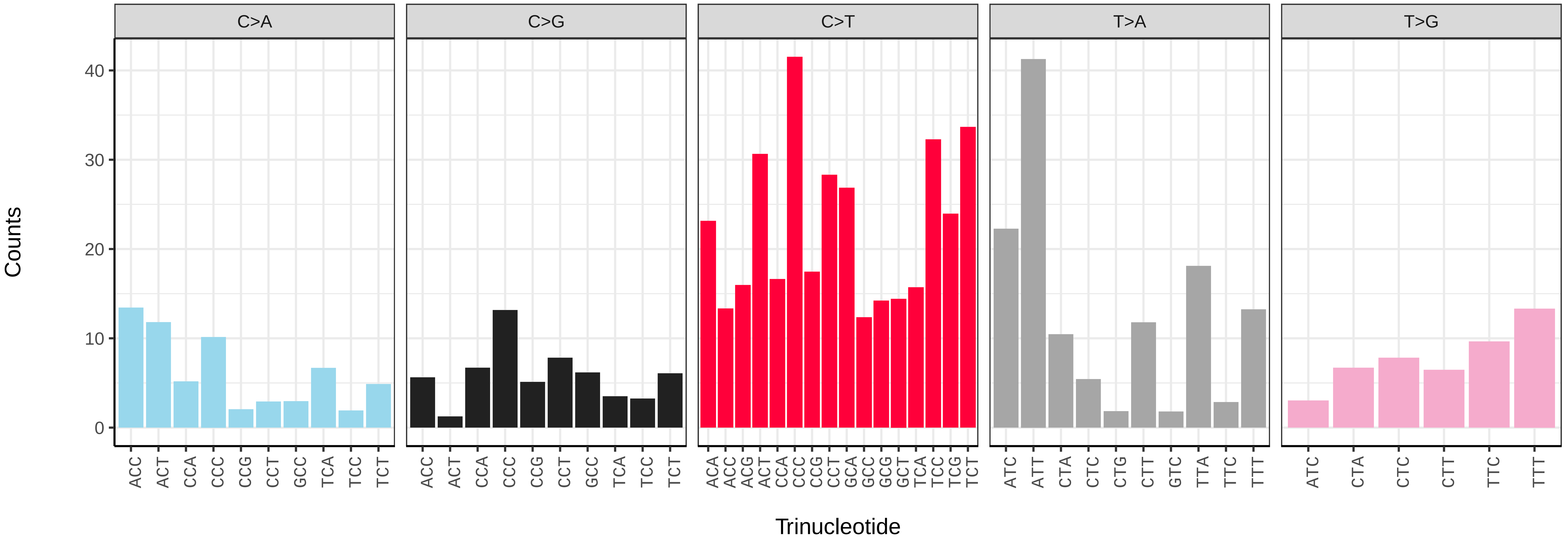

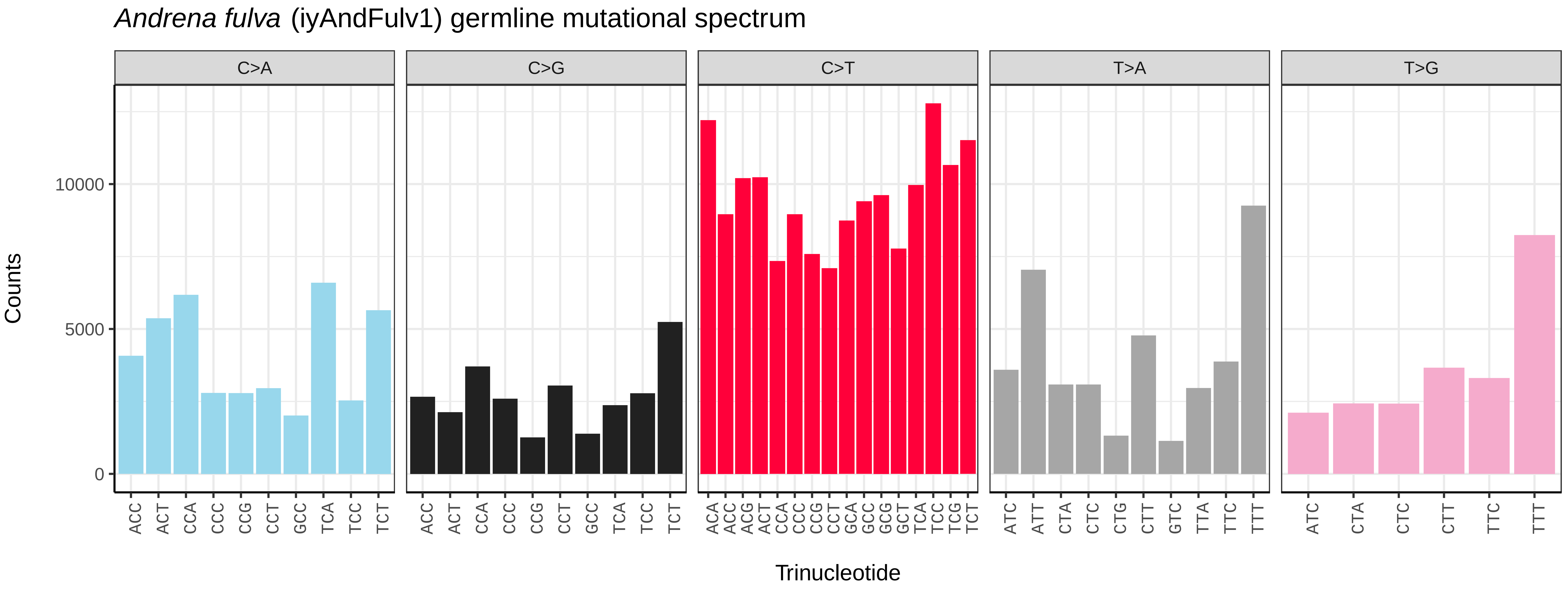
*Andrena fulva* (iyAndFulv1) germline mutational spectrum
C>A
C>G
C>T
T>A
T>G
Counts
10000
5000
0
ACC
ACT
Trinucleotide

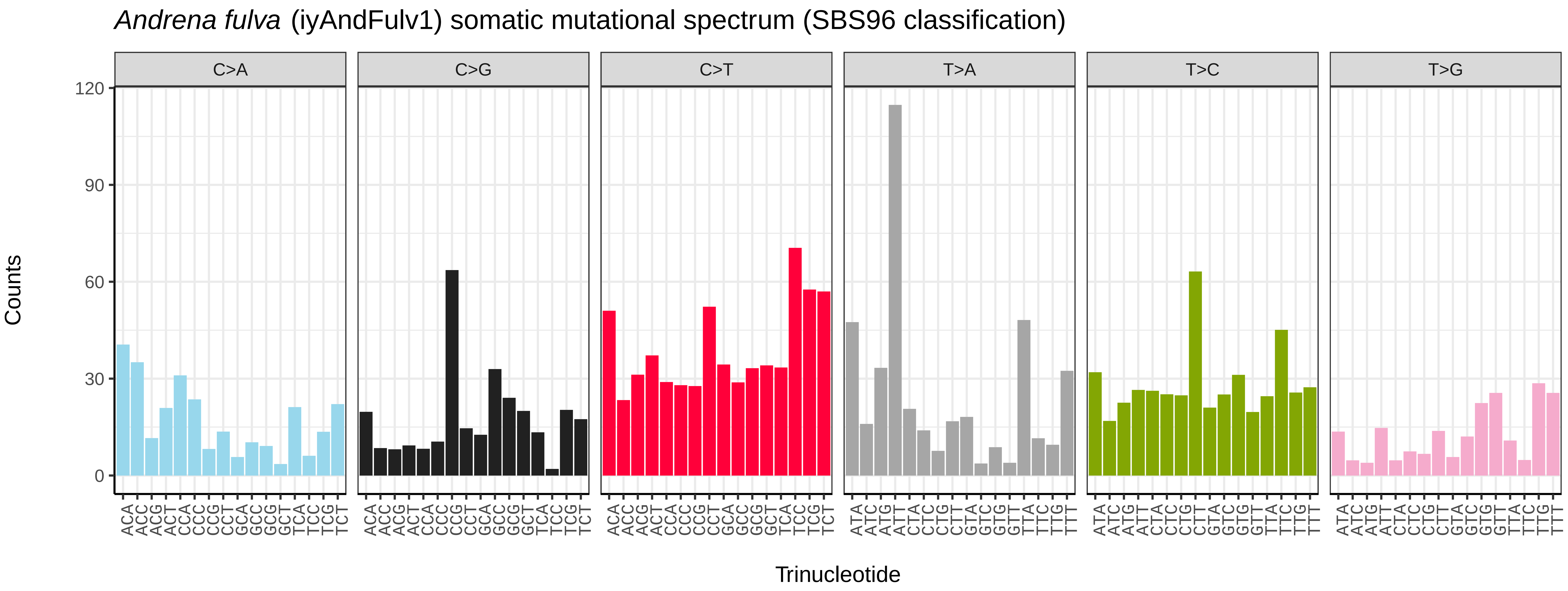
*Andrena fulva* (iyAndFulv1) somatic mutational spectrum (SBS96 classification)
C>A
C>G
C>T
T>A
T>C
T>G
Counts
120
90
60
30
0
Trinucleotide

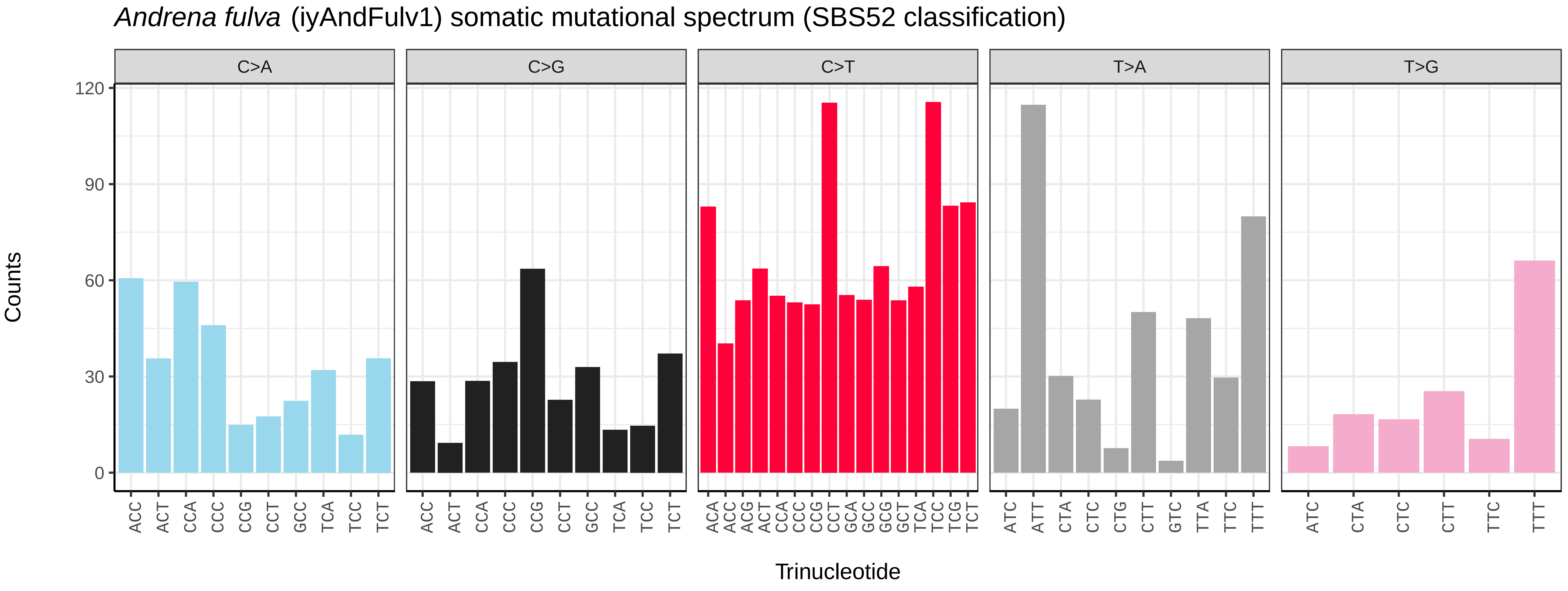
*Andrena fulva* (iyAndFulv1) somatic mutational spectrum (SBS52 classification)
C>A
C>G
C>T
T>A
T>G
Counts
120
90
60
30
0
ACC
ACT
Trinucleotide

*Andrena haemorrhoa* (iyAndHaem1) germline mutational spectrum

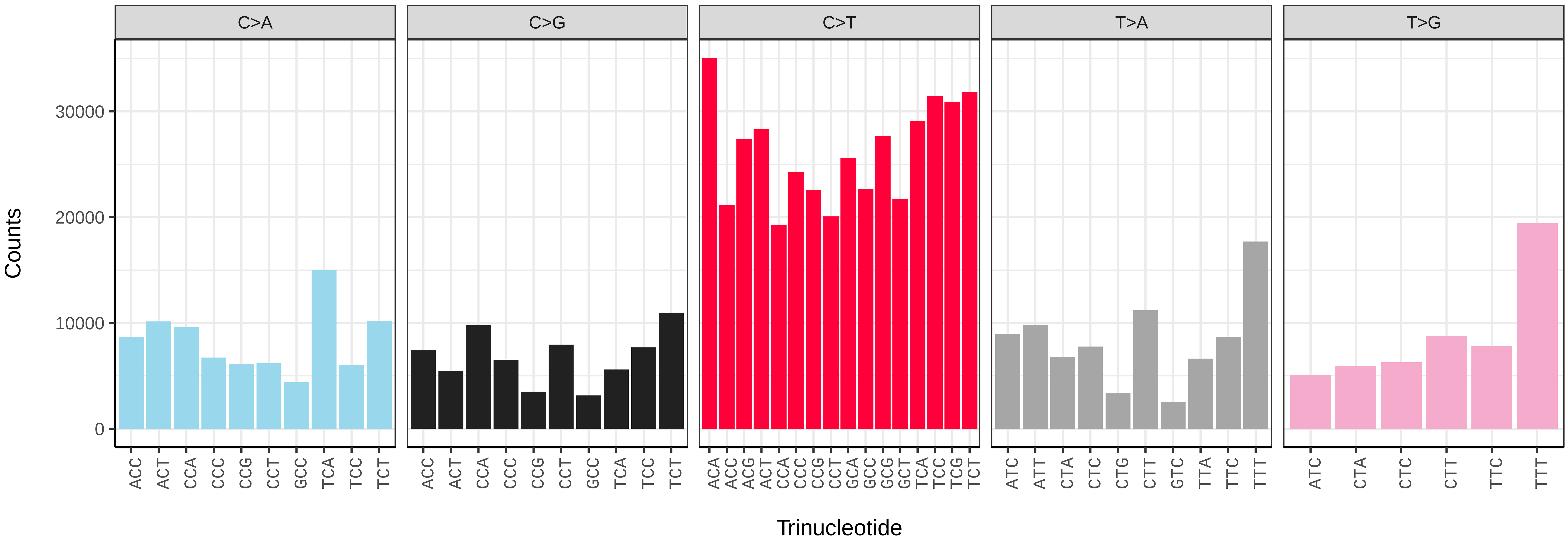


*Andrena haemorrhoa* (iyAndHaem1) somatic mutational spectrum (SBS96 classification)

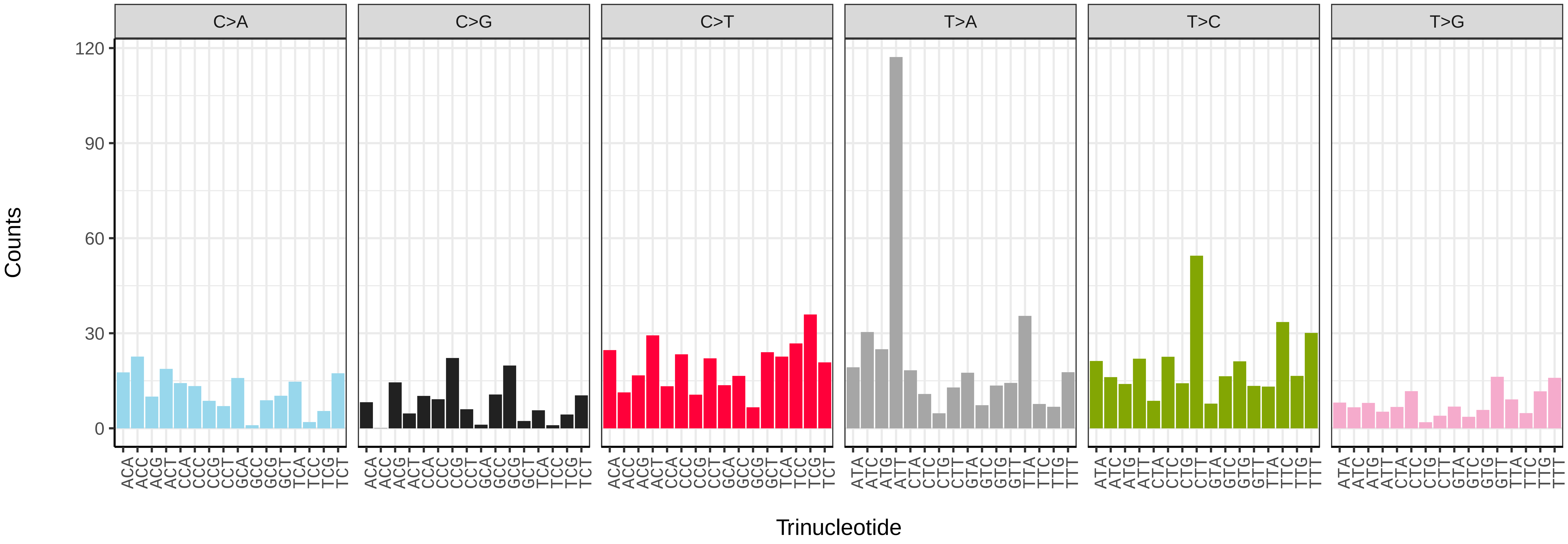


*Andrena haemorrhoa* (iyAndHaem1) somatic mutational spectrum (SBS52 classification)

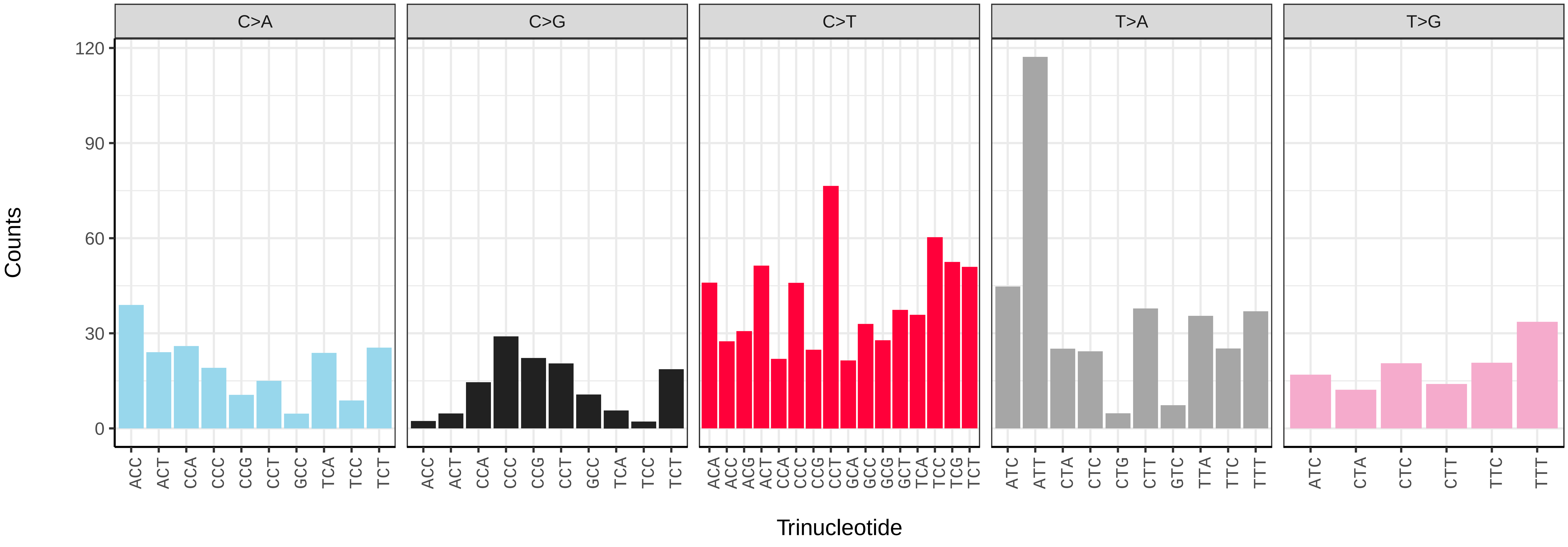

*Andrena hattorfiana* (iyAndHatt1) germline mutational spectrum

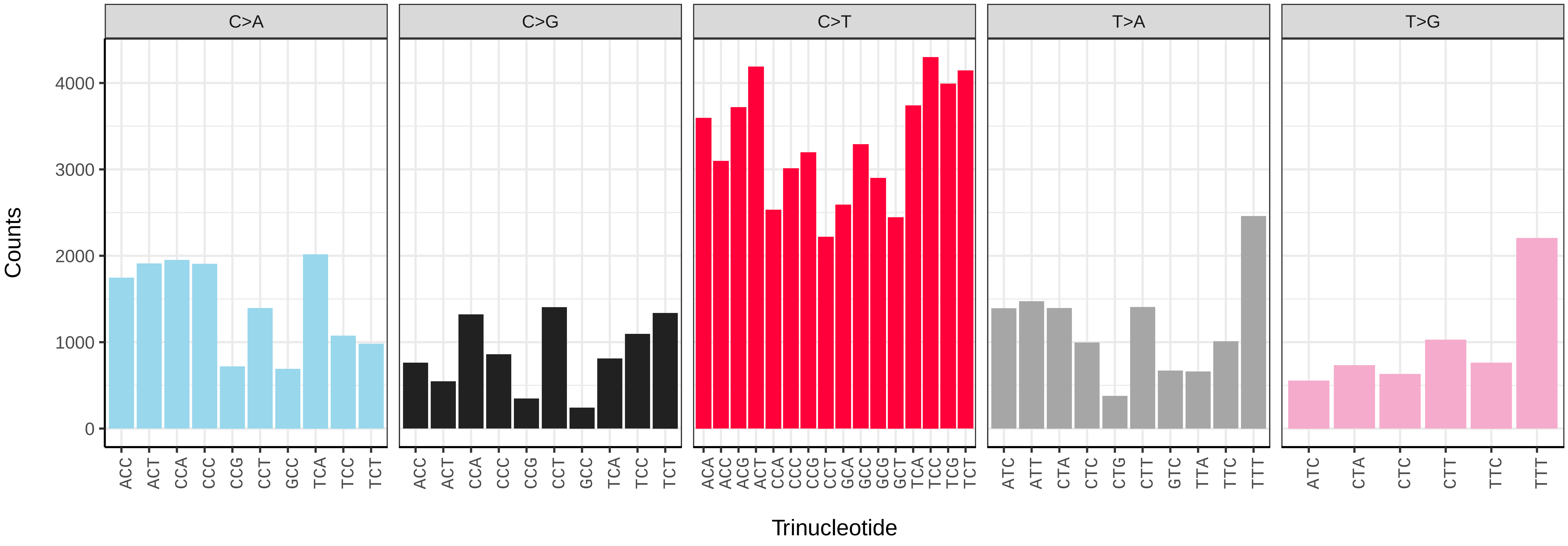


*Andrena hattorfiana* (iyAndHatt1) somatic mutational spectrum (SBS96 classification)

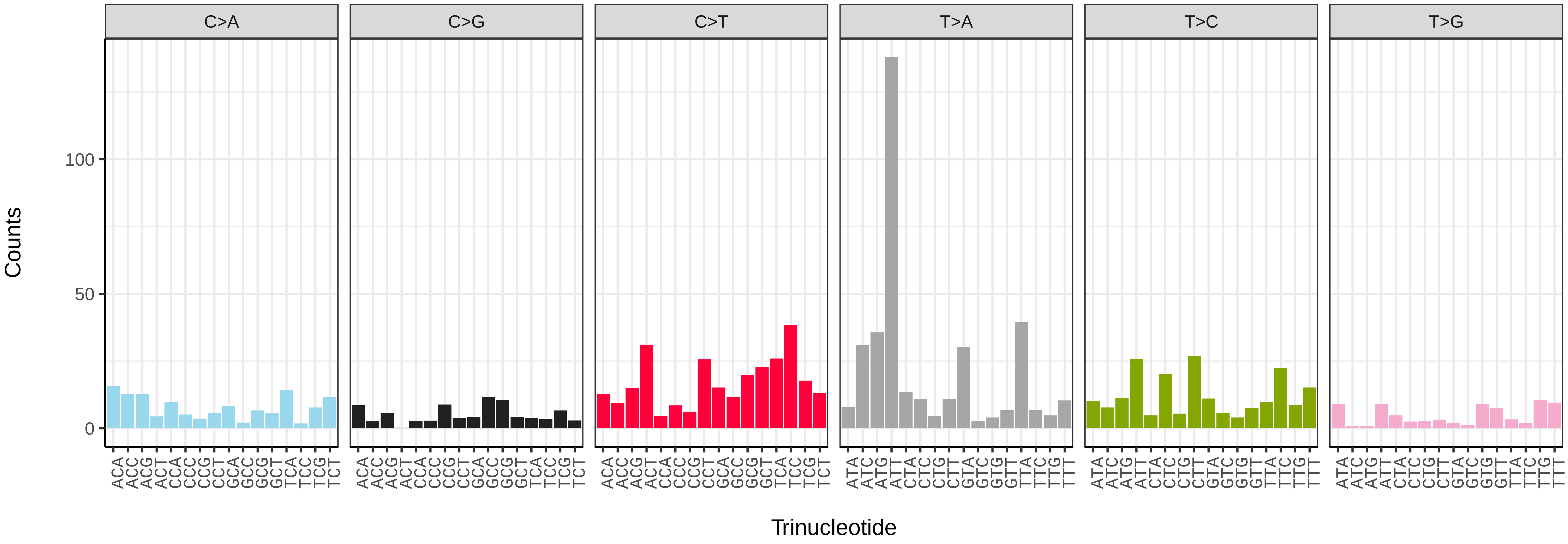


*Andrena hattorfiana* (iyAndHatt1) somatic mutational spectrum (SBS52 classification)

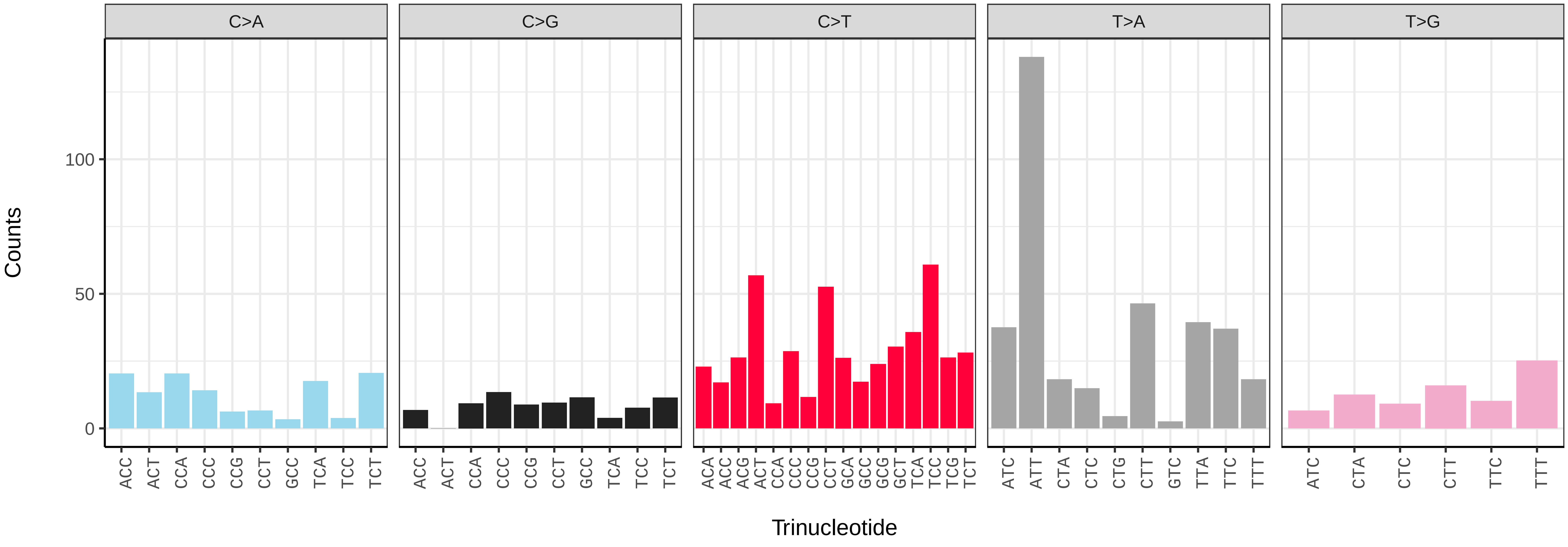

*Andrena minutula* (iyAndMinu1) germline mutational spectrum

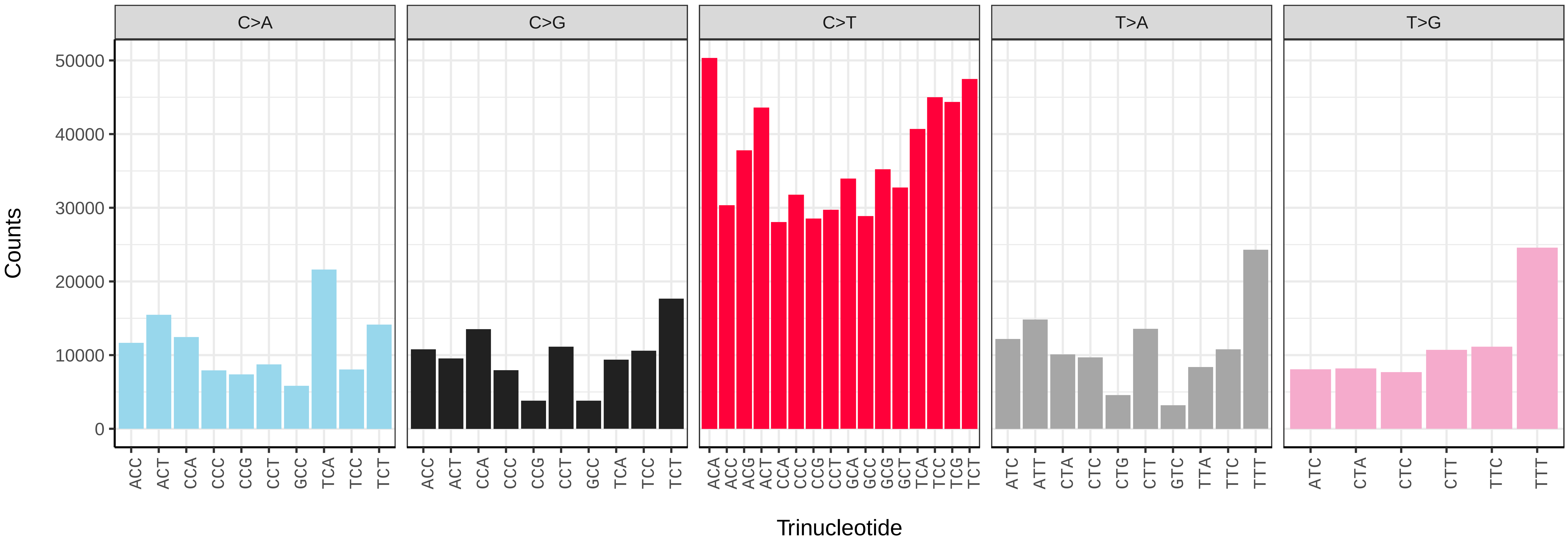


*Andrena minutula* (iyAndMinu1) somatic mutational spectrum (SBS96 classification)

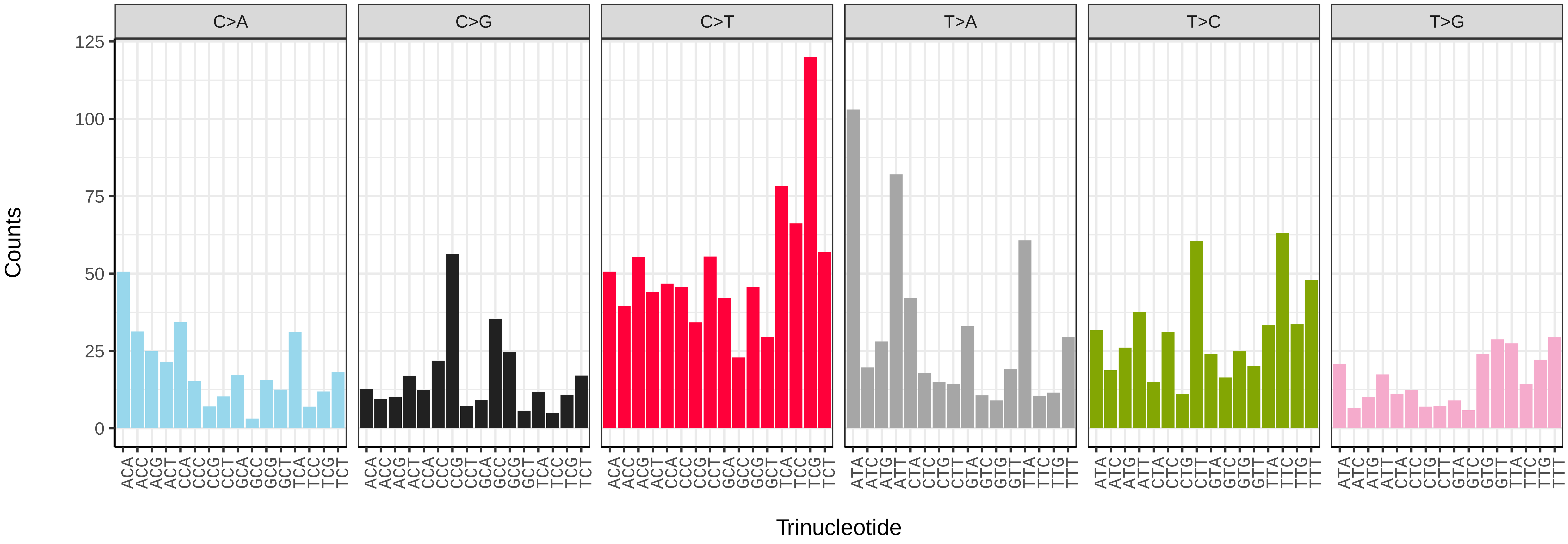


*Andrena minutula* (iyAndMinu1) somatic mutational spectrum (SBS52 classification)

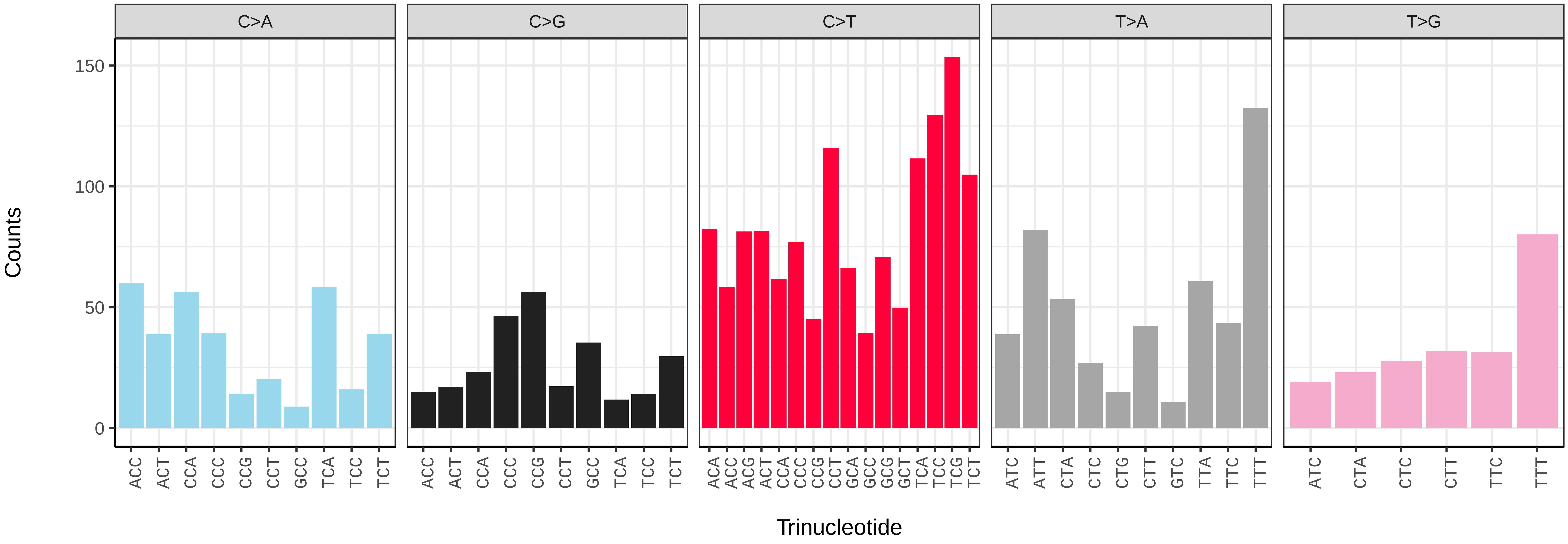

*Anoplius nigerrimus* (iyAnoNige1) germline mutational spectrum

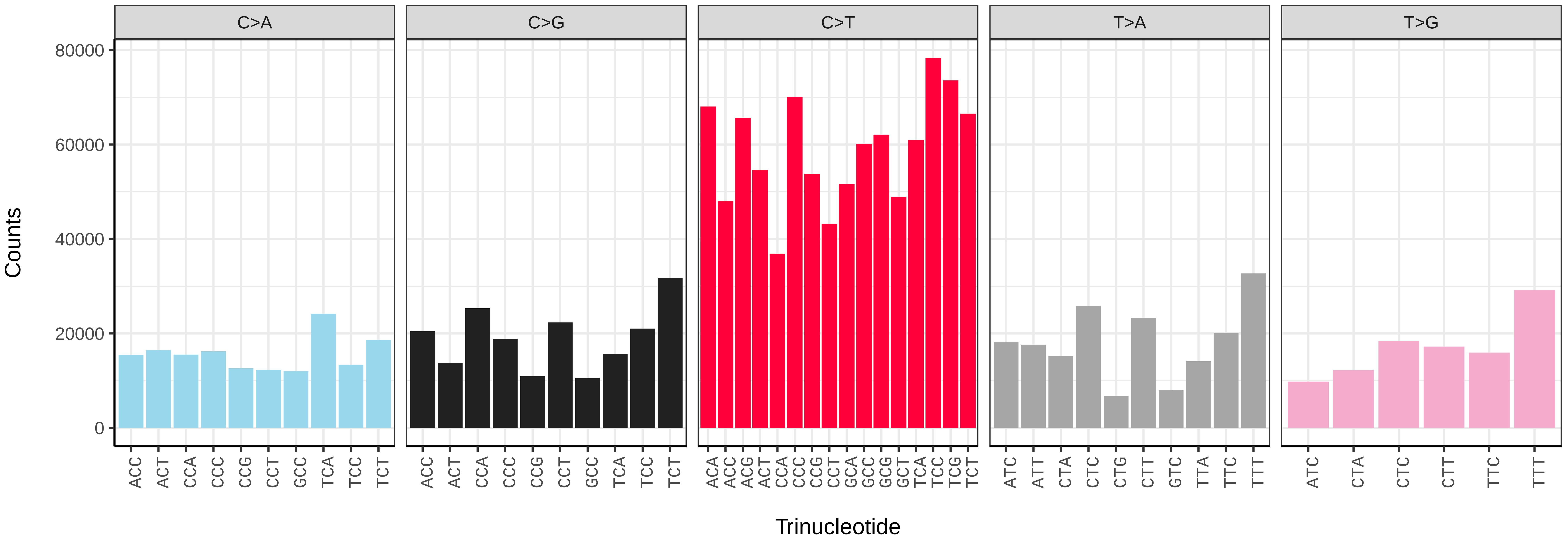


*Anoplius nigerrimus* (iyAnoNige1) somatic mutational spectrum (SBS96 classification)

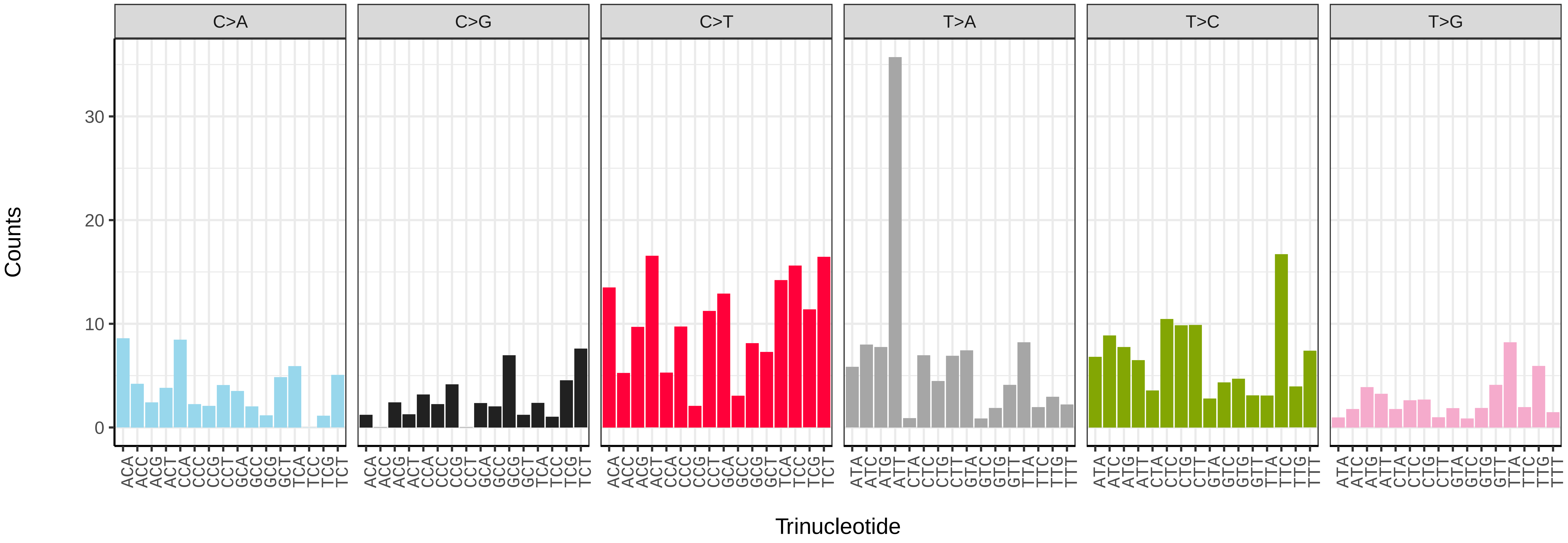


*Anoplius nigerrimus* (iyAnoNige1) somatic mutational spectrum (SBS52 classification)

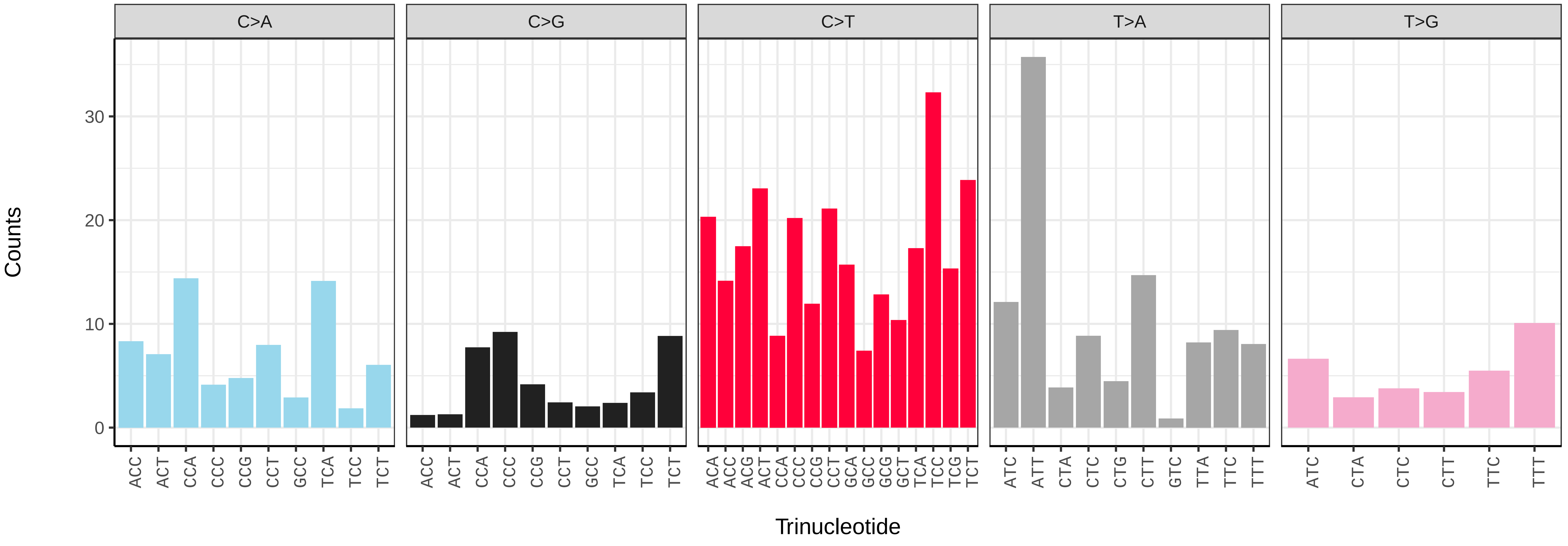

*Athalia rosae* (iyAthRosa1) germline mutational spectrum

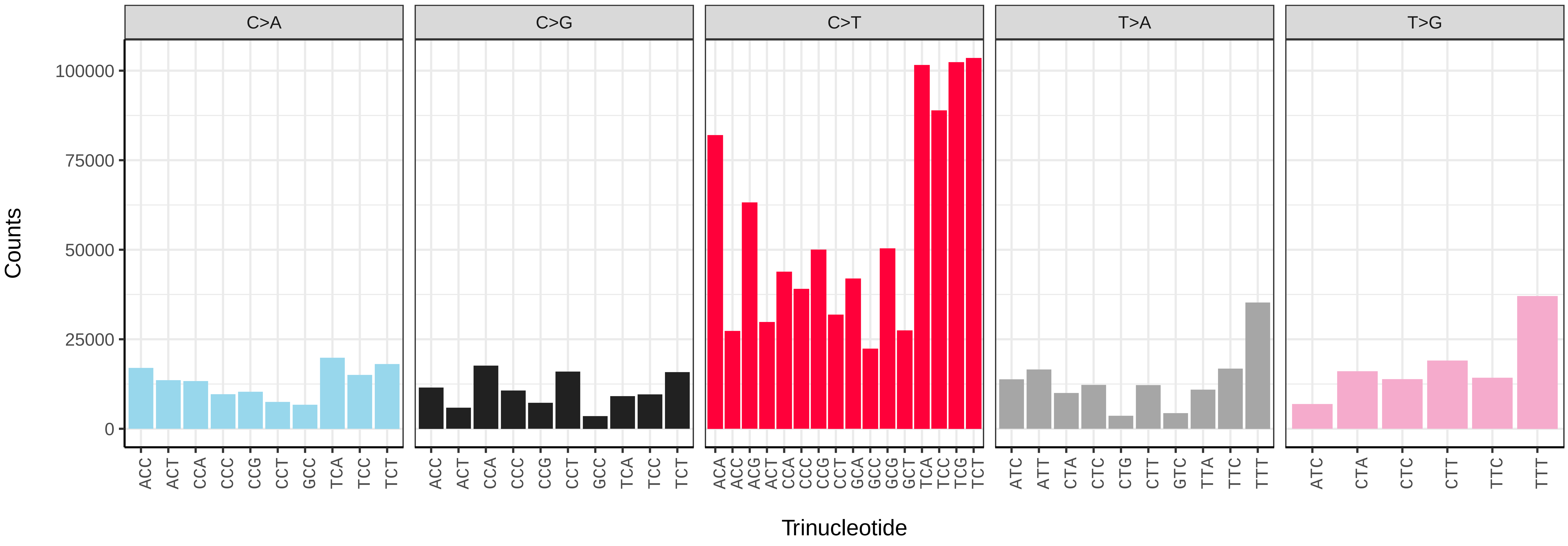


*Athalia rosae* (iyAthRosa1) somatic mutational spectrum (SBS96 classification)

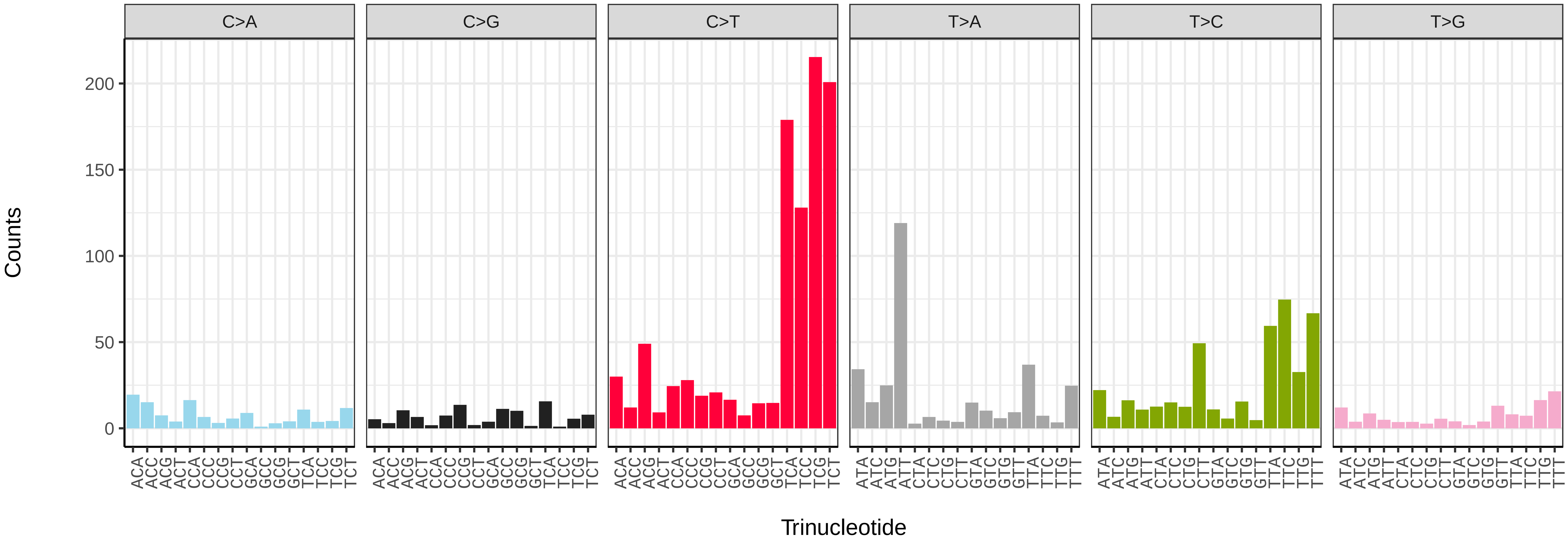


*Athalia rosae* (iyAthRosa1) somatic mutational spectrum (SBS52 classification)

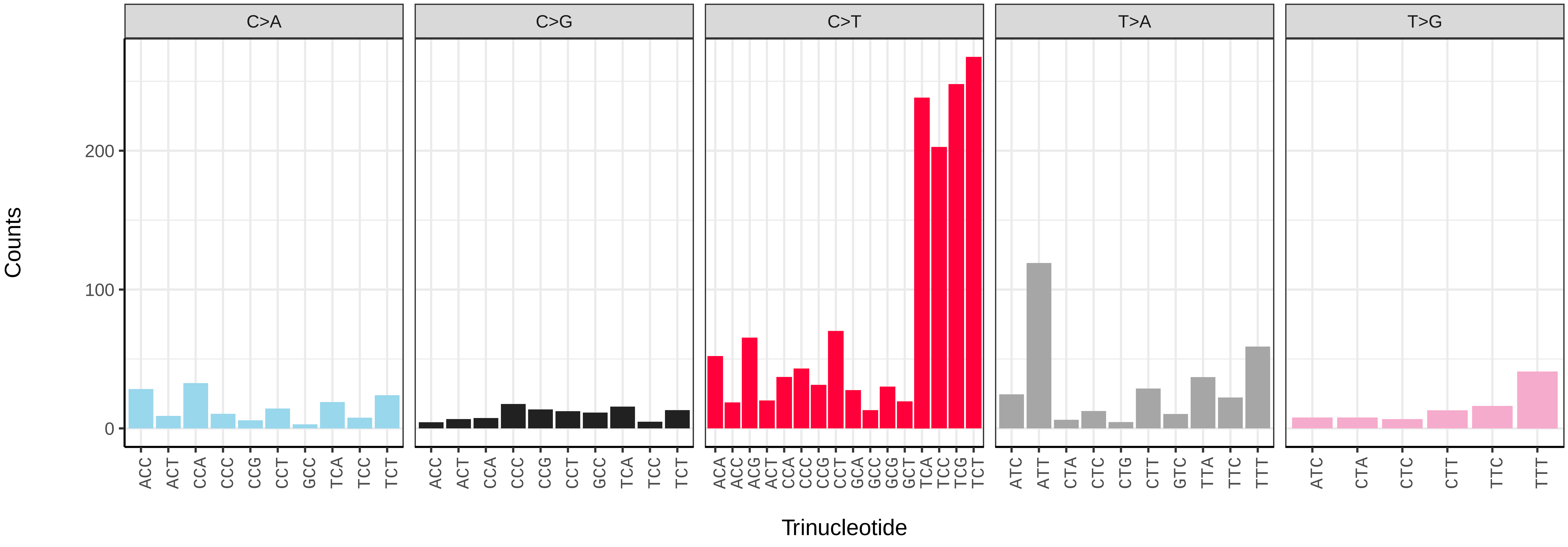

*Athalia rosae* (iyAthRosa1.iyAthRosa2) germline mutational spectrum

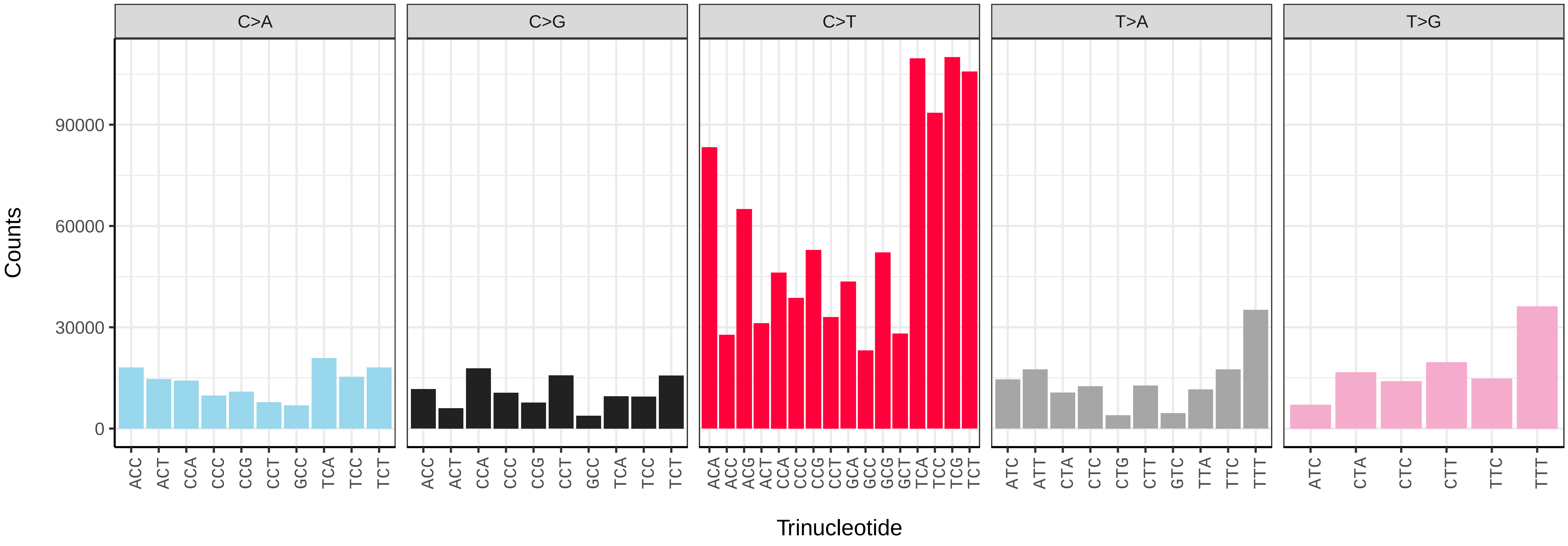


*Athalia rosae* (iyAthRosa1.iyAthRosa2) somatic mutational spectrum (SBS96 classification)

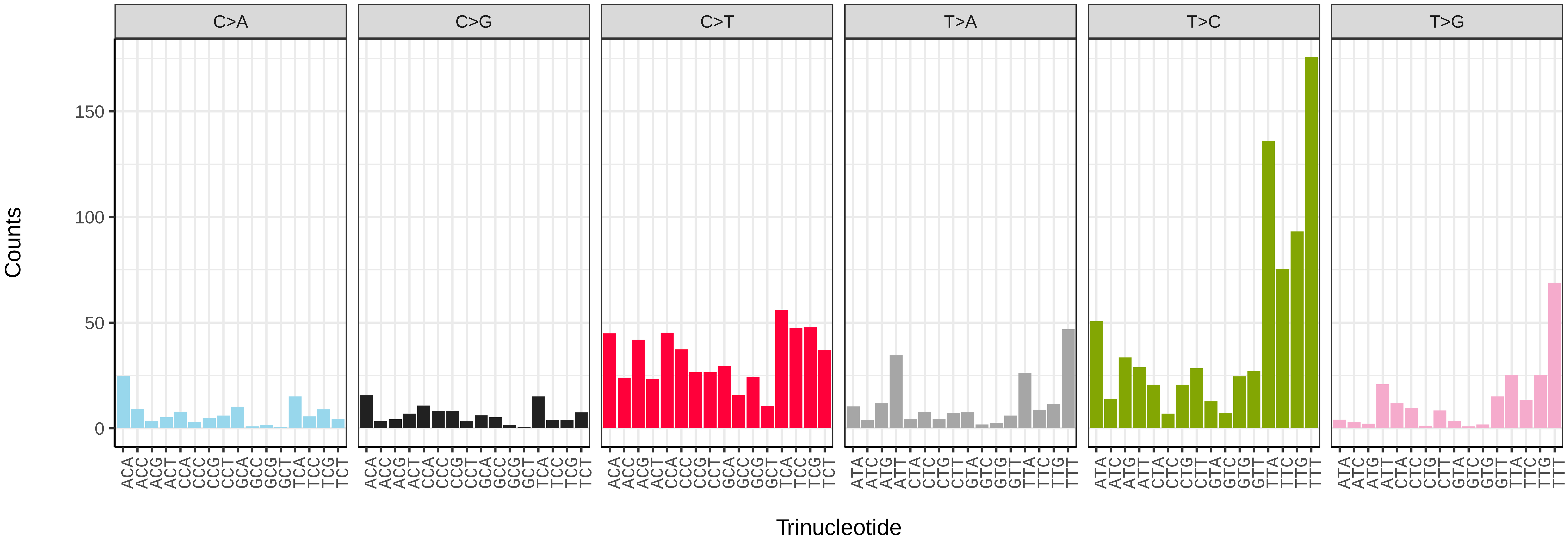


*Athalia rosae* (iyAthRosa1.iyAthRosa2) somatic mutational spectrum (SBS52 classification)

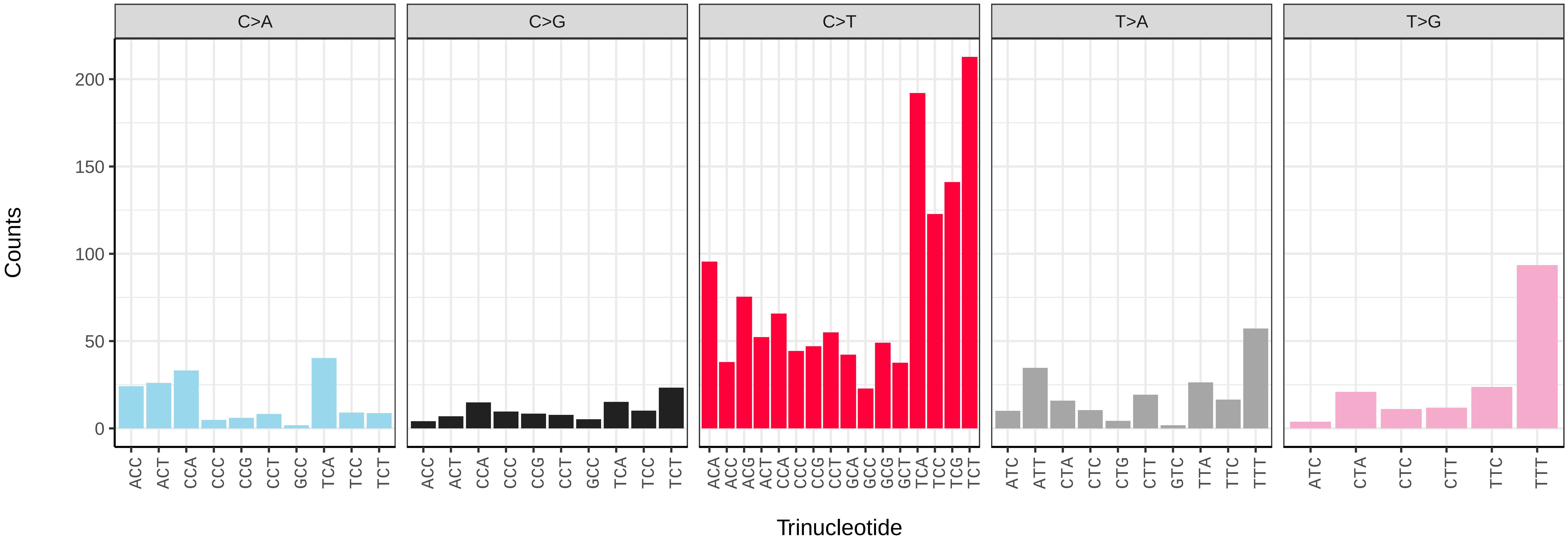

*Bombus hortorum* (iyBomHort1) germline mutational spectrum

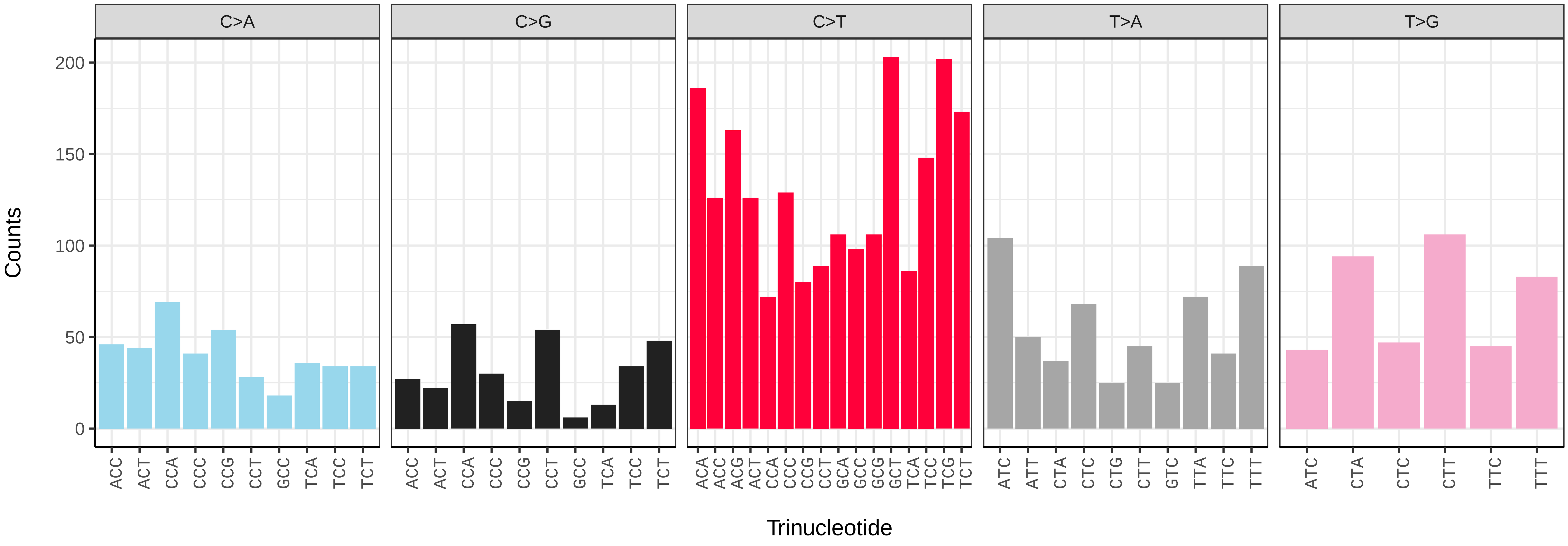


*Bombus hortorum* (iyBomHort1) somatic mutational spectrum (SBS96 classification)

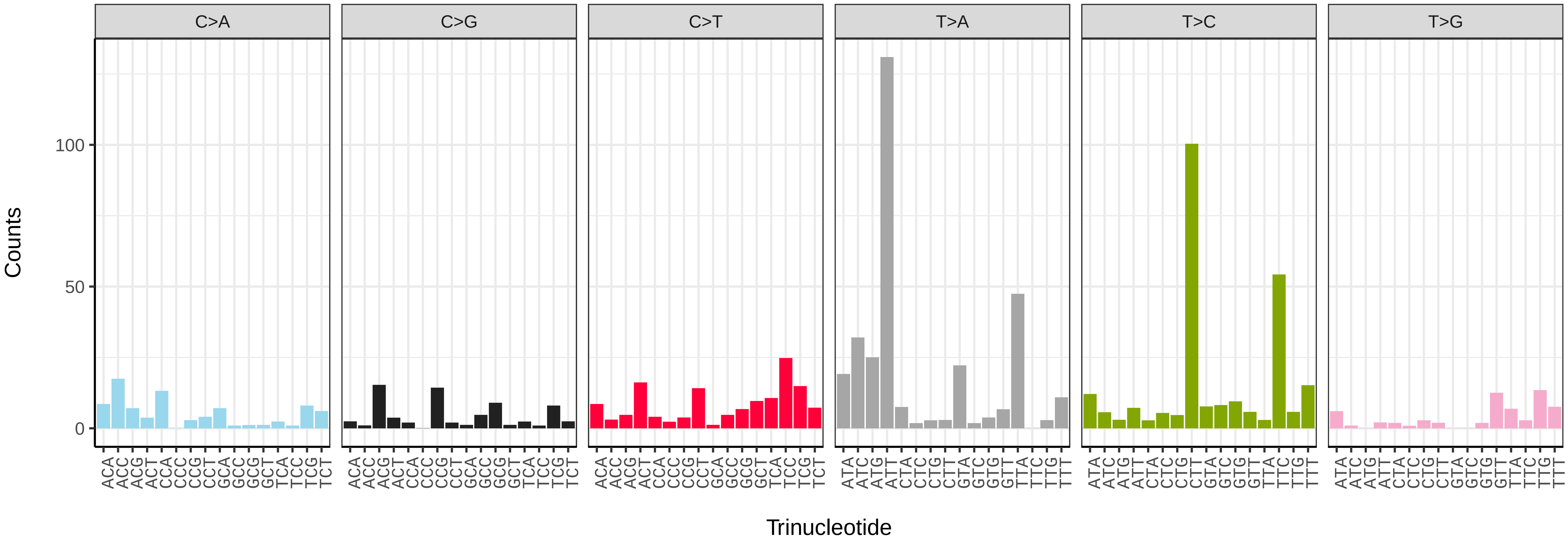


*Bombus hortorum* (iyBomHort1) somatic mutational spectrum (SBS52 classification)

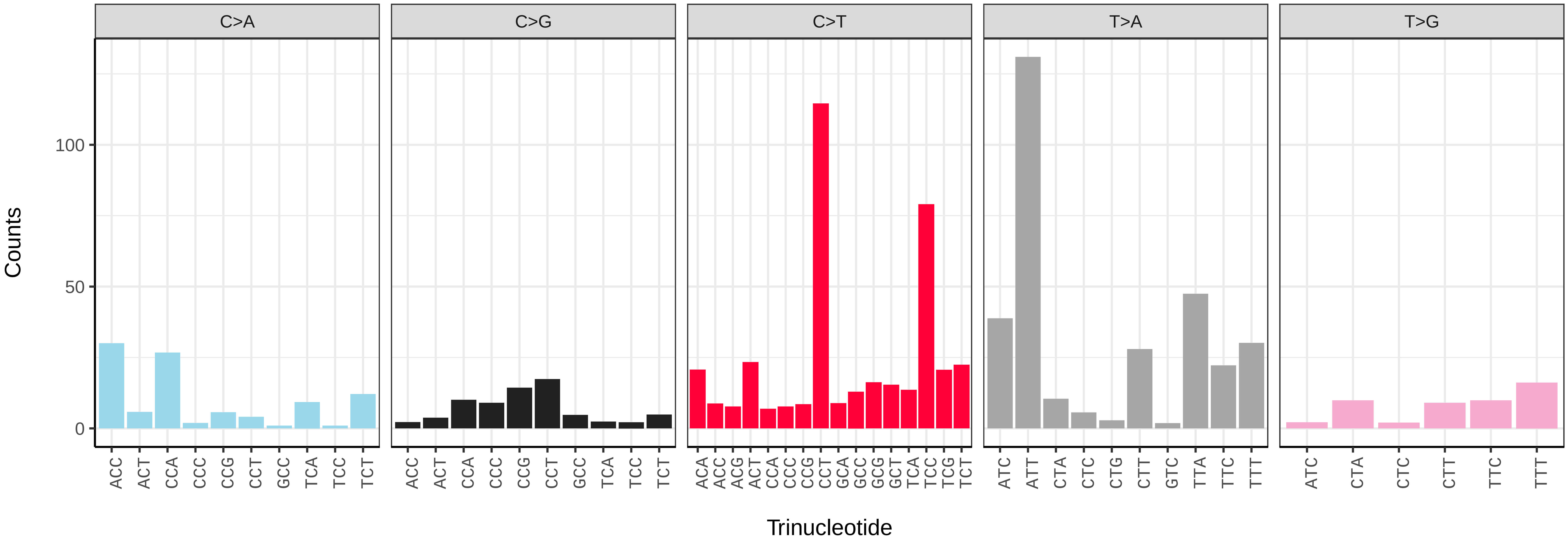

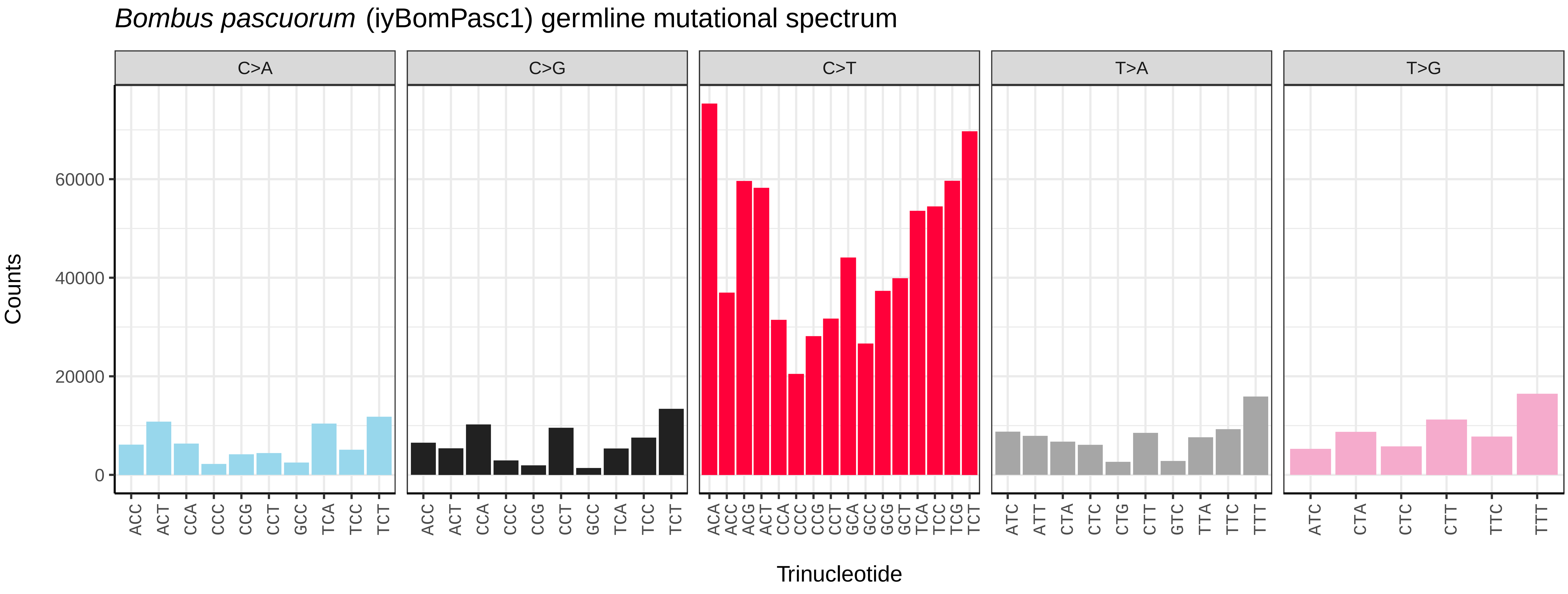
*Bombus pascuorum* (iyBomPasc1) germline mutational spectrum
C>A
C>G
C>T
T>A
T>G
Counts
0
20000
40000
60000
ACC ACT CCA CCC CCG CCT GCC TCA TCC TCT
ACC ACT CCA CCC CCG CCT GCC TCA TCC TCT
ACA ACC ACG ACT CCA CCC CCG CCT GCA GCC GCG GCT TCA TCC TCG TCT
ATC ATT CTA CTC CTG CTT GTC TTA TTC TTT
ATC CTA CTC CTT TTC TTT
Trinucleotide

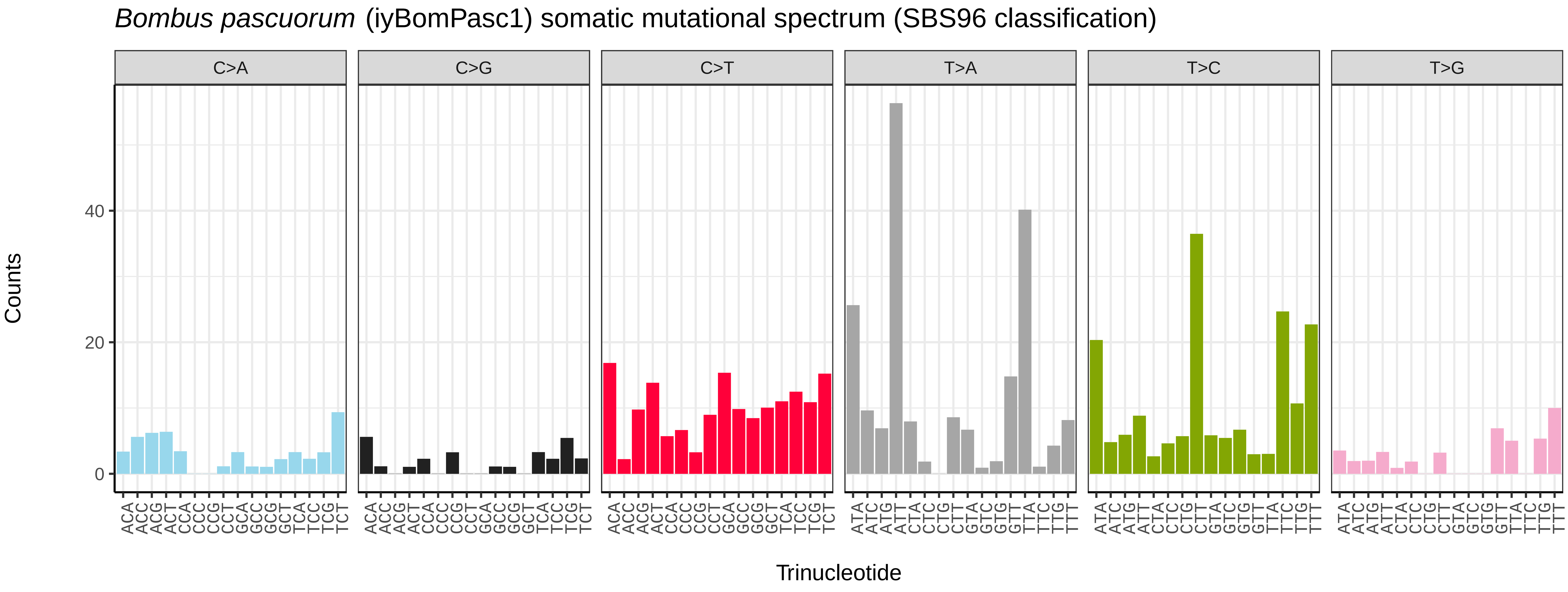
*Bombus pascuorum* (iyBomPasc1) somatic mutational spectrum (SBS96 classification)
C>A
C>G
C>T
T>A
T>C
T>G
Counts
0
20
40
ACA ACC ACG ACT CCA CCC CCG CCT GCA GCC GCG GCT TCA TCC TCG TCT
ATA ATC ATG ATT CTA CTC CTG CTT GTA GTC GTG GTT TTA TTC TTG TTT
Trinucleotide

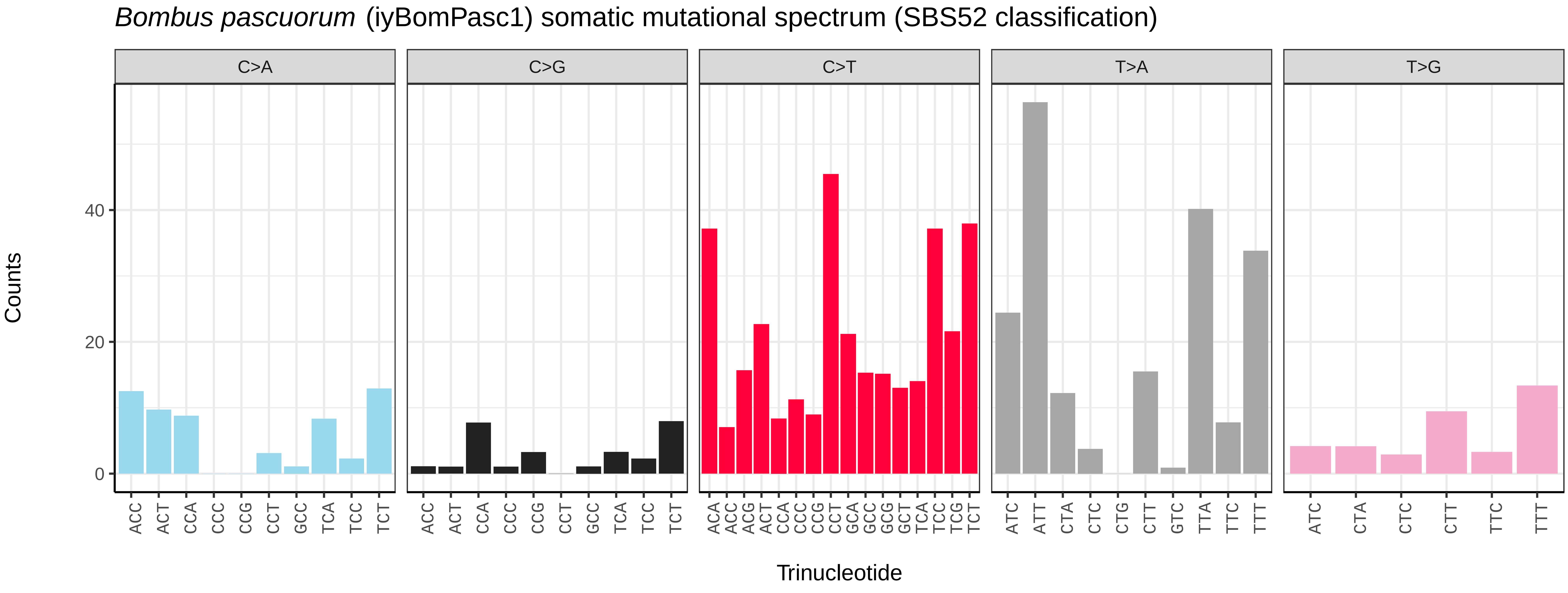
*Bombus pascuorum* (iyBomPasc1) somatic mutational spectrum (SBS52 classification)
C>A
C>G
C>T
T>A
T>G
Counts
0
20
40
ACC ACT CCA CCC CCG CCT GCC TCA TCC TCT
ACC ACT CCA CCC CCG CCT GCC TCA TCC TCT
ACA ACC ACG ACT CCA CCC CCG CCT GCA GCC GCG GCT TCA TCC TCG TCT
ATC ATT CTA CTC CTG CTT GTC TTA TTC TTT
ATC CTA CTC CTT TTC TTT
Trinucleotide

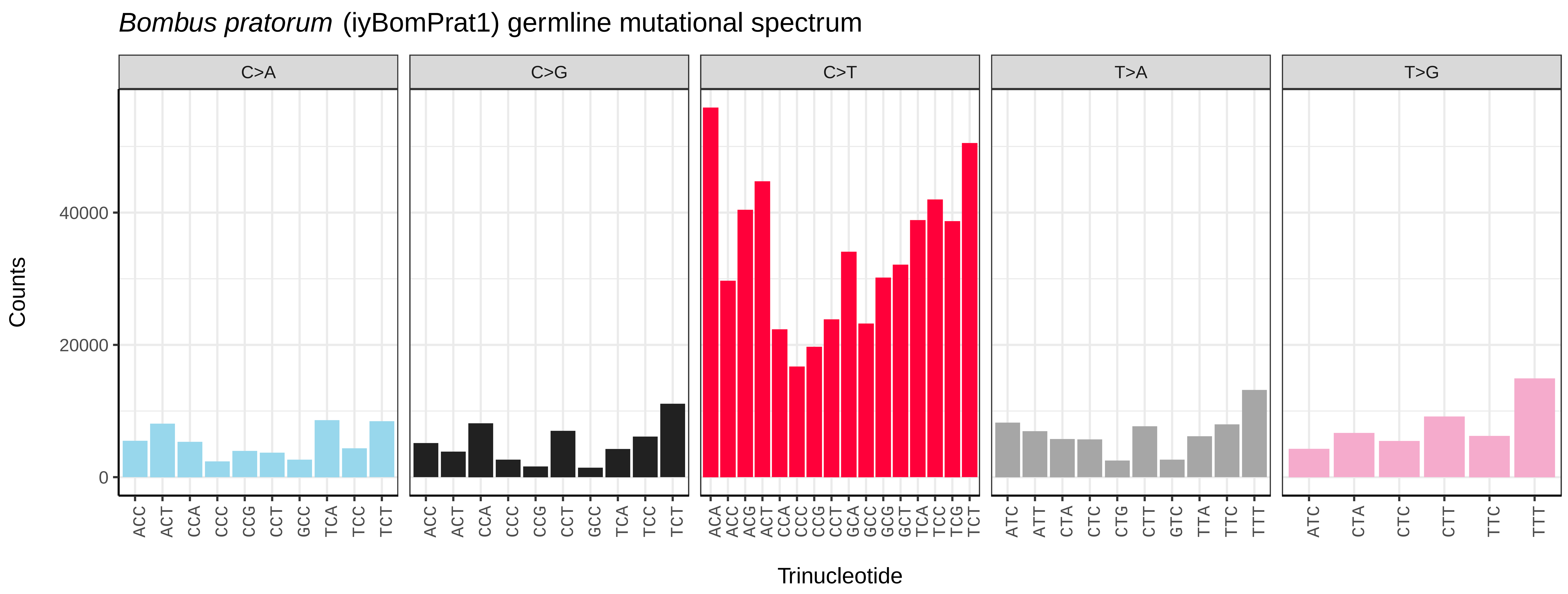
Bombus pratorum (iyBomPrat1) germline mutational spectrum
C>A
C>G
C>T
T>A
T>G
Counts
0
20000
40000
Trinucleotide

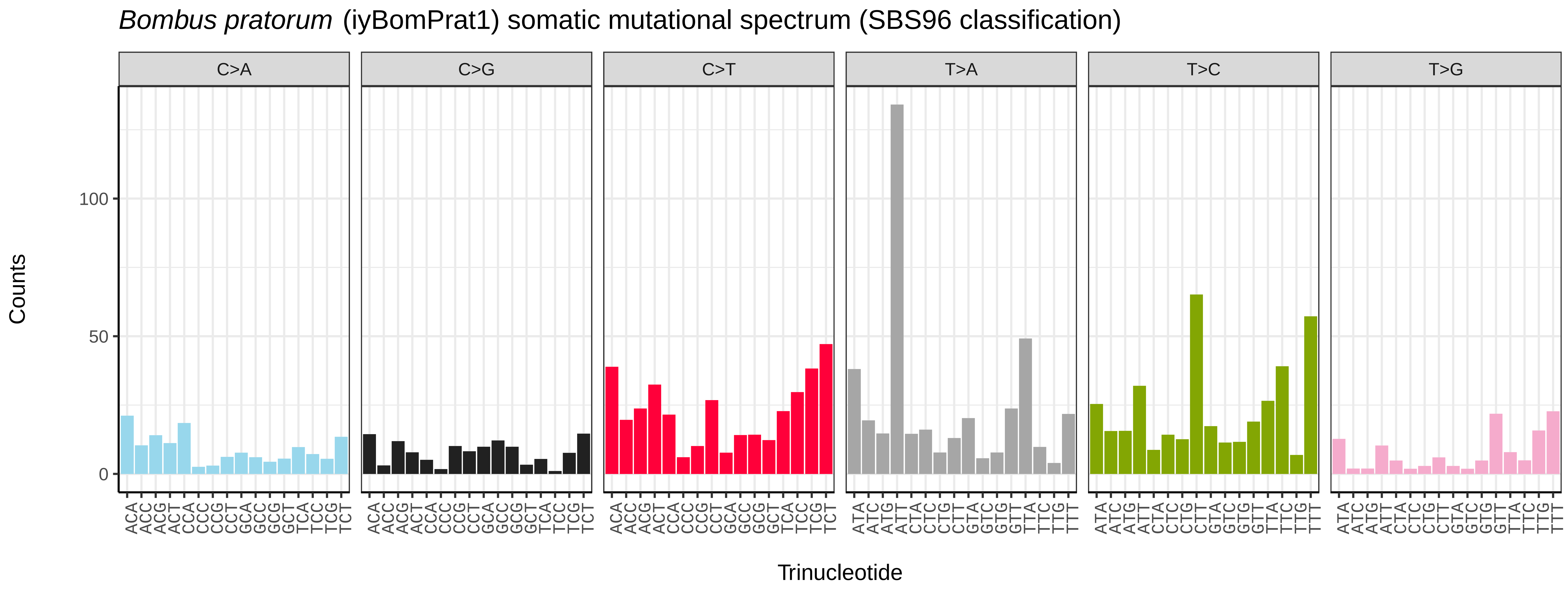
Bombus pratorum (iyBomPrat1) somatic mutational spectrum (SBS96 classification)
C>A
C>G
C>T
T>A
T>C
T>G
Counts
0
50
100
Trinucleotide

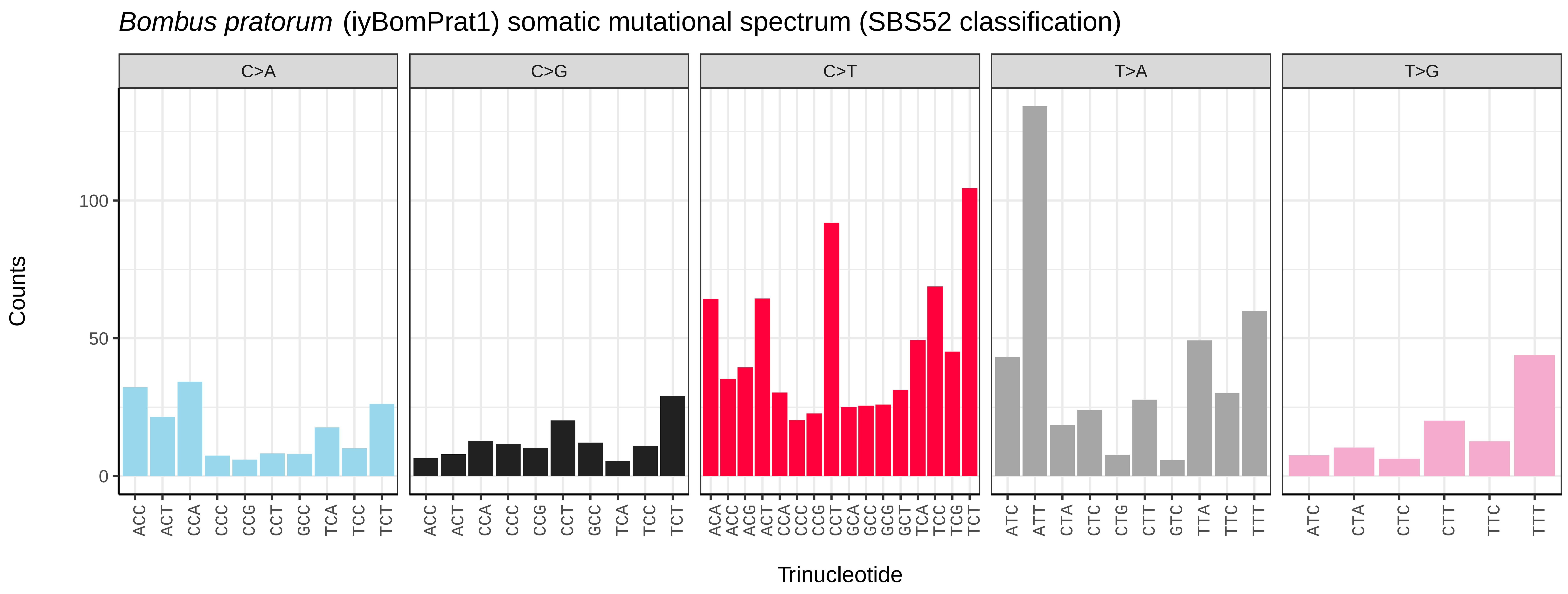
Bombus pratorum (iyBomPrat1) somatic mutational spectrum (SBS52 classification)
C>A
C>G
C>T
T>A
T>G
Counts
0
50
100
Trinucleotide

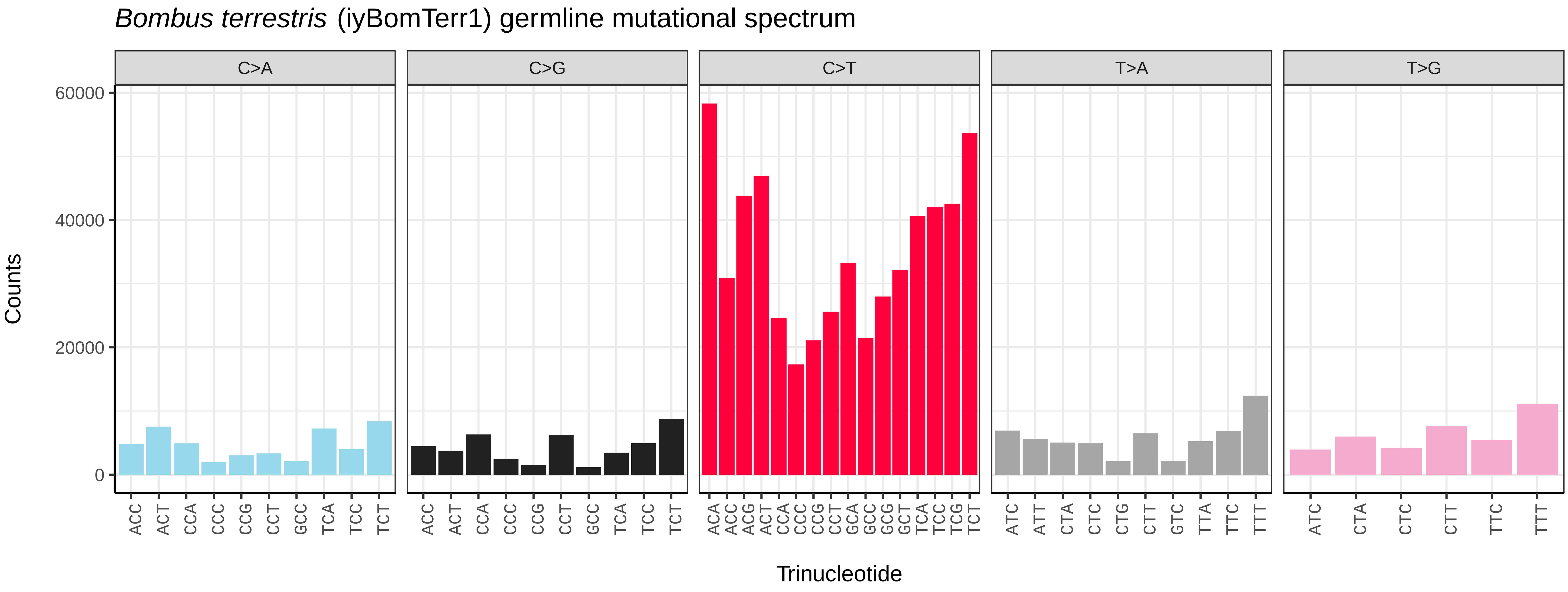
Bombus terrestris (iyBomTerr1) germline mutational spectrum
C>A
C>G
C>T
T>A
T>G
Counts
60000
40000
20000
0
ACC ACT CCA CCC CCG CCT GCC TCA TCC TCT
ACA ACC ACG ACT CCA CCC CCG CCT GCA GCC GCG GCT TCA TCC TCG TCT
ATC ATT CTA CTC CTG CTT GTC TTA TTC TTT
ATC CTA CTC CTT TTC TTT
Trinucleotide

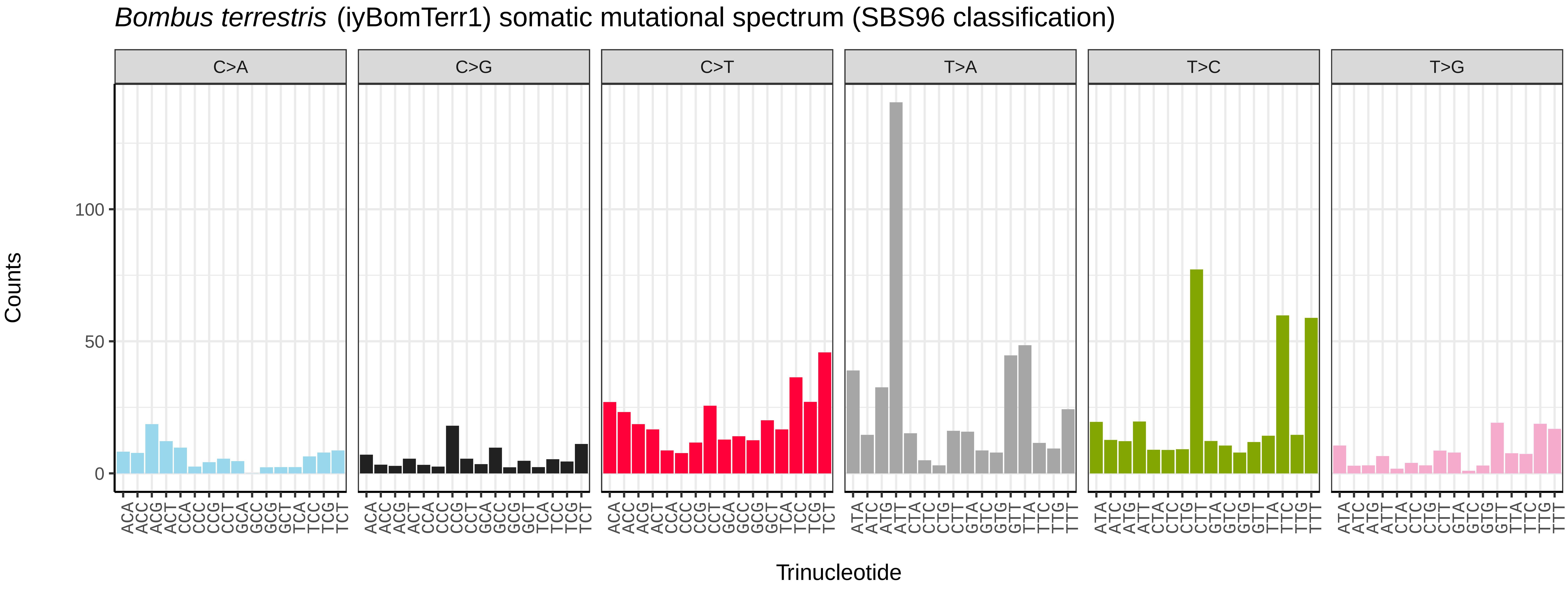
Bombus terrestris (iyBomTerr1) somatic mutational spectrum (SBS96 classification)
C>A
C>G
C>T
T>A
T>C
T>G
Counts
100
50
0
ACA ACC ACG ACT CCA CCC CCG CCT GCA GCC GCG GCT TCA TCC TCG TCT
ATA ATC ATG ATT CTA CTC CTG CTT GTA GTC GTG GTT TTA TTC TTG TTT
Trinucleotide

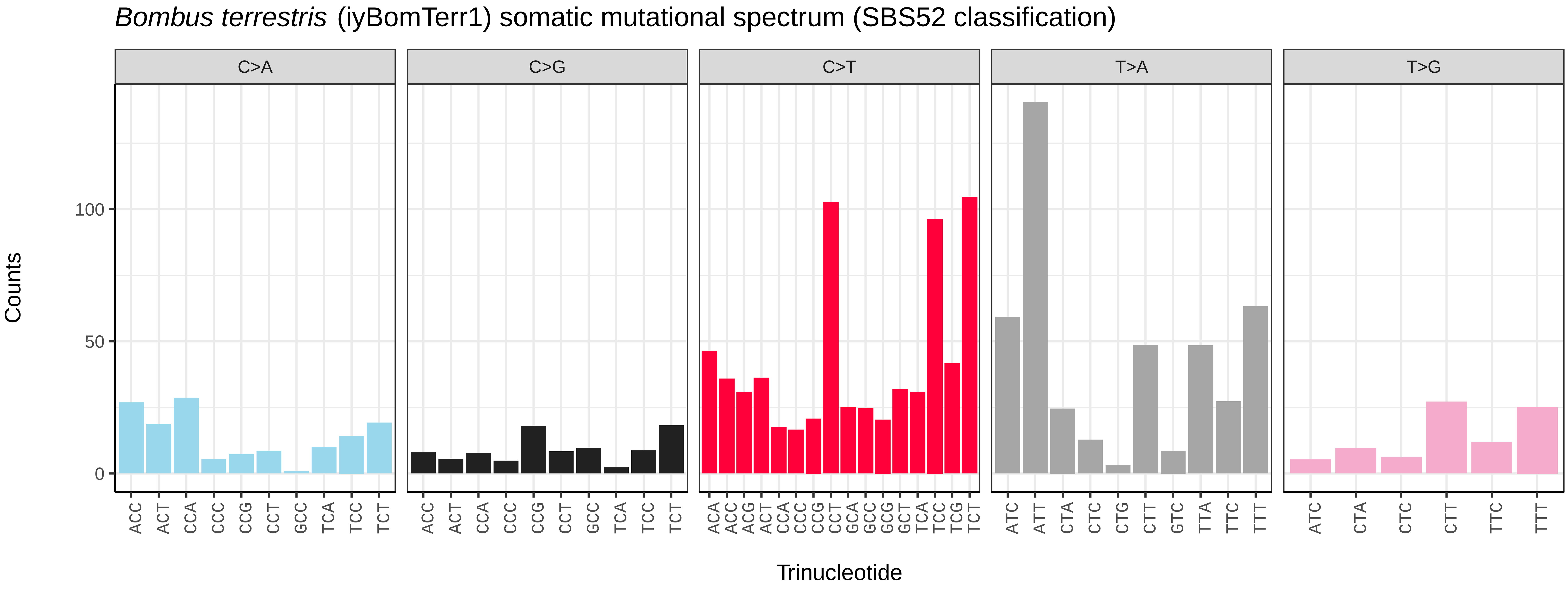
Bombus terrestris (iyBomTerr1) somatic mutational spectrum (SBS52 classification)
C>A
C>G
C>T
T>A
T>G
Counts
100
50
0
ACC ACT CCA CCC CCG CCT GCC TCA TCC TCT
ACA ACC ACG ACT CCA CCC CCG CCT GCA GCC GCG GCT TCA TCC TCG TCT
ATC ATT CTA CTC CTG CTT GTC TTA TTC TTT
ATC CTA CTC CTT TTC TTT
Trinucleotide

*Buathra laborator* (iyBuaLabo1) germline mutational spectrum

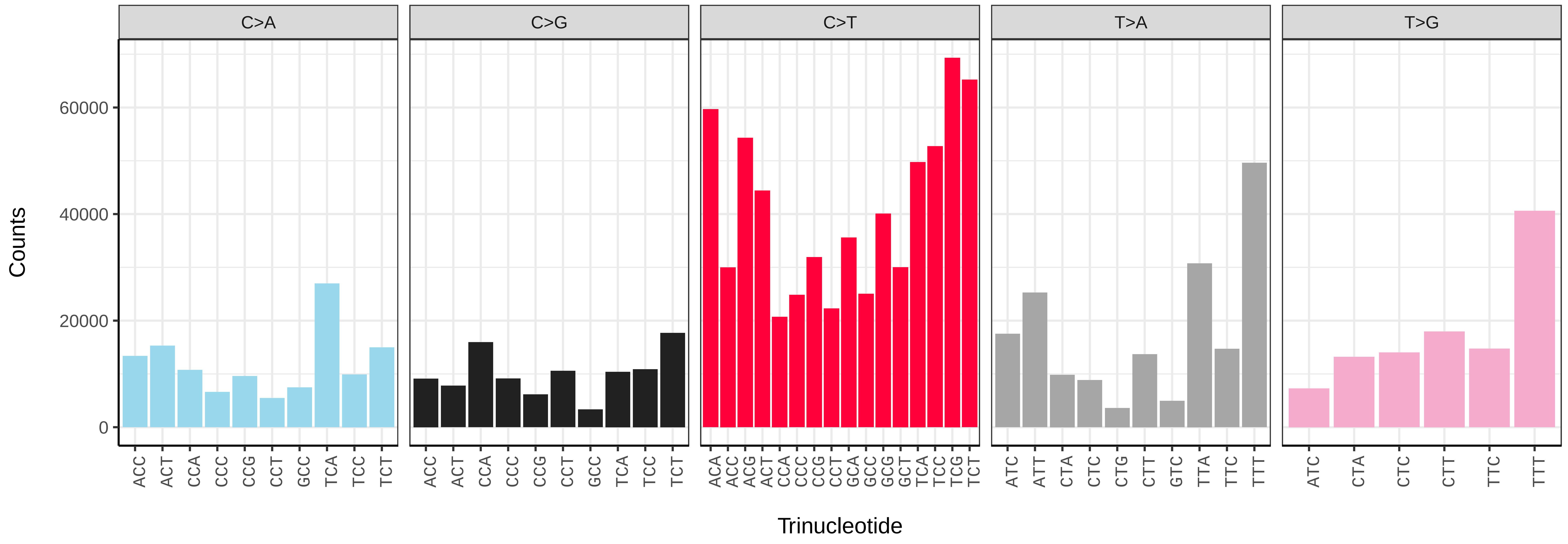


*Buathra laborator* (iyBuaLabo1) somatic mutational spectrum (SBS96 classification)

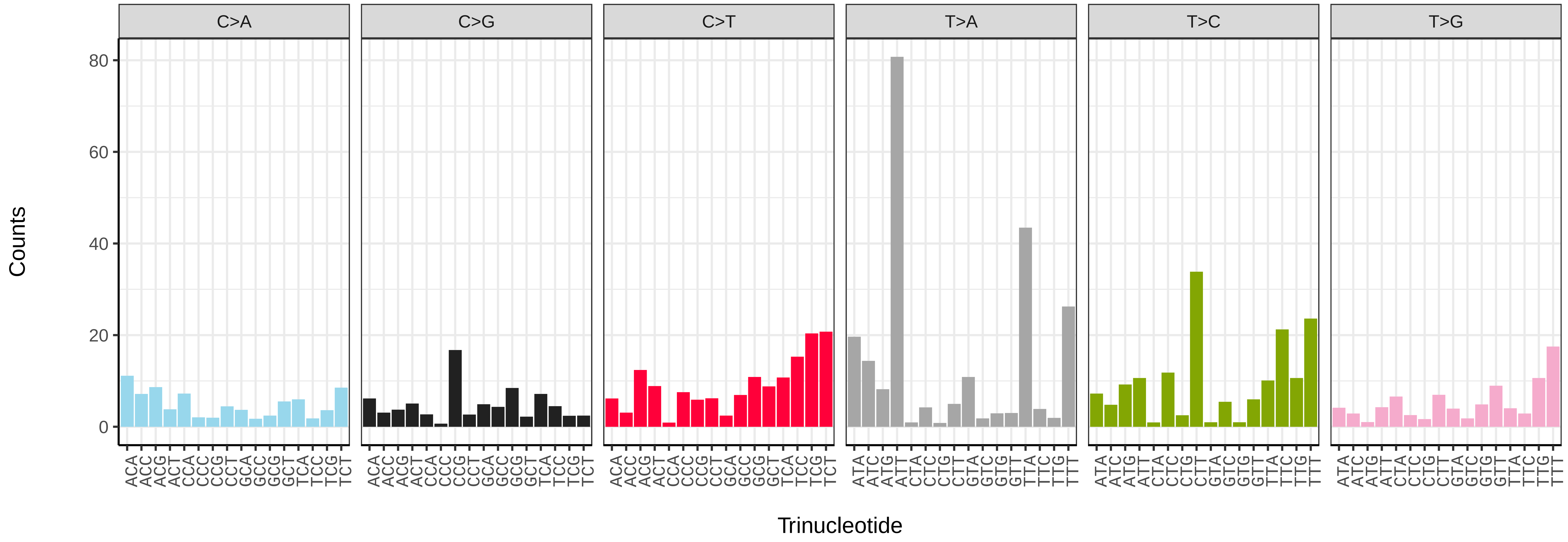


*Buathra laborator* (iyBuaLabo1) somatic mutational spectrum (SBS52 classification)

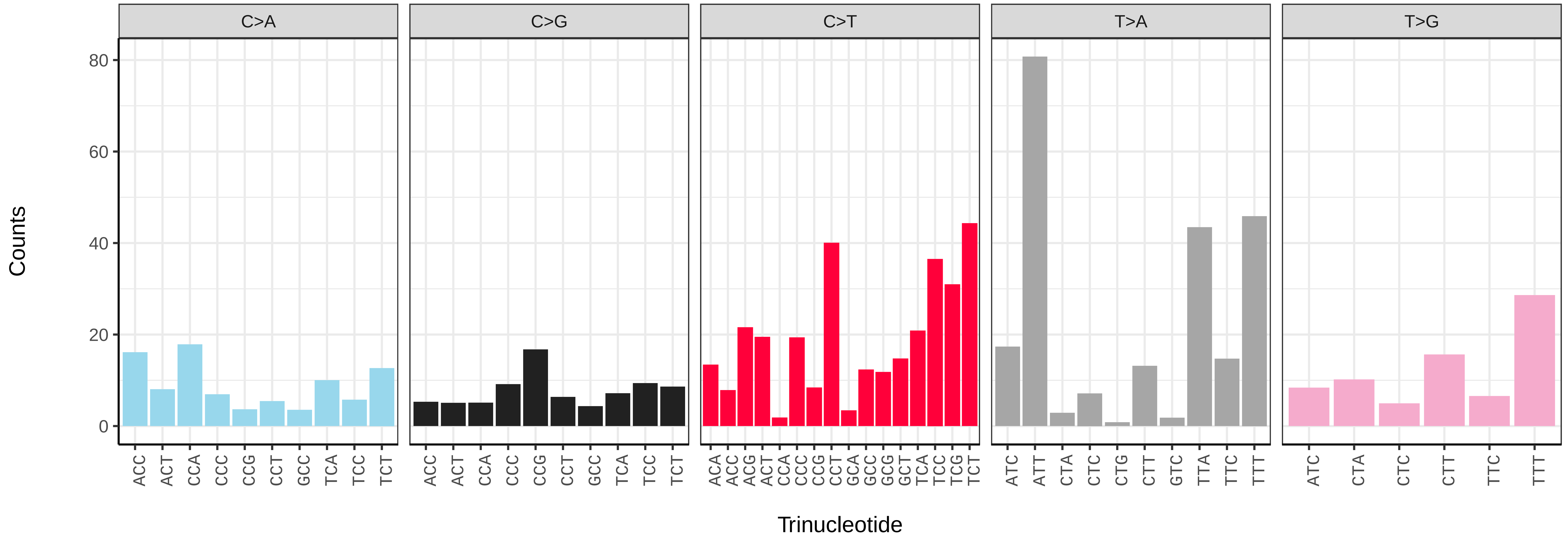

*Campoletis raptor* (iyCamRapt1) germline mutational spectrum

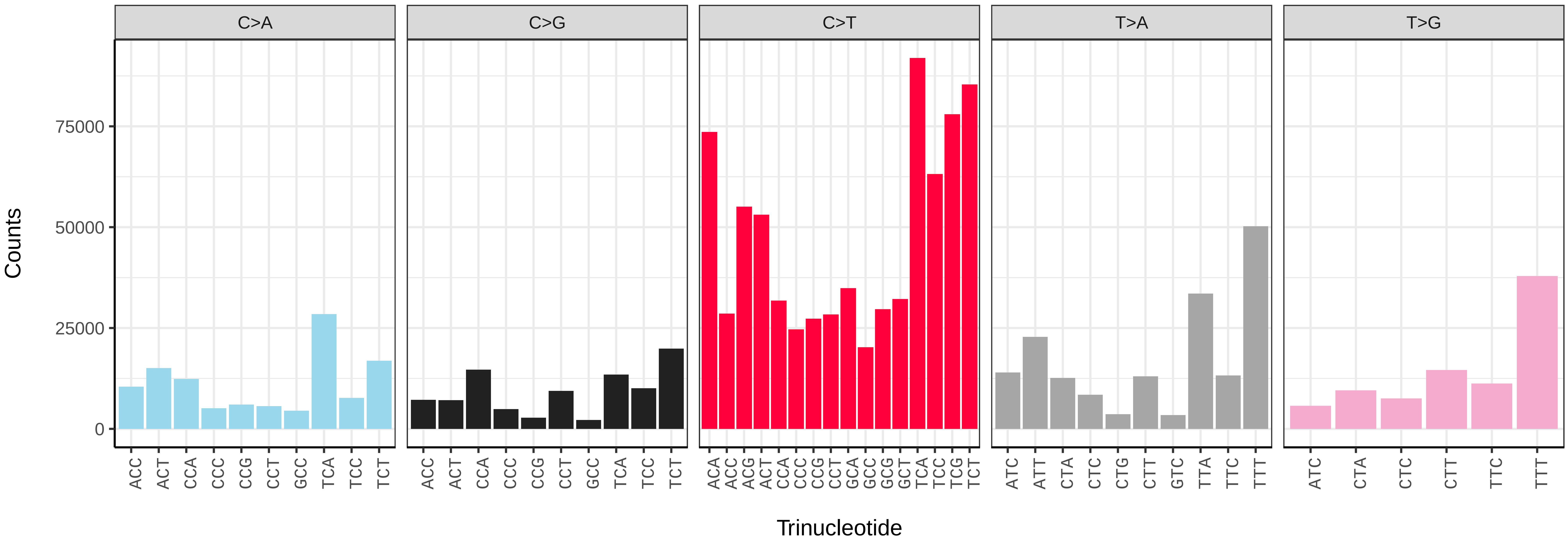


*Campoletis raptor* (iyCamRapt1) somatic mutational spectrum (SBS96 classification)

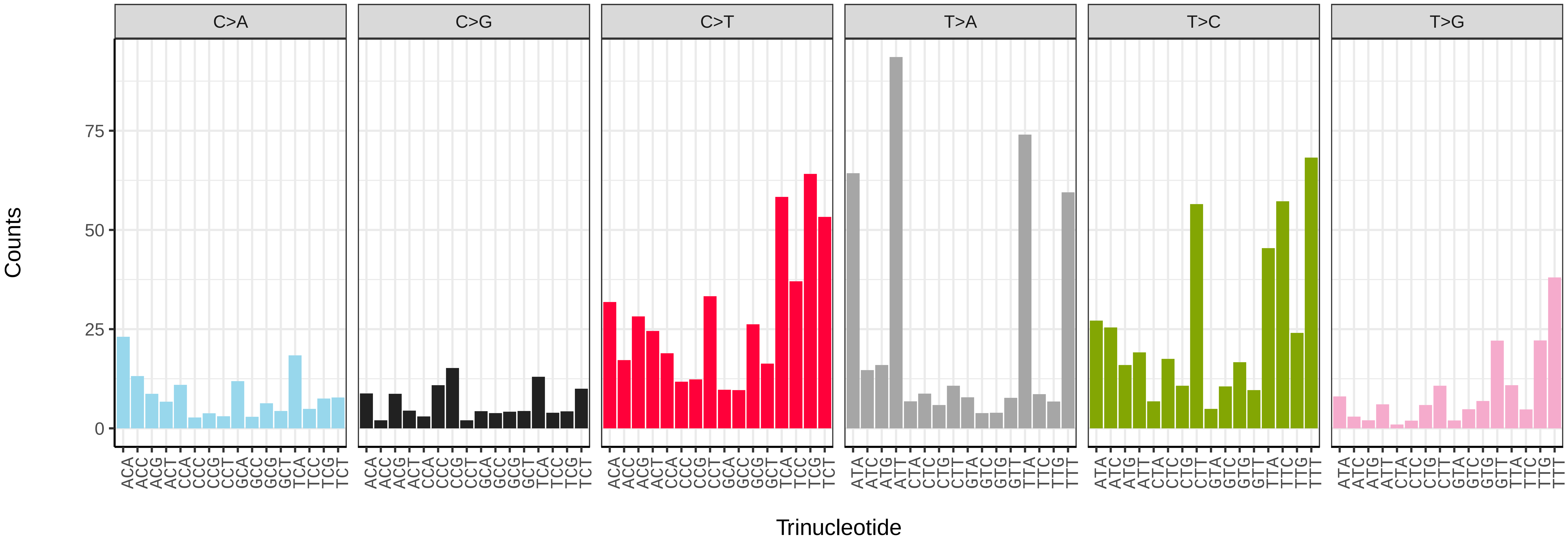


*Campoletis raptor* (iyCamRapt1) somatic mutational spectrum (SBS52 classification)

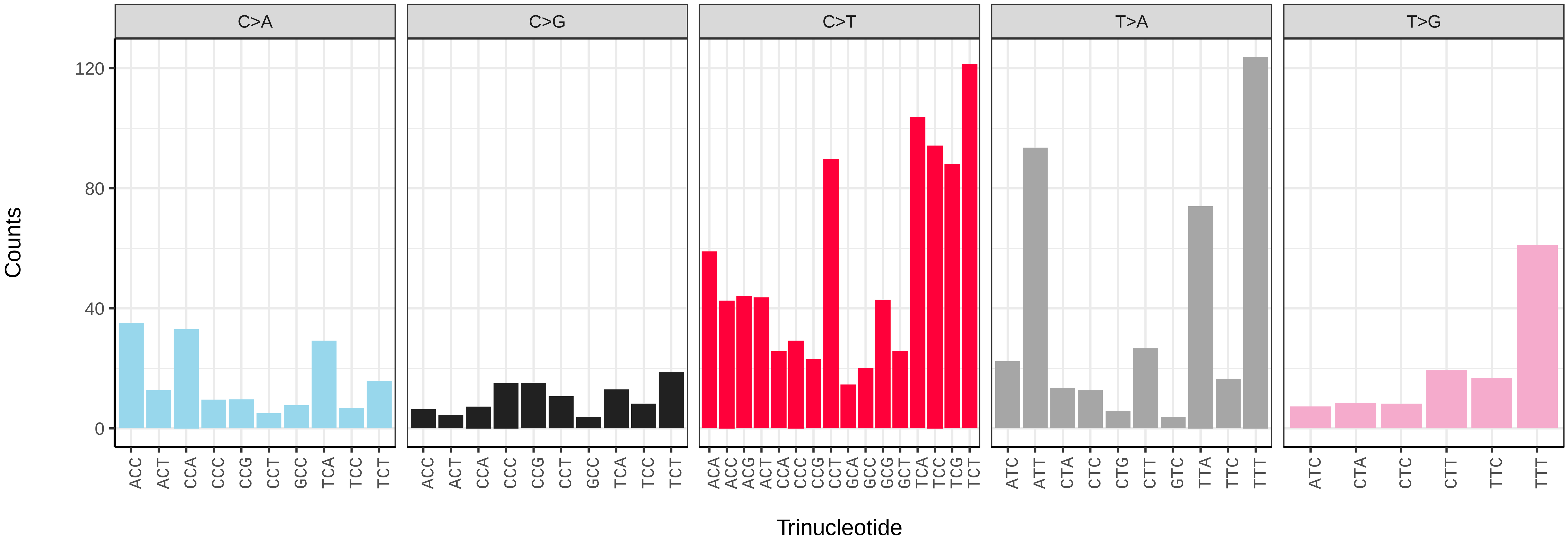

*Cerceris rybyensis* (iyCerRyby1) germline mutational spectrum

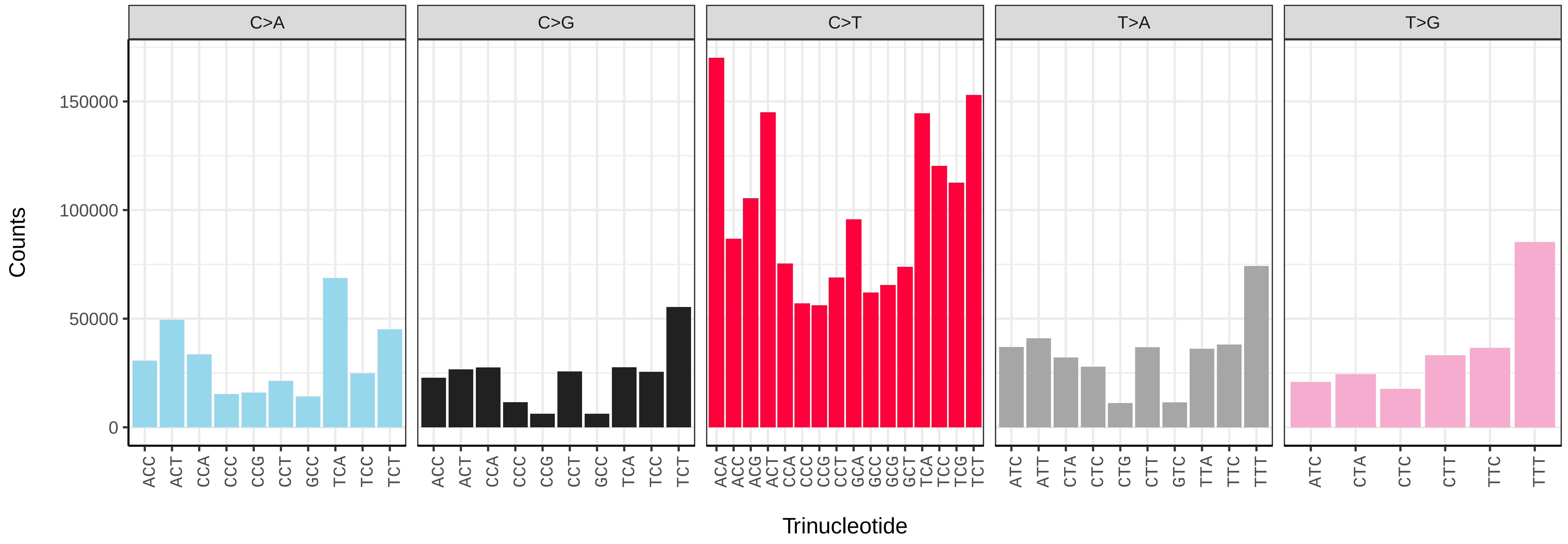


*Cerceris rybyensis* (iyCerRyby1) somatic mutational spectrum (SBS96 classification)

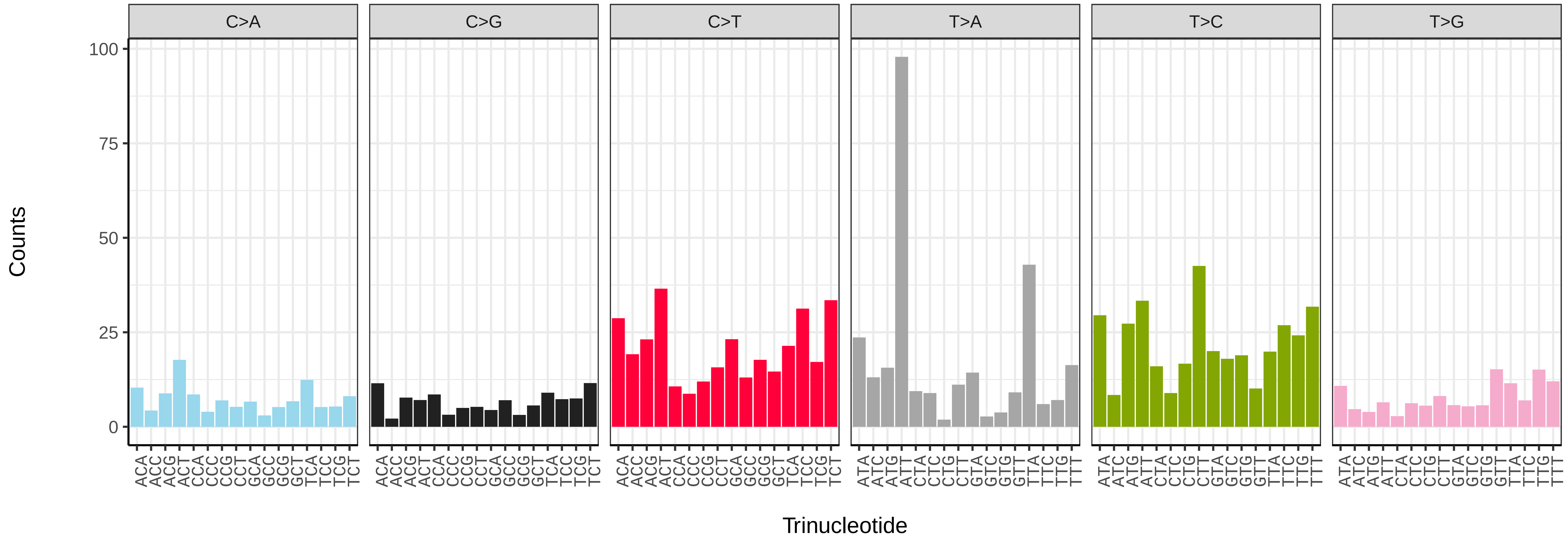


*Cerceris rybyensis* (iyCerRyby1) somatic mutational spectrum (SBS52 classification)

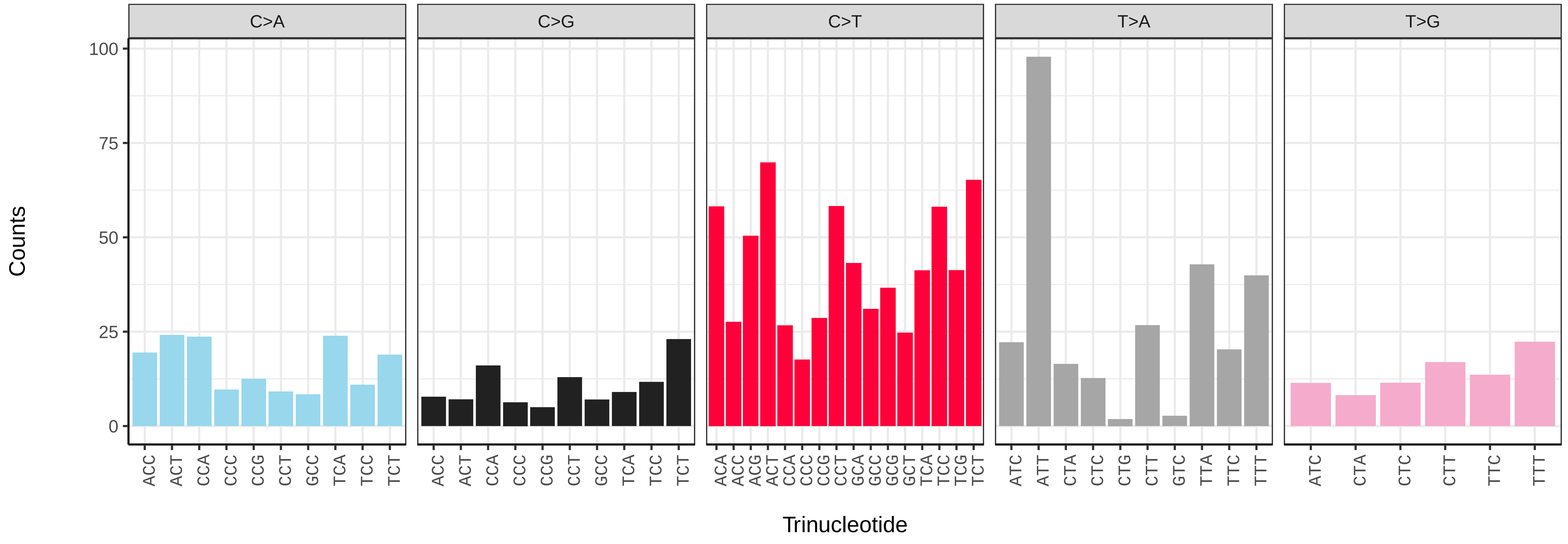

*Clistopyga incitator* (iyCliInci1) germline mutational spectrum

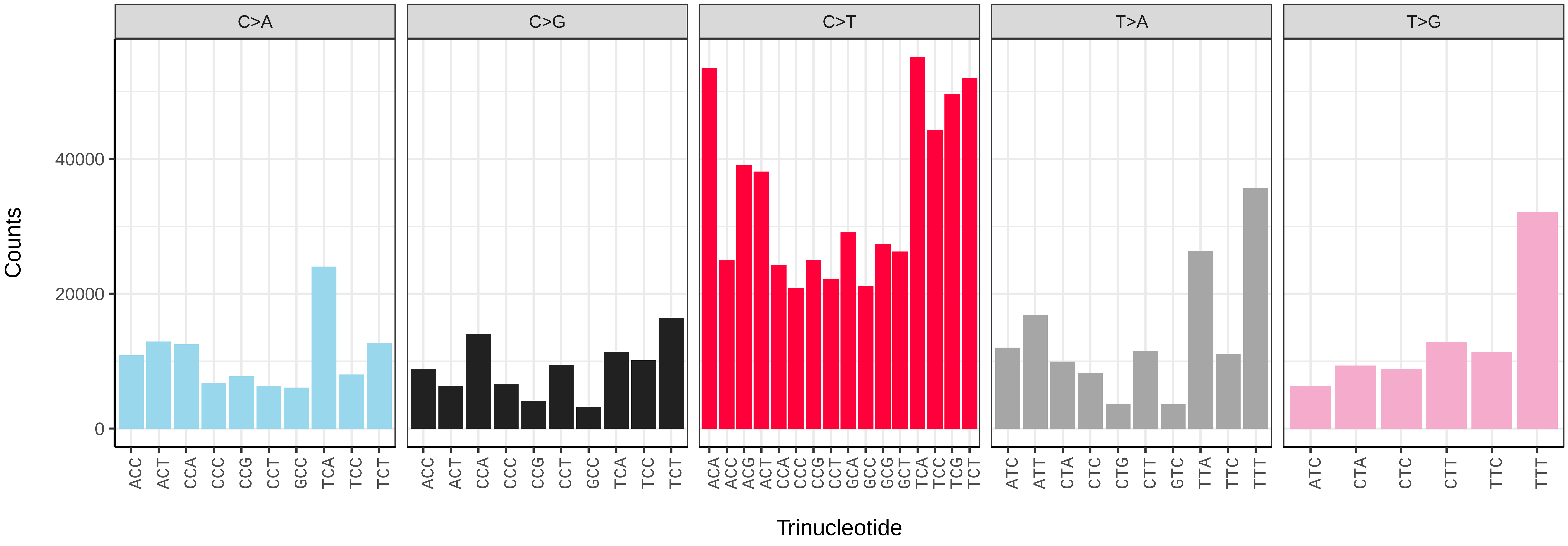


*Clistopyga incitator* (iyCliInci1) somatic mutational spectrum (SBS96 classification)

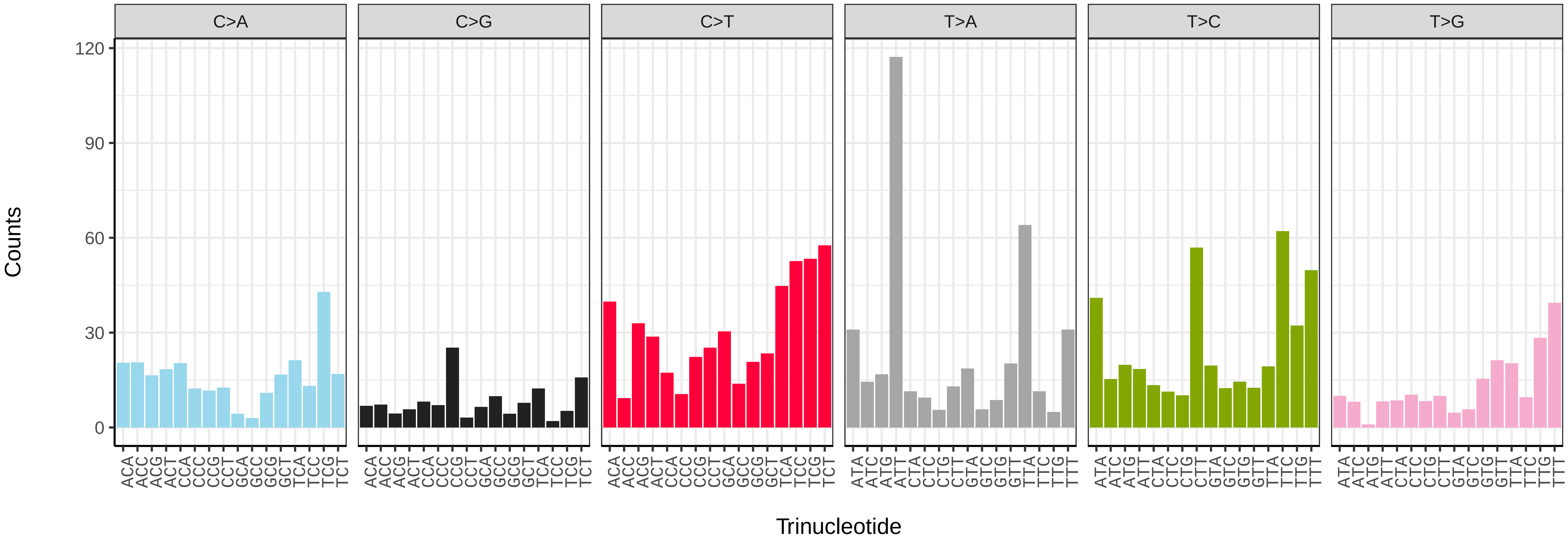


*Clistopyga incitator* (iyCliInci1) somatic mutational spectrum (SBS52 classification)

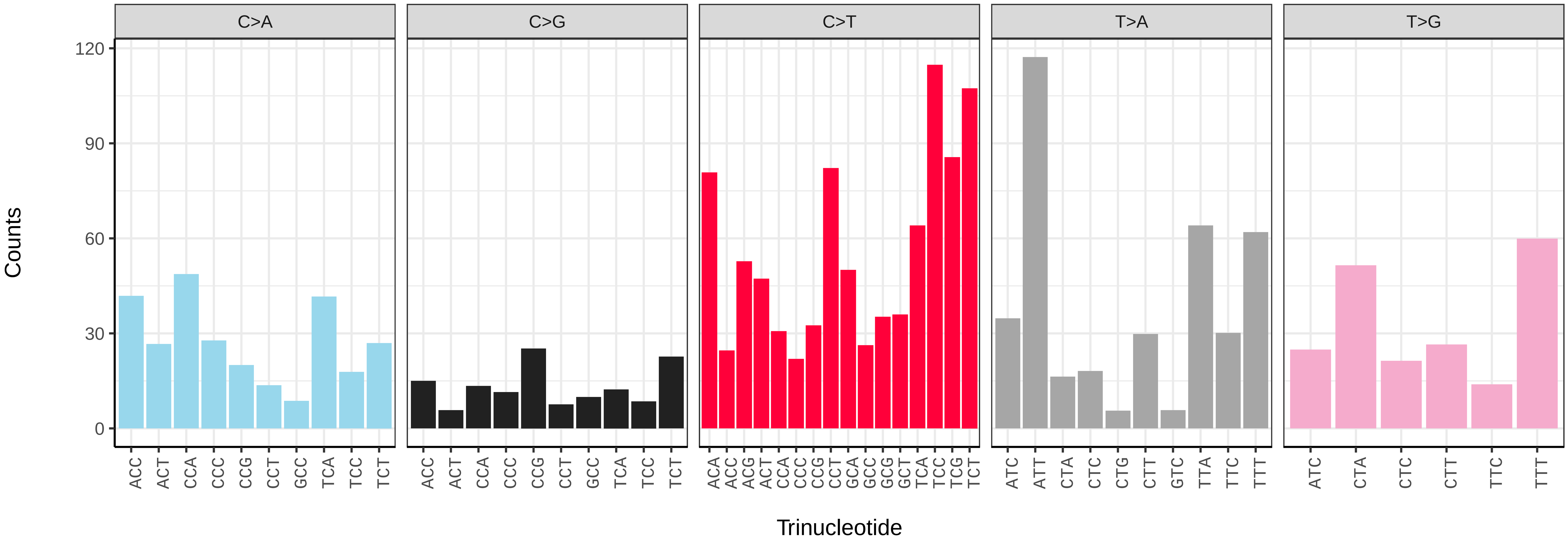

*Coelioxys conoideus* (iyCoeConi1) germline mutational spectrum

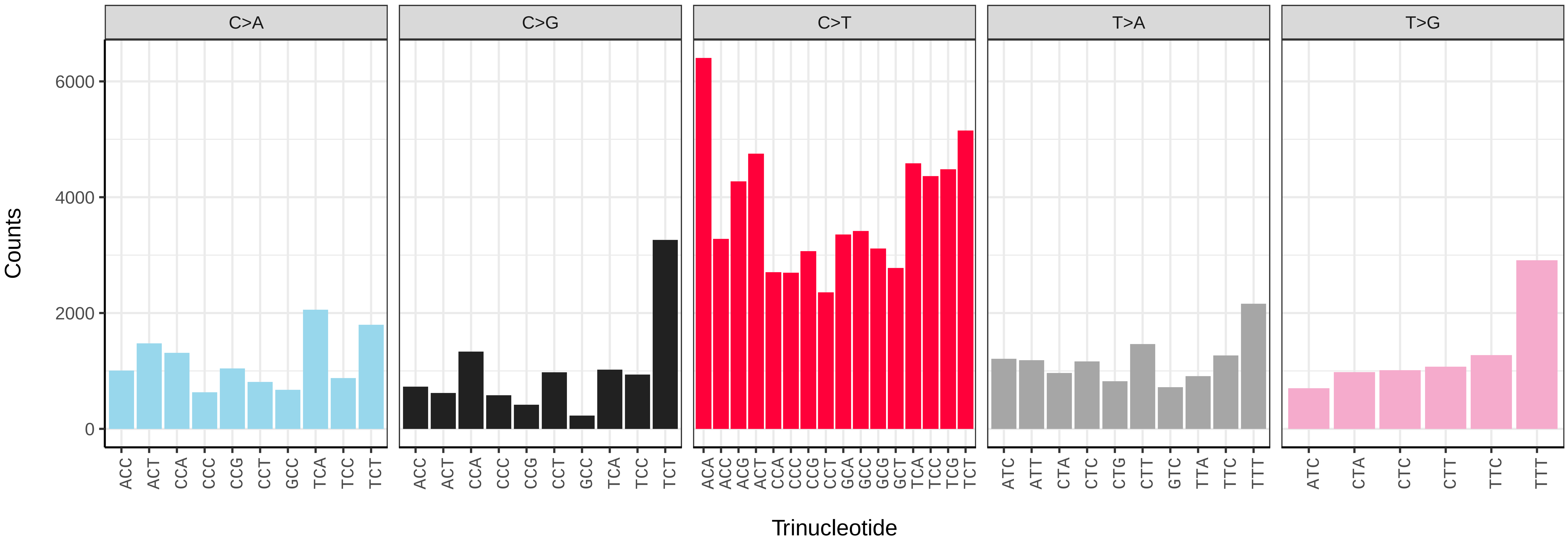


*Coelioxys conoideus* (iyCoeConi1) somatic mutational spectrum (SBS96 classification)

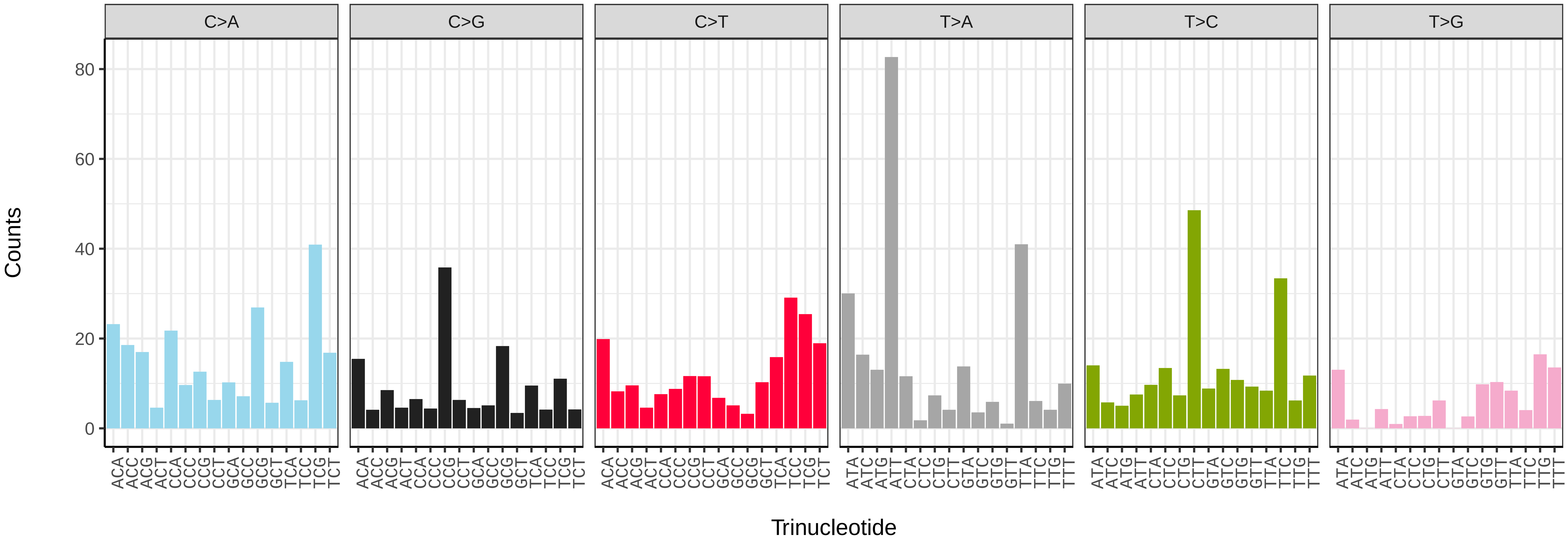


*Coelioxys conoideus* (iyCoeConi1) somatic mutational spectrum (SBS52 classification)

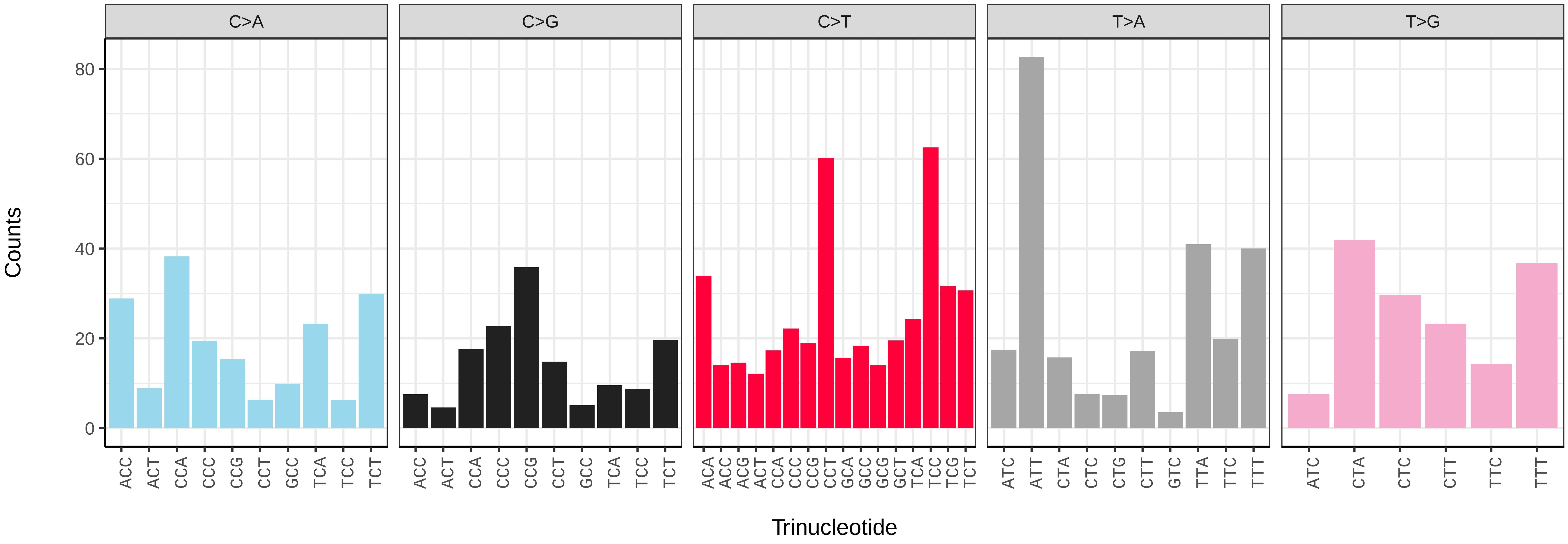

*Dolichovespula media* (iyDolMedi1) germline mutational spectrum

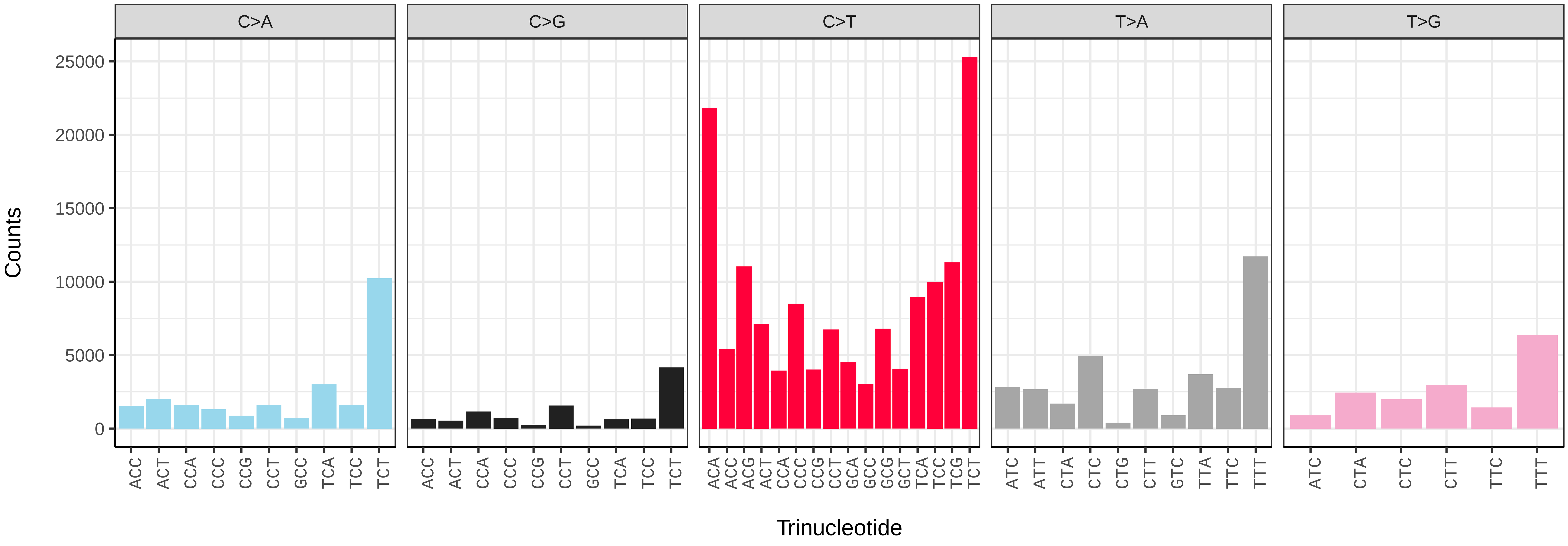


*Dolichovespula media* (iyDolMedi1) somatic mutational spectrum (SBS96 classification)

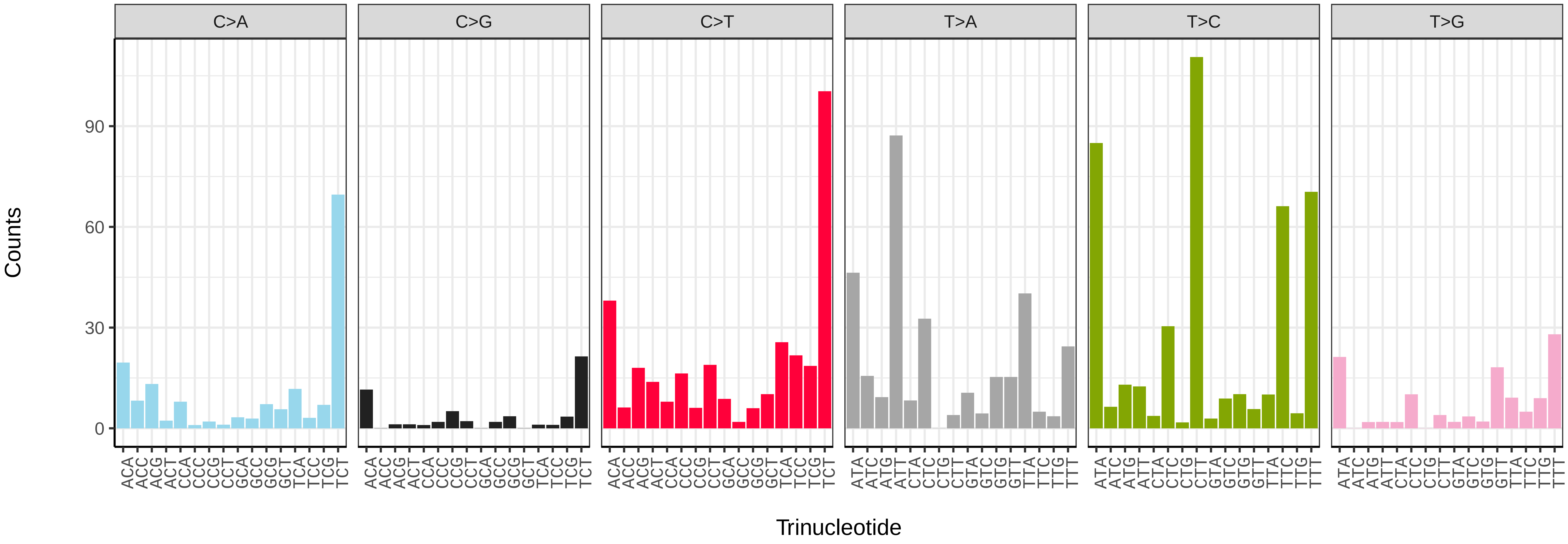


*Dolichovespula media* (iyDolMedi1) somatic mutational spectrum (SBS52 classification)

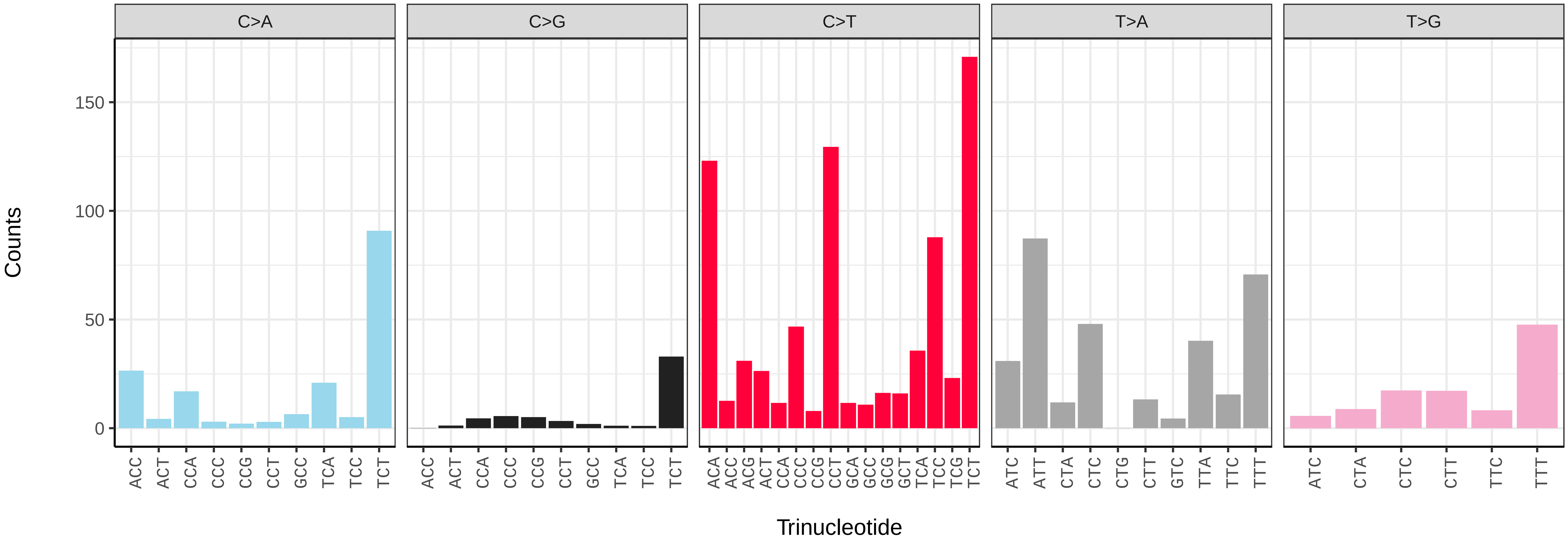

*Ectemnius continuus* (iyEctCont1) germline mutational spectrum

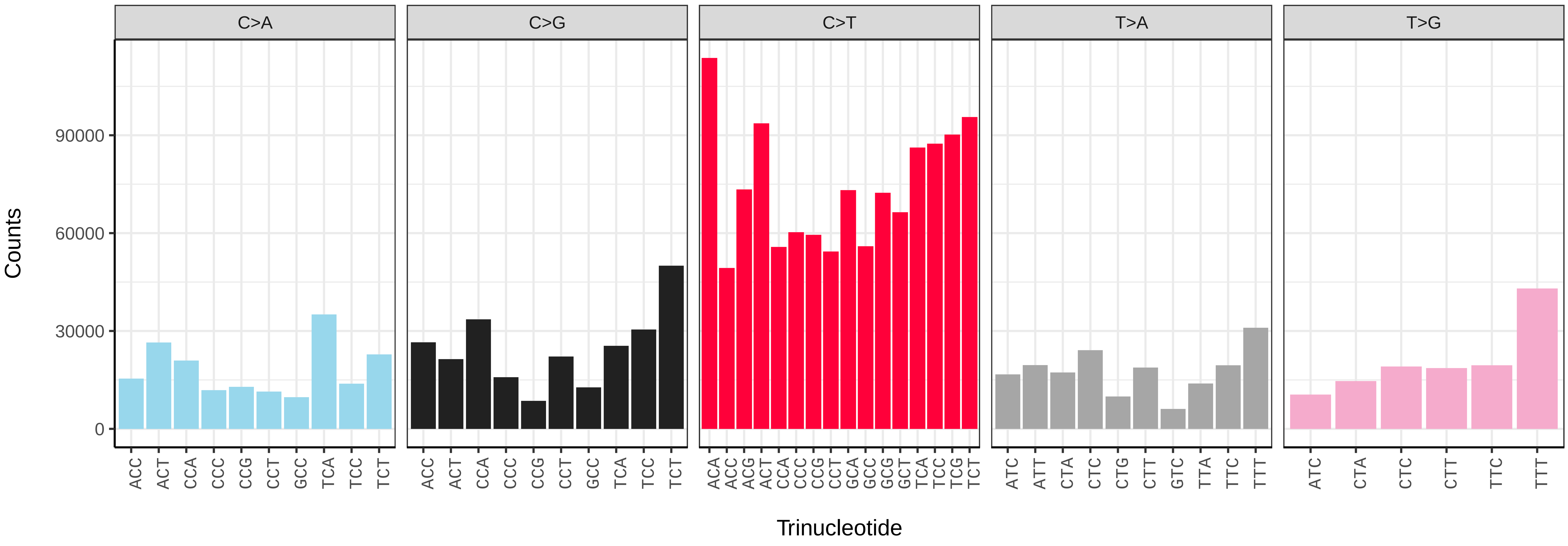


*Ectemnius continuus* (iyEctCont1) somatic mutational spectrum (SBS96 classification)

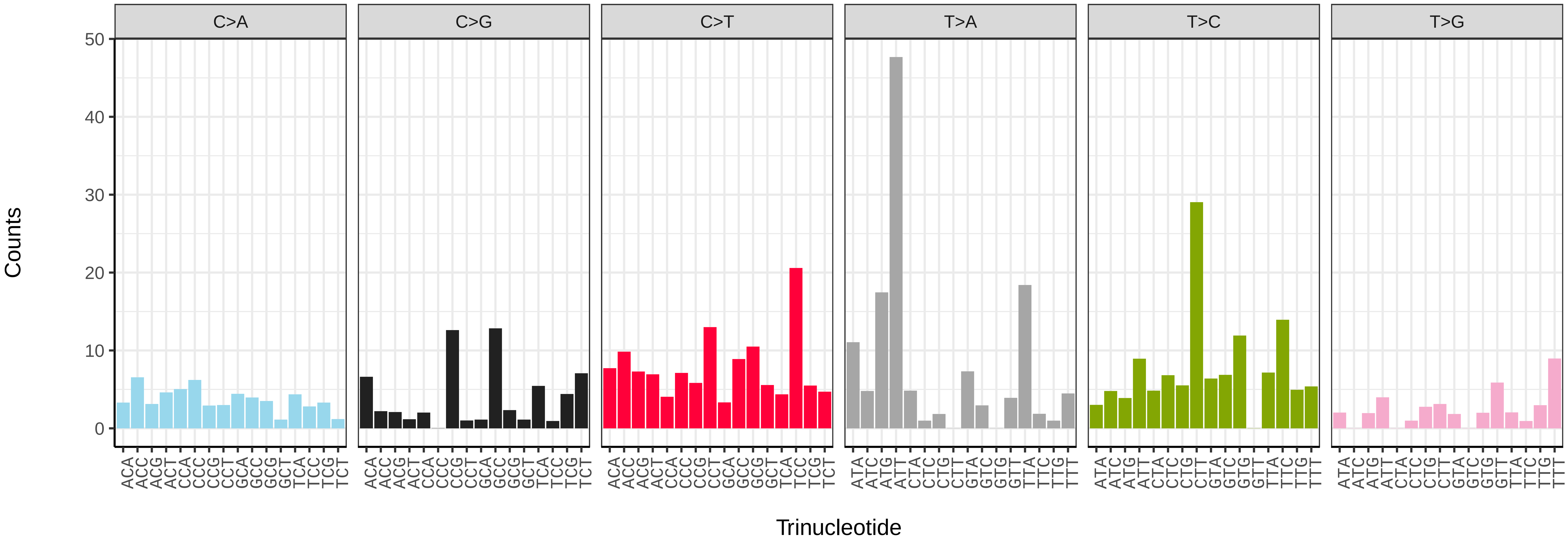


*Ectemnius continuus* (iyEctCont1) somatic mutational spectrum (SBS52 classification)

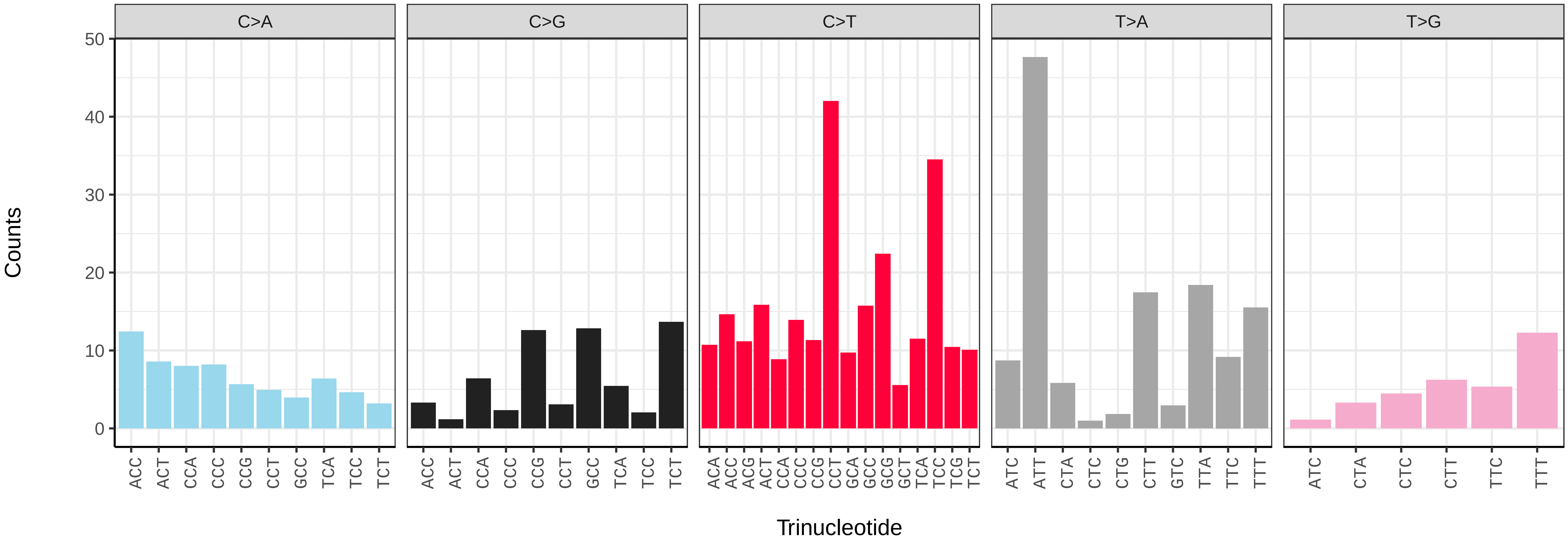

*Ectemnius lituratus* (iyEctLitu1) germline mutational spectrum

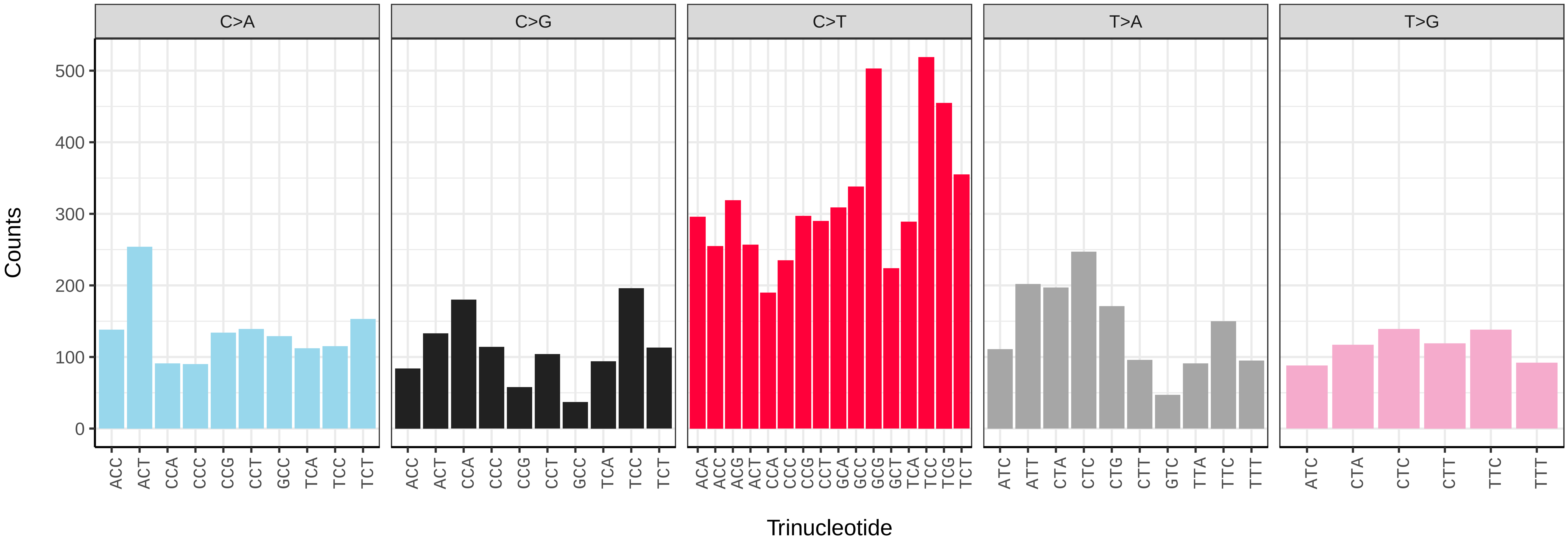


*Ectemnius lituratus* (iyEctLitu1) somatic mutational spectrum (SBS96 classification)

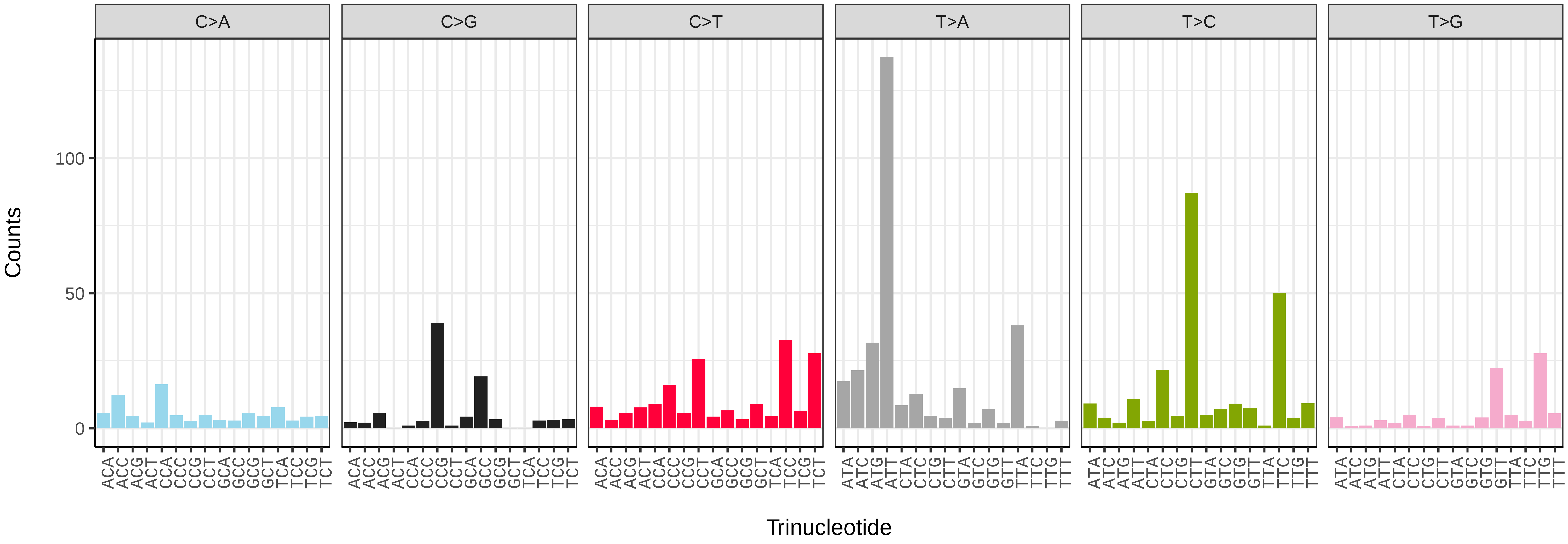


*Ectemnius lituratus* (iyEctLitu1) somatic mutational spectrum (SBS52 classification)

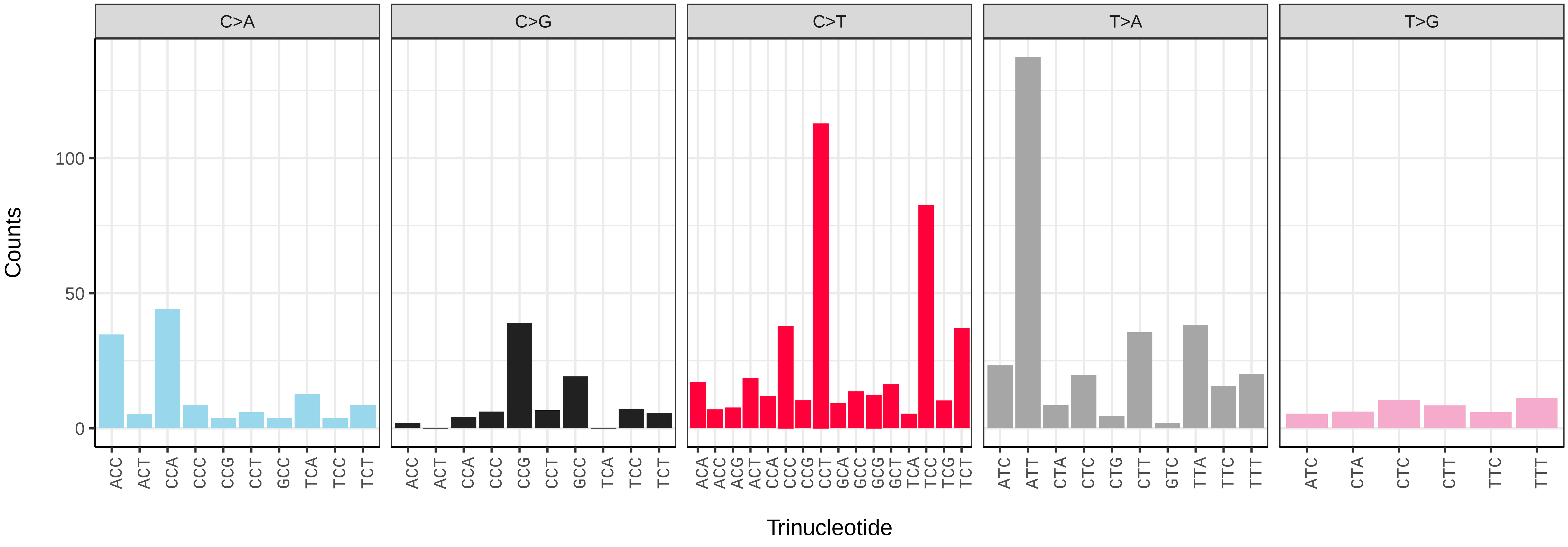

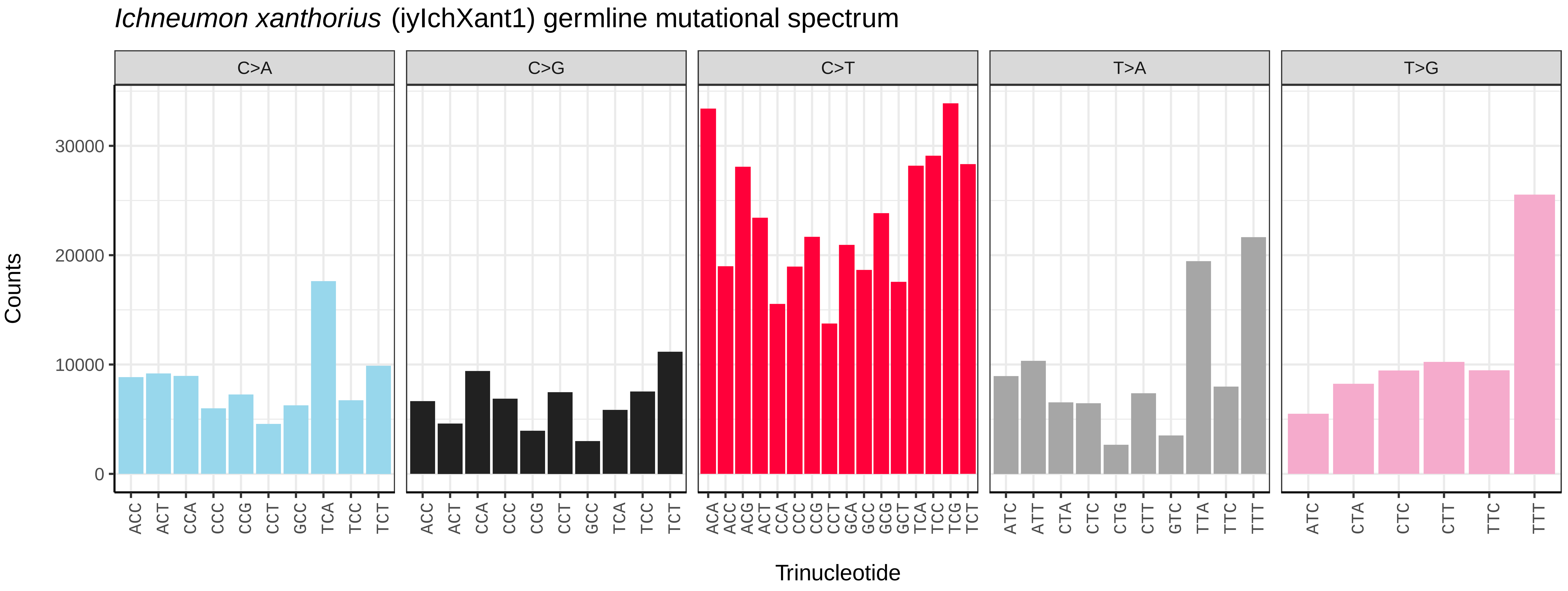
*Ichneumon xanthorius* (iyIchXant1) germline mutational spectrum
C>A
C>G
C>T
T>A
T>G
Counts
30000
20000
10000
0
ACC ACT CCA CCC CCG CCT GCC TCA TCC TCT
ACC ACT CCA CCC CCG CCT GCC TCA TCC TCT
ACA ACC ACG ACT CCA CCC CCG CCT GCA GCC GCG GCT TCA TCC TCG TCT
ATC ATT CTA CTC CTG CTT GTC TTA TTC TTT
ATC CTA CTC CTT TTC TTT
Trinucleotide

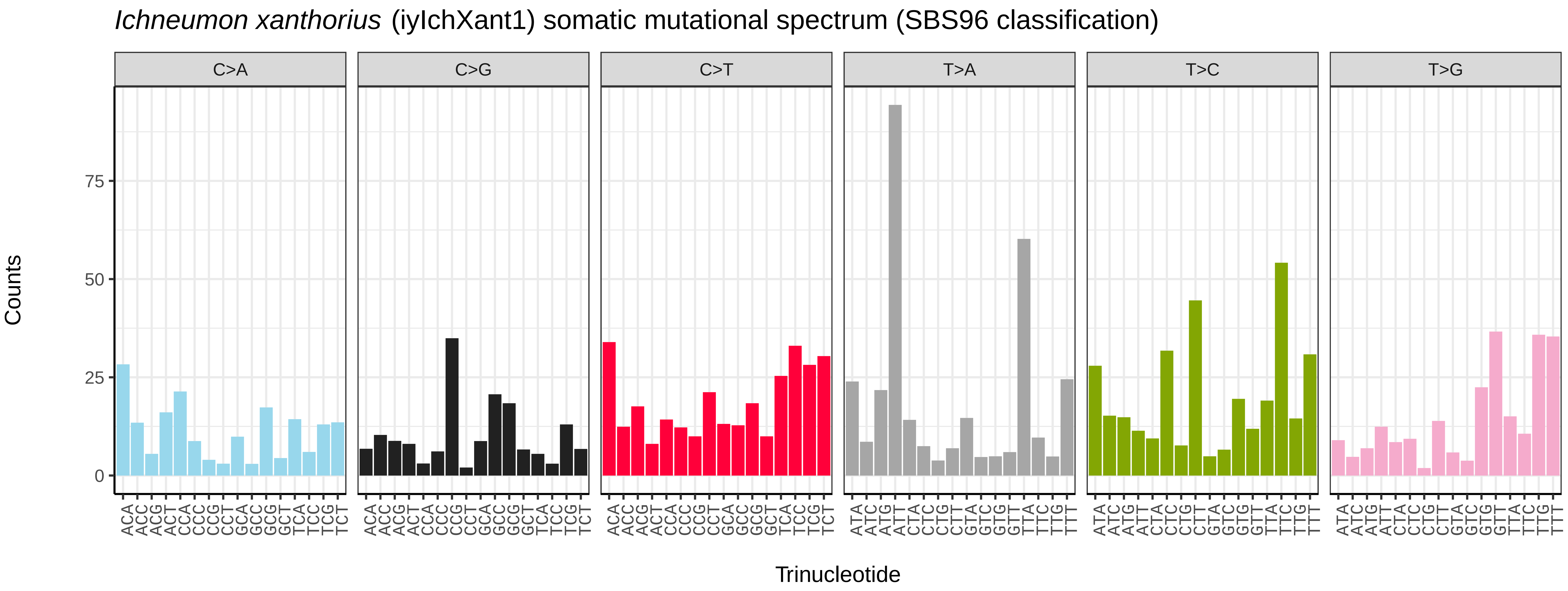
*Ichneumon xanthorius* (iyIchXant1) somatic mutational spectrum (SBS96 classification)
C>A
C>G
C>T
T>A
T>C
T>G
Counts
75
50
25
0
ACA ACC ACG ACT CCA CCC CCG CCT GCA GCC GCG GCT TCA TCC TCG TCT
ATA ATC ATG ATT CTA CTC CTG CTT GTA GTC GTG GTT TTA TTC TTG TTT
Trinucleotide

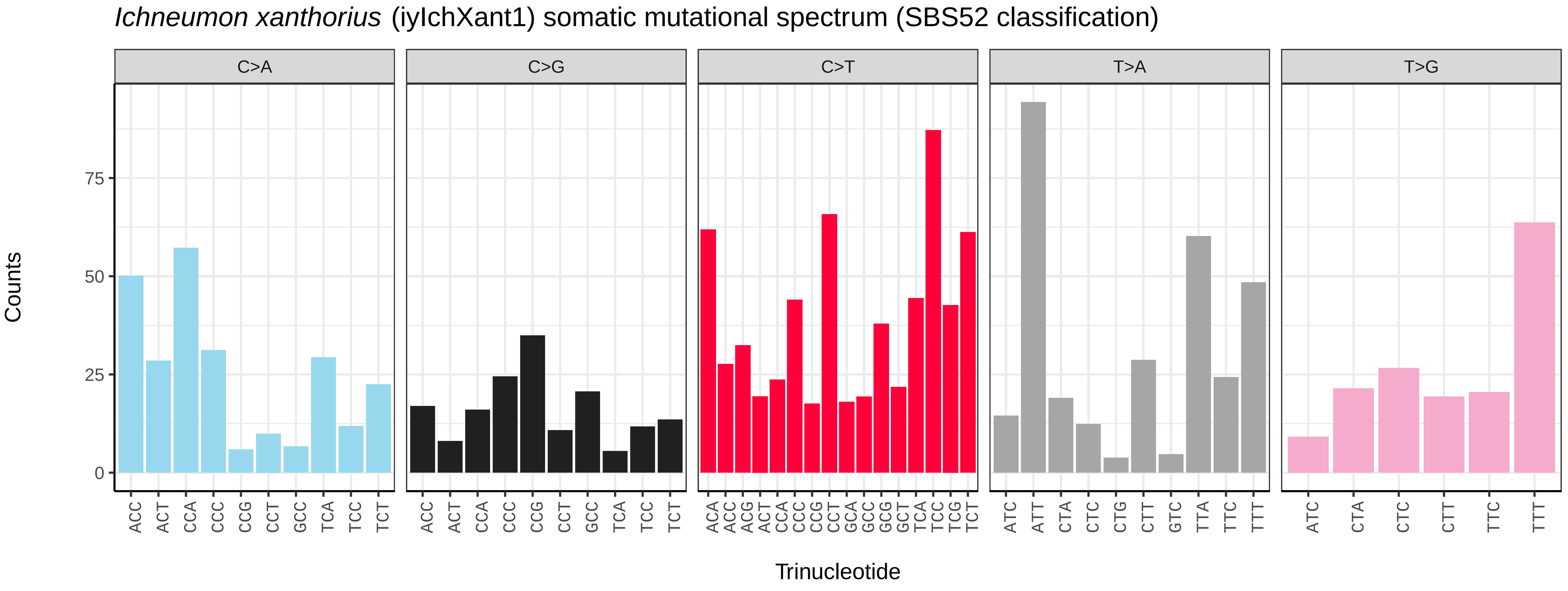
*Ichneumon xanthorius* (iyIchXant1) somatic mutational spectrum (SBS52 classification)
C>A
C>G
C>T
T>A
T>G
Counts
75
50
25
0
ACC ACT CCA CCC CCG CCT GCC TCA TCC TCT
ACC ACT CCA CCC CCG CCT GCC TCA TCC TCT
ACA ACC ACG ACT CCA CCC CCG CCT GCA GCC GCG GCT TCA TCC TCG TCT
ATC ATT CTA CTC CTG CTT GTC TTA TTC TTT
ATC CTA CTC CTT TTC TTT
Trinucleotide

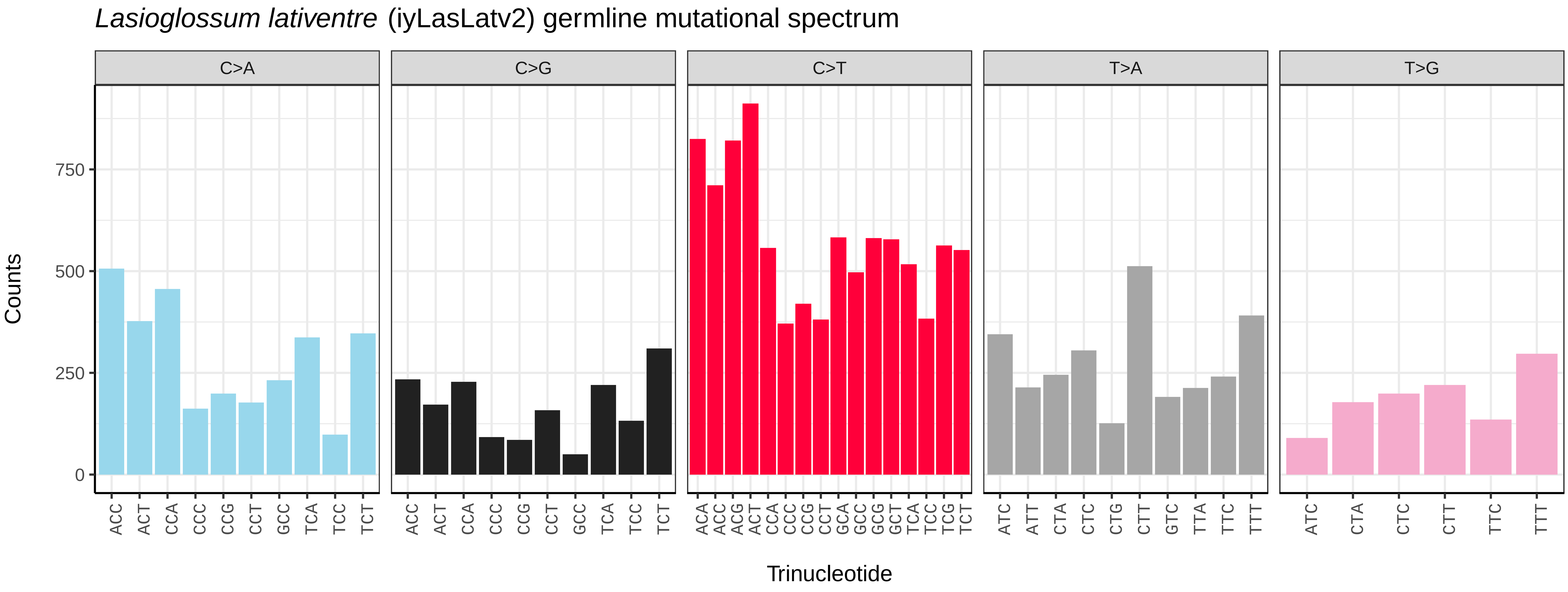

*Lasioglossum lativentre* (iyLasLatv2) germline mutational spectrum
C>A
C>G
C>T
T>A
T>G
Counts
750
500
250
0
ACC ACT CCA CCC CCG CCT GCC TCA TCC TCT
ACC ACT CCA CCC CCG CCT GCC TCA TCC TCT
ACA ACC ACG ACT CCA CCC CCG CCT GCA GCC GCG GCT TCA TCC TCG TCT
ATC ATT CTA CTC CTG CTT GTC TTA TTC TTT
ATC CTA CTC CTT TTC TTT
Trinucleotide


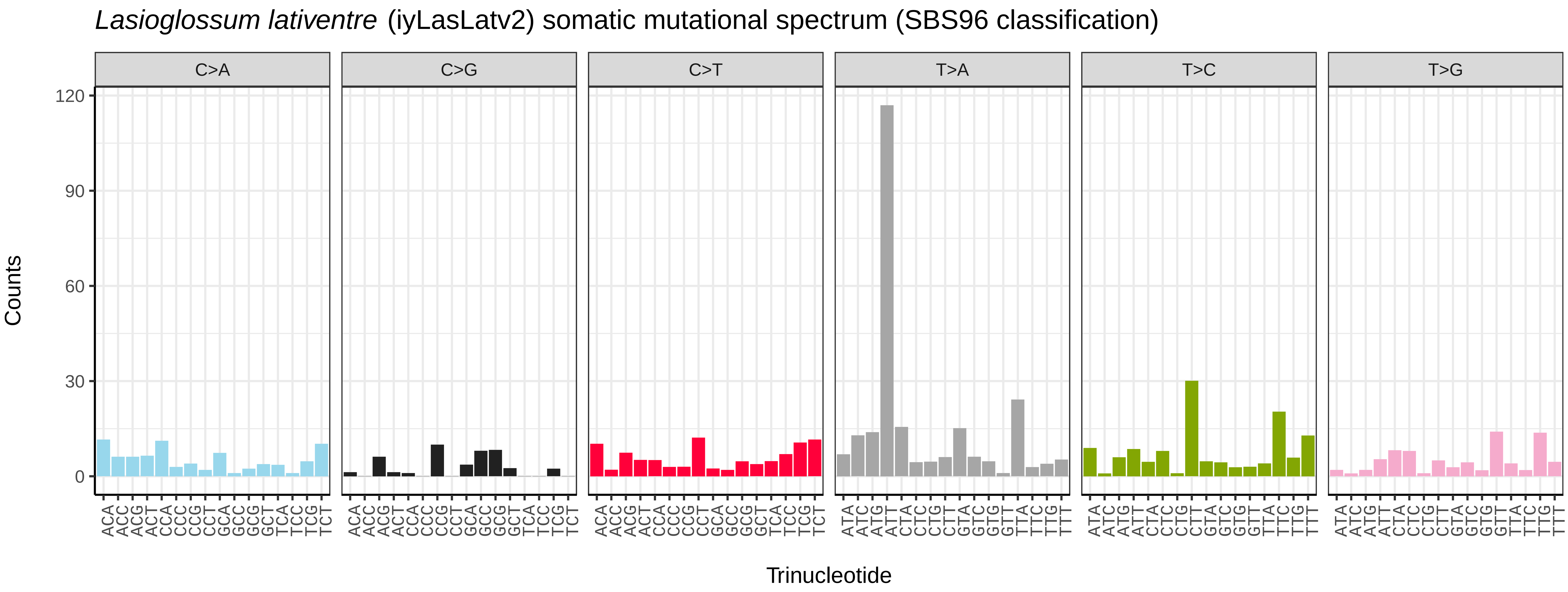

*Lasioglossum lativentre* (iyLasLatv2) somatic mutational spectrum (SBS96 classification)
C>A
C>G
C>T
T>A
T>C
T>G
Counts
120
90
60
30
0
Trinucleotide


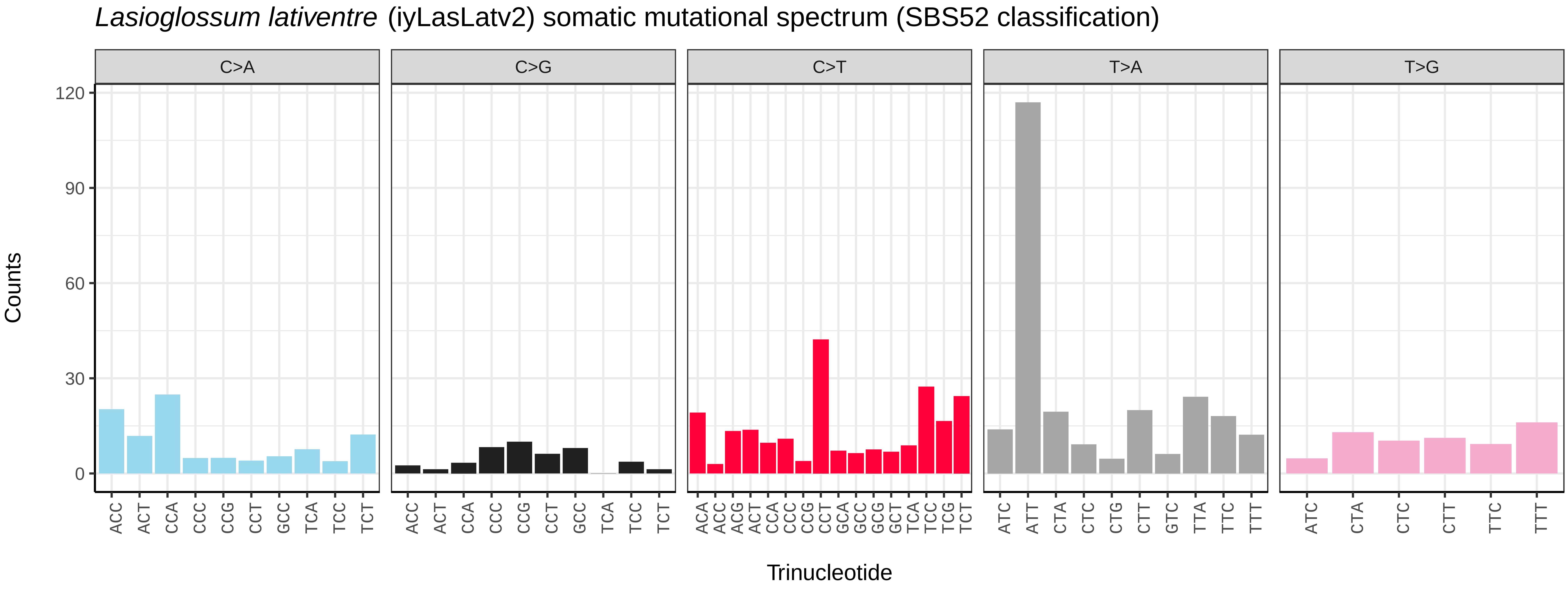

*Lasioglossum lativentre* (iyLasLatv2) somatic mutational spectrum (SBS52 classification)
C>A
C>G
C>T
T>A
T>G
Counts
120
90
60
30
0
ACC ACT CCA CCC CCG CCT GCC TCA TCC TCT
ACC ACT CCA CCC CCG CCT GCC TCA TCC TCT
ACA ACC ACG ACT CCA CCC CCG CCT GCA GCC GCG GCT TCA TCC TCG TCT
ATC ATT CTA CTC CTG CTT GTC TTA TTC TTT
ATC CTA CTC CTT TTC TTT
Trinucleotide

*Lasioglossum morio* (iyLasMori1) germline mutational spectrum

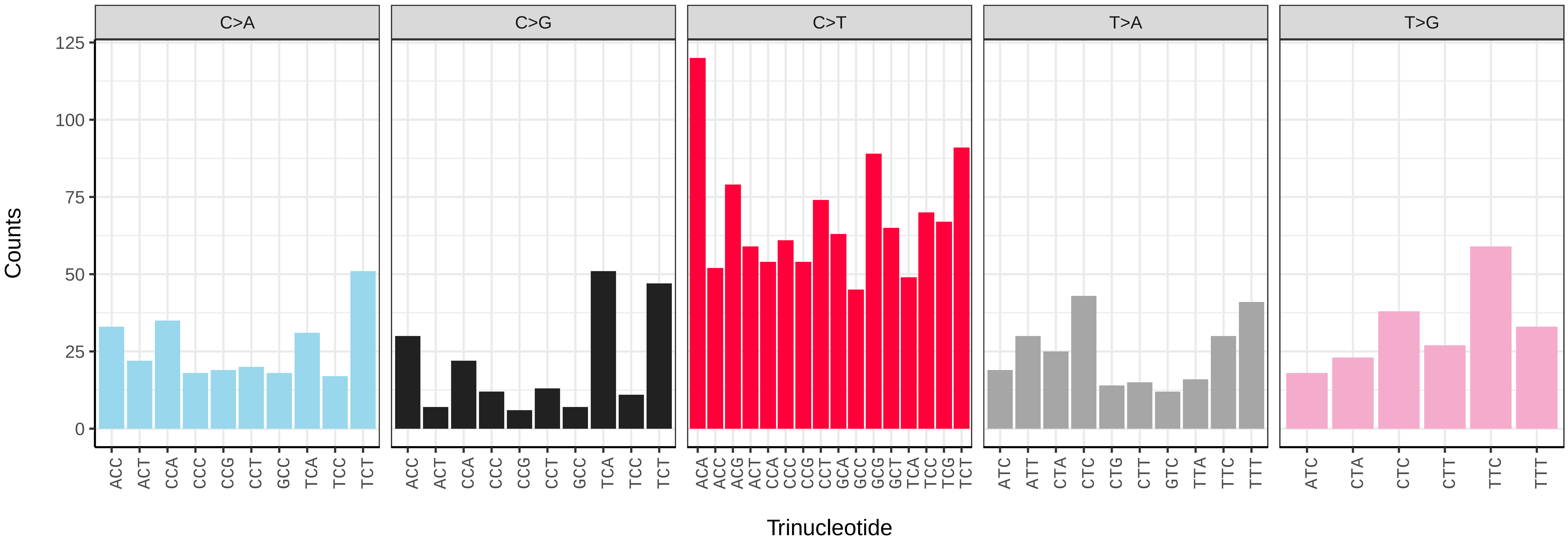


*Lasioglossum morio* (iyLasMori1) somatic mutational spectrum (SBS96 classification)

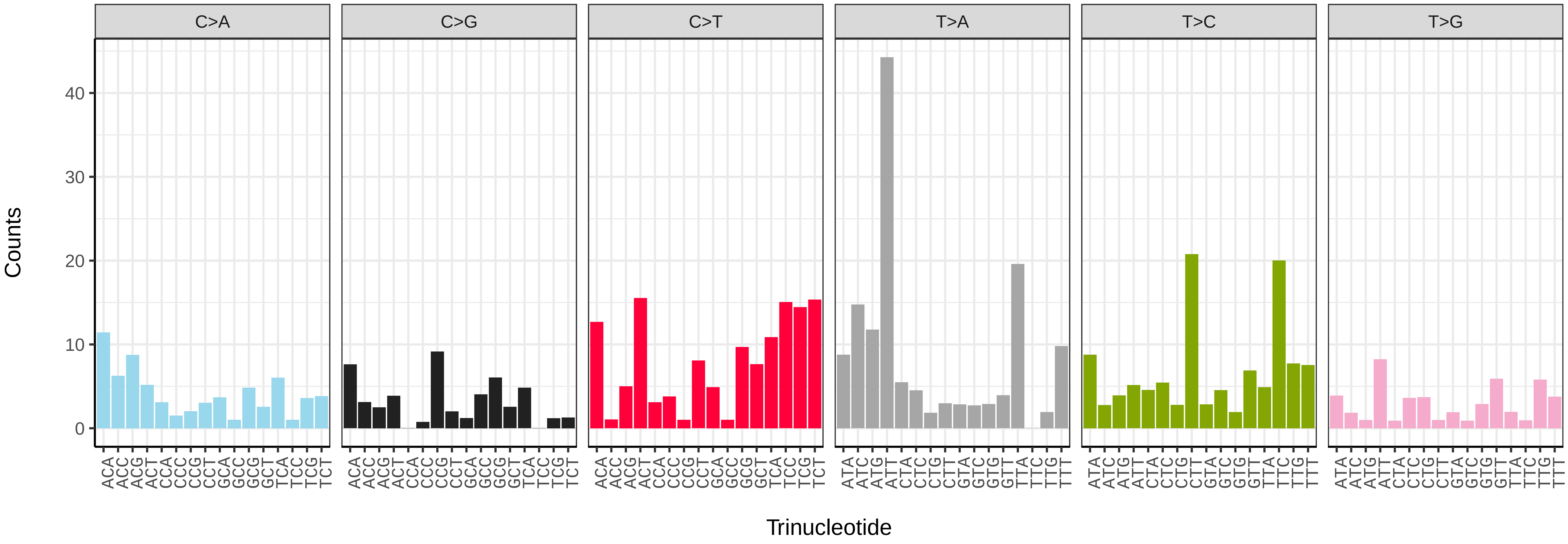


*Lasioglossum morio* (iyLasMori1) somatic mutational spectrum (SBS52 classification)

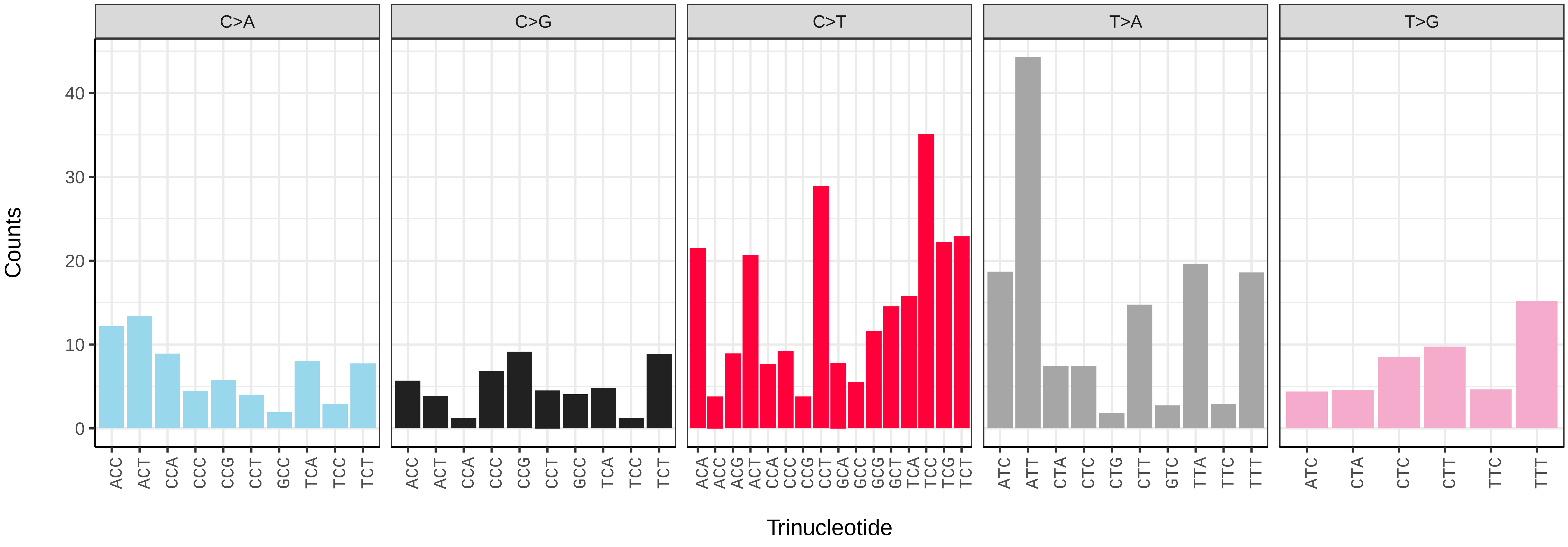

*Lasioglossum pauxillum* (iyLasPaux1) germline mutational spectrum

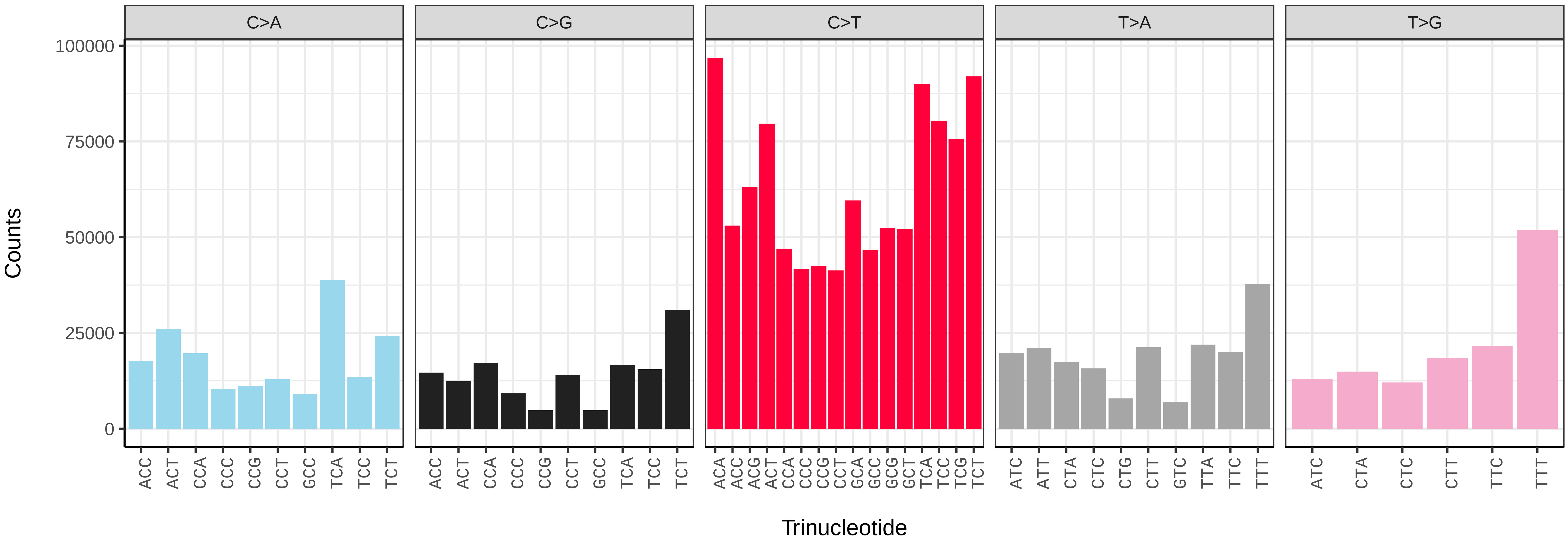


*Lasioglossum pauxillum* (iyLasPaux1) somatic mutational spectrum (SBS96 classification)

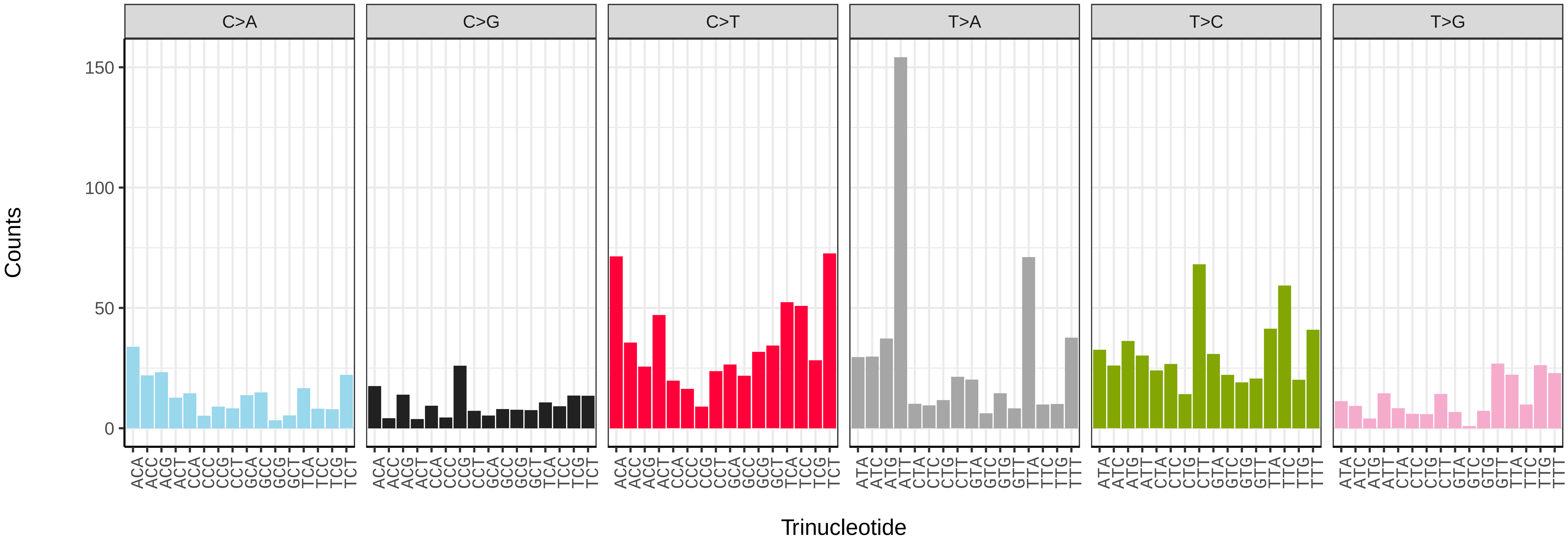


*Lasioglossum pauxillum* (iyLasPaux1) somatic mutational spectrum (SBS52 classification)

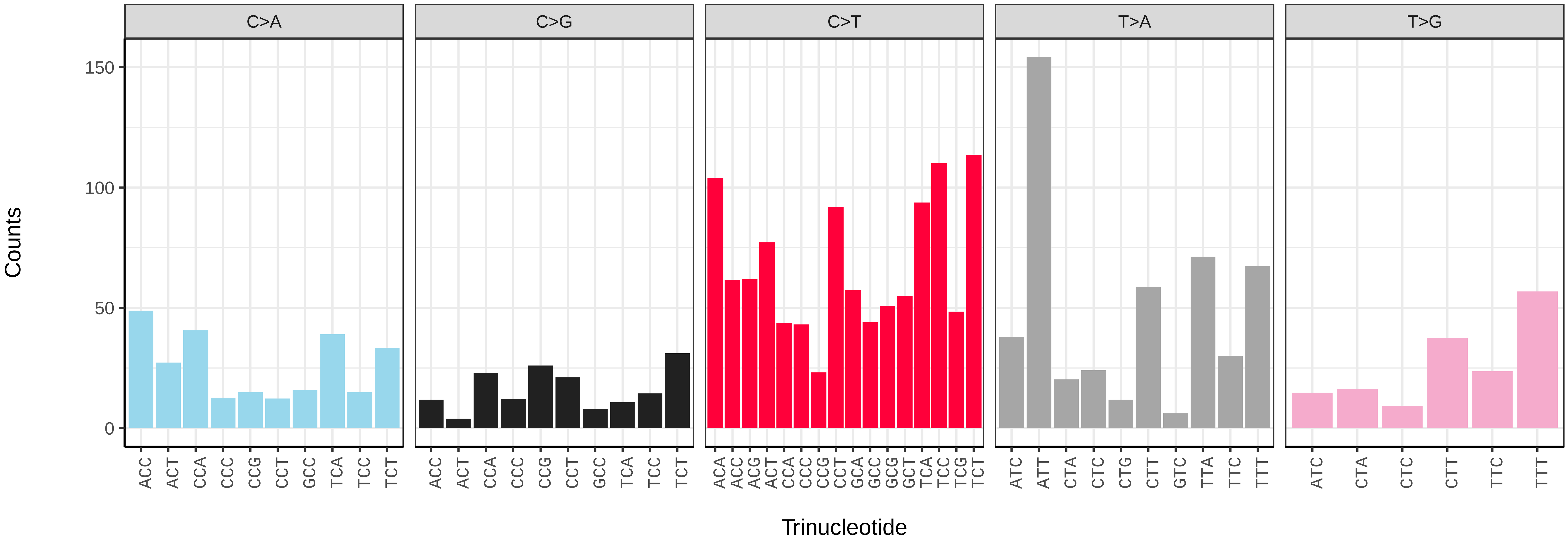

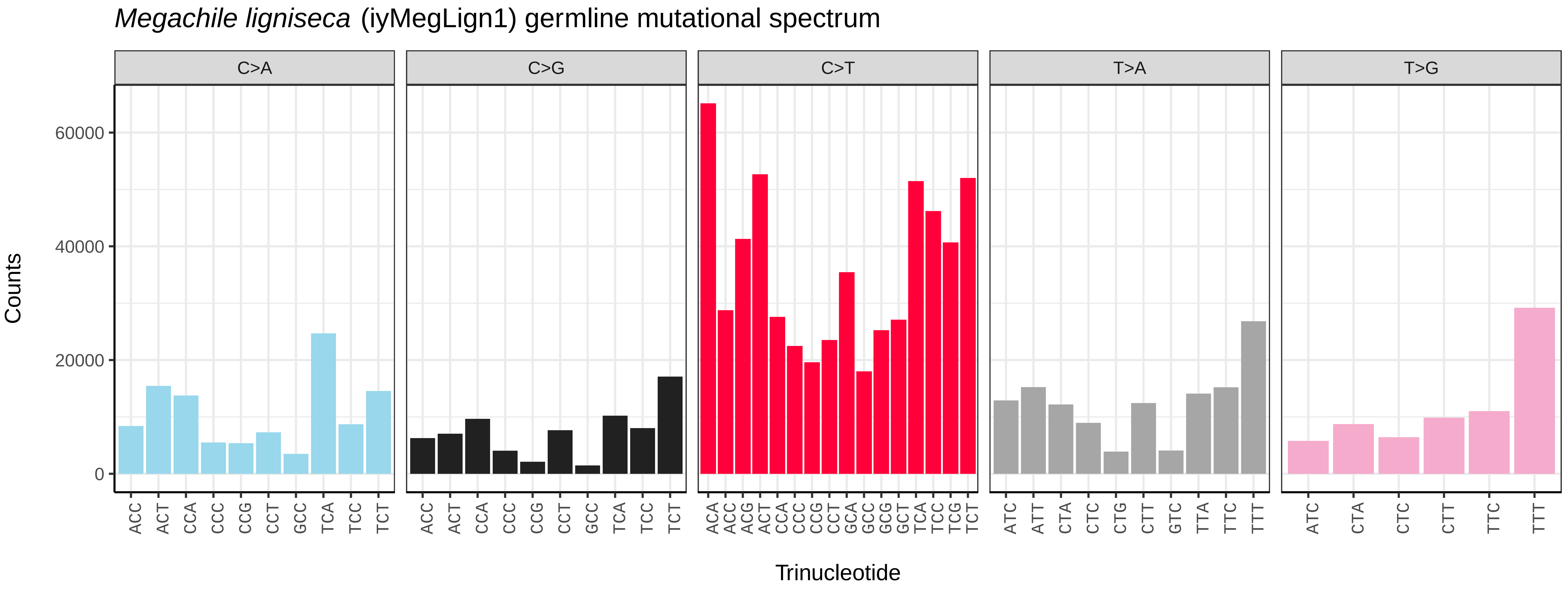
*Megachile ligniseca* (iyMegLign1) germline mutational spectrum
C>A
C>G
C>T
T>A
T>G
Counts
0
20000
40000
60000
Trinucleotide

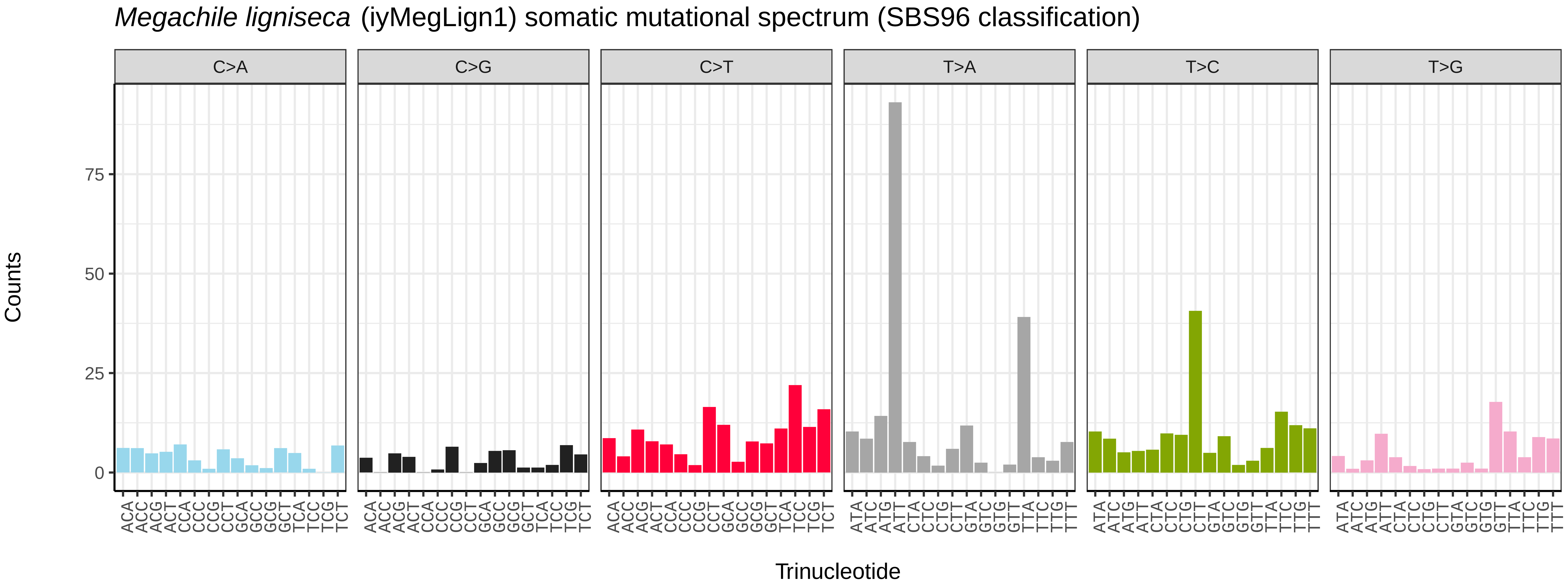
*Megachile ligniseca* (iyMegLign1) somatic mutational spectrum (SBS96 classification)
C>A
C>G
C>T
T>A
T>C
T>G
Counts
0
25
50
75
Trinucleotide

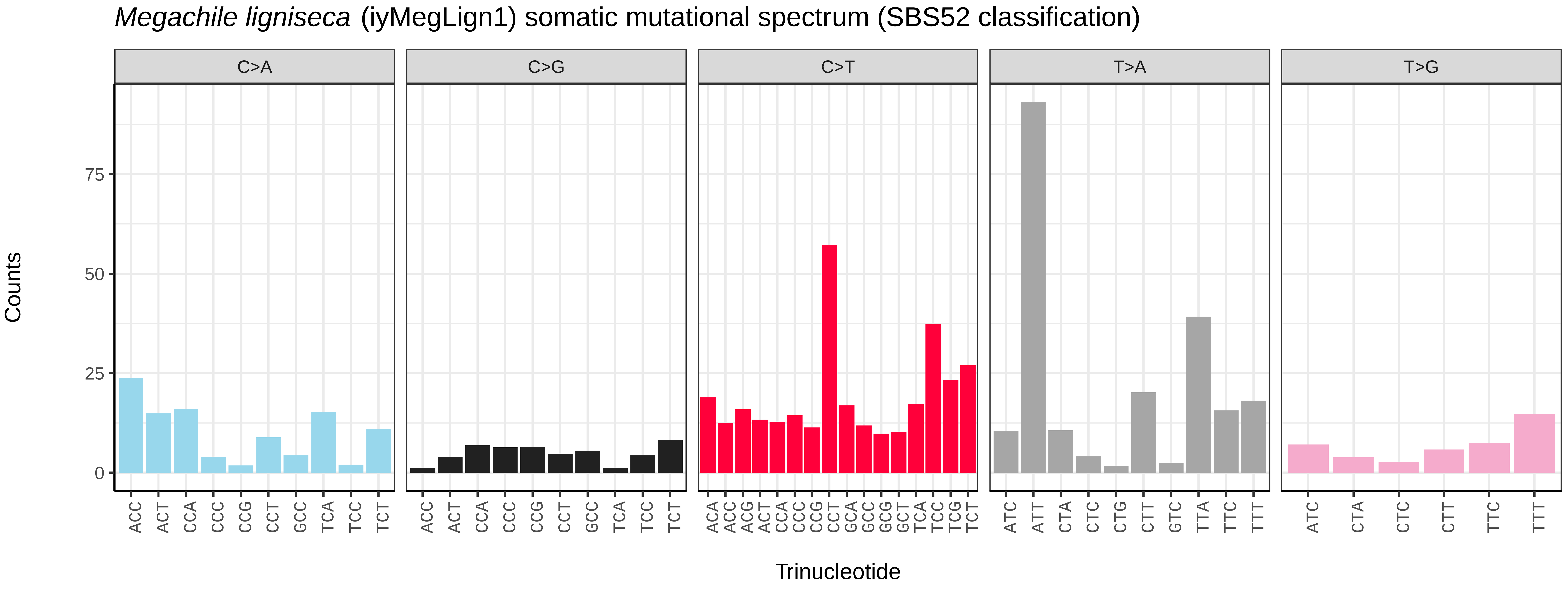
*Megachile ligniseca* (iyMegLign1) somatic mutational spectrum (SBS52 classification)
C>A
C>G
C>T
T>A
T>G
Counts
0
25
50
75
Trinucleotide

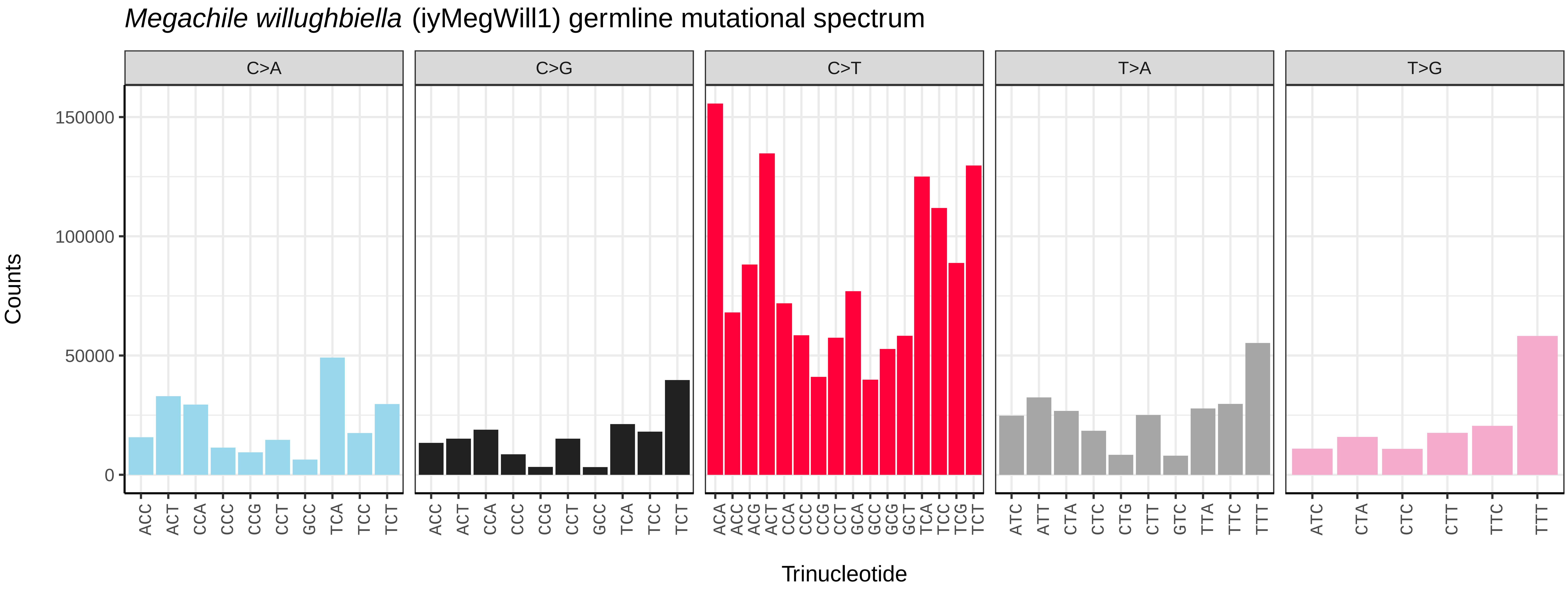
*Megachile willughbiella* (iyMegWill1) germline mutational spectrum
C>A
C>G
C>T
T>A
T>G
Counts
0
50000
100000
150000
ACC ACT CCA CCC CCG CCT GCC TCA TCC TCT
ACC ACT CCA CCC CCG CCT GCC TCA TCC TCT
ACA ACC ACG ACT CCA CCC CCG CCT GCA GCC GCG GCT TCA TCC TCG TCT
ATC ATT CTA CTC CTG CTT GTC TTA TTC TTT
ATC CTA CTC CTT TTC TTT
Trinucleotide

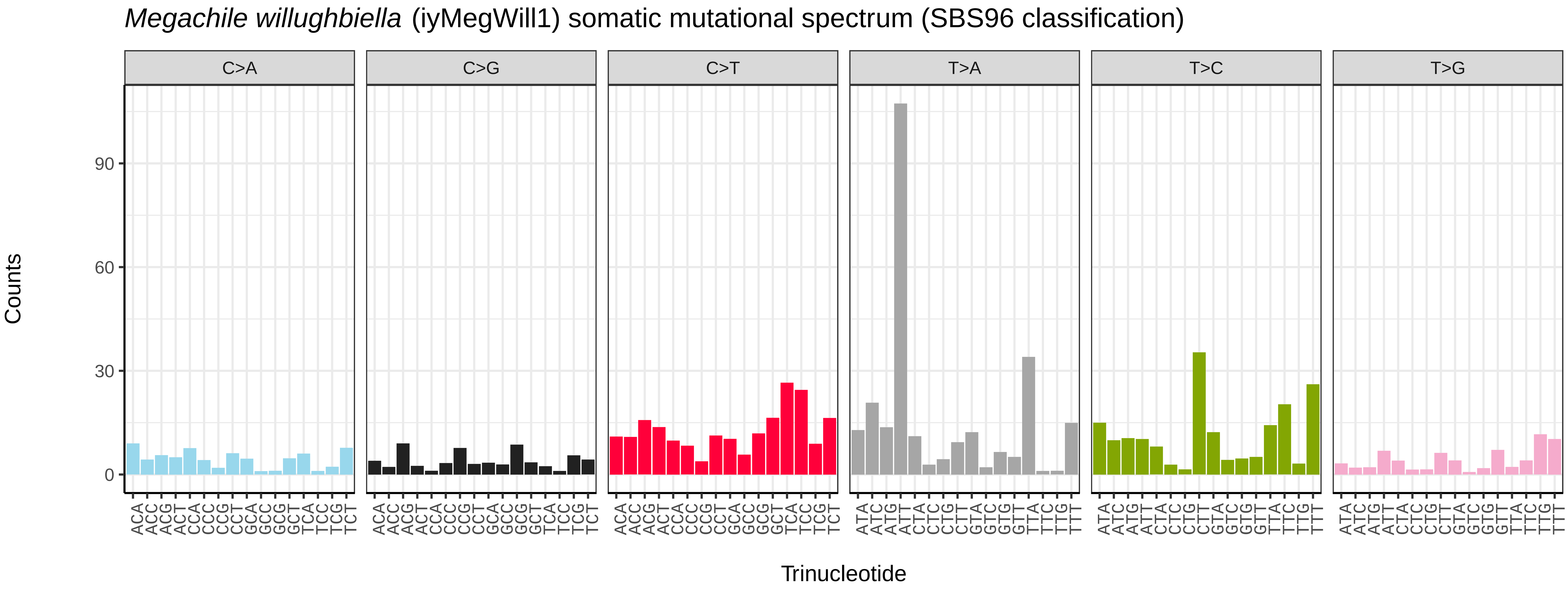
*Megachile willughbiella* (iyMegWill1) somatic mutational spectrum (SBS96 classification)
C>A
C>G
C>T
T>A
T>C
T>G
Counts
0
30
60
90
ACA ACC ACG ACT CCA CCC CCG CCT GCA GCC GCG GCT TCA TCC TCG TCT
ATA ATC ATG ATT CTA CTC CTG CTT GTA GTC GTG GTT TTA TTC TTG TTT
Trinucleotide

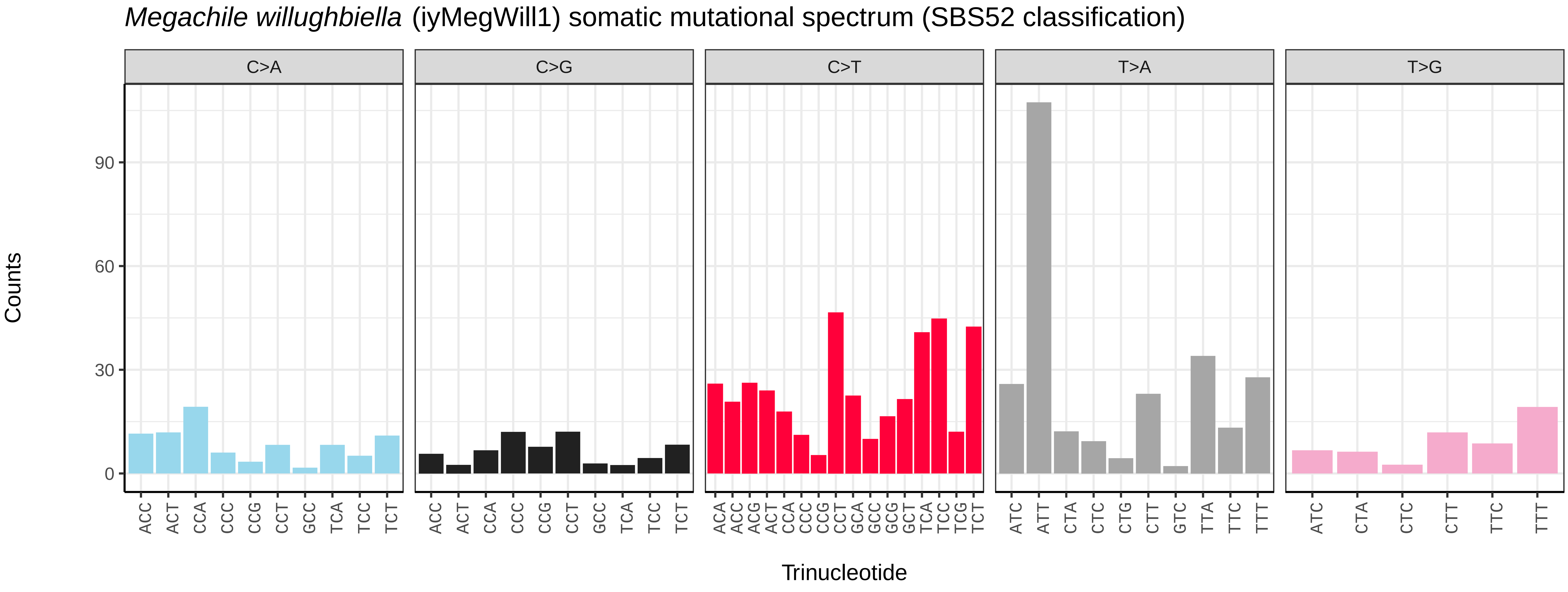
*Megachile willughbiella* (iyMegWill1) somatic mutational spectrum (SBS52 classification)
C>A
C>G
C>T
T>A
T>G
Counts
0
30
60
90
ACC ACT CCA CCC CCG CCT GCC TCA TCC TCT
ACC ACT CCA CCC CCG CCT GCC TCA TCC TCT
ACA ACC ACG ACT CCA CCC CCG CCT GCA GCC GCG GCT TCA TCC TCG TCT
ATC ATT CTA CTC CTG CTT GTC TTA TTC TTT
ATC CTA CTC CTT TTC TTT
Trinucleotide

*Mimumesa dahlbomi* (iyMimDahl1) germline mutational spectrum

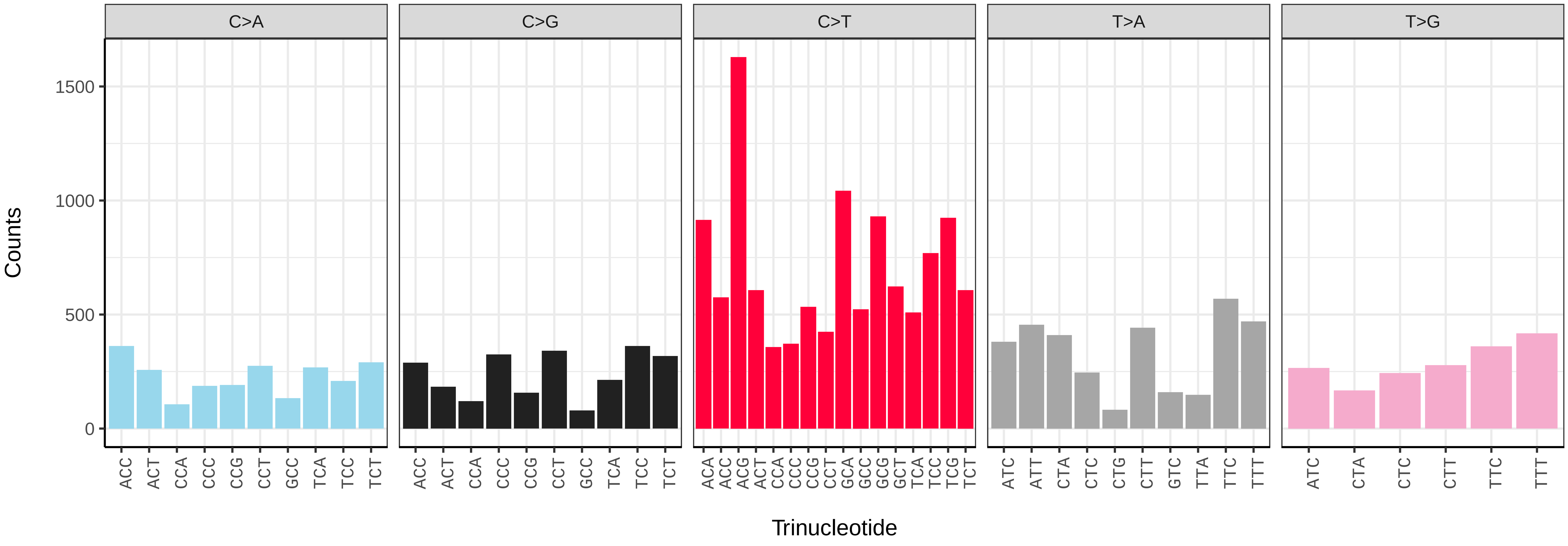


*Mimumesa dahlbomi* (iyMimDahl1) somatic mutational spectrum (SBS96 classification)

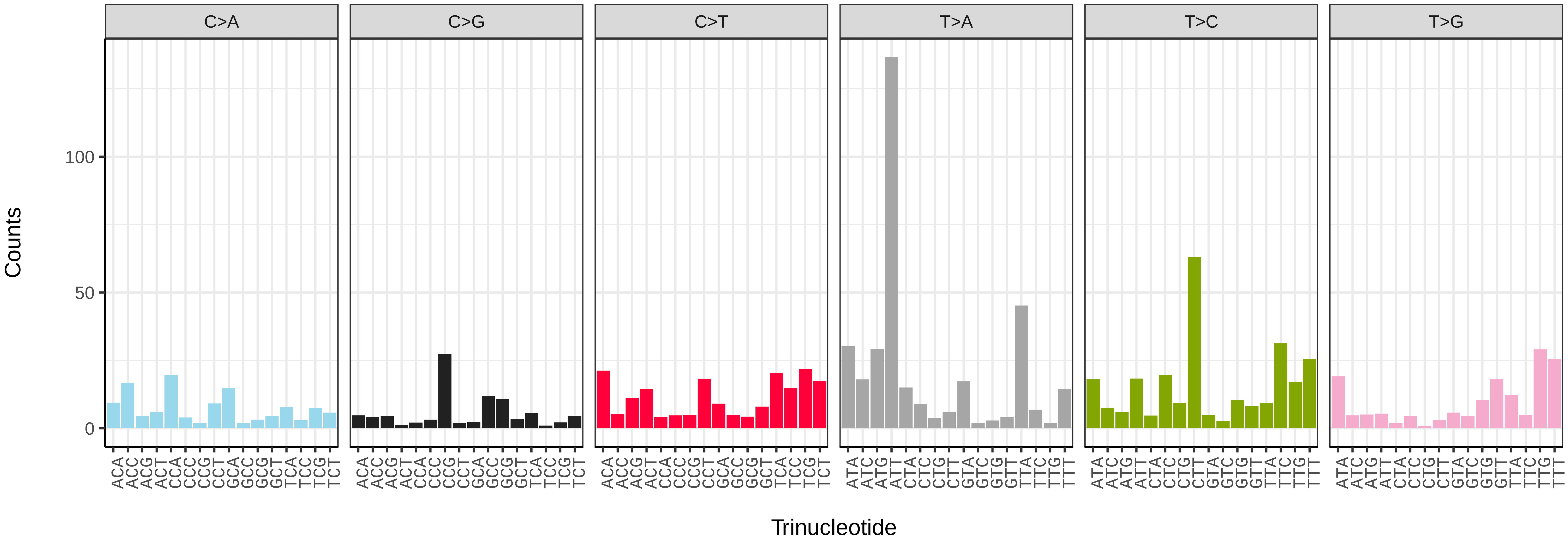


*Mimumesa dahlbomi* (iyMimDahl1) somatic mutational spectrum (SBS52 classification)

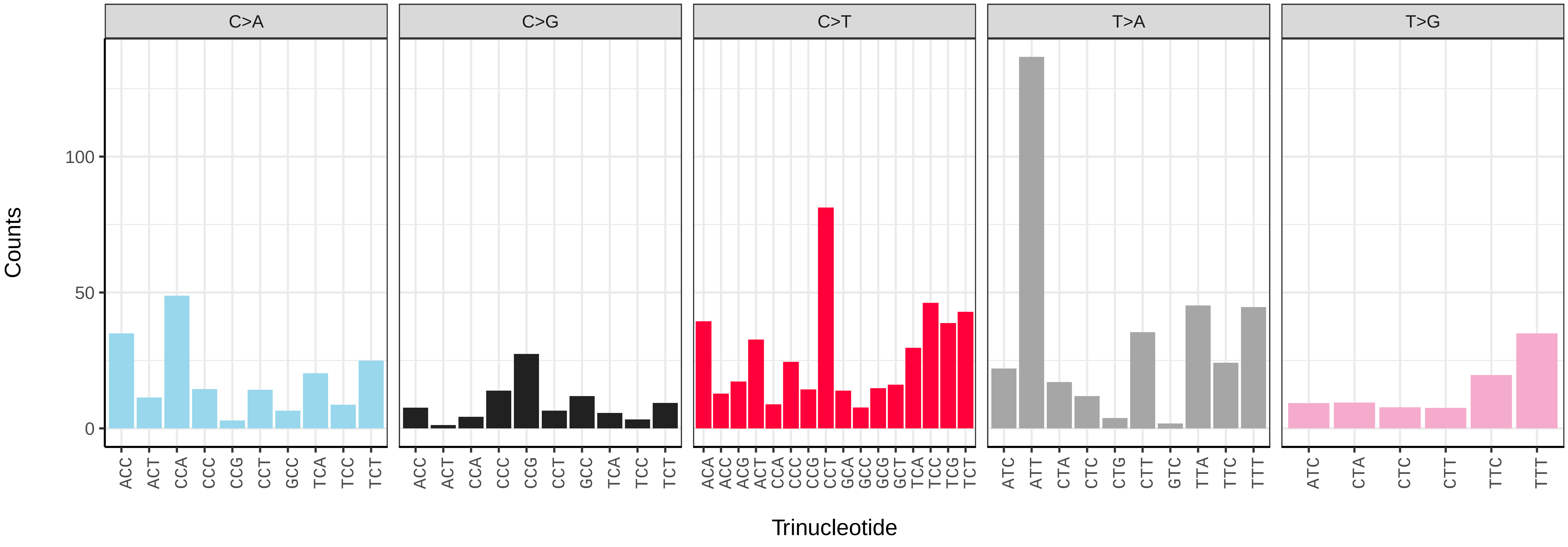

*Netelia fuscicornis* (iyNetDila1) germline mutational spectrum

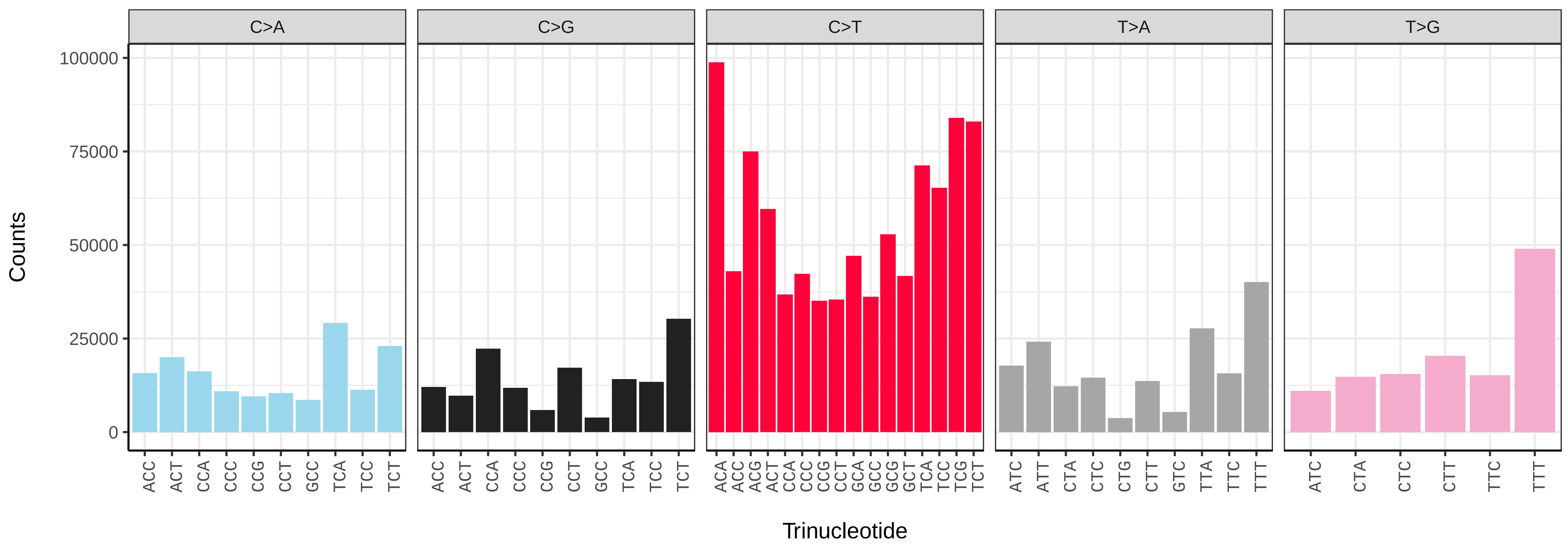


*Netelia fuscicornis* (iyNetDila1) somatic mutational spectrum (SBS96 classification)

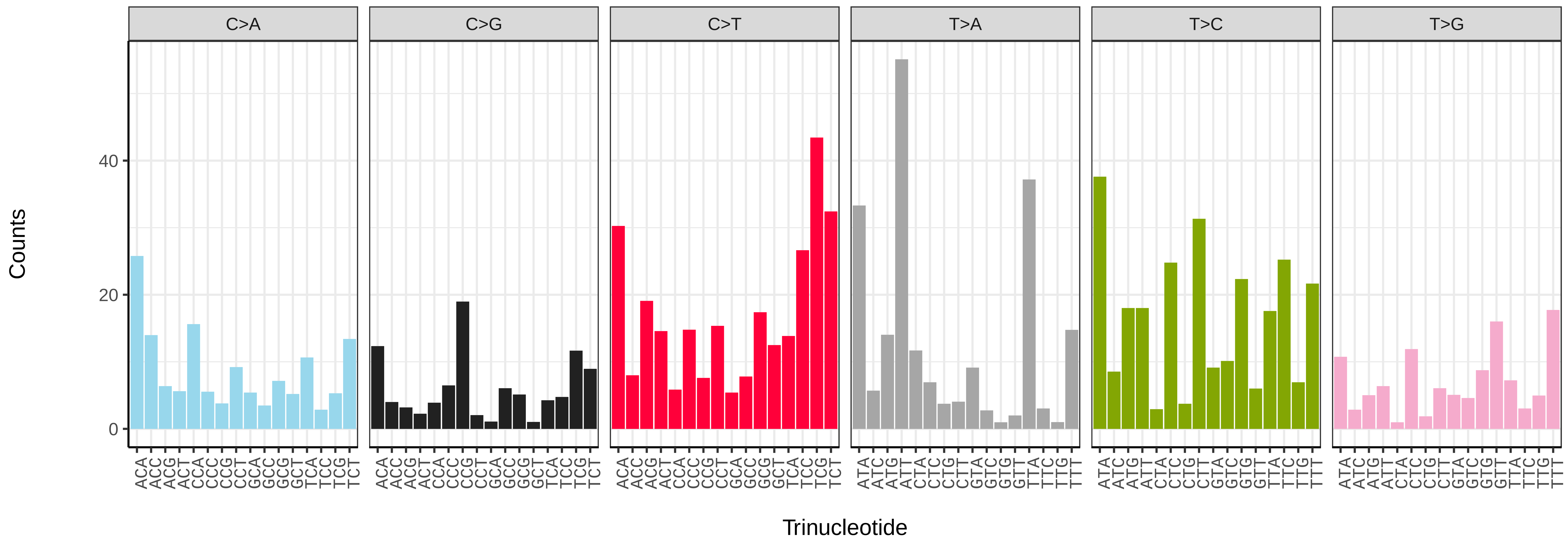


*Netelia fuscicornis* (iyNetDila1) somatic mutational spectrum (SBS52 classification)

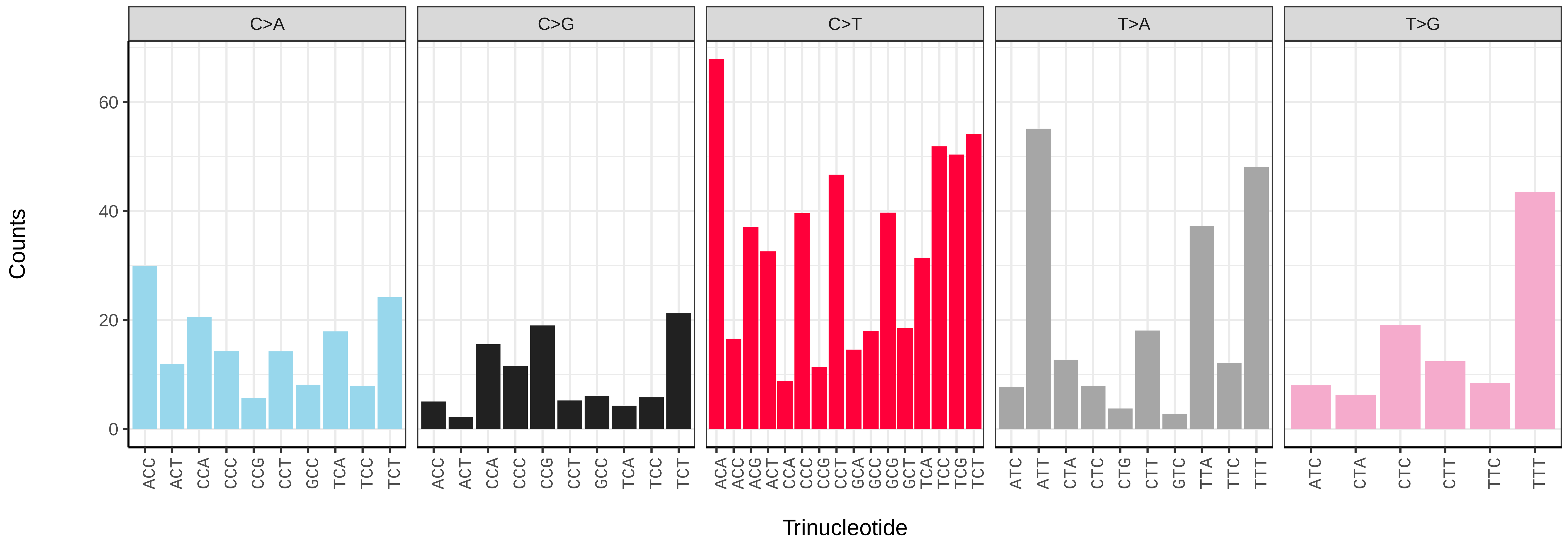

*Nomada fabriciana* (iyNomFabr1) germline mutational spectrum

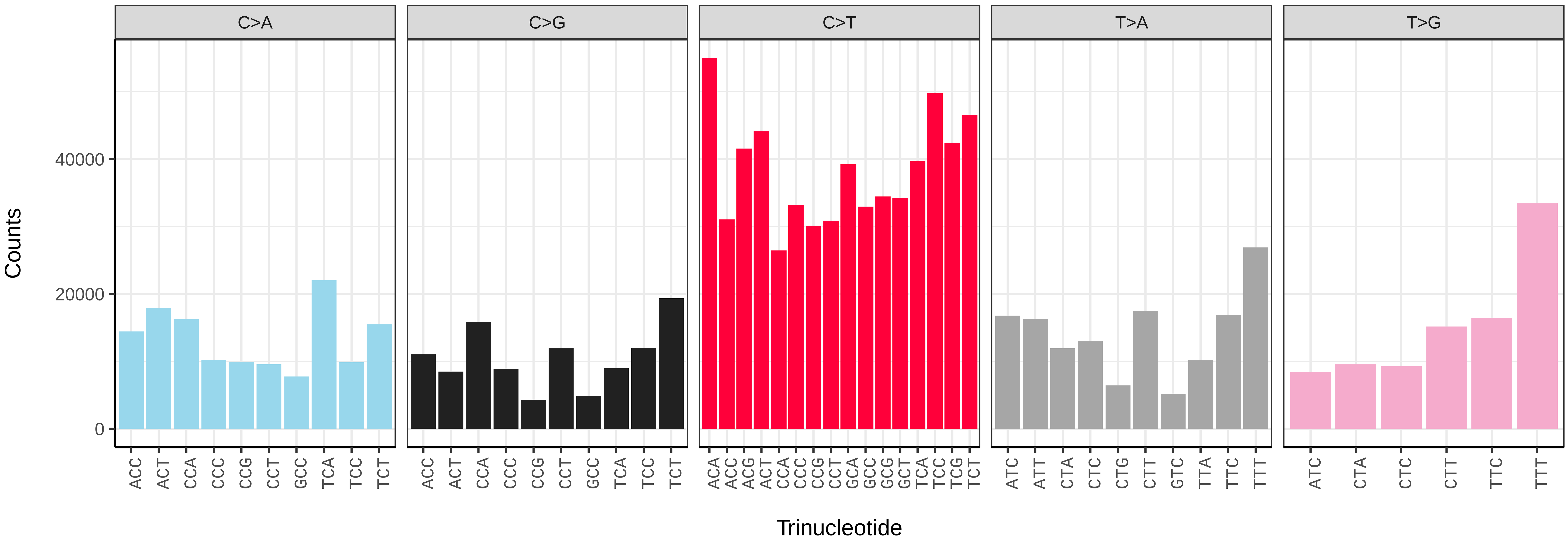


*Nomada fabriciana* (iyNomFabr1) somatic mutational spectrum (SBS96 classification)

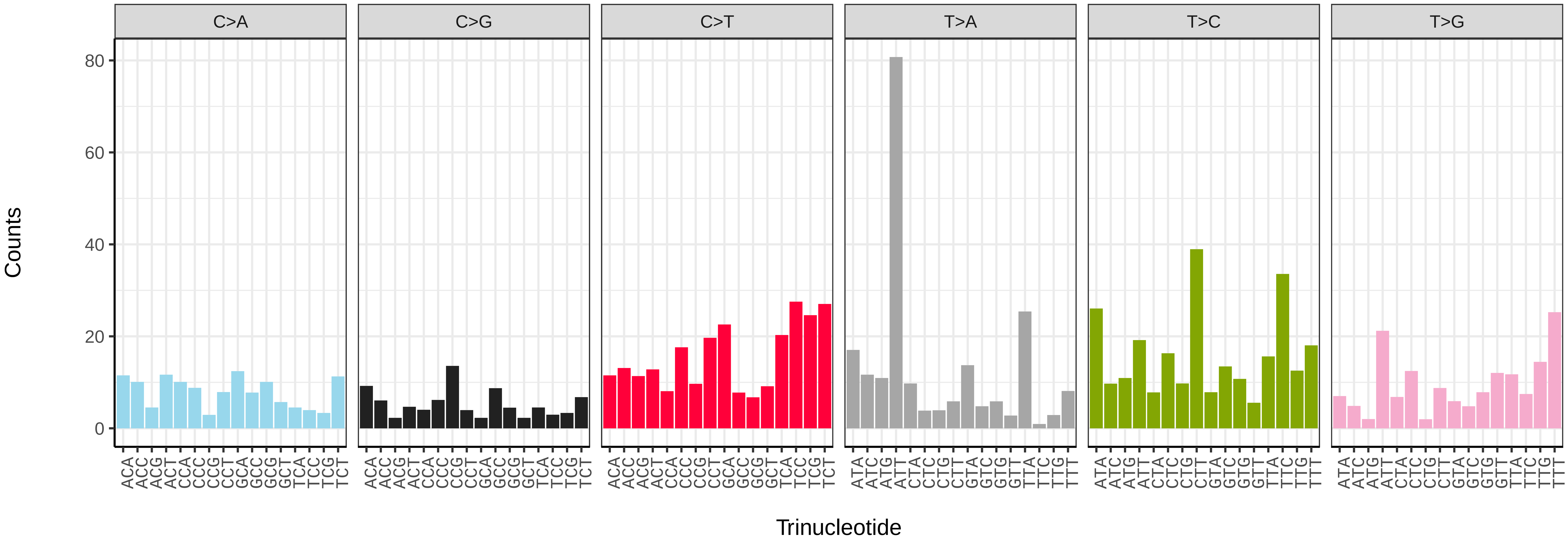


*Nomada fabriciana* (iyNomFabr1) somatic mutational spectrum (SBS52 classification)

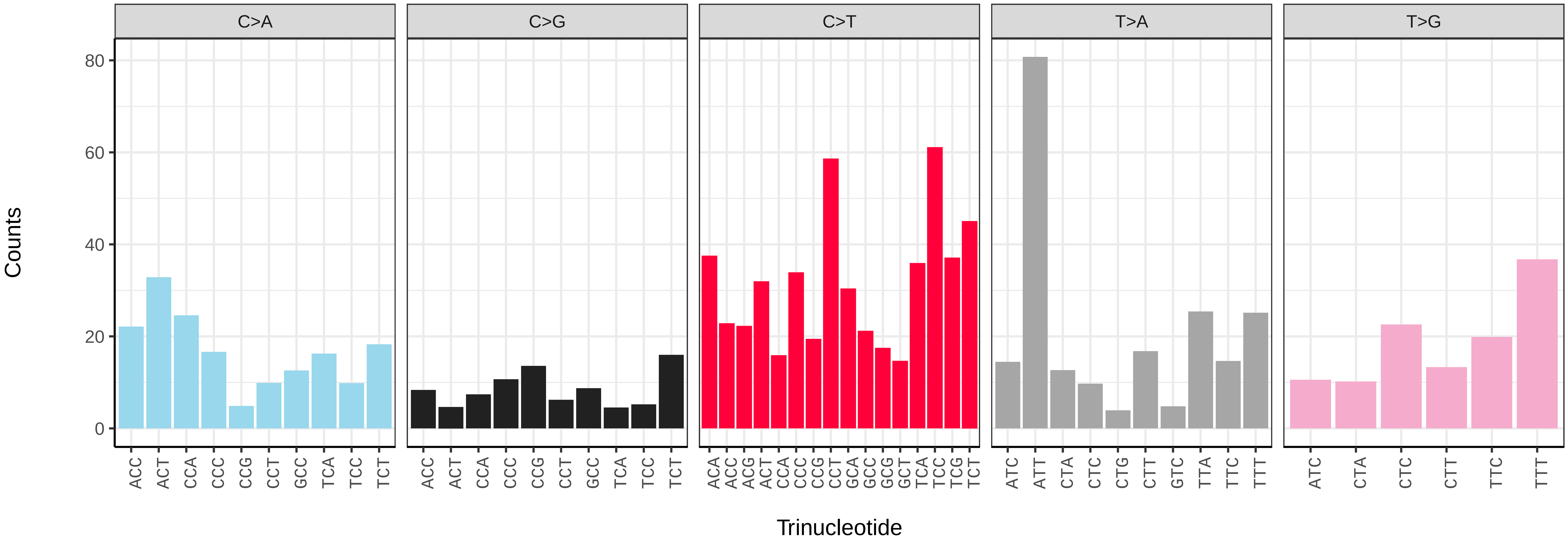

*Nomada fucata* (iyNomFuca1) germline mutational spectrum

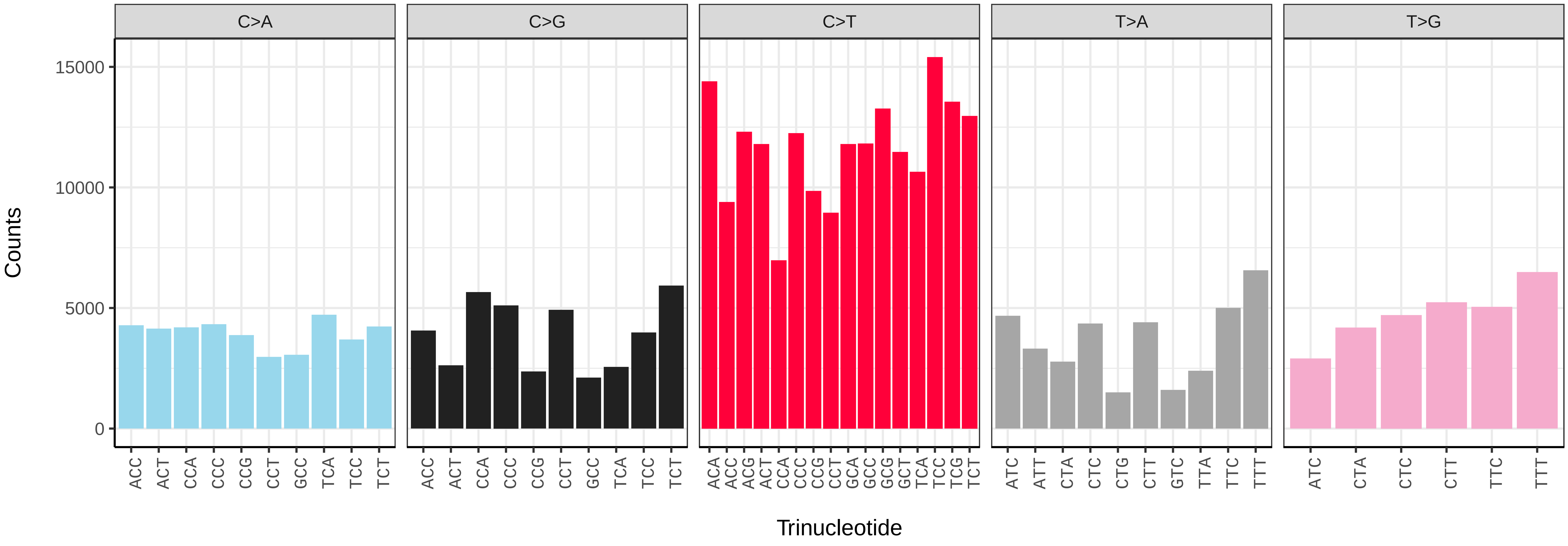


*Nomada fucata* (iyNomFuca1) somatic mutational spectrum (SBS96 classification)

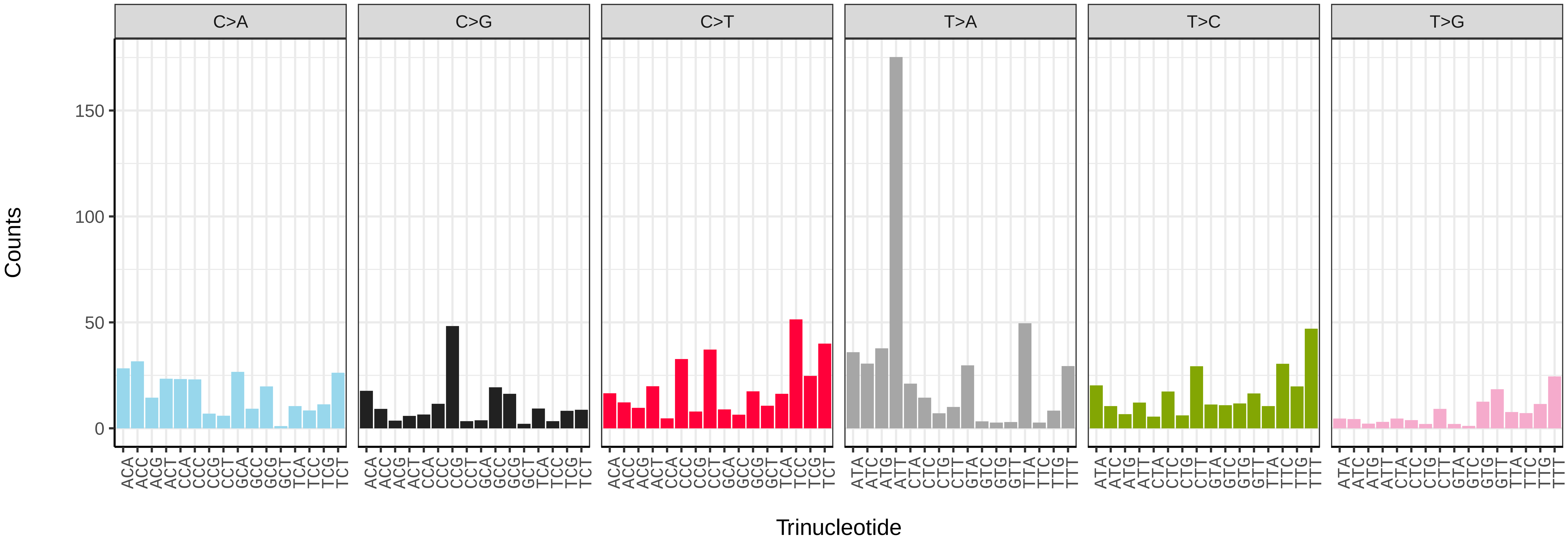


*Nomada fucata* (iyNomFuca1) somatic mutational spectrum (SBS52 classification)

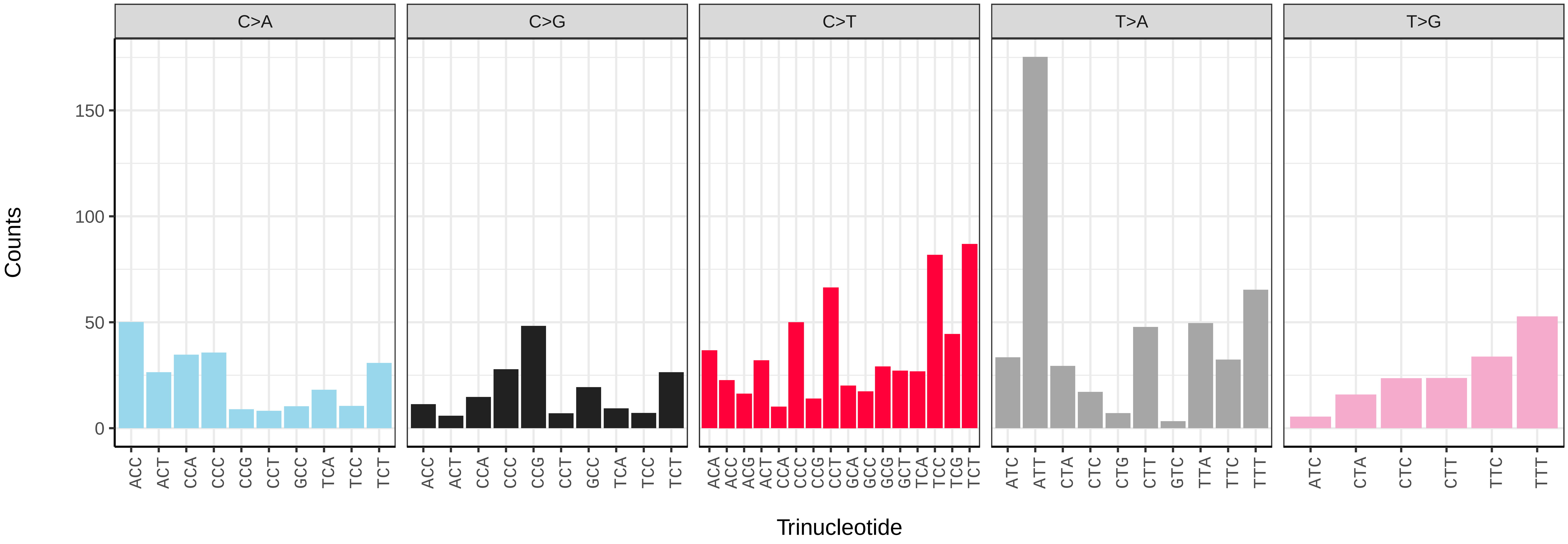

*Nysson spinosus* (iyNysSpin1) germline mutational spectrum

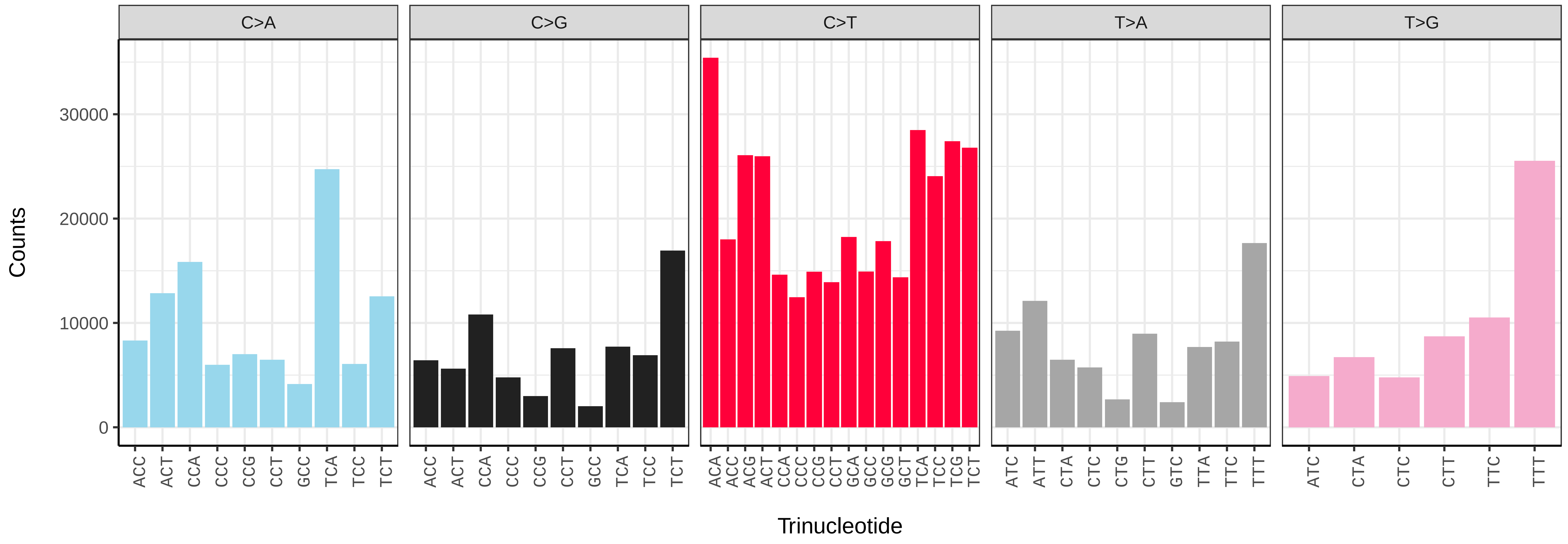


*Nysson spinosus* (iyNysSpin1) somatic mutational spectrum (SBS96 classification)

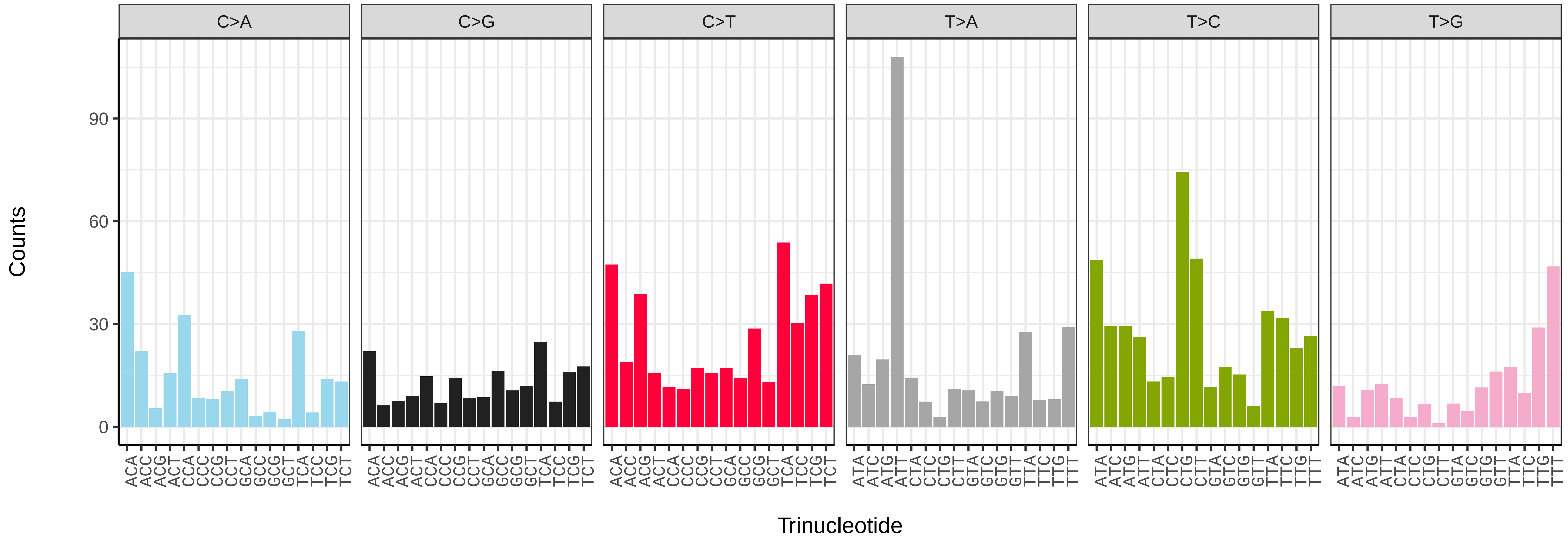


*Nysson spinosus* (iyNysSpin1) somatic mutational spectrum (SBS52 classification)

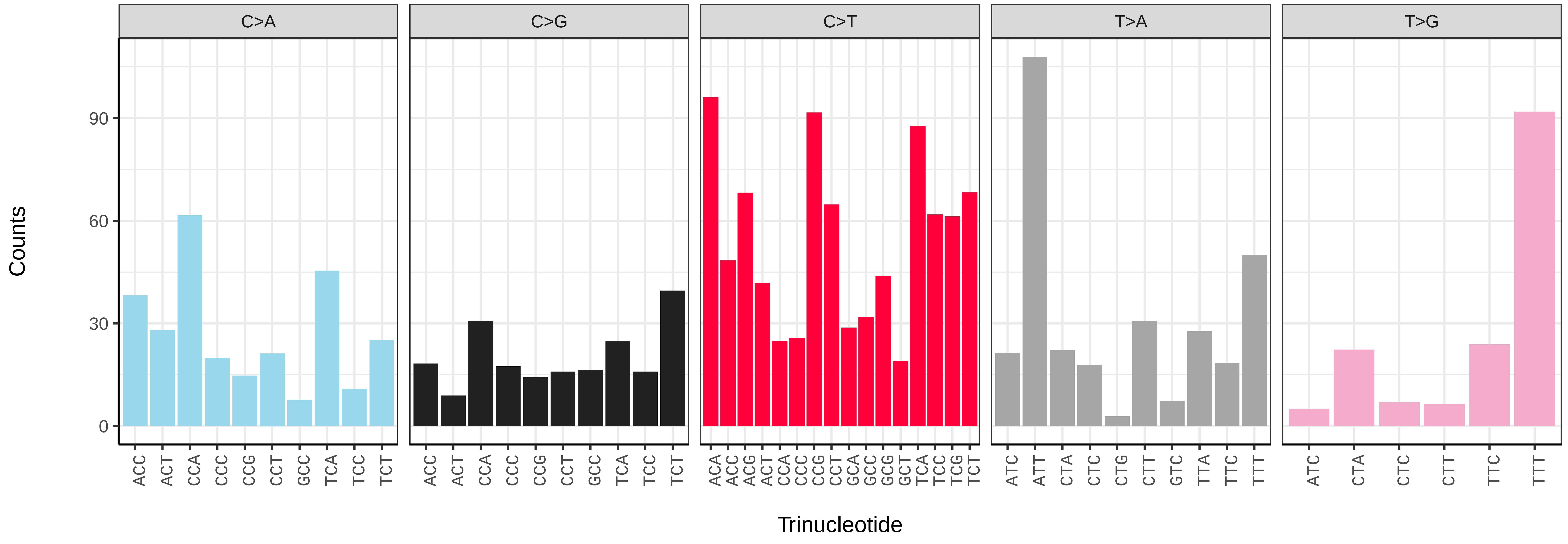

*Ophion slaviceki* (iyOphLute1) germline mutational spectrum

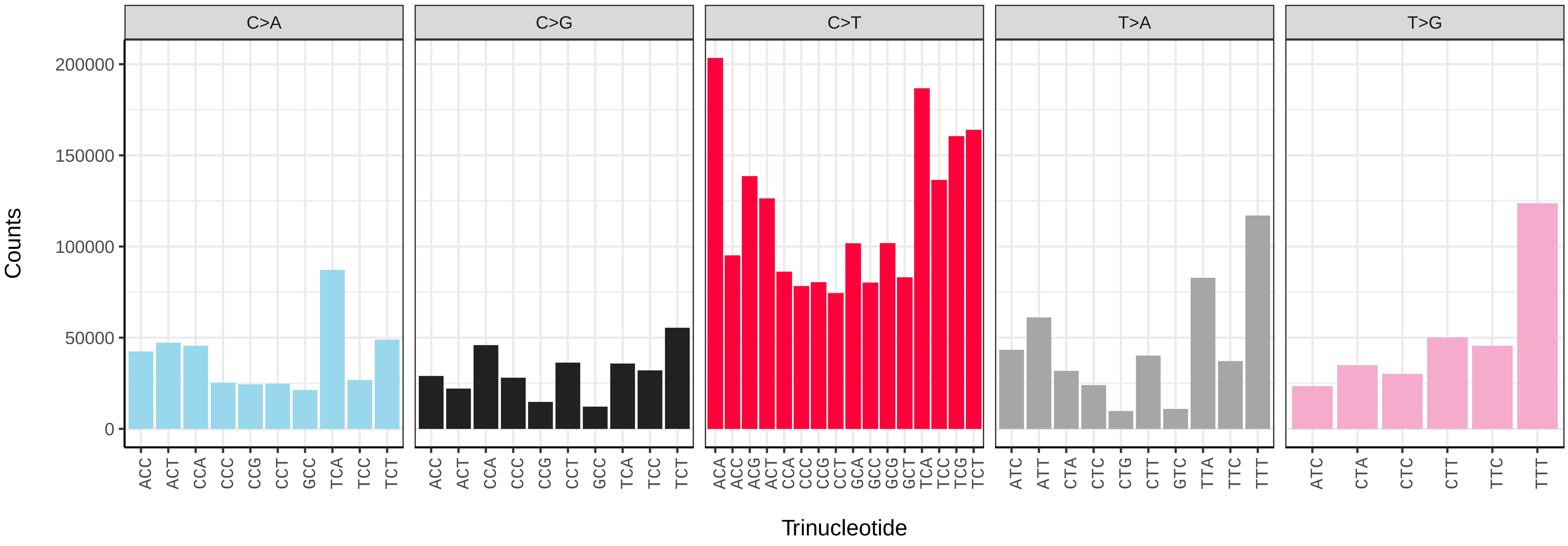


*Ophion slaviceki* (iyOphLute1) somatic mutational spectrum (SBS96 classification)

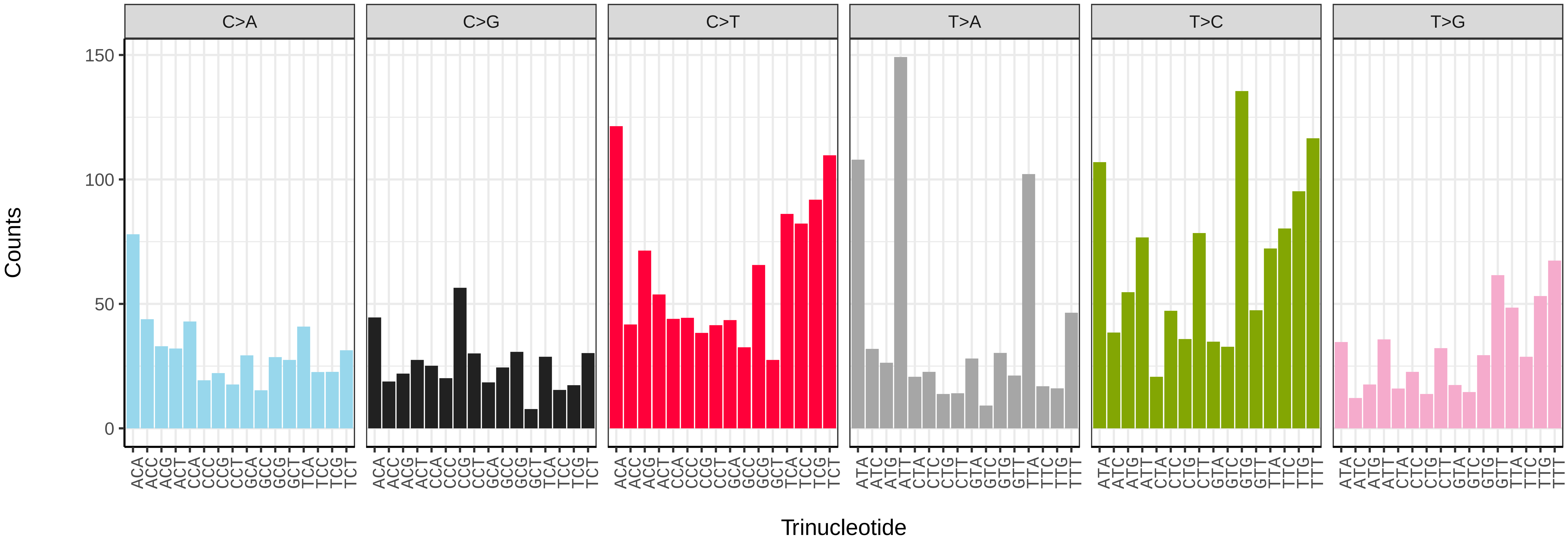


*Ophion slaviceki* (iyOphLute1) somatic mutational spectrum (SBS52 classification)

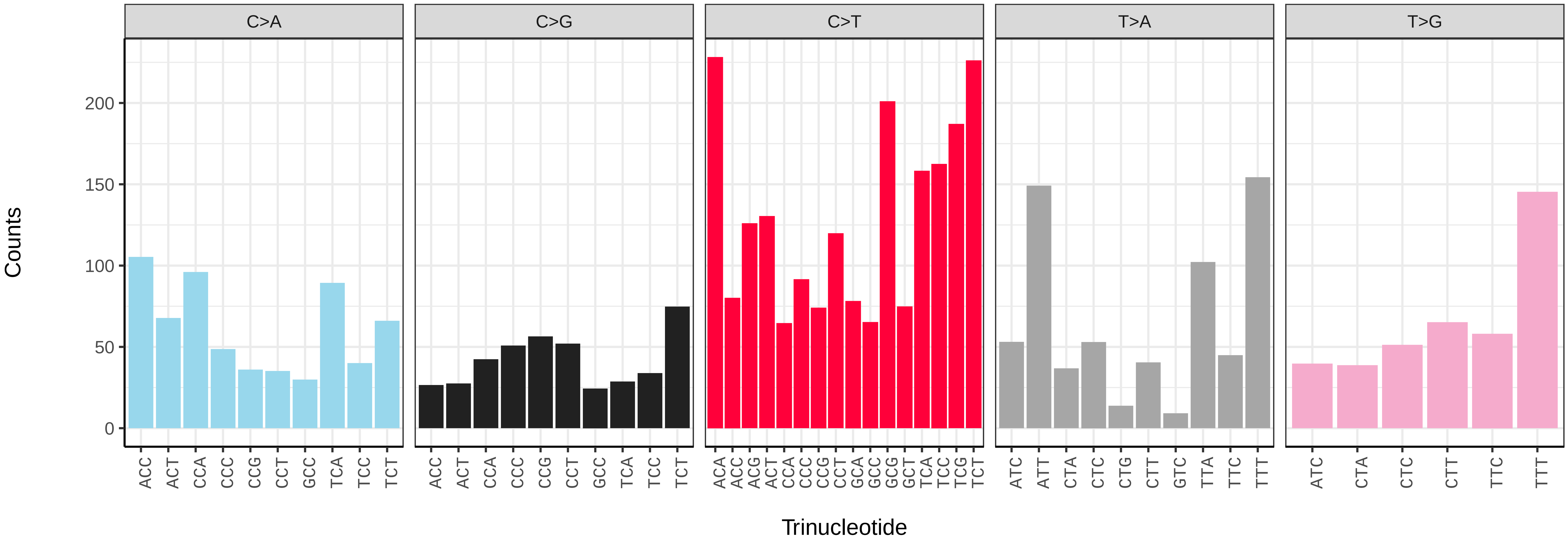

*Pemphredon lugubris* (iyPemLugu1) germline mutational spectrum

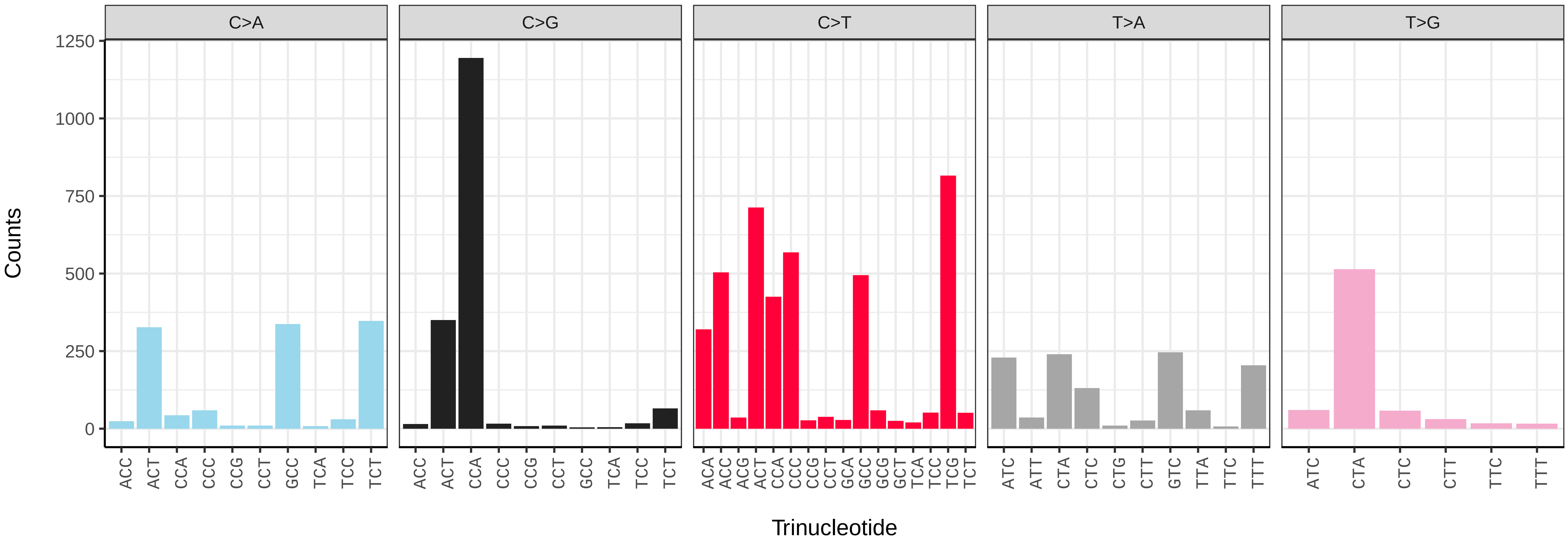


*Pemphredon lugubris* (iyPemLugu1) somatic mutational spectrum (SBS96 classification)

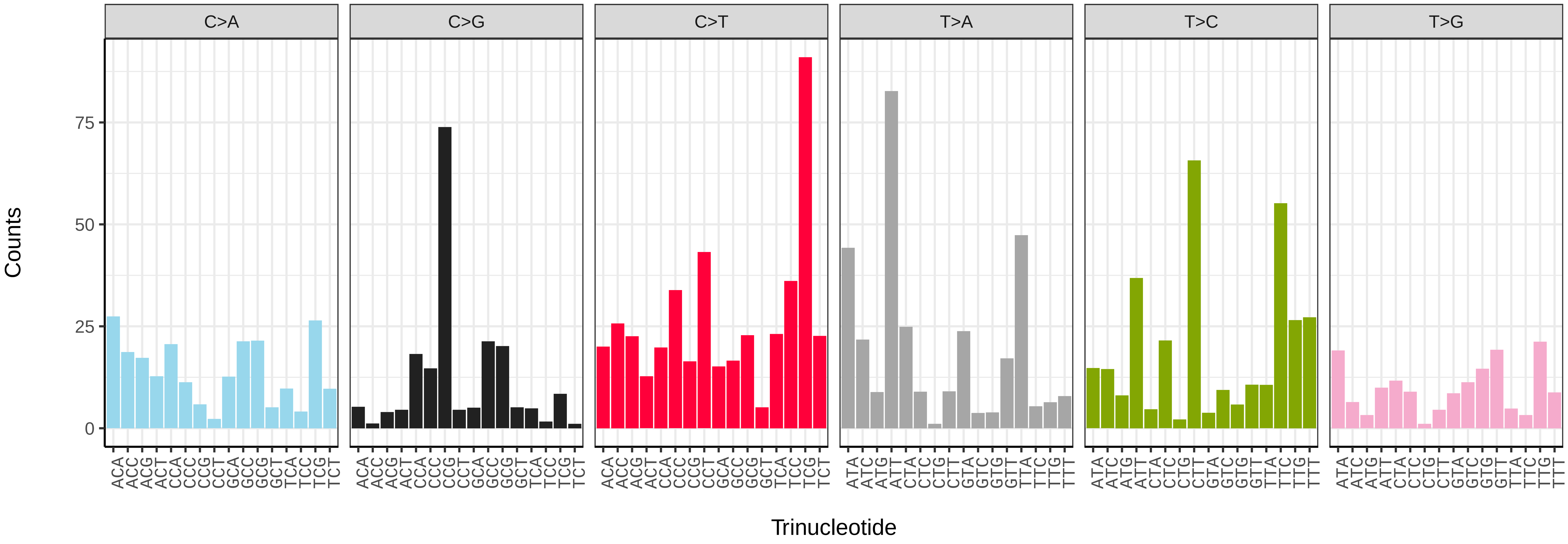


*Pemphredon lugubris* (iyPemLugu1) somatic mutational spectrum (SBS52 classification)

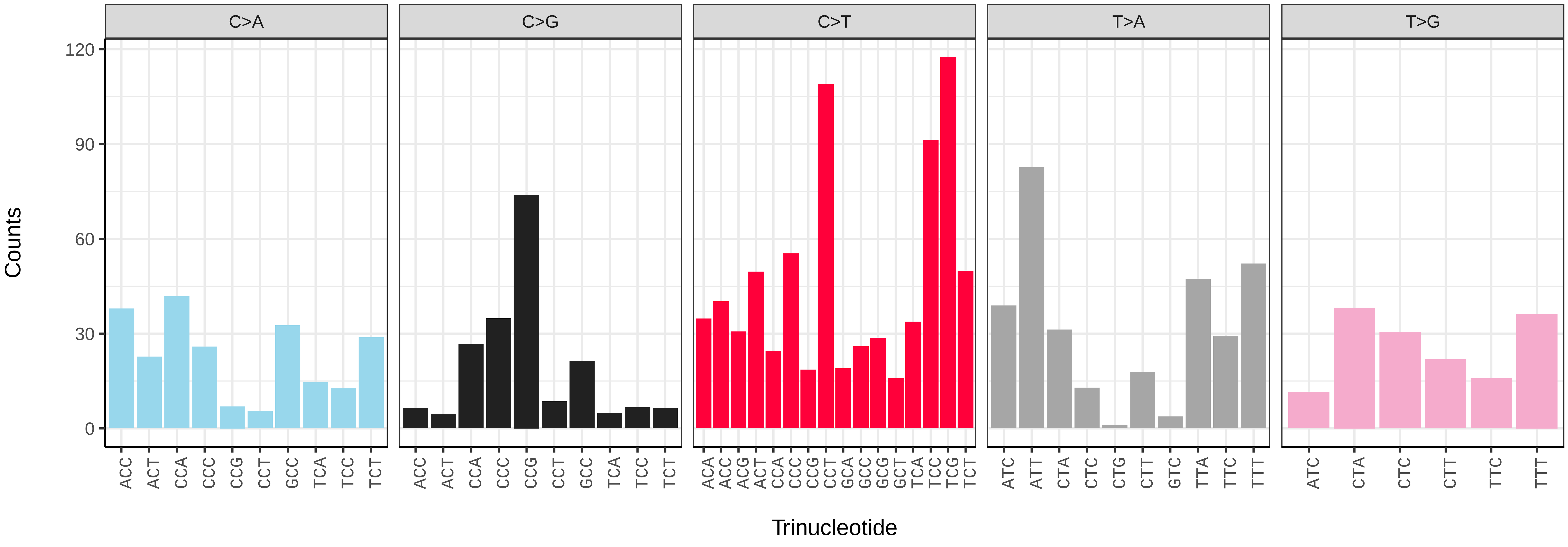

*Rhogogaster chlorosoma* (iyRhoChlo1) germline mutational spectrum

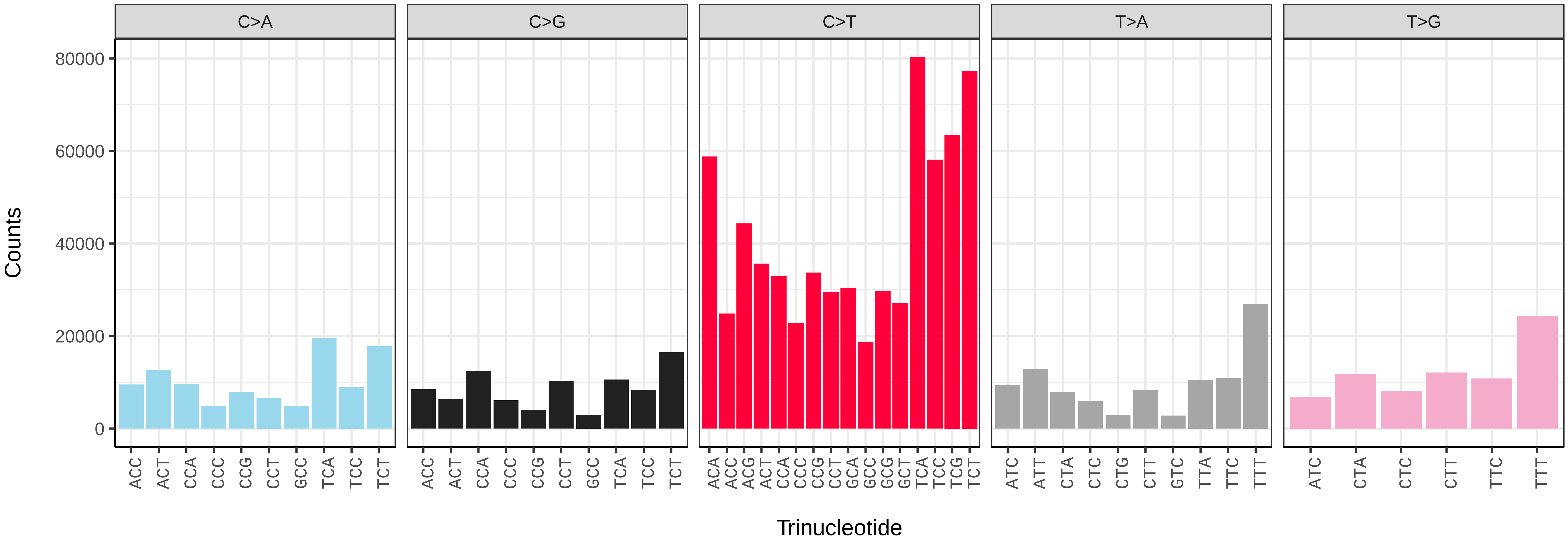


*Rhogogaster chlorosoma* (iyRhoChlo1) somatic mutational spectrum (SBS96 classification)

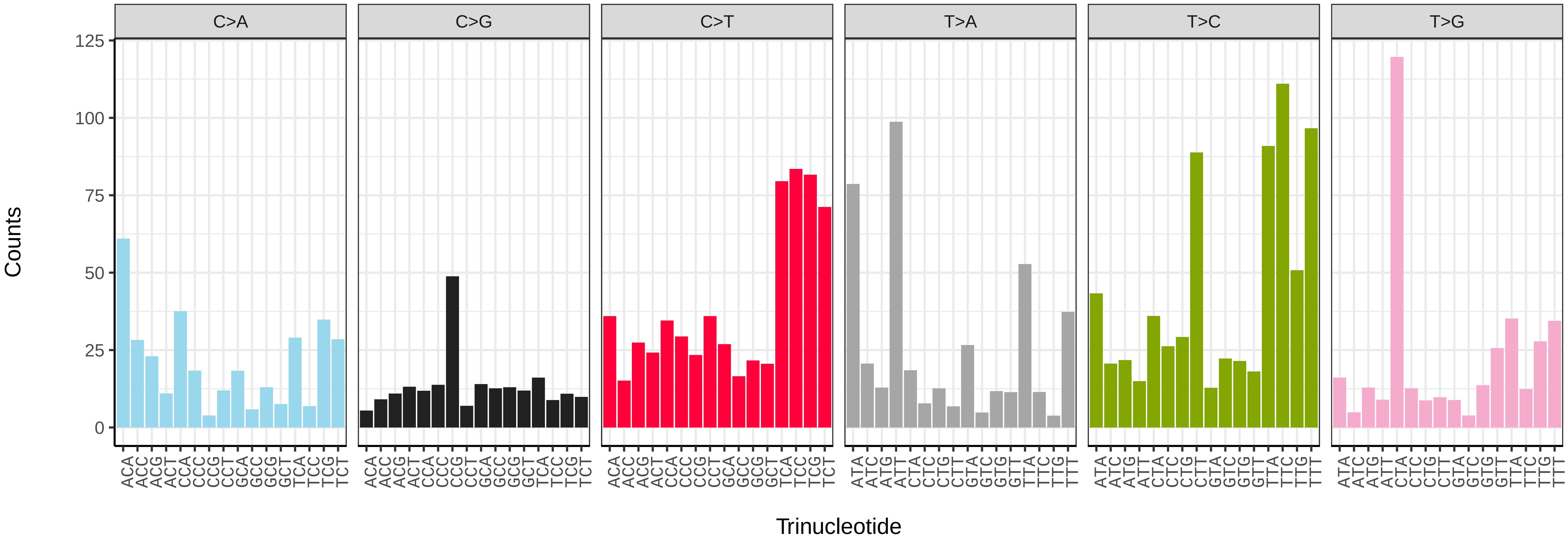


*Rhogogaster chlorosoma* (iyRhoChlo1) somatic mutational spectrum (SBS52 classification)

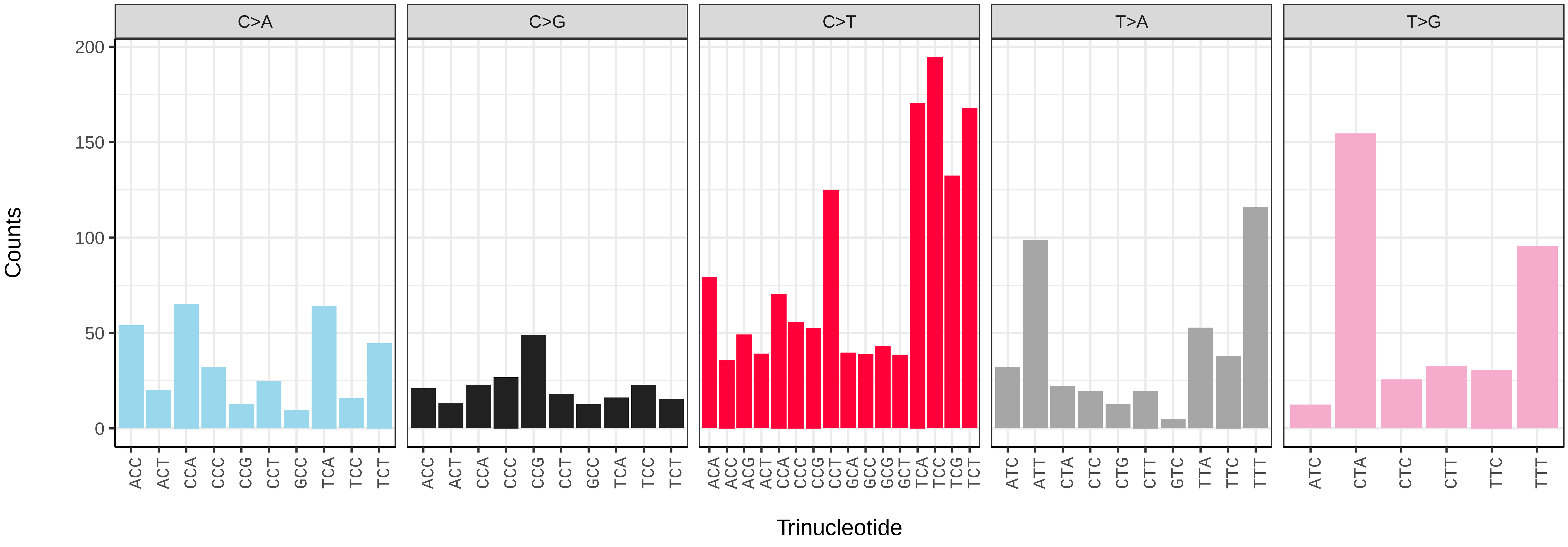

*Stelis phaeoptera* (iyStePhae1) germline mutational spectrum

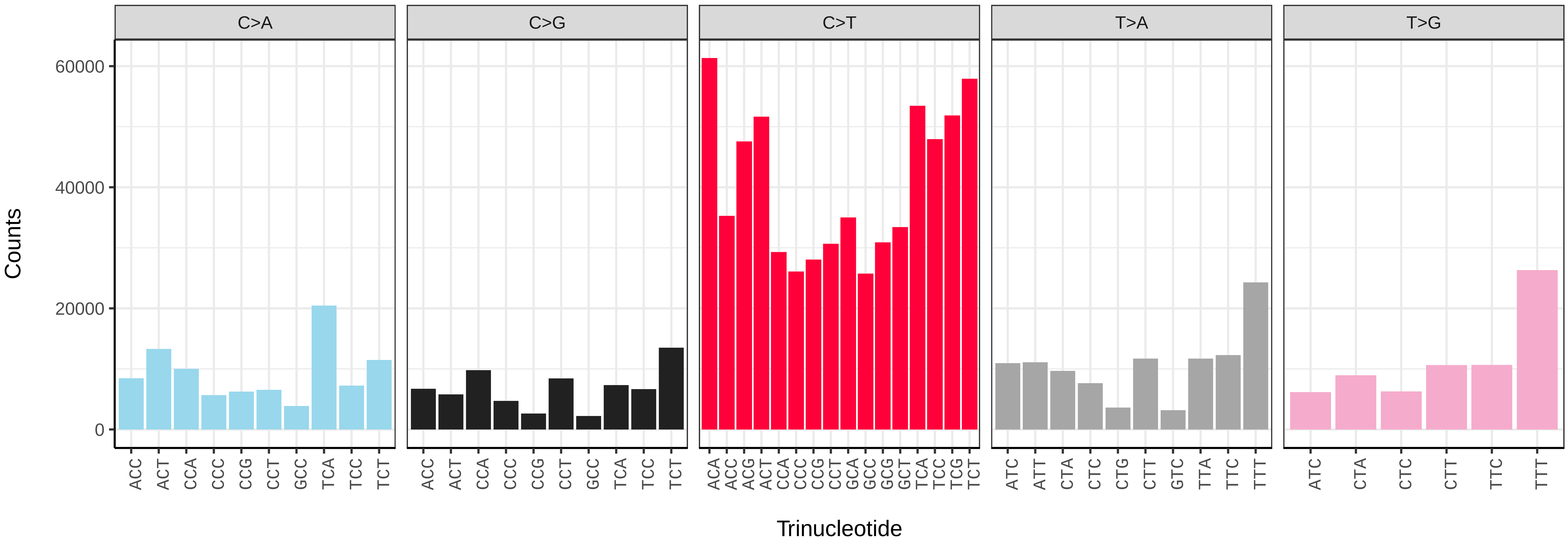


*Stelis phaeoptera* (iyStePhae1) somatic mutational spectrum (SBS96 classification)

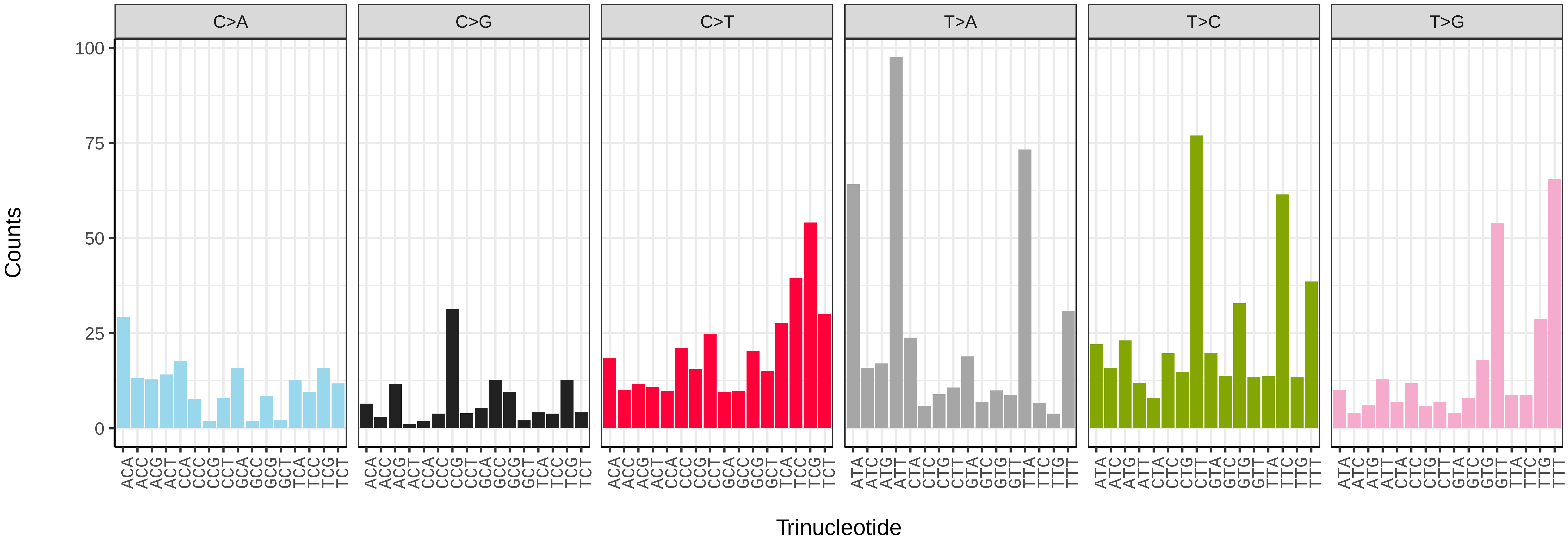


*Stelis phaeoptera* (iyStePhae1) somatic mutational spectrum (SBS52 classification)

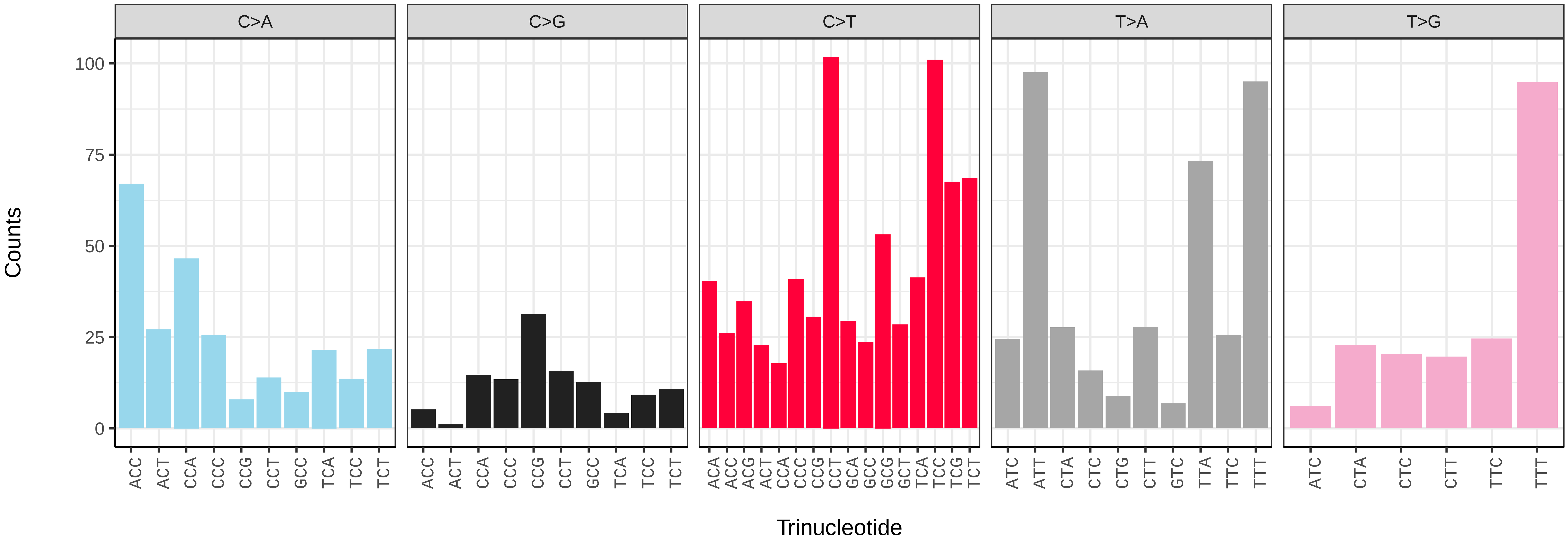

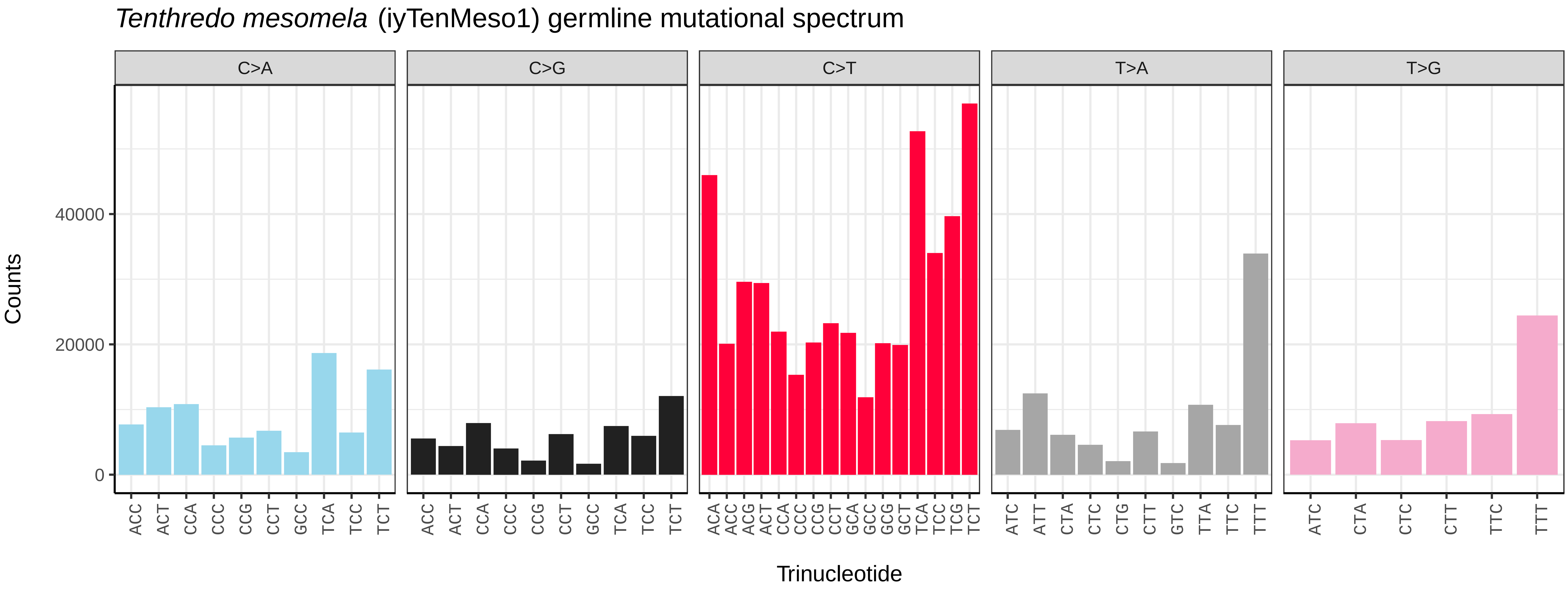
Tenthredo mesomela (iyTenMeso1) germline mutational spectrum
C>A
C>G
C>T
T>A
T>G
Counts
0
20000
40000
ACC ACT CCA CCC CCG CCT GCC TCA TCC TCT
ACA ACC ACG ACT CCA CCC CCG CCT GCA GCC GCG GCT TCA TCC TCG TCT
ATC ATT CTA CTC CTG CTT GTC TTA TTC TTT
ATC CTA CTC CTT TTC TTT
Trinucleotide

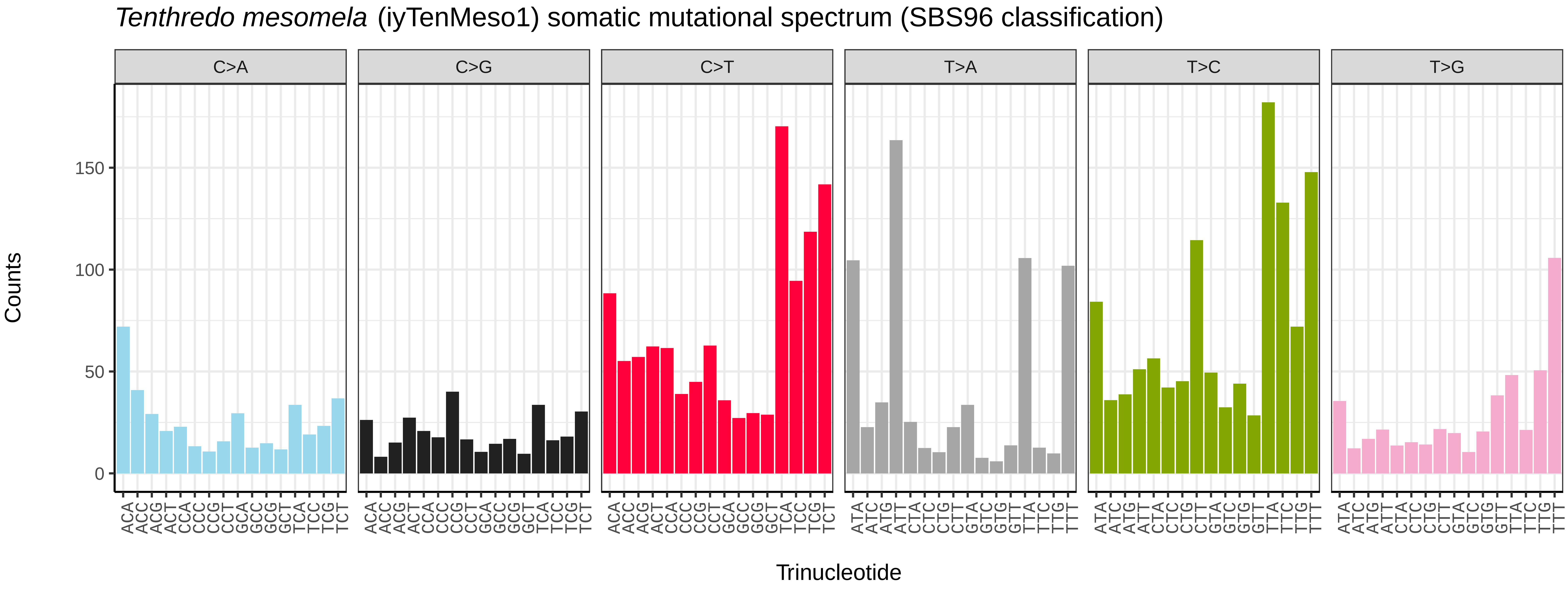
Tenthredo mesomela (iyTenMeso1) somatic mutational spectrum (SBS96 classification)
C>A
C>G
C>T
T>A
T>C
T>G
Counts
0
50
100
150
ACA ACC ACG ACT CCA CCC CCG CCT GCA GCC GCG GCT TCA TCC TCG TCT
ATA ATC ATG ATT CTA CTC CTG CTT GTA GTC GTG GTT TTA TTC TTG TTT
Trinucleotide

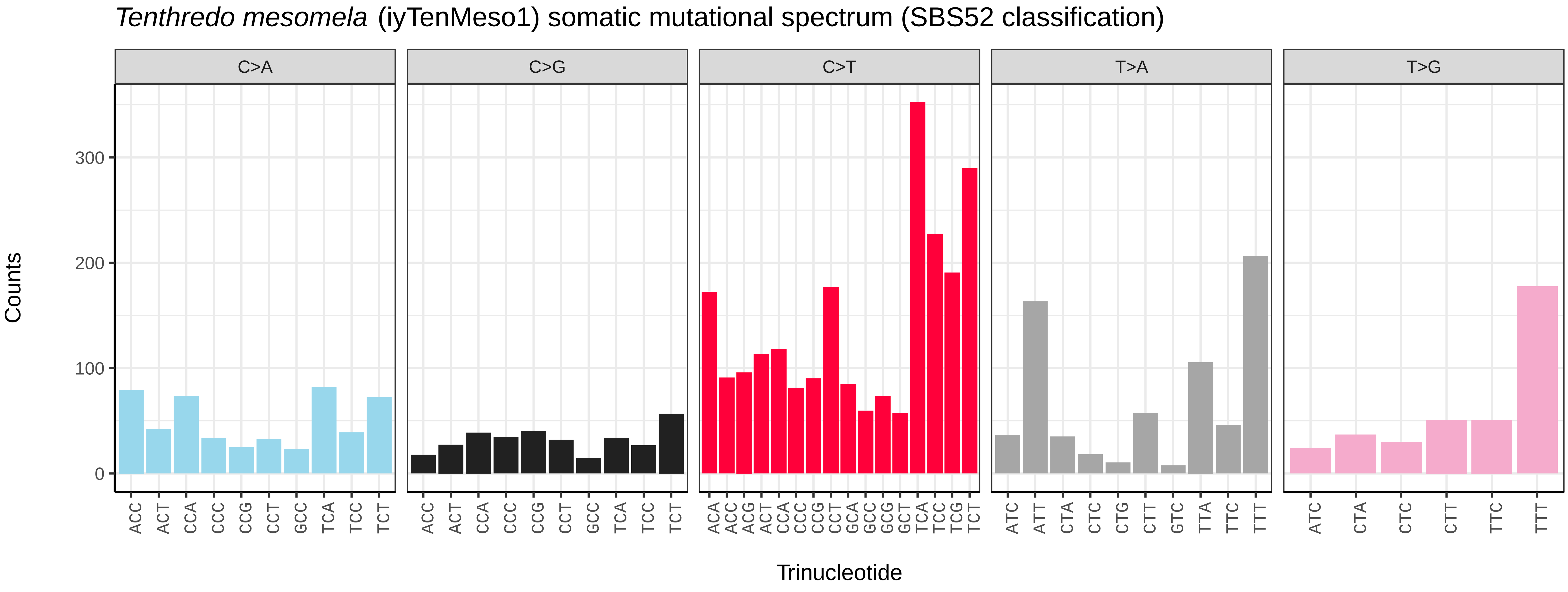
Tenthredo mesomela (iyTenMeso1) somatic mutational spectrum (SBS52 classification)
C>A
C>G
C>T
T>A
T>G
Counts
0
100
200
300
ACC ACT CCA CCC CCG CCT GCC TCA TCC TCT
ACA ACC ACG ACT CCA CCC CCG CCT GCA GCC GCG GCT TCA TCC TCG TCT
ATC ATT CTA CTC CTG CTT GTC TTA TTC TTT
ATC CTA CTC CTT TTC TTT
Trinucleotide

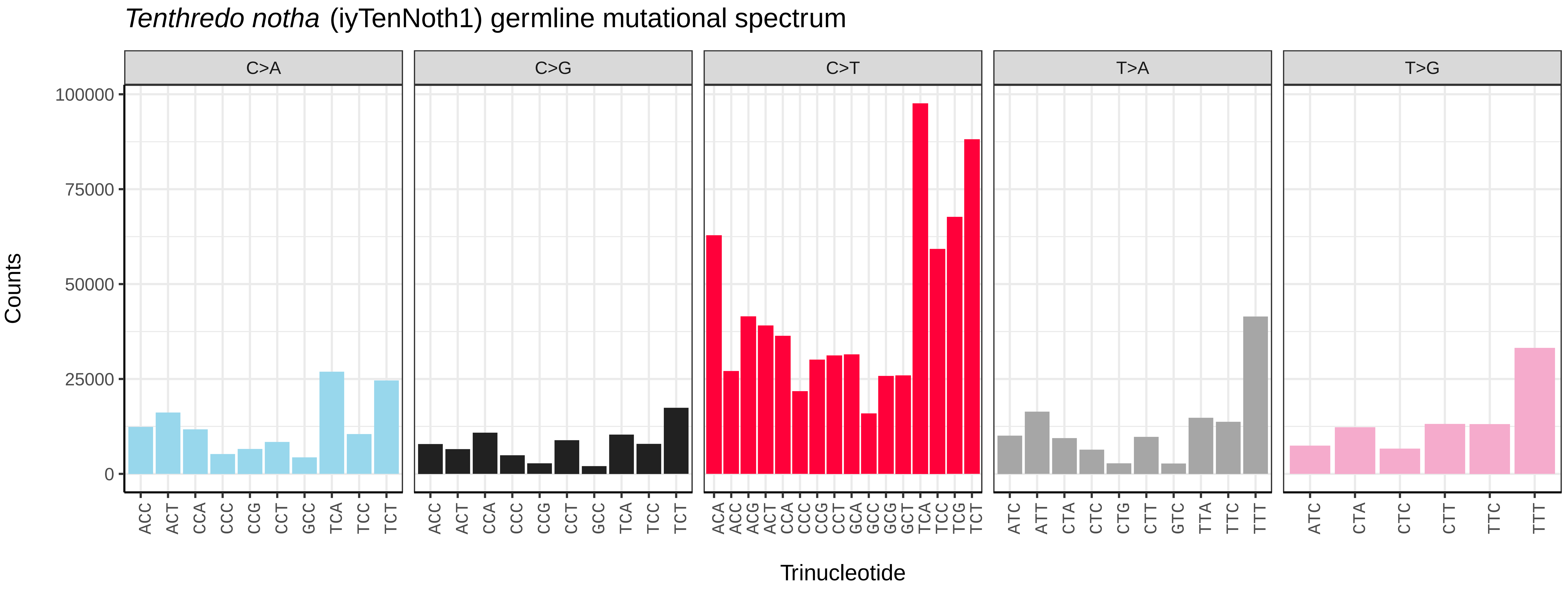
*Tenthredo notha* (iyTenNoth1) germline mutational spectrum
C>A
C>G
C>T
T>A
T>G
Counts
0
25000
50000
75000
100000
ACC ACT CCA CCC CCG CCT GCC TCA TCC TCT
ACA ACC ACG ACT CCA CCC CCG CCT GCA GCC GCG GCT TCA TCC TCG TCT
ATC ATT CTA CTC CTG CTT GTC TTA TTC TTT
ATC CTA CTC CTT TTC TTT
Trinucleotide

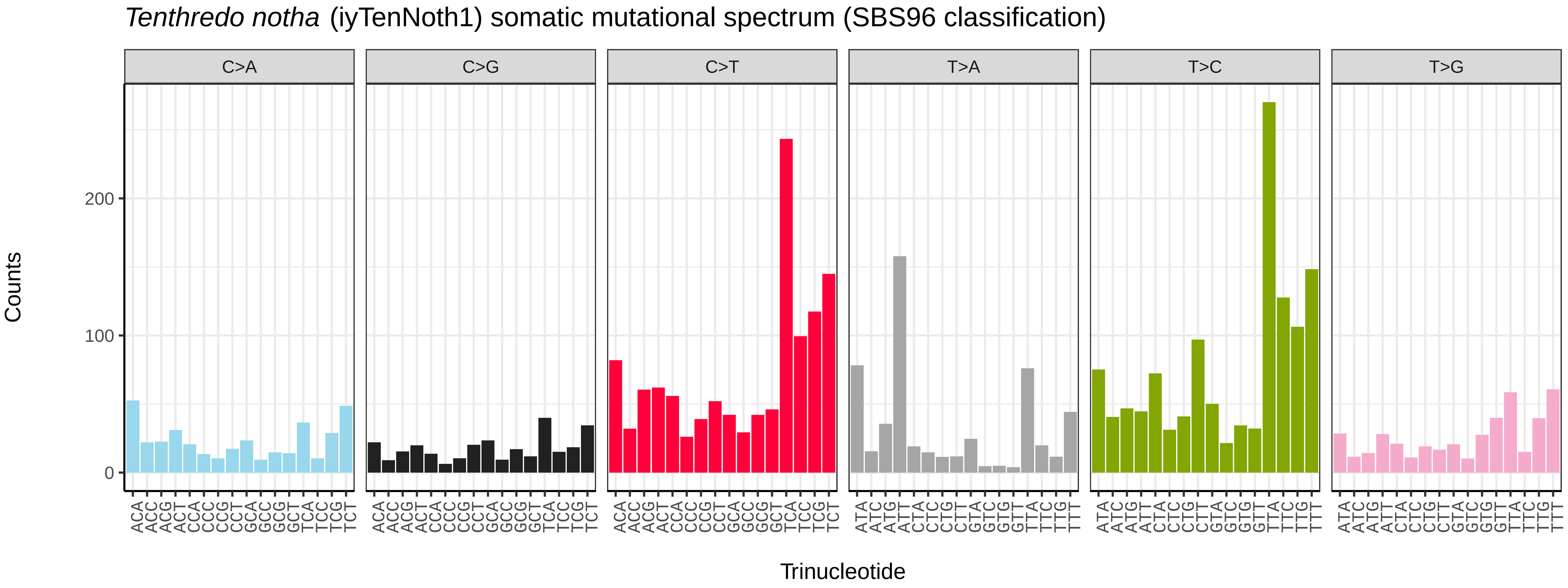
*Tenthredo notha* (iyTenNoth1) somatic mutational spectrum (SBS96 classification)
C>A
C>G
C>T
T>A
T>C
T>G
Counts
0
100
200
ACA ACC ACG ACT CCA CCC CCG CCT GCA GCC GCG GCT TCA TCC TCG TCT
ATA ATC ATG ATT CTA CTC CTG CTT GTA GTC GTG GTT TTA TTC TTG TTT
Trinucleotide

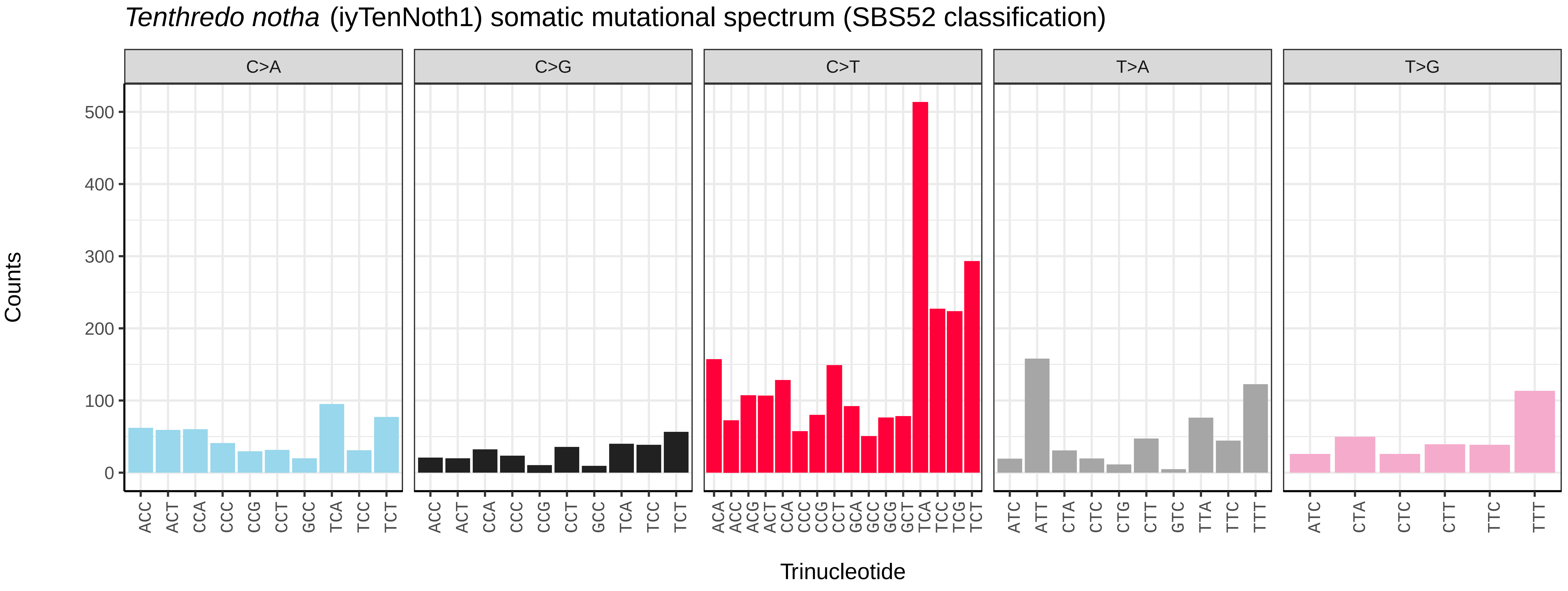
*Tenthredo notha* (iyTenNoth1) somatic mutational spectrum (SBS52 classification)
C>A
C>G
C>T
T>A
T>G
Counts
0
100
200
300
400
500
ACC ACT CCA CCC CCG CCT GCC TCA TCC TCT
ACA ACC ACG ACT CCA CCC CCG CCT GCA GCC GCG GCT TCA TCC TCG TCT
ATC ATT CTA CTC CTG CTT GTC TTA TTC TTT
ATC CTA CTC CTT TTC TTT
Trinucleotide

*Tiphia femorata* (iyTipFemo1) germline mutational spectrum

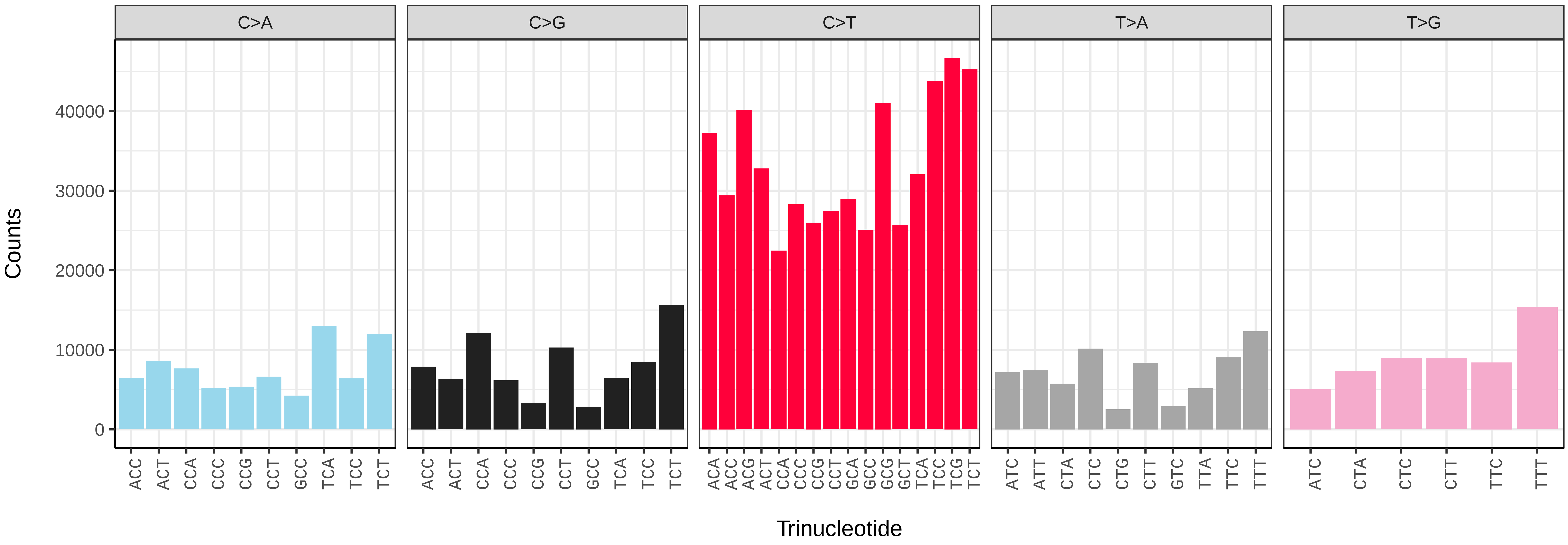


*Tiphia femorata* (iyTipFemo1) somatic mutational spectrum (SBS96 classification)

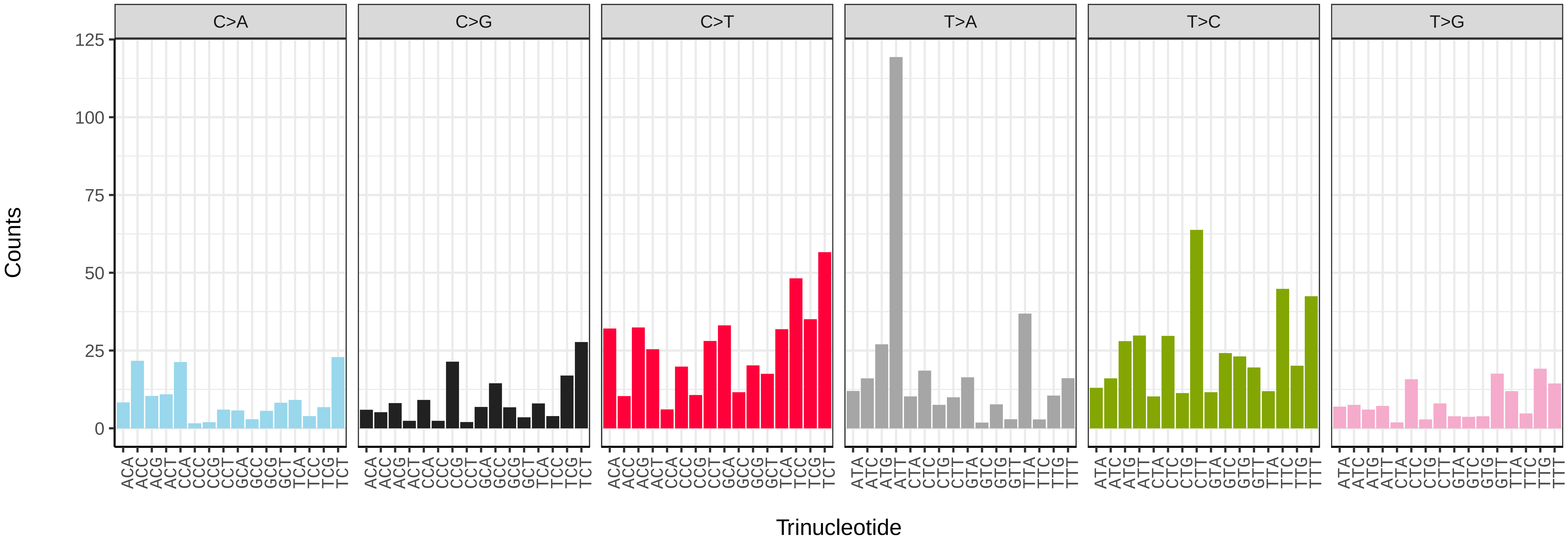


*Tiphia femorata* (iyTipFemo1) somatic mutational spectrum (SBS52 classification)

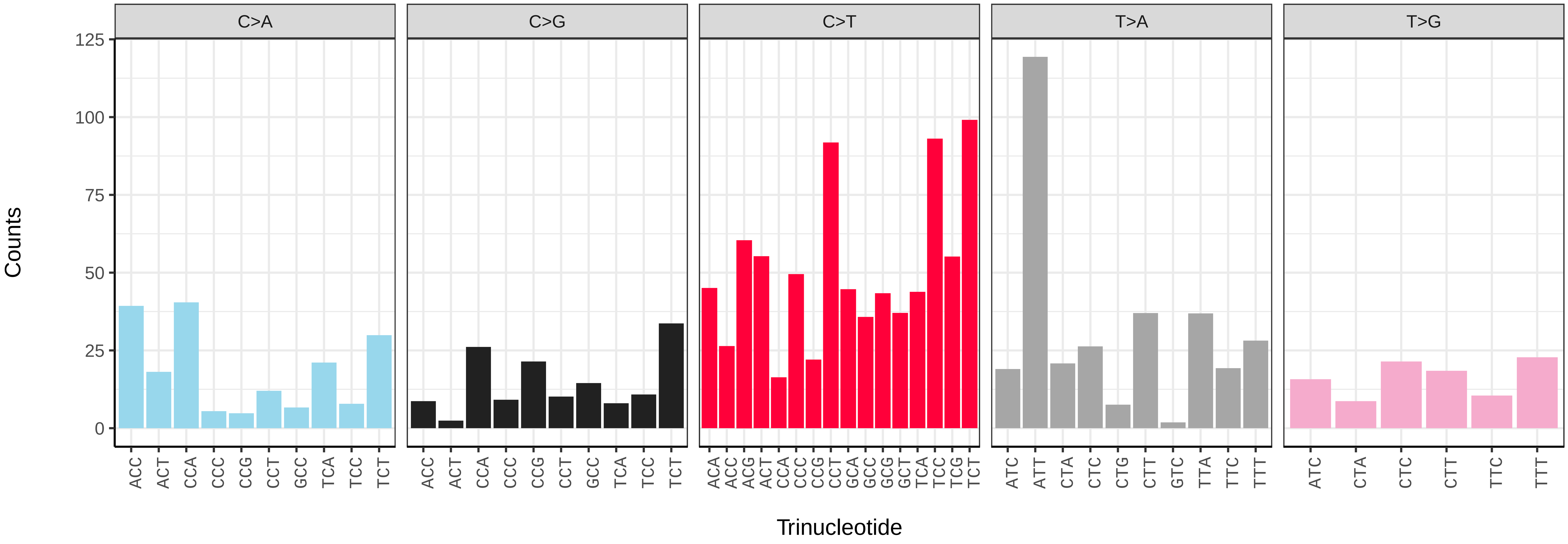

*Vespa crabro* (iyVesCrab1) germline mutational spectrum

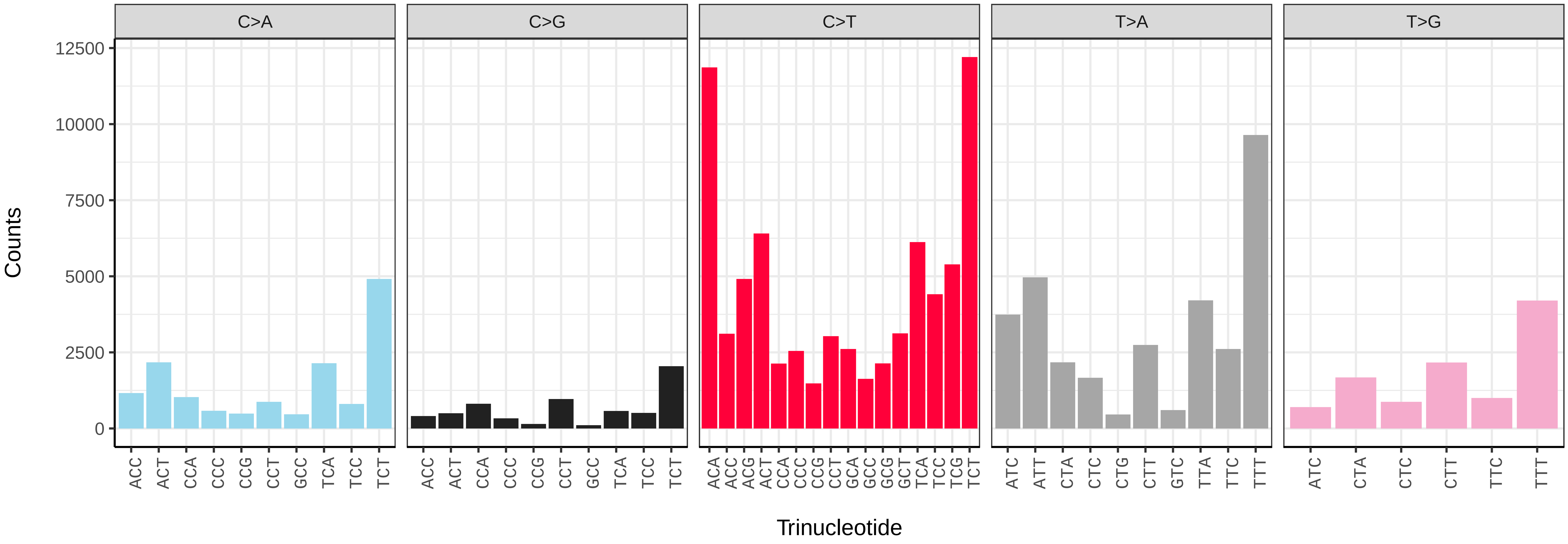


*Vespa crabro* (iyVesCrab1) somatic mutational spectrum (SBS96 classification)

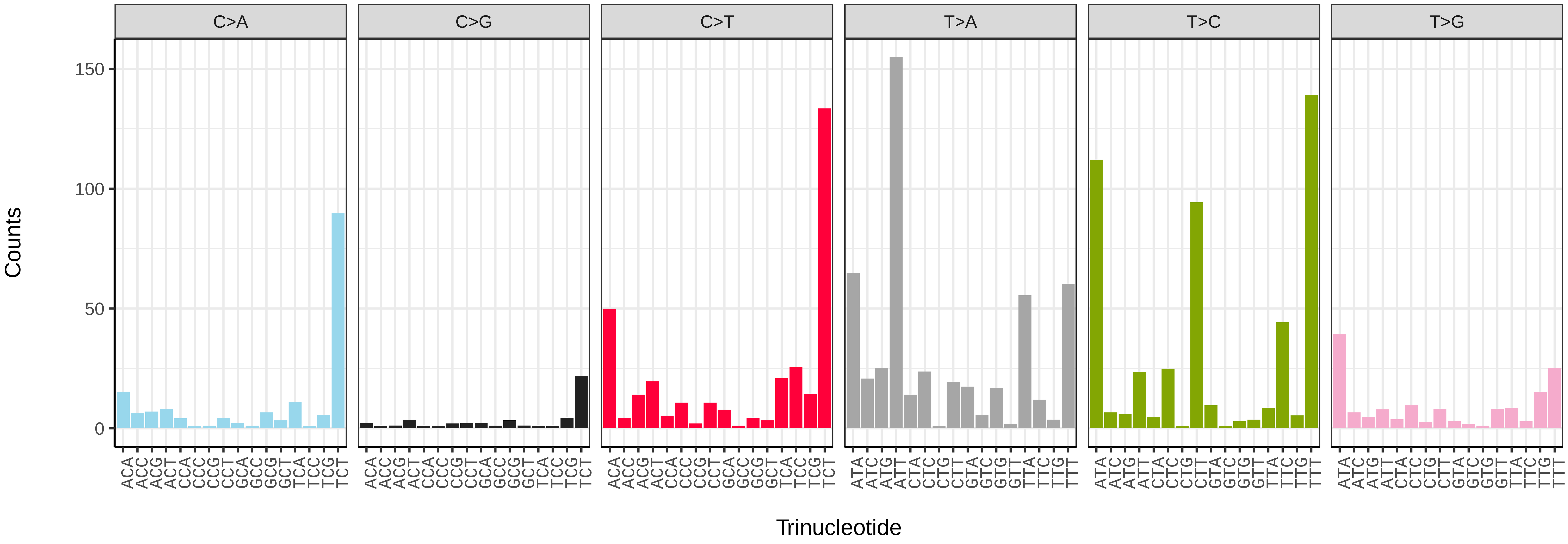


*Vespa crabro* (iyVesCrab1) somatic mutational spectrum (SBS52 classification)

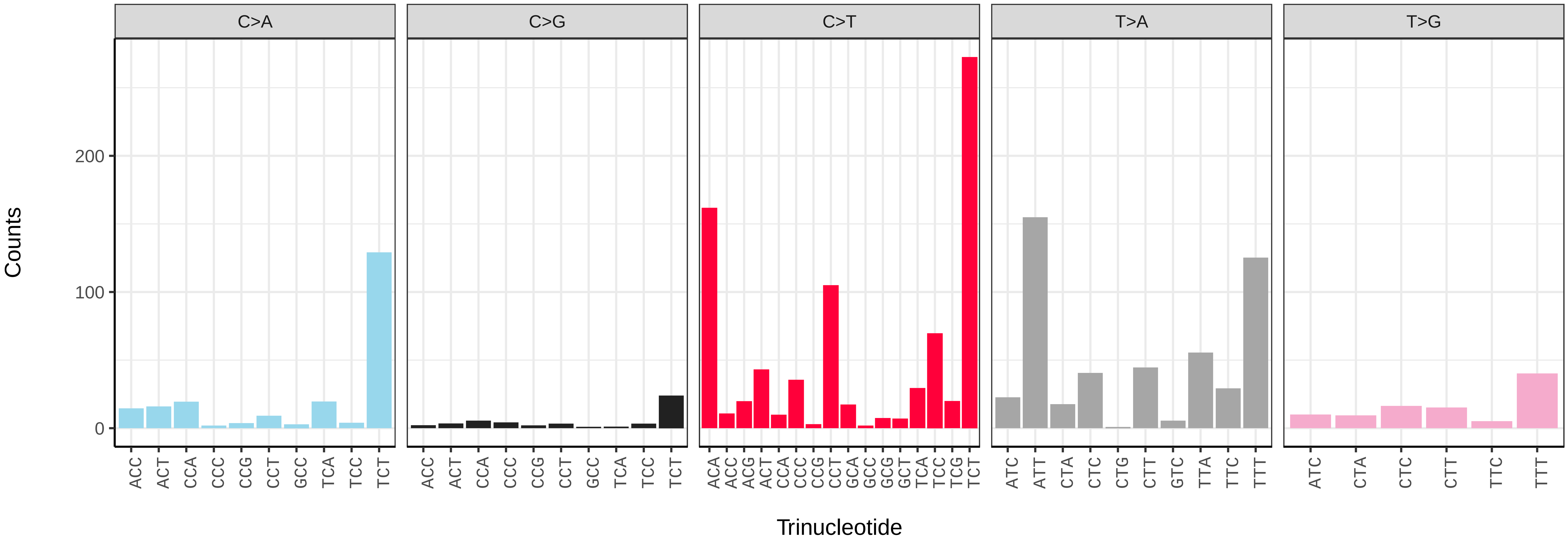

*Vespula germanica* (iyVesGerm1) germline mutational spectrum

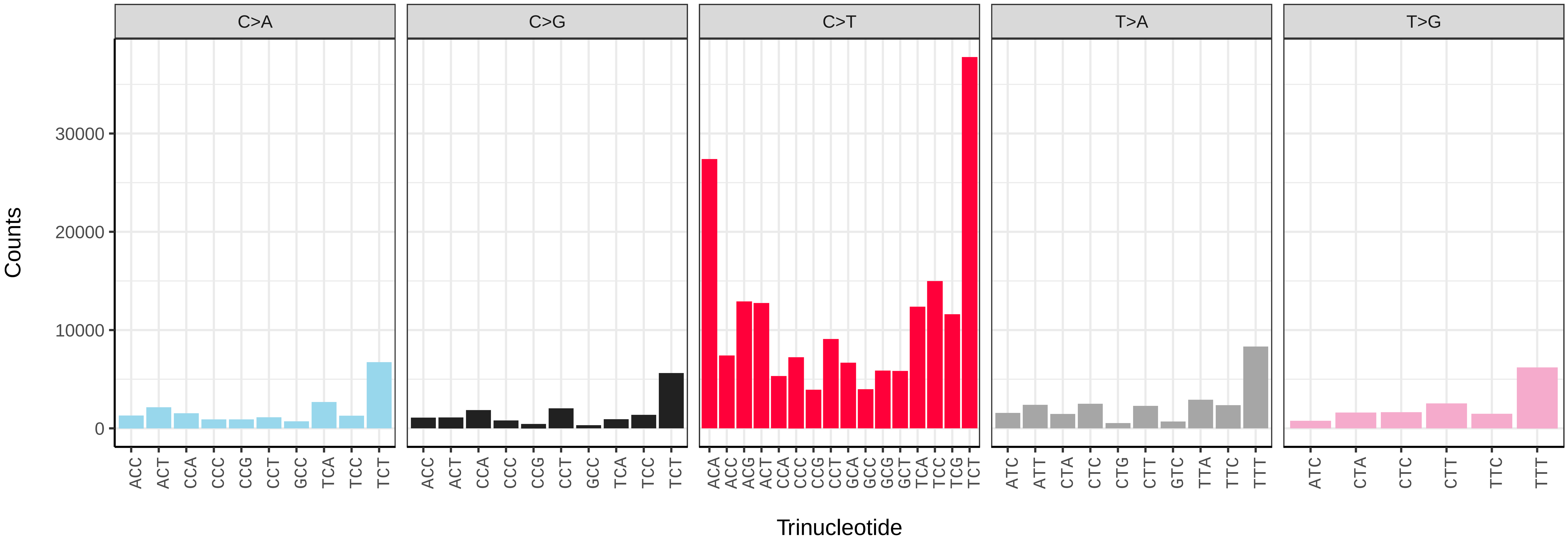


*Vespula germanica* (iyVesGerm1) somatic mutational spectrum (SBS96 classification)

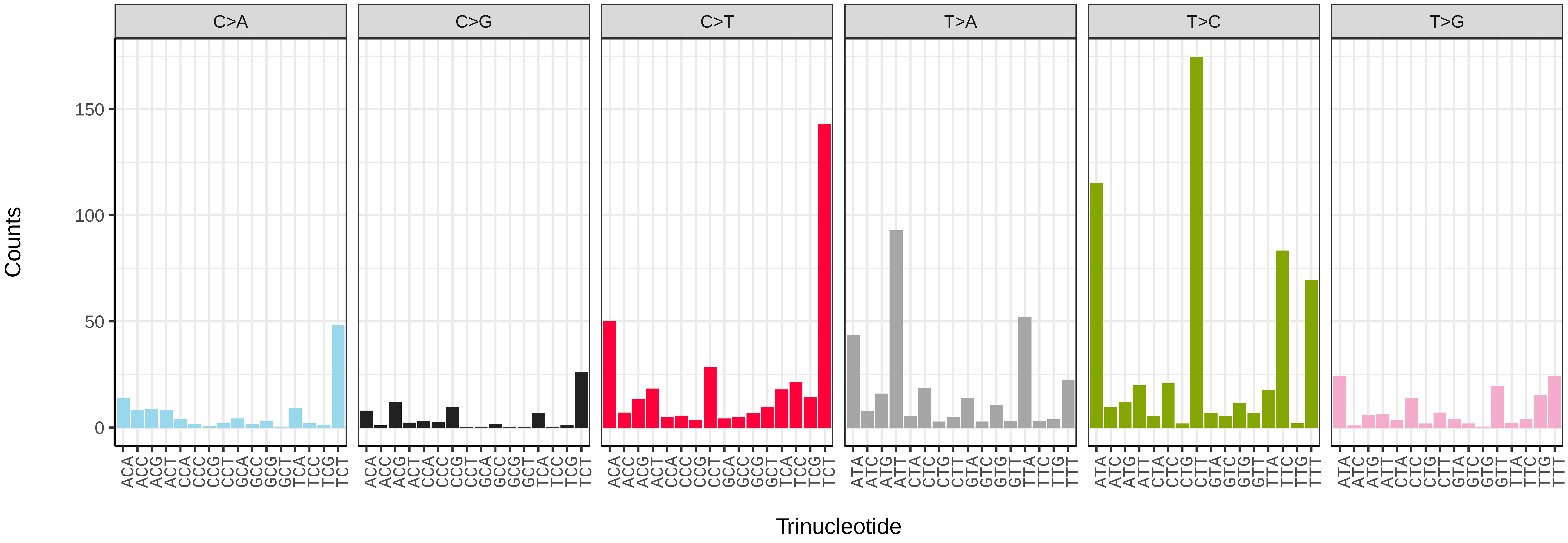


*Vespula germanica* (iyVesGerm1) somatic mutational spectrum (SBS52 classification)

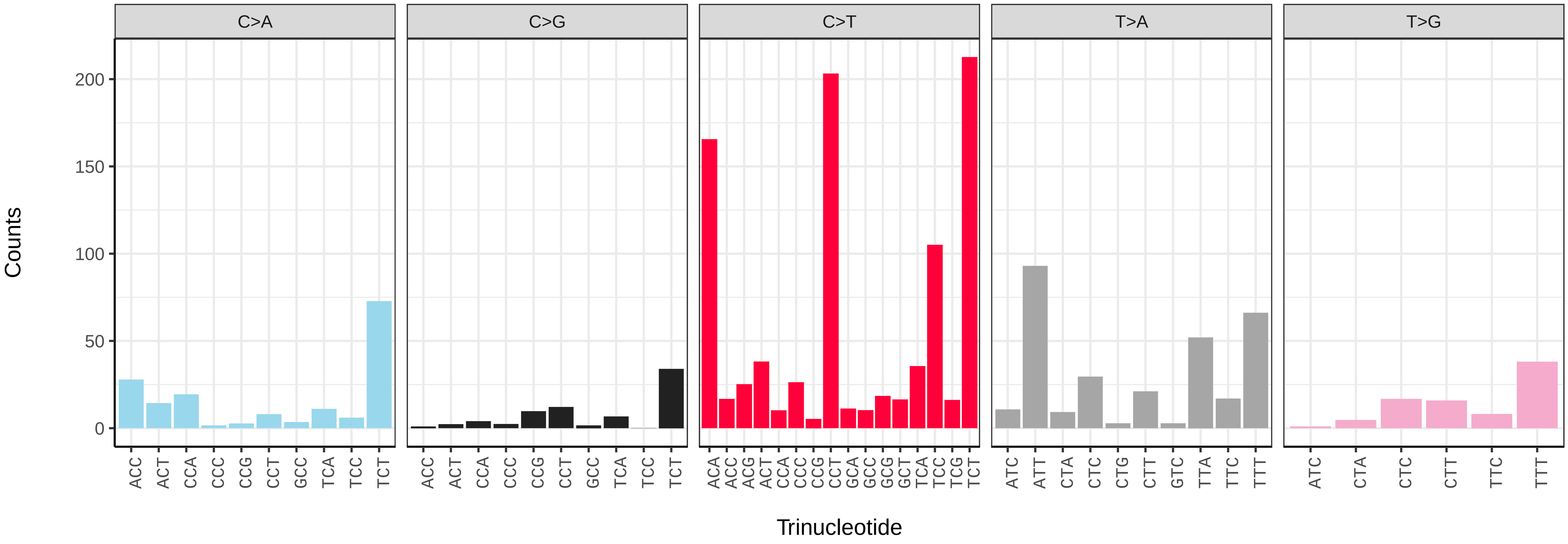

*Vespula vulgaris* (iyVesVulg1) germline mutational spectrum

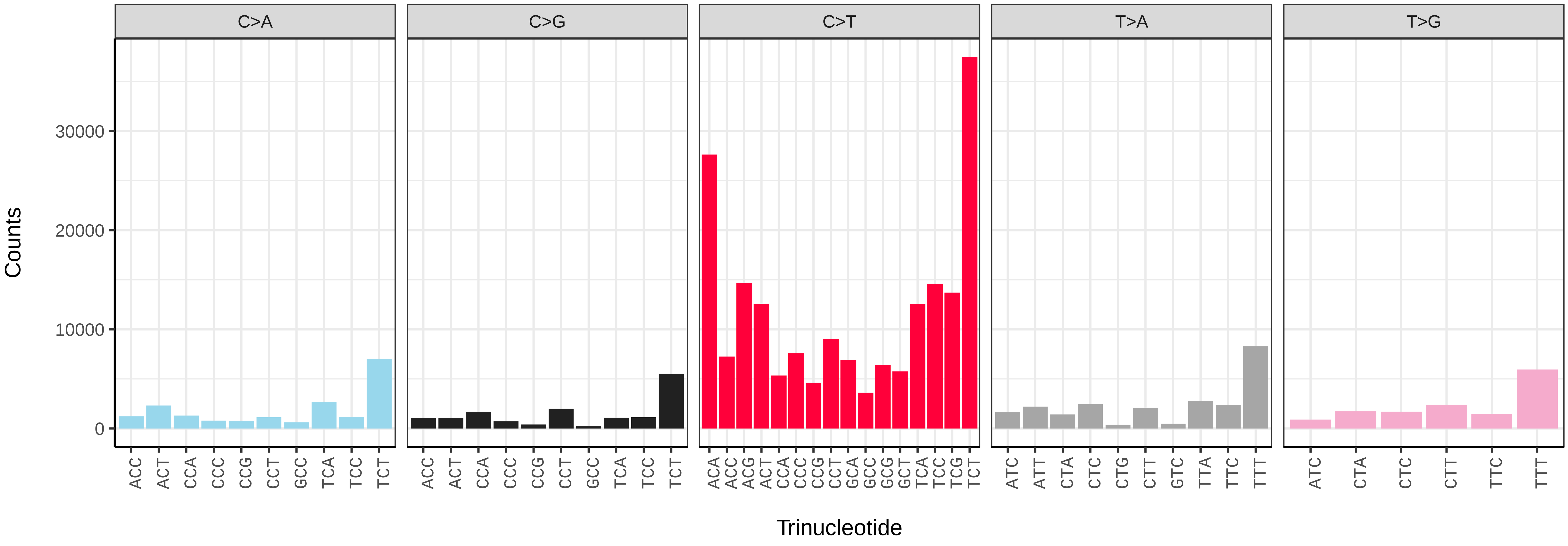


*Vespula vulgaris* (iyVesVulg1) somatic mutational spectrum (SBS96 classification)

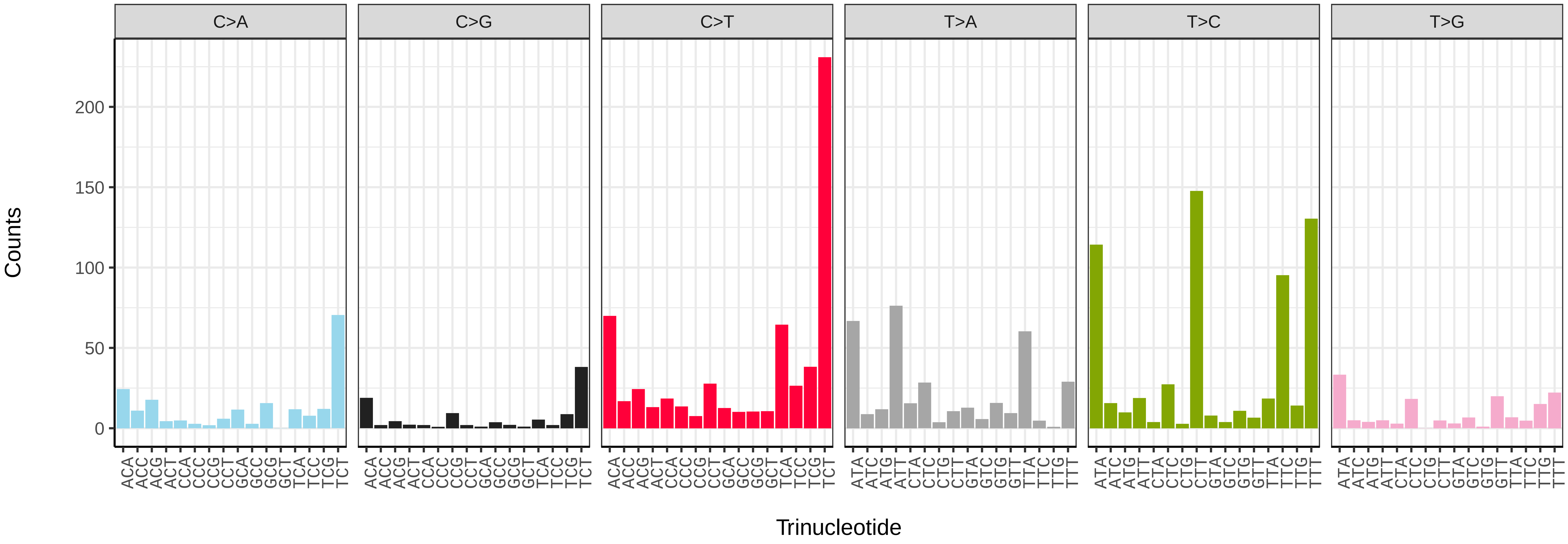


*Vespula vulgaris* (iyVesVulg1) somatic mutational spectrum (SBS52 classification)

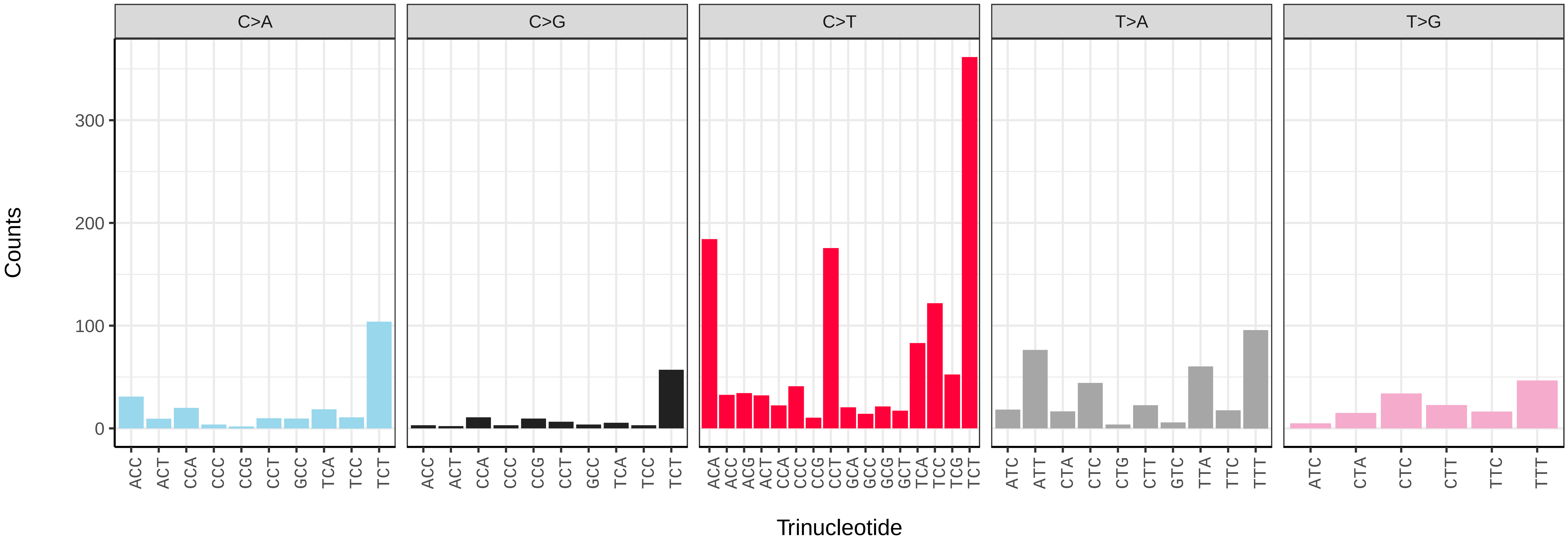

*Vespula vulgaris* (iyVesVulg1.iyVesVulg2) germline mutational spectrum

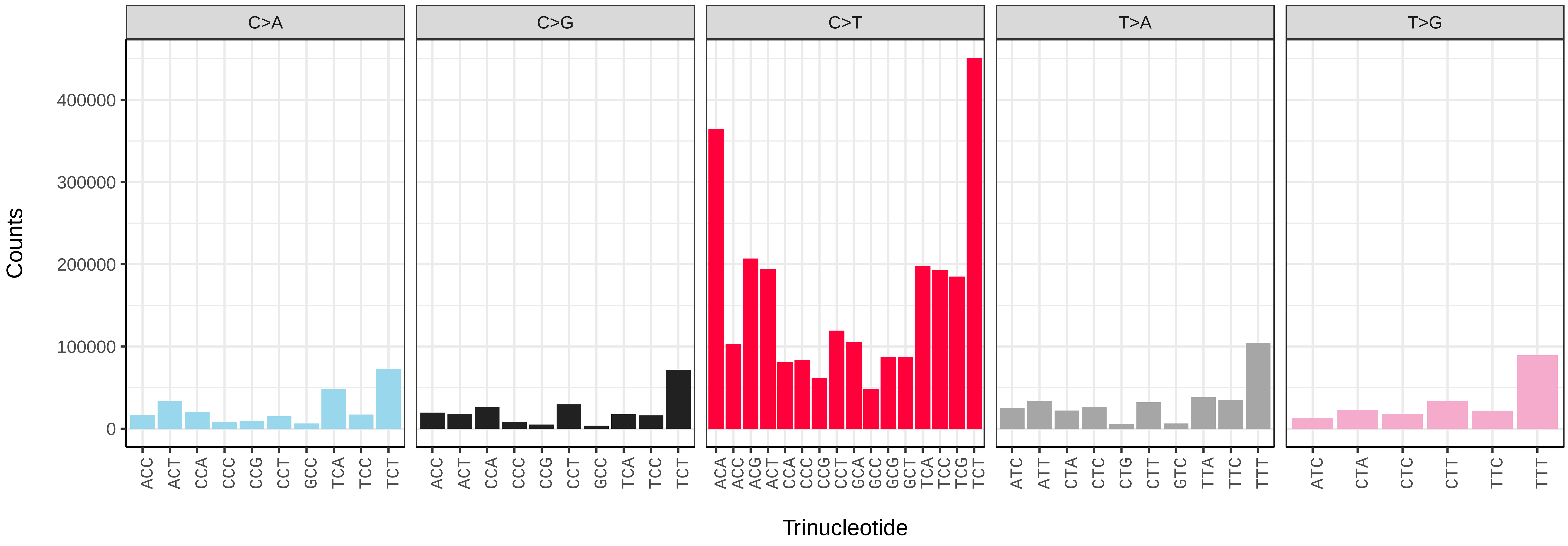


*Vespula vulgaris* (iyVesVulg1.iyVesVulg2) somatic mutational spectrum (SBS96 classification)

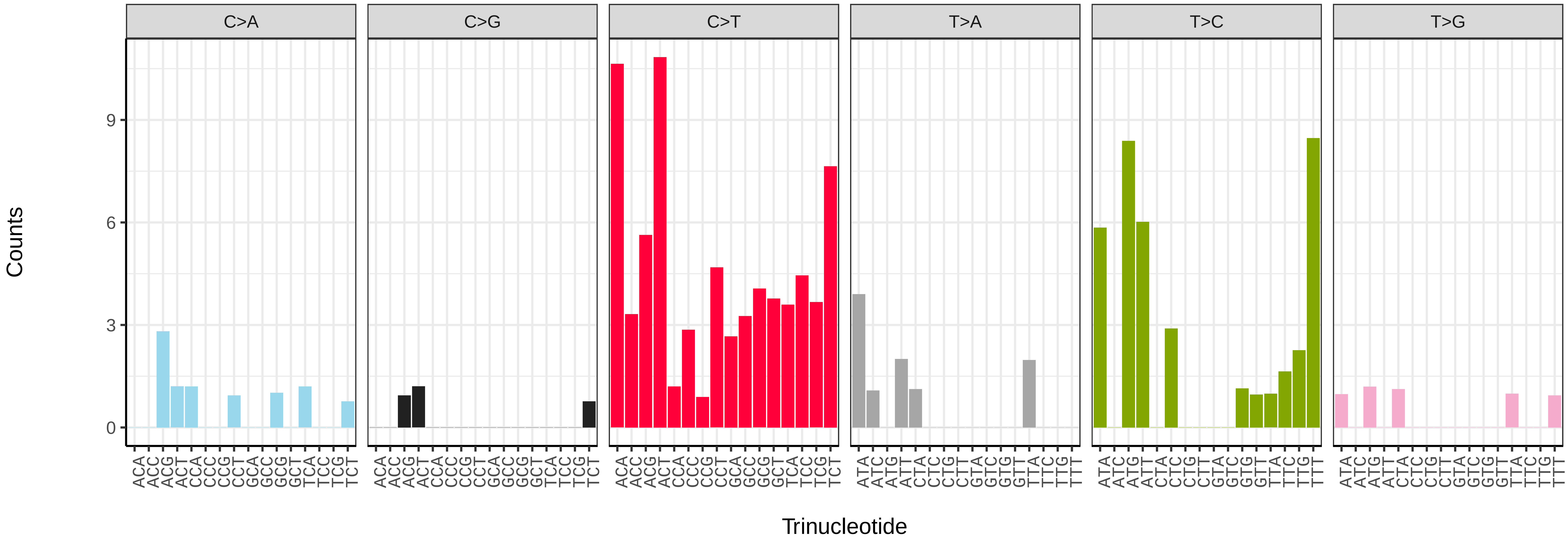


*Vespula vulgaris* (iyVesVulg1.iyVesVulg2) somatic mutational spectrum (SBS52 classification)

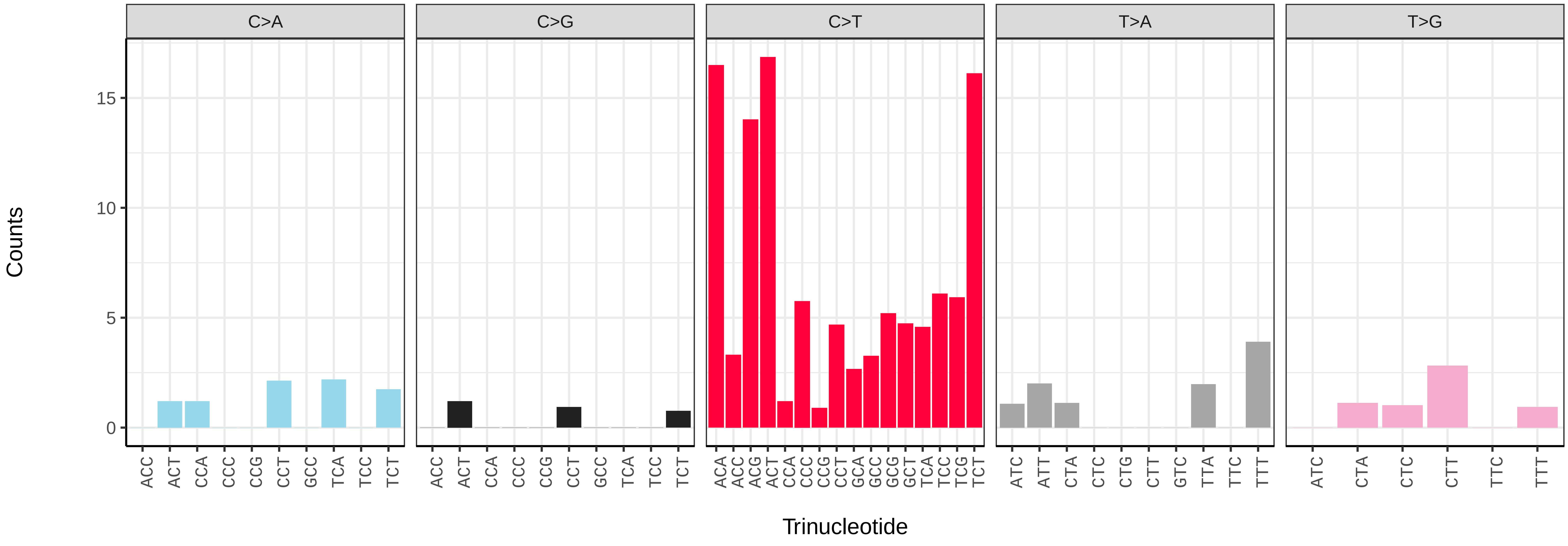

*Vespula vulgaris* (iyVesVulg1.iyVesVulg3) germline mutational spectrum

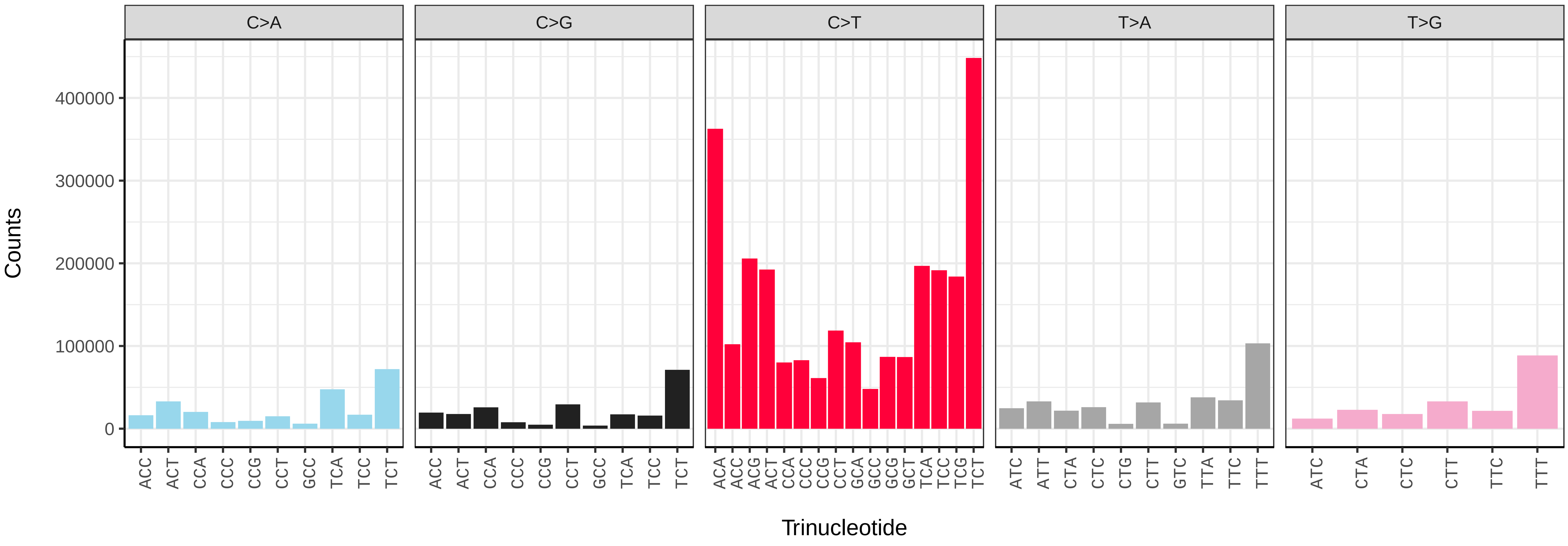


*Vespula vulgaris* (iyVesVulg1.iyVesVulg3) somatic mutational spectrum (SBS96 classification)

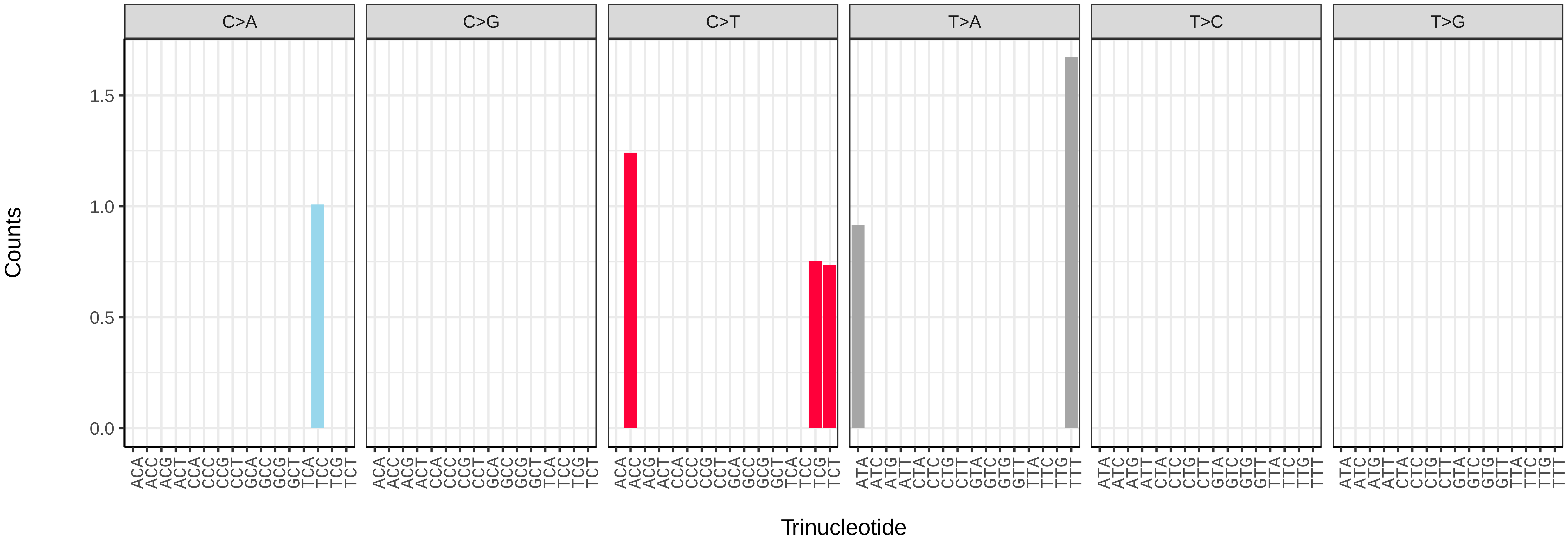


*Vespula vulgaris* (iyVesVulg1.iyVesVulg3) somatic mutational spectrum (SBS52 classification)

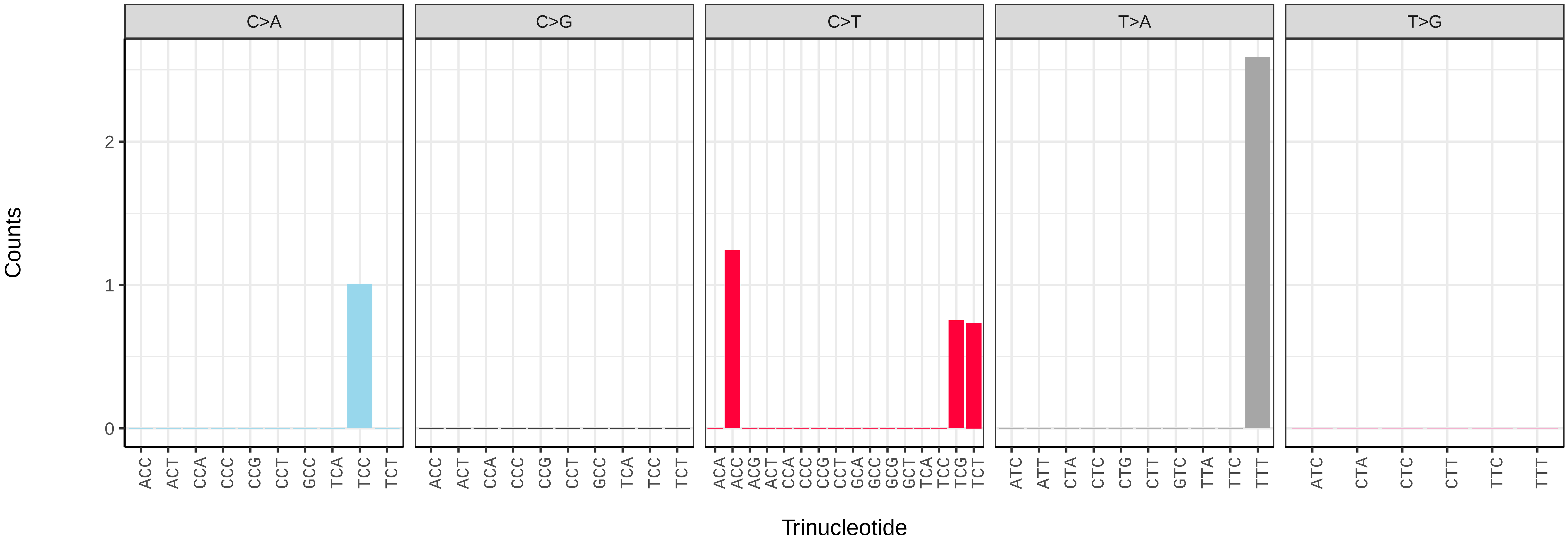

*Vespula vulgaris* (iyVesVulg1.iyVesVulg5) germline mutational spectrum

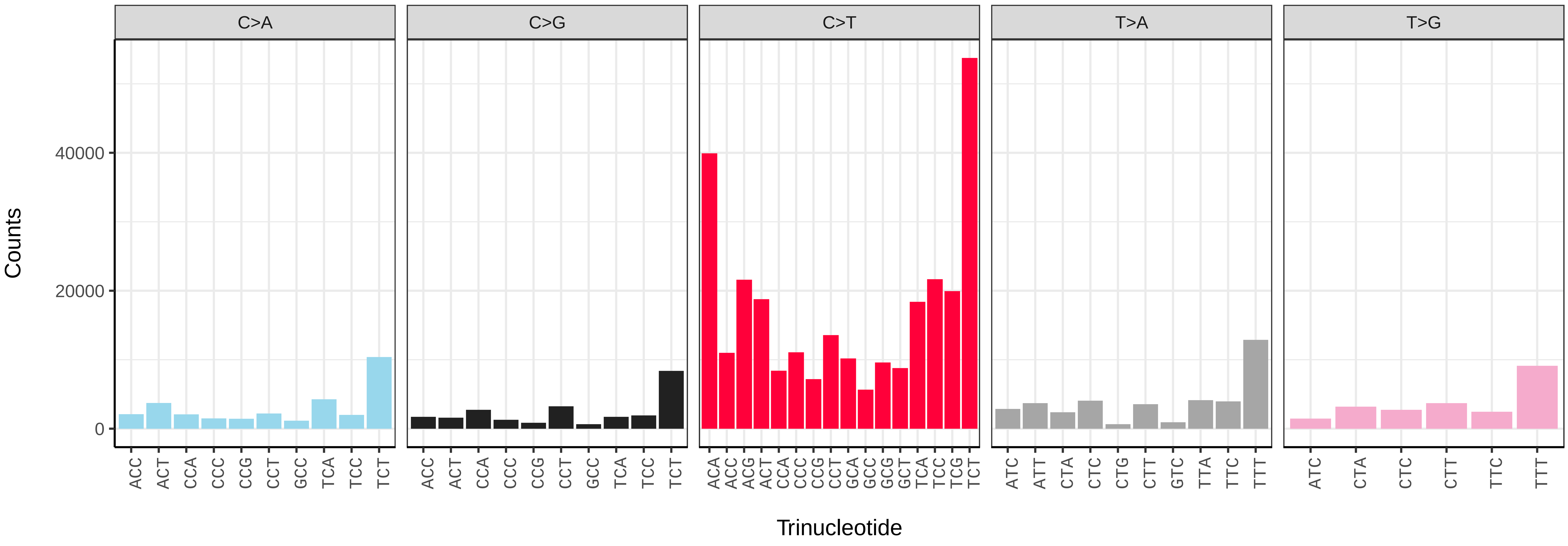


*Vespula vulgaris* (iyVesVulg1.iyVesVulg5) somatic mutational spectrum (SBS96 classification)

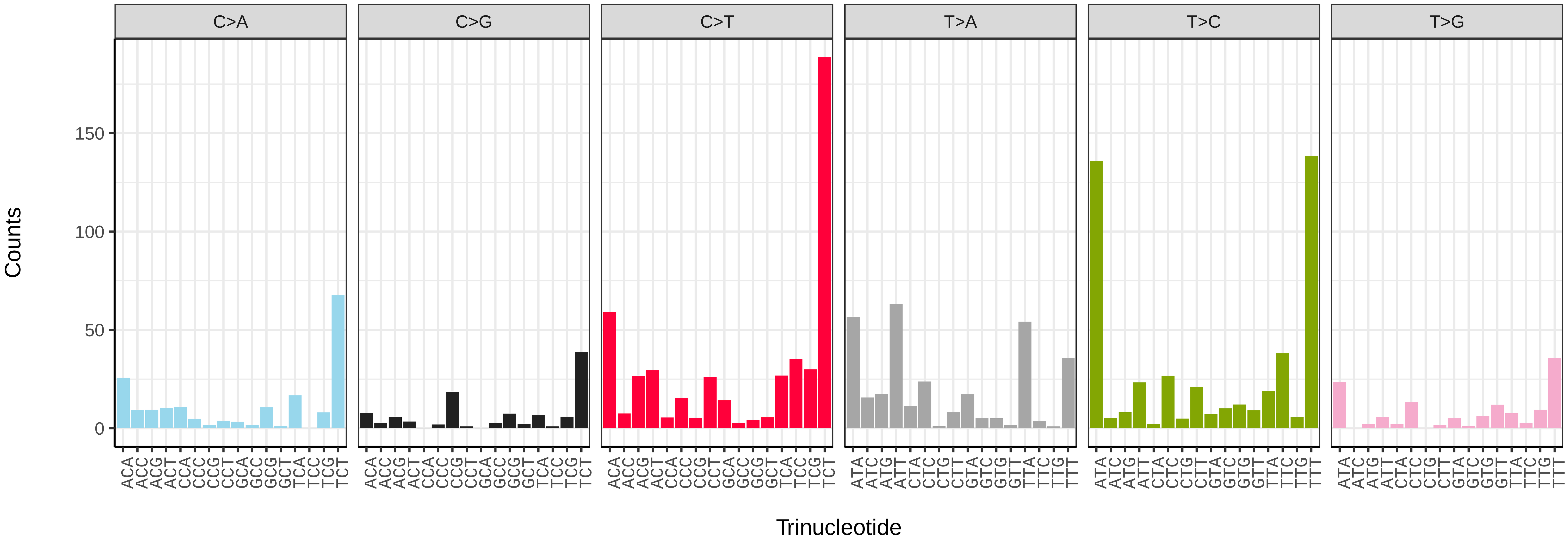


*Vespula vulgaris* (iyVesVulg1.iyVesVulg5) somatic mutational spectrum (SBS52 classification)

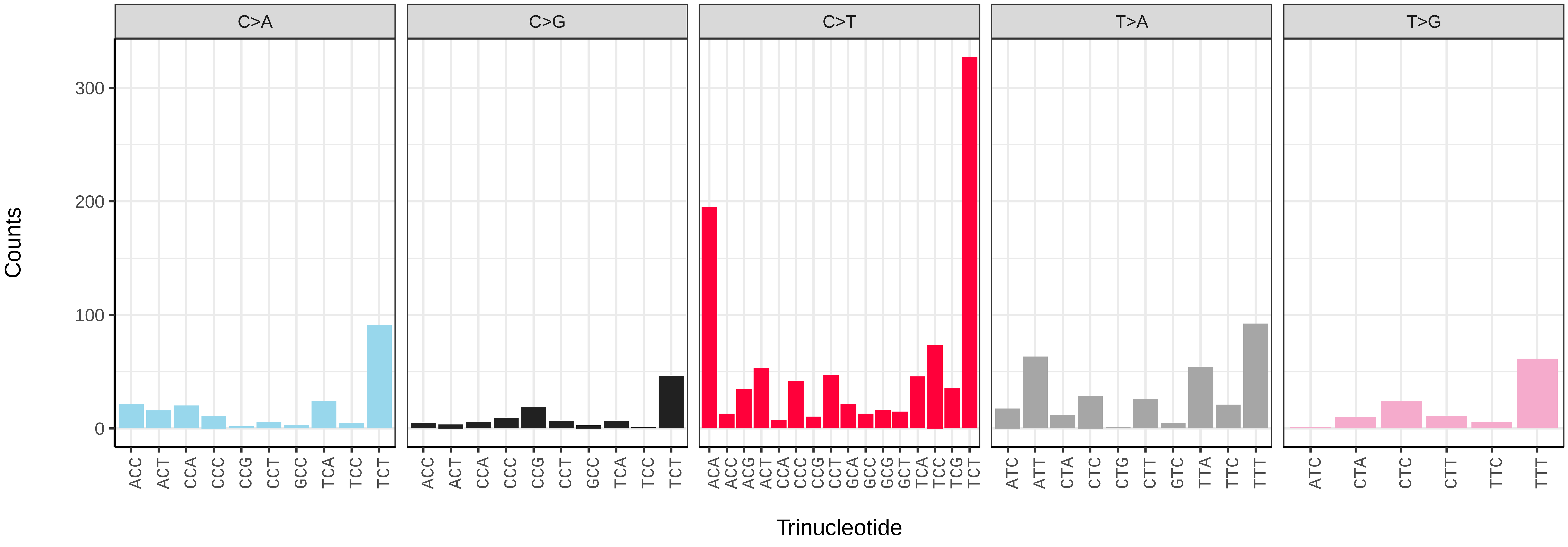

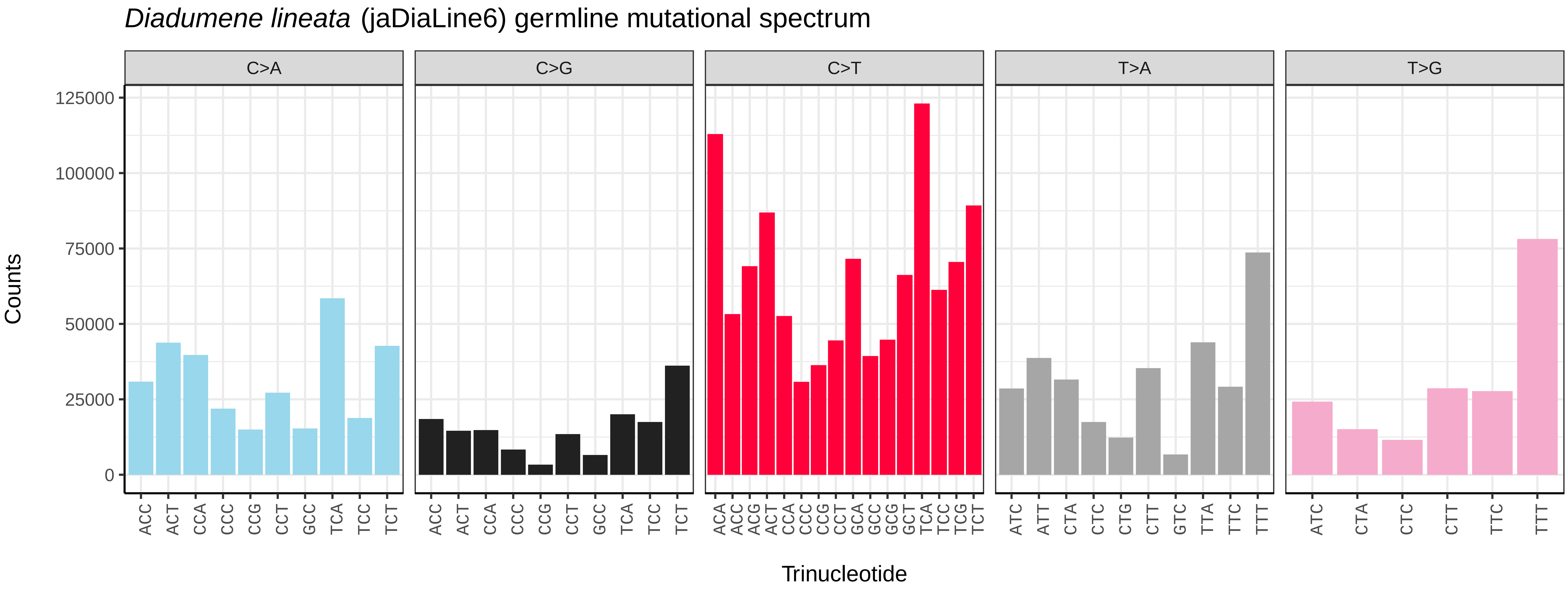
Diadumene lineata (jaDiaLine6) germline mutational spectrum
C>A
C>G
C>T
T>A
T>G
Counts
Trinucleotide
0
25000
50000
75000
100000
125000

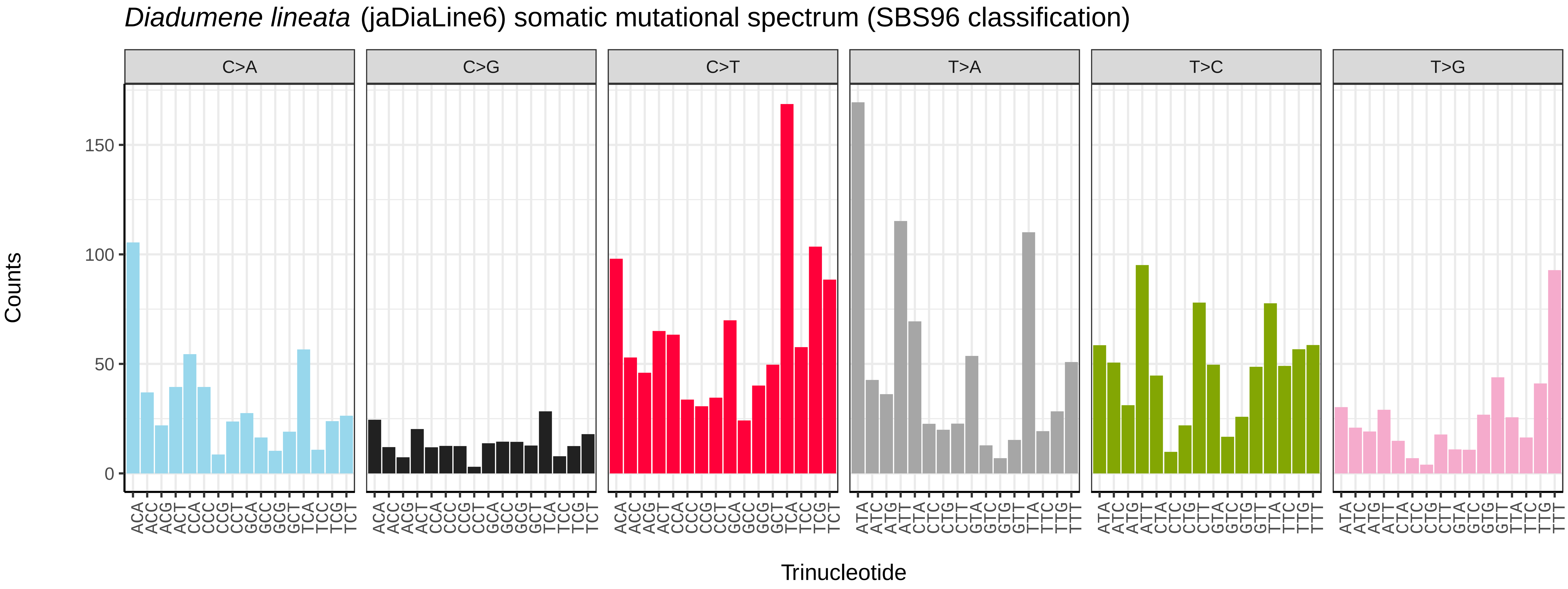
Diadumene lineata (jaDiaLine6) somatic mutational spectrum (SBS96 classification)
C>A
C>G
C>T
T>A
T>C
T>G
Counts
Trinucleotide
0
50
100
150

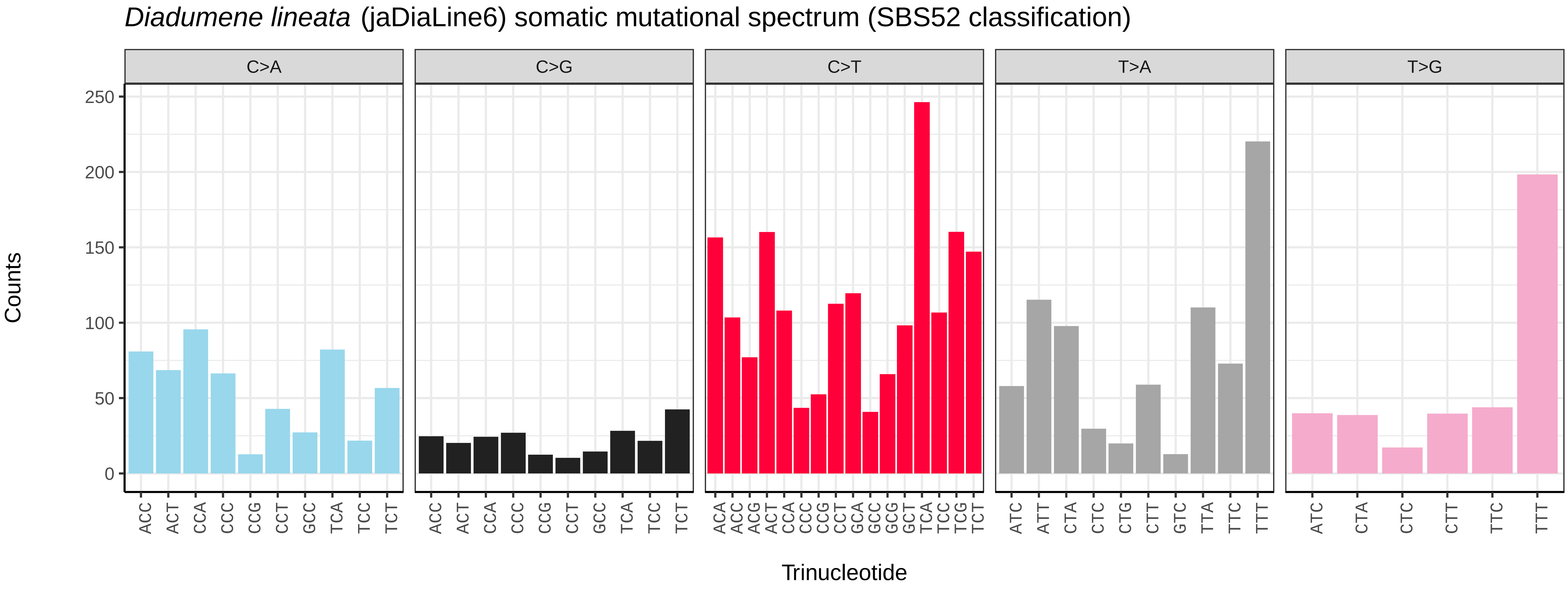
Diadumene lineata (jaDiaLine6) somatic mutational spectrum (SBS52 classification)
C>A
C>G
C>T
T>A
T>G
Counts
Trinucleotide
0
50
100
150
200
250

*Metridium senile* (jaMetSeni4) germline mutational spectrum

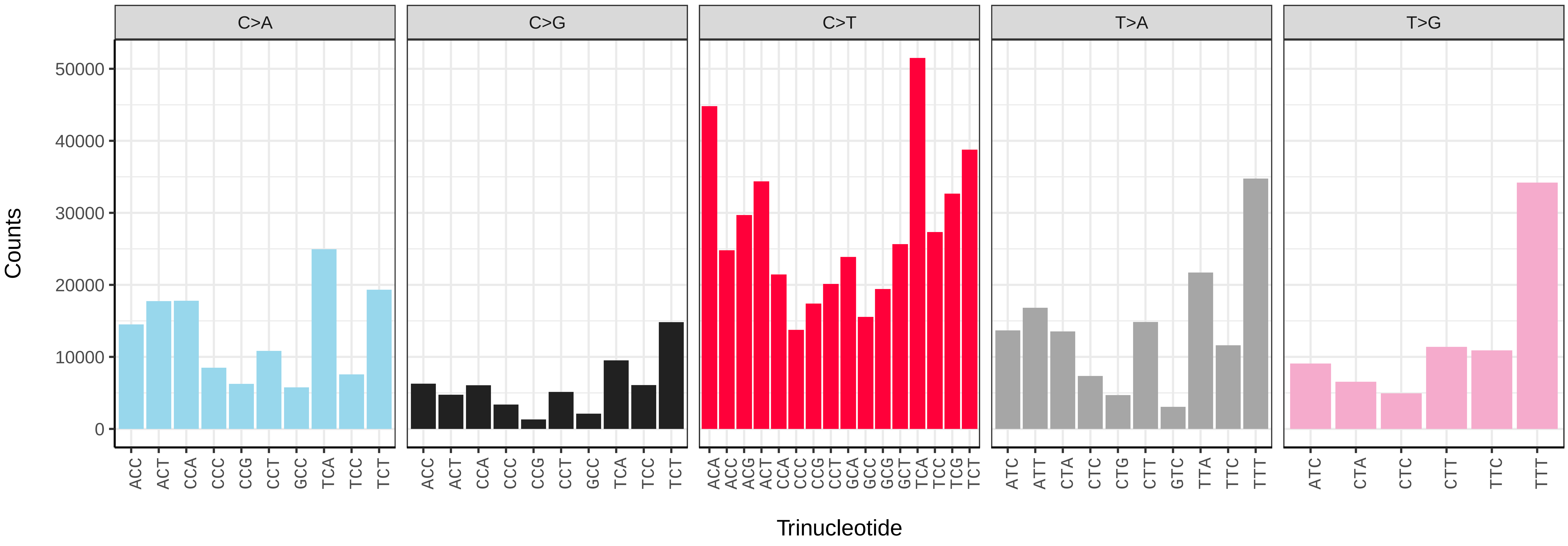


*Metridium senile* (jaMetSeni4) somatic mutational spectrum (SBS96 classification)

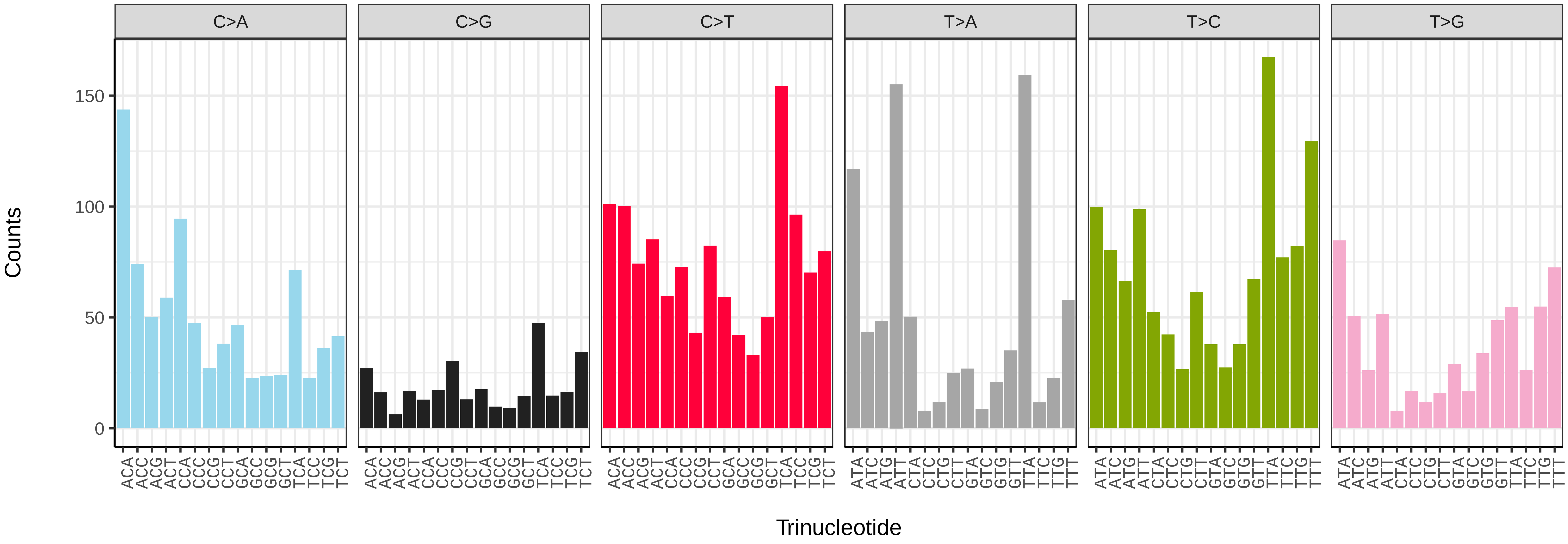


*Metridium senile* (jaMetSeni4) somatic mutational spectrum (SBS52 classification)

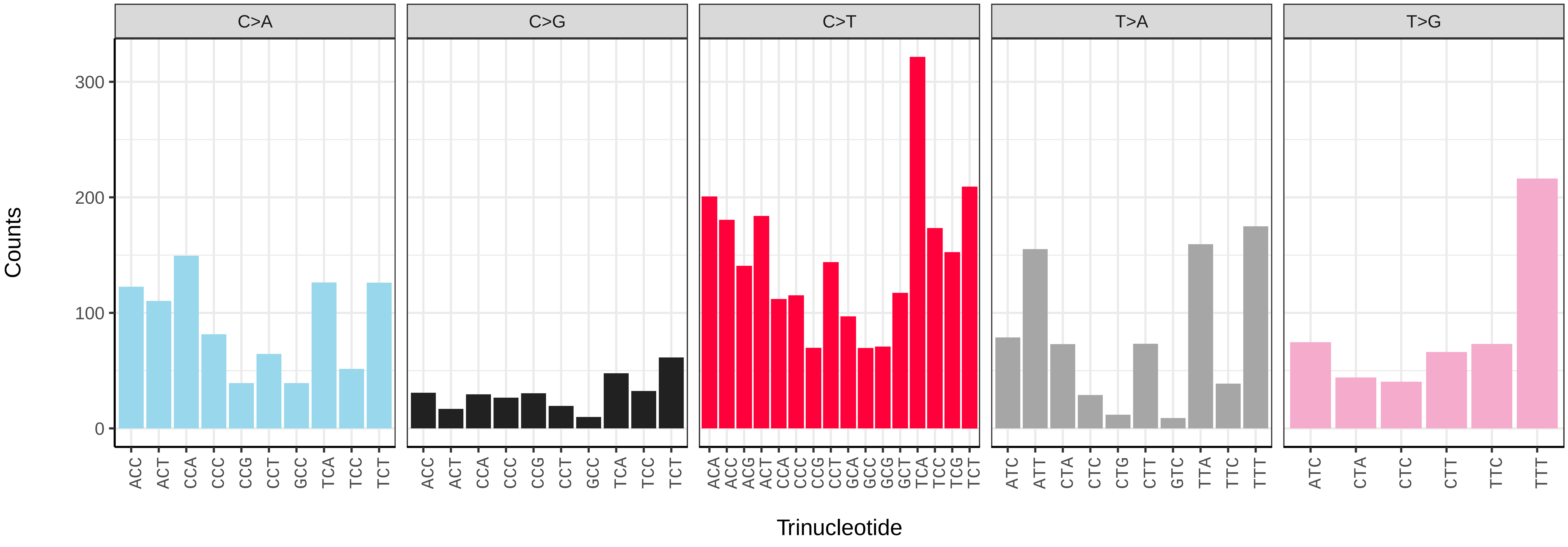

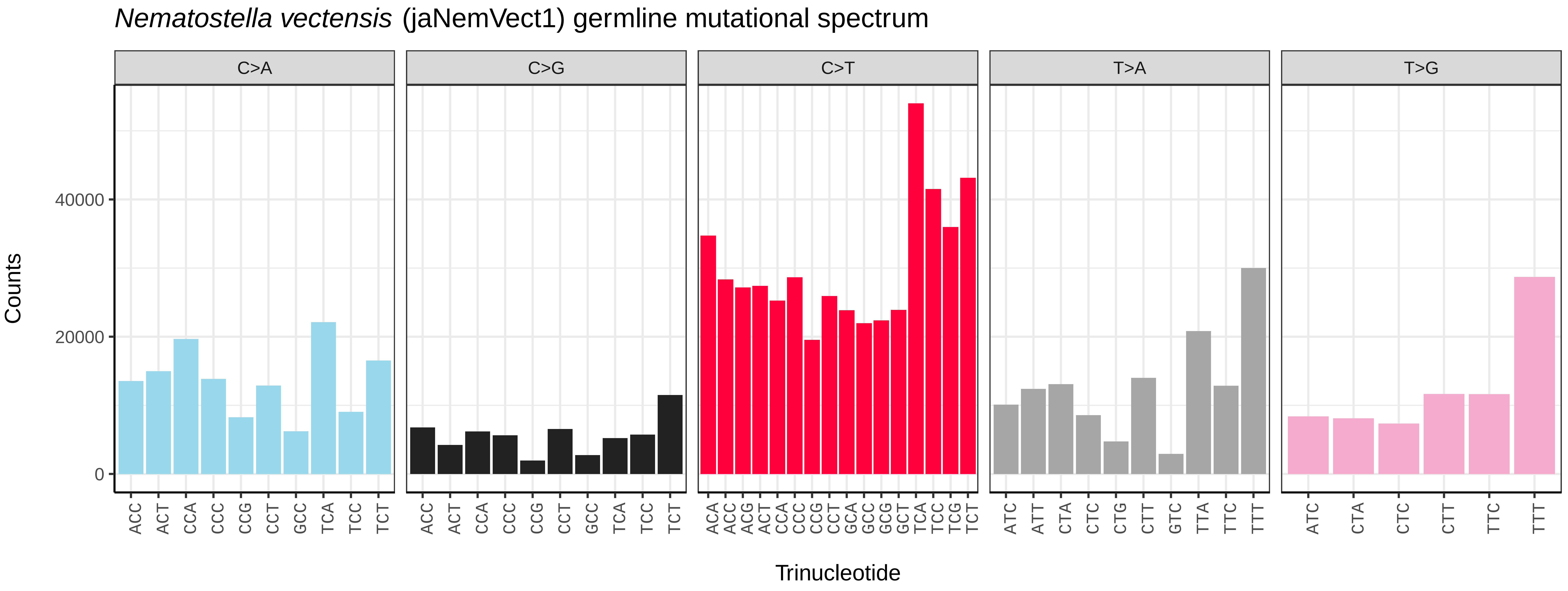
Nematostella vectensis (jaNemVect1) germline mutational spectrum
C>A
C>G
C>T
T>A
T>G
Counts
0
20000
40000
Trinucleotide
ACC ACT CCA CCC CCG CCT GCC TCA TCC TCT
ACA ACC ACG ACT CCA CCC CCG CCT GCA GCC GCG GCT TCA TCC TCG TCT
ATC ATT CTA CTC CTG CTT GTC TTA TTC TTT
ATC CTA CTC CTT TTC TTT

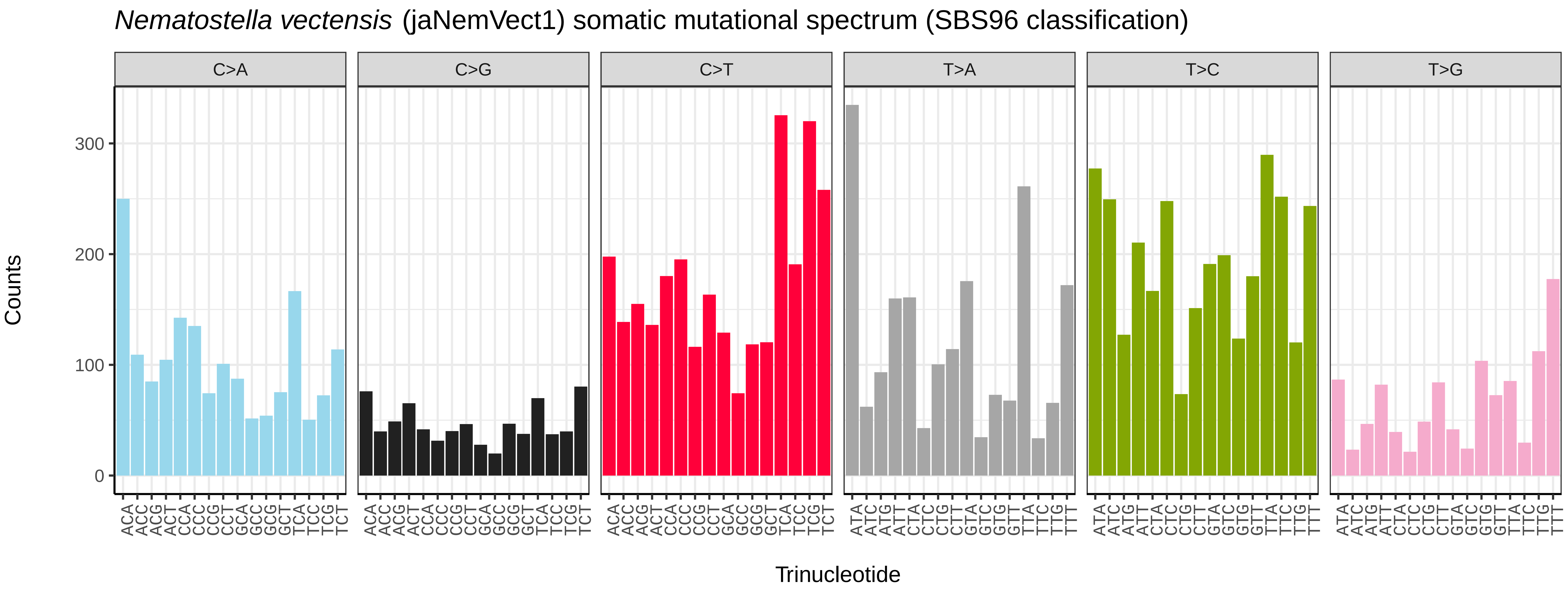
Nematostella vectensis (jaNemVect1) somatic mutational spectrum (SBS96 classification)
C>A
C>G
C>T
T>A
T>C
T>G
Counts
0
100
200
300
Trinucleotide
ACA ACC ACG ACT CCA CCC CCG CCT GCA GCC GCG GCT TCA TCC TCG TCT
ATA ATC ATG ATT CTA CTC CTG CTT GTA GTC GTG GTT TTA TTC TTG TTT

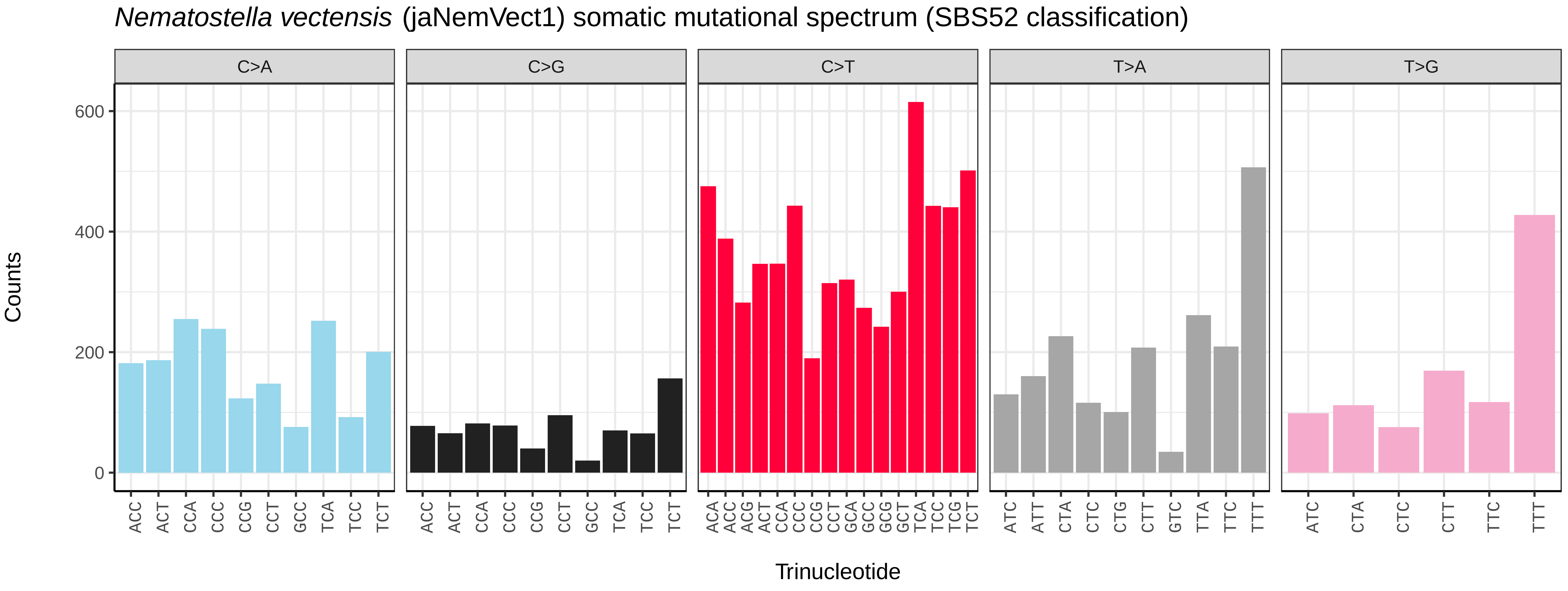
Nematostella vectensis (jaNemVect1) somatic mutational spectrum (SBS52 classification)
C>A
C>G
C>T
T>A
T>G
Counts
0
200
400
600
Trinucleotide
ACC ACT CCA CCC CCG CCT GCC TCA TCC TCT
ACA ACC ACG ACT CCA CCC CCG CCT GCA GCC GCG GCT TCA TCC TCG TCT
ATC ATT CTA CTC CTG CTT GTC TTA TTC TTT
ATC CTA CTC CTT TTC TTT

*Candelabrum cocksii* (jhCanCock6) germline mutational spectrum

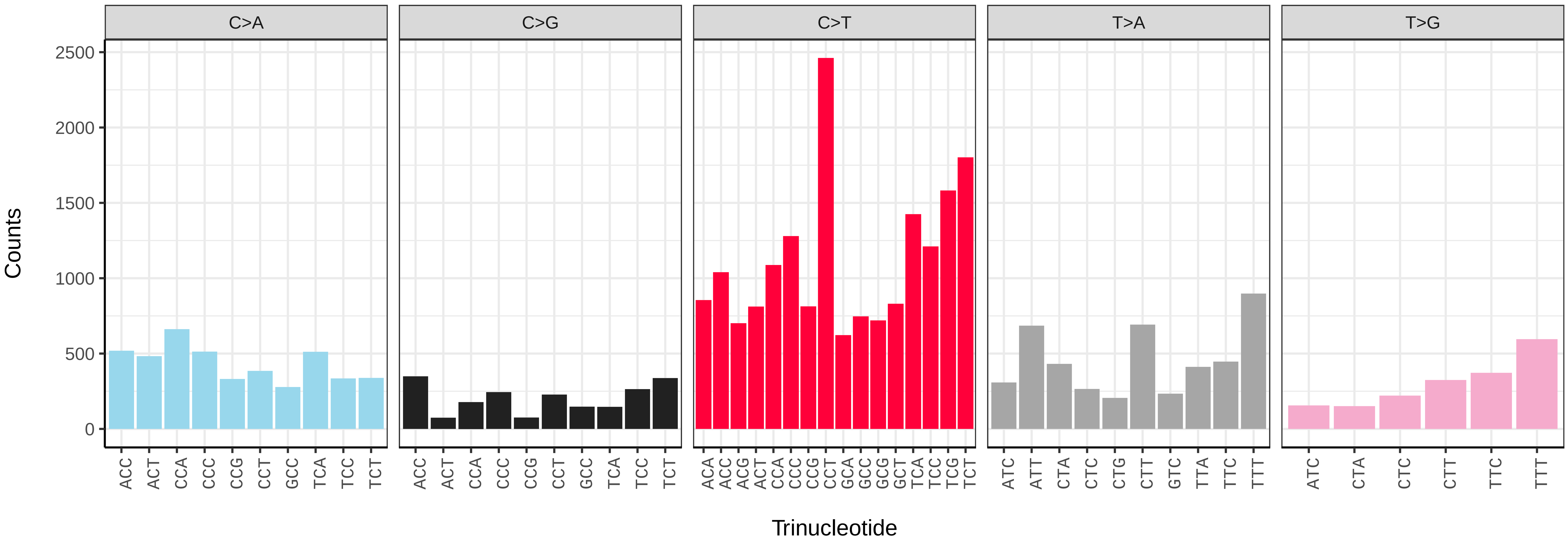


*Candelabrum cocksii* (jhCanCock6) somatic mutational spectrum (SBS96 classification)

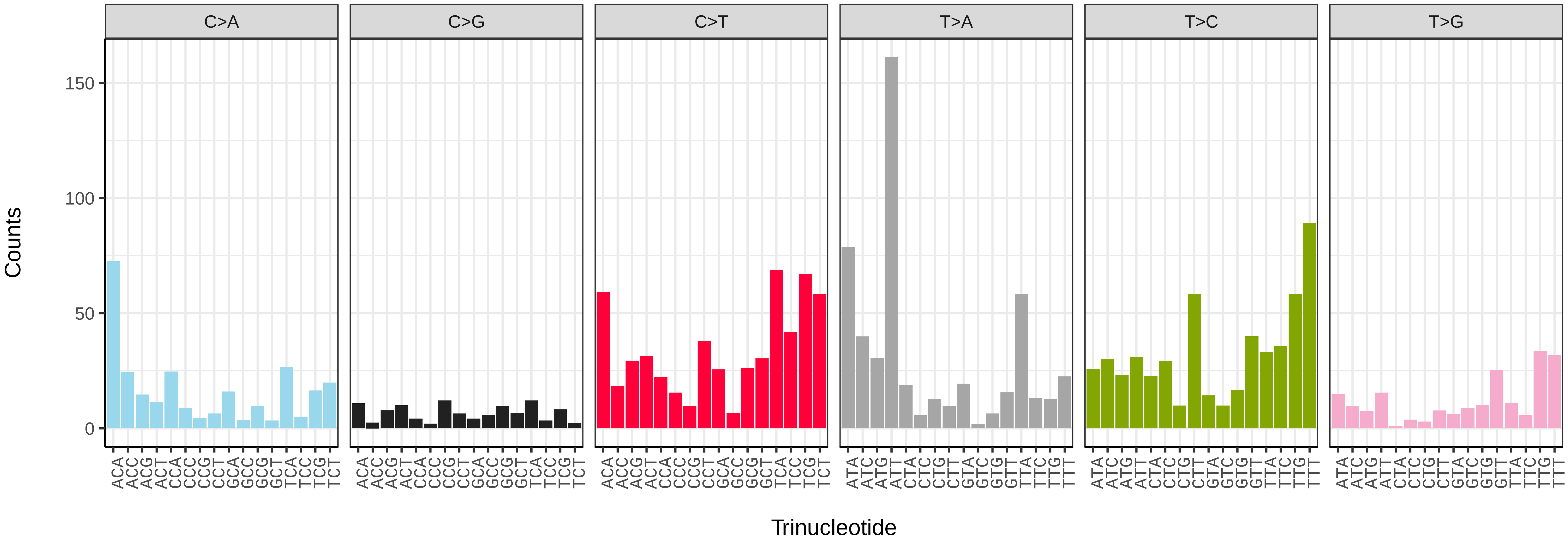


*Candelabrum cocksii* (jhCanCock6) somatic mutational spectrum (SBS52 classification)

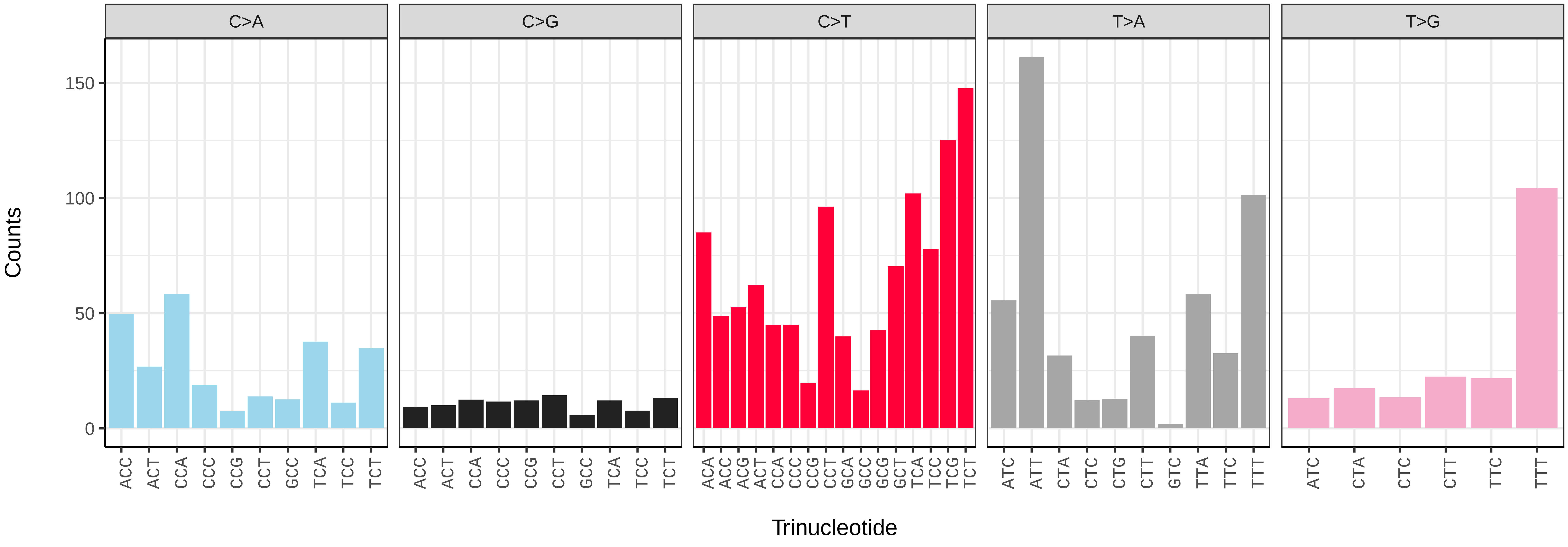

*Haliclystus octoradiatus* (jrHalOcto1) germline mutational spectrum

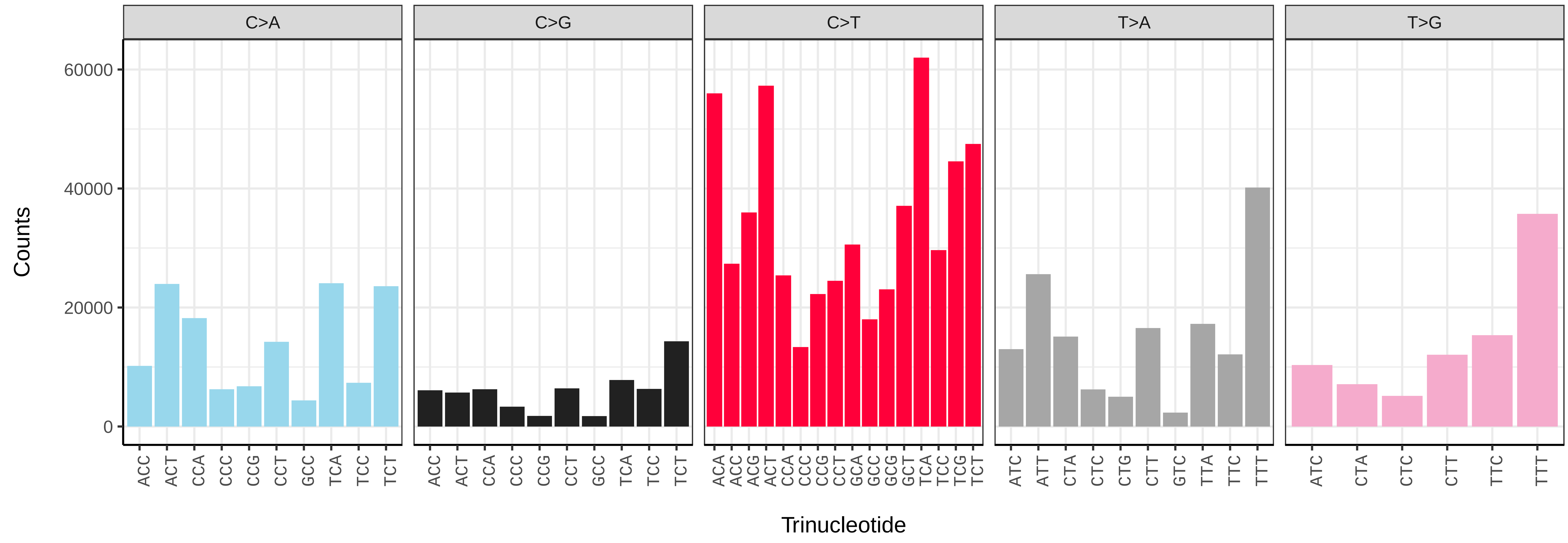


*Haliclystus octoradiatus* (jrHalOcto1) somatic mutational spectrum (SBS96 classification)

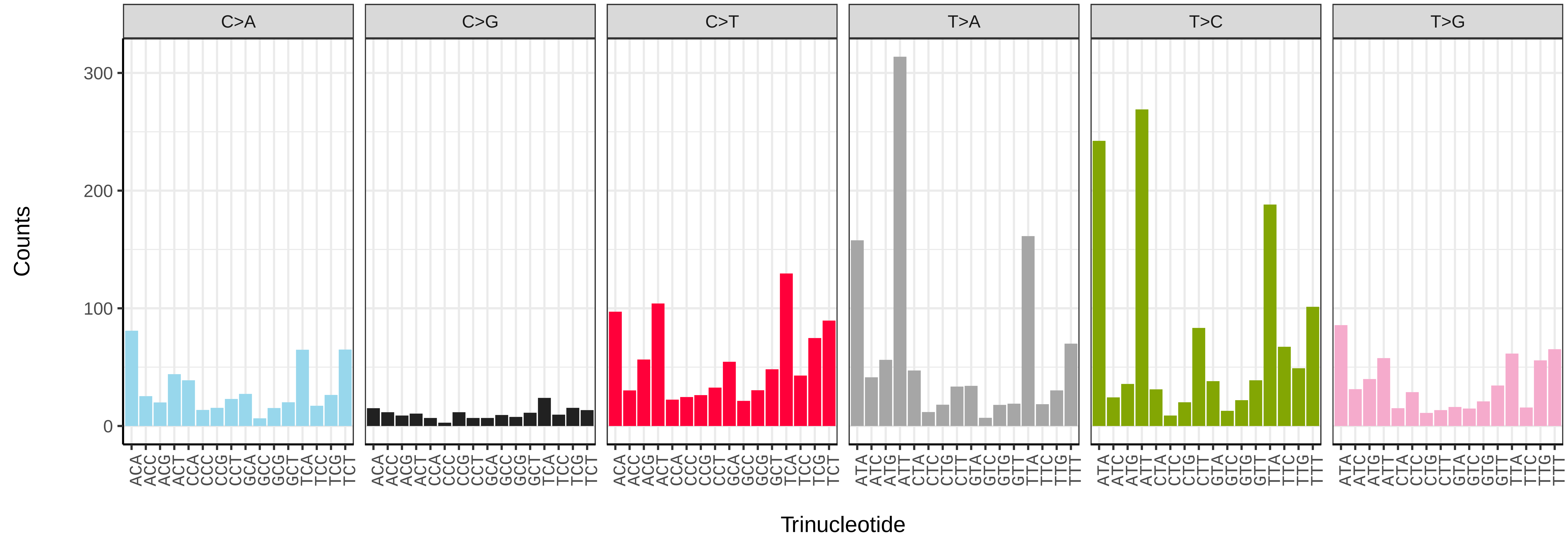


*Haliclystus octoradiatus* (jrHalOcto1) somatic mutational spectrum (SBS52 classification)

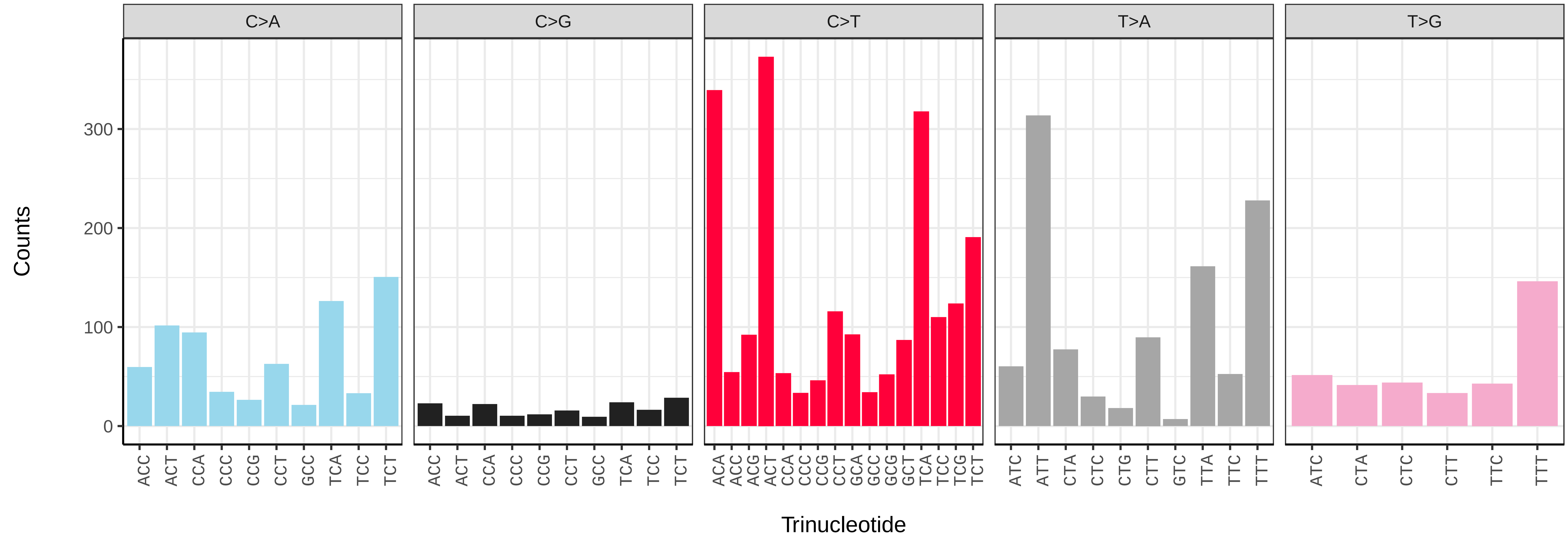

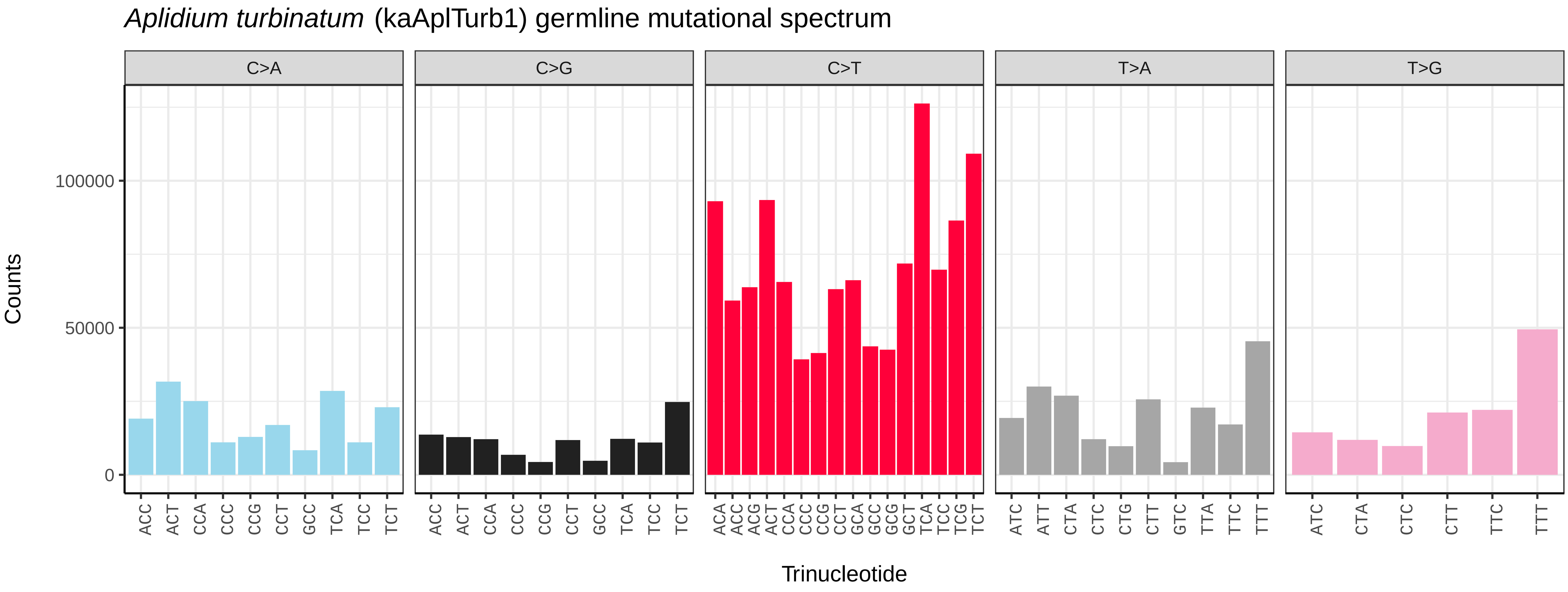
Aplidium turbinatum (kaAplTurb1) germline mutational spectrum
C>A
C>G
C>T
T>A
T>G
Counts
0
50000
100000
ACC ACT CCA CCC CCG CCT GCC TCA TCC TCT
ACC ACT CCA CCC CCG CCT GCC TCA TCC TCT
ACA ACC ACG ACT CCA CCC CCG CCT GCA GCC GCG GCT TCA TCC TCG TCT
ATC ATT CTA CTC CTG CTT GTC TTA TTC TTT
ATC CTA CTC CTT TTC TTT
Trinucleotide

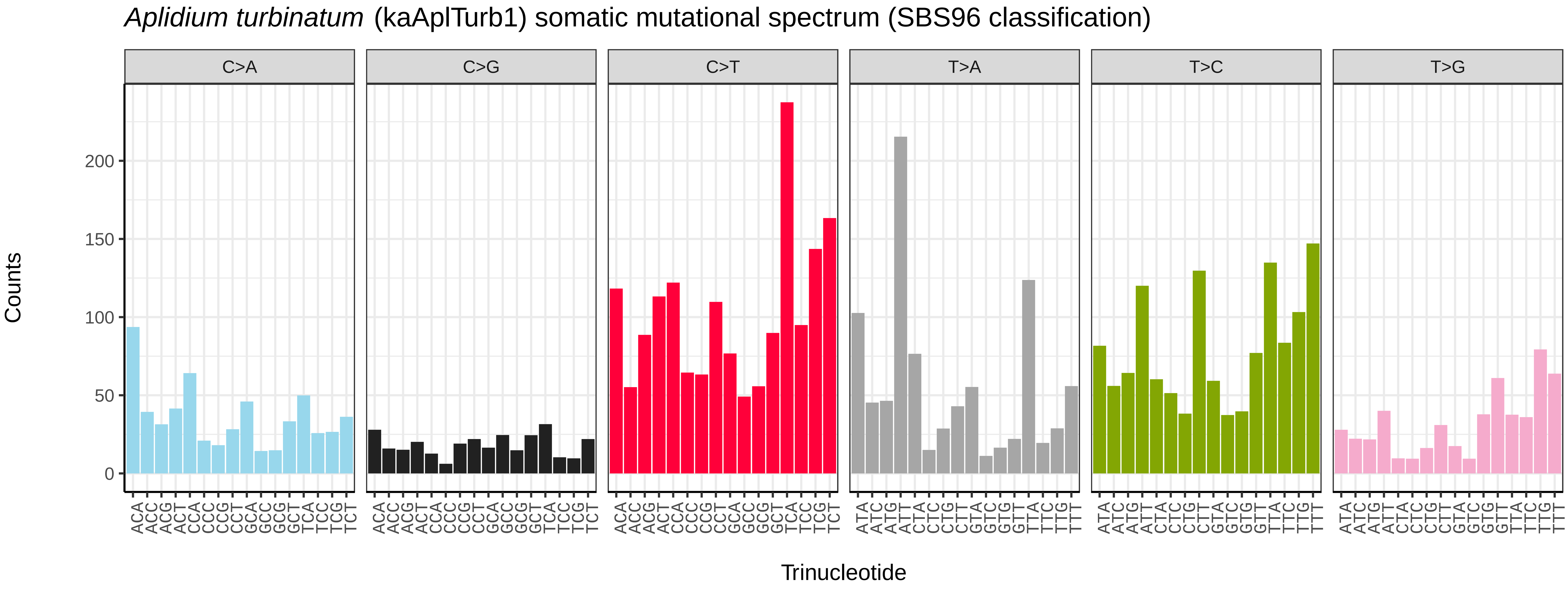
Aplidium turbinatum (kaAplTurb1) somatic mutational spectrum (SBS96 classification)
C>A
C>G
C>T
T>A
T>C
T>G
Counts
0
50
100
150
200
ACA ACC ACG ACT CCA CCC CCG CCT GCA GCC GCG GCT TCA TCC TCG TCT
ACA ACC ACG ACT CCA CCC CCG CCT GCA GCC GCG GCT TCA TCC TCG TCT
ACA ACC ACG ACT CCA CCC CCG CCT GCA GCC GCG GCT TCA TCC TCG TCT
ATA ATC ATG ATT CTA CTC CTG CTT GTA GTC GTG GTT TTA TTC TTG TTT
ATA ATC ATG ATT CTA CTC CTG CTT GTA GTC GTG GTT TTA TTC TTG TTT
ATA ATC ATG ATT CTA CTC CTG CTT GTA GTC GTG GTT TTA TTC TTG TTT
Trinucleotide

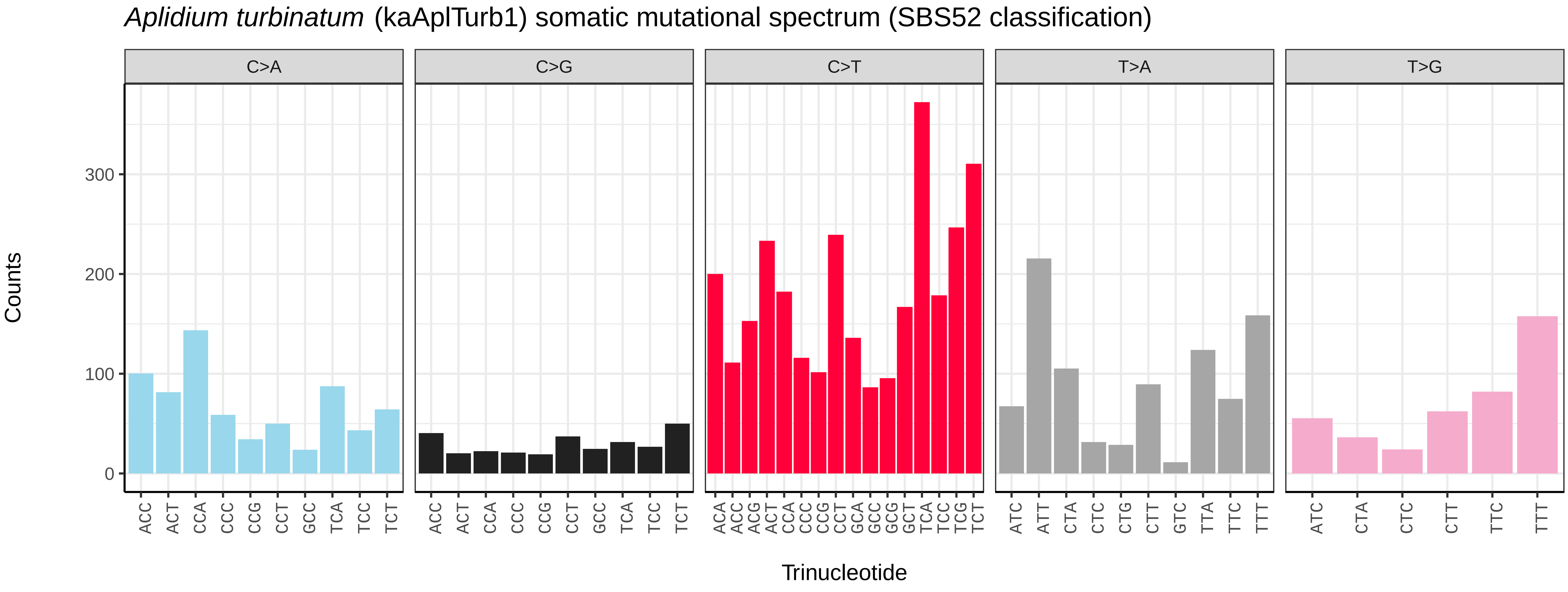
Aplidium turbinatum (kaAplTurb1) somatic mutational spectrum (SBS52 classification)
C>A
C>G
C>T
T>A
T>G
Counts
0
100
200
300
ACC ACT CCA CCC CCG CCT GCC TCA TCC TCT
ACC ACT CCA CCC CCG CCT GCC TCA TCC TCT
ACA ACC ACG ACT CCA CCC CCG CCT GCA GCC GCG GCT TCA TCC TCG TCT
ATC ATT CTA CTC CTG CTT GTC TTA TTC TTT
ATC CTA CTC CTT TTC TTT
Trinucleotide

*Ascidia mentula* (kaAscMent1) germline mutational spectrum

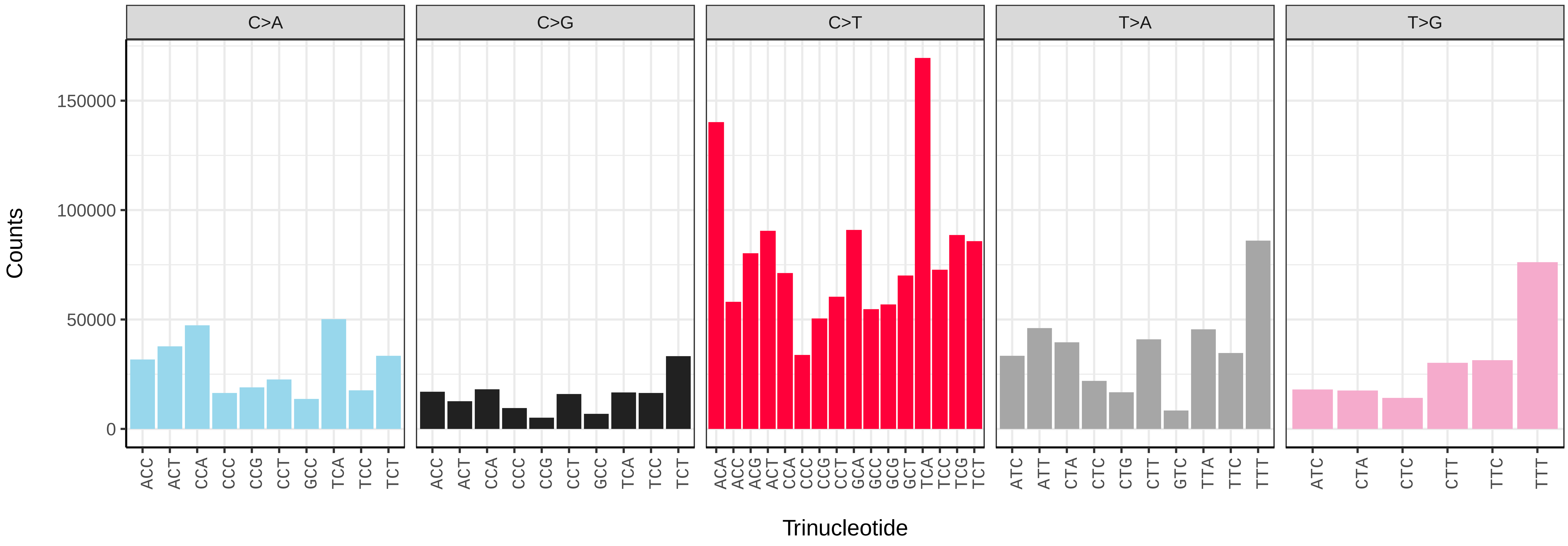


*Ascidia mentula* (kaAscMent1) somatic mutational spectrum (SBS96 classification)

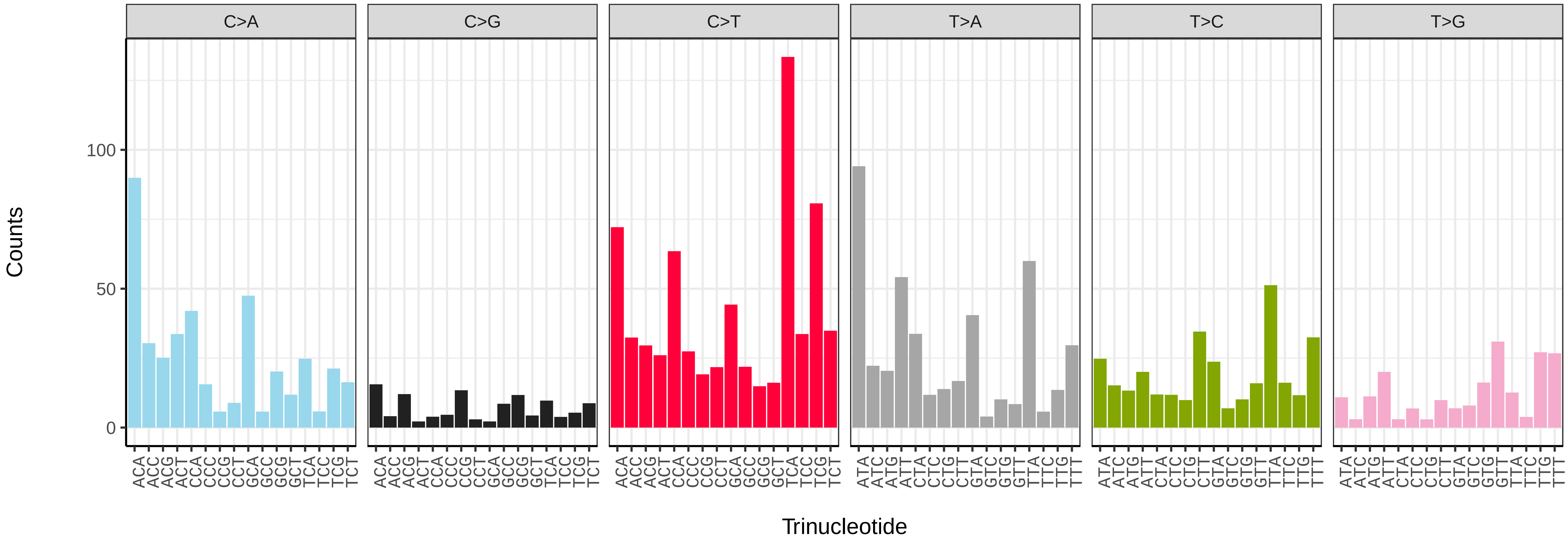


*Ascidia mentula* (kaAscMent1) somatic mutational spectrum (SBS52 classification)

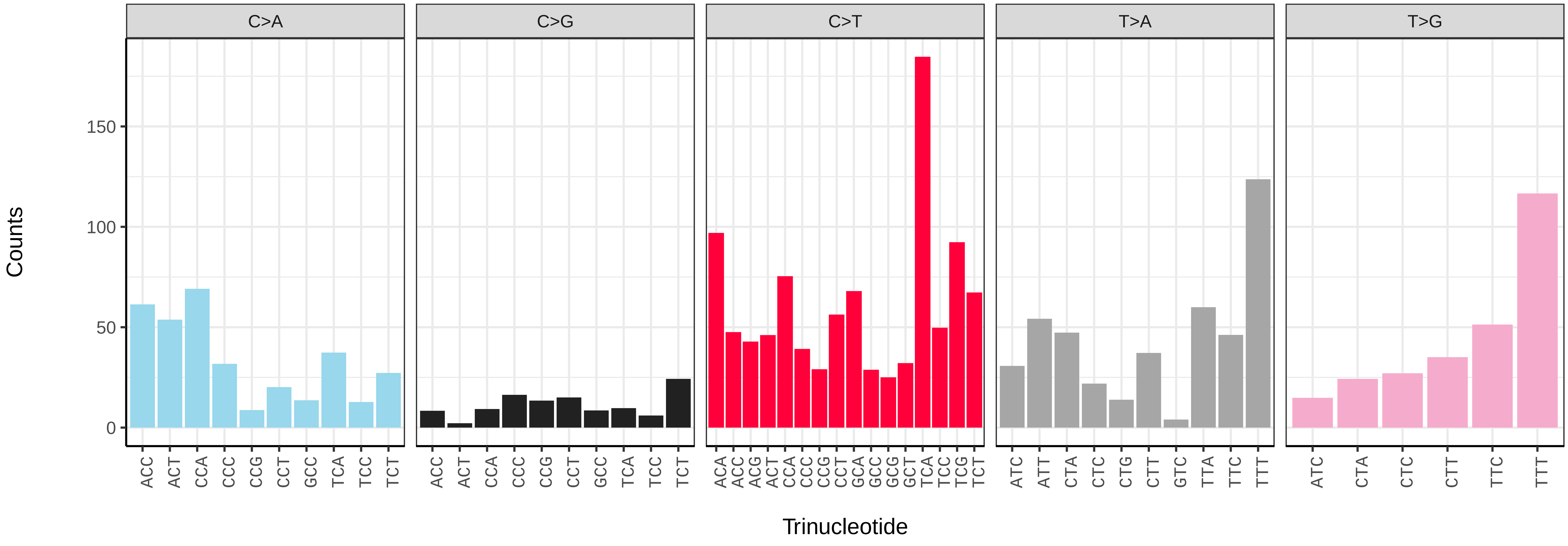

*Clavelina lepadiformis* (kaClaLepa1) germline mutational spectrum

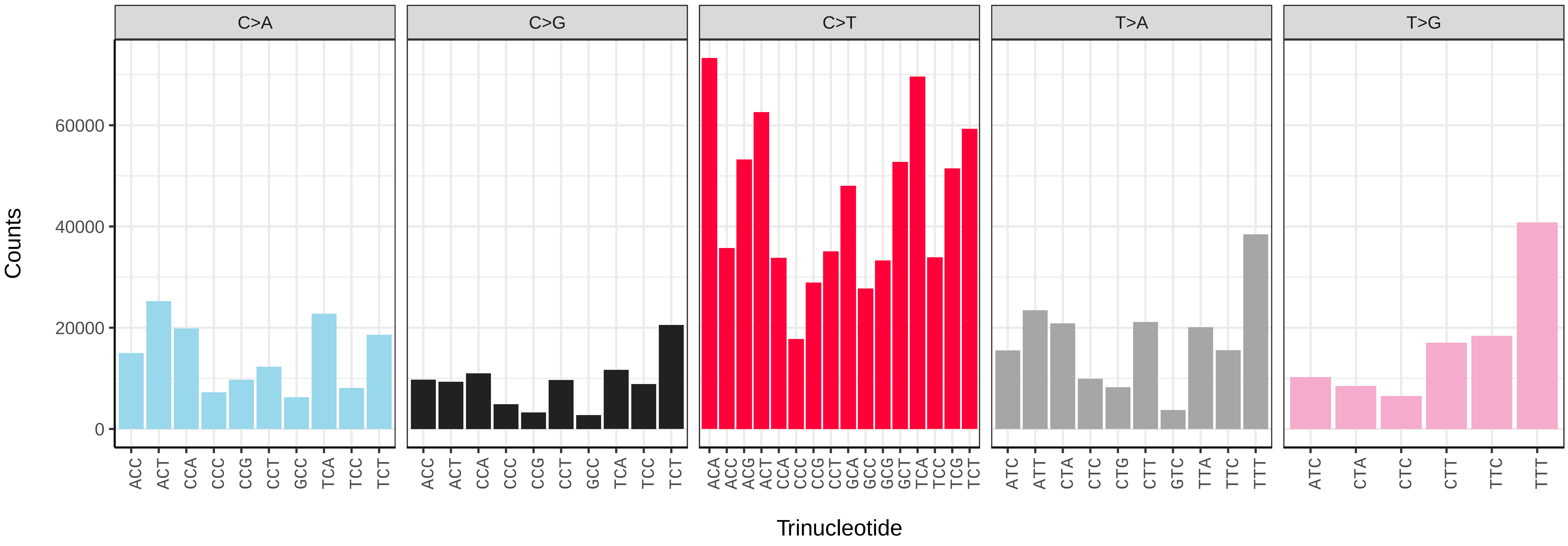


*Clavelina lepadiformis* (kaClaLepa1) somatic mutational spectrum (SBS96 classification)

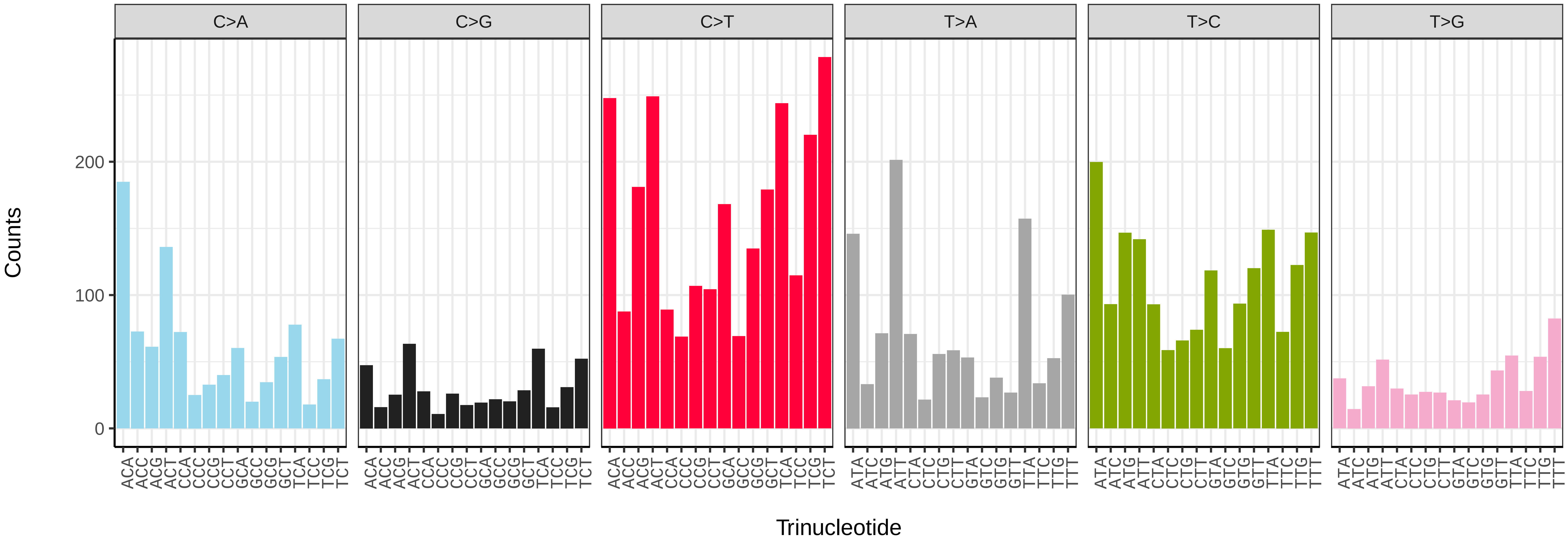


*Clavelina lepadiformis* (kaClaLepa1) somatic mutational spectrum (SBS52 classification)

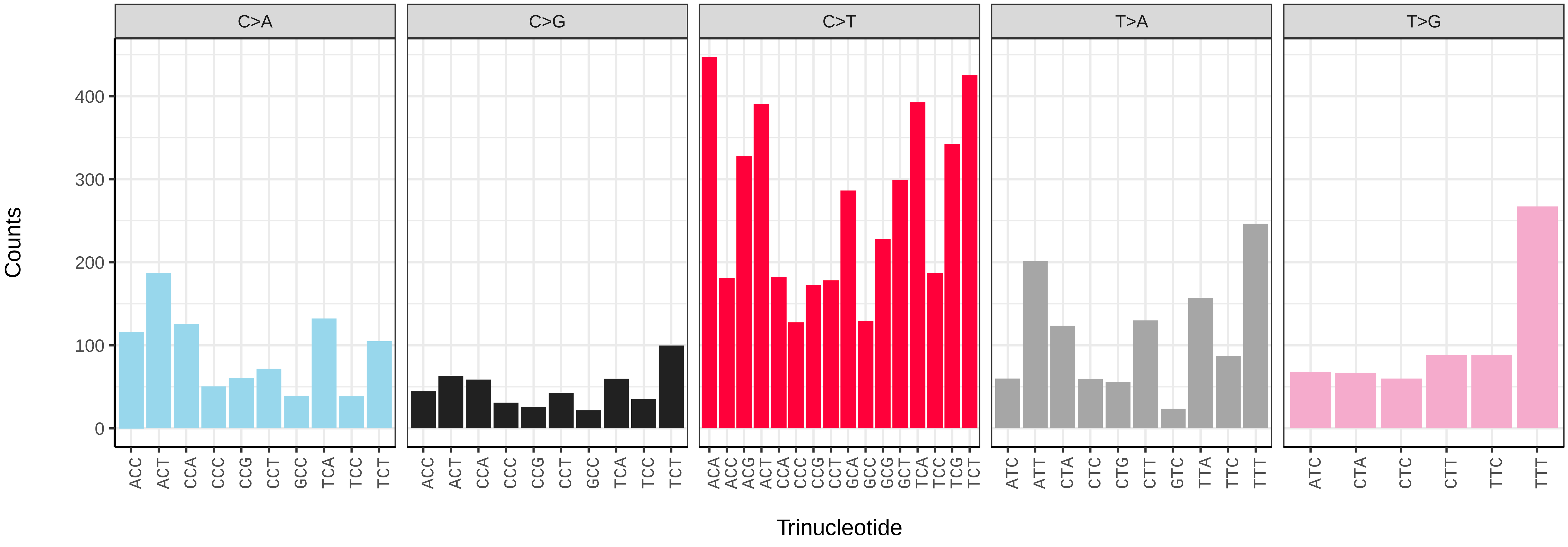

*Lampetra fluviatilis* (kcLamFluv1) germline mutational spectrum

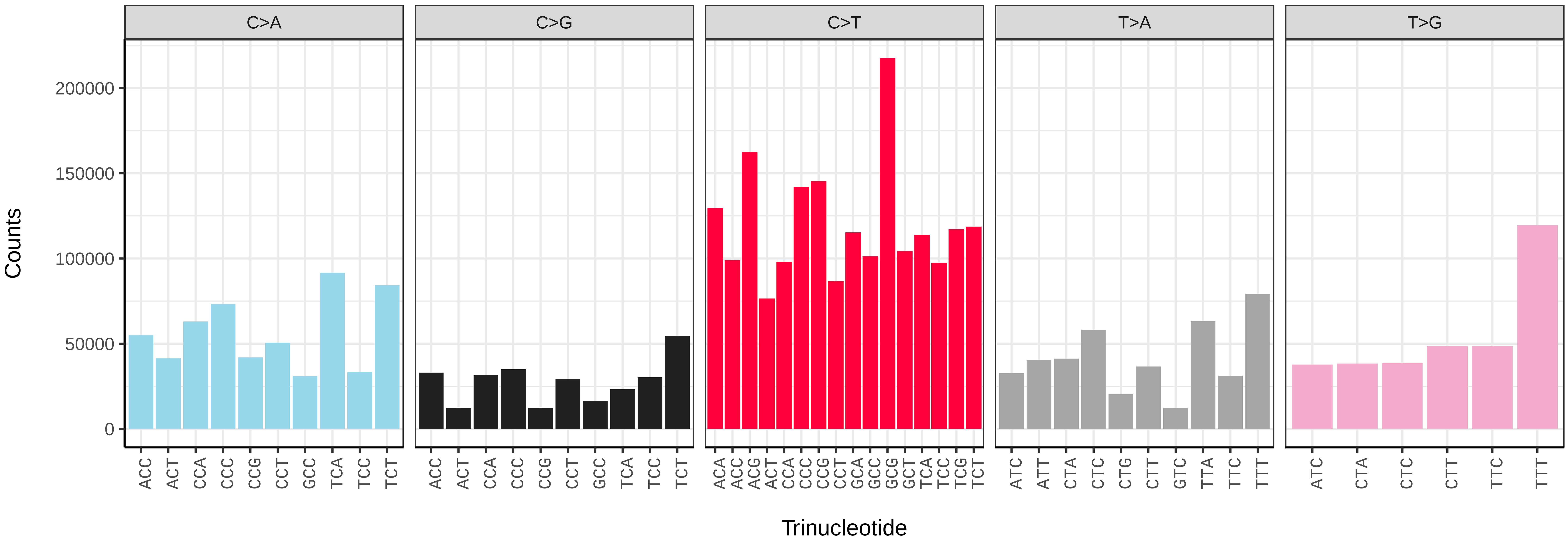


*Lampetra fluviatilis* (kcLamFluv1) somatic mutational spectrum (SBS96 classification)

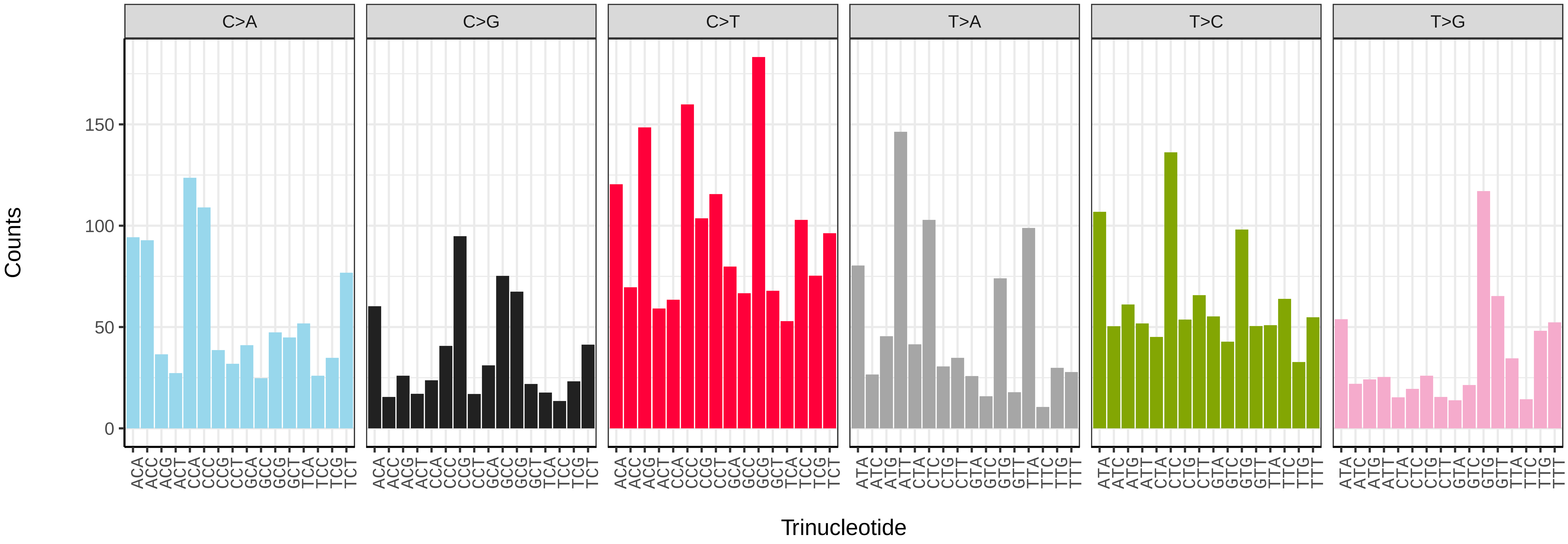


*Lampetra fluviatilis* (kcLamFluv1) somatic mutational spectrum (SBS52 classification)

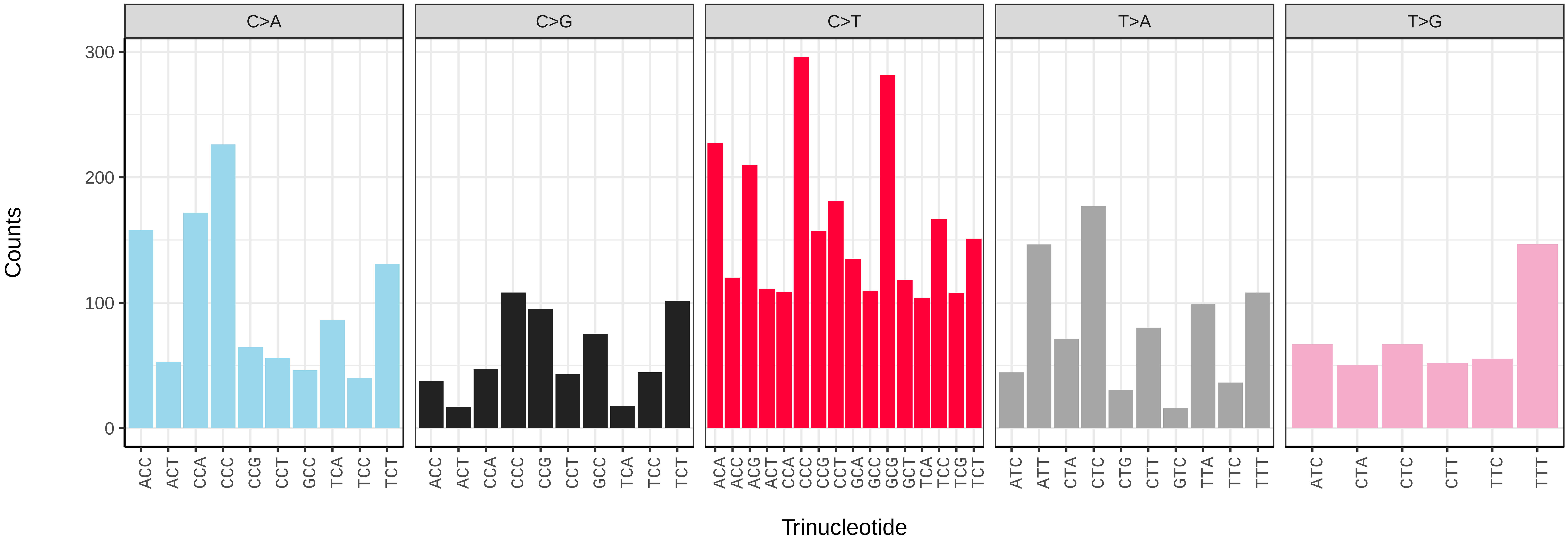

*Triglochin palustris* (laTriPals1) germline mutational spectrum

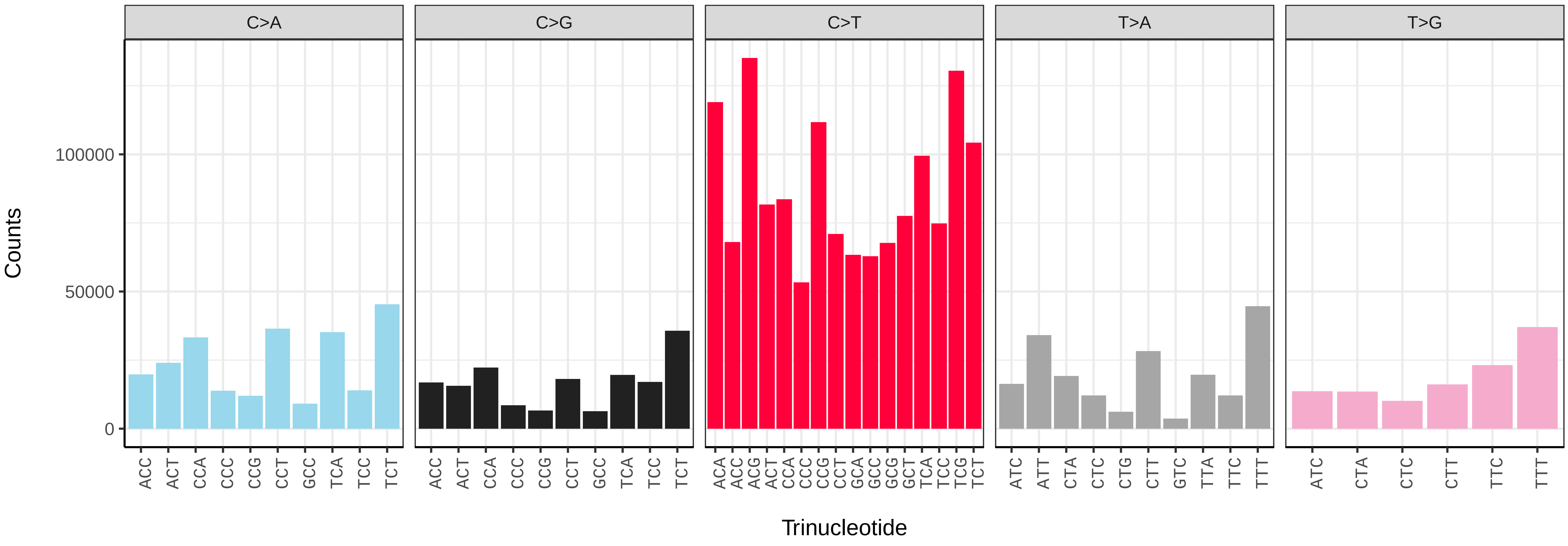


*Triglochin palustris* (laTriPals1) somatic mutational spectrum (SBS96 classification)

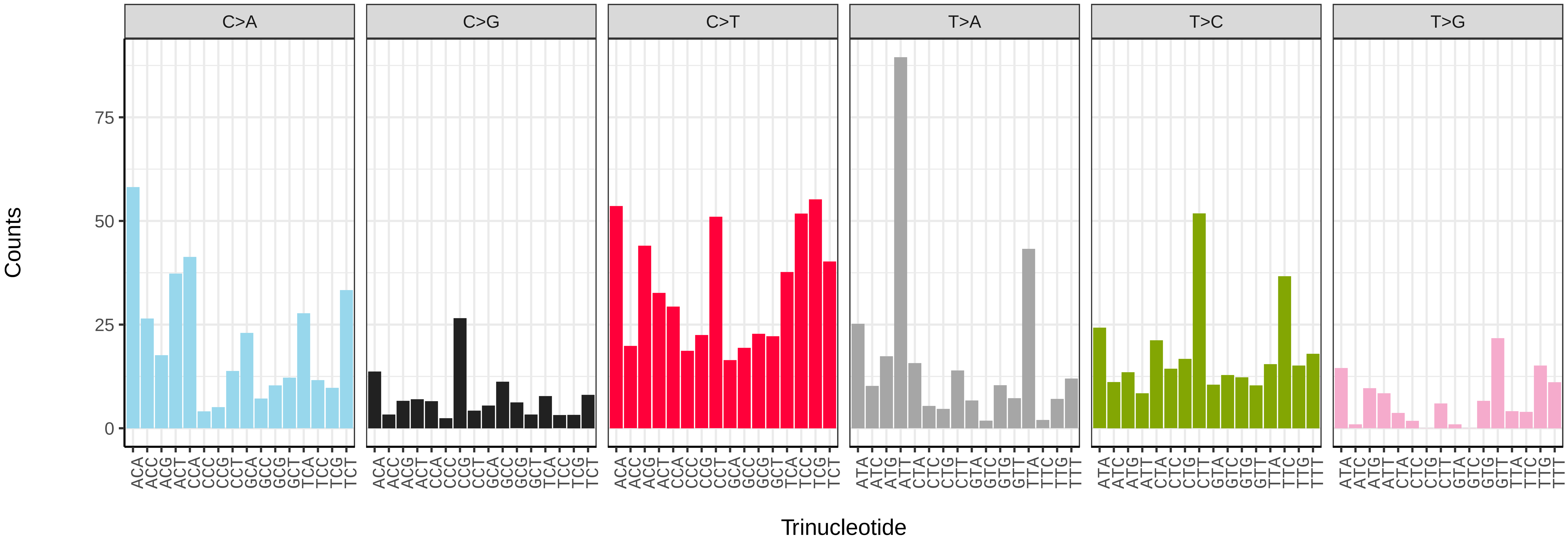


*Triglochin palustris* (laTriPals1) somatic mutational spectrum (SBS52 classification)

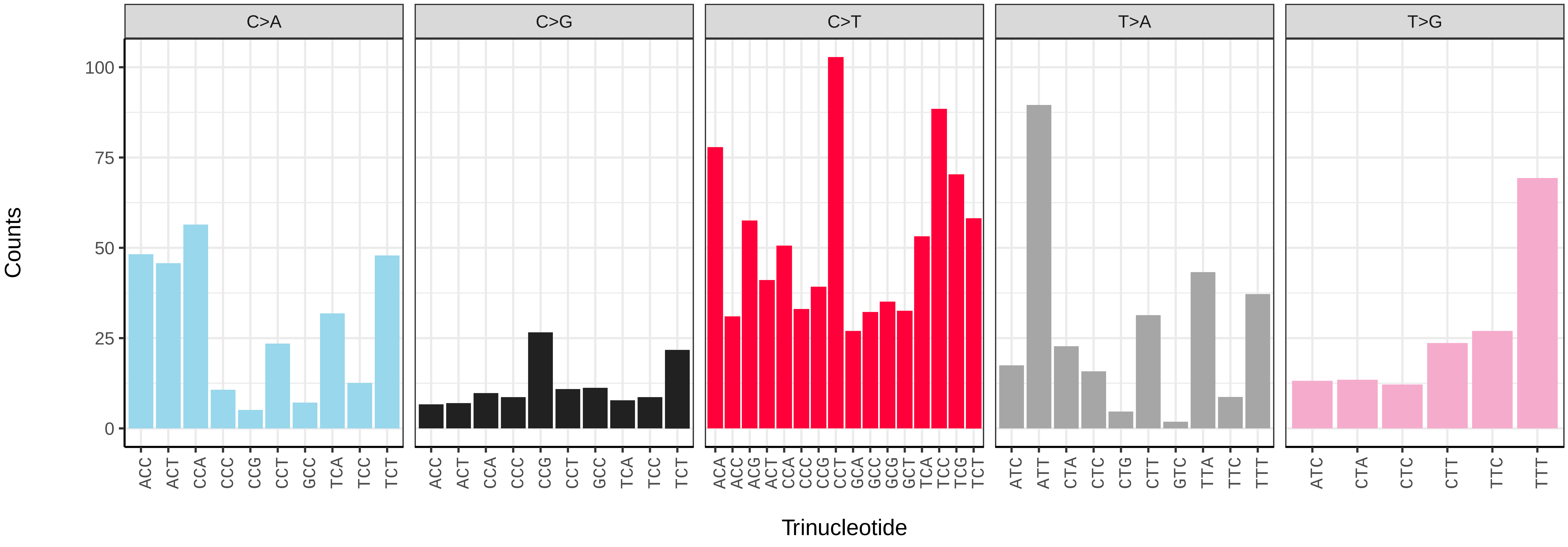

*Bromus sterilis* (lpBroSter1) germline mutational spectrum

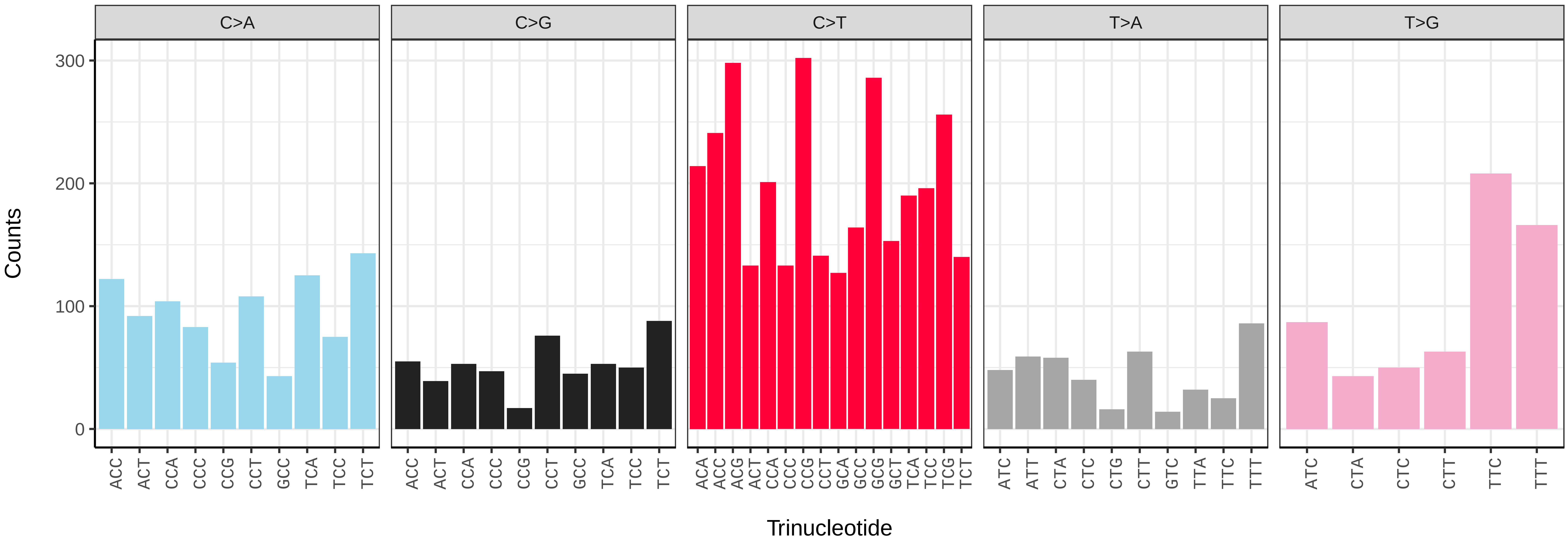


*Bromus sterilis* (lpBroSter1) somatic mutational spectrum (SBS96 classification)

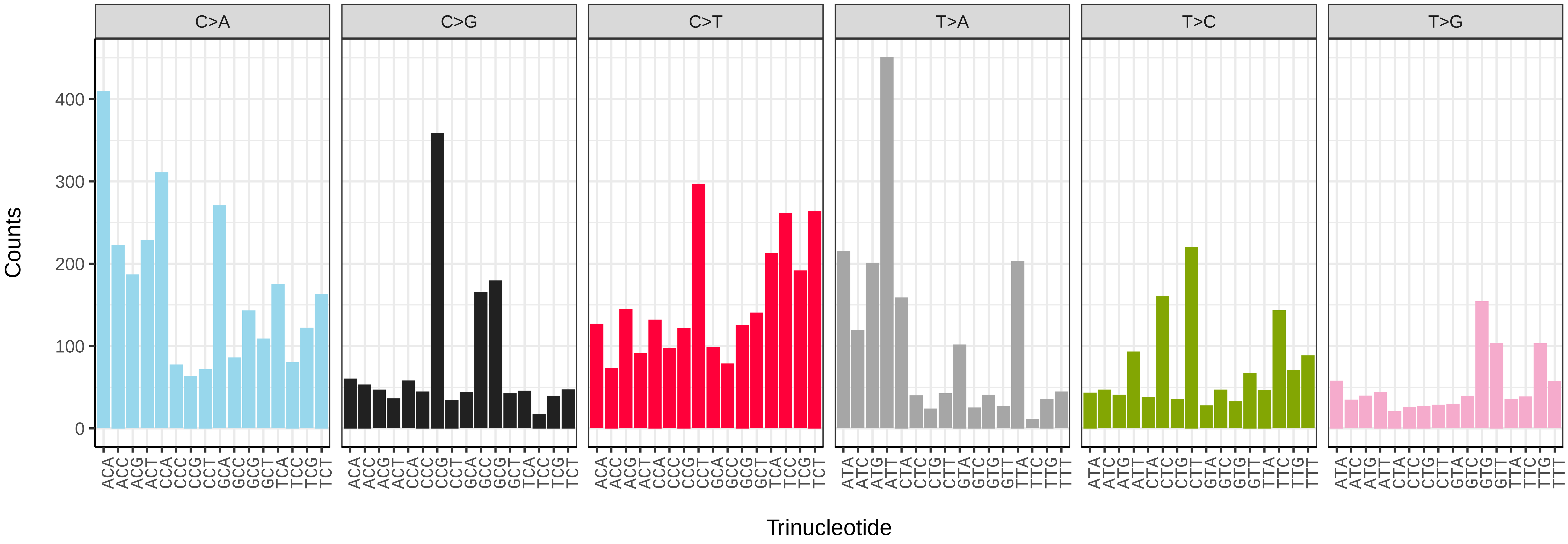


*Bromus sterilis* (lpBroSter1) somatic mutational spectrum (SBS52 classification)

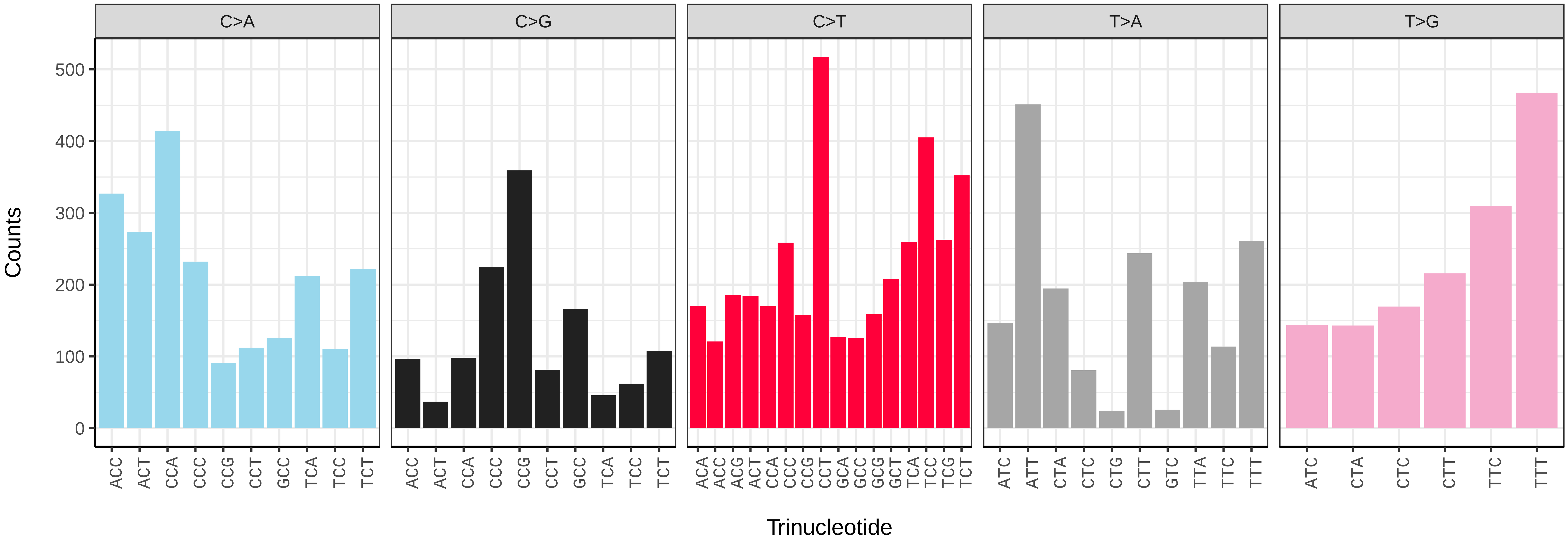

*Carex arenaria* (lpCarAren1) germline mutational spectrum

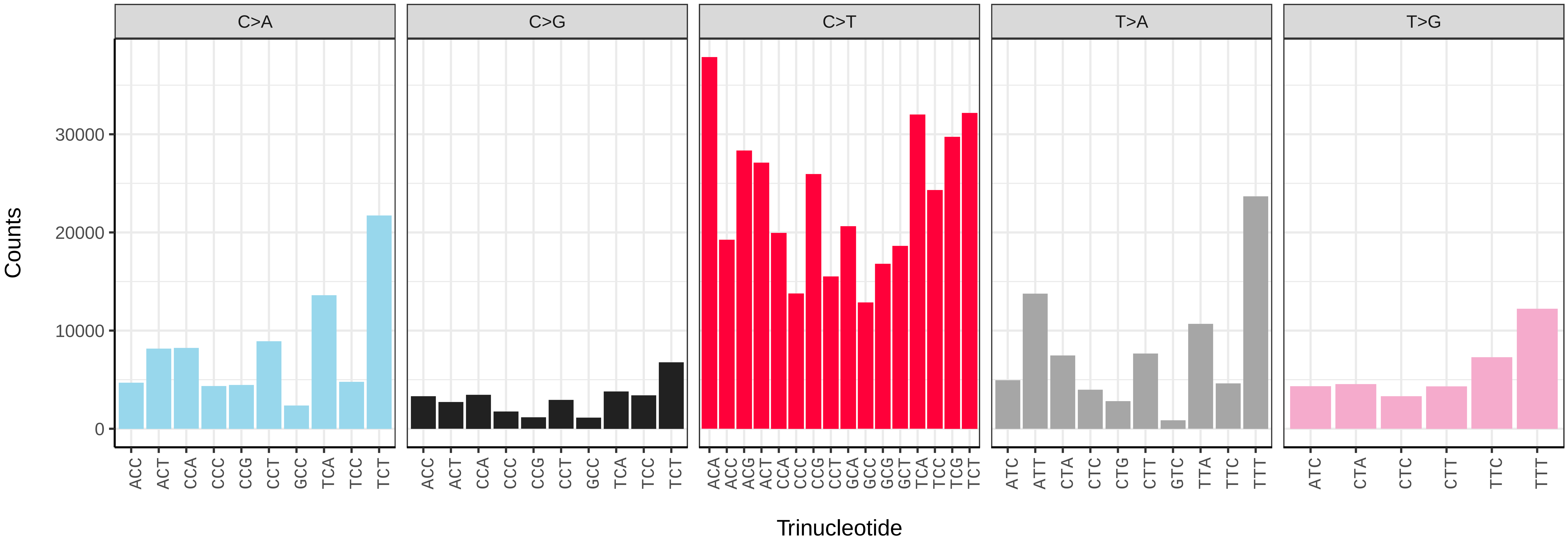


*Carex arenaria* (lpCarAren1) somatic mutational spectrum (SBS96 classification)

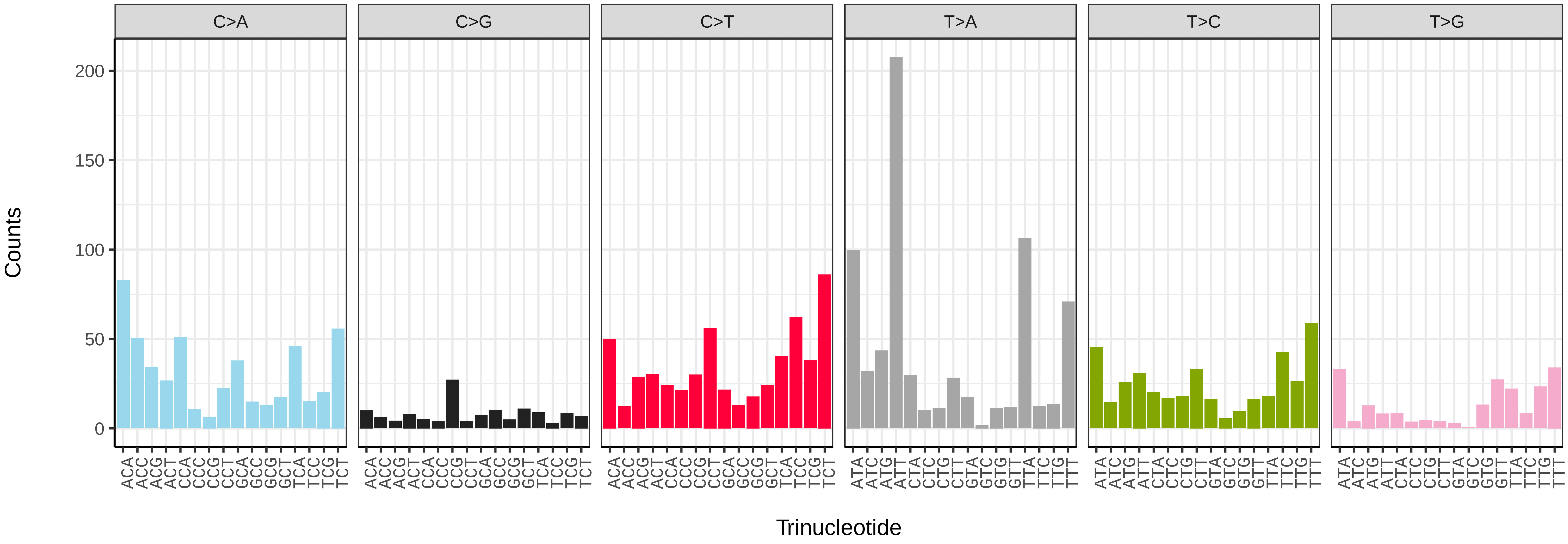


*Carex arenaria* (lpCarAren1) somatic mutational spectrum (SBS52 classification)

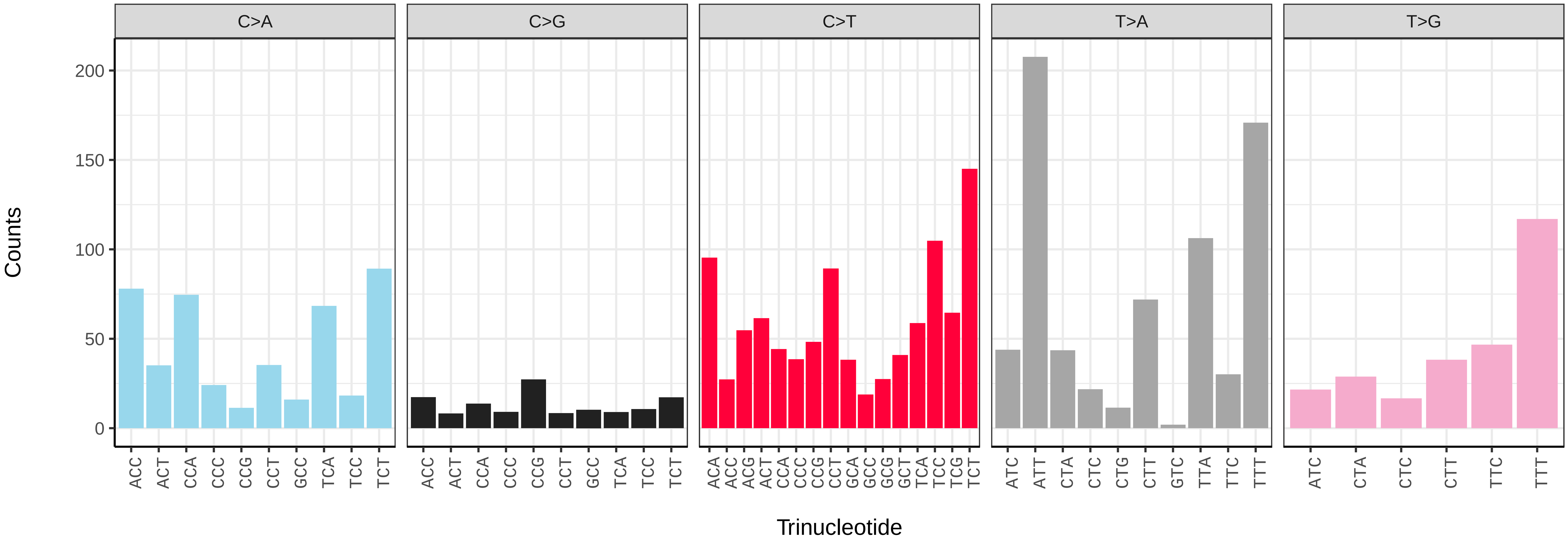

*Carex depauperata* (lpCarDepa1) germline mutational spectrum

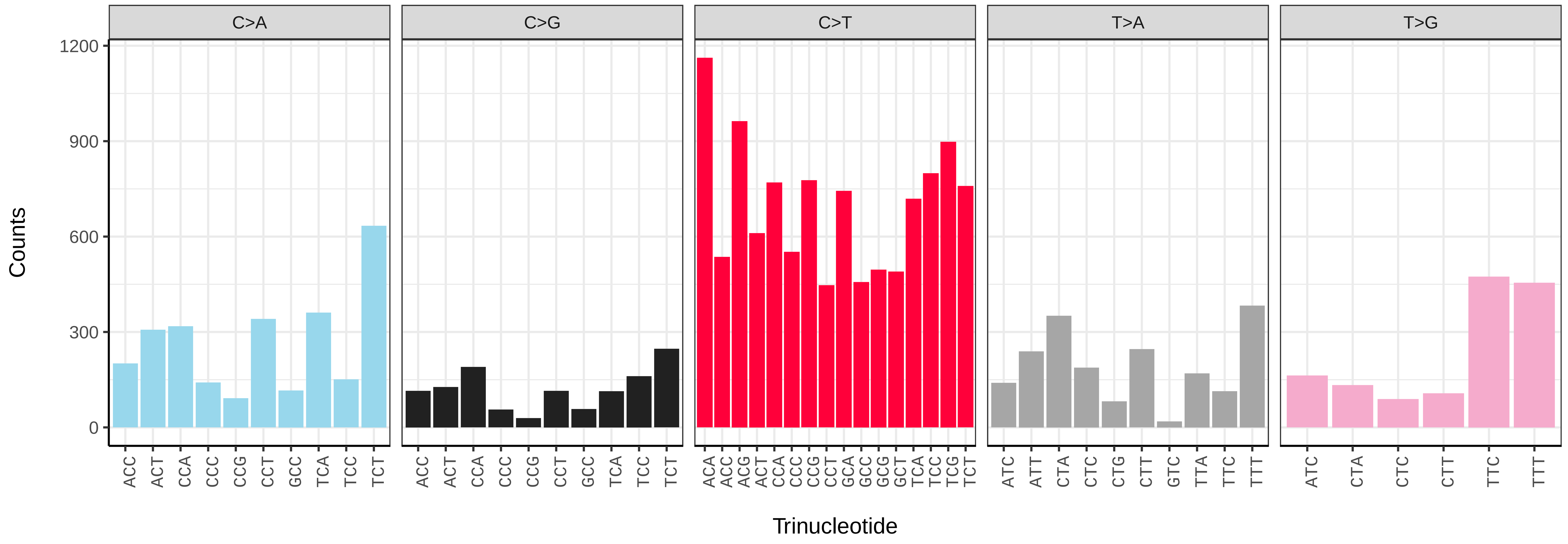


*Carex depauperata* (lpCarDepa1) somatic mutational spectrum (SBS96 classification)

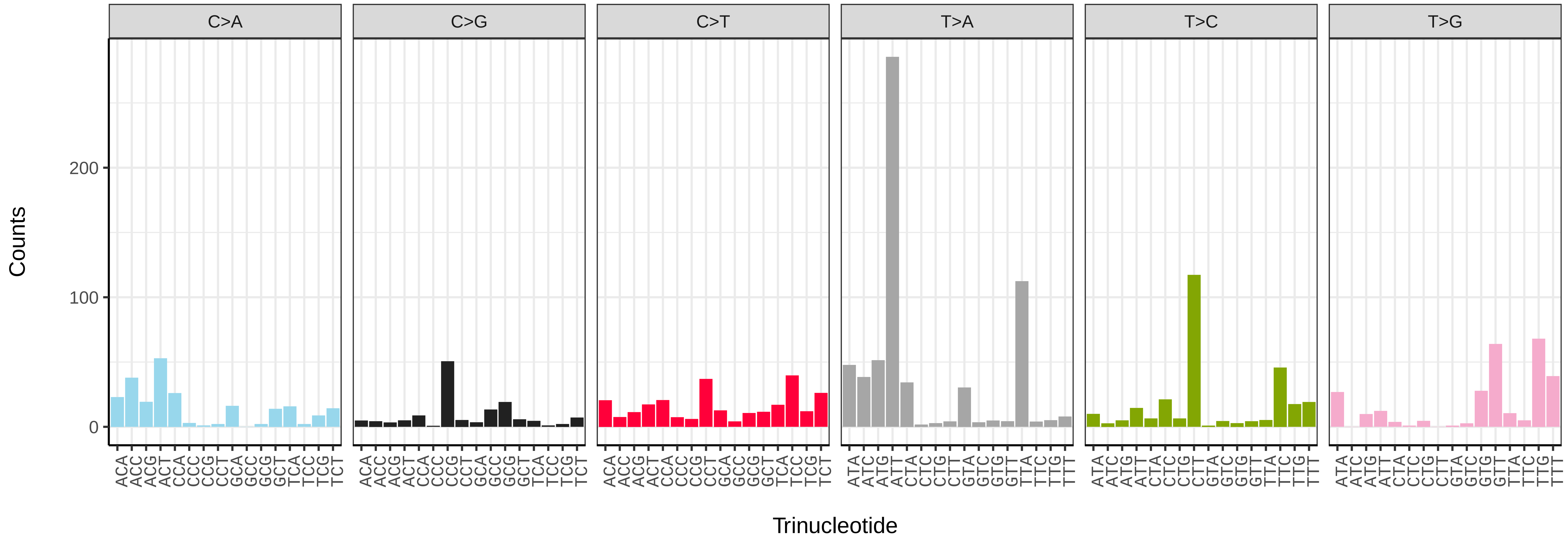


*Carex depauperata* (lpCarDepa1) somatic mutational spectrum (SBS52 classification)

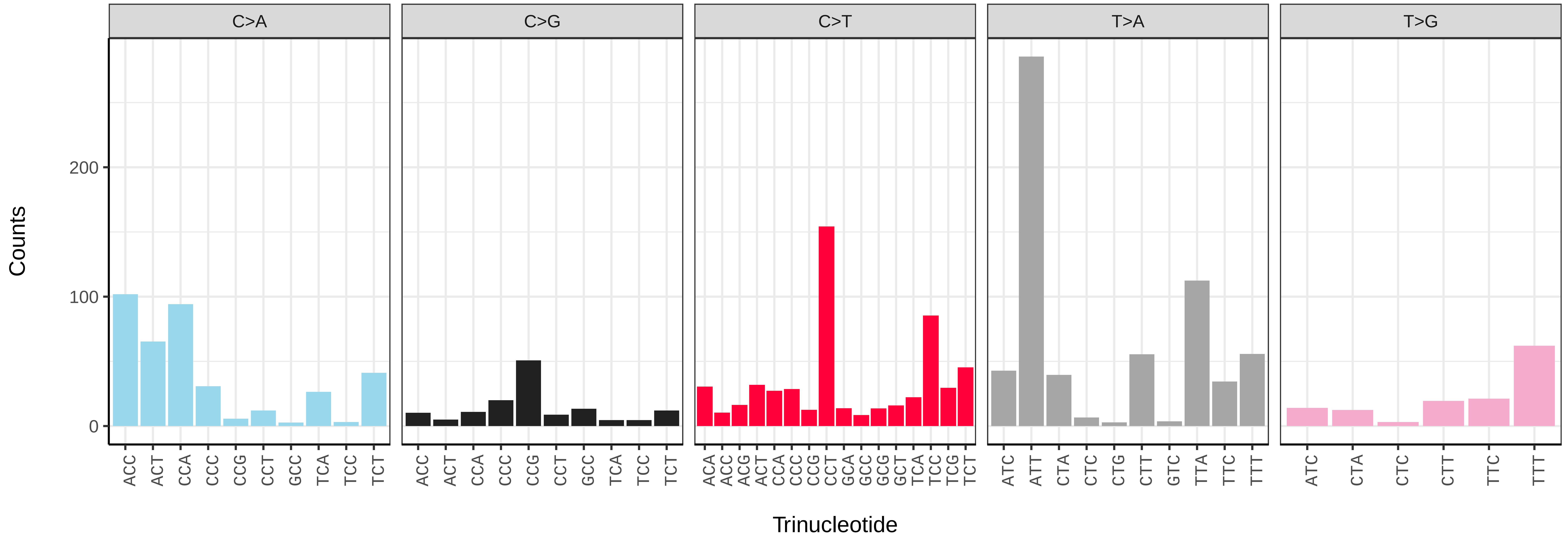

*Carex divulsa* (lpCarDivu1) germline mutational spectrum

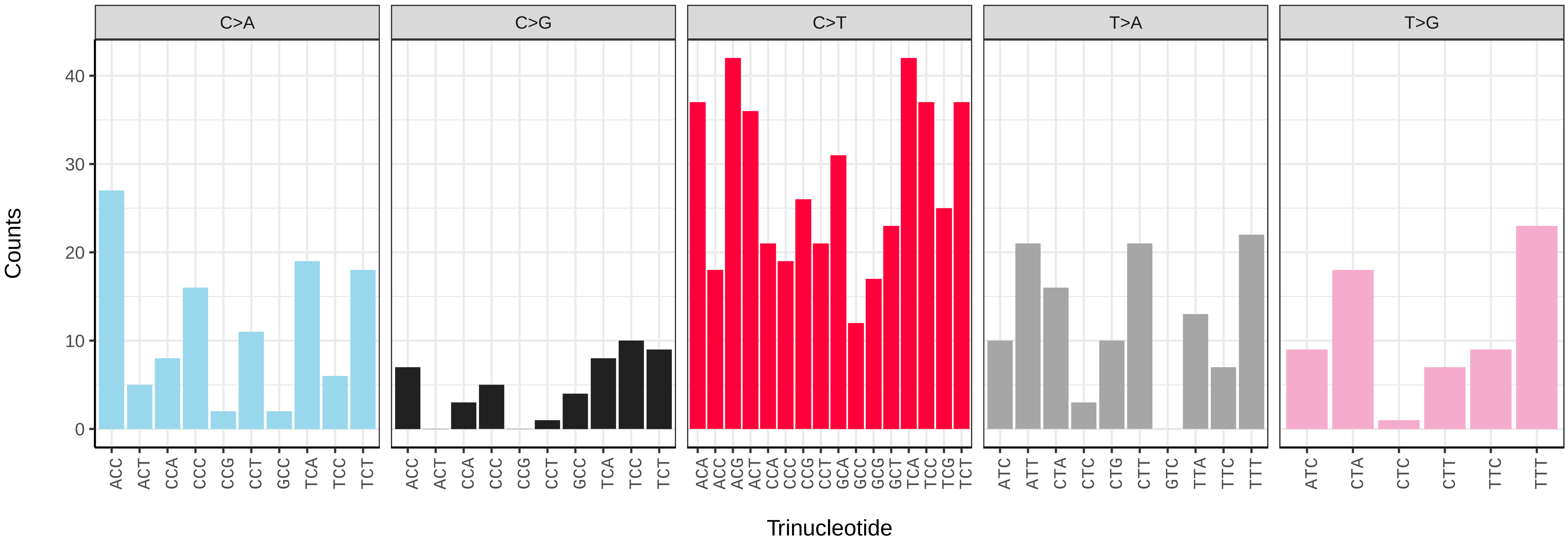


*Carex divulsa* (lpCarDivu1) somatic mutational spectrum (SBS96 classification)

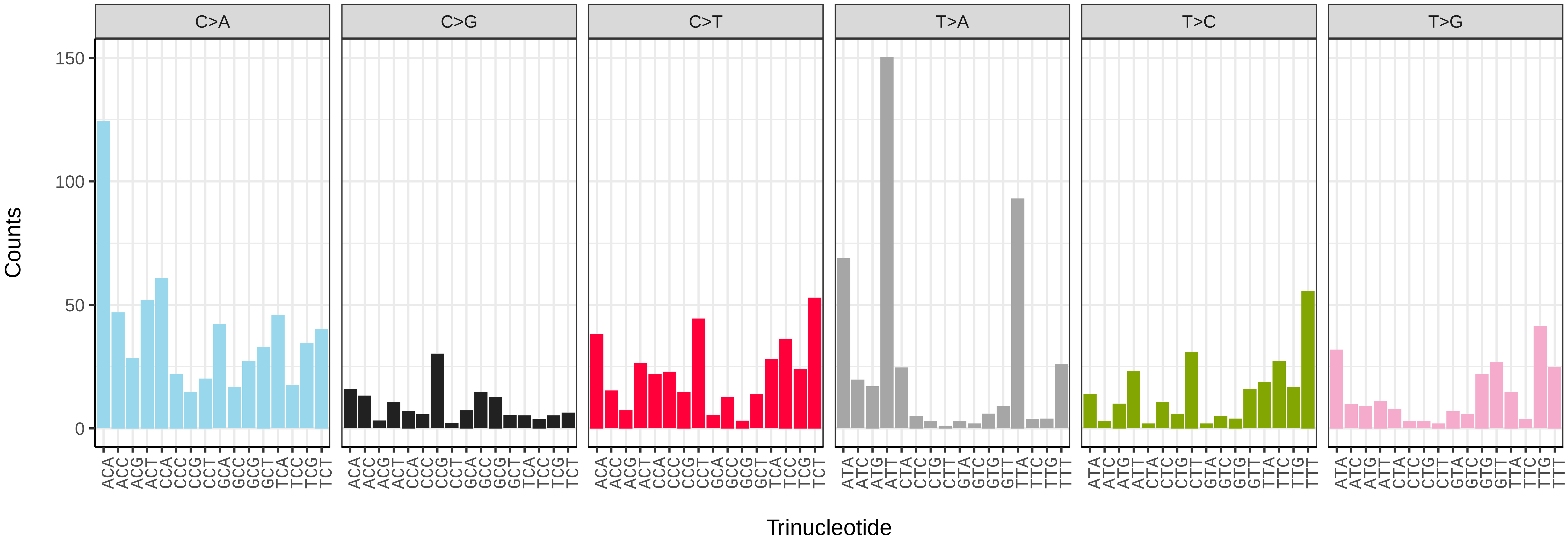


*Carex divulsa* (lpCarDivu1) somatic mutational spectrum (SBS52 classification)

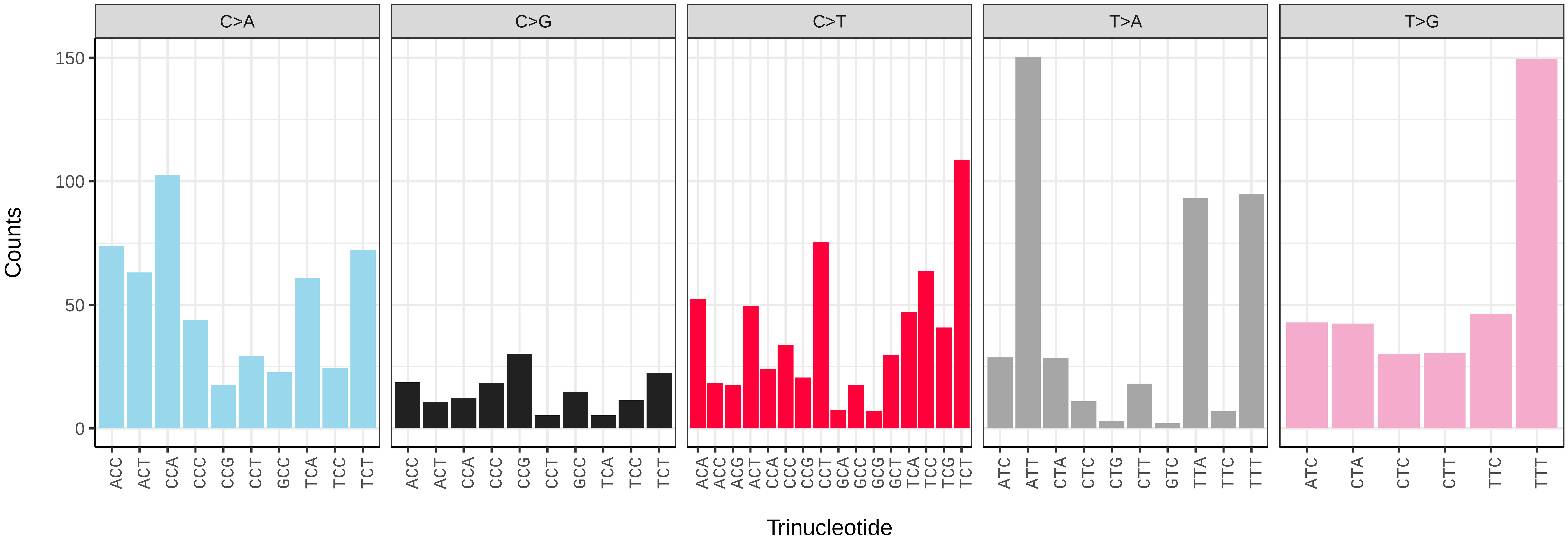

*Carex pendula* (lpCarPend1) germline mutational spectrum

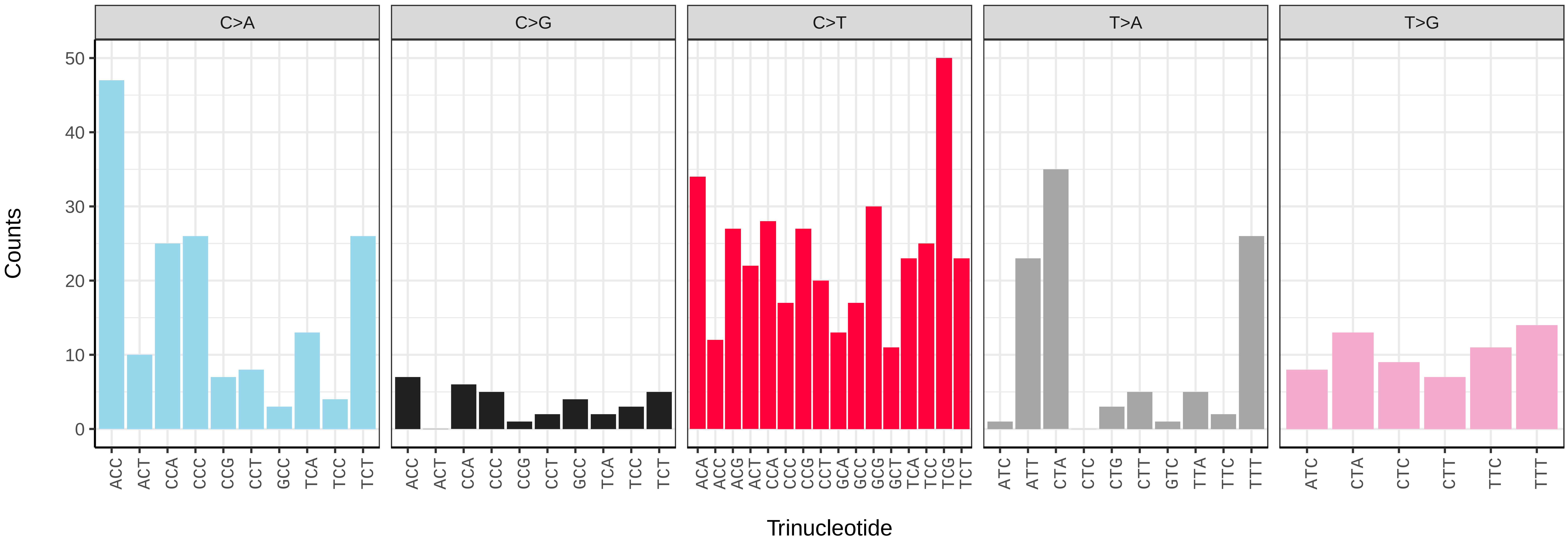


*Carex pendula* (lpCarPend1) somatic mutational spectrum (SBS96 classification)

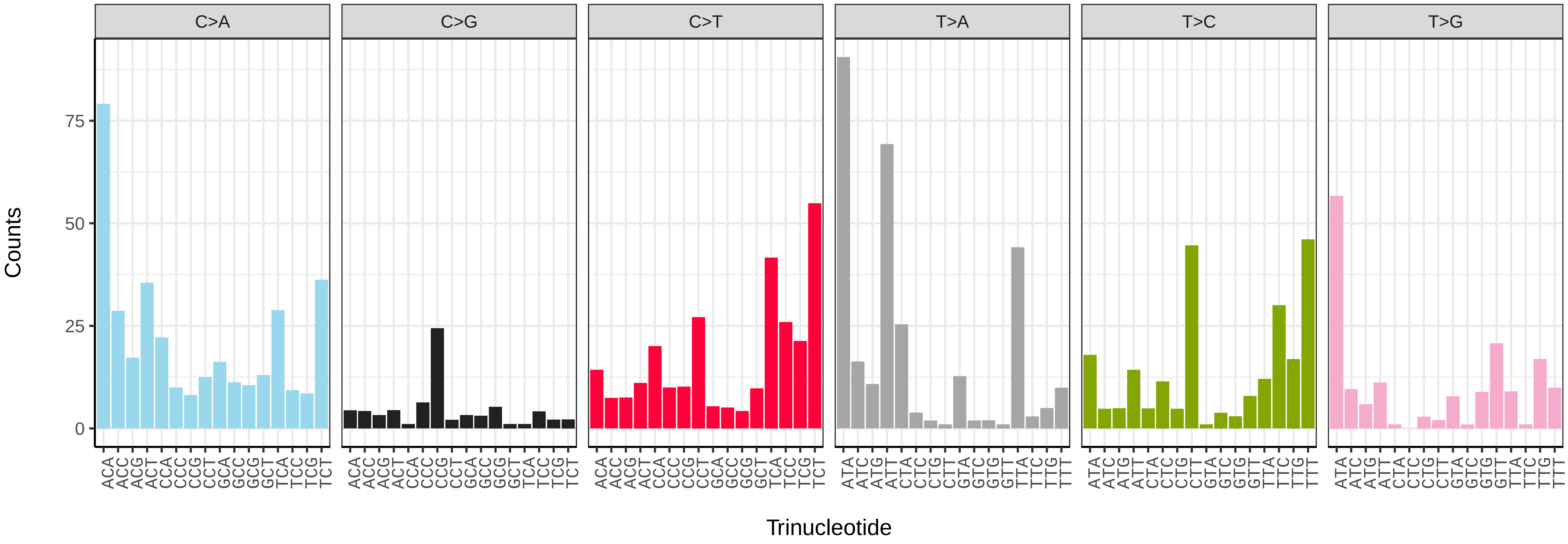


*Carex pendula* (lpCarPend1) somatic mutational spectrum (SBS52 classification)

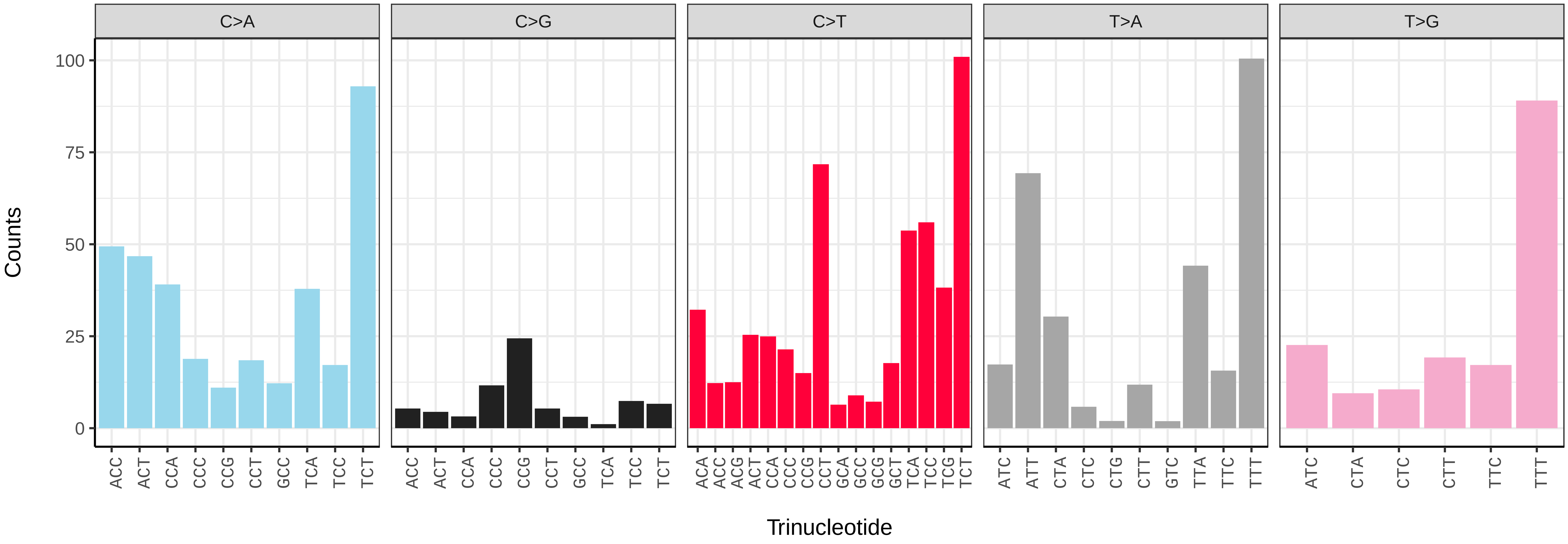

*Carex rostrata* (lpCarRost1) germline mutational spectrum

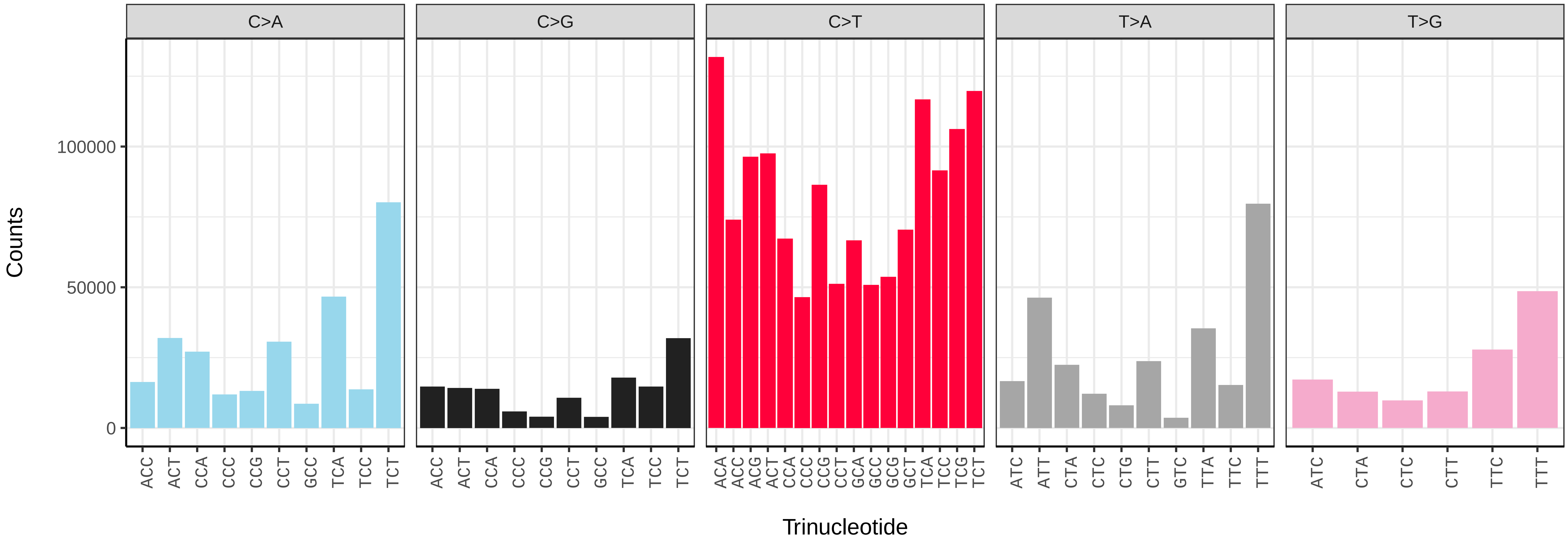


*Carex rostrata* (lpCarRost1) somatic mutational spectrum (SBS96 classification)

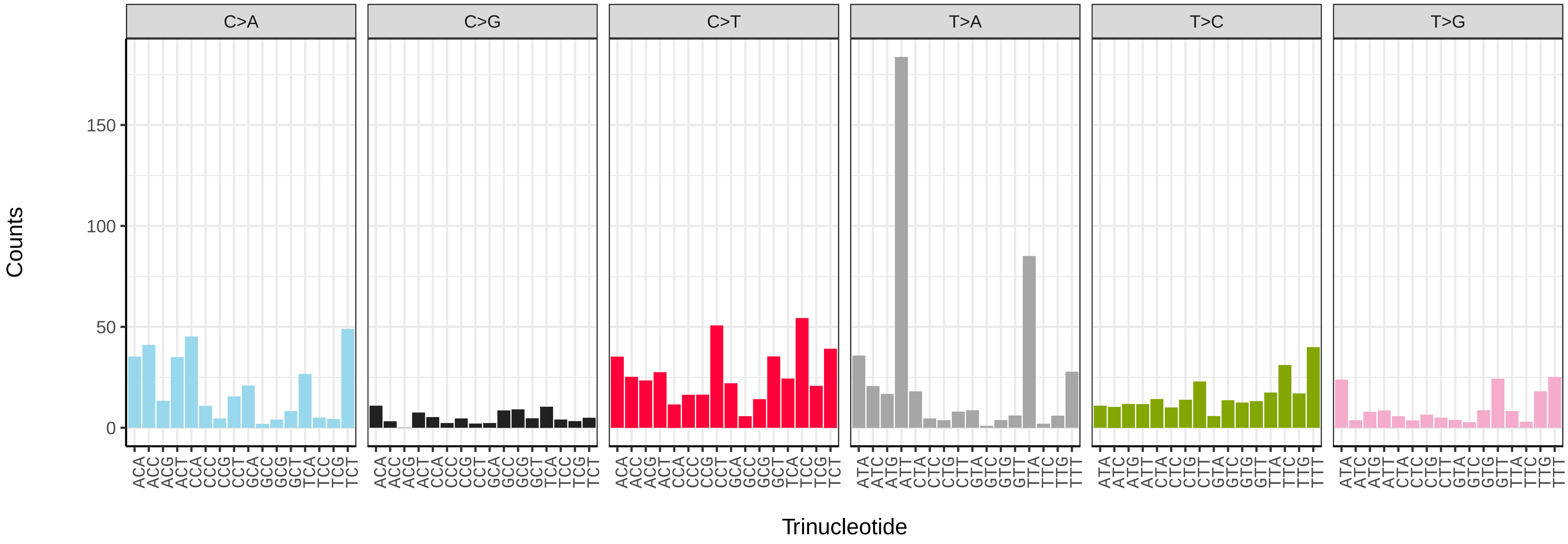


*Carex rostrata* (lpCarRost1) somatic mutational spectrum (SBS52 classification)

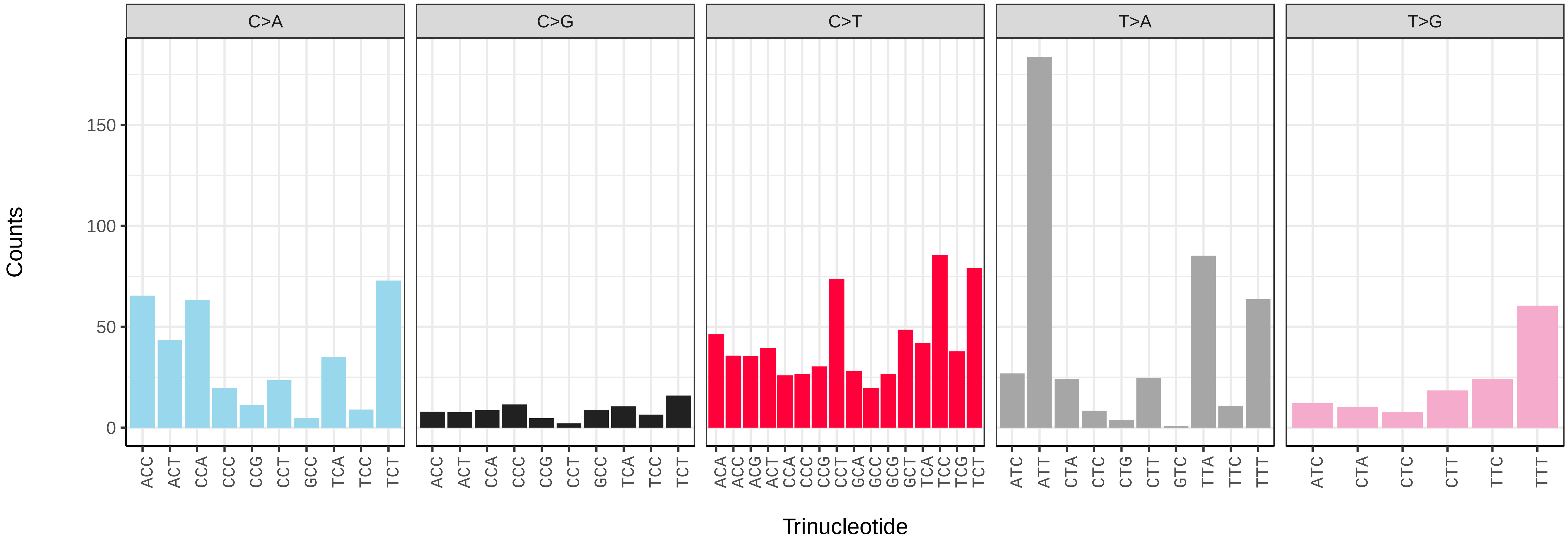

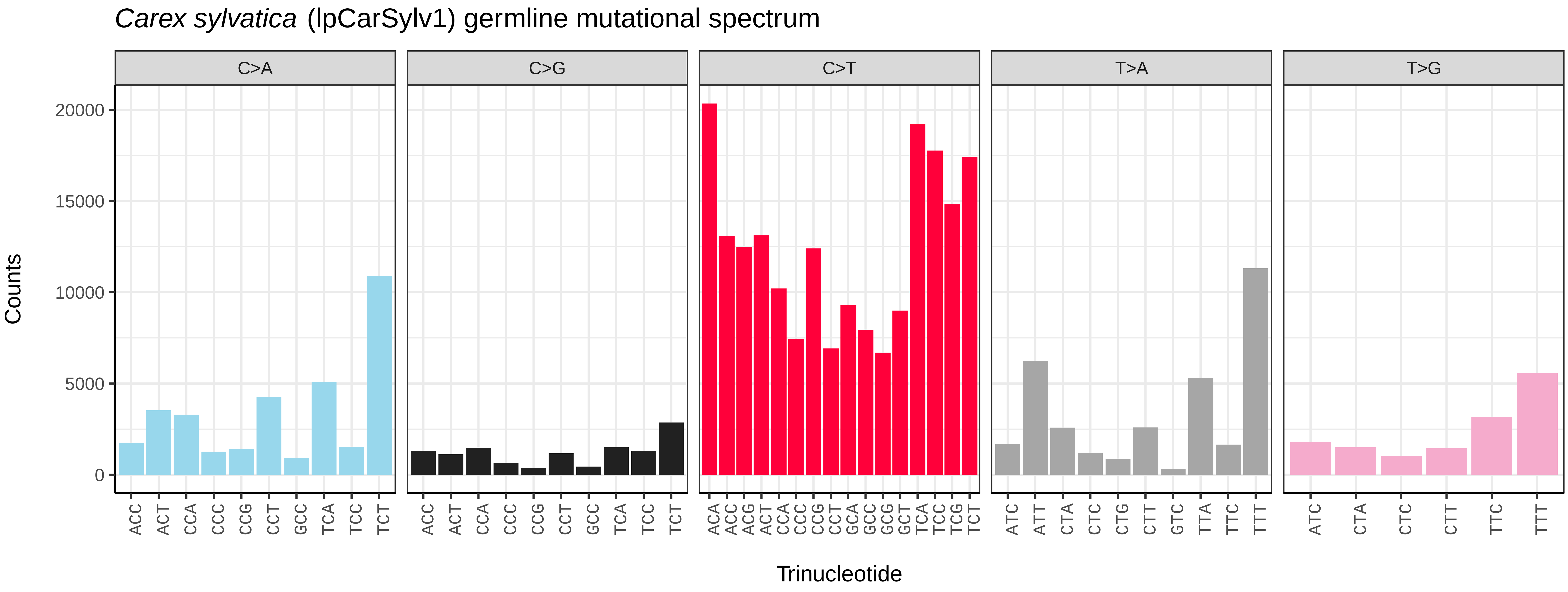
Carex sylvatica (lpCarSylv1) germline mutational spectrum
C>A
C>G
C>T
T>A
T>G
Counts
20000
15000
10000
5000
0
ACC ACT CCA CCC CCG CCT GCC TCA TCC TCT
ACC ACT CCA CCC CCG CCT GCC TCA TCC TCT
ACA ACC ACG ACT CCA CCC CCG CCT GCA GCC GCG GCT TCA TCC TCG TCT
ATC ATT CTA CTC CTG CTT GTC TTA TTC TTT
ATC CTA CTC CTT TTC TTT
Trinucleotide

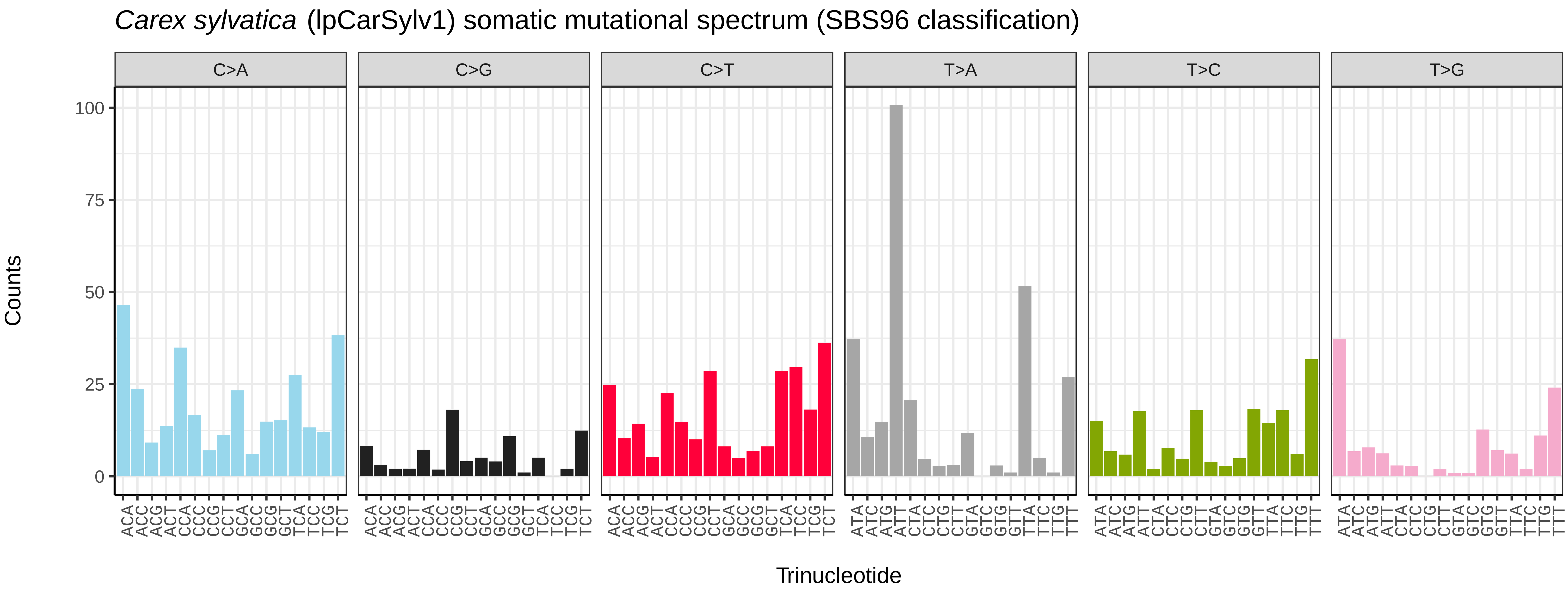
Carex sylvatica (lpCarSylv1) somatic mutational spectrum (SBS96 classification)
C>A
C>G
C>T
T>A
T>C
T>G
Counts
100
75
50
25
0
ACA ACC ACG ACT CCA CCC CCG CCT GCA GCC GCG GCT TCA TCC TCG TCT
ACA ACC ACG ACT CCA CCC CCG CCT GCA GCC GCG GCT TCA TCC TCG TCT
ACA ACC ACG ACT CCA CCC CCG CCT GCA GCC GCG GCT TCA TCC TCG TCT
ATA ATC ATG ATT CTA CTC CTG CTT GTA GTC GTG GTT TTA TTC TTG TTT
ATA ATC ATG ATT CTA CTC CTG CTT GTA GTC GTG GTT TTA TTC TTG TTT
ATA ATC ATG ATT CTA CTC CTG CTT GTA GTC GTG GTT TTA TTC TTG TTT
Trinucleotide

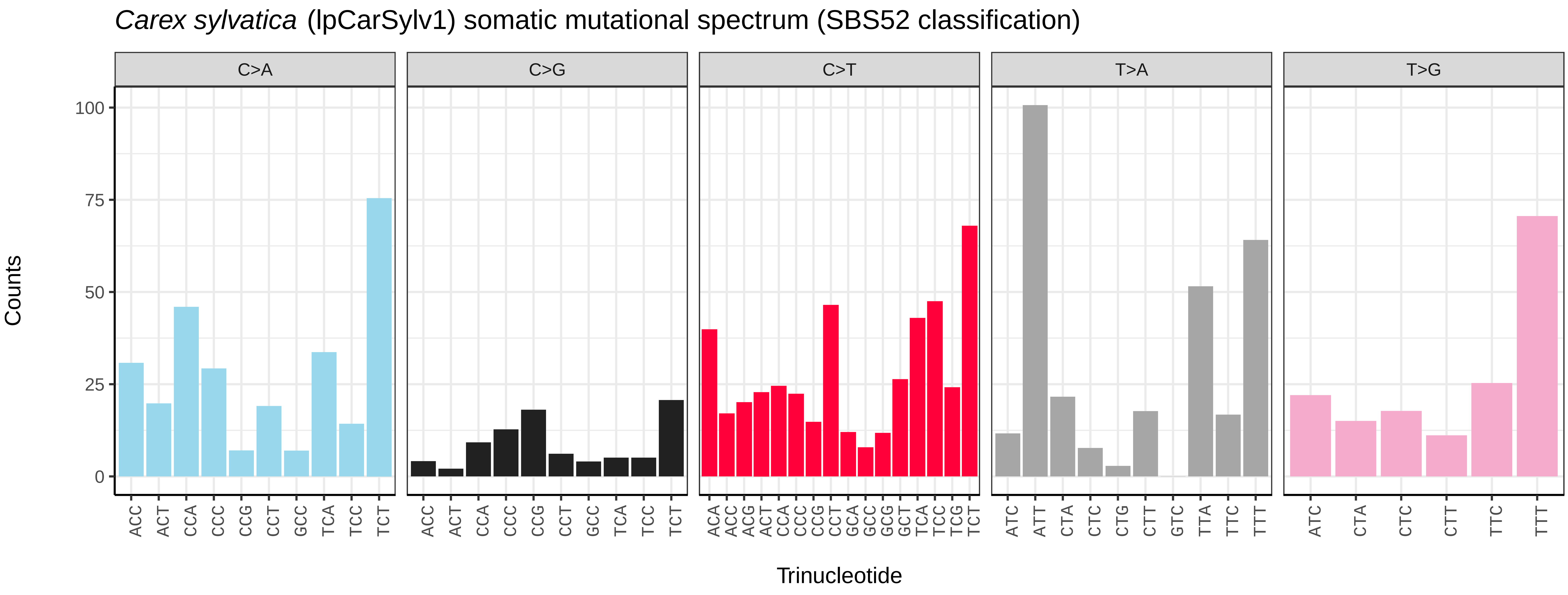
Carex sylvatica (lpCarSylv1) somatic mutational spectrum (SBS52 classification)
C>A
C>G
C>T
T>A
T>G
Counts
100
75
50
25
0
ACC ACT CCA CCC CCG CCT GCC TCA TCC TCT
ACC ACT CCA CCC CCG CCT GCC TCA TCC TCT
ACA ACC ACG ACT CCA CCC CCG CCT GCA GCC GCG GCT TCA TCC TCG TCT
ATC ATT CTA CTC CTG CTT GTC TTA TTC TTT
ATC CTA CTC CTT TTC TTT
Trinucleotide

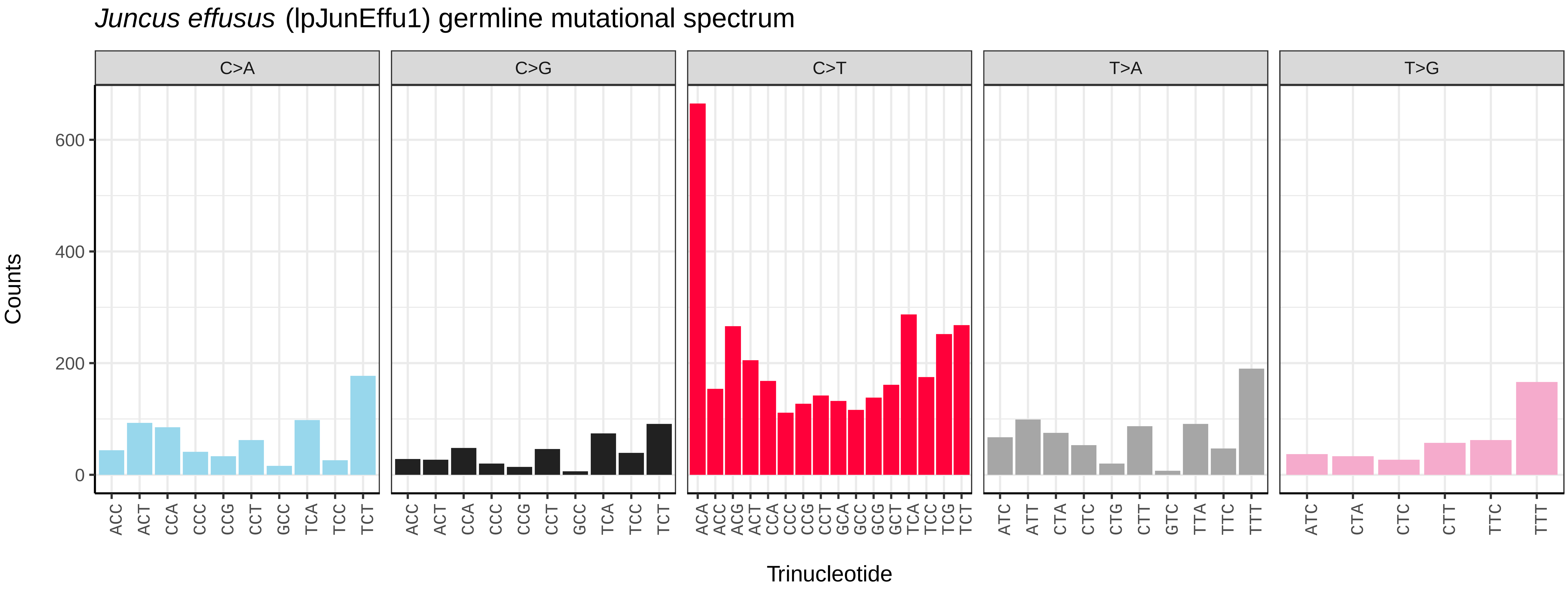
Juncus effusus (lpJunEffu1) germline mutational spectrum
C>A
C>G
C>T
T>A
T>G
Counts
0
200
400
600
Trinucleotide

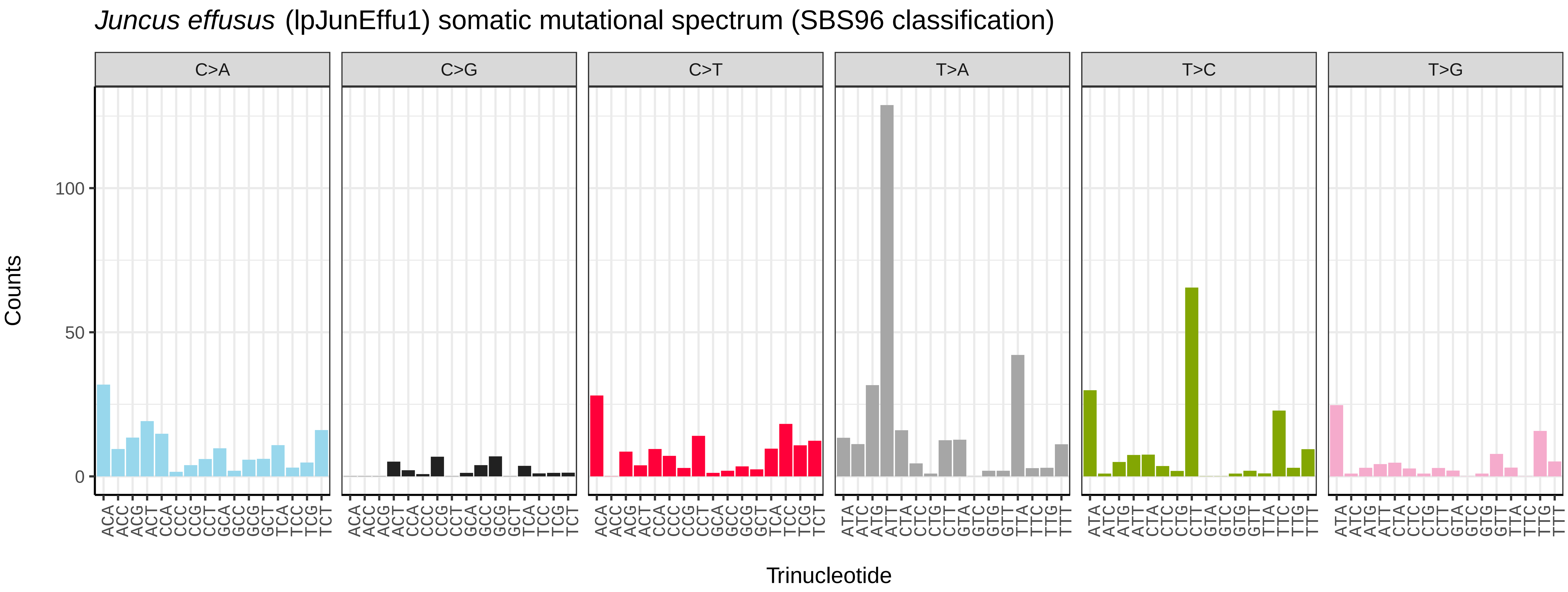
Juncus effusus (lpJunEffu1) somatic mutational spectrum (SBS96 classification)
C>A
C>G
C>T
T>A
T>C
T>G
Counts
0
50
100
Trinucleotide

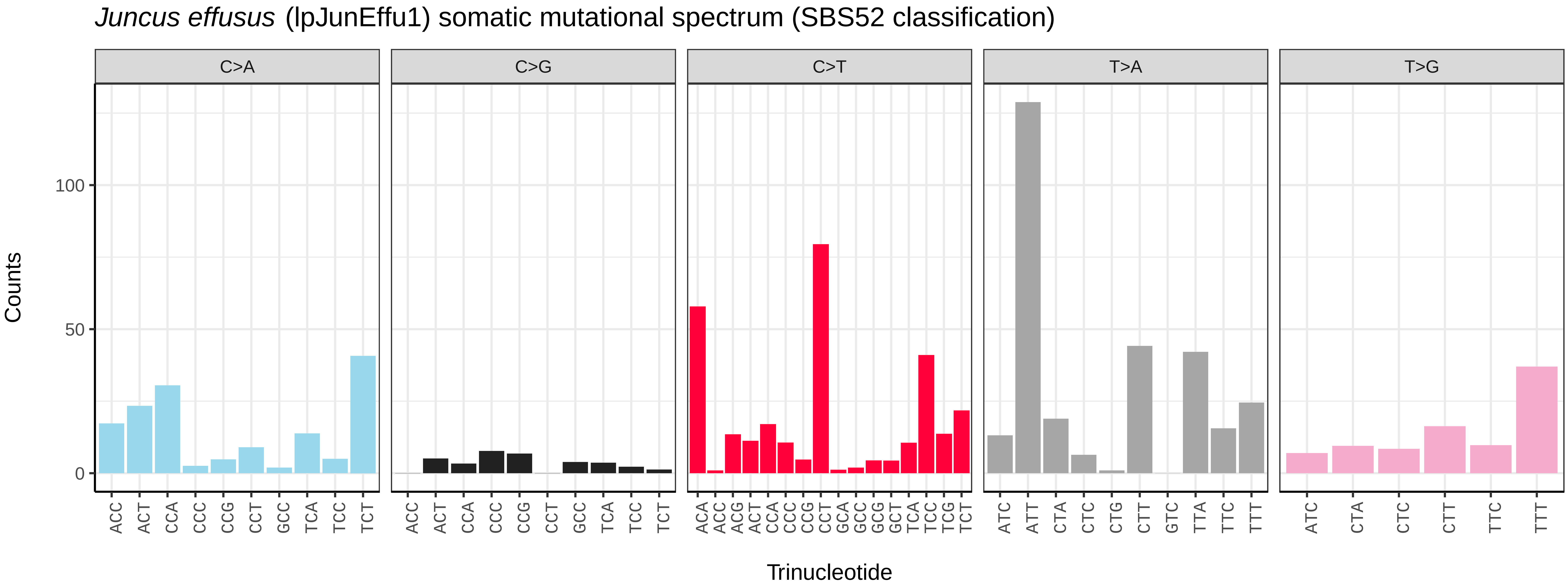
Juncus effusus (lpJunEffu1) somatic mutational spectrum (SBS52 classification)
C>A
C>G
C>T
T>A
T>G
Counts
0
50
100
Trinucleotide

*Luzula sylvatica* (lpLuzSylv1) germline mutational spectrum

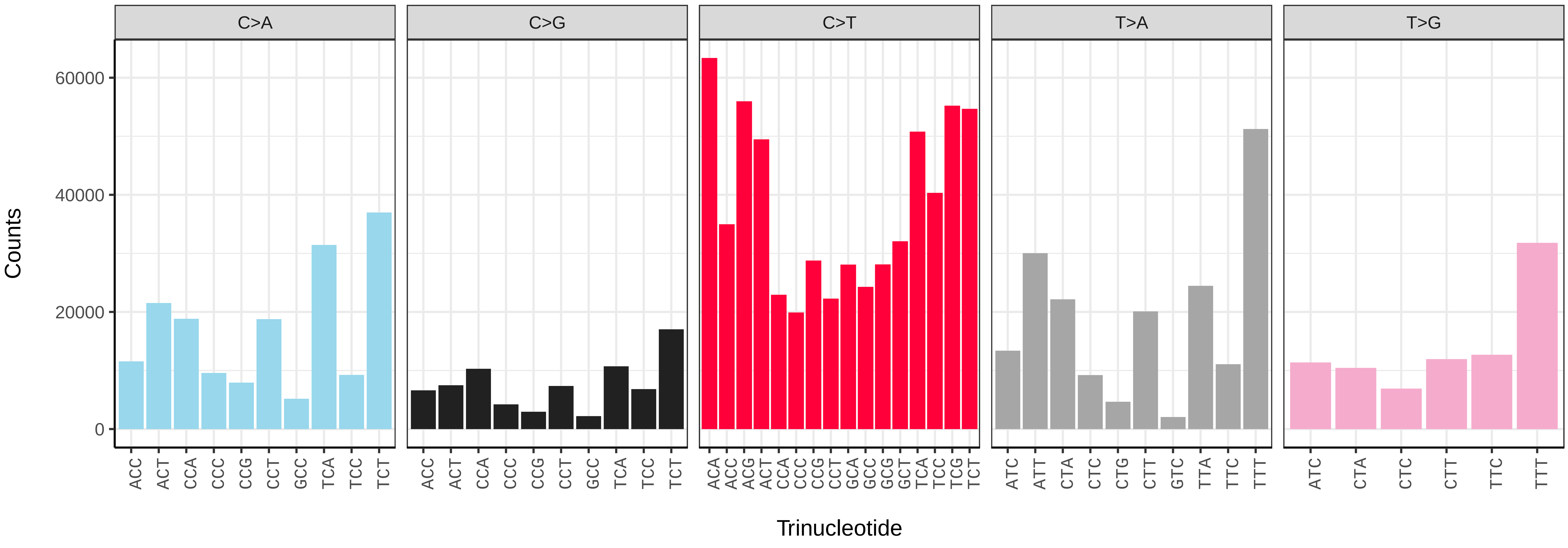


*Luzula sylvatica* (lpLuzSylv1) somatic mutational spectrum (SBS96 classification)

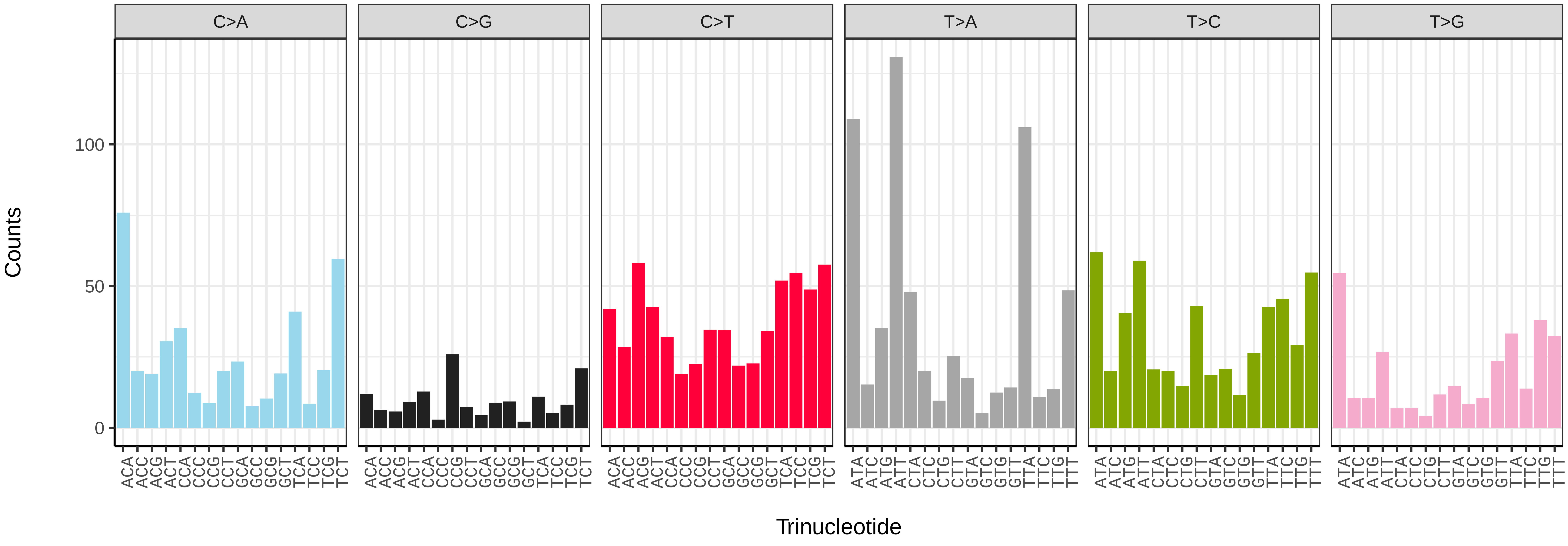


*Luzula sylvatica* (lpLuzSylv1) somatic mutational spectrum (SBS52 classification)

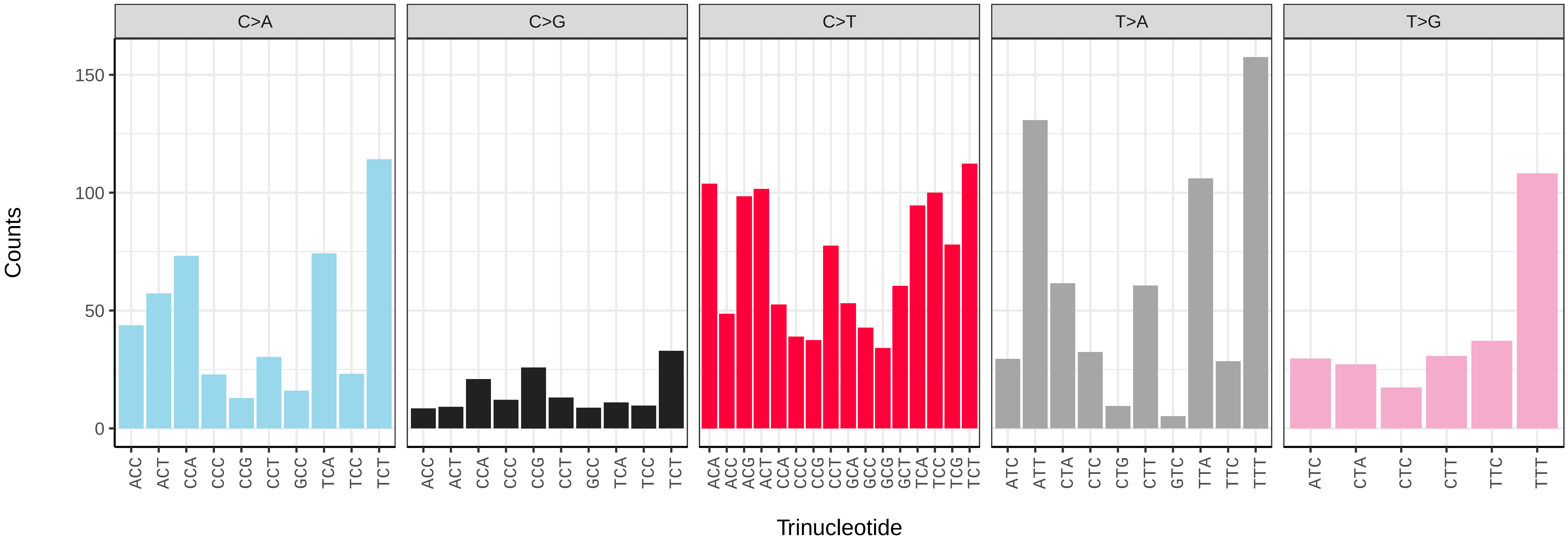

*Phragmites australis* (lpPhrAust1) germline mutational spectrum

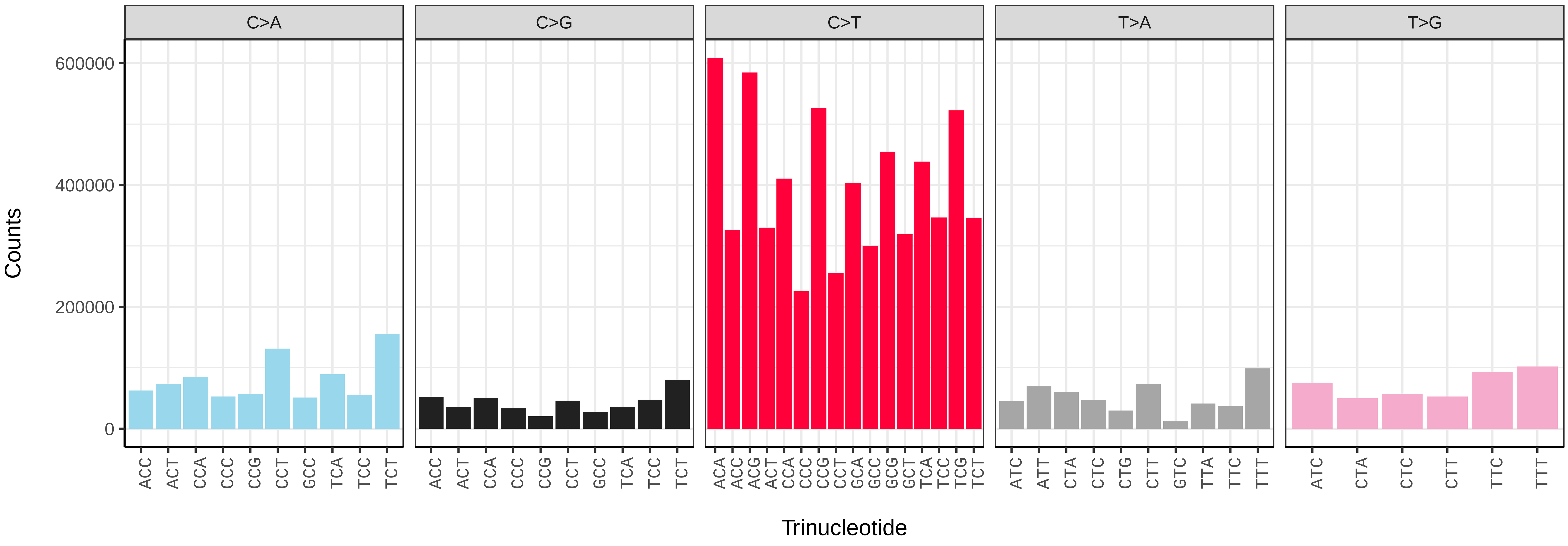


*Phragmites australis* (lpPhrAust1) somatic mutational spectrum (SBS96 classification)

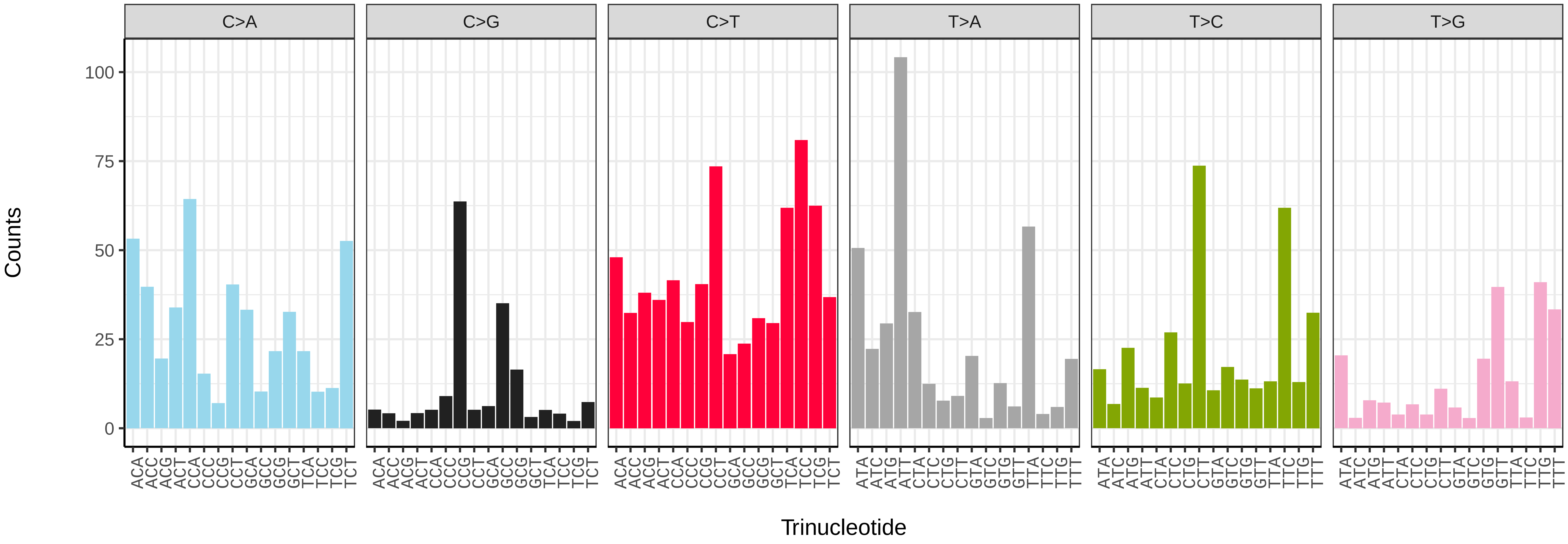


*Phragmites australis* (lpPhrAust1) somatic mutational spectrum (SBS52 classification)

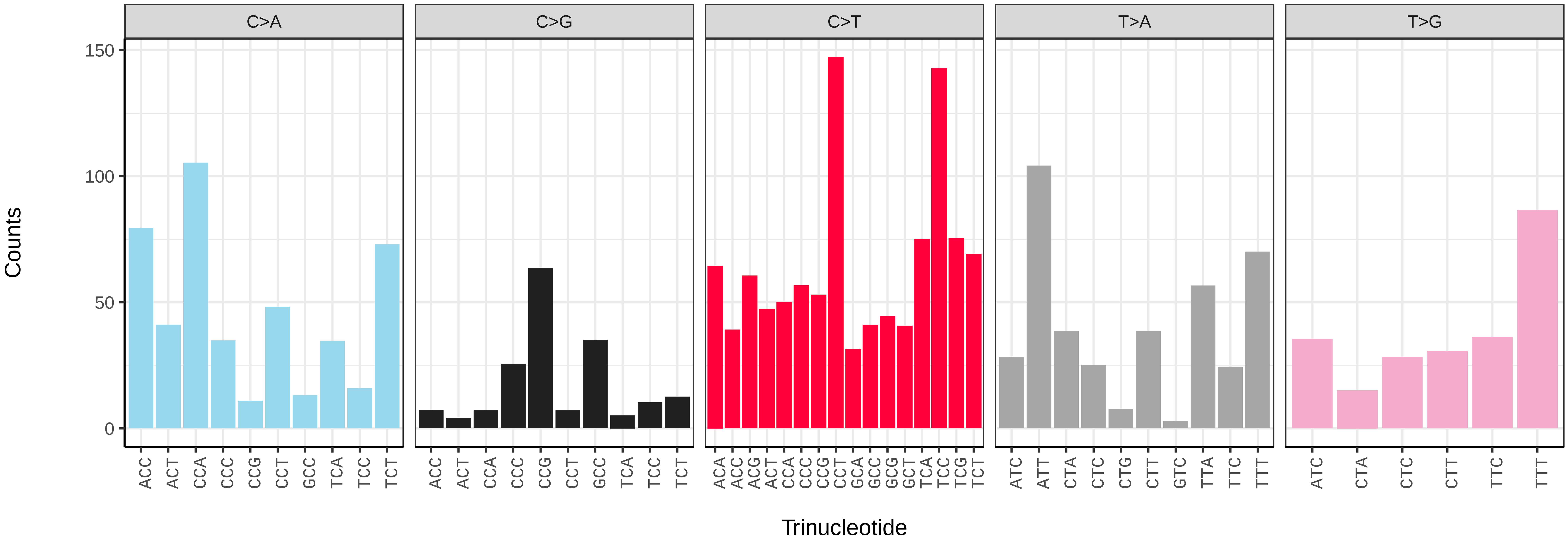

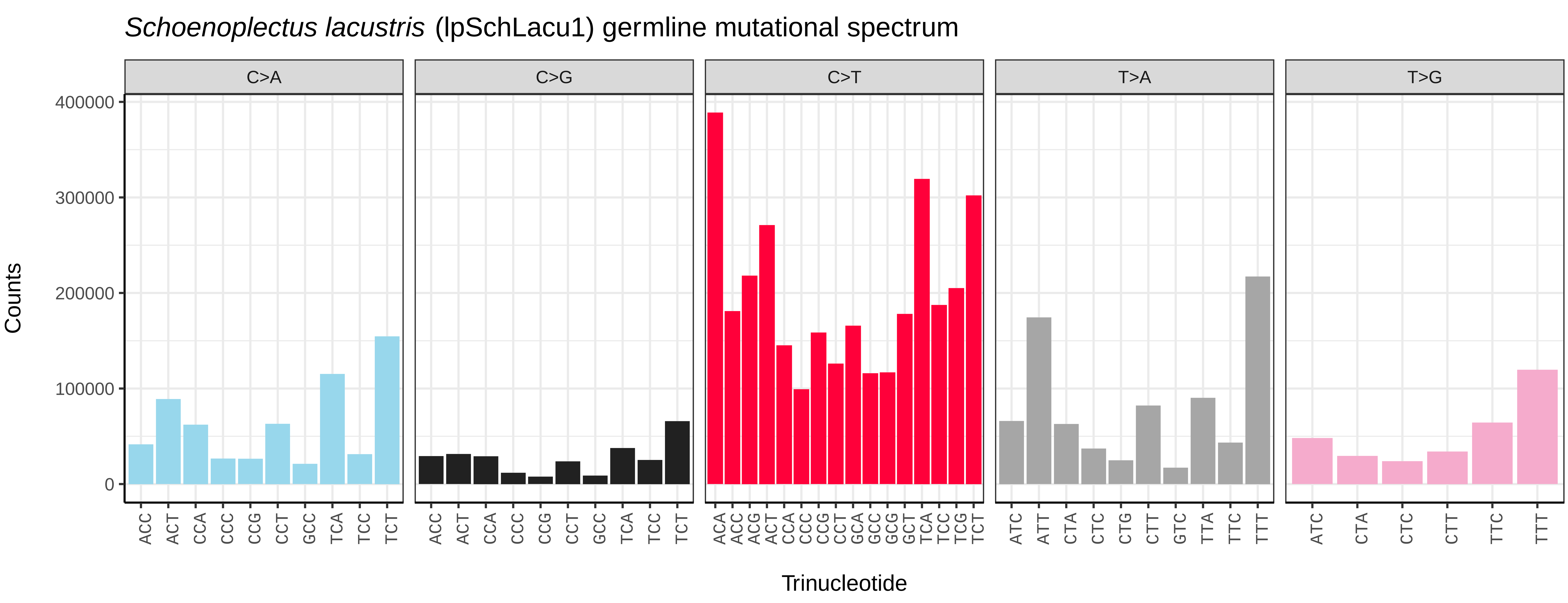
*Schoenoplectus lacustris* (lpSchLacu1) germline mutational spectrum
C>A
C>G
C>T
T>A
T>G
Counts
400000
300000
200000
100000
0
ACC ACT CCA CCC CCG CCT GCC TCA TCC TCT
ACA ACC ACG ACT CCA CCC CCG CCT GCA GCC GCG GCT TCA TCC TCG TCT
ATC ATT CTA CTC CTG CTT GTC TTA TTC TTT
ATC CTA CTC CTT TTC TTT
Trinucleotide

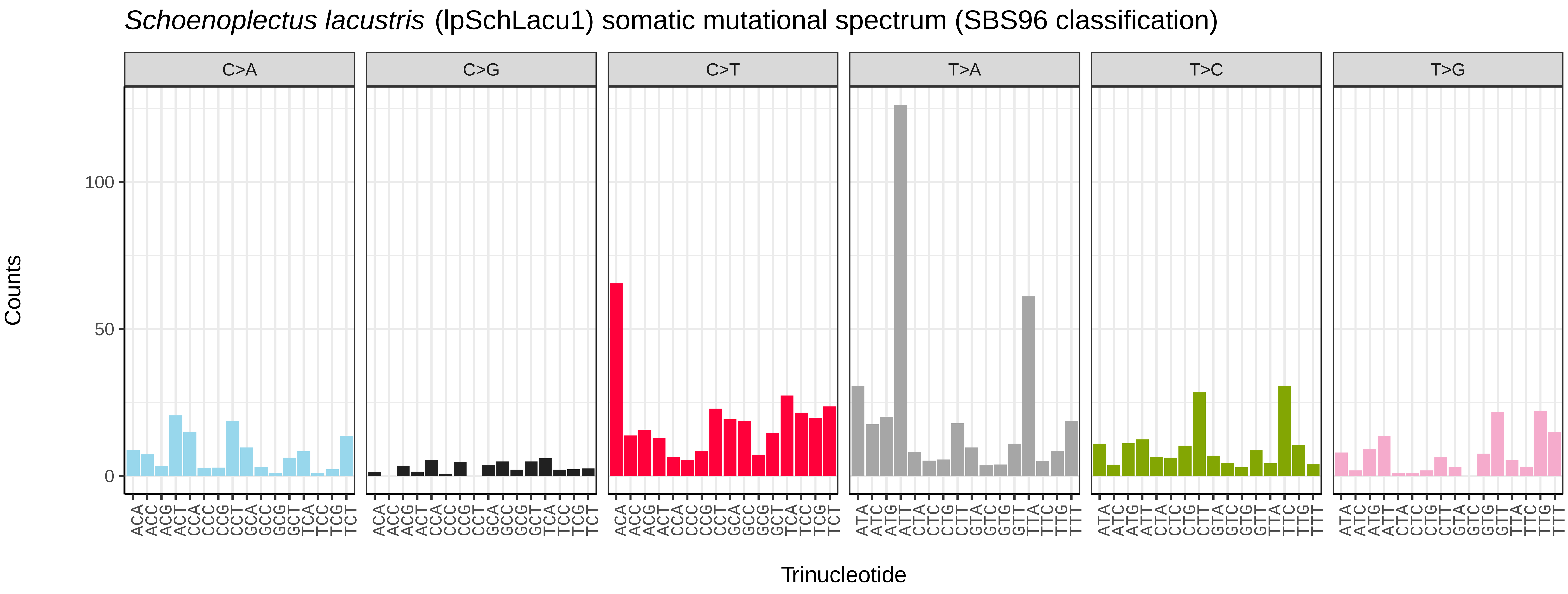
*Schoenoplectus lacustris* (lpSchLacu1) somatic mutational spectrum (SBS96 classification)
C>A
C>G
C>T
T>A
T>C
T>G
Counts
100
50
0
ACA ACC ACG ACT CCA CCC CCG CCT GCA GCC GCG GCT TCA TCC TCG TCT
ATA ATC ATG ATT CTA CTC CTG CTT GTA GTC GTG GTT TTA TTC TTG TTT
Trinucleotide

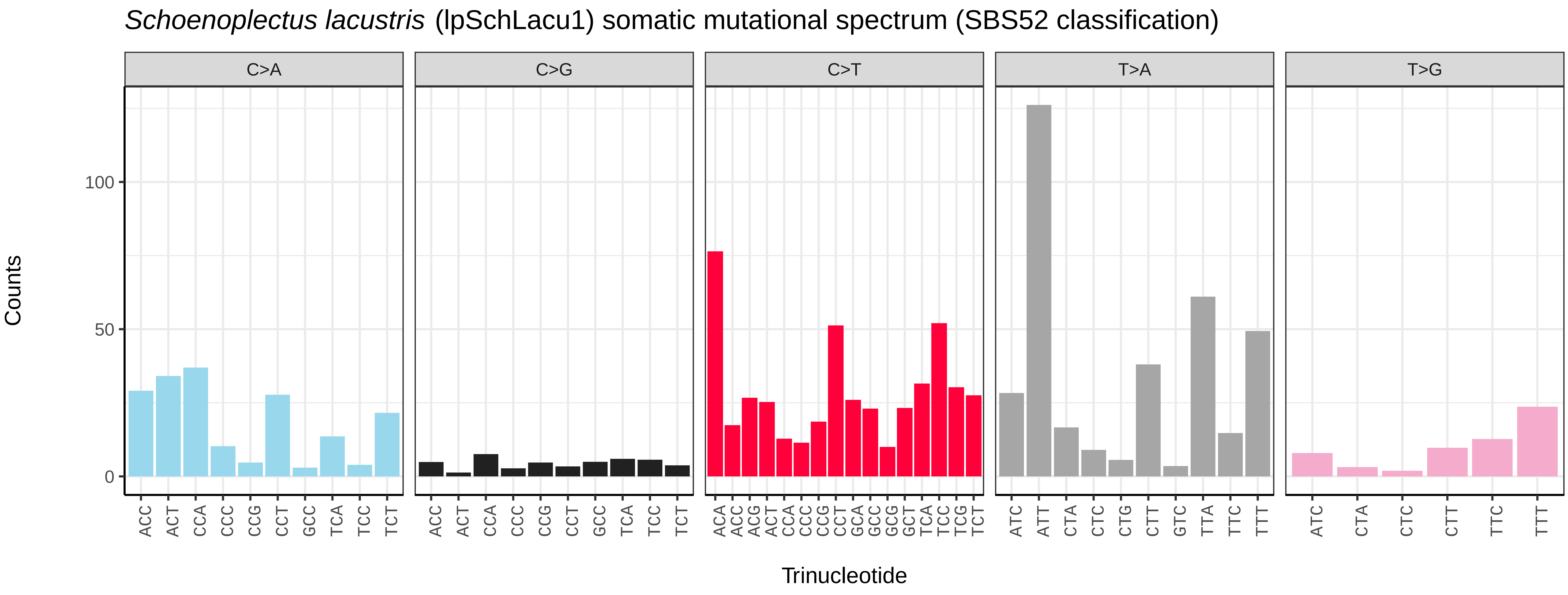
*Schoenoplectus lacustris* (lpSchLacu1) somatic mutational spectrum (SBS52 classification)
C>A
C>G
C>T
T>A
T>G
Counts
100
50
0
ACC ACT CCA CCC CCG CCT GCC TCA TCC TCT
ACA ACC ACG ACT CCA CCC CCG CCT GCA GCC GCG GCT TCA TCC TCG TCT
ATC ATT CTA CTC CTG CTT GTC TTA TTC TTT
ATC CTA CTC CTT TTC TTT
Trinucleotide

*Schoenoplectus triqueter* (lpSchTriq1) germline mutational spectrum

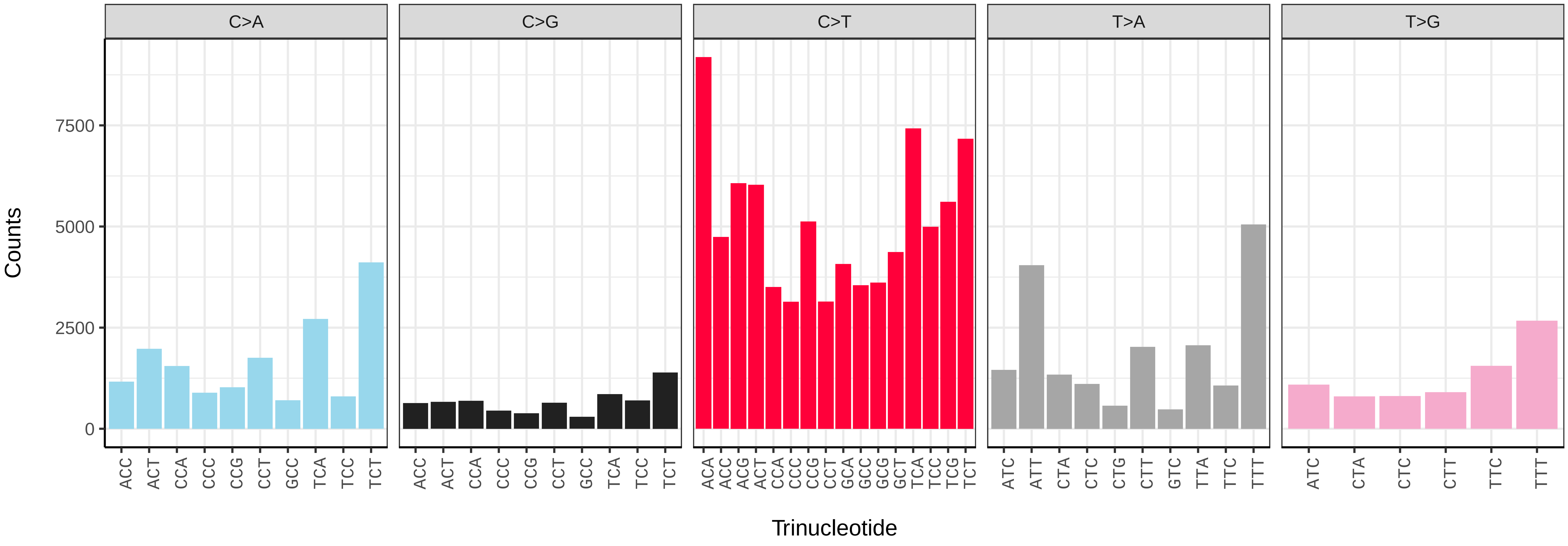


*Schoenoplectus triqueter* (lpSchTriq1) somatic mutational spectrum (SBS96 classification)

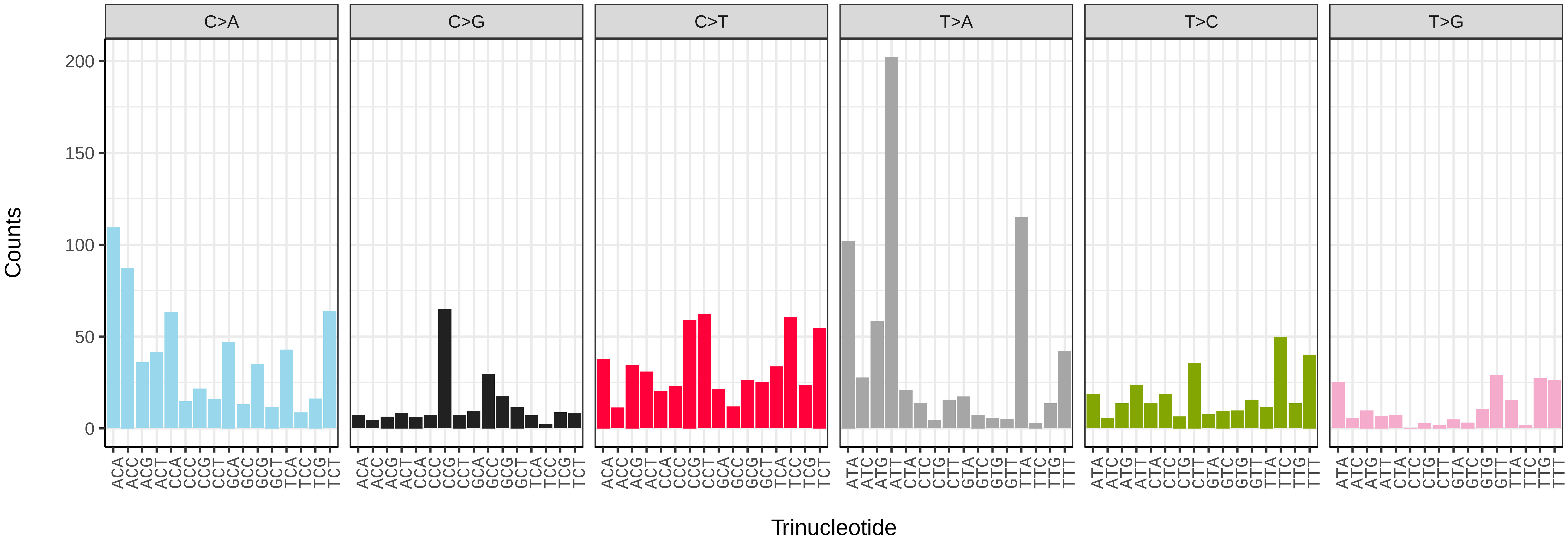


*Schoenoplectus triqueter* (lpSchTriq1) somatic mutational spectrum (SBS52 classification)

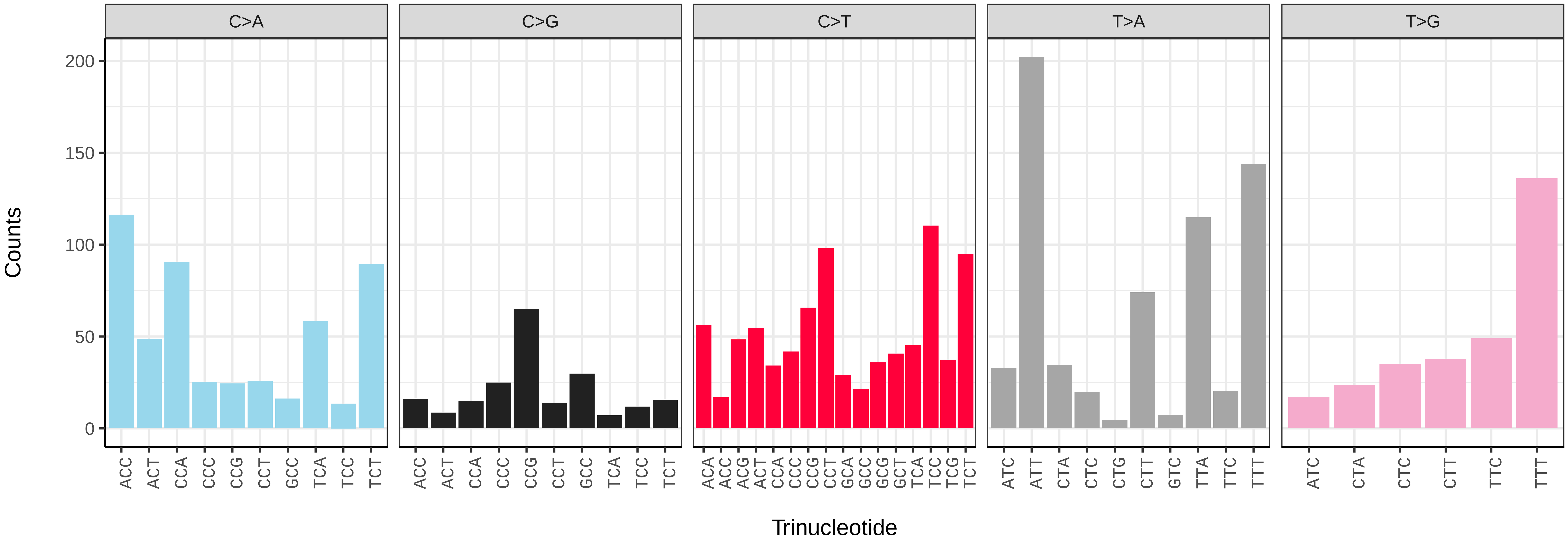

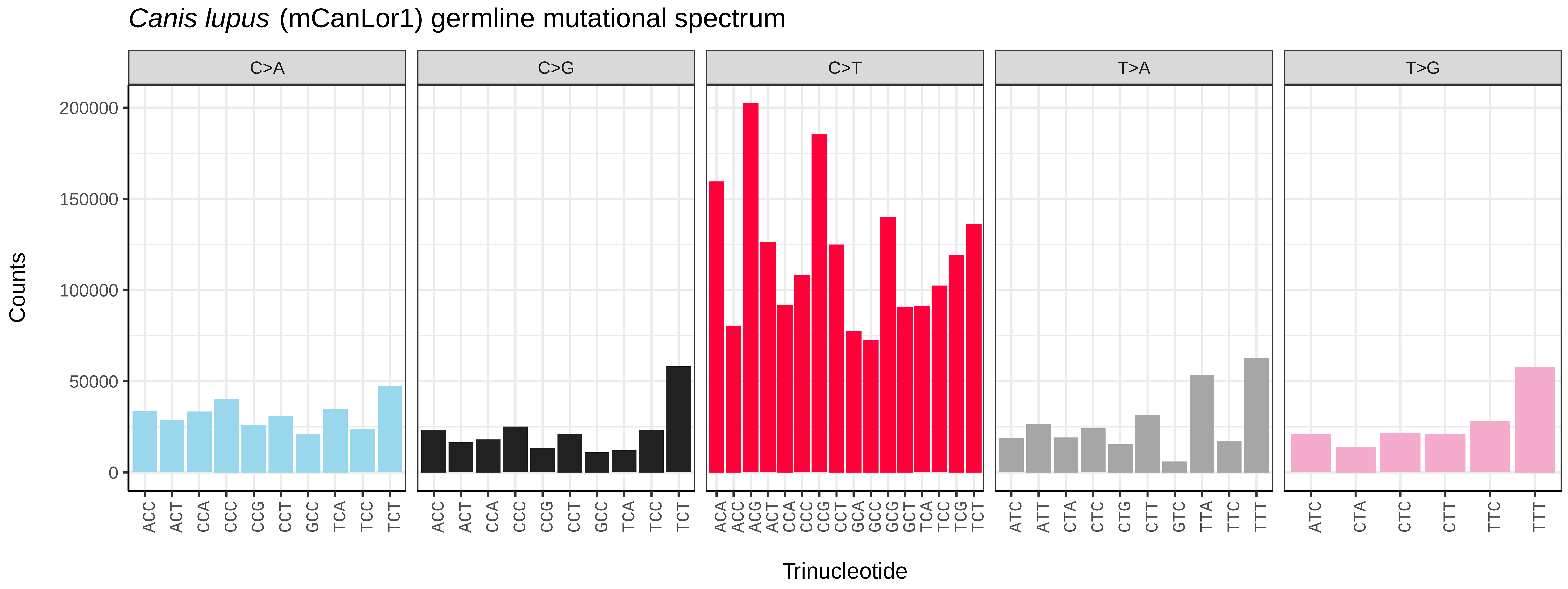
Canis lupus (mCanLor1) germline mutational spectrum
C>A
C>G
C>T
T>A
T>G
Counts
200000
150000
100000
50000
0
Trinucleotide

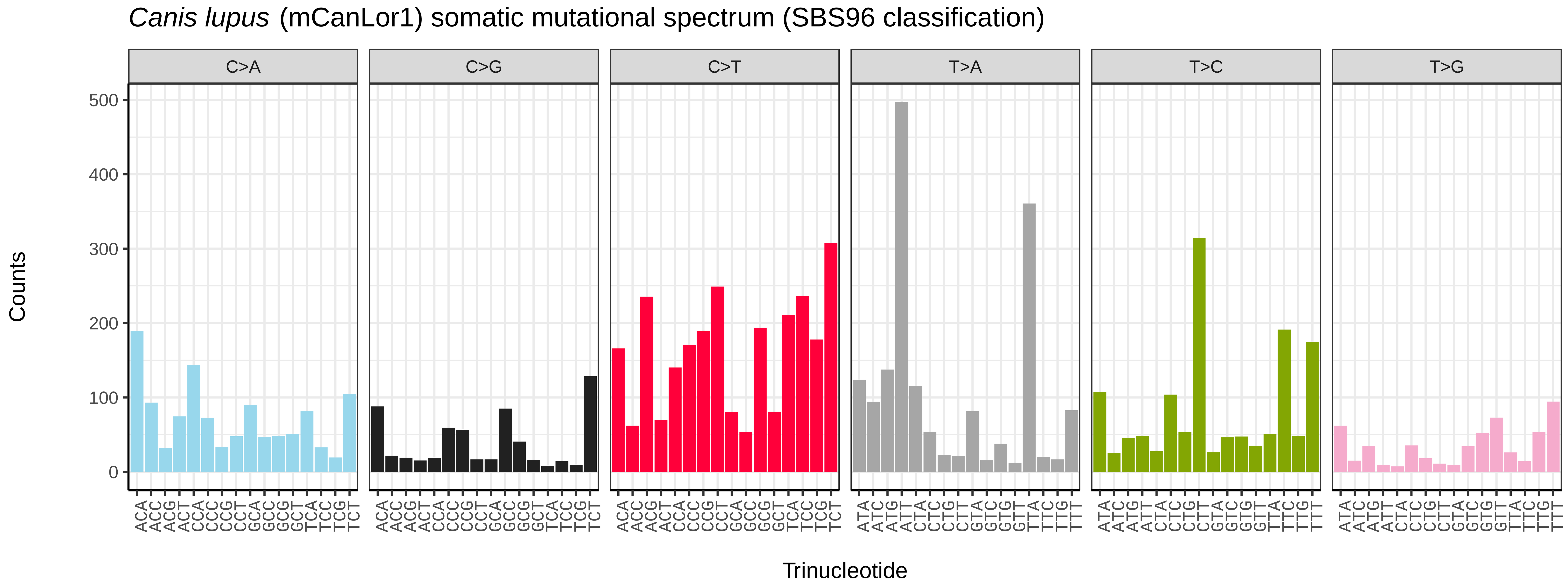
Canis lupus (mCanLor1) somatic mutational spectrum (SBS96 classification)
C>A
C>G
C>T
T>A
T>C
T>G
Counts
500
400
300
200
100
0
Trinucleotide

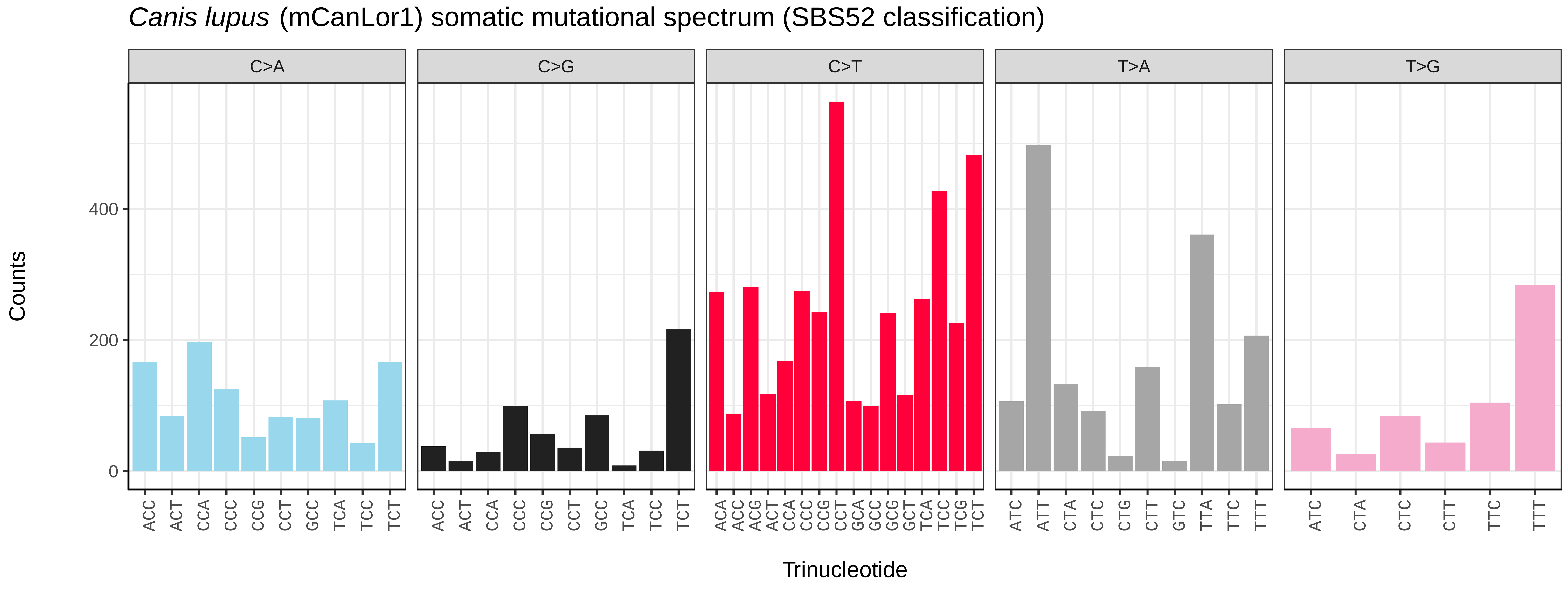
Canis lupus (mCanLor1) somatic mutational spectrum (SBS52 classification)
C>A
C>G
C>T
T>A
T>G
Counts
400
200
0
Trinucleotide

*Cervus elaphus* (mCerEla1) germline mutational spectrum

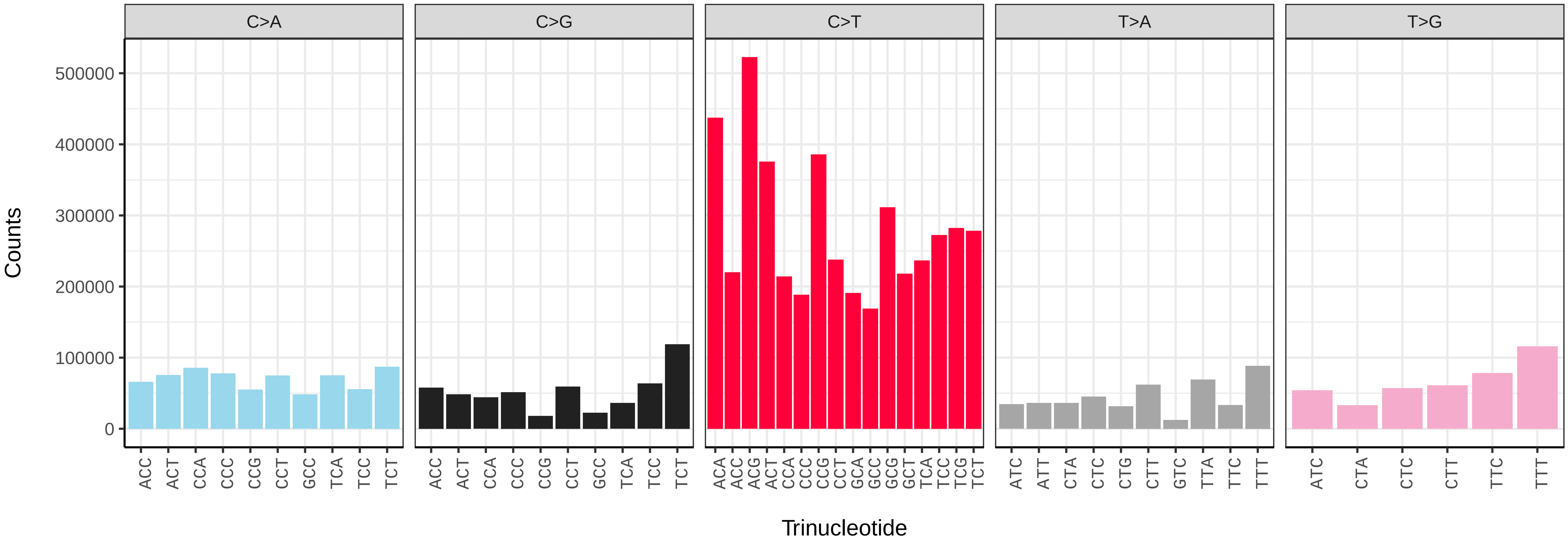


*Cervus elaphus* (mCerEla1) somatic mutational spectrum (SBS96 classification)

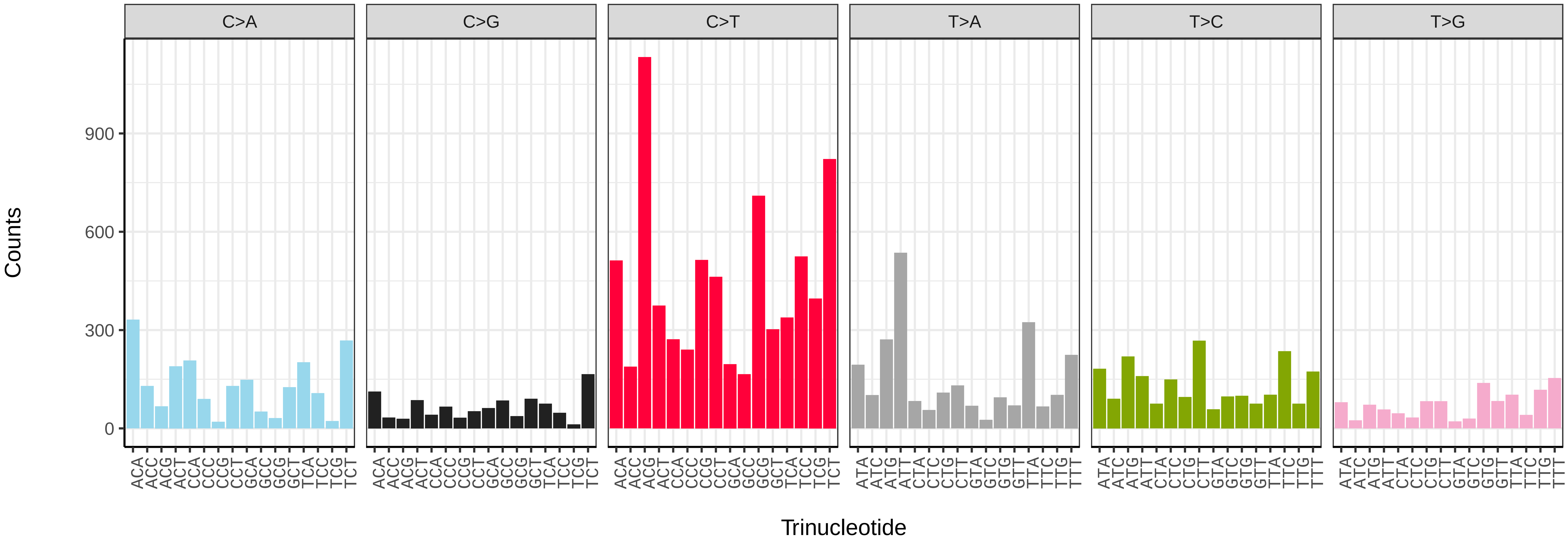


*Cervus elaphus* (mCerEla1) somatic mutational spectrum (SBS52 classification)

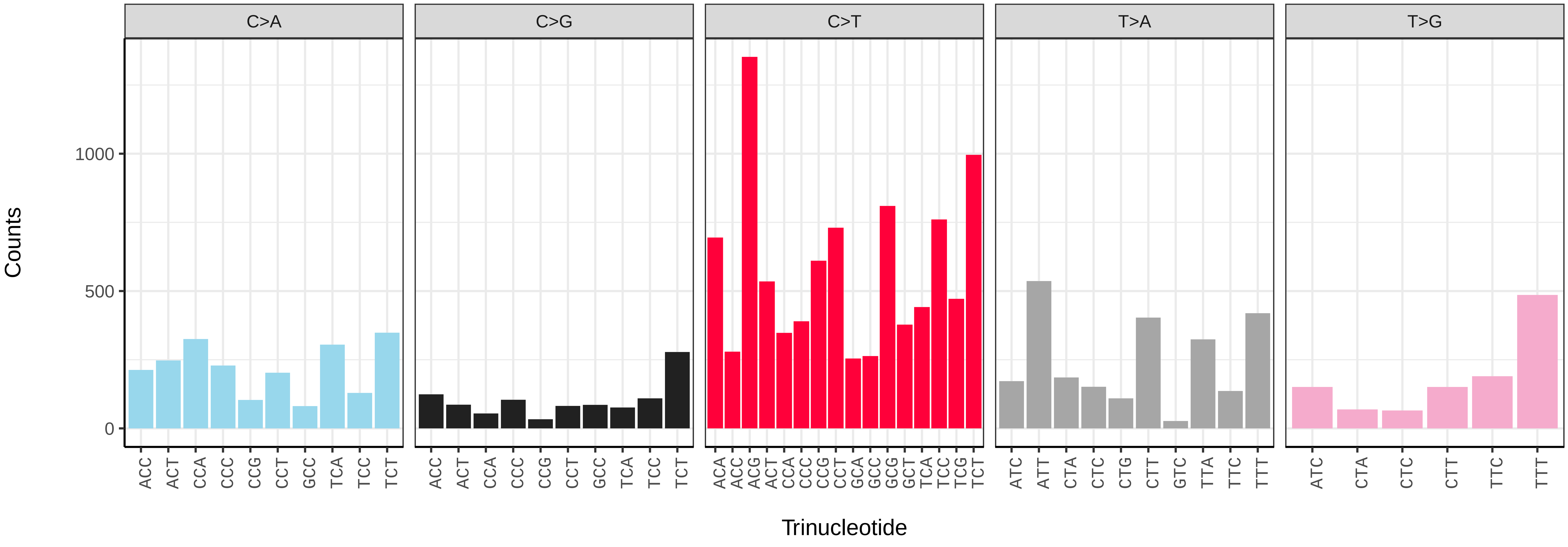

*Delphinus delphis* (mDelDel1) germline mutational spectrum

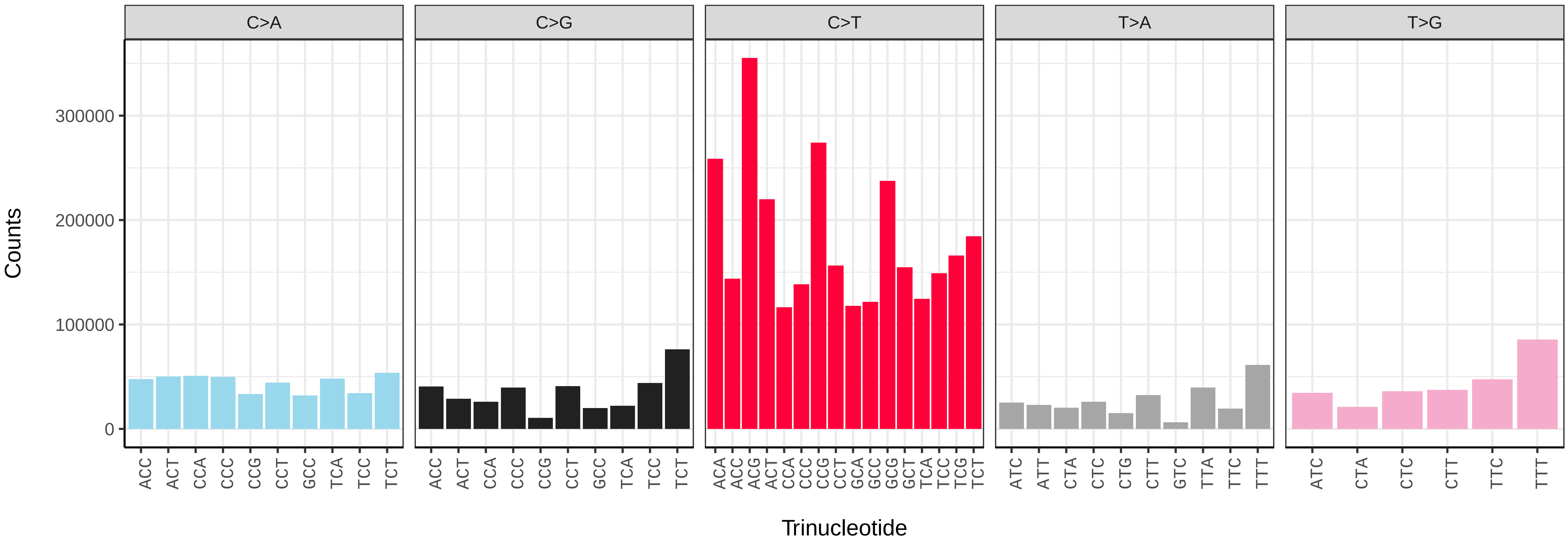


*Delphinus delphis* (mDelDel1) somatic mutational spectrum (SBS96 classification)

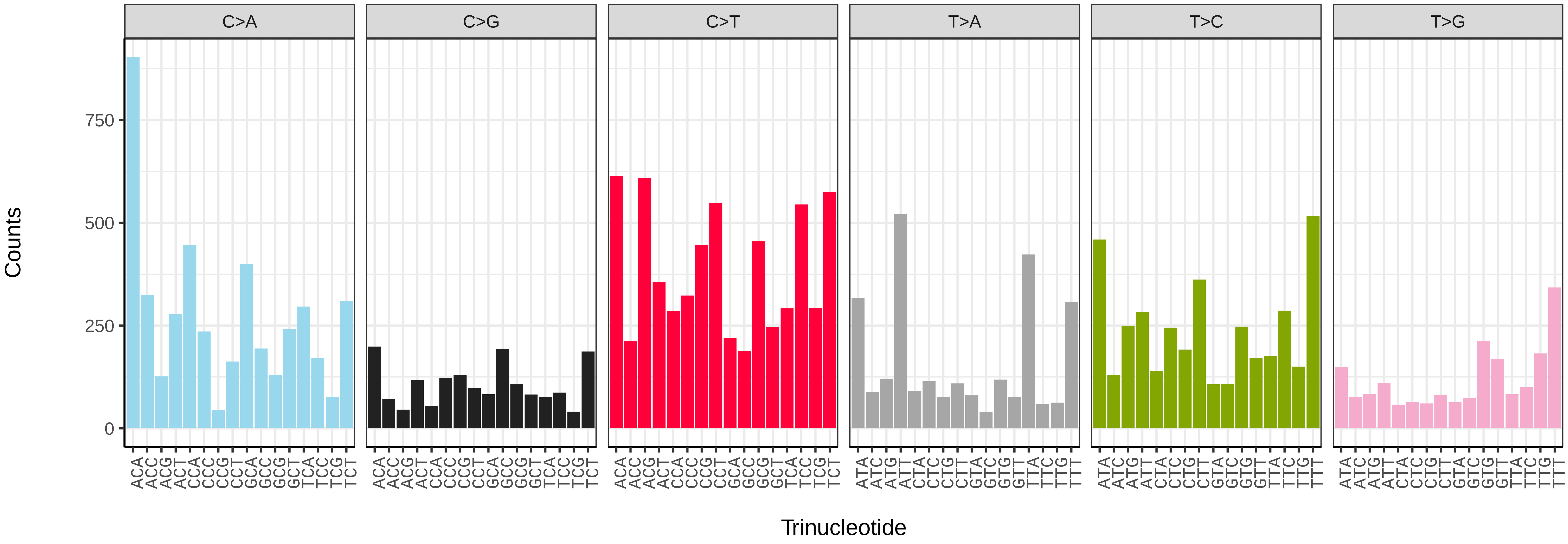


*Delphinus delphis* (mDelDel1) somatic mutational spectrum (SBS52 classification)

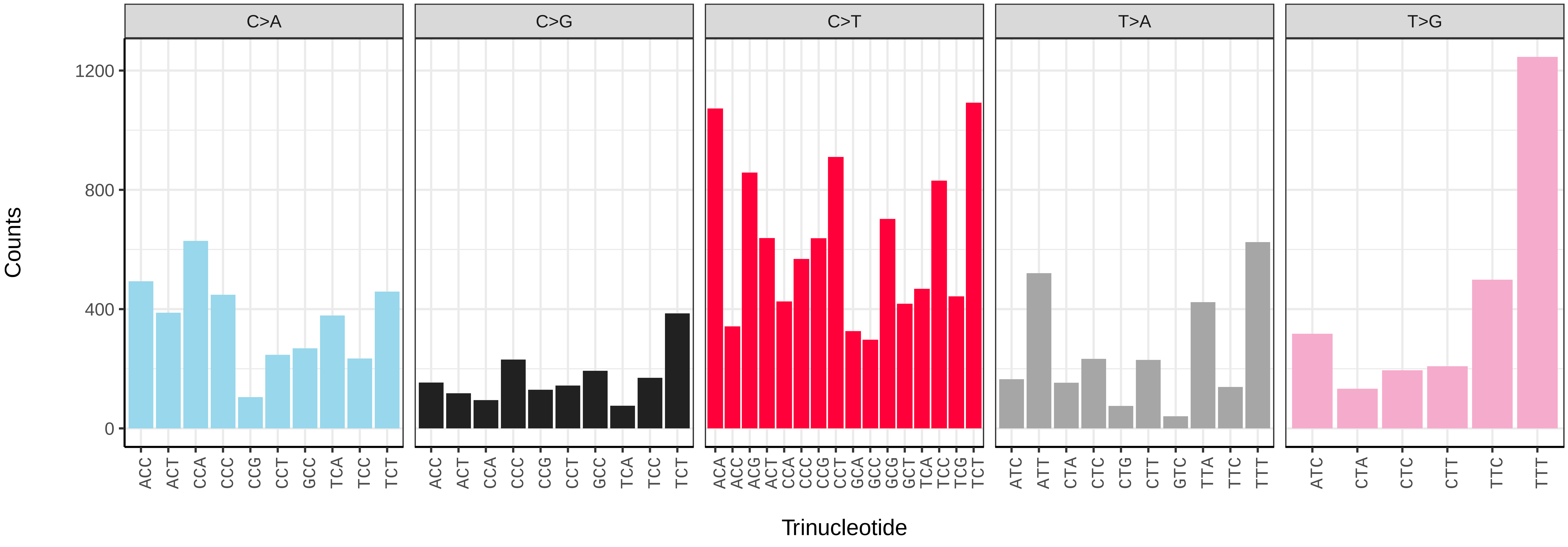

*Eptesicus nilssonii* (mEptNil1) germline mutational spectrum

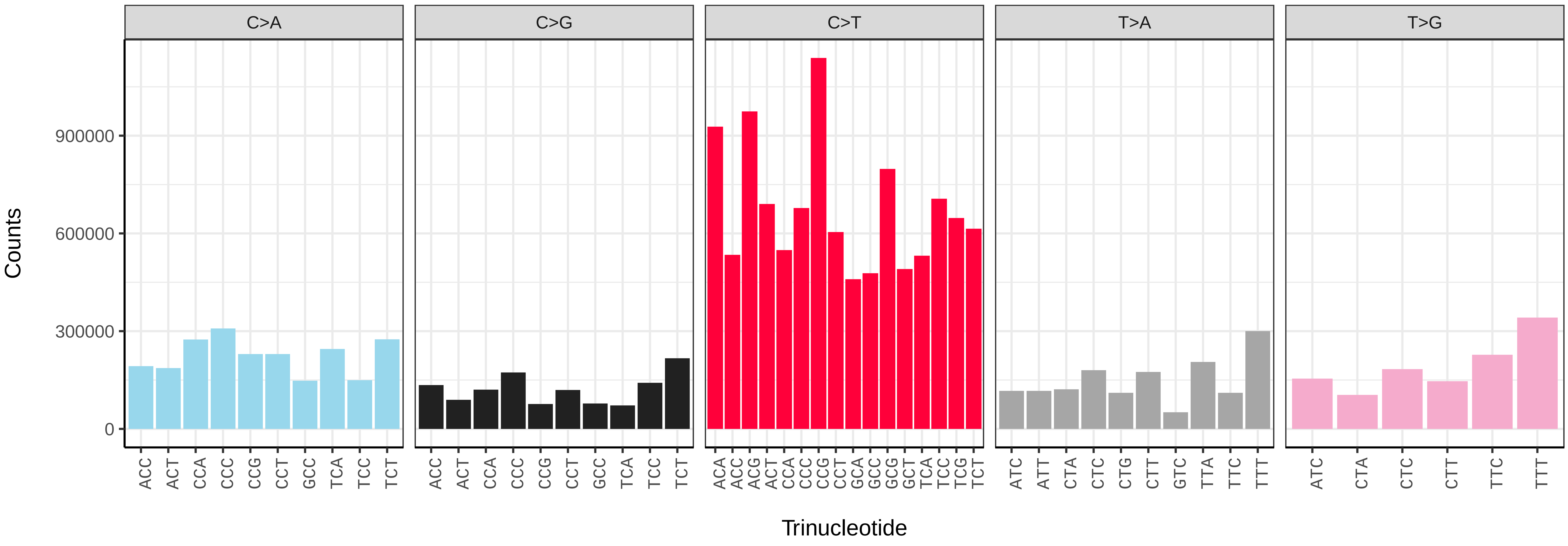


*Eptesicus nilssonii* (mEptNil1) somatic mutational spectrum (SBS96 classification)

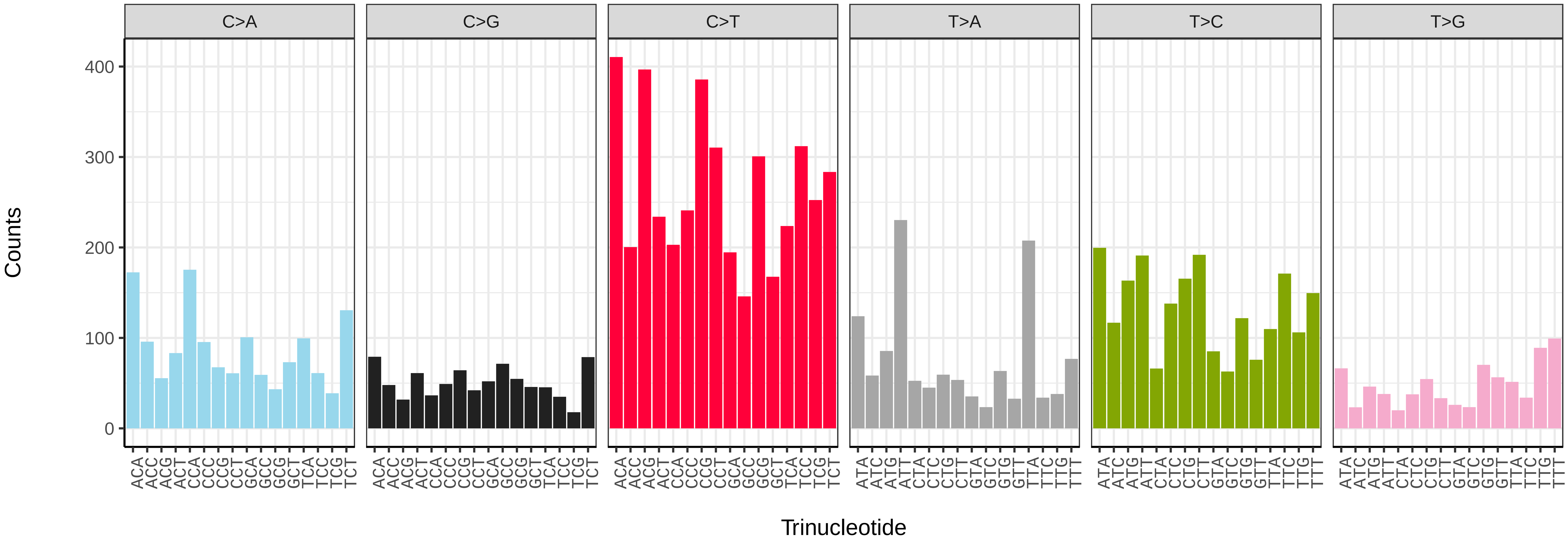


*Eptesicus nilssonii* (mEptNil1) somatic mutational spectrum (SBS52 classification)

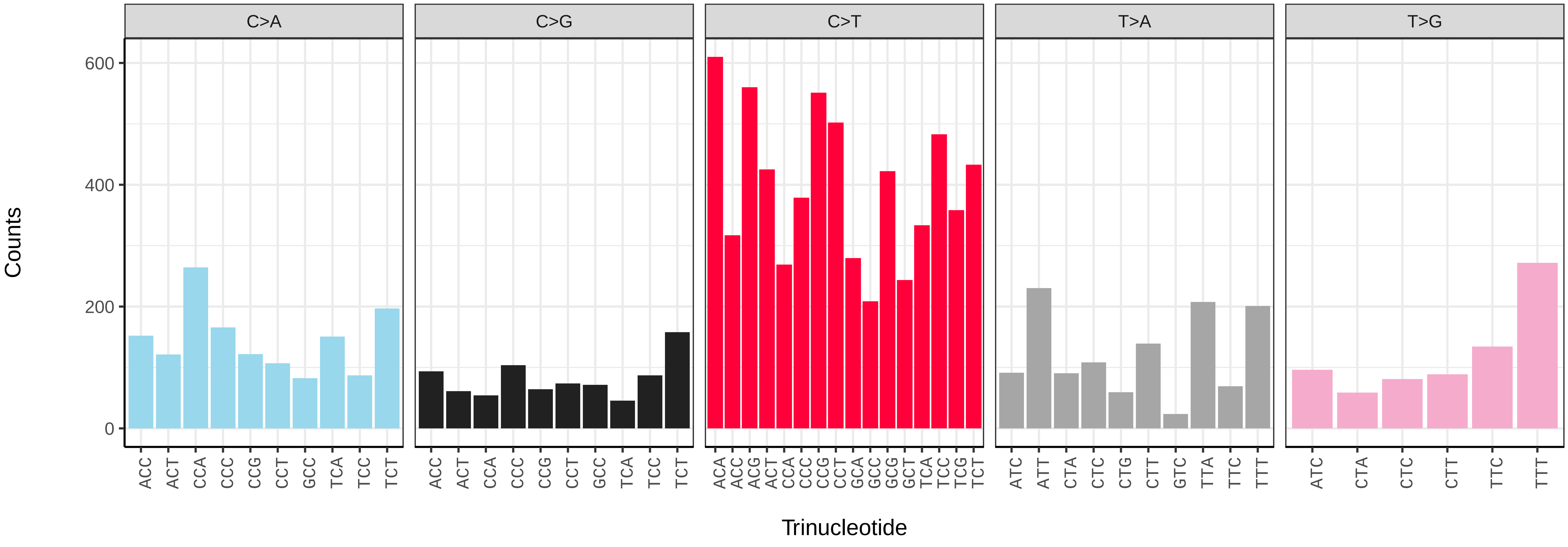

*Globicephala melas* (mGloMel1) germline mutational spectrum

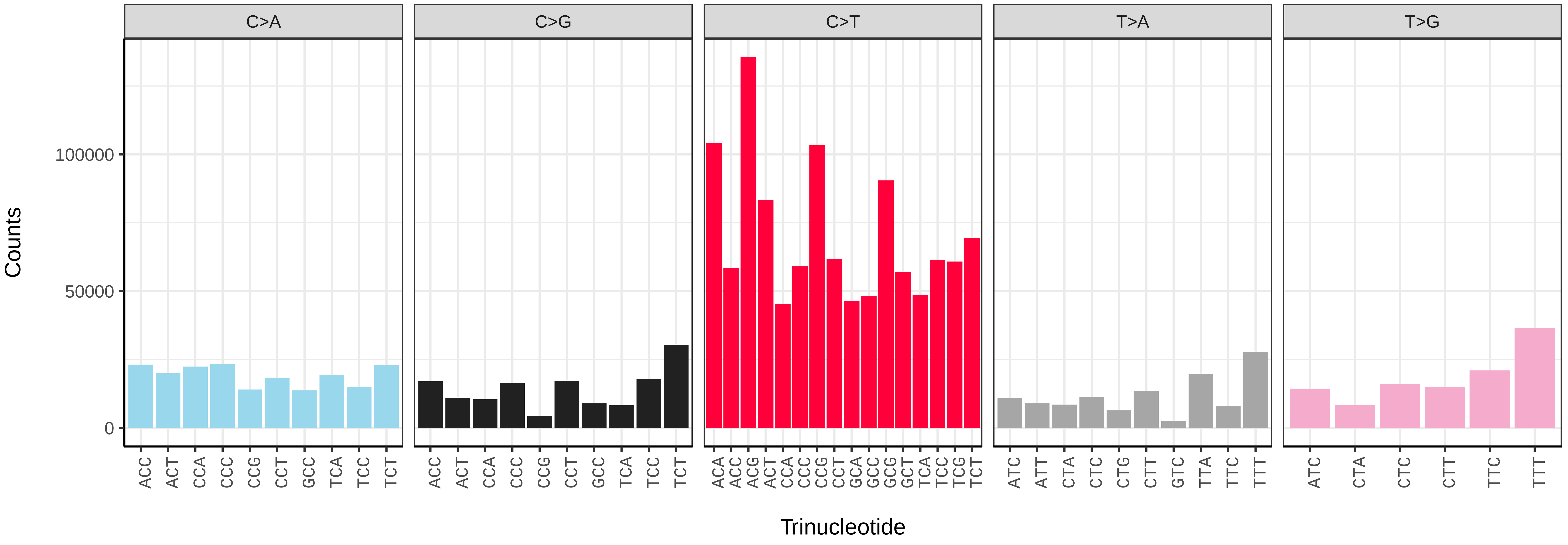


*Globicephala melas* (mGloMel1) somatic mutational spectrum (SBS96 classification)

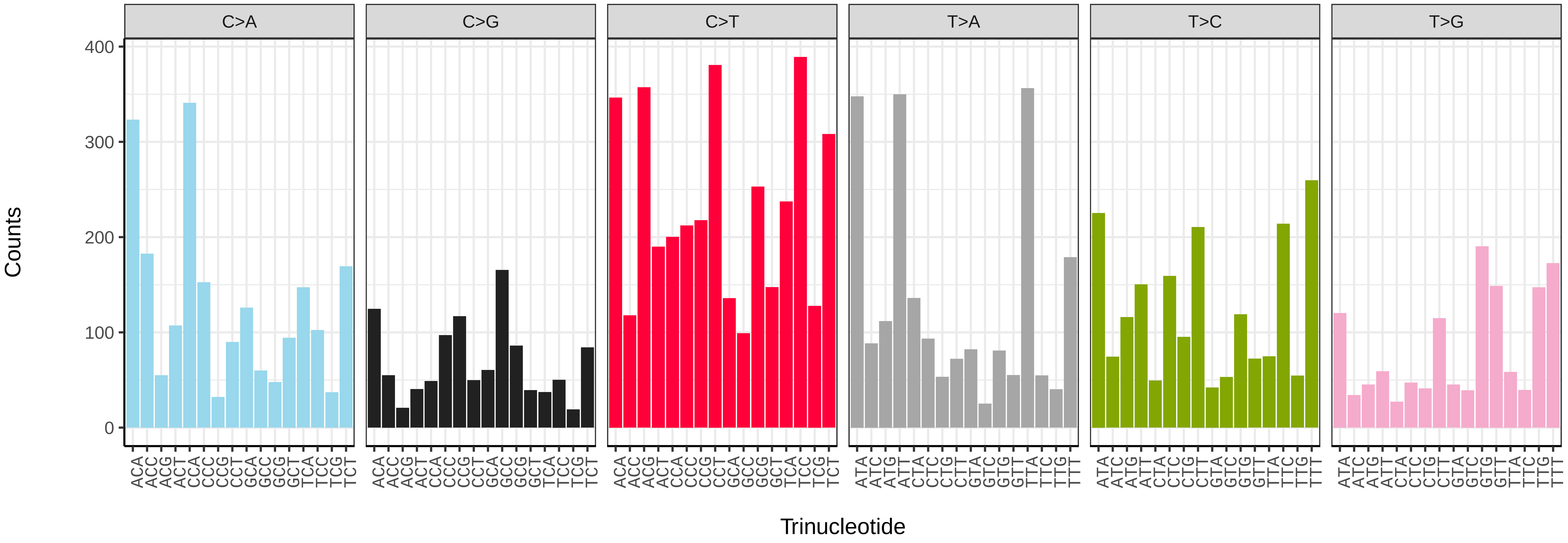


*Globicephala melas* (mGloMel1) somatic mutational spectrum (SBS52 classification)

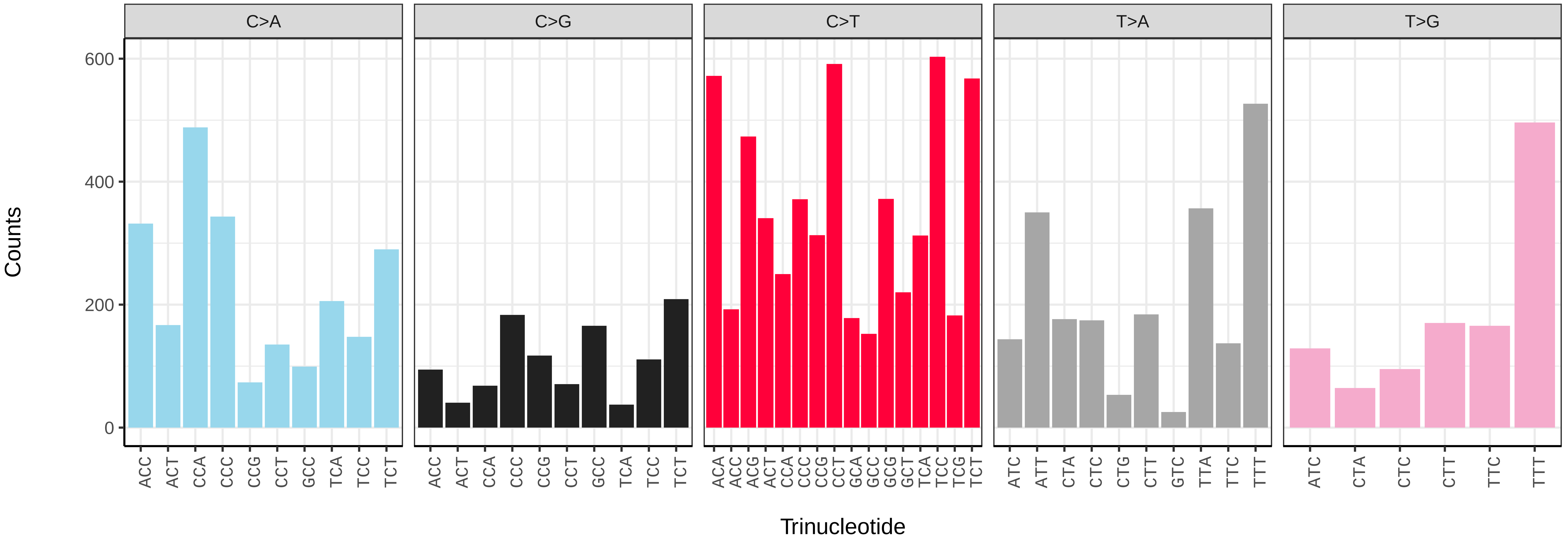

*Hyperoodon ampullatus* (mHypAmp2) germline mutational spectrum

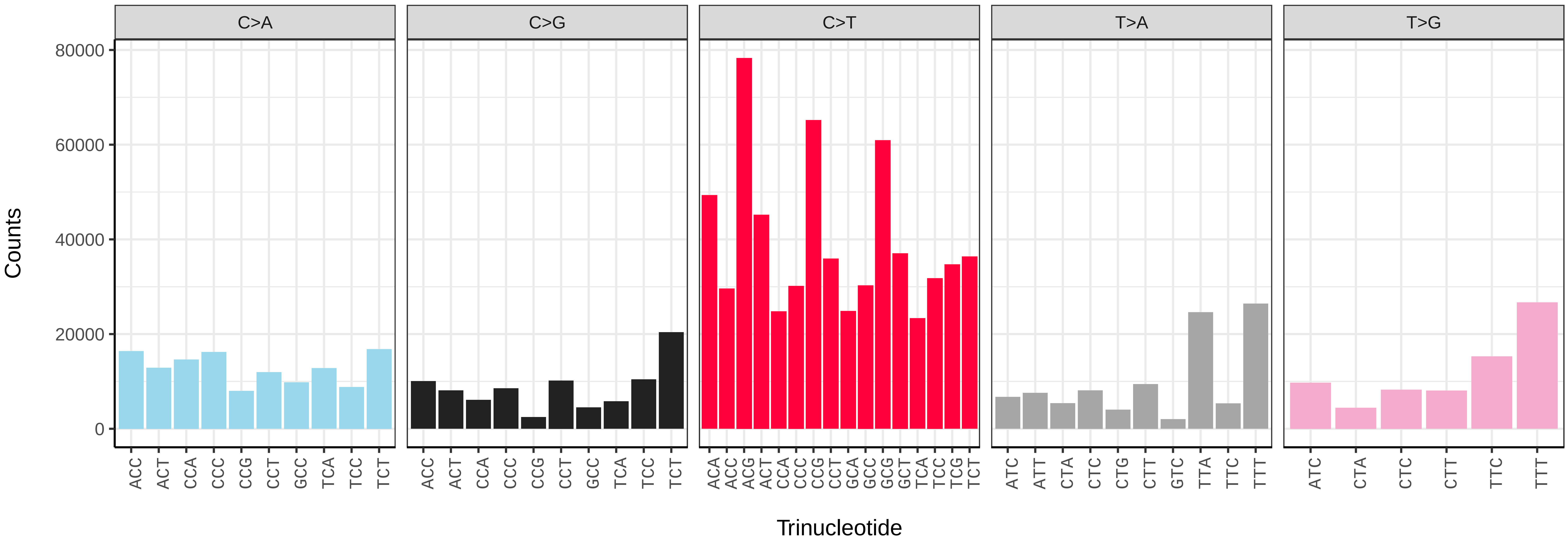


*Hyperoodon ampullatus* (mHypAmp2) somatic mutational spectrum (SBS96 classification)

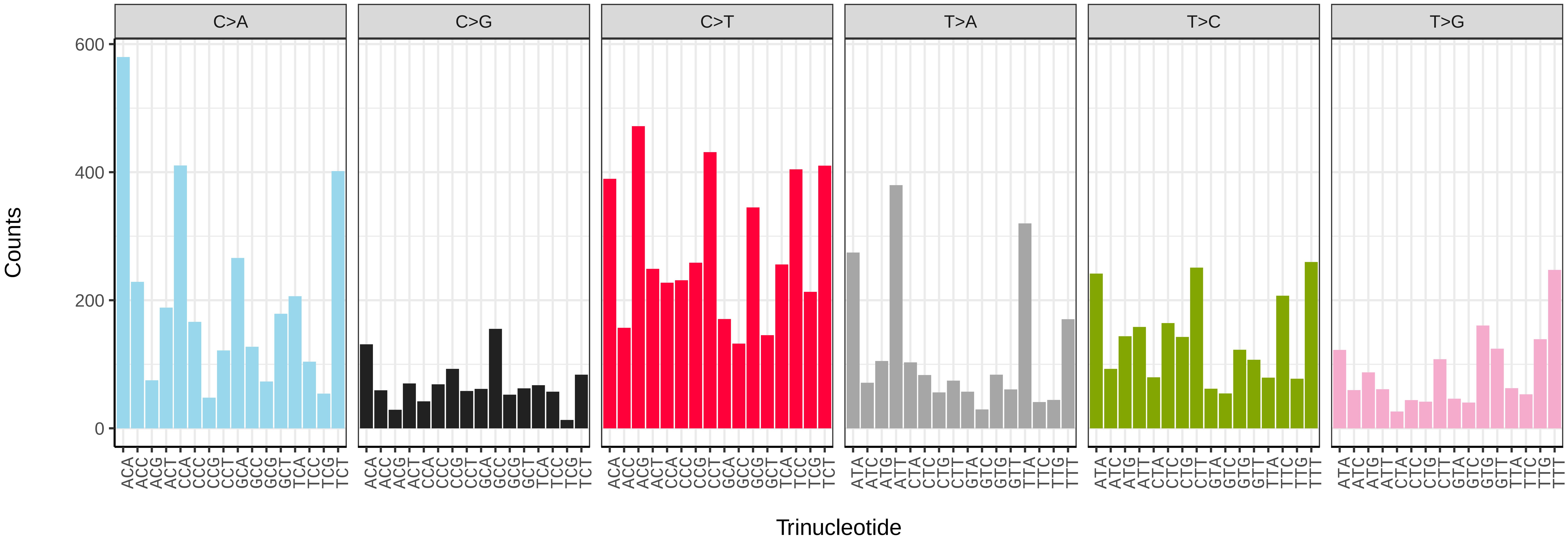


*Hyperoodon ampullatus* (mHypAmp2) somatic mutational spectrum (SBS52 classification)

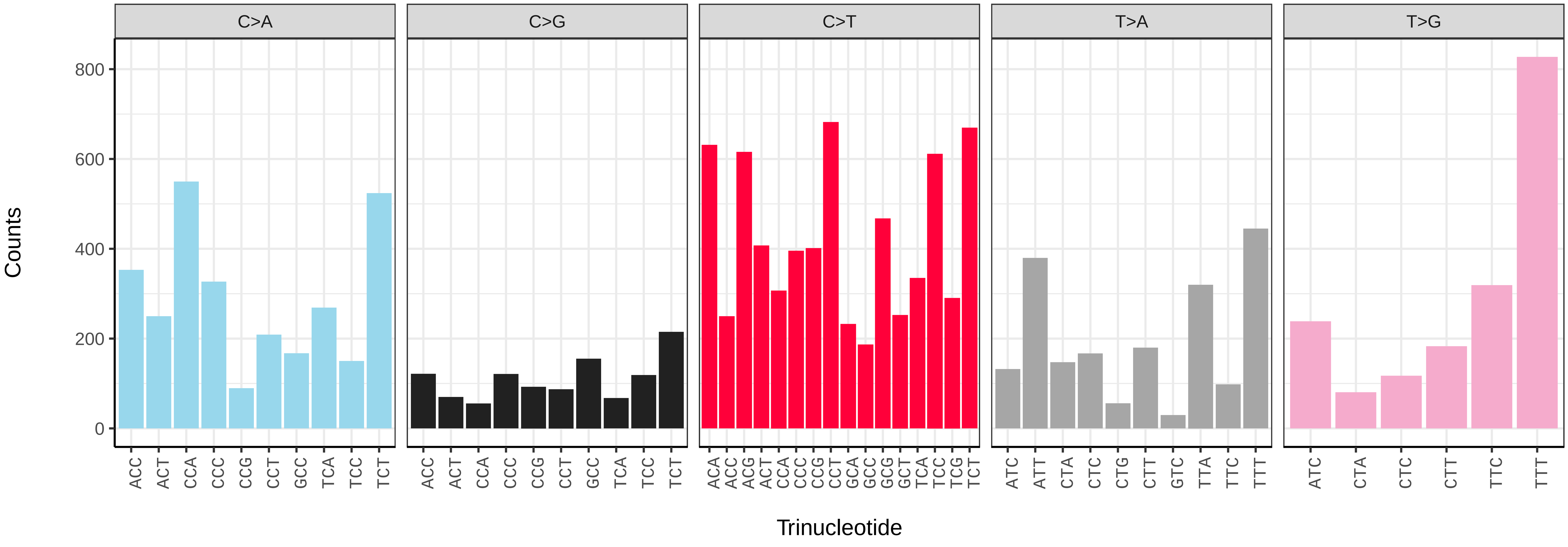

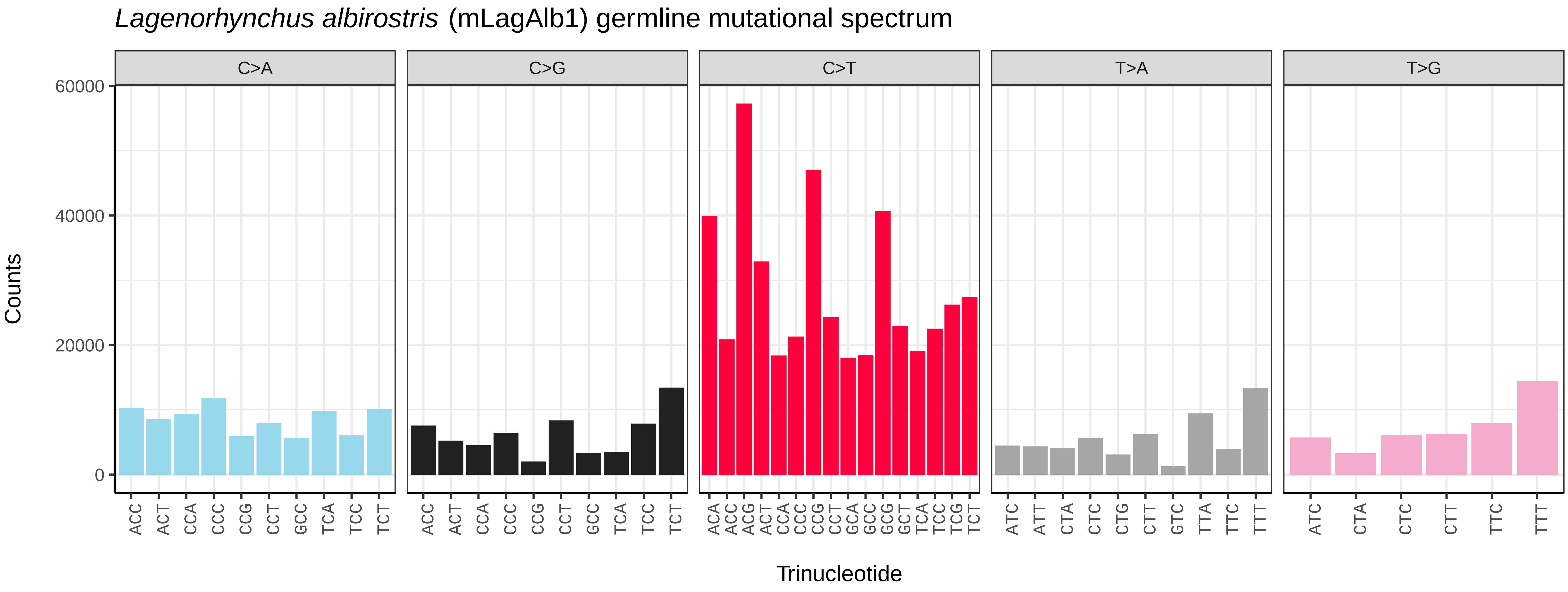
*Lagenorhynchus albirostris* (mLagAlb1) germline mutational spectrum
C>A
C>G
C>T
T>A
T>G
Counts
0
20000
40000
60000
ACC ACT CCA CCC CCG CCT GCC TCA TCC TCT
ACA ACC ACG ACT CCA CCC CCG CCT GCA GCC GCG GCT TCA TCC TCG TCT
ATC ATT CTA CTC CTG CTT GTC TTA TTC TTT
ATC CTA CTC CTT TTC TTT
Trinucleotide

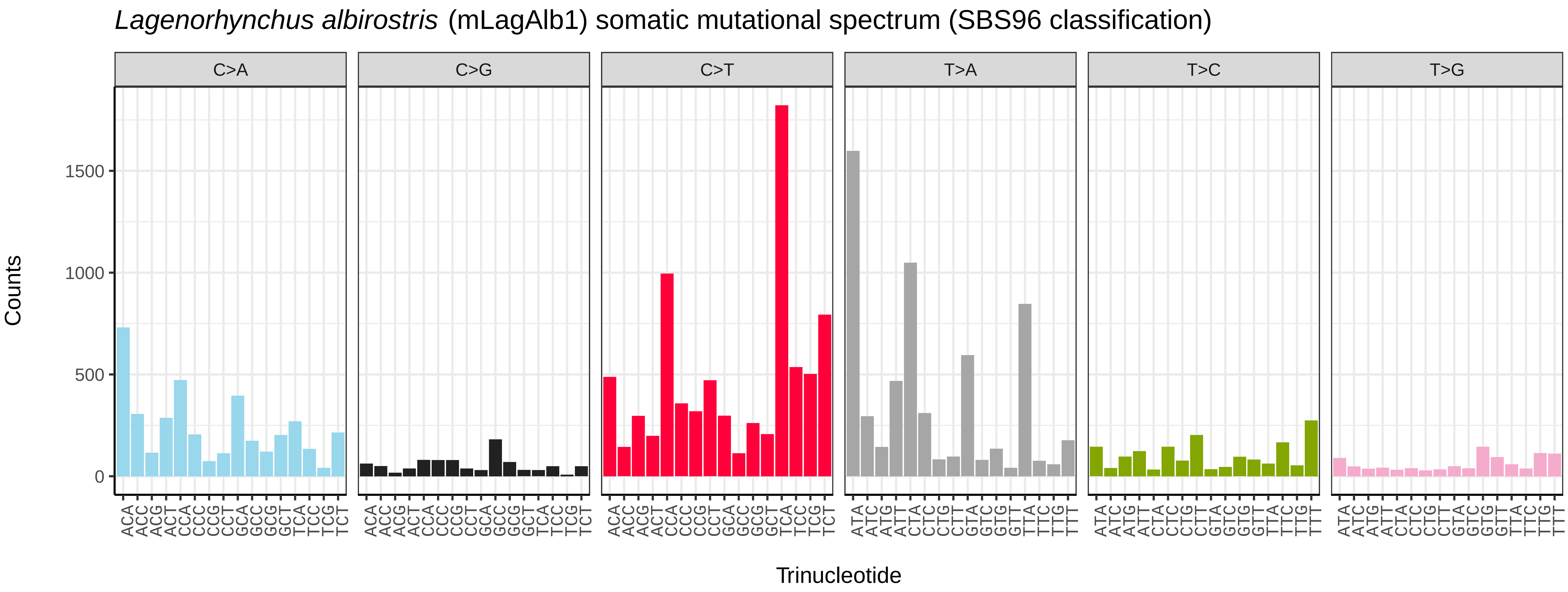
*Lagenorhynchus albirostris* (mLagAlb1) somatic mutational spectrum (SBS96 classification)
C>A
C>G
C>T
T>A
T>C
T>G
Counts
0
500
1000
1500
ACA ACC ACG ACT CCA CCC CCG CCT GCA GCC GCG GCT TCA TCC TCG TCT
ATA ATC ATG ATT CTA CTC CTG CTT GTA GTC GTG GTT TTA TTC TTG TTT
Trinucleotide

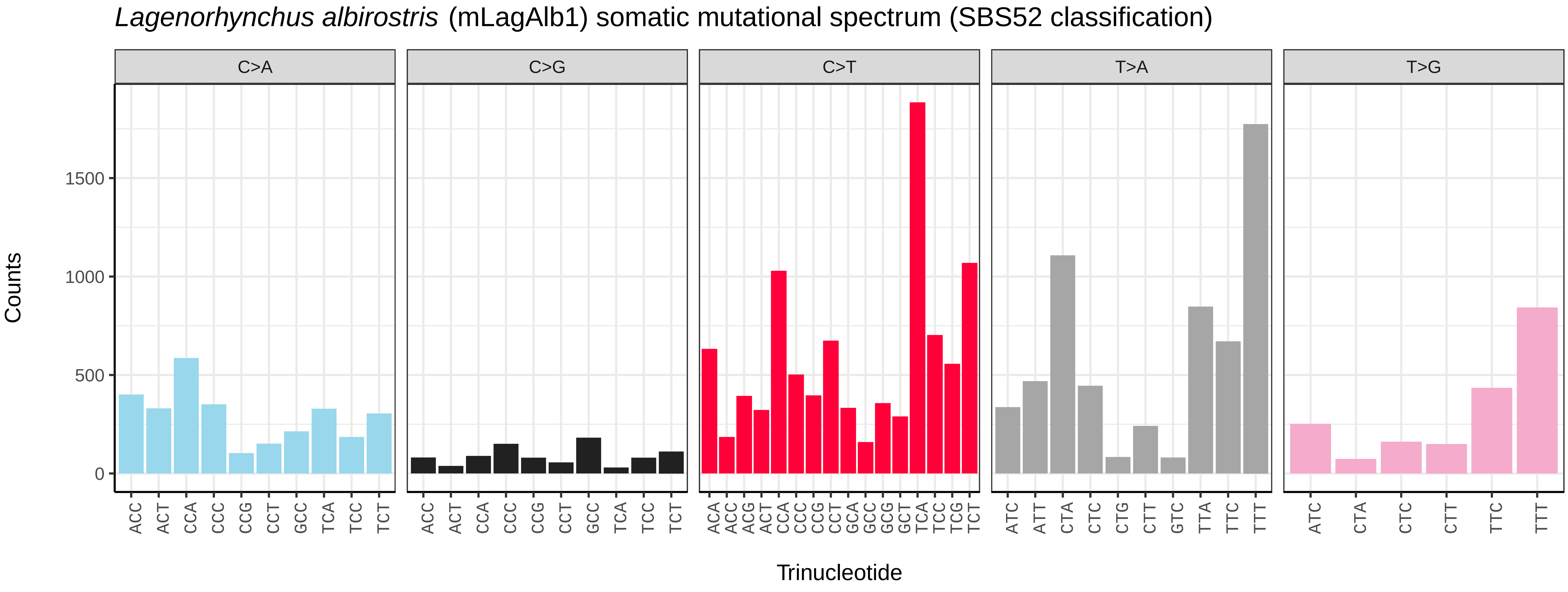
*Lagenorhynchus albirostris* (mLagAlb1) somatic mutational spectrum (SBS52 classification)
C>A
C>G
C>T
T>A
T>G
Counts
0
500
1000
1500
ACC ACT CCA CCC CCG CCT GCC TCA TCC TCT
ACA ACC ACG ACT CCA CCC CCG CCT GCA GCC GCG GCT TCA TCC TCG TCT
ATC ATT CTA CTC CTG CTT GTC TTA TTC TTT
ATC CTA CTC CTT TTC TTT
Trinucleotide

*Martes martes* (mMarMar1) germline mutational spectrum

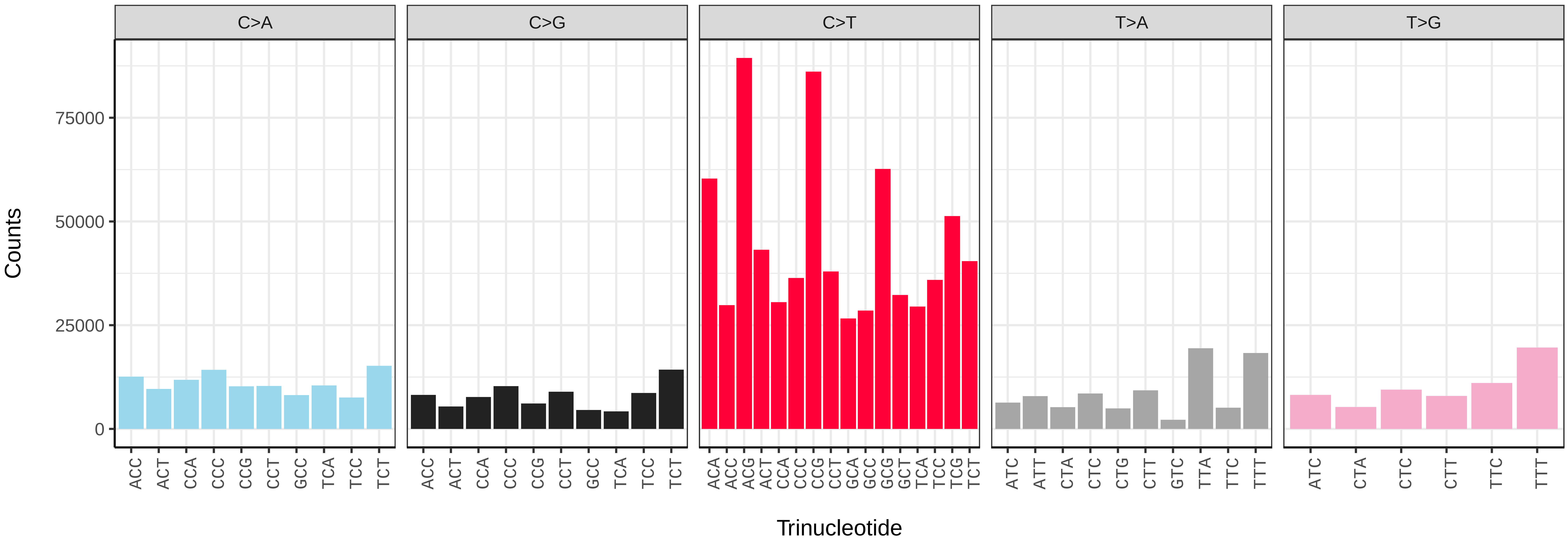


*Martes martes* (mMarMar1) somatic mutational spectrum (SBS96 classification)

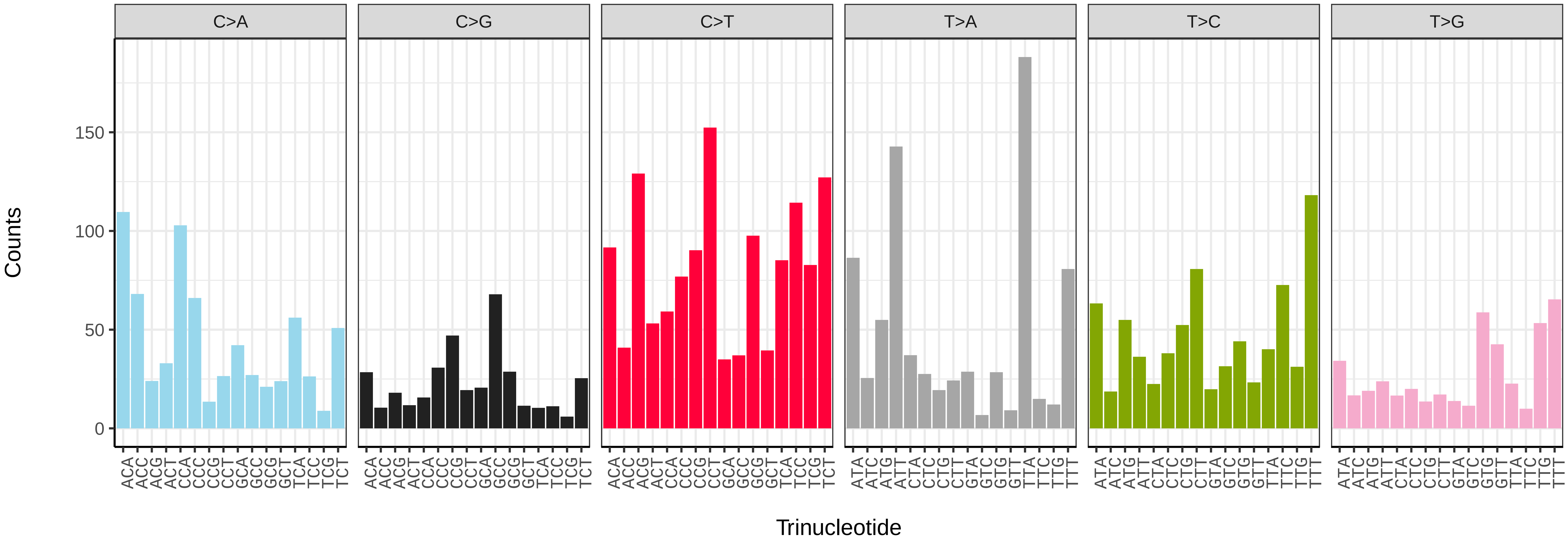


*Martes martes* (mMarMar1) somatic mutational spectrum (SBS52 classification)

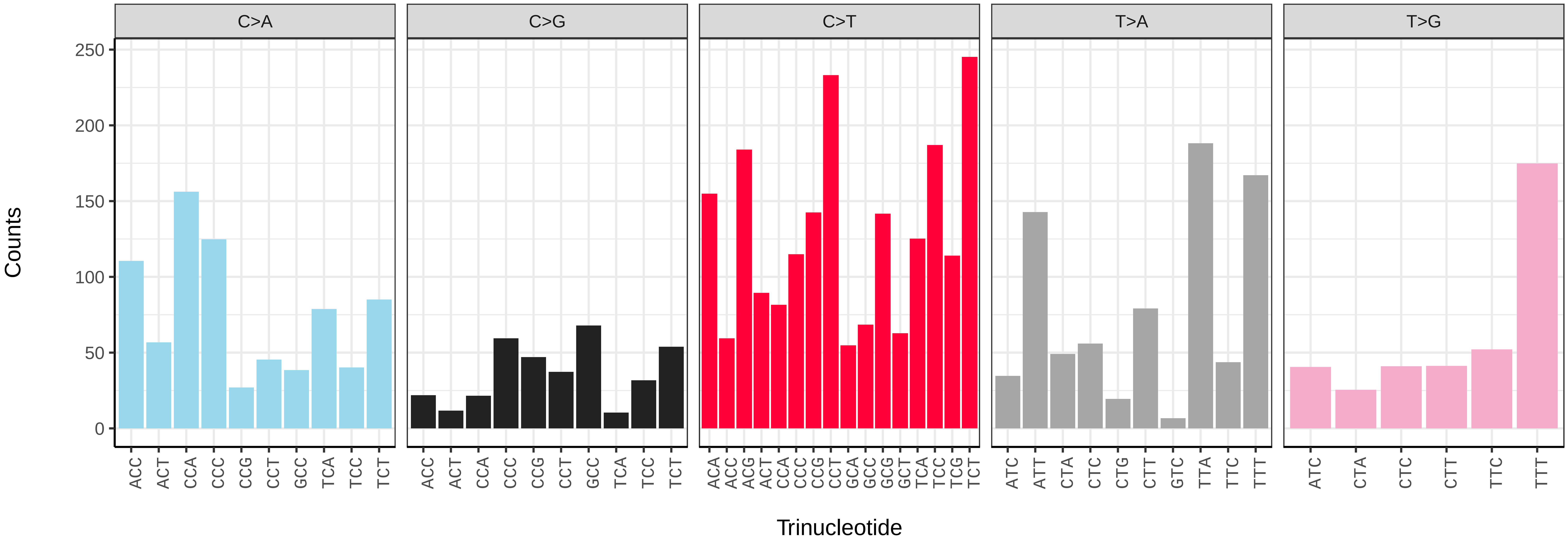

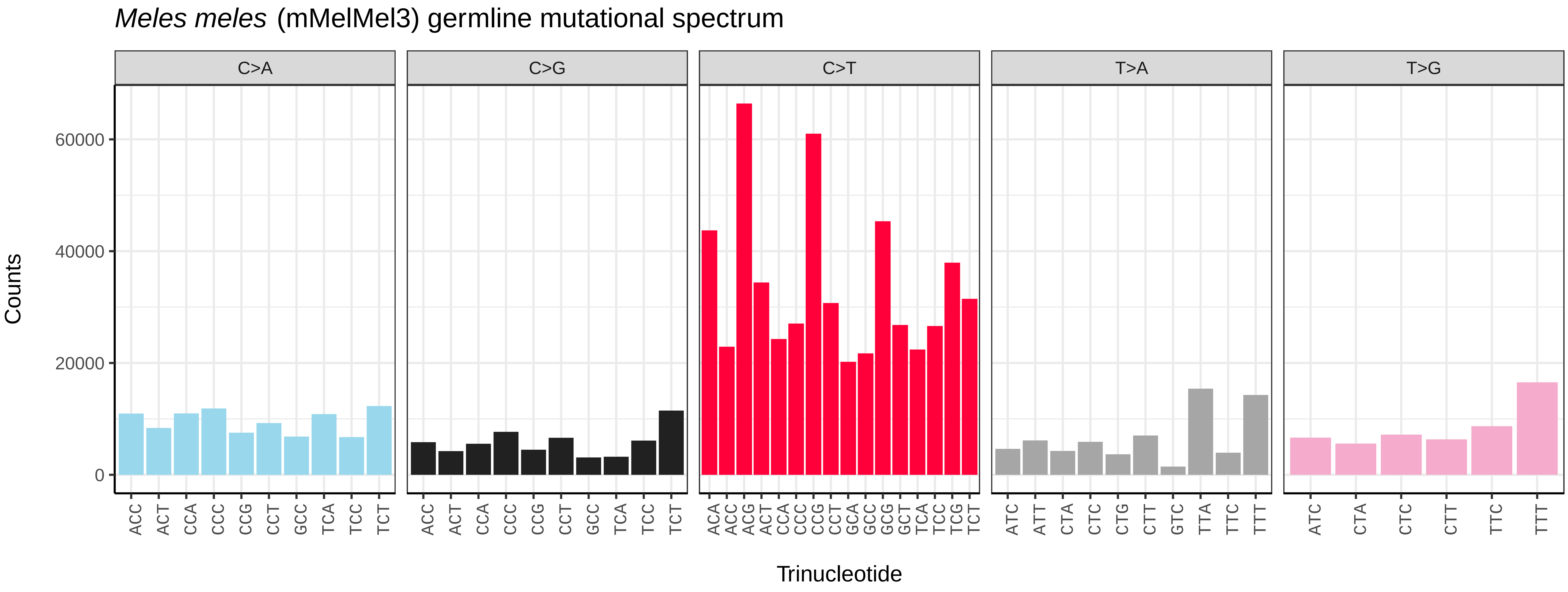
*Meles meles* (mMelMel3) germline mutational spectrum
C>A
C>G
C>T
T>A
T>G
60000
40000
20000
0
Counts
ACC ACT CCA CCC CCG CCT GCC TCA TCC TCT
ACC ACT CCA CCC CCG CCT GCC TCA TCC TCT
ACA ACC ACG ACT CCA CCC CCG CCT GCA GCC GCG GCT TCA TCC TCG TCT
ATC ATT CTA CTC CTG CTT GTC TTA TTC TTT
ATC CTA CTC CTT TTC TTT
Trinucleotide

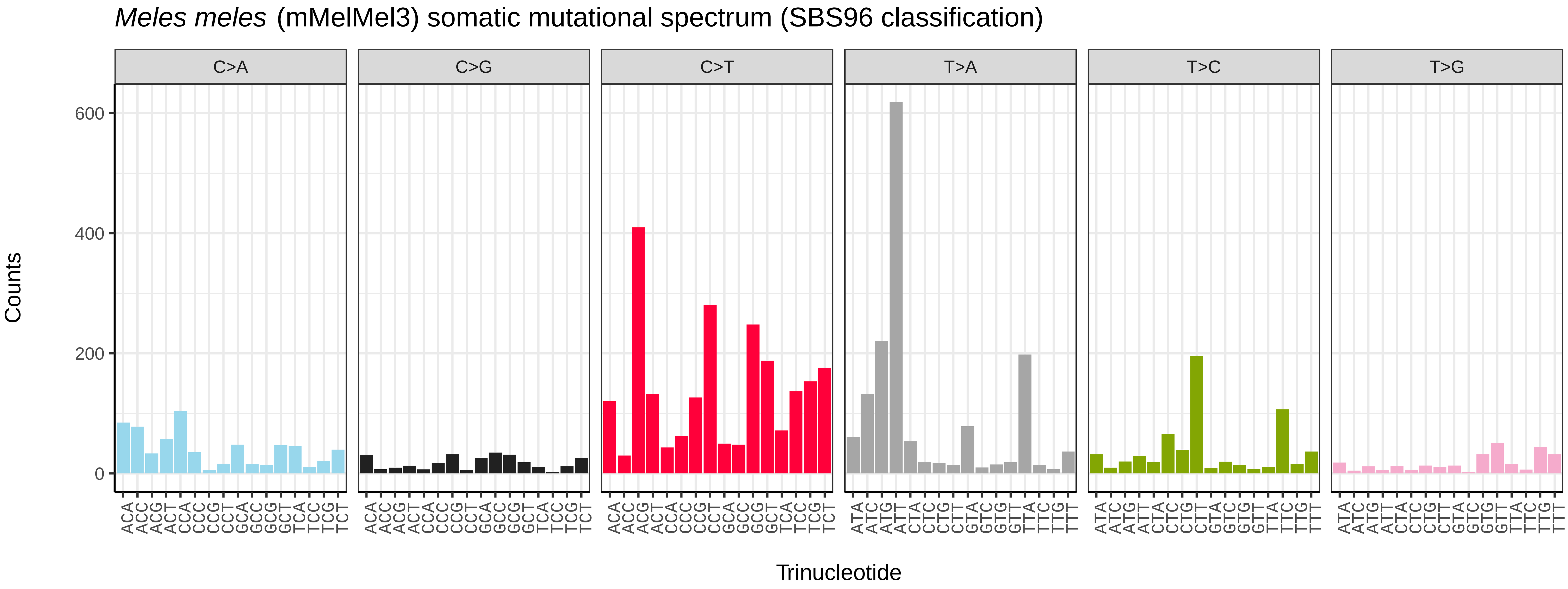
*Meles meles* (mMelMel3) somatic mutational spectrum (SBS96 classification)
C>A
C>G
C>T
T>A
T>C
T>G
600
400
200
0
Counts
Trinucleotide

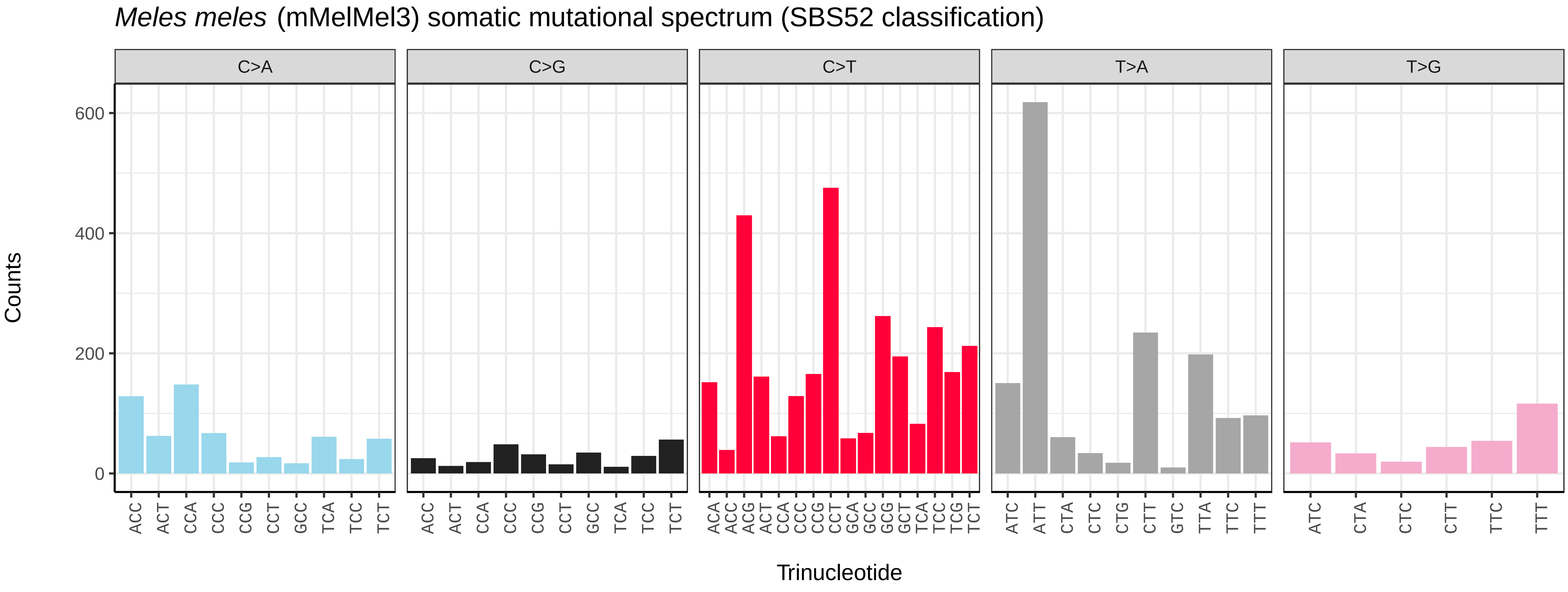
*Meles meles* (mMelMel3) somatic mutational spectrum (SBS52 classification)
C>A
C>G
C>T
T>A
T>G
600
400
200
0
Counts
ACC ACT CCA CCC CCG CCT GCC TCA TCC TCT
ACC ACT CCA CCC CCG CCT GCC TCA TCC TCT
ACA ACC ACG ACT CCA CCC CCG CCT GCA GCC GCG GCT TCA TCC TCG TCT
ATC ATT CTA CTC CTG CTT GTC TTA TTC TTT
ATC CTA CTC CTT TTC TTT
Trinucleotide

*Mus musculus* (mMusMuc1) germline mutational spectrum

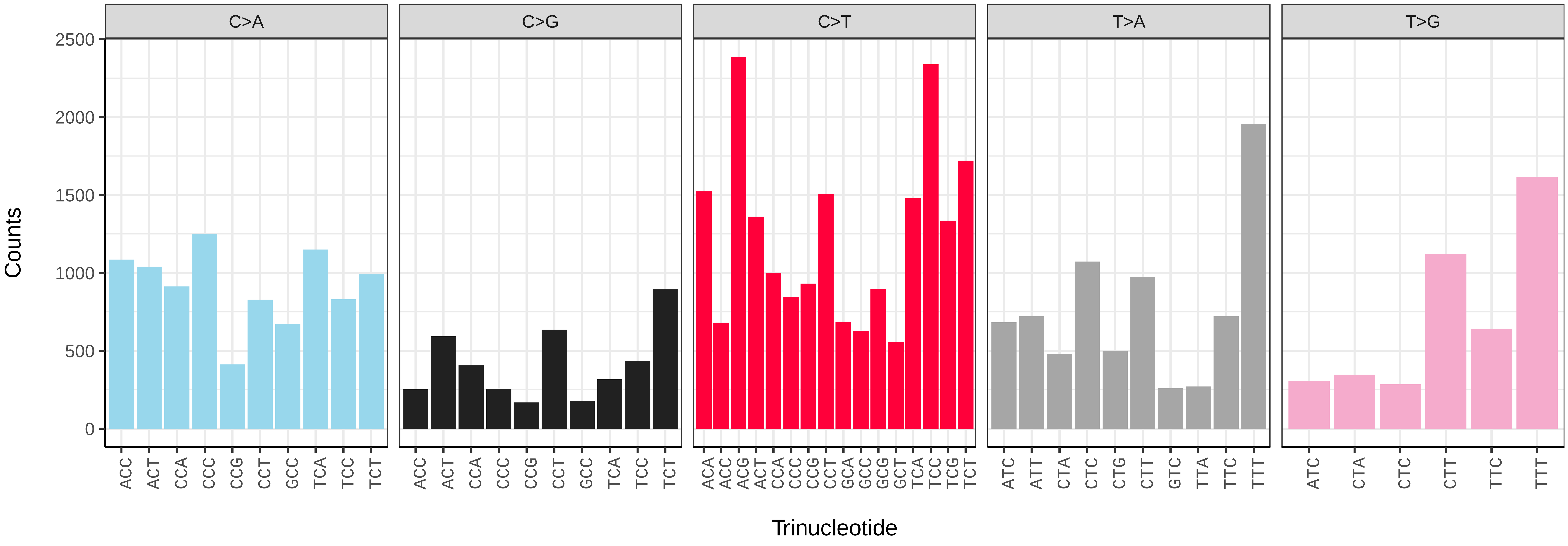


*Mus musculus* (mMusMuc1) somatic mutational spectrum (SBS96 classification)

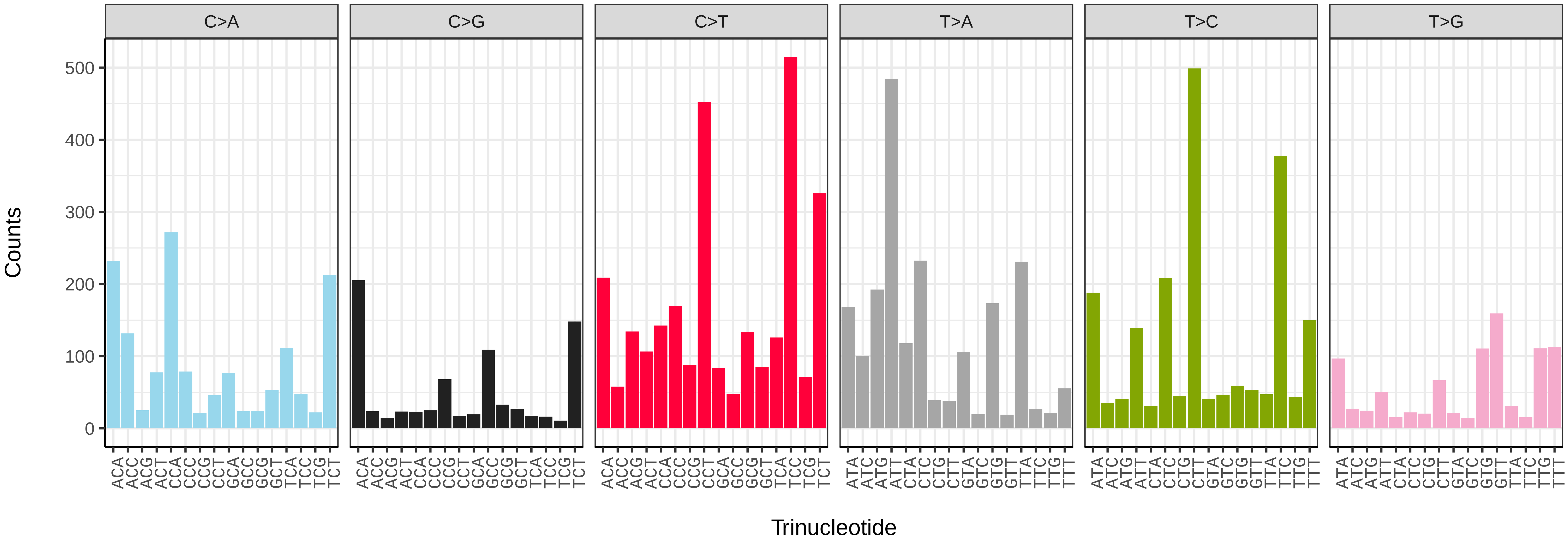


*Mus musculus* (mMusMuc1) somatic mutational spectrum (SBS52 classification)

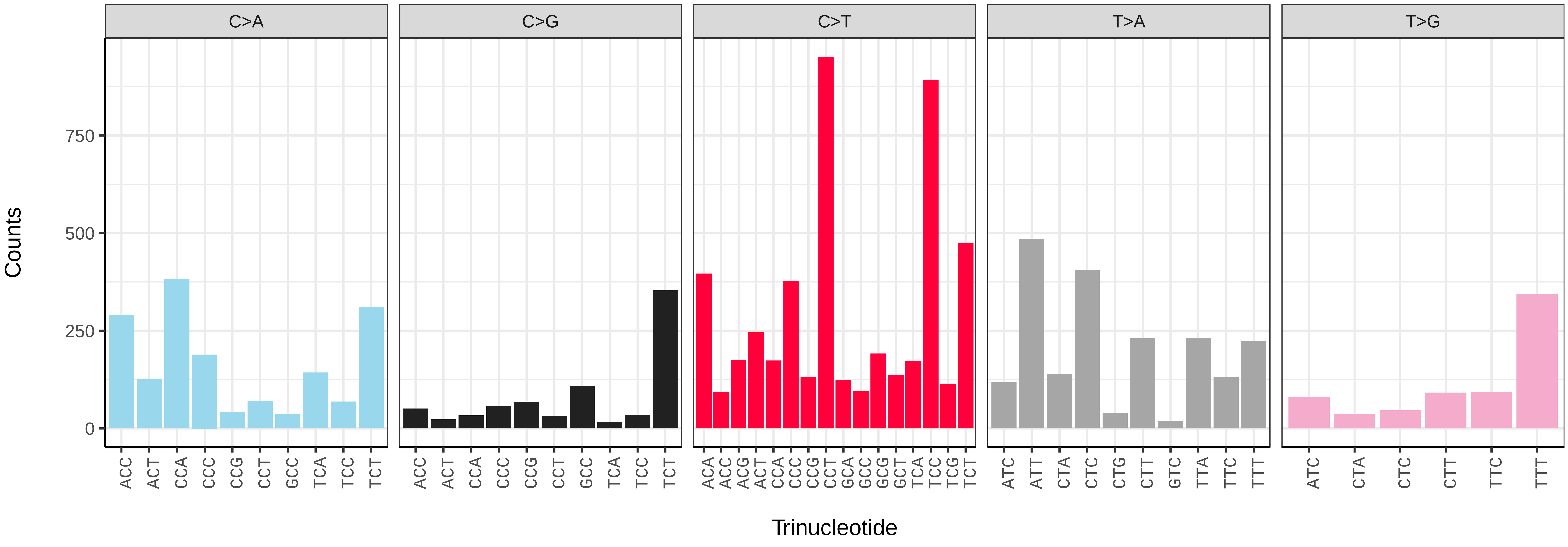

*Orcinus orca* (mOrcOrc1) germline mutational spectrum

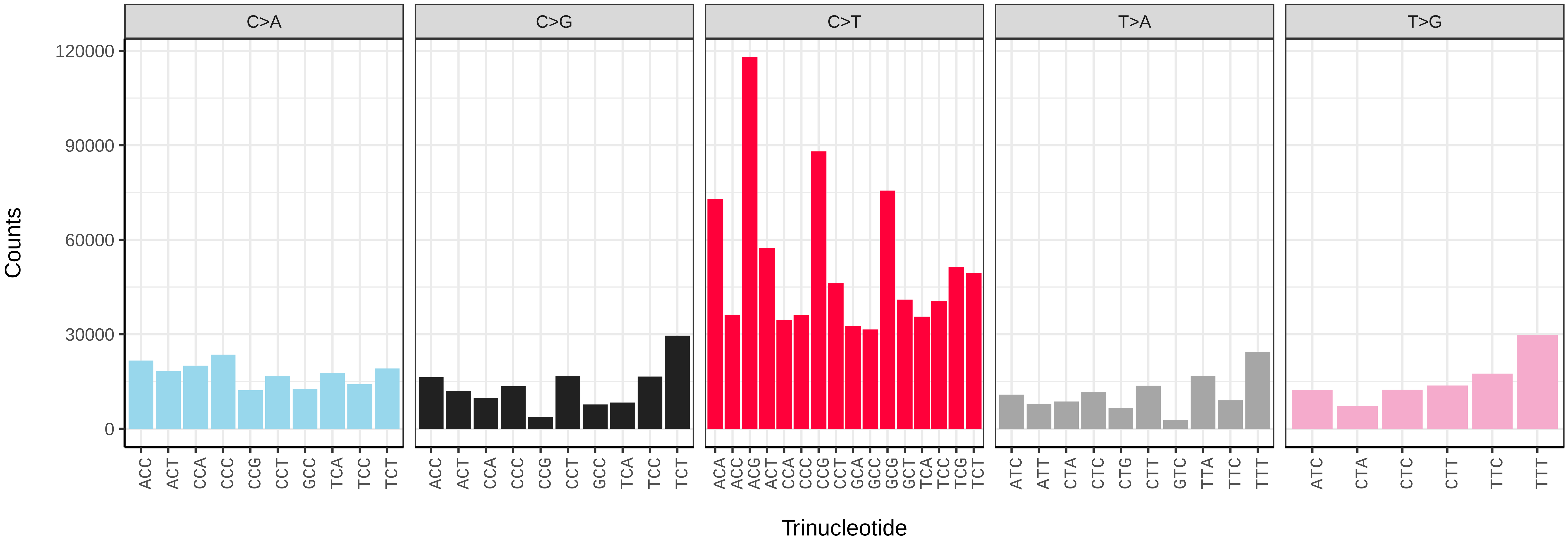


*Orcinus orca* (mOrcOrc1) somatic mutational spectrum (SBS96 classification)

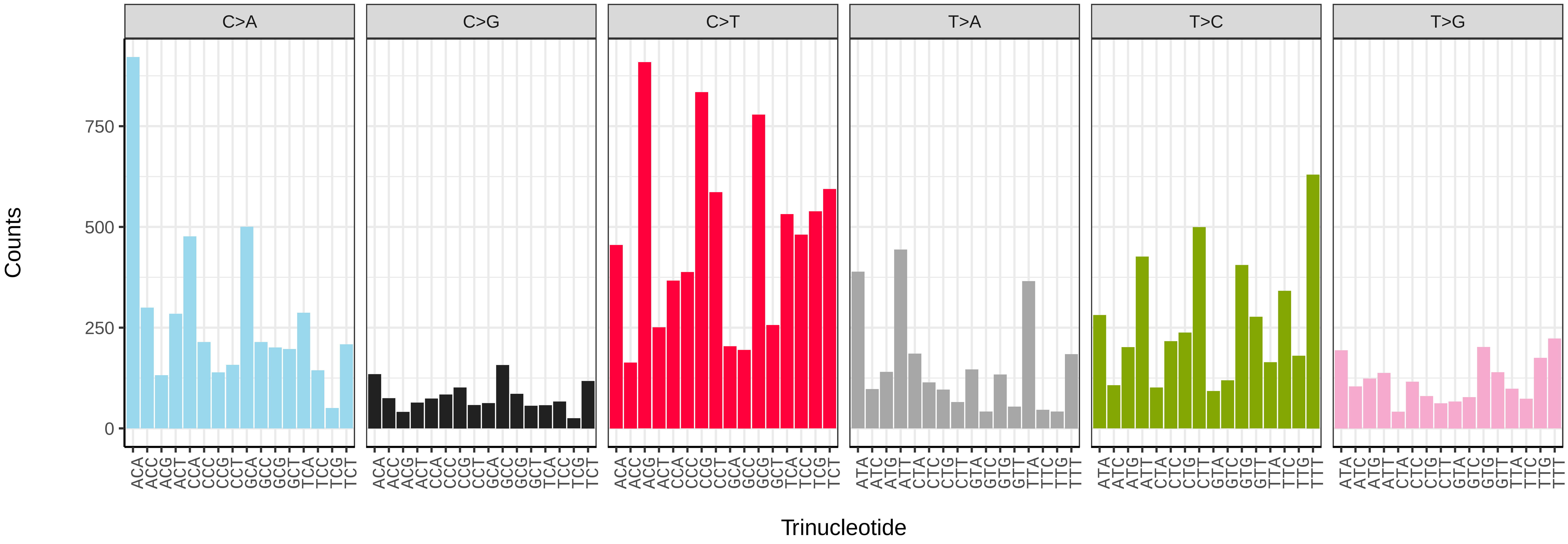


*Orcinus orca* (mOrcOrc1) somatic mutational spectrum (SBS52 classification)

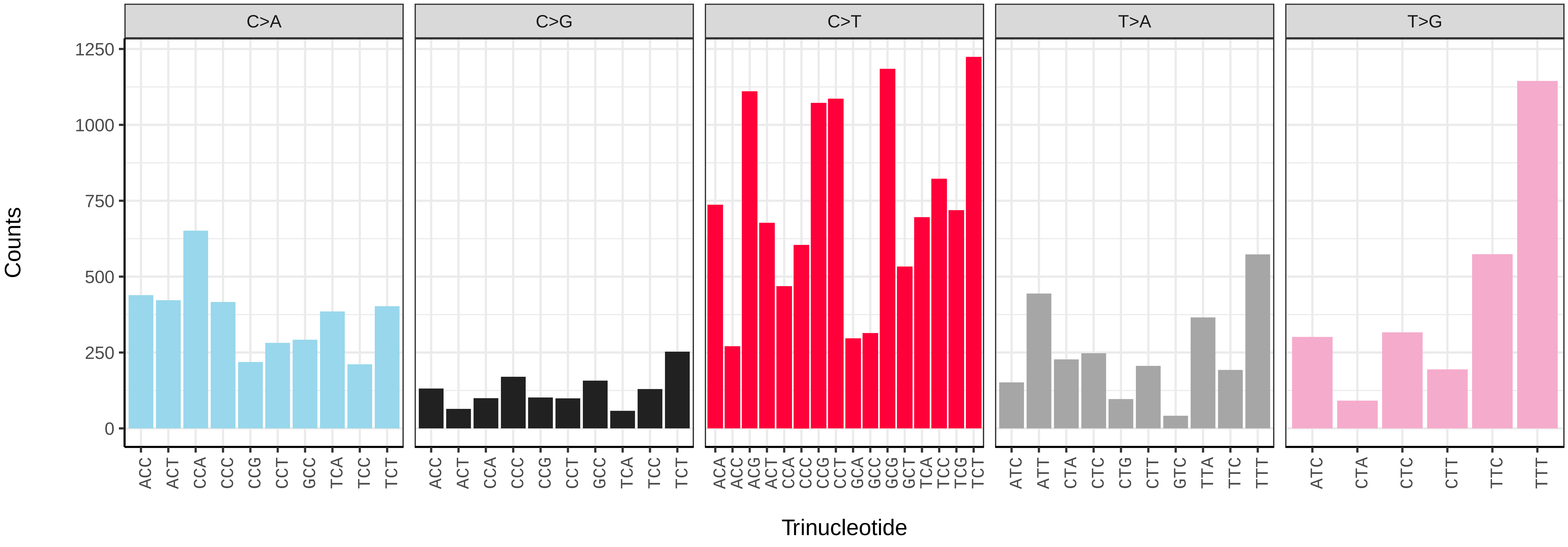

*Pipistrellus pygmaeus* (mPipPyg2) germline mutational spectrum

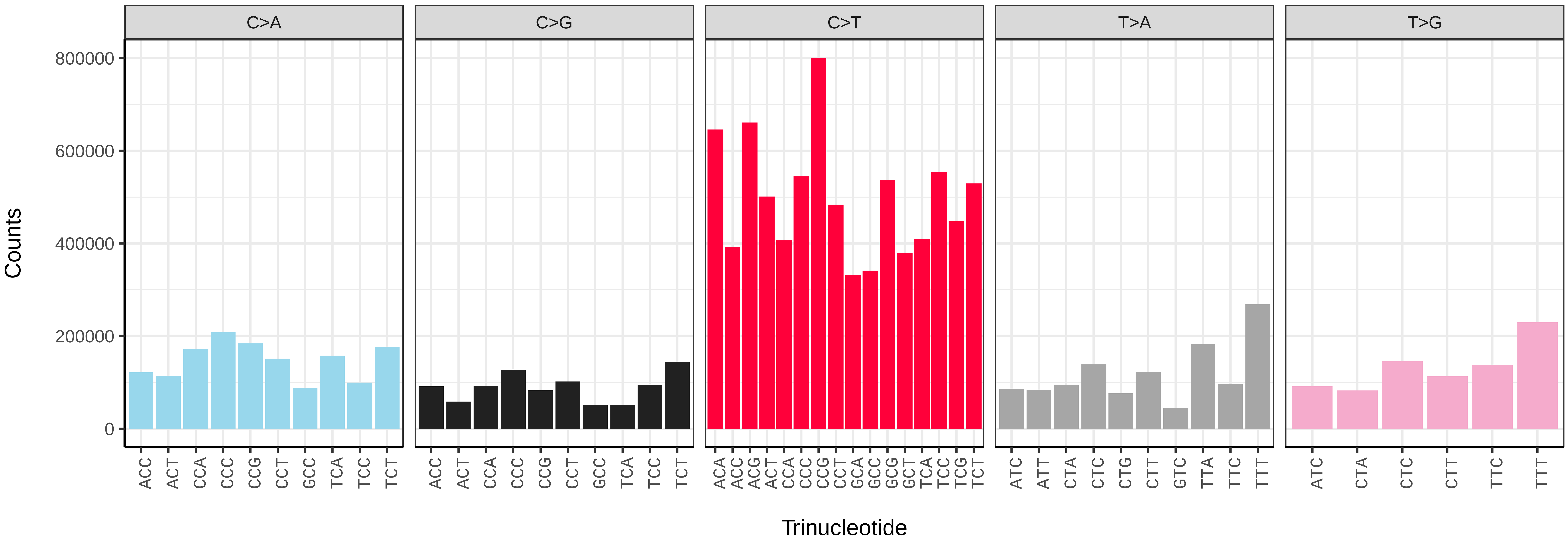


*Pipistrellus pygmaeus* (mPipPyg2) somatic mutational spectrum (SBS96 classification)

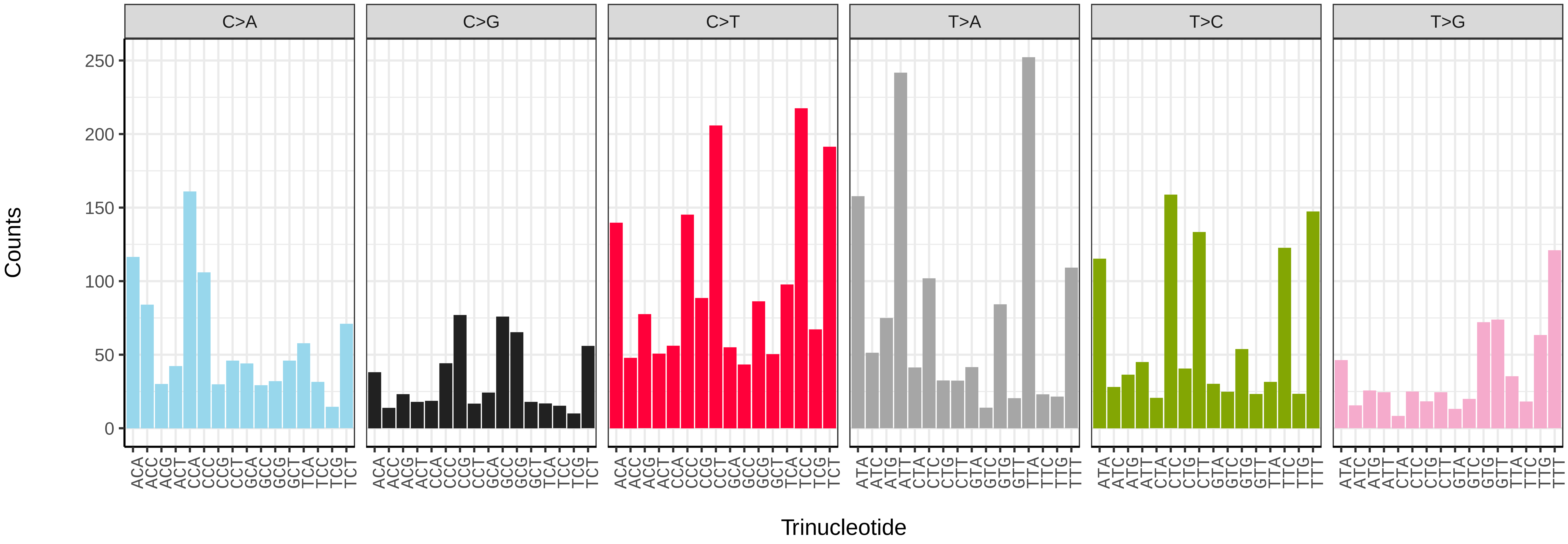


*Pipistrellus pygmaeus* (mPipPyg2) somatic mutational spectrum (SBS52 classification)

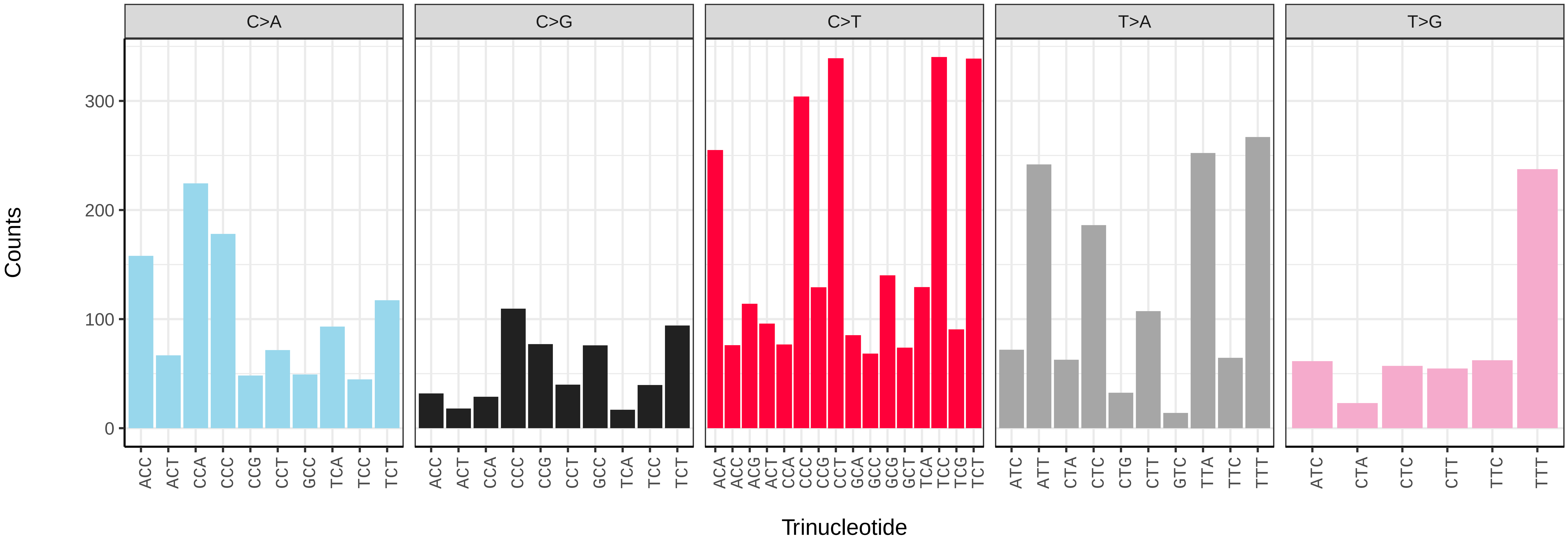

*Plecotus auritus* (mPleAur1) germline mutational spectrum

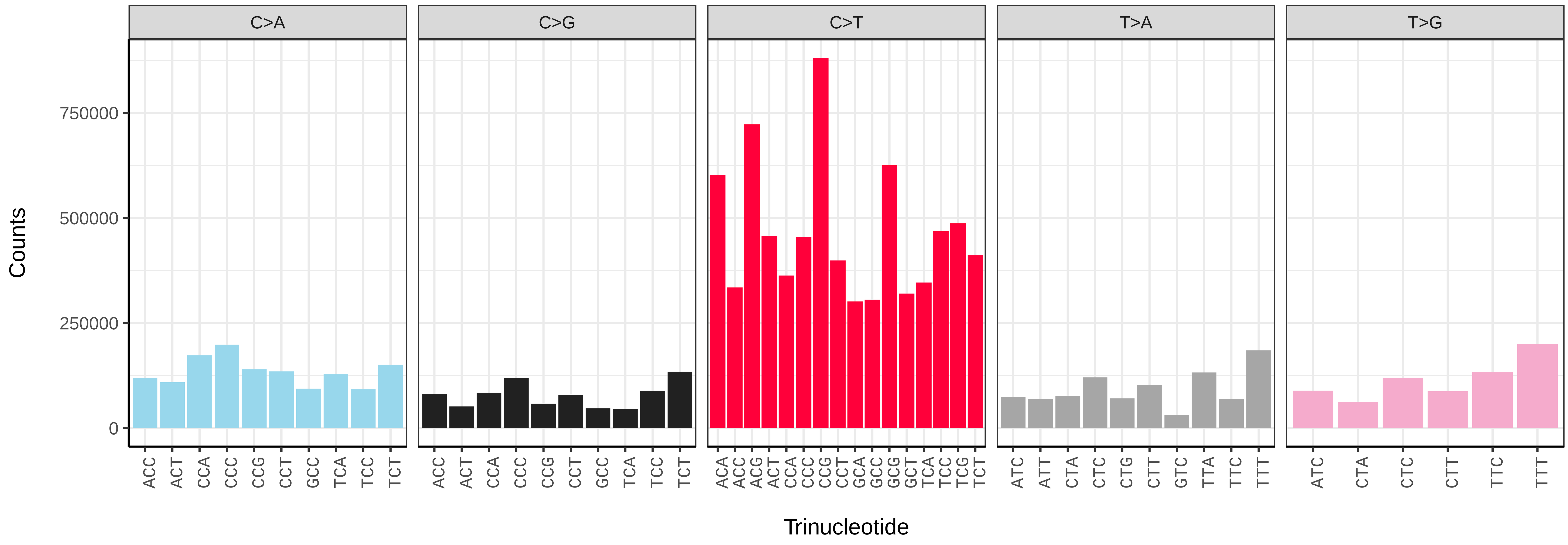


*Plecotus auritus* (mPleAur1) somatic mutational spectrum (SBS96 classification)

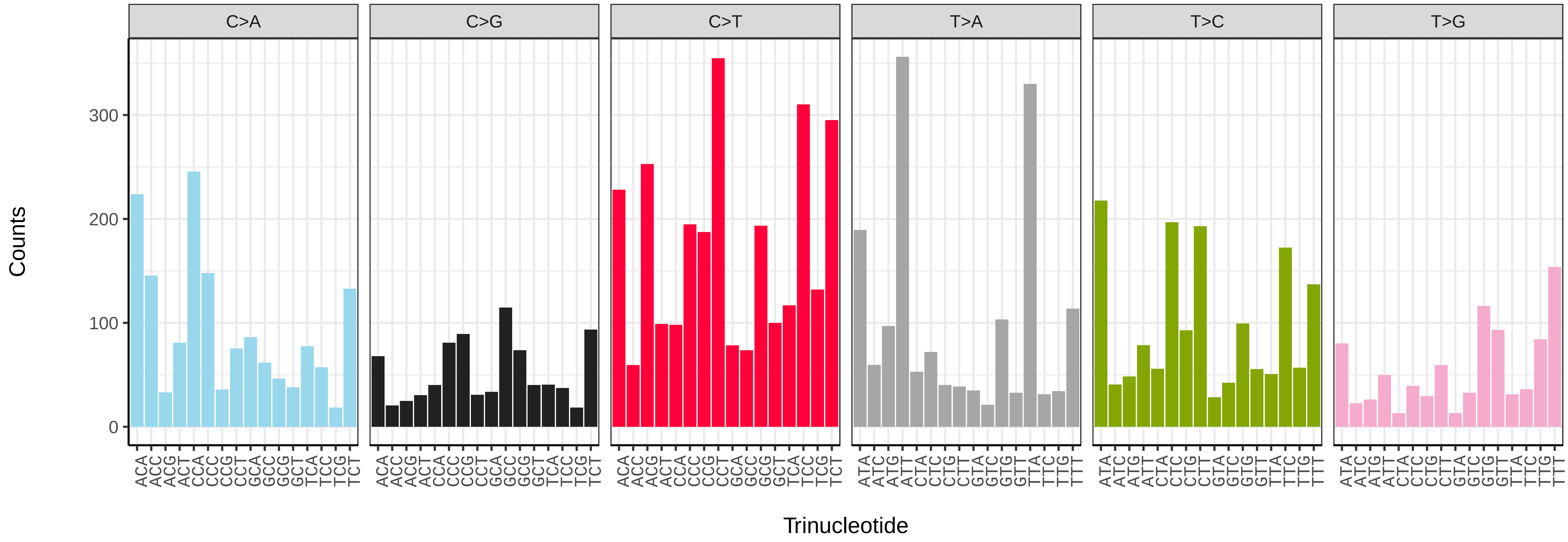


*Plecotus auritus* (mPleAur1) somatic mutational spectrum (SBS52 classification)

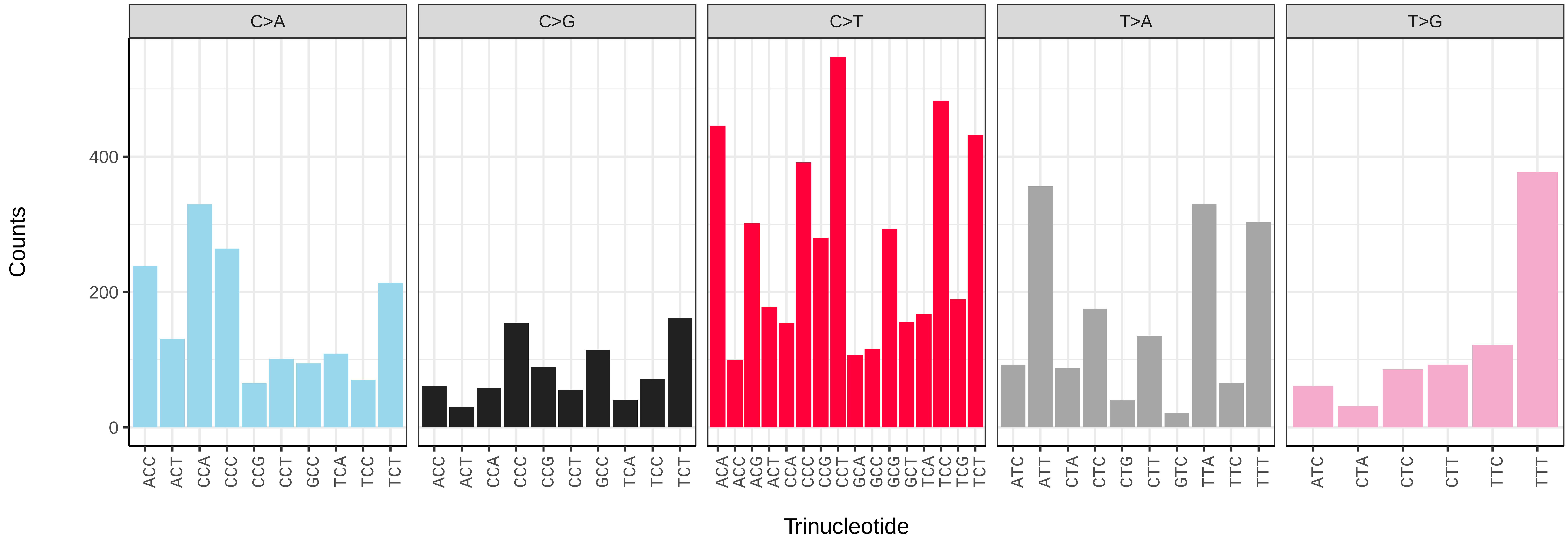

*Stenella coeruleoalba* (mSteCoe1) germline mutational spectrum

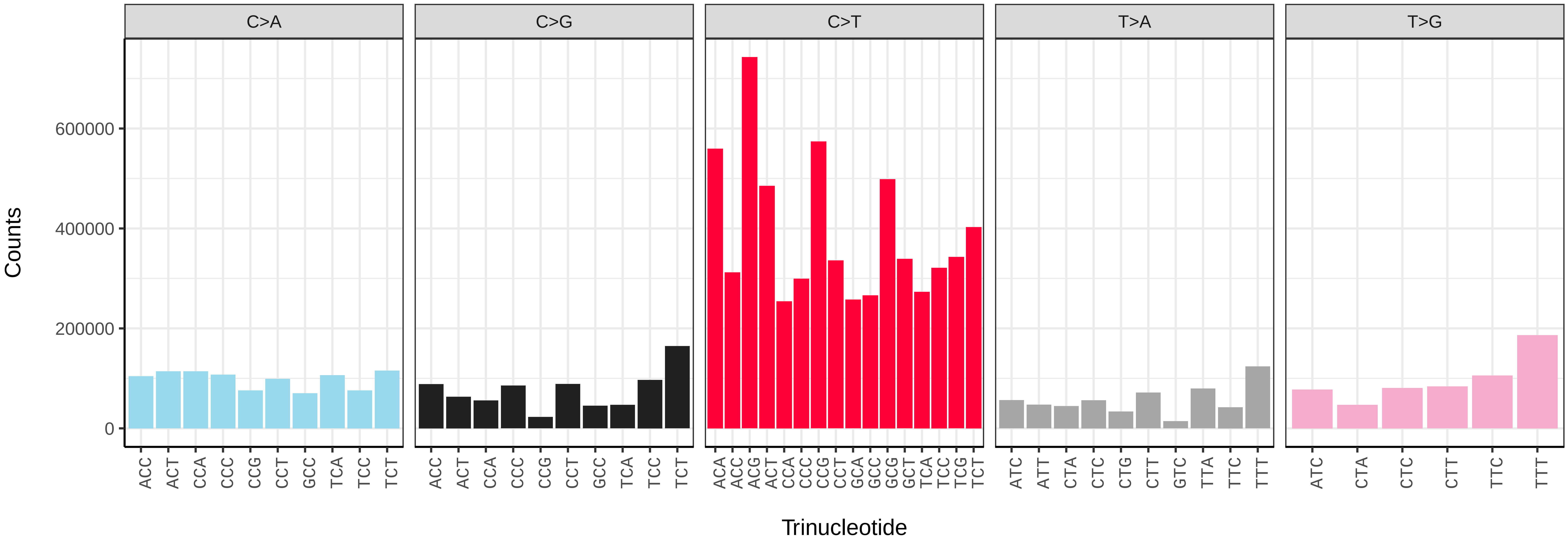


*Stenella coeruleoalba* (mSteCoe1) somatic mutational spectrum (SBS96 classification)

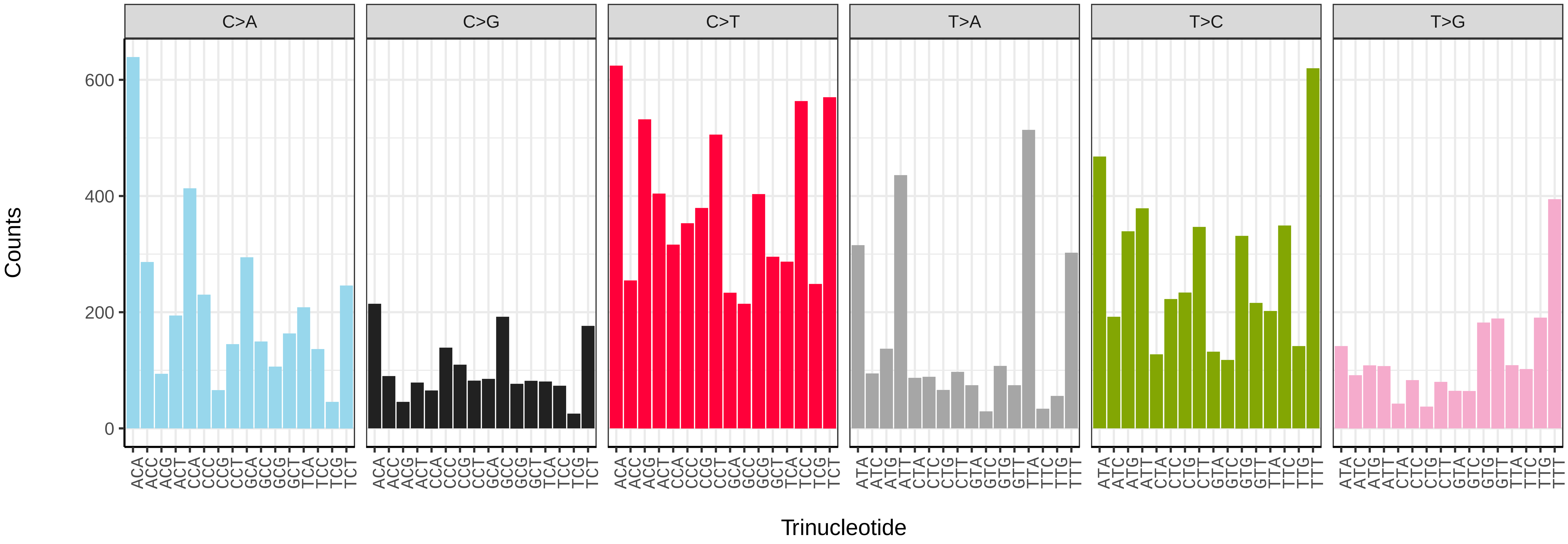


*Stenella coeruleoalba* (mSteCoe1) somatic mutational spectrum (SBS52 classification)

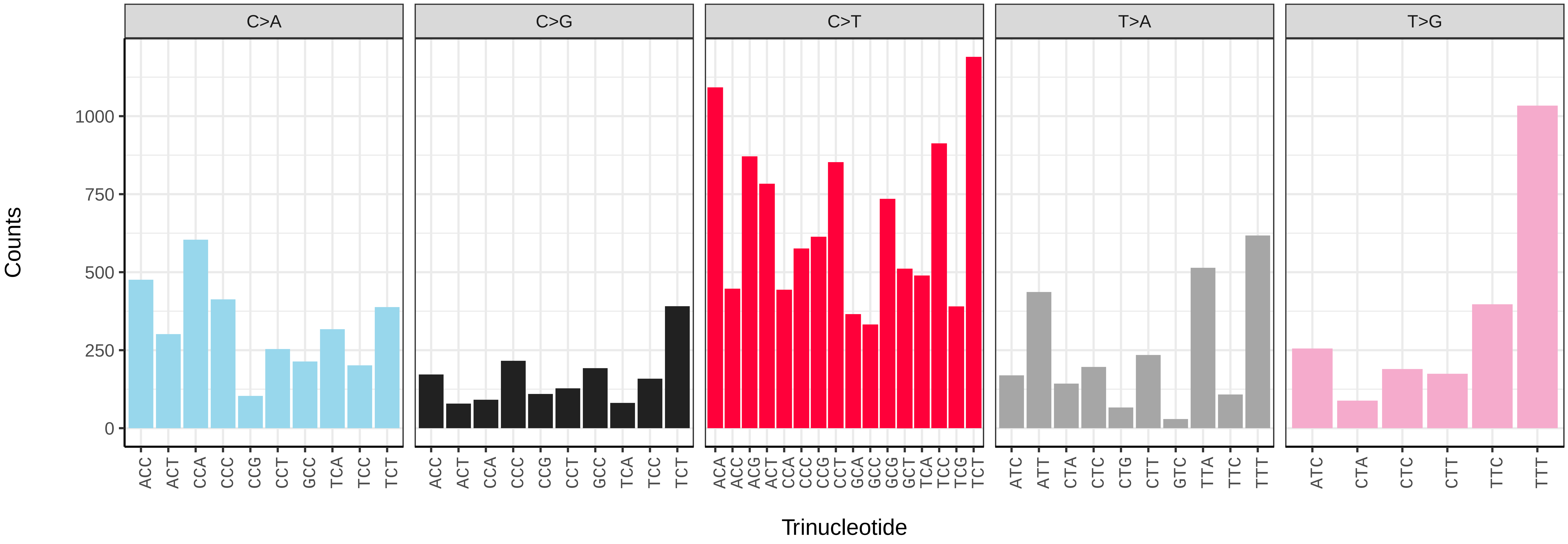

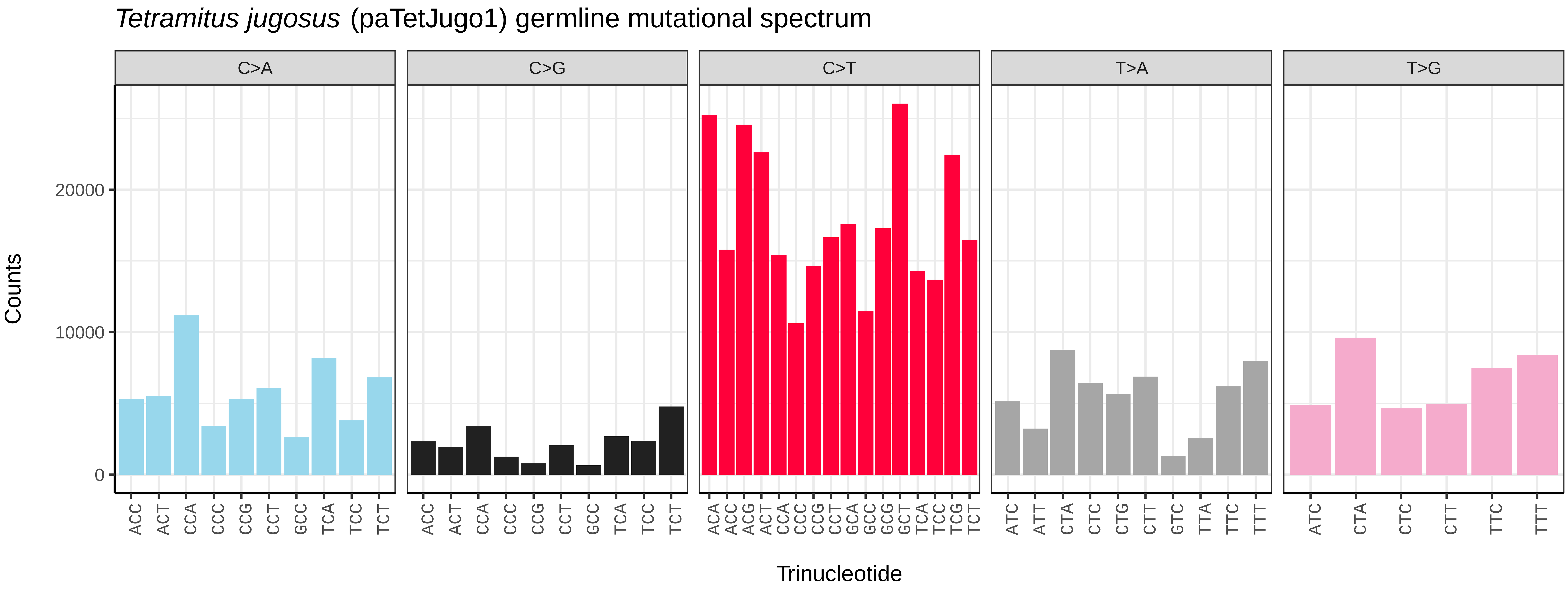
*Tetramitus jugosus* (paTetJugo1) germline mutational spectrum
C>A
C>G
C>T
T>A
T>G
Counts
0
10000
20000
ACC ACT CCA CCC CCG CCT GCC TCA TCC TCT
ACC ACT CCA CCC CCG CCT GCC TCA TCC TCT
ACA ACC ACG ACT CCA CCC CCG CCT GCA GCC GCG GCT TCA TCC TCG TCT
ATC ATT CTA CTC CTG CTT GTC TTA TTC TTT
ATC CTA CTC CTT TTC TTT
Trinucleotide

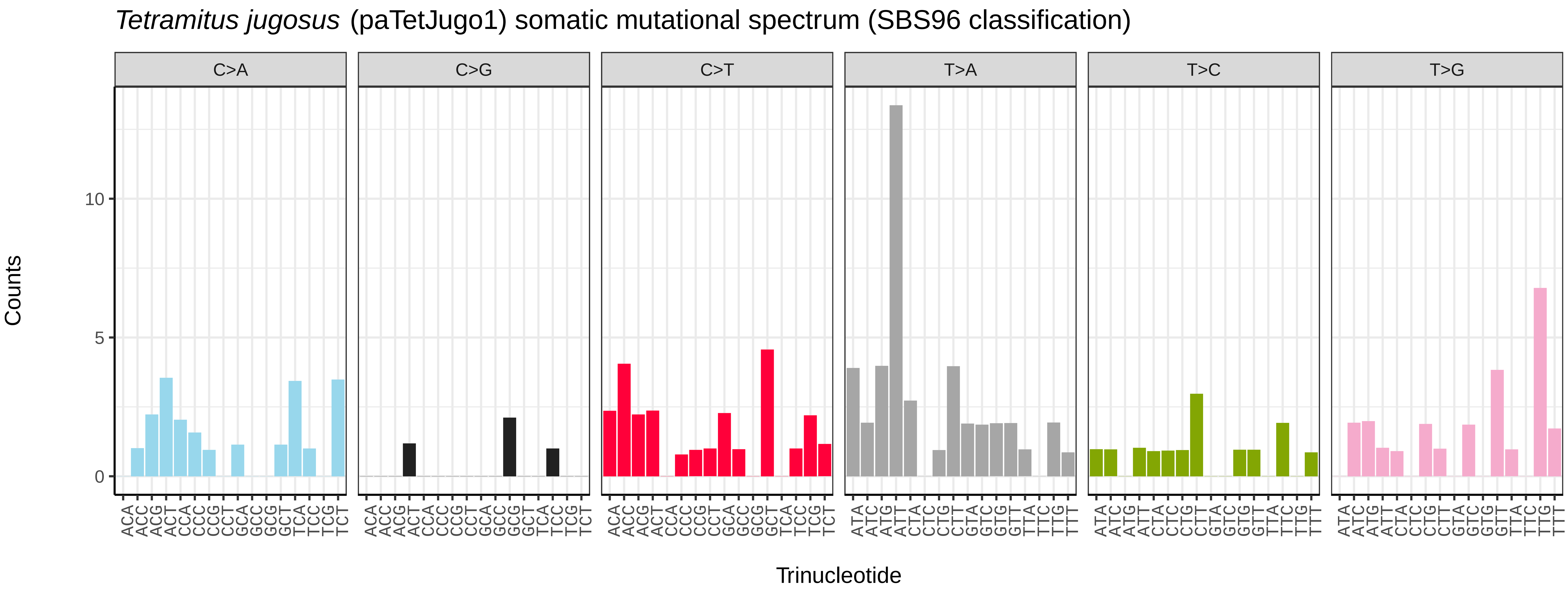
*Tetramitus jugosus* (paTetJugo1) somatic mutational spectrum (SBS96 classification)
C>A
C>G
C>T
T>A
T>C
T>G
Counts
0
5
10
ACA ACC ACG ACT CCA CCC CCG CCT GCA GCC GCG GCT TCA TCC TCG TCT
ACA ACC ACG ACT CCA CCC CCG CCT GCA GCC GCG GCT TCA TCC TCG TCT
ACA ACC ACG ACT CCA CCC CCG CCT GCA GCC GCG GCT TCA TCC TCG TCT
ATA ATC ATG ATT CTA CTC CTG CTT GTA GTC GTG GTT TTA TTC TTG TTT
ATA ATC ATG ATT CTA CTC CTG CTT GTA GTC GTG GTT TTA TTC TTG TTT
ATA ATC ATG ATT CTA CTC CTG CTT GTA GTC GTG GTT TTA TTC TTG TTT
Trinucleotide

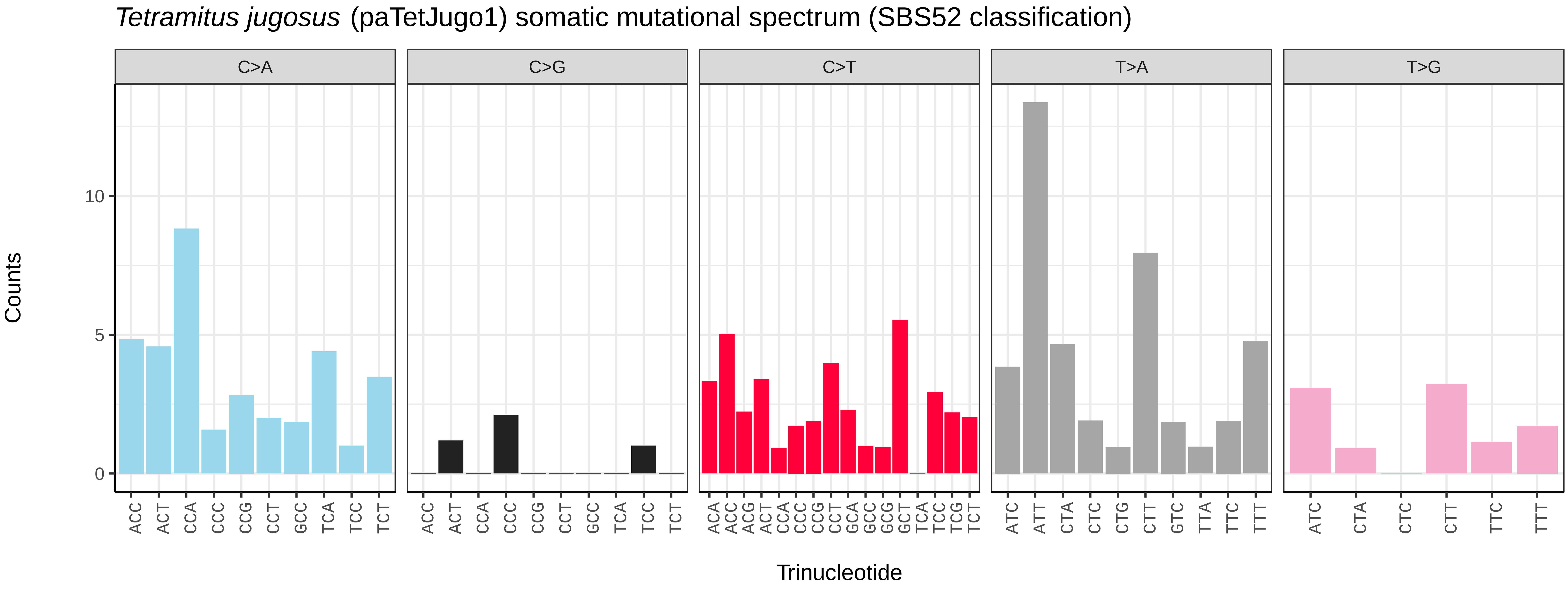
*Tetramitus jugosus* (paTetJugo1) somatic mutational spectrum (SBS52 classification)
C>A
C>G
C>T
T>A
T>G
Counts
0
5
10
ACC ACT CCA CCC CCG CCT GCC TCA TCC TCT
ACC ACT CCA CCC CCG CCT GCC TCA TCC TCT
ACA ACC ACG ACT CCA CCC CCG CCT GCA GCC GCG GCT TCA TCC TCG TCT
ATC ATT CTA CTC CTG CTT GTC TTA TTC TTT
ATC CTA CTC CTT TTC TTT
Trinucleotide

*Eimeria praecox* (pxEimPrae1) germline mutational spectrum

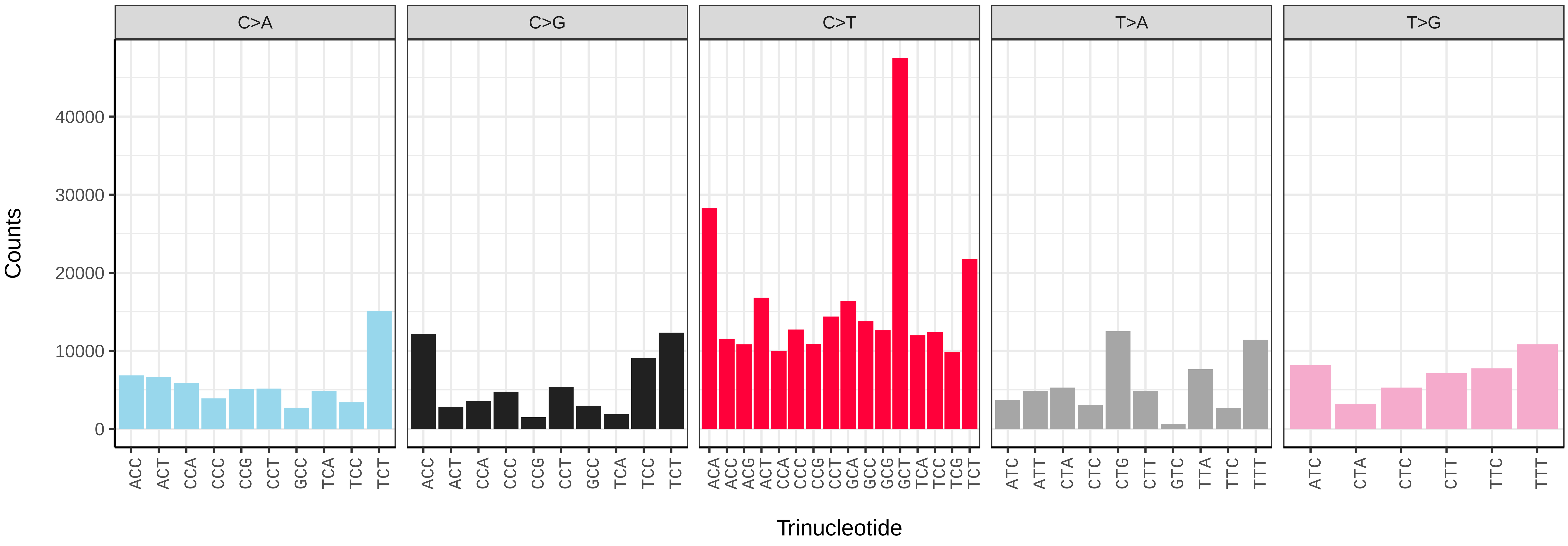


*Eimeria praecox* (pxEimPrae1) somatic mutational spectrum (SBS96 classification)

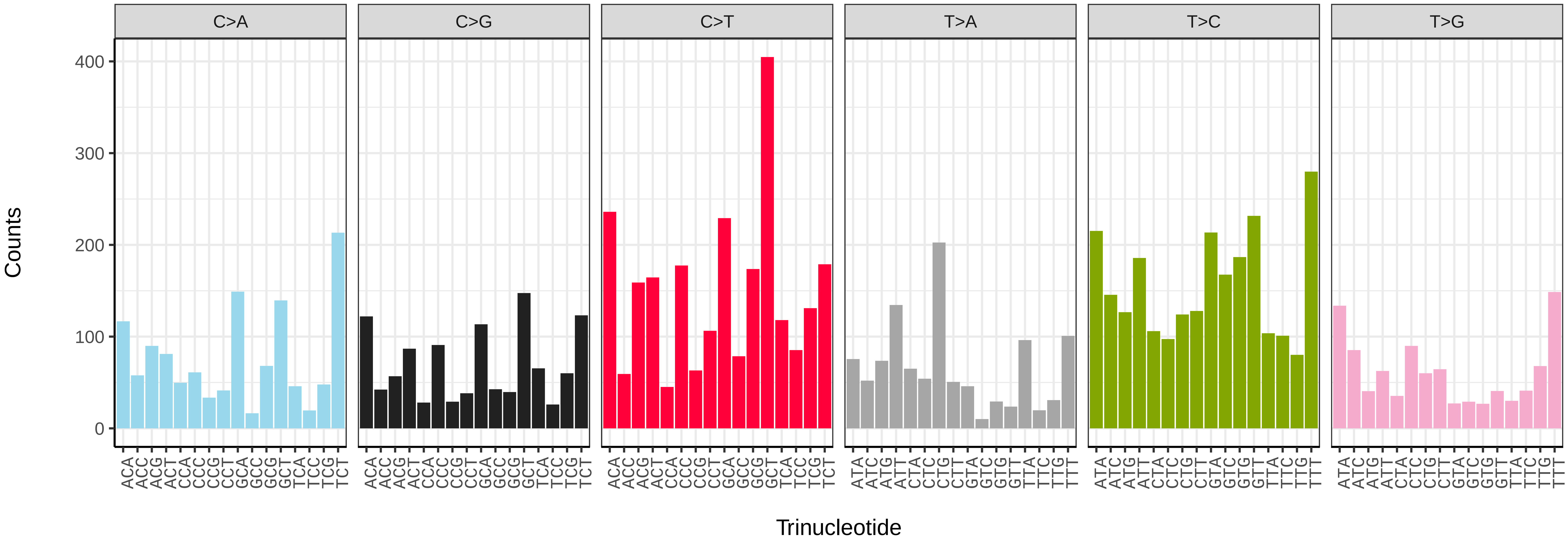


*Eimeria praecox* (pxEimPrae1) somatic mutational spectrum (SBS52 classification)

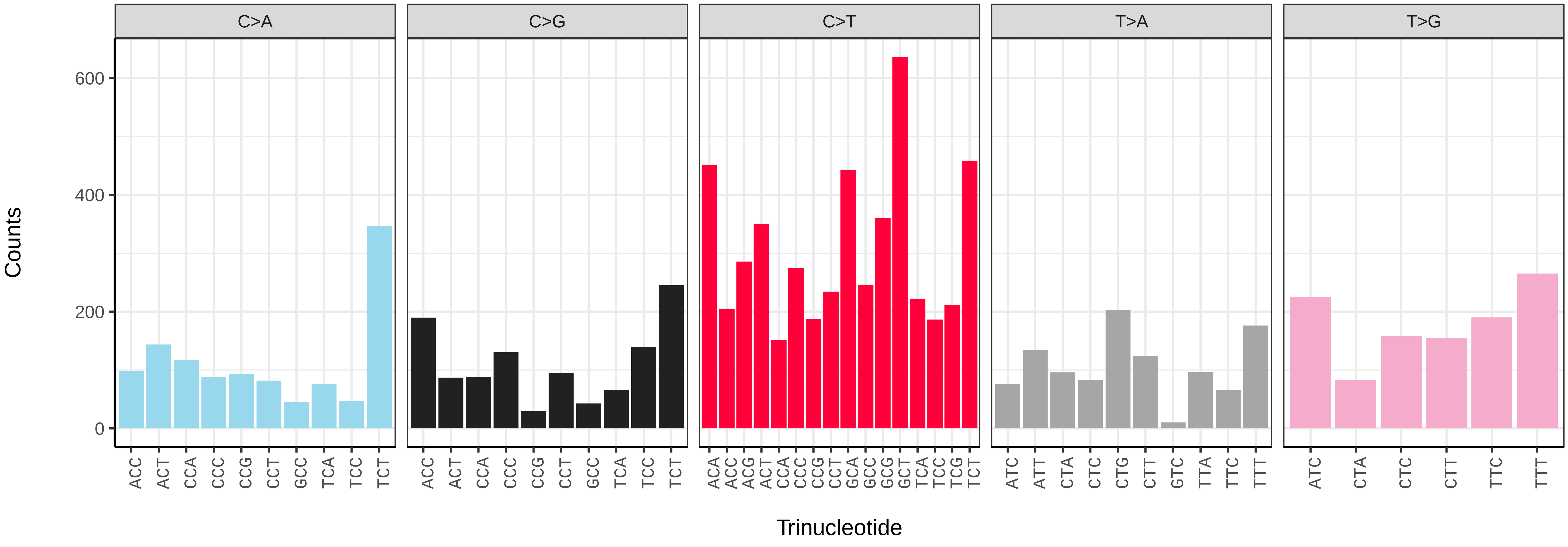

*Strigamia acuminata* (qcStrAcum1) germline mutational spectrum

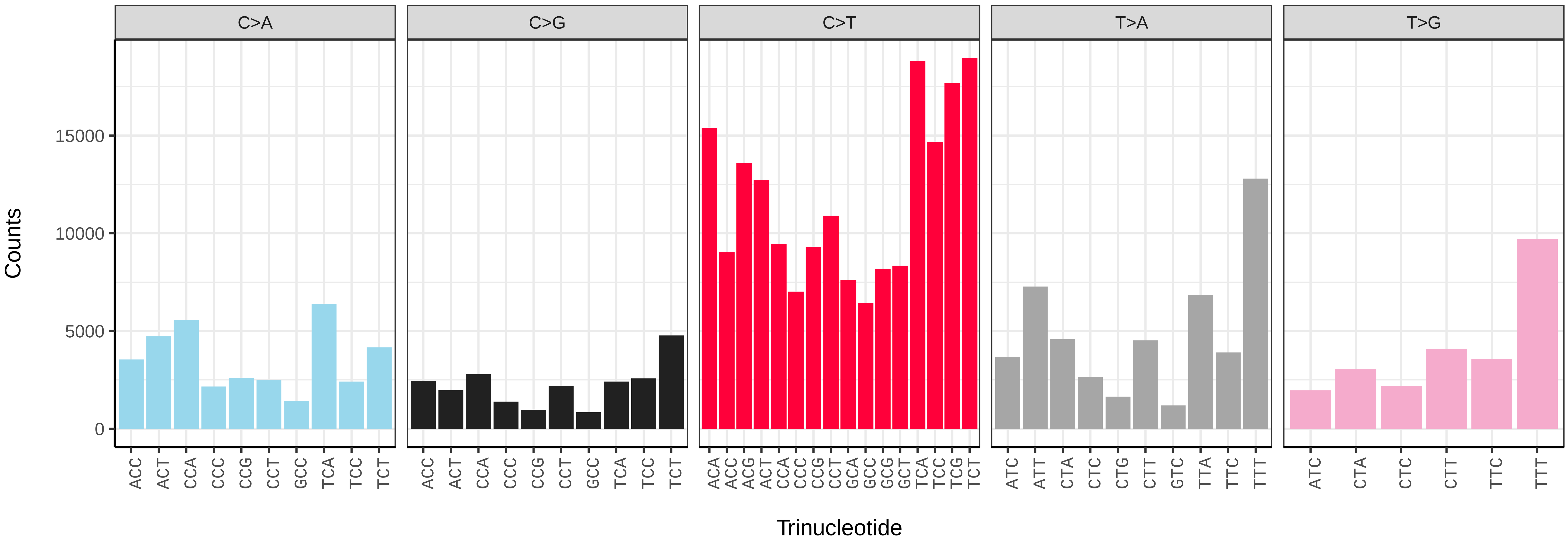


*Strigamia acuminata* (qcStrAcum1) somatic mutational spectrum (SBS96 classification)

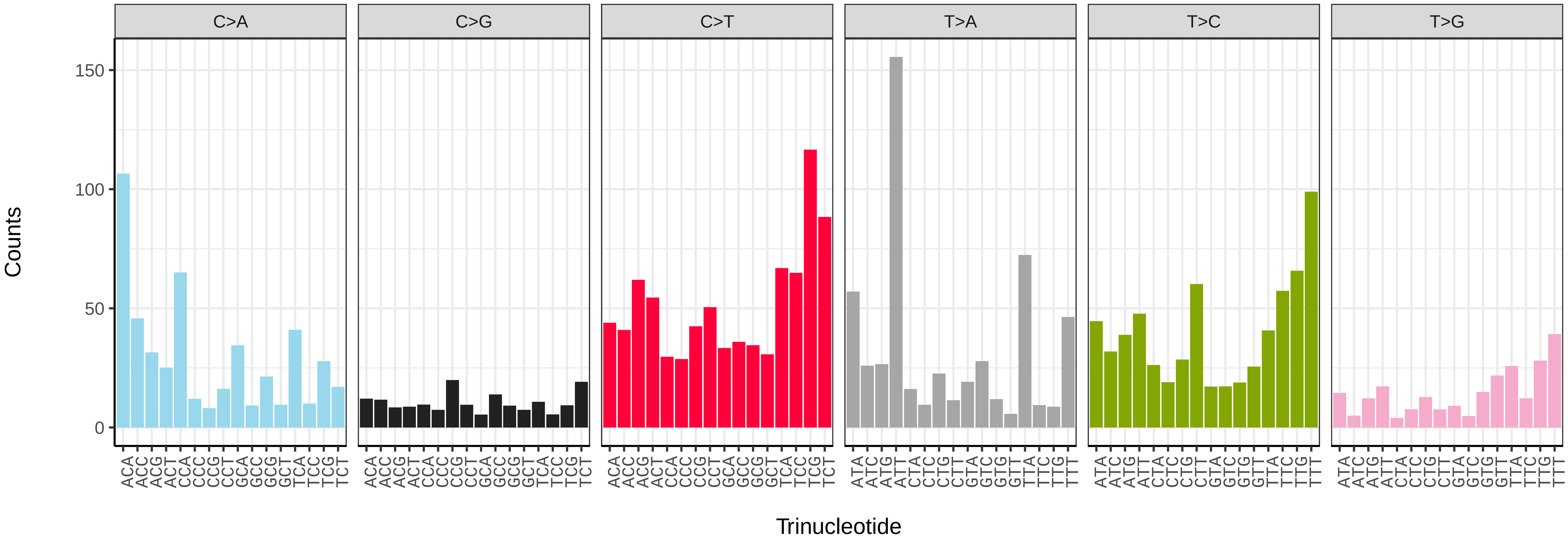


*Strigamia acuminata* (qcStrAcum1) somatic mutational spectrum (SBS52 classification)

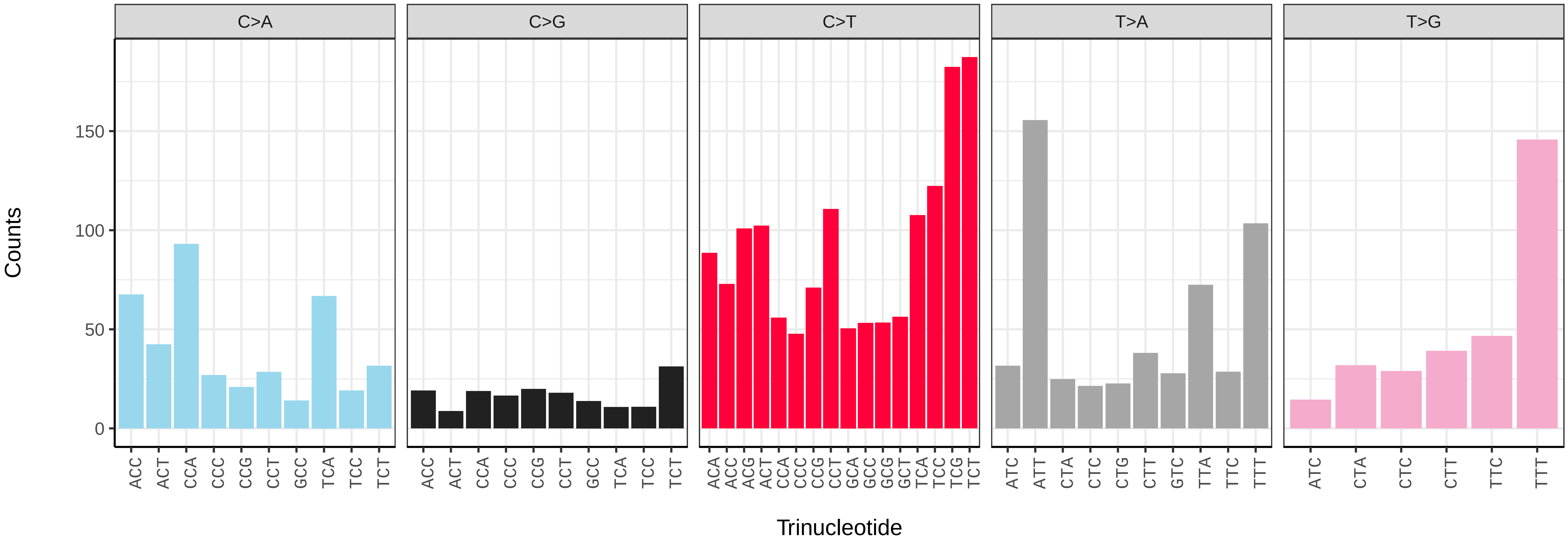

*Lepeophtheirus nordmannii* (qhLepNord3) germline mutational spectrum

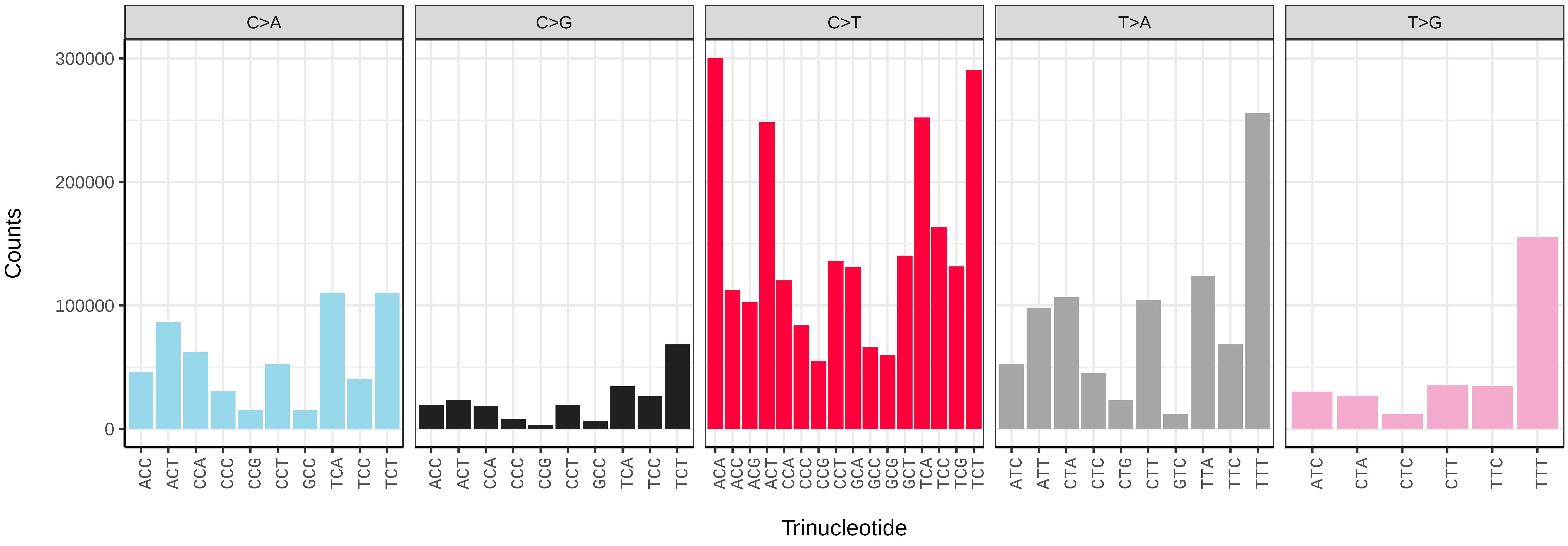


*Lepeophtheirus nordmannii* (qhLepNord3) somatic mutational spectrum (SBS96 classification)

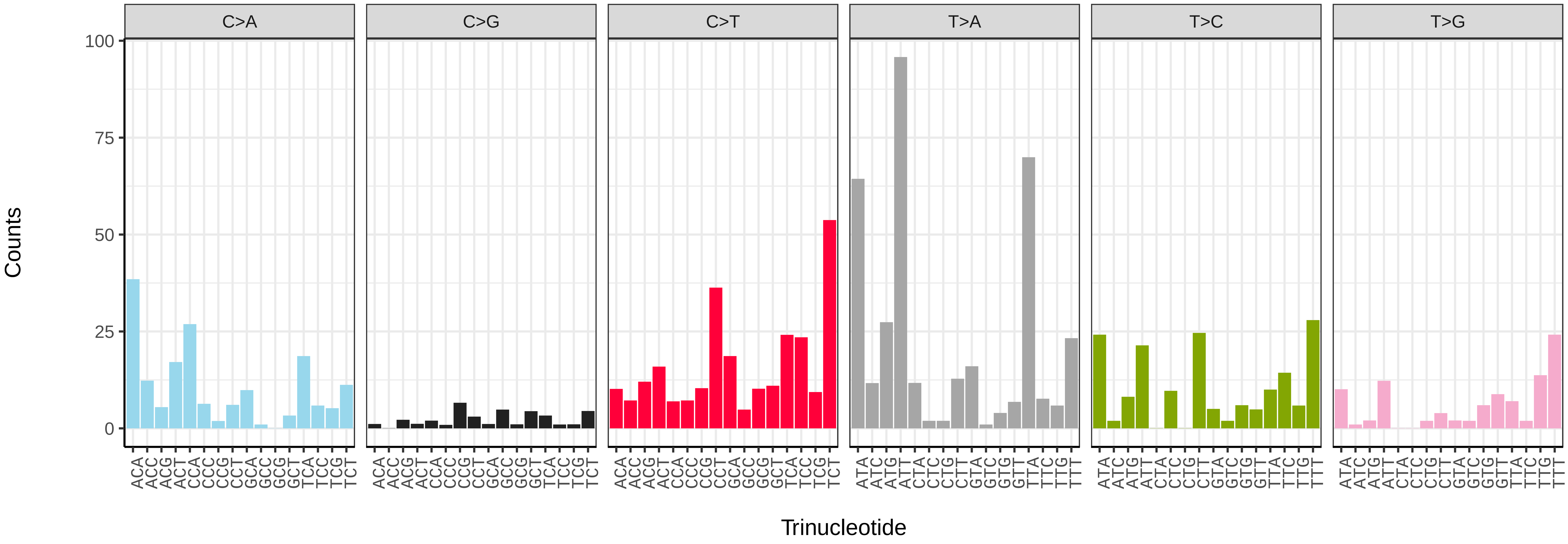


*Lepeophtheirus nordmannii* (qhLepNord3) somatic mutational spectrum (SBS52 classification)

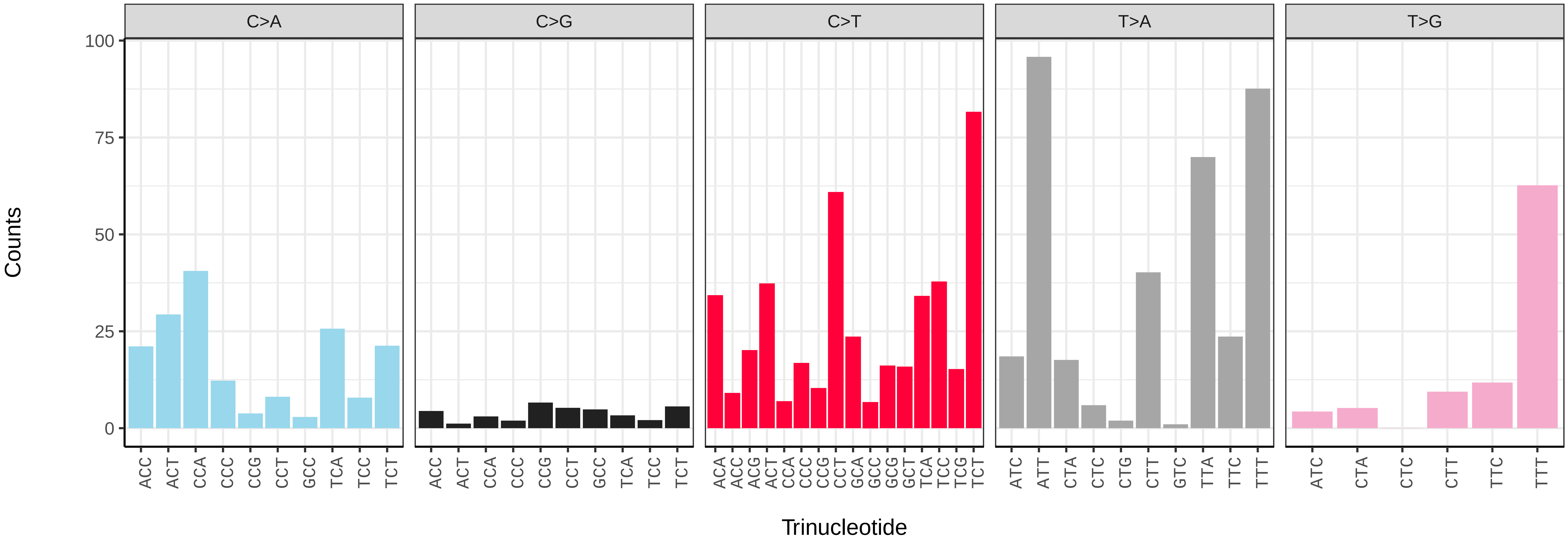

*Amaurobius ferox* (qqAmaFero1) germline mutational spectrum

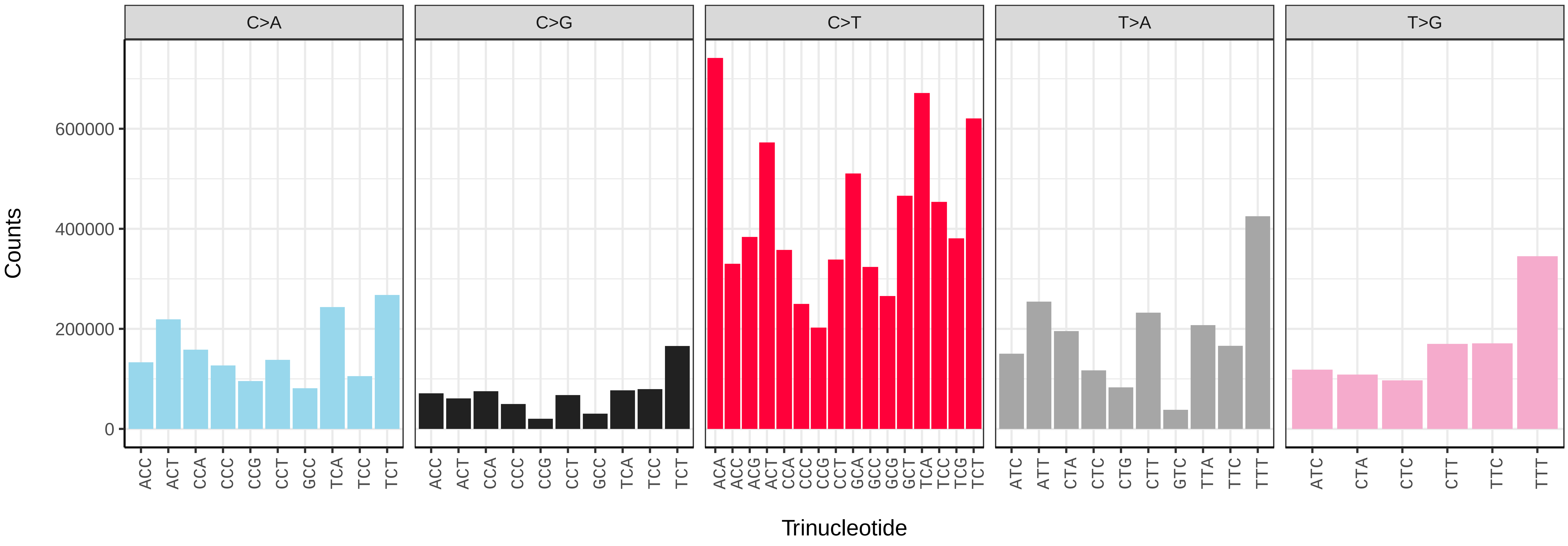


*Amaurobius ferox* (qqAmaFero1) somatic mutational spectrum (SBS96 classification)

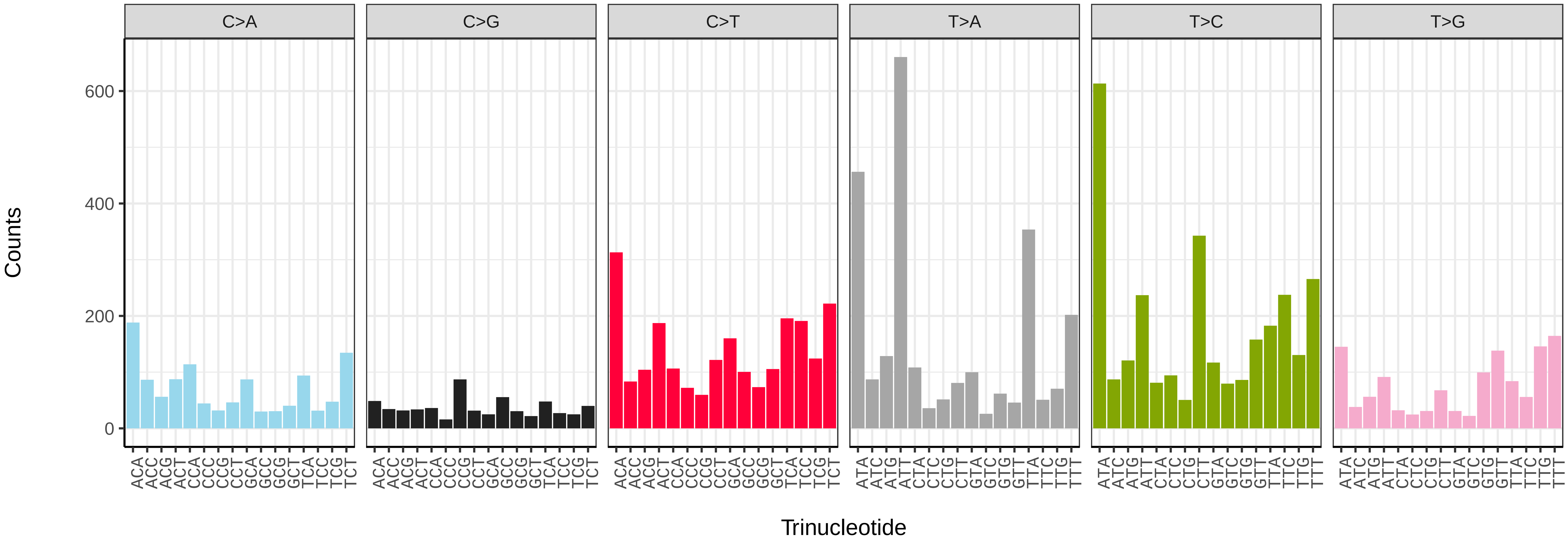


*Amaurobius ferox* (qqAmaFero1) somatic mutational spectrum (SBS52 classification)

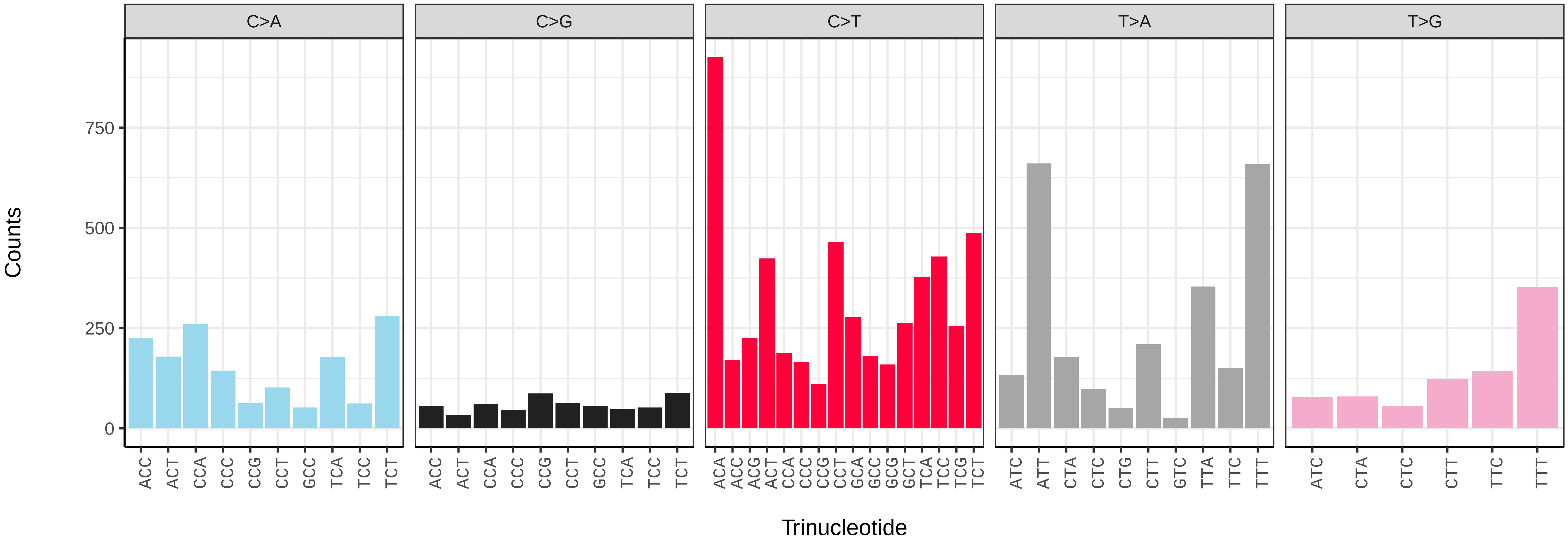

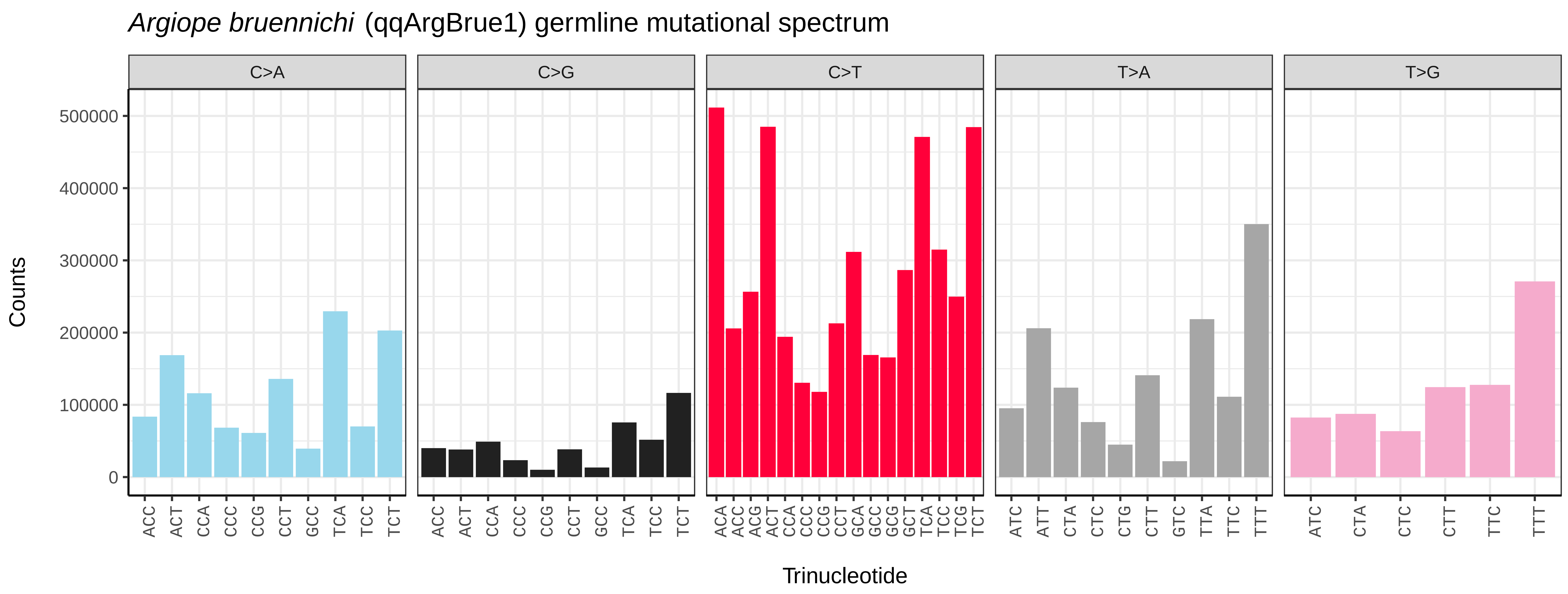
Argiope bruennichi (qqArgBrue1) germline mutational spectrum
C>A
C>G
C>T
T>A
T>G
Counts
500000
400000
300000
200000
100000
0
ACC ACT CCA CCC CCG CCT GCC TCA TCC TCT
ACA ACC ACG ACT CCA CCC CCG CCT GCA GCC GCG GCT TCA TCC TCG TCT
ATC ATT CTA CTC CTG CTT GTC TTA TTC TTT
ATC CTA CTC CTT TTC TTT
Trinucleotide

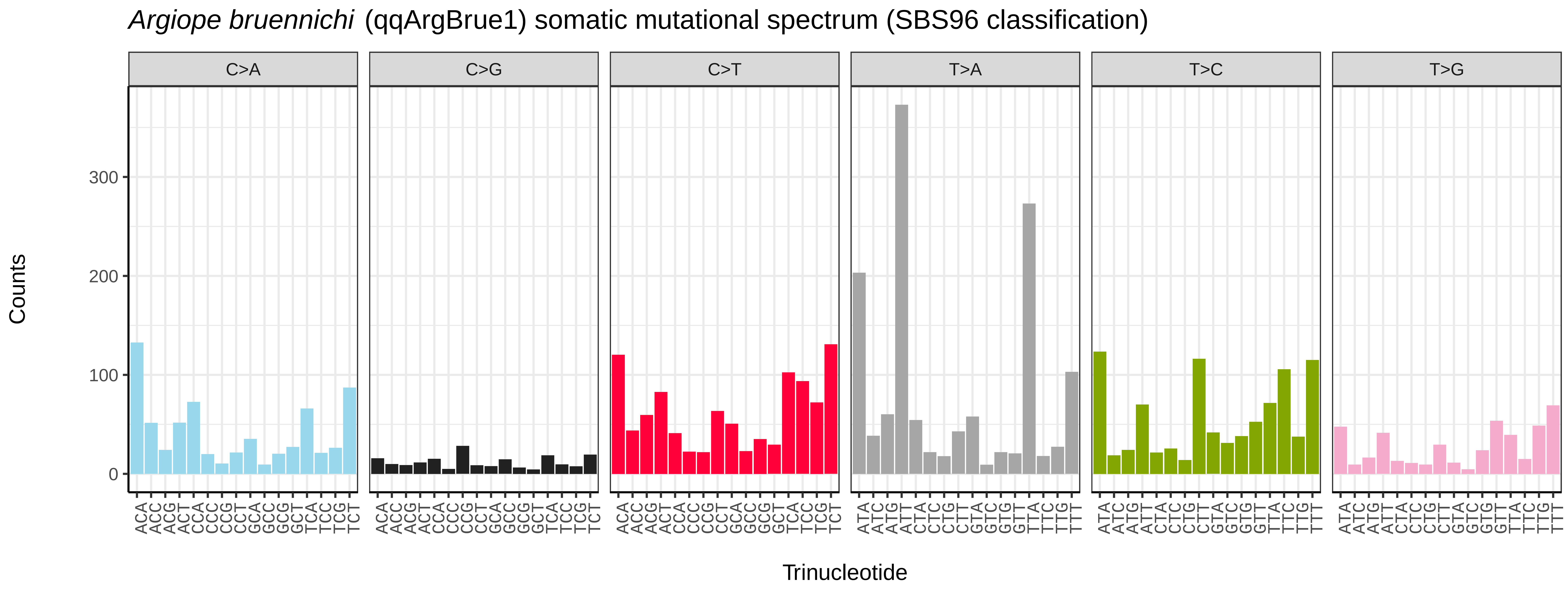
Argiope bruennichi (qqArgBrue1) somatic mutational spectrum (SBS96 classification)
C>A
C>G
C>T
T>A
T>C
T>G
Counts
300
200
100
0
ACA ACC ACG ACT CCA CCC CCG CCT GCA GCC GCG GCT TCA TCC TCG TCT
ATA ATC ATG ATT CTA CTC CTG CTT GTA GTC GTG GTT TTA TTC TTG TTT
Trinucleotide

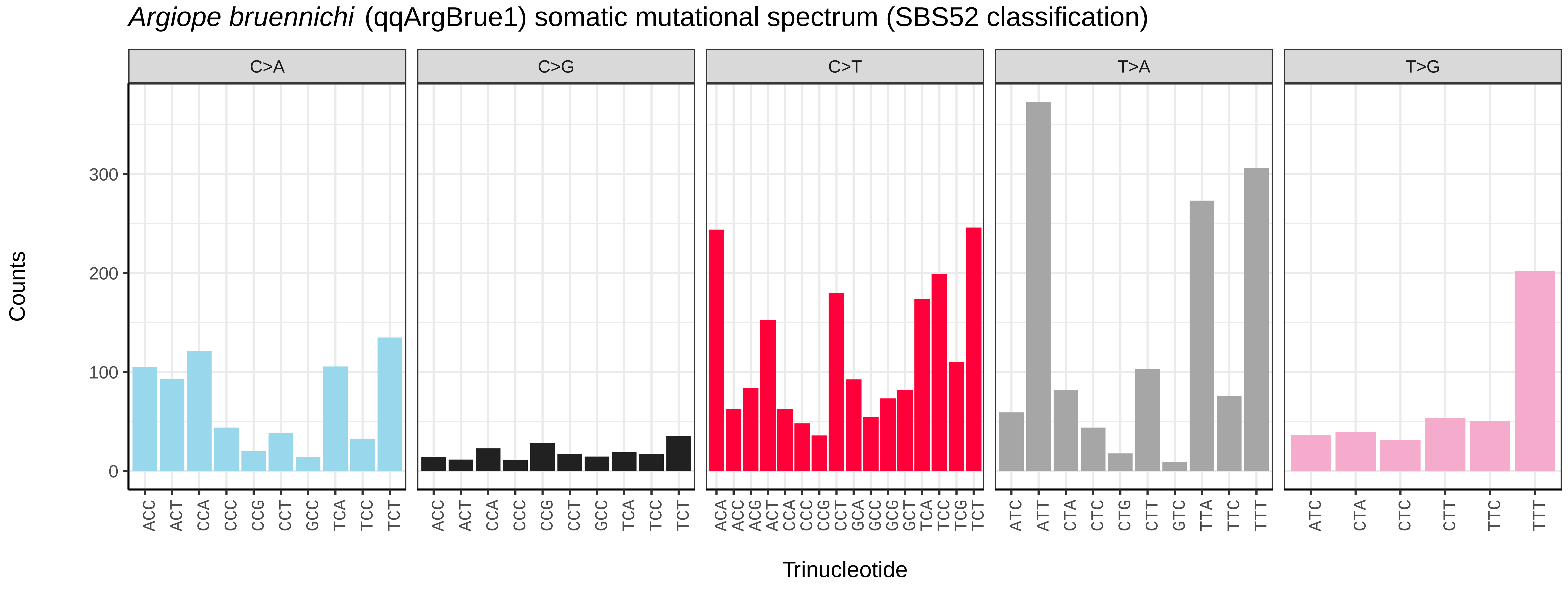
Argiope bruennichi (qqArgBrue1) somatic mutational spectrum (SBS52 classification)
C>A
C>G
C>T
T>A
T>G
Counts
300
200
100
0
ACC ACT CCA CCC CCG CCT GCC TCA TCC TCT
ACA ACC ACG ACT CCA CCC CCG CCT GCA GCC GCG GCT TCA TCC TCG TCT
ATC ATT CTA CTC CTG CTT GTC TTA TTC TTT
ATC CTA CTC CTT TTC TTT
Trinucleotide

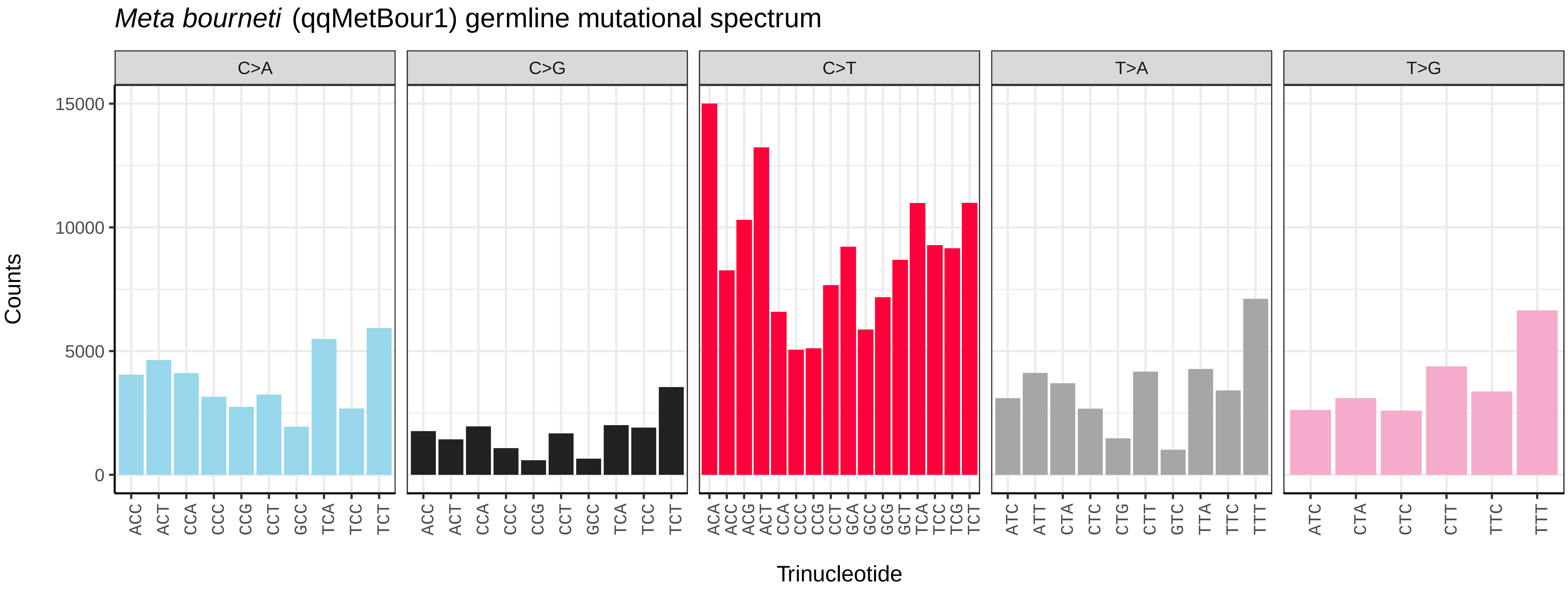
*Meta bourneti* (qqMetBour1) germline mutational spectrum
C>A
C>G
C>T
T>A
T>G
Counts
15000
10000
5000
0
ACC ACT CCA CCC CCG CCT GCC TCA TCC TCT
ACA ACC ACG ACT CCA CCC CCG CCT GCA GCC GCG GCT TCA TCC TCG TCT
ATC ATT CTA CTC CTG CTT GTC TTA TTC TTT
ATC CTA CTC CTT TTC TTT
Trinucleotide

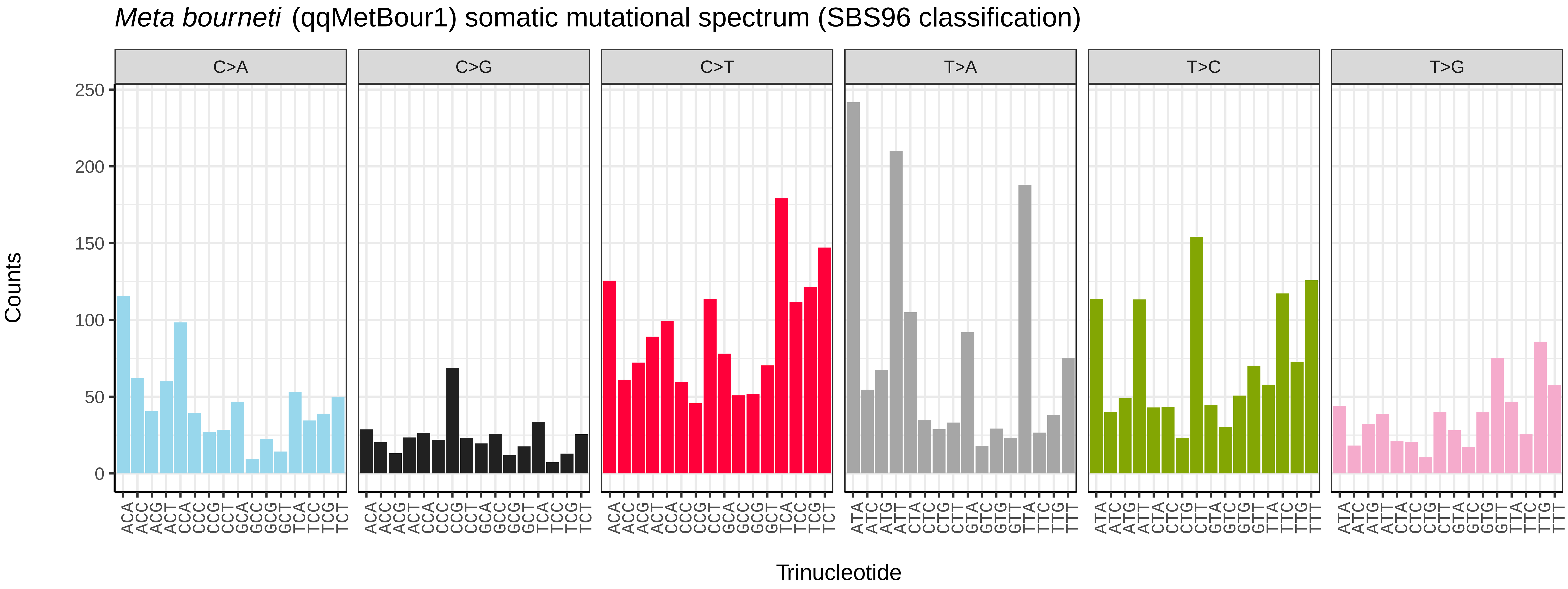
*Meta bourneti* (qqMetBour1) somatic mutational spectrum (SBS96 classification)
C>A
C>G
C>T
T>A
T>C
T>G
Counts
250
200
150
100
50
0
ACA ACC ACG ACT CCA CCC CCG CCT GCA GCC GCG GCT TCA TCC TCG TCT
ATA ATC ATG ATT CTA CTC CTG CTT GTA GTC GTG GTT TTA TTC TTG TTT
Trinucleotide

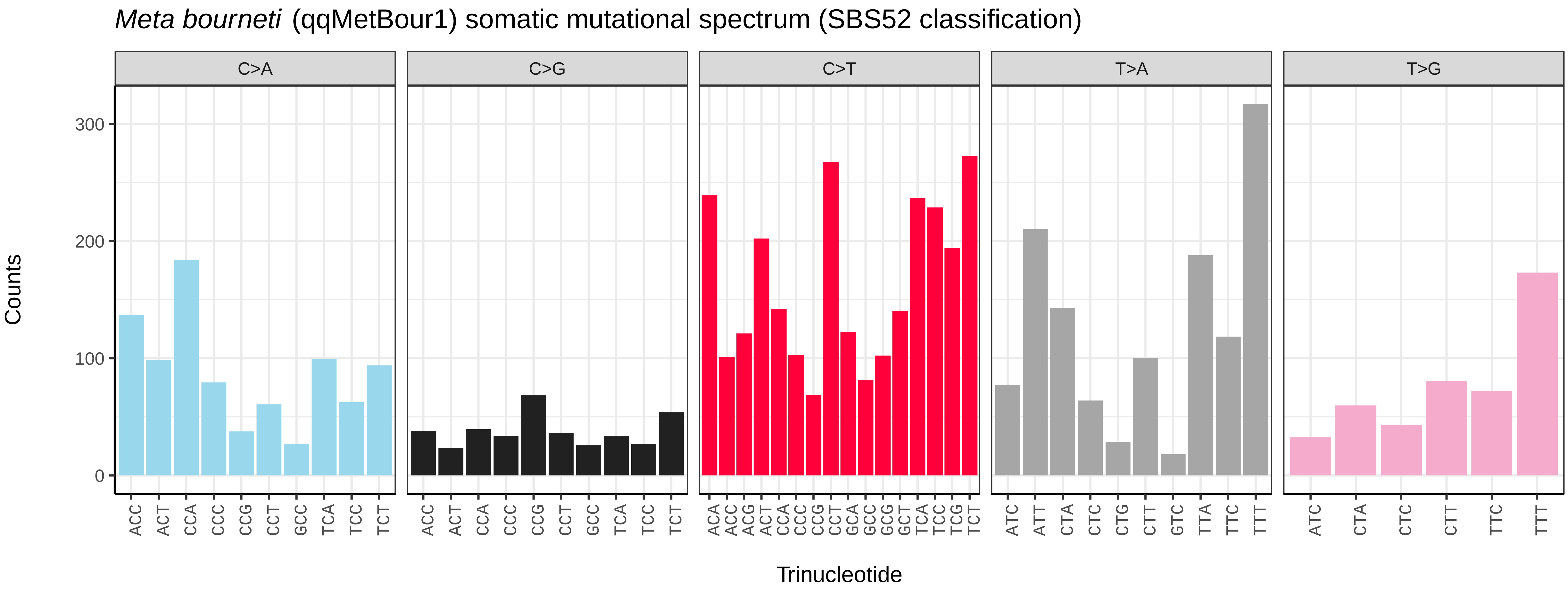
*Meta bourneti* (qqMetBour1) somatic mutational spectrum (SBS52 classification)
C>A
C>G
C>T
T>A
T>G
Counts
300
200
100
0
ACC ACT CCA CCC CCG CCT GCC TCA TCC TCT
ACA ACC ACG ACT CCA CCC CCG CCT GCA GCC GCG GCT TCA TCC TCG TCT
ATC ATT CTA CTC CTG CTT GTC TTA TTC TTT
ATC CTA CTC CTT TTC TTT
Trinucleotide

*Odiellus spinosus* (qqOdiSpin1) germline mutational spectrum

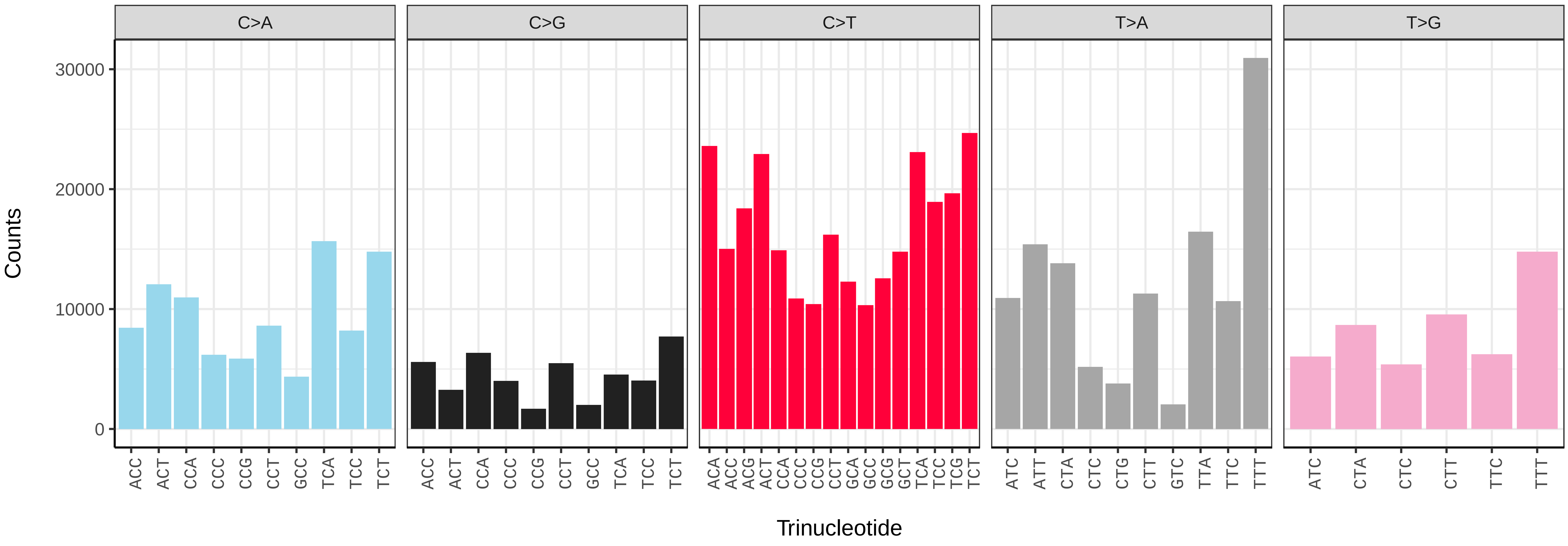


*Odiellus spinosus* (qqOdiSpin1) somatic mutational spectrum (SBS96 classification)

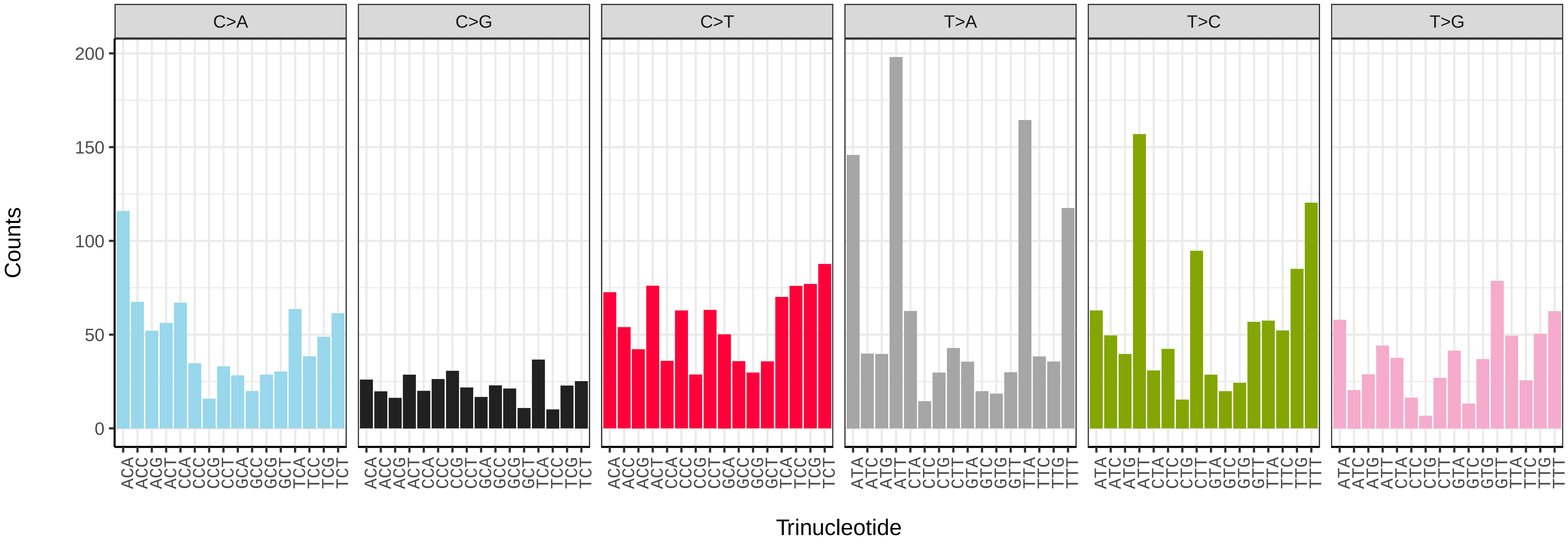


*Odiellus spinosus* (qqOdiSpin1) somatic mutational spectrum (SBS52 classification)

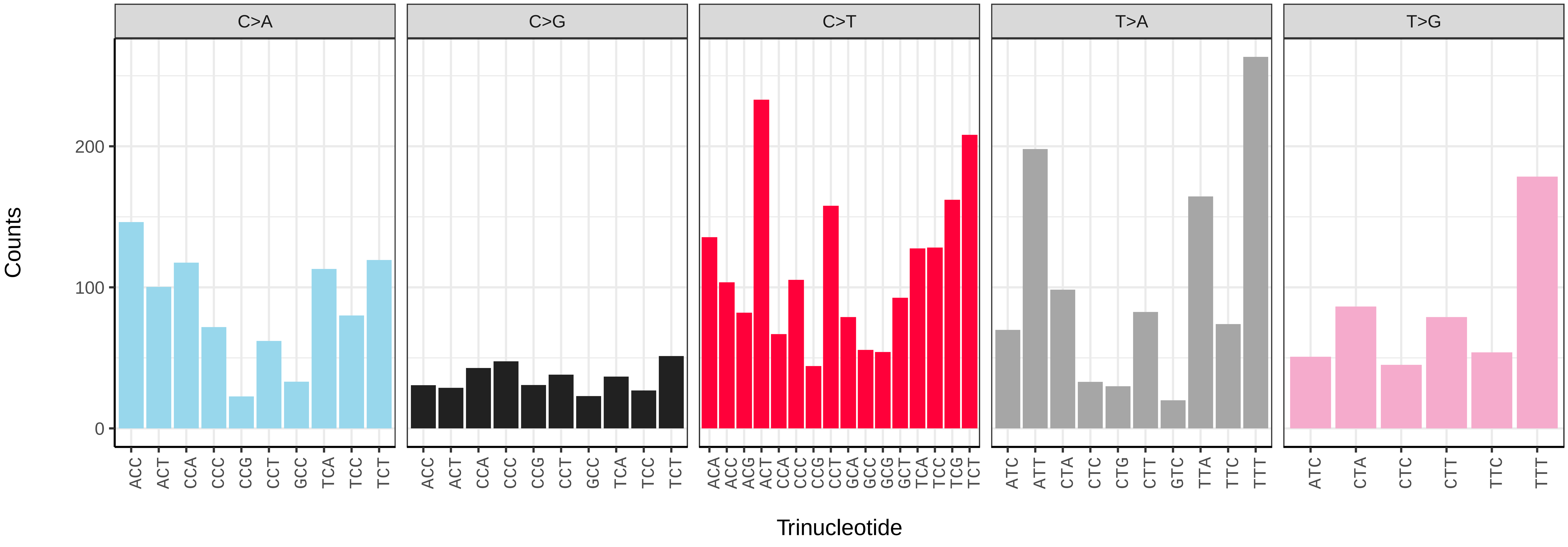

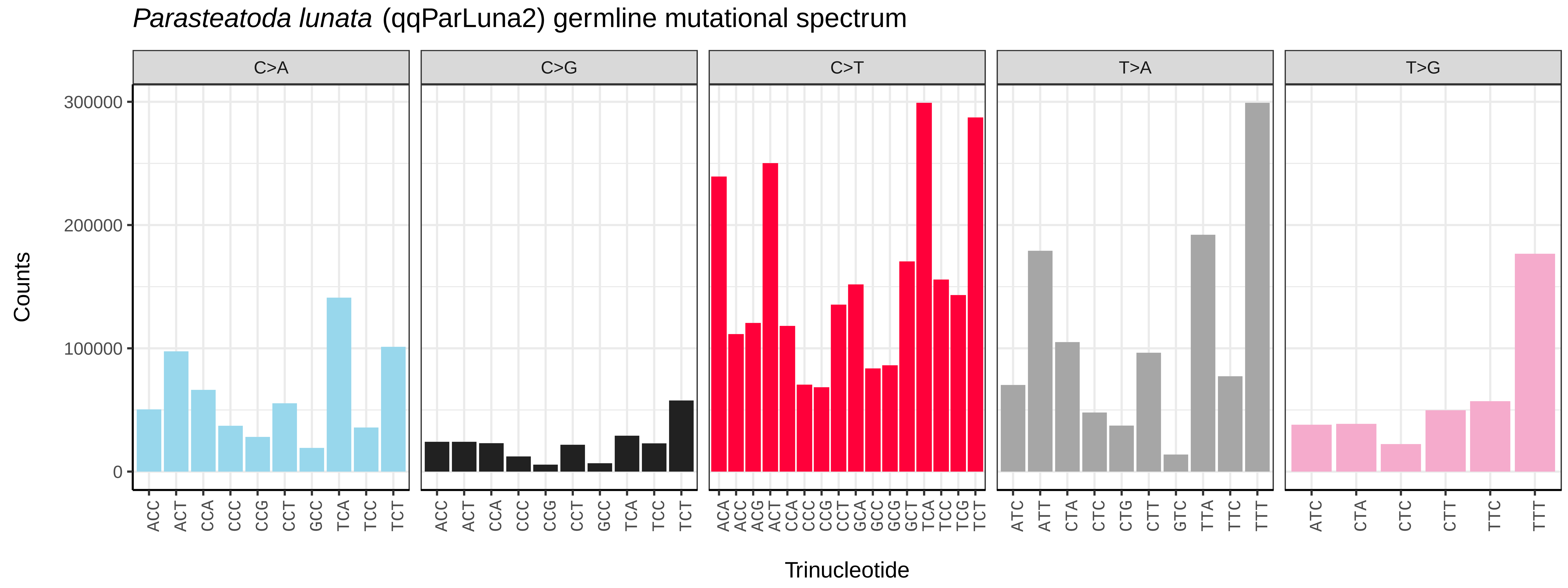

Parasteatoda lunata (qqParLuna2) germline mutational spectrum
C>A
C>G
C>T
T>A
T>G
Counts
Trinucleotide


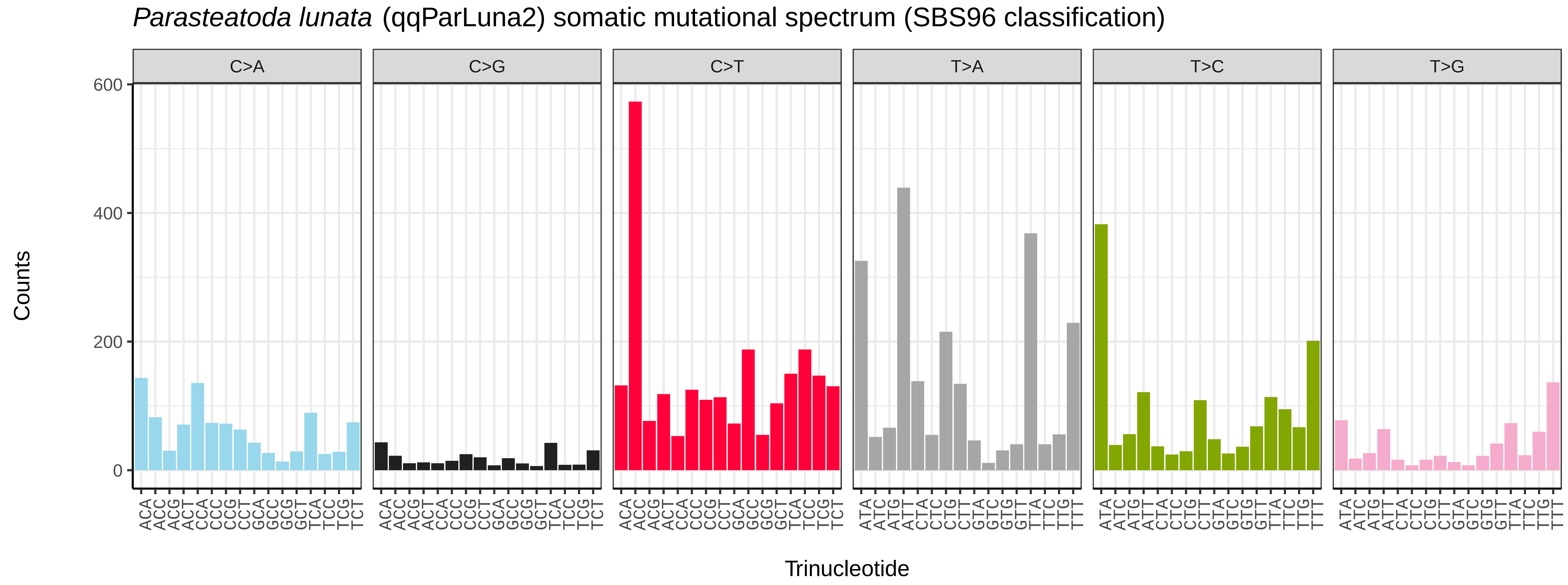

Parasteatoda lunata (qqParLuna2) somatic mutational spectrum (SBS96 classification)
C>A
C>G
C>T
T>A
T>C
T>G
Counts
Trinucleotide


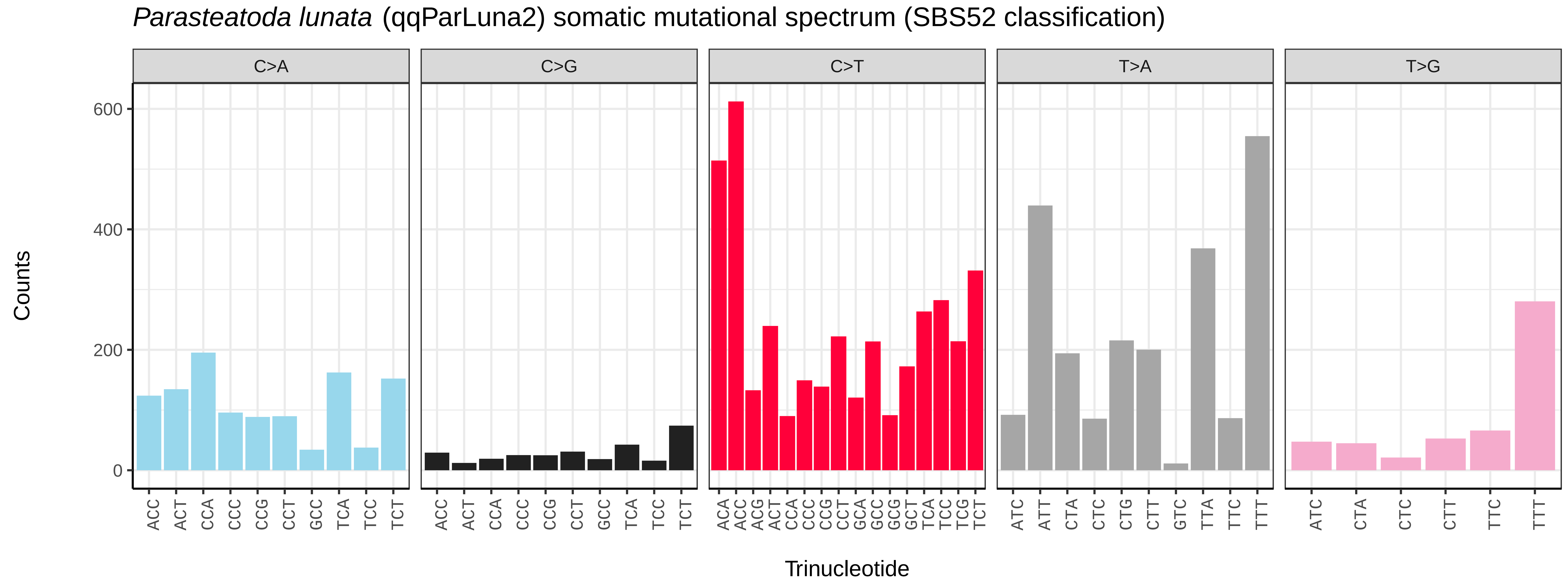

Parasteatoda lunata (qqParLuna2) somatic mutational spectrum (SBS52 classification)
C>A
C>G
C>T
T>A
T>G
Counts
Trinucleotide

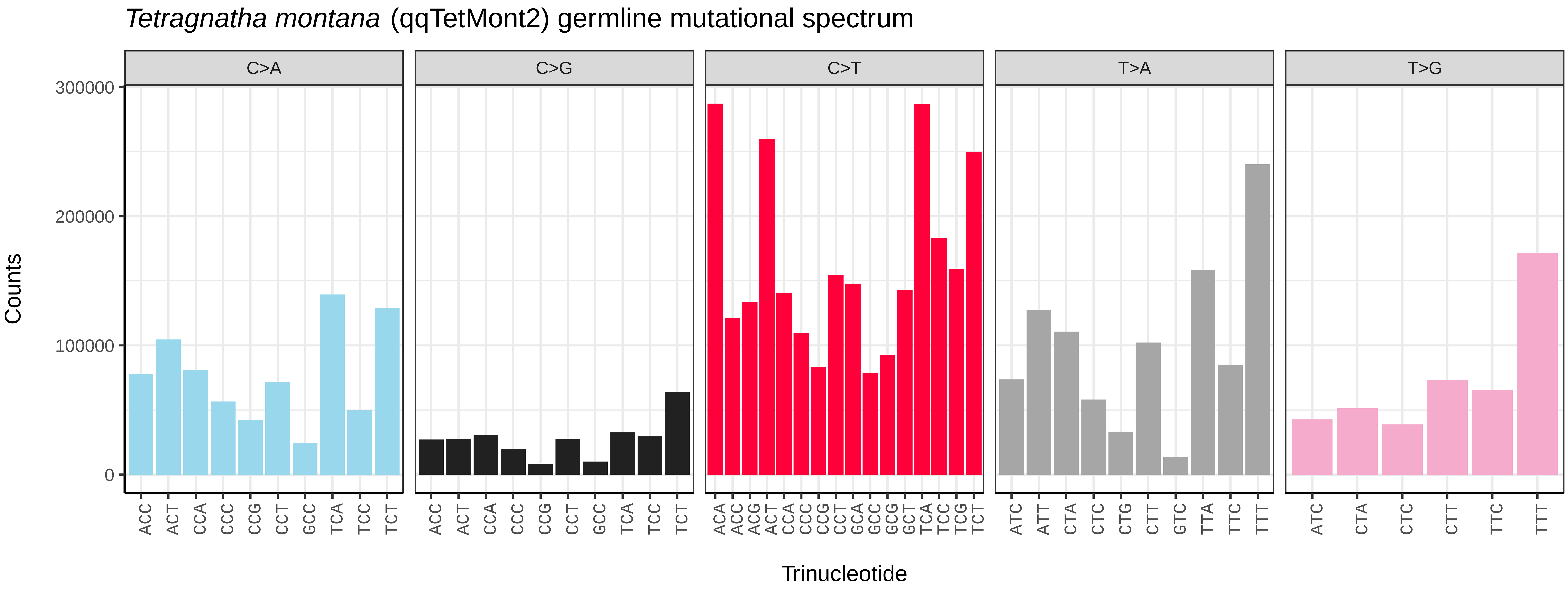
*Tetragnatha montana* (qqTetMont2) germline mutational spectrum
C>A
C>G
C>T
T>A
T>G
Counts
300000
200000
100000
0
ACC ACT CCA CCC CCG CCT GCC TCA TCC TCT
ACA ACC ACG ACT CCA CCC CCG CCT GCA GCC GCG GCT TCA TCC TCG TCT
ATC ATT CTA CTC CTG CTT GTC TTA TTC TTT
ATC CTA CTC CTT TTC TTT
Trinucleotide

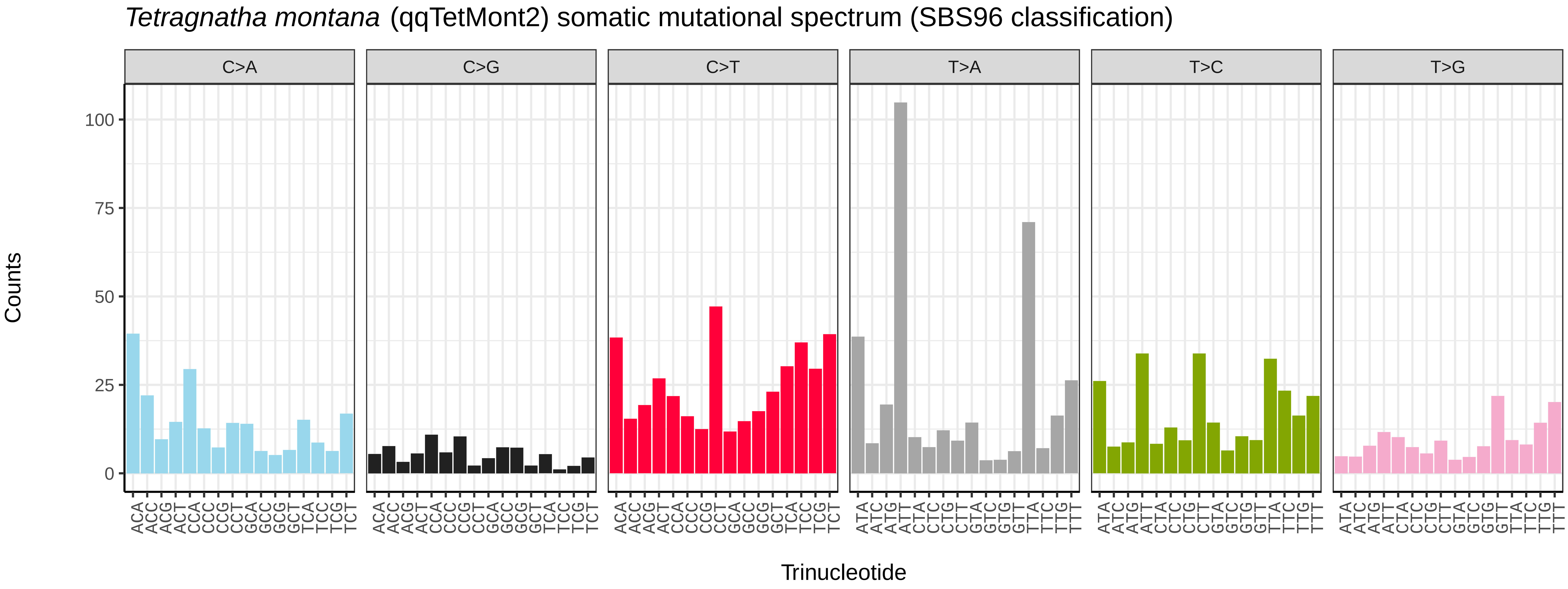
*Tetragnatha montana* (qqTetMont2) somatic mutational spectrum (SBS96 classification)
C>A
C>G
C>T
T>A
T>C
T>G
Counts
100
75
50
25
0
ACA ACC ACG ACT CCA CCC CCG CCT GCA GCC GCG GCT TCA TCC TCG TCT
ATA ATC ATG ATT CTA CTC CTG CTT GTA GTC GTG GTT TTA TTC TTG TTT
Trinucleotide

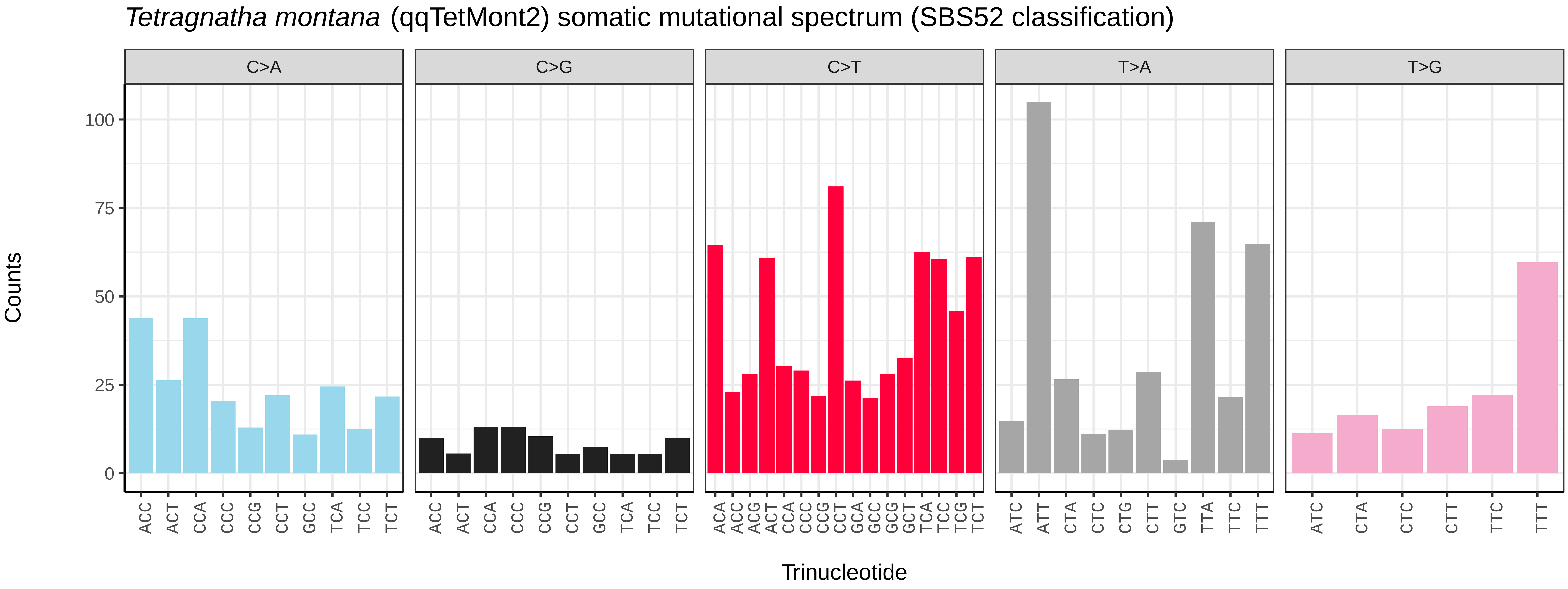
*Tetragnatha montana* (qqTetMont2) somatic mutational spectrum (SBS52 classification)
C>A
C>G
C>T
T>A
T>G
Counts
100
75
50
25
0
ACC ACT CCA CCC CCG CCT GCC TCA TCC TCT
ACA ACC ACG ACT CCA CCC CCG CCT GCA GCC GCG GCT TCA TCC TCG TCT
ATC ATT CTA CTC CTG CTT GTC TTA TTC TTT
ATC CTA CTC CTT TTC TTT
Trinucleotide

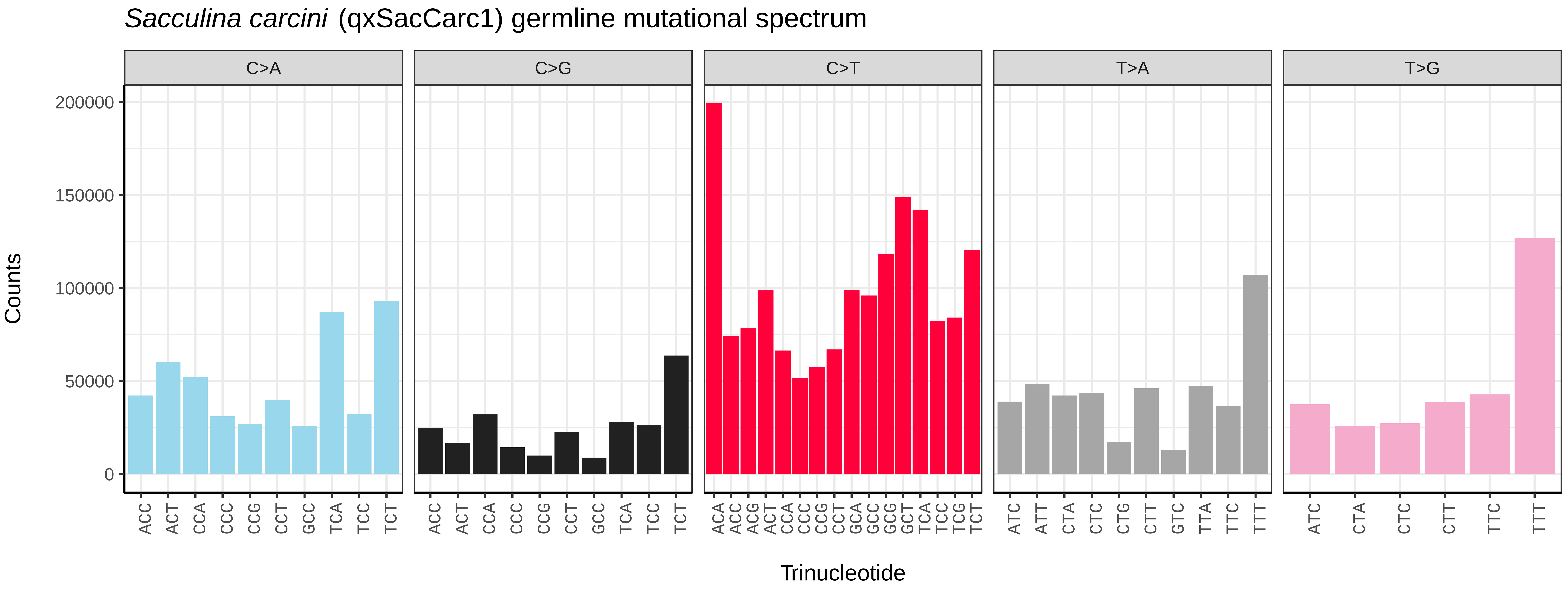
Sacculina carcini (qxSacCarc1) germline mutational spectrum
C>A
C>G
C>T
T>A
T>G
Counts
200000
150000
100000
50000
0
ACC ACT CCA CCC CCG CCT GCC TCA TCC TCT
ACC ACT CCA CCC CCG CCT GCC TCA TCC TCT
ACA ACC ACG ACT CCA CCC CCG CCT GCA GCC GCG GCT TCA TCC TCG TCT
ATC ATT CTA CTC CTG CTT GTC TTA TTC TTT
ATC CTA CTC CTT TTC TTT
Trinucleotide

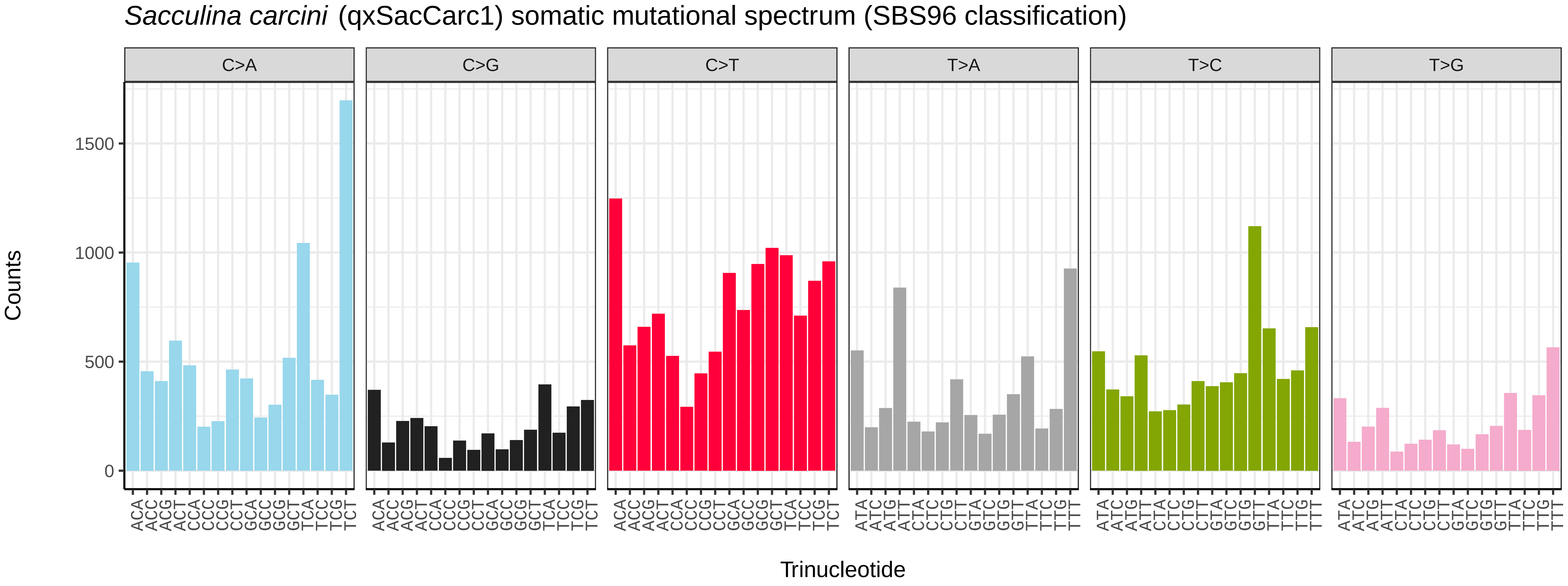
Sacculina carcini (qxSacCarc1) somatic mutational spectrum (SBS96 classification)
C>A
C>G
C>T
T>A
T>C
T>G
Counts
1500
1000
500
0
ACA ACC ACG ACT CCA CCC CCG CCT GCA GCC GCG GCT TCA TCC TCG TCT
ATA ATC ATG ATT CTA CTC CTG CTT GTA GTC GTG GTT TTA TTC TTG TTT
Trinucleotide

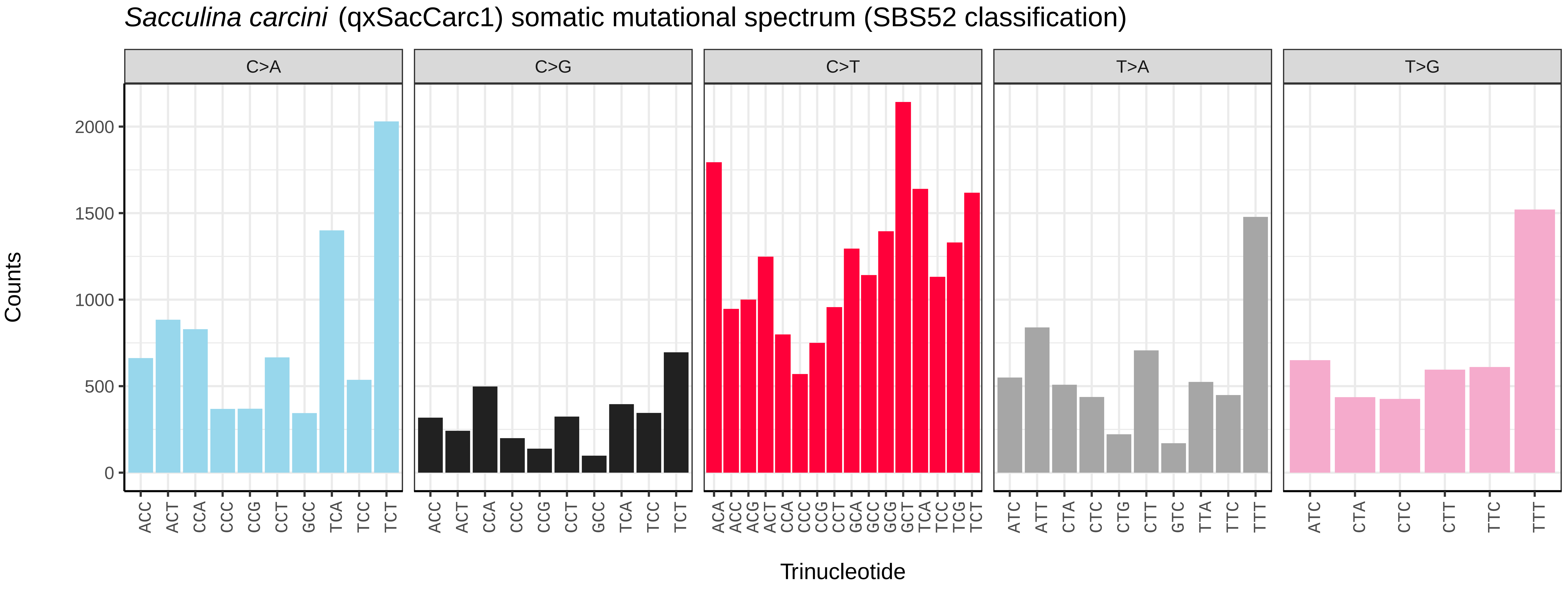
Sacculina carcini (qxSacCarc1) somatic mutational spectrum (SBS52 classification)
C>A
C>G
C>T
T>A
T>G
Counts
2000
1500
1000
500
0
ACC ACT CCA CCC CCG CCT GCC TCA TCC TCT
ACC ACT CCA CCC CCG CCT GCC TCA TCC TCT
ACA ACC ACG ACT CCA CCC CCG CCT GCA GCC GCG GCT TCA TCC TCG TCT
ATC ATT CTA CTC CTG CTT GTC TTA TTC TTT
ATC CTA CTC CTT TTC TTT
Trinucleotide

*Zootoca vivipara* (rZooViv1) germline mutational spectrum

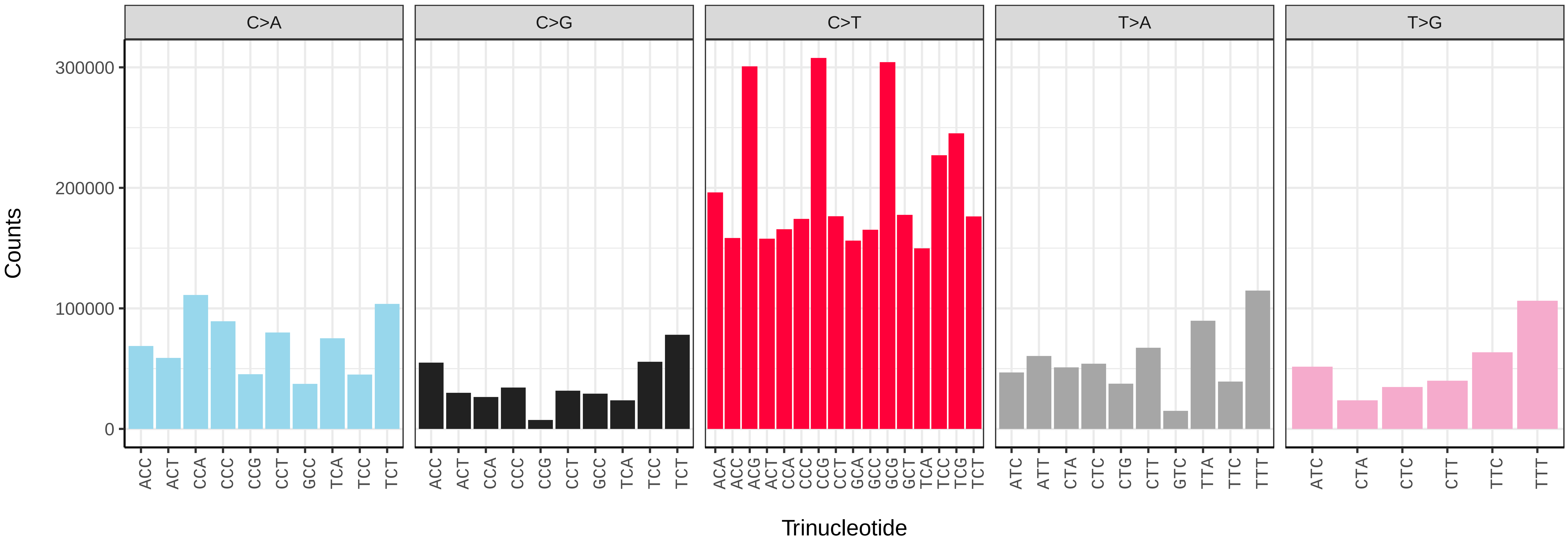


*Zootoca vivipara* (rZooViv1) somatic mutational spectrum (SBS96 classification)

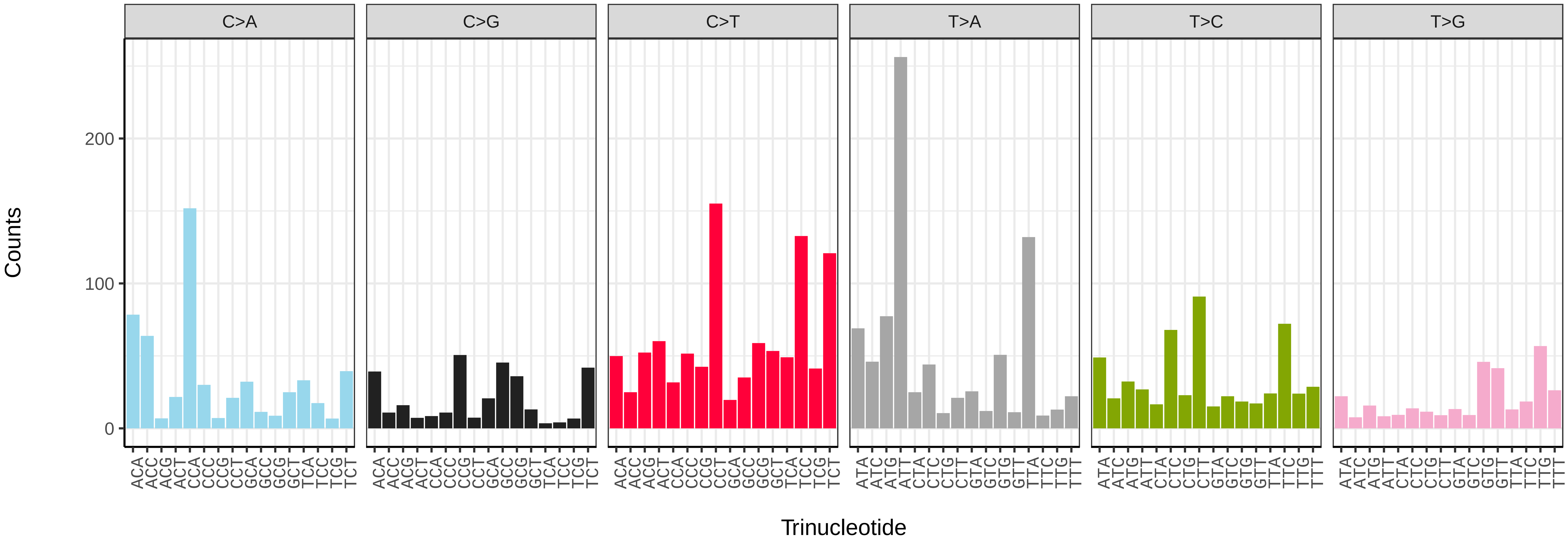


*Zootoca vivipara* (rZooViv1) somatic mutational spectrum (SBS52 classification)

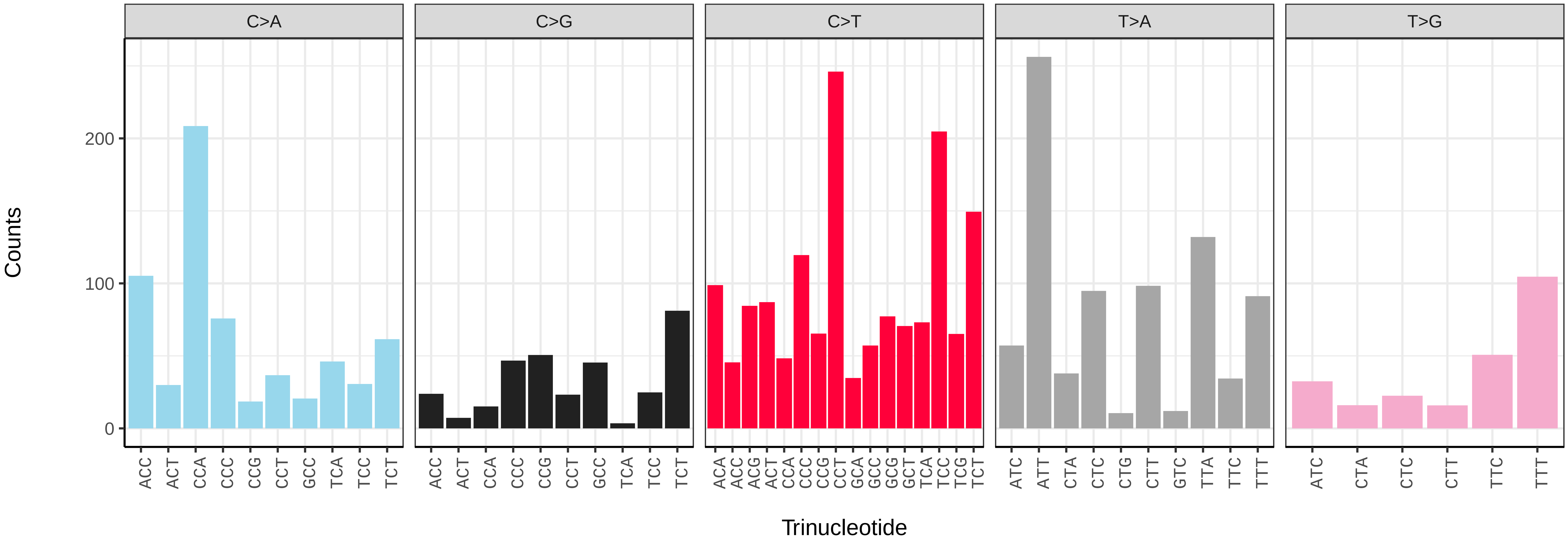

*Raja brachyura* (sRajBra1) germline mutational spectrum

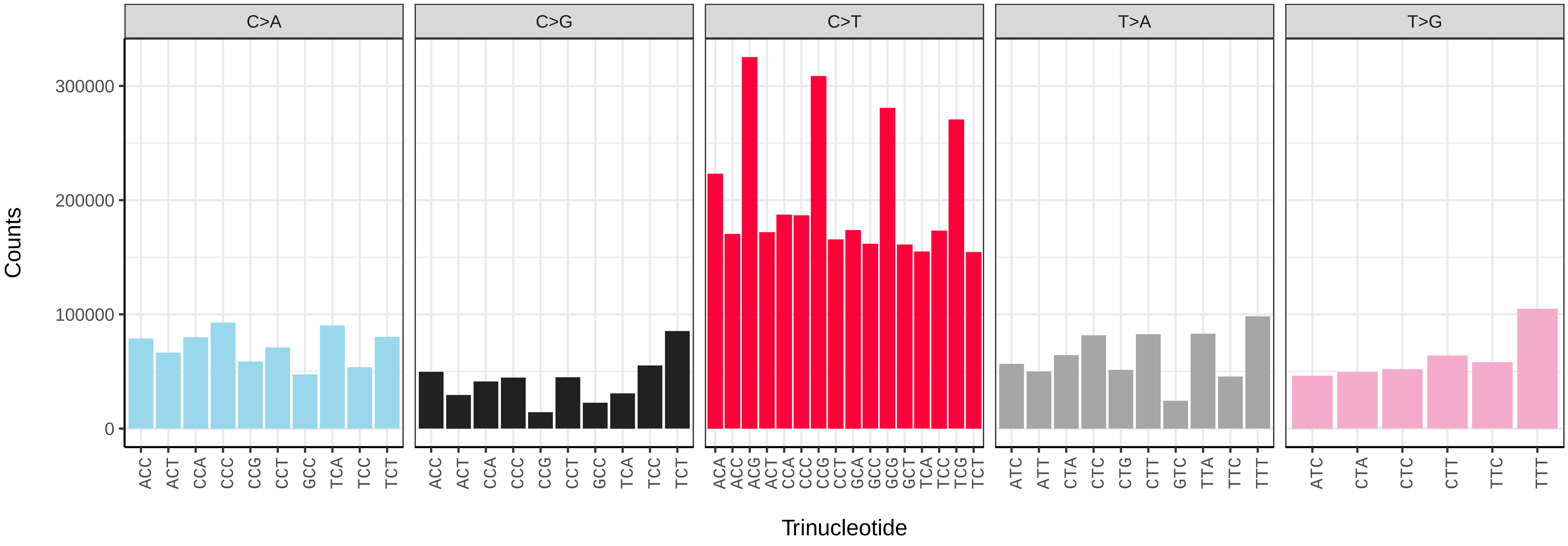


*Raja brachyura* (sRajBra1) somatic mutational spectrum (SBS96 classification)

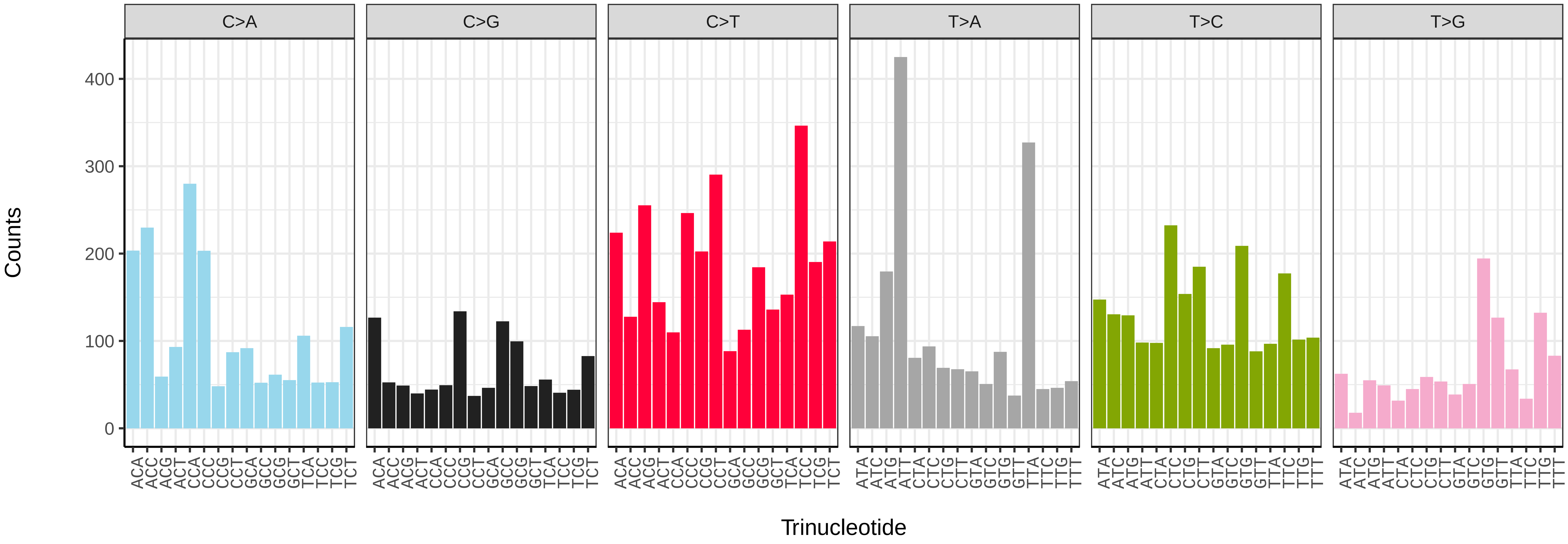


*Raja brachyura* (sRajBra1) somatic mutational spectrum (SBS52 classification)

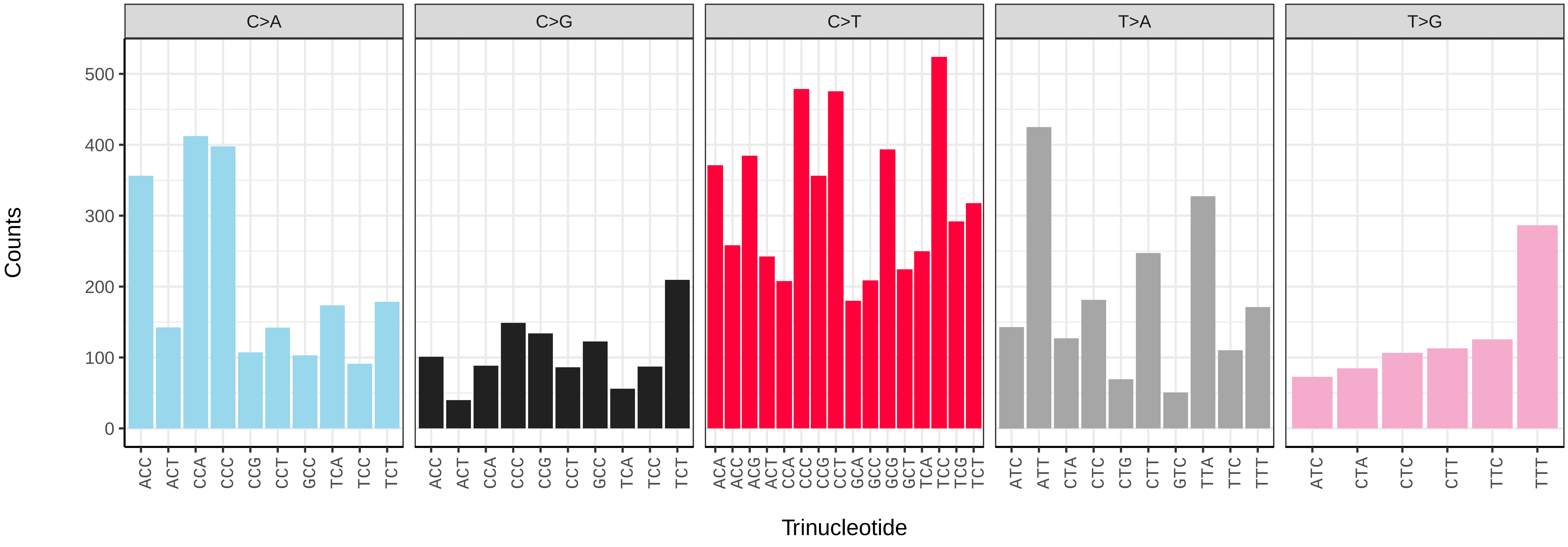

*Lineus longissimus* (tnLinLong1) germline mutational spectrum

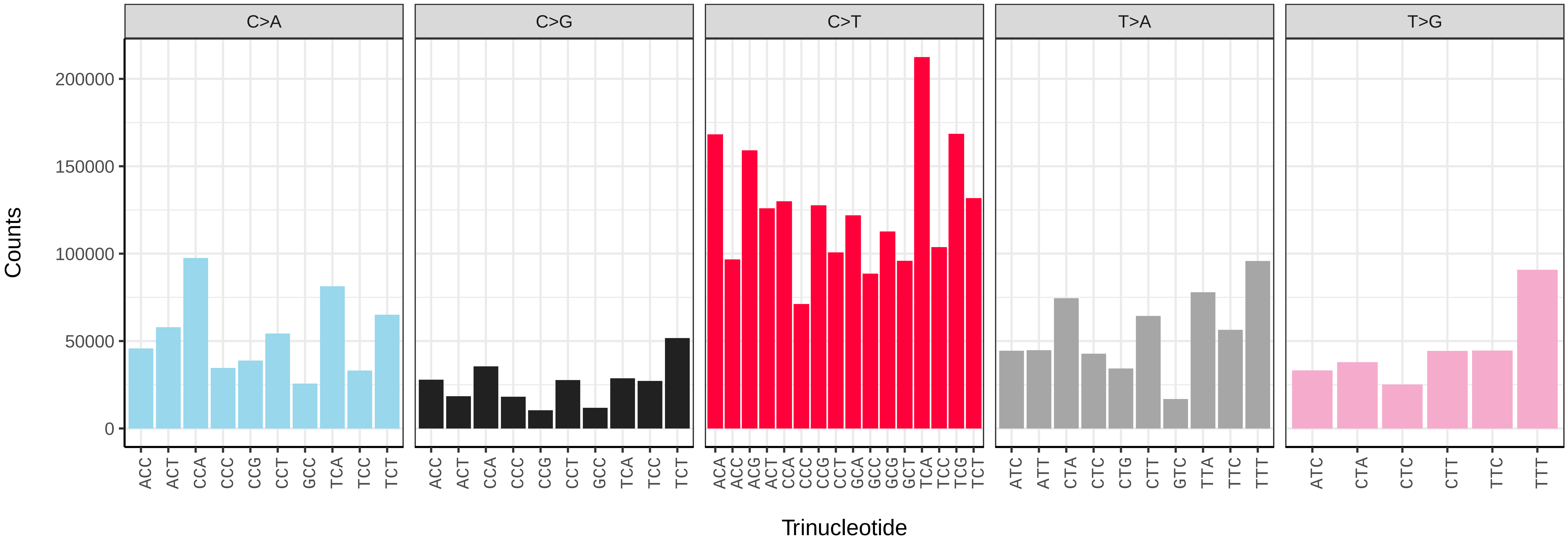


*Lineus longissimus* (tnLinLong1) somatic mutational spectrum (SBS96 classification)

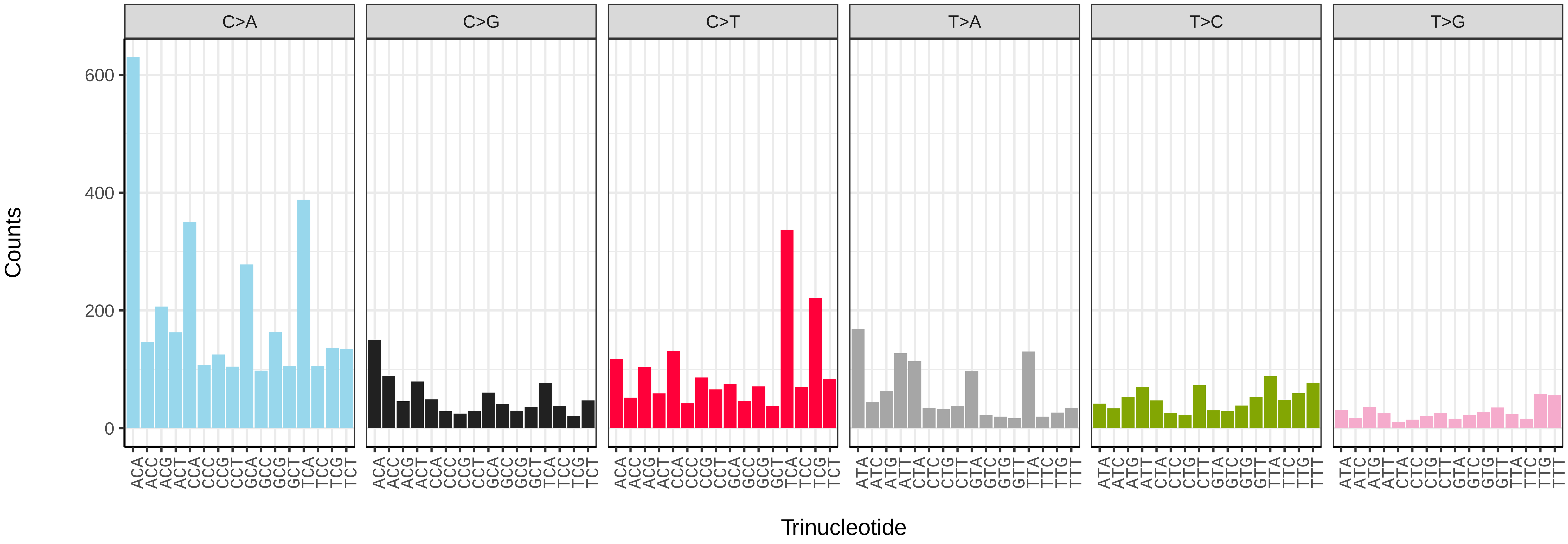


*Lineus longissimus* (tnLinLong1) somatic mutational spectrum (SBS52 classification)

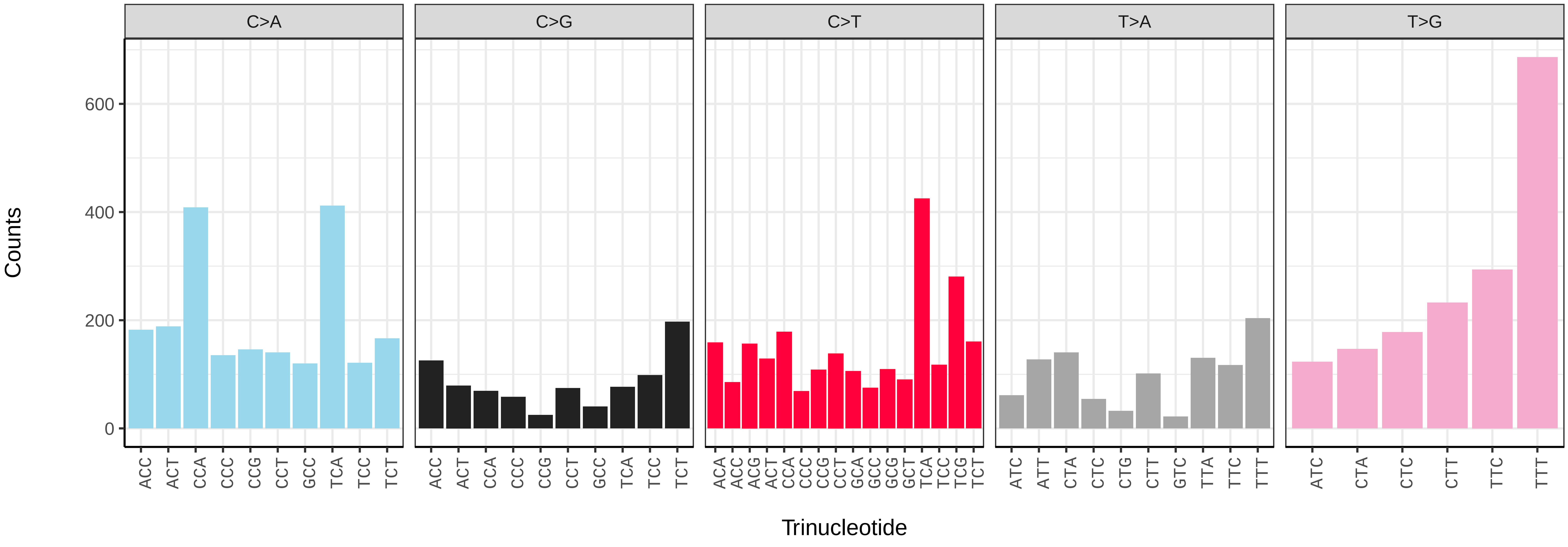

*Bugula neritina* (tzBugNeri2) germline mutational spectrum

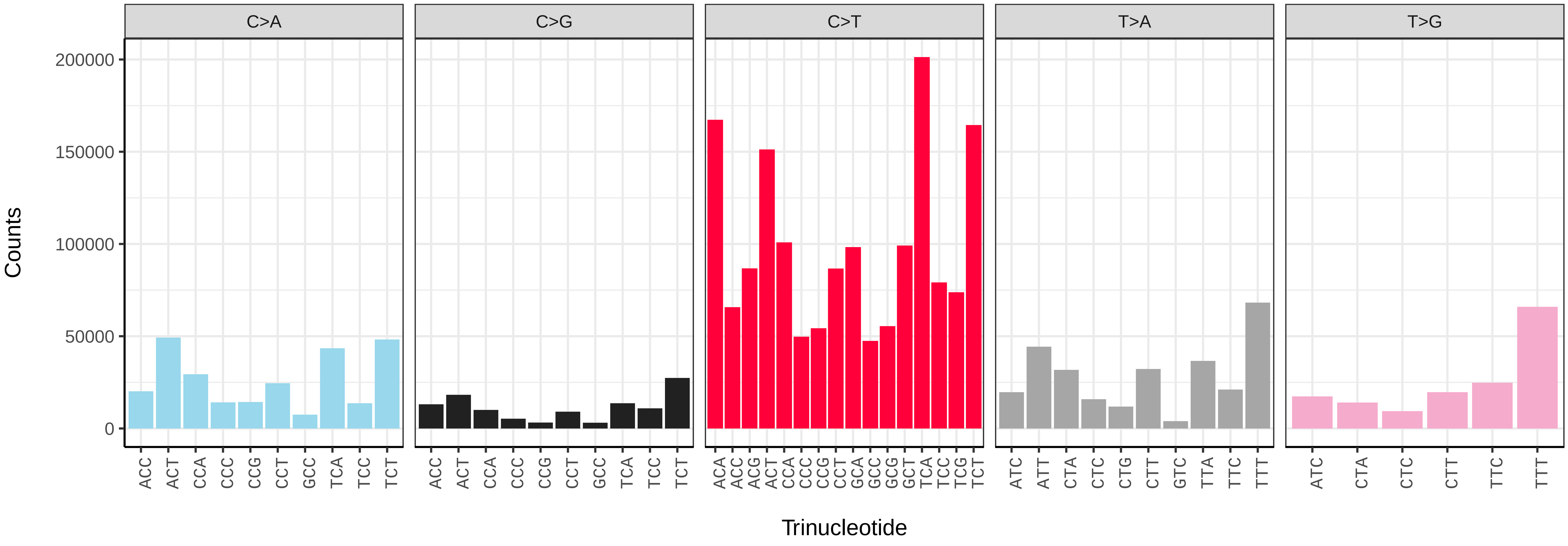


*Bugula neritina* (tzBugNeri2) somatic mutational spectrum (SBS96 classification)

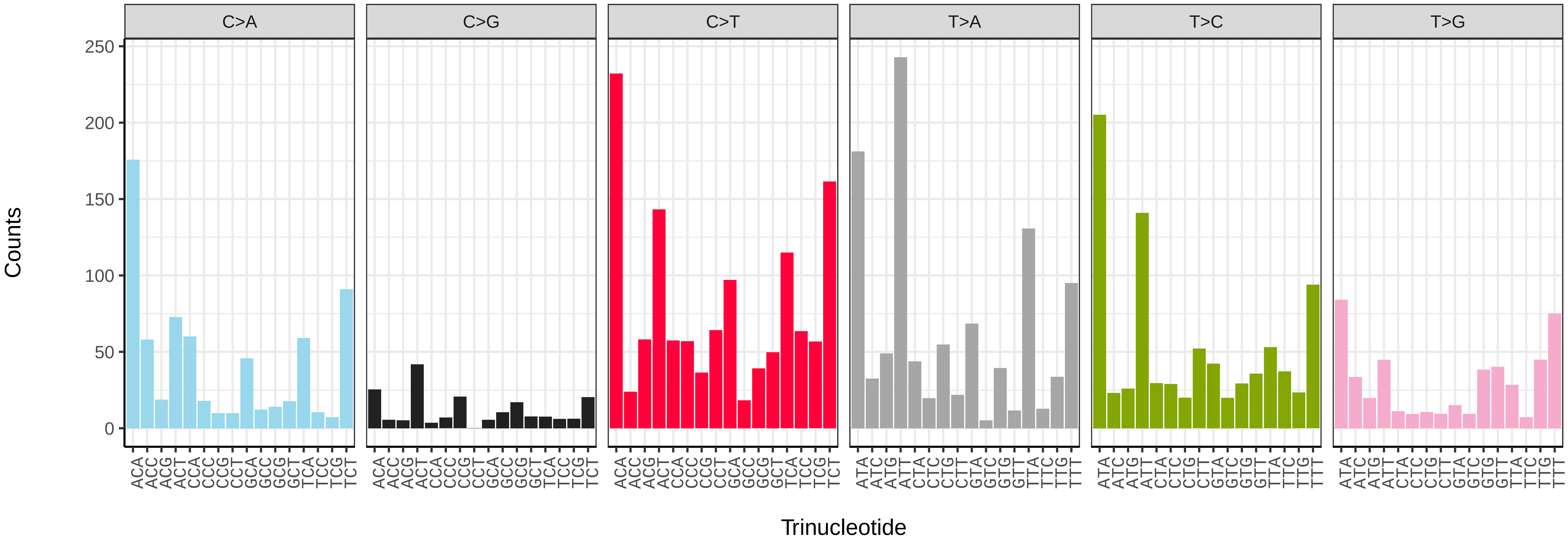


*Bugula neritina* (tzBugNeri2) somatic mutational spectrum (SBS52 classification)

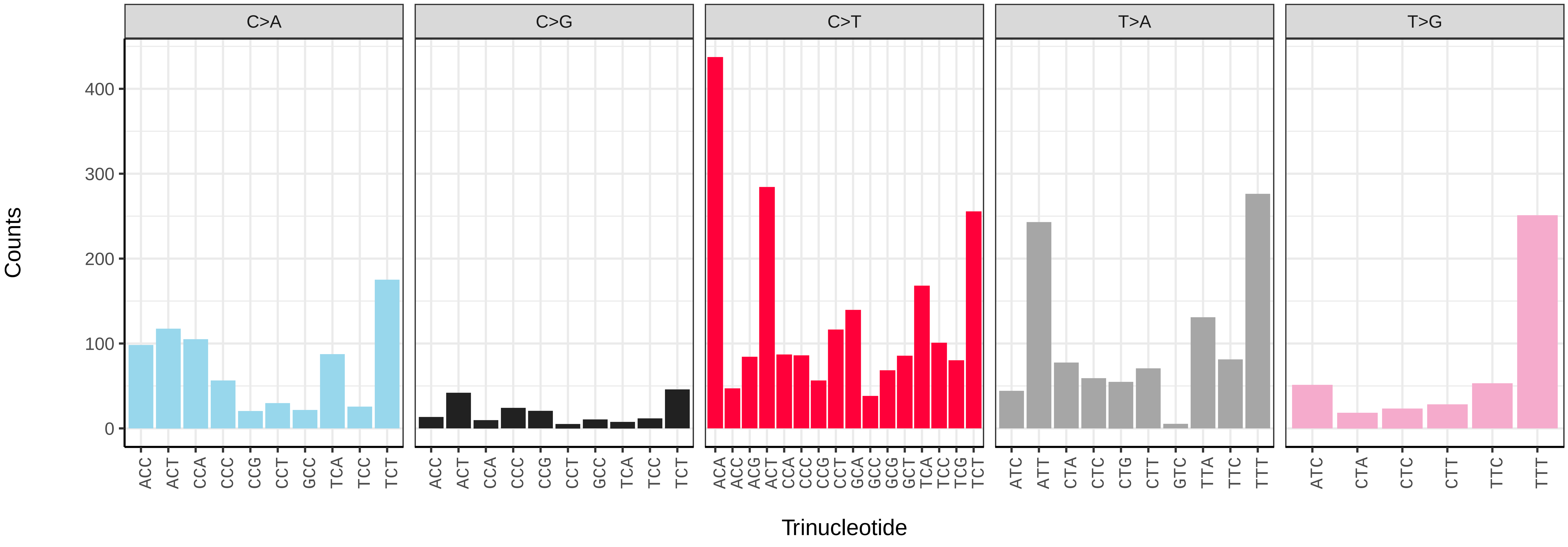

*Bugulina stolonifera* (tzBugStol2) germline mutational spectrum

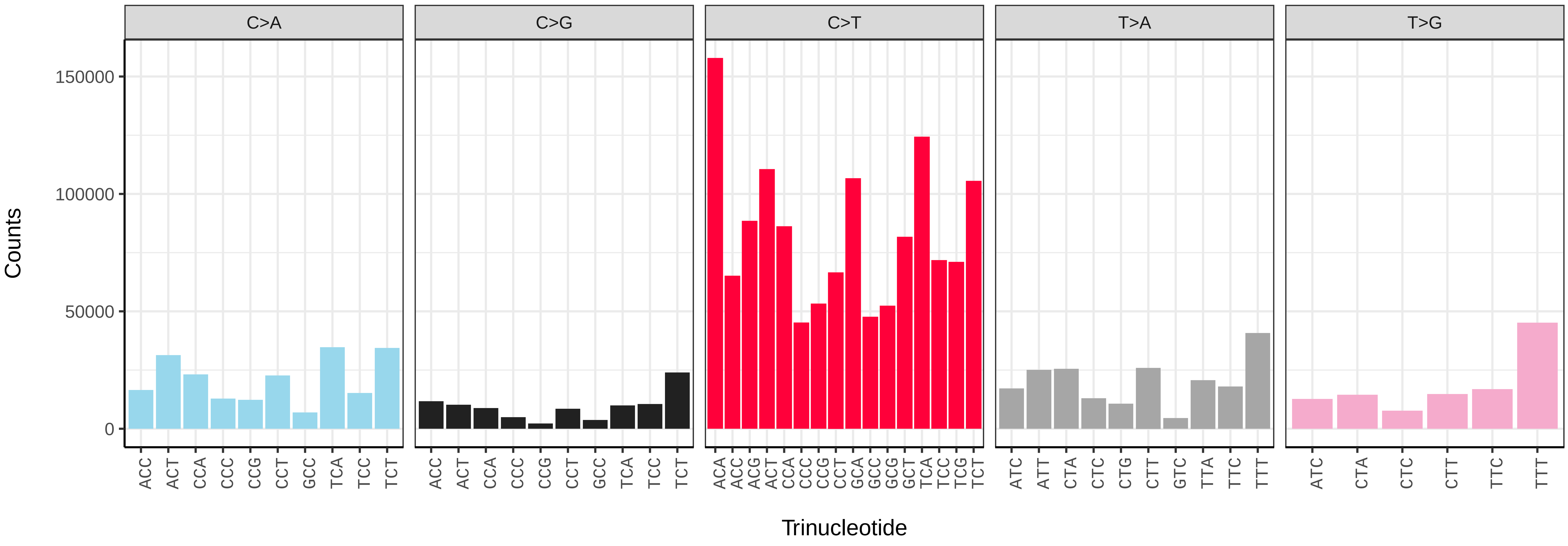


*Bugulina stolonifera* (tzBugStol2) somatic mutational spectrum (SBS96 classification)

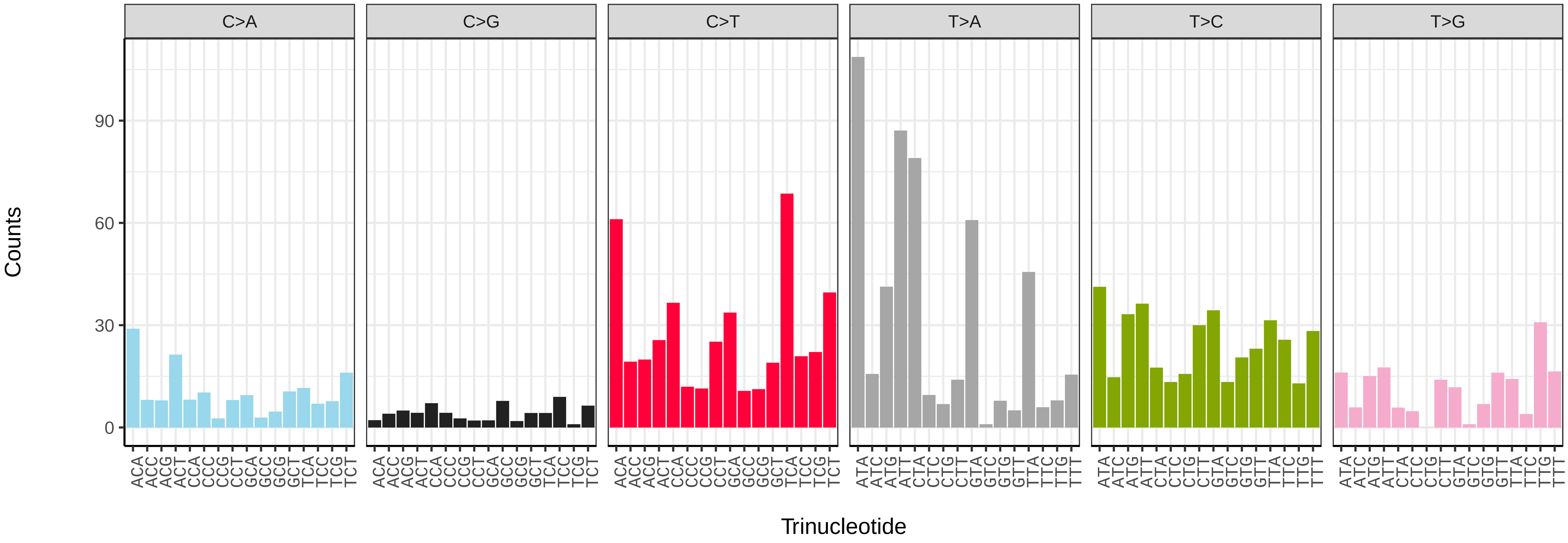


*Bugulina stolonifera* (tzBugStol2) somatic mutational spectrum (SBS52 classification)

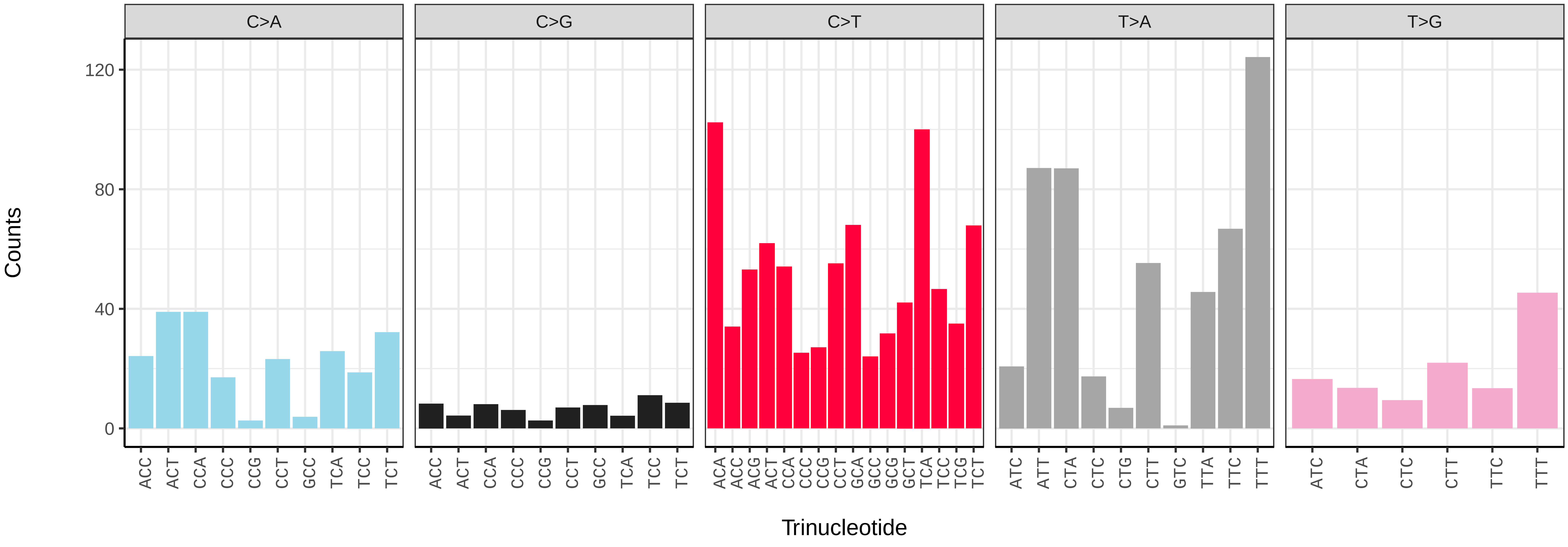

*Cryptosula pallasiana* (tzCryPall5) germline mutational spectrum

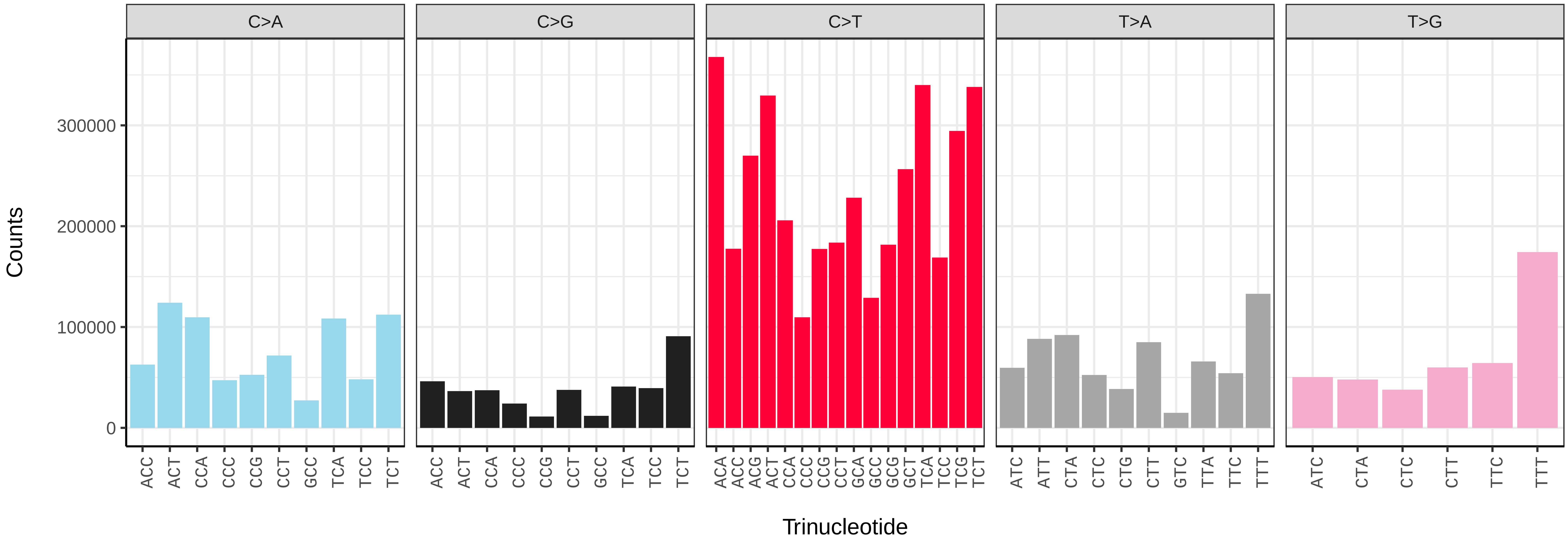


*Cryptosula pallasiana* (tzCryPall5) somatic mutational spectrum (SBS96 classification)

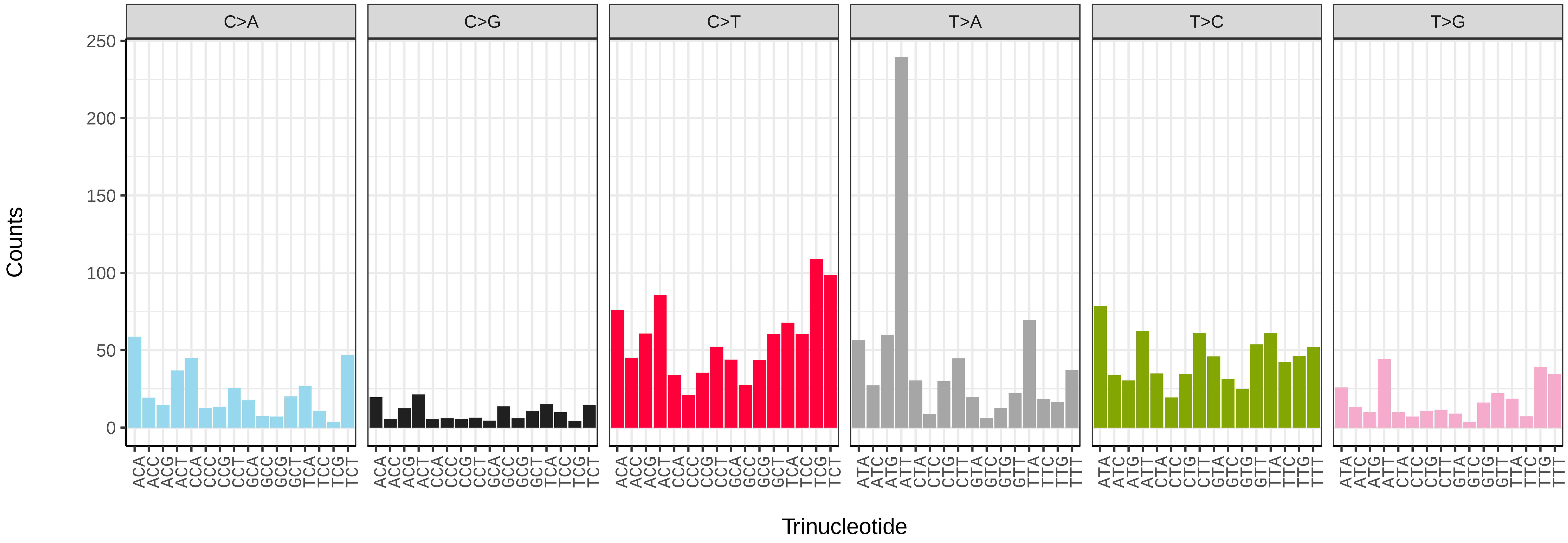


*Cryptosula pallasiana* (tzCryPall5) somatic mutational spectrum (SBS52 classification)

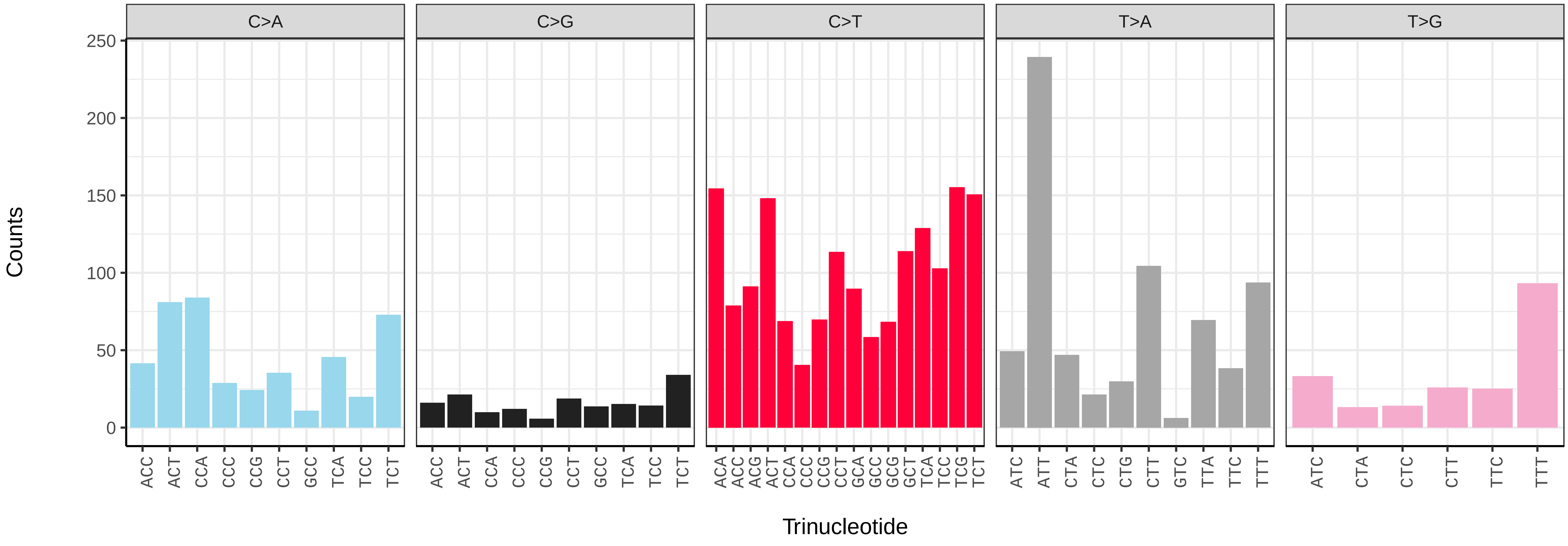

*Membranipora membranacea* (tzMemMemb1) germline mutational spectrum

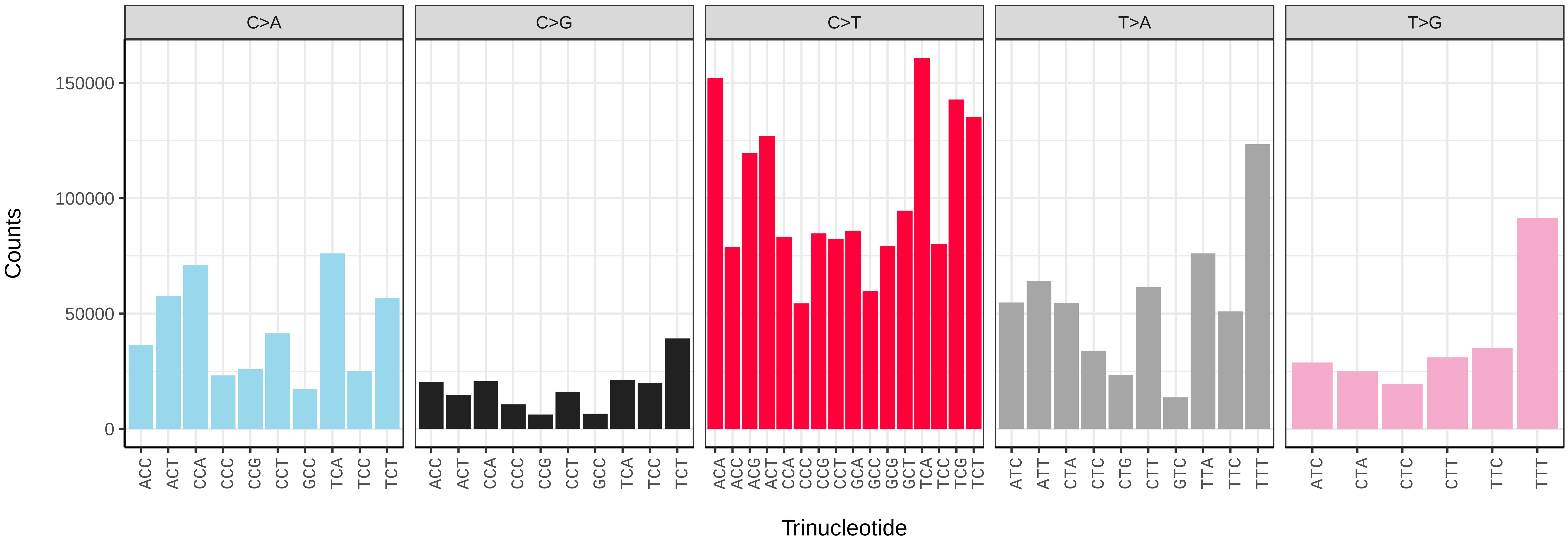


*Membranipora membranacea* (tzMemMemb1) somatic mutational spectrum (SBS96 classification)

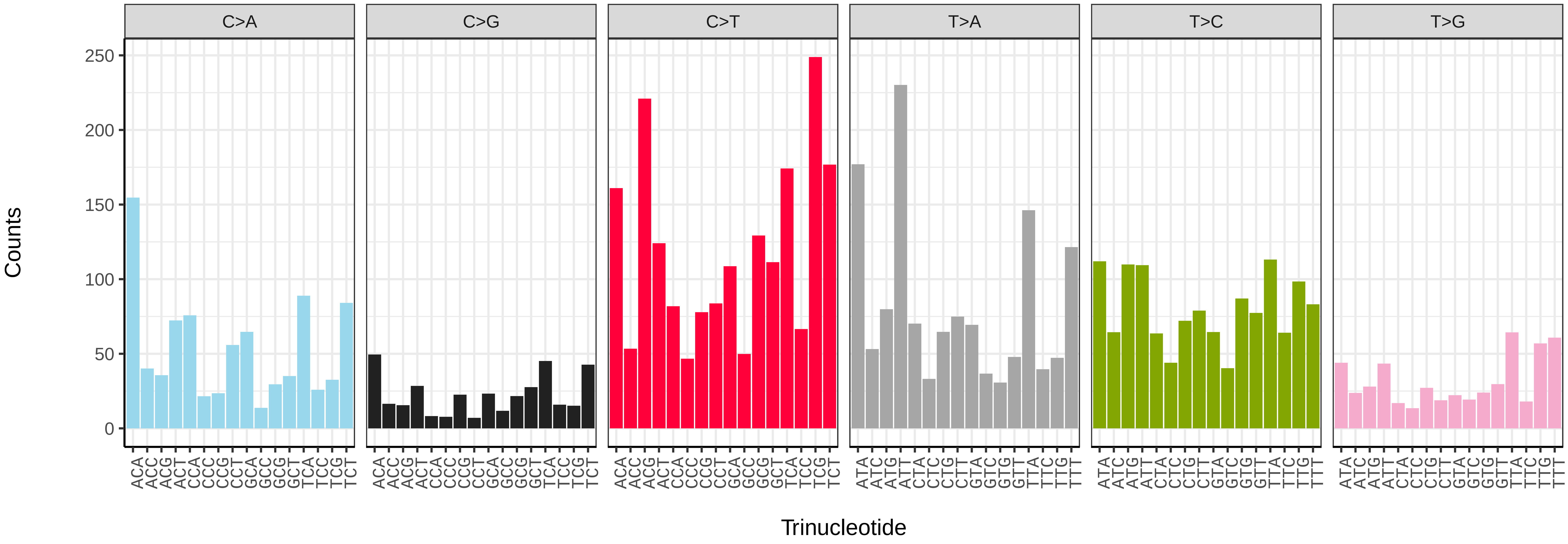


*Membranipora membranacea* (tzMemMemb1) somatic mutational spectrum (SBS52 classification)

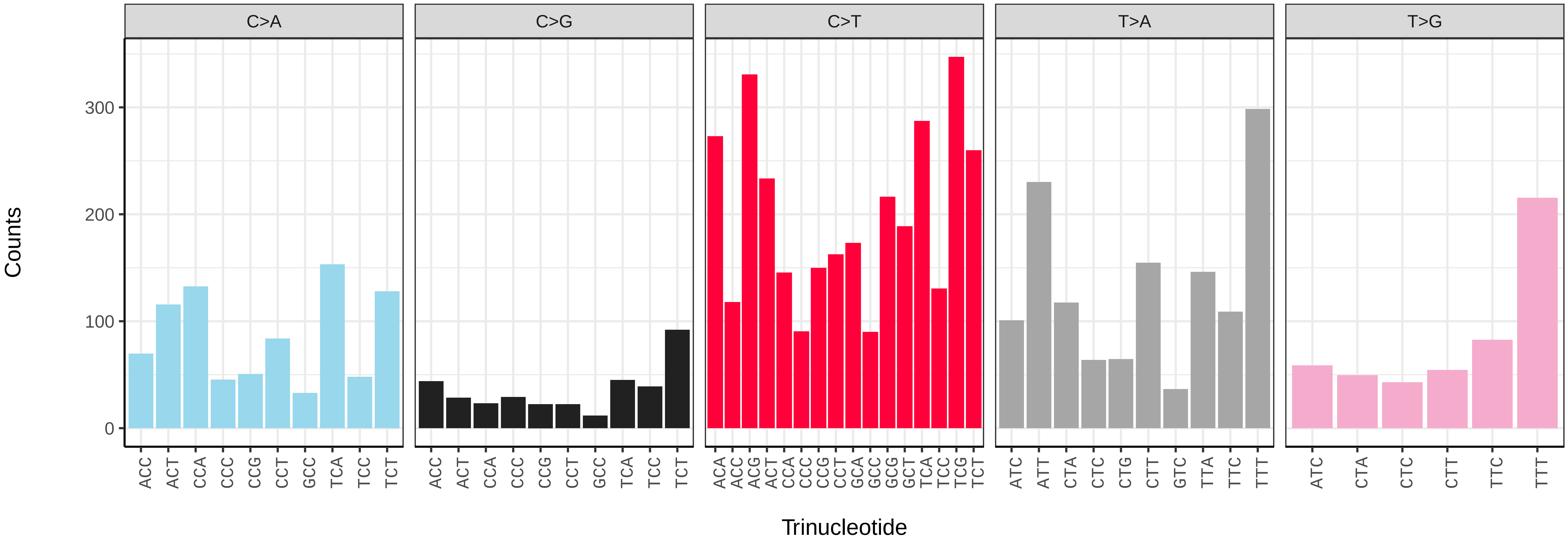

*Watersipora subatra* (tzWatSuba1) germline mutational spectrum

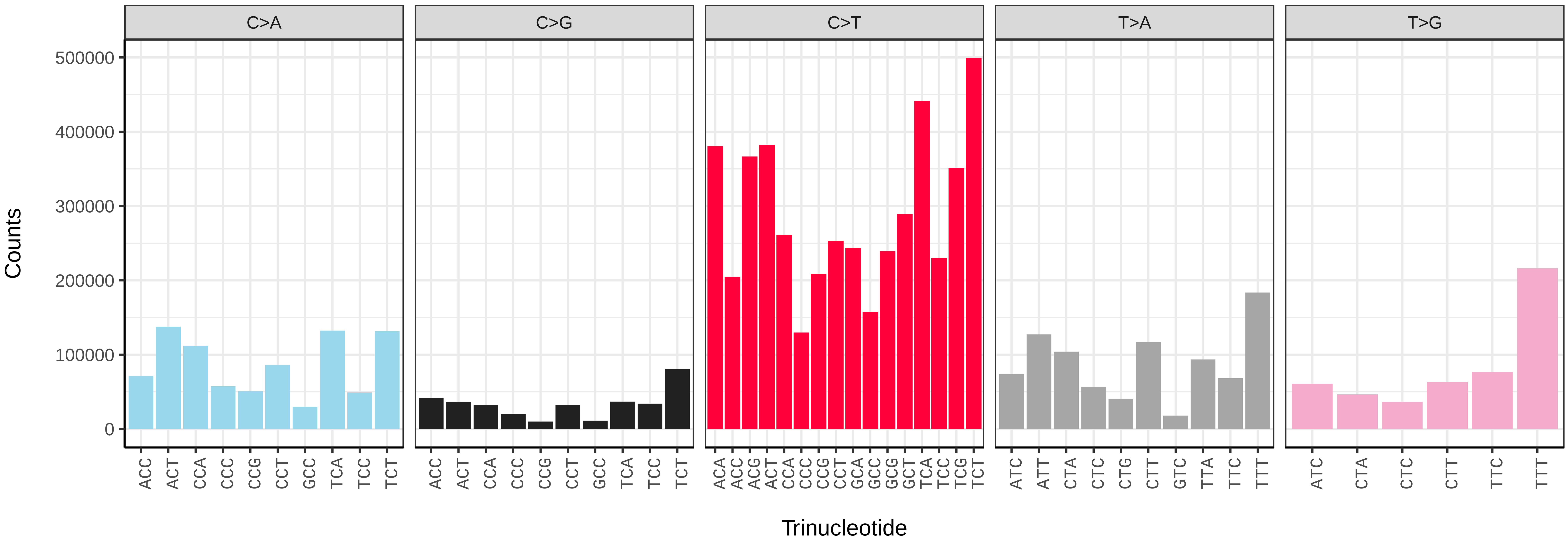


*Watersipora subatra* (tzWatSuba1) somatic mutational spectrum (SBS96 classification)

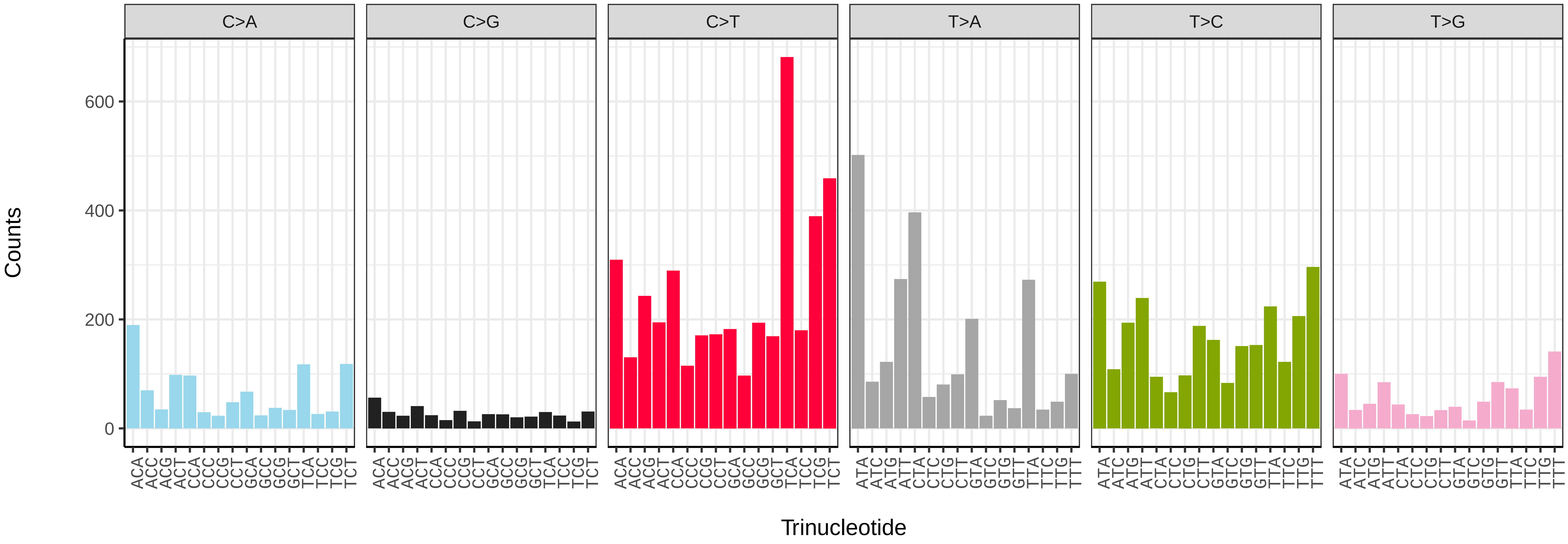


*Watersipora subatra* (tzWatSuba1) somatic mutational spectrum (SBS52 classification)

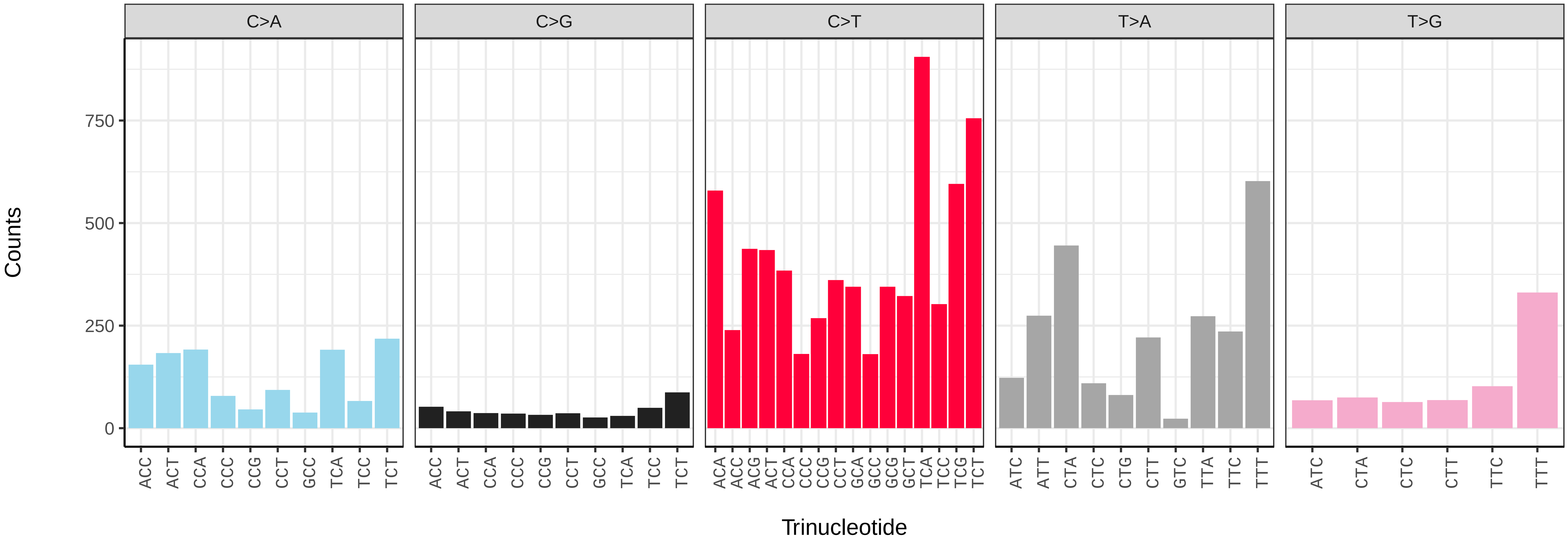

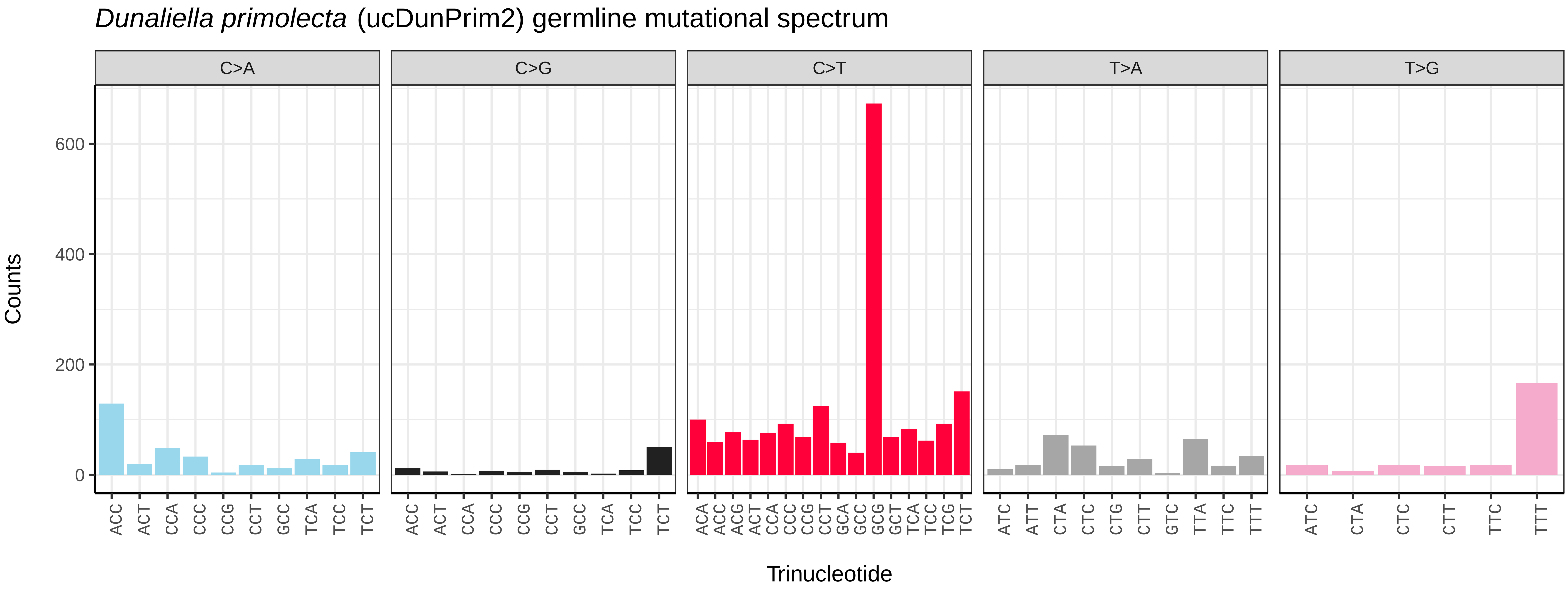
*Dunaliella primolecta* (ucDunPrim2) germline mutational spectrum
C>A
C>G
C>T
T>A
T>G
Counts
600
400
200
0
Trinucleotide

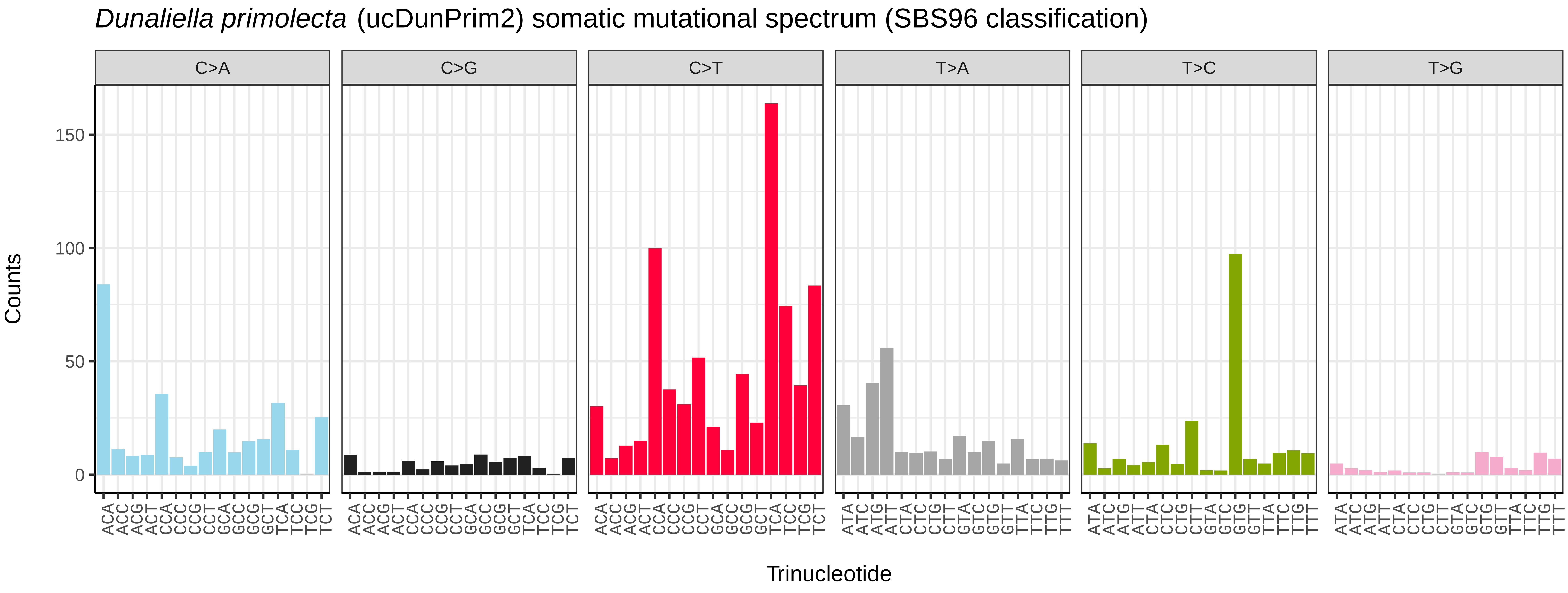
*Dunaliella primolecta* (ucDunPrim2) somatic mutational spectrum (SBS96 classification)
C>A
C>G
C>T
T>A
T>C
T>G
Counts
150
100
50
0
Trinucleotide

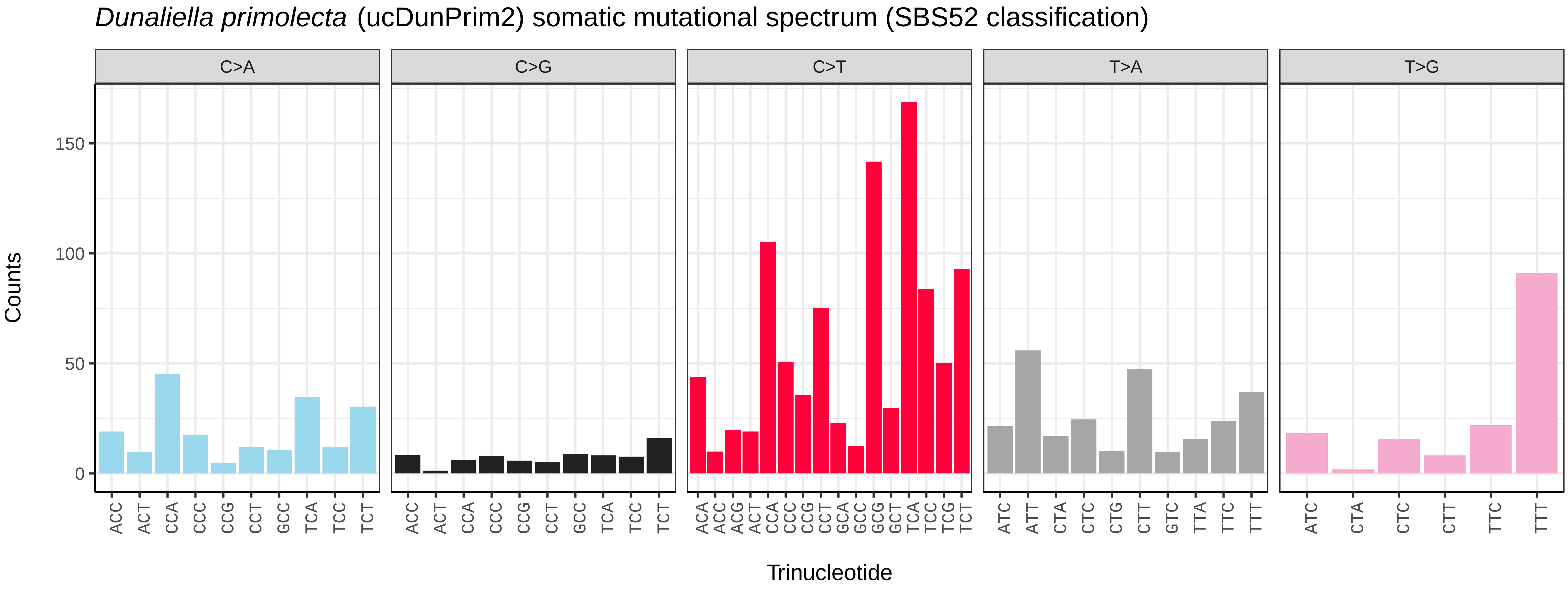
*Dunaliella primolecta* (ucDunPrim2) somatic mutational spectrum (SBS52 classification)
C>A
C>G
C>T
T>A
T>G
Counts
150
100
50
0
Trinucleotide

*Marvania coccoides* (ucMarCocc1) germline mutational spectrum

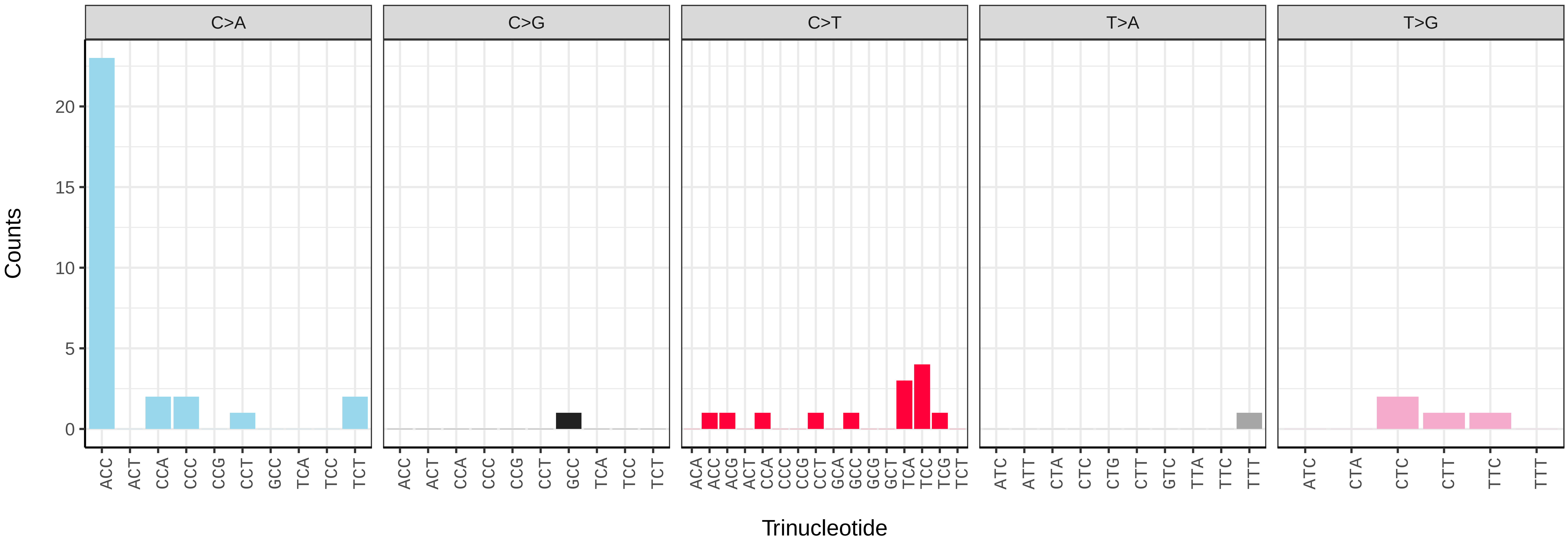


*Marvania coccoides* (ucMarCocc1) somatic mutational spectrum (SBS96 classification)

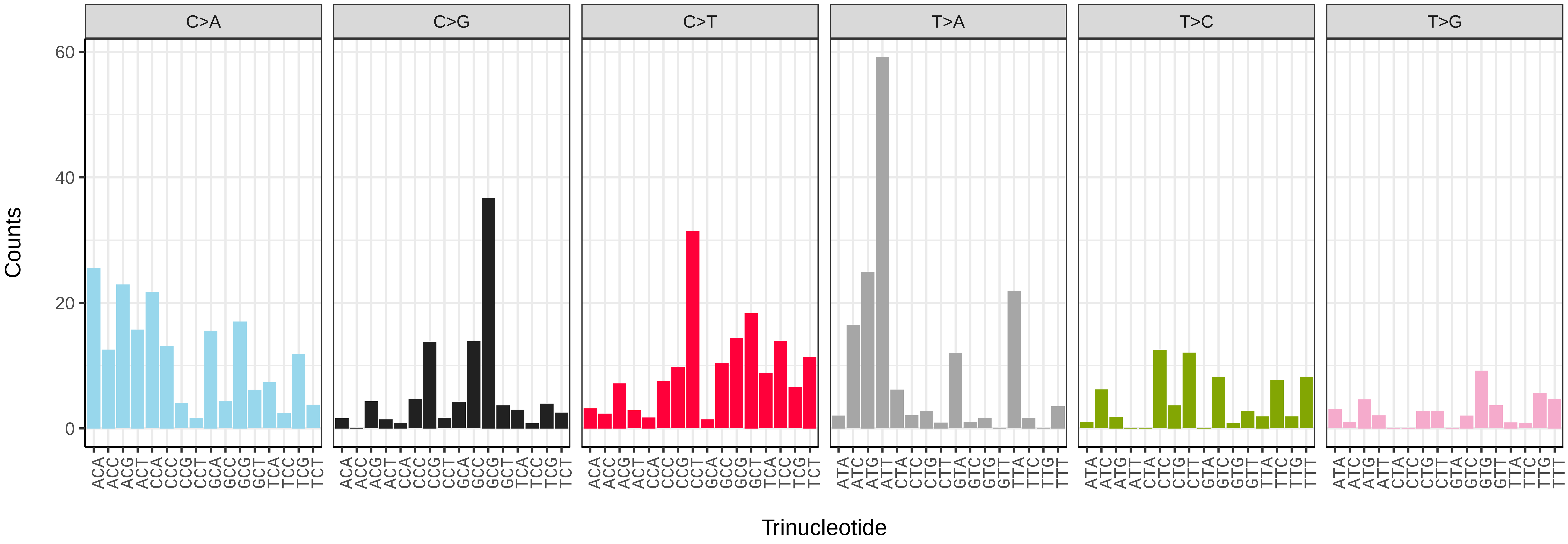


*Marvania coccoides* (ucMarCocc1) somatic mutational spectrum (SBS52 classification)

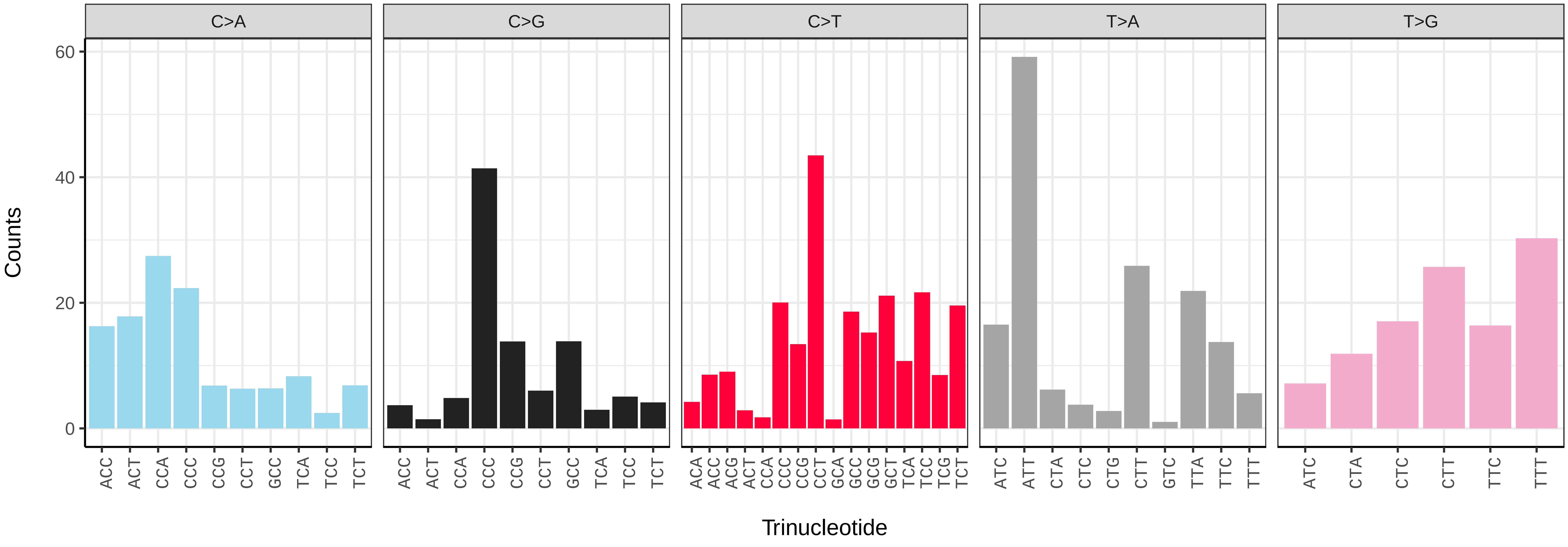

*Pycnococcus provasolii* (ucPycProv1) germline mutational spectrum

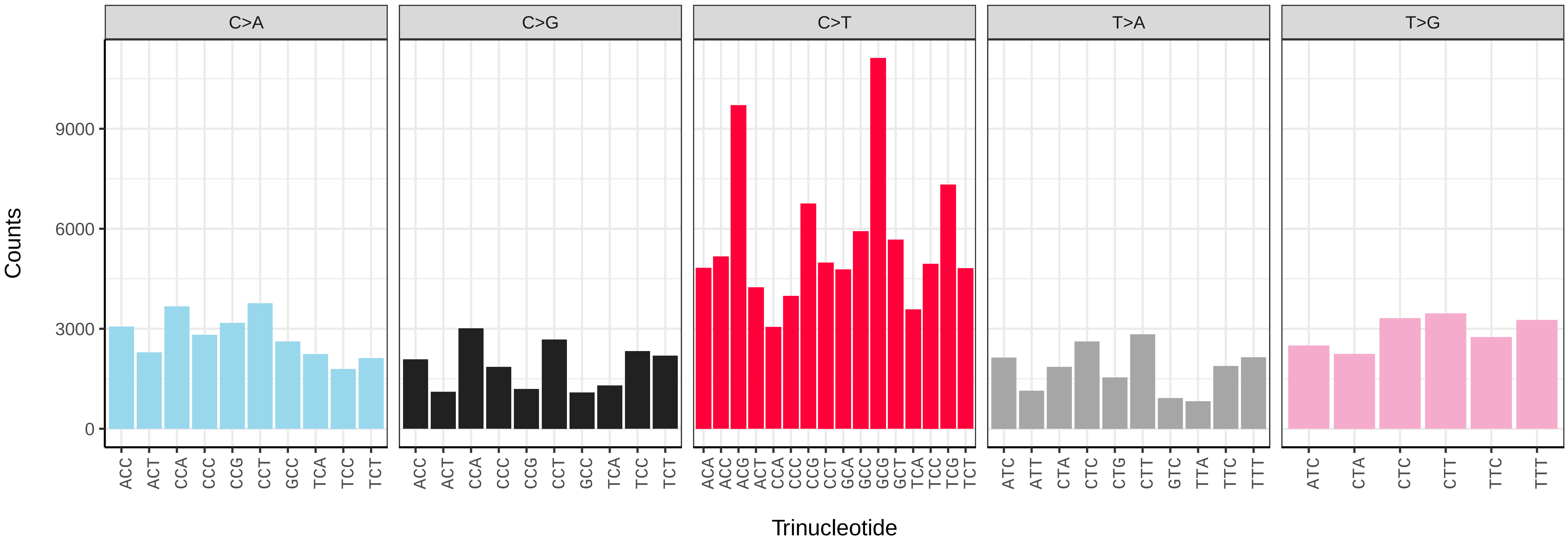


*Pycnococcus provasolii* (ucPycProv1) somatic mutational spectrum (SBS96 classification)

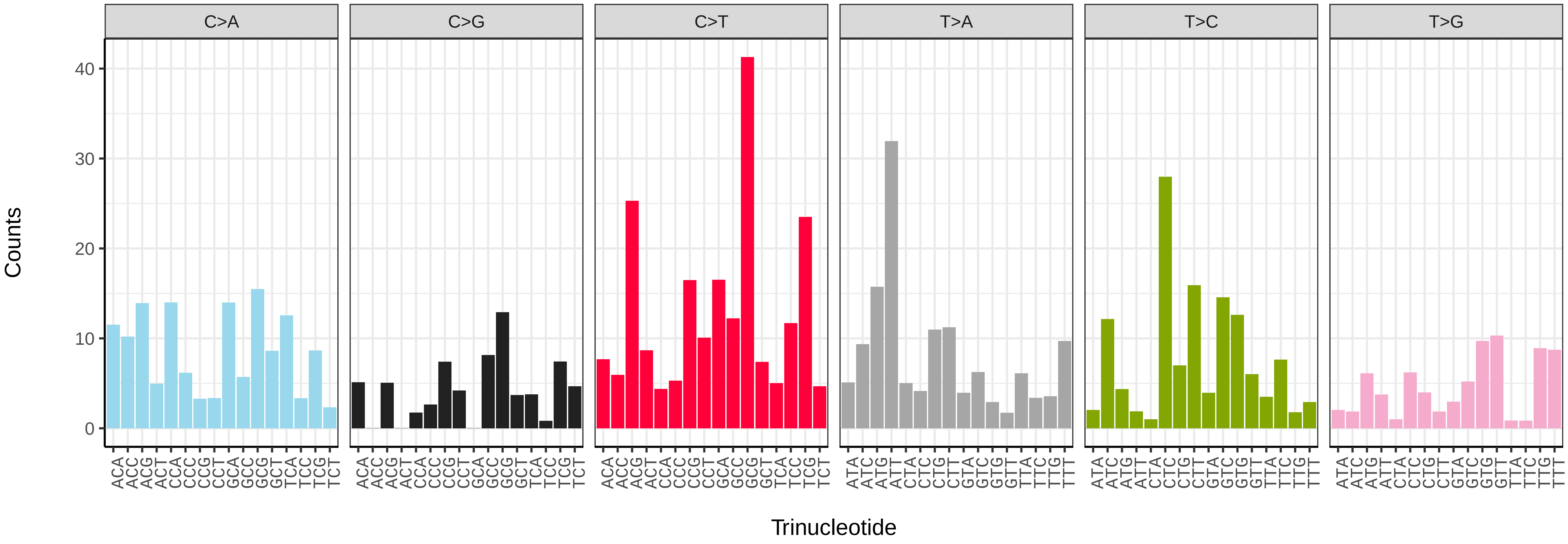


*Pycnococcus provasolii* (ucPycProv1) somatic mutational spectrum (SBS52 classification)

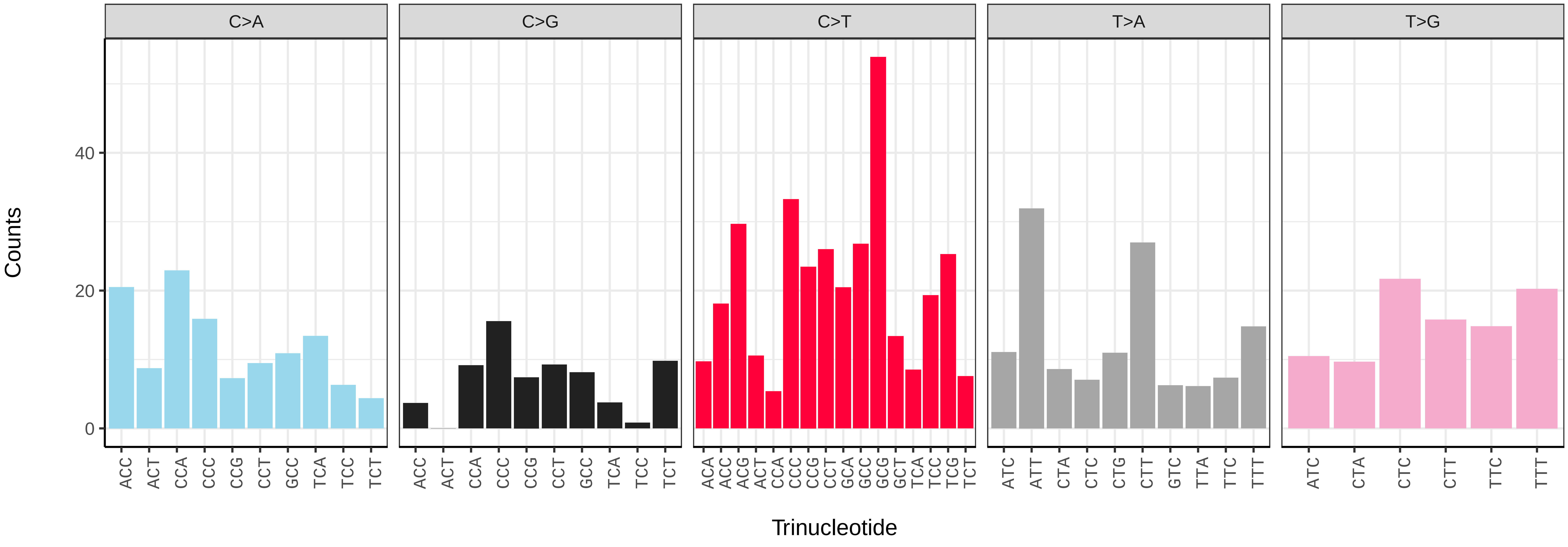

*Aporrectodea icterica* (wcApoIcte1) germline mutational spectrum

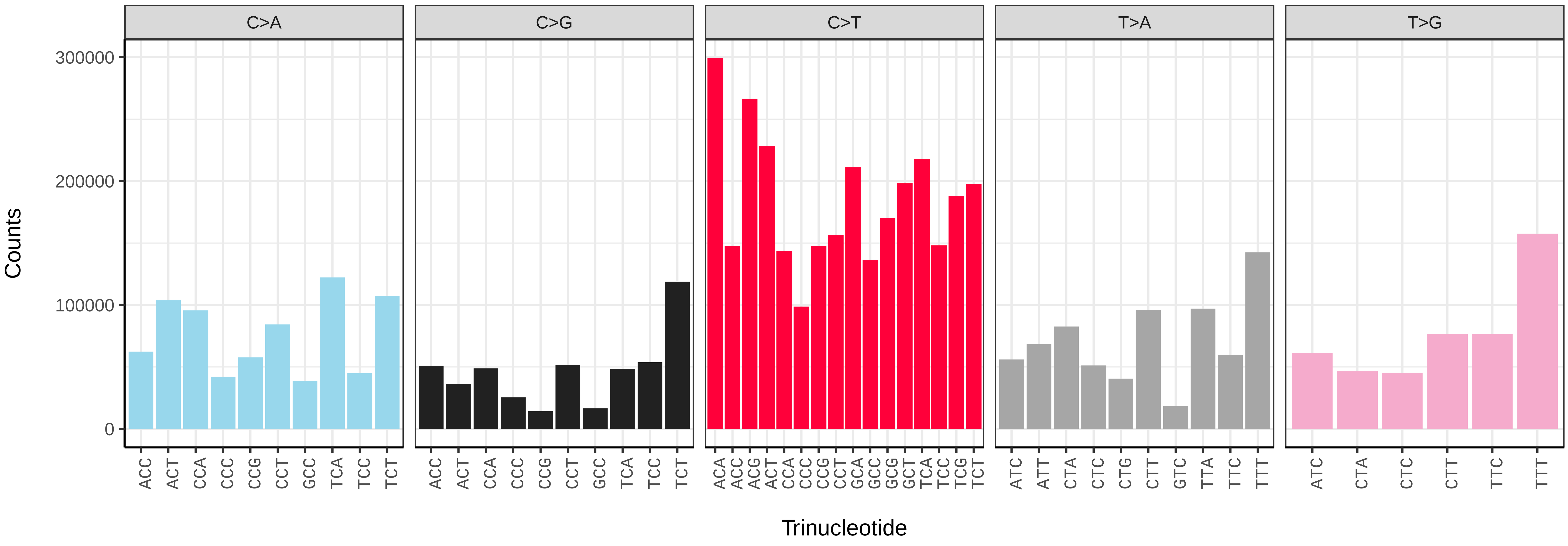


*Aporrectodea icterica* (wcApoIcte1) somatic mutational spectrum (SBS96 classification)

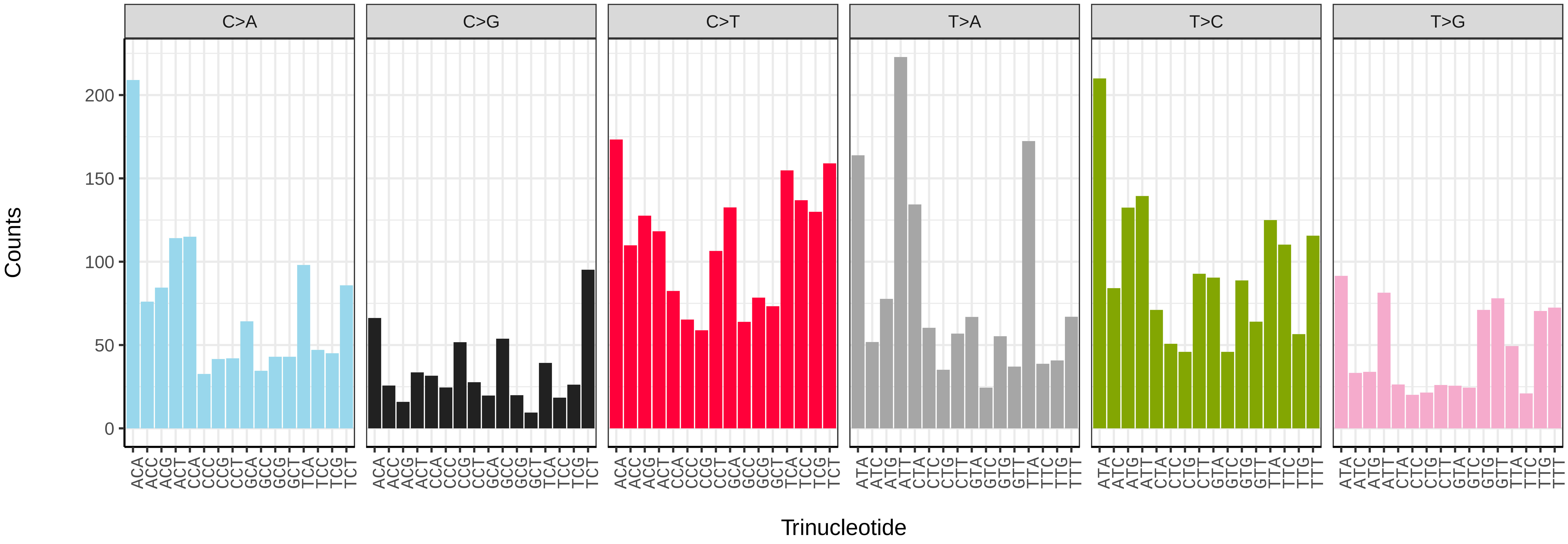


*Aporrectodea icterica* (wcApoIcte1) somatic mutational spectrum (SBS52 classification)

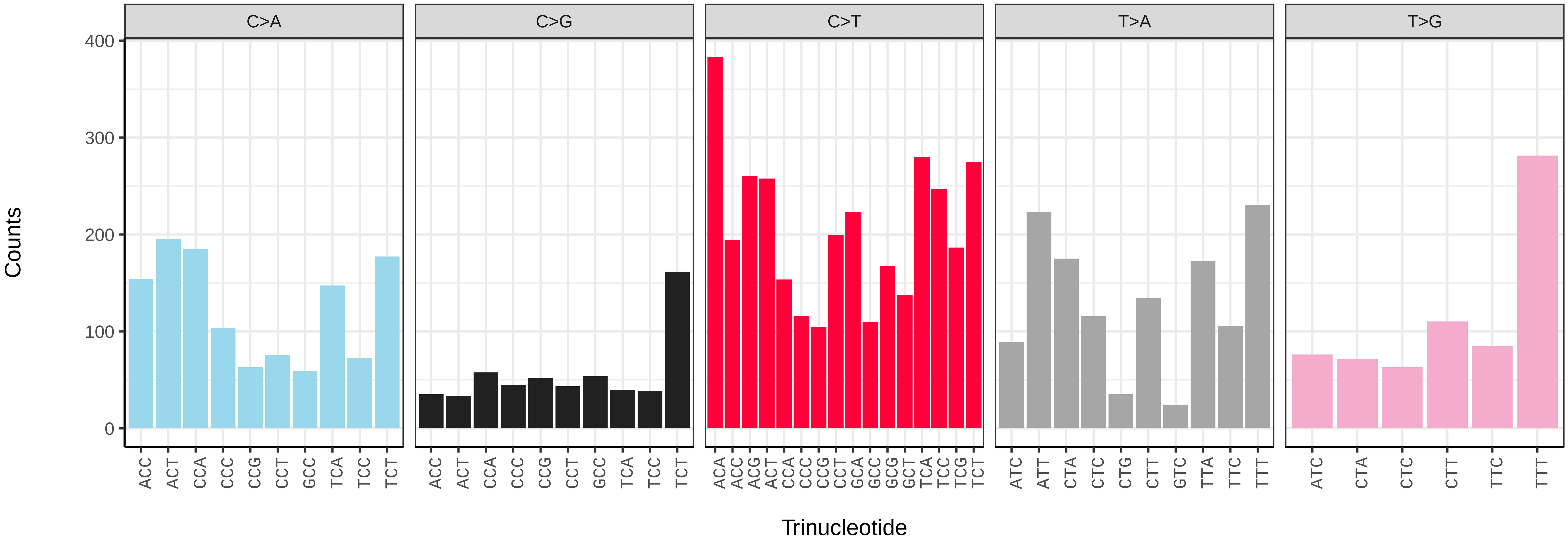

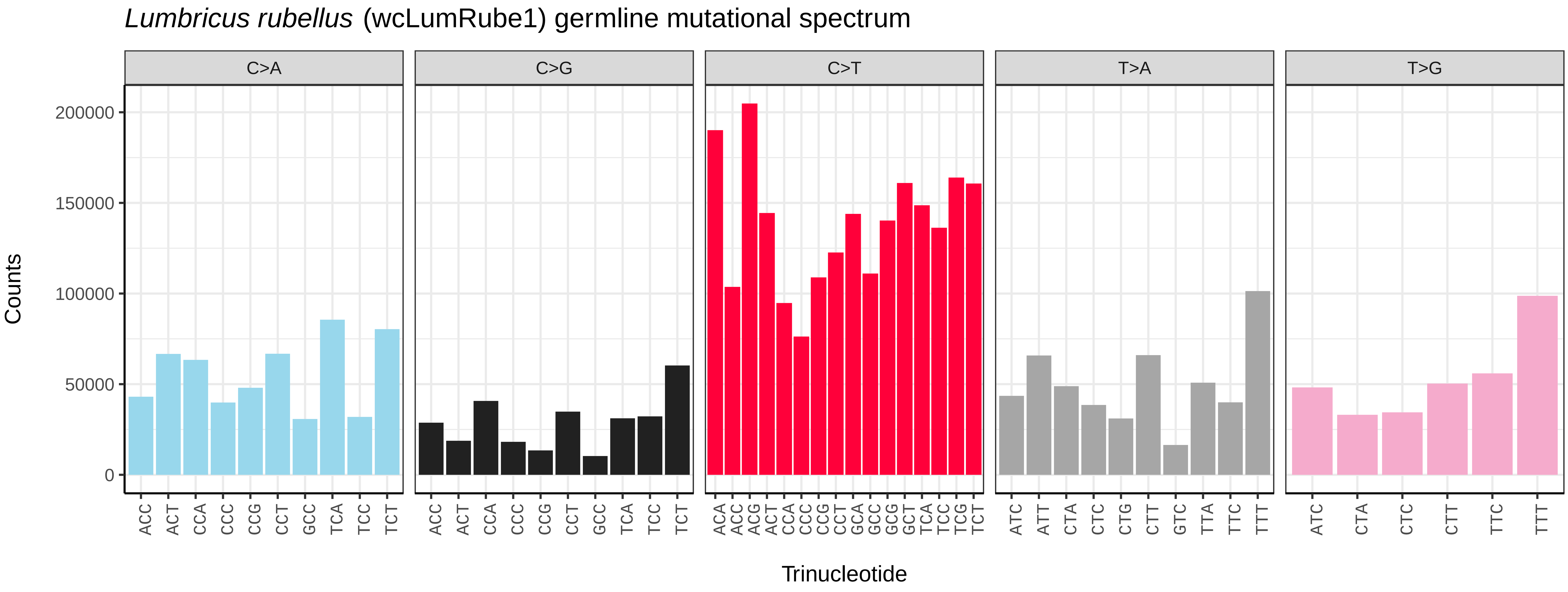
Lumbricus rubellus (wcLumRube1) germline mutational spectrum
C>A
C>G
C>T
T>A
T>G
Counts
200000
150000
100000
50000
0
ACC ACT CCA CCC CCG CCT GCC TCA TCC TCT
ACC ACT CCA CCC CCG CCT GCC TCA TCC TCT
ACA ACC ACG ACT CCA CCC CCG CCT GCA GCC GCG GCT TCA TCC TCG TCT
ATC ATT CTA CTC CTG CTT GTC TTA TTC TTT
ATC CTA CTC CTT TTC TTT
Trinucleotide

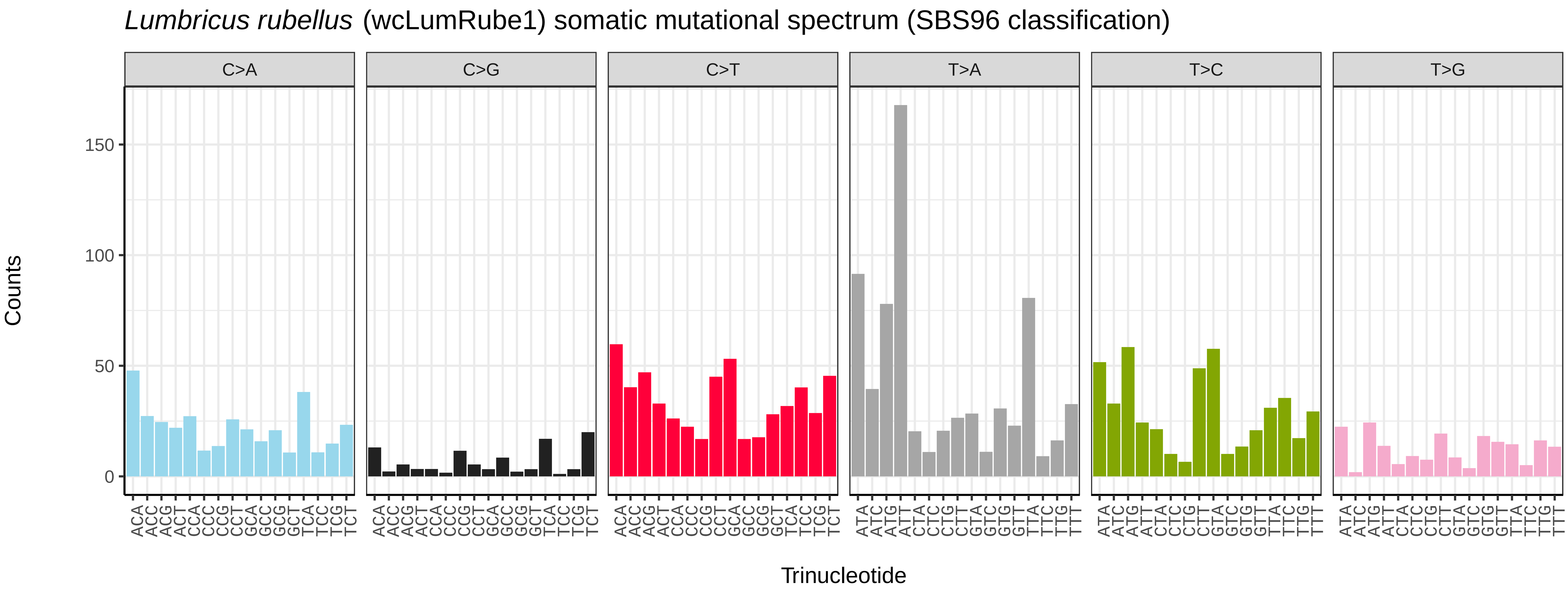
Lumbricus rubellus (wcLumRube1) somatic mutational spectrum (SBS96 classification)
C>A
C>G
C>T
T>A
T>C
T>G
Counts
150
100
50
0
ACA ACC ACG ACT CCA CCC CCG CCT GCA GCC GCG GCT TCA TCC TCG TCT
ACA ACC ACG ACT CCA CCC CCG CCT GCA GCC GCG GCT TCA TCC TCG TCT
ACA ACC ACG ACT CCA CCC CCG CCT GCA GCC GCG GCT TCA TCC TCG TCT
ATA ATC ATG ATT CTA CTC CTG CTT GTA GTC GTG GTT TTA TTC TTG TTT
ATA ATC ATG ATT CTA CTC CTG CTT GTA GTC GTG GTT TTA TTC TTG TTT
ATA ATC ATG ATT CTA CTC CTG CTT GTA GTC GTG GTT TTA TTC TTG TTT
Trinucleotide

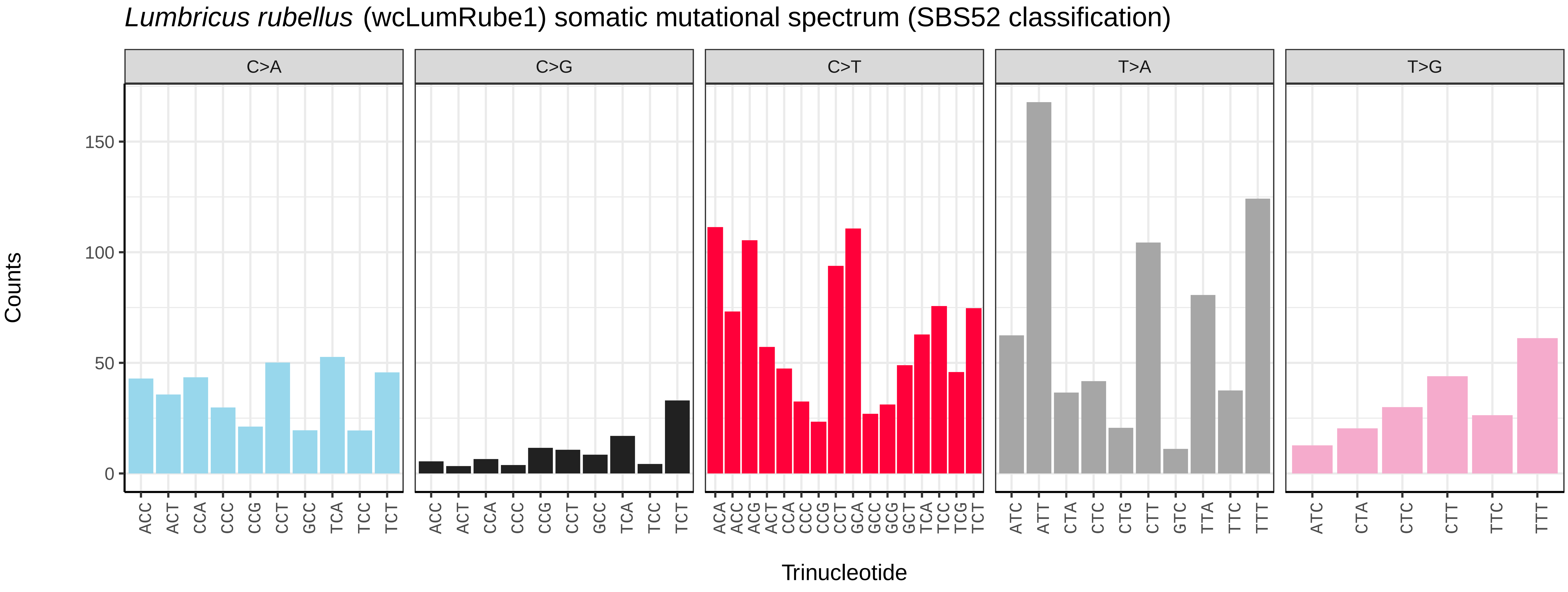
Lumbricus rubellus (wcLumRube1) somatic mutational spectrum (SBS52 classification)
C>A
C>G
C>T
T>A
T>G
Counts
150
100
50
0
ACC ACT CCA CCC CCG CCT GCC TCA TCC TCT
ACC ACT CCA CCC CCG CCT GCC TCA TCC TCT
ACA ACC ACG ACT CCA CCC CCG CCT GCA GCC GCG GCT TCA TCC TCG TCT
ATC ATT CTA CTC CTG CTT GTC TTA TTC TTT
ATC CTA CTC CTT TTC TTT
Trinucleotide

*Lumbricus terrestris* (wcLumTerr1) germline mutational spectrum

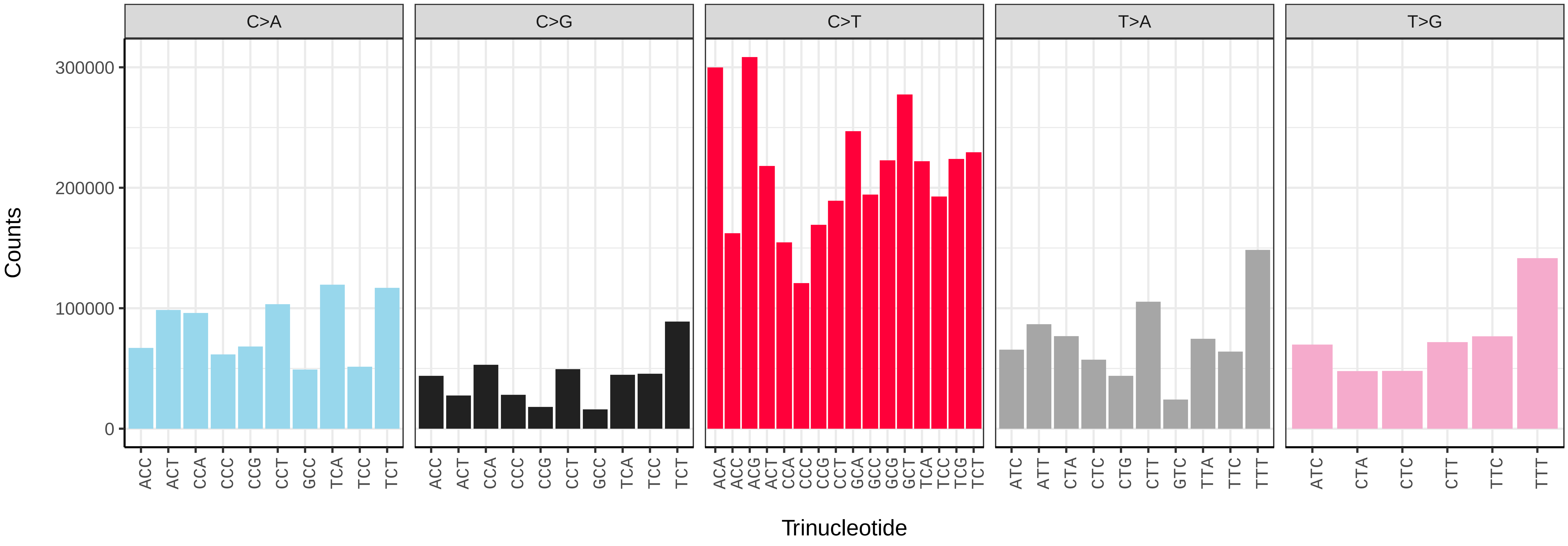


*Lumbricus terrestris* (wcLumTerr1) somatic mutational spectrum (SBS96 classification)

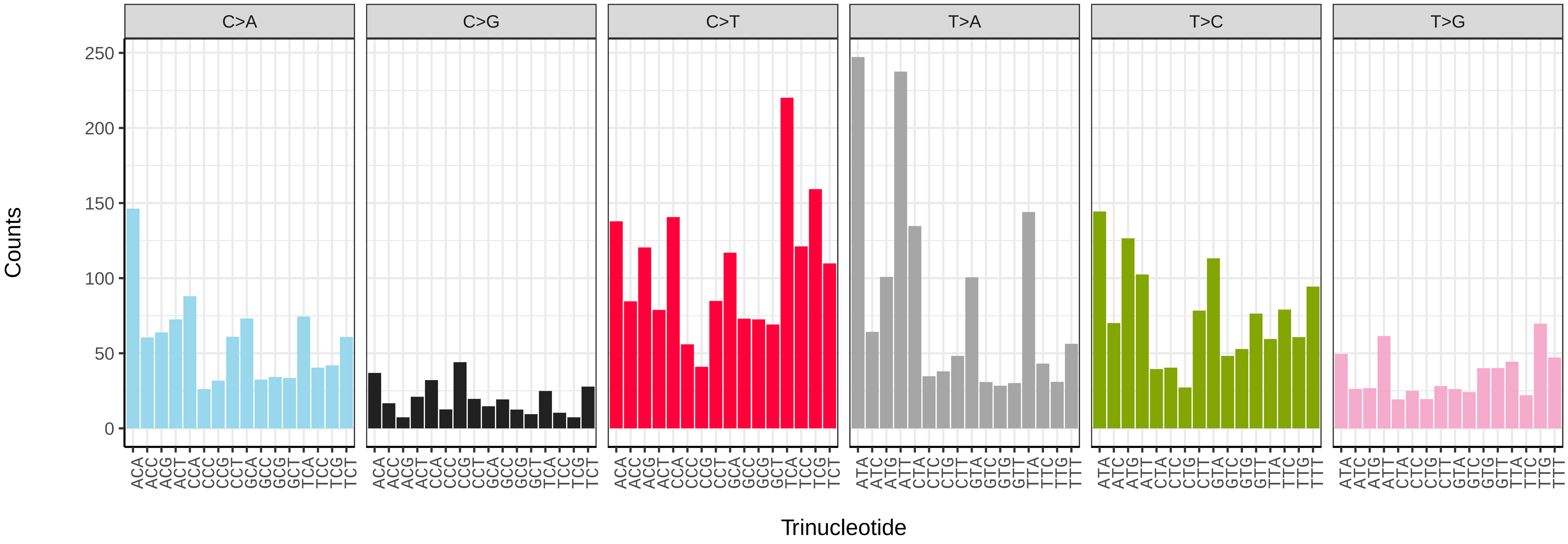


*Lumbricus terrestris* (wcLumTerr1) somatic mutational spectrum (SBS52 classification)

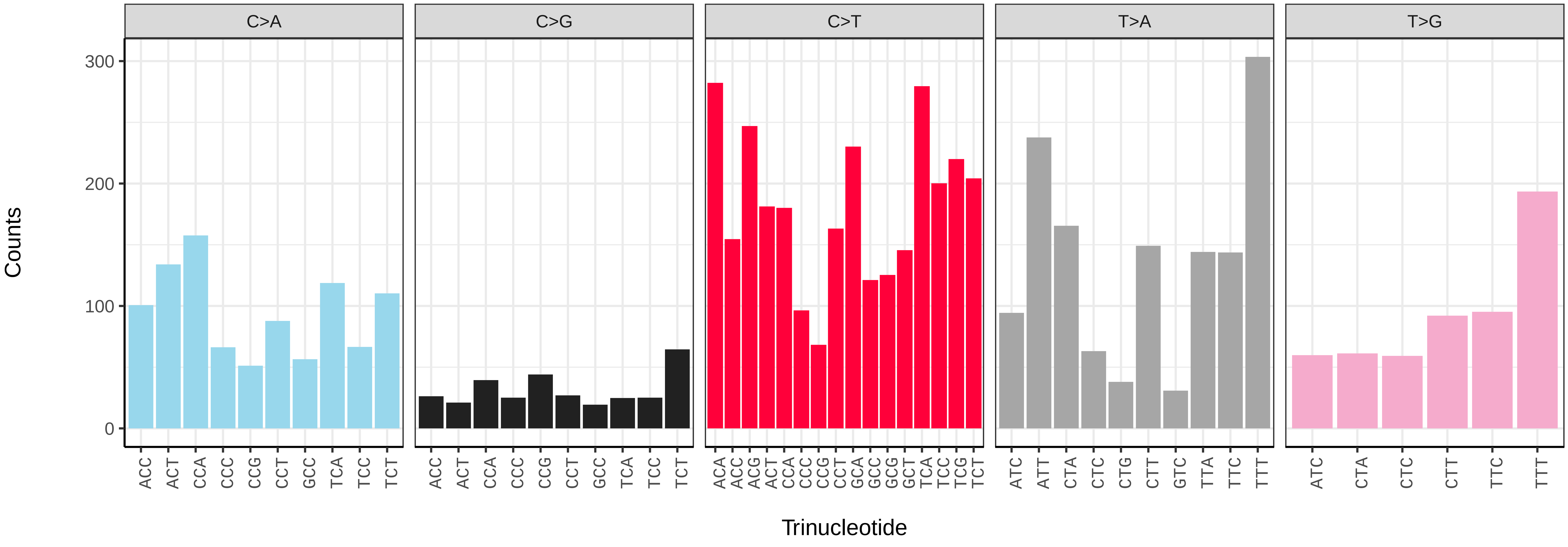

*Magelona johnstoni* (wdMagJohn1) germline mutational spectrum

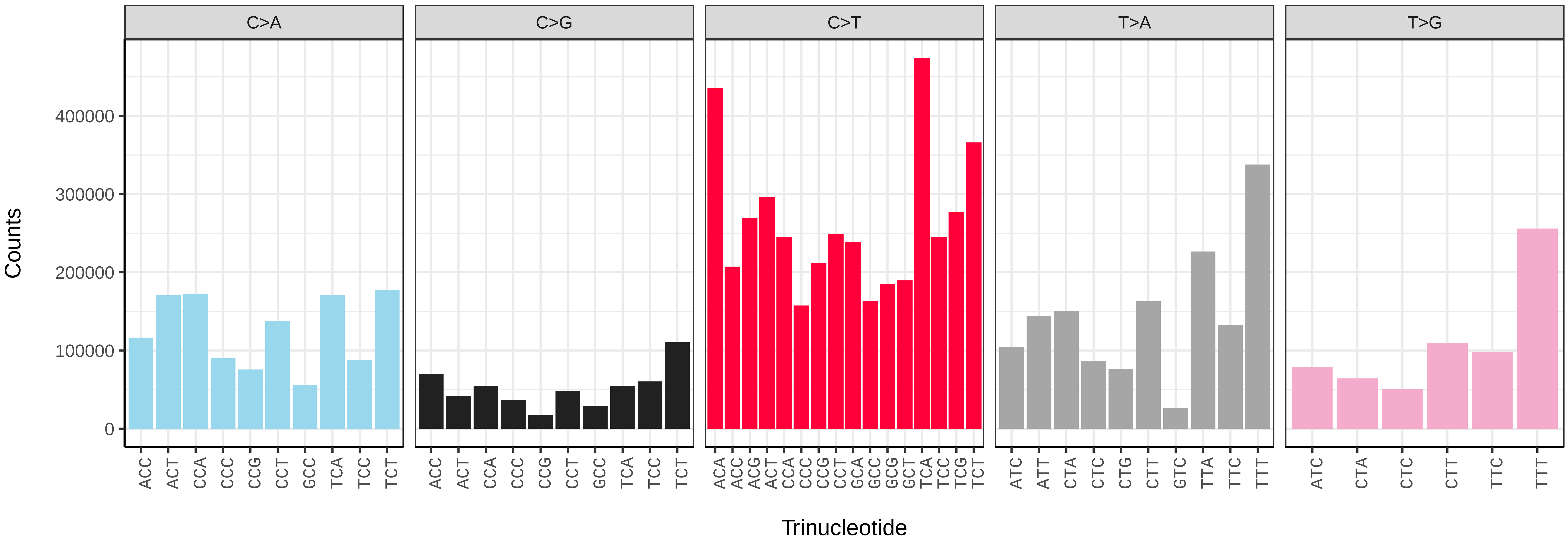


*Magelona johnstoni* (wdMagJohn1) somatic mutational spectrum (SBS96 classification)

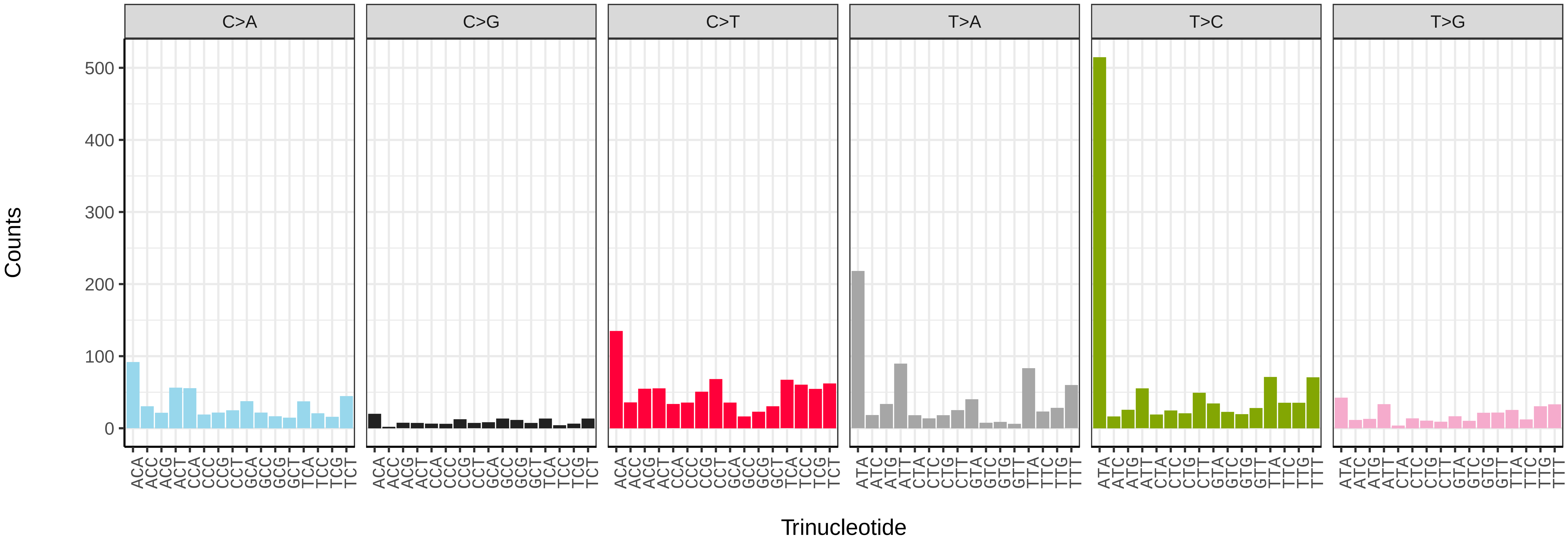


*Magelona johnstoni* (wdMagJohn1) somatic mutational spectrum (SBS52 classification)

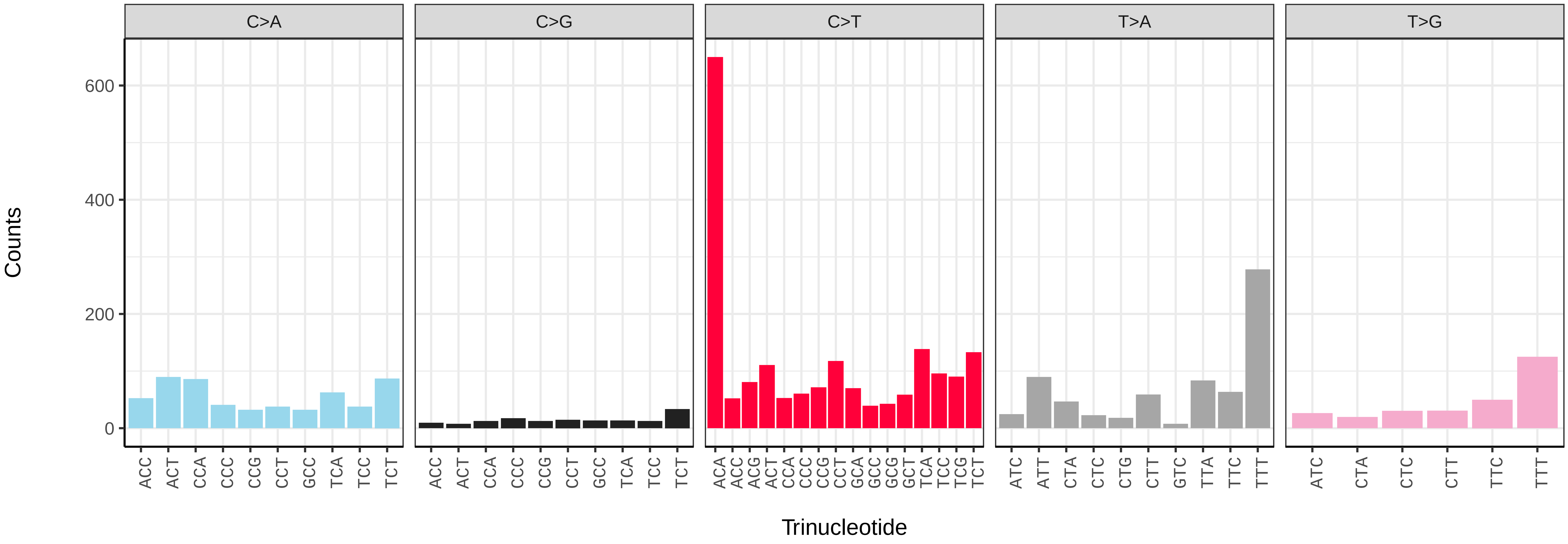

*Erpobdella octoculata* (wkErpOcto2) germline mutational spectrum

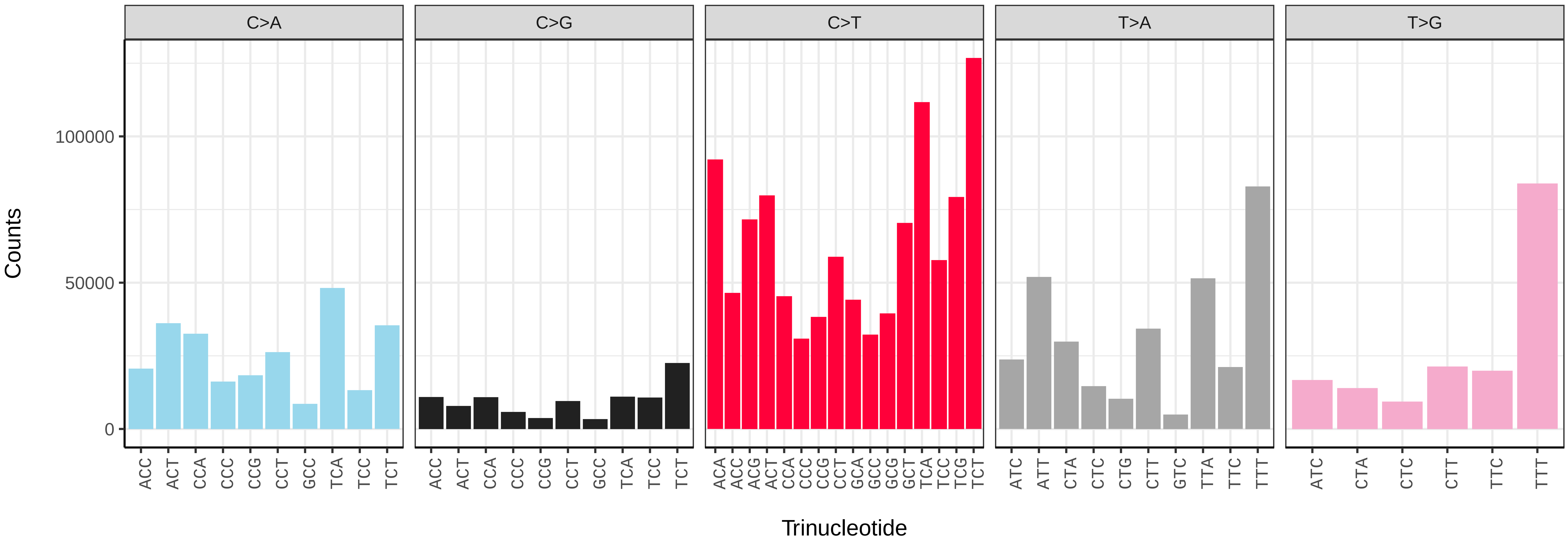


*Erpobdella octoculata* (wkErpOcto2) somatic mutational spectrum (SBS96 classification)

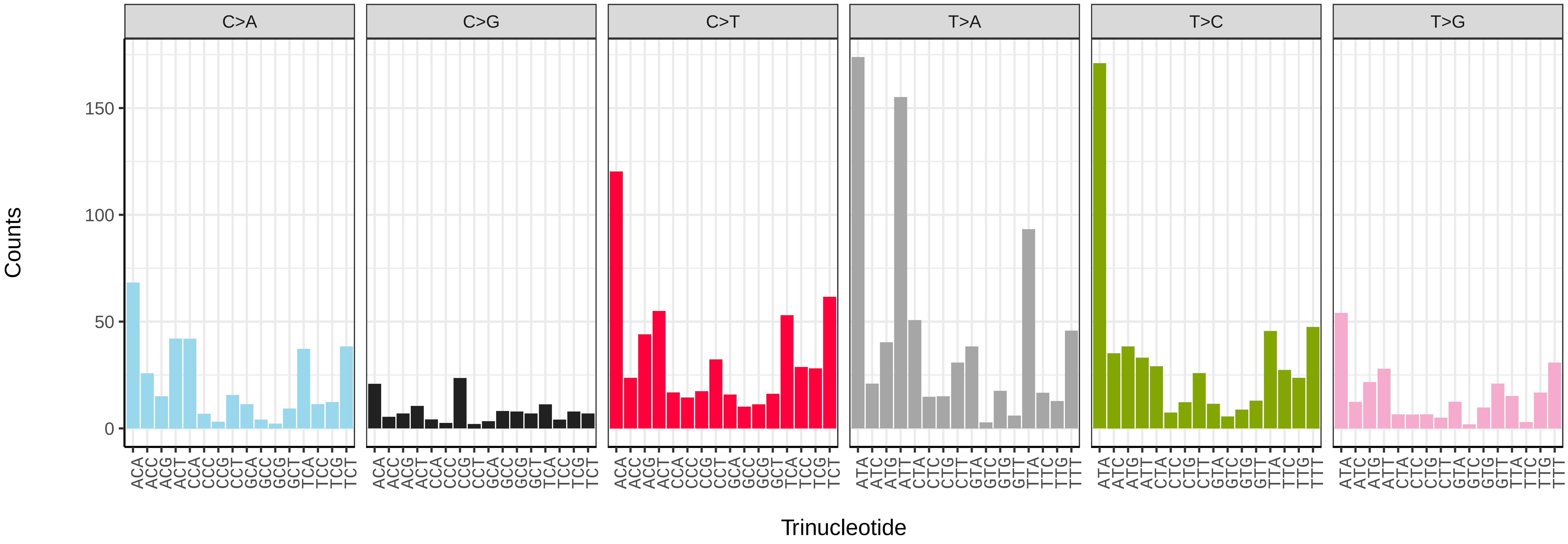


*Erpobdella octoculata* (wkErpOcto2) somatic mutational spectrum (SBS52 classification)

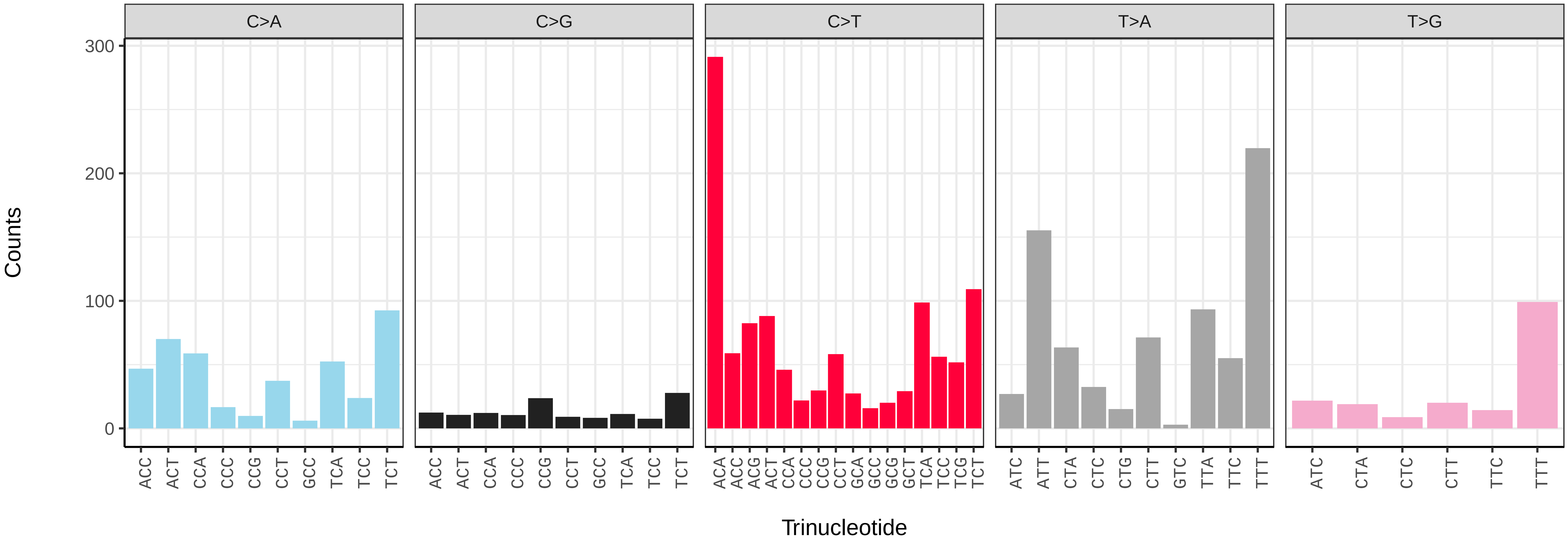

*Acholoe squamosa* (wpAchSqua1) germline mutational spectrum

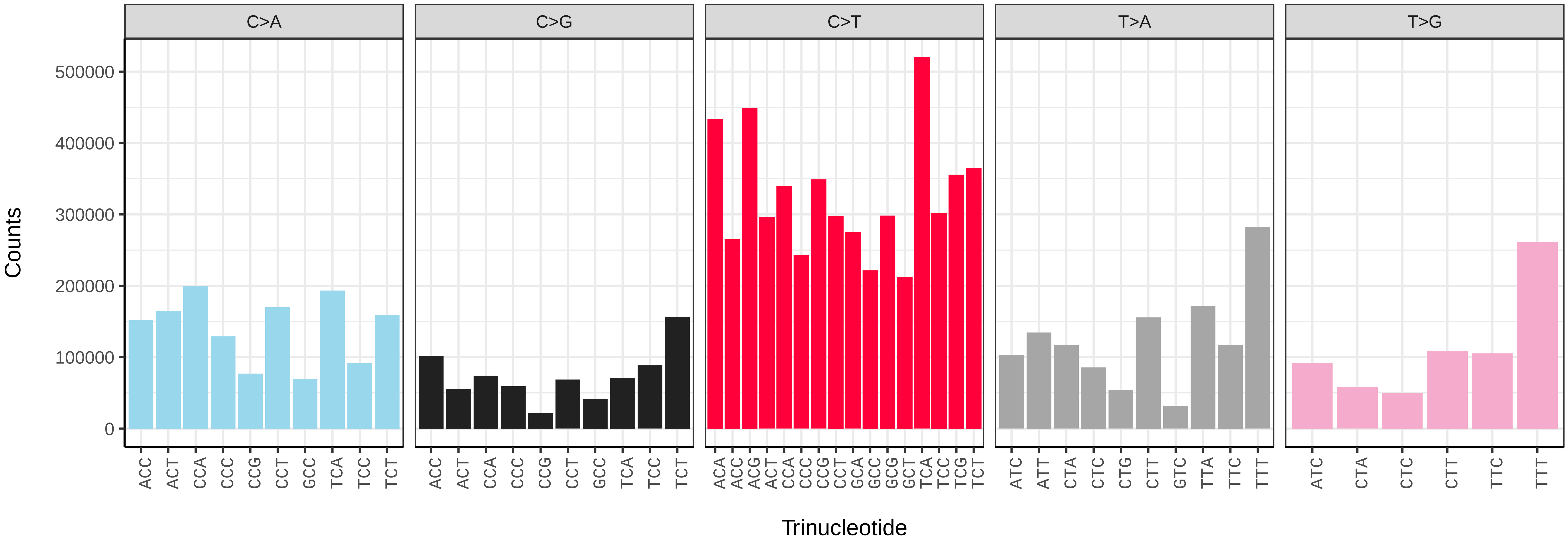


*Acholoe squamosa* (wpAchSqua1) somatic mutational spectrum (SBS96 classification)

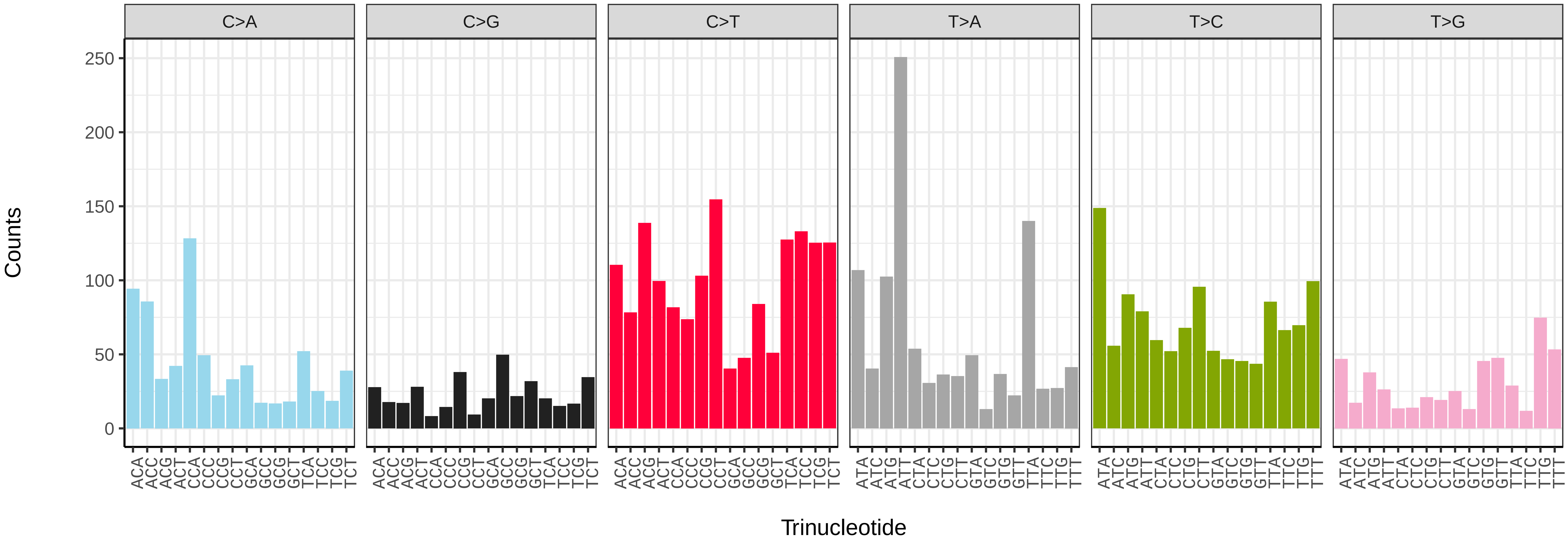


*Acholoe squamosa* (wpAchSqua1) somatic mutational spectrum (SBS52 classification)

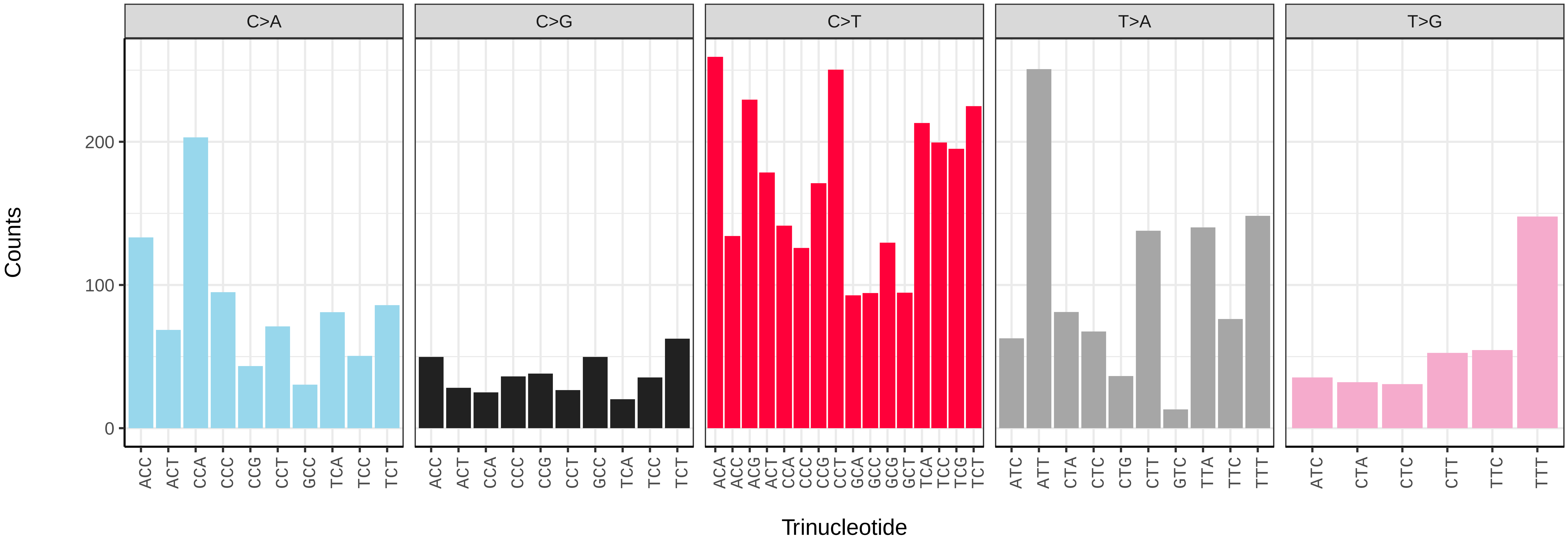

*Alentia gelatinosa* (wpAleGela1) germline mutational spectrum

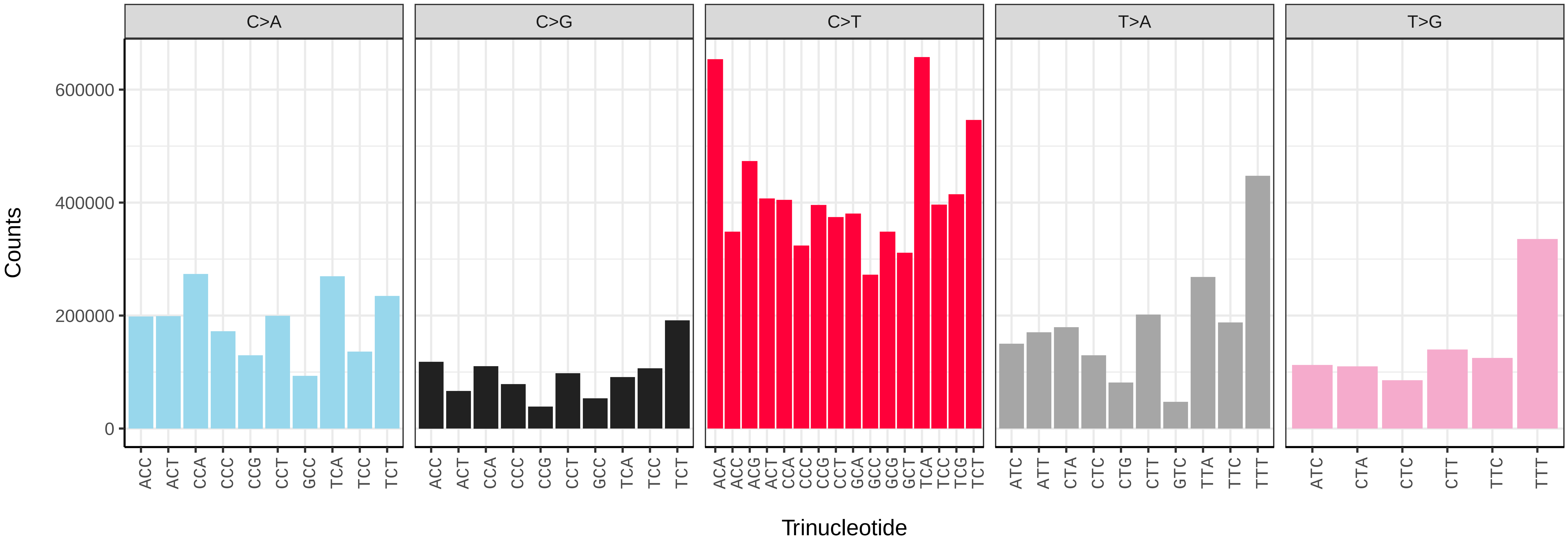


*Alentia gelatinosa* (wpAleGela1) somatic mutational spectrum (SBS96 classification)

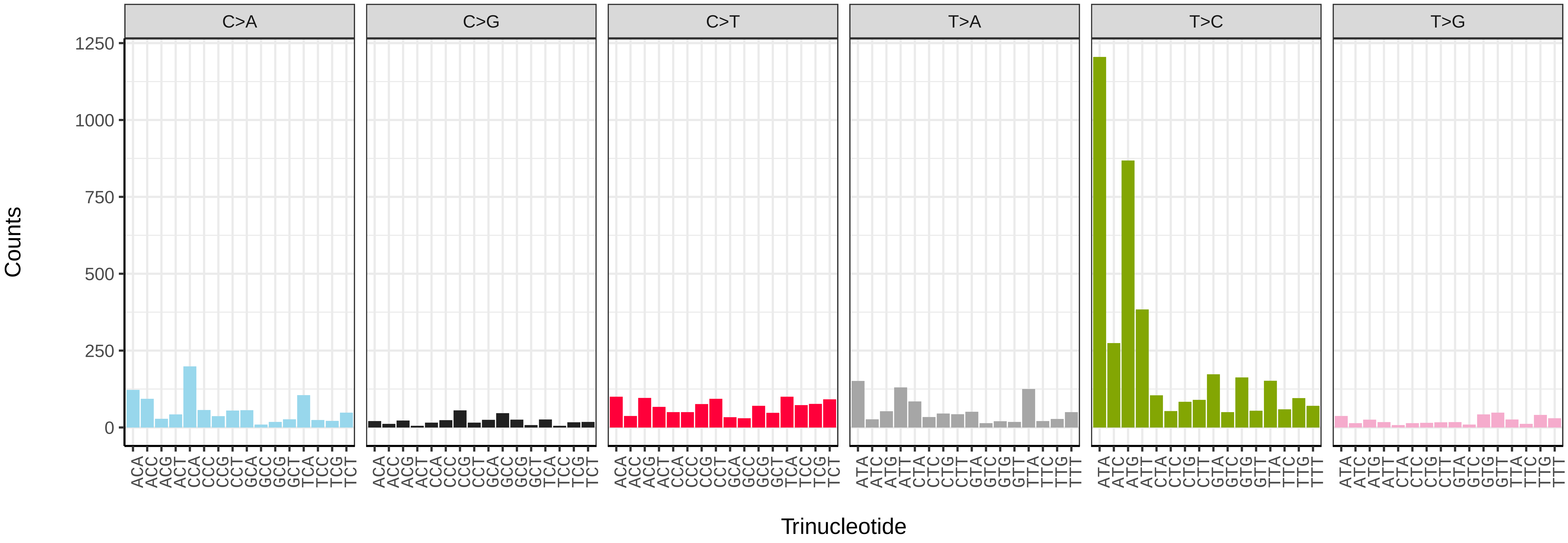


*Alentia gelatinosa* (wpAleGela1) somatic mutational spectrum (SBS52 classification)

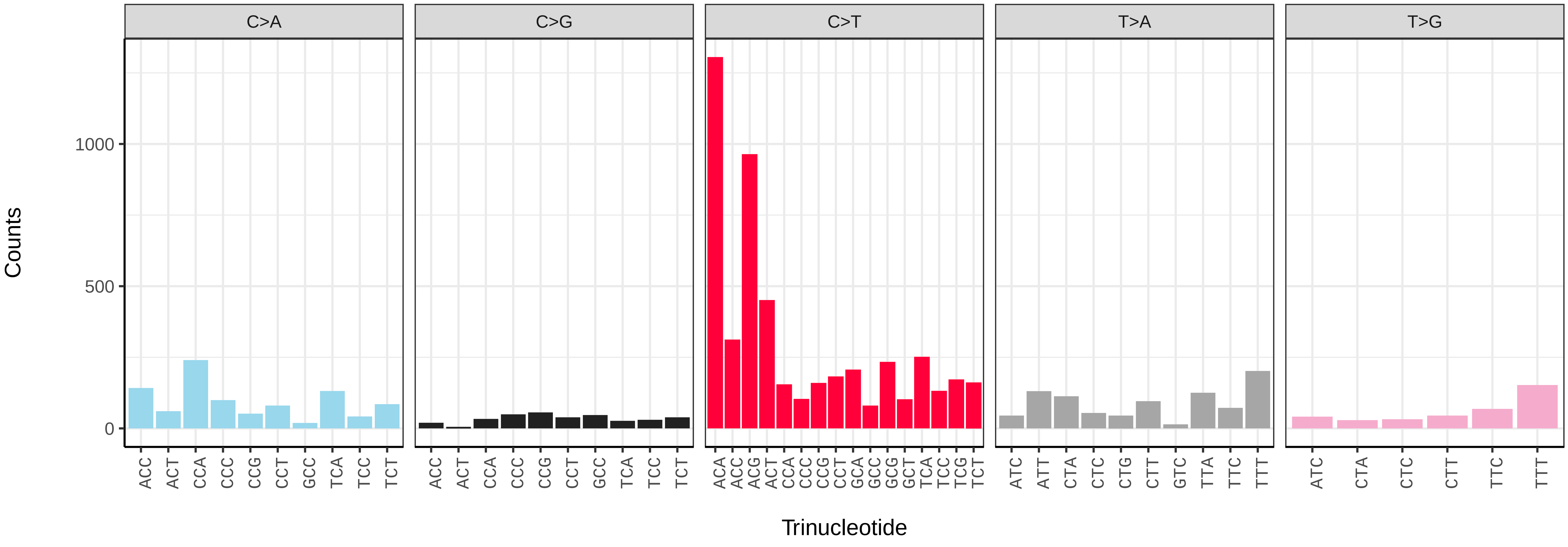

*Alitta virens* (wpAliVire1) germline mutational spectrum

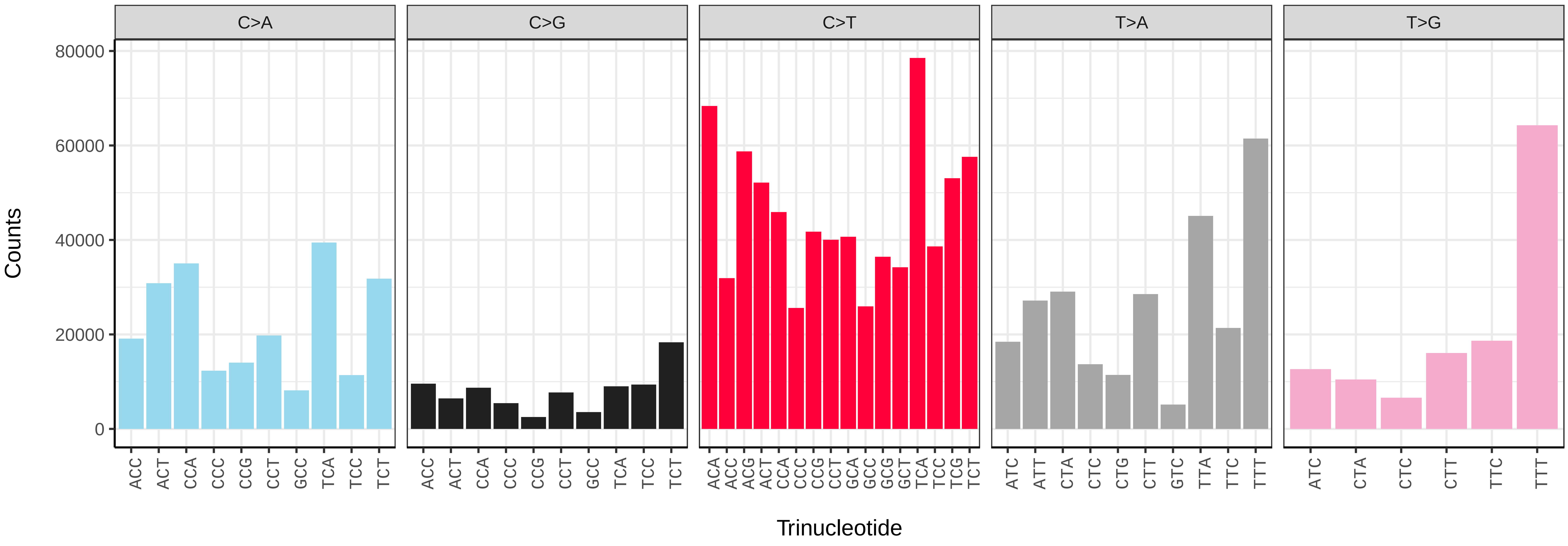


*Alitta virens* (wpAliVire1) somatic mutational spectrum (SBS96 classification)

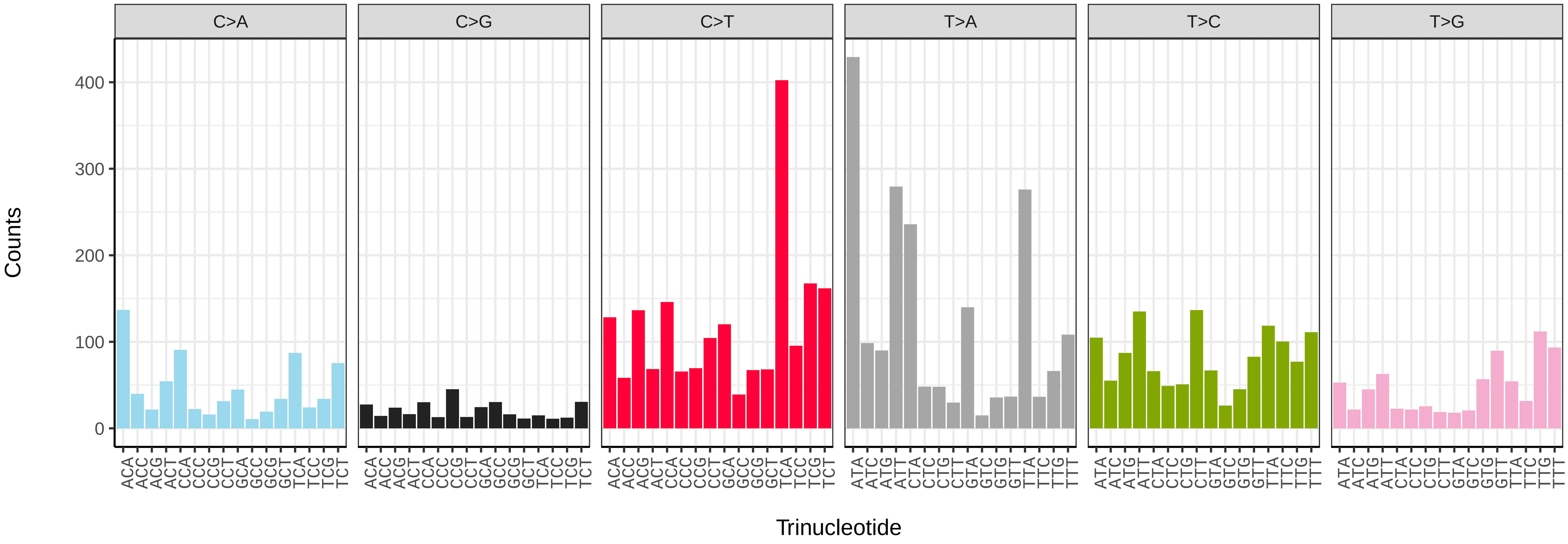


*Alitta virens* (wpAliVire1) somatic mutational spectrum (SBS52 classification)

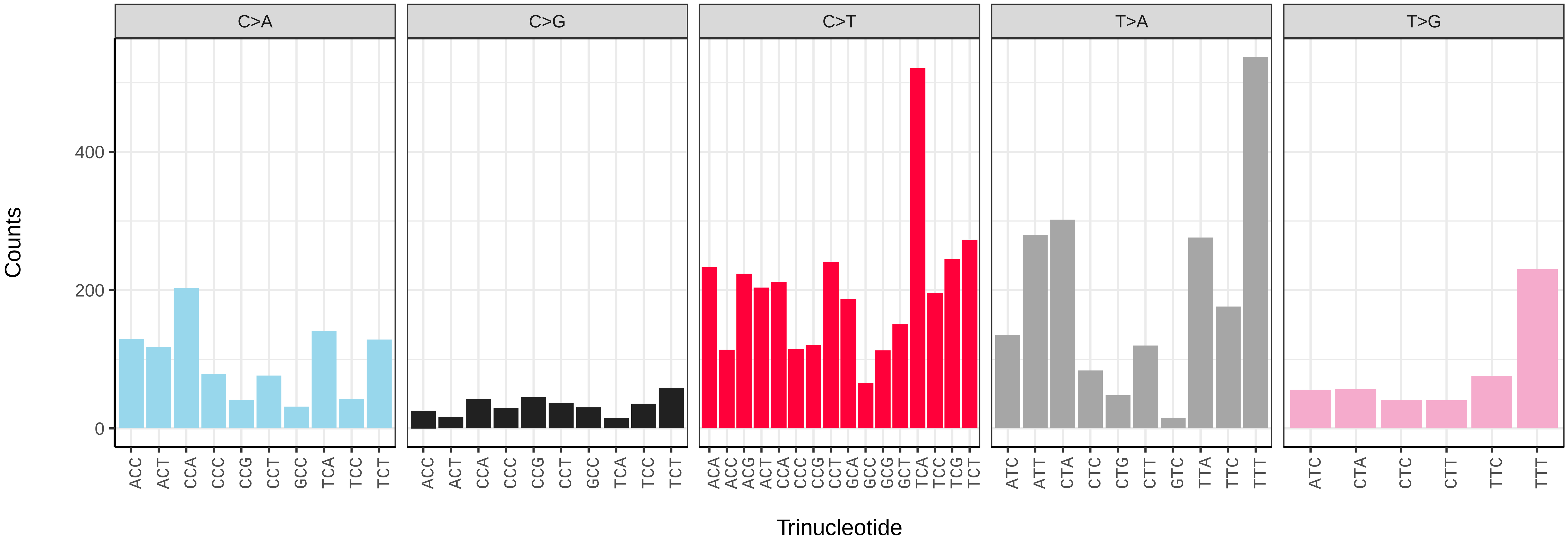

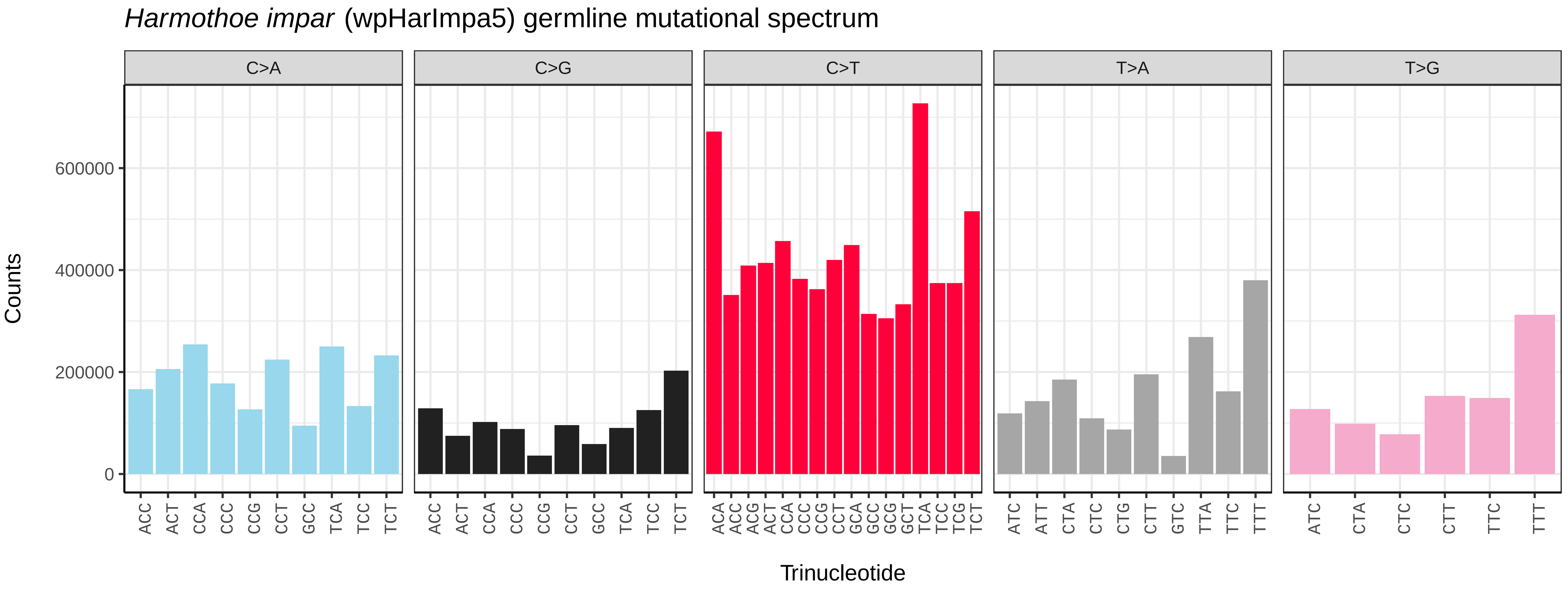
*Harmothoe impar* (wpHarImpa5) germline mutational spectrum
C>A
C>G
C>T
T>A
T>G
Counts
0
200000
400000
600000
ACC ACT CCA CCC CCG CCT GCC TCA TCC TCT
ACC ACT CCA CCC CCG CCT GCC TCA TCC TCT
ACA ACC ACG ACT CCA CCC CCG CCT GCA GCC GCG GCT TCA TCC TCG TCT
ATC ATT CTA CTC CTG CTT GTC TTA TTC TTT
ATC CTA CTC CTT TTC TTT
Trinucleotide

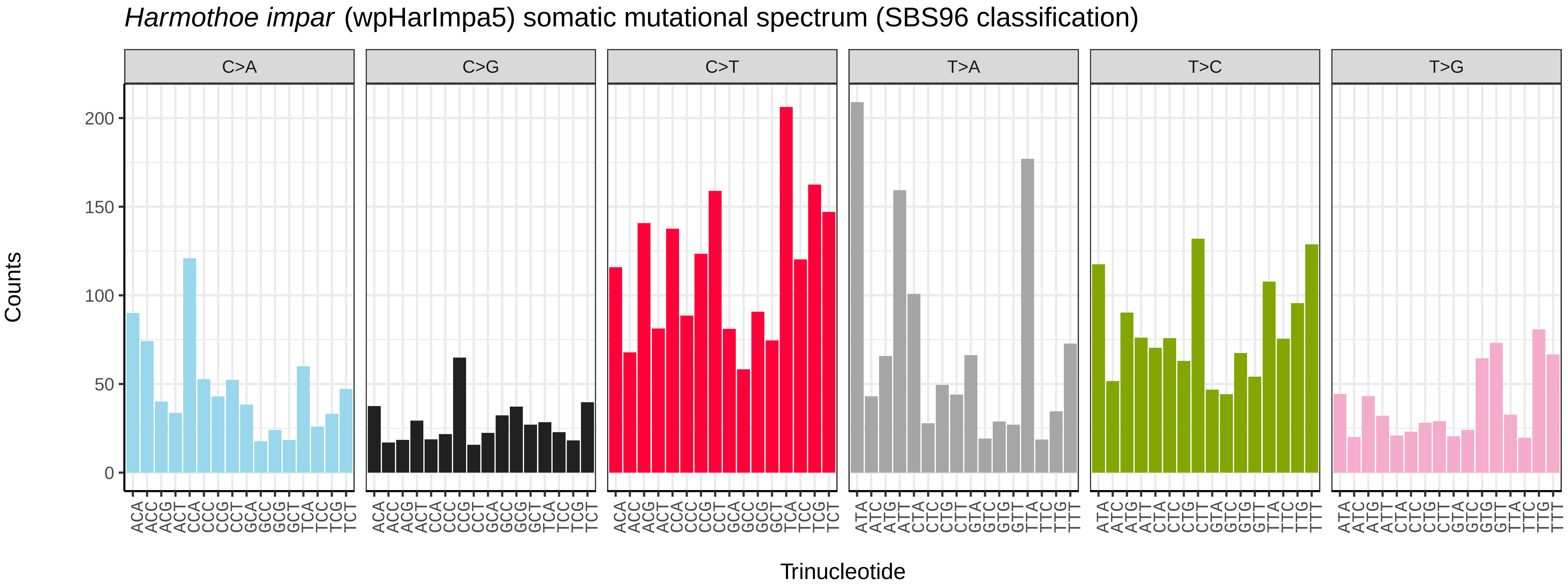
*Harmothoe impar* (wpHarImpa5) somatic mutational spectrum (SBS96 classification)
C>A
C>G
C>T
T>A
T>C
T>G
Counts
0
50
100
150
200
ACA ACC ACG ACT CCA CCC CCG CCT GCA GCC GCG GCT TCA TCC TCG TCT
ACA ACC ACG ACT CCA CCC CCG CCT GCA GCC GCG GCT TCA TCC TCG TCT
ACA ACC ACG ACT CCA CCC CCG CCT GCA GCC GCG GCT TCA TCC TCG TCT
ATA ATC ATG ATT CTA CTC CTG CTT GTA GTC GTG GTT TTA TTC TTG TTT
ATA ATC ATG ATT CTA CTC CTG CTT GTA GTC GTG GTT TTA TTC TTG TTT
ATA ATC ATG ATT CTA CTC CTG CTT GTA GTC GTG GTT TTA TTC TTG TTT
Trinucleotide

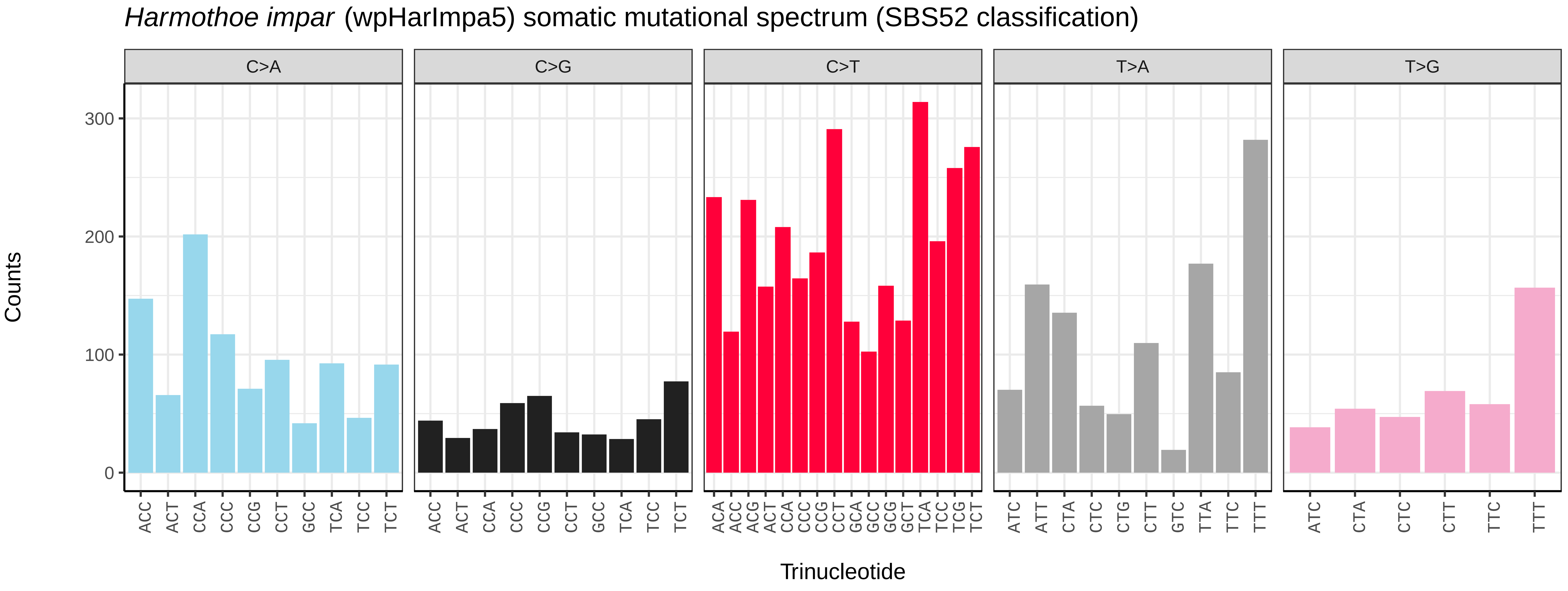
*Harmothoe impar* (wpHarImpa5) somatic mutational spectrum (SBS52 classification)
C>A
C>G
C>T
T>A
T>G
Counts
0
100
200
300
ACC ACT CCA CCC CCG CCT GCC TCA TCC TCT
ACC ACT CCA CCC CCG CCT GCC TCA TCC TCT
ACA ACC ACG ACT CCA CCC CCG CCT GCA GCC GCG GCT TCA TCC TCG TCT
ATC ATT CTA CTC CTG CTT GTC TTA TTC TTT
ATC CTA CTC CTT TTC TTT
Trinucleotide

*Sthenelais limicola* (wpSthLimi1) germline mutational spectrum

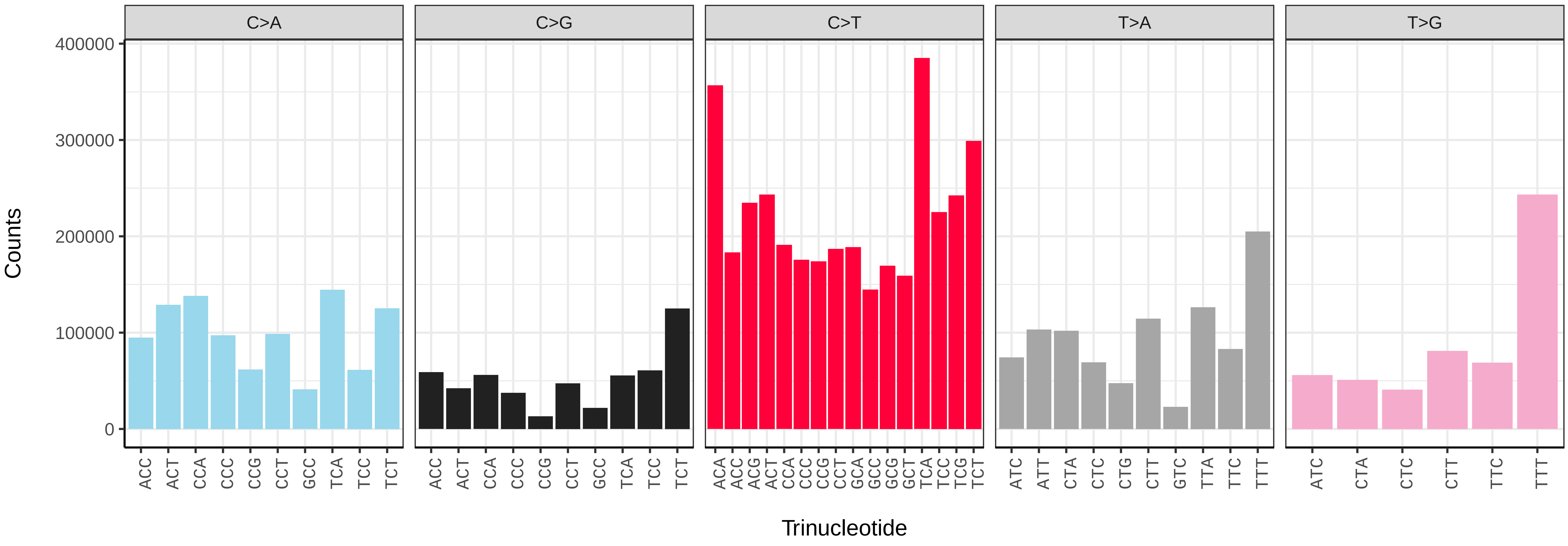


*Sthenelais limicola* (wpSthLimi1) somatic mutational spectrum (SBS96 classification)

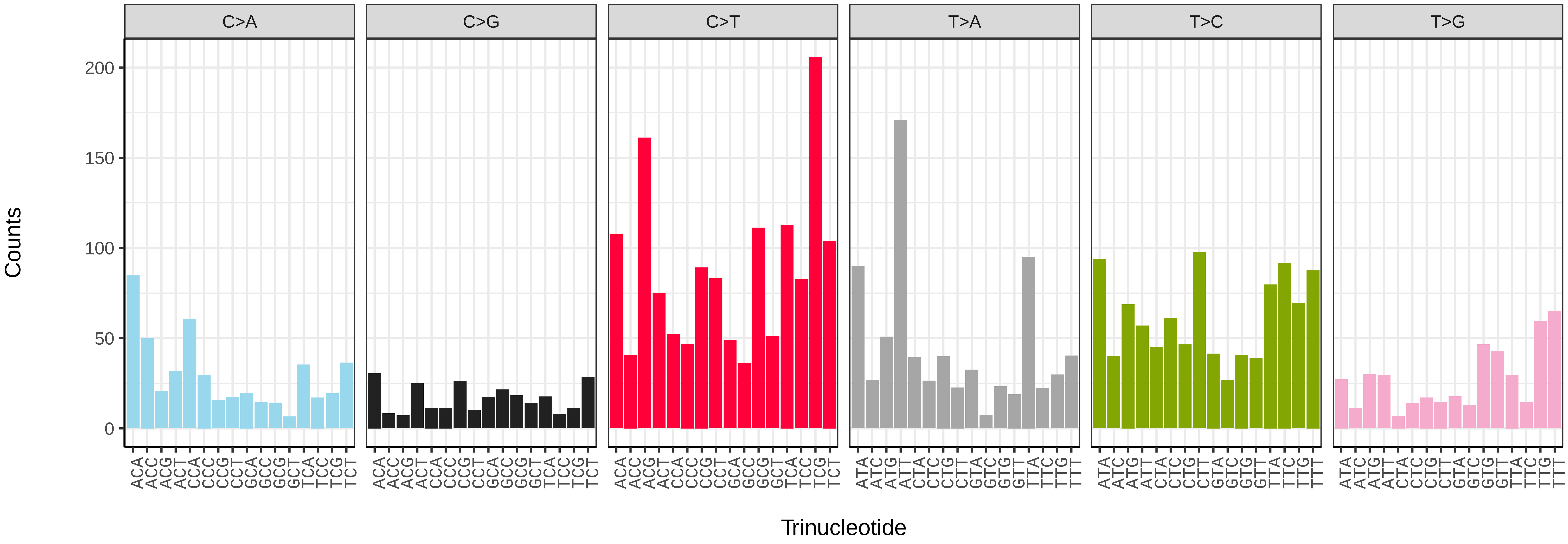


*Sthenelais limicola* (wpSthLimi1) somatic mutational spectrum (SBS52 classification)

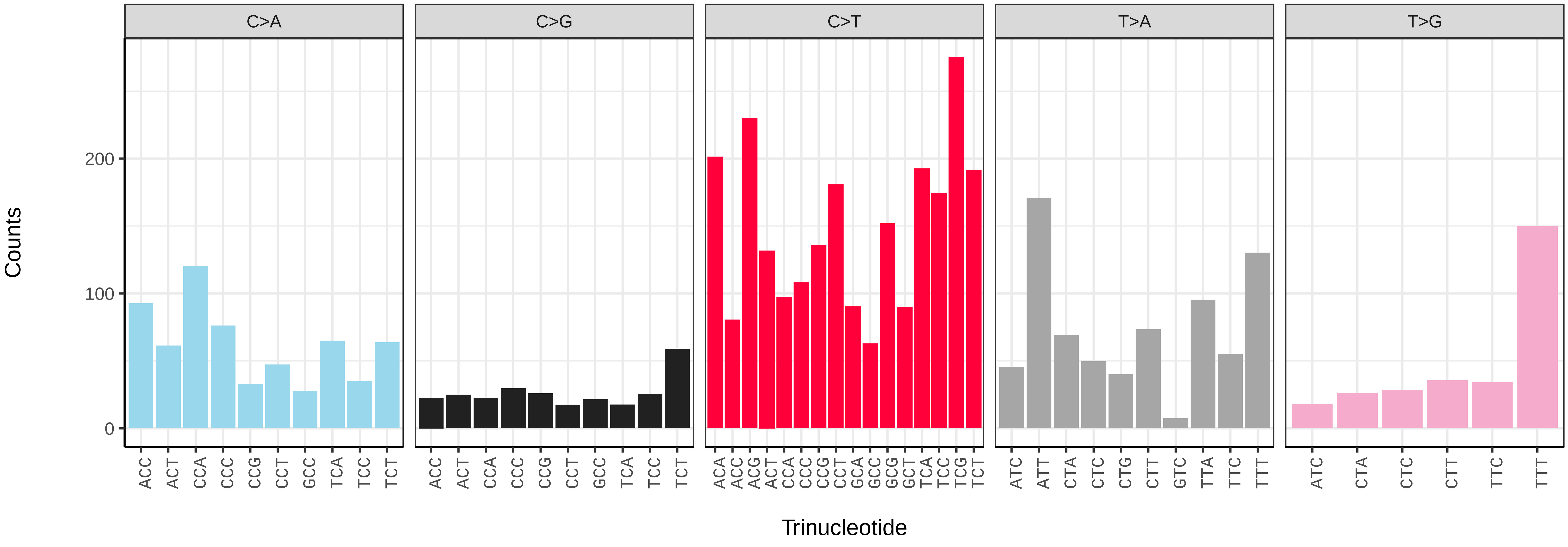

*Piscicola geometra* (wrPisGeom1) germline mutational spectrum

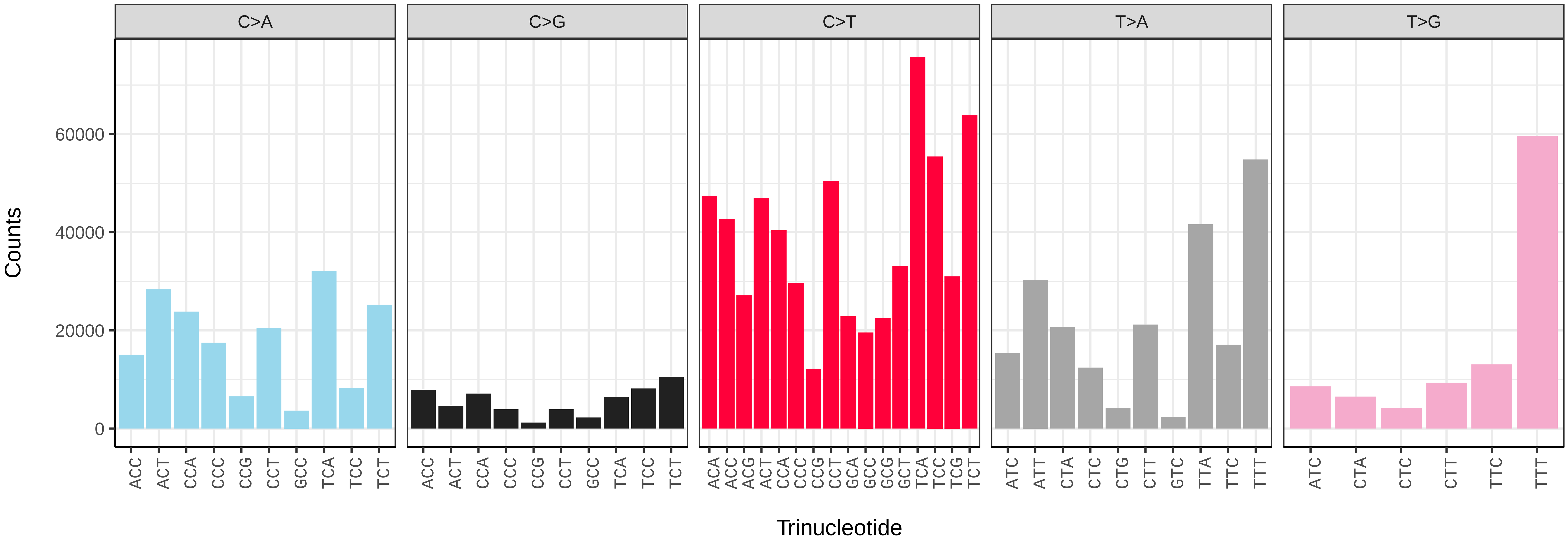


*Piscicola geometra* (wrPisGeom1) somatic mutational spectrum (SBS96 classification)

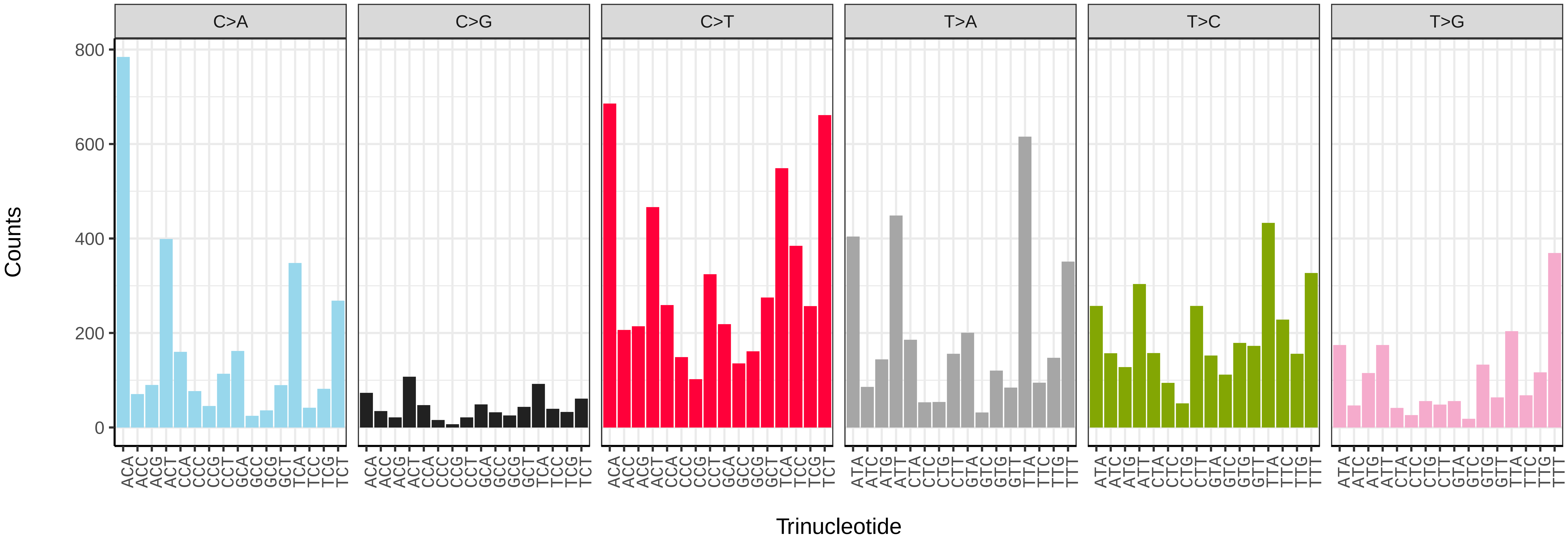


*Piscicola geometra* (wrPisGeom1) somatic mutational spectrum (SBS52 classification)

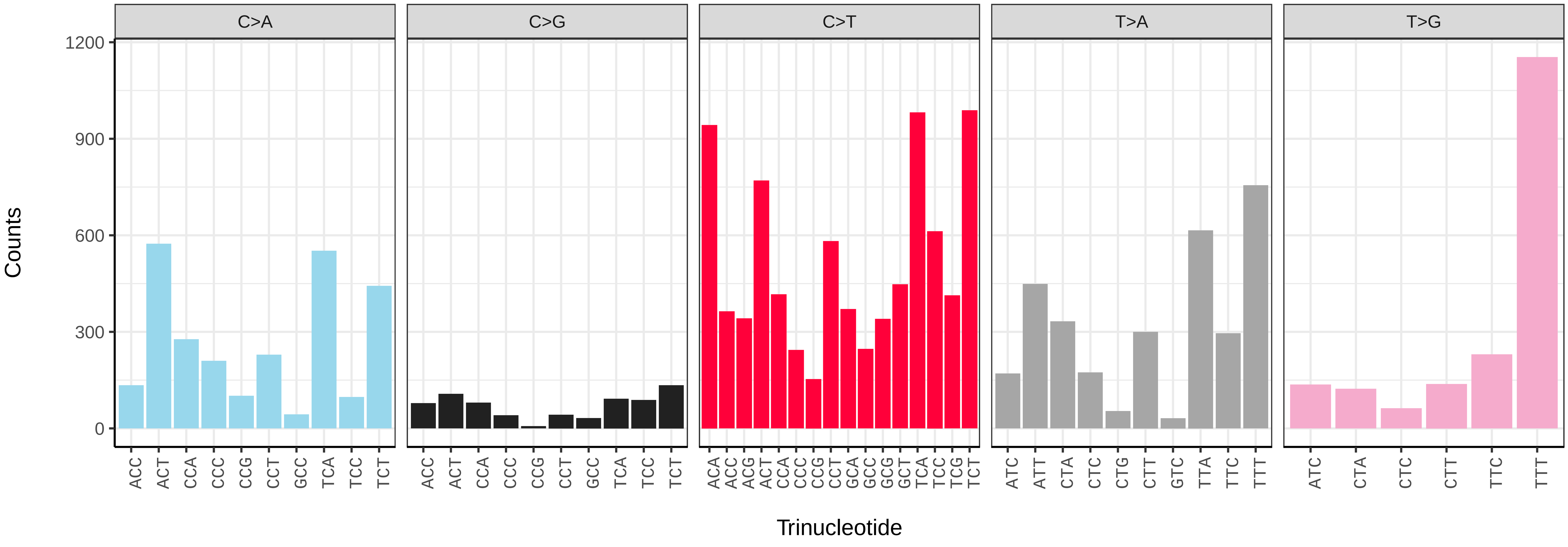

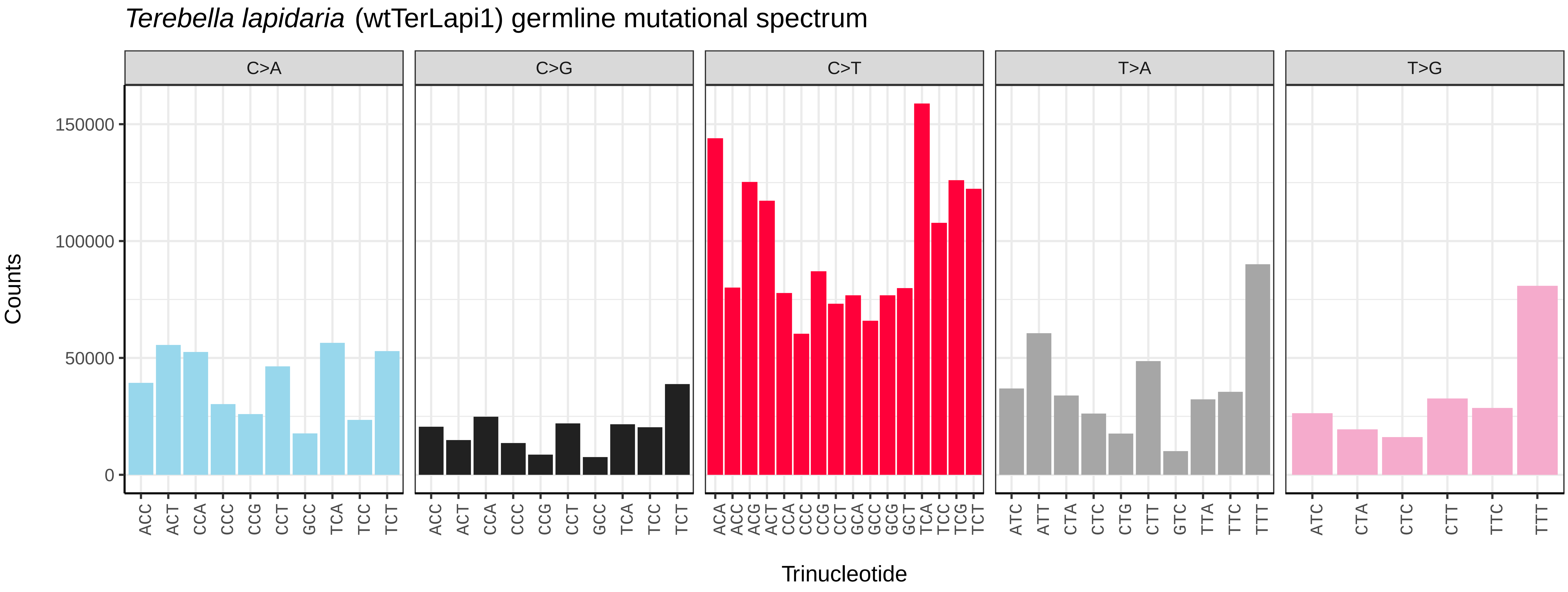
*Terebella lapidaria* (wtTerLapi1) germline mutational spectrum
C>A
C>G
C>T
T>A
T>G
Counts
150000
100000
50000
0
Trinucleotide

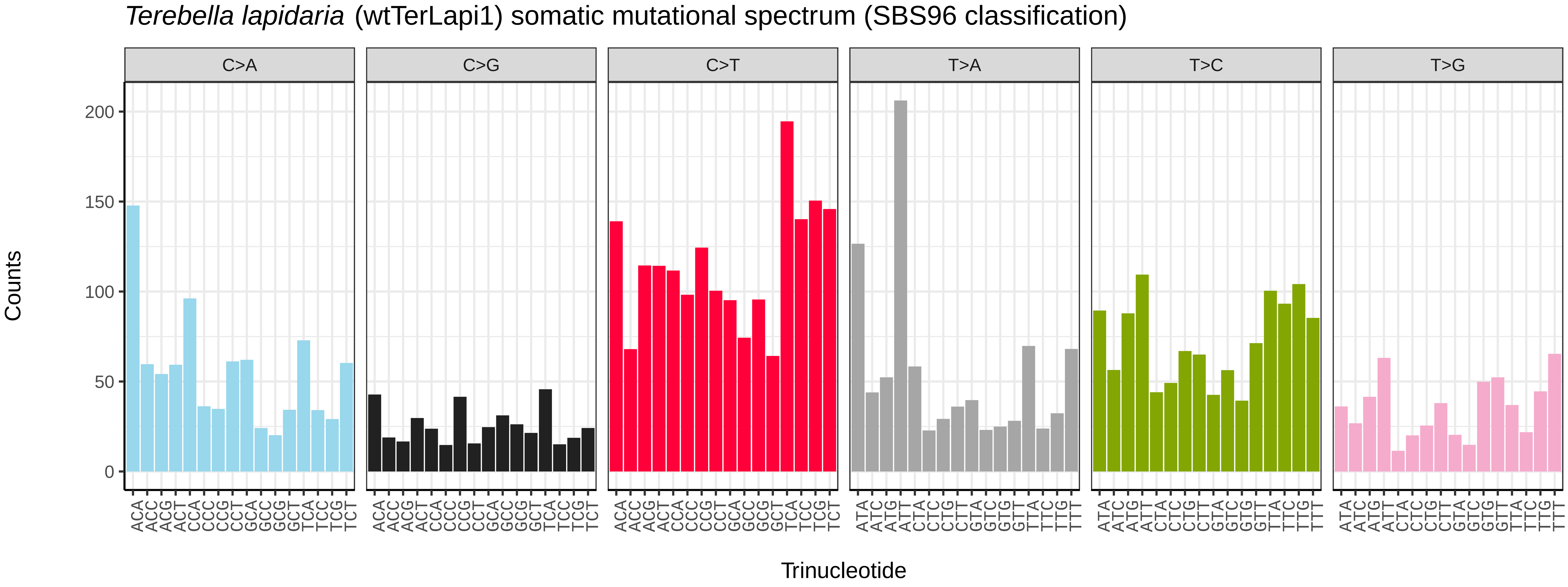
*Terebella lapidaria* (wtTerLapi1) somatic mutational spectrum (SBS96 classification)
C>A
C>G
C>T
T>A
T>C
T>G
Counts
200
150
100
50
0
Trinucleotide

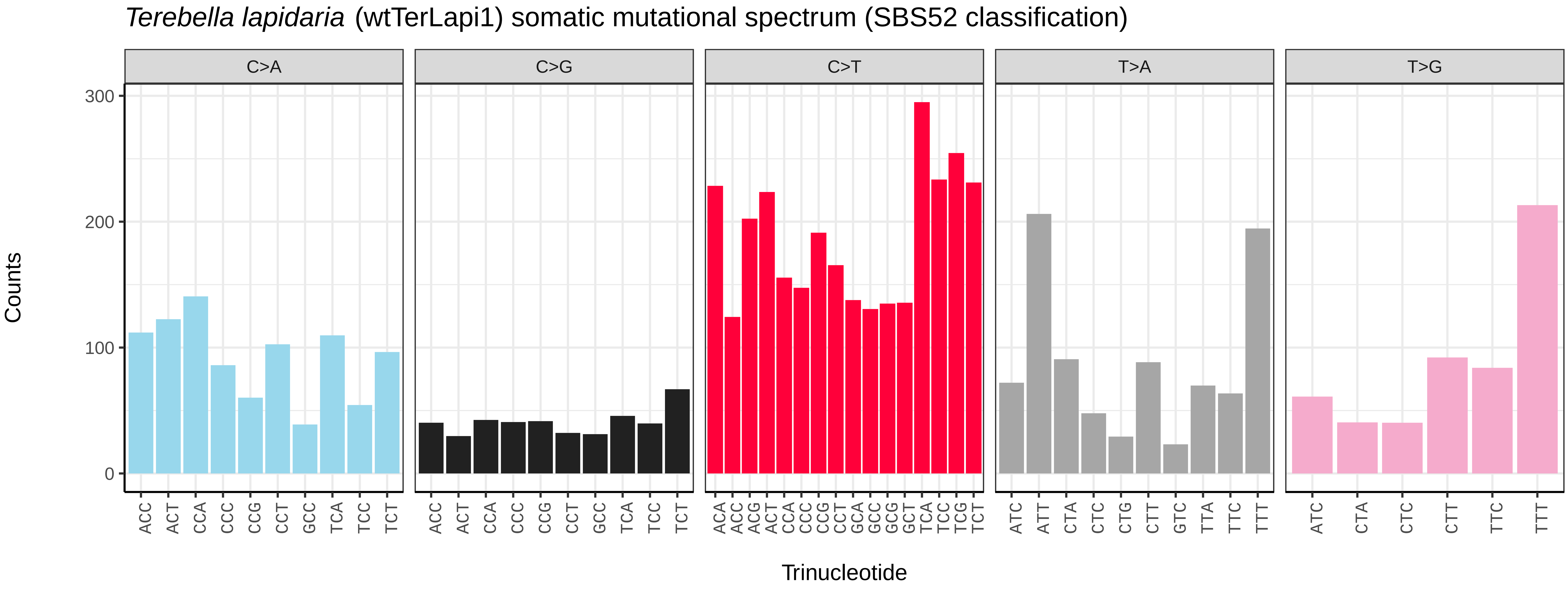
*Terebella lapidaria* (wtTerLapi1) somatic mutational spectrum (SBS52 classification)
C>A
C>G
C>T
T>A
T>G
Counts
300
200
100
0
Trinucleotide

*Arcuatula senhousia* (xbArcSenh1) germline mutational spectrum

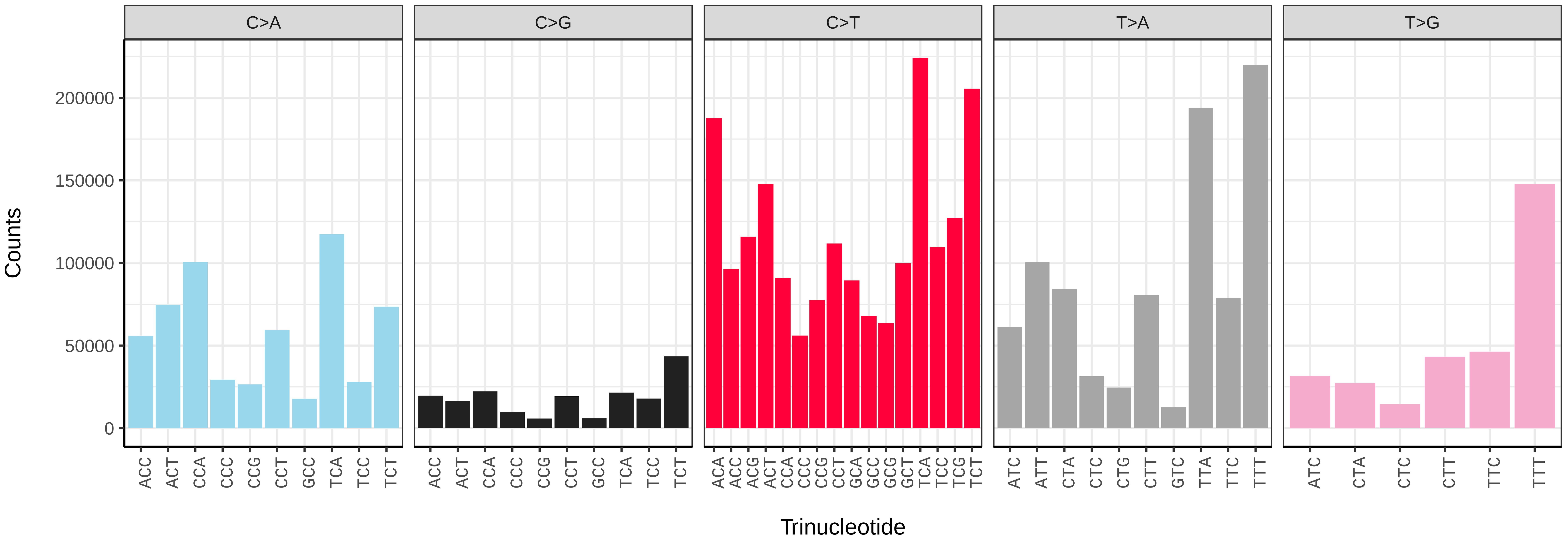


*Arcuatula senhousia* (xbArcSenh1) somatic mutational spectrum (SBS96 classification)

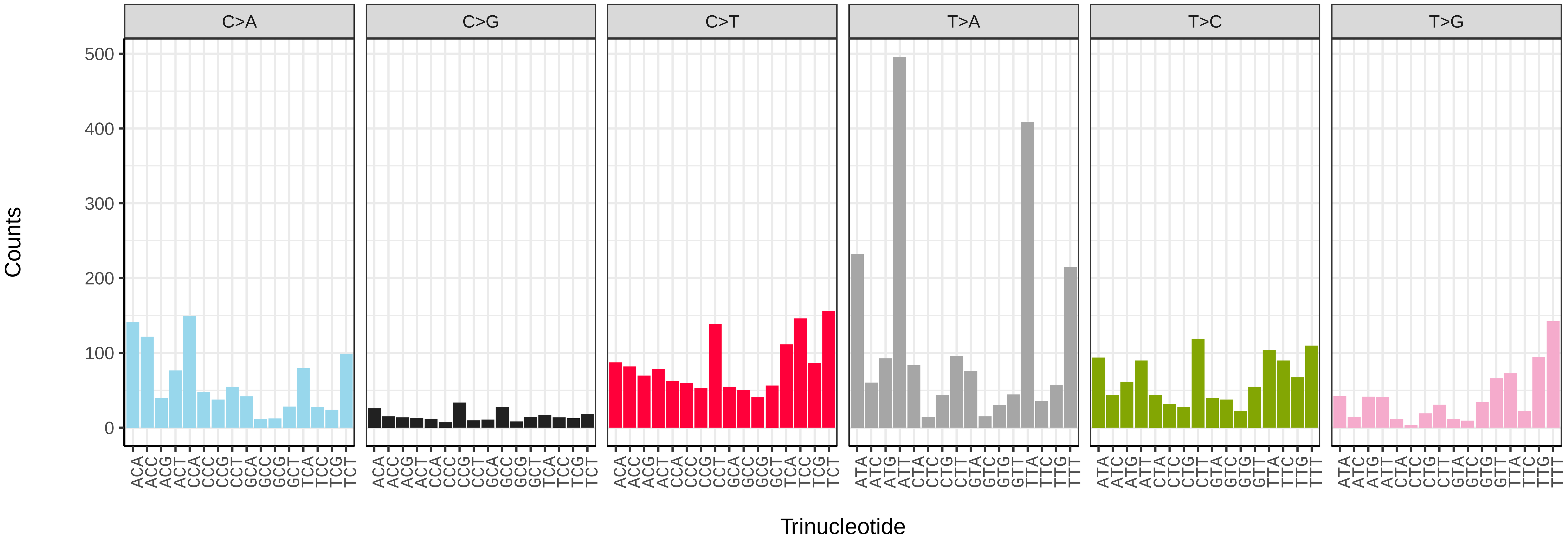


*Arcuatula senhousia* (xbArcSenh1) somatic mutational spectrum (SBS52 classification)

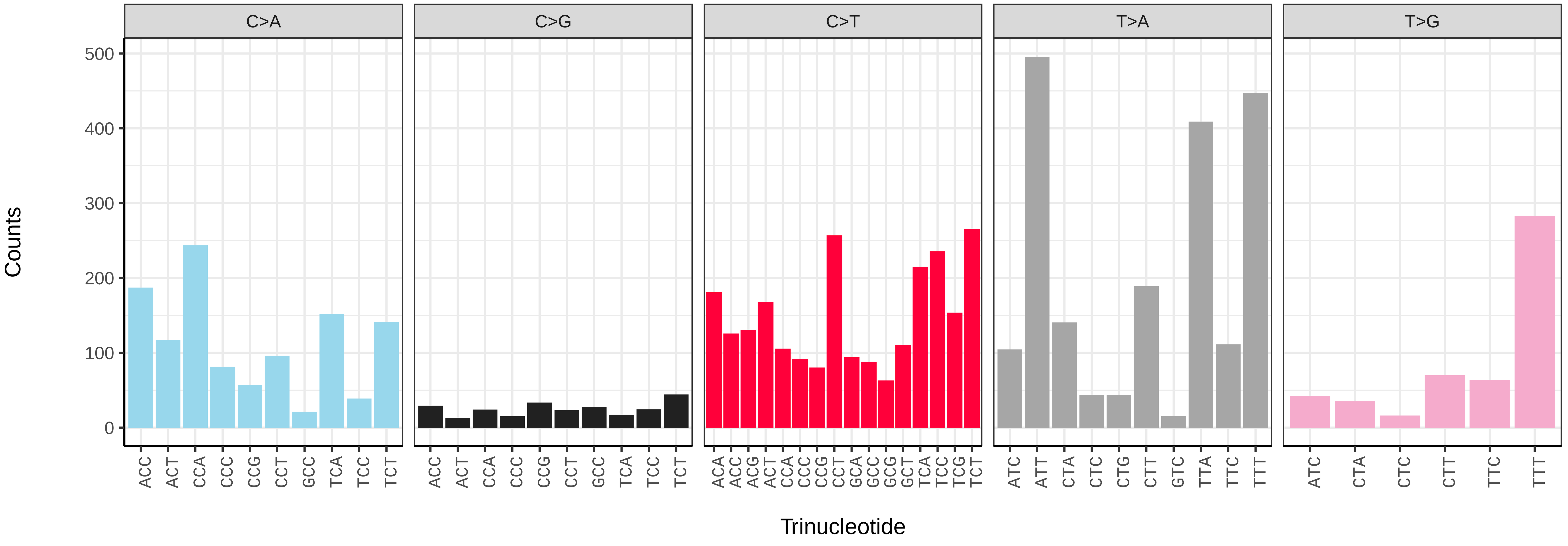

*Azorinus chamasolen* (xbAzoCham1) germline mutational spectrum

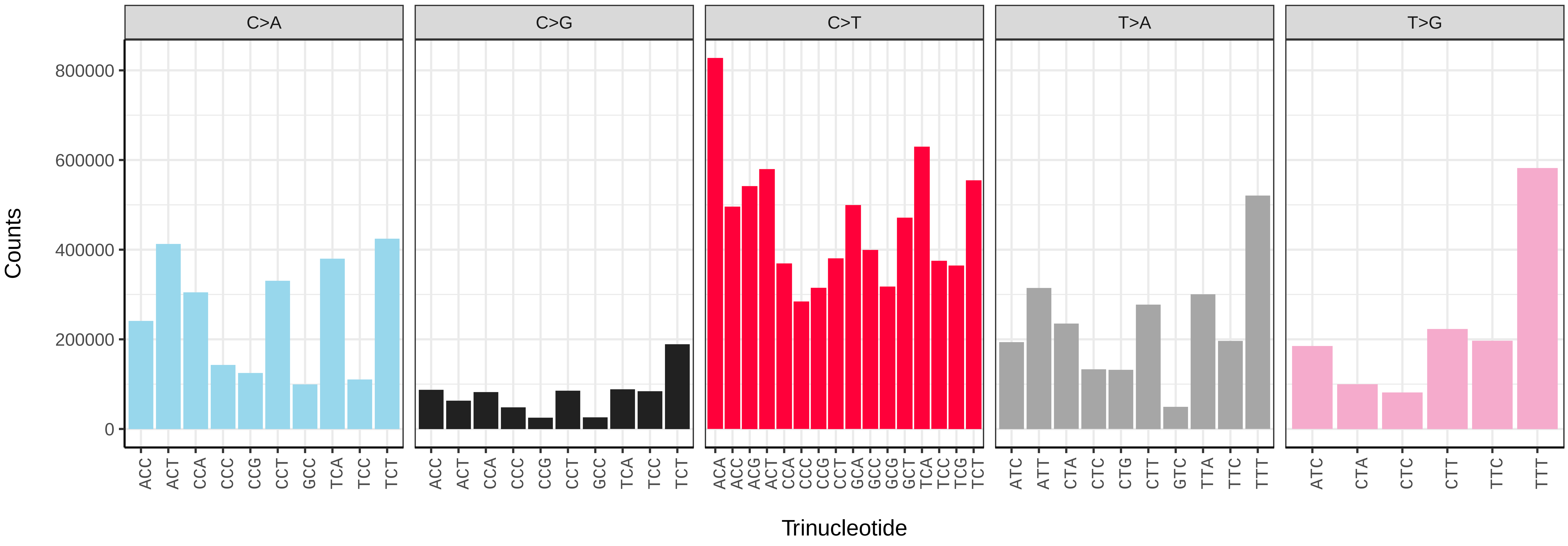


*Azorinus chamasolen* (xbAzoCham1) somatic mutational spectrum (SBS96 classification)

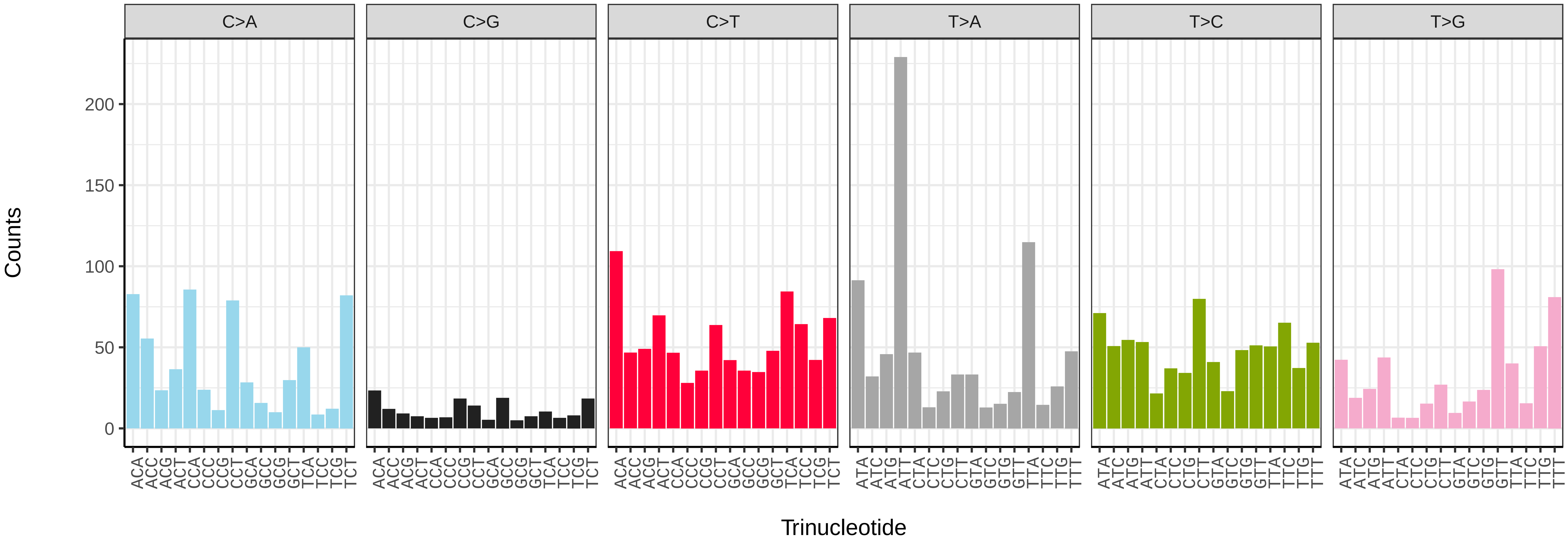


*Azorinus chamasolen* (xbAzoCham1) somatic mutational spectrum (SBS52 classification)

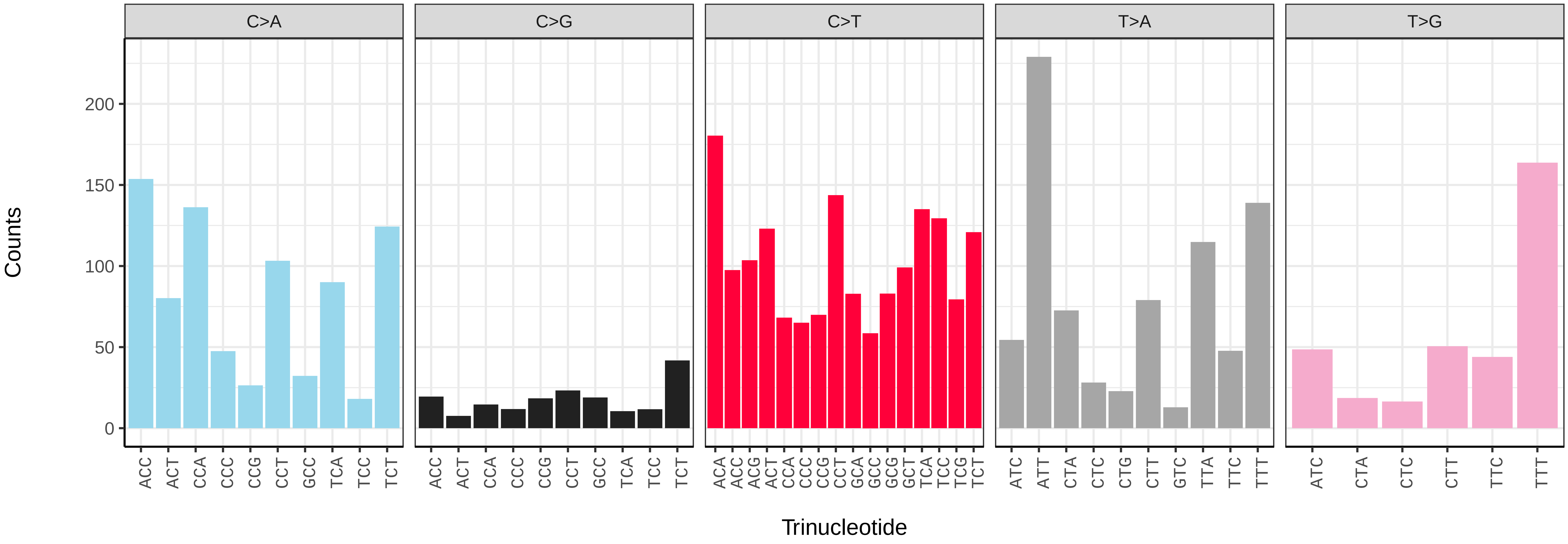

*Cerastoderma edule* (xbCerEdul1) germline mutational spectrum

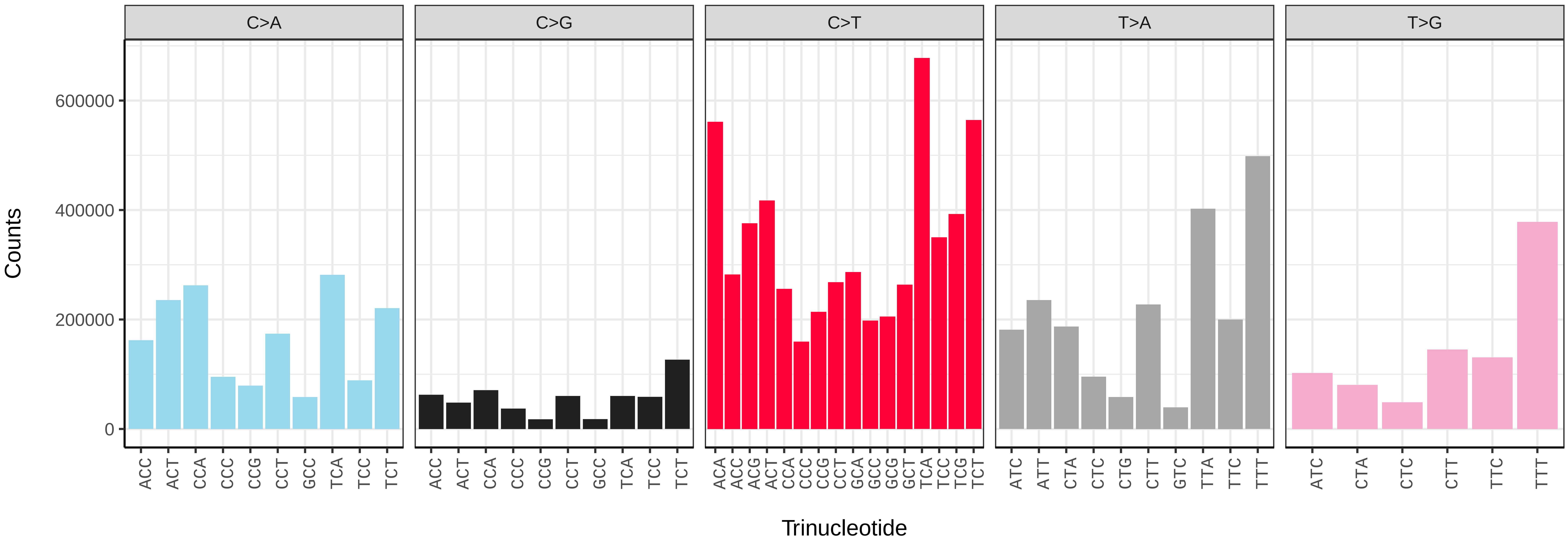


*Cerastoderma edule* (xbCerEdul1) somatic mutational spectrum (SBS96 classification)

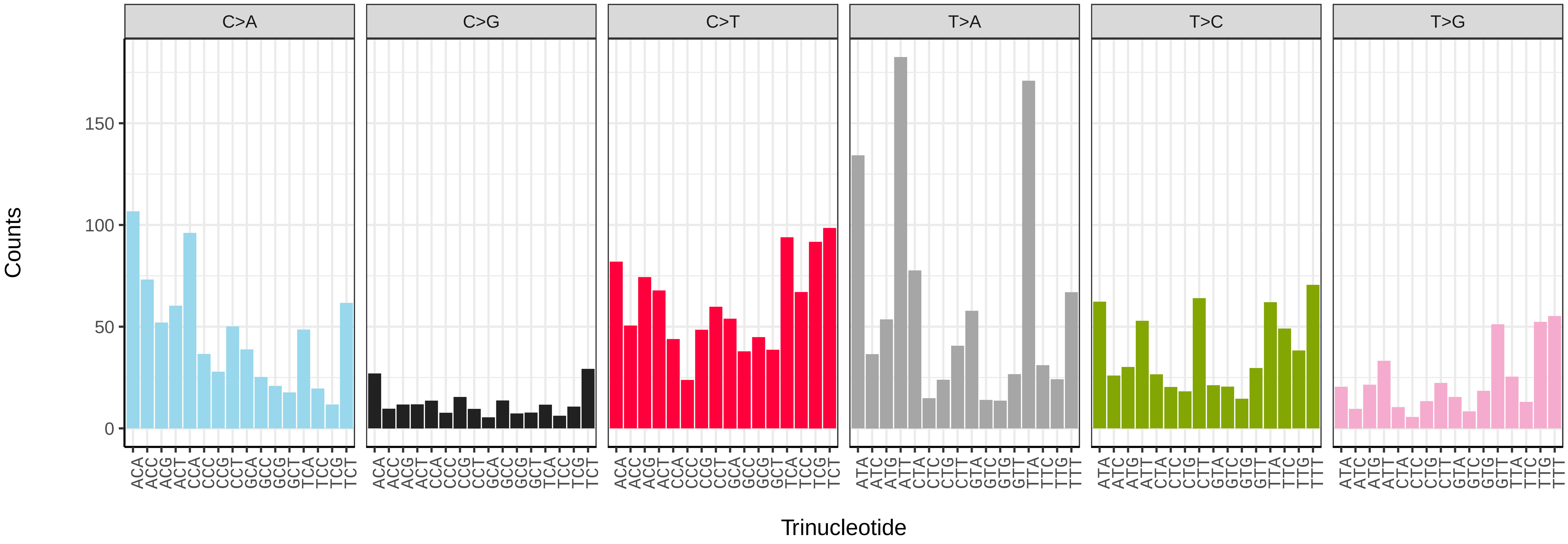


*Cerastoderma edule* (xbCerEdul1) somatic mutational spectrum (SBS52 classification)

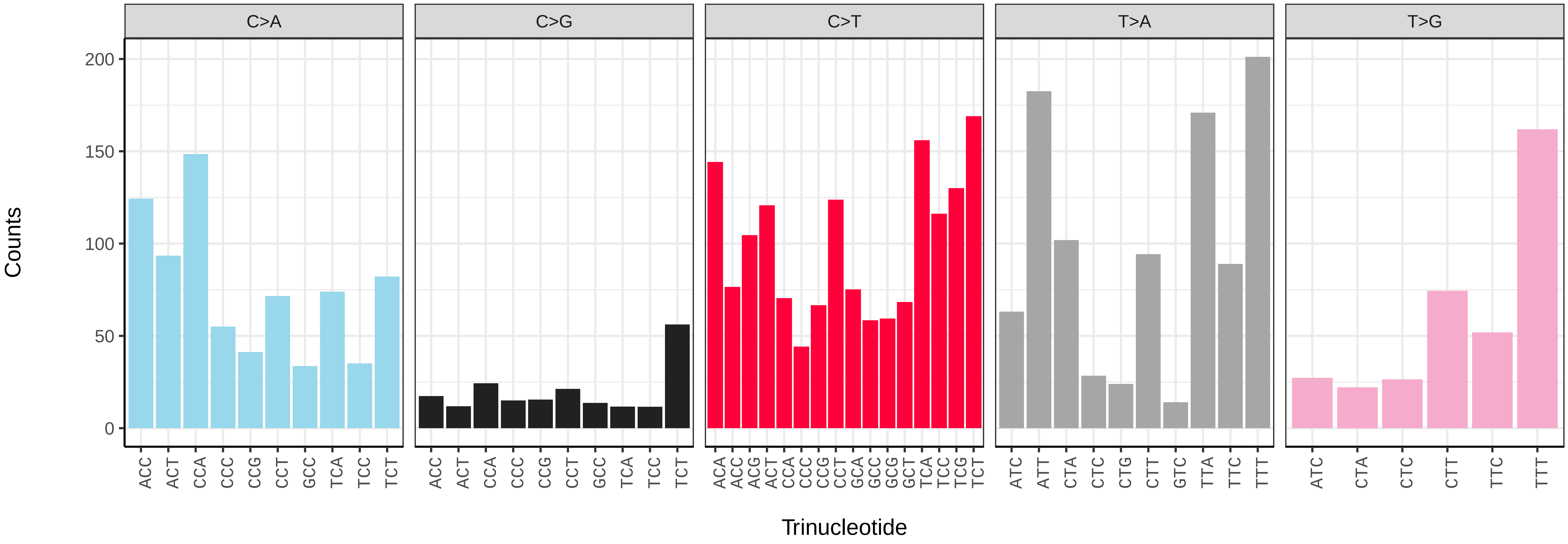

*Gari tellinella* (xbGarTell2) germline mutational spectrum

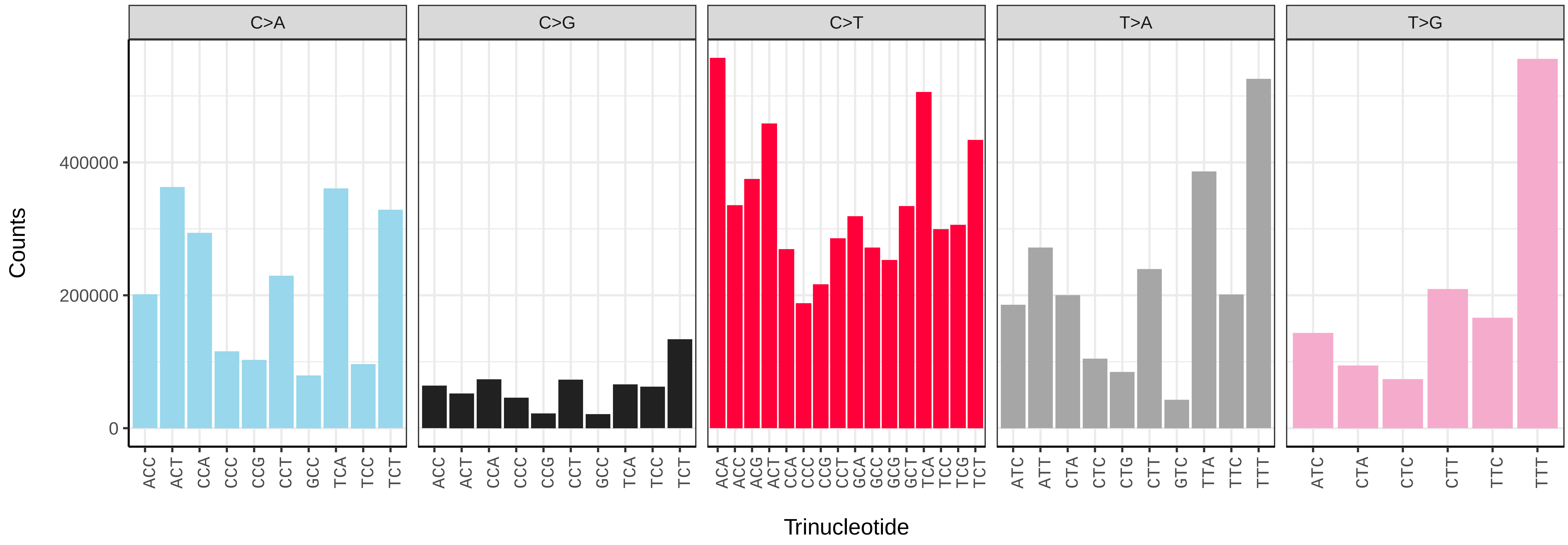


*Gari tellinella* (xbGarTell2) somatic mutational spectrum (SBS96 classification)

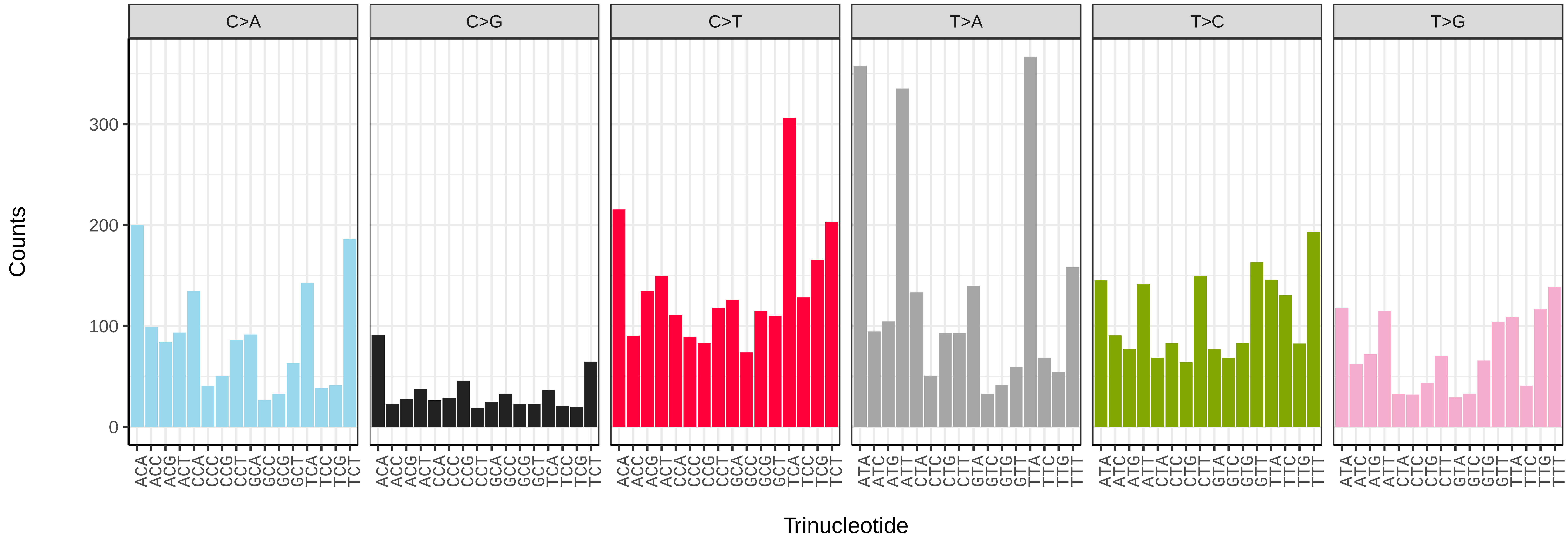


*Gari tellinella* (xbGarTell2) somatic mutational spectrum (SBS52 classification)

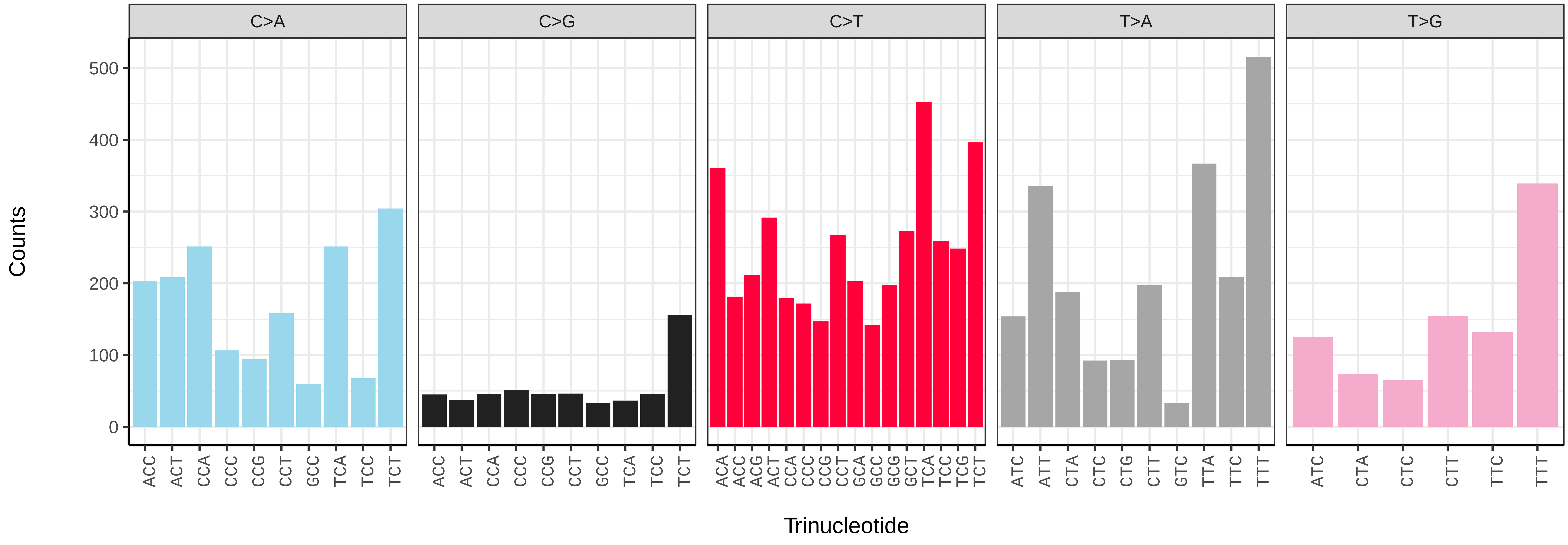

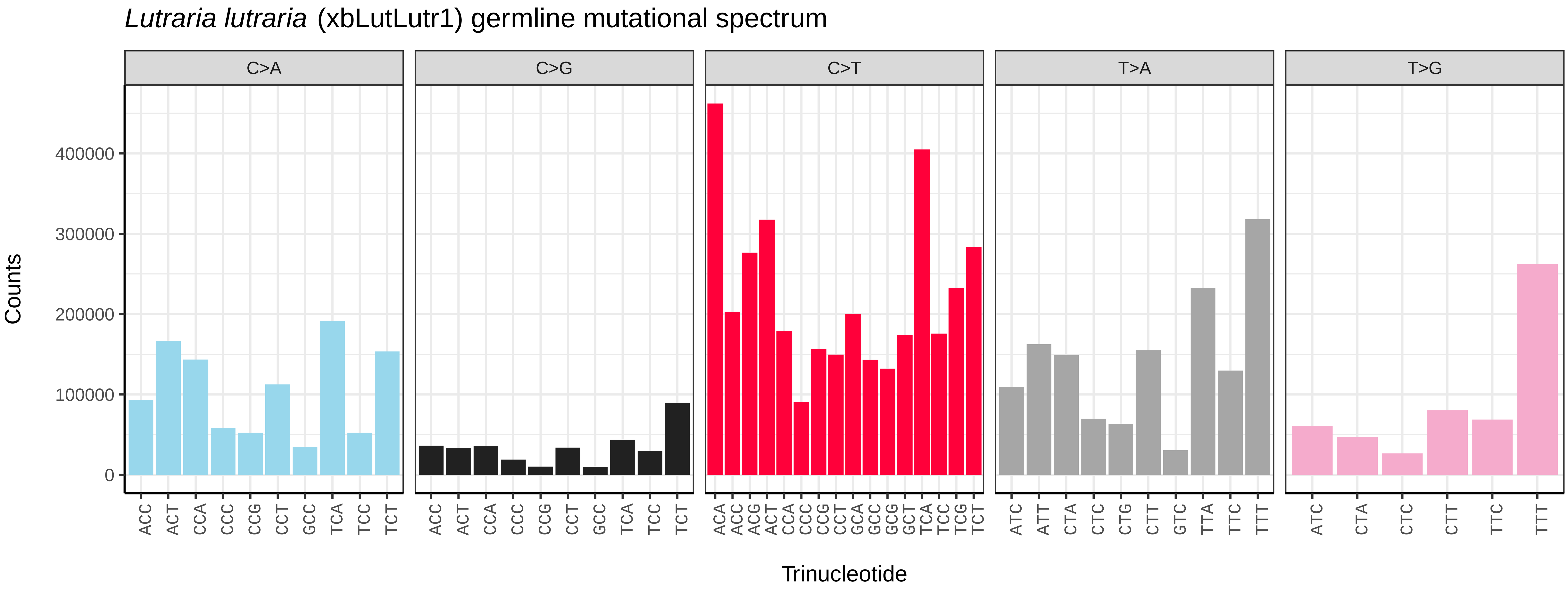
*Lutraria lutraria* (xbLutLutr1) germline mutational spectrum
C>A
C>G
C>T
T>A
T>G
Counts
Trinucleotide

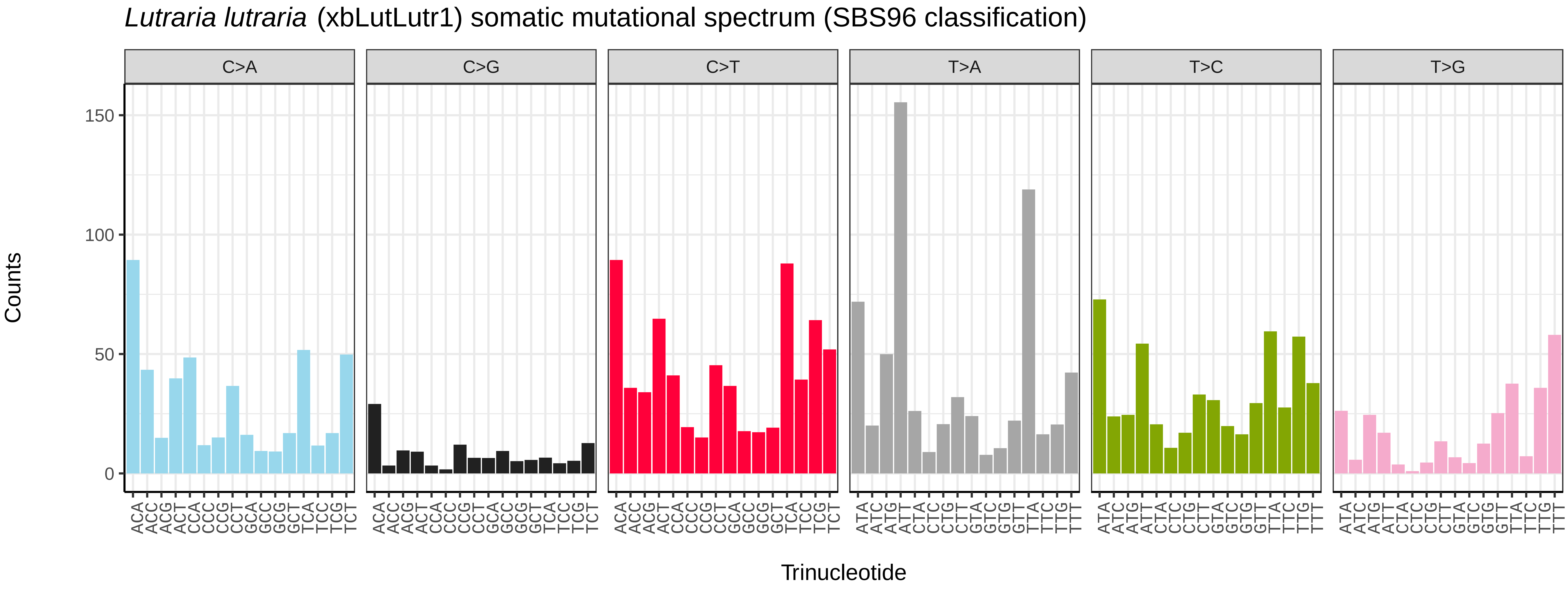
*Lutraria lutraria* (xbLutLutr1) somatic mutational spectrum (SBS96 classification)
C>A
C>G
C>T
T>A
T>C
T>G
Counts
Trinucleotide

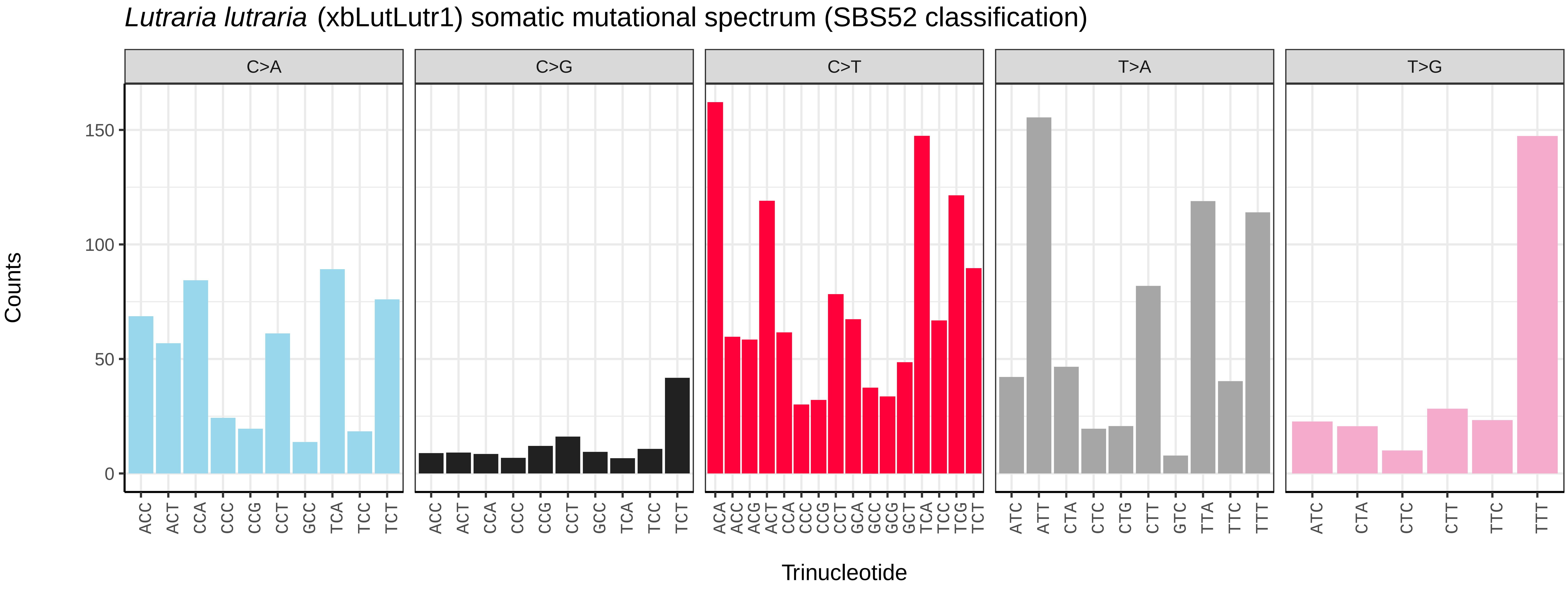
*Lutraria lutraria* (xbLutLutr1) somatic mutational spectrum (SBS52 classification)
C>A
C>G
C>T
T>A
T>G
Counts
Trinucleotide

*Magallana gigas* (xbMagGiga1) germline mutational spectrum

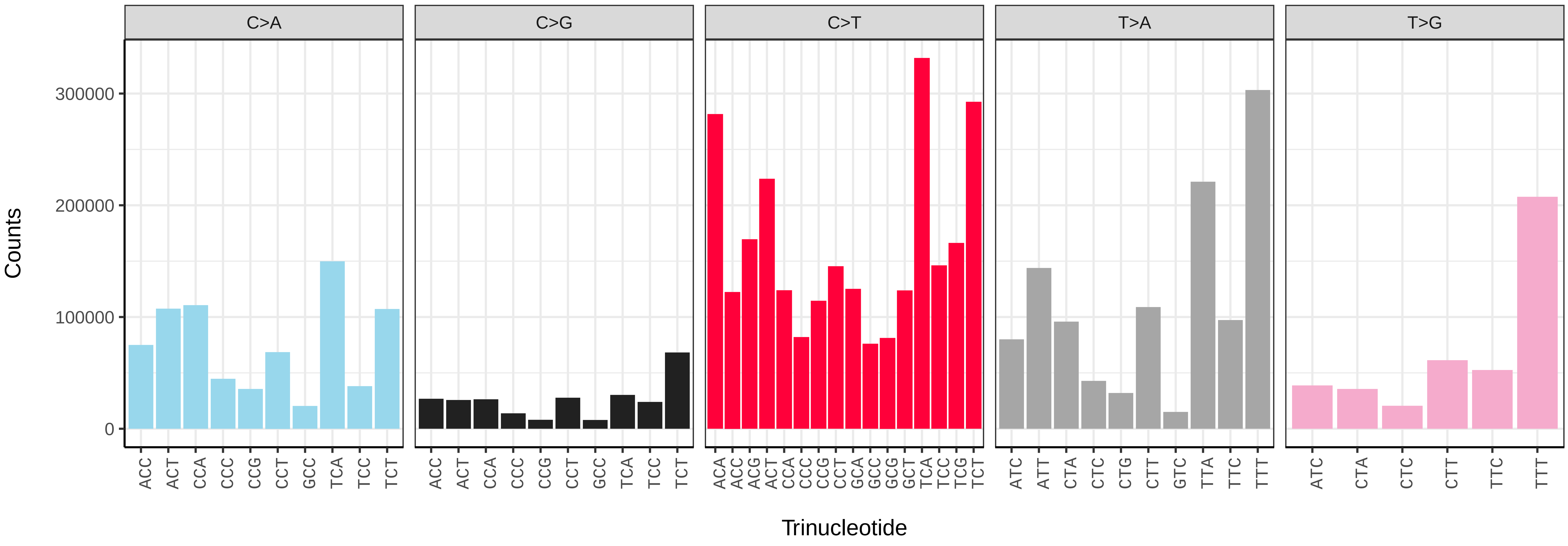


*Magallana gigas* (xbMagGiga1) somatic mutational spectrum (SBS96 classification)

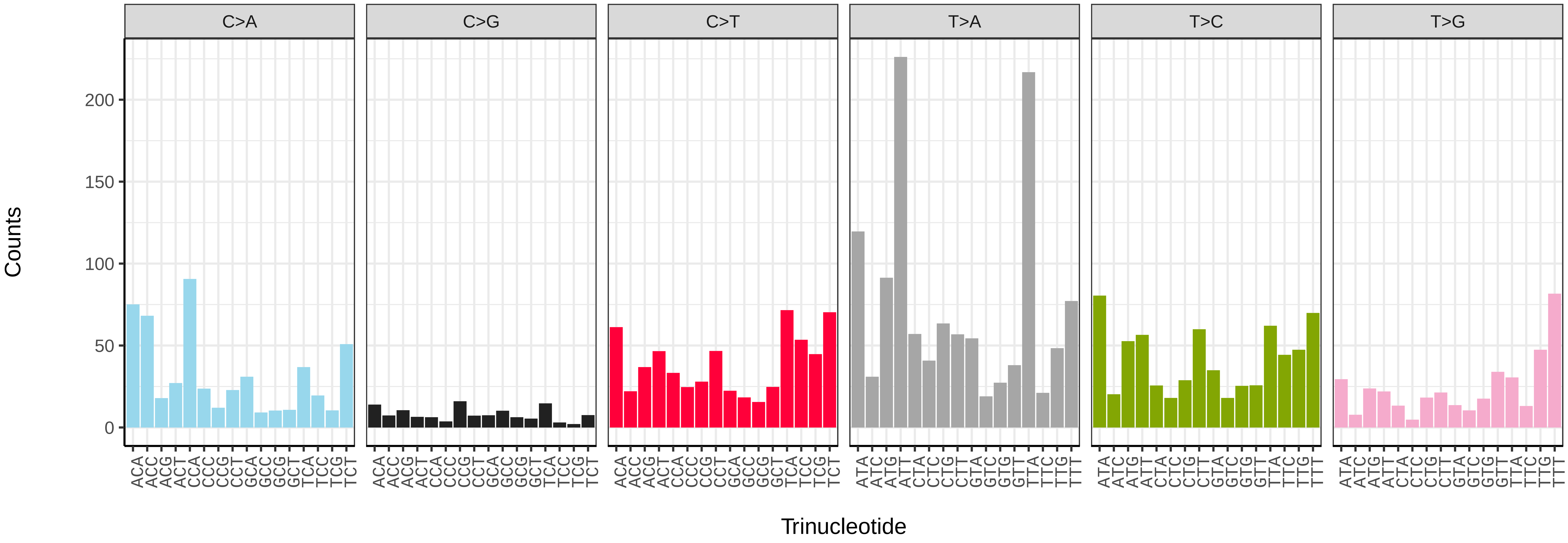


*Magallana gigas* (xbMagGiga1) somatic mutational spectrum (SBS52 classification)

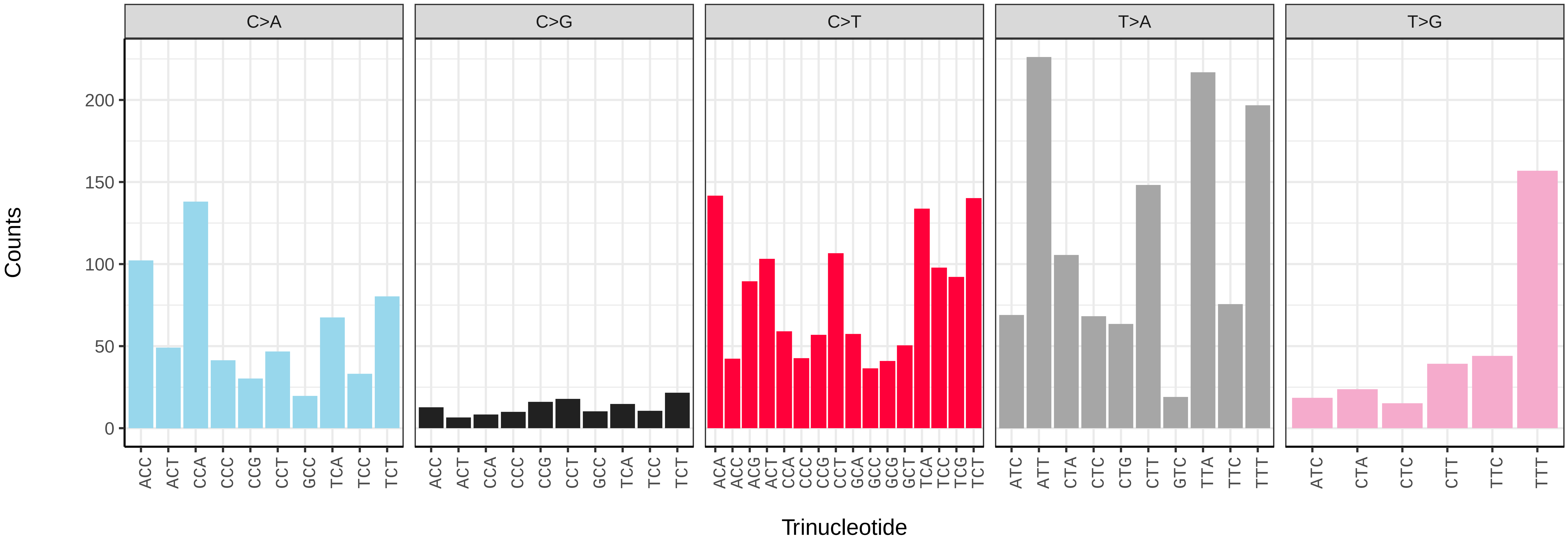

*Mimachlamys varia* (xbMimVari1) germline mutational spectrum

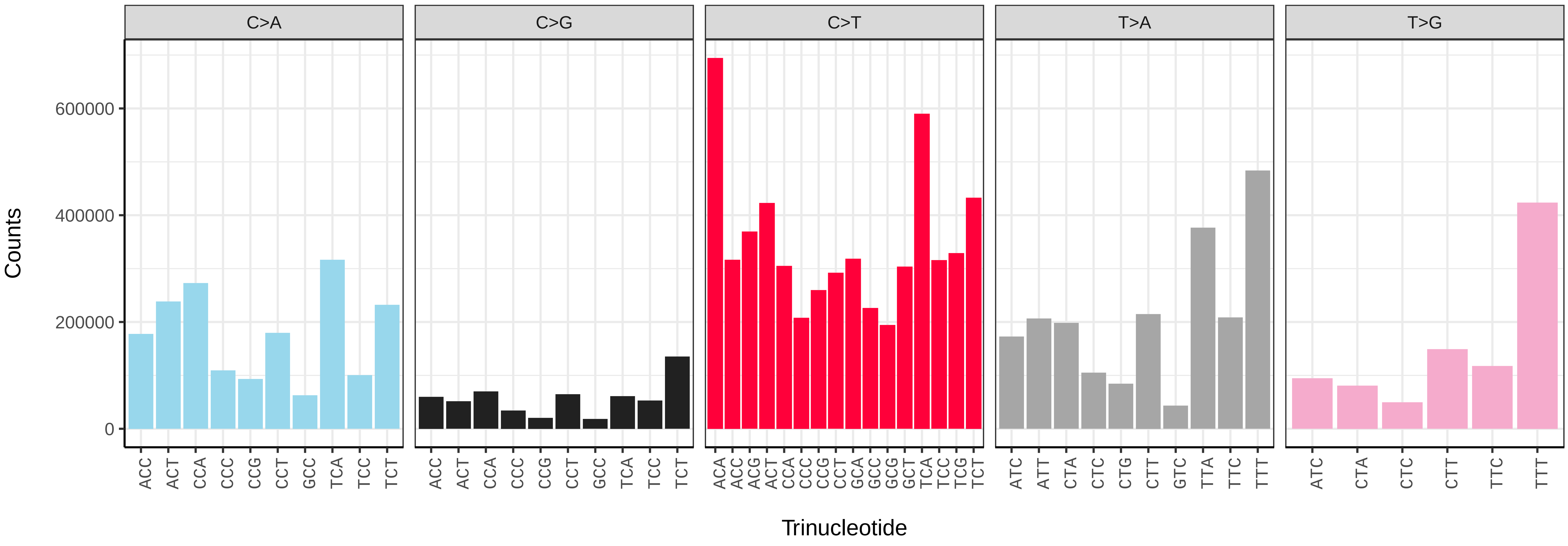


*Mimachlamys varia* (xbMimVari1) somatic mutational spectrum (SBS96 classification)

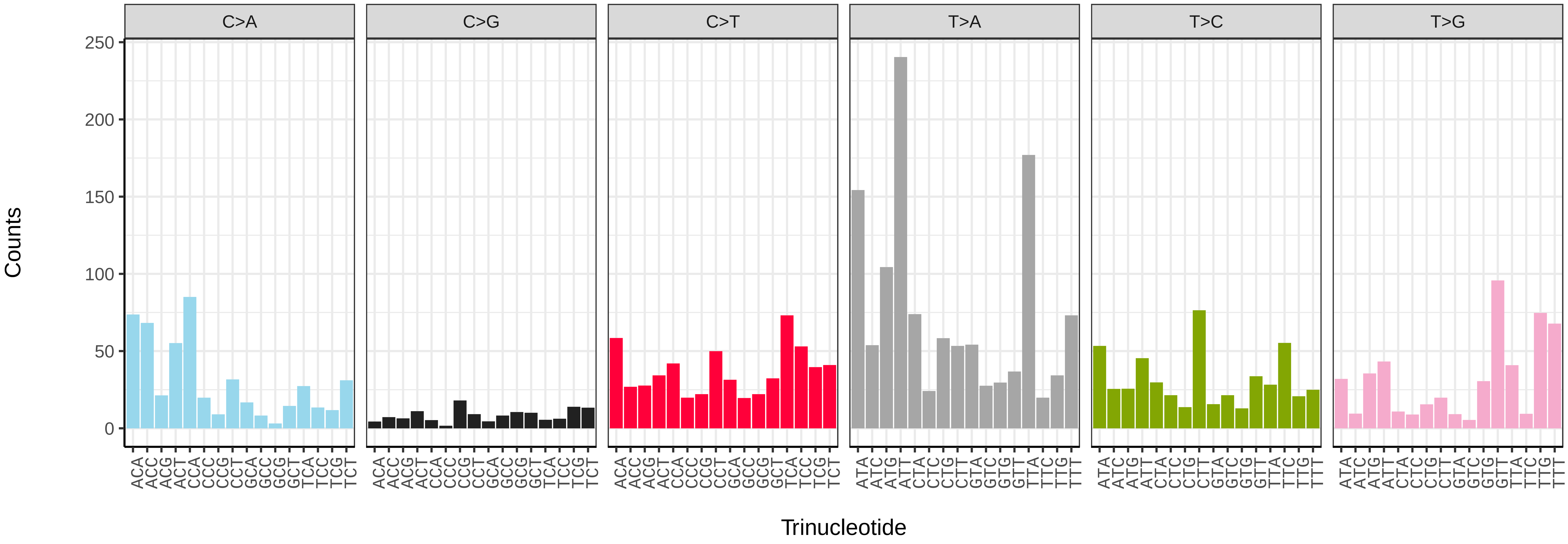


*Mimachlamys varia* (xbMimVari1) somatic mutational spectrum (SBS52 classification)

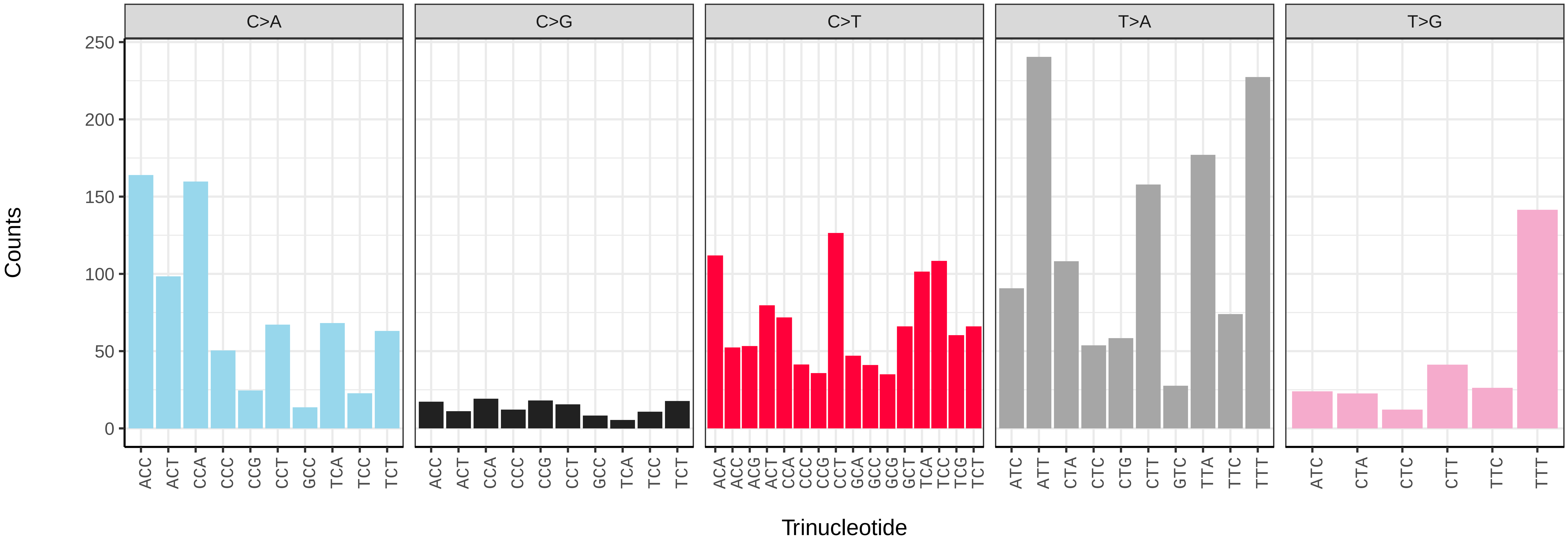

*Mytilus edulis* (xbMytEdul2) germline mutational spectrum

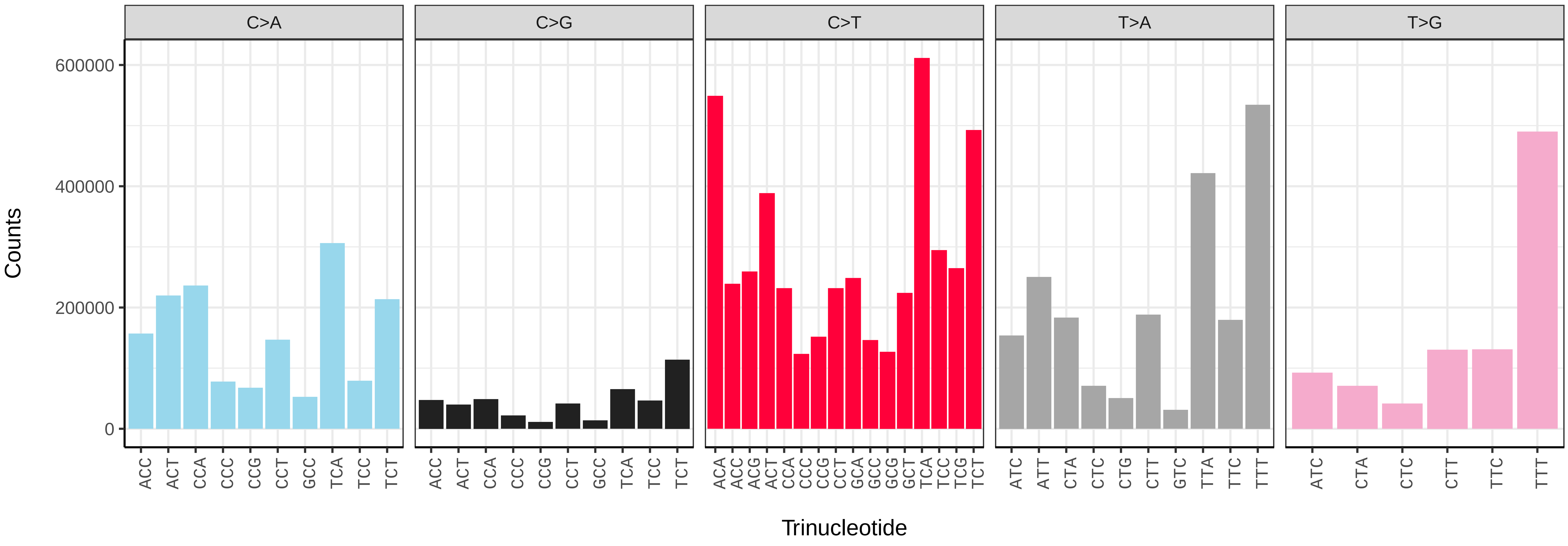


*Mytilus edulis* (xbMytEdul2) somatic mutational spectrum (SBS96 classification)

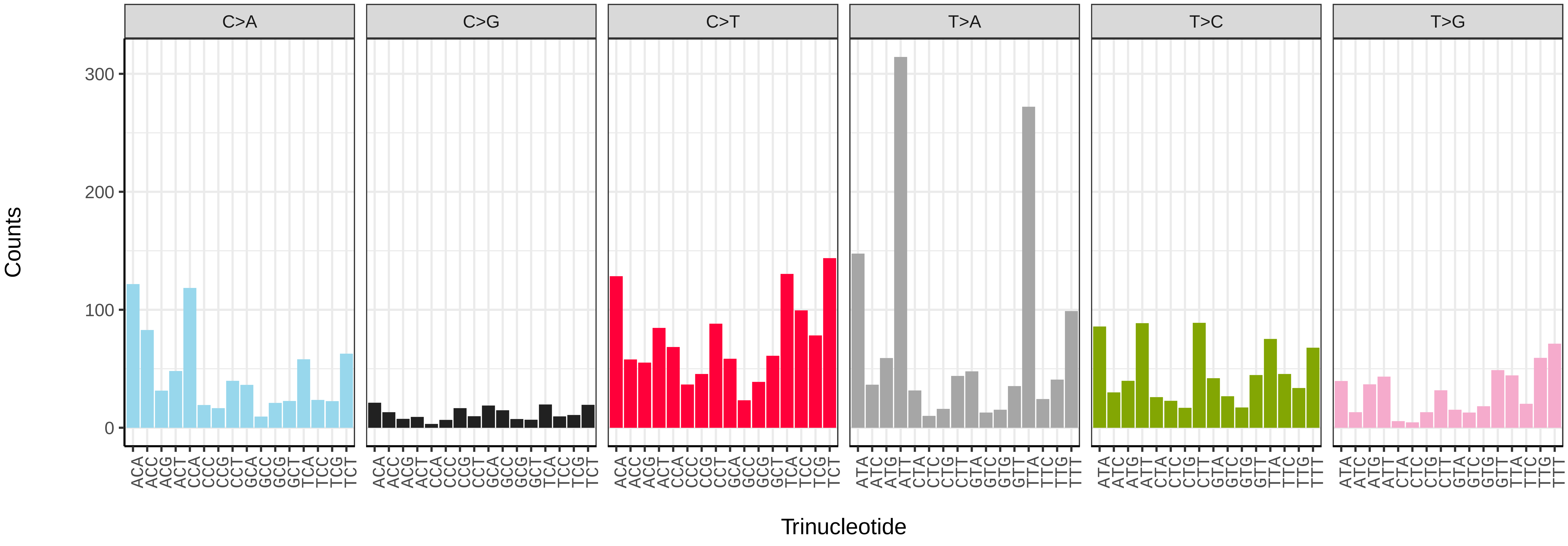


*Mytilus edulis* (xbMytEdul2) somatic mutational spectrum (SBS52 classification)

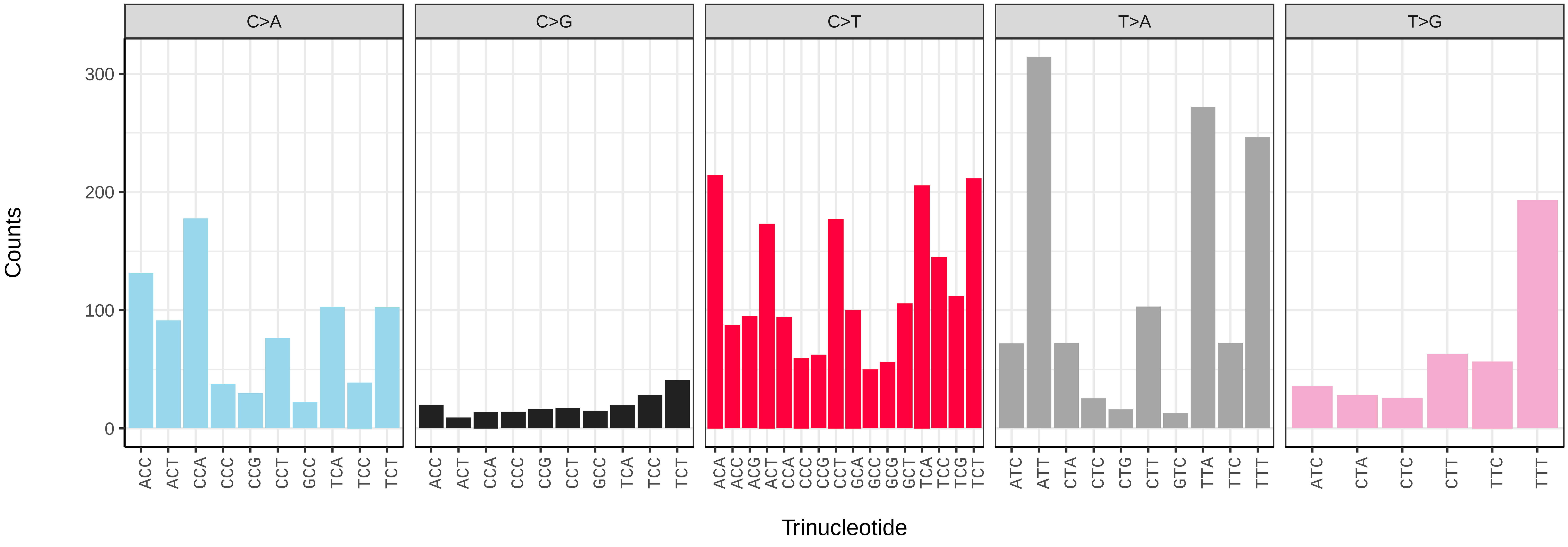

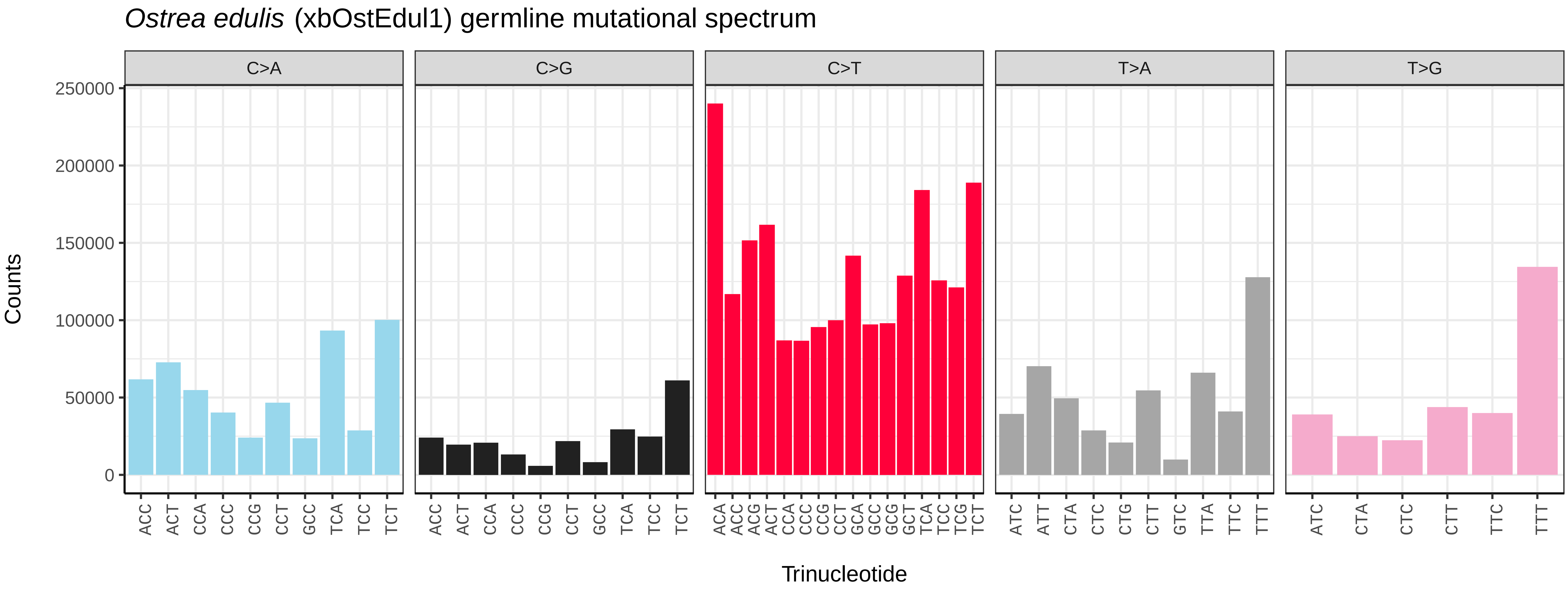
*Ostrea edulis* (xbOstEdul1) germline mutational spectrum
C>A
C>G
C>T
T>A
T>G
Counts
0
50000
100000
150000
200000
250000
Trinucleotide

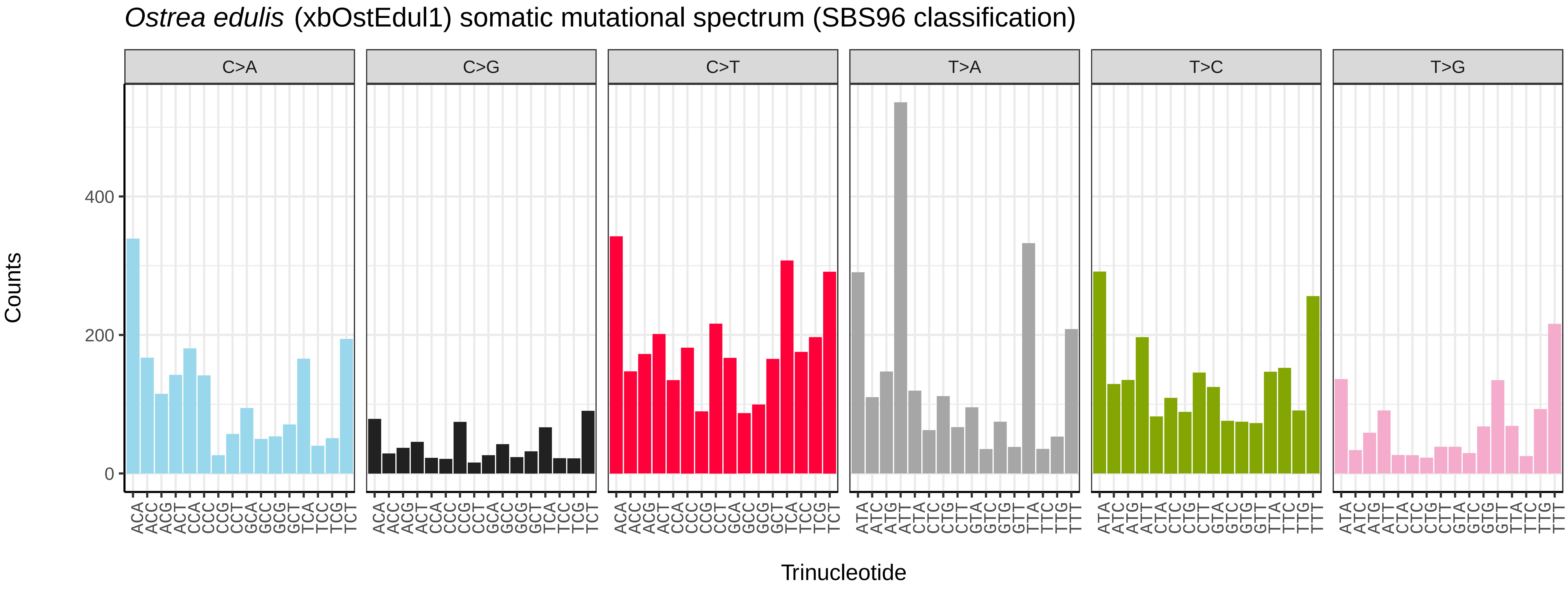
*Ostrea edulis* (xbOstEdul1) somatic mutational spectrum (SBS96 classification)
C>A
C>G
C>T
T>A
T>C
T>G
Counts
0
200
400
Trinucleotide

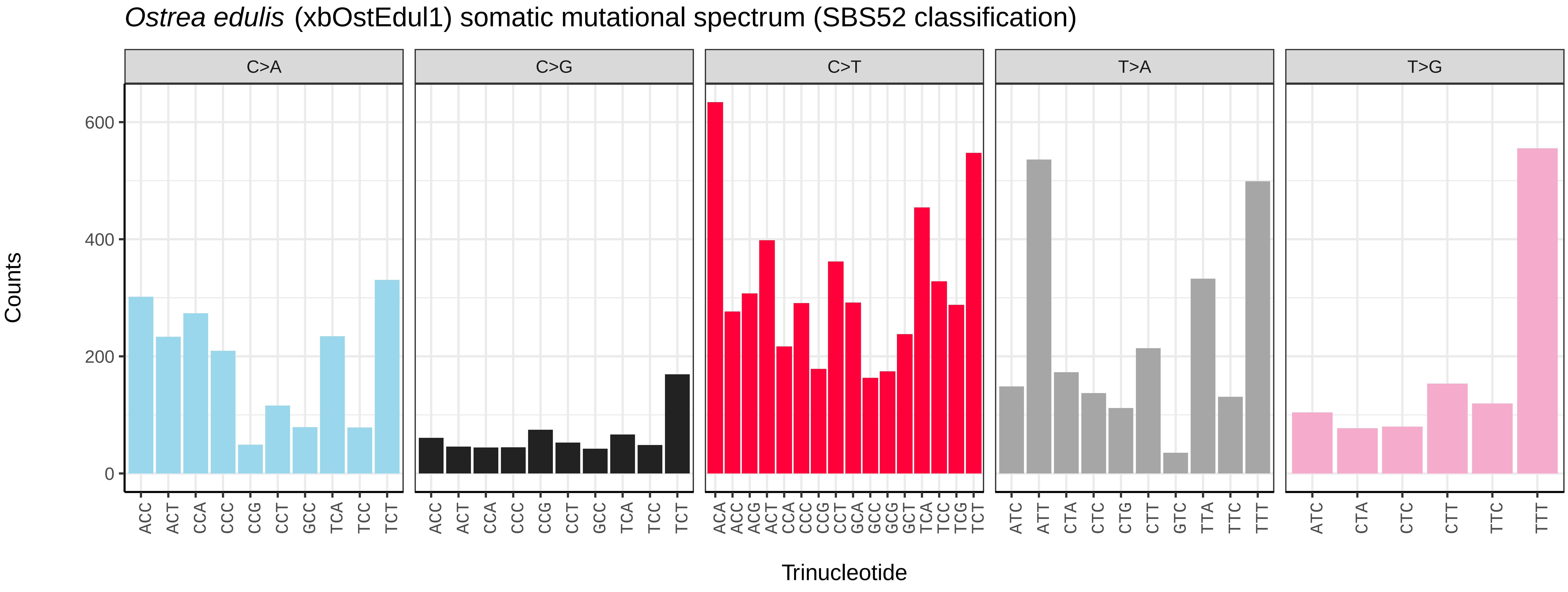
*Ostrea edulis* (xbOstEdul1) somatic mutational spectrum (SBS52 classification)
C>A
C>G
C>T
T>A
T>G
Counts
0
200
400
600
Trinucleotide

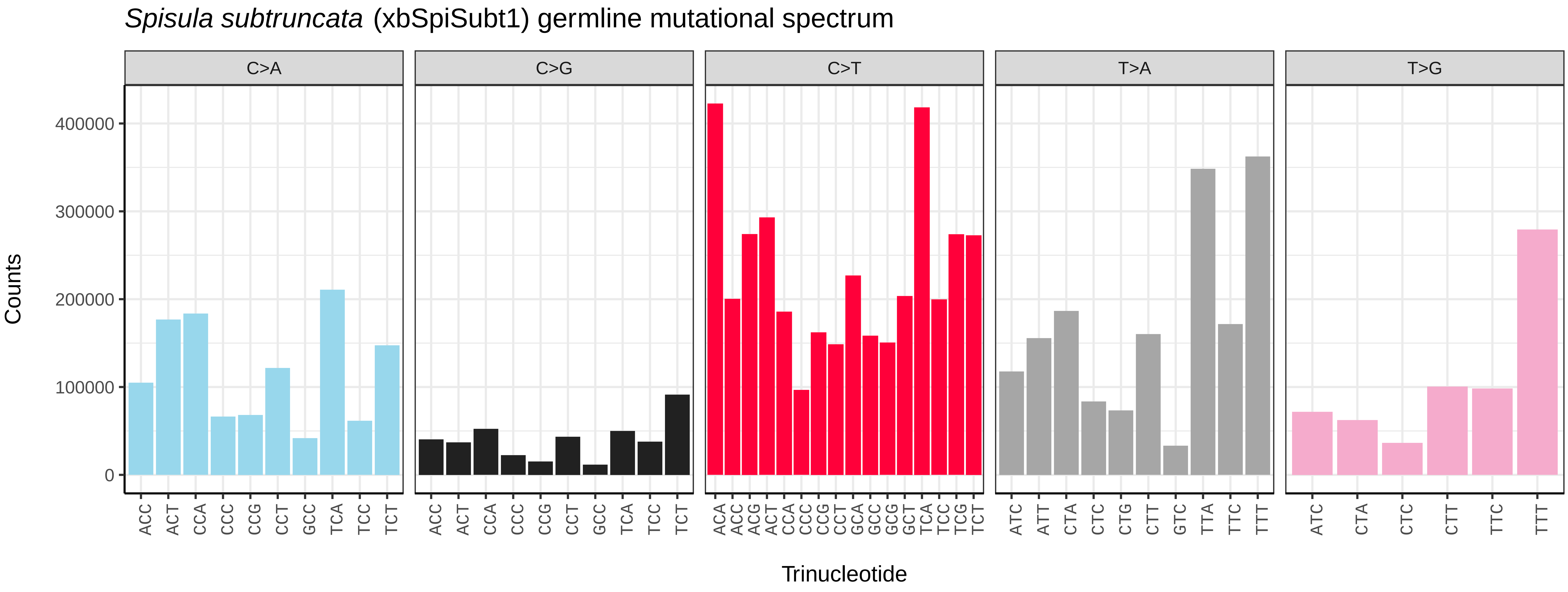
*Spisula subtruncata* (xbSpiSubt1) germline mutational spectrum
C>A
C>G
C>T
T>A
T>G
Counts
400000
300000
200000
100000
0
ACC ACT CCA CCC CCG CCT GCC TCA TCC TCT
ACA ACC ACG ACT CCA CCC CCG CCT GCA GCC GCG GCT TCA TCC TCG TCT
ATC ATT CTA CTC CTG CTT GTC TTA TTC TTT
ATC CTA CTC CTT TTC TTT
Trinucleotide

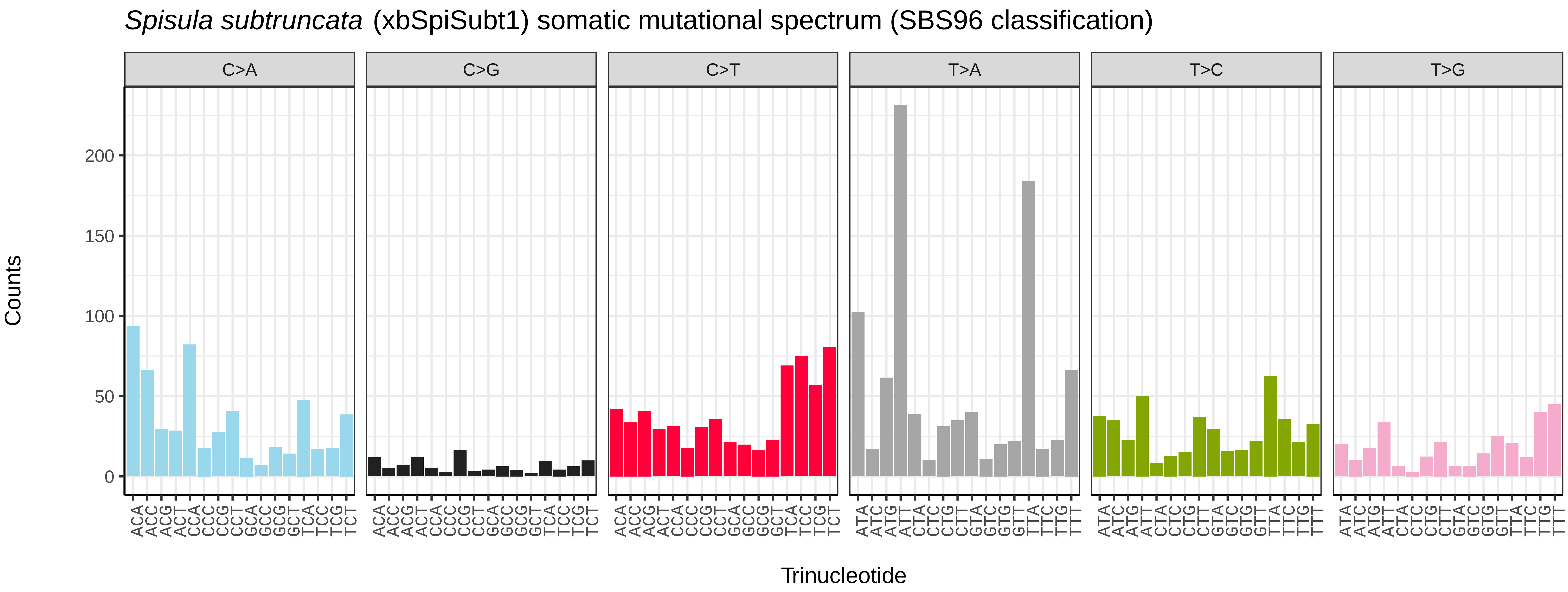
*Spisula subtruncata* (xbSpiSubt1) somatic mutational spectrum (SBS96 classification)
C>A
C>G
C>T
T>A
T>C
T>G
Counts
200
150
100
50
0
ACA ACC ACG ACT CCA CCC CCG CCT GCA GCC GCG GCT TCA TCC TCG TCT
ATA ATC ATG ATT CTA CTC CTG CTT GTA GTC GTG GTT TTA TTC TTG TTT
Trinucleotide

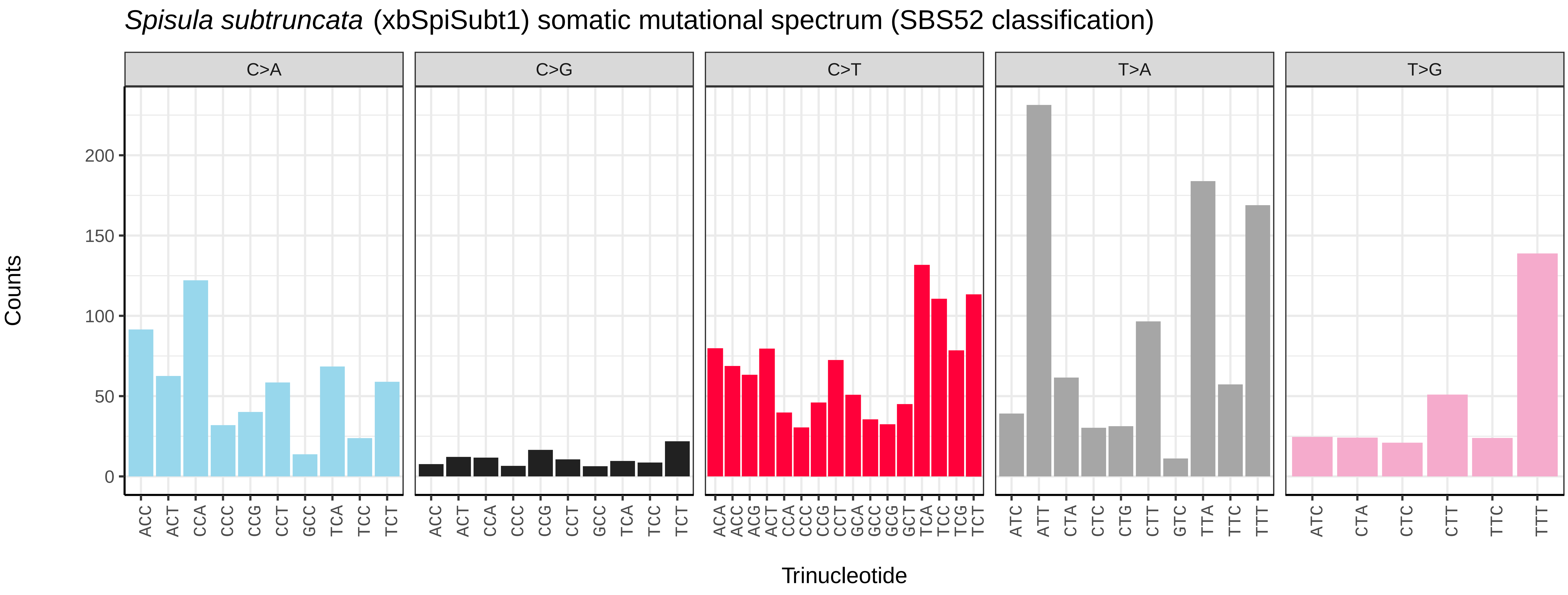
*Spisula subtruncata* (xbSpiSubt1) somatic mutational spectrum (SBS52 classification)
C>A
C>G
C>T
T>A
T>G
Counts
200
150
100
50
0
ACC ACT CCA CCC CCG CCT GCC TCA TCC TCT
ACA ACC ACG ACT CCA CCC CCG CCT GCA GCC GCG GCT TCA TCC TCG TCT
ATC ATT CTA CTC CTG CTT GTC TTA TTC TTT
ATC CTA CTC CTT TTC TTT
Trinucleotide

*Anisus vortex* (xgAniVort1) germline mutational spectrum

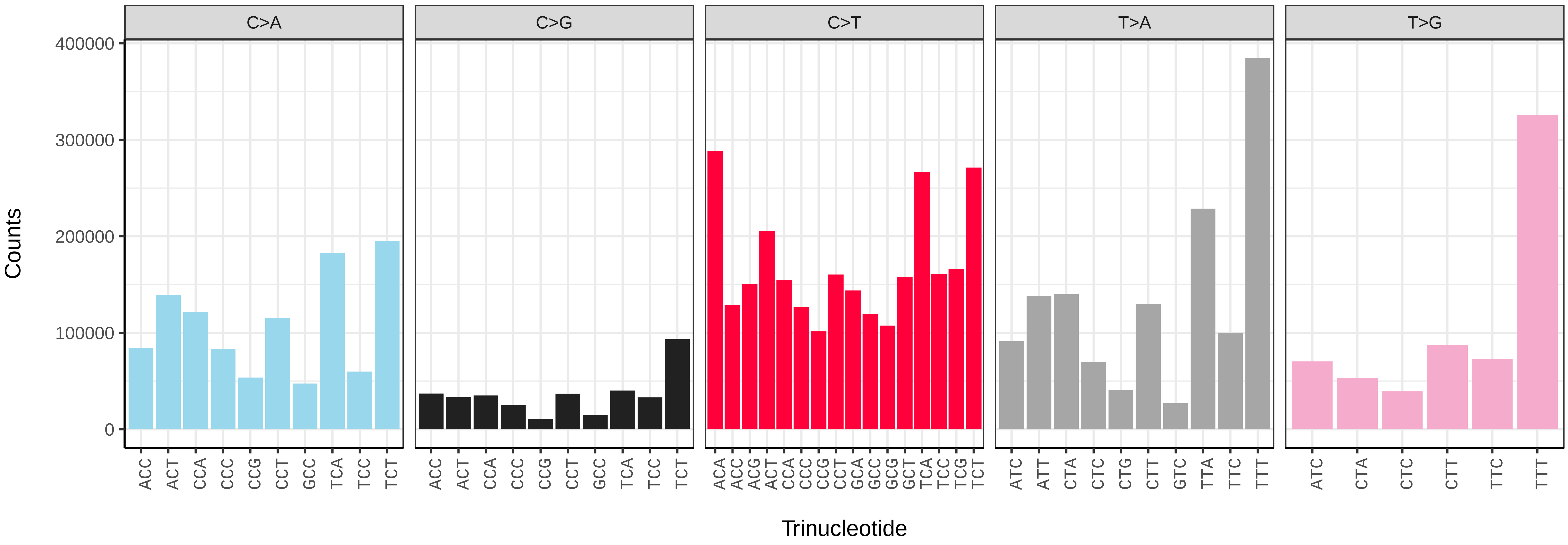


*Anisus vortex* (xgAniVort1) somatic mutational spectrum (SBS96 classification)

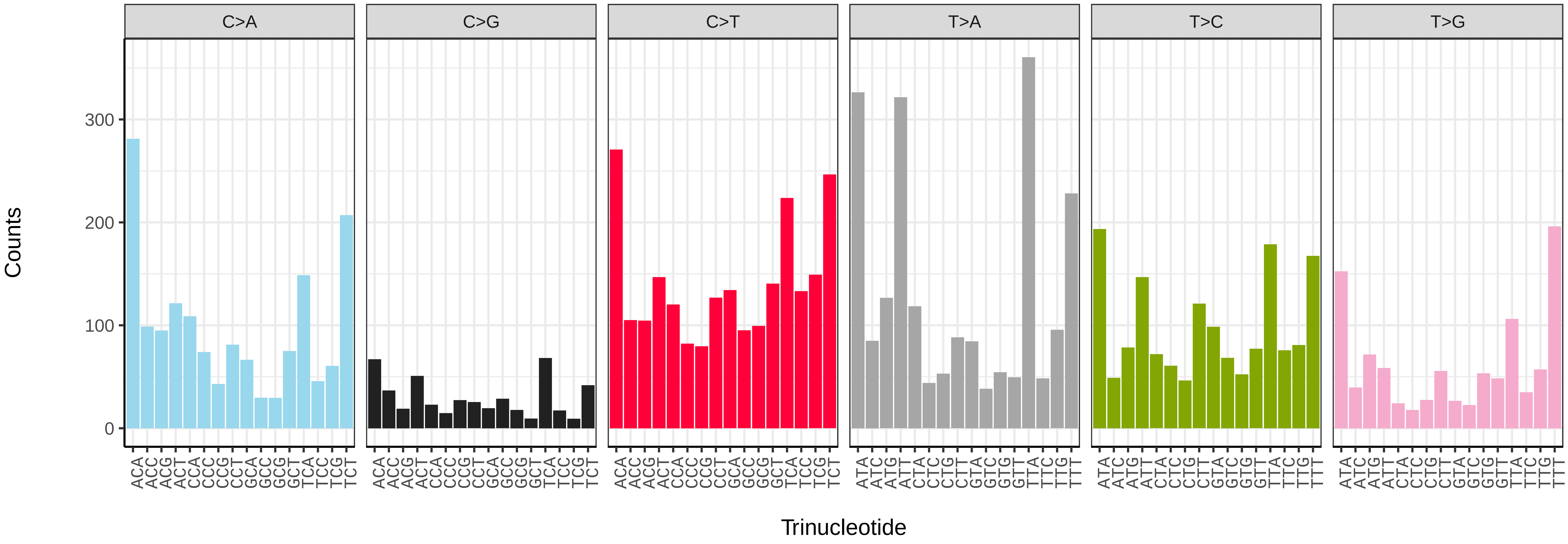


*Anisus vortex* (xgAniVort1) somatic mutational spectrum (SBS52 classification)

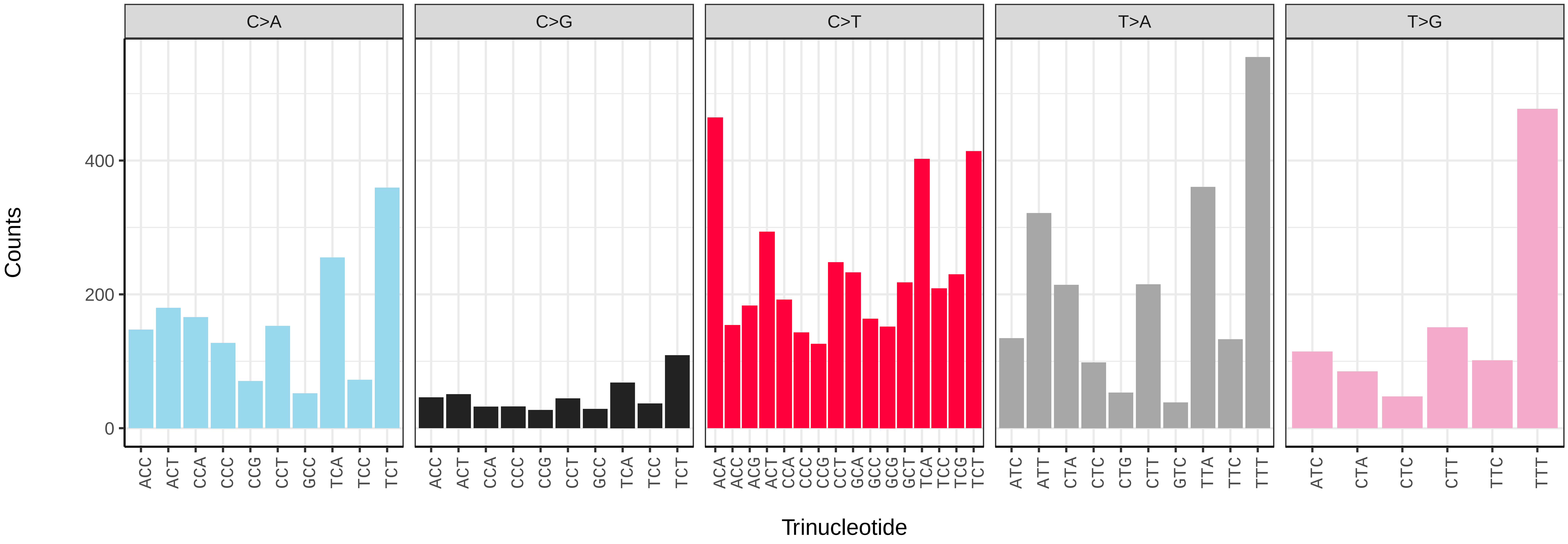

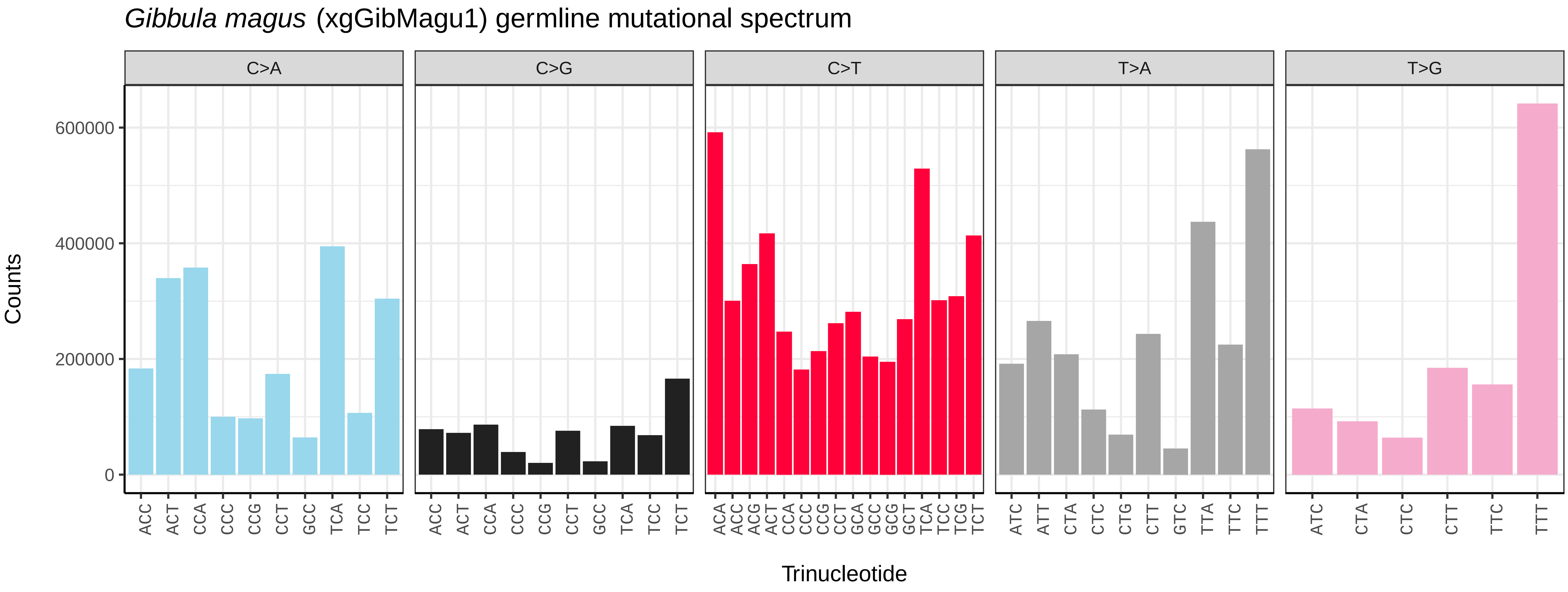
Gibbula magus (xgGibMagu1) germline mutational spectrum
C>A
C>G
C>T
T>A
T>G
Counts
0
200000
400000
600000
Trinucleotide

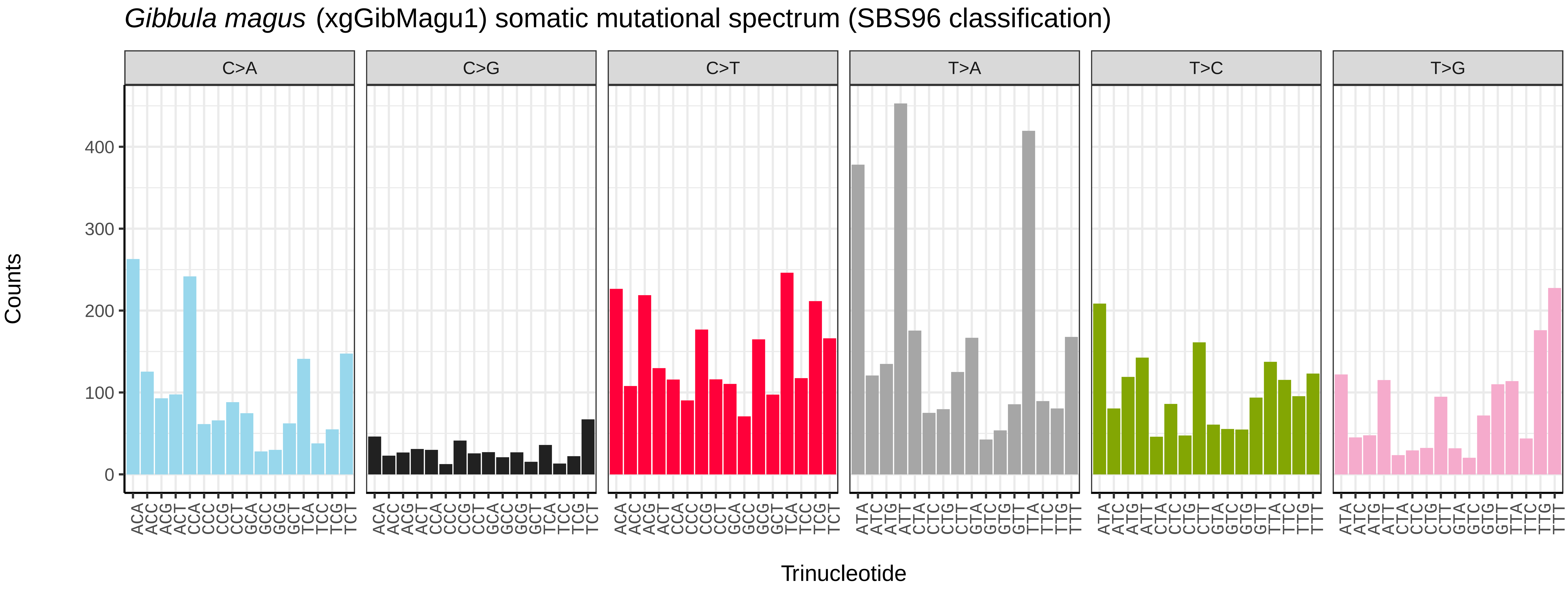
Gibbula magus (xgGibMagu1) somatic mutational spectrum (SBS96 classification)
C>A
C>G
C>T
T>A
T>C
T>G
Counts
0
100
200
300
400
Trinucleotide

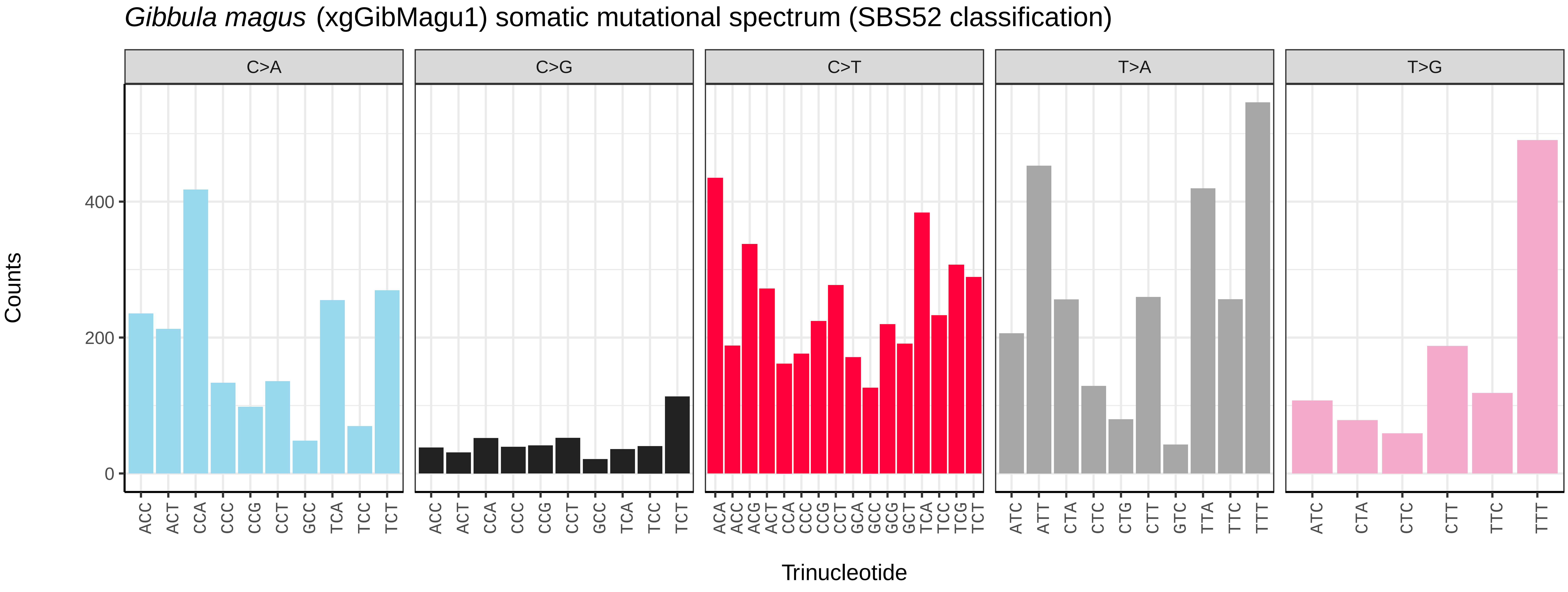
Gibbula magus (xgGibMagu1) somatic mutational spectrum (SBS52 classification)
C>A
C>G
C>T
T>A
T>G
Counts
0
200
400
Trinucleotide

*Onchidella celtica* (xgOncCelt3) germline mutational spectrum

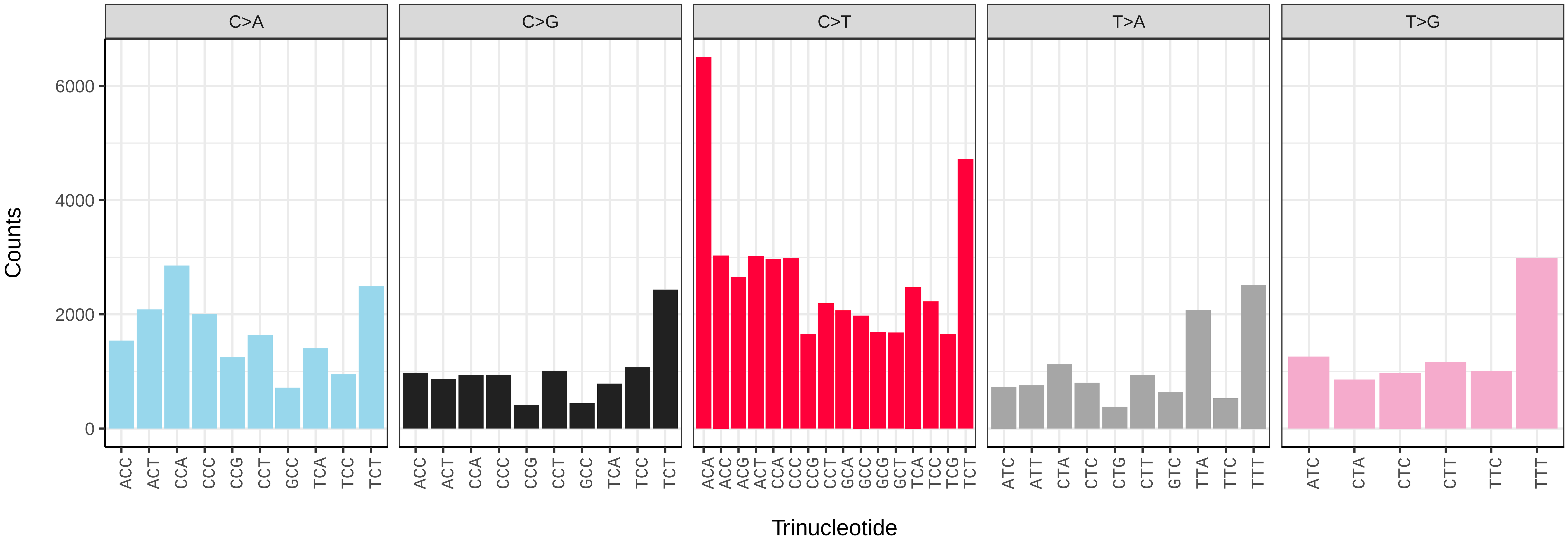


*Onchidella celtica* (xgOncCelt3) somatic mutational spectrum (SBS96 classification)

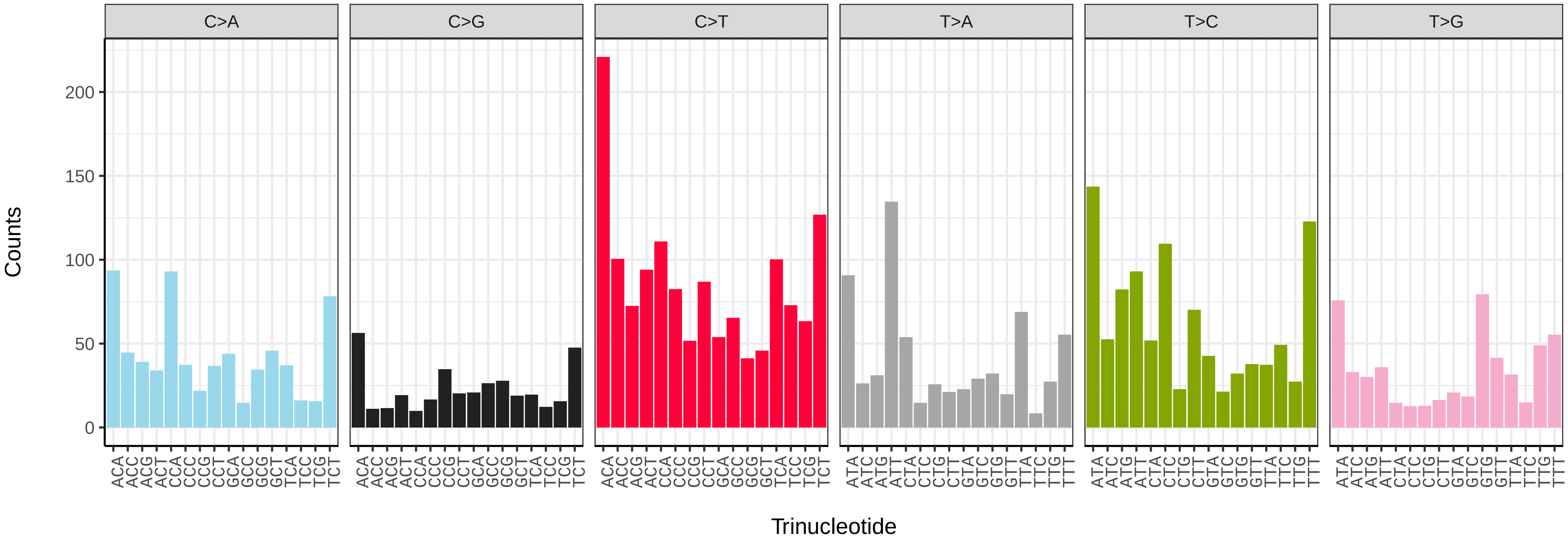


*Onchidella celtica* (xgOncCelt3) somatic mutational spectrum (SBS52 classification)

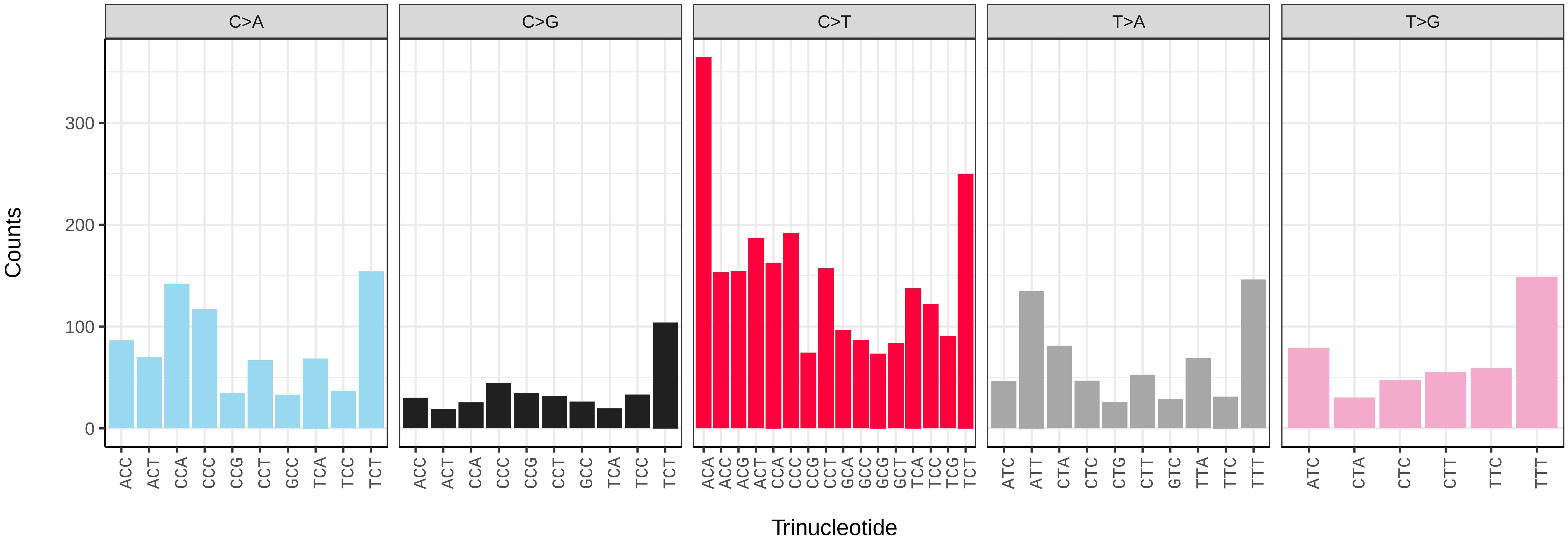

*Patella depressa* (xgPatDepr1) germline mutational spectrum

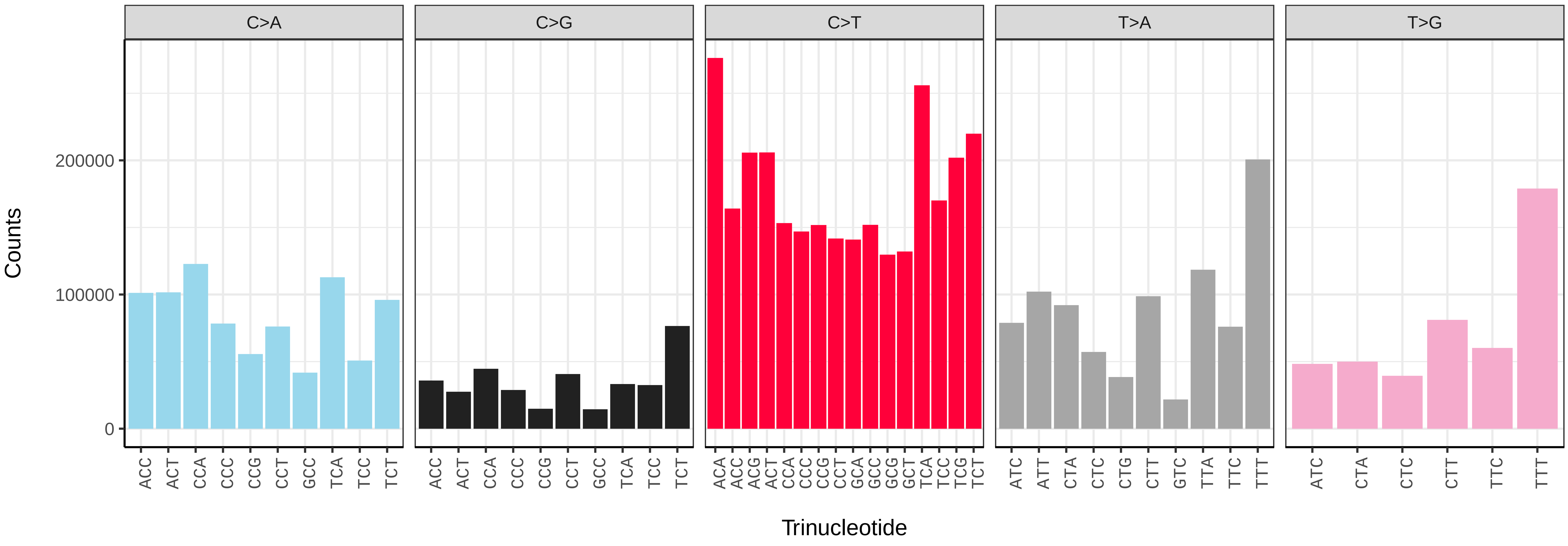


*Patella depressa* (xgPatDepr1) somatic mutational spectrum (SBS96 classification)

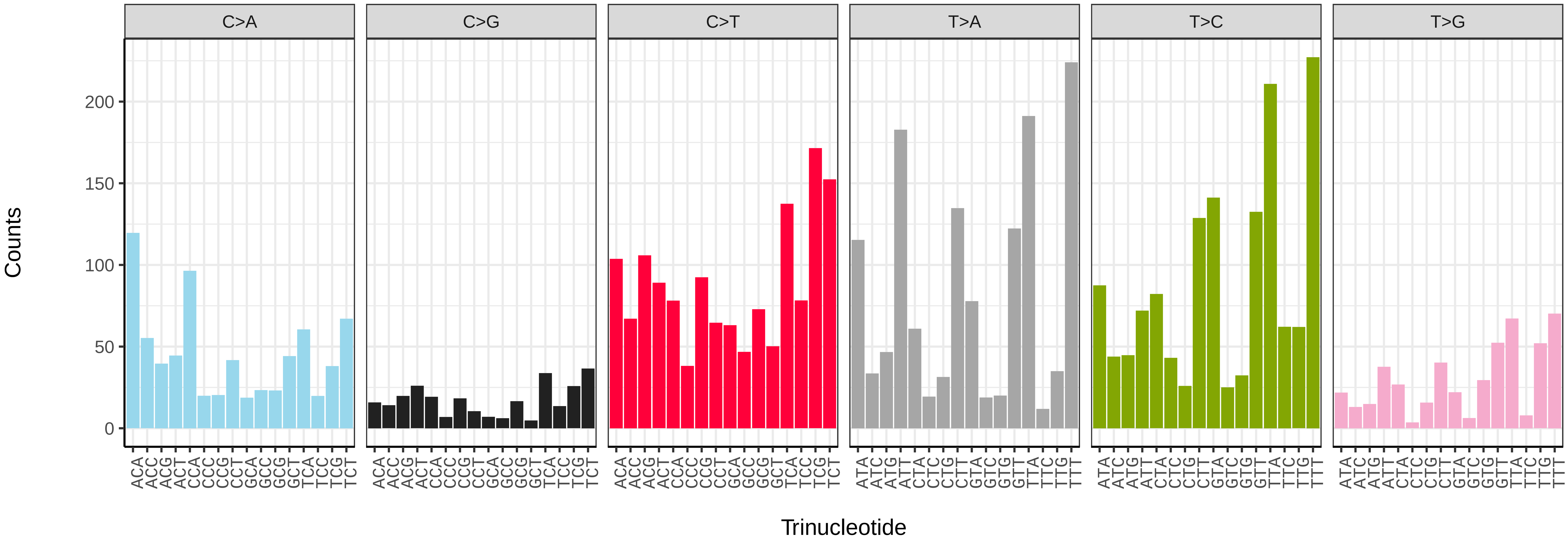


*Patella depressa* (xgPatDepr1) somatic mutational spectrum (SBS52 classification)

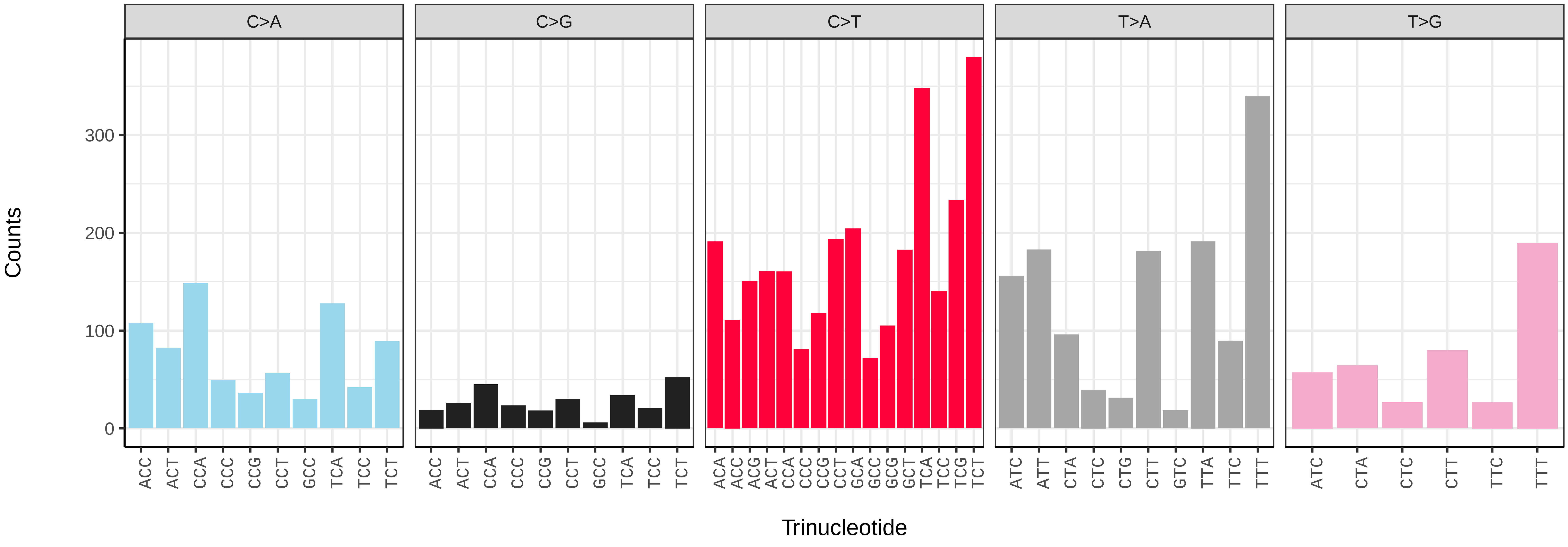

*Patella pellucida* (xgPatPell1) germline mutational spectrum

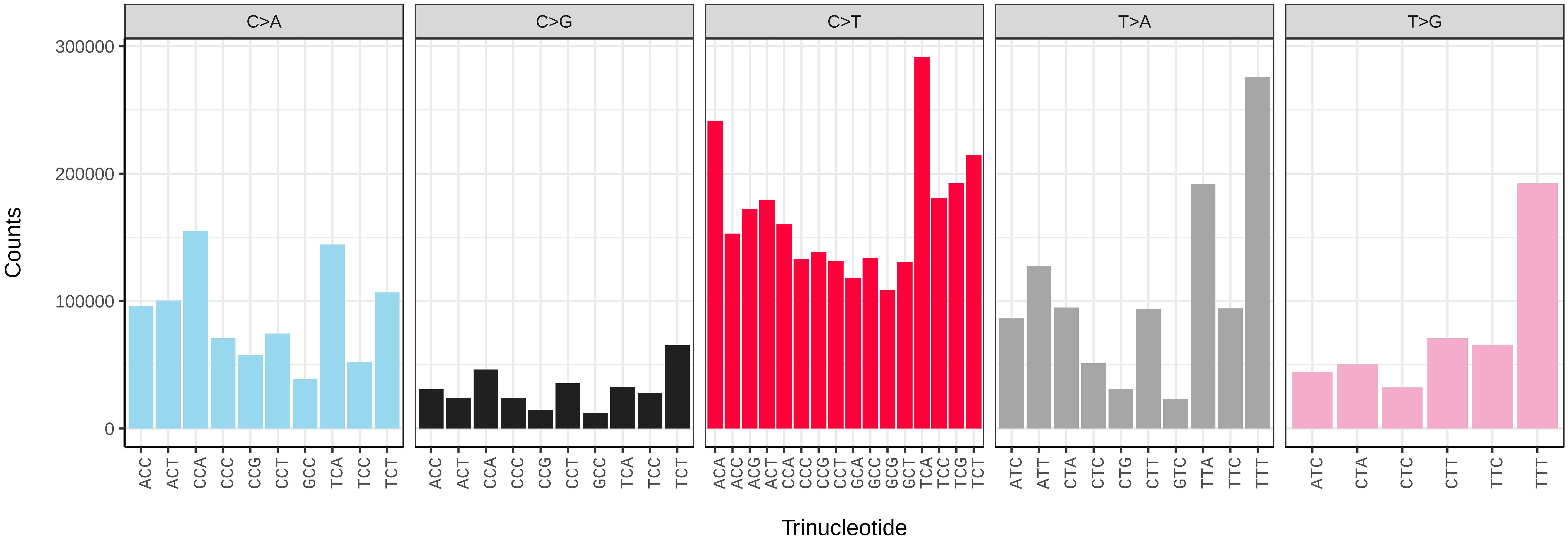


*Patella pellucida* (xgPatPell1) somatic mutational spectrum (SBS96 classification)

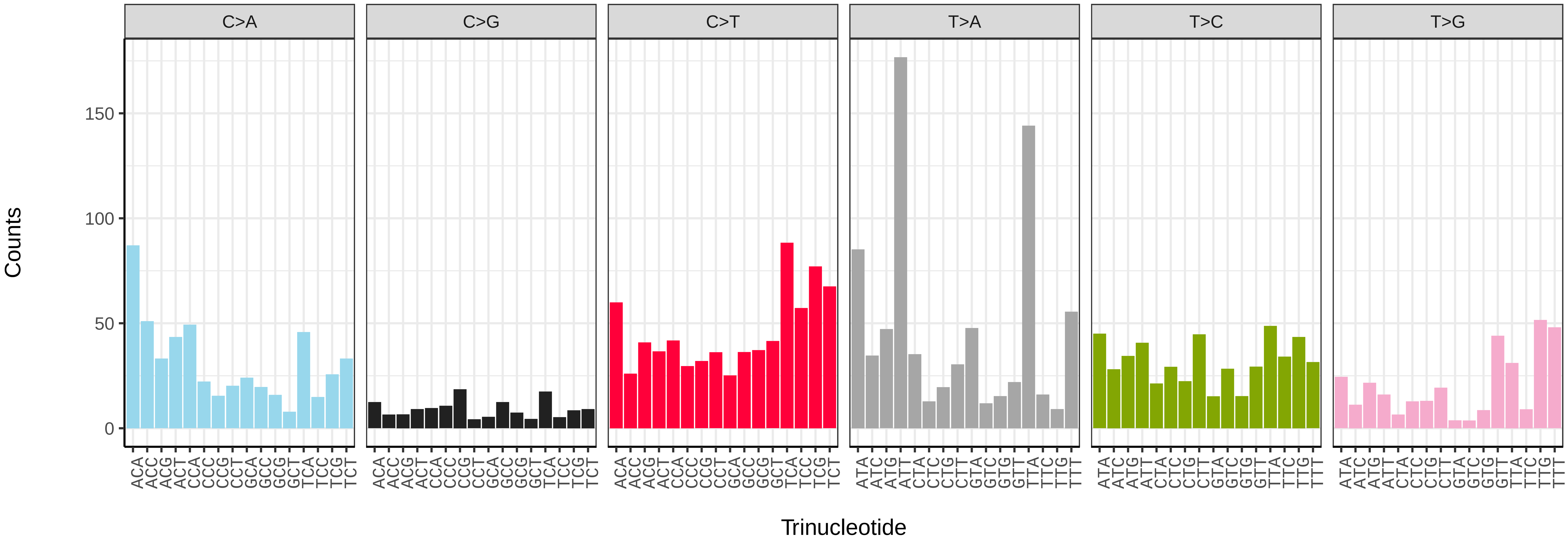


*Patella pellucida* (xgPatPell1) somatic mutational spectrum (SBS52 classification)

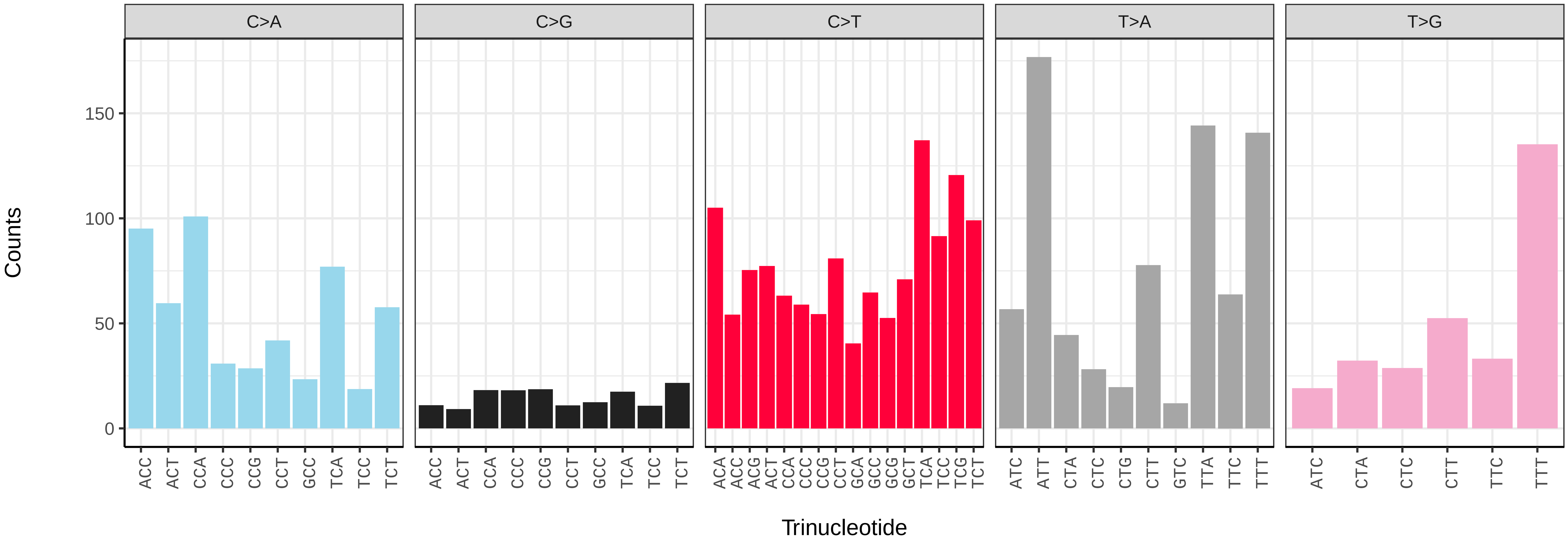

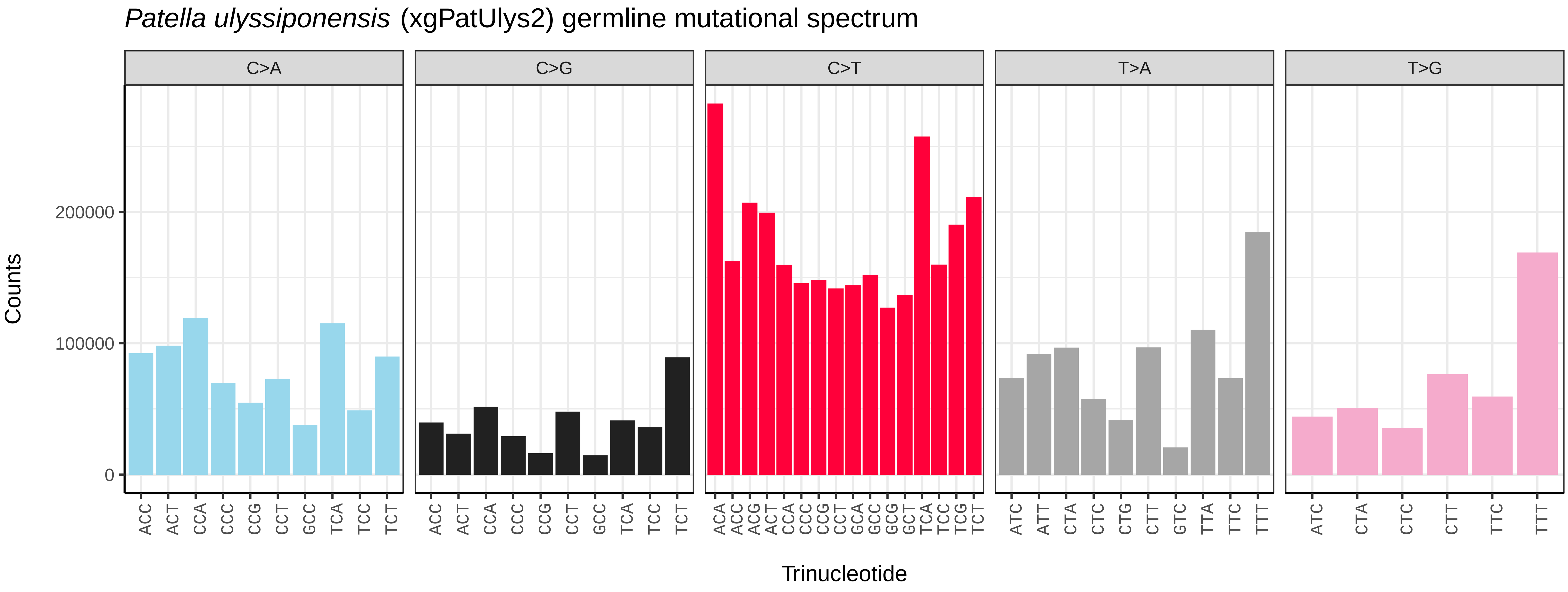

Patella ulyssiponensis (xgPatUlys2) germline mutational spectrum
C>A
C>G
C>T
T>A
T>G
Counts
200000
100000
0
Trinucleotide


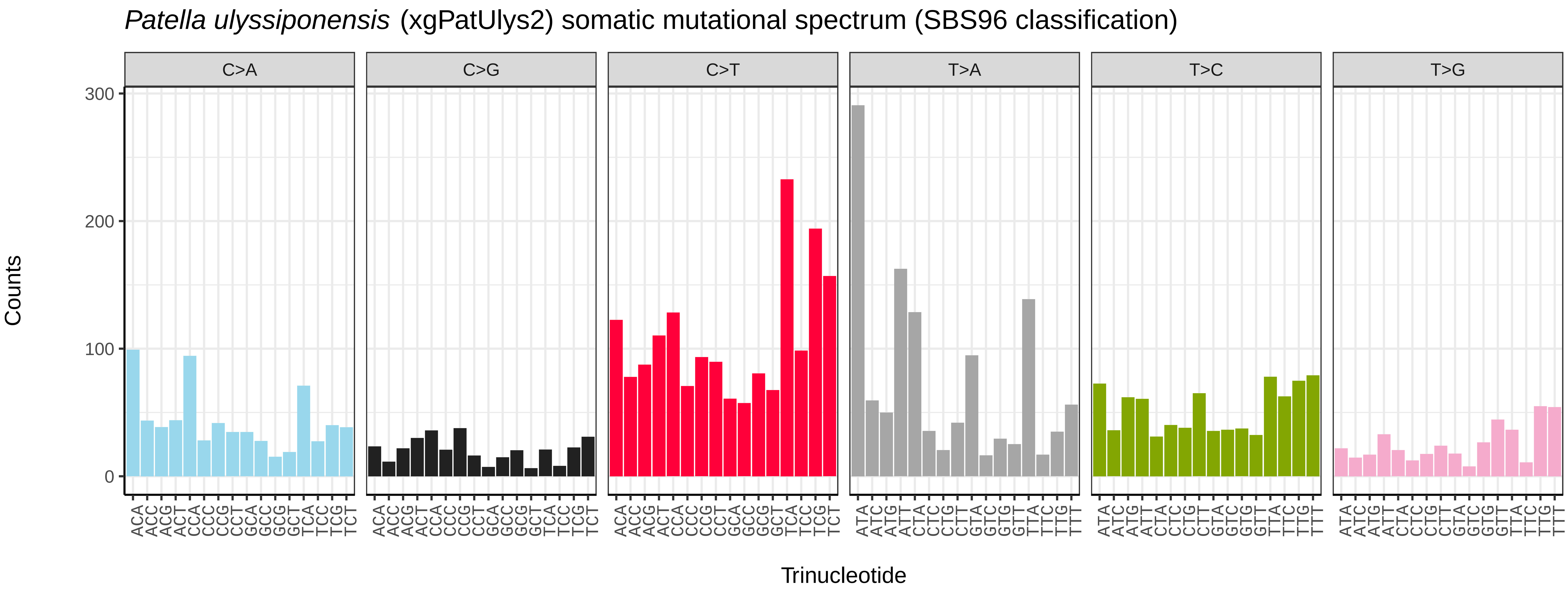

Patella ulyssiponensis (xgPatUlys2) somatic mutational spectrum (SBS96 classification)
C>A
C>G
C>T
T>A
T>C
T>G
Counts
300
200
100
0
Trinucleotide


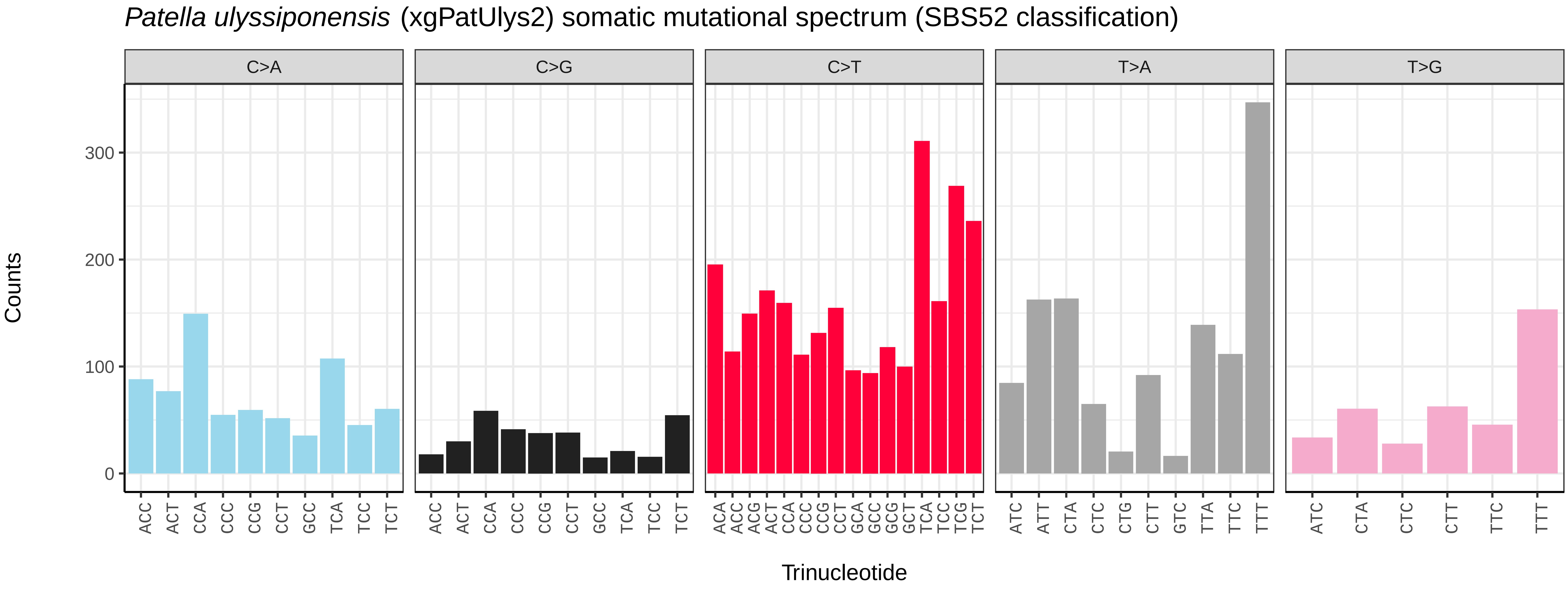

Patella ulyssiponensis (xgPatUlys2) somatic mutational spectrum (SBS52 classification)
C>A
C>G
C>T
T>A
T>G
Counts
300
200
100
0
Trinucleotide

*Patella vulgata* (xgPatVulg1) germline mutational spectrum

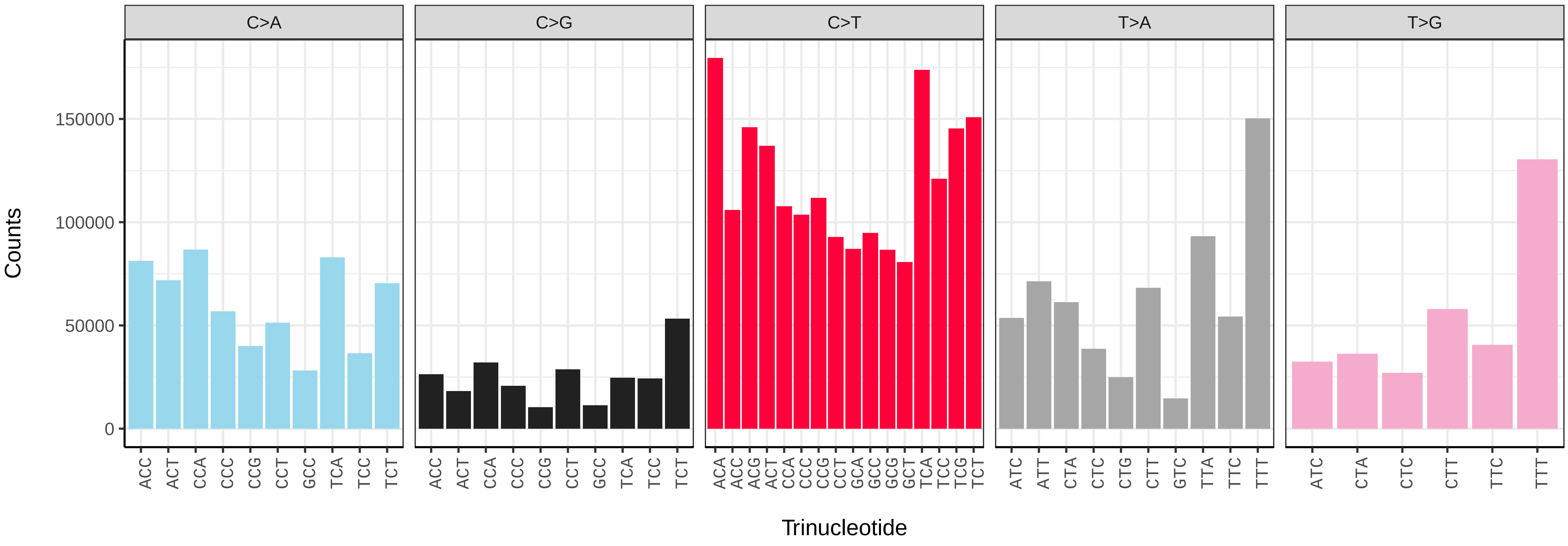


*Patella vulgata* (xgPatVulg1) somatic mutational spectrum (SBS96 classification)

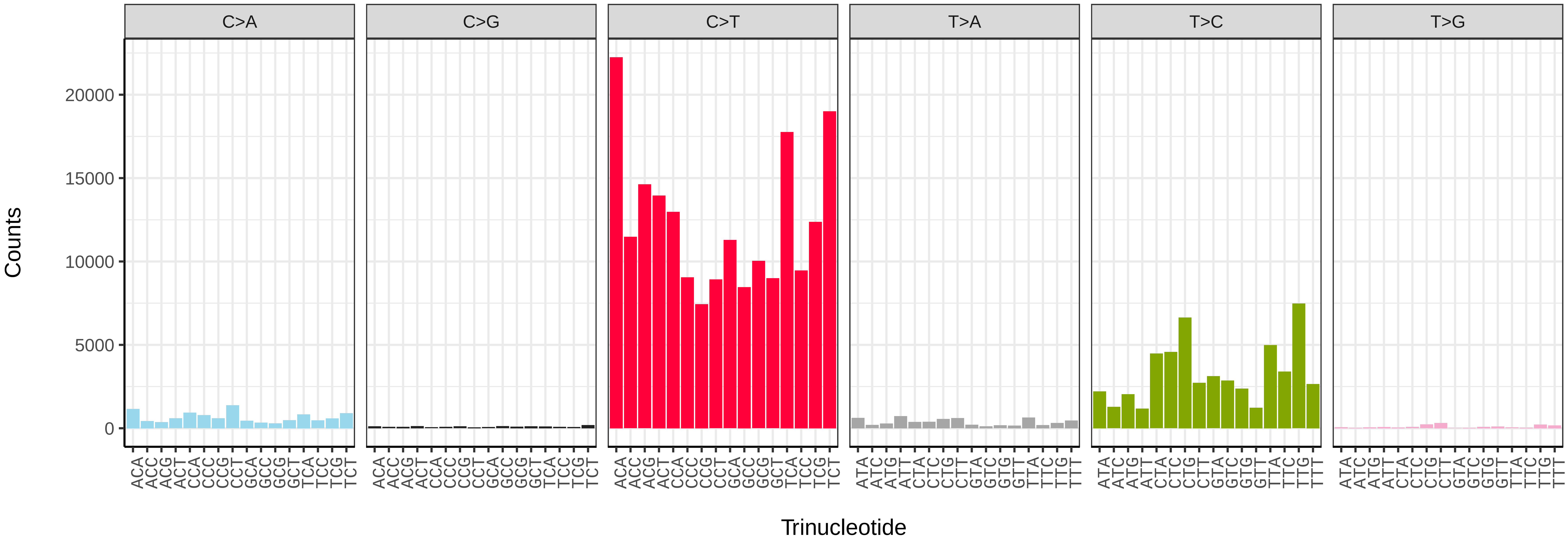


*Patella vulgata* (xgPatVulg1) somatic mutational spectrum (SBS52 classification)

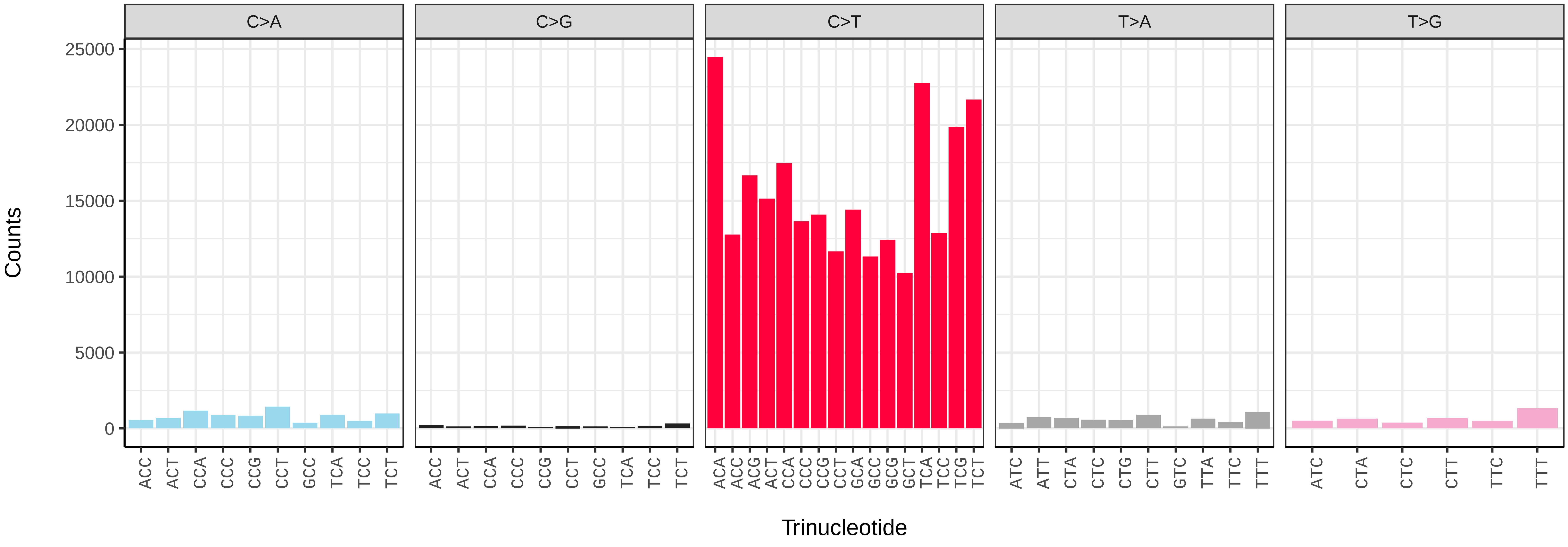

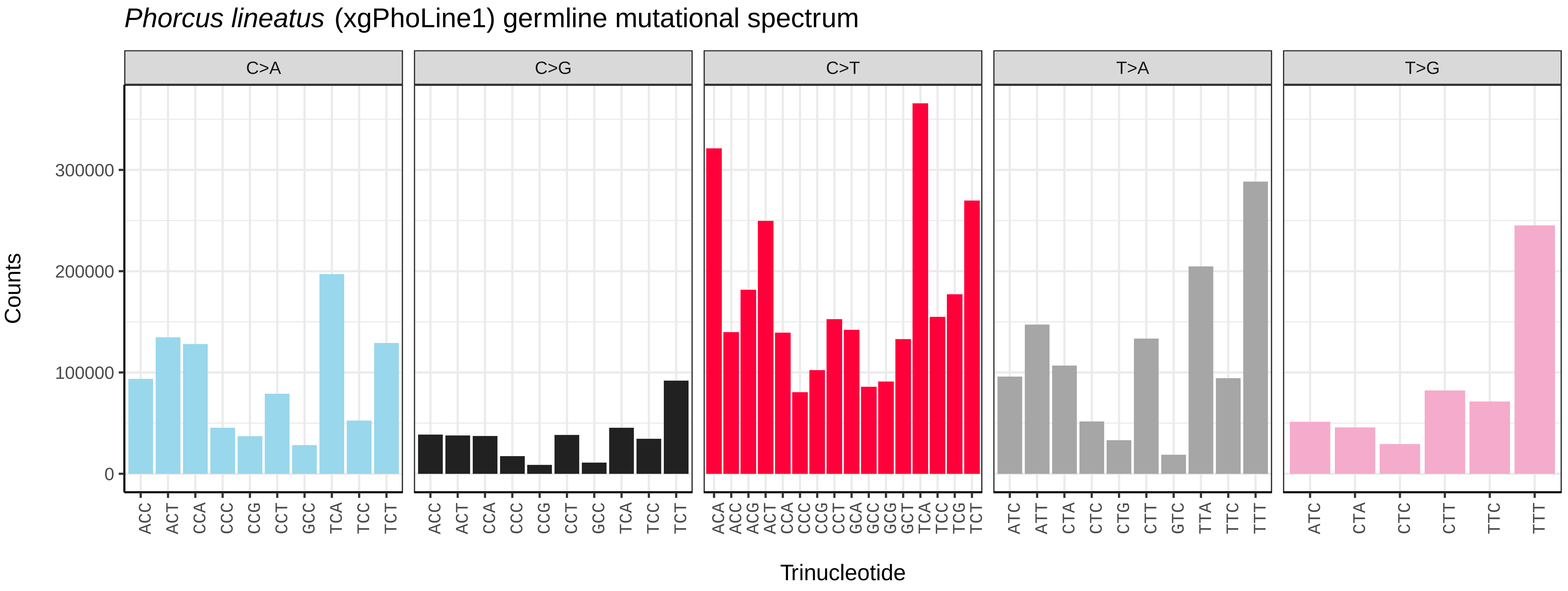
*Phorcus lineatus* (xgPhoLine1) germline mutational spectrum
C>A
C>G
C>T
T>A
T>G
Counts
0
100000
200000
300000
Trinucleotide
ACC ACT CCA CCC CCG CCT GCC TCA TCC TCT
ACA ACC ACG ACT CCA CCC CCG CCT GCA GCC GCG GCT TCA TCC TCG TCT
ATC ATT CTA CTC CTG CTT GTC TTA TTC TTT
ATC CTA CTC CTT TTC TTT

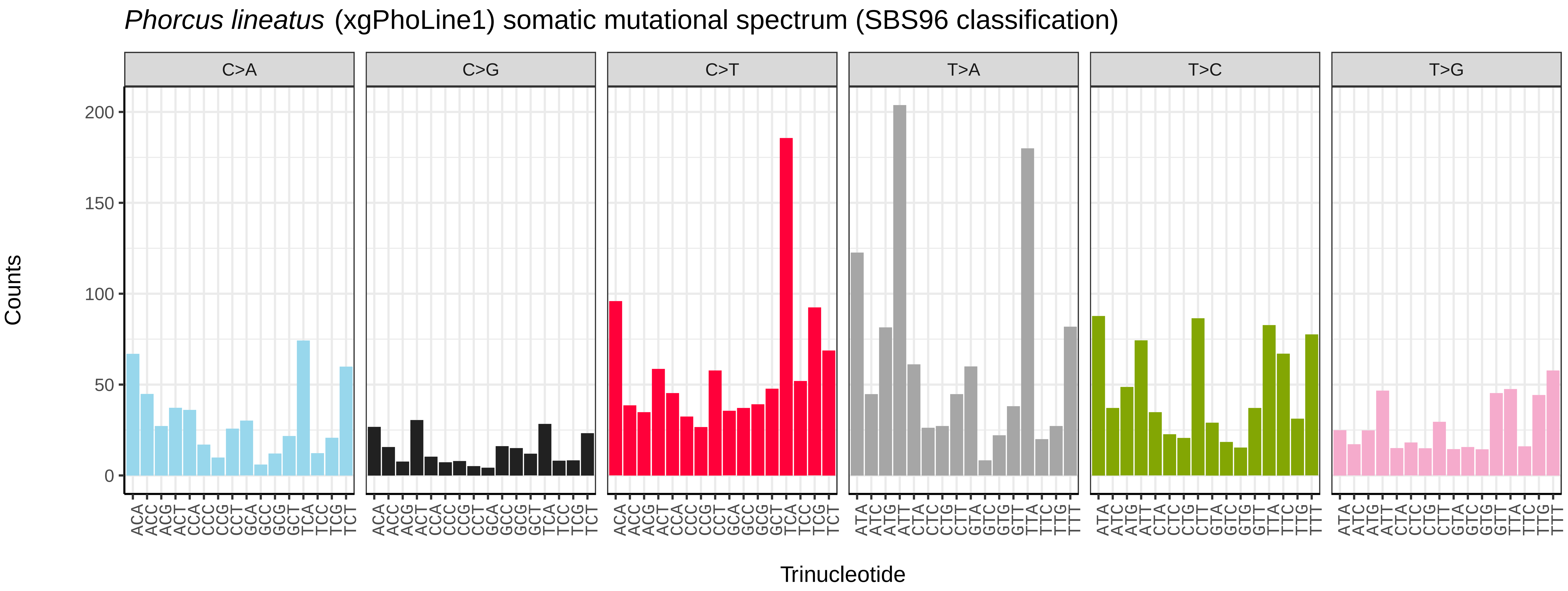
*Phorcus lineatus* (xgPhoLine1) somatic mutational spectrum (SBS96 classification)
C>A
C>G
C>T
T>A
T>C
T>G
Counts
0
50
100
150
200
Trinucleotide
ACA ACC ACG ACT CCA CCC CCG CCT GCA GCC GCG GCT TCA TCC TCG TCT
ATA ATC ATG ATT CTA CTC CTG CTT GTA GTC GTG GTT TTA TTC TTG TTT

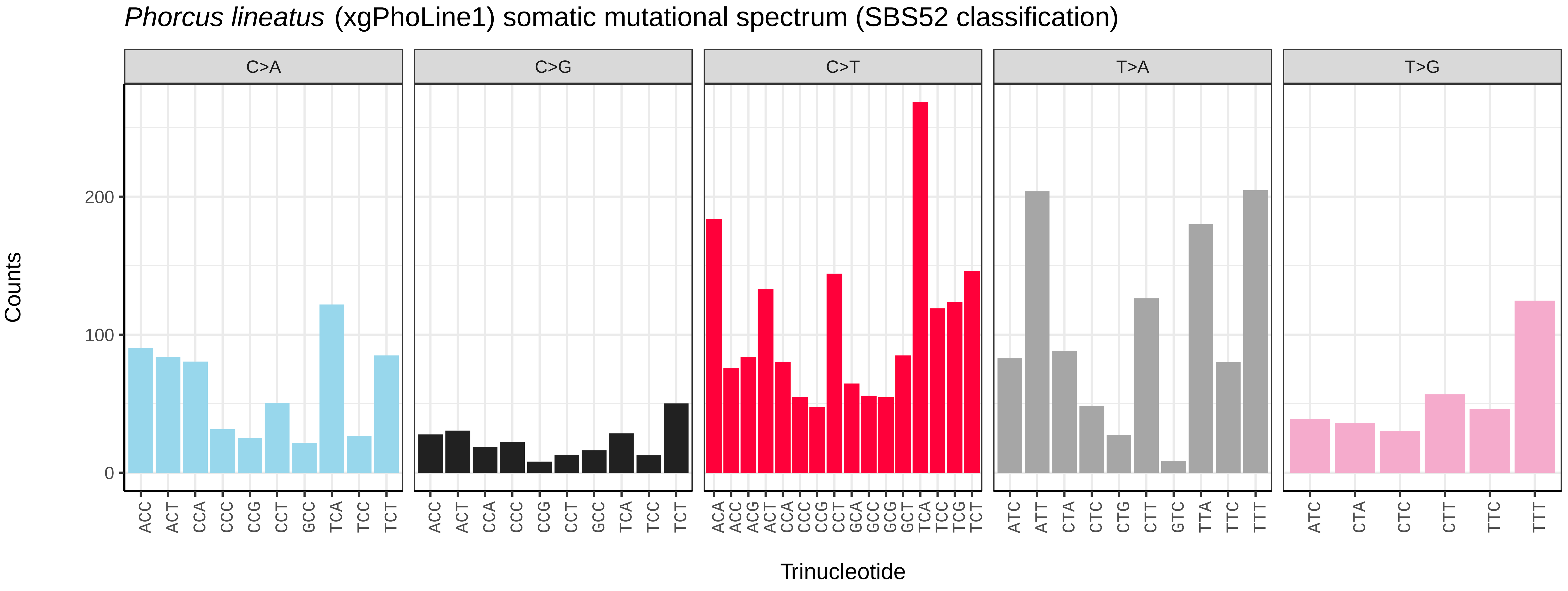
*Phorcus lineatus* (xgPhoLine1) somatic mutational spectrum (SBS52 classification)
C>A
C>G
C>T
T>A
T>G
Counts
0
100
200
Trinucleotide
ACC ACT CCA CCC CCG CCT GCC TCA TCC TCT
ACA ACC ACG ACT CCA CCC CCG CCT GCA GCC GCG GCT TCA TCC TCG TCT
ATC ATT CTA CTC CTG CTT GTC TTA TTC TTT
ATC CTA CTC CTT TTC TTT

*Phorcus lineatus* (xgPhoLine1.PHORD0020b) germline mutational spectrum

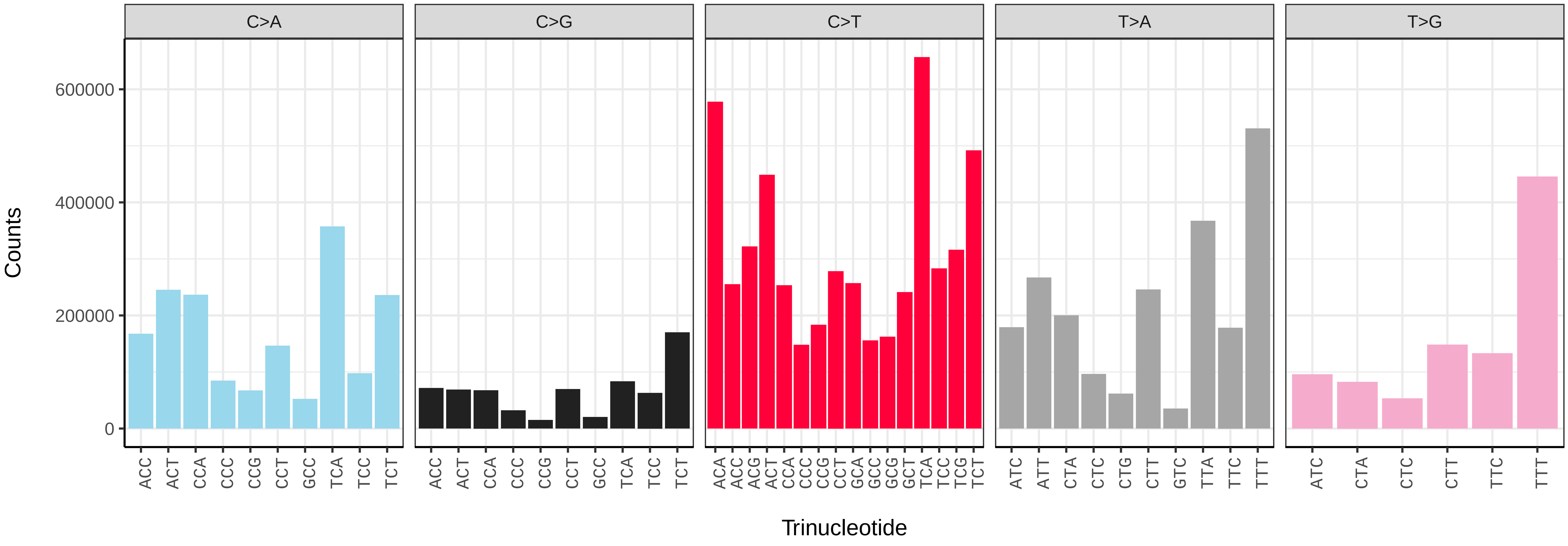


*Phorcus lineatus* (xgPhoLine1.PHORD0020b) somatic mutational spectrum (SBS96 classification)

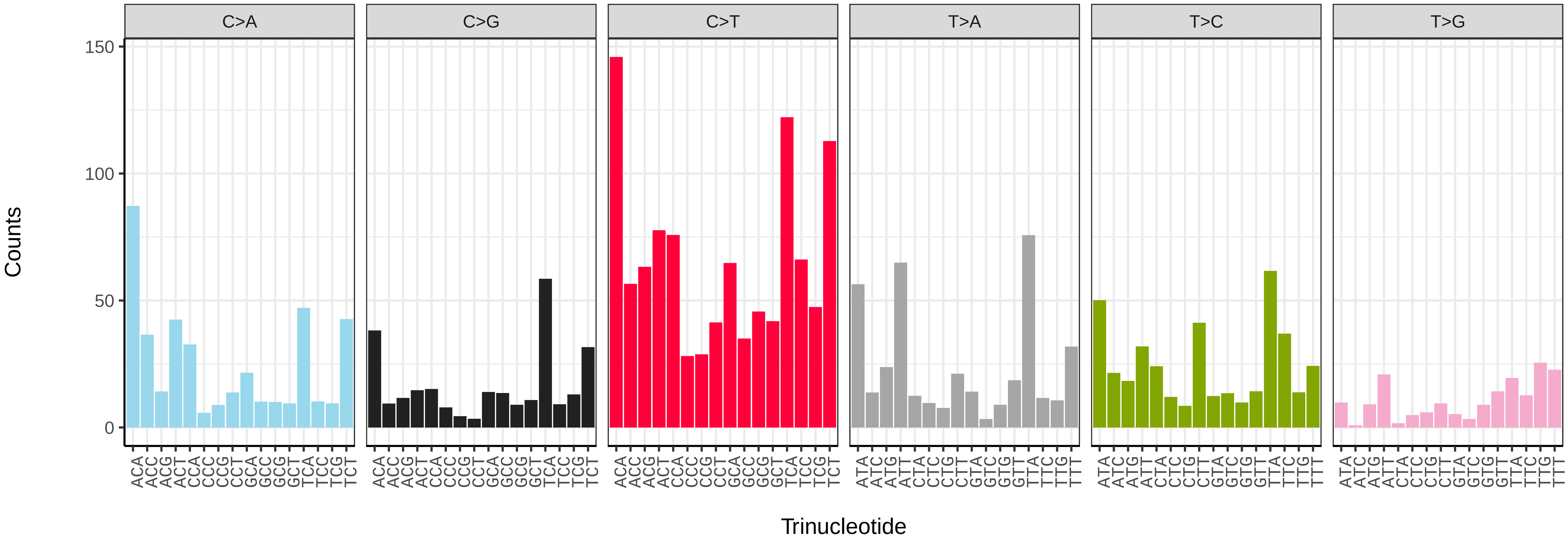


*Phorcus lineatus* (xgPhoLine1.PHORD0020b) somatic mutational spectrum (SBS52 classification)

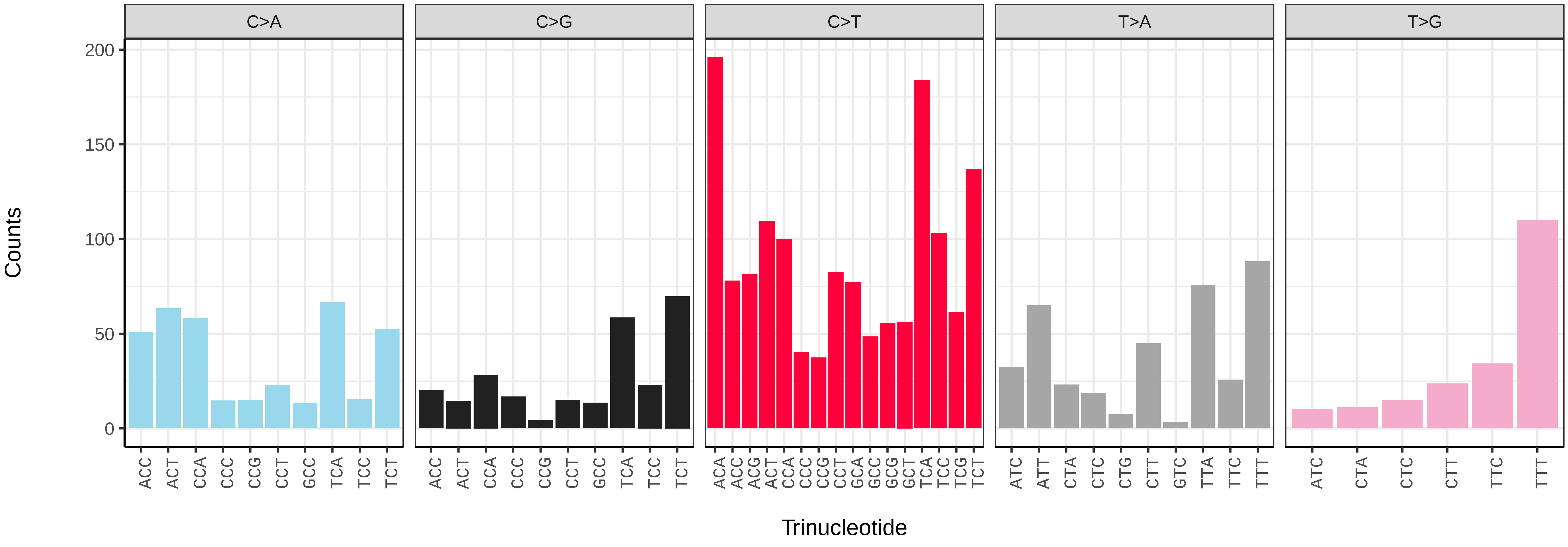

*Phorcus lineatus* (xgPhoLine1.PHORD0021b) germline mutational spectrum

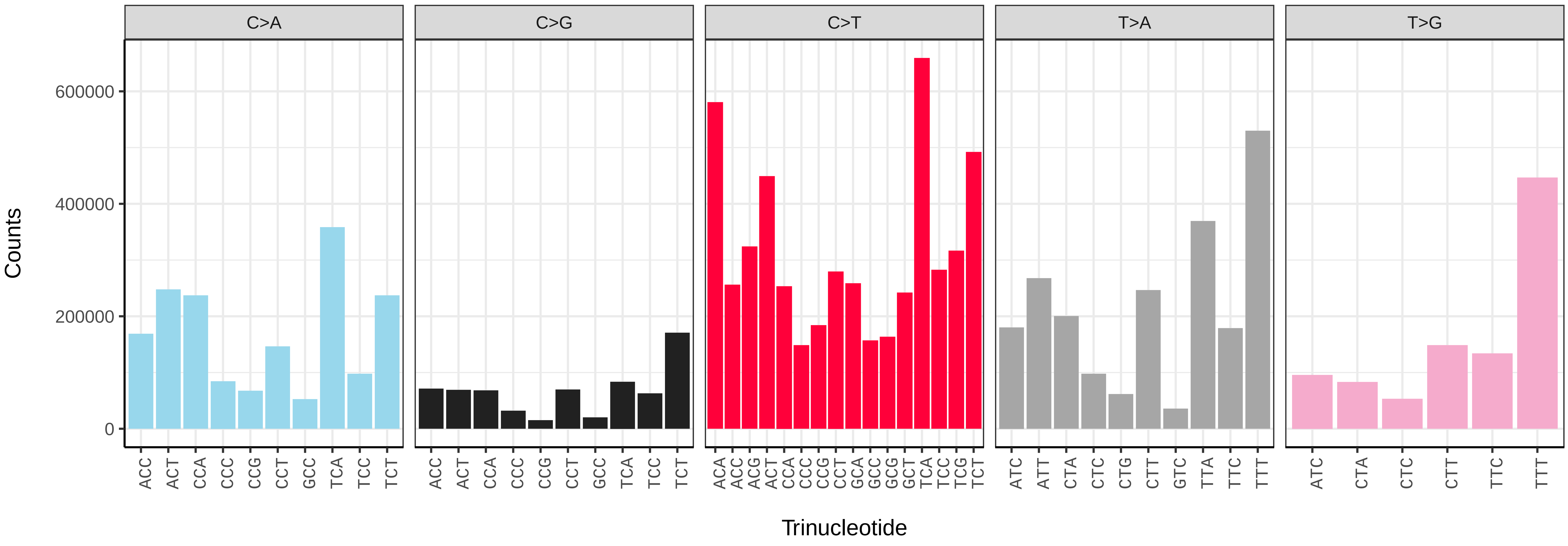


*Phorcus lineatus* (xgPhoLine1.PHORD0021b) somatic mutational spectrum (SBS96 classification)

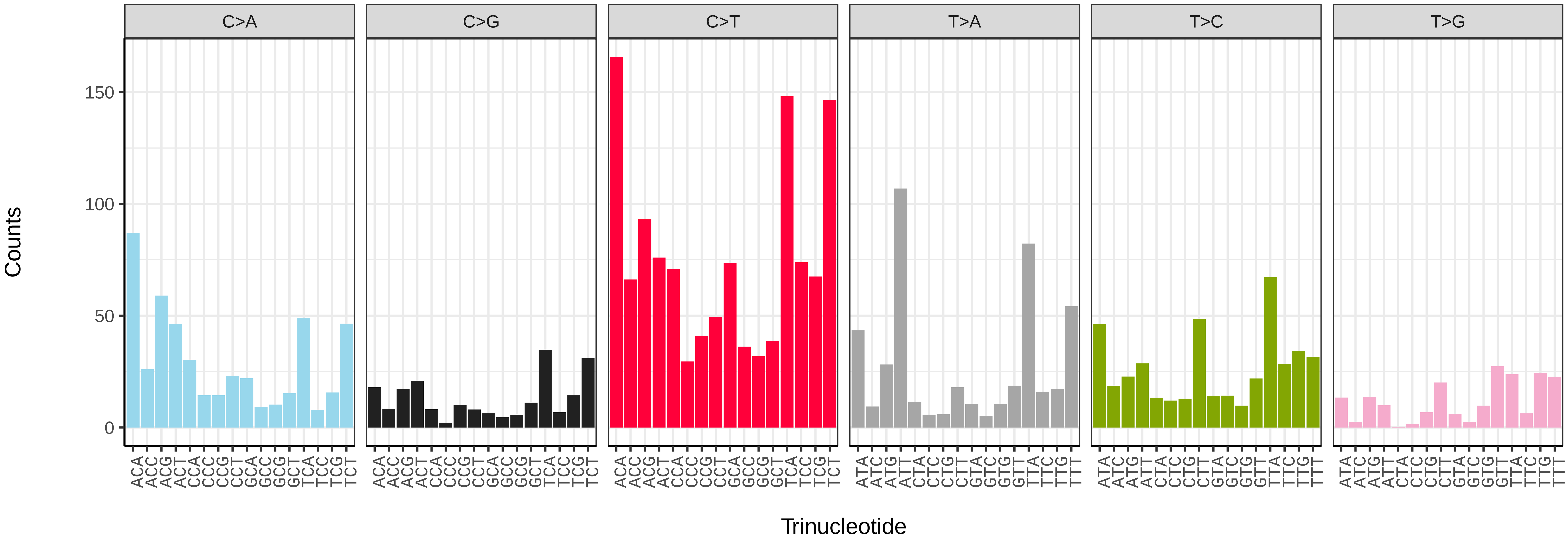


*Phorcus lineatus* (xgPhoLine1.PHORD0021b) somatic mutational spectrum (SBS52 classification)

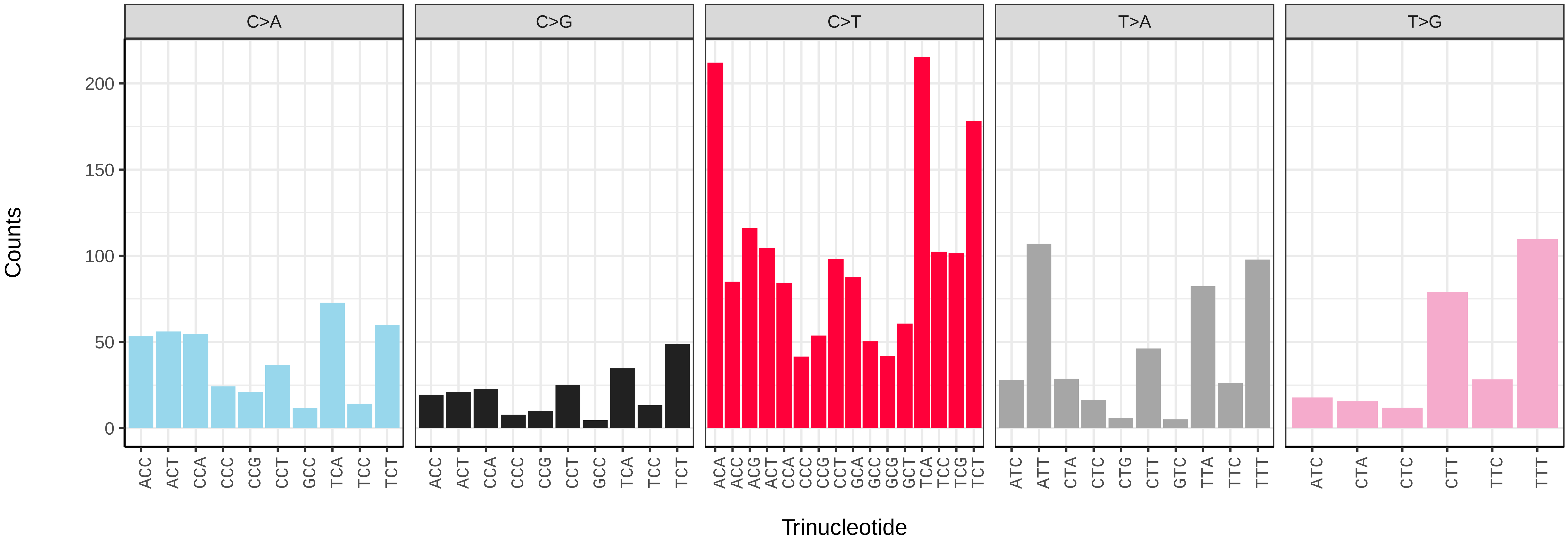

*Phorcus lineatus* (xgPhoLine1.PHORD0023b) germline mutational spectrum

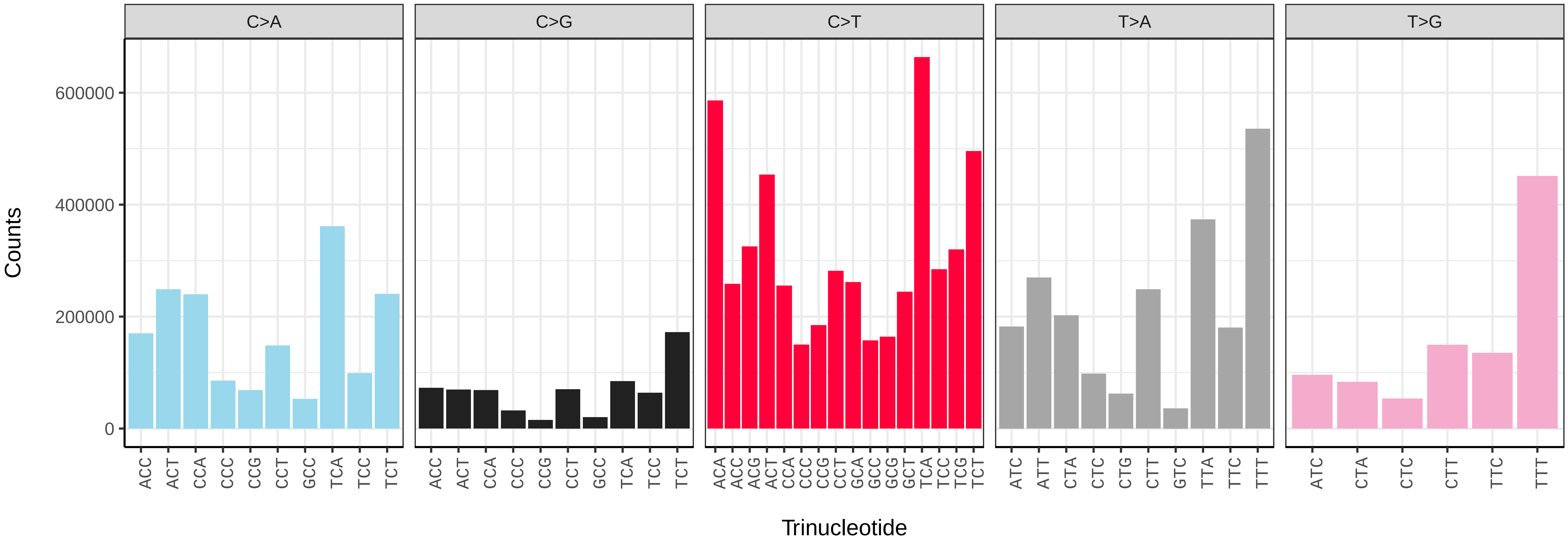


*Phorcus lineatus* (xgPhoLine1.PHORD0023b) somatic mutational spectrum (SBS96 classification)

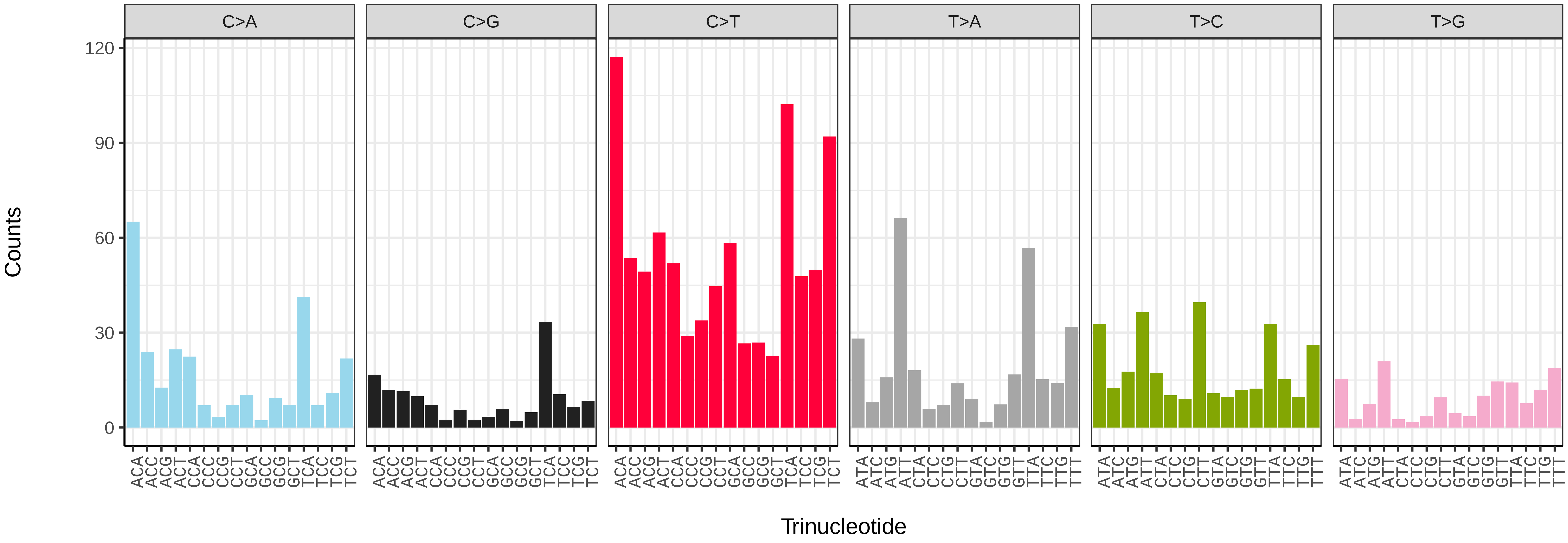


*Phorcus lineatus* (xgPhoLine1.PHORD0023b) somatic mutational spectrum (SBS52 classification)

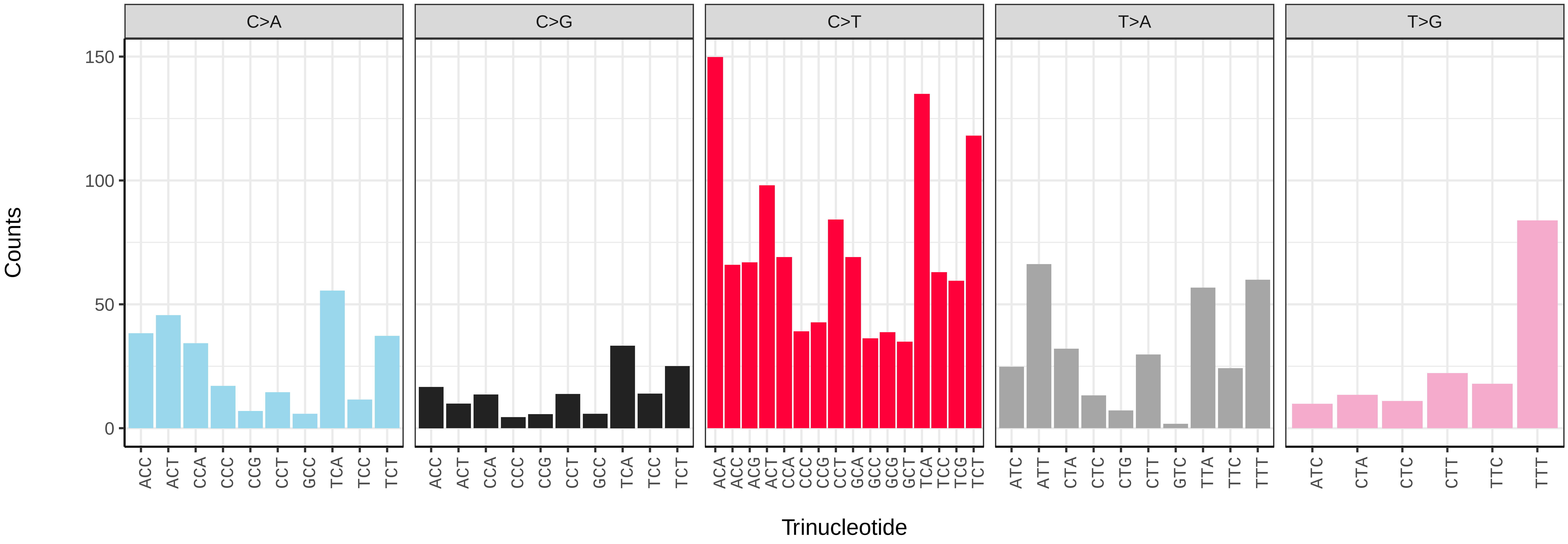

*Phorcus lineatus* (xgPhoLine1.PHORD0024b) germline mutational spectrum

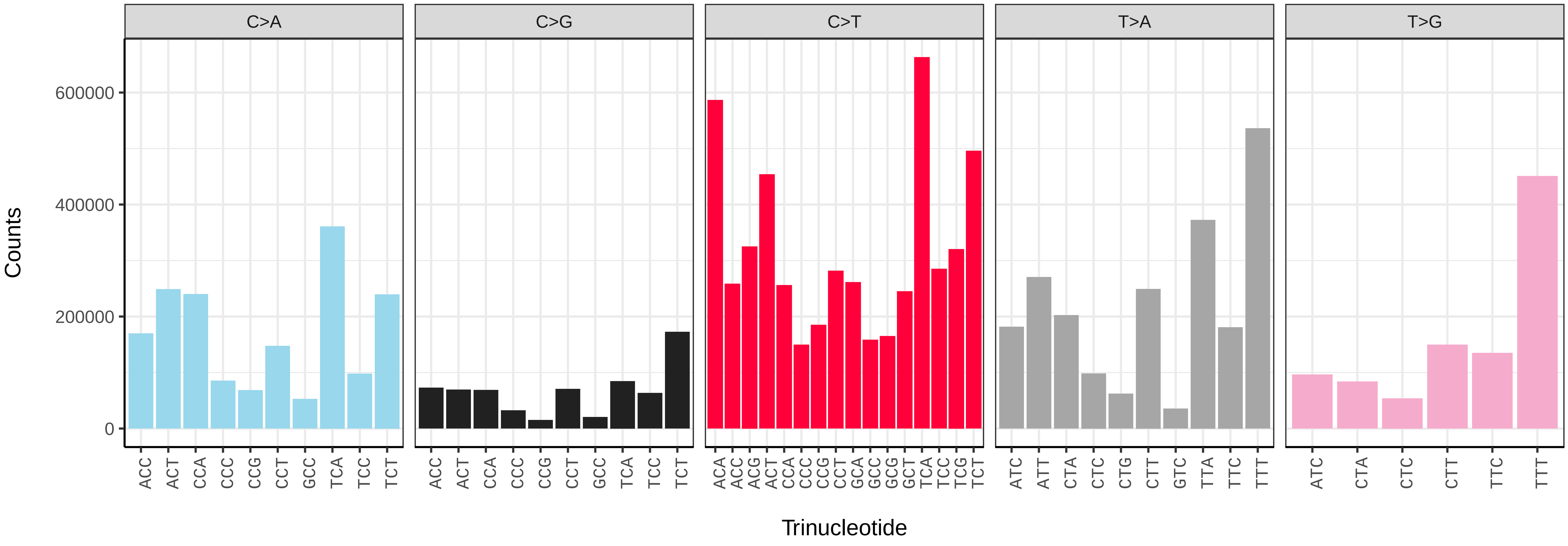


*Phorcus lineatus* (xgPhoLine1.PHORD0024b) somatic mutational spectrum (SBS96 classification)

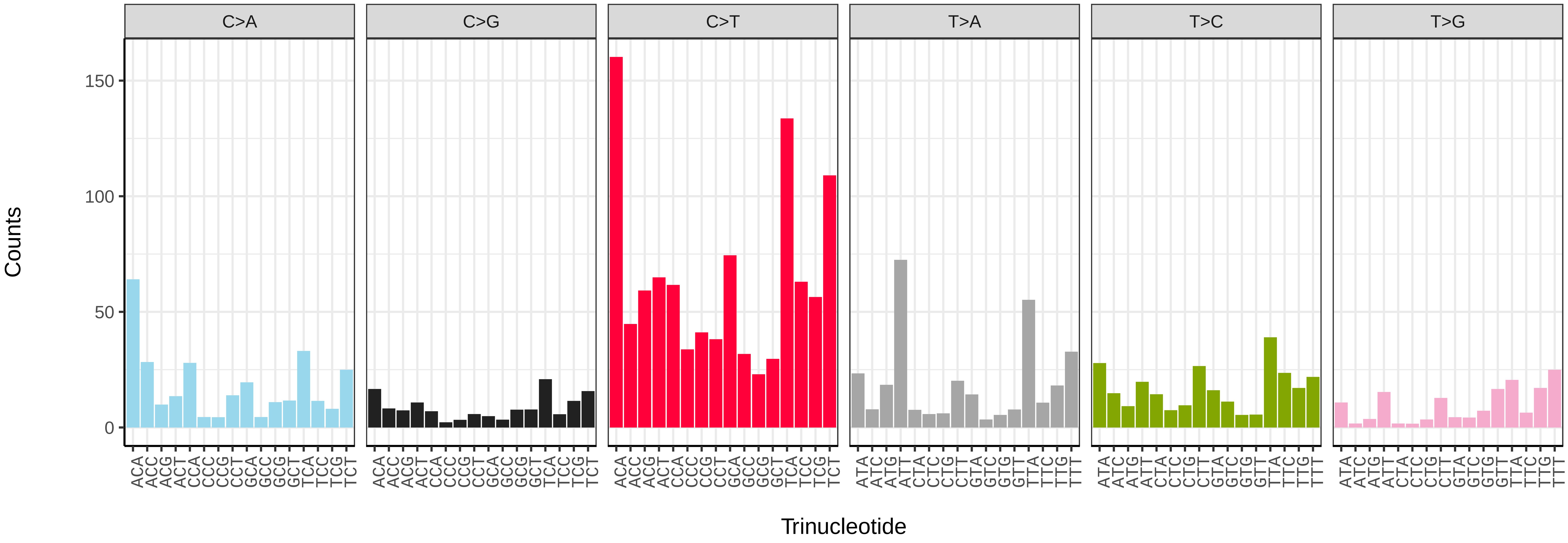


*Phorcus lineatus* (xgPhoLine1.PHORD0024b) somatic mutational spectrum (SBS52 classification)

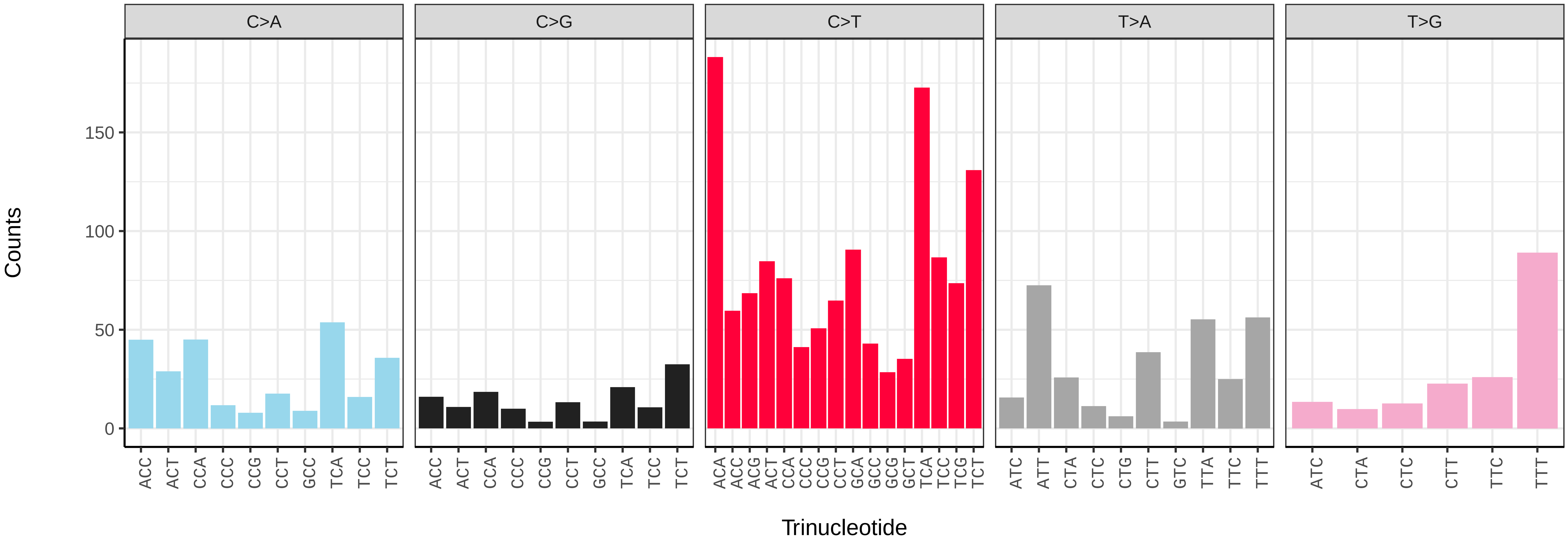

*Phorcus lineatus* (xgPhoLine1.PHORD0030b) germline mutational spectrum

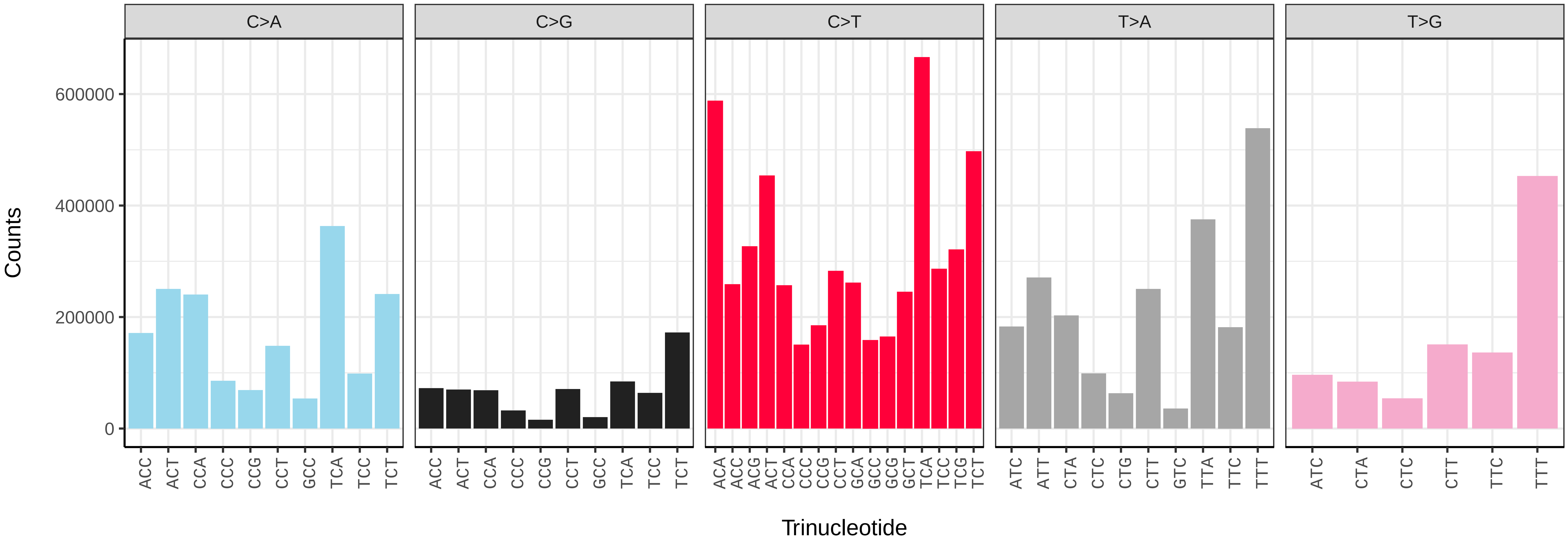


*Phorcus lineatus* (xgPhoLine1.PHORD0030b) somatic mutational spectrum (SBS96 classification)

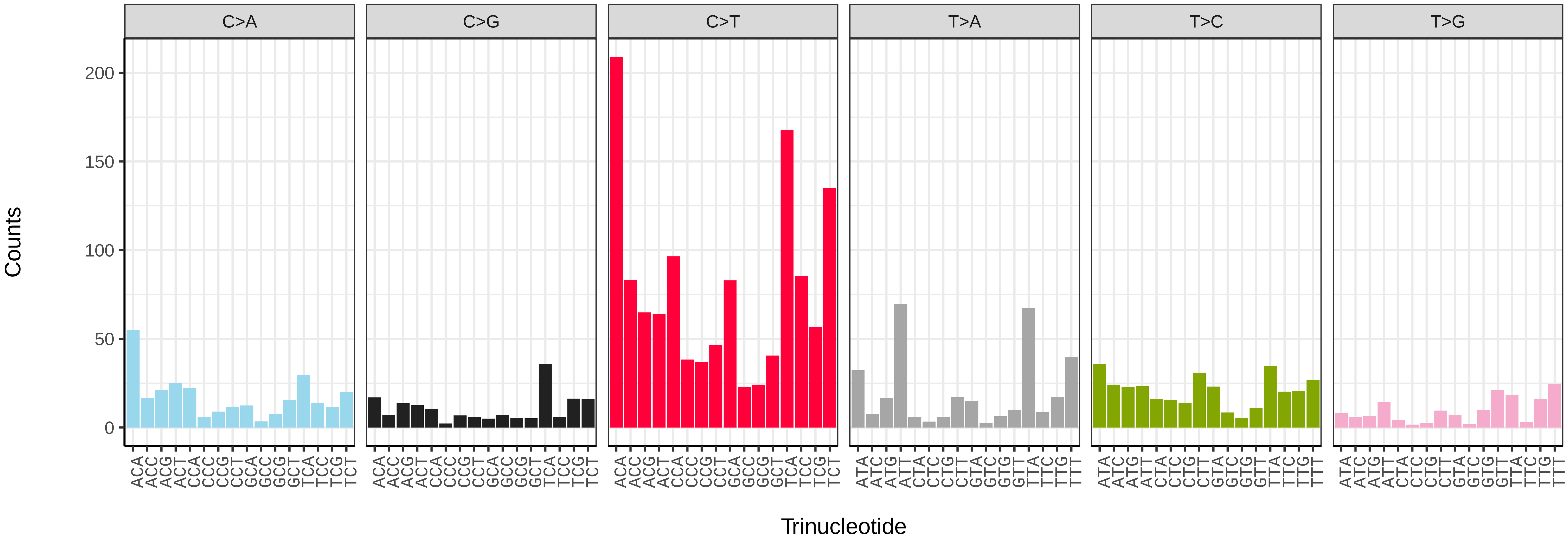


*Phorcus lineatus* (xgPhoLine1.PHORD0030b) somatic mutational spectrum (SBS52 classification)

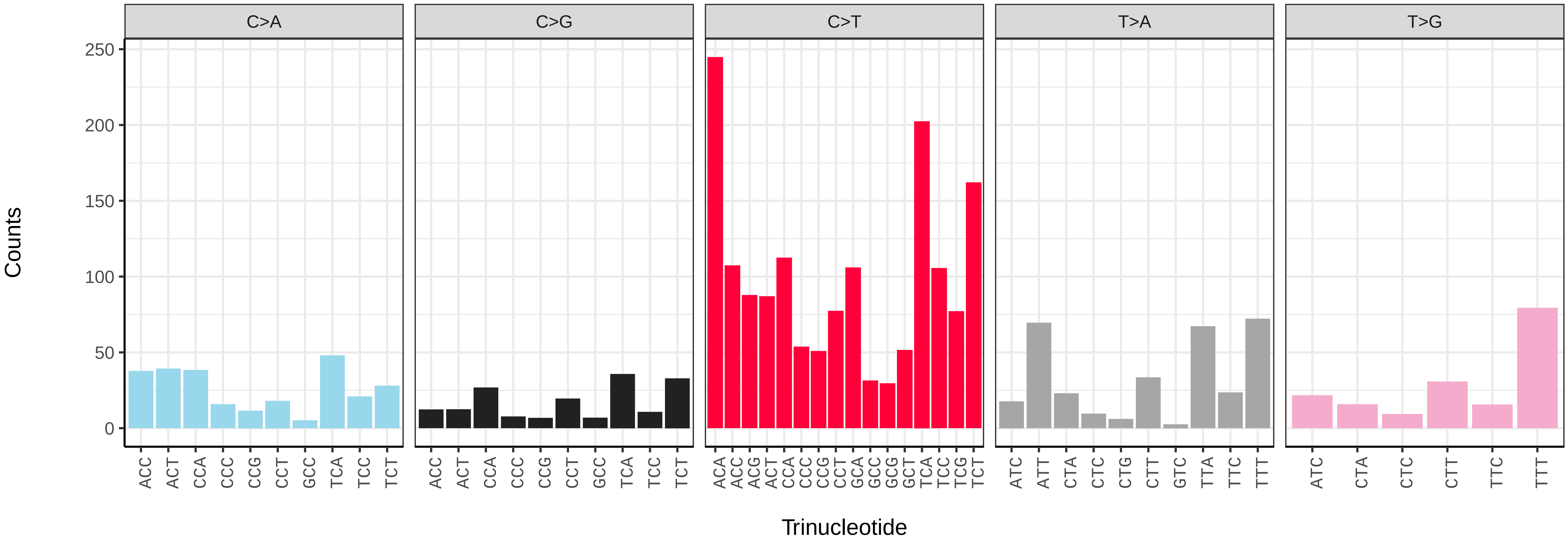

*Phorcus lineatus* (xgPhoLine1.PHORD0031b) germline mutational spectrum

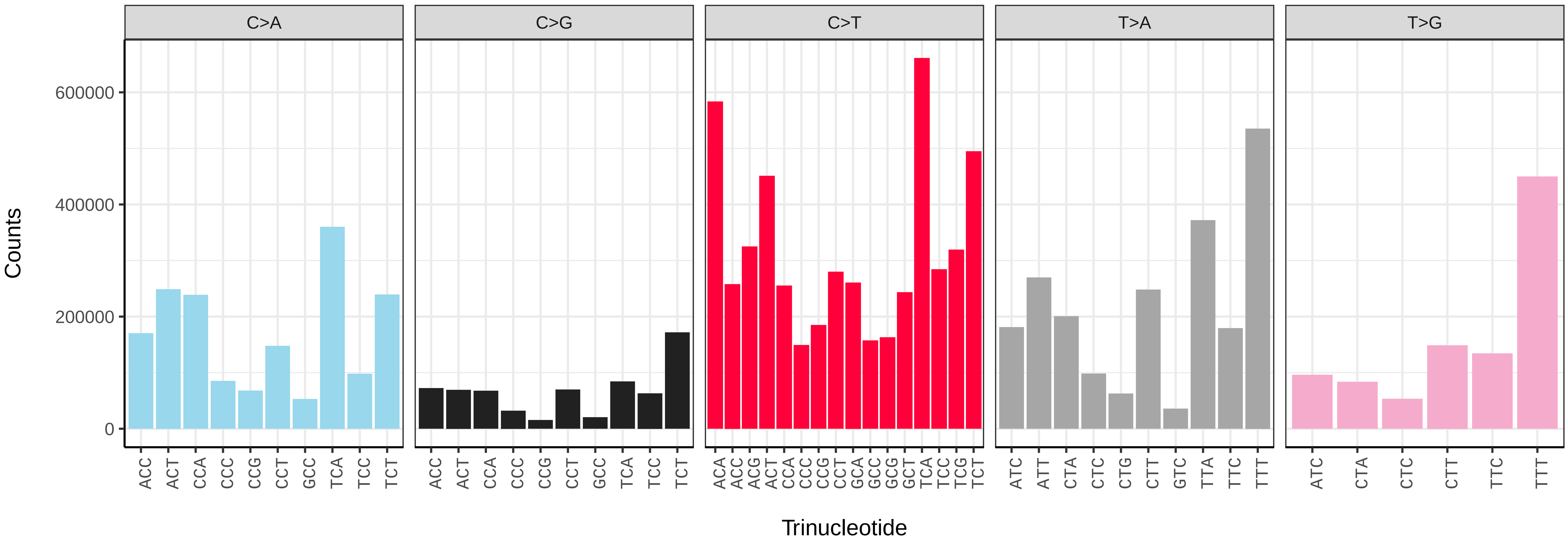


*Phorcus lineatus* (xgPhoLine1.PHORD0031b) somatic mutational spectrum (SBS96 classification)

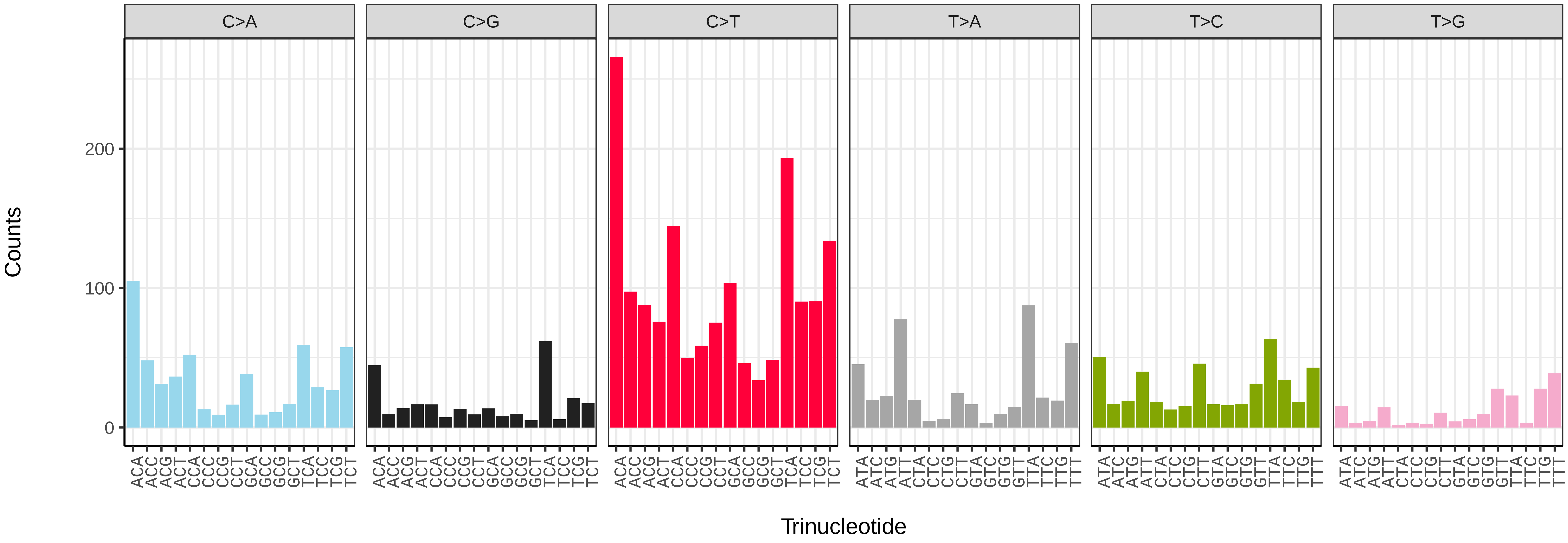


*Phorcus lineatus* (xgPhoLine1.PHORD0031b) somatic mutational spectrum (SBS52 classification)

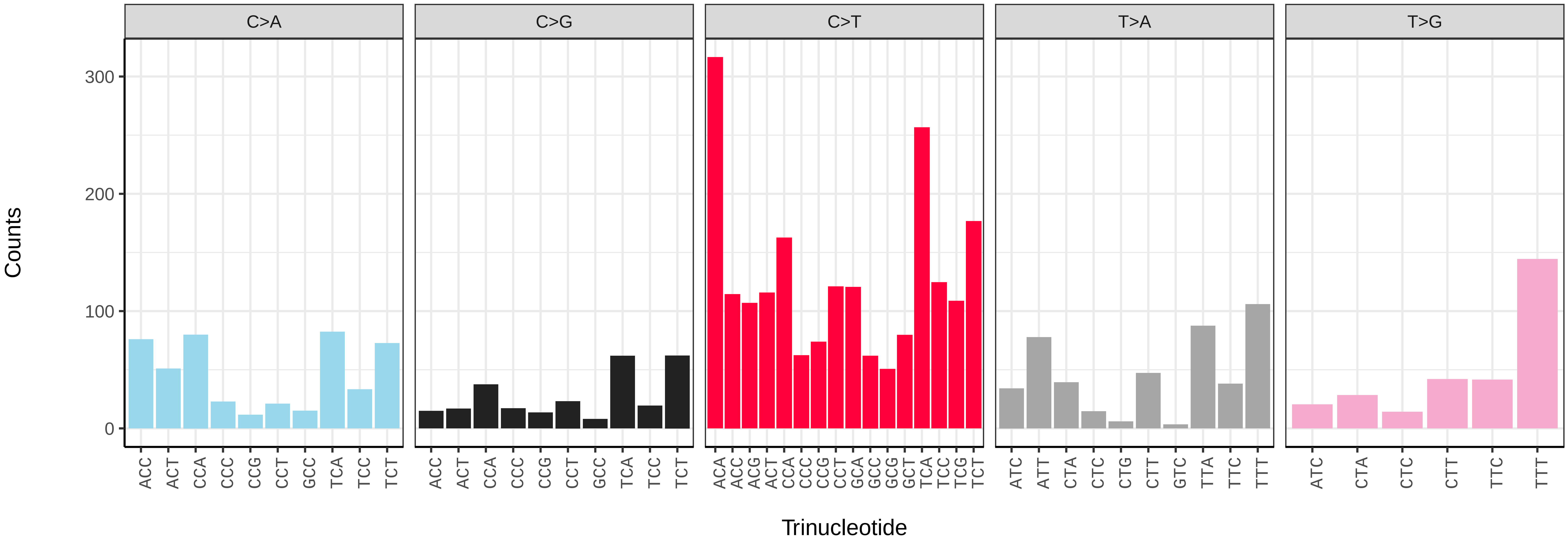

*Phorcus lineatus* (xgPhoLine1.PHORD0034b) germline mutational spectrum

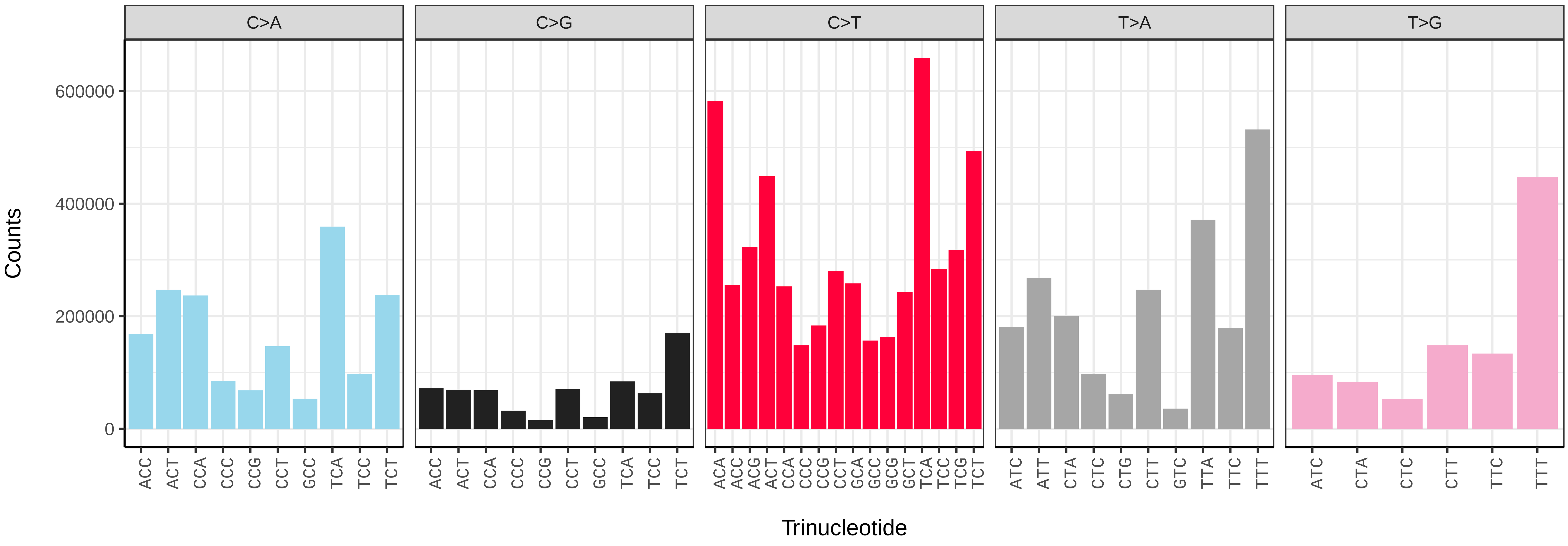


*Phorcus lineatus* (xgPhoLine1.PHORD0034b) somatic mutational spectrum (SBS96 classification)

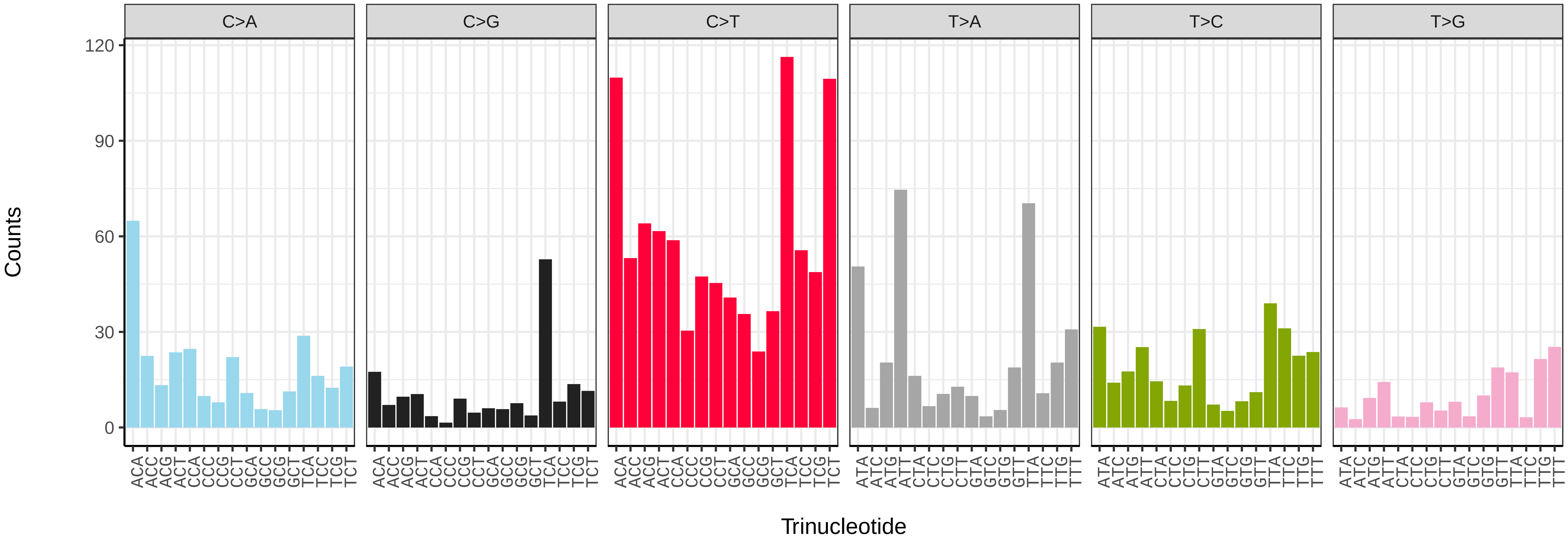


*Phorcus lineatus* (xgPhoLine1.PHORD0034b) somatic mutational spectrum (SBS52 classification)

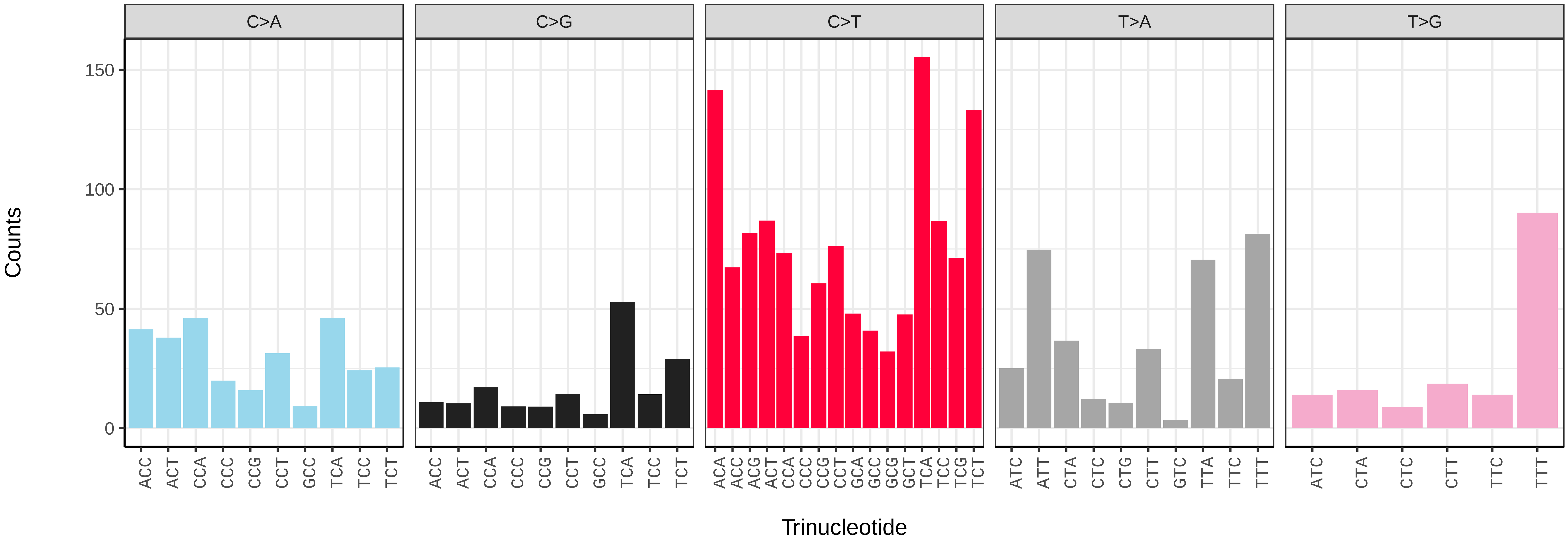

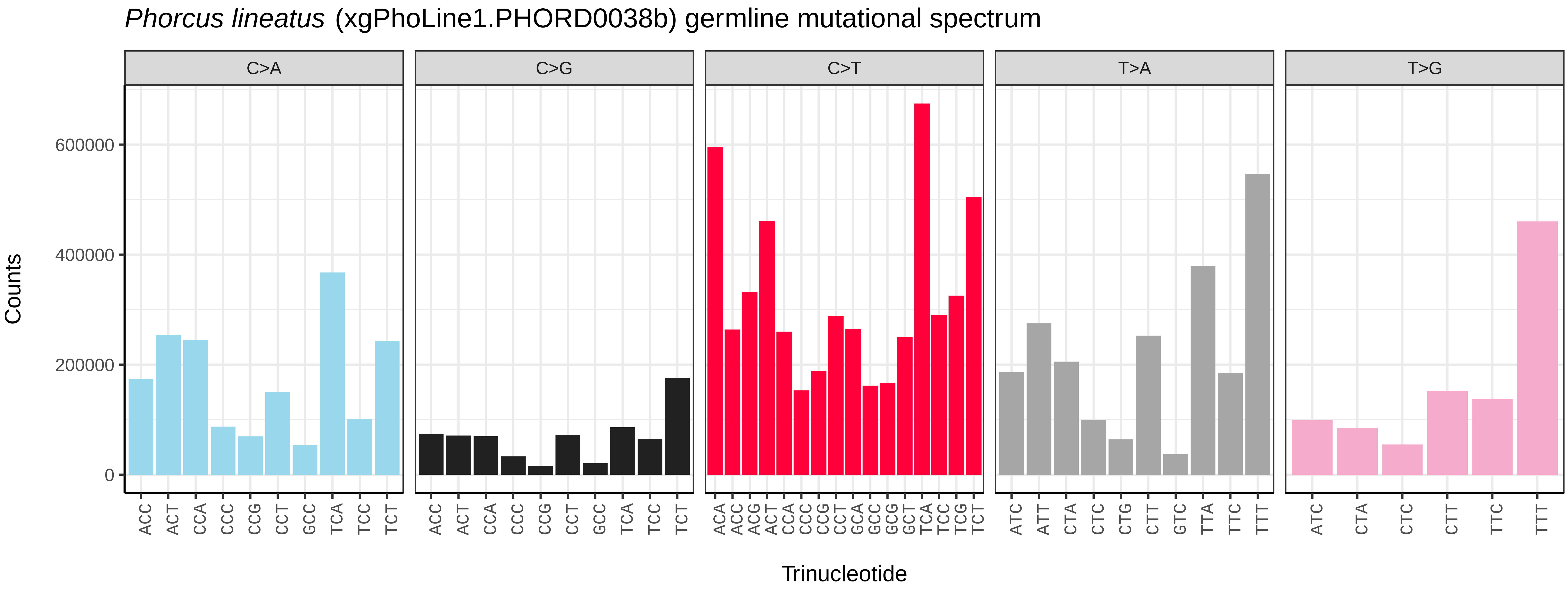
*Phorcus lineatus* (xgPhoLine1.PHORD0038b) germline mutational spectrum
C>A
C>G
C>T
T>A
T>G
Counts
0
200000
400000
600000
Trinucleotide

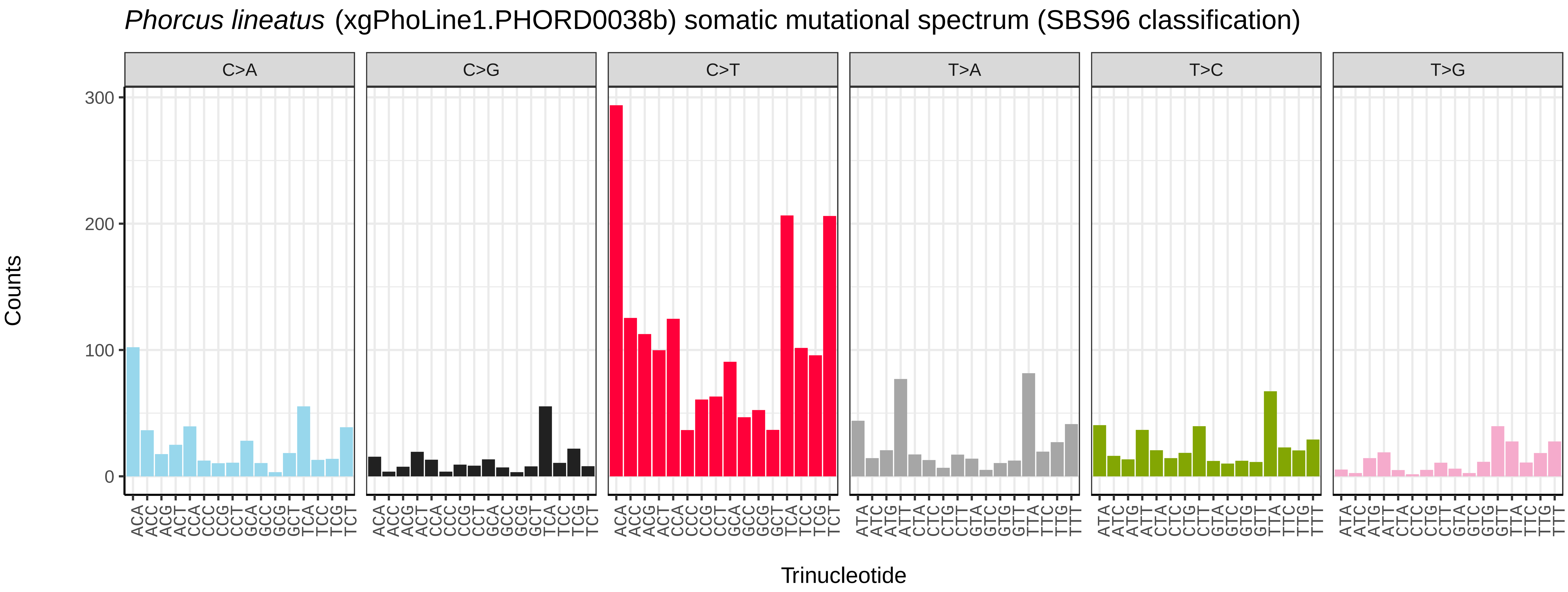
*Phorcus lineatus* (xgPhoLine1.PHORD0038b) somatic mutational spectrum (SBS96 classification)
C>A
C>G
C>T
T>A
T>C
T>G
Counts
0
100
200
300
Trinucleotide

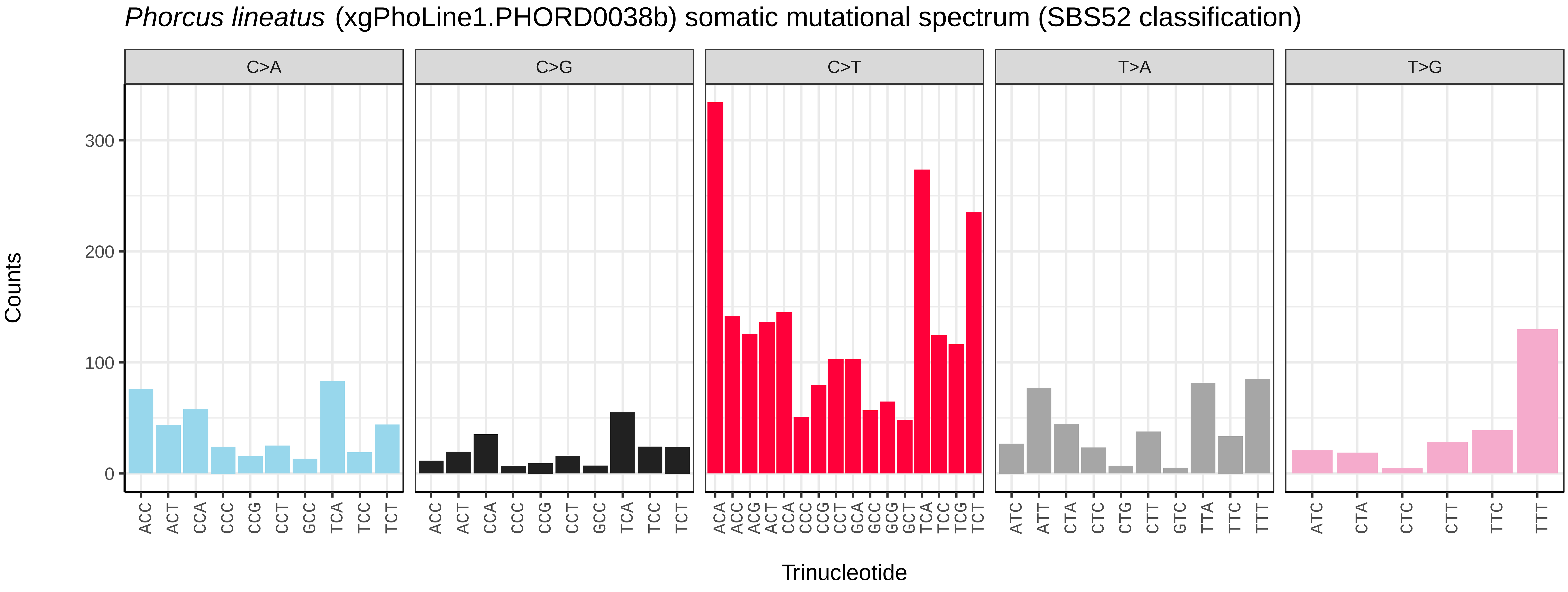
*Phorcus lineatus* (xgPhoLine1.PHORD0038b) somatic mutational spectrum (SBS52 classification)
C>A
C>G
C>T
T>A
T>G
Counts
0
100
200
300
Trinucleotide

*Phorcus lineatus* (xgPhoLine1.PHORD0051b) germline mutational spectrum

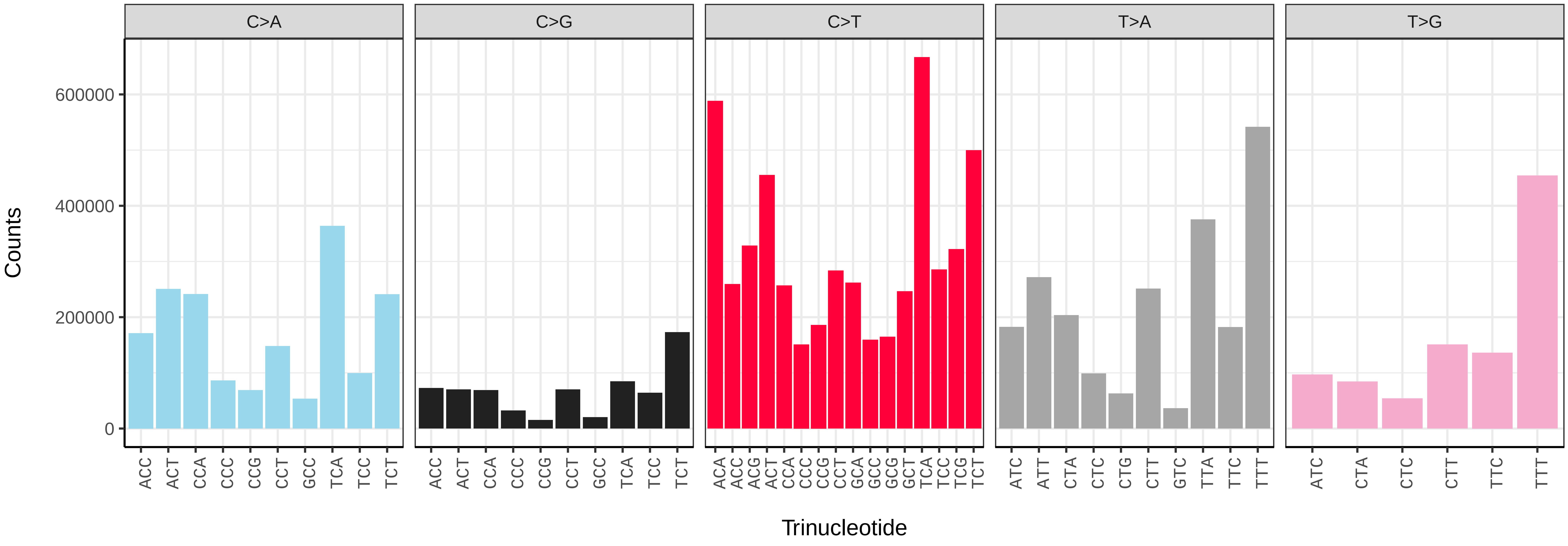


*Phorcus lineatus* (xgPhoLine1.PHORD0051b) somatic mutational spectrum (SBS96 classification)

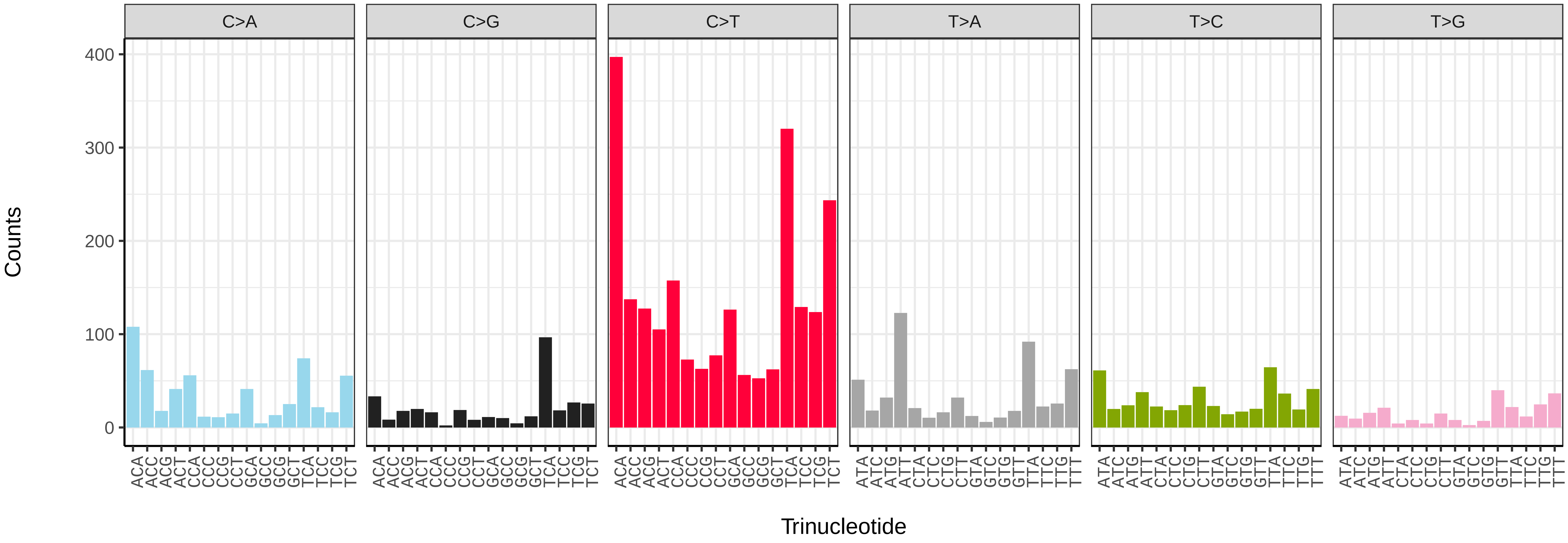


*Phorcus lineatus* (xgPhoLine1.PHORD0051b) somatic mutational spectrum (SBS52 classification)

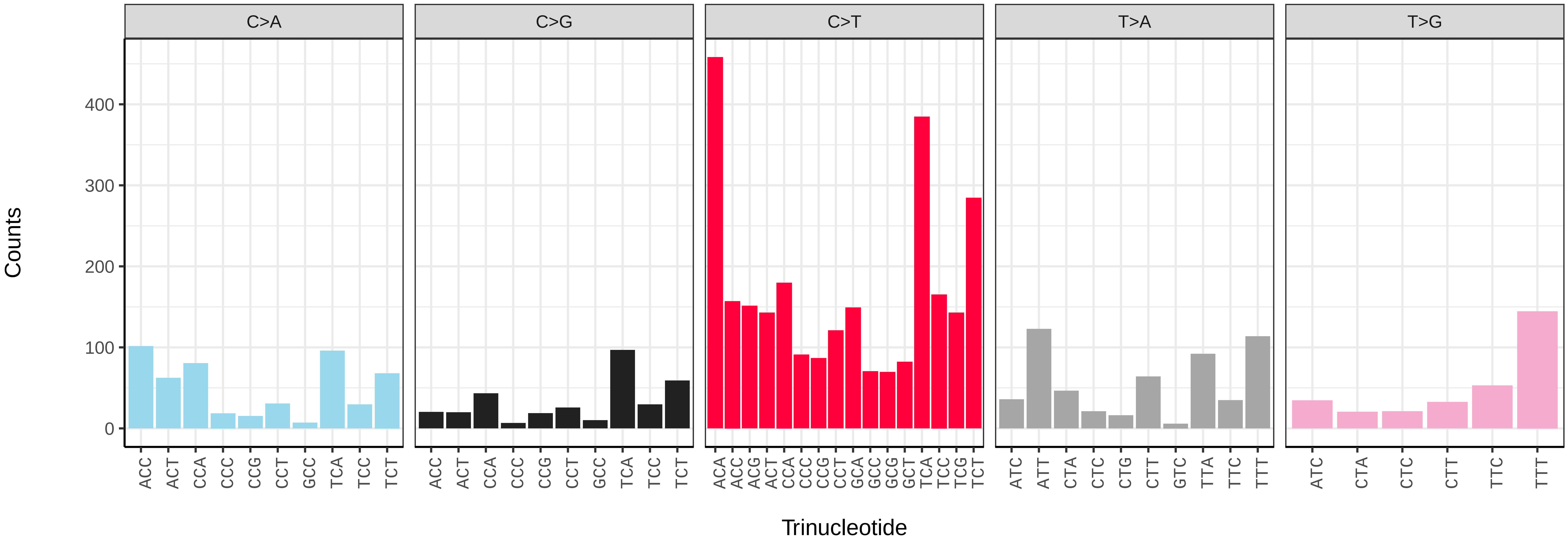

*Phorcus lineatus* (xgPhoLine1.PHORD0058b) germline mutational spectrum

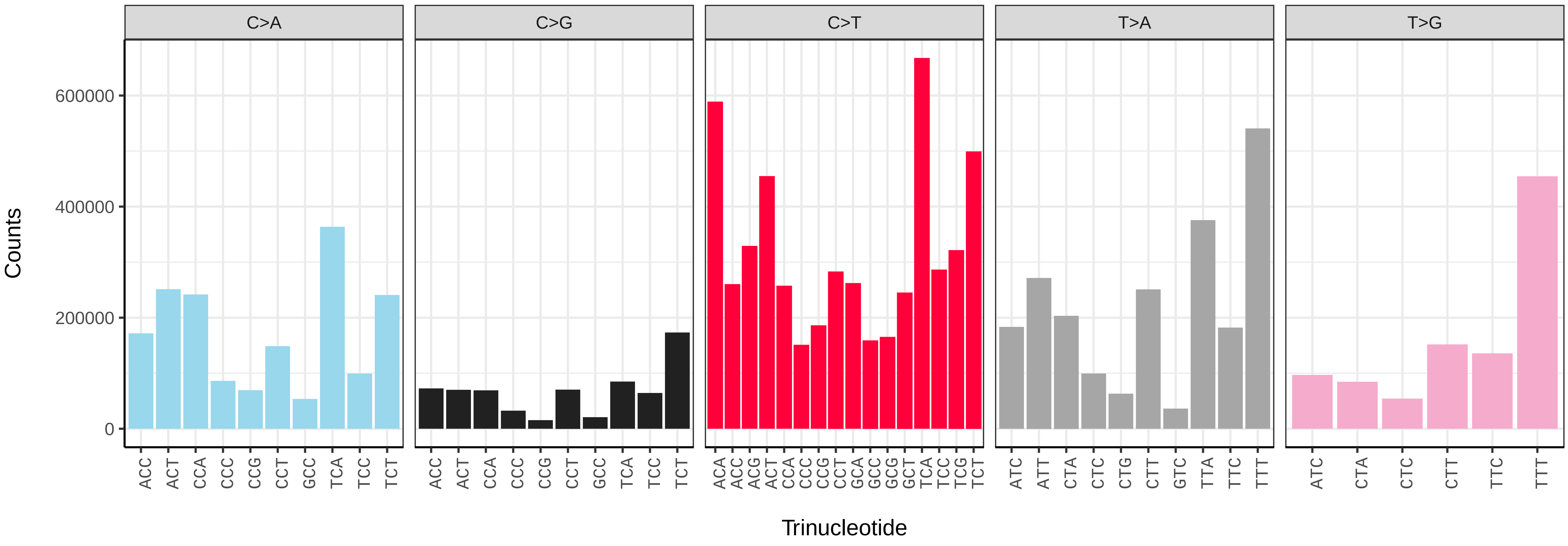


*Phorcus lineatus* (xgPhoLine1.PHORD0058b) somatic mutational spectrum (SBS96 classification)

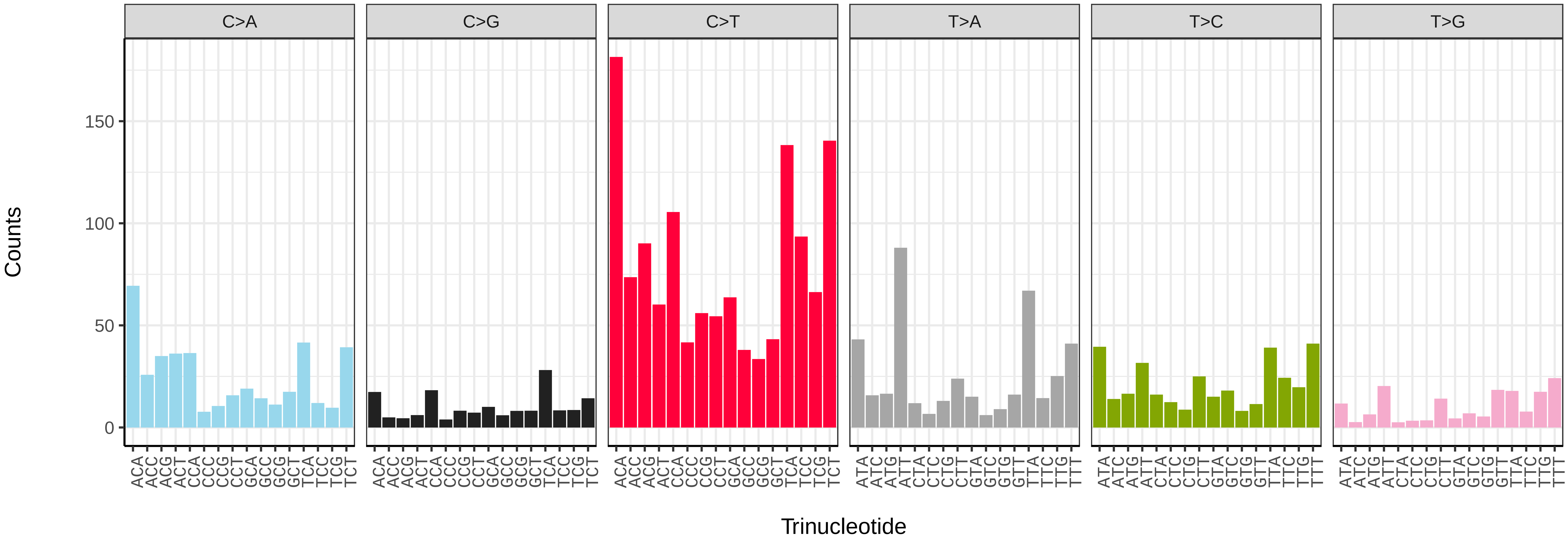


*Phorcus lineatus* (xgPhoLine1.PHORD0058b) somatic mutational spectrum (SBS52 classification)

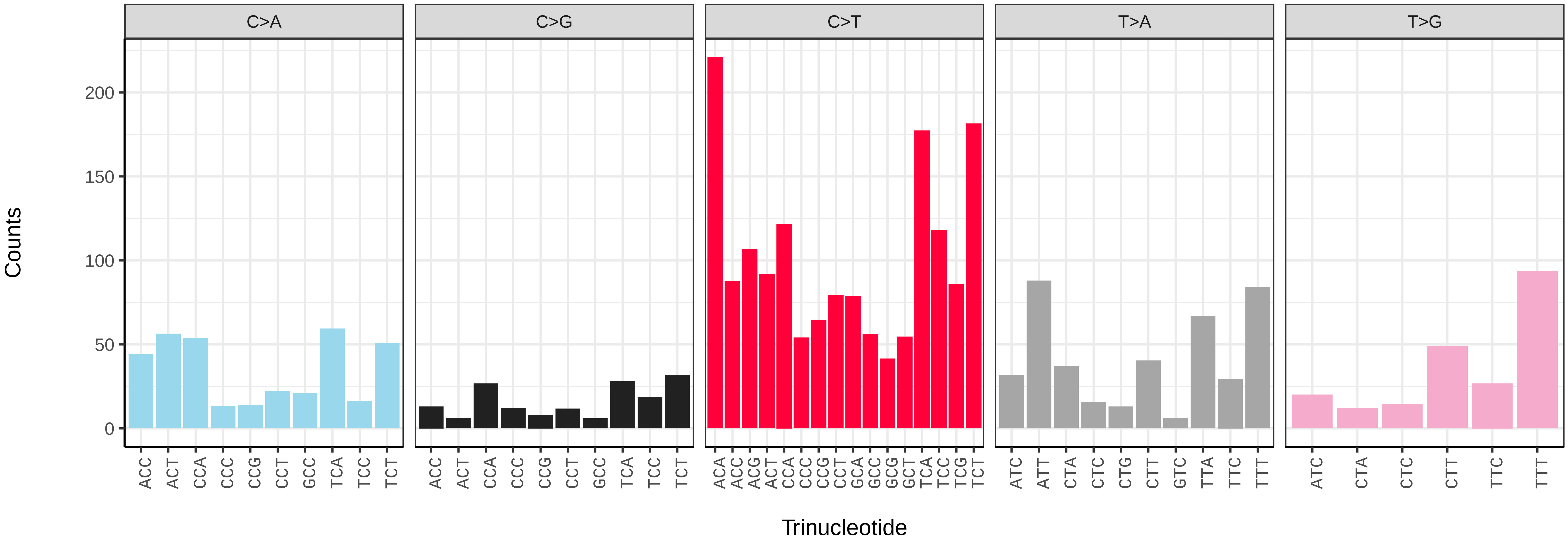

*Phorcus lineatus* (xgPhoLine1.PHORD0060b) germline mutational spectrum

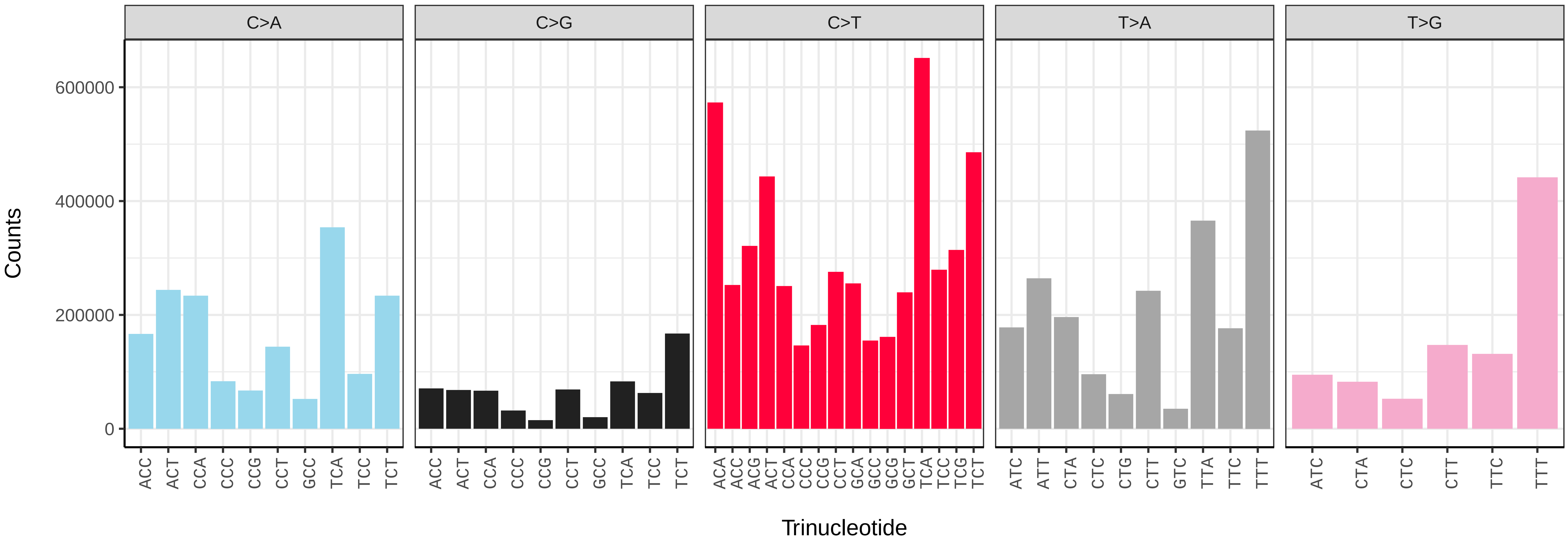


*Phorcus lineatus* (xgPhoLine1.PHORD0060b) somatic mutational spectrum (SBS96 classification)

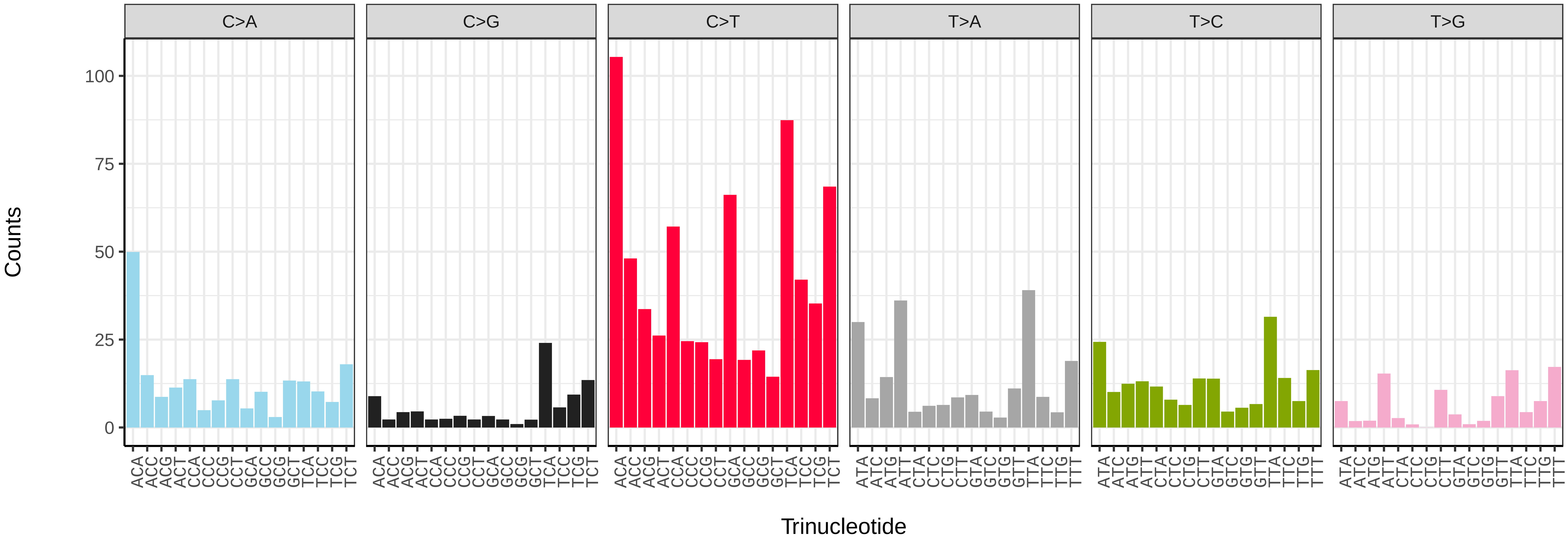


*Phorcus lineatus* (xgPhoLine1.PHORD0060b) somatic mutational spectrum (SBS52 classification)

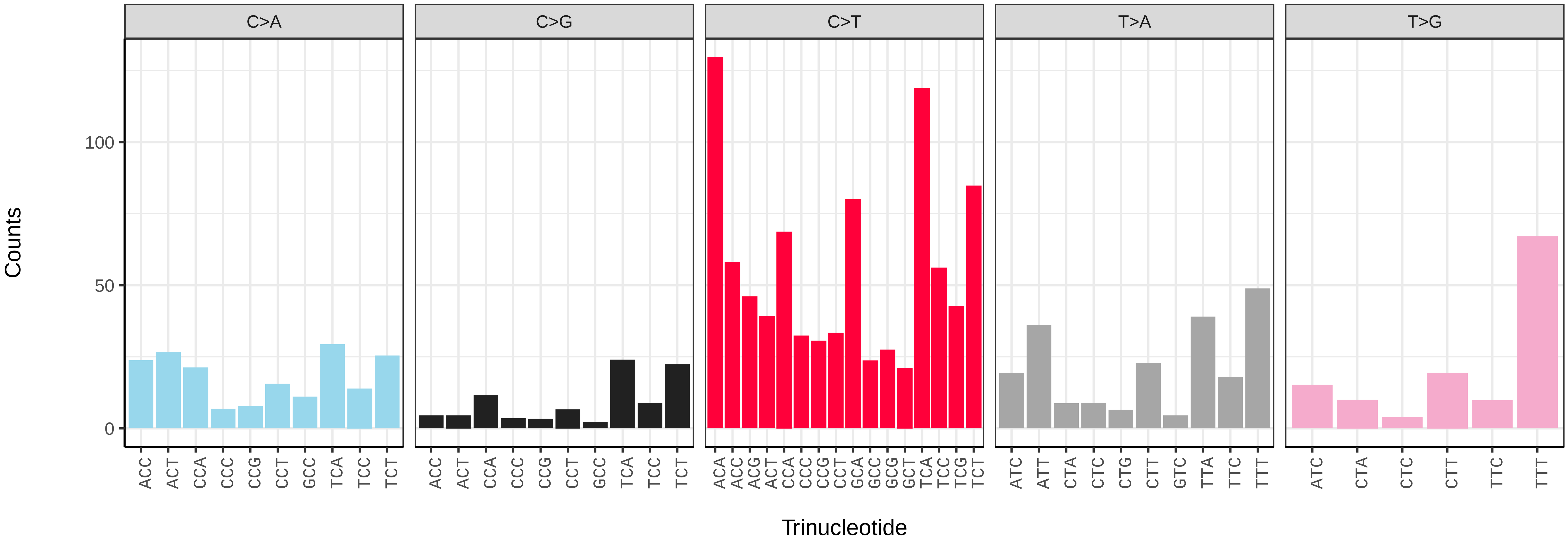

*Phorcus lineatus* (xgPhoLine1.xgPhoLine2) germline mutational spectrum

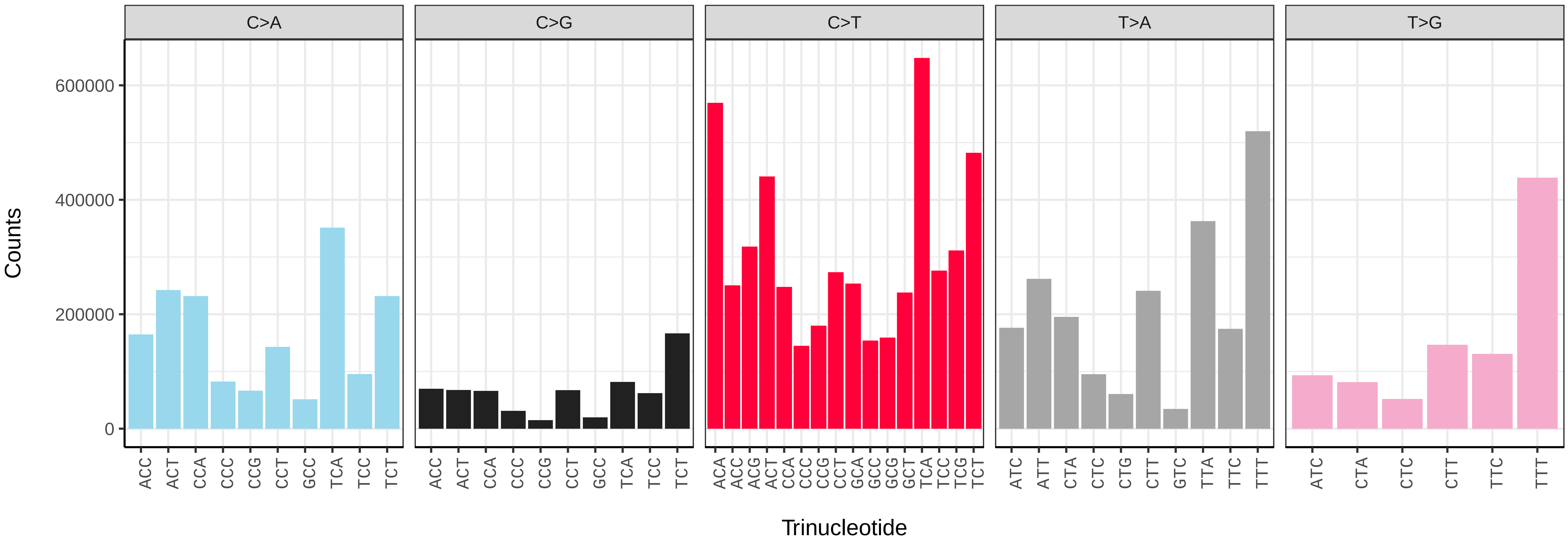


*Phorcus lineatus* (xgPhoLine1.xgPhoLine2) somatic mutational spectrum (SBS96 classification)

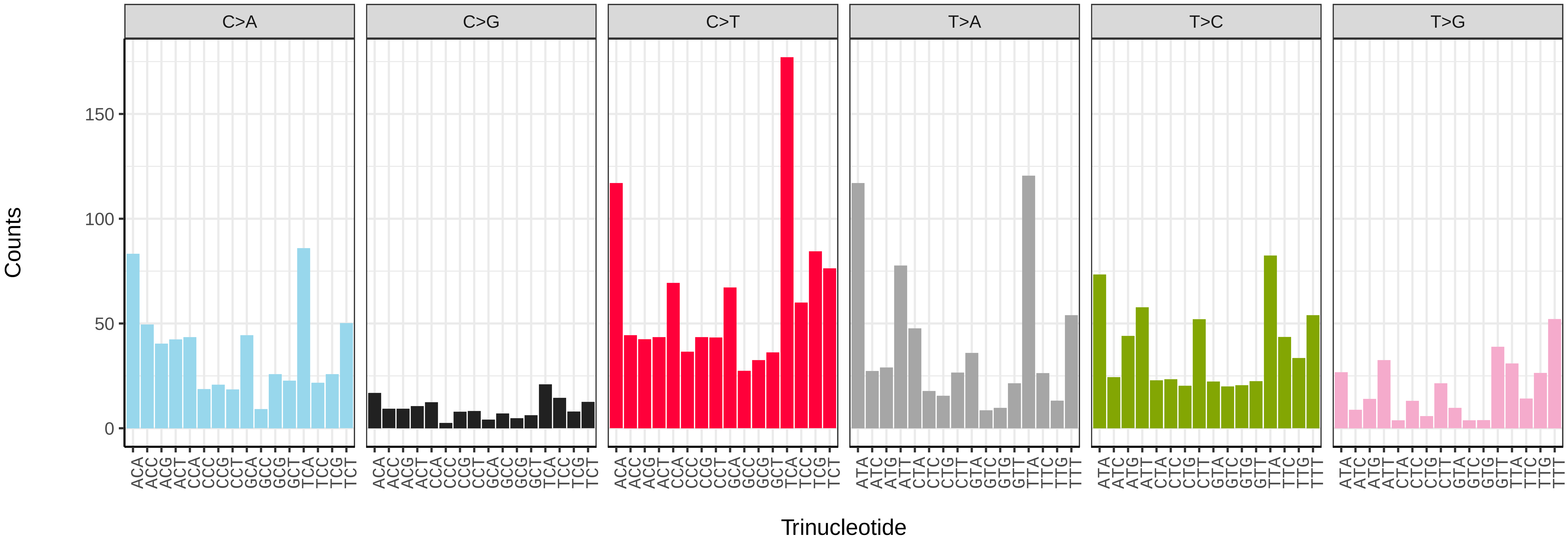


*Phorcus lineatus* (xgPhoLine1.xgPhoLine2) somatic mutational spectrum (SBS52 classification)

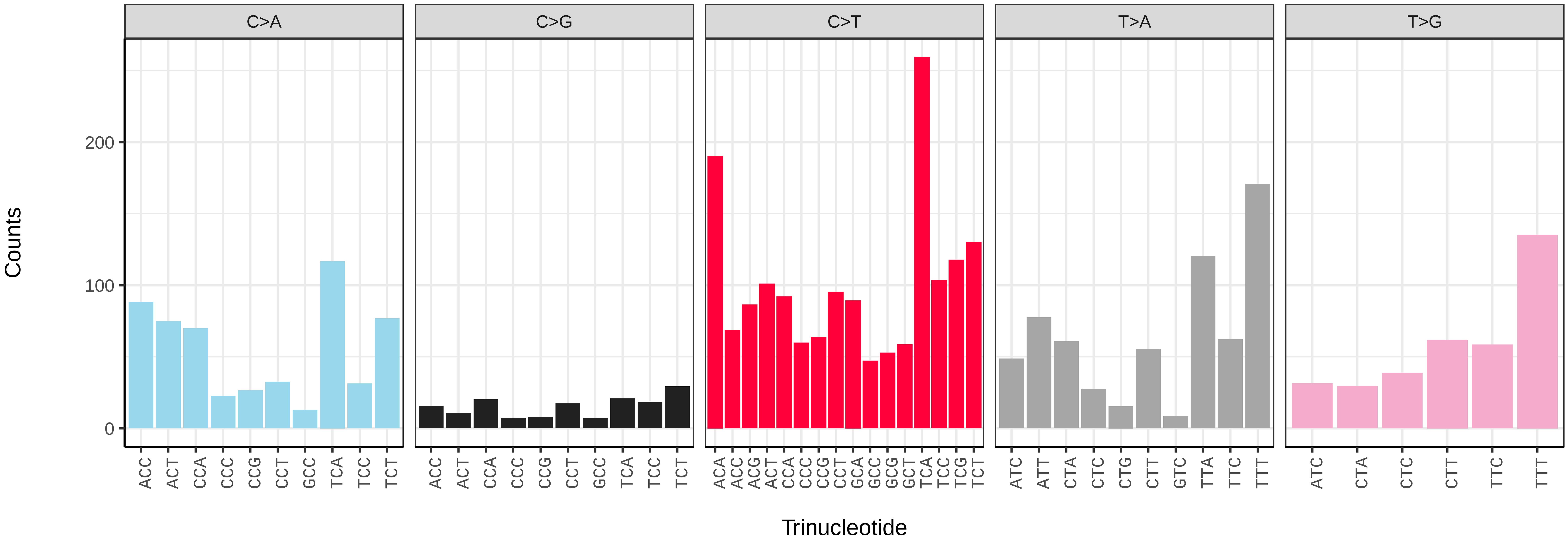

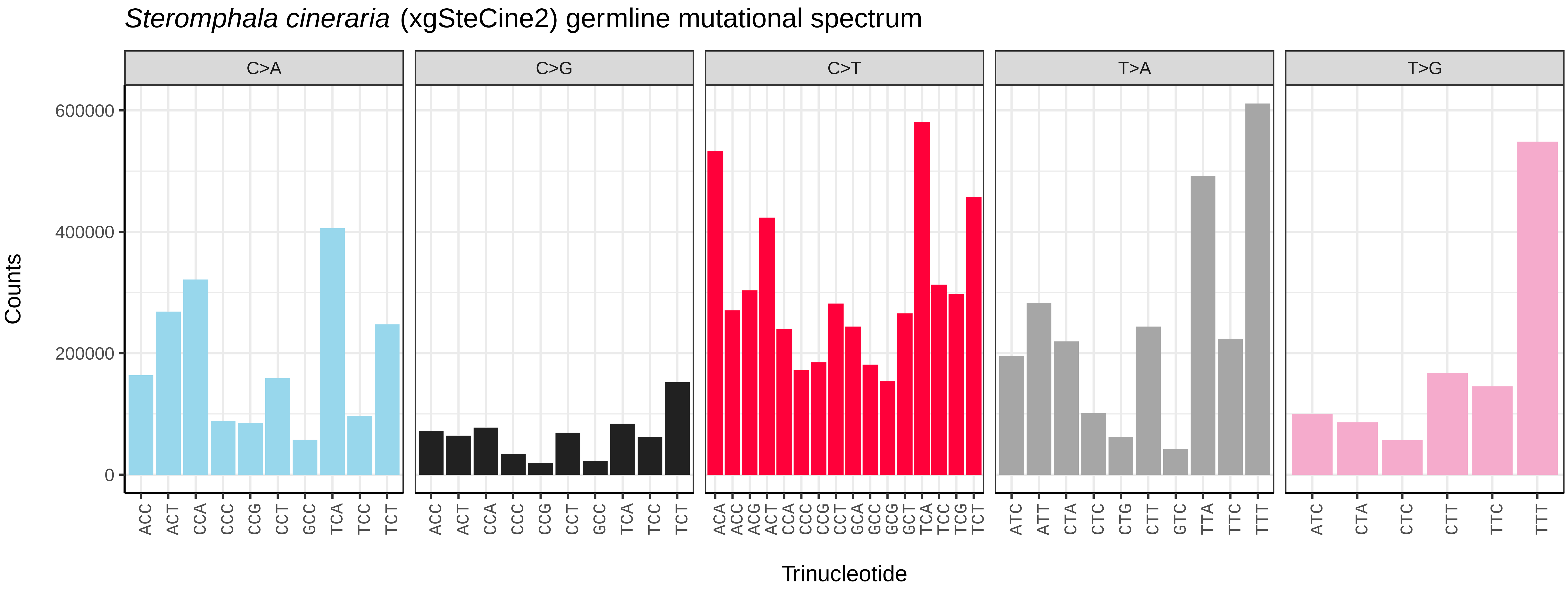
*Steromphala cineraria* (xgSteCine2) germline mutational spectrum
C>A
C>G
C>T
T>A
T>G
Counts
0
200000
400000
600000
Trinucleotide

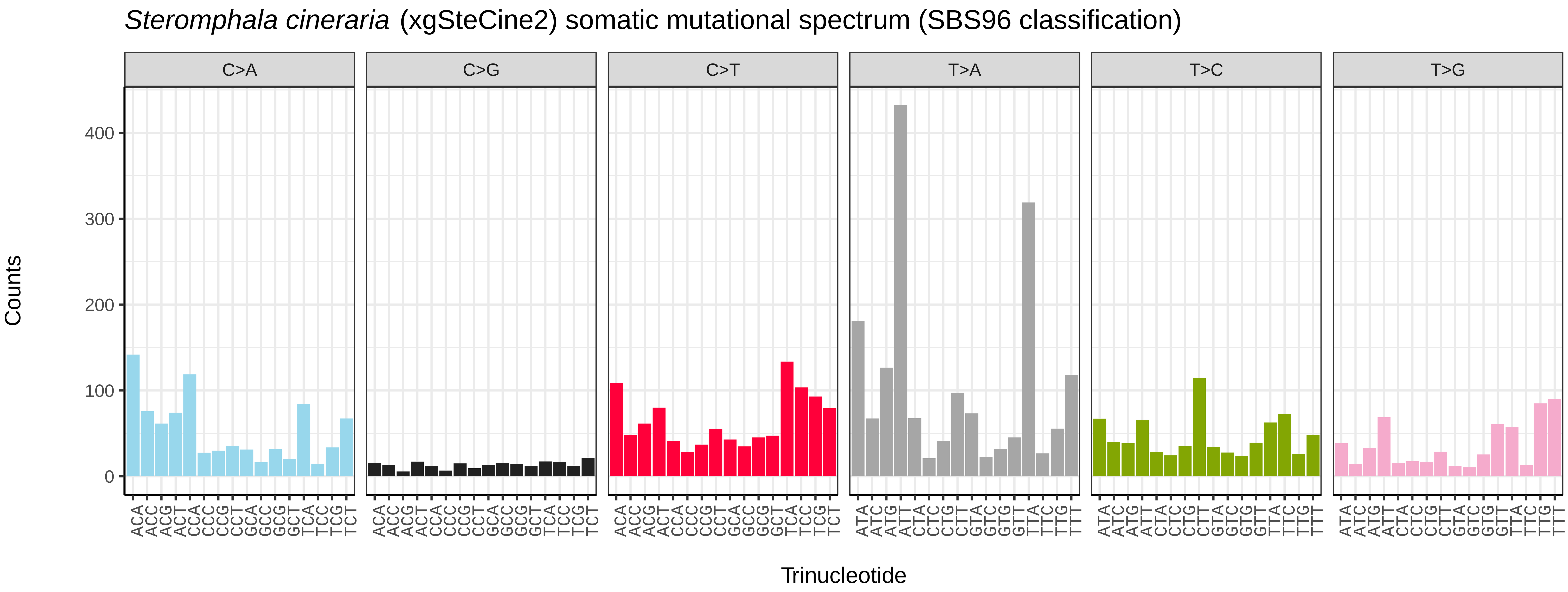
*Steromphala cineraria* (xgSteCine2) somatic mutational spectrum (SBS96 classification)
C>A
C>G
C>T
T>A
T>C
T>G
Counts
0
100
200
300
400
Trinucleotide

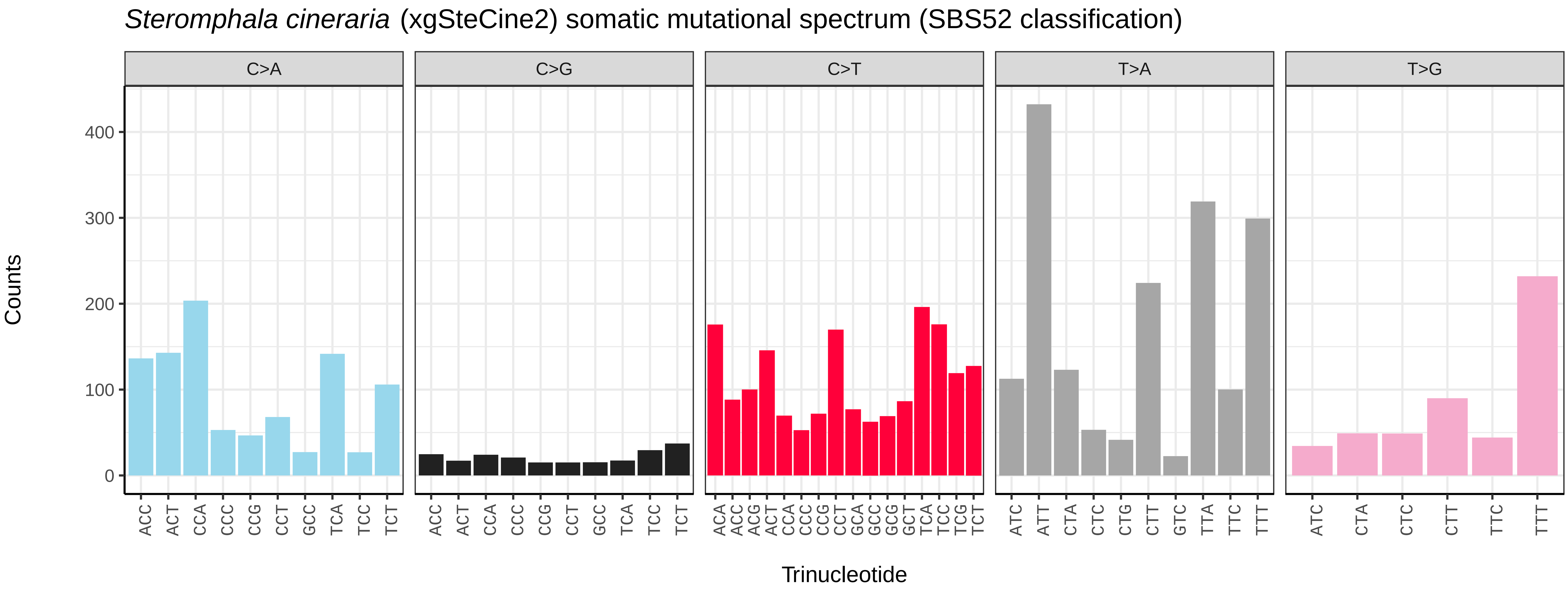
*Steromphala cineraria* (xgSteCine2) somatic mutational spectrum (SBS52 classification)
C>A
C>G
C>T
T>A
T>G
Counts
0
100
200
300
400
Trinucleotide

*Lepidochitona cinerea* (xpLepCine2) germline mutational spectrum

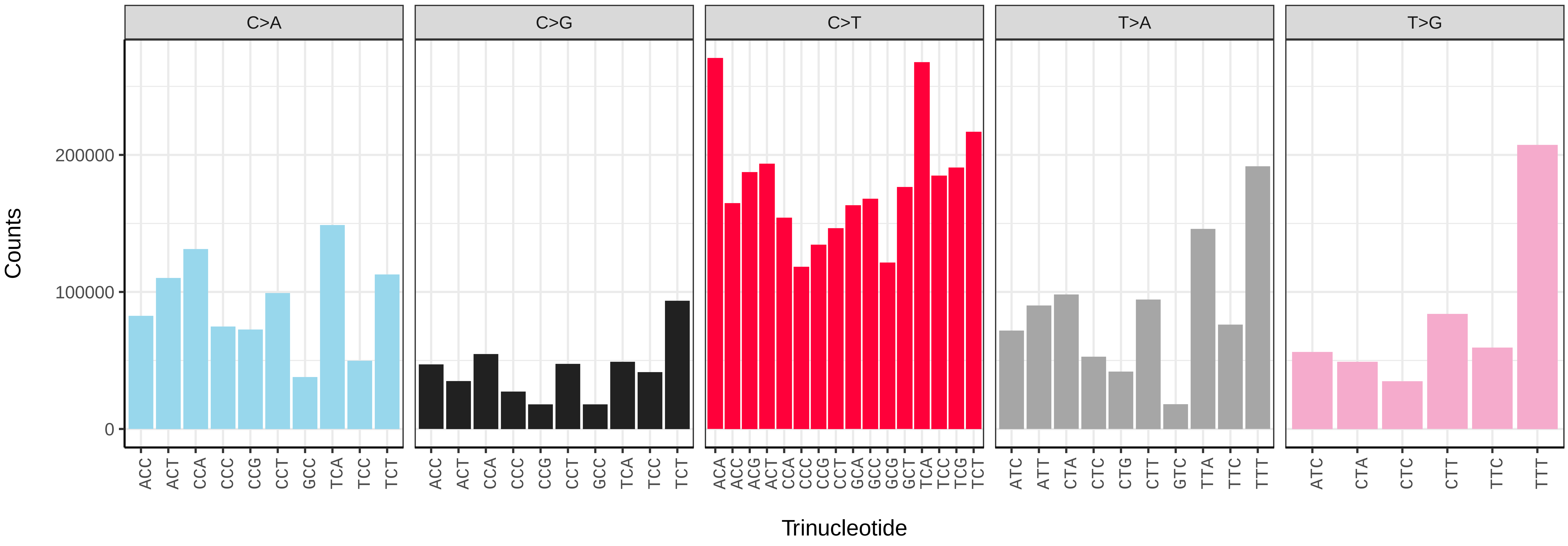


*Lepidochitona cinerea* (xpLepCine2) somatic mutational spectrum (SBS96 classification)

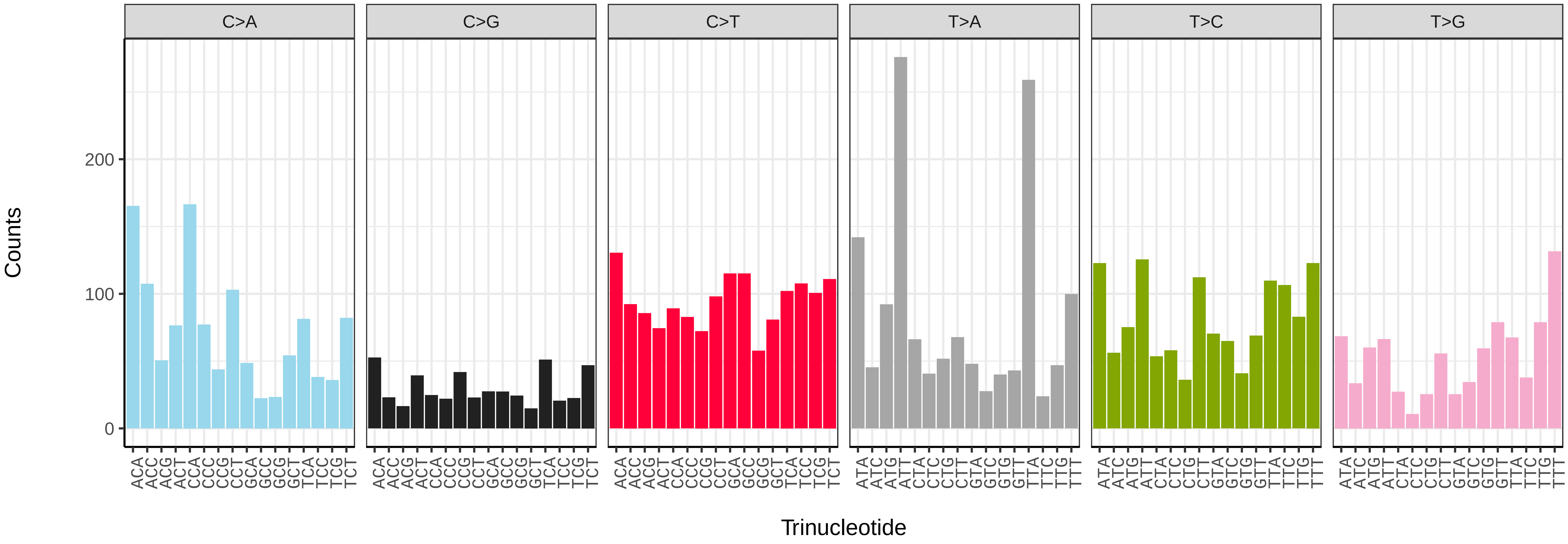


*Lepidochitona cinerea* (xpLepCine2) somatic mutational spectrum (SBS52 classification)

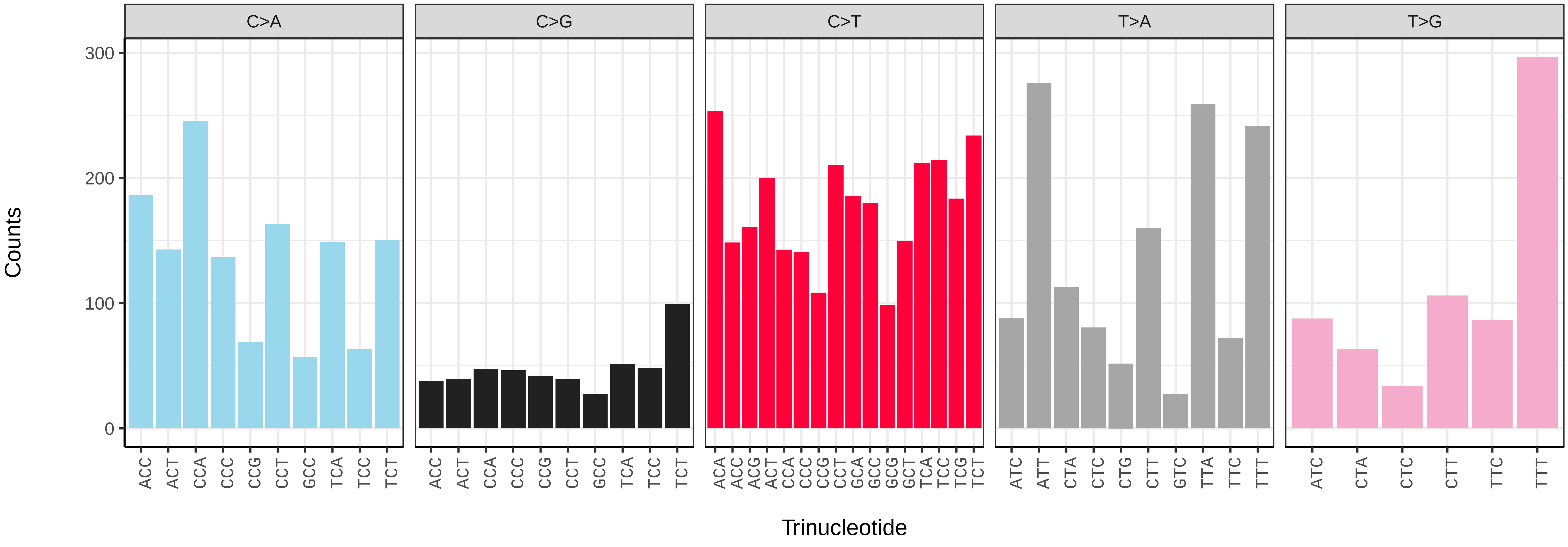
